# Analysis of SARS-CoV-2 Mutations Over Time Reveals Increasing Prevalence of Variants in the Spike Protein and RNA-Dependent RNA Polymerase

**DOI:** 10.1101/2021.03.05.433666

**Authors:** William M. Showers, Sonia M. Leach, Katerina Kechris, Michael Strong

## Abstract

Amid the ongoing COVID-19 pandemic, it has become increasingly important to monitor the mutations that arise in the SARS-CoV-2 virus, to prepare public health strategies and guide the further development of vaccines and therapeutics. The spike (S) protein and the proteins comprising the RNA-Dependent RNA Polymerase (RdRP) are key vaccine and drug targets, respectively, making mutation surveillance of these proteins of great importance.

Full protein sequences for the spike proteins and RNA-dependent RNA polymerase proteins were downloaded from the GISAID database, aligned, and the variants identified. Polymorphisms in the protein sequence were investigated at the protein structural level and examined longitudinally in order to identify sequence and strain variants that are emerging over time. Our analysis revealed a group of variants in the spike protein and the polymerase complex that appeared in August, and account for around five percent of the genomes analyzed up to the last week of October. A structural analysis also facilitated investigation of several unique variants in the receptor binding domain and the N-terminal domain of the spike protein, with high-frequency mutations occurring more commonly in these regions. The identification of new variants emphasizes the need for further study on the effects of these mutations and the implications of their increased prevalence, particularly as these mutations may impact vaccine or therapeutic efficacy.

## Introduction

The global pandemic of Coronavirus Respiratory Disease 2019 (COVID-19), caused by Severe Acute Respiratory Syndrome Coronavirus 2 (SARS-CoV-2), has caused significant disruption to public health and economic activity worldwide. As of February 15, 2021 there have been over 108 million confirmed cases of COVID-19 and 2.4 million deaths worldwide.^1^ To date, a number of COVID-19 vaccines have been developed and approved for use, including mRNA vaccines of the spike protein from Pfizer and Moderna.^2^ The speed of development for the vaccine is unprecedented and has been aided by the rapid and broad availability of viral genomic data. High throughput genomic analysis of SARS-CoV-2 strains has been greatly facilitated by databases, such as the Global Initiative on Sharing all Influenza Data (GISAID).^3^ GISAID was initially created after the global spread of the H5N1 avian flu, to break down barriers to data sharing, enabling users to share and analyze data in a timely manner, and allowing users to access unpublished genomic data under the conditions of a data use agreement that protects the intellectual property rights of data contributors.^3^ Many studies have leveraged the GISAID information to examine the emergence of genetic variants,^4–6^ to understand how the SARS-CoV-2 genomes are evolving over time and geographic locations, and to assess mutation rates.

The genome of SARS-CoV-2 consists of 14 open reading frames (ORFs) that encode 27 proteins.^7^ Four of the ORFs encode structural proteins, and are named with one letter corresponding to the name of the structural protein produced: E (envelope protein), N (nucleocapsid protein), M (membrane protein), and S (spike protein).^8^ The spike protein has been shown to be a key factor in viral entry, and binds to the human angiotensin-converting enzyme 2 (ACE2),^9^ resulting in the fusion with the host cell membrane.^10^ In addition to the four structural proteins, there are 16 non-structural proteins (Nsps) encoded by open reading frame 1ab.^7^ The non-structural proteins Nsp7, Nsp8, and Nsp12 have been found to form the viral RNA-dependent RNA polymerase (RdRP) complex, and each of the Nsps forming the RdRP complex must be present for the replication of viral genomic RNA to occur.^11^ Nsp12 contains the active site in which the antiviral drug remdesivir binds,^11^ making this protein of great importance for variant surveillance. Here we describe a comprehensive analysis of the amino acid variants in the spike protein and the RdRP complex from the beginning of the pandemic to October 31, 2020. We identify and document changes in the prevalence of variants over time, and examine the variants within the context of protein structures to identify patterns of variation that may impact host-pathogen interactions, as well as vaccine and therapeutic efficacy.

## Methods

### Raw Data Download

All protein sequences submitted to GISAID by November 14^th^ were downloaded in a single FASTA file. The file was pre-processed to amino acid format, with one entry for each protein in every sequence. Sequence headers contained metadata including the protein, the accession ID of the sequence, the date of collection, and the geographic location. A second file containing extended metadata was also downloaded; the file was formatted as a table with one row per sequence. The reference genome used in our analysis was the Severe Acute Respiratory Syndrome Coronavirus 2 Isolate WIV04 (WIV04), sequenced in Wuhan, China on December 30^th^, 2019.^12^ The raw FASTA file was split by protein into 27 files using a Python script in Jupyter Notebook (version 6.1.4),^13^ and each protein was processed separately through all subsequent steps.

### Filtering of Sequences

Sequences were filtered in Python using the Biopython SeqIO module.^14^ In order to reduce potential incomplete sequences and lower quality sequences, all sequences that were ten or more codons shorter or longer than the reference sequence were eliminated from the analysis, along with sequences containing more than 0.1% ambiguous (“X”) codons. The number of sequences for each protein remaining after filtering is listed in **Table 1**.

**Table 1:** Number of sequences remaining after each step in the analysis. The number of sequences clustered and aligned reflects the number of sequences in the cumulative analysis and structural visualizations.

|  | **Spike** | **Nsp7** | **Nsp8** | **Nsp12** |
| --- | --- | --- | --- | --- |
| Sequences Downloaded (November 14th, 2020) | 196,674 | 196,191 | 196,192 | 196,314 |
| Sequences Passing Filter Criteria | 155,118 | 193,786 | 192,168 | 178,648 |
| Sequences Clustered and Aligned | 150,069 | 193,713 | 192,032 | 177,643 |
| Clustered Sequences Linked to Metadata | 146,740 | 189,712 | 188,088 | 173,978 |
| Metadata-Sequence Pairs Included in Time Series Analysis | 139,835 | 181,856 | 180,432 | 166,617 |

### Sequence Dereplication

In order to streamline our computational pipeline, identical sequences were condensed into clusters using USEARCH (version 11.0.667).^15^ Clusters, representing unique sequences, were written out to a FASTA file with the ID of the cluster and the number of sequences in the cluster. Concurrently, a separate file for cluster information was created that linked the metadata for all sequences in each cluster to the cluster ID. Clusters of size one were not included in the analysis due to the low abundance and possibility of these clusters reflecting errors in sequencing rather than true variation.

### Sequence Alignment

Clustal Omega (version 1.2.4)^16,17^ was used to align clustered sequences for comparison with the reference sequence. Clustal Omega was selected based on the balance between alignment quality and speed. The option “--full” was specified to perform a full distance matrix calculation to improve alignment accuracy. Two iterations of alignment were performed to optimize both alignment quality and computational efficiency.^16^ Full distance matrix calculations were applied to the extra iteration using “--full-iter”, and the output was stored in FASTA format.

### Parsing of Multiple Sequence Alignment

A Python script was developed in Jupyter notebook to automatically parse the aligned sequences for variants given the ID of the cluster containing the reference sequence, which was determined by searching for “WIV04” in the cluster information file using RStudio (version 1.3.1093).^18^ The Python script scanned through the other clusters (**Supplementary Figure S1**), comparing each codon with the corresponding codon of the reference cluster. When variants were discovered, the program determined the type of mutation, and functions were run accordingly to store the position of the variant relative to the reference, the ID and size of the variant cluster, and the identity of the codon in the reference and variant clusters. Based on this information, a code was computed for the variant based on the nomenclature recommended by the Human Genome Variation Society (HGVS).^19^ This information was then stored for each mutation observed, as the variant events dataset. Deletions spanning multiple codons were recorded as a single event, and insertions at the beginning and end of the sequences were named as extensions with the format <Position of the first or last codon>ext<Identity of codon(s) inserted>. The variant events dataset was then grouped by position and the sum of the cluster sizes was taken to compute the total number of sequences with a mutation at each position, and a similar operation was performed to compute the total number of sequences containing each unique variant.

### Three-dimensional Visualization of Frequently Mutated Sites

Structures of the spike protein and the RNA-dependent RNA polymerase (RdRP) complex were downloaded from the Protein Data Bank (PDB)^20^ and visualized using PyMOL.^21^ For the spike protein, two structures were downloaded: PDB ID 6VSB,^22^ which shows the whole spike protein with one receptor binding domain in the up conformation, and PDB ID 6M17,^23^ which shows the receptor binding domain of the spike protein in contact with the ACE2 receptor. For the RdRP complex, the structure PDB ID 7BV2^24^ was used. The variants by position dataset was used to color each position in the structures by the frequency of variation using a log-10 scale. In a separate visualization, the spike protein and the RdRP complex were colored by domain and all positions mutated in more than 100 sequences were highlighted.

### Frequency of Mutations in Key Residues

The variants by code dataset was filtered by residue number to determine the frequency and identity of variants in key regions of the genome, such as the receptor-binding domain (RBD), the furin cleavage site,^25^ and superantigen motifs^26^ in the spike protein. For the RdRP complex, the binding site of remdesivir^11^ was analyzed.

### Time Series Analysis of Variants

To determine changes in the prevalence of variants over time, the variant events dataset was linked with the extended metadata according to the process outlined in **Supplementary Figure S2**. The GISAID accession ID for each sequence was extracted from the cluster information dataset using RStudio, yielding a dataframe mapping sequence accession IDs to the corresponding cluster ID. This dataframe was merged with the extended metadata on the common accession ID column, yielding a dataset listing the metadata for each sequence, along with the ID of the representative cluster. A dataframe mapping the cluster ID to the variants observed in the cluster was created from the variant events dataset, and this dataframe was merged with the metadata with cluster IDs to yield a dataset giving the metadata for each sequence, along with the variants observed. Subsets were then taken based on the collection date of the samples: weekly time intervals beginning on Sunday, January 5^th^, 2020 and ending on Saturday, October 31^st^, 2020, were used. The frequency of each unique variant was obtained for each week and stored in a separate dataset along with the total number of sequences analyzed; variant counts were then divided by the total number of sequences to give the percentage of sequences with each unique variant by week. Subsets were also taken by continent to perform a region-specific analysis. Sequences with no defined day of collection were excluded from analysis, as well as sequences with metadata entries that could not be linked to cluster IDs. There were only 53 sequences analyzed with a collection date later than October 31^st^, 2020: these were also excluded from analysis. The number of sequences included in the time series dataset is given in **Table 1**.

### Visualization of Variant Trends

The Python package Matplotlib^27,28^ was used to visualize trends in spike variants over time. The top ten most common variants were selected from the percentage table, and a line plot showing the prevalence of each mutation over time was created. To analyze trends in less common variants, a heatmap was used: all variants with a prevalence greater than or equal to 2 percent in at least one week were included. A color map for the heatmap was defined using a log-10 scale, with 0.10% as the lower bound for coloring cells. A histogram was generated to show the number of sequences represented in each week.

## Results

### Spike Protein

#### Time Series Analysis of Variants

The global prevalence of the top ten most common variants by week of sample collection is shown in **Figure 1A**, and the prevalence of the top ten most common variants on each continent are shown in **Figure 1 B-G**. The most common variant was the substitution D614G, which quickly became prevalent after its appearance in mid-January 2020. The variant was observed in more than 50% of sequences collected worldwide by the week of March 1^st^, and in more than 90% of sequences collected by the week of April 26^th^. D614G quickly became the dominant variant on all continents, though its rate of establishment was much lower in Asia (**Figure 1B**). D614G reached 90% prevalence in Asia during the week of June 14^th^. The N-terminal domain (NTD) substitution A222V and the signal peptide substitution L18F have also gained in prevalence since their appearance in late July and early August, respectively. These variants appear to be common in Europe (**Figure 1C**), though A222V has been observed to an increasing extent in Asia and Oceania (**Supplementary Figure S3A**), and L18F and A222V have been sporadically observed in North America (**Supplementary Figure S3B**). The receptor binding domain (RBD) substitution S477N appeared in June and reached peak prevalence during week of July 19^th^, appearing in 44% of sequences worldwide before decreasing in prevalence to 4.3% by the week of October 18^th^. An influx of sequences from Oceania was observed during this time (**Figure 1H**), and S477N was observed in more than 90% sequences from Oceania between the week of July 15^th^ and the week of August 16th (**Figure 1D**). S477N has also been observed in Europe with increasing frequency, but not on other continents (**Supplementary Figure S3C**). The RBD substitution N439K and the NTD deletion H69_V70del have slowly increased in prevalence since August and are present in 3.2 and 3.7 percent of the sequences collected worldwide on the week of October 18^th^, respectively. Both variants were in the top ten most common variants from Europe but were not consistently observed on other continents (**Supplementary Figure S3 D-E**). The signal peptide substitution L5F was observed on all continents at low prevalence (**Supplementary Figure S3F**).

**Figure 1A-H:**
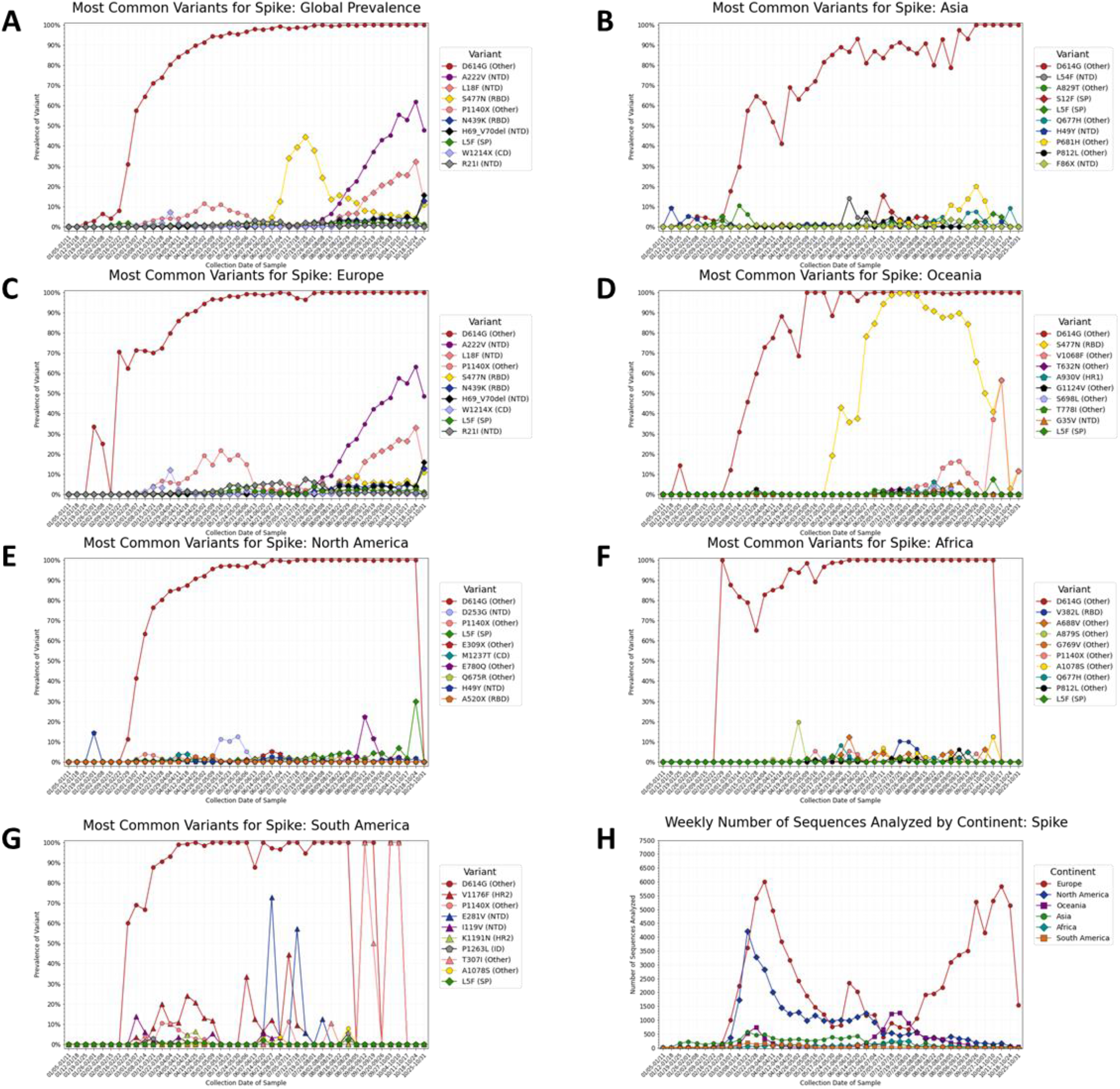
Prevalence of the top ten most common variants for the spike protein, by collection date in one-week intervals beginning on January 5th, 2020 and ending on October 31^st^, 2020. Prevalence of the ten most common variants **A)** worldwide, **B)** in Asia, **C)** in Europe, **D)** in Oceania, **E)** in North America, **F)** in Africa, and **G)** in South America. Variants that appear within the top ten most prevalent variants on multiple continents are given the same color and shape in every graph. The variant D614G quickly gained prevalence after its appearance in mid-January, occurring in 50% of the sequences sequenced by early march and 90% by early May. The variants A222V and L18F are increasing in prevalence in Europe and are not observed in other continents as of October 31, 2020. The spike in global prevalence of S477N appears to be driven by an influx of sequences from Oceania, where it appears in the majority of samples sequenced between June and September. The signal peptide substitution L5F was observed at low prevalence on all continents. N439K, which binds ACE2 with increased affinity and decreases the neutralization capability of some antibodies,^5^ have seen slight increases in prevalence in Europe since September. **H)** Number of complete spike protein sequences analyzed per week, by continent. Europe has contributed the majority of sequences from September to the end of October, 2020. Lower sample sizes in regions may explain the sudden shifts in prevalence seen in (A-G).

The data for North America (**Figure 1E**) suggests that there are no highly prevalent variants besides D614G as of October 31^st^, but the limited deposition of full-length spike sequences to GISAID since July (**Figure 1H, Supplementary Figure 4**) limits the conclusions that can be drawn from the latter months of the analysis. Data from Africa and South America (**Figure 1 F-G**) shows several variants unique to both regions that have appeared then later decreased in prevalence, but the sample size from these continents is very low (**Supplementary Figure 4**). Considerable changes in the prevalence of several mutations are observed in the last week of this analysis (October 25^th^ to October 31^st^), though there are only 1,500 sequences included in this week compared with 5,000 in preceding weeks (**Figure 1H**).

**Figure 2** shows time series trends for all variants present in at least two percent of samples worldwide from any given week. Of the 33 variants that met this criterion, 13 were in the NTD, five were in the RBD, two each were in the signal peptide, the intracellular domain, the cytoplasmic domain, and heptad repeat 2; and seven were outside of a named domain. Time series analyses of variants occurring within NTD (**Supplementary Figure S5**) and the RBD (**Supplementary Figure S6**) show that the N-terminal domain contains more variants than the RBD that are consistently present and increasing in prevalence with time. Four additional variants became more prevalent since late August (S98F, T723I, A626S, and E583D),^29^ while other variants emerged and then later disappeared. Two heptad repeat 2 residues, D1163Y and G1167V appear to increase in prevalence at the same time, near the end of August. Out of the 910 sequences containing either D1163Y, G1167V, or both, 802 contain both variants (**Supplementary Table S2**). The same trend is observed for the N-terminal domain residues A262S and P272L, and these variants occur together in 687 out of 1,084 sequences containing either A262S, P272L, or both. Some of the fluctuations in amino acid variations may reflect the geographic trends of data deposited to GISAID. Region-specific heatmaps showing all variants present in at least two percent of samples on each continent are shown in **Supplementary Figures S7-S12**, and a complete prevalence table for all variants is given in **Supplementary Table 1 A-G**.

**Figure 2:**
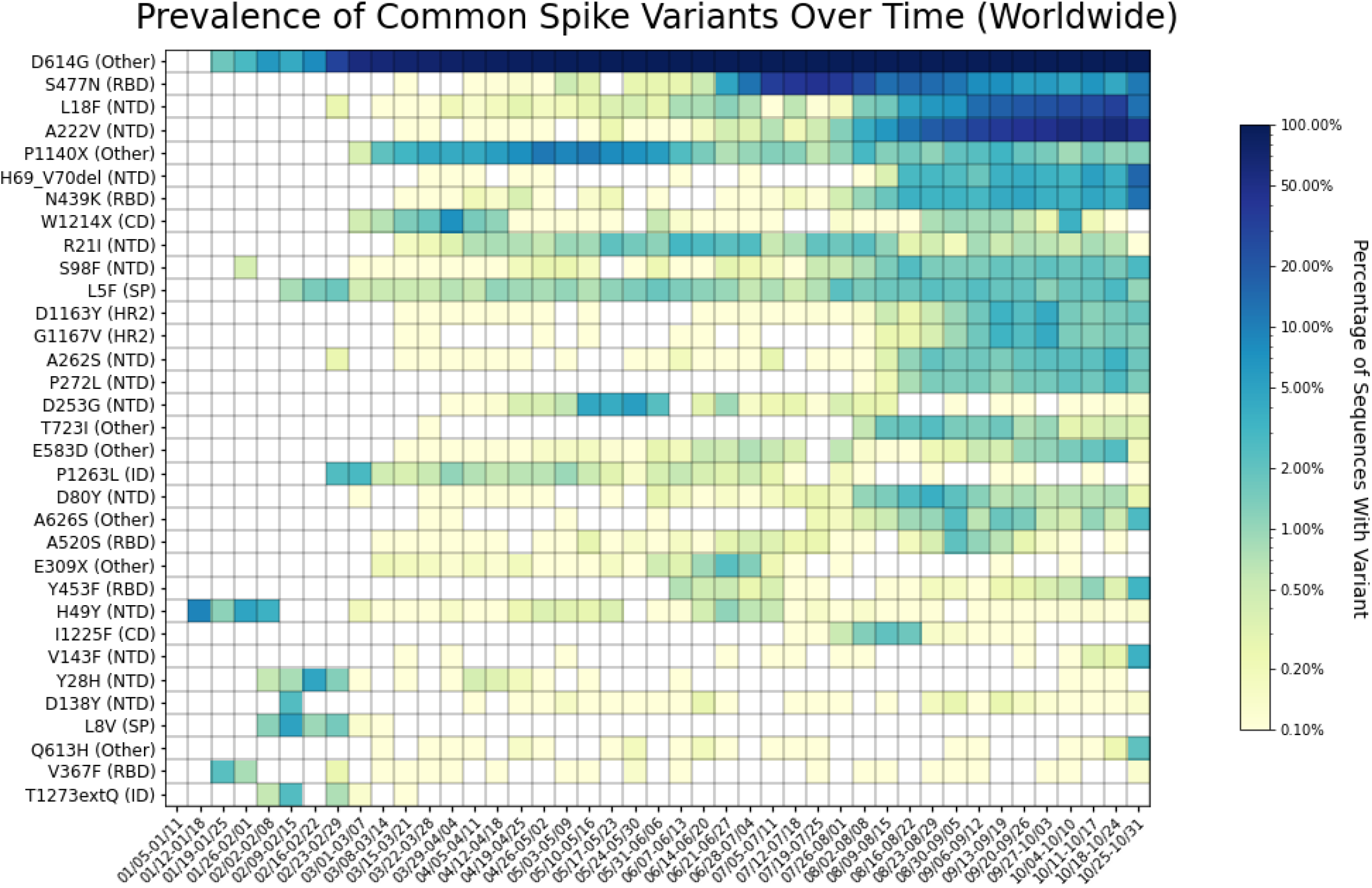
Heatmap of all spike protein variants observed in two percent or more of the genomes sequenced, for at least one week. Variants are listed on the y-axis, with the row “zero mutations in spike” showing the percentage of genomes sequenced with spike protein sequences identical to that of the reference genome. Parentheses indicate the spike protein domain in which each variant appears, according to the domain ranges specified in Huang et al. 2020.^29^ Key for domain abbreviations: SP=signal peptide, NTD=N-terminal domain, RBD=receptor-binding domain, FP=fusion peptide, HR1=heptad repeat 1, HR2=heptad repeat 2, CD=cytoplasmic domain, ID=intracellular domain. The heatmap is colored based on a log-10 scale, with prevalence values of zero colored in white, and values less than or equal to 0.10% colored with the lightest shade. Time on the x-axis is categorized by week of collection date, beginning on January 5^th^, 2020, and ending on October 31^st^, 2020. The heatmap reveals several variants that have increased in prevalence since August, as well as others that have appeared and have subsequently disappeared. Fifteen out of the 33 variants observed in at least two percent of samples are in the N-terminal domain, five out of 33 are in the receptor-binding domain, two each are in the signal peptide, the intracellular domain, the cytoplasmic domain, and heptad repeat 2, and seven are in unspecified domains (other).

#### Structural Visualization of Variants

We utilized the structural visualization program PyMOL to examine the frequency of variation at each position in the spike protein sequence and structure (**Figure 3 A-D**). The N-terminal domain and the receptor binding domain are mutated more frequently than other domains (**Figure 3A-B**). Of the 1,735 unique variants observed in the spike protein across all collection dates, 595 were observed in the N-terminal domain and 220 were observed in the receptor binding domain. Heptad repeats 1 and 2 contained 62 and 69 unique variants, respectively, the intracellular domain contained 63 variants, and the cytoplasmic domain contained 35 unique variants. The fusion peptide contained 25 variants, none of which were observed in more than 100 sequences. 636 variants were in regions of the spike protein not classified within a domain.

**Figure 3 A-D:**
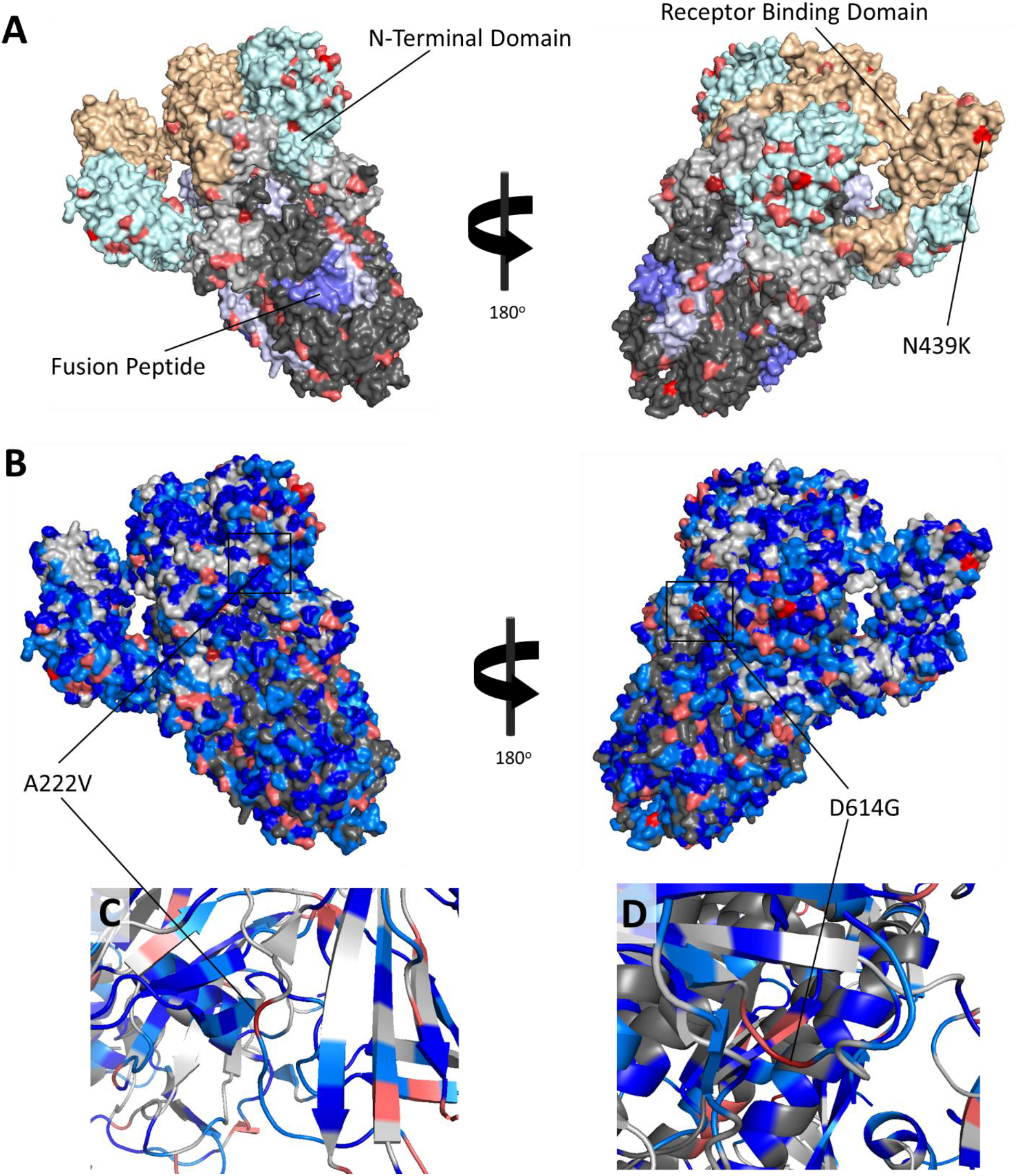
Structure of spike protein (PBD ID: 6VSB) with common amino acid variants labeled. **A)** Spike protein domains are shown with residues mutated in 100 or more sequences highlighted. The N-terminal domain is shown in light cyan, the receptor binding domain is shown in light orange, heptad repeat 1 is shown in light blue, and the fusion peptide is shown in blue. Residues in the S1 subunit with no classified domain are shown in light grey, and unclassified residues in the S2 subunit are shown in dark grey. Residues with a variant frequency of more than 100, 1,000, and 10,000 are shown in pink, red, and dark red, respectively. Variants are observed in more residues in the N-terminal domain and the receptor binding domain compared to the S2 domains, and higher-frequency variants appear to be concentrated in these domains. **B)** All residues with variants are shown. The color-coding for variants appearing in at least 100 sequences is the same as in A) and B); light blue is used for residues with 10-100 variants, and deep blue is used for variant frequencies of 2-10. Residues with zero variants are shown based on subunit colors specified in A) and B). **C)** A ribbon diagram visualizing the secondary structure of the A222V variant and surrounding residues. A222V is located on a loop region, as is the case with D614G **(D)**. Despite its distance from the receptor binding domain, D614G has been observed to alter the conformational state of the receptor binding domain by altering the conformation in the region surrounding codon 614, acting as a hinge. ^30^ PDB structures used: PDB ID: 6VSB.^22^

A comparison of the secondary structure of A222V (**Figure 3C**) to that of D614G (**Figure 3D**) shows that both variants occur in a loop region. A ribbon diagram of the secondary structures of A222V (**Figure 3C**), which quickly became more prevalent in Europe between mid-July and October 31^st^, shows that the variant occurs in a loop region, like D614G (**Figure 3D**). D614G has been shown to alter the conformational state of the receptor binding domain through a hinge mechanism involving its loop structure,^30^ and it is possible that A222V may have a similar effect on the conformational state.

The interface between the receptor binding domain of the spike protein and ACE2 is shown in **Figure 4** (PDB ID: 6M17).^31^ Three residues that directly contact ACE2 had more than 100 instances of variation: Y453, N501, and F486. The residues Y453 and F486 were shown to have pi-pi stacking interactions with the ACE2 receptor. Y453 was observed as mutating to a phenylalanine residue in 346 sequences, and F486 was observed to mutate to leucine in 140 sequences. N501 forms two polar contacts with ACE2 residues; it was observed to mutate to a tyrosine residue in 156 sequences. N439 and S477 do not contact the ACE2 receptor directly but in the vicinity of the binding site. S477N was observed in 9,931 sequences, and N439K was observed in 2,025 sequences.

**Figure 4:**
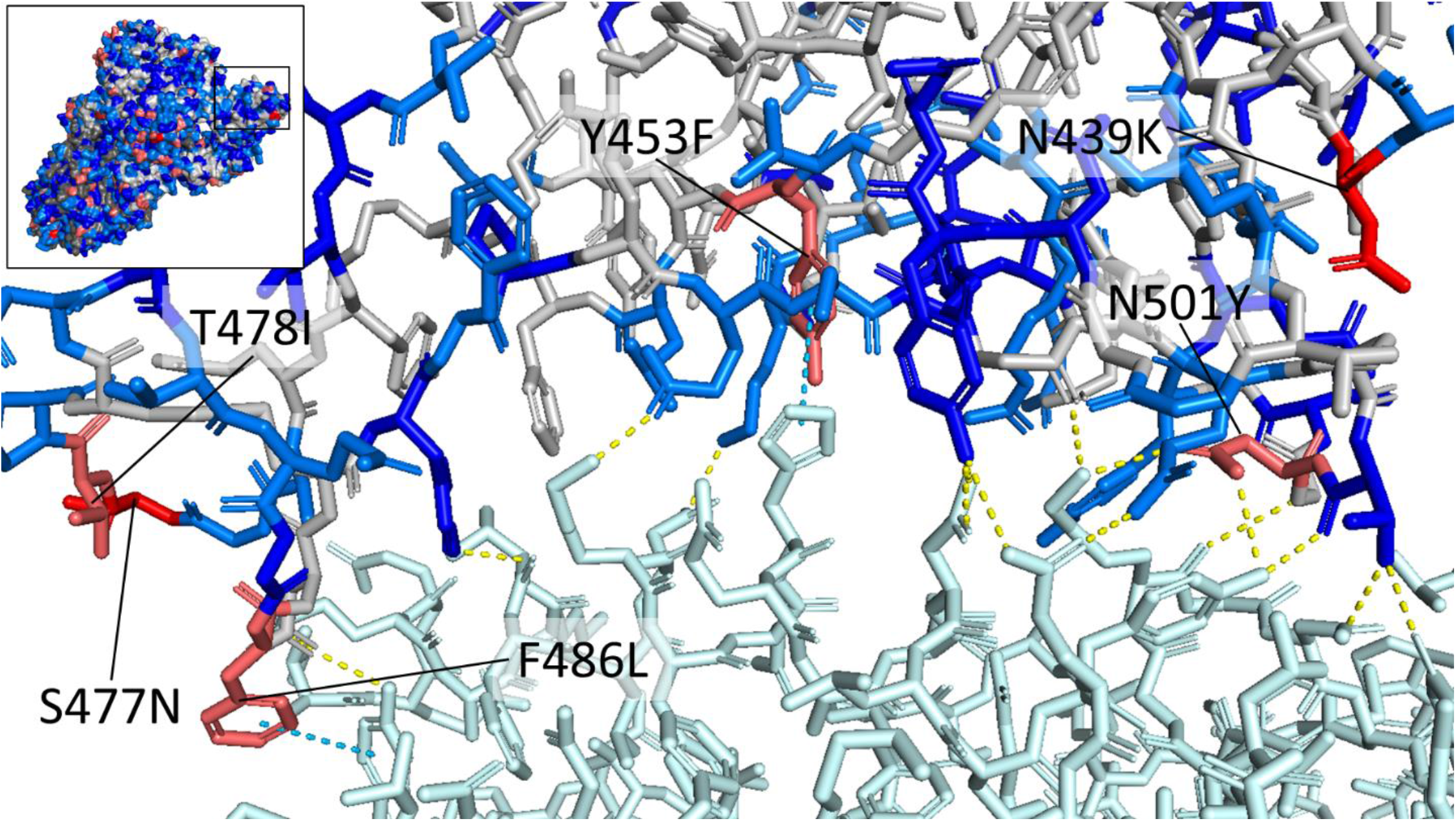
Structure of the interface between the spike receptor binding domain (RBD) and the human ACE2 receptor (top left shows the location of the RBD on the spike protein). The ACE2 receptor is shown in light cyan, and the spike residues are colored according to the frequency of variation: red is used for residues with variants in more than 1000 sequences, pink is used for variant frequencies of 100-1000, light blue for frequencies of 10-100, blue for frequencies of 2-10, and grey for residues with no observed variants. Residues mutated in more than 100 sequences are labeled with the one-letter codes of the reference residue and the most commonly observed variant. Direct contacts are shown with dotted lines: hydrogen bonds and salt bridges are shown in yellow, and pi-pi stacking interactions are shown in cyan. The residues Y453 and F486 form pi-pi stacking interactions with ACE2 residues: the variant Y453F would result in the loss of a hydroxyl group, and the variant F486L would disrupt the pi-pi stacking interaction due to the loss of the aromatic group in phenylalanine. N439 does not directly contact the receptor binding domain, though it is believed that the variant lysine residue, which is positively charged, could form a salt bridge with the negatively charged ACE2 residue E329, increasing binding affinity of the RBD to ACE2.^5^ PDB ID: 6M17^23^ (RBD-ACE2 interface), PDB ID: 6VSB^22^ (whole spike protein on upper left).

#### Furin Cleavage Site

SARS-CoV-2 contains an insertion not found in SARS-CoV that creates an additional cleavage site for the human protease furin.^25,32^ The furin cleavage site has been shown to strongly influence host mortality rates of other RNA-based viruses such as influenza^33^, and furin cleavage sites similar to those of SARS-CoV-2 have been observed in MERS-CoV.^34^ The furin cleavage site occurs between residues N679 and R685^25^; these residues were analyzed for the frequency and type of variants present. A high degree of conservation was observed at the furin cleavage site, with zero variants in R685 and less than 20 variants observed for residues S680, R682, and R683. P681 was mutated in 233 sequences to several different codons: the proline was substituted with a histidine in 144 sequences, with a leucine in 67 sequences, with an arginine in 12 sequences, and with a serine in seven sequences.

#### Superantigen Mimicry

Other spike protein motifs of interest include those that function as superantigens. Superantigens are proteins that stimulate the receptors of T lymphocytes (T cells) by binding at either the alpha or the beta variable chains,^35^ resulting in an overproduction of cytokines that leads to hyperinflammation and toxic shock syndrome.^26^ The symptoms of multisystem inflammatory syndrome in children (MIS-C) infected with SARS-CoV-2 are similar to those of toxic shock syndrome,^26^ suggesting that sequence patterns similar to superantigens exist in SARS-CoV-2 proteins. A sequence motif between the residues E661 and R685 was discovered that is capable of binding both the alpha and beta variable chains in T-cells, with residues S680 through R683 shown to directly contact the T-cell receptor.^26^ This motif overlaps with the furin cleavage site. The sequence that directly contacts the T-cell receptor is the same sequence that is cleaved by furin. A high degree of conservation was observed for residues in the superantigen motif: all residues except for Q675, Q677, and P681 contained variants in less than 100 sequences. A time series analysis of variants in the superantigen motif (**Supplementary Figure S13**) reveals five variants out of 44 total that are present in at least 0.5% of samples from any given week (Q675H, Q675R, Q677H, P681H, P681L). P681H appears to be gradually increasing in prevalence over time, while Q675H and Q677H appear stable in prevalence. Q675R and P681L appear to have increased in prevalence, peaked, and later disappeared. A deletion of the furin cleavage site (N679_S686del) was observed in 0.402% of sequences from the week of January 26^th^ and 0.045% of samples from the week of May 31^st^; the deletion was not observed in any other samples.

#### Emergent SARS-CoV-2 Strains

The time series trends of variants specific to the B.1.1.7, B.1.351, and P.1 strains first detected in the UK, South Africa, and Brazil, respectively, are shown in **Figure 5 A-C** with updated data current as of January 23^rd^, 2021. The ten variants associated with B.1.1.7 have very quickly gained prevalence worldwide since their appearance in mid-September, appearing in more than half of sequences collected worldwide during the week of January 17^th^. Variants associated with B.1.1.7 appear to be propagating at a greater rate than those in B.1.351 and P.1. The RBD variant N484K is observed in B.1.351 and P.1, and N501K is observed in all three strains.

**Figure 5 A-C:**
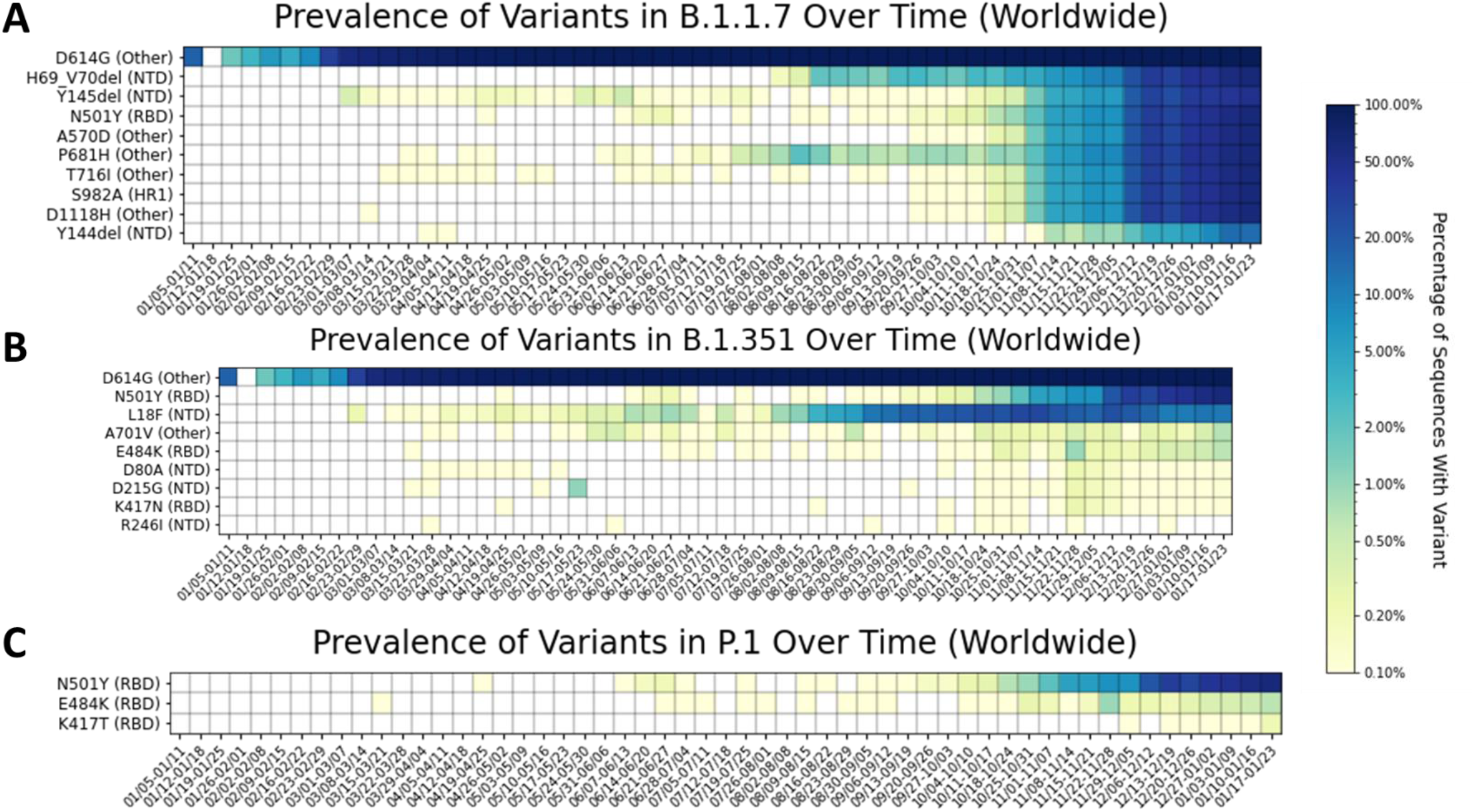
Heatmap of spike protein variants observed in the **A)** B.1.1.7 strain first detected in the UK, **B)** B.1.351 strain first detected in South Africa, and **C)** P.1 strain first detected in Brazil. Variants are listed on the y-axis, and time is categorized by week of collection date on the x-axis, beginning on January 5^th^, 2020, and ending on January 23^rd^, 2021. Parentheses give the domain in which each variant appears, according to the domain ranges specified in Huang et al. 2020.^29^ Key for domain abbreviations: SP=signal peptide, NTD=N-terminal domain, RBD=receptor-binding domain, FP=fusion peptide, HR1=heptad repeat 1, HR2=heptad repeat 2, CD=cytoplasmic domain, ID=intracellular domain. The heatmap is colored based on a log-10 scale, with prevalence values of zero colored in white, and values less than or equal to 0.10% colored with the lightest shade. **A)** The ten variants associated with the B.1.1.7 strain appeared together in mid-September and have quickly increased in prevalence. The variants associated are now observed in more than 50% of sequences worldwide. (Y144del sometimes appears as Y145del due to difficulties with identical adjacent amino acids in the alignment software). **B)** The variants associated with B.1.351 appeared together in mid-October and have increased in prevalence over time in samples worldwide, but to a lesser extent than the variants in B.1.1.7. **C)** The variants associated with P.1 appeared together in mid-December. The RBD variant E484K is also observed in B.1.351, and the RBD variant N501Y is also observed in B.1.1.7.

### RNA-Dependent RNA Polymerase (RdRP) Complex

#### Time Series Trends of Variants in Nsp12

The Nsp12: P323L variant appeared in late January 2020 and was present in 50% of the sequences by early March, and 90% of the sequences by late April (**Figure 6A**). The trend of predominance for Nsp12: P323L was observed on all continents (**Figure 6 B-G**), but the rate of establishment was lower in Asia than in other continents, as with spike: D614G. Other variants have emerged and are relatively rare, present in 3.5-4.0% of samples collected weekly. Variants that are increasing in prevalence globally include the Nsp12 variants V776L, E254D, A656S, and V720I, and the Nsp7 variants S25L and M75I. Time series trends in Europe (**Figure 6B**) match global trends. All the top ten global variants worldwide are seen in the top ten for Europe except for Nsp7: S25L, and several Nsp12 variants are present in 5.0%-10.0% of samples. This may be due to the predominance of samples from Europe since August 2020 (**Supplementary Figure S14 A-F**). In Asia (**Figure 6C**), the Nsp12 beta-hairpin substitution A97V appeared in late January and was observed in 49.7% of sequences collected during the week of April 12^th^. Afterward, the variant decreased in prevalence and was observed in between 5-25% of samples. The variant was observed in no samples during and after the week of September 13^th^. The Nsp7 variant S25L, which exists on the interface between Nsp7 and Nsp8, was observed in both Asia (**Figure 6C**) and North America (**Figure 6D**). Nsp7: S25L was first observed in North America during the week of March 1^st^, and in Asia during the week of March 8^th^. The prevalence of the variant in Asia peaked at 45.6% of sequences during the week of June 28^th^ and then declined in prevalence. After the week of September 13^th^, no sequences were observed with the variant. In North America, the variant is consistently observed in 1-7 % of sequences. Nsp7: S25L is rare on other continents: it is observed in less than 0.5% of sequences from Oceania, and less than 0.1% of sequences from Africa and Europe. In South America Nsp7: S25L was observed in three out of 16 sequences (18.75%) collected during the week of May 17^th^, though the low sample size from this continent makes it difficult to determine from this observation if the variant is common on the continent. In Africa (**Figure 6E**), the Nsp8 variant Y138H appears consistently in 20-40% of sequences since late June. The Nsp12 thumb subdomain variant G823S is also highly prevalent in samples from the week of August 16^th^, and the sharp increase in prevalence in this variant is observed at the same time as a decrease in the prevalence of Nsp12: P323L. In Oceania (**Figure 6F**), there are no stable variants in the RdRP complex besides Nsp12: P323L. Several variants, such as the Nsp12 variants A97V, K718N, and N215S, and the Nsp8 variant L9F, emerged, peaked in prevalence between 5-25%, and later decreased to less than 4% prevalence. However, one variant appeared recently and may be increasing in prevalence: the Nsp12 finger subdomain mutation A656S appeared in late September and is present in 5.26% of samples collected during the week of October 18^th^ (**Supplementary Table 2F**). In South America (**Figure 6G**), the Nsp12 thumb subdomain variant P918L was observed in 72.5% of samples collected during the week of June 21^st^ and 56.7% of sequences collected during the week of July 12^th^. This amino acid residue is near M924, which is involved in RNA binding.

**Figure 6 A-G:**
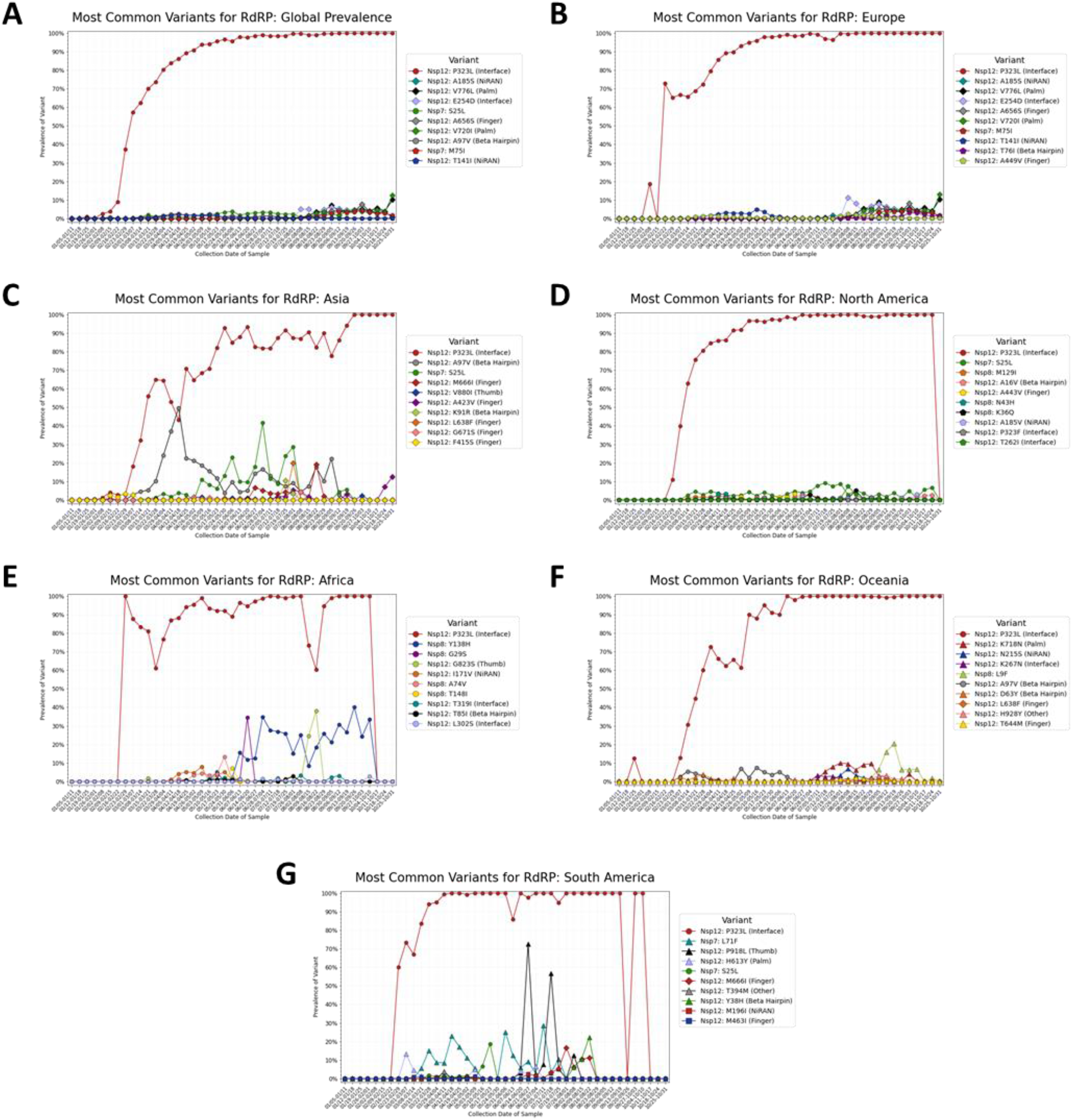
Prevalence of the top ten most common variants for the constituent proteins of the RNA-dependent RNA polymerase (RdRP) complex, by collection date in one-week intervals beginning on January 5^th^, 2020 and ending on October 31^st^, 2020.The top ten variants **A)** worldwide, **B)** in Europe, **C)** in Asia, **D)** in North America, **E)** in Africa, **F)** in Oceania, and **G)** in South America, are shown. Variants that appear within the top ten most prevalent variants on multiple continents are given the same color and shape in every graph.

A heatmap of time series data for variants in the RdRP complex is shown in **Figure 7**. The Nsp12 variants A185S and V776L appear to increase in prevalence together, suggesting a correlation between these variants. This is supported by the table of variant combinations for each cluster (**Supplementary Tables S4-S6**), which shows V776L occurring together with A185 in 2,652 sequences, and V776L and A185S occurring separately in 11 and 34 sequences, respectively. Relatively low sample sizes on some continents (**Supplementary Figure 14 B, D, F**) limit the conclusions that can be drawn from the time series data. **Supplementary Figures S15-S20** show regional heatmaps of all variants present in at least two percent of samples on each continent, and **Supplementary Table S3 A-G** shows complete prevalence data of all variants.

**Figure 7:**
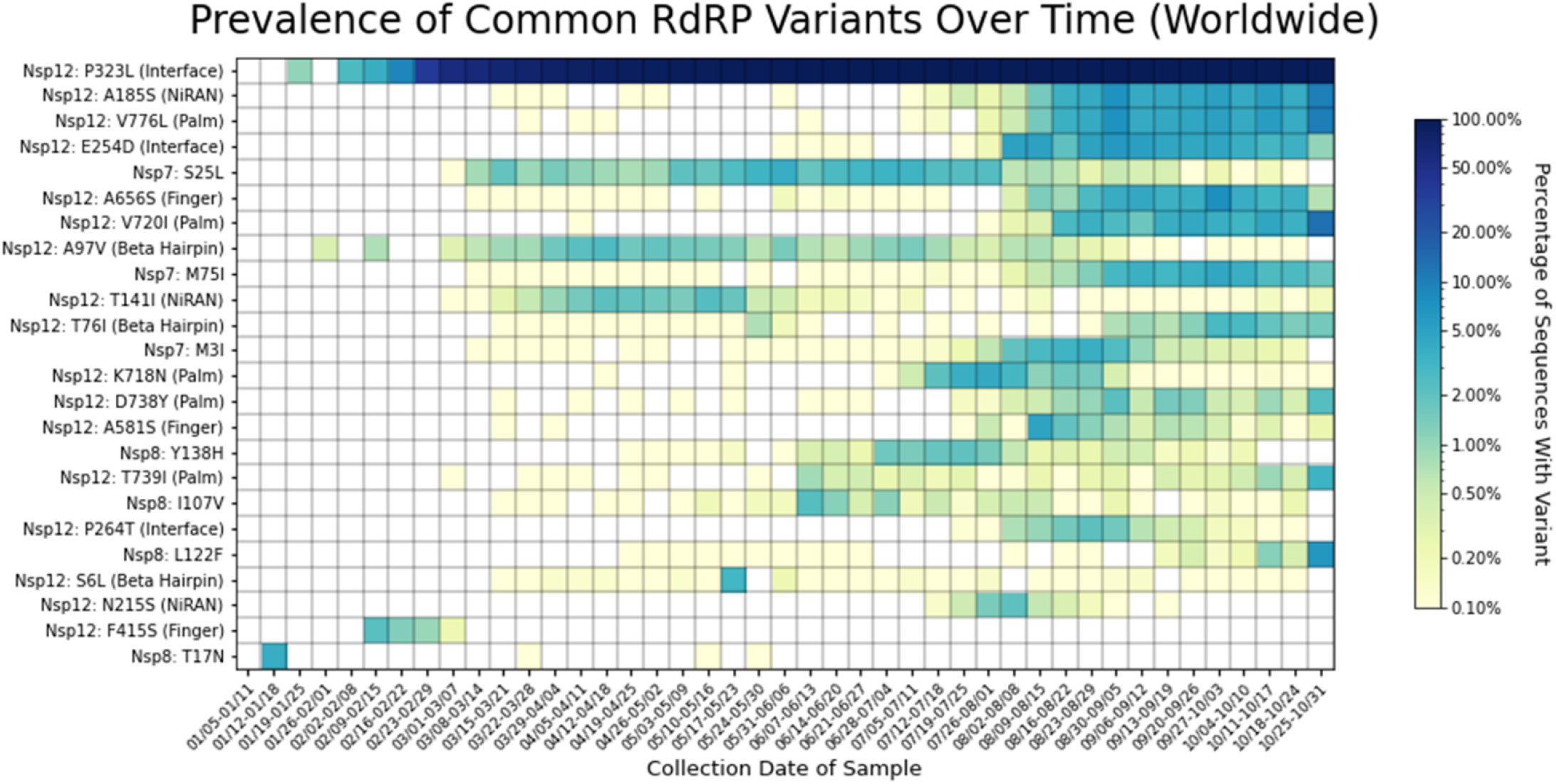
Heatmap of all RdRP complex variants observed in two percent or more of genomes sequenced for at least one week. Variants are listed on the y-axis, and parentheses give the domain in which each variant appears, according to the domain ranges specified in Yin et al. 2020^11^. The heatmap is colored based on a log-10 scale, with prevalence values of zero colored in white, and values less than or equal to 0.10% colored with the lightest shade. Time on the x-axis is categorized by week of collection date, beginning on January 5th, 2020, and ending on October 31st, 2020. P323L quickly increased in prevalence after its appearance in mid-January. The Nsp12 variants A185S, V776L, E254D, A656S, V720I, and T76I, as well as the Nsp7 variant M75I appear to be increasing in prevalence over time since early August. Common variants are observed in all domains except for the thumb subdomain of Nsp12. Of the 24 variants observed in at least two percent of genomes in any week, five are in the palm subdomain of Nsp12; three each are in the finger subdomain, the interface domain, the NiRAN domain, and the beta hairpin of Nsp12; zero are in the thumb subdomain of Nsp12, four are in Nsp8, three are in Nsp7.

#### Structural Visualization of Variants in RdRP Complex

325 unique variants were observed in Nsp8, 79 unique variants were observed in Nsp7, and 1,072 unique variants were observed in Nsp12 (**Figure 8A**). Of the 1,072 variants in Nsp12, 259 were observed in the finger subdomain, 177 were observed in the beta hairpin, 165 were observed in the palm subdomain, 161 were observed in the NiRAN domain, 133 were observed in the interface, and 118 were observed in the thumb subdomain. Fifty-nine variants occurred in an uncharacterized region of Nsp12.

**Figure 8 A-D:**
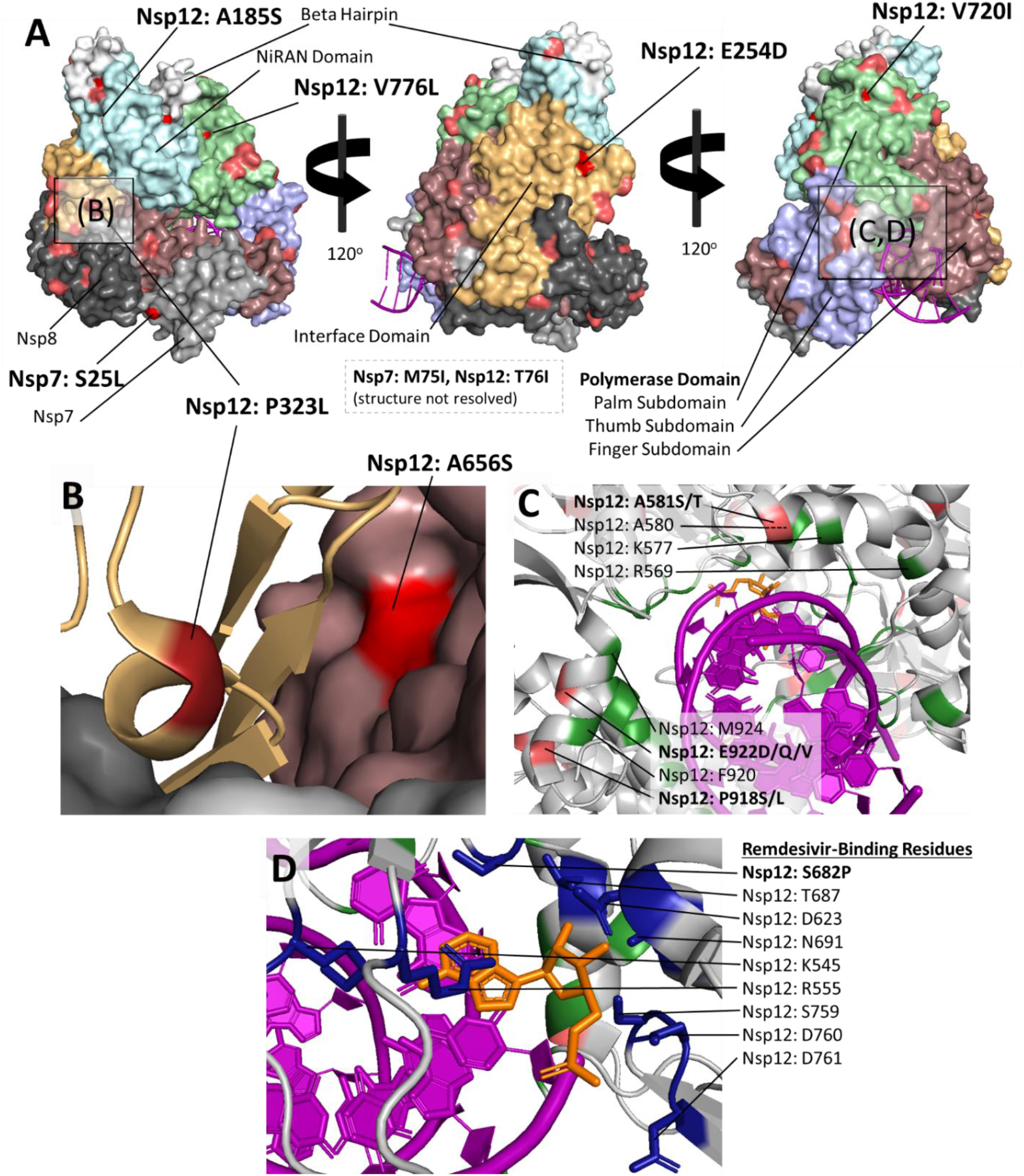
Structural visualization of variants in the RNA-Dependent RNA Polymerase (RdRP) complex. **A)** RdRP complex colored by domain, with high-frequency variants highlighted. Nsp7 and Nsp8 are colored in grey and dark grey, respectively. Nsp12 is colored and labeled by domain. Residues with more than 100 variants are highlighted according to the number of sequences with a variant at that position (worldwide, all collection dates). Dark red is used for residues with variants in10,000+ sequences, red is used for variant frequencies of 1000-10000, and pink is used for variant frequencies of 100-1000. Nsp7 has few variants compared with Nsp8 and Nsp12, and the interface and NiRAN domain of Nsp12 also appear to have relatively few variants compared with other domains. The common variant Nsp7: S25L exists at the interface between Nsp7 and Nsp8. **B)** A ribbon diagram of the highly prevalent Nsp12: P323L variant. The variant exists within a small alpha helix surrounded by loop regions. Proline helps maintain a tight helical geometry, and substitution of this variant may disrupt the secondary structure. The common Nsp12 variant A656S is also shown. **C)** Ribbon diagram of RNA-binding residues of Nsp12 (shown in green). Remdesivir is shown in orange, and variants are colored according to the scheme in **(A)**. There are no RNA-binding residues with variation in more than 100 sequences, though variation is observed in residues within two alpha-helix regions containing RNA-binding residues (labeled). **D)** Ribbon diagram of remdesivir-binding site with side chains of remdesivir-binding residues (dark blue). Variation was observed in only one remdesivir-binding residue, S862, which is mutated to proline in 2 sequences. PDB structures used: PDB ID: 7BV2.^24^

#### Secondary Structure of P323

The Nsp12 residue P323 occurs in a small alpha helix region within a loop secondary structure (**Figure 8B**). This residue is not known to directly bind residues in neighboring domains, though it is near residue A656, which contacts the interface domain and is mutated to a serine in 2,010 sequences across all time points (1.132%). A656 has also been observed to mutate to a threonine in 54 sequences, and to a valine in 13 sequences. A656S has been observed in increasing frequency since August, while A656T and A656V have not increased in prevalence with time (**Supplementary Figures S21-S22**).

#### RNA-Binding Residues

RNA-binding residues of in the finger, thumb, and palm subdomains were highly conserved (**Figure 8C**). Out of the 41 residues of Nsp12 known to bind RNA^11^, only seven were mutated in at least two sequences and no residues were mutated in more than 6 sequences (0.0034% of all sequences analyzed).

#### Variation in the Binding Site for Remdesivir

Remdesivir blocks viral replication by binding to the following residues in Nsp12: K545, S682, R555, T687, S759, N691, D623, D760, and D761.^11^ Remdesivir-binding residues are shown in **Figure 8D**. Of these nine amino acid residues, variants were observed for only one: S862, which mutated to proline in two sequences worldwide.

## Discussion

This study analyzed viral genomic data made available through GISAID and used structural information to identify trends in variant representation over time and to examine the structural features of variants. The study validated previous findings of the predominance of spike: D614G and Nsp12: P323L in the population^4^ while uncovering recent increases in the prevalence of new mutations in Nsp12 and the spike protein. Trends in the emergence of new variants vary by geographic area.

The very high prevalence of the D614G mutant in the sequences sampled reflects studies documenting increased viral counts in-vitro^4,36^ as well as higher viral loads in infected individuals.^4^ D614G has also been observed to co-occur with Nsp12: P323L^4^, the most common mutation observed in Nsp12. For both the spike protein and Nsp12, new variants appeared in late July to early August and have since steadily increased in prevalence. Most mutations generally have neutral or deleterious effects on protein function^37^; these variants would not be expected to increase in prevalence over time unless they occur with a variant that confers an increase in fitness, replication, or transmission potential. The increasing prevalence of new variants in the population suggests that these variants provide an evolutionary advantage, though follow-up studies *in-vitro* or *in-vivo* are required to definitively determine the effects of these variants.

The high conservation observed in the RNA-binding residues of the RdRP indicates that there is no evidence of remdesivir resistant mutations as of October 31, 2020. Continued surveillance of contact residues for remdesivir^11^ and other drug candidates targeting the RdRP^38^ is necessary to effectively respond to potential drug resistance in the future.

The N-terminal domain and the receptor binding domain (RBD) of the spike protein were observed to be variable, which may have implications for monoclonal antibody treatments that target these domains.^39–41^ The RBD variant N439K, which has been shown to result in antibody evasion,^5,42^ is becoming more common over time. Several variants observed in the N-terminal domain and the signal peptide (L18F, Y144del, and D253G) have also been shown to decrease the effectiveness of antibodies.^43^ It is essential to continue monitoring these variants and other variants in the RBD shown to escape antibodies^42,44^ to ensure that vaccines and monoclonal antibody treatments against SARS-CoV-2 remain effective against evolving and globally circulating variant strains.

In the case of antibody cocktails, the chance of any single mutation rendering the treatment ineffective is low. In some cases, antibody cocktails contain multiple antibodies that recognize different regions of the RBD. Studies on the Regeneron cocktail demonstrate enduring effectiveness despite the presence of individual escape mutations.^45^ Vaccines also result in the production of multiple antibodies targeting different regions of the spike and other proteins, and are likely to remain effective, even when encountering single escape mutations. Strains with many variants relative to the reference may be more difficult to protect against with current vaccines designed against the original SARS-CoV-2 strain, as has been observed with the South African SARS-CoV-2 variant B.1.351.^46^

The analysis in this study is limited in some aspects, since the files used and downloaded for this study consist only of amino acid sequences, without the accompanying nucleotide information. Amino acid data can robustly capture information on non-synonymous mutations, but analyses such as codon bias^47^ cannot be performed without nucleotide information. The lack of nucleotide information also limits the ability of alignment programs to distinguish between two possible deletions in regions where two or more identical amino acids are adjacent to one another; this causes the spike variant Y144del to be identified as Y145del in some sequences of our analysis. In addition, the identity of ambiguous “X” codons cannot be fully elucidated without nucleotide data. Despite stringent filtering of sequences, ambiguous codons were still observed in some sequences.

Due to the lack of accompanying clinical data posted to the GISAID platform, it is possible that some samples may represent longitudinal samples taken from the same patient across different time points. If this is the case, rare variants in these samples may be over-represented. The time series data may also be influenced by reporting bias. A sudden influx of samples from a region where a variant is present may over-represent that variant, and variants may appear to become less common due to decreasing reporting of sequences from regions where they are common. European variants are likely over-represented in the time series data from August onward due to the very high number of samples collected in this region relative to others in the same time interval. Without extensive sequencing of samples worldwide, it is likely that there are functionally significant variants present that have not yet been discovered.

Future directions for study include segmenting the data by location as well as time to characterize regional trends in variation. The analyses performed in this study can be repeated for other SARS-CoV-2 proteins, and differing trends in variation by protein could be determined by normalizing variant frequencies to the length of each protein. Also, correlation analysis of variants can be performed to see which mutations happen concurrently, verified by identifying variants that co-occur on specific viral protein sequences. This would allow for preemptive identification of new strains consisting of multiple variants, which can be evaluated in follow-up studies for functional implications.

## Supporting information

Supplementary Table 1 - Variant Prevalence for Spike Protein

Supplementary Table 2 - Combination of Variants Observed in Spike Protein

Supplementary Table 3 - Variant Prevalence for RNA Polymerase

Supplementary Table 4 - Combination of Variants Observed in Nsp7

Supplementary Table 5 - Combination of Variants Observed in Nsp8

Supplementary Table 6 - Combination of Variants Observed in Nsp12

Supplementary Table 7 - Acknowledgement of GISAID Data Contributors

## Acknowledgements

We gratefully acknowledge the authors of the submitting laboratories to the GISAID portal, as well as the authors from the originating laboratories who obtained the sequences. The entire analysis is based on data from the GISAID portal (**Supplementary Figure S7**). Many thanks to our colleagues, Cody Glickman, Elaine Epperson, Nabeeh Hasan, and Jo Hendrix, in the Strong Laboratory for their input and feedback. We also wish to thank Tzu Phang, Lauren Vanderlinden, Fuyong Xing for helpful discussions and training. WS thanks the countless faculty and peers that have served as role models and furthered his learning. MS acknowledges funding from the Colorado Advanced Industries grant program.

