## Supplementary Figures for "Analysis of SARS-CoV-2 Mutations Over Time Reveals Increasing Prevalence of Variants in the Spike Protein and RNA-Dependent RNA Polymerase"

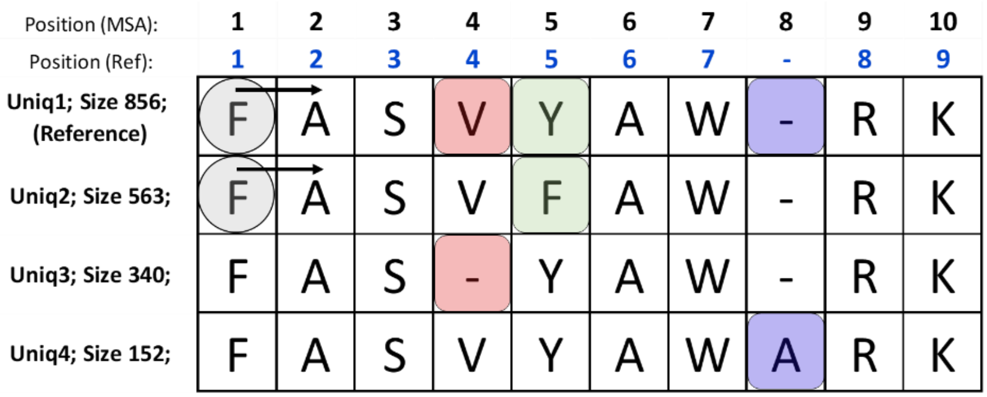


***Supplementary*** **Figure *S*1:** Schematic of the script used for multiple sequence alignment processing. The Biopython module creates a MultipleSeqAlignment object from the aligned sequences, which can be indexed by row and column in the same manner as a *pandas D*ata*F*rame. Each *residue* in the sequence has a position (black text) for indexing; due to the presence of insertions relative to the reference, the position in the MultipleSeqAlignment object does not correspond to the position of the *amino acid residue* relative to the reference sequence, which is required to describe the mutations observed. After a reference cluster is defined (Uniq1 in this example) the position of each *residue relative to* the reference (blue text) is computed*. Comparison of each sequence with the reference then proceeds from left to right beginning with the first residue of the first non-reference sequence (as designated by grey circles and arrows). Comparison continues* until a substitution (green), deletion (red), or insertion (blue) is observed. Each variant type triggers a separate function, which stores the code of the variant along with information on the position of the variant and the cluster in which it was discovered.

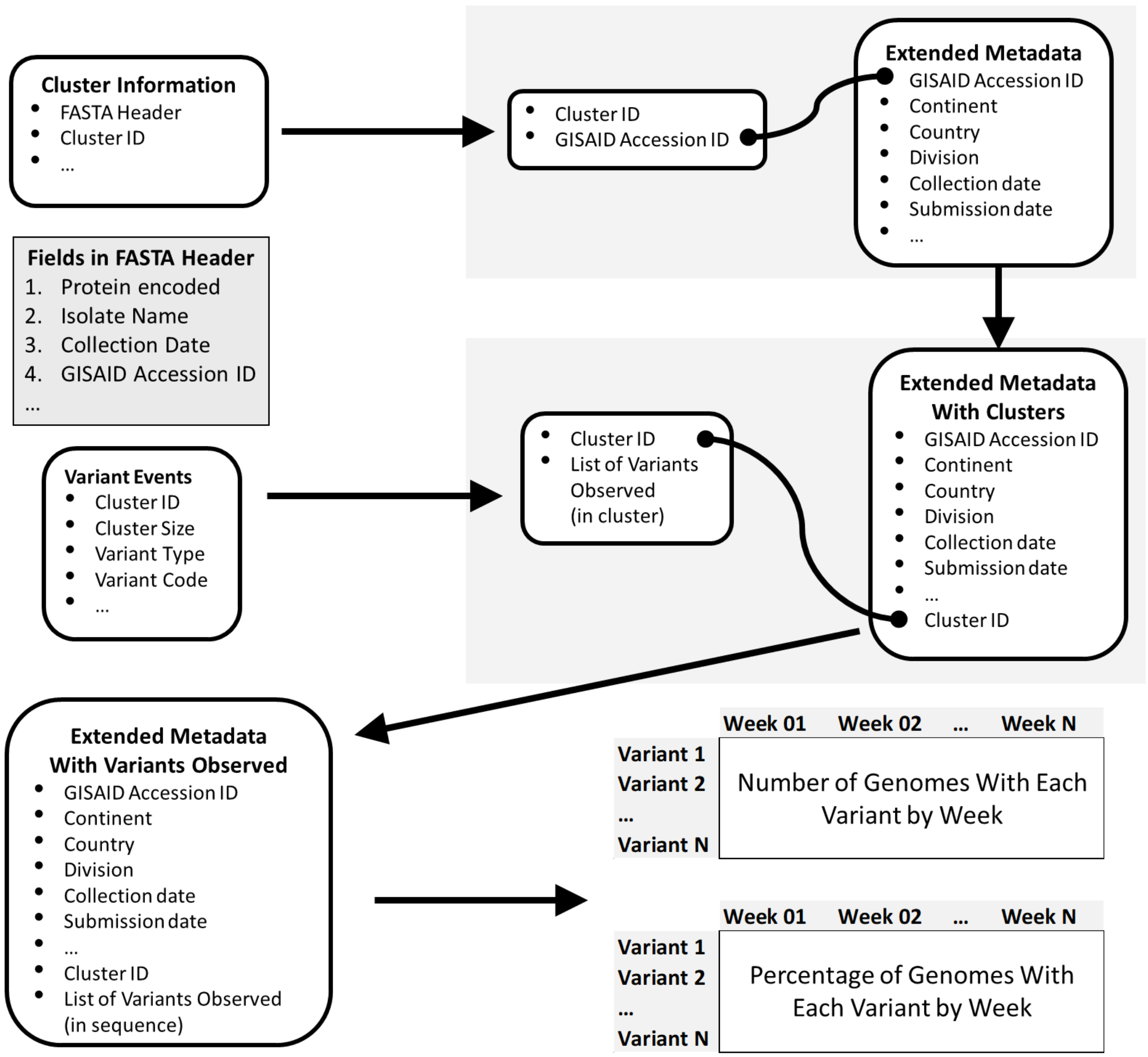


**Supplementary Figure S2:** Process for creation of time series data. Each dataset is represented by a rounded rectangle, and column names are given in bulleted lists. The ellipsis indicates the presence of additional columns not involved in the analysis. Related datasets to be merged are boxed in grey, with the common column between the two datasets indicated by the rounded arrow. Beginning with the cluster information generated by USEARCH, the GISAID accession ID is extracted from the FASTA header and added as a new column, while dropping all other columns besides the cluster ID. The resulting dataset is merged with the GISAID extended metadata using the common cluster ID column, yielding sequence metadata with each sequence mapped to its corresponding cluster. The variant events data produced from multiple sequence alignment parsing was then looped through to yield a dataset that maps each cluster to a list of all variants observed in the cluster, and this dataset was merged with the metadata with the cluster ID column to yield a dataset that maps each sequence to a list of the variants observed in that sequence. This dataset was then segmented by week of sample collection, and the total number of sequences in each week was recorded along with the number of sequences from that week containing each variant. The resulting data table (lower right) contained a row for each unique variant and a column for each week, with weekly frequency values of each mutation. The values in this table were divided by the total number of sequences for each week to yield a table giving the percentage of each variant over time.

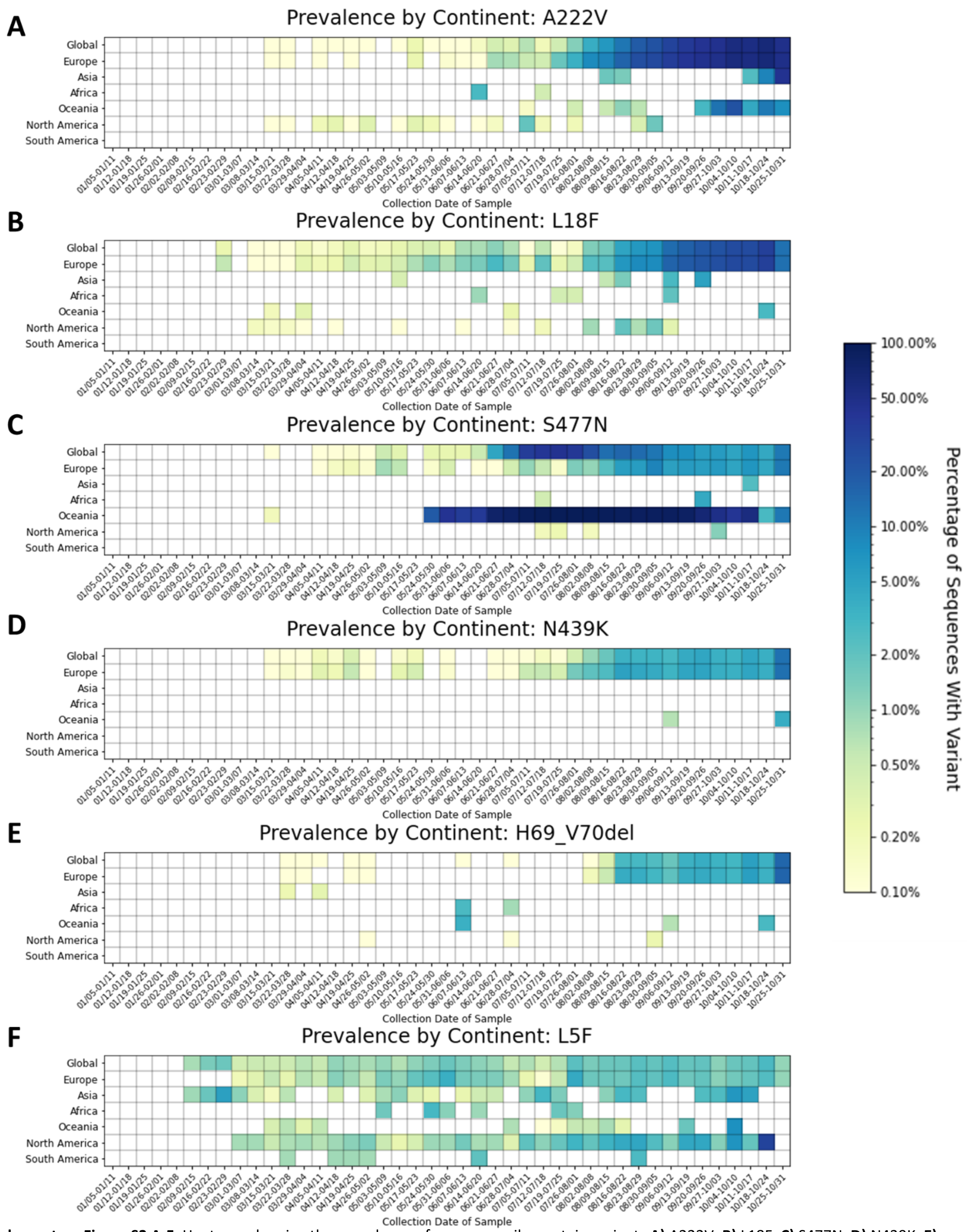


**Supplementary Figure S3 A-F:** Heatmap showing the prevalence of common spike protein variants **A)** A222V, **B)** L18F, **C)** S477N, **D)** N439K, **E)** H69_V70del, and **F)** L5F, on each continent and worldwide. The heatmap is colored based on a log-10 scale, with prevalence values of zero colored in white, and values less than or equal to 0.10% colored with the lightest shade. Time on the x-axis is categorized by week of collection date, beginning on January 5th, 2020, and ending on October 31st, 2020.

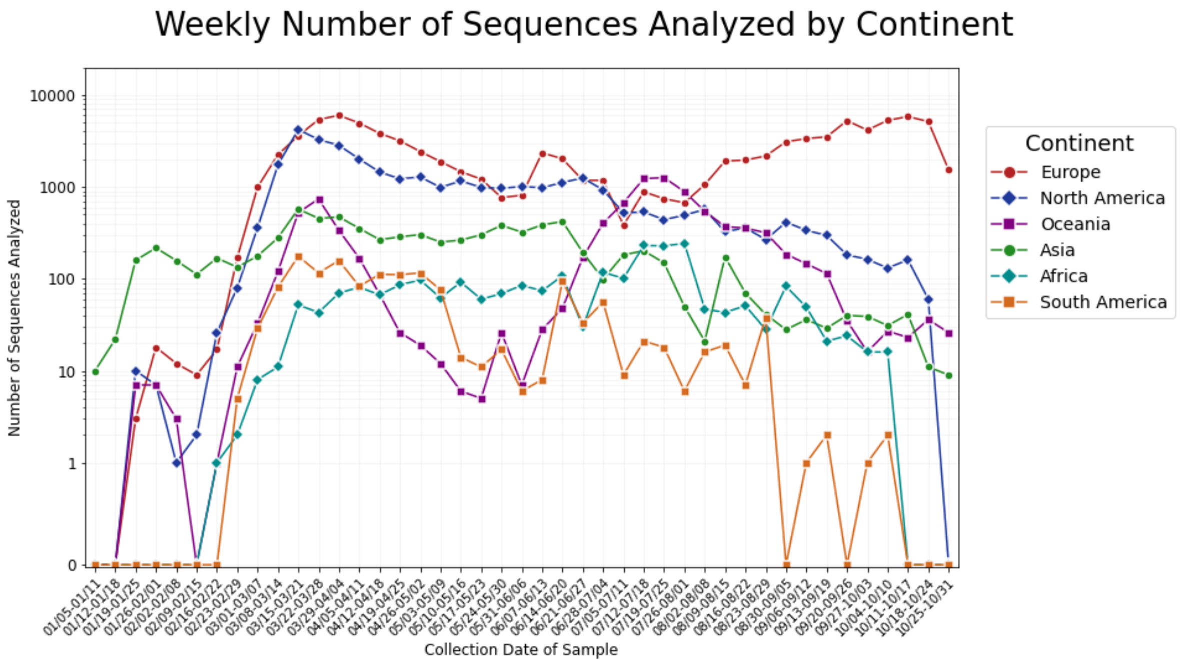


**Supplementary Figure S4:** Number of sequences analyzed for the spike protein, segmented by continent and week of collection date. A log-10 scale is used for the y-axis, and time on the x-axis is categorized by week of collection date, beginning on January 5th, 2020, and ending on October 31st, 2020.

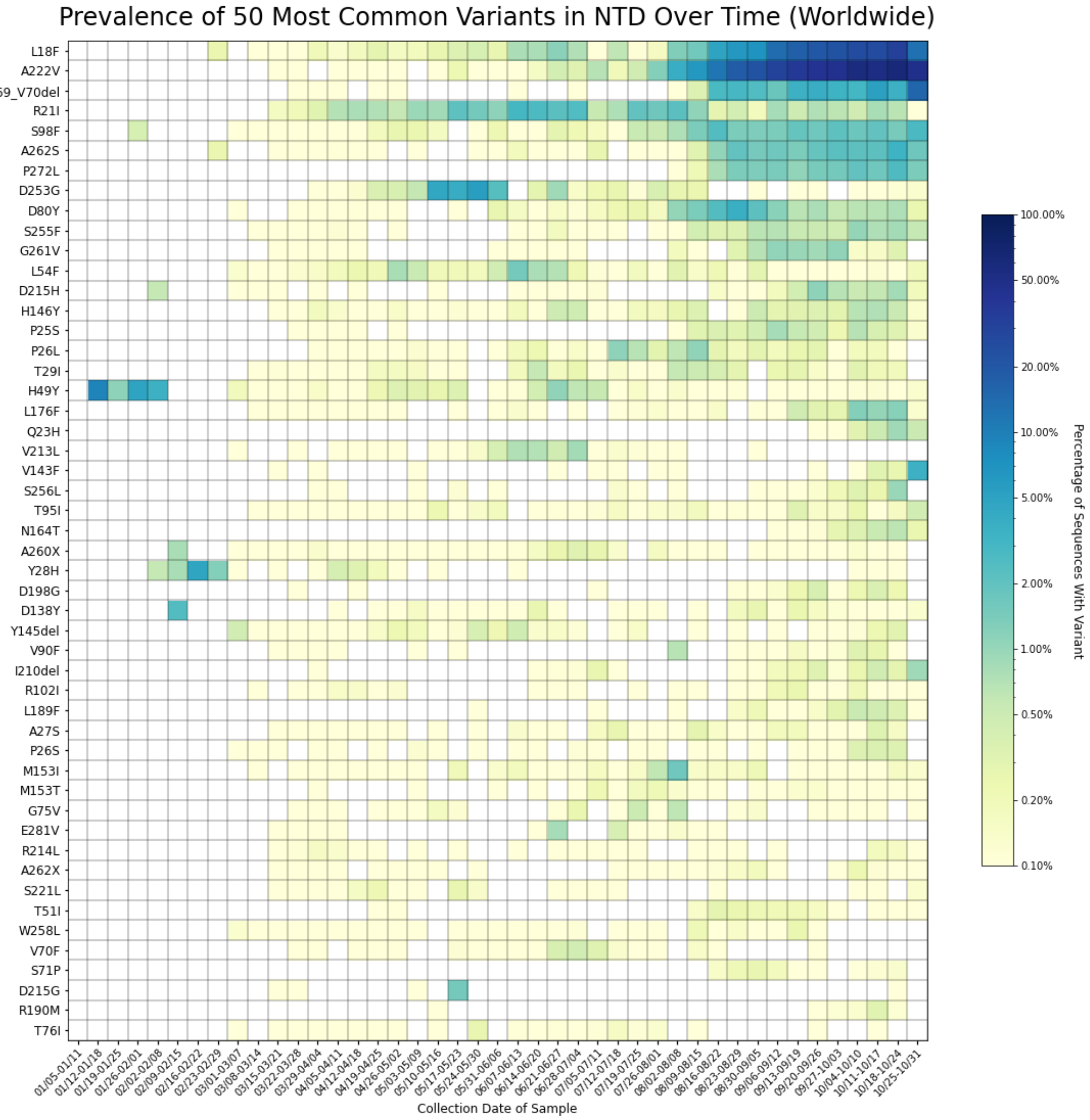


**Supplementary Figure S5:** Heatmap showing the top 50 most common variants within the N-Terminal Domain (NTD) of the spike protein (residues 14 to 305). The heatmap is colored based on a log-10 scale, with prevalence values of zero colored in white, and values less than or equal to 0.10% colored with the lightest shade. Time on the x-axis is categorized by week of collection date, beginning on January 5th, 2020, and ending on October 31st, 2020.

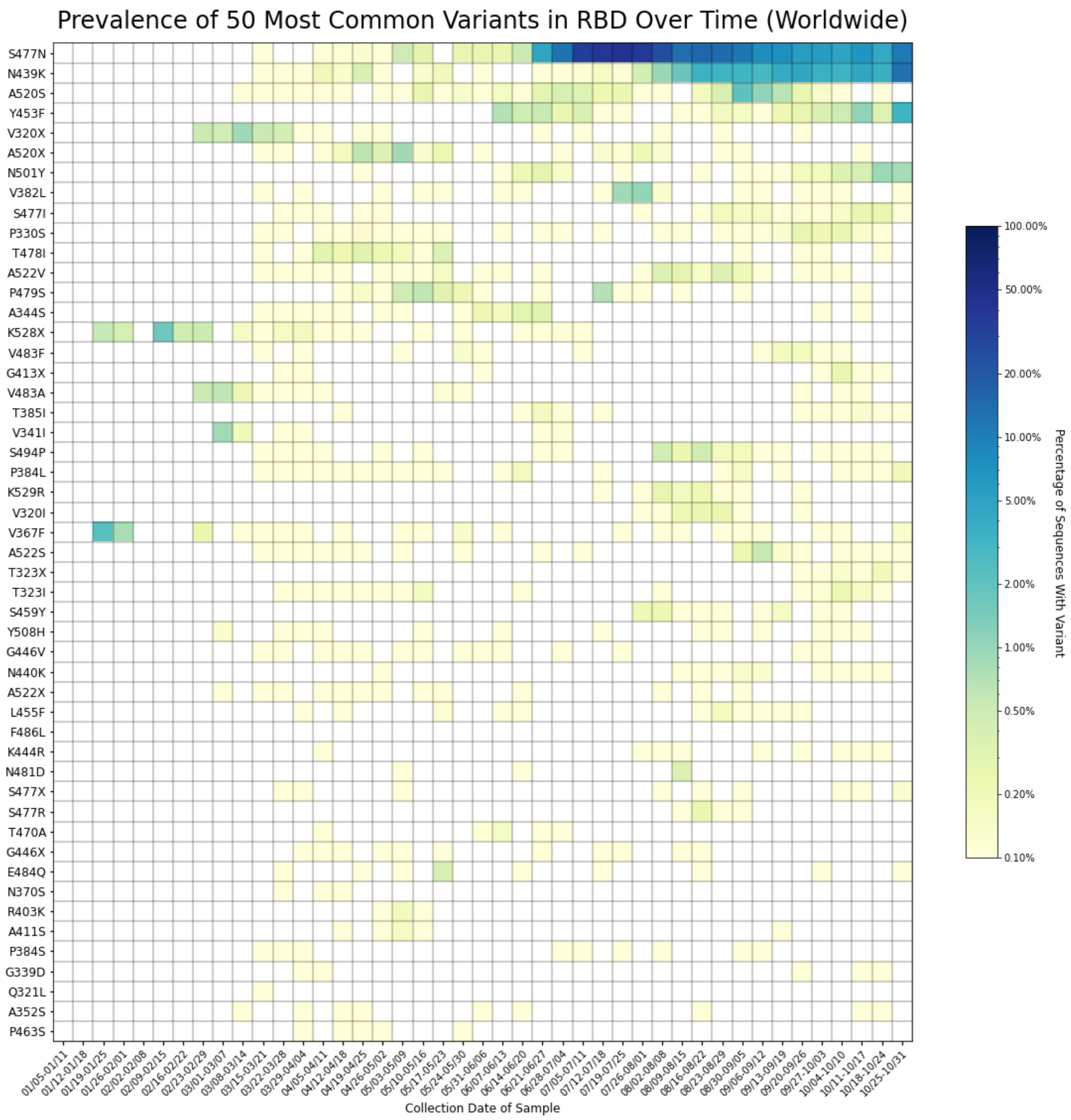


**Supplementary Figure S6:** Heatmap showing the top 50 most common variants within the Receptor Binding Domain (RBD) of the spike protein (residues 319 to 541). The heatmap is colored based on a log-10 scale, with prevalence values of zero colored in white, and values less than or equal to 0.10% colored with the lightest shade. Time on the x-axis is categorized by week of collection date, beginning on January 5th, 2020, and ending on October 31st, 2020.

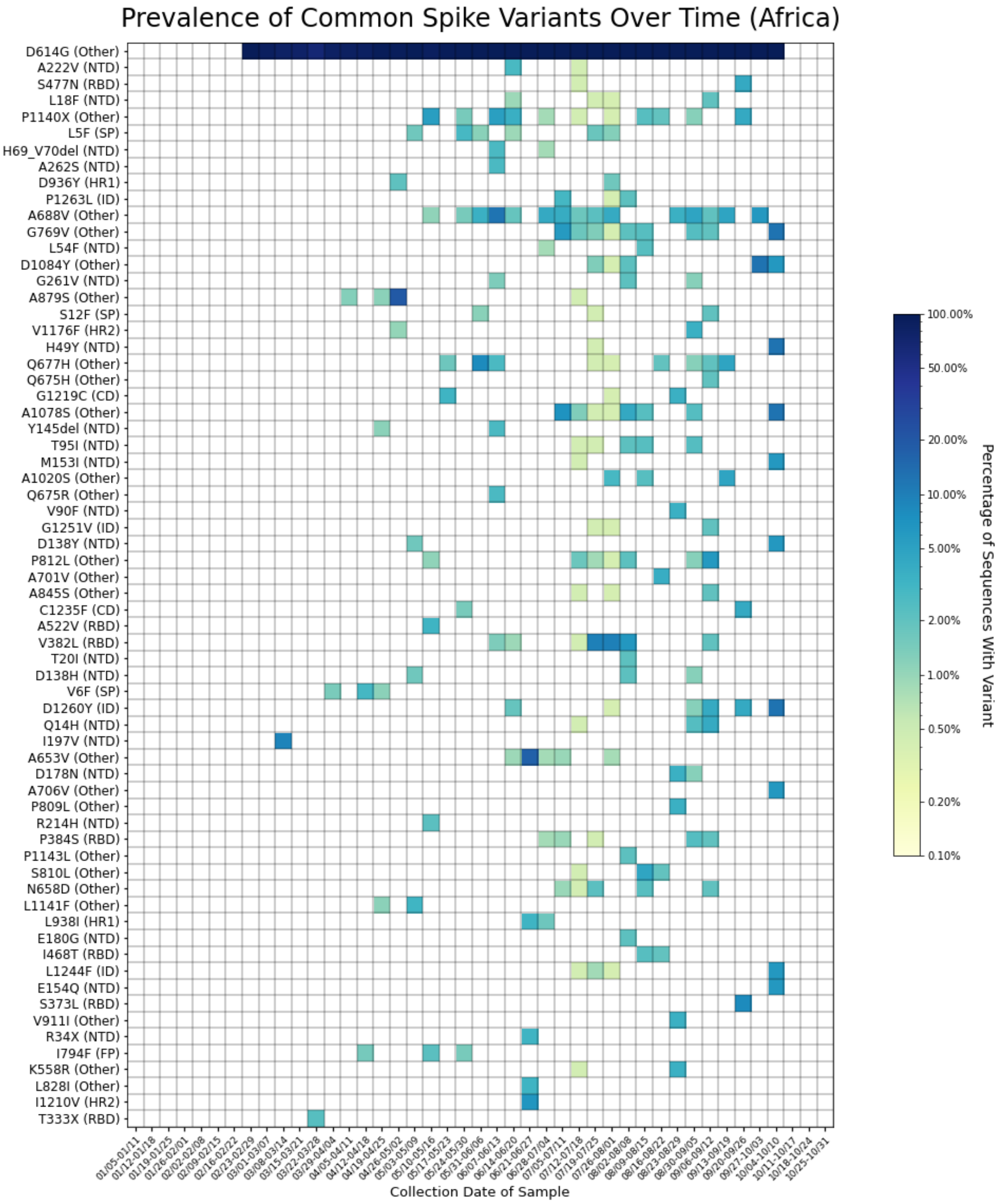


**Supplementary Figure S7:** Heatmap showing time series trends for all variants detected in at least two percent of sequences from Africa in any given week. The heatmap is colored based on a log-10 scale, with prevalence values of zero colored in white, and values less than or equal to 0.10% colored with the lightest shade.

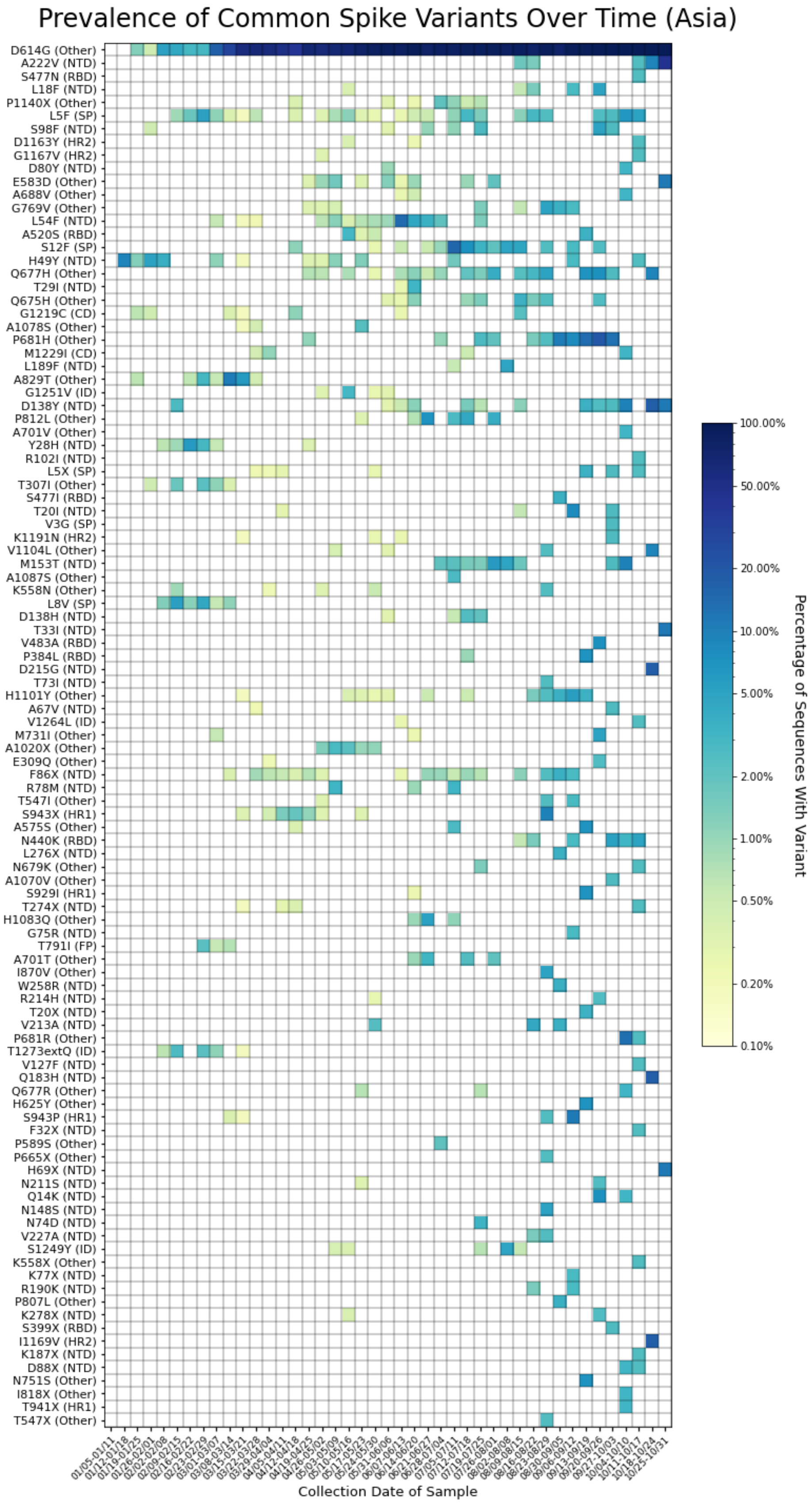


**Supplementary Figure S8:** Heatmap showing time series trends for all variants detected in at least two percent of sequences from Asia in any given week. The heatmap is colored based on a log-10 scale, with prevalence values of zero colored in white, and values less than or equal to 0.10% colored with the lightest shade.

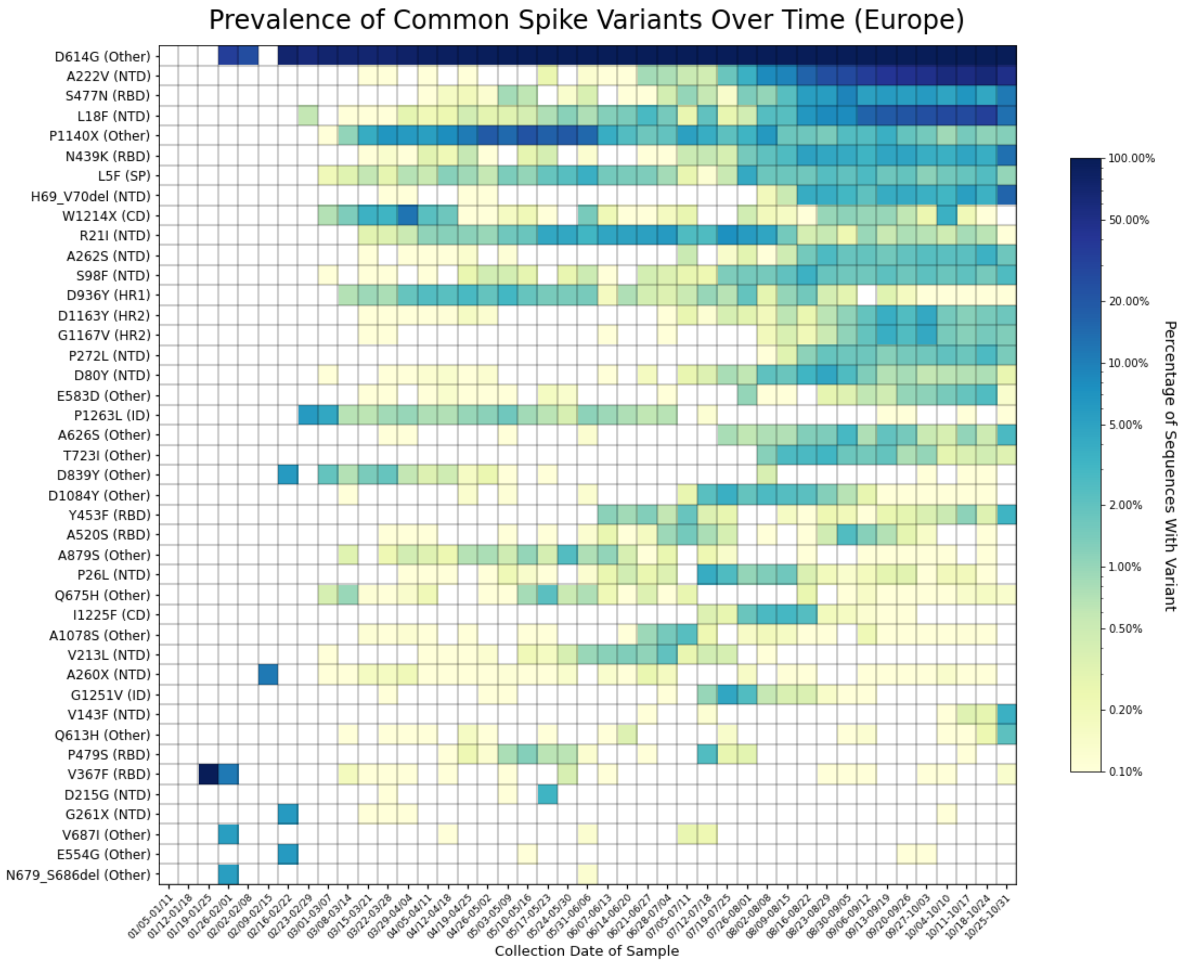


**Supplementary Figure S9:** Heatmap showing time series trends for all variants detected in at least two percent of sequences from Europe in any given week. The heatmap is colored based on a log-10 scale, with prevalence values of zero colored in white, and values less than or equal to 0.10% colored with the lightest shade.

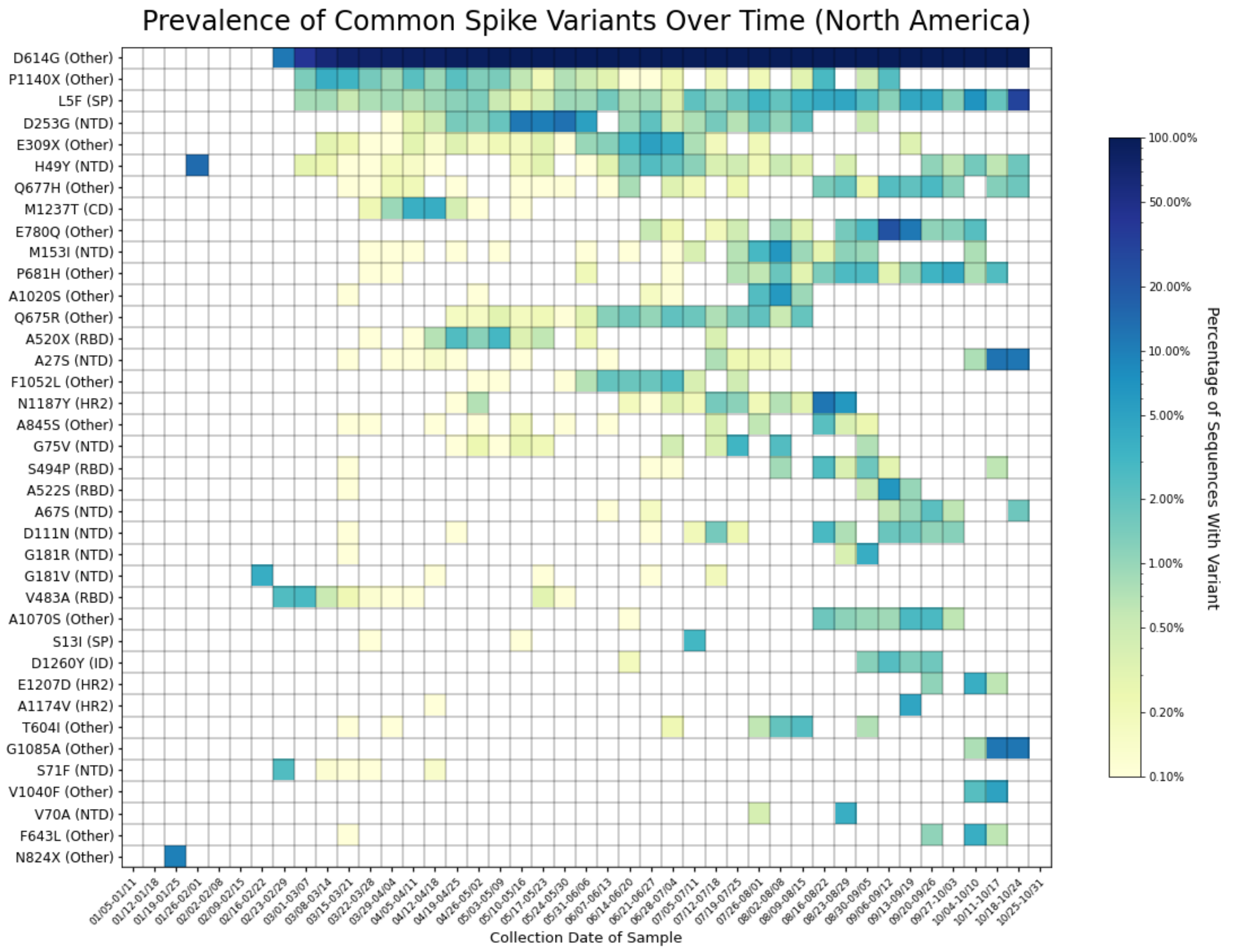


**Supplementary Figure S10:** Heatmap showing time series trends for all variants detected in at least two percent of sequences from North America in any given week. The heatmap is colored based on a log-10 scale, with prevalence values of zero colored in white, and values less than or equal to 0.10% colored with the lightest shade.

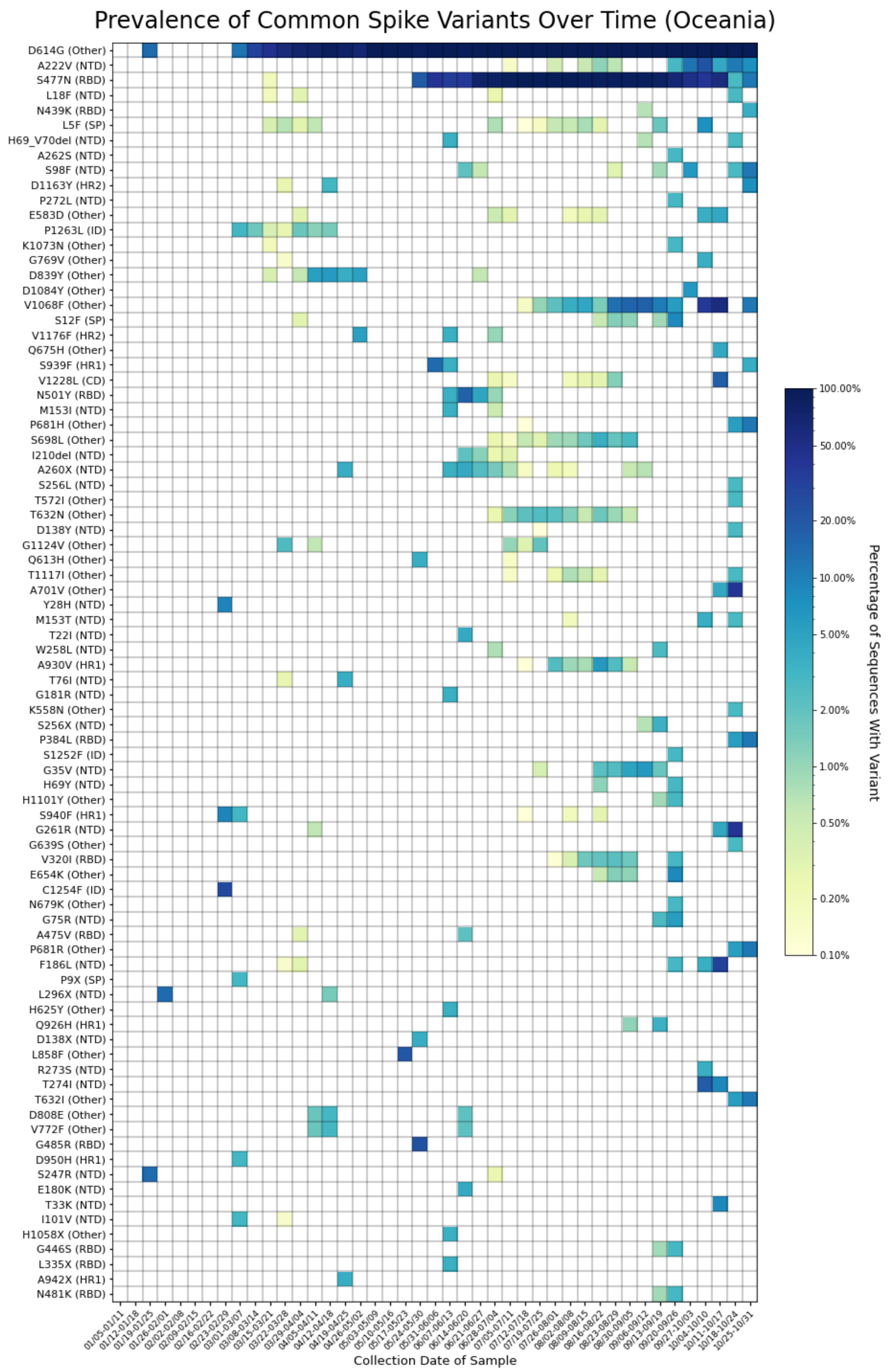


**Supplementary Figure S11:** Heatmap showing time series trends for all variants detected in at least two percent of sequences from Oceania in any given week. The heatmap is colored based on a log-10 scale, with prevalence values of zero colored in white, and values less than or equal to 0.10% colored with the lightest shade.

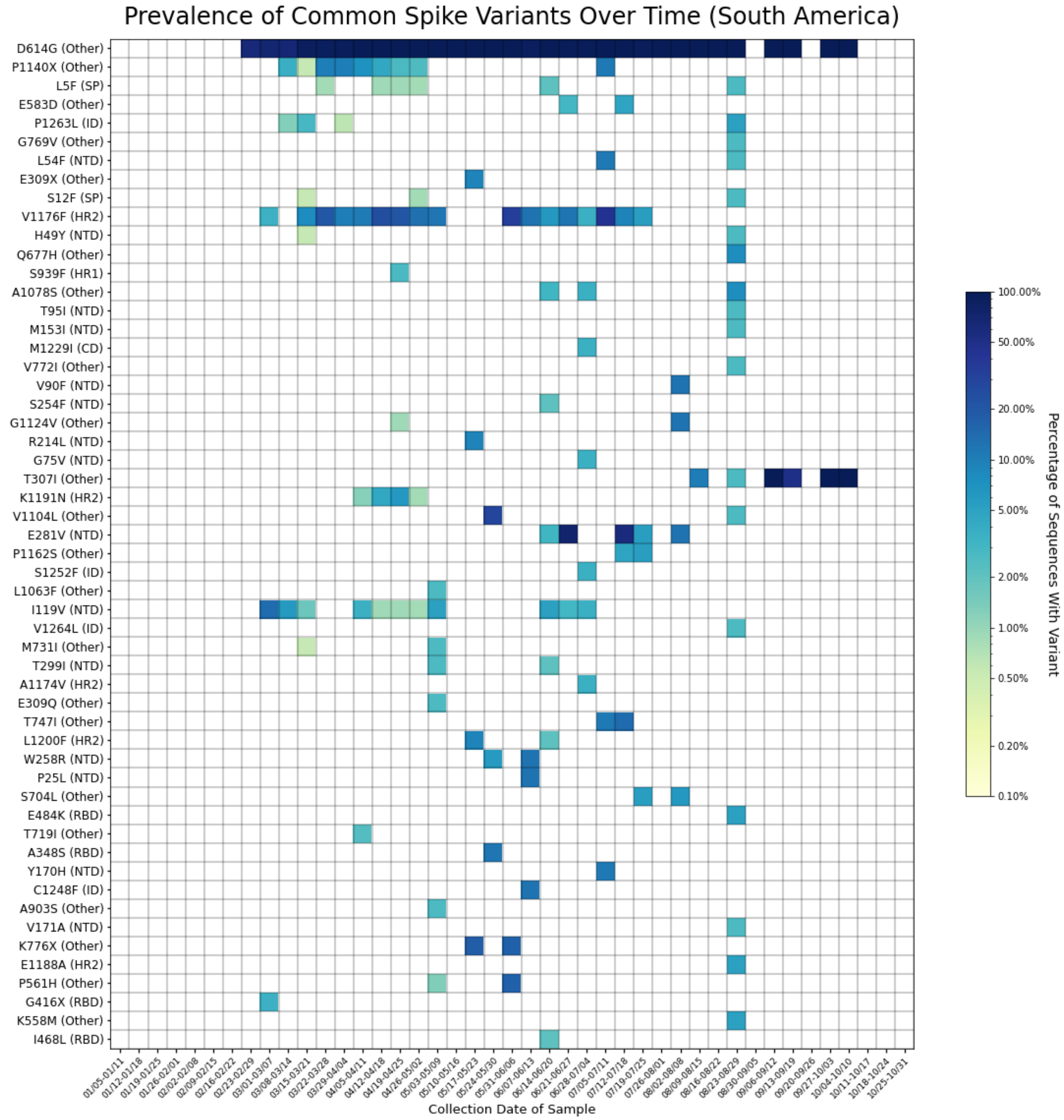


**Supplementary Figure S12:** Heatmap showing time series trends for all variants detected in at least two percent of sequences from South America in any given week. The heatmap is colored based on a log-10 scale, with prevalence values of zero colored in white, and values less than or equal to 0.10% colored with the lightest shade.

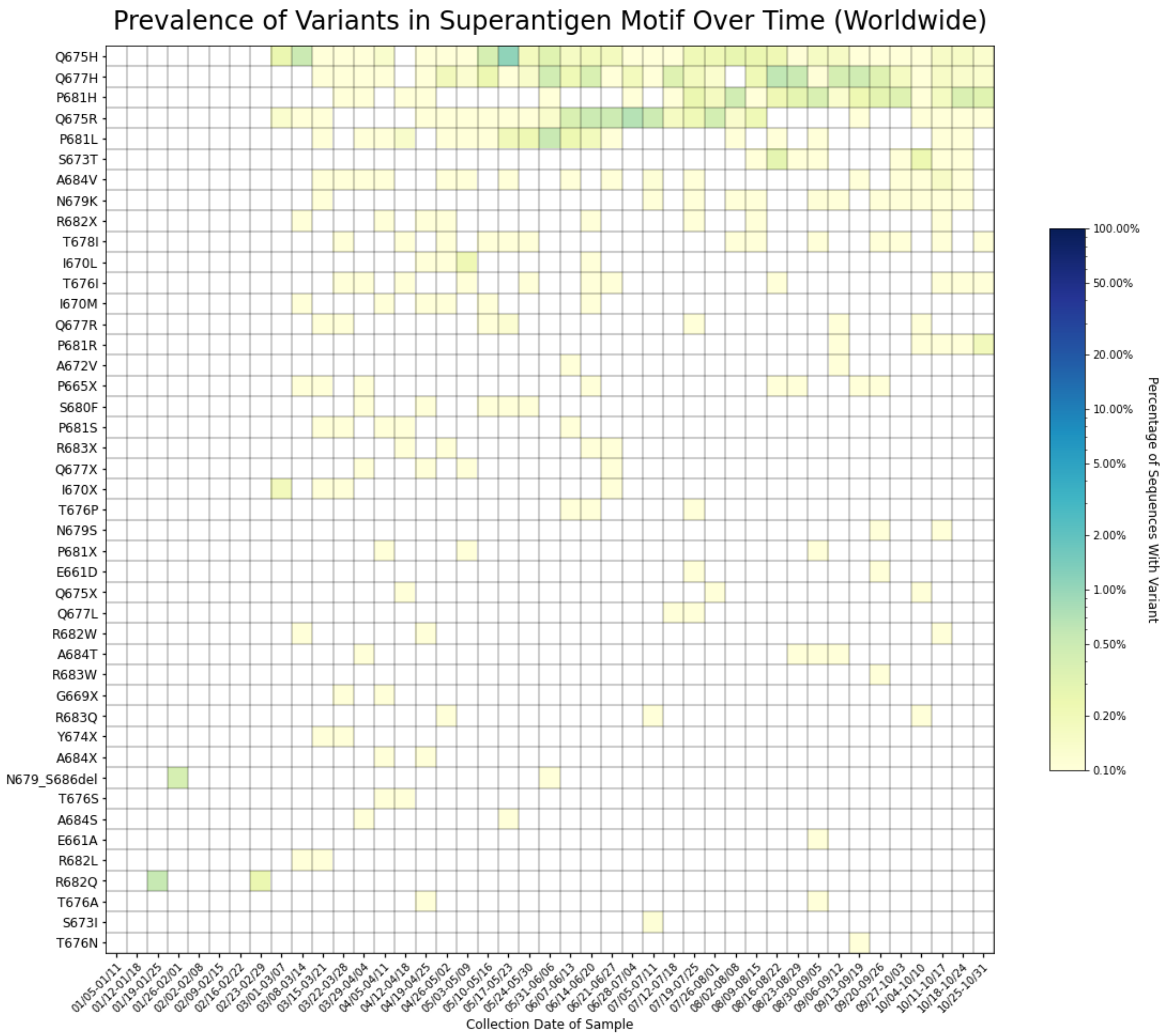


**Supplementary Figure S13:** Heatmap showing all variants within the superantigen motif of the spike protein (residues 661 to 685). The heatmap is colored based on a log-10 scale, with prevalence values of zero colored in white, and values less than or equal to 0.10% colored with the lightest shade.

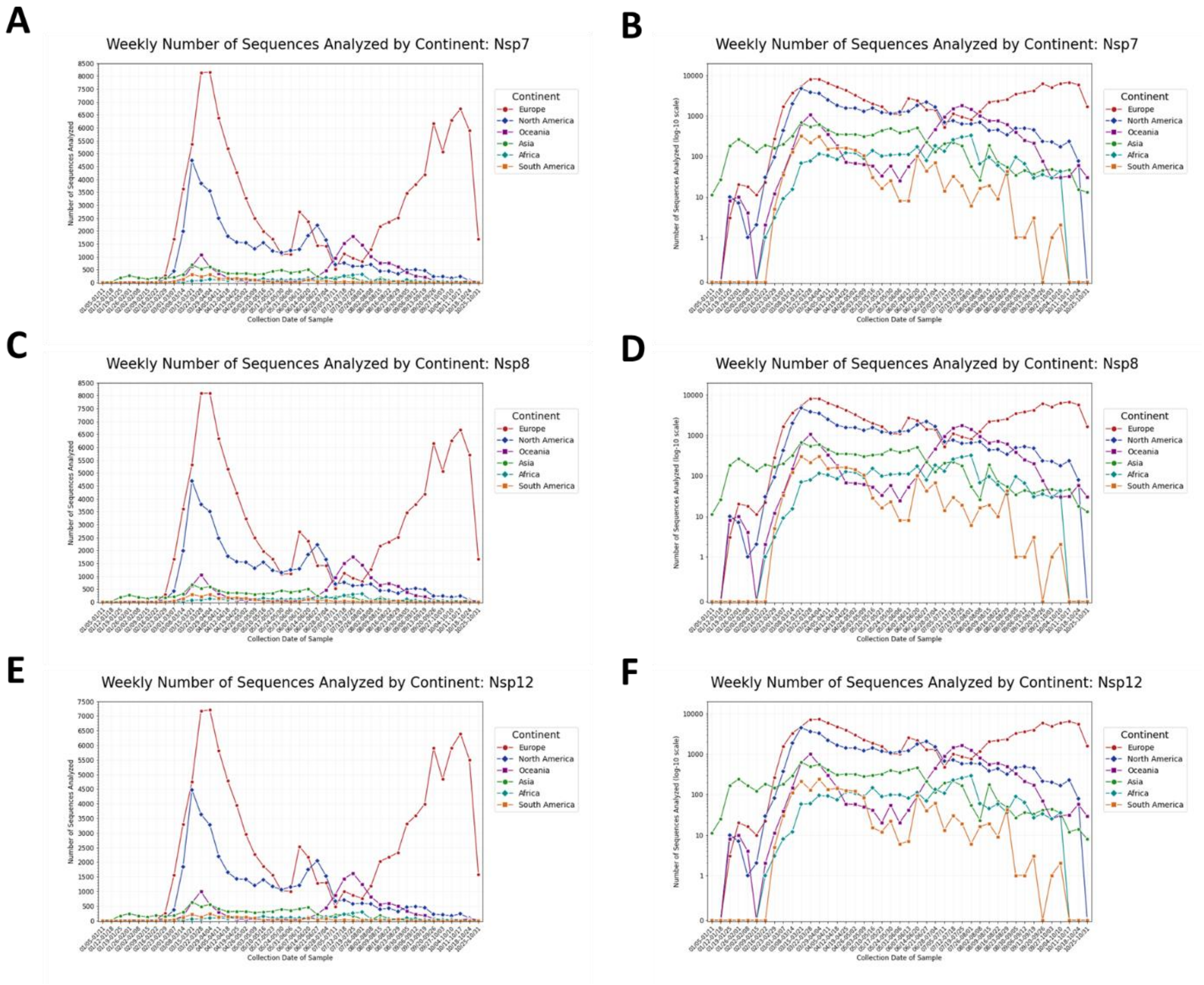


**Supplementary Figure S14 A-F:** Number of sequences in each week of the analysis by continent for the constituent proteins of the RNA-dependent RNA polymerase complex. Sequence counts for Nsp7 (**A-B**), Nsp8 (**C-D**), and Nsp12 (**E-F**) are displayed on both a linear and a log y-axis scale.

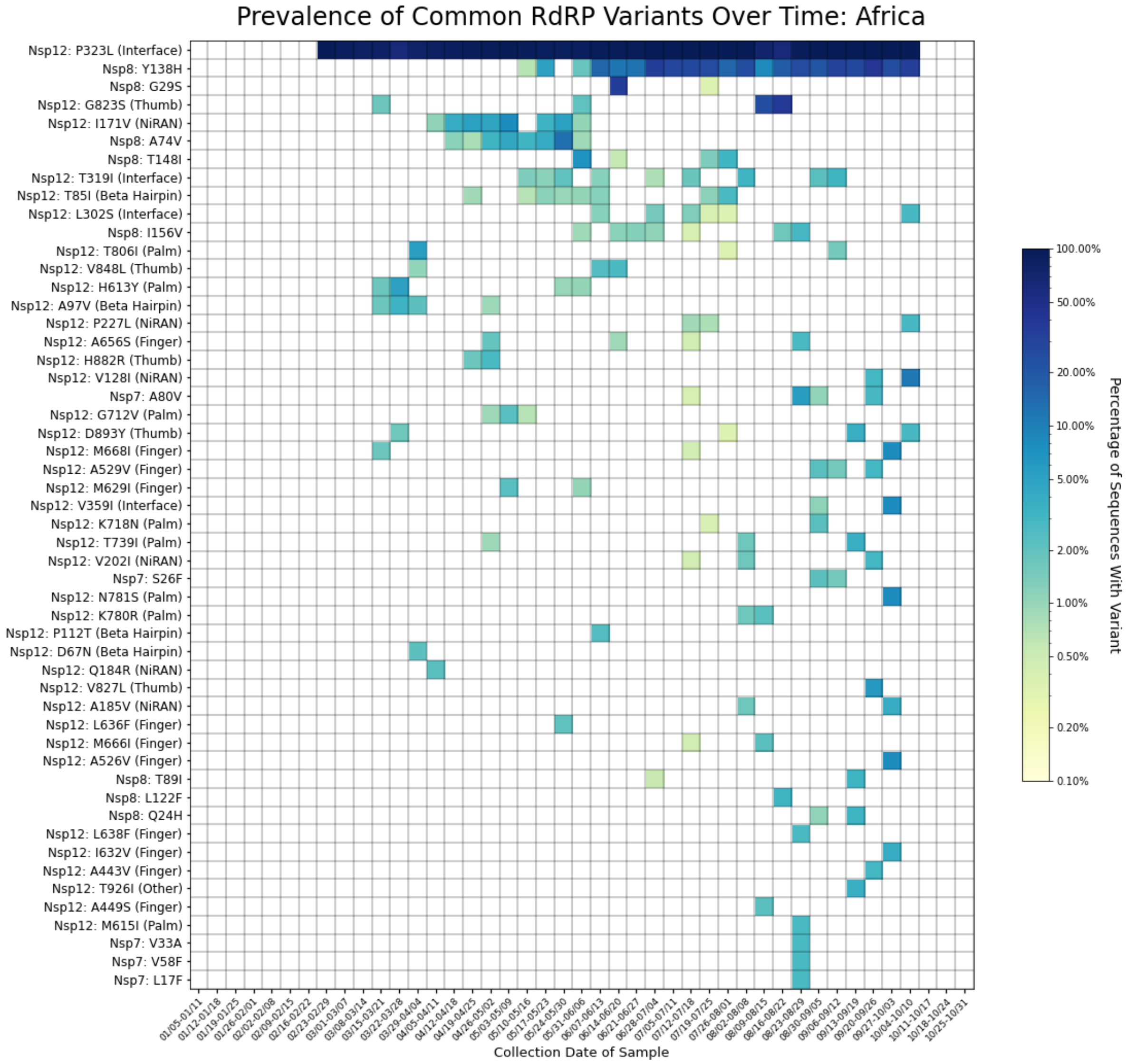


**Supplementary Figure S15:** Heatmap showing time series trends for all variants detected in at least two percent of sequences from Africa in any given week. The heatmap is colored based on a log-10 scale, with prevalence values of zero colored in white, and values less than or equal to 0.10% colored with the lightest shade.

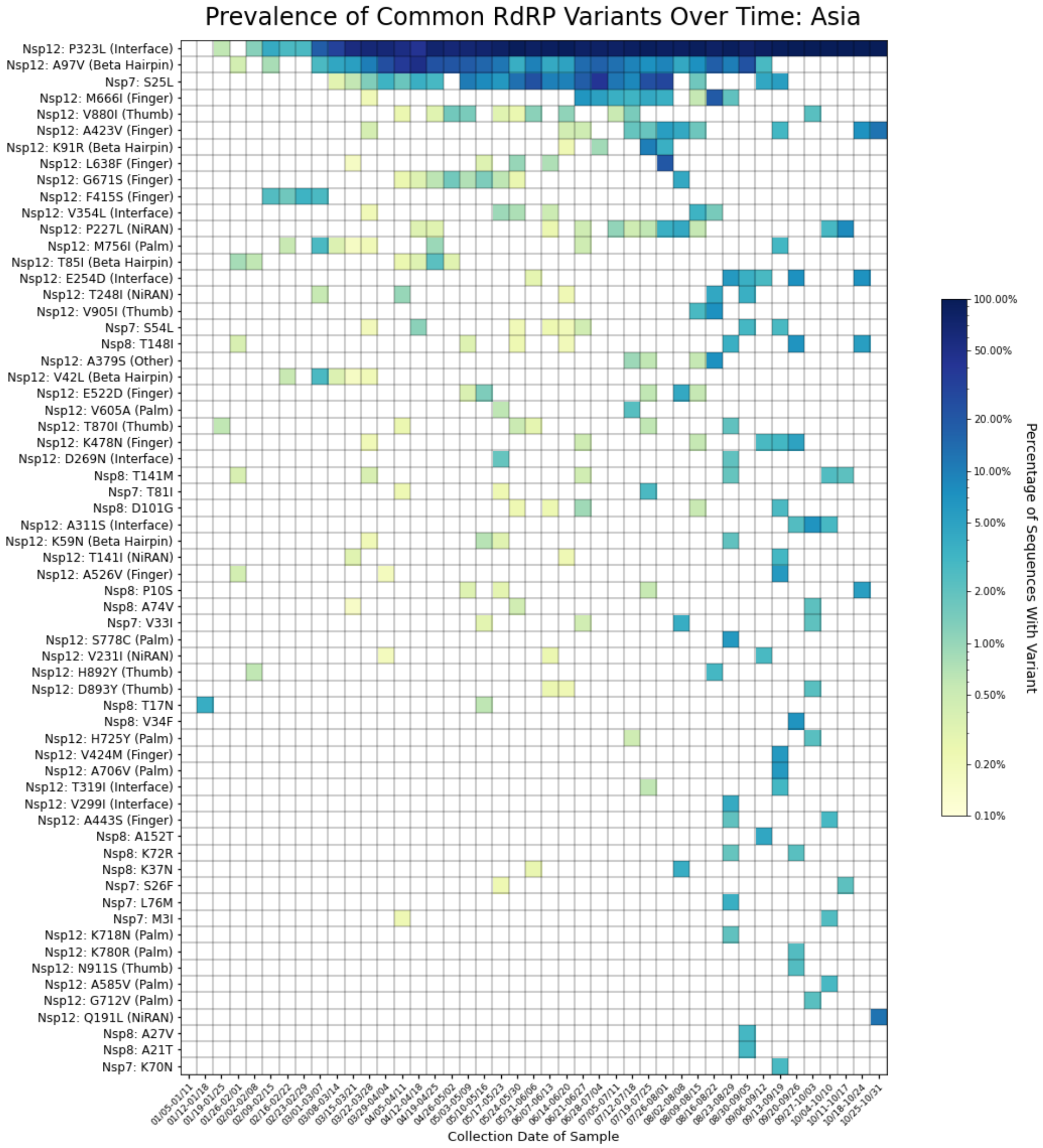


**Supplementary Figure S16:** Heatmap showing time series trends for all variants detected in at least two percent of sequences from Asia in any given week. The heatmap is colored based on a log-10 scale, with prevalence values of zero colored in white, and values less than or equal to 0.10% colored with the lightest shade.

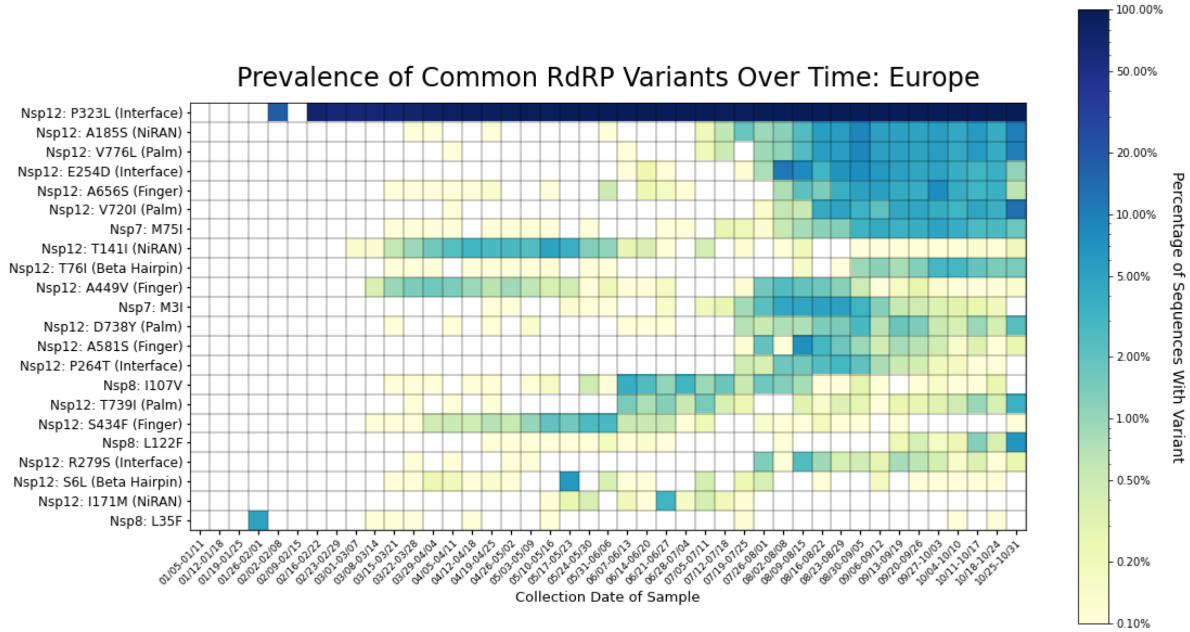


**Supplementary Figure S17:** Heatmap showing time series trends for all variants detected in at least two percent of sequences from Europe in any given week. The heatmap is colored based on a log-10 scale, with prevalence values of zero colored in white, and values less than or equal to 0.10% colored with the lightest shade.

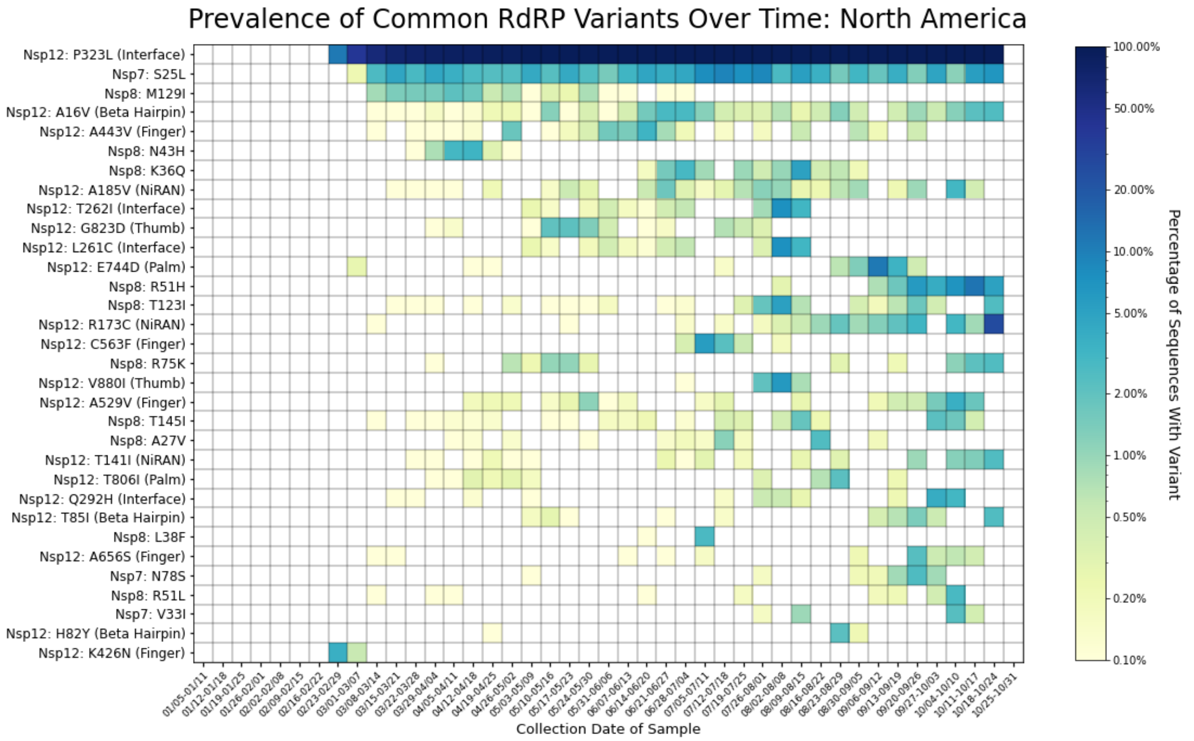


**Supplementary Figure S18:** Heatmap showing time series trends for all variants detected in at least two percent of sequences from North America in any given week. The heatmap is colored based on a log-10 scale, with prevalence values of zero colored in white, and values less than or equal to 0.10% colored with the lightest shade.

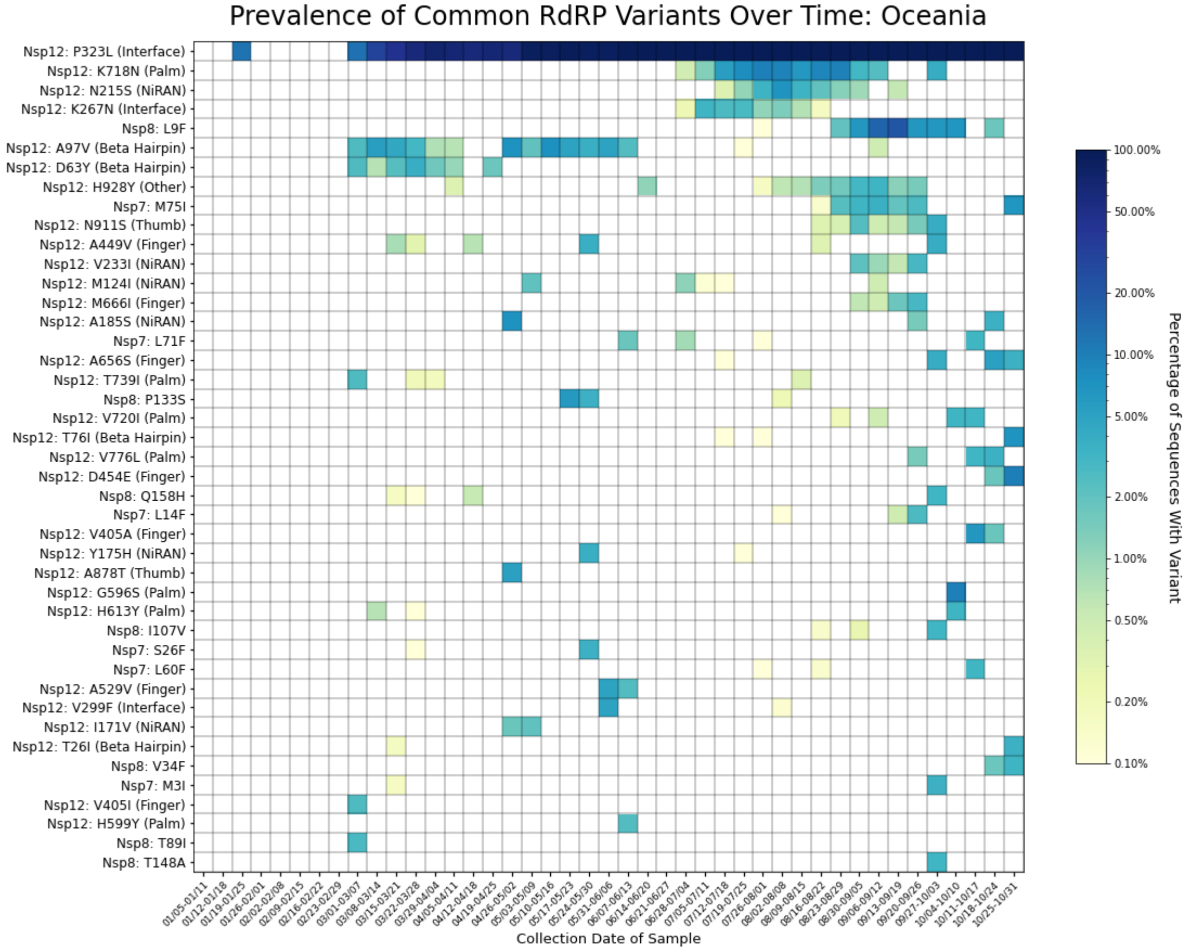


**Supplementary Figure S19:** Heatmap showing time series trends for all variants detected in at least two percent of sequences from Oceania in any given week. The heatmap is colored based on a log-10 scale, with prevalence values of zero colored in white, and values less than or equal to 0.10% colored with the lightest shade.

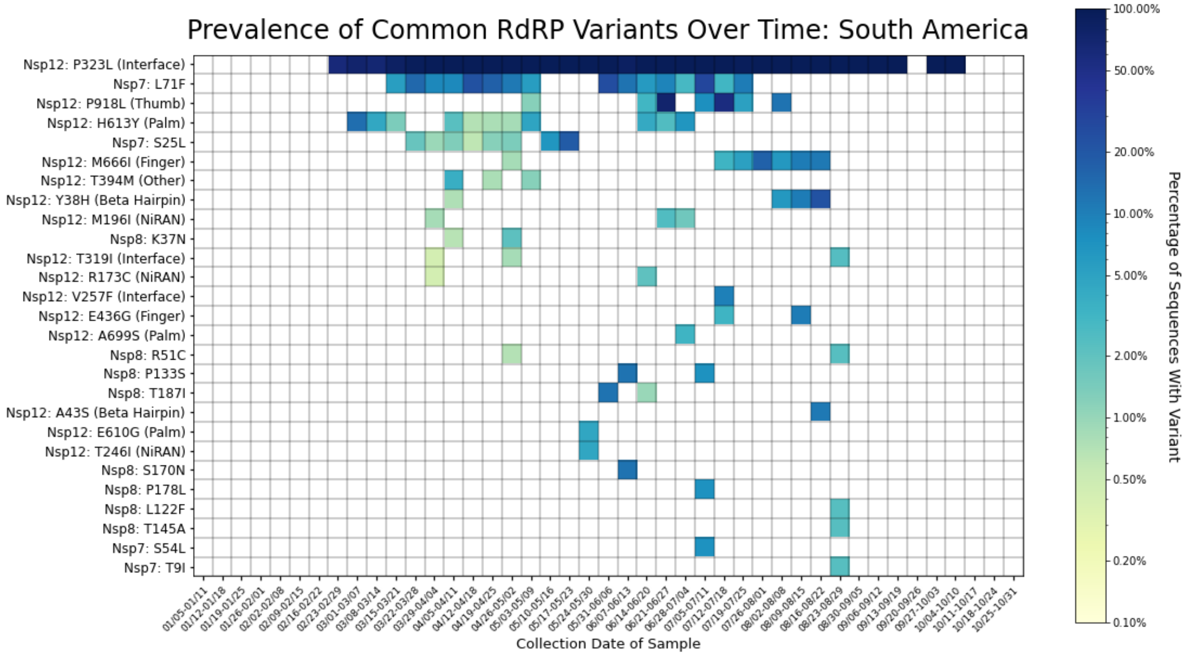


**Supplementary Figure S20:** Heatmap showing time series trends for all variants detected in at least two percent of sequences from South America in any given week. The heatmap is colored based on a log-10 scale, with prevalence values of zero colored in white, and values less than or equal to 0.10% colored with the lightest shade.

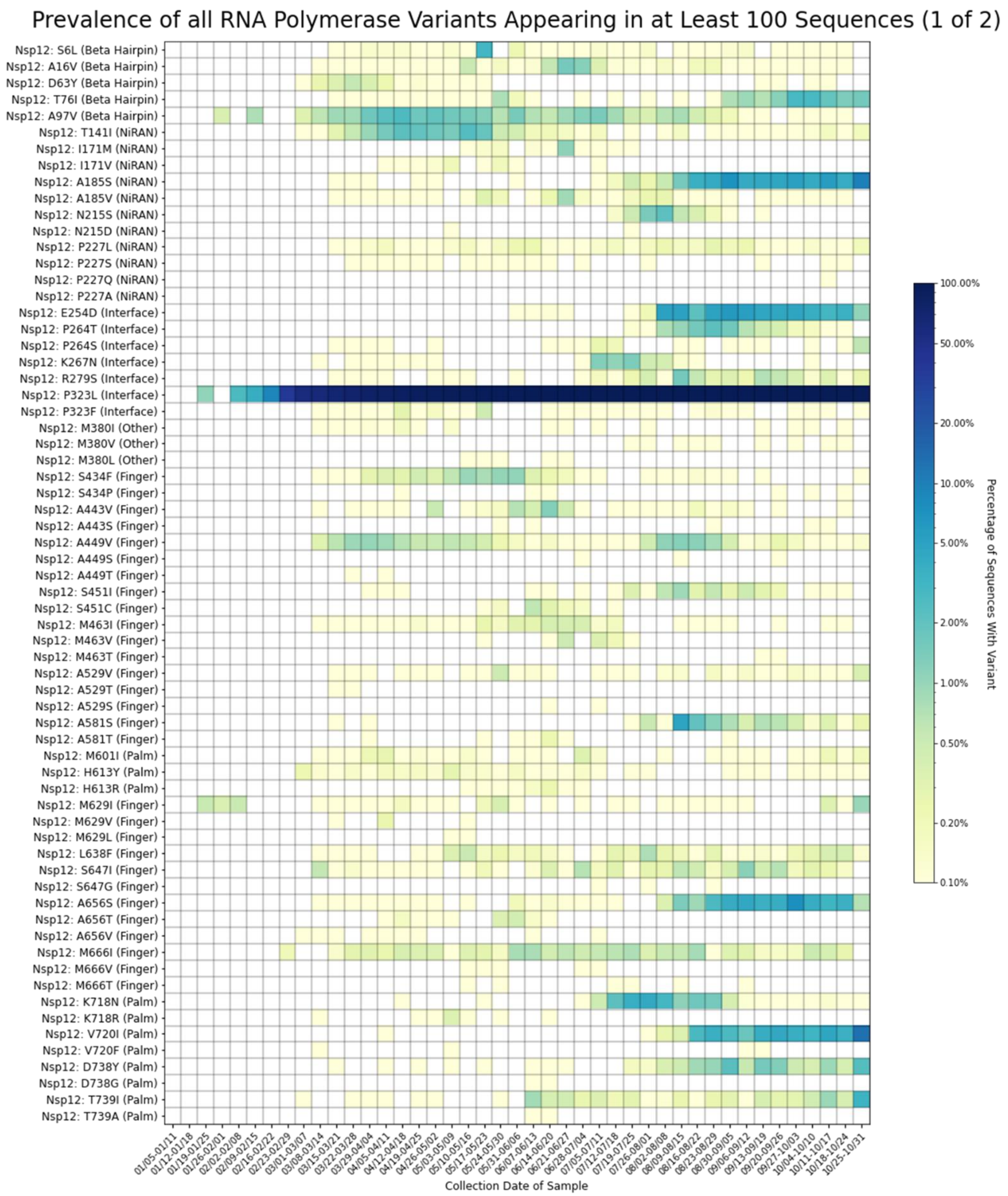


**Supplementary Figure S21:** Heatmap showing time series trends for all amino acid residues mutated in more than 100 sequences across all time points (1 of 2). The heatmap is colored based on a log-10 scale, with prevalence values of zero colored in white, and values less than or equal to 0.10% colored with the lightest shade.

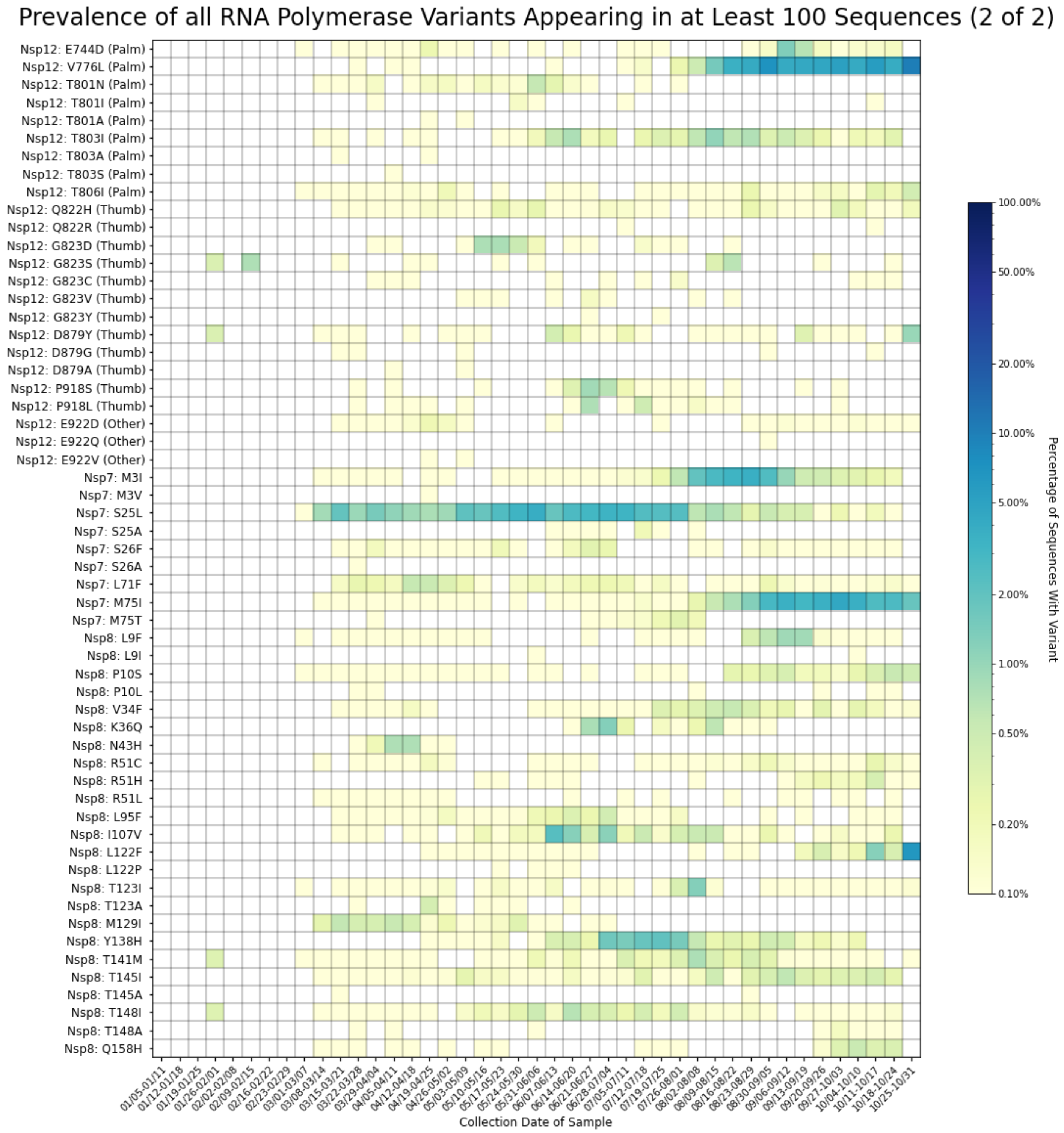


**Supplementary Figure S22:** Heatmap showing time series trends for all amino acid residues mutated in more than 100 sequences across all time points (2 of 2). The heatmap is colored based on a log-10 scale, with prevalence values of zero colored in white, and values less than or equal to 0.10% colored with the lightest shade.
