## Supplementary Table 7 - Acknowledgement of GISAID Data Contributors for "Analysis of SARS-CoV-2 Mutations Over Time Reveals Increasing Prevalence of Variants in the Spike Protein and RNA-Dependent RNA Polymerase"

We gratefully acknowledge the following Authors from the Originating laboratories responsible for obtaining the specimens, as well as the Submitting laboratories where the genome data were generated and shared via GISAID, on which this research is based.

All Submitters of data may be contacted directly via www.gisaid.org

Authors are sorted alphabetically.

| Accession ID | Originating Laboratory | Submitting Laboratory | Authors |
|---|---|---|---|
| EPI_ISL_414477 | The National Institute of Public Health Center for Epidemiology and Microbiology | State Veterinary Institute Prague | Alexander Nagy, Oldrich Bartos, Helena Jirincova, Klara Labska, Ludmila Novakova, Olga Storkanova, Dusan Trnka, Jaromira Vecerova |
| EPI_ISL_414577, EPI_ISL_414578 | Hospital de Talca, Chile | Instituto de Salud Publica de Chile | Andrés E. Castillo, Bárbara Parra, Paz Tapia, Alejandra Acevedo, Jaime Lagos, Winston Andrade, Loredana Arata, Gabriel Leal, Gisselle Barra, Carolina Tambley, Javier Tognarelli, Patricia Bustos, Soledad Ulloa, Rodrigo Fasce, Jorge Fernández. |
| EPI_ISL_414579 | Clinica Alemana de Santiago, Chile | Instituto de Salud Publica de Chile | Andrés E. Castillo, Bárbara Parra, Paz Tapia, Alejandra Acevedo, Jaime Lagos, Winston Andrade, Loredana Arata, Gabriel Leal, Gisselle Barra, Carolina Tambley, Javier Tognarelli, Patricia Bustos, Soledad Ulloa, Rodrigo Fasce, Jorge Fernández. |
| EPI_ISL_414580 | Clinica Santa Maria, Santiago, Chile | Instituto de Salud Publica de Chile | Andrés E. Castillo, Bárbara Parra, Paz Tapia, Alejandra Acevedo, Jaime Lagos, Winston Andrade, Loredana Arata, Gabriel Leal, Gisselle Barra, Carolina Tambley, Javier Tognarelli, Patricia Bustos, Soledad Ulloa, Rodrigo Fasce, Jorge Fernández. |
| EPI_ISL_415710 | WHO National Influenza Centre Russian Federation | WHO National Influenza Centre Russian Federation | Andrey Komissarov, Artem Fadeev, Anna Ivanova, Daria Danilenko |
| EPI_ISL_416482 | R. G. Lugar Center for Public Health Research, National Center for Disease Control and Public Health (NCDC) of Georgia. | R. G. Lugar Center for Public Health Research, National Center for Disease Control and Public Health (NCDC) of Georgia. | Adam Kotorashvili, Marine Murtskhvaladze, Nato Kotaria, Ann Machablishvili, Lela Sabadze, Mari Gavashelidze, Ana Papkiauri, Meri Pantsulaia, Gvantsa Brachveli, Tata Imnadze, Tamar Jashiashvili, Tea Tevdoradze, Ketevan Sidamonidze, Ekaterine Khmaladze, Ekaterine Zhghenti, Roena Sukhiashvili, Mariam Zakalashvili, Lela Urushadze, Magda Dgebuadze, Giorgi Tomashvili, Davit Tsaguria, Ekaterine Zangaladze, Nino Berishvili, Gvantsa Chanturia, Maia Alkhazashvili, Irma Burjanadze, Anna Kasradze, Khatuna Zakhashvili, Paata Imnadze, Amiran Gamkrelidze. |
| EPI_ISL_416488 | ViroGenetics - BSL3 Laboratory of Virology; Human Genome Variation Research Group & Genomics Centre MCB; Bioinformatics Research Group Department of Virology | ViroGenetics - BSL3 Laboratory of Virology; Human Genome Variation Research Group & Genomics Centre MCB; Bioinformatics Research Group Department of Virology | Aleksandra Milewska, Ewelina Popiech, Agata Jarosz, Adrianna Klajmon, Kamila Marszaek, Katarzyna Pancer, Magdalena Rzeczkowska, Tomasz Wokowicz, Katarzyna Zacharczuk, Agnieszka Koakowska-Kulesza, Natalia Wolaniuk, Ewelina Hallman-Szeliska, Pawe P abaj, Wojciech Branicki, Krzysztof Pyr |
| EPI_ISL_416521 | Public Health Laboratory | Public Health Laboratory, Saudi CDC | Albarrag, A |
| EPI_ISL_416522 | Public Health Laboratory, Saudi CDC | Public Health Laboratory, Saudi CDC | Albarrag,A |
| EPI_ISL_417064 | Prince of Wales Hospital | Hong Kong Department of Health | Alan K.L. Tsang, Peter C.W. Yip, Edman T.K. Lam, Rickjason C.W. Chan, Dominic N.C. Tsang |
| EPI_ISL_417186 | National Institute for Communicable Diseases of the National Health Laboratory Service | National Institute for Communicable Diseases of the National Health Laboratory Service | Allam M, Kwenda S, van Heusden P, Khumalo Z, Mohale T, Subramoney K, von Gottberg, A, Ismail A, Bhiman JN |
| EPI_ISL_419654, EPI_ISL_419655, EPI_ISL_419656, EPI_ISL_419657, EPI_ISL_419658, EPI_ISL_419659, EPI_ISL_419660, EPI_ISL_419661, EPI_ISL_419662, EPI_ISL_419663, EPI_ISL_419664, EPI_ISL_419665, EPI_ISL_419666, EPI_ISL_419667, EPI_ISL_419668, EPI_ISL_419669, EPI_ISL_419670, EPI_ISL_419671, EPI_ISL_419672, EPI_ISL_419673, EPI_ISL_419674 | | | |
| see above | Center for Virology, Medical University of Vienna | Bergthaler laboratory, CeMM Research Center for Molecular Medicine of the Austrian Academy of Sciences | Alexandra Popa, Benedikt Agerer, Henrique Colaco, Lukas Endler, Jakob-Wendelin Genger, Alexander Lercher, Mark Smyth, Thomas Penz, Michael Schuster, Judith Aberle, Stephan Aberle, Elisabeth Puchhammer-Stöckl, Christoph Bock, Andreas Bergthaler |
| EPI_ISL_420080 | WHO National Influenza Centre Russian Federation | WHO National Influenza Centre Russian Federation | Andrey Komissarov, Artem Fadeev, Anna Ivanova, Daria Danilenko |
| EPI_ISL_420081 | WHO National Influenza Centre Russian Federation | WHO National Influenza Centre Russian Federation | Andrey Komissarov, Artem Fadeev, Mariia Sergeeva, Anna Ivanova, Daria Danilenko |
| EPI_ISL_421275 | Russian State Collection of Viruses | Pathogenic Microorganisms Variability Laboratory | Alexey Shchetinin, Maria Nikiforova, Nadezhda Kuznetsova, Ekaterina Aksenova, Marina Kunda, Natalia Ryzhova, Olga Voronina, Inna Dolzhikova, Daria Grousova, Andrey Botikov, Denis Logunov, Alexander Gintsburg, Vladimir Gushchin |
| EPI_ISL_421348, EPI_ISL_421349, EPI_ISL_421350, EPI_ISL_421351, EPI_ISL_421352, EPI_ISL_421353, EPI_ISL_421354, EPI_ISL_421355, EPI_ISL_421356, EPI_ISL_421357, EPI_ISL_421358, EPI_ISL_421359, EPI_ISL_421360, EPI_ISL_421361, EPI_ISL_421362, EPI_ISL_421363, EPI_ISL_421364, EPI_ISL_421365, EPI_ISL_421366, EPI_ISL_421367, EPI_ISL_421368, EPI_ISL_421369, EPI_ISL_421370, EPI_ISL_421371, EPI_ISL_421372, EPI_ISL_421373, EPI_ISL_421374, EPI_ISL_421375, EPI_ISL_421376, EPI_ISL_421377, EPI_ISL_421378, EPI_ISL_421379, EPI_ISL_421380, EPI_ISL_421381, EPI_ISL_421382, EPI_ISL_421383, EPI_ISL_421384, EPI_ISL_421385, EPI_ISL_421386, EPI_ISL_421387, EPI_ISL_421388, EPI_ISL_421389, EPI_ISL_421390, EPI_ISL_421391, EPI_ISL_421392, EPI_ISL_421393, EPI_ISL_421394, EPI_ISL_421395, EPI_ISL_421396, EPI_ISL_421397, EPI_ISL_421398, EPI_ISL_421399, EPI_ISL_421400, EPI_ISL_421401, EPI_ISL_421402, EPI_ISL_421403, EPI_ISL_421404, EPI_ISL_421405, EPI_ISL_421406, EPI_ISL_421407, EPI_ISL_421408, EPI_ISL_421409, EPI_ISL_421410, EPI_ISL_421411, EPI_ISL_421412, EPI_ISL_421413, EPI_ISL_421414, EPI_ISL_421415, EPI_ISL_421416, EPI_ISL_421417, EPI_ISL_421418, EPI_ISL_421419, EPI_ISL_421420, EPI_ISL_421421, EPI_ISL_421422, EPI_ISL_421423, EPI_ISL_421424, EPI_ISL_421425, EPI_ISL_421426, EPI_ISL_421427, EPI_ISL_421428, EPI_ISL_421429, EPI_ISL_421430, EPI_ISL_421431, EPI_ISL_421432, EPI_ISL_421433, EPI_ISL_421434, EPI_ISL_421435, EPI_ISL_421593, EPI_ISL_421594, EPI_ISL_421595, EPI_ISL_421596, EPI_ISL_421597, EPI_ISL_421598, EPI_ISL_421599, EPI_ISL_421600, EPI_ISL_421601, EPI_ISL_421602, EPI_ISL_421603, EPI_ISL_421604, EPI_ISL_421605, EPI_ISL_421606, EPI_ISL_421607, EPI_ISL_421608, EPI_ISL_421609, EPI_ISL_421610, EPI_ISL_421611, EPI_ISL_421612, EPI_ISL_421613, EPI_ISL_421614, EPI_ISL_421615, EPI_ISL_421616, EPI_ISL_421617, EPI_ISL_421618, EPI_ISL_421619, EPI_ISL_421620, EPI_ISL_421621, EPI_ISL_421622, EPI_ISL_421623, EPI_ISL_421624, EPI_ISL_421625, EPI_ISL_421626, EPI_ISL_421627, EPI_ISL_421628, EPI_ISL_421629, EPI_ISL_421630, EPI_ISL_421631, EPI_ISL_421632, EPI_ISL_421633, EPI_ISL_421634, EPI_ISL_421635 | | | |
| see above | MSHS Clinical Microbiology Laboratories | MSHS Pathogen Surveillance Program | Ana S. Gonzalez-Reiche, Mitchell Sullivan, Ajay Obla, Gopi Patel, Emilia Sordillo, Melissa Gitman, Alberto Paniz-mondolfi, Matthew Hernandez, Shelcie Fabre, Jose Polanco, Zenab Khan, Bremy Albuquerque, Jayeeta Dutta, Juan Soto, Shwetha Sridhar Hara, Ying-Chih Wang, Melissa Smith, Robert Sebra, Lisa Miorin, Wen-chun Liu, Randy Albrecht, Judith Aberg, Florian Krammer, Adolfo Garcia-Sarstre, Viviana Simon, Harm van Bakel |
| EPI_ISL_422437, EPI_ISL_422438 | ULSS9 Distretto di Bussolengo | Istituto Zooprofilattico Sperimentale delle Venezie | Adelaide Milani, Alessia Schivo, Annalisa Salviato, Erika Giorgia Quaranta, Ambra Pastori, Bianca Zecchin, Alice Fusaro, Isabella Monne, Calogero Terregino, Antonia Ricci |
| EPI_ISL_422488, EPI_ISL_422489, EPI_ISL_422490, EPI_ISL_422491, EPI_ISL_422492, EPI_ISL_422493, EPI_ISL_422494, EPI_ISL_422495, EPI_ISL_422496, EPI_ISL_422497, EPI_ISL_422498, EPI_ISL_422499, EPI_ISL_422500, EPI_ISL_422501, EPI_ISL_422502, EPI_ISL_422503, EPI_ISL_422504, EPI_ISL_422505, EPI_ISL_422506, EPI_ISL_422507, EPI_ISL_422508, EPI_ISL_422509, EPI_ISL_422510, EPI_ISL_422511, EPI_ISL_422512, EPI_ISL_422513, EPI_ISL_422514, EPI_ISL_422515, EPI_ISL_422516, EPI_ISL_422517, EPI_ISL_422518, EPI_ISL_422519, EPI_ISL_422520, EPI_ISL_422521, EPI_ISL_422522, EPI_ISL_422523, EPI_ISL_422524, EPI_ISL_422525, EPI_ISL_422526, EPI_ISL_422527, EPI_ISL_422528, EPI_ISL_422529, EPI_ISL_422530, EPI_ISL_422531, EPI_ISL_422532, EPI_ISL_422533, EPI_ISL_422534, EPI_ISL_422535, EPI_ISL_422536, EPI_ISL_422537, EPI_ISL_422538, EPI_ISL_422539, EPI_ISL_422540, EPI_ISL_422541, EPI_ISL_422542, EPI_ISL_422543, EPI_ISL_422544, EPI_ISL_422545, EPI_ISL_422546, EPI_ISL_422547, EPI_ISL_422548, EPI_ISL_422549, EPI_ISL_422550, EPI_ISL_422551, EPI_ISL_422552, EPI_ISL_422553, EPI_ISL_422554, EPI_ISL_422555, EPI_ISL_422556, EPI_ISL_422557, EPI_ISL_422558, EPI_ISL_422559, EPI_ISL_422560, EPI_ISL_422561, EPI_ISL_422562 | | | |
| see above | MSHS Clinical Microbiology Laboratories | MSHS Pathogen Surveillance Program | Ana S. Gonzalez-Reiche, Mitchell Sullivan, Ajay Obla, Gopi Patel, Emilia Sordillo, Melissa Gitman, Alberto Paniz-mondolfi, Matthew Hernandez, Shelcie Fabre, Jose Polanco, Zenab Khan, Bremy Albuquerque, Jayeeta Dutta, Juan Soto, Shwetha Sridhar Hara, Ying-Chih Wang, Melissa Smith, Robert Sebra, Lisa Miorin, Wen-chun Liu, Randy Albrecht, Judith Aberg, Florian Krammer, Adolfo Garcia-Sarstre, Viviana Simon, Harm van Bakel |
| EPI_ISL_422636 | The National Institute of Public Health Center for Epidemiology and Microbiology | State Veterinary Institute Prague | Alexander Nagy, Helena Jirincova, Klara Labska, Ludmila Novakova, Olga Storkanova, Dusan Trnka, Jaromira Vecerova |
| EPI_ISL_424673 | Instituto de Diagnostico y Referencia Epidemiologicos | Instituto de Diagnostico y Referencia Epidemiologicos | Adnan Araiza Rodríguez, Edgar Mendieta Condado,Fabiola Garcés Ayala, Gisela Barrera Badillo, Irma López Martínez, Lucia Hernández Rivas, Celia Boukadida, Santiago Avila Ríos, Mario Mújica Sánchez, José Arturo Martínez Orozco,Eduardo Becerril Vargas, Joel Armando Vázquez Pérez, Victor Hugo Borja Aburto, Concepción Grajales Muñiz, Cesar Raúl González Bonilla, Carolina González Torres, Francisco Javier Gaytán Cervantes, José Esteban Muñoz Medina, Guillermo M. Ruiz-Palacios, Pilar Ramos Cervantes, Violeta Ibarra Gonzalez, Fernando Ledesma Barrientos, Luis Alberto García Andrade, Alfredo Ponce de León Garduño, Blanca Taboada, Alejandro Sánchez, Pavel Isa, Ricardo Grande, Gloria Vázquez, Francisco Pulido, Carlos F. Arias,José Ernesto Ramírez González |
| EPI_ISL_425177 | Public Health Ontario | Public Health Agency of Canada - National Microbiology Laboratory | Amrit S. Boese, Nikesh Tailor, Anders Leung, Joshua Quick, Shari Tyson, Morag Graham, Jonathan Audet, Natalie Knox, Darwyn Kobasa |
| EPI_ISL_425647, EPI_ISL_425648, EPI_ISL_425649, EPI_ISL_425650, EPI_ISL_425651, EPI_ISL_425652, EPI_ISL_425653, EPI_ISL_425654, EPI_ISL_425655, EPI_ISL_425656, EPI_ISL_425657, EPI_ISL_425658, EPI_ISL_425659, EPI_ISL_425660, EPI_ISL_425661, EPI_ISL_425662, EPI_ISL_425663, EPI_ISL_425664, EPI_ISL_425665, EPI_ISL_425666, EPI_ISL_425667, EPI_ISL_425668, EPI_ISL_425669, EPI_ISL_425670, EPI_ISL_425671, EPI_ISL_425672, EPI_ISL_425673, EPI_ISL_425674, EPI_ISL_425675, EPI_ISL_425676, EPI_ISL_425677, EPI_ISL_425678, EPI_ISL_425679, EPI_ISL_425680, EPI_ISL_425681, EPI_ISL_425682, EPI_ISL_425683, EPI_ISL_425684, EPI_ISL_425685, EPI_ISL_425686, EPI_ISL_425687, EPI_ISL_425688, EPI_ISL_425689, EPI_ISL_425690, EPI_ISL_425691, EPI_ISL_425692, EPI_ISL_425693, EPI_ISL_425694, EPI_ISL_425695, EPI_ISL_425696, EPI_ISL_425697, EPI_ISL_425698, EPI_ISL_425699, EPI_ISL_425700, | | | |

| | | | |
|---|---|---|---|
| EPI_ISL_425701, EPI_ISL_425702, EPI_ISL_425703, EPI_ISL_425704, EPI_ISL_425705, EPI_ISL_425706, EPI_ISL_425707, EPI_ISL_425708, EPI_ISL_425709, EPI_ISL_425710, EPI_ISL_425711, EPI_ISL_425712, EPI_ISL_425713, EPI_ISL_425714, EPI_ISL_425715, EPI_ISL_425716, EPI_ISL_425717, EPI_ISL_425718, EPI_ISL_425719, EPI_ISL_425720, EPI_ISL_425721, EPI_ISL_425722, EPI_ISL_425723, EPI_ISL_425724, EPI_ISL_425725, EPI_ISL_425726, EPI_ISL_425727, EPI_ISL_425728, EPI_ISL_425729, EPI_ISL_425730, EPI_ISL_425731, EPI_ISL_425732, EPI_ISL_425733, EPI_ISL_425734, EPI_ISL_425735, EPI_ISL_425736, EPI_ISL_425737, EPI_ISL_425738, EPI_ISL_425739, EPI_ISL_425740, EPI_ISL_425741, EPI_ISL_425742, EPI_ISL_425743, EPI_ISL_425744, EPI_ISL_425745, EPI_ISL_425746, EPI_ISL_425747, EPI_ISL_425748, EPI_ISL_425749, EPI_ISL_425750, EPI_ISL_425751, EPI_ISL_425752, EPI_ISL_425753, EPI_ISL_425754, EPI_ISL_425755, EPI_ISL_425756, EPI_ISL_425757, EPI_ISL_425758, EPI_ISL_425759, EPI_ISL_425760, EPI_ISL_425761, EPI_ISL_425762, EPI_ISL_425763, EPI_ISL_425764, EPI_ISL_425765, EPI_ISL_425766, EPI_ISL_425767, EPI_ISL_425768, EPI_ISL_425769, EPI_ISL_425770, EPI_ISL_425771, EPI_ISL_425772, EPI_ISL_425773, EPI_ISL_425774, EPI_ISL_425775, EPI_ISL_425776, EPI_ISL_425777, EPI_ISL_425778, EPI_ISL_425779, EPI_ISL_425780, EPI_ISL_425781, EPI_ISL_425782, EPI_ISL_425783, EPI_ISL_425784, EPI_ISL_425785, EPI_ISL_425786, EPI_ISL_425787, EPI_ISL_425788, EPI_ISL_425789, EPI_ISL_425790, EPI_ISL_425791, EPI_ISL_425792, EPI_ISL_425793, EPI_ISL_425794, EPI_ISL_425795, EPI_ISL_425796, EPI_ISL_425797, EPI_ISL_425798, EPI_ISL_425799, EPI_ISL_425800, EPI_ISL_425801, EPI_ISL_425802, EPI_ISL_425803, EPI_ISL_425804, EPI_ISL_425805, EPI_ISL_425806, EPI_ISL_425807, EPI_ISL_425808, EPI_ISL_425809, EPI_ISL_425810, EPI_ISL_425811, EPI_ISL_425812, EPI_ISL_425813, EPI_ISL_425814, EPI_ISL_425815, EPI_ISL_425816, EPI_ISL_425817, EPI_ISL_425818 | | | |
| see above | West of Scotland Specialist Virology Centre, NHSGGC / MRC-University of Glasgow Centre for Virus Research | COVID-19 Genomics UK (COG-UK) Consortium | Ana da Silva Filipe, Kathy Smollett, Stephen Carmichael, Natasha Johnson, Daniel Mair, Lily Tong, Jenna Nichols; Sarah McDonald; Richard Orton, Joseph Hughes, Sreenu Vattipally, David L Robertson; Kathy Li, Natasha Jesudason, Rajiv Shah, James Shepherd, Antonia Ho, Emma Thomson; Alasdair MacLean, Rory Gunson. |
| EPI_ISL_426627, EPI_ISL_426628 | Ochsner Health | BioInfoExperts, LLC | Amy Feehan, David Nolan, Rebecca Rose, Susanna Lamers, Sissy Cross, Julia-Garcia-Diaz, Tong Yang, Luke Caruso, David Moraga Amador, Wayra Navia, Lydia Von Borstel, Xiao Hui Zhou |
| EPI_ISL_427307, EPI_ISL_427308, EPI_ISL_427309, EPI_ISL_427310, EPI_ISL_427311, EPI_ISL_427312, EPI_ISL_427313, EPI_ISL_427314, EPI_ISL_427315, EPI_ISL_427316, EPI_ISL_427317, EPI_ISL_427318, EPI_ISL_427319, EPI_ISL_427320, EPI_ISL_427321, EPI_ISL_427322, EPI_ISL_427323, EPI_ISL_427324, EPI_ISL_427325, EPI_ISL_427326, EPI_ISL_427327, EPI_ISL_427328, EPI_ISL_427329, EPI_ISL_427330, EPI_ISL_427331, EPI_ISL_427332, EPI_ISL_427333, EPI_ISL_427334, EPI_ISL_427335, EPI_ISL_427336, EPI_ISL_427337, EPI_ISL_427338, EPI_ISL_427339 | | | |
| see above | WHO National Influenza Centre Russian Federation | WHO National Influenza Centre Russian Federation | Andrey Komissarov, Artem Fadeev, Mariia Sergeeva, Anna Ivanova, Daria Danilenko |
| EPI_ISL_427404, EPI_ISL_427405, EPI_ISL_427406, EPI_ISL_427407, EPI_ISL_427408, EPI_ISL_427409, EPI_ISL_427410, EPI_ISL_427411, EPI_ISL_427412, EPI_ISL_427414, EPI_ISL_427415, EPI_ISL_427416, EPI_ISL_427417, EPI_ISL_427418, EPI_ISL_427419, EPI_ISL_427420 | | | |
| see above | Ministry of Public Health (MoPH) | Biomedical Research Center (BRC) | Abdullatif Al-Khal, Muna A. S. Al-Maslamani, Ajaeb D. M. H. Al-Nabet, Peter V. Coyle, Einas A. E. Al-Kuwari, Nourah B. M. Younes, Hamad E. Al-Romaihi, Salih Al-Marri, Mohammed Al-Thani, Fatiha M. Benslimane, Heba A. Al-Khatib, Sonia Boughattas, Hadi M. Yassine, Asmaa A. Al-Thani. |
| EPI_ISL_427815 | WHO National Influenza Centre Russian Federation | WHO National Influenza Centre Russian Federation | Andrey Komissarov, Artem Fadeev, Anna Ivanova, Daria Danilenko |
| EPI_ISL_429076, EPI_ISL_429078, EPI_ISL_429079, EPI_ISL_429080, EPI_ISL_429081, EPI_ISL_429084, EPI_ISL_429086, EPI_ISL_429094, EPI_ISL_429095, EPI_ISL_429099, EPI_ISL_429100, EPI_ISL_429101, EPI_ISL_429102, EPI_ISL_429103, EPI_ISL_429104, EPI_ISL_429105 | | | |
| see above | The First Affiliated Hospital of Guangzhou Medical University | BGI-shenzhen & The First Affiliated Hospital of Guangzhou Medical University | |
| EPI_ISL_430067, EPI_ISL_430068, EPI_ISL_430069, EPI_ISL_430070, EPI_ISL_430071, EPI_ISL_430072, EPI_ISL_430073, EPI_ISL_430074, EPI_ISL_430075, EPI_ISL_430076, EPI_ISL_430077, EPI_ISL_430078, EPI_ISL_430079, EPI_ISL_430080, EPI_ISL_430081, EPI_ISL_430082, EPI_ISL_430083, EPI_ISL_430084, EPI_ISL_430085, EPI_ISL_430086, EPI_ISL_430087, EPI_ISL_430088, EPI_ISL_430089, EPI_ISL_430090, EPI_ISL_430091, EPI_ISL_430092, EPI_ISL_430093, EPI_ISL_430094, EPI_ISL_430095, EPI_ISL_430096, EPI_ISL_430097, EPI_ISL_430098, EPI_ISL_430099, EPI_ISL_430100, EPI_ISL_430101, EPI_ISL_430102, EPI_ISL_430103, EPI_ISL_430104, EPI_ISL_430105, EPI_ISL_430106, EPI_ISL_430107, EPI_ISL_430108, EPI_ISL_430109, EPI_ISL_430110, EPI_ISL_430111, EPI_ISL_430112 | | | |
| see above | WHO National Influenza Centre Russian Federation | WHO National Influenza Centre Russian Federation | Andrey Komissarov, Artem Fadeev, Mariia Sergeeva, Anna Ivanova, Daria Danilenko |
| EPI_ISL_430297 | National Institute for Communicable Diseases of the National Health Laboratory Service | National Institute for Communicable Diseases of the National Health Laboratory Service | Allam M, Kwenda S, van Heusden P, Khumalo Z, Mohale T, Subramoney K, von Gottberg, A, Ismail A, Bhiman JN |
| EPI_ISL_433268, EPI_ISL_433269, EPI_ISL_433270, EPI_ISL_433271, EPI_ISL_433272, EPI_ISL_433273, EPI_ISL_433274, EPI_ISL_433275, EPI_ISL_433276, EPI_ISL_433277, EPI_ISL_433278, EPI_ISL_433279, EPI_ISL_433280, EPI_ISL_433281, EPI_ISL_433282, EPI_ISL_433283, EPI_ISL_433284, EPI_ISL_433285, EPI_ISL_433286, EPI_ISL_433287, EPI_ISL_433288, EPI_ISL_433289, EPI_ISL_433290, EPI_ISL_433291, EPI_ISL_433292, EPI_ISL_433293, EPI_ISL_433294, EPI_ISL_433295, EPI_ISL_433296, EPI_ISL_433297, EPI_ISL_433298, EPI_ISL_433299, EPI_ISL_433300, EPI_ISL_433301, EPI_ISL_433302, EPI_ISL_433303, EPI_ISL_433304, EPI_ISL_433305, EPI_ISL_433306, EPI_ISL_433307, EPI_ISL_433308, EPI_ISL_433309, EPI_ISL_433310, EPI_ISL_433311, EPI_ISL_433312, EPI_ISL_433313, EPI_ISL_433314, EPI_ISL_433315, EPI_ISL_433316, EPI_ISL_433317, EPI_ISL_433318, EPI_ISL_433319, EPI_ISL_433320, EPI_ISL_433321, EPI_ISL_433322, EPI_ISL_433323, EPI_ISL_433324, EPI_ISL_433325, EPI_ISL_433326, EPI_ISL_433327, EPI_ISL_433328, EPI_ISL_433329, EPI_ISL_433330, EPI_ISL_433331, EPI_ISL_433332, EPI_ISL_433333, EPI_ISL_433334, EPI_ISL_433335, EPI_ISL_433336, EPI_ISL_433337, EPI_ISL_433338, EPI_ISL_433339, EPI_ISL_433340, EPI_ISL_433341, EPI_ISL_433342, EPI_ISL_433343, EPI_ISL_433344, EPI_ISL_433345, EPI_ISL_433346, EPI_ISL_433347, EPI_ISL_433348, EPI_ISL_433349, EPI_ISL_433350, EPI_ISL_433351, EPI_ISL_433352, EPI_ISL_433353, EPI_ISL_433354, EPI_ISL_433355, EPI_ISL_433356, EPI_ISL_433357, EPI_ISL_433358, EPI_ISL_433359, EPI_ISL_433360, EPI_ISL_433361, EPI_ISL_433362, EPI_ISL_433363, EPI_ISL_433364, EPI_ISL_433365, EPI_ISL_433366, EPI_ISL_433367, EPI_ISL_433368, EPI_ISL_433369, EPI_ISL_433370, EPI_ISL_433371, EPI_ISL_433372, EPI_ISL_433373, EPI_ISL_433498, EPI_ISL_433499, EPI_ISL_433500, EPI_ISL_433501, EPI_ISL_433502, EPI_ISL_433503, EPI_ISL_433504, EPI_ISL_433505, EPI_ISL_433506, EPI_ISL_433507, EPI_ISL_433508, EPI_ISL_433509, EPI_ISL_433510, EPI_ISL_433511, EPI_ISL_433512, EPI_ISL_433513, EPI_ISL_433514, EPI_ISL_433515, EPI_ISL_433516, EPI_ISL_433517, EPI_ISL_433518, EPI_ISL_433519, EPI_ISL_433520, EPI_ISL_433521, EPI_ISL_433522, EPI_ISL_433523, EPI_ISL_433524, EPI_ISL_433525, EPI_ISL_433526, EPI_ISL_433527, EPI_ISL_433528, EPI_ISL_433529, EPI_ISL_433530, EPI_ISL_433531, EPI_ISL_433532, EPI_ISL_433533, EPI_ISL_433534, EPI_ISL_433535, EPI_ISL_433536, EPI_ISL_433537, EPI_ISL_433538, EPI_ISL_433539, EPI_ISL_433540, EPI_ISL_433541, EPI_ISL_433542, EPI_ISL_433543, EPI_ISL_433544, EPI_ISL_433545, EPI_ISL_433546, EPI_ISL_433547, EPI_ISL_433548, EPI_ISL_433549, EPI_ISL_433550, EPI_ISL_433551, EPI_ISL_433552, EPI_ISL_433553, EPI_ISL_433554, EPI_ISL_433555, EPI_ISL_433556, EPI_ISL_433557, EPI_ISL_433558, EPI_ISL_433559, EPI_ISL_433560, EPI_ISL_433561, EPI_ISL_433562, EPI_ISL_433563, EPI_ISL_433564, EPI_ISL_433565, EPI_ISL_433566, EPI_ISL_433567, EPI_ISL_433568, EPI_ISL_433569, EPI_ISL_433570, EPI_ISL_433571, EPI_ISL_433572, EPI_ISL_433573, EPI_ISL_433574, EPI_ISL_433575, EPI_ISL_433576, EPI_ISL_433577, EPI_ISL_433578, EPI_ISL_433579, EPI_ISL_433580, EPI_ISL_433581, EPI_ISL_433582, EPI_ISL_433583, EPI_ISL_433584, EPI_ISL_433585, EPI_ISL_433586, EPI_ISL_433587, EPI_ISL_433588, EPI_ISL_433589, EPI_ISL_433590, EPI_ISL_433591, EPI_ISL_433592, EPI_ISL_433593, EPI_ISL_433594, EPI_ISL_433595, EPI_ISL_433596, EPI_ISL_433597, EPI_ISL_433598, EPI_ISL_433599, EPI_ISL_433600, EPI_ISL_433601, EPI_ISL_433602, EPI_ISL_433603, EPI_ISL_433604, EPI_ISL_433605, EPI_ISL_433606, EPI_ISL_433607, EPI_ISL_433608, EPI_ISL_433609, EPI_ISL_433610, EPI_ISL_433611, EPI_ISL_433612, EPI_ISL_433613, EPI_ISL_433614, EPI_ISL_433615, EPI_ISL_433616, EPI_ISL_433617, EPI_ISL_433618, EPI_ISL_433619, EPI_ISL_433620, EPI_ISL_433621, EPI_ISL_433622, EPI_ISL_433623, EPI_ISL_433624, EPI_ISL_433625, EPI_ISL_433626, EPI_ISL_433627, EPI_ISL_433628, EPI_ISL_433629, EPI_ISL_433630, EPI_ISL_433631, EPI_ISL_433632, EPI_ISL_433633, EPI_ISL_433634, EPI_ISL_433635, EPI_ISL_433636, EPI_ISL_433637, EPI_ISL_433638, EPI_ISL_433639, EPI_ISL_433640, EPI_ISL_433641, EPI_ISL_433642, EPI_ISL_433643, EPI_ISL_433644, EPI_ISL_433645, EPI_ISL_433646, EPI_ISL_433647, EPI_ISL_433648, EPI_ISL_433649, EPI_ISL_433650, EPI_ISL_433651, EPI_ISL_433652, EPI_ISL_433653, EPI_ISL_433654, EPI_ISL_433655, EPI_ISL_433656, EPI_ISL_433657, EPI_ISL_433658, EPI_ISL_433659, EPI_ISL_433660, EPI_ISL_433661, EPI_ISL_433662, EPI_ISL_433663, EPI_ISL_433664, EPI_ISL_433665 | | | |
| see above | West of Scotland Specialist Virology Centre, NHSGGC / MRC-University of Glasgow Centre for Virus Research | COVID-19 Genomics UK (COG-UK) Consortium | Ana da Silva Filipe, Natasha Johnson, Kathy Smollett, Daniel Mair, Stephen Carmichael, Lily Tong, Jenna Nichols, Elihu Aranday-Cortes, Kirstyn Brunker, Yasmin Parr, Kyriaki Nomikou; Sarah McDonald, Marc Niebel, Patawee Asamaphan; Richard Orton, Joseph Hughes, Sreenu Vattipally, David L Robertson; Alasdair MacLean, Rory Gunson; Kathy Li, Natasha Jesudason, Rajiv Shah, James Shepherd, Antonia Ho, Emma Thomson |
| EPI_ISL_434517, EPI_ISL_434518, EPI_ISL_434519, EPI_ISL_434520, EPI_ISL_434521, EPI_ISL_434522, EPI_ISL_434523, EPI_ISL_434524, EPI_ISL_434525, EPI_ISL_434526, EPI_ISL_434527, EPI_ISL_434528, EPI_ISL_434529, EPI_ISL_434530, EPI_ISL_434531, EPI_ISL_434532 | | | |
| see above | Robert Garry lab | Andersen lab at Scripps Research | Allison Smither, Gilberto Sabino-Santos, Patricia Snarski, Lilia Melnik, Antoinette Bell, Kaylynn Genemaras, Arnaud Drouin, Dahlene Fusco, Robert Garry with SEARCH Alliance San Diego |
| EPI_ISL_434572 | The National Institute of Public Health Center for Epidemiology and Microbiology | The National Institute of Public Health Center for Epidemiology and Microbiology | Alexander Nagy, Helena Jirincova, Ludmila Novakova, Dusan Trnka, Jaromira Vecerova |
| EPI_ISL_434653, EPI_ISL_434656 | Trollbackens VC | The Public Health Agency of Sweden | Amelie Holmqvist, Oskar Karlsson Lindsjo, Maria Lind Karlberg, Anna-Malin Linde, Olov Svartstrom, Anna Risberg, Theresa Enkirch, Mia Brytting, Karin Tegmark-Wisell |
| EPI_ISL_435058, EPI_ISL_435059 | National Institute for Communicable Diseases of the National Health Laboratory Service | National Institute for Communicable Diseases of the National Health Laboratory Service | Allam M, Kwenda S, van Heusden P, Khumalo Z, Mohale T, Subramoney K, von Gottberg, A, Ismail A, Bhiman JN |
| EPI_ISL_435119 | Mohammed Bin Rashid University of Medicine and Health Sciences | Al Jalila Children's Hospital | Ahmad Abou Tayoun, Tom Loney, Hamda Khansaheb, Sathishkumar Ramaswamy, Divinlal Harilal, Zulfa Omar Deesi, Rupa Murthy Varghese, Hanan Al Suwaidi, Abdulmajeed Alkhaja, Mohammed Uddin, Rifat Hamoudi, Rabih Halwani, Abiola Catherine Senok, Qutayba Hamid, Norbert Nowotny, Alawi Alsheikh-Ali |
| EPI_ISL_435120, EPI_ISL_435121, EPI_ISL_435122, EPI_ISL_435123, EPI_ISL_435124, EPI_ISL_435125, EPI_ISL_435126, EPI_ISL_435127, EPI_ISL_435128, EPI_ISL_435129, EPI_ISL_435130, EPI_ISL_435131, EPI_ISL_435132, EPI_ISL_435133, EPI_ISL_435134, EPI_ISL_435135, EPI_ISL_435136, EPI_ISL_435137, EPI_ISL_435138, EPI_ISL_435139, EPI_ISL_435140, EPI_ISL_435141, EPI_ISL_435142, EPI_ISL_435143 | | | |
| see above | Mohammed Bin Rashid University of Medicine and Health Sciences | Al Jalila Genomics Center | Ahmad Abou Tayoun, Tom Loney, Hamda Khansaheb, Sathishkumar Ramaswamy, Divinlal Harilal, Zulfa Omar Deesi, Rupa Murthy Varghese, Hanan Al Suwaidi, Abdulmajeed Alkhaja, Mohammed Uddin, Rifat Hamoudi, Rabih Halwani, Abiola Catherine Senok, Qutayba Hamid, Norbert Nowotny, Alawi Alsheikh-Ali |
| EPI_ISL_435445, EPI_ISL_435446, EPI_ISL_435447, EPI_ISL_435448, EPI_ISL_435449, EPI_ISL_435450, EPI_ISL_435451, EPI_ISL_435452, EPI_ISL_435453, EPI_ISL_435454, EPI_ISL_435455, EPI_ISL_435456, EPI_ISL_435457, EPI_ISL_435458, EPI_ISL_435459, EPI_ISL_435460, EPI_ISL_435461, EPI_ISL_435462, EPI_ISL_435463, EPI_ISL_435464, EPI_ISL_435465, EPI_ISL_435466, EPI_ISL_435467, EPI_ISL_435468, EPI_ISL_435469, EPI_ISL_435470, EPI_ISL_435471, EPI_ISL_435472 | | | |
| see above | Robert Garry lab | Andersen lab at Scripps Research | Allison Smither, Gilberto Sabino-Santos, Patricia Snarski, Lilia Melnik, Antoinette Bell, Kaylynn Genemaras, Arnaud Drouin, Dahlene Fusco, Robert Garry with SEARCH Alliance San Diego |
| EPI_ISL_436097 | Prince Charles Hospital | Public Health Virology Laboratory, Forensics and Scientific Services, Queensland Health | Alyssa Pyke, Neelima Nair, Natalie Simpson, Lisa Leckie, Jamie McMahon, Jean Barcelon, Amanda De Jong, Sean Moody, Doris Genge, Glen Hewitson, Peter Burtonclay, Judy Northill, Ian Maxwell Mackay, Carmel Taylor, Bixing Huang, David Warrilow, Mitchell Finger, Peter Moore, Sarah Wheatley, Sonja Hall-Mendelin, Andrew Van Den Hurk, Elisabeth Gamez, Inga Sultana and Frederick Moore |
| EPI_ISL_436098 | Royal Brisbane and Women's Hospital | Public Health Virology Laboratory, Forensic and Scientific Services, Queensland Health | Alyssa Pyke, Neelima Nair, Natalie Simpson, Lisa Leckie, Jamie McMahon, Jean Barcelon, Amanda De Jong, Sean Moody, Doris Genge, Glen Hewitson, Peter Burtonclay, Judy Northill, Ian Maxwell Mackay, Carmel Taylor, Bixing Huang, David Warrilow, Mitchell Finger, Peter Moore, Sarah Wheatley, Sonja Hall-Mendelin, Andrew Van Den Hurk, Elisabeth Gamez, Inga Sultana and Frederick Moore |

| | | | |
|---|---|---|---|
| EPI_ISL_436100 | TSGH-CP molecular lab | TSGH-CP molecular lab | Cherng-Lih Perng, Ming-Jr Jian, Chih-Kai Chang, Jung-Chung Lin, Kuo-Ming Yeh, Chien-Wen Chen, Sheng-Kang Chiu, Hsing-Yi Chung, Shih-Hung Tsai, Kuo-Sheng Hung, Tien-Yao Chang, Feng-Yee Chang, Hung-Sheng Shang |
| EPI_ISL_436715, EPI_ISL_436716, EPI_ISL_436717 | Genomics and Computational Biology Lab, Scientific Research Institute of Physical-Chemical Medicine, FMBA of Russia | Genomics and Computational Biology Lab, Scientific Research Institute of Physical-Chemical Medicine, FMBA of Russia | A. Pavlenko, O. Guskova, K. Klimina, V. Veselovsky, A. Manolov, D. Fedorov, V. Govorun and E. Ilina |
| EPI_ISL_436939, EPI_ISL_436940, EPI_ISL_436941, EPI_ISL_436942, EPI_ISL_436943, EPI_ISL_436944, EPI_ISL_436945, EPI_ISL_436946, EPI_ISL_436947, EPI_ISL_436948, EPI_ISL_436949, EPI_ISL_436950, EPI_ISL_436951, EPI_ISL_436952, EPI_ISL_436953, EPI_ISL_436954, EPI_ISL_436955, EPI_ISL_436956, EPI_ISL_436957, EPI_ISL_436958, EPI_ISL_436959, EPI_ISL_436960, EPI_ISL_436961 | | | |
| see above | Ochsner Health | Bioinfoexperts, LLC | Amy Feehan, David J. Nolan, Rebecca Rose, Sissy Cross, David Moraga Amador, Tong Yang, Luke Caruso, Wayra Navia, Lydia Von Borstel, Xiao Hui Zhou, Julia-Garcia-Diaz, Susanna L. Lamers |
| EPI_ISL_437438 | Department of MicroBiology, Government Medical College, Surat | Gujarat Biotechnology Research Centre | Amit Kanani, Akanksha Verma, Nitin Savaliya, Raghawendra Kumar, Dinesh Kumar, Zuber Saiyed, Dipa Kinariwala, Disha Patel, Binita Aring, Neeta Khandelwal, Geeta Vaghela, Sonia Barve, Bhavesh Modi, Kairavi Joshi, Gaurishankar Shrimali, Nidhi Sood, Pranay Shah, R D Dixit, Snehal Bagatharia, Kamlesh J Upadhyay, Ramesh Pandit, Tejas Shah, Ankit Hinsu, Pritesh Sabara, Apurvasinh Puvar, Janvi Raval, Monika Gandhi, Pinal Trivedi, Maharshi Pandya, Neelam Nathani, Chaitanya Joshi, Madhvi Joshi |
| EPI_ISL_437439 | Department of MicroBiology, Government Medical College, Surat | Gujarat Biotechnology Research Centre | Akanksha Verma, Nitin Savaliya, Raghawendra Kumar, Dinesh Kumar, Zuber Saiyed, Dipa Kinariwala, Disha Patel, Binita Aring, Neeta Khandelwal, Geeta Vaghela, Sonia Barve, Bhavesh Modi, Kairavi Joshi, Gaurishankar Shrimali, Nidhi Sood, Pranay Shah, R D Dixit, Snehal Bagatharia, Kamlesh J Upadhyay, Ramesh Pandit, Tejas Shah, Ankit Hinsu, Pritesh Sabara, Apurvasinh Puvar, Janvi Raval, Monika Gandhi, Pinal Trivedi, Maharshi Pandya, Amit Kanani, Armi Chaudhari, Chaitanya Joshi, Madhvi Joshi |
| EPI_ISL_437519 | The National Institute of Public Health Center for Epidemiology and Microbiology | The National Institute of Public Health Center for Epidemiology and Microbiology | Alexander Nagy, Helena Jirincova, Ludmila Novakova, Dusan Trnka, Jaromira Vecerova |
| EPI_ISL_437540, EPI_ISL_437541, EPI_ISL_437542, EPI_ISL_437544, EPI_ISL_437545, EPI_ISL_437546, EPI_ISL_437547, EPI_ISL_437548 | Robert Garry lab | Andersen lab at Scripps Research | Allison Smither, Gilberto Sabino-Santos, Patricia Snarski, Lilia Melnik, Antoinette Bell, Kaylynn Genemaras, Arnaud Drouin, Dahlene Fusco, Robert Garry with SEARCH Alliance San Diego |
| EPI_ISL_437913, EPI_ISL_437914, EPI_ISL_437915, EPI_ISL_437916, EPI_ISL_437917, EPI_ISL_437918, EPI_ISL_437919, EPI_ISL_437920, EPI_ISL_437921, EPI_ISL_437922, EPI_ISL_437923, EPI_ISL_437924, EPI_ISL_437925, EPI_ISL_437926, EPI_ISL_437927, EPI_ISL_437928, EPI_ISL_437929, EPI_ISL_437930, EPI_ISL_437931, EPI_ISL_437932 | | | |
| see above | Institut für Virologie am Department für Hygiene, Mikrobiologie und Public Health | Bergthaler laboratory, CeMM Research Center for Molecular Medicine of the Austrian Academy of Sciences | Alexandra Popa, Benedikt Agerer, Henrique Colaco, Lukas Endler, Jakob-Wendelin Genger, Alexander Lercher, Mark Smyth, Thomas Penz, Michael Schuster, Jan Laine, Martin Senekowitsch, Judith Aberle, Stephan Aberle, Elisabeth Puchhammer-Stoeckl, Manfred Nairz, Guenter Weiss, Wegene Borena, Dorothee von Laer, Christoph Bock, Andreas Bergthaler |
| EPI_ISL_437933, EPI_ISL_437934, EPI_ISL_437935, EPI_ISL_437936, EPI_ISL_437937, EPI_ISL_437938, EPI_ISL_437939, EPI_ISL_437940, EPI_ISL_437941, EPI_ISL_437942, EPI_ISL_437943, EPI_ISL_437944, EPI_ISL_437945, EPI_ISL_437946, EPI_ISL_437947, EPI_ISL_437948, EPI_ISL_437949, EPI_ISL_437950, EPI_ISL_437951, EPI_ISL_437952, EPI_ISL_437953, EPI_ISL_437954, EPI_ISL_437955, EPI_ISL_437956, EPI_ISL_437957, EPI_ISL_437958, EPI_ISL_437959, EPI_ISL_437960, EPI_ISL_437961, EPI_ISL_437962, EPI_ISL_437963, EPI_ISL_437964, EPI_ISL_437965, EPI_ISL_437966, EPI_ISL_437967, EPI_ISL_437968, EPI_ISL_437969, EPI_ISL_437970, EPI_ISL_437971, EPI_ISL_437972, EPI_ISL_437973 | | | |
| see above | Universitaetsklinik für Innere Medizin II Innsbruck | Bergthaler laboratory, CeMM Research Center for Molecular Medicine of the Austrian Academy of Sciences | Alexandra Popa, Benedikt Agerer, Henrique Colaco, Lukas Endler, Jakob-Wendelin Genger, Alexander Lercher, Mark Smyth, Thomas Penz, Michael Schuster, Jan Laine, Martin Senekowitsch, Judith Aberle, Stephan Aberle, Elisabeth Puchhammer-Stoeckl, Manfred Nairz, Guenter Weiss, Wegene Borena, Dorothee von Laer, Christoph Bock, Andreas Bergthaler |
| EPI_ISL_437974, EPI_ISL_437975, EPI_ISL_437976, EPI_ISL_437977, EPI_ISL_437978, EPI_ISL_437979, EPI_ISL_437980, EPI_ISL_437981, EPI_ISL_437982, EPI_ISL_437983, EPI_ISL_437984, EPI_ISL_437985, EPI_ISL_437986, EPI_ISL_437987, EPI_ISL_437988, EPI_ISL_437989, EPI_ISL_437990, EPI_ISL_437991, EPI_ISL_437992 | | | |
| see above | Institut für Virologie am Department für Hygiene, Mikrobiologie und Public Health | Bergthaler laboratory, CeMM Research Center for Molecular Medicine of the Austrian Academy of Sciences | Alexandra Popa, Benedikt Agerer, Henrique Colaco, Lukas Endler, Jakob-Wendelin Genger, Alexander Lercher, Mark Smyth, Thomas Penz, Michael Schuster, Jan Laine, Martin Senekowitsch, Judith Aberle, Stephan Aberle, Elisabeth Puchhammer-Stoeckl, Manfred Nairz, Guenter Weiss, Wegene Borena, Dorothee von Laer, Christoph Bock, Andreas Bergthaler |
| EPI_ISL_437993, EPI_ISL_437994, EPI_ISL_437995, EPI_ISL_437996, EPI_ISL_437997, EPI_ISL_437998, EPI_ISL_437999, EPI_ISL_438000, EPI_ISL_438001, EPI_ISL_438002, EPI_ISL_438003, EPI_ISL_438004, EPI_ISL_438005, EPI_ISL_438006, EPI_ISL_438007, EPI_ISL_438008, EPI_ISL_438009, EPI_ISL_438010, EPI_ISL_438011, EPI_ISL_438012, EPI_ISL_438013, EPI_ISL_438014, EPI_ISL_438015, EPI_ISL_438016, EPI_ISL_438017, EPI_ISL_438018, EPI_ISL_438019, EPI_ISL_438020, EPI_ISL_438021, EPI_ISL_438022, EPI_ISL_438023, EPI_ISL_438024, EPI_ISL_438025, EPI_ISL_438026, EPI_ISL_438027, EPI_ISL_438028, EPI_ISL_438029, EPI_ISL_438030, EPI_ISL_438031, EPI_ISL_438032, EPI_ISL_438033, EPI_ISL_438034, EPI_ISL_438035, EPI_ISL_438036, EPI_ISL_438037, EPI_ISL_438038, EPI_ISL_438039, EPI_ISL_438040, EPI_ISL_438041, EPI_ISL_438042, EPI_ISL_438043, EPI_ISL_438044, EPI_ISL_438045, EPI_ISL_438046, EPI_ISL_438047, EPI_ISL_438048, EPI_ISL_438049, EPI_ISL_438050, EPI_ISL_438051, EPI_ISL_438052, EPI_ISL_438053, EPI_ISL_438054, EPI_ISL_438055, EPI_ISL_438056, EPI_ISL_438057, EPI_ISL_438058, EPI_ISL_438059, EPI_ISL_438060, EPI_ISL_438061, EPI_ISL_438062, EPI_ISL_438063, EPI_ISL_438064, EPI_ISL_438065, EPI_ISL_438066, EPI_ISL_438067, EPI_ISL_438068, EPI_ISL_438069, EPI_ISL_438070, EPI_ISL_438071, EPI_ISL_438072, EPI_ISL_438073, EPI_ISL_438074, EPI_ISL_438075, EPI_ISL_438076, EPI_ISL_438077, EPI_ISL_438078, EPI_ISL_438079, EPI_ISL_438080, EPI_ISL_438081, EPI_ISL_438082, EPI_ISL_438083, EPI_ISL_438084, EPI_ISL_438085, EPI_ISL_438086, EPI_ISL_438087, EPI_ISL_438088, EPI_ISL_438089, EPI_ISL_438090, EPI_ISL_438091, EPI_ISL_438092, EPI_ISL_438093, EPI_ISL_438094, EPI_ISL_438095, EPI_ISL_438096, EPI_ISL_438097, EPI_ISL_438098, EPI_ISL_438099, EPI_ISL_438100, EPI_ISL_438101, EPI_ISL_438102, EPI_ISL_438103, EPI_ISL_438104, EPI_ISL_438105, EPI_ISL_438106, EPI_ISL_438107, EPI_ISL_438108, EPI_ISL_438109, EPI_ISL_438110, EPI_ISL_438111, EPI_ISL_438112, EPI_ISL_438113, EPI_ISL_438114, EPI_ISL_438115, EPI_ISL_438116, EPI_ISL_438117, EPI_ISL_438118, EPI_ISL_438119, EPI_ISL_438120, EPI_ISL_438121, EPI_ISL_438122, EPI_ISL_438123, EPI_ISL_438124, EPI_ISL_438125, EPI_ISL_438126, EPI_ISL_438127, EPI_ISL_438128 | | | |
| see above | Center for Virology, Medical University of Vienna | Bergthaler laboratory, CeMM Research Center for Molecular Medicine of the Austrian Academy of Sciences | Alexandra Popa, Benedikt Agerer, Henrique Colaco, Lukas Endler, Jakob-Wendelin Genger, Alexander Lercher, Mark Smyth, Thomas Penz, Michael Schuster, Jan Laine, Martin Senekowitsch, Judith Aberle, Stephan Aberle, Elisabeth Puchhammer-Stoeckl, Manfred Nairz, Guenter Weiss, Wegene Borena, Dorothee von Laer, Christoph Bock, Andreas Bergthaler |
| EPI_ISL_438749, EPI_ISL_438750, EPI_ISL_438751, EPI_ISL_438752, EPI_ISL_438753, EPI_ISL_438754, EPI_ISL_438755, EPI_ISL_438756, EPI_ISL_438757, EPI_ISL_438758, EPI_ISL_438759, EPI_ISL_438760, EPI_ISL_438761, EPI_ISL_438762, EPI_ISL_438763, EPI_ISL_438764, EPI_ISL_438765, EPI_ISL_438766, EPI_ISL_438767, EPI_ISL_438768, EPI_ISL_438769, EPI_ISL_438770, EPI_ISL_438771, EPI_ISL_438772, EPI_ISL_438773, EPI_ISL_438774, EPI_ISL_438775, EPI_ISL_438776, EPI_ISL_438777, EPI_ISL_438778, EPI_ISL_438779, EPI_ISL_438780, EPI_ISL_438781, EPI_ISL_438782, EPI_ISL_438783, EPI_ISL_438784, EPI_ISL_438785, EPI_ISL_438786, EPI_ISL_438787, EPI_ISL_438788, EPI_ISL_438789, EPI_ISL_438790, EPI_ISL_438791, EPI_ISL_438792, EPI_ISL_438793, EPI_ISL_438794, EPI_ISL_438795, EPI_ISL_438796, EPI_ISL_438797, EPI_ISL_438798, EPI_ISL_438799, EPI_ISL_438800, EPI_ISL_438801, EPI_ISL_438802, EPI_ISL_438803, EPI_ISL_438804, EPI_ISL_438805, EPI_ISL_438806, EPI_ISL_438807, EPI_ISL_438808, EPI_ISL_438809, EPI_ISL_438810, EPI_ISL_438811, EPI_ISL_438812, EPI_ISL_438813, EPI_ISL_438814, EPI_ISL_438815, EPI_ISL_438816, EPI_ISL_438817, EPI_ISL_438818, EPI_ISL_438819, EPI_ISL_438820, EPI_ISL_438821, EPI_ISL_438822, EPI_ISL_438823, EPI_ISL_438824, EPI_ISL_438825, EPI_ISL_438826, EPI_ISL_438827, EPI_ISL_438828, EPI_ISL_438829, EPI_ISL_438830, EPI_ISL_438831, EPI_ISL_438832, EPI_ISL_438833, EPI_ISL_438834, EPI_ISL_438835, EPI_ISL_438836, EPI_ISL_438837, EPI_ISL_438838, EPI_ISL_438839, EPI_ISL_438840, EPI_ISL_438841, EPI_ISL_438842, EPI_ISL_438843, EPI_ISL_438844, EPI_ISL_438845, EPI_ISL_438846, EPI_ISL_438847, EPI_ISL_438848, EPI_ISL_438849, EPI_ISL_438850, EPI_ISL_438851, EPI_ISL_438852, EPI_ISL_438853, EPI_ISL_438854, EPI_ISL_438855, EPI_ISL_438856, EPI_ISL_438857, EPI_ISL_438858, EPI_ISL_438859, EPI_ISL_438860, EPI_ISL_438861, EPI_ISL_438862, EPI_ISL_438863, EPI_ISL_438864, EPI_ISL_438865, EPI_ISL_438866, EPI_ISL_438867, EPI_ISL_438868, EPI_ISL_438869, EPI_ISL_438870, EPI_ISL_438871, EPI_ISL_438872, EPI_ISL_438873, EPI_ISL_438874, EPI_ISL_438875, EPI_ISL_438876, EPI_ISL_438877, EPI_ISL_438878, EPI_ISL_438879, EPI_ISL_438880, EPI_ISL_438881, EPI_ISL_438882, EPI_ISL_438883, EPI_ISL_438884, EPI_ISL_438885, EPI_ISL_438886, EPI_ISL_438887, EPI_ISL_438888, EPI_ISL_438889, EPI_ISL_438890, EPI_ISL_438891, EPI_ISL_438892, EPI_ISL_438893, EPI_ISL_438894, EPI_ISL_438895, EPI_ISL_438896, EPI_ISL_438897, EPI_ISL_438898, EPI_ISL_438899, EPI_ISL_438900, EPI_ISL_438901, EPI_ISL_438902, EPI_ISL_438903, EPI_ISL_438904, EPI_ISL_438905, EPI_ISL_438906, EPI_ISL_438907, EPI_ISL_438908, EPI_ISL_438909, EPI_ISL_438910, EPI_ISL_438911, EPI_ISL_438912, EPI_ISL_438913, EPI_ISL_438914, EPI_ISL_438915, EPI_ISL_438916, EPI_ISL_438917, EPI_ISL_438918, EPI_ISL_438919, EPI_ISL_438920, EPI_ISL_438921, EPI_ISL_438922, EPI_ISL_438923, EPI_ISL_438924, EPI_ISL_438925, EPI_ISL_438926, EPI_ISL_438927, EPI_ISL_438928, EPI_ISL_438929, EPI_ISL_438930, EPI_ISL_438931, EPI_ISL_438932, EPI_ISL_438933, EPI_ISL_438934, EPI_ISL_438935, EPI_ISL_438936, EPI_ISL_438937, EPI_ISL_438938, EPI_ISL_438939, EPI_ISL_438940, EPI_ISL_438941, EPI_ISL_438942, EPI_ISL_438943, EPI_ISL_438944, EPI_ISL_438945, EPI_ISL_438946, EPI_ISL_438973, EPI_ISL_438974, EPI_ISL_438975, EPI_ISL_438976, EPI_ISL_438977, EPI_ISL_438978, EPI_ISL_438979, EPI_ISL_438980, EPI_ISL_438981, EPI_ISL_438982, EPI_ISL_438983, EPI_ISL_438984, EPI_ISL_438985, EPI_ISL_438986, EPI_ISL_438987, EPI_ISL_438988, EPI_ISL_438989, EPI_ISL_438990, EPI_ISL_438991, EPI_ISL_438992, EPI_ISL_438993, EPI_ISL_438994, EPI_ISL_438995, EPI_ISL_438996, EPI_ISL_438997, EPI_ISL_438998, EPI_ISL_438999, EPI_ISL_439000, EPI_ISL_439001, EPI_ISL_439002, EPI_ISL_439003, EPI_ISL_439004, EPI_ISL_439005, EPI_ISL_439006, EPI_ISL_439007, EPI_ISL_439008, EPI_ISL_439009, EPI_ISL_439010, EPI_ISL_439011, EPI_ISL_439012, EPI_ISL_439013, EPI_ISL_439014, EPI_ISL_439015, EPI_ISL_439016, EPI_ISL_439017, EPI_ISL_439018, EPI_ISL_439019, EPI_ISL_439020, EPI_ISL_439021, EPI_ISL_439022, EPI_ISL_439023, EPI_ISL_439024, EPI_ISL_439025, EPI_ISL_439026, EPI_ISL_439027, EPI_ISL_439028, EPI_ISL_439029, EPI_ISL_439030, EPI_ISL_439031, EPI_ISL_439032, EPI_ISL_439033, EPI_ISL_439034, EPI_ISL_439035, EPI_ISL_439036, EPI_ISL_439037, EPI_ISL_439038, EPI_ISL_439039, EPI_ISL_439040, EPI_ISL_439041, EPI_ISL_439042, EPI_ISL_439043, EPI_ISL_439044, EPI_ISL_439045, EPI_ISL_439046, EPI_ISL_439047, EPI_ISL_439048, EPI_ISL_439049, EPI_ISL_439050, EPI_ISL_439051, EPI_ISL_439052, EPI_ISL_439053, EPI_ISL_439054, EPI_ISL_439055, EPI_ISL_439056, EPI_ISL_439057, EPI_ISL_439058, EPI_ISL_439059, EPI_ISL_439060, EPI_ISL_439061, EPI_ISL_439062, EPI_ISL_439063, EPI_ISL_439064, EPI_ISL_439065, EPI_ISL_439066, EPI_ISL_439067, EPI_ISL_439068, EPI_ISL_439069, EPI_ISL_439070, EPI_ISL_439071, EPI_ISL_439072, EPI_ISL_439073, EPI_ISL_439074, EPI_ISL_439075, EPI_ISL_439076, EPI_ISL_439077, EPI_ISL_439078, EPI_ISL_439079, EPI_ISL_439080, EPI_ISL_439081, EPI_ISL_439082, EPI_ISL_439083, EPI_ISL_439084, EPI_ISL_439085, EPI_ISL_439086, EPI_ISL_439087, EPI_ISL_439088, EPI_ISL_439089, EPI_ISL_439090, EPI_ISL_439091, EPI_ISL_439092, EPI_ISL_439093, EPI_ISL_439094, EPI_ISL_439095, EPI_ISL_439096, EPI_ISL_439097, EPI_ISL_439098, EPI_ISL_439099, EPI_ISL_439100, EPI_ISL_439101, EPI_ISL_439102, EPI_ISL_439103, EPI_ISL_439104, EPI_ISL_439105, EPI_ISL_439106, EPI_ISL_439107, EPI_ISL_439108, EPI_ISL_439109, EPI_ISL_439110, EPI_ISL_439111, EPI_ISL_439112, EPI_ISL_439113, EPI_ISL_439114, EPI_ISL_439115, EPI_ISL_439116, EPI_ISL_439117, EPI_ISL_439118, EPI_ISL_439119, EPI_ISL_439120, EPI_ISL_439121, EPI_ISL_439122, EPI_ISL_439123, EPI_ISL_439124, EPI_ISL_439125, EPI_ISL_439126, EPI_ISL_439127, EPI_ISL_439128, EPI_ISL_439129, EPI_ISL_439130, EPI_ISL_439131, EPI_ISL_439132, EPI_ISL_439133, EPI_ISL_439134, EPI_ISL_439135, EPI_ISL_439136, EPI_ISL_439137, EPI_ISL_439138, EPI_ISL_439139, EPI_ISL_439140, EPI_ISL_439141, EPI_ISL_439142, EPI_ISL_439143 | | | |
| see above | West of Scotland Specialist Virology Centre, NHSGGC / MRC-University of Glasgow Centre for Virus Research | COVID-19 Genomics UK (COG-UK) Consortium | Ana da Silva Filipe, Natasha Johnson, Kathy Smollett, Daniel Mair, Stephen Carmichael, Lily Tong, Jenna Nichols, Elihu Aranday-Cortes, Kirstyn Brunker, Yasmin Parr, Kyriaki Nomikou; Sarah McDonald, Marc Niebel, Patawee Asamaphan; Richard Orton, Joseph Hughes, Sreenu Vattipally, David L Robertson; Alasdair MacLean, Rory Gunson; Kathy Li, Natasha Jesudason, Rajiv Shah, James Shepherd, Antonia Ho, Emma Thomson |
| EPI_ISL_444468 | B.J. Medical College and Civil hospital | Gujarat Biotechnology Research Centre | Amit Kanani, Akanksha Verma, Nitin Savaliya, Raghawendra Kumar, Dinesh Kumar, Zuber Saiyed, Dipa Kinariwala, Disha Patel, Binita Aring, Neeta Khandelwal, Geeta Vaghela, Sonia Barve, Bhavesh Modi, Kairavi Joshi, Gaurishankar Shrimali, Nidhi Sood, Pranay Shah, R D Dixit, Snehal Bagatharia, Kamlesh J Upadhyay, Ramesh Pandit, Tejas Shah, Ankit Hinsu, Pritesh Sabara, Apurvasinh Puvar, Janvi Raval, Monika Gandhi, Pinal Trivedi, Maharshi Pandya, Nidhi Patel, Chaitanya Joshi, Madhvi Joshi |

| | | | |
|---|---|---|---|
| EPI_ISL_444469 | B.J. Medical College and Civil hospital | Gujarat Biotechnology Research Centre | Akanksha Verma, Nitin Savaliya, Raghawendra Kumar, Dinesh Kumar, Zuber Saiyed, Dipa Kinariwala, Disha Patel, Binita Aring, Neeta Khandelwal, Geeta Vaghela, Sonia Barve, Bhavesh Modi, Kairavi Joshi, Gaurishankar Shrimali, Nidhi Sood, Pranay Shah, R D Dixit, Snehal Bagatharia, Kamlesh J Upadhyay, Ramesh Pandit, Tejas Shah, Ankit Hinsu, Pritesh Sabara, Apurvasinh Puvar, Janvi Raval, Monika Gandhi, Pinal Trivedi, Maharshi Pandya, Amit Kanani, Priti Pandita, Chaitanya Joshi, Madhvi Joshi |
| EPI_ISL_444518 | unknown | Molecular Infectious Disease | Anderson,B.P., Rosenthal,S.H., Gerasimova,A., Kagan,R.M. and Owen,R. |
| EPI_ISL_445088 | Human Genetic Research Center, Kawsar Biotech Company | Human Genetic Research Center, Kawsar Biotech Company | Abbasalipour Bashash,M., Khosravi,M.A., Zeinali,S., Keshvar,Y., Sabeghi,S., Jadaliha,M. and Yazdani,R. |
| EPI_ISL_445119, EPI_ISL_445120, EPI_ISL_445121, EPI_ISL_445122, EPI_ISL_445123, EPI_ISL_445124, EPI_ISL_445125, EPI_ISL_445126, EPI_ISL_445127, EPI_ISL_445128, EPI_ISL_445129, EPI_ISL_445130, EPI_ISL_445131, EPI_ISL_445132, EPI_ISL_445133, EPI_ISL_445134, EPI_ISL_445135, EPI_ISL_445136, EPI_ISL_445137, EPI_ISL_445138, EPI_ISL_445139, EPI_ISL_445140, EPI_ISL_445141, EPI_ISL_445142, EPI_ISL_445143, EPI_ISL_445144, EPI_ISL_445145, EPI_ISL_445146, EPI_ISL_445147, EPI_ISL_445148, EPI_ISL_445149, EPI_ISL_445150, EPI_ISL_445151, EPI_ISL_445152, EPI_ISL_445153, EPI_ISL_445154, EPI_ISL_445155, EPI_ISL_445156, EPI_ISL_445157, EPI_ISL_445159, EPI_ISL_445160, EPI_ISL_445161, EPI_ISL_445162, EPI_ISL_445163 | | | |
| see above | Robert Garry lab | Andersen lab at Scripps Research | Allison Smither, Gilberto Sabino-Santos, Patricia Snarski, Lilia Melnik, Antoinette Bell, Kaylynn Genemaras, Arnaud Drouin, Dahlene Fusco, Robert Garry with SEARCH Alliance San Diego |
| EPI_ISL_445233 | Vardcentralen Brinken | The Public Health Agency of Sweden | Agnes Wigh, Oskar Karlsson Lindsjo, Maria Lind Karlberg, Anna-Malin Linde, Olov Svartstrom, Anna Risberg, Theresa Enkirch, Mia Brytting, Karin Tegmark-Wisell |
| EPI_ISL_445245 | CLINICA ALEMANA DE SANTIAGO S.A. | Instituto de Salud Publica de Chile | Andrés E Castillo, Bárbara Parra,Paz Tapia, Jaime Lagos, Loredana Arata, Alejandra Acevedo, Winston Andrade, Gabriel Leal, Carolina Tambley, Patricia Bustos, Rodrigo Fasce, Jorge Fernandez |
| EPI_ISL_445246 | HOSPITAL PUERTO MONTT | Instituto de Salud Publica de Chile | Andrés E Castillo, Bárbara Parra,Paz Tapia, Jaime Lagos, Loredana Arata, Alejandra Acevedo, Winston Andrade, Gabriel Leal, Carolina Tambley, Patricia Bustos, Rodrigo Fasce, Jorge Fernandez |
| EPI_ISL_445247 | UNIVERSIDAD DE LOS ANDES | Instituto de Salud Publica de Chile | Andrés E Castillo, Bárbara Parra,Paz Tapia, Jaime Lagos, Loredana Arata, Alejandra Acevedo, Winston Andrade, Gabriel Leal, Carolina Tambley, Patricia Bustos, Rodrigo Fasce, Jorge Fernandez |
| EPI_ISL_445248 | CLINICA ALEMANA DE SANTIAGO S.A. | Instituto de Salud Publica de Chile | Andrés E Castillo, Bárbara Parra,Paz Tapia, Jaime Lagos, Loredana Arata, Alejandra Acevedo, Winston Andrade, Gabriel Leal, Carolina Tambley, Patricia Bustos, Rodrigo Fasce, Jorge Fernandez |
| EPI_ISL_445249 | CLINICA SANTA MARIA S.A. | Instituto de Salud Publica de Chile | Andrés E Castillo, Bárbara Parra,Paz Tapia, Jaime Lagos, Loredana Arata, Alejandra Acevedo, Winston Andrade, Gabriel Leal, Carolina Tambley, Patricia Bustos, Rodrigo Fasce, Jorge Fernandez |
| EPI_ISL_445250 | CLINICA ALEMANA DE SANTIAGO S.A. | Instituto de Salud Publica de Chile | Andrés E Castillo, Bárbara Parra,Paz Tapia, Jaime Lagos, Loredana Arata, Alejandra Acevedo, Winston Andrade, Gabriel Leal, Carolina Tambley, Patricia Bustos, Rodrigo Fasce, Jorge Fernandez |
| EPI_ISL_445251 | HOSPITAL DE CARABINEROS | Instituto de Salud Publica de Chile | Andrés E Castillo, Bárbara Parra,Paz Tapia, Jaime Lagos, Loredana Arata, Alejandra Acevedo, Winston Andrade, Gabriel Leal, Carolina Tambley, Patricia Bustos, Rodrigo Fasce, Jorge Fernandez |
| EPI_ISL_445252 | PONTIFICIA U. CATOLICA FAC. MEDICINA | Instituto de Salud Publica de Chile | Andrés E Castillo, Bárbara Parra,Paz Tapia, Jaime Lagos, Loredana Arata, Alejandra Acevedo, Winston Andrade, Gabriel Leal, Carolina Tambley, Patricia Bustos, Rodrigo Fasce, Jorge Fernandez |
| EPI_ISL_445253, EPI_ISL_445254, EPI_ISL_445255 | CLINICA ALEMANA DE SANTIAGO S.A. | Instituto de Salud Publica de Chile | Andrés E Castillo, Bárbara Parra,Paz Tapia, Jaime Lagos, Loredana Arata, Alejandra Acevedo, Winston Andrade, Gabriel Leal, Carolina Tambley, Patricia Bustos, Rodrigo Fasce, Jorge Fernandez |
| EPI_ISL_445256 | CLINICA LAS CONDES S.A. | Instituto de Salud Publica de Chile | Andrés E Castillo, Bárbara Parra,Paz Tapia, Jaime Lagos, Loredana Arata, Alejandra Acevedo, Winston Andrade, Gabriel Leal, Carolina Tambley, Patricia Bustos, Rodrigo Fasce, Jorge Fernandez |
| EPI_ISL_445257 | CLINICA TABANCURA | Instituto de Salud Publica de Chile | Andrés E Castillo, Bárbara Parra,Paz Tapia, Jaime Lagos, Loredana Arata, Alejandra Acevedo, Winston Andrade, Gabriel Leal, Carolina Tambley, Patricia Bustos, Rodrigo Fasce, Jorge Fernandez |
| EPI_ISL_445258 | CLINICA ALEMANA DE SANTIAGO S.A. | Instituto de Salud Publica de Chile | Andrés E Castillo, Bárbara Parra,Paz Tapia, Jaime Lagos, Loredana Arata, Alejandra Acevedo, Winston Andrade, Gabriel Leal, Carolina Tambley, Patricia Bustos, Rodrigo Fasce, Jorge Fernandez |
| EPI_ISL_445259 | CLINICA LAS CONDES S.A. | Instituto de Salud Publica de Chile | Andrés E Castillo, Bárbara Parra,Paz Tapia, Jaime Lagos, Loredana Arata, Alejandra Acevedo, Winston Andrade, Gabriel Leal, Carolina Tambley, Patricia Bustos, Rodrigo Fasce, Jorge Fernandez |
| EPI_ISL_445260 | CLINICA ALEMANA DE SANTIAGO S.A. | Instituto de Salud Publica de Chile | Andrés E Castillo, Bárbara Parra,Paz Tapia, Jaime Lagos, Loredana Arata, Alejandra Acevedo, Winston Andrade, Gabriel Leal, Carolina Tambley, Patricia Bustos, Rodrigo Fasce, Jorge Fernandez |
| EPI_ISL_445261 | INTEGRAMEDICA LAB. CLINICO LTDA. | Instituto de Salud Publica de Chile | Andrés E Castillo, Bárbara Parra,Paz Tapia, Jaime Lagos, Loredana Arata, Alejandra Acevedo, Winston Andrade, Gabriel Leal, Carolina Tambley, Patricia Bustos, Rodrigo Fasce, Jorge Fernandez |
| EPI_ISL_445262 | CLINICA TABANCURA | Instituto de Salud Publica de Chile | Andrés E Castillo, Bárbara Parra,Paz Tapia, Jaime Lagos, Loredana Arata, Alejandra Acevedo, Winston Andrade, Gabriel Leal, Carolina Tambley, Patricia Bustos, Rodrigo Fasce, Jorge Fernandez |
| EPI_ISL_445263 | MEGASALUD SPA. | Instituto de Salud Publica de Chile | Andrés E Castillo, Bárbara Parra,Paz Tapia, Jaime Lagos, Loredana Arata, Alejandra Acevedo, Winston Andrade, Gabriel Leal, Carolina Tambley, Patricia Bustos, Rodrigo Fasce, Jorge Fernandez |
| EPI_ISL_445264 | CLINICA REDSALUD VITACURA. | Instituto de Salud Publica de Chile | Andrés E Castillo, Bárbara Parra,Paz Tapia, Jaime Lagos, Loredana Arata, Alejandra Acevedo, Winston Andrade, Gabriel Leal, Carolina Tambley, Patricia Bustos, Rodrigo Fasce, Jorge Fernandez |
| EPI_ISL_445265 | PONTIFICIA UNIVERSIDAD CATOLICA DE CHILE | Instituto de Salud Publica de Chile | Andrés E Castillo, Bárbara Parra,Paz Tapia, Jaime Lagos, Loredana Arata, Alejandra Acevedo, Winston Andrade, Gabriel Leal, Carolina Tambley, Patricia Bustos, Rodrigo Fasce, Jorge Fernandez |
| EPI_ISL_445266, EPI_ISL_445267 | CENTRO ONCOLOGICO DEL NORTE | Instituto de Salud Publica de Chile | Andrés E Castillo, Bárbara Parra,Paz Tapia, Jaime Lagos, Loredana Arata, Alejandra Acevedo, Winston Andrade, Gabriel Leal, Carolina Tambley, Patricia Bustos, Rodrigo Fasce, Jorge Fernandez |
| EPI_ISL_445268, EPI_ISL_445269 | HOSPITAL REG.LAUTARO NAVARRO AVARIA | Instituto de Salud Publica de Chile | Andrés E Castillo, Bárbara Parra,Paz Tapia, Jaime Lagos, Loredana Arata, Alejandra Acevedo, Winston Andrade, Gabriel Leal, Carolina Tambley, Patricia Bustos, Rodrigo Fasce, Jorge Fernandez |
| EPI_ISL_445270 | HOSPITAL DR.HERNAN HENRIQUEZ ARAVENA | Instituto de Salud Publica de Chile | Andrés E Castillo, Bárbara Parra,Paz Tapia, Jaime Lagos, Loredana Arata, Alejandra Acevedo, Winston Andrade, Gabriel Leal, Carolina Tambley, Patricia Bustos, Rodrigo Fasce, Jorge Fernandez |
| EPI_ISL_445272 | CLINICA CIUDAD DEL MAR | Instituto de Salud Publica de Chile | Andrés E Castillo, Bárbara Parra,Paz Tapia, Jaime Lagos, Loredana Arata, Alejandra Acevedo, Winston Andrade, Gabriel Leal, Carolina Tambley, Patricia Bustos, Rodrigo Fasce, Jorge Fernandez |
| EPI_ISL_445273, EPI_ISL_445274 | LABORATORIO TORRE MEDICA LTDA. | Instituto de Salud Publica de Chile | Andrés E Castillo, Bárbara Parra,Paz Tapia, Jaime Lagos, Loredana Arata, Alejandra Acevedo, Winston Andrade, Gabriel Leal, Carolina Tambley, Patricia Bustos, Rodrigo Fasce, Jorge Fernandez |
| EPI_ISL_445275, EPI_ISL_445276 | HOSPITAL CLINICO FUSAT | Instituto de Salud Publica de Chile | Andrés E Castillo, Bárbara Parra,Paz Tapia, Jaime Lagos, Loredana Arata, Alejandra Acevedo, Winston Andrade, Gabriel Leal, Carolina Tambley, Patricia Bustos, Rodrigo Fasce, Jorge Fernandez |
| EPI_ISL_445277 | FUNDACION DE SALUD EL TENIENTE | Instituto de Salud Publica de Chile | Andrés E Castillo, Bárbara Parra,Paz Tapia, Jaime Lagos, Loredana Arata, Alejandra Acevedo, Winston Andrade, Gabriel Leal, Carolina Tambley, Patricia Bustos, Rodrigo Fasce, Jorge Fernandez |
| EPI_ISL_445278 | LABORATORIO TORRE MEDICA LTDA. | Instituto de Salud Publica de Chile | Andrés E Castillo, Bárbara Parra,Paz Tapia, Jaime Lagos, Loredana Arata, Alejandra Acevedo, Winston Andrade, Gabriel Leal, Carolina Tambley, Patricia Bustos, Rodrigo Fasce, Jorge Fernandez |
| EPI_ISL_445279 | LABORATORIO INMUNOLAB SPA | Instituto de Salud Publica de Chile | Andrés E Castillo, Bárbara Parra,Paz Tapia, Jaime Lagos, Loredana Arata, Alejandra Acevedo, Winston Andrade, Gabriel Leal, Carolina Tambley, Patricia Bustos, Rodrigo Fasce, Jorge Fernandez |
| EPI_ISL_445280 | HOSPITAL REG.LAUTARO NAVARRO AVARIA | Instituto de Salud Publica de Chile | Andrés E Castillo, Bárbara Parra,Paz Tapia, Jaime Lagos, Loredana Arata, Alejandra Acevedo, Winston Andrade, Gabriel Leal, Carolina Tambley, Patricia Bustos, Rodrigo Fasce, Jorge Fernandez |

| | | | |
|---|---|---|---|
| EPI_ISL_445281 | HOSPITAL CLINICO DEL SUR | Instituto de Salud Publica de Chile | Andrés E Castillo, Bárbara Parra,Paz Tapia, Jaime Lagos, Loredana Arata, Alejandra Acevedo, Winston Andrade, Gabriel Leal, Carolina Tambley, Patricia Bustos, Rodrigo Fasce, Jorge Fernandez |
| EPI_ISL_445282, EPI_ISL_445283 | CLINICA MAGALLANES S.A. | Instituto de Salud Publica de Chile | Andrés E Castillo, Bárbara Parra,Paz Tapia, Jaime Lagos, Loredana Arata, Alejandra Acevedo, Winston Andrade, Gabriel Leal, Carolina Tambley, Patricia Bustos, Rodrigo Fasce, Jorge Fernandez |
| EPI_ISL_445284 | HOSPITAL REG.LAUTARO NAVARRO AVARIA | Instituto de Salud Publica de Chile | Andrés E Castillo, Bárbara Parra,Paz Tapia, Jaime Lagos, Loredana Arata, Alejandra Acevedo, Winston Andrade, Gabriel Leal, Carolina Tambley, Patricia Bustos, Rodrigo Fasce, Jorge Fernandez |
| EPI_ISL_445285 | CLINICA CIUDAD DEL MAR | Instituto de Salud Publica de Chile | Andrés E Castillo, Bárbara Parra,Paz Tapia, Jaime Lagos, Loredana Arata, Alejandra Acevedo, Winston Andrade, Gabriel Leal, Carolina Tambley, Patricia Bustos, Rodrigo Fasce, Jorge Fernandez |
| EPI_ISL_445286 | HOSPITAL HANGA ROA | Instituto de Salud Publica de Chile | Andrés E Castillo, Bárbara Parra,Paz Tapia, Jaime Lagos, Loredana Arata, Alejandra Acevedo, Winston Andrade, Gabriel Leal, Carolina Tambley, Patricia Bustos, Rodrigo Fasce, Jorge Fernandez |
| EPI_ISL_445287 | CLINICA CIUDAD DEL MAR | Instituto de Salud Publica de Chile | Andrés E Castillo, Bárbara Parra,Paz Tapia, Jaime Lagos, Loredana Arata, Alejandra Acevedo, Winston Andrade, Gabriel Leal, Carolina Tambley, Patricia Bustos, Rodrigo Fasce, Jorge Fernandez |
| EPI_ISL_445288 | HOSPITAL REG.LAUTARO NAVARRO AVARIA | Instituto de Salud Publica de Chile | Andrés E Castillo, Bárbara Parra,Paz Tapia, Jaime Lagos, Loredana Arata, Alejandra Acevedo, Winston Andrade, Gabriel Leal, Carolina Tambley, Patricia Bustos, Rodrigo Fasce, Jorge Fernandez |
| EPI_ISL_445289 | HOSPITAL NAVAL PUERTO WILLIAMS | Instituto de Salud Publica de Chile | Andrés E Castillo, Bárbara Parra,Paz Tapia, Jaime Lagos, Loredana Arata, Alejandra Acevedo, Winston Andrade, Gabriel Leal, Carolina Tambley, Patricia Bustos, Rodrigo Fasce, Jorge Fernandez |
| EPI_ISL_445290, EPI_ISL_445291, EPI_ISL_445292 | CLINICA MAGALLANES S.A. | Instituto de Salud Publica de Chile | Andrés E Castillo, Bárbara Parra,Paz Tapia, Jaime Lagos, Loredana Arata, Alejandra Acevedo, Winston Andrade, Gabriel Leal, Carolina Tambley, Patricia Bustos, Rodrigo Fasce, Jorge Fernandez |
| EPI_ISL_445293, EPI_ISL_445294, EPI_ISL_445295 | HOSPITAL REG.LAUTARO NAVARRO AVARIA | Instituto de Salud Publica de Chile | Andrés E Castillo, Bárbara Parra,Paz Tapia, Jaime Lagos, Loredana Arata, Alejandra Acevedo, Winston Andrade, Gabriel Leal, Carolina Tambley, Patricia Bustos, Rodrigo Fasce, Jorge Fernandez |
| EPI_ISL_445296 | HOSPITAL REGIONAL DE COYHAIQUE | Instituto de Salud Publica de Chile | Andrés E Castillo, Bárbara Parra,Paz Tapia, Jaime Lagos, Loredana Arata, Alejandra Acevedo, Winston Andrade, Gabriel Leal, Carolina Tambley, Patricia Bustos, Rodrigo Fasce, Jorge Fernandez |
| EPI_ISL_445297 | CLINICA INTEGRAL S.A. | Instituto de Salud Publica de Chile | Andrés E Castillo, Bárbara Parra,Paz Tapia, Jaime Lagos, Loredana Arata, Alejandra Acevedo, Winston Andrade, Gabriel Leal, Carolina Tambley, Patricia Bustos, Rodrigo Fasce, Jorge Fernandez |
| EPI_ISL_445298 | HOSPITAL DE SAN FERNANDO | Instituto de Salud Publica de Chile | Andrés E Castillo, Bárbara Parra,Paz Tapia, Jaime Lagos, Loredana Arata, Alejandra Acevedo, Winston Andrade, Gabriel Leal, Carolina Tambley, Patricia Bustos, Rodrigo Fasce, Jorge Fernandez |
| EPI_ISL_445299 | CLINICA MAGALLANES S.A. | Instituto de Salud Publica de Chile | Andrés E Castillo, Bárbara Parra,Paz Tapia, Jaime Lagos, Loredana Arata, Alejandra Acevedo, Winston Andrade, Gabriel Leal, Carolina Tambley, Patricia Bustos, Rodrigo Fasce, Jorge Fernandez |
| EPI_ISL_445300 | HOSPITAL DE RANCAGUA | Instituto de Salud Publica de Chile | Andrés E Castillo, Bárbara Parra,Paz Tapia, Jaime Lagos, Loredana Arata, Alejandra Acevedo, Winston Andrade, Gabriel Leal, Carolina Tambley, Patricia Bustos, Rodrigo Fasce, Jorge Fernandez |
| EPI_ISL_445301 | HOSPITAL NAVAL PUERTO WILLIAMS | Instituto de Salud Publica de Chile | Andrés E Castillo, Bárbara Parra,Paz Tapia, Jaime Lagos, Loredana Arata, Alejandra Acevedo, Winston Andrade, Gabriel Leal, Carolina Tambley, Patricia Bustos, Rodrigo Fasce, Jorge Fernandez |
| EPI_ISL_445302 | INSTITUTO MEDICO LEGAL | Instituto de Salud Publica de Chile | Andrés E Castillo, Bárbara Parra,Paz Tapia, Jaime Lagos, Loredana Arata, Alejandra Acevedo, Winston Andrade, Gabriel Leal, Carolina Tambley, Patricia Bustos, Rodrigo Fasce, Jorge Fernandez |
| EPI_ISL_445303 | CTRO.DE SALUD FAMILIAR DR. RAUL YAZIGI | Instituto de Salud Publica de Chile | Andrés E Castillo, Bárbara Parra,Paz Tapia, Jaime Lagos, Loredana Arata, Alejandra Acevedo, Winston Andrade, Gabriel Leal, Carolina Tambley, Patricia Bustos, Rodrigo Fasce, Jorge Fernandez |
| EPI_ISL_445304 | HOSPITAL SAN JUAN DE DIOS | Instituto de Salud Publica de Chile | Andrés E Castillo, Bárbara Parra,Paz Tapia, Jaime Lagos, Loredana Arata, Alejandra Acevedo, Winston Andrade, Gabriel Leal, Carolina Tambley, Patricia Bustos, Rodrigo Fasce, Jorge Fernandez |
| EPI_ISL_445305 | HOSPITAL DE CARABINEROS | Instituto de Salud Publica de Chile | Andrés E Castillo, Bárbara Parra,Paz Tapia, Jaime Lagos, Loredana Arata, Alejandra Acevedo, Winston Andrade, Gabriel Leal, Carolina Tambley, Patricia Bustos, Rodrigo Fasce, Jorge Fernandez |
| EPI_ISL_445306 | CLINICA UC SAN CARLOS DE APOQUINDO | Instituto de Salud Publica de Chile | Andrés E Castillo, Bárbara Parra,Paz Tapia, Jaime Lagos, Loredana Arata, Alejandra Acevedo, Winston Andrade, Gabriel Leal, Carolina Tambley, Patricia Bustos, Rodrigo Fasce, Jorge Fernandez |
| EPI_ISL_445307 | HOSPITAL SAN JOSE DE MAIPO | Instituto de Salud Publica de Chile | Andrés E Castillo, Bárbara Parra,Paz Tapia, Jaime Lagos, Loredana Arata, Alejandra Acevedo, Winston Andrade, Gabriel Leal, Carolina Tambley, Patricia Bustos, Rodrigo Fasce, Jorge Fernandez |
| EPI_ISL_445308 | HOSPITAL EL CARMEN DR.LUIS VALENTIN F. | Instituto de Salud Publica de Chile | Andrés E Castillo, Bárbara Parra,Paz Tapia, Jaime Lagos, Loredana Arata, Alejandra Acevedo, Winston Andrade, Gabriel Leal, Carolina Tambley, Patricia Bustos, Rodrigo Fasce, Jorge Fernandez |
| EPI_ISL_445309 | MUTUAL DE SEGURIDAD C.CH.C. | Instituto de Salud Publica de Chile | Andrés E Castillo, Bárbara Parra,Paz Tapia, Jaime Lagos, Loredana Arata, Alejandra Acevedo, Winston Andrade, Gabriel Leal, Carolina Tambley, Patricia Bustos, Rodrigo Fasce, Jorge Fernandez |
| EPI_ISL_445310 | HOSPITAL DR.SOTERO DEL RIO | Instituto de Salud Publica de Chile | Andrés E Castillo, Bárbara Parra,Paz Tapia, Jaime Lagos, Loredana Arata, Alejandra Acevedo, Winston Andrade, Gabriel Leal, Carolina Tambley, Patricia Bustos, Rodrigo Fasce, Jorge Fernandez |
| EPI_ISL_445311 | MEGASALUD S.A. | Instituto de Salud Publica de Chile | Andrés E Castillo, Bárbara Parra,Paz Tapia, Jaime Lagos, Loredana Arata, Alejandra Acevedo, Winston Andrade, Gabriel Leal, Carolina Tambley, Patricia Bustos, Rodrigo Fasce, Jorge Fernandez |
| EPI_ISL_445312 | CLINICA UC SAN CARLOS DE APOQUINDO | Instituto de Salud Publica de Chile | Andrés E Castillo, Bárbara Parra,Paz Tapia, Jaime Lagos, Loredana Arata, Alejandra Acevedo, Winston Andrade, Gabriel Leal, Carolina Tambley, Patricia Bustos, Rodrigo Fasce, Jorge Fernandez |
| EPI_ISL_445313, EPI_ISL_445314 | HOSPITAL EL CARMEN DR.LUIS VALENTIN F. | Instituto de Salud Publica de Chile | Andrés E Castillo, Bárbara Parra,Paz Tapia, Jaime Lagos, Loredana Arata, Alejandra Acevedo, Winston Andrade, Gabriel Leal, Carolina Tambley, Patricia Bustos, Rodrigo Fasce, Jorge Fernandez |
| EPI_ISL_445315 | CLINICA UC SAN CARLOS DE APOQUINDO | Instituto de Salud Publica de Chile | Andrés E Castillo, Bárbara Parra,Paz Tapia, Jaime Lagos, Loredana Arata, Alejandra Acevedo, Winston Andrade, Gabriel Leal, Carolina Tambley, Patricia Bustos, Rodrigo Fasce, Jorge Fernandez |
| EPI_ISL_445316 | CESFAM BALMACEDA DE RENCA | Instituto de Salud Publica de Chile | Andrés E Castillo, Bárbara Parra,Paz Tapia, Jaime Lagos, Loredana Arata, Alejandra Acevedo, Winston Andrade, Gabriel Leal, Carolina Tambley, Patricia Bustos, Rodrigo Fasce, Jorge Fernandez |
| EPI_ISL_445317 | UNIV.DE CHILE HOSP.CLINICO | Instituto de Salud Publica de Chile | Andrés E Castillo, Bárbara Parra,Paz Tapia, Jaime Lagos, Loredana Arata, Alejandra Acevedo, Winston Andrade, Gabriel Leal, Carolina Tambley, Patricia Bustos, Rodrigo Fasce, Jorge Fernandez |
| EPI_ISL_445318 | C.DE SALUD FAMILIAR PABLO NERUDA | Instituto de Salud Publica de Chile | Andrés E Castillo, Bárbara Parra,Paz Tapia, Jaime Lagos, Loredana Arata, Alejandra Acevedo, Winston Andrade, Gabriel Leal, Carolina Tambley, Patricia Bustos, Rodrigo Fasce, Jorge Fernandez |
| EPI_ISL_445319 | HOSPITAL FELIX BULNES | Instituto de Salud Publica de Chile | Andrés E Castillo, Bárbara Parra,Paz Tapia, Jaime Lagos, Loredana Arata, Alejandra Acevedo, Winston Andrade, Gabriel Leal, Carolina Tambley, Patricia Bustos, Rodrigo Fasce, Jorge Fernandez |
| EPI_ISL_445320 | C.C.SALUD FAMILIAR PADRE FELIX DONOSO G. | Instituto de Salud Publica de Chile | Andrés E Castillo, Bárbara Parra,Paz Tapia, Jaime Lagos, Loredana Arata, Alejandra Acevedo, Winston Andrade, Gabriel Leal, Carolina Tambley, Patricia Bustos, Rodrigo Fasce, Jorge Fernandez |
| EPI_ISL_445321 | HOSPITAL FELIX BULNES | Instituto de Salud Publica de Chile | Andrés E Castillo, Bárbara Parra,Paz Tapia, Jaime Lagos, Loredana Arata, Alejandra Acevedo, Winston Andrade, Gabriel Leal, Carolina Tambley, Patricia Bustos, Rodrigo Fasce, Jorge Fernandez |

| | | | |
|---|---|---|---|
| EPI_ISL_445322 | UNIV.DE CHILE HOSP.CLINICO | Instituto de Salud Publica de Chile | Andrés E Castillo, Bárbara Parra,Paz Tapia, Jaime Lagos, Loredana Arata, Alejandra Acevedo, Winston Andrade, Gabriel Leal, Carolina Tambley, Patricia Bustos, Rodrigo Fasce, Jorge Fernandez |
| EPI_ISL_445323 | HOSP.ENFERMEDADES INFECCIOSAS | Instituto de Salud Publica de Chile | Andrés E Castillo, Bárbara Parra,Paz Tapia, Jaime Lagos, Loredana Arata, Alejandra Acevedo, Winston Andrade, Gabriel Leal, Carolina Tambley, Patricia Bustos, Rodrigo Fasce, Jorge Fernandez |
| EPI_ISL_445324 | INTEGRAMEDICA S.A | Instituto de Salud Publica de Chile | Andrés E Castillo, Bárbara Parra,Paz Tapia, Jaime Lagos, Loredana Arata, Alejandra Acevedo, Winston Andrade, Gabriel Leal, Carolina Tambley, Patricia Bustos, Rodrigo Fasce, Jorge Fernandez |
| EPI_ISL_445325 | HOSPITAL DR.SOTERO DEL RIO | Instituto de Salud Publica de Chile | Andrés E Castillo, Bárbara Parra,Paz Tapia, Jaime Lagos, Loredana Arata, Alejandra Acevedo, Winston Andrade, Gabriel Leal, Carolina Tambley, Patricia Bustos, Rodrigo Fasce, Jorge Fernandez |
| EPI_ISL_445326 | ASISTENCIA PUBLICA DR.ALEJANDRO DEL RIO | Instituto de Salud Publica de Chile | Andrés E Castillo, Bárbara Parra,Paz Tapia, Jaime Lagos, Loredana Arata, Alejandra Acevedo, Winston Andrade, Gabriel Leal, Carolina Tambley, Patricia Bustos, Rodrigo Fasce, Jorge Fernandez |
| EPI_ISL_445327 | HOSPITAL PADRE HURTADO | Instituto de Salud Publica de Chile | Andrés E Castillo, Bárbara Parra,Paz Tapia, Jaime Lagos, Loredana Arata, Alejandra Acevedo, Winston Andrade, Gabriel Leal, Carolina Tambley, Patricia Bustos, Rodrigo Fasce, Jorge Fernandez |
| EPI_ISL_445328 | MEGASALUD S.A. | Instituto de Salud Publica de Chile | Andrés E Castillo, Bárbara Parra,Paz Tapia, Jaime Lagos, Loredana Arata, Alejandra Acevedo, Winston Andrade, Gabriel Leal, Carolina Tambley, Patricia Bustos, Rodrigo Fasce, Jorge Fernandez |
| EPI_ISL_445329 | HOSPITAL DR.SOTERO DEL RIO | Instituto de Salud Publica de Chile | Andrés E Castillo, Bárbara Parra,Paz Tapia, Jaime Lagos, Loredana Arata, Alejandra Acevedo, Winston Andrade, Gabriel Leal, Carolina Tambley, Patricia Bustos, Rodrigo Fasce, Jorge Fernandez |
| EPI_ISL_445330 | CLINICA VESPUCIO S. A. | Instituto de Salud Publica de Chile | Andrés E Castillo, Bárbara Parra,Paz Tapia, Jaime Lagos, Loredana Arata, Alejandra Acevedo, Winston Andrade, Gabriel Leal, Carolina Tambley, Patricia Bustos, Rodrigo Fasce, Jorge Fernandez |
| EPI_ISL_445333 | LABORATORIO CLINICA UNIVERSITARIA DE CONCEPCION | Instituto de Salud Publica de Chile | Andrés E Castillo, Bárbara Parra,Paz Tapia, Jaime Lagos, Loredana Arata, Alejandra Acevedo, Winston Andrade, Gabriel Leal, Carolina Tambley, Patricia Bustos, Rodrigo Fasce, Jorge Fernandez |
| EPI_ISL_445334 | CLINICA UNIVERSITARIA DE PUERTO MONTT S.A. | Instituto de Salud Publica de Chile | Andrés E Castillo, Bárbara Parra,Paz Tapia, Jaime Lagos, Loredana Arata, Alejandra Acevedo, Winston Andrade, Gabriel Leal, Carolina Tambley, Patricia Bustos, Rodrigo Fasce, Jorge Fernandez |
| EPI_ISL_445335 | HOSPITAL DE CALBUCO | Instituto de Salud Publica de Chile | Andrés E Castillo, Bárbara Parra,Paz Tapia, Jaime Lagos, Loredana Arata, Alejandra Acevedo, Winston Andrade, Gabriel Leal, Carolina Tambley, Patricia Bustos, Rodrigo Fasce, Jorge Fernandez |
| EPI_ISL_445336 | HOSPITAL LAS HIGUERAS DE TALCAHUANO | Instituto de Salud Publica de Chile | Andrés E Castillo, Bárbara Parra,Paz Tapia, Jaime Lagos, Loredana Arata, Alejandra Acevedo, Winston Andrade, Gabriel Leal, Carolina Tambley, Patricia Bustos, Rodrigo Fasce, Jorge Fernandez |
| EPI_ISL_445337 | HOSPITAL HANGA ROA | Instituto de Salud Publica de Chile | Andrés E Castillo, Bárbara Parra,Paz Tapia, Jaime Lagos, Loredana Arata, Alejandra Acevedo, Winston Andrade, Gabriel Leal, Carolina Tambley, Patricia Bustos, Rodrigo Fasce, Jorge Fernandez |
| EPI_ISL_445338, EPI_ISL_445339, EPI_ISL_445340, EPI_ISL_445341, EPI_ISL_445342, EPI_ISL_445343, EPI_ISL_445344, EPI_ISL_445345, EPI_ISL_445346, EPI_ISL_445347, EPI_ISL_445348 | | | |
| see above | HOSPITAL DR.HERNAN HENRIQUEZ ARAVENA | Instituto de Salud Publica de Chile | Andrés E Castillo, Bárbara Parra,Paz Tapia, Jaime Lagos, Loredana Arata, Alejandra Acevedo, Winston Andrade, Gabriel Leal, Carolina Tambley, Patricia Bustos, Rodrigo Fasce, Jorge Fernandez |
| EPI_ISL_445349, EPI_ISL_445350, EPI_ISL_445351 | HOSPITAL SAN JUAN DE DIOS | Instituto de Salud Publica de Chile | Andrés E Castillo, Bárbara Parra,Paz Tapia, Jaime Lagos, Loredana Arata, Alejandra Acevedo, Winston Andrade, Gabriel Leal, Carolina Tambley, Patricia Bustos, Rodrigo Fasce, Jorge Fernandez |
| EPI_ISL_445352 | HOSPITAL DEL PROFESOR | Instituto de Salud Publica de Chile | Andrés E Castillo, Bárbara Parra,Paz Tapia, Jaime Lagos, Loredana Arata, Alejandra Acevedo, Winston Andrade, Gabriel Leal, Carolina Tambley, Patricia Bustos, Rodrigo Fasce, Jorge Fernandez |
| EPI_ISL_445353 | HOSPITAL PADRE HURTADO | Instituto de Salud Publica de Chile | Andrés E Castillo, Bárbara Parra,Paz Tapia, Jaime Lagos, Loredana Arata, Alejandra Acevedo, Winston Andrade, Gabriel Leal, Carolina Tambley, Patricia Bustos, Rodrigo Fasce, Jorge Fernandez |
| EPI_ISL_445354 | HOSPITAL DE CARABINEROS | Instituto de Salud Publica de Chile | Andrés E Castillo, Bárbara Parra,Paz Tapia, Jaime Lagos, Loredana Arata, Alejandra Acevedo, Winston Andrade, Gabriel Leal, Carolina Tambley, Patricia Bustos, Rodrigo Fasce, Jorge Fernandez |
| EPI_ISL_445355 | MUTUAL DE SEGURIDAD C.CH.C. | Instituto de Salud Publica de Chile | Andrés E Castillo, Bárbara Parra,Paz Tapia, Jaime Lagos, Loredana Arata, Alejandra Acevedo, Winston Andrade, Gabriel Leal, Carolina Tambley, Patricia Bustos, Rodrigo Fasce, Jorge Fernandez |
| EPI_ISL_445356 | PONTIFICIA U. CATOLICA SERV. LABORATORIO | Instituto de Salud Publica de Chile | Andrés E Castillo, Bárbara Parra,Paz Tapia, Jaime Lagos, Loredana Arata, Alejandra Acevedo, Winston Andrade, Gabriel Leal, Carolina Tambley, Patricia Bustos, Rodrigo Fasce, Jorge Fernandez |
| EPI_ISL_445357 | INTEGRAMEDICA CENTROS MEDICOS S.A. | Instituto de Salud Publica de Chile | Andrés E Castillo, Bárbara Parra,Paz Tapia, Jaime Lagos, Loredana Arata, Alejandra Acevedo, Winston Andrade, Gabriel Leal, Carolina Tambley, Patricia Bustos, Rodrigo Fasce, Jorge Fernandez |
| EPI_ISL_445358 | MEGASALUD SPA. | Instituto de Salud Publica de Chile | Andrés E Castillo, Bárbara Parra,Paz Tapia, Jaime Lagos, Loredana Arata, Alejandra Acevedo, Winston Andrade, Gabriel Leal, Carolina Tambley, Patricia Bustos, Rodrigo Fasce, Jorge Fernandez |
| EPI_ISL_445359 | HOSP. SANTIAGO ORIENTE DR. LUIS TISNE B. | Instituto de Salud Publica de Chile | Andrés E Castillo, Bárbara Parra,Paz Tapia, Jaime Lagos, Loredana Arata, Alejandra Acevedo, Winston Andrade, Gabriel Leal, Carolina Tambley, Patricia Bustos, Rodrigo Fasce, Jorge Fernandez |
| EPI_ISL_445360 | HOSPITAL DEL PROFESOR | Instituto de Salud Publica de Chile | Andrés E Castillo, Bárbara Parra,Paz Tapia, Jaime Lagos, Loredana Arata, Alejandra Acevedo, Winston Andrade, Gabriel Leal, Carolina Tambley, Patricia Bustos, Rodrigo Fasce, Jorge Fernandez |
| EPI_ISL_445361 | CLINICA UC SAN CARLOS DE APOQUINDO | Instituto de Salud Publica de Chile | Andrés E Castillo, Bárbara Parra,Paz Tapia, Jaime Lagos, Loredana Arata, Alejandra Acevedo, Winston Andrade, Gabriel Leal, Carolina Tambley, Patricia Bustos, Rodrigo Fasce, Jorge Fernandez |
| EPI_ISL_445363 | ASISTENCIA PUBLICA DR.ALEJANDRO DEL RIO | Instituto de Salud Publica de Chile | Andrés E Castillo, Bárbara Parra,Paz Tapia, Jaime Lagos, Loredana Arata, Alejandra Acevedo, Winston Andrade, Gabriel Leal, Carolina Tambley, Patricia Bustos, Rodrigo Fasce, Jorge Fernandez |
| EPI_ISL_445364 | HOSPITAL EL CARMEN DR.LUIS VALENTIN F. | Instituto de Salud Publica de Chile | Andrés E Castillo, Bárbara Parra,Paz Tapia, Jaime Lagos, Loredana Arata, Alejandra Acevedo, Winston Andrade, Gabriel Leal, Carolina Tambley, Patricia Bustos, Rodrigo Fasce, Jorge Fernandez |
| EPI_ISL_445365, EPI_ISL_445366 | HOSPITAL DR.SOTERO DEL RIO | Instituto de Salud Publica de Chile | Andrés E Castillo, Bárbara Parra,Paz Tapia, Jaime Lagos, Loredana Arata, Alejandra Acevedo, Winston Andrade, Gabriel Leal, Carolina Tambley, Patricia Bustos, Rodrigo Fasce, Jorge Fernandez |
| EPI_ISL_445367 | ASISTENCIA PUBLICA DR.ALEJANDRO DEL RIO | Instituto de Salud Publica de Chile | Andrés E Castillo, Bárbara Parra,Paz Tapia, Jaime Lagos, Loredana Arata, Alejandra Acevedo, Winston Andrade, Gabriel Leal, Carolina Tambley, Patricia Bustos, Rodrigo Fasce, Jorge Fernandez |
| EPI_ISL_445368 | HOSPITAL DEL PROFESOR | Instituto de Salud Publica de Chile | Andrés E Castillo, Bárbara Parra,Paz Tapia, Jaime Lagos, Loredana Arata, Alejandra Acevedo, Winston Andrade, Gabriel Leal, Carolina Tambley, Patricia Bustos, Rodrigo Fasce, Jorge Fernandez |
| EPI_ISL_445369, EPI_ISL_445370 | HOSPITAL DE CARABINEROS | Instituto de Salud Publica de Chile | Andrés E Castillo, Bárbara Parra,Paz Tapia, Jaime Lagos, Loredana Arata, Alejandra Acevedo, Winston Andrade, Gabriel Leal, Carolina Tambley, Patricia Bustos, Rodrigo Fasce, Jorge Fernandez |
| EPI_ISL_445371 | HOSPITAL DR.SOTERO DEL RIO | Instituto de Salud Publica de Chile | Andrés E Castillo, Bárbara Parra,Paz Tapia, Jaime Lagos, Loredana Arata, Alejandra Acevedo, Winston Andrade, Gabriel Leal, Carolina Tambley, Patricia Bustos, Rodrigo Fasce, Jorge Fernandez |
| EPI_ISL_445372 | HOSPITAL FF.AA. "CIRUJANO C. GUZMAN | Instituto de Salud Publica de Chile | Andrés E Castillo, Bárbara Parra,Paz Tapia, Jaime Lagos, Loredana Arata, Alejandra Acevedo, Winston Andrade, Gabriel Leal, Carolina Tambley, Patricia Bustos, Rodrigo Fasce, Jorge Fernandez |
| EPI_ISL_445373, EPI_ISL_445374, | HOSPITAL SAN JUAN DE DIOS | Instituto de Salud Publica de Chile | Andrés E Castillo, Bárbara Parra,Paz Tapia, Jaime Lagos, Loredana Arata, Alejandra Acevedo, Winston Andrade, Gabriel Leal, Carolina Tambley, Patricia |

| | | | |
|---|---|---|---|
| EPI_ISL_445375, EPI_ISL_445376, EPI_ISL_445377 | | | Bustos, Rodrigo Fasce, Jorge Fernandez |
| EPI_ISL_445378 | HOSPITAL DE BULNES | Instituto de Salud Publica de Chile | Andrés E Castillo, Bárbara Parra,Paz Tapia, Jaime Lagos, Loredana Arata, Alejandra Acevedo, Winston Andrade, Gabriel Leal, Carolina Tambley, Patricia Bustos, Rodrigo Fasce, Jorge Fernandez |
| EPI_ISL_445379 | IMALAB- HOSPITAL FACH | Instituto de Salud Publica de Chile | Andrés E Castillo, Bárbara Parra,Paz Tapia, Jaime Lagos, Loredana Arata, Alejandra Acevedo, Winston Andrade, Gabriel Leal, Carolina Tambley, Patricia Bustos, Rodrigo Fasce, Jorge Fernandez |
| EPI_ISL_447040 | B.J. Medical College and Civil hospital | Gujarat Biotechnology Research Centre | Amit Kanani, Akanksha Verma, Nitin Savaliya, Raghawendra Kumar, Dinesh Kumar, Zuber Saiyed, Dipa Kinariwala, Disha Patel, Binita Aring, Neeta Khandelwal, Geeta Vaghela, Sonia Barve, Bhavesh Modi, Kairavi Joshi, Gaurishankar Shrimali, Nidhi Sood, Pranay Shah, R D Dixit, Snehal Bagatharia, Kamlesh J Upadhyay, Ramesh Pandit, Tejas Shah, Ankit Hinsu, Pritesh Sabara, Apurvasinh Puvar, Janvi Raval, Monika Gandhi, Pinal Trivedi, Maharshi Pandya, Anjali Rajwar, Chaitanya Joshi, Madhvi Joshi |
| EPI_ISL_447041 | B.J. Medical College and Civil hospital | Gujarat Biotechnology Research Centre | Akanksha Verma, Nitin Savaliya, Raghawendra Kumar, Dinesh Kumar, Zuber Saiyed, Dipa Kinariwala, Disha Patel, Binita Aring, Neeta Khandelwal, Geeta Vaghela, Sonia Barve, Bhavesh Modi, Kairavi Joshi, Gaurishankar Shrimali, Nidhi Sood, Pranay Shah, R D Dixit, Snehal Bagatharia, Kamlesh J Upadhyay, Ramesh Pandit, Tejas Shah, Ankit Hinsu, Pritesh Sabara, Apurvasinh Puvar, Janvi Raval, Monika Gandhi, Pinal Trivedi, Maharshi Pandya, Amit Kanani, Sharmistha Majumdar, Chaitanya Joshi, Madhvi Joshi |
| EPI_ISL_447119 | HOSPITAL DR.HERNAN HENRIQUEZ ARAVENA | Instituto de Salud Publica de Chile | Andrés E Castillo, Bárbara Parra,Paz Tapia, Jaime Lagos, Loredana Arata, Alejandra Acevedo, Winston Andrade, Gabriel Leal, Carolina Tambley, Patricia Bustos, Rodrigo Fasce, Jorge Fernandez |
| EPI_ISL_447549 | GMERS Medical College and Hospital, Gandhinagar | Gujarat Biotechnology Research Centre | Amit Kanani, Akanksha Verma, Nitin Savaliya, Raghawendra Kumar, Dinesh Kumar, Zuber Saiyed, Dipa Kinariwala, Disha Patel, Binita Aring, Neeta Khandelwal, Geeta Vaghela, Sonia Barve, Bhavesh Modi, Kairavi Joshi, Gaurishankar Shrimali, Nidhi Sood, Pranay Shah, R D Dixit, Snehal Bagatharia, Kamlesh J Upadhyay, Ramesh Pandit, Tejas Shah, Ankit Hinsu, Pritesh Sabara, Apurvasinh Puvar, Janvi Raval, Monika Gandhi, Pinal Trivedi, Maharshi Pandya, Neelam Nathani, Chaitanya Joshi, Madhvi Joshi |
| EPI_ISL_447550 | GMERS Medical College and Hospital, Gandhinagar | Gujarat Biotechnology Research Centre | Akanksha Verma, Nitin Savaliya, Raghawendra Kumar, Dinesh Kumar, Zuber Saiyed, Dipa Kinariwala, Disha Patel, Binita Aring, Neeta Khandelwal, Geeta Vaghela, Sonia Barve, Bhavesh Modi, Kairavi Joshi, Gaurishankar Shrimali, Nidhi Sood, Pranay Shah, R D Dixit, Snehal Bagatharia, Kamlesh J Upadhyay, Ramesh Pandit, Tejas Shah, Ankit Hinsu, Pritesh Sabara, Apurvasinh Puvar, Janvi Raval, Monika Gandhi, Pinal Trivedi, Maharshi Pandya, Amit Kanani, Armi Chaudhari, Chaitanya Joshi, Madhvi Joshi |
| EPI_ISL_448116, EPI_ISL_448117, EPI_ISL_448118, EPI_ISL_448119, EPI_ISL_448120, EPI_ISL_448121, EPI_ISL_448122, EPI_ISL_448123, EPI_ISL_448124, EPI_ISL_448125, EPI_ISL_448126, EPI_ISL_448127, EPI_ISL_448128, EPI_ISL_448129, EPI_ISL_448130, EPI_ISL_448131, EPI_ISL_448132, EPI_ISL_448133, EPI_ISL_448134, EPI_ISL_448135, EPI_ISL_448136, EPI_ISL_448137, EPI_ISL_448138, EPI_ISL_448139, EPI_ISL_448140, EPI_ISL_448141, EPI_ISL_448142, EPI_ISL_448143, EPI_ISL_448144, EPI_ISL_448145, EPI_ISL_448146, EPI_ISL_448147, EPI_ISL_448148, EPI_ISL_448149, EPI_ISL_448150, EPI_ISL_448151, EPI_ISL_448152, EPI_ISL_448153, EPI_ISL_448154, EPI_ISL_448155, EPI_ISL_448156, EPI_ISL_448157, EPI_ISL_448158, EPI_ISL_448159, EPI_ISL_448160, EPI_ISL_448161, EPI_ISL_448162, EPI_ISL_448163, EPI_ISL_448164, EPI_ISL_448165, EPI_ISL_448166, EPI_ISL_448167, EPI_ISL_448168, EPI_ISL_448169, EPI_ISL_448170, EPI_ISL_448171, EPI_ISL_448172, EPI_ISL_448173, EPI_ISL_448174, EPI_ISL_448175, EPI_ISL_448176, EPI_ISL_448177, EPI_ISL_448178, EPI_ISL_448179, EPI_ISL_448180, EPI_ISL_448181, EPI_ISL_448182, EPI_ISL_448183, EPI_ISL_448184, EPI_ISL_448185, EPI_ISL_448186, EPI_ISL_448187, EPI_ISL_448188, EPI_ISL_448189, EPI_ISL_448190, EPI_ISL_448191, EPI_ISL_448192, EPI_ISL_448193, EPI_ISL_448194, EPI_ISL_448195, EPI_ISL_448196, EPI_ISL_448197, EPI_ISL_448198, EPI_ISL_448199, EPI_ISL_448200, EPI_ISL_448201, EPI_ISL_448202, EPI_ISL_448203, EPI_ISL_448204, EPI_ISL_448205, EPI_ISL_448206, EPI_ISL_448207, EPI_ISL_448208, EPI_ISL_448209, EPI_ISL_448210, EPI_ISL_448211, EPI_ISL_448212, EPI_ISL_448213, EPI_ISL_448214, EPI_ISL_448215, EPI_ISL_448216, EPI_ISL_448217, EPI_ISL_448218, EPI_ISL_448219, EPI_ISL_448220, EPI_ISL_448221, EPI_ISL_449176, EPI_ISL_449177, EPI_ISL_449178, EPI_ISL_449179, EPI_ISL_449180, EPI_ISL_449181, EPI_ISL_449182, EPI_ISL_449183, EPI_ISL_449184, EPI_ISL_449185, EPI_ISL_449186, EPI_ISL_449187, EPI_ISL_449188, EPI_ISL_449189, EPI_ISL_449190, EPI_ISL_449191, EPI_ISL_449192, EPI_ISL_449193, EPI_ISL_449194, EPI_ISL_449195, EPI_ISL_449196, EPI_ISL_449197, EPI_ISL_449198, EPI_ISL_449199, EPI_ISL_449200, EPI_ISL_449201, EPI_ISL_449202, EPI_ISL_449203, EPI_ISL_449204, EPI_ISL_449205, EPI_ISL_449206, EPI_ISL_449207, EPI_ISL_449208, EPI_ISL_449209, EPI_ISL_449210, EPI_ISL_449211, EPI_ISL_449212, EPI_ISL_449213, EPI_ISL_449214, EPI_ISL_449215, EPI_ISL_449216, EPI_ISL_449217, EPI_ISL_449218, EPI_ISL_449219, EPI_ISL_449220, EPI_ISL_449221, EPI_ISL_449222, EPI_ISL_449223, EPI_ISL_449224, EPI_ISL_449225, EPI_ISL_449226, EPI_ISL_449227, EPI_ISL_449228, EPI_ISL_449229, EPI_ISL_449230, EPI_ISL_449231, EPI_ISL_449232, EPI_ISL_449233, EPI_ISL_449234, EPI_ISL_449235, EPI_ISL_449236, EPI_ISL_449237, EPI_ISL_449238, EPI_ISL_449239, EPI_ISL_449240, EPI_ISL_449241, EPI_ISL_449242, EPI_ISL_449243, EPI_ISL_449244, EPI_ISL_449245, EPI_ISL_449246, EPI_ISL_449247, EPI_ISL_449248, EPI_ISL_449249, EPI_ISL_449250, EPI_ISL_449251, EPI_ISL_449252, EPI_ISL_449253, EPI_ISL_449254, EPI_ISL_449255, EPI_ISL_449256, EPI_ISL_449257, EPI_ISL_449258 | | | |
| see above | West of Scotland Specialist Virology Centre, NHSGGC / MRC-University of Glasgow Centre for Virus Research | COVID-19 Genomics UK (COG-UK) Consortium | Ana da Silva Filipe, Natasha Johnson, Kathy Smollett, Daniel Mair, Stephen Carmichael, Lily Tong, Jenna Nichols, Elihu Aranday-Cortes, Kirstyn Brunker, Yasmin Parr, Kyriaki Nomikou, Sarah McDonald, Marc Niebel, Patawee Asamaphan, Richard Orton, Joseph Hughes, Sreenu Vattipally, David L Robertson, Alasdair MacLean, Rory Gunson, Kathy Li, Natasha Jesudason, Rajiv Shah, James Shepherd, Antonia Ho, Emma Thomson |
| EPI_ISL_449800 | HOSPITAL SAN JUAN DE DIOS | Instituto de Salud Publica de Chile | Andrés E Castillo, Bárbara Parra,Paz Tapia, Jaime Lagos, Loredana Arata, Alejandra Acevedo, Winston Andrade, Gabriel Leal, Carolina Tambley, Patricia Bustos, Rodrigo Fasce, Jorge Fernandez |
| EPI_ISL_450004, EPI_ISL_450005, EPI_ISL_450006, EPI_ISL_450007, EPI_ISL_450008, EPI_ISL_450009, EPI_ISL_450010, EPI_ISL_450011, EPI_ISL_450012, EPI_ISL_450013, EPI_ISL_450014, EPI_ISL_450015, EPI_ISL_450016, EPI_ISL_450017, EPI_ISL_450018, EPI_ISL_450019, EPI_ISL_450020, EPI_ISL_450021, EPI_ISL_450022, EPI_ISL_450023, EPI_ISL_450024, EPI_ISL_450025, EPI_ISL_450026, EPI_ISL_450027, EPI_ISL_450028, EPI_ISL_450029, EPI_ISL_450030, EPI_ISL_450031, EPI_ISL_450032, EPI_ISL_450033, EPI_ISL_450034, EPI_ISL_450035, EPI_ISL_450036, EPI_ISL_450037, EPI_ISL_450038, EPI_ISL_450039, EPI_ISL_450040, EPI_ISL_450041, EPI_ISL_450042, EPI_ISL_450043, EPI_ISL_450044, EPI_ISL_450045, EPI_ISL_450046, EPI_ISL_450047, EPI_ISL_450048, EPI_ISL_450049, EPI_ISL_450050, EPI_ISL_450051, EPI_ISL_450052, EPI_ISL_450053, EPI_ISL_450054, EPI_ISL_450055, EPI_ISL_450056, EPI_ISL_450057, EPI_ISL_450058, EPI_ISL_450059, EPI_ISL_450060, EPI_ISL_450061, EPI_ISL_450062, EPI_ISL_450063, EPI_ISL_450064, EPI_ISL_450065, EPI_ISL_450066, EPI_ISL_450067, EPI_ISL_450068, EPI_ISL_450069, EPI_ISL_450070, EPI_ISL_450071, EPI_ISL_450072, EPI_ISL_450073, EPI_ISL_450074, EPI_ISL_450075, EPI_ISL_450076, EPI_ISL_450077, EPI_ISL_450078, EPI_ISL_450079, EPI_ISL_450080, EPI_ISL_450081, EPI_ISL_450082, EPI_ISL_450083, EPI_ISL_450084, EPI_ISL_450085, EPI_ISL_450086, EPI_ISL_450087, EPI_ISL_450088, EPI_ISL_450089, EPI_ISL_450090, EPI_ISL_450091, EPI_ISL_450092, EPI_ISL_450093, EPI_ISL_450094, EPI_ISL_450095, EPI_ISL_450096, EPI_ISL_450097, EPI_ISL_450098, EPI_ISL_450099, EPI_ISL_450100, EPI_ISL_450101, EPI_ISL_450102, EPI_ISL_450103, EPI_ISL_450104, EPI_ISL_450105, EPI_ISL_450106, EPI_ISL_450107, EPI_ISL_450108, EPI_ISL_450109, EPI_ISL_450110, EPI_ISL_450111, EPI_ISL_450112, EPI_ISL_450113, EPI_ISL_450114, EPI_ISL_450115, EPI_ISL_450116, EPI_ISL_450117, EPI_ISL_450118, EPI_ISL_450119, EPI_ISL_450120, EPI_ISL_450121, EPI_ISL_450122, EPI_ISL_450123, EPI_ISL_450124, EPI_ISL_450125, EPI_ISL_450126, EPI_ISL_450127, EPI_ISL_450128, EPI_ISL_450129, EPI_ISL_450130, EPI_ISL_450131, EPI_ISL_450132, EPI_ISL_450133, EPI_ISL_450134, EPI_ISL_450135, EPI_ISL_450136, EPI_ISL_450137, EPI_ISL_450138, EPI_ISL_450139, EPI_ISL_450140, EPI_ISL_450141, EPI_ISL_450142, EPI_ISL_450143, EPI_ISL_450144, EPI_ISL_450145, EPI_ISL_450146, EPI_ISL_450147, EPI_ISL_450148, EPI_ISL_450149, EPI_ISL_450150, EPI_ISL_450151, EPI_ISL_450152, EPI_ISL_450153, EPI_ISL_450154, EPI_ISL_450155, EPI_ISL_450156, EPI_ISL_450157, EPI_ISL_450158, EPI_ISL_450159, EPI_ISL_450160, EPI_ISL_450161, EPI_ISL_450162, EPI_ISL_450163, EPI_ISL_450164, EPI_ISL_450165, EPI_ISL_450166, EPI_ISL_450167, EPI_ISL_450168, EPI_ISL_450169, EPI_ISL_450170 | | | |
| see above | MSHS Clinical Microbiology Laboratories | MSHS Pathogen Surveillance Program | Ana S. Gonzalez-Reiche, Mitchell Sullivan, Ajay Obla, Gopi Patel, Emilia Sordillo, Melissa Gitman, Alberto Paniz-mondolfi, Matthew Hernandez, Shelcie Fabre, Jose Polanco, Zenab Khan, Bremy Alburquerque, Jayeeta Dutta, Juan Soto, Shwetha Sridhar Hara, Ying-Chih Wang, Melissa Smith, Robert Sebra, Lisa Miorin, Wen-chun Liu, Randy Albrecht, Judith Aberg, Florian Krammer, Adolfo Garcia-Sastre, Viviana Simon, Harm van Bakel |
| EPI_ISL_450173, EPI_ISL_450174, EPI_ISL_450175, EPI_ISL_450176, EPI_ISL_450177, EPI_ISL_450178, EPI_ISL_450179, EPI_ISL_450180, EPI_ISL_450181, EPI_ISL_450182, EPI_ISL_450183, EPI_ISL_450184, EPI_ISL_450185, EPI_ISL_450231 | | | |
| see above | Robert Garry lab | Andersen lab at Scripps Research | Allison Smither, Gilberto Sabino-Santos, Patricia Snarski, Lilia Melnik, Antoinette Bell, Kaylynn Genemaras, Arnaud Drouin, Dahlene Fusco, Robert Garry with SEARCH Alliance San Diego |
| EPI_ISL_450241, EPI_ISL_450242, EPI_ISL_450243, EPI_ISL_450244, EPI_ISL_450245, EPI_ISL_450246, EPI_ISL_450247, EPI_ISL_450248, EPI_ISL_450249, EPI_ISL_450250, EPI_ISL_450251, EPI_ISL_450252, EPI_ISL_450253, EPI_ISL_450254, EPI_ISL_450255, EPI_ISL_450256, EPI_ISL_450257, EPI_ISL_450258, EPI_ISL_450259, EPI_ISL_450260, EPI_ISL_450261, EPI_ISL_450262, EPI_ISL_450263, EPI_ISL_450264, EPI_ISL_450265, EPI_ISL_450266, EPI_ISL_450267, EPI_ISL_450268, EPI_ISL_450269, EPI_ISL_450270, EPI_ISL_450271, EPI_ISL_450272, EPI_ISL_450273, EPI_ISL_450274, EPI_ISL_450275, EPI_ISL_450276, EPI_ISL_450277, EPI_ISL_450278, EPI_ISL_450279, EPI_ISL_450280, EPI_ISL_450281, EPI_ISL_450282, EPI_ISL_450283, EPI_ISL_450284, EPI_ISL_450285, EPI_ISL_450286, EPI_ISL_450287, EPI_ISL_450288, EPI_ISL_450289, EPI_ISL_450290, EPI_ISL_450291, EPI_ISL_450292, EPI_ISL_450293 | | | |
| see above | WHO National Influenza Centre Russian Federation | WHO National Influenza Centre Russian Federation | Andrey Komissarov, Artem Fadeev, Mariia Sergeeva, Anna Ivanova, Tamila Musaeva, Ksenia Komissarova, Mariia Timofeeva, Veronica Eder, Mariia Pisareva, Daria Danilenko |
| EPI_ISL_450296, EPI_ISL_450297, EPI_ISL_450298, EPI_ISL_450299, EPI_ISL_450300, EPI_ISL_450301 | National Institute for Communicable Diseases of the National Health Laboratory Service | National Institute for Communicable Diseases of the National Health Laboratory Service | Allam M, Ismail A, Khumalo Z, Kwenda S, van Heusden P, Mtshali P, Mnyameni F, Mohale T, Subramoney K, Bhiman JN |
| EPI_ISL_450346, EPI_ISL_450347, EPI_ISL_450348, EPI_ISL_450349, EPI_ISL_450350, EPI_ISL_450351, EPI_ISL_450352 | St.Olavs hospital/NTNU | Institute of Genomics Core Facility, University of Tartu | Aleksandr Ianevski, Tuuli Reisberg, Janne-Fossum Malmring, Svein Arne Nordbø, Denis Kainov |
| EPI_ISL_450495 | National Institute for Communicable Diseases of the National Health Laboratory Service | National Institute for Communicable Diseases of the National Health Laboratory Service | Allam M, Ismail A, Khumalo Z, Kwenda S, van Heusden P, Mtshali P, Mnyameni F, Mohale T, Subramoney K, Bhiman JN |
| EPI_ISL_450831, EPI_ISL_450833 | Wetterhalsan | The Public Health Agency of Sweden | Anders Tengblad, Anna-Malin Linde, Maria Lind Karlberg, Oskar Karlsson Lindsjo, Olov Svartstrom, Anna Risberg, Theresa Enkirch, Mia Brytting, Karin Tegmark-Wisell |
| EPI_ISL_451156 | Government Medical College, Vadodara | Gujarat Biotechnology Research Centre | Amit Kanani, Nidhi Patel, Nitin Savaliya, Raghawendra Kumar, Dinesh Kumar, Zuber Saiyed, Komal Patel, Labdhi Pandya, Snehal Bagatharia, Ramesh Pandit, Tejas Shah, Ankit Hinsu, Pritesh Sabara, Apurvasinh Puvar, Janvi Raval, Zarna Patel, Monika Gandhi, Pinal Trivedi, Maharshi Pandya, Manish Pattani, Tanuja Javadekar , Bhavesh Modi, Gaurishankar Shrimali, R D Dixit, A M Kadri, Bhavya Jindal, Chaitanya Joshi, Madhvi Joshi |
| EPI_ISL_452181, EPI_ISL_452182, | ULSS9 Distretto di Bussolengo | Istituto Zooprofilattico Sperimentale delle Venezie | Adelaide Milani, Alessia Schivo, Annalisa Salviato, Erika Giorgia Quaranta, Ambra Pastori, Bianca Zecchin, Alice Fusaro, Isabella Monne, Calogero |

| | | | |
|---|---|---|---|
| EPI_ISL_452183, EPI_ISL_452184, EPI_ISL_452185, EPI_ISL_452186, EPI_ISL_452187, EPI_ISL_452188, EPI_ISL_452189 | | | Terregino, Antonia Ricci |
| EPI_ISL_452190, EPI_ISL_452191 | ULSS9 Distretto di San Bonifacio | Istituto Zooprofilattico Sperimentale delle Venezie | Adelaide Milani, Alessia Schivo, Annalisa Salviato, Erika Giorgia Quaranta, Ambra Pastori, Bianca Zecchin, Alice Fusaro, Isabella Monne, Calogero Terregino, Antonia Ricci |
| EPI_ISL_452237, EPI_ISL_452240 | Huddinge VC | The Public Health Agency of Sweden | Anders Johansson, Anna-Malin Linde, Maria Lind Karlberg, Oskar Karlsson Lindsjo, Olov Svartstrom, Anna Risberg, Theresa Enkirch, Mia Brytting, Karin Tegmark-Wisell |
| EPI_ISL_452787, EPI_ISL_452788, EPI_ISL_452789, EPI_ISL_452790, EPI_ISL_452791, EPI_ISL_452792, EPI_ISL_452793, EPI_ISL_452794, EPI_ISL_452795 | ICAR-National Institute of High Security Animal Diseases | ICAR-National Institute of High Security Animal Diseases | Anamika Mishra, Ashutosh Aasdev, Sandeep Bhatia, Harshad Murugkar, Chakradhar Tosh, Niranjan Mishra, Shanmugasundaram Nagarajan, Katherukamem Rajukumar, Richa Sood, G Venkatesh, Atul Kumar Pateriya, Manoj Kumar, Shashi Bhushan Sudhakar, Fateh Singh, Sethil Kumar D, Senmannan Kalaiyarasu, Pradeep Gandhale, Naveen Kumar, Chandan Kumar Dubey, Sushil Tripathi, Sandeep Kumar Jhade, Meghna Tripathi, Suman Kumari Shah, Pushpendra Singh, Pushpendra Namdeo, Suman Mishra, Rupal Singh, Vishnupriya Patil, Dipesh Kumar Nayak, Vijendra Pal Singh, Ashwin Ashok Raut |
| EPI_ISL_453006, EPI_ISL_453007, EPI_ISL_453008, EPI_ISL_453009, EPI_ISL_453010, EPI_ISL_453011, EPI_ISL_453012, EPI_ISL_453013, EPI_ISL_453014, EPI_ISL_453015, EPI_ISL_453016, EPI_ISL_453017, EPI_ISL_453018, EPI_ISL_453019, EPI_ISL_453020, EPI_ISL_453021, EPI_ISL_453022, EPI_ISL_453023, EPI_ISL_453024, EPI_ISL_453025, EPI_ISL_453026, EPI_ISL_453027, EPI_ISL_453028, EPI_ISL_453029, EPI_ISL_453030, EPI_ISL_453031, EPI_ISL_453032, EPI_ISL_453033, EPI_ISL_453034, EPI_ISL_453035, EPI_ISL_453036, EPI_ISL_453037, EPI_ISL_453038, EPI_ISL_453039, EPI_ISL_453040, EPI_ISL_453041, EPI_ISL_453042, EPI_ISL_453043, EPI_ISL_453044, EPI_ISL_453045, EPI_ISL_453046, EPI_ISL_453047, EPI_ISL_453048, EPI_ISL_453049, EPI_ISL_453050, EPI_ISL_453051, EPI_ISL_453052, EPI_ISL_453053, EPI_ISL_453054, EPI_ISL_453055, EPI_ISL_453056, EPI_ISL_453057, EPI_ISL_453058, EPI_ISL_453059, EPI_ISL_453060, EPI_ISL_453061, EPI_ISL_453062, EPI_ISL_453063, EPI_ISL_453064, EPI_ISL_453065, EPI_ISL_453066, EPI_ISL_453067, EPI_ISL_453068, EPI_ISL_453069, EPI_ISL_453070, EPI_ISL_453071, EPI_ISL_453072, EPI_ISL_453073, EPI_ISL_453074, EPI_ISL_453075, EPI_ISL_453076, EPI_ISL_453077, EPI_ISL_453078, EPI_ISL_453079, EPI_ISL_453080, EPI_ISL_453081, EPI_ISL_453082, EPI_ISL_453083, EPI_ISL_453084, EPI_ISL_453085, EPI_ISL_453086, EPI_ISL_453087, EPI_ISL_453088, EPI_ISL_453089, EPI_ISL_453090, EPI_ISL_453091, EPI_ISL_453092, EPI_ISL_453093, EPI_ISL_453094, EPI_ISL_453095, EPI_ISL_453096, EPI_ISL_453097, EPI_ISL_453098 | | | |
| see above | West of Scotland Specialist Virology Centre, NHSGGC / MRC-University of Glasgow Centre for Virus Research | COVID-19 Genomics UK (COG-UK) Consortium | Ana da Silva Filipe, Natasha Johnson, Kathy Smollett, Daniel Mair, Stephen Carmichael, Lily Tong, Jenna Nichols, Elihu Aranday-Cortes, Kirstyn Brunker, Yasmin Parr, Kyriaki Nomikou; Sarah McDonald, Marc Niebel, Patawee Asamaphan; Richard Orton, Joseph Hughes, Sreenu Vattipally, David L Robertson; Alasdair MacLean, Rory Gunson; Kathy Li, Natasha Jesudason, Rajiv Shah, James Shepherd, Antonia Ho, Emma Thomson |
| EPI_ISL_454497, EPI_ISL_454498, EPI_ISL_454499, EPI_ISL_454500, EPI_ISL_454501, EPI_ISL_454502, EPI_ISL_454503, EPI_ISL_454504, EPI_ISL_454505, EPI_ISL_454506, EPI_ISL_454507, EPI_ISL_454508, EPI_ISL_454509, EPI_ISL_454510, EPI_ISL_454511, EPI_ISL_454512, EPI_ISL_454513, EPI_ISL_454514, EPI_ISL_454515, EPI_ISL_454516, EPI_ISL_454517, EPI_ISL_454518, EPI_ISL_454519, EPI_ISL_454520 | | | |
| see above | RSE "National Center for Biotechnology" | RSE "National Center for Biotechnology" | Alexandr Shevtsov, Ilyas Akhmetollayev, Viktoriya Lutsay, Asylulan Amirgazin, Askar Abdaliyev, Akbota Rakhmetova, Zabira Aushakhmetova, Ruslan Kalendar, Yerlan Ramankulov |
| EPI_ISL_454571 | National Center of Expertise | National Center for Expertise, National Center for Biotechnology, Kazakhstan | Abdaliyev Askar, Shevtsov Alexandr, Akhmetollayev Ilyas, Kalendar Ruslan, Rakhmetova Akbota, , Lutsay Viktoriya, Amirgazin Asylulan, Aushakhmetova Zabira, Ramankulov Yerlan |
| EPI_ISL_454572 | National Center of Expertise | National Center for Expertise, Kazakhstan National Center for Biotechnology, Kazakhstan | Abdaliyev Askar, Shevtsov Alexandr, Akhmetollayev Ilyas, Kalendar Ruslan, Rakhmetova Akbota, , Lutsay Viktoriya, Amirgazin Asylulan, Aushakhmetova Zabira, Ramankulov Yerlan |
| EPI_ISL_454575 | Laboratory of virology, National Center of Expertise | Laboratory of molecular-genetic research, National Center for Expertise, Kazakhstan National Center for Biotechnology, Kazakhstan | Abdaliyev Askar, Shevtsov Alexandr, Akhmetollayev Ilyas, Kalendar Ruslan, Rakhmetova Akbota, , Lutsay Viktoriya, Amirgazin Asylulan, Aushakhmetova Zabira, Ramankulov Yerlan |
| EPI_ISL_454576, EPI_ISL_454577, EPI_ISL_454579, EPI_ISL_454580, EPI_ISL_454582 | Laboratory of virology, National Center of Expertise | Laboratory of molecular-genetic research, National Center of Expertise, Kazakhstan National Center for Biotechnology, Kazakhstan | Abdaliyev Askar, Shevtsov Alexandr, Akhmetollayev Ilyas, Kalendar Ruslan, Rakhmetova Akbota, , Lutsay Viktoriya, Amirgazin Asylulan, Aushakhmetova Zabira, Ramankulov Yerlan |
| EPI_ISL_454584, EPI_ISL_454585, EPI_ISL_454586 | Laboratory of virology, National Center of Expertise | Laboratory of molecular-genetic research, National Center for Expertise, Kazakhstan National Center for Biotechnology, Kazakhstan | Abdaliyev Askar, Shevtsov Alexandr, Akhmetollayev Ilyas, Kalendar Ruslan, Rakhmetova Akbota, , Lutsay Viktoriya, Amirgazin Asylulan, Aushakhmetova Zabira, Ramankulov Yerlan |
| EPI_ISL_454587 | Laboratory of virology, National Center of Expertise | Laboratory of molecular-genetic research, National Center of Expertise, Kazakhstan National Center for Biotechnology, Kazakhstan | Abdaliyev Askar, Shevtsov Alexandr, Akhmetollayev Ilyas, Kalendar Ruslan, Rakhmetova Akbota, , Lutsay Viktoriya, Amirgazin Asylulan, Aushakhmetova Zabira, Ramankulov Yerlan |
| EPI_ISL_454589 | Laboratory of virology, National Center of Expertise | Laboratory of molecular-genetic research, National Center for Expertise, Kazakhstan National Center for Biotechnology, Kazakhstan | Abdaliyev Askar, Shevtsov Alexandr, Akhmetollayev Ilyas, Kalendar Ruslan, Rakhmetova Akbota, , Lutsay Viktoriya, Amirgazin Asylulan, Aushakhmetova Zabira, Ramankulov Yerlan |
| EPI_ISL_454590, EPI_ISL_454591, EPI_ISL_454593, EPI_ISL_454594 | Laboratory of virology, National Center of Expertise | Laboratory of molecular-genetic research, National Center of Expertise, Kazakhstan National Center for Biotechnology, Kazakhstan | Abdaliyev Askar, Shevtsov Alexandr, Akhmetollayev Ilyas, Kalendar Ruslan, Rakhmetova Akbota, , Lutsay Viktoriya, Amirgazin Asylulan, Aushakhmetova Zabira, Ramankulov Yerlan |
| EPI_ISL_454596, EPI_ISL_454597 | Laboratory of virology, National Center of Expertise | Laboratory of molecular-genetic research, National Center for Expertise, Kazakhstan National Center for Biotechnology, Kazakhstan | Abdaliyev Askar, Shevtsov Alexandr, Akhmetollayev Ilyas, Kalendar Ruslan, Rakhmetova Akbota, , Lutsay Viktoriya, Amirgazin Asylulan, Aushakhmetova Zabira, Ramankulov Yerlan |
| EPI_ISL_454598 | Laboratory of virology, National Center of Expertise | Laboratory of molecular-genetic research, National Center of Expertise, Kazakhstan National Center for Biotechnology, Kazakhstan | Abdaliyev Askar, Shevtsov Alexandr, Akhmetollayev Ilyas, Kalendar Ruslan, Rakhmetova Akbota, , Lutsay Viktoriya, Amirgazin Asylulan, Aushakhmetova Zabira, Ramankulov Yerlan |
| EPI_ISL_454599, EPI_ISL_454600 | Laboratory of virology, National Center of Expertise | Laboratory of molecular-genetic research, National Center for Expertise, Kazakhstan National Center for Biotechnology, Kazakhstan | Abdaliyev Askar, Shevtsov Alexandr, Akhmetollayev Ilyas, Kalendar Ruslan, Rakhmetova Akbota, , Lutsay Viktoriya, Amirgazin Asylulan, Aushakhmetova Zabira, Ramankulov Yerlan |
| EPI_ISL_454601, EPI_ISL_454603 | Laboratory of virology, National Center of Expertise | Laboratory of molecular-genetic research, National Center of Expertise, Kazakhstan National Center for Biotechnology, Kazakhstan | Abdaliyev Askar, Shevtsov Alexandr, Akhmetollayev Ilyas, Kalendar Ruslan, Rakhmetova Akbota, , Lutsay Viktoriya, Amirgazin Asylulan, Aushakhmetova Zabira, Ramankulov Yerlan |
| EPI_ISL_454604 | Laboratory of virology, National Center of Expertise | Laboratory of molecular-genetic research, National Center for Expertise, Kazakhstan National Center for Biotechnology, Kazakhstan | Abdaliyev Askar, Shevtsov Alexandr, Akhmetollayev Ilyas, Kalendar Ruslan, Rakhmetova Akbota, , Lutsay Viktoriya, Amirgazin Asylulan, Aushakhmetova Zabira, Ramankulov Yerlan |
| EPI_ISL_454690 | Emory Molecular Diagnostics Laboratory, Emory Healthcare | Piantadosi Lab, Emory Department of Pathology | Ahmed Babiker, Anne Piantadosi |
| EPI_ISL_454691, EPI_ISL_454692, EPI_ISL_454693 | Quest Diagnostics | Quest Diagnostics | Anderson,B.P., Rosenthal,S.H., Gerasimova,A., Kagan,R.M. and Owen, R. |
| EPI_ISL_454750, EPI_ISL_454751, EPI_ISL_454752, EPI_ISL_454753, EPI_ISL_454754, EPI_ISL_454755, EPI_ISL_454756, EPI_ISL_454757, EPI_ISL_454758, EPI_ISL_454759, EPI_ISL_454760, EPI_ISL_454761, EPI_ISL_454762, EPI_ISL_454763, EPI_ISL_454764, EPI_ISL_454765, EPI_ISL_454766, EPI_ISL_454767, EPI_ISL_454769, EPI_ISL_454770, EPI_ISL_454771, EPI_ISL_454772, EPI_ISL_454773, EPI_ISL_454774, EPI_ISL_454775, EPI_ISL_454776, EPI_ISL_454777, EPI_ISL_454778, EPI_ISL_454779, EPI_ISL_454780, EPI_ISL_454781, EPI_ISL_454782, EPI_ISL_454783, EPI_ISL_454784, EPI_ISL_454785, EPI_ISL_454786, EPI_ISL_454787, EPI_ISL_454788, EPI_ISL_454789, EPI_ISL_454790, EPI_ISL_454791, EPI_ISL_454792, EPI_ISL_454793, EPI_ISL_454794 | | | |
| see above | Dutch COVID-19 response team | National Institute for Public Health and the Environment (RIVM) | Adam Meijer, Harry Vennema, Jeroen Cremer, Sharon van den Brink, Pieter Overduin, Florian Zwagemaker, Dennis Schmitz, Chantal Reusken, on behalf of the national COVID-19 response team |
| EPI_ISL_455105 | Wetterhalsan | The Public Health Agency of Sweden | Anders Tengblad, Anna-Malin Linde, Maria Lind Karlberg, Oskar Karlsson Lindsjo, Olov Svartstrom, Anna Risberg, Theresa Enkirch, Mia Brytting, Karin Tegmark-Wisell |
| EPI_ISL_455355, EPI_ISL_455356, | Emory Molecular Diagnostics Laboratory, Emory Healthcare | Piantadosi Lab, Emory Department of Pathology | Ahmed Babiker, Anne Piantadosi |

| | | | |
|---|---|---|---|
| EPI_ISL_455358, EPI_ISL_455359, EPI_ISL_455360, EPI_ISL_455361 | | | |
| EPI_ISL_455420, EPI_ISL_455458, EPI_ISL_455459 | National Institute of Laboratory Medicine and Referral Center | Genomic Research Lab, BCSIR | Abu Sayeed Mohammad Mahmud, Mohammad Samir Uzzaman, Eshrar Osman, Md. Ahasan Habib, Shahina Akhter, Tanjina Akhter Banu, Barna Goswami, Iffat Jahan, Tasnim Nafisa, Md. Maruf Ahmed Molla, MahmudaYeasmin, Sheikh Md. Selim Al Din, Utpal Chandra Ray, Md. Salim Khan |
| EPI_ISL_456756, EPI_ISL_456757, EPI_ISL_456758, EPI_ISL_456759, EPI_ISL_456760, EPI_ISL_456761, EPI_ISL_456762, EPI_ISL_456763, EPI_ISL_456764, EPI_ISL_456765, EPI_ISL_456766, EPI_ISL_456767, EPI_ISL_456768, EPI_ISL_456769, EPI_ISL_456770, EPI_ISL_456771, EPI_ISL_456772, EPI_ISL_456773, EPI_ISL_456774, EPI_ISL_456775, EPI_ISL_456776, EPI_ISL_456777, EPI_ISL_456778, EPI_ISL_456779, EPI_ISL_456780, EPI_ISL_456781, EPI_ISL_456782, EPI_ISL_456783, EPI_ISL_456784, EPI_ISL_456785, EPI_ISL_456786, EPI_ISL_456787, EPI_ISL_456788, EPI_ISL_456789, EPI_ISL_456790, EPI_ISL_456791, EPI_ISL_456792, EPI_ISL_456793, EPI_ISL_456794, EPI_ISL_456795, EPI_ISL_456796, EPI_ISL_456797, EPI_ISL_456798, EPI_ISL_456799, EPI_ISL_456800, EPI_ISL_456801, EPI_ISL_456802, EPI_ISL_456803, EPI_ISL_456804, EPI_ISL_456805, EPI_ISL_456806, EPI_ISL_456807, EPI_ISL_456808, EPI_ISL_456809, EPI_ISL_456810, EPI_ISL_456811, EPI_ISL_456812, EPI_ISL_456813, EPI_ISL_456814, EPI_ISL_456815, EPI_ISL_456816, EPI_ISL_456817, EPI_ISL_456818, EPI_ISL_456819, EPI_ISL_456820, EPI_ISL_456821, EPI_ISL_456822, EPI_ISL_456823, EPI_ISL_456824, EPI_ISL_456825, EPI_ISL_456826, EPI_ISL_456827, EPI_ISL_456828, EPI_ISL_456829, EPI_ISL_456830, EPI_ISL_456831, EPI_ISL_456832, EPI_ISL_456833, EPI_ISL_456834, EPI_ISL_456835, EPI_ISL_456836, EPI_ISL_456837, EPI_ISL_456838, EPI_ISL_456839, EPI_ISL_456840, EPI_ISL_456841, EPI_ISL_456842, EPI_ISL_456843, EPI_ISL_456844, EPI_ISL_456845, EPI_ISL_456846, EPI_ISL_456847, EPI_ISL_456848, EPI_ISL_456849, EPI_ISL_456850, EPI_ISL_456851, EPI_ISL_456852, EPI_ISL_456853, EPI_ISL_456854, EPI_ISL_456855, EPI_ISL_456856, EPI_ISL_456857, EPI_ISL_456858, EPI_ISL_456859, EPI_ISL_456860, EPI_ISL_456861, EPI_ISL_456862, EPI_ISL_456863, EPI_ISL_456864, EPI_ISL_456865, EPI_ISL_456866, EPI_ISL_456867, EPI_ISL_456868, EPI_ISL_456869, EPI_ISL_456870, EPI_ISL_456871, EPI_ISL_456872, EPI_ISL_456873, EPI_ISL_456874, EPI_ISL_456875, EPI_ISL_456876, EPI_ISL_456877, EPI_ISL_456878, EPI_ISL_456879, EPI_ISL_456880, EPI_ISL_456881, EPI_ISL_456882, EPI_ISL_456883, EPI_ISL_456884, EPI_ISL_456885, EPI_ISL_456886, EPI_ISL_456887, EPI_ISL_456888, EPI_ISL_456889 | | | |
| see above | West of Scotland Specialist Virology Centre, NHSGGC / MRC-University of Glasgow Centre for Virus Research | COVID-19 Genomics UK (COG-UK) Consortium | Ana da Silva Filipe, Natasha Johnson, Kathy Smollett, Daniel Mair, Stephen Carmichael, Lily Tong, Jenna Nichols, Elihu Aranday-Cortes, Kirstyn Brunker, Yasmin Parr, Kyriaki Nomikou; Sarah McDonald, Marc Niebel, Patawee Asamaphan; Richard Orton, Joseph Hughes, Sreenu Vattipally, David L Robertson; Alasdair MacLean, Rory Gunson; Kathy Li, Natasha Jesudason, Rajiv Shah, James Shepherd, Antonia Ho, Emma Thomson |
| EPI_ISL_458079 | Mitra Keluarga Hospital Kenjeran | Institute of Tropical Disease, Universitas Airlangga | Aldise M Nastri, Jezzy R Dewantari, Rima R Prasetya, Krisnoadi Rahardjo, Anastasia W Jefuna, Gatot Soegiarto, Laksmi Wulandari, Retno A Setyoningrum, Resti Yudhawati, Yohko K Shimizu, Mitsuhiro Nishimura, Yasuko Mori, Soetjipto, Kazufumi Shimizu, Maria I Lusida |
| EPI_ISL_458092 | B.J. Medical College and Civil hospital | Gujarat Biotechnology Research Centre | Amit Kanani, Nidhi Patel, Nitin Savaliya, Raghawendra Kumar, Dinesh Kumar, Zuber Saiyed, Komal Patel, Labdhi Pandya, Snehal Bagatharia, Dhaval Vaghela, Ramesh Patel, Pranay Shah, Kamlesh J Upadhyay, Ramesh Pandit, Tejas Shah, Ankit Hinsu, Pritesh Sabara, Apurvasinh Puvar, Janvi Raval, Zarna Patel, Monika Gandhi, Pinal Trivedi, Maharshi Pandya, Camellia Chakraborty, Bhavesh Modi, Gaurishankar Shrimali, R D Dixit, A M Kadri, Umang Mishra, Chaitanya Joshi, Madhvi Joshi |
| EPI_ISL_458113 | Gujarat Biotechnology Research Centre | Gujarat Biotechnology Research Centre | Amit Kanani, Nidhi Patel, Nitin Savaliya, Raghawendra Kumar, Dinesh Kumar, Zuber Saiyed, Komal Patel, Labdhi Pandya, Snehal Bagatharia, Ramesh Pandit, Tejas Shah, Ankit Hinsu, Pritesh Sabara, Apurvasinh Puvar, Janvi Raval, Zarna Patel, Monika Gandhi, Pinal Trivedi, Maharshi Pandya, Afzal Ansari, Bhavesh Modi, Gaurishankar Shrimali, R D Dixit, A M Kadri, Umang Mishra, Chaitanya Joshi, Madhvi Joshi, , , , , |
| EPI_ISL_458236 | Hospital Mexico | Charité Virology-University of Costa Rica | Andres Moreira-Soto, Eugenia Corrales-Aguilar, Teresita Somogyi, Jan Felix Drexler |
| EPI_ISL_459095, EPI_ISL_459096, EPI_ISL_459097, EPI_ISL_459098, EPI_ISL_459099, EPI_ISL_459100, EPI_ISL_459101, EPI_ISL_459102, EPI_ISL_459103, EPI_ISL_459104, EPI_ISL_459105, EPI_ISL_459106, EPI_ISL_459107 | | | |
| see above | NHSGGC West of Scotland Specialist Virology Centre / MRC-University of Glasgow Centre for Virus Research | Wellcome Sanger Institute for the COVID-19 Genomics UK (COG-UK) consortium | Ana da Silva Filipe, Natasha Johnson, Kathy Smollett, Daniel Mair, Stephen Carmichael, Lily Tong, Jenna Nichols, Elihu Aranday-Cortes, Kirstyn Brunker, Yasmin Parr, Kyriaki Nomikou; Sarah McDonald, Marc Niebel, Patawee Asamaphan; Richard Orton, Joseph Hughes, Sreenu Vattipally, David L Robertson; Alasdair MacLean, Rory Gunson; Kathy Li, Natasha Jesudason, Rajiv Shah, James Shepherd, Antonia Ho, Alice Broos, Emma Thomson and Alex Alderton, Roberto Amato, Sonia Goncalves, Ewan Harrison, David K. Jackson, Ian Johnston, Dominic Kwiatkowski, Cordelia Langford, John Sillitoe on behalf of the Wellcome Sanger Institute COVID-19 Surveillance Team (http://www.sanger.ac.uk/covid-team) |
| EPI_ISL_459108 | NHSGGC West of Scotland Specialist Virology Centre / MRC-University of Glasgow Centre for Virus Research | Wellcome Sanger Institute for the COVID-19 Genomics UK (COG-UK) Consortium | Ana da Silva Filipe, Natasha Johnson, Kathy Smollett, Daniel Mair, Stephen Carmichael, Lily Tong, Jenna Nichols, Elihu Aranday-Cortes, Kirstyn Brunker, Yasmin Parr, Kyriaki Nomikou; Sarah McDonald, Marc Niebel, Patawee Asamaphan; Richard Orton, Joseph Hughes, Sreenu Vattipally, David L Robertson; Alasdair MacLean, Rory Gunson; Kathy Li, Natasha Jesudason, Rajiv Shah, James Shepherd, Antonia Ho, Alice Broos, Emma Thomson and Alex Alderton, Roberto Amato, Sonia Goncalves, Ewan Harrison, David K. Jackson, Ian Johnston, Dominic Kwiatkowski, Cordelia Langford, John Sillitoe on behalf of the Wellcome Sanger Institute COVID-19 Surveillance Team |
| EPI_ISL_459109, EPI_ISL_459110, EPI_ISL_459111, EPI_ISL_459112, EPI_ISL_459113, EPI_ISL_459114, EPI_ISL_459115, EPI_ISL_459116, EPI_ISL_459117, EPI_ISL_459118, EPI_ISL_459119, EPI_ISL_459120, EPI_ISL_459121, EPI_ISL_459122, EPI_ISL_459123, EPI_ISL_459124, EPI_ISL_459125, EPI_ISL_459126, EPI_ISL_459127, EPI_ISL_459128, EPI_ISL_459129, EPI_ISL_459130, EPI_ISL_459131, EPI_ISL_459132, EPI_ISL_459133, EPI_ISL_459134, EPI_ISL_459135, EPI_ISL_459136, EPI_ISL_459137, EPI_ISL_459138, EPI_ISL_459139, EPI_ISL_459140, EPI_ISL_459141, EPI_ISL_459142, EPI_ISL_459143, EPI_ISL_459144, EPI_ISL_459145, EPI_ISL_459146, EPI_ISL_459147, EPI_ISL_459148, EPI_ISL_459149, EPI_ISL_459150, EPI_ISL_459151, EPI_ISL_459152, EPI_ISL_459153, EPI_ISL_459154, EPI_ISL_459155, EPI_ISL_459156, EPI_ISL_459157, EPI_ISL_459158, EPI_ISL_459159, EPI_ISL_459160, EPI_ISL_459161, EPI_ISL_459162, EPI_ISL_459163, EPI_ISL_459164, EPI_ISL_459165, EPI_ISL_459505, EPI_ISL_459506, EPI_ISL_459507, EPI_ISL_459508, EPI_ISL_459509, EPI_ISL_459510, EPI_ISL_459511, EPI_ISL_459512, EPI_ISL_459513, EPI_ISL_459514, EPI_ISL_459515, EPI_ISL_459516, EPI_ISL_459517, EPI_ISL_459518, EPI_ISL_459519, EPI_ISL_459520, EPI_ISL_459521, EPI_ISL_459522, EPI_ISL_459523, EPI_ISL_459524, EPI_ISL_459525, EPI_ISL_459526, EPI_ISL_459527, EPI_ISL_459528, EPI_ISL_459529, EPI_ISL_459530, EPI_ISL_459531, EPI_ISL_459532, EPI_ISL_459533, EPI_ISL_459534, EPI_ISL_459535, EPI_ISL_459536, EPI_ISL_459537, EPI_ISL_459538, EPI_ISL_459539, EPI_ISL_459540, EPI_ISL_459541, EPI_ISL_459542, EPI_ISL_459543, EPI_ISL_459544, EPI_ISL_459545, EPI_ISL_459546, EPI_ISL_459547, EPI_ISL_459548, EPI_ISL_459549, EPI_ISL_459550, EPI_ISL_459551, EPI_ISL_459552, EPI_ISL_459553, EPI_ISL_459554, EPI_ISL_459555, EPI_ISL_459556, EPI_ISL_459557, EPI_ISL_459558, EPI_ISL_459559, EPI_ISL_459560, EPI_ISL_459561, EPI_ISL_459562, EPI_ISL_459563, EPI_ISL_459564, EPI_ISL_459565, EPI_ISL_459566, EPI_ISL_459567, EPI_ISL_459568, EPI_ISL_459569, EPI_ISL_459570, EPI_ISL_459571, EPI_ISL_459572, EPI_ISL_459573, EPI_ISL_459574, EPI_ISL_459575, EPI_ISL_459576, EPI_ISL_459577, EPI_ISL_459578, EPI_ISL_459579, EPI_ISL_459580, EPI_ISL_459581, EPI_ISL_459582, EPI_ISL_459583, EPI_ISL_459584, EPI_ISL_459585, EPI_ISL_459586, EPI_ISL_459587, EPI_ISL_459588, EPI_ISL_459589, EPI_ISL_459590, EPI_ISL_459591, EPI_ISL_459592, EPI_ISL_459593, EPI_ISL_459594, EPI_ISL_459595, EPI_ISL_459596, EPI_ISL_459597, EPI_ISL_459598, EPI_ISL_459599, EPI_ISL_459600, EPI_ISL_459601, EPI_ISL_459602, EPI_ISL_459603, EPI_ISL_459604, EPI_ISL_459605, EPI_ISL_459606, EPI_ISL_459607, EPI_ISL_459608, EPI_ISL_459609, EPI_ISL_459610, EPI_ISL_459611, EPI_ISL_459612, EPI_ISL_459613, EPI_ISL_459614, EPI_ISL_459615, EPI_ISL_459616, EPI_ISL_459617, EPI_ISL_459618, EPI_ISL_459619, EPI_ISL_459620, EPI_ISL_459621, EPI_ISL_459622, EPI_ISL_459623, EPI_ISL_459624, EPI_ISL_459625, EPI_ISL_459626, EPI_ISL_459627, EPI_ISL_459628, EPI_ISL_459629, EPI_ISL_459630, EPI_ISL_459631, EPI_ISL_459632, EPI_ISL_459633, EPI_ISL_459634, EPI_ISL_459635, EPI_ISL_459636, EPI_ISL_459637, EPI_ISL_459638, EPI_ISL_459639, EPI_ISL_459640, EPI_ISL_459641, EPI_ISL_459642, EPI_ISL_459643, EPI_ISL_459644, EPI_ISL_459645, EPI_ISL_459646, EPI_ISL_459647, EPI_ISL_459648, EPI_ISL_459649, EPI_ISL_459650, EPI_ISL_459651, EPI_ISL_459652, EPI_ISL_459653, EPI_ISL_459654, EPI_ISL_459655, EPI_ISL_459656, EPI_ISL_459657, EPI_ISL_459658, EPI_ISL_459659, EPI_ISL_459660, EPI_ISL_459661, EPI_ISL_459662, EPI_ISL_459663, EPI_ISL_459664, EPI_ISL_459665, EPI_ISL_459666, EPI_ISL_459667, EPI_ISL_459668, EPI_ISL_459669, EPI_ISL_459670, EPI_ISL_459671, EPI_ISL_459672, EPI_ISL_459673, EPI_ISL_459674, EPI_ISL_459675, EPI_ISL_459676, EPI_ISL_459677, EPI_ISL_459678, EPI_ISL_459679, EPI_ISL_459680, EPI_ISL_459681, EPI_ISL_459682, EPI_ISL_459683, EPI_ISL_459684, EPI_ISL_459685, EPI_ISL_459686, EPI_ISL_459687, EPI_ISL_459688, EPI_ISL_459689, EPI_ISL_459690, EPI_ISL_459691, EPI_ISL_459692, EPI_ISL_459693, EPI_ISL_459694, EPI_ISL_459695, EPI_ISL_459696, EPI_ISL_459697, EPI_ISL_459698, EPI_ISL_459699, EPI_ISL_459700, EPI_ISL_459701, EPI_ISL_459702, EPI_ISL_459703, EPI_ISL_459704, EPI_ISL_459705, EPI_ISL_459706, EPI_ISL_459707, EPI_ISL_459708, EPI_ISL_459709, EPI_ISL_459710, EPI_ISL_459711, EPI_ISL_459712, EPI_ISL_459713, EPI_ISL_459714, EPI_ISL_459715, EPI_ISL_459716, EPI_ISL_459717, EPI_ISL_459718, EPI_ISL_459719, EPI_ISL_459720, EPI_ISL_459721, EPI_ISL_459722, EPI_ISL_459723, EPI_ISL_459724 | | | |
| see above | NHSGGC West of Scotland Specialist Virology Centre / MRC-University of Glasgow Centre for Virus Research | Wellcome Sanger Institute for the COVID-19 Genomics UK (COG-UK) consortium | Ana da Silva Filipe, Natasha Johnson, Kathy Smollett, Daniel Mair, Stephen Carmichael, Lily Tong, Jenna Nichols, Elihu Aranday-Cortes, Kirstyn Brunker, Yasmin Parr, Kyriaki Nomikou; Sarah McDonald, Marc Niebel, Patawee Asamaphan; Richard Orton, Joseph Hughes, Sreenu Vattipally, David L Robertson; Alasdair MacLean, Rory Gunson; Kathy Li, Natasha Jesudason, Rajiv Shah, James Shepherd, Antonia Ho, Alice Broos, Emma Thomson and Alex Alderton, Roberto Amato, Sonia Goncalves, Ewan Harrison, David K. Jackson, Ian Johnston, Dominic Kwiatkowski, Cordelia Langford, John Sillitoe on behalf of the Wellcome Sanger Institute COVID-19 Surveillance Team (http://www.sanger.ac.uk/covid-team) |
| EPI_ISL_461486 | B.J. Medical College and Civil hospital | Gujarat Biotechnology Research Centre | Akanksha Verma, Pranay Shah, Kamlesh J Upadhyay, Tejas Shah, Ankit Hinsu, Pritesh Sabara, Apurvasinh Puvar, Janvi Raval, Zarna Patel, Monika Gandhi, Pinal Trivedi, Maharshi Pandya, Nidhi Patel, Nitin Savaliya, Raghawendra Kumar, Dinesh Kumar, Zuber Saiyed, Komal Patel, Labdhi Pandya, Snehal Bagatharia, R D Dixit, A M Kadri, Harsh Bakshi, Chaitanya Joshi, Madhvi Joshi, |
| EPI_ISL_461490 | B.J. Medical College and Civil hospital | Gujarat Biotechnology Research Centre | Afzal Ansari, Pritesh Sabara, Apurvasinh Puvar, Janvi Raval, Zarna Patel, Monika Gandhi, Pinal Trivedi, Maharshi Pandya, Nidhi Patel, Nitin Savaliya, Raghawendra Kumar, Dinesh Kumar, Zuber Saiyed, Komal Patel, Labdhi Pandya, Snehal Bagatharia, Pranay Shah, Kamlesh J Upadhyay, Tejas Shah, Ankit Hinsu, R D Dixit, A M Kadri, Harsh Bakshi, Chaitanya Joshi, Madhvi Joshi, |
| EPI_ISL_461499 | B.J. Medical College and Civil hospital | Gujarat Biotechnology Research Centre | Akanksha Verma, Raghawendra Kumar, Dinesh Kumar, Zuber Saiyed, Komal Patel, Labdhi Pandya, Snehal Bagatharia, Pranay Shah, Kamlesh J Upadhyay, Tejas Shah, Ankit Hinsu, Pritesh Sabara, Apurvasinh Puvar, Janvi Raval, Zarna Patel, Monika Gandhi, Pinal Trivedi, Maharshi Pandya, Nidhi Patel, Nitin Savaliya, R D Dixit, A M Kadri, Harsh Bakshi, Chaitanya Joshi, Madhvi Joshi, |
| EPI_ISL_461503 | B.J. Medical College and Civil hospital | Gujarat Biotechnology Research Centre | Afzal Ansari, Labdhi Pandya, Snehal Bagatharia, Pranay Shah, Kamlesh J Upadhyay, Tejas Shah, Ankit Hinsu, Pritesh Sabara, Apurvasinh Puvar, Janvi Raval, Zarna Patel, Monika Gandhi, Pinal Trivedi, Maharshi Pandya, Nidhi Patel, Nitin Savaliya, Raghawendra Kumar, Dinesh Kumar, Zuber Saiyed, Komal Patel, R D Dixit, A M Kadri, Harsh Bakshi, Chaitanya Joshi, Madhvi Joshi, |
| EPI_ISL_461589, EPI_ISL_461590, EPI_ISL_461591, EPI_ISL_461592, EPI_ISL_461593, EPI_ISL_461594, EPI_ISL_461595, EPI_ISL_461596, EPI_ISL_461597, EPI_ISL_461598, EPI_ISL_461599, EPI_ISL_461600, EPI_ISL_461601, EPI_ISL_461602, EPI_ISL_461603, EPI_ISL_461604, EPI_ISL_461605, EPI_ISL_461606, EPI_ISL_461607, EPI_ISL_461608, EPI_ISL_461609, EPI_ISL_461610, EPI_ISL_461611, EPI_ISL_461612, EPI_ISL_461613, EPI_ISL_461614, EPI_ISL_461615, EPI_ISL_461616, EPI_ISL_461617, EPI_ISL_461618, EPI_ISL_461619, EPI_ISL_461620, EPI_ISL_461621, EPI_ISL_461622, EPI_ISL_461623, EPI_ISL_461624, EPI_ISL_461625, EPI_ISL_461626, EPI_ISL_461627, EPI_ISL_461628, EPI_ISL_461629, EPI_ISL_461630, EPI_ISL_461631, EPI_ISL_461632, EPI_ISL_461633, EPI_ISL_461634, EPI_ISL_461635, EPI_ISL_461636, EPI_ISL_461637, EPI_ISL_461638, EPI_ISL_461639, EPI_ISL_461640, EPI_ISL_461641, EPI_ISL_461642, EPI_ISL_461643, EPI_ISL_461644, EPI_ISL_461645, EPI_ISL_461646, EPI_ISL_461647, EPI_ISL_461648, EPI_ISL_461649, EPI_ISL_461650, EPI_ISL_461651, EPI_ISL_461652, EPI_ISL_461653, EPI_ISL_461654, EPI_ISL_461655, EPI_ISL_461656, EPI_ISL_461657, EPI_ISL_461658, EPI_ISL_461659, EPI_ISL_461660, EPI_ISL_461661, EPI_ISL_461662, EPI_ISL_461663, EPI_ISL_461664, EPI_ISL_461665, EPI_ISL_461666, EPI_ISL_461667, EPI_ISL_461668, EPI_ISL_461669, EPI_ISL_461670, EPI_ISL_461671, EPI_ISL_461672, EPI_ISL_461673, EPI_ISL_461674, EPI_ISL_461675, EPI_ISL_461676, EPI_ISL_461677, EPI_ISL_461678, EPI_ISL_461679, EPI_ISL_461680, EPI_ISL_461681, EPI_ISL_461682, EPI_ISL_461683, EPI_ISL_461684, EPI_ISL_461685, EPI_ISL_461686, EPI_ISL_461687, EPI_ISL_461688, EPI_ISL_461689, EPI_ISL_461690, EPI_ISL_461691, EPI_ISL_461692, EPI_ISL_461693, EPI_ISL_461694, EPI_ISL_461695, EPI_ISL_461696, EPI_ISL_461697, EPI_ISL_461698, EPI_ISL_461699, EPI_ISL_461700, EPI_ISL_461701, EPI_ISL_461702, EPI_ISL_461703, EPI_ISL_461704, EPI_ISL_461705 | | | |
| see above | West of Scotland Specialist Virology Centre, NHSGGC / MRC-University of Glasgow Centre for Virus Research | COVID-19 Genomics UK (COG-UK) Consortium | Ana da Silva Filipe, Natasha Johnson, Kathy Smollett, Daniel Mair, Stephen Carmichael, Lily Tong, Jenna Nichols, Elihu Aranday-Cortes, Kirstyn Brunker, Yasmin Parr, Kyriaki Nomikou; Sarah McDonald, Marc Niebel, Patawee Asamaphan; Richard Orton, Joseph Hughes, Sreenu Vattipally, David L Robertson; |

| | | | |
|---|---|---|---|
| | | | Alasdair MacLean, Rory Gunson; Kathy Li, Natasha Jesudason, Rajiv Shah, James Shepherd, Antonia Ho, Emma Thomson |
| EPI_ISL_462093, EPI_ISL_462094, EPI_ISL_462095, EPI_ISL_462096, EPI_ISL_462097, EPI_ISL_462098 | National Institute of Laboratory Medicine and Referral Center | Genomic Research Lab, BCSIR | Abu Sayeed Mohammad Mahmud, Mohammad Samir Uzzaman, Eshrar Osman, Md. Ahasan Habib, Tanjina Akhter Banu, Shahina Akter, Barna Goswami, Iffat Jahan, Md. Saddam Hossain, Tasnim Nafisa, Md. Maruf Ahmed Molla, Mahmuda Yeasmin, Asish Kumar Ghosh, Bayzid Bin Monir, Arifa Akram, Sheikh Md. Selim Al Din, Salek Ahmed Sajib, Utpal Chandra Ray, Md. Salim Khan |
| EPI_ISL_463740 | Mohammed Bin Rashid University of Medicine and Health Sciences | Al Jalila Genomics Center | Ahmad Abou Tayoun, Tom Loney, Hamda Khansaheb, Sathishkumar Ramaswamy, Divinlal Harilal, Zulfa Omar Deesi, Rupa Murthy Varghese, Hanan Al Suwaidi, Abdulmajeed Alkhaja, Mohammed Uddin, Rifat Hamoudi, Rabih Halwani, Abiola Catherine Senok, Qutayba Hamid, Norbert Nowotny, Alawi Alsheikh-Ali |
| EPI_ISL_463970, EPI_ISL_463971, EPI_ISL_463972, EPI_ISL_463973, EPI_ISL_463974, EPI_ISL_463975, EPI_ISL_463976, EPI_ISL_463977, EPI_ISL_463978, EPI_ISL_463979, EPI_ISL_463980, EPI_ISL_463981, EPI_ISL_463982, EPI_ISL_463983, EPI_ISL_463984, EPI_ISL_463985, EPI_ISL_463986, EPI_ISL_463988, EPI_ISL_463989, EPI_ISL_463990, EPI_ISL_463991, EPI_ISL_463992, EPI_ISL_463993, EPI_ISL_463994 | | | |
| see above | Toronto Invasive Bacterial Diseases Network | McMaster University | Allison McGeer, Patryk Aftanas, Angel Li, Kuganya Nirmalarajah, Samira Mubareka, Andrew G. McArthur |
| EPI_ISL_466626, EPI_ISL_466627, EPI_ISL_466628, EPI_ISL_466629, EPI_ISL_466630, EPI_ISL_466636, EPI_ISL_466637, EPI_ISL_466638, EPI_ISL_466639, EPI_ISL_466644, EPI_ISL_466645, EPI_ISL_466649, EPI_ISL_466650, EPI_ISL_466686, EPI_ISL_466687, EPI_ISL_466688, EPI_ISL_466689, EPI_ISL_466690, EPI_ISL_466691, EPI_ISL_466693, EPI_ISL_466694 | | | |
| see above | National Institute of Laboratory Medicine and Referral Center | Genomic Research Lab, BCSIR | Abu Sayeed Mohammad Mahmud, Mohammad Samir Uzzaman, Eshrar Osman, Md. Ahasan Habib, Shahina Akter, Tanjina Akhter Banu, Md. Murshed Hasan Sarker, Iffat Jahan, Barna Goswami, Md. Saddam Hossain, Tasnim Nafisa, Md. Maruf Ahmed Molla, Mahmuda Yeasmin, Asish Kumar Ghosh, Arifa Akram, A. K. M. Shamsuzzaman, Sheikh Md. Selim Al Din, Utpal Chandra Ray, Salek Ahmed Sajib, Md. Salim Khan |
| EPI_ISL_467811, EPI_ISL_467812, EPI_ISL_467813, EPI_ISL_467814, EPI_ISL_467815, EPI_ISL_467816, EPI_ISL_467817, EPI_ISL_467818, EPI_ISL_467819, EPI_ISL_467820, EPI_ISL_467821, EPI_ISL_467822, EPI_ISL_467823, EPI_ISL_467824, EPI_ISL_467825, EPI_ISL_467826, EPI_ISL_467827, EPI_ISL_467828, EPI_ISL_467829, EPI_ISL_467830, EPI_ISL_467831, EPI_ISL_467832, EPI_ISL_467833, EPI_ISL_467834, EPI_ISL_467835, EPI_ISL_467836, EPI_ISL_467837, EPI_ISL_467838, EPI_ISL_467839, EPI_ISL_467840, EPI_ISL_467841, EPI_ISL_467842, EPI_ISL_467843, EPI_ISL_467844, EPI_ISL_467845, EPI_ISL_467846, EPI_ISL_467847, EPI_ISL_467848, EPI_ISL_467849, EPI_ISL_467850, EPI_ISL_467851, EPI_ISL_467852, EPI_ISL_467853, EPI_ISL_467854, EPI_ISL_467855, EPI_ISL_467856, EPI_ISL_467857, EPI_ISL_467858, EPI_ISL_467859, EPI_ISL_467860, EPI_ISL_467861, EPI_ISL_467862, EPI_ISL_467863, EPI_ISL_467864, EPI_ISL_467865, EPI_ISL_467866, EPI_ISL_467867, EPI_ISL_467868, EPI_ISL_467869, EPI_ISL_467870, EPI_ISL_467871, EPI_ISL_467872, EPI_ISL_467873, EPI_ISL_467874, EPI_ISL_467875, EPI_ISL_467876, EPI_ISL_467877, EPI_ISL_467878, EPI_ISL_467879, EPI_ISL_467880, EPI_ISL_467881, EPI_ISL_467882, EPI_ISL_467883, EPI_ISL_467884, EPI_ISL_467885, EPI_ISL_467886, EPI_ISL_467887, EPI_ISL_467888, EPI_ISL_467889, EPI_ISL_467890, EPI_ISL_467891, EPI_ISL_467892, EPI_ISL_467893, EPI_ISL_467894, EPI_ISL_467895, EPI_ISL_467896, EPI_ISL_467897, EPI_ISL_467898, EPI_ISL_467899, EPI_ISL_467900, EPI_ISL_467901, EPI_ISL_467902, EPI_ISL_467903, EPI_ISL_467904, EPI_ISL_467905, EPI_ISL_467906, EPI_ISL_467907, EPI_ISL_467908, EPI_ISL_467909, EPI_ISL_467910, EPI_ISL_467911, EPI_ISL_467912, EPI_ISL_467913, EPI_ISL_467914, EPI_ISL_467915, EPI_ISL_467916, EPI_ISL_467917, EPI_ISL_467918, EPI_ISL_467919, EPI_ISL_467920, EPI_ISL_467921, EPI_ISL_467922, EPI_ISL_467923, EPI_ISL_467924, EPI_ISL_467925, EPI_ISL_467926, EPI_ISL_467927, EPI_ISL_468560, EPI_ISL_468561, EPI_ISL_468562, EPI_ISL_468563, EPI_ISL_468564, EPI_ISL_468565, EPI_ISL_468566, EPI_ISL_468567, EPI_ISL_468568, EPI_ISL_468569, EPI_ISL_468570, EPI_ISL_468571, EPI_ISL_468572, EPI_ISL_468573, EPI_ISL_468574, EPI_ISL_468575, EPI_ISL_468576, EPI_ISL_468577, EPI_ISL_468578, EPI_ISL_468579, EPI_ISL_468580, EPI_ISL_468581, EPI_ISL_468582, EPI_ISL_468583, EPI_ISL_468584, EPI_ISL_468585, EPI_ISL_468586, EPI_ISL_468587, EPI_ISL_468588, EPI_ISL_468589, EPI_ISL_468590 | | | |
| see above | Quest Diagnostics | Quest Diagnostics | Anderson,B.P., Rosenthal,S.H., Gerasimova,A., Kagan,R.M. and Owen, R. |
| EPI_ISL_469276, EPI_ISL_469277, EPI_ISL_469278, EPI_ISL_469279, EPI_ISL_469280, EPI_ISL_469281 | Mohammed Bin Rashid University of Medicine and Health Sciences | Al Jalila Genomics Center | Ahmad Abou Tayoun, Tom Loney, Hamda Khansaheb, Sathishkumar Ramaswamy, Divinlal Harilal, Zulfa Omar Deesi, Rupa Murthy Varghese, Hanan Al Suwaidi, Abdulmajeed Alkhaja, Mohammed Uddin, Rifat Hamoudi, Rabih Halwani, Abiola Catherine Senok, Qutayba Hamid, Norbert Nowotny, Alawi Alsheikh-Ali |
| EPI_ISL_469300 | National Institute of Laboratory Medicine and Referral Center | Genomic Research Lab, BCSIR | Abu Sayeed Mohammad Mahmud, Mohammad Samir Uzzaman, Eshrar Osman, Md. Ahasan Habib, Shahina Akter, Tanjina Akhter Banu, Md. Murshed Hasan Sarkar, Barna Goswami, Iffat Jahan, Md. Saddam Hossain, Tasnim Nafisa, Md. Maruf Ahmed Molla, Mahmuda Yeasmin, Asish Kumar Ghosh, Bayzid Bin Monir, A. K. M. Shamsuzzaman, Sheikh Md. Selim Al Din, Utpal Chandra Ray, Salek Ahmed Sajib, Md. Salim Khan |
| EPI_ISL_469930, EPI_ISL_469931, EPI_ISL_469932, EPI_ISL_469933, EPI_ISL_469934, EPI_ISL_469935, EPI_ISL_469936, EPI_ISL_469937, EPI_ISL_469938, EPI_ISL_469939, EPI_ISL_469940, EPI_ISL_469941, EPI_ISL_469942, EPI_ISL_469943, EPI_ISL_469944, EPI_ISL_469945, EPI_ISL_469946, EPI_ISL_469947, EPI_ISL_469948, EPI_ISL_469949, EPI_ISL_469950, EPI_ISL_469951, EPI_ISL_469952, EPI_ISL_469953, EPI_ISL_469954, EPI_ISL_469955, EPI_ISL_469956, EPI_ISL_469957, EPI_ISL_469958, EPI_ISL_469959, EPI_ISL_469960, EPI_ISL_469961, EPI_ISL_469962, EPI_ISL_469963, EPI_ISL_469964, EPI_ISL_469965, EPI_ISL_469966, EPI_ISL_469967, EPI_ISL_469968, EPI_ISL_469969, EPI_ISL_469970, EPI_ISL_469971, EPI_ISL_469972, EPI_ISL_469973, EPI_ISL_469974, EPI_ISL_469975, EPI_ISL_469976, EPI_ISL_469977, EPI_ISL_469978, EPI_ISL_469979, EPI_ISL_469980, EPI_ISL_469981, EPI_ISL_469982, EPI_ISL_469983, EPI_ISL_469984, EPI_ISL_469985, EPI_ISL_469986, EPI_ISL_469987, EPI_ISL_469988, EPI_ISL_469989, EPI_ISL_469990, EPI_ISL_469991, EPI_ISL_469992, EPI_ISL_469993, EPI_ISL_469994, EPI_ISL_469995, EPI_ISL_469996, EPI_ISL_469997, EPI_ISL_469998, EPI_ISL_469999, EPI_ISL_470000, EPI_ISL_470001, EPI_ISL_470002, EPI_ISL_470003, EPI_ISL_470004, EPI_ISL_470005, EPI_ISL_470006, EPI_ISL_470007, EPI_ISL_470008, EPI_ISL_470009, EPI_ISL_470010, EPI_ISL_470011, EPI_ISL_470012 | | | |
| see above | NHSGGC West of Scotland Specialist Virology Centre / MRC-University of Glasgow Centre for Virus Research | Wellcome Sanger Institute for the COVID-19 Genomics UK (COG-UK) consortium | Ana da Silva Filipe, Natasha Johnson, Kathy Smollett, Daniel Mair, Stephen Carmichael, Lily Tong, Jenna Nichols, Elihu Aranday-Cortes, Kirstyn Brunker, Yasmin Parr, Kyriaki Nomikou; Sarah McDonald, Marc Niebel, Patawee Asamaphan; Richard Orton, Joseph Hughes, Sreenu Vattipally, David L Robertson; Alasdair MacLean, Rory Gunson; Kathy Li, Natasha Jesudason, Rajiv Shah, James Shepherd, Antonia Ho, Alice Broos, Emma Thomson and Alex Alderton, Roberto Amato, Sonia Goncalves, Ewan Harrison, David K. Jackson, Ian Johnston, Dominic Kwiatkowski, Cordelia Langford, John Sillitoe on behalf of the Wellcome Sanger Institute COVID-19 Surveillance Team (http://www.sanger.ac.uk/covid-team) |
| EPI_ISL_470568, EPI_ISL_470569, EPI_ISL_470570, EPI_ISL_470571, EPI_ISL_470572, EPI_ISL_470573, EPI_ISL_470574, EPI_ISL_470575, EPI_ISL_470576, EPI_ISL_470577, EPI_ISL_470578, EPI_ISL_470579, EPI_ISL_470580, EPI_ISL_470581, EPI_ISL_470582, EPI_ISL_470583, EPI_ISL_470584, EPI_ISL_470585, EPI_ISL_470586, EPI_ISL_470587, EPI_ISL_470588 | | | |
| see above | Hermes Pardini | Bioinformatics Laboratory / LNCC | Alexandra Gerber, Ana Paula Guimarães, Luiz Gonzaga Paula de Almeida, Ronaldo da Silva Francisco Junior, Mariane Talon, Filipe Romero, Átila Duque Rossi, Terezinha Marta Pereira, working group UFRJ, Jaqueline Goes de Jesus, Ingra Morales Claro, Ester Cerdeira Sabino, Nuno Rodrigues Faria, CADDE-group, Laboratorio Hermes Pardini, Laboratorio Simile, working group UFMG, Amilcar Tanuri, Carolina Voloch, Renato Santana Aguiar e Ana Tereza Vasconcelos |
| EPI_ISL_470589, EPI_ISL_470590, EPI_ISL_470591, EPI_ISL_470592, EPI_ISL_470593, EPI_ISL_470594, EPI_ISL_470595, EPI_ISL_470596, EPI_ISL_470597 | Simile | Bioinformatics Laboratory / LNCC | Alexandra Gerber, Ana Paula Guimarães, Luiz Gonzaga Paula de Almeida, Ronaldo da Silva Francisco Junior, Mariane Talon, Filipe Romero, Átila Duque Rossi, Terezinha Marta Pereira, working group UFRJ, Jaqueline Goes de Jesus, Ingra Morales Claro, Ester Cerdeira Sabino, Nuno Rodrigues Faria, CADDE-group, Laboratorio Hermes Pardini, Laboratorio Simile, working group UFMG, Amilcar Tanuri, Carolina Voloch, Renato Santana Aguiar e Ana Tereza Vasconcelos |
| EPI_ISL_470598, EPI_ISL_470599, EPI_ISL_470600, EPI_ISL_470601, EPI_ISL_470602, EPI_ISL_470603, EPI_ISL_470604, EPI_ISL_470605, EPI_ISL_470606, EPI_ISL_470607, EPI_ISL_470608, EPI_ISL_470609, EPI_ISL_470610, EPI_ISL_470611, EPI_ISL_470612, EPI_ISL_470613, EPI_ISL_470614 | | | |
| see above | Hermes Pardini | Bioinformatics Laboratory / LNCC | Alexandra Gerber, Ana Paula Guimarães, Luiz Gonzaga Paula de Almeida, Ronaldo da Silva Francisco Junior, Mariane Talon, Filipe Romero, Átila Duque Rossi, Terezinha Marta Pereira, working group UFRJ, Jaqueline Goes de Jesus, Ingra Morales Claro, Ester Cerdeira Sabino, Nuno Rodrigues Faria, CADDE-group, Laboratorio Hermes Pardini, Laboratorio Simile, working group UFMG, Amilcar Tanuri, Carolina Voloch, Renato Santana Aguiar e Ana Tereza Vasconcelos |
| EPI_ISL_470615, EPI_ISL_470616, EPI_ISL_470617, EPI_ISL_470618, EPI_ISL_470619, EPI_ISL_470620, EPI_ISL_470621, EPI_ISL_470622, EPI_ISL_470623, EPI_ISL_470624, EPI_ISL_470625, EPI_ISL_470626, EPI_ISL_470627, EPI_ISL_470628, EPI_ISL_470629, EPI_ISL_470630, EPI_ISL_470631, EPI_ISL_470632, EPI_ISL_470633, EPI_ISL_470634, EPI_ISL_470635, EPI_ISL_470636, EPI_ISL_470637, EPI_ISL_470638, EPI_ISL_470639, EPI_ISL_470640, EPI_ISL_470641, EPI_ISL_470642, EPI_ISL_470643, EPI_ISL_470644, EPI_ISL_470645, EPI_ISL_470646, EPI_ISL_470647, EPI_ISL_470648, EPI_ISL_470649, EPI_ISL_470650 | | | |
| see above | Laboratorio de Virologia Molecular / UFRJ | Bioinformatics Laboratory / LNCC | Alexandra Gerber, Ana Paula Guimarães, Luiz Gonzaga Paula de Almeida, Ronaldo da Silva Francisco Junior, Mariane Talon, Filipe Romero, Átila Duque Rossi, Terezinha Marta Pereira, working group UFRJ, Jaqueline Goes de Jesus, Ingra Morales Claro, Ester Cerdeira Sabino, Nuno Rodrigues Faria, CADDE-group, Laboratorio Hermes Pardini, Laboratorio Simile, working group UFMG, Amilcar Tanuri, Carolina Voloch, Renato Santana Aguiar e Ana Tereza Vasconcelos |
| EPI_ISL_470651, EPI_ISL_470652, EPI_ISL_470653, EPI_ISL_470654, EPI_ISL_470655 | Hermes Pardini | Bioinformatics Laboratory / LNCC | Alexandra Gerber, Ana Paula Guimarães, Luiz Gonzaga Paula de Almeida, Ronaldo da Silva Francisco Junior, Mariane Talon, Filipe Romero, Átila Duque Rossi, Terezinha Marta Pereira, working group UFRJ, Jaqueline Goes de Jesus, Ingra Morales Claro, Ester Cerdeira Sabino, Nuno Rodrigues Faria, CADDE-group, Laboratorio Hermes Pardini, Laboratorio Simile, working group UFMG, Amilcar Tanuri, Carolina Voloch, Renato Santana Aguiar e Ana Tereza Vasconcelos |
| EPI_ISL_470878, EPI_ISL_470879, EPI_ISL_470880 | National Institute for Communicable Diseases of the National Health Laboratory Service | National Institute for Communicable Diseases of the National Health Laboratory Service | Allam M, Ismail A, Khumalo Z, Kwenda S, van Heusden P, Mtshali P, Mnyameni F, Mohale T, Subramoney K, Bhiman JN |
| EPI_ISL_470896 | Russian State Collection of Viruses | Pathogenic Microorganisms Variability Laboratory | Alexey Shchetinin, Maria Nikiforova, Elena Shidlovskaya, Nadezhda Kuznetsova, Inna Dolzhikova, Daria Grousova, Andrey Botikov, Denis Logunov, Alexander Gintsburg, Vladimir Gushchin |
| EPI_ISL_470897, EPI_ISL_470898, EPI_ISL_470899 | Pathogenic Microorganisms Variability Laboratory | Pathogenic Microorganisms Variability Laboratory | Alexey Shchetinin, Maria Nikiforova, Elena Shidlovskaya, Nadezhda Kuznetsova, Andrey Botikov, Alexander Gintsburg, Vladimir Gushchin |
| EPI_ISL_470900, EPI_ISL_470901, EPI_ISL_470902 | Influenza etiology and epidemiology laboratory | Pathogenic Microorganisms Variability Laboratory | Alexey Shchetinin, Maria Nikiforova, Elena Shidlovskaya, Nadezhda Kuznetsova, Vladimir Gushchin, Inna Dolzhikova, Daria Grousova, Andrey Botikov, Denis Logunov, Kirill Krasnoslobotsev, Svetlana Trushakova, Elena Burtseva, Ludmila Kolobukhina, Svetlana Smetanina, Alexander Gintsburg |

| | | | |
|---|---|---|---|
| EPI_ISL_470903, EPI_ISL_470904 | Influenza etiology and epidemiology laboratory | Pathogenic Microorganisms Variability Laboratory | Alexey Shchetinin, Maria Nikiforova, Elena Shidlovskaya, Nadezhda Kuznetsova, Vladimir Gushchin, Inna Dolzhikova, Daria Grousova, Andrey Botikov, Denis Logunov, Anna Ignatjeva, Evgeniya Mukasheva, Elena Burtseva, Ludmila Kolobukhina, Svetlana Smetanina, Alexander Gintsburg |
| EPI_ISL_473508, EPI_ISL_473509, EPI_ISL_473510, EPI_ISL_473511, EPI_ISL_473512, EPI_ISL_473513, EPI_ISL_473514, EPI_ISL_473515, EPI_ISL_473516, EPI_ISL_473517, EPI_ISL_473518, EPI_ISL_473519, EPI_ISL_473520, EPI_ISL_473521, EPI_ISL_473522, EPI_ISL_473523, EPI_ISL_473524, EPI_ISL_473525, EPI_ISL_473526, EPI_ISL_473527, EPI_ISL_473528, EPI_ISL_473529, EPI_ISL_473530, EPI_ISL_473531, EPI_ISL_473532, EPI_ISL_473533, EPI_ISL_473534, EPI_ISL_473535, EPI_ISL_473536, EPI_ISL_473537, EPI_ISL_473538, EPI_ISL_473539, EPI_ISL_473540, EPI_ISL_473541, EPI_ISL_473542, EPI_ISL_473543, EPI_ISL_473544, EPI_ISL_473545, EPI_ISL_473546, EPI_ISL_473547, EPI_ISL_473548, EPI_ISL_473549, EPI_ISL_473550, EPI_ISL_473551, EPI_ISL_473552, EPI_ISL_473553, EPI_ISL_473554, EPI_ISL_473555, EPI_ISL_473556, EPI_ISL_473557, EPI_ISL_473558, EPI_ISL_473559, EPI_ISL_473560, EPI_ISL_473561, EPI_ISL_473562, EPI_ISL_473563, EPI_ISL_473564, EPI_ISL_473565, EPI_ISL_473566, EPI_ISL_473567, EPI_ISL_473568, EPI_ISL_473569, EPI_ISL_473570, EPI_ISL_473571, EPI_ISL_473572, EPI_ISL_473573, EPI_ISL_473574, EPI_ISL_473575, EPI_ISL_473576, EPI_ISL_473577, EPI_ISL_473578, EPI_ISL_473579, EPI_ISL_473580, EPI_ISL_473581, EPI_ISL_473582, EPI_ISL_473583, EPI_ISL_473584, EPI_ISL_473585, EPI_ISL_473586, EPI_ISL_473587, EPI_ISL_473588, EPI_ISL_473589, EPI_ISL_473590, EPI_ISL_473591, EPI_ISL_473592, EPI_ISL_473593, EPI_ISL_473594, EPI_ISL_473595, EPI_ISL_473596, EPI_ISL_473597, EPI_ISL_473598, EPI_ISL_473599, EPI_ISL_473600, EPI_ISL_473601, EPI_ISL_473602, EPI_ISL_473603, EPI_ISL_473604, EPI_ISL_473605, EPI_ISL_473606, EPI_ISL_473607, EPI_ISL_473608, EPI_ISL_473609, EPI_ISL_473610, EPI_ISL_473611, EPI_ISL_473612, EPI_ISL_473613, EPI_ISL_473614, EPI_ISL_473615, EPI_ISL_473616, EPI_ISL_473617, EPI_ISL_473618, EPI_ISL_473619, EPI_ISL_473620, EPI_ISL_473621, EPI_ISL_473622, EPI_ISL_473623, EPI_ISL_473624, EPI_ISL_473625, EPI_ISL_473626, EPI_ISL_473627, EPI_ISL_473628, EPI_ISL_473629, EPI_ISL_473630, EPI_ISL_473631, EPI_ISL_473632, EPI_ISL_473633, EPI_ISL_473634, EPI_ISL_473635, EPI_ISL_473636, EPI_ISL_473637, EPI_ISL_473638, EPI_ISL_473639, EPI_ISL_473640, EPI_ISL_473641, EPI_ISL_473642, EPI_ISL_473643, EPI_ISL_473644, EPI_ISL_473645, EPI_ISL_473646, EPI_ISL_473647, EPI_ISL_473648, EPI_ISL_473649, EPI_ISL_473650, EPI_ISL_473651, EPI_ISL_473652, EPI_ISL_473653, EPI_ISL_473654, EPI_ISL_473655, EPI_ISL_473656, EPI_ISL_473657, EPI_ISL_473658, EPI_ISL_473659, EPI_ISL_473660, EPI_ISL_473661, EPI_ISL_473662, EPI_ISL_473663, EPI_ISL_473664, EPI_ISL_473665, EPI_ISL_473666, EPI_ISL_473667, EPI_ISL_473668, EPI_ISL_473669, EPI_ISL_473670, EPI_ISL_473671, EPI_ISL_473672, EPI_ISL_473673, EPI_ISL_473674, EPI_ISL_473675, EPI_ISL_473676, EPI_ISL_473677, EPI_ISL_473678, EPI_ISL_473679, EPI_ISL_473680, EPI_ISL_473681, EPI_ISL_473682, EPI_ISL_473683, EPI_ISL_473684, EPI_ISL_473685, EPI_ISL_473686, EPI_ISL_473687, EPI_ISL_473688, EPI_ISL_473689, EPI_ISL_473690, EPI_ISL_473691, EPI_ISL_473692, EPI_ISL_473693, EPI_ISL_473694, EPI_ISL_473695, EPI_ISL_473696, EPI_ISL_473697, EPI_ISL_473698, EPI_ISL_473699, EPI_ISL_473700, EPI_ISL_473701, EPI_ISL_473702, EPI_ISL_473703, EPI_ISL_473704, EPI_ISL_473705, EPI_ISL_473706, EPI_ISL_473707, EPI_ISL_473708, EPI_ISL_473709, EPI_ISL_473710, EPI_ISL_473711, EPI_ISL_473712, EPI_ISL_473713, EPI_ISL_473714, EPI_ISL_473715, EPI_ISL_473716, EPI_ISL_473717, EPI_ISL_473718, EPI_ISL_473719, EPI_ISL_473720, EPI_ISL_473721, EPI_ISL_473722, EPI_ISL_473723, EPI_ISL_473724, EPI_ISL_473725, EPI_ISL_473726, EPI_ISL_473727, EPI_ISL_473728, EPI_ISL_473729, EPI_ISL_473730, EPI_ISL_473731, EPI_ISL_473732, EPI_ISL_473733, EPI_ISL_473734, EPI_ISL_473735, EPI_ISL_473736, EPI_ISL_473737, EPI_ISL_473738, EPI_ISL_473739, EPI_ISL_473740, EPI_ISL_473741, EPI_ISL_473742, EPI_ISL_473743, EPI_ISL_473744, EPI_ISL_473745, EPI_ISL_473746, EPI_ISL_473747, EPI_ISL_473748, EPI_ISL_473749, EPI_ISL_473750, EPI_ISL_473751, EPI_ISL_473752, EPI_ISL_473753, EPI_ISL_473754, EPI_ISL_473755, EPI_ISL_473756, EPI_ISL_473757, EPI_ISL_473758, EPI_ISL_473759, EPI_ISL_473760, EPI_ISL_473761, EPI_ISL_473762, EPI_ISL_473763, EPI_ISL_473764, EPI_ISL_473765, EPI_ISL_473766, EPI_ISL_473767, EPI_ISL_473768, EPI_ISL_473769, EPI_ISL_473770, EPI_ISL_473771, EPI_ISL_473772, EPI_ISL_473773, EPI_ISL_473774, EPI_ISL_473775, EPI_ISL_473776, EPI_ISL_473777, EPI_ISL_473778, EPI_ISL_473779, EPI_ISL_473780, EPI_ISL_473781, EPI_ISL_473782 | | | |
| see above | West of Scotland Specialist Virology Centre, NHSGGC / MRC-University of Glasgow Centre for Virus Research | COVID-19 Genomics UK (COG-UK) Consortium | Ana da Silva Filipe, Natasha Johnson, Kathy Smollett, Daniel Mair, Stephen Carmichael, Lily Tong, Jenna Nichols, Elihu Aranday-Cortes, Kirstyn Brunker, Yasmin Parr, Alice Broos, Kyriaki Nomikou; Sarah McDonald, Marc Niebel, Patawee Asamaphan; Richard Orton, Joseph Hughes, Sreenu Vattipally, David L Robertson; Alasdair MacLean, Rory Gunson; Kathy Li, Natasha Jesudason, Rajiv Shah, James Shepherd, Antonia Ho, Emma Thomson |
| EPI_ISL_475045 | Department of MicroBiology, Government Medical College, Surat | Gujarat Biotechnology Research Centre | Amit gamit, Pritesh Sabara, Apurvasinh Puvar, Janvi Raval, Zarna Patel, Monika Gandhi, Pinal Trivedi, Maharshi Pandya, Nidhi Patel, Nitin Savaliya, Raghawendra Kumar, Dinesh Kumar, Zuber Saiyed, Komal Patel, Labdhi Pandya, Snehal Bagatharia, Naresh Chauhan, Summaiya Mullan, Akanksha Verma, R D Dixit, A M Kadri, Harsh Bakshi, Chaitanya Joshi, Madhvi Joshi |
| EPI_ISL_475170, EPI_ISL_475171, EPI_ISL_475172, EPI_ISL_475173, EPI_ISL_475238 | National Institute of Laboratory Medicine and Referral Center | Genomic Research Lab, BCSIR | Abu Sayeed Mohammad Mahmud, Mohammad Samir Uzzaman, Eshrar Osman, Md. Ahasan Habib, Shahina Akter, Tanjina Akhter Banu, Md. Murshed Hasan Sarkar, Barna Goswami, Iffat Jahan, Md. Saddam Hossain, Tasnim Nafisa, Md. Maruf Ahmed Molla, Mahmuda Yeasmin, Asish Kumar Ghosh, Bayzid Bin Monir, A. K. M. Shamsuzzaman, Sheikh Md. Selim Al Din, Utpal Chandra Ray, Salek Ahmed Sajib, Md. Salim Khan |
| EPI_ISL_475272, EPI_ISL_475274, EPI_ISL_475275, EPI_ISL_475276, EPI_ISL_475277, EPI_ISL_475279, EPI_ISL_475280, EPI_ISL_475281, EPI_ISL_475282, EPI_ISL_475283, EPI_ISL_475285, EPI_ISL_475286, EPI_ISL_475287, EPI_ISL_475288, EPI_ISL_475289, EPI_ISL_475291, EPI_ISL_475292, EPI_ISL_475293, EPI_ISL_475295, EPI_ISL_475296, EPI_ISL_475297, EPI_ISL_475298, EPI_ISL_475299, EPI_ISL_475300, EPI_ISL_475301, EPI_ISL_475302, EPI_ISL_475303, EPI_ISL_475306, EPI_ISL_475307, EPI_ISL_475308, EPI_ISL_475309, EPI_ISL_475310, EPI_ISL_475311, EPI_ISL_475312, EPI_ISL_475313, EPI_ISL_475314, EPI_ISL_475315, EPI_ISL_475316, EPI_ISL_475319, EPI_ISL_475321, EPI_ISL_475322, EPI_ISL_475323, EPI_ISL_475324, EPI_ISL_475325, EPI_ISL_475326, EPI_ISL_475327, EPI_ISL_475328, EPI_ISL_475329, EPI_ISL_475330, EPI_ISL_475331, EPI_ISL_475332, EPI_ISL_475333, EPI_ISL_475334, EPI_ISL_475335, EPI_ISL_475336, EPI_ISL_475337, EPI_ISL_475339, EPI_ISL_475341 | | | |
| see above | Centre for Enzyme Innovation, University of Portsmouth / Translational Research Laboratory, Portsmouth Hospitals NHS Trust | COVID-19 Genomics UK (COG-UK) Consortium | Angela Beckett,Yann Bourgeois,Garry Scarlett,Sharon Glaysher,Scott Elliott,Kelly Bicknell,Robert Impey,Allyson Lloyd,Sarah Wyllie,Ethan Butcher,Anoop Chauhan,Samuel Robson |
| EPI_ISL_475570 | Genome Center | Genome Center | A. S. M. Rubayet- Ul- Alam, Ovinu Kibria Islam, Md. Shazid Hasan, Hassan M. Al-Emran, Shireen Nigar, Selina Akter, Pravas Chandra Roy, Md. Tanvir Islam, Shovon Lal Sarkar, M. Shaminur Rahman, M. Rafiul Islam, Habiba Ibnat, Md Nur Kabidul Azam, Chakrabortty Atonu, Proshanto Kumar Das, Md. Hasan al Pramanik, Md. Zannat Ali, Shohanur Rahaman, Md. Aminul Islam, Ashok Kumar, Md. Nazmul Hasan, Md. Iqbal Kabir Jahid, Md. Anwar Hossain |
| EPI_ISL_475760, EPI_ISL_475761 | National Institute of Laboratory Medicine and Referral Center | Genomic Research Lab, BCSIR | Abu Sayeed Mohammad Mahmud, Mohammad Samir Uzzaman, Eshrar Osman, Md. Ahasan Habib, Shahina Akter, Tanjina Akhter Banu, Md. Murshed Hasan Sarkar, Barna Goswami, Iffat Jahan, Md. Saddam Hossain, Tasnim Nafisa, Md. Maruf Ahmed Molla, Mahmuda Yeasmin, Asish Kumar Ghosh, Arifa Akram, A. K. M. Shamsuzzaman, Sheikh Md. Selim Al Din, Utpal Chandra Ray, Salek Ahmed Sajib, Md. Salim Khan |
| EPI_ISL_475763 | Institut für Virologie am Department für Hygiene, Mikrobiologie und Public Health | Bergthaler laboratory, CeMM Research Center for Molecular Medicine of the Austrian Academy of Sciences | Alexandra Popa, Benedikt Agerer, Henrique Colaco, Lukas Endler, Jakob-Wendelin Genger, Alexander Lercher, Mark Smyth, Thomas Penz, Michael Schuster, Jan Laine, Martin Senekowitsch, Judith Aberle, Stephan Aberle, Peter Hufnagl, Daniela Schmid, Franz Allerberger, Elisabeth Puchhammer-Stoeckl, Manfred Nairz, Guenter Weiss, Gregor Hörmann, Kinga Rigler-Hohenwarter, Rainer Gattringer, Wegene Borena, Dorothee von Laer, Christoph Bock, Andreas Bergthaler |
| EPI_ISL_475764, EPI_ISL_475765, EPI_ISL_475766, EPI_ISL_475767, EPI_ISL_475768 | Universitaetsklinik für Innere Medizin II Innsbruck | Bergthaler laboratory, CeMM Research Center for Molecular Medicine of the Austrian Academy of Sciences | Alexandra Popa, Benedikt Agerer, Henrique Colaco, Lukas Endler, Jakob-Wendelin Genger, Alexander Lercher, Mark Smyth, Thomas Penz, Michael Schuster, Jan Laine, Martin Senekowitsch, Judith Aberle, Stephan Aberle, Peter Hufnagl, Daniela Schmid, Franz Allerberger, Elisabeth Puchhammer-Stoeckl, Manfred Nairz, Guenter Weiss, Gregor Hörmann, Kinga Rigler-Hohenwarter, Rainer Gattringer, Wegene Borena, Dorothee von Laer, Christoph Bock, Andreas Bergthaler |
| EPI_ISL_475769 | Institut für Virologie am Department für Hygiene, Mikrobiologie und Public Health | Bergthaler laboratory, CeMM Research Center for Molecular Medicine of the Austrian Academy of Sciences | Alexandra Popa, Benedikt Agerer, Henrique Colaco, Lukas Endler, Jakob-Wendelin Genger, Alexander Lercher, Mark Smyth, Thomas Penz, Michael Schuster, Jan Laine, Martin Senekowitsch, Judith Aberle, Stephan Aberle, Peter Hufnagl, Daniela Schmid, Franz Allerberger, Elisabeth Puchhammer-Stoeckl, Manfred Nairz, Guenter Weiss, Gregor Hörmann, Kinga Rigler-Hohenwarter, Rainer Gattringer, Wegene Borena, Dorothee von Laer, Christoph Bock, Andreas Bergthaler |
| EPI_ISL_475770, EPI_ISL_475771, EPI_ISL_475772, EPI_ISL_475773, EPI_ISL_475774, EPI_ISL_475775, EPI_ISL_475776, EPI_ISL_475777, EPI_ISL_475778, EPI_ISL_475779, EPI_ISL_475780, EPI_ISL_475781, EPI_ISL_475782, EPI_ISL_475783, EPI_ISL_475784, EPI_ISL_475785, EPI_ISL_475786, EPI_ISL_475787, EPI_ISL_475788, EPI_ISL_475789, EPI_ISL_475790, EPI_ISL_475791, EPI_ISL_475792, EPI_ISL_475793, EPI_ISL_475794, EPI_ISL_475795, EPI_ISL_475796, EPI_ISL_475797, EPI_ISL_475798, EPI_ISL_475799, EPI_ISL_475800, EPI_ISL_475801, EPI_ISL_475802, EPI_ISL_475803, EPI_ISL_475804, EPI_ISL_475805, EPI_ISL_475806, EPI_ISL_475807, EPI_ISL_475808, EPI_ISL_475809, EPI_ISL_475810, EPI_ISL_475811, EPI_ISL_475812 | | | |
| see above | Center for Virology, Medical University of Vienna | Bergthaler laboratory, CeMM Research Center for Molecular Medicine of the Austrian Academy of Sciences | Alexandra Popa, Benedikt Agerer, Henrique Colaco, Lukas Endler, Jakob-Wendelin Genger, Alexander Lercher, Mark Smyth, Thomas Penz, Michael Schuster, Jan Laine, Martin Senekowitsch, Judith Aberle, Stephan Aberle, Peter Hufnagl, Daniela Schmid, Franz Allerberger, Elisabeth Puchhammer-Stoeckl, Manfred Nairz, Guenter Weiss, Gregor Hörmann, Kinga Rigler-Hohenwarter, Rainer Gattringer, Wegene Borena, Dorothee von Laer, Christoph Bock, Andreas Bergthaler |
| EPI_ISL_475813, EPI_ISL_475814, EPI_ISL_475815, EPI_ISL_475816, EPI_ISL_475817, EPI_ISL_475818, EPI_ISL_475819, EPI_ISL_475820, EPI_ISL_475821, EPI_ISL_475822, EPI_ISL_475823, EPI_ISL_475824, EPI_ISL_475825, EPI_ISL_475826, EPI_ISL_475827, EPI_ISL_475828, EPI_ISL_475829 | | | |
| see above | Institut für Virologie am Department für Hygiene, Mikrobiologie und Public Health | Bergthaler laboratory, CeMM Research Center for Molecular Medicine of the Austrian Academy of Sciences | Alexandra Popa, Benedikt Agerer, Henrique Colaco, Lukas Endler, Jakob-Wendelin Genger, Alexander Lercher, Mark Smyth, Thomas Penz, Michael Schuster, Jan Laine, Martin Senekowitsch, Judith Aberle, Stephan Aberle, Peter Hufnagl, Daniela Schmid, Franz Allerberger, Elisabeth Puchhammer-Stoeckl, Manfred Nairz, Guenter Weiss, Gregor Hörmann, Kinga Rigler-Hohenwarter, Rainer Gattringer, Wegene Borena, Dorothee von Laer, Christoph Bock, Andreas Bergthaler |
| EPI_ISL_475830, EPI_ISL_475831, EPI_ISL_475832, EPI_ISL_475833, EPI_ISL_475834, EPI_ISL_475835, EPI_ISL_475836, EPI_ISL_475837, EPI_ISL_475838, EPI_ISL_475839, EPI_ISL_475840, EPI_ISL_475841, EPI_ISL_475842, EPI_ISL_475843, EPI_ISL_475844, EPI_ISL_475845, EPI_ISL_475846, EPI_ISL_475847, EPI_ISL_475848, EPI_ISL_475849, EPI_ISL_475850, EPI_ISL_475851, EPI_ISL_475852, EPI_ISL_475853, EPI_ISL_475854, EPI_ISL_475855, EPI_ISL_475856, EPI_ISL_475857, EPI_ISL_475858, EPI_ISL_475859, EPI_ISL_475860, EPI_ISL_475861, EPI_ISL_475862, EPI_ISL_475863, EPI_ISL_475864, EPI_ISL_475865, EPI_ISL_475866, EPI_ISL_475867, EPI_ISL_475868, EPI_ISL_475869, EPI_ISL_475870, EPI_ISL_475871, EPI_ISL_475872, EPI_ISL_475873, EPI_ISL_475874, EPI_ISL_475875, EPI_ISL_475876, EPI_ISL_475877, EPI_ISL_475878, EPI_ISL_475879, EPI_ISL_475880, EPI_ISL_475881, EPI_ISL_475882, EPI_ISL_475883, EPI_ISL_475884, EPI_ISL_475885, EPI_ISL_475886 | | | |
| see above | Austrian Agency for Health and Food Safety (AGES) | Bergthaler laboratory, CeMM Research Center for Molecular Medicine of the Austrian Academy of Sciences | Alexandra Popa, Benedikt Agerer, Henrique Colaco, Lukas Endler, Jakob-Wendelin Genger, Alexander Lercher, Mark Smyth, Thomas Penz, Michael Schuster, Jan Laine, Martin Senekowitsch, Judith Aberle, Stephan Aberle, Peter Hufnagl, Daniela Schmid, Franz Allerberger, Elisabeth Puchhammer-Stoeckl, Manfred Nairz, Guenter Weiss, Gregor Hörmann, Kinga Rigler-Hohenwarter, Rainer Gattringer, Wegene Borena, Dorothee von Laer, Christoph Bock, Andreas Bergthaler |
| EPI_ISL_475887, EPI_ISL_475888, EPI_ISL_475889, EPI_ISL_475890, EPI_ISL_475891, EPI_ISL_475892, EPI_ISL_475893, EPI_ISL_475894, EPI_ISL_475895, EPI_ISL_475896, EPI_ISL_475897, EPI_ISL_475898, EPI_ISL_475899, EPI_ISL_475900, EPI_ISL_475901, EPI_ISL_475902, EPI_ISL_475903, EPI_ISL_475904, EPI_ISL_475905, EPI_ISL_475906, EPI_ISL_475907, EPI_ISL_475908, EPI_ISL_475909 | | | |
| see above | Zentralinstitut für medizinische und chemische Labordiagnostik, Universitätskliniken Insbruck | Bergthaler laboratory, CeMM Research Center for Molecular Medicine of the Austrian Academy of Sciences | Alexandra Popa, Benedikt Agerer, Henrique Colaco, Lukas Endler, Jakob-Wendelin Genger, Alexander Lercher, Mark Smyth, Thomas Penz, Michael Schuster, Jan Laine, Martin Senekowitsch, Judith Aberle, Stephan Aberle, Peter Hufnagl, Daniela Schmid, Franz Allerberger, Elisabeth Puchhammer-Stoeckl, Manfred Nairz, Guenter Weiss, Gregor Hörmann, Kinga Rigler-Hohenwarter, Rainer Gattringer, Wegene Borena, Dorothee von Laer, Christoph Bock, Andreas Bergthaler |

| | | | |
|---|---|---|---|
| EPI_ISL_475910, EPI_ISL_475911, EPI_ISL_475912, EPI_ISL_475913, EPI_ISL_475914, EPI_ISL_475915 | Klinikum Wels-Grieskirchen | Bergthaler laboratory, CeMM Research Center for Molecular Medicine of the Austrian Academy of Sciences | Alexandra Popa, Benedikt Agerer, Henrique Colaco, Lukas Endler, Jakob-Wendelin Genger, Alexander Lercher, Mark Smyth, Thomas Penz, Michael Schuster, Jan Laine, Martin Senekowitsch, Judith Aberle, Stephan Aberle, Peter Hufnagl, Daniela Schmid, Franz Allerberger, Elisabeth Puchhammer-Stoeckl, Manfred Nairz, Guenter Weiss, Gregor Hörmann, Kinga Rigler-Hohenwarter, Rainer Gattringer, Wegene Borena, Dorothee von Laer, Christoph Bock, Andreas Bergthaler |
| EPI_ISL_475916, EPI_ISL_475917, EPI_ISL_475918, EPI_ISL_475919, EPI_ISL_475920, EPI_ISL_475921, EPI_ISL_475922, EPI_ISL_475923, EPI_ISL_475924, EPI_ISL_475925, EPI_ISL_475926, EPI_ISL_475927, EPI_ISL_475928 | | | |
| see above | Institut für Virologie am Department für Hygiene, Mikrobiologie und Public Health | Bergthaler laboratory, CeMM Research Center for Molecular Medicine of the Austrian Academy of Sciences | Alexandra Popa, Benedikt Agerer, Henrique Colaco, Lukas Endler, Jakob-Wendelin Genger, Alexander Lercher, Mark Smyth, Thomas Penz, Michael Schuster, Jan Laine, Martin Senekowitsch, Judith Aberle, Stephan Aberle, Peter Hufnagl, Daniela Schmid, Franz Allerberger, Elisabeth Puchhammer-Stoeckl, Manfred Nairz, Guenter Weiss, Gregor Hörmann, Kinga Rigler-Hohenwarter, Rainer Gattringer, Wegene Borena, Dorothee von Laer, Christoph Bock, Andreas Bergthaler |
| EPI_ISL_475929, EPI_ISL_475930, EPI_ISL_475931, EPI_ISL_475932, EPI_ISL_475933, EPI_ISL_475934, EPI_ISL_475935, EPI_ISL_475936 | Universitaetsklinik für Innere Medizin II Innsbruck | Bergthaler laboratory, CeMM Research Center for Molecular Medicine of the Austrian Academy of Sciences | Alexandra Popa, Benedikt Agerer, Henrique Colaco, Lukas Endler, Jakob-Wendelin Genger, Alexander Lercher, Mark Smyth, Thomas Penz, Michael Schuster, Jan Laine, Martin Senekowitsch, Judith Aberle, Stephan Aberle, Peter Hufnagl, Daniela Schmid, Franz Allerberger, Elisabeth Puchhammer-Stoeckl, Manfred Nairz, Guenter Weiss, Gregor Hörmann, Kinga Rigler-Hohenwarter, Rainer Gattringer, Wegene Borena, Dorothee von Laer, Christoph Bock, Andreas Bergthaler |
| EPI_ISL_476565 | Hospital de Pediatría "Prof. Dr. Juan P Garrahan" | Héritas | Andrea Mangano, Maria Florencia Fernandez, Dalmacio Pereyra, Roberta Crespo, Mauricio Grisolia, Cristian Rohr, Fabian Fay, Martin Vazquez |
| EPI_ISL_476702, EPI_ISL_476703, EPI_ISL_476704 | Incubadora Venezolana de Ciencia, Venezuela | Incubadora Venezolana de Ciencia, Venezuela / Instituto Nacional de Salud, Bogotá, Colombia / Grupo de Investigaciones Microbiológicas-UR (GIMUR), Departamento de Biología, Facultad de Ciencias Naturales, Universidad del Rosario, Bogotá, Colombia / Icahn School of Medicine at Mount Sinai, New York, USA | Alberto Paniz-Mondolfi, Marina Muñoz, Luis Perez-Garcia, Lourdes Delgado, Carolina Florez, Sergio Gomez, Angelica Rico, Lisseth Pardo, Esther C. Barros, Carolina Hernández, Jesús E. Jaimes, Aníbal A. Teherán, Ana S. Gonzalez-Reiche, Matthew M. Hernandez, Emilia Mia Sordillo, Viviana Simon, Harm van Bakel, Juan David Ramírez |
| EPI_ISL_476868 | Banas Medical College and Research Institute | Gujarat Biotechnology Research Centre | Afzal Ansari, Nikha Trivedi, Radhika Khara, Sunil R Joshi, Viren s Doshi, Apurvasinh Puvar, Janvi Raval, Zarna Patel, Monika Gandhi, Pinal Trivedi, Maharshi Pandya, Nidhi Patel, Nitin Savaliya, Raghawendra Kumar, Dinesh Kumar, Zuber Saiyed, Komal Patel, Labdhi Pandya, R D Dixit, A M Kadri, Harsh Bakshi, Chaitanya Joshi, Madhvi Joshi |
| EPI_ISL_476872 | Department of MicroBiology, Government Medical College, Surat | Gujarat Biotechnology Research Centre | Amit gamit, Apurvasinh Puvar, Janvi Raval, Zarna Patel, Monika Gandhi, Pinal Trivedi, Maharshi Pandya, Nidhi Patel, Nitin Savaliya, Raghawendra Kumar, Dinesh Kumar, Zuber Saiyed, Komal Patel, Labdhi Pandya, Afzal Ansari, Nikha Trivedi, Naresh Chauhan, Summaiya Mullan, R D Dixit, A M Kadri, Harsh Bakshi, Chaitanya Joshi, Madhvi Joshi |
| EPI_ISL_477820, EPI_ISL_477821, EPI_ISL_477822, EPI_ISL_477823, EPI_ISL_477824, EPI_ISL_477825, EPI_ISL_477826, EPI_ISL_477827, EPI_ISL_477828, EPI_ISL_477829, EPI_ISL_477830, EPI_ISL_477831, EPI_ISL_477832, EPI_ISL_477833, EPI_ISL_477834, EPI_ISL_477835, EPI_ISL_477836, EPI_ISL_477837, EPI_ISL_477838, EPI_ISL_477839, EPI_ISL_477840, EPI_ISL_477841, EPI_ISL_477842, EPI_ISL_477843, EPI_ISL_477844, EPI_ISL_477845, EPI_ISL_477846, EPI_ISL_477847, EPI_ISL_477848, EPI_ISL_477849, EPI_ISL_477850, EPI_ISL_477851, EPI_ISL_477852, EPI_ISL_477853, EPI_ISL_477854, EPI_ISL_477855, EPI_ISL_477856, EPI_ISL_477857, EPI_ISL_477858, EPI_ISL_477859, EPI_ISL_477860, EPI_ISL_477861, EPI_ISL_477862, EPI_ISL_477863, EPI_ISL_477864, EPI_ISL_477865, EPI_ISL_477866, EPI_ISL_477867, EPI_ISL_477868, EPI_ISL_477869, EPI_ISL_477870, EPI_ISL_477871, EPI_ISL_477872, EPI_ISL_477873, EPI_ISL_477874, EPI_ISL_477875, EPI_ISL_477876, EPI_ISL_477877, EPI_ISL_477878, EPI_ISL_477879, EPI_ISL_477880, EPI_ISL_477881, EPI_ISL_477882, EPI_ISL_477883, EPI_ISL_477884, EPI_ISL_477885, EPI_ISL_477886, EPI_ISL_477887, EPI_ISL_477888, EPI_ISL_477889, EPI_ISL_477890, EPI_ISL_477891, EPI_ISL_477892, EPI_ISL_477893, EPI_ISL_477894, EPI_ISL_477895, EPI_ISL_477896, EPI_ISL_477897, EPI_ISL_477898, EPI_ISL_477899, EPI_ISL_477900, EPI_ISL_477901, EPI_ISL_477902, EPI_ISL_477903, EPI_ISL_477904, EPI_ISL_477905, EPI_ISL_477906, EPI_ISL_477907, EPI_ISL_477908, EPI_ISL_477909, EPI_ISL_477910, EPI_ISL_477911, EPI_ISL_477912, EPI_ISL_477913, EPI_ISL_477914, EPI_ISL_477915, EPI_ISL_477916, EPI_ISL_477917, EPI_ISL_477918, EPI_ISL_477919, EPI_ISL_477920, EPI_ISL_477921, EPI_ISL_477922, EPI_ISL_477923, EPI_ISL_477924, EPI_ISL_477925, EPI_ISL_477926, EPI_ISL_477927, EPI_ISL_477928, EPI_ISL_477929, EPI_ISL_477930, EPI_ISL_477931, EPI_ISL_477932, EPI_ISL_477933, EPI_ISL_477934, EPI_ISL_477935, EPI_ISL_477936, EPI_ISL_477937, EPI_ISL_477938, EPI_ISL_477939, EPI_ISL_477940, EPI_ISL_477941, EPI_ISL_477942, EPI_ISL_477943, EPI_ISL_477944, EPI_ISL_477945, EPI_ISL_477946, EPI_ISL_477947, EPI_ISL_477948, EPI_ISL_477949, EPI_ISL_477950, EPI_ISL_477951, EPI_ISL_477952, EPI_ISL_477953, EPI_ISL_477954, EPI_ISL_477955, EPI_ISL_477956, EPI_ISL_477957, EPI_ISL_477958, EPI_ISL_477959, EPI_ISL_477960, EPI_ISL_477961, EPI_ISL_477962, EPI_ISL_477963, EPI_ISL_477964, EPI_ISL_477965, EPI_ISL_477966, EPI_ISL_477967, EPI_ISL_477968, EPI_ISL_477969, EPI_ISL_477970, EPI_ISL_477971, EPI_ISL_477972, EPI_ISL_477973, EPI_ISL_477974, EPI_ISL_477975, EPI_ISL_477976, EPI_ISL_477977, EPI_ISL_477978, EPI_ISL_477979, EPI_ISL_477980, EPI_ISL_477981, EPI_ISL_477982, EPI_ISL_477983, EPI_ISL_477984, EPI_ISL_477985, EPI_ISL_477986, EPI_ISL_477987, EPI_ISL_477988, EPI_ISL_477989, EPI_ISL_477990, EPI_ISL_477991, EPI_ISL_477992, EPI_ISL_477993, EPI_ISL_477994, EPI_ISL_477995, EPI_ISL_477996, EPI_ISL_477997, EPI_ISL_477998, EPI_ISL_477999, EPI_ISL_478000, EPI_ISL_478001, EPI_ISL_478002, EPI_ISL_478003, EPI_ISL_478004, EPI_ISL_478005, EPI_ISL_478006, EPI_ISL_478007, EPI_ISL_478008, EPI_ISL_478009, EPI_ISL_478010, EPI_ISL_478011, EPI_ISL_478012, EPI_ISL_478013, EPI_ISL_478014, EPI_ISL_478015, EPI_ISL_478016, EPI_ISL_478017, EPI_ISL_478018, EPI_ISL_478019, EPI_ISL_478020, EPI_ISL_478021, EPI_ISL_478022, EPI_ISL_478023, EPI_ISL_478024, EPI_ISL_478025, EPI_ISL_478026, EPI_ISL_478027, EPI_ISL_478028, EPI_ISL_478029, EPI_ISL_478030, EPI_ISL_478031, EPI_ISL_478032, EPI_ISL_478033, EPI_ISL_478034, EPI_ISL_478035, EPI_ISL_478036, EPI_ISL_478037, EPI_ISL_478038, EPI_ISL_478039, EPI_ISL_478040, EPI_ISL_478041, EPI_ISL_478042, EPI_ISL_478043, EPI_ISL_478044, EPI_ISL_478045, EPI_ISL_478046, EPI_ISL_478047, EPI_ISL_478048, EPI_ISL_478049, EPI_ISL_478050, EPI_ISL_478051, EPI_ISL_478052, EPI_ISL_478053, EPI_ISL_478054, EPI_ISL_478055, EPI_ISL_478056, EPI_ISL_478057, EPI_ISL_478058, EPI_ISL_478059, EPI_ISL_478060, EPI_ISL_478061, EPI_ISL_478062, EPI_ISL_478063, EPI_ISL_478064, EPI_ISL_478065, EPI_ISL_478066, EPI_ISL_478067, EPI_ISL_478068, EPI_ISL_478069, EPI_ISL_478070, EPI_ISL_478071, EPI_ISL_478072, EPI_ISL_478073, EPI_ISL_478074, EPI_ISL_478075, EPI_ISL_478076, EPI_ISL_478077, EPI_ISL_478078, EPI_ISL_478079, EPI_ISL_478080, EPI_ISL_478081, EPI_ISL_478082, EPI_ISL_478083, EPI_ISL_478084, EPI_ISL_478085, EPI_ISL_478086, EPI_ISL_478087, EPI_ISL_478088, EPI_ISL_478089, EPI_ISL_478090, EPI_ISL_478091, EPI_ISL_478092, EPI_ISL_478093, EPI_ISL_478094, EPI_ISL_478095, EPI_ISL_478096, EPI_ISL_478097, EPI_ISL_478098, EPI_ISL_478099, EPI_ISL_478100, EPI_ISL_478101, EPI_ISL_478102, EPI_ISL_478103, EPI_ISL_478104, EPI_ISL_478105, EPI_ISL_478106, EPI_ISL_478107, EPI_ISL_478108, EPI_ISL_478109, EPI_ISL_478110, EPI_ISL_478111, EPI_ISL_478112, EPI_ISL_478113, EPI_ISL_478114, EPI_ISL_478115, EPI_ISL_478116, EPI_ISL_478117, EPI_ISL_478118, EPI_ISL_478119, EPI_ISL_478120, EPI_ISL_478121, EPI_ISL_478122, EPI_ISL_478123, EPI_ISL_478124, EPI_ISL_478125, EPI_ISL_478126, EPI_ISL_478127, EPI_ISL_478128, EPI_ISL_478129, EPI_ISL_478130, EPI_ISL_478131, EPI_ISL_478132, EPI_ISL_478133, EPI_ISL_478134, EPI_ISL_478135, EPI_ISL_478136, EPI_ISL_478137, EPI_ISL_478138, EPI_ISL_478139, EPI_ISL_478140, EPI_ISL_478141, EPI_ISL_478142, EPI_ISL_478143, EPI_ISL_478144, EPI_ISL_478145, EPI_ISL_478146, EPI_ISL_478147, EPI_ISL_478148, EPI_ISL_478149, EPI_ISL_478150, EPI_ISL_478151, EPI_ISL_478152, EPI_ISL_478153, EPI_ISL_478154, EPI_ISL_478155, EPI_ISL_478156, EPI_ISL_478157, EPI_ISL_478158, EPI_ISL_478159, EPI_ISL_478160, EPI_ISL_478161, EPI_ISL_478162 | | | |
| see above | West of Scotland Specialist Virology Centre, NHSGGC / MRC-University of Glasgow Centre for Virus Research | COVID-19 Genomics UK (COG-UK) Consortium | Ana da Silva Filipe, Natasha Johnson, Kathy Smollett, Daniel Mair, Stephen Carmichael, Lily Tong, Jenna Nichols, Elihu Aranday-Cortes, Kirstyn Brunker, Yasmin Parr, Alice Broos, Kyriaki Nomikou; Sarah McDonald, Marc Niebel, Patawee Asamaphan; Richard Orton, Joseph Hughes, Sreenu Vattipally, David L Robertson; Alasdair MacLean, Rory Gunson; Kathy Li, Natasha Jesudason, Rajiv Shah, James Shepherd, Antonia Ho, Emma Thomson |
| EPI_ISL_479175, EPI_ISL_479176, EPI_ISL_479177, EPI_ISL_479178, EPI_ISL_479179, EPI_ISL_479180, EPI_ISL_479181, EPI_ISL_479183, EPI_ISL_479184, EPI_ISL_479185, EPI_ISL_479186, EPI_ISL_479187, EPI_ISL_479188, EPI_ISL_479189, EPI_ISL_479190, EPI_ISL_479191, EPI_ISL_479192, EPI_ISL_479193, EPI_ISL_479194 | | | |
| see above | Centre for Enzyme Innovation, University of Portsmouth / Translational Research Laboratory, Portsmouth Hospitals NHS Trust | COVID-19 Genomics UK (COG-UK) Consortium | Angela Beckett,Yann Bourgeois,Garry Scarlett,Sharon Glaysher,Scott Elliott,Kelly Bicknell,Robert Impey,Allyson Lloyd,Sarah Wyllie,Ethan Butcher,Anoop Chauhan,Samuel Robson |
| EPI_ISL_480312, EPI_ISL_480313, EPI_ISL_480314 | Hospital Mexico | Charité Virology-University of Costa Rica | Andres Moreira-Soto, Eugenia Corrales-Aguilar, Ignacio Postigo-Hidalgo, Teresita Somogyi, Jan Felix Drexler |
| EPI_ISL_480315, EPI_ISL_480316, EPI_ISL_480317, EPI_ISL_480318, EPI_ISL_480319, EPI_ISL_480320 | Hospital Clínica Bíblica | Charité Virology-University of Costa Rica | Andres Moreira-Soto, Eugenia Corrales-Aguilar, Ignacio Postigo-Hidalgo, Karla Sofía Gutiérrez, Jan Felix Drexler |
| EPI_ISL_480321 | Laboratorio Clínico San José | Charité Virology-University of Costa Rica | Andres Moreira-Soto, Eugenia Corrales-Aguilar, Ignacio Postigo-Hidalgo, Hugo Núñez Navas, Jan Felix Drexler |
| EPI_ISL_480322, EPI_ISL_480324, EPI_ISL_480325, EPI_ISL_480326, EPI_ISL_480327 | Hospital Nacional de Niños | Charité Virology-University of Costa Rica | Andres Moreira-Soto, Eugenia Corrales-Aguilar, Ignacio Postigo-Hidalgo, Cristian Pérez Corrales, Andrei Montero Bonilla, Jan Felix Drexler |
| EPI_ISL_480328 | Laboratorio LABIN | Charité Virology-University of Costa Rica | Andres Moreira-Soto, Eugenia Corrales-Aguilar, Ignacio Postigo-Hidalgo, Ignacio Soto Pacheco, Jan Felix Drexler |
| EPI_ISL_480446, EPI_ISL_480447, EPI_ISL_480448, EPI_ISL_480449, EPI_ISL_480450 | National Institute of Laboratory Medicine and Referral Center | Genomic Research Lab, BCSIR | Abu Sayeed Mohammad Mahmud, Mohammad Samir Uzzaman, Eshrar Osman, Md. Ahasan Habib, Shahina Akter, Tanjina Akhter Banu, Md. Murshed Hasan Sarkar, Barna Goswami, Iffat Jahan, Md. Saddam Hossain, Tasnim Nafisa, Md. Maruf Ahmed Molla, Mahmuda Yeasmin, Asish Kumar Ghosh, Shahjahan Siddike, A. K. M. Shamsuzzaman, Sheikh Md. Selim Al Din, Utpal Chandra Ray, Salek Ahmed Sajib, Md. Salim Khan |
| EPI_ISL_480961, EPI_ISL_480975, EPI_ISL_480981, EPI_ISL_480989, EPI_ISL_480995, EPI_ISL_481003, EPI_ISL_481017, EPI_ISL_481025, EPI_ISL_481029, EPI_ISL_481034, EPI_ISL_481035 | | | |
| see above | ISGlobal, Institut de Salut Global de Barcelona | SeqCOVID-SPAIN consortium/IBV(CSIC) | Alfredo Mayor, Alberto L Garcia-Basteiro, Carlota Dobaño, Gemma Moncunill, Pau Cisteró and SeqCOVID-SPAIN consortium |
| EPI_ISL_481380 | Department for Virology, Molecular Biology and Genome Research, R. G. Lugar Center for Public Health Research, National Center for Disease Control and Public Health (NCDC) of Georgia. | Department for Virology, Molecular Biology and Genome Research, R. G. Lugar Center for Public Health Research, National Center for Disease Control and Public Health (NCDC) of Georgia. | Ana Papkiauri, Tata Imnadze, Giorgi Tomashvili, Meri Pantsulaia, Gvantsa Brachveli, Gvantsa Chanturia, Ann Machablishvili, Nato Kotaria, Marine Murtskhvaladze, Lela Sabadze, Mari Gavashelidze, Tamar Jashiashvili, Tea Tevdoradze, Ketevan Sidamonidze, Ekaterine Khmaladze, Ekaterine Zhghenti, Roena Sukhiashvili, Mariam Zakalashvili, Lela Urushadze, Magda Dgebuadze, Davit Tsaguria, Ekaterine Zangaladze, Nino Berishvili, Adam Kotorashvili, Maia Alkhazashvili, Irma Burjanadze, Anna Kasradze, Khatuna Zakhashvili, Paata Imnadze, Amiran Gamkrelidze. |
| EPI_ISL_482057, EPI_ISL_482058, EPI_ISL_482059, EPI_ISL_482060, EPI_ISL_482061, EPI_ISL_482062, EPI_ISL_482063, EPI_ISL_482064, EPI_ISL_482065, EPI_ISL_482066, EPI_ISL_482067, EPI_ISL_482068 | | | |
| see above | The Department of Microbiology, Torbay and South Devon NHS Foundation Trust | Wellcome Sanger Institute for the COVID-19 Genomics UK (COG-UK) consortium | Amy Hurd, Sophie Lloyd, Anthony Mogridge, Jack Howe, Helen Brown, Gary Booth, Mel Brown, Cheryl Bailiss, Michelle Harrison and Alex Alderton, Roberto Amato, Sonia Goncalves, Ewan Harrison, David K. Jackson, Ian Johnston, Dominic Kwiatkowski, Cordelia Langford, John Sillitoe on behalf of the |

| | | | |
|---|---|---|---|
| | | | Wellcome Sanger Institute COVID-19 Surveillance Team (http://www.sanger.ac.uk/covid-team) |
| EPI_ISL_482069, EPI_ISL_482071, EPI_ISL_482072, EPI_ISL_482073, EPI_ISL_482074, EPI_ISL_482076, EPI_ISL_482078, EPI_ISL_482079, EPI_ISL_482080, EPI_ISL_482081, EPI_ISL_482083, EPI_ISL_482084, EPI_ISL_482085, EPI_ISL_482086, EPI_ISL_482087, EPI_ISL_482088, EPI_ISL_482089, EPI_ISL_482090, EPI_ISL_482092, EPI_ISL_482093, EPI_ISL_482094, EPI_ISL_482095, EPI_ISL_482098, EPI_ISL_482099, EPI_ISL_482100, EPI_ISL_482103, EPI_ISL_482104, EPI_ISL_482106, EPI_ISL_482107, EPI_ISL_482108, EPI_ISL_482109, EPI_ISL_482110, EPI_ISL_482111, EPI_ISL_482112, EPI_ISL_482113, EPI_ISL_482114, EPI_ISL_482116, EPI_ISL_482117, EPI_ISL_482118, EPI_ISL_482119 | | | |
| see above | Microbiology Department, Hereford County Hospital | Wellcome Sanger Institute for the COVID-19 Genomics UK (COG-UK) consortium | Alison Johnson, Venkat Sivaprakasam, Fenella Halstead, Jane Thomas, Wendy Hogsden, Samantha Lamb and Alex Alderton, Roberto Amato, Sonia Goncalves, Ewan Harrison, David K. Jackson, Ian Johnston, Dominic Kwiatkowski, Cordelia Langford, John Sillitoe on behalf of the Wellcome Sanger Institute COVID-19 Surveillance Team (http://www.sanger.ac.uk/covid-team) |
| EPI_ISL_482120, EPI_ISL_482121, EPI_ISL_482122, EPI_ISL_482123, EPI_ISL_482125, EPI_ISL_482126, EPI_ISL_482127, EPI_ISL_482128, EPI_ISL_482129, EPI_ISL_482130 | The Department of Microbiology, Torbay and South Devon NHS Foundation Trust | Wellcome Sanger Institute for the COVID-19 Genomics UK (COG-UK) consortium | Amy Hurd, Sophie Lloyd, Anthony Mogridge, Jack Howe, Helen Brown, Gary Booth, Mel Brown, Cheryl Bailiss, Michelle Harrison and Alex Alderton, Roberto Amato, Sonia Goncalves, Ewan Harrison, David K. Jackson, Ian Johnston, Dominic Kwiatkowski, Cordelia Langford, John Sillitoe on behalf of the Wellcome Sanger Institute COVID-19 Surveillance Team (http://www.sanger.ac.uk/covid-team) |
| EPI_ISL_482292, EPI_ISL_482293, EPI_ISL_482294, EPI_ISL_482295, EPI_ISL_482296, EPI_ISL_482297, EPI_ISL_482298, EPI_ISL_482299, EPI_ISL_482300, EPI_ISL_482301, EPI_ISL_482302, EPI_ISL_482303, EPI_ISL_482304, EPI_ISL_482305, EPI_ISL_482306, EPI_ISL_482307, EPI_ISL_482308, EPI_ISL_482309, EPI_ISL_482310, EPI_ISL_482311, EPI_ISL_482312, EPI_ISL_482313, EPI_ISL_482314, EPI_ISL_482315, EPI_ISL_482316, EPI_ISL_482317, EPI_ISL_482318, EPI_ISL_482319, EPI_ISL_482320, EPI_ISL_482321, EPI_ISL_482322, EPI_ISL_482323, EPI_ISL_482324, EPI_ISL_482325, EPI_ISL_482326, EPI_ISL_482327, EPI_ISL_482328, EPI_ISL_482329, EPI_ISL_482330, EPI_ISL_482331, EPI_ISL_482332, EPI_ISL_482333, EPI_ISL_482334, EPI_ISL_482335, EPI_ISL_482336, EPI_ISL_482337, EPI_ISL_482338, EPI_ISL_482339, EPI_ISL_482340, EPI_ISL_482341, EPI_ISL_482342, EPI_ISL_482343, EPI_ISL_482344, EPI_ISL_482345, EPI_ISL_482346, EPI_ISL_482347, EPI_ISL_482348, EPI_ISL_482349, EPI_ISL_482350, EPI_ISL_482351, EPI_ISL_482352, EPI_ISL_482353, EPI_ISL_482354, EPI_ISL_482355, EPI_ISL_482356, EPI_ISL_482357, EPI_ISL_482358, EPI_ISL_482359, EPI_ISL_482360, EPI_ISL_482361, EPI_ISL_482362, EPI_ISL_482363, EPI_ISL_482364, EPI_ISL_482365, EPI_ISL_482366, EPI_ISL_482367, EPI_ISL_482368, EPI_ISL_482369, EPI_ISL_482370, EPI_ISL_482371, EPI_ISL_482372, EPI_ISL_482373, EPI_ISL_482374, EPI_ISL_482375, EPI_ISL_482376, EPI_ISL_482377, EPI_ISL_482378, EPI_ISL_482379, EPI_ISL_482380, EPI_ISL_482381, EPI_ISL_482382, EPI_ISL_482383, EPI_ISL_482384, EPI_ISL_482385, EPI_ISL_482386, EPI_ISL_482387, EPI_ISL_482388, EPI_ISL_482389, EPI_ISL_482390, EPI_ISL_482391, EPI_ISL_482392, EPI_ISL_482393, EPI_ISL_482394, EPI_ISL_482395, EPI_ISL_482396, EPI_ISL_482397, EPI_ISL_482398, EPI_ISL_482399, EPI_ISL_482400, EPI_ISL_482401, EPI_ISL_482402, EPI_ISL_482403, EPI_ISL_482404, EPI_ISL_482405, EPI_ISL_482406, EPI_ISL_482407, EPI_ISL_482408, EPI_ISL_482409, EPI_ISL_482410, EPI_ISL_482411, EPI_ISL_482412, EPI_ISL_482413, EPI_ISL_482414, EPI_ISL_482415, EPI_ISL_482416, EPI_ISL_482417, EPI_ISL_482418, EPI_ISL_482419, EPI_ISL_482420, EPI_ISL_482421, EPI_ISL_482422, EPI_ISL_482423, EPI_ISL_482424, EPI_ISL_482425, EPI_ISL_482426, EPI_ISL_482427, EPI_ISL_482428, EPI_ISL_482429, EPI_ISL_482430, EPI_ISL_482431, EPI_ISL_482432, EPI_ISL_482433, EPI_ISL_482434, EPI_ISL_482435, EPI_ISL_482436, EPI_ISL_482437, EPI_ISL_482438, EPI_ISL_482439, EPI_ISL_482440, EPI_ISL_482441, EPI_ISL_482442, EPI_ISL_482443, EPI_ISL_482444, EPI_ISL_482445, EPI_ISL_482446, EPI_ISL_482447, EPI_ISL_482448, EPI_ISL_482449, EPI_ISL_482450, EPI_ISL_482451, EPI_ISL_482452, EPI_ISL_482453, EPI_ISL_482454, EPI_ISL_482455, EPI_ISL_482456, EPI_ISL_482457, EPI_ISL_482458, EPI_ISL_482459, EPI_ISL_482460, EPI_ISL_482461, EPI_ISL_482462, EPI_ISL_482463, EPI_ISL_482464, EPI_ISL_482465, EPI_ISL_482466, EPI_ISL_482467 | | | |
| see above | Providence St. Joseph Health Molecular Genomics Laboratory | Providence St. Joseph Health Molecular Genomics Laboratory | Alexa K Dowdell, Brian D Piening, Fred L Robinson, Carlo B Bifulco, Mary Campbell |
| EPI_ISL_482485, EPI_ISL_482486, EPI_ISL_482487, EPI_ISL_482700, EPI_ISL_482701 | National Institute of Laboratory Medicine and Referral Center | Genomic Research Lab, BCSIR | Abu Sayeed Mohammad Mahmud, Mohammad Samir Uzzaman, Eshrar Osman, Md. Ahasan Habib, Shahina Akter, Tanjina Akhter Banu, Md. Murshed Hasan Sarkar, Barna Goswami, Iffat Jahan, Md. Saddam Hossain, Tasnim Nafisa, Md. Maruf Ahmed Molla, Mahmuda Yeasmin, Asish Kumar Ghosh, Shahjahan Siddike, A. K. M. Shamsuzzaman, Sheikh Md. Selim Al Din, Utpal Chandra Ray, Salek Ahmed Sajib, Md. Salim Khan |
| EPI_ISL_483398, EPI_ISL_483399, EPI_ISL_483400, EPI_ISL_483401, EPI_ISL_483402 | UC San Diego Center for Advanced Laboratory Medicine | Andersen lab at Scripps Research | Allison Smither, Gilberto Sabino-Santos, Patricia Snarski, Lilia Melnik, Antoinette Bell, Kaylynn Genemaras, Arnaud Drouin, Dahlene Fusco, Robert Garry with SEARCH Alliance San Diego |
| EPI_ISL_483693, EPI_ISL_483694, EPI_ISL_483695, EPI_ISL_483699, EPI_ISL_483700, EPI_ISL_483703, EPI_ISL_483705, EPI_ISL_483707, EPI_ISL_483710 | National Institute of Laboratory Medicine and Referral Center | Genomic Research Lab, BCSIR | Abu Sayeed Mohammad Mahmud, Mohammad Samir Uzzaman, Eshrar Osman, Md. Ahasan Habib, Shahina Akter, Tanjina Akhter Banu, Md. Murshed Hasan Sarkar, Barna Goswami, Iffat Jahan, Md. Saddam Hossain, Tasnim Nafisa, Md. Maruf Ahmed Molla, Mahmuda Yeasmin, Asish Kumar Ghosh, A. K. M. Shamsuzzaman, Sheikh Md. Selim Al Din, Utpal Chandra Ray, Salek Ahmed Sajib, Md. Salim Khan |
| EPI_ISL_483822 | Government Medical College, Vadodara | Gujarat Biotechnology Research Centre | Afzal Ansari, Nikha Trivedi, Meenakshi Shah, Neena Doshi, Varsha Godbole, Apurvasinh Puvar, Janvi Raval, Zarna Patel, Monika Gandhi, Pinal Trivedi, Maharshi Pandya, Nidhi Patel, Nitin Savaliya, Raghawendra Kumar, Dinesh Kumar, Zuber Saiyed, Komal Patel, Labdhi Pandya, R D Dixit, A M Kadri, Harsh Bakshi, Chaitanya Joshi, Madhvi Joshi |
| EPI_ISL_483839 | Pandit Deendayal Upadhyay Government Medical College, Rajkot | Gujarat Biotechnology Research Centre | Afzal Ansari, Nikha Trivedi, Gauravi Dhruv, Arti Trivedi, Apurvasinh Puvar, Janvi Raval, Zarna Patel, Monika Gandhi, Pinal Trivedi, Maharshi Pandya, Nidhi Patel, Nitin Savaliya, Raghawendra Kumar, Dinesh Kumar, Zuber Saiyed, Komal Patel, Labdhi Pandya, R D Dixit, A M Kadri, Harsh Bakshi, Chaitanya Joshi, Madhvi Joshi |
| EPI_ISL_483856 | Department of Microbiology, Government Medical College, Surat | Gujarat Biotechnology Research Centre | Afzal Ansari, Nikha Trivedi, Naresh Chauhan, Summaiya Mullan, Amit gamit, Apurvasinh Puvar, Janvi Raval, Zarna Patel, Monika Gandhi, Pinal Trivedi, Maharshi Pandya, Nidhi Patel, Nitin Savaliya, Raghawendra Kumar, Dinesh Kumar, Zuber Saiyed, Komal Patel, Labdhi Pandya, R D Dixit, A M Kadri, Harsh Bakshi, Chaitanya Joshi, Madhvi Joshi |
| EPI_ISL_483860 | Department of Microbiology, Government Medical College, Surat | Gujarat Biotechnology Research Centre | Amit gamit, Apurvasinh Puvar, Janvi Raval, Zarna Patel, Monika Gandhi, Pinal Trivedi, Maharshi Pandya, Nidhi Patel, Nitin Savaliya, Raghawendra Kumar, Dinesh Kumar, Zuber Saiyed, Komal Patel, Labdhi Pandya, Afzal Ansari, Nikha Trivedi, Naresh Chauhan, Summaiya Mullan, R D Dixit, A M Kadri, Harsh Bakshi, Chaitanya Joshi, Madhvi Joshi |
| EPI_ISL_483874 | Department of Microbiology, Government Medical College, Surat | Gujarat Biotechnology Research Centre | Afzal Ansari, Nikha Trivedi, Naresh Chauhan, Summaiya Mullan, Amit gamit, Apurvasinh Puvar, Janvi Raval, Zarna Patel, Monika Gandhi, Pinal Trivedi, Maharshi Pandya, Nidhi Patel, Nitin Savaliya, Raghawendra Kumar, Dinesh Kumar, Zuber Saiyed, Komal Patel, Labdhi Pandya, R D Dixit, A M Kadri, Harsh Bakshi, Chaitanya Joshi, Madhvi Joshi |
| EPI_ISL_483878 | Department of Microbiology, Government Medical College, Surat | Gujarat Biotechnology Research Centre | Amit gamit, Apurvasinh Puvar, Janvi Raval, Zarna Patel, Monika Gandhi, Pinal Trivedi, Maharshi Pandya, Nidhi Patel, Nitin Savaliya, Raghawendra Kumar, Dinesh Kumar, Zuber Saiyed, Komal Patel, Labdhi Pandya, Afzal Ansari, Nikha Trivedi, Naresh Chauhan, Summaiya Mullan, R D Dixit, A M Kadri, Harsh Bakshi, Chaitanya Joshi, Madhvi Joshi |
| EPI_ISL_484407, EPI_ISL_484410, EPI_ISL_484411, EPI_ISL_484412, EPI_ISL_484414, EPI_ISL_484415, EPI_ISL_484416, EPI_ISL_484417, EPI_ISL_484419, EPI_ISL_484421, EPI_ISL_484422, EPI_ISL_484423, EPI_ISL_484424, EPI_ISL_484425, EPI_ISL_484427, EPI_ISL_484428, EPI_ISL_484429 | | | |
| see above | Centre for Enzyme Innovation, University of Portsmouth / Translational Research Laboratory, Portsmouth Hospitals NHS Trust | COVID-19 Genomics UK (COG-UK) Consortium | Angela Beckett,Yann Bourgeois,Garry Scarlett,Sharon Glaysher,Scott Elliott,Kelly Bicknell,Robert Impey,Allyson Lloyd,Sarah Wyllie,Ethan Butcher,Anoop Chauhan,Samuel Robson |
| EPI_ISL_484519, EPI_ISL_484520, EPI_ISL_484521, EPI_ISL_484522, EPI_ISL_484523, EPI_ISL_484524, EPI_ISL_484525, EPI_ISL_484526, EPI_ISL_484527, EPI_ISL_484528, EPI_ISL_484529, EPI_ISL_484530, EPI_ISL_484531, EPI_ISL_484532, EPI_ISL_484533, EPI_ISL_484534, EPI_ISL_484535, EPI_ISL_484536, EPI_ISL_484537, EPI_ISL_484538, EPI_ISL_484539, EPI_ISL_484540, EPI_ISL_484541, EPI_ISL_484542, EPI_ISL_484543, EPI_ISL_484544, EPI_ISL_484545, EPI_ISL_484546, EPI_ISL_484547, EPI_ISL_484548, EPI_ISL_484549, EPI_ISL_484550, EPI_ISL_484551, EPI_ISL_484552, EPI_ISL_484553, EPI_ISL_484554, EPI_ISL_484555, EPI_ISL_484556, EPI_ISL_484557, EPI_ISL_484558, EPI_ISL_484559, EPI_ISL_484560, EPI_ISL_484561, EPI_ISL_484562, EPI_ISL_484563, EPI_ISL_484564, EPI_ISL_484565, EPI_ISL_484566, EPI_ISL_484567, EPI_ISL_484568, EPI_ISL_484569, EPI_ISL_484570, EPI_ISL_484571, EPI_ISL_484572, EPI_ISL_484573, EPI_ISL_484574, EPI_ISL_484575, EPI_ISL_484576, EPI_ISL_484577, EPI_ISL_484578, EPI_ISL_484579, EPI_ISL_484580, EPI_ISL_484581, EPI_ISL_484582, EPI_ISL_484583, EPI_ISL_484584, EPI_ISL_484585, EPI_ISL_484586, EPI_ISL_484587, EPI_ISL_484588, EPI_ISL_484589, EPI_ISL_484590, EPI_ISL_484591, EPI_ISL_484592, EPI_ISL_484593, EPI_ISL_484594, EPI_ISL_484595, EPI_ISL_484596, EPI_ISL_484597, EPI_ISL_484598, EPI_ISL_484599, EPI_ISL_484600, EPI_ISL_484601, EPI_ISL_484602, EPI_ISL_484603, EPI_ISL_484604, EPI_ISL_484605, EPI_ISL_484606, EPI_ISL_484607, EPI_ISL_484608, EPI_ISL_484609, EPI_ISL_484610, EPI_ISL_484611, EPI_ISL_484612, EPI_ISL_484613, EPI_ISL_484614, EPI_ISL_484615, EPI_ISL_484616, EPI_ISL_484617, EPI_ISL_484618, EPI_ISL_484619, EPI_ISL_484620, EPI_ISL_484621, EPI_ISL_484622, EPI_ISL_484623, EPI_ISL_484624, EPI_ISL_484625, EPI_ISL_484626, EPI_ISL_484627, EPI_ISL_484628, EPI_ISL_484629, EPI_ISL_484630, EPI_ISL_484631, EPI_ISL_484632, EPI_ISL_484633, EPI_ISL_484634, EPI_ISL_484635, EPI_ISL_484636, EPI_ISL_484637, EPI_ISL_484638, EPI_ISL_484639, EPI_ISL_484640, EPI_ISL_484641, EPI_ISL_484642, EPI_ISL_484643, EPI_ISL_484644, EPI_ISL_484645, EPI_ISL_484646, EPI_ISL_484647, EPI_ISL_484648, EPI_ISL_484649, EPI_ISL_484650, EPI_ISL_484651, EPI_ISL_484652, EPI_ISL_484653, EPI_ISL_484654, EPI_ISL_484655, EPI_ISL_484656, EPI_ISL_484657, EPI_ISL_484658, EPI_ISL_484659, EPI_ISL_484660, EPI_ISL_484661, EPI_ISL_484662, EPI_ISL_484663, EPI_ISL_484664, EPI_ISL_484665, EPI_ISL_484666, EPI_ISL_484667, EPI_ISL_484668, EPI_ISL_484669, EPI_ISL_484670, EPI_ISL_484671, EPI_ISL_484672, EPI_ISL_484673, EPI_ISL_484674, EPI_ISL_484675, EPI_ISL_484676, EPI_ISL_484677, EPI_ISL_484678, EPI_ISL_484679 | | | |
| see above | West of Scotland Specialist Virology Centre, NHSGGC / MRC-University of Glasgow Centre for Virus Research | COVID-19 Genomics UK (COG-UK) Consortium | Ana da Silva Filipe, Natasha Johnson, Kathy Smollett, Daniel Mair, Stephen Carmichael, Lily Tong, Jenna Nichols, Elihu Aranday-Cortes, Kirstyn Brunker, Yasmin Parr, Alice Broos, Kyriaki Nomikou; Sarah McDonald, Marc Niebel, Patawee Asamaphan; Richard Orton, Joseph Hughes, Sreenu Vattipally, David L Robertson; Alasdair MacLean, Rory Gunson; Kathy Li, Natasha Jesudason, Rajiv Shah, James Shepherd, Antonia Ho, Emma Thomson |
| EPI_ISL_486427 | unknown | Clinical Laboratory, Hospital Israelita Albert Einstein | Amgarteb,D., Malta,F., Guedes,R.L., Santana,R.A., de Menezes,F.G., Mangueira,C.L. and Pinho,J.R. |
| EPI_ISL_486830, EPI_ISL_486831 | Providence St. Joseph Health Molecular Genomics Laboratory | Providence St. Joseph Health Molecular Genomics Laboratory | Alexa K Dowdell, Brian D Piening, Fred L Robinson, Carlo B Bifulco, Mary Campbell |
| EPI_ISL_486888 | National Influenza Center, Bahrain | National Influenza Center, Bahrain | AlWasti,H., Altaif,Z., Zaed,A., Shehab,F. |
| EPI_ISL_486889 | National Influenza Center, Bahrain | National Influenza Center, Bahrain | Altaif,Z., AlWasti,H., Shehab,F., Zaed,A. |
| EPI_ISL_487270 | unknown | Communicable Disease Laboratory, Public Health Directorate | AlWasti,H., AlTaif,Z., Zaed,A., Shehab,F. |
| EPI_ISL_487272 | unknown | Communicable Disease Laboratory, Public Health Directorate | Altaif,z., AlWasti,H., Shehab,F., Zaed,A. |
| EPI_ISL_487274 | unknown | Communicable Disease Laboratory, Public Health Directorate | AlWasti,H., AlTaif,Z., Zaed,A., Shehab,F. |

| | | | |
|---|---|---|---|
| EPI_ISL_487386, EPI_ISL_487392, EPI_ISL_487393, EPI_ISL_487394, EPI_ISL_487395, EPI_ISL_487396 | National Institute of Laboratory Medicine and Referral Center | Genomic Research Lab, BCSIR | Abu Sayeed Mohammad Mahmud, Mohammad Samir Uzzaman, Eshrar Osman, Md. Ahasan Habib, Shahina Akter, Tanjina Akhter Banu, Md. Murshed Hasan Sarkar, Barna Goswami, Iffat Jahan, Md. Saddam Hossain, Tasnim Nafisa, Md. Maruf Ahmed Molla, Mahmuda Yeasmin, Asish Kumar Ghosh, A. K. M. Shamsuzzaman, Sheikh Md. Selim Al Din, Utpal Chandra Ray, Salek Ahmed Sajib, Md. Salim Khan |
| EPI_ISL_488837, EPI_ISL_488838, EPI_ISL_488839, EPI_ISL_488841, EPI_ISL_488842, EPI_ISL_488843, EPI_ISL_488844, EPI_ISL_488846, EPI_ISL_488848, EPI_ISL_488849, EPI_ISL_488850, EPI_ISL_488851, EPI_ISL_488852, EPI_ISL_488857, EPI_ISL_488858, EPI_ISL_488859, EPI_ISL_488860, EPI_ISL_488862, EPI_ISL_488865, EPI_ISL_488866, EPI_ISL_488867, EPI_ISL_488868, EPI_ISL_488869, EPI_ISL_488870, EPI_ISL_488871, EPI_ISL_488873 | | | |
| see above | Microbiology Department, Hereford County Hospital | Wellcome Sanger Institute for the COVID-19 Genomics UK (COG-UK) consortium | Alison Johnson, Venkat Sivaprakasam, Fenella Halstead, Jane Thomas, Wendy Hogsden, Samantha Lamb and Alex Alderton, Roberto Amato, Sonia Goncalves, Ewan Harrison, David K. Jackson, Ian Johnston, Dominic Kwiatkowski, Cordelia Langford, John Sillitoe on behalf of the Wellcome Sanger Institute COVID-19 Surveillance Team (http://www.sanger.ac.uk/covid-team) |
| EPI_ISL_489380, EPI_ISL_489391, EPI_ISL_489392, EPI_ISL_489393, EPI_ISL_489396, EPI_ISL_489401, EPI_ISL_489407, EPI_ISL_489408, EPI_ISL_489413, EPI_ISL_489418, EPI_ISL_489429, EPI_ISL_489442, EPI_ISL_489449, EPI_ISL_489455, EPI_ISL_489470, EPI_ISL_489480, EPI_ISL_489489, EPI_ISL_489523, EPI_ISL_489530, EPI_ISL_489536, EPI_ISL_489537, EPI_ISL_489543, EPI_ISL_489547, EPI_ISL_489567, EPI_ISL_489578, EPI_ISL_489579, EPI_ISL_489580, EPI_ISL_489582, EPI_ISL_489583, EPI_ISL_489584, EPI_ISL_489585, EPI_ISL_489586, EPI_ISL_489587, EPI_ISL_489588, EPI_ISL_489589, EPI_ISL_489590, EPI_ISL_489591, EPI_ISL_489592, EPI_ISL_489593, EPI_ISL_489594, EPI_ISL_489595, EPI_ISL_489596, EPI_ISL_489597, EPI_ISL_489598, EPI_ISL_489599, EPI_ISL_489600, EPI_ISL_489601, EPI_ISL_489602, EPI_ISL_489603, EPI_ISL_489604, EPI_ISL_489605, EPI_ISL_489606, EPI_ISL_489607, EPI_ISL_489608, EPI_ISL_489609, EPI_ISL_489610, EPI_ISL_489611, EPI_ISL_489612, EPI_ISL_489613, EPI_ISL_489614, EPI_ISL_489615, EPI_ISL_489616, EPI_ISL_489617, EPI_ISL_489618, EPI_ISL_489619, EPI_ISL_489620, EPI_ISL_489621, EPI_ISL_489622, EPI_ISL_489623, EPI_ISL_489624, EPI_ISL_489625, EPI_ISL_489626, EPI_ISL_489627, EPI_ISL_489628, EPI_ISL_489629, EPI_ISL_489630, EPI_ISL_489631, EPI_ISL_489632, EPI_ISL_489633, EPI_ISL_489634, EPI_ISL_489635, EPI_ISL_489636, EPI_ISL_489637, EPI_ISL_489638, EPI_ISL_489639, EPI_ISL_489640, EPI_ISL_489641, EPI_ISL_489642, EPI_ISL_489643, EPI_ISL_489644, EPI_ISL_489645, EPI_ISL_489646, EPI_ISL_489647, EPI_ISL_489648, EPI_ISL_489649, EPI_ISL_489650, EPI_ISL_489651, EPI_ISL_489652, EPI_ISL_489653, EPI_ISL_489654, EPI_ISL_489655, EPI_ISL_489656, EPI_ISL_489657, EPI_ISL_489658, EPI_ISL_489659, EPI_ISL_489660, EPI_ISL_489661, EPI_ISL_489662, EPI_ISL_489663, EPI_ISL_489664, EPI_ISL_489665, EPI_ISL_489666, EPI_ISL_489667, EPI_ISL_489668, EPI_ISL_489669, EPI_ISL_489670, EPI_ISL_489671, EPI_ISL_489672, EPI_ISL_489673, EPI_ISL_489674, EPI_ISL_489675, EPI_ISL_489676, EPI_ISL_489677, EPI_ISL_489678, EPI_ISL_489679, EPI_ISL_489680, EPI_ISL_489681, EPI_ISL_489682, EPI_ISL_489683, EPI_ISL_489684, EPI_ISL_489685, EPI_ISL_489686, EPI_ISL_489687, EPI_ISL_489688, EPI_ISL_489689, EPI_ISL_489690, EPI_ISL_489691, EPI_ISL_489692, EPI_ISL_489693, EPI_ISL_489694, EPI_ISL_489695, EPI_ISL_489696, EPI_ISL_489697, EPI_ISL_489698, EPI_ISL_489699, EPI_ISL_489700, EPI_ISL_489701, EPI_ISL_489702, EPI_ISL_489703, EPI_ISL_489704, EPI_ISL_489705, EPI_ISL_489706 | | | |
| see above | NHSGGC West of Scotland Specialist Virology Centre / MRC-University of Glasgow Centre for Virus Research | Wellcome Sanger Institute for the COVID-19 Genomics UK (COG-UK) consortium | Ana da Silva Filipe, Natasha Johnson, Kathy Smollett, Daniel Mair, Stephen Carmichael, Lily Tong, Jenna Nichols, Elihu Aranday-Cortes, Kirstyn Brunker, Yasmin Parr, Kyriaki Nomikou; Sarah McDonald, Marc Niebel, Patawee Asamaphan; Richard Orton, Joseph Hughes, Sreenu Vattipally, David L Robertson; Alasdair MacLean, Rory Gunson; Kathy Li, Natasha Jesudason, Rajiv Shah, James Shepherd, Antonia Ho, Alice Broos, Emma Thomson and Alex Alderton, Roberto Amato, Sonia Goncalves, Ewan Harrison, David K. Jackson, Ian Johnston, Dominic Kwiatkowski, Cordelia Langford, John Sillitoe on behalf of the Wellcome Sanger Institute COVID-19 Surveillance Team (http://www.sanger.ac.uk/covid-team) |
| EPI_ISL_489996, EPI_ISL_489997, EPI_ISL_489998, EPI_ISL_489999, EPI_ISL_490000, EPI_ISL_490001, EPI_ISL_490002, EPI_ISL_490003, EPI_ISL_490004, EPI_ISL_490005, EPI_ISL_490006, EPI_ISL_490007, EPI_ISL_490008, EPI_ISL_490009, EPI_ISL_490010, EPI_ISL_490011, EPI_ISL_490012 | | | |
| see above | King Fahad Medical City | King Fahad Medical City | Alosaimi,B., Naeem,A., Alghoraibi,M., Enani,M. |
| EPI_ISL_490167, EPI_ISL_490168 | National Institute of Laboratory Medicine and Referral Center | Genomic Research Lab, BCSIR | Abu Sayeed Mohammad Mahmud, Mohammad Samir Uzzaman, Eshrar Osman, Md. Ahasan Habib, Shahina Akter, Tanjina Akhter Banu, Md. Murshed Hasan Sarkar, Barna Goswami, Iffat Jahan, Md. Saddam Hossain, Tasnim Nafisa, Md. Maruf Ahmed Molla, Mahmuda Yeasmin, Asish Kumar Ghosh, A. K. M. Shamsuzzaman, Sheikh Md. Selim Al Din, Utpal Chandra Ray, Salek Ahmed Sajib, Md. Salim Khan |
| EPI_ISL_490201 | National Influenza Centre of INSPI-Ecuador | INSPI - Charité | Alfredo Bruno Caicedo, Domenica de Mora Coloma, Andres Moreira-Soto, Anna-Lena Sander, Nina Krause, Maritza Olmedo,Denisses Portugal, Manuel Gonzalez, Silvia Salgado, Alberto Orlando, Alexandra Usiña, Juan Carlos Zeballos,Jan Felix Drexler |
| EPI_ISL_490255, EPI_ISL_490256, EPI_ISL_490257, EPI_ISL_490258, EPI_ISL_490259, EPI_ISL_490260, EPI_ISL_490261, EPI_ISL_490262, EPI_ISL_490263, EPI_ISL_490264, EPI_ISL_490265, EPI_ISL_490266, EPI_ISL_490267, EPI_ISL_490268, EPI_ISL_490269, EPI_ISL_490270, EPI_ISL_490271, EPI_ISL_490272, EPI_ISL_490273, EPI_ISL_490274, EPI_ISL_490275, EPI_ISL_490276, EPI_ISL_490277, EPI_ISL_490278, EPI_ISL_490279, EPI_ISL_490280, EPI_ISL_490281, EPI_ISL_490282, EPI_ISL_490283, EPI_ISL_490284, EPI_ISL_490285, EPI_ISL_490286, EPI_ISL_490287, EPI_ISL_490288, EPI_ISL_490289, EPI_ISL_490290, EPI_ISL_490291, EPI_ISL_490292, EPI_ISL_490293, EPI_ISL_490294, EPI_ISL_490295, EPI_ISL_490296, EPI_ISL_490297, EPI_ISL_490298, EPI_ISL_490299, EPI_ISL_490300, EPI_ISL_490301, EPI_ISL_490302, EPI_ISL_490303, EPI_ISL_490304, EPI_ISL_490305, EPI_ISL_490306, EPI_ISL_490307, EPI_ISL_490308, EPI_ISL_490309, EPI_ISL_490310, EPI_ISL_490311, EPI_ISL_490312, EPI_ISL_490313, EPI_ISL_490314 | | | |
| see above | National Institute for Communicable Diseases of the National Health Laboratory Service | National Institute for Communicable Diseases of the National Health Laboratory Service | Allam M, Ismail A, Khumalo Z, Kwenda S, Mtshali P, Mnyameni F, Mohale T, Subramoney K, Bhiman JN |
| EPI_ISL_490330, EPI_ISL_490646, EPI_ISL_490647, EPI_ISL_490648, EPI_ISL_490649, EPI_ISL_490650, EPI_ISL_490651, EPI_ISL_490652, EPI_ISL_490653, EPI_ISL_490654, EPI_ISL_490655, EPI_ISL_490656, EPI_ISL_490657, EPI_ISL_490658, EPI_ISL_490659, EPI_ISL_490660, EPI_ISL_490661, EPI_ISL_490662, EPI_ISL_490663, EPI_ISL_490664, EPI_ISL_490665, EPI_ISL_490666, EPI_ISL_490667, EPI_ISL_490668, EPI_ISL_490669, EPI_ISL_490670, EPI_ISL_490671, EPI_ISL_490672, EPI_ISL_490673, EPI_ISL_490674, EPI_ISL_490675, EPI_ISL_490676, EPI_ISL_490677, EPI_ISL_490678, EPI_ISL_490679, EPI_ISL_490680, EPI_ISL_490681, EPI_ISL_490682, EPI_ISL_490683, EPI_ISL_490684, EPI_ISL_490685, EPI_ISL_490686, EPI_ISL_490687, EPI_ISL_490688, EPI_ISL_490689, EPI_ISL_490690, EPI_ISL_490691, EPI_ISL_490692, EPI_ISL_490693, EPI_ISL_490694, EPI_ISL_490695, EPI_ISL_490696, EPI_ISL_490697, EPI_ISL_490698, EPI_ISL_490699, EPI_ISL_490700, EPI_ISL_490701, EPI_ISL_490702, EPI_ISL_490703, EPI_ISL_490704, EPI_ISL_490705, EPI_ISL_490706, EPI_ISL_490707, EPI_ISL_490708, EPI_ISL_490709 | | | |
| see above | West of Scotland Specialist Virology Centre, NHSGGC / MRC-University of Glasgow Centre for Virus Research | COVID-19 Genomics UK (COG-UK) Consortium | Ana da Silva Filipe, Natasha Johnson, Kathy Smollett, Daniel Mair, Stephen Carmichael, Lily Tong, Jenna Nichols, Elihu Aranday-Cortes, Kirstyn Brunker, Yasmin Parr, Alice Broos, Kyriaki Nomikou; Sarah McDonald, Marc Niebel, Patawee Asamaphan; Richard Orton, Joseph Hughes, Sreenu Vattipally, David L Robertson; Alasdair MacLean, Rory Gunson; Kathy Li, Natasha Jesudason, Rajiv Shah, James Shepherd, Antonia Ho, Emma Thomson |
| EPI_ISL_491146, EPI_ISL_491147, EPI_ISL_491148, EPI_ISL_491149, EPI_ISL_491150, EPI_ISL_491151, EPI_ISL_491152, EPI_ISL_491153, EPI_ISL_491154, EPI_ISL_491155, EPI_ISL_491156, EPI_ISL_491157, EPI_ISL_491158 | | | |
| see above | Oman-National Influenza Center | Biotechnology & OMICs Laboratory | Abdul Latif Khan, Samira Al-Mahruqi, Ahmed Al-Harrasi, Samiha Al-Kharusi, Adil Khan, Ahmed Al-Rawahi, Sajjad Asaf, Amina Al-Jardani, Hanan Al-Kindi, Intisar Al-Shukri, Ahlam Al-Amri, Aisha Al-Amri, Aisha Al-Busaidi, Adil Al-Wahaibi, Seif Al-Abri. |
| EPI_ISL_491944, EPI_ISL_491945, EPI_ISL_491946, EPI_ISL_491947, EPI_ISL_491948, EPI_ISL_491949, EPI_ISL_491950, EPI_ISL_491951, EPI_ISL_491952, EPI_ISL_491953, EPI_ISL_491954 | | | |
| see above | Instituto Nacional de Investigación en Salud Pública - INSPI | INSPI - Charité | Alfredo Bruno Caicedo, Domenica de Mora Coloma, Andres Moreira-Soto, Anna-Lena Sander, Nina Krause, Maritza Olmedo,Denisses Portugal, Manuel Gonzalez, Silvia Salgado, Alberto Orlando, Alexandra Usiña, Juan Carlos Zeballos, Jan Felix Drexler |
| EPI_ISL_493741, EPI_ISL_493742, EPI_ISL_493743, EPI_ISL_493744, EPI_ISL_493745, EPI_ISL_493746, EPI_ISL_493747, EPI_ISL_493748, EPI_ISL_493749, EPI_ISL_493750, EPI_ISL_493751, EPI_ISL_493752, EPI_ISL_493753, EPI_ISL_493754, EPI_ISL_493755, EPI_ISL_493756, EPI_ISL_493757, EPI_ISL_493758, EPI_ISL_493759, EPI_ISL_493760, EPI_ISL_493761, EPI_ISL_493762, EPI_ISL_493763, EPI_ISL_493764, EPI_ISL_493765, EPI_ISL_493766, EPI_ISL_493767, EPI_ISL_493768, EPI_ISL_493769, EPI_ISL_493770, EPI_ISL_493771, EPI_ISL_493772, EPI_ISL_493773, EPI_ISL_493774, EPI_ISL_493775, EPI_ISL_493776, EPI_ISL_493777, EPI_ISL_493778, EPI_ISL_493779, EPI_ISL_493780, EPI_ISL_493781, EPI_ISL_493782, EPI_ISL_493783, EPI_ISL_493784, EPI_ISL_493785, EPI_ISL_493786, EPI_ISL_493787, EPI_ISL_493788, EPI_ISL_493789, EPI_ISL_493790, EPI_ISL_493791, EPI_ISL_493792, EPI_ISL_493793, EPI_ISL_493794, EPI_ISL_493795, EPI_ISL_493796, EPI_ISL_493797, EPI_ISL_493798, EPI_ISL_493799, EPI_ISL_493800, EPI_ISL_493801, EPI_ISL_493802, EPI_ISL_493803, EPI_ISL_493804, EPI_ISL_493805, EPI_ISL_493806, EPI_ISL_493807, EPI_ISL_493808, EPI_ISL_493809, EPI_ISL_493810, EPI_ISL_493811, EPI_ISL_493812, EPI_ISL_493813, EPI_ISL_493814, EPI_ISL_493815, EPI_ISL_493816, EPI_ISL_493817, EPI_ISL_493818, EPI_ISL_493819, EPI_ISL_493820, EPI_ISL_493821, EPI_ISL_493822, EPI_ISL_493823, EPI_ISL_493824, EPI_ISL_493825, EPI_ISL_493826, EPI_ISL_493827, EPI_ISL_493828, EPI_ISL_493829, EPI_ISL_493830, EPI_ISL_493831, EPI_ISL_493832, EPI_ISL_493833, EPI_ISL_493834, EPI_ISL_493835, EPI_ISL_493836, EPI_ISL_493837, EPI_ISL_493838, EPI_ISL_493839, EPI_ISL_493840, EPI_ISL_493841, EPI_ISL_493842, EPI_ISL_493843, EPI_ISL_493844, EPI_ISL_493845, EPI_ISL_493846, EPI_ISL_493847, EPI_ISL_493848, EPI_ISL_493849, EPI_ISL_493850, EPI_ISL_493851, EPI_ISL_493852, EPI_ISL_493853, EPI_ISL_493854, EPI_ISL_493855, EPI_ISL_493856, EPI_ISL_493857, EPI_ISL_493858, EPI_ISL_493859, EPI_ISL_493860, EPI_ISL_493861, EPI_ISL_493862, EPI_ISL_493863, EPI_ISL_493864, EPI_ISL_493865, EPI_ISL_493866, EPI_ISL_493867, EPI_ISL_493868, EPI_ISL_493869, EPI_ISL_493870, EPI_ISL_493871, EPI_ISL_493872, EPI_ISL_493873, EPI_ISL_493874, EPI_ISL_493875, EPI_ISL_493876, EPI_ISL_493877, EPI_ISL_493878, EPI_ISL_493879, EPI_ISL_493880, EPI_ISL_493881, EPI_ISL_493882, EPI_ISL_493883, EPI_ISL_493884, EPI_ISL_493885, EPI_ISL_493886, EPI_ISL_493887, EPI_ISL_493888, EPI_ISL_493889, EPI_ISL_493890, EPI_ISL_493891, EPI_ISL_493892, EPI_ISL_493893, EPI_ISL_493894, EPI_ISL_493895, EPI_ISL_493896, EPI_ISL_493897 | | | |
| see above | West of Scotland Specialist Virology Centre, NHSGGC / MRC-University of Glasgow Centre for Virus Research | COVID-19 Genomics UK (COG-UK) Consortium | Ana da Silva Filipe, Natasha Johnson, Kathy Smollett, Daniel Mair, Stephen Carmichael, Lily Tong, Jenna Nichols, Elihu Aranday-Cortes, Kirstyn Brunker, Yasmin Parr, Alice Broos, Kyriaki Nomikou; Sarah McDonald, Marc Niebel, Patawee Asamaphan; Richard Orton, Joseph Hughes, Sreenu Vattipally, David L Robertson; Alasdair MacLean, Rory Gunson; Kathy Li, Natasha Jesudason, Rajiv Shah, James Shepherd, Antonia Ho, Emma Thomson |
| EPI_ISL_494504, EPI_ISL_494505, EPI_ISL_494506, EPI_ISL_494507, EPI_ISL_494508, EPI_ISL_494509, EPI_ISL_494510, EPI_ISL_494511, EPI_ISL_494512, EPI_ISL_494513, EPI_ISL_494514, EPI_ISL_494515, EPI_ISL_494516, EPI_ISL_494517, EPI_ISL_494518, EPI_ISL_494519, EPI_ISL_494520, EPI_ISL_494521, EPI_ISL_494522, EPI_ISL_494523, EPI_ISL_494524, EPI_ISL_494525, EPI_ISL_494526, EPI_ISL_494527, EPI_ISL_494528, EPI_ISL_494529, EPI_ISL_494530, EPI_ISL_494531, EPI_ISL_494532, EPI_ISL_494533, EPI_ISL_494534, EPI_ISL_494535, EPI_ISL_494536, EPI_ISL_494537, EPI_ISL_494538, EPI_ISL_494539, EPI_ISL_494540, EPI_ISL_494541, EPI_ISL_494542, EPI_ISL_494543, EPI_ISL_494544, EPI_ISL_494545, EPI_ISL_494546, EPI_ISL_494547, EPI_ISL_494548, EPI_ISL_494549, EPI_ISL_494550, EPI_ISL_494551, EPI_ISL_494552 | | | |
| see above | Quest Diagnostics | Quest Diagnostics | Anderson,B.P., Rosenthal,S.H., Gerasimova,A., Kagan,R.M. and Owen, R. |
| EPI_ISL_495028, EPI_ISL_495046 | GMERS Medical College & Hospital, Gotri, Vadodara | Gujarat Biotechnology Research Centre | Afzal Ansari, Nikha Trivedi, Meenakshi Shah, Neena Doshi, Varsha Godbole, Apurvasinh Puvar, Janvi Raval, Zarna Patel, Monika Gandhi, Pinal Trivedi, Maharshi Pandya, Nidhi Patel, Nitin Savaliya, Raghawendra Kumar, Dinesh Kumar, Zuber Saiyed, Komal Patel, Labdhi Pandya, R D Dixit, A M Kadri, Harsh Bakshi, Chaitanya Joshi, Madhvi Joshi |
| EPI_ISL_495064 | GMERS Medical College & Hospital, Himmatnagar | Gujarat Biotechnology Research Centre | Afzal Ansari, Nikha Trivedi, Himanshu Khatri, Mayur Gandhi, Apurvasinh Puvar, Janvi Raval, Zarna Patel, Monika Gandhi, Pinal Trivedi, Maharshi Pandya, Nidhi Patel, Nitin Savaliya, Raghawendra Kumar, Dinesh Kumar, Zuber Saiyed, Komal Patel, Labdhi Pandya, R D Dixit, A M Kadri, Harsh Bakshi, Chaitanya Joshi, Madhvi Joshi, |
| EPI_ISL_495079 | Department of MicroBiology, Government Medical College, Surat | Gujarat Biotechnology Research Centre | Afzal Ansari, Nikha Trivedi, Naresh Chauhan, Summaiya Mullan, Amit gamit, Apurvasinh Puvar, Janvi Raval, Zarna Patel, Monika Gandhi, Pinal Trivedi, Maharshi Pandya, Nidhi Patel, Nitin Savaliya, Raghawendra Kumar, Dinesh Kumar, Zuber Saiyed, Komal Patel, Labdhi Pandya, R D Dixit, A M Kadri, Harsh Bakshi, Chaitanya Joshi, Madhvi Joshi |
| EPI_ISL_496482 | Dept. Infectious, Tropical Diseases & Microbiology, IRCCS Sacro Cuore Don Calabria Hospital | 1) Dept. Infectious, Tropical Diseases & Microbiology, IRCCS Sacro Cuore Don Calabria Hospital; 2) Centro Piattaforme | 1) Antonio Mori, Michela Deiana, Elena Pomari, Chiara Piubelli; 2) Monica Castellucci and Francesca Griggio; 3) Giovanni Malerba |

| | | | |
|---|---|---|---|
| | | Tecnologiche, University of Verona; 3) Dept. Neurosciences, Biomedicine and Movement Sciences, University of Verona. | |
| EPI_ISL_498551, EPI_ISL_498552, EPI_ISL_498554, EPI_ISL_498556 | Lebanese American University | Lebanese American University | Abi Habib,W., Abdallah,J., El Shesheny,R., Mokhbat,J., Webby,R.J., Goldstein,J. and Kayali,G. |
| EPI_ISL_498808, EPI_ISL_498809, EPI_ISL_498811, EPI_ISL_498814, EPI_ISL_498815, EPI_ISL_498816, EPI_ISL_498817, EPI_ISL_498818, EPI_ISL_498830, EPI_ISL_498892, EPI_ISL_498932, EPI_ISL_498933 | | | |
| see above | National Institute of Laboratory Medicine and Referral Center | Genomic Research Lab, BCSIR | Abu Sayeed Mohammad Mahmud, Mohammad Samir Uzzaman, Eshrar Osman, Md. Ahasan Habib, Shahina Akter, Tanjina Akhter Banu, Md. Murshed Hasan Sarkar, Barna Goswami, Iffat Jahan, Md. Saddam Hossain, Tasnim Nafisa, Md. Maruf Ahmed Molla, Mahmuda Yeasmin, Asish Kumar Ghosh, A. K. M. Shamsuzzaman, Sheikh Md. Selim Al Din, Utpal Chandra Ray, Salek Ahmed Sajib, Md. Salim Khan |
| EPI_ISL_499271, EPI_ISL_499272, EPI_ISL_499274, EPI_ISL_499278, EPI_ISL_499282, EPI_ISL_499283, EPI_ISL_499284, EPI_ISL_499285, EPI_ISL_499288, EPI_ISL_499289, EPI_ISL_499294, EPI_ISL_499295, EPI_ISL_499296, EPI_ISL_499297, EPI_ISL_499298, EPI_ISL_499300, EPI_ISL_499302, EPI_ISL_499303, EPI_ISL_499304, EPI_ISL_499306, EPI_ISL_499307, EPI_ISL_499308, EPI_ISL_499309, EPI_ISL_499310, EPI_ISL_499311, EPI_ISL_499312, EPI_ISL_499314, EPI_ISL_499315, EPI_ISL_499316, EPI_ISL_499317, EPI_ISL_499318, EPI_ISL_499321, EPI_ISL_499322, EPI_ISL_499323, EPI_ISL_499324, EPI_ISL_499325, EPI_ISL_499326, EPI_ISL_499327, EPI_ISL_499328, EPI_ISL_499329 | | | |
| see above | Centre for Enzyme Innovation, University of Portsmouth / Translational Research Laboratory, Portsmouth Hospitals NHS Trust | COVID-19 Genomics UK (COG-UK) Consortium | Angela Beckett,Yann Bourgeois,Garry Scarlett,Sharon Glaysher,Scott Elliott,Kelly Bicknell,Robert Impey,Allyson Lloyd,Sarah Wyllie,Ethan Butcher,Anoop Chauhan,Samuel Robson |
| EPI_ISL_499352, EPI_ISL_499353, EPI_ISL_499354 | West of Scotland Specialist Virology Centre, NHSGGC / MRC-University of Glasgow Centre for Virus Research | COVID-19 Genomics UK (COG-UK) Consortium | Ana da Silva Filipe, Natasha Johnson, Kathy Smollett, Daniel Mair, Stephen Carmichael, Lily Tong, Jenna Nichols, Elihu Aranday-Cortes, Kirstyn Brunker, Yasmin Parr, Alice Broos, Kyriaki Nomikou; Sarah McDonald, Marc Niebel, Patawee Asamaphan; Richard Orton, Joseph Hughes, Sreenu Vattipally, David L Robertson; Alasdair MacLean, Rory Gunson; Kathy Li, Natasha Jesudason, Rajiv Shah, James Shepherd, Antonia Ho, Emma Thomson |
| EPI_ISL_500864 | Instituto de Pesquisas Biomédicas, Hospital Naval Marcílio Dias | Bioinformatics Laboratory / LNCC | Andressa Rangel de Oliveira Lima, Luiz Gonzaga Paula de Almeida, Alexandra Lehmkuhl Gerber, Marlon Daniel Lima Tonin, Mônica Gomes Monteiro da R. Santos, Claudio Alberto Mule Monteiro, Átila Duque Rossi, Isabel de Medeiros Magalhaes de As, Luana Ferreira Martins de Toledo, Amilcar Tanuri, Albina Luciana da Silva Freitas, Fernanda Conceição Silva do Amorim, Shana Priscila Coutinho Barroso, Cynthia Chester Cardoso, Carolina Moreira Voloch, Ana Tereza Vasconcelos |
| EPI_ISL_501633, EPI_ISL_501634, EPI_ISL_501635 | NHSGGC West of Scotland Specialist Virology Centre / MRC-University of Glasgow Centre for Virus Research | Wellcome Sanger Institute for the COVID-19 Genomics UK (COG-UK) consortium | Ana da Silva Filipe, Natasha Johnson, Kathy Smollett, Daniel Mair, Stephen Carmichael, Lily Tong, Jenna Nichols, Elihu Aranday-Cortes, Kirstyn Brunker, Yasmin Parr, Kyriaki Nomikou; Sarah McDonald, Marc Niebel, Patawee Asamaphan; Richard Orton, Joseph Hughes, Sreenu Vattipally, David L Robertson; Alasdair MacLean, Rory Gunson; Kathy Li, Natasha Jesudason, Rajiv Shah, James Shepherd, Antonia Ho, Alice Broos, Emma Thomson and Alex Alderton, Roberto Amato, Sonia Goncalves, Ewan Harrison, David K. Jackson, Ian Johnston, Dominic Kwiatkowski, Cordelia Langford, John Sillitoe on behalf of the Wellcome Sanger Institute COVID-19 Surveillance Team (http://www.sanger.ac.uk/covid-team) |
| EPI_ISL_504137, EPI_ISL_504176, EPI_ISL_504177, EPI_ISL_504178, EPI_ISL_504179, EPI_ISL_504180, EPI_ISL_504181, EPI_ISL_504182, EPI_ISL_504183, EPI_ISL_504184 | National Institute of Laboratory Medicine and Referral Center | Genomic Research Lab, BCSIR | Abu Sayeed Mohammad Mahmud, Mohammad Samir Uzzaman, Eshrar Osman, Md. Ahasan Habib, Shahina Akter, Tanjina Akhter Banu, Md. Murshed Hasan Sarkar, Barna Goswami, Iffat Jahan, Md. Saddam Hossain, Tarannum Taznin, Tasnim Nafisa, Md. Maruf Ahmed Molla, Mahmuda Yeasmin, Asish Kumar Ghosh, A. K. M. Shamsuzzaman, Sheikh Md. Selim Al Din, Utpal Chandra Ray, Salek Ahmed Sajib, Md. Salim Khan |
| EPI_ISL_504186, EPI_ISL_504187, EPI_ISL_504188, EPI_ISL_504189, EPI_ISL_504190, EPI_ISL_504191, EPI_ISL_504192, EPI_ISL_504193, EPI_ISL_504194, EPI_ISL_504195, EPI_ISL_504196, EPI_ISL_504197, EPI_ISL_504198, EPI_ISL_504199, EPI_ISL_504200, EPI_ISL_504201, EPI_ISL_504202, EPI_ISL_504203, EPI_ISL_504204, EPI_ISL_504205, EPI_ISL_504206, EPI_ISL_504207, EPI_ISL_504208, EPI_ISL_504209, EPI_ISL_504210, EPI_ISL_504211, EPI_ISL_504212, EPI_ISL_504213, EPI_ISL_504214, EPI_ISL_504215, EPI_ISL_504216, EPI_ISL_504217, EPI_ISL_504218, EPI_ISL_504219, EPI_ISL_504220, EPI_ISL_504221, EPI_ISL_504222, EPI_ISL_504223, EPI_ISL_504224, EPI_ISL_504225, EPI_ISL_504226, EPI_ISL_504227, EPI_ISL_504228, EPI_ISL_504229, EPI_ISL_504230, EPI_ISL_504231, EPI_ISL_504232, EPI_ISL_504233, EPI_ISL_504234, EPI_ISL_504235, EPI_ISL_504236, EPI_ISL_504237, EPI_ISL_504238, EPI_ISL_504239, EPI_ISL_504240, EPI_ISL_504241, EPI_ISL_504242, EPI_ISL_504243, EPI_ISL_504244 | | | |
| see above | National Institute for Communicable Diseases of the National Health Laboratory Service | National Institute for Communicable Diseases of the National Health Laboratory Service | Allam M, Ismail A, Khumalo Z, Kwenda S, Mtshali P, Mnyameni F, Mohale T, Bhiman JN |
| EPI_ISL_507010, EPI_ISL_507011, EPI_ISL_507012, EPI_ISL_507013, EPI_ISL_507014, EPI_ISL_507015, EPI_ISL_507016, EPI_ISL_507017, EPI_ISL_507018, EPI_ISL_507019, EPI_ISL_507020, EPI_ISL_507021, EPI_ISL_507022, EPI_ISL_507023, EPI_ISL_507024, EPI_ISL_507025, EPI_ISL_507026, EPI_ISL_507027, EPI_ISL_507028, EPI_ISL_507029, EPI_ISL_507030, EPI_ISL_507031, EPI_ISL_507032, EPI_ISL_507033, EPI_ISL_507034, EPI_ISL_507035, EPI_ISL_507036, EPI_ISL_507037, EPI_ISL_507038 | | | |
| see above | unknown | Infectious Diseases Research, King Abdullah International Medical Research Center (KAIMRC) | Alghoribi,M.F. |
| EPI_ISL_507131, EPI_ISL_507132, EPI_ISL_507133, EPI_ISL_507134, EPI_ISL_507135, EPI_ISL_507137, EPI_ISL_507139, EPI_ISL_507140, EPI_ISL_507142, EPI_ISL_507143, EPI_ISL_507146, EPI_ISL_507147, EPI_ISL_507148, EPI_ISL_507149, EPI_ISL_507150, EPI_ISL_507151, EPI_ISL_507152, EPI_ISL_507153, EPI_ISL_507154 | | | |
| see above | Centre for Enzyme Innovation, University of Portsmouth / Translational Research Laboratory, Portsmouth Hospitals NHS Trust | COVID-19 Genomics UK (COG-UK) Consortium | Angela Beckett,Yann Bourgeois,Garry Scarlett,Sharon Glaysher,Scott Elliott,Kelly Bicknell,Robert Impey,Allyson Lloyd,Sarah Wyllie,Ethan Butcher,Anoop Chauhan,Samuel Robson |
| EPI_ISL_507206, EPI_ISL_507207, EPI_ISL_507208, EPI_ISL_507209, EPI_ISL_507210, EPI_ISL_507211, EPI_ISL_507212, EPI_ISL_507213, EPI_ISL_507214, EPI_ISL_507215 | Department of Experimental Modeling and Pathogenesis of Infectious Diseases | WHO National Influenza Centre Russian Federation | Andrey Komissarov, Artem Fadeev, Mariia Sergeeva, Anna Ivanova, Daria Danilenko |
| EPI_ISL_507224, EPI_ISL_507225, EPI_ISL_507226, EPI_ISL_507227, EPI_ISL_507229, EPI_ISL_507230, EPI_ISL_507231, EPI_ISL_507232, EPI_ISL_507233, EPI_ISL_507234, EPI_ISL_507235, EPI_ISL_507236, EPI_ISL_507237, EPI_ISL_507238, EPI_ISL_507239, EPI_ISL_507240, EPI_ISL_507241, EPI_ISL_507242, EPI_ISL_507243, EPI_ISL_507244, EPI_ISL_507245, EPI_ISL_507246, EPI_ISL_507247, EPI_ISL_507248, EPI_ISL_507249, EPI_ISL_507250, EPI_ISL_507251, EPI_ISL_507252, EPI_ISL_507253, EPI_ISL_507254, EPI_ISL_507255, EPI_ISL_507256, EPI_ISL_507257, EPI_ISL_507258, EPI_ISL_507259, EPI_ISL_507260, EPI_ISL_507261, EPI_ISL_507262, EPI_ISL_507263, EPI_ISL_507264, EPI_ISL_507265, EPI_ISL_507266, EPI_ISL_507267, EPI_ISL_507268, EPI_ISL_507269, EPI_ISL_507270, EPI_ISL_507271, EPI_ISL_507272, EPI_ISL_507273, EPI_ISL_507274, EPI_ISL_507275, EPI_ISL_507276, EPI_ISL_507277, EPI_ISL_507278, EPI_ISL_507279, EPI_ISL_507280, EPI_ISL_507281, EPI_ISL_507282, EPI_ISL_507283, EPI_ISL_507284, EPI_ISL_507285, EPI_ISL_507286, EPI_ISL_507287, EPI_ISL_507288, EPI_ISL_507289, EPI_ISL_507290, EPI_ISL_507291, EPI_ISL_507293, EPI_ISL_507294, EPI_ISL_507295, EPI_ISL_507296 | | | |
| see above | WHO National Influenza Centre Russian Federation | WHO National Influenza Centre Russian Federation | Andrey Komissarov, Artem Fadeev, Mariia Sergeeva, Anna Ivanova, Daria Danilenko |
| EPI_ISL_508615, EPI_ISL_508616, EPI_ISL_508617, EPI_ISL_508618, EPI_ISL_508619, EPI_ISL_508620, EPI_ISL_508621, EPI_ISL_508622, EPI_ISL_508623, EPI_ISL_508624, EPI_ISL_508625, EPI_ISL_508626, EPI_ISL_508627, EPI_ISL_508628, EPI_ISL_508629, EPI_ISL_508630, EPI_ISL_508631, EPI_ISL_508632, EPI_ISL_508633, EPI_ISL_508634, EPI_ISL_508635, EPI_ISL_508636, EPI_ISL_508637, EPI_ISL_508638, EPI_ISL_508639, EPI_ISL_508640, EPI_ISL_508641, EPI_ISL_508642, EPI_ISL_508643, EPI_ISL_508644, EPI_ISL_508645, EPI_ISL_508646, EPI_ISL_508647, EPI_ISL_508648, EPI_ISL_508649, EPI_ISL_508650, EPI_ISL_508651, EPI_ISL_508652, EPI_ISL_508653, EPI_ISL_508654, EPI_ISL_508655, EPI_ISL_508656, EPI_ISL_508657, EPI_ISL_508658, EPI_ISL_508659, EPI_ISL_508660, EPI_ISL_508661, EPI_ISL_508662, EPI_ISL_508663, EPI_ISL_508664, EPI_ISL_508665, EPI_ISL_508666, EPI_ISL_508667, EPI_ISL_508668, EPI_ISL_508669, EPI_ISL_508670, EPI_ISL_508671, EPI_ISL_508672, EPI_ISL_508673, EPI_ISL_508674, EPI_ISL_508675, EPI_ISL_508676, EPI_ISL_508677, EPI_ISL_508678, EPI_ISL_508679, EPI_ISL_508680, EPI_ISL_508681, EPI_ISL_508682, EPI_ISL_508683, EPI_ISL_508684, EPI_ISL_508685, EPI_ISL_508686 | | | |
| see above | Departamento de Microbiología, CDB, Hospital Clínic, Barcelona | SeqCOVID-SPAIN consortium/IBV(CSIC) | Andrea Vergara, Mikel Martínez, Elisa Rubio, Jéssica Navero, Aida Peiró and SeqCOVID-SPAIN consortium |
| EPI_ISL_508687, EPI_ISL_508688, EPI_ISL_508689, EPI_ISL_508690, EPI_ISL_508691, EPI_ISL_508692, EPI_ISL_508693, EPI_ISL_508694, EPI_ISL_508695, EPI_ISL_508696, EPI_ISL_508697, EPI_ISL_508698, EPI_ISL_508699, EPI_ISL_508700, EPI_ISL_508701, EPI_ISL_508702, EPI_ISL_508703, EPI_ISL_508704 | | | |
| see above | Institut für Virologie und Epidemiologie der Viruskrankheiten, Universitätsklinikum Tübingen | NGS Competence Center Tübingen, Institut für Medizinische Mikrobiologie und Hygiene, Universitätsklinikum Tübingen | Angel Angelov |
| EPI_ISL_510528 | Communicable Disease Laboratory, Public Health Directorate | Communicable Disease Laboratory, Public Health Directorate | Al Wasti,H. and AlTaif,Z. |
| EPI_ISL_512068 | Sardar Vallabhbhai Patel Institute of Medical Sciences & Research | Gujarat Biotechnology Research Centre | Afzal Ansari, Nikha Trivedi, Pranay Shah, Kamlesh J Upadhyay, Sanjay Kapadia, Apurvasinh Puvar, Janvi Raval, Zarna Patel, Monika Gandhi, Pinal Trivedi, Maharshi Pandya, Nidhi Patel, Nitin Savaliya, Raghawendra Kumar, Dinesh Kumar, Zuber Saiyed, Komal Patel, Labdhi Pandya, R D Dixit, A M Kadri, Harsh Bakshi, Chaitanya Joshi, Madhvi Joshi |
| EPI_ISL_512395, EPI_ISL_512396, EPI_ISL_512400, EPI_ISL_512407, EPI_ISL_512409, EPI_ISL_512410, EPI_ISL_512411, EPI_ISL_512412, EPI_ISL_512415, EPI_ISL_512416, EPI_ISL_512417 | | | |
| see above | Centre for Enzyme Innovation, University of Portsmouth / Translational Research Laboratory, Portsmouth Hospitals NHS Trust | COVID-19 Genomics UK (COG-UK) Consortium | Angela Beckett,Yann Bourgeois,Garry Scarlett,Sharon Glaysher,Scott Elliott,Kelly Bicknell,Robert Impey,Allyson Lloyd,Sarah Wyllie,Ethan Butcher,Anoop Chauhan,Samuel Robson |
| EPI_ISL_512436, EPI_ISL_512437, EPI_ISL_512438, EPI_ISL_512439, EPI_ISL_512440, EPI_ISL_512441, EPI_ISL_512442, EPI_ISL_512443, EPI_ISL_512444, EPI_ISL_512445, EPI_ISL_512446, EPI_ISL_512447, EPI_ISL_512448, EPI_ISL_512449, EPI_ISL_512450, EPI_ISL_512451, EPI_ISL_512452, EPI_ISL_512453, | | | |

| | | | |
|---|---|---|---|
| EPI_ISL_512454, EPI_ISL_512455, EPI_ISL_512456, EPI_ISL_512457, EPI_ISL_512458, EPI_ISL_512459, EPI_ISL_512460, EPI_ISL_512461, EPI_ISL_512462, EPI_ISL_512463, EPI_ISL_512464, EPI_ISL_512465, EPI_ISL_512466, EPI_ISL_512467, EPI_ISL_512468, EPI_ISL_512469, EPI_ISL_512470, EPI_ISL_512471, EPI_ISL_512472, EPI_ISL_512473, EPI_ISL_512474, EPI_ISL_512475, EPI_ISL_512476, EPI_ISL_512477, EPI_ISL_512478, EPI_ISL_512479, EPI_ISL_512480 | | | |
| see above | West of Scotland Specialist Virology Centre, NHSGGC / MRC-University of Glasgow Centre for Virus Research | COVID-19 Genomics UK (COG-UK) Consortium | Ana da Silva Filipe, Natasha Johnson, Kathy Smollett, Daniel Mair, Stephen Carmichael, Lily Tong, Jenna Nichols, Elihu Aranday-Cortes, Kirstyn Brunker, Yasmin Parr, Alice Broos, Kyriaki Nomikou; Sarah McDonald, Marc Niebel, Patawee Asamaphan; Richard Orton, Joseph Hughes, Sreenu Vattipally, David L Robertson; Alasdair MacLean, Rory Gunson; Kathy Li, Natasha Jesudason, Rajiv Shah, James Shepherd, Antonia Ho, Emma Thomson |
| EPI_ISL_512774 | Genomic Research Lab, BCSIR | Bangladesh Council of Scientific and Industrial Research | Abu Sayeed Mohammad Mahmud, Mohammad Samir Uzzaman, Eshrar Osman, Md. Ahasan Habib, Shahina Akter, Tanjina Akhter Banu, Md. Murshed Hasan Sarkar, Barna Goswami, Iffat Jahan, Md. Saddam Hossain, Tasnim Nafisa, Md. Maruf Ahmed Molla, Mahmuda Yeasmin, Asish Kumar Ghosh, A. K. M. Shamsuzzaman, Sheikh Md. Selim Al Din, Utpal Chandra Ray, Salek Ahmed Sajib, Md. Salim Khan |
| EPI_ISL_512989, EPI_ISL_512990, EPI_ISL_512991, EPI_ISL_512992, EPI_ISL_512993, EPI_ISL_512994, EPI_ISL_512995, EPI_ISL_512996, EPI_ISL_512997, EPI_ISL_512998, EPI_ISL_512999, EPI_ISL_513000, EPI_ISL_513001 | | | |
| see above | Pathogen Genomics Lab King Abdullah University of Science and Technology(KAUST) | Pathogen Genomics Lab King Abdullah University of Science and Technology(KAUST) | Amit Kumar Subudhi, Rahul P Salunke, Sara Mfarrej, Sharif Hala, Fadwa Alofi, Fathia Ben Rached, Afrah Alsomali, Asim Khogeer, Nashwa Al-khotani, Raeece Naeem, Anwar Hashem, Naif Almontashiri, Arnab Pain |
| EPI_ISL_513002, EPI_ISL_513003, EPI_ISL_513004, EPI_ISL_513005, EPI_ISL_513006, EPI_ISL_513007, EPI_ISL_513008, EPI_ISL_513009, EPI_ISL_513010, EPI_ISL_513011, EPI_ISL_513012, EPI_ISL_513013, EPI_ISL_513014, EPI_ISL_513015, EPI_ISL_513016, EPI_ISL_513017, EPI_ISL_513018, EPI_ISL_513019, EPI_ISL_513020, EPI_ISL_513021, EPI_ISL_513022, EPI_ISL_513023, EPI_ISL_513024, EPI_ISL_513025, EPI_ISL_513026 | | | |
| see above | Pathogen Genomics Lab King Abdullah University of Science and Technology(KAUST) | Pathogen Genomics Lab King Abdullah University of Science and Technology(KAUST) | Afrah Alsomali, Fathia Ben Rached, Raeece Naeem, Sharif Hala,Rahul P Salunke, Amanda Ooi, Luke Esau, Sara Mfarrej, Amit Kumar Subudhi, Fadwa Alofi, Asim Khogeer, Kahled Algithami, Anwar Hashem, Naif Almontashiri, Arnab Pain |
| EPI_ISL_513175, EPI_ISL_513176, EPI_ISL_513177, EPI_ISL_513178, EPI_ISL_513179, EPI_ISL_513180, EPI_ISL_513181, EPI_ISL_513182, EPI_ISL_513183, EPI_ISL_513184, EPI_ISL_513185, EPI_ISL_513186, EPI_ISL_513187, EPI_ISL_513188, EPI_ISL_513189 | | | |
| see above | Pathogen Genomics Lab King Abdullah University of Science and Technology(KAUST) | Pathogen Genomics Lab King Abdullah University of Science and Technology(KAUST) | Amit Kumar Subudhi, Rahul P Salunke, Sara Mfarrej, Sharif Hala, Fadwa Alofi, Fathia Ben Rached, Afrah Alsomali, Asim Khogeer, Nashwa Al-khotani, Raeece Naeem, Anwar Hashem, Naif Almontashiri, Arnab Pain |
| EPI_ISL_513190, EPI_ISL_513191, EPI_ISL_513192, EPI_ISL_513193, EPI_ISL_513194, EPI_ISL_513195, EPI_ISL_513196, EPI_ISL_513197 | Pathogen Genomics Lab King Abdullah University of Science and Technology(KAUST) | Pathogen Genomics Lab King Abdullah University of Science and Technology(KAUST) | Afrah Alsomali, Fathia Ben Rached, Raeece Naeem, Sharif Hala,Rahul P Salunke, Amanda Ooi, Luke Esau, Sara Mfarrej, Amit Kumar Subudhi, Fadwa Alofi, Asim Khogeer, Kahled Algithami, Anwar Hashem, Naif Almontashiri, Arnab Pain |
| EPI_ISL_513227, EPI_ISL_513228, EPI_ISL_513229, EPI_ISL_513230, EPI_ISL_513231, EPI_ISL_513232, EPI_ISL_513233, EPI_ISL_513234, EPI_ISL_513235, EPI_ISL_513236, EPI_ISL_513237, EPI_ISL_513238, EPI_ISL_513239, EPI_ISL_513240, EPI_ISL_513241, EPI_ISL_513242, EPI_ISL_513243, EPI_ISL_513244, EPI_ISL_513245, EPI_ISL_513246, EPI_ISL_513247, EPI_ISL_513248 | | | |
| see above | Pathogen Genomics Lab King Abdullah University of Science and Technology(KAUST) | Pathogen Genomics Lab King Abdullah University of Science and Technology(KAUST) | Amit Kumar Subudhi, Rahul P Salunke, Sara Mfarrej, Sharif Hala, Fadwa Alofi, Fathia Ben Rached, Afrah Alsomali, Asim Khogeer, Nashwa Al-khotani, Raeece Naeem, Anwar Hashem, Naif Almontashiri, Arnab Pain |
| EPI_ISL_513249, EPI_ISL_513250, EPI_ISL_513251, EPI_ISL_513252, EPI_ISL_513253, EPI_ISL_513254, EPI_ISL_513256, EPI_ISL_513257, EPI_ISL_513258, EPI_ISL_513259, EPI_ISL_513260, EPI_ISL_513261, EPI_ISL_513262, EPI_ISL_513263, EPI_ISL_513264 | | | |
| see above | Pathogen Genomics Lab King Abdullah University of Science and Technology(KAUST) | Pathogen Genomics Lab King Abdullah University of Science and Technology(KAUST) | Afrah Alsomali, Fathia Ben Rached, Raeece Naeem, Sharif Hala,Rahul P Salunke, Amanda Ooi, Luke Esau, Sara Mfarrej, Amit Kumar Subudhi, Fadwa Alofi, Asim Khogeer, Kahled Algithami, Anwar Hashem, Naif Almontashiri, Arnab Pain |
| EPI_ISL_514248, EPI_ISL_514249, EPI_ISL_514250, EPI_ISL_514251, EPI_ISL_514252 | National Institute of Laboratory Medicine and Referral Center | Genomic Research Lab, BCSIR | Abu Sayeed Mohammad Mahmud, Mohammad Samir Uzzaman, Eshrar Osman, Md. Ahashan Habib, Shahina Akter, Tanjina Akhter Banu, Md. Murshed Hasan Sarkar, Barna Goswami, Iffat Jahan, Md. Saddam Hossain, Tasnim Nafisa, Md. Maruf Ahmed Molla, Mahmuda Yeasmin, Asish Kumar Ghosh, A. K. M. Shamsuzzaman, Sheikh Md. Selim Al Din, Utpal Chandra Ray, Salek Ahmed Sajib, Md. Salim Khan |
| EPI_ISL_514253 | Advanced Biotechnology Laboratory | Genomic Research Lab, BCSIR | Abu Sayeed Mohammad Mahmud, Mohammad Samir Uzzaman, Eshrar Osman, Hossain Uddin Shekhar, M. Aftab Uddin, Md. Bayejid Hosen, Eunus Ali, Md. Ahashan Habib, Shahina Akter, Tanjina Akhter Banu, Md. Murshed Hasan Sarkar, Barna Goswami, Iffat Jahan, Md. Saddam Hossain, Utpal Chandra Ray, Salek Ahmed Sajib, Md. Salim Khan |
| EPI_ISL_514355, EPI_ISL_514356, EPI_ISL_514357, EPI_ISL_514358, EPI_ISL_514359, EPI_ISL_514360, EPI_ISL_514361, EPI_ISL_514362, EPI_ISL_514363, EPI_ISL_514364, EPI_ISL_514365, EPI_ISL_514366, EPI_ISL_514367, EPI_ISL_514368, EPI_ISL_514369, EPI_ISL_514370, EPI_ISL_514371, EPI_ISL_514372, EPI_ISL_514373, EPI_ISL_514374, EPI_ISL_514375, EPI_ISL_514376, EPI_ISL_514377, EPI_ISL_514378, EPI_ISL_514379, EPI_ISL_514380, EPI_ISL_514381, EPI_ISL_514382, EPI_ISL_514383, EPI_ISL_514384, EPI_ISL_514385, EPI_ISL_514386, EPI_ISL_514387, EPI_ISL_514388, EPI_ISL_514389, EPI_ISL_514390, EPI_ISL_514391, EPI_ISL_514392, EPI_ISL_514393, EPI_ISL_514394, EPI_ISL_514395, EPI_ISL_514396, EPI_ISL_514397, EPI_ISL_514398, EPI_ISL_514399, EPI_ISL_514400, EPI_ISL_514401, EPI_ISL_514402, EPI_ISL_514403, EPI_ISL_514404, EPI_ISL_514405, EPI_ISL_514406, EPI_ISL_514407, EPI_ISL_514408, EPI_ISL_514409, EPI_ISL_514410, EPI_ISL_514411, EPI_ISL_514412, EPI_ISL_514413, EPI_ISL_514414, EPI_ISL_514415, EPI_ISL_514416, EPI_ISL_514417, EPI_ISL_514418, EPI_ISL_514419, EPI_ISL_514420, EPI_ISL_514421, EPI_ISL_514422, EPI_ISL_514423, EPI_ISL_514424 | | | |
| see above | National Institute for Communicable Diseases of the National Health Laboratory Service | National Institute for Communicable Diseases of the National Health Laboratory Service | Allam M, Ismail A, Khumalo Z, Kwenda S, Mtshali P, Mnyameni F, Mohale T, Bhiman JN |
| EPI_ISL_514457, EPI_ISL_514459, EPI_ISL_514460, EPI_ISL_514461, EPI_ISL_514462, EPI_ISL_514463, EPI_ISL_514465, EPI_ISL_514466, EPI_ISL_514468, EPI_ISL_514469, EPI_ISL_514472, EPI_ISL_514473, EPI_ISL_514474, EPI_ISL_514475, EPI_ISL_514476, EPI_ISL_514477, EPI_ISL_514478, EPI_ISL_514479, EPI_ISL_514480, EPI_ISL_514482, EPI_ISL_514483, EPI_ISL_514484, EPI_ISL_514485, EPI_ISL_514486, EPI_ISL_514490, EPI_ISL_514505, EPI_ISL_514508, EPI_ISL_514510 | | | |
| see above | Centre for Enzyme Innovation, University of Portsmouth / Translational Research Laboratory, Portsmouth Hospitals NHS Trust | COVID-19 Genomics UK (COG-UK) Consortium | Angela Beckett,Yann Bourgeois,Garry Scarlett,Sharon Glaysher,Scott Elliott,Kelly Bicknell,Robert Impey,Allyson Lloyd,Sarah Wyllie,Ethan Butcher,Anoop Chauhan,Samuel Robson |
| EPI_ISL_514514, EPI_ISL_514515 | West of Scotland Specialist Virology Centre, NHSGGC / MRC-University of Glasgow Centre for Virus Research | COVID-19 Genomics UK (COG-UK) Consortium | Ana da Silva Filipe, Natasha Johnson, Kathy Smollett, Daniel Mair, Stephen Carmichael, Lily Tong, Jenna Nichols, Elihu Aranday-Cortes, Kirstyn Brunker, Yasmin Parr, Alice Broos, Kyriaki Nomikou; Sarah McDonald, Marc Niebel, Patawee Asamaphan; Richard Orton, Joseph Hughes, Sreenu Vattipally, David L Robertson; Alasdair MacLean, Rory Gunson; Kathy Li, Natasha Jesudason, Rajiv Shah, James Shepherd, Antonia Ho, Emma Thomson |
| EPI_ISL_514587, EPI_ISL_514605 | B.J. Medical College and Civil hospital, Ahmedabad | Gujarat Biotechnology Research Centre | Afzal Ansari, Nikha Trivedi, Pranay Shah, Kamlesh J Upadhyay, Sanjay Kapadia, Apurvasinh Puvar, Janvi Raval, Zarna Patel, Monika Gandhi, Pinal Trivedi, Maharshi Pandya, Nidhi Patel, Nitin Savaliya, Raghawendra Kumar, Dinesh Kumar, Zuber Saiyed, Komal Patel, Labdhi Pandya, R D Dixit, A M Kadri, Harsh Bakshi, Chaitanya Joshi, Madhvi Joshi |
| EPI_ISL_515133, EPI_ISL_515134, EPI_ISL_515135, EPI_ISL_515136, EPI_ISL_515137, EPI_ISL_515138, EPI_ISL_515139, EPI_ISL_515140, EPI_ISL_515141, EPI_ISL_515142, EPI_ISL_515143, EPI_ISL_515144, EPI_ISL_515145, EPI_ISL_515146, EPI_ISL_515147, EPI_ISL_515148, EPI_ISL_515149, EPI_ISL_515150, EPI_ISL_515151, EPI_ISL_515152, EPI_ISL_515153, EPI_ISL_515154, EPI_ISL_515155, EPI_ISL_515156, EPI_ISL_515157, EPI_ISL_515158, EPI_ISL_515159, EPI_ISL_515160, EPI_ISL_515161, EPI_ISL_515162, EPI_ISL_515163, EPI_ISL_515164, EPI_ISL_515165, EPI_ISL_515166, EPI_ISL_515167, EPI_ISL_515168, EPI_ISL_515169, EPI_ISL_515170, EPI_ISL_515171, EPI_ISL_515172, EPI_ISL_515174, EPI_ISL_515176, EPI_ISL_515177, EPI_ISL_515178, EPI_ISL_515179, EPI_ISL_515180 | | | |
| see above | National Institute for Communicable Diseases of the National Health Laboratory Service | National Institute for Communicable Diseases of the National Health Laboratory Service | Allam M, Ismail A, Khumalo Z, Kwenda S, Mtshali P, Mnyameni F, Mohale T, Bhiman JN |
| EPI_ISL_516748, EPI_ISL_516749, EPI_ISL_516750, EPI_ISL_516751, EPI_ISL_516752, EPI_ISL_516753, EPI_ISL_516754, EPI_ISL_516755, EPI_ISL_516756, EPI_ISL_516757, EPI_ISL_516758, EPI_ISL_516759, EPI_ISL_516760, EPI_ISL_516761, EPI_ISL_516762, EPI_ISL_516763, EPI_ISL_516764, EPI_ISL_516765, EPI_ISL_516766, EPI_ISL_516767, EPI_ISL_516768, EPI_ISL_516769, EPI_ISL_516770, EPI_ISL_516771, EPI_ISL_516772, EPI_ISL_516773, EPI_ISL_516774, EPI_ISL_516775, EPI_ISL_516776, EPI_ISL_516777, EPI_ISL_516778, EPI_ISL_516779, EPI_ISL_516780, EPI_ISL_516781, EPI_ISL_516782 | | | |
| see above | van Bakel Laboratory, Genetics and Genomics Sciences, Icahn School of Medicine at Mount Sinai | van Bakel Laboratory, Genetics and Genomics Sciences, Icahn School of Medicine at Mount Sinai | Andrew G. Letizia, Irene Ramos, Ajay Obla, Carl Goforth, Dawn Weir, Yongchao Ge, Marcas M. Bamman, Jayeeta Dutta, Ethan Ellis, Luis Estrella, Mary-Catherine George, Ana S. Gonzalez-Reiche, Darnell Graham, Adriana van de Guchte, Ramiro Gutierrez, Franca Jones, Aspasia Kalomoiri, Rhonda Lizewski, Stephen Lizewski, Jan Marayag, Nada Marjanovic, Eugene V. Millar, Venugopalan Nair, German Nudelman, Edgar Nunez, Brian Pike, James Regeimbal, Stas Rirak , Ernesto Santa Ana, Rachel S. Gelernter Sealfon, Robert Sebra, Mark Simons, Alessandra Soares-Schanoski, Michael Termini, Sindhu Vangeti, Carlos Williams, Harm van Bakel, Stuart C. Sealfon |
| EPI_ISL_517512, EPI_ISL_517513, EPI_ISL_517517, EPI_ISL_517519, EPI_ISL_517522, EPI_ISL_517526, EPI_ISL_517529, EPI_ISL_517532, EPI_ISL_517535, EPI_ISL_517536, EPI_ISL_517537, EPI_ISL_517540 | | | |
| see above | Centre for Enzyme Innovation, University of Portsmouth / Translational Research Laboratory, Portsmouth Hospitals NHS Trust | COVID-19 Genomics UK (COG-UK) Consortium | Angela Beckett,Yann Bourgeois,Garry Scarlett,Sharon Glaysher,Scott Elliott,Kelly Bicknell,Robert Impey,Allyson Lloyd,Sarah Wyllie,Ethan Butcher,Anoop Chauhan,Samuel Robson |
| EPI_ISL_517580, EPI_ISL_517581, EPI_ISL_517582 | West of Scotland Specialist Virology Centre, NHSGGC / MRC-University of Glasgow Centre for Virus Research | COVID-19 Genomics UK (COG-UK) Consortium | Ana da Silva Filipe, Natasha Johnson, Kathy Smollett, Daniel Mair, Stephen Carmichael, Lily Tong, Jenna Nichols, Elihu Aranday-Cortes, Kirstyn Brunker, Yasmin Parr, Alice Broos, Kyriaki Nomikou; Sarah McDonald, Marc Niebel, Patawee Asamaphan; Richard Orton, Joseph Hughes, Sreenu Vattipally, David L Robertson; Alasdair MacLean, Rory Gunson; Kathy Li, Natasha Jesudason, Rajiv Shah, James Shepherd, Antonia Ho, Emma Thomson |
| EPI_ISL_518824, EPI_ISL_518846, EPI_ISL_518847, EPI_ISL_518848, | Oman-National Influenza Center | Biotechnology & OMICs Laboratory, Natural & Medical Sciences Research Center, University of Nizwa | Abdul Latif Khan, Samira Al-Mahruqi, Ahmed Al-Harrasi, Samiha Al-Kharusi, Adil Khan, Ahmed Al-Rawahi, Sajjad Asaf, Amina Al-Jardani, Hanan Al-Kindi, Intisar Al-Shukri, Ahlam Al-Amri, Aisha Al-Amri, Aisha Al-Busaidi, Adil Al-Wahaibi, Seif Al-Abri. |

| | | | |
|---|---|---|---|
| EPI_ISL_518849, EPI_ISL_518850, EPI_ISL_518851, EPI_ISL_518852, EPI_ISL_518853, EPI_ISL_518854 | | | |
| EPI_ISL_520663, EPI_ISL_520664, EPI_ISL_520665, EPI_ISL_520666, EPI_ISL_520667, EPI_ISL_520668, EPI_ISL_520669, EPI_ISL_520670, EPI_ISL_520671, EPI_ISL_520672, EPI_ISL_520673, EPI_ISL_520674, EPI_ISL_520675, EPI_ISL_520676, EPI_ISL_520677, EPI_ISL_520678, EPI_ISL_520679, EPI_ISL_520680, EPI_ISL_520681, EPI_ISL_520682, EPI_ISL_520683, EPI_ISL_520684, EPI_ISL_520685, EPI_ISL_520686, EPI_ISL_520687, EPI_ISL_520688, EPI_ISL_520689, EPI_ISL_520690, EPI_ISL_520691, EPI_ISL_520692, EPI_ISL_520693, EPI_ISL_520694, EPI_ISL_520695, EPI_ISL_520696, EPI_ISL_520697, EPI_ISL_520698, EPI_ISL_520699, EPI_ISL_520700, EPI_ISL_520701, EPI_ISL_520702, EPI_ISL_520703, EPI_ISL_520704, EPI_ISL_520705, EPI_ISL_520706, EPI_ISL_520707, EPI_ISL_520708, EPI_ISL_520709, EPI_ISL_520710, EPI_ISL_520711, EPI_ISL_520712, EPI_ISL_520713, EPI_ISL_520714, EPI_ISL_520715, EPI_ISL_520716, EPI_ISL_520717, EPI_ISL_520718, EPI_ISL_520719, EPI_ISL_520720, EPI_ISL_520721, EPI_ISL_520722, EPI_ISL_520723, EPI_ISL_520724, EPI_ISL_520725, EPI_ISL_520726, EPI_ISL_520727, EPI_ISL_520728, EPI_ISL_520729, EPI_ISL_520730, EPI_ISL_520731, EPI_ISL_520732, EPI_ISL_520733, EPI_ISL_520734, EPI_ISL_520735, EPI_ISL_520736, EPI_ISL_520737, EPI_ISL_520738, EPI_ISL_520739, EPI_ISL_520740, EPI_ISL_520741, EPI_ISL_520742, EPI_ISL_520743, EPI_ISL_520744 | | | |
| see above | Mohammed Bin Rashid University of Medicine and Health Sciences | Al Jalila Genomics Center | Ahmad Abou Tayoun, Tom Loney, Hamda Khansaheb, Sathishkumar Ramaswamy, Divinlal Harilal, Zulfa Omar Deesi, Rupa Murthy Varghese, Hanan Al Suwaidi, Abdulmajeed Alkhaja, Mohammed Uddin, Rifat Hamoudi, Rabih Halwani, Abiola Catherine Senok, Qutayba Hamid, Norbert Nowotny, Alawi Alsheikh-Ali |
| EPI_ISL_522855 | ULSS9 Distretto di Bussolengo | Istituto Zooprofilattico Sperimentale delle Venezie | Adelaide Milani, Alessia Schivo, Annalisa Salviato, Erika Giorgia Quaranta, Ambra Pastori, Bianca Zecchin, Alice Fusaro, Isabella Monne, Calogero Terregino, Antonia Ricci |
| EPI_ISL_522856 | ULSS9 Distretto di San Bonifacio | Istituto Zooprofilattico Sperimentale delle Venezie | Adelaide Milani, Alessia Schivo, Annalisa Salviato, Erika Giorgia Quaranta, Ambra Pastori, Bianca Zecchin, Alice Fusaro, Isabella Monne, Calogero Terregino, Antonia Ricci |
| EPI_ISL_522857 | ULSS9 Scaligera | Istituto Zooprofilattico Sperimentale delle Venezie | Adelaide Milani, Alessia Schivo, Annalisa Salviato, Erika Giorgia Quaranta, Ambra Pastori, Bianca Zecchin, Alice Fusaro, Isabella Monne, Calogero Terregino, Antonia Ricci |
| EPI_ISL_522858 | ULSS9 Distretto di San Bonifacio | Istituto Zooprofilattico Sperimentale delle Venezie | Adelaide Milani, Alessia Schivo, Annalisa Salviato, Erika Giorgia Quaranta, Ambra Pastori, Bianca Zecchin, Alice Fusaro, Isabella Monne, Calogero Terregino, Antonia Ricci |
| EPI_ISL_522859 | ULSS9 Scaligera | Istituto Zooprofilattico Sperimentale delle Venezie | Adelaide Milani, Alessia Schivo, Annalisa Salviato, Erika Giorgia Quaranta, Ambra Pastori, Bianca Zecchin, Alice Fusaro, Isabella Monne, Calogero Terregino, Antonia Ricci |
| EPI_ISL_522860, EPI_ISL_522861, EPI_ISL_522862, EPI_ISL_522863, EPI_ISL_522864, EPI_ISL_522865, EPI_ISL_522866, EPI_ISL_522867, EPI_ISL_522868 | ULSS9 Distretto di Bussolengo | Istituto Zooprofilattico Sperimentale delle Venezie | Adelaide Milani, Alessia Schivo, Annalisa Salviato, Erika Giorgia Quaranta, Ambra Pastori, Bianca Zecchin, Alice Fusaro, Isabella Monne, Calogero Terregino, Antonia Ricci |
| EPI_ISL_523952, EPI_ISL_523953 | Mohammed Bin Rashid University of Medicine and Health Sciences | Al Jalila Genomics Center | Ahmad Abou Tayoun, Tom Loney, Hamda Khansaheb, Sathishkumar Ramaswamy, Divinlal Harilal, Zulfa Omar Deesi, Rupa Murthy Varghese, Hanan Al Suwaidi, Abdulmajeed Alkhaja, Mohammed Uddin, Rifat Hamoudi, Rabih Halwani, Abiola Catherine Senok, Qutayba Hamid, Norbert Nowotny, Alawi Alsheikh-Ali |
| EPI_ISL_523995, EPI_ISL_523996, EPI_ISL_523997, EPI_ISL_523998, EPI_ISL_523999, EPI_ISL_524000, EPI_ISL_524001, EPI_ISL_524002, EPI_ISL_524003, EPI_ISL_524004, EPI_ISL_524005, EPI_ISL_524006, EPI_ISL_524007, EPI_ISL_524008, EPI_ISL_524009, EPI_ISL_524010, EPI_ISL_524011, EPI_ISL_524012, EPI_ISL_524013, EPI_ISL_524014, EPI_ISL_524015, EPI_ISL_524016, EPI_ISL_524017, EPI_ISL_524018, EPI_ISL_524019, EPI_ISL_524020, EPI_ISL_524021, EPI_ISL_524022, EPI_ISL_524023, EPI_ISL_524024, EPI_ISL_524025, EPI_ISL_524026, EPI_ISL_524027, EPI_ISL_524028, EPI_ISL_524029, EPI_ISL_524030, EPI_ISL_524031, EPI_ISL_524032, EPI_ISL_524033, EPI_ISL_524034, EPI_ISL_524035, EPI_ISL_524036, EPI_ISL_524037, EPI_ISL_524038, EPI_ISL_524039, EPI_ISL_524040, EPI_ISL_524041, EPI_ISL_524042, EPI_ISL_524043, EPI_ISL_524044, EPI_ISL_524045, EPI_ISL_524046, EPI_ISL_524047, EPI_ISL_524048, EPI_ISL_524049, EPI_ISL_524050, EPI_ISL_524051, EPI_ISL_524052 | | | |
| see above | WHO National Influenza Centre Russian Federation | WHO National Influenza Centre Russian Federation | Andrey Komissarov, Artem Fadeev, Mariia Sergeeva, Anna Ivanova, Daria Danilenko |
| EPI_ISL_524494, EPI_ISL_524496, EPI_ISL_524499, EPI_ISL_524500, EPI_ISL_524504, EPI_ISL_524507, EPI_ISL_524510, EPI_ISL_524511, EPI_ISL_524512, EPI_ISL_524513, EPI_ISL_524514, EPI_ISL_524516, EPI_ISL_524517, EPI_ISL_524518, EPI_ISL_524519, EPI_ISL_524520, EPI_ISL_524523, EPI_ISL_524524, EPI_ISL_524526, EPI_ISL_524527, EPI_ISL_524532, EPI_ISL_524533, EPI_ISL_524534, EPI_ISL_524535, EPI_ISL_524536, EPI_ISL_524539, EPI_ISL_524540, EPI_ISL_524542, EPI_ISL_524543, EPI_ISL_524545, EPI_ISL_524547, EPI_ISL_524549, EPI_ISL_524550, EPI_ISL_524552, EPI_ISL_524554, EPI_ISL_524555, EPI_ISL_524556, EPI_ISL_524557, EPI_ISL_524559, EPI_ISL_524561, EPI_ISL_524562, EPI_ISL_524563, EPI_ISL_524565, EPI_ISL_524566, EPI_ISL_524567, EPI_ISL_524569, EPI_ISL_524571, EPI_ISL_524572 | | | |
| see above | NHSGGC West of Scotland Specialist Virology Centre / MRC-University of Glasgow Centre for Virus Research | Wellcome Sanger Institute for the COVID-19 Genomics UK (COG-UK) consortium | Ana da Silva Filipe, Natasha Johnson, Kathy Smollett, Daniel Mair, Stephen Carmichael, Lily Tong, Jenna Nichols, Elihu Aranday-Cortes, Kirstyn Brunker, Yasmin Parr, Kyriaki Nomikou; Sarah McDonald, Marc Niebel, Patawee Asamaphan; Richard Orton, Joseph Hughes, Sreenu Vattipally, David L Robertson; Alasdair MacLean, Rory Gunson; Kathy Li, Natasha Jesudason, Rajiv Shah, James Shepherd, Antonia Ho, Alice Broos, Emma Thomson and Alex Alderton, Roberto Amato, Sonia Goncalves, Ewan Harrison, David K. Jackson, Ian Johnston, Dominic Kwiatkowski, Cordelia Langford, John Sillitoe on behalf of the Wellcome Sanger Institute COVID-19 Surveillance Team (http://www.sanger.ac.uk/covid-team) |
| EPI_ISL_524726 | B.J. Medical College and Civil hospital, Ahmedabad | Gujarat Biotechnology Research Centre | Afzal Ansari, Nikha Trivedi, Pranay Shah, Kamlesh J Upadhyay, Sanjay Kapadia, Apurvasinh Puvar, Janvi Raval, Zarna Patel, Monika Gandhi, Pinal Trivedi, Maharshi Pandya, Nidhi Patel, Nitin Savaliya, Raghawendra Kumar, Dinesh Kumar, Zuber Saiyed, Komal Patel, Labdhi Pandya, R D Dixit, A M Kadri, Harsh Bakshi, Chaitanya Joshi, Madhvi Joshi |
| EPI_ISL_524733 | GMERS Medical College and Hospital, Dharpur, Patan | Gujarat Biotechnology Research Centre | A N Parmar, Apurvasinh Puvar, Janvi Raval, Zarna Patel, Monika Gandhi, Pinal Trivedi, Maharshi Pandya, Nidhi Patel, Nitin Savaliya, Raghawendra Kumar, Dinesh Kumar, Zuber Saiyed, Komal Patel, Labdhi Pandya, Afzal Ansari, Nikha Trivedi, R D Dixit, A M Kadri, Harsh Bakshi, Chaitanya Joshi, Madhvi Joshi |
| EPI_ISL_524747 | Sir M P Shah Government Medical College, Jamnagar | Gujarat Biotechnology Research Centre | Afzal Ansari, Nikha Trivedi, Binita Aring, Apurvasinh Puvar, Janvi Raval, Zarna Patel, Monika Gandhi, Pinal Trivedi, Maharshi Pandya, Nidhi Patel, Nitin Savaliya, Raghawendra Kumar, Dinesh Kumar, Zuber Saiyed, Komal Patel, Labdhi Pandya, R D Dixit, A M Kadri, Harsh Bakshi, Chaitanya Joshi, Madhvi Joshi |
| EPI_ISL_524762 | GMERS Medical College and Hospital, Dharpur, Patan | Gujarat Biotechnology Research Centre | Afzal Ansari, Nikha Trivedi, A N Parmar, Apurvasinh Puvar, Janvi Raval, Zarna Patel, Monika Gandhi, Pinal Trivedi, Maharshi Pandya, Nidhi Patel, Nitin Savaliya, Raghawendra Kumar, Dinesh Kumar, Zuber Saiyed, Komal Patel, Labdhi Pandya, R D Dixit, A M Kadri, Harsh Bakshi, Chaitanya Joshi, Madhvi Joshi |
| EPI_ISL_524764 | GMERS Medical College and Hospital, Dharpur, Patan | Gujarat Biotechnology Research Centre | A N Parmar, Apurvasinh Puvar, Janvi Raval, Zarna Patel, Monika Gandhi, Pinal Trivedi, Maharshi Pandya, Nidhi Patel, Nitin Savaliya, Raghawendra Kumar, Dinesh Kumar, Zuber Saiyed, Komal Patel, Labdhi Pandya, Afzal Ansari, Nikha Trivedi, R D Dixit, A M Kadri, Harsh Bakshi, Chaitanya Joshi, Madhvi Joshi |
| EPI_ISL_526434, EPI_ISL_526435 | Centre for Enzyme Innovation, University of Portsmouth / Translational Research Laboratory, Portsmouth Hospitals NHS Trust | COVID-19 Genomics UK (COG-UK) Consortium | Angela Beckett,Yann Bourgeois,Garry Scarlett,Sharon Glaysher,Scott Elliott,Kelly Bicknell,Robert Impey,Allyson Lloyd,Sarah Wyllie,Ethan Butcher,Anoop Chauhan,Samuel Robson |
| EPI_ISL_526451, EPI_ISL_526452, EPI_ISL_526453, EPI_ISL_526454, EPI_ISL_526455, EPI_ISL_526456, EPI_ISL_526457, EPI_ISL_526458 | West of Scotland Specialist Virology Centre, NHSGGC / MRC-University of Glasgow Centre for Virus Research | COVID-19 Genomics UK (COG-UK) Consortium | Ana da Silva Filipe, Natasha Johnson, Kathy Smollett, Daniel Mair, Stephen Carmichael, Lily Tong, Jenna Nichols, Elihu Aranday-Cortes, Kirstyn Brunker, Yasmin Parr, Alice Broos, Kyriaki Nomikou; Sarah McDonald, Marc Niebel, Patawee Asamaphan; Richard Orton, Joseph Hughes, Sreenu Vattipally, David L Robertson; Alasdair MacLean, Rory Gunson; Kathy Li, Natasha Jesudason, Rajiv Shah, James Shepherd, Antonia Ho, Emma Thomson |
| EPI_ISL_528538 | Alsafar | Alsafar | Andreas Henschel, Gihan Elsir Ahmed Daw Elbait, Samuel Feng, Rifat, Ernesto Damiani, Guan Tay, Habiba Alsafar |
| EPI_ISL_528686, EPI_ISL_528687, EPI_ISL_528688, EPI_ISL_528689, EPI_ISL_528690, EPI_ISL_528691, EPI_ISL_528692, EPI_ISL_528693, EPI_ISL_528694, EPI_ISL_528695, EPI_ISL_528696, EPI_ISL_528697, EPI_ISL_528698, EPI_ISL_528699, EPI_ISL_528700, EPI_ISL_528701, EPI_ISL_528702, EPI_ISL_528703, EPI_ISL_528704, EPI_ISL_528705, EPI_ISL_528706, EPI_ISL_528707, EPI_ISL_528708, EPI_ISL_528709, EPI_ISL_528710, EPI_ISL_528711, EPI_ISL_528712, EPI_ISL_528713, EPI_ISL_528714, EPI_ISL_528715, EPI_ISL_528716, EPI_ISL_528717, EPI_ISL_528718, EPI_ISL_528719, EPI_ISL_528720, EPI_ISL_528721 | | | |
| see above | Alsafar - Khalifa University Abu Dhabi | Alsafar - Khalifa University Abu Dhabi | Andreas Henschel, Gihan Daw Elbait, Samuel Feng, Rifat Hamoudi, Ernesto Damiani, Guan Tay, Habiba Alsafar |
| EPI_ISL_528788, EPI_ISL_528789, EPI_ISL_528790, EPI_ISL_528791, EPI_ISL_528792, EPI_ISL_528793, EPI_ISL_528794, EPI_ISL_528795, EPI_ISL_528796, EPI_ISL_528797, EPI_ISL_528798, EPI_ISL_528799, EPI_ISL_528800, EPI_ISL_528801, EPI_ISL_528802, EPI_ISL_528803, EPI_ISL_528804, EPI_ISL_528805, EPI_ISL_528806, EPI_ISL_528807, EPI_ISL_528808 | | | |
| see above | Microbiology Department, Barking Havering and Redbridge University Hospitals NHS trust | Wellcome Sanger Institute for the COVID-19 Genomics UK (COG-UK) consortium | Amy Ash, Fatima Ali, Cherian Koshy and Alex Alderton, Roberto Amato, Sonia Goncalves, Ewan Harrison, David K. Jackson, Ian Johnston, Dominic Kwiatkowski, Cordelia Langford, John Sillitoe on behalf of the Wellcome Sanger Institute COVID-19 Surveillance Team (http://www.sanger.ac.uk/covid-team) |
| EPI_ISL_529162, EPI_ISL_529163, | Department of Immunology, The Scripps Research Institute | Andersen lab at Scripps Research | Allison Smither, Gilberto Sabino-Santos, Patricia Snarski, Lilia Melnik, Antoinette Bell, Kaylynn Genemaras, Arnaud Drouin, Dahlene Fusco, Robert Garry |

| | | | |
|---|---|---|---|
| EPI_ISL_529164, EPI_ISL_529165, EPI_ISL_529166 | | | with SEARCH Alliance San Diego |
| EPI_ISL_529315, EPI_ISL_529316, EPI_ISL_529319, EPI_ISL_529409 | West of Scotland Specialist Virology Centre, NHSGGC / MRC-University of Glasgow Centre for Virus Research | COVID-19 Genomics UK (COG-UK) Consortium | Ana da Silva Filipe, Natasha Johnson, Kathy Smollett, Daniel Mair, Stephen Carmichael, Lily Tong, Jenna Nichols, Elihu Aranday-Cortes, Kirstyn Brunker, Yasmin Parr, Alice Broos, Kyriaki Nomikou; Sarah McDonald, Marc Niebel, Patawee Asamaphan; Richard Orton, Joseph Hughes, Sreenu Vattipally, David L Robertson; Alasdair MacLean, Rory Gunson; Kathy Li, Natasha Jesudason, Rajiv Shah, James Shepherd, Antonia Ho, Emma Thomson |
| EPI_ISL_529491, EPI_ISL_529511, EPI_ISL_529668, EPI_ISL_529669, EPI_ISL_529671, EPI_ISL_529672, EPI_ISL_529673, EPI_ISL_529674 | Centre for Enzyme Innovation, University of Portsmouth / Translational Research Laboratory, Portsmouth Hospitals NHS Trust | COVID-19 Genomics UK (COG-UK) Consortium | Angela Beckett,Yann Bourgeois,Garry Scarlett,Sharon Glaysher,Scott Elliott,Kelly Bicknell,Robert Impey,Allyson Lloyd,Sarah Wyllie,Ethan Butcher,Anoop Chauhan,Samuel Robson |
| EPI_ISL_529684, EPI_ISL_529685, EPI_ISL_529686, EPI_ISL_529687, EPI_ISL_529688 | West of Scotland Specialist Virology Centre, NHSGGC / MRC-University of Glasgow Centre for Virus Research | COVID-19 Genomics UK (COG-UK) Consortium | Ana da Silva Filipe, Natasha Johnson, Kathy Smollett, Daniel Mair, Stephen Carmichael, Lily Tong, Jenna Nichols, Elihu Aranday-Cortes, Kirstyn Brunker, Yasmin Parr, Alice Broos, Kyriaki Nomikou; Sarah McDonald, Marc Niebel, Patawee Asamaphan; Richard Orton, Joseph Hughes, Sreenu Vattipally, David L Robertson; Alasdair MacLean, Rory Gunson; Kathy Li, Natasha Jesudason, Rajiv Shah, James Shepherd, Antonia Ho, Emma Thomson |
| EPI_ISL_529964 | RSUD Bangil Pasuruan | Institute of Tropical Disease, Universitas Airlangga | Aldise M Nastri, Jezzy R Dewantari, Rima R Prasetya, Krisnoadi Rahardjo, Arma Roosalina, Gatot Soegiarto, Laksmi Wulandari, Retno A Setyoningrum, Resti Yudhawati, Yohko K Shimizu, Mitsuhiro Nishimura, Yasuko Mori, Soetjipto, Kazufumi Shimizu, Maria I Lusida |
| EPI_ISL_529967 | RSUD Sidoarjo | Institute of Tropical Disease, Universitas Airlangga | Aldise M Nastri, Jezzy R Dewantari, Rima R Prasetya, Krisnoadi Rahardjo, Atok Irawan, Gatot Soegiarto, Laksmi Wulandari, Retno A Setyoningrum, Resti Yudhawati, Yohko K Shimizu, Mitsuhiro Nishimura, Yasuko Mori, Soetjipto, Kazufumi Shimizu, Maria I Lusida |
| EPI_ISL_530361, EPI_ISL_530366, EPI_ISL_530370, EPI_ISL_530377, EPI_ISL_530386, EPI_ISL_530388, EPI_ISL_530391, EPI_ISL_530398, EPI_ISL_530410, EPI_ISL_530429, EPI_ISL_530435, EPI_ISL_530440, EPI_ISL_530468, EPI_ISL_530480, EPI_ISL_530488, EPI_ISL_530489, EPI_ISL_530523, EPI_ISL_530528, EPI_ISL_530533, EPI_ISL_530535, EPI_ISL_530542, EPI_ISL_530553, EPI_ISL_530562, EPI_ISL_530596, EPI_ISL_530642, EPI_ISL_530659, EPI_ISL_530673, EPI_ISL_530682 | | | |
| see above | NHSGGC West of Scotland Specialist Virology Centre / MRC-University of Glasgow Centre for Virus Research | Wellcome Sanger Institute for the COVID-19 Genomics UK (COG-UK) consortium | Ana da Silva Filipe, Natasha Johnson, Kathy Smollett, Daniel Mair, Stephen Carmichael, Lily Tong, Jenna Nichols, Elihu Aranday-Cortes, Kirstyn Brunker, Yasmin Parr, Kyriaki Nomikou; Sarah McDonald, Marc Niebel, Patawee Asamaphan; Richard Orton, Joseph Hughes, Sreenu Vattipally, David L Robertson; Alasdair MacLean, Rory Gunson; Kathy Li, Natasha Jesudason, Rajiv Shah, James Shepherd, Antonia Ho, Alice Broos, Emma Thomson and Alex Alderton, Roberto Amato, Sonia Goncalves, Ewan Harrison, David K. Jackson, Ian Johnston, Dominic Kwiatkowski, Cordelia Langford, John Sillitoe |
| EPI_ISL_530689 | NHSGGC West of Scotland Specialist Virology Centre / MRC-University of Glasgow Centre for Virus Research | Wellcome Sanger Institute for the COVID-19 Genomics UK (COG-UK) consortium | Ana da Silva Filipe, Natasha Johnson, Kathy Smollett, Daniel Mair, Stephen Carmichael, Lily Tong, Jenna Nichols, Elihu Aranday-Cortes, Kirstyn Brunker, Yasmin Parr, Kyriaki Nomikou; Sarah McDonald, Marc Niebel, Patawee Asamaphan; Richard Orton, Joseph Hughes, Sreenu Vattipally, David L Robertson; Alasdair MacLean, Rory Gunson; Kathy Li, Natasha Jesudason, Rajiv Shah, James Shepherd, Antonia Ho, Alice Broos, Emma Thomson and Alex Alderton, Roberto Amato, Sonia Goncalves, Ewan Harrison, David K. Jackson, Ian Johnston, Dominic Kwiatkowski, Cordelia Langford, John Sillitoe on behalf of the Wellcome Sanger Institute COVID-19 Surveillance Team (http://www.sanger.ac.uk/covid-team) |
| EPI_ISL_530708, EPI_ISL_530722, EPI_ISL_530740, EPI_ISL_530743 | NHSGGC West of Scotland Specialist Virology Centre / MRC-University of Glasgow Centre for Virus Research | Wellcome Sanger Institute for the COVID-19 Genomics UK (COG-UK) consortium | Ana da Silva Filipe, Natasha Johnson, Kathy Smollett, Daniel Mair, Stephen Carmichael, Lily Tong, Jenna Nichols, Elihu Aranday-Cortes, Kirstyn Brunker, Yasmin Parr, Kyriaki Nomikou; Sarah McDonald, Marc Niebel, Patawee Asamaphan; Richard Orton, Joseph Hughes, Sreenu Vattipally, David L Robertson; Alasdair MacLean, Rory Gunson; Kathy Li, Natasha Jesudason, Rajiv Shah, James Shepherd, Antonia Ho, Alice Broos, Emma Thomson and Alex Alderton, Roberto Amato, Sonia Goncalves, Ewan Harrison, David K. Jackson, Ian Johnston, Dominic Kwiatkowski, Cordelia Langford, John Sillitoe |
| EPI_ISL_530746 | NHSGGC West of Scotland Specialist Virology Centre / MRC-University of Glasgow Centre for Virus Research | Wellcome Sanger Institute for the COVID-19 Genomics UK (COG-UK) consortium | Ana da Silva Filipe, Natasha Johnson, Kathy Smollett, Daniel Mair, Stephen Carmichael, Lily Tong, Jenna Nichols, Elihu Aranday-Cortes, Kirstyn Brunker, Yasmin Parr, Kyriaki Nomikou; Sarah McDonald, Marc Niebel, Patawee Asamaphan; Richard Orton, Joseph Hughes, Sreenu Vattipally, David L Robertson; Alasdair MacLean, Rory Gunson; Kathy Li, Natasha Jesudason, Rajiv Shah, James Shepherd, Antonia Ho, Alice Broos, Emma Thomson and Alex Alderton, Roberto Amato, Sonia Goncalves, Ewan Harrison, David K. Jackson, Ian Johnston, Dominic Kwiatkowski, Cordelia Langford, John Sillitoe on behalf of the Wellcome Sanger Institute COVID-19 Surveillance Team (http://www.sanger.ac.uk/covid-team) |
| EPI_ISL_530754, EPI_ISL_530764, EPI_ISL_530766, EPI_ISL_530767, EPI_ISL_530768, EPI_ISL_530826, EPI_ISL_530836, EPI_ISL_530845, EPI_ISL_530862, EPI_ISL_530871, EPI_ISL_530873, EPI_ISL_530904, EPI_ISL_530906, EPI_ISL_530923, EPI_ISL_530929, EPI_ISL_530953, EPI_ISL_530959, EPI_ISL_530961, EPI_ISL_530962, EPI_ISL_530973, EPI_ISL_530976, EPI_ISL_530979, EPI_ISL_530983, EPI_ISL_530985, EPI_ISL_530986, EPI_ISL_530990, EPI_ISL_530996, EPI_ISL_530997, EPI_ISL_531002, EPI_ISL_531004, EPI_ISL_531006, EPI_ISL_531017, EPI_ISL_531020, EPI_ISL_531032, EPI_ISL_531033, EPI_ISL_531038, EPI_ISL_531040, EPI_ISL_531045, EPI_ISL_531048, EPI_ISL_531049, EPI_ISL_531052, EPI_ISL_531054, EPI_ISL_531056, EPI_ISL_531057, EPI_ISL_531065, EPI_ISL_531079, EPI_ISL_531083, EPI_ISL_531087, EPI_ISL_531091, EPI_ISL_531096, EPI_ISL_531099, EPI_ISL_531100, EPI_ISL_531103, EPI_ISL_531104, EPI_ISL_531114, EPI_ISL_531116, EPI_ISL_531118, EPI_ISL_531123, EPI_ISL_531126, EPI_ISL_531130, EPI_ISL_531131, EPI_ISL_531142, EPI_ISL_531147, EPI_ISL_531154, EPI_ISL_531155, EPI_ISL_531156, EPI_ISL_531157, EPI_ISL_531159, EPI_ISL_531166, EPI_ISL_531174, EPI_ISL_531175, EPI_ISL_531188, EPI_ISL_531189, EPI_ISL_531198, EPI_ISL_531207, EPI_ISL_531212, EPI_ISL_531214, EPI_ISL_531223, EPI_ISL_531249, EPI_ISL_531266, EPI_ISL_531321, EPI_ISL_531330, EPI_ISL_531365, EPI_ISL_531374, EPI_ISL_531382, EPI_ISL_531412, EPI_ISL_531416, EPI_ISL_531420, EPI_ISL_531422, EPI_ISL_531432, EPI_ISL_531442, EPI_ISL_531464, EPI_ISL_531470, EPI_ISL_531473, EPI_ISL_531476, EPI_ISL_531477, EPI_ISL_531480, EPI_ISL_531483, EPI_ISL_531484, EPI_ISL_531485, EPI_ISL_531492, EPI_ISL_531495, EPI_ISL_531500, EPI_ISL_531504, EPI_ISL_531524, EPI_ISL_531546, EPI_ISL_531548, EPI_ISL_531557, EPI_ISL_531563, EPI_ISL_531564, EPI_ISL_531565, EPI_ISL_531569, EPI_ISL_531570, EPI_ISL_531575, EPI_ISL_531577, EPI_ISL_531583, EPI_ISL_531590, EPI_ISL_531595, EPI_ISL_531598, EPI_ISL_531599, EPI_ISL_531607, EPI_ISL_531609, EPI_ISL_531617, EPI_ISL_531618, EPI_ISL_531625, EPI_ISL_531631, EPI_ISL_531634, EPI_ISL_531638, EPI_ISL_531639, EPI_ISL_531649, EPI_ISL_531660, EPI_ISL_531668, EPI_ISL_531672, EPI_ISL_531677, EPI_ISL_531678, EPI_ISL_531687, EPI_ISL_531689, EPI_ISL_531690, EPI_ISL_531691, EPI_ISL_531702, EPI_ISL_531703, EPI_ISL_531714, EPI_ISL_531727, EPI_ISL_531730, EPI_ISL_531732, EPI_ISL_531743, EPI_ISL_531753, EPI_ISL_531754, EPI_ISL_531755, EPI_ISL_531758, EPI_ISL_531768, EPI_ISL_531779, EPI_ISL_531788, EPI_ISL_531792, EPI_ISL_531801, EPI_ISL_531802, EPI_ISL_531805, EPI_ISL_531814, EPI_ISL_531815, EPI_ISL_531821, EPI_ISL_531824, EPI_ISL_531832, EPI_ISL_531842, EPI_ISL_531851, EPI_ISL_531856, EPI_ISL_531857, EPI_ISL_531858, EPI_ISL_531862, EPI_ISL_531863, EPI_ISL_531865, EPI_ISL_531873, EPI_ISL_531877, EPI_ISL_531880, EPI_ISL_531881, EPI_ISL_531884, EPI_ISL_531887, EPI_ISL_531889, EPI_ISL_531890, EPI_ISL_531892, EPI_ISL_531894, EPI_ISL_531905, EPI_ISL_531912, EPI_ISL_531913, EPI_ISL_531918, EPI_ISL_531919, EPI_ISL_531925, EPI_ISL_531929, EPI_ISL_531932, EPI_ISL_531933, EPI_ISL_531935, EPI_ISL_531936, EPI_ISL_531943, EPI_ISL_531945, EPI_ISL_531951, EPI_ISL_531954, EPI_ISL_531963, EPI_ISL_531973, EPI_ISL_531978, EPI_ISL_531979, EPI_ISL_531982, EPI_ISL_531984, EPI_ISL_531987, EPI_ISL_531990, EPI_ISL_531995, EPI_ISL_532003, EPI_ISL_532004, EPI_ISL_532005, EPI_ISL_532007, EPI_ISL_532018, EPI_ISL_532024, EPI_ISL_532032, EPI_ISL_532038, EPI_ISL_532049, EPI_ISL_532053, EPI_ISL_532055, EPI_ISL_532057, EPI_ISL_532062, EPI_ISL_532070, EPI_ISL_532071, EPI_ISL_532076, EPI_ISL_532077, EPI_ISL_532078, EPI_ISL_532080, EPI_ISL_532081, EPI_ISL_532085, EPI_ISL_532093, EPI_ISL_532094, EPI_ISL_532098, EPI_ISL_532102, EPI_ISL_532105, EPI_ISL_532118, EPI_ISL_532123, EPI_ISL_532125, EPI_ISL_532129, EPI_ISL_532130, EPI_ISL_532131, EPI_ISL_532138, EPI_ISL_532153, EPI_ISL_532168, EPI_ISL_532178, EPI_ISL_532186, EPI_ISL_532192, EPI_ISL_532194, EPI_ISL_532195, EPI_ISL_532196, EPI_ISL_532197, EPI_ISL_532200, EPI_ISL_532202, EPI_ISL_532203, EPI_ISL_532210, EPI_ISL_532214, EPI_ISL_532219, EPI_ISL_532221, EPI_ISL_532235, EPI_ISL_532241, EPI_ISL_532244, EPI_ISL_532249, EPI_ISL_532258, EPI_ISL_532263, EPI_ISL_532271, EPI_ISL_532272, EPI_ISL_532275, EPI_ISL_532276, EPI_ISL_532278, EPI_ISL_532283, EPI_ISL_532288, EPI_ISL_532293, EPI_ISL_532301, EPI_ISL_532311, EPI_ISL_532316, EPI_ISL_532317, EPI_ISL_532321, EPI_ISL_532327, EPI_ISL_532328, EPI_ISL_532334, EPI_ISL_532335, EPI_ISL_532338, EPI_ISL_532340, EPI_ISL_532342, EPI_ISL_532347, EPI_ISL_532349, EPI_ISL_532351, EPI_ISL_532353, EPI_ISL_532355, EPI_ISL_532356, EPI_ISL_532363, EPI_ISL_532364, EPI_ISL_532379, EPI_ISL_532384, EPI_ISL_532388, EPI_ISL_532390, EPI_ISL_532394, EPI_ISL_532399, EPI_ISL_532402, EPI_ISL_532404, EPI_ISL_532414, EPI_ISL_532430, EPI_ISL_532431, EPI_ISL_532432, EPI_ISL_532438, EPI_ISL_532440, EPI_ISL_532444, EPI_ISL_532446, EPI_ISL_532452, EPI_ISL_532454, EPI_ISL_532456, EPI_ISL_532463, EPI_ISL_532464, EPI_ISL_532469, EPI_ISL_532472, EPI_ISL_532473, EPI_ISL_532480, EPI_ISL_532484, EPI_ISL_532490, EPI_ISL_532492, EPI_ISL_532493, EPI_ISL_532506, EPI_ISL_532510, EPI_ISL_532512, EPI_ISL_532514, EPI_ISL_532515, EPI_ISL_532518, EPI_ISL_532523, EPI_ISL_532531, EPI_ISL_532538, EPI_ISL_532540, EPI_ISL_532563, EPI_ISL_532564, EPI_ISL_532572, EPI_ISL_532582, EPI_ISL_532587, EPI_ISL_532594, EPI_ISL_532598, EPI_ISL_532620, EPI_ISL_532621, EPI_ISL_532625, EPI_ISL_532626, EPI_ISL_532633, EPI_ISL_532639, EPI_ISL_532648, EPI_ISL_532670, EPI_ISL_532697, EPI_ISL_532715, EPI_ISL_532733, EPI_ISL_532738, EPI_ISL_532744, EPI_ISL_532746, EPI_ISL_532750, EPI_ISL_532753, EPI_ISL_532755, EPI_ISL_532765, EPI_ISL_532767, EPI_ISL_532784, EPI_ISL_532786, EPI_ISL_532791, EPI_ISL_532811, EPI_ISL_532835, EPI_ISL_532838, EPI_ISL_532844, EPI_ISL_532858, EPI_ISL_532864, EPI_ISL_532865, EPI_ISL_532874, EPI_ISL_532875, EPI_ISL_532876, EPI_ISL_532878, EPI_ISL_532885, EPI_ISL_532886, EPI_ISL_532891, EPI_ISL_532894, EPI_ISL_532896, EPI_ISL_532905, EPI_ISL_532906, EPI_ISL_532915, EPI_ISL_532932, EPI_ISL_532933, EPI_ISL_532948, EPI_ISL_532962, EPI_ISL_532988, EPI_ISL_533000, EPI_ISL_533016, EPI_ISL_533027, EPI_ISL_533072, EPI_ISL_533228, EPI_ISL_533230, EPI_ISL_533231, EPI_ISL_533232, EPI_ISL_533236, EPI_ISL_533237, EPI_ISL_533241, EPI_ISL_533252, EPI_ISL_533256, EPI_ISL_533262, EPI_ISL_533270, EPI_ISL_533271, EPI_ISL_533283, EPI_ISL_533289, EPI_ISL_533298, EPI_ISL_533311, EPI_ISL_533343, EPI_ISL_533346, EPI_ISL_533350, EPI_ISL_533351, EPI_ISL_533355, EPI_ISL_533358, EPI_ISL_533365, EPI_ISL_533376, EPI_ISL_533394, EPI_ISL_533396, EPI_ISL_533406, EPI_ISL_533419, EPI_ISL_533421 | | | |
| see above | NHSGGC West of Scotland Specialist Virology Centre / MRC-University of Glasgow Centre for Virus Research | Wellcome Sanger Institute for the COVID-19 Genomics UK (COG-UK) consortium | Ana da Silva Filipe, Natasha Johnson, Kathy Smollett, Daniel Mair, Stephen Carmichael, Lily Tong, Jenna Nichols, Elihu Aranday-Cortes, Kirstyn Brunker, Yasmin Parr, Kyriaki Nomikou; Sarah McDonald, Marc Niebel, Patawee Asamaphan; Richard Orton, Joseph Hughes, Sreenu Vattipally, David L Robertson; Alasdair MacLean, Rory Gunson; Kathy Li, Natasha Jesudason, Rajiv Shah, James Shepherd, Antonia Ho, Alice Broos, Emma Thomson and Alex Alderton, Roberto Amato, Sonia Goncalves, Ewan Harrison, David K. Jackson, Ian Johnston, Dominic Kwiatkowski, Cordelia Langford, John Sillitoe |
| EPI_ISL_534365, EPI_ISL_534366, EPI_ISL_534367, EPI_ISL_534368, EPI_ISL_534369, EPI_ISL_534370, EPI_ISL_534371, EPI_ISL_534372, EPI_ISL_534373, EPI_ISL_534374, EPI_ISL_534375, EPI_ISL_534376, EPI_ISL_534377, EPI_ISL_534378, EPI_ISL_534379, EPI_ISL_534380, EPI_ISL_534381, EPI_ISL_534382, EPI_ISL_534383, EPI_ISL_534384, EPI_ISL_534385, EPI_ISL_534386, EPI_ISL_534387, EPI_ISL_534388, EPI_ISL_534389, EPI_ISL_534390, EPI_ISL_534391, EPI_ISL_534392, EPI_ISL_534393, EPI_ISL_534394, EPI_ISL_534396, EPI_ISL_534397, EPI_ISL_534398, EPI_ISL_534399, EPI_ISL_534400, EPI_ISL_534401, EPI_ISL_534402, EPI_ISL_534404, EPI_ISL_534405, EPI_ISL_534406, EPI_ISL_534407, EPI_ISL_534408, EPI_ISL_534409, EPI_ISL_534410, EPI_ISL_534411, EPI_ISL_534412, EPI_ISL_534413, EPI_ISL_534414, EPI_ISL_534415, EPI_ISL_534416, EPI_ISL_534417, EPI_ISL_534418, EPI_ISL_534419, EPI_ISL_534420, EPI_ISL_534421, EPI_ISL_534422, EPI_ISL_534423, EPI_ISL_534424, EPI_ISL_534425, EPI_ISL_534426, EPI_ISL_534427, EPI_ISL_534428, EPI_ISL_534429, EPI_ISL_534430, EPI_ISL_534431, EPI_ISL_534432, EPI_ISL_534433, EPI_ISL_534434, EPI_ISL_534435, EPI_ISL_534436, EPI_ISL_534437, EPI_ISL_534438, EPI_ISL_534439, EPI_ISL_534440, EPI_ISL_534441, EPI_ISL_534442, EPI_ISL_534443, EPI_ISL_534444, EPI_ISL_534445, EPI_ISL_534446, EPI_ISL_534447, EPI_ISL_534448, EPI_ISL_534449, EPI_ISL_534450, EPI_ISL_534451, EPI_ISL_534452, EPI_ISL_534453, EPI_ISL_534454, EPI_ISL_534455, EPI_ISL_534456, EPI_ISL_534457, EPI_ISL_534458, EPI_ISL_534459, EPI_ISL_534460, EPI_ISL_534461, EPI_ISL_534462, EPI_ISL_534463, EPI_ISL_534464, EPI_ISL_534465, EPI_ISL_534466, EPI_ISL_534467, EPI_ISL_534468, EPI_ISL_534469, EPI_ISL_534470, EPI_ISL_534471, EPI_ISL_534472, EPI_ISL_534473, EPI_ISL_534474, EPI_ISL_534475, EPI_ISL_534476, EPI_ISL_534477, EPI_ISL_534478, EPI_ISL_534479, EPI_ISL_534480, EPI_ISL_534481, EPI_ISL_534482, EPI_ISL_534483, EPI_ISL_534484, EPI_ISL_534485, EPI_ISL_534486, EPI_ISL_534487, EPI_ISL_534488, EPI_ISL_534489, EPI_ISL_534490, EPI_ISL_534491, EPI_ISL_534492, EPI_ISL_534493, EPI_ISL_534494, EPI_ISL_534495, EPI_ISL_534496, EPI_ISL_534497, EPI_ISL_534498, EPI_ISL_534499, EPI_ISL_534500, EPI_ISL_534501, EPI_ISL_534502, EPI_ISL_534503, EPI_ISL_534504, EPI_ISL_534505, EPI_ISL_534506, EPI_ISL_534507, EPI_ISL_534508, EPI_ISL_534509, EPI_ISL_534510, EPI_ISL_534511, EPI_ISL_534512, EPI_ISL_534513, EPI_ISL_534514, EPI_ISL_534515, EPI_ISL_534516, EPI_ISL_534517, EPI_ISL_534518, EPI_ISL_534519, EPI_ISL_534520, EPI_ISL_534521, EPI_ISL_534522, EPI_ISL_534523, EPI_ISL_534524, EPI_ISL_534525, EPI_ISL_534526, EPI_ISL_534527, EPI_ISL_534528, EPI_ISL_534529, EPI_ISL_534530, EPI_ISL_534531, EPI_ISL_534532, EPI_ISL_534533, EPI_ISL_534534, EPI_ISL_534535, EPI_ISL_534536, EPI_ISL_534537, EPI_ISL_534538, EPI_ISL_534539, EPI_ISL_534540, EPI_ISL_534541, EPI_ISL_534542, EPI_ISL_534543, EPI_ISL_534544, EPI_ISL_534545, EPI_ISL_534546, EPI_ISL_534547, EPI_ISL_534548, EPI_ISL_534549, EPI_ISL_534550, EPI_ISL_534551, EPI_ISL_534552, EPI_ISL_534553, EPI_ISL_534554, EPI_ISL_534555, EPI_ISL_534556, EPI_ISL_534557, EPI_ISL_534558, EPI_ISL_534559, EPI_ISL_534560, EPI_ISL_534561, EPI_ISL_534562, EPI_ISL_534563, EPI_ISL_534564, EPI_ISL_534565, EPI_ISL_534566, EPI_ISL_534567, EPI_ISL_534568, EPI_ISL_534569, EPI_ISL_534570, EPI_ISL_534571, EPI_ISL_534572, EPI_ISL_534573, EPI_ISL_534574, EPI_ISL_534575, EPI_ISL_534576, EPI_ISL_534577, EPI_ISL_534578, EPI_ISL_534579, EPI_ISL_534580, EPI_ISL_534581, EPI_ISL_534582, EPI_ISL_534583, EPI_ISL_534584, EPI_ISL_534585, EPI_ISL_534586, EPI_ISL_534587, EPI_ISL_534588, EPI_ISL_534589, EPI_ISL_534590, EPI_ISL_534591, EPI_ISL_534592, EPI_ISL_534593, EPI_ISL_534594, EPI_ISL_534595, EPI_ISL_534596, EPI_ISL_534597, EPI_ISL_534598, EPI_ISL_534599, EPI_ISL_534600, EPI_ISL_534601, EPI_ISL_534602, EPI_ISL_534603, EPI_ISL_534604, EPI_ISL_534605, EPI_ISL_534606, EPI_ISL_534607, EPI_ISL_534608, EPI_ISL_534609, EPI_ISL_534610, EPI_ISL_534611, EPI_ISL_534612, EPI_ISL_534613, EPI_ISL_534614, EPI_ISL_534615, EPI_ISL_534616, EPI_ISL_534617, EPI_ISL_534618, EPI_ISL_534619, EPI_ISL_534620, EPI_ISL_534621, EPI_ISL_534622, EPI_ISL_534623, EPI_ISL_534624, EPI_ISL_534625, EPI_ISL_534626, EPI_ISL_534627, EPI_ISL_534628, EPI_ISL_534629, EPI_ISL_534630, EPI_ISL_534631, EPI_ISL_534632, EPI_ISL_534633, EPI_ISL_534635, EPI_ISL_534636, EPI_ISL_534637, EPI_ISL_534638, EPI_ISL_534639, EPI_ISL_534640, EPI_ISL_534641, EPI_ISL_534642, EPI_ISL_534643, EPI_ISL_534644, EPI_ISL_534645, EPI_ISL_534646, EPI_ISL_534647, EPI_ISL_534648, EPI_ISL_534649, EPI_ISL_534650, EPI_ISL_534651, EPI_ISL_534652, EPI_ISL_534653, EPI_ISL_534654, EPI_ISL_534655, EPI_ISL_534656, EPI_ISL_534657, EPI_ISL_534658, EPI_ISL_534659, EPI_ISL_534660, EPI_ISL_534661, EPI_ISL_534662, EPI_ISL_534663, EPI_ISL_534664, EPI_ISL_534665, EPI_ISL_534666, EPI_ISL_534667, EPI_ISL_534668, EPI_ISL_534669, EPI_ISL_534670, EPI_ISL_534671, EPI_ISL_534672, EPI_ISL_534673, EPI_ISL_534674, EPI_ISL_534675, EPI_ISL_534676, EPI_ISL_534677, EPI_ISL_534678, EPI_ISL_534679, EPI_ISL_534680, EPI_ISL_534681, EPI_ISL_534682, EPI_ISL_534683, EPI_ISL_534684, EPI_ISL_534685, EPI_ISL_534686, EPI_ISL_534687, EPI_ISL_534688, EPI_ISL_534689, EPI_ISL_534690 | | | |

| | | | |
|---|---|---|---|
| see above | NHSGGC West of Scotland Specialist Virology Centre / MRC-University of Glasgow Centre for Virus Research | Wellcome Sanger Institute for the COVID-19 Genomics UK (COG-UK) consortium | Ana da Silva Filipe, Natasha Johnson, Kathy Smollett, Daniel Mair, Stephen Carmichael, Lily Tong, Jenna Nichols, Elihu Aranday-Cortes, Kirstyn Brunker, Yasmin Parr, Kyriaki Nomikou; Sarah McDonald, Marc Niebel, Patawee Asamaphan; Richard Orton, Joseph Hughes, Sreenu Vattipally, David L Robertson; Alasdair MacLean, Rory Gunson; Kathy Li, Natasha Jesudason, Rajiv Shah, James Shepherd, Antonia Ho, Alice Broos, Emma Thomson and Alex Alderton, Roberto Amato, Sonia Goncalves, Ewan Harrison, David K. Jackson, Ian Johnston, Dominic Kwiatkowski, Cordelia Langford, John Sillitoe on behalf of the Wellcome Sanger Institute COVID-19 Surveillance Team (http://www.sanger.ac.uk/covid-team) |
| EPI_ISL_535024, EPI_ISL_535025, EPI_ISL_535027, EPI_ISL_535028, EPI_ISL_535029, EPI_ISL_535031, EPI_ISL_535032, EPI_ISL_535036, EPI_ISL_535038 | Centre for Enzyme Innovation, University of Portsmouth / Translational Research Laboratory, Portsmouth Hospitals NHS Trust | COVID-19 Genomics UK (COG-UK) Consortium | Angela Beckett,Yann Bourgeois,Garry Scarlett,Sharon Glaysher,Scott Elliott,Kelly Bicknell,Robert Impey,Allyson Lloyd,Sarah Wyllie,Ethan Butcher,Anoop Chauhan,Samuel Robson |
| EPI_ISL_535102, EPI_ISL_535103, EPI_ISL_535104, EPI_ISL_535105, EPI_ISL_535106, EPI_ISL_535107, EPI_ISL_535108, EPI_ISL_535109, EPI_ISL_535110, EPI_ISL_535111, EPI_ISL_535112, EPI_ISL_535113, EPI_ISL_535114, EPI_ISL_535115, EPI_ISL_535116, EPI_ISL_535117, EPI_ISL_535118, EPI_ISL_535119, EPI_ISL_535120, EPI_ISL_535121, EPI_ISL_535122, EPI_ISL_535123, EPI_ISL_535124, EPI_ISL_535125, EPI_ISL_535126, EPI_ISL_535127, EPI_ISL_535128, EPI_ISL_535129, EPI_ISL_535130, EPI_ISL_535131, EPI_ISL_535132, EPI_ISL_535133, EPI_ISL_535134, EPI_ISL_535135, EPI_ISL_535136, EPI_ISL_535137, EPI_ISL_535138, EPI_ISL_535139, EPI_ISL_535140, EPI_ISL_535141, EPI_ISL_535142, EPI_ISL_535143, EPI_ISL_535144, EPI_ISL_535145, EPI_ISL_535146, EPI_ISL_535147, EPI_ISL_535148, EPI_ISL_535149, EPI_ISL_535150, EPI_ISL_535151, EPI_ISL_535152, EPI_ISL_535153, EPI_ISL_535154, EPI_ISL_535155, EPI_ISL_535156, EPI_ISL_535157, EPI_ISL_535158, EPI_ISL_535159, EPI_ISL_535160, EPI_ISL_535161, EPI_ISL_535162, EPI_ISL_535163, EPI_ISL_535164, EPI_ISL_535165, EPI_ISL_535166, EPI_ISL_535167, EPI_ISL_535168, EPI_ISL_535169, EPI_ISL_535170, EPI_ISL_535171, EPI_ISL_535172, EPI_ISL_535173, EPI_ISL_535174, EPI_ISL_535175, EPI_ISL_535176, EPI_ISL_535177, EPI_ISL_535178, EPI_ISL_535179, EPI_ISL_535180, EPI_ISL_535181, EPI_ISL_535182 | | | |
| see above | West of Scotland Specialist Virology Centre, NHSGGC / MRC-University of Glasgow Centre for Virus Research | COVID-19 Genomics UK (COG-UK) Consortium | Ana da Silva Filipe, Natasha Johnson, Kathy Smollett, Daniel Mair, Stephen Carmichael, Lily Tong, Jenna Nichols, Elihu Aranday-Cortes, Yasmin Parr, Alice Broos, Kyriaki Nomikou; Sarah McDonald, Marc Niebel, Patawee Asamaphan; Richard Orton, Joseph Hughes, Sreenu Vattipally, David L Robertson; Alasdair MacLean, Rory Gunson; Kathy Li, Natasha Jesudason, Rajiv Shah, James Shepherd, Antonia Ho, Emma Thomson |
| EPI_ISL_537288, EPI_ISL_537289, EPI_ISL_537290, EPI_ISL_537291, EPI_ISL_537292, EPI_ISL_537293, EPI_ISL_537294, EPI_ISL_537295, EPI_ISL_537296, EPI_ISL_537297, EPI_ISL_537298, EPI_ISL_537299, EPI_ISL_537300, EPI_ISL_537301, EPI_ISL_537302, EPI_ISL_537303, EPI_ISL_537304, EPI_ISL_537305, EPI_ISL_537306, EPI_ISL_537307, EPI_ISL_537308, EPI_ISL_537309, EPI_ISL_537310, EPI_ISL_537311, EPI_ISL_537312, EPI_ISL_537313, EPI_ISL_537314, EPI_ISL_537315, EPI_ISL_537316, EPI_ISL_537317, EPI_ISL_537318, EPI_ISL_537319, EPI_ISL_537320, EPI_ISL_537321, EPI_ISL_537322, EPI_ISL_537323, EPI_ISL_537324, EPI_ISL_537325, EPI_ISL_537326, EPI_ISL_537327, EPI_ISL_537328, EPI_ISL_537329, EPI_ISL_537330, EPI_ISL_537331, EPI_ISL_537332, EPI_ISL_537333, EPI_ISL_537334, EPI_ISL_537335, EPI_ISL_537336, EPI_ISL_537337, EPI_ISL_537338, EPI_ISL_537339, EPI_ISL_537340, EPI_ISL_537341, EPI_ISL_537342, EPI_ISL_537343, EPI_ISL_537344, EPI_ISL_537345, EPI_ISL_537346, EPI_ISL_537347, EPI_ISL_537348, EPI_ISL_537349, EPI_ISL_537350, EPI_ISL_537351, EPI_ISL_537352, EPI_ISL_537353, EPI_ISL_537354, EPI_ISL_537355, EPI_ISL_537356, EPI_ISL_537357, EPI_ISL_537358, EPI_ISL_537359, EPI_ISL_537360, EPI_ISL_537361, EPI_ISL_537362, EPI_ISL_537363, EPI_ISL_537364, EPI_ISL_537365, EPI_ISL_537366, EPI_ISL_537367, EPI_ISL_537368, EPI_ISL_537369, EPI_ISL_537370, EPI_ISL_537371, EPI_ISL_537372, EPI_ISL_537373, EPI_ISL_537374, EPI_ISL_537375, EPI_ISL_537376, EPI_ISL_537377, EPI_ISL_537378, EPI_ISL_537379, EPI_ISL_537609, EPI_ISL_537610, EPI_ISL_537611, EPI_ISL_537612, EPI_ISL_537613, EPI_ISL_537614, EPI_ISL_537615, EPI_ISL_537616, EPI_ISL_537617, EPI_ISL_537618, EPI_ISL_537619, EPI_ISL_537620, EPI_ISL_537621, EPI_ISL_537622, EPI_ISL_537623, EPI_ISL_537624, EPI_ISL_537625, EPI_ISL_537626, EPI_ISL_537627, EPI_ISL_537628, EPI_ISL_537629, EPI_ISL_537630, EPI_ISL_537631, EPI_ISL_537632, EPI_ISL_537633, EPI_ISL_537634, EPI_ISL_537635, EPI_ISL_537636, EPI_ISL_537637, EPI_ISL_537638, EPI_ISL_537639, EPI_ISL_537640, EPI_ISL_537641, EPI_ISL_537642, EPI_ISL_537643, EPI_ISL_537644, EPI_ISL_537645, EPI_ISL_537646, EPI_ISL_537647, EPI_ISL_537648, EPI_ISL_537649, EPI_ISL_537650, EPI_ISL_537651, EPI_ISL_537652, EPI_ISL_537653, EPI_ISL_537654, EPI_ISL_537655, EPI_ISL_537656, EPI_ISL_537657, EPI_ISL_537658, EPI_ISL_537659, EPI_ISL_537660, EPI_ISL_537661, EPI_ISL_537662, EPI_ISL_537663, EPI_ISL_537664, EPI_ISL_537665, EPI_ISL_537666, EPI_ISL_537667, EPI_ISL_537668, EPI_ISL_537669, EPI_ISL_537670, EPI_ISL_537671, EPI_ISL_537672, EPI_ISL_537673, EPI_ISL_537674, EPI_ISL_537675, EPI_ISL_537676, EPI_ISL_537677, EPI_ISL_537678, EPI_ISL_537679, EPI_ISL_537680, EPI_ISL_537681, EPI_ISL_537682, EPI_ISL_537683, EPI_ISL_537684, EPI_ISL_537685, EPI_ISL_537686, EPI_ISL_537687, EPI_ISL_537688, EPI_ISL_537689, EPI_ISL_537690 | | | |
| see above | Universidad de León | SeqCOVID-SPAIN consortium/IBV(CSIC) | Ana Carvajal, Vicente Martín, Héctor Argüello, Juan M. Fregeneda, Tania Fernández-Villa, Antonio J. Molina and SeqCOVID-SPAIN consortium |
| EPI_ISL_539785, EPI_ISL_539786, EPI_ISL_539787, EPI_ISL_539788 | Institute of Microbiology, Universidad San Francisco de Quito | Institute of Microbiology, Universidad San Francisco de Quito | Andrea Macias, Belén Prado-Vivar, Sully Márquez, Juan José Guadalupe, Monica Becerra-Wong, Bernardo Gutiérrez, Verónica Barragán, Patricio Rojas-Silva, Gabriel Trueba, Michelle Grunauer, Paúl Cárdenas |
| EPI_ISL_539803 | Queen Mary Hospital | Hong Kong Department of Health | Alan K.L. Tsang, Peter C.W. Yip, Edman T.K. Lam, Rickjason C.W. Chan, Dominic N.C. Tsang |
| EPI_ISL_539804 | Yan Chai Hospital | Hong Kong Department of Health | Alan K.L. Tsang, Peter C.W. Yip, Edman T.K. Lam, Rickjason C.W. Chan, Dominic N.C. Tsang |
| EPI_ISL_539805 | Kwong Wah Hospital | Hong Kong Department of Health | Alan K.L. Tsang, Peter C.W. Yip, Edman T.K. Lam, Rickjason C.W. Chan, Dominic N.C. Tsang |
| EPI_ISL_539806, EPI_ISL_539807, EPI_ISL_539808 | Princess Margaret Hospital | Hong Kong Department of Health | Alan K.L. Tsang, Peter C.W. Yip, Edman T.K. Lam, Rickjason C.W. Chan, Dominic N.C. Tsang |
| EPI_ISL_539809, EPI_ISL_539810, EPI_ISL_539811, EPI_ISL_539812 | Asiaworld Expo Command Post | Hong Kong Department of Health | Alan K.L. Tsang, Peter C.W. Yip, Edman T.K. Lam, Rickjason C.W. Chan, Dominic N.C. Tsang |
| EPI_ISL_539813 | Prince of Wales Hospital | Hong Kong Department of Health | Alan K.L. Tsang, Peter C.W. Yip, Edman T.K. Lam, Rickjason C.W. Chan, Dominic N.C. Tsang |
| EPI_ISL_539814 | Tuen Mun Hospital | Hong Kong Department of Health | Alan K.L. Tsang, Peter C.W. Yip, Edman T.K. Lam, Rickjason C.W. Chan, Dominic N.C. Tsang |
| EPI_ISL_539815 | Queen Elizabeth Hospital | Hong Kong Department of Health | Alan K.L. Tsang, Peter C.W. Yip, Edman T.K. Lam, Rickjason C.W. Chan, Dominic N.C. Tsang |
| EPI_ISL_539816, EPI_ISL_539817 | Queen Mary Hospital | Hong Kong Department of Health | Alan K.L. Tsang, Peter C.W. Yip, Edman T.K. Lam, Rickjason C.W. Chan, Dominic N.C. Tsang |
| EPI_ISL_539818, EPI_ISL_539819 | Tuen Mun Hospital | Hong Kong Department of Health | Alan K.L. Tsang, Peter C.W. Yip, Edman T.K. Lam, Rickjason C.W. Chan, Dominic N.C. Tsang |
| EPI_ISL_539820 | Queen Mary Hospital | Hong Kong Department of Health | Alan K.L. Tsang, Peter C.W. Yip, Edman T.K. Lam, Rickjason C.W. Chan, Dominic N.C. Tsang |
| EPI_ISL_539821 | Asiaworld Expo Command Post | Hong Kong Department of Health | Alan K.L. Tsang, Peter C.W. Yip, Edman T.K. Lam, Rickjason C.W. Chan, Dominic N.C. Tsang |
| EPI_ISL_539822, EPI_ISL_539823 | Communicable Disease Branch | Hong Kong Department of Health | Alan K.L. Tsang, Peter C.W. Yip, Edman T.K. Lam, Rickjason C.W. Chan, Dominic N.C. Tsang |
| EPI_ISL_539850, EPI_ISL_539851 | Pok Oi Hospital | Hong Kong Department of Health | Alan K.L. Tsang, Peter C.W. Yip, Edman T.K. Lam, Rickjason C.W. Chan, Dominic N.C. Tsang |
| EPI_ISL_540775, EPI_ISL_540776, EPI_ISL_540777, EPI_ISL_540778, EPI_ISL_540779, EPI_ISL_540780, EPI_ISL_540781, EPI_ISL_540782, EPI_ISL_540783, EPI_ISL_540784, EPI_ISL_540785, EPI_ISL_540786, EPI_ISL_540787, EPI_ISL_540788, EPI_ISL_540789, EPI_ISL_540790, EPI_ISL_540791, EPI_ISL_540792, EPI_ISL_540793, EPI_ISL_540794, EPI_ISL_540795, EPI_ISL_540796, EPI_ISL_540797, EPI_ISL_540798, EPI_ISL_540799, EPI_ISL_540800, EPI_ISL_540801, EPI_ISL_540802, EPI_ISL_540803, EPI_ISL_540804, EPI_ISL_540805, EPI_ISL_540806, EPI_ISL_540807, EPI_ISL_540808, EPI_ISL_540809, EPI_ISL_540810, EPI_ISL_540811, EPI_ISL_540812, EPI_ISL_540813, EPI_ISL_540814, EPI_ISL_540815, EPI_ISL_540816, EPI_ISL_540817, EPI_ISL_540818, EPI_ISL_540819, EPI_ISL_540820, EPI_ISL_540821, EPI_ISL_540822 | | | |
| see above | West of Scotland Specialist Virology Centre, NHSGGC / MRC-University of Glasgow Centre for Virus Research | COVID-19 Genomics UK (COG-UK) Consortium | Ana da Silva Filipe, Natasha Johnson, Kathy Smollett, Daniel Mair, Stephen Carmichael, Lily Tong, Jenna Nichols, Elihu Aranday-Cortes, Kyriaki Nomikou; Sarah McDonald, Marc Niebel, Patawee Asamaphan; Richard Orton, Joseph Hughes, Sreenu Vattipally, David L Robertson; Alasdair MacLean, Rory Gunson; Kathy Li, Igor Starinskij, Natasha Jesudason, Rajiv Shah, James Shepherd, Antonia Ho, Emma Thomson |
| EPI_ISL_540823, EPI_ISL_540824, EPI_ISL_540825, EPI_ISL_540826, EPI_ISL_540827, EPI_ISL_540828, EPI_ISL_540829, EPI_ISL_540830, EPI_ISL_540831, EPI_ISL_540832, EPI_ISL_540833, EPI_ISL_540834, EPI_ISL_540835, EPI_ISL_540836, EPI_ISL_540837, EPI_ISL_540838, EPI_ISL_540839, EPI_ISL_540840, EPI_ISL_540841, EPI_ISL_540842, EPI_ISL_540843, EPI_ISL_540844, EPI_ISL_540845, EPI_ISL_540846, EPI_ISL_540847, EPI_ISL_540848, EPI_ISL_540849, EPI_ISL_540850, EPI_ISL_540851, EPI_ISL_540852, EPI_ISL_540853, EPI_ISL_540854, EPI_ISL_540855, EPI_ISL_540856, EPI_ISL_540857, EPI_ISL_540858, EPI_ISL_540859, EPI_ISL_540860, EPI_ISL_540861, EPI_ISL_540862, EPI_ISL_540863, EPI_ISL_540864, EPI_ISL_540865, EPI_ISL_540866, EPI_ISL_540867, EPI_ISL_540868, EPI_ISL_540869, EPI_ISL_540870 | | | |
| see above | Lighthouse Lab in Glasgow / MRC-University of Glasgow Centre for Virus Research | COVID-19 Genomics UK (COG-UK) Consortium | Ana da Silva Filipe, Natasha Johnson, Kathy Smollett, Daniel Mair, Stephen Carmichael, Lily Tong, Jenna Nichols, Elihu Aranday-Cortes, Kyriaki Nomikou; Sarah McDonald, Marc Niebel, Patawee Asamaphan; Harper VanSteenhouse, Yumi Kasai, David Gray, Carol Clugston, Anna Dominiczak; Alasdair MacLean, Rory Gunson; Richard Orton, Joseph Hughes, Sreenu Vattipally, David L Robertson; Sharif Shaaban, Matthew Holden; Kathy Li, Natasha Jesudason, Rajiv Shah, James Shepherd, Antonia Ho, Emma Thomson |
| EPI_ISL_541753, EPI_ISL_541756, EPI_ISL_541767, EPI_ISL_541773, EPI_ISL_541774 | Microbiology Department, Barking Havering and Redbridge University Hospitals NHS trust | Wellcome Sanger Institute for the COVID-19 Genomics UK (COG-UK) consortium | Amy Ash, Fatima Ali, Cherian Koshy and Alex Alderton, Roberto Amato, Sonia Goncalves, Ewan Harrison, David K. Jackson, Ian Johnston, Dominic Kwiatkowski, Cordelia Langford, John Sillitoe on behalf of the Wellcome Sanger Institute COVID-19 Surveillance Team |
| EPI_ISL_547445, EPI_ISL_547446, EPI_ISL_547447, EPI_ISL_547448, EPI_ISL_547449, EPI_ISL_547450, EPI_ISL_547451, EPI_ISL_547452, EPI_ISL_547453, EPI_ISL_547455, EPI_ISL_547456, EPI_ISL_547457, EPI_ISL_547458, EPI_ISL_547460, EPI_ISL_547461, EPI_ISL_547462, EPI_ISL_547463, EPI_ISL_547464, EPI_ISL_547465, EPI_ISL_547469, EPI_ISL_547470, EPI_ISL_547473, EPI_ISL_547474, EPI_ISL_547475, EPI_ISL_547476, EPI_ISL_547477, EPI_ISL_547478, EPI_ISL_547479, EPI_ISL_547480, EPI_ISL_547481, EPI_ISL_547482, EPI_ISL_547483, EPI_ISL_547484, EPI_ISL_547485, EPI_ISL_547486, EPI_ISL_547491, EPI_ISL_547492, EPI_ISL_547493, EPI_ISL_547494, EPI_ISL_547495, EPI_ISL_547496, EPI_ISL_547497, EPI_ISL_547498, EPI_ISL_547499, EPI_ISL_547501, EPI_ISL_547502, EPI_ISL_547503, EPI_ISL_547504, EPI_ISL_547505, EPI_ISL_547506, EPI_ISL_547507, EPI_ISL_547508, EPI_ISL_547509, EPI_ISL_547511, EPI_ISL_547513, EPI_ISL_547514, EPI_ISL_547515, EPI_ISL_547516, EPI_ISL_547517, EPI_ISL_547518, EPI_ISL_547519, EPI_ISL_547520, EPI_ISL_547521, EPI_ISL_547522, EPI_ISL_547524, EPI_ISL_547525, EPI_ISL_547526, EPI_ISL_547527, EPI_ISL_547528, EPI_ISL_547529, EPI_ISL_547531, EPI_ISL_547532, EPI_ISL_547533, EPI_ISL_547534, EPI_ISL_547536, EPI_ISL_547537, EPI_ISL_547538, EPI_ISL_547539, EPI_ISL_547541, EPI_ISL_547557, EPI_ISL_547558, EPI_ISL_547559, EPI_ISL_547560 | | | |
| see above | Dutch COVID-19 response team | National Institute for Public Health and the Environment (RIVM) | Adam Meijer, Harry Vennema, Jeroen Cremer, Sharon van den Brink, Bas van der Veer, AnneMarie van den Brandt, Florian Zwagemaker, Dennis Schmitz, Chantal Reusken, on behalf of the national COVID-19 response team |
| EPI_ISL_549374, EPI_ISL_549386, EPI_ISL_549388, EPI_ISL_549389, EPI_ISL_549390, EPI_ISL_549391, | Department of Pathology, University of Cambridge | COVID-19 Genomics UK (COG-UK) Consortium | Aminu S. Jahun, Yasmin Chaudhry, Grant Hall, Iliana Georgana, Myra Hosmillo, Martin D. Curran, Malte Pinckert, Surendra Parmar, Ian Goodfellow |

| | | | |
|---|---|---|---|
| EPI_ISL_549392, EPI_ISL_549393, EPI_ISL_549394 | | | |
| EPI_ISL_549472 | Centre for Enzyme Innovation, University of Portsmouth / Translational Research Laboratory, Portsmouth Hospitals NHS Trust | COVID-19 Genomics UK (COG-UK) Consortium | Angela Beckett,Yann Bourgeois,Garry Scarlett,Sharon Glaysher,Scott Elliott,Kelly Bicknell,Robert Impey,Allyson Lloyd,Sarah Wyllie,Ethan Butcher,Anoop Chauhan,Samuel Robson |
| EPI_ISL_559947, EPI_ISL_559948, EPI_ISL_559949, EPI_ISL_559950, EPI_ISL_559951, EPI_ISL_559952, EPI_ISL_559953, EPI_ISL_559954, EPI_ISL_559955, EPI_ISL_559956, EPI_ISL_559957, EPI_ISL_559958, EPI_ISL_559959, EPI_ISL_559960, EPI_ISL_559961, EPI_ISL_559962, EPI_ISL_559963, EPI_ISL_559964, EPI_ISL_559965, EPI_ISL_559966, EPI_ISL_559967, EPI_ISL_559968, EPI_ISL_559969, EPI_ISL_559970 | | | |
| see above | Lighthouse Lab in Glasgow / MRC-University of Glasgow Centre for Virus Research | COVID-19 Genomics UK (COG-UK) Consortium | Ana da Silva Filipe, Natasha Johnson, Kathy Smollett, Daniel Mair, Stephen Carmichael, Lily Tong, Jenna Nichols, Elihu Aranday-Cortes, Kyriaki Nomikou; Sarah McDonald, Marc Niebel, Patawee Asamaphan; Harper VanSteenhouse, Yumi Kasai, David Gray, Carol Clugston, Anna Dominiczak; Alasdair MacLean, Rory Gunson; Richard Orton, Joseph Hughes, Sreenu Vattipally, David L Robertson; Sharif Shaaban, Matthew Holden; Kathy Li, Natasha Jesudason, Rajiv Shah, James Shepherd, Antonia Ho, Emma Thomson |
| EPI_ISL_560308, EPI_ISL_560309, EPI_ISL_560310, EPI_ISL_560311, EPI_ISL_560312, EPI_ISL_560313, EPI_ISL_560314, EPI_ISL_560315, EPI_ISL_560316, EPI_ISL_560317 | UMMC-Health | WHO National Influenza Centre Russian Federation | Andrey Komissarov, Artem Fadeev, Anna Ivanova, Tatiana Platonova, Daria Danilenko |
| EPI_ISL_560992, EPI_ISL_560993, EPI_ISL_560994, EPI_ISL_560995, EPI_ISL_560996, EPI_ISL_560997, EPI_ISL_560998, EPI_ISL_560999, EPI_ISL_561000, EPI_ISL_561001, EPI_ISL_561002, EPI_ISL_561003, EPI_ISL_561004, EPI_ISL_561005, EPI_ISL_561006, EPI_ISL_561007, EPI_ISL_561008, EPI_ISL_561009, EPI_ISL_561010, EPI_ISL_561011, EPI_ISL_561012, EPI_ISL_561013, EPI_ISL_561014, EPI_ISL_561015, EPI_ISL_561016, EPI_ISL_561017, EPI_ISL_561018, EPI_ISL_561019, EPI_ISL_561020, EPI_ISL_561021, EPI_ISL_561022, EPI_ISL_561023, EPI_ISL_561024, EPI_ISL_561025, EPI_ISL_561026, EPI_ISL_561027, EPI_ISL_561028, EPI_ISL_561029, EPI_ISL_561030, EPI_ISL_561031, EPI_ISL_561032, EPI_ISL_561033, EPI_ISL_561034, EPI_ISL_561035, EPI_ISL_561036, EPI_ISL_561037, EPI_ISL_561038, EPI_ISL_561039, EPI_ISL_561040, EPI_ISL_561041, EPI_ISL_561042, EPI_ISL_561043, EPI_ISL_561044, EPI_ISL_561045, EPI_ISL_561046, EPI_ISL_561047, EPI_ISL_561048, EPI_ISL_561049, EPI_ISL_561050, EPI_ISL_561051, EPI_ISL_561052, EPI_ISL_561053, EPI_ISL_561054, EPI_ISL_561055, EPI_ISL_561056, EPI_ISL_561057, EPI_ISL_561058, EPI_ISL_561059, EPI_ISL_561060, EPI_ISL_561061, EPI_ISL_561062, EPI_ISL_561063, EPI_ISL_561064, EPI_ISL_561065, EPI_ISL_561066, EPI_ISL_561067, EPI_ISL_561068, EPI_ISL_561069, EPI_ISL_561070, EPI_ISL_561071, EPI_ISL_561072, EPI_ISL_561073, EPI_ISL_561074, EPI_ISL_561075, EPI_ISL_561076, EPI_ISL_561077, EPI_ISL_561078, EPI_ISL_561079, EPI_ISL_561080, EPI_ISL_561081, EPI_ISL_561082, EPI_ISL_561083, EPI_ISL_561084, EPI_ISL_561085, EPI_ISL_561086, EPI_ISL_561087, EPI_ISL_561088, EPI_ISL_561089, EPI_ISL_561090, EPI_ISL_561091, EPI_ISL_561092, EPI_ISL_561093, EPI_ISL_561094, EPI_ISL_561095, EPI_ISL_561096, EPI_ISL_561097, EPI_ISL_561098, EPI_ISL_561099, EPI_ISL_561100, EPI_ISL_561101, EPI_ISL_561102, EPI_ISL_561103, EPI_ISL_561104, EPI_ISL_561105, EPI_ISL_561106, EPI_ISL_561107, EPI_ISL_561108, EPI_ISL_561109, EPI_ISL_561110, EPI_ISL_561111, EPI_ISL_561112, EPI_ISL_561113, EPI_ISL_561114, EPI_ISL_561115, EPI_ISL_561116, EPI_ISL_561117, EPI_ISL_561118, EPI_ISL_561119, EPI_ISL_561120, EPI_ISL_561121, EPI_ISL_561122, EPI_ISL_561123, EPI_ISL_561124, EPI_ISL_561125, EPI_ISL_561126, EPI_ISL_561127, EPI_ISL_561128, EPI_ISL_561129, EPI_ISL_561130, EPI_ISL_561131, EPI_ISL_561132, EPI_ISL_561133, EPI_ISL_561134, EPI_ISL_561135, EPI_ISL_561136, EPI_ISL_561137, EPI_ISL_561138, EPI_ISL_561139, EPI_ISL_561140, EPI_ISL_561141, EPI_ISL_561142, EPI_ISL_561143, EPI_ISL_561144, EPI_ISL_561145, EPI_ISL_561146, EPI_ISL_561147, EPI_ISL_561148, EPI_ISL_561149, EPI_ISL_561150, EPI_ISL_561151, EPI_ISL_561152, EPI_ISL_561153, EPI_ISL_561154, EPI_ISL_561155, EPI_ISL_561156, EPI_ISL_561157, EPI_ISL_561158, EPI_ISL_561159, EPI_ISL_561160, EPI_ISL_561161, EPI_ISL_561162, EPI_ISL_561163, EPI_ISL_561164, EPI_ISL_561165, EPI_ISL_561166, EPI_ISL_561167, EPI_ISL_561168, EPI_ISL_561169, EPI_ISL_561170, EPI_ISL_561171, EPI_ISL_561172, EPI_ISL_561173, EPI_ISL_561174, EPI_ISL_561175, EPI_ISL_561176, EPI_ISL_561177, EPI_ISL_561178, EPI_ISL_561179, EPI_ISL_561180, EPI_ISL_561181, EPI_ISL_561182, EPI_ISL_561183, EPI_ISL_561184, EPI_ISL_561185, EPI_ISL_561186, EPI_ISL_561187, EPI_ISL_561188, EPI_ISL_561189, EPI_ISL_561190, EPI_ISL_561191, EPI_ISL_561192, EPI_ISL_561193, EPI_ISL_561194, EPI_ISL_561195, EPI_ISL_561196, EPI_ISL_561197, EPI_ISL_561198, EPI_ISL_561199, EPI_ISL_561200, EPI_ISL_561201, EPI_ISL_561202, EPI_ISL_561203, EPI_ISL_561204, EPI_ISL_561207, EPI_ISL_561209, EPI_ISL_561210, EPI_ISL_561211, EPI_ISL_561212, EPI_ISL_561213, EPI_ISL_561214, EPI_ISL_561215, EPI_ISL_561216, EPI_ISL_561217, EPI_ISL_561218, EPI_ISL_561219, EPI_ISL_561220, EPI_ISL_561221, EPI_ISL_561222, EPI_ISL_561223, EPI_ISL_561224, EPI_ISL_561225, EPI_ISL_561226, EPI_ISL_561227, EPI_ISL_561228, EPI_ISL_561229, EPI_ISL_561230, EPI_ISL_561234, EPI_ISL_561235, EPI_ISL_561236, EPI_ISL_561237, EPI_ISL_561238, EPI_ISL_561240, EPI_ISL_561241, EPI_ISL_561249, EPI_ISL_561252, EPI_ISL_561253, EPI_ISL_561255, EPI_ISL_561256, EPI_ISL_561258, EPI_ISL_561259, EPI_ISL_561260, EPI_ISL_561261, EPI_ISL_561262, EPI_ISL_561263, EPI_ISL_561264, EPI_ISL_561265, EPI_ISL_561266, EPI_ISL_561267, EPI_ISL_561268, EPI_ISL_561269, EPI_ISL_561272, EPI_ISL_561274, EPI_ISL_561276, EPI_ISL_561281, EPI_ISL_561283, EPI_ISL_561285, EPI_ISL_561288, EPI_ISL_561289, EPI_ISL_561290, EPI_ISL_561291, EPI_ISL_561292, EPI_ISL_561293, EPI_ISL_561294, EPI_ISL_561295, EPI_ISL_561297, EPI_ISL_561302, EPI_ISL_561304, EPI_ISL_561309, EPI_ISL_561310, EPI_ISL_561312, EPI_ISL_561315, EPI_ISL_561316, EPI_ISL_561319, EPI_ISL_561322, EPI_ISL_561325, EPI_ISL_561328 | | | |
| see above | MRCG at LSHTM Genomics lab | MRCG at LSHTM Genomics lab | Abdul Karim sesay, Abdoulie Kanteh, Jarra Manneh, Mariama Kujabi, Bakary Sanyang |
| EPI_ISL_566077, EPI_ISL_566078, EPI_ISL_566079, EPI_ISL_566080, EPI_ISL_566081, EPI_ISL_566082, EPI_ISL_566083 | Pathogenic Microorganisms Variability Laboratory | WHO National Influenza Centre Russian Federation | Andrey Komissarov, Artem Fadeev, Anna Ivanova, Kseniya Komissarova, Daria Danilenko, Dmitry Lioznov, Nadezhda Kuznetsova, Elena Shidlovskaya, Elizaveta Divisenko, Ekaterina Milashenko, Kirill Krasnoslobotsev, Evgeniya Mukasheva, Anna Ignatieva, Svetlana Trushakova, Alexey Shchetinin, Maria Nikiforova, Andrey Pochtovyy, Valeria Bacalin, Evgeny Usachev, Olga Burgasova, Ludmila Kolobukhina, Svetlana Smetanina, Elena Burtseva, Artem Tkachuk, Vladimir Gushchin, Alexander Gintsburg |
| EPI_ISL_572398 | Laboratory of Biology and Identification of Arboviruses | Pathogenic Microorganisms Variability Laboratory | Alexey Shchetinin, Maria Nikiforova, Andrei Siniavin, Victor Larichev, Alina Kozlova, Muhammad Saifullin, Alexey Prilipov, Vladimir Gushchin, Alexander Gintsburg |
| EPI_ISL_572415, EPI_ISL_572470 | West of Scotland Specialist Virology Centre, NHSGGC / MRC-University of Glasgow Centre for Virus Research | COVID-19 Genomics UK (COG-UK) Consortium | Ana da Silva Filipe, Natasha Johnson, Kathy Smollett, Daniel Mair, Stephen Carmichael, Lily Tong, Jenna Nichols, Elihu Aranday-Cortes, Kyriaki Nomikou; Sarah McDonald, Marc Niebel, Patawee Asamaphan; Richard Orton, Joseph Hughes, Sreenu Vattipally, David L Robertson; Alasdair MacLean, Rory Gunson; Kathy Li, Igor Starinskij, Natasha Jesudason, Rajiv Shah, James Shepherd, Antonia Ho, Emma Thomson |
| EPI_ISL_572471, EPI_ISL_572472, EPI_ISL_572484 | Department of Pathology, University of Cambridge | COVID-19 Genomics UK (COG-UK) Consortium | Aminu S. Jahun, Yasmin Chaudhry, Grant Hall, Iliana Georgana, Myra Hosmillo, Martin D. Curran, Malte Pinckert, Surendra Parmar, Ian Goodfellow |
| EPI_ISL_572518 | West of Scotland Specialist Virology Centre, NHSGGC / MRC-University of Glasgow Centre for Virus Research | COVID-19 Genomics UK (COG-UK) Consortium | Ana da Silva Filipe, Natasha Johnson, Kathy Smollett, Daniel Mair, Stephen Carmichael, Lily Tong, Jenna Nichols, Elihu Aranday-Cortes, Kyriaki Nomikou; Sarah McDonald, Marc Niebel, Patawee Asamaphan; Richard Orton, Joseph Hughes, Sreenu Vattipally, David L Robertson; Alasdair MacLean, Rory Gunson; Kathy Li, Igor Starinskij, Natasha Jesudason, Rajiv Shah, James Shepherd, Antonia Ho, Emma Thomson |
| EPI_ISL_572519 | Department of Pathology, University of Cambridge | COVID-19 Genomics UK (COG-UK) Consortium | Aminu S. Jahun, Yasmin Chaudhry, Grant Hall, Iliana Georgana, Myra Hosmillo, Martin D. Curran, Malte Pinckert, Surendra Parmar, Ian Goodfellow |
| EPI_ISL_572523 | West of Scotland Specialist Virology Centre, NHSGGC / MRC-University of Glasgow Centre for Virus Research | COVID-19 Genomics UK (COG-UK) Consortium | Ana da Silva Filipe, Natasha Johnson, Kathy Smollett, Daniel Mair, Stephen Carmichael, Lily Tong, Jenna Nichols, Elihu Aranday-Cortes, Kyriaki Nomikou; Sarah McDonald, Marc Niebel, Patawee Asamaphan; Richard Orton, Joseph Hughes, Sreenu Vattipally, David L Robertson; Alasdair MacLean, Rory Gunson; Kathy Li, Igor Starinskij, Natasha Jesudason, Rajiv Shah, James Shepherd, Antonia Ho, Emma Thomson |
| EPI_ISL_572524 | Department of Pathology, University of Cambridge | COVID-19 Genomics UK (COG-UK) Consortium | Aminu S. Jahun, Yasmin Chaudhry, Grant Hall, Iliana Georgana, Myra Hosmillo, Martin D. Curran, Malte Pinckert, Surendra Parmar, Ian Goodfellow |
| EPI_ISL_572528, EPI_ISL_572534 | West of Scotland Specialist Virology Centre, NHSGGC / MRC-University of Glasgow Centre for Virus Research | COVID-19 Genomics UK (COG-UK) Consortium | Ana da Silva Filipe, Natasha Johnson, Kathy Smollett, Daniel Mair, Stephen Carmichael, Lily Tong, Jenna Nichols, Elihu Aranday-Cortes, Kyriaki Nomikou; Sarah McDonald, Marc Niebel, Patawee Asamaphan; Richard Orton, Joseph Hughes, Sreenu Vattipally, David L Robertson; Alasdair MacLean, Rory Gunson; Kathy Li, Igor Starinskij, Natasha Jesudason, Rajiv Shah, James Shepherd, Antonia Ho, Emma Thomson |
| EPI_ISL_572541 | Department of Pathology, University of Cambridge | COVID-19 Genomics UK (COG-UK) Consortium | Aminu S. Jahun, Yasmin Chaudhry, Grant Hall, Iliana Georgana, Myra Hosmillo, Martin D. Curran, Malte Pinckert, Surendra Parmar, Ian Goodfellow |
| EPI_ISL_572585 | West of Scotland Specialist Virology Centre, NHSGGC / MRC-University of Glasgow Centre for Virus Research | COVID-19 Genomics UK (COG-UK) Consortium | Ana da Silva Filipe, Natasha Johnson, Kathy Smollett, Daniel Mair, Stephen Carmichael, Lily Tong, Jenna Nichols, Elihu Aranday-Cortes, Kyriaki Nomikou; Sarah McDonald, Marc Niebel, Patawee Asamaphan; Richard Orton, Joseph Hughes, Sreenu Vattipally, David L Robertson; Alasdair MacLean, Rory Gunson; Kathy Li, Igor Starinskij, Natasha Jesudason, Rajiv Shah, James Shepherd, Antonia Ho, Emma Thomson |
| EPI_ISL_572592, EPI_ISL_572616 | Department of Pathology, University of Cambridge | COVID-19 Genomics UK (COG-UK) Consortium | Aminu S. Jahun, Yasmin Chaudhry, Grant Hall, Iliana Georgana, Myra Hosmillo, Martin D. Curran, Malte Pinckert, Surendra Parmar, Ian Goodfellow |
| EPI_ISL_572620, EPI_ISL_572621, EPI_ISL_572622, EPI_ISL_572623, EPI_ISL_572624, EPI_ISL_572625, EPI_ISL_572626, EPI_ISL_572627, EPI_ISL_572628, EPI_ISL_572629, EPI_ISL_572630, EPI_ISL_572631, EPI_ISL_572632, EPI_ISL_572633, EPI_ISL_572634, EPI_ISL_572635, EPI_ISL_572636, EPI_ISL_572637, EPI_ISL_572638, EPI_ISL_572639, EPI_ISL_572640, EPI_ISL_572641 | | | |
| see above | West of Scotland Specialist Virology Centre, NHSGGC / MRC-University of Glasgow Centre for Virus Research | COVID-19 Genomics UK (COG-UK) Consortium | Ana da Silva Filipe, Natasha Johnson, Kathy Smollett, Daniel Mair, Stephen Carmichael, Lily Tong, Jenna Nichols, Elihu Aranday-Cortes, Kyriaki Nomikou; Sarah McDonald, Marc Niebel, Patawee Asamaphan; Richard Orton, Joseph Hughes, Sreenu Vattipally, David L Robertson; Alasdair MacLean, Rory Gunson; Kathy Li, Igor Starinskij, Natasha Jesudason, Rajiv Shah, James Shepherd, Antonia Ho, Emma Thomson |
| EPI_ISL_572642 | Department of Pathology, University of Cambridge | COVID-19 Genomics UK (COG-UK) Consortium | Aminu S. Jahun, Yasmin Chaudhry, Grant Hall, Iliana Georgana, Myra Hosmillo, Martin D. Curran, Malte Pinckert, Surendra Parmar, Ian Goodfellow |
| EPI_ISL_572686, EPI_ISL_572741, EPI_ISL_572746, EPI_ISL_572763, EPI_ISL_572765 | West of Scotland Specialist Virology Centre, NHSGGC / MRC-University of Glasgow Centre for Virus Research | COVID-19 Genomics UK (COG-UK) Consortium | Ana da Silva Filipe, Natasha Johnson, Kathy Smollett, Daniel Mair, Stephen Carmichael, Lily Tong, Jenna Nichols, Elihu Aranday-Cortes, Kyriaki Nomikou; Sarah McDonald, Marc Niebel, Patawee Asamaphan; Richard Orton, Joseph Hughes, Sreenu Vattipally, David L Robertson; Alasdair MacLean, Rory Gunson; Kathy Li, Igor Starinskij, Natasha Jesudason, Rajiv Shah, James Shepherd, Antonia Ho, Emma Thomson |
| EPI_ISL_572779 | Department of Pathology, University of Cambridge | COVID-19 Genomics UK (COG-UK) Consortium | Aminu S. Jahun, Yasmin Chaudhry, Grant Hall, Iliana Georgana, Myra Hosmillo, Martin D. Curran, Malte Pinckert, Surendra Parmar, Ian Goodfellow |
| EPI_ISL_572788 | West of Scotland Specialist Virology Centre, NHSGGC / MRC-University of Glasgow Centre for Virus Research | COVID-19 Genomics UK (COG-UK) Consortium | Ana da Silva Filipe, Natasha Johnson, Kathy Smollett, Daniel Mair, Stephen Carmichael, Lily Tong, Jenna Nichols, Elihu Aranday-Cortes, Kyriaki Nomikou; Sarah McDonald, Marc Niebel, Patawee Asamaphan; Richard Orton, Joseph Hughes, Sreenu Vattipally, David L Robertson; Alasdair MacLean, Rory Gunson; Kathy Li, Igor Starinskij, Natasha Jesudason, Rajiv Shah, James Shepherd, Antonia Ho, Emma Thomson |
| EPI_ISL_572792 | Department of Pathology, University of Cambridge | COVID-19 Genomics UK (COG-UK) Consortium | Aminu S. Jahun, Yasmin Chaudhry, Grant Hall, Iliana Georgana, Myra Hosmillo, Martin D. Curran, Malte Pinckert, Surendra Parmar, Ian Goodfellow |

| | | | |
|---|---|---|---|
| EPI_ISL_572794, EPI_ISL_572832 | West of Scotland Specialist Virology Centre, NHSGGC / MRC-University of Glasgow Centre for Virus Research | COVID-19 Genomics UK (COG-UK) Consortium | Ana da Silva Filipe, Natasha Johnson, Kathy Smollett, Daniel Mair, Stephen Carmichael, Lily Tong, Jenna Nichols, Elihu Aranday-Cortes, Kyriaki Nomikou; Sarah McDonald, Marc Niebel, Patawee Asamaphan; Richard Orton, Joseph Hughes, Sreenu Vattipally, David L Robertson; Alasdair MacLean, Rory Gunson; Kathy Li, Igor Starinskij, Natasha Jesudason, Rajiv Shah, James Shepherd, Antonia Ho, Emma Thomson |
| EPI_ISL_572851, EPI_ISL_572852, EPI_ISL_572862, EPI_ISL_572863, EPI_ISL_572864, EPI_ISL_572865, EPI_ISL_572866, EPI_ISL_572867, EPI_ISL_572868, EPI_ISL_573138, EPI_ISL_573139, EPI_ISL_573140, EPI_ISL_573141, EPI_ISL_573142, EPI_ISL_573143, EPI_ISL_573144, EPI_ISL_573145, EPI_ISL_573146, EPI_ISL_573147, EPI_ISL_573148, EPI_ISL_573149, EPI_ISL_573150, EPI_ISL_573151, EPI_ISL_573153 | | | |
| see above | Department of Pathology, University of Cambridge | COVID-19 Genomics UK (COG-UK) Consortium | Aminu S. Jahun, Yasmin Chaudhry, Grant Hall, Iliana Georgana, Myra Hosmillo, Martin D. Curran, Malte Pinckert, Surendra Parmar, Ian Goodfellow |
| EPI_ISL_573765, EPI_ISL_573766, EPI_ISL_573767, EPI_ISL_573768 | West of Scotland Specialist Virology Centre, NHSGGC / MRC-University of Glasgow Centre for Virus Research | COVID-19 Genomics UK (COG-UK) Consortium | Ana da Silva Filipe, Natasha Johnson, Kathy Smollett, Daniel Mair, Stephen Carmichael, Lily Tong, Jenna Nichols, Elihu Aranday-Cortes, Kyriaki Nomikou; Sarah McDonald, Marc Niebel, Patawee Asamaphan; Richard Orton, Joseph Hughes, Sreenu Vattipally, David L Robertson; Alasdair MacLean, Rory Gunson; Kathy Li, Igor Starinskij, Natasha Jesudason, Rajiv Shah, James Shepherd, Antonia Ho, Emma Thomson |
| EPI_ISL_573769 | Lighthouse Lab in Glasgow / MRC-University of Glasgow Centre for Virus Research | COVID-19 Genomics UK (COG-UK) Consortium | Ana da Silva Filipe, Natasha Johnson, Kathy Smollett, Daniel Mair, Stephen Carmichael, Lily Tong, Jenna Nichols, Elihu Aranday-Cortes, Kyriaki Nomikou; Sarah McDonald, Marc Niebel, Patawee Asamaphan; Harper VanSteenhouse, Yumi Kasai, David Gray, Carol Clugston, Anna Dominiczak; Alasdair MacLean, Rory Gunson; Richard Orton, Joseph Hughes, Sreenu Vattipally, David L Robertson; Sharif Shaaban, Matthew Holden; Kathy Li, Natasha Jesudason, Rajiv Shah, James Shepherd, Antonia Ho, Emma Thomson |
| EPI_ISL_573770, EPI_ISL_573771, EPI_ISL_573772, EPI_ISL_573773, EPI_ISL_573774, EPI_ISL_573775, EPI_ISL_573776, EPI_ISL_573777, EPI_ISL_573778, EPI_ISL_573779, EPI_ISL_573780, EPI_ISL_573781, EPI_ISL_573782, EPI_ISL_573783, EPI_ISL_573784, EPI_ISL_573785, EPI_ISL_573786, EPI_ISL_573787, EPI_ISL_573788, EPI_ISL_573789, EPI_ISL_573790, EPI_ISL_573791, EPI_ISL_573792, EPI_ISL_573793 | | | |
| see above | West of Scotland Specialist Virology Centre, NHSGGC / MRC-University of Glasgow Centre for Virus Research | COVID-19 Genomics UK (COG-UK) Consortium | Ana da Silva Filipe, Natasha Johnson, Kathy Smollett, Daniel Mair, Stephen Carmichael, Lily Tong, Jenna Nichols, Elihu Aranday-Cortes, Kyriaki Nomikou; Sarah McDonald, Marc Niebel, Patawee Asamaphan; Richard Orton, Joseph Hughes, Sreenu Vattipally, David L Robertson; Alasdair MacLean, Rory Gunson; Kathy Li, Igor Starinskij, Natasha Jesudason, Rajiv Shah, James Shepherd, Antonia Ho, Emma Thomson |
| EPI_ISL_573870 | Centre for Enzyme Innovation, University of Portsmouth / Translational Research Laboratory, Portsmouth Hospitals NHS Trust | COVID-19 Genomics UK (COG-UK) Consortium | Angela Beckett,Yann Bourgeois,Garry Scarlett,Sharon Glaysher,Scott Elliott,Kelly Bicknell,Robert Impey,Allyson Lloyd,Sarah Wyllie,Ethan Butcher,Anoop Chauhan,Samuel Robson |
| EPI_ISL_576146, EPI_ISL_576147, EPI_ISL_576148 | Department of Respiratory & Other Viral Infections of L.V. Gromashevsky Institute of Epidemiology & Infectious Diseases NAMS of Ukraine | Department of Respiratory & Other Viral Infections of L.V. Gromashevsky Institute of Epidemiology & Infectious Diseases NAMS of Ukraine, JSC "Farmak" | Alla Mironenko, Ihor Kravchuk, Liudmyla Bolotova, Larysa Radchenko, Nataliia Teteriuk |
| EPI_ISL_576149 | Department of Respiratory & Other Viral Infections of L.V. Gromashevsky Institute of Epidemiology & Infectious Diseases NAMS of Ukraine | Department of Respiratory & Other Viral Infections of L.V. Gromashevsky Institute of Epidemiology & Infectious Diseases NAMS of Ukraine, JSC "Farmak" | Alla Mironenko, Andriy Goy, Ihor Kravchuk, Liudmyla Bolotova, Larysa Radchenko, Nataliia Teteriuk |
| EPI_ISL_576897, EPI_ISL_576898, EPI_ISL_576899, EPI_ISL_576900, EPI_ISL_576901, EPI_ISL_576902, EPI_ISL_576906, EPI_ISL_576907, EPI_ISL_576908, EPI_ISL_576909, EPI_ISL_576910, EPI_ISL_576911, EPI_ISL_576912, EPI_ISL_576913, EPI_ISL_576914, EPI_ISL_576915, EPI_ISL_576916, EPI_ISL_576917, EPI_ISL_576918, EPI_ISL_576919 | | | |
| see above | Department of Pathology, University of Cambridge | COVID-19 Genomics UK (COG-UK) Consortium | Aminu S. Jahun, Yasmin Chaudhry, Grant Hall, Iliana Georgana, Myra Hosmillo, Martin D. Curran, Malte Pinckert, Surendra Parmar, Ian Goodfellow |
| EPI_ISL_577196, EPI_ISL_577197, EPI_ISL_577204, EPI_ISL_577205, EPI_ISL_577207, EPI_ISL_577208, EPI_ISL_577209, EPI_ISL_577214, EPI_ISL_577385, EPI_ISL_577386, EPI_ISL_577389, EPI_ISL_577390, EPI_ISL_577391, EPI_ISL_577395, EPI_ISL_577396, EPI_ISL_577397, EPI_ISL_577399, EPI_ISL_577400 | | | |
| see above | Centre for Enzyme Innovation, University of Portsmouth / Translational Research Laboratory, Portsmouth Hospitals NHS Trust | COVID-19 Genomics UK (COG-UK) Consortium | Angela Beckett,Yann Bourgeois,Garry Scarlett,Sharon Glaysher,Scott Elliott,Kelly Bicknell,Robert Impey,Allyson Lloyd,Sarah Wyllie,Ethan Butcher,Anoop Chauhan,Samuel Robson |
| EPI_ISL_581452, EPI_ISL_581453, EPI_ISL_581454 | Medizinische Klinik Innere Medizin I, Universitätsklinikum Tübingen | NGS Competence Center Tübingen, Institut für Medizinische Mikrobiologie und Hygiene, Universitätsklinikum Tübingen | Angel Angelov |
| EPI_ISL_581455 | Fondation Congolaise pour la recherche medicale (FCRM) | NGS Competence Center Tübingen, Institut für Medizinische Mikrobiologie und Hygiene, Universitätsklinikum Tübingen | Angel Angelov |
| EPI_ISL_581456, EPI_ISL_581457, EPI_ISL_581458, EPI_ISL_581459, EPI_ISL_581460, EPI_ISL_581461, EPI_ISL_581463, EPI_ISL_581464, EPI_ISL_581465, EPI_ISL_581466, EPI_ISL_581467, EPI_ISL_581468, EPI_ISL_581469, EPI_ISL_581470, EPI_ISL_581471, EPI_ISL_581473, EPI_ISL_581474, EPI_ISL_581475, EPI_ISL_581476, EPI_ISL_581477, EPI_ISL_581478, EPI_ISL_581479, EPI_ISL_581480, EPI_ISL_581481, EPI_ISL_581482, EPI_ISL_581483, EPI_ISL_581484, EPI_ISL_581485 | | | |
| see above | Medizinische Klinik Innere Medizin I, Universitätsklinikum Tübingen | NGS Competence Center Tübingen, Institut für Medizinische Mikrobiologie und Hygiene, Universitätsklinikum Tübingen | Angel Angelov |
| EPI_ISL_581487, EPI_ISL_581489, EPI_ISL_581490, EPI_ISL_581491, EPI_ISL_581492, EPI_ISL_581493 | Fondation Congolaise pour la recherche medicale (FCRM) | NGS Competence Center Tübingen, Institut für Medizinische Mikrobiologie und Hygiene, Universitätsklinikum Tübingen | Angel Angelov |
| EPI_ISL_582125, EPI_ISL_582126 | Sheikh Khalifa Medical City | Molecular Surveillance lab Sheikh Khalifa Medical City | Amirtharaj Francis, Sajeed Abdul, Hala Imambaccus, Sahar Almarzooqi, Hiba Saud, Stefan Weber |
| EPI_ISL_582509 | Department of Respiratory and other Viral Infections of L.V.Gromashevsky Institute of Epidemiology & Infectious Diseases NAMS of Ukraine | Department of Respiratory and other Viral Infections of L.V.Gromashevsky Institute of Epidemiology & Infectious Diseases NAMS of Ukraine, JSC "Farmak" | Alla Mironenko, Andriy Goy, Ihor Kravchuk, Ludmyla Bolotova, Larysa Radchenko, Nataliia Teteriuk |
| EPI_ISL_582510 | Department of Respiratory and other Viral Infections of L.V.Gromashevsky Institute of Epidemiology & Infectious Diseases NAMS of Ukraine | Department of Respiratory and other Viral Infections of L.V.Gromashevsky Institute of Epidemiology & Infectious Diseases NAMS of Ukrain, JSC "Farmak" | Alla Mironenko, Andriy Goy, Ihor Kravchuk, Ludmyla Bolotova, Larysa Radchenko, Nataliia Teteriuk |
| EPI_ISL_582511, EPI_ISL_582512, EPI_ISL_582513 | Department of Respiratory and other Viral Infections of L.V.Gromashevsky Institute of Epidemiology & Infectious Diseases NAMS of Ukrain | Department of Respiratory and other Viral Infections of L.V.Gromashevsky Institute of Epidemiology & Infectious Diseases NAMS of Ukrain, JSC "Farmak" | Alla Mironenko, Andriy Goy, Ihor Kravchuk, Ludmyla Bolotova, Larysa Radchenko, Nataliia Teteriuk |
| EPI_ISL_582608, EPI_ISL_582609, EPI_ISL_582610, EPI_ISL_582611, EPI_ISL_582612, EPI_ISL_582613, EPI_ISL_582614, EPI_ISL_582615, EPI_ISL_582616, EPI_ISL_582617, EPI_ISL_582618, EPI_ISL_582619, EPI_ISL_582620, EPI_ISL_582621, EPI_ISL_582622, EPI_ISL_582623, EPI_ISL_582624, EPI_ISL_582625, EPI_ISL_582626, EPI_ISL_582627, EPI_ISL_582628, EPI_ISL_582629, EPI_ISL_582630, EPI_ISL_582631, EPI_ISL_582632, EPI_ISL_582633, EPI_ISL_582634, EPI_ISL_582635, EPI_ISL_582636, EPI_ISL_582637, EPI_ISL_582638, EPI_ISL_582639, EPI_ISL_582640, EPI_ISL_582641, EPI_ISL_582642, EPI_ISL_582643, EPI_ISL_582644, EPI_ISL_582645, EPI_ISL_582646, EPI_ISL_582647, EPI_ISL_582648, EPI_ISL_582649, EPI_ISL_582650, EPI_ISL_582651, EPI_ISL_582652, EPI_ISL_582653, EPI_ISL_582654, EPI_ISL_582655, EPI_ISL_582656, EPI_ISL_582657, EPI_ISL_582658, EPI_ISL_582659, EPI_ISL_582660, EPI_ISL_582661, EPI_ISL_582662, EPI_ISL_582663, EPI_ISL_582664, EPI_ISL_582665, EPI_ISL_582666, EPI_ISL_582667, EPI_ISL_582668, EPI_ISL_582669, EPI_ISL_582670, EPI_ISL_582671, EPI_ISL_582672, EPI_ISL_582673, EPI_ISL_582674, EPI_ISL_582675, EPI_ISL_582676, EPI_ISL_582677, EPI_ISL_582678, EPI_ISL_582679, EPI_ISL_582680, EPI_ISL_582681, EPI_ISL_582682, EPI_ISL_582683, EPI_ISL_582684, EPI_ISL_582685, EPI_ISL_582686, EPI_ISL_582687, EPI_ISL_582688, EPI_ISL_582689, EPI_ISL_582690, EPI_ISL_582691 | | | |
| see above | Sheikh Khalifa Medical City | Molecular/Surveillance lab Sheikh Khalifa Medical City | Amirtharaj Francis, Sajeed Abdul, Hala Imambaccus, Sahar Almarzooqi, Hiba Saud, Stefan Weber |
| EPI_ISL_583557, EPI_ISL_583558, EPI_ISL_583559, EPI_ISL_583560, EPI_ISL_583561, EPI_ISL_583562 | Universitaetsklinik für Innere Medizin II Innsbruck | Bergthaler laboratory, CeMM Research Center for Molecular Medicine of the Austrian Academy of Sciences | Alexandra Popa, Benedikt Agerer, Henrique Colaco, Lukas Endler, Jakob-Wendelin Genger, Alexander Lercher, Mark Smyth, Thomas Penz, Michael Schuster, Jan Laine, Martin Senekowitsch, Judith Aberle, Stephan Aberle, Peter Hufnagl, Daniela Schmid, Franz Allerberger, Elisabeth Puchhammer-Stoeckl, Manfred Nairz, Guenter Weiss, Gregor Hörmann, Kinga Rigler-Hohenwarter, Rainer Gattringer, Wegene Borena, Dorothee von Laer, Gernot Walder, Peter Obrist, Christian Paar, Sabine Sussitz-Rack, Gunther Vogl, Adi Steinrigl, Christoph Bock, Andreas Bergthaler |
| EPI_ISL_583563, EPI_ISL_583564, EPI_ISL_583565, EPI_ISL_583566, EPI_ISL_583567, EPI_ISL_583568, EPI_ISL_583569, EPI_ISL_583570, EPI_ISL_583571 | Center for Virology, Medical University of Vienna | Bergthaler laboratory, CeMM Research Center for Molecular Medicine of the Austrian Academy of Sciences | Alexandra Popa, Benedikt Agerer, Henrique Colaco, Lukas Endler, Jakob-Wendelin Genger, Alexander Lercher, Mark Smyth, Thomas Penz, Michael Schuster, Jan Laine, Martin Senekowitsch, Judith Aberle, Stephan Aberle, Peter Hufnagl, Daniela Schmid, Franz Allerberger, Elisabeth Puchhammer-Stoeckl, Manfred Nairz, Guenter Weiss, Gregor Hörmann, Kinga Rigler-Hohenwarter, Rainer Gattringer, Wegene Borena, Dorothee von Laer, Gernot Walder, Peter Obrist, Christian Paar, Sabine Sussitz-Rack, Gunther Vogl, Adi Steinrigl, Christoph Bock, Andreas Bergthaler |
| EPI_ISL_583572 | Institut für Virologie am Department für Hygiene, Mikrobiologie und Public Health | Bergthaler laboratory, CeMM Research Center for Molecular Medicine of the Austrian Academy of Sciences | Alexandra Popa, Benedikt Agerer, Henrique Colaco, Lukas Endler, Jakob-Wendelin Genger, Alexander Lercher, Mark Smyth, Thomas Penz, Michael Schuster, Jan Laine, Martin Senekowitsch, Judith Aberle, Stephan Aberle, Peter Hufnagl, Daniela Schmid, Franz Allerberger, Elisabeth Puchhammer-Stoeckl, Manfred Nairz, Guenter Weiss, Gregor Hörmann, Kinga Rigler-Hohenwarter, Rainer Gattringer, Wegene Borena, Dorothee von Laer, Gernot Walder, Peter Obrist, Christian Paar, Sabine Sussitz-Rack, Gunther Vogl, Adi Steinrigl, Christoph Bock, Andreas Bergthaler |

| | | | |
|---|---|---|---|
| EPI_ISL_583573, EPI_ISL_583574, EPI_ISL_583575, EPI_ISL_583576, EPI_ISL_583577, EPI_ISL_583578 | Austrian Agency for Health and Food Safety (AGES) | Bergthaler laboratory, CeMM Research Center for Molecular Medicine of the Austrian Academy of Sciences | Alexandra Popa, Benedikt Agerer, Henrique Colaco, Lukas Endler, Jakob-Wendelin Genger, Alexander Lercher, Mark Smyth, Thomas Penz, Michael Schuster, Jan Laine, Martin Senekowitsch, Judith Aberle, Stephan Aberle, Peter Hufnagl, Daniela Schmid, Franz Allerberger, Elisabeth Puchhammer-Stoeckl, Manfred Nairz, Guenter Weiss, Gregor Hörmann, Kinga Rigler-Hohenwarter, Rainer Gattringer, Wegene Borena, Dorothee von Laer, Gernot Walder, Peter Obrist, Christian Paar, Sabine Sussitz-Rack, Gunther Vogl, Adi Steinrigl, Christoph Bock, Andreas Bergthaler |
| EPI_ISL_583579, EPI_ISL_583580, EPI_ISL_583581, EPI_ISL_583582, EPI_ISL_583583, EPI_ISL_583584, EPI_ISL_583585, EPI_ISL_583586, EPI_ISL_583587, EPI_ISL_583588, EPI_ISL_583589, EPI_ISL_583590, EPI_ISL_583591, EPI_ISL_583592, EPI_ISL_583593, EPI_ISL_583594, EPI_ISL_583595, EPI_ISL_583596, EPI_ISL_583597, EPI_ISL_583598 | | | |
| see above | Institute for Medical and Chemical Laboratory Diagnostics, Kepler Universitätsklinikum | Bergthaler laboratory, CeMM Research Center for Molecular Medicine of the Austrian Academy of Sciences | Alexandra Popa, Benedikt Agerer, Henrique Colaco, Lukas Endler, Jakob-Wendelin Genger, Alexander Lercher, Mark Smyth, Thomas Penz, Michael Schuster, Jan Laine, Martin Senekowitsch, Judith Aberle, Stephan Aberle, Peter Hufnagl, Daniela Schmid, Franz Allerberger, Elisabeth Puchhammer-Stoeckl, Manfred Nairz, Guenter Weiss, Gregor Hörmann, Kinga Rigler-Hohenwarter, Rainer Gattringer, Wegene Borena, Dorothee von Laer, Gernot Walder, Peter Obrist, Christian Paar, Sabine Sussitz-Rack, Gunther Vogl, Adi Steinrigl, Christoph Bock, Andreas Bergthaler |
| EPI_ISL_583599 | Zentralinstitut für medizinische und chemische Labordiagnostik, Universitätskliniken Insbruck | Bergthaler laboratory, CeMM Research Center for Molecular Medicine of the Austrian Academy of Sciences | Alexandra Popa, Benedikt Agerer, Henrique Colaco, Lukas Endler, Jakob-Wendelin Genger, Alexander Lercher, Mark Smyth, Thomas Penz, Michael Schuster, Jan Laine, Martin Senekowitsch, Judith Aberle, Stephan Aberle, Peter Hufnagl, Daniela Schmid, Franz Allerberger, Elisabeth Puchhammer-Stoeckl, Manfred Nairz, Guenter Weiss, Gregor Hörmann, Kinga Rigler-Hohenwarter, Rainer Gattringer, Wegene Borena, Dorothee von Laer, Gernot Walder, Peter Obrist, Christian Paar, Sabine Sussitz-Rack, Gunther Vogl, Adi Steinrigl, Christoph Bock, Andreas Bergthaler |
| EPI_ISL_583600, EPI_ISL_583601, EPI_ISL_583602, EPI_ISL_583603, EPI_ISL_583604, EPI_ISL_583605, EPI_ISL_583606, EPI_ISL_583607, EPI_ISL_583608, EPI_ISL_583609, EPI_ISL_583610, EPI_ISL_583611, EPI_ISL_583612, EPI_ISL_583613, EPI_ISL_583614, EPI_ISL_583615, EPI_ISL_583616 | | | |
| see above | Institut für Virologie am Department für Hygiene, Mikrobiologie und Public Health | Bergthaler laboratory, CeMM Research Center for Molecular Medicine of the Austrian Academy of Sciences | Alexandra Popa, Benedikt Agerer, Henrique Colaco, Lukas Endler, Jakob-Wendelin Genger, Alexander Lercher, Mark Smyth, Thomas Penz, Michael Schuster, Jan Laine, Martin Senekowitsch, Judith Aberle, Stephan Aberle, Peter Hufnagl, Daniela Schmid, Franz Allerberger, Elisabeth Puchhammer-Stoeckl, Manfred Nairz, Guenter Weiss, Gregor Hörmann, Kinga Rigler-Hohenwarter, Rainer Gattringer, Wegene Borena, Dorothee von Laer, Gernot Walder, Peter Obrist, Christian Paar, Sabine Sussitz-Rack, Gunther Vogl, Adi Steinrigl, Christoph Bock, Andreas Bergthaler |
| EPI_ISL_583617, EPI_ISL_583618 | Universitaetsklinik für Innere Medizin II Innsbruck | Bergthaler laboratory, CeMM Research Center for Molecular Medicine of the Austrian Academy of Sciences | Alexandra Popa, Benedikt Agerer, Henrique Colaco, Lukas Endler, Jakob-Wendelin Genger, Alexander Lercher, Mark Smyth, Thomas Penz, Michael Schuster, Jan Laine, Martin Senekowitsch, Judith Aberle, Stephan Aberle, Peter Hufnagl, Daniela Schmid, Franz Allerberger, Elisabeth Puchhammer-Stoeckl, Manfred Nairz, Guenter Weiss, Gregor Hörmann, Kinga Rigler-Hohenwarter, Rainer Gattringer, Wegene Borena, Dorothee von Laer, Gernot Walder, Peter Obrist, Christian Paar, Sabine Sussitz-Rack, Gunther Vogl, Adi Steinrigl, Christoph Bock, Andreas Bergthaler |
| EPI_ISL_583619, EPI_ISL_583620, EPI_ISL_583621, EPI_ISL_583622, EPI_ISL_583623, EPI_ISL_583624, EPI_ISL_583625, EPI_ISL_583626, EPI_ISL_583627, EPI_ISL_583628, EPI_ISL_583629 | | | |
| see above | Pathologie-Labor Dr. Obrist- Dr. Brunhuber | Bergthaler laboratory, CeMM Research Center for Molecular Medicine of the Austrian Academy of Sciences | Alexandra Popa, Benedikt Agerer, Henrique Colaco, Lukas Endler, Jakob-Wendelin Genger, Alexander Lercher, Mark Smyth, Thomas Penz, Michael Schuster, Jan Laine, Martin Senekowitsch, Judith Aberle, Stephan Aberle, Peter Hufnagl, Daniela Schmid, Franz Allerberger, Elisabeth Puchhammer-Stoeckl, Manfred Nairz, Guenter Weiss, Gregor Hörmann, Kinga Rigler-Hohenwarter, Rainer Gattringer, Wegene Borena, Dorothee von Laer, Gernot Walder, Peter Obrist, Christian Paar, Sabine Sussitz-Rack, Gunther Vogl, Adi Steinrigl, Christoph Bock, Andreas Bergthaler |
| EPI_ISL_583630, EPI_ISL_583631, EPI_ISL_583633, EPI_ISL_583634, EPI_ISL_583635, EPI_ISL_583636, EPI_ISL_583637, EPI_ISL_583638, EPI_ISL_583639, EPI_ISL_583640, EPI_ISL_583641, EPI_ISL_583642, EPI_ISL_583643, EPI_ISL_583644, EPI_ISL_583645, EPI_ISL_583646, EPI_ISL_583647, EPI_ISL_583648, EPI_ISL_583649, EPI_ISL_583650, EPI_ISL_583651, EPI_ISL_583652, EPI_ISL_583653, EPI_ISL_583654, EPI_ISL_583655, EPI_ISL_583656, EPI_ISL_583657, EPI_ISL_583658, EPI_ISL_583659, EPI_ISL_583660, EPI_ISL_583661, EPI_ISL_583662, EPI_ISL_583663, EPI_ISL_583664, EPI_ISL_583665, EPI_ISL_583666, EPI_ISL_583667, EPI_ISL_583668, EPI_ISL_583669, EPI_ISL_583670, EPI_ISL_583671, EPI_ISL_583672, EPI_ISL_583673, EPI_ISL_583674, EPI_ISL_583675, EPI_ISL_583676, EPI_ISL_583677, EPI_ISL_583678, EPI_ISL_583679, EPI_ISL_583680, EPI_ISL_583681, EPI_ISL_583682, EPI_ISL_583683, EPI_ISL_583684, EPI_ISL_583685, EPI_ISL_583686, EPI_ISL_583687, EPI_ISL_583688, EPI_ISL_583689, EPI_ISL_583690 | | | |
| see above | Austrian Agency for Health and Food Safety (AGES) | Bergthaler laboratory, CeMM Research Center for Molecular Medicine of the Austrian Academy of Sciences | Alexandra Popa, Benedikt Agerer, Henrique Colaco, Lukas Endler, Jakob-Wendelin Genger, Alexander Lercher, Mark Smyth, Thomas Penz, Michael Schuster, Jan Laine, Martin Senekowitsch, Judith Aberle, Stephan Aberle, Peter Hufnagl, Daniela Schmid, Franz Allerberger, Elisabeth Puchhammer-Stoeckl, Manfred Nairz, Guenter Weiss, Gregor Hörmann, Kinga Rigler-Hohenwarter, Rainer Gattringer, Wegene Borena, Dorothee von Laer, Gernot Walder, Peter Obrist, Christian Paar, Sabine Sussitz-Rack, Gunther Vogl, Adi Steinrigl, Christoph Bock, Andreas Bergthaler |
| EPI_ISL_583691 | Institut für Virologie am Department für Hygiene, Mikrobiologie und Public Health | Bergthaler laboratory, CeMM Research Center for Molecular Medicine of the Austrian Academy of Sciences | Alexandra Popa, Benedikt Agerer, Henrique Colaco, Lukas Endler, Jakob-Wendelin Genger, Alexander Lercher, Mark Smyth, Thomas Penz, Michael Schuster, Jan Laine, Martin Senekowitsch, Judith Aberle, Stephan Aberle, Peter Hufnagl, Daniela Schmid, Franz Allerberger, Elisabeth Puchhammer-Stoeckl, Manfred Nairz, Guenter Weiss, Gregor Hörmann, Kinga Rigler-Hohenwarter, Rainer Gattringer, Wegene Borena, Dorothee von Laer, Gernot Walder, Peter Obrist, Christian Paar, Sabine Sussitz-Rack, Gunther Vogl, Adi Steinrigl, Christoph Bock, Andreas Bergthaler |
| EPI_ISL_583692, EPI_ISL_583693, EPI_ISL_583694, EPI_ISL_583695, EPI_ISL_583696, EPI_ISL_583697, EPI_ISL_583698, EPI_ISL_583699, EPI_ISL_583700, EPI_ISL_583701, EPI_ISL_583702, EPI_ISL_583703, EPI_ISL_583704, EPI_ISL_583705, EPI_ISL_583706, EPI_ISL_583707, EPI_ISL_583708, EPI_ISL_583709, EPI_ISL_583710, EPI_ISL_583711, EPI_ISL_583712, EPI_ISL_583713, EPI_ISL_583714, EPI_ISL_583715, EPI_ISL_583716, EPI_ISL_583717, EPI_ISL_583718, EPI_ISL_583719, EPI_ISL_583720, EPI_ISL_583721, EPI_ISL_583722, EPI_ISL_583723, EPI_ISL_583724, EPI_ISL_583725, EPI_ISL_583726 | | | |
| see above | Center for Virology, Medical University of Vienna | Bergthaler laboratory, CeMM Research Center for Molecular Medicine of the Austrian Academy of Sciences | Alexandra Popa, Benedikt Agerer, Henrique Colaco, Lukas Endler, Jakob-Wendelin Genger, Alexander Lercher, Mark Smyth, Thomas Penz, Michael Schuster, Jan Laine, Martin Senekowitsch, Judith Aberle, Stephan Aberle, Peter Hufnagl, Daniela Schmid, Franz Allerberger, Elisabeth Puchhammer-Stoeckl, Manfred Nairz, Guenter Weiss, Gregor Hörmann, Kinga Rigler-Hohenwarter, Rainer Gattringer, Wegene Borena, Dorothee von Laer, Gernot Walder, Peter Obrist, Christian Paar, Sabine Sussitz-Rack, Gunther Vogl, Adi Steinrigl, Christoph Bock, Andreas Bergthaler |
| EPI_ISL_583727, EPI_ISL_583728, EPI_ISL_583729, EPI_ISL_583730, EPI_ISL_583731, EPI_ISL_583732, EPI_ISL_583733, EPI_ISL_583734, EPI_ISL_583735, EPI_ISL_583736, EPI_ISL_583737, EPI_ISL_583738, EPI_ISL_583739, EPI_ISL_583740, EPI_ISL_583741, EPI_ISL_583742, EPI_ISL_583743, EPI_ISL_583744, EPI_ISL_583745, EPI_ISL_583746, EPI_ISL_583747, EPI_ISL_583748, EPI_ISL_583749, EPI_ISL_583750, EPI_ISL_583751, EPI_ISL_583752, EPI_ISL_583753, EPI_ISL_583754, EPI_ISL_583755, EPI_ISL_583756, EPI_ISL_583757, EPI_ISL_583758, EPI_ISL_583759, EPI_ISL_583760, EPI_ISL_583761, EPI_ISL_583762, EPI_ISL_583763, EPI_ISL_583764, EPI_ISL_583765, EPI_ISL_583766, EPI_ISL_583767, EPI_ISL_583768, EPI_ISL_583769, EPI_ISL_583770, EPI_ISL_583771, EPI_ISL_583772, EPI_ISL_583773, EPI_ISL_583774, EPI_ISL_583775, EPI_ISL_583776, EPI_ISL_583777, EPI_ISL_583778, EPI_ISL_583779, EPI_ISL_583780, EPI_ISL_583781, EPI_ISL_583782, EPI_ISL_583783, EPI_ISL_583784, EPI_ISL_583785, EPI_ISL_583786, EPI_ISL_583787, EPI_ISL_583788, EPI_ISL_583789, EPI_ISL_583790, EPI_ISL_583791, EPI_ISL_583792, EPI_ISL_583793, EPI_ISL_583794, EPI_ISL_583795, EPI_ISL_583796, EPI_ISL_583797, EPI_ISL_583798, EPI_ISL_583799, EPI_ISL_583800, EPI_ISL_583801, EPI_ISL_583802, EPI_ISL_583803, EPI_ISL_583804, EPI_ISL_583805, EPI_ISL_583806, EPI_ISL_583807, EPI_ISL_583808, EPI_ISL_583809, EPI_ISL_583810, EPI_ISL_583811, EPI_ISL_583812, EPI_ISL_583813, EPI_ISL_583814, EPI_ISL_583815, EPI_ISL_583816, EPI_ISL_583817, EPI_ISL_583818, EPI_ISL_583819, EPI_ISL_583820, EPI_ISL_583821, EPI_ISL_583822, EPI_ISL_583823, EPI_ISL_583824, EPI_ISL_583825, EPI_ISL_583826, EPI_ISL_583827, EPI_ISL_583828, EPI_ISL_583829, EPI_ISL_583830, EPI_ISL_583831, EPI_ISL_583832, EPI_ISL_583833, EPI_ISL_583834, EPI_ISL_583835, EPI_ISL_583836, EPI_ISL_583837, EPI_ISL_583838, EPI_ISL_583839, EPI_ISL_583840, EPI_ISL_583841, EPI_ISL_583842, EPI_ISL_583843, EPI_ISL_583844, EPI_ISL_583845, EPI_ISL_583846, EPI_ISL_583847, EPI_ISL_583848, EPI_ISL_583849, EPI_ISL_583850, EPI_ISL_583851, EPI_ISL_583852 | | | |
| see above | Dr. Gernot Walder GmbH | Bergthaler laboratory, CeMM Research Center for Molecular Medicine of the Austrian Academy of Sciences | Alexandra Popa, Benedikt Agerer, Henrique Colaco, Lukas Endler, Jakob-Wendelin Genger, Alexander Lercher, Mark Smyth, Thomas Penz, Michael Schuster, Jan Laine, Martin Senekowitsch, Judith Aberle, Stephan Aberle, Peter Hufnagl, Daniela Schmid, Franz Allerberger, Elisabeth Puchhammer-Stoeckl, Manfred Nairz, Guenter Weiss, Gregor Hörmann, Kinga Rigler-Hohenwarter, Rainer Gattringer, Wegene Borena, Dorothee von Laer, Gernot Walder, Peter Obrist, Christian Paar, Sabine Sussitz-Rack, Gunther Vogl, Adi Steinrigl, Christoph Bock, Andreas Bergthaler |
| EPI_ISL_583853, EPI_ISL_583854, EPI_ISL_583855, EPI_ISL_583856, EPI_ISL_583857, EPI_ISL_583858, EPI_ISL_583859, EPI_ISL_583860, EPI_ISL_583861, EPI_ISL_583862, EPI_ISL_583863, EPI_ISL_583864, EPI_ISL_583865, EPI_ISL_583866, EPI_ISL_583867, EPI_ISL_583868 | | | |
| see above | Institute for Laboratory Diagnostics and Microbiology, Klinikum Klagenfurt am Worthersee | Bergthaler laboratory, CeMM Research Center for Molecular Medicine of the Austrian Academy of Sciences | Alexandra Popa, Benedikt Agerer, Henrique Colaco, Lukas Endler, Jakob-Wendelin Genger, Alexander Lercher, Mark Smyth, Thomas Penz, Michael Schuster, Jan Laine, Martin Senekowitsch, Judith Aberle, Stephan Aberle, Peter Hufnagl, Daniela Schmid, Franz Allerberger, Elisabeth Puchhammer-Stoeckl, Manfred Nairz, Guenter Weiss, Gregor Hörmann, Kinga Rigler-Hohenwarter, Rainer Gattringer, Wegene Borena, Dorothee von Laer, Gernot Walder, Peter Obrist, Christian Paar, Sabine Sussitz-Rack, Gunther Vogl, Adi Steinrigl, Christoph Bock, Andreas Bergthaler |
| EPI_ISL_583869, EPI_ISL_583870, EPI_ISL_583871, EPI_ISL_583872, EPI_ISL_583873, EPI_ISL_583874, EPI_ISL_583875, EPI_ISL_583876, EPI_ISL_583877, EPI_ISL_583878, EPI_ISL_583879, EPI_ISL_583880, EPI_ISL_583881, EPI_ISL_583882 | | | |
| see above | Center for Virology, Medical University of Vienna | Bergthaler laboratory, CeMM Research Center for Molecular Medicine of the Austrian Academy of Sciences | Alexandra Popa, Benedikt Agerer, Henrique Colaco, Lukas Endler, Jakob-Wendelin Genger, Alexander Lercher, Mark Smyth, Thomas Penz, Michael Schuster, Jan Laine, Martin Senekowitsch, Judith Aberle, Stephan Aberle, Peter Hufnagl, Daniela Schmid, Franz Allerberger, Elisabeth Puchhammer-Stoeckl, Manfred Nairz, Guenter Weiss, Gregor Hörmann, Kinga Rigler-Hohenwarter, Rainer Gattringer, Wegene Borena, Dorothee von Laer, Gernot Walder, Peter Obrist, Christian Paar, Sabine Sussitz-Rack, Gunther Vogl, Adi Steinrigl, Christoph Bock, Andreas Bergthaler |
| EPI_ISL_583883, EPI_ISL_583884, EPI_ISL_583885, EPI_ISL_583886, EPI_ISL_583887, EPI_ISL_583888, EPI_ISL_583889, EPI_ISL_583890, EPI_ISL_583891, EPI_ISL_583892 | Austrian Agency for Health and Food Safety (AGES) | Bergthaler laboratory, CeMM Research Center for Molecular Medicine of the Austrian Academy of Sciences | Alexandra Popa, Benedikt Agerer, Henrique Colaco, Lukas Endler, Jakob-Wendelin Genger, Alexander Lercher, Mark Smyth, Thomas Penz, Michael Schuster, Jan Laine, Martin Senekowitsch, Judith Aberle, Stephan Aberle, Peter Hufnagl, Daniela Schmid, Franz Allerberger, Elisabeth Puchhammer-Stoeckl, Manfred Nairz, Guenter Weiss, Gregor Hörmann, Kinga Rigler-Hohenwarter, Rainer Gattringer, Wegene Borena, Dorothee von Laer, Gernot Walder, Peter Obrist, Christian Paar, Sabine Sussitz-Rack, Gunther Vogl, Adi Steinrigl, Christoph Bock, Andreas Bergthaler |
| EPI_ISL_584283, EPI_ISL_584286, EPI_ISL_584287, EPI_ISL_584288, EPI_ISL_584289, EPI_ISL_584290, EPI_ISL_584291, EPI_ISL_584294, EPI_ISL_584295, EPI_ISL_584296, EPI_ISL_584297, EPI_ISL_584298, EPI_ISL_584299, EPI_ISL_584300, EPI_ISL_584301, EPI_ISL_584302, EPI_ISL_584304, EPI_ISL_584305, EPI_ISL_584306, EPI_ISL_584307, EPI_ISL_584309, EPI_ISL_584310, EPI_ISL_584311, EPI_ISL_584312, EPI_ISL_584313, EPI_ISL_584314, EPI_ISL_584316, EPI_ISL_584317, EPI_ISL_584318, EPI_ISL_584321, EPI_ISL_584322, EPI_ISL_584323 | | | |
| see above | Department of Pathology, University of Cambridge | COVID-19 Genomics UK (COG-UK) Consortium | Aminu S. Jahun, Yasmin Chaudhry, Grant Hall, Iliana Georgana, Myra Hosmillo, Martin D. Curran, Malte Pinckert, Surendra Parmar, Ian Goodfellow |
| EPI_ISL_585261, EPI_ISL_585262, EPI_ISL_585263, EPI_ISL_585264, EPI_ISL_585265, EPI_ISL_585266, EPI_ISL_585267, EPI_ISL_585268, EPI_ISL_585269, EPI_ISL_585270, EPI_ISL_585271, EPI_ISL_585272, EPI_ISL_585273, EPI_ISL_585274, EPI_ISL_585275, EPI_ISL_585276, EPI_ISL_585277, EPI_ISL_585278, | | | |

| | | | |
|---|---|---|---|
| EPI_ISL_585279, EPI_ISL_585280, EPI_ISL_585281, EPI_ISL_585282, EPI_ISL_585283, EPI_ISL_585284, EPI_ISL_585285, EPI_ISL_585286, EPI_ISL_585287, EPI_ISL_585288, EPI_ISL_585289, EPI_ISL_585290, EPI_ISL_585291, EPI_ISL_585292, EPI_ISL_585293, EPI_ISL_585294, EPI_ISL_585295, EPI_ISL_585296 | | | |
| see above | West of Scotland Specialist Virology Centre, NHSGGC / MRC-University of Glasgow Centre for Virus Research | COVID-19 Genomics UK (COG-UK) Consortium | Ana da Silva Filipe, Natasha Johnson, Kathy Smollett, Daniel Mair, Stephen Carmichael, Lily Tong, Jenna Nichols, Elihu Aranday-Cortes, Kyriaki Nomikou; Sarah McDonald, Marc Niebel, Patawee Asamaphan; Richard Orton, Joseph Hughes, Sreenu Vattipally, David L Robertson; Alasdair MacLean, Rory Gunson; Kathy Li, Igor Starinskij, Natasha Jesudason, Rajiv Shah, James Shepherd, Antonia Ho, Emma Thomson |
| EPI_ISL_585297, EPI_ISL_585298, EPI_ISL_585299, EPI_ISL_585300, EPI_ISL_585301, EPI_ISL_585302, EPI_ISL_585303, EPI_ISL_585304, EPI_ISL_585305, EPI_ISL_585306, EPI_ISL_585307, EPI_ISL_585308, EPI_ISL_585309, EPI_ISL_585310, EPI_ISL_585311, EPI_ISL_585312, EPI_ISL_585313, EPI_ISL_585314, EPI_ISL_585315, EPI_ISL_585316, EPI_ISL_585317, EPI_ISL_585318, EPI_ISL_585319, EPI_ISL_585320, EPI_ISL_585321, EPI_ISL_585322, EPI_ISL_585323, EPI_ISL_585324, EPI_ISL_585325, EPI_ISL_585326, EPI_ISL_585327, EPI_ISL_585328, EPI_ISL_585329, EPI_ISL_585330, EPI_ISL_585331, EPI_ISL_585332, EPI_ISL_585333, EPI_ISL_585334, EPI_ISL_585335, EPI_ISL_585336, EPI_ISL_585337, EPI_ISL_585338, EPI_ISL_585339, EPI_ISL_585340, EPI_ISL_585341, EPI_ISL_585342, EPI_ISL_585343, EPI_ISL_585344, EPI_ISL_585345, EPI_ISL_585346, EPI_ISL_585347, EPI_ISL_585348, EPI_ISL_585349, EPI_ISL_585350, EPI_ISL_585351, EPI_ISL_585352, EPI_ISL_585353, EPI_ISL_585354, EPI_ISL_585355, EPI_ISL_585356, EPI_ISL_585357, EPI_ISL_585358, EPI_ISL_585359, EPI_ISL_585360, EPI_ISL_585361, EPI_ISL_585362, EPI_ISL_585363, EPI_ISL_585364, EPI_ISL_585365, EPI_ISL_585366, EPI_ISL_585367, EPI_ISL_585368, EPI_ISL_585369, EPI_ISL_585370, EPI_ISL_585371, EPI_ISL_585372, EPI_ISL_585373, EPI_ISL_585374, EPI_ISL_585375, EPI_ISL_585376, EPI_ISL_585377, EPI_ISL_585378, EPI_ISL_585379, EPI_ISL_585380, EPI_ISL_585381, EPI_ISL_585382, EPI_ISL_585383, EPI_ISL_585384, EPI_ISL_585385, EPI_ISL_585386, EPI_ISL_585387, EPI_ISL_585388, EPI_ISL_585389, EPI_ISL_585390, EPI_ISL_585391, EPI_ISL_585392, EPI_ISL_585393, EPI_ISL_585394, EPI_ISL_585395, EPI_ISL_585396, EPI_ISL_585397, EPI_ISL_585398, EPI_ISL_585399, EPI_ISL_585400, EPI_ISL_585401, EPI_ISL_585402, EPI_ISL_585403, EPI_ISL_585404, EPI_ISL_585405, EPI_ISL_585406, EPI_ISL_585407, EPI_ISL_585408, EPI_ISL_585409, EPI_ISL_585410, EPI_ISL_585411, EPI_ISL_585412, EPI_ISL_585413, EPI_ISL_585414, EPI_ISL_585415, EPI_ISL_585416, EPI_ISL_585417, EPI_ISL_585418, EPI_ISL_585419, EPI_ISL_585420, EPI_ISL_585421 | | | |
| see above | Lighthouse Lab in Glasgow / MRC-University of Glasgow Centre for Virus Research | COVID-19 Genomics UK (COG-UK) Consortium | Ana da Silva Filipe, Natasha Johnson, Kathy Smollett, Daniel Mair, Stephen Carmichael, Lily Tong, Jenna Nichols, Elihu Aranday-Cortes, Kyriaki Nomikou; Sarah McDonald, Marc Niebel, Patawee Asamaphan; Harper VanSteenhouse, Yumi Kasai, David Gray, Carol Clugston, Anna Dominiczak; Alasdair MacLean, Rory Gunson; Richard Orton, Joseph Hughes, Sreenu Vattipally, David L Robertson; Sharif Shaaban, Matthew Holden; Kathy Li, Natasha Jesudason, Rajiv Shah, James Shepherd, Antonia Ho, Emma Thomson |
| EPI_ISL_585422, EPI_ISL_585423, EPI_ISL_585424, EPI_ISL_585425 | West of Scotland Specialist Virology Centre, NHSGGC / MRC-University of Glasgow Centre for Virus Research | COVID-19 Genomics UK (COG-UK) Consortium | Ana da Silva Filipe, Natasha Johnson, Kathy Smollett, Daniel Mair, Stephen Carmichael, Lily Tong, Jenna Nichols, Elihu Aranday-Cortes, Kyriaki Nomikou; Sarah McDonald, Marc Niebel, Patawee Asamaphan; Richard Orton, Joseph Hughes, Sreenu Vattipally, David L Robertson; Alasdair MacLean, Rory Gunson; Kathy Li, Igor Starinskij, Natasha Jesudason, Rajiv Shah, James Shepherd, Antonia Ho, Emma Thomson |
| EPI_ISL_586267, EPI_ISL_586268, EPI_ISL_586269, EPI_ISL_586270, EPI_ISL_586271, EPI_ISL_586272, EPI_ISL_586273, EPI_ISL_586274, EPI_ISL_586275, EPI_ISL_586276, EPI_ISL_586277, EPI_ISL_586278, EPI_ISL_586279, EPI_ISL_586280, EPI_ISL_586281, EPI_ISL_586282, EPI_ISL_586283, EPI_ISL_586284, EPI_ISL_586285, EPI_ISL_586286, EPI_ISL_586287, EPI_ISL_586288, EPI_ISL_586289, EPI_ISL_586290, EPI_ISL_586291, EPI_ISL_586292, EPI_ISL_586293, EPI_ISL_586294, EPI_ISL_586295, EPI_ISL_586296, EPI_ISL_586297, EPI_ISL_586298, EPI_ISL_586299, EPI_ISL_586300, EPI_ISL_586301, EPI_ISL_586302, EPI_ISL_586303, EPI_ISL_586304, EPI_ISL_586305, EPI_ISL_586306, EPI_ISL_586307, EPI_ISL_586308, EPI_ISL_586309, EPI_ISL_586310, EPI_ISL_586311, EPI_ISL_586312, EPI_ISL_586313, EPI_ISL_586314, EPI_ISL_586315, EPI_ISL_586316, EPI_ISL_586317, EPI_ISL_586318, EPI_ISL_586319, EPI_ISL_586320, EPI_ISL_586321, EPI_ISL_586322, EPI_ISL_586323, EPI_ISL_586324, EPI_ISL_586325, EPI_ISL_586326, EPI_ISL_586327, EPI_ISL_586328, EPI_ISL_586329, EPI_ISL_586330, EPI_ISL_586331, EPI_ISL_586332, EPI_ISL_586333, EPI_ISL_586334, EPI_ISL_586335, EPI_ISL_586336, EPI_ISL_586337, EPI_ISL_586338, EPI_ISL_586339, EPI_ISL_586340, EPI_ISL_586341, EPI_ISL_586342, EPI_ISL_586343, EPI_ISL_586344, EPI_ISL_586345, EPI_ISL_586346, EPI_ISL_586347, EPI_ISL_586348, EPI_ISL_586349, EPI_ISL_586350, EPI_ISL_586351, EPI_ISL_586352, EPI_ISL_586353, EPI_ISL_586354, EPI_ISL_586355, EPI_ISL_586356, EPI_ISL_586357, EPI_ISL_586358, EPI_ISL_586359, EPI_ISL_586360, EPI_ISL_586361, EPI_ISL_586362, EPI_ISL_586363, EPI_ISL_586364, EPI_ISL_586365, EPI_ISL_586366, EPI_ISL_586367, EPI_ISL_586368, EPI_ISL_586369, EPI_ISL_586370, EPI_ISL_586371, EPI_ISL_586372, EPI_ISL_586373, EPI_ISL_586374, EPI_ISL_586375, EPI_ISL_586376, EPI_ISL_586377, EPI_ISL_586378, EPI_ISL_586379, EPI_ISL_586380, EPI_ISL_586381, EPI_ISL_586382, EPI_ISL_586383, EPI_ISL_586384, EPI_ISL_586385, EPI_ISL_586386, EPI_ISL_586387, EPI_ISL_586388, EPI_ISL_586389, EPI_ISL_586390, EPI_ISL_586391, EPI_ISL_586392, EPI_ISL_586393, EPI_ISL_586394, EPI_ISL_586395, EPI_ISL_586396, EPI_ISL_586397, EPI_ISL_586398, EPI_ISL_586399, EPI_ISL_586400, EPI_ISL_586401, EPI_ISL_586402, EPI_ISL_586403, EPI_ISL_586404, EPI_ISL_586405, EPI_ISL_586406, EPI_ISL_586407, EPI_ISL_586408, EPI_ISL_586409, EPI_ISL_586410, EPI_ISL_586411, EPI_ISL_586412, EPI_ISL_586413, EPI_ISL_586414, EPI_ISL_586415, EPI_ISL_586416, EPI_ISL_586417, EPI_ISL_586418, EPI_ISL_586419, EPI_ISL_586420, EPI_ISL_586421, EPI_ISL_586422, EPI_ISL_586423, EPI_ISL_586424, EPI_ISL_586425, EPI_ISL_586426, EPI_ISL_586427, EPI_ISL_586428, EPI_ISL_586430, EPI_ISL_586431, EPI_ISL_586432, EPI_ISL_586433, EPI_ISL_586434, EPI_ISL_586435, EPI_ISL_586436, EPI_ISL_586437, EPI_ISL_586438, EPI_ISL_586440, EPI_ISL_586441, EPI_ISL_586442, EPI_ISL_586443, EPI_ISL_586444, EPI_ISL_586445, EPI_ISL_586446, EPI_ISL_586447, EPI_ISL_586448, EPI_ISL_586449, EPI_ISL_586452, EPI_ISL_586453, EPI_ISL_586455, EPI_ISL_586456, EPI_ISL_586458, EPI_ISL_586459, EPI_ISL_586460, EPI_ISL_586461, EPI_ISL_586462, EPI_ISL_586463, EPI_ISL_586466, EPI_ISL_586467, EPI_ISL_586468, EPI_ISL_586469, EPI_ISL_586470, EPI_ISL_586471, EPI_ISL_586472, EPI_ISL_586473, EPI_ISL_586474, EPI_ISL_586475, EPI_ISL_586476, EPI_ISL_586477, EPI_ISL_586478, EPI_ISL_586479, EPI_ISL_586480, EPI_ISL_586481, EPI_ISL_586482, EPI_ISL_586483, EPI_ISL_586484, EPI_ISL_586485, EPI_ISL_586486, EPI_ISL_586487, EPI_ISL_586488, EPI_ISL_586489, EPI_ISL_586490, EPI_ISL_586491, EPI_ISL_586492, EPI_ISL_586493, EPI_ISL_586494, EPI_ISL_586495, EPI_ISL_586496, EPI_ISL_586497, EPI_ISL_586498, EPI_ISL_586499, EPI_ISL_586500, EPI_ISL_586501, EPI_ISL_586502, EPI_ISL_586503, EPI_ISL_586504, EPI_ISL_586505, EPI_ISL_586506, EPI_ISL_586507, EPI_ISL_586508 | | | |
| see above | Toronto Invasive Bacterial Diseases Network | McMaster University | Allison McGeer, Patryk Aftanas, Hooman Derakhshani, Angel Li, Kuganya Nirmalarajah, Emily Panousis, Ahmed Draia, Jalees Nasir, Michael Surette, Samira Mubareka, Andrew G. McArthur |
| EPI_ISL_586522 | B.J. Medical College and Civil hospital, Ahmedabad | Gujarat Biotechnology Research Centre | Afzal Ansari, Nikha Trivedi, Pranay Shah, Kamlesh J Upadhyay, Sanjay Kapadia, Apurvasinh Puvar, Janvi Raval, Zarna Patel, Monika Gandhi, Pinal Trivedi, Maharshi Pandya, Nidhi Patel, Nitin Savaliya, Raghawendra Kumar, Dinesh Kumar, Zuber Saiyed, Komal Patel, Labdhi Pandya, R D Dixit, A M Kadri, Harsh Bakshi, Chaitanya Joshi, Madhvi Joshi |
| EPI_ISL_586544 | Government General Hospital, Jam Khambhaliya, Devbhoomi Dwarka | Gujarat Biotechnology Research Centre | Afzal Ansari, Nikha Trivedi, Harish Matani, Apurvasinh Puvar, Janvi Raval, Zarna Patel, Monika Gandhi, Pinal Trivedi, Maharshi Pandya, Nidhi Patel, Nitin Savaliya, Raghawendra Kumar, Dinesh Kumar, Zuber Saiyed, Komal Patel, Labdhi Pandya, R D Dixit, A M Kadri, Harsh Bakshi, Chaitanya Joshi, Madhvi Joshi |
| EPI_ISL_586561 | CCC,Veraval | Gujarat Biotechnology Research Centre | Afzal Ansari, Nikha Trivedi, Jignesh Parmar, Jitendra Bamrotia, Apurvasinh Puvar, Janvi Raval, Zarna Patel, Monika Gandhi, Pinal Trivedi, Maharshi Pandya, Nidhi Patel, Nitin Savaliya, Raghawendra Kumar, Dinesh Kumar, Zuber Saiyed, Komal Patel, Labdhi Pandya, R D Dixit, A M Kadri, Harsh Bakshi, Chaitanya Joshi, Madhvi Joshi |
| EPI_ISL_591100, EPI_ISL_591101, EPI_ISL_591102, EPI_ISL_591103, EPI_ISL_591104, EPI_ISL_591105, EPI_ISL_591106, EPI_ISL_591107, EPI_ISL_591108, EPI_ISL_591109, EPI_ISL_591110, EPI_ISL_591111, EPI_ISL_591112, EPI_ISL_591113, EPI_ISL_591114, EPI_ISL_591115, EPI_ISL_591116, EPI_ISL_591117, EPI_ISL_591118, EPI_ISL_591119, EPI_ISL_591120, EPI_ISL_591121, EPI_ISL_591122, EPI_ISL_591123, EPI_ISL_591124, EPI_ISL_591125, EPI_ISL_591126, EPI_ISL_591127, EPI_ISL_591128, EPI_ISL_591129, EPI_ISL_591130, EPI_ISL_591131, EPI_ISL_591132, EPI_ISL_591133, EPI_ISL_591134, EPI_ISL_591135, EPI_ISL_591136, EPI_ISL_591137, EPI_ISL_591138, EPI_ISL_591139, EPI_ISL_591140, EPI_ISL_591141, EPI_ISL_591143, EPI_ISL_591148, EPI_ISL_591150, EPI_ISL_591152, EPI_ISL_591154, EPI_ISL_591155, EPI_ISL_591156, EPI_ISL_591157, EPI_ISL_591158, EPI_ISL_591159, EPI_ISL_591160, EPI_ISL_591164, EPI_ISL_591165, EPI_ISL_591166, EPI_ISL_591168, EPI_ISL_591169, EPI_ISL_591170, EPI_ISL_591175, EPI_ISL_591176, EPI_ISL_591177, EPI_ISL_591178, EPI_ISL_591179, EPI_ISL_591180, EPI_ISL_591181, EPI_ISL_591182, EPI_ISL_591183, EPI_ISL_591184, EPI_ISL_591185, EPI_ISL_591186, EPI_ISL_591187, EPI_ISL_591188, EPI_ISL_591189, EPI_ISL_591190, EPI_ISL_591191, EPI_ISL_591192, EPI_ISL_591193, EPI_ISL_591194, EPI_ISL_591195, EPI_ISL_591196, EPI_ISL_591197, EPI_ISL_591198, EPI_ISL_591199, EPI_ISL_591200, EPI_ISL_591201, EPI_ISL_591202, EPI_ISL_591203, EPI_ISL_591204, EPI_ISL_591205, EPI_ISL_591206, EPI_ISL_591207, EPI_ISL_591208, EPI_ISL_591209, EPI_ISL_591210, EPI_ISL_591211, EPI_ISL_591212, EPI_ISL_591216, EPI_ISL_591220, EPI_ISL_591222, EPI_ISL_591223, EPI_ISL_591224, EPI_ISL_591225, EPI_ISL_591237, EPI_ISL_591238, EPI_ISL_591240, EPI_ISL_591245, EPI_ISL_591246, EPI_ISL_591247, EPI_ISL_591249, EPI_ISL_591253, EPI_ISL_591254, EPI_ISL_591255, EPI_ISL_591256, EPI_ISL_591257, EPI_ISL_591258, EPI_ISL_591259, EPI_ISL_591260, EPI_ISL_591261, EPI_ISL_591262, EPI_ISL_591263, EPI_ISL_591264, EPI_ISL_591265, EPI_ISL_591266, EPI_ISL_591267, EPI_ISL_591268, EPI_ISL_591269 | | | |
| see above | Toronto Invasive Bacterial Diseases Network | McMaster University | Allison McGeer, Patryk Aftanas, Hooman Derakhshani, Angel Li, Kuganya Nirmalarajah, Emily Panousis, Ahmed Draia, Jalees Nasir, Michael Surette, Samira Mubareka, Andrew G. McArthur |
| EPI_ISL_594645, EPI_ISL_594646, EPI_ISL_594647, EPI_ISL_594648, EPI_ISL_594649, EPI_ISL_594650, EPI_ISL_594651, EPI_ISL_594652, EPI_ISL_594653, EPI_ISL_594654, EPI_ISL_594655, EPI_ISL_594656, EPI_ISL_594657, EPI_ISL_594658, EPI_ISL_594659, EPI_ISL_594660, EPI_ISL_594661, EPI_ISL_594662, EPI_ISL_594663, EPI_ISL_594664, EPI_ISL_594665, EPI_ISL_594666, EPI_ISL_594667, EPI_ISL_594668, EPI_ISL_594669, EPI_ISL_594670, EPI_ISL_594671, EPI_ISL_594672, EPI_ISL_594673, EPI_ISL_594674, EPI_ISL_594675, EPI_ISL_594676, EPI_ISL_594677, EPI_ISL_594678, EPI_ISL_594679, EPI_ISL_594680, EPI_ISL_594681, EPI_ISL_594682, EPI_ISL_594683, EPI_ISL_594684, EPI_ISL_594685, EPI_ISL_594686, EPI_ISL_594687, EPI_ISL_594688, EPI_ISL_594689, EPI_ISL_594690, EPI_ISL_594691, EPI_ISL_594692, EPI_ISL_594693, EPI_ISL_594694, EPI_ISL_594695, EPI_ISL_594696, EPI_ISL_594697, EPI_ISL_594698, EPI_ISL_594699, EPI_ISL_594700, EPI_ISL_594701, EPI_ISL_594702, EPI_ISL_594703, EPI_ISL_594704, EPI_ISL_594705, EPI_ISL_594706, EPI_ISL_594707, EPI_ISL_594708, EPI_ISL_594709, EPI_ISL_594710, EPI_ISL_594711, EPI_ISL_594712, EPI_ISL_594713, EPI_ISL_594714, EPI_ISL_594715, EPI_ISL_594716, EPI_ISL_594717, EPI_ISL_594718, EPI_ISL_594719, EPI_ISL_594720, EPI_ISL_594721, EPI_ISL_594722, EPI_ISL_594723, EPI_ISL_594724, EPI_ISL_594725, EPI_ISL_594726, EPI_ISL_594727, EPI_ISL_594728, EPI_ISL_594729, EPI_ISL_594730, EPI_ISL_594731, EPI_ISL_594732, EPI_ISL_594733, EPI_ISL_594734, EPI_ISL_594735, EPI_ISL_594736, EPI_ISL_594737, EPI_ISL_594738, EPI_ISL_594739, EPI_ISL_594740, EPI_ISL_594741, EPI_ISL_594742, EPI_ISL_594743, EPI_ISL_594744, EPI_ISL_594745, EPI_ISL_594746, EPI_ISL_594747, EPI_ISL_594748, EPI_ISL_594749, EPI_ISL_594750, EPI_ISL_594751, EPI_ISL_594752, EPI_ISL_594753, EPI_ISL_594754, EPI_ISL_594755, EPI_ISL_594756, EPI_ISL_594757, EPI_ISL_594758, EPI_ISL_594759, EPI_ISL_594760, EPI_ISL_594761, EPI_ISL_594762, EPI_ISL_594763, EPI_ISL_594764, EPI_ISL_594765, EPI_ISL_594766, EPI_ISL_594767, EPI_ISL_594768, EPI_ISL_594769, EPI_ISL_594770, EPI_ISL_594771, EPI_ISL_594772, EPI_ISL_594773, EPI_ISL_594774, EPI_ISL_594775, EPI_ISL_594776, EPI_ISL_594777, EPI_ISL_594778, EPI_ISL_594779, EPI_ISL_594780, EPI_ISL_594781, EPI_ISL_594782, EPI_ISL_594783, EPI_ISL_594784, EPI_ISL_594785, EPI_ISL_594786, EPI_ISL_594787, EPI_ISL_594788, EPI_ISL_594789, EPI_ISL_594790, EPI_ISL_594791, EPI_ISL_594792, EPI_ISL_594793, EPI_ISL_594794, EPI_ISL_594795, EPI_ISL_594796, EPI_ISL_594797, EPI_ISL_594798, EPI_ISL_594799, EPI_ISL_594800, EPI_ISL_594801, EPI_ISL_594802, EPI_ISL_594803, EPI_ISL_594804, EPI_ISL_594805, EPI_ISL_594806, EPI_ISL_594807, EPI_ISL_594808, EPI_ISL_594809, EPI_ISL_594810, EPI_ISL_594811, EPI_ISL_594812, EPI_ISL_594813, EPI_ISL_594814, EPI_ISL_594815 | | | |
| see above | West of Scotland Specialist Virology Centre, NHSGGC / MRC-University of Glasgow Centre for Virus Research | COVID-19 Genomics UK (COG-UK) Consortium | Ana da Silva Filipe, Natasha Johnson, Kathy Smollett, Daniel Mair, Stephen Carmichael, Lily Tong, Jenna Nichols, Elihu Aranday-Cortes, Kyriaki Nomikou; Sarah McDonald, Marc Niebel, Patawee Asamaphan; Richard Orton, Joseph Hughes, Sreenu Vattipally, David L Robertson; Alasdair MacLean, Rory Gunson; Kathy Li, Igor Starinskij, Natasha Jesudason, Rajiv Shah, James Shepherd, Antonia Ho, Emma Thomson |
| EPI_ISL_595102, EPI_ISL_595105, EPI_ISL_595106, EPI_ISL_595109, EPI_ISL_595110, EPI_ISL_595112 | Centre for Enzyme Innovation, University of Portsmouth / Translational Research Laboratory, Portsmouth Hospitals NHS Trust | COVID-19 Genomics UK (COG-UK) Consortium | Angela Beckett,Yann Bourgeois,Garry Scarlett,Sharon Glaysher,Scott Elliott,Kelly Bicknell,Robert Impey,Allyson Lloyd,Sarah Wyllie,Ethan Butcher,Anoop Chauhan,Samuel Robson |
| EPI_ISL_596228 | WHO National Influenza Centre Russian Federation | WHO National Influenza Centre Russian Federation | Andrey Komissarov, Artem Fadeev, Anna Ivanova, Kseniya Komissarova, Dmitry Bazhenov, Daria Danilenko, Ksenia Safina, Elena Nabieva, Georgii Bazykin, Dmitry Lioznov |
| EPI_ISL_596229, EPI_ISL_596230, EPI_ISL_596231 | WHO National Influenza Centre Russian Federation | WHO National Influenza Centre Russian Federation | Andrey Komissarov, Artem Fadeev, Anna Ivanova, Kseniya Komissarova, Dmitry Bazhenov, Daria Danilenko |
| EPI_ISL_596232 | WHO National Influenza Centre Russian Federation | WHO National Influenza Centre Russian Federation | Andrey Komissarov, Artem Fadeev, Anna Ivanova, Mariia Sergeeva, Kseniya Komissarova, Dmitry Bazhenov, Daria Danilenko |
| EPI_ISL_596235, EPI_ISL_596236, EPI_ISL_596237, EPI_ISL_596238, EPI_ISL_596239, EPI_ISL_596240, EPI_ISL_596241, EPI_ISL_596242, EPI_ISL_596243, EPI_ISL_596244, EPI_ISL_596245, EPI_ISL_596246, EPI_ISL_596247, EPI_ISL_596248, EPI_ISL_596249 | | | |
| see above | WHO National Influenza Centre Russian Federation | WHO National Influenza Centre Russian Federation | Andrey Komissarov, Artem Fadeev, Anna Ivanova, Kseniya Komissarova, Dmitry Bazhenov, Daria Danilenko |
| EPI_ISL_596250, EPI_ISL_596251, EPI_ISL_596252, EPI_ISL_596253, EPI_ISL_596254, EPI_ISL_596255, EPI_ISL_596256, EPI_ISL_596257, EPI_ISL_596258, EPI_ISL_596259, EPI_ISL_596260, EPI_ISL_596261, EPI_ISL_596262, EPI_ISL_596263 | | | |
| see above | HELIX LCC | WHO National Influenza Centre Russian Federation | Andrey Komissarov, Artem Fadeev, Anna Ivanova, Kseniya Komissarova, Dmitry Bazhenov, Daria Danilenko |

| | | | |
|---|---|---|---|
| EPI_ISL_596264, EPI_ISL_596265 | WHO National Influenza Centre Russian Federation | WHO National Influenza Centre Russian Federation | Andrey Komissarov, Artem Fadeev, Anna Ivanova, Mariia Sergeeva, Kseniya Komissarova, Dmitry Bazhenov, Daria Danilenko |
| EPI_ISL_596266, EPI_ISL_596267 | WHO National Influenza Centre Russian Federation | WHO National Influenza Centre Russian Federation | Andrey Komissarov, Artem Fadeev, Anna Ivanova, Kseniya Komissarova, Dmitry Bazhenov, Daria Danilenko |
| EPI_ISL_596268, EPI_ISL_596269, EPI_ISL_596270, EPI_ISL_596271, EPI_ISL_596272, EPI_ISL_596273, EPI_ISL_596274, EPI_ISL_596275, EPI_ISL_596276, EPI_ISL_596277, EPI_ISL_596278, EPI_ISL_596279, EPI_ISL_596280, EPI_ISL_596281, EPI_ISL_596282, EPI_ISL_596283, EPI_ISL_596284, EPI_ISL_596285, EPI_ISL_596286, EPI_ISL_596287, EPI_ISL_596288, EPI_ISL_596289, EPI_ISL_596290, EPI_ISL_596291, EPI_ISL_596292, EPI_ISL_596293, EPI_ISL_596294, EPI_ISL_596295, EPI_ISL_596296, EPI_ISL_596297, EPI_ISL_596298, EPI_ISL_596299, EPI_ISL_596300, EPI_ISL_596301, EPI_ISL_596302, EPI_ISL_596303, EPI_ISL_596304, EPI_ISL_596305, EPI_ISL_596306, EPI_ISL_596307, EPI_ISL_596308, EPI_ISL_596309, EPI_ISL_596310, EPI_ISL_596311, EPI_ISL_596312, EPI_ISL_596313, EPI_ISL_596314, EPI_ISL_596315, EPI_ISL_596316, EPI_ISL_596317 | | | |
| see above | HELIX LCC | WHO National Influenza Centre Russian Federation | Andrey Komissarov, Artem Fadeev, Anna Ivanova, Kseniya Komissarova, Dmitry Bazhenov, Daria Danilenko |
| EPI_ISL_596318, EPI_ISL_596319, EPI_ISL_596320, EPI_ISL_596321, EPI_ISL_596322, EPI_ISL_596323, EPI_ISL_596324, EPI_ISL_596325, EPI_ISL_596326, EPI_ISL_596327, EPI_ISL_596328, EPI_ISL_596329, EPI_ISL_596330, EPI_ISL_596331, EPI_ISL_596332, EPI_ISL_596333, EPI_ISL_596334, EPI_ISL_596335, EPI_ISL_596336, EPI_ISL_596337, EPI_ISL_596338, EPI_ISL_596339, EPI_ISL_596340, EPI_ISL_596341, EPI_ISL_596342, EPI_ISL_596343, EPI_ISL_596344, EPI_ISL_596345, EPI_ISL_596346, EPI_ISL_596347, EPI_ISL_596348, EPI_ISL_596349, EPI_ISL_596350, EPI_ISL_596351, EPI_ISL_596352 | | | |
| see above | Pathogenic Microorganisms Variability Laboratory | WHO National Influenza Centre Russian Federation | Andrey Komissarov, Artem Fadeev, Anna Ivanova, Kseniya Komissarova, Dmitry Bazhenov, Daria Danilenko, Dmitry Lioznov, Nadezhda Kuznetsova, Elena Shidlovskaya, Elizaveta Divisenko, Ekaterina Milashenko, Kirill Krasnoslobotsev, Evgeniya Mukasheva, Anna Ignatieva, Svetlana Trushakova, Alexey Shchetinin, Maria Nikiforova, Andrey Pochtovyy, Valeria Bacalin, Evgeny Usachev, Olga Burgasova, Ludmila Kolobukhina, Svetlana Smetanina, Elena Burtseva, Artem Tkachuk, Vladimir Gushchin, Alexander Gintsburg |
| EPI_ISL_596353, EPI_ISL_596354, EPI_ISL_596355, EPI_ISL_596356 | WHO National Influenza Centre Russian Federation | WHO National Influenza Centre Russian Federation | Andrey Komissarov, Artem Fadeev, Anna Ivanova, Kseniya Komissarova, Dmitry Bazhenov, Daria Danilenko |
| EPI_ISL_596357, EPI_ISL_596358, EPI_ISL_596359, EPI_ISL_596360, EPI_ISL_596361, EPI_ISL_596362, EPI_ISL_596363, EPI_ISL_596364, EPI_ISL_596365, EPI_ISL_596366, EPI_ISL_596367, EPI_ISL_596368, EPI_ISL_596369, EPI_ISL_596370, EPI_ISL_596371, EPI_ISL_596372, EPI_ISL_596373, EPI_ISL_596374, EPI_ISL_596375, EPI_ISL_596376, EPI_ISL_596377 | | | |
| see above | HELIX LCC | WHO National Influenza Centre Russian Federation | Andrey Komissarov, Artem Fadeev, Anna Ivanova, Kseniya Komissarova, Dmitry Bazhenov, Daria Danilenko |
| EPI_ISL_602329, EPI_ISL_602330, EPI_ISL_602331, EPI_ISL_602332, EPI_ISL_602333, EPI_ISL_602334, EPI_ISL_602335, EPI_ISL_602336, EPI_ISL_602337, EPI_ISL_602338, EPI_ISL_602339, EPI_ISL_602340, EPI_ISL_602341, EPI_ISL_602342, EPI_ISL_602343, EPI_ISL_602344, EPI_ISL_602345, EPI_ISL_602346, EPI_ISL_602347, EPI_ISL_602348, EPI_ISL_602349, EPI_ISL_602350, EPI_ISL_602351, EPI_ISL_602352, EPI_ISL_602353, EPI_ISL_602354, EPI_ISL_602355, EPI_ISL_602356, EPI_ISL_602357, EPI_ISL_602358, EPI_ISL_602359, EPI_ISL_602360, EPI_ISL_602361, EPI_ISL_602362, EPI_ISL_602363, EPI_ISL_602364, EPI_ISL_602365, EPI_ISL_602366, EPI_ISL_602367, EPI_ISL_602368, EPI_ISL_602369, EPI_ISL_602370, EPI_ISL_602371, EPI_ISL_602372, EPI_ISL_602373, EPI_ISL_602374, EPI_ISL_602375, EPI_ISL_602376, EPI_ISL_602377, EPI_ISL_602378, EPI_ISL_602379, EPI_ISL_602380, EPI_ISL_602381, EPI_ISL_602382, EPI_ISL_602383, EPI_ISL_602384, EPI_ISL_602385, EPI_ISL_602386, EPI_ISL_602387, EPI_ISL_602388, EPI_ISL_602389, EPI_ISL_602390, EPI_ISL_602391, EPI_ISL_602392, EPI_ISL_602393, EPI_ISL_602394, EPI_ISL_602395, EPI_ISL_602396, EPI_ISL_602397, EPI_ISL_602398, EPI_ISL_602399, EPI_ISL_602400, EPI_ISL_602401, EPI_ISL_602402, EPI_ISL_602403, EPI_ISL_602404, EPI_ISL_602405, EPI_ISL_602406, EPI_ISL_602407, EPI_ISL_602408, EPI_ISL_602409, EPI_ISL_602410, EPI_ISL_602411 | | | |
| see above | HELIX LLC | WHO National Influenza Centre Russian Federation | Andrey Komissarov, Artem Fadeev, Kseniya Komissarova, Anna Ivanova, Dmitry Bazhenov, Daria Danilenko |
| EPI_ISL_602413, EPI_ISL_602414, EPI_ISL_602415, EPI_ISL_602416 | WHO National Influenza Centre Russian Federation | WHO National Influenza Centre Russian Federation | Andrey Komissarov, Artem Fadeev, Kseniya Komissarova, Anna Ivanova, Dmitry Bazhenov, Daria Danilenko |
| EPI_ISL_602417, EPI_ISL_602418, EPI_ISL_602419, EPI_ISL_602420, EPI_ISL_602421, EPI_ISL_602422, EPI_ISL_602423, EPI_ISL_602424, EPI_ISL_602425, EPI_ISL_602426, EPI_ISL_602427, EPI_ISL_602428, EPI_ISL_602429, EPI_ISL_602430, EPI_ISL_602431, EPI_ISL_602432, EPI_ISL_602433, EPI_ISL_602434, EPI_ISL_602435, EPI_ISL_602436, EPI_ISL_602437, EPI_ISL_602438, EPI_ISL_602439, EPI_ISL_602440, EPI_ISL_602441, EPI_ISL_602442, EPI_ISL_602443, EPI_ISL_602444, EPI_ISL_602445, EPI_ISL_602446, EPI_ISL_602447, EPI_ISL_602448, EPI_ISL_602449, EPI_ISL_602450, EPI_ISL_602451, EPI_ISL_602452, EPI_ISL_602453, EPI_ISL_602454, EPI_ISL_602455, EPI_ISL_602456, EPI_ISL_602457, EPI_ISL_602458, EPI_ISL_602459, EPI_ISL_602460, EPI_ISL_602461 | | | |
| see above | HELIX LLC | WHO National Influenza Centre Russian Federation | Andrey Komissarov, Artem Fadeev, Kseniya Komissarova, Anna Ivanova, Dmitry Bazhenov, Daria Danilenko |
| EPI_ISL_603221, EPI_ISL_603222 | National Institute of Laboratory Medicine and Referral Center | Genomic Research Lab, BCSIR | Abu Sayeed Mohammad Mahmud, Mohammad Samir Uzzaman, Eshrar Osman, Md. Ahashan Habib, Shahina Akter, Tanjina Akhter Banu, Md. Murshed Hasan Sarkar, Barna Goswami, Iffat Jahan, Md. Saddam Hossain, Tasnim Nafisa, Md. Maruf Ahmed Molla, Mahmuda Yeasmin, Asish Kumar Ghosh, A. K. M. Shamsuzzaman, Monira Parveen, Md. Masum Hossain Arif, Md. Salim Khan |
| EPI_ISL_611533, EPI_ISL_611547, EPI_ISL_611548 | West of Scotland Specialist Virology Centre, NHSGGC / MRC-University of Glasgow Centre for Virus Research | COVID-19 Genomics UK (COG-UK) Consortium | Ana da Silva Filipe, Natasha Johnson, Kathy Smollett, Daniel Mair, Stephen Carmichael, Lily Tong, Jenna Nichols, Elihu Aranday-Cortes, Kyriaki Nomikou; Sarah McDonald, Marc Niebel, Patawee Asamaphan; Richard Orton, Joseph Hughes, Sreenu Vattipally, David L Robertson; Alasdair MacLean, Rory Gunson; Kathy Li, Igor Starinskij, Natasha Jesudason, Rajiv Shah, James Shepherd, Antonia Ho, Emma Thomson |
| EPI_ISL_611593 | Department of Pathology, University of Cambridge | COVID-19 Genomics UK (COG-UK) Consortium | Aminu S. Jahun, Yasmin Chaudhry, Grant Hall, Iliana Georgana, Myra Hosmillo, Martin D. Curran, Malte Pinckert, Surendra Parmar, Ian Goodfellow |
| EPI_ISL_611610, EPI_ISL_611611, EPI_ISL_611614 | West of Scotland Specialist Virology Centre, NHSGGC / MRC-University of Glasgow Centre for Virus Research | COVID-19 Genomics UK (COG-UK) Consortium | Ana da Silva Filipe, Natasha Johnson, Kathy Smollett, Daniel Mair, Stephen Carmichael, Lily Tong, Jenna Nichols, Elihu Aranday-Cortes, Kyriaki Nomikou; Sarah McDonald, Marc Niebel, Patawee Asamaphan; Richard Orton, Joseph Hughes, Sreenu Vattipally, David L Robertson; Alasdair MacLean, Rory Gunson; Kathy Li, Igor Starinskij, Natasha Jesudason, Rajiv Shah, James Shepherd, Antonia Ho, Emma Thomson |
| EPI_ISL_611639 | Centre for Enzyme Innovation, University of Portsmouth / Translational Research Laboratory, Portsmouth Hospitals NHS Trust | COVID-19 Genomics UK (COG-UK) Consortium | Angela Beckett,Yann Bourgeois,Garry Scarlett,Sharon Glaysher,Scott Elliott,Kelly Bicknell,Robert Impey,Allyson Lloyd,Sarah Wyllie,Ethan Butcher,Anoop Chauhan,Samuel Robson |
| EPI_ISL_611660, EPI_ISL_611693, EPI_ISL_611708, EPI_ISL_611709, EPI_ISL_611710, EPI_ISL_611712, EPI_ISL_611713, EPI_ISL_611714, EPI_ISL_611715, EPI_ISL_611716, EPI_ISL_611717, EPI_ISL_611718, EPI_ISL_611719, EPI_ISL_611720 | | | |
| see above | West of Scotland Specialist Virology Centre, NHSGGC / MRC-University of Glasgow Centre for Virus Research | COVID-19 Genomics UK (COG-UK) Consortium | Ana da Silva Filipe, Natasha Johnson, Kathy Smollett, Daniel Mair, Stephen Carmichael, Lily Tong, Jenna Nichols, Elihu Aranday-Cortes, Kyriaki Nomikou; Sarah McDonald, Marc Niebel, Patawee Asamaphan; Richard Orton, Joseph Hughes, Sreenu Vattipally, David L Robertson; Alasdair MacLean, Rory Gunson; Kathy Li, Igor Starinskij, Natasha Jesudason, Rajiv Shah, James Shepherd, Antonia Ho, Emma Thomson |
| EPI_ISL_611751 | Department of Pathology, University of Cambridge | COVID-19 Genomics UK (COG-UK) Consortium | Aminu S. Jahun, Yasmin Chaudhry, Grant Hall, Iliana Georgana, Myra Hosmillo, Martin D. Curran, Malte Pinckert, Surendra Parmar, Ian Goodfellow |
| EPI_ISL_611754 | Centre for Enzyme Innovation, University of Portsmouth / Translational Research Laboratory, Portsmouth Hospitals NHS Trust | COVID-19 Genomics UK (COG-UK) Consortium | Angela Beckett,Yann Bourgeois,Garry Scarlett,Sharon Glaysher,Scott Elliott,Kelly Bicknell,Robert Impey,Allyson Lloyd,Sarah Wyllie,Ethan Butcher,Anoop Chauhan,Samuel Robson |
| EPI_ISL_611818, EPI_ISL_611820, EPI_ISL_611826, EPI_ISL_611848, EPI_ISL_611852, EPI_ISL_611866, EPI_ISL_611867, EPI_ISL_611870, EPI_ISL_611871, EPI_ISL_611872, EPI_ISL_611873, EPI_ISL_611874, EPI_ISL_611875, EPI_ISL_611876, EPI_ISL_611877, EPI_ISL_611878, EPI_ISL_611879, EPI_ISL_611880, EPI_ISL_611881, EPI_ISL_611882, EPI_ISL_611883, EPI_ISL_611884, EPI_ISL_611890 | | | |
| see above | West of Scotland Specialist Virology Centre, NHSGGC / MRC-University of Glasgow Centre for Virus Research | COVID-19 Genomics UK (COG-UK) Consortium | Ana da Silva Filipe, Natasha Johnson, Kathy Smollett, Daniel Mair, Stephen Carmichael, Lily Tong, Jenna Nichols, Elihu Aranday-Cortes, Kyriaki Nomikou; Sarah McDonald, Marc Niebel, Patawee Asamaphan; Richard Orton, Joseph Hughes, Sreenu Vattipally, David L Robertson; Alasdair MacLean, Rory Gunson; Kathy Li, Igor Starinskij, Natasha Jesudason, Rajiv Shah, James Shepherd, Antonia Ho, Emma Thomson |
| EPI_ISL_611927, EPI_ISL_611928, EPI_ISL_611933 | Centre for Enzyme Innovation, University of Portsmouth / Translational Research Laboratory, Portsmouth Hospitals NHS Trust | COVID-19 Genomics UK (COG-UK) Consortium | Angela Beckett,Yann Bourgeois,Garry Scarlett,Sharon Glaysher,Scott Elliott,Kelly Bicknell,Robert Impey,Allyson Lloyd,Sarah Wyllie,Ethan Butcher,Anoop Chauhan,Samuel Robson |
| EPI_ISL_611938 | Department of Pathology, University of Cambridge | COVID-19 Genomics UK (COG-UK) Consortium | Aminu S. Jahun, Yasmin Chaudhry, Grant Hall, Iliana Georgana, Myra Hosmillo, Martin D. Curran, Malte Pinckert, Surendra Parmar, Ian Goodfellow |
| EPI_ISL_611957, EPI_ISL_611958 | Centre for Enzyme Innovation, University of Portsmouth / Translational Research Laboratory, Portsmouth Hospitals NHS Trust | COVID-19 Genomics UK (COG-UK) Consortium | Angela Beckett,Yann Bourgeois,Garry Scarlett,Sharon Glaysher,Scott Elliott,Kelly Bicknell,Robert Impey,Allyson Lloyd,Sarah Wyllie,Ethan Butcher,Anoop Chauhan,Samuel Robson |
| EPI_ISL_611964, EPI_ISL_611965, EPI_ISL_611966, EPI_ISL_611967 | Department of Pathology, University of Cambridge | COVID-19 Genomics UK (COG-UK) Consortium | Aminu S. Jahun, Yasmin Chaudhry, Grant Hall, Iliana Georgana, Myra Hosmillo, Martin D. Curran, Malte Pinckert, Surendra Parmar, Ian Goodfellow |
| EPI_ISL_611971 | Centre for Enzyme Innovation, University of Portsmouth / Translational Research Laboratory, Portsmouth Hospitals NHS Trust | COVID-19 Genomics UK (COG-UK) Consortium | Angela Beckett,Yann Bourgeois,Garry Scarlett,Sharon Glaysher,Scott Elliott,Kelly Bicknell,Robert Impey,Allyson Lloyd,Sarah Wyllie,Ethan Butcher,Anoop Chauhan,Samuel Robson |
| EPI_ISL_611990 | West of Scotland Specialist Virology Centre, NHSGGC / MRC-University of Glasgow Centre for Virus Research | COVID-19 Genomics UK (COG-UK) Consortium | Ana da Silva Filipe, Natasha Johnson, Kathy Smollett, Daniel Mair, Stephen Carmichael, Lily Tong, Jenna Nichols, Elihu Aranday-Cortes, Kyriaki Nomikou; Sarah McDonald, Marc Niebel, Patawee Asamaphan; Richard Orton, Joseph Hughes, Sreenu Vattipally, David L Robertson; Alasdair MacLean, Rory Gunson; Kathy Li, Igor Starinskij, Natasha Jesudason, Rajiv Shah, James Shepherd, Antonia Ho, Emma Thomson |
| EPI_ISL_612007, EPI_ISL_612034 | Department of Pathology, University of Cambridge | COVID-19 Genomics UK (COG-UK) Consortium | Aminu S. Jahun, Yasmin Chaudhry, Grant Hall, Iliana Georgana, Myra Hosmillo, Martin D. Curran, Malte Pinckert, Surendra Parmar, Ian Goodfellow |

| | | | |
|---|---|---|---|
| EPI_ISL_612063 | West of Scotland Specialist Virology Centre, NHSGGC / MRC-University of Glasgow Centre for Virus Research | COVID-19 Genomics UK (COG-UK) Consortium | Ana da Silva Filipe, Natasha Johnson, Kathy Smollett, Daniel Mair, Stephen Carmichael, Lily Tong, Jenna Nichols, Elihu Aranday-Cortes, Kyriaki Nomikou; Sarah McDonald, Marc Niebel, Patawee Asamaphan; Richard Orton, Joseph Hughes, Sreenu Vattipally, David L Robertson; Alasdair MacLean, Rory Gunson; Kathy Li, Igor Starinskij, Natasha Jesudason, Rajiv Shah, James Shepherd, Antonia Ho, Emma Thomson |
| EPI_ISL_612099, EPI_ISL_612106 | Department of Pathology, University of Cambridge | COVID-19 Genomics UK (COG-UK) Consortium | Aminu S. Jahun, Yasmin Chaudhry, Grant Hall, Iliana Georgana, Myra Hosmillo, Martin D. Curran, Malte Pinckert, Surendra Parmar, Ian Goodfellow |
| EPI_ISL_612113 | West of Scotland Specialist Virology Centre, NHSGGC / MRC-University of Glasgow Centre for Virus Research | COVID-19 Genomics UK (COG-UK) Consortium | Ana da Silva Filipe, Natasha Johnson, Kathy Smollett, Daniel Mair, Stephen Carmichael, Lily Tong, Jenna Nichols, Elihu Aranday-Cortes, Kyriaki Nomikou; Sarah McDonald, Marc Niebel, Patawee Asamaphan; Richard Orton, Joseph Hughes, Sreenu Vattipally, David L Robertson; Alasdair MacLean, Rory Gunson; Kathy Li, Igor Starinskij, Natasha Jesudason, Rajiv Shah, James Shepherd, Antonia Ho, Emma Thomson |
| EPI_ISL_612123 | Department of Pathology, University of Cambridge | COVID-19 Genomics UK (COG-UK) Consortium | Aminu S. Jahun, Yasmin Chaudhry, Grant Hall, Iliana Georgana, Myra Hosmillo, Martin D. Curran, Malte Pinckert, Surendra Parmar, Ian Goodfellow |
| EPI_ISL_612124, EPI_ISL_612125 | West of Scotland Specialist Virology Centre, NHSGGC / MRC-University of Glasgow Centre for Virus Research | COVID-19 Genomics UK (COG-UK) Consortium | Ana da Silva Filipe, Natasha Johnson, Kathy Smollett, Daniel Mair, Stephen Carmichael, Lily Tong, Jenna Nichols, Elihu Aranday-Cortes, Kyriaki Nomikou; Sarah McDonald, Marc Niebel, Patawee Asamaphan; Richard Orton, Joseph Hughes, Sreenu Vattipally, David L Robertson; Alasdair MacLean, Rory Gunson; Kathy Li, Igor Starinskij, Natasha Jesudason, Rajiv Shah, James Shepherd, Antonia Ho, Emma Thomson |
| EPI_ISL_612186, EPI_ISL_612262, EPI_ISL_612263, EPI_ISL_612264, EPI_ISL_612265, EPI_ISL_612266, EPI_ISL_612267, EPI_ISL_612268, EPI_ISL_612269, EPI_ISL_612270, EPI_ISL_612271, EPI_ISL_612272, EPI_ISL_612273, EPI_ISL_612274, EPI_ISL_612275, EPI_ISL_612276, EPI_ISL_612277, EPI_ISL_612278, EPI_ISL_612279, EPI_ISL_612280, EPI_ISL_612281, EPI_ISL_612282, EPI_ISL_612283 | | | |
| see above | Department of Pathology, University of Cambridge | COVID-19 Genomics UK (COG-UK) Consortium | Aminu S. Jahun, Yasmin Chaudhry, Grant Hall, Iliana Georgana, Myra Hosmillo, Martin D. Curran, Malte Pinckert, Surendra Parmar, Ian Goodfellow |
| EPI_ISL_612284, EPI_ISL_612285, EPI_ISL_612286, EPI_ISL_612287, EPI_ISL_612288, EPI_ISL_612289, EPI_ISL_612290, EPI_ISL_612291, EPI_ISL_612292, EPI_ISL_612293, EPI_ISL_612294, EPI_ISL_612295, EPI_ISL_612296, EPI_ISL_612297, EPI_ISL_612298, EPI_ISL_612299, EPI_ISL_612300, EPI_ISL_612301, EPI_ISL_612302, EPI_ISL_612303, EPI_ISL_612304, EPI_ISL_612305, EPI_ISL_612306, EPI_ISL_612307, EPI_ISL_612308, EPI_ISL_612309, EPI_ISL_612310, EPI_ISL_612311, EPI_ISL_612312, EPI_ISL_612313 | | | |
| see above | West of Scotland Specialist Virology Centre, NHSGGC / MRC-University of Glasgow Centre for Virus Research | COVID-19 Genomics UK (COG-UK) Consortium | Ana da Silva Filipe, Natasha Johnson, Kathy Smollett, Daniel Mair, Stephen Carmichael, Lily Tong, Jenna Nichols, Elihu Aranday-Cortes, Kyriaki Nomikou; Sarah McDonald, Marc Niebel, Patawee Asamaphan; Richard Orton, Joseph Hughes, Sreenu Vattipally, David L Robertson; Alasdair MacLean, Rory Gunson; Kathy Li, Igor Starinskij, Natasha Jesudason, Rajiv Shah, James Shepherd, Antonia Ho, Emma Thomson |
| EPI_ISL_613282, EPI_ISL_613283, EPI_ISL_613284, EPI_ISL_613286, EPI_ISL_613287, EPI_ISL_613288, EPI_ISL_613289, EPI_ISL_613290, EPI_ISL_613291, EPI_ISL_613292, EPI_ISL_613293 | | | |
| see above | Centre for Enzyme Innovation, University of Portsmouth / Translational Research Laboratory, Portsmouth Hospitals NHS Trust | COVID-19 Genomics UK (COG-UK) Consortium | Angela Beckett,Yann Bourgeois,Garry Scarlett,Sharon Glaysher,Scott Elliott,Kelly Bicknell,Robert Impey,Allyson Lloyd,Sarah Wyllie,Ethan Butcher,Anoop Chauhan,Samuel Robson |
| EPI_ISL_613457 | QUARANTINE CAMP | Hong Kong Department of Health | Alan K.L. Tsang, Peter C.W. Yip, Edman T.K. Lam, Rickjason C.W. Chan, Dominic N.C. Tsang |
| EPI_ISL_622894, EPI_ISL_622895, EPI_ISL_622896, EPI_ISL_622897, EPI_ISL_622898, EPI_ISL_622899, EPI_ISL_622900, EPI_ISL_622901, EPI_ISL_622902, EPI_ISL_622903, EPI_ISL_622904, EPI_ISL_622905, EPI_ISL_622906, EPI_ISL_622907, EPI_ISL_622908, EPI_ISL_622909, EPI_ISL_622910, EPI_ISL_622911, EPI_ISL_622912, EPI_ISL_622913, EPI_ISL_622914, EPI_ISL_622915, EPI_ISL_622916, EPI_ISL_622917, EPI_ISL_622918, EPI_ISL_622919, EPI_ISL_622920, EPI_ISL_622921, EPI_ISL_622922, EPI_ISL_622923, EPI_ISL_622924, EPI_ISL_622925, EPI_ISL_622926, EPI_ISL_622927, EPI_ISL_622928, EPI_ISL_622929, EPI_ISL_622930, EPI_ISL_622931, EPI_ISL_622932, EPI_ISL_622933, EPI_ISL_622934, EPI_ISL_622935, EPI_ISL_622936, EPI_ISL_622937, EPI_ISL_622938, EPI_ISL_622939, EPI_ISL_622940, EPI_ISL_622941 | | | |
| see above | National Institute for Communicable Diseases of the National Health Laboratory Service | National Institute for Communicable Diseases of the National Health Laboratory Service | Allam M, Ismail A, Khumalo Z, Kwenda S, Mtshali P, Mnyameni F, Mohale T, Subramoney K, Bhiman JN |
| EPI_ISL_622942 | National Health Laboratory Service | National Institute for Communicable Diseases of the National Health Laboratory Service | Allam M, Ismail A, Khumalo Z, Kwenda S, Mtshali P, Mnyameni F, Mohale T, Subramoney K, Bhiman JN |
| EPI_ISL_622943 | Lancet Laboratories | National Institute for Communicable Diseases of the National Health Laboratory Service | Allam M, Ismail A, Khumalo Z, Kwenda S, Mtshali P, Mnyameni F, Mohale T, Subramoney K, Bhiman JN |
| EPI_ISL_622944, EPI_ISL_622945 | National Health Laboratory Service | National Institute for Communicable Diseases of the National Health Laboratory Service | Allam M, Ismail A, Khumalo Z, Kwenda S, Mtshali P, Mnyameni F, Mohale T, Subramoney K, Bhiman JN |
| EPI_ISL_622946, EPI_ISL_622947, EPI_ISL_622948, EPI_ISL_622949, EPI_ISL_622950, EPI_ISL_622951, EPI_ISL_622952, EPI_ISL_622953 | Lancet Laboratories | National Institute for Communicable Diseases of the National Health Laboratory Service | Allam M, Ismail A, Khumalo Z, Kwenda S, Mtshali P, Mnyameni F, Mohale T, Subramoney K, Bhiman JN |
| EPI_ISL_622954 | National Health Laboratory Service | National Institute for Communicable Diseases of the National Health Laboratory Service | Allam M, Ismail A, Khumalo Z, Kwenda S, Mtshali P, Mnyameni F, Mohale T, Subramoney K, Bhiman JN |
| EPI_ISL_622955, EPI_ISL_622957 | Lancet Laboratories | National Institute for Communicable Diseases of the National Health Laboratory Service | Allam M, Ismail A, Khumalo Z, Kwenda S, Mtshali P, Mnyameni F, Mohale T, Subramoney K, Bhiman JN |
| EPI_ISL_622958 | National Health Laboratory Service | National Institute for Communicable Diseases of the National Health Laboratory Service | Allam M, Ismail A, Khumalo Z, Kwenda S, Mtshali P, Mnyameni F, Mohale T, Subramoney K, Bhiman JN |
| EPI_ISL_622959, EPI_ISL_622960, EPI_ISL_622961, EPI_ISL_622962, EPI_ISL_622963, EPI_ISL_622964, EPI_ISL_622965, EPI_ISL_622966, EPI_ISL_622967, EPI_ISL_622968, EPI_ISL_622969 | | | |
| see above | Lancet Laboratories | National Institute for Communicable Diseases of the National Health Laboratory Service | Allam M, Ismail A, Khumalo Z, Kwenda S, Mtshali P, Mnyameni F, Mohale T, Subramoney K, Bhiman JN |
| EPI_ISL_622970, EPI_ISL_622971, EPI_ISL_622972, EPI_ISL_622973, EPI_ISL_622974, EPI_ISL_622975, EPI_ISL_622976, EPI_ISL_622977 | National Health Laboratory Service | National Institute for Communicable Diseases of the National Health Laboratory Service | Allam M, Ismail A, Khumalo Z, Kwenda S, Mtshali P, Mnyameni F, Mohale T, Subramoney K, Bhiman JN |
| EPI_ISL_622978 | Lancet Laboratories | National Institute for Communicable Diseases of the National Health Laboratory Service | Allam M, Ismail A, Khumalo Z, Kwenda S, Mtshali P, Mnyameni F, Mohale T, Subramoney K, Bhiman JN |
| EPI_ISL_622979, EPI_ISL_622980, EPI_ISL_622981, EPI_ISL_622982 | National Health Laboratory Service | National Institute for Communicable Diseases of the National Health Laboratory Service | Allam M, Ismail A, Khumalo Z, Kwenda S, Mtshali P, Mnyameni F, Mohale T, Subramoney K, Bhiman JN |
| EPI_ISL_622983 | Lancet Laboratories | National Institute for Communicable Diseases of the National Health Laboratory Service | Allam M, Ismail A, Khumalo Z, Kwenda S, Mtshali P, Mnyameni F, Mohale T, Subramoney K, Bhiman JN |
| EPI_ISL_622984 | National Health Laboratory Service | National Institute for Communicable Diseases of the National Health Laboratory Service | Allam M, Ismail A, Khumalo Z, Kwenda S, Mtshali P, Mnyameni F, Mohale T, Subramoney K, Bhiman JN |
| EPI_ISL_622985, EPI_ISL_622986 | Lancet Laboratories | National Institute for Communicable Diseases of the National Health Laboratory Service | Allam M, Ismail A, Khumalo Z, Kwenda S, Mtshali P, Mnyameni F, Mohale T, Subramoney K, Bhiman JN |
| EPI_ISL_622987 | National Health Laboratory Service | National Institute for Communicable Diseases of the National Health Laboratory Service | Allam M, Ismail A, Khumalo Z, Kwenda S, Mtshali P, Mnyameni F, Mohale T, Subramoney K, Bhiman JN |
| EPI_ISL_622988 | Lancet Laboratories | National Institute for Communicable Diseases of the National Health Laboratory Service | Allam M, Ismail A, Khumalo Z, Kwenda S, Mtshali P, Mnyameni F, Mohale T, Subramoney K, Bhiman JN |
| EPI_ISL_622989 | National Health Laboratory Service | National Institute for Communicable Diseases of the National Health Laboratory Service | Allam M, Ismail A, Khumalo Z, Kwenda S, Mtshali P, Mnyameni F, Mohale T, Subramoney K, Bhiman JN |
| EPI_ISL_622990 | Lancet Laboratories | National Institute for Communicable Diseases of the National Health Laboratory Service | Allam M, Ismail A, Khumalo Z, Kwenda S, Mtshali P, Mnyameni F, Mohale T, Subramoney K, Bhiman JN |
| EPI_ISL_622991, EPI_ISL_622992, | National Health Laboratory Service | National Institute for Communicable Diseases of the National | Allam M, Ismail A, Khumalo Z, Kwenda S, Mtshali P, Mnyameni F, Mohale T, Subramoney K, Bhiman JN |

| | | | |
|---|---|---|---|
| EPI_ISL_622993 | | Health Laboratory Service | |
| EPI_ISL_622994 | Lancet Laboratories | National Institute for Communicable Diseases of the National Health Laboratory Service | Allam M, Ismail A, Khumalo Z, Kwenda S, Mtshali P, Mnyameni F, Mohale T, Subramoney K, Bhiman JN |
| EPI_ISL_622995, EPI_ISL_622996, EPI_ISL_622997, EPI_ISL_622998, EPI_ISL_622999, EPI_ISL_623001, EPI_ISL_623002, EPI_ISL_623003, EPI_ISL_623004, EPI_ISL_623005, EPI_ISL_623007, EPI_ISL_623008, EPI_ISL_623009, EPI_ISL_623010, EPI_ISL_623011, EPI_ISL_623012, EPI_ISL_623013, EPI_ISL_623014, EPI_ISL_623015, EPI_ISL_623016, EPI_ISL_623017, EPI_ISL_623018, EPI_ISL_623019, EPI_ISL_623020, EPI_ISL_623021, EPI_ISL_623022, EPI_ISL_623023, EPI_ISL_623026, EPI_ISL_623027, EPI_ISL_623029, EPI_ISL_623030, EPI_ISL_623031, EPI_ISL_623032, EPI_ISL_623033, EPI_ISL_623034, EPI_ISL_623035, EPI_ISL_623036, EPI_ISL_623037, EPI_ISL_623038, EPI_ISL_623039, EPI_ISL_623040, EPI_ISL_623041, EPI_ISL_623043, EPI_ISL_623044, EPI_ISL_623045, EPI_ISL_623046, EPI_ISL_623047, EPI_ISL_623048, EPI_ISL_623049, EPI_ISL_623050, EPI_ISL_623051, EPI_ISL_623053, EPI_ISL_623054, EPI_ISL_623055, EPI_ISL_623056, EPI_ISL_623057, EPI_ISL_623060, EPI_ISL_623062 | | | |
| see above | National Health Laboratory Service | National Institute for Communicable Diseases of the National Health Laboratory Service | Allam M, Ismail A, Khumalo Z, Kwenda S, Mtshali P, Mnyameni F, Mohale T, Subramoney K, Bhiman JN |
| EPI_ISL_623064 | Lancet Laboratories | National Institute for Communicable Diseases of the National Health Laboratory Service | Allam M, Ismail A, Khumalo Z, Kwenda S, Mtshali P, Mnyameni F, Mohale T, Subramoney K, Bhiman JN |
| EPI_ISL_623065, EPI_ISL_623066 | National Health Laboratory Service | National Institute for Communicable Diseases of the National Health Laboratory Service | Allam M, Ismail A, Khumalo Z, Kwenda S, Mtshali P, Mnyameni F, Mohale T, Subramoney K, Bhiman JN |
| EPI_ISL_623068 | Lancet Laboratories | National Institute for Communicable Diseases of the National Health Laboratory Service | Allam M, Ismail A, Khumalo Z, Kwenda S, Mtshali P, Mnyameni F, Mohale T, Subramoney K, Bhiman JN |
| EPI_ISL_623069, EPI_ISL_623070, EPI_ISL_623071, EPI_ISL_623072 | National Health Laboratory Service | National Institute for Communicable Diseases of the National Health Laboratory Service | Allam M, Ismail A, Khumalo Z, Kwenda S, Mtshali P, Mnyameni F, Mohale T, Subramoney K, Bhiman JN |
| EPI_ISL_623073 | Lancet Laboratories | National Institute for Communicable Diseases of the National Health Laboratory Service | Allam M, Ismail A, Khumalo Z, Kwenda S, Mtshali P, Mnyameni F, Mohale T, Subramoney K, Bhiman JN |
| EPI_ISL_626687, EPI_ISL_626691, EPI_ISL_626692, EPI_ISL_626749, EPI_ISL_626758, EPI_ISL_626812, EPI_ISL_626818, EPI_ISL_626822, EPI_ISL_626825, EPI_ISL_626826, EPI_ISL_626831, EPI_ISL_626835, EPI_ISL_626837, EPI_ISL_626838, EPI_ISL_626839, EPI_ISL_626840, EPI_ISL_626841, EPI_ISL_626842, EPI_ISL_626843, EPI_ISL_626844, EPI_ISL_626845, EPI_ISL_626846, EPI_ISL_626847, EPI_ISL_626848, EPI_ISL_626849, EPI_ISL_626851, EPI_ISL_626855, EPI_ISL_626873, EPI_ISL_626897, EPI_ISL_626904, EPI_ISL_626917, EPI_ISL_626927, EPI_ISL_626933, EPI_ISL_626934, EPI_ISL_626935, EPI_ISL_626952, EPI_ISL_626957, EPI_ISL_626958, EPI_ISL_626962, EPI_ISL_626963, EPI_ISL_626966, EPI_ISL_626967, EPI_ISL_626968, EPI_ISL_626969, EPI_ISL_626970, EPI_ISL_626971, EPI_ISL_626972, EPI_ISL_626973, EPI_ISL_626974, EPI_ISL_626975, EPI_ISL_626976, EPI_ISL_626977, EPI_ISL_626978, EPI_ISL_626979, EPI_ISL_626980, EPI_ISL_626981, EPI_ISL_626982, EPI_ISL_626983, EPI_ISL_626984, EPI_ISL_626985, EPI_ISL_626986, EPI_ISL_626987, EPI_ISL_626988, EPI_ISL_626989, EPI_ISL_626990, EPI_ISL_626991, EPI_ISL_626992, EPI_ISL_626995, EPI_ISL_627014, EPI_ISL_627030, EPI_ISL_627040, EPI_ISL_627045, EPI_ISL_627049, EPI_ISL_627063, EPI_ISL_627117, EPI_ISL_627165, EPI_ISL_627173, EPI_ISL_627195, EPI_ISL_627200, EPI_ISL_627201, EPI_ISL_627202, EPI_ISL_627208, EPI_ISL_627209, EPI_ISL_627329, EPI_ISL_627330, EPI_ISL_627331, EPI_ISL_627332, EPI_ISL_627333, EPI_ISL_627334, EPI_ISL_627335, EPI_ISL_627336, EPI_ISL_627337, EPI_ISL_627338, EPI_ISL_627339, EPI_ISL_627340, EPI_ISL_627341, EPI_ISL_627342, EPI_ISL_627343, EPI_ISL_627344, EPI_ISL_627345, EPI_ISL_627346, EPI_ISL_627347, EPI_ISL_627348, EPI_ISL_627349, EPI_ISL_627350, EPI_ISL_627351, EPI_ISL_627352, EPI_ISL_627353, EPI_ISL_627354, EPI_ISL_627355, EPI_ISL_627356, EPI_ISL_627357, EPI_ISL_627358, EPI_ISL_627359, EPI_ISL_627360, EPI_ISL_627361, EPI_ISL_627362, EPI_ISL_627363, EPI_ISL_627364, EPI_ISL_627365, EPI_ISL_627366, EPI_ISL_627367, EPI_ISL_627368, EPI_ISL_627369, EPI_ISL_627370, EPI_ISL_627371, EPI_ISL_627372, EPI_ISL_627373, EPI_ISL_627374, EPI_ISL_627375, EPI_ISL_627376, EPI_ISL_627377, EPI_ISL_627378, EPI_ISL_627379, EPI_ISL_627380, EPI_ISL_627381, EPI_ISL_627382, EPI_ISL_627383, EPI_ISL_627384, EPI_ISL_627385, EPI_ISL_627386, EPI_ISL_627387, EPI_ISL_627388, EPI_ISL_627389, EPI_ISL_627390, EPI_ISL_627391, EPI_ISL_627392, EPI_ISL_627393, EPI_ISL_627394, EPI_ISL_627395, EPI_ISL_627396, EPI_ISL_627397, EPI_ISL_627398, EPI_ISL_627399, EPI_ISL_627400, EPI_ISL_627401, EPI_ISL_627402, EPI_ISL_627403, EPI_ISL_627404, EPI_ISL_627405, EPI_ISL_627406, EPI_ISL_627407, EPI_ISL_627408, EPI_ISL_627409, EPI_ISL_627410, EPI_ISL_627411, EPI_ISL_627412, EPI_ISL_627413, EPI_ISL_627414, EPI_ISL_627415, EPI_ISL_627416, EPI_ISL_627417, EPI_ISL_627418, EPI_ISL_627419, EPI_ISL_627420, EPI_ISL_627421, EPI_ISL_627422, EPI_ISL_627423 | | | |
| see above | West of Scotland Specialist Virology Centre, NHSGGC / MRC-University of Glasgow Centre for Virus Research | COVID-19 Genomics UK (COG-UK) Consortium | Ana da Silva Filipe, Natasha Johnson, Kathy Smollett, Daniel Mair, Stephen Carmichael, Lily Tong, Jenna Nichols, Elihu Aranday-Cortes, Kyriaki Nomikou; Sarah McDonald, Marc Niebel, Patawee Asamaphan; Richard Orton, Joseph Hughes, Sreenu Vattipally, David L Robertson; Alasdair MacLean, Rory Gunson; Kathy Li, Igor Starinskij, Natasha Jesudason, Rajiv Shah, James Shepherd, Antonia Ho, Emma Thomson |
| EPI_ISL_628312, EPI_ISL_628313, EPI_ISL_628314, EPI_ISL_628315, EPI_ISL_628316, EPI_ISL_628317, EPI_ISL_628318, EPI_ISL_628319 | Centre for Enzyme Innovation, University of Portsmouth / Translational Research Laboratory, Portsmouth Hospitals NHS Trust | COVID-19 Genomics UK (COG-UK) Consortium | Angela Beckett,Yann Bourgeois,Garry Scarlett,Sharon Glaysher,Scott Elliott,Kelly Bicknell,Robert Impey,Allyson Lloyd,Sarah Wyllie,Ethan Butcher,Anoop Chauhan,Samuel Robson |
| EPI_ISL_629000 | West of Scotland Specialist Virology Centre, NHSGGC | MRC-University of Glasgow Centre for Virus Research | Ana da Silva Filipe, Kathy Smollett, Stephen Carmichael, Natasha Johnson, Daniel Mair, Lily Tong, Jenna Nichols; Sarah McDonald; Richard Orton, Joseph Hughes, Sreenu Vattipally, David L Robertson; Kathy Li, Natasha Jesudason, Rajiv Shah, James Shepherd, Antonia Ho, Emma Thomson; Alasdair MacLean, Rory Gunson. |
| EPI_ISL_632242, EPI_ISL_632243, EPI_ISL_632244, EPI_ISL_632245, EPI_ISL_632246, EPI_ISL_632247, EPI_ISL_632248, EPI_ISL_632249 | NHSGGC West of Scotland Specialist Virology Centre | MRC-University of Glasgow Centre for Virus Research | Ana da Silva Filipe, Natasha Johnson, Kathy Smollett, Daniel Mair, Stephen Carmichael, Lily Tong, Jenna Nichols, Kyriaki Nomikou; Richard Orton, Joseph Hughes, Sreenu Vattipally, David L Robertson; Alasdair MacLean, Rory Gunson; Kathy Li, Natasha Jesudason, Rajiv Shah, James Shepherd, Antonia Ho, Emma Thomson; Alex Alderton, Roberto Amato, Sonia Goncalves, Ewan Harrison, David K. Jackson, Ian Johnston, Dominic Kwiatkowski, Cordelia Langford, John Sillitoe on behalf of the Wellcome Sanger Institute COVID-19 Surveillance Team (http://www.sanger.ac.uk/covid-team) |
| EPI_ISL_632250, EPI_ISL_632251, EPI_ISL_632252, EPI_ISL_632253, EPI_ISL_632254, EPI_ISL_632255 | Communicable Disease Laboratory, Public Health Directorate | Communicable Disease Laboratory, Public Health Directorate | AlWasti,H., Altaif,Z., AlHujairi,Z., AlAbbas,Z. |
| EPI_ISL_632256, EPI_ISL_632257, EPI_ISL_632258 | Communicable Disease Laboratory, Public Health Directorate | Communicable Disease Laboratory, Public Health Directorate | AlHujairi,Z., Altaif,Z., AlWasti,H., AlAbbas,Z. |
| EPI_ISL_632259, EPI_ISL_632260 | Communicable Disease Laboratory, Public Health Directorate | Communicable Disease Laboratory, Public Health Directorate | AlTaif,Z., AlWasti,H., AlHujairi,Z., AlAbbas,Z. |
| EPI_ISL_632261, EPI_ISL_632262, EPI_ISL_632263, EPI_ISL_632264, EPI_ISL_632265, EPI_ISL_632266, EPI_ISL_632267, EPI_ISL_632268, EPI_ISL_632269, EPI_ISL_632270, EPI_ISL_632271, EPI_ISL_632272, EPI_ISL_632273, EPI_ISL_632274, EPI_ISL_632275, EPI_ISL_632276, EPI_ISL_632277, EPI_ISL_632278, EPI_ISL_632279, EPI_ISL_632280, EPI_ISL_632281, EPI_ISL_632282, EPI_ISL_632283, EPI_ISL_632284, EPI_ISL_632285 | | | |
| see above | Communicable Disease Laboratory, Public Health Directorate | Communicable Disease Laboratory, Public Health Directorate | AlWasti,H., AlTaif,Z., AlHujairi,Z., AlAbbas,Z. |
| EPI_ISL_632287 | Environmental and Global Health, University of Florida | Environmental and Global Health, University of Florida | Alam,M.M., Lauzardo,M., Elbadry,M.A., Stephenson,C.J., Gibson,J.C., Morris,J.G. and Lednicky,J.A. |
| EPI_ISL_632899, EPI_ISL_632900, EPI_ISL_632901, EPI_ISL_632902, EPI_ISL_632903 | Communicable Disease Laboratory, Public Health Directorate | Communicable Disease Laboratory, Public Health Directorate | AlAbbas,Z., Altaif,Z., AlWasti,H., Alhujairi,Z. |
| EPI_ISL_632904 | Communicable Disease Laboratory, Public Health Directorate | Communicable Disease Laboratory, Public Health Directorate | AlTaif,Z., AlHujairi,Z., AlWasti,H., AlAbbas,Z. |
| EPI_ISL_632905 | Communicable Disease Laboratory, Public Health Directorate | Communicable Disease Laboratory, Public Health Directorate | AlHujairi,Z., Altaif,Z., AlWasti,H., AlAbbas,Z. |
| EPI_ISL_632906, EPI_ISL_632907 | Communicable Disease Laboratory, Public Health Directorate | Communicable Disease Laboratory, Public Health Directorate | AlAbbas,Z., Altaif,Z., AlWasti,H., Alhujairi,Z. |
| EPI_ISL_632908 | Genomic Sciences, Rehman Medical Institute | Genomic Sciences, Rehman Medical Institute | Ali,J., Afridi,U.K., Haider,S.A., Sabiha,B., Jan,H. and Jehanzeb,V. |

We gratefully acknowledge the following Authors from the Originating laboratories responsible for obtaining the specimens, as well as the Submitting laboratories where the genome data were generated and shared via GISAID, on which this research is based.

All Submitters of data may be contacted directly via www.gisaid.org

Authors are sorted alphabetically.

| Accession ID | Originating Laboratory | Submitting Laboratory | Authors |
|---|---|---|---|
| EPI_ISL_402132 | Wuhan Jinyintan Hospital | Hubei Provincial Center for Disease Control and Prevention | Bin Fang, Xiang Li, Xiao Yu, Linlin Liu, Bo Yang, Faxian Zhan, Guojun Ye, Xixiang Huo, Junqiang Xu, Bo Yu, Kun Cai, Jing Li, Yongzhong Jiang. |
| EPI_ISL_406034, EPI_ISL_406036 | California Department of Public Health | Pathogen Discovery, Respiratory Viruses Branch, Division of Viral Diseases, Centers for Dieases Control and Prevention | Anna Uehara, Krista Queen, Ying Tao, Yan Li, Clinton R. Paden, Jing Zhang, Xiaoyan Lu, Brian Lynch, Senthil Kumar K. Sakthivel, Brett L. Whitaker, Shifaq Kamili, Lijuan Wang, Janna' R. Murray, Susan I. Gerber, Stephen Lindstrom, Suxiang Tong |
| EPI_ISL_407894, EPI_ISL_407896 | Pathology Queensland | Public Health Virology Laboratory | Ben Huang, Alyssa Pyke, Amanda De Jong, Andrew Van Den Hurk, Carmel Taylor, David Warrilow, Doris Genge, Elisabeth Gamez, Glen Hewitson, Ian Maxwell Mackay, Inga Sultana, Jamie McMahon, Jean Barcelon, Judy Northill, Mitchell Finger, Natalie Simpson, Neelima Nair, Peter Burtonclay, Peter Moore, Sarah Wheatley, Sean Moody, Sonja Hall-Mendelin, Timothy Gardam, and Frederick Moore. |
| EPI_ISL_407976 | KU Leuven, Clinical and Epidemiological Virology | KU Leuven, Clinical and Epidemiological Virology | Bert Vanmechelen, Elke Wollants, Annabel Rector, Els Keyaerts, Lies Laenen, Marc Van Ranst, and Piet Maes |
| EPI_ISL_410486 | CNR Virus des Infections Respiratoires - France SUD | CNR Virus des Infections Respiratoires - France SUD | Bal, Antonin; Destras, Gregory; Gaymard, Alexandre; Bouscambert-Duchamp, Maude; Cheynet, Valérie; Brengel-Pesce, Karen; Morfin-Sherpa, Florence; Valette, Martine; Josset, Laurence; Lina, Bruno. |
| EPI_ISL_410717, EPI_ISL_410718 | Pathology Queensland | Public Health Virology Laboratory | Ben Huang, Alyssa Pyke, Amanda De Jong, Andrew Van Den Hurk, Carmel Taylor, David Warrilow, Doris Genge, Elisabeth Gamez, Glen Hewitson, Ian Maxwell Mackay, Inga Sultana, Jamie McMahon, Jean Barcelon, Judy Northill, Mitchell Finger, Natalie Simpson, Neelima Nair, Peter Burtonclay, Peter Moore, Sarah Wheatley, Sean Moody, Sonja Hall-Mendelin, Timothy Gardam, and Frederick Moore. |
| EPI_ISL_411951 | Unit for Laboratory Development and Technology Transfer, Public Health Agency of Sweden | Unit for Laboratory Development and Technology Transfer, Public Health Agency of Sweden | Bengner,M., Palmerus,M., Lindsjo,O., Lind Karlberg,M., Monteil,V., Appelberg,S., Brave,A., Muradrasoli,S. and Tegmark-Wisell,K. |
| EPI_ISL_412459 | Jingzhou Center for Disease Control and Prevention | Hubei Provincial Center for Disease Control and Prevention | Bin Fang, Xiang Li, Xiao Yu, Linlin Liu, Bo Yang, Faxian Zhan, Guojun Ye, Xixiang Huo, Junqiang Xu, Bo Yu, Kun Cai, Jing Li, Maoyi Chen,Jie Hu, Chunlin Mao, Yongzhong Jiang. |
| EPI_ISL_412898, EPI_ISL_412899, EPI_ISL_412900 | Wuhan Jinyintan Hospital | Hubei Provincial Center for Disease Control and Prevention | Bin Fang, Xiang Li, Xiao Yu, Linlin Liu, Bo Yang, Faxian Zhan, Guojun Ye, Xixiang Huo, Junqiang Xu, Bo Yu, Kun Cai, Jing Li, Yongzhong Jiang. |
| EPI_ISL_412978 | The Central Hospital Of Wuhan | Hubei Provincial Center for Disease Control and Prevention | Bin Fang, Xiang Li, Xiao Yu, Linlin Liu, Bo Yang, Faxian Zhan, Guojun Ye, Xixiang Huo, Junqiang Xu, Bo Yu, Kun Cai, Jing Li, Yongzhong Jiang. |
| EPI_ISL_412979, EPI_ISL_412980 | Union Hospital of Tongji Medical College, Huazhong University of Science and Technology | Hubei Provincial Center for Disease Control and Prevention | Bin Fang, Xiang Li, Xiao Yu, Linlin Liu, Bo Yang, Faxian Zhan, Guojun Ye, Xixiang Huo, Junqiang Xu, Bo Yu, Kun Cai, Jing Li, Yongzhong Jiang. |
| EPI_ISL_412981 | CR&WISCO GENERAL HOSPITAL | Hubei Provincial Center for Disease Control and Prevention | Bin Fang, Xiang Li, Xiao Yu, Linlin Liu, Bo Yang, Faxian Zhan, Guojun Ye, Xixiang Huo, Junqiang Xu, Bo Yu, Kun Cai, Jing Li, Yongzhong Jiang. |
| EPI_ISL_412982 | Wuhan Lung Hospital | Hubei Provincial Center for Disease Control and Prevention | Bin Fang, Xiang Li, Xiao Yu, Linlin Liu, Bo Yang, Faxian Zhan, Guojun Ye, Xixiang Huo, Junqiang Xu, Bo Yu, Kun Cai, Jing Li, Yongzhong Jiang. |
| EPI_ISL_412983 | Tianmen Center for Disease Control and Prevention | Hubei Provincial Center for Disease Control and Prevention | Bin Fang, Xiang Li, Xiao Yu, Linlin Liu, Bo Yang, Faxian Zhan, Guojun Ye, Xixiang Huo, Junqiang Xu, Bo Yu, Kun Cai, Jing Li, YiFa Zhu, Yangyang Tao,Xierong Li,Yongzhong Jiang. |
| EPI_ISL_413606, EPI_ISL_413607, EPI_ISL_413608, EPI_ISL_413609, EPI_ISL_413610, EPI_ISL_413611 | unknown | Pathogen Discovery, Respiratory Viruses Branch, Division of Viral Diseases, Centers for Dieases Control and Prevention | Anna Uehara, Ying Tao, Clinton R. Paden, Krista Queen, Jing Zhang, Yan Li, Mary S. Keckler, Alison S Laufer Halpin, Haibin Wang, Jasmine Padilla, Justin Lee, Christopher A. Elkins, Susan I. Gerber, Suxiang Tong |
| EPI_ISL_414414 | Pathology Queensland | Public Health Virology Laboratory | Bixing Huang, Alyssa Pyke, Amanda De Jong, Andrew Van Den Hurk, Carmel Taylor, David Warrilow, Doris Genge, Elisabeth Gamez, Glen Hewitson, Ian Maxwell Mackay, Inga Sultana, Jamie McMahon, Jean Barcelon, Judy Northill, Mitchell Finger, Natalie Simpson, Neelima Nair, Peter Burtonclay, Peter Moore, Sarah Wheatley, Sean Moody, Sonja Hall-Mendelin, Timothy Gardam, and Frederick Moore |
| EPI_ISL_415153 | KU Leuven, Clinical and Epidemiological Virology | KU Leuven, Clinical and Epidemiological Virology | Bert Vanmechelen, Joan Marti-Carreras, Tony Wawina, Marc Van Ranst, Piet Maes |
| EPI_ISL_415154 | KU Leuven, Clincal and Epidemiological Virology | KU Leuven, Clincal and Epidemiological Virology | Bert Vanmechelen, Joan Marti-Carreras, Tony Wawina, Marc Van Ranst, Piet Maes. |
| EPI_ISL_415155 | KU Leuven, Clinical and Epidemiological Virology | KU Leuven, Clinical and Epidemiological Virology | Bert Vanmechelen, Joan Marti-Carreras, Tony Wawina, Marc Van Ranst, Piet Maes |
| EPI_ISL_415156, EPI_ISL_415157, EPI_ISL_415158, EPI_ISL_415159 | KU Leuven, Clinical and Epidemiological Virology | KU Leuven, Clinical and Epidemiological Virology | Bert Vanmechelen, Joan Marti-Carreras, Tony Wawina, Piet Maes |
| EPI_ISL_416467, EPI_ISL_416468, EPI_ISL_416469, EPI_ISL_416470, EPI_ISL_416471, EPI_ISL_416472, EPI_ISL_416475, EPI_ISL_416476 | KU Leuven, Clinical and Epidemiological Virology | KU Leuven, Clinical and Epidemiological Virology | Bert Vanmechelen, Tony Wawina, Joan Marti-Carreras, Piet Maes |
| EPI_ISL_416480 | R. G. Lugar Center for Public Health Research, National Center for Disease Control and Public Health (NCDC) of Georgia. | R. G. Lugar Center for Public Health Research, National Center for Disease Control and Public Health (NCDC) of Georgia. | Ann Machablishvili, Nato Kotaria, Marine Murtskhvaladze, Lela Sabadze, Mari Gavashelidze, Ana Papkiauri, Meri Pantsulaia, Gvantsa Brachveli, Tata Imnadze, Tamar Jashiashvili, Tea Tevdoradze, Ketevan Sidamonidze, Ekaterine Khmaladze, Ekaterine Zhghenti, Roena Sukhiashvili, Mariam Zakalashvili, Lela Urushadze, Magda Dgebuadze, Giorgi Tomashvili, Davit Tsaguria, Ekaterine Zangaladze, Nino Berishvili, Gvantsa Chanturia, Adam Kotorashvili, Maia Alkhazashvili, Irma Burjanadze, Anna Kasradze, Khatuna Zakhashvili, Paata Imnadze, Amiran Gamkrelidze. |
| EPI_ISL_416745, EPI_ISL_416746 | CNR Virus des Infections Respiratoires - France SUD | CNR Virus des Infections Respiratoires - France SUD | Bal, Antonin; Destras, Gregory; Gaymard, Alexandre; Bouscambert-Duchamp, Maude; Cheynet, Valérie; Brengel-Pesce, Karen; Morfin-Sherpa, Florence; Valette, Martine; Josset, Laurence; Lina, Bruno. |
| EPI_ISL_416747, EPI_ISL_416748 | Institut des Agents Infectieux (IAI) Hospices Civils de Lyon | CNR Virus des Infections Respiratoires - France SUD | Bal, Antonin; Destras, Gregory; Gaymard, Alexandre; Bouscambert-Duchamp, Maude; Cheynet, Valérie; Brengel-Pesce, Karen; Morfin-Sherpa, Florence; Valette, Martine; Josset, Laurence; Lina, Bruno. |
| EPI_ISL_416749 | Centre Hospitalier de Valence | CNR Virus des Infections Respiratoires - France SUD | Bal, Antonin; Destras, Gregory; Gaymard, Alexandre; Bouscambert-Duchamp, Maude; Cheynet, Valérie; Brengel-Pesce, Karen; Morfin-Sherpa, Florence; Valette, Martine; Josset, Laurence; Lina, Bruno. |
| EPI_ISL_416750 | Institut des Agents Infectieux (IAI) Hospices Civils de Lyon | CNR Virus des Infections Respiratoires - France SUD | Bal, Antonin; Destras, Gregory; Gaymard, Alexandre; Bouscambert-Duchamp, Maude; Cheynet, Valérie; Brengel-Pesce, Karen; Morfin-Sherpa, Florence; Valette, Martine; Josset, Laurence; Lina, Bruno. |
| EPI_ISL_416751, EPI_ISL_416752 | CHU Gabriel Montpied | CNR Virus des Infections Respiratoires - France SUD | Bal, Antonin; Destras, Gregory; Gaymard, Alexandre; Bouscambert-Duchamp, Maude; Cheynet, Valérie; Brengel-Pesce, Karen; Morfin-Sherpa, Florence; Valette, Martine; Josset, Laurence; Lina, Bruno. |
| EPI_ISL_416753, EPI_ISL_416754, EPI_ISL_416756 | Institut des Agents Infectieux (IAI) Hospices Civils de Lyon | CNR Virus des Infections Respiratoires - France SUD | Bal, Antonin; Destras, Gregory; Gaymard, Alexandre; Bouscambert-Duchamp, Maude; Cheynet, Valérie; Brengel-Pesce, Karen; Morfin-Sherpa, Florence; Valette, Martine; Josset, Laurence; Lina, Bruno. |
| EPI_ISL_416757 | Centre Hospitalier de Bourg en Bresse | CNR Virus des Infections Respiratoires - France SUD | Bal, Antonin; Destras, Gregory; Gaymard, Alexandre; Bouscambert-Duchamp, Maude; Cheynet, Valérie; Brengel-Pesce, Karen; Morfin-Sherpa, Florence; Valette, Martine; Josset, Laurence; Lina, Bruno. |
| EPI_ISL_416758 | Institut des Agents Infectieux (IAI) Hospices Civils de Lyon | CNR Virus des Infections Respiratoires - France SUD | Bal, Antonin; Destras, Gregory; Gaymard, Alexandre; Bouscambert-Duchamp, Maude; Cheynet, Valérie; Brengel-Pesce, Karen; Morfin-Sherpa, Florence; Valette, Martine; Josset, Laurence; Lina, Bruno. |

| | | | |
|---|---|---|---|
| EPI_ISL_417031 | Pathology Queensland | Public Health Virology Laboratory | Bixing Huang, Alyssa Pyke, Amanda De Jong, Andrew Van Den Hurk, Carmel Taylor, David Warrilow, Doris Genge, Elisabeth Gamez, Glen Hewitson, Ian Maxwell Mackay, Inga Sultana, Jamie McMahon, Jean Barcelon, Judy Northill, Mitchell Finger, Natalie Simpson, Neelima Nair, Peter Burtonclay, Peter Moore, Sarah Wheatley, Sean Moody, Sonja Hall-Mendelin, Timothy Gardam, and Frederick Moore |
| EPI_ISL_417032 | Rockhampton Base Hospital | Public Health Virology Laboratory | Bixing Huang, Alyssa Pyke, Amanda De Jong, Andrew Van Den Hurk, Carmel Taylor, David Warrilow, Doris Genge, Elisabeth Gamez, Glen Hewitson, Ian Maxwell Mackay, Inga Sultana, Jamie McMahon, Jean Barcelon, Judy Northill, Mitchell Finger, Natalie Simpson, Neelima Nair, Peter Burtonclay, Peter Moore, Sarah Wheatley, Sean Moody, Sonja Hall-Mendelin, Timothy Gardam, and Frederick Moore |
| EPI_ISL_417033 | Sullivan Nicolaides Pathology | Public Health Virology Laboratory | Bixing Huang, Alyssa Pyke, Amanda De Jong, Andrew Van Den Hurk, Carmel Taylor, David Warrilow, Doris Genge, Elisabeth Gamez, Glen Hewitson, Ian Maxwell Mackay, Inga Sultana, Jamie McMahon, Jean Barcelon, Judy Northill, Mitchell Finger, Natalie Simpson, Neelima Nair, Peter Burtonclay, Peter Moore, Sarah Wheatley, Sean Moody, Sonja Hall-Mendelin, Timothy Gardam, and Frederick Moore |
| EPI_ISL_417333, EPI_ISL_417334, EPI_ISL_417335, EPI_ISL_417336, EPI_ISL_417337 | Institut des Agents Infectieux (IAI), Hospices Civils de Lyon | CNR Virus des Infections Respiratoires - France SUD | Antonin Bal, Gregory Destras, Gwendolyne Burfin, Solenne Brun, Carine Moustaud, Raphaelle Lamy, Alexandre Gaymard, Maude Bouscambert-Duchamp, Florence Morfin-Sherpa, Martine Valette, Laurence Josset, Bruno Lina |
| EPI_ISL_417338 | Centre Hospitalier de Macon | CNR Virus des Infections Respiratoires - France SUD | Antonin Bal, Gregory Destras, Gwendolyne Burfin, Solenne Brun, Carine Moustaud, Raphaelle Lamy, Alexandre Gaymard, Maude Bouscambert-Duchamp, Florence Morfin-Sherpa, Martine Valette, Laurence Josset, Bruno Lina |
| EPI_ISL_417339 | Institut des Agents Infectieux (IAI), Hospices Civils de Lyon | CNR Virus des Infections Respiratoires - France SUD | Antonin Bal, Gregory Destras, Gwendolyne Burfin, Solenne Brun, Carine Moustaud, Raphaelle Lamy, Alexandre Gaymard, Maude Bouscambert-Duchamp, Florence Morfin-Sherpa, Martine Valette, Laurence Josset, Bruno Lina |
| EPI_ISL_417340 | Centre Hospitalier de Bourg en Bresse | CNR Virus des Infections Respiratoires - France SUD | Antonin Bal, Gregory Destras, Gwendolyne Burfin, Solenne Brun, Carine Moustaud, Raphaelle Lamy, Alexandre Gaymard, Maude Bouscambert-Duchamp, Florence Morfin-Sherpa, Martine Valette, Laurence Josset, Bruno Lina |
| EPI_ISL_417389, EPI_ISL_417406 | Centre for Infectious Diseases and Microbiology Public Health | NSW Health Pathology - Institute of Clinical Pathology and Medical Research; Westmead Hospital; University of Sydney | Arnott A, Sadsad R, Carter I, Rahman H, Holmes EC, O'Sullivan MV, Sintchenko V, Chen SC, Maddocks S, Kok J, Dwyer DE, Rockett R, Eden J-S, Lam C, Gray K, Timms V and Gall M for the 2019-nCoV Study Group |
| EPI_ISL_417491 | Virology Laboratory, Department of Biomedical Sciences and Public Health, University Politecnica delle Marche | Virology and Legal Medicine Laboratories, Department of Biomedical Sciences and Public Health, University Politecnica delle Marche | Bagnarelli,P., Caucci,S., Di Sante,L., Menzo,S., Alessandrini,F., Onofri,V., Turchi,C., Tagliabracci,A. |
| EPI_ISL_417526, EPI_ISL_417527, EPI_ISL_417528, EPI_ISL_417529, EPI_ISL_417530, EPI_ISL_417531, EPI_ISL_417532, EPI_ISL_417533, EPI_ISL_417534 | Laboratoire Nationale de Santé, Microbiology, Virology | Laboratoire Nationale de Santé, Microbiology, Epidemiology and Microbial Genomics | Anke Wienecke-Baldacchino, Ardashel Latsuzbaia, Jessica Tapp, Catherine Ragimbeau, Guillaume Fournier, Tamir Abdelrahman, Trung Nguyen Nguyen, Joel Mossong |
| EPI_ISL_418412 | Centre Hospitalier des Vals d'Ardeche | CNR Virus des Infections Respiratoires - France SUD | Antonin Bal, Gregory Destras, Gwendolyne Burfin, Solenne Brun, Carine Moustaud, Raphaelle Lamy, Alexandre Gaymard, Maude Bouscambert-Duchamp, Florence Morfin-Sherpa, Martine Valette, Bruno Lina, Laurence Josset |
| EPI_ISL_418413 | Centre Hospitalier de Macon | CNR Virus des Infections Respiratoires - France SUD | Antonin Bal, Gregory Destras, Gwendolyne Burfin, Solenne Brun, Carine Moustaud, Raphaelle Lamy, Alexandre Gaymard, Maude Bouscambert-Duchamp, Florence Morfin-Sherpa, Martine Valette, Bruno Lina, Laurence Josset |
| EPI_ISL_418414, EPI_ISL_418415 | Centre Hospitalier de Valence | CNR Virus des Infections Respiratoires - France SUD | Antonin Bal, Gregory Destras, Gwendolyne Burfin, Solenne Brun, Carine Moustaud, Raphaelle Lamy, Alexandre Gaymard, Maude Bouscambert-Duchamp, Florence Morfin-Sherpa, Martine Valette, Bruno Lina, Laurence Josset |
| EPI_ISL_418416 | GH Les Portes du Sud | CNR Virus des Infections Respiratoires - France SUD | Antonin Bal, Gregory Destras, Gwendolyne Burfin, Solenne Brun, Carine Moustaud, Raphaelle Lamy, Alexandre Gaymard, Maude Bouscambert-Duchamp, Florence Morfin-Sherpa, Martine Valette, Bruno Lina, Laurence Josset |
| EPI_ISL_418417 | Centre Hospitalier de Valence | CNR Virus des Infections Respiratoires - France SUD | Antonin Bal, Gregory Destras, Gwendolyne Burfin, Solenne Brun, Carine Moustaud, Raphaelle Lamy, Alexandre Gaymard, Maude Bouscambert-Duchamp, Florence Morfin-Sherpa, Martine Valette, Bruno Lina, Laurence Josset |
| EPI_ISL_418418, EPI_ISL_418419 | Centre Hospitalier Saint Joseph Saint Luc | CNR Virus des Infections Respiratoires - France SUD | Antonin Bal, Gregory Destras, Gwendolyne Burfin, Solenne Brun, Carine Moustaud, Raphaelle Lamy, Alexandre Gaymard, Maude Bouscambert-Duchamp, Florence Morfin-Sherpa, Martine Valette, Bruno Lina, Laurence Josset |
| EPI_ISL_418420, EPI_ISL_418421, EPI_ISL_418422, EPI_ISL_418423, EPI_ISL_418424, EPI_ISL_418425 | Institut des Agents Infectieux (IAI), Hospices Civils de Lyon | CNR Virus des Infections Respiratoires - France SUD | Antonin Bal, Gregory Destras, Gwendolyne Burfin, Solenne Brun, Carine Moustaud, Raphaelle Lamy, Alexandre Gaymard, Maude Bouscambert-Duchamp, Florence Morfin-Sherpa, Martine Valette, Bruno Lina, Laurence Josset |
| EPI_ISL_418426 | Centre Hospitalier de Bourg en Bresse | CNR Virus des Infections Respiratoires - France SUD | Antonin Bal, Gregory Destras, Gwendolyne Burfin, Solenne Brun, Carine Moustaud, Raphaelle Lamy, Alexandre Gaymard, Maude Bouscambert-Duchamp, Florence Morfin-Sherpa, Martine Valette, Bruno Lina, Laurence Josset |
| EPI_ISL_418427 | Hopital Privé de l'Est Lyonnais | CNR Virus des Infections Respiratoires - France SUD | Antonin Bal, Gregory Destras, Gwendolyne Burfin, Solenne Brun, Carine Moustaud, Raphaelle Lamy, Alexandre Gaymard, Maude Bouscambert-Duchamp, Florence Morfin-Sherpa, Martine Valette, Bruno Lina, Laurence Josset |
| EPI_ISL_418428 | Centre Hospitalier Lucien Hussel | CNR Virus des Infections Respiratoires - France SUD | Antonin Bal, Gregory Destras, Gwendolyne Burfin, Solenne Brun, Carine Moustaud, Raphaelle Lamy, Alexandre Gaymard, Maude Bouscambert-Duchamp, Florence Morfin-Sherpa, Martine Valette, Bruno Lina, Laurence Josset |
| EPI_ISL_418429, EPI_ISL_418430, EPI_ISL_418431, EPI_ISL_418432 | Institut des Agents Infectieux (IAI), Hospices Civils de Lyon | CNR Virus des Infections Respiratoires - France SUD | Antonin Bal, Gregory Destras, Gwendolyne Burfin, Solenne Brun, Carine Moustaud, Raphaelle Lamy, Alexandre Gaymard, Maude Bouscambert-Duchamp, Florence Morfin-Sherpa, Martine Valette, Bruno Lina, Laurence Josset |
| EPI_ISL_418792, EPI_ISL_418793 | KU Leuven, Clinical and Epidemiological Virology | KU Leuven, Clinical and Epidemiological Virology | Bert Vanmechelen, Tony Wawina, Joan Marti-Carreras, Piet Maes |
| EPI_ISL_418794, EPI_ISL_418795, EPI_ISL_418796, EPI_ISL_418797, EPI_ISL_418798 | KU Leuven, Clinical and Epidemiological Virology | KU Leuven, Clinical and Epidemiological Virology | Bert Vanmechelen, Joan Marti-Carreras, Tony Wawina, Piet Maes |
| EPI_ISL_418799 | Mater Pathology | Public Health Virology Laboratory | Bixing Huang, Alyssa Pyke, Amanda De Jong, Andrew Van Den Hurk, Carmel Taylor, David Warrilow, Doris Genge, Elisabeth Gamez, Glen Hewitson, Ian Maxwell Mackay, Inga Sultana, Jamie McMahon, Jean Barcelon, Judy Northill, Mitchell Finger, Natalie Simpson, Neelima Nair, Peter Burtonclay, Peter Moore, Sarah Wheatley, Sean Moody, Sonja Hall-Mendelin, Timothy Gardam, and Frederick Moore |
| EPI_ISL_418800 | KU Leuven, Clinical and Epidemiological Virology | KU Leuven, Clinical and Epidemiological Virology | Bert Vanmechelen, Joan Marti-Carreras, Tony Wawina, Piet Maes |
| EPI_ISL_418801 | Mater Pathology | Public Health Virology Laboratory | Bixing Huang, Alyssa Pyke, Amanda De Jong, Andrew Van Den Hurk, Carmel Taylor, David Warrilow, Doris Genge, Elisabeth Gamez, Glen Hewitson, Ian Maxwell Mackay, Inga Sultana, Jamie McMahon, Jean Barcelon, Judy Northill, Mitchell Finger, Natalie Simpson, Neelima Nair, Peter Burtonclay, Peter Moore, Sarah Wheatley, Sean Moody, Sonja Hall-Mendelin, Timothy Gardam, and Frederick Moore |
| EPI_ISL_418802, EPI_ISL_418803, EPI_ISL_418804 | Pathology Queensland | Public Health Virology Laboratory | Bixing Huang, Alyssa Pyke, Amanda De Jong, Andrew Van Den Hurk, Carmel Taylor, David Warrilow, Doris Genge, Elisabeth Gamez, Glen Hewitson, Ian Maxwell Mackay, Inga Sultana, Jamie McMahon, Jean Barcelon, Judy Northill, Mitchell Finger, Natalie Simpson, Neelima Nair, Peter Burtonclay, Peter Moore, Sarah Wheatley, Sean Moody, Sonja Hall-Mendelin, Timothy Gardam, and Frederick Moore |
| EPI_ISL_418805, EPI_ISL_418806 | KU Leuven, Clinical and Epidemiological Virology | KU Leuven, Clinical and Epidemiological Virology | Bert Vanmechelen, Joan Marti-Carreras, Tony Wawina, Piet Maes |
| EPI_ISL_418807, EPI_ISL_418808 | Pathology Queensland | Public Health Virology Laboratory | Bixing Huang, Alyssa Pyke, Amanda De Jong, Andrew Van Den Hurk, Carmel Taylor, David Warrilow, Doris Genge, Elisabeth Gamez, Glen Hewitson, Ian Maxwell Mackay, Inga Sultana, Jamie McMahon, Jean Barcelon, Judy Northill, Mitchell Finger, Natalie Simpson, Neelima Nair, Peter Burtonclay, Peter Moore, Sarah Wheatley, Sean Moody, Sonja Hall-Mendelin, Timothy Gardam, and Frederick Moore |
| EPI_ISL_418811 | Dr. Georges-L.-Dumont University Hospital Centre | National Microbiology Laboratory | Anna Majer, Shari Tyson, Grace Seo, Philip Mabon, Natalie Knox, Morag Graham, Richard Garceau, Guillaume Desnoyers, Nathalie Bastien, Yan Li, Matthew Gilmour, Timothy Booth |
| EPI_ISL_418812, EPI_ISL_418813 | Cadham Provincial Laboratory | National Microbiology Laboratory | Anna Majer, Shari Tyson, Grace Seo, Philip Mabon, Natalie Knox, Morag Graham, Paul Van Caeseele, Jared Bullard, David Alexander, Kerry Dust, Nathalie Bastien, Yan Li, Matthew Gilmour, Timothy Booth |

| | | | |
|---|---|---|---|
| EPI_ISL_418814 | Queen Elizabeth II Health Science Centre | National Microbiology Laboratory | Anna Majer, Shari Tyson, Grace Seo, Philip Mabon, Natalie Knox, Morag Graham, Todd Hatchette, Jason LeBlanc, Nathalie Bastien, Yan Li, Matthew Gilmour, Timothy Booth |
| EPI_ISL_418863, EPI_ISL_418981, EPI_ISL_418982, EPI_ISL_418983, EPI_ISL_418984, EPI_ISL_418985, EPI_ISL_418986, EPI_ISL_418987 | KU Leuven, Clinical and Epidemiological Virology | KU Leuven, Clinical and Epidemiological Virology | Bert Vanmechelen, Joan Marti-Carreras, Tony Wawina, Piet Maes |
| EPI_ISL_418988 | Institute information KU Leuven, Clinical and Epidemiological Virology | Institute information KU Leuven, Clinical and Epidemiological Virology | Bert Vanmechelen, Joan Marti-Carreras, Tony Wawina, Piet Maes |
| EPI_ISL_418989 | KU Leuven, Clinical and Epidemiological Virology | KU Leuven, Clinical and Epidemiological Virology | Bert Vanmechelen, Joan Marti-Carreras, Tony Wawina, Piet Maes |
| EPI_ISL_419168 | Centre Hospitalier de Valence | CNR Virus des Infections Respiratoires - France SUD | Antonin Bal, Gregory Destras, Gwendolyne Burfin, Solenne Brun, Carine Moustaud, Raphaelle Lamy, Alexandre Gaymard, Maude Bouscambert-Duchamp, Florence Morfin-Sherpa, Martine Valette, Bruno Lina, Laurence Josset |
| EPI_ISL_419169, EPI_ISL_419170, EPI_ISL_419171, EPI_ISL_419172, EPI_ISL_419173 | Institut des Agents Infectieux (IAI), Hospices Civils de Lyon | CNR Virus des Infections Respiratoires - France SUD | Antonin Bal, Gregory Destras, Gwendolyne Burfin, Solenne Brun, Carine Moustaud, Raphaelle Lamy, Alexandre Gaymard, Maude Bouscambert-Duchamp, Florence Morfin-Sherpa, Martine Valette, Bruno Lina, Laurence Josset |
| EPI_ISL_419174, EPI_ISL_419175, EPI_ISL_419176 | Centre Hospitalier de Macon | CNR Virus des Infections Respiratoires - France SUD | Antonin Bal, Gregory Destras, Gwendolyne Burfin, Solenne Brun, Carine Moustaud, Raphaelle Lamy, Alexandre Gaymard, Maude Bouscambert-Duchamp, Florence Morfin-Sherpa, Martine Valette, Bruno Lina, Laurence Josset |
| EPI_ISL_419177, EPI_ISL_419178, EPI_ISL_419179, EPI_ISL_419180, EPI_ISL_419181, EPI_ISL_419182 | Institut des Agents Infectieux (IAI), Hospices Civils de Lyon | CNR Virus des Infections Respiratoires - France SUD | Antonin Bal, Gregory Destras, Gwendolyne Burfin, Solenne Brun, Carine Moustaud, Raphaelle Lamy, Alexandre Gaymard, Maude Bouscambert-Duchamp, Florence Morfin-Sherpa, Martine Valette, Bruno Lina, Laurence Josset |
| EPI_ISL_419183 | Centre Hospitalier de Bourg en Bresse | CNR Virus des Infections Respiratoires - France SUD | Antonin Bal, Gregory Destras, Gwendolyne Burfin, Solenne Brun, Carine Moustaud, Raphaelle Lamy, Alexandre Gaymard, Maude Bouscambert-Duchamp, Florence Morfin-Sherpa, Martine Valette, Bruno Lina, Laurence Josset |
| EPI_ISL_419184 | Institut des Agents Infectieux (IAI), Hospices Civils de Lyon | CNR Virus des Infections Respiratoires - France SUD | Antonin Bal, Gregory Destras, Gwendolyne Burfin, Solenne Brun, Carine Moustaud, Raphaelle Lamy, Alexandre Gaymard, Maude Bouscambert-Duchamp, Florence Morfin-Sherpa, Martine Valette, Bruno Lina, Laurence Josset |
| EPI_ISL_419185, EPI_ISL_419186 | Centre Hospitalier de Bourg en Bresse | CNR Virus des Infections Respiratoires - France SUD | Antonin Bal, Gregory Destras, Gwendolyne Burfin, Solenne Brun, Carine Moustaud, Raphaelle Lamy, Alexandre Gaymard, Maude Bouscambert-Duchamp, Florence Morfin-Sherpa, Martine Valette, Bruno Lina, Laurence Josset |
| EPI_ISL_419187, EPI_ISL_419188 | Centre Hospitalier de Macon | CNR Virus des Infections Respiratoires - France SUD | Antonin Bal, Gregory Destras, Gwendolyne Burfin, Solenne Brun, Carine Moustaud, Raphaelle Lamy, Alexandre Gaymard, Maude Bouscambert-Duchamp, Florence Morfin-Sherpa, Martine Valette, Bruno Lina, Laurence Josset |
| EPI_ISL_419254 | INMI Lazzaro Spallanzani IRCCS | Laboratory of Virology, INMI Lazzaro Spallanzani IRCCS | Barbara Bartolini, Martina Rueca, Francesco Messina, Cesare E. M. Gruber, Emanuela Giombini, Maria R. Capobianchi, Fabrizio Carletti, Francesca Colavita, Concetta Castilletti, Eleonora Lalle, Daniele Lapa, Giuseppe Ippolito. |
| EPI_ISL_419255 | INMI Lazzaro Spallanzani IRCCS | INMI Lazzaro Spallanzani IRCCS | Antonino Di Caro, Cesare E. M. Gruber, Martina Rueca, Barbara Bartolini, Francesco Messina, Emanuela Giombini, Maria R. Capobianchi, Fabrizio Carletti, Francesca Colavita, Concetta Castilletti, Eleonora Lalle, Daniele Lapa, Giuseppe Ippolito. |
| EPI_ISL_419559, EPI_ISL_419560 | FL Bureau of Public Health Laboratories-Tampa | Pathogen Discovery, Respiratory Viruses Branch, Division of Viral Diseases, Centers for Disease Control and Prevention | Anna Uehara, Ying Tao, Jing Zhang, Krista Queen, Clinton R. Paden, Yan Li, Haibin Wang, Jasmine Padilla, Justin Lee, Suxiang Tong |
| EPI_ISL_419562, EPI_ISL_419563, EPI_ISL_419564, EPI_ISL_419565, EPI_ISL_419566, EPI_ISL_419567, EPI_ISL_419568, EPI_ISL_419569, EPI_ISL_419570, EPI_ISL_419571, EPI_ISL_419572, EPI_ISL_419573, EPI_ISL_419574, EPI_ISL_419575, EPI_ISL_419576, EPI_ISL_419577, EPI_ISL_419578, EPI_ISL_419579, EPI_ISL_419580, EPI_ISL_419581, EPI_ISL_419582, EPI_ISL_419583, EPI_ISL_419584, EPI_ISL_419585, EPI_ISL_419586, EPI_ISL_419587, EPI_ISL_419588, EPI_ISL_419589, EPI_ISL_419590, EPI_ISL_419591, EPI_ISL_419592, EPI_ISL_419593, EPI_ISL_419594, EPI_ISL_419595, EPI_ISL_419596, EPI_ISL_419597, EPI_ISL_419598, EPI_ISL_419599, EPI_ISL_419600, EPI_ISL_419601, EPI_ISL_419602, EPI_ISL_419603, EPI_ISL_419604, EPI_ISL_419605, EPI_ISL_419606, EPI_ISL_419607 | | | |
| see above | Laboratoire National de Santé, Microbiology, Virology | Laboratoire National de Santé, Microbiology, Epidemiology and Microbial Genomics | Anke Wienecke-Baldacchino, Ardashel Latsuzbaia, Jessica Tapp, Catherine Ragimbeau, Guillaume Fournier, Tamir Abdelrahman, Trung Nguyen Nguyen, Joel Mossong |
| EPI_ISL_420113 | Servicio de Microbiología. Consorcio Hospital General Universitario de Valencia | Sequencing and Bioinformatics Service and Molecular Epidemiology Research Group. FISABIO-Public Health | Beatriz Beamud, Lidia Ruiz Roldan, Marta Pla Diaz, Neris Garcia-Gonzalez, Loreto Ferrús Abad, Inma Galán Vendrell, Paula Ruiz-Hueso, Mariana Reyes-Prieto, Vicente Soriano Chirona, Maria Alma Bracho, Maria Dolores Ocete, LlÚcia Martínez-PriegoGriselda De Marco, , Concepcion Gimeno, Giuseppe D'Auria, Fernando Gonzalez-Candelas |
| EPI_ISL_420598, EPI_ISL_420599, EPI_ISL_420600 | Servicio Virosis Respiratorias-Departamento Virología-INEI | Instituto Nacional Enfermedades Infecciosas C.G.Malbran | Baumeister E., Avaro M., Benedetti E., Russo M., Dattero ME, Pontoriero A., Cisterna D., Molina V., Perandones C., Tuduri E., Lorenzo F., Poklepovich T., Campos J. |
| EPI_ISL_420604, EPI_ISL_420605, EPI_ISL_420606, EPI_ISL_420607, EPI_ISL_420608, EPI_ISL_420609, EPI_ISL_420610, EPI_ISL_420611 | Institut des Agents Infectieux (IAI), Hospices Civils de Lyon | CNR Virus des Infections Respiratoires - France SUD | Antonin Bal, Gregory Destras, Gwendolyne Burfin, Solenne Brun, Carine Moustaud, Raphaelle Lamy, Alexandre Gaymard, Maude Bouscambert-Duchamp, Florence Morfin-Sherpa, Martine Valette, Bruno Lina, Laurence Josset |
| EPI_ISL_420612, EPI_ISL_420613, EPI_ISL_420614 | Centre Hospitalier de Macon | CNR Virus des Infections Respiratoires - France SUD | Antonin Bal, Gregory Destras, Gwendolyne Burfin, Solenne Brun, Carine Moustaud, Raphaelle Lamy, Alexandre Gaymard, Maude Bouscambert-Duchamp, Florence Morfin-Sherpa, Martine Valette, Bruno Lina, Laurence Josset |
| EPI_ISL_420615, EPI_ISL_420616 | Institut des Agents Infectieux (IAI), Hospices Civils de Lyon | CNR Virus des Infections Respiratoires - France SUD | Antonin Bal, Gregory Destras, Gwendolyne Burfin, Solenne Brun, Carine Moustaud, Raphaelle Lamy, Alexandre Gaymard, Maude Bouscambert-Duchamp, Florence Morfin-Sherpa, Martine Valette, Bruno Lina, Laurence Josset |
| EPI_ISL_420617 | Centre Hospitalier Saint Joseph Saint Luc | CNR Virus des Infections Respiratoires - France SUD | Antonin Bal, Gregory Destras, Gwendolyne Burfin, Solenne Brun, Carine Moustaud, Raphaelle Lamy, Alexandre Gaymard, Maude Bouscambert-Duchamp, Florence Morfin-Sherpa, Martine Valette, Bruno Lina, Laurence Josset |
| EPI_ISL_420618, EPI_ISL_420619 | Institut des Agents Infectieux (IAI), Hospices Civils de Lyon | CNR Virus des Infections Respiratoires - France SUD | Antonin Bal, Gregory Destras, Gwendolyne Burfin, Solenne Brun, Carine Moustaud, Raphaelle Lamy, Alexandre Gaymard, Maude Bouscambert-Duchamp, Florence Morfin-Sherpa, Martine Valette, Bruno Lina, Laurence Josset |
| EPI_ISL_420620 | Centre Hospitalier de Bourg en Bresse | CNR Virus des Infections Respiratoires - France SUD | Antonin Bal, Gregory Destras, Gwendolyne Burfin, Solenne Brun, Carine Moustaud, Raphaelle Lamy, Alexandre Gaymard, Maude Bouscambert-Duchamp, Florence Morfin-Sherpa, Martine Valette, Bruno Lina, Laurence Josset |
| EPI_ISL_420621, EPI_ISL_420622, EPI_ISL_420623, EPI_ISL_420624, EPI_ISL_420625 | Institut des Agents Infectieux (IAI), Hospices Civils de Lyon | CNR Virus des Infections Respiratoires - France SUD | Antonin Bal, Gregory Destras, Gwendolyne Burfin, Solenne Brun, Carine Moustaud, Raphaelle Lamy, Alexandre Gaymard, Maude Bouscambert-Duchamp, Florence Morfin-Sherpa, Martine Valette, Bruno Lina, Laurence Josset |
| EPI_ISL_420878, EPI_ISL_420879 | Mater Pathology | Public Health Virology Laboratory | Bixing Huang, Alyssa Pyke, Amanda De Jong, Andrew Van Den Hurk, Carmel Taylor, David Warrilow, Doris Genge, Elisabeth Gamez, Glen Hewitson, Ian Maxwell Mackay, Inga Sultana, Jamie McMahon, Jean Barcelon, Judy Northill, Mitchell Finger, Natalie Simpson, Neelima Nair, Peter Burtonclay, Peter Moore, Sarah Wheatley, Sean Moody, Sonja Hall-Mendelin, Timothy Gardam, and Frederick Moore |
| EPI_ISL_421636 | Pathology North | Public Health Virology Laboratory | Bixing Huang, Alyssa Pyke, Amanda De Jong, Andrew Van Den Hurk, Carmel Taylor, David Warrilow, Doris Genge, Elisabeth Gamez, Glen Hewitson, Ian Maxwell Mackay, Inga Sultana, Jamie McMahon, Jean Barcelon, Judy Northill, Mitchell Finger, Natalie Simpson, Neelima Nair, Peter Burtonclay, Peter Moore, Sarah Wheatley, Sean Moody, Sonja Hall-Mendelin, Timothy Gardam, and Frederick Moore |
| EPI_ISL_421734, EPI_ISL_421735, EPI_ISL_421736, EPI_ISL_421737, EPI_ISL_421738, EPI_ISL_421739, EPI_ISL_421740, EPI_ISL_421741, EPI_ISL_421742, EPI_ISL_421743, EPI_ISL_421744, EPI_ISL_421745, EPI_ISL_421746, EPI_ISL_421747, EPI_ISL_421748, EPI_ISL_421749, EPI_ISL_421750, EPI_ISL_421751, EPI_ISL_421752, EPI_ISL_421753, EPI_ISL_421754, EPI_ISL_421755, EPI_ISL_421756, EPI_ISL_421757, EPI_ISL_421758, EPI_ISL_421759, EPI_ISL_421760, EPI_ISL_421761, EPI_ISL_421762, EPI_ISL_421763 | | | |
| see above | Laboratoire National de Sante, Microbiology, Virology | Laboratoire National de Sante, Microbiology, Epidemiology and Microbial Genomics | Anke Wienecke-Baldacchino, Ardashel Latsuzbaia, Jessica Tapp, Catherine Ragimbeau, Guillaume Fournier, Tamir Abdelrahman, Trung Nguyen Nguyen, Joel Mossong |
| EPI_ISL_422563 | Institute of Microbiology Universidad San Francisco de Quito | Institute of Microbiology Universidad San Francisco de Quito | Belen Prado-Vivar, Sully Marquez, Juan Jose Guadalupe, Bernardo Gutierrez, Francisco Mora, Juan Gaviria, Alejandra Ramones, Franklin Espinoza, Edison Ligña, Jorge Reyes, Patricio Rojas-Silva, Veronica Barragan, Gabriel Trueba, Michelle Grunauer, Paul Cardenas |

| | | | |
|---|---|---|---|
| EPI_ISL_422566, EPI_ISL_422567, EPI_ISL_422568, EPI_ISL_422569, EPI_ISL_422570, EPI_ISL_422571, EPI_ISL_422572, EPI_ISL_422573, EPI_ISL_422574, EPI_ISL_422575, EPI_ISL_422576, EPI_ISL_422577, EPI_ISL_422578, EPI_ISL_422579, EPI_ISL_422580, EPI_ISL_422581, EPI_ISL_422582, EPI_ISL_422583, EPI_ISL_422584, EPI_ISL_422585, EPI_ISL_422586, EPI_ISL_422587, EPI_ISL_422588, EPI_ISL_422589, EPI_ISL_422590, EPI_ISL_422591, EPI_ISL_422592, EPI_ISL_422593, EPI_ISL_422594, EPI_ISL_422595, EPI_ISL_422596, EPI_ISL_422597, EPI_ISL_422598, EPI_ISL_422599, EPI_ISL_422600, EPI_ISL_422601, EPI_ISL_422602, EPI_ISL_422603, EPI_ISL_422604, EPI_ISL_422605, EPI_ISL_422606, EPI_ISL_422607, EPI_ISL_422608, EPI_ISL_422609, EPI_ISL_422610, EPI_ISL_422611, EPI_ISL_422612, EPI_ISL_422613, EPI_ISL_422614, EPI_ISL_422615, EPI_ISL_422616, EPI_ISL_422617, EPI_ISL_422618, EPI_ISL_422619, EPI_ISL_422620, EPI_ISL_422621, EPI_ISL_422622, EPI_ISL_422623, EPI_ISL_422624, EPI_ISL_422625, EPI_ISL_422626, EPI_ISL_422627, EPI_ISL_422628, EPI_ISL_422629, EPI_ISL_422630, EPI_ISL_422631, EPI_ISL_422632, EPI_ISL_422633, EPI_ISL_422634, EPI_ISL_422635, EPI_ISL_422637, EPI_ISL_422638, EPI_ISL_422639, EPI_ISL_422640, EPI_ISL_422641, EPI_ISL_422642, EPI_ISL_422643, EPI_ISL_422644, EPI_ISL_422645, EPI_ISL_422646, EPI_ISL_422647, EPI_ISL_422648, EPI_ISL_422649, EPI_ISL_422650, EPI_ISL_422651, EPI_ISL_422652, EPI_ISL_422653, EPI_ISL_422654, EPI_ISL_422655, EPI_ISL_422656, EPI_ISL_422657, EPI_ISL_422658, EPI_ISL_422659, EPI_ISL_422660, EPI_ISL_422661, EPI_ISL_422662, EPI_ISL_422663, EPI_ISL_422664, EPI_ISL_422665, EPI_ISL_422666, EPI_ISL_422667, EPI_ISL_422668, EPI_ISL_422669, EPI_ISL_422670, EPI_ISL_422671, EPI_ISL_422672, EPI_ISL_422673, EPI_ISL_422674, EPI_ISL_422675, EPI_ISL_422676, EPI_ISL_422677, EPI_ISL_422678, EPI_ISL_422679, EPI_ISL_422680, EPI_ISL_422681, EPI_ISL_422682, EPI_ISL_422683, EPI_ISL_422684, EPI_ISL_422685, EPI_ISL_422686, EPI_ISL_422687, EPI_ISL_422688, EPI_ISL_422689, EPI_ISL_422690, EPI_ISL_422691, EPI_ISL_422692, EPI_ISL_422693, EPI_ISL_422694, EPI_ISL_422695, EPI_ISL_422696, EPI_ISL_422697, EPI_ISL_422698, EPI_ISL_422699, EPI_ISL_422700, EPI_ISL_422701, EPI_ISL_422702, EPI_ISL_422703, EPI_ISL_422704, EPI_ISL_422705, EPI_ISL_422706, EPI_ISL_422707, EPI_ISL_422708, EPI_ISL_422709, EPI_ISL_422710, EPI_ISL_422711, EPI_ISL_422712, EPI_ISL_422713, EPI_ISL_422714, EPI_ISL_422715, EPI_ISL_422716, EPI_ISL_422717, EPI_ISL_422718, EPI_ISL_422719, EPI_ISL_422720, EPI_ISL_422721, EPI_ISL_422722, EPI_ISL_422723, EPI_ISL_422724, EPI_ISL_422725, EPI_ISL_422726, EPI_ISL_422727, EPI_ISL_422728, EPI_ISL_422729, EPI_ISL_422730, EPI_ISL_422731, EPI_ISL_422732, EPI_ISL_422733, EPI_ISL_422734, EPI_ISL_422735, EPI_ISL_422736, EPI_ISL_422737, EPI_ISL_422738, EPI_ISL_422739, EPI_ISL_422740, EPI_ISL_422741, EPI_ISL_422742, EPI_ISL_422743, EPI_ISL_422744, EPI_ISL_422745, EPI_ISL_422746, EPI_ISL_422747, EPI_ISL_422748, EPI_ISL_422749, EPI_ISL_422750, EPI_ISL_422751, EPI_ISL_422752, EPI_ISL_422753, EPI_ISL_422754, EPI_ISL_422755, EPI_ISL_422756, EPI_ISL_422757, EPI_ISL_422758, EPI_ISL_422759, EPI_ISL_422760, EPI_ISL_422761, EPI_ISL_422762, EPI_ISL_422763, EPI_ISL_422764, EPI_ISL_422765, EPI_ISL_422766, EPI_ISL_422767, EPI_ISL_422768, EPI_ISL_422769, EPI_ISL_422770, EPI_ISL_422771, EPI_ISL_422772, EPI_ISL_422773, EPI_ISL_422774, EPI_ISL_422775, EPI_ISL_422776, EPI_ISL_422777, EPI_ISL_422778, EPI_ISL_422779, EPI_ISL_422780, EPI_ISL_422781, EPI_ISL_422782, EPI_ISL_422783, EPI_ISL_422784, EPI_ISL_422785, EPI_ISL_422786, EPI_ISL_422787, EPI_ISL_422788, EPI_ISL_422789, EPI_ISL_422790, EPI_ISL_422791, EPI_ISL_422792, EPI_ISL_422793, EPI_ISL_422794, EPI_ISL_422795, EPI_ISL_422796, EPI_ISL_422797, EPI_ISL_422798, EPI_ISL_422799, EPI_ISL_422800, EPI_ISL_422801, EPI_ISL_422802, EPI_ISL_422803, EPI_ISL_422804, EPI_ISL_422805, EPI_ISL_422806, EPI_ISL_422807, EPI_ISL_422808, EPI_ISL_422809, EPI_ISL_422810, EPI_ISL_422811, EPI_ISL_422812, EPI_ISL_422813, EPI_ISL_422814, EPI_ISL_422815, EPI_ISL_422816, EPI_ISL_422817, EPI_ISL_422818, EPI_ISL_422819, EPI_ISL_422820, EPI_ISL_422821, EPI_ISL_422822, EPI_ISL_422823, EPI_ISL_422824, EPI_ISL_422825, EPI_ISL_422826, EPI_ISL_422827, EPI_ISL_422828, EPI_ISL_422829, EPI_ISL_422830, EPI_ISL_422831, EPI_ISL_422832, EPI_ISL_422833, EPI_ISL_422834, EPI_ISL_422835, EPI_ISL_422836, EPI_ISL_422837, EPI_ISL_422838, EPI_ISL_422839, EPI_ISL_422840, EPI_ISL_422841, EPI_ISL_422842, EPI_ISL_422843, EPI_ISL_422844, EPI_ISL_422845, EPI_ISL_422846, EPI_ISL_422847, EPI_ISL_422848, EPI_ISL_422849, EPI_ISL_422850, EPI_ISL_422851, EPI_ISL_422852, EPI_ISL_422853, EPI_ISL_422854, EPI_ISL_422855, EPI_ISL_422856, EPI_ISL_422857, EPI_ISL_422858, EPI_ISL_422859, EPI_ISL_422860, EPI_ISL_422861, EPI_ISL_422862, EPI_ISL_422863, EPI_ISL_422864, EPI_ISL_422865, EPI_ISL_422866, EPI_ISL_422867, EPI_ISL_422868, EPI_ISL_422869, EPI_ISL_422870, EPI_ISL_422871, EPI_ISL_422872, EPI_ISL_422873, EPI_ISL_422874, EPI_ISL_422875, EPI_ISL_422876, EPI_ISL_422877, EPI_ISL_422878, EPI_ISL_422879, EPI_ISL_422880, EPI_ISL_422881, EPI_ISL_422885, EPI_ISL_422886, EPI_ISL_422887, EPI_ISL_422888, EPI_ISL_422889, EPI_ISL_422890, EPI_ISL_422891, EPI_ISL_422892, EPI_ISL_422893, EPI_ISL_422894, EPI_ISL_422895, EPI_ISL_422896, EPI_ISL_422897, EPI_ISL_422898, EPI_ISL_422899, EPI_ISL_422901, EPI_ISL_422902, EPI_ISL_422903, EPI_ISL_422904, EPI_ISL_422905, EPI_ISL_422906, EPI_ISL_422907, EPI_ISL_422908, EPI_ISL_422909, EPI_ISL_422910, EPI_ISL_422911, EPI_ISL_422912, EPI_ISL_422913, EPI_ISL_422914, EPI_ISL_422915, EPI_ISL_422916, EPI_ISL_422917, EPI_ISL_422918, EPI_ISL_422919, EPI_ISL_422920, EPI_ISL_422921, EPI_ISL_422922, EPI_ISL_422923, EPI_ISL_422924, EPI_ISL_422925, EPI_ISL_422926, EPI_ISL_422927, EPI_ISL_422928, EPI_ISL_422929, EPI_ISL_422930, EPI_ISL_422931, EPI_ISL_422932, EPI_ISL_422933, EPI_ISL_422934, EPI_ISL_422935, EPI_ISL_422936, EPI_ISL_422937, EPI_ISL_422938, EPI_ISL_422939, EPI_ISL_422940, EPI_ISL_422941, EPI_ISL_422942, EPI_ISL_422943, EPI_ISL_422944, EPI_ISL_422945, EPI_ISL_422946, EPI_ISL_422947, EPI_ISL_422948, EPI_ISL_422949, EPI_ISL_422950, EPI_ISL_422951, EPI_ISL_422952, EPI_ISL_422953, EPI_ISL_422954, EPI_ISL_422955, EPI_ISL_422956, EPI_ISL_422957, EPI_ISL_422958, EPI_ISL_422959, EPI_ISL_422960, EPI_ISL_423034 | | | |
| see above | Dutch COVID-19 response team | Erasmus Medical Center | Bas Oude Munnink, David Nieuwenhuijse, Reina Sikkema, Claudia Schapendonk, Irina Chestakova, Anne van der Linden, Theo Bestebroer, Stefan van Nieuwkoop, Mark Pronk, Pascal Lexmond, Corien Swaan, Manon Haverkate, Madelief Mollers, Mart Stein, Sandra Kengne Kamga Mobou, Jeroen van Kampen, Jolanda Voermans, Aura Timen, Corine GeurtsvanKessel, Annemiek van der Eijk, Richard Molenkamp, Marion Koopmans, on behalf of the Dutch national COVID-19 response team. |
| EPI_ISL_424981, EPI_ISL_424982, EPI_ISL_424984, EPI_ISL_424985, EPI_ISL_424986, EPI_ISL_424988, EPI_ISL_424989, EPI_ISL_424990, EPI_ISL_424992 | Bureau of Laboratories, Michigan Department of Health and Human Services | Michigan Department of Health and Human Services | Blankenship HM, Riner D, Soehnlen MK |
| EPI_ISL_425181, EPI_ISL_425195 | Servicio de Microbiología. Consorcio Hospital General Universitario de Valencia | Sequencing and Bioinformatics Service and Molecular Epidemiology Research Group. FISABIO-Public Health | Beatriz Beamud, Lidia Ruiz Roldan, Marta Pla Diaz, Neris Garcia-Gonzalez, Inma Galán Vendrell, Sandra Carbo, Loreto Ferrús Abad, Paula Ruiz-Hueso, Mariana Reyes-Prieto, Vicente Soriano Chirona, Ivan Ansari, David Navarro, Maria Alma Bracho, Griselda De Marco, Llúcia Martínez-Priego, Giuseppe D'Auria, Fernando Gonzalez-Candelas |
| EPI_ISL_425206 | Servicio de Microbiologia. Hospital Clinico Universitario de Valencia | Sequencing and Bioinformatics Service and Molecular Epidemiology Research Group. FISABIO-Public Health | Beatriz Beamud, Lidia Ruiz Roldan, Marta Pla Diaz,Neris Garcia-Gonzalez, Loreto Ferrús Abad, Maria Dolores Ocete, Inma Galán Vendrell, Paula Ruiz-Hueso, Mariana Reyes-Prieto, Vicente Soriano Chirona, Maria Alma Bracho, Griselda De Marco, Llúcia Martínez-Priego, Concepcion Gimeno, Giuseppe D'Auria, Fernando Gonzalez-Candelas |
| EPI_ISL_425215 | Servicio de Microbiología. Consorcio Hospital General Universitario de Valencia | Sequencing and Bioinformatics Service and Molecular Epidemiology Research Group. FISABIO-Public Health | Beatriz Beamud, Lidia Ruiz Roldan, Marta Pla Diaz, Neris Garcia-Gonzalez, Inma Galán Vendrell, Sandra Carbo, Loreto Ferrús Abad, Paula Ruiz-Hueso, Mariana Reyes-Prieto, Vicente Soriano Chirona, Ivan Ansari, David Navarro, Maria Alma Bracho, Griselda De Marco, Llúcia Martínez-Priego, Giuseppe D'Auria, Fernando Gonzalez-Candelas |
| EPI_ISL_426416 | CT-Dr. Katherine A. Kelley State Public Health Lab | Pathogen Discovery, Respiratory Viruses Branch, Division of Viral Diseases, Centers for Disease Control and Prevention | Anna Uehara, Yan Li, Krista Queen, Clinton R. Paden, Rachel Marine, Ying Tao, Jing Zhang, Haibin Wang, Mary S. Keckler, Alison S. Laufer Halpin, Christopher A. Elkins, Suxiang Tong |
| EPI_ISL_426417 | GA Department of Public Health Laboratory | Pathogen Discovery, Respiratory Viruses Branch, Division of Viral Diseases, Centers for Disease Control and Prevention | Anna Uehara, Yan Li, Krista Queen, Clinton R. Paden, Rachel Marine, Ying Tao, Jing Zhang, Haibin Wang, Mary S. Keckler, Alison S. Laufer Halpin, Christopher A. Elkins, Suxiang Tong |
| EPI_ISL_426420, EPI_ISL_426421 | HI Dept. of Health, State Laboratories Division | Pathogen Discovery, Respiratory Viruses Branch, Division of Viral Diseases, Centers for Disease Control and Prevention | Anna Uehara, Yan Li, Krista Queen, Clinton R. Paden, Rachel Marine, Ying Tao, Jing Zhang, Haibin Wang, Mary S. Keckler, Alison S. Laufer Halpin, Christopher A. Elkins, Suxiang Tong |
| EPI_ISL_427043 | Laboratory of Microbiology, Medical School, National and Kapodistrian University of Athens | Laboratory of Biology, Department of Medicine, Democritus University of Thrace | Bampali,M., Dovrolis,N., Gatzidou,E., Froukala,E., Stavropoulou,A., Veletza,S., Tsakris,A., Spanakis,N. and Karakasiliotis,I. |
| EPI_ISL_427648 | Centre for Infectious Diseases and Microbiology Public Health | NSW Health Pathology - Institute of Clinical Pathology and Medical Research; Westmead Hospital; University of Sydney | Bachmann N, Rockett R, Lam C, Gray K, Timms V, Gall M, Arnott A, Sadsad R, Draper J, Sim E, Carter I, Holmes EC, O'Sullivan MV, Byun R, Sintchenko V, Chen SC, Eden JS, Maddocks S, Kok J, Propenko M, Sorrell T, Chang S, Basile K, Dwyer DE for the 2019-nCoV Study Group |
| EPI_ISL_427651 | Centre for Infectious Diseases and Microbiology Public Health | NSW Health Pathology - Institute of Clinical Pathology and Medical Research; Westmead Hospital; University of Sydney | Arnott A, Sadsad R, Draper J, Sim E, Bachmann N, Rockett R, Lam C, Gray K, Timms V, Gall M, Carter I, Holmes EC, O'Sullivan MV, Byun R, Sintchenko V, Chen SC, Eden JS, Maddocks S, Kok J, Propenko M, Sorrell T, Chang S, Basile K, Dwyer DE for the 2019-nCoV Study Group |
| EPI_ISL_427656, EPI_ISL_427659, EPI_ISL_427666 | Centre for Infectious Diseases and Microbiology Public Health | NSW Health Pathology - Institute of Clinical Pathology and Medical Research; Westmead Hospital; University of Sydney | Bachmann N, Rockett R, Lam C, Gray K, Timms V, Gall M, Arnott A, Sadsad R, Draper J, Sim E, Carter I, Holmes EC, O'Sullivan MV, Byun R, Sintchenko V, Chen SC, Eden JS, Maddocks S, Kok J, Propenko M, Sorrell T, Chang S, Basile K, Dwyer DE for the 2019-nCoV Study Group |
| EPI_ISL_427670 | Centre for Infectious Diseases and Microbiology Public Health | NSW Health Pathology - Institute of Clinical Pathology and Medical Research; Westmead Hospital; University of Sydney | Arnott A, Sadsad R, Draper J, Sim E, Bachmann N, Rockett R, Lam C, Gray K, Timms V, Gall M, Carter I, Holmes EC, O'Sullivan MV, Byun R, Sintchenko V, Chen SC, Eden JS, Maddocks S, Kok J, Propenko M, Sorrell T, Chang S, Basile K, Dwyer DE for the 2019-nCoV Study Group |
| EPI_ISL_427671, EPI_ISL_427675 | Centre for Infectious Diseases and Microbiology Public Health | NSW Health Pathology - Institute of Clinical Pathology and Medical Research; Westmead Hospital; University of Sydney | Bachmann N, Rockett R, Lam C, Gray K, Timms V, Gall M, Arnott A, Sadsad R, Draper J, Sim E, Carter I, Holmes EC, O'Sullivan MV, Byun R, Sintchenko V, Chen SC, Eden JS, Maddocks S, Kok J, Propenko M, Sorrell T, Chang S, Basile K, Dwyer DE for the 2019-nCoV Study Group |
| EPI_ISL_427683, EPI_ISL_427689, EPI_ISL_427693, EPI_ISL_427694, EPI_ISL_427702, EPI_ISL_427707 | Centre for Infectious Diseases and Microbiology Public Health | NSW Health Pathology - Institute of Clinical Pathology and Medical Research; Westmead Hospital; University of Sydney | Arnott A, Sadsad R, Draper J, Sim E, Bachmann N, Rockett R, Lam C, Gray K, Timms V, Gall M, Carter I, Holmes EC, O'Sullivan MV, Byun R, Sintchenko V, Chen SC, Eden JS, Maddocks S, Kok J, Propenko M, Sorrell T, Chang S, Basile K, Dwyer DE for the 2019-nCoV Study Group |
| EPI_ISL_427713 | Centre for Infectious Diseases and Microbiology Public Health | NSW Health Pathology - Institute of Clinical Pathology and Medical Research; Westmead Hospital; University of Sydney | Bachmann N, Rockett R, Lam C, Gray K, Timms V, Gall M, Arnott A, Sadsad R, Draper J, Sim E, Carter I, Holmes EC, O'Sullivan MV, Byun R, Sintchenko V, Chen SC, Eden JS, Maddocks S, Kok J, Propenko M, Sorrell T, Chang S, Basile K, Dwyer DE for the 2019-nCoV Study Group |
| EPI_ISL_427735, EPI_ISL_427739 | Centre for Infectious Diseases and Microbiology Public Health | NSW Health Pathology - Institute of Clinical Pathology and Medical Research; Westmead Hospital; University of Sydney | Arnott A, Sadsad R, Draper J, Sim E, Bachmann N, Rockett R, Lam C, Gray K, Timms V, Gall M, Carter I, Holmes EC, O'Sullivan MV, Byun R, Sintchenko V, Chen SC, Eden JS, Maddocks S, Kok J, Propenko M, Sorrell T, Chang S, Basile K, Dwyer DE for the 2019-nCoV Study Group |
| EPI_ISL_427744, EPI_ISL_427749 | Centre for Infectious Diseases and Microbiology Public Health | NSW Health Pathology - Institute of Clinical Pathology and Medical Research; Westmead Hospital; University of Sydney | Bachmann N, Rockett R, Lam C, Gray K, Timms V, Gall M, Arnott A, Sadsad R, Draper J, Sim E, Carter I, Holmes EC, O'Sullivan MV, Byun R, Sintchenko V, Chen SC, Eden JS, Maddocks S, Kok J, Propenko M, Sorrell T, Chang S, Basile K, Dwyer DE for the 2019-nCoV Study Group |
| EPI_ISL_427751, EPI_ISL_427753 | Centre for Infectious Diseases and Microbiology Public Health | NSW Health Pathology - Institute of Clinical Pathology and Medical Research; Westmead Hospital; University of Sydney | Arnott A, Sadsad R, Draper J, Sim E, Bachmann N, Rockett R, Lam C, Gray K, Timms V, Gall M, Carter I, Holmes EC, O'Sullivan MV, Byun R, Sintchenko V, Chen SC, Eden JS, Maddocks S, Kok J, Propenko M, Sorrell T, Chang S, Basile K, Dwyer DE for the 2019-nCoV Study Group |
| EPI_ISL_427758 | Centre for Infectious Diseases and Microbiology Public Health | NSW Health Pathology - Institute of Clinical Pathology and Medical Research; Westmead Hospital; University of Sydney | Bachmann N, Rockett R, Lam C, Gray K, Timms V, Gall M, Arnott A, Sadsad R, Draper J, Sim E, Carter I, Holmes EC, O'Sullivan MV, Byun R, Sintchenko V, Chen SC, Eden JS, Maddocks S, Kok J, Propenko M, Sorrell T, Chang S, Basile K, Dwyer DE for the 2019-nCoV Study Group |
| EPI_ISL_427761 | Centre for Infectious Diseases and Microbiology Public Health | NSW Health Pathology - Institute of Clinical Pathology and Medical Research; Westmead Hospital; University of Sydney | Arnott A, Sadsad R, Draper J, Sim E, Bachmann N, Rockett R, Lam C, Gray K, Timms V, Gall M, Carter I, Holmes EC, O'Sullivan MV, Byun R, Sintchenko V, Chen SC, Eden JS, Maddocks S, Kok J, Propenko M, Sorrell T, Chang S, Basile K, Dwyer DE for the 2019-nCoV Study Group |
| EPI_ISL_427762 | Centre for Infectious Diseases and Microbiology Public Health | NSW Health Pathology - Institute of Clinical Pathology and Medical Research; Westmead Hospital; University of Sydney | Bachmann N, Rockett R, Lam C, Gray K, Timms V, Gall M, Arnott A, Sadsad R, Draper J, Sim E, Carter I, Holmes EC, O'Sullivan MV, Byun R, Sintchenko V, Chen SC, Eden JS, Maddocks S, Kok J, Propenko M, Sorrell T, Chang S, Basile K, Dwyer DE for the 2019-nCoV Study Group |
| EPI_ISL_427770, EPI_ISL_427774 | Centre for Infectious Diseases and Microbiology Public Health | NSW Health Pathology - Institute of Clinical Pathology and | Arnott A, Sadsad R, Draper J, Sim E, Bachmann N, Rockett R, Lam C, Gray K, Timms V, Gall M, Carter I, Holmes EC, O'Sullivan MV, Byun R, Sintchenko |

| | | | |
|---|---|---|---|
| | | Medical Research; Westmead Hospital; University of Sydney | V, Chen SC, Eden JS, Maddocks S, Kok J, Propenko M, Sorrell T, Chang S, Basile K, Dwyer DE for the 2019-nCoV Study Group |
| EPI_ISL_427778, EPI_ISL_427787, EPI_ISL_427797 | Centre for Infectious Diseases and Microbiology Public Health | NSW Health Pathology - Institute of Clinical Pathology and Medical Research; Westmead Hospital; University of Sydney | Bachmann N, Rockett R, Lam C, Gray K, Timms V, Gall M, Arnott A, Sadsad R, Draper J, Sim E, Carter I, Holmes EC, O'Sullivan MV, Byun R, Sintchenko V, Chen SC, Eden JS, Maddocks S, Kok J, Propenko M, Sorrell T, Chang S, Basile K, Dwyer DE for the 2019-nCoV Study Group |
| EPI_ISL_427804 | Centre for Infectious Diseases and Microbiology Public Health | NSW Health Pathology - Institute of Clinical Pathology and Medical Research; Westmead Hospital; University of Sydney | Arnott A, Sadsad R, Draper J, Sim E, Bachmann N, Rockett R, Lam C, Gray K, Timms V, Gall M, Carter I, Holmes EC, O'Sullivan MV, Byun R, Sintchenko V, Chen SC, Eden JS, Maddocks S, Kok J, Propenko M, Sorrell T, Chang S, Basile K, Dwyer DE for the 2019-nCoV Study Group |
| EPI_ISL_427805 | Centre for Infectious Diseases and Microbiology Public Health | NSW Health Pathology - Institute of Clinical Pathology and Medical Research; Westmead Hospital; University of Sydney | Bachmann N, Rockett R, Lam C, Gray K, Timms V, Gall M, Arnott A, Sadsad R, Draper J, Sim E, Carter I, Holmes EC, O'Sullivan MV, Byun R, Sintchenko V, Chen SC, Eden JS, Maddocks S, Kok J, Propenko M, Sorrell T, Chang S, Basile K, Dwyer DE for the 2019-nCoV Study Group |
| EPI_ISL_428939, EPI_ISL_428940, EPI_ISL_428941, EPI_ISL_428942, EPI_ISL_428943, EPI_ISL_428944, EPI_ISL_428945, EPI_ISL_428946, EPI_ISL_428947, EPI_ISL_428948, EPI_ISL_428949, EPI_ISL_428950, EPI_ISL_428951, EPI_ISL_428952, EPI_ISL_428953, EPI_ISL_428954, EPI_ISL_428955, EPI_ISL_428956, EPI_ISL_428957, EPI_ISL_428958, EPI_ISL_428959, EPI_ISL_428960, EPI_ISL_428961, EPI_ISL_428962 | | | |
| see above | Laboratoire National de Sante, Microbiology, Virology | Laboratoire National de Sante, Microbiology, Epidemiology and Microbial Genomics | Anke Wienecke-Baldacchino, Ardashel Latsuzbaia, Jessica Tapp, Catherine Ragimbeau, Guillaume Fournier, Tamir Abdelrahman, Trung Nguyen Nguyen, Joel Mossong |
| EPI_ISL_429115, EPI_ISL_429116, EPI_ISL_429117, EPI_ISL_429118, EPI_ISL_429119, EPI_ISL_429120, EPI_ISL_429121, EPI_ISL_429122, EPI_ISL_429123, EPI_ISL_429124, EPI_ISL_429125 | | | |
| see above | Klinisk mikrobiologi och vardhygien Halmstad | The Public Health Agency of Sweden | Arne Kotz, Olov Svartstrom, Maria Lind Karlberg, Anna-Malin Linde, Oskar Karlsson Lindsjo, Anna Risberg, Shaman Muradrasoli, Karin Tegmark-Wisell |
| EPI_ISL_429706, EPI_ISL_429707, EPI_ISL_429708, EPI_ISL_429709, EPI_ISL_429710, EPI_ISL_429711, EPI_ISL_429712, EPI_ISL_429713, EPI_ISL_429714, EPI_ISL_429715, EPI_ISL_429716, EPI_ISL_429717, EPI_ISL_429718, EPI_ISL_429719, EPI_ISL_429720, EPI_ISL_429721, EPI_ISL_429722, EPI_ISL_429723, EPI_ISL_429724, EPI_ISL_429725, EPI_ISL_429726, EPI_ISL_429727, EPI_ISL_429728, EPI_ISL_429729, EPI_ISL_429730, EPI_ISL_429731, EPI_ISL_429732, EPI_ISL_429733, EPI_ISL_429734, EPI_ISL_429735, EPI_ISL_429736, EPI_ISL_429737, EPI_ISL_429738, EPI_ISL_429739, EPI_ISL_429740, EPI_ISL_429741, EPI_ISL_429742, EPI_ISL_429743, EPI_ISL_429744, EPI_ISL_429745, EPI_ISL_429746, EPI_ISL_429747, EPI_ISL_429748, EPI_ISL_429749, EPI_ISL_429750, EPI_ISL_429751, EPI_ISL_429752, EPI_ISL_429753, EPI_ISL_429754, EPI_ISL_429755, EPI_ISL_429756, EPI_ISL_429757, EPI_ISL_429758, EPI_ISL_429759, EPI_ISL_429760, EPI_ISL_429761, EPI_ISL_429762, EPI_ISL_429763, EPI_ISL_429764, EPI_ISL_429765, EPI_ISL_429766, EPI_ISL_429767, EPI_ISL_429768, EPI_ISL_429769, EPI_ISL_429770, EPI_ISL_429771, EPI_ISL_429772, EPI_ISL_429773, EPI_ISL_429774, EPI_ISL_429775, EPI_ISL_429776, EPI_ISL_429777, EPI_ISL_429778, EPI_ISL_429779, EPI_ISL_429780, EPI_ISL_429781, EPI_ISL_429782, EPI_ISL_429783, EPI_ISL_429784, EPI_ISL_429785, EPI_ISL_429786, EPI_ISL_429787, EPI_ISL_429788, EPI_ISL_429789, EPI_ISL_429790, EPI_ISL_429791, EPI_ISL_429793, EPI_ISL_429794, EPI_ISL_429795, EPI_ISL_429796, EPI_ISL_429797, EPI_ISL_429798, EPI_ISL_429799, EPI_ISL_429800 | | | |
| see above | Laboratoire National de Sante, Microbiology, Virology | Laboratoire National de Sante, Microbiology, Epidemiology and Microbial Genomics | Anke Wienecke-Baldacchino, Ardashel Latsuzbaia, Jessica Tapp, Catherine Ragimbeau, Guillaume Fournier, Tamir Abdelrahman, Trung Nguyen Nguyen, Joel Mossong |
| EPI_ISL_429806 | Dr. Georges-L.-Dumont University Hospital Centre | National Microbiology Laboratory | Anna Majer, Shari Tyson, Grace Seo, Kristyn Burak, Philip Mabon, Elsie Grudeski, Rhiannon Huzarewich, Russell Mandes, Jennifer Tanner, Natalie Knox, Morag Graham, Gary Van Domselaar, Richard Garceau, Guillaume Desnoyers, Nathalie Bastien, Yan Li, Timothy Booth, Matthew Gilmour |
| EPI_ISL_429807 | Cadham Provincial Laboratory | National Microbiology Laboratory | Anna Majer, Shari Tyson, Grace Seo, Kristyn Burak, Philip Mabon, Elsie Grudeski, Rhiannon Huzarewich, Russell Mandes, Jennifer Tanner, Natalie Knox, Morag Graham, Gary Van Domselaar, Paul Van Caeseele, Jared Bullard, David Alexander, Kerry Dust, Nathalie Bastien, Yan Li, Timothy Booth, Matthew Gilmour |
| EPI_ISL_429811 | Dr. Georges-L.-Dumont University Hospital Centre | National Microbiology Laboratory | Anna Majer, Shari Tyson, Grace Seo, Kristyn Burak, Philip Mabon, Elsie Grudeski, Rhiannon Huzarewich, Russell Mandes, Jennifer Tanner, Natalie Knox, Morag Graham, Gary Van Domselaar, Richard Garceau, Guillaume Desnoyers, Nathalie Bastien, Yan Li, Timothy Booth, Matthew Gilmour |
| EPI_ISL_429812, EPI_ISL_429813, EPI_ISL_429814 | Queen Elizabeth II Health Science Centre | National Microbiology Laboratory | Anna Majer, Shari Tyson, Grace Seo, Kristyn Burak, Philip Mabon, Elsie Grudeski, Rhiannon Huzarewich, Russell Mandes, Jennifer Tanner, Natalie Knox, Morag Graham, Gary Van Domselaar, Todd Hatchette, Jason LeBlanc, Nathalie Bastien, Yan Li, Timothy Booth, Matthew Gilmour |
| EPI_ISL_429815 | Public Health Laboratory | National Microbiology Laboratory | Anna Majer, Shari Tyson, Grace Seo, Kristyn Burak, Philip Mabon, Elsie Grudeski, Rhiannon Huzarewich, Russell Mandes, Jennifer Tanner, Natalie Knox, Morag Graham, Gary Van Domselaar, Robert Needle, Yang Yu, Adel Malek, Laura Gilbert, George Zahariadis, Nathalie Bastien, Yan Li, Timothy Booth, Matthew Gilmour |
| EPI_ISL_429816 | Dr. Georges-L.-Dumont University Hospital Centre | National Microbiology Laboratory | Anna Majer, Shari Tyson, Grace Seo, Kristyn Burak, Philip Mabon, Elsie Grudeski, Rhiannon Huzarewich, Russell Mandes, Jennifer Tanner, Natalie Knox, Morag Graham, Gary Van Domselaar, Richard Garceau, Guillaume Desnoyers, Nathalie Bastien, Yan Li, Timothy Booth, Matthew Gilmour |
| EPI_ISL_429817, EPI_ISL_429818, EPI_ISL_429819 | Cadham Provincial Laboratory | National Microbiology Laboratory | Anna Majer, Shari Tyson, Grace Seo, Kristyn Burak, Philip Mabon, Elsie Grudeski, Rhiannon Huzarewich, Russell Mandes, Jennifer Tanner, Natalie Knox, Morag Graham, Gary Van Domselaar, Paul Van Caeseele, Jared Bullard, David Alexander, Kerry Dust, Nathalie Bastien, Yan Li, Timothy Booth, Matthew Gilmour |
| EPI_ISL_429820 | Cadham Provincial Laboratory | National Microbiology Laboratory | Anna Majer, Shari Tyson, Grace Seo, Kristyn Burak, Philip Mabon, Elsie Grudeski, Rhiannon Huzarewich, Russell Mandez, Jennifer Tanner, Natalie Knox, Morag Graham, Gary Van Domselaar, Paul Van Caeseele, Jared Bullard, David Alexander, Kerry Dust, Nathalie Bastien, Yan Li, Timothy Booth, Matthew Gilmour |
| EPI_ISL_430464, EPI_ISL_430465, EPI_ISL_430466, EPI_ISL_430467, EPI_ISL_430468 | ICMR-National Institute of Cholera and Enteric Diseases | National Institute of Biomedical Genomics | Arindam Maitra, Mamta Chawla Sarkar, Sreedhar Chinnaswamy, Hasina Banu, Ananya Chatterjee, Shanta Dutta, Saumitra Das |
| EPI_ISL_430846 | General Intensive Care Unit, Raymond Poincaré Hospital (AP-HP), Lab Inflammation & Infection, U1173 University Paris Saclay-UVSQ/INSERM, Garches, France. | Institut Pasteur, Laboratory for Urgent Response to biological Threats | Annane Djillali, Vanhomwegen Jessica, Caro Valérie, Manuguerra Jean-Claude |
| EPI_ISL_430848, EPI_ISL_430849, EPI_ISL_430850, EPI_ISL_430851, EPI_ISL_430852, EPI_ISL_430853, EPI_ISL_430854, EPI_ISL_430855 | Klinisk mikrobiologi och vardhygien Halmstad | The Public Health Agency of Sweden | Arne Kotz, Oskar Karlsson Lindsjo, Maria Lind Karlberg, Anna-Malin Linde, Olov Svartstrom, Anna Risberg, Shaman Muradrasoli, Karin Tegmark-Wisell |
| EPI_ISL_434487, EPI_ISL_434488, EPI_ISL_434489, EPI_ISL_434490, EPI_ISL_434491, EPI_ISL_434492, EPI_ISL_434493, EPI_ISL_434494, EPI_ISL_434495, EPI_ISL_434496, EPI_ISL_434497, EPI_ISL_434498, EPI_ISL_434499, EPI_ISL_434500, EPI_ISL_434501, EPI_ISL_434502, EPI_ISL_434503, EPI_ISL_434504, EPI_ISL_434505, EPI_ISL_434506, EPI_ISL_434507, EPI_ISL_434508, EPI_ISL_434509, EPI_ISL_434510, EPI_ISL_434511, EPI_ISL_434512, EPI_ISL_434513, EPI_ISL_434514, EPI_ISL_434515 | | | |
| see above | Laboratoire National de Sante, Microbiology, Virology | Laboratoire National de Sante, Microbiology, Epidemiology and Microbial Genomics | Anke Wienecke-Baldacchino, Ardashel Latsuzbaia, Jessica Tapp, Catherine Ragimbeau, Guillaume Fournier, Tamir Abdelrahman, Trung Nguyen Nguyen, Joel Mossong |
| EPI_ISL_434643, EPI_ISL_434655 | Uppsala Narakut Aleris | The Public Health Agency of Sweden | Annika Nilsson, Oskar Karlsson Lindsjo, Maria Lind Karlberg, Anna-Malin Linde, Olov Svartstrom, Anna Risberg, Theresa Enkirch, Mia Brytting, Karin Tegmark-Wisell |
| EPI_ISL_434661, EPI_ISL_434662, EPI_ISL_434664 | Omtanken Grimmered | The Public Health Agency of Sweden | Bernd Sengpiel, Oskar Karlsson Lindsjo, Maria Lind Karlberg, Anna-Malin Linde, Olov Svartstrom, Anna Risberg, Theresa Enkirch, Mia Brytting, Karin Tegmark-Wisell |
| EPI_ISL_434665 | Uppsala Narakut Aleris | The Public Health Agency of Sweden | Annika Nilsson, Oskar Karlsson Lindsjo, Maria Lind Karlberg, Anna-Malin Linde, Olov Svartstrom, Anna Risberg, Theresa Enkirch, Mia Brytting, Karin Tegmark-Wisell |
| EPI_ISL_434673 | Omtanken Grimmered | The Public Health Agency of Sweden | Bernd Sengpiel, Oskar Karlsson Lindsjo, Maria Lind Karlberg, Anna-Malin Linde, Olov Svartstrom, Anna Risberg, Theresa Enkirch, Mia Brytting, Karin Tegmark-Wisell |
| EPI_ISL_435050 | B.J. Medical College and Civil hospital | Gujarat Biotechnology Research Centre | Ankit Hinsu, Pritesh Sabara, Apurvasinh Puvar, Janvi Raval, Monika Gandhi, Pinal Trivedi, Maharshi Pandya, Amit Kanani, Akanksha Verma, Nitin Savaliya, Raghawendra Kumar, Dinesh Kumar, Zuber Saiyed, Dipa Kinariwala, Disha Patel, Binita Aring, Geeta Vaghela, Sonia Barve, Bhavesh Modi, Kairavi Joshi, Gaurishankar Shrimali, Nidhi Sood, Pranay Shah, R D Dixit, Snehal Bagatharia, Kamlesh J Upadhyay, Ramesh Pandit, Tejas Shah, Dipeshwari Shewale, Chaitanya Joshi, Madhvi Joshi |
| EPI_ISL_435052 | B.J. Medical College and Civil hospital | Gujarat Biotechnology Research Centre | Apurvasinh Puvar, Janvi Raval, Monika Gandhi, Pinal Trivedi, Maharshi Pandya, Amit Kanani, Akanksha Verma, Nitin Savaliya, Raghawendra Kumar, Dinesh Kumar, Zuber Saiyed, Dipa Kinariwala, Disha Patel, Binita Aring, Geeta Vaghela, Sonia Barve, Bhavesh Modi, Kairavi Joshi, Gaurishankar Shrimali, Nidhi Sood, Pranay Shah, R D Dixit, Snehal Bagatharia, Kamlesh J Upadhyay, Ramesh Pandit, Tejas Shah, Ankit Hinsu, Pritesh Sabara, Pooja P Doshi, Chaitanya Joshi, Madhvi Joshi |
| EPI_ISL_436197, EPI_ISL_436205, EPI_ISL_436213, EPI_ISL_436221, | Servicio de Microbiología. Consorcio Hospital General Universitario de Valencia | Sequencing and Bioinformatics Service and Molecular Epidemiology Research Group. FISABIO-Public Health | Beatriz Beamud, Lidia Ruiz Roldan, Marta Pla Diaz,Neris Garcia-Gonzalez, Loreto Ferrús Abad, Maria Dolores Ocete, Inma Galán Vendrell, Paula Ruiz-Hueso, Mariana Reyes-Prieto, Vicente Soriano Chirona, Maria Alma Bracho, Griselda De Marco, Llúcia Martínez-Priego, Concepcion Gimeno, |

| | | | |
|---|---|---|---|
| EPI_ISL_436228, EPI_ISL_436272 | | | Giuseppe D'Auria, Fernando Gonzalez-Candelas |
| EPI_ISL_436280 | Servicio de Microbiologia. Hospital Clinico Universitario de Valencia | Sequencing and Bioinformatics Service and Molecular Epidemiology Research Group. FISABIO-Public Health | Beatriz Beamud, Lidia Ruiz Roldan, Marta Pla Diaz, Neris Garcia-Gonzalez, Inma Galán Vendrell, Sandra Carbo, Loreto Ferrús Abad, Paula Ruiz-Hueso, Mariana Reyes-Prieto, Vicente Soriano Chirona, Ivan Ansari, David Navarro, Maria Alma Bracho, Griselda De Marco, Llúcia Martínez-Priego, Giuseppe D'Auria, Fernando Gonzalez-Candelas |
| EPI_ISL_436294 | Servicio de Microbiologia. Hospital Clinico Universitario de Valencia | Sequencing and Bioinformatics Service and Molecular Epidemiology Research Group. FISABIO-Public Health | Beatriz Beamud, Lidia Ruiz Roldan, Marta Pla Diaz, Neris Garcia-Gonzalez, Inma Galán Vendrell, Sandra Carbo, Loreto Ferrús Abad, Paula Ruiz-Hueso, Mariana Reyes-Prieto, Vicente Soriano Chirona, Ivan Ansari, David Navarro, Maria Alma Bracho, Griselda De Marco, Llúcia Martínez-Priego, Giuseppe D'Auria, Fernando Gonzalez-Candelas |
| EPI_ISL_436300 | Servicio de Microbiologia. Hospital Clinico Universitario de Valencia | Sequencing and Bioinformatics Service and Molecular Epidemiology Research Group. FISABIO-Public Health | Beatriz Beamud, Lidia Ruiz Roldan, Marta Pla Diaz, Neris Garcia-Gonzalez, Inma Galán Vendrell, Sandra Carbo, Loreto Ferrús Abad, Paula Ruiz-Hueso, Mariana Reyes-Prieto, Vicente Soriano Chirona, Ivan Ansari, David Navarro, Maria Alma Bracho, Griselda De Marco, Llúcia Martínez-Priego, Giuseppe D'Auria, Fernando Gonzalez-Candelas |
| EPI_ISL_436314 | Servicio de Microbiologia. Hospital Clinico Universitario de Valencia | Sequencing and Bioinformatics Service and Molecular Epidemiology Research Group. FISABIO-Public Health | Beatriz Beamud, Lidia Ruiz Roldan, Marta Pla Diaz, Neris Garcia-Gonzalez, Inma Galán Vendrell, Sandra Carbo, Loreto Ferrús Abad, Paula Ruiz-Hueso, Mariana Reyes-Prieto, Vicente Soriano Chirona, Ivan Ansari, David Navarro, Maria Alma Bracho, Griselda De Marco, Llúcia Martínez-Priego, Giuseppe D'Auria, Fernando Gonzalez-Candelas |
| EPI_ISL_436320 | Servicio de Microbiologia. Hospital Clinico Universitario de Valencia | Sequencing and Bioinformatics Service and Molecular Epidemiology Research Group. FISABIO-Public Health | Beatriz Beamud, Lidia Ruiz Roldan, Marta Pla Diaz, Neris Garcia-Gonzalez, Inma Galán Vendrell, Sandra Carbo, Loreto Ferrús Abad, Paula Ruiz-Hueso, Mariana Reyes-Prieto, Vicente Soriano Chirona, Ivan Ansari, David Navarro, Maria Alma Bracho, Griselda De Marco, Llúcia Martínez-Priego, Giuseppe D'Auria, Fernando Gonzalez-Candelas |
| EPI_ISL_436334 | Servicio de Microbiologia. Hospital Clinico Universitario de Valencia | Sequencing and Bioinformatics Service and Molecular Epidemiology Research Group. FISABIO-Public Health | Beatriz Beamud, Lidia Ruiz Roldan, Marta Pla Diaz, Neris Garcia-Gonzalez, Inma Galán Vendrell, Sandra Carbo, Loreto Ferrús Abad, Paula Ruiz-Hueso, Mariana Reyes-Prieto, Vicente Soriano Chirona, Ivan Ansari, David Navarro, Maria Alma Bracho, Griselda De Marco, Llúcia Martínez-Priego, Giuseppe D'Auria, Fernando Gonzalez-Candelas |
| EPI_ISL_436340 | Servicio de Microbiologia. Hospital Clinico Universitario de Valencia | Sequencing and Bioinformatics Service and Molecular Epidemiology Research Group. FISABIO-Public Health | Beatriz Beamud, Lidia Ruiz Roldan, Marta Pla Diaz, Neris Garcia-Gonzalez, Inma Galán Vendrell, Sandra Carbo, Loreto Ferrús Abad, Paula Ruiz-Hueso, Mariana Reyes-Prieto, Vicente Soriano Chirona, Ivan Ansari, David Navarro, Maria Alma Bracho, Griselda De Marco, Llúcia Martínez-Priego, Giuseppe D'Auria, Fernando Gonzalez-Candelas |
| EPI_ISL_436354 | Servicio de Microbiologia. Hospital Clinico Universitario de Valencia | Sequencing and Bioinformatics Service and Molecular Epidemiology Research Group. FISABIO-Public Health | Beatriz Beamud, Lidia Ruiz Roldan, Marta Pla Diaz, Neris Garcia-Gonzalez, Inma Galán Vendrell, Sandra Carbo, Loreto Ferrús Abad, Paula Ruiz-Hueso, Mariana Reyes-Prieto, Vicente Soriano Chirona, Ivan Ansari, David Navarro, Maria Alma Bracho, Griselda De Marco, Llúcia Martínez-Priego, Giuseppe D'Auria, Fernando Gonzalez-Candelas |
| EPI_ISL_436364 | Servicio de Microbiologia. Hospital Clinico Universitario de Valencia | Sequencing and Bioinformatics Service and Molecular Epidemiology Research Group. FISABIO-Public Health | Beatriz Beamud, Lidia Ruiz Roldan, Marta Pla Diaz, Neris Garcia-Gonzalez, Inma Galán Vendrell, Sandra Carbo, Loreto Ferrús Abad, Paula Ruiz-Hueso, Mariana Reyes-Prieto, Vicente Soriano Chirona, Ivan Ansari, David Navarro, Maria Alma Bracho, Griselda De Marco, Llúcia Martínez-Priego, Giuseppe D'Auria, Fernando Gonzalez-Candelas |
| EPI_ISL_436385, EPI_ISL_436392 | Servicio de Microbiología. Consorcio Hospital General Universitario de Valencia | Sequencing and Bioinformatics Service and Molecular Epidemiology Research Group. FISABIO-Public Health | Beatriz Beamud, Lidia Ruiz Roldan, Marta Pla Diaz,Neris Garcia-Gonzalez, Loreto Ferrús Abad, Maria Dolores Ocete, Inma Galán Vendrell, Paula Ruiz-Hueso, Mariana Reyes-Prieto, Vicente Soriano Chirona, Maria Alma Bracho, Griselda De Marco, Llúcia Martínez-Priego, Concepcion Gimeno, Giuseppe D'Auria, Fernando Gonzalez-Candelas |
| EPI_ISL_436405 | Servicio de Microbiologia. Hospital Clinico Universitario de Valencia | Sequencing and Bioinformatics Service and Molecular Epidemiology Research Group. FISABIO-Public Health | Beatriz Beamud, Lidia Ruiz Roldan, Marta Pla Diaz, Neris Garcia-Gonzalez, Inma Galán Vendrell, Sandra Carbo, Loreto Ferrús Abad, Paula Ruiz-Hueso, Mariana Reyes-Prieto, Vicente Soriano Chirona, Ivan Ansari, David Navarro, Maria Alma Bracho, Griselda De Marco, Llúcia Martínez-Priego, Giuseppe D'Auria, Fernando Gonzalez-Candelas |
| EPI_ISL_436800, EPI_ISL_436801, EPI_ISL_436802, EPI_ISL_436803, EPI_ISL_436804, EPI_ISL_436805, EPI_ISL_436806, EPI_ISL_436807, EPI_ISL_436808, EPI_ISL_436809, EPI_ISL_436810, EPI_ISL_436811, EPI_ISL_436812, EPI_ISL_436813, EPI_ISL_436814, EPI_ISL_436815, EPI_ISL_436816, EPI_ISL_436817, EPI_ISL_436818, EPI_ISL_436819, EPI_ISL_436820, EPI_ISL_436821, EPI_ISL_436822, EPI_ISL_436823, EPI_ISL_436824, EPI_ISL_436825, EPI_ISL_436826, EPI_ISL_436827, EPI_ISL_436828, EPI_ISL_436829, EPI_ISL_436830, EPI_ISL_436831, EPI_ISL_436832, EPI_ISL_436833, EPI_ISL_436834, EPI_ISL_436835, EPI_ISL_436836, EPI_ISL_436837, EPI_ISL_436838, EPI_ISL_436839, EPI_ISL_436840, EPI_ISL_436841, EPI_ISL_436842, EPI_ISL_436843, EPI_ISL_436844, EPI_ISL_436845, EPI_ISL_436846, EPI_ISL_436847, EPI_ISL_436848, EPI_ISL_436849, EPI_ISL_436850, EPI_ISL_436851, EPI_ISL_436852, EPI_ISL_436853, EPI_ISL_436854, EPI_ISL_436855, EPI_ISL_436856, EPI_ISL_436857, EPI_ISL_436858, EPI_ISL_436859, EPI_ISL_436860, EPI_ISL_436861, EPI_ISL_436862, EPI_ISL_436863, EPI_ISL_436864, EPI_ISL_436865, EPI_ISL_436866, EPI_ISL_436867, EPI_ISL_436868, EPI_ISL_436869, EPI_ISL_436870, EPI_ISL_436871, EPI_ISL_436872, EPI_ISL_436873, EPI_ISL_436874, EPI_ISL_436875, EPI_ISL_436876, EPI_ISL_436877, EPI_ISL_436878, EPI_ISL_436879, EPI_ISL_436880, EPI_ISL_436881, EPI_ISL_436882, EPI_ISL_436883, EPI_ISL_436884, EPI_ISL_436885, EPI_ISL_436886, EPI_ISL_436887, EPI_ISL_436888, EPI_ISL_436889, EPI_ISL_436890, EPI_ISL_437097, EPI_ISL_437098, EPI_ISL_437099, EPI_ISL_437100, EPI_ISL_437101, EPI_ISL_437102, EPI_ISL_437103, EPI_ISL_437104, EPI_ISL_437105, EPI_ISL_437106, EPI_ISL_437107, EPI_ISL_437108, EPI_ISL_437109, EPI_ISL_437110, EPI_ISL_437111, EPI_ISL_437112, EPI_ISL_437113, EPI_ISL_437114, EPI_ISL_437115, EPI_ISL_437116, EPI_ISL_437117, EPI_ISL_437118, EPI_ISL_437119, EPI_ISL_437120, EPI_ISL_437121, EPI_ISL_437122, EPI_ISL_437123, EPI_ISL_437124, EPI_ISL_437125, EPI_ISL_437126, EPI_ISL_437127, EPI_ISL_437128, EPI_ISL_437129, EPI_ISL_437130, EPI_ISL_437131, EPI_ISL_437132, EPI_ISL_437133, EPI_ISL_437134, EPI_ISL_437135, EPI_ISL_437136, EPI_ISL_437137, EPI_ISL_437138, EPI_ISL_437139, EPI_ISL_437140, EPI_ISL_437141, EPI_ISL_437142, EPI_ISL_437143, EPI_ISL_437144, EPI_ISL_437145, EPI_ISL_437146, EPI_ISL_437147, EPI_ISL_437148, EPI_ISL_437149, EPI_ISL_437150, EPI_ISL_437151, EPI_ISL_437152, EPI_ISL_437153, EPI_ISL_437154, EPI_ISL_437155, EPI_ISL_437156, EPI_ISL_437157, EPI_ISL_437158, EPI_ISL_437159, EPI_ISL_437160, EPI_ISL_437161, EPI_ISL_437162, EPI_ISL_437163, EPI_ISL_437164, EPI_ISL_437165, EPI_ISL_437166, EPI_ISL_437167, EPI_ISL_437168, EPI_ISL_437169, EPI_ISL_437170, EPI_ISL_437171, EPI_ISL_437172, EPI_ISL_437173, EPI_ISL_437174, EPI_ISL_437175, EPI_ISL_437176, EPI_ISL_437177, EPI_ISL_437178, EPI_ISL_437179, EPI_ISL_437180, EPI_ISL_437181, EPI_ISL_437182, EPI_ISL_437183, EPI_ISL_437184, EPI_ISL_437185, EPI_ISL_437186 | | | |
| see above | Michigan Department of Health and Human Services, Bureau of Laboratories | Michigan Department of Health and Human Services, Bureau of Laboratories | Blankenship HM, Riner D, Soehnlen MK |
| EPI_ISL_437433 | Bozeman Health Deaconess Hospital | Wiedenheft lab, Montana State University | Artem Nemudryi, Anna Nemudraia, Kevin Surya, Tanner Wiegand, Murat Buyukyoruk, Royce Wilkinson, Blake Wiedenheft |
| EPI_ISL_437446 | B.J. Medical College and Civil hospital | Gujarat Biotechnology Research Centre | Binita Aring, Neeta Khandelwal, Geeta Vaghela, Sonia Barve, Bhavesh Modi, Kairavi Joshi, Gaurishankar Shrimali, Nidhi Sood, Pranay Shah, R D Dixit, Snehal Bagatharia, Kamlesh J Upadhyay, Ramesh Pandit, Tejas Shah, Ankit Hinsu, Pritesh Sabara, Apurvasinh Puvar, Janvi Raval, Monika Gandhi, Pinal Trivedi, Maharshi Pandya, Amit Kanani, Akanksha Verma, Nitin Savaliya, Raghawendra Kumar, Dinesh Kumar, Zuber Saiyed, Dipa Kinariwala, Disha Patel, Priti Pandita, Chaitanya Joshi, Madhvi Joshi |
| EPI_ISL_437450 | B.J. Medical College and Civil hospital | Gujarat Biotechnology Research Centre | Bhavesh Modi, Kairavi Joshi, Gaurishankar Shrimali, Nidhi Sood, Pranay Shah, R D Dixit, Snehal Bagatharia, Kamlesh J Upadhyay, Ramesh Pandit, Tejas Shah, Ankit Hinsu, Pritesh Sabara, Apurvasinh Puvar, Janvi Raval, Monika Gandhi, Pinal Trivedi, Maharshi Pandya, Amit Kanani, Akanksha Verma, Nitin Savaliya, Raghawendra Kumar, Dinesh Kumar, Zuber Saiyed, Dipa Kinariwala, Disha Patel, Binita Aring, Neeta Khandelwal, Geeta Vaghela, Sonia Barve, Armi Chaudhari, Chaitanya Joshi, Madhvi Joshi |
| EPI_ISL_437536, EPI_ISL_437537, EPI_ISL_437538, EPI_ISL_437539 | ICMR-National Institute of Cholera and Enteric Diseases | National Institute of Biomedical Genomics | Arindam Maitra, Mamta Chawla Sarkar, Sreedhar Chinnaswamy, Hasina Banu, Ananya Chatterjee, Shanta Dutta, Saumitra Das |
| EPI_ISL_444461 | B.J. Medical College and Civil hospital | Gujarat Biotechnology Research Centre | Ankit Hinsu, Pritesh Sabara, Apurvasinh Puvar, Janvi Raval, Monika Gandhi, Pinal Trivedi, Maharshi Pandya, Amit Kanani, Akanksha Verma, Nitin Savaliya, Raghawendra Kumar, Dinesh Kumar, Zuber Saiyed, Dipa Kinariwala, Disha Patel, Binita Aring, Neeta Khandelwal, Geeta Vaghela, Sonia Barve, Bhavesh Modi, Kairavi Joshi, Gaurishankar Shrimali, Nidhi Sood, Pranay Shah, R D Dixit, Snehal Bagatharia, Kamlesh J Upadhyay, Ramesh Pandit, Neelam Nathani, Chaitanya Joshi, Madhvi Joshi, Tejas Shah |
| EPI_ISL_444463 | B.J. Medical College and Civil hospital | Gujarat Biotechnology Research Centre | Apurvasinh Puvar, Janvi Raval, Monika Gandhi, Pinal Trivedi, Maharshi Pandya, Amit Kanani, Akanksha Verma, Nitin Savaliya, Raghawendra Kumar, Dinesh Kumar, Zuber Saiyed, Dipa Kinariwala, Disha Patel, Binita Aring, Neeta Khandelwal, Geeta Vaghela, Sonia Barve, Bhavesh Modi, Kairavi Joshi, Gaurishankar Shrimali, Nidhi Sood, Pranay Shah, R D Dixit, Snehal Bagatharia, Kamlesh J Upadhyay, Ramesh Pandit, Tejas Shah, Ankit Hinsu, Pritesh Sabara, Bhavya Jindal, Chaitanya Joshi, Madhvi Joshi |
| EPI_ISL_444476 | B.J. Medical College and Civil hospital | Gujarat Biotechnology Research Centre | Binita Aring, Neeta Khandelwal, Geeta Vaghela, Sonia Barve, Bhavesh Modi, Kairavi Joshi, Gaurishankar Shrimali, Nidhi Sood, Pranay Shah, R D Dixit, Snehal Bagatharia, Kamlesh J Upadhyay, Ramesh Pandit, Tejas Shah, Ankit Hinsu, Pritesh Sabara, Apurvasinh Puvar, Janvi Raval, Monika Gandhi, Pinal Trivedi, Maharshi Pandya, Amit Kanani, Akanksha Verma, Nitin Savaliya, Raghawendra Kumar, Dinesh Kumar, Zuber Saiyed, Dipa Kinariwala, Disha Patel, Chaitanya Joshi, Madhvi Joshi, Dipeshwari Shewale |
| EPI_ISL_444480 | B.J. Medical College and Civil hospital | Gujarat Biotechnology Research Centre | Bhavesh Modi, Kairavi Joshi, Gaurishankar Shrimali, Nidhi Sood, Pranay Shah, R D Dixit, Snehal Bagatharia, Kamlesh J Upadhyay, Ramesh Pandit, Tejas Shah, Ankit Hinsu, Pritesh Sabara, Apurvasinh Puvar, Janvi Raval, Monika Gandhi, Pinal Trivedi, Maharshi Pandya, Amit Kanani, Akanksha Verma, Nitin Savaliya, Raghawendra Kumar, Dinesh Kumar, Zuber Saiyed, Dipa Kinariwala, Disha Patel, Binita Aring, Neeta Khandelwal, Geeta Vaghela, Sonia Barve, Priti Pandita, Chaitanya Joshi, Madhvi Joshi |
| EPI_ISL_444611 | Pathology Queensland | Public Health Virology Laboratory | Bixing Huang, Alyssa Pyke, Amanda De Jong, Andrew Van Den Hurk, Carmel Taylor, David Warrilow, Doris Genge, Elisabeth Gamez, Glen Hewitson, Ian Maxwell Mackay, Inga Sultana, Jamie McMahon, Jean Barcelon, Judy Northill, Mitchell Finger, Natalie Simpson, Neelima Nair, Peter Burtonclay, Peter |

| | | | |
|---|---|---|---|
| | | | Moore, Sarah Wheatley, Sean Moody, Sonja Hall-Mendelin, Timothy Gardam, and Frederick Moore |
| EPI_ISL_444612 | QML Pathology | Public Health Virology Laboratory | Bixing Huang, Alyssa Pyke, Amanda De Jong, Andrew Van Den Hurk, Carmel Taylor, David Warrilow, Doris Genge, Elisabeth Gamez, Glen Hewitson, Ian Maxwell Mackay, Inga Sultana, Jamie McMahon, Jean Barcelon, Judy Northill, Mitchell Finger, Natalie Simpson, Neelima Nair, Peter Burtonclay, Peter Moore, Sarah Wheatley, Sean Moody, Sonja Hall-Mendelin, Timothy Gardam, and Frederick Moore |
| EPI_ISL_444793 | Pathology Queensland | Public Health Virology Laboratory | Bixing Huang, Alyssa Pyke, Amanda De Jong, Andrew Van Den Hurk, Carmel Taylor, David Warrilow, Doris Genge, Elisabeth Gamez, Glen Hewitson, Ian Maxwell Mackay, Inga Sultana, Jamie McMahon, Jean Barcelon, Judy Northill, Mitchell Finger, Natalie Simpson, Neelima Nair, Peter Burtonclay, Peter Moore, Sarah Wheatley, Sean Moody, Sonja Hall-Mendelin, Timothy Gardam, and Frederick Moore |
| EPI_ISL_444794 | Cairns Hospital | Public Health Virology Laboratory | Bixing Huang, Alyssa Pyke, Amanda De Jong, Andrew Van Den Hurk, Carmel Taylor, David Warrilow, Doris Genge, Elisabeth Gamez, Glen Hewitson, Ian Maxwell Mackay, Inga Sultana, Jamie McMahon, Jean Barcelon, Judy Northill, Mitchell Finger, Natalie Simpson, Neelima Nair, Peter Burtonclay, Peter Moore, Sarah Wheatley, Sean Moody, Sonja Hall-Mendelin, Timothy Gardam, and Frederick Moore |
| EPI_ISL_445053, EPI_ISL_445054, EPI_ISL_445055, EPI_ISL_445056, EPI_ISL_445057, EPI_ISL_445058, EPI_ISL_445059, EPI_ISL_445060, EPI_ISL_445061, EPI_ISL_445062, EPI_ISL_445063, EPI_ISL_445064, EPI_ISL_445065, EPI_ISL_445066, EPI_ISL_445067, EPI_ISL_445068, EPI_ISL_445069, EPI_ISL_445070, EPI_ISL_445071, EPI_ISL_445072, EPI_ISL_445073, EPI_ISL_445074, EPI_ISL_445075, EPI_ISL_445076 | | | |
| see above | Laboratoire National de Sante, Microbiology, Virology | Laboratoire National de Sante, Microbiology, Epidemiology and Microbial Genomics | Anke Wienecke-Baldacchino, Ardashel Latsuzbaia, Jessica Tapp, Catherine Ragimbeau, Guillaume Fournier, Tamir Abdelrahman, Trung Nguyen Nguyen, Joel Mossong |
| EPI_ISL_445219 | Universidad del Valle, Laboratorio de Microbiologia, VIREM | Universidad del Valle, Universidad Nacional de Colombia-Sede Palmira, International Center for Tropical Agriculture | Beatriz Parra, Diana López-Alvarez, Wilmer J. Cuellar |
| EPI_ISL_445227, EPI_ISL_445230, EPI_ISL_445231, EPI_ISL_445239 | Uppsala Narakut Aleris | The Public Health Agency of Sweden | Annika Nilsson, Oskar Karlsson Lindsjo, Maria Lind Karlberg, Anna-Malin Linde, Olov Svartstrom, Anna Risberg, Theresa Enkirch, Mia Brytting, Karin Tegmark-Wisell |
| EPI_ISL_447033 | B.J. Medical College and Civil hospital | Gujarat Biotechnology Research Centre | Ankit Hinsu, Pritesh Sabara, Apurvasinh Puvar, Janvi Raval, Monika Gandhi, Pinal Trivedi, Maharshi Pandya, Amit Kanani, Akanksha Verma, Nitin Savaliya, Raghawendra Kumar, Dinesh Kumar, Zuber Saiyed, Dipa Kinariwala, Disha Patel, Binita Aring, Neeta Khandelwal, Geeta Vaghela, Sonia Barve, Bhavesh Modi, Kairavi Joshi, Gaurishankar Shrimali, Nidhi Sood, Pranay Shah, R D Dixit, Snehal Bagatharia, Kamlesh J Upadhyay, Ramesh Pandit, Tejas Shah, Nidhi Patel, Chaitanya Joshi, Madhvi Joshi |
| EPI_ISL_447035 | B.J. Medical College and Civil hospital | Gujarat Biotechnology Research Centre | Apurvasinh Puvar, Janvi Raval, Monika Gandhi, Pinal Trivedi, Maharshi Pandya, Amit Kanani, Akanksha Verma, Nitin Savaliya, Raghawendra Kumar, Dinesh Kumar, Zuber Saiyed, Dipa Kinariwala, Disha Patel, Binita Aring, Neeta Khandelwal, Geeta Vaghela, Sonia Barve, Bhavesh Modi, Kairavi Joshi, Gaurishankar Shrimali, Nidhi Sood, Pranay Shah, R D Dixit, Snehal Bagatharia, Kamlesh J Upadhyay, Ramesh Pandit, Tejas Shah, Ankit Hinsu, Pritesh Sabara, Neha Rajpara, Chaitanya Joshi, Madhvi Joshi |
| EPI_ISL_447048 | GMERS Medical College and Hospital, Gandhinagar | Gujarat Biotechnology Research Centre | Binita Aring, Neeta Khandelwal, Geeta Vaghela, Sonia Barve, Bhavesh Modi, Kairavi Joshi, Gaurishankar Shrimali, Nidhi Sood, Pranay Shah, R D Dixit, Snehal Bagatharia, Kamlesh J Upadhyay, Ramesh Pandit, Tejas Shah, Ankit Hinsu, Pritesh Sabara, Apurvasinh Puvar, Janvi Raval, Monika Gandhi, Pinal Trivedi, Maharshi Pandya, Amit Kanani, Akanksha Verma, Nitin Savaliya, Raghawendra Kumar, Dinesh Kumar, Zuber Saiyed, Dipa Kinariwala, Disha Patel, Armi Chaudhari, Chaitanya Joshi, Madhvi Joshi |
| EPI_ISL_447052 | GMERS Medical College and Hospital, Gandhinagar | Gujarat Biotechnology Research Centre | Bhavesh Modi, Kairavi Joshi, Gaurishankar Shrimali, Nidhi Sood, Pranay Shah, R D Dixit, Snehal Bagatharia, Kamlesh J Upadhyay, Ramesh Pandit, Tejas Shah, Ankit Hinsu, Pritesh Sabara, Apurvasinh Puvar, Janvi Raval, Monika Gandhi, Pinal Trivedi, Maharshi Pandya, Amit Kanani, Akanksha Verma, Nitin Savaliya, Raghawendra Kumar, Dinesh Kumar, Zuber Saiyed, Dipa Kinariwala, Disha Patel, Binita Aring, Neeta Khandelwal, Geeta Vaghela, Sonia Barve, Sharmistha Majumdar, Chaitanya Joshi, Madhvi Joshi |
| EPI_ISL_447057, EPI_ISL_447058, EPI_ISL_447059, EPI_ISL_447060, EPI_ISL_447061, EPI_ISL_447062, EPI_ISL_447063, EPI_ISL_447064, EPI_ISL_447065, EPI_ISL_447066, EPI_ISL_447067, EPI_ISL_447068, EPI_ISL_447069, EPI_ISL_447070, EPI_ISL_447071, EPI_ISL_447072, EPI_ISL_447073, EPI_ISL_447074, EPI_ISL_447075, EPI_ISL_447076, EPI_ISL_447077, EPI_ISL_447078, EPI_ISL_447079, EPI_ISL_447080, EPI_ISL_447081, EPI_ISL_447082, EPI_ISL_447083, EPI_ISL_447084, EPI_ISL_447085, EPI_ISL_447086, EPI_ISL_447087, EPI_ISL_447088, EPI_ISL_447089, EPI_ISL_447090, EPI_ISL_447091, EPI_ISL_447092, EPI_ISL_447093, EPI_ISL_447094, EPI_ISL_447095, EPI_ISL_447096, EPI_ISL_447097, EPI_ISL_447098, EPI_ISL_447099, EPI_ISL_447100, EPI_ISL_447101, EPI_ISL_447102, EPI_ISL_447103, EPI_ISL_447104, EPI_ISL_447105, EPI_ISL_447106, EPI_ISL_447107, EPI_ISL_447108, EPI_ISL_447109, EPI_ISL_447110, EPI_ISL_447111, EPI_ISL_447112, EPI_ISL_447113, EPI_ISL_447114, EPI_ISL_447115, EPI_ISL_447116, EPI_ISL_447117, EPI_ISL_447118, EPI_ISL_447163, EPI_ISL_447164, EPI_ISL_447165, EPI_ISL_447166, EPI_ISL_447167, EPI_ISL_447168, EPI_ISL_447169, EPI_ISL_447170, EPI_ISL_447171, EPI_ISL_447172, EPI_ISL_447173, EPI_ISL_447174, EPI_ISL_447175, EPI_ISL_447176, EPI_ISL_447177, EPI_ISL_447178, EPI_ISL_447179, EPI_ISL_447180, EPI_ISL_447181, EPI_ISL_447182, EPI_ISL_447183, EPI_ISL_447184, EPI_ISL_447185, EPI_ISL_447186, EPI_ISL_447187, EPI_ISL_447188, EPI_ISL_447189, EPI_ISL_447190, EPI_ISL_447191, EPI_ISL_447192, EPI_ISL_447193, EPI_ISL_447194, EPI_ISL_447195, EPI_ISL_447196, EPI_ISL_447197, EPI_ISL_447198, EPI_ISL_447199, EPI_ISL_447200, EPI_ISL_447201, EPI_ISL_447202, EPI_ISL_447203, EPI_ISL_447204, EPI_ISL_447205, EPI_ISL_447206, EPI_ISL_447207, EPI_ISL_447208, EPI_ISL_447209, EPI_ISL_447210, EPI_ISL_447211, EPI_ISL_447212, EPI_ISL_447213, EPI_ISL_447214, EPI_ISL_447215, EPI_ISL_447216, EPI_ISL_447217, EPI_ISL_447218, EPI_ISL_447219, EPI_ISL_447220, EPI_ISL_447221, EPI_ISL_447222, EPI_ISL_447223, EPI_ISL_447224, EPI_ISL_447225, EPI_ISL_447226, EPI_ISL_447227, EPI_ISL_447228, EPI_ISL_447229 | | | |
| see above | Michigan Department of Health and Human Services, Bureau of Laboratories | Michigan Department of Health and Human Services, Bureau of Laboratories | Blankenship HM, Riner D, Soehnlen MK |
| EPI_ISL_447542 | Gujarat Biotechnology Research Centre | Gujarat Biotechnology Research Centre | Ankit Hinsu, Pritesh Sabara, Apurvasinh Puvar, Janvi Raval, Monika Gandhi, Pinal Trivedi, Maharshi Pandya, Amit Kanani, Akanksha Verma, Nitin Savaliya, Raghawendra Kumar, Dinesh Kumar, Zuber Saiyed, Dipa Kinariwala, Disha Patel, Binita Aring, Neeta Khandelwal, Geeta Vaghela, Sonia Barve, Bhavesh Modi, Kairavi Joshi, Gaurishankar Shrimali, Nidhi Sood, Pranay Shah, R D Dixit, Snehal Bagatharia, Kamlesh J Upadhyay, Ramesh Pandit, Tejas Shah, Dipeshwari Shewale, Chaitanya Joshi, Madhvi Joshi |
| EPI_ISL_447544 | Gujarat Biotechnology Research Centre | Gujarat Biotechnology Research Centre | Apurvasinh Puvar, Janvi Raval, Monika Gandhi, Pinal Trivedi, Maharshi Pandya, Amit Kanani, Akanksha Verma, Nitin Savaliya, Raghawendra Kumar, Dinesh Kumar, Zuber Saiyed, Dipa Kinariwala, Disha Patel, Binita Aring, Neeta Khandelwal, Geeta Vaghela, Sonia Barve, Bhavesh Modi, Kairavi Joshi, Gaurishankar Shrimali, Nidhi Sood, Pranay Shah, R D Dixit, Snehal Bagatharia, Kamlesh J Upadhyay, Ramesh Pandit, Tejas Shah, Ankit Hinsu, Pritesh Sabara, Pooja P Doshi, Chaitanya Joshi, Madhvi Joshi |
| EPI_ISL_447594 | Caloundra Hospital | Public Health Virology Laboratory | Bixing Huang, Alyssa Pyke, Amanda De Jong, Andrew Van Den Hurk, Carmel Taylor, David Warrilow, Doris Genge, Elisabeth Gamez, Glen Hewitson, Ian Maxwell Mackay, Inga Sultana, Jamie McMahon, Jean Barcelon, Judy Northill, Mitchell Finger, Natalie Simpson, Neelima Nair, Peter Burtonclay, Peter Moore, Sarah Wheatley, Sean Moody, Sonja Hall-Mendelin, Timothy Gardam, and Frederick Moore |
| EPI_ISL_447595 | Pathology Queensland, Sunshine Coast University Hospital | Public Health Virology Laboratory | Bixing Huang, Alyssa Pyke, Amanda De Jong, Andrew Van Den Hurk, Carmel Taylor, David Warrilow, Doris Genge, Elisabeth Gamez, Glen Hewitson, Ian Maxwell Mackay, Inga Sultana, Jamie McMahon, Jean Barcelon, Judy Northill, Mitchell Finger, Natalie Simpson, Neelima Nair, Peter Burtonclay, Peter Moore, Sarah Wheatley, Sean Moody, Sonja Hall-Mendelin, Timothy Gardam, and Frederick Moore |
| EPI_ISL_450198, EPI_ISL_450199, EPI_ISL_450200, EPI_ISL_450203, EPI_ISL_450204, EPI_ISL_450207, EPI_ISL_450208, EPI_ISL_450209 | Department of Virology | Department of Virology | Boehmer,M.M., Buchholz,U., Corman,V.M., Hoch,M., Katz,K., Marosevic,D.V., Boehm,S., Woudenberg,T., Ackermann,N., Konrad,R., Eberle,U., Treis,B., Dangel,A., Bengs,K., Fingerle,V., Berger,A., Hoermansdorfer,S., Ippisch,S., Wicklein,B., Grahl,A., Poertner,K., Muller,N., Zeitlmann,N., Boender,T.S., Cai,W., Reich,A., an der Heiden,M., Rexroth,U., Hamouda,O., Schneider,J., Veith,T., Muehlemann,B., Woelfel,R., Antwerpen,M., Walter,M., Protzer,U., Liebl,B., Haas,W., Sing,A., Drosten,C., Zapf,A., Jones,T.C. |
| EPI_ISL_450445, EPI_ISL_450446, EPI_ISL_450447, EPI_ISL_450448, EPI_ISL_450449, EPI_ISL_450450, EPI_ISL_450451, EPI_ISL_450452, EPI_ISL_450453, EPI_ISL_450454, EPI_ISL_450455, EPI_ISL_450456, EPI_ISL_450457, EPI_ISL_450458, EPI_ISL_450459, EPI_ISL_450460, EPI_ISL_450461, EPI_ISL_450462, EPI_ISL_450463, EPI_ISL_450464, EPI_ISL_450465, EPI_ISL_450466, EPI_ISL_450467, EPI_ISL_450468, EPI_ISL_450469, EPI_ISL_450470, EPI_ISL_450471, EPI_ISL_450472, EPI_ISL_450473, EPI_ISL_450474, EPI_ISL_450475, EPI_ISL_450476, EPI_ISL_450477, EPI_ISL_450478, EPI_ISL_450479, EPI_ISL_450480, EPI_ISL_450481 | | | |
| see above | Stanford clinical virology lab | Chan-Zuckerberg Biohub | Benjamin Pinksy, Katharine Walter, Victoria N. Parikh, John Gorzynski, Hannah N. DeJong, Matthew T. Wheeler, Jason Andrews, Manuel Rivas, Carlos Bustamante, Euan Ashley, with CZB Cliahub Consortium |
| EPI_ISL_450570, EPI_ISL_450571, EPI_ISL_450572, EPI_ISL_450573, EPI_ISL_450574, EPI_ISL_450575, EPI_ISL_450576, EPI_ISL_450577, EPI_ISL_450578, EPI_ISL_450579, EPI_ISL_450580, EPI_ISL_450581, EPI_ISL_450582, EPI_ISL_450583, EPI_ISL_450584, EPI_ISL_450585, EPI_ISL_450586, EPI_ISL_450587, EPI_ISL_450588, EPI_ISL_450589, EPI_ISL_450590, EPI_ISL_450591, EPI_ISL_450592, EPI_ISL_450593, EPI_ISL_450594, EPI_ISL_450595, EPI_ISL_450596, EPI_ISL_450597, EPI_ISL_450598, EPI_ISL_450599, EPI_ISL_450600 | | | |
| see above | Michigan Department of Health and Human Services, Bureau of Laboratories | Michigan Department of Health and Human Services, Bureau of Laboratories | Blankenship HM; Riner D; Soehnlen MK |
| EPI_ISL_450601, EPI_ISL_450602, EPI_ISL_450603, EPI_ISL_450604, EPI_ISL_450605, EPI_ISL_450606, EPI_ISL_450607, EPI_ISL_450608, EPI_ISL_450609, EPI_ISL_450610, EPI_ISL_450611, EPI_ISL_450612, EPI_ISL_450613, EPI_ISL_450614, EPI_ISL_450615, EPI_ISL_450616, EPI_ISL_450617, EPI_ISL_450618, EPI_ISL_450619, EPI_ISL_450620, EPI_ISL_450621, EPI_ISL_450622, EPI_ISL_450623, EPI_ISL_450624, EPI_ISL_450625, EPI_ISL_450626, EPI_ISL_450627, EPI_ISL_450628, EPI_ISL_450629, EPI_ISL_450630, EPI_ISL_450631, EPI_ISL_450632, EPI_ISL_450633, EPI_ISL_450634, EPI_ISL_450635, EPI_ISL_450636, EPI_ISL_450637 | | | |

| | | | |
|---|---|---|---|
| see above | Michigan Department of Health and Human Services, Bureau of Laboratories | Michigan Department of Health and Human Services, Bureau of Laboratories | Blankenship HM, Riner D, Soehnlen MK |
| EPI_ISL_450789 | Pandit Deendayal Upadhyay Government Medical College, Rajkot | Gujarat Biotechnology Research Centre | Ankit Hinsu, Pritesh Sabara, Apurvasinh Puvar, Janvi Raval, Zarna Patel, Monika Gandhi, Pinal Trivedi, Maharshi Pandya, Amit Kanani, Nidhi Patel, Nitin Savaliya, Raghawendra Kumar, Dinesh Kumar, Zuber Saiyed, Komal Patel, Labdhi Pandya, Snehal Bagatharia, Prakash Modi, Sejul Antala, Manish Pattani, Ramesh Pandit, Tejas Shah, Bhavesh Modi, Gaurishankar Shrimali, R D Dixit, A M Kadri, Siddhant Kumar, Chaitanya Joshi, Madhvi Joshi |
| EPI_ISL_450812 | Uppsala Narakut Aleris | The Public Health Agency of Sweden | Annika Nilsson, Anna-Malin Linde, Maria Lind Karlberg, Oskar Karlsson Lindsjo, Olov Svartstrom, Anna Risberg, Theresa Enkirch, Mia Brytting, Karin Tegmark-Wisell |
| EPI_ISL_450819 | Hovas Askim Familjelakare och BVC | The Public Health Agency of Sweden | Anna Wendel, Anna-Malin Linde, Maria Lind Karlberg, Oskar Karlsson Lindsjo, Olov Svartstrom, Anna Risberg, Theresa Enkirch, Mia Brytting, Karin Tegmark-Wisell |
| EPI_ISL_450826, EPI_ISL_450827, EPI_ISL_450828 | Uppsala Narakut Aleris | The Public Health Agency of Sweden | Annika Nilsson, Anna-Malin Linde, Maria Lind Karlberg, Oskar Karlsson Lindsjo, Olov Svartstrom, Anna Risberg, Theresa Enkirch, Mia Brytting, Karin Tegmark-Wisell |
| EPI_ISL_450834 | Klinisk mikrobiologi och vardhygien Halmstad | The Public Health Agency of Sweden | Arne Kotz, Anna-Malin Linde, Maria Lind Karlberg, Oskar Karlsson Lindsjo, Olov Svartstrom, Anna Risberg, Theresa Enkirch, Mia Brytting, Karin Tegmark-Wisell |
| EPI_ISL_450836, EPI_ISL_450837 | Laboratoriemedicin | The Public Health Agency of Sweden | Anna-Malin Linde, Maria Lind Karlberg, Oskar Karlsson Lindsjo, Olov Svartstrom, Anna Risberg, Theresa Enkirch, Mia Brytting, Karin Tegmark-Wisell |
| EPI_ISL_451076 | West China Hospital of Sichuan University | State Key Laboratory of Biotherapy of Sichuan University | Baowen Du, Minjin Wang, Chao Tanga, Chuan Chena, Yongzhao Zhou, Mingxia Yu, Han-Cheng Wei, Weimin Li, Jing-wen Lin, Jia Geng, Binwu Ying, Lu Chen |
| EPI_ISL_451306 | Molecular Virology Unit, Fondazione IRCCS Policlinico San Matteo , Pavia | Laboratory of Virology, INMI Lazzaro Spallanzani IRCCS | Antonio Piralla, Fausto Baldanti, Martina Rueca, Antonino Di Caro, Maria R. Capobianchi, Cesare E.M. Gruber, Barbara Bartolini |
| EPI_ISL_451308 | Molecular Virology Unit, Fondazione IRCCS Policlinico San Matteo , Pavia | Laboratory of Virology, INMI Lazzaro Spallanzani IRCCS | Antonio Piralla, Fausto Baldanti, Maria R. Capobianchi, Cesare E.M. Gruber, Martina Rueca, Barbara Bartolini, Antonino Di Caro |
| EPI_ISL_451312, EPI_ISL_451313, EPI_ISL_451314, EPI_ISL_451315, EPI_ISL_451316, EPI_ISL_451317, EPI_ISL_451318, EPI_ISL_451319, EPI_ISL_451320, EPI_ISL_451321, EPI_ISL_451322, EPI_ISL_451323, EPI_ISL_451324, EPI_ISL_451325, EPI_ISL_451326, EPI_ISL_451327, EPI_ISL_451328, EPI_ISL_451329, EPI_ISL_451330, EPI_ISL_451331, EPI_ISL_451332, EPI_ISL_451333, EPI_ISL_451334, EPI_ISL_451335, EPI_ISL_451336, EPI_ISL_451337, EPI_ISL_451338, EPI_ISL_451339, EPI_ISL_451340, EPI_ISL_451341, EPI_ISL_451342, EPI_ISL_451343, EPI_ISL_451344, EPI_ISL_451345, EPI_ISL_451346, EPI_ISL_451347, EPI_ISL_451348, EPI_ISL_451349, EPI_ISL_451350, EPI_ISL_451351, EPI_ISL_451352, EPI_ISL_451353, EPI_ISL_451354, EPI_ISL_451355, EPI_ISL_451356, EPI_ISL_451357, EPI_ISL_451358, EPI_ISL_451359, EPI_ISL_451360, EPI_ISL_451361, EPI_ISL_451362, EPI_ISL_451363, EPI_ISL_451364, EPI_ISL_451365, EPI_ISL_451366, EPI_ISL_451367, EPI_ISL_451368, EPI_ISL_451369, EPI_ISL_451370, EPI_ISL_451371, EPI_ISL_451372, EPI_ISL_451373, EPI_ISL_451374, EPI_ISL_451375, EPI_ISL_451376, EPI_ISL_451378, EPI_ISL_451379, EPI_ISL_451380, EPI_ISL_451381, EPI_ISL_451382, EPI_ISL_451383, EPI_ISL_451384, EPI_ISL_451385, EPI_ISL_451386, EPI_ISL_451387, EPI_ISL_451388, EPI_ISL_451389, EPI_ISL_451390, EPI_ISL_451391, EPI_ISL_451392, EPI_ISL_451393, EPI_ISL_451394, EPI_ISL_451395, EPI_ISL_451396, EPI_ISL_451397, EPI_ISL_451398, EPI_ISL_451399 | | | |
| see above | West China Hospital of Sichuan University | State Key Laboratory of Biotherapy of Sichuan University | Baowen Du, Minjin Wang, Chao Tang, Chuan Chen, Yongzhao Zhou, Mingxia Yu, Hancheng Wei, Weimin Li, Jing-wen Lin, Jia Geng, Binwu Ying, Lu Chen |
| EPI_ISL_451666 | M.P Shah Government Medocal college Jamnagar | Gujarat Biotechnology Research Centre | Binita Aring, Janvi Raval, Zarna Patel, Monika Gandhi, Pinal Trivedi, Maharshi Pandya, Amit Kanani, Nidhi Patel, Nitin Savaliya, Raghawendra Kumar, Dinesh Kumar, Zuber Saiyed, Komal Patel, Labdhi Pandya, Snehal Bagatharia, Ramesh Pandit, Tejas Shah, Ankit Hinsu, Pritesh Sabara, Apurvasinh Puvar, Bhavesh Modi, Gaurishankar Shrimali, R D Dixit, A M Kadri, Akanksha Verma, Chaitanya Joshi, Madhvi Joshi, |
| EPI_ISL_452137, EPI_ISL_452138 | VI-US Virgin Islands Department of Health | Pathogen Discovery, Respiratory Viruses Branch, Division of Viral Diseases, Centers for Disease Control and Prevention | Anna Uehara, Yan Li, Anna Montmayeur, Ying Tao, Krista Queen, Jing Zhang, Clinton R. Paden, Rachel Marine, Haibin Wang, Bettina Bankamp, Zachary Weiner, Suxiang Tong |
| EPI_ISL_452262, EPI_ISL_452263, EPI_ISL_452264, EPI_ISL_452265, EPI_ISL_452266, EPI_ISL_452267, EPI_ISL_452268, EPI_ISL_452269, EPI_ISL_452270, EPI_ISL_452271, EPI_ISL_452272, EPI_ISL_452273, EPI_ISL_452274, EPI_ISL_452275, EPI_ISL_452276, EPI_ISL_452277, EPI_ISL_452278, EPI_ISL_452279, EPI_ISL_452280, EPI_ISL_452281, EPI_ISL_452282, EPI_ISL_452283, EPI_ISL_452284, EPI_ISL_452285, EPI_ISL_452286, EPI_ISL_452287, EPI_ISL_452288, EPI_ISL_452289, EPI_ISL_452290, EPI_ISL_452291, EPI_ISL_452292, EPI_ISL_452293, EPI_ISL_452294, EPI_ISL_452295, EPI_ISL_452296, EPI_ISL_452297, EPI_ISL_452298, EPI_ISL_452299, EPI_ISL_452300, EPI_ISL_452301, EPI_ISL_452302, EPI_ISL_452303, EPI_ISL_452304, EPI_ISL_452305, EPI_ISL_452306, EPI_ISL_452307, EPI_ISL_452308, EPI_ISL_452309, EPI_ISL_452310, EPI_ISL_452311, EPI_ISL_452312, EPI_ISL_452313, EPI_ISL_452314, EPI_ISL_452315, EPI_ISL_452316, EPI_ISL_452317, EPI_ISL_452318, EPI_ISL_452319, EPI_ISL_452320, EPI_ISL_452321, EPI_ISL_452322, EPI_ISL_452323, EPI_ISL_452324, EPI_ISL_452325, EPI_ISL_452326 | | | |
| see above | Michigan Department of Health and Human Services, Bureau of Laboratories | Michigan Department of Health and Human Services, Bureau of Laboratories | Blankenship HM, Riner D, Soehnlen MK |
| EPI_ISL_453665 | Centre for Enzyme Innovation, University of Portsmouth / Translational Research Laboratory, Portsmouth Hospitals NHS Trust | University of Portsmouth | Angela Beckett,,Yann Bourgeois,,Garry Scarlett,,Sharon Glaysher,,Scott Elliott,,Kelly Bicknell,,Robert Impey,,Allyson Lloyd,,Sarah Wyllie,,Ethan Butcher,,Anoop Chauhan,,Samuel Robson |
| EPI_ISL_453666, EPI_ISL_453667, EPI_ISL_453668, EPI_ISL_453669, EPI_ISL_453670, EPI_ISL_453671, EPI_ISL_453672, EPI_ISL_453673, EPI_ISL_453674, EPI_ISL_453675, EPI_ISL_453676, EPI_ISL_453677, EPI_ISL_453678, EPI_ISL_453679, EPI_ISL_453680, EPI_ISL_453681, EPI_ISL_453682, EPI_ISL_453683, EPI_ISL_453684, EPI_ISL_453685, EPI_ISL_453686, EPI_ISL_453687, EPI_ISL_453688, EPI_ISL_453689, EPI_ISL_453690, EPI_ISL_453691, EPI_ISL_453692, EPI_ISL_453693, EPI_ISL_453694, EPI_ISL_453695 | | | |
| see above | Centre for Enzyme Innovation, University of Portsmouth / Translational Research Laboratory, Portsmouth Hospitals NHS Trust | COVID-19 Genomics UK (COG-UK) Consortium | Angela Beckett,,Yann Bourgeois,,Garry Scarlett,,Sharon Glaysher,,Scott Elliott,,Kelly Bicknell,,Robert Impey,,Allyson Lloyd,,Sarah Wyllie,,Ethan Butcher,,Anoop Chauhan,,Samuel Robson |
| EPI_ISL_454436, EPI_ISL_454437, EPI_ISL_454438, EPI_ISL_454439, EPI_ISL_454440, EPI_ISL_454441, EPI_ISL_454442, EPI_ISL_454443, EPI_ISL_454444, EPI_ISL_454445, EPI_ISL_454446 | | | |
| see above | Halmstad klinisk mikrobiologi | The Public Health Agency of Sweden | Anna-Malin Linde, Maria Lind Karlberg, Mattias Haukland, Reza Advani, Olov Svartstrom, Oskar Karlsson Lindsjo, Petra Edquist, Shamam Muradrasoli, Anna Risberg, Karin Tegmark-Wisell |
| EPI_ISL_454447, EPI_ISL_454448, EPI_ISL_454449, EPI_ISL_454450, EPI_ISL_454451, EPI_ISL_454452, EPI_ISL_454453, EPI_ISL_454454, EPI_ISL_454455, EPI_ISL_454456, EPI_ISL_454457, EPI_ISL_454458, EPI_ISL_454459, EPI_ISL_454460, EPI_ISL_454461, EPI_ISL_454462, EPI_ISL_454463, EPI_ISL_454464, EPI_ISL_454465, EPI_ISL_454466, EPI_ISL_454467, EPI_ISL_454468, EPI_ISL_454469, EPI_ISL_454470, EPI_ISL_454471, EPI_ISL_454472, EPI_ISL_454473, EPI_ISL_454474, EPI_ISL_454475, EPI_ISL_454476, EPI_ISL_454477, EPI_ISL_454478, EPI_ISL_454479, EPI_ISL_454480, EPI_ISL_454481, EPI_ISL_454482, EPI_ISL_454483, EPI_ISL_454484, EPI_ISL_454485, EPI_ISL_454486, EPI_ISL_454487, EPI_ISL_454488, EPI_ISL_454489, EPI_ISL_454490, EPI_ISL_454491, EPI_ISL_454492, EPI_ISL_454493 | | | |
| see above | Karolinska Universitetslaboratoriet | The Public Health Agency of Sweden | Anna-Malin Linde, Maria Lind Karlberg, Mattias Haukland, Reza Advani, Olov Svartstrom, Oskar Karlsson Lindsjo, Petra Edquist, Shamam Muradrasoli, Anna Risberg, Karin Tegmark-Wisell |
| EPI_ISL_454494, EPI_ISL_454495, EPI_ISL_454496 | Skovde/Unilabs | The Public Health Agency of Sweden | Anna-Malin Linde, Maria Lind Karlberg, Mattias Haukland, Reza Advani, Olov Svartstrom, Oskar Karlsson Lindsjo, Petra Edquist, Shamam Muradrasoli, Anna Risberg, Karin Tegmark-Wisell |
| EPI_ISL_454868, EPI_ISL_454869, EPI_ISL_454870, EPI_ISL_454871, EPI_ISL_454872, EPI_ISL_454873, EPI_ISL_454874, EPI_ISL_454875, EPI_ISL_454876, EPI_ISL_454877, EPI_ISL_454878, EPI_ISL_454879, EPI_ISL_454880, EPI_ISL_454881, EPI_ISL_454882, EPI_ISL_454883, EPI_ISL_454884, EPI_ISL_454885, EPI_ISL_454886, EPI_ISL_454887, EPI_ISL_454888, EPI_ISL_454889, EPI_ISL_454890, EPI_ISL_454891, EPI_ISL_454892, EPI_ISL_454893, EPI_ISL_454894, EPI_ISL_454895, EPI_ISL_454896, EPI_ISL_454897, EPI_ISL_454898, EPI_ISL_454899, EPI_ISL_454900, EPI_ISL_454901, EPI_ISL_454902 | | | |
| see above | Karolinska Universitetslaboratoriet | The Public Health Agency of Sweden | Anna-Malin Linde, Maria Lind Karlberg, Mattias Haukland, Reza Advani, Olov Svartstrom, Oskar Karlsson Lindsjo, Petra Edquist, Shamam Muradrasoli, Anna Risberg, Karin Tegmark-Wisell |
| EPI_ISL_454903 | Klinisk mikrobiologi, UAS | The Public Health Agency of Sweden | Anna-Malin Linde, Maria Lind Karlberg, Mattias Haukland, Reza Advani, Olov Svartstrom, Oskar Karlsson Lindsjo, Petra Edquist, Shamam Muradrasoli, Anna Risberg, Karin Tegmark-Wisell |
| EPI_ISL_454904, EPI_ISL_454905, EPI_ISL_454906, EPI_ISL_454907, EPI_ISL_454908, EPI_ISL_454909, EPI_ISL_454910, EPI_ISL_454911, EPI_ISL_454912, EPI_ISL_454913, EPI_ISL_454914, EPI_ISL_454915, EPI_ISL_454916, EPI_ISL_454917, EPI_ISL_454918, EPI_ISL_454919, EPI_ISL_454920, EPI_ISL_454921, EPI_ISL_454922, EPI_ISL_454923, EPI_ISL_454924, EPI_ISL_454925, EPI_ISL_454926, EPI_ISL_454927, EPI_ISL_454928, EPI_ISL_454929, EPI_ISL_454930, EPI_ISL_454931, EPI_ISL_454932, EPI_ISL_454933, EPI_ISL_454934, EPI_ISL_454935, EPI_ISL_454936, EPI_ISL_454937, EPI_ISL_454938, EPI_ISL_454939, EPI_ISL_454940, EPI_ISL_454941, EPI_ISL_454942, EPI_ISL_454943, EPI_ISL_454944, EPI_ISL_454945, EPI_ISL_454946, EPI_ISL_454947, EPI_ISL_454948, EPI_ISL_454949, EPI_ISL_454950, EPI_ISL_454951, EPI_ISL_454952, EPI_ISL_454953, EPI_ISL_454954, EPI_ISL_454955, EPI_ISL_454956, EPI_ISL_454957, EPI_ISL_454958, EPI_ISL_454959, EPI_ISL_454960, EPI_ISL_454961, EPI_ISL_454962, EPI_ISL_454963, EPI_ISL_454964, EPI_ISL_454965, EPI_ISL_454966, EPI_ISL_454967, EPI_ISL_454968, EPI_ISL_454969, EPI_ISL_454970, EPI_ISL_454971, EPI_ISL_454972, EPI_ISL_454973, EPI_ISL_454974, EPI_ISL_454975, EPI_ISL_454976, EPI_ISL_454977, EPI_ISL_454978, EPI_ISL_454979, EPI_ISL_454980, EPI_ISL_454981, EPI_ISL_454982, EPI_ISL_454983, EPI_ISL_454984, EPI_ISL_454985, EPI_ISL_454986, EPI_ISL_454987, EPI_ISL_454988, EPI_ISL_454989, EPI_ISL_454990, EPI_ISL_454991, EPI_ISL_454992, EPI_ISL_454993, EPI_ISL_454994, EPI_ISL_454995, EPI_ISL_454996, EPI_ISL_454997, EPI_ISL_454998, EPI_ISL_454999, EPI_ISL_455000, EPI_ISL_455001, EPI_ISL_455002, EPI_ISL_455003, EPI_ISL_455004, EPI_ISL_455005, EPI_ISL_455006, EPI_ISL_455007, EPI_ISL_455008, EPI_ISL_455009, EPI_ISL_455010, EPI_ISL_455011, EPI_ISL_455012, EPI_ISL_455013, EPI_ISL_455014 | | | |
| see above | Wuhan Chain Medical Labs (CMLabs) | State Key Laboratory of Biotherapy of Sichuan University | Baowen Du, Minjin Wang, Chao Tang, Chuan Chen, Yongzhao Zhou, Mingxia Yu, Hancheng Wei, Weimin Li, Jing-wen Lin, Jia Geng, Binwu Ying, Lu Chen |
| EPI_ISL_455021 | Government Medical College, Vadodara | Gujarat Biotechnology Research Centre | Ankit Hinsu, Pritesh Sabara, Apurvasinh Puvar, Janvi Raval, Zarna Patel, Monika Gandhi, Pinal Trivedi, Maharshi Pandya, Amit Kanani, Nidhi Patel, Nitin Savaliya, Raghawendra Kumar, Dinesh Kumar, Zuber Saiyed, Komal Patel, Labdhi Pandya, Snehal Bagatharia, Tanuja Javadekar , R N Daveshwar, |

| | | | |
|---|---|---|---|
| | | | Ramesh Pandit, Tejas Shah, Camellia Chakraborty, Bhavesh Modi, Gaurishankar Shrimali, R D Dixit, A M Kadri, Umang Mishra, Chaitanya Joshi, Madhvi Joshi, |
| EPI_ISL_455023 | Government Medical College, Vadodara | Gujarat Biotechnology Research Centre | Apurvasinh Puvar, Janvi Raval, Zarna Patel, Monika Gandhi, Pinal Trivedi, Maharshi Pandya, Amit Kanani, Nidhi Patel, Nitin Savaliya, Raghawendra Kumar, Dinesh Kumar, Zuber Saiyed, Komal Patel, Labdhi Pandya, Snehal Bagatharia, Tanuja Javadekar , R N Daveshwar, Ramesh Pandit, Tejas Shah, Ankit Hinsu, Pritesh Sabara, Priyanka P Vatsa, Bhavesh Modi, Gaurishankar Shrimali, R D Dixit, A M Kadri, Umang Mishra, Chaitanya Joshi, Madhvi Joshi, |
| EPI_ISL_455100 | Orsa VC | The Public Health Agency of Sweden | Anna-Karin Lundqvist, Anna-Malin Linde, Maria Lind Karlberg, Oskar Karlsson Lindsjo, Olov Svartstrom, Anna Risberg, Theresa Enkirch, Mia Brytting, Karin Tegmark-Wisell |
| EPI_ISL_455111 | Olof Norrby | The Public Health Agency of Sweden | Bla Kustens halsocentral, Anna-Malin Linde, Maria Lind Karlberg, Oskar Karlsson Lindsjo, Olov Svartstrom, Anna Risberg, Theresa Enkirch, Mia Brytting, Karin Tegmark-Wisell |
| EPI_ISL_455112, EPI_ISL_455113, EPI_ISL_455114, EPI_ISL_455115, EPI_ISL_455116, EPI_ISL_455117, EPI_ISL_455118, EPI_ISL_455119, EPI_ISL_455120, EPI_ISL_455121, EPI_ISL_455122, EPI_ISL_455123, EPI_ISL_455124, EPI_ISL_455125, EPI_ISL_455126, EPI_ISL_455127, EPI_ISL_455128, EPI_ISL_455129, EPI_ISL_455130, EPI_ISL_455131, EPI_ISL_455132, EPI_ISL_455133, EPI_ISL_455134, EPI_ISL_455135, EPI_ISL_455136, EPI_ISL_455137, EPI_ISL_455138, EPI_ISL_455139, EPI_ISL_455140, EPI_ISL_455141, EPI_ISL_455142, EPI_ISL_455143, EPI_ISL_455144, EPI_ISL_455145, EPI_ISL_455146, EPI_ISL_455147, EPI_ISL_455148, EPI_ISL_455149, EPI_ISL_455150, EPI_ISL_455151, EPI_ISL_455152, EPI_ISL_455153, EPI_ISL_455154, EPI_ISL_455155, EPI_ISL_455156, EPI_ISL_455157, EPI_ISL_455158, EPI_ISL_455159, EPI_ISL_455160, EPI_ISL_455161, EPI_ISL_455162, EPI_ISL_455163, EPI_ISL_455164, EPI_ISL_455165, EPI_ISL_455166, EPI_ISL_455167, EPI_ISL_455168, EPI_ISL_455169, EPI_ISL_455170, EPI_ISL_455171, EPI_ISL_455172, EPI_ISL_455173, EPI_ISL_455174, EPI_ISL_455175, EPI_ISL_455176, EPI_ISL_455177, EPI_ISL_455178, EPI_ISL_455179, EPI_ISL_455180, EPI_ISL_455194, EPI_ISL_455195, EPI_ISL_455196, EPI_ISL_455197, EPI_ISL_455198, EPI_ISL_455199, EPI_ISL_455200, EPI_ISL_455202, EPI_ISL_455203, EPI_ISL_455204, EPI_ISL_455205, EPI_ISL_455206, EPI_ISL_455207, EPI_ISL_455208, EPI_ISL_455209, EPI_ISL_455210, EPI_ISL_455211, EPI_ISL_455212, EPI_ISL_455213, EPI_ISL_455214, EPI_ISL_455215, EPI_ISL_455216, EPI_ISL_455217, EPI_ISL_455218, EPI_ISL_455219, EPI_ISL_455220, EPI_ISL_455221, EPI_ISL_455222, EPI_ISL_455226, EPI_ISL_455227, EPI_ISL_455228, EPI_ISL_455229, EPI_ISL_455230, EPI_ISL_455231, EPI_ISL_455232, EPI_ISL_455233, EPI_ISL_455234, EPI_ISL_455235, EPI_ISL_455236, EPI_ISL_455237, EPI_ISL_455238, EPI_ISL_455239, EPI_ISL_455240, EPI_ISL_455241, EPI_ISL_455242, EPI_ISL_455243, EPI_ISL_455244, EPI_ISL_455245, EPI_ISL_455246, EPI_ISL_455247, EPI_ISL_455248, EPI_ISL_455249, EPI_ISL_455250, EPI_ISL_455251, EPI_ISL_455252, EPI_ISL_455253, EPI_ISL_455254, EPI_ISL_455255, EPI_ISL_455256, EPI_ISL_455257, EPI_ISL_455258, EPI_ISL_455261, EPI_ISL_455263, EPI_ISL_455264, EPI_ISL_455265, EPI_ISL_455266, EPI_ISL_455267, EPI_ISL_455268, EPI_ISL_455269, EPI_ISL_455270, EPI_ISL_455271, EPI_ISL_455272, EPI_ISL_455273, EPI_ISL_455274, EPI_ISL_455275, EPI_ISL_455276, EPI_ISL_455277, EPI_ISL_455278, EPI_ISL_455279, EPI_ISL_455280, EPI_ISL_455281, EPI_ISL_455282, EPI_ISL_455283, EPI_ISL_455284, EPI_ISL_455285, EPI_ISL_455286, EPI_ISL_455287, EPI_ISL_455288, EPI_ISL_455289, EPI_ISL_455290, EPI_ISL_455291, EPI_ISL_455292, EPI_ISL_455293, EPI_ISL_455294, EPI_ISL_455295, EPI_ISL_455296, EPI_ISL_455297, EPI_ISL_455298, EPI_ISL_455299, EPI_ISL_455300, EPI_ISL_455301, EPI_ISL_455302, EPI_ISL_455303, EPI_ISL_455304, EPI_ISL_455305, EPI_ISL_455306, EPI_ISL_455307 | | | |
| see above | Dutch COVID-19 response team | Erasmus Medical Center | Bas Oude Munnink, David Nieuwenhuijse, Reina Sikkema, Claudia Schapendonk, Irina Chestakova, Anne van der Linden, Theo Bestebroer, Stefan van Nieuwkoop, Mark Pronk, Pascal Lexmond, Corien Swaan, Manon Haverkate, Madelief Mollers, Mart Stein, Sandra Kengne Kamga Mobou, Jeroen van Kampen, Jolanda Voermans, Aura Timen, Corine GeurtsvanKessel, Annemiek van der Eijk, Richard Molenkamp, Marion Koopmans, on behalf of the Dutch national COVID-19 response team. |
| EPI_ISL_455363, EPI_ISL_455364, EPI_ISL_455365, EPI_ISL_455366, EPI_ISL_455367, EPI_ISL_455368, EPI_ISL_455369, EPI_ISL_455370, EPI_ISL_455371, EPI_ISL_455372, EPI_ISL_455373, EPI_ISL_455374, EPI_ISL_455375, EPI_ISL_455376, EPI_ISL_455377, EPI_ISL_455378, EPI_ISL_455379, EPI_ISL_455380, EPI_ISL_455381, EPI_ISL_455382, EPI_ISL_455383, EPI_ISL_455384, EPI_ISL_455385, EPI_ISL_455386, EPI_ISL_455387, EPI_ISL_455388, EPI_ISL_455389, EPI_ISL_455390, EPI_ISL_455391, EPI_ISL_455392, EPI_ISL_455393, EPI_ISL_455394, EPI_ISL_455395, EPI_ISL_455396, EPI_ISL_455397, EPI_ISL_455398, EPI_ISL_455399, EPI_ISL_455400, EPI_ISL_455401, EPI_ISL_455402, EPI_ISL_455403, EPI_ISL_455404, EPI_ISL_455405, EPI_ISL_455406, EPI_ISL_455407, EPI_ISL_455408, EPI_ISL_455409, EPI_ISL_455410, EPI_ISL_455411 | | | |
| see above | Wuhan Chain Medical Labs (CMLabs) | State Key Laboratory of Biotherapy of Sichuan University | Baowen Du, Minjin Wang, Chao Tang, Chuan Chen, Yongzhao Zhou, Mingxia Yu, Hancheng Wei, Weimin Li, Jing-wen Lin, Jia Geng, Binwu Ying, Lu Chen |
| EPI_ISL_455437, EPI_ISL_455438 | Instituto de Diagnostico y Referencia Epidemiologicos (INDRE) | Instituto de Diagnostico y Referencia Epidemiologicos (INDRE) | Araiza-Rodriguez Adnan, Garces-Ayala Fabiola. Ramirez-Gonzalez Ernesto, Mendieta-Condado Edgar, Rodriguez-Maldonado Abril, Wong-Arambula Claudia, Barrera-Badillo Gisela, Hernandez-Rivas Lucia, Lopez-Martinez Irma, Taboada Ramírez Blanca. |
| EPI_ISL_455595, EPI_ISL_455596, EPI_ISL_455597, EPI_ISL_455598, EPI_ISL_455599, EPI_ISL_455600, EPI_ISL_455601, EPI_ISL_455602, EPI_ISL_455603 | SA Pathology | VPRL | Beard, MR., Van Der Hoek, K., Lim, C.K., Leong, L.E.X., Coldbeck-Shackley, R., Shue, B., Kirby, E., Merrett, J., Llamas, B. |
| EPI_ISL_455640, EPI_ISL_455641, EPI_ISL_455642, EPI_ISL_455643, EPI_ISL_455644, EPI_ISL_455645, EPI_ISL_455646, EPI_ISL_455647, EPI_ISL_455648, EPI_ISL_455649, EPI_ISL_455650, EPI_ISL_455651, EPI_ISL_455652, EPI_ISL_455653, EPI_ISL_455654, EPI_ISL_455655, EPI_ISL_455656, EPI_ISL_455657, EPI_ISL_455658, EPI_ISL_455659, EPI_ISL_455660, EPI_ISL_455661, EPI_ISL_455662, EPI_ISL_455663, EPI_ISL_455664, EPI_ISL_455665, EPI_ISL_455666, EPI_ISL_455667, EPI_ISL_455668, EPI_ISL_455669, EPI_ISL_455670, EPI_ISL_455671, EPI_ISL_455672, EPI_ISL_455673, EPI_ISL_455674, EPI_ISL_455675, EPI_ISL_455676, EPI_ISL_455677, EPI_ISL_455678, EPI_ISL_455679 | | | |
| see above | ICMR-National Institute of Cholera and Enteric Diseases | National Institute of Biomedical Genomics | Arindam Maitra, Mamta Chawla Sarkar, Sreedhar Chinnaswamy, Hasina Banu, Ananya Chatterjee, Shanta Dutta, Saumitra Das |
| EPI_ISL_455840, EPI_ISL_455841 | Orebro klinisk mikrobiologi | The Public Health Agency of Sweden | Anna-Malin Linde, Maria Lind Karlberg, Mattias Haukland, Reza Advani, Olov Svartstrom, Oskar Karlsson Lindsjo, Petra Edquist, Shamam Muradrasoli, Anna Risberg, Karin Tegmark-Wisell |
| EPI_ISL_455842, EPI_ISL_455843, EPI_ISL_455844, EPI_ISL_455845 | Skovde/Unilabs | The Public Health Agency of Sweden | Anna-Malin Linde, Maria Lind Karlberg, Mattias Haukland, Reza Advani, Olov Svartstrom, Oskar Karlsson Lindsjo, Petra Edquist, Shamam Muradrasoli, Anna Risberg, Karin Tegmark-Wisell |
| EPI_ISL_455846, EPI_ISL_455847, EPI_ISL_455848, EPI_ISL_455849, EPI_ISL_455850, EPI_ISL_455851, EPI_ISL_455852, EPI_ISL_455853, EPI_ISL_455854, EPI_ISL_455855, EPI_ISL_455856, EPI_ISL_455857, EPI_ISL_455858, EPI_ISL_455859, EPI_ISL_455860, EPI_ISL_455861, EPI_ISL_455862, EPI_ISL_455863, EPI_ISL_455864, EPI_ISL_455865, EPI_ISL_455866, EPI_ISL_455867, EPI_ISL_455868, EPI_ISL_455869, EPI_ISL_455870, EPI_ISL_455871, EPI_ISL_455872, EPI_ISL_455873, EPI_ISL_455874, EPI_ISL_455875, EPI_ISL_455876, EPI_ISL_455877, EPI_ISL_455878, EPI_ISL_455879, EPI_ISL_455880, EPI_ISL_455881, EPI_ISL_455882, EPI_ISL_455883, EPI_ISL_455884, EPI_ISL_455885, EPI_ISL_455886, EPI_ISL_455887, EPI_ISL_455888, EPI_ISL_455889, EPI_ISL_455890, EPI_ISL_455891, EPI_ISL_455892, EPI_ISL_455893, EPI_ISL_455894, EPI_ISL_455895, EPI_ISL_455896, EPI_ISL_455897, EPI_ISL_455898, EPI_ISL_455899, EPI_ISL_455900, EPI_ISL_455901 | | | |
| see above | Karolinska Universitetslaboratoriet | The Public Health Agency of Sweden | Anna-Malin Linde, Maria Lind Karlberg, Mattias Haukland, Reza Advani, Olov Svartstrom, Oskar Karlsson Lindsjo, Petra Edquist, Shamam Muradrasoli, Anna Risberg, Karin Tegmark-Wisell |
| EPI_ISL_455902 | Halmstad klinisk mikrobiologi | The Public Health Agency of Sweden | Anna-Malin Linde, Maria Lind Karlberg, Mattias Haukland, Reza Advani, Olov Svartstrom, Oskar Karlsson Lindsjo, Petra Edquist, Shamam Muradrasoli, Anna Risberg, Karin Tegmark-Wisell |
| EPI_ISL_455903 | Folkhalsomyndigheten | The Public Health Agency of Sweden | Anna-Malin Linde, Maria Lind Karlberg, Mattias Haukland, Reza Advani, Olov Svartstrom, Oskar Karlsson Lindsjo, Petra Edquist, Shamam Muradrasoli, Anna Risberg, Karin Tegmark-Wisell |
| EPI_ISL_455904, EPI_ISL_455905, EPI_ISL_455906, EPI_ISL_455907 | Klinisk mikrobiologi, UAS | The Public Health Agency of Sweden | Anna-Malin Linde, Maria Lind Karlberg, Mattias Haukland, Reza Advani, Olov Svartstrom, Oskar Karlsson Lindsjo, Petra Edquist, Shamam Muradrasoli, Anna Risberg, Karin Tegmark-Wisell |
| EPI_ISL_457028, EPI_ISL_457029, EPI_ISL_457031, EPI_ISL_457032, EPI_ISL_457033, EPI_ISL_457034, EPI_ISL_457035, EPI_ISL_457036, EPI_ISL_457037, EPI_ISL_457038, EPI_ISL_457039, EPI_ISL_457040, EPI_ISL_457041, EPI_ISL_457042, EPI_ISL_457043, EPI_ISL_457044, EPI_ISL_457045, EPI_ISL_457046, EPI_ISL_457047, EPI_ISL_457048, EPI_ISL_457049, EPI_ISL_457050, EPI_ISL_457051, EPI_ISL_457052, EPI_ISL_457053, EPI_ISL_457054, EPI_ISL_457055, EPI_ISL_457056, EPI_ISL_457057, EPI_ISL_457058, EPI_ISL_457059, EPI_ISL_457060, EPI_ISL_457061, EPI_ISL_457062, EPI_ISL_457063, EPI_ISL_457064, EPI_ISL_457065, EPI_ISL_457066, EPI_ISL_457067, EPI_ISL_457068, EPI_ISL_457069, EPI_ISL_457070, EPI_ISL_457071, EPI_ISL_457072, EPI_ISL_457073, EPI_ISL_457074, EPI_ISL_457075, EPI_ISL_457076, EPI_ISL_457077, EPI_ISL_457078, EPI_ISL_457079, EPI_ISL_457080, EPI_ISL_457081, EPI_ISL_457082, EPI_ISL_457083, EPI_ISL_457084, EPI_ISL_457085, EPI_ISL_457086, EPI_ISL_457087, EPI_ISL_457088, EPI_ISL_457089, EPI_ISL_457090, EPI_ISL_457091, EPI_ISL_457092, EPI_ISL_457093, EPI_ISL_457094, EPI_ISL_457095, EPI_ISL_457096, EPI_ISL_457097, EPI_ISL_457098, EPI_ISL_457099, EPI_ISL_457100, EPI_ISL_457101, EPI_ISL_457102, EPI_ISL_457103, EPI_ISL_457104, EPI_ISL_457105, EPI_ISL_457106, EPI_ISL_457107, EPI_ISL_457108, EPI_ISL_457109, EPI_ISL_457110, EPI_ISL_457111, EPI_ISL_457112, EPI_ISL_457113, EPI_ISL_457114, EPI_ISL_457115, EPI_ISL_457116, EPI_ISL_457117, EPI_ISL_457118, EPI_ISL_457119, EPI_ISL_457120, EPI_ISL_457121, EPI_ISL_457122, EPI_ISL_457123, EPI_ISL_457124, EPI_ISL_457125, EPI_ISL_457126, EPI_ISL_457127, EPI_ISL_457128, EPI_ISL_457129, EPI_ISL_457130, EPI_ISL_457131, EPI_ISL_457132, EPI_ISL_457133, EPI_ISL_457134, EPI_ISL_457135, EPI_ISL_457136, EPI_ISL_457138, EPI_ISL_457139, EPI_ISL_457140, EPI_ISL_457141, EPI_ISL_457142, EPI_ISL_457143, EPI_ISL_457144, EPI_ISL_457145, EPI_ISL_457146, EPI_ISL_457147, EPI_ISL_457149, EPI_ISL_457150, EPI_ISL_457151, EPI_ISL_457153, EPI_ISL_457154, EPI_ISL_457155, EPI_ISL_457156, EPI_ISL_457157, EPI_ISL_457158, EPI_ISL_457159, EPI_ISL_457160, EPI_ISL_457161, EPI_ISL_457162, EPI_ISL_457163, EPI_ISL_457164, EPI_ISL_457165, EPI_ISL_457166, EPI_ISL_457167, EPI_ISL_457168, EPI_ISL_457169, EPI_ISL_457170, EPI_ISL_457171, EPI_ISL_457172, EPI_ISL_457173, EPI_ISL_457174, EPI_ISL_457175, EPI_ISL_457176, EPI_ISL_457177, EPI_ISL_457178, EPI_ISL_457179, EPI_ISL_457180, EPI_ISL_457181, EPI_ISL_457182, EPI_ISL_457183, EPI_ISL_457184, EPI_ISL_457185, EPI_ISL_457186, EPI_ISL_457187, EPI_ISL_457188, EPI_ISL_457189, EPI_ISL_457190, EPI_ISL_457191, EPI_ISL_457192, EPI_ISL_457193, EPI_ISL_457194, EPI_ISL_457195, EPI_ISL_457196, EPI_ISL_457197, EPI_ISL_457198, EPI_ISL_457199, EPI_ISL_457200, EPI_ISL_457201, EPI_ISL_457202, EPI_ISL_457203, EPI_ISL_457204, EPI_ISL_457205, EPI_ISL_457206, EPI_ISL_457207, EPI_ISL_457208, EPI_ISL_457209, EPI_ISL_457210, EPI_ISL_457211, EPI_ISL_457212, EPI_ISL_457213, EPI_ISL_457214, EPI_ISL_457215, EPI_ISL_457216, EPI_ISL_457217, EPI_ISL_457218, EPI_ISL_457219, EPI_ISL_457220, EPI_ISL_457221, EPI_ISL_457222, EPI_ISL_457223, EPI_ISL_457224, EPI_ISL_457225, EPI_ISL_457226, EPI_ISL_457227, EPI_ISL_457228, EPI_ISL_457229, EPI_ISL_457230, EPI_ISL_457231, EPI_ISL_457232, EPI_ISL_457233, EPI_ISL_457234, EPI_ISL_457235, EPI_ISL_457236, EPI_ISL_457237, EPI_ISL_457238, EPI_ISL_457239, EPI_ISL_457240, EPI_ISL_457241, EPI_ISL_457242, EPI_ISL_457243, EPI_ISL_457244, EPI_ISL_457245, EPI_ISL_457246, EPI_ISL_457247, EPI_ISL_457248, EPI_ISL_457249, EPI_ISL_457250, EPI_ISL_457251, EPI_ISL_457252, EPI_ISL_457253, EPI_ISL_457254, EPI_ISL_457255, EPI_ISL_457256, EPI_ISL_457257, EPI_ISL_457258 | | | |
| see above | University of Exeter | COVID-19 Genomics UK (COG-UK) Consortium | Ben Temperton,Aaron Jeffries,Michelle Michelsen,Joanna Warwick-Dugdale,Audrey Farbos,Robyn Manley,Stephen Michell,Jane Masoli |
| EPI_ISL_458105 | Gujarat Biotechnology Research Centre | Gujarat Biotechnology Research Centre | Ankit Hinsu, Pritesh Sabara, Apurvasinh Puvar, Janvi Raval, Zarna Patel, Monika Gandhi, Pinal Trivedi, Maharshi Pandya, Amit Kanani, Nidhi Patel, Nitin Savaliya, Raghawendra Kumar, Dinesh Kumar, Zuber Saiyed, Komal Patel, Labdhi Pandya, Snehal Bagatharia, Ramesh Pandit, Tejas Shah, Camellia Chakraborty, Bhavesh Modi, Gaurishankar Shrimali, R D Dixit, A M Kadri, Umang Mishra, Chaitanya Joshi, Madhvi Joshi, , , , , |
| EPI_ISL_458107 | Gujarat Biotechnology Research Centre | Gujarat Biotechnology Research Centre | Apurvasinh Puvar, Janvi Raval, Zarna Patel, Monika Gandhi, Pinal Trivedi, Maharshi Pandya, Amit Kanani, Nidhi Patel, Nitin Savaliya, Raghawendra Kumar, Dinesh Kumar, Zuber Saiyed, Komal Patel, Labdhi Pandya, Snehal Bagatharia, Ramesh Pandit, Tejas Shah, Ankit Hinsu, Pritesh Sabara, Priyanka P Vatsa, Bhavesh Modi, Gaurishankar Shrimali, R D Dixit, A M Kadri, Umang Mishra, Chaitanya Joshi, Madhvi Joshi, , , , , |
| EPI_ISL_459893, EPI_ISL_459894, EPI_ISL_459895, EPI_ISL_459896, EPI_ISL_459897, EPI_ISL_459898, EPI_ISL_459899, EPI_ISL_459900, EPI_ISL_459901, EPI_ISL_459902, EPI_ISL_459903, EPI_ISL_459904, EPI_ISL_459905, EPI_ISL_459906 | | | |

| | | | |
|---|---|---|---|
| see above | Laboratoire National de Sante, Microbiology, Virology | Laboratoire National de Sante, Microbiology, Epidemiology and Microbial Genomics | Anke Wienecke-Baldacchino, Jessica Tapp, Guillaume Fournier, Tamir Abdelrahman, Trung Nguyen Nguyen, Catherine Ragimbeau |
| EPI_ISL_459992, EPI_ISL_459993, EPI_ISL_459994, EPI_ISL_459995, EPI_ISL_459996, EPI_ISL_459997, EPI_ISL_459998, EPI_ISL_459999, EPI_ISL_460000, EPI_ISL_460001, EPI_ISL_460002, EPI_ISL_460003, EPI_ISL_460004, EPI_ISL_460005, EPI_ISL_460006, EPI_ISL_460007, EPI_ISL_460008, EPI_ISL_460009, EPI_ISL_460010, EPI_ISL_460011, EPI_ISL_460012, EPI_ISL_460013, EPI_ISL_460014, EPI_ISL_460015, EPI_ISL_460016, EPI_ISL_460017, EPI_ISL_460018, EPI_ISL_460019, EPI_ISL_460020, EPI_ISL_460021, EPI_ISL_460022, EPI_ISL_460023, EPI_ISL_460024, EPI_ISL_460025, EPI_ISL_460026, EPI_ISL_460027, EPI_ISL_460028, EPI_ISL_460029, EPI_ISL_460030, EPI_ISL_460031, EPI_ISL_460032, EPI_ISL_460033, EPI_ISL_460034, EPI_ISL_460035, EPI_ISL_460036, EPI_ISL_460037, EPI_ISL_460038, EPI_ISL_460039, EPI_ISL_460040, EPI_ISL_460041, EPI_ISL_460042, EPI_ISL_460043 | | | |
| see above | Michigan Department of Health and Human Services, Bureau of Laboratories | Michigan Department of Health and Human Services, Bureau of Laboratories | Blankenship HM, Riner D, Soehnlen MK |
| EPI_ISL_460079 | Molecular Virology Unit, Fondazione IRCCS Policlinico San Matteo , Pavia | Laboratory of Virology, INMI Lazzaro Spallanzani IRCCS | Barbara Bartolini, Cesare E.M. Gruber, Maria R. Capobianchi, Martina Rueca, Antonio Piralla, Fausto Baldanti, Antonino Di Caro |
| EPI_ISL_460080 | Molecular Virology Unit, Fondazione IRCCS Policlinico San Matteo , Pavia | Laboratory of Virology, INMI Lazzaro Spallanzani IRCCS | Antonio Piralla, Barbara Bartolini, Fausto Baldanti, Martina Rueca, Antonino Di Caro, Cesare E.M. Gruber, Maria R. Capobianchi |
| EPI_ISL_460089 | Molecular Virology Unit, Fondazione IRCCS Policlinico San Matteo , Pavia | Laboratory of Virology, INMI Lazzaro Spallanzani IRCCS | Antonino Di Caro, Barbara Bartolini, Martina Rueca, Cesare E.M. Gruber, Antonio Piralla, Fausto Baldanti, Maria R. Capobianchi |
| EPI_ISL_460090 | Molecular Virology Unit, Fondazione IRCCS Policlinico San Matteo , Pavia | Laboratory of Virology, INMI Lazzaro Spallanzani IRCCS | Antonio Piralla, Cesare E.M. Gruber, Antonino Di Caro, Maria R. Capobianchi, Martina Rueca, Barbara Bartolini, Fausto Baldanti |
| EPI_ISL_460091 | Molecular Virology Unit, Fondazione IRCCS Policlinico San Matteo , Pavia | Laboratory of Virology, INMI Lazzaro Spallanzani IRCCS | Antonino Di Caro, Antonio Piralla, Martina Rueca, Fausto Baldanti, Barbara Bartolini, Maria R. Capobianchi, Cesare E.M. Gruber |
| EPI_ISL_460094 | Molecular Virology Unit, Fondazione IRCCS Policlinico San Matteo , Pavia | Laboratory of Virology, INMI Lazzaro Spallanzani IRCCS | Barbara Bartolini, Maria R. Capobianchi, Antonino Di Caro, Antonio Piralla, Cesare E.M. Gruber, Martina Rueca, Fausto Baldanti |
| EPI_ISL_460095 | Molecular Virology Unit, Fondazione IRCCS Policlinico San Matteo , Pavia | Laboratory of Virology, INMI Lazzaro Spallanzani IRCCS | Barbara Bartolini, Antonino Di Caro, Fausto Baldanti, Cesare E.M. Gruber, Maria R. Capobianchi, Martina Rueca, Antonio Piralla |
| EPI_ISL_460554, EPI_ISL_460555, EPI_ISL_460556, EPI_ISL_460557, EPI_ISL_460558, EPI_ISL_460559, EPI_ISL_460560, EPI_ISL_460561, EPI_ISL_460562, EPI_ISL_460563, EPI_ISL_460564, EPI_ISL_460565, EPI_ISL_460566, EPI_ISL_460567, EPI_ISL_460568, EPI_ISL_460569, EPI_ISL_460570, EPI_ISL_460571, EPI_ISL_460572, EPI_ISL_460573, EPI_ISL_460574, EPI_ISL_460575, EPI_ISL_460576, EPI_ISL_460577, EPI_ISL_460578, EPI_ISL_460579, EPI_ISL_460580, EPI_ISL_460581, EPI_ISL_460582, EPI_ISL_460583, EPI_ISL_460584, EPI_ISL_460585, EPI_ISL_460586, EPI_ISL_460587, EPI_ISL_460588, EPI_ISL_460589, EPI_ISL_460590, EPI_ISL_460591, EPI_ISL_460592, EPI_ISL_460593, EPI_ISL_460594, EPI_ISL_460595, EPI_ISL_460596, EPI_ISL_460597, EPI_ISL_460598, EPI_ISL_460599, EPI_ISL_460600, EPI_ISL_460601, EPI_ISL_460602 | | | |
| see above | Michigan Department of Health and Human Services, Bureau of Laboratories | Michigan Department of Health and Human Services, Bureau of Laboratories | Blankenship HM, Riner D, Soehnlen MK |
| EPI_ISL_460636, EPI_ISL_460637, EPI_ISL_460638, EPI_ISL_460639, EPI_ISL_460640, EPI_ISL_460641, EPI_ISL_460642, EPI_ISL_460643, EPI_ISL_460644, EPI_ISL_460645, EPI_ISL_460646, EPI_ISL_460647, EPI_ISL_460648, EPI_ISL_460649, EPI_ISL_460650, EPI_ISL_460651, EPI_ISL_460652, EPI_ISL_460653, EPI_ISL_460654, EPI_ISL_460655, EPI_ISL_460656, EPI_ISL_460657, EPI_ISL_460658, EPI_ISL_460659, EPI_ISL_460660, EPI_ISL_460661, EPI_ISL_460662, EPI_ISL_460663, EPI_ISL_460664, EPI_ISL_460665, EPI_ISL_460666, EPI_ISL_460667, EPI_ISL_460668, EPI_ISL_460669, EPI_ISL_460670, EPI_ISL_460671, EPI_ISL_460672, EPI_ISL_460673, EPI_ISL_460674, EPI_ISL_460675, EPI_ISL_460676, EPI_ISL_460677, EPI_ISL_460678, EPI_ISL_460679, EPI_ISL_460680, EPI_ISL_460681, EPI_ISL_460682, EPI_ISL_460683, EPI_ISL_460684, EPI_ISL_460685, EPI_ISL_460686, EPI_ISL_460687, EPI_ISL_460688, EPI_ISL_460689, EPI_ISL_460690, EPI_ISL_460691, EPI_ISL_460692, EPI_ISL_460693, EPI_ISL_460694, EPI_ISL_460695, EPI_ISL_460696, EPI_ISL_460697, EPI_ISL_460698, EPI_ISL_460699, EPI_ISL_460700, EPI_ISL_460701, EPI_ISL_460702, EPI_ISL_460703, EPI_ISL_460704, EPI_ISL_460705, EPI_ISL_460714, EPI_ISL_460715, EPI_ISL_460716, EPI_ISL_460717, EPI_ISL_460718, EPI_ISL_460719, EPI_ISL_460720, EPI_ISL_460721, EPI_ISL_460723, EPI_ISL_460724, EPI_ISL_460725, EPI_ISL_460726, EPI_ISL_460727, EPI_ISL_460728, EPI_ISL_460729, EPI_ISL_460730, EPI_ISL_460731, EPI_ISL_460732, EPI_ISL_460733, EPI_ISL_460734, EPI_ISL_460735, EPI_ISL_460736, EPI_ISL_460737, EPI_ISL_460738, EPI_ISL_460753, EPI_ISL_460754, EPI_ISL_460755, EPI_ISL_460756, EPI_ISL_460757, EPI_ISL_460758, EPI_ISL_460759, EPI_ISL_460760, EPI_ISL_460761, EPI_ISL_460762, EPI_ISL_460763, EPI_ISL_460764, EPI_ISL_460765, EPI_ISL_460766, EPI_ISL_460767, EPI_ISL_460768, EPI_ISL_460770, EPI_ISL_460771, EPI_ISL_460773, EPI_ISL_460774, EPI_ISL_460775, EPI_ISL_460776, EPI_ISL_460777, EPI_ISL_460778, EPI_ISL_460779, EPI_ISL_460780, EPI_ISL_460781, EPI_ISL_460782, EPI_ISL_460783, EPI_ISL_460784, EPI_ISL_460785, EPI_ISL_460786, EPI_ISL_460787, EPI_ISL_460788, EPI_ISL_460789, EPI_ISL_460790, EPI_ISL_460791, EPI_ISL_460792, EPI_ISL_460793, EPI_ISL_460794, EPI_ISL_460795, EPI_ISL_460796, EPI_ISL_460797, EPI_ISL_460798, EPI_ISL_460799, EPI_ISL_460800, EPI_ISL_460801, EPI_ISL_460802, EPI_ISL_460803, EPI_ISL_460804, EPI_ISL_460805, EPI_ISL_460806, EPI_ISL_460807, EPI_ISL_460808, EPI_ISL_460809, EPI_ISL_460810, EPI_ISL_460811, EPI_ISL_460812, EPI_ISL_460813, EPI_ISL_460814, EPI_ISL_460815, EPI_ISL_460816, EPI_ISL_460817, EPI_ISL_460818, EPI_ISL_460819, EPI_ISL_460820, EPI_ISL_460821, EPI_ISL_460823, EPI_ISL_460824, EPI_ISL_460825, EPI_ISL_460826, EPI_ISL_460827, EPI_ISL_460828, EPI_ISL_460829, EPI_ISL_460830, EPI_ISL_460831, EPI_ISL_460832, EPI_ISL_460833, EPI_ISL_460834, EPI_ISL_460835, EPI_ISL_460836, EPI_ISL_460837, EPI_ISL_460838, EPI_ISL_460839, EPI_ISL_460840, EPI_ISL_460841, EPI_ISL_460842, EPI_ISL_460843, EPI_ISL_460844, EPI_ISL_460845, EPI_ISL_460846, EPI_ISL_460847, EPI_ISL_460848, EPI_ISL_460849, EPI_ISL_460850, EPI_ISL_460851, EPI_ISL_460852, EPI_ISL_460853, EPI_ISL_460854, EPI_ISL_460855, EPI_ISL_460856, EPI_ISL_460857, EPI_ISL_460858, EPI_ISL_460859, EPI_ISL_460860, EPI_ISL_460861, EPI_ISL_460862, EPI_ISL_460863, EPI_ISL_460864, EPI_ISL_460865, EPI_ISL_460866, EPI_ISL_460867, EPI_ISL_460868, EPI_ISL_460869, EPI_ISL_460870, EPI_ISL_460871, EPI_ISL_460872, EPI_ISL_460873, EPI_ISL_460874, EPI_ISL_460875, EPI_ISL_460876, EPI_ISL_460877, EPI_ISL_460878, EPI_ISL_460879, EPI_ISL_460880, EPI_ISL_460881, EPI_ISL_460882, EPI_ISL_460883, EPI_ISL_460884, EPI_ISL_460885, EPI_ISL_460886, EPI_ISL_460887, EPI_ISL_460888, EPI_ISL_460889, EPI_ISL_460890, EPI_ISL_460892, EPI_ISL_460893, EPI_ISL_460894, EPI_ISL_460895, EPI_ISL_460896, EPI_ISL_460897, EPI_ISL_460898, EPI_ISL_460899, EPI_ISL_460900, EPI_ISL_460901, EPI_ISL_460902, EPI_ISL_460903, EPI_ISL_460904, EPI_ISL_460905, EPI_ISL_460906, EPI_ISL_460907, EPI_ISL_460908, EPI_ISL_460909, EPI_ISL_460910, EPI_ISL_460911, EPI_ISL_460912, EPI_ISL_460913, EPI_ISL_460914, EPI_ISL_460915, EPI_ISL_460916, EPI_ISL_460917, EPI_ISL_460918, EPI_ISL_460919, EPI_ISL_460920, EPI_ISL_460921, EPI_ISL_460922, EPI_ISL_460923, EPI_ISL_460924, EPI_ISL_460925, EPI_ISL_460926, EPI_ISL_460927, EPI_ISL_460928, EPI_ISL_460929, EPI_ISL_460930, EPI_ISL_460931, EPI_ISL_460932, EPI_ISL_460933, EPI_ISL_460934, EPI_ISL_460935, EPI_ISL_460936, EPI_ISL_460937, EPI_ISL_460938, EPI_ISL_460939, EPI_ISL_460940, EPI_ISL_460941, EPI_ISL_460942, EPI_ISL_460943, EPI_ISL_460944, EPI_ISL_460945, EPI_ISL_460946, EPI_ISL_460947, EPI_ISL_460948, EPI_ISL_460949, EPI_ISL_460950, EPI_ISL_460951, EPI_ISL_460952, EPI_ISL_460953, EPI_ISL_460954, EPI_ISL_460955, EPI_ISL_460956, EPI_ISL_460957, EPI_ISL_460958, EPI_ISL_460959, EPI_ISL_460960, EPI_ISL_460961, EPI_ISL_460962, EPI_ISL_460963, EPI_ISL_460964, EPI_ISL_460965, EPI_ISL_460966, EPI_ISL_460967, EPI_ISL_460968, EPI_ISL_460969, EPI_ISL_460970, EPI_ISL_460971, EPI_ISL_460972, EPI_ISL_460973, EPI_ISL_460974, EPI_ISL_460975, EPI_ISL_460976, EPI_ISL_460977, EPI_ISL_460978, EPI_ISL_460979, EPI_ISL_460980, EPI_ISL_460981, EPI_ISL_460982, EPI_ISL_460983, EPI_ISL_460984, EPI_ISL_460985, EPI_ISL_460986, EPI_ISL_460987, EPI_ISL_460988, EPI_ISL_460989, EPI_ISL_460990, EPI_ISL_460991, EPI_ISL_460992, EPI_ISL_460993, EPI_ISL_460994, EPI_ISL_460995, EPI_ISL_460996, EPI_ISL_460997, EPI_ISL_460998, EPI_ISL_460999, EPI_ISL_461000, EPI_ISL_461001, EPI_ISL_461002, EPI_ISL_461003, EPI_ISL_461004, EPI_ISL_461005, EPI_ISL_461006, EPI_ISL_461007, EPI_ISL_461008, EPI_ISL_461009, EPI_ISL_461010, EPI_ISL_461011, EPI_ISL_461012, EPI_ISL_461013, EPI_ISL_461014, EPI_ISL_461015, EPI_ISL_461016, EPI_ISL_461017, EPI_ISL_461018, EPI_ISL_461019, EPI_ISL_461020, EPI_ISL_461021, EPI_ISL_461022, EPI_ISL_461023, EPI_ISL_461024, EPI_ISL_461025, EPI_ISL_461026, EPI_ISL_461027, EPI_ISL_461028, EPI_ISL_461029, EPI_ISL_461030, EPI_ISL_461031, EPI_ISL_461032, EPI_ISL_461033, EPI_ISL_461034, EPI_ISL_461035, EPI_ISL_461036, EPI_ISL_461037, EPI_ISL_461038, EPI_ISL_461039, EPI_ISL_461040, EPI_ISL_461041, EPI_ISL_461042, EPI_ISL_461043, EPI_ISL_461044, EPI_ISL_461045, EPI_ISL_461046, EPI_ISL_461047, EPI_ISL_461048, EPI_ISL_461049, EPI_ISL_461050, EPI_ISL_461061, EPI_ISL_461062, EPI_ISL_461063, EPI_ISL_461064, EPI_ISL_461065, EPI_ISL_461066, EPI_ISL_461067, EPI_ISL_461068, EPI_ISL_461069, EPI_ISL_461070, EPI_ISL_461071, EPI_ISL_461072, EPI_ISL_461073, EPI_ISL_461074, EPI_ISL_461075, EPI_ISL_461076, EPI_ISL_461077, EPI_ISL_461078, EPI_ISL_461079, EPI_ISL_461080, EPI_ISL_461081, EPI_ISL_461082, EPI_ISL_461083, EPI_ISL_461084, EPI_ISL_461085, EPI_ISL_461086, EPI_ISL_461087, EPI_ISL_461101, EPI_ISL_461102, EPI_ISL_461103, EPI_ISL_461104, EPI_ISL_461105, EPI_ISL_461106, EPI_ISL_461107, EPI_ISL_461108, EPI_ISL_461109, EPI_ISL_461110, EPI_ISL_461111, EPI_ISL_461113, EPI_ISL_461114, EPI_ISL_461115, EPI_ISL_461116, EPI_ISL_461117, EPI_ISL_461118, EPI_ISL_461119, EPI_ISL_461120, EPI_ISL_461121, EPI_ISL_461122, EPI_ISL_461123, EPI_ISL_461124, EPI_ISL_461125, EPI_ISL_461126, EPI_ISL_461127, EPI_ISL_461128, EPI_ISL_461129, EPI_ISL_461130, EPI_ISL_461131, EPI_ISL_461132, EPI_ISL_461133, EPI_ISL_461134, EPI_ISL_461135, EPI_ISL_461136, EPI_ISL_461137, EPI_ISL_461138, EPI_ISL_461139, EPI_ISL_461140, EPI_ISL_461141, EPI_ISL_461142, EPI_ISL_461143, EPI_ISL_461144, EPI_ISL_461145, EPI_ISL_461146, EPI_ISL_461147, EPI_ISL_461148, EPI_ISL_461149, EPI_ISL_461150, EPI_ISL_461151, EPI_ISL_461152, EPI_ISL_461153, EPI_ISL_461154, EPI_ISL_461155, EPI_ISL_461156, EPI_ISL_461157, EPI_ISL_461158, EPI_ISL_461159, EPI_ISL_461161, EPI_ISL_461163, EPI_ISL_461164, EPI_ISL_461165, EPI_ISL_461166, EPI_ISL_461167, EPI_ISL_461172, EPI_ISL_461177, EPI_ISL_461179, EPI_ISL_461181, EPI_ISL_461182, EPI_ISL_461183, EPI_ISL_461184, EPI_ISL_461185, EPI_ISL_461186, EPI_ISL_461187, EPI_ISL_461188, EPI_ISL_461189, EPI_ISL_461193, EPI_ISL_461194, EPI_ISL_461195, EPI_ISL_461196, EPI_ISL_461197, EPI_ISL_461198, EPI_ISL_461199, EPI_ISL_461201, EPI_ISL_461202, EPI_ISL_461204, EPI_ISL_461205, EPI_ISL_461206, EPI_ISL_461207, EPI_ISL_461208, EPI_ISL_461209, EPI_ISL_461210, EPI_ISL_461211, EPI_ISL_461212, EPI_ISL_461213, EPI_ISL_461214, EPI_ISL_461215, EPI_ISL_461216, EPI_ISL_461217, EPI_ISL_461218, EPI_ISL_461219, EPI_ISL_461220, EPI_ISL_461221, EPI_ISL_461222, EPI_ISL_461223, EPI_ISL_461224, EPI_ISL_461225, EPI_ISL_461226, EPI_ISL_461227, EPI_ISL_461228, EPI_ISL_461229, EPI_ISL_461230, EPI_ISL_461231, EPI_ISL_461232, EPI_ISL_461234, EPI_ISL_461235, EPI_ISL_461236, EPI_ISL_461237, EPI_ISL_461238, EPI_ISL_461239, EPI_ISL_461240, EPI_ISL_461241, EPI_ISL_461242, EPI_ISL_461243, EPI_ISL_461244, EPI_ISL_461245, EPI_ISL_461246, EPI_ISL_461247, EPI_ISL_461248, EPI_ISL_461249, EPI_ISL_461250, EPI_ISL_461251, EPI_ISL_461252, EPI_ISL_461253, EPI_ISL_461254, EPI_ISL_461255, EPI_ISL_461256, EPI_ISL_461257, EPI_ISL_461258, EPI_ISL_461259, EPI_ISL_461260, EPI_ISL_461261, EPI_ISL_461262, EPI_ISL_461263, EPI_ISL_461264, EPI_ISL_461265, EPI_ISL_461266, EPI_ISL_461267, EPI_ISL_461268, EPI_ISL_461269, EPI_ISL_461270, EPI_ISL_461271, EPI_ISL_461272, EPI_ISL_461273, EPI_ISL_461274, EPI_ISL_461275, EPI_ISL_461276, EPI_ISL_461277, EPI_ISL_461278, EPI_ISL_461279, EPI_ISL_461280, EPI_ISL_461281, EPI_ISL_461282, EPI_ISL_461283, EPI_ISL_461284, EPI_ISL_461285, EPI_ISL_461286, EPI_ISL_461287, EPI_ISL_461288, EPI_ISL_461289, EPI_ISL_461290, EPI_ISL_461291, EPI_ISL_461292, EPI_ISL_461293, EPI_ISL_461294, EPI_ISL_461295, EPI_ISL_461296, EPI_ISL_461297, EPI_ISL_461298, EPI_ISL_461299, EPI_ISL_461300, EPI_ISL_461301, EPI_ISL_461302, EPI_ISL_461303, EPI_ISL_461304, EPI_ISL_461305, EPI_ISL_461306, EPI_ISL_461307, EPI_ISL_461308, EPI_ISL_461309, EPI_ISL_461310, EPI_ISL_461311, EPI_ISL_461312, EPI_ISL_461313, EPI_ISL_461314, EPI_ISL_461315, EPI_ISL_461316, EPI_ISL_461317, EPI_ISL_461318, EPI_ISL_461319, EPI_ISL_461320, EPI_ISL_461321, EPI_ISL_461322, EPI_ISL_461323, EPI_ISL_461324, EPI_ISL_461325, EPI_ISL_461326, EPI_ISL_461327, EPI_ISL_461328, EPI_ISL_461329, EPI_ISL_461330, EPI_ISL_461331, EPI_ISL_461332, EPI_ISL_461333, EPI_ISL_461334, EPI_ISL_461335, EPI_ISL_461336, EPI_ISL_461337, EPI_ISL_461338, EPI_ISL_461339, EPI_ISL_461340, EPI_ISL_461341, EPI_ISL_461342, EPI_ISL_461343, EPI_ISL_461344, EPI_ISL_461345, EPI_ISL_461346, EPI_ISL_461347, EPI_ISL_461348, EPI_ISL_461349, EPI_ISL_461350, EPI_ISL_461351, EPI_ISL_461352, EPI_ISL_461353, EPI_ISL_461354, EPI_ISL_461355, EPI_ISL_461356, EPI_ISL_461357, EPI_ISL_461358, EPI_ISL_461359, EPI_ISL_461360, EPI_ISL_461361, EPI_ISL_461362, EPI_ISL_461363, EPI_ISL_461364, EPI_ISL_461365, EPI_ISL_461366, EPI_ISL_461367, EPI_ISL_461368, EPI_ISL_461369, EPI_ISL_461370, EPI_ISL_461371, EPI_ISL_461372, EPI_ISL_461373, EPI_ISL_461375, EPI_ISL_461376, EPI_ISL_461377, EPI_ISL_461378, EPI_ISL_461379, EPI_ISL_461380, EPI_ISL_461381, EPI_ISL_461383, EPI_ISL_461384, EPI_ISL_461385, EPI_ISL_461386, EPI_ISL_461387, EPI_ISL_461388, EPI_ISL_461389, EPI_ISL_461390, EPI_ISL_461391, EPI_ISL_461394, EPI_ISL_461396, EPI_ISL_461397, EPI_ISL_461398 | | | |
| see above | Dutch COVID-19 response team | Erasmus Medical Center | Bas Oude Munnink, David Nieuwenhuijse, Reina Sikkema, Claudia Schapendonk, Irina Chestakova, Anne van der Linden, Theo Bestebroer, Stefan van Nieuwkoop, Mark Pronk, Pascal Lexmond, Corien Swaan, Manon Haverkate, Madelief Mollers, Mart Stein, Sandra Kengne Kamga Mobou, Jeroen van Kampen, Jolanda Voermans, Aura Timen, Corine GeurtsvanKessel, Annemiek van der Eijk, Richard Molenkamp, Marion Koopmans, on behalf of the Dutch national COVID-19 response team. |
| EPI_ISL_461480 | Government Medical College, Vadodara | Gujarat Biotechnology Research Centre | Armi Chaudhari, Raghawendra Kumar, Dinesh Kumar, Zuber Saiyed, Komal Patel, Labdhi Pandya, Snehal Bagatharia, Tanuja Javadekar , R N Daveshwar, Tejas Shah, Ankit Hinsu, Pritesh Sabara, Apurvasinh Puvar, Janvi Raval, Zarna Patel, Monika Gandhi, Pinal Trivedi, Maharshi Pandya, Nidhi Patel, Nitin Savaliya, R D Dixit, A M Kadri, Harsh Bakshi, Chaitanya Joshi, Madhvi Joshi, |
| EPI_ISL_461481 | Pandit Deendayal Upadhyay Government Medical College, Rajkot | Gujarat Biotechnology Research Centre | Bhavya Jindal, Dinesh Kumar, Zuber Saiyed, Komal Patel, Labdhi Pandya, Snehal Bagatharia, Prakash Modi, Sejul Antala, Manish Pattani, Tejas Shah, Ankit Hinsu, Pritesh Sabara, Apurvasinh Puvar, Janvi Raval, Zarna Patel, Monika Gandhi, Pinal Trivedi, Maharshi Pandya, Nidhi Patel, Nitin Savaliya, Raghawendra Kumar, R D Dixit, A M Kadri, Harsh Bakshi, Chaitanya Joshi, Madhvi Joshi |
| EPI_ISL_461482 | Pandit Deendayal Upadhyay Government Medical College, Rajkot | Gujarat Biotechnology Research Centre | Anjali Rajwar, Zuber Saiyed, Komal Patel, Labdhi Pandya, Snehal Bagatharia, Prakash Modi, Sejul Antala, Manish Pattani, Tejas Shah, Ankit Hinsu, Pritesh Sabara, Apurvasinh Puvar, Janvi Raval, Zarna Patel, Monika Gandhi, Pinal Trivedi, Maharshi Pandya, Nidhi Patel, Nitin Savaliya, Raghawendra Kumar, Dinesh Kumar, R D Dixit, A M Kadri, Harsh Bakshi, Chaitanya Joshi, Madhvi Joshi |
| EPI_ISL_461493 | B.J. Medical College and Civil hospital | Gujarat Biotechnology Research Centre | Armi Chaudhari, Zarna Patel, Monika Gandhi, Pinal Trivedi, Maharshi Pandya, Nidhi Patel, Nitin Savaliya, Raghawendra Kumar, Dinesh Kumar, Zuber |

| | | | |
|---|---|---|---|
| | | | Saiyed, Komal Patel, Labdhi Pandya, Snehal Bagatharia, Pranay Shah, Kamlesh J Upadhyay, Tejas Shah, Ankit Hinsu, Pritesh Sabara, Apurvasinh Puvar, Janvi Raval, R D Dixit, A M Kadri, Harsh Bakshi, Chaitanya Joshi, Madhvi Joshi, |
| EPI_ISL_461494 | B.J. Medical College and Civil hospital | Gujarat Biotechnology Research Centre | Bhavya Jindal, Monika Gandhi, Pinal Trivedi, Maharshi Pandya, Nidhi Patel, Nitin Savaliya, Raghawendra Kumar, Dinesh Kumar, Zuber Saiyed, Komal Patel, Labdhi Pandya, Snehal Bagatharia, Pranay Shah, Kamlesh J Upadhyay, Tejas Shah, Ankit Hinsu, Pritesh Sabara, Apurvasinh Puvar, Janvi Raval, Zarna Patel, R D Dixit, A M Kadri, Harsh Bakshi, Chaitanya Joshi, Madhvi Joshi, |
| EPI_ISL_461495 | B.J. Medical College and Civil hospital | Gujarat Biotechnology Research Centre | Anjali Rajwar, Pinal Trivedi, Maharshi Pandya, Nidhi Patel, Nitin Savaliya, Raghawendra Kumar, Dinesh Kumar, Zuber Saiyed, Komal Patel, Labdhi Pandya, Snehal Bagatharia, Pranay Shah, Kamlesh J Upadhyay, Tejas Shah, Ankit Hinsu, Pritesh Sabara, Apurvasinh Puvar, Janvi Raval, Zarna Patel, Monika Gandhi, R D Dixit, A M Kadri, Harsh Bakshi, Chaitanya Joshi, Madhvi Joshi, |
| EPI_ISL_461970, EPI_ISL_461971, EPI_ISL_461972, EPI_ISL_461973, EPI_ISL_461974, EPI_ISL_461975, EPI_ISL_461976, EPI_ISL_461977, EPI_ISL_461978, EPI_ISL_461979, EPI_ISL_461980, EPI_ISL_461981, EPI_ISL_461983, EPI_ISL_461984, EPI_ISL_461985, EPI_ISL_461986, EPI_ISL_461987, EPI_ISL_461988, EPI_ISL_461989, EPI_ISL_461990, EPI_ISL_461991, EPI_ISL_461992, EPI_ISL_461993, EPI_ISL_461994, EPI_ISL_461995, EPI_ISL_461996, EPI_ISL_461997 | | | |
| see above | Centre for Enzyme Innovation, University of Portsmouth / Translational Research Laboratory, Portsmouth Hospitals NHS Trust | COVID-19 Genomics UK (COG-UK) Consortium | Angela Beckett,Yann Bourgeois,Garry Scarlett,Sharon Glaysher,Scott Elliott,Kelly Bicknell,Robert Impey,Allyson Lloyd,Sarah Wyllie,Ethan Butcher,Anoop Chauhan,Samuel Robson |
| EPI_ISL_462090 | National Institute of Laboratory Medicine and Referral Center | Genomic Research Lab, BCSIR | Barna Goswami, Abu Sayeed Mohammad Mahmud, Mohammad Samir Uzzaman, Eshrar Osman, Md. Ahasan Habib, Shahina Akter, Tanjina Akhter Banu, Iffat Jahan, Md. Saddam Hossain, Tasnim Nafisa, Md. Maruf Ahmed Molla, Mahmuda Yeasmin, Asish Kumar Ghos, Bayzid Bin Monir, Arifa Akram, Sheikh Md. Selim Al Din, Salek Ahmed Sajib, Utpal Chandra Ray, Md. Salim Khan |
| EPI_ISL_462636, EPI_ISL_462637, EPI_ISL_462638, EPI_ISL_462639, EPI_ISL_462640, EPI_ISL_462641, EPI_ISL_462642, EPI_ISL_462643, EPI_ISL_462644, EPI_ISL_462645, EPI_ISL_462646, EPI_ISL_462647, EPI_ISL_462648, EPI_ISL_462649, EPI_ISL_462650, EPI_ISL_462651, EPI_ISL_462652, EPI_ISL_462653, EPI_ISL_462654, EPI_ISL_462655, EPI_ISL_462656, EPI_ISL_462657, EPI_ISL_462658, EPI_ISL_462659, EPI_ISL_462660, EPI_ISL_462661, EPI_ISL_462662, EPI_ISL_462663, EPI_ISL_462664, EPI_ISL_462665, EPI_ISL_462666, EPI_ISL_462667, EPI_ISL_462668, EPI_ISL_462669, EPI_ISL_462670, EPI_ISL_462671, EPI_ISL_462672, EPI_ISL_462673, EPI_ISL_462674, EPI_ISL_462675, EPI_ISL_462676, EPI_ISL_462677, EPI_ISL_462678, EPI_ISL_462679, EPI_ISL_462680, EPI_ISL_462681, EPI_ISL_462682, EPI_ISL_462683, EPI_ISL_462684, EPI_ISL_462685, EPI_ISL_462686, EPI_ISL_462687, EPI_ISL_462688, EPI_ISL_462689, EPI_ISL_462690, EPI_ISL_462691, EPI_ISL_462692, EPI_ISL_462693, EPI_ISL_462694, EPI_ISL_462695, EPI_ISL_462696, EPI_ISL_462697, EPI_ISL_462698, EPI_ISL_462699, EPI_ISL_462700, EPI_ISL_462701, EPI_ISL_462702, EPI_ISL_462703, EPI_ISL_462704, EPI_ISL_462705, EPI_ISL_462706, EPI_ISL_462707, EPI_ISL_462708, EPI_ISL_462709, EPI_ISL_462710, EPI_ISL_462711, EPI_ISL_462712, EPI_ISL_462713, EPI_ISL_462714, EPI_ISL_462715, EPI_ISL_462716, EPI_ISL_462717, EPI_ISL_462718, EPI_ISL_462719, EPI_ISL_462720, EPI_ISL_462721, EPI_ISL_462722, EPI_ISL_462723, EPI_ISL_462724, EPI_ISL_462725, EPI_ISL_462726, EPI_ISL_462727, EPI_ISL_462728, EPI_ISL_462729, EPI_ISL_462730, EPI_ISL_462731, EPI_ISL_462732, EPI_ISL_462733, EPI_ISL_462734, EPI_ISL_462735, EPI_ISL_462736 | | | |
| see above | Michigan Department of Health and Human Services, Bureau of Laboratories | Michigan Department of Health and Human Services, Bureau of Laboratories | Blankenship HM, Riner D, Soehnlen MK |
| EPI_ISL_463008 | Institute of Molecular Virology, University Münster | Institute of Molecular Virology, University Münster | Angeles Mecate Zambrano, Linda Brunotte, Stephan Ludwig, Joachim Kühn, Alexander Mellmann |
| EPI_ISL_464165, EPI_ISL_464166 | National Institute of Laboratory Medicine and Referral Center | Genomic Research Lab, BCSIR | Barna Goswami, Abu Sayeed Mohammad Mahmud, Mohammad Samir Uzzaman, Eshrar Osman, Md. Ahasan Habib, Shahina Akter, Tanjina Akhter Banu, Md. Murshed Hasan Sarker, Iffat Jahan, Md. Saddam Hossain, Tasnim Nafisa, Md. Maruf Ahmed Molla, Mahmuda Yeasmin, Asish Kumar Ghosh, Arifa Akram, A. K. M. Shamsuzzaman, Sheikh Md. Selim Al Din, Utpal Chandra Ray, Salek Ahmed Sajib, Md. Salim Khan |
| EPI_ISL_466839 | National Genomics Core-Center for DNA Fingerprinting and Diagnostics | National Genomics Core- Center for DNA Fingerprinting and Diagnostics (NGC-CDFD)- DBT's PAN-INDIA-1000 Genome consortium | Bala Pratyusha, Vinay Donipadi, G Shashikanth, Amrita Bhattacherjee, Rajeshree Sanyal, Raju Kumar, Ajay Kumar Chaudhary, Akash Chinchole, Brahmaji Sontyana, C. Arun Kumar, R HARINARAYANAN, RASHNA BHANDARI, MURALI DHARAN BASHYAM, DEBASHIS MITRA, DIVYA VASHISHT, ASHWIN DALAL |
| EPI_ISL_466840, EPI_ISL_466841, EPI_ISL_466842, EPI_ISL_466843, EPI_ISL_466844 | National Genomics Core-Center for DNA Fingerprinting and Diagnostics | National Genomics Core- Center for DNA Fingerprinting and Diagnostics (NGC-CDFD)- DBT's PAN-INDIA-1000 Genome consortium | Bala Pratyusha, Vinay Donipadi, G Shashikanth, Amrita Bhattacherjee, Nalini Raghunathan, Rajeshree Sanyal, Raju Kumar, Ajay Kumar Chaudhary, Akash Chinchole, Brahmaji Sontyana, C. Arun Kumar, R Harinarayanan, Rashna Bhandari, Murali Dharan Bashyam, Debashis Mitra, Divya Vashisht, Ashwin Dalal |
| EPI_ISL_466845, EPI_ISL_466846, EPI_ISL_466847 | National Genomics Core-Center for DNA Fingerprinting and Diagnostics | National Genomics Core- Center for DNA Fingerprinting and Diagnostics (NGC-CDFD)- DBT's PAN-INDIA-1000 Genome consortium | Bala Pratyusha, Vinay Donipadi, G Shashikanth, Amrita Bhattacherjee, Chandra Shekhar V, Chilakala Gangi Reddy, Chinthakindi KrishnaPrasad, Edurugatla Dinesh, Guru Raja, Hilal Ahmad Reshi, R HARINARAYANAN, RASHNA BHANDARI, MURALI DHARAN BASHYAM, DEBASHIS MITRA, DIVYA VASHISHT, ASHWIN DALAL |
| EPI_ISL_466848, EPI_ISL_466849, EPI_ISL_466850, EPI_ISL_466851, EPI_ISL_466852 | National Genomics Core-Center for DNA Fingerprinting and Diagnostics | National Genomics Core- Center for DNA Fingerprinting and Diagnostics (NGC-CDFD)- DBT's PAN-INDIA-1000 Genome consortium | Bala Pratyusha, Vinay Donipadi, G Shashikanth, Amrita Bhattacherjee, J. Mallikarjun, K. Viswakalyan, Kaisar Ahmad Lone, Kausika Kumar Malik, N. Sudheer, Neeraj Kumar, R HARINARAYANAN, RASHNA BHANDARI, MURALI DHARAN BASHYAM, DEBASHIS MITRA, DIVYA VASHISHT, ASHWIN DALAL |
| EPI_ISL_466853, EPI_ISL_466854, EPI_ISL_466855, EPI_ISL_466856, EPI_ISL_466857 | National Genomics Core-Center for DNA Fingerprinting and Diagnostics | National Genomics Core- Center for DNA Fingerprinting and Diagnostics (NGC-CDFD)- DBT's PAN-INDIA-1000 Genome consortium | Bala Pratyusha, Vinay Donipadi, G Shashikanth, Amrita Bhattacherjee, Niteen Pathak, Pradipta Hore, Rahul Baroi, Sayantan Goswami, Shaffiqu T S, Shalini Aricthota, R HARINARAYANAN, RASHNA BHANDARI, MURALI DHARAN BASHYAM, DEBASHIS MITRA, DIVYA VASHISHT, ASHWIN DALAL |
| EPI_ISL_466858, EPI_ISL_466859, EPI_ISL_466860, EPI_ISL_466861, EPI_ISL_466862 | National Genomics Core-Center for DNA Fingerprinting and Diagnostics | National Genomics Core- Center for DNA Fingerprinting and Diagnostics (NGC-CDFD)- DBT's PAN-INDIA-1000 Genome consortium | Bala Pratyusha, Vinay Donipadi, G Shashikanth, Amrita Bhattacherjee, Sobhan Babu, SPR Prasad, Yogesh Patidar, Arjita Jaiswal, Arpita Singh, Devanshi Gupta, R HARINARAYANAN, RASHNA BHANDARI, MURALI DHARAN BASHYAM, DEBASHIS MITRA, DIVYA VASHISHT, ASHWIN DALAL |
| EPI_ISL_466863, EPI_ISL_466864, EPI_ISL_466865, EPI_ISL_466866, EPI_ISL_466867 | National Genomics Core-Center for DNA Fingerprinting and Diagnostics | National Genomics Core- Center for DNA Fingerprinting and Diagnostics (NGC-CDFD)- DBT's PAN-INDIA-1000 Genome consortium | Bala Pratyusha, Vinay Donipadi, G Shashikanth, Amrita Bhattacherjee, Romila Moirangthem, Sanjana Sarkar, Shivani Yadav, Shubhra Ganguli, Suchitra Upreti, Swathi Chodisetty , R HARINARAYANAN, RASHNA BHANDARI, MURALI DHARAN BASHYAM, DEBASHIS MITRA, DIVYA VASHISHT, ASHWIN DALAL |
| EPI_ISL_466868, EPI_ISL_466869, EPI_ISL_466870, EPI_ISL_466871, EPI_ISL_466872 | National Genomics Core-Center for DNA Fingerprinting and Diagnostics | National Genomics Core- Center for DNA Fingerprinting and Diagnostics (NGC-CDFD)- DBT's PAN-INDIA-1000 Genome consortium | Bala Pratyusha, Vinay Donipadi, G Shashikanth, Amrita Bhattacherjee, Vani Singh, Shubhra Ganguli, Suchitra Upreti, Swathi Chodisetty , Vani Singh , R HARINARAYANAN, RASHNA BHANDARI, MURALI DHARAN BASHYAM, DEBASHIS MITRA, DIVYA VASHISHT, ASHWIN DALAL |
| EPI_ISL_467031 | GMERS Medical College and Hospital, Gandhinagar | Gujarat Biotechnology Research Centre | Bhavesh Modi, Bharti Rajani, Tejas Shah, Ankit Hinsu, Pritesh Sabara, Apurvasinh Puvar, Janvi Raval, Zarna Patel, Monika Gandhi, Pinal Trivedi, Maharshi Pandya, Nidhi Patel, Nitin Savaliya, Raghawendra Kumar, Dinesh Kumar, Zuber Saiyed, Komal Patel, Labdhi Pandya, Snehal Bagatharia, Seema Bhatt, Gaurishankar Shrimali, Pooja P Doshi, R D Dixit, A M Kadri, Harsh Bakshi, Chaitanya Joshi, Madhvi Joshi |
| EPI_ISL_467032 | GMERS Medical College and Hospital, Gandhinagar | Gujarat Biotechnology Research Centre | Bharti Rajani, Tejas Shah, Ankit Hinsu, Pritesh Sabara, Apurvasinh Puvar, Janvi Raval, Zarna Patel, Monika Gandhi, Pinal Trivedi, Maharshi Pandya, Nidhi Patel, Nitin Savaliya, Raghawendra Kumar, Dinesh Kumar, Zuber Saiyed, Komal Patel, Labdhi Pandya, Snehal Bagatharia, Seema Bhatt, Gaurishankar Shrimali, Bhavesh Modi, Akanksha Verma, R D Dixit, A M Kadri, Harsh Bakshi, Chaitanya Joshi, Madhvi Joshi |
| EPI_ISL_467034 | GMERS Medical College and Hospital, Gandhinagar | Gujarat Biotechnology Research Centre | Ankit Hinsu, Pritesh Sabara, Apurvasinh Puvar, Janvi Raval, Zarna Patel, Monika Gandhi, Pinal Trivedi, Maharshi Pandya, Nidhi Patel, Nitin Savaliya, Raghawendra Kumar, Dinesh Kumar, Zuber Saiyed, Komal Patel, Labdhi Pandya, Snehal Bagatharia, Seema Bhatt, Gaurishankar Shrimali, Bhavesh Modi, Bharti Rajani, Tejas Shah, Pragya Sharma, R D Dixit, A M Kadri, Harsh Bakshi, Chaitanya Joshi, Madhvi Joshi |
| EPI_ISL_467036 | GMERS Medical College and Hospital, Gandhinagar | Gujarat Biotechnology Research Centre | Apurvasinh Puvar, Janvi Raval, Zarna Patel, Monika Gandhi, Pinal Trivedi, Maharshi Pandya, Nidhi Patel, Nitin Savaliya, Raghawendra Kumar, Dinesh Kumar, Zuber Saiyed, Komal Patel, Labdhi Pandya, Snehal Bagatharia, Seema Bhatt, Gaurishankar Shrimali, Bhavesh Modi, Bharti Rajani, Tejas Shah, Ankit Hinsu, Pritesh Sabara, Afzal Ansari, R D Dixit, A M Kadri, Harsh Bakshi, Chaitanya Joshi, Madhvi Joshi |
| EPI_ISL_468765, EPI_ISL_468766, EPI_ISL_468767, EPI_ISL_468768, EPI_ISL_468769, EPI_ISL_468770, EPI_ISL_468771, EPI_ISL_468772, EPI_ISL_468773, EPI_ISL_468774, EPI_ISL_468775, EPI_ISL_468776, EPI_ISL_468777, EPI_ISL_468778, EPI_ISL_468779, EPI_ISL_468780, EPI_ISL_468781, EPI_ISL_468782, EPI_ISL_468783, EPI_ISL_468784, EPI_ISL_468785, EPI_ISL_468786, EPI_ISL_468787, EPI_ISL_468788, EPI_ISL_468789, EPI_ISL_468790, EPI_ISL_468791, EPI_ISL_468792, EPI_ISL_468793, EPI_ISL_468794, EPI_ISL_468795, EPI_ISL_468796, EPI_ISL_468797, EPI_ISL_468798, EPI_ISL_468799, EPI_ISL_468800, EPI_ISL_468801, EPI_ISL_468802, EPI_ISL_468803, EPI_ISL_468804, EPI_ISL_468805, EPI_ISL_468806, EPI_ISL_468807, EPI_ISL_468808, EPI_ISL_468809, EPI_ISL_468810, EPI_ISL_468811, EPI_ISL_468812, EPI_ISL_468813, EPI_ISL_468814, EPI_ISL_468815, EPI_ISL_468816, EPI_ISL_468817, EPI_ISL_468818, EPI_ISL_468819, EPI_ISL_468820, EPI_ISL_468821, EPI_ISL_468822, EPI_ISL_468823, EPI_ISL_468824, EPI_ISL_468825, EPI_ISL_468826, EPI_ISL_468827, EPI_ISL_468828, EPI_ISL_468829, EPI_ISL_468830, EPI_ISL_468831, EPI_ISL_468832, EPI_ISL_468833, EPI_ISL_468834, EPI_ISL_468835, EPI_ISL_468836, EPI_ISL_468837, EPI_ISL_468838, EPI_ISL_468839, EPI_ISL_468840, EPI_ISL_468841, EPI_ISL_468842, EPI_ISL_468843, EPI_ISL_468844, EPI_ISL_468845, EPI_ISL_468846, EPI_ISL_468847, EPI_ISL_468848, EPI_ISL_468849, EPI_ISL_468850, EPI_ISL_468851, EPI_ISL_468852, EPI_ISL_468853, EPI_ISL_468854, EPI_ISL_468855, EPI_ISL_468856, EPI_ISL_468857, EPI_ISL_468858, EPI_ISL_468859 | | | |
| see above | Servicio de Microbiología, Hospital Miguel Servet, Zaragoza | SeqCOVID-SPAIN consortium/IBV(CSIC) | Antonio Rezusta López, Alexander Tristancho Baró, Ana Milagro, Yolanda Gracia Grataloup, Nieves Martínez Cameo and SeqCOVID-SPAIN consortium |
| EPI_ISL_469025 | B.J. Medical College and Civil hospital | Gujarat Biotechnology Research Centre | Ankit Hinsu, Pritesh Sabara, Apurvasinh Puvar, Janvi Raval, Zarna Patel, Monika Gandhi, Pinal Trivedi, Maharshi Pandya, Nidhi Patel, Nitin Savaliya, Raghawendra Kumar, Dinesh Kumar, Zuber Saiyed, Komal Patel, Labdhi Pandya, Snehal Bagatharia, Pranay Shah, Kamlesh J Upadhyay, Nirav Mungalpara, Tejas Shah, Pragya Sharma, R D Dixit, A M Kadri, Harsh Bakshi, Chaitanya Joshi, Madhvi Joshi |
| EPI_ISL_469027 | B.J. Medical College and Civil hospital | Gujarat Biotechnology Research Centre | Apurvasinh Puvar, Janvi Raval, Zarna Patel, Monika Gandhi, Pinal Trivedi, Maharshi Pandya, Nidhi Patel, Nitin Savaliya, Raghawendra Kumar, Dinesh |

| | | | |
|---|---|---|---|
| | | | Kumar, Zuber Saiyed, Komal Patel, Labdhi Pandya, Snehal Bagatharia, Pranay Shah, Kamlesh J Upadhyay, Nirav Mungalpara, Tejas Shah, Ankit Hinsu, Pritesh Sabara, Afzal Ansari, R D Dixit, A M Kadri, Harsh Bakshi, Chaitanya Joshi, Madhvi Joshi |
| EPI_ISL_469047 | Dr. N. D. Desai Medical College & Hospital | Gujarat Biotechnology Research Centre | Ankit Hinsu, Pritesh Sabara, Apurvasinh Puvar, Janvi Raval, Zarna Patel, Monika Gandhi, Pinal Trivedi, Maharshi Pandya, Nidhi Patel, Nitin Savaliya, Raghawendra Kumar, Dinesh Kumar, Zuber Saiyed, Komal Patel, Labdhi Pandya, Snehal Bagatharia, J G Buch, Jigar Gusani, Supreet Prabhu, Tejas Shah, Akanksha Verma, R D Dixit, A M Kadri, Harsh Bakshi, Chaitanya Joshi, Madhvi Joshi |
| EPI_ISL_469224, EPI_ISL_469225, EPI_ISL_469226, EPI_ISL_469227, EPI_ISL_469228, EPI_ISL_469229, EPI_ISL_469230, EPI_ISL_469231, EPI_ISL_469232, EPI_ISL_469233, EPI_ISL_469234, EPI_ISL_469235, EPI_ISL_469236, EPI_ISL_469237, EPI_ISL_469238, EPI_ISL_469239, EPI_ISL_469240 | | | |
| see above | Public Health Laboratory | National Microbiology Laboratory | Anna Majer, Shari Tyson, Grace Seo, Kristyn Burak, Philip Mabon, Elsie Grudeski, Rhiannon Huzarewich, Russell Mandes, Jennifer Tanner, Natalie Knox, Morag Graham, Gary Van Domselaar, Robert Needle, Yang Yu, Adel Malek, Laura Gilbert, George Zahariadis, Nathalie Bastien, Yan Li, Timothy Booth, Matthew Gilmour |
| EPI_ISL_469241, EPI_ISL_469242, EPI_ISL_469243, EPI_ISL_469244, EPI_ISL_469245, EPI_ISL_469246, EPI_ISL_469247, EPI_ISL_469248, EPI_ISL_469249, EPI_ISL_469250, EPI_ISL_469251, EPI_ISL_469252 | | | |
| see above | Special Infectious Agents Unit | Special Infectious Agents Unit | Azhar,E.I., Hassan,A.M., Tolah,A.M., Uthman,N.A., Al-Sobahy,T.L., Farraj,S.A., El-Kafrawy,S.A. |
| EPI_ISL_469297 | National Institute of Laboratory Medicine and Referral Center | Genomic Research Lab, BCSIR | Barna Goswami, Abu Sayeed Mohammad Mahmud, Mohammad Samir Uzzaman, Eshrar Osman, Md. Ahasan Habib, Shahina Akter, Tanjina Akhter Banu, Md. Murshed Hasan Sarkar, Iffat Jahan, Md. Saddam Hossain, Tasnim Nafisa, Md. Maruf Ahmed Molla, Mahmuda Yeasmin, Asish Kumar Ghosh, Bayzid Bin Monir, A. K. M. Shamsuzzaman, Sheikh Md. Selim Al Din, Utpal Chandra Ray, Salek Ahmed Sajib, Md. Salim Khan |
| EPI_ISL_471268 | Hospital IESS Babahoyo | Institute of Microbiology, Universidad San Francisco de Quito | Belén Prado-Vivar, Sully Márquez, Juan José Guadalupe, Bernardo Gutiérrez, Francisco Cordova, Ninfa Henriquez, Killen Briones-Zamora, Killen Briones-Claudette, Verónica Barragán, Patricio Rojas-Silva, Gabriel Trueba, Michelle Grunauer, Paúl Cárdenas |
| EPI_ISL_471676, EPI_ISL_471677, EPI_ISL_471678, EPI_ISL_471679, EPI_ISL_471680, EPI_ISL_471681, EPI_ISL_471682, EPI_ISL_471683, EPI_ISL_471684, EPI_ISL_471685, EPI_ISL_471686, EPI_ISL_471687, EPI_ISL_471688, EPI_ISL_471689, EPI_ISL_471690, EPI_ISL_471691, EPI_ISL_471692, EPI_ISL_471693, EPI_ISL_471694, EPI_ISL_471695, EPI_ISL_471696, EPI_ISL_471697, EPI_ISL_471698, EPI_ISL_471699, EPI_ISL_471700, EPI_ISL_471701, EPI_ISL_471702, EPI_ISL_471703, EPI_ISL_471704, EPI_ISL_471705, EPI_ISL_471706, EPI_ISL_471707, EPI_ISL_471708, EPI_ISL_471709, EPI_ISL_471710, EPI_ISL_471711, EPI_ISL_471712, EPI_ISL_471713, EPI_ISL_471714, EPI_ISL_471715, EPI_ISL_471716, EPI_ISL_471717, EPI_ISL_471718, EPI_ISL_471719, EPI_ISL_471720, EPI_ISL_471721, EPI_ISL_471722, EPI_ISL_471723, EPI_ISL_471724, EPI_ISL_471725, EPI_ISL_471726, EPI_ISL_471727, EPI_ISL_471728, EPI_ISL_471729, EPI_ISL_471730, EPI_ISL_471731, EPI_ISL_471732, EPI_ISL_471733, EPI_ISL_471734, EPI_ISL_471735, EPI_ISL_471736, EPI_ISL_471737, EPI_ISL_471738, EPI_ISL_471739, EPI_ISL_471740, EPI_ISL_471741, EPI_ISL_471742, EPI_ISL_471743, EPI_ISL_471744, EPI_ISL_471745, EPI_ISL_471746, EPI_ISL_471747, EPI_ISL_471748, EPI_ISL_471749, EPI_ISL_471750, EPI_ISL_471751, EPI_ISL_471752, EPI_ISL_471753, EPI_ISL_471754, EPI_ISL_471755, EPI_ISL_471756, EPI_ISL_471757, EPI_ISL_471758, EPI_ISL_471759, EPI_ISL_471760, EPI_ISL_471761, EPI_ISL_471762, EPI_ISL_471763, EPI_ISL_471764, EPI_ISL_471765, EPI_ISL_471766, EPI_ISL_471767, EPI_ISL_471768, EPI_ISL_471769, EPI_ISL_471770, EPI_ISL_471771, EPI_ISL_471772, EPI_ISL_471773, EPI_ISL_471774, EPI_ISL_471775, EPI_ISL_471776, EPI_ISL_471777, EPI_ISL_471778, EPI_ISL_471779, EPI_ISL_471780, EPI_ISL_471781, EPI_ISL_471782, EPI_ISL_471783, EPI_ISL_471784, EPI_ISL_471785, EPI_ISL_471786, EPI_ISL_471787, EPI_ISL_471788, EPI_ISL_471789, EPI_ISL_471790, EPI_ISL_471791, EPI_ISL_471792, EPI_ISL_471793, EPI_ISL_471794, EPI_ISL_471795, EPI_ISL_471796, EPI_ISL_471797, EPI_ISL_471798, EPI_ISL_471799, EPI_ISL_471800, EPI_ISL_471801, EPI_ISL_471802, EPI_ISL_471803, EPI_ISL_471804, EPI_ISL_471805, EPI_ISL_471806, EPI_ISL_471807, EPI_ISL_471808, EPI_ISL_471809, EPI_ISL_471810, EPI_ISL_471811, EPI_ISL_471812, EPI_ISL_471813, EPI_ISL_471814, EPI_ISL_471815, EPI_ISL_471816, EPI_ISL_471817, EPI_ISL_471818, EPI_ISL_471819, EPI_ISL_471820, EPI_ISL_471821, EPI_ISL_471822, EPI_ISL_471823, EPI_ISL_471824, EPI_ISL_471825, EPI_ISL_471826, EPI_ISL_471827, EPI_ISL_471828, EPI_ISL_471829, EPI_ISL_471830, EPI_ISL_471831, EPI_ISL_471832, EPI_ISL_471833, EPI_ISL_471834, EPI_ISL_471835, EPI_ISL_471836, EPI_ISL_471837, EPI_ISL_471838, EPI_ISL_471839, EPI_ISL_471840, EPI_ISL_471841, EPI_ISL_471842, EPI_ISL_471843, EPI_ISL_471844, EPI_ISL_471845, EPI_ISL_471846, EPI_ISL_471847, EPI_ISL_471848, EPI_ISL_471849, EPI_ISL_471850, EPI_ISL_471851, EPI_ISL_471852, EPI_ISL_471853, EPI_ISL_471854, EPI_ISL_471855, EPI_ISL_471856, EPI_ISL_471857, EPI_ISL_471858, EPI_ISL_471859, EPI_ISL_471860, EPI_ISL_471861, EPI_ISL_471862, EPI_ISL_471863, EPI_ISL_471864, EPI_ISL_471865, EPI_ISL_471866, EPI_ISL_471867, EPI_ISL_471868, EPI_ISL_471869, EPI_ISL_471870, EPI_ISL_471871, EPI_ISL_471872, EPI_ISL_471873, EPI_ISL_471874, EPI_ISL_471875, EPI_ISL_471876, EPI_ISL_471877, EPI_ISL_471878, EPI_ISL_471879, EPI_ISL_471880, EPI_ISL_471881, EPI_ISL_471882, EPI_ISL_471883, EPI_ISL_471884, EPI_ISL_471885, EPI_ISL_471886, EPI_ISL_471887, EPI_ISL_471888, EPI_ISL_471889, EPI_ISL_471890, EPI_ISL_471891, EPI_ISL_471892, EPI_ISL_471893, EPI_ISL_471894, EPI_ISL_471895, EPI_ISL_471896, EPI_ISL_471897, EPI_ISL_471898, EPI_ISL_471899, EPI_ISL_471900, EPI_ISL_471901, EPI_ISL_471902, EPI_ISL_471903, EPI_ISL_471904, EPI_ISL_471905, EPI_ISL_471906, EPI_ISL_471907, EPI_ISL_471908, EPI_ISL_471909, EPI_ISL_471910 | | | |
| see above | Michigan Department of Health and Human Services, Bureau of Laboratories | Michigan Department of Health and Human Services, Bureau of Laboratories | Blankenship HM, Riner D, Soehnlen MK |
| EPI_ISL_471911, EPI_ISL_471912, EPI_ISL_471913, EPI_ISL_471914, EPI_ISL_471915, EPI_ISL_471916, EPI_ISL_471917, EPI_ISL_471918, EPI_ISL_471919, EPI_ISL_471920, EPI_ISL_471921, EPI_ISL_471922, EPI_ISL_471924, EPI_ISL_471925, EPI_ISL_471926, EPI_ISL_471927, EPI_ISL_471928, EPI_ISL_471929, EPI_ISL_471930, EPI_ISL_471931, EPI_ISL_471932, EPI_ISL_471933, EPI_ISL_471934, EPI_ISL_471935, EPI_ISL_471936, EPI_ISL_471937, EPI_ISL_471938, EPI_ISL_471939, EPI_ISL_471940, EPI_ISL_471941, EPI_ISL_471942, EPI_ISL_471943, EPI_ISL_471944, EPI_ISL_471945, EPI_ISL_471946, EPI_ISL_471947, EPI_ISL_471948, EPI_ISL_471949, EPI_ISL_471950, EPI_ISL_471951, EPI_ISL_471952, EPI_ISL_471953, EPI_ISL_471954, EPI_ISL_471955, EPI_ISL_471956, EPI_ISL_471957, EPI_ISL_471958, EPI_ISL_471959, EPI_ISL_471960, EPI_ISL_471961, EPI_ISL_471962, EPI_ISL_471963, EPI_ISL_471964, EPI_ISL_471965, EPI_ISL_471966, EPI_ISL_471967, EPI_ISL_471968, EPI_ISL_471969, EPI_ISL_471970, EPI_ISL_471971, EPI_ISL_471972, EPI_ISL_471973, EPI_ISL_471974, EPI_ISL_471975, EPI_ISL_471976, EPI_ISL_471977, EPI_ISL_471978, EPI_ISL_471979, EPI_ISL_471980, EPI_ISL_471981, EPI_ISL_471982, EPI_ISL_471983, EPI_ISL_471984, EPI_ISL_471985, EPI_ISL_471986, EPI_ISL_471987, EPI_ISL_471988, EPI_ISL_471989, EPI_ISL_471990, EPI_ISL_471991, EPI_ISL_471992 | | | |
| see above | University of Exeter | COVID-19 Genomics UK (COG-UK) Consortium | Ben Temperton,Aaron Jeffries,Michelle Michelsen,Joanna Warwick-Dugdale,Audrey Farbos,Robyn Manley,Stephen Michell,Jane Masoli |
| EPI_ISL_475029 | Banas Medical College and Research Institute | Gujarat Biotechnology Research Centre | Apurvasinh Puvar, Janvi Raval, Zarna Patel, Monika Gandhi, Pinal Trivedi, Maharshi Pandya, Nidhi Patel, Nitin Savaliya, Raghawendra Kumar, Dinesh Kumar, Zuber Saiyed, Komal Patel, Labdhi Pandya, Snehal Bagatharia, Radhika Khara, Sunil R Joshi, Viren s Doshi, Pritesh Sabara, Neelam Nathani, R D Dixit, A M Kadri, Harsh Bakshi, Chaitanya Joshi, Madhvi Joshi |
| EPI_ISL_475047 | GMERS Medical College & Hospital | Gujarat Biotechnology Research Centre | Apurvasinh Puvar, Janvi Raval, Zarna Patel, Monika Gandhi, Pinal Trivedi, Maharshi Pandya, Nidhi Patel, Nitin Savaliya, Raghawendra Kumar, Dinesh Kumar, Zuber Saiyed, Komal Patel, Labdhi Pandya, Snehal Bagatharia, Meenakshi Shah, Neena Doshi, Varsha Godbole, Pritesh Sabara, Pragya Sharma, R D Dixit, A M Kadri, Harsh Bakshi, Chaitanya Joshi, Madhvi Joshi |
| EPI_ISL_475058 | GAIMS & G K General Hospital | Gujarat Biotechnology Research Centre | Babulal Babhoria, Hitesh Assudani, Komal Patel, Labdhi Pandya, Snehal Bagatharia, Pritesh Sabara, Apurvasinh Puvar, Janvi Raval, Zarna Patel, Monika Gandhi, Pinal Trivedi, Maharshi Pandya, Nidhi Patel, Nitin Savaliya, Raghawendra Kumar, Dinesh Kumar, Zuber Saiyed, Pragya Sharma, R D Dixit, A M Kadri, Harsh Bakshi, Chaitanya Joshi, Madhvi Joshi |
| EPI_ISL_475167 | National Institute of Laboratory Medicine and Referral Center | Genomic Research Lab, BCSIR | Barna Goswami, Abu Sayeed Mohammad Mahmud, Mohammad Samir Uzzaman, Eshrar Osman, Md. Ahasan Habib, Shahina Akter, Tanjina Akhter Banu, Md. Murshed Hasan Sarkar, Iffat Jahan, Md. Saddam Hossain, Tasnim Nafisa, Md. Maruf Ahmed Molla, Mahmuda Yeasmin, Asish Kumar Ghosh, Bayzid Bin Monir, A. K. M. Shamsuzzaman, Sheikh Md. Selim Al Din, Utpal Chandra Ray, Salek Ahmed Sajib, Md. Salim Khan |
| EPI_ISL_475239, EPI_ISL_475240, EPI_ISL_475241, EPI_ISL_475242, EPI_ISL_475243, EPI_ISL_475244, EPI_ISL_475245, EPI_ISL_475246, EPI_ISL_475247, EPI_ISL_475248, EPI_ISL_475249, EPI_ISL_475250, EPI_ISL_475251, EPI_ISL_475252, EPI_ISL_475253, EPI_ISL_475254, EPI_ISL_475255, EPI_ISL_475256, EPI_ISL_475257, EPI_ISL_475258, EPI_ISL_475259, EPI_ISL_475260, EPI_ISL_475261, EPI_ISL_475262, EPI_ISL_475263, EPI_ISL_475264, EPI_ISL_475265, EPI_ISL_475267, EPI_ISL_475268, EPI_ISL_475269, EPI_ISL_475270, EPI_ISL_475271, EPI_ISL_475273, EPI_ISL_475278, EPI_ISL_475284, EPI_ISL_475290, EPI_ISL_475317, EPI_ISL_475318, EPI_ISL_475320, EPI_ISL_475338, EPI_ISL_475340 | | | |
| see above | Centre for Enzyme Innovation, University of Portsmouth / Translational Research Laboratory, Portsmouth Hospitals NHS Trust | COVID-19 Genomics UK (COG-UK) Consortium | Angela Beckett,Yann Bourgeois,Garry Scarlett,Sharon Glaysher,Scott Elliott,Kelly Bicknell,Robert Impey,Allyson Lloyd,Sarah Wyllie,Ethan Butcher,Anoop Chauhan,Samuel Robson |
| EPI_ISL_475757 | National Institute of Laboratory Medicine and Referral Center | Genomic Research Lab, BCSIR | Barna Goswami, Abu Sayeed Mohammad Mahmud, Mohammad Samir Uzzaman, Eshrar Osman, Md. Ahasan Habib, Shahina Akter, Tanjina Akhter Banu, Md. Murshed Hasan Sarkar, Iffat Jahan, Md. Saddam Hossain, Tasnim Nafisa, Md. Maruf Ahmed Molla, Mahmuda Yeasmin, Asish Kumar Ghosh, Arifa Akram, A. K. M. Shamsuzzaman, Sheikh Md. Selim Al Din, Utpal Chandra Ray, Salek Ahmed Sajib, Md. Salim Khan |
| EPI_ISL_476027, EPI_ISL_476028, EPI_ISL_476029, EPI_ISL_476030, EPI_ISL_476031, EPI_ISL_476032, EPI_ISL_476033, EPI_ISL_476034, EPI_ISL_476035, EPI_ISL_476036, EPI_ISL_476037, EPI_ISL_476038, EPI_ISL_476039, EPI_ISL_476040, EPI_ISL_476041, EPI_ISL_476042, EPI_ISL_476043, EPI_ISL_476044, EPI_ISL_476045, EPI_ISL_476046, EPI_ISL_476047, EPI_ISL_476048, EPI_ISL_476049, EPI_ISL_476050, EPI_ISL_476051, EPI_ISL_476052, EPI_ISL_476053, EPI_ISL_476054, EPI_ISL_476055, EPI_ISL_476056, EPI_ISL_476057, EPI_ISL_476058, EPI_ISL_476059, EPI_ISL_476060, EPI_ISL_476061, EPI_ISL_476062, EPI_ISL_476063, EPI_ISL_476064, EPI_ISL_476065, EPI_ISL_476066 | | | |
| see above | Michigan Department of Health and Human Services, Bureau of Laboratories | Michigan Department of Health and Human Services, Bureau of Laboratories | Blankenship HM, Riner D, Soehnlen MK |
| EPI_ISL_476768, EPI_ISL_476769, EPI_ISL_476770, EPI_ISL_476771, EPI_ISL_476772, EPI_ISL_476773, EPI_ISL_476774, EPI_ISL_476775, EPI_ISL_476776, EPI_ISL_476777, EPI_ISL_476778, EPI_ISL_476779, EPI_ISL_476780, EPI_ISL_476781, EPI_ISL_476782, EPI_ISL_476783, EPI_ISL_476784, EPI_ISL_476785, EPI_ISL_476786, EPI_ISL_476787, EPI_ISL_476788, EPI_ISL_476789, EPI_ISL_476790, EPI_ISL_476791, EPI_ISL_476792, EPI_ISL_476793, EPI_ISL_476794 | | | |
| see above | Stanford clinical virology lab | Chan-Zuckerberg Biohub | Benjamin Pinksy, Katharine Walter, Victoria N. Parikh, John Gorzynski, Hannah N. DeJong, Matthew T. Wheeler, Jason Andrews, Manuel Rivas, Carlos Bustamante, Euan Ashley, with CZB Cliahub Consortium |
| EPI_ISL_476855 | GMERS Medical College & Hospital, Gotri, Vadodara | Gujarat Biotechnology Research Centre | Apurvasinh Puvar, Janvi Raval, Zarna Patel, Monika Gandhi, Pinal Trivedi, Maharshi Pandya, Nidhi Patel, Nitin Savaliya, Raghawendra Kumar, Dinesh Kumar, Zuber Saiyed, Komal Patel, Labdhi Pandya, Afzal Ansari, Nikha Trivedi, Meenakshi Shah, Neena Doshi, Varsha Godbole, R D Dixit, A M Kadri, Harsh Bakshi, Chaitanya Joshi, Madhvi Joshi |
| EPI_ISL_476873 | Department of MicroBiology, Government Medical College, Surat | Gujarat Biotechnology Research Centre | Apurvasinh Puvar, Janvi Raval, Zarna Patel, Monika Gandhi, Pinal Trivedi, Maharshi Pandya, Nidhi Patel, Nitin Savaliya, Raghawendra Kumar, Dinesh Kumar, Zuber Saiyed, Komal Patel, Labdhi Pandya, Afzal Ansari, Nikha Trivedi, Naresh Chauhan, Summaiya Mullan, Amit gamit, R D Dixit, A M Kadri, Harsh Bakshi, Chaitanya Joshi, Madhvi Joshi |
| EPI_ISL_477014 | Institute of Microbiology, Universidad San Francisco de Quito | Institute of Microbiology, Universidad San Francisco de Quito | Belen Prado-Vivar, Sully Marquez, Juan Jose Guadalupe, Monica Becerra-Wong, Carla Torres, Bernardo Gutierrez, Francisco Mora, Juan Gaviria, |

| | | | |
|---|---|---|---|
| | | | Alejandra Ramones, Franklin Espinoza, Edison Ligña, Jorge Reyes, Patricio Rojas-Silva, Veronica Barragan, Gabriel Trueba, Michelle Grunauer, Paul Cardenas |
| EPI_ISL_478274, EPI_ISL_478275, EPI_ISL_478276, EPI_ISL_478277, EPI_ISL_478278, EPI_ISL_478279, EPI_ISL_478281, EPI_ISL_478282, EPI_ISL_478283, EPI_ISL_478284, EPI_ISL_478285, EPI_ISL_478286, EPI_ISL_478287, EPI_ISL_478288 | | | |
| see above | University of Exeter | COVID-19 Genomics UK (COG-UK) Consortium | Ben Temperton,Aaron Jeffries,Michelle Michelsen,Joanna Warwick-Dugdale,Audrey Farbos,Robyn Manley,Stephen Michell,Jane Masoli |
| EPI_ISL_479625, EPI_ISL_479626, EPI_ISL_479627, EPI_ISL_479628, EPI_ISL_479629, EPI_ISL_479630, EPI_ISL_479631, EPI_ISL_479632, EPI_ISL_479633, EPI_ISL_479634, EPI_ISL_479635, EPI_ISL_479636, EPI_ISL_479637, EPI_ISL_479638, EPI_ISL_479639, EPI_ISL_479640, EPI_ISL_479641, EPI_ISL_479642, EPI_ISL_479643, EPI_ISL_479644, EPI_ISL_479645, EPI_ISL_479646, EPI_ISL_479647, EPI_ISL_479648, EPI_ISL_479649, EPI_ISL_479650, EPI_ISL_479651 | | | |
| see above | Dr. Georges-L.-Dumont University Hospital Centre | National Microbiology Laboratory | Anna Majer, Shari Tyson, Grace Seo, Kristyn Burak, Philip Mabon, Elsie Grudeski, Rhiannon Huzarewich, Russell Mandes, Jennifer Tanner, Natalie Knox, Morag Graham, Gary Van Domselaar, Richard Garceau, Guillaume Desnoyers, Nathalie Bastien, Yan Li, Timothy Booth |
| EPI_ISL_480441, EPI_ISL_480442 | National Institute of Laboratory Medicine and Referral Center | Bangladesh Council of Scientific and Industrial Research | Barna Goswami, Abu Sayeed Mohammad Mahmud, Mohammad Samir Uzzaman, Eshrar Osman, Md. Ahasan Habib, Shahina Akter, Tanjina Akhter Banu, Md. Murshed Hasan Sarkar, Iffat Jahan, Md. Saddam Hossain, Tasnim Nafisa, Md. Maruf Ahmed Molla, Mahmuda Yeasmin, Asish Kumar Ghosh, Shahjahan Siddike, A. K. M. Shamsuzzaman, Sheikh Md. Selim Al Din, Utpal Chandra Ray, Salek Ahmed Sajib, Md. Salim Khan |
| EPI_ISL_481244, EPI_ISL_481245, EPI_ISL_481246, EPI_ISL_481247, EPI_ISL_481248 | Hospital IESS Babahoyo | Institute of Microbiology, Universidad San Francisco de Quito | Belén Prado-Vivar, Sully Márquez, Juan José Guadalupe, Monica Becerra-Wong, Carla Torres, Bernardo Gutiérrez, Francisco Cordova, Ninfa Henriquez, Killen Briones-Zamora, Killen Briones-Claudette, Verónica Barragán, Patricio Rojas-Silva, Gabriel Trueba, Michelle Grunauer, Paúl Cárdenas |
| EPI_ISL_481742, EPI_ISL_481743, EPI_ISL_481744, EPI_ISL_481745, EPI_ISL_481746, EPI_ISL_481747, EPI_ISL_481748, EPI_ISL_481749, EPI_ISL_481750, EPI_ISL_481751, EPI_ISL_481752, EPI_ISL_481753, EPI_ISL_481754, EPI_ISL_481755, EPI_ISL_481756, EPI_ISL_481757, EPI_ISL_481758 | | | |
| see above | Dr. Georges-L.-Dumont University Hospital Centre | National Microbiology Laboratory | Anna Majer, Shari Tyson, Grace Seo, Kristyn Burak, Philip Mabon, Elsie Grudeski, Rhiannon Huzarewich, Russell Mandes, Jennifer Tanner, Natalie Knox, Morag Graham, Gary Van Domselaar, Richard Garceau, Guillaume Desnoyers, Nathalie Bastien, Yan Li, Timothy Booth |
| EPI_ISL_482469, EPI_ISL_482470 | Queen Elizabeth II Health Science Centre | National Microbiology Laboratory | Anna Majer, Shari Tyson, Grace Seo, Kristyn Burak, Philip Mabon, Elsie Grudeski, Rhiannon Huzarewich, Russell Mandes, Jennifer Tanner, Natalie Knox, Morag Graham, Gary Van Domselaar, Todd Hatchette, Jason LeBlanc, Nathalie Bastien, Yan Li, Timothy Booth |
| EPI_ISL_482471, EPI_ISL_482472, EPI_ISL_482473 | Dr. Georges-L.-Dumont University Hospital Centre | National Microbiology Laboratory | Anna Majer, Shari Tyson, Grace Seo, Kristyn Burak, Philip Mabon, Elsie Grudeski, Rhiannon Huzarewich, Russell Mandes, Jennifer Tanner, Natalie Knox, Morag Graham, Gary Van Domselaar, Richard Garceau, Guillaume Desnoyers, Nathalie Bastien, Yan Li, Timothy Booth |
| EPI_ISL_482474, EPI_ISL_482475 | Cadham Provincial Laboratory | National Microbiology Laboratory | Anna Majer, Shari Tyson, Grace Seo, Kristyn Burak, Philip Mabon, Elsie Grudeski, Rhiannon Huzarewich, Russell Mandes, Jennifer Tanner, Natalie Knox, Morag Graham, Gary Van Domselaar, Paul Van Caeseele, Jared Bullard, David Alexander, Kerry Dust, Nathalie Bastien, Yan Li, Timothy Booth, |
| EPI_ISL_482476, EPI_ISL_482477 | Queen Elizabeth II Health Science Centre | National Microbiology Laboratory | Anna Majer, Shari Tyson, Grace Seo, Kristyn Burak, Philip Mabon, Elsie Grudeski, Rhiannon Huzarewich, Russell Mandes, Jennifer Tanner, Natalie Knox, Morag Graham, Gary Van Domselaar, Todd Hatchette, Jason LeBlanc, Nathalie Bastien, Yan Li, Timothy Booth |
| EPI_ISL_482478 | Cadham Provincial Laboratory | National Microbiology Laboratory | Anna Majer, Shari Tyson, Grace Seo, Kristyn Burak, Philip Mabon, Elsie Grudeski, Rhiannon Huzarewich, Russell Mandes, Jennifer Tanner, Natalie Knox, Morag Graham, Gary Van Domselaar, Paul Van Caeseele, Jared Bullard, David Alexander, Kerry Dust, Nathalie Bastien, Yan Li, Timothy Booth, |
| EPI_ISL_482479 | Public Health Laboratory | National Microbiology Laboratory | Anna Majer, Shari Tyson, Grace Seo, Kristyn Burak, Philip Mabon, Elsie Grudeski, Rhiannon Huzarewich, Russell Mandes, Jennifer Tanner, Natalie Knox, Morag Graham, Gary Van Domselaar, Robert Needle, Yang Yu, Adel Malek, Laura Gilbert, George Zahariadis, Nathalie Bastien, Yan Li, Timothy Booth |
| EPI_ISL_482480, EPI_ISL_482481, EPI_ISL_482482, EPI_ISL_482483, EPI_ISL_482484 | Cadham Provincial Laboratory | National Microbiology Laboratory | Anna Majer, Shari Tyson, Grace Seo, Kristyn Burak, Philip Mabon, Elsie Grudeski, Rhiannon Huzarewich, Russell Mandes, Jennifer Tanner, Natalie Knox, Morag Graham, Gary Van Domselaar, Paul Van Caeseele, Jared Bullard, David Alexander, Kerry Dust, Nathalie Bastien, Yan Li, Timothy Booth, |
| EPI_ISL_483542, EPI_ISL_483543, EPI_ISL_483544, EPI_ISL_483545, EPI_ISL_483546, EPI_ISL_483547, EPI_ISL_483548, EPI_ISL_483549, EPI_ISL_483550, EPI_ISL_483551, EPI_ISL_483552, EPI_ISL_483553, EPI_ISL_483554, EPI_ISL_483555, EPI_ISL_483556, EPI_ISL_483557, EPI_ISL_483558, EPI_ISL_483559, EPI_ISL_483560, EPI_ISL_483561, EPI_ISL_483562, EPI_ISL_483563, EPI_ISL_483564, EPI_ISL_483565 | | | |
| see above | Kingdom of Bahrein Ministry of Health | Erasmus Medical Center | Bas Oude Munnink, David Nieuwenhuijse, Reina Sikkema, Fatema, Ebrahim Shehad, Amjad Ghanem Mohamed, Hashmeya Al Wasti, Claudia Schapendonk, Irina Chestakova, Anne van der Linden, Theo Bestebroer, Stefan van Nieuwkoop, Mark Pronk, Pascal Lexmond, Richard Molenkamp, Marion Koopmans, on behalf of the Dutch national COVID-19 response team. |
| EPI_ISL_483629, EPI_ISL_483630 | National Institute of Laboratory Medicine and Referral Center | Genomic Research Lab, BCSIR | Asish Kumar Ghosh, Abu Sayeed Mohammad Mahmud, Mohammad Samir Uzzaman, Eshrar Osman, Md. Ahasan Habib, Shahina Akter, Tanjina Akhter Banu, Md. Murshed Hasan Sarkar, Barna Goswami, Iffat Jahan, Md. Saddam Hossain, Tasnim Nafisa, Md. Maruf Ahmed Molla, Mahmuda Yeasmin, A. K. M. Shamsuzzaman, Sheikh Md. Selim Al Din, Utpal Chandra Ray, Salek Ahmed Sajib, Md. Salim Khan |
| EPI_ISL_483638, EPI_ISL_483639, EPI_ISL_483640 | Kingdom of Bahrein Ministry of Health | Erasmus Medical Center | Bas Oude Munnink, David Nieuwenhuijse, Reina Sikkema, Fatema, Ebrahim Shehad, Amjad Ghanem Mohamed, Hashmeya Al Wasti, Claudia Schapendonk, Irina Chestakova, Anne van der Linden, Theo Bestebroer, Stefan van Nieuwkoop, Mark Pronk, Pascal Lexmond, Richard Molenkamp, Marion Koopmans, on behalf of the Dutch national COVID-19 response team. |
| EPI_ISL_483641, EPI_ISL_483642 | National Institute of Laboratory Medicine and Referral Center | Genomic Research Lab, BCSIR | Barna Goswami, Abu Sayeed Mohammad Mahmud, Mohammad Samir Uzzaman, Eshrar Osman, Md. Ahasan Habib, Shahina Akter, Tanjina Akhter Banu, Md. Murshed Hasan Sarkar, Iffat Jahan, Md. Saddam Hossain, Tasnim Nafisa, Md. Maruf Ahmed Molla, Mahmuda Yeasmin, Asish Kumar Ghosh, A. K. M. Shamsuzzaman, Sheikh Md. Selim Al Din, Utpal Chandra Ray, Salek Ahmed Sajib, Md. Salim Khan |
| EPI_ISL_483826 | GMERS Medical College Himmatnagar | Gujarat Biotechnology Research Centre | Apurvasinh Puvar, Janvi Raval, Zarna Patel, Monika Gandhi, Pinal Trivedi, Maharshi Pandya, Nidhi Patel, Nitin Savaliya, Raghawendra Kumar, Dinesh Kumar, Zuber Saiyed, Komal Patel, Labdhi Pandya, Afzal Ansari, Nikha Trivedi, Himanshu Khatri, Mayur Gandhi, R D Dixit, A M Kadri, Harsh Bakshi, Chaitanya Joshi, Madhvi Joshi |
| EPI_ISL_483842 | Pandit Deendayal Upadhyay Government Medical College, Rajkot | Gujarat Biotechnology Research Centre | Arti Trivedi, Apurvasinh Puvar, Janvi Raval, Zarna Patel, Monika Gandhi, Pinal Trivedi, Maharshi Pandya, Nidhi Patel, Nitin Savaliya, Raghawendra Kumar, Dinesh Kumar, Zuber Saiyed, Komal Patel, Labdhi Pandya, Afzal Ansari, Nikha Trivedi, Gauravi Dhruv, R D Dixit, A M Kadri, Harsh Bakshi, Chaitanya Joshi, Madhvi Joshi |
| EPI_ISL_483843 | Pandit Deendayal Upadhyay Government Medical College, Rajkot | Gujarat Biotechnology Research Centre | Apurvasinh Puvar, Janvi Raval, Zarna Patel, Monika Gandhi, Pinal Trivedi, Maharshi Pandya, Nidhi Patel, Nitin Savaliya, Raghawendra Kumar, Dinesh Kumar, Zuber Saiyed, Komal Patel, Labdhi Pandya, Afzal Ansari, Nikha Trivedi, Gauravi Dhruv, Arti Trivedi, R D Dixit, A M Kadri, Harsh Bakshi, Chaitanya Joshi, Madhvi Joshi |
| EPI_ISL_483861, EPI_ISL_483879 | Department of Microbiology, Government Medical College, Surat | Gujarat Biotechnology Research Centre | Apurvasinh Puvar, Janvi Raval, Zarna Patel, Monika Gandhi, Pinal Trivedi, Maharshi Pandya, Nidhi Patel, Nitin Savaliya, Raghawendra Kumar, Dinesh Kumar, Zuber Saiyed, Komal Patel, Labdhi Pandya, Afzal Ansari, Nikha Trivedi, Naresh Chauhan, Summaiya Mullan, Amit gamit, R D Dixit, A M Kadri, Harsh Bakshi, Chaitanya Joshi, Madhvi Joshi |
| EPI_ISL_484408, EPI_ISL_484413, EPI_ISL_484418, EPI_ISL_484420, EPI_ISL_484426, EPI_ISL_484430, EPI_ISL_484431, EPI_ISL_484432 | Centre for Enzyme Innovation, University of Portsmouth / Translational Research Laboratory, Portsmouth Hospitals NHS Trust | COVID-19 Genomics UK (COG-UK) Consortium | Angela Beckett,Yann Bourgeois,Garry Scarlett,Sharon Glaysher,Scott Elliott,Kelly Bicknell,Robert Impey,Allyson Lloyd,Sarah Wyllie,Ethan Butcher,Anoop Chauhan,Samuel Robson |
| EPI_ISL_485809 | Institut für Virologie und Epidemiologie der Viruskrankheiten, Universitätsklinikum Tübingen | NGS Competence Center Tübingen, Institut für Medizinische Mikrobiologie und Hygiene, Universitätsklinikum Tübingen | Angelov at al. |
| EPI_ISL_485810, EPI_ISL_485811, EPI_ISL_485812, EPI_ISL_485813 | Institut für Virologie und Epidemiologie der Viruskrankheiten, Universitätsklinikum Tübingen | NGS Competence Center Tübingen, Institut für Medizinische Mikrobiologie und Hygiene, Universitätsklinikum Tübingen | Angelov et al. |
| EPI_ISL_486842, EPI_ISL_486843, EPI_ISL_486844 | Institute of Microbiology, Universidad San Francisco de Quito | Institute of Microbiology, Universidad San Francisco de Quito | Belén Prado-Vivar, Sully Márquez, Juan José Guadalupe, Monica Becerra-Wong, Carla Torres, Bernardo Gutiérrez, Fausto Maldonado, Geovanny Carzola, Verónica Barragán, Patricio Rojas-Silva, Gabriel Trueba, Michelle Grunauer, Paúl Cárdenas |
| EPI_ISL_486845, EPI_ISL_486846, EPI_ISL_486847, EPI_ISL_486848, EPI_ISL_486849, EPI_ISL_486850, EPI_ISL_486851 | Institute of Microbiology, Universidad San Francisco de Quito | Institute of Microbiology, Universidad San Francisco de Quito | Belén Prado-Vivar, Sully Márquez, Juan José Guadalupe, Monica Becerra-Wong, Carla Torres, Bernardo Gutiérrez, Jonathan Araujo, Verónica Barragán, Patricio Rojas-Silva, Gabriel Trueba, Michelle Grunauer, Paúl Cárdenas |
| EPI_ISL_487368, EPI_ISL_487371, | National Institute of Laboratory Medicine and Referral Center | Genomic Research Lab, BCSIR | Barna Goswami, Abu Sayeed Mohammad Mahmud, Mohammad Samir Uzzaman, Eshrar Osman, Md. Ahasan Habib, Shahina Akter, Tanjina Akhter Banu, |

| | | | |
|---|---|---|---|
| EPI_ISL_490144 | | | Md. Murshed Hasan Sarkar, Iffat Jahan, Md. Saddam Hossain, Tasnim Nafisa, Md. Maruf Ahmed Molla, Mahmuda Yeasmin, Asish Kumar Ghosh, A. K. M. Shamsuzzaman, Sheikh Md. Selim Al Din, Utpal Chandra Ray, Salek Ahmed Sajib, Md. Salim Khan |
| EPI_ISL_491932 | Institute of Microbiology, Universidad San Francisco de Quito | Institute of Microbiology, Universidad San Francisco de Quito | Belén Prado-Vivar, Sully Márquez, Juan José Guadalupe, Monica Becerra-Wong, Bernardo Gutiérrez, Carlos Mena, Nabih Dahik, Verónica Barragán, Patricio Rojas-Silva, Gabriel Trueba, Michelle Grunauer, Paúl Cárdenas |
| EPI_ISL_491936 | Institute of Microbiology, Universidad San Francisco de Quito | Institute of Microbiology, Universidad San Francisco de Quito | Belén Prado-Vivar, Sully Márquez, Juan José Guadalupe, Monica Becerra-Wong, Bernardo Gutiérrez, Carlos Guerrero, Verónica Barragán, Patricio Rojas-Silva, Gabriel Trueba, Michelle Grunauer, Paúl Cárdenas |
| EPI_ISL_491937, EPI_ISL_491938 | Institute of Microbiology, Universidad San Francisco de Quito | Institute of Microbiology, Universidad San Francisco de Quito | Belén Prado-Vivar, Sully Márquez, Juan José Guadalupe, Monica Becerra-Wong, Bernardo Gutiérrez, Rosario Erazo, Verónica Barragán, Patricio Rojas-Silva, Gabriel Trueba, Michelle Grunauer, Paúl Cárdenas |
| EPI_ISL_491939, EPI_ISL_491940 | Institute of Microbiology, Universidad San Francisco de Quito | Institute of Microbiology, Universidad San Francisco de Quito | Belén Prado-Vivar, Sully Márquez, Juan José Guadalupe, Monica Becerra-Wong, Bernardo Gutiérrez, Carlos Mena, Nabih Dahik, Verónica Barragán, Patricio Rojas-Silva, Gabriel Trueba, Michelle Grunauer, Paúl Cárdenas |
| EPI_ISL_492032 | Instituto de Biologia do Exército | Laboratório Metabolismo Macromolecular FirminoTorres de Castro, Instituto de Biofísica Carlos Chagas Filho, Universidade Federal do Rio de Janeiro | Bianca Catarina Azevedo Cabral, Aline Rosa Vianna de Souza , Marcos Dornelas-Ribeiro, Tatiana LS Nogueira, Nádia Vaez Gonçalves da Cruz, Caleb GM Santos, Elizabeth Valentin, Marcio da Costa Cipitelli, Virginia Sara Grancieri do Amaral, Rodrigo Soares de Moura Neto, Clarissa Damaso, Rosane Silva |
| EPI_ISL_492033 | Instituto de Biologia do Exército | Laboratório Metabolismo Macromolecular FirminoTorres de Castro, Instituto de Biofísica Carlos Chagas Filho, Universidade Federal do Rio de Janeiro | Bianca Catarina Azevedo Cabral, Aline Rosa Vianna de Souza, Caleb GM Santos, Marcos Dornelas-Ribeiro, Tatiana LS Nogueira, Nádia Vaez Gonçalves da Cruz, Elizabeth Valentin, Marcio da Costa Cipitelli, Virginia Sara Grancieri do Amaral, Rodrigo Soares de Moura Neto, Clarissa Damaso, Rosane Silva |
| EPI_ISL_492034 | Instituto de Biologia do Exército | Laboratório Metabolismo Macromolecular FirminoTorres de Castro, Instituto de Biofísica Carlos Chagas Filho, Universidade Federal do Rio de Janeiro | Bianca Catarina Azevedo Cabral, Aline Rosa Vianna de Souza, Nádia Vaez Gonçalves da Cruz, Caleb GM Santos, Marcos Dornelas-Ribeiro, Tatiana LS Nogueira, Elizabeth Valentin, Marcio da Costa Cipitelli, Virginia Sara Grancieri do Amaral, Rodrigo Soares de Moura Neto, Clarissa Damaso, Rosane Silva |
| EPI_ISL_492035 | Instituto de Biologia do Exército | Laboratório Metabolismo Macromolecular FirminoTorres de Castro, Instituto de Biofísica Carlos Chagas Filho, Universidade Federal do Rio de Janeiro | Bianca Catarina Azevedo Cabral, Aline Rosa Vianna de Souza, Tatiana LS Nogueira, Nádia Vaez Gonçalves da Cruz, Caleb GM Santos, Marcos Dornelas-Ribeiro, Elizabeth Valentin, Marcio da Costa Cipitelli, Virginia Sara Grancieri do Amaral, Rodrigo Soares de Moura Neto, Clarissa Damaso, Rosane Silva |
| EPI_ISL_492036 | Instituto de Biologia do Exército | Laboratório Metabolismo Macromolecular FirminoTorres de Castro, Instituto de Biofísica Carlos Chagas Filho, Universidade Federal do Rio de Janeiro | Bianca Catarina Azevedo Cabral, Aline Rosa Vianna de Souza , Marcos Dornelas-Ribeiro, Tatiana LS Nogueira, Nádia Vaez Gonçalves da Cruz, Caleb GM Santos, Elizabeth Valentin, Marcio da Costa Cipitelli, Virginia Sara Grancieri do Amaral, Rodrigo Soares de Moura Neto, Clarissa Damaso, Rosane Silva |
| EPI_ISL_492037 | Instituto de Biologia do Exército | Laboratório Metabolismo Macromolecular FirminoTorres de Castro, Instituto de Biofísica Carlos Chagas Filho, Universidade Federal do Rio de Janeiro | Bianca Catarina Azevedo Cabral, Aline Rosa Vianna de Souza, Caleb GM Santos, Marcos Dornelas-Ribeiro, Tatiana LS Nogueira, Nádia Vaez Gonçalves da Cruz, Elizabeth Valentin, Marcio da Costa Cipitelli, Virginia Sara Grancieri do Amaral, Rodrigo Soares de Moura Neto, Clarissa Damaso, Rosane Silva |
| EPI_ISL_492038 | Instituto de Biologia do Exército | Laboratório Metabolismo Macromolecular FirminoTorres de Castro, Instituto de Biofísica Carlos Chagas Filho, Universidade Federal do Rio de Janeiro | Bianca Catarina Azevedo Cabral, Aline Rosa Vianna de Souza, Nádia Vaez Gonçalves da Cruz, Caleb GM Santos, Marcos Dornelas-Ribeiro, Tatiana LS Nogueira, Elizabeth Valentin, Marcio da Costa Cipitelli, Virginia Sara Grancieri do Amaral, Rodrigo Soares de Moura Neto, Clarissa Damaso, Rosane Silva |
| EPI_ISL_492039 | Instituto de Biologia do Exército | Laboratório Metabolismo Macromolecular FirminoTorres de Castro, Instituto de Biofísica Carlos Chagas Filho, Universidade Federal do Rio de Janeiro | Bianca Catarina Azevedo Cabral, Aline Rosa Vianna de Souza, Tatiana LS Nogueira, Nádia Vaez Gonçalves da Cruz, Caleb GM Santos, Marcos Dornelas-Ribeiro, Elizabeth Valentin, Marcio da Costa Cipitelli, Virginia Sara Grancieri do Amaral, Rodrigo Soares de Moura Neto, Clarissa Damaso, Rosane Silva |
| EPI_ISL_492040 | Instituto de Biologia do Exército | Laboratório Metabolismo Macromolecular FirminoTorres de Castro, Instituto de Biofísica Carlos Chagas Filho, Universidade Federal do Rio de Janeiro | Bianca Catarina Azevedo Cabral, Aline Rosa Vianna de Souza , Marcos Dornelas-Ribeiro, Tatiana LS Nogueira, Nádia Vaez Gonçalves da Cruz, Caleb GM Santos, Elizabeth Valentin, Marcio da Costa Cipitelli, Virginia Sara Grancieri do Amaral, Rodrigo Soares de Moura Neto, Clarissa Damaso, Rosane Silva |
| EPI_ISL_492041 | Instituto de Biologia do Exército | Laboratório Metabolismo Macromolecular FirminoTorres de Castro, Instituto de Biofísica Carlos Chagas Filho, Universidade Federal do Rio de Janeiro | Bianca Catarina Azevedo Cabral, Aline Rosa Vianna de Souza, Caleb GM Santos, Marcos Dornelas-Ribeiro, Tatiana LS Nogueira, Nádia Vaez Gonçalves da Cruz, Elizabeth Valentin, Marcio da Costa Cipitelli, Virginia Sara Grancieri do Amaral, Rodrigo Soares de Moura Neto, Clarissa Damaso, Rosane Silva |
| EPI_ISL_492042 | Instituto de Biologia do Exército | Laboratório Metabolismo Macromolecular FirminoTorres de Castro, Instituto de Biofísica Carlos Chagas Filho, Universidade Federal do Rio de Janeiro | Bianca Catarina Azevedo Cabral, Aline Rosa Vianna de Souza, Nádia Vaez Gonçalves da Cruz, Caleb GM Santos, Marcos Dornelas-Ribeiro, Tatiana LS Nogueira, Elizabeth Valentin, Marcio da Costa Cipitelli, Virginia Sara Grancieri do Amaral, Rodrigo Soares de Moura Neto, Clarissa Damaso, Rosane Silva |
| EPI_ISL_492043 | Instituto de Biologia do Exército | Laboratório Metabolismo Macromolecular FirminoTorres de Castro, Instituto de Biofísica Carlos Chagas Filho, Universidade Federal do Rio de Janeiro | Bianca Catarina Azevedo Cabral, Aline Rosa Vianna de Souza, Tatiana LS Nogueira, Nádia Vaez Gonçalves da Cruz, Caleb GM Santos, Marcos Dornelas-Ribeiro, Elizabeth Valentin, Marcio da Costa Cipitelli, Virginia Sara Grancieri do Amaral, Rodrigo Soares de Moura Neto, Clarissa Damaso, Rosane Silva |
| EPI_ISL_492044 | Instituto de Biologia do Exército | Laboratório Metabolismo Macromolecular FirminoTorres de Castro, Instituto de Biofísica Carlos Chagas Filho, Universidade Federal do Rio de Janeiro | Bianca Catarina Azevedo Cabral, Aline Rosa Vianna de Souza , Marcos Dornelas-Ribeiro, Tatiana LS Nogueira, Nádia Vaez Gonçalves da Cruz, Caleb GM Santos, Elizabeth Valentin, Marcio da Costa Cipitelli, Virginia Sara Grancieri do Amaral, Rodrigo Soares de Moura Neto, Clarissa Damaso, Rosane Silva |
| EPI_ISL_492045 | Instituto de Biologia do Exército | Laboratório Metabolismo Macromolecular FirminoTorres de Castro, Instituto de Biofísica Carlos Chagas Filho, Universidade Federal do Rio de Janeiro | Bianca Catarina Azevedo Cabral, Aline Rosa Vianna de Souza, Caleb GM Santos, Marcos Dornelas-Ribeiro, Tatiana LS Nogueira, Nádia Vaez Gonçalves da Cruz, Elizabeth Valentin, Marcio da Costa Cipitelli, Virginia Sara Grancieri do Amaral, Rodrigo Soares de Moura Neto, Clarissa Damaso, Rosane Silva |
| EPI_ISL_492046 | Instituto de Biologia do Exército | Laboratório Metabolismo Macromolecular FirminoTorres de Castro, Instituto de Biofísica Carlos Chagas Filho, Universidade Federal do Rio de Janeiro | Bianca Catarina Azevedo Cabral, Aline Rosa Vianna de Souza, Nádia Vaez Gonçalves da Cruz, Caleb GM Santos, Marcos Dornelas-Ribeiro, Tatiana LS Nogueira, Elizabeth Valentin, Marcio da Costa Cipitelli, Virginia Sara Grancieri do Amaral, Rodrigo Soares de Moura Neto, Clarissa Damaso, Rosane Silva |
| EPI_ISL_492047 | Instituto de Biologia do Exército | Laboratório Metabolismo Macromolecular FirminoTorres de Castro, Instituto de Biofísica Carlos Chagas Filho, Universidade Federal do Rio de Janeiro | Bianca Catarina Azevedo Cabral, Aline Rosa Vianna de Souza, Tatiana LS Nogueira, Nádia Vaez Gonçalves da Cruz, Caleb GM Santos, Marcos Dornelas-Ribeiro, Elizabeth Valentin, Marcio da Costa Cipitelli, Virginia Sara Grancieri do Amaral, Rodrigo Soares de Moura Neto, Clarissa Damaso, Rosane Silva |
| EPI_ISL_492048 | Instituto de Biologia do Exército | Laboratório Metabolismo Macromolecular FirminoTorres de Castro, Instituto de Biofísica Carlos Chagas Filho, Universidade Federal do Rio de Janeiro | Bianca Catarina Azevedo Cabral, Aline Rosa Vianna de Souza , Marcos Dornelas-Ribeiro, Tatiana LS Nogueira, Nádia Vaez Gonçalves da Cruz, Caleb GM Santos, Elizabeth Valentin, Marcio da Costa Cipitelli, Virginia Sara Grancieri do Amaral, Rodrigo Soares de Moura Neto, Clarissa Damaso, Rosane Silva |
| EPI_ISL_492980, EPI_ISL_492981, EPI_ISL_492982, EPI_ISL_492983, EPI_ISL_492984, EPI_ISL_492985, EPI_ISL_492986, EPI_ISL_492987 | IRCCS Sacro Cuore Don Calabria Hospital, Department of Infectious, Tropical Diseases & Microbiology | University of Verona, Department of Biotechnology | Antonio Mori, Michela Deiana, Elena Pomari, Chiara Piubelli; Giulia Lopatriello, Luca Marcolungo, Cristina Beltrami, Chiara Degli Esposti, Emanuela Cosentino, Massimo Delledonne |
| EPI_ISL_493329 | INMI Lazzaro Spallanzani IRCCS | INMI Lazzaro Spallanzani IRCCS | Barbara Bartolini, Martina Rueca, Cesare E.M. Gruber, Francesco Messina, Antonino Di Caro, Maria R. Capobianchi |
| EPI_ISL_493333 | Istituto Zooprofilattico Sperimentale del Mezzogiorno | INMI Lazzaro Spallanzani IRCCS | Barbara Bartolini, Martina Rueca, Cesare E.M. Gruber, Francesco Messina, Antonino Di Caro, Giovanna Fusco, Maurizio Viscardi, Giorgia Borriello, Maria R. Capobianchi |
| EPI_ISL_493615, EPI_ISL_493616, EPI_ISL_493617, EPI_ISL_493618, EPI_ISL_493619, EPI_ISL_493620, EPI_ISL_493621, EPI_ISL_493622, EPI_ISL_493623, EPI_ISL_493625, EPI_ISL_493626, EPI_ISL_493627, EPI_ISL_493628, EPI_ISL_493629, EPI_ISL_493630, EPI_ISL_493631, EPI_ISL_493632, EPI_ISL_493633, EPI_ISL_493634, EPI_ISL_493635, EPI_ISL_493636, EPI_ISL_493637, EPI_ISL_493638, EPI_ISL_493639, EPI_ISL_493640, EPI_ISL_493641, EPI_ISL_493642, EPI_ISL_493643, EPI_ISL_493644, EPI_ISL_493645, EPI_ISL_493646, EPI_ISL_493647, EPI_ISL_493648, EPI_ISL_493649, EPI_ISL_493650, EPI_ISL_493651, EPI_ISL_493652, EPI_ISL_493653, EPI_ISL_493654, EPI_ISL_493655, EPI_ISL_493656, EPI_ISL_493657, EPI_ISL_493658, EPI_ISL_493659, EPI_ISL_493660, EPI_ISL_493661, EPI_ISL_493662, EPI_ISL_493663, EPI_ISL_493664, EPI_ISL_493665 | | | |
| see above | Centre for Enzyme Innovation, University of Portsmouth / Translational Research Laboratory, Portsmouth Hospitals NHS Trust | COVID-19 Genomics UK (COG-UK) Consortium | Angela Beckett,Yann Bourgeois,Garry Scarlett,Sharon Glaysher,Scott Elliott,Kelly Bicknell,Robert Impey,Allyson Lloyd,Sarah Wyllie,Ethan Butcher,Anoop Chauhan,Samuel Robson |

| | | | |
|---|---|---|---|
| EPI_ISL_495033, EPI_ISL_495051 | GMERS Medical College & Hospital, Gotri, Vadodara | Gujarat Biotechnology Research Centre | Apurvasinh Puvar, Janvi Raval, Zarna Patel, Monika Gandhi, Pinal Trivedi, Maharshi Pandya, Nidhi Patel, Nitin Savaliya, Raghawendra Kumar, Dinesh Kumar, Zuber Saiyed, Komal Patel, Labdhi Pandya, Afzal Ansari, Nikha Trivedi, Meenakshi Shah, Neena Doshi, Varsha Godbole, R D Dixit, A M Kadri, Harsh Bakshi, Chaitanya Joshi, Madhvi Joshi |
| EPI_ISL_495066 | GAIMS & G K General Hospital | Gujarat Biotechnology Research Centre | Babulal Babhoria, Hitesh Assudani, Apurvasinh Puvar, Janvi Raval, Zarna Patel, Monika Gandhi, Pinal Trivedi, Maharshi Pandya, Nidhi Patel, Nitin Savaliya, Raghawendra Kumar, Dinesh Kumar, Zuber Saiyed, Komal Patel, Labdhi Pandya, Afzal Ansari, Nikha Trivedi, R D Dixit, A M Kadri, Harsh Bakshi, Chaitanya Joshi, Madhvi Joshi, |
| EPI_ISL_495070 | Department of MicroBiology, Government Medical College, Surat | Gujarat Biotechnology Research Centre | Apurvasinh Puvar, Janvi Raval, Zarna Patel, Monika Gandhi, Pinal Trivedi, Maharshi Pandya, Nidhi Patel, Nitin Savaliya, Raghawendra Kumar, Dinesh Kumar, Zuber Saiyed, Komal Patel, Labdhi Pandya, Afzal Ansari, Nikha Trivedi, Naresh Chauhan, Summaiya Mullan, Amit gamit, R D Dixit, A M Kadri, Harsh Bakshi, Chaitanya Joshi, Madhvi Joshi |
| EPI_ISL_498267, EPI_ISL_498268 | National Institute of Laboratory Medicine and Referral Center | Genomic Research Lab, BCSIR | Barna Goswami, Abu Sayeed Mohammad Mahmud, Mohammad Samir Uzzaman, Eshrar Osman, Md. Ahasan Habib, Shahina Akter, Tanjina Akhter Banu, Md. Murshed Hasan Sarkar, Iffat Jahan, Md. Saddam Hossain, Tasnim Nafisa, Md. Maruf Ahmed Molla, Mahmuda Yeasmin, Asish Kumar Ghosh, A. K. M. Shamsuzzaman, Sheikh Md. Selim Al Din, Utpal Chandra Ray, Salek Ahmed Sajib, Md. Salim Khan |
| EPI_ISL_498549, EPI_ISL_498550 | National Institute of Laboratory Medicine and Referral Center | Genomic Research Lab, BCSIR | Asish Kumar Ghosh, Abu Sayeed Mohammad Mahmud, Mohammad Samir Uzzaman, Eshrar Osman, Md. Ahasan Habib, Shahina Akter, Tanjina Akhter Banu, Md. Murshed Hasan Sarkar, Barna Goswami, Iffat Jahan, Md. Saddam Hossain, Tasnim Nafisa, Md. Maruf Ahmed Molla, Mahmuda Yeasmin, A. K. M. Shamsuzzaman, Sheikh Md. Selim Al Din, Utpal Chandra Ray, Salek Ahmed Sajib, Md. Salim Khan |
| EPI_ISL_499270, EPI_ISL_499273, EPI_ISL_499275, EPI_ISL_499276, EPI_ISL_499277, EPI_ISL_499279, EPI_ISL_499280, EPI_ISL_499281, EPI_ISL_499286, EPI_ISL_499287, EPI_ISL_499290, EPI_ISL_499291, EPI_ISL_499292, EPI_ISL_499293, EPI_ISL_499299, EPI_ISL_499301, EPI_ISL_499305, EPI_ISL_499313, EPI_ISL_499319, EPI_ISL_499320 | | | |
| see above | Centre for Enzyme Innovation, University of Portsmouth / Translational Research Laboratory, Portsmouth Hospitals NHS Trust | COVID-19 Genomics UK (COG-UK) Consortium | Angela Beckett,Yann Bourgeois,Garry Scarlett,Sharon Glaysher,Scott Elliott,Kelly Bicknell,Robert Impey,Allyson Lloyd,Sarah Wyllie,Ethan Butcher,Anoop Chauhan,Samuel Robson |
| EPI_ISL_500287, EPI_ISL_500288, EPI_ISL_500290, EPI_ISL_500291, EPI_ISL_500336, EPI_ISL_500337, EPI_ISL_500338, EPI_ISL_500339, EPI_ISL_500340, EPI_ISL_500341, EPI_ISL_500343, EPI_ISL_500344, EPI_ISL_500345, EPI_ISL_500346, EPI_ISL_500347, EPI_ISL_500349, EPI_ISL_500350, EPI_ISL_500351, EPI_ISL_500352, EPI_ISL_500353, EPI_ISL_500354, EPI_ISL_500355, EPI_ISL_500356, EPI_ISL_500357, EPI_ISL_500359, EPI_ISL_500360, EPI_ISL_500361, EPI_ISL_500362, EPI_ISL_500363, EPI_ISL_500364, EPI_ISL_500365, EPI_ISL_500366, EPI_ISL_500367, EPI_ISL_500368 | | | |
| see above | Servicio de Microbiología, Hospital Miguel Servet, Zaragoza | SeqCOVID-SPAIN consortium/IBV(CSIC) | Antonio Rezusta López, Alexander Tristancho Baró, Ana Milagro, Yolanda Gracia Grataloup, Nieves Martínez Cameo and SeqCOVID-SPAIN consortium |
| EPI_ISL_507136, EPI_ISL_507138, EPI_ISL_507141, EPI_ISL_507144, EPI_ISL_507145, EPI_ISL_507155, EPI_ISL_507156, EPI_ISL_507157 | Centre for Enzyme Innovation, University of Portsmouth / Translational Research Laboratory, Portsmouth Hospitals NHS Trust | COVID-19 Genomics UK (COG-UK) Consortium | Angela Beckett,Yann Bourgeois,Garry Scarlett,Sharon Glaysher,Scott Elliott,Kelly Bicknell,Robert Impey,Allyson Lloyd,Sarah Wyllie,Ethan Butcher,Anoop Chauhan,Samuel Robson |
| EPI_ISL_507415, EPI_ISL_507416, EPI_ISL_507417, EPI_ISL_507418, EPI_ISL_507419, EPI_ISL_507420, EPI_ISL_507421, EPI_ISL_507422, EPI_ISL_507423, EPI_ISL_507424, EPI_ISL_507425, EPI_ISL_507426, EPI_ISL_507427, EPI_ISL_507428, EPI_ISL_507429, EPI_ISL_507430, EPI_ISL_507431, EPI_ISL_507432, EPI_ISL_507433, EPI_ISL_507434, EPI_ISL_507435, EPI_ISL_507436, EPI_ISL_507437, EPI_ISL_507438, EPI_ISL_507439, EPI_ISL_507440, EPI_ISL_507441, EPI_ISL_507442, EPI_ISL_507443, EPI_ISL_507444, EPI_ISL_507445, EPI_ISL_507446, EPI_ISL_507447, EPI_ISL_507448, EPI_ISL_507449, EPI_ISL_507450, EPI_ISL_507451, EPI_ISL_507452, EPI_ISL_507453, EPI_ISL_507454, EPI_ISL_507455, EPI_ISL_507456, EPI_ISL_507457, EPI_ISL_507458, EPI_ISL_507459, EPI_ISL_507460, EPI_ISL_507461, EPI_ISL_507462, EPI_ISL_507463, EPI_ISL_507464, EPI_ISL_507465, EPI_ISL_507466, EPI_ISL_507467, EPI_ISL_507468, EPI_ISL_507469, EPI_ISL_507470, EPI_ISL_507471, EPI_ISL_507472, EPI_ISL_507473, EPI_ISL_507474, EPI_ISL_507475, EPI_ISL_507476, EPI_ISL_507477, EPI_ISL_507478, EPI_ISL_507479, EPI_ISL_507480, EPI_ISL_507481, EPI_ISL_507482, EPI_ISL_507483, EPI_ISL_507484, EPI_ISL_507485, EPI_ISL_507486, EPI_ISL_507487, EPI_ISL_507488, EPI_ISL_507489, EPI_ISL_507490, EPI_ISL_507491, EPI_ISL_507492, EPI_ISL_507493, EPI_ISL_507494, EPI_ISL_507495, EPI_ISL_507496, EPI_ISL_507497, EPI_ISL_507498, EPI_ISL_507499, EPI_ISL_507500, EPI_ISL_507501, EPI_ISL_507502, EPI_ISL_507503, EPI_ISL_507504, EPI_ISL_507505, EPI_ISL_507506, EPI_ISL_507507, EPI_ISL_507508, EPI_ISL_507509, EPI_ISL_507510, EPI_ISL_507511, EPI_ISL_507512, EPI_ISL_507513, EPI_ISL_507514, EPI_ISL_507515, EPI_ISL_507516, EPI_ISL_507517, EPI_ISL_507518, EPI_ISL_507519, EPI_ISL_507520, EPI_ISL_507521, EPI_ISL_507522, EPI_ISL_507523, EPI_ISL_507524, EPI_ISL_507525, EPI_ISL_507526, EPI_ISL_507527, EPI_ISL_507528, EPI_ISL_507529, EPI_ISL_507530, EPI_ISL_507531, EPI_ISL_507532, EPI_ISL_507533, EPI_ISL_507534, EPI_ISL_507535, EPI_ISL_507536, EPI_ISL_507537, EPI_ISL_507538, EPI_ISL_507539, EPI_ISL_507540, EPI_ISL_507541, EPI_ISL_507542, EPI_ISL_507543, EPI_ISL_507544, EPI_ISL_507545, EPI_ISL_507546, EPI_ISL_507547, EPI_ISL_507548, EPI_ISL_507549, EPI_ISL_507550, EPI_ISL_507551, EPI_ISL_507552, EPI_ISL_507553, EPI_ISL_507554, EPI_ISL_507555, EPI_ISL_507556, EPI_ISL_507557, EPI_ISL_507558, EPI_ISL_507559, EPI_ISL_507560, EPI_ISL_507561, EPI_ISL_507562, EPI_ISL_507563, EPI_ISL_507564, EPI_ISL_507565, EPI_ISL_507566, EPI_ISL_507567, EPI_ISL_507568, EPI_ISL_507569, EPI_ISL_507570, EPI_ISL_507571, EPI_ISL_507572, EPI_ISL_507573, EPI_ISL_507574, EPI_ISL_507575, EPI_ISL_507576, EPI_ISL_507577, EPI_ISL_507578, EPI_ISL_507579, EPI_ISL_507580, EPI_ISL_507581, EPI_ISL_507582, EPI_ISL_507583, EPI_ISL_507584, EPI_ISL_507585, EPI_ISL_507586, EPI_ISL_507587, EPI_ISL_507588, EPI_ISL_507589, EPI_ISL_507590, EPI_ISL_507591, EPI_ISL_507592, EPI_ISL_507593, EPI_ISL_507594, EPI_ISL_507595, EPI_ISL_507596, EPI_ISL_507597, EPI_ISL_507598, EPI_ISL_507599, EPI_ISL_507600, EPI_ISL_507601, EPI_ISL_507602, EPI_ISL_507603, EPI_ISL_507604, EPI_ISL_507605, EPI_ISL_507606, EPI_ISL_507607, EPI_ISL_507608, EPI_ISL_507609, EPI_ISL_507610, EPI_ISL_507611, EPI_ISL_507612, EPI_ISL_507613, EPI_ISL_507614, EPI_ISL_507615, EPI_ISL_507616, EPI_ISL_507617, EPI_ISL_507618, EPI_ISL_507619, EPI_ISL_507620, EPI_ISL_507621, EPI_ISL_507622, EPI_ISL_507623, EPI_ISL_507624, EPI_ISL_507625, EPI_ISL_507626, EPI_ISL_507627, EPI_ISL_507628, EPI_ISL_507629, EPI_ISL_507630, EPI_ISL_507631, EPI_ISL_507632, EPI_ISL_507633, EPI_ISL_507634, EPI_ISL_507635, EPI_ISL_507636, EPI_ISL_507637, EPI_ISL_507638, EPI_ISL_507639, EPI_ISL_507640, EPI_ISL_507641, EPI_ISL_507642, EPI_ISL_507643, EPI_ISL_507644, EPI_ISL_507645, EPI_ISL_507646, EPI_ISL_507647, EPI_ISL_507648, EPI_ISL_507649, EPI_ISL_507650, EPI_ISL_507651, EPI_ISL_507652, EPI_ISL_507653, EPI_ISL_507654, EPI_ISL_507655, EPI_ISL_507656, EPI_ISL_507657, EPI_ISL_507658, EPI_ISL_507659, EPI_ISL_507660, EPI_ISL_507661, EPI_ISL_507662, EPI_ISL_507663, EPI_ISL_507664, EPI_ISL_507665, EPI_ISL_507666, EPI_ISL_507667, EPI_ISL_507668, EPI_ISL_507669, EPI_ISL_507670, EPI_ISL_507671, EPI_ISL_507672, EPI_ISL_507673, EPI_ISL_507674, EPI_ISL_507675, EPI_ISL_507676, EPI_ISL_507677, EPI_ISL_507678, EPI_ISL_507679, EPI_ISL_507680, EPI_ISL_507681, EPI_ISL_507682, EPI_ISL_507683, EPI_ISL_507684, EPI_ISL_507685, EPI_ISL_507686, EPI_ISL_507687, EPI_ISL_507688, EPI_ISL_507689, EPI_ISL_507690, EPI_ISL_507691, EPI_ISL_507692, EPI_ISL_507693, EPI_ISL_507694, EPI_ISL_507695, EPI_ISL_507696, EPI_ISL_507697, EPI_ISL_507698, EPI_ISL_507699, EPI_ISL_507700, EPI_ISL_507701, EPI_ISL_507702, EPI_ISL_507703, EPI_ISL_507704, EPI_ISL_507705, EPI_ISL_507706, EPI_ISL_507707, EPI_ISL_507708, EPI_ISL_507709, EPI_ISL_507710, EPI_ISL_507711, EPI_ISL_507712, EPI_ISL_507713, EPI_ISL_507714, EPI_ISL_507715, EPI_ISL_507716, EPI_ISL_507717, EPI_ISL_507718, EPI_ISL_507719, EPI_ISL_507720, EPI_ISL_507721, EPI_ISL_507722, EPI_ISL_507723, EPI_ISL_507724, EPI_ISL_507725, EPI_ISL_507726, EPI_ISL_507727, EPI_ISL_507728, EPI_ISL_507729, EPI_ISL_507730, EPI_ISL_507731, EPI_ISL_507732, EPI_ISL_507733, EPI_ISL_507734, EPI_ISL_507735, EPI_ISL_507736, EPI_ISL_507737, EPI_ISL_507738, EPI_ISL_507739, EPI_ISL_507740, EPI_ISL_507741, EPI_ISL_507742, EPI_ISL_507743, EPI_ISL_507744, EPI_ISL_507745, EPI_ISL_507746, EPI_ISL_507747, EPI_ISL_507748, EPI_ISL_507749, EPI_ISL_507750, EPI_ISL_507751, EPI_ISL_507752, EPI_ISL_507753, EPI_ISL_507754, EPI_ISL_507755, EPI_ISL_507756, EPI_ISL_507757, EPI_ISL_507758, EPI_ISL_507759, EPI_ISL_507760, EPI_ISL_507761, EPI_ISL_507762, EPI_ISL_507763, EPI_ISL_507764, EPI_ISL_507765, EPI_ISL_507766, EPI_ISL_507767, EPI_ISL_507768, EPI_ISL_507769, EPI_ISL_507770, EPI_ISL_507771, EPI_ISL_507772, EPI_ISL_507773, EPI_ISL_507774, EPI_ISL_507775, EPI_ISL_507776, EPI_ISL_507777, EPI_ISL_507778, EPI_ISL_507779, EPI_ISL_507780, EPI_ISL_507781, EPI_ISL_507782, EPI_ISL_507783, EPI_ISL_507784, EPI_ISL_507785, EPI_ISL_507786, EPI_ISL_507787, EPI_ISL_507788, EPI_ISL_507789, EPI_ISL_507790, EPI_ISL_507791, EPI_ISL_507792, EPI_ISL_507793, EPI_ISL_507794, EPI_ISL_507795, EPI_ISL_507796, EPI_ISL_507797, EPI_ISL_507798, EPI_ISL_507799, EPI_ISL_507800, EPI_ISL_507801, EPI_ISL_507802, EPI_ISL_507803, EPI_ISL_507804, EPI_ISL_507805, EPI_ISL_507806, EPI_ISL_507807, EPI_ISL_507808, EPI_ISL_507809, EPI_ISL_507810, EPI_ISL_507811, EPI_ISL_507812, EPI_ISL_507813, EPI_ISL_507814, EPI_ISL_507815, EPI_ISL_507816, EPI_ISL_507817, EPI_ISL_507818, EPI_ISL_507819, EPI_ISL_507820, EPI_ISL_507821, EPI_ISL_507822, EPI_ISL_507823, EPI_ISL_507824, EPI_ISL_507825, EPI_ISL_507826, EPI_ISL_507827, EPI_ISL_507828, EPI_ISL_507829, EPI_ISL_507830, EPI_ISL_507831, EPI_ISL_507832, EPI_ISL_507833, EPI_ISL_507834, EPI_ISL_507835, EPI_ISL_507836, EPI_ISL_507837, EPI_ISL_507838, EPI_ISL_507839, EPI_ISL_507840, EPI_ISL_507841, EPI_ISL_507842, EPI_ISL_507843, EPI_ISL_507844, EPI_ISL_507845, EPI_ISL_507846, EPI_ISL_507847, EPI_ISL_507848, EPI_ISL_507849, EPI_ISL_507850, EPI_ISL_507851, EPI_ISL_507852, EPI_ISL_507853, EPI_ISL_507854, EPI_ISL_507855, EPI_ISL_507856, EPI_ISL_507857, EPI_ISL_507858, EPI_ISL_507859, EPI_ISL_507860, EPI_ISL_507861, EPI_ISL_507862, EPI_ISL_507863, EPI_ISL_507864, EPI_ISL_507865, EPI_ISL_507866, EPI_ISL_507867, EPI_ISL_507868, EPI_ISL_507869, EPI_ISL_507870, EPI_ISL_507871, EPI_ISL_507872, EPI_ISL_507873, EPI_ISL_507874, EPI_ISL_507875, EPI_ISL_507876, EPI_ISL_507877, EPI_ISL_507878, EPI_ISL_507879, EPI_ISL_507880, EPI_ISL_507881, EPI_ISL_507882, EPI_ISL_507883, EPI_ISL_507884, EPI_ISL_507885, EPI_ISL_507886, EPI_ISL_507887, EPI_ISL_507888, EPI_ISL_507889, EPI_ISL_507890, EPI_ISL_507891, EPI_ISL_507892, EPI_ISL_507893, EPI_ISL_507894, EPI_ISL_507895, EPI_ISL_507896, EPI_ISL_507897, EPI_ISL_507898, EPI_ISL_507899, EPI_ISL_507900, EPI_ISL_507901, EPI_ISL_507902, EPI_ISL_507903, EPI_ISL_507904, EPI_ISL_507905, EPI_ISL_507906, EPI_ISL_507907, EPI_ISL_507908, EPI_ISL_507909, EPI_ISL_507910, EPI_ISL_507911, EPI_ISL_507912, EPI_ISL_507913, EPI_ISL_507914, EPI_ISL_507915, EPI_ISL_507916, EPI_ISL_507917, EPI_ISL_507918, EPI_ISL_507919, EPI_ISL_507920, EPI_ISL_507921, EPI_ISL_507922, EPI_ISL_507923, EPI_ISL_507924, EPI_ISL_507925, EPI_ISL_507926, EPI_ISL_507927, EPI_ISL_507928, EPI_ISL_507929, EPI_ISL_507930, EPI_ISL_507931, EPI_ISL_507932, EPI_ISL_507933 | | | |
| see above | Michigan Department of Health and Human Services, Bureau of Laboratories | Michigan Department of Health and Human Services, Bureau of Laboratories | Blankenship HM, Riner D, Soehnlen MK |
| EPI_ISL_508156, EPI_ISL_508157, EPI_ISL_508158, EPI_ISL_508159, EPI_ISL_508160, EPI_ISL_508161, EPI_ISL_508162, EPI_ISL_508163, EPI_ISL_508164, EPI_ISL_508165, EPI_ISL_508166, EPI_ISL_508167, EPI_ISL_508168, EPI_ISL_508169, EPI_ISL_508170, EPI_ISL_508171, EPI_ISL_508172, EPI_ISL_508173, EPI_ISL_508174, EPI_ISL_508175, EPI_ISL_508176, EPI_ISL_508177, EPI_ISL_508178, EPI_ISL_508179, EPI_ISL_508180, EPI_ISL_508181, EPI_ISL_508182, EPI_ISL_508183, EPI_ISL_508184, EPI_ISL_508185, EPI_ISL_508186, EPI_ISL_508187, EPI_ISL_508188, EPI_ISL_508189, EPI_ISL_508190, EPI_ISL_508191, EPI_ISL_508192, EPI_ISL_508193, EPI_ISL_508194, EPI_ISL_508195, EPI_ISL_508196, EPI_ISL_508197, EPI_ISL_508198, EPI_ISL_508199, EPI_ISL_508200, EPI_ISL_508201, EPI_ISL_508202, EPI_ISL_508203, EPI_ISL_508204, EPI_ISL_508205, EPI_ISL_508206 | | | |
| see above | All india institute of Medical Sciences Rishikesh | National Institute of Biomedical Genomics | Arindam Maitra, Deepjyoti Kalita, Amit Mangla, Ravi Kant, Saumitra Das |
| EPI_ISL_508207, EPI_ISL_508208, EPI_ISL_508209, EPI_ISL_508210, EPI_ISL_508211, EPI_ISL_508212, EPI_ISL_508213, EPI_ISL_508214, EPI_ISL_508215, EPI_ISL_508216, EPI_ISL_508217, EPI_ISL_508218, EPI_ISL_508219, EPI_ISL_508220, EPI_ISL_508221, EPI_ISL_508222, EPI_ISL_508223, EPI_ISL_508224, EPI_ISL_508225, EPI_ISL_508226, EPI_ISL_508227, EPI_ISL_508228, EPI_ISL_508229, EPI_ISL_508230, EPI_ISL_508231, EPI_ISL_508232, EPI_ISL_508233, EPI_ISL_508234, EPI_ISL_508235, EPI_ISL_508236, EPI_ISL_508237, EPI_ISL_508238, EPI_ISL_508239, EPI_ISL_508240, EPI_ISL_508241, EPI_ISL_508242, EPI_ISL_508243, EPI_ISL_508244, EPI_ISL_508245, EPI_ISL_508246, EPI_ISL_508247, EPI_ISL_508248, EPI_ISL_508249, EPI_ISL_508250, EPI_ISL_508251, EPI_ISL_508252, EPI_ISL_508253, EPI_ISL_508254, EPI_ISL_508255, EPI_ISL_508256, EPI_ISL_508257, EPI_ISL_508258, EPI_ISL_508259, EPI_ISL_508260, EPI_ISL_508261, EPI_ISL_508262, EPI_ISL_508263, EPI_ISL_508264, EPI_ISL_508265, EPI_ISL_508266, EPI_ISL_508267, EPI_ISL_508268, EPI_ISL_508269, EPI_ISL_508270, EPI_ISL_508271, EPI_ISL_508272, EPI_ISL_508273, EPI_ISL_508274, EPI_ISL_508275, EPI_ISL_508276, EPI_ISL_508277, EPI_ISL_508278, EPI_ISL_508279, EPI_ISL_508280, EPI_ISL_508281, EPI_ISL_508282, EPI_ISL_508283, EPI_ISL_508284, EPI_ISL_508285, EPI_ISL_508286 | | | |
| see above | Government Medical College | National Institute of Biomedical Genomics | Arindam Maitra, Jyoti Iravane, Dhaval Khatri, Maitrik Dave, Saumitra Das |
| EPI_ISL_508287, EPI_ISL_508288, EPI_ISL_508289, EPI_ISL_508290, EPI_ISL_508291, EPI_ISL_508292, EPI_ISL_508293, EPI_ISL_508294, EPI_ISL_508295, EPI_ISL_508296, EPI_ISL_508297, EPI_ISL_508298, EPI_ISL_508299, EPI_ISL_508300, EPI_ISL_508301, EPI_ISL_508302, EPI_ISL_508303, EPI_ISL_508304, EPI_ISL_508305, EPI_ISL_508306, EPI_ISL_508307, EPI_ISL_508308, EPI_ISL_508309, EPI_ISL_508310, EPI_ISL_508311, EPI_ISL_508312, EPI_ISL_508313, EPI_ISL_508314, EPI_ISL_508315, EPI_ISL_508316, EPI_ISL_508317, EPI_ISL_508318, EPI_ISL_508319, EPI_ISL_508320, EPI_ISL_508321, EPI_ISL_508322, EPI_ISL_508323, EPI_ISL_508324, EPI_ISL_508325, EPI_ISL_508326, EPI_ISL_508327, EPI_ISL_508328, EPI_ISL_508329, EPI_ISL_508330, EPI_ISL_508331, EPI_ISL_508332, EPI_ISL_508333, EPI_ISL_508334, EPI_ISL_508335, EPI_ISL_508336, EPI_ISL_508337 | | | |
| see above | Indian Institute of Science | National Institute of Biomedical Genomics | Arindam Maitra, Bharath K Sundararaj, Harsha Raheja, N. Srinivasan, Deepak K Saini, Amit Singh, Saumitra Das |
| EPI_ISL_508338, EPI_ISL_508339, EPI_ISL_508340, EPI_ISL_508341, EPI_ISL_508342, EPI_ISL_508343, EPI_ISL_508344, EPI_ISL_508345, EPI_ISL_508346, EPI_ISL_508347, EPI_ISL_508348, EPI_ISL_508349, EPI_ISL_508350, EPI_ISL_508351, EPI_ISL_508352, EPI_ISL_508353, EPI_ISL_508354, EPI_ISL_508355, EPI_ISL_508356, EPI_ISL_508357, EPI_ISL_508358, EPI_ISL_508359, EPI_ISL_508360, EPI_ISL_508361, EPI_ISL_508362, EPI_ISL_508363, EPI_ISL_508364, EPI_ISL_508365, EPI_ISL_508366, EPI_ISL_508367, EPI_ISL_508368, EPI_ISL_508369, EPI_ISL_508370, EPI_ISL_508371, EPI_ISL_508372, EPI_ISL_508373, EPI_ISL_508374, EPI_ISL_508375, EPI_ISL_508376, EPI_ISL_508377, EPI_ISL_508378, EPI_ISL_508379, EPI_ISL_508380, EPI_ISL_508381, EPI_ISL_508382, EPI_ISL_508383, EPI_ISL_508384, EPI_ISL_508385, EPI_ISL_508386, EPI_ISL_508387, EPI_ISL_508388, EPI_ISL_508389, EPI_ISL_508390, EPI_ISL_508391, EPI_ISL_508392, EPI_ISL_508393, EPI_ISL_508394, EPI_ISL_508395, EPI_ISL_508396, EPI_ISL_508397, EPI_ISL_508398, EPI_ISL_508399, EPI_ISL_508400, EPI_ISL_508401, EPI_ISL_508402, EPI_ISL_508403, EPI_ISL_508404, EPI_ISL_508405, EPI_ISL_508406, EPI_ISL_508407, EPI_ISL_508408, EPI_ISL_508409, EPI_ISL_508410, EPI_ISL_508411, EPI_ISL_508412, EPI_ISL_508413, EPI_ISL_508414 | | | |
| see above | Institute of Post Graduate Medical Education & Research | National Institute of Biomedical Genomics | Arindam Maitra, Aritra Biswas, Jayeeta Haldar, Raja Ray, Monimoy Banerjee, Saumitra Das |

| | | | |
|---|---|---|---|
| EPI_ISL_508415, EPI_ISL_508416, EPI_ISL_508417, EPI_ISL_508418, EPI_ISL_508419, EPI_ISL_508420, EPI_ISL_508421, EPI_ISL_508422 | Maulana Azad Medical College | National Institute of Biomedical Genomics | Arindam Maitra, Sonal Saxena, Vikas Manchanda, Oves Siddiqui, Saumitra Das |
| EPI_ISL_508423, EPI_ISL_508424, EPI_ISL_508425, EPI_ISL_508426, EPI_ISL_508427, EPI_ISL_508428, EPI_ISL_508429, EPI_ISL_508430, EPI_ISL_508431, EPI_ISL_508432, EPI_ISL_508433, EPI_ISL_508434, EPI_ISL_508435, EPI_ISL_508436, EPI_ISL_508437, EPI_ISL_508438, EPI_ISL_508439, EPI_ISL_508440 | | | |
| see above | Mahatma Gandhi Institute of Medical Sciences | National Institute of Biomedical Genomics | Arindam Maitra, Vijayshri Deotale, Rahul Narang, Deepashri Maraskolhe, Saumitra Das |
| EPI_ISL_508441, EPI_ISL_508442, EPI_ISL_508443, EPI_ISL_508444, EPI_ISL_508445, EPI_ISL_508446, EPI_ISL_508447, EPI_ISL_508448, EPI_ISL_508449, EPI_ISL_508450, EPI_ISL_508451, EPI_ISL_508452, EPI_ISL_508453, EPI_ISL_508454, EPI_ISL_508455, EPI_ISL_508456, EPI_ISL_508457, EPI_ISL_508458, EPI_ISL_508459, EPI_ISL_508460, EPI_ISL_508461, EPI_ISL_508462, EPI_ISL_508463, EPI_ISL_508464, EPI_ISL_508465, EPI_ISL_508466, EPI_ISL_508467, EPI_ISL_508468, EPI_ISL_508469, EPI_ISL_508470, EPI_ISL_508471, EPI_ISL_508472, EPI_ISL_508473, EPI_ISL_508474, EPI_ISL_508475, EPI_ISL_508476, EPI_ISL_508477, EPI_ISL_508478, EPI_ISL_508479, EPI_ISL_508480, EPI_ISL_508481, EPI_ISL_508482, EPI_ISL_508483, EPI_ISL_508484, EPI_ISL_508485, EPI_ISL_508486, EPI_ISL_508487, EPI_ISL_508488 | | | |
| see above | ICMR-National Institute of Cholera and Enteric Diseases | National Institute of Biomedical Genomics | Arindam Maitra, Mamta Chawla Sarkar, Sreedhar Chinnaswamy, Hasina Banu, Ananya Chatterjee, Shanta Dutta, Saumitra Das |
| EPI_ISL_508489, EPI_ISL_508490, EPI_ISL_508491, EPI_ISL_508492, EPI_ISL_508493, EPI_ISL_508494, EPI_ISL_508495, EPI_ISL_508496, EPI_ISL_508497, EPI_ISL_508498, EPI_ISL_508499, EPI_ISL_508500, EPI_ISL_508501, EPI_ISL_508502, EPI_ISL_508503, EPI_ISL_508504, EPI_ISL_508505, EPI_ISL_508506, EPI_ISL_508507, EPI_ISL_508508, EPI_ISL_508509 | | | |
| see above | Translational Health Science and Technology Institute | National Institute of Biomedical Genomics | Arindam Maitra, Guruprasad Medigeshi, Sharanabasava Patil, Anbalagan Ananthraj, Madhu Pareek, Imran Khan, Gagandeep Kang, Saumitra Das |
| EPI_ISL_508871, EPI_ISL_508872, EPI_ISL_508873, EPI_ISL_508874 | Institut des Agents Infectieux (IAI), Hospices Civils de Lyon | CNR Virus des Infections Respiratoires - France SUD | Antonin Bal, Gregory Destras, Gwendolyne Burfin, Solenne Brun, Carine Moustaud, Raphaelle Lamy, Alexandre Gaymard, Maude Bouscambert-Duchamp, Florence Morfin-Sherpa, Martine Valette, Bruno Lina, Laurence Josset |
| EPI_ISL_508875, EPI_ISL_508876 | Centre Hospitalier de Macon | CNR Virus des Infections Respiratoires - France SUD | Antonin Bal, Gregory Destras, Gwendolyne Burfin, Solenne Brun, Carine Moustaud, Raphaelle Lamy, Alexandre Gaymard, Maude Bouscambert-Duchamp, Florence Morfin-Sherpa, Martine Valette, Bruno Lina, Laurence Josset |
| EPI_ISL_508877 | Institut des Agents Infectieux (IAI), Hospices Civils de Lyon | CNR Virus des Infections Respiratoires - France SUD | Antonin Bal, Gregory Destras, Gwendolyne Burfin, Solenne Brun, Carine Moustaud, Raphaelle Lamy, Alexandre Gaymard, Maude Bouscambert-Duchamp, Florence Morfin-Sherpa, Martine Valette, Bruno Lina, Laurence Josset |
| EPI_ISL_508878 | GH Les Portes du Sud | CNR Virus des Infections Respiratoires - France SUD | Antonin Bal, Gregory Destras, Gwendolyne Burfin, Solenne Brun, Carine Moustaud, Raphaelle Lamy, Alexandre Gaymard, Maude Bouscambert-Duchamp, Florence Morfin-Sherpa, Martine Valette, Bruno Lina, Laurence Josset |
| EPI_ISL_508879, EPI_ISL_508880 | Centre Hospitalier Saint Joseph Saint Luc | CNR Virus des Infections Respiratoires - France SUD | Antonin Bal, Gregory Destras, Gwendolyne Burfin, Solenne Brun, Carine Moustaud, Raphaelle Lamy, Alexandre Gaymard, Maude Bouscambert-Duchamp, Florence Morfin-Sherpa, Martine Valette, Bruno Lina, Laurence Josset |
| EPI_ISL_508881 | Centre Hospitalier de Valence | CNR Virus des Infections Respiratoires - France SUD | Antonin Bal, Gregory Destras, Gwendolyne Burfin, Solenne Brun, Carine Moustaud, Raphaelle Lamy, Alexandre Gaymard, Maude Bouscambert-Duchamp, Florence Morfin-Sherpa, Martine Valette, Bruno Lina, Laurence Josset |
| EPI_ISL_508882, EPI_ISL_508883, EPI_ISL_508884, EPI_ISL_508885, EPI_ISL_508886, EPI_ISL_508887, EPI_ISL_508888, EPI_ISL_508889, EPI_ISL_508890, EPI_ISL_508891, EPI_ISL_508892, EPI_ISL_508893, EPI_ISL_508894, EPI_ISL_508895, EPI_ISL_508896, EPI_ISL_508897, EPI_ISL_508898, EPI_ISL_508899, EPI_ISL_508900, EPI_ISL_508901, EPI_ISL_508902, EPI_ISL_508903, EPI_ISL_508904, EPI_ISL_508905, EPI_ISL_508906, EPI_ISL_508907, EPI_ISL_508908, EPI_ISL_508909, EPI_ISL_508910, EPI_ISL_508911 | | | |
| see above | Institut des Agents Infectieux (IAI), Hospices Civils de Lyon | CNR Virus des Infections Respiratoires - France SUD | Antonin Bal, Gregory Destras, Gwendolyne Burfin, Solenne Brun, Carine Moustaud, Raphaelle Lamy, Alexandre Gaymard, Maude Bouscambert-Duchamp, Florence Morfin-Sherpa, Martine Valette, Bruno Lina, Laurence Josset |
| EPI_ISL_508912, EPI_ISL_508913, EPI_ISL_508914, EPI_ISL_508915, EPI_ISL_508916, EPI_ISL_508917, EPI_ISL_508918, EPI_ISL_508919, EPI_ISL_508920, EPI_ISL_508921, EPI_ISL_508922, EPI_ISL_508923, EPI_ISL_508924, EPI_ISL_508925, EPI_ISL_508926, EPI_ISL_508927, EPI_ISL_508928, EPI_ISL_508929, EPI_ISL_508930 | | | |
| see above | CNR Virus des Infections Respiratoires - France SUD | CNR Virus des Infections Respiratoires - France SUD | Antonin Bal, Gregory Destras, Gwendolyne Burfin, Solenne Brun, Carine Moustaud, Raphaelle Lamy, Alexandre Gaymard, Maude Bouscambert-Duchamp, Florence Morfin-Sherpa, Martine Valette, Bruno Lina, Laurence Josset |
| EPI_ISL_508931 | L'Hôpital Nord-Ouest Tarare-Grandris | CNR Virus des Infections Respiratoires - France SUD | Antonin Bal, Gregory Destras, Gwendolyne Burfin, Solenne Brun, Carine Moustaud, Raphaelle Lamy, Alexandre Gaymard, Maude Bouscambert-Duchamp, Florence Morfin-Sherpa, Martine Valette, Bruno Lina, Laurence Josset |
| EPI_ISL_508932 | Centre Hospitalier de Villefranche | CNR Virus des Infections Respiratoires - France SUD | Antonin Bal, Gregory Destras, Gwendolyne Burfin, Solenne Brun, Carine Moustaud, Raphaelle Lamy, Alexandre Gaymard, Maude Bouscambert-Duchamp, Florence Morfin-Sherpa, Martine Valette, Bruno Lina, Laurence Josset |
| EPI_ISL_508933 | CNR Virus des Infections Respiratoires - France SUD | CNR Virus des Infections Respiratoires - France SUD | Antonin Bal, Gregory Destras, Gwendolyne Burfin, Solenne Brun, Carine Moustaud, Raphaelle Lamy, Alexandre Gaymard, Maude Bouscambert-Duchamp, Florence Morfin-Sherpa, Martine Valette, Bruno Lina, Laurence Josset |
| EPI_ISL_508934, EPI_ISL_508935, EPI_ISL_508936 | Centre hospitalier Métropole Savoie | CNR Virus des Infections Respiratoires - France SUD | Antonin Bal, Gregory Destras, Gwendolyne Burfin, Solenne Brun, Carine Moustaud, Raphaelle Lamy, Alexandre Gaymard, Maude Bouscambert-Duchamp, Florence Morfin-Sherpa, Martine Valette, Bruno Lina, Laurence Josset |
| EPI_ISL_508937 | CNR Virus des Infections Respiratoires - France SUD | CNR Virus des Infections Respiratoires - France SUD | Antonin Bal, Gregory Destras, Gwendolyne Burfin, Solenne Brun, Carine Moustaud, Raphaelle Lamy, Alexandre Gaymard, Maude Bouscambert-Duchamp, Florence Morfin-Sherpa, Martine Valette, Bruno Lina, Laurence Josset |
| EPI_ISL_508938 | Centre Hospitalier de Bourg en Bresse | CNR Virus des Infections Respiratoires - France SUD | Antonin Bal, Gregory Destras, Gwendolyne Burfin, Solenne Brun, Carine Moustaud, Raphaelle Lamy, Alexandre Gaymard, Maude Bouscambert-Duchamp, Florence Morfin-Sherpa, Martine Valette, Bruno Lina, Laurence Josset |
| EPI_ISL_508939, EPI_ISL_508940 | CNR Virus des Infections Respiratoires - France SUD | CNR Virus des Infections Respiratoires - France SUD | Antonin Bal, Gregory Destras, Gwendolyne Burfin, Solenne Brun, Carine Moustaud, Raphaelle Lamy, Alexandre Gaymard, Maude Bouscambert-Duchamp, Florence Morfin-Sherpa, Martine Valette, Bruno Lina, Laurence Josset |
| EPI_ISL_508941 | Centre Hospitalier de Macon | CNR Virus des Infections Respiratoires - France SUD | Antonin Bal, Gregory Destras, Gwendolyne Burfin, Solenne Brun, Carine Moustaud, Raphaelle Lamy, Alexandre Gaymard, Maude Bouscambert-Duchamp, Florence Morfin-Sherpa, Martine Valette, Bruno Lina, Laurence Josset |
| EPI_ISL_508942 | CNR Virus des Infections Respiratoires - France SUD | CNR Virus des Infections Respiratoires - France SUD | Antonin Bal, Gregory Destras, Gwendolyne Burfin, Solenne Brun, Carine Moustaud, Raphaelle Lamy, Alexandre Gaymard, Maude Bouscambert-Duchamp, Florence Morfin-Sherpa, Martine Valette, Bruno Lina, Laurence Josset |
| EPI_ISL_508943 | Centre Hospitalier de Macon | CNR Virus des Infections Respiratoires - France SUD | Antonin Bal, Gregory Destras, Gwendolyne Burfin, Solenne Brun, Carine Moustaud, Raphaelle Lamy, Alexandre Gaymard, Maude Bouscambert-Duchamp, Florence Morfin-Sherpa, Martine Valette, Bruno Lina, Laurence Josset |
| EPI_ISL_508944 | Centre Hospitalier de Bourg en Bresse | CNR Virus des Infections Respiratoires - France SUD | Antonin Bal, Gregory Destras, Gwendolyne Burfin, Solenne Brun, Carine Moustaud, Raphaelle Lamy, Alexandre Gaymard, Maude Bouscambert-Duchamp, Florence Morfin-Sherpa, Martine Valette, Bruno Lina, Laurence Josset |
| EPI_ISL_508945 | CNR Virus des Infections Respiratoires - France SUD | CNR Virus des Infections Respiratoires - France SUD | Antonin Bal, Gregory Destras, Gwendolyne Burfin, Solenne Brun, Carine Moustaud, Raphaelle Lamy, Alexandre Gaymard, Maude Bouscambert-Duchamp, Florence Morfin-Sherpa, Martine Valette, Bruno Lina, Laurence Josset |
| EPI_ISL_508946 | Centre Hospitalier de Macon | CNR Virus des Infections Respiratoires - France SUD | Antonin Bal, Gregory Destras, Gwendolyne Burfin, Solenne Brun, Carine Moustaud, Raphaelle Lamy, Alexandre Gaymard, Maude Bouscambert-Duchamp, Florence Morfin-Sherpa, Martine Valette, Bruno Lina, Laurence Josset |
| EPI_ISL_508947 | Centre Hospitalier Lucien Hussel | CNR Virus des Infections Respiratoires - France SUD | Antonin Bal, Gregory Destras, Gwendolyne Burfin, Solenne Brun, Carine Moustaud, Raphaelle Lamy, Alexandre Gaymard, Maude Bouscambert-Duchamp, Florence Morfin-Sherpa, Martine Valette, Bruno Lina, Laurence Josset |
| EPI_ISL_508948 | CNR Virus des Infections Respiratoires - France SUD | CNR Virus des Infections Respiratoires - France SUD | Antonin Bal, Gregory Destras, Gwendolyne Burfin, Solenne Brun, Carine Moustaud, Raphaelle Lamy, Alexandre Gaymard, Maude Bouscambert-Duchamp, Florence Morfin-Sherpa, Martine Valette, Bruno Lina, Laurence Josset |
| EPI_ISL_508949, EPI_ISL_508950 | Centre Hospitalier de Macon | CNR Virus des Infections Respiratoires - France SUD | Antonin Bal, Gregory Destras, Gwendolyne Burfin, Solenne Brun, Carine Moustaud, Raphaelle Lamy, Alexandre Gaymard, Maude Bouscambert-Duchamp, Florence Morfin-Sherpa, Martine Valette, Bruno Lina, Laurence Josset |
| EPI_ISL_508951 | CNR Virus des Infections Respiratoires - France SUD | CNR Virus des Infections Respiratoires - France SUD | Antonin Bal, Gregory Destras, Gwendolyne Burfin, Solenne Brun, Carine Moustaud, Raphaelle Lamy, Alexandre Gaymard, Maude Bouscambert-Duchamp, Florence Morfin-Sherpa, Martine Valette, Bruno Lina, Laurence Josset |
| EPI_ISL_508952 | Centre Hospitalier de Macon | CNR Virus des Infections Respiratoires - France SUD | Antonin Bal, Gregory Destras, Gwendolyne Burfin, Solenne Brun, Carine Moustaud, Raphaelle Lamy, Alexandre Gaymard, Maude Bouscambert-Duchamp, Florence Morfin-Sherpa, Martine Valette, Bruno Lina, Laurence Josset |
| EPI_ISL_508953, EPI_ISL_508954, | CNR Virus des Infections Respiratoires - France SUD | CNR Virus des Infections Respiratoires - France SUD | Antonin Bal, Gregory Destras, Gwendolyne Burfin, Solenne Brun, Carine Moustaud, Raphaelle Lamy, Alexandre Gaymard, Maude |

| | | | |
|---|---|---|---|
| EPI_ISL_508955, EPI_ISL_508956, EPI_ISL_508957 | | | Bouscambert-Duchamp, Florence Morfin-Sherpa, Martine Valette, Bruno Lina, Laurence Josset |
| EPI_ISL_508958 | Centre Hospitalier Alpes Leman | CNR Virus des Infections Respiratoires - France SUD | Antonin Bal, Gregory Destras, Gwendolyne Burfin, Solenne Brun, Carine Moustaud, Raphaelle Lamy, Alexandre Gaymard, Maude Bouscambert-Duchamp, Florence Morfin-Sherpa, Martine Valette, Bruno Lina, Laurence Josset |
| EPI_ISL_508959, EPI_ISL_508960 | Centre Hospitalier Saint Joseph Saint Luc | CNR Virus des Infections Respiratoires - France SUD | Antonin Bal, Gregory Destras, Gwendolyne Burfin, Solenne Brun, Carine Moustaud, Raphaelle Lamy, Alexandre Gaymard, Maude Bouscambert-Duchamp, Florence Morfin-Sherpa, Martine Valette, Bruno Lina, Laurence Josset |
| EPI_ISL_508961, EPI_ISL_508962, EPI_ISL_508963, EPI_ISL_508964, EPI_ISL_508965 | CNR Virus des Infections Respiratoires - France SUD | CNR Virus des Infections Respiratoires - France SUD | Antonin Bal, Gregory Destras, Gwendolyne Burfin, Solenne Brun, Carine Moustaud, Raphaelle Lamy, Alexandre Gaymard, Maude Bouscambert-Duchamp, Florence Morfin-Sherpa, Martine Valette, Bruno Lina, Laurence Josset |
| EPI_ISL_508966 | Centre Hospitalier Pierre Oudot | CNR Virus des Infections Respiratoires - France SUD | Antonin Bal, Gregory Destras, Gwendolyne Burfin, Solenne Brun, Carine Moustaud, Raphaelle Lamy, Alexandre Gaymard, Maude Bouscambert-Duchamp, Florence Morfin-Sherpa, Martine Valette, Bruno Lina, Laurence Josset |
| EPI_ISL_508967 | CNR Virus des Infections Respiratoires - France SUD | CNR Virus des Infections Respiratoires - France SUD | Antonin Bal, Gregory Destras, Gwendolyne Burfin, Solenne Brun, Carine Moustaud, Raphaelle Lamy, Alexandre Gaymard, Maude Bouscambert-Duchamp, Florence Morfin-Sherpa, Martine Valette, Bruno Lina, Laurence Josset |
| EPI_ISL_508968 | Centre Hospitalier de Bourg en Bresse | CNR Virus des Infections Respiratoires - France SUD | Antonin Bal, Gregory Destras, Gwendolyne Burfin, Solenne Brun, Carine Moustaud, Raphaelle Lamy, Alexandre Gaymard, Maude Bouscambert-Duchamp, Florence Morfin-Sherpa, Martine Valette, Bruno Lina, Laurence Josset |
| EPI_ISL_508969, EPI_ISL_508970, EPI_ISL_508971, EPI_ISL_508972, EPI_ISL_508973, EPI_ISL_508974 | CNR Virus des Infections Respiratoires - France SUD | CNR Virus des Infections Respiratoires - France SUD | Antonin Bal, Gregory Destras, Gwendolyne Burfin, Solenne Brun, Carine Moustaud, Raphaelle Lamy, Alexandre Gaymard, Maude Bouscambert-Duchamp, Florence Morfin-Sherpa, Martine Valette, Bruno Lina, Laurence Josset |
| EPI_ISL_508975 | Centre Hospitalier Pierre Oudot | CNR Virus des Infections Respiratoires - France SUD | Antonin Bal, Gregory Destras, Gwendolyne Burfin, Solenne Brun, Carine Moustaud, Raphaelle Lamy, Alexandre Gaymard, Maude Bouscambert-Duchamp, Florence Morfin-Sherpa, Martine Valette, Bruno Lina, Laurence Josset |
| EPI_ISL_508976, EPI_ISL_508977 | CNR Virus des Infections Respiratoires - France SUD | CNR Virus des Infections Respiratoires - France SUD | Antonin Bal, Gregory Destras, Gwendolyne Burfin, Solenne Brun, Carine Moustaud, Raphaelle Lamy, Alexandre Gaymard, Maude Bouscambert-Duchamp, Florence Morfin-Sherpa, Martine Valette, Bruno Lina, Laurence Josset |
| EPI_ISL_508978 | Centre Hospitalier de Villefranche | CNR Virus des Infections Respiratoires - France SUD | Antonin Bal, Gregory Destras, Gwendolyne Burfin, Solenne Brun, Carine Moustaud, Raphaelle Lamy, Alexandre Gaymard, Maude Bouscambert-Duchamp, Florence Morfin-Sherpa, Martine Valette, Bruno Lina, Laurence Josset |
| EPI_ISL_508979, EPI_ISL_508980, EPI_ISL_508981, EPI_ISL_508982, EPI_ISL_508983, EPI_ISL_508984, EPI_ISL_508985 | CNR Virus des Infections Respiratoires - France SUD | CNR Virus des Infections Respiratoires - France SUD | Antonin Bal, Gregory Destras, Gwendolyne Burfin, Solenne Brun, Carine Moustaud, Raphaelle Lamy, Alexandre Gaymard, Maude Bouscambert-Duchamp, Florence Morfin-Sherpa, Martine Valette, Bruno Lina, Laurence Josset |
| EPI_ISL_508986, EPI_ISL_508987, EPI_ISL_508988, EPI_ISL_508989 | CNR Virus des Infections Respiratoires - France SUD | CNR Virus des Infections Respiratoires - France SUD | Antonin Bal, Gregory Destras, Gwendolyne Burfin, Solenne Brun, Alexandre Gaymard, Maude Bouscambert-Duchamp, Florence Morfin-Sherpa, Martine Valette, Bruno Lina, Laurence Josset |
| EPI_ISL_508990, EPI_ISL_508991, EPI_ISL_508992, EPI_ISL_508993, EPI_ISL_508994, EPI_ISL_508995, EPI_ISL_508996, EPI_ISL_508997 | CNR Virus des Infections Respiratoires - France SUD | CNR Virus des Infections Respiratoires - France SUD | Antonin Bal, Gregory Destras, Gwendolyne Burfin, Solenne Brun, Carine Moustaud, Raphaelle Lamy, Alexandre Gaymard, Maude Bouscambert-Duchamp, Florence Morfin-Sherpa, Martine Valette, Bruno Lina, Laurence Josset |
| EPI_ISL_508998 | Centre Hospitalier Pierre Oudot | CNR Virus des Infections Respiratoires - France SUD | Antonin Bal, Gregory Destras, Gwendolyne Burfin, Solenne Brun, Carine Moustaud, Raphaelle Lamy, Alexandre Gaymard, Maude Bouscambert-Duchamp, Florence Morfin-Sherpa, Martine Valette, Bruno Lina, Laurence Josset |
| EPI_ISL_508999, EPI_ISL_509000, EPI_ISL_509001, EPI_ISL_509002 | CNR Virus des Infections Respiratoires - France SUD | CNR Virus des Infections Respiratoires - France SUD | Antonin Bal, Gregory Destras, Gwendolyne Burfin, Solenne Brun, Carine Moustaud, Raphaelle Lamy, Alexandre Gaymard, Maude Bouscambert-Duchamp, Florence Morfin-Sherpa, Martine Valette, Bruno Lina, Laurence Josset |
| EPI_ISL_509003, EPI_ISL_509004 | GH Les Portes du Sud | CNR Virus des Infections Respiratoires - France SUD | Antonin Bal, Gregory Destras, Gwendolyne Burfin, Solenne Brun, Carine Moustaud, Raphaelle Lamy, Alexandre Gaymard, Maude Bouscambert-Duchamp, Florence Morfin-Sherpa, Martine Valette, Bruno Lina, Laurence Josset |
| EPI_ISL_509005 | Centre Hospitalier de Macon | CNR Virus des Infections Respiratoires - France SUD | Antonin Bal, Gregory Destras, Gwendolyne Burfin, Solenne Brun, Carine Moustaud, Raphaelle Lamy, Alexandre Gaymard, Maude Bouscambert-Duchamp, Florence Morfin-Sherpa, Martine Valette, Bruno Lina, Laurence Josset |
| EPI_ISL_509006 | Centre Hospitalier de Villefranche | CNR Virus des Infections Respiratoires - France SUD | Antonin Bal, Gregory Destras, Gwendolyne Burfin, Solenne Brun, Carine Moustaud, Raphaelle Lamy, Alexandre Gaymard, Maude Bouscambert-Duchamp, Florence Morfin-Sherpa, Martine Valette, Bruno Lina, Laurence Josset |
| EPI_ISL_509007, EPI_ISL_509008, EPI_ISL_509009, EPI_ISL_509010, EPI_ISL_509011 | Institut des Agents Infectieux (IAI), Hospices Civils de Lyon | CNR Virus des Infections Respiratoires - France SUD | Antonin Bal, Gregory Destras, Gwendolyne Burfin, Solenne Brun, Carine Moustaud, Raphaelle Lamy, Alexandre Gaymard, Maude Bouscambert-Duchamp, Florence Morfin-Sherpa, Martine Valette, Bruno Lina, Laurence Josset |
| EPI_ISL_509012 | Centre Hospitalier Alpes Leman | CNR Virus des Infections Respiratoires - France SUD | Antonin Bal, Gregory Destras, Gwendolyne Burfin, Solenne Brun, Carine Moustaud, Raphaelle Lamy, Alexandre Gaymard, Maude Bouscambert-Duchamp, Florence Morfin-Sherpa, Martine Valette, Bruno Lina, Laurence Josset |
| EPI_ISL_509013, EPI_ISL_509014 | Institut des Agents Infectieux (IAI), Hospices Civils de Lyon | CNR Virus des Infections Respiratoires - France SUD | Antonin Bal, Gregory Destras, Gwendolyne Burfin, Solenne Brun, Carine Moustaud, Raphaelle Lamy, Alexandre Gaymard, Maude Bouscambert-Duchamp, Florence Morfin-Sherpa, Martine Valette, Bruno Lina, Laurence Josset |
| EPI_ISL_509015 | Centre Hospitalier de Villefranche | CNR Virus des Infections Respiratoires - France SUD | Antonin Bal, Gregory Destras, Gwendolyne Burfin, Solenne Brun, Carine Moustaud, Raphaelle Lamy, Alexandre Gaymard, Maude Bouscambert-Duchamp, Florence Morfin-Sherpa, Martine Valette, Bruno Lina, Laurence Josset |
| EPI_ISL_509016 | Centre Hospitalier du Haut-Bugey | CNR Virus des Infections Respiratoires - France SUD | Antonin Bal, Gregory Destras, Gwendolyne Burfin, Solenne Brun, Carine Moustaud, Raphaelle Lamy, Alexandre Gaymard, Maude Bouscambert-Duchamp, Florence Morfin-Sherpa, Martine Valette, Bruno Lina, Laurence Josset |
| EPI_ISL_510064, EPI_ISL_510065, EPI_ISL_510066, EPI_ISL_510067, EPI_ISL_510068, EPI_ISL_510069, EPI_ISL_510070, EPI_ISL_510071, EPI_ISL_510072, EPI_ISL_510073, EPI_ISL_510074, EPI_ISL_510075, EPI_ISL_510076, EPI_ISL_510077 | | | |
| see above | Instituto de Investigaciones Biomédicas de Barcelona (CSIC), Hospital Clinic i Provincial de Barcelona, Instituto de Biomedicina de Valencia (CSIC), Hospital de Sant Pau | SeqCOVID-SPAIN consortium/IBV(CSIC) | Anna M. Planas, Mª Angeles Marcos, Miguel J. Martínez, Andrea Vergara, Alex Soriano, Jordi Pérez Tur, Israel Fernández Cadenas and SeqCOVID-SPAIN consortium |
| EPI_ISL_510333, EPI_ISL_510334, EPI_ISL_510335, EPI_ISL_510336, EPI_ISL_510337, EPI_ISL_510338, EPI_ISL_510339, EPI_ISL_510340, EPI_ISL_510341, EPI_ISL_510342, EPI_ISL_510343, EPI_ISL_510344, EPI_ISL_510345, EPI_ISL_510346, EPI_ISL_510347, EPI_ISL_510348, EPI_ISL_510349, EPI_ISL_510350, EPI_ISL_510351, EPI_ISL_510352, EPI_ISL_510353, EPI_ISL_510354, EPI_ISL_510355, EPI_ISL_510356, EPI_ISL_510357, EPI_ISL_510358, EPI_ISL_510359, EPI_ISL_510360, EPI_ISL_510361, EPI_ISL_510362, EPI_ISL_510363, EPI_ISL_510364, EPI_ISL_510365, EPI_ISL_510366, EPI_ISL_510367, EPI_ISL_510368, EPI_ISL_510369, EPI_ISL_510370, EPI_ISL_510371, EPI_ISL_510372, EPI_ISL_510373, EPI_ISL_510374, EPI_ISL_510375, EPI_ISL_510376, EPI_ISL_510377, EPI_ISL_510378, EPI_ISL_510379, EPI_ISL_510380, EPI_ISL_510381, EPI_ISL_510382, EPI_ISL_510383, EPI_ISL_510384, EPI_ISL_510385, EPI_ISL_510386, EPI_ISL_510387, EPI_ISL_510388, EPI_ISL_510389, EPI_ISL_510390 | | | |
| see above | Servicio de Microbiología, Hospital Miguel Servet, Zaragoza | SeqCOVID-SPAIN consortium/IBV(CSIC) | Antonio Rezusta López, Alexander Tristancho Baró, Ana Milagro, Yolanda Gracia Grataloup, Nieves Martínez Cameo and SeqCOVID-SPAIN consortium |
| EPI_ISL_510461, EPI_ISL_510462, EPI_ISL_510463, EPI_ISL_510464, EPI_ISL_510465, EPI_ISL_510466 | Instituto de Investigaciones Biomédicas de Barcelona (CSIC), Hospital Clinic i Provincial de Barcelona, Instituto de Biomedicina de Valencia (CSIC), Hospital de Sant Pau | SeqCOVID-SPAIN consortium/IBV(CSIC) | Anna M. Planas, Mª Angeles Marcos, Miguel J. Martínez, Andrea Vergara, Alex Soriano, Jordi Pérez Tur, Israel Fernández Cadenas and SeqCOVID-SPAIN consortium |
| EPI_ISL_510527 | Unite des virus emergents, UMR190 | Unite des virus emergents, UMR190 | Baronti, C., Piorkowski,G., Coutard,B., Charrel,R. and de Lamballerie,X. |
| EPI_ISL_511899 | ICMR-National Institute of Cholera and Enteric Diseases | National Institute of Biomedical Genomics - DBT's PAN-INDIA 1000 SARS--CoV-2 RNA Genome Sequencing Consortium | Arindam Maitra, Mamta Chawla Sarkar, Sreedhar Chinnaswamy, Hasina Banu, Ananya Chatterjee, Shanta Dutta, Saumitra Das |
| EPI_ISL_511900, EPI_ISL_511901, EPI_ISL_511902, EPI_ISL_511903, | Institute of Post Graduate Medical Education & Research | National Institute of Biomedical Genomics - DBT's PAN-INDIA 1000 SARS--CoV-2 RNA Genome Sequencing Consortium | Arindam Maitra, Aritra Biswas, Jayeeta Haldar, Raja Ray, Monimoy Banerjee, Saumitra Das |

| | | | |
|---|---|---|---|
| EPI_ISL_511904, EPI_ISL_511905, EPI_ISL_511906, EPI_ISL_511907 | | | |
| EPI_ISL_511908, EPI_ISL_511909, EPI_ISL_511910, EPI_ISL_511911, EPI_ISL_511912, EPI_ISL_511913, EPI_ISL_511914, EPI_ISL_511915, EPI_ISL_511916, EPI_ISL_511917, EPI_ISL_511918, EPI_ISL_511919, EPI_ISL_511920, EPI_ISL_511921, EPI_ISL_511922 | | | |
| see above | All india institute of Medical Sciences Rishikesh | National Institute of Biomedical Genomics - DBT's PAN-INDIA 1000 SARS--CoV-2 RNA Genome Sequencing Consortium | Arindam Maitra, Deepjyoti Kalita, Amit Mangla, Ravi Kant, Saumitra Das |
| EPI_ISL_511923, EPI_ISL_511924, EPI_ISL_511925, EPI_ISL_511926, EPI_ISL_511927, EPI_ISL_511928, EPI_ISL_511929 | Mahatma Gandhi Institute of Medical Sciences | National Institute of Biomedical Genomics - DBT's PAN-INDIA 1000 SARS--CoV-2 RNA Genome Sequencing Consortium | Arindam Maitra, Vijayshri Deotale, Rahul Narang, Deepashri Maraskolhe, Saumitra Das |
| EPI_ISL_511930, EPI_ISL_511931, EPI_ISL_511932, EPI_ISL_511933, EPI_ISL_511934, EPI_ISL_511935, EPI_ISL_511936, EPI_ISL_511937, EPI_ISL_511938, EPI_ISL_511939, EPI_ISL_511940, EPI_ISL_511941 | | | |
| see above | Government Medical College | National Institute of Biomedical Genomics - DBT's PAN-INDIA 1000 SARS--CoV-2 RNA Genome Sequencing Consortium | Arindam Maitra, Jyoti Iravane, Dhaval Khatri, Maitrik Dave, Saumitra Das |
| EPI_ISL_511942, EPI_ISL_511943, EPI_ISL_511944 | Translational Health Science and Technology Institute | National Institute of Biomedical Genomics - DBT's PAN-INDIA 1000 SARS--CoV-2 RNA Genome Sequencing Consortium | Arindam Maitra, Guruprasad Medigeshi, Sharanabasava Patil, Anbalagan Ananthraj, Madhu Pareek, Imran Khan, Gagandeep Kang, Saumitra Das |
| EPI_ISL_511945, EPI_ISL_511946, EPI_ISL_511947, EPI_ISL_511948, EPI_ISL_511949, EPI_ISL_511950, EPI_ISL_511951, EPI_ISL_511952, EPI_ISL_511953, EPI_ISL_511954 | Indian Institute of Science | National Institute of Biomedical Genomics - DBT's PAN-INDIA 1000 SARS--CoV-2 RNA Genome Sequencing Consortium | Arindam Maitra, Bharath K Sundararaj, Harsha Raheja, N. Srinivasan, Deepak K Saini, Amit Singh, Saumitra Das |
| EPI_ISL_512073 | Saikrishna Hospital, Mehsana | Gujarat Biotechnology Research Centre | Apurvasinh Puvar, Janvi Raval, Zarna Patel, Monika Gandhi, Pinal Trivedi, Maharshi Pandya, Nidhi Patel, Nitin Savaliya, Raghawendra Kumar, Dinesh Kumar, Zuber Saiyed, Komal Patel, Labdhi Pandya, Afzal Ansari, Nikha Trivedi, Harshadbhai Parmar, R D Dixit, A M Kadri, Harsh Bakshi, Chaitanya Joshi, Madhvi Joshi |
| EPI_ISL_512336, EPI_ISL_512337, EPI_ISL_512338, EPI_ISL_512339, EPI_ISL_512340, EPI_ISL_512341 | University of Exeter | COVID-19 Genomics UK (COG-UK) Consortium | Ben Temperton,Aaron Jeffries,Michelle Michelsen,Joanna Warwick-Dugdale,Audrey Farbos,Robyn Manley,Stephen Michell,Jane Masoli |
| EPI_ISL_512384, EPI_ISL_512385, EPI_ISL_512386, EPI_ISL_512387, EPI_ISL_512388, EPI_ISL_512392, EPI_ISL_512393, EPI_ISL_512394, EPI_ISL_512397, EPI_ISL_512398, EPI_ISL_512399, EPI_ISL_512401, EPI_ISL_512402, EPI_ISL_512403, EPI_ISL_512404, EPI_ISL_512405, EPI_ISL_512406, EPI_ISL_512408, EPI_ISL_512413, EPI_ISL_512414, EPI_ISL_512418, EPI_ISL_512419, EPI_ISL_512420, EPI_ISL_512421, EPI_ISL_512422, EPI_ISL_512423, EPI_ISL_512424, EPI_ISL_512425, EPI_ISL_512426, EPI_ISL_512427, EPI_ISL_512428, EPI_ISL_512429, EPI_ISL_512430, EPI_ISL_512431, EPI_ISL_512432, EPI_ISL_512433, EPI_ISL_512434, EPI_ISL_512435 | | | |
| see above | Centre for Enzyme Innovation, University of Portsmouth / Translational Research Laboratory, Portsmouth Hospitals NHS Trust | COVID-19 Genomics UK (COG-UK) Consortium | Angela Beckett,Yann Bourgeois,Garry Scarlett,Sharon Glaysher,Scott Elliott,Kelly Bicknell,Robert Impey,Allyson Lloyd,Sarah Wyllie,Ethan Butcher,Anoop Chauhan,Samuel Robson |
| EPI_ISL_514228, EPI_ISL_514229, EPI_ISL_514230 | National Institute of Laboratory Medicine and Referral Center | Genomic Research Lab, BCSIR | Barna Goswami, Abu Sayeed Mohammad Mahmud, Mohammad Samir Uzzaman, Eshrar Osman, Md. Ahashan Habib, Shahina Akter, Tanjina Akhter Banu, Md. Murshed Hasan Sarkar, Iffat Jahan, Md. Saddam Hossain, Tasnim Nafisa, Md. Maruf Ahmed Molla, Mahmuda Yeasmin, Asish Kumar Ghosh, A. K. M. Shamsuzzaman, Sheikh Md. Selim Al Din, Utpal Chandra Ray, Salek Ahmed Sajib, Md. Salim Khan |
| EPI_ISL_514455, EPI_ISL_514456, EPI_ISL_514464, EPI_ISL_514470, EPI_ISL_514471, EPI_ISL_514481, EPI_ISL_514487, EPI_ISL_514488, EPI_ISL_514489, EPI_ISL_514491, EPI_ISL_514492, EPI_ISL_514493, EPI_ISL_514494, EPI_ISL_514495, EPI_ISL_514496, EPI_ISL_514497, EPI_ISL_514498, EPI_ISL_514499, EPI_ISL_514500, EPI_ISL_514501, EPI_ISL_514502, EPI_ISL_514503, EPI_ISL_514504, EPI_ISL_514506, EPI_ISL_514507, EPI_ISL_514509, EPI_ISL_514511, EPI_ISL_514512, EPI_ISL_514513 | | | |
| see above | Centre for Enzyme Innovation, University of Portsmouth / Translational Research Laboratory, Portsmouth Hospitals NHS Trust | COVID-19 Genomics UK (COG-UK) Consortium | Angela Beckett,Yann Bourgeois,Garry Scarlett,Sharon Glaysher,Scott Elliott,Kelly Bicknell,Robert Impey,Allyson Lloyd,Sarah Wyllie,Ethan Butcher,Anoop Chauhan,Samuel Robson |
| EPI_ISL_514592 | B.J. Medical College and Civil hospital, Ahmedabad | Gujarat Biotechnology Research Centre | Apurvasinh Puvar, Janvi Raval, Zarna Patel, Monika Gandhi, Pinal Trivedi, Maharshi Pandya, Nidhi Patel, Nitin Savaliya, Raghawendra Kumar, Dinesh Kumar, Zuber Saiyed, Komal Patel, Labdhi Pandya, Afzal Ansari, Nikha Trivedi, Pranay Shah, Kamlesh J Upadhyay, Sanjay Kapadia, R D Dixit, A M Kadri, Harsh Bakshi, Chaitanya Joshi, Madhvi Joshi |
| EPI_ISL_514750 | Pirogov Russian National Research Medical University | Pirogov Russian National Research Medical University, Research and Development | Blagodatskikh, K.A. |
| EPI_ISL_515181, EPI_ISL_515182, EPI_ISL_515183, EPI_ISL_515184, EPI_ISL_515247, EPI_ISL_515248, EPI_ISL_515249 | Kumasi Centre for Collaborative Research in Tropical Medicine, Kumasi. | Institute of Virology, Charité - Universitätsmedizin Berlin | Augustina Sylverken, Philip El-Duah, Michael Owusu, Julia Schneider, Richmond Yeboah, Richmond Gorman, Eric Adu, Sherihane Aryeetey, Jesse Addo Asamoah,Jörn Beheim-Schwarzbach, Victor Max Corman, Christian Drosten, Richard Phillips. |
| EPI_ISL_516079, EPI_ISL_516080, EPI_ISL_516081, EPI_ISL_516082, EPI_ISL_516083, EPI_ISL_516084, EPI_ISL_516085, EPI_ISL_516086, EPI_ISL_516087, EPI_ISL_516088 | Biomedical Sciences and Public Health, Polytechnic University of Marche | Biomedical Sciences and Public Health, Polytechnic University of Marche | Bagnarelli,P., Caucci,S., Di Sante,L., Menzo,S., Alessandrini,F., Onofri,V., Turchi,C., Melchionda,F., Tagliabracci,A. |
| EPI_ISL_516194, EPI_ISL_516195, EPI_ISL_516196, EPI_ISL_516197, EPI_ISL_516198, EPI_ISL_516199, EPI_ISL_516200, EPI_ISL_516201, EPI_ISL_516202, EPI_ISL_516203, EPI_ISL_516204, EPI_ISL_516205, EPI_ISL_516206, EPI_ISL_516207, EPI_ISL_516208, EPI_ISL_516209, EPI_ISL_516210, EPI_ISL_516211, EPI_ISL_516212, EPI_ISL_516213, EPI_ISL_516214, EPI_ISL_516215, EPI_ISL_516216, EPI_ISL_516217, EPI_ISL_516218, EPI_ISL_516219, EPI_ISL_516220, EPI_ISL_516221, EPI_ISL_516222, EPI_ISL_516223, EPI_ISL_516224, EPI_ISL_516225, EPI_ISL_516226, EPI_ISL_516227, EPI_ISL_516228, EPI_ISL_516229, EPI_ISL_516230, EPI_ISL_516231, EPI_ISL_516232, EPI_ISL_516233, EPI_ISL_516234, EPI_ISL_516235, EPI_ISL_516236, EPI_ISL_516237, EPI_ISL_516238, EPI_ISL_516239, EPI_ISL_516240, EPI_ISL_516241, EPI_ISL_516242, EPI_ISL_516243, EPI_ISL_516244, EPI_ISL_516245, EPI_ISL_516246, EPI_ISL_516247, EPI_ISL_516248, EPI_ISL_516249, EPI_ISL_516250, EPI_ISL_516251, EPI_ISL_516252, EPI_ISL_516253, EPI_ISL_516254, EPI_ISL_516255, EPI_ISL_516256, EPI_ISL_516257, EPI_ISL_516258, EPI_ISL_516259, EPI_ISL_516260, EPI_ISL_516261, EPI_ISL_516262, EPI_ISL_516263, EPI_ISL_516264, EPI_ISL_516265, EPI_ISL_516266, EPI_ISL_516267, EPI_ISL_516268, EPI_ISL_516269, EPI_ISL_516270, EPI_ISL_516271, EPI_ISL_516272, EPI_ISL_516273, EPI_ISL_516274, EPI_ISL_516275, EPI_ISL_516276, EPI_ISL_516277, EPI_ISL_516278, EPI_ISL_516279, EPI_ISL_516280, EPI_ISL_516281, EPI_ISL_516282, EPI_ISL_516283, EPI_ISL_516284, EPI_ISL_516285, EPI_ISL_516286, EPI_ISL_516287, EPI_ISL_516288, EPI_ISL_516289, EPI_ISL_516290, EPI_ISL_516291, EPI_ISL_516292, EPI_ISL_516293, EPI_ISL_516294, EPI_ISL_516295, EPI_ISL_516296, EPI_ISL_516297, EPI_ISL_516298, EPI_ISL_516299, EPI_ISL_516300, EPI_ISL_516301, EPI_ISL_516302, EPI_ISL_516303, EPI_ISL_516304, EPI_ISL_516305, EPI_ISL_516306, EPI_ISL_516307, EPI_ISL_516308, EPI_ISL_516309, EPI_ISL_516310, EPI_ISL_516311, EPI_ISL_516312, EPI_ISL_516313, EPI_ISL_516314, EPI_ISL_516315, EPI_ISL_516316, EPI_ISL_516317, EPI_ISL_516318, EPI_ISL_516319, EPI_ISL_516320, EPI_ISL_516321, EPI_ISL_516322, EPI_ISL_516323, EPI_ISL_516324, EPI_ISL_516325, EPI_ISL_516326, EPI_ISL_516327, EPI_ISL_516328, EPI_ISL_516329, EPI_ISL_516330, EPI_ISL_516331, EPI_ISL_516332, EPI_ISL_516333, EPI_ISL_516334, EPI_ISL_516335, EPI_ISL_516336, EPI_ISL_516337, EPI_ISL_516338, EPI_ISL_516339, EPI_ISL_516340, EPI_ISL_516341, EPI_ISL_516342, EPI_ISL_516343, EPI_ISL_516344, EPI_ISL_516345, EPI_ISL_516346, EPI_ISL_516347, EPI_ISL_516348, EPI_ISL_516349, EPI_ISL_516350, EPI_ISL_516351, EPI_ISL_516352, EPI_ISL_516353, EPI_ISL_516354, EPI_ISL_516355, EPI_ISL_516356, EPI_ISL_516357, EPI_ISL_516358, EPI_ISL_516359, EPI_ISL_516360, EPI_ISL_516361, EPI_ISL_516362, EPI_ISL_516363, EPI_ISL_516364, EPI_ISL_516365, EPI_ISL_516366, EPI_ISL_516367, EPI_ISL_516368, EPI_ISL_516369, EPI_ISL_516370, EPI_ISL_516371, EPI_ISL_516372, EPI_ISL_516373, EPI_ISL_516374, EPI_ISL_516375, EPI_ISL_516376, EPI_ISL_516377, EPI_ISL_516378, EPI_ISL_516379, EPI_ISL_516380, EPI_ISL_516381, EPI_ISL_516382, EPI_ISL_516383, EPI_ISL_516384, EPI_ISL_516385, EPI_ISL_516386, EPI_ISL_516387, EPI_ISL_516388, EPI_ISL_516389, EPI_ISL_516390, EPI_ISL_516391, EPI_ISL_516392, EPI_ISL_516393, EPI_ISL_516394, EPI_ISL_516395, EPI_ISL_516396, EPI_ISL_516397, EPI_ISL_516398, EPI_ISL_516399, EPI_ISL_516400, EPI_ISL_516401, EPI_ISL_516402, EPI_ISL_516403, EPI_ISL_516404, EPI_ISL_516405, EPI_ISL_516406, EPI_ISL_516407, EPI_ISL_516408, EPI_ISL_516409, EPI_ISL_516410, EPI_ISL_516411, EPI_ISL_516412 | | | |
| see above | Michigan Department of Health and Human Services, Bureau of Laboratories | Michigan Department of Health and Human Services, Bureau of Laboratories | Blankenship HM, Riner D, Soehnlen MK |
| EPI_ISL_517510, EPI_ISL_517511, EPI_ISL_517514, EPI_ISL_517515, EPI_ISL_517516, EPI_ISL_517518, EPI_ISL_517520, EPI_ISL_517521, EPI_ISL_517523, EPI_ISL_517524, EPI_ISL_517525, EPI_ISL_517527, EPI_ISL_517528, EPI_ISL_517530, EPI_ISL_517533, EPI_ISL_517534, EPI_ISL_517538, EPI_ISL_517539, EPI_ISL_517541 | | | |
| see above | Centre for Enzyme Innovation, University of Portsmouth / Translational Research Laboratory, Portsmouth Hospitals NHS Trust | COVID-19 Genomics UK (COG-UK) Consortium | Angela Beckett,Yann Bourgeois,Garry Scarlett,Sharon Glaysher,Scott Elliott,Kelly Bicknell,Robert Impey,Allyson Lloyd,Sarah Wyllie,Ethan Butcher,Anoop Chauhan,Samuel Robson |
| EPI_ISL_517616, EPI_ISL_517621, EPI_ISL_517622, EPI_ISL_517623, EPI_ISL_517624, EPI_ISL_517625, EPI_ISL_517626, EPI_ISL_517627, EPI_ISL_517628, EPI_ISL_517629, EPI_ISL_517630, EPI_ISL_517631, EPI_ISL_517632, EPI_ISL_517633, EPI_ISL_517634, EPI_ISL_517635, EPI_ISL_517636, EPI_ISL_517637, EPI_ISL_517638, EPI_ISL_517639, EPI_ISL_517640, EPI_ISL_517641, EPI_ISL_517642, EPI_ISL_517643, EPI_ISL_517644, EPI_ISL_517645, EPI_ISL_517646, EPI_ISL_517647, EPI_ISL_517648, EPI_ISL_517649, EPI_ISL_517650, EPI_ISL_517651, EPI_ISL_517652, EPI_ISL_517653, EPI_ISL_517654, EPI_ISL_517655, EPI_ISL_517656, EPI_ISL_517657, EPI_ISL_517658, EPI_ISL_517659, EPI_ISL_517660, EPI_ISL_517661, EPI_ISL_517662 | | | |

| | | | |
|---|---|---|---|
| see above | Academic Hospital Paramaribo | Erasmus Medical Center | Bas Oude Munnink, Dion Gajadin, Ed Ijzerman, Emmanuelle Munger, Gary Gummels, Ingrid Krishnadath, Lycke Woittiez, Marion Koopmans, Mireille Van de Veer, Princes Wongsowidjojo, Radjesh Ori, Rohma Banwari, Stephen Vreden |
| EPI_ISL_518031, EPI_ISL_518032 | Indian Institute of Science | National Institute of Biomedical Genomics - DBT's PAN-INDIA 1000 SARS--CoV-2 RNA Genome Sequencing Consortium | Arindam Maitra, Bharath K Sundararaj, Harsha Raheja, N. Srinivasan, Deepak K Saini, Amit Singh, Saumitra Das |
| EPI_ISL_518799, EPI_ISL_518800, EPI_ISL_518801, EPI_ISL_518802, EPI_ISL_518803, EPI_ISL_518804, EPI_ISL_518805, EPI_ISL_518806, EPI_ISL_518807, EPI_ISL_518808, EPI_ISL_518809, EPI_ISL_518810, EPI_ISL_518811, EPI_ISL_518812, EPI_ISL_518813, EPI_ISL_518814, EPI_ISL_518815, EPI_ISL_518816, EPI_ISL_518817, EPI_ISL_518818 | | | |
| see above | Academic Hospital Paramaribo | Erasmus Medical Center | Bas Oude Munnink, Dion Gajadin, Ed Ijzerman, Emmanuelle Munger, Gary Gummels, Ingrid Krishnadath, Lycke Woittiez, Marion Koopmans, Mireille Van de Veer, Princes Wongsowidjojo, Radjesh Ori, Rohma Banwari, Stephen Vreden |
| EPI_ISL_518819 | Qadr Hospital, Tangerang, Banten | Biosafety Level-3 Laboratory, Indonesian Institute of Sciences (LIPI) | Anik Budhi Dharmayanthi, Syam Budi Iryanto, Andri Wardiana, Anggia Prasetyoputri, Isa Nuryana, Ade Andriani, Ario Betha Juanssilfero, Asep M Ridwanuloh, Ahmad Fathoni, Rifki Sadikin, Ratih Asmana Ningrum, Wien Kusharyoto, Puspita Lisdiyanti |
| EPI_ISL_518820 | Klinik Apotek Dein, Jakarta, Indonesia | Biosafety Level-3 Laboratory, Indonesian Institute of Sciences (LIPI) | Anik Budhi Dharmayanthi, Syam Budi Iryanto, Andri Wardiana, Anggia Prasetyoputri, Isa Nuryana, Ade Andriani, Ario Betha Juanssilfero, Asep M Ridwanuloh, Ahmad Fathoni, Rifki Sadikin, Ratih Asmana Ningrum, Wien Kusharyoto, Puspita Lisdiyanti |
| EPI_ISL_522437 | Mahatma Gandhi Institute of Medical Sciences | National Institute of Biomedical Genomics - DBT's PAN-INDIA 1000 SARS--CoV-2 RNA Genome Sequencing Consortium | Arindam Maitra, Vijayshri Deotale, Rahul Narang, Deepashri Maraskolhe, Saumitra Das |
| EPI_ISL_523121, EPI_ISL_523122, EPI_ISL_523123, EPI_ISL_523124, EPI_ISL_523125, EPI_ISL_523126, EPI_ISL_523127, EPI_ISL_523128, EPI_ISL_523129, EPI_ISL_523130, EPI_ISL_523131, EPI_ISL_523132, EPI_ISL_523133, EPI_ISL_523134, EPI_ISL_523135, EPI_ISL_523136, EPI_ISL_523137, EPI_ISL_523138, EPI_ISL_523139, EPI_ISL_523140, EPI_ISL_523141, EPI_ISL_523142, EPI_ISL_523143, EPI_ISL_523144, EPI_ISL_523145, EPI_ISL_523146, EPI_ISL_523147, EPI_ISL_523148, EPI_ISL_523149, EPI_ISL_523150, EPI_ISL_523151, EPI_ISL_523152, EPI_ISL_523153, EPI_ISL_523154, EPI_ISL_523155, EPI_ISL_523156, EPI_ISL_523157, EPI_ISL_523158, EPI_ISL_523159, EPI_ISL_523160, EPI_ISL_523161, EPI_ISL_523162, EPI_ISL_523163, EPI_ISL_523164, EPI_ISL_523165, EPI_ISL_523166, EPI_ISL_523167, EPI_ISL_523168, EPI_ISL_523169, EPI_ISL_523170, EPI_ISL_523171, EPI_ISL_523172, EPI_ISL_523173, EPI_ISL_523174, EPI_ISL_523175, EPI_ISL_523176, EPI_ISL_523177, EPI_ISL_523178, EPI_ISL_523179, EPI_ISL_523180, EPI_ISL_523181, EPI_ISL_523182, EPI_ISL_523183, EPI_ISL_523184, EPI_ISL_523185, EPI_ISL_523186, EPI_ISL_523187, EPI_ISL_523188, EPI_ISL_523189, EPI_ISL_523190, EPI_ISL_523191, EPI_ISL_523192, EPI_ISL_523193, EPI_ISL_523194, EPI_ISL_523195, EPI_ISL_523196, EPI_ISL_523197, EPI_ISL_523198, EPI_ISL_523199, EPI_ISL_523200, EPI_ISL_523201, EPI_ISL_523202, EPI_ISL_523203, EPI_ISL_523204, EPI_ISL_523205, EPI_ISL_523206, EPI_ISL_523207, EPI_ISL_523208, EPI_ISL_523209, EPI_ISL_523210, EPI_ISL_523211, EPI_ISL_523212, EPI_ISL_523235, EPI_ISL_523236, EPI_ISL_523237, EPI_ISL_523238, EPI_ISL_523239, EPI_ISL_523240, EPI_ISL_523241, EPI_ISL_523242, EPI_ISL_523243, EPI_ISL_523244, EPI_ISL_523245, EPI_ISL_523246, EPI_ISL_523247, EPI_ISL_523248, EPI_ISL_523249, EPI_ISL_523250, EPI_ISL_523251, EPI_ISL_523252, EPI_ISL_523253, EPI_ISL_523254, EPI_ISL_523255, EPI_ISL_523256, EPI_ISL_523257, EPI_ISL_523258, EPI_ISL_523259, EPI_ISL_523260, EPI_ISL_523261, EPI_ISL_523262, EPI_ISL_523263, EPI_ISL_523264, EPI_ISL_523265, EPI_ISL_523266, EPI_ISL_523267, EPI_ISL_523268, EPI_ISL_523269, EPI_ISL_523270, EPI_ISL_523271, EPI_ISL_523272, EPI_ISL_523273, EPI_ISL_523275, EPI_ISL_523276, EPI_ISL_523277, EPI_ISL_523278, EPI_ISL_523279, EPI_ISL_523280, EPI_ISL_523281, EPI_ISL_523282, EPI_ISL_523283, EPI_ISL_523284, EPI_ISL_523285, EPI_ISL_523286, EPI_ISL_523287, EPI_ISL_523288, EPI_ISL_523289, EPI_ISL_523290, EPI_ISL_523291, EPI_ISL_523301, EPI_ISL_523302, EPI_ISL_523303, EPI_ISL_523304, EPI_ISL_523305, EPI_ISL_523306, EPI_ISL_523307, EPI_ISL_523308, EPI_ISL_523309, EPI_ISL_523332, EPI_ISL_523334, EPI_ISL_523335, EPI_ISL_523336, EPI_ISL_523337, EPI_ISL_523338, EPI_ISL_523339, EPI_ISL_523340, EPI_ISL_523341, EPI_ISL_523342, EPI_ISL_523343, EPI_ISL_523344, EPI_ISL_523345, EPI_ISL_523346, EPI_ISL_523347, EPI_ISL_523348, EPI_ISL_523349, EPI_ISL_523350, EPI_ISL_523351, EPI_ISL_523352, EPI_ISL_523353, EPI_ISL_523354, EPI_ISL_523355, EPI_ISL_523356, EPI_ISL_523357, EPI_ISL_523358, EPI_ISL_523359, EPI_ISL_523360, EPI_ISL_523361, EPI_ISL_523362, EPI_ISL_523363, EPI_ISL_523364, EPI_ISL_523365, EPI_ISL_523366, EPI_ISL_523367, EPI_ISL_523368, EPI_ISL_523369, EPI_ISL_523370, EPI_ISL_523371, EPI_ISL_523372, EPI_ISL_523373, EPI_ISL_523374, EPI_ISL_523375, EPI_ISL_523376, EPI_ISL_523377, EPI_ISL_523378, EPI_ISL_523379, EPI_ISL_523380, EPI_ISL_523381, EPI_ISL_523382, EPI_ISL_523383, EPI_ISL_523384, EPI_ISL_523385, EPI_ISL_523401, EPI_ISL_523402, EPI_ISL_523403, EPI_ISL_523404, EPI_ISL_523405, EPI_ISL_523406, EPI_ISL_523407, EPI_ISL_523408, EPI_ISL_523409, EPI_ISL_523410, EPI_ISL_523411, EPI_ISL_523412, EPI_ISL_523413, EPI_ISL_523414, EPI_ISL_523415, EPI_ISL_523416, EPI_ISL_523417, EPI_ISL_523418, EPI_ISL_523419, EPI_ISL_523420, EPI_ISL_523421, EPI_ISL_523422, EPI_ISL_523423, EPI_ISL_523424, EPI_ISL_523425, EPI_ISL_523426, EPI_ISL_523427, EPI_ISL_523428, EPI_ISL_523429, EPI_ISL_523430, EPI_ISL_523431, EPI_ISL_523432, EPI_ISL_523433, EPI_ISL_523434, EPI_ISL_523435, EPI_ISL_523436, EPI_ISL_523437, EPI_ISL_523438, EPI_ISL_523439, EPI_ISL_523440, EPI_ISL_523441, EPI_ISL_523442, EPI_ISL_523443, EPI_ISL_523444, EPI_ISL_523445, EPI_ISL_523446, EPI_ISL_523447, EPI_ISL_523448, EPI_ISL_523449, EPI_ISL_523450, EPI_ISL_523451, EPI_ISL_523452, EPI_ISL_523453, EPI_ISL_523454, EPI_ISL_523455, EPI_ISL_523456, EPI_ISL_523457, EPI_ISL_523458, EPI_ISL_523459, EPI_ISL_523460, EPI_ISL_523461, EPI_ISL_523462, EPI_ISL_523463, EPI_ISL_523464, EPI_ISL_523465, EPI_ISL_523466, EPI_ISL_523467, EPI_ISL_523468, EPI_ISL_523469, EPI_ISL_523470, EPI_ISL_523471, EPI_ISL_523472, EPI_ISL_523473, EPI_ISL_523474, EPI_ISL_523475, EPI_ISL_523476, EPI_ISL_523477, EPI_ISL_523478, EPI_ISL_523479, EPI_ISL_523480, EPI_ISL_523481, EPI_ISL_523482, EPI_ISL_523483, EPI_ISL_523484, EPI_ISL_523485, EPI_ISL_523486, EPI_ISL_523487, EPI_ISL_523488, EPI_ISL_523489, EPI_ISL_523490, EPI_ISL_523491, EPI_ISL_523492, EPI_ISL_523493, EPI_ISL_523494, EPI_ISL_523495, EPI_ISL_523496, EPI_ISL_523497, EPI_ISL_523498, EPI_ISL_523499, EPI_ISL_523501, EPI_ISL_523502, EPI_ISL_523503, EPI_ISL_523504, EPI_ISL_523505, EPI_ISL_523506, EPI_ISL_523507, EPI_ISL_523508, EPI_ISL_523509, EPI_ISL_523510, EPI_ISL_523511, EPI_ISL_523512, EPI_ISL_523513, EPI_ISL_523514, EPI_ISL_523515, EPI_ISL_523516, EPI_ISL_523517, EPI_ISL_523522, EPI_ISL_523523, EPI_ISL_523524, EPI_ISL_523525, EPI_ISL_523526, EPI_ISL_523528, EPI_ISL_523529, EPI_ISL_523530, EPI_ISL_523531, EPI_ISL_523532, EPI_ISL_523533, EPI_ISL_523534, EPI_ISL_523535, EPI_ISL_523536, EPI_ISL_523537, EPI_ISL_523538, EPI_ISL_523539, EPI_ISL_523540, EPI_ISL_523541, EPI_ISL_523542, EPI_ISL_523543, EPI_ISL_523544, EPI_ISL_523545, EPI_ISL_523546, EPI_ISL_523547, EPI_ISL_523548, EPI_ISL_523549, EPI_ISL_523550, EPI_ISL_523551, EPI_ISL_523552, EPI_ISL_523553, EPI_ISL_523554, EPI_ISL_523555, EPI_ISL_523556, EPI_ISL_523557, EPI_ISL_523558, EPI_ISL_523568, EPI_ISL_523569, EPI_ISL_523570, EPI_ISL_523571, EPI_ISL_523572, EPI_ISL_523573, EPI_ISL_523574, EPI_ISL_523575, EPI_ISL_523576, EPI_ISL_523577, EPI_ISL_523578, EPI_ISL_523595, EPI_ISL_523596, EPI_ISL_523597, EPI_ISL_523598, EPI_ISL_523599, EPI_ISL_523600, EPI_ISL_523601, EPI_ISL_523602, EPI_ISL_523603, EPI_ISL_523604, EPI_ISL_523605, EPI_ISL_523606, EPI_ISL_523607, EPI_ISL_523608, EPI_ISL_523609, EPI_ISL_523610, EPI_ISL_523611, EPI_ISL_523613, EPI_ISL_523614, EPI_ISL_523615, EPI_ISL_523616, EPI_ISL_523617, EPI_ISL_523618, EPI_ISL_523619, EPI_ISL_523620, EPI_ISL_523621, EPI_ISL_523622, EPI_ISL_523623, EPI_ISL_523624, EPI_ISL_523625, EPI_ISL_523626, EPI_ISL_523627, EPI_ISL_523628, EPI_ISL_523629, EPI_ISL_523630, EPI_ISL_523631, EPI_ISL_523632, EPI_ISL_523633, EPI_ISL_523634, EPI_ISL_523635, EPI_ISL_523636, EPI_ISL_523637, EPI_ISL_523638, EPI_ISL_523639, EPI_ISL_523640, EPI_ISL_523641, EPI_ISL_523642, EPI_ISL_523643, EPI_ISL_523644, EPI_ISL_523645, EPI_ISL_523646, EPI_ISL_523647, EPI_ISL_523648, EPI_ISL_523649, EPI_ISL_523650, EPI_ISL_523651, EPI_ISL_523652, EPI_ISL_523653, EPI_ISL_523654, EPI_ISL_523655, EPI_ISL_523656, EPI_ISL_523657, EPI_ISL_523659, EPI_ISL_523660, EPI_ISL_523661, EPI_ISL_523662, EPI_ISL_523663, EPI_ISL_523664, EPI_ISL_523665, EPI_ISL_523666, EPI_ISL_523667, EPI_ISL_523668, EPI_ISL_523669, EPI_ISL_523670, EPI_ISL_523671, EPI_ISL_523672, EPI_ISL_523673, EPI_ISL_523674, EPI_ISL_523675, EPI_ISL_523676, EPI_ISL_523677, EPI_ISL_523678, EPI_ISL_523679, EPI_ISL_523680, EPI_ISL_523681, EPI_ISL_523682, EPI_ISL_523683, EPI_ISL_523684, EPI_ISL_523685, EPI_ISL_523686, EPI_ISL_523687, EPI_ISL_523688, EPI_ISL_523689, EPI_ISL_523690, EPI_ISL_523691, EPI_ISL_523692, EPI_ISL_523693, EPI_ISL_523694, EPI_ISL_523695, EPI_ISL_523696, EPI_ISL_523697, EPI_ISL_523698, EPI_ISL_523699, EPI_ISL_523700, EPI_ISL_523701, EPI_ISL_523702, EPI_ISL_523703, EPI_ISL_523704, EPI_ISL_523705, EPI_ISL_523706, EPI_ISL_523707, EPI_ISL_523708, EPI_ISL_523709, EPI_ISL_523710, EPI_ISL_523711, EPI_ISL_523712, EPI_ISL_523713, EPI_ISL_523714, EPI_ISL_523715, EPI_ISL_523716, EPI_ISL_523717, EPI_ISL_523718, EPI_ISL_523719, EPI_ISL_523720, EPI_ISL_523721, EPI_ISL_523722, EPI_ISL_523723, EPI_ISL_523724, EPI_ISL_523725, EPI_ISL_523726, EPI_ISL_523727, EPI_ISL_523728, EPI_ISL_523729, EPI_ISL_523730, EPI_ISL_523731, EPI_ISL_523732, EPI_ISL_523733, EPI_ISL_523734, EPI_ISL_523735, EPI_ISL_523736, EPI_ISL_523737, EPI_ISL_523738, EPI_ISL_523739, EPI_ISL_523740, EPI_ISL_523741, EPI_ISL_523742, EPI_ISL_523743, EPI_ISL_523744, EPI_ISL_523745, EPI_ISL_523746, EPI_ISL_523747, EPI_ISL_523748, EPI_ISL_523749, EPI_ISL_523750, EPI_ISL_523751, EPI_ISL_523752, EPI_ISL_523753, EPI_ISL_523754, EPI_ISL_523755, EPI_ISL_523756, EPI_ISL_523757, EPI_ISL_523758, EPI_ISL_523759, EPI_ISL_523760, EPI_ISL_523761, EPI_ISL_523762, EPI_ISL_523763, EPI_ISL_523764, EPI_ISL_523765, EPI_ISL_523766, EPI_ISL_523767, EPI_ISL_523768, EPI_ISL_523769, EPI_ISL_523770, EPI_ISL_523771, EPI_ISL_523772, EPI_ISL_523773, EPI_ISL_523774, EPI_ISL_523775, EPI_ISL_523776, EPI_ISL_523777, EPI_ISL_523778, EPI_ISL_523779, EPI_ISL_523780, EPI_ISL_523781, EPI_ISL_523782, EPI_ISL_523783, EPI_ISL_523784, EPI_ISL_523785, EPI_ISL_523786, EPI_ISL_523787, EPI_ISL_523788, EPI_ISL_523789, EPI_ISL_523790, EPI_ISL_523791, EPI_ISL_523792, EPI_ISL_523793, EPI_ISL_523794, EPI_ISL_523795, EPI_ISL_523796, EPI_ISL_523797, EPI_ISL_523798, EPI_ISL_523799, EPI_ISL_523800, EPI_ISL_523801, EPI_ISL_523802, EPI_ISL_523803, EPI_ISL_523804, EPI_ISL_523805, EPI_ISL_523806, EPI_ISL_523807, EPI_ISL_523808, EPI_ISL_523809 | | | |
| see above | Dutch COVID-19 response team | Erasmus Medical Center | Bas Oude Munnink, David Nieuwenhuijse, Reina Sikkema, Claudia Schapendonk, Irina Chestakova, Anne van der Linden, Theo Bestebroer, Stefan van Nieuwkoop, Mark Pronk, Pascal Lexmond, Corien Swaan, Manon Haverkate, Madelief Mollers, Mart Stein, Sandra Kengne Kamga Mobou, Jeroen van Kampen, Jolanda Voermans, Aura Timen, Corine GeurtsvanKessel, Annemiek van der Eijk, Richard Molenkamp, Marion Koopmans, on behalf of the Dutch national COVID-19 response team. |
| EPI_ISL_524428, EPI_ISL_524429, EPI_ISL_524430, EPI_ISL_524431, EPI_ISL_524432 | Dirk Dittmer | Dirk Dittmer | Bailey,A.G., Caro-Vegas,C.P., Dittmer,D., Eason,A.B., Juarez,A., Landis,J.T., McNamara,R.P., Miller,M.B., Moorad,R., Pluta,L.J.,Seltzer,T.A., Thompson,C., Vahrson,W. and Villamor,F. |
| EPI_ISL_524713 | B.J. Medical College and Civil hospital, Ahmedabad | Gujarat Biotechnology Research Centre | Apurvasinh Puvar, Janvi Raval, Zarna Patel, Monika Gandhi, Pinal Trivedi, Maharshi Pandya, Nidhi Patel, Nitin Savaliya, Raghawendra Kumar, Dinesh Kumar, Zuber Saiyed, Komal Patel, Labdhi Pandya, Afzal Ansari, Nikha Trivedi, Pranay Shah, Kamlesh J Upadhyay, Sanjay Kapadia, R D Dixit, A M Kadri, Harsh Bakshi, Chaitanya Joshi, Madhvi Joshi |
| EPI_ISL_524734 | GMERS Medical College and Hospital, Dharpur, Patan | Gujarat Biotechnology Research Centre | Apurvasinh Puvar, Janvi Raval, Zarna Patel, Monika Gandhi, Pinal Trivedi, Maharshi Pandya, Nidhi Patel, Nitin Savaliya, Raghawendra Kumar, Dinesh Kumar, Zuber Saiyed, Komal Patel, Labdhi Pandya, Afzal Ansari, Nikha Trivedi, A N Parmar, R D Dixit, A M Kadri, Harsh Bakshi, Chaitanya Joshi, Madhvi Joshi |
| EPI_ISL_524746 | Sir M P Shah Government Medical College, Jamnagar | Gujarat Biotechnology Research Centre | Binita Aring, Afzal Ansari, Nikha Trivedi, Apurvasinh Puvar, Janvi Raval, Zarna Patel, Monika Gandhi, Pinal Trivedi, Maharshi Pandya, Nidhi Patel, Nitin Savaliya, Raghawendra Kumar, Dinesh Kumar, Zuber Saiyed, Komal Patel, Labdhi Pandya, R D Dixit, A M Kadri, Harsh Bakshi, Chaitanya Joshi, Madhvi Joshi |
| EPI_ISL_524749 | Sir M P Shah Government Medical College, Jamnagar | Gujarat Biotechnology Research Centre | Apurvasinh Puvar, Janvi Raval, Zarna Patel, Monika Gandhi, Pinal Trivedi, Maharshi Pandya, Nidhi Patel, Nitin Savaliya, Raghawendra Kumar, Dinesh Kumar, Zuber Saiyed, Komal Patel, Labdhi Pandya, Afzal Ansari, Nikha Trivedi, Binita Aring, R D Dixit, A M Kadri, Harsh Bakshi, Chaitanya Joshi, Madhvi Joshi |
| EPI_ISL_524765 | GMERS Medical College and Hospital, Dharpur, Patan | Gujarat Biotechnology Research Centre | Apurvasinh Puvar, Janvi Raval, Zarna Patel, Monika Gandhi, Pinal Trivedi, Maharshi Pandya, Nidhi Patel, Nitin Savaliya, Raghawendra Kumar, Dinesh Kumar, Zuber Saiyed, Komal Patel, Labdhi Pandya, Afzal Ansari, Nikha Trivedi, A N Parmar, R D Dixit, A M Kadri, Harsh Bakshi, Chaitanya Joshi, Madhvi Joshi |
| EPI_ISL_525536, EPI_ISL_525537, EPI_ISL_525538 | CNR Virus des Infections Respiratoires - France SUD | CNR Virus des Infections Respiratoires - France SUD | Antonin Bal, Gregory Destras, Gwendolyne Burfin, Solenne Brun, Alexandre Gaymard, Maude Bouscambert-Duchamp, Florence Morfin-Sherpa, Martine Valette, Bruno Lina, Laurence Josset |
| EPI_ISL_525539 | Centre Hospitalier de Bourg en Bresse | CNR Virus des Infections Respiratoires - France SUD | Antonin Bal, Gregory Destras, Gwendolyne Burfin, Solenne Brun, Alexandre Gaymard, Maude Bouscambert-Duchamp, Florence Morfin-Sherpa, Martine Valette, Bruno Lina, Laurence Josset |
| EPI_ISL_525540, EPI_ISL_525541, EPI_ISL_525542, EPI_ISL_525543 | CNR Virus des Infections Respiratoires - France SUD | CNR Virus des Infections Respiratoires - France SUD | Antonin Bal, Gregory Destras, Gwendolyne Burfin, Solenne Brun, Alexandre Gaymard, Maude Bouscambert-Duchamp, Florence Morfin-Sherpa, Martine Valette, Bruno Lina, Laurence Josset |
| EPI_ISL_526433, EPI_ISL_526436 | Centre for Enzyme Innovation, University of Portsmouth / Translational Research Laboratory, Portsmouth Hospitals NHS Trust | COVID-19 Genomics UK (COG-UK) Consortium | Angela Beckett,Yann Bourgeois,Garry Scarlett,Sharon Glaysher,Scott Elliott,Kelly Bicknell,Robert Impey,Allyson Lloyd,Sarah Wyllie,Ethan Butcher,Anoop Chauhan,Samuel Robson |

| | | | |
|---|---|---|---|
| EPI_ISL_527809, EPI_ISL_527810, EPI_ISL_527811 | Institute of Microbiology, Universidad San Francisco de Quito | Institute of Microbiology, Universidad San Francisco de Quito | Belén Prado-Vivar, Sully Márquez, Juan José Guadalupe, Monica Becerra-Wong, Bernardo Gutiérrez, Stephanie Arregui, Rene Bracho, Karina Barragan, Anita Garcia, Carlos Tobar, Verónica Barragán, Patricio Rojas-Silva, Gabriel Trueba, Michelle Grunauer, Paúl Cárdenas |
| EPI_ISL_527812, EPI_ISL_527813, EPI_ISL_527814, EPI_ISL_527815, EPI_ISL_527816, EPI_ISL_527817 | Institute of Microbiology, Universidad San Francisco de Quito | Institute of Microbiology, Universidad San Francisco de Quito | Belén Prado-Vivar, Sully Márquez, Juan José Guadalupe, Monica Becerra-Wong, Bernardo Gutiérrez, Khurram Mahbbob, Verónica Barragán, Patricio Rojas-Silva, Gabriel Trueba, Michelle Grunauer, Paúl Cárdenas |
| EPI_ISL_527822, EPI_ISL_527823, EPI_ISL_527824 | Texas Department of State Health Services | Texas Department of State Health Services | Bonnie Oh, Rashmi Tuladhar, Jenny Zhang, Maliha Rahman, Anita Pokharel, Myong Koag, Chun Wang, Rachel Lee, Grace Kubin |
| EPI_ISL_527861 | Hospital e Maternidade Celso Pierro | Instituto Adolfo Lutz, Interdiciplinary Procedures Center, Strategic Laboratory | Av. Dr. Arnaldo, 355 - Brazil, Cerqueira Cesar, São Paulo - SP, 01246-1301 |
| EPI_ISL_527866 | PS Municipal Dr Lauro Ribas Braga | Instituto Adolfo Lutz, Interdiciplinary Procedures Center, Strategic Laboratory | Av. Dr. Arnaldo, 355 - Brazil, Cerqueira Cesar, São Paulo - SP, 01246-1301 |
| EPI_ISL_528428, EPI_ISL_528429 | National Genomics Core-Center for DNA Fingerprinting and Diagnostics | National Genomics Core- Center for DNA Fingerprinting and Diagnostics (NGC-CDFD)- DBT's PAN-INDIA-1000 Genome consortium | Ashwin Dalal, Bala Pratyusha, Heena Shah, G Shashikanth, Vinay Donipadi, Neeraj Kumar, Niteen Pathak, Pradipta Hore, Rahul Baroi, Sayantan Goswami, Shaffiqu T S, Shalini Aricthota, Sobhan Babu, R Harinarayanan, Rashna Bhandari, Murali Dharan Bashyam, Debashish Mitra, Divya Vashisht |
| EPI_ISL_528596, EPI_ISL_528597, EPI_ISL_528598, EPI_ISL_528599, EPI_ISL_528600, EPI_ISL_528601, EPI_ISL_528602 | National Genomics Core-Center for DNA Fingerprinting and Diagnostics | National Genomics Core- Center for DNA Fingerprinting and Diagnostics (NGC-CDFD)- DBT's PAN-INDIA-1000 Genome consortium | Bala Pratyusha, Heena Shah, G Shashikanth, Vinay Donipadi, Edurugatla Dinesh, Guru Raja, Hilal Ahmad Reshi, J. Mallikarjun, K. Viswakalyan, Kaisar Ahmad Lone,Kausika Kumar Malik, N. Sudheer, R Harinarayanan, Rashna Bhandari, Murali Dharan Bashyam, Debashish Mitra, Divya Vashisht, Ashwin Dalal |
| EPI_ISL_528603, EPI_ISL_528604 | National Genomics Core-Center for DNA Fingerprinting and Diagnostics | National Genomics Core- Center for DNA Fingerprinting and Diagnostics (NGC-CDFD)- DBT's PAN-INDIA-1000 Genome consortium | Ashwin Dalal, Bala Pratyusha, Heena Shah, G Shashikanth, Vinay Donipadi, Neeraj Kumar, Niteen Pathak, Pradipta Hore, Rahul Baroi, Sayantan Goswami, Shaffiqu T S, Shalini Aricthota, Sobhan Babu, R Harinarayanan, Rashna Bhandari, Murali Dharan Bashyam, Debashish Mitra, Divya Vashisht |
| EPI_ISL_528605, EPI_ISL_528606, EPI_ISL_528607, EPI_ISL_528608 | National Genomics Core-Center for DNA Fingerprinting and Diagnostics | National Genomics Core- Center for DNA Fingerprinting and Diagnostics (NGC-CDFD)- DBT's PAN-INDIA-1000 Genome consortium | Ashwin Dalal, Bala Pratyusha, Heena Shah, G Shashikanth, Vinay Donipadi, K.Manohar, Madhumohan Rao,Neeraj Kumar, Niteen Pathak, Pradipta Hore, Rahul Baroi, Sayantan Goswami, Shaffiqu T S, Shalini Aricthota, Sobhan Babu, R Harinarayanan, Rashna Bhandari, Murali Dharan Bashyam, Debashish Mitra, Divya Vashisht |
| EPI_ISL_528748 | Dinkes Provinsi Jawa Barat | School of Life Sciences and Technology & School of Pharmacy-Institut Teknologi Bandung; Molecular Genetics Laboratory-Faculty of Medicine-Universitas Padjadjaran; Laboratorium Kesehatan Provinsi Jawa Barat | Azzania Fibriani, Catur Riani, Marselina Irasonia Tan, Yunia Sribudiani, Husna Nugrahapraja, Tarwadi, Ema Rahmawati, Savira Ekawardhani, Hesti Lina Wiraswati, Ryan Bayusantika Ristandi, Rifky Waluyajati Rachman, Cut Nur Cinthia Alamanda, Lia Faridah, Gusti Ayu Prani Pradani, Adelina Khristiani Rahayu, Hammam Riza, Sony Solistia Wirawan, Agung Eru Wibowo, Irvan Faizal |
| EPI_ISL_528753 | Dinkes Kota Bogor | School of Life Sciences and Technology & School of Pharmacy-Institut Teknologi Bandung; Molecular Genetics Laboratory-Faculty of Medicine-Universitas Padjadjaran; Laboratorium Kesehatan Provinsi Jawa Barat | Azzania Fibriani, Catur Riani, Marselina Irasonia Tan, Yunia Sribudiani, Husna Nugrahapraja, Tarwadi, Ema Rahmawati, Savira Ekawardhani, Hesti Lina Wiraswati, Ryan Bayusantika Ristandi, Rifky Waluyajati Rachman, Cut Nur Cinthia Alamanda, Lia Faridah, Davin H. E. Setiamarga, Rizki Mardian , Hammam Riza, Sony Solistia Wirawan, Agung Eru Wibowo, Irvan Faizal |
| EPI_ISL_529154 | Department of Immunology, The Scripps Research Institute | Andersen lab at Scripps Research | Basler, T., Shephard, J., Austin, B. with SEARCH Alliance San Diego |
| EPI_ISL_529169 | Michigan Department of Health and Human Services, Bureau of Laboratories | Michigan Department of Health and Human Services, Bureau of Laboratories | Blankenship,H.M., Riner,D., Soehnlen,M. |
| EPI_ISL_529272, EPI_ISL_529303, EPI_ISL_529341, EPI_ISL_529342, EPI_ISL_529343, EPI_ISL_529344, EPI_ISL_529345, EPI_ISL_529346, EPI_ISL_529347, EPI_ISL_529348, EPI_ISL_529349, EPI_ISL_529350, EPI_ISL_529351, EPI_ISL_529352, EPI_ISL_529373, EPI_ISL_529374, EPI_ISL_529375, EPI_ISL_529376, EPI_ISL_529377, EPI_ISL_529431, EPI_ISL_529432, EPI_ISL_529465, EPI_ISL_529481, EPI_ISL_529525, EPI_ISL_529670 | | | |
| see above | Centre for Enzyme Innovation, University of Portsmouth / Translational Research Laboratory, Portsmouth Hospitals NHS Trust | COVID-19 Genomics UK (COG-UK) Consortium | Angela Beckett,Yann Bourgeois,Garry Scarlett,Sharon Glaysher,Scott Elliott,Kelly Bicknell,Robert Impey,Allyson Lloyd,Sarah Wyllie,Ethan Butcher,Anoop Chauhan,Samuel Robson |
| EPI_ISL_529804, EPI_ISL_529805, EPI_ISL_529806, EPI_ISL_529807, EPI_ISL_529808, EPI_ISL_529809, EPI_ISL_529810, EPI_ISL_529811, EPI_ISL_529812, EPI_ISL_529813, EPI_ISL_529814, EPI_ISL_529815, EPI_ISL_529816, EPI_ISL_529817, EPI_ISL_529818, EPI_ISL_529819, EPI_ISL_529820, EPI_ISL_529821, EPI_ISL_529822, EPI_ISL_529823, EPI_ISL_529824, EPI_ISL_529825, EPI_ISL_529826, EPI_ISL_529827, EPI_ISL_529828, EPI_ISL_529829, EPI_ISL_529830, EPI_ISL_529831, EPI_ISL_529832, EPI_ISL_529833, EPI_ISL_529834, EPI_ISL_529835, EPI_ISL_529836, EPI_ISL_529837, EPI_ISL_529838, EPI_ISL_529839, EPI_ISL_529840, EPI_ISL_529841, EPI_ISL_529842, EPI_ISL_529843, EPI_ISL_529844, EPI_ISL_529845, EPI_ISL_529846, EPI_ISL_529847, EPI_ISL_529848, EPI_ISL_529849, EPI_ISL_529850, EPI_ISL_529851, EPI_ISL_529852, EPI_ISL_529853, EPI_ISL_529854, EPI_ISL_529855, EPI_ISL_529856, EPI_ISL_529857, EPI_ISL_529858, EPI_ISL_529859, EPI_ISL_529860, EPI_ISL_529861, EPI_ISL_529862, EPI_ISL_529863, EPI_ISL_529864, EPI_ISL_529865, EPI_ISL_529866, EPI_ISL_529867, EPI_ISL_529868, EPI_ISL_529869, EPI_ISL_529870, EPI_ISL_529871, EPI_ISL_529872, EPI_ISL_529873, EPI_ISL_529874, EPI_ISL_529875, EPI_ISL_529876, EPI_ISL_529877, EPI_ISL_529878, EPI_ISL_529879, EPI_ISL_529880, EPI_ISL_529881, EPI_ISL_529882, EPI_ISL_529883, EPI_ISL_529884, EPI_ISL_529885, EPI_ISL_529886, EPI_ISL_529887, EPI_ISL_529888, EPI_ISL_529889, EPI_ISL_529890, EPI_ISL_529891, EPI_ISL_529892, EPI_ISL_529893, EPI_ISL_529894, EPI_ISL_529895, EPI_ISL_529896, EPI_ISL_529897, EPI_ISL_529898, EPI_ISL_529899, EPI_ISL_529900, EPI_ISL_529901, EPI_ISL_529902, EPI_ISL_529903, EPI_ISL_529904, EPI_ISL_529905, EPI_ISL_529906, EPI_ISL_529907 | | | |
| see above | Michigan Department of Health and Human Services, Bureau of Laboratories | Michigan Department of Health and Human Services, Bureau of Laboratories | Blankenship HM, Riner D, Soehnlen MK |
| EPI_ISL_534224 | Universitetssjukhuset i Linkoping | The Public Health Agency of Sweden | Anna-Malin Linde, Maria Lind Karlberg, Mattias Haukland, Reza Advani, Olov Svartstrom, Oskar Karlsson Lindsjo, Sandra Broddesson, Petra Edquist, Mia Brytting, Anna Risberg, Karin Tegmark-Wisell |
| EPI_ISL_534225, EPI_ISL_534226, EPI_ISL_534227, EPI_ISL_534228, EPI_ISL_534229 | Skanes universitetssjukhus Lund | The Public Health Agency of Sweden | Anna-Malin Linde, Maria Lind Karlberg, Mattias Haukland, Reza Advani, Olov Svartstrom, Oskar Karlsson Lindsjo, Sandra Broddesson, Petra Edquist, Mia Brytting, Anna Risberg, Karin Tegmark-Wisell |
| EPI_ISL_534230, EPI_ISL_534231, EPI_ISL_534232, EPI_ISL_534233 | Capio S:t Gorans sjukhus | The Public Health Agency of Sweden | Anna-Malin Linde, Maria Lind Karlberg, Mattias Haukland, Reza Advani, Olov Svartstrom, Oskar Karlsson Lindsjo, Sandra Broddesson, Petra Edquist, Mia Brytting, Anna Risberg, Karin Tegmark-Wisell |
| EPI_ISL_534234, EPI_ISL_534235, EPI_ISL_534236 | Karolinska universitetslaboratoriet SOLNA | The Public Health Agency of Sweden | Anna-Malin Linde, Maria Lind Karlberg, Mattias Haukland, Reza Advani, Olov Svartstrom, Oskar Karlsson Lindsjo, Sandra Broddesson, Petra Edquist, Mia Brytting, Anna Risberg, Karin Tegmark-Wisell |
| EPI_ISL_534237, EPI_ISL_534238 | Lanssjukhuset Kalmar | The Public Health Agency of Sweden | Anna-Malin Linde, Maria Lind Karlberg, Mattias Haukland, Reza Advani, Olov Svartstrom, Oskar Karlsson Lindsjo, Sandra Broddesson, Petra Edquist, Mia Brytting, Anna Risberg, Karin Tegmark-Wisell |
| EPI_ISL_534239, EPI_ISL_534240, EPI_ISL_534241, EPI_ISL_534242, EPI_ISL_534243 | Norra Alvsborgs lanssjukhus | The Public Health Agency of Sweden | Anna-Malin Linde, Maria Lind Karlberg, Mattias Haukland, Reza Advani, Olov Svartstrom, Oskar Karlsson Lindsjo, Sandra Broddesson, Petra Edquist, Mia Brytting, Anna Risberg, Karin Tegmark-Wisell |
| EPI_ISL_534244 | Sundsvalls sjukhus | The Public Health Agency of Sweden | Anna-Malin Linde, Maria Lind Karlberg, Mattias Haukland, Reza Advani, Olov Svartstrom, Oskar Karlsson Lindsjo, Sandra Broddesson, Petra Edquist, Mia Brytting, Anna Risberg, Karin Tegmark-Wisell |
| EPI_ISL_534245 | Kliniskt mikrobiologiska laboratoriet | The Public Health Agency of Sweden | Anna-Malin Linde, Maria Lind Karlberg, Mattias Haukland, Reza Advani, Olov Svartstrom, Oskar Karlsson Lindsjo, Sandra Broddesson, Petra Edquist, Mia Brytting, Anna Risberg, Karin Tegmark-Wisell |
| EPI_ISL_534246, EPI_ISL_534247 | Universitetssjukhuset i Linkoping | The Public Health Agency of Sweden | Anna-Malin Linde, Maria Lind Karlberg, Mattias Haukland, Reza Advani, Olov Svartstrom, Oskar Karlsson Lindsjo, Sandra Broddesson, Petra Edquist, Mia Brytting, Anna Risberg, Karin Tegmark-Wisell |
| EPI_ISL_534248 | Laboratoriemedicin Vasternorrland | The Public Health Agency of Sweden | Anna-Malin Linde, Maria Lind Karlberg, Mattias Haukland, Reza Advani, Olov Svartstrom, Oskar Karlsson Lindsjo, Sandra Broddesson, Petra Edquist, Mia Brytting, Anna Risberg, Karin Tegmark-Wisell |
| EPI_ISL_534249, EPI_ISL_534250, EPI_ISL_534251 | Gavle Sjukhus | The Public Health Agency of Sweden | Anna-Malin Linde, Maria Lind Karlberg, Mattias Haukland, Reza Advani, Olov Svartstrom, Oskar Karlsson Lindsjo, Sandra Broddesson, Petra Edquist, Mia Brytting, Anna Risberg, Karin Tegmark-Wisell |
| EPI_ISL_534252, EPI_ISL_534253, | Laboratoriemedicin Vasternorrland | The Public Health Agency of Sweden | Anna-Malin Linde, Maria Lind Karlberg, Mattias Haukland, Reza Advani, Olov Svartstrom, Oskar Karlsson Lindsjo, Sandra Broddesson, Petra Edquist, Mia |

| | | | |
|---|---|---|---|
| EPI_ISL_534254, EPI_ISL_534255 | | | Brytting, Anna Risberg, Karin Tegmark-Wisell |
| EPI_ISL_534256 | Klinisk mikrobiologi, Laboratoriemedicin Gavleborg | The Public Health Agency of Sweden | Anna-Malin Linde, Maria Lind Karlberg, Mattias Haukland, Reza Advani, Olov Svartstrom, Oskar Karlsson Lindsjo, Sandra Broddesson, Petra Edquist, Mia Brytting, Anna Risberg, Karin Tegmark-Wisell |
| EPI_ISL_534257, EPI_ISL_534258 | Laboratoriemedicin Vasternorrland | The Public Health Agency of Sweden | Anna-Malin Linde, Maria Lind Karlberg, Mattias Haukland, Reza Advani, Olov Svartstrom, Oskar Karlsson Lindsjo, Sandra Broddesson, Petra Edquist, Mia Brytting, Anna Risberg, Karin Tegmark-Wisell |
| EPI_ISL_534259 | Karolinska universitetslaboratoriet | The Public Health Agency of Sweden | Anna-Malin Linde, Maria Lind Karlberg, Mattias Haukland, Reza Advani, Olov Svartstrom, Oskar Karlsson Lindsjo, Sandra Broddesson, Petra Edquist, Mia Brytting, Anna Risberg, Karin Tegmark-Wisell |
| EPI_ISL_535030, EPI_ISL_535033, EPI_ISL_535034, EPI_ISL_535035, EPI_ISL_535037, EPI_ISL_535039, EPI_ISL_535040, EPI_ISL_535041, EPI_ISL_535042 | Centre for Enzyme Innovation, University of Portsmouth / Translational Research Laboratory, Portsmouth Hospitals NHS Trust | COVID-19 Genomics UK (COG-UK) Consortium | Angela Beckett,Yann Bourgeois,Garry Scarlett,Sharon Glaysher,Scott Elliott,Kelly Bicknell,Robert Impey,Allyson Lloyd,Sarah Wyllie,Ethan Butcher,Anoop Chauhan,Samuel Robson |
| EPI_ISL_537380, EPI_ISL_537381 | Complejo Hospitalario Universitario de Vigo | SeqCOVID-SPAIN consortium/IBV(CSIC) | Benito Regueiro and SeqCOVID-SPAIN consortium |
| EPI_ISL_537729, EPI_ISL_537730, EPI_ISL_537731, EPI_ISL_537732, EPI_ISL_537733, EPI_ISL_537734, EPI_ISL_537735, EPI_ISL_537736, EPI_ISL_537737, EPI_ISL_537738, EPI_ISL_537739, EPI_ISL_537740, EPI_ISL_537741, EPI_ISL_537742, EPI_ISL_537743, EPI_ISL_537744, EPI_ISL_537745, EPI_ISL_537746, EPI_ISL_537747, EPI_ISL_537748, EPI_ISL_537749, EPI_ISL_537750, EPI_ISL_537751, EPI_ISL_537752, EPI_ISL_537753, EPI_ISL_537754, EPI_ISL_537755, EPI_ISL_537756, EPI_ISL_537757, EPI_ISL_537758, EPI_ISL_537759, EPI_ISL_537760, EPI_ISL_537761, EPI_ISL_537762, EPI_ISL_537763, EPI_ISL_537764, EPI_ISL_537765, EPI_ISL_537766, EPI_ISL_537767, EPI_ISL_537768, EPI_ISL_537769, EPI_ISL_537770, EPI_ISL_537771, EPI_ISL_537772, EPI_ISL_537773, EPI_ISL_537774, EPI_ISL_537775, EPI_ISL_537776, EPI_ISL_537777, EPI_ISL_537778, EPI_ISL_537779, EPI_ISL_537780, EPI_ISL_537781, EPI_ISL_537782, EPI_ISL_537783, EPI_ISL_537784, EPI_ISL_537785, EPI_ISL_537786, EPI_ISL_537787 | | | |
| see above | Servicio de Microbiología, Hospital Miguel Servet, Zaragoza | SeqCOVID-SPAIN consortium/IBV(CSIC) | Antonio Rezusta López, Alexander Tristancho Baró, Ana Milagro, Yolanda Gracia Grataloup, Nieves Martínez Cameo and SeqCOVID-SPAIN consortium |
| EPI_ISL_538302 | Texas Department of State Health Services | Texas Department of State Health Services | Bonnie Oh, Rashmi Tuladhar, Jenny Zhang, Maliha Rahman, Anita Pokharel, Myong Koag, Chun Wang, Rachel Lee, Grace Kubin |
| EPI_ISL_539789, EPI_ISL_539790, EPI_ISL_539791, EPI_ISL_539792, EPI_ISL_539793 | Institute of Microbiology, Universidad San Francisco de Quito | Institute of Microbiology, Universidad San Francisco de Quito | Belén Prado-Vivar, Sully Márquez, Juan José Guadalupe, Monica Becerra-Wong, Bernardo Gutiérrez, Ligia Briceño, Nabih Dahik, Verónica Barragán, Patricio Rojas-Silva, Gabriel Trueba, Michelle Grunauer, Paúl Cárdenas |
| EPI_ISL_539881 | Kungsbacka Narakut | The Public Health Agency of Sweden | Anna-Malin Linde, Maria Lind Karlberg, Oskar Karlsson Lindsjo, Olov Svartstrom, Mattias Haukland, Reza Advani, Sandra Broddesson, Anna Risberg, Theresa Enkirch, Mia Brytting, Karin Tegmark-Wisell |
| EPI_ISL_539882 | Narhalsan Olskroken VC | The Public Health Agency of Sweden | Anna-Malin Linde, Maria Lind Karlberg, Oskar Karlsson Lindsjo, Olov Svartstrom, Mattias Haukland, Reza Advani, Sandra Broddesson, Anna Risberg, Theresa Enkirch, Mia Brytting, Karin Tegmark-Wisell |
| EPI_ISL_539883, EPI_ISL_539884 | Omtanken Grimmered | The Public Health Agency of Sweden | Anna-Malin Linde, Maria Lind Karlberg, Oskar Karlsson Lindsjo, Olov Svartstrom, Mattias Haukland, Reza Advani, Sandra Broddesson, Anna Risberg, Theresa Enkirch, Mia Brytting, Karin Tegmark-Wisell |
| EPI_ISL_539885 | Narhalsan Fjallbacka VC | The Public Health Agency of Sweden | Anna-Malin Linde, Maria Lind Karlberg, Oskar Karlsson Lindsjo, Olov Svartstrom, Mattias Haukland, Reza Advani, Sandra Broddesson, Anna Risberg, Theresa Enkirch, Mia Brytting, Karin Tegmark-Wisell |
| EPI_ISL_539886 | Barnakuten | The Public Health Agency of Sweden | Anna-Malin Linde, Maria Lind Karlberg, Oskar Karlsson Lindsjo, Olov Svartstrom, Mattias Haukland, Reza Advani, Sandra Broddesson, Anna Risberg, Theresa Enkirch, Mia Brytting, Karin Tegmark-Wisell |
| EPI_ISL_540579, EPI_ISL_540582, EPI_ISL_540583 | University of Exeter | COVID-19 Genomics UK (COG-UK) Consortium | Ben Temperton,Aaron Jeffries,Michelle Michelsen,Joanna Warwick-Dugdale,Audrey Farbos,Robyn Manley,Stephen Michell,Jane Masoli |
| EPI_ISL_541008, EPI_ISL_541009, EPI_ISL_541010, EPI_ISL_541011, EPI_ISL_541012, EPI_ISL_541013, EPI_ISL_541014, EPI_ISL_541015, EPI_ISL_541016, EPI_ISL_541017, EPI_ISL_541018, EPI_ISL_541019, EPI_ISL_541020, EPI_ISL_541021, EPI_ISL_541022, EPI_ISL_541023, EPI_ISL_541024, EPI_ISL_541025, EPI_ISL_541026, EPI_ISL_541027, EPI_ISL_541028, EPI_ISL_541029, EPI_ISL_541030, EPI_ISL_541031, EPI_ISL_541032, EPI_ISL_541033, EPI_ISL_541034, EPI_ISL_541035, EPI_ISL_541036 | | | |
| see above | Servicio de Microbiología, Hospital Miguel Servet, Zaragoza | SeqCOVID-SPAIN consortium/Institute of Biomedicine of Valencia, IBV-CSIC | Antonio Rezusta López, Alexander Tristancho Baró, Ana Milagro, Yolanda Gracia Grataloup, Nieves Martínez Cameo and SeqCOVID-SPAIN consortium |
| EPI_ISL_548245, EPI_ISL_548246 | Skovde/Unilabs | The Public Health Agency of Sweden | Anna-Malin Linde, Maria Lind Karlberg, Mattias Haukland, Reza Advani, Olov Svartstrom, Oskar Karlsson Lindsjo, Sandra Broddesson, Petra Edquist, Mia Brytting, Anna Risberg, Karin Tegmark-Wisell |
| EPI_ISL_548247 | Halmstad klinisk mikrobiologi | The Public Health Agency of Sweden | Anna-Malin Linde, Maria Lind Karlberg, Mattias Haukland, Reza Advani, Olov Svartstrom, Oskar Karlsson Lindsjo, Sandra Broddesson, Petra Edquist, Mia Brytting, Anna Risberg, Karin Tegmark-Wisell |
| EPI_ISL_548248 | Ostersund klinisk mikrobiologi | The Public Health Agency of Sweden | Anna-Malin Linde, Maria Lind Karlberg, Mattias Haukland, Reza Advani, Olov Svartstrom, Oskar Karlsson Lindsjo, Sandra Broddesson, Petra Edquist, Mia Brytting, Anna Risberg, Karin Tegmark-Wisell |
| EPI_ISL_548249 | Halmstad klinisk mikrobiologi | The Public Health Agency of Sweden | Anna-Malin Linde, Maria Lind Karlberg, Mattias Haukland, Reza Advani, Olov Svartstrom, Oskar Karlsson Lindsjo, Sandra Broddesson, Petra Edquist, Mia Brytting, Anna Risberg, Karin Tegmark-Wisell |
| EPI_ISL_548250, EPI_ISL_548251 | Klinisk mikrobiologi NAL Trollhattan | The Public Health Agency of Sweden | Anna-Malin Linde, Maria Lind Karlberg, Mattias Haukland, Reza Advani, Olov Svartstrom, Oskar Karlsson Lindsjo, Sandra Broddesson, Petra Edquist, Mia Brytting, Anna Risberg, Karin Tegmark-Wisell |
| EPI_ISL_548252 | Klinisk mikrobiologi centralsjukhuset Karlstad | The Public Health Agency of Sweden | Anna-Malin Linde, Maria Lind Karlberg, Mattias Haukland, Reza Advani, Olov Svartstrom, Oskar Karlsson Lindsjo, Sandra Broddesson, Petra Edquist, Mia Brytting, Anna Risberg, Karin Tegmark-Wisell |
| EPI_ISL_548253, EPI_ISL_548254 | Klinisk mikrobiologi NAL Trollhattan | The Public Health Agency of Sweden | Anna-Malin Linde, Maria Lind Karlberg, Mattias Haukland, Reza Advani, Olov Svartstrom, Oskar Karlsson Lindsjo, Sandra Broddesson, Petra Edquist, Mia Brytting, Anna Risberg, Karin Tegmark-Wisell |
| EPI_ISL_548255 | Lanssjukhuset Kalmar | The Public Health Agency of Sweden | Anna-Malin Linde, Maria Lind Karlberg, Mattias Haukland, Reza Advani, Olov Svartstrom, Oskar Karlsson Lindsjo, Sandra Broddesson, Petra Edquist, Mia Brytting, Anna Risberg, Karin Tegmark-Wisell |
| EPI_ISL_548256 | Centralsjukhuset | The Public Health Agency of Sweden | Anna-Malin Linde, Maria Lind Karlberg, Mattias Haukland, Reza Advani, Olov Svartstrom, Oskar Karlsson Lindsjo, Sandra Broddesson, Petra Edquist, Mia Brytting, Anna Risberg, Karin Tegmark-Wisell |
| EPI_ISL_548257, EPI_ISL_548258 | Karolinska universitetslaboratoriet | The Public Health Agency of Sweden | Anna-Malin Linde, Maria Lind Karlberg, Mattias Haukland, Reza Advani, Olov Svartstrom, Oskar Karlsson Lindsjo, Sandra Broddesson, Petra Edquist, Mia Brytting, Anna Risberg, Karin Tegmark-Wisell |
| EPI_ISL_549341, EPI_ISL_549343, EPI_ISL_549360, EPI_ISL_549363, EPI_ISL_549364, EPI_ISL_549366, EPI_ISL_549460, EPI_ISL_549461, EPI_ISL_549462, EPI_ISL_549463, EPI_ISL_549464, EPI_ISL_549465, EPI_ISL_549466, EPI_ISL_549467, EPI_ISL_549468, EPI_ISL_549469, EPI_ISL_549470, EPI_ISL_549471, EPI_ISL_549473, EPI_ISL_549474, EPI_ISL_549475, EPI_ISL_549476, EPI_ISL_559983 | | | |
| see above | Centre for Enzyme Innovation, University of Portsmouth / Translational Research Laboratory, Portsmouth Hospitals NHS Trust | COVID-19 Genomics UK (COG-UK) Consortium | Angela Beckett,Yann Bourgeois,Garry Scarlett,Sharon Glaysher,Scott Elliott,Kelly Bicknell,Robert Impey,Allyson Lloyd,Sarah Wyllie,Ethan Butcher,Anoop Chauhan,Samuel Robson |
| EPI_ISL_560407 | Istituto Zooprofilattico Sperimentale del Mezzogiorno | INMI Lazzaro Spallanzani IRCCS | Barbara Bartolini, Cesare E.M. Gruber, Martina Rueca, Francesco Messina, Antonino Di Caro, Giovanna Fusco, Maurizio Viscardi, Giorgia Borriello, Sergio Brandi, Maria R. Capobianchi |
| EPI_ISL_560972 | Orebro klinisk mikrobiologi | The Public Health Agency of Sweden | Anna-Malin Linde, Maria Lind Karlberg, Mattias Haukland, Reza Advani, Olov Svartstrom, Oskar Karlsson Lindsjo, Sandra Broddesson, Petra Edquist, Mia Brytting, Anna Risberg, Karin Tegmark-Wisell |
| EPI_ISL_560973 | Klinisk mikrobiologi NAL Trollhattan | The Public Health Agency of Sweden | Anna-Malin Linde, Maria Lind Karlberg, Mattias Haukland, Reza Advani, Olov Svartstrom, Oskar Karlsson Lindsjo, Sandra Broddesson, Petra Edquist, Mia Brytting, Anna Risberg, Karin Tegmark-Wisell |
| EPI_ISL_560974 | Karolinska Universitetslaboratoriet | The Public Health Agency of Sweden | Anna-Malin Linde, Maria Lind Karlberg, Mattias Haukland, Reza Advani, Olov Svartstrom, Oskar Karlsson Lindsjo, Sandra Broddesson, Petra Edquist, Mia Brytting, Anna Risberg, Karin Tegmark-Wisell |

| | | | |
|---|---|---|---|
| EPI_ISL_560975 | Klinisk mikrobiologi centralsjukhuset Karlstad | The Public Health Agency of Sweden | Anna-Malin Linde, Maria Lind Karlberg, Mattias Haukland, Reza Advani, Olov Svartstrom, Oskar Karlsson Lindsjo, Sandra Broddesson, Petra Edquist, Mia Brytting, Anna Risberg, Karin Tegmark-Wisell |
| EPI_ISL_560976 | Klinsisk mikrobiologi Linkoping | The Public Health Agency of Sweden | Anna-Malin Linde, Maria Lind Karlberg, Mattias Haukland, Reza Advani, Olov Svartstrom, Oskar Karlsson Lindsjo, Sandra Broddesson, Petra Edquist, Mia Brytting, Anna Risberg, Karin Tegmark-Wisell |
| EPI_ISL_560977, EPI_ISL_560978, EPI_ISL_560979 | Universitetssjukhuset i Linkoping | The Public Health Agency of Sweden | Anna-Malin Linde, Maria Lind Karlberg, Mattias Haukland, Reza Advani, Olov Svartstrom, Oskar Karlsson Lindsjo, Sandra Broddesson, Petra Edquist, Mia Brytting, Anna Risberg, Karin Tegmark-Wisell |
| EPI_ISL_560980 | Skanes universitetssjukhus Lund | The Public Health Agency of Sweden | Anna-Malin Linde, Maria Lind Karlberg, Mattias Haukland, Reza Advani, Olov Svartstrom, Oskar Karlsson Lindsjo, Sandra Broddesson, Petra Edquist, Mia Brytting, Anna Risberg, Karin Tegmark-Wisell |
| EPI_ISL_560981 | Capio S:t Gorans sjukhus | The Public Health Agency of Sweden | Anna-Malin Linde, Maria Lind Karlberg, Mattias Haukland, Reza Advani, Olov Svartstrom, Oskar Karlsson Lindsjo, Sandra Broddesson, Petra Edquist, Mia Brytting, Anna Risberg, Karin Tegmark-Wisell |
| EPI_ISL_560982, EPI_ISL_560983, EPI_ISL_560984, EPI_ISL_560985 | Karolinska universitetslaboratoriet SOLNA | The Public Health Agency of Sweden | Anna-Malin Linde, Maria Lind Karlberg, Mattias Haukland, Reza Advani, Olov Svartstrom, Oskar Karlsson Lindsjo, Sandra Broddesson, Petra Edquist, Mia Brytting, Anna Risberg, Karin Tegmark-Wisell |
| EPI_ISL_560986 | Sundsvalls sjukhus | The Public Health Agency of Sweden | Anna-Malin Linde, Maria Lind Karlberg, Mattias Haukland, Reza Advani, Olov Svartstrom, Oskar Karlsson Lindsjo, Sandra Broddesson, Petra Edquist, Mia Brytting, Anna Risberg, Karin Tegmark-Wisell |
| EPI_ISL_560987 | Kliniskt mikrobiologiska laboratoriet | The Public Health Agency of Sweden | Anna-Malin Linde, Maria Lind Karlberg, Mattias Haukland, Reza Advani, Olov Svartstrom, Oskar Karlsson Lindsjo, Sandra Broddesson, Petra Edquist, Mia Brytting, Anna Risberg, Karin Tegmark-Wisell |
| EPI_ISL_560988 | Klinisk mikrobiologi, Laboratoriemedicin Gavleborg | The Public Health Agency of Sweden | Anna-Malin Linde, Maria Lind Karlberg, Mattias Haukland, Reza Advani, Olov Svartstrom, Oskar Karlsson Lindsjo, Sandra Broddesson, Petra Edquist, Mia Brytting, Anna Risberg, Karin Tegmark-Wisell |
| EPI_ISL_560989, EPI_ISL_560990 | Karolinska universitetslaboratoriet | The Public Health Agency of Sweden | Anna-Malin Linde, Maria Lind Karlberg, Mattias Haukland, Reza Advani, Olov Svartstrom, Oskar Karlsson Lindsjo, Sandra Broddesson, Petra Edquist, Mia Brytting, Anna Risberg, Karin Tegmark-Wisell |
| EPI_ISL_565833, EPI_ISL_565834, EPI_ISL_565835, EPI_ISL_565836, EPI_ISL_565837, EPI_ISL_565838, EPI_ISL_565839, EPI_ISL_565840, EPI_ISL_565841, EPI_ISL_565842, EPI_ISL_565843, EPI_ISL_565844, EPI_ISL_565845, EPI_ISL_565846, EPI_ISL_565847, EPI_ISL_565848, EPI_ISL_565849, EPI_ISL_565850, EPI_ISL_565851, EPI_ISL_565852, EPI_ISL_565853, EPI_ISL_565854, EPI_ISL_565855, EPI_ISL_565856, EPI_ISL_565857, EPI_ISL_565858, EPI_ISL_565859, EPI_ISL_565860, EPI_ISL_565861, EPI_ISL_565862, EPI_ISL_565863, EPI_ISL_565864, EPI_ISL_565865, EPI_ISL_565866, EPI_ISL_565867, EPI_ISL_565868, EPI_ISL_565869, EPI_ISL_565870, EPI_ISL_565871, EPI_ISL_565872, EPI_ISL_565873, EPI_ISL_565874, EPI_ISL_565875, EPI_ISL_565876, EPI_ISL_565877, EPI_ISL_565878, EPI_ISL_565879, EPI_ISL_565880, EPI_ISL_565881, EPI_ISL_565882, EPI_ISL_565883, EPI_ISL_565884, EPI_ISL_565885, EPI_ISL_565886, EPI_ISL_565887, EPI_ISL_565888, EPI_ISL_565889, EPI_ISL_565890, EPI_ISL_565891, EPI_ISL_565892, EPI_ISL_565893, EPI_ISL_565894, EPI_ISL_565895, EPI_ISL_565896, EPI_ISL_565897, EPI_ISL_565898, EPI_ISL_565899, EPI_ISL_565900, EPI_ISL_565901, EPI_ISL_565902, EPI_ISL_565903, EPI_ISL_565904, EPI_ISL_565905, EPI_ISL_565906, EPI_ISL_565907, EPI_ISL_565926, EPI_ISL_565927, EPI_ISL_565928, EPI_ISL_565929, EPI_ISL_565930, EPI_ISL_565931, EPI_ISL_565932, EPI_ISL_565933, EPI_ISL_565934, EPI_ISL_565935, EPI_ISL_565936, EPI_ISL_565937, EPI_ISL_565938, EPI_ISL_565939, EPI_ISL_565940, EPI_ISL_565941, EPI_ISL_565942, EPI_ISL_565943, EPI_ISL_565944, EPI_ISL_565945, EPI_ISL_565946, EPI_ISL_565947, EPI_ISL_565948, EPI_ISL_565949, EPI_ISL_565950, EPI_ISL_565951, EPI_ISL_565952, EPI_ISL_565953, EPI_ISL_565954, EPI_ISL_565955, EPI_ISL_565956, EPI_ISL_565957, EPI_ISL_565958, EPI_ISL_565959, EPI_ISL_565960, EPI_ISL_565961, EPI_ISL_565962, EPI_ISL_565963, EPI_ISL_565964, EPI_ISL_565965, EPI_ISL_565966, EPI_ISL_565967, EPI_ISL_565968, EPI_ISL_565969, EPI_ISL_565970, EPI_ISL_565971, EPI_ISL_565972, EPI_ISL_565973, EPI_ISL_565974, EPI_ISL_565975, EPI_ISL_565976, EPI_ISL_565977, EPI_ISL_565978, EPI_ISL_565979, EPI_ISL_565980, EPI_ISL_565981, EPI_ISL_565982, EPI_ISL_565983, EPI_ISL_565984, EPI_ISL_565985, EPI_ISL_565986, EPI_ISL_565987, EPI_ISL_565988, EPI_ISL_565989, EPI_ISL_565990, EPI_ISL_565991, EPI_ISL_565992, EPI_ISL_565993, EPI_ISL_565994, EPI_ISL_565995, EPI_ISL_565996, EPI_ISL_565997, EPI_ISL_565998, EPI_ISL_565999, EPI_ISL_566000, EPI_ISL_566001, EPI_ISL_566002, EPI_ISL_566003, EPI_ISL_566004, EPI_ISL_566005, EPI_ISL_566006, EPI_ISL_566007, EPI_ISL_566008, EPI_ISL_566009, EPI_ISL_566010, EPI_ISL_566011, EPI_ISL_566012, EPI_ISL_566013, EPI_ISL_566014, EPI_ISL_566015, EPI_ISL_566016, EPI_ISL_566017, EPI_ISL_566018, EPI_ISL_566019, EPI_ISL_566020, EPI_ISL_566021, EPI_ISL_566022, EPI_ISL_566023, EPI_ISL_566024, EPI_ISL_566025, EPI_ISL_566026, EPI_ISL_566027, EPI_ISL_566028, EPI_ISL_566029, EPI_ISL_566030, EPI_ISL_566031, EPI_ISL_566032, EPI_ISL_566033, EPI_ISL_566034, EPI_ISL_566035 | | | |
| see above | Michigan Department of Health and Human Services, Bureau of Laboratories | Michigan Department of Health and Human Services, Bureau of Laboratories | Blankenship HM, Riner D, Soehnlen MK |
| EPI_ISL_568877, EPI_ISL_568878, EPI_ISL_568879, EPI_ISL_568880, EPI_ISL_568881, EPI_ISL_568882, EPI_ISL_568883, EPI_ISL_568884, EPI_ISL_568885, EPI_ISL_568886, EPI_ISL_568887, EPI_ISL_568888, EPI_ISL_568889, EPI_ISL_568890, EPI_ISL_568891, EPI_ISL_568892, EPI_ISL_568893, EPI_ISL_568894, EPI_ISL_568895, EPI_ISL_568896, EPI_ISL_568897, EPI_ISL_568898, EPI_ISL_568899, EPI_ISL_568900, EPI_ISL_568901, EPI_ISL_568902, EPI_ISL_568903, EPI_ISL_568904, EPI_ISL_568905, EPI_ISL_568906, EPI_ISL_568907, EPI_ISL_568908, EPI_ISL_568909, EPI_ISL_568910, EPI_ISL_568911, EPI_ISL_568912, EPI_ISL_568913, EPI_ISL_568914, EPI_ISL_568915, EPI_ISL_568916, EPI_ISL_568917, EPI_ISL_568918, EPI_ISL_568919, EPI_ISL_568920, EPI_ISL_568921, EPI_ISL_568922, EPI_ISL_568923, EPI_ISL_568924, EPI_ISL_568925, EPI_ISL_568926, EPI_ISL_568927, EPI_ISL_568928, EPI_ISL_568929, EPI_ISL_568930, EPI_ISL_568931, EPI_ISL_568932, EPI_ISL_568933, EPI_ISL_568934, EPI_ISL_568935, EPI_ISL_568936, EPI_ISL_568937, EPI_ISL_568938, EPI_ISL_568939, EPI_ISL_568940, EPI_ISL_568941, EPI_ISL_568942, EPI_ISL_568943, EPI_ISL_568944, EPI_ISL_568945, EPI_ISL_568946, EPI_ISL_568947, EPI_ISL_568948, EPI_ISL_568949, EPI_ISL_568950, EPI_ISL_568951, EPI_ISL_568952, EPI_ISL_568953, EPI_ISL_568954, EPI_ISL_568955, EPI_ISL_568956, EPI_ISL_568957, EPI_ISL_568958, EPI_ISL_568959, EPI_ISL_568960, EPI_ISL_568961, EPI_ISL_568962, EPI_ISL_568963, EPI_ISL_568964, EPI_ISL_568965, EPI_ISL_568966, EPI_ISL_568967, EPI_ISL_568968, EPI_ISL_568969, EPI_ISL_568970, EPI_ISL_568971, EPI_ISL_568972, EPI_ISL_568973, EPI_ISL_568974, EPI_ISL_568975, EPI_ISL_568976, EPI_ISL_568977, EPI_ISL_568978, EPI_ISL_568979, EPI_ISL_568980, EPI_ISL_568981, EPI_ISL_568982, EPI_ISL_568983, EPI_ISL_568984, EPI_ISL_568985, EPI_ISL_568986, EPI_ISL_568987, EPI_ISL_568988, EPI_ISL_568989, EPI_ISL_568990, EPI_ISL_568991, EPI_ISL_568992, EPI_ISL_568993, EPI_ISL_568994, EPI_ISL_568995, EPI_ISL_568996, EPI_ISL_568997, EPI_ISL_568998, EPI_ISL_568999, EPI_ISL_569000, EPI_ISL_569001, EPI_ISL_569002, EPI_ISL_569003, EPI_ISL_569004, EPI_ISL_569005, EPI_ISL_569006, EPI_ISL_569007, EPI_ISL_569008, EPI_ISL_569009, EPI_ISL_569010, EPI_ISL_569011, EPI_ISL_569012, EPI_ISL_569013, EPI_ISL_569014, EPI_ISL_569015, EPI_ISL_569016, EPI_ISL_569017, EPI_ISL_569018, EPI_ISL_569019, EPI_ISL_569020, EPI_ISL_569021, EPI_ISL_569022, EPI_ISL_569023, EPI_ISL_569024, EPI_ISL_569025, EPI_ISL_569026, EPI_ISL_569027, EPI_ISL_569028, EPI_ISL_569029, EPI_ISL_569030, EPI_ISL_569031, EPI_ISL_569032, EPI_ISL_569033, EPI_ISL_569034, EPI_ISL_569035, EPI_ISL_569036, EPI_ISL_569037, EPI_ISL_569038, EPI_ISL_569039, EPI_ISL_569040, EPI_ISL_569041, EPI_ISL_569042, EPI_ISL_569043, EPI_ISL_569044, EPI_ISL_569045, EPI_ISL_569046, EPI_ISL_569047, EPI_ISL_569048, EPI_ISL_569049, EPI_ISL_569050, EPI_ISL_569051, EPI_ISL_569052, EPI_ISL_569053, EPI_ISL_569054, EPI_ISL_569055, EPI_ISL_569056, EPI_ISL_569057, EPI_ISL_569058, EPI_ISL_569059, EPI_ISL_569060, EPI_ISL_569061, EPI_ISL_569062, EPI_ISL_569063, EPI_ISL_569064, EPI_ISL_569065, EPI_ISL_569066, EPI_ISL_569067, EPI_ISL_569068, EPI_ISL_569069, EPI_ISL_569070, EPI_ISL_569071, EPI_ISL_569072, EPI_ISL_569073, EPI_ISL_569074, EPI_ISL_569075, EPI_ISL_569076, EPI_ISL_569077, EPI_ISL_569078, EPI_ISL_569079, EPI_ISL_569080, EPI_ISL_569081, EPI_ISL_569082, EPI_ISL_569083, EPI_ISL_569084, EPI_ISL_569085, EPI_ISL_569086, EPI_ISL_569087, EPI_ISL_569088, EPI_ISL_569089, EPI_ISL_569090, EPI_ISL_569091, EPI_ISL_569092, EPI_ISL_569093, EPI_ISL_569094, EPI_ISL_569095, EPI_ISL_569096, EPI_ISL_569097, EPI_ISL_569098, EPI_ISL_569099, EPI_ISL_569100, EPI_ISL_569101, EPI_ISL_569102, EPI_ISL_569103, EPI_ISL_569104, EPI_ISL_569105, EPI_ISL_569106, EPI_ISL_569107, EPI_ISL_569108, EPI_ISL_569109, EPI_ISL_569110, EPI_ISL_569111, EPI_ISL_569112, EPI_ISL_569113, EPI_ISL_569114, EPI_ISL_569115, EPI_ISL_569116, EPI_ISL_569117, EPI_ISL_569118, EPI_ISL_569119, EPI_ISL_569120, EPI_ISL_569121, EPI_ISL_569122, EPI_ISL_569123, EPI_ISL_569124, EPI_ISL_569125, EPI_ISL_569126, EPI_ISL_569127, EPI_ISL_569128, EPI_ISL_569129, EPI_ISL_569130, EPI_ISL_569131, EPI_ISL_569132, EPI_ISL_569133, EPI_ISL_569134, EPI_ISL_569135, EPI_ISL_569136, EPI_ISL_569137, EPI_ISL_569138, EPI_ISL_569139, EPI_ISL_569140, EPI_ISL_569141, EPI_ISL_569142, EPI_ISL_569143, EPI_ISL_569144, EPI_ISL_569145, EPI_ISL_569146, EPI_ISL_569147, EPI_ISL_569148, EPI_ISL_569149, EPI_ISL_569150, EPI_ISL_569151, EPI_ISL_569152, EPI_ISL_569153, EPI_ISL_569154, EPI_ISL_569155, EPI_ISL_569156, EPI_ISL_569157, EPI_ISL_569158, EPI_ISL_569159, EPI_ISL_569160, EPI_ISL_569161, EPI_ISL_569162, EPI_ISL_569163, EPI_ISL_569164, EPI_ISL_569165, EPI_ISL_569166, EPI_ISL_569167, EPI_ISL_569168, EPI_ISL_569169, EPI_ISL_569170, EPI_ISL_569171, EPI_ISL_569172, EPI_ISL_569173, EPI_ISL_569174, EPI_ISL_569175, EPI_ISL_569176, EPI_ISL_569177, EPI_ISL_569178, EPI_ISL_569179, EPI_ISL_569180, EPI_ISL_569181, EPI_ISL_569182, EPI_ISL_569183, EPI_ISL_569184, EPI_ISL_569185, EPI_ISL_569186, EPI_ISL_569187, EPI_ISL_569188, EPI_ISL_569189, EPI_ISL_569190, EPI_ISL_569191, EPI_ISL_569192, EPI_ISL_569193, EPI_ISL_569194, EPI_ISL_569195, EPI_ISL_569196, EPI_ISL_569197, EPI_ISL_569198, EPI_ISL_569199, EPI_ISL_569200, EPI_ISL_569201, EPI_ISL_569202, EPI_ISL_569203, EPI_ISL_569204, EPI_ISL_569205, EPI_ISL_569206, EPI_ISL_569207, EPI_ISL_569208, EPI_ISL_569209, EPI_ISL_569210, EPI_ISL_569211, EPI_ISL_569212, EPI_ISL_569213, EPI_ISL_569214, EPI_ISL_569215, EPI_ISL_569216, EPI_ISL_569217, EPI_ISL_569218, EPI_ISL_569219, EPI_ISL_569220, EPI_ISL_569221, EPI_ISL_569222, EPI_ISL_569223, EPI_ISL_569224, EPI_ISL_569225, EPI_ISL_569226, EPI_ISL_569227, EPI_ISL_569228, EPI_ISL_569229, EPI_ISL_569230, EPI_ISL_569231, EPI_ISL_569232, EPI_ISL_569233, EPI_ISL_569234, EPI_ISL_569235, EPI_ISL_569236, EPI_ISL_569237, EPI_ISL_569238, EPI_ISL_569239, EPI_ISL_569240, EPI_ISL_569241, EPI_ISL_569242, EPI_ISL_569243, EPI_ISL_569244, EPI_ISL_569245, EPI_ISL_569246, EPI_ISL_569247, EPI_ISL_569248, EPI_ISL_569249, EPI_ISL_569250, EPI_ISL_569251, EPI_ISL_569252, EPI_ISL_569253, EPI_ISL_569254, EPI_ISL_569255, EPI_ISL_569256, EPI_ISL_569257, EPI_ISL_569258, EPI_ISL_569259, EPI_ISL_569260, EPI_ISL_569261, EPI_ISL_569262, EPI_ISL_569263, EPI_ISL_569264, EPI_ISL_569265, EPI_ISL_569266, EPI_ISL_569267, EPI_ISL_569268, EPI_ISL_569269, EPI_ISL_569270, EPI_ISL_569271, EPI_ISL_569272, EPI_ISL_569273, EPI_ISL_569274, EPI_ISL_569275, EPI_ISL_569276, EPI_ISL_569277, EPI_ISL_569278, EPI_ISL_569279, EPI_ISL_569280, EPI_ISL_569281, EPI_ISL_569282, EPI_ISL_569283, EPI_ISL_569284, EPI_ISL_569285, EPI_ISL_569286, EPI_ISL_569287, EPI_ISL_569288, EPI_ISL_569289, EPI_ISL_569290, EPI_ISL_569291, EPI_ISL_569292, EPI_ISL_569293, EPI_ISL_569294, EPI_ISL_569295, EPI_ISL_569296, EPI_ISL_569297, EPI_ISL_569298, EPI_ISL_569299, EPI_ISL_569300, EPI_ISL_569301, EPI_ISL_569302, EPI_ISL_569303, EPI_ISL_569304, EPI_ISL_569305, EPI_ISL_569306, EPI_ISL_569307, EPI_ISL_569308, EPI_ISL_569309, EPI_ISL_569310, EPI_ISL_569311, EPI_ISL_569312, EPI_ISL_569313, EPI_ISL_569314, EPI_ISL_569315, EPI_ISL_569316, EPI_ISL_569317, EPI_ISL_569318, EPI_ISL_569319, EPI_ISL_569320, EPI_ISL_569321, EPI_ISL_569322, EPI_ISL_569323, EPI_ISL_569324, EPI_ISL_569325, EPI_ISL_569326, EPI_ISL_569327, EPI_ISL_569328, EPI_ISL_569329, EPI_ISL_569330, EPI_ISL_569331, EPI_ISL_569332, EPI_ISL_569333, EPI_ISL_569334, EPI_ISL_569335, EPI_ISL_569336, EPI_ISL_569337, EPI_ISL_569338, EPI_ISL_569339, EPI_ISL_569340, EPI_ISL_569341, EPI_ISL_569342, EPI_ISL_569343, EPI_ISL_569344, EPI_ISL_569345, EPI_ISL_569346, EPI_ISL_569347, EPI_ISL_569348, EPI_ISL_569349, EPI_ISL_569350, EPI_ISL_569351, EPI_ISL_569352, EPI_ISL_569353, EPI_ISL_569354, EPI_ISL_569355, EPI_ISL_569356, EPI_ISL_569357, EPI_ISL_569358, EPI_ISL_569359, EPI_ISL_569360, EPI_ISL_569361, EPI_ISL_569362, EPI_ISL_569363, EPI_ISL_569364, EPI_ISL_569365, EPI_ISL_569366, EPI_ISL_569367, EPI_ISL_569368, EPI_ISL_569369, EPI_ISL_569370, EPI_ISL_569371, EPI_ISL_569372, EPI_ISL_569373, EPI_ISL_569374, EPI_ISL_569375, EPI_ISL_569376, EPI_ISL_569377, EPI_ISL_569378, EPI_ISL_569379, EPI_ISL_569380, EPI_ISL_569381, EPI_ISL_569382, EPI_ISL_569383, EPI_ISL_569384, EPI_ISL_569385, EPI_ISL_569386, EPI_ISL_569387, EPI_ISL_569388, EPI_ISL_569389, EPI_ISL_569390, EPI_ISL_569391, EPI_ISL_569392, EPI_ISL_569393, EPI_ISL_569394, EPI_ISL_569395, EPI_ISL_569396, EPI_ISL_569397, EPI_ISL_569398, EPI_ISL_569399, EPI_ISL_569400, EPI_ISL_569401, EPI_ISL_569402, EPI_ISL_569403, EPI_ISL_569404, EPI_ISL_569405, EPI_ISL_569406, EPI_ISL_569407, EPI_ISL_569408, EPI_ISL_569409, EPI_ISL_569410, EPI_ISL_569411, EPI_ISL_569412, EPI_ISL_569413, EPI_ISL_569414, EPI_ISL_569415, EPI_ISL_569416, EPI_ISL_569417, EPI_ISL_569418, EPI_ISL_569419, EPI_ISL_569420, EPI_ISL_569421, EPI_ISL_569422, EPI_ISL_569423, EPI_ISL_569424, EPI_ISL_569425, EPI_ISL_569426, EPI_ISL_569427, EPI_ISL_569428, EPI_ISL_569429, EPI_ISL_569430, EPI_ISL_569431, EPI_ISL_569432, EPI_ISL_569433, EPI_ISL_569434, EPI_ISL_569435, EPI_ISL_569436, EPI_ISL_569437, EPI_ISL_569438, EPI_ISL_569439, EPI_ISL_569440, EPI_ISL_569441, EPI_ISL_569442, EPI_ISL_569443, EPI_ISL_569444, EPI_ISL_569445, EPI_ISL_569446, EPI_ISL_569447, EPI_ISL_569448, EPI_ISL_569449, EPI_ISL_569450, EPI_ISL_569451, EPI_ISL_569452, EPI_ISL_569453, EPI_ISL_569454, EPI_ISL_569455, EPI_ISL_569456, EPI_ISL_569457, EPI_ISL_569458, EPI_ISL_569459, EPI_ISL_569460, EPI_ISL_569461, EPI_ISL_569462, EPI_ISL_569463, EPI_ISL_569464, EPI_ISL_569465, EPI_ISL_569466, EPI_ISL_569467, EPI_ISL_569468, EPI_ISL_569469, EPI_ISL_569470, EPI_ISL_569471, EPI_ISL_569472, EPI_ISL_569473, EPI_ISL_569474, EPI_ISL_569475, EPI_ISL_569476, EPI_ISL_569477, EPI_ISL_569478, EPI_ISL_569479, EPI_ISL_569480, EPI_ISL_569481, EPI_ISL_569482, EPI_ISL_569483, EPI_ISL_569484, EPI_ISL_569485, EPI_ISL_569486, EPI_ISL_569487, EPI_ISL_569488, EPI_ISL_569489, EPI_ISL_569490, EPI_ISL_569491, EPI_ISL_569492, EPI_ISL_569493, EPI_ISL_569494, EPI_ISL_569495, EPI_ISL_569496, EPI_ISL_569497, EPI_ISL_569498, EPI_ISL_569499, EPI_ISL_569500, EPI_ISL_569501, EPI_ISL_569502, EPI_ISL_569503, EPI_ISL_569504, EPI_ISL_569505, EPI_ISL_569506, EPI_ISL_569507, EPI_ISL_569508, EPI_ISL_569509, EPI_ISL_569510, EPI_ISL_569511, EPI_ISL_569512, EPI_ISL_569513, EPI_ISL_569514, EPI_ISL_569515, EPI_ISL_569516, EPI_ISL_569517, EPI_ISL_569518, EPI_ISL_569519, EPI_ISL_569520, EPI_ISL_569521, EPI_ISL_569522, EPI_ISL_569523, EPI_ISL_569524, EPI_ISL_569525, EPI_ISL_569526, EPI_ISL_569527, EPI_ISL_569528, EPI_ISL_569529, EPI_ISL_569530, EPI_ISL_569531, EPI_ISL_569532, EPI_ISL_569533, EPI_ISL_569534, EPI_ISL_569535, EPI_ISL_569536, EPI_ISL_569537, EPI_ISL_569538, EPI_ISL_569539, EPI_ISL_569540, EPI_ISL_569541, EPI_ISL_569542, EPI_ISL_569543, EPI_ISL_569544, EPI_ISL_569545, EPI_ISL_569546, EPI_ISL_569547, EPI_ISL_569548, EPI_ISL_569549, EPI_ISL_569550, EPI_ISL_569551, EPI_ISL_569552, EPI_ISL_569553, EPI_ISL_569554, EPI_ISL_569555, EPI_ISL_569556, EPI_ISL_569557, EPI_ISL_569558, EPI_ISL_569559, EPI_ISL_569560, EPI_ISL_569561, EPI_ISL_569562, EPI_ISL_569563, EPI_ISL_569564, EPI_ISL_569565, EPI_ISL_569566, EPI_ISL_569567, EPI_ISL_569568, EPI_ISL_569569, EPI_ISL_569570, EPI_ISL_569571, EPI_ISL_569572, EPI_ISL_569573, EPI_ISL_569574, EPI_ISL_569575, EPI_ISL_569576, EPI_ISL_569577, EPI_ISL_569578, EPI_ISL_569579, EPI_ISL_569580, EPI_ISL_569581, EPI_ISL_569582, EPI_ISL_569583, EPI_ISL_569584, EPI_ISL_569585, EPI_ISL_569586, EPI_ISL_569587, EPI_ISL_569588, EPI_ISL_569589, EPI_ISL_569590, EPI_ISL_569591, EPI_ISL_569592, EPI_ISL_569593, EPI_ISL_569594, EPI_ISL_569595, EPI_ISL_569596, EPI_ISL_569597, EPI_ISL_569598, EPI_ISL_569599, EPI_ISL_569600, EPI_ISL_569601, EPI_ISL_569602, EPI_ISL_569603, EPI_ISL_569604, EPI_ISL_569605, EPI_ISL_569606, EPI_ISL_569607, EPI_ISL_569608 | | | |
| see above | MEPHI, Aix Marseille University | MEPHI, Aix Marseille University | Anthony LEVASSEUR |
| EPI_ISL_569735, EPI_ISL_569736, EPI_ISL_569737, EPI_ISL_569738, EPI_ISL_569739, EPI_ISL_569740, EPI_ISL_569741, EPI_ISL_569742, EPI_ISL_569743, EPI_ISL_569744, EPI_ISL_569745, EPI_ISL_569746, EPI_ISL_569747, EPI_ISL_569748, EPI_ISL_569749, EPI_ISL_569750, EPI_ISL_569751, EPI_ISL_569752, EPI_ISL_569753, EPI_ISL_569754, EPI_ISL_569755, EPI_ISL_569756, EPI_ISL_569757, EPI_ISL_569758, EPI_ISL_569759, EPI_ISL_569760, EPI_ISL_569761, EPI_ISL_569762, EPI_ISL_569763, EPI_ISL_569764, EPI_ISL_569765, EPI_ISL_569766, EPI_ISL_569767, EPI_ISL_569768, EPI_ISL_569769, EPI_ISL_569770, EPI_ISL_569771, EPI_ISL_569772, EPI_ISL_569773, EPI_ISL_569774, EPI_ISL_569775, EPI_ISL_569776, EPI_ISL_569777, EPI_ISL_569778, EPI_ISL_569779, EPI_ISL_569780, EPI_ISL_569781, EPI_ISL_569782, EPI_ISL_569783, EPI_ISL_569784, EPI_ISL_569785, EPI_ISL_569786, EPI_ISL_569787, EPI_ISL_569788, EPI_ISL_569789, EPI_ISL_569790, EPI_ISL_569791, EPI_ISL_569792, EPI_ISL_569793, EPI_ISL_569794, EPI_ISL_569795, EPI_ISL_569796, EPI_ISL_569797, EPI_ISL_569798, EPI_ISL_569799, EPI_ISL_569800, EPI_ISL_569801, EPI_ISL_569802, EPI_ISL_569803, EPI_ISL_569804, EPI_ISL_569805, EPI_ISL_569806, EPI_ISL_569807, EPI_ISL_569808, EPI_ISL_569809, EPI_ISL_569810, EPI_ISL_569811, EPI_ISL_569812, EPI_ISL_569813, EPI_ISL_569814, EPI_ISL_569815, EPI_ISL_569816, EPI_ISL_569817, EPI_ISL_569818, EPI_ISL_569819, EPI_ISL_569820, EPI_ISL_569821, EPI_ISL_569822, EPI_ISL_569823, EPI_ISL_569824, | | | |

| | | | |
|---|---|---|---|
| EPI_ISL_569825, EPI_ISL_569826, EPI_ISL_569827, EPI_ISL_569828, EPI_ISL_569829, EPI_ISL_569830, EPI_ISL_569831, EPI_ISL_569832, EPI_ISL_569833, EPI_ISL_569834, EPI_ISL_569835, EPI_ISL_569836, EPI_ISL_569837, EPI_ISL_569838, EPI_ISL_569839, EPI_ISL_569840, EPI_ISL_569841, EPI_ISL_569842, EPI_ISL_569843, EPI_ISL_569844, EPI_ISL_569845, EPI_ISL_569846, EPI_ISL_569847, EPI_ISL_569848, EPI_ISL_569849, EPI_ISL_569850, EPI_ISL_569851, EPI_ISL_569852, EPI_ISL_569853, EPI_ISL_569854, EPI_ISL_569855, EPI_ISL_569856, EPI_ISL_569857 | | | |
| see above | Omsk Research Institute of Natural Focal Infections | WHO National Influenza Centre Russian Federation | Artem Fadeev, Ekaterina Gradoboeva, Ekaterina Savkina, Daria Nashatyreva, Elena Poleshchuk, Aleksei Vasilenko, Valery Yakimenko, Andrey Komissarov |
| EPI_ISL_572715, EPI_ISL_572716, EPI_ISL_572721, EPI_ISL_573253, EPI_ISL_573254, EPI_ISL_573255, EPI_ISL_573256, EPI_ISL_573257, EPI_ISL_573258, EPI_ISL_573259, EPI_ISL_573260, EPI_ISL_573261, EPI_ISL_573262, EPI_ISL_573263, EPI_ISL_573264, EPI_ISL_573265 | | | |
| see above | Centre for Enzyme Innovation, University of Portsmouth / Translational Research Laboratory, Portsmouth Hospitals NHS Trust | COVID-19 Genomics UK (COG-UK) Consortium | Angela Beckett,Yann Bourgeois,Garry Scarlett,Sharon Glaysher,Scott Elliott,Kelly Bicknell,Robert Impey,Allyson Lloyd,Sarah Wyllie,Ethan Butcher,Anoop Chauhan,Samuel Robson |
| EPI_ISL_574431 | Hospital IESS Babahoyo | Institute of Microbiology, Universidad San Francisco de Quito | Belén Prado-Vivar, Sully Márquez, Juan José Guadalupe, Monica Becerra-Wong, Fernanda Zurita, Bernardo Gutiérrez, Francisco Cordova, Ninfa Henriquez, Killen Briones-Zamora, Killen Briones-Claudette, Verónica Barragán, Patricio Rojas-Silva, Gabriel Trueba, Michelle Grunauer, Paúl Cárdenas |
| EPI_ISL_574779, EPI_ISL_574780, EPI_ISL_574781, EPI_ISL_574786, EPI_ISL_574787, EPI_ISL_574788, EPI_ISL_574789 | Dutch COVID-19 response team | Erasmus Medical Center | Bas Oude Munnink, Reina Sikkema, David Nieuwenhuijse, Irina Chestakova, Anne van der Linden, Marjan Boter, Emmanuelle Munger, Corine GeurtsvanKessel, Annemiek van der Eijk, Richard Molenkamp, Marion Koopmans, on behalf of the Dutch national COVID-19 response team. |
| EPI_ISL_576117, EPI_ISL_576118 | Laboratory, The Bio Arte Limited | Laboratory, The Bio Arte Limited | Biazzo,M., Madeddu,S., Santoro,F., Pinzauti,D. |
| EPI_ISL_576119 | Laboratory, The Bio Arte Limited | Laboratory, The Bio Arte Limited | Biazzo,M., Madeddu,S., Santoro,F., Pinzauti,D. |
| EPI_ISL_576120, EPI_ISL_576121, EPI_ISL_576122, EPI_ISL_576123, EPI_ISL_576124, EPI_ISL_576125, EPI_ISL_576126 | Laboratory, The Bio Arte Limited | Laboratory, The Bio Arte Limited | Biazzo,M., Madeddu,S., Santoro,F., Pinzauti,D. |
| EPI_ISL_576198, EPI_ISL_576199, EPI_ISL_576200, EPI_ISL_576201, EPI_ISL_576202, EPI_ISL_576203, EPI_ISL_576204, EPI_ISL_576205, EPI_ISL_576206, EPI_ISL_576207, EPI_ISL_576208, EPI_ISL_576209, EPI_ISL_576210, EPI_ISL_576211, EPI_ISL_576212, EPI_ISL_576213, EPI_ISL_576214, EPI_ISL_576215, EPI_ISL_576216, EPI_ISL_576217, EPI_ISL_576218, EPI_ISL_576219, EPI_ISL_576220, EPI_ISL_576221 | | | |
| see above | Michigan Department of Health and Human Services, Bureau of Laboratories | Michigan Department of Health and Human Services, Bureau of Laboratories | Blankenship HM, Riner D, Soehnlen MK |
| EPI_ISL_577040, EPI_ISL_577041, EPI_ISL_577042, EPI_ISL_577043, EPI_ISL_577044, EPI_ISL_577045, EPI_ISL_577046, EPI_ISL_577047, EPI_ISL_577048, EPI_ISL_577049, EPI_ISL_577050, EPI_ISL_577051, EPI_ISL_577052, EPI_ISL_577053, EPI_ISL_577054, EPI_ISL_577055, EPI_ISL_577056, EPI_ISL_577057, EPI_ISL_577058, EPI_ISL_577059, EPI_ISL_577060, EPI_ISL_577061, EPI_ISL_577062, EPI_ISL_577063, EPI_ISL_577064, EPI_ISL_577065, EPI_ISL_577066, EPI_ISL_577067, EPI_ISL_577193, EPI_ISL_577194, EPI_ISL_577195, EPI_ISL_577198, EPI_ISL_577199, EPI_ISL_577200, EPI_ISL_577201, EPI_ISL_577202, EPI_ISL_577203, EPI_ISL_577210, EPI_ISL_577211, EPI_ISL_577212, EPI_ISL_577213, EPI_ISL_577215, EPI_ISL_577216, EPI_ISL_577217, EPI_ISL_577218, EPI_ISL_577219 | | | |
| see above | Centre for Enzyme Innovation, University of Portsmouth / Translational Research Laboratory, Portsmouth Hospitals NHS Trust | COVID-19 Genomics UK (COG-UK) Consortium | Angela Beckett,Yann Bourgeois,Garry Scarlett,Sharon Glaysher,Scott Elliott,Kelly Bicknell,Robert Impey,Allyson Lloyd,Sarah Wyllie,Ethan Butcher,Anoop Chauhan,Samuel Robson |
| EPI_ISL_577220, EPI_ISL_577221, EPI_ISL_577222, EPI_ISL_577223, EPI_ISL_577224, EPI_ISL_577225, EPI_ISL_577226, EPI_ISL_577227, EPI_ISL_577228, EPI_ISL_577229, EPI_ISL_577230, EPI_ISL_577231, EPI_ISL_577232, EPI_ISL_577233, EPI_ISL_577234, EPI_ISL_577235, EPI_ISL_577236, EPI_ISL_577237, EPI_ISL_577238, EPI_ISL_577239, EPI_ISL_577240, EPI_ISL_577241, EPI_ISL_577242, EPI_ISL_577243, EPI_ISL_577244, EPI_ISL_577245, EPI_ISL_577246, EPI_ISL_577247, EPI_ISL_577248, EPI_ISL_577249, EPI_ISL_577250, EPI_ISL_577251, EPI_ISL_577252, EPI_ISL_577253, EPI_ISL_577254, EPI_ISL_577255, EPI_ISL_577256, EPI_ISL_577257, EPI_ISL_577258, EPI_ISL_577259, EPI_ISL_577260, EPI_ISL_577261, EPI_ISL_577262, EPI_ISL_577263, EPI_ISL_577264, EPI_ISL_577265, EPI_ISL_577266, EPI_ISL_577267, EPI_ISL_577268, EPI_ISL_577269, EPI_ISL_577270, EPI_ISL_577271, EPI_ISL_577272, EPI_ISL_577273, EPI_ISL_577274, EPI_ISL_577275, EPI_ISL_577276, EPI_ISL_577277, EPI_ISL_577278, EPI_ISL_577279, EPI_ISL_577280, EPI_ISL_577281, EPI_ISL_577282, EPI_ISL_577283, EPI_ISL_577284, EPI_ISL_577285, EPI_ISL_577286, EPI_ISL_577287, EPI_ISL_577288, EPI_ISL_577289, EPI_ISL_577290, EPI_ISL_577291, EPI_ISL_577292, EPI_ISL_577293, EPI_ISL_577294, EPI_ISL_577295, EPI_ISL_577296, EPI_ISL_577297, EPI_ISL_577298 | | | |
| see above | University of Exeter | COVID-19 Genomics UK (COG-UK) Consortium | Ben Temperton,Aaron Jeffries,Michelle Michelsen,Joanna Warwick-Dugdale,Audrey Farbos,Robyn Manley,Stephen Michell,Jane Masoli |
| EPI_ISL_577299, EPI_ISL_577300, EPI_ISL_577301, EPI_ISL_577302, EPI_ISL_577303, EPI_ISL_577304, EPI_ISL_577305, EPI_ISL_577306, EPI_ISL_577307, EPI_ISL_577308, EPI_ISL_577309, EPI_ISL_577310, EPI_ISL_577311, EPI_ISL_577312, EPI_ISL_577313, EPI_ISL_577314, EPI_ISL_577315, EPI_ISL_577316, EPI_ISL_577317, EPI_ISL_577318, EPI_ISL_577319, EPI_ISL_577320, EPI_ISL_577321, EPI_ISL_577322, EPI_ISL_577323, EPI_ISL_577324, EPI_ISL_577325, EPI_ISL_577326, EPI_ISL_577327, EPI_ISL_577328, EPI_ISL_577329, EPI_ISL_577330, EPI_ISL_577331, EPI_ISL_577332, EPI_ISL_577333, EPI_ISL_577334, EPI_ISL_577335, EPI_ISL_577336, EPI_ISL_577337, EPI_ISL_577338, EPI_ISL_577339, EPI_ISL_577340, EPI_ISL_577341, EPI_ISL_577342, EPI_ISL_577343, EPI_ISL_577344, EPI_ISL_577345 | | | |
| see above | Centre for Enzyme Innovation, University of Portsmouth / Translational Research Laboratory, Portsmouth Hospitals NHS Trust | COVID-19 Genomics UK (COG-UK) Consortium | Angela Beckett,Yann Bourgeois,Garry Scarlett,Sharon Glaysher,Scott Elliott,Kelly Bicknell,Robert Impey,Allyson Lloyd,Sarah Wyllie,Ethan Butcher,Anoop Chauhan,Samuel Robson |
| EPI_ISL_577346, EPI_ISL_577359, EPI_ISL_577360, EPI_ISL_577361, EPI_ISL_577362, EPI_ISL_577363, EPI_ISL_577364, EPI_ISL_577365, EPI_ISL_577366, EPI_ISL_577367, EPI_ISL_577368, EPI_ISL_577369, EPI_ISL_577370, EPI_ISL_577371, EPI_ISL_577372, EPI_ISL_577373, EPI_ISL_577374, EPI_ISL_577375, EPI_ISL_577376, EPI_ISL_577377, EPI_ISL_577378, EPI_ISL_577379, EPI_ISL_577380 | | | |
| see above | University of Exeter | COVID-19 Genomics UK (COG-UK) Consortium | Ben Temperton,Aaron Jeffries,Michelle Michelsen,Joanna Warwick-Dugdale,Audrey Farbos,Robyn Manley,Stephen Michell,Jane Masoli |
| EPI_ISL_577381, EPI_ISL_577382, EPI_ISL_577387, EPI_ISL_577388, EPI_ISL_577392, EPI_ISL_577393, EPI_ISL_577394, EPI_ISL_577398 | Centre for Enzyme Innovation, University of Portsmouth / Translational Research Laboratory, Portsmouth Hospitals NHS Trust | COVID-19 Genomics UK (COG-UK) Consortium | Angela Beckett,Yann Bourgeois,Garry Scarlett,Sharon Glaysher,Scott Elliott,Kelly Bicknell,Robert Impey,Allyson Lloyd,Sarah Wyllie,Ethan Butcher,Anoop Chauhan,Samuel Robson |
| EPI_ISL_577544 | University of Exeter | COVID-19 Genomics UK (COG-UK) Consortium | Ben Temperton,Aaron Jeffries,Michelle Michelsen,Joanna Warwick-Dugdale,Audrey Farbos,Robyn Manley,Stephen Michell,Jane Masoli |
| EPI_ISL_577545, EPI_ISL_577546 | Centre for Enzyme Innovation, University of Portsmouth / Translational Research Laboratory, Portsmouth Hospitals NHS Trust | COVID-19 Genomics UK (COG-UK) Consortium | Angela Beckett,Yann Bourgeois,Garry Scarlett,Sharon Glaysher,Scott Elliott,Kelly Bicknell,Robert Impey,Allyson Lloyd,Sarah Wyllie,Ethan Butcher,Anoop Chauhan,Samuel Robson |
| EPI_ISL_577547, EPI_ISL_577548, EPI_ISL_577549, EPI_ISL_577550, EPI_ISL_577551, EPI_ISL_577552, EPI_ISL_577553, EPI_ISL_577554, EPI_ISL_577555, EPI_ISL_577556, EPI_ISL_577557, EPI_ISL_577558, EPI_ISL_577559, EPI_ISL_577560, EPI_ISL_577561, EPI_ISL_577562, EPI_ISL_577563, EPI_ISL_577564, EPI_ISL_577565, EPI_ISL_577566, EPI_ISL_577567, EPI_ISL_577568, EPI_ISL_577569, EPI_ISL_577570, EPI_ISL_577571, EPI_ISL_577572, EPI_ISL_577573, EPI_ISL_577574, EPI_ISL_577575, EPI_ISL_577576, EPI_ISL_577577, EPI_ISL_577578, EPI_ISL_577579, EPI_ISL_577580, EPI_ISL_577581, EPI_ISL_577582, EPI_ISL_577583, EPI_ISL_577584, EPI_ISL_577585, EPI_ISL_577586, EPI_ISL_577587, EPI_ISL_577588, EPI_ISL_577589, EPI_ISL_577590, EPI_ISL_577591, EPI_ISL_577592, EPI_ISL_577593, EPI_ISL_577594, EPI_ISL_577595, EPI_ISL_577596 | | | |
| see above | Michigan Department of Health and Human Services, Bureau of Laboratories | Michigan Department of Health and Human Services, Bureau of Laboratories | Blankenship HM, Riner D, Soehnlen MK |
| EPI_ISL_577743, EPI_ISL_577744, EPI_ISL_577745, EPI_ISL_577746, EPI_ISL_577747, EPI_ISL_577829, EPI_ISL_577830, EPI_ISL_577831, EPI_ISL_577832, EPI_ISL_577833, EPI_ISL_577834, EPI_ISL_577835, EPI_ISL_577836, EPI_ISL_577837, EPI_ISL_577838, EPI_ISL_577839, EPI_ISL_577840, EPI_ISL_577841, EPI_ISL_577842, EPI_ISL_577843, EPI_ISL_577844, EPI_ISL_577847, EPI_ISL_577848, EPI_ISL_577849, EPI_ISL_577850, EPI_ISL_577853, EPI_ISL_577854, EPI_ISL_577855, EPI_ISL_577856, EPI_ISL_577857, EPI_ISL_577858, EPI_ISL_577859, EPI_ISL_577860, EPI_ISL_577861, EPI_ISL_577862, EPI_ISL_577863, EPI_ISL_577864, EPI_ISL_577865, EPI_ISL_577866, EPI_ISL_577872, EPI_ISL_577873, EPI_ISL_577874, EPI_ISL_577875, EPI_ISL_577876, EPI_ISL_577877, EPI_ISL_577878, EPI_ISL_577879, EPI_ISL_577880, EPI_ISL_577881, EPI_ISL_577882, EPI_ISL_577886, EPI_ISL_577887, EPI_ISL_577888, EPI_ISL_577889, EPI_ISL_577890, EPI_ISL_577891, EPI_ISL_577892, EPI_ISL_577893, EPI_ISL_577894, EPI_ISL_577895, EPI_ISL_577896, EPI_ISL_577897, EPI_ISL_577898, EPI_ISL_577899, EPI_ISL_577900, EPI_ISL_577901, EPI_ISL_577902, EPI_ISL_577903, EPI_ISL_577904, EPI_ISL_577905, EPI_ISL_577908, EPI_ISL_577909, EPI_ISL_577910, EPI_ISL_577911, EPI_ISL_577912, EPI_ISL_577913, EPI_ISL_577914, EPI_ISL_577915, EPI_ISL_577916, EPI_ISL_577926, EPI_ISL_577927, EPI_ISL_577928, EPI_ISL_577929, EPI_ISL_577930, EPI_ISL_577931, EPI_ISL_577932, EPI_ISL_577933, EPI_ISL_577934, EPI_ISL_577935, EPI_ISL_577936, EPI_ISL_577937, EPI_ISL_577938, EPI_ISL_577939, EPI_ISL_577940, EPI_ISL_577941, EPI_ISL_577942, EPI_ISL_577943, EPI_ISL_577944, EPI_ISL_577948, EPI_ISL_577949, EPI_ISL_577950, EPI_ISL_577951, EPI_ISL_577952, EPI_ISL_577953, EPI_ISL_577954, EPI_ISL_577964, EPI_ISL_577965, EPI_ISL_577966, EPI_ISL_577967, EPI_ISL_577968, EPI_ISL_577969, EPI_ISL_577970, EPI_ISL_577971, EPI_ISL_577972, EPI_ISL_577973, EPI_ISL_577974, EPI_ISL_577975, EPI_ISL_577976, EPI_ISL_577977, EPI_ISL_577978, EPI_ISL_577979, EPI_ISL_577980, EPI_ISL_577981, EPI_ISL_577982, EPI_ISL_577983, EPI_ISL_577984, EPI_ISL_577985, EPI_ISL_577986, EPI_ISL_577987, EPI_ISL_577988, EPI_ISL_577989, EPI_ISL_577990, EPI_ISL_577991, EPI_ISL_577992, EPI_ISL_577993, EPI_ISL_577994, EPI_ISL_577995, EPI_ISL_577996, EPI_ISL_577997, EPI_ISL_577998, EPI_ISL_577999, EPI_ISL_578000, EPI_ISL_578004, EPI_ISL_578005, EPI_ISL_578006, EPI_ISL_578007, EPI_ISL_578008, EPI_ISL_578009, EPI_ISL_578010, EPI_ISL_578011, EPI_ISL_578012, EPI_ISL_578013, EPI_ISL_578014, EPI_ISL_578015, EPI_ISL_578016, EPI_ISL_578017, EPI_ISL_578018, EPI_ISL_578019, EPI_ISL_578020, EPI_ISL_578021, EPI_ISL_578022, EPI_ISL_578023, EPI_ISL_578024, EPI_ISL_578025, EPI_ISL_578026, EPI_ISL_578027, EPI_ISL_578028, EPI_ISL_578029, EPI_ISL_578030, EPI_ISL_578031, EPI_ISL_578032, EPI_ISL_578033, EPI_ISL_578034, EPI_ISL_578035, EPI_ISL_578036, EPI_ISL_578037, EPI_ISL_578038, EPI_ISL_578039, EPI_ISL_578040, EPI_ISL_578041, EPI_ISL_578042, EPI_ISL_578043, EPI_ISL_578044, EPI_ISL_578045, EPI_ISL_578046, EPI_ISL_578047, EPI_ISL_578048, EPI_ISL_578049, EPI_ISL_578050, EPI_ISL_578051, EPI_ISL_578052, EPI_ISL_578053, EPI_ISL_578054, EPI_ISL_578055, EPI_ISL_578056, EPI_ISL_578057, EPI_ISL_578058, EPI_ISL_578059, EPI_ISL_578060, EPI_ISL_578061, EPI_ISL_578062, EPI_ISL_578063, EPI_ISL_578064, EPI_ISL_578065, EPI_ISL_578066, EPI_ISL_578067, EPI_ISL_578068, EPI_ISL_578069, EPI_ISL_578070, EPI_ISL_578071, EPI_ISL_578073, EPI_ISL_578074, EPI_ISL_578075, EPI_ISL_578076, EPI_ISL_578077, EPI_ISL_578078, EPI_ISL_578079 | | | |
| see above | Dutch COVID-19 response team | Erasmus Medical Center | Bas Oude Munnink, Reina Sikkema, David Nieuwenhuijse, Irina Chestakova, Anne van der Linden, Marjan Boter, Emmanuelle Munger, Corine GeurtsvanKessel, Annemiek van der Eijk, Richard Molenkamp, Marion Koopmans, on behalf of the Dutch national COVID-19 response team. |
| EPI_ISL_578176, EPI_ISL_578177 | CNR Virus des Infections Respiratoires - France SUD | CNR Virus des Infections Respiratoires - France SUD | Antonin Bal, Gregory Queromes, Emilie Frobert, Gregory Destras, Gwendolyne Burfin, Solenne Brun, Alexandre Gaymard, Maude Bouscambert-Duchamp, |

| | | | |
|---|---|---|---|
| | | | Florence Morfin-Sherpa, Martine Valette, Bruno Lina, Laurence Josset |
| EPI_ISL_579515, EPI_ISL_579516, EPI_ISL_579517, EPI_ISL_579518, EPI_ISL_579519, EPI_ISL_579520, EPI_ISL_579521, EPI_ISL_579522, EPI_ISL_579523, EPI_ISL_579524, EPI_ISL_579525, EPI_ISL_579526, EPI_ISL_579527, EPI_ISL_579528, EPI_ISL_579529, EPI_ISL_579530, EPI_ISL_579531, EPI_ISL_579532, EPI_ISL_579533, EPI_ISL_579534, EPI_ISL_579535, EPI_ISL_579536, EPI_ISL_579537, EPI_ISL_579538, EPI_ISL_579539, EPI_ISL_579540, EPI_ISL_579541, EPI_ISL_579542, EPI_ISL_579543, EPI_ISL_579544, EPI_ISL_579545, EPI_ISL_579546, EPI_ISL_579547, EPI_ISL_579548, EPI_ISL_579549, EPI_ISL_579550, EPI_ISL_579551, EPI_ISL_579552, EPI_ISL_579553, EPI_ISL_579554, EPI_ISL_579555, EPI_ISL_579556, EPI_ISL_579557, EPI_ISL_579558, EPI_ISL_579559, EPI_ISL_579560, EPI_ISL_579561, EPI_ISL_579562, EPI_ISL_579563, EPI_ISL_579564, EPI_ISL_579565, EPI_ISL_579566, EPI_ISL_579567, EPI_ISL_579568, EPI_ISL_579569, EPI_ISL_579570, EPI_ISL_579571, EPI_ISL_579572, EPI_ISL_579573, EPI_ISL_579574, EPI_ISL_579575, EPI_ISL_579576, EPI_ISL_579577, EPI_ISL_579578, EPI_ISL_579579, EPI_ISL_579580, EPI_ISL_579581, EPI_ISL_579582, EPI_ISL_579583, EPI_ISL_579584, EPI_ISL_579585, EPI_ISL_579586, EPI_ISL_579587, EPI_ISL_579588, EPI_ISL_579589, EPI_ISL_579590, EPI_ISL_579591, EPI_ISL_579592, EPI_ISL_579593, EPI_ISL_579594, EPI_ISL_579595, EPI_ISL_579596, EPI_ISL_579597, EPI_ISL_579598, EPI_ISL_579599, EPI_ISL_579600, EPI_ISL_579601, EPI_ISL_579602, EPI_ISL_579603, EPI_ISL_579604, EPI_ISL_579605, EPI_ISL_579606, EPI_ISL_579607, EPI_ISL_579608, EPI_ISL_579609, EPI_ISL_579610, EPI_ISL_579611, EPI_ISL_579612, EPI_ISL_579613, EPI_ISL_579614, EPI_ISL_579615, EPI_ISL_579616, EPI_ISL_579617, EPI_ISL_579618, EPI_ISL_579619, EPI_ISL_579620, EPI_ISL_579621, EPI_ISL_579622, EPI_ISL_579623, EPI_ISL_579624, EPI_ISL_579625, EPI_ISL_579626, EPI_ISL_579627, EPI_ISL_579628, EPI_ISL_579629, EPI_ISL_579630, EPI_ISL_579631, EPI_ISL_579632, EPI_ISL_579633, EPI_ISL_579634, EPI_ISL_579635, EPI_ISL_579636, EPI_ISL_579637, EPI_ISL_579638, EPI_ISL_579639, EPI_ISL_579640, EPI_ISL_579641, EPI_ISL_579642, EPI_ISL_579643, EPI_ISL_579644, EPI_ISL_579645, EPI_ISL_579646, EPI_ISL_579647, EPI_ISL_579648, EPI_ISL_579649, EPI_ISL_579650, EPI_ISL_579651, EPI_ISL_579652, EPI_ISL_579653, EPI_ISL_579654, EPI_ISL_579655, EPI_ISL_579656, EPI_ISL_579657, EPI_ISL_579658, EPI_ISL_579659, EPI_ISL_579660, EPI_ISL_579661, EPI_ISL_579662, EPI_ISL_579663, EPI_ISL_579664, EPI_ISL_579665, EPI_ISL_579666, EPI_ISL_579667, EPI_ISL_579668, EPI_ISL_579669, EPI_ISL_579670, EPI_ISL_579671, EPI_ISL_579672, EPI_ISL_579673, EPI_ISL_579674, EPI_ISL_579675, EPI_ISL_579676, EPI_ISL_579677, EPI_ISL_579678, EPI_ISL_579679, EPI_ISL_579680, EPI_ISL_579681, EPI_ISL_579682, EPI_ISL_579683, EPI_ISL_579684, EPI_ISL_579685, EPI_ISL_579686, EPI_ISL_579687, EPI_ISL_579688, EPI_ISL_579689, EPI_ISL_579690, EPI_ISL_579691, EPI_ISL_579692, EPI_ISL_579693, EPI_ISL_579694, EPI_ISL_579695, EPI_ISL_579696, EPI_ISL_579697, EPI_ISL_579698, EPI_ISL_579699, EPI_ISL_579700, EPI_ISL_579701, EPI_ISL_579702, EPI_ISL_579703, EPI_ISL_579704, EPI_ISL_579705, EPI_ISL_579706, EPI_ISL_579707, EPI_ISL_579708, EPI_ISL_579709, EPI_ISL_579710, EPI_ISL_579711, EPI_ISL_579712, EPI_ISL_579713, EPI_ISL_579714, EPI_ISL_579715, EPI_ISL_579716, EPI_ISL_579717, EPI_ISL_579718, EPI_ISL_579719, EPI_ISL_579720, EPI_ISL_579721, EPI_ISL_579722, EPI_ISL_579723, EPI_ISL_579724, EPI_ISL_579725, EPI_ISL_579726, EPI_ISL_579727, EPI_ISL_579728, EPI_ISL_579729, EPI_ISL_579730, EPI_ISL_579731, EPI_ISL_579732, EPI_ISL_579733, EPI_ISL_579734, EPI_ISL_579735, EPI_ISL_579736, EPI_ISL_579737, EPI_ISL_579738, EPI_ISL_579739, EPI_ISL_579740, EPI_ISL_579741, EPI_ISL_579742, EPI_ISL_579743, EPI_ISL_579744, EPI_ISL_579745, EPI_ISL_579746, EPI_ISL_579747, EPI_ISL_579748, EPI_ISL_579749, EPI_ISL_579750, EPI_ISL_579751, EPI_ISL_579752, EPI_ISL_579753, EPI_ISL_579754, EPI_ISL_579755, EPI_ISL_579756, EPI_ISL_579757, EPI_ISL_579758 | | | |
| see above | QEII Health Sciences Centre | National Microbiology Laboratory (NML) | Anna Majer, Shari Tyson, Grace Seo, Philip Mabon, Darian Hole, Elsie Grudeski, Rhiannon Huzarewich, Russell Mandes, Anneliese Landgraff, Jennifer Tanner, Natalie Knox, Morag Graham, Gary Van Domselaar, Todd Hatchette, Jason LeBlanc, Nathalie Bastien, Yan Li, Timothy Booth, CanCOGeN's metadata curation team, Public Health Agency of Canada's CanCOGeN team |
| EPI_ISL_582110, EPI_ISL_582111, EPI_ISL_582112, EPI_ISL_582113, EPI_ISL_582114, EPI_ISL_582115, EPI_ISL_582116, EPI_ISL_582117, EPI_ISL_582118, EPI_ISL_582119, EPI_ISL_582120 | | | |
| see above | CNR Virus des Infections Respiratoires - France SUD | CNR Virus des Infections Respiratoires - France SUD | Antonin Bal, Gregory Destras, Gwendolyne Burfin, Hadrien Règue, Alexandre Gaymard, Maude Bouscambert-Duchamp, Florence Morfin-Sherpa, Martine Valette, Bruno Lina, Laurence Josset |
| EPI_ISL_582121 | Centre Hospitalier de Bourg en Bresse | CNR Virus des Infections Respiratoires - France SUD | Antonin Bal, Gregory Destras, Gwendolyne Burfin, Hadrien Règue, Alexandre Gaymard, Maude Bouscambert-Duchamp, Florence Morfin-Sherpa, Martine Valette, Bruno Lina, Laurence Josset |
| EPI_ISL_582122 | CNR Virus des Infections Respiratoires - France SUD | CNR Virus des Infections Respiratoires - France SUD | Antonin Bal, Gregory Destras, Gwendolyne Burfin, Hadrien Règue, Alexandre Gaymard, Maude Bouscambert-Duchamp, Florence Morfin-Sherpa, Martine Valette, Bruno Lina, Laurence Josset |
| EPI_ISL_582242, EPI_ISL_582243, EPI_ISL_582244, EPI_ISL_582245, EPI_ISL_582246, EPI_ISL_582247, EPI_ISL_582248, EPI_ISL_582249, EPI_ISL_582251, EPI_ISL_582252, EPI_ISL_582253, EPI_ISL_582254, EPI_ISL_582255, EPI_ISL_582256, EPI_ISL_582257, EPI_ISL_582258, EPI_ISL_582259, EPI_ISL_582260, EPI_ISL_582261, EPI_ISL_582262, EPI_ISL_582263, EPI_ISL_582264, EPI_ISL_582265, EPI_ISL_582266, EPI_ISL_582267, EPI_ISL_582268, EPI_ISL_582269, EPI_ISL_582270, EPI_ISL_582271, EPI_ISL_582272, EPI_ISL_582273, EPI_ISL_582274, EPI_ISL_582275, EPI_ISL_582276, EPI_ISL_582277, EPI_ISL_582278, EPI_ISL_582279, EPI_ISL_582280, EPI_ISL_582281, EPI_ISL_582282, EPI_ISL_582283, EPI_ISL_582284, EPI_ISL_582285, EPI_ISL_582286, EPI_ISL_582288, EPI_ISL_582289, EPI_ISL_582290, EPI_ISL_582291, EPI_ISL_582292, EPI_ISL_582293, EPI_ISL_582294, EPI_ISL_582295, EPI_ISL_582296, EPI_ISL_582297, EPI_ISL_582298, EPI_ISL_582299, EPI_ISL_582300, EPI_ISL_582301, EPI_ISL_582302, EPI_ISL_582303, EPI_ISL_582304, EPI_ISL_582305, EPI_ISL_582306, EPI_ISL_582308, EPI_ISL_582311, EPI_ISL_582312, EPI_ISL_582313, EPI_ISL_582314, EPI_ISL_582315, EPI_ISL_582316, EPI_ISL_582317, EPI_ISL_582318, EPI_ISL_582319, EPI_ISL_582320, EPI_ISL_582321, EPI_ISL_582322, EPI_ISL_582323, EPI_ISL_582324, EPI_ISL_582325, EPI_ISL_582326, EPI_ISL_582327, EPI_ISL_582328, EPI_ISL_582329, EPI_ISL_582330, EPI_ISL_582331, EPI_ISL_582332, EPI_ISL_582333, EPI_ISL_582334, EPI_ISL_582335, EPI_ISL_582336, EPI_ISL_582337, EPI_ISL_582338, EPI_ISL_582339, EPI_ISL_582340, EPI_ISL_582341, EPI_ISL_582342, EPI_ISL_582343, EPI_ISL_582345, EPI_ISL_582346, EPI_ISL_582347, EPI_ISL_582349, EPI_ISL_582350, EPI_ISL_582351, EPI_ISL_582352, EPI_ISL_582353, EPI_ISL_582354, EPI_ISL_582355, EPI_ISL_582356, EPI_ISL_582357, EPI_ISL_582358, EPI_ISL_582359, EPI_ISL_582360, EPI_ISL_582361, EPI_ISL_582362, EPI_ISL_582363, EPI_ISL_582364, EPI_ISL_582365, EPI_ISL_582366, EPI_ISL_582367, EPI_ISL_582368, EPI_ISL_582369, EPI_ISL_582370, EPI_ISL_582371, EPI_ISL_582372, EPI_ISL_582373, EPI_ISL_582374, EPI_ISL_582375, EPI_ISL_582376, EPI_ISL_582377, EPI_ISL_582378, EPI_ISL_582379, EPI_ISL_582380, EPI_ISL_582381, EPI_ISL_582382, EPI_ISL_582383, EPI_ISL_582384, EPI_ISL_582385, EPI_ISL_582386, EPI_ISL_582387, EPI_ISL_582388, EPI_ISL_582389, EPI_ISL_582390, EPI_ISL_582391, EPI_ISL_582392, EPI_ISL_582393, EPI_ISL_582394, EPI_ISL_582395, EPI_ISL_582396, EPI_ISL_582397, EPI_ISL_582399, EPI_ISL_582400, EPI_ISL_582401, EPI_ISL_582402, EPI_ISL_582403, EPI_ISL_582404, EPI_ISL_582405, EPI_ISL_582406, EPI_ISL_582407, EPI_ISL_582408, EPI_ISL_582409, EPI_ISL_582410, EPI_ISL_582411, EPI_ISL_582412, EPI_ISL_582413, EPI_ISL_582414, EPI_ISL_582416, EPI_ISL_582417, EPI_ISL_582418, EPI_ISL_582419, EPI_ISL_582420, EPI_ISL_582422, EPI_ISL_582423, EPI_ISL_582424, EPI_ISL_582425, EPI_ISL_582426, EPI_ISL_582427, EPI_ISL_582428, EPI_ISL_582429, EPI_ISL_582430, EPI_ISL_582431, EPI_ISL_582432, EPI_ISL_582433, EPI_ISL_582434, EPI_ISL_582435, EPI_ISL_582436, EPI_ISL_582437, EPI_ISL_582438, EPI_ISL_582439, EPI_ISL_582440, EPI_ISL_582441, EPI_ISL_582442, EPI_ISL_582443, EPI_ISL_582445, EPI_ISL_582446, EPI_ISL_582447, EPI_ISL_582448, EPI_ISL_582449, EPI_ISL_582450, EPI_ISL_582451, EPI_ISL_582452, EPI_ISL_582453, EPI_ISL_582454, EPI_ISL_582455, EPI_ISL_582456, EPI_ISL_582457, EPI_ISL_582458, EPI_ISL_582459, EPI_ISL_582460, EPI_ISL_582461, EPI_ISL_582462, EPI_ISL_582463, EPI_ISL_582464, EPI_ISL_582465, EPI_ISL_582466, EPI_ISL_582467, EPI_ISL_582468, EPI_ISL_582469, EPI_ISL_582470, EPI_ISL_582471, EPI_ISL_582472, EPI_ISL_582473, EPI_ISL_582474, EPI_ISL_582475, EPI_ISL_582476, EPI_ISL_582477, EPI_ISL_582478, EPI_ISL_582479, EPI_ISL_582480, EPI_ISL_582481, EPI_ISL_582482, EPI_ISL_582483, EPI_ISL_582484, EPI_ISL_582485, EPI_ISL_582486, EPI_ISL_582487, EPI_ISL_582488, EPI_ISL_582489, EPI_ISL_582490, EPI_ISL_582491, EPI_ISL_582492, EPI_ISL_582493, EPI_ISL_582494, EPI_ISL_582495, EPI_ISL_582496, EPI_ISL_582497, EPI_ISL_582498, EPI_ISL_582499, EPI_ISL_582500, EPI_ISL_582501, EPI_ISL_582502, EPI_ISL_582503, EPI_ISL_582504, EPI_ISL_582505, EPI_ISL_582506, EPI_ISL_582507 | | | |
| see above | Cadham Provincial Laboratory | National Microbiology Laboratory (NML) | Anna Majer, Shari Tyson, Grace Seo, Philip Mabon, Elsie Grudeski, Rhiannon Huzarewich, Russell Mandes, Anneliese Landgraff, Jennifer Tanner, Natalie Knox, Morag Graham, Gary Van Domselaar, Paul Van Caeseele, Jared Bullard, David Alexander, Kerry Dust, Nathalie Bastien, Yan Li, Timothy Booth, Darian Hole, Madison Chapel, CanCOGeN's metadata curation team, Public Health Agency of Canada CanCOGeN team |
| EPI_ISL_582508 | CNR Virus des Infections Respiratoires - France SUD | CNR Virus des Infections Respiratoires - France SUD | Antonin Bal, Gregory Destras, Gwendolyne Burfin, Hadrien Règue, Alexandre Gaymard, Maude Bouscambert-Duchamp, Florence Morfin-Sherpa, Martine Valette, Bruno Lina, Laurence Josset |
| EPI_ISL_582771, EPI_ISL_582772, EPI_ISL_582773, EPI_ISL_582774, EPI_ISL_582775 | Klinsisk mikrobiologi Linkoping | The Public Health Agency of Sweden | Anna-Malin Linde, Maria Lind Karlberg, Mattias Haukland, Reza Advani, Olov Svartstrom, Oskar Karlsson Lindsjo, Sandra Broddesson, Petra Edquist, Mia Brytting, Anna Risberg, Karin Tegmark-Wisell |
| EPI_ISL_582776, EPI_ISL_582777, EPI_ISL_582778, EPI_ISL_582779, EPI_ISL_582780, EPI_ISL_582781, EPI_ISL_582782, EPI_ISL_582783, EPI_ISL_582784 | Uppsala klinisk mikrobiologi | The Public Health Agency of Sweden | Anna-Malin Linde, Maria Lind Karlberg, Mattias Haukland, Reza Advani, Olov Svartstrom, Oskar Karlsson Lindsjo, Sandra Broddesson, Petra Edquist, Mia Brytting, Anna Risberg, Karin Tegmark-Wisell |
| EPI_ISL_582785, EPI_ISL_582786, EPI_ISL_582787, EPI_ISL_582788, EPI_ISL_582789, EPI_ISL_582791, EPI_ISL_582792, EPI_ISL_582793 | Orebro klinisk mikrobiologi | The Public Health Agency of Sweden | Anna-Malin Linde, Maria Lind Karlberg, Mattias Haukland, Reza Advani, Olov Svartstrom, Oskar Karlsson Lindsjo, Sandra Broddesson, Petra Edquist, Mia Brytting, Anna Risberg, Karin Tegmark-Wisell |
| EPI_ISL_582794, EPI_ISL_582795, EPI_ISL_582796 | Unilabs Eskilstuna | The Public Health Agency of Sweden | Anna-Malin Linde, Maria Lind Karlberg, Mattias Haukland, Reza Advani, Olov Svartstrom, Oskar Karlsson Lindsjo, Sandra Broddesson, Petra Edquist, Mia Brytting, Anna Risberg, Karin Tegmark-Wisell |
| EPI_ISL_582797, EPI_ISL_582798 | Klinisk mikrobiologi Vasternorrland | The Public Health Agency of Sweden | Anna-Malin Linde, Maria Lind Karlberg, Mattias Haukland, Reza Advani, Olov Svartstrom, Oskar Karlsson Lindsjo, Sandra Broddesson, Petra Edquist, Mia Brytting, Anna Risberg, Karin Tegmark-Wisell |
| EPI_ISL_582799 | Stockholm_KUL Solna | The Public Health Agency of Sweden | Anna-Malin Linde, Maria Lind Karlberg, Mattias Haukland, Reza Advani, Olov Svartstrom, Oskar Karlsson Lindsjo, Sandra Broddesson, Petra Edquist, Mia Brytting, Anna Risberg, Karin Tegmark-Wisell |
| EPI_ISL_582800 | Klinisk mikrobiologi Vasternorrland | The Public Health Agency of Sweden | Anna-Malin Linde, Maria Lind Karlberg, Mattias Haukland, Reza Advani, Olov Svartstrom, Oskar Karlsson Lindsjo, Sandra Broddesson, Petra Edquist, Mia Brytting, Anna Risberg, Karin Tegmark-Wisell |
| EPI_ISL_582801 | Gavle klinisk mikrobiologi | The Public Health Agency of Sweden | Anna-Malin Linde, Maria Lind Karlberg, Mattias Haukland, Reza Advani, Olov Svartstrom, Oskar Karlsson Lindsjo, Sandra Broddesson, Petra Edquist, Mia Brytting, Anna Risberg, Karin Tegmark-Wisell |
| EPI_ISL_582803, EPI_ISL_582804, EPI_ISL_582805 | Halmstad klinisk mikrobiologi | The Public Health Agency of Sweden | Anna-Malin Linde, Maria Lind Karlberg, Mattias Haukland, Reza Advani, Olov Svartstrom, Oskar Karlsson Lindsjo, Sandra Broddesson, Petra Edquist, Mia Brytting, Anna Risberg, Karin Tegmark-Wisell |
| EPI_ISL_582806, EPI_ISL_582807 | Klinisk mikrobiologi SAS Boras | The Public Health Agency of Sweden | Anna-Malin Linde, Maria Lind Karlberg, Mattias Haukland, Reza Advani, Olov Svartstrom, Oskar Karlsson Lindsjo, Sandra Broddesson, Petra Edquist, Mia Brytting, Anna Risberg, Karin Tegmark-Wisell |
| EPI_ISL_582808 | Klinisk Mikrobiologi | The Public Health Agency of Sweden | Anna-Malin Linde, Maria Lind Karlberg, Mattias Haukland, Reza Advani, Olov Svartstrom, Oskar Karlsson Lindsjo, Sandra Broddesson, Petra Edquist, Mia Brytting, Anna Risberg, Karin Tegmark-Wisell |
| EPI_ISL_582809, EPI_ISL_582834 | Gavle klinisk mikrobiologi | The Public Health Agency of Sweden | Anna-Malin Linde, Maria Lind Karlberg, Mattias Haukland, Reza Advani, Olov Svartstrom, Oskar Karlsson Lindsjo, Sandra Broddesson, Petra Edquist, Mia Brytting, Anna Risberg, Karin Tegmark-Wisell |
| EPI_ISL_582835, EPI_ISL_582836, | Klinisk mikrobiologi Vasternorrland | The Public Health Agency of Sweden | Anna-Malin Linde, Maria Lind Karlberg, Mattias Haukland, Reza Advani, Olov Svartstrom, Oskar Karlsson Lindsjo, Sandra Broddesson, Petra Edquist, Mia |

| | | | |
|---|---|---|---|
| EPI_ISL_582837, EPI_ISL_582838 | | | Brytting, Anna Risberg, Karin Tegmark-Wisell |
| EPI_ISL_582839 | Stockholm_KUL Solna | The Public Health Agency of Sweden | Anna-Malin Linde, Maria Lind Karlberg, Mattias Haukland, Reza Advani, Olov Svartstrom, Oskar Karlsson Lindsjo, Sandra Broddesson, Petra Edquist, Mia Brytting, Anna Risberg, Karin Tegmark-Wisell |
| EPI_ISL_582840 | Narhalsan Fjallbacka VC | The Public Health Agency of Sweden | Anna-Malin Linde, Maria Lind Karlberg, Mattias Haukland, Reza Advani, Olov Svartstrom, Oskar Karlsson Lindsjo, Sandra Broddesson, Petra Edquist, Mia Brytting, Anna Risberg, Karin Tegmark-Wisell |
| EPI_ISL_582841 | Orebro klinisk mikrobiologi | The Public Health Agency of Sweden | Anna-Malin Linde, Maria Lind Karlberg, Mattias Haukland, Reza Advani, Olov Svartstrom, Oskar Karlsson Lindsjo, Sandra Broddesson, Petra Edquist, Mia Brytting, Anna Risberg, Karin Tegmark-Wisell |
| EPI_ISL_583506, EPI_ISL_583507, EPI_ISL_583508, EPI_ISL_583509, EPI_ISL_583510, EPI_ISL_583511, EPI_ISL_583512, EPI_ISL_583513, EPI_ISL_583514, EPI_ISL_583515, EPI_ISL_583516, EPI_ISL_583517, EPI_ISL_583518, EPI_ISL_583519, EPI_ISL_583520, EPI_ISL_583521, EPI_ISL_583522, EPI_ISL_583523, EPI_ISL_583524, EPI_ISL_583525, EPI_ISL_583526, EPI_ISL_583527, EPI_ISL_583528, EPI_ISL_583529, EPI_ISL_583530, EPI_ISL_583531, EPI_ISL_583532, EPI_ISL_583533, EPI_ISL_583534 | | | |
| see above | Michigan Department of Health and Human Services, Bureau of Laboratories | Michigan Department of Health and Human Services, Bureau of Laboratories | Blankenship HM, Riner D, Soehnlen MK |
| EPI_ISL_584168, EPI_ISL_584169, EPI_ISL_584170, EPI_ISL_584171, EPI_ISL_584172, EPI_ISL_584173, EPI_ISL_584174, EPI_ISL_584175, EPI_ISL_584176, EPI_ISL_584177, EPI_ISL_584178, EPI_ISL_584179, EPI_ISL_584180, EPI_ISL_584181, EPI_ISL_584182, EPI_ISL_584183, EPI_ISL_584184, EPI_ISL_584185, EPI_ISL_584186, EPI_ISL_584187, EPI_ISL_584193, EPI_ISL_584194, EPI_ISL_584195, EPI_ISL_584200, EPI_ISL_584201, EPI_ISL_584202, EPI_ISL_584203, EPI_ISL_584204, EPI_ISL_584205, EPI_ISL_584206, EPI_ISL_584212, EPI_ISL_584213, EPI_ISL_584214, EPI_ISL_584215, EPI_ISL_584216, EPI_ISL_584809, EPI_ISL_584810 | | | |
| see above | Centre for Enzyme Innovation, University of Portsmouth / Translational Research Laboratory, Portsmouth Hospitals NHS Trust | COVID-19 Genomics UK (COG-UK) Consortium | Angela Beckett,Yann Bourgeois,Garry Scarlett,Sharon Glaysher,Scott Elliott,Kelly Bicknell,Robert Impey,Allyson Lloyd,Sarah Wyllie,Ethan Butcher,Anoop Chauhan,Samuel Robson |
| EPI_ISL_584883, EPI_ISL_584884, EPI_ISL_584885, EPI_ISL_584886, EPI_ISL_584887, EPI_ISL_584888, EPI_ISL_584889, EPI_ISL_584890, EPI_ISL_584891, EPI_ISL_584892, EPI_ISL_584893, EPI_ISL_584894, EPI_ISL_584895, EPI_ISL_584896, EPI_ISL_584897, EPI_ISL_584898, EPI_ISL_584899, EPI_ISL_584900, EPI_ISL_584901, EPI_ISL_584902, EPI_ISL_584903, EPI_ISL_584904, EPI_ISL_584905, EPI_ISL_585533, EPI_ISL_585534, EPI_ISL_585535, EPI_ISL_585536, EPI_ISL_585537, EPI_ISL_585538, EPI_ISL_585539, EPI_ISL_585540, EPI_ISL_585541, EPI_ISL_585542, EPI_ISL_585543, EPI_ISL_585544, EPI_ISL_585545, EPI_ISL_585546, EPI_ISL_585547, EPI_ISL_585548, EPI_ISL_585549, EPI_ISL_585550, EPI_ISL_585551, EPI_ISL_585552, EPI_ISL_585553, EPI_ISL_585554, EPI_ISL_585555, EPI_ISL_585556, EPI_ISL_585557, EPI_ISL_585558, EPI_ISL_585559, EPI_ISL_585560, EPI_ISL_585561, EPI_ISL_585562, EPI_ISL_585563, EPI_ISL_585564, EPI_ISL_585565, EPI_ISL_585566, EPI_ISL_585567, EPI_ISL_585568, EPI_ISL_585569, EPI_ISL_585570, EPI_ISL_585571, EPI_ISL_585572, EPI_ISL_585573, EPI_ISL_585574, EPI_ISL_585575, EPI_ISL_585576, EPI_ISL_585577, EPI_ISL_585578, EPI_ISL_585579, EPI_ISL_585580, EPI_ISL_585581, EPI_ISL_585582 | | | |
| see above | University of Exeter | COVID-19 Genomics UK (COG-UK) Consortium | Ben Temperton,Aaron Jeffries,Michelle Michelsen,Joanna Warwick-Dugdale,Audrey Farbos,Robyn Manley,Stephen Michell,Jane Masoli |
| EPI_ISL_586527 | B.J. Medical College and Civil hospital, Ahmedabad | Gujarat Biotechnology Research Centre | Apurvasinh Puvar, Janvi Raval, Zarna Patel, Monika Gandhi, Pinal Trivedi, Maharshi Pandya, Nidhi Patel, Nitin Savaliya, Raghawendra Kumar, Dinesh Kumar, Zuber Saiyed, Komal Patel, Labdhi Pandya, Afzal Ansari, Nikha Trivedi, Pranay Shah, Kamlesh J Upadhyay, Sanjay Kapadia, R D Dixit, A M Kadri, Harsh Bakshi, Chaitanya Joshi, Madhvi Joshi |
| EPI_ISL_586532 | General Hospital,Chhotaudepur | Gujarat Biotechnology Research Centre | Arti Thakur, Dharak Pandya, Maharshi Pandya, Nidhi Patel, Nitin Savaliya, Raghawendra Kumar, Dinesh Kumar, Zuber Saiyed, Komal Patel, Labdhi Pandya, Afzal Ansari, Nikha Trivedi, Apurvasinh Puvar, Janvi Raval, Zarna Patel, Monika Gandhi, Pinal Trivedi, Yogesh Parmar, R D Dixit, A M Kadri, Harsh Bakshi, Chaitanya Joshi, Madhvi Joshi |
| EPI_ISL_586563 | GMERS GMERS Medical College and Hospital, Junagadh | Gujarat Biotechnology Research Centre | Bhavesh Bagda , Ravi Dedaniya, Apurvasinh Puvar, Janvi Raval, Zarna Patel, Monika Gandhi, Pinal Trivedi, Maharshi Pandya, Nidhi Patel, Nitin Savaliya, Raghawendra Kumar, Dinesh Kumar, Zuber Saiyed, Komal Patel, Labdhi Pandya, Afzal Ansari, Nikha Trivedi, R D Dixit, A M Kadri, Harsh Bakshi, Chaitanya Joshi, Madhvi Joshi |
| EPI_ISL_586565 | GMERS GMERS Medical College and Hospital, Junagadh | Gujarat Biotechnology Research Centre | Apurvasinh Puvar, Janvi Raval, Zarna Patel, Monika Gandhi, Pinal Trivedi, Maharshi Pandya, Nidhi Patel, Nitin Savaliya, Raghawendra Kumar, Dinesh Kumar, Zuber Saiyed, Komal Patel, Labdhi Pandya, Afzal Ansari, Nikha Trivedi, Bhavesh Bagda , Ravi Dedaniya, R D Dixit, A M Kadri, Harsh Bakshi, Chaitanya Joshi, Madhvi Joshi |
| EPI_ISL_590690 | CCC,Veraval | Gujarat Biotechnology Research Centre | Apurvasinh Puvar, Janvi Raval, Zarna Patel, Monika Gandhi, Pinal Trivedi, Maharshi Pandya, Nidhi Patel, Nitin Savaliya, Raghawendra Kumar, Dinesh Kumar, Zuber Saiyed, Komal Patel, Labdhi Pandya, Afzal Ansari, Nikha Trivedi, Jignesh Parmar, Jitendra Bamrotia, R D Dixit, A M Kadri, Harsh Bakshi, Chaitanya Joshi, Madhvi Joshi |
| EPI_ISL_590694 | INMI Lazzaro Spallanzani IRCCS | INMI Lazzaro Spallanzani IRCCS | Barbara Bartolini, Martina Rueca, Francesco Messina, Cesare E.M. Gruber, Emanuela Giombini, Beatrice Valli, Eleonora Lalle, Simone Lanini, Francesco Vairo, Maria R. Capobianchi, Antonino Di Caro |
| EPI_ISL_590698 | INMI Lazzaro Spallanzani IRCCS | INMI Lazzaro Spallanzani IRCCS | Barbara Bartolini, Francesco Messina, Cesare E.M. Gruber, Martina Rueca, Emanuela Giombini, Beatrice Valli, Eleonora Lalle, Simone Lanini, Francesco Vairo, Maria R. Capobianchi, Antonino Di Caro |
| EPI_ISL_594148 | Halmstad klinisk mikrobiologi | The Public Health Agency of Sweden | Anna-Malin Linde, Maria Lind Karlberg, Mattias Haukland, Reza Advani, Olov Svartstrom, Oskar Karlsson Lindsjo, Sandra Broddesson, Petra Edquist, Mia Brytting, Anna Risberg, Karin Tegmark-Wisell |
| EPI_ISL_594149, EPI_ISL_594150, EPI_ISL_594151, EPI_ISL_594152, EPI_ISL_594153 | Klinsisk mikrobiologi Linkoping | The Public Health Agency of Sweden | Anna-Malin Linde, Maria Lind Karlberg, Mattias Haukland, Reza Advani, Olov Svartstrom, Oskar Karlsson Lindsjo, Sandra Broddesson, Petra Edquist, Mia Brytting, Anna Risberg, Karin Tegmark-Wisell |
| EPI_ISL_594154 | Unilabs Eskilstuna | The Public Health Agency of Sweden | Anna-Malin Linde, Maria Lind Karlberg, Mattias Haukland, Reza Advani, Olov Svartstrom, Oskar Karlsson Lindsjo, Sandra Broddesson, Petra Edquist, Mia Brytting, Anna Risberg, Karin Tegmark-Wisell |
| EPI_ISL_594190, EPI_ISL_594191, EPI_ISL_594192, EPI_ISL_594193, EPI_ISL_594194, EPI_ISL_594195, EPI_ISL_594196, EPI_ISL_594197, EPI_ISL_594198, EPI_ISL_594199, EPI_ISL_594200, EPI_ISL_594201, EPI_ISL_594202, EPI_ISL_594203, EPI_ISL_594204, EPI_ISL_594205, EPI_ISL_594206, EPI_ISL_594207, EPI_ISL_594208, EPI_ISL_594209, EPI_ISL_594210, EPI_ISL_594211, EPI_ISL_594212, EPI_ISL_594213, EPI_ISL_594214, EPI_ISL_594215, EPI_ISL_594216, EPI_ISL_594217, EPI_ISL_594218 | | | |
| see above | Michigan Department of Health and Human Services, Bureau of Laboratories | Michigan Department of Health and Human Services, Bureau of Laboratories | Blankenship HM, Riner D, Soehnlen MK |
| EPI_ISL_595100, EPI_ISL_595101, EPI_ISL_595103, EPI_ISL_595104, EPI_ISL_595107, EPI_ISL_595108, EPI_ISL_595111, EPI_ISL_595113, EPI_ISL_595114 | Centre for Enzyme Innovation, University of Portsmouth / Translational Research Laboratory, Portsmouth Hospitals NHS Trust | COVID-19 Genomics UK (COG-UK) Consortium | Angela Beckett,Yann Bourgeois,Garry Scarlett,Sharon Glaysher,Scott Elliott,Kelly Bicknell,Robert Impey,Allyson Lloyd,Sarah Wyllie,Ethan Butcher,Anoop Chauhan,Samuel Robson |
| EPI_ISL_603142 | INMI Lazzaro Spallanzani IRCCS | INMI Lazzaro Spallanzani IRCCS | Barbara Bartolini, Cesare E.M. Gruber, Francesco Messina, Martina Rueca, Simone Lanini, Emanuela Giombini, Maria R. Capobianchi, Antonino Di Caro |
| EPI_ISL_603150 | INMI Lazzaro Spallanzani IRCCS | INMI Lazzaro Spallanzani IRCCS | Barbara Bartolini, Francesco Messina, Martina Rueca, Cesare E.M. Gruber, Simone Lanini, Emanuela Giombini, Patrizia Massarelli, Maria R. Capobianchi, Antonino Di Caro |
| EPI_ISL_603152 | INMI Lazzaro Spallanzani IRCCS | INMI Lazzaro Spallanzani IRCCS | Barbara Bartolini, Cesare E.M. Gruber, Francesco Messina, Martina Rueca, Emanuela Giombini, Simone Lanini, Patrizia Massarelli, Antonino Di Caro, Maria R. Capobianchi |
| EPI_ISL_603156 | INMI Lazzaro Spallanzani IRCCS | INMI Lazzaro Spallanzani IRCCS | Barbara Bartolini, Francesco Messina, Cesare E.M. Gruber, Martina Rueca, Emanuela Giombini, Simone Lanini, Maria R. Capobianchi, Antonino Di Caro |
| EPI_ISL_603158 | INMI Lazzaro Spallanzani IRCCS | INMI Lazzaro Spallanzani IRCCS | Barbara Bartolini, Cesare E.M. Gruber, Francesco Messina, Martina Rueca, Emanuela Giombini, Simone Lanini, Fulvia Pimpinelli, Maria R. Capobianchi, Antonino Di Caro |
| EPI_ISL_603159 | INMI Lazzaro Spallanzani IRCCS | INMI Lazzaro Spallanzani IRCCS | Barbara Bartolini, Francesco Messina, Cesare E.M. Gruber, Martina Rueca, Simone Lanini, Emanuela Giombini, Fulvia Pimpinelli, Maria R. Capobianchi, Antonino Di Caro |
| EPI_ISL_603160 | INMI Lazzaro Spallanzani IRCCS | INMI Lazzaro Spallanzani IRCCS | Barbara Bartolini, Simone Lanini, Emanuela Giombini, Francesco Messina, Cesare E.M. Gruber, Martina Rueca, Fulvia Pimpinelli, Maria R. Capobianchi, Antonino Di Caro |
| EPI_ISL_603171 | INMI Lazzaro Spallanzani IRCCS | INMI Lazzaro Spallanzani IRCCS | Barbara Bartolini, Cesare E.M. Gruber, Francesco Messina, Martina Rueca, Emanuela Giombini, Simone Lanini, Mariarosa Gaudio, Antonino Di Caro, Maria R. Capobianchi |

| | | | |
|---|---|---|---|
| EPI_ISL_603173 | INMI Lazzaro Spallanzani IRCCS | INMI Lazzaro Spallanzani IRCCS | Barbara Bartolini, Cesare E.M. Gruber, Martina Rueca, Francesco Messina, Simone Lanini, Emanuela Giombini, Giuseppina Cappiello, Maria R. Capobianchi, Antonino Di Caro |
| EPI_ISL_603175 | INMI Lazzaro Spallanzani IRCCS | INMI Lazzaro Spallanzani IRCCS | Barbara Bartolini, Cesare E.M. Gruber, Simone Lanini, Emanuela Giombini, Francesco Messina, Martina Rueca, Mariarosa Gaudio, Antonino Di Caro, Maria R. Capobianchi |
| EPI_ISL_603177 | INMI Lazzaro Spallanzani IRCCS | INMI Lazzaro Spallanzani IRCCS | Barbara Bartolini, Francesco Messina, Emanuela Giombini, Martina Rueca, Cesare E.M. Gruber, Simone Lanini, Maria R. Capobianchi, Antonino Di Caro |
| EPI_ISL_603180 | INMI Lazzaro Spallanzani IRCCS | INMI Lazzaro Spallanzani IRCCS | Barbara Bartolini, Cesare E.M. Gruber, Francesco Messina, Simone Lanini, Martina Rueca, Emanuela Giombini, Maria R. Capobianchi, Antonino Di Caro |
| EPI_ISL_603183 | INMI Lazzaro Spallanzani IRCCS | INMI Lazzaro Spallanzani IRCCS | Barbara Bartolini, Francesco Messina, Martina Rueca, Cesare E.M. Gruber, Emanuela Giombini, Simone Lanini, Maria R. Capobianchi, Antonino Di Caro |
| EPI_ISL_603242, EPI_ISL_603243 | National Institute of Laboratory Medicine and Referral Center | Genomic Research Lab, BCSIR | Barna Goswami, Abu Sayeed Mohammad Mahmud, Mohammad Samir Uzzaman, Eshrar Osman, Md. Ahashan Habib, Shahina Akter, Tanjina Akhter Banu, Md. Murshed Hasan Sarkar, Iffat Jahan, Md. Saddam Hossain, Tasnim Nafisa, Md. Maruf Ahmed Molla, Mahmuda Yeasmin, Asish Kumar Ghosh, A. K. M. Shamsuzzaman, Monira Parveen, Md. Masum Hossain Arif, Md. Salim Khan |
| EPI_ISL_609990 | INMI Lazzaro Spallanzani IRCCS | INMI Lazzaro Spallanzani IRCCS | B Bartolini, C.E.M Gruber, M Rueca, F Messina, E Giombini, MR Capobianchi, A Di Caro |
| EPI_ISL_609999 | INMI Lazzaro Spallanzani IRCCS | INMI Lazzaro Spallanzani IRCCS | B Bartolini, M Rueca, C.E.M Gruber, F Messina, E Giombini, MR Capobianchi, A Di Caro |
| EPI_ISL_611526, EPI_ISL_611532, EPI_ISL_611535, EPI_ISL_611537, EPI_ISL_611538 | Centre for Enzyme Innovation, University of Portsmouth / Translational Research Laboratory, Portsmouth Hospitals NHS Trust | COVID-19 Genomics UK (COG-UK) Consortium | Angela Beckett,Yann Bourgeois,Garry Scarlett,Sharon Glaysher,Scott Elliott,Kelly Bicknell,Robert Impey,Allyson Lloyd,Sarah Wyllie,Ethan Butcher,Anoop Chauhan,Samuel Robson |
| EPI_ISL_611539, EPI_ISL_611540, EPI_ISL_611551, EPI_ISL_611552 | University of Exeter | COVID-19 Genomics UK (COG-UK) Consortium | Ben Temperton,Aaron Jeffries,Michelle Michelsen,Joanna Warwick-Dugdale,Audrey Farbos,Robyn Manley,Stephen Michell,Jane Masoli |
| EPI_ISL_611555, EPI_ISL_611557, EPI_ISL_611585, EPI_ISL_611589, EPI_ISL_611590, EPI_ISL_611592 | Centre for Enzyme Innovation, University of Portsmouth / Translational Research Laboratory, Portsmouth Hospitals NHS Trust | COVID-19 Genomics UK (COG-UK) Consortium | Angela Beckett,Yann Bourgeois,Garry Scarlett,Sharon Glaysher,Scott Elliott,Kelly Bicknell,Robert Impey,Allyson Lloyd,Sarah Wyllie,Ethan Butcher,Anoop Chauhan,Samuel Robson |
| EPI_ISL_611599, EPI_ISL_611607, EPI_ISL_611615, EPI_ISL_611616, EPI_ISL_611617, EPI_ISL_611618, EPI_ISL_611619, EPI_ISL_611620 | University of Exeter | COVID-19 Genomics UK (COG-UK) Consortium | Ben Temperton,Aaron Jeffries,Michelle Michelsen,Joanna Warwick-Dugdale,Audrey Farbos,Robyn Manley,Stephen Michell,Jane Masoli |
| EPI_ISL_611638 | Centre for Enzyme Innovation, University of Portsmouth / Translational Research Laboratory, Portsmouth Hospitals NHS Trust | COVID-19 Genomics UK (COG-UK) Consortium | Angela Beckett,Yann Bourgeois,Garry Scarlett,Sharon Glaysher,Scott Elliott,Kelly Bicknell,Robert Impey,Allyson Lloyd,Sarah Wyllie,Ethan Butcher,Anoop Chauhan,Samuel Robson |
| EPI_ISL_611641 | University of Exeter | COVID-19 Genomics UK (COG-UK) Consortium | Ben Temperton,Aaron Jeffries,Michelle Michelsen,Joanna Warwick-Dugdale,Audrey Farbos,Robyn Manley,Stephen Michell,Jane Masoli |
| EPI_ISL_611689 | Centre for Enzyme Innovation, University of Portsmouth / Translational Research Laboratory, Portsmouth Hospitals NHS Trust | COVID-19 Genomics UK (COG-UK) Consortium | Angela Beckett,Yann Bourgeois,Garry Scarlett,Sharon Glaysher,Scott Elliott,Kelly Bicknell,Robert Impey,Allyson Lloyd,Sarah Wyllie,Ethan Butcher,Anoop Chauhan,Samuel Robson |
| EPI_ISL_611701, EPI_ISL_611702, EPI_ISL_611703 | University of Exeter | COVID-19 Genomics UK (COG-UK) Consortium | Ben Temperton,Aaron Jeffries,Michelle Michelsen,Joanna Warwick-Dugdale,Audrey Farbos,Robyn Manley,Stephen Michell,Jane Masoli |
| EPI_ISL_611755, EPI_ISL_611761, EPI_ISL_611763, EPI_ISL_611771, EPI_ISL_611772, EPI_ISL_611773 | Centre for Enzyme Innovation, University of Portsmouth / Translational Research Laboratory, Portsmouth Hospitals NHS Trust | COVID-19 Genomics UK (COG-UK) Consortium | Angela Beckett,Yann Bourgeois,Garry Scarlett,Sharon Glaysher,Scott Elliott,Kelly Bicknell,Robert Impey,Allyson Lloyd,Sarah Wyllie,Ethan Butcher,Anoop Chauhan,Samuel Robson |
| EPI_ISL_611836, EPI_ISL_611837, EPI_ISL_611838 | University of Exeter | COVID-19 Genomics UK (COG-UK) Consortium | Ben Temperton,Aaron Jeffries,Michelle Michelsen,Joanna Warwick-Dugdale,Audrey Farbos,Robyn Manley,Stephen Michell,Jane Masoli |
| EPI_ISL_611841 | Centre for Enzyme Innovation, University of Portsmouth / Translational Research Laboratory, Portsmouth Hospitals NHS Trust | COVID-19 Genomics UK (COG-UK) Consortium | Angela Beckett,Yann Bourgeois,Garry Scarlett,Sharon Glaysher,Scott Elliott,Kelly Bicknell,Robert Impey,Allyson Lloyd,Sarah Wyllie,Ethan Butcher,Anoop Chauhan,Samuel Robson |
| EPI_ISL_611843, EPI_ISL_611844 | University of Exeter | COVID-19 Genomics UK (COG-UK) Consortium | Ben Temperton,Aaron Jeffries,Michelle Michelsen,Joanna Warwick-Dugdale,Audrey Farbos,Robyn Manley,Stephen Michell,Jane Masoli |
| EPI_ISL_611916, EPI_ISL_611925, EPI_ISL_611934 | Centre for Enzyme Innovation, University of Portsmouth / Translational Research Laboratory, Portsmouth Hospitals NHS Trust | COVID-19 Genomics UK (COG-UK) Consortium | Angela Beckett,Yann Bourgeois,Garry Scarlett,Sharon Glaysher,Scott Elliott,Kelly Bicknell,Robert Impey,Allyson Lloyd,Sarah Wyllie,Ethan Butcher,Anoop Chauhan,Samuel Robson |
| EPI_ISL_611939, EPI_ISL_611940 | University of Exeter | COVID-19 Genomics UK (COG-UK) Consortium | Ben Temperton,Aaron Jeffries,Michelle Michelsen,Joanna Warwick-Dugdale,Audrey Farbos,Robyn Manley,Stephen Michell,Jane Masoli |
| EPI_ISL_611959, EPI_ISL_611960, EPI_ISL_611962 | Centre for Enzyme Innovation, University of Portsmouth / Translational Research Laboratory, Portsmouth Hospitals NHS Trust | COVID-19 Genomics UK (COG-UK) Consortium | Angela Beckett,Yann Bourgeois,Garry Scarlett,Sharon Glaysher,Scott Elliott,Kelly Bicknell,Robert Impey,Allyson Lloyd,Sarah Wyllie,Ethan Butcher,Anoop Chauhan,Samuel Robson |
| EPI_ISL_611976 | University of Exeter | COVID-19 Genomics UK (COG-UK) Consortium | Ben Temperton,Aaron Jeffries,Michelle Michelsen,Joanna Warwick-Dugdale,Audrey Farbos,Robyn Manley,Stephen Michell,Jane Masoli |
| EPI_ISL_611983 | Centre for Enzyme Innovation, University of Portsmouth / Translational Research Laboratory, Portsmouth Hospitals NHS Trust | COVID-19 Genomics UK (COG-UK) Consortium | Angela Beckett,Yann Bourgeois,Garry Scarlett,Sharon Glaysher,Scott Elliott,Kelly Bicknell,Robert Impey,Allyson Lloyd,Sarah Wyllie,Ethan Butcher,Anoop Chauhan,Samuel Robson |
| EPI_ISL_611987 | University of Exeter | COVID-19 Genomics UK (COG-UK) Consortium | Ben Temperton,Aaron Jeffries,Michelle Michelsen,Joanna Warwick-Dugdale,Audrey Farbos,Robyn Manley,Stephen Michell,Jane Masoli |
| EPI_ISL_611998 | Centre for Enzyme Innovation, University of Portsmouth / Translational Research Laboratory, Portsmouth Hospitals NHS Trust | COVID-19 Genomics UK (COG-UK) Consortium | Angela Beckett,Yann Bourgeois,Garry Scarlett,Sharon Glaysher,Scott Elliott,Kelly Bicknell,Robert Impey,Allyson Lloyd,Sarah Wyllie,Ethan Butcher,Anoop Chauhan,Samuel Robson |
| EPI_ISL_612056, EPI_ISL_612057 | University of Exeter | COVID-19 Genomics UK (COG-UK) Consortium | Ben Temperton,Aaron Jeffries,Michelle Michelsen,Joanna Warwick-Dugdale,Audrey Farbos,Robyn Manley,Stephen Michell,Jane Masoli |
| EPI_ISL_612068, EPI_ISL_612069, EPI_ISL_612082, EPI_ISL_612089, EPI_ISL_612092, EPI_ISL_612094 | Centre for Enzyme Innovation, University of Portsmouth / Translational Research Laboratory, Portsmouth Hospitals NHS Trust | COVID-19 Genomics UK (COG-UK) Consortium | Angela Beckett,Yann Bourgeois,Garry Scarlett,Sharon Glaysher,Scott Elliott,Kelly Bicknell,Robert Impey,Allyson Lloyd,Sarah Wyllie,Ethan Butcher,Anoop Chauhan,Samuel Robson |
| EPI_ISL_612095, EPI_ISL_612096, EPI_ISL_612118, EPI_ISL_612130, EPI_ISL_612375, EPI_ISL_612376, EPI_ISL_612377, EPI_ISL_612378, EPI_ISL_612379, EPI_ISL_612380, EPI_ISL_612381, EPI_ISL_612382, EPI_ISL_612383, EPI_ISL_612384, EPI_ISL_612385, EPI_ISL_612386, EPI_ISL_612387, EPI_ISL_612388, EPI_ISL_612389, EPI_ISL_612390, EPI_ISL_612391, EPI_ISL_612392, EPI_ISL_612393, EPI_ISL_612395, EPI_ISL_612396, EPI_ISL_612397, EPI_ISL_612398, EPI_ISL_612399, EPI_ISL_612400 | | | |
| see above | University of Exeter | COVID-19 Genomics UK (COG-UK) Consortium | Ben Temperton,Aaron Jeffries,Michelle Michelsen,Joanna Warwick-Dugdale,Audrey Farbos,Robyn Manley,Stephen Michell,Jane Masoli |
| EPI_ISL_613285, EPI_ISL_613294, EPI_ISL_613295, EPI_ISL_613296, EPI_ISL_613297, EPI_ISL_613298, EPI_ISL_613302, EPI_ISL_613303, EPI_ISL_613304, EPI_ISL_613305, EPI_ISL_613306 | | | |
| see above | Centre for Enzyme Innovation, University of Portsmouth / Translational Research Laboratory, Portsmouth Hospitals NHS Trust | COVID-19 Genomics UK (COG-UK) Consortium | Angela Beckett,Yann Bourgeois,Garry Scarlett,Sharon Glaysher,Scott Elliott,Kelly Bicknell,Robert Impey,Allyson Lloyd,Sarah Wyllie,Ethan Butcher,Anoop Chauhan,Samuel Robson |
| EPI_ISL_614158, EPI_ISL_614159, EPI_ISL_614160, EPI_ISL_614161, EPI_ISL_614162, EPI_ISL_614163, EPI_ISL_614164, EPI_ISL_614165, EPI_ISL_614166, EPI_ISL_614167, EPI_ISL_614168, EPI_ISL_614169, EPI_ISL_614170, EPI_ISL_614171, EPI_ISL_614172, EPI_ISL_614173, EPI_ISL_614174, EPI_ISL_614175, EPI_ISL_614176, EPI_ISL_614177, EPI_ISL_614178, EPI_ISL_614179, EPI_ISL_614180, EPI_ISL_614181, EPI_ISL_614182, EPI_ISL_614183, EPI_ISL_614184, EPI_ISL_614185, EPI_ISL_614186, EPI_ISL_614187, EPI_ISL_614188, EPI_ISL_614189, EPI_ISL_614190, EPI_ISL_614191, EPI_ISL_614192, EPI_ISL_614193, EPI_ISL_614194, EPI_ISL_614195, EPI_ISL_614196, EPI_ISL_614197, EPI_ISL_614198, EPI_ISL_614199, EPI_ISL_614200, EPI_ISL_614201, EPI_ISL_614202, EPI_ISL_614203, EPI_ISL_614204, EPI_ISL_614205, EPI_ISL_614206, EPI_ISL_614207, EPI_ISL_614208, EPI_ISL_614209, EPI_ISL_614210, EPI_ISL_614211, | | | |

| | | | |
|---|---|---|---|
| EPI_ISL_614212, EPI_ISL_614213, EPI_ISL_614214, EPI_ISL_614215, EPI_ISL_614216, EPI_ISL_614217, EPI_ISL_614218, EPI_ISL_614219, EPI_ISL_614220, EPI_ISL_614221, EPI_ISL_614222, EPI_ISL_614223, EPI_ISL_614224, EPI_ISL_614225, EPI_ISL_614226, EPI_ISL_614227, EPI_ISL_614228, EPI_ISL_614229, EPI_ISL_614230, EPI_ISL_614231, EPI_ISL_614232, EPI_ISL_614233, EPI_ISL_614234, EPI_ISL_614235, EPI_ISL_614236, EPI_ISL_614237, EPI_ISL_614238, EPI_ISL_614239, EPI_ISL_614240, EPI_ISL_614241, EPI_ISL_614242, EPI_ISL_614243, EPI_ISL_614244, EPI_ISL_614245, EPI_ISL_614246 | | | |
| see above | Michigan Department of Health and Human Services, Bureau of Laboratories | Michigan Department of Health and Human Services, Bureau of Laboratories | Blankenship HM, Riner D, Soehnlen MK |
| EPI_ISL_615076, EPI_ISL_615077, EPI_ISL_615078 | Uppsala klinisk mikrobiologi | The Public Health Agency of Sweden | Anna-Malin Linde, Maria Lind Karlberg, Mattias Haukland, Reza Advani, Olov Svartstrom, Oskar Karlsson Lindsjo, Sandra Broddesson, Petra Edquist, Mia Brytting, Anna Risberg, Karin Tegmark-Wisell |
| EPI_ISL_615079 | Orebro klinisk mikrobiologi | The Public Health Agency of Sweden | Anna-Malin Linde, Maria Lind Karlberg, Mattias Haukland, Reza Advani, Olov Svartstrom, Oskar Karlsson Lindsjo, Sandra Broddesson, Petra Edquist, Mia Brytting, Anna Risberg, Karin Tegmark-Wisell |
| EPI_ISL_615080, EPI_ISL_615081, EPI_ISL_615082, EPI_ISL_615083, EPI_ISL_615084, EPI_ISL_615085 | Klinisk mikrobiologi, Skanes universitetssjukhus, Lund | The Public Health Agency of Sweden | Anna-Malin Linde, Maria Lind Karlberg, Mattias Haukland, Reza Advani, Olov Svartstrom, Oskar Karlsson Lindsjo, Sandra Broddesson, Petra Edquist, Mia Brytting, Anna Risberg, Karin Tegmark-Wisell |
| EPI_ISL_615086, EPI_ISL_615087, EPI_ISL_615088, EPI_ISL_615089 | Karolinska Universitetslaboratoriet | The Public Health Agency of Sweden | Anna-Malin Linde, Maria Lind Karlberg, Mattias Haukland, Reza Advani, Olov Svartstrom, Oskar Karlsson Lindsjo, Sandra Broddesson, Petra Edquist, Mia Brytting, Anna Risberg, Karin Tegmark-Wisell |
| EPI_ISL_615090, EPI_ISL_615091 | Klinisk mikrobiologi NAL Trollhattan | The Public Health Agency of Sweden | Anna-Malin Linde, Maria Lind Karlberg, Mattias Haukland, Reza Advani, Olov Svartstrom, Oskar Karlsson Lindsjo, Sandra Broddesson, Petra Edquist, Mia Brytting, Anna Risberg, Karin Tegmark-Wisell |
| EPI_ISL_615092, EPI_ISL_615093, EPI_ISL_615094, EPI_ISL_615095, EPI_ISL_615096 | Umea klinisk mikrobiologi | The Public Health Agency of Sweden | Anna-Malin Linde, Maria Lind Karlberg, Mattias Haukland, Reza Advani, Olov Svartstrom, Oskar Karlsson Lindsjo, Sandra Broddesson, Petra Edquist, Mia Brytting, Anna Risberg, Karin Tegmark-Wisell |
| EPI_ISL_615097, EPI_ISL_615098, EPI_ISL_615099, EPI_ISL_615100 | Klinisk mikrobiologi Lanssjukhuset Ryhov, Jonkoping | The Public Health Agency of Sweden | Anna-Malin Linde, Maria Lind Karlberg, Mattias Haukland, Reza Advani, Olov Svartstrom, Oskar Karlsson Lindsjo, Sandra Broddesson, Petra Edquist, Mia Brytting, Anna Risberg, Karin Tegmark-Wisell |
| EPI_ISL_615101, EPI_ISL_615102, EPI_ISL_615103 | Gavle klinisk mikrobiologi | The Public Health Agency of Sweden | Anna-Malin Linde, Maria Lind Karlberg, Mattias Haukland, Reza Advani, Olov Svartstrom, Oskar Karlsson Lindsjo, Sandra Broddesson, Petra Edquist, Mia Brytting, Anna Risberg, Karin Tegmark-Wisell |
| EPI_ISL_615104, EPI_ISL_615105, EPI_ISL_615106, EPI_ISL_615107 | Halmstad klinisk mikrobiologi | The Public Health Agency of Sweden | Anna-Malin Linde, Maria Lind Karlberg, Mattias Haukland, Reza Advani, Olov Svartstrom, Oskar Karlsson Lindsjo, Sandra Broddesson, Petra Edquist, Mia Brytting, Anna Risberg, Karin Tegmark-Wisell |
| EPI_ISL_615108 | Klinisk Mikrobiologi | The Public Health Agency of Sweden | Anna-Malin Linde, Maria Lind Karlberg, Mattias Haukland, Reza Advani, Olov Svartstrom, Oskar Karlsson Lindsjo, Sandra Broddesson, Petra Edquist, Mia Brytting, Anna Risberg, Karin Tegmark-Wisell |
| EPI_ISL_615109 | Klinisk mikrobiologi NAL Trollhattan | The Public Health Agency of Sweden | Anna-Malin Linde, Maria Lind Karlberg, Mattias Haukland, Reza Advani, Olov Svartstrom, Oskar Karlsson Lindsjo, Sandra Broddesson, Petra Edquist, Mia Brytting, Anna Risberg, Karin Tegmark-Wisell |
| EPI_ISL_615110 | Klinsisk mikrobiologi Linkoping | The Public Health Agency of Sweden | Anna-Malin Linde, Maria Lind Karlberg, Mattias Haukland, Reza Advani, Olov Svartstrom, Oskar Karlsson Lindsjo, Sandra Broddesson, Petra Edquist, Mia Brytting, Anna Risberg, Karin Tegmark-Wisell |
| EPI_ISL_615111, EPI_ISL_615112, EPI_ISL_615113, EPI_ISL_615114 | Orebro klinisk mikrobiologi | The Public Health Agency of Sweden | Anna-Malin Linde, Maria Lind Karlberg, Mattias Haukland, Reza Advani, Olov Svartstrom, Oskar Karlsson Lindsjo, Sandra Broddesson, Petra Edquist, Mia Brytting, Anna Risberg, Karin Tegmark-Wisell |
| EPI_ISL_615115, EPI_ISL_615116 | Skovde/Unilabs | The Public Health Agency of Sweden | Anna-Malin Linde, Maria Lind Karlberg, Mattias Haukland, Reza Advani, Olov Svartstrom, Oskar Karlsson Lindsjo, Sandra Broddesson, Petra Edquist, Mia Brytting, Anna Risberg, Karin Tegmark-Wisell |
| EPI_ISL_615117, EPI_ISL_615118, EPI_ISL_615119 | Klinsisk mikrobiologi Linkoping | The Public Health Agency of Sweden | Anna-Malin Linde, Maria Lind Karlberg, Mattias Haukland, Reza Advani, Olov Svartstrom, Oskar Karlsson Lindsjo, Sandra Broddesson, Petra Edquist, Mia Brytting, Anna Risberg, Karin Tegmark-Wisell |
| EPI_ISL_615120 | Orebro klinisk mikrobiologi | The Public Health Agency of Sweden | Anna-Malin Linde, Maria Lind Karlberg, Mattias Haukland, Reza Advani, Olov Svartstrom, Oskar Karlsson Lindsjo, Sandra Broddesson, Petra Edquist, Mia Brytting, Anna Risberg, Karin Tegmark-Wisell |
| EPI_ISL_623076, EPI_ISL_623077, EPI_ISL_623078, EPI_ISL_623079 | Uppsala klinisk mikrobiologi | The Public Health Agency of Sweden | Anna-Malin Linde, Maria Lind Karlberg, Mattias Haukland, Reza Advani, Olov Svartstrom, Oskar Karlsson Lindsjo, Sandra Broddesson, Petra Edquist, Mia Brytting, Anna Risberg, Karin Tegmark-Wisell |
| EPI_ISL_623080, EPI_ISL_623081, EPI_ISL_623082, EPI_ISL_623083, EPI_ISL_623084, EPI_ISL_623085 | Klinisk mikrobiologi, Skanes universitetssjukhus, Lund | The Public Health Agency of Sweden | Anna-Malin Linde, Maria Lind Karlberg, Mattias Haukland, Reza Advani, Olov Svartstrom, Oskar Karlsson Lindsjo, Sandra Broddesson, Petra Edquist, Mia Brytting, Anna Risberg, Karin Tegmark-Wisell |
| EPI_ISL_623086 | Halmstad klinisk mikrobiologi | The Public Health Agency of Sweden | Anna-Malin Linde, Maria Lind Karlberg, Mattias Haukland, Reza Advani, Olov Svartstrom, Oskar Karlsson Lindsjo, Sandra Broddesson, Petra Edquist, Mia Brytting, Anna Risberg, Karin Tegmark-Wisell |
| EPI_ISL_623087, EPI_ISL_623088 | Klinisk mikrobiologi NAL Trollhattan | The Public Health Agency of Sweden | Anna-Malin Linde, Maria Lind Karlberg, Mattias Haukland, Reza Advani, Olov Svartstrom, Oskar Karlsson Lindsjo, Sandra Broddesson, Petra Edquist, Mia Brytting, Anna Risberg, Karin Tegmark-Wisell |
| EPI_ISL_623089 | Klinisk mikrobiologi Linkoping | The Public Health Agency of Sweden | Anna-Malin Linde, Maria Lind Karlberg, Mattias Haukland, Reza Advani, Olov Svartstrom, Oskar Karlsson Lindsjo, Sandra Broddesson, Petra Edquist, Mia Brytting, Anna Risberg, Karin Tegmark-Wisell |
| EPI_ISL_623090 | Klinisk mikrobiologi Vasternorrland | The Public Health Agency of Sweden | Anna-Malin Linde, Maria Lind Karlberg, Mattias Haukland, Reza Advani, Olov Svartstrom, Oskar Karlsson Lindsjo, Sandra Broddesson, Petra Edquist, Mia Brytting, Anna Risberg, Karin Tegmark-Wisell |
| EPI_ISL_623091 | Kalmar klinisk mikrobiologi | The Public Health Agency of Sweden | Anna-Malin Linde, Maria Lind Karlberg, Mattias Haukland, Reza Advani, Olov Svartstrom, Oskar Karlsson Lindsjo, Sandra Broddesson, Petra Edquist, Mia Brytting, Anna Risberg, Karin Tegmark-Wisell |
| EPI_ISL_623092, EPI_ISL_623093, EPI_ISL_623094 | Klinisk mikrobiologi Lanssjukhuset Ryhov, Jonkoping | The Public Health Agency of Sweden | Anna-Malin Linde, Maria Lind Karlberg, Mattias Haukland, Reza Advani, Olov Svartstrom, Oskar Karlsson Lindsjo, Sandra Broddesson, Petra Edquist, Mia Brytting, Anna Risberg, Karin Tegmark-Wisell |
| EPI_ISL_623095 | Gavle klinisk mikrobiologi | The Public Health Agency of Sweden | Anna-Malin Linde, Maria Lind Karlberg, Mattias Haukland, Reza Advani, Olov Svartstrom, Oskar Karlsson Lindsjo, Sandra Broddesson, Petra Edquist, Mia Brytting, Anna Risberg, Karin Tegmark-Wisell |
| EPI_ISL_623098, EPI_ISL_623099, EPI_ISL_623100, EPI_ISL_623101, EPI_ISL_623102 | CNR Virus des Infections Respiratoires - France SUD | CNR Virus des Infections Respiratoires - France SUD | Antonin Bal, Gregory Destras, Gwendolyne Burfin, Hadrien Règue, Alexandre Gaymard, Maude Bouscambert-Duchamp, Florence Morfin-Sherpa, Martine Valette, Bruno Lina, Laurence Josset |
| EPI_ISL_626662, EPI_ISL_626666, EPI_ISL_626668, EPI_ISL_626669, EPI_ISL_626671, EPI_ISL_626672, EPI_ISL_626673, EPI_ISL_626676 | University of Exeter | COVID-19 Genomics UK (COG-UK) Consortium | Ben Temperton,Aaron Jeffries,Michelle Michelsen,Joanna Warwick-Dugdale,Audrey Farbos,Robyn Manley,Stephen Michell,Jane Masoli |
| EPI_ISL_626677, EPI_ISL_626678 | Centre for Enzyme Innovation, University of Portsmouth / Translational Research Laboratory, Portsmouth Hospitals NHS Trust | COVID-19 Genomics UK (COG-UK) Consortium | Angela Beckett,Yann Bourgeois,Garry Scarlett,Sharon Glaysher,Scott Elliott,Kelly Bicknell,Robert Impey,Allyson Lloyd,Sarah Wyllie,Ethan Butcher,Anoop Chauhan,Samuel Robson |
| EPI_ISL_626682, EPI_ISL_626683 | University of Exeter | COVID-19 Genomics UK (COG-UK) Consortium | Ben Temperton,Aaron Jeffries,Michelle Michelsen,Joanna Warwick-Dugdale,Audrey Farbos,Robyn Manley,Stephen Michell,Jane Masoli |
| EPI_ISL_626684 | Centre for Enzyme Innovation, University of Portsmouth / Translational Research Laboratory, Portsmouth Hospitals NHS Trust | COVID-19 Genomics UK (COG-UK) Consortium | Angela Beckett,Yann Bourgeois,Garry Scarlett,Sharon Glaysher,Scott Elliott,Kelly Bicknell,Robert Impey,Allyson Lloyd,Sarah Wyllie,Ethan Butcher,Anoop Chauhan,Samuel Robson |
| EPI_ISL_626685, EPI_ISL_626686 | University of Exeter | COVID-19 Genomics UK (COG-UK) Consortium | Ben Temperton,Aaron Jeffries,Michelle Michelsen,Joanna Warwick-Dugdale,Audrey Farbos,Robyn Manley,Stephen Michell,Jane Masoli |

| | | | |
|---|---|---|---|
| EPI_ISL_626689, EPI_ISL_626690 | Centre for Enzyme Innovation, University of Portsmouth / Translational Research Laboratory, Portsmouth Hospitals NHS Trust | COVID-19 Genomics UK (COG-UK) Consortium | Angela Beckett,Yann Bourgeois,Garry Scarlett,Sharon Glaysher,Scott Elliott,Kelly Bicknell,Robert Impey,Allyson Lloyd,Sarah Wyllie,Ethan Butcher,Anoop Chauhan,Samuel Robson |
| EPI_ISL_626713, EPI_ISL_626715, EPI_ISL_626716, EPI_ISL_626717, EPI_ISL_626718, EPI_ISL_626719, EPI_ISL_626720, EPI_ISL_626721, EPI_ISL_626722, EPI_ISL_626723, EPI_ISL_626724, EPI_ISL_626725, EPI_ISL_626726, EPI_ISL_626727, EPI_ISL_626728, EPI_ISL_626729, EPI_ISL_626730, EPI_ISL_626731 | | | |
| see above | University of Exeter | COVID-19 Genomics UK (COG-UK) Consortium | Ben Temperton,Aaron Jeffries,Michelle Michelsen,Joanna Warwick-Dugdale,Audrey Farbos,Robyn Manley,Stephen Michell,Jane Masoli |
| EPI_ISL_626738 | Centre for Enzyme Innovation, University of Portsmouth / Translational Research Laboratory, Portsmouth Hospitals NHS Trust | COVID-19 Genomics UK (COG-UK) Consortium | Angela Beckett,Yann Bourgeois,Garry Scarlett,Sharon Glaysher,Scott Elliott,Kelly Bicknell,Robert Impey,Allyson Lloyd,Sarah Wyllie,Ethan Butcher,Anoop Chauhan,Samuel Robson |
| EPI_ISL_626772, EPI_ISL_626773, EPI_ISL_626774, EPI_ISL_626775, EPI_ISL_626776, EPI_ISL_626777 | University of Exeter | COVID-19 Genomics UK (COG-UK) Consortium | Ben Temperton,Aaron Jeffries,Michelle Michelsen,Joanna Warwick-Dugdale,Audrey Farbos,Robyn Manley,Stephen Michell,Jane Masoli |
| EPI_ISL_626778 | Centre for Enzyme Innovation, University of Portsmouth / Translational Research Laboratory, Portsmouth Hospitals NHS Trust | COVID-19 Genomics UK (COG-UK) Consortium | Angela Beckett,Yann Bourgeois,Garry Scarlett,Sharon Glaysher,Scott Elliott,Kelly Bicknell,Robert Impey,Allyson Lloyd,Sarah Wyllie,Ethan Butcher,Anoop Chauhan,Samuel Robson |
| EPI_ISL_626785 | University of Exeter | COVID-19 Genomics UK (COG-UK) Consortium | Ben Temperton,Aaron Jeffries,Michelle Michelsen,Joanna Warwick-Dugdale,Audrey Farbos,Robyn Manley,Stephen Michell,Jane Masoli |
| EPI_ISL_626792 | Centre for Enzyme Innovation, University of Portsmouth / Translational Research Laboratory, Portsmouth Hospitals NHS Trust | COVID-19 Genomics UK (COG-UK) Consortium | Angela Beckett,Yann Bourgeois,Garry Scarlett,Sharon Glaysher,Scott Elliott,Kelly Bicknell,Robert Impey,Allyson Lloyd,Sarah Wyllie,Ethan Butcher,Anoop Chauhan,Samuel Robson |
| EPI_ISL_626793, EPI_ISL_626794, EPI_ISL_626797, EPI_ISL_626798, EPI_ISL_626799, EPI_ISL_626800, EPI_ISL_626801, EPI_ISL_626802, EPI_ISL_626803, EPI_ISL_626804, EPI_ISL_626805, EPI_ISL_626806, EPI_ISL_626810, EPI_ISL_626813 | | | |
| see above | University of Exeter | COVID-19 Genomics UK (COG-UK) Consortium | Ben Temperton,Aaron Jeffries,Michelle Michelsen,Joanna Warwick-Dugdale,Audrey Farbos,Robyn Manley,Stephen Michell,Jane Masoli |
| EPI_ISL_626878, EPI_ISL_626879, EPI_ISL_626880 | Centre for Enzyme Innovation, University of Portsmouth / Translational Research Laboratory, Portsmouth Hospitals NHS Trust | COVID-19 Genomics UK (COG-UK) Consortium | Angela Beckett,Yann Bourgeois,Garry Scarlett,Sharon Glaysher,Scott Elliott,Kelly Bicknell,Robert Impey,Allyson Lloyd,Sarah Wyllie,Ethan Butcher,Anoop Chauhan,Samuel Robson |
| EPI_ISL_626882, EPI_ISL_626883, EPI_ISL_626884, EPI_ISL_626885, EPI_ISL_626886, EPI_ISL_626887, EPI_ISL_626888, EPI_ISL_626889, EPI_ISL_626890, EPI_ISL_626891 | University of Exeter | COVID-19 Genomics UK (COG-UK) Consortium | Ben Temperton,Aaron Jeffries,Michelle Michelsen,Joanna Warwick-Dugdale,Audrey Farbos,Robyn Manley,Stephen Michell,Jane Masoli |
| EPI_ISL_626895 | Centre for Enzyme Innovation, University of Portsmouth / Translational Research Laboratory, Portsmouth Hospitals NHS Trust | COVID-19 Genomics UK (COG-UK) Consortium | Angela Beckett,Yann Bourgeois,Garry Scarlett,Sharon Glaysher,Scott Elliott,Kelly Bicknell,Robert Impey,Allyson Lloyd,Sarah Wyllie,Ethan Butcher,Anoop Chauhan,Samuel Robson |
| EPI_ISL_626910, EPI_ISL_626925 | University of Exeter | COVID-19 Genomics UK (COG-UK) Consortium | Ben Temperton,Aaron Jeffries,Michelle Michelsen,Joanna Warwick-Dugdale,Audrey Farbos,Robyn Manley,Stephen Michell,Jane Masoli |
| EPI_ISL_626936 | Centre for Enzyme Innovation, University of Portsmouth / Translational Research Laboratory, Portsmouth Hospitals NHS Trust | COVID-19 Genomics UK (COG-UK) Consortium | Angela Beckett,Yann Bourgeois,Garry Scarlett,Sharon Glaysher,Scott Elliott,Kelly Bicknell,Robert Impey,Allyson Lloyd,Sarah Wyllie,Ethan Butcher,Anoop Chauhan,Samuel Robson |
| EPI_ISL_626937, EPI_ISL_626939, EPI_ISL_626940, EPI_ISL_626941, EPI_ISL_626942, EPI_ISL_626943, EPI_ISL_626944, EPI_ISL_626945, EPI_ISL_626946, EPI_ISL_627031, EPI_ISL_627052 | | | |
| see above | University of Exeter | COVID-19 Genomics UK (COG-UK) Consortium | Ben Temperton,Aaron Jeffries,Michelle Michelsen,Joanna Warwick-Dugdale,Audrey Farbos,Robyn Manley,Stephen Michell,Jane Masoli |
| EPI_ISL_627056, EPI_ISL_627057 | Centre for Enzyme Innovation, University of Portsmouth / Translational Research Laboratory, Portsmouth Hospitals NHS Trust | COVID-19 Genomics UK (COG-UK) Consortium | Angela Beckett,Yann Bourgeois,Garry Scarlett,Sharon Glaysher,Scott Elliott,Kelly Bicknell,Robert Impey,Allyson Lloyd,Sarah Wyllie,Ethan Butcher,Anoop Chauhan,Samuel Robson |
| EPI_ISL_627058, EPI_ISL_627059, EPI_ISL_627060, EPI_ISL_627061, EPI_ISL_627062, EPI_ISL_627064, EPI_ISL_627065, EPI_ISL_627067, EPI_ISL_627102, EPI_ISL_627111, EPI_ISL_627112 | | | |
| see above | University of Exeter | COVID-19 Genomics UK (COG-UK) Consortium | Ben Temperton,Aaron Jeffries,Michelle Michelsen,Joanna Warwick-Dugdale,Audrey Farbos,Robyn Manley,Stephen Michell,Jane Masoli |
| EPI_ISL_627116 | Centre for Enzyme Innovation, University of Portsmouth / Translational Research Laboratory, Portsmouth Hospitals NHS Trust | COVID-19 Genomics UK (COG-UK) Consortium | Angela Beckett,Yann Bourgeois,Garry Scarlett,Sharon Glaysher,Scott Elliott,Kelly Bicknell,Robert Impey,Allyson Lloyd,Sarah Wyllie,Ethan Butcher,Anoop Chauhan,Samuel Robson |
| EPI_ISL_627118 | University of Exeter | COVID-19 Genomics UK (COG-UK) Consortium | Ben Temperton,Aaron Jeffries,Michelle Michelsen,Joanna Warwick-Dugdale,Audrey Farbos,Robyn Manley,Stephen Michell,Jane Masoli |
| EPI_ISL_627128 | Centre for Enzyme Innovation, University of Portsmouth / Translational Research Laboratory, Portsmouth Hospitals NHS Trust | COVID-19 Genomics UK (COG-UK) Consortium | Angela Beckett,Yann Bourgeois,Garry Scarlett,Sharon Glaysher,Scott Elliott,Kelly Bicknell,Robert Impey,Allyson Lloyd,Sarah Wyllie,Ethan Butcher,Anoop Chauhan,Samuel Robson |
| EPI_ISL_627143 | University of Exeter | COVID-19 Genomics UK (COG-UK) Consortium | Ben Temperton,Aaron Jeffries,Michelle Michelsen,Joanna Warwick-Dugdale,Audrey Farbos,Robyn Manley,Stephen Michell,Jane Masoli |
| EPI_ISL_627144 | Centre for Enzyme Innovation, University of Portsmouth / Translational Research Laboratory, Portsmouth Hospitals NHS Trust | COVID-19 Genomics UK (COG-UK) Consortium | Angela Beckett,Yann Bourgeois,Garry Scarlett,Sharon Glaysher,Scott Elliott,Kelly Bicknell,Robert Impey,Allyson Lloyd,Sarah Wyllie,Ethan Butcher,Anoop Chauhan,Samuel Robson |
| EPI_ISL_627145, EPI_ISL_627149 | University of Exeter | COVID-19 Genomics UK (COG-UK) Consortium | Ben Temperton,Aaron Jeffries,Michelle Michelsen,Joanna Warwick-Dugdale,Audrey Farbos,Robyn Manley,Stephen Michell,Jane Masoli |
| EPI_ISL_627150 | Centre for Enzyme Innovation, University of Portsmouth / Translational Research Laboratory, Portsmouth Hospitals NHS Trust | COVID-19 Genomics UK (COG-UK) Consortium | Angela Beckett,Yann Bourgeois,Garry Scarlett,Sharon Glaysher,Scott Elliott,Kelly Bicknell,Robert Impey,Allyson Lloyd,Sarah Wyllie,Ethan Butcher,Anoop Chauhan,Samuel Robson |
| EPI_ISL_627159, EPI_ISL_627163, EPI_ISL_627166, EPI_ISL_627182, EPI_ISL_627186, EPI_ISL_627189, EPI_ISL_627190, EPI_ISL_627194, EPI_ISL_627196, EPI_ISL_627210, EPI_ISL_627291, EPI_ISL_627311, EPI_ISL_627312, EPI_ISL_627314, EPI_ISL_627315, EPI_ISL_627316, EPI_ISL_627317, EPI_ISL_627318, EPI_ISL_627319, EPI_ISL_627320, EPI_ISL_627321, EPI_ISL_627322, EPI_ISL_627323, EPI_ISL_627324, EPI_ISL_627325, EPI_ISL_627326, EPI_ISL_627327, EPI_ISL_627328, EPI_ISL_627424, EPI_ISL_627425, EPI_ISL_627426, EPI_ISL_627427, EPI_ISL_627428, EPI_ISL_627429, EPI_ISL_627430, EPI_ISL_627431, EPI_ISL_627432, EPI_ISL_627433, EPI_ISL_627434, EPI_ISL_627435, EPI_ISL_627436, EPI_ISL_627437, EPI_ISL_627438, EPI_ISL_627439, EPI_ISL_627440, EPI_ISL_627441, EPI_ISL_627444 | | | |
| see above | University of Exeter | COVID-19 Genomics UK (COG-UK) Consortium | Ben Temperton,Aaron Jeffries,Michelle Michelsen,Joanna Warwick-Dugdale,Audrey Farbos,Robyn Manley,Stephen Michell,Jane Masoli |
| EPI_ISL_628320, EPI_ISL_628321, EPI_ISL_628322, EPI_ISL_628323, EPI_ISL_628324, EPI_ISL_628325, EPI_ISL_628326, EPI_ISL_628327, EPI_ISL_628328, EPI_ISL_628329, EPI_ISL_628330, EPI_ISL_628331, EPI_ISL_628332, EPI_ISL_628333 | | | |
| see above | Centre for Enzyme Innovation, University of Portsmouth / Translational Research Laboratory, Portsmouth Hospitals NHS Trust | COVID-19 Genomics UK (COG-UK) Consortium | Angela Beckett,Yann Bourgeois,Garry Scarlett,Sharon Glaysher,Scott Elliott,Kelly Bicknell,Robert Impey,Allyson Lloyd,Sarah Wyllie,Ethan Butcher,Anoop Chauhan,Samuel Robson |
| EPI_ISL_629011, EPI_ISL_629012, EPI_ISL_629013, EPI_ISL_629014, EPI_ISL_629015, EPI_ISL_629016, EPI_ISL_629017, EPI_ISL_629018, EPI_ISL_629019, EPI_ISL_629020, EPI_ISL_629021, EPI_ISL_629022, EPI_ISL_629023, EPI_ISL_629024, EPI_ISL_629025, EPI_ISL_629026, EPI_ISL_629027, EPI_ISL_629028 | | | |
| see above | Centro de Biotecnología Vegetal, Universidad Andrés Bello, Center for Genome Regulation | Center for Mathematical Modeling and Center for Genome Regulation. Santiago, Chile | Bastias M, Sanhueza D, Travisany D, Allende ML, Maass A, González M, Bustos F, Arriagada G, Montecino, M, Orellana A, Castro E, Meneses C. |
| EPI_ISL_629031, EPI_ISL_629032, EPI_ISL_629033, EPI_ISL_629034 | Omsk Research Institute of Natural Focal Infections | WHO National Influenza Centre Russian Federation | Artem Fadeev, Ekaterina Gradoboeva, Ekaterina Savkina, Daria Nashatyreva, Elena Poleshchuk, Aleksei Vasilenko, Valery Yakimenko, Andrey Komissarov |

EPI_ISL_629081, EPI_ISL_629082, EPI_ISL_629083, EPI_ISL_629084, EPI_ISL_629085, EPI_ISL_629086, EPI_ISL_629087, EPI_ISL_629088, EPI_ISL_629089, EPI_ISL_629090, EPI_ISL_629091, EPI_ISL_629092, EPI_ISL_629093, EPI_ISL_629094, EPI_ISL_629095, EPI_ISL_629096, EPI_ISL_629097, EPI_ISL_629098, EPI_ISL_629099, EPI_ISL_629100, EPI_ISL_629101, EPI_ISL_629102, EPI_ISL_629103, EPI_ISL_629104, EPI_ISL_629105, EPI_ISL_629106, EPI_ISL_629107, EPI_ISL_629108

| | | | |
|---|---|---|---|
| see above | Laboratoire du Centre Hospitalier Annecy Genevois | CNR Virus des Infections Respiratoires - France SUD | Antonin Bal, Gregory Destras, Gwendolyne Burfin, Hadrien Règue, Quentin Semanas, Martine Valette, Bruno Lina, Hélène Petitprez, Bruno Chanzy, Laurence Josset |

EPI_ISL_632321, EPI_ISL_632322, EPI_ISL_632323, EPI_ISL_632325, EPI_ISL_632326, EPI_ISL_632327, EPI_ISL_632328, EPI_ISL_632329, EPI_ISL_632330, EPI_ISL_632331, EPI_ISL_632332, EPI_ISL_632333, EPI_ISL_632335, EPI_ISL_632336, EPI_ISL_632337, EPI_ISL_632338, EPI_ISL_632339, EPI_ISL_632340, EPI_ISL_632341, EPI_ISL_632342, EPI_ISL_632343, EPI_ISL_632344, EPI_ISL_632345, EPI_ISL_632346, EPI_ISL_632347, EPI_ISL_632349, EPI_ISL_632350, EPI_ISL_632352, EPI_ISL_632354, EPI_ISL_632355, EPI_ISL_632356, EPI_ISL_632359, EPI_ISL_632360, EPI_ISL_632361, EPI_ISL_632362, EPI_ISL_632363, EPI_ISL_632364, EPI_ISL_632366, EPI_ISL_632367, EPI_ISL_632368, EPI_ISL_632369, EPI_ISL_632370, EPI_ISL_632371, EPI_ISL_632372, EPI_ISL_632373, EPI_ISL_632374, EPI_ISL_632375, EPI_ISL_632376, EPI_ISL_632377, EPI_ISL_632378, EPI_ISL_632379, EPI_ISL_632381, EPI_ISL_632383, EPI_ISL_632386, EPI_ISL_632387, EPI_ISL_632388, EPI_ISL_632389, EPI_ISL_632390, EPI_ISL_632391, EPI_ISL_632392, EPI_ISL_632393, EPI_ISL_632394, EPI_ISL_632395, EPI_ISL_632397, EPI_ISL_632398, EPI_ISL_632399, EPI_ISL_632400, EPI_ISL_632401, EPI_ISL_632402, EPI_ISL_632403, EPI_ISL_632404, EPI_ISL_632405, EPI_ISL_632406, EPI_ISL_632407, EPI_ISL_632408, EPI_ISL_632409, EPI_ISL_632410, EPI_ISL_632411, EPI_ISL_632412, EPI_ISL_632413, EPI_ISL_632414, EPI_ISL_632415, EPI_ISL_632416, EPI_ISL_632417, EPI_ISL_632418, EPI_ISL_632419, EPI_ISL_632421, EPI_ISL_632422, EPI_ISL_632423, EPI_ISL_632424, EPI_ISL_632425, EPI_ISL_632426, EPI_ISL_632427, EPI_ISL_632440, EPI_ISL_632441, EPI_ISL_632442, EPI_ISL_632443, EPI_ISL_632444, EPI_ISL_632445, EPI_ISL_632446, EPI_ISL_632447, EPI_ISL_632448, EPI_ISL_632449, EPI_ISL_632460, EPI_ISL_632461, EPI_ISL_632480, EPI_ISL_632481, EPI_ISL_632482, EPI_ISL_632493, EPI_ISL_632494, EPI_ISL_632495, EPI_ISL_632496, EPI_ISL_632497, EPI_ISL_632498, EPI_ISL_632499, EPI_ISL_632500, EPI_ISL_632501, EPI_ISL_632502, EPI_ISL_632503, EPI_ISL_632504, EPI_ISL_632505, EPI_ISL_632506, EPI_ISL_632507, EPI_ISL_632508, EPI_ISL_632509, EPI_ISL_632510, EPI_ISL_632511, EPI_ISL_632512, EPI_ISL_632513, EPI_ISL_632514, EPI_ISL_632515, EPI_ISL_632516, EPI_ISL_632517, EPI_ISL_632518, EPI_ISL_632519, EPI_ISL_632520, EPI_ISL_632521, EPI_ISL_632522, EPI_ISL_632523, EPI_ISL_632524, EPI_ISL_632525, EPI_ISL_632526, EPI_ISL_632527, EPI_ISL_632528, EPI_ISL_632529, EPI_ISL_632530, EPI_ISL_632531, EPI_ISL_632532, EPI_ISL_632533, EPI_ISL_632534, EPI_ISL_632535, EPI_ISL_632536, EPI_ISL_632537, EPI_ISL_632538, EPI_ISL_632539, EPI_ISL_632540, EPI_ISL_632541, EPI_ISL_632542, EPI_ISL_632543, EPI_ISL_632544, EPI_ISL_632545, EPI_ISL_632546, EPI_ISL_632547, EPI_ISL_632548, EPI_ISL_632549, EPI_ISL_632550, EPI_ISL_632551, EPI_ISL_632552, EPI_ISL_632553, EPI_ISL_632554, EPI_ISL_632555, EPI_ISL_632556, EPI_ISL_632557, EPI_ISL_632558, EPI_ISL_632559, EPI_ISL_632560, EPI_ISL_632561, EPI_ISL_632562, EPI_ISL_632563, EPI_ISL_632564, EPI_ISL_632565, EPI_ISL_632566, EPI_ISL_632567, EPI_ISL_632568, EPI_ISL_632569, EPI_ISL_632570, EPI_ISL_632571, EPI_ISL_632572, EPI_ISL_632573, EPI_ISL_632574, EPI_ISL_632575, EPI_ISL_632576, EPI_ISL_632577, EPI_ISL_632578, EPI_ISL_632579, EPI_ISL_632580, EPI_ISL_632581, EPI_ISL_632582, EPI_ISL_632583, EPI_ISL_632584, EPI_ISL_632585, EPI_ISL_632586, EPI_ISL_632587, EPI_ISL_632588, EPI_ISL_632589, EPI_ISL_632590, EPI_ISL_632591, EPI_ISL_632592, EPI_ISL_632593, EPI_ISL_632594, EPI_ISL_632595, EPI_ISL_632596, EPI_ISL_632597, EPI_ISL_632598, EPI_ISL_632599, EPI_ISL_632600, EPI_ISL_632601, EPI_ISL_632602, EPI_ISL_632603, EPI_ISL_632604, EPI_ISL_632605, EPI_ISL_632606, EPI_ISL_632607, EPI_ISL_632608, EPI_ISL_632609, EPI_ISL_632610, EPI_ISL_632611, EPI_ISL_632612, EPI_ISL_632613, EPI_ISL_632614, EPI_ISL_632615, EPI_ISL_632616, EPI_ISL_632617, EPI_ISL_632618, EPI_ISL_632619, EPI_ISL_632620, EPI_ISL_632621, EPI_ISL_632622, EPI_ISL_632623, EPI_ISL_632624, EPI_ISL_632625, EPI_ISL_632626, EPI_ISL_632627, EPI_ISL_632628, EPI_ISL_632629, EPI_ISL_632630, EPI_ISL_632631, EPI_ISL_632632, EPI_ISL_632633, EPI_ISL_632634, EPI_ISL_632635, EPI_ISL_632636, EPI_ISL_632637, EPI_ISL_632638, EPI_ISL_632639, EPI_ISL_632640, EPI_ISL_632641, EPI_ISL_632642, EPI_ISL_632643, EPI_ISL_632644, EPI_ISL_632645, EPI_ISL_632646, EPI_ISL_632647, EPI_ISL_632648, EPI_ISL_632649, EPI_ISL_632650, EPI_ISL_632651, EPI_ISL_632652, EPI_ISL_632653, EPI_ISL_632654, EPI_ISL_632655, EPI_ISL_632656, EPI_ISL_632657, EPI_ISL_632658, EPI_ISL_632659, EPI_ISL_632660, EPI_ISL_632661, EPI_ISL_632662, EPI_ISL_632663, EPI_ISL_632664, EPI_ISL_632665, EPI_ISL_632666, EPI_ISL_632667, EPI_ISL_632668, EPI_ISL_632669, EPI_ISL_632670, EPI_ISL_632671, EPI_ISL_632672, EPI_ISL_632673, EPI_ISL_632674, EPI_ISL_632675, EPI_ISL_632676, EPI_ISL_632677, EPI_ISL_632678, EPI_ISL_632679, EPI_ISL_632680, EPI_ISL_632681, EPI_ISL_632682, EPI_ISL_632683, EPI_ISL_632684, EPI_ISL_632685, EPI_ISL_632686, EPI_ISL_632687, EPI_ISL_632688, EPI_ISL_632689, EPI_ISL_632690, EPI_ISL_632691, EPI_ISL_632692, EPI_ISL_632693, EPI_ISL_632694, EPI_ISL_632695, EPI_ISL_632696, EPI_ISL_632697, EPI_ISL_632698, EPI_ISL_632699, EPI_ISL_632700, EPI_ISL_632701, EPI_ISL_632702, EPI_ISL_632703, EPI_ISL_632704, EPI_ISL_632705, EPI_ISL_632706, EPI_ISL_632707, EPI_ISL_632708, EPI_ISL_632709, EPI_ISL_632710, EPI_ISL_632711, EPI_ISL_632712, EPI_ISL_632713, EPI_ISL_632714, EPI_ISL_632715, EPI_ISL_632716, EPI_ISL_632717, EPI_ISL_632718, EPI_ISL_632719, EPI_ISL_632720, EPI_ISL_632721, EPI_ISL_632722, EPI_ISL_632723, EPI_ISL_632724, EPI_ISL_632725, EPI_ISL_632726, EPI_ISL_632727, EPI_ISL_632728, EPI_ISL_632729, EPI_ISL_632730, EPI_ISL_632731, EPI_ISL_632732, EPI_ISL_632733, EPI_ISL_632734, EPI_ISL_632735, EPI_ISL_632736, EPI_ISL_632737, EPI_ISL_632738, EPI_ISL_632739, EPI_ISL_632740, EPI_ISL_632741, EPI_ISL_632742, EPI_ISL_632743, EPI_ISL_632744, EPI_ISL_632745, EPI_ISL_632746, EPI_ISL_632747, EPI_ISL_632748, EPI_ISL_632749, EPI_ISL_632750, EPI_ISL_632751, EPI_ISL_632752, EPI_ISL_632753, EPI_ISL_632754, EPI_ISL_632755, EPI_ISL_632756, EPI_ISL_632757, EPI_ISL_632758, EPI_ISL_632759, EPI_ISL_632760, EPI_ISL_632761, EPI_ISL_632762, EPI_ISL_632763, EPI_ISL_632764, EPI_ISL_632765, EPI_ISL_632766, EPI_ISL_632767, EPI_ISL_632768, EPI_ISL_632769, EPI_ISL_632770, EPI_ISL_632771, EPI_ISL_632772, EPI_ISL_632773, EPI_ISL_632774, EPI_ISL_632775, EPI_ISL_632776, EPI_ISL_632777, EPI_ISL_632778, EPI_ISL_632779, EPI_ISL_632780, EPI_ISL_632781, EPI_ISL_632782, EPI_ISL_632783, EPI_ISL_632784, EPI_ISL_632785, EPI_ISL_632786, EPI_ISL_632787, EPI_ISL_632788, EPI_ISL_632789, EPI_ISL_632790, EPI_ISL_632791, EPI_ISL_632792, EPI_ISL_632793, EPI_ISL_632794, EPI_ISL_632795

| | | | |
|---|---|---|---|
| see above | Dutch COVID-19 response team | Erasmus Medical Center | Bas Oude Munnink, David Nieuwenhuijse, Reina Sikkema, Claudia Schapendonk, Irina Chestakova, Anne van der Linden, Theo Bestebroer, Stefan van Nieuwkoop, Mark Pronk, Pascal Lexmond, Corien Swaan, Manon Haverkate, Madelief Mollers, Mart Stein, Sandra Kengne Kamga Mobou, Jeroen van Kampen, Jolanda Voermans, Aura Timen, Corine GeurtsvanKessel, Annemiek van der Eijk, Richard Molenkamp, Marion Koopmans, on behalf of the Dutch national COVID-19 response team. |

EPI_ISL_632909, EPI_ISL_632910, EPI_ISL_632911, EPI_ISL_632912, EPI_ISL_632913, EPI_ISL_632914, EPI_ISL_632915, EPI_ISL_632916, EPI_ISL_632917, EPI_ISL_632918, EPI_ISL_632919, EPI_ISL_632920, EPI_ISL_632921, EPI_ISL_632922, EPI_ISL_632923, EPI_ISL_632924, EPI_ISL_632925, EPI_ISL_632926, EPI_ISL_632927, EPI_ISL_632928, EPI_ISL_632929, EPI_ISL_632930, EPI_ISL_632931, EPI_ISL_632932, EPI_ISL_632933

| | | | |
|---|---|---|---|
| see above | Cadham Provincial laboratory | Cadham Provincial laboratory | Anna Majer, Shari Tyson, Grace Seo, Philip Mabon, Elsie Grudeski, Rhiannon Huzarewich, Russell Mandes, Anneliese Landgraff, Jennifer Tanner, Natalie Knox, Morag Graham, Gary Van Domselaar, Paul Van Caeseele, Jared Bullard, David Alexander, Kerry Dust, Nathalie Bastien, Yan Li, Timothy Booth, Darian Hole, Madison Chapel, CanCOGeN's metadata curation team, Public Health Agency of Canada CanCOGeN team |

We gratefully acknowledge the following Authors from the Originating laboratories responsible for obtaining the specimens, as well as the Submitting laboratories where the genome data were generated and shared via GISAID, on which this research is based.

All Submitters of data may be contacted directly via www.gisaid.org

Authors are sorted alphabetically.

| Accession ID | Originating Laboratory | Submitting Laboratory | Authors |
| --- | --- | --- | --- |
| EPI_ISL_406844 | Monash Medical Centre | Collaboration between the University of Melbourne at The Peter Doherty Institute for Infection and Immunity, and the Victorian Infectious Disease Reference Laboratory | Caly,L., Seemann,T., Schultz,M., Druce,J. and Taiaroa,G |
| EPI_ISL_415787 | Laboratorio de Referencia Nacional de Virus Respiratorio. Instituto Nacional de Salud. Peru | Laboratorio de Referencia Nacional de Biotecnologia y Biologia Molecular.Instituto Nacional de Salud.Peru | Carlos Padilla Rojas, Priscila Lope Pari, Karolyn Vega Chozo, Johanna Balbuena Torres, Omar Caceres Rey, Hemri Bailon Calderon, Maribel Huaringa Nuñez, Nancy Rojas Serrano |
| EPI_ISL_416410, EPI_ISL_416411, EPI_ISL_416412, EPI_ISL_416413, EPI_ISL_416414, EPI_ISL_416415 | Victorian Infectious Diseases Reference Laboratory (VIDRL) | Victorian Infectious Diseases Reference Laboratory and Microbiological Diagnostic Unit Public Health Laboratory, Doherty Institute | Caly L., Seemann T., Schultz M., Druce J., Taiaroa, G. |
| EPI_ISL_416514, EPI_ISL_416515, EPI_ISL_416516, EPI_ISL_416517, EPI_ISL_416518 | Victorian Infectious Diseases Reference Laboratory (VIDRL) | Victorian Infectious Diseases Reference Laboratory and Microbiological Diagnostic Unit Public Health Laboratory, Doherty Institute | Caly L., Seemann T., Schultz M., Taiaroa, G., Druce J. |
| EPI_ISL_417391, EPI_ISL_417408 | Centre for Infectious Diseases and Microbiology Public Health | NSW Health Pathology - Institute of Clinical Pathology and Medical Research; Westmead Hospital; University of Sydney | Carter I, Rahman H, Holmes EC, O'Sullivan MV, Sintchenko V, Chen SC, Maddocks S, Kok J, Dwyer DE, Rockett R, Eden J-S, Lam C, Gray K, Timms V, Gall M, Arnott A and Sadsad R for the 2019-nCoV Study Group |
| EPI_ISL_417432 | UCD National Virus Reference Laboratory | Public Health Wales Microbiology Cardiff | Catherine Dempsey, Olive Murphy, Michael Prentice, Cillian F De Gascun, |
| EPI_ISL_418269 | Microbiology and Immunology department, Pasteur institute in Ho Chi Minh city | Microbiology and Immunology department, Pasteur institute in Ho Chi Minh city | Cao,T.M., Nguyen,H.T., Pham,H.T.T., Vu,N.P.H., Dao,M.H., Huynh,L.T.K., Nguyen,L.T., Nguyen,N.T., Nguyen,T.T.N., Nguyen,A.H., Luong,Q.C., Nguyen,T.V., Tran,K.C., Pham,Q.D., Tran,T., Hoang,C.Q., Nguyen,T.T., Le,H.Q., Phung,T.M., Vo,T.N.A., Nguyen,S.N., Pham,D.T., Phan,L.T. and Nguyen,T.V. |
| EPI_ISL_419733, EPI_ISL_419734, EPI_ISL_419735, EPI_ISL_419736, EPI_ISL_419737, EPI_ISL_419738, EPI_ISL_419739, EPI_ISL_419740, EPI_ISL_419741, EPI_ISL_419742, EPI_ISL_419743, EPI_ISL_419744, EPI_ISL_419745, EPI_ISL_419746, EPI_ISL_419747, EPI_ISL_419748, EPI_ISL_419749, EPI_ISL_419750, EPI_ISL_419751, EPI_ISL_419752, EPI_ISL_419753, EPI_ISL_419754, EPI_ISL_419755, EPI_ISL_419756, EPI_ISL_419757, EPI_ISL_419758, EPI_ISL_419759, EPI_ISL_419760, EPI_ISL_419761, EPI_ISL_419762, EPI_ISL_419763, EPI_ISL_419764, EPI_ISL_419765, EPI_ISL_419766, EPI_ISL_419767, EPI_ISL_419768, EPI_ISL_419769, EPI_ISL_419770, EPI_ISL_419771, EPI_ISL_419772, EPI_ISL_419773, EPI_ISL_419774, EPI_ISL_419775, EPI_ISL_419776, EPI_ISL_419777, EPI_ISL_419778, EPI_ISL_419779, EPI_ISL_419780, EPI_ISL_419781, EPI_ISL_419782, EPI_ISL_419783, EPI_ISL_419784, EPI_ISL_419785, EPI_ISL_419786, EPI_ISL_419787, EPI_ISL_419788, EPI_ISL_419789, EPI_ISL_419790, EPI_ISL_419791, EPI_ISL_419792, EPI_ISL_419793, EPI_ISL_419794, EPI_ISL_419795, EPI_ISL_419796, EPI_ISL_419797, EPI_ISL_419798, EPI_ISL_419799, EPI_ISL_419800, EPI_ISL_419801, EPI_ISL_419802, EPI_ISL_419803, EPI_ISL_419804, EPI_ISL_419805, EPI_ISL_419806, EPI_ISL_419807, EPI_ISL_419808, EPI_ISL_419809, EPI_ISL_419810, EPI_ISL_419811, EPI_ISL_419812, EPI_ISL_419813, EPI_ISL_419814, EPI_ISL_419815, EPI_ISL_419816, EPI_ISL_419817, EPI_ISL_419818, EPI_ISL_419819, EPI_ISL_419820, EPI_ISL_419821, EPI_ISL_419822, EPI_ISL_419826, EPI_ISL_419834, EPI_ISL_419836, EPI_ISL_419837, EPI_ISL_419838, EPI_ISL_419839, EPI_ISL_419840, EPI_ISL_419841, EPI_ISL_419842, EPI_ISL_419843, EPI_ISL_419844, EPI_ISL_419845, EPI_ISL_419846, EPI_ISL_419847, EPI_ISL_419848, EPI_ISL_419849, EPI_ISL_419850, EPI_ISL_419851, EPI_ISL_419852, EPI_ISL_419853, EPI_ISL_419854, EPI_ISL_419855, EPI_ISL_419856, EPI_ISL_419857, EPI_ISL_419858, EPI_ISL_419859, EPI_ISL_419860, EPI_ISL_419861, EPI_ISL_419862, EPI_ISL_419863, EPI_ISL_419864, EPI_ISL_419865, EPI_ISL_419866, EPI_ISL_419867, EPI_ISL_419868, EPI_ISL_419869, EPI_ISL_419870, EPI_ISL_419871, EPI_ISL_419872, EPI_ISL_419873, EPI_ISL_419874, EPI_ISL_419875, EPI_ISL_419876, EPI_ISL_419877, EPI_ISL_419878, EPI_ISL_419879, EPI_ISL_419880, EPI_ISL_419881, EPI_ISL_419882, EPI_ISL_419883, EPI_ISL_419884, EPI_ISL_419885, EPI_ISL_419886, EPI_ISL_419887, EPI_ISL_419888, EPI_ISL_419889, EPI_ISL_419890, EPI_ISL_419891, EPI_ISL_419892, EPI_ISL_419893, EPI_ISL_419894, EPI_ISL_419895, EPI_ISL_419896, EPI_ISL_419897, EPI_ISL_419898, EPI_ISL_419899, EPI_ISL_419900, EPI_ISL_419901, EPI_ISL_419902, EPI_ISL_419903, EPI_ISL_419904, EPI_ISL_419905, EPI_ISL_419906, EPI_ISL_419907, EPI_ISL_419908, EPI_ISL_419909, EPI_ISL_419910, EPI_ISL_419911, EPI_ISL_419912, EPI_ISL_419913, EPI_ISL_419914, EPI_ISL_419915, EPI_ISL_419916, EPI_ISL_419917, EPI_ISL_419918, EPI_ISL_419919, EPI_ISL_419920, EPI_ISL_419921, EPI_ISL_419922, EPI_ISL_419923, EPI_ISL_419924, EPI_ISL_419925, EPI_ISL_419926, EPI_ISL_419927, EPI_ISL_419928, EPI_ISL_419929, EPI_ISL_419930, EPI_ISL_419931, EPI_ISL_419932, EPI_ISL_419933, EPI_ISL_419934, EPI_ISL_419935, EPI_ISL_419936, EPI_ISL_419937, EPI_ISL_419938, EPI_ISL_419939, EPI_ISL_419940, EPI_ISL_419941, EPI_ISL_419942, EPI_ISL_419943, EPI_ISL_419944, EPI_ISL_419945, EPI_ISL_419946, EPI_ISL_419947, EPI_ISL_419948, EPI_ISL_419949, EPI_ISL_419950, EPI_ISL_419951, EPI_ISL_419952, EPI_ISL_419953, EPI_ISL_419954, EPI_ISL_419955, EPI_ISL_419956, EPI_ISL_419957, EPI_ISL_419958, EPI_ISL_419959, EPI_ISL_419960, EPI_ISL_419961, EPI_ISL_419962, EPI_ISL_419963, EPI_ISL_419964, EPI_ISL_419965, EPI_ISL_419966, EPI_ISL_419967, EPI_ISL_419968, EPI_ISL_419969, EPI_ISL_419970, EPI_ISL_419971, EPI_ISL_419972, EPI_ISL_419973, EPI_ISL_419974, EPI_ISL_419975, EPI_ISL_419976, EPI_ISL_419977, EPI_ISL_419978, EPI_ISL_419979, EPI_ISL_419982, EPI_ISL_419983, EPI_ISL_419984, EPI_ISL_419985, EPI_ISL_419986, EPI_ISL_419987, EPI_ISL_419988, EPI_ISL_419989, EPI_ISL_419990, EPI_ISL_419991, EPI_ISL_419992, EPI_ISL_419993, EPI_ISL_419994, EPI_ISL_419995, EPI_ISL_419996, EPI_ISL_419997, EPI_ISL_419998, EPI_ISL_420004, EPI_ISL_420005, EPI_ISL_420036 | | | |
| see above | Victorian Infectious Diseases Reference Laboratory (VIDRL) | Victorian Infectious Diseases Reference Laboratory and Microbiological Diagnostic Unit Public Health Laboratory, Doherty Institute | Caly L., Seemann T., Sait, M., Schultz M., Druce J., Sherry, N. |
| EPI_ISL_426484 | n/a | Worobey Lab on behalf of the Arizona COVID-19 Genomics Union | Brendan B. Larsen, Megan Folkerts, Krystal Sheridan, Ashlyn Pfeiffer, Danielle Yasquez, Hayley Yaglom, Darrin Lemmer, Jolene Bowers,Evan Bolyen, Jason W. Sahl, Nicholas A. Bokulich, J. Gregory Caporaso, Crystal Hepp, Jason Ladner,David M. Engelthaler, Paul Keim, Michael Worobey |
| EPI_ISL_426637, EPI_ISL_426638, EPI_ISL_426639, EPI_ISL_426640, EPI_ISL_426641, EPI_ISL_426642, EPI_ISL_426643, EPI_ISL_426644, EPI_ISL_426645, EPI_ISL_426646, EPI_ISL_426647, EPI_ISL_426648, EPI_ISL_426649, EPI_ISL_426650, EPI_ISL_426651, EPI_ISL_426652, EPI_ISL_426653, EPI_ISL_426654, EPI_ISL_426655, EPI_ISL_426656, EPI_ISL_426657, EPI_ISL_426658, EPI_ISL_426659, EPI_ISL_426660, EPI_ISL_426661, EPI_ISL_426662, EPI_ISL_426663, EPI_ISL_426664, EPI_ISL_426665, EPI_ISL_426666, EPI_ISL_426667, EPI_ISL_426668, EPI_ISL_426669, EPI_ISL_426670, EPI_ISL_426671, EPI_ISL_426672, EPI_ISL_426673, EPI_ISL_426674, EPI_ISL_426675, EPI_ISL_426676, EPI_ISL_426677, EPI_ISL_426678, EPI_ISL_426679, EPI_ISL_426680, EPI_ISL_426681, EPI_ISL_426682, EPI_ISL_426683, EPI_ISL_426684, EPI_ISL_426685, EPI_ISL_426686, EPI_ISL_426687, EPI_ISL_426688, EPI_ISL_426689, EPI_ISL_426690, EPI_ISL_426691, EPI_ISL_426692, EPI_ISL_426693, EPI_ISL_426694, EPI_ISL_426695, EPI_ISL_426696, EPI_ISL_426697, EPI_ISL_426698, EPI_ISL_426699, EPI_ISL_426700, EPI_ISL_426701, EPI_ISL_426702, EPI_ISL_426703, EPI_ISL_426704, EPI_ISL_426705, EPI_ISL_426706, EPI_ISL_426707, EPI_ISL_426708, EPI_ISL_426709, EPI_ISL_426710, EPI_ISL_426711, EPI_ISL_426712, EPI_ISL_426713, EPI_ISL_426714, EPI_ISL_426715, EPI_ISL_426716, EPI_ISL_426717, EPI_ISL_426718, EPI_ISL_426719, EPI_ISL_426720, EPI_ISL_426721, EPI_ISL_426722, EPI_ISL_426723, EPI_ISL_426724, EPI_ISL_426725, EPI_ISL_426726, EPI_ISL_426727, EPI_ISL_426728, EPI_ISL_426729, EPI_ISL_426730, EPI_ISL_426731, EPI_ISL_426732, EPI_ISL_426733, EPI_ISL_426734, EPI_ISL_426735, EPI_ISL_426736, EPI_ISL_426737, EPI_ISL_426738, EPI_ISL_426739, EPI_ISL_426740, EPI_ISL_426741, EPI_ISL_426742, EPI_ISL_426743, EPI_ISL_426744, EPI_ISL_426745, EPI_ISL_426746, EPI_ISL_426747, EPI_ISL_426748, EPI_ISL_426749, EPI_ISL_426750, EPI_ISL_426751, EPI_ISL_426752, EPI_ISL_426753, EPI_ISL_426754, EPI_ISL_426755, EPI_ISL_426756, EPI_ISL_426757, EPI_ISL_426758, EPI_ISL_426759, EPI_ISL_426760, EPI_ISL_426761, EPI_ISL_426762, EPI_ISL_426763, EPI_ISL_426764, EPI_ISL_426765, EPI_ISL_426766, EPI_ISL_426767, EPI_ISL_426768, EPI_ISL_426769, EPI_ISL_426770, EPI_ISL_426771, EPI_ISL_426772, EPI_ISL_426773, EPI_ISL_426774, EPI_ISL_426775, EPI_ISL_426776, EPI_ISL_426777, EPI_ISL_426778, EPI_ISL_426779, EPI_ISL_426780, EPI_ISL_426781, EPI_ISL_426782, EPI_ISL_426783, EPI_ISL_426784, EPI_ISL_426785, EPI_ISL_426786, EPI_ISL_426787, EPI_ISL_426788, EPI_ISL_426789, EPI_ISL_426790, EPI_ISL_426791, EPI_ISL_426792, EPI_ISL_426793, EPI_ISL_426794, EPI_ISL_426795, EPI_ISL_426796, EPI_ISL_426797, EPI_ISL_426798, EPI_ISL_426799, EPI_ISL_426800, EPI_ISL_426801, EPI_ISL_426802, EPI_ISL_426803, EPI_ISL_426804, EPI_ISL_426805, EPI_ISL_426806, EPI_ISL_426807, EPI_ISL_426808, EPI_ISL_426809, EPI_ISL_426810, EPI_ISL_426811, EPI_ISL_426812, EPI_ISL_426813, EPI_ISL_426814, EPI_ISL_426815, EPI_ISL_426816, EPI_ISL_426817, EPI_ISL_426818, EPI_ISL_426819, EPI_ISL_426820, EPI_ISL_426821, EPI_ISL_426822, EPI_ISL_426823, EPI_ISL_426824, EPI_ISL_426825, EPI_ISL_426826, EPI_ISL_426827, EPI_ISL_426828, EPI_ISL_426829, EPI_ISL_426830, EPI_ISL_426831, EPI_ISL_426832, EPI_ISL_426833, EPI_ISL_426834, EPI_ISL_426835, EPI_ISL_426836, EPI_ISL_426837, EPI_ISL_426838, EPI_ISL_426839, EPI_ISL_426840, EPI_ISL_426841, EPI_ISL_426842, EPI_ISL_426843, EPI_ISL_426844, EPI_ISL_426845, EPI_ISL_426846, EPI_ISL_426847, EPI_ISL_426848, EPI_ISL_426849, EPI_ISL_426850, EPI_ISL_426851, EPI_ISL_426852, EPI_ISL_426853, EPI_ISL_426854, EPI_ISL_426855, EPI_ISL_426856, EPI_ISL_426857, EPI_ISL_426858, EPI_ISL_426859, EPI_ISL_426860, EPI_ISL_426861, EPI_ISL_426862, EPI_ISL_426863, EPI_ISL_426864, EPI_ISL_426865, EPI_ISL_426866, EPI_ISL_426867, EPI_ISL_426869, EPI_ISL_426870, EPI_ISL_426871, EPI_ISL_426872, EPI_ISL_426873, EPI_ISL_426874, EPI_ISL_426875, EPI_ISL_426876, EPI_ISL_426877, EPI_ISL_426878, EPI_ISL_426879, EPI_ISL_426880, EPI_ISL_426881, EPI_ISL_426882, EPI_ISL_426923, EPI_ISL_426924, EPI_ISL_426925, EPI_ISL_426926, EPI_ISL_426927, EPI_ISL_426928, EPI_ISL_426929, EPI_ISL_426930, EPI_ISL_426931, EPI_ISL_426932, EPI_ISL_426933, EPI_ISL_426934, EPI_ISL_426935, EPI_ISL_426936, EPI_ISL_426937, EPI_ISL_426938, EPI_ISL_426939, EPI_ISL_426940, EPI_ISL_426941, EPI_ISL_426942, EPI_ISL_426943, EPI_ISL_426944, EPI_ISL_426945, EPI_ISL_426946, EPI_ISL_426947, EPI_ISL_426948, EPI_ISL_426949, EPI_ISL_426950, EPI_ISL_426951, EPI_ISL_426952, EPI_ISL_426953, EPI_ISL_426954, EPI_ISL_426955, EPI_ISL_426956, EPI_ISL_426957, EPI_ISL_426958, EPI_ISL_426959, EPI_ISL_426960, EPI_ISL_426961, EPI_ISL_426962, EPI_ISL_426963, EPI_ISL_426964, EPI_ISL_426965, EPI_ISL_426966, EPI_ISL_426967, EPI_ISL_426968, EPI_ISL_426969, EPI_ISL_426970, EPI_ISL_426971, EPI_ISL_426972, EPI_ISL_426973, EPI_ISL_426974, EPI_ISL_426975, EPI_ISL_426976, EPI_ISL_426977, EPI_ISL_426978, EPI_ISL_426979, EPI_ISL_426980, EPI_ISL_426981, EPI_ISL_426982, EPI_ISL_426983, EPI_ISL_426984, EPI_ISL_426985, EPI_ISL_426986, EPI_ISL_426987, EPI_ISL_426988, EPI_ISL_426989, EPI_ISL_426990, EPI_ISL_426991, EPI_ISL_426992, EPI_ISL_426993, EPI_ISL_426994, EPI_ISL_426995, EPI_ISL_426996, EPI_ISL_426997, EPI_ISL_426998, EPI_ISL_426999, EPI_ISL_427000, EPI_ISL_427001, EPI_ISL_427002, EPI_ISL_427003, EPI_ISL_427004, EPI_ISL_427005, EPI_ISL_427006, EPI_ISL_427007, EPI_ISL_427008, EPI_ISL_427009, EPI_ISL_427010, EPI_ISL_427011, EPI_ISL_427012, EPI_ISL_427013, EPI_ISL_427014, EPI_ISL_427015, EPI_ISL_427016, EPI_ISL_427017, EPI_ISL_427018, EPI_ISL_427019, EPI_ISL_427020, EPI_ISL_427022, EPI_ISL_427023, EPI_ISL_427024, EPI_ISL_427025, EPI_ISL_427026, EPI_ISL_427027, EPI_ISL_427028, EPI_ISL_427029, EPI_ISL_427030, EPI_ISL_427031, EPI_ISL_427032, EPI_ISL_427033, EPI_ISL_427034, EPI_ISL_427035, EPI_ISL_427036, EPI_ISL_427037, EPI_ISL_427038, EPI_ISL_427039, EPI_ISL_427040, EPI_ISL_427041, EPI_ISL_427042, EPI_ISL_427044, EPI_ISL_427045, EPI_ISL_427046, EPI_ISL_427047, EPI_ISL_427048, EPI_ISL_427049, EPI_ISL_427050, EPI_ISL_427051, EPI_ISL_427052, EPI_ISL_427053, EPI_ISL_427079, EPI_ISL_427080, EPI_ISL_427081, EPI_ISL_427082, EPI_ISL_427083, EPI_ISL_427084, EPI_ISL_427085, EPI_ISL_427086, EPI_ISL_427087, EPI_ISL_427088, EPI_ISL_427089, EPI_ISL_427090, EPI_ISL_427091, EPI_ISL_427092, EPI_ISL_427093, EPI_ISL_427094, EPI_ISL_427095, EPI_ISL_427096, EPI_ISL_427097, EPI_ISL_427098, EPI_ISL_427099, EPI_ISL_427100, EPI_ISL_427101, EPI_ISL_427102, EPI_ISL_427103, EPI_ISL_427104, EPI_ISL_427105, EPI_ISL_427106, EPI_ISL_427107, EPI_ISL_427108, EPI_ISL_427109, EPI_ISL_427110, EPI_ISL_427111, EPI_ISL_427112, EPI_ISL_427113, EPI_ISL_427114, EPI_ISL_427115, EPI_ISL_427116, EPI_ISL_427117, EPI_ISL_427118, EPI_ISL_427119, EPI_ISL_427120, EPI_ISL_427121, EPI_ISL_427122, EPI_ISL_427123, EPI_ISL_427124, EPI_ISL_427125, EPI_ISL_427126, EPI_ISL_427127, EPI_ISL_427128, EPI_ISL_427129, EPI_ISL_427130, EPI_ISL_427131, EPI_ISL_427132, EPI_ISL_427134, EPI_ISL_427135, EPI_ISL_427136, EPI_ISL_427137, EPI_ISL_427138, EPI_ISL_427139, EPI_ISL_427140, EPI_ISL_427141, EPI_ISL_427142, EPI_ISL_427143, EPI_ISL_427144, EPI_ISL_427145, EPI_ISL_427146, EPI_ISL_427147, EPI_ISL_427150, EPI_ISL_427151, EPI_ISL_427152, EPI_ISL_427153, EPI_ISL_427154, EPI_ISL_427155, EPI_ISL_427156, EPI_ISL_427157, EPI_ISL_427158, EPI_ISL_427159, EPI_ISL_427160 | | | |
| see above | Victorian Infectious Diseases Reference Laboratory (VIDRL) | Microbiological Diagnostic Unit Public Health Laboratory and Victorian Infectious Diseases Reference Laboratory, Doherty Institute | Caly L., Seemann T., Sait, M., Schultz M., Druce J., Sherry, N. |
| EPI_ISL_428440, EPI_ISL_428441, EPI_ISL_428442, EPI_ISL_428443, EPI_ISL_428444, EPI_ISL_428445, EPI_ISL_428446, EPI_ISL_428447, EPI_ISL_428448, EPI_ISL_428449, EPI_ISL_428450, EPI_ISL_428451, EPI_ISL_428452, EPI_ISL_428453, EPI_ISL_428454, EPI_ISL_428455, EPI_ISL_428456, EPI_ISL_428457, EPI_ISL_428458, EPI_ISL_428459, EPI_ISL_428460, EPI_ISL_428461, EPI_ISL_428462, EPI_ISL_428463, EPI_ISL_428464, EPI_ISL_428465, EPI_ISL_428466, EPI_ISL_428467, EPI_ISL_428468, EPI_ISL_428469, EPI_ISL_428470, EPI_ISL_428471, EPI_ISL_428472, EPI_ISL_428473, EPI_ISL_428474, EPI_ISL_428475, EPI_ISL_428476, EPI_ISL_428477, EPI_ISL_428478 | | | |

| | | | |
|---|---|---|---|
| see above | Guangdong Provincial Center for Diseases Control and Prevention;Guangdong Provincial Institute of Public Health | School of Public Health, The University of Hong Kong | Bosheng Li, Haogao Gu, Lijun Liang, Zhencui Li, Hui-Ling Yen, Yao Hu, Yingchao Song , Hanri Zeng, Tie Song, Jie Wu, Leo L.M. Poon |
| EPI_ISL_428990, EPI_ISL_428991, EPI_ISL_428992, EPI_ISL_428993, EPI_ISL_428994, EPI_ISL_428995, EPI_ISL_428996, EPI_ISL_428997, EPI_ISL_428998, EPI_ISL_428999, EPI_ISL_429000, EPI_ISL_429001, EPI_ISL_429002, EPI_ISL_429003, EPI_ISL_429004, EPI_ISL_429005, EPI_ISL_429006, EPI_ISL_429007, EPI_ISL_429008, EPI_ISL_429009, EPI_ISL_429010, EPI_ISL_429011, EPI_ISL_429012, EPI_ISL_429013, EPI_ISL_429014, EPI_ISL_429015, EPI_ISL_429016, EPI_ISL_429017, EPI_ISL_429018, EPI_ISL_429019, EPI_ISL_429020, EPI_ISL_429021, EPI_ISL_429022, EPI_ISL_429023, EPI_ISL_429024, EPI_ISL_429025, EPI_ISL_429026, EPI_ISL_429027, EPI_ISL_429028, EPI_ISL_429029, EPI_ISL_429030, EPI_ISL_429031, EPI_ISL_429032, EPI_ISL_429033, EPI_ISL_429034, EPI_ISL_429035, EPI_ISL_429036, EPI_ISL_429037, EPI_ISL_429038, EPI_ISL_429039, EPI_ISL_429040, EPI_ISL_429041, EPI_ISL_429042, EPI_ISL_429043, EPI_ISL_429044, EPI_ISL_429045, EPI_ISL_429046, EPI_ISL_429047, EPI_ISL_429048, EPI_ISL_429049, EPI_ISL_429050, EPI_ISL_429051, EPI_ISL_429052, EPI_ISL_429053, EPI_ISL_429054, EPI_ISL_429055, EPI_ISL_429056, EPI_ISL_429057, EPI_ISL_429058, EPI_ISL_429059, EPI_ISL_429060, EPI_ISL_429061, EPI_ISL_429062, EPI_ISL_429063, EPI_ISL_429064, EPI_ISL_429065, EPI_ISL_429066, EPI_ISL_429067, EPI_ISL_429068, EPI_ISL_429069, EPI_ISL_429070, EPI_ISL_429071, EPI_ISL_429072, EPI_ISL_429073 | | | |
| see above | UCSF Clinical Microbiology Laboratory | Chan-Zuckerberg Biohub | CZB Cliahub Consortium |
| EPI_ISL_430473, EPI_ISL_430474, EPI_ISL_430475, EPI_ISL_430476, EPI_ISL_430477, EPI_ISL_430478, EPI_ISL_430479, EPI_ISL_430480, EPI_ISL_430481, EPI_ISL_430482, EPI_ISL_430483, EPI_ISL_430484, EPI_ISL_430485, EPI_ISL_430486, EPI_ISL_430487, EPI_ISL_430488, EPI_ISL_430489, EPI_ISL_430490, EPI_ISL_430491, EPI_ISL_430492, EPI_ISL_430493, EPI_ISL_430498, EPI_ISL_430499, EPI_ISL_430500, EPI_ISL_430501, EPI_ISL_430502, EPI_ISL_430503, EPI_ISL_430504, EPI_ISL_430505, EPI_ISL_430506, EPI_ISL_430507, EPI_ISL_430508, EPI_ISL_430509, EPI_ISL_430510, EPI_ISL_430511, EPI_ISL_430512, EPI_ISL_430513, EPI_ISL_430514, EPI_ISL_430515, EPI_ISL_430516, EPI_ISL_430517, EPI_ISL_430518, EPI_ISL_430519, EPI_ISL_430520, EPI_ISL_430521, EPI_ISL_430522, EPI_ISL_430523, EPI_ISL_430524, EPI_ISL_430525, EPI_ISL_430526, EPI_ISL_430527, EPI_ISL_430528, EPI_ISL_430529, EPI_ISL_430530, EPI_ISL_430531, EPI_ISL_430532, EPI_ISL_430533, EPI_ISL_430534, EPI_ISL_430535, EPI_ISL_430536, EPI_ISL_430537, EPI_ISL_430538, EPI_ISL_430539, EPI_ISL_430540, EPI_ISL_430542, EPI_ISL_430543, EPI_ISL_430544, EPI_ISL_430545, EPI_ISL_430546, EPI_ISL_430547, EPI_ISL_430548, EPI_ISL_430549, EPI_ISL_430550, EPI_ISL_430551, EPI_ISL_430552, EPI_ISL_430553, EPI_ISL_430554, EPI_ISL_430555, EPI_ISL_430556, EPI_ISL_430557, EPI_ISL_430558, EPI_ISL_430559, EPI_ISL_430560, EPI_ISL_430561, EPI_ISL_430562, EPI_ISL_430563, EPI_ISL_430564, EPI_ISL_430565, EPI_ISL_430566, EPI_ISL_430567, EPI_ISL_430569, EPI_ISL_430570, EPI_ISL_430571, EPI_ISL_430572, EPI_ISL_430573, EPI_ISL_430574, EPI_ISL_430575, EPI_ISL_430576, EPI_ISL_430577, EPI_ISL_430578, EPI_ISL_430579, EPI_ISL_430580, EPI_ISL_430581, EPI_ISL_430582, EPI_ISL_430583, EPI_ISL_430584, EPI_ISL_430585, EPI_ISL_430586, EPI_ISL_430587, EPI_ISL_430588, EPI_ISL_430589, EPI_ISL_430590, EPI_ISL_430591, EPI_ISL_430592, EPI_ISL_430593, EPI_ISL_430594, EPI_ISL_430595, EPI_ISL_430596, EPI_ISL_430597, EPI_ISL_430598, EPI_ISL_430599, EPI_ISL_430600, EPI_ISL_430601, EPI_ISL_430602, EPI_ISL_430603, EPI_ISL_430604, EPI_ISL_430605, EPI_ISL_430606, EPI_ISL_430607, EPI_ISL_430608, EPI_ISL_430609, EPI_ISL_430610, EPI_ISL_430611, EPI_ISL_430612, EPI_ISL_430613, EPI_ISL_430614, EPI_ISL_430615, EPI_ISL_430616, EPI_ISL_430617, EPI_ISL_430618, EPI_ISL_430619, EPI_ISL_430620, EPI_ISL_430621, EPI_ISL_430622, EPI_ISL_430623, EPI_ISL_430624, EPI_ISL_430625, EPI_ISL_430627, EPI_ISL_430628, EPI_ISL_430629, EPI_ISL_430630, EPI_ISL_430638, EPI_ISL_430688, EPI_ISL_430689, EPI_ISL_430690, EPI_ISL_430691, EPI_ISL_430692, EPI_ISL_430693, EPI_ISL_430694, EPI_ISL_430695, EPI_ISL_430696, EPI_ISL_430697, EPI_ISL_430698, EPI_ISL_430700, EPI_ISL_430701, EPI_ISL_430702, EPI_ISL_430703, EPI_ISL_430704, EPI_ISL_430705, EPI_ISL_430706, EPI_ISL_430707, EPI_ISL_430708, EPI_ISL_430709, EPI_ISL_430710, EPI_ISL_430711, EPI_ISL_430712, EPI_ISL_430713, EPI_ISL_430714 | | | |
| see above | Victorian Infectious Diseases Reference Laboratory (VIDRL) | Microbiological Diagnostic Unit Public Health Laboratory and Victorian Infectious Diseases Reference Laboratory, The Peter Doherty Institute for Infection and Immunity | Caly L., Seemann T., Sait, M., Schultz M., Druce J., Sherry, N. |
| EPI_ISL_430791, EPI_ISL_430792 | UCSF Clinical Microbiology Laboratory | Chan-Zuckerberg Biohub | CZB Cliahub Consortium |
| EPI_ISL_431833 | National Institutes of Health, University of the Philippines Manila | Philippine Genome Center, University of the Philippines System | Carlo M. Lapid, Francis A. Tablizo, Benedict A. Maralit, Jan Michael C. Yap, Raul V. Destura, Marissa M. Alejandria, El King D. Morado, Joshua Gregor A. Dizon, Jo-Hannah S. Llames, Shiela Mae M. Araiza, Kris P. Punayan, Kristianne Arielle D. Gabriel, Shebna Rose D. Fabilloren, Shana F. Genavia, Jarvin E. Nipales, Alessandra C. Sanchez, Haifa L.Gaza, Joy Ann Petronio-Santos, Julius Aaron Mejia, Maribell Dollete, Sonia Salamat, Christina Tan, Bernard Demot, John Mark Velasco, Eva Maria Cutiongco-de la Paz, and Cynthia P. Saloma |
| EPI_ISL_434555, EPI_ISL_434556, EPI_ISL_434557, EPI_ISL_434558 | National Institutes of Health, University of the Philippines Manila | Philippine Genome Center | Carlo M. Lapid, Francis A. Tablizo, Benedict A. Maralit, Jan Michael C. Yap, Raul V. Destura, Marissa M. Alejandria, El King D. Morado, Joshua Gregor A. Dizon, Jo-Hannah S. Llames, Shiela Mae M. Araiza, Kris P. Punayan, Kristianne Arielle D. Gabriel, Shebna Rose D. Fabilloren, Shana F. Genavia, Jarvin E. Nipales, Alessandra C. Sanchez, Haifa L.Gaza, Joy Ann Petronio-Santos, Julius Aaron Mejia, Maribell Dollete, Sonia Salamat, Christina Tan, Bernard Demot, John Mark Velasco, Eva Maria Cutiongco-de la Paz, and Cynthia P. Saloma |
| EPI_ISL_434666 | Hornefors Halsocentral | The Public Health Agency of Sweden | Camilla Eiback, Oskar Karlsson Lindsjo, Maria Lind Karlberg, Anna-Malin Linde, Olov Svartstrom, Anna Risberg, Theresa Enkirch, Mia Brytting, Karin Tegmark-Wisell |
| EPI_ISL_434669 | Lakargruppen | The Public Health Agency of Sweden | Boris Klanger, Oskar Karlsson Lindsjo, Maria Lind Karlberg, Anna-Malin Linde, Olov Svartstrom, Anna Risberg, Theresa Enkirch, Mia Brytting, Karin Tegmark-Wisell |
| EPI_ISL_436111, EPI_ISL_436112, EPI_ISL_436113, EPI_ISL_436114, EPI_ISL_436115, EPI_ISL_436116, EPI_ISL_436117, EPI_ISL_436119, EPI_ISL_436120, EPI_ISL_436121, EPI_ISL_436122, EPI_ISL_436123, EPI_ISL_436124, EPI_ISL_436125, EPI_ISL_436126, EPI_ISL_436127, EPI_ISL_436128, EPI_ISL_436129, EPI_ISL_436130, EPI_ISL_436131, EPI_ISL_436132 | | | |
| see above | Victorian Infectious Diseases Reference Laboratory (VIDRL) | Microbiological Diagnostic Unit Public Health Laboratory and Victorian Infectious Diseases Reference Laboratory, Doherty Institute | Caly L., Seemann T., Sait, M., Schultz M., Druce J., Sherry, N. |
| EPI_ISL_436641, EPI_ISL_436642, EPI_ISL_436643, EPI_ISL_436644, EPI_ISL_436645, EPI_ISL_436646, EPI_ISL_436647, EPI_ISL_436648, EPI_ISL_436649, EPI_ISL_436650, EPI_ISL_436651, EPI_ISL_436652, EPI_ISL_436653, EPI_ISL_436654, EPI_ISL_436655, EPI_ISL_436656, EPI_ISL_436657, EPI_ISL_436658, EPI_ISL_436659, EPI_ISL_436660, EPI_ISL_436661, EPI_ISL_436662, EPI_ISL_436663, EPI_ISL_436664, EPI_ISL_436665, EPI_ISL_436666, EPI_ISL_436667, EPI_ISL_436668, EPI_ISL_436669, EPI_ISL_436670, EPI_ISL_436671, EPI_ISL_436672, EPI_ISL_436673, EPI_ISL_436674, EPI_ISL_436675, EPI_ISL_436676, EPI_ISL_436677, EPI_ISL_436678, EPI_ISL_436679, EPI_ISL_436680, EPI_ISL_436681, EPI_ISL_436682, EPI_ISL_436683 | | | |
| see above | County of Santa Clara Public Health Department | Chan-Zuckerberg Biohub | CZB Cliahub Consortium |
| EPI_ISL_436688, EPI_ISL_436689 | Victorian Infectious Diseases Reference Laboratory (VIDRL) | Microbiological Diagnostic Unit Public Health Laboratory and Victorian Infectious Diseases Reference Laboratory, The Peter Doherty Institute for Infection & Immunity | Caly L., Seemann T., Sait, M., Schultz M., Druce J., Sherry, N. |
| EPI_ISL_437043, EPI_ISL_437044, EPI_ISL_437045, EPI_ISL_437046, EPI_ISL_437047, EPI_ISL_437048, EPI_ISL_437049, EPI_ISL_437050, EPI_ISL_437051, EPI_ISL_437052, EPI_ISL_437053, EPI_ISL_437054, EPI_ISL_437055, EPI_ISL_437056, EPI_ISL_437057, EPI_ISL_437058, EPI_ISL_437059, EPI_ISL_437060, EPI_ISL_437061, EPI_ISL_437062, EPI_ISL_437063, EPI_ISL_437064, EPI_ISL_437065, EPI_ISL_437066, EPI_ISL_437067, EPI_ISL_437068, EPI_ISL_437069, EPI_ISL_437070, EPI_ISL_437071, EPI_ISL_437072, EPI_ISL_437073, EPI_ISL_437074, EPI_ISL_437075, EPI_ISL_437076, EPI_ISL_437077, EPI_ISL_437078, EPI_ISL_437079, EPI_ISL_437080, EPI_ISL_437081, EPI_ISL_437082, EPI_ISL_437083, EPI_ISL_437084, EPI_ISL_437085, EPI_ISL_437086, EPI_ISL_437087, EPI_ISL_437088 | | | |
| see above | County of Santa Clara Public Health | Chan-Zuckerberg Biohub | CZB Cliahub Consortium |
| EPI_ISL_437520, EPI_ISL_437521, EPI_ISL_437522, EPI_ISL_437523, EPI_ISL_437524, EPI_ISL_437525, EPI_ISL_437526, EPI_ISL_437527, EPI_ISL_437528, EPI_ISL_437529, EPI_ISL_437530, EPI_ISL_437531, EPI_ISL_437532, EPI_ISL_437533, EPI_ISL_437534, EPI_ISL_437535 | | | |
| see above | OHSU Lab Services Molecular Microbiology Lab | Oregon SARS-CoV-2 Genome Sequencing Center | Brendan L. O'Connell, Ruth V. Nichols, Alec J. Hirsch, Guang Fan, Daniel N. Streblow, William B. Messer, Andrew C. Adey, Benjamin N. Bimber, Brian J. O'Roak |
| EPI_ISL_444023, EPI_ISL_444024, EPI_ISL_444025, EPI_ISL_444026 | County of Santa Clara Public Health | Chan-Zuckerberg Biohub | CZB Cliahub Consortium |
| EPI_ISL_444051, EPI_ISL_444052, EPI_ISL_444053, EPI_ISL_444054, EPI_ISL_444055, EPI_ISL_444056, EPI_ISL_444057, EPI_ISL_444058, EPI_ISL_444059, EPI_ISL_444060, EPI_ISL_444061, EPI_ISL_444062, EPI_ISL_444063, EPI_ISL_444064, EPI_ISL_444065, EPI_ISL_444066, EPI_ISL_444067, EPI_ISL_444068, EPI_ISL_444069, EPI_ISL_444070, EPI_ISL_444071, EPI_ISL_444072, EPI_ISL_444073, EPI_ISL_444074, EPI_ISL_444075, EPI_ISL_444076, EPI_ISL_444077, EPI_ISL_444078, EPI_ISL_445169, EPI_ISL_445170, EPI_ISL_445171, EPI_ISL_445172, EPI_ISL_445173, EPI_ISL_445174, EPI_ISL_445175, EPI_ISL_445176, EPI_ISL_445177, EPI_ISL_445178, EPI_ISL_445179, EPI_ISL_445180, EPI_ISL_445181, EPI_ISL_445182 | | | |
| see above | UCSF Clinical Microbiology Laboratory | Chan-Zuckerberg Biohub | CZB Cliahub Consortium |
| EPI_ISL_447887, EPI_ISL_447888, EPI_ISL_447889, EPI_ISL_447890, EPI_ISL_447891, EPI_ISL_447892, EPI_ISL_447893, EPI_ISL_447894, EPI_ISL_447895, EPI_ISL_447896 | University of California, Davis | Chan-Zuckerberg Biohub | CZB Cliahub Consortium |
| EPI_ISL_450484, EPI_ISL_450485, EPI_ISL_450486, EPI_ISL_450487 | unknown | Data Science | Carroll,T.D., Tran,N.K., Cohen,S.H., Miller,C.J. |
| EPI_ISL_451486 | Australian Clinical Labs | NSW Health Pathology - Institute of Clinical Pathology and Medical Research; Westmead Hospital; University of Sydney | CIDM-PH et al. |
| EPI_ISL_451487, EPI_ISL_451488 | Pathology North - NSW Health Pathology | NSW Health Pathology - Institute of Clinical Pathology and Medical Research; Westmead Hospital; University of Sydney | CIDM-PH et al. |
| EPI_ISL_451489 | Laverty Pathology | NSW Health Pathology - Institute of Clinical Pathology and Medical Research; Westmead Hospital; University of Sydney | CIDM-PH et al. |
| EPI_ISL_451490, EPI_ISL_451491, EPI_ISL_451492, EPI_ISL_451493, EPI_ISL_451494, EPI_ISL_451495, EPI_ISL_451496, EPI_ISL_451497, EPI_ISL_451498, EPI_ISL_451499, EPI_ISL_451500, EPI_ISL_451501, EPI_ISL_451502, EPI_ISL_451503, EPI_ISL_451504, EPI_ISL_451505, EPI_ISL_451506, EPI_ISL_451507, EPI_ISL_451508, EPI_ISL_451509, EPI_ISL_451510, EPI_ISL_451511, EPI_ISL_451512, EPI_ISL_451513, EPI_ISL_451514, EPI_ISL_451515, EPI_ISL_451516 | | | |

| | | | |
|---|---|---|---|
| see above | Pathology West - NSW Health Pathology | NSW Health Pathology - Institute of Clinical Pathology and Medical Research; Westmead Hospital; University of Sydney | CIDM-PH et al. |
| EPI_ISL_451517, EPI_ISL_451518, EPI_ISL_451519, EPI_ISL_451521 | South Eastern Area Laboratory Services | NSW Health Pathology - Institute of Clinical Pathology and Medical Research; Westmead Hospital; University of Sydney | CIDM-PH et al. |
| EPI_ISL_451522, EPI_ISL_451523, EPI_ISL_451524, EPI_ISL_451525, EPI_ISL_451526, EPI_ISL_451527, EPI_ISL_451528, EPI_ISL_451529 | Pathology West - NSW Health Pathology | NSW Health Pathology - Institute of Clinical Pathology and Medical Research; Westmead Hospital; University of Sydney | CIDM-PH et al. |
| EPI_ISL_451530, EPI_ISL_451531 | Pathology Sydney South West - NSW Health Pathology | NSW Health Pathology - Institute of Clinical Pathology and Medical Research; Westmead Hospital; University of Sydney | CIDM-PH et al. |
| EPI_ISL_451532 | Pathology West - NSW Health Pathology | NSW Health Pathology - Institute of Clinical Pathology and Medical Research; Westmead Hospital; University of Sydney | CIDM-PH et al. |
| EPI_ISL_451533 | Pathology Sydney South West - NSW Health Pathology | NSW Health Pathology - Institute of Clinical Pathology and Medical Research; Westmead Hospital; University of Sydney | CIDM-PH et al. |
| EPI_ISL_451534 | Medlab Pathology | NSW Health Pathology - Institute of Clinical Pathology and Medical Research; Westmead Hospital; University of Sydney | CIDM-PH et al. |
| EPI_ISL_451535 | Pathology West - NSW Health Pathology | NSW Health Pathology - Institute of Clinical Pathology and Medical Research; Westmead Hospital; University of Sydney | CIDM-PH et al. |
| EPI_ISL_451536 | Pathology Sydney South West - NSW Health Pathology | NSW Health Pathology - Institute of Clinical Pathology and Medical Research; Westmead Hospital; University of Sydney | CIDM-PH et al. |
| EPI_ISL_451537, EPI_ISL_451538, EPI_ISL_451539 | Pathology West - NSW Health Pathology | NSW Health Pathology - Institute of Clinical Pathology and Medical Research; Westmead Hospital; University of Sydney | CIDM-PH et al. |
| EPI_ISL_451540 | ACT pathology | NSW Health Pathology - Institute of Clinical Pathology and Medical Research; Westmead Hospital; University of Sydney | CIDM-PH et al. |
| EPI_ISL_451541 | Pathology West - NSW Health Pathology | NSW Health Pathology - Institute of Clinical Pathology and Medical Research; Westmead Hospital; University of Sydney | CIDM-PH et al. |
| EPI_ISL_451542, EPI_ISL_451543, EPI_ISL_451544 | Pathology Sydney South West - NSW Health Pathology | NSW Health Pathology - Institute of Clinical Pathology and Medical Research; Westmead Hospital; University of Sydney | CIDM-PH et al. |
| EPI_ISL_451545 | Laverty Pathology | NSW Health Pathology - Institute of Clinical Pathology and Medical Research; Westmead Hospital; University of Sydney | CIDM-PH et al. |
| EPI_ISL_451546, EPI_ISL_451547 | Pathology West - NSW Health Pathology | NSW Health Pathology - Institute of Clinical Pathology and Medical Research; Westmead Hospital; University of Sydney | CIDM-PH et al. |
| EPI_ISL_451548 | Childrens Hospital Westmead | NSW Health Pathology - Institute of Clinical Pathology and Medical Research; Westmead Hospital; University of Sydney | CIDM-PH et al. |
| EPI_ISL_451549 | Pathology Sydney South West - NSW Health Pathology | NSW Health Pathology - Institute of Clinical Pathology and Medical Research; Westmead Hospital; University of Sydney | CIDM-PH et al. |
| EPI_ISL_451550, EPI_ISL_451551 | Pathology West - NSW Health Pathology | NSW Health Pathology - Institute of Clinical Pathology and Medical Research; Westmead Hospital; University of Sydney | CIDM-PH et al. |
| EPI_ISL_451552 | Pathology North - NSW Health Pathology | NSW Health Pathology - Institute of Clinical Pathology and Medical Research; Westmead Hospital; University of Sydney | CIDM-PH et al. |
| EPI_ISL_451553, EPI_ISL_451554, EPI_ISL_451555 | Medlab Pathology | NSW Health Pathology - Institute of Clinical Pathology and Medical Research; Westmead Hospital; University of Sydney | CIDM-PH et al. |
| EPI_ISL_451556, EPI_ISL_451557 | Pathology West - NSW Health Pathology | NSW Health Pathology - Institute of Clinical Pathology and Medical Research; Westmead Hospital; University of Sydney | CIDM-PH et al. |
| EPI_ISL_451558 | Medlab Pathology | NSW Health Pathology - Institute of Clinical Pathology and Medical Research; Westmead Hospital; University of Sydney | CIDM-PH et al. |
| EPI_ISL_451559, EPI_ISL_451560, EPI_ISL_451561, EPI_ISL_451562, EPI_ISL_451563, EPI_ISL_451564, EPI_ISL_451565 | Pathology Sydney South West - NSW Health Pathology | NSW Health Pathology - Institute of Clinical Pathology and Medical Research; Westmead Hospital; University of Sydney | CIDM-PH et al. |
| EPI_ISL_451566 | Pathology West - NSW Health Pathology | NSW Health Pathology - Institute of Clinical Pathology and Medical Research; Westmead Hospital; University of Sydney | CIDM-PH et al. |
| EPI_ISL_451567 | Medlab Pathology | NSW Health Pathology - Institute of Clinical Pathology and Medical Research; Westmead Hospital; University of Sydney | CIDM-PH et al. |
| EPI_ISL_451568, EPI_ISL_451569, EPI_ISL_451570, EPI_ISL_451571 | Pathology West - NSW Health Pathology | NSW Health Pathology - Institute of Clinical Pathology and Medical Research; Westmead Hospital; University of Sydney | CIDM-PH et al. |
| EPI_ISL_451572 | Laverty Pathology | NSW Health Pathology - Institute of Clinical Pathology and Medical Research; Westmead Hospital; University of Sydney | CIDM-PH et al. |
| EPI_ISL_451573 | Pathology West - NSW Health Pathology | NSW Health Pathology - Institute of Clinical Pathology and Medical Research; Westmead Hospital; University of Sydney | CIDM-PH et al. |
| EPI_ISL_451574 | Australian Clinical Labs | NSW Health Pathology - Institute of Clinical Pathology and Medical Research; Westmead Hospital; University of Sydney | CIDM-PH et al. |
| EPI_ISL_451575, EPI_ISL_451576, EPI_ISL_451577 | Pathology West - NSW Health Pathology | NSW Health Pathology - Institute of Clinical Pathology and Medical Research; Westmead Hospital; University of Sydney | CIDM-PH et al. |
| EPI_ISL_451578 | Pathology Sydney South West - NSW Health Pathology | NSW Health Pathology - Institute of Clinical Pathology and Medical Research; Westmead Hospital; University of Sydney | CIDM-PH et al. |
| EPI_ISL_451579, EPI_ISL_451580 | Pathology West - NSW Health Pathology | NSW Health Pathology - Institute of Clinical Pathology and Medical Research; Westmead Hospital; University of Sydney | CIDM-PH et al. |
| EPI_ISL_451581 | Pathology Sydney South West - NSW Health Pathology | NSW Health Pathology - Institute of Clinical Pathology and Medical Research; Westmead Hospital; University of Sydney | CIDM-PH et al. |
| EPI_ISL_451582 | Pathology West - NSW Health Pathology | NSW Health Pathology - Institute of Clinical Pathology and | CIDM-PH et al. |

| | | | |
|---|---|---|---|
| | | Medical Research; Westmead Hospital; University of Sydney | |
| EPI_ISL_451583 | Medlab Pathology | NSW Health Pathology - Institute of Clinical Pathology and Medical Research; Westmead Hospital; University of Sydney | CIDM-PH et al. |
| EPI_ISL_451584, EPI_ISL_451585, EPI_ISL_451586 | Pathology West - NSW Health Pathology | NSW Health Pathology - Institute of Clinical Pathology and Medical Research; Westmead Hospital; University of Sydney | CIDM-PH et al. |
| EPI_ISL_451587 | Pathology North Hunter- NSW Health Pathology | NSW Health Pathology - Institute of Clinical Pathology and Medical Research; Westmead Hospital; University of Sydney | CIDM-PH et al. |
| EPI_ISL_451588 | ACT pathology | NSW Health Pathology - Institute of Clinical Pathology and Medical Research; Westmead Hospital; University of Sydney | CIDM-PH et al. |
| EPI_ISL_451589 | Pathology North Hunter- NSW Health Pathology | NSW Health Pathology - Institute of Clinical Pathology and Medical Research; Westmead Hospital; University of Sydney | CIDM-PH et al. |
| EPI_ISL_451590, EPI_ISL_451591, EPI_ISL_451592, EPI_ISL_451593 | ACT pathology | NSW Health Pathology - Institute of Clinical Pathology and Medical Research; Westmead Hospital; University of Sydney | CIDM-PH et al. |
| EPI_ISL_451594 | Childrens Hospital Westmead | NSW Health Pathology - Institute of Clinical Pathology and Medical Research; Westmead Hospital; University of Sydney | CIDM-PH et al. |
| EPI_ISL_451595, EPI_ISL_451596 | ACT pathology | NSW Health Pathology - Institute of Clinical Pathology and Medical Research; Westmead Hospital; University of Sydney | CIDM-PH et al. |
| EPI_ISL_451597 | Medlab Pathology | NSW Health Pathology - Institute of Clinical Pathology and Medical Research; Westmead Hospital; University of Sydney | CIDM-PH et al. |
| EPI_ISL_451598 | ACT pathology | NSW Health Pathology - Institute of Clinical Pathology and Medical Research; Westmead Hospital; University of Sydney | CIDM-PH et al. |
| EPI_ISL_451599 | Australian Clinical Labs | NSW Health Pathology - Institute of Clinical Pathology and Medical Research; Westmead Hospital; University of Sydney | CIDM-PH et al. |
| EPI_ISL_451600 | Pathology North Hunter- NSW Health Pathology | NSW Health Pathology - Institute of Clinical Pathology and Medical Research; Westmead Hospital; University of Sydney | CIDM-PH et al. |
| EPI_ISL_451601 | Douglas Hanly Moir Pathology | NSW Health Pathology - Institute of Clinical Pathology and Medical Research; Westmead Hospital; University of Sydney | CIDM-PH et al. |
| EPI_ISL_451602 | Laverty Pathology | NSW Health Pathology - Institute of Clinical Pathology and Medical Research; Westmead Hospital; University of Sydney | CIDM-PH et al. |
| EPI_ISL_451603 | Pathology Sydney South West - NSW Health Pathology | NSW Health Pathology - Institute of Clinical Pathology and Medical Research; Westmead Hospital; University of Sydney | CIDM-PH et al. |
| EPI_ISL_451604 | Pathology North - NSW Health Pathology | NSW Health Pathology - Institute of Clinical Pathology and Medical Research; Westmead Hospital; University of Sydney | CIDM-PH et al. |
| EPI_ISL_451605 | Childrens Hospital Westmead | NSW Health Pathology - Institute of Clinical Pathology and Medical Research; Westmead Hospital; University of Sydney | CIDM-PH et al. |
| EPI_ISL_451606, EPI_ISL_451607 | Pathology West - NSW Health Pathology | NSW Health Pathology - Institute of Clinical Pathology and Medical Research; Westmead Hospital; University of Sydney | CIDM-PH et al. |
| EPI_ISL_451608 | Pathology Sydney South West - NSW Health Pathology | NSW Health Pathology - Institute of Clinical Pathology and Medical Research; Westmead Hospital; University of Sydney | CIDM-PH et al. |
| EPI_ISL_451609 | Laverty Pathology | NSW Health Pathology - Institute of Clinical Pathology and Medical Research; Westmead Hospital; University of Sydney | CIDM-PH et al. |
| EPI_ISL_451610 | Medlab Pathology | NSW Health Pathology - Institute of Clinical Pathology and Medical Research; Westmead Hospital; University of Sydney | CIDM-PH et al. |
| EPI_ISL_451611 | Laverty Pathology | NSW Health Pathology - Institute of Clinical Pathology and Medical Research; Westmead Hospital; University of Sydney | CIDM-PH et al. |
| EPI_ISL_451612 | ACT pathology | NSW Health Pathology - Institute of Clinical Pathology and Medical Research; Westmead Hospital; University of Sydney | CIDM-PH et al. |
| EPI_ISL_451613, EPI_ISL_451614, EPI_ISL_451615, EPI_ISL_451616, EPI_ISL_451617, EPI_ISL_451618, EPI_ISL_451619, EPI_ISL_451620, EPI_ISL_451621, EPI_ISL_451622, EPI_ISL_451623, EPI_ISL_451624, EPI_ISL_451625, EPI_ISL_451626, EPI_ISL_451627, EPI_ISL_451628, EPI_ISL_451629, EPI_ISL_451630 | | | |
| see above | Pathology West - NSW Health Pathology | NSW Health Pathology - Institute of Clinical Pathology and Medical Research; Westmead Hospital; University of Sydney | CIDM-PH et al. |
| EPI_ISL_451631 | Laverty Pathology | NSW Health Pathology - Institute of Clinical Pathology and Medical Research; Westmead Hospital; University of Sydney | CIDM-PH et al. |
| EPI_ISL_451632, EPI_ISL_451633, EPI_ISL_451634, EPI_ISL_451635, EPI_ISL_451636 | South Eastern Area Laboratory Services | NSW Health Pathology - Institute of Clinical Pathology and Medical Research; Westmead Hospital; University of Sydney | CIDM-PH et al. |
| EPI_ISL_451637, EPI_ISL_451638, EPI_ISL_451639, EPI_ISL_451640 | Pathology West - NSW Health Pathology | NSW Health Pathology - Institute of Clinical Pathology and Medical Research; Westmead Hospital; University of Sydney | CIDM-PH et al. |
| EPI_ISL_451641 | Laverty Pathology | NSW Health Pathology - Institute of Clinical Pathology and Medical Research; Westmead Hospital; University of Sydney | CIDM-PH et al. |
| EPI_ISL_451642, EPI_ISL_451643 | Pathology Sydney South West - NSW Health Pathology | NSW Health Pathology - Institute of Clinical Pathology and Medical Research; Westmead Hospital; University of Sydney | CIDM-PH et al. |
| EPI_ISL_453813, EPI_ISL_453814, EPI_ISL_453815, EPI_ISL_453816, EPI_ISL_453817, EPI_ISL_453818, EPI_ISL_453819, EPI_ISL_453820, EPI_ISL_453821, EPI_ISL_453822, EPI_ISL_453823, EPI_ISL_453824, EPI_ISL_453825, EPI_ISL_453826, EPI_ISL_453827, EPI_ISL_453828, EPI_ISL_453829, EPI_ISL_453830, EPI_ISL_453831, EPI_ISL_453832, EPI_ISL_453833, EPI_ISL_453834, EPI_ISL_453835, EPI_ISL_453836, EPI_ISL_453837, EPI_ISL_453838, EPI_ISL_453839, EPI_ISL_453840, EPI_ISL_453841, EPI_ISL_453842, EPI_ISL_453843, EPI_ISL_453844, EPI_ISL_453845, EPI_ISL_453846, EPI_ISL_453847, EPI_ISL_453848, EPI_ISL_453849, EPI_ISL_453850, EPI_ISL_453851, EPI_ISL_453852, EPI_ISL_453853, EPI_ISL_453854, EPI_ISL_453855, EPI_ISL_453856, EPI_ISL_453857, EPI_ISL_453858, EPI_ISL_453859, EPI_ISL_453860, EPI_ISL_453861, EPI_ISL_453862, EPI_ISL_453863, EPI_ISL_453864, EPI_ISL_453865, EPI_ISL_453866, EPI_ISL_453867, EPI_ISL_453868, EPI_ISL_453869, EPI_ISL_453870, EPI_ISL_453871, EPI_ISL_453872, EPI_ISL_453873, EPI_ISL_453874, EPI_ISL_453875, EPI_ISL_453876, EPI_ISL_453877, EPI_ISL_453878, EPI_ISL_453879, EPI_ISL_453880, EPI_ISL_453881, EPI_ISL_453882, EPI_ISL_453883, EPI_ISL_453884, EPI_ISL_453885, EPI_ISL_453886, EPI_ISL_453887, EPI_ISL_453888, EPI_ISL_453889, EPI_ISL_453890, EPI_ISL_453891, EPI_ISL_453892, EPI_ISL_453893, EPI_ISL_453894, EPI_ISL_453895, EPI_ISL_453896, EPI_ISL_453897, EPI_ISL_453898, EPI_ISL_453899, EPI_ISL_453900, EPI_ISL_453901, EPI_ISL_453902, EPI_ISL_453903, EPI_ISL_453904, EPI_ISL_453905, EPI_ISL_453906, EPI_ISL_453907, EPI_ISL_453908, EPI_ISL_453909, EPI_ISL_453910, EPI_ISL_453911, EPI_ISL_453912, EPI_ISL_453913, EPI_ISL_453914, EPI_ISL_453915, EPI_ISL_453916, EPI_ISL_453917, EPI_ISL_453918, EPI_ISL_453919, EPI_ISL_453920, EPI_ISL_453921, EPI_ISL_453922, EPI_ISL_453923, EPI_ISL_453924, EPI_ISL_453925, EPI_ISL_453926, EPI_ISL_453927, EPI_ISL_453928, EPI_ISL_453929, EPI_ISL_453930, EPI_ISL_453931, EPI_ISL_453932, EPI_ISL_453933, EPI_ISL_453934, EPI_ISL_453935, EPI_ISL_453936, EPI_ISL_453937, EPI_ISL_453938, EPI_ISL_453939, EPI_ISL_453940, EPI_ISL_453941, EPI_ISL_453942, EPI_ISL_453943, EPI_ISL_453944, EPI_ISL_453945, EPI_ISL_453946, EPI_ISL_453947, EPI_ISL_453948, EPI_ISL_453949, EPI_ISL_453950, EPI_ISL_453951, EPI_ISL_453952, EPI_ISL_453953, EPI_ISL_453954, EPI_ISL_453955, EPI_ISL_453956, EPI_ISL_453957, EPI_ISL_453958, EPI_ISL_453959, EPI_ISL_453960, EPI_ISL_453961, EPI_ISL_453962, EPI_ISL_453963, EPI_ISL_453964, EPI_ISL_453965, EPI_ISL_453966, EPI_ISL_453967, EPI_ISL_453968, EPI_ISL_453969, EPI_ISL_453970, EPI_ISL_453971, EPI_ISL_453972, EPI_ISL_453973, EPI_ISL_453974, EPI_ISL_453975, EPI_ISL_453976, EPI_ISL_453977, EPI_ISL_453978, EPI_ISL_453979, EPI_ISL_453980, EPI_ISL_453981, EPI_ISL_453982, EPI_ISL_453983, EPI_ISL_453984, EPI_ISL_453985, EPI_ISL_453986, EPI_ISL_453987, EPI_ISL_453988, EPI_ISL_453989, EPI_ISL_453990, EPI_ISL_453991, EPI_ISL_453992, EPI_ISL_453993, EPI_ISL_453994, EPI_ISL_453995, EPI_ISL_453996, EPI_ISL_453997, EPI_ISL_453998, EPI_ISL_453999, EPI_ISL_454000, EPI_ISL_454001, EPI_ISL_454002, EPI_ISL_454003, EPI_ISL_454004, EPI_ISL_454005, EPI_ISL_454006, EPI_ISL_454007, EPI_ISL_454008, EPI_ISL_454009, EPI_ISL_454010, | | | |

| | | | |
|---|---|---|---|
| EPI_ISL_454011, EPI_ISL_454012, EPI_ISL_454013, EPI_ISL_454014, EPI_ISL_454015, EPI_ISL_454016, EPI_ISL_454017, EPI_ISL_454018, EPI_ISL_454019, EPI_ISL_454020, EPI_ISL_454021, EPI_ISL_454022, EPI_ISL_454023, EPI_ISL_454024, EPI_ISL_454025, EPI_ISL_454026, EPI_ISL_454027, EPI_ISL_454028, EPI_ISL_454029, EPI_ISL_454030, EPI_ISL_454031, EPI_ISL_454032, EPI_ISL_454033, EPI_ISL_454034, EPI_ISL_454035, EPI_ISL_454036, EPI_ISL_454037, EPI_ISL_454038, EPI_ISL_454039, EPI_ISL_454040, EPI_ISL_454041, EPI_ISL_454042, EPI_ISL_454043, EPI_ISL_454044, EPI_ISL_454045, EPI_ISL_454046, EPI_ISL_454047, EPI_ISL_454048, EPI_ISL_454049, EPI_ISL_454050, EPI_ISL_454051, EPI_ISL_454052, EPI_ISL_454053, EPI_ISL_454054, EPI_ISL_454055, EPI_ISL_454056, EPI_ISL_454057, EPI_ISL_454058, EPI_ISL_454059, EPI_ISL_454060, EPI_ISL_454061, EPI_ISL_454062, EPI_ISL_454063, EPI_ISL_454064, EPI_ISL_454065, EPI_ISL_454066, EPI_ISL_454067, EPI_ISL_454068, EPI_ISL_454069, EPI_ISL_454070, EPI_ISL_454071, EPI_ISL_454072, EPI_ISL_454073, EPI_ISL_454074, EPI_ISL_454075, EPI_ISL_454076, EPI_ISL_454077, EPI_ISL_454078, EPI_ISL_454079, EPI_ISL_454080, EPI_ISL_454081, EPI_ISL_454082, EPI_ISL_454083, EPI_ISL_454084, EPI_ISL_454085, EPI_ISL_454086, EPI_ISL_454087, EPI_ISL_454088, EPI_ISL_454089, EPI_ISL_454090, EPI_ISL_454091, EPI_ISL_454092, EPI_ISL_454093, EPI_ISL_454094, EPI_ISL_454095, EPI_ISL_454096, EPI_ISL_454097, EPI_ISL_454098, EPI_ISL_454099, EPI_ISL_454100, EPI_ISL_454101, EPI_ISL_454102, EPI_ISL_454103, EPI_ISL_454104, EPI_ISL_454105, EPI_ISL_454106, EPI_ISL_454107, EPI_ISL_454108, EPI_ISL_454109, EPI_ISL_454110, EPI_ISL_454111, EPI_ISL_454112, EPI_ISL_454113, EPI_ISL_454114, EPI_ISL_454115, EPI_ISL_454116, EPI_ISL_454117, EPI_ISL_454118, EPI_ISL_454119, EPI_ISL_454120, EPI_ISL_454121, EPI_ISL_454122, EPI_ISL_454123, EPI_ISL_454124, EPI_ISL_454125, EPI_ISL_454126, EPI_ISL_454127, EPI_ISL_454128, EPI_ISL_454129, EPI_ISL_454130, EPI_ISL_454131, EPI_ISL_454132, EPI_ISL_454133, EPI_ISL_454134, EPI_ISL_454135, EPI_ISL_454136, EPI_ISL_454137, EPI_ISL_454138, EPI_ISL_454139, EPI_ISL_454140, EPI_ISL_454141, EPI_ISL_454142, EPI_ISL_454143, EPI_ISL_454144, EPI_ISL_454145, EPI_ISL_454146, EPI_ISL_454147, EPI_ISL_454148, EPI_ISL_454149, EPI_ISL_454150, EPI_ISL_454151, EPI_ISL_454152, EPI_ISL_454153, EPI_ISL_454154, EPI_ISL_454155, EPI_ISL_454156, EPI_ISL_454157, EPI_ISL_454158, EPI_ISL_454159, EPI_ISL_454160, EPI_ISL_454161, EPI_ISL_454162, EPI_ISL_454163, EPI_ISL_454164, EPI_ISL_454165, EPI_ISL_454166, EPI_ISL_454167, EPI_ISL_454168, EPI_ISL_454169, EPI_ISL_454170, EPI_ISL_454171, EPI_ISL_454172, EPI_ISL_454173, EPI_ISL_454174, EPI_ISL_454175, EPI_ISL_454176, EPI_ISL_454177, EPI_ISL_454178, EPI_ISL_454179, EPI_ISL_454180, EPI_ISL_454181, EPI_ISL_454182, EPI_ISL_454183, EPI_ISL_454184, EPI_ISL_454185, EPI_ISL_454186, EPI_ISL_454187, EPI_ISL_454188, EPI_ISL_454189, EPI_ISL_454190, EPI_ISL_454191, EPI_ISL_454192, EPI_ISL_454193, EPI_ISL_454194, EPI_ISL_454195, EPI_ISL_454196, EPI_ISL_454197, EPI_ISL_454198, EPI_ISL_454199, EPI_ISL_454200, EPI_ISL_454201, EPI_ISL_454202, EPI_ISL_454203, EPI_ISL_454204, EPI_ISL_454205, EPI_ISL_454206, EPI_ISL_454207, EPI_ISL_454208, EPI_ISL_454209, EPI_ISL_454210, EPI_ISL_454211, EPI_ISL_454212, EPI_ISL_454213, EPI_ISL_454214, EPI_ISL_454215, EPI_ISL_454216, EPI_ISL_454217, EPI_ISL_454218, EPI_ISL_454219, EPI_ISL_454220, EPI_ISL_454221, EPI_ISL_454222, EPI_ISL_454223, EPI_ISL_454224, EPI_ISL_454225, EPI_ISL_454226, EPI_ISL_454227, EPI_ISL_454228, EPI_ISL_454229, EPI_ISL_454230, EPI_ISL_454231, EPI_ISL_454232, EPI_ISL_454233, EPI_ISL_454234, EPI_ISL_454235, EPI_ISL_454236, EPI_ISL_454237, EPI_ISL_454238, EPI_ISL_454239, EPI_ISL_454240, EPI_ISL_454241, EPI_ISL_454242, EPI_ISL_454243, EPI_ISL_454244, EPI_ISL_454245, EPI_ISL_454246, EPI_ISL_454247, EPI_ISL_454248, EPI_ISL_454249, EPI_ISL_454250, EPI_ISL_454251, EPI_ISL_454252, EPI_ISL_454253, EPI_ISL_454254, EPI_ISL_454255, EPI_ISL_454256, EPI_ISL_454257, EPI_ISL_454258, EPI_ISL_454259, EPI_ISL_454260, EPI_ISL_454261, EPI_ISL_454262, EPI_ISL_454263, EPI_ISL_454264, EPI_ISL_454265, EPI_ISL_454266, EPI_ISL_454267, EPI_ISL_454268, EPI_ISL_454269, EPI_ISL_454270, EPI_ISL_454271, EPI_ISL_454272, EPI_ISL_454273, EPI_ISL_454274, EPI_ISL_454275, EPI_ISL_454276, EPI_ISL_454277, EPI_ISL_454278, EPI_ISL_454279, EPI_ISL_454280, EPI_ISL_454281, EPI_ISL_454282, EPI_ISL_454283, EPI_ISL_454284, EPI_ISL_454285, EPI_ISL_454286, EPI_ISL_454287, EPI_ISL_454288, EPI_ISL_454289, EPI_ISL_454290, EPI_ISL_454291, EPI_ISL_454292, EPI_ISL_454293, EPI_ISL_454294, EPI_ISL_454295, EPI_ISL_454296, EPI_ISL_454297, EPI_ISL_454298, EPI_ISL_454299, EPI_ISL_454300, EPI_ISL_454301, EPI_ISL_454302, EPI_ISL_454303, EPI_ISL_454304, EPI_ISL_454305, EPI_ISL_454306, EPI_ISL_454307, EPI_ISL_454308, EPI_ISL_454309, EPI_ISL_454310, EPI_ISL_454311, EPI_ISL_454312, EPI_ISL_454313, EPI_ISL_454314, EPI_ISL_454315, EPI_ISL_454316, EPI_ISL_454317, EPI_ISL_454318, EPI_ISL_454319, EPI_ISL_454320, EPI_ISL_454321, EPI_ISL_454322, EPI_ISL_454323, EPI_ISL_454324, EPI_ISL_454325, EPI_ISL_454326, EPI_ISL_454327, EPI_ISL_454328, EPI_ISL_454329, EPI_ISL_454330, EPI_ISL_454331, EPI_ISL_454332, EPI_ISL_454333, EPI_ISL_454334, EPI_ISL_454335, EPI_ISL_454336, EPI_ISL_454337, EPI_ISL_454338, EPI_ISL_454339, EPI_ISL_454340, EPI_ISL_454341, EPI_ISL_454342, EPI_ISL_454343, EPI_ISL_454344, EPI_ISL_454345, EPI_ISL_454346, EPI_ISL_454347, EPI_ISL_454348, EPI_ISL_454349, EPI_ISL_454350, EPI_ISL_454351, EPI_ISL_454352 | | | |
| see above | unknown | Instituto Nacional de Saude (INSA) | Borges et al |
| EPI_ISL_454607, EPI_ISL_454608, EPI_ISL_454609, EPI_ISL_454610, EPI_ISL_454611, EPI_ISL_454612, EPI_ISL_454613 | Alameda County Public Health Lab | Chan-Zuckerberg Biohub | CZB Cliahub Consortium |
| EPI_ISL_454614, EPI_ISL_454615, EPI_ISL_454616, EPI_ISL_454617, EPI_ISL_454618, EPI_ISL_454619, EPI_ISL_454620, EPI_ISL_454621, EPI_ISL_454622, EPI_ISL_454623, EPI_ISL_454624, EPI_ISL_454625, EPI_ISL_454626, EPI_ISL_454627, EPI_ISL_454628, EPI_ISL_454629, EPI_ISL_454630, EPI_ISL_454631, EPI_ISL_454632, EPI_ISL_454633, EPI_ISL_454634 | | | |
| see above | UCSF Clinical Microbiology Laboratory | Chan-Zuckerberg Biohub | CZB Cliahub Consortium |
| EPI_ISL_454635 | County Of San Luis Obispo Public Health Laboratory | Chan-Zuckerberg Biohub | CZB Cliahub Consortium |
| EPI_ISL_454636, EPI_ISL_454637, EPI_ISL_454638, EPI_ISL_454639, EPI_ISL_454640, EPI_ISL_454641 | Humboldt County Public Health Laboratory | Chan-Zuckerberg Biohub | CZB Cliahub Consortium |
| EPI_ISL_454653, EPI_ISL_454654, EPI_ISL_454655, EPI_ISL_454656, EPI_ISL_454657, EPI_ISL_454658, EPI_ISL_454659, EPI_ISL_454661, EPI_ISL_454662, EPI_ISL_454663, EPI_ISL_454664, EPI_ISL_454665, EPI_ISL_454666, EPI_ISL_454667, EPI_ISL_454668, EPI_ISL_454669, EPI_ISL_454670, EPI_ISL_454672, EPI_ISL_454673, EPI_ISL_454674, EPI_ISL_454675, EPI_ISL_454676, EPI_ISL_454677, EPI_ISL_454678, EPI_ISL_454679, EPI_ISL_454680, EPI_ISL_454681, EPI_ISL_454682, EPI_ISL_454683, EPI_ISL_454684, EPI_ISL_454685, EPI_ISL_454686, EPI_ISL_454687, EPI_ISL_454688, EPI_ISL_454689 | | | |
| see above | County of Santa Clara Public Health Department | Chan-Zuckerberg Biohub | CZB Cliahub Consortium |
| EPI_ISL_455028, EPI_ISL_455029, EPI_ISL_455030, EPI_ISL_455031, EPI_ISL_455032, EPI_ISL_455033, EPI_ISL_455034, EPI_ISL_455035 | Pathology West - NSW Health Pathology | NSW Health Pathology - Institute of Clinical Pathology and Medical Research; Westmead Hospital; University of Sydney | CIDM-PH et al. |
| EPI_ISL_455036 | South Eastern Area Laboratory Services | NSW Health Pathology - Institute of Clinical Pathology and Medical Research; Westmead Hospital; University of Sydney | CIDM-PH et al. |
| EPI_ISL_455037 | Pathology Sydney South West - NSW Health Pathology | NSW Health Pathology - Institute of Clinical Pathology and Medical Research; Westmead Hospital; University of Sydney | CIDM-PH et al. |
| EPI_ISL_455038, EPI_ISL_455039, EPI_ISL_455040 | Pathology West - NSW Health Pathology | NSW Health Pathology - Institute of Clinical Pathology and Medical Research; Westmead Hospital; University of Sydney | CIDM-PH et al. |
| EPI_ISL_455041 | Laverty Pathology | NSW Health Pathology - Institute of Clinical Pathology and Medical Research; Westmead Hospital; University of Sydney | CIDM-PH et al. |
| EPI_ISL_455042, EPI_ISL_455043 | ACT Pathology | NSW Health Pathology - Institute of Clinical Pathology and Medical Research; Westmead Hospital; University of Sydney | CIDM-PH et al. |
| EPI_ISL_455044, EPI_ISL_455045, EPI_ISL_455046, EPI_ISL_455047 | Pathology West - NSW Health Pathology | NSW Health Pathology - Institute of Clinical Pathology and Medical Research; Westmead Hospital; University of Sydney | CIDM-PH et al. |
| EPI_ISL_455048 | Laverty Pathology | NSW Health Pathology - Institute of Clinical Pathology and Medical Research; Westmead Hospital; University of Sydney | CIDM-PH et al. |
| EPI_ISL_455049 | Pathology West - NSW Health Pathology | NSW Health Pathology - Institute of Clinical Pathology and Medical Research; Westmead Hospital; University of Sydney | CIDM-PH et al. |
| EPI_ISL_455050 | ACT Pathology | NSW Health Pathology - Institute of Clinical Pathology and Medical Research; Westmead Hospital; University of Sydney | CIDM-PH et al. |
| EPI_ISL_455051, EPI_ISL_455052, EPI_ISL_455053 | Douglas Hanly Moir Pathology | NSW Health Pathology - Institute of Clinical Pathology and Medical Research; Westmead Hospital; University of Sydney | CIDM-PH et al. |
| EPI_ISL_455054 | South Eastern Area Laboratory Services | NSW Health Pathology - Institute of Clinical Pathology and Medical Research; Westmead Hospital; University of Sydney | CIDM-PH et al. |
| EPI_ISL_455055, EPI_ISL_455056, EPI_ISL_455057, EPI_ISL_455058, EPI_ISL_455059, EPI_ISL_455060, EPI_ISL_455061, EPI_ISL_455062 | Pathology West - NSW Health Pathology | NSW Health Pathology - Institute of Clinical Pathology and Medical Research; Westmead Hospital; University of Sydney | CIDM-PH et al. |
| EPI_ISL_455063 | Laverty Pathology | NSW Health Pathology - Institute of Clinical Pathology and Medical Research; Westmead Hospital; University of Sydney | CIDM-PH et al. |
| EPI_ISL_455064 | Pathology West - NSW Health Pathology | NSW Health Pathology - Institute of Clinical Pathology and Medical Research; Westmead Hospital; University of Sydney | CIDM-PH et al. |
| EPI_ISL_455065 | Douglas Hanly Moir Pathology | NSW Health Pathology - Institute of Clinical Pathology and Medical Research; Westmead Hospital; University of Sydney | CIDM-PH et al. |
| EPI_ISL_455066, EPI_ISL_455067 | Pathology West - NSW Health Pathology | NSW Health Pathology - Institute of Clinical Pathology and Medical Research; Westmead Hospital; University of Sydney | CIDM-PH et al. |
| EPI_ISL_455068 | Childrens Hospital Westmead | NSW Health Pathology - Institute of Clinical Pathology and Medical Research; Westmead Hospital; University of Sydney | CIDM-PH et al. |

| | | | |
|---|---|---|---|
| EPI_ISL_455069, EPI_ISL_455070 | Pathology West - NSW Health Pathology | NSW Health Pathology - Institute of Clinical Pathology and Medical Research; Westmead Hospital; University of Sydney | CIDM-PH et al. |
| EPI_ISL_455071 | Pathology Sydney South West - NSW Health Pathology | NSW Health Pathology - Institute of Clinical Pathology and Medical Research; Westmead Hospital; University of Sydney | CIDM-PH et al. |
| EPI_ISL_455072, EPI_ISL_455073 | Sullivan Nicolaides Pathology | NSW Health Pathology - Institute of Clinical Pathology and Medical Research; Westmead Hospital; University of Sydney | CIDM-PH et al. |
| EPI_ISL_455074 | ACT Pathology | NSW Health Pathology - Institute of Clinical Pathology and Medical Research; Westmead Hospital; University of Sydney | CIDM-PH et al. |
| EPI_ISL_455075, EPI_ISL_455076, EPI_ISL_455077, EPI_ISL_455078, EPI_ISL_455079, EPI_ISL_455080, EPI_ISL_455081, EPI_ISL_455082, EPI_ISL_455083, EPI_ISL_455084, EPI_ISL_455085, EPI_ISL_455086, EPI_ISL_455087, EPI_ISL_455088, EPI_ISL_455089, EPI_ISL_455090, EPI_ISL_455091, EPI_ISL_455092, EPI_ISL_455093 | | | |
| see above | South Eastern Area Laboratory Services | NSW Health Pathology - Institute of Clinical Pathology and Medical Research; Westmead Hospital; University of Sydney | CIDM-PH et al. |
| EPI_ISL_455094 | Pathology Sydney South West - NSW Health Pathology | NSW Health Pathology - Institute of Clinical Pathology and Medical Research; Westmead Hospital; University of Sydney | CIDM-PH et al. |
| EPI_ISL_455095 | Pathology West - NSW Health Pathology | NSW Health Pathology - Institute of Clinical Pathology and Medical Research; Westmead Hospital; University of Sydney | CIDM-PH et al. |
| EPI_ISL_455096, EPI_ISL_455097, EPI_ISL_455098, EPI_ISL_455099 | South Eastern Area Laboratory Services | NSW Health Pathology - Institute of Clinical Pathology and Medical Research; Westmead Hospital; University of Sydney | CIDM-PH et al. |
| EPI_ISL_455625, EPI_ISL_455626, EPI_ISL_455627 | unknown | Instituto Nacional de Saude (INSA) | Borges et al |
| EPI_ISL_456410, EPI_ISL_456411, EPI_ISL_456412, EPI_ISL_456413, EPI_ISL_456414, EPI_ISL_456415, EPI_ISL_456416, EPI_ISL_456417, EPI_ISL_456419, EPI_ISL_456420, EPI_ISL_456421, EPI_ISL_456422, EPI_ISL_456423, EPI_ISL_456424, EPI_ISL_456425, EPI_ISL_456426, EPI_ISL_456427, EPI_ISL_456428, EPI_ISL_456429, EPI_ISL_456430, EPI_ISL_456431, EPI_ISL_456432, EPI_ISL_456433, EPI_ISL_456434, EPI_ISL_456435, EPI_ISL_456436, EPI_ISL_456438, EPI_ISL_456440, EPI_ISL_456441, EPI_ISL_456442, EPI_ISL_456443, EPI_ISL_456444, EPI_ISL_456445, EPI_ISL_456446, EPI_ISL_456447, EPI_ISL_456448, EPI_ISL_456449, EPI_ISL_456450, EPI_ISL_456451, EPI_ISL_456452, EPI_ISL_456453, EPI_ISL_456454, EPI_ISL_456455, EPI_ISL_456456, EPI_ISL_456457, EPI_ISL_456458, EPI_ISL_456459, EPI_ISL_456460, EPI_ISL_456461, EPI_ISL_456462, EPI_ISL_456463, EPI_ISL_456464, EPI_ISL_456465, EPI_ISL_456466, EPI_ISL_456467, EPI_ISL_456468, EPI_ISL_456469, EPI_ISL_456470, EPI_ISL_456471, EPI_ISL_456472, EPI_ISL_456473, EPI_ISL_456474, EPI_ISL_456475, EPI_ISL_456476, EPI_ISL_456477, EPI_ISL_456480, EPI_ISL_456481, EPI_ISL_456482, EPI_ISL_456483, EPI_ISL_456484, EPI_ISL_456485, EPI_ISL_456486, EPI_ISL_456487, EPI_ISL_456488, EPI_ISL_456489, EPI_ISL_456490, EPI_ISL_456491, EPI_ISL_456492, EPI_ISL_456499, EPI_ISL_456500, EPI_ISL_456502, EPI_ISL_456503, EPI_ISL_456504, EPI_ISL_456505, EPI_ISL_456508, EPI_ISL_456509, EPI_ISL_456510, EPI_ISL_456511, EPI_ISL_456512, EPI_ISL_456513, EPI_ISL_456514, EPI_ISL_456515, EPI_ISL_456516, EPI_ISL_456517, EPI_ISL_456519, EPI_ISL_456520, EPI_ISL_456521, EPI_ISL_456522, EPI_ISL_456523, EPI_ISL_456524, EPI_ISL_456525, EPI_ISL_456526, EPI_ISL_456527, EPI_ISL_456528, EPI_ISL_456529, EPI_ISL_456530, EPI_ISL_456531, EPI_ISL_456532, EPI_ISL_456533, EPI_ISL_456534, EPI_ISL_456535, EPI_ISL_456536, EPI_ISL_456537, EPI_ISL_456538, EPI_ISL_456539, EPI_ISL_456540, EPI_ISL_456541, EPI_ISL_456542, EPI_ISL_456543, EPI_ISL_456544, EPI_ISL_456545, EPI_ISL_456546, EPI_ISL_456547, EPI_ISL_456548, EPI_ISL_456549, EPI_ISL_456550, EPI_ISL_456551, EPI_ISL_456552, EPI_ISL_456553, EPI_ISL_456554, EPI_ISL_456555, EPI_ISL_456557, EPI_ISL_456558, EPI_ISL_456559, EPI_ISL_456560, EPI_ISL_456561, EPI_ISL_456562, EPI_ISL_456563, EPI_ISL_456564, EPI_ISL_456565, EPI_ISL_456566, EPI_ISL_456567, EPI_ISL_456568, EPI_ISL_456569, EPI_ISL_456571, EPI_ISL_456572, EPI_ISL_456573, EPI_ISL_456574, EPI_ISL_456575, EPI_ISL_456576, EPI_ISL_456577, EPI_ISL_456578, EPI_ISL_456579, EPI_ISL_456580, EPI_ISL_456581, EPI_ISL_456582, EPI_ISL_456583, EPI_ISL_456584, EPI_ISL_456585, EPI_ISL_456587, EPI_ISL_456588, EPI_ISL_456589, EPI_ISL_456590, EPI_ISL_456591, EPI_ISL_456592, EPI_ISL_456593, EPI_ISL_456613, EPI_ISL_456614, EPI_ISL_456615, EPI_ISL_456616, EPI_ISL_456617, EPI_ISL_456618, EPI_ISL_456619, EPI_ISL_456620, EPI_ISL_456621, EPI_ISL_456622, EPI_ISL_456625, EPI_ISL_456626, EPI_ISL_456628, EPI_ISL_456629, EPI_ISL_456630, EPI_ISL_456631, EPI_ISL_456632, EPI_ISL_456633, EPI_ISL_456634, EPI_ISL_456635, EPI_ISL_456636, EPI_ISL_456637, EPI_ISL_456638, EPI_ISL_456640, EPI_ISL_456641, EPI_ISL_456642, EPI_ISL_456643, EPI_ISL_456644, EPI_ISL_456645, EPI_ISL_456646, EPI_ISL_456647, EPI_ISL_456648, EPI_ISL_456649, EPI_ISL_456650, EPI_ISL_456651, EPI_ISL_456652, EPI_ISL_456653, EPI_ISL_456654, EPI_ISL_456655 | | | |
| see above | Victorian Infectious Diseases Reference Laboratory (VIDRL) | Microbiological Diagnostic Unit Public Health Laboratory and Victorian Infectious Diseases Reference Laboratory, Doherty Institute | Caly L., Seemann T., Sait, M., Schultz M., Druce J., Sherry, N. |
| EPI_ISL_467055, EPI_ISL_467056, EPI_ISL_467057, EPI_ISL_467058 | Servicio de Microbiología, Hospital Universitario Son Espases | SeqCOVID-SPAIN consortium/IBV(CSIC) | Carla López-Causapé, Jordi Reina y Antonio Oliver and SeqCOVID-SPAIN consortium |
| EPI_ISL_468067 | Microbiology and Immunology, The Peter Doherty Institute for Infection and Immunity | Microbiology and Immunology, The Peter Doherty Institute for Infection and Immunity | Caly,L., Seemann,T., Sait,M., Schultz,M.B., Druce,J., Sherry,N., Meumann,E., Soares da Silva,E., Dolores de Jesus da Costa,M., Salles de Sousa,A., Jayanti Pereira Tilman,A., Antonia da Costa,E., Barreto,I., Marr,I., Wapling,J., Francis,J., Ximenes,J., Canisia,D., Freeman,K., Dakh,F., Douglas,N. and Baird,R. |
| EPI_ISL_468080, EPI_ISL_468081, EPI_ISL_468082, EPI_ISL_468083, EPI_ISL_468084, EPI_ISL_468085, EPI_ISL_468086, EPI_ISL_468087, EPI_ISL_468088, EPI_ISL_468089, EPI_ISL_468090, EPI_ISL_468091, EPI_ISL_468092, EPI_ISL_468093, EPI_ISL_468094, EPI_ISL_468095, EPI_ISL_468096, EPI_ISL_468097, EPI_ISL_468098, EPI_ISL_468099, EPI_ISL_468100, EPI_ISL_468101, EPI_ISL_468102, EPI_ISL_468103, EPI_ISL_468104, EPI_ISL_468105, EPI_ISL_468106, EPI_ISL_468107, EPI_ISL_468108, EPI_ISL_468109, EPI_ISL_468110, EPI_ISL_468111, EPI_ISL_468112, EPI_ISL_468113, EPI_ISL_468114, EPI_ISL_468115, EPI_ISL_468116, EPI_ISL_468117, EPI_ISL_468118, EPI_ISL_468119, EPI_ISL_468120, EPI_ISL_468121, EPI_ISL_468122, EPI_ISL_468123, EPI_ISL_468124, EPI_ISL_468125, EPI_ISL_468126, EPI_ISL_468127, EPI_ISL_468128, EPI_ISL_468129 | | | |
| see above | OHSU Lab Services Molecular Microbiology Lab | Oregon SARS-CoV-2 Genome Sequencing Center | Brendan L. O'Connell, Ruth V. Nichols, Alec J. Hirsch, Guang Fan, Daniel N. Streblow, William B. Messer, Andrew C. Adey, Benjamin N. Bimber, Brian J. O'Roak |
| EPI_ISL_468345, EPI_ISL_468346, EPI_ISL_468347, EPI_ISL_468348, EPI_ISL_468349, EPI_ISL_468350, EPI_ISL_468351, EPI_ISL_468352, EPI_ISL_468353, EPI_ISL_468354, EPI_ISL_468355, EPI_ISL_468356 | | | |
| see above | County of Santa Clara Public Health Department | Chan-Zuckerberg Biohub | CZB Cliahub Consortium |
| EPI_ISL_468357, EPI_ISL_468358, EPI_ISL_468359, EPI_ISL_468360, EPI_ISL_468361, EPI_ISL_468362, EPI_ISL_468363, EPI_ISL_468364, EPI_ISL_468365, EPI_ISL_468366, EPI_ISL_468367, EPI_ISL_468368, EPI_ISL_468369, EPI_ISL_468370, EPI_ISL_468371, EPI_ISL_468372, EPI_ISL_468373, EPI_ISL_468374, EPI_ISL_468375, EPI_ISL_468376, EPI_ISL_468377, EPI_ISL_468378, EPI_ISL_468379, EPI_ISL_468380, EPI_ISL_468381, EPI_ISL_468382, EPI_ISL_468383, EPI_ISL_468384, EPI_ISL_468385, EPI_ISL_468386, EPI_ISL_468387 | | | |
| see above | Alameda County Public Health Lab | Chan-Zuckerberg Biohub | CZB Cliahub Consortium |
| EPI_ISL_468388, EPI_ISL_468389, EPI_ISL_468390, EPI_ISL_468391, EPI_ISL_468392, EPI_ISL_468393, EPI_ISL_468394, EPI_ISL_468395, EPI_ISL_468396, EPI_ISL_468397, EPI_ISL_468398, EPI_ISL_468399, EPI_ISL_468400, EPI_ISL_468401, EPI_ISL_468402, EPI_ISL_468403, EPI_ISL_468404, EPI_ISL_468405, EPI_ISL_468406, EPI_ISL_468407, EPI_ISL_468408, EPI_ISL_468409, EPI_ISL_468410, EPI_ISL_468411, EPI_ISL_468412, EPI_ISL_468413, EPI_ISL_468414, EPI_ISL_468415, EPI_ISL_468416, EPI_ISL_468417, EPI_ISL_468418, EPI_ISL_468419, EPI_ISL_468420, EPI_ISL_468421, EPI_ISL_468422, EPI_ISL_468423, EPI_ISL_468424, EPI_ISL_468425, EPI_ISL_468426, EPI_ISL_468427, EPI_ISL_468428, EPI_ISL_468429, EPI_ISL_468430, EPI_ISL_468431, EPI_ISL_468432, EPI_ISL_468433, EPI_ISL_468434, EPI_ISL_468435, EPI_ISL_468436, EPI_ISL_468437 | | | |
| see above | County of San Luis Obispo Public Health Laboratory | Chan-Zuckerberg Biohub | CZB Cliahub Consortium |
| EPI_ISL_468438, EPI_ISL_468439, EPI_ISL_468440, EPI_ISL_468441, EPI_ISL_468442, EPI_ISL_468443, EPI_ISL_468444, EPI_ISL_468445, EPI_ISL_468446, EPI_ISL_468447, EPI_ISL_468448, EPI_ISL_468449, EPI_ISL_468450, EPI_ISL_468451, EPI_ISL_468452, EPI_ISL_468453, EPI_ISL_468454, EPI_ISL_468455, EPI_ISL_468456, EPI_ISL_468457, EPI_ISL_468458, EPI_ISL_468459, EPI_ISL_468460, EPI_ISL_468461 | | | |
| see above | Humboldt County Public Health Laboratory | Chan-Zuckerberg Biohub | CZB Cliahub Consortium |
| EPI_ISL_468462, EPI_ISL_468463, EPI_ISL_468464, EPI_ISL_468465, EPI_ISL_468466, EPI_ISL_468467, EPI_ISL_468468, EPI_ISL_468469, EPI_ISL_468470, EPI_ISL_468471, EPI_ISL_468472, EPI_ISL_468473, EPI_ISL_468474, EPI_ISL_468475, EPI_ISL_468476, EPI_ISL_468477, EPI_ISL_468478, EPI_ISL_468479, EPI_ISL_468480, EPI_ISL_468481, EPI_ISL_468482, EPI_ISL_468483, EPI_ISL_468484, EPI_ISL_468485, EPI_ISL_468486, EPI_ISL_468487, EPI_ISL_468488, EPI_ISL_468489, EPI_ISL_468490, EPI_ISL_468491, EPI_ISL_468492, EPI_ISL_468493, EPI_ISL_468494, EPI_ISL_468495, EPI_ISL_468496, EPI_ISL_468497, EPI_ISL_468498, EPI_ISL_468499, EPI_ISL_468500, EPI_ISL_468501, EPI_ISL_468502, EPI_ISL_468503, EPI_ISL_468504, EPI_ISL_468505 | | | |
| see above | Ventura County Public Health Lab | Chan-Zuckerberg Biohub | CZB Cliahub Consortium |
| EPI_ISL_468506, EPI_ISL_468507, EPI_ISL_468508, EPI_ISL_468509, EPI_ISL_468510, EPI_ISL_468511, EPI_ISL_468512, EPI_ISL_468513, EPI_ISL_468514, EPI_ISL_468515, EPI_ISL_468516, EPI_ISL_468517, EPI_ISL_468518, EPI_ISL_468519, EPI_ISL_468520, EPI_ISL_468521, EPI_ISL_468522, EPI_ISL_468523, EPI_ISL_468524, EPI_ISL_468525, EPI_ISL_468526, EPI_ISL_468527, EPI_ISL_468528, EPI_ISL_468529, EPI_ISL_468530, EPI_ISL_468531, EPI_ISL_468532, EPI_ISL_468533, EPI_ISL_468534, EPI_ISL_468535, EPI_ISL_468536, EPI_ISL_468537, EPI_ISL_468538, EPI_ISL_468539, EPI_ISL_468540, EPI_ISL_468541, EPI_ISL_468542, EPI_ISL_468543, EPI_ISL_468544, EPI_ISL_468545, EPI_ISL_468546, EPI_ISL_468547, EPI_ISL_468548, EPI_ISL_468549, EPI_ISL_468550, EPI_ISL_468551, EPI_ISL_468552, EPI_ISL_468553, EPI_ISL_468554, EPI_ISL_468555, EPI_ISL_468556, EPI_ISL_468557, EPI_ISL_468558, EPI_ISL_468559 | | | |
| see above | San Joaquin County Public Health Lab | Chan-Zuckerberg Biohub | CZB Cliahub Consortium |
| EPI_ISL_468592, EPI_ISL_468593, EPI_ISL_468594, EPI_ISL_468595, EPI_ISL_468596, EPI_ISL_468597, EPI_ISL_468598, EPI_ISL_468599, EPI_ISL_468600, EPI_ISL_468601, EPI_ISL_468602, EPI_ISL_468603, EPI_ISL_468604, EPI_ISL_468605, EPI_ISL_468606, EPI_ISL_468607, EPI_ISL_468608, EPI_ISL_468609, EPI_ISL_468610, EPI_ISL_468611, EPI_ISL_468612, EPI_ISL_468613, EPI_ISL_468614 | | | |
| see above | Orange County Public Health Lab | Chan-Zuckerberg Biohub | CZB Cliahub Consortium |
| EPI_ISL_468615, EPI_ISL_468616, EPI_ISL_468617, EPI_ISL_468618, EPI_ISL_468619, EPI_ISL_468620, EPI_ISL_468621, EPI_ISL_468622, EPI_ISL_468623, EPI_ISL_468624, EPI_ISL_468625, EPI_ISL_468626, EPI_ISL_468627, EPI_ISL_468628, EPI_ISL_468629, EPI_ISL_468630, EPI_ISL_468631, EPI_ISL_468632, EPI_ISL_468633, EPI_ISL_468634, EPI_ISL_468635, EPI_ISL_468636, EPI_ISL_468637, EPI_ISL_468638, EPI_ISL_468639, EPI_ISL_468640, EPI_ISL_468641, EPI_ISL_468642, EPI_ISL_468643, EPI_ISL_468644, EPI_ISL_468645, EPI_ISL_468646, EPI_ISL_468647, EPI_ISL_468648, EPI_ISL_468649, EPI_ISL_468650, EPI_ISL_468651, EPI_ISL_468652, EPI_ISL_468653, EPI_ISL_468654, EPI_ISL_468655 | | | |
| see above | Contra Costa Public Health Lab | Chan-Zuckerberg Biohub | CZB Cliahub Consortium |

| | | | |
|---|---|---|---|
| EPI_ISL_468952, EPI_ISL_468953, EPI_ISL_468954, EPI_ISL_468955, EPI_ISL_468956, EPI_ISL_468957, EPI_ISL_468958, EPI_ISL_468959, EPI_ISL_468960, EPI_ISL_468961, EPI_ISL_468962, EPI_ISL_468963, EPI_ISL_468964, EPI_ISL_468965, EPI_ISL_468966, EPI_ISL_468967, EPI_ISL_468968, EPI_ISL_468969, EPI_ISL_468970, EPI_ISL_468971, EPI_ISL_468972, EPI_ISL_468973, EPI_ISL_468974, EPI_ISL_468975, EPI_ISL_468976, EPI_ISL_468977, EPI_ISL_468978, EPI_ISL_468979, EPI_ISL_468980, EPI_ISL_468981, EPI_ISL_468982, EPI_ISL_468983, EPI_ISL_468984, EPI_ISL_468985, EPI_ISL_468986, EPI_ISL_468987, EPI_ISL_468988, EPI_ISL_468989, EPI_ISL_468990, EPI_ISL_468991, EPI_ISL_468992, EPI_ISL_468993, EPI_ISL_468994, EPI_ISL_468995, EPI_ISL_468996, EPI_ISL_468997, EPI_ISL_468998, EPI_ISL_468999, EPI_ISL_469000, EPI_ISL_469001, EPI_ISL_469002, EPI_ISL_469003, EPI_ISL_469004, EPI_ISL_469005, EPI_ISL_469006, EPI_ISL_469007, EPI_ISL_469008, EPI_ISL_469009, EPI_ISL_469010, EPI_ISL_469011, EPI_ISL_469012, EPI_ISL_469013, EPI_ISL_469014, EPI_ISL_469015 | | | |
| see above | Servicio de Microbiología, Hospital Universitario Son Espases | SeqCOVID-SPAIN consortium/IBV(CSIC) | Carla López-Causapé, Jordi Reina, Antonio Oliver and SeqCOVID-SPAIN consortium |
| EPI_ISL_471471 | Victorian Infectious Diseases Reference Laboratory | Victorian Infectious Diseases Reference Laboratory | Caly,L., Seemann,T., Sait,M., Schultz,M.B., Druce,J., Sherry,N. Meumann,E., Soares da Silva,E., Dolores de Jesus da Costa,M., Salles de Sousa,A., Jayanti Pereira Tilman,A., Antonia da Costa,E., Barreto,I., Marr,I., Wapling,J., Francis,J., Ximenes,J., Canisia,D., Freeman,K., Dakh,F., Douglas,N. and Baird,R. |
| EPI_ISL_478669, EPI_ISL_478670 | unknown | Microbiology, Koc University | Can,F., Ozer,B., Nurtop,E., Dogan,O. |
| EPI_ISL_478673, EPI_ISL_478674 | Pathology North - Royal North Shore Hospital - NSW Health Pathology | NSW Health Pathology - Institute of Clinical Pathology and Medical Research; Westmead Hospital; University of Sydney | CIDM-PH et al. |
| EPI_ISL_478675, EPI_ISL_478676, EPI_ISL_478677, EPI_ISL_478678, EPI_ISL_478679, EPI_ISL_478680, EPI_ISL_478681, EPI_ISL_478682 | Sydney South West Pathology Service (SSWPS) - Liverpool Hospital - NSW Health Pathology | NSW Health Pathology - Institute of Clinical Pathology and Medical Research; Westmead Hospital; University of Sydney | CIDM-PH et al. |
| EPI_ISL_478683, EPI_ISL_478684, EPI_ISL_478685, EPI_ISL_478686, EPI_ISL_478687, EPI_ISL_478688, EPI_ISL_478689, EPI_ISL_478690, EPI_ISL_478691, EPI_ISL_478692, EPI_ISL_478693, EPI_ISL_478694, EPI_ISL_478695, EPI_ISL_478696, EPI_ISL_478697, EPI_ISL_478698, EPI_ISL_478699, EPI_ISL_478700, EPI_ISL_478701, EPI_ISL_478702, EPI_ISL_478703, EPI_ISL_478704, EPI_ISL_478705 | | | |
| see above | South Eastern Area Laboratory Services (SEALS) | NSW Health Pathology - Institute of Clinical Pathology and Medical Research; Westmead Hospital; University of Sydney | CIDM-PH et al. |
| EPI_ISL_478706, EPI_ISL_478707 | Sydney South West Pathology Service (SSWPS) - Liverpool Hospital - NSW Health Pathology | NSW Health Pathology - Institute of Clinical Pathology and Medical Research; Westmead Hospital; University of Sydney | CIDM-PH et al. |
| EPI_ISL_478708, EPI_ISL_478709, EPI_ISL_478710, EPI_ISL_478711 | South Eastern Area Laboratory Services (SEALS) | NSW Health Pathology - Institute of Clinical Pathology and Medical Research; Westmead Hospital; University of Sydney | CIDM-PH et al. |
| EPI_ISL_478712, EPI_ISL_478713, EPI_ISL_478714, EPI_ISL_478715, EPI_ISL_478716 | Sydney South West Pathology Service (SSWPS) - Liverpool Hospital - NSW Health Pathology | NSW Health Pathology - Institute of Clinical Pathology and Medical Research; Westmead Hospital; University of Sydney | CIDM-PH et al. |
| EPI_ISL_478717, EPI_ISL_478718 | Sydney South West Pathology Service (SSWPS) - Concord Repatriation General Hospital - NSW Health Pathology | NSW Health Pathology - Institute of Clinical Pathology and Medical Research; Westmead Hospital; University of Sydney | CIDM-PH et al. |
| EPI_ISL_480563, EPI_ISL_480564, EPI_ISL_480565, EPI_ISL_480566, EPI_ISL_480567, EPI_ISL_480568, EPI_ISL_480571, EPI_ISL_480577, EPI_ISL_480579, EPI_ISL_480581, EPI_ISL_480582, EPI_ISL_480583, EPI_ISL_480584, EPI_ISL_480586, EPI_ISL_480588, EPI_ISL_480589, EPI_ISL_480590, EPI_ISL_480591, EPI_ISL_480592, EPI_ISL_480593, EPI_ISL_480594, EPI_ISL_480598, EPI_ISL_480599, EPI_ISL_480604, EPI_ISL_480605, EPI_ISL_480606, EPI_ISL_480607, EPI_ISL_480608, EPI_ISL_480609, EPI_ISL_480611, EPI_ISL_480621, EPI_ISL_480622, EPI_ISL_480624, EPI_ISL_480625, EPI_ISL_480626, EPI_ISL_480627, EPI_ISL_480628, EPI_ISL_480629, EPI_ISL_480630, EPI_ISL_480631, EPI_ISL_480632, EPI_ISL_480633, EPI_ISL_480634, EPI_ISL_480635, EPI_ISL_480636, EPI_ISL_480637, EPI_ISL_480638, EPI_ISL_480639, EPI_ISL_480640, EPI_ISL_480641, EPI_ISL_480642, EPI_ISL_480643, EPI_ISL_480644, EPI_ISL_480645, EPI_ISL_480646, EPI_ISL_480647, EPI_ISL_480648, EPI_ISL_480649, EPI_ISL_480650, EPI_ISL_480651, EPI_ISL_480652, EPI_ISL_480653, EPI_ISL_480654, EPI_ISL_480655, EPI_ISL_480656, EPI_ISL_480657, EPI_ISL_480659, EPI_ISL_480660, EPI_ISL_480661, EPI_ISL_480662, EPI_ISL_480663, EPI_ISL_480664, EPI_ISL_480665, EPI_ISL_480666, EPI_ISL_480667, EPI_ISL_480668, EPI_ISL_480669, EPI_ISL_480670, EPI_ISL_480671, EPI_ISL_480672, EPI_ISL_480673, EPI_ISL_480674, EPI_ISL_480675, EPI_ISL_480676, EPI_ISL_480677, EPI_ISL_480678, EPI_ISL_480679, EPI_ISL_480680, EPI_ISL_480681, EPI_ISL_480682, EPI_ISL_480683, EPI_ISL_480684, EPI_ISL_480685, EPI_ISL_480686, EPI_ISL_480688, EPI_ISL_480689, EPI_ISL_480698, EPI_ISL_480699, EPI_ISL_480700, EPI_ISL_480701, EPI_ISL_480702, EPI_ISL_480703, EPI_ISL_480704, EPI_ISL_480706, EPI_ISL_480707, EPI_ISL_480708, EPI_ISL_480709, EPI_ISL_480710, EPI_ISL_480711, EPI_ISL_480712, EPI_ISL_480713, EPI_ISL_480714, EPI_ISL_480715, EPI_ISL_480716, EPI_ISL_480717, EPI_ISL_480718, EPI_ISL_480719, EPI_ISL_480721, EPI_ISL_480722, EPI_ISL_480723, EPI_ISL_480724, EPI_ISL_480725, EPI_ISL_480726, EPI_ISL_480727, EPI_ISL_480728, EPI_ISL_480729, EPI_ISL_480730, EPI_ISL_480731, EPI_ISL_480732, EPI_ISL_480733, EPI_ISL_480734, EPI_ISL_480735, EPI_ISL_480736, EPI_ISL_480737, EPI_ISL_480738, EPI_ISL_480739, EPI_ISL_480740, EPI_ISL_480741, EPI_ISL_480742, EPI_ISL_480743, EPI_ISL_480744, EPI_ISL_480764, EPI_ISL_480765, EPI_ISL_480766, EPI_ISL_480778, EPI_ISL_480779, EPI_ISL_480780, EPI_ISL_480781 | | | |
| see above | Victorian Infectious Diseases Reference Laboratory (VIDRL) | VIDRL and MDU-PHL | Caly L., Seemann T., Sait, M., Schultz M., Druce J., Sherry, N. |
| EPI_ISL_482468 | Laboratorio de Referencia Nacional de Virus Respiratorios. Instituto Nacional de Salud Peru | Laboratorio de Referencia Nacional de Biotecnologia y Biologia Molecular. Instituto Nacional de Salud Peru | Carlos Padilla Rojas, Priscila Lope Pari, Karolyn Vega Chozo, Johanna Balbuena Torres, Omar Caceres Rey, Henri Bailon Calderon, Maribel Huaringa Nuñez, Nancy Rojas Serrano |
| EPI_ISL_486115, EPI_ISL_486116, EPI_ISL_486117, EPI_ISL_486118, EPI_ISL_486119 | County of Santa Clara Public Health Department | Chan-Zuckerberg Biohub | CZB Cliahub Consortium |
| EPI_ISL_486120, EPI_ISL_486121, EPI_ISL_486122, EPI_ISL_486123, EPI_ISL_486124, EPI_ISL_486125, EPI_ISL_486126, EPI_ISL_486127, EPI_ISL_486128, EPI_ISL_486129, EPI_ISL_486130, EPI_ISL_486131, EPI_ISL_486132, EPI_ISL_486133, EPI_ISL_486134, EPI_ISL_486136, EPI_ISL_486137, EPI_ISL_486138, EPI_ISL_486139, EPI_ISL_486140, EPI_ISL_486141, EPI_ISL_486142, EPI_ISL_486143, EPI_ISL_486144, EPI_ISL_486145, EPI_ISL_486146, EPI_ISL_486147, EPI_ISL_486148, EPI_ISL_486149, EPI_ISL_486150, EPI_ISL_486151, EPI_ISL_486152, EPI_ISL_486153, EPI_ISL_486154, EPI_ISL_486155, EPI_ISL_486156, EPI_ISL_486157, EPI_ISL_486158, EPI_ISL_486159, EPI_ISL_486160, EPI_ISL_486161, EPI_ISL_486162, EPI_ISL_486163, EPI_ISL_486164, EPI_ISL_486165, EPI_ISL_486166, EPI_ISL_486167, EPI_ISL_486168, EPI_ISL_486169, EPI_ISL_486170, EPI_ISL_486171, EPI_ISL_486172, EPI_ISL_486173, EPI_ISL_486174, EPI_ISL_486176, EPI_ISL_486177, EPI_ISL_486178, EPI_ISL_486179, EPI_ISL_486180, EPI_ISL_486181, EPI_ISL_486182, EPI_ISL_486183, EPI_ISL_486184, EPI_ISL_486185, EPI_ISL_486186, EPI_ISL_486187, EPI_ISL_486188, EPI_ISL_486189, EPI_ISL_486190, EPI_ISL_486191, EPI_ISL_486192, EPI_ISL_486193, EPI_ISL_486194, EPI_ISL_486195, EPI_ISL_486196, EPI_ISL_486197, EPI_ISL_486198, EPI_ISL_486199, EPI_ISL_486200, EPI_ISL_486201, EPI_ISL_486202, EPI_ISL_486203, EPI_ISL_486204, EPI_ISL_486205, EPI_ISL_486206, EPI_ISL_486207, EPI_ISL_486208, EPI_ISL_486209, EPI_ISL_486210, EPI_ISL_486211, EPI_ISL_486212, EPI_ISL_486213, EPI_ISL_486214, EPI_ISL_486215, EPI_ISL_486216, EPI_ISL_486217, EPI_ISL_486218, EPI_ISL_486219, EPI_ISL_486220, EPI_ISL_486221, EPI_ISL_486222, EPI_ISL_486223, EPI_ISL_486224, EPI_ISL_486225, EPI_ISL_486226, EPI_ISL_486227, EPI_ISL_486228, EPI_ISL_486229, EPI_ISL_486230, EPI_ISL_486231, EPI_ISL_486232, EPI_ISL_486233, EPI_ISL_486234, EPI_ISL_486235, EPI_ISL_486236, EPI_ISL_486237, EPI_ISL_486238, EPI_ISL_486239, EPI_ISL_486240, EPI_ISL_486241, EPI_ISL_486242, EPI_ISL_486243, EPI_ISL_486244, EPI_ISL_486245, EPI_ISL_486246, EPI_ISL_486247, EPI_ISL_486248, EPI_ISL_486249, EPI_ISL_486250, EPI_ISL_486251, EPI_ISL_486252, EPI_ISL_486253, EPI_ISL_486254, EPI_ISL_486255, EPI_ISL_486256, EPI_ISL_486257, EPI_ISL_486258, EPI_ISL_486259, EPI_ISL_486260, EPI_ISL_486261, EPI_ISL_486262, EPI_ISL_486263, EPI_ISL_486264, EPI_ISL_486265, EPI_ISL_486266, EPI_ISL_486267, EPI_ISL_486268, EPI_ISL_486269, EPI_ISL_486270, EPI_ISL_486271, EPI_ISL_486272, EPI_ISL_486273, EPI_ISL_486274, EPI_ISL_486275, EPI_ISL_486276, EPI_ISL_486277, EPI_ISL_486278, EPI_ISL_486279 | | | |
| see above | Orange County Public Health Laboratory | Chan-Zuckerberg Biohub | CZB Cliahub Consortium |
| EPI_ISL_486280, EPI_ISL_486281, EPI_ISL_486282, EPI_ISL_486283, EPI_ISL_486284, EPI_ISL_486285, EPI_ISL_486286 | Humboldt County Public Health Laboratory | Chan-Zuckerberg Biohub | CZB Cliahub Consortium |
| EPI_ISL_486287, EPI_ISL_486288, EPI_ISL_486289, EPI_ISL_486290, EPI_ISL_486291, EPI_ISL_486292, EPI_ISL_486293, EPI_ISL_486294, EPI_ISL_486295, EPI_ISL_486296, EPI_ISL_486297, EPI_ISL_486298, EPI_ISL_486299, EPI_ISL_486300, EPI_ISL_486301, EPI_ISL_486302, EPI_ISL_486303, EPI_ISL_486304, EPI_ISL_486305, EPI_ISL_486306, EPI_ISL_486307, EPI_ISL_486308, EPI_ISL_486309, EPI_ISL_486310, EPI_ISL_486311, EPI_ISL_486312, EPI_ISL_486313, EPI_ISL_486314, EPI_ISL_486315, EPI_ISL_486316, EPI_ISL_486317, EPI_ISL_486318, EPI_ISL_486319, EPI_ISL_486320, EPI_ISL_486321, EPI_ISL_486322, EPI_ISL_486323, EPI_ISL_486324, EPI_ISL_486325, EPI_ISL_486326, EPI_ISL_486327, EPI_ISL_486328, EPI_ISL_486329, EPI_ISL_486330, EPI_ISL_486331, EPI_ISL_486332, EPI_ISL_486333, EPI_ISL_486334, EPI_ISL_486335, EPI_ISL_486336, EPI_ISL_486337, EPI_ISL_486338 | | | |
| see above | San Joaquin County Public Health Lab | Chan-Zuckerberg Biohub | CZB Cliahub Consortium |
| EPI_ISL_486339, EPI_ISL_486340, EPI_ISL_486341, EPI_ISL_486342, EPI_ISL_486343, EPI_ISL_486344, EPI_ISL_486345, EPI_ISL_486346, EPI_ISL_486347, EPI_ISL_486348, EPI_ISL_486349, EPI_ISL_486350, EPI_ISL_486351, EPI_ISL_486352, EPI_ISL_486353, EPI_ISL_486354, EPI_ISL_486355, EPI_ISL_486356, EPI_ISL_486357, EPI_ISL_486358, EPI_ISL_486359, EPI_ISL_486360, EPI_ISL_486361, EPI_ISL_486362, EPI_ISL_486363, EPI_ISL_486364, EPI_ISL_486365 | | | |
| see above | UCSF Clinical Microbiology Laboratory | Chan-Zuckerberg Biohub | CZB Cliahub Consortium |
| EPI_ISL_487269 | Laboratorio de Referencia Nacional de Virus Respiratorio. Instituto Nacional de Salud. | Laboratorio de Referencia Nacional de Biotecnología y Biología Molecular. Instituto Nacional de Salud. | Carlos Padilla Rojas, Karolyn Chozo Vega, Priscila Lope Pari, Omar Caceres Rey, Marco Galarza Perez, Maribel Huaringa Nuñez, Johanna Balbuena Torrez, Henri Bailon Calderon, Nancy Rojas Serrano |
| EPI_ISL_487271 | unknown | MDU-PHL, The Peter Doherty Institute for Infection and Immunity | Caly,L., Seemann,T., Sait,M., Schultz,M.B., Sherry,N., Meumann,E., Baird,R., Leong,L., Lim,C.K., Turra,M., Bastian,I., Higgins,G., Soares da Silva,E., Dolores de Jesus da Costa,M., Salles de Sousa,A., Jayanti Pereira Tilman,A., Antonia da Costa,E., Baretto,I., Marr,I., Wapling,J., Francis,J., Ximenes,J., Canisia,D., Freeman,K., Dakh,F., Douglas,N. |
| EPI_ISL_489836 | Laboratorio de Referencia Nacional de Virus Respiratorio. Instituto Nacional de Salud Perú | Laboratorio de Referencia Nacional de Biotecnología y Biología Molecular. Instituto Nacional de Salud Perú | Carlos Padilla Rojas, Karolyn Chozo Vega, Priscila Lope Pari, Omar Caceres Rey, Marco Galarza Perez, Maribel Huaringa Nuñez, Johanna Balbuena Torres, Henri Bailon Calderon, Nancy Rojas Serrano |
| EPI_ISL_489837, EPI_ISL_489838 | Laboratorio de Referencia Nacional de Virus Respiratorio. Instituto Nacional de Salud Perú | Laboratorio de Referencia Nacional de Biotecnología y Biología Molecular. Instituto Nacional de Salud Perú | Carlos Padilla Rojas, Karolyn Chozo Vega, Priscila Lope Pari, Omar Caceres Rey, Marco Galarza Perez, Maribel Huaringa Nuñez, Johanna Balbuena Torrez, Henri Bailon Calderon, Nancy Rojas Serrano |
| EPI_ISL_489839, EPI_ISL_489987, | Laboratorio de Referencia Nacional de Virus Respiratorio. | Laboratorio de Referencia Nacional de Biotecnología y | Carlos Padilla Rojas, Karolyn Chozo Vega, Priscila Lope Pari, Omar Caceres Rey, Marco Galarza Perez, Maribel Huaringa Nuñez, Johanna Balbuena |

| | | | |
|---|---|---|---|
| EPI_ISL_489988, EPI_ISL_489989, EPI_ISL_489990 | Instituto Nacional de Salud Perú | Biología Molecular. Instituto Nacional de Salud Perú | Torres, Henri Bailon Calderon, Nancy Rojas Serrano. |
| EPI_ISL_490017, EPI_ISL_490018, EPI_ISL_490019 | South Eastern Area Laboratory Services (SEALS) | NSW Health Pathology - Institute of Clinical Pathology and Medical Research; Westmead Hospital; University of Sydney | CIDM-PH et al. |
| EPI_ISL_490020 | Sydney South West Pathology Service (SSWPS) - Liverpool Hospital - NSW Health Pathology | NSW Health Pathology - Institute of Clinical Pathology and Medical Research; Westmead Hospital; University of Sydney | CIDM-PH et al. |
| EPI_ISL_490021, EPI_ISL_490022, EPI_ISL_490023, EPI_ISL_490024, EPI_ISL_490025, EPI_ISL_490026, EPI_ISL_490027, EPI_ISL_490028, EPI_ISL_490029, EPI_ISL_490030, EPI_ISL_490031, EPI_ISL_490032, EPI_ISL_490033, EPI_ISL_490034, EPI_ISL_490035 | | | |
| see above | South Eastern Area Laboratory Services (SEALS) | NSW Health Pathology - Institute of Clinical Pathology and Medical Research; Westmead Hospital; University of Sydney | CIDM-PH et al. |
| EPI_ISL_490036 | Pathology West - NSW Health Pathology | NSW Health Pathology - Institute of Clinical Pathology and Medical Research; Westmead Hospital; University of Sydney | CIDM-PH et al. |
| EPI_ISL_490037 | Sydney South West Pathology Service (SSWPS) - Liverpool Hospital - NSW Health Pathology | NSW Health Pathology - Institute of Clinical Pathology and Medical Research; Westmead Hospital; University of Sydney | CIDM-PH et al. |
| EPI_ISL_490038, EPI_ISL_490039 | Pathology West - NSW Health Pathology | NSW Health Pathology - Institute of Clinical Pathology and Medical Research; Westmead Hospital; University of Sydney | CIDM-PH et al. |
| EPI_ISL_490040, EPI_ISL_490041, EPI_ISL_490042 | South Eastern Area Laboratory Services (SEALS) | NSW Health Pathology - Institute of Clinical Pathology and Medical Research; Westmead Hospital; University of Sydney | CIDM-PH et al. |
| EPI_ISL_490043 | Pathology North Hunter- NSW Health Pathology | NSW Health Pathology - Institute of Clinical Pathology and Medical Research; Westmead Hospital; University of Sydney | CIDM-PH et al. |
| EPI_ISL_490044 | Pathology West - NSW Health Pathology | NSW Health Pathology - Institute of Clinical Pathology and Medical Research; Westmead Hospital; University of Sydney | CIDM-PH et al. |
| EPI_ISL_490045 | South Eastern Area Laboratory Services (SEALS) | NSW Health Pathology - Institute of Clinical Pathology and Medical Research; Westmead Hospital; University of Sydney | CIDM-PH et al. |
| EPI_ISL_490046 | Pathology West - NSW Health Pathology | NSW Health Pathology - Institute of Clinical Pathology and Medical Research; Westmead Hospital; University of Sydney | CIDM-PH et al. |
| EPI_ISL_490105 | South Eastern Area Laboratory Services (SEALS) | NSW Health Pathology - Institute of Clinical Pathology and Medical Research; Westmead Hospital; University of Sydney | CIDM-PH et al. |
| EPI_ISL_490209, EPI_ISL_490315, EPI_ISL_490316 | Laboratorio de Referencia Nacional de Virus Respiratorio. Instituto Nacional de Salud Perú | Laboratorio de Referencia Nacional de Biotecnología y Biología Molecular. Instituto Nacional de Salud Perú | Carlos Padilla Rojas, Karolyn Chozo Vega, Priscila Lope Pari, Omar Caceres Rey, Marco Galarza Perez, Maribel Huaringa Nuñez, Johanna Balbuena Torres, Henri Bailon Calderon, Nancy Rojas Serrano |
| EPI_ISL_490975, EPI_ISL_490976 | Laboratorio de Referencia Nacional de Virus Respiratorio. Instituto Nacional de Salud Perú | Laboratorio de Referencia Nacional de Biotecnología y Biología Molecular. Instituto Nacional de Salud Perú | Carlos Padilla Rojas, Karolyn Chozo Vega, Priscila Lope Pari, Omar Caceres Rey, Marco Galarza Perez, Maribel Huaringa Nuñez, Johanna Balbuena Torres, Henri Bailon Calderon, Nancy Rojas Serrano. |
| EPI_ISL_491172 | Laboratorio de Referencia Nacional de Virus Respiratorio. Instituto Nacional de Salud Perú | Laboratorio de Referencia Nacional de Biotecnología y Biología Molecular. Instituto Nacional de Salud Perú. | Carlos Padilla Rojas, Karolyn Chozo Vega, Priscila Lope Pari, Omar Caceres Rey, Marco Galarza Perez, Maribel Huaringa Nuñez, Johanna Balbuena Torres, Henri Bailon Calderon, Nancy Rojas Serrano |
| EPI_ISL_491427 | Laboratorio de Referencia Nacional de Virus Respiratorio. Instituto Nacional de Salud Perú | Laboratorio de Referencia Nacional de Biotecnología y Biología Molecular. Instituto Nacional de Salud Perú | Carlos Padilla Rojas, Karolyn Chozo Vega, Priscila Lope Pari, Omar Caceres Rey, Marco Galarza Perez, Maribel Huaringa Nuñez, Johanna Balbuena Torres, Henri Bailon Calderon, Nancy Rojas Serrano. |
| EPI_ISL_491428 | Laboratorio de Referencia Nacional de Virus Respiratorio. Instituto Nacional de Salud Perú | Laboratorio de Referencia Nacional de Biotecnología y Biología Molecular. Instituto Nacional de Salud Perú | Carlos Padilla Rojas, Karolyn Vega Chozo, Priscila Lope Pari, Omar Caceres Rey, Marco Galarza Perez, Maribel Huaringa Nuñez, Johanna Balbuena Torres, Henri Bailon Calderon, Nancy Rojas Serrano. |
| EPI_ISL_491429 | Laboratorio de Referencia Nacional de Virus Respiratorio. Instituto Nacional de Salud Perú | Laboratorio de Referencia Nacional de Biotecnología y Biología Molecular. Instituto Nacional de Salud Perú | Carlos Padilla Rojas, Karolyn Vega Chozo, Priscila Lope Pari, Omar Caceres Rey, Marco Galarza Perez, Maribel Huaringa Nuñez, Johanna Balbuena Torrez, Henri Bailon Calderon, Nancy Rojas Serrano |
| EPI_ISL_491430 | Laboratorio de Referencia Nacional de Virus Respiratorio. Instituto Nacional de Salud Perú | Laboratorio de Referencia Nacional de Biotecnología y Biología Molecular. Instituto Nacional de Salud Perú | Carlos Padilla Rojas, Karolyn Vega Chozo, Priscila Lope Pari, Omar Caceres Rey, Marco Galarza Perez, Maribel Huaringa Nuñez, Johanna Balbuena Torres, Henri Bailon Calderon, Nancy Rojas Serrano. |
| EPI_ISL_491431 | Laboratorio de Referencia Nacional de Virus Respiratorio. Instituto Nacional de Salud. Perú | Laboratorio de Referencia Nacional de Biotecnología y Biología Molecular. Instituto Nacional de Salud.Perú | Carlos Padilla Rojas, Karolyn Vega Chozo, Priscila Lope Pari, Omar Caceres Rey, Marco Galarza Perez, Maribel Huaringa Nuñez, Johanna Balbuena Torres, Henri Bailon Calderon, Nancy Rojas Serrano. |
| EPI_ISL_491432 | Laboratorio de Referencia Nacional de Virus Respiratorio. Instituto Nacional de Salud.Perú | Laboratorio de Referencia Nacional de Biotecnología y Biología Molecular. Instituto Nacional de Salud.Perú | Carlos Padilla Rojas, Karolyn Vega Chozo, Priscila Lope Pari, Omar Caceres Rey, Marco Galarza Perez, Maribel Huaringa Nuñez, Johanna Balbuena Torres, Henri Bailon Calderon, Nancy Rojas Serrano. |
| EPI_ISL_491433, EPI_ISL_491434, EPI_ISL_491435 | Laboratorio de Referencia Nacional de Virus Respiratorio. Instituto Nacional de Salud Perú | Laboratorio de Referencia Nacional de Biotecnología y Biología Molecular. Instituto Nacional de Salud Perú | Carlos Padilla Rojas, Karolyn Vega Chozo, Priscila Lope Pari, Omar Caceres Rey, Marco Galarza Perez, Maribel Huaringa Nuñez, Johanna Balbuena Torres, Henri Bailon Calderon, Nancy Rojas Serrano. |
| EPI_ISL_491436 | Laboratorio de Referencia Nacional de Virus Respiratorio. Instituto Nacional de Salud Perú | Laboratorio de Referencia Nacional de Biotecnología y Biología Molecular. Instituto Nacional de Salud Perú | Carlos Padilla Rojas, Karolyn Vega Chozo, Priscila Lope Pari, Omar Caceres Rey, Marco Galarza Perez, Maribel Huaringa Nuñez, Johanna Balbuena Torres, Henri Bailon Calderon, Nancy Rojas Serrano |
| EPI_ISL_491458, EPI_ISL_491459, EPI_ISL_491460, EPI_ISL_491461, EPI_ISL_491462 | Laboratorio de Referencia Nacional de Virus Respiratorio. Instituto Nacional de Salud Perú | Laboratorio de Referencia Nacional de Biotecnología y Biología Molecular. Instituto Nacional de Salud Perú | Carlos Padilla Rojas, Karolyn Vega Chozo, Priscila Lope Pari, Omar Caceres Rey, Marco Galarza Perez, Maribel Huaringa Nuñez, Johanna Balbuena Torrez, Henri Bailon Calderon, Nancy Rojas Serrano |
| EPI_ISL_491463 | Laboratorio de Referencia Nacional de Virus Respiratorio. Instituto Nacional de Salud Perú | Laboratorio de Referencia Nacional de Biotecnología y Biología Molecular. Instituto Nacional de Salud Perú | Carlos Padilla Rojas, Karolyn Vega Chozo, Priscila Lope Pari, Omar Caceres Rey, Marco Galarza Perez, Maribel Huaringa Nuñez, Johanna Balbuena Torres, Henri Bailon Calderon, Nancy Rojas Serrano |
| EPI_ISL_491464 | Laboratorio de Referencia Nacional de Virus Respiratorio. Instituto Nacional de Salud Perú | Laboratorio de Referencia Nacional de Biotecnología y Biología Molecular. Instituto Nacional de Salud Perú | Carlos Padilla Rojas, Karolyn Vega Chozo, Priscila Lope Pari, Omar Caceres Rey, Marco Galarza Perez, Maribel Huaringa Nuñez, Johanna Balbuena Torrez, Henri Bailon Calderon, Nancy Rojas Serrano |
| EPI_ISL_498748, EPI_ISL_498749, EPI_ISL_498750 | Pathology West - NSW Health Pathology | NSW Health Pathology - Institute of Clinical Pathology and Medical Research; Westmead Hospital; University of Sydney | CIDM-PH et al. |
| EPI_ISL_498751 | South Eastern Area Laboratory Services (SEALS) | NSW Health Pathology - Institute of Clinical Pathology and Medical Research; Westmead Hospital; University of Sydney | CIDM-PH et al. |
| EPI_ISL_498761, EPI_ISL_498762 | Pathology West - NSW Health Pathology | NSW Health Pathology - Institute of Clinical Pathology and Medical Research; Westmead Hospital; University of Sydney | CIDM-PH et al. |
| EPI_ISL_498763, EPI_ISL_498764 | Sydney South West Pathology Service (SSWPS) - Liverpool Hospital - NSW Health Pathology | NSW Health Pathology - Institute of Clinical Pathology and Medical Research; Westmead Hospital; University of Sydney | CIDM-PH et al. |
| EPI_ISL_498765 | Sydney South West Pathology Service (SSWPS) - Concord Repatriation General Hospital - NSW Health Pathology | NSW Health Pathology - Institute of Clinical Pathology and Medical Research; Westmead Hospital; University of Sydney | CIDM-PH et al. |
| EPI_ISL_498766, EPI_ISL_498767 | Pathology West - NSW Health Pathology | NSW Health Pathology - Institute of Clinical Pathology and Medical Research; Westmead Hospital; University of Sydney | CIDM-PH et al. |
| EPI_ISL_498768 | South Eastern Area Laboratory Services (SEALS) | NSW Health Pathology - Institute of Clinical Pathology and Medical Research; Westmead Hospital; University of Sydney | CIDM-PH et al. |

| | | | |
|---|---|---|---|
| EPI_ISL_498769, EPI_ISL_498770, EPI_ISL_498771, EPI_ISL_498772 | Laverty Pathology | NSW Health Pathology - Institute of Clinical Pathology and Medical Research; Westmead Hospital; University of Sydney | CIDM-PH et al. |
| EPI_ISL_498773 | Douglass Hanly Moir Pathology | NSW Health Pathology - Institute of Clinical Pathology and Medical Research; Westmead Hospital; University of Sydney | CIDM-PH et al. |
| EPI_ISL_498774, EPI_ISL_498775 | Histopath | NSW Health Pathology - Institute of Clinical Pathology and Medical Research; Westmead Hospital; University of Sydney | CIDM-PH et al. |
| EPI_ISL_498776, EPI_ISL_498777 | Pathology West - NSW Health Pathology | NSW Health Pathology - Institute of Clinical Pathology and Medical Research; Westmead Hospital; University of Sydney | CIDM-PH et al. |
| EPI_ISL_498778 | Sydney South West Pathology Service (SSWPS) - Liverpool Hospital - NSW Health Pathology | NSW Health Pathology - Institute of Clinical Pathology and Medical Research; Westmead Hospital; University of Sydney | CIDM-PH et al. |
| EPI_ISL_498779, EPI_ISL_498780 | Pathology West - NSW Health Pathology | NSW Health Pathology - Institute of Clinical Pathology and Medical Research; Westmead Hospital; University of Sydney | CIDM-PH et al. |
| EPI_ISL_498781, EPI_ISL_498782 | Sydney South West Pathology Service (SSWPS) - Liverpool Hospital - NSW Health Pathology | NSW Health Pathology - Institute of Clinical Pathology and Medical Research; Westmead Hospital; University of Sydney | CIDM-PH et al. |
| EPI_ISL_509059, EPI_ISL_509060, EPI_ISL_509061, EPI_ISL_509062, EPI_ISL_509063, EPI_ISL_509064, EPI_ISL_509065, EPI_ISL_509066, EPI_ISL_509067, EPI_ISL_509068, EPI_ISL_509069, EPI_ISL_509070, EPI_ISL_509071, EPI_ISL_509072, EPI_ISL_509073, EPI_ISL_509074, EPI_ISL_509075, EPI_ISL_509076, EPI_ISL_509077, EPI_ISL_509078, EPI_ISL_509079, EPI_ISL_509080, EPI_ISL_509081, EPI_ISL_509082, EPI_ISL_509083, EPI_ISL_509084, EPI_ISL_509085, EPI_ISL_509086, EPI_ISL_509087, EPI_ISL_509088, EPI_ISL_509089, EPI_ISL_509090, EPI_ISL_509091, EPI_ISL_509092, EPI_ISL_509093, EPI_ISL_509094, EPI_ISL_509095, EPI_ISL_509096, EPI_ISL_509097, EPI_ISL_509098, EPI_ISL_509099, EPI_ISL_509100, EPI_ISL_509101, EPI_ISL_509102, EPI_ISL_509103, EPI_ISL_509104, EPI_ISL_509105, EPI_ISL_509106, EPI_ISL_509107, EPI_ISL_509108, EPI_ISL_509109, EPI_ISL_509110, EPI_ISL_509111, EPI_ISL_509112, EPI_ISL_509113, EPI_ISL_509114, EPI_ISL_509115, EPI_ISL_509116, EPI_ISL_509117, EPI_ISL_509118, EPI_ISL_509119, EPI_ISL_509120, EPI_ISL_509121, EPI_ISL_509122, EPI_ISL_509123, EPI_ISL_509124, EPI_ISL_509125, EPI_ISL_509126, EPI_ISL_509127, EPI_ISL_509128, EPI_ISL_509129, EPI_ISL_509130, EPI_ISL_509131, EPI_ISL_509132, EPI_ISL_509133, EPI_ISL_509134, EPI_ISL_509135, EPI_ISL_509136, EPI_ISL_509137, EPI_ISL_509138, EPI_ISL_509139, EPI_ISL_509140, EPI_ISL_509141, EPI_ISL_509142, EPI_ISL_509143, EPI_ISL_509144, EPI_ISL_509145, EPI_ISL_509146, EPI_ISL_509147, EPI_ISL_509148, EPI_ISL_509149, EPI_ISL_509150, EPI_ISL_509151, EPI_ISL_509152, EPI_ISL_509153, EPI_ISL_509154, EPI_ISL_509155, EPI_ISL_509156, EPI_ISL_509157, EPI_ISL_509158, EPI_ISL_509159, EPI_ISL_509160, EPI_ISL_509161, EPI_ISL_509162, EPI_ISL_509163, EPI_ISL_509164, EPI_ISL_509165, EPI_ISL_509166, EPI_ISL_509167, EPI_ISL_509168, EPI_ISL_509169, EPI_ISL_509170, EPI_ISL_509171, EPI_ISL_509172, EPI_ISL_509173, EPI_ISL_509174, EPI_ISL_509175, EPI_ISL_509176, EPI_ISL_509177, EPI_ISL_509178, EPI_ISL_509179, EPI_ISL_509180, EPI_ISL_509181, EPI_ISL_509182, EPI_ISL_509183, EPI_ISL_509184, EPI_ISL_509185, EPI_ISL_509186, EPI_ISL_509187, EPI_ISL_509188, EPI_ISL_509189, EPI_ISL_509190, EPI_ISL_509191, EPI_ISL_509192, EPI_ISL_509193, EPI_ISL_509194, EPI_ISL_509195, EPI_ISL_509196, EPI_ISL_509197, EPI_ISL_509198, EPI_ISL_509199, EPI_ISL_509200, EPI_ISL_509201, EPI_ISL_509202, EPI_ISL_509203, EPI_ISL_509204, EPI_ISL_509205, EPI_ISL_509206, EPI_ISL_509207, EPI_ISL_509208, EPI_ISL_509209, EPI_ISL_509210, EPI_ISL_509211, EPI_ISL_509212, EPI_ISL_509213, EPI_ISL_509214 | | | |
| see above | OHSU Lab Services Molecular Microbiology Lab | Oregon SARS-CoV-2 Genome Sequencing Center | Brendan L. O'Connell, Ruth V. Nichols, Sally B. Grindstaff, Alec J. Hirsch, Guang Fan, Daniel N. Streblow, William B. Messer, Andrew C. Adey, Benjamin N. Bimber, Brian J. O'Roak |
| EPI_ISL_510889, EPI_ISL_510890, EPI_ISL_510891, EPI_ISL_510892, EPI_ISL_510893, EPI_ISL_510894, EPI_ISL_510895, EPI_ISL_510896, EPI_ISL_510897, EPI_ISL_510898, EPI_ISL_510899, EPI_ISL_510900, EPI_ISL_510901, EPI_ISL_510902, EPI_ISL_510903, EPI_ISL_510904, EPI_ISL_510905, EPI_ISL_510906, EPI_ISL_510907, EPI_ISL_510908, EPI_ISL_510909, EPI_ISL_510910, EPI_ISL_510911, EPI_ISL_510912, EPI_ISL_510913, EPI_ISL_510914, EPI_ISL_510915, EPI_ISL_510916, EPI_ISL_510917, EPI_ISL_510918, EPI_ISL_510919, EPI_ISL_510920, EPI_ISL_510921, EPI_ISL_510922, EPI_ISL_510923, EPI_ISL_510924, EPI_ISL_510925, EPI_ISL_510926, EPI_ISL_510927, EPI_ISL_510928, EPI_ISL_510929, EPI_ISL_510930, EPI_ISL_510931, EPI_ISL_510932, EPI_ISL_510933, EPI_ISL_510934, EPI_ISL_510935, EPI_ISL_510936, EPI_ISL_510937, EPI_ISL_510938, EPI_ISL_510939, EPI_ISL_510940, EPI_ISL_510941, EPI_ISL_510942, EPI_ISL_510943, EPI_ISL_510944, EPI_ISL_510945, EPI_ISL_510946, EPI_ISL_510947, EPI_ISL_510948, EPI_ISL_510949, EPI_ISL_510950, EPI_ISL_510951, EPI_ISL_510952, EPI_ISL_510953, EPI_ISL_510954, EPI_ISL_510955, EPI_ISL_510956, EPI_ISL_510957, EPI_ISL_510958, EPI_ISL_510959, EPI_ISL_510960, EPI_ISL_510961, EPI_ISL_510962, EPI_ISL_510963, EPI_ISL_510964, EPI_ISL_510965, EPI_ISL_510966, EPI_ISL_510967, EPI_ISL_510968, EPI_ISL_510969, EPI_ISL_510970, EPI_ISL_510971, EPI_ISL_510972, EPI_ISL_510973, EPI_ISL_510974, EPI_ISL_510975, EPI_ISL_510976, EPI_ISL_510977, EPI_ISL_510978, EPI_ISL_510979, EPI_ISL_510980, EPI_ISL_510981, EPI_ISL_510982, EPI_ISL_510983, EPI_ISL_510984, EPI_ISL_510985, EPI_ISL_510986, EPI_ISL_510987, EPI_ISL_510988, EPI_ISL_510989, EPI_ISL_510990, EPI_ISL_510991, EPI_ISL_510992, EPI_ISL_510993, EPI_ISL_510994, EPI_ISL_510995, EPI_ISL_510996, EPI_ISL_510997, EPI_ISL_510998, EPI_ISL_510999, EPI_ISL_511000, EPI_ISL_511001, EPI_ISL_511002, EPI_ISL_511003, EPI_ISL_511004, EPI_ISL_511005, EPI_ISL_511006, EPI_ISL_511007, EPI_ISL_511008, EPI_ISL_511009, EPI_ISL_511010, EPI_ISL_511011, EPI_ISL_511012, EPI_ISL_511013, EPI_ISL_511014, EPI_ISL_511015, EPI_ISL_511016, EPI_ISL_511017, EPI_ISL_511018, EPI_ISL_511019, EPI_ISL_511020, EPI_ISL_511021, EPI_ISL_511022, EPI_ISL_511023, EPI_ISL_511024, EPI_ISL_511025, EPI_ISL_511026, EPI_ISL_511027, EPI_ISL_511028, EPI_ISL_511029, EPI_ISL_511030, EPI_ISL_511031, EPI_ISL_511032, EPI_ISL_511033, EPI_ISL_511034, EPI_ISL_511035, EPI_ISL_511036, EPI_ISL_511037, EPI_ISL_511038, EPI_ISL_511039, EPI_ISL_511040, EPI_ISL_511041, EPI_ISL_511042, EPI_ISL_511043, EPI_ISL_511044, EPI_ISL_511045, EPI_ISL_511046, EPI_ISL_511047, EPI_ISL_511048, EPI_ISL_511049, EPI_ISL_511050, EPI_ISL_511051, EPI_ISL_511052, EPI_ISL_511053, EPI_ISL_511054, EPI_ISL_511055, EPI_ISL_511056, EPI_ISL_511057, EPI_ISL_511058, EPI_ISL_511059, EPI_ISL_511060, EPI_ISL_511061, EPI_ISL_511062, EPI_ISL_511063, EPI_ISL_511064, EPI_ISL_511065, EPI_ISL_511066, EPI_ISL_511067, EPI_ISL_511068, EPI_ISL_511069, EPI_ISL_511070, EPI_ISL_511071, EPI_ISL_511072, EPI_ISL_511073, EPI_ISL_511074, EPI_ISL_511075, EPI_ISL_511076, EPI_ISL_511077, EPI_ISL_511078, EPI_ISL_511079, EPI_ISL_511080, EPI_ISL_511081, EPI_ISL_511082, EPI_ISL_511083, EPI_ISL_511084, EPI_ISL_511085, EPI_ISL_511086, EPI_ISL_511087, EPI_ISL_511088, EPI_ISL_511089, EPI_ISL_511090, EPI_ISL_511091, EPI_ISL_511092, EPI_ISL_511093, EPI_ISL_511094, EPI_ISL_511095, EPI_ISL_511096, EPI_ISL_511097, EPI_ISL_511098, EPI_ISL_511099, EPI_ISL_511100, EPI_ISL_511101, EPI_ISL_511102, EPI_ISL_511103, EPI_ISL_511104, EPI_ISL_511105, EPI_ISL_511106, EPI_ISL_511107, EPI_ISL_511108, EPI_ISL_511109, EPI_ISL_511110, EPI_ISL_511111, EPI_ISL_511112, EPI_ISL_511113, EPI_ISL_511114, EPI_ISL_511115, EPI_ISL_511116, EPI_ISL_511117, EPI_ISL_511118, EPI_ISL_511119, EPI_ISL_511120, EPI_ISL_511121, EPI_ISL_511122, EPI_ISL_511123, EPI_ISL_511124, EPI_ISL_511125, EPI_ISL_511126, EPI_ISL_511127, EPI_ISL_511128, EPI_ISL_511129, EPI_ISL_511130, EPI_ISL_511131, EPI_ISL_511132, EPI_ISL_511133, EPI_ISL_511134, EPI_ISL_511135, EPI_ISL_511136, EPI_ISL_511137, EPI_ISL_511138, EPI_ISL_511139, EPI_ISL_511140, EPI_ISL_511141, EPI_ISL_511142, EPI_ISL_511143, EPI_ISL_511144, EPI_ISL_511145, EPI_ISL_511146, EPI_ISL_511147, EPI_ISL_511148, EPI_ISL_511149, EPI_ISL_511150, EPI_ISL_511151, EPI_ISL_511152, EPI_ISL_511153, EPI_ISL_511154, EPI_ISL_511155, EPI_ISL_511156, EPI_ISL_511157, EPI_ISL_511158, EPI_ISL_511159, EPI_ISL_511160, EPI_ISL_511161, EPI_ISL_511162, EPI_ISL_511163, EPI_ISL_511164, EPI_ISL_511165, EPI_ISL_511166, EPI_ISL_511167, EPI_ISL_511168, EPI_ISL_511169, EPI_ISL_511170, EPI_ISL_511171, EPI_ISL_511172, EPI_ISL_511173, EPI_ISL_511174, EPI_ISL_511175, EPI_ISL_511176, EPI_ISL_511177, EPI_ISL_511178, EPI_ISL_511179, EPI_ISL_511180, EPI_ISL_511181 | | | |
| see above | Instituto Nacional de Saude (INSA) | Instituto Nacional de Saude (INSA) | Borges et al |
| EPI_ISL_511182, EPI_ISL_511183, EPI_ISL_511184, EPI_ISL_511185, EPI_ISL_511186, EPI_ISL_511187, EPI_ISL_511188 | Instituto Nacional de Saude (INSA) and Instituto Gulbenkian de Ciencia (IGC) | Instituto Nacional de Saude (INSA) and Instituto Gulbenkian de Ciencia (IGC) | Borges et al |
| EPI_ISL_511189 | Instituto Nacional de Saude (INSA) | Instituto Nacional de Saude (INSA) | Borges et al |
| EPI_ISL_511190, EPI_ISL_511191, EPI_ISL_511192, EPI_ISL_511193, EPI_ISL_511194, EPI_ISL_511195, EPI_ISL_511196, EPI_ISL_511197, EPI_ISL_511198, EPI_ISL_511199, EPI_ISL_511200, EPI_ISL_511201, EPI_ISL_511202, EPI_ISL_511203, EPI_ISL_511204, EPI_ISL_511205, EPI_ISL_511206, EPI_ISL_511207, EPI_ISL_511208, EPI_ISL_511209, EPI_ISL_511210, EPI_ISL_511211, EPI_ISL_511212, EPI_ISL_511213, EPI_ISL_511214, EPI_ISL_511215, EPI_ISL_511216, EPI_ISL_511217, EPI_ISL_511218, EPI_ISL_511219, EPI_ISL_511220, EPI_ISL_511221, EPI_ISL_511222, EPI_ISL_511223, EPI_ISL_511224, EPI_ISL_511225, EPI_ISL_511226, EPI_ISL_511227, EPI_ISL_511228, EPI_ISL_511229, EPI_ISL_511230, EPI_ISL_511231, EPI_ISL_511232, EPI_ISL_511233, EPI_ISL_511234, EPI_ISL_511235, EPI_ISL_511236, EPI_ISL_511237, EPI_ISL_511238, EPI_ISL_511239, EPI_ISL_511240, EPI_ISL_511241, EPI_ISL_511242, EPI_ISL_511243, EPI_ISL_511244, EPI_ISL_511245, EPI_ISL_511246, EPI_ISL_511247, EPI_ISL_511248, EPI_ISL_511249, EPI_ISL_511250, EPI_ISL_511251, EPI_ISL_511252, EPI_ISL_511253, EPI_ISL_511254, EPI_ISL_511255, EPI_ISL_511256, EPI_ISL_511257, EPI_ISL_511258, EPI_ISL_511259, EPI_ISL_511260, EPI_ISL_511261, EPI_ISL_511262, EPI_ISL_511263, EPI_ISL_511264, EPI_ISL_511265, EPI_ISL_511266, EPI_ISL_511267, EPI_ISL_511268, EPI_ISL_511269, EPI_ISL_511270, EPI_ISL_511271, EPI_ISL_511272, EPI_ISL_511273, EPI_ISL_511274, EPI_ISL_511275, EPI_ISL_511276, EPI_ISL_511277, EPI_ISL_511278, EPI_ISL_511279, EPI_ISL_511280, EPI_ISL_511281, EPI_ISL_511282, EPI_ISL_511283, EPI_ISL_511284, EPI_ISL_511285, EPI_ISL_511286, EPI_ISL_511287, EPI_ISL_511288, EPI_ISL_511289, EPI_ISL_511290, EPI_ISL_511291, EPI_ISL_511292, EPI_ISL_511293, EPI_ISL_511294, EPI_ISL_511295, EPI_ISL_511296, EPI_ISL_511297, EPI_ISL_511298, EPI_ISL_511299, EPI_ISL_511300, EPI_ISL_511301, EPI_ISL_511302, EPI_ISL_511303, EPI_ISL_511304, EPI_ISL_511305, EPI_ISL_511306, EPI_ISL_511307, EPI_ISL_511308, EPI_ISL_511309, EPI_ISL_511310, EPI_ISL_511311, EPI_ISL_511312, EPI_ISL_511313, EPI_ISL_511314, EPI_ISL_511315 | | | |
| see above | Instituto Nacional de Saude (INSA) and Instituto Gulbenkian de Ciencia (IGC) | Instituto Nacional de Saude (INSA) and Instituto Gulbenkian de Ciencia (IGC) | Borges et al |
| EPI_ISL_511316, EPI_ISL_511317, EPI_ISL_511318, EPI_ISL_511319, EPI_ISL_511320, EPI_ISL_511321, EPI_ISL_511322, EPI_ISL_511323, EPI_ISL_511324, EPI_ISL_511325, EPI_ISL_511326, EPI_ISL_511327, EPI_ISL_511328, EPI_ISL_511329, EPI_ISL_511330, EPI_ISL_511331, EPI_ISL_511332, EPI_ISL_511333, EPI_ISL_511334, EPI_ISL_511335, EPI_ISL_511336, EPI_ISL_511337, EPI_ISL_511338, EPI_ISL_511339, EPI_ISL_511340, EPI_ISL_511341, EPI_ISL_511342, EPI_ISL_511343, EPI_ISL_511344, EPI_ISL_511345, EPI_ISL_511346, EPI_ISL_511347, EPI_ISL_511348, EPI_ISL_511349, EPI_ISL_511350, EPI_ISL_511351, EPI_ISL_511352, EPI_ISL_511353, EPI_ISL_511354, EPI_ISL_511355, EPI_ISL_511356, EPI_ISL_511357, EPI_ISL_511358, EPI_ISL_511359, EPI_ISL_511360, EPI_ISL_511361, EPI_ISL_511362, EPI_ISL_511363, EPI_ISL_511364, EPI_ISL_511365, EPI_ISL_511366, EPI_ISL_511367, EPI_ISL_511368, EPI_ISL_511369, EPI_ISL_511370, EPI_ISL_511371, EPI_ISL_511372, EPI_ISL_511373, EPI_ISL_511374, EPI_ISL_511375, EPI_ISL_511376, EPI_ISL_511377, EPI_ISL_511378, EPI_ISL_511379, EPI_ISL_511380, EPI_ISL_511381, EPI_ISL_511382, EPI_ISL_511383, EPI_ISL_511384, EPI_ISL_511385, EPI_ISL_511386, EPI_ISL_511387, EPI_ISL_511388, EPI_ISL_511389, EPI_ISL_511390, EPI_ISL_511391, EPI_ISL_511392, EPI_ISL_511393, EPI_ISL_511394, EPI_ISL_511395, EPI_ISL_511396, EPI_ISL_511397, EPI_ISL_511398, EPI_ISL_511399, EPI_ISL_511400, EPI_ISL_511401, EPI_ISL_511402, EPI_ISL_511403, EPI_ISL_511404, EPI_ISL_511405, EPI_ISL_511406, EPI_ISL_511407, EPI_ISL_511408, EPI_ISL_511409, EPI_ISL_511410, EPI_ISL_511411, EPI_ISL_511412, EPI_ISL_511413, EPI_ISL_511414, EPI_ISL_511415, EPI_ISL_511416, EPI_ISL_511417, EPI_ISL_511418, EPI_ISL_511419, EPI_ISL_511420, EPI_ISL_511421, EPI_ISL_511422, EPI_ISL_511423, EPI_ISL_511424, EPI_ISL_511425, EPI_ISL_511426, EPI_ISL_511427, EPI_ISL_511428, EPI_ISL_511429, EPI_ISL_511430, EPI_ISL_511431, EPI_ISL_511432, EPI_ISL_511433, EPI_ISL_511434, EPI_ISL_511435, EPI_ISL_511436, EPI_ISL_511437, EPI_ISL_511438, EPI_ISL_511439, EPI_ISL_511440, EPI_ISL_511441, EPI_ISL_511442, EPI_ISL_511443, EPI_ISL_511444, EPI_ISL_511445, EPI_ISL_511446, EPI_ISL_511447, EPI_ISL_511448, EPI_ISL_511449, EPI_ISL_511450, EPI_ISL_511451, EPI_ISL_511452, EPI_ISL_511453, EPI_ISL_511454, EPI_ISL_511455, EPI_ISL_511456, EPI_ISL_511457, EPI_ISL_511458, EPI_ISL_511459, EPI_ISL_511460, EPI_ISL_511461, EPI_ISL_511462, EPI_ISL_511463, EPI_ISL_511464, EPI_ISL_511465, EPI_ISL_511466, EPI_ISL_511467, EPI_ISL_511468, EPI_ISL_511469, EPI_ISL_511470, EPI_ISL_511471, EPI_ISL_511472, EPI_ISL_511473, EPI_ISL_511474, EPI_ISL_511475, EPI_ISL_511476, EPI_ISL_511477, EPI_ISL_511478, EPI_ISL_511479, EPI_ISL_511480, EPI_ISL_511481, EPI_ISL_511482, EPI_ISL_511483 | | | |
| see above | Instituto Nacional de Saude (INSA) | Instituto Nacional de Saude (INSA) | Borges et al |
| EPI_ISL_511484, EPI_ISL_511485, EPI_ISL_511486, EPI_ISL_511487, EPI_ISL_511488, EPI_ISL_511489, EPI_ISL_511490, EPI_ISL_511491, EPI_ISL_511492, EPI_ISL_511493, EPI_ISL_511494, EPI_ISL_511495, EPI_ISL_511496, EPI_ISL_511497, EPI_ISL_511498, EPI_ISL_511499, EPI_ISL_511500, EPI_ISL_511501, EPI_ISL_511502, EPI_ISL_511503, EPI_ISL_511504, EPI_ISL_511505, EPI_ISL_511506, EPI_ISL_511507, EPI_ISL_511508 | | | |
| see above | Instituto Nacional de Saude (INSA) | Instituto Nacional de Saude (INSA) and Instituto Gulbenkian de Ciencia (IGC) | Borges et al |
| EPI_ISL_511509, EPI_ISL_511510, EPI_ISL_511511, EPI_ISL_511512, EPI_ISL_511513, EPI_ISL_511514, EPI_ISL_511515, EPI_ISL_511516, EPI_ISL_511517, EPI_ISL_511518, EPI_ISL_511519, EPI_ISL_511520, EPI_ISL_511521, EPI_ISL_511522, EPI_ISL_511523, EPI_ISL_511524, EPI_ISL_511525, EPI_ISL_511526, EPI_ISL_511527, EPI_ISL_511528, EPI_ISL_511529, EPI_ISL_511530, EPI_ISL_511531, EPI_ISL_511532, EPI_ISL_511533, EPI_ISL_511534, EPI_ISL_511535, EPI_ISL_511536, EPI_ISL_511537, EPI_ISL_511538, EPI_ISL_511539, EPI_ISL_511540, EPI_ISL_511541, EPI_ISL_511542, EPI_ISL_511543, EPI_ISL_511544, EPI_ISL_511545, EPI_ISL_511546, EPI_ISL_511547, EPI_ISL_511548, EPI_ISL_511549, EPI_ISL_511550, EPI_ISL_511551, EPI_ISL_511552, EPI_ISL_511553, EPI_ISL_511554, EPI_ISL_511555, EPI_ISL_511556, EPI_ISL_511557, EPI_ISL_511558, EPI_ISL_511559, EPI_ISL_511560, EPI_ISL_511561, EPI_ISL_511562, EPI_ISL_511563, EPI_ISL_511564, EPI_ISL_511565, EPI_ISL_511566, EPI_ISL_511567, EPI_ISL_511568, EPI_ISL_511569, EPI_ISL_511570, EPI_ISL_511571, EPI_ISL_511572, EPI_ISL_511573, EPI_ISL_511574, EPI_ISL_511575, EPI_ISL_511576 | | | |
| see above | Instituto Nacional de Saude (INSA) | Instituto Nacional de Saude (INSA) | Borges et al |

| | | | |
|---|---|---|---|
| EPI_ISL_511577, EPI_ISL_511578 | Hospital de Braga | Instituto Nacional de Saude (INSA) | Borges et al |
| EPI_ISL_511579, EPI_ISL_511580, EPI_ISL_511581, EPI_ISL_511582, EPI_ISL_511583, EPI_ISL_511584, EPI_ISL_511585, EPI_ISL_511586, EPI_ISL_511587, EPI_ISL_511588, EPI_ISL_511589, EPI_ISL_511590, EPI_ISL_511591, EPI_ISL_511592, EPI_ISL_511593, EPI_ISL_511594, EPI_ISL_511595, EPI_ISL_511596, EPI_ISL_511597, EPI_ISL_511598, EPI_ISL_511599, EPI_ISL_511600, EPI_ISL_511601, EPI_ISL_511602, EPI_ISL_511603, EPI_ISL_511604, EPI_ISL_511605, EPI_ISL_511606, EPI_ISL_511607, EPI_ISL_511608, EPI_ISL_511609, EPI_ISL_511610, EPI_ISL_511611, EPI_ISL_511612, EPI_ISL_511613, EPI_ISL_511614, EPI_ISL_511615, EPI_ISL_511616, EPI_ISL_511617, EPI_ISL_511618, EPI_ISL_511619, EPI_ISL_511620, EPI_ISL_511621, EPI_ISL_511622, EPI_ISL_511623, EPI_ISL_511624, EPI_ISL_511625, EPI_ISL_511626, EPI_ISL_511627, EPI_ISL_511628, EPI_ISL_511629, EPI_ISL_511630, EPI_ISL_511631, EPI_ISL_511632, EPI_ISL_511633, EPI_ISL_511634, EPI_ISL_511635, EPI_ISL_511636, EPI_ISL_511637, EPI_ISL_511638, EPI_ISL_511639, EPI_ISL_511640, EPI_ISL_511641, EPI_ISL_511642, EPI_ISL_511643 | | | |
| see above | Instituto Nacional de Saude (INSA) | Instituto Nacional de Saude (INSA) | Borges et al |
| EPI_ISL_511644 | Life and Health Sciences Research Institute, University of Minho, Braga | Instituto Nacional de Saude (INSA) | Borges et al |
| EPI_ISL_511645, EPI_ISL_511646, EPI_ISL_511647, EPI_ISL_511648, EPI_ISL_511649, EPI_ISL_511650, EPI_ISL_511651, EPI_ISL_511652, EPI_ISL_511653, EPI_ISL_511654, EPI_ISL_511655, EPI_ISL_511656, EPI_ISL_511657, EPI_ISL_511658, EPI_ISL_511659, EPI_ISL_511660, EPI_ISL_511661, EPI_ISL_511662, EPI_ISL_511663, EPI_ISL_511664, EPI_ISL_511665, EPI_ISL_511666, EPI_ISL_511667, EPI_ISL_511668, EPI_ISL_511669, EPI_ISL_511670, EPI_ISL_511671, EPI_ISL_511672, EPI_ISL_511673, EPI_ISL_511674, EPI_ISL_511675, EPI_ISL_511676, EPI_ISL_511677, EPI_ISL_511678, EPI_ISL_511679, EPI_ISL_511680, EPI_ISL_511681, EPI_ISL_511682, EPI_ISL_511683, EPI_ISL_511684, EPI_ISL_511685, EPI_ISL_511686, EPI_ISL_511687, EPI_ISL_511688, EPI_ISL_511689, EPI_ISL_511690, EPI_ISL_511691, EPI_ISL_511692, EPI_ISL_511693, EPI_ISL_511694, EPI_ISL_511695, EPI_ISL_511696, EPI_ISL_511697, EPI_ISL_511698, EPI_ISL_511699, EPI_ISL_511700, EPI_ISL_511701, EPI_ISL_511702, EPI_ISL_511703, EPI_ISL_511704, EPI_ISL_511705, EPI_ISL_511706, EPI_ISL_511707, EPI_ISL_511708, EPI_ISL_511709, EPI_ISL_511710, EPI_ISL_511711, EPI_ISL_511712, EPI_ISL_511713, EPI_ISL_511714, EPI_ISL_511715, EPI_ISL_511716, EPI_ISL_511717, EPI_ISL_511718, EPI_ISL_511719, EPI_ISL_511720, EPI_ISL_511721, EPI_ISL_511722, EPI_ISL_511723, EPI_ISL_511724, EPI_ISL_511725, EPI_ISL_511726, EPI_ISL_511727, EPI_ISL_511728, EPI_ISL_511729, EPI_ISL_511730, EPI_ISL_511731, EPI_ISL_511732, EPI_ISL_511733, EPI_ISL_511734, EPI_ISL_511735, EPI_ISL_511736, EPI_ISL_511737, EPI_ISL_511738, EPI_ISL_511739, EPI_ISL_511740, EPI_ISL_511741, EPI_ISL_511742, EPI_ISL_511743, EPI_ISL_511744, EPI_ISL_511745, EPI_ISL_511746, EPI_ISL_511747, EPI_ISL_511748, EPI_ISL_511749, EPI_ISL_511750 | | | |
| see above | Instituto Nacional de Saude (INSA) | Instituto Nacional de Saude (INSA) | Borges et al |
| EPI_ISL_511751, EPI_ISL_511752, EPI_ISL_511753, EPI_ISL_511754, EPI_ISL_511755, EPI_ISL_511756, EPI_ISL_511757, EPI_ISL_511758, EPI_ISL_511759, EPI_ISL_511760, EPI_ISL_511761, EPI_ISL_511762, EPI_ISL_511763, EPI_ISL_511764, EPI_ISL_511765, EPI_ISL_511766, EPI_ISL_511767, EPI_ISL_511768, EPI_ISL_511769, EPI_ISL_511770, EPI_ISL_511771 | | | |
| see above | Instituto Nacional de Saude (INSA) | Instituto Nacional de Saude (INSA) and Instituto Gulbenkian de Ciencia (IGC) | Borges et al |
| EPI_ISL_512775, EPI_ISL_512776, EPI_ISL_512777, EPI_ISL_512778, EPI_ISL_512779, EPI_ISL_512780, EPI_ISL_512781, EPI_ISL_512782, EPI_ISL_512783, EPI_ISL_512784 | Utah Public Health Laboratory, Utah Public Health Laboratory Infectious Disease submission group | Utah Public Health Laboratory, Utah Public Health Laboratory Infectious Disease submission group | Butz,H.A., Young,E.L., Oakeson,K. |
| EPI_ISL_513315, EPI_ISL_513316, EPI_ISL_513317 | South Eastern Area Laboratory Services (SEALS) | NSW Health Pathology - Institute of Clinical Pathology and Medical Research; Westmead Hospital; University of Sydney | CIDM-PH et al. |
| EPI_ISL_513318, EPI_ISL_513319, EPI_ISL_513320 | Histopath | NSW Health Pathology - Institute of Clinical Pathology and Medical Research; Westmead Hospital; University of Sydney | CIDM-PH et al. |
| EPI_ISL_513321, EPI_ISL_513322 | Pathology West - NSW Health Pathology | NSW Health Pathology - Institute of Clinical Pathology and Medical Research; Westmead Hospital; University of Sydney | CIDM-PH et al. |
| EPI_ISL_513323 | Sydney South West Pathology Service (SSWPS) - Liverpool Hospital - NSW Health Pathology | NSW Health Pathology - Institute of Clinical Pathology and Medical Research; Westmead Hospital; University of Sydney | CIDM-PH et al. |
| EPI_ISL_513324, EPI_ISL_513325, EPI_ISL_513326 | Pathology West - NSW Health Pathology | NSW Health Pathology - Institute of Clinical Pathology and Medical Research; Westmead Hospital; University of Sydney | CIDM-PH et al. |
| EPI_ISL_513327, EPI_ISL_513328, EPI_ISL_513329 | Sydney South West Pathology Service (SSWPS) - Liverpool Hospital - NSW Health Pathology | NSW Health Pathology - Institute of Clinical Pathology and Medical Research; Westmead Hospital; University of Sydney | CIDM-PH et al. |
| EPI_ISL_513330 | St Vincent's Pathology (SydPath) | NSW Health Pathology - Institute of Clinical Pathology and Medical Research; Westmead Hospital; University of Sydney | CIDM-PH et al. |
| EPI_ISL_513331 | Australian Clinical Labs | NSW Health Pathology - Institute of Clinical Pathology and Medical Research; Westmead Hospital; University of Sydney | CIDM-PH et al. |
| EPI_ISL_513332 | Pathology West - NSW Health Pathology | NSW Health Pathology - Institute of Clinical Pathology and Medical Research; Westmead Hospital; University of Sydney | CIDM-PH et al. |
| EPI_ISL_513333, EPI_ISL_513334 | Sydney South West Pathology Service (SSWPS) - Liverpool Hospital - NSW Health Pathology | NSW Health Pathology - Institute of Clinical Pathology and Medical Research; Westmead Hospital; University of Sydney | CIDM-PH et al. |
| EPI_ISL_513335 | Pathology North - Hunter - NSW Health Pathology | NSW Health Pathology - Institute of Clinical Pathology and Medical Research; Westmead Hospital; University of Sydney | CIDM-PH et al. |
| EPI_ISL_513336, EPI_ISL_513337 | Sydney South West Pathology Service (SSWPS) - Liverpool Hospital - NSW Health Pathology | NSW Health Pathology - Institute of Clinical Pathology and Medical Research; Westmead Hospital; University of Sydney | CIDM-PH et al. |
| EPI_ISL_513338, EPI_ISL_513339 | Pathology West - NSW Health Pathology | NSW Health Pathology - Institute of Clinical Pathology and Medical Research; Westmead Hospital; University of Sydney | CIDM-PH et al. |
| EPI_ISL_513340 | Austech Medical Laboratories | NSW Health Pathology - Institute of Clinical Pathology and Medical Research; Westmead Hospital; University of Sydney | CIDM-PH et al. |
| EPI_ISL_513341 | Medlab Pathology | NSW Health Pathology - Institute of Clinical Pathology and Medical Research; Westmead Hospital; University of Sydney | CIDM-PH et al. |
| EPI_ISL_513342 | Pathology West - NSW Health Pathology | NSW Health Pathology - Institute of Clinical Pathology and Medical Research; Westmead Hospital; University of Sydney | CIDM-PH et al. |
| EPI_ISL_513343, EPI_ISL_513344 | Children Westmead Hospital | NSW Health Pathology - Institute of Clinical Pathology and Medical Research; Westmead Hospital; University of Sydney | CIDM-PH et al. |
| EPI_ISL_513345, EPI_ISL_513346 | Pathology West - NSW Health Pathology | NSW Health Pathology - Institute of Clinical Pathology and Medical Research; Westmead Hospital; University of Sydney | CIDM-PH et al. |
| EPI_ISL_513347 | St Vincent's Pathology (SydPath) | NSW Health Pathology - Institute of Clinical Pathology and Medical Research; Westmead Hospital; University of Sydney | CIDM-PH et al. |
| EPI_ISL_513348, EPI_ISL_513349 | 4Cyte Pathology | NSW Health Pathology - Institute of Clinical Pathology and Medical Research; Westmead Hospital; University of Sydney | CIDM-PH et al. |
| EPI_ISL_513350, EPI_ISL_513351 | Pathology West - NSW Health Pathology | NSW Health Pathology - Institute of Clinical Pathology and Medical Research; Westmead Hospital; University of Sydney | CIDM-PH et al. |
| EPI_ISL_513352 | South Eastern Area Laboratory Services (SEALS) | NSW Health Pathology - Institute of Clinical Pathology and Medical Research; Westmead Hospital; University of Sydney | CIDM-PH et al. |
| EPI_ISL_513353, EPI_ISL_513354, EPI_ISL_513355, EPI_ISL_513356 | St Vincent's Pathology (SydPath) | NSW Health Pathology - Institute of Clinical Pathology and Medical Research; Westmead Hospital; University of Sydney | CIDM-PH et al. |
| EPI_ISL_513357 | Douglas Hanly Moir | NSW Health Pathology - Institute of Clinical Pathology and Medical Research; Westmead Hospital; University of Sydney | CIDM-PH et al. |

| | | | |
|---|---|---|---|
| EPI_ISL_513358, EPI_ISL_513359 | Pathology West - NSW Health Pathology | NSW Health Pathology - Institute of Clinical Pathology and Medical Research; Westmead Hospital; University of Sydney | CIDM-PH et al. |
| EPI_ISL_513360 | Pathology North - Hunter - NSW Health Pathology | NSW Health Pathology - Institute of Clinical Pathology and Medical Research; Westmead Hospital; University of Sydney | CIDM-PH et al. |
| EPI_ISL_513361, EPI_ISL_513362, EPI_ISL_513363, EPI_ISL_513364, EPI_ISL_513365 | St Vincent's Pathology (SydPath) | NSW Health Pathology - Institute of Clinical Pathology and Medical Research; Westmead Hospital; University of Sydney | CIDM-PH et al. |
| EPI_ISL_513366, EPI_ISL_513367 | Sydney South West Pathology Service (SSWPS) - Liverpool Hospital - NSW Health Pathology | NSW Health Pathology - Institute of Clinical Pathology and Medical Research; Westmead Hospital; University of Sydney | CIDM-PH et al. |
| EPI_ISL_513368 | Pathology North - Hunter - NSW Health Pathology | NSW Health Pathology - Institute of Clinical Pathology and Medical Research; Westmead Hospital; University of Sydney | CIDM-PH et al. |
| EPI_ISL_513369, EPI_ISL_513370, EPI_ISL_513371 | Pathology West - NSW Health Pathology | NSW Health Pathology - Institute of Clinical Pathology and Medical Research; Westmead Hospital; University of Sydney | CIDM-PH et al. |
| EPI_ISL_513372 | St Vincent's Pathology (SydPath) | NSW Health Pathology - Institute of Clinical Pathology and Medical Research; Westmead Hospital; University of Sydney | CIDM-PH et al. |
| EPI_ISL_513373, EPI_ISL_513374 | Sydney South West Pathology Service (SSWPS) - Liverpool Hospital - NSW Health Pathology | NSW Health Pathology - Institute of Clinical Pathology and Medical Research; Westmead Hospital; University of Sydney | CIDM-PH et al. |
| EPI_ISL_513375, EPI_ISL_513376 | Pathology North - Royal North Shore Hospital - NSW Health Pathology | NSW Health Pathology - Institute of Clinical Pathology and Medical Research; Westmead Hospital; University of Sydney | CIDM-PH et al. |
| EPI_ISL_513377 | South Eastern Area Laboratory Services (SEALS) | NSW Health Pathology - Institute of Clinical Pathology and Medical Research; Westmead Hospital; University of Sydney | CIDM-PH et al. |
| EPI_ISL_513378 | Pathology West - NSW Health Pathology | NSW Health Pathology - Institute of Clinical Pathology and Medical Research; Westmead Hospital; University of Sydney | CIDM-PH et al. |
| EPI_ISL_513379 | St Vincent's Pathology (SydPath) | NSW Health Pathology - Institute of Clinical Pathology and Medical Research; Westmead Hospital; University of Sydney | CIDM-PH et al. |
| EPI_ISL_513380 | Pathology West - NSW Health Pathology | NSW Health Pathology - Institute of Clinical Pathology and Medical Research; Westmead Hospital; University of Sydney | CIDM-PH et al. |
| EPI_ISL_513381, EPI_ISL_513382, EPI_ISL_513383 | Sydney South West Pathology Service (SSWPS) - Liverpool Hospital - NSW Health Pathology | NSW Health Pathology - Institute of Clinical Pathology and Medical Research; Westmead Hospital; University of Sydney | CIDM-PH et al. |
| EPI_ISL_513384, EPI_ISL_513385 | St Vincent's Pathology (SydPath) | NSW Health Pathology - Institute of Clinical Pathology and Medical Research; Westmead Hospital; University of Sydney | CIDM-PH et al. |
| EPI_ISL_513386, EPI_ISL_513387, EPI_ISL_513388 | Pathology North - NSW Health Pathology | NSW Health Pathology - Institute of Clinical Pathology and Medical Research; Westmead Hospital; University of Sydney | CIDM-PH et al. |
| EPI_ISL_513389 | Douglas Hanly Moir | NSW Health Pathology - Institute of Clinical Pathology and Medical Research; Westmead Hospital; University of Sydney | CIDM-PH et al. |
| EPI_ISL_513390, EPI_ISL_513391, EPI_ISL_513392, EPI_ISL_513393, EPI_ISL_513394, EPI_ISL_513395 | Pathology West - NSW Health Pathology | NSW Health Pathology - Institute of Clinical Pathology and Medical Research; Westmead Hospital; University of Sydney | CIDM-PH et al. |
| EPI_ISL_513396, EPI_ISL_513397, EPI_ISL_513398, EPI_ISL_513399 | St Vincent's Pathology (SydPath) | NSW Health Pathology - Institute of Clinical Pathology and Medical Research; Westmead Hospital; University of Sydney | CIDM-PH et al. |
| EPI_ISL_513400 | Sydney South West Pathology Service (SSWPS) - Royal Prince Alfred Hospital - NSW Health Pathology | NSW Health Pathology - Institute of Clinical Pathology and Medical Research; Westmead Hospital; University of Sydney | CIDM-PH et al. |
| EPI_ISL_513401 | Sydney South West Pathology Service (SSWPS) - Liverpool Hospital - NSW Health Pathology | NSW Health Pathology - Institute of Clinical Pathology and Medical Research; Westmead Hospital; University of Sydney | CIDM-PH et al. |
| EPI_ISL_513402 | 4Cyte Pathology | NSW Health Pathology - Institute of Clinical Pathology and Medical Research; Westmead Hospital; University of Sydney | CIDM-PH et al. |
| EPI_ISL_513403, EPI_ISL_513404, EPI_ISL_513405, EPI_ISL_513406, EPI_ISL_513407, EPI_ISL_513408 | Laverty Pathology | NSW Health Pathology - Institute of Clinical Pathology and Medical Research; Westmead Hospital; University of Sydney | CIDM-PH et al. |
| EPI_ISL_513409, EPI_ISL_513410, EPI_ISL_513411, EPI_ISL_513412 | Sydney South West Pathology Service (SSWPS) - Liverpool Hospital - NSW Health Pathology | NSW Health Pathology - Institute of Clinical Pathology and Medical Research; Westmead Hospital; University of Sydney | CIDM-PH et al. |
| EPI_ISL_513413 | 4Cyte Pathology | NSW Health Pathology - Institute of Clinical Pathology and Medical Research; Westmead Hospital; University of Sydney | CIDM-PH et al. |
| EPI_ISL_513707, EPI_ISL_513708, EPI_ISL_513709, EPI_ISL_513710, EPI_ISL_513711, EPI_ISL_513712, EPI_ISL_513713, EPI_ISL_513714, EPI_ISL_513715, EPI_ISL_513716, EPI_ISL_513717, EPI_ISL_513718, EPI_ISL_513719, EPI_ISL_513720, EPI_ISL_513721, EPI_ISL_513722, EPI_ISL_513723, EPI_ISL_513724, EPI_ISL_513725, EPI_ISL_513726, EPI_ISL_513727, EPI_ISL_513728, EPI_ISL_513729, EPI_ISL_513730, EPI_ISL_513731, EPI_ISL_513732, EPI_ISL_513733, EPI_ISL_513734, EPI_ISL_513735, EPI_ISL_513736, EPI_ISL_513737, EPI_ISL_513738, EPI_ISL_513739, EPI_ISL_513740, EPI_ISL_513741, EPI_ISL_513742, EPI_ISL_513743, EPI_ISL_513744, EPI_ISL_513745, EPI_ISL_513746, EPI_ISL_513747, EPI_ISL_513748, EPI_ISL_513749, EPI_ISL_513750, EPI_ISL_513751, EPI_ISL_513752, EPI_ISL_513753, EPI_ISL_513754, EPI_ISL_513755, EPI_ISL_513756, EPI_ISL_513757, EPI_ISL_513758, EPI_ISL_513759, EPI_ISL_513760, EPI_ISL_513761, EPI_ISL_513762, EPI_ISL_513763, EPI_ISL_513764, EPI_ISL_513765, EPI_ISL_513766, EPI_ISL_513767, EPI_ISL_513768, EPI_ISL_513769, EPI_ISL_513770, EPI_ISL_513771, EPI_ISL_513772 | | | |
| see above | Orange County Public Health Lab | Chan-Zuckerberg Biohub | CZB Cliahub Consortium |
| EPI_ISL_513773, EPI_ISL_513774, EPI_ISL_513775, EPI_ISL_513776, EPI_ISL_513777, EPI_ISL_513778, EPI_ISL_513779, EPI_ISL_513780, EPI_ISL_513781, EPI_ISL_513782, EPI_ISL_513783, EPI_ISL_513784, EPI_ISL_513785, EPI_ISL_513786, EPI_ISL_513787, EPI_ISL_513788 | | | |
| see above | County of Santa Clara Public Health Department | Chan-Zuckerberg Biohub | CZB Cliahub Consortium |
| EPI_ISL_513789, EPI_ISL_513790, EPI_ISL_513791, EPI_ISL_513792, EPI_ISL_513793, EPI_ISL_513794, EPI_ISL_513795, EPI_ISL_513796, EPI_ISL_513797, EPI_ISL_513798, EPI_ISL_513799, EPI_ISL_513800, EPI_ISL_513801, EPI_ISL_513802, EPI_ISL_513803, EPI_ISL_513804, EPI_ISL_513806, EPI_ISL_513807, EPI_ISL_513808, EPI_ISL_513809, EPI_ISL_513810, EPI_ISL_513811, EPI_ISL_513812, EPI_ISL_513813, EPI_ISL_513814, EPI_ISL_513815, EPI_ISL_513816, EPI_ISL_513817, EPI_ISL_513818, EPI_ISL_513819, EPI_ISL_513820, EPI_ISL_513821, EPI_ISL_513822, EPI_ISL_513823, EPI_ISL_513824, EPI_ISL_513825, EPI_ISL_513826, EPI_ISL_513827, EPI_ISL_513828, EPI_ISL_513829, EPI_ISL_513830, EPI_ISL_513831, EPI_ISL_513832, EPI_ISL_513833, EPI_ISL_513834, EPI_ISL_513835, EPI_ISL_513836, EPI_ISL_513837, EPI_ISL_513838, EPI_ISL_513839 | | | |
| see above | Orange County Public Health Laboratory | Chan-Zuckerberg Biohub | CZB Cliahub Consortium |
| EPI_ISL_513840, EPI_ISL_513841, EPI_ISL_513842, EPI_ISL_513843, EPI_ISL_513844, EPI_ISL_513845, EPI_ISL_513846, EPI_ISL_513847, EPI_ISL_513848, EPI_ISL_513849, EPI_ISL_513850, EPI_ISL_513851, EPI_ISL_513852, EPI_ISL_513853, EPI_ISL_513854, EPI_ISL_513855 | | | |
| see above | Humboldt County Public Health Laboratory | Chan-Zuckerberg Biohub | CZB Cliahub Consortium |
| EPI_ISL_513856, EPI_ISL_513857, EPI_ISL_513858, EPI_ISL_513859, EPI_ISL_513860, EPI_ISL_513861, EPI_ISL_513862, EPI_ISL_513863, EPI_ISL_513864, EPI_ISL_513865, EPI_ISL_513866, EPI_ISL_513867, EPI_ISL_513868, EPI_ISL_513869, EPI_ISL_513870, EPI_ISL_513871, EPI_ISL_513872, EPI_ISL_513873, EPI_ISL_513874, EPI_ISL_513875, EPI_ISL_513876, EPI_ISL_513877, EPI_ISL_513878, EPI_ISL_513879, EPI_ISL_513880, EPI_ISL_513881, EPI_ISL_513882 | | | |
| see above | San Francisco Public Health Laboratory | Chan-Zuckerberg Biohub | CZB Cliahub Consortium |
| EPI_ISL_513883, EPI_ISL_513884, EPI_ISL_513885, EPI_ISL_513886, EPI_ISL_513887, EPI_ISL_513888, EPI_ISL_513889, EPI_ISL_513890, EPI_ISL_513891, EPI_ISL_513892, EPI_ISL_513893, EPI_ISL_513894, EPI_ISL_513895, EPI_ISL_513896, EPI_ISL_513897, EPI_ISL_513898, EPI_ISL_513899, EPI_ISL_513900, | | | |

| | | | |
|---|---|---|---|
| EPI_ISL_513901, EPI_ISL_513902, EPI_ISL_513903, EPI_ISL_513904, EPI_ISL_513905, EPI_ISL_513906, EPI_ISL_513907, EPI_ISL_513908, EPI_ISL_513909, EPI_ISL_513910 | | | |
| see above | UCSF Clinical Microbiology Laboratory | Chan-Zuckerberg Biohub | CZB Cliahub Consortium |
| EPI_ISL_513941, EPI_ISL_513942 | Utah Public Health Laboratory, Utah Public Health Laboratory Infectious Disease submission group | Utah Public Health Laboratory, Utah Public Health Laboratory Infectious Disease submission group | Butz,H.A., Young,E.L. and Oakeson,K. |
| EPI_ISL_514226 | Laboratorio de Referencia Nacional de Virus Respiratorio. Instituto Nacional de Salud Perú | Laboratorio de Referencia Nacional de Biotecnología y Biología Molecular. Instituto Nacional de Salud Perú | Carlos Padilla Rojas, Karolyn Vega Chozo, Priscila Lope Pari, Omar Caceres Rey, Marco Galarza Perez, Maribel Huaringa Nuñez, Johanna Balbuena Torrez, Henri Bailon Calderon, Nancy Rojas Serrano |
| EPI_ISL_514227 | Laboratorio de Referencia Nacional de Virus Respiratorio. Instituto Nacional de Salud. Peru | Laboratorio de Referencia Nacional de Biotecnologia y Biologia Molecular.Instituto Nacional de Salud.Peru | Carlos Padilla Rojas, Karolyn Vega Chozo, Priscila Lope Pari, Omar Caceres Rey, Marco Galarza Perez, Maribel Huaringa Nuñez, Johanna Balbuena Torrez, Henri Bailon Calderon, Nancy Rojas Serrano |
| EPI_ISL_514264, EPI_ISL_514265 | Laboratorio de Referencia Nacional de Virus Respiratorio. Instituto Nacional de Salud. Perú | Laboratorio de Referencia Nacional de Biotecnologia y Biologia Molecular.Instituto Nacional de Salud.Perú | Carlos Padilla Rojas, Karolyn Vega Chozo, Priscila Lope Pari, Omar Caceres Rey, Marco Galarza Perez, Maribel Huaringa Nuñez, Johanna Balbuena Torrez, Henri Bailon Calderon, Nancy Rojas Serrano |
| EPI_ISL_514317 | Laboratorio de Referencia Nacional de Virus Respiratorio. Instituto Nacional de Salud Perú | Laboratorio de Referencia Nacional de Biotecnología y Biología Molecular. Instituto Nacional de Salud Perú | Carlos Padilla Rojas, Karolyn Vega Chozo, Priscila Lope Pari, Omar Caceres Rey, Marco Galarza Perez, Maribel Huaringa Nuñez, Johanna Balbuena Torres, Henri Bailon Calderon, Nancy Rojas Serrano. |
| EPI_ISL_514338, EPI_ISL_514339, EPI_ISL_514340, EPI_ISL_514341, EPI_ISL_514342 | Laboratorio de Referencia Nacional de Virus Respiratorio. Instituto Nacional de Salud Perú | Laboratorio de Referencia Nacional de Biotecnología y Biología Molecular. Instituto Nacional de Salud Perú | Carlos Padilla Rojas, Karolyn Vega Chozo, Priscila Lope Pari, Omar Caceres Rey, Marco Galarza Perez, Maribel Huaringa Nuñez, Johanna Balbuena Torres, Henri Bailon Calderon, Nancy Rojas Serrano. |
| EPI_ISL_515894, EPI_ISL_515895, EPI_ISL_515896, EPI_ISL_515897, EPI_ISL_515898, EPI_ISL_515899, EPI_ISL_515900, EPI_ISL_515901, EPI_ISL_515902, EPI_ISL_515903, EPI_ISL_515904, EPI_ISL_515905, EPI_ISL_515906, EPI_ISL_515907, EPI_ISL_515908, EPI_ISL_515909, EPI_ISL_515910, EPI_ISL_515911, EPI_ISL_515912, EPI_ISL_515913, EPI_ISL_515914, EPI_ISL_515915, EPI_ISL_515916, EPI_ISL_515917, EPI_ISL_515918, EPI_ISL_515919, EPI_ISL_515920, EPI_ISL_515921, EPI_ISL_515922, EPI_ISL_515923, EPI_ISL_515924, EPI_ISL_515925, EPI_ISL_515926, EPI_ISL_515927, EPI_ISL_515928, EPI_ISL_515929 | | | |
| see above | California Department of Public Health | California Department of Public Health | CDPH IDLB COVIDNet |
| EPI_ISL_516626, EPI_ISL_516628, EPI_ISL_516646, EPI_ISL_516647, EPI_ISL_516653, EPI_ISL_516654, EPI_ISL_516685, EPI_ISL_516686, EPI_ISL_516721, EPI_ISL_516722 | Laboratorio de Referencia Nacional de Virus Respiratorio. Centro Nacional de Salud Publica. Instituto Nacional de Salud Peru. | Laboratorio de Referencia Nacional de Biotecnología y Biología Molecular. Centro Nacional de Salud Publica. Instituto Nacional de Salud Peru. | Carlos Padilla Rojas, Karolyn Vega Chozo, Priscila Lope Pari, Omar Caceres Rey, Marco Galarza Perez, Maribel Huaringa Nuñez, Johanna Balbuena Torres, Henri Bailon Calderon, Nancy Rojas Serrano. |
| EPI_ISL_516723, EPI_ISL_516724, EPI_ISL_516725, EPI_ISL_516726, EPI_ISL_516727, EPI_ISL_516728, EPI_ISL_516729, EPI_ISL_516730, EPI_ISL_516731, EPI_ISL_516732, EPI_ISL_516733, EPI_ISL_516734, EPI_ISL_516735, EPI_ISL_516736, EPI_ISL_516737, EPI_ISL_516738, EPI_ISL_516739, EPI_ISL_516740, EPI_ISL_516741, EPI_ISL_516742, EPI_ISL_516743, EPI_ISL_516744, EPI_ISL_516745, EPI_ISL_516746, EPI_ISL_516747 | | | |
| see above | UCSF Clinical Microbiology Laboratory | Chan-Zuckerberg Biohub | CZB Cliahub Consortium |
| EPI_ISL_516987, EPI_ISL_517531, EPI_ISL_517686, EPI_ISL_517687, EPI_ISL_517713 | Laboratorio de Referencia Nacional de Virus Respiratorio. Centro Nacional de Salud Publica. Instituto Nacional de Salud Peru. | Laboratorio de Referencia Nacional de Biotecnología y Biología Molecular. Centro Nacional de Salud Publica. Instituto Nacional de Salud Peru. | Carlos Padilla Rojas, Karolyn Vega Chozo, Priscila Lope Pari, Omar Caceres Rey, Marco Galarza Perez, Maribel Huaringa Nuñez, Johanna Balbuena Torres, Henri Bailon Calderon, Nancy Rojas Serrano. |
| EPI_ISL_517770 | Laboratorio de Referencia Nacional de Virus Respiratorio. Centro Nacional de Salud Publica. Instituto Nacional de Salud Peru | Laboratorio de Referencia Nacional de Biotecnología y Biología Molecular. Centro Nacional de Salud Publica. Instituto Nacional de Salud Peru. | Carlos Padilla Rojas, Karolyn Vega Chozo, Priscila Lope Pari, Omar Caceres Rey, Marco Galarza Perez, Maribel Huaringa Nuñez, Johanna Balbuena Torres, Henri Bailon Calderon, Nancy Rojas Serrano. |
| EPI_ISL_517958 | Laboratorio de Referencia Nacional de Virus Respiratorio. Centro Nacional de Salud Publica. Instituto Nacional de Salud Peru. | Laboratorio de Referencia Nacional de Biotecnología y Biología Molecular. Centro Nacional de Salud Publica. Instituto Nacional de Salud Peru. | Carlos Padilla Rojas, Karolyn Vega Chozo, Priscila Lope Pari, Omar Caceres Rey, Marco Galarza Perez, Maribel Huaringa Nuñez, Johanna Balbuena Torres, Henri Bailon Calderon, Nancy Rojas Serrano. |
| EPI_ISL_517959 | Laboratorio de Referencia Nacional de Virus Respiratorio. Centro Nacional de Salud Publica. Instituto Nacional de Salud Peru | Laboratorio de Referencia Nacional de Biotecnología y Biología Molecular. Centro Nacional de Salud Publica. Instituto Nacional de Salud Peru. | Carlos Padilla Rojas, Karolyn Vega Chozo, Priscila Lope Pari, Omar Caceres Rey, Marco Galarza Perez, Maribel Huaringa Nuñez, Johanna Balbuena Torres, Henri Bailon Calderon, Nancy Rojas Serrano. |
| EPI_ISL_518081, EPI_ISL_518082, EPI_ISL_518083, EPI_ISL_518084, EPI_ISL_518085, EPI_ISL_518086, EPI_ISL_518087, EPI_ISL_518088, EPI_ISL_518089, EPI_ISL_518090, EPI_ISL_518091, EPI_ISL_518092, EPI_ISL_518098, EPI_ISL_518099, EPI_ISL_518100, EPI_ISL_518101, EPI_ISL_518102, EPI_ISL_518103, EPI_ISL_518104, EPI_ISL_518105, EPI_ISL_518106, EPI_ISL_518107, EPI_ISL_518108, EPI_ISL_518109, EPI_ISL_518110, EPI_ISL_518111, EPI_ISL_518112, EPI_ISL_518113, EPI_ISL_518114, EPI_ISL_518115, EPI_ISL_518116, EPI_ISL_518117, EPI_ISL_518119, EPI_ISL_518120, EPI_ISL_518121, EPI_ISL_518122, EPI_ISL_518123, EPI_ISL_518124, EPI_ISL_518125, EPI_ISL_518126, EPI_ISL_518127, EPI_ISL_518128, EPI_ISL_518129, EPI_ISL_518130, EPI_ISL_518131, EPI_ISL_518132, EPI_ISL_518133, EPI_ISL_518134, EPI_ISL_518135, EPI_ISL_518136, EPI_ISL_518137, EPI_ISL_518138, EPI_ISL_518139, EPI_ISL_518140, EPI_ISL_518141, EPI_ISL_518142, EPI_ISL_518143, EPI_ISL_518144, EPI_ISL_518145, EPI_ISL_518146, EPI_ISL_518147, EPI_ISL_518148, EPI_ISL_518149, EPI_ISL_518150, EPI_ISL_518151, EPI_ISL_518152, EPI_ISL_518153, EPI_ISL_518154, EPI_ISL_518155, EPI_ISL_518156, EPI_ISL_518157, EPI_ISL_518158, EPI_ISL_518159, EPI_ISL_518160, EPI_ISL_518161, EPI_ISL_518162, EPI_ISL_518163, EPI_ISL_518164, EPI_ISL_518165, EPI_ISL_518166, EPI_ISL_518167, EPI_ISL_518168, EPI_ISL_518169, EPI_ISL_518170, EPI_ISL_518171, EPI_ISL_518172, EPI_ISL_518173, EPI_ISL_518174, EPI_ISL_518175, EPI_ISL_518176, EPI_ISL_518177, EPI_ISL_518178, EPI_ISL_518179, EPI_ISL_518180, EPI_ISL_518181, EPI_ISL_518182, EPI_ISL_518183, EPI_ISL_518184, EPI_ISL_518185, EPI_ISL_518186, EPI_ISL_518187, EPI_ISL_518188, EPI_ISL_518189, EPI_ISL_518190, EPI_ISL_518191, EPI_ISL_518192, EPI_ISL_518193, EPI_ISL_518194, EPI_ISL_518195, EPI_ISL_518196, EPI_ISL_518197, EPI_ISL_518198, EPI_ISL_518199, EPI_ISL_518200, EPI_ISL_518201, EPI_ISL_518202, EPI_ISL_518203, EPI_ISL_518204, EPI_ISL_518205, EPI_ISL_518206, EPI_ISL_518207, EPI_ISL_518208, EPI_ISL_518209, EPI_ISL_518210, EPI_ISL_518211, EPI_ISL_518212, EPI_ISL_518213, EPI_ISL_518214, EPI_ISL_518215, EPI_ISL_518216, EPI_ISL_518217, EPI_ISL_518218, EPI_ISL_518219, EPI_ISL_518220, EPI_ISL_518221, EPI_ISL_518222, EPI_ISL_518223, EPI_ISL_518224, EPI_ISL_518225, EPI_ISL_518226, EPI_ISL_518227, EPI_ISL_518228, EPI_ISL_518229, EPI_ISL_518230, EPI_ISL_518239, EPI_ISL_518240, EPI_ISL_518244, EPI_ISL_518245, EPI_ISL_518246, EPI_ISL_518247, EPI_ISL_518248, EPI_ISL_518249, EPI_ISL_518388, EPI_ISL_518389, EPI_ISL_518390, EPI_ISL_518391, EPI_ISL_518392, EPI_ISL_518393, EPI_ISL_518394, EPI_ISL_518395, EPI_ISL_518396, EPI_ISL_518397, EPI_ISL_518398, EPI_ISL_518399, EPI_ISL_518400, EPI_ISL_518401, EPI_ISL_518424, EPI_ISL_518425, EPI_ISL_518426, EPI_ISL_518427, EPI_ISL_518428, EPI_ISL_518429, EPI_ISL_518430, EPI_ISL_518431, EPI_ISL_518432, EPI_ISL_518433, EPI_ISL_518434, EPI_ISL_518435, EPI_ISL_518436, EPI_ISL_518437, EPI_ISL_518438, EPI_ISL_518439, EPI_ISL_518440, EPI_ISL_518441, EPI_ISL_518442, EPI_ISL_518443, EPI_ISL_518444, EPI_ISL_518445, EPI_ISL_518446, EPI_ISL_518447, EPI_ISL_518448, EPI_ISL_518449, EPI_ISL_518450, EPI_ISL_518451, EPI_ISL_518452, EPI_ISL_518453, EPI_ISL_518454, EPI_ISL_518455, EPI_ISL_518456, EPI_ISL_518457, EPI_ISL_518458, EPI_ISL_518459, EPI_ISL_518460, EPI_ISL_518461, EPI_ISL_518462, EPI_ISL_518463, EPI_ISL_518464, EPI_ISL_518465, EPI_ISL_518466, EPI_ISL_518467, EPI_ISL_518468, EPI_ISL_518469, EPI_ISL_518470, EPI_ISL_518471, EPI_ISL_518723, EPI_ISL_518976, EPI_ISL_518981, EPI_ISL_518982, EPI_ISL_518983, EPI_ISL_518984, EPI_ISL_518985, EPI_ISL_518986, EPI_ISL_518987, EPI_ISL_518988, EPI_ISL_518989, EPI_ISL_518990, EPI_ISL_518991, EPI_ISL_518992, EPI_ISL_518993, EPI_ISL_518994, EPI_ISL_518995, EPI_ISL_518996, EPI_ISL_518997, EPI_ISL_518998, EPI_ISL_518999, EPI_ISL_519000, EPI_ISL_519001, EPI_ISL_519002, EPI_ISL_519003, EPI_ISL_519004, EPI_ISL_519005, EPI_ISL_519006, EPI_ISL_519007, EPI_ISL_519008, EPI_ISL_519009, EPI_ISL_519010, EPI_ISL_519011, EPI_ISL_519012, EPI_ISL_519013, EPI_ISL_519014, EPI_ISL_519015, EPI_ISL_519016, EPI_ISL_519017, EPI_ISL_519018, EPI_ISL_519019, EPI_ISL_519020, EPI_ISL_519021, EPI_ISL_519022, EPI_ISL_519023, EPI_ISL_519025, EPI_ISL_519026, EPI_ISL_519027, EPI_ISL_519028, EPI_ISL_519029, EPI_ISL_519030, EPI_ISL_519031, EPI_ISL_519032, EPI_ISL_519033, EPI_ISL_519034, EPI_ISL_519035, EPI_ISL_519036, EPI_ISL_519037, EPI_ISL_519038, EPI_ISL_519039, EPI_ISL_519040, EPI_ISL_519041, EPI_ISL_519042, EPI_ISL_519043, EPI_ISL_519044, EPI_ISL_519045, EPI_ISL_519046, EPI_ISL_519048, EPI_ISL_519049, EPI_ISL_519050, EPI_ISL_519051, EPI_ISL_519256, EPI_ISL_519257, EPI_ISL_519258, EPI_ISL_519259, EPI_ISL_519260, EPI_ISL_519261, EPI_ISL_519262, EPI_ISL_519263, EPI_ISL_519264, EPI_ISL_519265, EPI_ISL_519266, EPI_ISL_519267, EPI_ISL_519268, EPI_ISL_519269, EPI_ISL_519270, EPI_ISL_519271, EPI_ISL_519272, EPI_ISL_519273, EPI_ISL_519274, EPI_ISL_519275, EPI_ISL_519276, EPI_ISL_519277, EPI_ISL_519278, EPI_ISL_519279, EPI_ISL_519280, EPI_ISL_519281, EPI_ISL_519282, EPI_ISL_519283, EPI_ISL_519284, EPI_ISL_519285, EPI_ISL_519286, EPI_ISL_519287, EPI_ISL_519288, EPI_ISL_519289, EPI_ISL_519290, EPI_ISL_519291, EPI_ISL_519292, EPI_ISL_519293, EPI_ISL_519294, EPI_ISL_519295, EPI_ISL_519296, EPI_ISL_519297, EPI_ISL_519298, EPI_ISL_519299, EPI_ISL_519300, EPI_ISL_519301, EPI_ISL_519302, EPI_ISL_519303, EPI_ISL_519304, EPI_ISL_519305, EPI_ISL_519306, EPI_ISL_519307, EPI_ISL_519308, EPI_ISL_519309, EPI_ISL_519310, EPI_ISL_519311, EPI_ISL_519312, EPI_ISL_519313, EPI_ISL_519314, EPI_ISL_519315, EPI_ISL_519316, EPI_ISL_519317, EPI_ISL_519318, EPI_ISL_519319, EPI_ISL_519320, EPI_ISL_519321, EPI_ISL_519322, EPI_ISL_519323, EPI_ISL_519324, EPI_ISL_519325, EPI_ISL_519326, EPI_ISL_519385, EPI_ISL_519519, EPI_ISL_519520, EPI_ISL_519521, EPI_ISL_519948, EPI_ISL_519949, EPI_ISL_519950, EPI_ISL_520181, EPI_ISL_520182, EPI_ISL_520183, EPI_ISL_520184, EPI_ISL_520185, EPI_ISL_520186, EPI_ISL_520187, EPI_ISL_520188, EPI_ISL_520189, EPI_ISL_520190, EPI_ISL_520191, EPI_ISL_520192, EPI_ISL_520193, EPI_ISL_520194, EPI_ISL_520195, EPI_ISL_520196, EPI_ISL_520197, EPI_ISL_520198, EPI_ISL_520199, EPI_ISL_520200, EPI_ISL_520202, EPI_ISL_520203, EPI_ISL_520204, EPI_ISL_520205, EPI_ISL_520206, EPI_ISL_520207, EPI_ISL_520208, EPI_ISL_520209, EPI_ISL_520210, EPI_ISL_520211, EPI_ISL_520212, EPI_ISL_520213, EPI_ISL_520214, EPI_ISL_520249, EPI_ISL_520250, EPI_ISL_520251, EPI_ISL_520252, EPI_ISL_520253, EPI_ISL_520254, EPI_ISL_520255, EPI_ISL_520256, EPI_ISL_520257, EPI_ISL_520258, EPI_ISL_520259, EPI_ISL_520260, EPI_ISL_520261, EPI_ISL_520262, EPI_ISL_520263, EPI_ISL_520264, EPI_ISL_520265, EPI_ISL_520266, EPI_ISL_520267, EPI_ISL_520268, EPI_ISL_520269, EPI_ISL_520270, EPI_ISL_520271, EPI_ISL_520272, EPI_ISL_520273, EPI_ISL_520274, EPI_ISL_520275, EPI_ISL_520276, EPI_ISL_520277, EPI_ISL_520278, EPI_ISL_520279, EPI_ISL_520280, EPI_ISL_520281, EPI_ISL_520282, EPI_ISL_520283, EPI_ISL_520284, EPI_ISL_520285, EPI_ISL_520286, EPI_ISL_520287, EPI_ISL_520288, EPI_ISL_520289, EPI_ISL_520290, EPI_ISL_520291, EPI_ISL_520292, EPI_ISL_520293, EPI_ISL_520294, EPI_ISL_520295, EPI_ISL_520296, EPI_ISL_520297, EPI_ISL_520298, EPI_ISL_520299, EPI_ISL_520300, EPI_ISL_520301, EPI_ISL_520302, EPI_ISL_520303, EPI_ISL_520304, EPI_ISL_520305, EPI_ISL_520306, EPI_ISL_520307, EPI_ISL_520309, EPI_ISL_520310, EPI_ISL_520311, EPI_ISL_520312, EPI_ISL_520313, EPI_ISL_520315, EPI_ISL_520316, EPI_ISL_520317, EPI_ISL_520318, EPI_ISL_520319, EPI_ISL_520320, EPI_ISL_520321, EPI_ISL_520322, EPI_ISL_520323, EPI_ISL_520324, EPI_ISL_520325, EPI_ISL_520326, EPI_ISL_520327, EPI_ISL_520328, EPI_ISL_520329, EPI_ISL_520331, EPI_ISL_520332, EPI_ISL_520333, EPI_ISL_520334, EPI_ISL_520335, EPI_ISL_520336, EPI_ISL_520337, EPI_ISL_520338, EPI_ISL_520339, EPI_ISL_520340, EPI_ISL_520341, EPI_ISL_520342, EPI_ISL_520343, EPI_ISL_520344, EPI_ISL_520345, EPI_ISL_520346, EPI_ISL_520347, EPI_ISL_520348, EPI_ISL_520349, EPI_ISL_520350, EPI_ISL_520351, EPI_ISL_520352, EPI_ISL_520353, EPI_ISL_520354, EPI_ISL_520355, EPI_ISL_520356, EPI_ISL_520357, EPI_ISL_520358, EPI_ISL_520359, EPI_ISL_520360, EPI_ISL_520361, EPI_ISL_520362, EPI_ISL_520363, EPI_ISL_520364, EPI_ISL_520365, EPI_ISL_520366, EPI_ISL_520367, EPI_ISL_520368, EPI_ISL_520369, EPI_ISL_520370, EPI_ISL_520371, EPI_ISL_520372, EPI_ISL_520373, EPI_ISL_520374, EPI_ISL_520375, EPI_ISL_520376, EPI_ISL_520377, EPI_ISL_520378, EPI_ISL_520379, EPI_ISL_520380, EPI_ISL_520381, EPI_ISL_520382, EPI_ISL_520383, EPI_ISL_520384, EPI_ISL_520385, EPI_ISL_520386, EPI_ISL_520387, EPI_ISL_520388, EPI_ISL_520389, EPI_ISL_520390, EPI_ISL_520391, EPI_ISL_520745, EPI_ISL_520746, EPI_ISL_520747, EPI_ISL_520748, EPI_ISL_520749, EPI_ISL_520750, EPI_ISL_520751, EPI_ISL_520752, EPI_ISL_520753, EPI_ISL_520754, EPI_ISL_520755, EPI_ISL_520756, EPI_ISL_520757, EPI_ISL_520758, EPI_ISL_520759, EPI_ISL_520760, EPI_ISL_520761, EPI_ISL_520762, EPI_ISL_520763, EPI_ISL_520764, EPI_ISL_520791, EPI_ISL_520792, EPI_ISL_520793, EPI_ISL_520794, EPI_ISL_520795, EPI_ISL_520796, EPI_ISL_520797, EPI_ISL_520798, EPI_ISL_520799, EPI_ISL_520800, EPI_ISL_520801, EPI_ISL_520802, EPI_ISL_520803, EPI_ISL_520804, EPI_ISL_520805, EPI_ISL_520806, EPI_ISL_520807, EPI_ISL_520808, EPI_ISL_520809, EPI_ISL_520810, EPI_ISL_520811, EPI_ISL_520812, EPI_ISL_520813, EPI_ISL_520814, EPI_ISL_520815, EPI_ISL_520816, EPI_ISL_520817, EPI_ISL_520818, EPI_ISL_520819, EPI_ISL_520820, EPI_ISL_520821, EPI_ISL_520822, EPI_ISL_520823, EPI_ISL_520824, EPI_ISL_520825, EPI_ISL_520826, EPI_ISL_520827, EPI_ISL_520828, EPI_ISL_520829, EPI_ISL_520830, EPI_ISL_520831, EPI_ISL_520832, EPI_ISL_520833, EPI_ISL_520834, EPI_ISL_520835, EPI_ISL_520836, EPI_ISL_520837, EPI_ISL_520838, EPI_ISL_520840, EPI_ISL_520841, EPI_ISL_520842, EPI_ISL_520843, EPI_ISL_520864, EPI_ISL_520865, EPI_ISL_520866, EPI_ISL_520867, EPI_ISL_520868, EPI_ISL_520869, EPI_ISL_520870, EPI_ISL_520871, EPI_ISL_520872, EPI_ISL_520873, EPI_ISL_520874, EPI_ISL_520875, EPI_ISL_520876, EPI_ISL_520877, EPI_ISL_520878, EPI_ISL_520879, EPI_ISL_520880, EPI_ISL_520881, EPI_ISL_520882, EPI_ISL_520883, EPI_ISL_520884, EPI_ISL_520885, EPI_ISL_520886, EPI_ISL_520887, EPI_ISL_520888, EPI_ISL_520889, EPI_ISL_520890, EPI_ISL_520891, EPI_ISL_520892, EPI_ISL_520893, EPI_ISL_520894, EPI_ISL_520895, EPI_ISL_520896, EPI_ISL_520897, EPI_ISL_520898, EPI_ISL_520899, EPI_ISL_520900, EPI_ISL_520901, EPI_ISL_520902, EPI_ISL_520903, EPI_ISL_520904, EPI_ISL_520905, EPI_ISL_520906, EPI_ISL_520907, EPI_ISL_520908, EPI_ISL_520909, EPI_ISL_520910, EPI_ISL_520911, EPI_ISL_520912, EPI_ISL_520913, EPI_ISL_520914, EPI_ISL_520915, EPI_ISL_520916, EPI_ISL_520917, EPI_ISL_520918, EPI_ISL_520919, EPI_ISL_520920, EPI_ISL_520921, EPI_ISL_520922, EPI_ISL_520923, EPI_ISL_520924, EPI_ISL_520925, EPI_ISL_520926, EPI_ISL_520927, EPI_ISL_520928, EPI_ISL_521102, EPI_ISL_521103, EPI_ISL_521104, EPI_ISL_521105, EPI_ISL_521106, EPI_ISL_521107, EPI_ISL_521108, EPI_ISL_521109, EPI_ISL_521110, EPI_ISL_521111, EPI_ISL_521112, EPI_ISL_521113, EPI_ISL_521114, EPI_ISL_521115, EPI_ISL_521116, EPI_ISL_521117, EPI_ISL_521118, EPI_ISL_521119, EPI_ISL_521120, EPI_ISL_521121, EPI_ISL_521122, EPI_ISL_521123, EPI_ISL_521124, EPI_ISL_521125, EPI_ISL_521126, EPI_ISL_521127, EPI_ISL_521128, EPI_ISL_521129, EPI_ISL_521130, EPI_ISL_521131, EPI_ISL_521132, EPI_ISL_521133, EPI_ISL_521134, EPI_ISL_521135, EPI_ISL_521136, EPI_ISL_521137, EPI_ISL_521138, EPI_ISL_521139, EPI_ISL_521140, EPI_ISL_521141, EPI_ISL_521142, EPI_ISL_521143, EPI_ISL_521144, EPI_ISL_521145, EPI_ISL_521146, EPI_ISL_521147, EPI_ISL_521148, EPI_ISL_521149, EPI_ISL_521150, EPI_ISL_521151, EPI_ISL_521152, EPI_ISL_521270, EPI_ISL_521271, EPI_ISL_521272, EPI_ISL_521273, EPI_ISL_521274, EPI_ISL_521275, EPI_ISL_521276, EPI_ISL_521277, EPI_ISL_521315, EPI_ISL_521316, EPI_ISL_521317, EPI_ISL_521318, EPI_ISL_521319, EPI_ISL_521320, | | | |

| | | | |
|---|---|---|---|
| EPI_ISL_521321, EPI_ISL_521322, EPI_ISL_521323, EPI_ISL_521324, EPI_ISL_521325, EPI_ISL_521326, EPI_ISL_521327, EPI_ISL_521328, EPI_ISL_521329, EPI_ISL_521330, EPI_ISL_521331, EPI_ISL_521332, EPI_ISL_521333, EPI_ISL_521334, EPI_ISL_521335, EPI_ISL_521337, EPI_ISL_521338, EPI_ISL_521339, EPI_ISL_521340, EPI_ISL_521341, EPI_ISL_521342, EPI_ISL_521343, EPI_ISL_521344, EPI_ISL_521345, EPI_ISL_521346, EPI_ISL_521347, EPI_ISL_521348, EPI_ISL_521349, EPI_ISL_521350, EPI_ISL_521351, EPI_ISL_521352, EPI_ISL_521353, EPI_ISL_521354, EPI_ISL_521355, EPI_ISL_521356, EPI_ISL_521357, EPI_ISL_521358, EPI_ISL_521359, EPI_ISL_521360, EPI_ISL_521361, EPI_ISL_521362, EPI_ISL_521363, EPI_ISL_521364, EPI_ISL_521365, EPI_ISL_521366, EPI_ISL_521367, EPI_ISL_521368, EPI_ISL_521369, EPI_ISL_521370, EPI_ISL_521371, EPI_ISL_521372, EPI_ISL_521373, EPI_ISL_521374, EPI_ISL_521375, EPI_ISL_521376, EPI_ISL_521377, EPI_ISL_521378, EPI_ISL_521379, EPI_ISL_521380, EPI_ISL_521381, EPI_ISL_521382, EPI_ISL_521383, EPI_ISL_521384, EPI_ISL_521385, EPI_ISL_521386, EPI_ISL_521387, EPI_ISL_521388, EPI_ISL_521389, EPI_ISL_521390, EPI_ISL_521391, EPI_ISL_521392, EPI_ISL_521393, EPI_ISL_521394, EPI_ISL_521395, EPI_ISL_521396, EPI_ISL_521397, EPI_ISL_521398, EPI_ISL_521399, EPI_ISL_521400, EPI_ISL_521401, EPI_ISL_521402, EPI_ISL_521403, EPI_ISL_521404, EPI_ISL_521405, EPI_ISL_521406, EPI_ISL_521407, EPI_ISL_521408, EPI_ISL_521410, EPI_ISL_521411, EPI_ISL_521412, EPI_ISL_521414, EPI_ISL_521415, EPI_ISL_521416, EPI_ISL_521417, EPI_ISL_521418, EPI_ISL_521419, EPI_ISL_521420, EPI_ISL_521421, EPI_ISL_521422, EPI_ISL_521423, EPI_ISL_521424, EPI_ISL_521425, EPI_ISL_521426, EPI_ISL_521427, EPI_ISL_521428, EPI_ISL_521429, EPI_ISL_521430, EPI_ISL_521431, EPI_ISL_521432, EPI_ISL_521433, EPI_ISL_521434, EPI_ISL_521435, EPI_ISL_521436, EPI_ISL_521437, EPI_ISL_521438, EPI_ISL_521439, EPI_ISL_521440, EPI_ISL_521441, EPI_ISL_521442, EPI_ISL_521443, EPI_ISL_521444, EPI_ISL_521445, EPI_ISL_521446, EPI_ISL_521447, EPI_ISL_521448, EPI_ISL_521449, EPI_ISL_521463, EPI_ISL_521464, EPI_ISL_521465, EPI_ISL_521466, EPI_ISL_521467, EPI_ISL_521468, EPI_ISL_521469, EPI_ISL_521470, EPI_ISL_521471, EPI_ISL_521472, EPI_ISL_521473, EPI_ISL_521474, EPI_ISL_521475, EPI_ISL_521476, EPI_ISL_521477, EPI_ISL_521478, EPI_ISL_521479, EPI_ISL_521480, EPI_ISL_521481, EPI_ISL_521482, EPI_ISL_521483, EPI_ISL_521484, EPI_ISL_521485, EPI_ISL_521486, EPI_ISL_521487, EPI_ISL_521488, EPI_ISL_521489, EPI_ISL_521490, EPI_ISL_521491, EPI_ISL_521492, EPI_ISL_521493, EPI_ISL_521494, EPI_ISL_521495, EPI_ISL_521496, EPI_ISL_521497, EPI_ISL_521498, EPI_ISL_521499, EPI_ISL_521500, EPI_ISL_521501, EPI_ISL_521502, EPI_ISL_521503, EPI_ISL_521504, EPI_ISL_521505, EPI_ISL_521506, EPI_ISL_521507, EPI_ISL_521508, EPI_ISL_521509, EPI_ISL_521510, EPI_ISL_521511, EPI_ISL_521512, EPI_ISL_521513, EPI_ISL_521514, EPI_ISL_521515, EPI_ISL_521516, EPI_ISL_521517, EPI_ISL_521518, EPI_ISL_521519, EPI_ISL_521520, EPI_ISL_521521, EPI_ISL_521522, EPI_ISL_521523, EPI_ISL_521524, EPI_ISL_521525, EPI_ISL_521526, EPI_ISL_521527, EPI_ISL_521528, EPI_ISL_521529, EPI_ISL_521530, EPI_ISL_521531, EPI_ISL_521532, EPI_ISL_521533, EPI_ISL_521534, EPI_ISL_521535, EPI_ISL_521536, EPI_ISL_521537, EPI_ISL_521538, EPI_ISL_521539, EPI_ISL_521540, EPI_ISL_521541, EPI_ISL_521542, EPI_ISL_521543, EPI_ISL_521544, EPI_ISL_521545, EPI_ISL_521546, EPI_ISL_521547, EPI_ISL_521548, EPI_ISL_521549, EPI_ISL_521550, EPI_ISL_521551, EPI_ISL_521552, EPI_ISL_521553, EPI_ISL_521554, EPI_ISL_521555, EPI_ISL_521556, EPI_ISL_521557, EPI_ISL_521558, EPI_ISL_521559, EPI_ISL_521560, EPI_ISL_521561, EPI_ISL_521840, EPI_ISL_521841, EPI_ISL_521842, EPI_ISL_521843, EPI_ISL_521844, EPI_ISL_521845, EPI_ISL_521846, EPI_ISL_521847, EPI_ISL_521848, EPI_ISL_521849, EPI_ISL_521850, EPI_ISL_521851, EPI_ISL_521852, EPI_ISL_521853, EPI_ISL_521854, EPI_ISL_521855, EPI_ISL_521856, EPI_ISL_521857, EPI_ISL_521858, EPI_ISL_521859, EPI_ISL_521864, EPI_ISL_521865, EPI_ISL_521866, EPI_ISL_521867, EPI_ISL_521868, EPI_ISL_521870, EPI_ISL_521871, EPI_ISL_521872, EPI_ISL_521873, EPI_ISL_521874, EPI_ISL_521875, EPI_ISL_521876, EPI_ISL_521877, EPI_ISL_521878, EPI_ISL_521879, EPI_ISL_521880, EPI_ISL_521881, EPI_ISL_521882, EPI_ISL_521883, EPI_ISL_521884, EPI_ISL_521885, EPI_ISL_521886, EPI_ISL_521889, EPI_ISL_521890, EPI_ISL_521891, EPI_ISL_521892, EPI_ISL_521893, EPI_ISL_521894, EPI_ISL_521895, EPI_ISL_521896, EPI_ISL_521897, EPI_ISL_521898, EPI_ISL_521899, EPI_ISL_521900, EPI_ISL_521901, EPI_ISL_521902, EPI_ISL_521903, EPI_ISL_521904, EPI_ISL_521905, EPI_ISL_521912, EPI_ISL_521913, EPI_ISL_521914, EPI_ISL_521915, EPI_ISL_521916, EPI_ISL_521917, EPI_ISL_521918, EPI_ISL_521919, EPI_ISL_521920, EPI_ISL_521921, EPI_ISL_521922, EPI_ISL_521924, EPI_ISL_521925, EPI_ISL_521945, EPI_ISL_521946, EPI_ISL_521947, EPI_ISL_521948, EPI_ISL_521949, EPI_ISL_521950, EPI_ISL_521951, EPI_ISL_521952, EPI_ISL_521953, EPI_ISL_521954, EPI_ISL_521955, EPI_ISL_521956, EPI_ISL_521957, EPI_ISL_521958, EPI_ISL_521959, EPI_ISL_521960, EPI_ISL_521961, EPI_ISL_521963, EPI_ISL_521964, EPI_ISL_521965, EPI_ISL_521966, EPI_ISL_521967, EPI_ISL_521968, EPI_ISL_521969, EPI_ISL_521970, EPI_ISL_521971, EPI_ISL_521975, EPI_ISL_521976, EPI_ISL_521977, EPI_ISL_521978, EPI_ISL_521980, EPI_ISL_521981, EPI_ISL_521982, EPI_ISL_521983, EPI_ISL_521984, EPI_ISL_521985, EPI_ISL_521986, EPI_ISL_521987, EPI_ISL_521988, EPI_ISL_522055, EPI_ISL_522056, EPI_ISL_522057, EPI_ISL_522058, EPI_ISL_522059, EPI_ISL_522060, EPI_ISL_522061, EPI_ISL_522062, EPI_ISL_522063, EPI_ISL_522064, EPI_ISL_522065, EPI_ISL_522066, EPI_ISL_522067, EPI_ISL_522068, EPI_ISL_522069, EPI_ISL_522070, EPI_ISL_522071, EPI_ISL_522072, EPI_ISL_522073, EPI_ISL_522074, EPI_ISL_522075, EPI_ISL_522076, EPI_ISL_522077, EPI_ISL_522078, EPI_ISL_522079, EPI_ISL_522080, EPI_ISL_522081, EPI_ISL_522082, EPI_ISL_522083, EPI_ISL_522084, EPI_ISL_522085, EPI_ISL_522086, EPI_ISL_522087, EPI_ISL_522088, EPI_ISL_522089, EPI_ISL_522091, EPI_ISL_522092, EPI_ISL_522093, EPI_ISL_522094, EPI_ISL_522095, EPI_ISL_522097, EPI_ISL_522098, EPI_ISL_522099, EPI_ISL_522100, EPI_ISL_522101, EPI_ISL_522102, EPI_ISL_522103, EPI_ISL_522104, EPI_ISL_522105, EPI_ISL_522106, EPI_ISL_522107, EPI_ISL_522108, EPI_ISL_522109, EPI_ISL_522110, EPI_ISL_522119, EPI_ISL_522120, EPI_ISL_522121, EPI_ISL_522122, EPI_ISL_522123, EPI_ISL_522124, EPI_ISL_522125, EPI_ISL_522126, EPI_ISL_522127, EPI_ISL_522128, EPI_ISL_522129, EPI_ISL_522130, EPI_ISL_522131, EPI_ISL_522132, EPI_ISL_522133, EPI_ISL_522134, EPI_ISL_522135, EPI_ISL_522136, EPI_ISL_522137, EPI_ISL_522138, EPI_ISL_522139, EPI_ISL_522140, EPI_ISL_522141, EPI_ISL_522142, EPI_ISL_522143, EPI_ISL_522144, EPI_ISL_522145, EPI_ISL_522146, EPI_ISL_522147, EPI_ISL_522148, EPI_ISL_522149, EPI_ISL_522150, EPI_ISL_522151, EPI_ISL_522152, EPI_ISL_522153, EPI_ISL_522154, EPI_ISL_522155, EPI_ISL_522156, EPI_ISL_522157, EPI_ISL_522158, EPI_ISL_522159, EPI_ISL_522160, EPI_ISL_522161, EPI_ISL_522162, EPI_ISL_522163, EPI_ISL_522164, EPI_ISL_522165, EPI_ISL_522166, EPI_ISL_522167, EPI_ISL_522168, EPI_ISL_522169, EPI_ISL_522170, EPI_ISL_522171, EPI_ISL_522172, EPI_ISL_522173, EPI_ISL_522174, EPI_ISL_522175, EPI_ISL_522176, EPI_ISL_522177, EPI_ISL_522178, EPI_ISL_522179, EPI_ISL_522180, EPI_ISL_522181, EPI_ISL_522182, EPI_ISL_522183, EPI_ISL_522184, EPI_ISL_522185, EPI_ISL_522186, EPI_ISL_522187, EPI_ISL_522233, EPI_ISL_522234, EPI_ISL_522235, EPI_ISL_522236, EPI_ISL_522237, EPI_ISL_522238, EPI_ISL_522239, EPI_ISL_522240, EPI_ISL_522241, EPI_ISL_522242 | | | |
| see above | Victorian Infectious Diseases Reference Laboratory (VIDRL) | VIDRL and MDU-PHL | Caly L., Seemann T., Sait, M., Schultz M., Druce J., Sherry, N. |
| EPI_ISL_523810, EPI_ISL_523954, EPI_ISL_523979, EPI_ISL_523994, EPI_ISL_524471, EPI_ISL_524473, EPI_ISL_525206 | Laboratorio de Referencia Nacional de Virus Respiratorio. Centro Nacional de Salud Publica. Instituto Nacional de Salud Peru. | Laboratorio de Referencia Nacional de Biotecnología y Biología Molecular. Centro Nacional de Salud Publica. Instituto Nacional de Salud Peru. | Carlos Padilla Rojas, Karolyn Vega Chozo, Priscila Lope Pari, Omar Caceres Rey, Marco Galarza Perez, Maribel Huaringa Nuñez, Johanna Balbuena Torres, Henri Bailon Calderon, Nancy Rojas Serrano. |
| EPI_ISL_525802 | OHSU Lab Services Molecular Microbiology Lab | Oregon SARS-CoV-2 Genome Sequencing Center | Brendan L. O'Connell, Ruth V. Nichols, Alec J. Hirsch, Guang Fan, Daniel N. Streblow, William B. Messer, Andrew C. Adey, Benjamin N. Bimber, Brian J. O'Roak |
| EPI_ISL_525803 | OHSU Lab Services Molecular Microbiology Lab | Ginkgo Bioworks Clinical Laboratory | Brendan L. O'Connell, Ruth V. Nichols, Alec J. Hirsch, Guang Fan, Daniel N. Streblow, Malaika Mckenzie-Bennett, James McGann, Jim Griffin, Keith Robison, Alex Plocik, Becky Schilling, Rebecca Littlefield, Michelle Spencer, Birgitte Simen, William B. Messer, Andrew C. Adey, Benjamin N. Bimber, Brian J. O'Roak |
| EPI_ISL_525804, EPI_ISL_525805, EPI_ISL_525806, EPI_ISL_525807 | OHSU Lab Services Molecular Microbiology Lab | Oregon SARS-CoV-2 Genome Sequencing Center | Brendan L. O'Connell, Ruth V. Nichols, Alec J. Hirsch, Guang Fan, Daniel N. Streblow, William B. Messer, Andrew C. Adey, Benjamin N. Bimber, Brian J. O'Roak |
| EPI_ISL_525808, EPI_ISL_525809, EPI_ISL_525810, EPI_ISL_525811, EPI_ISL_525812, EPI_ISL_525813, EPI_ISL_525814, EPI_ISL_525815, EPI_ISL_525816, EPI_ISL_525817, EPI_ISL_525818, EPI_ISL_525820, EPI_ISL_525821, EPI_ISL_525822, EPI_ISL_525823, EPI_ISL_525824 | | | |
| see above | OHSU Lab Services Molecular Microbiology Lab | Ginkgo Bioworks Clinical Laboratory | Brendan L. O'Connell, Ruth V. Nichols, Alec J. Hirsch, Guang Fan, Daniel N. Streblow, Malaika Mckenzie-Bennett, James McGann, Jim Griffin, Keith Robison, Alex Plocik, Becky Schilling, Rebecca Littlefield, Michelle Spencer, Birgitte Simen, William B. Messer, Andrew C. Adey, Benjamin N. Bimber, Brian J. O'Roak |
| EPI_ISL_525825 | OHSU Lab Services Molecular Microbiology Lab | Oregon SARS-CoV-2 Genome Sequencing Center | Brendan L. O'Connell, Ruth V. Nichols, Alec J. Hirsch, Guang Fan, Daniel N. Streblow, William B. Messer, Andrew C. Adey, Benjamin N. Bimber, Brian J. O'Roak |
| EPI_ISL_525826 | OHSU Lab Services Molecular Microbiology Lab | Ginkgo Bioworks Clinical Laboratory | Brendan L. O'Connell, Ruth V. Nichols, Alec J. Hirsch, Guang Fan, Daniel N. Streblow, Malaika Mckenzie-Bennett, James McGann, Jim Griffin, Keith Robison, Alex Plocik, Becky Schilling, Rebecca Littlefield, Michelle Spencer, Birgitte Simen, William B. Messer, Andrew C. Adey, Benjamin N. Bimber, Brian J. O'Roak |
| EPI_ISL_525827 | OHSU Lab Services Molecular Microbiology Lab | Oregon SARS-CoV-2 Genome Sequencing Center | Brendan L. O'Connell, Ruth V. Nichols, Alec J. Hirsch, Guang Fan, Daniel N. Streblow, William B. Messer, Andrew C. Adey, Benjamin N. Bimber, Brian J. O'Roak |
| EPI_ISL_525828, EPI_ISL_525829, EPI_ISL_525830, EPI_ISL_525831, EPI_ISL_525832 | OHSU Lab Services Molecular Microbiology Lab | Ginkgo Bioworks Clinical Laboratory | Brendan L. O'Connell, Ruth V. Nichols, Alec J. Hirsch, Guang Fan, Daniel N. Streblow, Malaika Mckenzie-Bennett, James McGann, Jim Griffin, Keith Robison, Alex Plocik, Becky Schilling, Rebecca Littlefield, Michelle Spencer, Birgitte Simen, William B. Messer, Andrew C. Adey, Benjamin N. Bimber, Brian J. O'Roak |
| EPI_ISL_525833 | OHSU Lab Services Molecular Microbiology Lab | Oregon SARS-CoV-2 Genome Sequencing Center | Brendan L. O'Connell, Ruth V. Nichols, Alec J. Hirsch, Guang Fan, Daniel N. Streblow, William B. Messer, Andrew C. Adey, Benjamin N. Bimber, Brian J. O'Roak |
| EPI_ISL_525834, EPI_ISL_525835, EPI_ISL_525836, EPI_ISL_525837, EPI_ISL_525838, EPI_ISL_525839, EPI_ISL_525840 | OHSU Lab Services Molecular Microbiology Lab | Ginkgo Bioworks Clinical Laboratory | Brendan L. O'Connell, Ruth V. Nichols, Alec J. Hirsch, Guang Fan, Daniel N. Streblow, Malaika Mckenzie-Bennett, James McGann, Jim Griffin, Keith Robison, Alex Plocik, Becky Schilling, Rebecca Littlefield, Michelle Spencer, Birgitte Simen, William B. Messer, Andrew C. Adey, Benjamin N. Bimber, Brian J. O'Roak |
| EPI_ISL_525841 | OHSU Lab Services Molecular Microbiology Lab | Oregon SARS-CoV-2 Genome Sequencing Center | Brendan L. O'Connell, Ruth V. Nichols, Alec J. Hirsch, Guang Fan, Daniel N. Streblow, William B. Messer, Andrew C. Adey, Benjamin N. Bimber, Brian J. O'Roak |
| EPI_ISL_525842, EPI_ISL_525843, EPI_ISL_525844, EPI_ISL_525845, EPI_ISL_525846 | OHSU Lab Services Molecular Microbiology Lab | Ginkgo Bioworks Clinical Laboratory | Brendan L. O'Connell, Ruth V. Nichols, Alec J. Hirsch, Guang Fan, Daniel N. Streblow, Malaika Mckenzie-Bennett, James McGann, Jim Griffin, Keith Robison, Alex Plocik, Becky Schilling, Rebecca Littlefield, Michelle Spencer, Birgitte Simen, William B. Messer, Andrew C. Adey, Benjamin N. Bimber, Brian J. O'Roak |
| EPI_ISL_525847 | OHSU Lab Services Molecular Microbiology Lab | Oregon SARS-CoV-2 Genome Sequencing Center | Brendan L. O'Connell, Ruth V. Nichols, Alec J. Hirsch, Guang Fan, Daniel N. Streblow, William B. Messer, Andrew C. Adey, Benjamin N. Bimber, Brian J. O'Roak |
| EPI_ISL_525848 | OHSU Lab Services Molecular Microbiology Lab | Ginkgo Bioworks Clinical Laboratory | Brendan L. O'Connell, Ruth V. Nichols, Alec J. Hirsch, Guang Fan, Daniel N. Streblow, Malaika Mckenzie-Bennett, James McGann, Jim Griffin, Keith Robison, Alex Plocik, Becky Schilling, Rebecca Littlefield, Michelle Spencer, Birgitte Simen, William B. Messer, Andrew C. Adey, Benjamin N. Bimber, Brian J. O'Roak |
| EPI_ISL_525849 | OHSU Lab Services Molecular Microbiology Lab | Oregon SARS-CoV-2 Genome Sequencing Center | Brendan L. O'Connell, Ruth V. Nichols, Alec J. Hirsch, Guang Fan, Daniel N. Streblow, William B. Messer, Andrew C. Adey, Benjamin N. Bimber, Brian J. O'Roak |
| EPI_ISL_525850 | OHSU Lab Services Molecular Microbiology Lab | Ginkgo Bioworks Clinical Laboratory | Brendan L. O'Connell, Ruth V. Nichols, Alec J. Hirsch, Guang Fan, Daniel N. Streblow, Malaika Mckenzie-Bennett, James McGann, Jim Griffin, Keith Robison, Alex Plocik, Becky Schilling, Rebecca Littlefield, Michelle Spencer, Birgitte Simen, William B. Messer, Andrew C. Adey, Benjamin N. Bimber, Brian J. O'Roak |
| EPI_ISL_525851 | OHSU Lab Services Molecular Microbiology Lab | Oregon SARS-CoV-2 Genome Sequencing Center | Brendan L. O'Connell, Ruth V. Nichols, Alec J. Hirsch, Guang Fan, Daniel N. Streblow, William B. Messer, Andrew C. Adey, Benjamin N. Bimber, Brian J. O'Roak |
| EPI_ISL_525852, EPI_ISL_525853, | OHSU Lab Services Molecular Microbiology Lab | Ginkgo Bioworks Clinical Laboratory | Brendan L. O'Connell, Ruth V. Nichols, Alec J. Hirsch, Guang Fan, Daniel N. Streblow, Malaika Mckenzie-Bennett, James McGann, Jim Griffin, Keith |

| | | | |
|---|---|---|---|
| EPI_ISL_525854, EPI_ISL_525855 | | | Robison, Alex Plocik, Becky Schilling, Rebecca Littlefield, Michelle Spencer, Birgitte Simen, William B. Messer, Andrew C. Adey, Benjamin N. Bimber, Brian J. O'Roak |
| EPI_ISL_525856, EPI_ISL_525857, EPI_ISL_525858 | OHSU Lab Services Molecular Microbiology Lab | Oregon SARS-CoV-2 Genome Sequencing Center | Brendan L. O'Connell, Ruth V. Nichols, Alec J. Hirsch, Guang Fan, Daniel N. Streblow, William B. Messer, Andrew C. Adey, Benjamin N. Bimber, Brian J. O'Roak |
| EPI_ISL_525859 | OHSU Lab Services Molecular Microbiology Lab | Ginkgo Bioworks Clinical Laboratory | Brendan L. O'Connell, Ruth V. Nichols, Alec J. Hirsch, Guang Fan, Daniel N. Streblow, Malaika Mckenzie-Bennett, James McGann, Jim Griffin, Keith Robison, Alex Plocik, Becky Schilling, Rebecca Littlefield, Michelle Spencer, Birgitte Simen, William B. Messer, Andrew C. Adey, Benjamin N. Bimber, Brian J. O'Roak |
| EPI_ISL_525860 | OHSU Lab Services Molecular Microbiology Lab | Oregon SARS-CoV-2 Genome Sequencing Center | Brendan L. O'Connell, Ruth V. Nichols, Alec J. Hirsch, Guang Fan, Daniel N. Streblow, William B. Messer, Andrew C. Adey, Benjamin N. Bimber, Brian J. O'Roak |
| EPI_ISL_525861 | OHSU Lab Services Molecular Microbiology Lab | Ginkgo Bioworks Clinical Laboratory | Brendan L. O'Connell, Ruth V. Nichols, Alec J. Hirsch, Guang Fan, Daniel N. Streblow, Malaika Mckenzie-Bennett, James McGann, Jim Griffin, Keith Robison, Alex Plocik, Becky Schilling, Rebecca Littlefield, Michelle Spencer, Birgitte Simen, William B. Messer, Andrew C. Adey, Benjamin N. Bimber, Brian J. O'Roak |
| EPI_ISL_525862 | OHSU Lab Services Molecular Microbiology Lab | Oregon SARS-CoV-2 Genome Sequencing Center | Brendan L. O'Connell, Ruth V. Nichols, Alec J. Hirsch, Guang Fan, Daniel N. Streblow, William B. Messer, Andrew C. Adey, Benjamin N. Bimber, Brian J. O'Roak |
| EPI_ISL_525863, EPI_ISL_525864 | OHSU Lab Services Molecular Microbiology Lab | Ginkgo Bioworks Clinical Laboratory | Brendan L. O'Connell, Ruth V. Nichols, Alec J. Hirsch, Guang Fan, Daniel N. Streblow, Malaika Mckenzie-Bennett, James McGann, Jim Griffin, Keith Robison, Alex Plocik, Becky Schilling, Rebecca Littlefield, Michelle Spencer, Birgitte Simen, William B. Messer, Andrew C. Adey, Benjamin N. Bimber, Brian J. O'Roak |
| EPI_ISL_525865 | OHSU Lab Services Molecular Microbiology Lab | Oregon SARS-CoV-2 Genome Sequencing Center | Brendan L. O'Connell, Ruth V. Nichols, Alec J. Hirsch, Guang Fan, Daniel N. Streblow, William B. Messer, Andrew C. Adey, Benjamin N. Bimber, Brian J. O'Roak |
| EPI_ISL_525866, EPI_ISL_525867 | OHSU Lab Services Molecular Microbiology Lab | Ginkgo Bioworks Clinical Laboratory | Brendan L. O'Connell, Ruth V. Nichols, Alec J. Hirsch, Guang Fan, Daniel N. Streblow, Malaika Mckenzie-Bennett, James McGann, Jim Griffin, Keith Robison, Alex Plocik, Becky Schilling, Rebecca Littlefield, Michelle Spencer, Birgitte Simen, William B. Messer, Andrew C. Adey, Benjamin N. Bimber, Brian J. O'Roak |
| EPI_ISL_525868 | OHSU Lab Services Molecular Microbiology Lab | Oregon SARS-CoV-2 Genome Sequencing Center | Brendan L. O'Connell, Ruth V. Nichols, Alec J. Hirsch, Guang Fan, Daniel N. Streblow, William B. Messer, Andrew C. Adey, Benjamin N. Bimber, Brian J. O'Roak |
| EPI_ISL_525869 | OHSU Lab Services Molecular Microbiology Lab | Ginkgo Bioworks Clinical Laboratory | Brendan L. O'Connell, Ruth V. Nichols, Alec J. Hirsch, Guang Fan, Daniel N. Streblow, Malaika Mckenzie-Bennett, James McGann, Jim Griffin, Keith Robison, Alex Plocik, Becky Schilling, Rebecca Littlefield, Michelle Spencer, Birgitte Simen, William B. Messer, Andrew C. Adey, Benjamin N. Bimber, Brian J. O'Roak |
| EPI_ISL_525870 | OHSU Lab Services Molecular Microbiology Lab | Oregon SARS-CoV-2 Genome Sequencing Center | Brendan L. O'Connell, Ruth V. Nichols, Alec J. Hirsch, Guang Fan, Daniel N. Streblow, William B. Messer, Andrew C. Adey, Benjamin N. Bimber, Brian J. O'Roak |
| EPI_ISL_525871, EPI_ISL_525872, EPI_ISL_525873 | OHSU Lab Services Molecular Microbiology Lab | Ginkgo Bioworks Clinical Laboratory | Brendan L. O'Connell, Ruth V. Nichols, Alec J. Hirsch, Guang Fan, Daniel N. Streblow, Malaika Mckenzie-Bennett, James McGann, Jim Griffin, Keith Robison, Alex Plocik, Becky Schilling, Rebecca Littlefield, Michelle Spencer, Birgitte Simen, William B. Messer, Andrew C. Adey, Benjamin N. Bimber, Brian J. O'Roak |
| EPI_ISL_525874 | OHSU Lab Services Molecular Microbiology Lab | Oregon SARS-CoV-2 Genome Sequencing Center | Brendan L. O'Connell, Ruth V. Nichols, Alec J. Hirsch, Guang Fan, Daniel N. Streblow, William B. Messer, Andrew C. Adey, Benjamin N. Bimber, Brian J. O'Roak |
| EPI_ISL_525875 | OHSU Lab Services Molecular Microbiology Lab | Ginkgo Bioworks Clinical Laboratory | Brendan L. O'Connell, Ruth V. Nichols, Alec J. Hirsch, Guang Fan, Daniel N. Streblow, Malaika Mckenzie-Bennett, James McGann, Jim Griffin, Keith Robison, Alex Plocik, Becky Schilling, Rebecca Littlefield, Michelle Spencer, Birgitte Simen, William B. Messer, Andrew C. Adey, Benjamin N. Bimber, Brian J. O'Roak |
| EPI_ISL_525876 | OHSU Lab Services Molecular Microbiology Lab | Oregon SARS-CoV-2 Genome Sequencing Center | Brendan L. O'Connell, Ruth V. Nichols, Alec J. Hirsch, Guang Fan, Daniel N. Streblow, William B. Messer, Andrew C. Adey, Benjamin N. Bimber, Brian J. O'Roak |
| EPI_ISL_525877 | OHSU Lab Services Molecular Microbiology Lab | Ginkgo Bioworks Clinical Laboratory | Brendan L. O'Connell, Ruth V. Nichols, Alec J. Hirsch, Guang Fan, Daniel N. Streblow, Malaika Mckenzie-Bennett, James McGann, Jim Griffin, Keith Robison, Alex Plocik, Becky Schilling, Rebecca Littlefield, Michelle Spencer, Birgitte Simen, William B. Messer, Andrew C. Adey, Benjamin N. Bimber, Brian J. O'Roak |
| EPI_ISL_525878 | OHSU Lab Services Molecular Microbiology Lab | Oregon SARS-CoV-2 Genome Sequencing Center | Brendan L. O'Connell, Ruth V. Nichols, Alec J. Hirsch, Guang Fan, Daniel N. Streblow, William B. Messer, Andrew C. Adey, Benjamin N. Bimber, Brian J. O'Roak |
| EPI_ISL_525879 | OHSU Lab Services Molecular Microbiology Lab | Ginkgo Bioworks Clinical Laboratory | Brendan L. O'Connell, Ruth V. Nichols, Alec J. Hirsch, Guang Fan, Daniel N. Streblow, Malaika Mckenzie-Bennett, James McGann, Jim Griffin, Keith Robison, Alex Plocik, Becky Schilling, Rebecca Littlefield, Michelle Spencer, Birgitte Simen, William B. Messer, Andrew C. Adey, Benjamin N. Bimber, Brian J. O'Roak |
| EPI_ISL_525880, EPI_ISL_525881 | OHSU Lab Services Molecular Microbiology Lab | Oregon SARS-CoV-2 Genome Sequencing Center | Brendan L. O'Connell, Ruth V. Nichols, Alec J. Hirsch, Guang Fan, Daniel N. Streblow, William B. Messer, Andrew C. Adey, Benjamin N. Bimber, Brian J. O'Roak |
| EPI_ISL_525882 | OHSU Lab Services Molecular Microbiology Lab | Ginkgo Bioworks Clinical Laboratory | Brendan L. O'Connell, Ruth V. Nichols, Alec J. Hirsch, Guang Fan, Daniel N. Streblow, Malaika Mckenzie-Bennett, James McGann, Jim Griffin, Keith Robison, Alex Plocik, Becky Schilling, Rebecca Littlefield, Michelle Spencer, Birgitte Simen, William B. Messer, Andrew C. Adey, Benjamin N. Bimber, Brian J. O'Roak |
| EPI_ISL_525883 | OHSU Lab Services Molecular Microbiology Lab | Oregon SARS-CoV-2 Genome Sequencing Center | Brendan L. O'Connell, Ruth V. Nichols, Alec J. Hirsch, Guang Fan, Daniel N. Streblow, William B. Messer, Andrew C. Adey, Benjamin N. Bimber, Brian J. O'Roak |
| EPI_ISL_525884 | OHSU Lab Services Molecular Microbiology Lab | Ginkgo Bioworks Clinical Laboratory | Brendan L. O'Connell, Ruth V. Nichols, Alec J. Hirsch, Guang Fan, Daniel N. Streblow, Malaika Mckenzie-Bennett, James McGann, Jim Griffin, Keith Robison, Alex Plocik, Becky Schilling, Rebecca Littlefield, Michelle Spencer, Birgitte Simen, William B. Messer, Andrew C. Adey, Benjamin N. Bimber, Brian J. O'Roak |
| EPI_ISL_525885 | OHSU Lab Services Molecular Microbiology Lab | Oregon SARS-CoV-2 Genome Sequencing Center | Brendan L. O'Connell, Ruth V. Nichols, Alec J. Hirsch, Guang Fan, Daniel N. Streblow, William B. Messer, Andrew C. Adey, Benjamin N. Bimber, Brian J. O'Roak |
| EPI_ISL_525886, EPI_ISL_525887, EPI_ISL_525888, EPI_ISL_525889, EPI_ISL_525890 | OHSU Lab Services Molecular Microbiology Lab | Ginkgo Bioworks Clinical Laboratory | Brendan L. O'Connell, Ruth V. Nichols, Alec J. Hirsch, Guang Fan, Daniel N. Streblow, Malaika Mckenzie-Bennett, James McGann, Jim Griffin, Keith Robison, Alex Plocik, Becky Schilling, Rebecca Littlefield, Michelle Spencer, Birgitte Simen, William B. Messer, Andrew C. Adey, Benjamin N. Bimber, Brian J. O'Roak |
| EPI_ISL_525891, EPI_ISL_525892, EPI_ISL_525893, EPI_ISL_525894, EPI_ISL_525895, EPI_ISL_525896, EPI_ISL_525897, EPI_ISL_525898, EPI_ISL_525899, EPI_ISL_525900, EPI_ISL_525901, EPI_ISL_525902, EPI_ISL_525903, EPI_ISL_525904, EPI_ISL_525905, EPI_ISL_525906, EPI_ISL_525907, EPI_ISL_525908, EPI_ISL_525909, EPI_ISL_525910, EPI_ISL_525911, EPI_ISL_525912, EPI_ISL_525913, EPI_ISL_525914, EPI_ISL_525915, EPI_ISL_525916, EPI_ISL_525917, EPI_ISL_525918, EPI_ISL_525919, EPI_ISL_525920, EPI_ISL_525921, EPI_ISL_525922, EPI_ISL_525923, EPI_ISL_525924, EPI_ISL_525925, EPI_ISL_525926, EPI_ISL_525927, EPI_ISL_525928, EPI_ISL_525929, EPI_ISL_525930, EPI_ISL_525931, EPI_ISL_525932, EPI_ISL_525933, EPI_ISL_525934, EPI_ISL_525935, EPI_ISL_525936, EPI_ISL_525937, EPI_ISL_525938, EPI_ISL_525939, EPI_ISL_525940, EPI_ISL_525941, EPI_ISL_525942, EPI_ISL_525943, EPI_ISL_525944, EPI_ISL_525945, EPI_ISL_525946, EPI_ISL_525947, EPI_ISL_525948, EPI_ISL_525949, EPI_ISL_525950, EPI_ISL_525951, EPI_ISL_525952, EPI_ISL_525953, EPI_ISL_525954, EPI_ISL_525955, EPI_ISL_525956, EPI_ISL_525957, EPI_ISL_525958, EPI_ISL_525959, EPI_ISL_525960, EPI_ISL_525961, EPI_ISL_525962, EPI_ISL_525963, EPI_ISL_525964, EPI_ISL_525965, EPI_ISL_525966, EPI_ISL_525967, EPI_ISL_525968, EPI_ISL_525969, EPI_ISL_525970, EPI_ISL_525971, EPI_ISL_525972, EPI_ISL_525973, EPI_ISL_525974, EPI_ISL_525975, EPI_ISL_525976, EPI_ISL_525977, EPI_ISL_525978, EPI_ISL_525979, EPI_ISL_525980, EPI_ISL_525981, EPI_ISL_525982, EPI_ISL_525983, EPI_ISL_525984, EPI_ISL_525985, EPI_ISL_525986, EPI_ISL_525987, EPI_ISL_525988, EPI_ISL_525989, EPI_ISL_525990, EPI_ISL_525991, EPI_ISL_525992, EPI_ISL_525993, EPI_ISL_525994, EPI_ISL_525995, EPI_ISL_525996, EPI_ISL_525997, EPI_ISL_525998, EPI_ISL_525999, EPI_ISL_526000, EPI_ISL_526001, EPI_ISL_526002, EPI_ISL_526003, EPI_ISL_526004, EPI_ISL_526005, EPI_ISL_526006, EPI_ISL_526007, EPI_ISL_526008, EPI_ISL_526009, EPI_ISL_526010, EPI_ISL_526011, EPI_ISL_526012, EPI_ISL_526013, EPI_ISL_526014, EPI_ISL_526015, EPI_ISL_526016, EPI_ISL_526017, EPI_ISL_526018, EPI_ISL_526019, EPI_ISL_526020, EPI_ISL_526021, EPI_ISL_526022, EPI_ISL_526023, EPI_ISL_526024, EPI_ISL_526025, EPI_ISL_526026, EPI_ISL_526027, EPI_ISL_526028, EPI_ISL_526029, EPI_ISL_526030, EPI_ISL_526031, EPI_ISL_526032, EPI_ISL_526033, EPI_ISL_526034, EPI_ISL_526035, EPI_ISL_526036, EPI_ISL_526037, EPI_ISL_526038, EPI_ISL_526039, EPI_ISL_526040, EPI_ISL_526041, EPI_ISL_526042, EPI_ISL_526043, EPI_ISL_526044, EPI_ISL_526045, EPI_ISL_526046, EPI_ISL_526047, EPI_ISL_526048, EPI_ISL_526049, EPI_ISL_526050, EPI_ISL_526051, EPI_ISL_526052, EPI_ISL_526053, EPI_ISL_526054, EPI_ISL_526055, EPI_ISL_526056, EPI_ISL_526057, EPI_ISL_526058, EPI_ISL_526059, EPI_ISL_526060, EPI_ISL_526061, EPI_ISL_526062, EPI_ISL_526063, EPI_ISL_526064, EPI_ISL_526065, EPI_ISL_526066, EPI_ISL_526067, EPI_ISL_526068, EPI_ISL_526069, EPI_ISL_526070, EPI_ISL_526071, EPI_ISL_526072, EPI_ISL_526073, EPI_ISL_526074, EPI_ISL_526075, EPI_ISL_526076, EPI_ISL_526077, EPI_ISL_526078, EPI_ISL_526079, EPI_ISL_526080, EPI_ISL_526081, EPI_ISL_526082, EPI_ISL_526083, EPI_ISL_526084, EPI_ISL_526085, EPI_ISL_526086 | | | |
| see above | OHSU Lab Services Molecular Microbiology Lab | Oregon SARS-CoV-2 Genome Sequencing Center | Brendan L. O'Connell, Ruth V. Nichols, Alec J. Hirsch, Guang Fan, Daniel N. Streblow, William B. Messer, Andrew C. Adey, Benjamin N. Bimber, Brian J. |

| | | | |
|---|---|---|---|
| | | | O'Roak |
| EPI_ISL_526087, EPI_ISL_526088, EPI_ISL_526089, EPI_ISL_526090 | OHSU Lab Services Molecular Microbiology Lab | Ginkgo Bioworks Clinical Laboratory | Brendan L. O'Connell, Ruth V. Nichols, Alec J. Hirsch, Guang Fan, Daniel N. Streblow, Malaika Mckenzie-Bennett, James McGann, Jim Griffin, Keith Robison, Alex Plocik, Becky Schilling, Rebecca Littlefield, Michelle Spencer, Birgitte Simen, William B. Messer, Andrew C. Adey, Benjamin N. Bimber, Brian J. O'Roak |
| EPI_ISL_526091 | OHSU Lab Services Molecular Microbiology Lab | Oregon SARS-CoV-2 Genome Sequencing Center | Brendan L. O'Connell, Ruth V. Nichols, Alec J. Hirsch, Guang Fan, Daniel N. Streblow, William B. Messer, Andrew C. Adey, Benjamin N. Bimber, Brian J. O'Roak |
| EPI_ISL_526092, EPI_ISL_526093 | OHSU Lab Services Molecular Microbiology Lab | Ginkgo Bioworks Clinical Laboratory | Brendan L. O'Connell, Ruth V. Nichols, Alec J. Hirsch, Guang Fan, Daniel N. Streblow, Malaika Mckenzie-Bennett, James McGann, Jim Griffin, Keith Robison, Alex Plocik, Becky Schilling, Rebecca Littlefield, Michelle Spencer, Birgitte Simen, William B. Messer, Andrew C. Adey, Benjamin N. Bimber, Brian J. O'Roak |
| EPI_ISL_526094, EPI_ISL_526095, EPI_ISL_526096 | OHSU Lab Services Molecular Microbiology Lab | Oregon SARS-CoV-2 Genome Sequencing Center | Brendan L. O'Connell, Ruth V. Nichols, Alec J. Hirsch, Guang Fan, Daniel N. Streblow, William B. Messer, Andrew C. Adey, Benjamin N. Bimber, Brian J. O'Roak |
| EPI_ISL_526097 | OHSU Lab Services Molecular Microbiology Lab | Ginkgo Bioworks Clinical Laboratory | Brendan L. O'Connell, Ruth V. Nichols, Alec J. Hirsch, Guang Fan, Daniel N. Streblow, Malaika Mckenzie-Bennett, James McGann, Jim Griffin, Keith Robison, Alex Plocik, Becky Schilling, Rebecca Littlefield, Michelle Spencer, Birgitte Simen, William B. Messer, Andrew C. Adey, Benjamin N. Bimber, |
| EPI_ISL_526098, EPI_ISL_526099, EPI_ISL_526100, EPI_ISL_526101, EPI_ISL_526102, EPI_ISL_526103, EPI_ISL_526104, EPI_ISL_526105, EPI_ISL_526106, EPI_ISL_526107, EPI_ISL_526108, EPI_ISL_526109, EPI_ISL_526110, EPI_ISL_526111, EPI_ISL_526112, EPI_ISL_526113, EPI_ISL_526114 | | | |
| see above | OHSU Lab Services Molecular Microbiology Lab | Oregon SARS-CoV-2 Genome Sequencing Center | Brendan L. O'Connell, Ruth V. Nichols, Alec J. Hirsch, Guang Fan, Daniel N. Streblow, William B. Messer, Andrew C. Adey, Benjamin N. Bimber, Brian J. O'Roak |
| EPI_ISL_526115, EPI_ISL_526116, EPI_ISL_526117, EPI_ISL_526118 | Pathology West - NSW Health Pathology | NSW Health Pathology - Institute of Clinical Pathology and Medical Research; Westmead Hospital; University of Sydney | CIDM-PH et al. |
| EPI_ISL_526119, EPI_ISL_526120 | Sydney South West Pathology Service (SSWPS) - Liverpool Hospital - NSW Health Pathology | NSW Health Pathology - Institute of Clinical Pathology and Medical Research; Westmead Hospital; University of Sydney | CIDM-PH et al. |
| EPI_ISL_526121, EPI_ISL_526122, EPI_ISL_526123 | St Vincent's Pathology (SydPath) | NSW Health Pathology - Institute of Clinical Pathology and Medical Research; Westmead Hospital; University of Sydney | CIDM-PH et al. |
| EPI_ISL_526124 | Sydney South West Pathology Service (SSWPS) - Liverpool Hospital - NSW Health Pathology | NSW Health Pathology - Institute of Clinical Pathology and Medical Research; Westmead Hospital; University of Sydney | CIDM-PH et al. |
| EPI_ISL_526125 | Sydney South West Pathology Service (SSWPS) - Concord Repatriation General Hospital - NSW Health Pathology | NSW Health Pathology - Institute of Clinical Pathology and Medical Research; Westmead Hospital; University of Sydney | CIDM-PH et al. |
| EPI_ISL_526126, EPI_ISL_526127 | Sydney South West Pathology Service (SSWPS) - Liverpool Hospital - NSW Health Pathology | NSW Health Pathology - Institute of Clinical Pathology and Medical Research; Westmead Hospital; University of Sydney | CIDM-PH et al. |
| EPI_ISL_526128 | Pathology North - Royal North Shore Hospital - NSW Health Pathology | NSW Health Pathology - Institute of Clinical Pathology and Medical Research; Westmead Hospital; University of Sydney | CIDM-PH et al. |
| EPI_ISL_526129 | The Children's Hospital at Westmead | NSW Health Pathology - Institute of Clinical Pathology and Medical Research; Westmead Hospital; University of Sydney | CIDM-PH et al. |
| EPI_ISL_526130 | Pathology North - Royal North Shore Hospital - NSW Health Pathology | NSW Health Pathology - Institute of Clinical Pathology and Medical Research; Westmead Hospital; University of Sydney | CIDM-PH et al. |
| EPI_ISL_526131 | Sydney South West Pathology Service (SSWPS) - Liverpool Hospital - NSW Health Pathology | NSW Health Pathology - Institute of Clinical Pathology and Medical Research; Westmead Hospital; University of Sydney | CIDM-PH et al. |
| EPI_ISL_526132, EPI_ISL_526133 | Pathology North - Royal North Shore Hospital - NSW Health Pathology | NSW Health Pathology - Institute of Clinical Pathology and Medical Research; Westmead Hospital; University of Sydney | CIDM-PH et al. |
| EPI_ISL_526134, EPI_ISL_526135, EPI_ISL_526136, EPI_ISL_526137 | Sydney South West Pathology Service (SSWPS) - Royal Prince Alfred Hospital - NSW Health Pathology | NSW Health Pathology - Institute of Clinical Pathology and Medical Research; Westmead Hospital; University of Sydney | CIDM-PH et al. |
| EPI_ISL_526138, EPI_ISL_526139, EPI_ISL_526140, EPI_ISL_526141 | Pathology West - NSW Health Pathology | NSW Health Pathology - Institute of Clinical Pathology and Medical Research; Westmead Hospital; University of Sydney | CIDM-PH et al. |
| EPI_ISL_526142 | South Eastern Area Laboratory Services (SEALS) | NSW Health Pathology - Institute of Clinical Pathology and Medical Research; Westmead Hospital; University of Sydney | CIDM-PH et al. |
| EPI_ISL_526143 | Laverty Pathology | NSW Health Pathology - Institute of Clinical Pathology and Medical Research; Westmead Hospital; University of Sydney | CIDM-PH et al. |
| EPI_ISL_526144, EPI_ISL_526145, EPI_ISL_526146, EPI_ISL_526147, EPI_ISL_526148, EPI_ISL_526149 | South Eastern Area Laboratory Services (SEALS) | NSW Health Pathology - Institute of Clinical Pathology and Medical Research; Westmead Hospital; University of Sydney | CIDM-PH et al. |
| EPI_ISL_526150, EPI_ISL_526151 | Pathology West - NSW Health Pathology | NSW Health Pathology - Institute of Clinical Pathology and Medical Research; Westmead Hospital; University of Sydney | CIDM-PH et al. |
| EPI_ISL_526152 | Pathology North - Hunter - NSW Health Pathology | NSW Health Pathology - Institute of Clinical Pathology and Medical Research; Westmead Hospital; University of Sydney | CIDM-PH et al. |
| EPI_ISL_526153 | Douglass Hanly Moir Pathology | NSW Health Pathology - Institute of Clinical Pathology and Medical Research; Westmead Hospital; University of Sydney | CIDM-PH et al. |
| EPI_ISL_526154, EPI_ISL_526155 | Pathology North - Royal North Shore Hospital - NSW Health Pathology | NSW Health Pathology - Institute of Clinical Pathology and Medical Research; Westmead Hospital; University of Sydney | CIDM-PH et al. |
| EPI_ISL_526156 | Pathology West - NSW Health Pathology | NSW Health Pathology - Institute of Clinical Pathology and Medical Research; Westmead Hospital; University of Sydney | CIDM-PH et al. |
| EPI_ISL_526157, EPI_ISL_526158, EPI_ISL_526159, EPI_ISL_526160 | Sydney South West Pathology Service (SSWPS) - Liverpool Hospital - NSW Health Pathology | NSW Health Pathology - Institute of Clinical Pathology and Medical Research; Westmead Hospital; University of Sydney | CIDM-PH et al. |
| EPI_ISL_526161, EPI_ISL_526162, EPI_ISL_526163, EPI_ISL_526164, EPI_ISL_526165 | Pathology West - NSW Health Pathology | NSW Health Pathology - Institute of Clinical Pathology and Medical Research; Westmead Hospital; University of Sydney | CIDM-PH et al. |
| EPI_ISL_526166, EPI_ISL_526167, EPI_ISL_526168, EPI_ISL_526169 | Austech Medical Laboratories | NSW Health Pathology - Institute of Clinical Pathology and Medical Research; Westmead Hospital; University of Sydney | CIDM-PH et al. |
| EPI_ISL_526170 | Pathology North - Royal North Shore Hospital - NSW Health Pathology | NSW Health Pathology - Institute of Clinical Pathology and Medical Research; Westmead Hospital; University of Sydney | CIDM-PH et al. |
| EPI_ISL_526171 | St Vincent's Pathology (SydPath) | NSW Health Pathology - Institute of Clinical Pathology and Medical Research; Westmead Hospital; University of Sydney | CIDM-PH et al. |

| | | | |
|---|---|---|---|
| EPI_ISL_526172, EPI_ISL_526173, EPI_ISL_526174, EPI_ISL_526175, EPI_ISL_526176 | Sydney South West Pathology Service (SSWPS) - Liverpool Hospital - NSW Health Pathology | NSW Health Pathology - Institute of Clinical Pathology and Medical Research; Westmead Hospital; University of Sydney | CIDM-PH et al. |
| EPI_ISL_526177, EPI_ISL_526178, EPI_ISL_526179, EPI_ISL_526180, EPI_ISL_526181, EPI_ISL_526182, EPI_ISL_526183, EPI_ISL_526184, EPI_ISL_526185 | St Vincent's Pathology (SydPath) | NSW Health Pathology - Institute of Clinical Pathology and Medical Research; Westmead Hospital; University of Sydney | CIDM-PH et al. |
| EPI_ISL_526186 | 4Cyte Pathology | NSW Health Pathology - Institute of Clinical Pathology and Medical Research; Westmead Hospital; University of Sydney | CIDM-PH et al. |
| EPI_ISL_526187, EPI_ISL_526188, EPI_ISL_526189, EPI_ISL_526190 | St Vincent's Pathology (SydPath) | NSW Health Pathology - Institute of Clinical Pathology and Medical Research; Westmead Hospital; University of Sydney | CIDM-PH et al. |
| EPI_ISL_526191 | South Eastern Area Laboratory Services (SEALS) | NSW Health Pathology - Institute of Clinical Pathology and Medical Research; Westmead Hospital; University of Sydney | CIDM-PH et al. |
| EPI_ISL_526192 | Medlab Pathology | NSW Health Pathology - Institute of Clinical Pathology and Medical Research; Westmead Hospital; University of Sydney | CIDM-PH et al. |
| EPI_ISL_526193 | Sydney South West Pathology Service (SSWPS) - Liverpool Hospital - NSW Health Pathology | NSW Health Pathology - Institute of Clinical Pathology and Medical Research; Westmead Hospital; University of Sydney | CIDM-PH et al. |
| EPI_ISL_526194, EPI_ISL_526195 | St Vincent's Pathology (SydPath) | NSW Health Pathology - Institute of Clinical Pathology and Medical Research; Westmead Hospital; University of Sydney | CIDM-PH et al. |
| EPI_ISL_526196 | Sydney South West Pathology Service (SSWPS) - Royal Prince Alfred Hospital - NSW Health Pathology | NSW Health Pathology - Institute of Clinical Pathology and Medical Research; Westmead Hospital; University of Sydney | CIDM-PH et al. |
| EPI_ISL_526197, EPI_ISL_526198 | Sydney South West Pathology Service (SSWPS) - Liverpool Hospital - NSW Health Pathology | NSW Health Pathology - Institute of Clinical Pathology and Medical Research; Westmead Hospital; University of Sydney | CIDM-PH et al. |
| EPI_ISL_526199 | Pathology West - NSW Health Pathology | NSW Health Pathology - Institute of Clinical Pathology and Medical Research; Westmead Hospital; University of Sydney | CIDM-PH et al. |
| EPI_ISL_526200 | St Vincent's Pathology (SydPath) | NSW Health Pathology - Institute of Clinical Pathology and Medical Research; Westmead Hospital; University of Sydney | CIDM-PH et al. |
| EPI_ISL_526201 | Sydney South West Pathology Service (SSWPS) - Royal Prince Alfred Hospital - NSW Health Pathology | NSW Health Pathology - Institute of Clinical Pathology and Medical Research; Westmead Hospital; University of Sydney | CIDM-PH et al. |
| EPI_ISL_526202, EPI_ISL_526203, EPI_ISL_526204 | Laverty Pathology | NSW Health Pathology - Institute of Clinical Pathology and Medical Research; Westmead Hospital; University of Sydney | CIDM-PH et al. |
| EPI_ISL_526205 | The Children's Hospital at Westmead | NSW Health Pathology - Institute of Clinical Pathology and Medical Research; Westmead Hospital; University of Sydney | CIDM-PH et al. |
| EPI_ISL_526206 | Sydney South West Pathology Service (SSWPS) - Liverpool Hospital - NSW Health Pathology | NSW Health Pathology - Institute of Clinical Pathology and Medical Research; Westmead Hospital; University of Sydney | CIDM-PH et al. |
| EPI_ISL_526207, EPI_ISL_526208, EPI_ISL_526209 | Sydney South West Pathology Service (SSWPS) - Royal Prince Alfred Hospital - NSW Health Pathology | NSW Health Pathology - Institute of Clinical Pathology and Medical Research; Westmead Hospital; University of Sydney | CIDM-PH et al. |
| EPI_ISL_526210, EPI_ISL_526211 | Sydney South West Pathology Service (SSWPS) - Liverpool Hospital - NSW Health Pathology | NSW Health Pathology - Institute of Clinical Pathology and Medical Research; Westmead Hospital; University of Sydney | CIDM-PH et al. |
| EPI_ISL_526212 | Australian Clinical Labs (formerly Healthscope Pathology) | NSW Health Pathology - Institute of Clinical Pathology and Medical Research; Westmead Hospital; University of Sydney | CIDM-PH et al. |
| EPI_ISL_526213, EPI_ISL_526214 | Douglass Hanly Moir Pathology | NSW Health Pathology - Institute of Clinical Pathology and Medical Research; Westmead Hospital; University of Sydney | CIDM-PH et al. |
| EPI_ISL_526551, EPI_ISL_526552 | OHSU Lab Services Molecular Microbiology Lab | Oregon SARS-CoV-2 Genome Sequencing Center | Brendan L. O'Connell, Ruth V. Nichols, Alec J. Hirsch, Guang Fan, Daniel N. Streblow, William B. Messer, Andrew C. Adey, Benjamin N. Bimber, Brian J. O'Roak |
| EPI_ISL_526689 | Pathology West - NSW Health Pathology | NSW Health Pathology - Institute of Clinical Pathology and Medical Research; Westmead Hospital; University of Sydney | CIDM-PH et al. |
| EPI_ISL_526697 | South Eastern Area Laboratory Services (SEALS) | NSW Health Pathology - Institute of Clinical Pathology and Medical Research; Westmead Hospital; University of Sydney | CIDM-PH et al. |
| EPI_ISL_527787, EPI_ISL_527789 | Laboratorio de Referencia Nacional de Virus Respiratorio. Centro Nacional de Salud Publica. Instituto Nacional de Salud Peru. | Laboratorio de Referencia Nacional de Biotecnología y Biología Molecular. Centro Nacional de Salud Publica. Instituto Nacional de Salud Peru. | Carlos Padilla Rojas, Karolyn Vega Chozo, Priscila Lope Pari, Omar Caceres Rey, Marco Galarza Perez, Maribel Huaringa Nuñez, Johanna Balbuena Torres, Henri Bailon Calderon, Nancy Rojas Serrano. |
| EPI_ISL_527820, EPI_ISL_527821, EPI_ISL_527825, EPI_ISL_527826, EPI_ISL_527827, EPI_ISL_527828, EPI_ISL_527829, EPI_ISL_527830, EPI_ISL_527831, EPI_ISL_527832, EPI_ISL_527833, EPI_ISL_527834, EPI_ISL_527835, EPI_ISL_527836, EPI_ISL_527837, EPI_ISL_527838, EPI_ISL_527839, EPI_ISL_527840, EPI_ISL_527841, EPI_ISL_527842, EPI_ISL_527843, EPI_ISL_527844, EPI_ISL_527845, EPI_ISL_527846, EPI_ISL_527847, EPI_ISL_527848, EPI_ISL_527849, EPI_ISL_527850, EPI_ISL_527851, EPI_ISL_527852, EPI_ISL_527853, EPI_ISL_527854, EPI_ISL_527855 | | | |
| see above | Texas Department of State Health Services | Texas Department of State Health Services | Bonnie Oh, Rashmi Tuladhar, Jenny Zhang, Maliha Rahman, Anita Pokharel, Myong Koag, Chun Wang, Rachel Lee, Grace Kubin |
| EPI_ISL_535650, EPI_ISL_535652, EPI_ISL_535654, EPI_ISL_535658, EPI_ISL_535659, EPI_ISL_535661 | AR Dept. of Health-Public Health Lab | Pathogen Discovery, Respiratory Viruses Branch, Division of Viral Diseases, Centers for Disease Control and Prevention | Brian Lynch, Yan Li, Jing Zhang, Ying Tao, Krista Queen, Anna Uehara, Clinton R. Paden, Rachel Marine, Haibin Wang, Suxiang Tong |
| EPI_ISL_535675, EPI_ISL_535701, EPI_ISL_535708 | CDPH, Microbial Diseases Laboratory | Pathogen Discovery, Respiratory Viruses Branch, Division of Viral Diseases, Centers for Disease Control and Prevention | Brian Lynch, Yan Li, Jing Zhang, Ying Tao, Krista Queen, Anna Uehara, Clinton R. Paden, Rachel Marine, Haibin Wang, Suxiang Tong |
| EPI_ISL_538275, EPI_ISL_538276, EPI_ISL_538277, EPI_ISL_538278, EPI_ISL_538279, EPI_ISL_538280, EPI_ISL_538281, EPI_ISL_538282, EPI_ISL_538283, EPI_ISL_538284, EPI_ISL_538285, EPI_ISL_538286, EPI_ISL_538287, EPI_ISL_538288, EPI_ISL_538289, EPI_ISL_538290, EPI_ISL_538291, EPI_ISL_538292, EPI_ISL_538293, EPI_ISL_538294, EPI_ISL_538295, EPI_ISL_538296, EPI_ISL_538297, EPI_ISL_538298, EPI_ISL_538299, EPI_ISL_538300, EPI_ISL_538301, EPI_ISL_538303, EPI_ISL_538304, EPI_ISL_538305, EPI_ISL_538306, EPI_ISL_538307, EPI_ISL_538308, EPI_ISL_538309, EPI_ISL_538310, EPI_ISL_538311, EPI_ISL_538312, EPI_ISL_538313, EPI_ISL_538314, EPI_ISL_538315, EPI_ISL_538316, EPI_ISL_538317, EPI_ISL_538318, EPI_ISL_538319, EPI_ISL_538320 | | | |
| see above | Texas Department of State Health Services | Texas Department of State Health Services | Bonnie Oh, Rashmi Tuladhar, Jenny Zhang, Maliha Rahman, Anita Pokharel, Myong Koag, Chun Wang, Rachel Lee, Grace Kubin |
| EPI_ISL_541949, EPI_ISL_541950, EPI_ISL_541951, EPI_ISL_541952, EPI_ISL_541953, EPI_ISL_541954, EPI_ISL_541955, EPI_ISL_541956, EPI_ISL_541957, EPI_ISL_541958, EPI_ISL_541959, EPI_ISL_541960, EPI_ISL_541961, EPI_ISL_541962, EPI_ISL_541963, EPI_ISL_541964, EPI_ISL_541965, EPI_ISL_541966, EPI_ISL_541967, EPI_ISL_541968, EPI_ISL_541969 | | | |
| see above | Servicio de Microbiología, Hospital Universitario Son Espases | SeqCOVID-SPAIN consortium/IBV(CSIC) | Carla López-Causapé, Jordi Reina, Antonio Oliver and SeqCOVID-SPAIN consortium |
| EPI_ISL_544955 | Pathology West - NSW Health Pathology | NSW Health Pathology - Institute of Clinical Pathology and Medical Research; Westmead Hospital; University of Sydney | CIDM-PH et al. |
| EPI_ISL_544956 | Sydney South West Pathology Service (SSWPS) - Concord | NSW Health Pathology - Institute of Clinical Pathology and | CIDM-PH et al. |

| | | | |
|---|---|---|---|
| | Repatriation General Hospital - NSW Health Pathology | Medical Research; Westmead Hospital; University of Sydney | |
| EPI_ISL_544957 | Laverty Pathology | NSW Health Pathology - Institute of Clinical Pathology and Medical Research; Westmead Hospital; University of Sydney | CIDM-PH et al. |
| EPI_ISL_544958, EPI_ISL_544959 | Sydney South West Pathology Service (SSWPS) - Liverpool Hospital - NSW Health Pathology | NSW Health Pathology - Institute of Clinical Pathology and Medical Research; Westmead Hospital; University of Sydney | CIDM-PH et al. |
| EPI_ISL_544960 | Histopath | NSW Health Pathology - Institute of Clinical Pathology and Medical Research; Westmead Hospital; University of Sydney | CIDM-PH et al. |
| EPI_ISL_544961 | St Vincent's Pathology (SydPath) | NSW Health Pathology - Institute of Clinical Pathology and Medical Research; Westmead Hospital; University of Sydney | CIDM-PH et al. |
| EPI_ISL_544962 | Laverty Pathology | NSW Health Pathology - Institute of Clinical Pathology and Medical Research; Westmead Hospital; University of Sydney | CIDM-PH et al. |
| EPI_ISL_544963 | The Children's Hospital at Westmead | NSW Health Pathology - Institute of Clinical Pathology and Medical Research; Westmead Hospital; University of Sydney | CIDM-PH et al. |
| EPI_ISL_544964, EPI_ISL_544965 | Sydney South West Pathology Service (SSWPS) - Liverpool Hospital - NSW Health Pathology | NSW Health Pathology - Institute of Clinical Pathology and Medical Research; Westmead Hospital; University of Sydney | CIDM-PH et al. |
| EPI_ISL_544966, EPI_ISL_544967 | Australian Clinical Labs | NSW Health Pathology - Institute of Clinical Pathology and Medical Research; Westmead Hospital; University of Sydney | CIDM-PH et al. |
| EPI_ISL_544968 | St Vincent's Pathology (SydPath) | NSW Health Pathology - Institute of Clinical Pathology and Medical Research; Westmead Hospital; University of Sydney | CIDM-PH et al. |
| EPI_ISL_544969 | Pathology North - Royal North Shore Hospital - NSW Health Pathology | NSW Health Pathology - Institute of Clinical Pathology and Medical Research; Westmead Hospital; University of Sydney | CIDM-PH et al. |
| EPI_ISL_544970 | South Eastern Area Laboratory Services (SEALS) | NSW Health Pathology - Institute of Clinical Pathology and Medical Research; Westmead Hospital; University of Sydney | CIDM-PH et al. |
| EPI_ISL_544971, EPI_ISL_544972, EPI_ISL_544973 | St Vincent's Pathology (SydPath) | NSW Health Pathology - Institute of Clinical Pathology and Medical Research; Westmead Hospital; University of Sydney | CIDM-PH et al. |
| EPI_ISL_544974 | Sydney South West Pathology Service (SSWPS) - Royal Prince Alfred Hospital - NSW Health Pathology | NSW Health Pathology - Institute of Clinical Pathology and Medical Research; Westmead Hospital; University of Sydney | CIDM-PH et al. |
| EPI_ISL_544975 | Histopath | NSW Health Pathology - Institute of Clinical Pathology and Medical Research; Westmead Hospital; University of Sydney | CIDM-PH et al. |
| EPI_ISL_544976 | Pathology North - Royal North Shore Hospital - NSW Health Pathology | NSW Health Pathology - Institute of Clinical Pathology and Medical Research; Westmead Hospital; University of Sydney | CIDM-PH et al. |
| EPI_ISL_544977, EPI_ISL_544978, EPI_ISL_544979, EPI_ISL_544980, EPI_ISL_544981 | St Vincent's Pathology (SydPath) | NSW Health Pathology - Institute of Clinical Pathology and Medical Research; Westmead Hospital; University of Sydney | CIDM-PH et al. |
| EPI_ISL_544982 | Pathology North - Royal North Shore Hospital - NSW Health Pathology | NSW Health Pathology - Institute of Clinical Pathology and Medical Research; Westmead Hospital; University of Sydney | CIDM-PH et al. |
| EPI_ISL_544983, EPI_ISL_544984 | South Eastern Area Laboratory Services (SEALS) | NSW Health Pathology - Institute of Clinical Pathology and Medical Research; Westmead Hospital; University of Sydney | CIDM-PH et al. |
| EPI_ISL_544985 | Sydney South West Pathology Service (SSWPS) - Liverpool Hospital - NSW Health Pathology | NSW Health Pathology - Institute of Clinical Pathology and Medical Research; Westmead Hospital; University of Sydney | CIDM-PH et al. |
| EPI_ISL_544986 | Pathology West - NSW Health Pathology | NSW Health Pathology - Institute of Clinical Pathology and Medical Research; Westmead Hospital; University of Sydney | CIDM-PH et al. |
| EPI_ISL_544987 | Histopath | NSW Health Pathology - Institute of Clinical Pathology and Medical Research; Westmead Hospital; University of Sydney | CIDM-PH et al. |
| EPI_ISL_544988 | Australian Clinical Labs | NSW Health Pathology - Institute of Clinical Pathology and Medical Research; Westmead Hospital; University of Sydney | CIDM-PH et al. |
| EPI_ISL_544989, EPI_ISL_544990 | Pathology West - NSW Health Pathology | NSW Health Pathology - Institute of Clinical Pathology and Medical Research; Westmead Hospital; University of Sydney | CIDM-PH et al. |
| EPI_ISL_544991 | Pathology North - Royal North Shore Hospital - NSW Health Pathology | NSW Health Pathology - Institute of Clinical Pathology and Medical Research; Westmead Hospital; University of Sydney | CIDM-PH et al. |
| EPI_ISL_544992, EPI_ISL_544993, EPI_ISL_544994, EPI_ISL_544995, EPI_ISL_544996, EPI_ISL_544997, EPI_ISL_544998, EPI_ISL_544999, EPI_ISL_545000, EPI_ISL_545001 | St Vincent's Pathology (SydPath) | NSW Health Pathology - Institute of Clinical Pathology and Medical Research; Westmead Hospital; University of Sydney | CIDM-PH et al. |
| EPI_ISL_545002 | The Children's Hospital at Westmead | NSW Health Pathology - Institute of Clinical Pathology and Medical Research; Westmead Hospital; University of Sydney | CIDM-PH et al. |
| EPI_ISL_545003 | Sydney South West Pathology Service (SSWPS) - Liverpool Hospital - NSW Health Pathology | NSW Health Pathology - Institute of Clinical Pathology and Medical Research; Westmead Hospital; University of Sydney | CIDM-PH et al. |
| EPI_ISL_545004, EPI_ISL_545005 | Sydney South West Pathology Service (SSWPS) - Royal Prince Alfred Hospital - NSW Health Pathology | NSW Health Pathology - Institute of Clinical Pathology and Medical Research; Westmead Hospital; University of Sydney | CIDM-PH et al. |
| EPI_ISL_545006 | Pathology West - NSW Health Pathology | NSW Health Pathology - Institute of Clinical Pathology and Medical Research; Westmead Hospital; University of Sydney | CIDM-PH et al. |
| EPI_ISL_545007 | Histopath | NSW Health Pathology - Institute of Clinical Pathology and Medical Research; Westmead Hospital; University of Sydney | CIDM-PH et al. |
| EPI_ISL_545008 | Pathology North - Royal North Shore Hospital - NSW Health Pathology | NSW Health Pathology - Institute of Clinical Pathology and Medical Research; Westmead Hospital; University of Sydney | CIDM-PH et al. |
| EPI_ISL_545009, EPI_ISL_545010, EPI_ISL_545011, EPI_ISL_545012, EPI_ISL_545013, EPI_ISL_545014, EPI_ISL_545015, EPI_ISL_545016 | South Eastern Area Laboratory Services (SEALS) | NSW Health Pathology - Institute of Clinical Pathology and Medical Research; Westmead Hospital; University of Sydney | CIDM-PH et al. |

| | | | |
|---|---|---|---|
| EPI_ISL_545017, EPI_ISL_545018, EPI_ISL_545019, EPI_ISL_545020, EPI_ISL_545021, EPI_ISL_545022, EPI_ISL_545023 | Sydney South West Pathology Service (SSWPS) - Royal Prince Alfred Hospital - NSW Health Pathology | NSW Health Pathology - Institute of Clinical Pathology and Medical Research; Westmead Hospital; University of Sydney | CIDM-PH et al. |
| EPI_ISL_545024 | Australian Clinical Labs | NSW Health Pathology - Institute of Clinical Pathology and Medical Research; Westmead Hospital; University of Sydney | CIDM-PH et al. |
| EPI_ISL_545025 | Sydney South West Pathology Service (SSWPS) - Liverpool Hospital - NSW Health Pathology | NSW Health Pathology - Institute of Clinical Pathology and Medical Research; Westmead Hospital; University of Sydney | CIDM-PH et al. |
| EPI_ISL_545026 | St Vincent's Pathology (SydPath) | NSW Health Pathology - Institute of Clinical Pathology and Medical Research; Westmead Hospital; University of Sydney | CIDM-PH et al. |
| EPI_ISL_545027 | Douglass Hanly Moir Pathology | NSW Health Pathology - Institute of Clinical Pathology and Medical Research; Westmead Hospital; University of Sydney | CIDM-PH et al. |
| EPI_ISL_545028 | Sydney South West Pathology Service (SSWPS) - Royal Prince Alfred Hospital - NSW Health Pathology | NSW Health Pathology - Institute of Clinical Pathology and Medical Research; Westmead Hospital; University of Sydney | CIDM-PH et al. |
| EPI_ISL_545029 | Pathology West - NSW Health Pathology | NSW Health Pathology - Institute of Clinical Pathology and Medical Research; Westmead Hospital; University of Sydney | CIDM-PH et al. |
| EPI_ISL_545030 | St Vincent's Pathology (SydPath) | NSW Health Pathology - Institute of Clinical Pathology and Medical Research; Westmead Hospital; University of Sydney | CIDM-PH et al. |
| EPI_ISL_545031 | South Eastern Area Laboratory Services (SEALS) | NSW Health Pathology - Institute of Clinical Pathology and Medical Research; Westmead Hospital; University of Sydney | CIDM-PH et al. |
| EPI_ISL_545032, EPI_ISL_545033 | Pathology West - NSW Health Pathology | NSW Health Pathology - Institute of Clinical Pathology and Medical Research; Westmead Hospital; University of Sydney | CIDM-PH et al. |
| EPI_ISL_545034, EPI_ISL_545035, EPI_ISL_545036 | Histopath | NSW Health Pathology - Institute of Clinical Pathology and Medical Research; Westmead Hospital; University of Sydney | CIDM-PH et al. |
| EPI_ISL_545037 | Sydney South West Pathology Service (SSWPS) - Concord Repatriation General Hospital - NSW Health Pathology | NSW Health Pathology - Institute of Clinical Pathology and Medical Research; Westmead Hospital; University of Sydney | CIDM-PH et al. |
| EPI_ISL_545038 | Pathology West - NSW Health Pathology | NSW Health Pathology - Institute of Clinical Pathology and Medical Research; Westmead Hospital; University of Sydney | CIDM-PH et al. |
| EPI_ISL_545039 | Sydney South West Pathology Service (SSWPS) - Royal Prince Alfred Hospital - NSW Health Pathology | NSW Health Pathology - Institute of Clinical Pathology and Medical Research; Westmead Hospital; University of Sydney | CIDM-PH et al. |
| EPI_ISL_545040 | Laverty Pathology | NSW Health Pathology - Institute of Clinical Pathology and Medical Research; Westmead Hospital; University of Sydney | CIDM-PH et al. |
| EPI_ISL_545041 | Sydney South West Pathology Service (SSWPS) - Royal Prince Alfred Hospital - NSW Health Pathology | NSW Health Pathology - Institute of Clinical Pathology and Medical Research; Westmead Hospital; University of Sydney | CIDM-PH et al. |
| EPI_ISL_545042 | Sydney South West Pathology Service (SSWPS) - Liverpool Hospital - NSW Health Pathology | NSW Health Pathology - Institute of Clinical Pathology and Medical Research; Westmead Hospital; University of Sydney | CIDM-PH et al. |
| EPI_ISL_545043 | Sydney South West Pathology Service (SSWPS) - Royal Prince Alfred Hospital - NSW Health Pathology | NSW Health Pathology - Institute of Clinical Pathology and Medical Research; Westmead Hospital; University of Sydney | CIDM-PH et al. |
| EPI_ISL_545044 | Sydney South West Pathology Service (SSWPS) - Concord Repatriation General Hospital - NSW Health Pathology | NSW Health Pathology - Institute of Clinical Pathology and Medical Research; Westmead Hospital; University of Sydney | CIDM-PH et al. |
| EPI_ISL_545045 | Sydney South West Pathology Service (SSWPS) - Royal Prince Alfred Hospital - NSW Health Pathology | NSW Health Pathology - Institute of Clinical Pathology and Medical Research; Westmead Hospital; University of Sydney | CIDM-PH et al. |
| EPI_ISL_545046, EPI_ISL_545047 | Pathology West - NSW Health Pathology | NSW Health Pathology - Institute of Clinical Pathology and Medical Research; Westmead Hospital; University of Sydney | CIDM-PH et al. |
| EPI_ISL_545048 | South Eastern Area Laboratory Services (SEALS) | NSW Health Pathology - Institute of Clinical Pathology and Medical Research; Westmead Hospital; University of Sydney | CIDM-PH et al. |
| EPI_ISL_545049 | Sydney South West Pathology Service (SSWPS) - Concord Repatriation General Hospital - NSW Health Pathology | NSW Health Pathology - Institute of Clinical Pathology and Medical Research; Westmead Hospital; University of Sydney | CIDM-PH et al. |
| EPI_ISL_545050 | Pathology West - NSW Health Pathology | NSW Health Pathology - Institute of Clinical Pathology and Medical Research; Westmead Hospital; University of Sydney | CIDM-PH et al. |
| EPI_ISL_547594 | Sydney South West Pathology Service (SSWPS) - Liverpool Hospital - NSW Health Pathology | NSW Health Pathology - Institute of Clinical Pathology and Medical Research; Westmead Hospital; University of Sydney | CIDM-PH et al. |
| EPI_ISL_547595, EPI_ISL_547596 | St Vincent's Pathology (SydPath) | NSW Health Pathology - Institute of Clinical Pathology and Medical Research; Westmead Hospital; University of Sydney | CIDM-PH et al. |
| EPI_ISL_547597 | Australian Clinical Labs | NSW Health Pathology - Institute of Clinical Pathology and Medical Research; Westmead Hospital; University of Sydney | CIDM-PH et al. |
| EPI_ISL_547598 | Sydney South West Pathology Service (SSWPS) - Concord Repatriation General Hospital - NSW Health Pathology | NSW Health Pathology - Institute of Clinical Pathology and Medical Research; Westmead Hospital; University of Sydney | CIDM-PH et al. |
| EPI_ISL_547599 | Sydney South West Pathology Service (SSWPS) - Royal Prince Alfred Hospital - NSW Health Pathology | NSW Health Pathology - Institute of Clinical Pathology and Medical Research; Westmead Hospital; University of Sydney | CIDM-PH et al. |
| EPI_ISL_547600 | Histopath | NSW Health Pathology - Institute of Clinical Pathology and Medical Research; Westmead Hospital; University of Sydney | CIDM-PH et al. |
| EPI_ISL_547601 | South Eastern Area Laboratory Services (SEALS) | NSW Health Pathology - Institute of Clinical Pathology and Medical Research; Westmead Hospital; University of Sydney | CIDM-PH et al. |
| EPI_ISL_548264 | County of Santa Clara Public Health Department | Chan-Zuckerberg Biohub | CZB Cliahub Consortium |
| EPI_ISL_548265 | County of San Luis Obispo Public Health Laboratory | Chan-Zuckerberg Biohub | CZB Cliahub Consortium |
| EPI_ISL_548266, EPI_ISL_548267 | County of Santa Clara Public Health Department | Chan-Zuckerberg Biohub | CZB Cliahub Consortium |
| EPI_ISL_548268, EPI_ISL_548269 | County of San Luis Obispo Public Health Laboratory | Chan-Zuckerberg Biohub | CZB Cliahub Consortium |
| EPI_ISL_548270, EPI_ISL_548271, EPI_ISL_548272, EPI_ISL_548273, | Orange County Public Health Laboratory | Chan-Zuckerberg Biohub | CZB Cliahub Consortium |

| | | | |
|---|---|---|---|
| EPI_ISL_548274, EPI_ISL_548275, EPI_ISL_548276 | | | |
| EPI_ISL_548277 | County of Santa Clara Public Health Department | Chan-Zuckerberg Biohub | CZB Cliahub Consortium |
| EPI_ISL_548278 | Orange County Public Health Laboratory | Chan-Zuckerberg Biohub | CZB Cliahub Consortium |
| EPI_ISL_548279 | County of Santa Clara Public Health Department | Chan-Zuckerberg Biohub | CZB Cliahub Consortium |
| EPI_ISL_548280, EPI_ISL_548281 | County of San Luis Obispo Public Health Laboratory | Chan-Zuckerberg Biohub | CZB Cliahub Consortium |
| EPI_ISL_548282 | County of Santa Clara Public Health Department | Chan-Zuckerberg Biohub | CZB Cliahub Consortium |
| EPI_ISL_548283 | Orange County Public Health Laboratory | Chan-Zuckerberg Biohub | CZB Cliahub Consortium |
| EPI_ISL_548284 | County of San Luis Obispo Public Health Laboratory | Chan-Zuckerberg Biohub | CZB Cliahub Consortium |
| EPI_ISL_548285 | Orange County Public Health Laboratory | Chan-Zuckerberg Biohub | CZB Cliahub Consortium |
| EPI_ISL_548286 | County of San Luis Obispo Public Health Laboratory | Chan-Zuckerberg Biohub | CZB Cliahub Consortium |
| EPI_ISL_548287 | Orange County Public Health Laboratory | Chan-Zuckerberg Biohub | CZB Cliahub Consortium |
| EPI_ISL_548288 | County of San Luis Obispo Public Health Laboratory | Chan-Zuckerberg Biohub | CZB Cliahub Consortium |
| EPI_ISL_548289 | Orange County Public Health Laboratory | Chan-Zuckerberg Biohub | CZB Cliahub Consortium |
| EPI_ISL_548290 | County of Santa Clara Public Health Department | Chan-Zuckerberg Biohub | CZB Cliahub Consortium |
| EPI_ISL_548291 | Orange County Public Health Laboratory | Chan-Zuckerberg Biohub | CZB Cliahub Consortium |
| EPI_ISL_548292 | County of San Luis Obispo Public Health Laboratory | Chan-Zuckerberg Biohub | CZB Cliahub Consortium |
| EPI_ISL_548293 | Orange County Public Health Laboratory | Chan-Zuckerberg Biohub | CZB Cliahub Consortium |
| EPI_ISL_548294, EPI_ISL_548295 | County of San Luis Obispo Public Health Laboratory | Chan-Zuckerberg Biohub | CZB Cliahub Consortium |
| EPI_ISL_548296, EPI_ISL_548297, EPI_ISL_548298 | Orange County Public Health Laboratory | Chan-Zuckerberg Biohub | CZB Cliahub Consortium |
| EPI_ISL_548299 | County of Santa Clara Public Health Department | Chan-Zuckerberg Biohub | CZB Cliahub Consortium |
| EPI_ISL_548300, EPI_ISL_548301 | Orange County Public Health Laboratory | Chan-Zuckerberg Biohub | CZB Cliahub Consortium |
| EPI_ISL_548302, EPI_ISL_548303 | County of Santa Clara Public Health Department | Chan-Zuckerberg Biohub | CZB Cliahub Consortium |
| EPI_ISL_548304 | County of San Luis Obispo Public Health Laboratory | Chan-Zuckerberg Biohub | CZB Cliahub Consortium |
| EPI_ISL_548305, EPI_ISL_548306 | Orange County Public Health Laboratory | Chan-Zuckerberg Biohub | CZB Cliahub Consortium |
| EPI_ISL_548307 | County of Santa Clara Public Health Department | Chan-Zuckerberg Biohub | CZB Cliahub Consortium |
| EPI_ISL_548308, EPI_ISL_548309 | Orange County Public Health Laboratory | Chan-Zuckerberg Biohub | CZB Cliahub Consortium |
| EPI_ISL_548310 | County of Santa Clara Public Health Department | Chan-Zuckerberg Biohub | CZB Cliahub Consortium |
| EPI_ISL_548311 | County of San Luis Obispo Public Health Laboratory | Chan-Zuckerberg Biohub | CZB Cliahub Consortium |
| EPI_ISL_548312, EPI_ISL_548313 | Orange County Public Health Laboratory | Chan-Zuckerberg Biohub | CZB Cliahub Consortium |
| EPI_ISL_548314 | County of Santa Clara Public Health Department | Chan-Zuckerberg Biohub | CZB Cliahub Consortium |
| EPI_ISL_548315 | Orange County Public Health Laboratory | Chan-Zuckerberg Biohub | CZB Cliahub Consortium |
| EPI_ISL_548316 | County of San Luis Obispo Public Health Laboratory | Chan-Zuckerberg Biohub | CZB Cliahub Consortium |
| EPI_ISL_548317, EPI_ISL_548318 | Orange County Public Health Laboratory | Chan-Zuckerberg Biohub | CZB Cliahub Consortium |
| EPI_ISL_548319, EPI_ISL_548320, EPI_ISL_548321 | County of Santa Clara Public Health Department | Chan-Zuckerberg Biohub | CZB Cliahub Consortium |
| EPI_ISL_548322 | County of San Luis Obispo Public Health Laboratory | Chan-Zuckerberg Biohub | CZB Cliahub Consortium |
| EPI_ISL_548323 | Orange County Public Health Laboratory | Chan-Zuckerberg Biohub | CZB Cliahub Consortium |
| EPI_ISL_548324 | County of Santa Clara Public Health Department | Chan-Zuckerberg Biohub | CZB Cliahub Consortium |
| EPI_ISL_548325 | County of San Luis Obispo Public Health Laboratory | Chan-Zuckerberg Biohub | CZB Cliahub Consortium |
| EPI_ISL_548326 | Orange County Public Health Laboratory | Chan-Zuckerberg Biohub | CZB Cliahub Consortium |
| EPI_ISL_548327 | County of Santa Clara Public Health Department | Chan-Zuckerberg Biohub | CZB Cliahub Consortium |
| EPI_ISL_548328 | Orange County Public Health Laboratory | Chan-Zuckerberg Biohub | CZB Cliahub Consortium |
| EPI_ISL_548329, EPI_ISL_548330 | County of San Luis Obispo Public Health Laboratory | Chan-Zuckerberg Biohub | CZB Cliahub Consortium |
| EPI_ISL_548331 | County of Santa Clara Public Health Department | Chan-Zuckerberg Biohub | CZB Cliahub Consortium |
| EPI_ISL_548332 | Orange County Public Health Laboratory | Chan-Zuckerberg Biohub | CZB Cliahub Consortium |
| EPI_ISL_548333 | County of Santa Clara Public Health Department | Chan-Zuckerberg Biohub | CZB Cliahub Consortium |
| EPI_ISL_548334 | Orange County Public Health Laboratory | Chan-Zuckerberg Biohub | CZB Cliahub Consortium |
| EPI_ISL_548335, EPI_ISL_548336 | County of San Luis Obispo Public Health Laboratory | Chan-Zuckerberg Biohub | CZB Cliahub Consortium |
| EPI_ISL_548337, EPI_ISL_548338, EPI_ISL_548339, EPI_ISL_548340, EPI_ISL_548341, EPI_ISL_548342, EPI_ISL_548343, EPI_ISL_548344, EPI_ISL_548345 | County of Santa Clara Public Health Department | Chan-Zuckerberg Biohub | CZB Cliahub Consortium |
| EPI_ISL_548346 | Ventura County Public Health Lab | Chan-Zuckerberg Biohub | CZB Cliahub Consortium |
| EPI_ISL_548347 | County of San Luis Obispo Public Health Laboratory | Chan-Zuckerberg Biohub | CZB Cliahub Consortium |
| EPI_ISL_548348, EPI_ISL_548349 | Ventura County Public Health Lab | Chan-Zuckerberg Biohub | CZB Cliahub Consortium |
| EPI_ISL_548350, EPI_ISL_548351 | County of San Luis Obispo Public Health Laboratory | Chan-Zuckerberg Biohub | CZB Cliahub Consortium |
| EPI_ISL_548352, EPI_ISL_548353, EPI_ISL_548354, EPI_ISL_548355, EPI_ISL_548356 | Ventura County Public Health Lab | Chan-Zuckerberg Biohub | CZB Cliahub Consortium |

| | | | |
|---|---|---|---|
| EPI_ISL_548357 | County of San Luis Obispo Public Health Laboratory | Chan-Zuckerberg Biohub | CZB Cliahub Consortium |
| EPI_ISL_548358 | Ventura County Public Health Lab | Chan-Zuckerberg Biohub | CZB Cliahub Consortium |
| EPI_ISL_548359, EPI_ISL_548360, EPI_ISL_548361 | County of San Luis Obispo Public Health Laboratory | Chan-Zuckerberg Biohub | CZB Cliahub Consortium |
| EPI_ISL_548362 | Ventura County Public Health Lab | Chan-Zuckerberg Biohub | CZB Cliahub Consortium |
| EPI_ISL_548363 | Tuolumne County Public Health | Chan-Zuckerberg Biohub | CZB Cliahub Consortium |
| EPI_ISL_548364 | Ventura County Public Health Lab | Chan-Zuckerberg Biohub | CZB Cliahub Consortium |
| EPI_ISL_548365, EPI_ISL_548366 | County of San Luis Obispo Public Health Laboratory | Chan-Zuckerberg Biohub | CZB Cliahub Consortium |
| EPI_ISL_548367, EPI_ISL_548368 | Ventura County Public Health Lab | Chan-Zuckerberg Biohub | CZB Cliahub Consortium |
| EPI_ISL_548369, EPI_ISL_548370 | County of San Luis Obispo Public Health Laboratory | Chan-Zuckerberg Biohub | CZB Cliahub Consortium |
| EPI_ISL_548371 | County of Santa Clara Public Health Department | Chan-Zuckerberg Biohub | CZB Cliahub Consortium |
| EPI_ISL_548372 | Ventura County Public Health Lab | Chan-Zuckerberg Biohub | CZB Cliahub Consortium |
| EPI_ISL_548373 | Orange County Public Health Laboratory | Chan-Zuckerberg Biohub | CZB Cliahub Consortium |
| EPI_ISL_548374 | University of California, Davis | Chan-Zuckerberg Biohub | CZB Cliahub Consortium |
| EPI_ISL_548375 | Orange County Public Health Laboratory | Chan-Zuckerberg Biohub | CZB Cliahub Consortium |
| EPI_ISL_548376 | Ventura County Public Health Lab | Chan-Zuckerberg Biohub | CZB Cliahub Consortium |
| EPI_ISL_548377, EPI_ISL_548378 | Orange County Public Health Laboratory | Chan-Zuckerberg Biohub | CZB Cliahub Consortium |
| EPI_ISL_548379 | County of Santa Clara Public Health Department | Chan-Zuckerberg Biohub | CZB Cliahub Consortium |
| EPI_ISL_548380 | University of California, Davis | Chan-Zuckerberg Biohub | CZB Cliahub Consortium |
| EPI_ISL_548381 | Orange County Public Health Laboratory | Chan-Zuckerberg Biohub | CZB Cliahub Consortium |
| EPI_ISL_548382 | University of California, Davis | Chan-Zuckerberg Biohub | CZB Cliahub Consortium |
| EPI_ISL_548383 | Orange County Public Health Laboratory | Chan-Zuckerberg Biohub | CZB Cliahub Consortium |
| EPI_ISL_548384, EPI_ISL_548385 | County of Santa Clara Public Health Department | Chan-Zuckerberg Biohub | CZB Cliahub Consortium |
| EPI_ISL_548386, EPI_ISL_548387 | Orange County Public Health Laboratory | Chan-Zuckerberg Biohub | CZB Cliahub Consortium |
| EPI_ISL_548388 | County of Santa Clara Public Health Department | Chan-Zuckerberg Biohub | CZB Cliahub Consortium |
| EPI_ISL_548389 | Orange County Public Health Laboratory | Chan-Zuckerberg Biohub | CZB Cliahub Consortium |
| EPI_ISL_548390 | Ventura County Public Health Lab | Chan-Zuckerberg Biohub | CZB Cliahub Consortium |
| EPI_ISL_548391 | County of San Luis Obispo Public Health Laboratory | Chan-Zuckerberg Biohub | CZB Cliahub Consortium |
| EPI_ISL_548392 | County of Santa Clara Public Health Department | Chan-Zuckerberg Biohub | CZB Cliahub Consortium |
| EPI_ISL_548393 | Orange County Public Health Laboratory | Chan-Zuckerberg Biohub | CZB Cliahub Consortium |
| EPI_ISL_548394, EPI_ISL_548395, EPI_ISL_548396 | County of Santa Clara Public Health Department | Chan-Zuckerberg Biohub | CZB Cliahub Consortium |
| EPI_ISL_548397 | University of California, Davis | Chan-Zuckerberg Biohub | CZB Cliahub Consortium |
| EPI_ISL_548398 | Ventura County Public Health Lab | Chan-Zuckerberg Biohub | CZB Cliahub Consortium |
| EPI_ISL_548399 | County of Santa Clara Public Health Department | Chan-Zuckerberg Biohub | CZB Cliahub Consortium |
| EPI_ISL_548400 | Orange County Public Health Laboratory | Chan-Zuckerberg Biohub | CZB Cliahub Consortium |
| EPI_ISL_548401 | County of Santa Clara Public Health Department | Chan-Zuckerberg Biohub | CZB Cliahub Consortium |
| EPI_ISL_548402 | Ventura County Public Health Lab | Chan-Zuckerberg Biohub | CZB Cliahub Consortium |
| EPI_ISL_548403 | County of Santa Clara Public Health Department | Chan-Zuckerberg Biohub | CZB Cliahub Consortium |
| EPI_ISL_548404, EPI_ISL_548405 | Orange County Public Health Laboratory | Chan-Zuckerberg Biohub | CZB Cliahub Consortium |
| EPI_ISL_548406 | Ventura County Public Health Lab | Chan-Zuckerberg Biohub | CZB Cliahub Consortium |
| EPI_ISL_548407 | County of Santa Clara Public Health Department | Chan-Zuckerberg Biohub | CZB Cliahub Consortium |
| EPI_ISL_548408 | Orange County Public Health Laboratory | Chan-Zuckerberg Biohub | CZB Cliahub Consortium |
| EPI_ISL_548409, EPI_ISL_548410 | Ventura County Public Health Lab | Chan-Zuckerberg Biohub | CZB Cliahub Consortium |
| EPI_ISL_548411, EPI_ISL_548412, EPI_ISL_548413 | Orange County Public Health Laboratory | Chan-Zuckerberg Biohub | CZB Cliahub Consortium |
| EPI_ISL_548414 | University of California, Davis | Chan-Zuckerberg Biohub | CZB Cliahub Consortium |
| EPI_ISL_548415 | County of Santa Clara Public Health Department | Chan-Zuckerberg Biohub | CZB Cliahub Consortium |
| EPI_ISL_548416 | Orange County Public Health Laboratory | Chan-Zuckerberg Biohub | CZB Cliahub Consortium |
| EPI_ISL_548417 | County of Santa Clara Public Health Department | Chan-Zuckerberg Biohub | CZB Cliahub Consortium |
| EPI_ISL_548418 | Orange County Public Health Laboratory | Chan-Zuckerberg Biohub | CZB Cliahub Consortium |
| EPI_ISL_548419, EPI_ISL_548420, EPI_ISL_548421, EPI_ISL_548422, EPI_ISL_548423 | County of Santa Clara Public Health Department | Chan-Zuckerberg Biohub | CZB Cliahub Consortium |
| EPI_ISL_548424 | Orange County Public Health Laboratory | Chan-Zuckerberg Biohub | CZB Cliahub Consortium |
| EPI_ISL_548425 | County of San Luis Obispo Public Health Laboratory | Chan-Zuckerberg Biohub | CZB Cliahub Consortium |
| EPI_ISL_548426 | County of Santa Clara Public Health Department | Chan-Zuckerberg Biohub | CZB Cliahub Consortium |
| EPI_ISL_548427 | Ventura County Public Health Lab | Chan-Zuckerberg Biohub | CZB Cliahub Consortium |
| EPI_ISL_548428 | County of San Luis Obispo Public Health Laboratory | Chan-Zuckerberg Biohub | CZB Cliahub Consortium |
| EPI_ISL_548429 | County of Santa Clara Public Health Department | Chan-Zuckerberg Biohub | CZB Cliahub Consortium |

| | | | |
|---|---|---|---|
| EPI_ISL_548430 | Ventura County Public Health Lab | Chan-Zuckerberg Biohub | CZB Cliahub Consortium |
| EPI_ISL_548431 | Orange County Public Health Laboratory | Chan-Zuckerberg Biohub | CZB Cliahub Consortium |
| EPI_ISL_548432, EPI_ISL_548433, EPI_ISL_548434 | County of Santa Clara Public Health Department | Chan-Zuckerberg Biohub | CZB Cliahub Consortium |
| EPI_ISL_548435 | University of California, Davis | Chan-Zuckerberg Biohub | CZB Cliahub Consortium |
| EPI_ISL_548436 | County of San Luis Obispo Public Health Laboratory | Chan-Zuckerberg Biohub | CZB Cliahub Consortium |
| EPI_ISL_548437 | County of Santa Clara Public Health Department | Chan-Zuckerberg Biohub | CZB Cliahub Consortium |
| EPI_ISL_548438, EPI_ISL_548439, EPI_ISL_548440, EPI_ISL_548441 | Orange County Public Health Laboratory | Chan-Zuckerberg Biohub | CZB Cliahub Consortium |
| EPI_ISL_548442, EPI_ISL_548443 | Ventura County Public Health Lab | Chan-Zuckerberg Biohub | CZB Cliahub Consortium |
| EPI_ISL_548444, EPI_ISL_548445 | Orange County Public Health Laboratory | Chan-Zuckerberg Biohub | CZB Cliahub Consortium |
| EPI_ISL_548446 | County of Santa Clara Public Health Department | Chan-Zuckerberg Biohub | CZB Cliahub Consortium |
| EPI_ISL_548447 | University of California, Davis | Chan-Zuckerberg Biohub | CZB Cliahub Consortium |
| EPI_ISL_548448 | County of Santa Clara Public Health Department | Chan-Zuckerberg Biohub | CZB Cliahub Consortium |
| EPI_ISL_548449 | Orange County Public Health Laboratory | Chan-Zuckerberg Biohub | CZB Cliahub Consortium |
| EPI_ISL_548450 | Ventura County Public Health Lab | Chan-Zuckerberg Biohub | CZB Cliahub Consortium |
| EPI_ISL_548451 | County of Santa Clara Public Health Department | Chan-Zuckerberg Biohub | CZB Cliahub Consortium |
| EPI_ISL_548452 | University of California, Davis | Chan-Zuckerberg Biohub | CZB Cliahub Consortium |
| EPI_ISL_548453 | County of San Luis Obispo Public Health Laboratory | Chan-Zuckerberg Biohub | CZB Cliahub Consortium |
| EPI_ISL_548454 | Orange County Public Health Laboratory | Chan-Zuckerberg Biohub | CZB Cliahub Consortium |
| EPI_ISL_548455 | Ventura County Public Health Lab | Chan-Zuckerberg Biohub | CZB Cliahub Consortium |
| EPI_ISL_548456 | Orange County Public Health Laboratory | Chan-Zuckerberg Biohub | CZB Cliahub Consortium |
| EPI_ISL_548457 | County of San Luis Obispo Public Health Laboratory | Chan-Zuckerberg Biohub | CZB Cliahub Consortium |
| EPI_ISL_548458 | University of California, Davis | Chan-Zuckerberg Biohub | CZB Cliahub Consortium |
| EPI_ISL_548459 | Ventura County Public Health Lab | Chan-Zuckerberg Biohub | CZB Cliahub Consortium |
| EPI_ISL_548460 | Orange County Public Health Laboratory | Chan-Zuckerberg Biohub | CZB Cliahub Consortium |
| EPI_ISL_548461 | Ventura County Public Health Lab | Chan-Zuckerberg Biohub | CZB Cliahub Consortium |
| EPI_ISL_548462, EPI_ISL_548463, EPI_ISL_548464, EPI_ISL_548465 | Orange County Public Health Laboratory | Chan-Zuckerberg Biohub | CZB Cliahub Consortium |
| EPI_ISL_548466 | County of Santa Clara Public Health Department | Chan-Zuckerberg Biohub | CZB Cliahub Consortium |
| EPI_ISL_548467 | Orange County Public Health Laboratory | Chan-Zuckerberg Biohub | CZB Cliahub Consortium |
| EPI_ISL_548468 | County of Santa Clara Public Health Department | Chan-Zuckerberg Biohub | CZB Cliahub Consortium |
| EPI_ISL_548469 | Orange County Public Health Laboratory | Chan-Zuckerberg Biohub | CZB Cliahub Consortium |
| EPI_ISL_548470 | County of Santa Clara Public Health Department | Chan-Zuckerberg Biohub | CZB Cliahub Consortium |
| EPI_ISL_548471 | Ventura County Public Health Lab | Chan-Zuckerberg Biohub | CZB Cliahub Consortium |
| EPI_ISL_548472 | County of Santa Clara Public Health Department | Chan-Zuckerberg Biohub | CZB Cliahub Consortium |
| EPI_ISL_548473 | Ventura County Public Health Lab | Chan-Zuckerberg Biohub | CZB Cliahub Consortium |
| EPI_ISL_548474 | University of California, Davis | Chan-Zuckerberg Biohub | CZB Cliahub Consortium |
| EPI_ISL_548475, EPI_ISL_548476 | Ventura County Public Health Lab | Chan-Zuckerberg Biohub | CZB Cliahub Consortium |
| EPI_ISL_548477 | University of California, Davis | Chan-Zuckerberg Biohub | CZB Cliahub Consortium |
| EPI_ISL_548478 | County of Santa Clara Public Health Department | Chan-Zuckerberg Biohub | CZB Cliahub Consortium |
| EPI_ISL_548479 | University of California, Davis | Chan-Zuckerberg Biohub | CZB Cliahub Consortium |
| EPI_ISL_548480, EPI_ISL_548481 | Ventura County Public Health Lab | Chan-Zuckerberg Biohub | CZB Cliahub Consortium |
| EPI_ISL_548482 | County of Santa Clara Public Health Department | Chan-Zuckerberg Biohub | CZB Cliahub Consortium |
| EPI_ISL_548483 | Ventura County Public Health Lab | Chan-Zuckerberg Biohub | CZB Cliahub Consortium |
| EPI_ISL_548484, EPI_ISL_548485 | Orange County Public Health Laboratory | Chan-Zuckerberg Biohub | CZB Cliahub Consortium |
| EPI_ISL_548486 | University of California, Davis | Chan-Zuckerberg Biohub | CZB Cliahub Consortium |
| EPI_ISL_548487 | Ventura County Public Health Lab | Chan-Zuckerberg Biohub | CZB Cliahub Consortium |
| EPI_ISL_548488 | Orange County Public Health Laboratory | Chan-Zuckerberg Biohub | CZB Cliahub Consortium |
| EPI_ISL_548489 | Ventura County Public Health Lab | Chan-Zuckerberg Biohub | CZB Cliahub Consortium |
| EPI_ISL_548490 | University of California, Davis | Chan-Zuckerberg Biohub | CZB Cliahub Consortium |
| EPI_ISL_548491, EPI_ISL_548492 | Contra Costa Public Health Lab | Chan-Zuckerberg Biohub | CZB Cliahub Consortium |
| EPI_ISL_548493 | County of San Luis Obispo Public Health Laboratory | Chan-Zuckerberg Biohub | CZB Cliahub Consortium |
| EPI_ISL_548494 | County of Santa Clara Public Health Department | Chan-Zuckerberg Biohub | CZB Cliahub Consortium |
| EPI_ISL_548495 | Contra Costa Public Health Lab | Chan-Zuckerberg Biohub | CZB Cliahub Consortium |
| EPI_ISL_548496, EPI_ISL_548497, EPI_ISL_548498 | County of Santa Clara Public Health Department | Chan-Zuckerberg Biohub | CZB Cliahub Consortium |
| EPI_ISL_548499 | Contra Costa Public Health Lab | Chan-Zuckerberg Biohub | CZB Cliahub Consortium |
| EPI_ISL_548500 | County of Santa Clara Public Health Department | Chan-Zuckerberg Biohub | CZB Cliahub Consortium |
| EPI_ISL_548501 | Contra Costa Public Health Lab | Chan-Zuckerberg Biohub | CZB Cliahub Consortium |

| | | | |
|---|---|---|---|
| EPI_ISL_548502 | County of San Luis Obispo Public Health Laboratory | Chan-Zuckerberg Biohub | CZB Cliahub Consortium |
| EPI_ISL_548503 | County of Santa Clara Public Health Department | Chan-Zuckerberg Biohub | CZB Cliahub Consortium |
| EPI_ISL_548504 | County of San Luis Obispo Public Health Laboratory | Chan-Zuckerberg Biohub | CZB Cliahub Consortium |
| EPI_ISL_548505 | UCSF Clinical Microbiology Laboratory | Chan-Zuckerberg Biohub | CZB Cliahub Consortium |
| EPI_ISL_548506, EPI_ISL_548507, EPI_ISL_548508 | Contra Costa Public Health Lab | Chan-Zuckerberg Biohub | CZB Cliahub Consortium |
| EPI_ISL_548509 | County of Santa Clara Public Health Department | Chan-Zuckerberg Biohub | CZB Cliahub Consortium |
| EPI_ISL_548510 | Contra Costa Public Health Lab | Chan-Zuckerberg Biohub | CZB Cliahub Consortium |
| EPI_ISL_548511, EPI_ISL_548512 | County of Santa Clara Public Health Department | Chan-Zuckerberg Biohub | CZB Cliahub Consortium |
| EPI_ISL_548513 | UCSF Clinical Microbiology Laboratory | Chan-Zuckerberg Biohub | CZB Cliahub Consortium |
| EPI_ISL_548514 | County of Santa Clara Public Health Department | Chan-Zuckerberg Biohub | CZB Cliahub Consortium |
| EPI_ISL_548515, EPI_ISL_548516 | Contra Costa Public Health Lab | Chan-Zuckerberg Biohub | CZB Cliahub Consortium |
| EPI_ISL_548517 | County of San Luis Obispo Public Health Laboratory | Chan-Zuckerberg Biohub | CZB Cliahub Consortium |
| EPI_ISL_548518 | County of Santa Clara Public Health Department | Chan-Zuckerberg Biohub | CZB Cliahub Consortium |
| EPI_ISL_548519 | County of San Luis Obispo Public Health Laboratory | Chan-Zuckerberg Biohub | CZB Cliahub Consortium |
| EPI_ISL_548520 | Contra Costa Public Health Lab | Chan-Zuckerberg Biohub | CZB Cliahub Consortium |
| EPI_ISL_548521 | County of San Luis Obispo Public Health Laboratory | Chan-Zuckerberg Biohub | CZB Cliahub Consortium |
| EPI_ISL_548522 | UCSF Clinical Microbiology Laboratory | Chan-Zuckerberg Biohub | CZB Cliahub Consortium |
| EPI_ISL_548523 | County of Santa Clara Public Health Department | Chan-Zuckerberg Biohub | CZB Cliahub Consortium |
| EPI_ISL_548524 | Contra Costa Public Health Lab | Chan-Zuckerberg Biohub | CZB Cliahub Consortium |
| EPI_ISL_548525, EPI_ISL_548526 | County of Santa Clara Public Health Department | Chan-Zuckerberg Biohub | CZB Cliahub Consortium |
| EPI_ISL_548527, EPI_ISL_548528 | Contra Costa Public Health Lab | Chan-Zuckerberg Biohub | CZB Cliahub Consortium |
| EPI_ISL_548529, EPI_ISL_548530 | County of Santa Clara Public Health Department | Chan-Zuckerberg Biohub | CZB Cliahub Consortium |
| EPI_ISL_548531 | Contra Costa Public Health Lab | Chan-Zuckerberg Biohub | CZB Cliahub Consortium |
| EPI_ISL_548532 | County of San Luis Obispo Public Health Laboratory | Chan-Zuckerberg Biohub | CZB Cliahub Consortium |
| EPI_ISL_548533 | County of Santa Clara Public Health Department | Chan-Zuckerberg Biohub | CZB Cliahub Consortium |
| EPI_ISL_548534, EPI_ISL_548535 | Contra Costa Public Health Lab | Chan-Zuckerberg Biohub | CZB Cliahub Consortium |
| EPI_ISL_548536 | County of Santa Clara Public Health Department | Chan-Zuckerberg Biohub | CZB Cliahub Consortium |
| EPI_ISL_548537, EPI_ISL_548538, EPI_ISL_548539, EPI_ISL_548540 | Contra Costa Public Health Lab | Chan-Zuckerberg Biohub | CZB Cliahub Consortium |
| EPI_ISL_548541 | County of Santa Clara Public Health Department | Chan-Zuckerberg Biohub | CZB Cliahub Consortium |
| EPI_ISL_548542 | County of San Luis Obispo Public Health Laboratory | Chan-Zuckerberg Biohub | CZB Cliahub Consortium |
| EPI_ISL_548543 | County of Santa Clara Public Health Department | Chan-Zuckerberg Biohub | CZB Cliahub Consortium |
| EPI_ISL_548544 | County of San Luis Obispo Public Health Laboratory | Chan-Zuckerberg Biohub | CZB Cliahub Consortium |
| EPI_ISL_548545 | Contra Costa Public Health Lab | Chan-Zuckerberg Biohub | CZB Cliahub Consortium |
| EPI_ISL_548546, EPI_ISL_548547 | County of Santa Clara Public Health Department | Chan-Zuckerberg Biohub | CZB Cliahub Consortium |
| EPI_ISL_548548, EPI_ISL_548549 | Contra Costa Public Health Lab | Chan-Zuckerberg Biohub | CZB Cliahub Consortium |
| EPI_ISL_548550, EPI_ISL_548551 | County of Santa Clara Public Health Department | Chan-Zuckerberg Biohub | CZB Cliahub Consortium |
| EPI_ISL_548552 | Contra Costa Public Health Lab | Chan-Zuckerberg Biohub | CZB Cliahub Consortium |
| EPI_ISL_548553 | County of Santa Clara Public Health Department | Chan-Zuckerberg Biohub | CZB Cliahub Consortium |
| EPI_ISL_548554 | Contra Costa Public Health Lab | Chan-Zuckerberg Biohub | CZB Cliahub Consortium |
| EPI_ISL_548555 | UCSF Clinical Microbiology Laboratory | Chan-Zuckerberg Biohub | CZB Cliahub Consortium |
| EPI_ISL_548556, EPI_ISL_548557, EPI_ISL_548558 | Contra Costa Public Health Lab | Chan-Zuckerberg Biohub | CZB Cliahub Consortium |
| EPI_ISL_548559 | County of Santa Clara Public Health Department | Chan-Zuckerberg Biohub | CZB Cliahub Consortium |
| EPI_ISL_548560 | County of San Luis Obispo Public Health Laboratory | Chan-Zuckerberg Biohub | CZB Cliahub Consortium |
| EPI_ISL_548561 | Contra Costa Public Health Lab | Chan-Zuckerberg Biohub | CZB Cliahub Consortium |
| EPI_ISL_548562, EPI_ISL_548563 | County of Santa Clara Public Health Department | Chan-Zuckerberg Biohub | CZB Cliahub Consortium |
| EPI_ISL_548564, EPI_ISL_548565, EPI_ISL_548566, EPI_ISL_548567 | Contra Costa Public Health Lab | Chan-Zuckerberg Biohub | CZB Cliahub Consortium |
| EPI_ISL_548568 | County of Santa Clara Public Health Department | Chan-Zuckerberg Biohub | CZB Cliahub Consortium |
| EPI_ISL_548569 | Contra Costa Public Health Lab | Chan-Zuckerberg Biohub | CZB Cliahub Consortium |
| EPI_ISL_548570 | County of Santa Clara Public Health Department | Chan-Zuckerberg Biohub | CZB Cliahub Consortium |
| EPI_ISL_548571 | UCSF Clinical Microbiology Laboratory | Chan-Zuckerberg Biohub | CZB Cliahub Consortium |
| EPI_ISL_548572, EPI_ISL_548573, EPI_ISL_548574, EPI_ISL_548575 | Contra Costa Public Health Lab | Chan-Zuckerberg Biohub | CZB Cliahub Consortium |
| EPI_ISL_548576 | County of Santa Clara Public Health Department | Chan-Zuckerberg Biohub | CZB Cliahub Consortium |
| EPI_ISL_548577 | Contra Costa Public Health Lab | Chan-Zuckerberg Biohub | CZB Cliahub Consortium |
| EPI_ISL_548578 | County of Santa Clara Public Health Department | Chan-Zuckerberg Biohub | CZB Cliahub Consortium |
| EPI_ISL_548579 | Contra Costa Public Health Lab | Chan-Zuckerberg Biohub | CZB Cliahub Consortium |

| | | | |
|---|---|---|---|
| EPI_ISL_548580, EPI_ISL_548581 | Orange County Public Health Laboratory | Chan-Zuckerberg Biohub | CZB Cliahub Consortium |
| EPI_ISL_548582, EPI_ISL_548583, EPI_ISL_548584 | County of Santa Clara Public Health Department | Chan-Zuckerberg Biohub | CZB Cliahub Consortium |
| EPI_ISL_548585, EPI_ISL_548586, EPI_ISL_548587 | Orange County Public Health Laboratory | Chan-Zuckerberg Biohub | CZB Cliahub Consortium |
| EPI_ISL_548588 | County of Santa Clara Public Health Department | Chan-Zuckerberg Biohub | CZB Cliahub Consortium |
| EPI_ISL_548589, EPI_ISL_548590, EPI_ISL_548591, EPI_ISL_548592 | Orange County Public Health Laboratory | Chan-Zuckerberg Biohub | CZB Cliahub Consortium |
| EPI_ISL_548593 | County of Santa Clara Public Health Department | Chan-Zuckerberg Biohub | CZB Cliahub Consortium |
| EPI_ISL_548594 | Orange County Public Health Laboratory | Chan-Zuckerberg Biohub | CZB Cliahub Consortium |
| EPI_ISL_548595, EPI_ISL_548596, EPI_ISL_548597, EPI_ISL_548598 | County of Santa Clara Public Health Department | Chan-Zuckerberg Biohub | CZB Cliahub Consortium |
| EPI_ISL_548599 | Tuolumne County Public Health | Chan-Zuckerberg Biohub | CZB Cliahub Consortium |
| EPI_ISL_548600 | County of Santa Clara Public Health Department | Chan-Zuckerberg Biohub | CZB Cliahub Consortium |
| EPI_ISL_548601 | Orange County Public Health Laboratory | Chan-Zuckerberg Biohub | CZB Cliahub Consortium |
| EPI_ISL_548602 | County of Santa Clara Public Health Department | Chan-Zuckerberg Biohub | CZB Cliahub Consortium |
| EPI_ISL_548603 | County of San Luis Obispo Public Health Laboratory | Chan-Zuckerberg Biohub | CZB Cliahub Consortium |
| EPI_ISL_548604, EPI_ISL_548605, EPI_ISL_548606, EPI_ISL_548607 | County of Santa Clara Public Health Department | Chan-Zuckerberg Biohub | CZB Cliahub Consortium |
| EPI_ISL_548608, EPI_ISL_548609 | Orange County Public Health Laboratory | Chan-Zuckerberg Biohub | CZB Cliahub Consortium |
| EPI_ISL_548610 | County of Santa Clara Public Health Department | Chan-Zuckerberg Biohub | CZB Cliahub Consortium |
| EPI_ISL_548611 | Orange County Public Health Laboratory | Chan-Zuckerberg Biohub | CZB Cliahub Consortium |
| EPI_ISL_548612 | County of Santa Clara Public Health Department | Chan-Zuckerberg Biohub | CZB Cliahub Consortium |
| EPI_ISL_548613, EPI_ISL_548614, EPI_ISL_548615 | Orange County Public Health Laboratory | Chan-Zuckerberg Biohub | CZB Cliahub Consortium |
| EPI_ISL_548616, EPI_ISL_548617, EPI_ISL_548618, EPI_ISL_548619 | County of Santa Clara Public Health Department | Chan-Zuckerberg Biohub | CZB Cliahub Consortium |
| EPI_ISL_548620 | Orange County Public Health Laboratory | Chan-Zuckerberg Biohub | CZB Cliahub Consortium |
| EPI_ISL_548621, EPI_ISL_548622 | County of Santa Clara Public Health Department | Chan-Zuckerberg Biohub | CZB Cliahub Consortium |
| EPI_ISL_548623 | Orange County Public Health Laboratory | Chan-Zuckerberg Biohub | CZB Cliahub Consortium |
| EPI_ISL_548624 | County of Santa Clara Public Health Department | Chan-Zuckerberg Biohub | CZB Cliahub Consortium |
| EPI_ISL_548625 | Orange County Public Health Laboratory | Chan-Zuckerberg Biohub | CZB Cliahub Consortium |
| EPI_ISL_548626 | Tuolumne County Public Health | Chan-Zuckerberg Biohub | CZB Cliahub Consortium |
| EPI_ISL_548627, EPI_ISL_548628 | Orange County Public Health Laboratory | Chan-Zuckerberg Biohub | CZB Cliahub Consortium |
| EPI_ISL_548629, EPI_ISL_548630 | County of Santa Clara Public Health Department | Chan-Zuckerberg Biohub | CZB Cliahub Consortium |
| EPI_ISL_548631 | County of San Luis Obispo Public Health Laboratory | Chan-Zuckerberg Biohub | CZB Cliahub Consortium |
| EPI_ISL_548632, EPI_ISL_548633 | County of Santa Clara Public Health Department | Chan-Zuckerberg Biohub | CZB Cliahub Consortium |
| EPI_ISL_548634, EPI_ISL_548635 | Orange County Public Health Laboratory | Chan-Zuckerberg Biohub | CZB Cliahub Consortium |
| EPI_ISL_548636 | County of Santa Clara Public Health Department | Chan-Zuckerberg Biohub | CZB Cliahub Consortium |
| EPI_ISL_548637, EPI_ISL_548638 | Orange County Public Health Laboratory | Chan-Zuckerberg Biohub | CZB Cliahub Consortium |
| EPI_ISL_548639 | County of Santa Clara Public Health Department | Chan-Zuckerberg Biohub | CZB Cliahub Consortium |
| EPI_ISL_548640, EPI_ISL_548641 | Orange County Public Health Laboratory | Chan-Zuckerberg Biohub | CZB Cliahub Consortium |
| EPI_ISL_548642, EPI_ISL_548643 | County of Santa Clara Public Health Department | Chan-Zuckerberg Biohub | CZB Cliahub Consortium |
| EPI_ISL_548644 | County of San Luis Obispo Public Health Laboratory | Chan-Zuckerberg Biohub | CZB Cliahub Consortium |
| EPI_ISL_548645 | County of Santa Clara Public Health Department | Chan-Zuckerberg Biohub | CZB Cliahub Consortium |
| EPI_ISL_548646 | Orange County Public Health Laboratory | Chan-Zuckerberg Biohub | CZB Cliahub Consortium |
| EPI_ISL_548647 | County of Santa Clara Public Health Department | Chan-Zuckerberg Biohub | CZB Cliahub Consortium |
| EPI_ISL_548648 | Orange County Public Health Laboratory | Chan-Zuckerberg Biohub | CZB Cliahub Consortium |
| EPI_ISL_548649, EPI_ISL_548650 | County of Santa Clara Public Health Department | Chan-Zuckerberg Biohub | CZB Cliahub Consortium |
| EPI_ISL_548651 | Orange County Public Health Laboratory | Chan-Zuckerberg Biohub | CZB Cliahub Consortium |
| EPI_ISL_548652 | County of Santa Clara Public Health Department | Chan-Zuckerberg Biohub | CZB Cliahub Consortium |
| EPI_ISL_548653, EPI_ISL_548654 | Orange County Public Health Laboratory | Chan-Zuckerberg Biohub | CZB Cliahub Consortium |
| EPI_ISL_548655, EPI_ISL_548656, EPI_ISL_548657, EPI_ISL_548658, EPI_ISL_548659 | County of Santa Clara Public Health Department | Chan-Zuckerberg Biohub | CZB Cliahub Consortium |
| EPI_ISL_548660 | Orange County Public Health Laboratory | Chan-Zuckerberg Biohub | CZB Cliahub Consortium |
| EPI_ISL_548661 | County of Santa Clara Public Health Department | Chan-Zuckerberg Biohub | CZB Cliahub Consortium |
| EPI_ISL_548662 | Orange County Public Health Laboratory | Chan-Zuckerberg Biohub | CZB Cliahub Consortium |
| EPI_ISL_548663 | County of San Luis Obispo Public Health Laboratory | Chan-Zuckerberg Biohub | CZB Cliahub Consortium |
| EPI_ISL_548664 | County of Santa Clara Public Health Department | Chan-Zuckerberg Biohub | CZB Cliahub Consortium |
| EPI_ISL_548665, EPI_ISL_548666, EPI_ISL_548667, EPI_ISL_548668 | Orange County Public Health Laboratory | Chan-Zuckerberg Biohub | CZB Cliahub Consortium |

| | | | |
|---|---|---|---|
| EPI_ISL_548669, EPI_ISL_548670 | County of Santa Clara Public Health Department | Chan-Zuckerberg Biohub | CZB Cliahub Consortium |
| EPI_ISL_548671 | County of San Luis Obispo Public Health Laboratory | Chan-Zuckerberg Biohub | CZB Cliahub Consortium |
| EPI_ISL_548672, EPI_ISL_548673 | County of Santa Clara Public Health Department | Chan-Zuckerberg Biohub | CZB Cliahub Consortium |
| EPI_ISL_548674, EPI_ISL_548675 | Orange County Public Health Laboratory | Chan-Zuckerberg Biohub | CZB Cliahub Consortium |
| EPI_ISL_548676 | County of Santa Clara Public Health Department | Chan-Zuckerberg Biohub | CZB Cliahub Consortium |
| EPI_ISL_548677 | Orange County Public Health Laboratory | Chan-Zuckerberg Biohub | CZB Cliahub Consortium |
| EPI_ISL_548678, EPI_ISL_548679 | County of Santa Clara Public Health Department | Chan-Zuckerberg Biohub | CZB Cliahub Consortium |
| EPI_ISL_561380, EPI_ISL_561388, EPI_ISL_561392, EPI_ISL_561397, EPI_ISL_561403, EPI_ISL_561404, EPI_ISL_561407, EPI_ISL_561409, EPI_ISL_561414, EPI_ISL_561416, EPI_ISL_561420, EPI_ISL_561425, EPI_ISL_561426, EPI_ISL_561428, EPI_ISL_561433, EPI_ISL_561434, EPI_ISL_561440, EPI_ISL_561450, EPI_ISL_561453, EPI_ISL_561458, EPI_ISL_561462, EPI_ISL_561465, EPI_ISL_561468, EPI_ISL_561491, EPI_ISL_561498, EPI_ISL_561508, EPI_ISL_561514, EPI_ISL_561515, EPI_ISL_561517, EPI_ISL_561519, EPI_ISL_561523, EPI_ISL_561532, EPI_ISL_561533, EPI_ISL_561535, EPI_ISL_561541, EPI_ISL_561544, EPI_ISL_561559, EPI_ISL_561565, EPI_ISL_561578, EPI_ISL_561582, EPI_ISL_561584, EPI_ISL_561586, EPI_ISL_561590, EPI_ISL_561591, EPI_ISL_561612, EPI_ISL_561613, EPI_ISL_561614, EPI_ISL_561621, EPI_ISL_561622, EPI_ISL_561629, EPI_ISL_561651, EPI_ISL_561653, EPI_ISL_561655, EPI_ISL_561656, EPI_ISL_561657, EPI_ISL_561661, EPI_ISL_561673, EPI_ISL_561679, EPI_ISL_561682, EPI_ISL_561686, EPI_ISL_561690, EPI_ISL_561697, EPI_ISL_561707, EPI_ISL_561711, EPI_ISL_561715, EPI_ISL_561721, EPI_ISL_561728, EPI_ISL_561741, EPI_ISL_561750, EPI_ISL_561752, EPI_ISL_561764, EPI_ISL_561765, EPI_ISL_561777, EPI_ISL_561778, EPI_ISL_561787, EPI_ISL_561788, EPI_ISL_561791, EPI_ISL_561793, EPI_ISL_561799, EPI_ISL_561803, EPI_ISL_561808, EPI_ISL_561816, EPI_ISL_561825, EPI_ISL_561840, EPI_ISL_561843, EPI_ISL_561844, EPI_ISL_561852, EPI_ISL_561856, EPI_ISL_561858, EPI_ISL_561863, EPI_ISL_561866, EPI_ISL_561879, EPI_ISL_561893, EPI_ISL_561894, EPI_ISL_561935, EPI_ISL_561936, EPI_ISL_561946, EPI_ISL_561957, EPI_ISL_561975, EPI_ISL_561986, EPI_ISL_561987, EPI_ISL_561988, EPI_ISL_561991, EPI_ISL_561994, EPI_ISL_562011, EPI_ISL_562012, EPI_ISL_562016, EPI_ISL_562021, EPI_ISL_562023, EPI_ISL_562033, EPI_ISL_562048, EPI_ISL_562055, EPI_ISL_562056, EPI_ISL_562057, EPI_ISL_562066, EPI_ISL_562067, EPI_ISL_562069, EPI_ISL_562076, EPI_ISL_562080, EPI_ISL_562081, EPI_ISL_562085, EPI_ISL_562104, EPI_ISL_562105, EPI_ISL_562106, EPI_ISL_562109, EPI_ISL_562110, EPI_ISL_562115, EPI_ISL_562123, EPI_ISL_562136, EPI_ISL_562138, EPI_ISL_562152, EPI_ISL_562155, EPI_ISL_562158, EPI_ISL_562163, EPI_ISL_562164, EPI_ISL_562166, EPI_ISL_562175, EPI_ISL_562179, EPI_ISL_562189, EPI_ISL_562194, EPI_ISL_562200, EPI_ISL_562202, EPI_ISL_562208, EPI_ISL_562209, EPI_ISL_562211, EPI_ISL_562213, EPI_ISL_562216, EPI_ISL_562217, EPI_ISL_562224, EPI_ISL_562225, EPI_ISL_562231, EPI_ISL_562237, EPI_ISL_562250, EPI_ISL_562258, EPI_ISL_562259, EPI_ISL_562261, EPI_ISL_562264, EPI_ISL_562267, EPI_ISL_562272, EPI_ISL_562273, EPI_ISL_562274, EPI_ISL_562280, EPI_ISL_562282, EPI_ISL_562286, EPI_ISL_562288, EPI_ISL_562289, EPI_ISL_562290, EPI_ISL_562294, EPI_ISL_562296, EPI_ISL_562297, EPI_ISL_562308, EPI_ISL_562317, EPI_ISL_562322, EPI_ISL_562326, EPI_ISL_562328, EPI_ISL_562331, EPI_ISL_562333, EPI_ISL_562335, EPI_ISL_562342, EPI_ISL_562348, EPI_ISL_562351, EPI_ISL_562354, EPI_ISL_562357, EPI_ISL_562363, EPI_ISL_562366, EPI_ISL_562376, EPI_ISL_562379, EPI_ISL_562381, EPI_ISL_562390, EPI_ISL_562393, EPI_ISL_562396, EPI_ISL_562406, EPI_ISL_562410, EPI_ISL_562412, EPI_ISL_562414, EPI_ISL_562415, EPI_ISL_562430, EPI_ISL_562431, EPI_ISL_562435, EPI_ISL_562437, EPI_ISL_562440, EPI_ISL_562441, EPI_ISL_562442, EPI_ISL_562445, EPI_ISL_562460, EPI_ISL_562463, EPI_ISL_562466, EPI_ISL_562483, EPI_ISL_562484, EPI_ISL_562488, EPI_ISL_562502, EPI_ISL_562503, EPI_ISL_562509, EPI_ISL_562513, EPI_ISL_562517, EPI_ISL_562524, EPI_ISL_562526, EPI_ISL_562531, EPI_ISL_562535, EPI_ISL_562536, EPI_ISL_562538, EPI_ISL_562539, EPI_ISL_562540, EPI_ISL_562542, EPI_ISL_562545, EPI_ISL_562546, EPI_ISL_562548, EPI_ISL_562552, EPI_ISL_562557, EPI_ISL_562558, EPI_ISL_562559, EPI_ISL_562585, EPI_ISL_562587, EPI_ISL_562588, EPI_ISL_562589, EPI_ISL_562593, EPI_ISL_562595, EPI_ISL_562628, EPI_ISL_562640, EPI_ISL_562645, EPI_ISL_562646, EPI_ISL_562658, EPI_ISL_562659, EPI_ISL_562669, EPI_ISL_562671, EPI_ISL_562696, EPI_ISL_562697, EPI_ISL_562700, EPI_ISL_562705, EPI_ISL_562754, EPI_ISL_562791, EPI_ISL_562792, EPI_ISL_562803, EPI_ISL_562804, EPI_ISL_562805, EPI_ISL_562806, EPI_ISL_562807, EPI_ISL_562808, EPI_ISL_562809, EPI_ISL_562838, EPI_ISL_562839, EPI_ISL_562840, EPI_ISL_562841, EPI_ISL_562842, EPI_ISL_562843, EPI_ISL_562844, EPI_ISL_562845, EPI_ISL_562910, EPI_ISL_562911, EPI_ISL_562912, EPI_ISL_562913, EPI_ISL_562924, EPI_ISL_562929, EPI_ISL_562936, EPI_ISL_562946, EPI_ISL_562951, EPI_ISL_562961, EPI_ISL_563000, EPI_ISL_563131, EPI_ISL_563132, EPI_ISL_563133, EPI_ISL_563136, EPI_ISL_563137, EPI_ISL_563138, EPI_ISL_563139, EPI_ISL_563140, EPI_ISL_563141, EPI_ISL_563142, EPI_ISL_563143, EPI_ISL_563144, EPI_ISL_563407, EPI_ISL_563408, EPI_ISL_563409, EPI_ISL_563410, EPI_ISL_563411, EPI_ISL_563412, EPI_ISL_563413, EPI_ISL_563414, EPI_ISL_563415, EPI_ISL_563416, EPI_ISL_563417, EPI_ISL_563418, EPI_ISL_563419, EPI_ISL_563420, EPI_ISL_563421, EPI_ISL_563422, EPI_ISL_563423, EPI_ISL_563497, EPI_ISL_563498, EPI_ISL_563499, EPI_ISL_563551, EPI_ISL_563609, EPI_ISL_563721, EPI_ISL_563722, EPI_ISL_563723, EPI_ISL_563790, EPI_ISL_563792, EPI_ISL_563808, EPI_ISL_563811, EPI_ISL_563817, EPI_ISL_563820, EPI_ISL_563832, EPI_ISL_563840, EPI_ISL_563866, EPI_ISL_563880, EPI_ISL_563925, EPI_ISL_563938, EPI_ISL_563973, EPI_ISL_563974, EPI_ISL_563975, EPI_ISL_563977, EPI_ISL_563978, EPI_ISL_563987, EPI_ISL_563997, EPI_ISL_563998, EPI_ISL_563999, EPI_ISL_564000, EPI_ISL_564001, EPI_ISL_564002, EPI_ISL_564003, EPI_ISL_564004, EPI_ISL_564005, EPI_ISL_564006, EPI_ISL_564007, EPI_ISL_564008, EPI_ISL_564009, EPI_ISL_564010, EPI_ISL_564011, EPI_ISL_564012, EPI_ISL_564013, EPI_ISL_564015, EPI_ISL_564016, EPI_ISL_564017, EPI_ISL_564018, EPI_ISL_564019, EPI_ISL_564020, EPI_ISL_564021, EPI_ISL_564022, EPI_ISL_564023, EPI_ISL_564024, EPI_ISL_564025, EPI_ISL_564026, EPI_ISL_564027, EPI_ISL_564028, EPI_ISL_564029, EPI_ISL_564030, EPI_ISL_564031, EPI_ISL_564032, EPI_ISL_564033, EPI_ISL_564034, EPI_ISL_564035, EPI_ISL_564036, EPI_ISL_564037, EPI_ISL_564038, EPI_ISL_564039, EPI_ISL_564040, EPI_ISL_564041, EPI_ISL_564042, EPI_ISL_564043, EPI_ISL_564044, EPI_ISL_564045, EPI_ISL_564046, EPI_ISL_564047, EPI_ISL_564048, EPI_ISL_564049, EPI_ISL_564055, EPI_ISL_564056, EPI_ISL_564057, EPI_ISL_564058, EPI_ISL_564059, EPI_ISL_564060, EPI_ISL_564061, EPI_ISL_564062, EPI_ISL_564063, EPI_ISL_564064, EPI_ISL_564065, EPI_ISL_564066, EPI_ISL_564067, EPI_ISL_564068, EPI_ISL_564069, EPI_ISL_564070, EPI_ISL_564071, EPI_ISL_564072, EPI_ISL_564074, EPI_ISL_564076, EPI_ISL_564077, EPI_ISL_564078, EPI_ISL_564079, EPI_ISL_564080, EPI_ISL_564081, EPI_ISL_564082, EPI_ISL_564083, EPI_ISL_564084, EPI_ISL_564085, EPI_ISL_564086, EPI_ISL_564087, EPI_ISL_564088, EPI_ISL_564089, EPI_ISL_564090, EPI_ISL_564091, EPI_ISL_564092, EPI_ISL_564093, EPI_ISL_564094, EPI_ISL_564095, EPI_ISL_564096, EPI_ISL_564097, EPI_ISL_564098, EPI_ISL_564099, EPI_ISL_564100, EPI_ISL_564101, EPI_ISL_564102, EPI_ISL_564103, EPI_ISL_564104, EPI_ISL_564105, EPI_ISL_564106, EPI_ISL_564107, EPI_ISL_564108, EPI_ISL_564109, EPI_ISL_564110, EPI_ISL_564180, EPI_ISL_564181, EPI_ISL_564182, EPI_ISL_564183, EPI_ISL_564184, EPI_ISL_564185, EPI_ISL_564186, EPI_ISL_564187, EPI_ISL_564188, EPI_ISL_564189, EPI_ISL_564190, EPI_ISL_564191, EPI_ISL_564192, EPI_ISL_564193, EPI_ISL_564194, EPI_ISL_564195, EPI_ISL_564196, EPI_ISL_564197, EPI_ISL_564198, EPI_ISL_564199, EPI_ISL_564200, EPI_ISL_564202, EPI_ISL_564203, EPI_ISL_564204, EPI_ISL_564205, EPI_ISL_564206, EPI_ISL_564207, EPI_ISL_564208, EPI_ISL_564209, EPI_ISL_564210, EPI_ISL_564211, EPI_ISL_564212, EPI_ISL_564213, EPI_ISL_564214, EPI_ISL_564215, EPI_ISL_564216, EPI_ISL_564217, EPI_ISL_564218, EPI_ISL_564219, EPI_ISL_564220, EPI_ISL_564221, EPI_ISL_564222, EPI_ISL_564223, EPI_ISL_564224, EPI_ISL_564225, EPI_ISL_564226, EPI_ISL_564227, EPI_ISL_564228, EPI_ISL_564229, EPI_ISL_564230, EPI_ISL_564231, EPI_ISL_564232, EPI_ISL_564233, EPI_ISL_564234, EPI_ISL_564235, EPI_ISL_564236, EPI_ISL_564237, EPI_ISL_564238, EPI_ISL_564239, EPI_ISL_564240, EPI_ISL_564241, EPI_ISL_564242, EPI_ISL_564243, EPI_ISL_564244, EPI_ISL_564245, EPI_ISL_564246, EPI_ISL_564247, EPI_ISL_564248, EPI_ISL_564249, EPI_ISL_564250, EPI_ISL_564251, EPI_ISL_564252, EPI_ISL_564253, EPI_ISL_564254, EPI_ISL_564270, EPI_ISL_564318, EPI_ISL_564319, EPI_ISL_564320, EPI_ISL_564321, EPI_ISL_564322, EPI_ISL_564323, EPI_ISL_564324, EPI_ISL_564325, EPI_ISL_564326, EPI_ISL_564327, EPI_ISL_564328, EPI_ISL_564329, EPI_ISL_564330, EPI_ISL_564331, EPI_ISL_564332, EPI_ISL_564333, EPI_ISL_564334, EPI_ISL_564335, EPI_ISL_564336, EPI_ISL_564337, EPI_ISL_564338, EPI_ISL_564339, EPI_ISL_564340, EPI_ISL_564341, EPI_ISL_564342, EPI_ISL_564343, EPI_ISL_564344, EPI_ISL_564345, EPI_ISL_564346, EPI_ISL_564347, EPI_ISL_564348, EPI_ISL_564349, EPI_ISL_564350, EPI_ISL_564353, EPI_ISL_564354, EPI_ISL_564355, EPI_ISL_564356, EPI_ISL_564357, EPI_ISL_564358, EPI_ISL_564359, EPI_ISL_564360, EPI_ISL_564361, EPI_ISL_564362, EPI_ISL_564363, EPI_ISL_564364, EPI_ISL_564365, EPI_ISL_564366, EPI_ISL_564367, EPI_ISL_564368, EPI_ISL_564369, EPI_ISL_564370, EPI_ISL_564371, EPI_ISL_564372, EPI_ISL_564373, EPI_ISL_564374, EPI_ISL_564375, EPI_ISL_564376, EPI_ISL_564377, EPI_ISL_564378, EPI_ISL_564379, EPI_ISL_564380, EPI_ISL_564381, EPI_ISL_564382, EPI_ISL_564383, EPI_ISL_564384, EPI_ISL_564385, EPI_ISL_564386, EPI_ISL_564387, EPI_ISL_564388, EPI_ISL_564389, EPI_ISL_564390, EPI_ISL_564391, EPI_ISL_564392, EPI_ISL_564393, EPI_ISL_564394, EPI_ISL_564395, EPI_ISL_564396, EPI_ISL_564397, EPI_ISL_564404, EPI_ISL_564544, EPI_ISL_564546, EPI_ISL_564547, EPI_ISL_564548, EPI_ISL_564549, EPI_ISL_564550, EPI_ISL_564551, EPI_ISL_564552, EPI_ISL_564553, EPI_ISL_564554, EPI_ISL_564555, EPI_ISL_564556, EPI_ISL_564557, EPI_ISL_564558, EPI_ISL_564559, EPI_ISL_564560, EPI_ISL_564561, EPI_ISL_564562, EPI_ISL_564563, EPI_ISL_564564, EPI_ISL_565136, EPI_ISL_565137, EPI_ISL_565138, EPI_ISL_565139, EPI_ISL_565140, EPI_ISL_565141, EPI_ISL_565142, EPI_ISL_565143, EPI_ISL_565144, EPI_ISL_565145, EPI_ISL_565146, EPI_ISL_565147, EPI_ISL_565148, EPI_ISL_565149, EPI_ISL_565150, EPI_ISL_565151, EPI_ISL_565152, EPI_ISL_565153, EPI_ISL_565154, EPI_ISL_565155, EPI_ISL_565156, EPI_ISL_565157, EPI_ISL_565158, EPI_ISL_565159, EPI_ISL_565160, EPI_ISL_565161, EPI_ISL_565162, EPI_ISL_565163, EPI_ISL_565164, EPI_ISL_565165, EPI_ISL_565166, EPI_ISL_565167, EPI_ISL_565168, EPI_ISL_565169, EPI_ISL_565170, EPI_ISL_565171, EPI_ISL_565172, EPI_ISL_565173, EPI_ISL_565174, EPI_ISL_565175, EPI_ISL_565176, EPI_ISL_565177, EPI_ISL_565178, EPI_ISL_565179, EPI_ISL_565180, EPI_ISL_565181, EPI_ISL_565182, EPI_ISL_565183, EPI_ISL_565184, EPI_ISL_565185, EPI_ISL_565186, EPI_ISL_565187, EPI_ISL_565188, EPI_ISL_565189, EPI_ISL_565190, EPI_ISL_565191, EPI_ISL_565192, EPI_ISL_565193, EPI_ISL_565194, EPI_ISL_565195, EPI_ISL_565196, EPI_ISL_565197, EPI_ISL_565198, EPI_ISL_565199, EPI_ISL_565200, EPI_ISL_565201, EPI_ISL_565202, EPI_ISL_565203, EPI_ISL_565204, EPI_ISL_565205, EPI_ISL_565206, EPI_ISL_565207, EPI_ISL_565208, EPI_ISL_565209, EPI_ISL_565210, EPI_ISL_565211, EPI_ISL_565212, EPI_ISL_565213, EPI_ISL_565214, EPI_ISL_565215, EPI_ISL_565216, EPI_ISL_565217, EPI_ISL_565218, EPI_ISL_565219, EPI_ISL_565220, EPI_ISL_565221, EPI_ISL_565222, EPI_ISL_565382, EPI_ISL_565385, EPI_ISL_565394, EPI_ISL_565395, EPI_ISL_565396, EPI_ISL_565399, EPI_ISL_565401, EPI_ISL_565402, EPI_ISL_565403, EPI_ISL_565404, EPI_ISL_565409, EPI_ISL_565415, EPI_ISL_565459, EPI_ISL_565461, EPI_ISL_565463, EPI_ISL_565464, EPI_ISL_565465, EPI_ISL_565466, EPI_ISL_565467, EPI_ISL_565469, EPI_ISL_565509, EPI_ISL_565730, EPI_ISL_565731, EPI_ISL_565732, EPI_ISL_565733, EPI_ISL_565734, EPI_ISL_565735, EPI_ISL_565736, EPI_ISL_565737, EPI_ISL_565738, EPI_ISL_565739, EPI_ISL_565740, EPI_ISL_565741, EPI_ISL_565742 | | | |
| see above | Victorian Infectious Diseases Reference Laboratory (VIDRL) | VIDRL and MDU-PHL | Caly, L., Seemann, T., Sait, M., Schultz, M. B., Druce J., Sherry, N. |
| EPI_ISL_577740, EPI_ISL_577741, EPI_ISL_577742 | Institute of Virology, Biomedical Research Center of the Slovak Academy of Sciences, Bratislava | Faculty of Natural Sciences, Comenius University, Bratislava | Broa Brejová, Viktória Hodorová, Kristína Boršová, Viktória abanová, Dominika Friová, Sabina Fumaová Havlíková, Juraj Kopáek, Martina Liková, ubomíra Lukáiková, Martina Neboháová, Monika Sláviková, Edita Staroová, Elena Tichá, Tomáš Vina, Jozef Nosek, Boris Klempa |
| EPI_ISL_582514, EPI_ISL_582515 | unknown | Instituto Nacional de Saude (INSA) and Instituto Gulbenkian de Ciencia (IGC) | Borges et al |
| EPI_ISL_582516, EPI_ISL_582517, EPI_ISL_582518, EPI_ISL_582519, EPI_ISL_582520, EPI_ISL_582521 | unknown | Instituto Nacional de Saude (INSA) | Borges et al |
| EPI_ISL_582522, EPI_ISL_582523 | unknown | Instituto Nacional de Saude (INSA) and Instituto Gulbenkian de Ciencia (IGC) | Borges et al |
| EPI_ISL_582851, EPI_ISL_582852, EPI_ISL_582853, EPI_ISL_582854, EPI_ISL_582855, EPI_ISL_582856, EPI_ISL_582857, EPI_ISL_582858, EPI_ISL_582859, EPI_ISL_582860, EPI_ISL_582861, EPI_ISL_582862, EPI_ISL_582863, EPI_ISL_582864, EPI_ISL_582865, EPI_ISL_582866, EPI_ISL_582867, EPI_ISL_582868, EPI_ISL_582869, EPI_ISL_582870, EPI_ISL_582871, EPI_ISL_582872, EPI_ISL_582873, EPI_ISL_582874, EPI_ISL_582875, EPI_ISL_582876, EPI_ISL_582877, EPI_ISL_582878, EPI_ISL_582879, EPI_ISL_582880, EPI_ISL_582881, EPI_ISL_582882, EPI_ISL_582883, EPI_ISL_582884, EPI_ISL_582885, EPI_ISL_582886, EPI_ISL_582887, EPI_ISL_582888, EPI_ISL_582889, EPI_ISL_582890, EPI_ISL_582891, EPI_ISL_582892, EPI_ISL_582893, EPI_ISL_582894, EPI_ISL_582895, EPI_ISL_582896, EPI_ISL_582897, EPI_ISL_582898, EPI_ISL_582899, EPI_ISL_582900, EPI_ISL_582901, EPI_ISL_582902, EPI_ISL_582903, EPI_ISL_582904, EPI_ISL_582905, EPI_ISL_582906, EPI_ISL_582907, EPI_ISL_582908, EPI_ISL_582909, EPI_ISL_582910, EPI_ISL_582911, EPI_ISL_582912, EPI_ISL_582913, EPI_ISL_582914, EPI_ISL_582915, EPI_ISL_582916, EPI_ISL_582917, EPI_ISL_582918, EPI_ISL_582919, EPI_ISL_582921, EPI_ISL_582922, EPI_ISL_582923, EPI_ISL_582924, EPI_ISL_582925, EPI_ISL_582926, EPI_ISL_582927, EPI_ISL_582928, EPI_ISL_582929, EPI_ISL_582930, EPI_ISL_582931, EPI_ISL_582932, EPI_ISL_582933, EPI_ISL_582934, EPI_ISL_582935, EPI_ISL_582936, EPI_ISL_582937, EPI_ISL_582938, EPI_ISL_582939, EPI_ISL_582940, EPI_ISL_582941, EPI_ISL_582942, EPI_ISL_582943, EPI_ISL_582944, EPI_ISL_582945, EPI_ISL_582946, EPI_ISL_582947, EPI_ISL_582948, EPI_ISL_582949, EPI_ISL_582950, EPI_ISL_582951, EPI_ISL_582952, EPI_ISL_582953, EPI_ISL_582954, EPI_ISL_582955, EPI_ISL_582956, EPI_ISL_582957, EPI_ISL_582958 | | | |
| see above | County of Santa Clara Public Health Department | Chan-Zuckerberg Biohub | CZB Cliahub Consortium |
| EPI_ISL_582959, EPI_ISL_582960, EPI_ISL_582961, EPI_ISL_582962, EPI_ISL_582963, EPI_ISL_582964, EPI_ISL_582965, EPI_ISL_582966, EPI_ISL_582967, EPI_ISL_582968, EPI_ISL_582969, EPI_ISL_582970, EPI_ISL_582971, EPI_ISL_582972, EPI_ISL_582973, EPI_ISL_582974 | | | |
| see above | San Luis Obispo Public Health Department | Chan-Zuckerberg Biohub | CZB Cliahub Consortium |
| EPI_ISL_582975, EPI_ISL_582976, EPI_ISL_582977, EPI_ISL_582978, EPI_ISL_582979, EPI_ISL_582980, EPI_ISL_582981, EPI_ISL_582982, EPI_ISL_582983, EPI_ISL_582984, EPI_ISL_582985, EPI_ISL_582986, EPI_ISL_582987, EPI_ISL_582988, EPI_ISL_582989, EPI_ISL_582990, EPI_ISL_582991, EPI_ISL_582992, EPI_ISL_582993, EPI_ISL_582994, EPI_ISL_582995, EPI_ISL_582996, EPI_ISL_582997, EPI_ISL_582998, EPI_ISL_582999, EPI_ISL_583000, EPI_ISL_583001, EPI_ISL_583002, EPI_ISL_583003, EPI_ISL_583004, EPI_ISL_583005, EPI_ISL_583006, EPI_ISL_583007, EPI_ISL_583008, EPI_ISL_583009, EPI_ISL_583010, EPI_ISL_583011, EPI_ISL_583012, EPI_ISL_583013, EPI_ISL_583014, EPI_ISL_583015, EPI_ISL_583016, EPI_ISL_583017, EPI_ISL_583018, EPI_ISL_583019, EPI_ISL_583020, EPI_ISL_583021, EPI_ISL_583022, EPI_ISL_583023, EPI_ISL_583024, EPI_ISL_583025, EPI_ISL_583026, EPI_ISL_583027, EPI_ISL_583028, EPI_ISL_583029, EPI_ISL_583030, EPI_ISL_583031, EPI_ISL_583032, EPI_ISL_583033, EPI_ISL_583034, EPI_ISL_583035, EPI_ISL_583036, EPI_ISL_583037, EPI_ISL_583038, EPI_ISL_583039, EPI_ISL_583040, EPI_ISL_583041, EPI_ISL_583042, EPI_ISL_583043, EPI_ISL_583044, EPI_ISL_583045 | | | |
| see above | Orange County Public Health Lab | Chan-Zuckerberg Biohub | CZB Cliahub Consortium |
| EPI_ISL_583046, EPI_ISL_583047, EPI_ISL_583048, EPI_ISL_583049, EPI_ISL_583050, EPI_ISL_583051, EPI_ISL_583052, EPI_ISL_583053, EPI_ISL_583054, EPI_ISL_583055, EPI_ISL_583056, EPI_ISL_583057, EPI_ISL_583059, EPI_ISL_583060, EPI_ISL_583061, EPI_ISL_583062, EPI_ISL_583063, EPI_ISL_583064, EPI_ISL_583065, EPI_ISL_583066, EPI_ISL_583067, EPI_ISL_583068, EPI_ISL_583069, EPI_ISL_583070, EPI_ISL_583071, EPI_ISL_583072, EPI_ISL_583073, EPI_ISL_583074, EPI_ISL_583075, EPI_ISL_583076, EPI_ISL_583077, EPI_ISL_583078, EPI_ISL_583079, EPI_ISL_583080, EPI_ISL_583081, EPI_ISL_583082, EPI_ISL_583083, EPI_ISL_583084, EPI_ISL_583085, EPI_ISL_583086, EPI_ISL_583087, EPI_ISL_583088, EPI_ISL_583089, EPI_ISL_583090, EPI_ISL_583091, EPI_ISL_583092, EPI_ISL_583093, EPI_ISL_583094, EPI_ISL_583095, EPI_ISL_583096, EPI_ISL_583097, EPI_ISL_583098, EPI_ISL_583099, EPI_ISL_583100, EPI_ISL_583101, EPI_ISL_583102, EPI_ISL_583103, EPI_ISL_583104, EPI_ISL_583105, EPI_ISL_583106, EPI_ISL_583107, EPI_ISL_583108, EPI_ISL_583109, EPI_ISL_583110, EPI_ISL_583111, EPI_ISL_583112, EPI_ISL_583113, EPI_ISL_583114, EPI_ISL_583115, EPI_ISL_583116, EPI_ISL_583117, EPI_ISL_583118, | | | |

| | | | |
|---|---|---|---|
| EPI_ISL_583119, EPI_ISL_583120, EPI_ISL_583121, EPI_ISL_583122, EPI_ISL_583123, EPI_ISL_583124, EPI_ISL_583125, EPI_ISL_583126, EPI_ISL_583127, EPI_ISL_583128, EPI_ISL_583129, EPI_ISL_583130, EPI_ISL_583131, EPI_ISL_583132, EPI_ISL_583133, EPI_ISL_583134, EPI_ISL_583135, EPI_ISL_583136, EPI_ISL_583137, EPI_ISL_583138, EPI_ISL_583139, EPI_ISL_583140, EPI_ISL_583141, EPI_ISL_583142, EPI_ISL_583143, EPI_ISL_583144, EPI_ISL_583145, EPI_ISL_583146, EPI_ISL_583147, EPI_ISL_583148, EPI_ISL_583149, EPI_ISL_583150, EPI_ISL_583151, EPI_ISL_583152, EPI_ISL_583153, EPI_ISL_583154, EPI_ISL_583155, EPI_ISL_583156, EPI_ISL_583157, EPI_ISL_583158, EPI_ISL_583159, EPI_ISL_583160, EPI_ISL_583161, EPI_ISL_583162, EPI_ISL_583163, EPI_ISL_583164, EPI_ISL_583165, EPI_ISL_583166, EPI_ISL_583167, EPI_ISL_583168, EPI_ISL_583169, EPI_ISL_583170 | | | |
| see above | Humboldt County Public Health Laboratory | Chan-Zuckerberg Biohub | CZB Cliahub Consortium |
| EPI_ISL_583171, EPI_ISL_583172 | Tulare County Public Health Lab | Chan-Zuckerberg Biohub | CZB Cliahub Consortium |
| EPI_ISL_583173, EPI_ISL_583174, EPI_ISL_583175, EPI_ISL_583176, EPI_ISL_583177, EPI_ISL_583178, EPI_ISL_583179, EPI_ISL_583180, EPI_ISL_583181, EPI_ISL_583182, EPI_ISL_583183, EPI_ISL_583184, EPI_ISL_583185, EPI_ISL_583186, EPI_ISL_583187, EPI_ISL_583188, EPI_ISL_583189 | | | |
| see above | San Bernardino County Public Health Lab | Chan-Zuckerberg Biohub | CZB Cliahub Consortium |
| EPI_ISL_583190, EPI_ISL_583191, EPI_ISL_583192, EPI_ISL_583193, EPI_ISL_583194, EPI_ISL_583195, EPI_ISL_583196, EPI_ISL_583197 | San Francisco Public Health Laboratory | Chan-Zuckerberg Biohub | CZB Cliahub Consortium |
| EPI_ISL_583198 | Marin County Public Health Department | Chan-Zuckerberg Biohub | CZB Cliahub Consortium |
| EPI_ISL_583199, EPI_ISL_583200, EPI_ISL_583201, EPI_ISL_583202, EPI_ISL_583203, EPI_ISL_583204, EPI_ISL_583205, EPI_ISL_583206, EPI_ISL_583207, EPI_ISL_583208, EPI_ISL_583209, EPI_ISL_583210, EPI_ISL_583211, EPI_ISL_583212, EPI_ISL_583213, EPI_ISL_583214, EPI_ISL_583215, EPI_ISL_583216, EPI_ISL_583217, EPI_ISL_583218, EPI_ISL_583219, EPI_ISL_583220, EPI_ISL_583221, EPI_ISL_583222, EPI_ISL_583223, EPI_ISL_583224, EPI_ISL_583225, EPI_ISL_583226, EPI_ISL_583227, EPI_ISL_583228, EPI_ISL_583229, EPI_ISL_583230, EPI_ISL_583231 | | | |
| see above | UCSF Clinical Microbiology Laboratory | Chan-Zuckerberg Biohub | CZB Cliahub Consortium |
| EPI_ISL_583487, EPI_ISL_583488 | Institute of Virology, Biomedical Research Center of the Slovak Academy of Sciences, Bratislava | Faculty of Natural Sciences, Comenius University, Bratislava | Broa Brejová, Viktória Hodorová, Kristína Boršová, Viktória abanová, Dominika Friová, Sabina Fumaová Havlíková, Juraj Kopáek, Martina Liková, ubomíra Lukáiková, Martina Neboháová, Monika Sláviková, Edita Staroová, Elena Tichá, Tomáš Vina, Jozef Nosek, Boris Klempa |
| EPI_ISL_586106, EPI_ISL_586107, EPI_ISL_586112, EPI_ISL_586115, EPI_ISL_586119, EPI_ISL_586120, EPI_ISL_586124, EPI_ISL_586126, EPI_ISL_586131, EPI_ISL_586134, EPI_ISL_586138, EPI_ISL_586140, EPI_ISL_586141, EPI_ISL_586143, EPI_ISL_586144, EPI_ISL_586146, EPI_ISL_586148, EPI_ISL_586151, EPI_ISL_586160, EPI_ISL_586161, EPI_ISL_586166, EPI_ISL_586169, EPI_ISL_586170, EPI_ISL_586173, EPI_ISL_586174, EPI_ISL_586175, EPI_ISL_586181, EPI_ISL_586183, EPI_ISL_586185, EPI_ISL_586186, EPI_ISL_586187, EPI_ISL_586189, EPI_ISL_586192, EPI_ISL_586195, EPI_ISL_586196, EPI_ISL_586197, EPI_ISL_586201, EPI_ISL_586208, EPI_ISL_586209, EPI_ISL_586212, EPI_ISL_586215, EPI_ISL_586216, EPI_ISL_586218, EPI_ISL_586219, EPI_ISL_586222, EPI_ISL_586223, EPI_ISL_586226, EPI_ISL_586228, EPI_ISL_586233, EPI_ISL_586239 | | | |
| see above | Wales Specialist Virology Centre Sequencing lab: Pathogen Genomics Unit | COVID-19 Genomics UK (COG-UK) Consortium | Catherine Moore, Johnathan Evans, Laura Gifford, Malorie Perry, Simon Cottrell, Angela Marchbank, Alec Birchley, Alexander Adams, Amy Gaskin, Bree Gatica-Wilcox, Jason Coombes, Joel Southgate, Lauren Gilbert, Lee Graham, Nicole Pacchiarini, Sara Kumziene-Summerhayes, Sarah Taylor, Sophie Jones, Sara Rey, Matthew Bull, Joanne Watkins, Sally Corden, Tom Connor |
| EPI_ISL_591336 | Dipartimento di Biotecnologie Mediche, University of Siena | Dipartimento di Biotecnologie Mediche, University of Siena | COVID |
| EPI_ISL_591484 | Australian Clinical Labs | NSW Health Pathology - Institute of Clinical Pathology and Medical Research; Westmead Hospital; University of Sydney | CIDM-PH et al. |
| EPI_ISL_591485, EPI_ISL_591486 | Histopath | NSW Health Pathology - Institute of Clinical Pathology and Medical Research; Westmead Hospital; University of Sydney | CIDM-PH et al. |
| EPI_ISL_591487, EPI_ISL_591488, EPI_ISL_591489, EPI_ISL_591490, EPI_ISL_591491 | Laverty Pathology | NSW Health Pathology - Institute of Clinical Pathology and Medical Research; Westmead Hospital; University of Sydney | CIDM-PH et al. |
| EPI_ISL_591492 | Medlab Pathology | NSW Health Pathology - Institute of Clinical Pathology and Medical Research; Westmead Hospital; University of Sydney | CIDM-PH et al. |
| EPI_ISL_591493 | Pathology North - Royal North Shore Hospital - NSW Health Pathology | NSW Health Pathology - Institute of Clinical Pathology and Medical Research; Westmead Hospital; University of Sydney | CIDM-PH et al. |
| EPI_ISL_591494, EPI_ISL_591495, EPI_ISL_591496, EPI_ISL_591497, EPI_ISL_591498, EPI_ISL_591499, EPI_ISL_591500, EPI_ISL_591501 | Pathology West - NSW Health Pathology | NSW Health Pathology - Institute of Clinical Pathology and Medical Research; Westmead Hospital; University of Sydney | CIDM-PH et al. |
| EPI_ISL_591502, EPI_ISL_591503, EPI_ISL_591504 | South Eastern Area Laboratory Services (SEALS) | NSW Health Pathology - Institute of Clinical Pathology and Medical Research; Westmead Hospital; University of Sydney | CIDM-PH et al. |
| EPI_ISL_591505, EPI_ISL_591506 | St Vincent's Pathology (SydPath) | NSW Health Pathology - Institute of Clinical Pathology and Medical Research; Westmead Hospital; University of Sydney | CIDM-PH et al. |
| EPI_ISL_591507 | Sydney South West Pathology Service (SSWPS) - Concord Repatriation General Hospital - NSW Health Pathology | NSW Health Pathology - Institute of Clinical Pathology and Medical Research; Westmead Hospital; University of Sydney | CIDM-PH et al. |
| EPI_ISL_591508, EPI_ISL_591509, EPI_ISL_591510, EPI_ISL_591511, EPI_ISL_591512, EPI_ISL_591513, EPI_ISL_591514, EPI_ISL_591515, EPI_ISL_591516 | Sydney South West Pathology Service (SSWPS) - Liverpool Hospital - NSW Health Pathology | NSW Health Pathology - Institute of Clinical Pathology and Medical Research; Westmead Hospital; University of Sydney | CIDM-PH et al. |
| EPI_ISL_591517 | Sydney South West Pathology Service (SSWPS) - Royal Prince Alfred Hospital - NSW Health Pathology | NSW Health Pathology - Institute of Clinical Pathology and Medical Research; Westmead Hospital; University of Sydney | CIDM-PH et al. |
| EPI_ISL_591518 | The Children's Hospital at Westmead | NSW Health Pathology - Institute of Clinical Pathology and Medical Research; Westmead Hospital; University of Sydney | CIDM-PH et al. |
| EPI_ISL_591558, EPI_ISL_591562, EPI_ISL_591565, EPI_ISL_591571, EPI_ISL_591573, EPI_ISL_591574, EPI_ISL_591578, EPI_ISL_591580, EPI_ISL_591584, EPI_ISL_591585, EPI_ISL_591587, EPI_ISL_591590, EPI_ISL_591591, EPI_ISL_591592, EPI_ISL_591595, EPI_ISL_591603, EPI_ISL_591609, EPI_ISL_591615, EPI_ISL_591619, EPI_ISL_591621, EPI_ISL_591623, EPI_ISL_591626, EPI_ISL_591638, EPI_ISL_591640, EPI_ISL_591643, EPI_ISL_591645, EPI_ISL_591652, EPI_ISL_591653, EPI_ISL_591654, EPI_ISL_591656, EPI_ISL_591657, EPI_ISL_591659, EPI_ISL_591666, EPI_ISL_591667, EPI_ISL_591669, EPI_ISL_591670, EPI_ISL_591672, EPI_ISL_591679, EPI_ISL_591681, EPI_ISL_591684, EPI_ISL_591688, EPI_ISL_591698, EPI_ISL_591700, EPI_ISL_591703, EPI_ISL_591704, EPI_ISL_591706, EPI_ISL_591707, EPI_ISL_591708, EPI_ISL_591715, EPI_ISL_591721, EPI_ISL_591723, EPI_ISL_591725, EPI_ISL_591726, EPI_ISL_591729, EPI_ISL_591734, EPI_ISL_591737, EPI_ISL_591743, EPI_ISL_591744, EPI_ISL_591746, EPI_ISL_591754, EPI_ISL_591755, EPI_ISL_591758, EPI_ISL_591760, EPI_ISL_591761, EPI_ISL_591763, EPI_ISL_591767, EPI_ISL_591771, EPI_ISL_591772, EPI_ISL_591773, EPI_ISL_591775, EPI_ISL_591778, EPI_ISL_591779, EPI_ISL_591782, EPI_ISL_591783, EPI_ISL_591786, EPI_ISL_591791, EPI_ISL_591792, EPI_ISL_591799, EPI_ISL_591800, EPI_ISL_591807, EPI_ISL_591822, EPI_ISL_591824, EPI_ISL_591825, EPI_ISL_591849, EPI_ISL_591850, EPI_ISL_591851, EPI_ISL_591852, EPI_ISL_591854, EPI_ISL_591856, EPI_ISL_591857, EPI_ISL_591858, EPI_ISL_591859, EPI_ISL_591860, EPI_ISL_591861, EPI_ISL_591862, EPI_ISL_591863, EPI_ISL_591864, EPI_ISL_591865, EPI_ISL_591866, EPI_ISL_591867, EPI_ISL_591868, EPI_ISL_591869, EPI_ISL_591870, EPI_ISL_591871, EPI_ISL_591872, EPI_ISL_591873, EPI_ISL_591874, EPI_ISL_591875, EPI_ISL_591876, EPI_ISL_591878, EPI_ISL_591879, EPI_ISL_591880, EPI_ISL_591881, EPI_ISL_591882, EPI_ISL_591883, EPI_ISL_591884, EPI_ISL_591885, EPI_ISL_591887, EPI_ISL_591888, EPI_ISL_591889, EPI_ISL_591890, EPI_ISL_591891, EPI_ISL_591892, EPI_ISL_591893, EPI_ISL_591894, EPI_ISL_591895, EPI_ISL_591896, EPI_ISL_591897, EPI_ISL_591898, EPI_ISL_591899, EPI_ISL_591900, EPI_ISL_591901, EPI_ISL_591904, EPI_ISL_591907, EPI_ISL_591908, EPI_ISL_591909, EPI_ISL_591912, EPI_ISL_591917, EPI_ISL_591919, EPI_ISL_591925, EPI_ISL_591928, EPI_ISL_591932, EPI_ISL_591940, EPI_ISL_591941, EPI_ISL_591942, EPI_ISL_591943, EPI_ISL_591946, EPI_ISL_591947, EPI_ISL_591948, EPI_ISL_591949, EPI_ISL_591950, EPI_ISL_591951, EPI_ISL_591952, EPI_ISL_591953, EPI_ISL_591954, EPI_ISL_591955, EPI_ISL_591956, EPI_ISL_591957, EPI_ISL_591959, EPI_ISL_592136, EPI_ISL_592137, EPI_ISL_592138, EPI_ISL_592139, EPI_ISL_592142, EPI_ISL_592143, EPI_ISL_592144, EPI_ISL_592145, EPI_ISL_592146, EPI_ISL_592147, EPI_ISL_592148, EPI_ISL_592149, EPI_ISL_592150, EPI_ISL_592151, EPI_ISL_592152, EPI_ISL_592154, EPI_ISL_592155, EPI_ISL_592156, EPI_ISL_592157, EPI_ISL_592158, EPI_ISL_592159, EPI_ISL_592160, EPI_ISL_592161, EPI_ISL_592162, EPI_ISL_592163, EPI_ISL_592164, EPI_ISL_592165, EPI_ISL_592166, EPI_ISL_592167, EPI_ISL_592168, EPI_ISL_592169, EPI_ISL_592170, EPI_ISL_592171, EPI_ISL_592175, EPI_ISL_592178, EPI_ISL_592179, EPI_ISL_592180, EPI_ISL_592181, EPI_ISL_592182, EPI_ISL_592183, EPI_ISL_592184, EPI_ISL_592185, EPI_ISL_592188, EPI_ISL_592189, EPI_ISL_592190, EPI_ISL_592191, EPI_ISL_592192, EPI_ISL_592193, EPI_ISL_592194, EPI_ISL_592195, EPI_ISL_592485, EPI_ISL_592486, EPI_ISL_592488, EPI_ISL_592489, EPI_ISL_592490, EPI_ISL_592491, EPI_ISL_592492, EPI_ISL_592493, EPI_ISL_592494, EPI_ISL_592495, EPI_ISL_592496, EPI_ISL_592497, EPI_ISL_592498, EPI_ISL_592499, EPI_ISL_592500, EPI_ISL_592501, EPI_ISL_592502, EPI_ISL_592503, EPI_ISL_592504, EPI_ISL_592505, EPI_ISL_592506, EPI_ISL_592507, EPI_ISL_592509, EPI_ISL_592511, EPI_ISL_592512, EPI_ISL_592514, EPI_ISL_592515, EPI_ISL_592516, EPI_ISL_592537, EPI_ISL_592548, EPI_ISL_592564, EPI_ISL_592568, EPI_ISL_592577, EPI_ISL_592590, EPI_ISL_592597, EPI_ISL_592600, EPI_ISL_592616, EPI_ISL_592660, EPI_ISL_592661, EPI_ISL_592668, EPI_ISL_592716, EPI_ISL_592720, EPI_ISL_592730, EPI_ISL_592770, EPI_ISL_592771, EPI_ISL_592781, EPI_ISL_592783, EPI_ISL_592790, EPI_ISL_592795, EPI_ISL_592804, EPI_ISL_592814, EPI_ISL_592820, EPI_ISL_592821, EPI_ISL_592824, EPI_ISL_592828, EPI_ISL_592836, EPI_ISL_592837, EPI_ISL_592858, EPI_ISL_592859, EPI_ISL_592860, EPI_ISL_592862, EPI_ISL_592863, EPI_ISL_592864, EPI_ISL_592865, EPI_ISL_592878, EPI_ISL_592879, EPI_ISL_592880, EPI_ISL_592881, EPI_ISL_592958, EPI_ISL_592969, EPI_ISL_592970, EPI_ISL_592971, EPI_ISL_592972, EPI_ISL_592973, EPI_ISL_592974, EPI_ISL_592975, EPI_ISL_592976, EPI_ISL_592977, EPI_ISL_592978, EPI_ISL_592979, EPI_ISL_592980, EPI_ISL_592981, EPI_ISL_592982, EPI_ISL_592983, EPI_ISL_592984, EPI_ISL_592985, EPI_ISL_593003, EPI_ISL_593004, EPI_ISL_593005, EPI_ISL_593006, EPI_ISL_593007, EPI_ISL_593033, EPI_ISL_593035, EPI_ISL_593048, EPI_ISL_593125, EPI_ISL_593126, EPI_ISL_593127, EPI_ISL_593128, EPI_ISL_593215, EPI_ISL_593216, EPI_ISL_593256, EPI_ISL_593411, EPI_ISL_593412, EPI_ISL_593476, EPI_ISL_593477 | | | |
| see above | Victorian Infectious Diseases Reference Laboratory (VIDRL) | VIDRL and MDU-PHL | Caly L., Seemann T., Sait, M., Schultz, M. B., Druce J., Sherry, N. |
| EPI_ISL_593646, EPI_ISL_593647, | 4Cyte Pathology | NSW Health Pathology - Institute of Clinical Pathology and | CIDM-PH et al. |

| | | | |
|---|---|---|---|
| EPI_ISL_593648 | | Medical Research; Westmead Hospital; University of Sydney | |
| EPI_ISL_593649 | Austech Medical Laboratories | NSW Health Pathology - Institute of Clinical Pathology and Medical Research; Westmead Hospital; University of Sydney | CIDM-PH et al. |
| EPI_ISL_593650, EPI_ISL_593651 | Douglass Hanly Moir Pathology | NSW Health Pathology - Institute of Clinical Pathology and Medical Research; Westmead Hospital; University of Sydney | CIDM-PH et al. |
| EPI_ISL_593652 | Histopath | NSW Health Pathology - Institute of Clinical Pathology and Medical Research; Westmead Hospital; University of Sydney | CIDM-PH et al. |
| EPI_ISL_593653 | Laverty Pathology | NSW Health Pathology - Institute of Clinical Pathology and Medical Research; Westmead Hospital; University of Sydney | CIDM-PH et al. |
| EPI_ISL_593654 | Medlab Pathology | NSW Health Pathology - Institute of Clinical Pathology and Medical Research; Westmead Hospital; University of Sydney | CIDM-PH et al. |
| EPI_ISL_593655, EPI_ISL_593656, EPI_ISL_593657, EPI_ISL_593658 | Pathology North - Hunter - NSW Health Pathology | NSW Health Pathology - Institute of Clinical Pathology and Medical Research; Westmead Hospital; University of Sydney | CIDM-PH et al. |
| EPI_ISL_593659 | Pathology North - Royal North Shore Hospital - NSW Health Pathology | NSW Health Pathology - Institute of Clinical Pathology and Medical Research; Westmead Hospital; University of Sydney | CIDM-PH et al. |
| EPI_ISL_593660, EPI_ISL_593661, EPI_ISL_593662, EPI_ISL_593663, EPI_ISL_593664, EPI_ISL_593665, EPI_ISL_593666, EPI_ISL_593667, EPI_ISL_593668, EPI_ISL_593669, EPI_ISL_593670, EPI_ISL_593671, EPI_ISL_593672, EPI_ISL_593673, EPI_ISL_593674, EPI_ISL_593675, EPI_ISL_593676, EPI_ISL_593677, EPI_ISL_593678, EPI_ISL_593679, EPI_ISL_593680, EPI_ISL_593681, EPI_ISL_593682 | | | |
| see above | Pathology West - NSW Health Pathology | NSW Health Pathology - Institute of Clinical Pathology and Medical Research; Westmead Hospital; University of Sydney | CIDM-PH et al. |
| EPI_ISL_593683, EPI_ISL_593684, EPI_ISL_593685, EPI_ISL_593686, EPI_ISL_593687, EPI_ISL_593688, EPI_ISL_593689, EPI_ISL_593690, EPI_ISL_593691, EPI_ISL_593692, EPI_ISL_593693, EPI_ISL_593694, EPI_ISL_593695, EPI_ISL_593696, EPI_ISL_593697, EPI_ISL_593698, EPI_ISL_593699, EPI_ISL_593700, EPI_ISL_593701, EPI_ISL_593702, EPI_ISL_593703, EPI_ISL_593704, EPI_ISL_593705, EPI_ISL_593706, EPI_ISL_593707, EPI_ISL_593708, EPI_ISL_593709, EPI_ISL_593710, EPI_ISL_593711, EPI_ISL_593712, EPI_ISL_593713, EPI_ISL_593714, EPI_ISL_593715, EPI_ISL_593716, EPI_ISL_593717, EPI_ISL_593718, EPI_ISL_593719, EPI_ISL_593720, EPI_ISL_593721, EPI_ISL_593722, EPI_ISL_593723, EPI_ISL_593724, EPI_ISL_593725, EPI_ISL_593726, EPI_ISL_593727, EPI_ISL_593728, EPI_ISL_593729, EPI_ISL_593730, EPI_ISL_593731, EPI_ISL_593732, EPI_ISL_593733, EPI_ISL_593734, EPI_ISL_593735, EPI_ISL_593736, EPI_ISL_593737, EPI_ISL_593738, EPI_ISL_593739, EPI_ISL_593740, EPI_ISL_593741, EPI_ISL_593742, EPI_ISL_593743, EPI_ISL_593744, EPI_ISL_593745, EPI_ISL_593746, EPI_ISL_593747, EPI_ISL_593748, EPI_ISL_593749, EPI_ISL_593750, EPI_ISL_593751, EPI_ISL_593752, EPI_ISL_593753, EPI_ISL_593754 | | | |
| see above | South Eastern Area Laboratory Services (SEALS) | NSW Health Pathology - Institute of Clinical Pathology and Medical Research; Westmead Hospital; University of Sydney | CIDM-PH et al. |
| EPI_ISL_593755, EPI_ISL_593756, EPI_ISL_593757, EPI_ISL_593758, EPI_ISL_593759, EPI_ISL_593760, EPI_ISL_593761, EPI_ISL_593762, EPI_ISL_593763, EPI_ISL_593764, EPI_ISL_593765 | | | |
| see above | Sydney South West Pathology Service (SSWPS) - Liverpool Hospital - NSW Health Pathology | NSW Health Pathology - Institute of Clinical Pathology and Medical Research; Westmead Hospital; University of Sydney | CIDM-PH et al. |
| EPI_ISL_593766, EPI_ISL_593767, EPI_ISL_593768 | Sydney South West Pathology Service (SSWPS) - Royal Prince Alfred Hospital - NSW Health Pathology | NSW Health Pathology - Institute of Clinical Pathology and Medical Research; Westmead Hospital; University of Sydney | CIDM-PH et al. |
| EPI_ISL_593769 | The Children's Hospital at Westmead | NSW Health Pathology - Institute of Clinical Pathology and Medical Research; Westmead Hospital; University of Sydney | CIDM-PH et al. |
| EPI_ISL_593770 | Pathology West - NSW Health Pathology | NSW Health Pathology - Institute of Clinical Pathology and Medical Research; Westmead Hospital; University of Sydney | CIDM-PH et al. |
| EPI_ISL_593771 | South Eastern Area Laboratory Services (SEALS) | NSW Health Pathology - Institute of Clinical Pathology and Medical Research; Westmead Hospital; University of Sydney | CIDM-PH et al. |
| EPI_ISL_594134, EPI_ISL_594135, EPI_ISL_594136, EPI_ISL_594137, EPI_ISL_594138, EPI_ISL_594139, EPI_ISL_594140, EPI_ISL_594141, EPI_ISL_594142, EPI_ISL_594143, EPI_ISL_594144, EPI_ISL_594145, EPI_ISL_594146, EPI_ISL_594147 | | | |
| see above | MDU-PHL, The Peter Doherty Institute for Infection and Immunity | MDU-PHL, The Peter Doherty Institute for Infection and Immunity | Caly,L., Seemann,T., Sait,M.L., Schultz,M.B., Druce,J., Sherry,N.L. |
| EPI_ISL_595329, EPI_ISL_595330, EPI_ISL_595331, EPI_ISL_595332, EPI_ISL_595333, EPI_ISL_595336, EPI_ISL_595337, EPI_ISL_595339, EPI_ISL_595340, EPI_ISL_595341, EPI_ISL_595343, EPI_ISL_595344, EPI_ISL_595345, EPI_ISL_595346, EPI_ISL_595349, EPI_ISL_595350, EPI_ISL_595352, EPI_ISL_595355, EPI_ISL_595356, EPI_ISL_595358, EPI_ISL_595359, EPI_ISL_595361, EPI_ISL_595363, EPI_ISL_595364, EPI_ISL_595365, EPI_ISL_595369, EPI_ISL_595370, EPI_ISL_595371, EPI_ISL_595372, EPI_ISL_595374, EPI_ISL_595375, EPI_ISL_595376, EPI_ISL_595377, EPI_ISL_595378, EPI_ISL_595380, EPI_ISL_595381, EPI_ISL_595382, EPI_ISL_595383, EPI_ISL_595385, EPI_ISL_595386, EPI_ISL_595387, EPI_ISL_595388, EPI_ISL_595389, EPI_ISL_595390, EPI_ISL_595391, EPI_ISL_595392, EPI_ISL_595393, EPI_ISL_595394, EPI_ISL_595396, EPI_ISL_595397, EPI_ISL_595400, EPI_ISL_595402, EPI_ISL_595403, EPI_ISL_595404, EPI_ISL_595405, EPI_ISL_595406, EPI_ISL_595407, EPI_ISL_595408, EPI_ISL_595409, EPI_ISL_595410, EPI_ISL_595412, EPI_ISL_595413, EPI_ISL_595414, EPI_ISL_595416, EPI_ISL_595417, EPI_ISL_595418, EPI_ISL_595419, EPI_ISL_595423, EPI_ISL_595424, EPI_ISL_595425, EPI_ISL_595426, EPI_ISL_595427, EPI_ISL_595428, EPI_ISL_595429, EPI_ISL_595430, EPI_ISL_595431, EPI_ISL_595432, EPI_ISL_595434, EPI_ISL_595435, EPI_ISL_595437, EPI_ISL_595439, EPI_ISL_595441, EPI_ISL_595443, EPI_ISL_595444, EPI_ISL_595447, EPI_ISL_595448, EPI_ISL_595449, EPI_ISL_595450, EPI_ISL_595451, EPI_ISL_595452, EPI_ISL_595453, EPI_ISL_595455, EPI_ISL_595457, EPI_ISL_595459, EPI_ISL_595460, EPI_ISL_595462, EPI_ISL_595463, EPI_ISL_595465, EPI_ISL_595466, EPI_ISL_595469, EPI_ISL_595470, EPI_ISL_595471, EPI_ISL_595473, EPI_ISL_595475, EPI_ISL_595476, EPI_ISL_595478, EPI_ISL_595481, EPI_ISL_595483, EPI_ISL_595484, EPI_ISL_595485, EPI_ISL_595487, EPI_ISL_595488, EPI_ISL_595490, EPI_ISL_595491, EPI_ISL_595492, EPI_ISL_595493, EPI_ISL_595494, EPI_ISL_595495, EPI_ISL_595496, EPI_ISL_595497, EPI_ISL_595498, EPI_ISL_595501, EPI_ISL_595502, EPI_ISL_595504, EPI_ISL_595505, EPI_ISL_595506, EPI_ISL_595510, EPI_ISL_595511, EPI_ISL_595512, EPI_ISL_595513, EPI_ISL_595514, EPI_ISL_595515, EPI_ISL_595518, EPI_ISL_595519, EPI_ISL_595520, EPI_ISL_595523, EPI_ISL_595525, EPI_ISL_595526, EPI_ISL_595527, EPI_ISL_595529, EPI_ISL_595530, EPI_ISL_595532, EPI_ISL_595534, EPI_ISL_595535, EPI_ISL_595536, EPI_ISL_595538, EPI_ISL_595540, EPI_ISL_595541, EPI_ISL_595542, EPI_ISL_595543, EPI_ISL_595544, EPI_ISL_595545, EPI_ISL_595546, EPI_ISL_595547, EPI_ISL_595548, EPI_ISL_595549, EPI_ISL_595550, EPI_ISL_595551, EPI_ISL_595552, EPI_ISL_595553, EPI_ISL_595554, EPI_ISL_595555, EPI_ISL_595556, EPI_ISL_595559, EPI_ISL_595561, EPI_ISL_595562, EPI_ISL_595563, EPI_ISL_595564, EPI_ISL_595565, EPI_ISL_595570, EPI_ISL_595571, EPI_ISL_595572, EPI_ISL_595573, EPI_ISL_595576, EPI_ISL_595577, EPI_ISL_595578, EPI_ISL_595579, EPI_ISL_595580, EPI_ISL_595581, EPI_ISL_595582, EPI_ISL_595584, EPI_ISL_595586, EPI_ISL_595587, EPI_ISL_595588, EPI_ISL_595589, EPI_ISL_595590, EPI_ISL_595592, EPI_ISL_595593, EPI_ISL_595594, EPI_ISL_595595, EPI_ISL_595596, EPI_ISL_595598, EPI_ISL_595599, EPI_ISL_595601, EPI_ISL_595603, EPI_ISL_595604, EPI_ISL_595605, EPI_ISL_595606, EPI_ISL_595607, EPI_ISL_595609, EPI_ISL_595610, EPI_ISL_595611, EPI_ISL_595612, EPI_ISL_595613, EPI_ISL_595614, EPI_ISL_595616 | | | |
| see above | Wales Specialist Virology Centre Sequencing lab: Pathogen Genomics Unit | COVID-19 Genomics UK (COG-UK) Consortium | Catherine Moore, Johnathan Evans, Laura Gifford, Malorie Perry, Simon Cottrell, Angela Marchbank, Alec Birchley, Alexander Adams, Amy Gaskin, Bree Gatica-Wilcox, Jason Coombes, Joel Southgate, Lauren Gilbert, Lee Graham, Nicole Pacchiarini, Sara Kumziene-Summerhayes, Sarah Taylor, Sophie Jones, Sara Rey, Matthew Bull, Joanne Watkins, Sally Corden, Tom Connor |
| EPI_ISL_609989 | INMI Lazzaro Spallanzani IRCCS | INMI Lazzaro Spallanzani IRCCS | C.E.M Gruber, B Bartolini, M Rueca, F Messina, E Giombini, A Di Caro, MR Capobianchi |
| EPI_ISL_609994 | INMI Lazzaro Spallanzani IRCCS | INMI Lazzaro Spallanzani IRCCS | C.E.M Gruber, F Messina, M Rueca, B Bartolini, E Giombini, MR Capobianchi, A Di Caro |
| EPI_ISL_611636, EPI_ISL_611683, EPI_ISL_611711, EPI_ISL_611792, EPI_ISL_611832, EPI_ISL_611887, EPI_ISL_611905, EPI_ISL_611926, EPI_ISL_611972, EPI_ISL_612117, EPI_ISL_612119, EPI_ISL_612122, EPI_ISL_612133, EPI_ISL_612136, EPI_ISL_612138, EPI_ISL_612139, EPI_ISL_612140, EPI_ISL_612142, EPI_ISL_612145, EPI_ISL_612147, EPI_ISL_612151, EPI_ISL_612160, EPI_ISL_612161, EPI_ISL_612162, EPI_ISL_612163, EPI_ISL_612164, EPI_ISL_612167, EPI_ISL_612168, EPI_ISL_612169, EPI_ISL_612170, EPI_ISL_612172, EPI_ISL_612180, EPI_ISL_612184, EPI_ISL_612193, EPI_ISL_612200, EPI_ISL_612201, EPI_ISL_612204, EPI_ISL_612205, EPI_ISL_612206, EPI_ISL_612464, EPI_ISL_612465, EPI_ISL_612472, EPI_ISL_612475, EPI_ISL_612480, EPI_ISL_612487, EPI_ISL_612498, EPI_ISL_612504, EPI_ISL_612509, EPI_ISL_612511, EPI_ISL_612516, EPI_ISL_612607, EPI_ISL_612608, EPI_ISL_612609, EPI_ISL_612610, EPI_ISL_612611, EPI_ISL_612612, EPI_ISL_612613, EPI_ISL_612615, EPI_ISL_612616, EPI_ISL_612617, EPI_ISL_612618, EPI_ISL_612621, EPI_ISL_612622, EPI_ISL_612623, EPI_ISL_612624, EPI_ISL_612628, EPI_ISL_612629, EPI_ISL_612630, EPI_ISL_612631, EPI_ISL_612632, EPI_ISL_612633, EPI_ISL_612635, EPI_ISL_612636, EPI_ISL_612637, EPI_ISL_612638, EPI_ISL_612639, EPI_ISL_612640, EPI_ISL_612641, EPI_ISL_612642, EPI_ISL_612643, EPI_ISL_612644, EPI_ISL_612645, EPI_ISL_612646, EPI_ISL_612647, EPI_ISL_612648, EPI_ISL_612649, EPI_ISL_612650, EPI_ISL_612651, EPI_ISL_612652, EPI_ISL_612653, EPI_ISL_612654, EPI_ISL_612655, EPI_ISL_612656, EPI_ISL_612657, EPI_ISL_612658, EPI_ISL_612659, EPI_ISL_612660, EPI_ISL_612661, EPI_ISL_612662, EPI_ISL_612663, EPI_ISL_612664, EPI_ISL_612665, EPI_ISL_612666, EPI_ISL_612667, EPI_ISL_612668, EPI_ISL_612669, EPI_ISL_612670, EPI_ISL_612671, EPI_ISL_612672, EPI_ISL_612673, EPI_ISL_612676, EPI_ISL_612677, EPI_ISL_612678, EPI_ISL_612679, EPI_ISL_612680, EPI_ISL_612681, EPI_ISL_612682, EPI_ISL_612683, EPI_ISL_612684, EPI_ISL_612685, EPI_ISL_612686, EPI_ISL_612687, EPI_ISL_612688, EPI_ISL_612690, EPI_ISL_612691, EPI_ISL_612692, EPI_ISL_612693, EPI_ISL_612694, EPI_ISL_612695, EPI_ISL_612696, EPI_ISL_612697, EPI_ISL_612698, EPI_ISL_612699, EPI_ISL_612700, EPI_ISL_612701, EPI_ISL_612702, EPI_ISL_612703, EPI_ISL_612704, EPI_ISL_612705, EPI_ISL_612706, EPI_ISL_612707, EPI_ISL_612708, EPI_ISL_612709, EPI_ISL_612710, EPI_ISL_612711, EPI_ISL_612712, EPI_ISL_612713, EPI_ISL_612714, EPI_ISL_612715, EPI_ISL_612717, EPI_ISL_612719, EPI_ISL_612720, EPI_ISL_612721, EPI_ISL_612724, EPI_ISL_612725, EPI_ISL_612726, EPI_ISL_612727, EPI_ISL_612728, EPI_ISL_612729, EPI_ISL_612730, EPI_ISL_612731, EPI_ISL_612732, EPI_ISL_612733, EPI_ISL_612734, EPI_ISL_612735, EPI_ISL_612736, EPI_ISL_612737, EPI_ISL_612738, EPI_ISL_612739, EPI_ISL_612741, EPI_ISL_612742, EPI_ISL_612743, EPI_ISL_612744, EPI_ISL_612745, EPI_ISL_612746, EPI_ISL_612747, EPI_ISL_612748, EPI_ISL_612749, EPI_ISL_612750, EPI_ISL_612751, EPI_ISL_612752, EPI_ISL_612753, EPI_ISL_612754, EPI_ISL_612755, EPI_ISL_612756, EPI_ISL_612757, EPI_ISL_612758, EPI_ISL_612759, EPI_ISL_612760, EPI_ISL_612761, EPI_ISL_612762, EPI_ISL_612763, EPI_ISL_612764, EPI_ISL_612766, EPI_ISL_612768, EPI_ISL_612769, EPI_ISL_612771, EPI_ISL_612772, EPI_ISL_612773, EPI_ISL_612774, EPI_ISL_612776, EPI_ISL_612777, EPI_ISL_612778, EPI_ISL_612779, EPI_ISL_612780, EPI_ISL_612781, EPI_ISL_612782, EPI_ISL_612783, EPI_ISL_612784, EPI_ISL_612785, EPI_ISL_612786, EPI_ISL_612787, EPI_ISL_612788, EPI_ISL_612791, EPI_ISL_612792, EPI_ISL_612793, EPI_ISL_612794, EPI_ISL_612795, EPI_ISL_612796, EPI_ISL_612797, EPI_ISL_612798, EPI_ISL_612799, EPI_ISL_612800, EPI_ISL_612801, EPI_ISL_612802, EPI_ISL_612803, EPI_ISL_612804, EPI_ISL_612805, EPI_ISL_612806, EPI_ISL_612808, EPI_ISL_612809, EPI_ISL_612810, EPI_ISL_612811, EPI_ISL_612812, EPI_ISL_612813, EPI_ISL_612814, EPI_ISL_612815, EPI_ISL_612816, EPI_ISL_612817, EPI_ISL_612818, EPI_ISL_612819, EPI_ISL_612820, EPI_ISL_612821, EPI_ISL_612822, EPI_ISL_612823, EPI_ISL_612824, EPI_ISL_612826, EPI_ISL_612827, EPI_ISL_612829, EPI_ISL_612830, EPI_ISL_612831, EPI_ISL_612832, EPI_ISL_612833, EPI_ISL_612834, EPI_ISL_612835, EPI_ISL_612836, EPI_ISL_612837, EPI_ISL_612838, EPI_ISL_612839, EPI_ISL_612841, EPI_ISL_612842, EPI_ISL_612843, EPI_ISL_612844, EPI_ISL_612846, EPI_ISL_612847, EPI_ISL_612850, EPI_ISL_612851, EPI_ISL_612852, EPI_ISL_612853, EPI_ISL_612854, EPI_ISL_612856, EPI_ISL_612857, EPI_ISL_612858, EPI_ISL_612859, EPI_ISL_612860, EPI_ISL_612861, EPI_ISL_612862, EPI_ISL_612863, EPI_ISL_612864, EPI_ISL_612865, EPI_ISL_612866, EPI_ISL_612869, EPI_ISL_612870, EPI_ISL_612871, EPI_ISL_612873, EPI_ISL_612874, EPI_ISL_612875, EPI_ISL_612876, EPI_ISL_612878, EPI_ISL_612903, EPI_ISL_612904, EPI_ISL_612905, EPI_ISL_612906, EPI_ISL_612907, EPI_ISL_612908, EPI_ISL_612923, EPI_ISL_612924, EPI_ISL_612925, EPI_ISL_612927, EPI_ISL_612928, EPI_ISL_612929, EPI_ISL_612930, EPI_ISL_612931, EPI_ISL_612932, EPI_ISL_612934, EPI_ISL_612935, EPI_ISL_612937, EPI_ISL_612938, EPI_ISL_612939, EPI_ISL_612941, EPI_ISL_612942, EPI_ISL_612943, EPI_ISL_612944, EPI_ISL_612945, EPI_ISL_612946, EPI_ISL_612947, EPI_ISL_612948, EPI_ISL_612949, EPI_ISL_612950, EPI_ISL_612951, EPI_ISL_612952, EPI_ISL_612953, EPI_ISL_612954, EPI_ISL_612955, EPI_ISL_612956, EPI_ISL_612958, EPI_ISL_612960, EPI_ISL_612961, EPI_ISL_612962, EPI_ISL_612963, EPI_ISL_612964, EPI_ISL_612965, EPI_ISL_612966, EPI_ISL_612967, EPI_ISL_612970, EPI_ISL_612971, EPI_ISL_612972, EPI_ISL_612973, EPI_ISL_612974, EPI_ISL_612977, EPI_ISL_612978, EPI_ISL_612979, | | | |

| | | | |
|---|---|---|---|
| EPI_ISL_612980, EPI_ISL_612982, EPI_ISL_612983, EPI_ISL_612984, EPI_ISL_612985, EPI_ISL_612986, EPI_ISL_612987, EPI_ISL_612988, EPI_ISL_612989, EPI_ISL_612990, EPI_ISL_612991, EPI_ISL_612992, EPI_ISL_612993, EPI_ISL_612994, EPI_ISL_612995, EPI_ISL_612996, EPI_ISL_612997, EPI_ISL_612998, EPI_ISL_613000, EPI_ISL_613001, EPI_ISL_613002, EPI_ISL_613003, EPI_ISL_613004, EPI_ISL_613005, EPI_ISL_613007, EPI_ISL_613008, EPI_ISL_613009, EPI_ISL_613011, EPI_ISL_613012, EPI_ISL_613016, EPI_ISL_613017, EPI_ISL_613018, EPI_ISL_613019, EPI_ISL_613109, EPI_ISL_613267, EPI_ISL_613268, EPI_ISL_613269, EPI_ISL_613270, EPI_ISL_613271, EPI_ISL_613272, EPI_ISL_613273, EPI_ISL_613274, EPI_ISL_613276, EPI_ISL_613277, EPI_ISL_613278, EPI_ISL_613279, EPI_ISL_613280, EPI_ISL_613281 | | | |
| see above | Wales Specialist Virology Centre Sequencing lab: Pathogen Genomics Unit | COVID-19 Genomics UK (COG-UK) Consortium | Catherine Moore, Johnathan Evans, Laura Gifford, Malorie Perry, Simon Cottrell, Angela Marchbank, Alec Birchley, Alexander Adams, Amy Gaskin, Bree Gatica-Wilcox, Jason Coombes, Joel Southgate, Lauren Gilbert, Lee Graham, Nicole Pacchiarini, Sara Kumziene-Summerhayes, Sarah Taylor, Sophie Jones, Sara Rey, Matthew Bull, Joanne Watkins, Sally Corden, Tom Connor |
| EPI_ISL_623104, EPI_ISL_623105 | Simile Medicina Diagnóstica | Bioinformatics Laboratory / LNCC | Carolina M Voloch, Ronaldo S Francisco Jr, Luiz G P de Almeida, Otavio J. Brustolini, Cynthia C Cardoso, Alexandra L Gerber, Ana Paula de C Guimarães, Diana Mariani, Covid19-UFRJ Workgroup, Luís Cristóvão Pôrto, Renato S Aguiar, Terezinha M P P Castiñeiras, Orlando C. Ferreira, Amilcar Tanuri, Ana Tereza R de Vasconcelos |
| EPI_ISL_623106, EPI_ISL_623107, EPI_ISL_623108, EPI_ISL_623109, EPI_ISL_623110, EPI_ISL_623111, EPI_ISL_623112, EPI_ISL_623113, EPI_ISL_623114, EPI_ISL_623115, EPI_ISL_623116, EPI_ISL_623117, EPI_ISL_623118, EPI_ISL_623119, EPI_ISL_623120, EPI_ISL_623121, EPI_ISL_623122, EPI_ISL_623123, EPI_ISL_623124, EPI_ISL_623125, EPI_ISL_623126, EPI_ISL_623127, EPI_ISL_623128, EPI_ISL_623129, EPI_ISL_623130, EPI_ISL_623131, EPI_ISL_623132, EPI_ISL_623133, EPI_ISL_623134, EPI_ISL_623135, EPI_ISL_623136, EPI_ISL_623137, EPI_ISL_623138, EPI_ISL_623139, EPI_ISL_623140, EPI_ISL_623141, EPI_ISL_623142, EPI_ISL_623143, EPI_ISL_623144, EPI_ISL_623145, EPI_ISL_623146, EPI_ISL_623147, EPI_ISL_623148, EPI_ISL_623149, EPI_ISL_623150, EPI_ISL_623151, EPI_ISL_623152, EPI_ISL_623153, EPI_ISL_623154, EPI_ISL_623155, EPI_ISL_623156, EPI_ISL_623157, EPI_ISL_623158, EPI_ISL_623159, EPI_ISL_623160, EPI_ISL_623161, EPI_ISL_623162, EPI_ISL_623163, EPI_ISL_623164, EPI_ISL_623165, EPI_ISL_623166, EPI_ISL_623167, EPI_ISL_623168, EPI_ISL_623169 | | | |
| see above | Laboratorio de Virologia Molecular / UFRJ | Bioinformatics Laboratory / LNCC | Carolina M Voloch, Ronaldo S Francisco Jr, Luiz G P de Almeida, Otavio J. Brustolini, Cynthia C Cardoso, Alexandra L Gerber, Ana Paula de C Guimarães, Diana Mariani, Covid19-UFRJ Workgroup, Luís Cristóvão Pôrto, Renato S Aguiar, Terezinha M P P Castiñeiras, Orlando C. Ferreira, Amilcar Tanuri, Ana Tereza R de Vasconcelos |
| EPI_ISL_625478, EPI_ISL_625479, EPI_ISL_625480, EPI_ISL_625481, EPI_ISL_625482, EPI_ISL_625483, EPI_ISL_625484, EPI_ISL_625485, EPI_ISL_625486, EPI_ISL_625487, EPI_ISL_625488, EPI_ISL_625489, EPI_ISL_625490, EPI_ISL_625491, EPI_ISL_625492, EPI_ISL_625493, EPI_ISL_625494, EPI_ISL_625495, EPI_ISL_625496, EPI_ISL_625497, EPI_ISL_625498, EPI_ISL_625499 | | | |
| see above | Santa Clara County Public Health Laboratory | Chan-Zuckerberg Biohub | CZB Cliahub Consortium |
| EPI_ISL_625500, EPI_ISL_625501, EPI_ISL_625502, EPI_ISL_625503, EPI_ISL_625504, EPI_ISL_625505, EPI_ISL_625506, EPI_ISL_625507, EPI_ISL_625508, EPI_ISL_625509, EPI_ISL_625510, EPI_ISL_625511, EPI_ISL_625512, EPI_ISL_625513, EPI_ISL_625514, EPI_ISL_625515, EPI_ISL_625516, EPI_ISL_625517, EPI_ISL_625518, EPI_ISL_625519, EPI_ISL_625520, EPI_ISL_625521, EPI_ISL_625522, EPI_ISL_625523, EPI_ISL_625524, EPI_ISL_625525, EPI_ISL_625526, EPI_ISL_625527, EPI_ISL_625528, EPI_ISL_625529, EPI_ISL_625530, EPI_ISL_625531, EPI_ISL_625532, EPI_ISL_625533, EPI_ISL_625534, EPI_ISL_625535, EPI_ISL_625536, EPI_ISL_625537, EPI_ISL_625538, EPI_ISL_625539, EPI_ISL_625540, EPI_ISL_625541, EPI_ISL_625542, EPI_ISL_625543, EPI_ISL_625544, EPI_ISL_625545, EPI_ISL_625546 | | | |
| see above | Alameda County Public Health Lab | Chan-Zuckerberg Biohub | CZB Cliahub Consortium |
| EPI_ISL_625547, EPI_ISL_625548, EPI_ISL_625549 | County of San Luis Obispo Public Health Laboratory | Chan-Zuckerberg Biohub | CZB Cliahub Consortium |
| EPI_ISL_625550, EPI_ISL_625551, EPI_ISL_625552, EPI_ISL_625553, EPI_ISL_625554, EPI_ISL_625555, EPI_ISL_625556, EPI_ISL_625557, EPI_ISL_625558 | UCSF Clinical Microbiology Laboratory | Chan-Zuckerberg Biohub | CZB Cliahub Consortium |
| EPI_ISL_625559, EPI_ISL_625560, EPI_ISL_625561, EPI_ISL_625562, EPI_ISL_625563, EPI_ISL_625564, EPI_ISL_625565, EPI_ISL_625566, EPI_ISL_625567, EPI_ISL_625568, EPI_ISL_625569, EPI_ISL_625570, EPI_ISL_625571, EPI_ISL_625572, EPI_ISL_625573, EPI_ISL_625574, EPI_ISL_625575, EPI_ISL_625576, EPI_ISL_625577, EPI_ISL_625578, EPI_ISL_625579, EPI_ISL_625580, EPI_ISL_625581, EPI_ISL_625582, EPI_ISL_625583, EPI_ISL_625584, EPI_ISL_625585, EPI_ISL_625586, EPI_ISL_625587, EPI_ISL_625588, EPI_ISL_625589, EPI_ISL_625590, EPI_ISL_625591, EPI_ISL_625592, EPI_ISL_625593 | | | |
| see above | NaN | Chan-Zuckerberg Biohub | CZB Cliahub Consortium |
| EPI_ISL_625594, EPI_ISL_625595, EPI_ISL_625596, EPI_ISL_625597, EPI_ISL_625598, EPI_ISL_625599, EPI_ISL_625600, EPI_ISL_625601, EPI_ISL_625602, EPI_ISL_625603, EPI_ISL_625604, EPI_ISL_625605, EPI_ISL_625606, EPI_ISL_625607, EPI_ISL_625608, EPI_ISL_625609, EPI_ISL_625610 | | | |
| see above | Santa Clara County Public Health Laboratory | Chan-Zuckerberg Biohub | CZB Cliahub Consortium |
| EPI_ISL_625611, EPI_ISL_625612, EPI_ISL_625613, EPI_ISL_625614, EPI_ISL_625615, EPI_ISL_625616, EPI_ISL_625617, EPI_ISL_625618, EPI_ISL_625619, EPI_ISL_625620, EPI_ISL_625621, EPI_ISL_625622 | | | |
| see above | Alameda County Public Health Lab | Chan-Zuckerberg Biohub | CZB Cliahub Consortium |
| EPI_ISL_625623, EPI_ISL_625624, EPI_ISL_625625, EPI_ISL_625626 | County of San Luis Obispo Public Health Laboratory | Chan-Zuckerberg Biohub | CZB Cliahub Consortium |
| EPI_ISL_625627, EPI_ISL_625628, EPI_ISL_625629, EPI_ISL_625630, EPI_ISL_625631, EPI_ISL_625632, EPI_ISL_625633, EPI_ISL_625634, EPI_ISL_625635, EPI_ISL_625636, EPI_ISL_625637, EPI_ISL_625638, EPI_ISL_625639, EPI_ISL_625640, EPI_ISL_625641, EPI_ISL_625642, EPI_ISL_625643, EPI_ISL_625644, EPI_ISL_625645, EPI_ISL_625647, EPI_ISL_625648, EPI_ISL_625649, EPI_ISL_625650, EPI_ISL_625651, EPI_ISL_625652, EPI_ISL_625653, EPI_ISL_625654, EPI_ISL_625655, EPI_ISL_625656, EPI_ISL_625657, EPI_ISL_625658, EPI_ISL_625659 | | | |
| see above | Orange County Public Health Lab | Chan-Zuckerberg Biohub | CZB Cliahub Consortium |
| EPI_ISL_625660 | San Joaquin County Public Health Lab | Chan-Zuckerberg Biohub | CZB Cliahub Consortium |
| EPI_ISL_625661, EPI_ISL_625662, EPI_ISL_625663, EPI_ISL_625664, EPI_ISL_625665, EPI_ISL_625666, EPI_ISL_625667, EPI_ISL_625668, EPI_ISL_625669, EPI_ISL_625670 | UCSF Clinical Microbiology Laboratory | Chan-Zuckerberg Biohub | CZB Cliahub Consortium |
| EPI_ISL_626833, EPI_ISL_626867, EPI_ISL_626869, EPI_ISL_626875, EPI_ISL_626959, EPI_ISL_627001, EPI_ISL_627011, EPI_ISL_627044, EPI_ISL_627055, EPI_ISL_627069, EPI_ISL_627078, EPI_ISL_627086, EPI_ISL_627087, EPI_ISL_627088, EPI_ISL_627089, EPI_ISL_627090, EPI_ISL_627091, EPI_ISL_627093, EPI_ISL_627094, EPI_ISL_627095, EPI_ISL_627096, EPI_ISL_627097, EPI_ISL_627205, EPI_ISL_627206, EPI_ISL_627207, EPI_ISL_627212, EPI_ISL_627217, EPI_ISL_627218, EPI_ISL_627219, EPI_ISL_627220, EPI_ISL_627222, EPI_ISL_627226, EPI_ISL_627229, EPI_ISL_627230, EPI_ISL_627232, EPI_ISL_627233, EPI_ISL_627234, EPI_ISL_627235, EPI_ISL_627236, EPI_ISL_627237, EPI_ISL_627238, EPI_ISL_627239, EPI_ISL_627240, EPI_ISL_627243, EPI_ISL_627244, EPI_ISL_627245, EPI_ISL_627247, EPI_ISL_627248, EPI_ISL_627249, EPI_ISL_627265, EPI_ISL_627266, EPI_ISL_627268, EPI_ISL_627269, EPI_ISL_627272, EPI_ISL_627273, EPI_ISL_627275, EPI_ISL_627276, EPI_ISL_627277, EPI_ISL_627278, EPI_ISL_627279, EPI_ISL_627296, EPI_ISL_627301, EPI_ISL_627307, EPI_ISL_627487, EPI_ISL_627488, EPI_ISL_627732, EPI_ISL_627733, EPI_ISL_627734, EPI_ISL_627736, EPI_ISL_627737, EPI_ISL_627740, EPI_ISL_627741, EPI_ISL_627742, EPI_ISL_627743, EPI_ISL_627745, EPI_ISL_627746, EPI_ISL_627752, EPI_ISL_627753, EPI_ISL_627754, EPI_ISL_627755, EPI_ISL_627756, EPI_ISL_627757, EPI_ISL_627759, EPI_ISL_627760, EPI_ISL_627761, EPI_ISL_627762, EPI_ISL_627764, EPI_ISL_627765, EPI_ISL_627766, EPI_ISL_627767, EPI_ISL_627768, EPI_ISL_627769, EPI_ISL_627770, EPI_ISL_627771, EPI_ISL_627772, EPI_ISL_627773, EPI_ISL_627774, EPI_ISL_627775, EPI_ISL_627776, EPI_ISL_627777, EPI_ISL_627778, EPI_ISL_627779, EPI_ISL_627780, EPI_ISL_627782, EPI_ISL_627783, EPI_ISL_627784, EPI_ISL_627785, EPI_ISL_627786, EPI_ISL_627787, EPI_ISL_627788, EPI_ISL_627789, EPI_ISL_627790, EPI_ISL_627791, EPI_ISL_627792, EPI_ISL_627793, EPI_ISL_627795, EPI_ISL_627796, EPI_ISL_627797, EPI_ISL_627798, EPI_ISL_627800, EPI_ISL_627801, EPI_ISL_627802, EPI_ISL_627803, EPI_ISL_627804, EPI_ISL_627805, EPI_ISL_627806, EPI_ISL_627807, EPI_ISL_627808, EPI_ISL_627809, EPI_ISL_627810, EPI_ISL_627812, EPI_ISL_627813, EPI_ISL_627814, EPI_ISL_627815, EPI_ISL_627816, EPI_ISL_627817, EPI_ISL_627818, EPI_ISL_627819, EPI_ISL_627820, EPI_ISL_627821, EPI_ISL_627822, EPI_ISL_627823, EPI_ISL_627825, EPI_ISL_627826, EPI_ISL_627827, EPI_ISL_627828, EPI_ISL_627829, EPI_ISL_627830, EPI_ISL_627831, EPI_ISL_627832, EPI_ISL_627833, EPI_ISL_627835, EPI_ISL_627836, EPI_ISL_627837, EPI_ISL_627838, EPI_ISL_627839, EPI_ISL_627840, EPI_ISL_627842, EPI_ISL_627845, EPI_ISL_627847, EPI_ISL_627849, EPI_ISL_627850, EPI_ISL_627852, EPI_ISL_627853, EPI_ISL_627854, EPI_ISL_627855, EPI_ISL_627856, EPI_ISL_627857, EPI_ISL_627858, EPI_ISL_627860, EPI_ISL_627861, EPI_ISL_627862, EPI_ISL_627864, EPI_ISL_627866, EPI_ISL_627867, EPI_ISL_627869, EPI_ISL_627870, EPI_ISL_627871, EPI_ISL_627872, EPI_ISL_627874, EPI_ISL_627875, EPI_ISL_627876, EPI_ISL_627877, EPI_ISL_627879, EPI_ISL_627881, EPI_ISL_627882, EPI_ISL_627883, EPI_ISL_627884, EPI_ISL_627885, EPI_ISL_627887, EPI_ISL_627888, EPI_ISL_627889, EPI_ISL_627891, EPI_ISL_627892, EPI_ISL_627893, EPI_ISL_627894, EPI_ISL_627895, EPI_ISL_627896, EPI_ISL_627897, EPI_ISL_627898, EPI_ISL_627899, EPI_ISL_627900, EPI_ISL_627901, EPI_ISL_627902, EPI_ISL_627903, EPI_ISL_627904, EPI_ISL_627905, EPI_ISL_627906, EPI_ISL_627907, EPI_ISL_627908, EPI_ISL_627909, EPI_ISL_627910, EPI_ISL_627911, EPI_ISL_627913, EPI_ISL_627914, EPI_ISL_627915, EPI_ISL_627916, EPI_ISL_627917, EPI_ISL_627918, EPI_ISL_627919, EPI_ISL_627920, EPI_ISL_627921, EPI_ISL_627922, EPI_ISL_627923, EPI_ISL_627924, EPI_ISL_627925, EPI_ISL_627926, EPI_ISL_627928, EPI_ISL_627929, EPI_ISL_627930, EPI_ISL_627931, EPI_ISL_627932, EPI_ISL_627933, EPI_ISL_627935, EPI_ISL_627937, EPI_ISL_627939, EPI_ISL_627940, EPI_ISL_627941, EPI_ISL_627942, EPI_ISL_627943, EPI_ISL_627945, EPI_ISL_627947, EPI_ISL_627948, EPI_ISL_627949, EPI_ISL_627950, EPI_ISL_627951, EPI_ISL_627952, EPI_ISL_627953, EPI_ISL_627954, EPI_ISL_627955, EPI_ISL_627957, EPI_ISL_627958, EPI_ISL_627959, EPI_ISL_627960, EPI_ISL_627961, EPI_ISL_627962, EPI_ISL_627963, EPI_ISL_627964, EPI_ISL_627967, EPI_ISL_627968, EPI_ISL_627970, EPI_ISL_627971, EPI_ISL_627972, EPI_ISL_627973, EPI_ISL_627975, EPI_ISL_627976, EPI_ISL_627977, EPI_ISL_627978, EPI_ISL_627980, EPI_ISL_627981, EPI_ISL_627982, EPI_ISL_627983, EPI_ISL_627984, EPI_ISL_627985, EPI_ISL_627987, EPI_ISL_627989, EPI_ISL_627990, EPI_ISL_627991, EPI_ISL_627992, EPI_ISL_627993, EPI_ISL_627994, EPI_ISL_627995, EPI_ISL_627996, EPI_ISL_627997, EPI_ISL_627998, EPI_ISL_627999, EPI_ISL_628000, EPI_ISL_628004, EPI_ISL_628005, EPI_ISL_628006, EPI_ISL_628007, EPI_ISL_628009, EPI_ISL_628010, EPI_ISL_628011, EPI_ISL_628012, EPI_ISL_628013, EPI_ISL_628014, EPI_ISL_628017, EPI_ISL_628018, EPI_ISL_628019, EPI_ISL_628020, EPI_ISL_628021, EPI_ISL_628022, EPI_ISL_628023, EPI_ISL_628024, EPI_ISL_628025, EPI_ISL_628028, EPI_ISL_628029, EPI_ISL_628030, EPI_ISL_628031, EPI_ISL_628032, EPI_ISL_628033, EPI_ISL_628035, EPI_ISL_628036, EPI_ISL_628037, EPI_ISL_628038, EPI_ISL_628039, EPI_ISL_628041, EPI_ISL_628042, EPI_ISL_628043, EPI_ISL_628044, EPI_ISL_628045, EPI_ISL_628046, EPI_ISL_628047, EPI_ISL_628048, EPI_ISL_628049, EPI_ISL_628050, EPI_ISL_628051, EPI_ISL_628052, EPI_ISL_628053, EPI_ISL_628054, EPI_ISL_628055, EPI_ISL_628056, EPI_ISL_628057, EPI_ISL_628058, EPI_ISL_628059, EPI_ISL_628060, EPI_ISL_628062, EPI_ISL_628063, EPI_ISL_628064, EPI_ISL_628065, EPI_ISL_628066, EPI_ISL_628067, EPI_ISL_628068, EPI_ISL_628070, EPI_ISL_628071, EPI_ISL_628072, EPI_ISL_628073, EPI_ISL_628074, EPI_ISL_628075, EPI_ISL_628077, EPI_ISL_628078, EPI_ISL_628079, EPI_ISL_628080, EPI_ISL_628081, EPI_ISL_628082, EPI_ISL_628083, EPI_ISL_628084, EPI_ISL_628085, EPI_ISL_628086, EPI_ISL_628087, EPI_ISL_628089, EPI_ISL_628090, EPI_ISL_628091, EPI_ISL_628093, EPI_ISL_628094, EPI_ISL_628095, EPI_ISL_628096, EPI_ISL_628098, EPI_ISL_628099, EPI_ISL_628100, EPI_ISL_628101, EPI_ISL_628103, EPI_ISL_628104, EPI_ISL_628105, EPI_ISL_628106, EPI_ISL_628108, EPI_ISL_628109, EPI_ISL_628110, EPI_ISL_628111, EPI_ISL_628112, EPI_ISL_628113, EPI_ISL_628114, EPI_ISL_628115, EPI_ISL_628116, EPI_ISL_628117, EPI_ISL_628118, EPI_ISL_628119, EPI_ISL_628120, EPI_ISL_628121, EPI_ISL_628122, EPI_ISL_628124, EPI_ISL_628125, EPI_ISL_628126, EPI_ISL_628127, EPI_ISL_628129, EPI_ISL_628130, EPI_ISL_628131, EPI_ISL_628132, EPI_ISL_628133, EPI_ISL_628135, EPI_ISL_628136, EPI_ISL_628137, EPI_ISL_628139, EPI_ISL_628140, EPI_ISL_628142, EPI_ISL_628143, EPI_ISL_628144, EPI_ISL_628145, EPI_ISL_628146, EPI_ISL_628147, EPI_ISL_628148, EPI_ISL_628149, EPI_ISL_628150, EPI_ISL_628151, EPI_ISL_628152, EPI_ISL_628153, EPI_ISL_628154, EPI_ISL_628155, EPI_ISL_628156, EPI_ISL_628157, EPI_ISL_628158, EPI_ISL_628159, EPI_ISL_628160, EPI_ISL_628161, EPI_ISL_628164, EPI_ISL_628165, EPI_ISL_628166, EPI_ISL_628167, EPI_ISL_628168, EPI_ISL_628169, EPI_ISL_628170, EPI_ISL_628171, EPI_ISL_628172, EPI_ISL_628173, EPI_ISL_628174, EPI_ISL_628175, EPI_ISL_628176, EPI_ISL_628177, EPI_ISL_628178, EPI_ISL_628179, EPI_ISL_628180, EPI_ISL_628181, EPI_ISL_628182, EPI_ISL_628184, EPI_ISL_628185, EPI_ISL_628186, EPI_ISL_628187, EPI_ISL_628188, EPI_ISL_628190, EPI_ISL_628191, EPI_ISL_628192, EPI_ISL_628193, EPI_ISL_628194, EPI_ISL_628195, EPI_ISL_628196, EPI_ISL_628197, EPI_ISL_628198, EPI_ISL_628199, EPI_ISL_628200, EPI_ISL_628201, EPI_ISL_628202, EPI_ISL_628203, EPI_ISL_628204, EPI_ISL_628206, EPI_ISL_628207, EPI_ISL_628208, EPI_ISL_628209, EPI_ISL_628210, EPI_ISL_628212, EPI_ISL_628213, EPI_ISL_628214, EPI_ISL_628215, EPI_ISL_628216, EPI_ISL_628217, EPI_ISL_628218, EPI_ISL_628219, EPI_ISL_628221, EPI_ISL_628222, EPI_ISL_628223, EPI_ISL_628224, EPI_ISL_628226, EPI_ISL_628227, EPI_ISL_628228, EPI_ISL_628229, EPI_ISL_628230, EPI_ISL_628231, EPI_ISL_628232, EPI_ISL_628233, EPI_ISL_628234, EPI_ISL_628235, EPI_ISL_628237, EPI_ISL_628238, EPI_ISL_628239, EPI_ISL_628240, EPI_ISL_628241, EPI_ISL_628242, EPI_ISL_628243, EPI_ISL_628244, EPI_ISL_628246, EPI_ISL_628247, EPI_ISL_628248, EPI_ISL_628249, EPI_ISL_628250, | | | |

EPI_ISL_628251, EPI_ISL_628252, EPI_ISL_628253, EPI_ISL_628254, EPI_ISL_628255, EPI_ISL_628256, EPI_ISL_628257, EPI_ISL_628258, EPI_ISL_628260, EPI_ISL_628261, EPI_ISL_628262, EPI_ISL_628265, EPI_ISL_628266, EPI_ISL_628268, EPI_ISL_628269, EPI_ISL_628270, EPI_ISL_628271, EPI_ISL_628274, EPI_ISL_628275, EPI_ISL_628276, EPI_ISL_628277, EPI_ISL_628278, EPI_ISL_628279, EPI_ISL_628281, EPI_ISL_628282, EPI_ISL_628283, EPI_ISL_628284, EPI_ISL_628285, EPI_ISL_628286, EPI_ISL_628287, EPI_ISL_628288, EPI_ISL_628289, EPI_ISL_628290, EPI_ISL_628291, EPI_ISL_628293, EPI_ISL_628294, EPI_ISL_628295, EPI_ISL_628296, EPI_ISL_628297, EPI_ISL_628298, EPI_ISL_628299, EPI_ISL_628300, EPI_ISL_628301, EPI_ISL_628302, EPI_ISL_628303, EPI_ISL_628304, EPI_ISL_628305, EPI_ISL_628306, EPI_ISL_628308, EPI_ISL_628310, EPI_ISL_628311

| | | | |
|---|---|---|---|
| see above | Wales Specialist Virology Centre Sequencing lab: Pathogen Genomics Unit | COVID-19 Genomics UK (COG-UK) Consortium | Catherine Moore, Johnathan Evans, Laura Gifford, Malorie Perry, Simon Cottrell, Angela Marchbank, Alec Birchley, Alexander Adams, Amy Gaskin, Bree Gatica-Wilcox, Jason Coombes, Joel Southgate, Lauren Gilbert, Lee Graham, Nicole Pacchiarini, Sara Kumziene-Summerhayes, Sarah Taylor, Sophie Jones, Sara Rey, Matthew Bull, Joanne Watkins, Sally Corden, Tom Connor |
| EPI_ISL_629001 | South Eastern Area Laboratory Services (SEALS) | NSW Health Pathology - Institute of Clinical Pathology and Medical Research; Westmead Hospital; University of Sydney | CIDM-PH et al. |
| EPI_ISL_629002 | Sydney South West Pathology Service (SSWPS) - Liverpool Hospital - NSW Health Pathology | NSW Health Pathology - Institute of Clinical Pathology and Medical Research; Westmead Hospital; University of Sydney | CIDM-PH et al. |
| EPI_ISL_629003, EPI_ISL_629004, EPI_ISL_629005, EPI_ISL_629006 | South Eastern Area Laboratory Services (SEALS) | NSW Health Pathology - Institute of Clinical Pathology and Medical Research; Westmead Hospital; University of Sydney | CIDM-PH et al. |
| EPI_ISL_629007, EPI_ISL_629008, EPI_ISL_629009 | Laverty Pathology | NSW Health Pathology - Institute of Clinical Pathology and Medical Research; Westmead Hospital; University of Sydney | CIDM-PH et al. |
| EPI_ISL_629010 | Sydney South West Pathology Service (SSWPS) - Royal Prince Alfred Hospital - NSW Health Pathology | NSW Health Pathology - Institute of Clinical Pathology and Medical Research; Westmead Hospital; University of Sydney | CIDM-PH et al. |
| EPI_ISL_629029, EPI_ISL_629030 | South Eastern Area Laboratory Services (SEALS) | NSW Health Pathology - Institute of Clinical Pathology and Medical Research; Westmead Hospital; University of Sydney | CIDM-PH et al. |

We gratefully acknowledge the following Authors from the Originating laboratories responsible for obtaining the specimens, as well as the Submitting laboratories where the genome data were generated and shared via GISAID, on which this research is based.

All Submitters of data may be contacted directly via www.gisaid.org

Authors are sorted alphabetically.

| Accession ID | Originating Laboratory | Submitting Laboratory | Authors |
|---|---|---|---|
| EPI_ISL_405839, EPI_ISL_406030 | The University of Hong Kong - Shenzhen Hospital | Li Ka Shing Faculty of Medicine, The University of Hong Kong | Chan,J.F.-W., Yuan,S., Kok,K.H., To,K.K.-W., Chu,H., Yang,J., Xing,F., Liu,J., Yip,C.C.-Y., Poon,R.W.-S., Tsai,H.W., Lo,S.K.-F., Chan,K.H., Poon,V.K.-M., Chan,W.M., Ip,J.D., Cai,J.P., Cheng,V.C.-C., Chen,H., Hui,C.K.-M. and Yuen,K.Y. |
| EPI_ISL_406716, EPI_ISL_406717 | State Key Laboratory of Virology, Wuhan University | State Key Laboratory of Virology, Wuhan University | Chen,L., Liu,W., Zhang,Q., Xu,K., Ye,G., Wu,W., Sun,Z., Liu,F., Wu,K., Mei,Y., Zhang,W., Chen,Y., Li,Y., Shi,M., Lan,K. and Liu,Y. |
| EPI_ISL_411060, EPI_ISL_411066 | Fujian Center for Disease Control and Prevention | Fujian Center for Disease Control and Prevention | Chen Wei, Zhang Yanhua, He Wenxiang, Weng Yuwei |
| EPI_ISL_412026 | Second Hospital of Anhui Medical University | Second Hospital of Anhui Medical University | Changtai Wang, Zhongping Liua, Zixiang Chen, Xin Huang, Mengyuan Xua, Tengfei He, Mengji Lu, Zhenhua Zhang |
| EPI_ISL_413017, EPI_ISL_413018 | Department of Microbiology, Institute for Viral Diseases, College of Medicine, Korea University | Department of Microbiology, Institute for Viral Diseases, College of Medicine, Korea University | Changmin Kang, Joon-Yong Bae, Jungmin Lee, Heedo Park, Juyoung Cho, Jeonghun Kim, Gee eun Lee, Cui Chunguang, Kyeong-ryeol Shin, Dong Min Kim, Jin Il Kim, Man-Seong Park |
| EPI_ISL_413513 | Division of Infectious Diseases, Department of Internal Medicine, Korea University College of Medicine | Department of Microbiology, Institute for Viral Diseases, College of Medicine, Korea University | Changmin Kang, Joon-Yong Bae, Jungmin Lee, Jin Gu Yoon, Heedo Park, Juyoung Cho, Jeonghun Kim, Gee Eun Lee, Cui Chunguang, Kyeong-ryeol Shin, Ji Yun Noh, Joon Young Song, Hee Jin Cheong, Woo Joo Kim, Jin Il Kim, Man-Seong Park |
| EPI_ISL_413514 | Department of Microbiology, Institute for Viral Diseases, College of Medicine, Korea University | Department of Microbiology, Institute for Viral Diseases, College of Medicine, Korea University | Changmin Kang, Joon-Yong Bae, Jungmin Lee, Jin Gu Yoon, Heedo Park, Juyoung Cho, Jeonghun Kim, Gee Eun Lee, Cui Chunguang, Kyeong-ryeol Shin, Ji Yun Noh, Joon Young Song, Hee Jin Cheong, Woo Joo Kim, Jin Il Kim, Man-Seong Park |
| EPI_ISL_413515 | Division of Infectious Diseases, Department of Internal Medicine, Korea University College of Medicine | Department of Microbiology, Institute for Viral Diseases, College of Medicine, Korea University | Changmin Kang, Joon-Yong Bae, Jungmin Lee, Jin Gu Yoon, Heedo Park, Juyoung Cho, Jeonghun Kim, Gee Eun Lee, Cui Chunguang, Kyeong-ryeol Shin, Ji Yun Noh, Joon Young Song, Hee Jin Cheong, Woo Joo Kim, Jin Il Kim, Man-Seong Park |
| EPI_ISL_413516 | Department of Microbiology, Institute for Viral Diseases, College of Medicine, Korea University | Department of Microbiology, Institute for Viral Diseases, College of Medicine, Korea University | Changmin Kang, Joon-Yong Bae, Jungmin Lee, Jin Gu Yoon, Heedo Park, Juyoung Cho, Jeonghun Kim, Gee Eun Lee, Cui Chunguang, Kyeong-ryeol Shin, Ji Yun Noh, Joon Young Song, Hee Jin Cheong, Woo Joo Kim, Jin Il Kim, Man-Seong Park |
| EPI_ISL_413555 | Wales Specialist Virology Centre | Public Health Wales Microbiology Cardiff | Catherine Moore, Cen Sabu, Joanne Watkins, Sally Corden, Tom Connor |
| EPI_ISL_413556 | Wales Specialist Virology Centre | Public Health Wales Microbiology Cardiff | Catherine Moore, Tim Jones, Joanne Watkins, Sally Corden, Tom Connor |
| EPI_ISL_414486, EPI_ISL_414488, EPI_ISL_415041, EPI_ISL_415435, EPI_ISL_415453, EPI_ISL_415536, EPI_ISL_415537, EPI_ISL_415538, EPI_ISL_415655, EPI_ISL_415656 | Wales Specialist Virology Centre | Public Health Wales Microbiology Cardiff | Catherine Moore, Joanne Watkins, Sally Corden, Tom Connor |
| EPI_ISL_415657 | Wales Specialist Virology Centre | Public Health Wales Microbiology Cardiff | Catherine Moore, Joanne watkins, Sally Corden, Tom Connor |
| EPI_ISL_415659, EPI_ISL_415916, EPI_ISL_415918, EPI_ISL_415919, EPI_ISL_415920, EPI_ISL_415977, EPI_ISL_415978, EPI_ISL_415991, EPI_ISL_416024, EPI_ISL_416026, EPI_ISL_416027 | | | |
| see above | Wales Specialist Virology Centre | Public Health Wales Microbiology Cardiff | Catherine Moore, Joanne Watkins, Sally Corden, Tom Connor |
| EPI_ISL_417199 | Department of Clinical Pathology, Pamela Youde Nethersole Eastern Hospital | Department of Health Technology and Informatics, Faculty of Health and Social Science, The Hong Kong Polytechnic University | Chong-Yee YAU, Hiu-Yin LAO, Ming-Pan CHOI, Kingsley King-Gee TAM, Lam-Kwong LEE, Barry Kin-Chung WONG, Alex Yat-Man HO, Kam-Tong Yip, Kwok-Cheung LUNG, Raymond Wai-To LIU, Eugene Yuk-Keung TSO, Wai-Shing LEUNG, Man-Chun CHAN, Yuk-Yung NG, Kit-Man SIN, Kitty Sau-Chun FUNG, Sandy Ka-Yee CHAU, Wing-Kin TO, Tak-Lun Que, David Ho-Keung SHUM, Shea Ping YIP, Wing Cheong YAM, Gilman Kit-Hang SIU |
| EPI_ISL_417396 | Centre for Infectious Diseases and Microbiology Public Health | NSW Health Pathology - Institute of Clinical Pathology and Medical Research; Westmead Hospital; University of Sydney | Chen SC, Maddocks S, Kok J, Dwyer DE, Rockett R, Eden J-S, Lam C, Gray K, Timms V, Gall M, Arnott A, Sadsad R, Carter I, Rahman H, Holmes EC, O'Sullivan MV and Sintchenko V for the 2019-nCoV Study Group |
| EPI_ISL_417922 | INMI Lazzaro Spallanzani IRCCS | Laboratory of Virology, INMI Lazzaro Spallanzani IRCCS | Cesare E. M. Gruber, Martina Rueca, Barbara Bartolini, Francesco Messina, Emanuela Giombini, Maria R. Capobianchi, Fabrizio Carletti, Francesca Colavita, Concetta Castilletti, Eleonora Lalle, Daniele Lapa, Giuseppe Ippolito. |
| EPI_ISL_418083, EPI_ISL_418084, EPI_ISL_418085, EPI_ISL_418086, EPI_ISL_418087, EPI_ISL_418088, EPI_ISL_418089, EPI_ISL_418090, EPI_ISL_418091, EPI_ISL_418092, EPI_ISL_418093, EPI_ISL_418094, EPI_ISL_418095, EPI_ISL_418096, EPI_ISL_418097, EPI_ISL_418098, EPI_ISL_418100, EPI_ISL_418101, EPI_ISL_418102, EPI_ISL_418103, EPI_ISL_418104, EPI_ISL_418105, EPI_ISL_418106, EPI_ISL_418107, EPI_ISL_418108, EPI_ISL_418109, EPI_ISL_418110, EPI_ISL_418111, EPI_ISL_418112, EPI_ISL_418113, EPI_ISL_418114, EPI_ISL_418115, EPI_ISL_418116, EPI_ISL_418117, EPI_ISL_418118, EPI_ISL_418119, EPI_ISL_418120, EPI_ISL_418121, EPI_ISL_418122, EPI_ISL_418123, EPI_ISL_418124, EPI_ISL_418125, EPI_ISL_418126, EPI_ISL_418127, EPI_ISL_418128, EPI_ISL_418129, EPI_ISL_418130, EPI_ISL_418131, EPI_ISL_418132, EPI_ISL_418133, EPI_ISL_418134, EPI_ISL_418135, EPI_ISL_418136, EPI_ISL_418137, EPI_ISL_418138, EPI_ISL_418139, EPI_ISL_418140, EPI_ISL_418141, EPI_ISL_418142, EPI_ISL_418143, EPI_ISL_418144, EPI_ISL_418145, EPI_ISL_418146, EPI_ISL_418147, EPI_ISL_418148, EPI_ISL_418149, EPI_ISL_418150, EPI_ISL_418151, EPI_ISL_418152, EPI_ISL_418153, EPI_ISL_418154, EPI_ISL_418155, EPI_ISL_418156, EPI_ISL_418157, EPI_ISL_418158, EPI_ISL_418159, EPI_ISL_418160, EPI_ISL_418161, EPI_ISL_418162, EPI_ISL_418163, EPI_ISL_418164, EPI_ISL_418165, EPI_ISL_419399, EPI_ISL_419400, EPI_ISL_419401, EPI_ISL_419402, EPI_ISL_419403, EPI_ISL_419404, EPI_ISL_419405, EPI_ISL_419406, EPI_ISL_419407, EPI_ISL_419408, EPI_ISL_419409, EPI_ISL_419410, EPI_ISL_419411, EPI_ISL_419412, EPI_ISL_419413, EPI_ISL_419414, EPI_ISL_419415, EPI_ISL_419416, EPI_ISL_419417, EPI_ISL_419418, EPI_ISL_419419, EPI_ISL_419420, EPI_ISL_419421, EPI_ISL_419422, EPI_ISL_419423, EPI_ISL_419424, EPI_ISL_419425, EPI_ISL_419426, EPI_ISL_419427, EPI_ISL_419428, EPI_ISL_419429, EPI_ISL_419430, EPI_ISL_419431, EPI_ISL_419432, EPI_ISL_419433, EPI_ISL_419434, EPI_ISL_419435, EPI_ISL_419436, EPI_ISL_419437, EPI_ISL_419438, EPI_ISL_419439, EPI_ISL_419440, EPI_ISL_419441, EPI_ISL_419442, EPI_ISL_419443, EPI_ISL_419444, EPI_ISL_419445, EPI_ISL_419446, EPI_ISL_419447, EPI_ISL_419448, EPI_ISL_419449, EPI_ISL_419450, EPI_ISL_419452, EPI_ISL_419453, EPI_ISL_419454, EPI_ISL_419455, EPI_ISL_419457, EPI_ISL_419458, EPI_ISL_419459, EPI_ISL_419460, EPI_ISL_419461, EPI_ISL_419462, EPI_ISL_419463, EPI_ISL_419464, EPI_ISL_419465, EPI_ISL_419466, EPI_ISL_419467, EPI_ISL_419468, EPI_ISL_419469, EPI_ISL_419470, EPI_ISL_419471, EPI_ISL_419472, EPI_ISL_419473, EPI_ISL_419474, EPI_ISL_419475, EPI_ISL_419476, EPI_ISL_419477, EPI_ISL_419478, EPI_ISL_419479, EPI_ISL_419480, EPI_ISL_419481, EPI_ISL_419482, EPI_ISL_419483, EPI_ISL_419484, EPI_ISL_419485, EPI_ISL_419486, EPI_ISL_419487, EPI_ISL_419488, EPI_ISL_419489, EPI_ISL_419490, EPI_ISL_419491, EPI_ISL_419492, EPI_ISL_419493, EPI_ISL_419494, EPI_ISL_419495, EPI_ISL_419496, EPI_ISL_419497, EPI_ISL_419498, EPI_ISL_419499, EPI_ISL_419500, EPI_ISL_419501, EPI_ISL_419502, EPI_ISL_419503, EPI_ISL_419504, EPI_ISL_419505, EPI_ISL_419506, EPI_ISL_419507, EPI_ISL_419508, EPI_ISL_419509, EPI_ISL_419510, EPI_ISL_419511 | | | |
| see above | Wales Specialist Virology Centre | Public Health Wales Microbiology Cardiff | Catherine Moore, Joanne Watkins, Sally Corden, Sara Rey, Matt Bull, Tom Connor |
| EPI_ISL_420303 | Alaska State Virology Laboratory | Alaska State Virology Laboratory | Chen, J |
| EPI_ISL_420304, EPI_ISL_420305, EPI_ISL_420306 | Alaska State Virology Laboratory | Alaska State Virology Laboratory | Chen, J. |
| EPI_ISL_420456 | PathWest Laboratory Medicine WA | PathWest Laboratory Medicine WA | Chisha Sikazwe, Jurissa Lang, Avram Levy, David Smith and David Speers |
| EPI_ISL_420531, EPI_ISL_420532, EPI_ISL_420533, EPI_ISL_420534, EPI_ISL_420536, EPI_ISL_420537, EPI_ISL_420538, EPI_ISL_420539 | Department of Microbiology, PathWest QEII Medical Centre | Department of Microbiology, PathWest QEII Medical Centre | Chisha Sikazwe, Jurissa Lang, Avram Levy, David Speers and David Smith |
| EPI_ISL_420855, EPI_ISL_420876, EPI_ISL_420877 | Geelong Centre for Emerging Infectious Diseases | Geelong Centre for Emerging Infectious Diseases | Chamings,A., Raj Bhatta T., Alexandersen S. |
| EPI_ISL_420913, EPI_ISL_420914, EPI_ISL_420915, EPI_ISL_420917, EPI_ISL_420918, EPI_ISL_420919, EPI_ISL_420920, EPI_ISL_420921, EPI_ISL_420922, EPI_ISL_420923, EPI_ISL_420924, EPI_ISL_420925, EPI_ISL_420926, EPI_ISL_420927, EPI_ISL_420928, EPI_ISL_420929, EPI_ISL_420930, EPI_ISL_420931, EPI_ISL_420932, EPI_ISL_420933, EPI_ISL_420934, EPI_ISL_420935, EPI_ISL_420936, EPI_ISL_420937, EPI_ISL_420938, EPI_ISL_420939, EPI_ISL_420940, EPI_ISL_420941, EPI_ISL_420942, EPI_ISL_420943, EPI_ISL_420944, EPI_ISL_420945, EPI_ISL_420946, EPI_ISL_420947, EPI_ISL_420948, EPI_ISL_420949, EPI_ISL_420950, EPI_ISL_420951, EPI_ISL_420952, EPI_ISL_420953, EPI_ISL_420954, EPI_ISL_420955, EPI_ISL_420956, EPI_ISL_420957, EPI_ISL_420958, EPI_ISL_420959, EPI_ISL_420960, EPI_ISL_420961, EPI_ISL_420962, EPI_ISL_420963, EPI_ISL_420964, EPI_ISL_420965, EPI_ISL_420966, EPI_ISL_420967, EPI_ISL_420968, EPI_ISL_420969, EPI_ISL_420970, EPI_ISL_420971, EPI_ISL_420972, EPI_ISL_420973, EPI_ISL_420974, EPI_ISL_420975, EPI_ISL_420976, EPI_ISL_420977, EPI_ISL_420978, EPI_ISL_420979, EPI_ISL_420980, EPI_ISL_420981, EPI_ISL_420982, EPI_ISL_420983, EPI_ISL_420984, EPI_ISL_420985, EPI_ISL_420986, EPI_ISL_420987, EPI_ISL_420988, EPI_ISL_420989, EPI_ISL_420990, EPI_ISL_420991, EPI_ISL_420992, EPI_ISL_420993, EPI_ISL_420994, EPI_ISL_420995, EPI_ISL_420996, EPI_ISL_420997, EPI_ISL_420998, EPI_ISL_420999, EPI_ISL_421000, EPI_ISL_421001, EPI_ISL_421002, EPI_ISL_421003, EPI_ISL_421004, EPI_ISL_421005, EPI_ISL_421006, EPI_ISL_421007, EPI_ISL_421008, EPI_ISL_421009, EPI_ISL_421010, EPI_ISL_421011 | | | |
| see above | Wales Specialist Virology Centre | Public Health Wales Microbiology Cardiff | Catherine Moore, Joanne Watkins, Sally Corden, Malorie Perry, Simon Cottrell Sara Rey, Matt Bull, Tom Connor |
| EPI_ISL_422016, EPI_ISL_422017, EPI_ISL_422018, EPI_ISL_422019, EPI_ISL_422020, EPI_ISL_422021, EPI_ISL_422022, EPI_ISL_422023, EPI_ISL_422024, EPI_ISL_422025, EPI_ISL_422026, EPI_ISL_422027, EPI_ISL_422028, EPI_ISL_422029, EPI_ISL_422030, EPI_ISL_422031, EPI_ISL_422032, EPI_ISL_422033, | | | |

| | | | |
|---|---|---|---|
| EPI_ISL_422034, EPI_ISL_422035, EPI_ISL_422036, EPI_ISL_422037, EPI_ISL_422038, EPI_ISL_422039, EPI_ISL_422040, EPI_ISL_422041, EPI_ISL_422042, EPI_ISL_422043, EPI_ISL_422044, EPI_ISL_422045, EPI_ISL_422046, EPI_ISL_422047, EPI_ISL_422048, EPI_ISL_422049, EPI_ISL_422050, EPI_ISL_422051, EPI_ISL_422052, EPI_ISL_422053, EPI_ISL_422054, EPI_ISL_422055, EPI_ISL_422056, EPI_ISL_422057, EPI_ISL_422058, EPI_ISL_422059, EPI_ISL_422060, EPI_ISL_422061, EPI_ISL_422062, EPI_ISL_422063, EPI_ISL_422064, EPI_ISL_422065, EPI_ISL_422066, EPI_ISL_422067, EPI_ISL_422068, EPI_ISL_422069, EPI_ISL_422070, EPI_ISL_422071, EPI_ISL_422072, EPI_ISL_422073, EPI_ISL_422074, EPI_ISL_422075, EPI_ISL_422076, EPI_ISL_422077, EPI_ISL_422078, EPI_ISL_422079, EPI_ISL_422080, EPI_ISL_422081, EPI_ISL_422082, EPI_ISL_422083, EPI_ISL_422084, EPI_ISL_422085, EPI_ISL_422086, EPI_ISL_422087, EPI_ISL_422088, EPI_ISL_422089, EPI_ISL_422090, EPI_ISL_422091, EPI_ISL_422092, EPI_ISL_422093, EPI_ISL_422094, EPI_ISL_422095, EPI_ISL_422096, EPI_ISL_422097, EPI_ISL_422098, EPI_ISL_422099, EPI_ISL_422100, EPI_ISL_422102, EPI_ISL_422103, EPI_ISL_422104, EPI_ISL_422105, EPI_ISL_422106, EPI_ISL_422107, EPI_ISL_422108, EPI_ISL_422109, EPI_ISL_422110, EPI_ISL_422111, EPI_ISL_422112, EPI_ISL_422113, EPI_ISL_422114, EPI_ISL_422115, EPI_ISL_422116, EPI_ISL_422117, EPI_ISL_422118, EPI_ISL_422119, EPI_ISL_422120, EPI_ISL_422121, EPI_ISL_422122, EPI_ISL_422123, EPI_ISL_422124, EPI_ISL_422125, EPI_ISL_422126, EPI_ISL_422127, EPI_ISL_422128, EPI_ISL_422129, EPI_ISL_422130, EPI_ISL_422131, EPI_ISL_422132, EPI_ISL_422133, EPI_ISL_422134, EPI_ISL_422135, EPI_ISL_422136, EPI_ISL_422137, EPI_ISL_422138, EPI_ISL_422139, EPI_ISL_422140, EPI_ISL_422141, EPI_ISL_422142, EPI_ISL_422143, EPI_ISL_422144, EPI_ISL_422145, EPI_ISL_422146, EPI_ISL_422147, EPI_ISL_422148, EPI_ISL_422149, EPI_ISL_422150, EPI_ISL_422151, EPI_ISL_422152, EPI_ISL_422153, EPI_ISL_422154, EPI_ISL_422155, EPI_ISL_422156, EPI_ISL_422157, EPI_ISL_422158, EPI_ISL_422159, EPI_ISL_422160, EPI_ISL_422161, EPI_ISL_422162, EPI_ISL_422163, EPI_ISL_422164, EPI_ISL_422165, EPI_ISL_422166, EPI_ISL_422167, EPI_ISL_422168, EPI_ISL_422169, EPI_ISL_422170, EPI_ISL_422171, EPI_ISL_422172, EPI_ISL_422173, EPI_ISL_422174, EPI_ISL_422175, EPI_ISL_422176, EPI_ISL_422177, EPI_ISL_422178, EPI_ISL_422179, EPI_ISL_422180, EPI_ISL_422181, EPI_ISL_422182, EPI_ISL_422183, EPI_ISL_422184, EPI_ISL_422185, EPI_ISL_422186, EPI_ISL_422187, EPI_ISL_422188, EPI_ISL_422189, EPI_ISL_422190, EPI_ISL_422191, EPI_ISL_422192, EPI_ISL_422193, EPI_ISL_422194, EPI_ISL_422195, EPI_ISL_422196, EPI_ISL_422197, EPI_ISL_422198, EPI_ISL_422199, EPI_ISL_422200, EPI_ISL_422201, EPI_ISL_422202, EPI_ISL_422203, EPI_ISL_422204, EPI_ISL_422205, EPI_ISL_422206, EPI_ISL_422207, EPI_ISL_422208, EPI_ISL_422209, EPI_ISL_422210, EPI_ISL_422211, EPI_ISL_422212, EPI_ISL_422213, EPI_ISL_422214, EPI_ISL_422215, EPI_ISL_422216, EPI_ISL_422217, EPI_ISL_422218, EPI_ISL_422219, EPI_ISL_422220, EPI_ISL_422221, EPI_ISL_422222, EPI_ISL_422223, EPI_ISL_422224, EPI_ISL_422225, EPI_ISL_422226, EPI_ISL_422227, EPI_ISL_422228, EPI_ISL_422229, EPI_ISL_422230, EPI_ISL_422231, EPI_ISL_422232, EPI_ISL_422233, EPI_ISL_422234, EPI_ISL_422235, EPI_ISL_422236, EPI_ISL_422237, EPI_ISL_422238, EPI_ISL_422239, EPI_ISL_422240, EPI_ISL_422241, EPI_ISL_422242, EPI_ISL_422243, EPI_ISL_422244, EPI_ISL_422245, EPI_ISL_422246, EPI_ISL_422247, EPI_ISL_422248, EPI_ISL_422249, EPI_ISL_422250, EPI_ISL_422251, EPI_ISL_422252, EPI_ISL_422253, EPI_ISL_422254, EPI_ISL_422255, EPI_ISL_422256, EPI_ISL_422257, EPI_ISL_422258, EPI_ISL_422259, EPI_ISL_422260, EPI_ISL_422261, EPI_ISL_422262, EPI_ISL_422263, EPI_ISL_422264, EPI_ISL_422265, EPI_ISL_422266, EPI_ISL_422267, EPI_ISL_422268, EPI_ISL_422269, EPI_ISL_422270, EPI_ISL_422271, EPI_ISL_422272, EPI_ISL_422273, EPI_ISL_422274, EPI_ISL_422275, EPI_ISL_422276, EPI_ISL_422277, EPI_ISL_422278, EPI_ISL_422279, EPI_ISL_422280, EPI_ISL_422281, EPI_ISL_422282, EPI_ISL_422283, EPI_ISL_422284, EPI_ISL_422285, EPI_ISL_422286, EPI_ISL_422287, EPI_ISL_422288, EPI_ISL_422289, EPI_ISL_422290, EPI_ISL_422291, EPI_ISL_422292, EPI_ISL_422293, EPI_ISL_422294, EPI_ISL_422295, EPI_ISL_422296, EPI_ISL_422297, EPI_ISL_422298, EPI_ISL_422299, EPI_ISL_422300, EPI_ISL_422301, EPI_ISL_422302, EPI_ISL_422303, EPI_ISL_422304, EPI_ISL_422305, EPI_ISL_422306, EPI_ISL_422307, EPI_ISL_422308, EPI_ISL_422309, EPI_ISL_422310, EPI_ISL_422311, EPI_ISL_422312, EPI_ISL_422313, EPI_ISL_422314, EPI_ISL_422315, EPI_ISL_422316, EPI_ISL_422317, EPI_ISL_422318, EPI_ISL_422319, EPI_ISL_422320, EPI_ISL_422321, EPI_ISL_422322, EPI_ISL_422323, EPI_ISL_422324, EPI_ISL_422325, EPI_ISL_422326, EPI_ISL_422327, EPI_ISL_422328, EPI_ISL_422329, EPI_ISL_422330, EPI_ISL_422331, EPI_ISL_422332, EPI_ISL_422333, EPI_ISL_422334, EPI_ISL_422335, EPI_ISL_422336, EPI_ISL_422337, EPI_ISL_422338, EPI_ISL_422339, EPI_ISL_422340, EPI_ISL_422341, EPI_ISL_422342, EPI_ISL_422343, EPI_ISL_422344, EPI_ISL_422345, EPI_ISL_422346, EPI_ISL_422347, EPI_ISL_422348, EPI_ISL_422349, EPI_ISL_422350, EPI_ISL_422351, EPI_ISL_422352, EPI_ISL_422353, EPI_ISL_422354, EPI_ISL_422355, EPI_ISL_422356, EPI_ISL_422357, EPI_ISL_422358, EPI_ISL_422359, EPI_ISL_422360, EPI_ISL_422361, EPI_ISL_422362, EPI_ISL_422363, EPI_ISL_422364, EPI_ISL_422365, EPI_ISL_422366, EPI_ISL_422368, EPI_ISL_422369, EPI_ISL_422370, EPI_ISL_422371, EPI_ISL_422372, EPI_ISL_422373, EPI_ISL_422374, EPI_ISL_422375, EPI_ISL_422376, EPI_ISL_422377 | | | |
| see above | Wales Specialist Virology Centre | Public Health Wales Microbiology Cardiff | Catherine Moore, Johnathan Evans, Malorie Perry, Simon Cottrell, Alec Birchley, Alexander Adams, Amy Gaskin, Bree Gatica-Wilcox, Jason Coombes, Lauren Gilbert, Lee Graham, Nicole Pacchiarini, Sara Kumziene-Summerhayes, Sarah Taylor, Sophie Jones, Sara Rey, Matthew Bull, Joanne Watkins, Sally Corden, Tom Connor |
| EPI_ISL_424346, EPI_ISL_424347 | Alaska State Virology Laboratory | Alaska State Virology Laboratory | Chen, J. |
| EPI_ISL_426476, EPI_ISL_426477, EPI_ISL_426478 | Microbial Genomics Laboratory, Institut Pasteur Montevideo | Microbial Genomics Laboratory, Institut Pasteur Montevideo, Uruguay | Cecilia Salazar, Florencia Díaz-Viraqué, Marianoel Pereira, Pilar Moreno, Gonzalo Moratorio, Gregorio Iraola |
| EPI_ISL_426479, EPI_ISL_426480 | Microbial Genomics Laboratory, Institut Pasteur Montevideo | Microbial Genomics Laboratory, Institut Pasteur Montevideo | Cecilia Salazar, Florencia Díaz-Viraqué, Marianoel Pereira, Pilar Moreno, Gonzalo Moratorio, Gregorio Iraola |
| EPI_ISL_426481, EPI_ISL_426482 | Microbial Genomics Laboratory, Institut Pasteur Montevideo, Uruguay | Microbial Genomics Laboratory, Institut Pasteur Montevideo | Cecilia Salazar, Florencia Díaz-Viraqué, Marianoel Pereira, Pilar Moreno, Gonzalo Moratorio, Gregorio Iraola |
| EPI_ISL_426584 | Microbial Genomics Laboratory, Institut Pasteur Montevideo, Uruguay | Microbial Genomics Laboratory, Institut Pasteur Montevideo, Uruguay | Cecilia Salazar, Florencia Díaz-Viraqué, Marianoel Pereira, Pilar Moreno, Gonzalo Moratorio, Gregorio Iraola |
| EPI_ISL_426629, EPI_ISL_426630 | TSGH-CP molecular lab | TSGH-CP molecular lab | Cherng-Lih Perng, Ming-Jr Jian, Chih-Kai Chang, Jung-Chung Lin, Kuo-Ming Yeh, Chien-Wen Chen, Sheng-Kang Chiu, Hsing-Yi Chung, Shih-Hung Tsai, Kuo-Sheng Hung, Feng-Yee Chang, Hung-Sheng Shang |
| EPI_ISL_426631, EPI_ISL_426632, EPI_ISL_427392, EPI_ISL_427393, EPI_ISL_427394, EPI_ISL_427395, EPI_ISL_427397, EPI_ISL_427398 | TSGH-CP molecular lab | TSGH-CP molecular lab | Cherng-Lih Perng, Ming-Jr Jian, Chih-Kai Chang, Jung-Chung Lin, Kuo-Ming Yeh, Chien-Wen Chen, Sheng-Kang Chiu, Hsing-Yi Chung, Shih-Hung Tsai, Kuo-Sheng Hung, Tien-Yao Chang, Feng-Yee Chang, Hung-Sheng Shang |
| EPI_ISL_427619, EPI_ISL_427620, EPI_ISL_427621, EPI_ISL_427622 | Alaska State Virology Laboratory | Alaska State Virology Laboratory | Chen, J. |
| EPI_ISL_428229, EPI_ISL_428230 | TSGH-CP molecular lab | TSGH-CP molecular lab | Cherng-Lih Perng, Ming-Jr Jian, Chih-Kai Chang, Jung-Chung Lin, Kuo-Ming Yeh, Chien-Wen Chen, Sheng-Kang Chiu, Hsing-Yi Chung, Shih-Hung Tsai, Kuo-Sheng Hung, Tien-Yao Chang, Feng-Yee Chang, Hung-Sheng Shang |
| EPI_ISL_428231 | TSGH-CP molecular lab | TSGH-CP molecular lab | Cherng-Lih Perng, Ming-Jr JIAN, Chih-Kai Chang, Jung-Chung Lin, Kuo-Ming Yeh, Chien-Wen Chen, Sheng-Kang Chiu, Hsing-Yi Chung, Shih-Hung Tsai, Kuo-Sheng Hung, Tien-Yao Chang, Feng-Yee Chang, Hung-Sheng Shang |
| EPI_ISL_428479, EPI_ISL_428481, EPI_ISL_428482, EPI_ISL_428483, EPI_ISL_428484, EPI_ISL_428485, EPI_ISL_428486, EPI_ISL_428487 | District Surveillence Unit | Department of Neurovirology, National Institute of Mental Health and Neuroscience (NIMHANS) | Chitra Pattabiraman, Vijayalakshmi Reddy, Harsha PK, Risha Rasheed, Shafeeq S Hameed, Manjunatha Venkataswamy, Anita Desai, Ravi Vasanthapuram |
| EPI_ISL_428670 | Centre for Dengue Research | Centre for Dengue Research | Chandima Jeewandara, Dinuka Ariyaratne, Laksiri Gomes, Deshni Jayathilaka, Ananda Wijewickrama, Eranga Narangoda, Damayanthi Idampitiya, Neelika Malaige |
| EPI_ISL_428671 | Centre for Dengue Research | Centre for Dengue Research | Chandima Jeewandara, Dinuka Ariyatane, Laksiri Gomes, Deshni Jayathilaka, Diyanath Ranasinghe, Ananda Wijewickrama, Eranga Narangoda, Damayanthi Tdampitiya, Neelika Malavige |
| EPI_ISL_428672, EPI_ISL_428673 | Centre for Dengue Research | Centre for Dengue Research | Chandima Jeewandara, Dinuka Ariyaratne, Laksiri Gomes, Deshni Jayathilaka, Diyanath Ranasinghe, Ananda Wijewickrama, Eranga Narangoda, Damayanthi Idampitiya, Neelika Malavige |
| EPI_ISL_429257 | Microbial Genomics Laboratory, Institut Pasteur Montevideo | Microbial Genomics Laboratory, Institut Pasteur Montevideo | Cecilia Salazar, Florencia Díaz-Viraqué, Marianoel Pereira, Pilar Moreno, Gonzalo Moratorio, Gregorio Iraola |
| EPI_ISL_430064, EPI_ISL_430065, EPI_ISL_430066 | Geelong Centre for Emerging Infectious Diseases | Geelong Centre for Emerging Infectious Diseases | Chamings A., Bhatta T.R., Alexandersen S. |
| EPI_ISL_431901, EPI_ISL_431902, EPI_ISL_431903, EPI_ISL_431904, EPI_ISL_431905, EPI_ISL_431906, EPI_ISL_431907, EPI_ISL_431908, EPI_ISL_431909, EPI_ISL_431910, EPI_ISL_431911, EPI_ISL_431912, EPI_ISL_431913, EPI_ISL_431914, EPI_ISL_431916, EPI_ISL_431917, EPI_ISL_431918, EPI_ISL_431919, EPI_ISL_431920, EPI_ISL_431921, EPI_ISL_431922, EPI_ISL_431923, EPI_ISL_431924, EPI_ISL_431925, EPI_ISL_431926, EPI_ISL_431927, EPI_ISL_431928, EPI_ISL_431929, EPI_ISL_431930, EPI_ISL_431931, EPI_ISL_431932, EPI_ISL_431933, EPI_ISL_431934, EPI_ISL_431935, EPI_ISL_431936, EPI_ISL_431937, EPI_ISL_431938, EPI_ISL_431939, EPI_ISL_431940, EPI_ISL_431941, EPI_ISL_431942, EPI_ISL_431943, EPI_ISL_431944, EPI_ISL_431945, EPI_ISL_431946, EPI_ISL_431947, EPI_ISL_431948, EPI_ISL_431949, EPI_ISL_431950, EPI_ISL_431951, EPI_ISL_431952, EPI_ISL_431953, EPI_ISL_431954, EPI_ISL_431955, EPI_ISL_431956, EPI_ISL_431957, EPI_ISL_431958, EPI_ISL_431959, EPI_ISL_431960, EPI_ISL_431961, EPI_ISL_431962, EPI_ISL_431963, EPI_ISL_431964, EPI_ISL_431965, EPI_ISL_431966, EPI_ISL_431967, EPI_ISL_431968, EPI_ISL_431971, EPI_ISL_431972, EPI_ISL_431973, EPI_ISL_431974, EPI_ISL_431975, EPI_ISL_431976, EPI_ISL_431977, EPI_ISL_431978, EPI_ISL_431980, EPI_ISL_431981, EPI_ISL_431982, EPI_ISL_431983, EPI_ISL_431984, EPI_ISL_431985, EPI_ISL_431986, EPI_ISL_431987, EPI_ISL_431988, EPI_ISL_431989, EPI_ISL_431990, EPI_ISL_431991, EPI_ISL_431992, EPI_ISL_431994, EPI_ISL_431995, EPI_ISL_431996, EPI_ISL_431997, EPI_ISL_431998, EPI_ISL_431999, EPI_ISL_432000, EPI_ISL_432001, EPI_ISL_432002, EPI_ISL_432003, EPI_ISL_432004, EPI_ISL_432005, EPI_ISL_432006, EPI_ISL_432007, EPI_ISL_432008, EPI_ISL_432009, EPI_ISL_432010, EPI_ISL_432011, EPI_ISL_432012, EPI_ISL_432013, EPI_ISL_432014, EPI_ISL_432015, EPI_ISL_432016, EPI_ISL_432017, EPI_ISL_432018, EPI_ISL_432019, EPI_ISL_432021, EPI_ISL_432022, EPI_ISL_432023, EPI_ISL_432024, EPI_ISL_432025, EPI_ISL_432026, EPI_ISL_432027, EPI_ISL_432028, EPI_ISL_432029, EPI_ISL_432030, EPI_ISL_432031, EPI_ISL_432032, EPI_ISL_432033, EPI_ISL_432034, EPI_ISL_432035, EPI_ISL_432036, EPI_ISL_432037, EPI_ISL_432038, EPI_ISL_432039, EPI_ISL_432040, EPI_ISL_432041, EPI_ISL_432042, EPI_ISL_432043, EPI_ISL_432044, EPI_ISL_432045, EPI_ISL_432046, EPI_ISL_432047, EPI_ISL_432048, EPI_ISL_432049, EPI_ISL_432050, EPI_ISL_432051, EPI_ISL_432052, EPI_ISL_432053, EPI_ISL_432054, EPI_ISL_432055, EPI_ISL_432056, EPI_ISL_432057, EPI_ISL_432058, EPI_ISL_432059, EPI_ISL_432062, EPI_ISL_432065, EPI_ISL_432066, EPI_ISL_432067, EPI_ISL_432068, EPI_ISL_432069, EPI_ISL_432070, EPI_ISL_432071, EPI_ISL_432072, EPI_ISL_432073, EPI_ISL_432074, EPI_ISL_432075, EPI_ISL_432077, EPI_ISL_432078, EPI_ISL_432079, EPI_ISL_432080, EPI_ISL_432081, EPI_ISL_432082, EPI_ISL_432083, EPI_ISL_432084, EPI_ISL_432085, EPI_ISL_432086, EPI_ISL_432088, EPI_ISL_432089, EPI_ISL_432090, EPI_ISL_432091, EPI_ISL_432092, EPI_ISL_432093, EPI_ISL_432094, EPI_ISL_432095, EPI_ISL_432096, EPI_ISL_432097, EPI_ISL_432098, EPI_ISL_432099, EPI_ISL_432100, EPI_ISL_432103, EPI_ISL_432105, EPI_ISL_432106, EPI_ISL_432107, EPI_ISL_432108, EPI_ISL_432109, EPI_ISL_432110, EPI_ISL_432111, EPI_ISL_432112, EPI_ISL_432113, EPI_ISL_432115, EPI_ISL_432116, EPI_ISL_432117, EPI_ISL_432118, EPI_ISL_432119, EPI_ISL_432120, EPI_ISL_432121, EPI_ISL_432122, EPI_ISL_432123, EPI_ISL_432124, EPI_ISL_432125, EPI_ISL_432126, EPI_ISL_432127, EPI_ISL_432128, EPI_ISL_432129, EPI_ISL_432130, EPI_ISL_432131, EPI_ISL_432132, EPI_ISL_432133, EPI_ISL_432134, EPI_ISL_432135, EPI_ISL_432136, EPI_ISL_432137, EPI_ISL_432138, EPI_ISL_432139, EPI_ISL_432140, EPI_ISL_432141, EPI_ISL_432142, EPI_ISL_432143, EPI_ISL_432144, EPI_ISL_432145, EPI_ISL_432146, EPI_ISL_432147, EPI_ISL_432148, EPI_ISL_432151, EPI_ISL_432152, EPI_ISL_432153, EPI_ISL_432154, EPI_ISL_432155, EPI_ISL_432156, EPI_ISL_432157, EPI_ISL_432158, EPI_ISL_432159, EPI_ISL_432160, EPI_ISL_432161, EPI_ISL_432162, EPI_ISL_432163, EPI_ISL_432165, EPI_ISL_432166, EPI_ISL_432167, EPI_ISL_432168, EPI_ISL_432169, EPI_ISL_432170, EPI_ISL_432171, EPI_ISL_432172, EPI_ISL_432173, EPI_ISL_432174, EPI_ISL_432175, EPI_ISL_432176, EPI_ISL_432177, EPI_ISL_432178, EPI_ISL_432179, EPI_ISL_432180, EPI_ISL_432181, EPI_ISL_432182, EPI_ISL_432183, EPI_ISL_432184, EPI_ISL_432185, EPI_ISL_432186, EPI_ISL_432187, EPI_ISL_432188, EPI_ISL_432189, EPI_ISL_432190, EPI_ISL_432191, EPI_ISL_432192, EPI_ISL_432193, EPI_ISL_432194, EPI_ISL_432195, EPI_ISL_432196, EPI_ISL_432197, EPI_ISL_432198, EPI_ISL_432199, EPI_ISL_432200, EPI_ISL_432201, EPI_ISL_432202, EPI_ISL_432203, EPI_ISL_432204, EPI_ISL_432205, EPI_ISL_432206, EPI_ISL_432207, EPI_ISL_432208, EPI_ISL_432209, EPI_ISL_432210, EPI_ISL_432211, EPI_ISL_432212, EPI_ISL_432213, EPI_ISL_432214, EPI_ISL_432215, EPI_ISL_432216, EPI_ISL_432217, EPI_ISL_432218, EPI_ISL_432219, EPI_ISL_432220, EPI_ISL_432221, EPI_ISL_432222, EPI_ISL_432223, EPI_ISL_432224, EPI_ISL_432225, EPI_ISL_432226, EPI_ISL_432227, EPI_ISL_432228, EPI_ISL_432229, EPI_ISL_432230, EPI_ISL_432231, EPI_ISL_432232, EPI_ISL_432233, EPI_ISL_432234, EPI_ISL_432235, EPI_ISL_432236, EPI_ISL_432237, EPI_ISL_432238, EPI_ISL_432239, EPI_ISL_432240, EPI_ISL_432241, EPI_ISL_432242, EPI_ISL_432243, EPI_ISL_432244, EPI_ISL_432245, EPI_ISL_432246, EPI_ISL_432247, EPI_ISL_432248, EPI_ISL_432249, EPI_ISL_432250, EPI_ISL_432251, EPI_ISL_432252, EPI_ISL_432253, EPI_ISL_432254, EPI_ISL_432255, EPI_ISL_432256, EPI_ISL_432257, EPI_ISL_432258, EPI_ISL_432259, EPI_ISL_432260, EPI_ISL_432261, EPI_ISL_432262, EPI_ISL_432263, EPI_ISL_432264, EPI_ISL_432265, EPI_ISL_432266, EPI_ISL_432267, EPI_ISL_432268, EPI_ISL_432269, EPI_ISL_432270, EPI_ISL_432271, EPI_ISL_432272, EPI_ISL_432273, EPI_ISL_432274, EPI_ISL_432275, EPI_ISL_432276, EPI_ISL_432277, EPI_ISL_432278, EPI_ISL_432279, EPI_ISL_432280, EPI_ISL_432281, EPI_ISL_432282, EPI_ISL_432283, EPI_ISL_432284, EPI_ISL_432285, EPI_ISL_432286, EPI_ISL_432287, EPI_ISL_432288, EPI_ISL_432289, EPI_ISL_432290, EPI_ISL_432291, EPI_ISL_432292, EPI_ISL_432293, EPI_ISL_432294, EPI_ISL_432295, EPI_ISL_432296, EPI_ISL_432297, EPI_ISL_432298, EPI_ISL_432299, EPI_ISL_432300, EPI_ISL_432301, EPI_ISL_432302, EPI_ISL_432303, EPI_ISL_432304, EPI_ISL_432305, EPI_ISL_432306, EPI_ISL_432307, EPI_ISL_432309, EPI_ISL_432310, EPI_ISL_432312, EPI_ISL_432313, EPI_ISL_432314, EPI_ISL_432315, EPI_ISL_432316, EPI_ISL_432317, EPI_ISL_432318, EPI_ISL_432319, EPI_ISL_432320, EPI_ISL_432321, EPI_ISL_432322, EPI_ISL_432323, EPI_ISL_432324, EPI_ISL_432325, EPI_ISL_432327, EPI_ISL_432328, EPI_ISL_432329, EPI_ISL_432330, EPI_ISL_432331, EPI_ISL_432332, EPI_ISL_432333, EPI_ISL_432334, EPI_ISL_432335, EPI_ISL_432337, | | | |

| | | | |
|---|---|---|---|
| EPI_ISL_432338, EPI_ISL_432339, EPI_ISL_432340, EPI_ISL_432341, EPI_ISL_432342, EPI_ISL_432343, EPI_ISL_432344, EPI_ISL_432345, EPI_ISL_432346, EPI_ISL_432347, EPI_ISL_432348, EPI_ISL_432349, EPI_ISL_432350, EPI_ISL_432351, EPI_ISL_432352, EPI_ISL_432353, EPI_ISL_432354, EPI_ISL_432355, EPI_ISL_432356, EPI_ISL_432357, EPI_ISL_432358, EPI_ISL_432359, EPI_ISL_432360, EPI_ISL_432361, EPI_ISL_432362, EPI_ISL_432363, EPI_ISL_432364, EPI_ISL_432365, EPI_ISL_432366, EPI_ISL_432367, EPI_ISL_432368, EPI_ISL_432369, EPI_ISL_432370, EPI_ISL_432371, EPI_ISL_432372, EPI_ISL_432373, EPI_ISL_432374, EPI_ISL_432377, EPI_ISL_432378, EPI_ISL_432379, EPI_ISL_432380, EPI_ISL_432381, EPI_ISL_432382, EPI_ISL_432383, EPI_ISL_432384, EPI_ISL_432385, EPI_ISL_432386, EPI_ISL_432387, EPI_ISL_432388, EPI_ISL_432389, EPI_ISL_432390, EPI_ISL_432391, EPI_ISL_432392, EPI_ISL_432393, EPI_ISL_432394, EPI_ISL_432395, EPI_ISL_432396, EPI_ISL_432397, EPI_ISL_432398, EPI_ISL_432399, EPI_ISL_432400, EPI_ISL_432401, EPI_ISL_432402, EPI_ISL_432403, EPI_ISL_432404, EPI_ISL_432407, EPI_ISL_432408, EPI_ISL_432409, EPI_ISL_432410, EPI_ISL_432411, EPI_ISL_432412, EPI_ISL_432413, EPI_ISL_432414, EPI_ISL_432416, EPI_ISL_432417, EPI_ISL_432418, EPI_ISL_432419, EPI_ISL_432420, EPI_ISL_432421, EPI_ISL_432422, EPI_ISL_432423, EPI_ISL_432424, EPI_ISL_432425, EPI_ISL_432426, EPI_ISL_432427, EPI_ISL_432428, EPI_ISL_432429, EPI_ISL_432430, EPI_ISL_432431, EPI_ISL_432432, EPI_ISL_432433, EPI_ISL_432434, EPI_ISL_432435, EPI_ISL_432436, EPI_ISL_432437, EPI_ISL_432438, EPI_ISL_432439, EPI_ISL_432440, EPI_ISL_432441, EPI_ISL_432442, EPI_ISL_432443, EPI_ISL_432444, EPI_ISL_432445, EPI_ISL_432446, EPI_ISL_432447, EPI_ISL_432448, EPI_ISL_432449, EPI_ISL_432450 | | | |
| see above | Wales Specialist Virology Centre | Public Health Wales Microbiology Cardiff | Catherine Moore, Johnathan Evans, Malorie Perry, Simon Cottrell, Alec Birchley, Alexander Adams, Amy Gaskin, Bree Gatica-Wilcox, Jason Coombes, Lauren Gilbert, Lee Graham, Nicole Pacchiarini, Sara Kumziene-Summerhayes, Sarah Taylor, Sophie Jones, Sara Rey, Matthew Bull, Joanne Watkins, Sally Corden, Tom Connor |
| EPI_ISL_434571 | Microbiology, The University of Hong Kong | Microbiology, The University of Hong Kong | Chan,J.F.W. and Yuen,K.-Y. |
| EPI_ISL_436099, EPI_ISL_436101, EPI_ISL_436102, EPI_ISL_436103, EPI_ISL_436104 | TSGH-CP molecular lab | TSGH-CP molecular lab | Cherng-Lih Perng, Ming-Jr JIAN, Chih-Kai Chang, Jung-Chung Lin, Kuo-Ming Yeh, Chien-Wen Chen, Sheng-Kang Chiu, Hsing-Yi Chung, Shih-Hung Tsai, Kuo-Sheng Hung, Tien-Yao Chang, Feng-Yee Chang, Hung-Sheng Shang |
| EPI_ISL_436105 | TSGH-CP molecular lab | TSGH-CP molecular lab | Cherng-Lih Perng, Ming-Jr Jian, Chih-Kai Chang, Jung-Chung Lin, Kuo-Ming Yeh, Chien-Wen Chen, Sheng-Kang Chiu, Hsing-Yi Chung, Shih-Hung Tsai, Kuo-Sheng Hung, Tien-Yao Chang, Feng-Yee Chang, Hung-Sheng Shang |
| EPI_ISL_436106, EPI_ISL_436107, EPI_ISL_436108 | TSGH-CP molecular lab | TSGH-CP molecular lab | Cherng-Lih Perng, Ming-Jr JIAN, Chih-Kai Chang, Jung-Chung Lin, Kuo-Ming Yeh, Chien-Wen Chen, Sheng-Kang Chiu, Hsing-Yi Chung, Shih-Hung Tsai, Kuo-Sheng Hung, Tien-Yao Chang, Feng-Yee Chang, Hung-Sheng Shang |
| EPI_ISL_436137, EPI_ISL_436138, EPI_ISL_436139, EPI_ISL_436140, EPI_ISL_436141, EPI_ISL_436156, EPI_ISL_436157 | District Surveillance Unit | Department of Neurovirology, National Institute of Mental Health and Neuroscience (NIMHANS) | Chitra Pattabiraman, Vijayalakshmi Reddy, Harsha PK, Risha Rasheed, Shafeeq S Hameed, Manjunatha Venkataswamy, Anita Desai, Ravi Vasanthapuram |
| EPI_ISL_437336 | TSGH-CP molecular lab, Division of Clinical Pathology, Department of Pathology | TSGH-CP molecular lab, Division of Clinical Pathology, Department of Pathology | Cherng-Lih Perng, Ming-Jr JIAN, Chih-Kai Chang, Jung-Chung Lin, Kuo-Ming Yeh, Chien-Wen Chen, Sheng-Kang Chiu, Hsing-Yi Chung, Shih-Hung Tsai, Kuo-Sheng Hung, Tien-Yao Chang, Feng-Yee Chang, Hung-Sheng Shang |
| EPI_ISL_445381, EPI_ISL_445382, EPI_ISL_445383, EPI_ISL_445384, EPI_ISL_445385, EPI_ISL_445386, EPI_ISL_445387, EPI_ISL_445388, EPI_ISL_445389, EPI_ISL_445390, EPI_ISL_445391, EPI_ISL_445392, EPI_ISL_445393, EPI_ISL_445394, EPI_ISL_445395, EPI_ISL_445396, EPI_ISL_445397, EPI_ISL_445398, EPI_ISL_445399, EPI_ISL_445400, EPI_ISL_445401, EPI_ISL_445402, EPI_ISL_445403, EPI_ISL_445404, EPI_ISL_445405, EPI_ISL_445406, EPI_ISL_445407, EPI_ISL_445408, EPI_ISL_445409, EPI_ISL_445410, EPI_ISL_445411, EPI_ISL_445412, EPI_ISL_445413, EPI_ISL_445414, EPI_ISL_445415, EPI_ISL_445416, EPI_ISL_445417, EPI_ISL_445418, EPI_ISL_445419, EPI_ISL_445420, EPI_ISL_445421, EPI_ISL_445422, EPI_ISL_445423, EPI_ISL_445424, EPI_ISL_445425, EPI_ISL_445426, EPI_ISL_445427, EPI_ISL_445428, EPI_ISL_445429, EPI_ISL_445430, EPI_ISL_445431, EPI_ISL_445432, EPI_ISL_445433, EPI_ISL_445434, EPI_ISL_445435, EPI_ISL_445436, EPI_ISL_445437, EPI_ISL_445438, EPI_ISL_445439, EPI_ISL_445440, EPI_ISL_445441, EPI_ISL_445442, EPI_ISL_445443, EPI_ISL_445444, EPI_ISL_445445, EPI_ISL_445446, EPI_ISL_445447, EPI_ISL_445448, EPI_ISL_445449, EPI_ISL_445450, EPI_ISL_445451, EPI_ISL_445452, EPI_ISL_445453, EPI_ISL_445454, EPI_ISL_445455, EPI_ISL_445456, EPI_ISL_445457, EPI_ISL_445458, EPI_ISL_445459, EPI_ISL_445460, EPI_ISL_445461, EPI_ISL_445462, EPI_ISL_445463, EPI_ISL_445464, EPI_ISL_445465, EPI_ISL_445466, EPI_ISL_445467, EPI_ISL_445468, EPI_ISL_445469, EPI_ISL_445470, EPI_ISL_445471, EPI_ISL_445472, EPI_ISL_445473, EPI_ISL_445474, EPI_ISL_445475, EPI_ISL_445476, EPI_ISL_445477, EPI_ISL_445478, EPI_ISL_445479, EPI_ISL_445480, EPI_ISL_445481, EPI_ISL_445482, EPI_ISL_445483, EPI_ISL_445484, EPI_ISL_445485, EPI_ISL_445486, EPI_ISL_445487, EPI_ISL_445488, EPI_ISL_445489, EPI_ISL_445490, EPI_ISL_445491, EPI_ISL_445492, EPI_ISL_445493, EPI_ISL_445494, EPI_ISL_445495, EPI_ISL_445496, EPI_ISL_445497, EPI_ISL_445498, EPI_ISL_445499, EPI_ISL_445500, EPI_ISL_445501, EPI_ISL_445502, EPI_ISL_445503, EPI_ISL_445504, EPI_ISL_445505, EPI_ISL_445506, EPI_ISL_445507, EPI_ISL_445508, EPI_ISL_445509, EPI_ISL_445510, EPI_ISL_445511, EPI_ISL_445512, EPI_ISL_445513, EPI_ISL_445514, EPI_ISL_445515, EPI_ISL_445516, EPI_ISL_445517, EPI_ISL_445518, EPI_ISL_445519, EPI_ISL_445520, EPI_ISL_445521, EPI_ISL_445522, EPI_ISL_445523, EPI_ISL_445524, EPI_ISL_445525, EPI_ISL_445526, EPI_ISL_445527, EPI_ISL_445528, EPI_ISL_445529, EPI_ISL_445530, EPI_ISL_445531, EPI_ISL_445533, EPI_ISL_445534, EPI_ISL_445535, EPI_ISL_445536, EPI_ISL_445537, EPI_ISL_445538, EPI_ISL_445539, EPI_ISL_445540, EPI_ISL_445541, EPI_ISL_445542, EPI_ISL_445543, EPI_ISL_445544, EPI_ISL_445545, EPI_ISL_445546, EPI_ISL_445547, EPI_ISL_445548, EPI_ISL_445549, EPI_ISL_445550, EPI_ISL_445551, EPI_ISL_445552, EPI_ISL_445553, EPI_ISL_445554, EPI_ISL_445555, EPI_ISL_445556, EPI_ISL_445557, EPI_ISL_445558, EPI_ISL_445559, EPI_ISL_445560, EPI_ISL_445561, EPI_ISL_445562, EPI_ISL_445563, EPI_ISL_445564, EPI_ISL_445565, EPI_ISL_445566, EPI_ISL_445567, EPI_ISL_445568, EPI_ISL_445569, EPI_ISL_445570, EPI_ISL_445571, EPI_ISL_445572, EPI_ISL_445573, EPI_ISL_445574, EPI_ISL_445575, EPI_ISL_445576, EPI_ISL_445577, EPI_ISL_445578, EPI_ISL_445579, EPI_ISL_445580, EPI_ISL_445581, EPI_ISL_445582, EPI_ISL_445583, EPI_ISL_445584, EPI_ISL_445585, EPI_ISL_445586, EPI_ISL_445587, EPI_ISL_445588, EPI_ISL_445589, EPI_ISL_445590, EPI_ISL_445591, EPI_ISL_445592, EPI_ISL_445593, EPI_ISL_445594, EPI_ISL_445595, EPI_ISL_445596, EPI_ISL_445597, EPI_ISL_445598, EPI_ISL_445599, EPI_ISL_445600, EPI_ISL_445601, EPI_ISL_445602, EPI_ISL_445603, EPI_ISL_445604, EPI_ISL_445605, EPI_ISL_445606, EPI_ISL_445607, EPI_ISL_445608, EPI_ISL_445609, EPI_ISL_445610, EPI_ISL_445611, EPI_ISL_445612, EPI_ISL_445613, EPI_ISL_445614, EPI_ISL_445615, EPI_ISL_445616, EPI_ISL_445617, EPI_ISL_445618, EPI_ISL_445619, EPI_ISL_445620, EPI_ISL_445621, EPI_ISL_445622, EPI_ISL_445623, EPI_ISL_445624, EPI_ISL_445625, EPI_ISL_445626, EPI_ISL_445627, EPI_ISL_445628, EPI_ISL_445629, EPI_ISL_445630, EPI_ISL_445631, EPI_ISL_445632, EPI_ISL_445633, EPI_ISL_445634, EPI_ISL_445635, EPI_ISL_445636, EPI_ISL_445637, EPI_ISL_445638, EPI_ISL_445639, EPI_ISL_445640, EPI_ISL_445641, EPI_ISL_445642, EPI_ISL_445643, EPI_ISL_445644, EPI_ISL_445645, EPI_ISL_445646, EPI_ISL_445647, EPI_ISL_445648, EPI_ISL_445649, EPI_ISL_445650, EPI_ISL_445651, EPI_ISL_445652, EPI_ISL_445653, EPI_ISL_445654, EPI_ISL_445655, EPI_ISL_445656, EPI_ISL_445657, EPI_ISL_445658, EPI_ISL_445659, EPI_ISL_445660, EPI_ISL_445661, EPI_ISL_445662, EPI_ISL_445663, EPI_ISL_445664, EPI_ISL_445665, EPI_ISL_445666, EPI_ISL_445667, EPI_ISL_445668, EPI_ISL_445669, EPI_ISL_445670, EPI_ISL_445671, EPI_ISL_445672, EPI_ISL_445673, EPI_ISL_445674, EPI_ISL_445675, EPI_ISL_445676, EPI_ISL_445677, EPI_ISL_445678, EPI_ISL_445679, EPI_ISL_445680, EPI_ISL_445681, EPI_ISL_445682, EPI_ISL_445683, EPI_ISL_445684, EPI_ISL_445685, EPI_ISL_445686, EPI_ISL_445687, EPI_ISL_445688, EPI_ISL_445689, EPI_ISL_445690, EPI_ISL_445691, EPI_ISL_445692, EPI_ISL_445693, EPI_ISL_445694, EPI_ISL_445695, EPI_ISL_445696, EPI_ISL_445697, EPI_ISL_445698, EPI_ISL_445699, EPI_ISL_445700, EPI_ISL_445701, EPI_ISL_445702, EPI_ISL_445703, EPI_ISL_445704, EPI_ISL_445705, EPI_ISL_445706, EPI_ISL_445707, EPI_ISL_445708, EPI_ISL_445709, EPI_ISL_445710, EPI_ISL_445711, EPI_ISL_445712, EPI_ISL_445713, EPI_ISL_445714, EPI_ISL_445715, EPI_ISL_445716, EPI_ISL_445717, EPI_ISL_445718, EPI_ISL_445719, EPI_ISL_445720, EPI_ISL_445721, EPI_ISL_445722, EPI_ISL_445723, EPI_ISL_445724, EPI_ISL_445725, EPI_ISL_445726, EPI_ISL_445727, EPI_ISL_445728, EPI_ISL_445729, EPI_ISL_445730, EPI_ISL_445731, EPI_ISL_445732, EPI_ISL_445733, EPI_ISL_445734, EPI_ISL_445735, EPI_ISL_445736, EPI_ISL_445737, EPI_ISL_445738, EPI_ISL_445739, EPI_ISL_445740, EPI_ISL_445741, EPI_ISL_445742, EPI_ISL_445743, EPI_ISL_445744, EPI_ISL_445745, EPI_ISL_445746, EPI_ISL_445747, EPI_ISL_445749, EPI_ISL_445750, EPI_ISL_445751, EPI_ISL_445752, EPI_ISL_445753, EPI_ISL_445754, EPI_ISL_445755, EPI_ISL_445756, EPI_ISL_445757, EPI_ISL_445758, EPI_ISL_445759, EPI_ISL_445760, EPI_ISL_445761, EPI_ISL_445762, EPI_ISL_445763, EPI_ISL_445764, EPI_ISL_445765, EPI_ISL_445766, EPI_ISL_445767, EPI_ISL_445768, EPI_ISL_445769, EPI_ISL_445770, EPI_ISL_445771, EPI_ISL_445772, EPI_ISL_445773, EPI_ISL_445774, EPI_ISL_445775, EPI_ISL_445776, EPI_ISL_445777, EPI_ISL_445778, EPI_ISL_445779, EPI_ISL_445780, EPI_ISL_445781, EPI_ISL_445783, EPI_ISL_445784, EPI_ISL_445785, EPI_ISL_445786, EPI_ISL_445787, EPI_ISL_445788, EPI_ISL_445789, EPI_ISL_445790, EPI_ISL_445791, EPI_ISL_445792, EPI_ISL_445793, EPI_ISL_445794, EPI_ISL_445795, EPI_ISL_445796, EPI_ISL_445797, EPI_ISL_445798, EPI_ISL_445799, EPI_ISL_445800, EPI_ISL_445801, EPI_ISL_445802, EPI_ISL_445803, EPI_ISL_445804, EPI_ISL_445805, EPI_ISL_445806, EPI_ISL_445807, EPI_ISL_445808, EPI_ISL_445809, EPI_ISL_445810, EPI_ISL_445811, EPI_ISL_445812, EPI_ISL_445813, EPI_ISL_445814, EPI_ISL_445815, EPI_ISL_445816, EPI_ISL_445817, EPI_ISL_445818, EPI_ISL_445819, EPI_ISL_445820, EPI_ISL_445821, EPI_ISL_445822, EPI_ISL_445823, EPI_ISL_445824, EPI_ISL_445825, EPI_ISL_445826, EPI_ISL_445827, EPI_ISL_445828, EPI_ISL_445829, EPI_ISL_445830, EPI_ISL_445831, EPI_ISL_445832, EPI_ISL_445833, EPI_ISL_445834, EPI_ISL_445835, EPI_ISL_445836, EPI_ISL_445837, EPI_ISL_445838, EPI_ISL_445839, EPI_ISL_445840, EPI_ISL_445841, EPI_ISL_445842, EPI_ISL_445843, EPI_ISL_445844, EPI_ISL_445845, EPI_ISL_445846, EPI_ISL_445847, EPI_ISL_445848, EPI_ISL_445849, EPI_ISL_445850, EPI_ISL_445851, EPI_ISL_445852, EPI_ISL_445853, EPI_ISL_445854, EPI_ISL_445855, EPI_ISL_445856, EPI_ISL_445857, EPI_ISL_445858, EPI_ISL_445859, EPI_ISL_445860, EPI_ISL_445861, EPI_ISL_445862, EPI_ISL_445863, EPI_ISL_445864, EPI_ISL_445865, EPI_ISL_445866, EPI_ISL_445867, EPI_ISL_445868, EPI_ISL_445869, EPI_ISL_445870, EPI_ISL_445871, EPI_ISL_445872, EPI_ISL_445873, EPI_ISL_445874, EPI_ISL_445875, EPI_ISL_445876, EPI_ISL_445877, EPI_ISL_445878, EPI_ISL_445879, EPI_ISL_445881, EPI_ISL_445882, EPI_ISL_445883, EPI_ISL_445884, EPI_ISL_445885, EPI_ISL_445886, EPI_ISL_445887, EPI_ISL_445888, EPI_ISL_445889, EPI_ISL_445890, EPI_ISL_445891, EPI_ISL_445892, EPI_ISL_445893, EPI_ISL_445894, EPI_ISL_445895, EPI_ISL_445896, EPI_ISL_445897, EPI_ISL_445898, EPI_ISL_445899, EPI_ISL_445900, EPI_ISL_445901, EPI_ISL_445902, EPI_ISL_445903, EPI_ISL_445904, EPI_ISL_445905, EPI_ISL_445906, EPI_ISL_445907, EPI_ISL_445908, EPI_ISL_445909, EPI_ISL_445910, EPI_ISL_445911, EPI_ISL_445912, EPI_ISL_445913, EPI_ISL_445914, EPI_ISL_445915, EPI_ISL_445916, EPI_ISL_445917, EPI_ISL_445918, EPI_ISL_445919, EPI_ISL_445920, EPI_ISL_445921, EPI_ISL_445922, EPI_ISL_445923, EPI_ISL_445924, EPI_ISL_445925, EPI_ISL_445926, EPI_ISL_445927, EPI_ISL_445928, EPI_ISL_445929, EPI_ISL_445930, EPI_ISL_445931, EPI_ISL_445932, EPI_ISL_445933, EPI_ISL_445934, EPI_ISL_445935, EPI_ISL_445936, EPI_ISL_445937, EPI_ISL_445938, EPI_ISL_445939, EPI_ISL_445940, EPI_ISL_445941, EPI_ISL_445942, EPI_ISL_445943, EPI_ISL_445944, EPI_ISL_445945, EPI_ISL_445946, EPI_ISL_445947, EPI_ISL_445948, EPI_ISL_445949, EPI_ISL_445950, EPI_ISL_445951, EPI_ISL_445952, EPI_ISL_445953, EPI_ISL_445954, EPI_ISL_445955, EPI_ISL_445956, EPI_ISL_445957, EPI_ISL_445958, EPI_ISL_445959, EPI_ISL_445960, EPI_ISL_445961, EPI_ISL_445962, EPI_ISL_445963, EPI_ISL_445964, EPI_ISL_445965, EPI_ISL_445966, EPI_ISL_445967, EPI_ISL_445968, EPI_ISL_445969, EPI_ISL_445970, EPI_ISL_445971, EPI_ISL_445972, EPI_ISL_445973, EPI_ISL_445974, EPI_ISL_445975, EPI_ISL_445976, EPI_ISL_445977, EPI_ISL_445978, EPI_ISL_445979, EPI_ISL_445980, EPI_ISL_445981, EPI_ISL_445982, EPI_ISL_445983, EPI_ISL_445984, EPI_ISL_445985, EPI_ISL_445986, EPI_ISL_445987, EPI_ISL_445988, EPI_ISL_445989, EPI_ISL_445990, EPI_ISL_445991, EPI_ISL_445992, EPI_ISL_445993, EPI_ISL_445994, EPI_ISL_445995, EPI_ISL_445996, EPI_ISL_445997, EPI_ISL_445998, EPI_ISL_445999, EPI_ISL_446000, EPI_ISL_446001, EPI_ISL_446002, EPI_ISL_446003, EPI_ISL_446004, EPI_ISL_446005, EPI_ISL_446006, EPI_ISL_446007, EPI_ISL_446008, EPI_ISL_446009, EPI_ISL_446010, EPI_ISL_446011, EPI_ISL_446012, EPI_ISL_446013, EPI_ISL_446014, EPI_ISL_446015, EPI_ISL_446016, EPI_ISL_446017, EPI_ISL_446018, EPI_ISL_446019, EPI_ISL_446020, EPI_ISL_446022, EPI_ISL_446023, EPI_ISL_446024, EPI_ISL_446025, EPI_ISL_446026, EPI_ISL_446027, EPI_ISL_446028, EPI_ISL_446029, EPI_ISL_446030, EPI_ISL_446031, EPI_ISL_446032, EPI_ISL_446033, EPI_ISL_446034, EPI_ISL_446035, EPI_ISL_446036, EPI_ISL_446037, EPI_ISL_446038, EPI_ISL_446039, EPI_ISL_446040, EPI_ISL_446041, EPI_ISL_446042, EPI_ISL_446043, EPI_ISL_446044, EPI_ISL_446045, EPI_ISL_446046, EPI_ISL_446047, EPI_ISL_446048, EPI_ISL_446049, EPI_ISL_446050, EPI_ISL_446051, EPI_ISL_446052, EPI_ISL_446053, EPI_ISL_446054, EPI_ISL_446055, EPI_ISL_446056, EPI_ISL_446057, EPI_ISL_446058, EPI_ISL_446059, EPI_ISL_446060, EPI_ISL_446061, EPI_ISL_446062, EPI_ISL_446063, EPI_ISL_446064, EPI_ISL_446065, EPI_ISL_446066, EPI_ISL_446067, EPI_ISL_446068, EPI_ISL_446069, EPI_ISL_446070, EPI_ISL_446071, EPI_ISL_446072, EPI_ISL_446073, EPI_ISL_446074, EPI_ISL_446075, EPI_ISL_446076, EPI_ISL_446077, EPI_ISL_446078, EPI_ISL_446079, EPI_ISL_446080, EPI_ISL_446081, EPI_ISL_446082, EPI_ISL_446083, EPI_ISL_446084, EPI_ISL_446085, EPI_ISL_446086, EPI_ISL_446087, EPI_ISL_446088, EPI_ISL_446089, EPI_ISL_446090, EPI_ISL_446091, EPI_ISL_446092, EPI_ISL_446093, EPI_ISL_446094, EPI_ISL_446096, EPI_ISL_446097, EPI_ISL_446098, EPI_ISL_446099, EPI_ISL_446100, EPI_ISL_446101, EPI_ISL_446102, EPI_ISL_446103, EPI_ISL_446104, EPI_ISL_446105, EPI_ISL_446106, EPI_ISL_446107, EPI_ISL_446108, EPI_ISL_446109, EPI_ISL_446110, EPI_ISL_446111, EPI_ISL_446112, EPI_ISL_446113, EPI_ISL_446114, EPI_ISL_446115, EPI_ISL_446116, EPI_ISL_446117, EPI_ISL_446118, EPI_ISL_446119, EPI_ISL_446120, EPI_ISL_446121, EPI_ISL_446122, EPI_ISL_446123, EPI_ISL_446124, EPI_ISL_446125, EPI_ISL_446126, EPI_ISL_446127, EPI_ISL_446128, EPI_ISL_446129, EPI_ISL_446130, EPI_ISL_446131, EPI_ISL_446132, EPI_ISL_446133, EPI_ISL_446134, EPI_ISL_446135, EPI_ISL_446136, EPI_ISL_446137, EPI_ISL_446138, EPI_ISL_446139, EPI_ISL_446140, EPI_ISL_446141, EPI_ISL_446142, EPI_ISL_446143, EPI_ISL_446144, EPI_ISL_446145, EPI_ISL_446146, EPI_ISL_446147, EPI_ISL_446148, EPI_ISL_446149, EPI_ISL_446150, EPI_ISL_446151, EPI_ISL_446152, EPI_ISL_446153, EPI_ISL_446154, EPI_ISL_446155, EPI_ISL_446156, EPI_ISL_446157, EPI_ISL_446158, EPI_ISL_446159, EPI_ISL_446160, EPI_ISL_446161, EPI_ISL_446162, EPI_ISL_446163, EPI_ISL_446164, EPI_ISL_446165, EPI_ISL_446166, EPI_ISL_446167, EPI_ISL_446168, EPI_ISL_446169, EPI_ISL_446170, EPI_ISL_446171, EPI_ISL_446172, EPI_ISL_446173, EPI_ISL_446174, EPI_ISL_446175, EPI_ISL_446176, EPI_ISL_446177, EPI_ISL_446178, EPI_ISL_446179, EPI_ISL_446180, EPI_ISL_446181, EPI_ISL_446182, EPI_ISL_446183, EPI_ISL_446184, EPI_ISL_446185, EPI_ISL_446186, EPI_ISL_446187, EPI_ISL_446188, EPI_ISL_446189, EPI_ISL_446190, EPI_ISL_446191, EPI_ISL_446192, EPI_ISL_446193, EPI_ISL_446194, EPI_ISL_446195, EPI_ISL_446196, EPI_ISL_446197, EPI_ISL_446198, EPI_ISL_446199, EPI_ISL_446200, EPI_ISL_446201, EPI_ISL_446202, EPI_ISL_446203, EPI_ISL_446204, EPI_ISL_446205, EPI_ISL_446206, EPI_ISL_446207, EPI_ISL_446208, EPI_ISL_446209, EPI_ISL_446210, EPI_ISL_446211, EPI_ISL_446212, EPI_ISL_446213, EPI_ISL_446214, EPI_ISL_446215, EPI_ISL_446216, EPI_ISL_446217, EPI_ISL_446218, EPI_ISL_446219, EPI_ISL_446220, EPI_ISL_446221, EPI_ISL_446222, EPI_ISL_446223, EPI_ISL_446224, EPI_ISL_446225, EPI_ISL_446226, EPI_ISL_446227, EPI_ISL_446228, EPI_ISL_446229, EPI_ISL_446230, EPI_ISL_446231, EPI_ISL_446232, EPI_ISL_446233, EPI_ISL_446234, EPI_ISL_446235, EPI_ISL_446236, EPI_ISL_446237, EPI_ISL_446238, EPI_ISL_446239, EPI_ISL_446240, EPI_ISL_446241, EPI_ISL_446242, EPI_ISL_446243, EPI_ISL_446244, EPI_ISL_446245, EPI_ISL_446246, EPI_ISL_446247, EPI_ISL_446248, EPI_ISL_446249, EPI_ISL_446250, EPI_ISL_446251, EPI_ISL_446252, EPI_ISL_446253, EPI_ISL_446254, EPI_ISL_446255, EPI_ISL_446256, EPI_ISL_446257, EPI_ISL_446258, EPI_ISL_446259, EPI_ISL_446260, EPI_ISL_446261, EPI_ISL_446262, EPI_ISL_446263, EPI_ISL_446264, EPI_ISL_446265, EPI_ISL_446266, EPI_ISL_446267, EPI_ISL_446268, EPI_ISL_446269, EPI_ISL_446270, EPI_ISL_446271, EPI_ISL_446272, EPI_ISL_446273, EPI_ISL_446274, EPI_ISL_446275, EPI_ISL_446276, EPI_ISL_446277, EPI_ISL_446278, EPI_ISL_446279, EPI_ISL_446280, EPI_ISL_446281, EPI_ISL_446282, EPI_ISL_446283, EPI_ISL_446284, EPI_ISL_446285, EPI_ISL_446286, EPI_ISL_446287, EPI_ISL_446288, EPI_ISL_446289, EPI_ISL_446290, EPI_ISL_446291, EPI_ISL_446292, EPI_ISL_446293, EPI_ISL_446294, EPI_ISL_446295, EPI_ISL_446296, EPI_ISL_446297, EPI_ISL_446298, EPI_ISL_446299, EPI_ISL_446300, EPI_ISL_446301, EPI_ISL_446302, EPI_ISL_446303, EPI_ISL_446304, EPI_ISL_446305, EPI_ISL_446306, EPI_ISL_446307, EPI_ISL_446308, EPI_ISL_446309, EPI_ISL_446310, EPI_ISL_446311, EPI_ISL_446312, EPI_ISL_446313, EPI_ISL_446314, EPI_ISL_446315, EPI_ISL_446316, EPI_ISL_446317, EPI_ISL_446318, EPI_ISL_446319, EPI_ISL_446320, EPI_ISL_446321, EPI_ISL_446322, EPI_ISL_446323, EPI_ISL_446324, EPI_ISL_446325, EPI_ISL_446326, EPI_ISL_446327, EPI_ISL_446328, EPI_ISL_446329, EPI_ISL_446330, EPI_ISL_446331, EPI_ISL_446332, EPI_ISL_446333, EPI_ISL_446334, EPI_ISL_446335, EPI_ISL_446336, EPI_ISL_446337, EPI_ISL_446338, EPI_ISL_446339, EPI_ISL_446340, EPI_ISL_446341, EPI_ISL_446342, EPI_ISL_446343, EPI_ISL_446344, EPI_ISL_446345, EPI_ISL_446346, EPI_ISL_446347, EPI_ISL_446348, EPI_ISL_446349, EPI_ISL_446350, EPI_ISL_446351, EPI_ISL_446352, EPI_ISL_446353, EPI_ISL_446354, EPI_ISL_446355, EPI_ISL_446356, EPI_ISL_446357, EPI_ISL_446359, EPI_ISL_446360, EPI_ISL_446361, EPI_ISL_446362, EPI_ISL_446363, EPI_ISL_446364, EPI_ISL_446365, EPI_ISL_446366, EPI_ISL_446367, EPI_ISL_446368, EPI_ISL_446369, EPI_ISL_446370, EPI_ISL_446371, EPI_ISL_446372, EPI_ISL_446373, EPI_ISL_446374, EPI_ISL_446375, EPI_ISL_446376, EPI_ISL_446377, EPI_ISL_446378, EPI_ISL_446379, EPI_ISL_446380, EPI_ISL_446381, EPI_ISL_446382, EPI_ISL_446383, EPI_ISL_446384, EPI_ISL_446385, EPI_ISL_446386, EPI_ISL_446387, EPI_ISL_446388, EPI_ISL_446389, EPI_ISL_446390, EPI_ISL_446391, EPI_ISL_446392, EPI_ISL_446393, EPI_ISL_446394, EPI_ISL_446395, EPI_ISL_446396, EPI_ISL_446397, EPI_ISL_446398, EPI_ISL_446399, EPI_ISL_446400, EPI_ISL_446401, EPI_ISL_446402, EPI_ISL_446403, EPI_ISL_446404, EPI_ISL_446405, EPI_ISL_446406, EPI_ISL_446407, EPI_ISL_446408, EPI_ISL_446409, EPI_ISL_446410, EPI_ISL_446411, EPI_ISL_446412, EPI_ISL_446413, EPI_ISL_446414, EPI_ISL_446415, EPI_ISL_446416, EPI_ISL_446417, EPI_ISL_446418, EPI_ISL_446419, EPI_ISL_446420, EPI_ISL_446421, EPI_ISL_446422, EPI_ISL_446423, EPI_ISL_446424, EPI_ISL_446425, EPI_ISL_446426, EPI_ISL_446427, EPI_ISL_446428, EPI_ISL_446429, EPI_ISL_446430, EPI_ISL_446431, EPI_ISL_446432, EPI_ISL_446433, EPI_ISL_446434, EPI_ISL_446435, EPI_ISL_446436, EPI_ISL_446437, EPI_ISL_446438, EPI_ISL_446439, EPI_ISL_446440, EPI_ISL_446441, EPI_ISL_446442, EPI_ISL_446443, EPI_ISL_446444, EPI_ISL_446445, EPI_ISL_446446, EPI_ISL_446447, EPI_ISL_446448, EPI_ISL_446449, EPI_ISL_446450, EPI_ISL_446451, EPI_ISL_446452, EPI_ISL_446453, EPI_ISL_446454, EPI_ISL_446455, EPI_ISL_446456, EPI_ISL_446457, EPI_ISL_446458, EPI_ISL_446459, EPI_ISL_446460, EPI_ISL_446461, EPI_ISL_446462, EPI_ISL_446463, EPI_ISL_446464, EPI_ISL_446465, EPI_ISL_446466, EPI_ISL_446467, | | | |

| | | | |
|---|---|---|---|
| EPI_ISL_446468, EPI_ISL_446469, EPI_ISL_446470, EPI_ISL_446471, EPI_ISL_446472, EPI_ISL_446473, EPI_ISL_446474, EPI_ISL_446475, EPI_ISL_446476, EPI_ISL_446477, EPI_ISL_446478, EPI_ISL_446479, EPI_ISL_446480, EPI_ISL_446481, EPI_ISL_446482, EPI_ISL_446483, EPI_ISL_446484, EPI_ISL_446485, EPI_ISL_446486, EPI_ISL_446487, EPI_ISL_446488, EPI_ISL_446489, EPI_ISL_446490, EPI_ISL_446491, EPI_ISL_446492, EPI_ISL_446493, EPI_ISL_446494, EPI_ISL_446495, EPI_ISL_446496, EPI_ISL_446497, EPI_ISL_446498, EPI_ISL_446499, EPI_ISL_446500, EPI_ISL_446501, EPI_ISL_446502, EPI_ISL_446503, EPI_ISL_446504, EPI_ISL_446505, EPI_ISL_446506, EPI_ISL_446507, EPI_ISL_446508, EPI_ISL_446509, EPI_ISL_446510, EPI_ISL_446511, EPI_ISL_446512, EPI_ISL_446513, EPI_ISL_446514, EPI_ISL_446515, EPI_ISL_446516, EPI_ISL_446517, EPI_ISL_446518, EPI_ISL_446519, EPI_ISL_446520, EPI_ISL_446521, EPI_ISL_446522, EPI_ISL_446523, EPI_ISL_446524, EPI_ISL_446525, EPI_ISL_446526, EPI_ISL_446527, EPI_ISL_446528, EPI_ISL_446529, EPI_ISL_446530, EPI_ISL_446531, EPI_ISL_446532, EPI_ISL_446533, EPI_ISL_446534, EPI_ISL_446535, EPI_ISL_446536, EPI_ISL_446537, EPI_ISL_446538, EPI_ISL_446539, EPI_ISL_446540, EPI_ISL_446541, EPI_ISL_446542, EPI_ISL_446543, EPI_ISL_446544, EPI_ISL_446545, EPI_ISL_446546, EPI_ISL_446547, EPI_ISL_446548, EPI_ISL_446549, EPI_ISL_446550, EPI_ISL_446551, EPI_ISL_446552, EPI_ISL_446553, EPI_ISL_446554, EPI_ISL_446555, EPI_ISL_446556, EPI_ISL_446557, EPI_ISL_446558, EPI_ISL_446559, EPI_ISL_446560, EPI_ISL_446561, EPI_ISL_446562, EPI_ISL_446563, EPI_ISL_446564, EPI_ISL_446565, EPI_ISL_446566, EPI_ISL_446567, EPI_ISL_446568, EPI_ISL_446569, EPI_ISL_446570, EPI_ISL_446571, EPI_ISL_446572, EPI_ISL_446573, EPI_ISL_446574, EPI_ISL_446575, EPI_ISL_446576, EPI_ISL_446577, EPI_ISL_446578, EPI_ISL_446579, EPI_ISL_446580, EPI_ISL_446581, EPI_ISL_446582, EPI_ISL_446583, EPI_ISL_446584, EPI_ISL_446585, EPI_ISL_446586, EPI_ISL_446587, EPI_ISL_446588, EPI_ISL_446589, EPI_ISL_446590, EPI_ISL_446591, EPI_ISL_446592, EPI_ISL_446593, EPI_ISL_446594, EPI_ISL_446595, EPI_ISL_446596, EPI_ISL_446597, EPI_ISL_446598, EPI_ISL_446599, EPI_ISL_446600, EPI_ISL_446601, EPI_ISL_446602, EPI_ISL_446603, EPI_ISL_446604, EPI_ISL_446605, EPI_ISL_446606, EPI_ISL_446607, EPI_ISL_446608, EPI_ISL_446609, EPI_ISL_446610, EPI_ISL_446611, EPI_ISL_446612, EPI_ISL_446613, EPI_ISL_446614, EPI_ISL_446615, EPI_ISL_446616, EPI_ISL_446617, EPI_ISL_446618, EPI_ISL_446619, EPI_ISL_446620, EPI_ISL_446621, EPI_ISL_446622, EPI_ISL_446623, EPI_ISL_446624, EPI_ISL_446625, EPI_ISL_446626, EPI_ISL_446627, EPI_ISL_446628, EPI_ISL_446629, EPI_ISL_446630, EPI_ISL_446631, EPI_ISL_446632, EPI_ISL_446633, EPI_ISL_446634, EPI_ISL_446635, EPI_ISL_446636, EPI_ISL_446637, EPI_ISL_446638, EPI_ISL_446639, EPI_ISL_446640, EPI_ISL_446641, EPI_ISL_446642, EPI_ISL_446643, EPI_ISL_446644, EPI_ISL_446645, EPI_ISL_446646, EPI_ISL_446647, EPI_ISL_446648, EPI_ISL_446649, EPI_ISL_446650, EPI_ISL_446651, EPI_ISL_446652, EPI_ISL_446653, EPI_ISL_446654, EPI_ISL_446655, EPI_ISL_446656, EPI_ISL_446657, EPI_ISL_446658, EPI_ISL_446659, EPI_ISL_446660, EPI_ISL_446661, EPI_ISL_446662, EPI_ISL_446663, EPI_ISL_446664, EPI_ISL_446665, EPI_ISL_446666, EPI_ISL_446667, EPI_ISL_446668, EPI_ISL_446669, EPI_ISL_446670, EPI_ISL_446671, EPI_ISL_446672, EPI_ISL_446673, EPI_ISL_446674, EPI_ISL_446675, EPI_ISL_446676, EPI_ISL_446677, EPI_ISL_446678, EPI_ISL_446679, EPI_ISL_446680, EPI_ISL_446681, EPI_ISL_446682, EPI_ISL_446683, EPI_ISL_446684, EPI_ISL_446685, EPI_ISL_446686, EPI_ISL_446687, EPI_ISL_446688, EPI_ISL_446689, EPI_ISL_446690, EPI_ISL_446691, EPI_ISL_446692, EPI_ISL_446693, EPI_ISL_446694, EPI_ISL_446695, EPI_ISL_446696, EPI_ISL_446697, EPI_ISL_446698, EPI_ISL_446699, EPI_ISL_446700, EPI_ISL_446701, EPI_ISL_446702, EPI_ISL_446703, EPI_ISL_446704, EPI_ISL_446705, EPI_ISL_446706, EPI_ISL_446707, EPI_ISL_446708, EPI_ISL_446709, EPI_ISL_446710, EPI_ISL_446711, EPI_ISL_446712, EPI_ISL_446713, EPI_ISL_446714, EPI_ISL_446715, EPI_ISL_446716, EPI_ISL_446717, EPI_ISL_446718, EPI_ISL_446719, EPI_ISL_446720, EPI_ISL_446721, EPI_ISL_446722, EPI_ISL_446723, EPI_ISL_446724, EPI_ISL_446725, EPI_ISL_446726, EPI_ISL_446727, EPI_ISL_446728, EPI_ISL_446729, EPI_ISL_446730, EPI_ISL_446731, EPI_ISL_446732, EPI_ISL_446733, EPI_ISL_446734, EPI_ISL_446735, EPI_ISL_446736, EPI_ISL_446737, EPI_ISL_446738, EPI_ISL_446739, EPI_ISL_446741, EPI_ISL_446742, EPI_ISL_446743, EPI_ISL_446744, EPI_ISL_446745, EPI_ISL_446746, EPI_ISL_446747, EPI_ISL_446748, EPI_ISL_446749, EPI_ISL_446750, EPI_ISL_446751, EPI_ISL_446752, EPI_ISL_446753, EPI_ISL_446754, EPI_ISL_446755, EPI_ISL_446756, EPI_ISL_446757, EPI_ISL_446758, EPI_ISL_446759, EPI_ISL_446760, EPI_ISL_446761, EPI_ISL_446762, EPI_ISL_446763, EPI_ISL_446764, EPI_ISL_446765, EPI_ISL_446766, EPI_ISL_446767, EPI_ISL_446768, EPI_ISL_446769, EPI_ISL_446770, EPI_ISL_446771, EPI_ISL_446772, EPI_ISL_446773, EPI_ISL_446774, EPI_ISL_446775, EPI_ISL_446776, EPI_ISL_446777, EPI_ISL_446778, EPI_ISL_446779, EPI_ISL_446780, EPI_ISL_446781, EPI_ISL_446782, EPI_ISL_446783, EPI_ISL_446784, EPI_ISL_446785, EPI_ISL_446786, EPI_ISL_446787, EPI_ISL_446788, EPI_ISL_446789, EPI_ISL_446790, EPI_ISL_446791, EPI_ISL_446792, EPI_ISL_446793, EPI_ISL_446794, EPI_ISL_446795, EPI_ISL_446796, EPI_ISL_446797, EPI_ISL_446798, EPI_ISL_446799, EPI_ISL_446800, EPI_ISL_446801, EPI_ISL_446802, EPI_ISL_446803, EPI_ISL_446804, EPI_ISL_446805, EPI_ISL_446806, EPI_ISL_446807, EPI_ISL_446808, EPI_ISL_446809, EPI_ISL_446810, EPI_ISL_446811, EPI_ISL_446812, EPI_ISL_446813, EPI_ISL_446814, EPI_ISL_446815, EPI_ISL_446816, EPI_ISL_446817, EPI_ISL_446818, EPI_ISL_446819, EPI_ISL_446820, EPI_ISL_446821, EPI_ISL_446822, EPI_ISL_446823, EPI_ISL_446824, EPI_ISL_446825, EPI_ISL_446826, EPI_ISL_446827, EPI_ISL_446828, EPI_ISL_446829, EPI_ISL_446830, EPI_ISL_446831, EPI_ISL_446832, EPI_ISL_446833, EPI_ISL_446834, EPI_ISL_446835, EPI_ISL_446836, EPI_ISL_446837, EPI_ISL_446838, EPI_ISL_446839, EPI_ISL_446840, EPI_ISL_446841, EPI_ISL_446842, EPI_ISL_446843, EPI_ISL_446844, EPI_ISL_446845, EPI_ISL_446846, EPI_ISL_446847, EPI_ISL_446848, EPI_ISL_446849, EPI_ISL_446850, EPI_ISL_446851, EPI_ISL_446852, EPI_ISL_446853, EPI_ISL_446854, EPI_ISL_446855, EPI_ISL_446856, EPI_ISL_446857, EPI_ISL_446858, EPI_ISL_446859, EPI_ISL_446860, EPI_ISL_446861, EPI_ISL_446862, EPI_ISL_446863, EPI_ISL_446864, EPI_ISL_446865, EPI_ISL_446866, EPI_ISL_446867, EPI_ISL_446868, EPI_ISL_446869, EPI_ISL_446870, EPI_ISL_446871, EPI_ISL_446872, EPI_ISL_446873, EPI_ISL_446874, EPI_ISL_446875, EPI_ISL_446876, EPI_ISL_446877, EPI_ISL_446878, EPI_ISL_446879, EPI_ISL_446880, EPI_ISL_446881, EPI_ISL_446882, EPI_ISL_446883, EPI_ISL_446884, EPI_ISL_446885, EPI_ISL_446886, EPI_ISL_446887, EPI_ISL_446888, EPI_ISL_446889, EPI_ISL_446890, EPI_ISL_446891, EPI_ISL_446892, EPI_ISL_446893, EPI_ISL_446894, EPI_ISL_446895, EPI_ISL_446896, EPI_ISL_446897, EPI_ISL_446898, EPI_ISL_446899, EPI_ISL_446900, EPI_ISL_446901, EPI_ISL_446902, EPI_ISL_446903, EPI_ISL_446904, EPI_ISL_446905, EPI_ISL_446906, EPI_ISL_446907, EPI_ISL_446908, EPI_ISL_446909, EPI_ISL_446910, EPI_ISL_446911, EPI_ISL_446912, EPI_ISL_446913, EPI_ISL_446914, EPI_ISL_446915, EPI_ISL_446916, EPI_ISL_446917, EPI_ISL_446918, EPI_ISL_446919, EPI_ISL_446920, EPI_ISL_446921, EPI_ISL_446922, EPI_ISL_446923, EPI_ISL_446924, EPI_ISL_446925, EPI_ISL_446926, EPI_ISL_446927, EPI_ISL_446928, EPI_ISL_446929, EPI_ISL_446930, EPI_ISL_446931, EPI_ISL_446932, EPI_ISL_446933, EPI_ISL_446934, EPI_ISL_446935, EPI_ISL_446936, EPI_ISL_446937, EPI_ISL_446938, EPI_ISL_446939, EPI_ISL_446940, EPI_ISL_446941, EPI_ISL_446942, EPI_ISL_446943, EPI_ISL_446944, EPI_ISL_446945, EPI_ISL_446946, EPI_ISL_446947, EPI_ISL_446948, EPI_ISL_446949, EPI_ISL_446950, EPI_ISL_446951, EPI_ISL_446952, EPI_ISL_446953, EPI_ISL_446954, EPI_ISL_446955, EPI_ISL_446956, EPI_ISL_446957, EPI_ISL_446958, EPI_ISL_446959, EPI_ISL_446960, EPI_ISL_446961, EPI_ISL_446962, EPI_ISL_446963, EPI_ISL_446964, EPI_ISL_446965, EPI_ISL_446966, EPI_ISL_446967, EPI_ISL_446968, EPI_ISL_446969, EPI_ISL_446970, EPI_ISL_446971, EPI_ISL_446972, EPI_ISL_446973, EPI_ISL_446974, EPI_ISL_446975, EPI_ISL_446976, EPI_ISL_446977, EPI_ISL_446978, EPI_ISL_446979, EPI_ISL_446980, EPI_ISL_446981, EPI_ISL_446982, EPI_ISL_446983, EPI_ISL_446984, EPI_ISL_446985, EPI_ISL_446986, EPI_ISL_446987, EPI_ISL_446988, EPI_ISL_446989, EPI_ISL_446990, EPI_ISL_446991, EPI_ISL_446992, EPI_ISL_446993, EPI_ISL_446994, EPI_ISL_446995 | | | |
| see above | Wales Specialist Virology Centre | Public Health Wales Microbiology Cardiff | Catherine Moore, Johnathan Evans, Laura Gifford, Malorie Perry, Simon Cottrell, Alec Birchley, Alexander Adams, Amy Gaskin, Bree Gatica-Wilcox, Jason Coombes, Lauren Gilbert, Lee Graham, Nicole Pacchiarini, Sara Kumziene-Summerhayes, Sarah Taylor, Sophie Jones, Sara Rey, Matthew Bull, Joanne Watkins, Sally Corden, Tom Connor |
| EPI_ISL_447252, EPI_ISL_447253, EPI_ISL_447254, EPI_ISL_447255, EPI_ISL_447256, EPI_ISL_447257, EPI_ISL_447591, EPI_ISL_447592, EPI_ISL_447593 | TSGH-CP molecular lab | TSGH-CP molecular lab | Cherng-Lih Perng, Ming-Jr JIAN, Chih-Kai Chang, Jung-Chung Lin, Kuo-Ming Yeh, Chien-Wen Chen, Sheng-Kang Chiu, Hsing-Yi Chung, Shih-Hung Tsai, Kuo-Sheng Hung, Tien-Yao Chang, Feng-Yee Chang, Hung-Sheng Shang |
| EPI_ISL_451298, EPI_ISL_451300, EPI_ISL_451302, EPI_ISL_451304 | Laboratory of Virology, INMI Lazzaro Spallanzani IRCCS | Laboratory of Virology, INMI Lazzaro Spallanzani IRCCS | Cesare E.M. Gruber, Martina Rueca, Barbara Bartolini, Francesco Messina, Antonino Di Caro, Maria R. Capobianchi, Giuseppe Ippolito |
| EPI_ISL_451694, EPI_ISL_451717, EPI_ISL_451759, EPI_ISL_451763, EPI_ISL_451788, EPI_ISL_451791, EPI_ISL_451832, EPI_ISL_451839 | Viollier AG | Department of Biosystems Science and Engineering, ETH Zürich | Christian Beisel, Sarah Nadeau, Ivan Topolsky, Pedro Ferreira, Philipp Jablonski, Susana Posada-Céspedes, Tobias Schär, Ina Nissen, Natascha Santacroce, Elodie Burcklen, Christiane Beckmann, Maurice Redondo, Olivier Kobel, Christoph Noppen, Sophie Seidel, Noemie Santamaria de Souza, Niko Beerenwinkel, Tanja Stadler |
| EPI_ISL_457726, EPI_ISL_457730, EPI_ISL_457733, EPI_ISL_458029 | TSGH-CP molecular lab | TSGH-CP molecular lab | Cherng-Lih Perng, Ming-Jr JIAN, Chih-Kai Chang, Jung-Chung Lin, Kuo-Ming Yeh, Chien-Wen Chen, Sheng-Kang Chiu, Hsing-Yi Chung, Shih-Hung Tsai, Kuo-Sheng Hung, Tien-Yao Chang, Feng-Yee Chang, Hung-Sheng Shang |
| EPI_ISL_458578, EPI_ISL_458579, EPI_ISL_458580, EPI_ISL_458581, EPI_ISL_458582, EPI_ISL_458583, EPI_ISL_458584, EPI_ISL_458585, EPI_ISL_458586, EPI_ISL_458587, EPI_ISL_458588, EPI_ISL_458589, EPI_ISL_458590, EPI_ISL_458591, EPI_ISL_458592, EPI_ISL_458593, EPI_ISL_458594, EPI_ISL_458595, EPI_ISL_458596, EPI_ISL_458597, EPI_ISL_458598, EPI_ISL_458599, EPI_ISL_458600, EPI_ISL_458601, EPI_ISL_458602, EPI_ISL_458603, EPI_ISL_458604, EPI_ISL_458605, EPI_ISL_458606, EPI_ISL_458607, EPI_ISL_458608, EPI_ISL_458609, EPI_ISL_458610, EPI_ISL_458611, EPI_ISL_458612, EPI_ISL_458613, EPI_ISL_458614, EPI_ISL_458615, EPI_ISL_458616, EPI_ISL_458617, EPI_ISL_458618, EPI_ISL_458619, EPI_ISL_458620, EPI_ISL_458621, EPI_ISL_458622, EPI_ISL_458623, EPI_ISL_458624, EPI_ISL_458625, EPI_ISL_458626, EPI_ISL_458627, EPI_ISL_458628, EPI_ISL_458629, EPI_ISL_458630, EPI_ISL_458631, EPI_ISL_458632, EPI_ISL_458633, EPI_ISL_458634, EPI_ISL_458635, EPI_ISL_458636, EPI_ISL_458637, EPI_ISL_458638, EPI_ISL_458639, EPI_ISL_458640, EPI_ISL_458641, EPI_ISL_458642, EPI_ISL_458643, EPI_ISL_458644, EPI_ISL_458645, EPI_ISL_458646, EPI_ISL_458647, EPI_ISL_458648, EPI_ISL_458649, EPI_ISL_458650, EPI_ISL_458651, EPI_ISL_458652, EPI_ISL_458653, EPI_ISL_458654, EPI_ISL_458655, EPI_ISL_458656, EPI_ISL_458657, EPI_ISL_458658, EPI_ISL_458659, EPI_ISL_458660, EPI_ISL_458661, EPI_ISL_458662, EPI_ISL_458663, EPI_ISL_458664, EPI_ISL_458665, EPI_ISL_458666, EPI_ISL_458667, EPI_ISL_458668, EPI_ISL_458669, EPI_ISL_458670, EPI_ISL_458671, EPI_ISL_458672, EPI_ISL_458673, EPI_ISL_458674, EPI_ISL_458675, EPI_ISL_458676, EPI_ISL_458677, EPI_ISL_458678, EPI_ISL_458679, EPI_ISL_458680, EPI_ISL_458681, EPI_ISL_458682, EPI_ISL_458683, EPI_ISL_458684, EPI_ISL_458685, EPI_ISL_458686, EPI_ISL_458687, EPI_ISL_458688, EPI_ISL_458689, EPI_ISL_458690, EPI_ISL_458691, EPI_ISL_458692, EPI_ISL_458693, EPI_ISL_458694, EPI_ISL_458695, EPI_ISL_458696, EPI_ISL_458697, EPI_ISL_458698, EPI_ISL_458699, EPI_ISL_458700, EPI_ISL_458701, EPI_ISL_458702, EPI_ISL_458703, EPI_ISL_458704, EPI_ISL_458705, EPI_ISL_458706, EPI_ISL_458707, EPI_ISL_458708, EPI_ISL_458709, EPI_ISL_458710, EPI_ISL_458711 | | | |
| see above | NU-OMICS DNA Sequencing research facility, Northumbria University | Wellcome Sanger Institute for the COVID-19 Genomics UK (COG-UK) consortium | Chris Duncan, Sheia Waugh, Shirelle Burton-Fanning, Gary Eltringham, Jennifer Collins, Brendan Payne, Yusri Taha, Emma Swindells, Jane Greenaway, Edward Barton, Garren Scott, Debra Padgett, Clive Graham, Sarah Essex, Steve Ligget, Paul Baker, Lynn Dover, Wen Yew, Gary Black, John Allan, Joshua Loh, Greg Young, Matthew Bashton, Andrew Nelson, Darren Smith and Alex Alderton, Roberto Amato, Sonia Goncalves, Ewan Harrison, David K. Jackson, Ian Johnston, Dominic Kwiatkowski, Cordelia Langford, John Sillitoe on behalf of the Wellcome Sanger Institute COVID-19 Surveillance Team (http://www.sanger.ac.uk/covid-team) |
| EPI_ISL_458712 | NU-OMICS DNA Sequencing research facility, Northumbria University | Wellcome Sanger Institute for the COVID-19 Genomics UK (COG-UK) Consortium | Chris Duncan, Sheia Waugh, Shirelle Burton-Fanning, Gary Eltringham, Jennifer Collins, Brendan Payne, Yusri Taha, Emma Swindells, Jane Greenaway, Edward Barton, Garren Scott, Debra Padgett, Clive Graham, Sarah Essex, Steve Ligget, Paul Baker, Lynn Dover, Wen Yew, Gary Black, John Allan, Joshua Loh, Greg Young, Matthew Bashton, Andrew Nelson, Darren Smith and Alex Alderton, Roberto Amato, Sonia Goncalves, Ewan Harrison, David K. Jackson, Ian Johnston, Dominic Kwiatkowski, Cordelia Langford, John Sillitoe on behalf of the Wellcome Sanger Institute COVID-19 Surveillance Team |
| EPI_ISL_458713, EPI_ISL_458714, EPI_ISL_458715, EPI_ISL_458716, EPI_ISL_458717, EPI_ISL_458718 | NU-OMICS DNA Sequencing research facility, Northumbria University | Wellcome Sanger Institute for the COVID-19 Genomics UK (COG-UK) consortium | Chris Duncan, Sheia Waugh, Shirelle Burton-Fanning, Gary Eltringham, Jennifer Collins, Brendan Payne, Yusri Taha, Emma Swindells, Jane Greenaway, Edward Barton, Garren Scott, Debra Padgett, Clive Graham, Sarah Essex, Steve Ligget, Paul Baker, Lynn Dover, Wen Yew, Gary Black, John Allan, Joshua Loh, Greg Young, Matthew Bashton, Andrew Nelson, Darren Smith and Alex Alderton, Roberto Amato, Sonia Goncalves, Ewan Harrison, David K. Jackson, Ian Johnston, Dominic Kwiatkowski, Cordelia Langford, John Sillitoe on behalf of the Wellcome Sanger Institute COVID-19 Surveillance Team (http://www.sanger.ac.uk/covid-team) |
| EPI_ISL_460085 | Molecular Virology Unit, Fondazione IRCCS Policlinico San Matteo , Pavia | Laboratory of Virology, INMI Lazzaro Spallanzani IRCCS | Cesare E.M. Gruber, Maria R. Capobianchi, Barbara Bartolini, Fausto Baldanti, Martina Rueca, Antonio Piralla, Antonino Di Caro |
| EPI_ISL_460087 | Molecular Virology Unit, Fondazione IRCCS Policlinico San Matteo , Pavia | Laboratory of Virology, INMI Lazzaro Spallanzani IRCCS | Cesare E.M. Gruber, Maria R. Capobianchi, Martina Rueca, Barbara Bartolini, Antonino Di Caro, Antonio Piralla, Fausto Baldanti |
| EPI_ISL_460092 | Molecular Virology Unit, Fondazione IRCCS Policlinico San Matteo , Pavia | Laboratory of Virology, INMI Lazzaro Spallanzani IRCCS | Cesare E.M. Gruber, Martina Rueca, Maria R. Capobianchi, Antonino Di Caro, Antonio Piralla, Barbara Bartolini, Fausto Baldanti |
| EPI_ISL_468247, EPI_ISL_468257 | Viollier AG | Department of Biosystems Science and Engineering, ETH Zürich | Christian Beisel, Sarah Nadeau, Ivan Topolsky, Pedro Ferreira, Philipp Jablonski, Susana Posada-Céspedes, Tobias Schär, Ina Nissen, Natascha Santacroce, Elodie Burcklen, Christiane Beckmann, Maurice Redondo, Olivier Kobel, Christoph Noppen, Sophie Seidel, Noemie Santamaria de Souza, Niko Beerenwinkel, Tanja Stadler |
| EPI_ISL_469302, EPI_ISL_469303, EPI_ISL_469304, EPI_ISL_469305, EPI_ISL_469306, EPI_ISL_469307, EPI_ISL_469308, EPI_ISL_469309, EPI_ISL_469310, EPI_ISL_469311, EPI_ISL_469312, EPI_ISL_469313, EPI_ISL_469314, EPI_ISL_469315, EPI_ISL_469316, EPI_ISL_469317, EPI_ISL_469318, EPI_ISL_469319, EPI_ISL_469320, EPI_ISL_469321, EPI_ISL_469322, EPI_ISL_469323, EPI_ISL_469324, EPI_ISL_469325, EPI_ISL_469326, EPI_ISL_469327, EPI_ISL_469328, EPI_ISL_469329, EPI_ISL_469330, EPI_ISL_469331, EPI_ISL_469332, EPI_ISL_469333, EPI_ISL_469334, EPI_ISL_469335, EPI_ISL_469336, EPI_ISL_469337, EPI_ISL_469338, EPI_ISL_469339, EPI_ISL_469340, EPI_ISL_469341, EPI_ISL_469342, EPI_ISL_469343, EPI_ISL_469344, EPI_ISL_469805, EPI_ISL_469915 | | | |

| | | | |
|---|---|---|---|
| see above | NU-OMICS DNA Sequencing research facility, Northumbria University | Wellcome Sanger Institute for the COVID-19 Genomics UK (COG-UK) consortium | Chris Duncan, Sheia Waugh, Shirelle Burton-Fanning, Gary Eltringham, Jennifer Collins, Brendan Payne, Yusri Taha, Emma Swindells, Jane Greenaway, Edward Barton, Garren Scott, Debra Padgett, Clive Graham, Sarah Essex, Steve Ligget, Paul Baker, Lynn Dover, Wen Yew, Gary Black, John Allan, Joshua Loh, Greg Young, Matthew Bashton, Andrew Nelson, Darren Smith and Alex Alderton, Roberto Amato, Sonia Goncalves, Ewan Harrison, David K. Jackson, Ian Johnston, Dominic Kwiatkowski, Cordelia Langford, John Sillitoe on behalf of the Wellcome Sanger Institute COVID-19 Surveillance Team (http://www.sanger.ac.uk/covid-team) |

EPI_ISL_470830, EPI_ISL_470831, EPI_ISL_470832, EPI_ISL_470833, EPI_ISL_470834, EPI_ISL_470835, EPI_ISL_470836, EPI_ISL_470839, EPI_ISL_470840, EPI_ISL_470841, EPI_ISL_470842, EPI_ISL_470843, EPI_ISL_470844, EPI_ISL_470845, EPI_ISL_470846, EPI_ISL_470847, EPI_ISL_470848, EPI_ISL_470849, EPI_ISL_470850, EPI_ISL_470851, EPI_ISL_470852, EPI_ISL_470853, EPI_ISL_470854, EPI_ISL_470855, EPI_ISL_470856, EPI_ISL_470857, EPI_ISL_470858, EPI_ISL_470859, EPI_ISL_470860, EPI_ISL_470861, EPI_ISL_470862, EPI_ISL_470863, EPI_ISL_470864, EPI_ISL_470865, EPI_ISL_470866, EPI_ISL_470867, EPI_ISL_470868, EPI_ISL_470869, EPI_ISL_470870, EPI_ISL_470871, EPI_ISL_470872, EPI_ISL_470873, EPI_ISL_470874, EPI_ISL_470875

| | | | |
|---|---|---|---|
| see above | PathWest Laboratory Medicine WA | PathWest Laboratory Medicine WA | Chisha Sikazwe, Jurissa Lang, Avram Levy, David Smith and David Speers |

EPI_ISL_472432, EPI_ISL_472433, EPI_ISL_472434, EPI_ISL_472435, EPI_ISL_472436, EPI_ISL_472437, EPI_ISL_472438, EPI_ISL_472439, EPI_ISL_472440, EPI_ISL_472441, EPI_ISL_472442, EPI_ISL_472443, EPI_ISL_472444, EPI_ISL_472445, EPI_ISL_472446, EPI_ISL_472447, EPI_ISL_472448, EPI_ISL_472449, EPI_ISL_472450, EPI_ISL_472451, EPI_ISL_472452, EPI_ISL_472453, EPI_ISL_472454, EPI_ISL_472455, EPI_ISL_472456, EPI_ISL_472457, EPI_ISL_472458, EPI_ISL_472459, EPI_ISL_472460, EPI_ISL_472461, EPI_ISL_472462, EPI_ISL_472463, EPI_ISL_472464, EPI_ISL_472465, EPI_ISL_472466, EPI_ISL_472467, EPI_ISL_472468, EPI_ISL_472469, EPI_ISL_472470, EPI_ISL_472471, EPI_ISL_472472, EPI_ISL_472473, EPI_ISL_472474, EPI_ISL_472475, EPI_ISL_472476, EPI_ISL_472477, EPI_ISL_472478, EPI_ISL_472479, EPI_ISL_472480, EPI_ISL_472481, EPI_ISL_472482, EPI_ISL_472483, EPI_ISL_472484, EPI_ISL_472485, EPI_ISL_472486, EPI_ISL_472487, EPI_ISL_472488, EPI_ISL_472489, EPI_ISL_472490, EPI_ISL_472491, EPI_ISL_472492, EPI_ISL_472493, EPI_ISL_472494, EPI_ISL_472495, EPI_ISL_472496, EPI_ISL_472497, EPI_ISL_472498, EPI_ISL_472499, EPI_ISL_472500, EPI_ISL_472501, EPI_ISL_472502, EPI_ISL_472503, EPI_ISL_472504, EPI_ISL_472505, EPI_ISL_472506, EPI_ISL_472507, EPI_ISL_472508, EPI_ISL_472509, EPI_ISL_472510, EPI_ISL_472511, EPI_ISL_472512, EPI_ISL_472513, EPI_ISL_472514, EPI_ISL_472515, EPI_ISL_472516, EPI_ISL_472517, EPI_ISL_472518, EPI_ISL_472519, EPI_ISL_472520, EPI_ISL_472521, EPI_ISL_472522, EPI_ISL_472523, EPI_ISL_472524, EPI_ISL_472525, EPI_ISL_472526, EPI_ISL_472527, EPI_ISL_472528, EPI_ISL_472529, EPI_ISL_472530, EPI_ISL_472531, EPI_ISL_472532, EPI_ISL_472533, EPI_ISL_472534, EPI_ISL_472535, EPI_ISL_472536, EPI_ISL_472537, EPI_ISL_472538, EPI_ISL_472539, EPI_ISL_472540, EPI_ISL_472541, EPI_ISL_472542, EPI_ISL_472543, EPI_ISL_472544, EPI_ISL_472545, EPI_ISL_472546, EPI_ISL_472547, EPI_ISL_472548, EPI_ISL_472549, EPI_ISL_472550, EPI_ISL_472551, EPI_ISL_472552, EPI_ISL_472553, EPI_ISL_472554, EPI_ISL_472555, EPI_ISL_472556, EPI_ISL_472557, EPI_ISL_472558, EPI_ISL_472559, EPI_ISL_472560, EPI_ISL_472561, EPI_ISL_472562, EPI_ISL_472563, EPI_ISL_472564, EPI_ISL_472565, EPI_ISL_472566, EPI_ISL_472567, EPI_ISL_472568, EPI_ISL_472569, EPI_ISL_472570, EPI_ISL_472571, EPI_ISL_472572, EPI_ISL_472573, EPI_ISL_472574, EPI_ISL_472575, EPI_ISL_472576, EPI_ISL_472577, EPI_ISL_472578, EPI_ISL_472579, EPI_ISL_472580, EPI_ISL_472581, EPI_ISL_472582, EPI_ISL_472583, EPI_ISL_472584, EPI_ISL_472585, EPI_ISL_472586, EPI_ISL_472587, EPI_ISL_472588, EPI_ISL_472589, EPI_ISL_472590, EPI_ISL_472591, EPI_ISL_472592, EPI_ISL_472593, EPI_ISL_472594, EPI_ISL_472595, EPI_ISL_472596, EPI_ISL_472597, EPI_ISL_472598, EPI_ISL_472600, EPI_ISL_472601, EPI_ISL_472602, EPI_ISL_472603, EPI_ISL_472604, EPI_ISL_472605, EPI_ISL_472606, EPI_ISL_472607, EPI_ISL_472608, EPI_ISL_472609, EPI_ISL_472610, EPI_ISL_472611, EPI_ISL_472612, EPI_ISL_472613, EPI_ISL_472614, EPI_ISL_472615, EPI_ISL_472616, EPI_ISL_472617, EPI_ISL_472618, EPI_ISL_472619, EPI_ISL_472620, EPI_ISL_472621, EPI_ISL_472622, EPI_ISL_472623, EPI_ISL_472624, EPI_ISL_472625, EPI_ISL_472626, EPI_ISL_472627, EPI_ISL_472628, EPI_ISL_472629, EPI_ISL_472630, EPI_ISL_472631, EPI_ISL_472632, EPI_ISL_472633, EPI_ISL_472634, EPI_ISL_472635, EPI_ISL_472636, EPI_ISL_472637, EPI_ISL_472638, EPI_ISL_472639, EPI_ISL_472640, EPI_ISL_472641, EPI_ISL_472642, EPI_ISL_472643, EPI_ISL_472644, EPI_ISL_472645, EPI_ISL_472646, EPI_ISL_472647, EPI_ISL_472648, EPI_ISL_472649, EPI_ISL_472650, EPI_ISL_472651, EPI_ISL_472652, EPI_ISL_472653, EPI_ISL_472654, EPI_ISL_472655, EPI_ISL_472656, EPI_ISL_472657, EPI_ISL_472658, EPI_ISL_472659, EPI_ISL_472660, EPI_ISL_472661, EPI_ISL_472662, EPI_ISL_472663, EPI_ISL_472664, EPI_ISL_472665, EPI_ISL_472666, EPI_ISL_472667, EPI_ISL_472668, EPI_ISL_472669, EPI_ISL_472670, EPI_ISL_472671, EPI_ISL_472672, EPI_ISL_472673, EPI_ISL_472674, EPI_ISL_472675, EPI_ISL_472676, EPI_ISL_472677, EPI_ISL_472678, EPI_ISL_472679, EPI_ISL_472680, EPI_ISL_472681, EPI_ISL_472682, EPI_ISL_472683, EPI_ISL_472684, EPI_ISL_472685, EPI_ISL_472686, EPI_ISL_472687, EPI_ISL_472688, EPI_ISL_472689, EPI_ISL_472690, EPI_ISL_472691, EPI_ISL_472692, EPI_ISL_472693, EPI_ISL_472694, EPI_ISL_472695, EPI_ISL_472696, EPI_ISL_472697, EPI_ISL_472698, EPI_ISL_472699, EPI_ISL_472700, EPI_ISL_472701, EPI_ISL_472702, EPI_ISL_472703, EPI_ISL_472704, EPI_ISL_472705, EPI_ISL_472706, EPI_ISL_472707, EPI_ISL_472708, EPI_ISL_472709, EPI_ISL_472710, EPI_ISL_472711, EPI_ISL_472712, EPI_ISL_472713, EPI_ISL_472714, EPI_ISL_472715, EPI_ISL_472716, EPI_ISL_472717, EPI_ISL_472718, EPI_ISL_472719, EPI_ISL_472720, EPI_ISL_472721, EPI_ISL_472722, EPI_ISL_472723, EPI_ISL_472724, EPI_ISL_472725, EPI_ISL_472726, EPI_ISL_472727, EPI_ISL_472728, EPI_ISL_472729, EPI_ISL_472730, EPI_ISL_472731, EPI_ISL_472732, EPI_ISL_472733, EPI_ISL_472734, EPI_ISL_472735, EPI_ISL_472736, EPI_ISL_472737, EPI_ISL_472738, EPI_ISL_472739, EPI_ISL_472740, EPI_ISL_472741, EPI_ISL_472742, EPI_ISL_472743, EPI_ISL_472744, EPI_ISL_472745, EPI_ISL_472746, EPI_ISL_472747, EPI_ISL_472748, EPI_ISL_472749, EPI_ISL_472750, EPI_ISL_472751, EPI_ISL_472752, EPI_ISL_472753, EPI_ISL_472754, EPI_ISL_472755, EPI_ISL_472756, EPI_ISL_472757, EPI_ISL_472758, EPI_ISL_472759, EPI_ISL_472760, EPI_ISL_472761, EPI_ISL_472762, EPI_ISL_472763, EPI_ISL_472764, EPI_ISL_472765, EPI_ISL_472766, EPI_ISL_472767, EPI_ISL_472768, EPI_ISL_472769, EPI_ISL_472770, EPI_ISL_472771, EPI_ISL_472772, EPI_ISL_472773, EPI_ISL_472774, EPI_ISL_472775, EPI_ISL_472776, EPI_ISL_472777, EPI_ISL_472778, EPI_ISL_472779, EPI_ISL_472780, EPI_ISL_472781, EPI_ISL_472782, EPI_ISL_472783, EPI_ISL_472784, EPI_ISL_472785, EPI_ISL_472786, EPI_ISL_472787, EPI_ISL_472788, EPI_ISL_472789, EPI_ISL_472790, EPI_ISL_472791, EPI_ISL_472792, EPI_ISL_472793, EPI_ISL_472794, EPI_ISL_472795, EPI_ISL_472796, EPI_ISL_472797, EPI_ISL_472798, EPI_ISL_472799, EPI_ISL_472800, EPI_ISL_472801, EPI_ISL_472802, EPI_ISL_472803, EPI_ISL_472804, EPI_ISL_472805, EPI_ISL_472806, EPI_ISL_472807, EPI_ISL_472808, EPI_ISL_472809, EPI_ISL_472810, EPI_ISL_472811, EPI_ISL_472812, EPI_ISL_472813, EPI_ISL_472814, EPI_ISL_472815, EPI_ISL_472816, EPI_ISL_472817, EPI_ISL_472818, EPI_ISL_472819, EPI_ISL_472820, EPI_ISL_472821, EPI_ISL_472822, EPI_ISL_472823, EPI_ISL_472824, EPI_ISL_472825, EPI_ISL_472826, EPI_ISL_472827, EPI_ISL_472828, EPI_ISL_472829, EPI_ISL_472830, EPI_ISL_472831, EPI_ISL_472832, EPI_ISL_472833, EPI_ISL_472834, EPI_ISL_472835, EPI_ISL_472836, EPI_ISL_472837, EPI_ISL_472838, EPI_ISL_472839, EPI_ISL_472840, EPI_ISL_472841, EPI_ISL_472842, EPI_ISL_472843, EPI_ISL_472844, EPI_ISL_472845, EPI_ISL_472846, EPI_ISL_472847, EPI_ISL_472848, EPI_ISL_472849, EPI_ISL_472850, EPI_ISL_472851, EPI_ISL_472852, EPI_ISL_472853, EPI_ISL_472854, EPI_ISL_472855, EPI_ISL_472856, EPI_ISL_472857, EPI_ISL_472858, EPI_ISL_472859, EPI_ISL_472860, EPI_ISL_472861, EPI_ISL_472862, EPI_ISL_472863, EPI_ISL_472864, EPI_ISL_472865, EPI_ISL_472866, EPI_ISL_472867, EPI_ISL_472868, EPI_ISL_472869, EPI_ISL_472870, EPI_ISL_472871, EPI_ISL_472872, EPI_ISL_472873, EPI_ISL_472874, EPI_ISL_472875, EPI_ISL_472876, EPI_ISL_472877, EPI_ISL_472878, EPI_ISL_472879, EPI_ISL_472880, EPI_ISL_472881, EPI_ISL_472882, EPI_ISL_472883, EPI_ISL_472884, EPI_ISL_472885, EPI_ISL_472886, EPI_ISL_472887, EPI_ISL_472888, EPI_ISL_472889, EPI_ISL_472890, EPI_ISL_472891, EPI_ISL_472892, EPI_ISL_472893, EPI_ISL_472894, EPI_ISL_472895, EPI_ISL_472896, EPI_ISL_472897, EPI_ISL_472898, EPI_ISL_472899, EPI_ISL_472900, EPI_ISL_472901, EPI_ISL_472902, EPI_ISL_472903, EPI_ISL_472904, EPI_ISL_472905, EPI_ISL_472906, EPI_ISL_472907, EPI_ISL_472908, EPI_ISL_472909, EPI_ISL_472910, EPI_ISL_472911, EPI_ISL_472912, EPI_ISL_472913, EPI_ISL_472914, EPI_ISL_472915, EPI_ISL_472916, EPI_ISL_472917, EPI_ISL_472918, EPI_ISL_472919, EPI_ISL_472920, EPI_ISL_472921, EPI_ISL_472922, EPI_ISL_472923, EPI_ISL_472924, EPI_ISL_472925, EPI_ISL_472926, EPI_ISL_472927, EPI_ISL_472928, EPI_ISL_472929, EPI_ISL_472930, EPI_ISL_472931, EPI_ISL_472932, EPI_ISL_472933, EPI_ISL_472934, EPI_ISL_472935, EPI_ISL_472937, EPI_ISL_472938, EPI_ISL_472939, EPI_ISL_472940, EPI_ISL_472941, EPI_ISL_472942, EPI_ISL_472943, EPI_ISL_472944, EPI_ISL_472946, EPI_ISL_472947, EPI_ISL_472948, EPI_ISL_472949, EPI_ISL_472950, EPI_ISL_472951, EPI_ISL_472952, EPI_ISL_472953, EPI_ISL_472954, EPI_ISL_472955, EPI_ISL_472956, EPI_ISL_472957, EPI_ISL_472958, EPI_ISL_472959, EPI_ISL_472960, EPI_ISL_472961, EPI_ISL_472962, EPI_ISL_472963, EPI_ISL_472964, EPI_ISL_472965, EPI_ISL_472966, EPI_ISL_472967, EPI_ISL_472968, EPI_ISL_472969, EPI_ISL_472970, EPI_ISL_472971, EPI_ISL_472972, EPI_ISL_472973, EPI_ISL_472974, EPI_ISL_472975, EPI_ISL_472976, EPI_ISL_472977, EPI_ISL_472978, EPI_ISL_472979, EPI_ISL_472980, EPI_ISL_472981, EPI_ISL_472982, EPI_ISL_472983, EPI_ISL_472984, EPI_ISL_472985, EPI_ISL_472986, EPI_ISL_472987, EPI_ISL_472988, EPI_ISL_472989, EPI_ISL_472990, EPI_ISL_472991, EPI_ISL_472992, EPI_ISL_472993, EPI_ISL_472994, EPI_ISL_472995, EPI_ISL_472996, EPI_ISL_472997, EPI_ISL_472998, EPI_ISL_472999, EPI_ISL_473000, EPI_ISL_473001, EPI_ISL_473002, EPI_ISL_473003, EPI_ISL_473004, EPI_ISL_473005, EPI_ISL_473006, EPI_ISL_473007, EPI_ISL_473008, EPI_ISL_473009, EPI_ISL_473010, EPI_ISL_473011, EPI_ISL_473012, EPI_ISL_473013, EPI_ISL_473014, EPI_ISL_473015, EPI_ISL_473016, EPI_ISL_473017, EPI_ISL_473018, EPI_ISL_473019, EPI_ISL_473020, EPI_ISL_473021, EPI_ISL_473022, EPI_ISL_473023, EPI_ISL_473024, EPI_ISL_473025, EPI_ISL_473026, EPI_ISL_473027, EPI_ISL_473028, EPI_ISL_473029, EPI_ISL_473030, EPI_ISL_473031, EPI_ISL_473032, EPI_ISL_473033, EPI_ISL_473034, EPI_ISL_473035, EPI_ISL_473036, EPI_ISL_473037, EPI_ISL_473038, EPI_ISL_473039, EPI_ISL_473040, EPI_ISL_473041, EPI_ISL_473042, EPI_ISL_473043, EPI_ISL_473044, EPI_ISL_473045, EPI_ISL_473046, EPI_ISL_473047, EPI_ISL_473048, EPI_ISL_473049, EPI_ISL_473050, EPI_ISL_473051, EPI_ISL_473052, EPI_ISL_473053, EPI_ISL_473054, EPI_ISL_473055, EPI_ISL_473056, EPI_ISL_473057, EPI_ISL_473058, EPI_ISL_473059, EPI_ISL_473060, EPI_ISL_473061, EPI_ISL_473062, EPI_ISL_473063, EPI_ISL_473064, EPI_ISL_473065, EPI_ISL_473066, EPI_ISL_473067, EPI_ISL_473068, EPI_ISL_473069, EPI_ISL_473070, EPI_ISL_473071, EPI_ISL_473072, EPI_ISL_473073, EPI_ISL_473074, EPI_ISL_473075, EPI_ISL_473076, EPI_ISL_473077, EPI_ISL_473078, EPI_ISL_473079, EPI_ISL_473080, EPI_ISL_473081, EPI_ISL_473082, EPI_ISL_473083, EPI_ISL_473084, EPI_ISL_473085, EPI_ISL_473086, EPI_ISL_473087, EPI_ISL_473088, EPI_ISL_473089, EPI_ISL_473090, EPI_ISL_473091, EPI_ISL_473092, EPI_ISL_473093, EPI_ISL_473094, EPI_ISL_473095, EPI_ISL_473096, EPI_ISL_473097, EPI_ISL_473098, EPI_ISL_473099, EPI_ISL_473100, EPI_ISL_473101, EPI_ISL_473102, EPI_ISL_473103, EPI_ISL_473104, EPI_ISL_473105, EPI_ISL_473106, EPI_ISL_473107, EPI_ISL_473108, EPI_ISL_473109, EPI_ISL_473110, EPI_ISL_473111, EPI_ISL_473112, EPI_ISL_473113, EPI_ISL_473114, EPI_ISL_473115, EPI_ISL_473116, EPI_ISL_473117, EPI_ISL_473118, EPI_ISL_473119, EPI_ISL_473120, EPI_ISL_473121, EPI_ISL_473122, EPI_ISL_473123, EPI_ISL_473124, EPI_ISL_473125, EPI_ISL_473126, EPI_ISL_473127, EPI_ISL_473128, EPI_ISL_473129, EPI_ISL_473130, EPI_ISL_473131, EPI_ISL_473132, EPI_ISL_473133, EPI_ISL_473134, EPI_ISL_473135, EPI_ISL_473136, EPI_ISL_473137, EPI_ISL_473138, EPI_ISL_473139, EPI_ISL_473140, EPI_ISL_473141, EPI_ISL_473142, EPI_ISL_473143, EPI_ISL_473144, EPI_ISL_473145, EPI_ISL_473146, EPI_ISL_473147, EPI_ISL_473148, EPI_ISL_473149, EPI_ISL_473150, EPI_ISL_473151, EPI_ISL_473152, EPI_ISL_473153, EPI_ISL_473154, EPI_ISL_473155, EPI_ISL_473156, EPI_ISL_473157, EPI_ISL_473158, EPI_ISL_473159, EPI_ISL_473160, EPI_ISL_473161, EPI_ISL_473162, EPI_ISL_473163, EPI_ISL_473164, EPI_ISL_473165, EPI_ISL_473166, EPI_ISL_473167, EPI_ISL_473168, EPI_ISL_473169, EPI_ISL_473170, EPI_ISL_473171, EPI_ISL_473172, EPI_ISL_473173, EPI_ISL_473174, EPI_ISL_473175, EPI_ISL_473176, EPI_ISL_473177, EPI_ISL_473178, EPI_ISL_473179, EPI_ISL_473180, EPI_ISL_473181, EPI_ISL_473182, EPI_ISL_473183, EPI_ISL_473184, EPI_ISL_473185, EPI_ISL_473186, EPI_ISL_473187, EPI_ISL_473188, EPI_ISL_473189, EPI_ISL_473190, EPI_ISL_473191, EPI_ISL_473192, EPI_ISL_473193, EPI_ISL_473194, EPI_ISL_473195, EPI_ISL_473196, EPI_ISL_473197, EPI_ISL_473198, EPI_ISL_473199, EPI_ISL_473200, EPI_ISL_473201, EPI_ISL_473202, EPI_ISL_473203, EPI_ISL_473204, EPI_ISL_473205, EPI_ISL_473206, EPI_ISL_473207, EPI_ISL_473208, EPI_ISL_473209, EPI_ISL_473210, EPI_ISL_473211, EPI_ISL_473212, EPI_ISL_473213, EPI_ISL_473214, EPI_ISL_473215, EPI_ISL_473216, EPI_ISL_473217, EPI_ISL_473218, EPI_ISL_473219, EPI_ISL_473220, EPI_ISL_473221, EPI_ISL_473222, EPI_ISL_473223, EPI_ISL_473224, EPI_ISL_473225, EPI_ISL_473226, EPI_ISL_473227, EPI_ISL_473228, EPI_ISL_473229, EPI_ISL_473230, EPI_ISL_473231, EPI_ISL_473232, EPI_ISL_473233, EPI_ISL_473234, EPI_ISL_473235, EPI_ISL_473236, EPI_ISL_473237, EPI_ISL_473238, EPI_ISL_473239, EPI_ISL_473240, EPI_ISL_473241, EPI_ISL_473242, EPI_ISL_473243, EPI_ISL_473244, EPI_ISL_473245, EPI_ISL_473246, EPI_ISL_473247, EPI_ISL_473248, EPI_ISL_473249, EPI_ISL_473250, EPI_ISL_473251, EPI_ISL_473252, EPI_ISL_473253, EPI_ISL_473254, EPI_ISL_473255, EPI_ISL_473256, EPI_ISL_473257, EPI_ISL_473258, EPI_ISL_473259, EPI_ISL_473260, EPI_ISL_473261, EPI_ISL_473262, EPI_ISL_473263, EPI_ISL_473264, EPI_ISL_473265, EPI_ISL_473266, EPI_ISL_473267, EPI_ISL_473268, EPI_ISL_473269, EPI_ISL_473270, EPI_ISL_473271, EPI_ISL_473272, EPI_ISL_473273, EPI_ISL_473274, EPI_ISL_473275, EPI_ISL_473276, EPI_ISL_473277, EPI_ISL_473278, EPI_ISL_473279, EPI_ISL_473280, EPI_ISL_473281, EPI_ISL_473282, EPI_ISL_473283, EPI_ISL_473284, EPI_ISL_473285, EPI_ISL_473286, EPI_ISL_473287, EPI_ISL_473288, EPI_ISL_473289, EPI_ISL_473290, EPI_ISL_473291, EPI_ISL_473292, EPI_ISL_473293, EPI_ISL_473294, EPI_ISL_473295, EPI_ISL_473296, EPI_ISL_473297, EPI_ISL_473298, EPI_ISL_473299, EPI_ISL_473300, EPI_ISL_473301, EPI_ISL_473302, EPI_ISL_473303, EPI_ISL_473304, EPI_ISL_473305, EPI_ISL_473956, EPI_ISL_473957, EPI_ISL_473958, EPI_ISL_473959, EPI_ISL_473960, EPI_ISL_473961, EPI_ISL_473962, EPI_ISL_473963, EPI_ISL_473964, EPI_ISL_473965, EPI_ISL_473966, EPI_ISL_473967, EPI_ISL_473968, EPI_ISL_473969, EPI_ISL_473970, EPI_ISL_473971, EPI_ISL_473972, EPI_ISL_473973, EPI_ISL_473974, EPI_ISL_473975, EPI_ISL_473976, EPI_ISL_473977, EPI_ISL_473978, EPI_ISL_473979, EPI_ISL_473980, EPI_ISL_473981, EPI_ISL_473982, EPI_ISL_473983, EPI_ISL_473984, EPI_ISL_473985, EPI_ISL_473986, EPI_ISL_473987, EPI_ISL_473988, EPI_ISL_473989, EPI_ISL_473990, EPI_ISL_473991, EPI_ISL_473992, EPI_ISL_473993, EPI_ISL_473994, EPI_ISL_473995, EPI_ISL_473996, EPI_ISL_473997, EPI_ISL_473998, EPI_ISL_473999, EPI_ISL_474000, EPI_ISL_474001, EPI_ISL_474002, EPI_ISL_474003, EPI_ISL_474004, EPI_ISL_474005, EPI_ISL_474006, EPI_ISL_474007, EPI_ISL_474008, EPI_ISL_474009, EPI_ISL_474010, EPI_ISL_474011, EPI_ISL_474012, EPI_ISL_474013, EPI_ISL_474014, EPI_ISL_474015, EPI_ISL_474016, EPI_ISL_474017, EPI_ISL_474018, EPI_ISL_474019, EPI_ISL_474020, EPI_ISL_474021, EPI_ISL_474022, EPI_ISL_474023, EPI_ISL_474024, EPI_ISL_474025, EPI_ISL_474026, EPI_ISL_474027, EPI_ISL_474028, EPI_ISL_474029, EPI_ISL_474030, EPI_ISL_474031, EPI_ISL_474032, EPI_ISL_474033, EPI_ISL_474034, EPI_ISL_474035, EPI_ISL_474036, EPI_ISL_474037, EPI_ISL_474038, EPI_ISL_474039, EPI_ISL_474040, EPI_ISL_474041, EPI_ISL_474042, EPI_ISL_474043, EPI_ISL_474044, EPI_ISL_474045, EPI_ISL_474046, EPI_ISL_474047, EPI_ISL_474048, EPI_ISL_474049, EPI_ISL_474050, EPI_ISL_474051, EPI_ISL_474052, EPI_ISL_474053, EPI_ISL_474054, EPI_ISL_474055, EPI_ISL_474056, EPI_ISL_474057, EPI_ISL_474058, EPI_ISL_474059, EPI_ISL_474060, EPI_ISL_474061, EPI_ISL_474062, EPI_ISL_474063, EPI_ISL_474064, EPI_ISL_474065, EPI_ISL_474066, EPI_ISL_474067, EPI_ISL_474068, EPI_ISL_474069, EPI_ISL_474070, EPI_ISL_474071, EPI_ISL_474072, EPI_ISL_474073, EPI_ISL_474074, EPI_ISL_474075, EPI_ISL_474076, EPI_ISL_474077, EPI_ISL_474078, EPI_ISL_474079, EPI_ISL_474080, EPI_ISL_474081, EPI_ISL_474082, EPI_ISL_474083, EPI_ISL_474084, EPI_ISL_474085, EPI_ISL_474086, EPI_ISL_474087, EPI_ISL_474088, EPI_ISL_474089, EPI_ISL_474090, EPI_ISL_474091, EPI_ISL_474092, EPI_ISL_474093, EPI_ISL_474094, EPI_ISL_474095, EPI_ISL_474096, EPI_ISL_474097, EPI_ISL_474098, EPI_ISL_474099, EPI_ISL_474100, EPI_ISL_474101, EPI_ISL_474102, EPI_ISL_474103, EPI_ISL_474104, EPI_ISL_474105, EPI_ISL_474106, EPI_ISL_474107, EPI_ISL_474108, EPI_ISL_474109, EPI_ISL_474110, EPI_ISL_474111, EPI_ISL_474112, EPI_ISL_474113, EPI_ISL_474114, EPI_ISL_474115, EPI_ISL_474116, EPI_ISL_474117, EPI_ISL_474118, EPI_ISL_474119, EPI_ISL_474120, EPI_ISL_474121, EPI_ISL_474122, EPI_ISL_474123, EPI_ISL_474124, EPI_ISL_474125, EPI_ISL_474126, EPI_ISL_474127, EPI_ISL_474128, EPI_ISL_474129, EPI_ISL_474130, EPI_ISL_474131, EPI_ISL_474132, EPI_ISL_474133, EPI_ISL_474134, EPI_ISL_474135, EPI_ISL_474136, EPI_ISL_474137, EPI_ISL_474138, EPI_ISL_474139, EPI_ISL_474140, EPI_ISL_474141, EPI_ISL_474142, EPI_ISL_474143, EPI_ISL_474144, EPI_ISL_474145, EPI_ISL_474146, EPI_ISL_474147, EPI_ISL_474148, EPI_ISL_474149, EPI_ISL_474150, EPI_ISL_474151, EPI_ISL_474152, EPI_ISL_474153, EPI_ISL_474154, EPI_ISL_474155, EPI_ISL_474156, EPI_ISL_474157, EPI_ISL_474158, EPI_ISL_474159, EPI_ISL_474160, EPI_ISL_474161, EPI_ISL_474162, EPI_ISL_474163, EPI_ISL_474164, EPI_ISL_474165, EPI_ISL_474166, EPI_ISL_474167, EPI_ISL_474168, EPI_ISL_474169, EPI_ISL_474170, EPI_ISL_474171, EPI_ISL_474172, EPI_ISL_474173, EPI_ISL_474174, EPI_ISL_474175, EPI_ISL_474176, EPI_ISL_474177, EPI_ISL_474178, EPI_ISL_474179, EPI_ISL_474180, EPI_ISL_474181, EPI_ISL_474182, EPI_ISL_474183, EPI_ISL_474184, EPI_ISL_474185, EPI_ISL_474186, EPI_ISL_474187, EPI_ISL_474188, EPI_ISL_474189, EPI_ISL_474190, EPI_ISL_474191, EPI_ISL_474192, EPI_ISL_474193, EPI_ISL_474194, EPI_ISL_474195, EPI_ISL_474196, EPI_ISL_474197, EPI_ISL_474198, EPI_ISL_474199, EPI_ISL_474200, EPI_ISL_474201, EPI_ISL_474202, EPI_ISL_474203, EPI_ISL_474204, EPI_ISL_474205, EPI_ISL_474206, EPI_ISL_474207, EPI_ISL_474208, EPI_ISL_474209, EPI_ISL_474210, EPI_ISL_474211, EPI_ISL_474212, EPI_ISL_474213, EPI_ISL_474214, EPI_ISL_474215, EPI_ISL_474216, EPI_ISL_474217, EPI_ISL_474218, EPI_ISL_474219, EPI_ISL_474220, EPI_ISL_474221, EPI_ISL_474223, EPI_ISL_474224, EPI_ISL_474225, EPI_ISL_474226, EPI_ISL_474227, EPI_ISL_474228, EPI_ISL_474229, EPI_ISL_474230, EPI_ISL_474231, EPI_ISL_474232, EPI_ISL_474233, EPI_ISL_474234, EPI_ISL_474235, EPI_ISL_474236, EPI_ISL_474237, EPI_ISL_474238, EPI_ISL_474239, EPI_ISL_474240, EPI_ISL_474241, EPI_ISL_474242, EPI_ISL_474243, EPI_ISL_474244, EPI_ISL_474245, EPI_ISL_474246, EPI_ISL_474247, EPI_ISL_474248, EPI_ISL_474249, EPI_ISL_474250, EPI_ISL_474251, EPI_ISL_474252, EPI_ISL_474253, EPI_ISL_474254, EPI_ISL_474255, EPI_ISL_474256, EPI_ISL_474257, EPI_ISL_474258, EPI_ISL_474259, EPI_ISL_474260, EPI_ISL_474261, EPI_ISL_474262, EPI_ISL_474263, EPI_ISL_474264, EPI_ISL_474265, EPI_ISL_474266, EPI_ISL_474267, EPI_ISL_474268, EPI_ISL_474269, EPI_ISL_474270, EPI_ISL_474271, EPI_ISL_474272, EPI_ISL_474273, EPI_ISL_474274, EPI_ISL_474275, EPI_ISL_474276, EPI_ISL_474277, EPI_ISL_474278, EPI_ISL_474279, EPI_ISL_474280, EPI_ISL_474281, EPI_ISL_474282, EPI_ISL_474283, EPI_ISL_474284, EPI_ISL_474285, EPI_ISL_474286, EPI_ISL_474287, EPI_ISL_474288, EPI_ISL_474289, EPI_ISL_474290, EPI_ISL_474291, EPI_ISL_474292, EPI_ISL_474293, EPI_ISL_474294, EPI_ISL_474295, EPI_ISL_474296, EPI_ISL_474297, EPI_ISL_474298, EPI_ISL_474299, EPI_ISL_474300, EPI_ISL_474301, EPI_ISL_474302, EPI_ISL_474303, EPI_ISL_474304, EPI_ISL_474305, EPI_ISL_474306, EPI_ISL_474307, EPI_ISL_474308, EPI_ISL_474309, EPI_ISL_474310, EPI_ISL_474311, EPI_ISL_474312, EPI_ISL_474313, EPI_ISL_474314, EPI_ISL_474315, EPI_ISL_474316, EPI_ISL_474317, EPI_ISL_474318, EPI_ISL_474319, EPI_ISL_474320, EPI_ISL_474321, EPI_ISL_474322, EPI_ISL_474323, EPI_ISL_474324, EPI_ISL_474325, EPI_ISL_474326, EPI_ISL_474327, EPI_ISL_474328, EPI_ISL_474329, EPI_ISL_474330, EPI_ISL_474331, EPI_ISL_474332, EPI_ISL_474333, EPI_ISL_474334, EPI_ISL_474335, EPI_ISL_474336, EPI_ISL_474337, EPI_ISL_474338, EPI_ISL_474339, EPI_ISL_474340, EPI_ISL_474341, EPI_ISL_474342, EPI_ISL_474343, EPI_ISL_474344, EPI_ISL_474345, EPI_ISL_474346, EPI_ISL_474347, EPI_ISL_474348, EPI_ISL_474349, EPI_ISL_474350, EPI_ISL_474351, EPI_ISL_474352, EPI_ISL_474353, EPI_ISL_474354, EPI_ISL_474355, EPI_ISL_474356, EPI_ISL_474357, EPI_ISL_474358, EPI_ISL_474359, EPI_ISL_474360, EPI_ISL_474361, EPI_ISL_474362, EPI_ISL_474363, EPI_ISL_474364, EPI_ISL_474365, EPI_ISL_474366, EPI_ISL_474367, EPI_ISL_474368, EPI_ISL_474369, EPI_ISL_474370, EPI_ISL_474371, EPI_ISL_474372, EPI_ISL_474373, EPI_ISL_474374, EPI_ISL_474375, EPI_ISL_474376, EPI_ISL_474377, EPI_ISL_474378, EPI_ISL_474379, EPI_ISL_474380, EPI_ISL_474381, EPI_ISL_474382, EPI_ISL_474383, EPI_ISL_474384, EPI_ISL_474385, EPI_ISL_474386, EPI_ISL_474387, EPI_ISL_474388, EPI_ISL_474389, EPI_ISL_474390, EPI_ISL_474391, EPI_ISL_474392, EPI_ISL_474393, EPI_ISL_474394, EPI_ISL_474395, EPI_ISL_474396, EPI_ISL_474397, EPI_ISL_474398, EPI_ISL_474399, EPI_ISL_474400, EPI_ISL_474401, EPI_ISL_474402, EPI_ISL_474403, EPI_ISL_474404, EPI_ISL_474405, EPI_ISL_474406, EPI_ISL_474407, EPI_ISL_474408, EPI_ISL_474409, EPI_ISL_474410, EPI_ISL_474411, EPI_ISL_474412, EPI_ISL_474413, EPI_ISL_474414, EPI_ISL_474415, EPI_ISL_474416, EPI_ISL_474417, EPI_ISL_474418, EPI_ISL_474419, EPI_ISL_474420, EPI_ISL_474421, EPI_ISL_474422, EPI_ISL_474423, EPI_ISL_474424, EPI_ISL_474425, EPI_ISL_474426, EPI_ISL_474427, EPI_ISL_474428, EPI_ISL_474429, EPI_ISL_474430, EPI_ISL_474431, EPI_ISL_474432, EPI_ISL_474433, EPI_ISL_474434, EPI_ISL_474435, EPI_ISL_474436, EPI_ISL_474437, EPI_ISL_474438, EPI_ISL_474439, EPI_ISL_474440, EPI_ISL_474441, EPI_ISL_474442, EPI_ISL_474443, EPI_ISL_474444, EPI_ISL_474445, EPI_ISL_474446, EPI_ISL_474447, EPI_ISL_474448, EPI_ISL_474449, EPI_ISL_474450, EPI_ISL_474451, EPI_ISL_474452, EPI_ISL_474453, EPI_ISL_474454, EPI_ISL_474455, EPI_ISL_474456, EPI_ISL_474457, EPI_ISL_474458, EPI_ISL_474459, EPI_ISL_474460, EPI_ISL_474461, EPI_ISL_474462, EPI_ISL_474463, EPI_ISL_474464, EPI_ISL_474465, EPI_ISL_474466, EPI_ISL_474467, EPI_ISL_474468, EPI_ISL_474469, EPI_ISL_474470, EPI_ISL_474471,

| | | | |
|---|---|---|---|
| EPI_ISL_474472, EPI_ISL_474473, EPI_ISL_474474, EPI_ISL_474475, EPI_ISL_474476, EPI_ISL_474477, EPI_ISL_474478, EPI_ISL_474479, EPI_ISL_474480, EPI_ISL_474481, EPI_ISL_474482, EPI_ISL_474483, EPI_ISL_474484, EPI_ISL_474485, EPI_ISL_474486, EPI_ISL_474487, EPI_ISL_474488, EPI_ISL_474489, EPI_ISL_474490, EPI_ISL_474491, EPI_ISL_474492, EPI_ISL_474493, EPI_ISL_474494, EPI_ISL_474495, EPI_ISL_474496, EPI_ISL_474497, EPI_ISL_474498, EPI_ISL_474499, EPI_ISL_474500, EPI_ISL_474501, EPI_ISL_474502, EPI_ISL_474503, EPI_ISL_474504, EPI_ISL_474505, EPI_ISL_474506, EPI_ISL_474507, EPI_ISL_474508, EPI_ISL_474509, EPI_ISL_474510, EPI_ISL_474511, EPI_ISL_474512, EPI_ISL_474513, EPI_ISL_474514, EPI_ISL_474515, EPI_ISL_474516, EPI_ISL_474517, EPI_ISL_474518, EPI_ISL_474519, EPI_ISL_474520, EPI_ISL_474521, EPI_ISL_474522, EPI_ISL_474523, EPI_ISL_474524, EPI_ISL_474525, EPI_ISL_474526, EPI_ISL_474527, EPI_ISL_474528, EPI_ISL_474529, EPI_ISL_474530, EPI_ISL_474531, EPI_ISL_474532, EPI_ISL_474533, EPI_ISL_474534, EPI_ISL_474535, EPI_ISL_474536, EPI_ISL_474537, EPI_ISL_474538, EPI_ISL_474539, EPI_ISL_474540, EPI_ISL_474541, EPI_ISL_474542, EPI_ISL_474543, EPI_ISL_474544, EPI_ISL_474545, EPI_ISL_474546, EPI_ISL_474547, EPI_ISL_474548, EPI_ISL_474549, EPI_ISL_474550, EPI_ISL_474551, EPI_ISL_474552, EPI_ISL_474553, EPI_ISL_474554, EPI_ISL_474555, EPI_ISL_474556, EPI_ISL_474557, EPI_ISL_474558, EPI_ISL_474559, EPI_ISL_474560, EPI_ISL_474561, EPI_ISL_474562, EPI_ISL_474563, EPI_ISL_474564, EPI_ISL_474565, EPI_ISL_474566, EPI_ISL_474567, EPI_ISL_474568, EPI_ISL_474569, EPI_ISL_474570, EPI_ISL_474571, EPI_ISL_474572, EPI_ISL_474573, EPI_ISL_474574, EPI_ISL_474575, EPI_ISL_474576, EPI_ISL_474577, EPI_ISL_474578, EPI_ISL_474579, EPI_ISL_474580, EPI_ISL_474581, EPI_ISL_474582, EPI_ISL_474583, EPI_ISL_474584, EPI_ISL_474585, EPI_ISL_474586, EPI_ISL_474587, EPI_ISL_474588, EPI_ISL_474589, EPI_ISL_474590, EPI_ISL_474591, EPI_ISL_474592, EPI_ISL_474593, EPI_ISL_474594, EPI_ISL_474595, EPI_ISL_474596, EPI_ISL_474597, EPI_ISL_474598, EPI_ISL_474599, EPI_ISL_474600, EPI_ISL_474601, EPI_ISL_474602, EPI_ISL_474603, EPI_ISL_474604, EPI_ISL_474605, EPI_ISL_474606, EPI_ISL_474607, EPI_ISL_474608, EPI_ISL_474609, EPI_ISL_474610, EPI_ISL_474611, EPI_ISL_474612, EPI_ISL_474613, EPI_ISL_474614, EPI_ISL_474615, EPI_ISL_474616, EPI_ISL_474617, EPI_ISL_474618, EPI_ISL_474619, EPI_ISL_474620, EPI_ISL_474621, EPI_ISL_474622, EPI_ISL_474623, EPI_ISL_474624, EPI_ISL_474625, EPI_ISL_474626, EPI_ISL_474627, EPI_ISL_474628, EPI_ISL_474629, EPI_ISL_474630, EPI_ISL_474631, EPI_ISL_474632, EPI_ISL_474633, EPI_ISL_474634, EPI_ISL_474635, EPI_ISL_474636, EPI_ISL_474637, EPI_ISL_474638, EPI_ISL_474639, EPI_ISL_474640, EPI_ISL_474641, EPI_ISL_474642, EPI_ISL_474643, EPI_ISL_474644, EPI_ISL_474645, EPI_ISL_474646, EPI_ISL_474647, EPI_ISL_474648, EPI_ISL_474649, EPI_ISL_474650, EPI_ISL_474651, EPI_ISL_474652, EPI_ISL_474653, EPI_ISL_474654, EPI_ISL_474655, EPI_ISL_474656, EPI_ISL_474657, EPI_ISL_474658, EPI_ISL_474659, EPI_ISL_474660, EPI_ISL_474661, EPI_ISL_474662, EPI_ISL_474663, EPI_ISL_474664, EPI_ISL_474665, EPI_ISL_474666, EPI_ISL_474667, EPI_ISL_474668, EPI_ISL_474669, EPI_ISL_474670, EPI_ISL_474671, EPI_ISL_474672, EPI_ISL_474673, EPI_ISL_474674, EPI_ISL_474675, EPI_ISL_474676, EPI_ISL_474677, EPI_ISL_474678, EPI_ISL_474679, EPI_ISL_474680, EPI_ISL_474681, EPI_ISL_474682, EPI_ISL_474683, EPI_ISL_474684, EPI_ISL_474685, EPI_ISL_474686, EPI_ISL_474687, EPI_ISL_474688, EPI_ISL_474689, EPI_ISL_474690, EPI_ISL_474691, EPI_ISL_474692, EPI_ISL_474693, EPI_ISL_474694, EPI_ISL_474695, EPI_ISL_474696, EPI_ISL_474697, EPI_ISL_474698, EPI_ISL_474699, EPI_ISL_474700, EPI_ISL_474701, EPI_ISL_474702, EPI_ISL_474703, EPI_ISL_474704, EPI_ISL_474705, EPI_ISL_474706, EPI_ISL_474707, EPI_ISL_474708, EPI_ISL_474709, EPI_ISL_474710, EPI_ISL_474711, EPI_ISL_474712, EPI_ISL_474713, EPI_ISL_474714, EPI_ISL_474715, EPI_ISL_474716, EPI_ISL_474717, EPI_ISL_474718, EPI_ISL_474719, EPI_ISL_474720, EPI_ISL_474721, EPI_ISL_474722, EPI_ISL_474723, EPI_ISL_474724, EPI_ISL_474725, EPI_ISL_474726, EPI_ISL_474727, EPI_ISL_474728, EPI_ISL_474729, EPI_ISL_474730, EPI_ISL_474731, EPI_ISL_474732, EPI_ISL_474733, EPI_ISL_474734, EPI_ISL_474735, EPI_ISL_474736, EPI_ISL_474737, EPI_ISL_474738, EPI_ISL_474739, EPI_ISL_474740, EPI_ISL_474741, EPI_ISL_474742, EPI_ISL_474743, EPI_ISL_474744, EPI_ISL_474745, EPI_ISL_474746, EPI_ISL_474747, EPI_ISL_474748, EPI_ISL_474749, EPI_ISL_474750, EPI_ISL_474751, EPI_ISL_474752, EPI_ISL_474753, EPI_ISL_474754, EPI_ISL_474755, EPI_ISL_474756, EPI_ISL_474757, EPI_ISL_474758, EPI_ISL_474759, EPI_ISL_474760, EPI_ISL_474761, EPI_ISL_474762, EPI_ISL_474763, EPI_ISL_474764, EPI_ISL_474765, EPI_ISL_474766, EPI_ISL_474767, EPI_ISL_474768, EPI_ISL_474769, EPI_ISL_474770, EPI_ISL_474771, EPI_ISL_474772, EPI_ISL_474773, EPI_ISL_474774, EPI_ISL_474775, EPI_ISL_474776, EPI_ISL_474777, EPI_ISL_474778, EPI_ISL_474779, EPI_ISL_474780, EPI_ISL_474781, EPI_ISL_474782, EPI_ISL_474783, EPI_ISL_474784, EPI_ISL_474785, EPI_ISL_474786, EPI_ISL_474787, EPI_ISL_474788, EPI_ISL_474789, EPI_ISL_474790, EPI_ISL_474791, EPI_ISL_474792, EPI_ISL_474793, EPI_ISL_474794, EPI_ISL_474795, EPI_ISL_474796 | | | |
| see above | Wales Specialist Virology Centre Sequencing lab: Pathogen Genomics Unit | COVID-19 Genomics UK (COG-UK) Consortium | Catherine Moore, Johnathan Evans, Laura Gifford, Malorie Perry, Simon Cottrell, Angela Marchbank, Alec Birchley, Alexander Adams, Amy Gaskin, Bree Gatica-Wilcox, Jason Coombes, Joel Southgate, Lauren Gilbert, Lee Graham, Nicole Pacchiarini, Sara Kumziene-Summerhayes, Sarah Taylor, Sophie Jones, Sara Rey, Matthew Bull, Joanne Watkins, Sally Corden, Tom Connor |
| EPI_ISL_476083, EPI_ISL_476095, EPI_ISL_476100, EPI_ISL_476116, EPI_ISL_476127 | Viollier AG | Department of Biosystems Science and Engineering, ETH Zürich | Christian Beisel, Sarah Nadeau, Ivan Topolsky, Pedro Ferreira, Philipp Jablonski, Susana Posada-Céspedes, Tobias Schär, Ina Nissen, Natascha Santacroce, Elodie Burcklen, Christiane Beckmann, Maurice Redondo, Olivier Kobel, Christoph Noppen, Sophie Seidel, Noemie Santamaria de Souza, Niko Beerenwinkel, Tanja Stadler |
| EPI_ISL_476795, EPI_ISL_476796, EPI_ISL_476797 | Department of Laboratory Medicine, Tan Tock Seng Hospital | Department of Laboratory Medicine, Tan Tock Seng Hospital | Chen YYC, Zair X, Li C, Tang WY, Maurer-Stroh S, Barkham TMS, Nagarajan N, Sessions OM |
| EPI_ISL_476805, EPI_ISL_476806, EPI_ISL_476807, EPI_ISL_476808, EPI_ISL_476809, EPI_ISL_476810, EPI_ISL_476811, EPI_ISL_476812 | Department of Laboratory Medicine Tan Tock Seng Hospital | Department of Laboratory Medicine Tan Tock Seng Hospital | Chen YYC, Zair X, Li C, Tang WY, Maurer-Stroh S, Barkham TMS, Nagarajan N, Sessions OM |
| EPI_ISL_476813, EPI_ISL_476814 | Department of Laboratory Medicine, Tan Tock Seng Hospital | Department of Laboratory Medicine, Tan Tock Seng Hospital | Chen YYC, Zair X, Li C, Tang WY, Maurer-Stroh S, Barkham TMS, Nagarajan N, Sessions OM |
| EPI_ISL_476815 | Department of Laboratory Medicine Tan Tock Seng Hospital | Department of Laboratory Medicine Tan Tock Seng Hospital | Chen YYC, Zair X, Li C, Tang WY, Maurer-Stroh S, Barkham TMS, Nagarajan N, Sessions OM |
| EPI_ISL_476816, EPI_ISL_476817, EPI_ISL_476818, EPI_ISL_476819, EPI_ISL_476820, EPI_ISL_476821 | Department of Laboratory Medicine, Tan Tock Seng Hospital | Department of Laboratory Medicine, Tan Tock Seng Hospital | Chen YYC, Zair X, Li C, Tang WY, Maurer-Stroh S, Barkham TMS, Nagarajan N, Sessions OM |
| EPI_ISL_477170 | Department of Laboratory, Medicine Tan Tock Seng Hospital | Department of Laboratory Medicine Tan Tock Seng Hospital | Chen YYC, Zair X, Li C, Tang WY, Maurer-Stroh S, Barkham TMS, Nagarajan N, Sessions OM |
| EPI_ISL_477171 | Department of Laboratory, Medicine Tan Tock Seng Hospital | Department of Laboratory, Medicine Tan Tock Seng Hospital | Chen YYC, Zair X, Li C, Tang WY, Maurer-Stroh S, Barkham TMS, Nagarajan N, Sessions OM |
| EPI_ISL_477172, EPI_ISL_477174, EPI_ISL_477175, EPI_ISL_477177, EPI_ISL_477178, EPI_ISL_477180, EPI_ISL_477182, EPI_ISL_477184, EPI_ISL_477187, EPI_ISL_477188, EPI_ISL_477189, EPI_ISL_477190, EPI_ISL_477191, EPI_ISL_477192 | | | |
| see above | Department of Laboratory Medicine Tan Tock Seng Hospital | Department of Laboratory Medicine Tan Tock Seng Hospital | Chen YYC, Zair X, Li C, Tang WY, Maurer-Stroh S, Barkham TMS, Nagarajan N, Sessions OM |
| EPI_ISL_479284, EPI_ISL_479285, EPI_ISL_479286, EPI_ISL_479287, EPI_ISL_479288, EPI_ISL_479289, EPI_ISL_479290, EPI_ISL_479291, EPI_ISL_479292, EPI_ISL_479293, EPI_ISL_479294, EPI_ISL_479295, EPI_ISL_479296, EPI_ISL_479297, EPI_ISL_479298, EPI_ISL_479299, EPI_ISL_479300, EPI_ISL_479301, EPI_ISL_479302, EPI_ISL_479303, EPI_ISL_479304, EPI_ISL_479305, EPI_ISL_479306, EPI_ISL_479307, EPI_ISL_479308, EPI_ISL_479309, EPI_ISL_479310, EPI_ISL_479311, EPI_ISL_479312, EPI_ISL_479313, EPI_ISL_479314, EPI_ISL_479315, EPI_ISL_479316, EPI_ISL_479317, EPI_ISL_479318, EPI_ISL_479319, EPI_ISL_479320, EPI_ISL_479321, EPI_ISL_479322, EPI_ISL_479323, EPI_ISL_479324, EPI_ISL_479325, EPI_ISL_479326, EPI_ISL_479327, EPI_ISL_479328, EPI_ISL_479329, EPI_ISL_479330, EPI_ISL_479331, EPI_ISL_479332, EPI_ISL_479333, EPI_ISL_479334, EPI_ISL_479335, EPI_ISL_479336, EPI_ISL_479337, EPI_ISL_479338, EPI_ISL_479339, EPI_ISL_479340, EPI_ISL_479341, EPI_ISL_479342, EPI_ISL_479343, EPI_ISL_479344, EPI_ISL_479345, EPI_ISL_479346, EPI_ISL_479347, EPI_ISL_479348, EPI_ISL_479349, EPI_ISL_479350, EPI_ISL_479351, EPI_ISL_479352, EPI_ISL_479353, EPI_ISL_479354, EPI_ISL_479355, EPI_ISL_479356, EPI_ISL_479357, EPI_ISL_479358, EPI_ISL_479359, EPI_ISL_479360, EPI_ISL_479361, EPI_ISL_479362, EPI_ISL_479363, EPI_ISL_479364, EPI_ISL_479365, EPI_ISL_479366, EPI_ISL_479367, EPI_ISL_479368, EPI_ISL_479369, EPI_ISL_479370, EPI_ISL_479371, EPI_ISL_479372, EPI_ISL_479373, EPI_ISL_479374, EPI_ISL_479375, EPI_ISL_479376, EPI_ISL_479377, EPI_ISL_479378, EPI_ISL_479379, EPI_ISL_479380, EPI_ISL_479381, EPI_ISL_479382, EPI_ISL_479383, EPI_ISL_479384, EPI_ISL_479385, EPI_ISL_479386, EPI_ISL_479387, EPI_ISL_479388, EPI_ISL_479389, EPI_ISL_479390, EPI_ISL_479391, EPI_ISL_479392, EPI_ISL_479393, EPI_ISL_479394, EPI_ISL_479395, EPI_ISL_479396, EPI_ISL_479397, EPI_ISL_479398, EPI_ISL_479399, EPI_ISL_479400, EPI_ISL_479401, EPI_ISL_479402, EPI_ISL_479403, EPI_ISL_479404, EPI_ISL_479405, EPI_ISL_479406, EPI_ISL_479407, EPI_ISL_479408, EPI_ISL_479409, EPI_ISL_479410, EPI_ISL_479411, EPI_ISL_479412, EPI_ISL_479413, EPI_ISL_479414, EPI_ISL_479415, EPI_ISL_479416, EPI_ISL_479417, EPI_ISL_479418, EPI_ISL_479419, EPI_ISL_479420, EPI_ISL_479421, EPI_ISL_479422, EPI_ISL_479423, EPI_ISL_479424, EPI_ISL_479425, EPI_ISL_479426, EPI_ISL_479427, EPI_ISL_479428, EPI_ISL_479429, EPI_ISL_479430, EPI_ISL_479431, EPI_ISL_479432, EPI_ISL_479433, EPI_ISL_479434, EPI_ISL_479435, EPI_ISL_479436, EPI_ISL_479437, EPI_ISL_479438, EPI_ISL_479439, EPI_ISL_479440, EPI_ISL_479441, EPI_ISL_479442, EPI_ISL_479443, EPI_ISL_479444, EPI_ISL_479445, EPI_ISL_479446, EPI_ISL_479447, EPI_ISL_479448, EPI_ISL_479449, EPI_ISL_479450, EPI_ISL_479451, EPI_ISL_479452, EPI_ISL_479453, EPI_ISL_479454, EPI_ISL_479455, EPI_ISL_479456, EPI_ISL_479457, EPI_ISL_479458, EPI_ISL_479459, EPI_ISL_479460, EPI_ISL_479461, EPI_ISL_479462, EPI_ISL_479463, EPI_ISL_479464, EPI_ISL_479465, EPI_ISL_479466, EPI_ISL_479467, EPI_ISL_479468, EPI_ISL_479469, EPI_ISL_479470, EPI_ISL_479471, EPI_ISL_479472, EPI_ISL_479473, EPI_ISL_479474, EPI_ISL_479475, EPI_ISL_479476, EPI_ISL_479477, EPI_ISL_479478, EPI_ISL_479479, EPI_ISL_479480, EPI_ISL_479481 | | | |
| see above | Wales Specialist Virology Centre Sequencing lab: Pathogen Genomics Unit | COVID-19 Genomics UK (COG-UK) Consortium | Catherine Moore, Johnathan Evans, Laura Gifford, Malorie Perry, Simon Cottrell, Angela Marchbank, Alec Birchley, Alexander Adams, Amy Gaskin, Bree Gatica-Wilcox, Jason Coombes, Joel Southgate, Lauren Gilbert, Lee Graham, Nicole Pacchiarini, Sara Kumziene-Summerhayes, Sarah Taylor, Sophie Jones, Sara Rey, Matthew Bull, Joanne Watkins, Sally Corden, Tom Connor |
| EPI_ISL_479482, EPI_ISL_479483, EPI_ISL_479484, EPI_ISL_479485, EPI_ISL_479486, EPI_ISL_479487, EPI_ISL_479488, EPI_ISL_479489, EPI_ISL_479490, EPI_ISL_479491, EPI_ISL_479492 | | | |
| see above | Department of Laboratory Medicine Tan Tock Seng Hospital | Department of Laboratory Medicine Tan Tock Seng Hospital | Chen YYC, Zair X, Li C, Tang WY, Maurer-Stroh S, Barkham TMS, Nagarajan N, Sessions OM |
| EPI_ISL_480331, EPI_ISL_480332, EPI_ISL_480333, EPI_ISL_480334, EPI_ISL_480335, EPI_ISL_480336, EPI_ISL_480337, EPI_ISL_480338, EPI_ISL_480339, EPI_ISL_480340, EPI_ISL_480341, EPI_ISL_480342, EPI_ISL_480343, EPI_ISL_480344, EPI_ISL_480345, EPI_ISL_480346, EPI_ISL_480347, EPI_ISL_480348 | | | |
| see above | Microbial Genomics Laboratory, Institut Pasteur de Montevideo | Microbial Genomics Laboratory, Institut Pasteur de Montevideo | Cecilia Salazar, Marianoel Pereira, Ignacio Ferrés, Gonzalo Moratorio, Pilar Moreno, Gregorio Iraola |
| EPI_ISL_483651, EPI_ISL_483652, EPI_ISL_483654, EPI_ISL_483660 | Viollier AG | Department of Biosystems Science and Engineering, ETH Zürich | Christian Beisel, Sarah Nadeau, Ivan Topolsky, Pedro Ferreira, Philipp Jablonski, Susana Posada-Céspedes, Tobias Schär, Ina Nissen, Natascha Santacroce, Elodie Burcklen, Christiane Beckmann, Maurice Redondo, Olivier Kobel, Christoph Noppen, Sophie Seidel, Noemie Santamaria de Souza, Niko Beerenwinkel, Tanja Stadler |
| EPI_ISL_483669, EPI_ISL_483671, EPI_ISL_483676, EPI_ISL_483678, EPI_ISL_483680, EPI_ISL_483681, EPI_ISL_483683 | University Hospital Zurich | Department of Biosystems Science and Engineering, ETH Zürich | Christian Beisel, Sarah Nadeau, Ivan Topolsky, Pedro Ferreira, Philipp Jablonski, Susana Posada-Céspedes, Tobias Schär, Ina Nissen, Natascha Santacroce, Elodie Burcklen, Julia Martinez-Gomez, Phil Cheng, Mitch Levesque, Philipp Bosshard, Niko Beerenwinkel, Tanja Stadler |
| EPI_ISL_483916, EPI_ISL_483917, EPI_ISL_483918, EPI_ISL_483919, EPI_ISL_483920, EPI_ISL_483921, EPI_ISL_483922, EPI_ISL_483923, EPI_ISL_483924, EPI_ISL_483925, EPI_ISL_483926, EPI_ISL_483927, EPI_ISL_483928, EPI_ISL_483929, EPI_ISL_483930, EPI_ISL_483931, EPI_ISL_483932, EPI_ISL_483933, EPI_ISL_483934, EPI_ISL_483935, EPI_ISL_483936, EPI_ISL_483937, EPI_ISL_483938, EPI_ISL_483939, EPI_ISL_483940, EPI_ISL_483941, EPI_ISL_483942, EPI_ISL_483943, EPI_ISL_483944, EPI_ISL_483945, EPI_ISL_483946, EPI_ISL_483947, EPI_ISL_483948, EPI_ISL_483949, EPI_ISL_483950, EPI_ISL_483951, EPI_ISL_483952, EPI_ISL_483953, EPI_ISL_483954, EPI_ISL_483955, EPI_ISL_483956, EPI_ISL_483957, EPI_ISL_483958, EPI_ISL_483959, EPI_ISL_483960, EPI_ISL_483961, EPI_ISL_483962, EPI_ISL_483963, EPI_ISL_483964, EPI_ISL_483965, EPI_ISL_483966, EPI_ISL_483967, EPI_ISL_483968, EPI_ISL_483969, EPI_ISL_483970, EPI_ISL_483971, EPI_ISL_483972, EPI_ISL_483973, EPI_ISL_483974, EPI_ISL_483975, EPI_ISL_483976, EPI_ISL_483977, EPI_ISL_483978, EPI_ISL_483979, EPI_ISL_483980, EPI_ISL_483981, EPI_ISL_483982, EPI_ISL_483983, EPI_ISL_483984, EPI_ISL_483985, EPI_ISL_483986, EPI_ISL_483987, EPI_ISL_483988, EPI_ISL_483989, EPI_ISL_483990, EPI_ISL_483991, EPI_ISL_483992, EPI_ISL_483993, EPI_ISL_483994, EPI_ISL_483995, EPI_ISL_483996, EPI_ISL_483997, EPI_ISL_483998, EPI_ISL_483999, EPI_ISL_484000, EPI_ISL_484001, EPI_ISL_484002, EPI_ISL_484003, EPI_ISL_484004, EPI_ISL_484005, EPI_ISL_484006, EPI_ISL_484007, EPI_ISL_484008, EPI_ISL_484009, EPI_ISL_484010, EPI_ISL_484011, EPI_ISL_484012, EPI_ISL_484013, EPI_ISL_484014, EPI_ISL_484015, EPI_ISL_484016, EPI_ISL_484017, EPI_ISL_484018, EPI_ISL_484019, EPI_ISL_484020, EPI_ISL_484021, EPI_ISL_484022, EPI_ISL_484023, EPI_ISL_484024, EPI_ISL_484025, EPI_ISL_484026, EPI_ISL_484027, EPI_ISL_484028, EPI_ISL_484029, EPI_ISL_484030, EPI_ISL_484031, EPI_ISL_484032, EPI_ISL_484033, EPI_ISL_484034, EPI_ISL_484035, EPI_ISL_484036, EPI_ISL_484037, EPI_ISL_484038, EPI_ISL_484039, EPI_ISL_484040, EPI_ISL_484041, EPI_ISL_484042, EPI_ISL_484043, EPI_ISL_484044, EPI_ISL_484045, EPI_ISL_484046, EPI_ISL_484047, EPI_ISL_484048, EPI_ISL_484049, EPI_ISL_484050, EPI_ISL_484051, EPI_ISL_484052, EPI_ISL_484053, EPI_ISL_484054, EPI_ISL_484055, EPI_ISL_484056, EPI_ISL_484057, EPI_ISL_484058, EPI_ISL_484059, EPI_ISL_484060, EPI_ISL_484061, EPI_ISL_484062, EPI_ISL_484063, EPI_ISL_484064, EPI_ISL_484065, EPI_ISL_484066, EPI_ISL_484067, EPI_ISL_484068, EPI_ISL_484069, EPI_ISL_484070, EPI_ISL_484071, EPI_ISL_484072, EPI_ISL_484073, EPI_ISL_484074, EPI_ISL_484075, EPI_ISL_484076, EPI_ISL_484077, EPI_ISL_484078, EPI_ISL_484079, EPI_ISL_484080, EPI_ISL_484081, EPI_ISL_484082, EPI_ISL_484083, EPI_ISL_484084, EPI_ISL_484085, EPI_ISL_484086, EPI_ISL_484087, EPI_ISL_484088, EPI_ISL_484089, EPI_ISL_484090, EPI_ISL_484091, EPI_ISL_484092, EPI_ISL_484093, EPI_ISL_484094, EPI_ISL_484095, EPI_ISL_484096, EPI_ISL_484097, EPI_ISL_484098, EPI_ISL_484099, EPI_ISL_484100, EPI_ISL_484101, EPI_ISL_484102, EPI_ISL_484103, EPI_ISL_484104, EPI_ISL_484105, EPI_ISL_484106, EPI_ISL_484107, EPI_ISL_484108, EPI_ISL_484109, EPI_ISL_484110, EPI_ISL_484111, EPI_ISL_484112, EPI_ISL_484113, | | | |

| | | | |
|---|---|---|---|
| EPI_ISL_484114, EPI_ISL_484115, EPI_ISL_484116, EPI_ISL_484117, EPI_ISL_484118, EPI_ISL_484119, EPI_ISL_484120, EPI_ISL_484121, EPI_ISL_484122, EPI_ISL_484123, EPI_ISL_484124, EPI_ISL_484125, EPI_ISL_484126, EPI_ISL_484127, EPI_ISL_484128, EPI_ISL_484129, EPI_ISL_484130, EPI_ISL_484131, EPI_ISL_484132, EPI_ISL_484133, EPI_ISL_484134, EPI_ISL_484135, EPI_ISL_484136, EPI_ISL_484137, EPI_ISL_484138, EPI_ISL_484139, EPI_ISL_484140, EPI_ISL_484141, EPI_ISL_484142, EPI_ISL_484143, EPI_ISL_484144, EPI_ISL_484145, EPI_ISL_484146, EPI_ISL_484147, EPI_ISL_484148, EPI_ISL_484149, EPI_ISL_484150, EPI_ISL_484151, EPI_ISL_484152, EPI_ISL_484153, EPI_ISL_484154, EPI_ISL_484155, EPI_ISL_484156, EPI_ISL_484157, EPI_ISL_484158, EPI_ISL_484159, EPI_ISL_484160, EPI_ISL_484161, EPI_ISL_484162, EPI_ISL_484163, EPI_ISL_484164, EPI_ISL_484165, EPI_ISL_484166, EPI_ISL_484167, EPI_ISL_484168, EPI_ISL_484169, EPI_ISL_484170, EPI_ISL_484171, EPI_ISL_484172, EPI_ISL_484173, EPI_ISL_484174, EPI_ISL_484175, EPI_ISL_484176, EPI_ISL_484177, EPI_ISL_484178, EPI_ISL_484179, EPI_ISL_484180, EPI_ISL_484181, EPI_ISL_484182, EPI_ISL_484183, EPI_ISL_484184, EPI_ISL_484185, EPI_ISL_484186, EPI_ISL_484187, EPI_ISL_484188, EPI_ISL_484189, EPI_ISL_484190, EPI_ISL_484191, EPI_ISL_484192, EPI_ISL_484193, EPI_ISL_484194, EPI_ISL_484195, EPI_ISL_484196, EPI_ISL_484197, EPI_ISL_484198, EPI_ISL_484199, EPI_ISL_484200, EPI_ISL_484201, EPI_ISL_484202, EPI_ISL_484203, EPI_ISL_484204, EPI_ISL_484205, EPI_ISL_484206, EPI_ISL_484207, EPI_ISL_484208, EPI_ISL_484209, EPI_ISL_484210, EPI_ISL_484211, EPI_ISL_484212, EPI_ISL_484213, EPI_ISL_484214, EPI_ISL_484215, EPI_ISL_484216, EPI_ISL_484217 | | | |
| see above | Centre for Clinical Infection and Diagnostics Research and Genomics Innovation Unit, Guy's and St. Thomas' NHS Trust | COVID-19 Genomics UK (COG-UK) Consortium | Chloe Fisher, Luke Snell, Penny Cliff, Rahul Batra, Jonathan Edgeworth, Ali Raza Awan |
| EPI_ISL_484685, EPI_ISL_484686, EPI_ISL_484687, EPI_ISL_484688, EPI_ISL_484689, EPI_ISL_484690, EPI_ISL_484691 | Wales Specialist Virology Centre Sequencing lab: Pathogen Genomics Unit | COVID-19 Genomics UK (COG-UK) Consortium | Catherine Moore, Johnathan Evans, Laura Gifford, Malorie Perry, Simon Cottrell, Angela Marchbank, Alec Birchley, Alexander Adams, Amy Gaskin, Bree Gatica-Wilcox, Jason Coombes, Joel Southgate, Lauren Gilbert, Lee Graham, Nicole Pacchiarini, Sara Kumziene-Summerhayes, Sarah Taylor, Sophie Jones, Sara Rey, Matthew Bull, Joanne Watkins, Sally Corden, Tom Connor |
| EPI_ISL_486382 | District Surveillence Unit | Department of Neurovirology, National Institute of Mental Health and Neuroscience (NIMHANS) | Chitra Pattabiraman, Vijayalakshmi Reddy, Harsha PK, Risha Rasheed, Shafeeq S Hameed, Manjunatha Venkataswamy, Anita Desai, Ravi Vasanthapuram |
| EPI_ISL_486383 | CV Raman Hospital | Department of Neurovirology, National Institute of Mental Health and Neuroscience (NIMHANS) | Chitra Pattabiraman, Vijayalakshmi Reddy, Harsha PK, Risha Rasheed, Shafeeq S Hameed, Manjunatha Venkataswamy, Anita Desai, Ravi Vasanthapuram |
| EPI_ISL_486385, EPI_ISL_486386 | Victoria Hospital | Department of Neurovirology, National Institute of Mental Health and Neuroscience (NIMHANS) | Chitra Pattabiraman, Vijayalakshmi Reddy, Harsha PK, Risha Rasheed, Shafeeq S Hameed, Manjunatha Venkataswamy, Anita Desai, Ravi Vasanthapuram |
| EPI_ISL_486387, EPI_ISL_486388, EPI_ISL_486389 | DH | Department of Neurovirology, National Institute of Mental Health and Neuroscience (NIMHANS) | Chitra Pattabiraman, Vijayalakshmi Reddy, Harsha PK, Risha Rasheed, Shafeeq S Hameed, Manjunatha Venkataswamy, Anita Desai, Ravi Vasanthapuram |
| EPI_ISL_486392 | Victoria Hospital | Department of Neurovirology, National Institute of Mental Health and Neuroscience (NIMHANS) | Chitra Pattabiraman, Vijayalakshmi Reddy, Harsha PK, Risha Rasheed, Shafeeq S Hameed, Manjunatha Venkataswamy, Anita Desai, Ravi Vasanthapuram |
| EPI_ISL_486394 | MIMS | Department of Neurovirology, National Institute of Mental Health and Neuroscience (NIMHANS) | Chitra Pattabiraman, Vijayalakshmi Reddy, Harsha PK, Risha Rasheed, Shafeeq S Hameed, Manjunatha Venkataswamy, Anita Desai, Ravi Vasanthapuram |
| EPI_ISL_486395 | BIMS | Department of Neurovirology, National Institute of Mental Health and Neuroscience (NIMHANS) | Chitra Pattabiraman, Vijayalakshmi Reddy, Harsha PK, Risha Rasheed, Shafeeq S Hameed, Manjunatha Venkataswamy, Anita Desai, Ravi Vasanthapuram |
| EPI_ISL_486396 | Jayanagar General Hospital to Victoria Hospital | Department of Neurovirology, National Institute of Mental Health and Neuroscience (NIMHANS) | Chitra Pattabiraman, Vijayalakshmi Reddy, Harsha PK, Risha Rasheed, Shafeeq S Hameed, Manjunatha Venkataswamy, Anita Desai, Ravi Vasanthapuram |
| EPI_ISL_486397 | KC General Hospital | Department of Neurovirology, National Institute of Mental Health and Neuroscience (NIMHANS) | Chitra Pattabiraman, Vijayalakshmi Reddy, Harsha PK, Risha Rasheed, Shafeeq S Hameed, Manjunatha Venkataswamy, Anita Desai, Ravi Vasanthapuram |
| EPI_ISL_486398, EPI_ISL_486399 | MIMS | Department of Neurovirology, National Institute of Mental Health and Neuroscience (NIMHANS) | Chitra Pattabiraman, Vijayalakshmi Reddy, Harsha PK, Risha Rasheed, Shafeeq S Hameed, Manjunatha Venkataswamy, Anita Desai, Ravi Vasanthapuram |
| EPI_ISL_486400 | Victoria Hospital | Department of Neurovirology, National Institute of Mental Health and Neuroscience (NIMHANS) | Chitra Pattabiraman, Vijayalakshmi Reddy, Harsha PK, Risha Rasheed, Shafeeq S Hameed, Manjunatha Venkataswamy, Anita Desai, Ravi Vasanthapuram |
| EPI_ISL_486401, EPI_ISL_486402, EPI_ISL_486403 | DH | Department of Neurovirology, National Institute of Mental Health and Neuroscience (NIMHANS) | Chitra Pattabiraman, Vijayalakshmi Reddy, Harsha PK, Risha Rasheed, Shafeeq S Hameed, Manjunatha Venkataswamy, Anita Desai, Ravi Vasanthapuram |
| EPI_ISL_486404 | Victoria Hospital | Department of Neurovirology, National Institute of Mental Health and Neuroscience (NIMHANS) | Chitra Pattabiraman, Vijayalakshmi Reddy, Harsha PK, Risha Rasheed, Shafeeq S Hameed, Manjunatha Venkataswamy, Anita Desai, Ravi Vasanthapuram |
| EPI_ISL_486405, EPI_ISL_486406, EPI_ISL_486407, EPI_ISL_486408, EPI_ISL_486409 | DH | Department of Neurovirology, National Institute of Mental Health and Neuroscience (NIMHANS) | Chitra Pattabiraman, Vijayalakshmi Reddy, Harsha PK, Risha Rasheed, Shafeeq S Hameed, Manjunatha Venkataswamy, Anita Desai, Ravi Vasanthapuram |
| EPI_ISL_486481, EPI_ISL_486499, EPI_ISL_486510, EPI_ISL_486515, EPI_ISL_486518, EPI_ISL_486527, EPI_ISL_486531, EPI_ISL_486535 | Viollier AG | Department of Biosystems Science and Engineering, ETH Zürich | Christian Beisel, Sarah Nadeau, Ivan Topolsky, Pedro Ferreira, Philipp Jablonski, Susana Posada-Céspedes, Tobias Schär, Ina Nissen, Natascha Santacroce, Elodie Burcklen, Christiane Beckmann, Maurice Redondo, Olivier Kobel, Christoph Noppen, Sophie Seidel, Noemie Santamaria de Souza, Niko Beerenwinkel, Tanja Stadler |
| EPI_ISL_486881 | CV Raman Hospital | Department of Neurovirology, National Institute of Mental Health and Neuroscience (NIMHANS) | Chitra Pattabiraman, Vijayalakshmi Reddy, Harsha PK, Risha Rasheed, Shafeeq S Hameed, Manjunatha Venkataswamy, Anita Desai, Ravi Vasanthapuram |
| EPI_ISL_488003, EPI_ISL_488004, EPI_ISL_488005, EPI_ISL_488006, EPI_ISL_488007, EPI_ISL_488008, EPI_ISL_488009, EPI_ISL_488010, EPI_ISL_488011, EPI_ISL_488012, EPI_ISL_488013, EPI_ISL_488014, EPI_ISL_488015, EPI_ISL_488016, EPI_ISL_488017, EPI_ISL_488018, EPI_ISL_488019, EPI_ISL_488020, EPI_ISL_488021, EPI_ISL_488022, EPI_ISL_488023, EPI_ISL_488024, EPI_ISL_488025, EPI_ISL_488026, EPI_ISL_488027, EPI_ISL_488028, EPI_ISL_488029, EPI_ISL_488030, EPI_ISL_488031, EPI_ISL_488032, EPI_ISL_488033, EPI_ISL_488034, EPI_ISL_488035, EPI_ISL_488036, EPI_ISL_488037, EPI_ISL_488038, EPI_ISL_488039, EPI_ISL_488040, EPI_ISL_488041, EPI_ISL_488042, EPI_ISL_488043, EPI_ISL_488044, EPI_ISL_488045, EPI_ISL_488046, EPI_ISL_488047, EPI_ISL_488048, EPI_ISL_488049, EPI_ISL_488050, EPI_ISL_488051, EPI_ISL_488052, EPI_ISL_488053, EPI_ISL_488054, EPI_ISL_488055, EPI_ISL_488056, EPI_ISL_488057, EPI_ISL_488058, EPI_ISL_488059, EPI_ISL_488060, EPI_ISL_488061, EPI_ISL_488062, EPI_ISL_488063, EPI_ISL_488064, EPI_ISL_488065, EPI_ISL_488066, EPI_ISL_488067, EPI_ISL_488068, EPI_ISL_488069, EPI_ISL_488070, EPI_ISL_488071, EPI_ISL_488072, EPI_ISL_488073, EPI_ISL_488074, EPI_ISL_488075, EPI_ISL_488076, EPI_ISL_488077, EPI_ISL_488078, EPI_ISL_488079, EPI_ISL_488080, EPI_ISL_488081, EPI_ISL_488082, EPI_ISL_488083, EPI_ISL_488084, EPI_ISL_488085, EPI_ISL_488086, EPI_ISL_488087, EPI_ISL_488088, EPI_ISL_488089, EPI_ISL_488090, EPI_ISL_488091, EPI_ISL_488092, EPI_ISL_488093, EPI_ISL_488094, EPI_ISL_488095, EPI_ISL_488096, EPI_ISL_488097, EPI_ISL_488098, EPI_ISL_488099, EPI_ISL_488100, EPI_ISL_488101, EPI_ISL_488102, EPI_ISL_488103, EPI_ISL_488104, EPI_ISL_488105, EPI_ISL_488106, EPI_ISL_488107, EPI_ISL_488108, EPI_ISL_488109, EPI_ISL_488110, EPI_ISL_488111, EPI_ISL_488112, EPI_ISL_488113, EPI_ISL_488114, EPI_ISL_488115, EPI_ISL_488116, EPI_ISL_488117, EPI_ISL_488118, EPI_ISL_488119, EPI_ISL_488120, EPI_ISL_488121, EPI_ISL_488122, EPI_ISL_488123, EPI_ISL_488124, EPI_ISL_488125, EPI_ISL_488126, EPI_ISL_488127, EPI_ISL_488128, EPI_ISL_488129, EPI_ISL_488130, EPI_ISL_488131, EPI_ISL_488132, EPI_ISL_488133, EPI_ISL_488134, EPI_ISL_488135, EPI_ISL_488136, EPI_ISL_488137, EPI_ISL_488138, EPI_ISL_488139, EPI_ISL_488140, EPI_ISL_488141, EPI_ISL_488142, EPI_ISL_488143, EPI_ISL_488144, EPI_ISL_488145, EPI_ISL_488146, EPI_ISL_488147, EPI_ISL_488148, EPI_ISL_488149, EPI_ISL_488150, EPI_ISL_488151, EPI_ISL_488152, EPI_ISL_488153, EPI_ISL_488154, EPI_ISL_488155, EPI_ISL_488156, EPI_ISL_488157, EPI_ISL_488158, EPI_ISL_488159, EPI_ISL_488160, EPI_ISL_488161, EPI_ISL_488162, EPI_ISL_488163, EPI_ISL_488164, EPI_ISL_488165, EPI_ISL_488166, EPI_ISL_488167, EPI_ISL_488168, EPI_ISL_488169, EPI_ISL_488170, EPI_ISL_488171, EPI_ISL_488172, EPI_ISL_488173, EPI_ISL_488174, EPI_ISL_488175, EPI_ISL_488176, EPI_ISL_488177, EPI_ISL_488178, EPI_ISL_488179, EPI_ISL_488180, EPI_ISL_488181, EPI_ISL_488182, EPI_ISL_488183, EPI_ISL_488184, EPI_ISL_488185, EPI_ISL_488186, EPI_ISL_488456, EPI_ISL_488457, EPI_ISL_488458, EPI_ISL_488459, EPI_ISL_488460, EPI_ISL_488461, EPI_ISL_488462, EPI_ISL_488463, EPI_ISL_488464, EPI_ISL_488465, EPI_ISL_488466, EPI_ISL_488467, EPI_ISL_488468, EPI_ISL_488469, EPI_ISL_488470, EPI_ISL_488471, EPI_ISL_488472, EPI_ISL_488473, EPI_ISL_488474, EPI_ISL_488475, EPI_ISL_488476, EPI_ISL_488477, EPI_ISL_488478, EPI_ISL_488479, EPI_ISL_488480, EPI_ISL_488481, EPI_ISL_488482, EPI_ISL_488483, EPI_ISL_488484, EPI_ISL_488485, EPI_ISL_488486, EPI_ISL_488487, EPI_ISL_488488, EPI_ISL_488489, EPI_ISL_488490, EPI_ISL_488491, EPI_ISL_488492, EPI_ISL_488493, EPI_ISL_488494, EPI_ISL_488495, EPI_ISL_488496, EPI_ISL_488497, EPI_ISL_488498, EPI_ISL_488499, EPI_ISL_488500, EPI_ISL_488501, EPI_ISL_488502, EPI_ISL_488503, EPI_ISL_488504, EPI_ISL_488505, EPI_ISL_488506, EPI_ISL_488507, EPI_ISL_488508, EPI_ISL_488509, EPI_ISL_488510, EPI_ISL_488511, EPI_ISL_488512, EPI_ISL_488513, EPI_ISL_488514, EPI_ISL_488515, EPI_ISL_488516, EPI_ISL_488517, EPI_ISL_488518, EPI_ISL_488519, EPI_ISL_488520, EPI_ISL_488521, EPI_ISL_488522, EPI_ISL_488523, EPI_ISL_488524, EPI_ISL_488525, EPI_ISL_488526, EPI_ISL_488527, EPI_ISL_488528, EPI_ISL_488529, EPI_ISL_488530, EPI_ISL_488531, EPI_ISL_488532, EPI_ISL_488533, EPI_ISL_488534, EPI_ISL_488535, EPI_ISL_488536, EPI_ISL_488537, EPI_ISL_488538, EPI_ISL_488539, EPI_ISL_488540, EPI_ISL_488541, EPI_ISL_488542, EPI_ISL_488543, EPI_ISL_488544, EPI_ISL_488545, EPI_ISL_488546, EPI_ISL_488547, EPI_ISL_488548, EPI_ISL_488549, EPI_ISL_488550, EPI_ISL_488551, EPI_ISL_488552, EPI_ISL_488553, EPI_ISL_488554, EPI_ISL_488555, EPI_ISL_488556, EPI_ISL_488557, EPI_ISL_488558, EPI_ISL_488559, EPI_ISL_488560, EPI_ISL_488561, EPI_ISL_488562, EPI_ISL_488563, EPI_ISL_488564, EPI_ISL_488565, EPI_ISL_488566, EPI_ISL_488567, EPI_ISL_488568, EPI_ISL_488569, EPI_ISL_488570, EPI_ISL_488571, EPI_ISL_488572, EPI_ISL_488573, EPI_ISL_488574, EPI_ISL_488575, EPI_ISL_488576, EPI_ISL_488577, EPI_ISL_488578, EPI_ISL_488579, EPI_ISL_488580, EPI_ISL_488581, EPI_ISL_488582, EPI_ISL_488583, EPI_ISL_488584, EPI_ISL_488585, EPI_ISL_488586, EPI_ISL_488587, EPI_ISL_488588, EPI_ISL_488589, EPI_ISL_488590, EPI_ISL_488591, EPI_ISL_488592, EPI_ISL_488593, EPI_ISL_488594, EPI_ISL_488595, EPI_ISL_488596, EPI_ISL_488597, EPI_ISL_488598, EPI_ISL_488599, EPI_ISL_488600, EPI_ISL_488601, EPI_ISL_488602, EPI_ISL_488603, EPI_ISL_488604, EPI_ISL_488605, EPI_ISL_488606, EPI_ISL_488607, EPI_ISL_488608, EPI_ISL_488609, EPI_ISL_488610, EPI_ISL_488611, EPI_ISL_488612, EPI_ISL_488613, EPI_ISL_488614, EPI_ISL_488615, EPI_ISL_488616, EPI_ISL_488617, EPI_ISL_488618, EPI_ISL_488619, EPI_ISL_488620, EPI_ISL_488621, EPI_ISL_488622, EPI_ISL_488623, EPI_ISL_488624, EPI_ISL_488625, EPI_ISL_488626, EPI_ISL_488627, EPI_ISL_488628, EPI_ISL_488629, EPI_ISL_488630, EPI_ISL_488631, EPI_ISL_488632, EPI_ISL_488633, EPI_ISL_488634, EPI_ISL_488635, EPI_ISL_488636, EPI_ISL_488637, EPI_ISL_488638, EPI_ISL_488639, EPI_ISL_488640, EPI_ISL_488641, EPI_ISL_488642, EPI_ISL_488643, EPI_ISL_488644, EPI_ISL_488645, EPI_ISL_488646, EPI_ISL_488647, EPI_ISL_488648, EPI_ISL_488649, EPI_ISL_488650, EPI_ISL_488651, EPI_ISL_488652, EPI_ISL_488653, EPI_ISL_488654, EPI_ISL_488655, EPI_ISL_488656, EPI_ISL_488657, EPI_ISL_488658, EPI_ISL_488659, EPI_ISL_488660, EPI_ISL_488661, EPI_ISL_488662, EPI_ISL_488663, EPI_ISL_488664, EPI_ISL_488665, EPI_ISL_488666, EPI_ISL_488667, EPI_ISL_488668, EPI_ISL_488669, EPI_ISL_488670, EPI_ISL_488671, EPI_ISL_488672, EPI_ISL_488673, EPI_ISL_488674, EPI_ISL_488675, EPI_ISL_488676, EPI_ISL_488677, EPI_ISL_488678, EPI_ISL_488679, EPI_ISL_488680, EPI_ISL_488681, EPI_ISL_488682, EPI_ISL_488683, EPI_ISL_488684, EPI_ISL_488685, EPI_ISL_488686, EPI_ISL_488687, EPI_ISL_488688, EPI_ISL_488689, EPI_ISL_488690, EPI_ISL_488691, EPI_ISL_488692, EPI_ISL_488693, EPI_ISL_488694, EPI_ISL_488695, EPI_ISL_488696, EPI_ISL_488697, EPI_ISL_488698, EPI_ISL_488699, EPI_ISL_488700, EPI_ISL_488701, EPI_ISL_488702, EPI_ISL_488703, EPI_ISL_488704, EPI_ISL_488705, EPI_ISL_488706, EPI_ISL_488707, EPI_ISL_488708, EPI_ISL_488709, EPI_ISL_488710, EPI_ISL_488711, EPI_ISL_488712, EPI_ISL_488713, EPI_ISL_488714, EPI_ISL_488715, EPI_ISL_488716, EPI_ISL_488717, EPI_ISL_488718, EPI_ISL_488719, EPI_ISL_488720, EPI_ISL_488721, EPI_ISL_488722, EPI_ISL_488723, EPI_ISL_488724, EPI_ISL_488725, EPI_ISL_488726, EPI_ISL_488727, EPI_ISL_488728, EPI_ISL_488729, EPI_ISL_488730, EPI_ISL_488731, EPI_ISL_488732, EPI_ISL_488733, EPI_ISL_488734, EPI_ISL_488735, EPI_ISL_488736, EPI_ISL_488737, EPI_ISL_488738, EPI_ISL_488739, EPI_ISL_488740, EPI_ISL_488741, EPI_ISL_488742, EPI_ISL_488743, EPI_ISL_488744, EPI_ISL_488745, EPI_ISL_488746, EPI_ISL_488747, EPI_ISL_488748, EPI_ISL_488749, EPI_ISL_488750, EPI_ISL_488751, EPI_ISL_488752, EPI_ISL_488753, EPI_ISL_488754, EPI_ISL_488755, EPI_ISL_488756, EPI_ISL_488757, EPI_ISL_488758, EPI_ISL_488759, EPI_ISL_488760, EPI_ISL_488761, EPI_ISL_488762, EPI_ISL_488763, EPI_ISL_488764, EPI_ISL_488765, EPI_ISL_488766, EPI_ISL_488767, EPI_ISL_488768, EPI_ISL_488769, EPI_ISL_488770, EPI_ISL_488771, EPI_ISL_488772, EPI_ISL_488773, EPI_ISL_488774, EPI_ISL_488775, EPI_ISL_488776, EPI_ISL_488777, EPI_ISL_488778, EPI_ISL_488779, EPI_ISL_488780, EPI_ISL_488781, EPI_ISL_488782, EPI_ISL_488783, EPI_ISL_488784, EPI_ISL_488785, EPI_ISL_488786, EPI_ISL_488787, EPI_ISL_488788, EPI_ISL_488789, EPI_ISL_488790, EPI_ISL_488791, EPI_ISL_488792, EPI_ISL_488793, EPI_ISL_488794, EPI_ISL_488795, EPI_ISL_488796, EPI_ISL_488797, EPI_ISL_488798, EPI_ISL_488799, EPI_ISL_488800, EPI_ISL_488801, EPI_ISL_488802, EPI_ISL_488803, EPI_ISL_488804, EPI_ISL_488805, EPI_ISL_488806, EPI_ISL_488807, EPI_ISL_488808, EPI_ISL_488809, EPI_ISL_488810, EPI_ISL_488811, EPI_ISL_488812, EPI_ISL_488813, EPI_ISL_488814, EPI_ISL_488815, EPI_ISL_488816, EPI_ISL_488817, EPI_ISL_488818, EPI_ISL_488819, EPI_ISL_488820, EPI_ISL_488821, EPI_ISL_488822, EPI_ISL_488823, EPI_ISL_488824, EPI_ISL_488825, EPI_ISL_488826, EPI_ISL_488827, EPI_ISL_488828, EPI_ISL_488829, | | | |

| | | | |
|---|---|---|---|
| EPI_ISL_488830, EPI_ISL_488831, EPI_ISL_488832, EPI_ISL_488833, EPI_ISL_488834, EPI_ISL_488835, EPI_ISL_488836, EPI_ISL_489064, EPI_ISL_489065, EPI_ISL_489066, EPI_ISL_489067, EPI_ISL_489068, EPI_ISL_489069, EPI_ISL_489070, EPI_ISL_489071, EPI_ISL_489072, EPI_ISL_489073, EPI_ISL_489074, EPI_ISL_489075, EPI_ISL_489076, EPI_ISL_489077, EPI_ISL_489078, EPI_ISL_489079, EPI_ISL_489080, EPI_ISL_489081, EPI_ISL_489082, EPI_ISL_489083, EPI_ISL_489084, EPI_ISL_489085, EPI_ISL_489086, EPI_ISL_489087, EPI_ISL_489088, EPI_ISL_489089, EPI_ISL_489090, EPI_ISL_489091, EPI_ISL_489092, EPI_ISL_489093, EPI_ISL_489094, EPI_ISL_489095, EPI_ISL_489096, EPI_ISL_489097, EPI_ISL_489098, EPI_ISL_489099, EPI_ISL_489100, EPI_ISL_489101, EPI_ISL_489102, EPI_ISL_489103, EPI_ISL_489104, EPI_ISL_489105, EPI_ISL_489106, EPI_ISL_489107, EPI_ISL_489108, EPI_ISL_489109, EPI_ISL_489110, EPI_ISL_489111, EPI_ISL_489112, EPI_ISL_489113, EPI_ISL_489114, EPI_ISL_489115, EPI_ISL_489116, EPI_ISL_489117, EPI_ISL_489118, EPI_ISL_489119, EPI_ISL_489121, EPI_ISL_489122, EPI_ISL_489123, EPI_ISL_489124, EPI_ISL_489125, EPI_ISL_489126, EPI_ISL_489127, EPI_ISL_489128, EPI_ISL_489129, EPI_ISL_489130, EPI_ISL_489131, EPI_ISL_489132, EPI_ISL_489133, EPI_ISL_489134, EPI_ISL_489135, EPI_ISL_489136, EPI_ISL_489137, EPI_ISL_489138, EPI_ISL_489139, EPI_ISL_489140, EPI_ISL_489141, EPI_ISL_489142, EPI_ISL_489143, EPI_ISL_489144, EPI_ISL_489145, EPI_ISL_489146, EPI_ISL_489147, EPI_ISL_489148, EPI_ISL_489149, EPI_ISL_489150, EPI_ISL_489151, EPI_ISL_489152, EPI_ISL_489153, EPI_ISL_489154, EPI_ISL_489155 | | | |
| see above | NU-OMICS DNA Sequencing research facility, Northumbria University | Wellcome Sanger Institute for the COVID-19 Genomics UK (COG-UK) consortium | Chris Duncan, Sheia Waugh, Shirelle Burton-Fanning, Gary Eltringham, Jennifer Collins, Brendan Payne, Yusri Taha, Emma Swindells, Jane Greenaway, Edward Barton, Garren Scott, Debra Padgett, Clive Graham, Sarah Essex, Steve Ligget, Paul Baker, Lynn Dover, Wen Yew, Gary Black, John Allan, Joshua Loh, Greg Young, Matthew Bashton, Andrew Nelson, Darren Smith and Alex Alderton, Roberto Amato, Sonia Goncalves, Ewan Harrison, David K. Jackson, Ian Johnston, Dominic Kwiatkowski, Cordelia Langford, John Sillitoe on behalf of the Wellcome Sanger Institute COVID-19 Surveillance Team (http://www.sanger.ac.uk/covid-team) |
| EPI_ISL_489961, EPI_ISL_489967 | Viollier AG | Department of Biosystems Science and Engineering, ETH Zürich | Christian Beisel, Sarah Nadeau, Ivan Topolsky, Pedro Ferreira, Philipp Jablonski, Susana Posada-Céspedes, Tobias Schär, Ina Nissen, Natascha Santacroce, Elodie Burcklen, Christiane Beckmann, Maurice Redondo, Olivier Kobel, Christoph Noppen, Sophie Seidel, Noemie Santamaria de Souza, Niko Beerenwinkel, Tanja Stadler |
| EPI_ISL_490710, EPI_ISL_490711, EPI_ISL_490712, EPI_ISL_490713, EPI_ISL_490714, EPI_ISL_490715, EPI_ISL_490716, EPI_ISL_490717, EPI_ISL_490718, EPI_ISL_490719, EPI_ISL_490720, EPI_ISL_490721, EPI_ISL_490722, EPI_ISL_490723, EPI_ISL_490724, EPI_ISL_490725, EPI_ISL_490726, EPI_ISL_490727, EPI_ISL_490728, EPI_ISL_490729, EPI_ISL_490730, EPI_ISL_490731, EPI_ISL_490732, EPI_ISL_490733, EPI_ISL_490734, EPI_ISL_490735, EPI_ISL_490736, EPI_ISL_490737, EPI_ISL_490738, EPI_ISL_490739, EPI_ISL_490740, EPI_ISL_490741, EPI_ISL_490742, EPI_ISL_490743, EPI_ISL_490744, EPI_ISL_490745, EPI_ISL_490746, EPI_ISL_490747, EPI_ISL_490748, EPI_ISL_490749, EPI_ISL_490750, EPI_ISL_490751, EPI_ISL_490752, EPI_ISL_490753, EPI_ISL_490754, EPI_ISL_490755, EPI_ISL_490756, EPI_ISL_490757, EPI_ISL_490758, EPI_ISL_490759, EPI_ISL_490760, EPI_ISL_490761, EPI_ISL_490762, EPI_ISL_490763, EPI_ISL_490764, EPI_ISL_490765, EPI_ISL_490766, EPI_ISL_490767, EPI_ISL_490768, EPI_ISL_490769, EPI_ISL_490770, EPI_ISL_490771, EPI_ISL_490772, EPI_ISL_490773, EPI_ISL_490774, EPI_ISL_490775, EPI_ISL_490776, EPI_ISL_490777, EPI_ISL_490778, EPI_ISL_490779, EPI_ISL_490780, EPI_ISL_490781, EPI_ISL_490782, EPI_ISL_490783, EPI_ISL_490784, EPI_ISL_490785, EPI_ISL_490786, EPI_ISL_490787, EPI_ISL_490788, EPI_ISL_490789, EPI_ISL_490790, EPI_ISL_490791, EPI_ISL_490792, EPI_ISL_490793, EPI_ISL_490794, EPI_ISL_490795, EPI_ISL_490796, EPI_ISL_490797, EPI_ISL_490798, EPI_ISL_490799, EPI_ISL_490800, EPI_ISL_490801, EPI_ISL_490802, EPI_ISL_490803, EPI_ISL_490804, EPI_ISL_490805, EPI_ISL_490806, EPI_ISL_490807, EPI_ISL_490808, EPI_ISL_490809, EPI_ISL_490810, EPI_ISL_490811, EPI_ISL_490812, EPI_ISL_490813, EPI_ISL_490814, EPI_ISL_490815, EPI_ISL_490816, EPI_ISL_490817, EPI_ISL_490818, EPI_ISL_490819, EPI_ISL_490820, EPI_ISL_490821, EPI_ISL_490822, EPI_ISL_490823, EPI_ISL_490824, EPI_ISL_490825, EPI_ISL_490826, EPI_ISL_490827, EPI_ISL_490828, EPI_ISL_490829, EPI_ISL_490830, EPI_ISL_490831, EPI_ISL_490832, EPI_ISL_490833, EPI_ISL_490834, EPI_ISL_490835, EPI_ISL_490836, EPI_ISL_490837, EPI_ISL_490838, EPI_ISL_490839, EPI_ISL_490840, EPI_ISL_490841, EPI_ISL_490842, EPI_ISL_490843, EPI_ISL_490844, EPI_ISL_490845, EPI_ISL_490846, EPI_ISL_490847, EPI_ISL_490848, EPI_ISL_490849, EPI_ISL_490850, EPI_ISL_490851, EPI_ISL_490852, EPI_ISL_490853, EPI_ISL_490854, EPI_ISL_490855, EPI_ISL_490856, EPI_ISL_490857, EPI_ISL_490858, EPI_ISL_490859, EPI_ISL_490860, EPI_ISL_490861, EPI_ISL_490862, EPI_ISL_490863, EPI_ISL_490864, EPI_ISL_490865, EPI_ISL_490866, EPI_ISL_490867 | | | |
| see above | Wales Specialist Virology Centre Sequencing lab: Pathogen Genomics Unit | COVID-19 Genomics UK (COG-UK) Consortium | Catherine Moore, Johnathan Evans, Laura Gifford, Malorie Perry, Simon Cottrell, Angela Marchbank, Alec Birchley, Alexander Adams, Amy Gaskin, Bree Gatica-Wilcox, Jason Coombes, Joel Southgate, Lauren Gilbert, Lee Graham, Nicole Pacchiarini, Sara Kumziene-Summerhayes, Sarah Taylor, Sophie Jones, Sara Rey, Matthew Bull, Joanne Watkins, Sally Corden, Tom Connor |
| EPI_ISL_491202, EPI_ISL_491203, EPI_ISL_491204, EPI_ISL_491205, EPI_ISL_491206, EPI_ISL_491207, EPI_ISL_491208, EPI_ISL_491209, EPI_ISL_491210, EPI_ISL_491211, EPI_ISL_491212, EPI_ISL_491213, EPI_ISL_491214, EPI_ISL_491215, EPI_ISL_491216, EPI_ISL_491217, EPI_ISL_491218, EPI_ISL_491219, EPI_ISL_491220, EPI_ISL_491221, EPI_ISL_491222, EPI_ISL_491223, EPI_ISL_491224, EPI_ISL_491225, EPI_ISL_491226, EPI_ISL_491227, EPI_ISL_491228, EPI_ISL_491229, EPI_ISL_491230 | | | |
| see above | Instituto Gulbenkian de Ciência | Instituto Gulbenkian de Ciência | Cathy Paulino, Joao Sobral, Susana Ladeiro, João Costa, Ricardo Leite |
| EPI_ISL_492049, EPI_ISL_492050, EPI_ISL_492051, EPI_ISL_492052, EPI_ISL_492053, EPI_ISL_492054, EPI_ISL_492055, EPI_ISL_492056, EPI_ISL_492057, EPI_ISL_492058, EPI_ISL_492059, EPI_ISL_492060, EPI_ISL_492061, EPI_ISL_492062, EPI_ISL_492063, EPI_ISL_492087 | | | |
| see above | Alaska State Virology Laboratory | Alaska State Virology Laboratory | Chen J et al with Pathogenomics group Dagdag R, Redlinger M, Milton E, George W, Kovalenko A, Drown DM, Bortz E |
| EPI_ISL_492207, EPI_ISL_492208, EPI_ISL_492209, EPI_ISL_492210, EPI_ISL_492450, EPI_ISL_492451, EPI_ISL_492452, EPI_ISL_492453, EPI_ISL_492454, EPI_ISL_492455, EPI_ISL_492456, EPI_ISL_492458, EPI_ISL_492459, EPI_ISL_492460, EPI_ISL_492461, EPI_ISL_492462, EPI_ISL_492463, EPI_ISL_492464, EPI_ISL_492465, EPI_ISL_492466, EPI_ISL_492468, EPI_ISL_492469, EPI_ISL_492470, EPI_ISL_492471, EPI_ISL_492472, EPI_ISL_492473, EPI_ISL_492474, EPI_ISL_492475, EPI_ISL_492476, EPI_ISL_492477, EPI_ISL_492478, EPI_ISL_492479, EPI_ISL_492480, EPI_ISL_492481, EPI_ISL_492482, EPI_ISL_492483, EPI_ISL_492484, EPI_ISL_492485, EPI_ISL_492486, EPI_ISL_492487, EPI_ISL_492488, EPI_ISL_492489, EPI_ISL_492490, EPI_ISL_492491, EPI_ISL_492492, EPI_ISL_492493, EPI_ISL_492494, EPI_ISL_492916, EPI_ISL_492917, EPI_ISL_492918, EPI_ISL_492919, EPI_ISL_492920, EPI_ISL_492921, EPI_ISL_492922, EPI_ISL_492923, EPI_ISL_492924, EPI_ISL_492925, EPI_ISL_492926, EPI_ISL_492927, EPI_ISL_492928, EPI_ISL_492930, EPI_ISL_492932, EPI_ISL_492933, EPI_ISL_492934, EPI_ISL_492935, EPI_ISL_492936, EPI_ISL_492937, EPI_ISL_492938, EPI_ISL_492939, EPI_ISL_492940, EPI_ISL_492941, EPI_ISL_492942, EPI_ISL_492943, EPI_ISL_492944, EPI_ISL_492945, EPI_ISL_492946, EPI_ISL_492947, EPI_ISL_492948, EPI_ISL_492949, EPI_ISL_492950, EPI_ISL_492951, EPI_ISL_492952, EPI_ISL_492953, EPI_ISL_492954, EPI_ISL_492955, EPI_ISL_492956, EPI_ISL_492957, EPI_ISL_492958, EPI_ISL_492959, EPI_ISL_492960, EPI_ISL_492961, EPI_ISL_492962, EPI_ISL_492963, EPI_ISL_492964, EPI_ISL_492965, EPI_ISL_492966, EPI_ISL_492967, EPI_ISL_492968, EPI_ISL_492969, EPI_ISL_492970, EPI_ISL_492971, EPI_ISL_492972, EPI_ISL_492973, EPI_ISL_492974, EPI_ISL_492975, EPI_ISL_492976, EPI_ISL_492977 | | | |
| see above | NU-OMICS DNA Sequencing research facility, Northumbria University | Wellcome Sanger Institute for the COVID-19 Genomics UK (COG-UK) consortium | Chris Duncan, Sheia Waugh, Shirelle Burton-Fanning, Gary Eltringham, Jennifer Collins, Brendan Payne, Yusri Taha, Emma Swindells, Jane Greenaway, Edward Barton, Garren Scott, Debra Padgett, Clive Graham, Sarah Essex, Steve Ligget, Paul Baker, Lynn Dover, Wen Yew, Gary Black, John Allan, Joshua Loh, Greg Young, Matthew Bashton, Andrew Nelson, Darren Smith and Alex Alderton, Roberto Amato, Sonia Goncalves, Ewan Harrison, David K. Jackson, Ian Johnston, Dominic Kwiatkowski, Cordelia Langford, John Sillitoe on behalf of the Wellcome Sanger Institute COVID-19 Surveillance Team (http://www.sanger.ac.uk/covid-team) |
| EPI_ISL_492978, EPI_ISL_492979 | Department of Laboratory Medicine Tan Tock Seng Hospital | Department of Laboratory Medicine Tan Tock Seng Hospital | Chen YYC, Zair X, Li C, Tang WY, Maurer-Stroh S, Barkham TMS, Nagarajan N, Sessions OM |
| EPI_ISL_493330 | INMI Lazzaro Spallanzani IRCCS | INMI Lazzaro Spallanzani IRCCS | Cesare E.M. Gruber, Martina Rueca, Barbara Bartolini, Francesco Messina, Maria R. Capobianchi, Antonino Di Caro |
| EPI_ISL_493332 | Istituto Zooprofilattico Sperimentale del Mezzogiorno | INMI Lazzaro Spallanzani IRCCS | Cesare E.M. Gruber, Martina Rueca, Barbara Bartolini, Francesco Messina, Antonino Di Caro, Giovanna Fusco, Maurizio Viscardi, Giorgia Borriello, Maria R. Capobianchi |
| EPI_ISL_493975, EPI_ISL_493976, EPI_ISL_493977, EPI_ISL_493979, EPI_ISL_493980, EPI_ISL_493982, EPI_ISL_493983, EPI_ISL_493984, EPI_ISL_493985, EPI_ISL_493986, EPI_ISL_493987, EPI_ISL_493988, EPI_ISL_493989, EPI_ISL_493990, EPI_ISL_493991, EPI_ISL_493992, EPI_ISL_493993, EPI_ISL_493994, EPI_ISL_493995, EPI_ISL_493996, EPI_ISL_493997, EPI_ISL_493998, EPI_ISL_493999, EPI_ISL_494000, EPI_ISL_494001, EPI_ISL_494002, EPI_ISL_494003, EPI_ISL_494004, EPI_ISL_494005, EPI_ISL_494006, EPI_ISL_494007, EPI_ISL_494008, EPI_ISL_494009, EPI_ISL_494010, EPI_ISL_494011, EPI_ISL_494012, EPI_ISL_494013, EPI_ISL_494014, EPI_ISL_494015, EPI_ISL_494016, EPI_ISL_494017, EPI_ISL_494018, EPI_ISL_494019, EPI_ISL_494020, EPI_ISL_494021, EPI_ISL_494022, EPI_ISL_494023, EPI_ISL_494024, EPI_ISL_494025, EPI_ISL_494026, EPI_ISL_494027, EPI_ISL_494028, EPI_ISL_494029, EPI_ISL_494030, EPI_ISL_494031, EPI_ISL_494032, EPI_ISL_494033, EPI_ISL_494034, EPI_ISL_494035, EPI_ISL_494036, EPI_ISL_494037, EPI_ISL_494038, EPI_ISL_494039, EPI_ISL_494040, EPI_ISL_494041, EPI_ISL_494042, EPI_ISL_494043, EPI_ISL_494044, EPI_ISL_494045, EPI_ISL_494046, EPI_ISL_494047, EPI_ISL_494048, EPI_ISL_494049, EPI_ISL_494050, EPI_ISL_494051, EPI_ISL_494052, EPI_ISL_494053, EPI_ISL_494054, EPI_ISL_494055, EPI_ISL_494056, EPI_ISL_494057, EPI_ISL_494058, EPI_ISL_494059, EPI_ISL_494060, EPI_ISL_494061, EPI_ISL_494062, EPI_ISL_494063, EPI_ISL_494064, EPI_ISL_494065, EPI_ISL_494066, EPI_ISL_494067, EPI_ISL_494068, EPI_ISL_494069, EPI_ISL_494070, EPI_ISL_494071, EPI_ISL_494073, EPI_ISL_494074, EPI_ISL_494075, EPI_ISL_494076, EPI_ISL_494077, EPI_ISL_494078, EPI_ISL_494079, EPI_ISL_494080, EPI_ISL_494081, EPI_ISL_494082, EPI_ISL_494083, EPI_ISL_494084, EPI_ISL_494085, EPI_ISL_494086, EPI_ISL_494087, EPI_ISL_494088, EPI_ISL_494089, EPI_ISL_494090, EPI_ISL_494091, EPI_ISL_494092, EPI_ISL_494093, EPI_ISL_494094, EPI_ISL_494095, EPI_ISL_494096, EPI_ISL_494097, EPI_ISL_494098, EPI_ISL_494099, EPI_ISL_494100, EPI_ISL_494101, EPI_ISL_494102, EPI_ISL_494103, EPI_ISL_494104, EPI_ISL_494105, EPI_ISL_494106, EPI_ISL_494107, EPI_ISL_494108, EPI_ISL_494109, EPI_ISL_494110, EPI_ISL_494111, EPI_ISL_494112, EPI_ISL_494113, EPI_ISL_494114, EPI_ISL_494115, EPI_ISL_494116, EPI_ISL_494117, EPI_ISL_494118, EPI_ISL_494119, EPI_ISL_494120, EPI_ISL_494121, EPI_ISL_494122, EPI_ISL_494123, EPI_ISL_494124, EPI_ISL_494125, EPI_ISL_494126, EPI_ISL_494127, EPI_ISL_494128, EPI_ISL_494129, EPI_ISL_494130, EPI_ISL_494131, EPI_ISL_494132, EPI_ISL_494133, EPI_ISL_494134, EPI_ISL_494135, EPI_ISL_494136, EPI_ISL_494137, EPI_ISL_494138, EPI_ISL_494139, EPI_ISL_494140, EPI_ISL_494141, EPI_ISL_494142, EPI_ISL_494143, EPI_ISL_494144, EPI_ISL_494145, EPI_ISL_494146, EPI_ISL_494147, EPI_ISL_494148, EPI_ISL_494149, EPI_ISL_494150, EPI_ISL_494151, EPI_ISL_494152, EPI_ISL_494153, EPI_ISL_494154, EPI_ISL_494155, EPI_ISL_494156, EPI_ISL_494157, EPI_ISL_494158, EPI_ISL_494159, EPI_ISL_494160, EPI_ISL_494161, EPI_ISL_494162, EPI_ISL_494163, EPI_ISL_494164, EPI_ISL_494165, EPI_ISL_494166, EPI_ISL_494167, EPI_ISL_494168, EPI_ISL_494169, EPI_ISL_494170, EPI_ISL_494171, EPI_ISL_494172, EPI_ISL_494173, EPI_ISL_494174, EPI_ISL_494175, EPI_ISL_494176, EPI_ISL_494177, EPI_ISL_494178, EPI_ISL_494179, EPI_ISL_494180, EPI_ISL_494181, EPI_ISL_494182, EPI_ISL_494183, EPI_ISL_494184, EPI_ISL_494185, EPI_ISL_494186, EPI_ISL_494187, EPI_ISL_494188, EPI_ISL_494189, EPI_ISL_494190, EPI_ISL_494191, EPI_ISL_494192, EPI_ISL_494194, EPI_ISL_494195, EPI_ISL_494196, EPI_ISL_494197, EPI_ISL_494198, EPI_ISL_494199, EPI_ISL_494200, EPI_ISL_494201, EPI_ISL_494202, EPI_ISL_494203, EPI_ISL_494204, EPI_ISL_494205, EPI_ISL_494206, EPI_ISL_494207, EPI_ISL_494208, EPI_ISL_494209, EPI_ISL_494210, EPI_ISL_494211, EPI_ISL_494212, EPI_ISL_494213, EPI_ISL_494214, EPI_ISL_494215, EPI_ISL_494216, EPI_ISL_494217, EPI_ISL_494218, EPI_ISL_494219, EPI_ISL_494220, EPI_ISL_494221, EPI_ISL_494222, EPI_ISL_494223, EPI_ISL_494224, EPI_ISL_494225, EPI_ISL_494226, EPI_ISL_494227, EPI_ISL_494228, EPI_ISL_494229, EPI_ISL_494230, EPI_ISL_494231, EPI_ISL_494232, EPI_ISL_494233, EPI_ISL_494234, EPI_ISL_494235, EPI_ISL_494236, EPI_ISL_494237, EPI_ISL_494238, EPI_ISL_494239, EPI_ISL_494240, EPI_ISL_494241, EPI_ISL_494242, EPI_ISL_494243, EPI_ISL_494244, EPI_ISL_494245, EPI_ISL_494246, EPI_ISL_494247, EPI_ISL_494248, EPI_ISL_494249, EPI_ISL_494250, EPI_ISL_494251, EPI_ISL_494252, EPI_ISL_494253, EPI_ISL_494254, EPI_ISL_494255, EPI_ISL_494256, EPI_ISL_494257, EPI_ISL_494258, EPI_ISL_494259, EPI_ISL_494260, EPI_ISL_494261, EPI_ISL_494262, EPI_ISL_494263, EPI_ISL_494264, EPI_ISL_494265, EPI_ISL_494266, EPI_ISL_494267, EPI_ISL_494268, EPI_ISL_494269, EPI_ISL_494270, EPI_ISL_494271, EPI_ISL_494272, EPI_ISL_494273, EPI_ISL_494274, EPI_ISL_494275, EPI_ISL_494276, EPI_ISL_494277, EPI_ISL_494278, EPI_ISL_494279, EPI_ISL_494280, EPI_ISL_494281, EPI_ISL_494282, EPI_ISL_494283, EPI_ISL_494284, EPI_ISL_494285, EPI_ISL_494286, EPI_ISL_494287, EPI_ISL_494288, EPI_ISL_494289, EPI_ISL_494290, EPI_ISL_494291, EPI_ISL_494292, EPI_ISL_494294, EPI_ISL_494295, EPI_ISL_494296, EPI_ISL_494297, EPI_ISL_494298, EPI_ISL_494299, EPI_ISL_494300, EPI_ISL_494301, EPI_ISL_494302, EPI_ISL_494303, EPI_ISL_494304, EPI_ISL_494305, EPI_ISL_494306, EPI_ISL_494307, EPI_ISL_494308, EPI_ISL_494309, EPI_ISL_494310, EPI_ISL_494311, EPI_ISL_494312, EPI_ISL_494313, EPI_ISL_494314, EPI_ISL_494315, EPI_ISL_494316, EPI_ISL_494317, EPI_ISL_494318, EPI_ISL_494319, EPI_ISL_494320, EPI_ISL_494321, EPI_ISL_494322, EPI_ISL_494323, EPI_ISL_494324, EPI_ISL_494325, EPI_ISL_494326, EPI_ISL_494327, EPI_ISL_494328, EPI_ISL_494329, EPI_ISL_494330, EPI_ISL_494331, EPI_ISL_494332, EPI_ISL_494333, EPI_ISL_494334, EPI_ISL_494335, EPI_ISL_494336, EPI_ISL_494337, EPI_ISL_494338, EPI_ISL_494339, EPI_ISL_494340, EPI_ISL_494341, EPI_ISL_494342, EPI_ISL_494343, EPI_ISL_494344, EPI_ISL_494345, EPI_ISL_494346, EPI_ISL_494347, EPI_ISL_494348, EPI_ISL_494349, EPI_ISL_494350, EPI_ISL_494351, EPI_ISL_494352, EPI_ISL_494353, EPI_ISL_494354, EPI_ISL_494355, EPI_ISL_494356, EPI_ISL_494357, EPI_ISL_494358, EPI_ISL_494359, EPI_ISL_494360, EPI_ISL_494361, EPI_ISL_494362, EPI_ISL_494363, EPI_ISL_494364, EPI_ISL_494365, EPI_ISL_494366, EPI_ISL_494367, EPI_ISL_494368, EPI_ISL_494369, EPI_ISL_494370, EPI_ISL_494371 | | | |
| see above | Wales Specialist Virology Centre Sequencing lab: Pathogen Genomics Unit | COVID-19 Genomics UK (COG-UK) Consortium | Catherine Moore, Johnathan Evans, Laura Gifford, Malorie Perry, Simon Cottrell, Angela Marchbank, Alec Birchley, Alexander Adams, Amy Gaskin, Bree Gatica-Wilcox, Jason Coombes, Joel Southgate, Lauren Gilbert, Lee Graham, Nicole Pacchiarini, Sara Kumziene-Summerhayes, Sarah Taylor, Sophie Jones, Sara Rey, Matthew Bull, Joanne Watkins, Sally Corden, Tom Connor |
| EPI_ISL_498627 | Viollier AG | Department of Biosystems Science and Engineering, ETH Zürich | Christian Beisel, Sarah Nadeau, Ivan Topolsky, Pedro Ferreira, Philipp Jablonski, Susana Posada-Céspedes, Tobias Schär, Ina Nissen, Natascha Santacroce, Elodie Burcklen, Christiane Beckmann, Maurice Redondo, Olivier Kobel, Christoph Noppen, Sophie Seidel, Noemie Santamaria de Souza, Niko Beerenwinkel, Tanja Stadler |
| EPI_ISL_499355, EPI_ISL_499356, EPI_ISL_499357, EPI_ISL_499358, EPI_ISL_499359, EPI_ISL_499360, EPI_ISL_499361, EPI_ISL_499362, EPI_ISL_499363, EPI_ISL_499364, EPI_ISL_499365, EPI_ISL_499366, EPI_ISL_499367, EPI_ISL_499368, EPI_ISL_499369, EPI_ISL_499370, EPI_ISL_499371, EPI_ISL_499372, EPI_ISL_499373, EPI_ISL_499374, EPI_ISL_499375, EPI_ISL_499376, EPI_ISL_499377, EPI_ISL_499378, EPI_ISL_499379, EPI_ISL_499380, EPI_ISL_499381, EPI_ISL_499382, EPI_ISL_499383, EPI_ISL_499384, EPI_ISL_499385, EPI_ISL_499386, EPI_ISL_499387, EPI_ISL_499388, EPI_ISL_499389, EPI_ISL_499390, EPI_ISL_499391, EPI_ISL_499392, EPI_ISL_499393, EPI_ISL_499394, EPI_ISL_499395, EPI_ISL_499396, EPI_ISL_499397, EPI_ISL_499398, EPI_ISL_499399, EPI_ISL_499400, EPI_ISL_499401, EPI_ISL_499402, EPI_ISL_499403, EPI_ISL_499404, EPI_ISL_499405, EPI_ISL_499406, EPI_ISL_499407, EPI_ISL_499408, | | | |

| | | | |
|---|---|---|---|
| EPI_ISL_499409, EPI_ISL_499410, EPI_ISL_499411, EPI_ISL_499412, EPI_ISL_499413, EPI_ISL_499414, EPI_ISL_499415, EPI_ISL_499416, EPI_ISL_499417, EPI_ISL_499418, EPI_ISL_499419, EPI_ISL_499420, EPI_ISL_499421, EPI_ISL_499422, EPI_ISL_499423, EPI_ISL_499424, EPI_ISL_499425, EPI_ISL_499426, EPI_ISL_499427, EPI_ISL_499428, EPI_ISL_499429, EPI_ISL_499430, EPI_ISL_499431, EPI_ISL_499432, EPI_ISL_499433, EPI_ISL_499434, EPI_ISL_499435, EPI_ISL_499436, EPI_ISL_499437, EPI_ISL_499438, EPI_ISL_499439, EPI_ISL_499440, EPI_ISL_499441, EPI_ISL_499442, EPI_ISL_499443, EPI_ISL_499444, EPI_ISL_499445, EPI_ISL_499446, EPI_ISL_499447, EPI_ISL_499448, EPI_ISL_499449, EPI_ISL_499450, EPI_ISL_499451, EPI_ISL_499452, EPI_ISL_499453, EPI_ISL_499454, EPI_ISL_499455, EPI_ISL_499456, EPI_ISL_499457, EPI_ISL_499458, EPI_ISL_499459 | | | |
| see above | Wales Specialist Virology Centre Sequencing lab: Pathogen Genomics Unit | COVID-19 Genomics UK (COG-UK) Consortium | Catherine Moore, Johnathan Evans, Laura Gifford, Malorie Perry, Simon Cottrell, Angela Marchbank, Alec Birchley, Alexander Adams, Amy Gaskin, Bree Gatica-Wilcox, Jason Coombes, Joel Southgate, Lauren Gilbert, Lee Graham, Nicole Pacchiarini, Sara Kumziene-Summerhayes, Sarah Taylor, Sophie Jones, Sara Rey, Matthew Bull, Joanne Watkins, Sally Corden, Tom Connor |
| EPI_ISL_500882, EPI_ISL_500890, EPI_ISL_500903, EPI_ISL_500905, EPI_ISL_500907, EPI_ISL_500908, EPI_ISL_500919, EPI_ISL_500923, EPI_ISL_500925, EPI_ISL_500932, EPI_ISL_500936, EPI_ISL_500938, EPI_ISL_500940 | | | |
| see above | Viollier AG | Department of Biosystems Science and Engineering, ETH Zürich | Christian Beisel, Sarah Nadeau, Ivan Topolsky, Pedro Ferreira, Philipp Jablonski, Susana Posada-Céspedes, Tobias Schär, Ina Nissen, Natascha Santacroce, Elodie Burcklen, Christiane Beckmann, Maurice Redondo, Olivier Kobel, Christoph Noppen, Sophie Seidel, Noemie Santamaria de Souza, Niko Beerenwinkel, Tanja Stadler |
| EPI_ISL_507002, EPI_ISL_507003, EPI_ISL_507004, EPI_ISL_507005, EPI_ISL_507006 | Department of Laboratory Medicine, Tan Tock Seng Hospital | Department of Laboratory Medicine, Tan Tock Seng Hospital | Chen YYC, Zair X, Li C, Tang WY, Maurer-Stroh S, Barkham TMS, Nagarajan N, Sessions OM |
| EPI_ISL_510692, EPI_ISL_510693, EPI_ISL_510695, EPI_ISL_510700, EPI_ISL_510709, EPI_ISL_510710, EPI_ISL_510721, EPI_ISL_510723, EPI_ISL_510728, EPI_ISL_510729, EPI_ISL_510742, EPI_ISL_510748, EPI_ISL_510763, EPI_ISL_510785, EPI_ISL_510809, EPI_ISL_511991, EPI_ISL_511997, EPI_ISL_512000, EPI_ISL_512001, EPI_ISL_512007, EPI_ISL_512012, EPI_ISL_512019, EPI_ISL_512020, EPI_ISL_512025, EPI_ISL_512026, EPI_ISL_512027, EPI_ISL_512038, EPI_ISL_512039, EPI_ISL_512040, EPI_ISL_512044, EPI_ISL_512048, EPI_ISL_512049 | | | |
| see above | Viollier AG | Department of Biosystems Science and Engineering, ETH Zürich | Christian Beisel, Sarah Nadeau, Ivan Topolsky, Pedro Ferreira, Philipp Jablonski, Susana Posada-Céspedes, Tobias Schär, Ina Nissen, Natascha Santacroce, Elodie Burcklen, Christiane Beckmann, Maurice Redondo, Olivier Kobel, Christoph Noppen, Sophie Seidel, Noemie Santamaria de Souza, Niko Beerenwinkel, Tanja Stadler |
| EPI_ISL_512135, EPI_ISL_512136, EPI_ISL_512137, EPI_ISL_512138, EPI_ISL_512139, EPI_ISL_512140, EPI_ISL_512141, EPI_ISL_512142, EPI_ISL_512143, EPI_ISL_512144, EPI_ISL_512145, EPI_ISL_512146, EPI_ISL_512147, EPI_ISL_512148, EPI_ISL_512149, EPI_ISL_512150, EPI_ISL_512151, EPI_ISL_512152, EPI_ISL_512153, EPI_ISL_512154, EPI_ISL_512155, EPI_ISL_512156, EPI_ISL_512157 | | | |
| see above | Alaska State Virology Laboratory | Alaska State Virology Laboratory | Chen J et al with Pathogenomics group Dagdag R, Redlinger M, Milton E, George W, Kovalenko A, Drown DM, Bortz E |
| EPI_ISL_512481, EPI_ISL_512482, EPI_ISL_512483, EPI_ISL_512484, EPI_ISL_512485, EPI_ISL_512486, EPI_ISL_512487, EPI_ISL_512488, EPI_ISL_512489, EPI_ISL_512490, EPI_ISL_512491, EPI_ISL_512492, EPI_ISL_512493, EPI_ISL_512494, EPI_ISL_512495, EPI_ISL_512496, EPI_ISL_512497, EPI_ISL_512498, EPI_ISL_512499, EPI_ISL_512500, EPI_ISL_512501, EPI_ISL_512502, EPI_ISL_512503, EPI_ISL_512504, EPI_ISL_512505, EPI_ISL_512506, EPI_ISL_512507, EPI_ISL_512508, EPI_ISL_512509, EPI_ISL_512510, EPI_ISL_512511, EPI_ISL_512512, EPI_ISL_512513, EPI_ISL_512514, EPI_ISL_512515, EPI_ISL_512516, EPI_ISL_512517, EPI_ISL_512518, EPI_ISL_512519, EPI_ISL_512520, EPI_ISL_512521, EPI_ISL_512522, EPI_ISL_512523, EPI_ISL_512524, EPI_ISL_512525, EPI_ISL_512526, EPI_ISL_512527, EPI_ISL_512528, EPI_ISL_512529, EPI_ISL_512530, EPI_ISL_512531, EPI_ISL_512532, EPI_ISL_512533, EPI_ISL_512534, EPI_ISL_512535, EPI_ISL_512536, EPI_ISL_512537, EPI_ISL_512538, EPI_ISL_512539, EPI_ISL_512540, EPI_ISL_512541, EPI_ISL_512542, EPI_ISL_512543, EPI_ISL_512544, EPI_ISL_512545, EPI_ISL_512546, EPI_ISL_514563, EPI_ISL_514564, EPI_ISL_514565, EPI_ISL_514566, EPI_ISL_514567, EPI_ISL_514568, EPI_ISL_514569, EPI_ISL_514570, EPI_ISL_514571, EPI_ISL_514572, EPI_ISL_514573, EPI_ISL_514574, EPI_ISL_514575, EPI_ISL_514576, EPI_ISL_514577, EPI_ISL_514578, EPI_ISL_514579 | | | |
| see above | Wales Specialist Virology Centre Sequencing lab: Pathogen Genomics Unit | COVID-19 Genomics UK (COG-UK) Consortium | Catherine Moore, Johnathan Evans, Laura Gifford, Malorie Perry, Simon Cottrell, Angela Marchbank, Alec Birchley, Alexander Adams, Amy Gaskin, Bree Gatica-Wilcox, Jason Coombes, Joel Southgate, Lauren Gilbert, Lee Graham, Nicole Pacchiarini, Sara Kumziene-Summerhayes, Sarah Taylor, Sophie Jones, Sara Rey, Matthew Bull, Joanne Watkins, Sally Corden, Tom Connor |
| EPI_ISL_515932 | DH | Department of Neurovirology, National Institute of Mental Health and Neuroscience (NIMHANS) | Chitra Pattabiraman,Vijayalakshmi Reddy, Harsha PK, Risha Rasheed, Pramada Prasad, Shafeeq S Hameed, Manjunatha Venkataswamy, Anita Desai, Ravi Vasanthapuram |
| EPI_ISL_515933 | BIMS | Department of Neurovirology, National Institute of Mental Health and Neuroscience (NIMHANS) | Chitra Pattabiraman,Vijayalakshmi Reddy, Harsha PK, Risha Rasheed, Pramada Prasad, Shafeeq S Hameed, Manjunatha Venkataswamy, Anita Desai, Ravi Vasanthapuram |
| EPI_ISL_515934, EPI_ISL_515935 | DH | Department of Neurovirology, National Institute of Mental Health and Neuroscience (NIMHANS) | Chitra Pattabiraman,Vijayalakshmi Reddy, Harsha PK, Risha Rasheed, Pramada Prasad, Shafeeq S Hameed, Manjunatha Venkataswamy, Anita Desai, Ravi Vasanthapuram |
| EPI_ISL_515937 | BIMS | Department of Neurovirology, National Institute of Mental Health and Neuroscience (NIMHANS) | Chitra Pattabiraman,Vijayalakshmi Reddy, Harsha PK, Risha Rasheed, Pramada Prasad, Shafeeq S Hameed, Manjunatha Venkataswamy, Anita Desai, Ravi Vasanthapuram |
| EPI_ISL_515938, EPI_ISL_515939, EPI_ISL_515941, EPI_ISL_515942 | CV RAMAN HOSPITAL | Department of Neurovirology, National Institute of Mental Health and Neuroscience (NIMHANS) | Chitra Pattabiraman,Vijayalakshmi Reddy, Harsha PK, Risha Rasheed, Pramada Prasad, Shafeeq S Hameed, Manjunatha Venkataswamy, Anita Desai, Ravi Vasanthapuram |
| EPI_ISL_515943, EPI_ISL_515944 | ESIC | Department of Neurovirology, National Institute of Mental Health and Neuroscience (NIMHANS) | Chitra Pattabiraman,Vijayalakshmi Reddy, Harsha PK, Risha Rasheed, Pramada Prasad, Shafeeq S Hameed, Manjunatha Venkataswamy, Anita Desai, Ravi Vasanthapuram |
| EPI_ISL_515945, EPI_ISL_515946, EPI_ISL_515947, EPI_ISL_515948, EPI_ISL_515949 | DH | Department of Neurovirology, National Institute of Mental Health and Neuroscience (NIMHANS) | Chitra Pattabiraman,Vijayalakshmi Reddy, Harsha PK, Risha Rasheed, Pramada Prasad, Shafeeq S Hameed, Manjunatha Venkataswamy, Anita Desai, Ravi Vasanthapuram |
| EPI_ISL_515950, EPI_ISL_515951, EPI_ISL_515952, EPI_ISL_515953 | VICTORIA HOSPITAL | Department of Neurovirology, National Institute of Mental Health and Neuroscience (NIMHANS) | Chitra Pattabiraman,Vijayalakshmi Reddy, Harsha PK, Risha Rasheed, Pramada Prasad, Shafeeq S Hameed, Manjunatha Venkataswamy, Anita Desai, Ravi Vasanthapuram |
| EPI_ISL_515954, EPI_ISL_515955 | DH | Department of Neurovirology, National Institute of Mental Health and Neuroscience (NIMHANS) | Chitra Pattabiraman,Vijayalakshmi Reddy, Harsha PK, Risha Rasheed, Pramada Prasad, Shafeeq S Hameed, Manjunatha Venkataswamy, Anita Desai, Ravi Vasanthapuram |
| EPI_ISL_515956 | SHEKAR HOSPITAL | Department of Neurovirology, National Institute of Mental Health and Neuroscience (NIMHANS) | Chitra Pattabiraman,Vijayalakshmi Reddy, Harsha PK, Risha Rasheed, Pramada Prasad, Shafeeq S Hameed, Manjunatha Venkataswamy, Anita Desai, Ravi Vasanthapuram |
| EPI_ISL_515957, EPI_ISL_515958, EPI_ISL_515959 | DH | Department of Neurovirology, National Institute of Mental Health and Neuroscience (NIMHANS) | Chitra Pattabiraman,Vijayalakshmi Reddy, Harsha PK, Risha Rasheed, Pramada Prasad, Shafeeq S Hameed, Manjunatha Venkataswamy, Anita Desai, Ravi Vasanthapuram |
| EPI_ISL_515960 | JGH | Department of Neurovirology, National Institute of Mental Health and Neuroscience (NIMHANS) | Chitra Pattabiraman,Vijayalakshmi Reddy, Harsha PK, Risha Rasheed, Pramada Prasad, Shafeeq S Hameed, Manjunatha Venkataswamy, Anita Desai, Ravi Vasanthapuram |
| EPI_ISL_515961, EPI_ISL_515962 | DH | Department of Neurovirology, National Institute of Mental Health and Neuroscience (NIMHANS) | Chitra Pattabiraman,Vijayalakshmi Reddy, Harsha PK, Risha Rasheed, Pramada Prasad, Shafeeq S Hameed, Manjunatha Venkataswamy, Anita Desai, Ravi Vasanthapuram |
| EPI_ISL_515963, EPI_ISL_515964 | VICTORIA HOSPITAL | Department of Neurovirology, National Institute of Mental Health and Neuroscience (NIMHANS) | Chitra Pattabiraman,Vijayalakshmi Reddy, Harsha PK, Risha Rasheed, Pramada Prasad, Shafeeq S Hameed, Manjunatha Venkataswamy, Anita Desai, Ravi Vasanthapuram |
| EPI_ISL_515965 | DH | Department of Neurovirology, National Institute of Mental Health and Neuroscience (NIMHANS) | Chitra Pattabiraman,Vijayalakshmi Reddy, Harsha PK, Risha Rasheed, Pramada Prasad, Shafeeq S Hameed, Manjunatha Venkataswamy, Anita Desai, Ravi Vasanthapuram |
| EPI_ISL_515966, EPI_ISL_515967 | VICTORIA HOSPITAL | Department of Neurovirology, National Institute of Mental Health and Neuroscience (NIMHANS) | Chitra Pattabiraman,Vijayalakshmi Reddy, Harsha PK, Risha Rasheed, Pramada Prasad, Shafeeq S Hameed, Manjunatha Venkataswamy, Anita Desai, Ravi Vasanthapuram |
| EPI_ISL_515968, EPI_ISL_515969, EPI_ISL_515970, EPI_ISL_515971, EPI_ISL_515972, EPI_ISL_515973 | MIMS | Department of Neurovirology, National Institute of Mental Health and Neuroscience (NIMHANS) | Chitra Pattabiraman,Vijayalakshmi Reddy, Harsha PK, Risha Rasheed, Pramada Prasad, Shafeeq S Hameed, Manjunatha Venkataswamy, Anita Desai, Ravi Vasanthapuram |
| EPI_ISL_516075 | BIMS | Department of Neurovirology, National Institute of Mental Health and Neuroscience (NIMHANS) | Chitra Pattabiraman,Vijayalakshmi Reddy, Harsha PK, Risha Rasheed, Pramada Prasad, Shafeeq S Hameed, Manjunatha Venkataswamy, Anita Desai, Ravi Vasanthapuram |
| EPI_ISL_516076, EPI_ISL_516077, EPI_ISL_516078 | VICTORIA HOSPITAL | Department of Neurovirology, National Institute of Mental Health and Neuroscience (NIMHANS) | Chitra Pattabiraman,Vijayalakshmi Reddy, Harsha PK, Risha Rasheed, Pramada Prasad, Shafeeq S Hameed, Manjunatha Venkataswamy, Anita Desai, Ravi Vasanthapuram |
| EPI_ISL_516552, EPI_ISL_516572, EPI_ISL_516574, EPI_ISL_516575, EPI_ISL_516581, EPI_ISL_516582, EPI_ISL_516600 | Viollier AG | Department of Biosystems Science and Engineering, ETH Zürich | Christian Beisel, Sarah Nadeau, Ivan Topolsky, Pedro Ferreira, Philipp Jablonski, Susana Posada-Céspedes, Tobias Schär, Ina Nissen, Natascha Santacroce, Elodie Burcklen, Christiane Beckmann, Maurice Redondo, Olivier Kobel, Christoph Noppen, Sophie Seidel, Noemie Santamaria de Souza, Niko Beerenwinkel, Tanja Stadler |

| | | | |
|---|---|---|---|
| EPI_ISL_516801, EPI_ISL_516802, EPI_ISL_516803, EPI_ISL_516804, EPI_ISL_516805 | Department of Laboratory Medicine, Tan Tock Seng Hospital | Department of Laboratory Medicine, Tan Tock Seng Hospital | Chen YYC, Zair X, Li C, Tang WY, Maurer-Stroh S, Barkham TMS, Nagarajan N, Sessions OM |
| EPI_ISL_517590, EPI_ISL_517591, EPI_ISL_517592, EPI_ISL_517593, EPI_ISL_517594, EPI_ISL_517595, EPI_ISL_517596, EPI_ISL_517597, EPI_ISL_517598, EPI_ISL_517599, EPI_ISL_517600, EPI_ISL_517601, EPI_ISL_517602, EPI_ISL_517603, EPI_ISL_517604, EPI_ISL_517605, EPI_ISL_517606, EPI_ISL_517607, EPI_ISL_517608, EPI_ISL_517609, EPI_ISL_517610 | | | |
| see above | Wales Specialist Virology Centre Sequencing lab: Pathogen Genomics Unit | COVID-19 Genomics UK (COG-UK) Consortium | Catherine Moore, Johnathan Evans, Laura Gifford, Malorie Perry, Simon Cottrell, Angela Marchbank, Alec Birchley, Alexander Adams, Amy Gaskin, Bree Gatica-Wilcox, Jason Coombes, Joel Southgate, Lauren Gilbert, Lee Graham, Nicole Pacchiarini, Sara Kumziene-Summerhayes, Sarah Taylor, Sophie Jones, Sara Rey, Matthew Bull, Joanne Watkins, Sally Corden, Tom Connor |
| EPI_ISL_523857, EPI_ISL_523878, EPI_ISL_523879, EPI_ISL_523885, EPI_ISL_523898, EPI_ISL_523899, EPI_ISL_523900, EPI_ISL_523903, EPI_ISL_523904, EPI_ISL_523908, EPI_ISL_523909, EPI_ISL_523926 | | | |
| see above | Viollier AG | Department of Biosystems Science and Engineering, ETH Zürich | Christian Beisel, Sarah Nadeau, Ivan Topolsky, Pedro Ferreira, Philipp Jablonski, Susana Posada-Céspedes, Tobias Schär, Ina Nissen, Natascha Santacroce, Elodie Burcklen, Christiane Beckmann, Maurice Redondo, Olivier Kobel, Christoph Noppen, Sophie Seidel, Noemie Santamaria de Souza, Niko Beerenwinkel, Tanja Stadler |
| EPI_ISL_525474 | Centre for Dengue Research | Centre for Dengue Research, USJ, SL | Chandima Jeewandara, Deshni Jayathilaka, Dinuka Ariyaratne, Laksiri Gomes, Diyanath Ranasinghe, Dinuka Guruge, Ruwan Wijayamuni, Gathsaurie Neelika Malavige |
| EPI_ISL_525476 | Centre for Dengue Research | Centre for Dengue Research | Chandima Jeewandara, Deshni Jayathilaka, Dinuka Ariyaratne, Laksiri Gomes, Diyanath Ranasinghe, Dinuka Guruge, Ruwan Wijayamuni, Gathsaurie Neelika Malavige |
| EPI_ISL_525478, EPI_ISL_525479, EPI_ISL_525481 | Centre for Dengue Research | Centre for Dengue Research | Chandima Jeewandara, Deshni Jayathilaka, Dinuka Ariyaratne, Laksiri Gomes, Diyanath Ranasinghe, Ananda Wijewickrama, Eranga Narangoda, Damayanthi Idampitiya, Gathsaurie Neelika Malavige |
| EPI_ISL_525486, EPI_ISL_525488, EPI_ISL_525489 | Centre for Dengue Research | Centre for Dengue Research | Chandima Jeewandara, Deshni Jayathilaka, Dinuka Ariyaratne, Laksiri Gomes, Diyanath Ranasinghe, Ananda Wijewickrama, Malika Karunaratne, Eranga Narangoda, Damayanthi Idampitiya, Gathsaurie Neelika Malavige |
| EPI_ISL_528485, EPI_ISL_528486, EPI_ISL_528487, EPI_ISL_528488, EPI_ISL_528489, EPI_ISL_528490, EPI_ISL_528491, EPI_ISL_528492, EPI_ISL_528493, EPI_ISL_528494, EPI_ISL_528495, EPI_ISL_528496, EPI_ISL_528497, EPI_ISL_528498, EPI_ISL_528499, EPI_ISL_528500, EPI_ISL_528501, EPI_ISL_528502, EPI_ISL_528503, EPI_ISL_528504, EPI_ISL_528505, EPI_ISL_528506, EPI_ISL_528507, EPI_ISL_528508, EPI_ISL_528509, EPI_ISL_528510, EPI_ISL_528511, EPI_ISL_528512, EPI_ISL_528513, EPI_ISL_528514, EPI_ISL_528515, EPI_ISL_528516, EPI_ISL_528517, EPI_ISL_528518, EPI_ISL_528519, EPI_ISL_528520, EPI_ISL_528521, EPI_ISL_528522, EPI_ISL_528523, EPI_ISL_528524, EPI_ISL_528525, EPI_ISL_528526, EPI_ISL_528527, EPI_ISL_528528, EPI_ISL_528529, EPI_ISL_528530, EPI_ISL_528531, EPI_ISL_528532, EPI_ISL_528533, EPI_ISL_528534, EPI_ISL_528535, EPI_ISL_528536, EPI_ISL_528537 | | | |
| see above | Alaska State Virology Laboratory | Alaska State Virology Laboratory | Chen J et al with Pathogenomics group Dagdag R, Redlinger M, Milton E, George W, Kovalenko A, Drown DM, Bortz E |
| EPI_ISL_528747 | Santo Borromeus Hospital | School of Pharmacy & School of Life Sciences and Technology - Institut Teknologi Bandung; Molecular Genetics Laboratory-Faculty of Medicine-Universitas Padjadjaran; Laboratorium Kesehatan Provinsi Jawa Barat | Catur Riani, Marselina Irasonia Tan, Yunia Sribudiani, Azzania Fibriani, Husna Nugrahapraja, Tarwadi, Ema Rahmawati, Savira Ekawardhani, Hesti Lina Wiraswati, Ryan Bayusantika Ristandi, Rifky Waluyajati Rachman, Cut Nur Cinthia Alamanda, Lia Faridah, Miftahul Faridl, Karimatu Khoirunnisa, Hammam Riza, Sony Solistia Wirawan, Agung Eru Wibowo, Irvan Faizal |
| EPI_ISL_528752 | Dr. H. A. Rotinsulu Lung Hospital | School of Pharmacy & School of Life Sciences and Technology - Institut Teknologi Bandung; Molecular Genetics Laboratory-Faculty of Medicine-Universitas Padjadjaran; Laboratorium Kesehatan Provinsi Jawa Barat | Catur Riani, Marselina Irasonia Tan, Yunia Sribudiani, Azzania Fibriani, Husna Nugrahapraja, Tarwadi, Ema Rahmawati, Savira Ekawardhani, Hesti Lina Wiraswati, Ryan Bayusantika Ristandi, Rifky Waluyajati Rachman, Cut Nur Cinthia Alamanda, Lia Faridah, Gusti Ayu Prani Pradani, Adelina Khristiani Rahayu, Hammam Riza, Sony Solistia Wirawan, Agung Eru Wibowo, Irvan Faizal |
| EPI_ISL_529226, EPI_ISL_529227, EPI_ISL_529228, EPI_ISL_529229, EPI_ISL_529230, EPI_ISL_529231, EPI_ISL_529232, EPI_ISL_529233, EPI_ISL_529253, EPI_ISL_529297, EPI_ISL_529418, EPI_ISL_529447, EPI_ISL_529505, EPI_ISL_529527, EPI_ISL_529697, EPI_ISL_529698, EPI_ISL_529699, EPI_ISL_529700, EPI_ISL_529701, EPI_ISL_529702, EPI_ISL_529703, EPI_ISL_529704, EPI_ISL_529705, EPI_ISL_529706, EPI_ISL_529707, EPI_ISL_529708, EPI_ISL_529709, EPI_ISL_529710, EPI_ISL_529711, EPI_ISL_529712, EPI_ISL_529713, EPI_ISL_529714, EPI_ISL_529715, EPI_ISL_529716, EPI_ISL_529717, EPI_ISL_535184, EPI_ISL_535185, EPI_ISL_535186, EPI_ISL_535187, EPI_ISL_535188, EPI_ISL_535189, EPI_ISL_535190, EPI_ISL_535191, EPI_ISL_535192, EPI_ISL_535193, EPI_ISL_535194, EPI_ISL_535195, EPI_ISL_535196, EPI_ISL_535197, EPI_ISL_535198, EPI_ISL_535199, EPI_ISL_535200, EPI_ISL_535201, EPI_ISL_535202, EPI_ISL_535203, EPI_ISL_535204, EPI_ISL_535205, EPI_ISL_535206, EPI_ISL_535207, EPI_ISL_535208, EPI_ISL_535209, EPI_ISL_535210, EPI_ISL_535211, EPI_ISL_535212, EPI_ISL_535213, EPI_ISL_535214, EPI_ISL_535215, EPI_ISL_535216, EPI_ISL_535217, EPI_ISL_535218, EPI_ISL_535219, EPI_ISL_535220, EPI_ISL_535221, EPI_ISL_535222, EPI_ISL_535223, EPI_ISL_535224, EPI_ISL_535225, EPI_ISL_535226, EPI_ISL_535227, EPI_ISL_535228, EPI_ISL_535229, EPI_ISL_535230, EPI_ISL_535231, EPI_ISL_535232, EPI_ISL_535233, EPI_ISL_535234, EPI_ISL_535235, EPI_ISL_535236, EPI_ISL_535237, EPI_ISL_535238, EPI_ISL_535239, EPI_ISL_535240, EPI_ISL_535241, EPI_ISL_535242, EPI_ISL_535243, EPI_ISL_535244, EPI_ISL_535245, EPI_ISL_535246, EPI_ISL_535247, EPI_ISL_535248, EPI_ISL_535249, EPI_ISL_535250, EPI_ISL_535251, EPI_ISL_535252, EPI_ISL_535253, EPI_ISL_535254, EPI_ISL_535255, EPI_ISL_535256, EPI_ISL_535257, EPI_ISL_535258, EPI_ISL_535259, EPI_ISL_535260, EPI_ISL_535261, EPI_ISL_535262 | | | |
| see above | Wales Specialist Virology Centre Sequencing lab: Pathogen Genomics Unit | COVID-19 Genomics UK (COG-UK) Consortium | Catherine Moore, Johnathan Evans, Laura Gifford, Malorie Perry, Simon Cottrell, Angela Marchbank, Alec Birchley, Alexander Adams, Amy Gaskin, Bree Gatica-Wilcox, Jason Coombes, Joel Southgate, Lauren Gilbert, Lee Graham, Nicole Pacchiarini, Sara Kumziene-Summerhayes, Sarah Taylor, Sophie Jones, Sara Rey, Matthew Bull, Joanne Watkins, Sally Corden, Tom Connor |
| EPI_ISL_535595, EPI_ISL_535609, EPI_ISL_535615, EPI_ISL_535623, EPI_ISL_535643, EPI_ISL_535645 | Viollier AG | Department of Biosystems Science and Engineering, ETH Zürich | Christian Beisel, Sarah Nadeau, Ivan Topolsky, Pedro Ferreira, Philipp Jablonski, Susana Posada-Céspedes, Tobias Schär, Ina Nissen, Natascha Santacroce, Elodie Burcklen, Christiane Beckmann, Maurice Redondo, Olivier Kobel, Christoph Noppen, Sophie Seidel, Noemie Santamaria de Souza, Niko Beerenwinkel, Tanja Stadler |
| EPI_ISL_538435, EPI_ISL_538436, EPI_ISL_538437, EPI_ISL_538438, EPI_ISL_538439, EPI_ISL_538440, EPI_ISL_538441, EPI_ISL_538442, EPI_ISL_538443, EPI_ISL_538444, EPI_ISL_538445, EPI_ISL_538446, EPI_ISL_538447, EPI_ISL_538448, EPI_ISL_538449, EPI_ISL_538450, EPI_ISL_538451, EPI_ISL_538452, EPI_ISL_538453, EPI_ISL_538454, EPI_ISL_538455, EPI_ISL_538456, EPI_ISL_538457, EPI_ISL_538458, EPI_ISL_538459, EPI_ISL_538460, EPI_ISL_538461, EPI_ISL_538462, EPI_ISL_538463, EPI_ISL_538464, EPI_ISL_538465, EPI_ISL_538466, EPI_ISL_538467, EPI_ISL_538468, EPI_ISL_538469, EPI_ISL_538470, EPI_ISL_538471, EPI_ISL_538472, EPI_ISL_538473, EPI_ISL_538474, EPI_ISL_538475, EPI_ISL_538476, EPI_ISL_538477, EPI_ISL_538478, EPI_ISL_538479, EPI_ISL_538480, EPI_ISL_538481, EPI_ISL_538482, EPI_ISL_538483, EPI_ISL_538484, EPI_ISL_538485, EPI_ISL_538486, EPI_ISL_538487, EPI_ISL_538488, EPI_ISL_538489, EPI_ISL_538490, EPI_ISL_538491, EPI_ISL_538492, EPI_ISL_538493, EPI_ISL_538494, EPI_ISL_538495, EPI_ISL_538496, EPI_ISL_538497 | | | |
| see above | Department of Laboratory Medicine, Tan Tock Seng Hospital | Department of Laboratory Medicine, Tan Tock Seng Hospital | Chen YYC, Zair X, Lim JX, Li C, Tang WY, Maurer-Stroh S, Barkham TMS, Nagarajan N, Sessions OM |
| EPI_ISL_538514, EPI_ISL_538515, EPI_ISL_538516, EPI_ISL_538517, EPI_ISL_538518, EPI_ISL_538519, EPI_ISL_538520, EPI_ISL_538521 | Infectious Diseases, North Carolina State Laboratory of Public Health COVID-19 Response Team | Infectious Diseases, North Carolina State Laboratory of Public Health COVID-19 Response Team | Chase,K. |
| EPI_ISL_538522, EPI_ISL_538523 | Infectious Diseases, North Carolina State Laboratory of Public Health COVID-19 Response Team | North Carolina State Laboratory of Public Health | Chase,K. |
| EPI_ISL_539344, EPI_ISL_539346, EPI_ISL_539348, EPI_ISL_539379, EPI_ISL_539382, EPI_ISL_539389, EPI_ISL_539410, EPI_ISL_539429, EPI_ISL_539430, EPI_ISL_539439, EPI_ISL_539446, EPI_ISL_539454, EPI_ISL_539473, EPI_ISL_539474 | | | |
| see above | Viollier AG | Department of Biosystems Science and Engineering, ETH Zürich | Christian Beisel, Sarah Nadeau, Ivan Topolsky, Pedro Ferreira, Philipp Jablonski, Susana Posada-Céspedes, Tobias Schär, Ina Nissen, Natascha Santacroce, Elodie Burcklen, Christiane Beckmann, Maurice Redondo, Olivier Kobel, Christoph Noppen, Sophie Seidel, Noemie Santamaria de Souza, Niko Beerenwinkel, Tanja Stadler |
| EPI_ISL_540897, EPI_ISL_540898, EPI_ISL_540899, EPI_ISL_540900, EPI_ISL_540901, EPI_ISL_540902, EPI_ISL_540903, EPI_ISL_540904, EPI_ISL_540905, EPI_ISL_540906, EPI_ISL_540907, EPI_ISL_540908, EPI_ISL_540909, EPI_ISL_540910, EPI_ISL_540911, EPI_ISL_540912, EPI_ISL_540913, EPI_ISL_540914, EPI_ISL_540915, EPI_ISL_540916, EPI_ISL_540917, EPI_ISL_540918, EPI_ISL_540919, EPI_ISL_540920 | | | |
| see above | Wales Specialist Virology Centre Sequencing lab: Pathogen Genomics Unit | COVID-19 Genomics UK (COG-UK) Consortium | Catherine Moore, Johnathan Evans, Laura Gifford, Malorie Perry, Simon Cottrell, Angela Marchbank, Alec Birchley, Alexander Adams, Amy Gaskin, Bree Gatica-Wilcox, Jason Coombes, Joel Southgate, Lauren Gilbert, Lee Graham, Nicole Pacchiarini, Sara Kumziene-Summerhayes, Sarah Taylor, Sophie Jones, Sara Rey, Matthew Bull, Joanne Watkins, Sally Corden, Tom Connor |
| EPI_ISL_541510, EPI_ISL_541512, EPI_ISL_541513, EPI_ISL_541516, EPI_ISL_541517, EPI_ISL_541519, EPI_ISL_541520, EPI_ISL_541521, EPI_ISL_541522, EPI_ISL_541523, EPI_ISL_541524, EPI_ISL_541525, EPI_ISL_541526, EPI_ISL_541527, EPI_ISL_541528, EPI_ISL_541529, EPI_ISL_541530, EPI_ISL_541531, EPI_ISL_541532, EPI_ISL_541533, EPI_ISL_541534, EPI_ISL_541536, EPI_ISL_541538, EPI_ISL_541539 | | | |
| see above | Viollier AG | Department of Biosystems Science and Engineering, ETH Zürich | Christian Beisel, Sarah Nadeau, Ivan Topolsky, Pedro Ferreira, Philipp Jablonski, Susana Posada-Céspedes, Tobias Schär, Ina Nissen, Natascha Santacroce, Elodie Burcklen, Christiane Beckmann, Maurice Redondo, Olivier Kobel, Christoph Noppen, Sophie Seidel, Noemie Santamaria de Souza, Niko Beerenwinkel, Tanja Stadler |
| EPI_ISL_560083, EPI_ISL_560084, EPI_ISL_560085, EPI_ISL_560086, EPI_ISL_560087, EPI_ISL_560088, EPI_ISL_560089, EPI_ISL_560090, EPI_ISL_560091, EPI_ISL_560092, EPI_ISL_560093, EPI_ISL_560094, EPI_ISL_560095, EPI_ISL_560096, EPI_ISL_560097, EPI_ISL_560098, EPI_ISL_560099, EPI_ISL_560100, EPI_ISL_560101, EPI_ISL_560102, EPI_ISL_560103, EPI_ISL_560104, EPI_ISL_560105, EPI_ISL_560106, EPI_ISL_560107, EPI_ISL_560108, EPI_ISL_560109, EPI_ISL_560110, EPI_ISL_560111, EPI_ISL_560112, EPI_ISL_560113, EPI_ISL_560114, EPI_ISL_560115, EPI_ISL_560116, EPI_ISL_560117, EPI_ISL_560118, EPI_ISL_560119, EPI_ISL_560120, EPI_ISL_560121, EPI_ISL_560122, EPI_ISL_560123, EPI_ISL_560124, EPI_ISL_560125, EPI_ISL_560126, EPI_ISL_560127, EPI_ISL_560128, EPI_ISL_560129, EPI_ISL_560130, EPI_ISL_560131, EPI_ISL_560132, EPI_ISL_560133, EPI_ISL_560134, EPI_ISL_560135, EPI_ISL_560136, EPI_ISL_560137, EPI_ISL_560138, EPI_ISL_560139, EPI_ISL_560140, EPI_ISL_560141, EPI_ISL_560142, EPI_ISL_560143, EPI_ISL_560144, EPI_ISL_560145, EPI_ISL_560146, EPI_ISL_560147, EPI_ISL_560148, EPI_ISL_560149, EPI_ISL_560150, EPI_ISL_560151, EPI_ISL_560152, EPI_ISL_560153, EPI_ISL_560154, | | | |

| | | | |
|---|---|---|---|
| EPI_ISL_560155, EPI_ISL_560156, EPI_ISL_560157, EPI_ISL_560158, EPI_ISL_560159, EPI_ISL_560160, EPI_ISL_560161, EPI_ISL_560162, EPI_ISL_560163, EPI_ISL_560164, EPI_ISL_560165, EPI_ISL_560166, EPI_ISL_560167, EPI_ISL_560168, EPI_ISL_560169, EPI_ISL_560170, EPI_ISL_560171, EPI_ISL_560172, EPI_ISL_560173, EPI_ISL_560174, EPI_ISL_560175, EPI_ISL_560176, EPI_ISL_560177, EPI_ISL_560178, EPI_ISL_560179, EPI_ISL_560180, EPI_ISL_560181, EPI_ISL_560182, EPI_ISL_560183, EPI_ISL_560184, EPI_ISL_560185, EPI_ISL_560186, EPI_ISL_560187, EPI_ISL_560188, EPI_ISL_560189, EPI_ISL_560190, EPI_ISL_560191, EPI_ISL_560192, EPI_ISL_560193, EPI_ISL_560194, EPI_ISL_560195, EPI_ISL_560196, EPI_ISL_560197, EPI_ISL_560198, EPI_ISL_560199, EPI_ISL_560200, EPI_ISL_560201, EPI_ISL_560202, EPI_ISL_560203, EPI_ISL_560204, EPI_ISL_560205, EPI_ISL_560206, EPI_ISL_560207, EPI_ISL_560208, EPI_ISL_560209, EPI_ISL_560210, EPI_ISL_560211, EPI_ISL_560212, EPI_ISL_560213, EPI_ISL_560214, EPI_ISL_560215, EPI_ISL_560216, EPI_ISL_560217, EPI_ISL_560218, EPI_ISL_560219, EPI_ISL_560220, EPI_ISL_560221, EPI_ISL_560222, EPI_ISL_560223, EPI_ISL_560224, EPI_ISL_560225, EPI_ISL_560226, EPI_ISL_560227, EPI_ISL_560228, EPI_ISL_560229, EPI_ISL_560230, EPI_ISL_560231, EPI_ISL_560232, EPI_ISL_560233, EPI_ISL_560234, EPI_ISL_560235, EPI_ISL_560236, EPI_ISL_560237, EPI_ISL_560238, EPI_ISL_560239, EPI_ISL_560240, EPI_ISL_560241, EPI_ISL_560242, EPI_ISL_560243, EPI_ISL_560244, EPI_ISL_560245, EPI_ISL_560246, EPI_ISL_560247, EPI_ISL_560248, EPI_ISL_560249, EPI_ISL_560250, EPI_ISL_560251, EPI_ISL_560252, EPI_ISL_560253, EPI_ISL_560254, EPI_ISL_560255, EPI_ISL_560256, EPI_ISL_560257, EPI_ISL_560258, EPI_ISL_560259, EPI_ISL_560260, EPI_ISL_560261, EPI_ISL_560262, EPI_ISL_560263, EPI_ISL_560264, EPI_ISL_560265, EPI_ISL_560266, EPI_ISL_560267, EPI_ISL_560268, EPI_ISL_560269, EPI_ISL_560270, EPI_ISL_560271, EPI_ISL_560272, EPI_ISL_560273, EPI_ISL_560274, EPI_ISL_560275, EPI_ISL_560276, EPI_ISL_560277, EPI_ISL_560278, EPI_ISL_560279, EPI_ISL_560280, EPI_ISL_560281, EPI_ISL_560282, EPI_ISL_560283, EPI_ISL_560284, EPI_ISL_560285, EPI_ISL_560286, EPI_ISL_560287, EPI_ISL_560288, EPI_ISL_560289, EPI_ISL_560290, EPI_ISL_560291, EPI_ISL_560292, EPI_ISL_560293, EPI_ISL_560294, EPI_ISL_560295, EPI_ISL_560296, EPI_ISL_560297, EPI_ISL_560298, EPI_ISL_560299, EPI_ISL_560300, EPI_ISL_560301, EPI_ISL_560302, EPI_ISL_560303, EPI_ISL_560304 | | | |
| see above | Wales Specialist Virology Centre Sequencing lab: Pathogen Genomics Unit | COVID-19 Genomics UK (COG-UK) Consortium | Catherine Moore, Johnathan Evans, Laura Gifford, Malorie Perry, Simon Cottrell, Angela Marchbank, Alec Birchley, Alexander Adams, Amy Gaskin, Bree Gatica-Wilcox, Jason Coombes, Joel Southgate, Lauren Gilbert, Lee Graham, Nicole Pacchiarini, Sara Kumziene-Summerhayes, Sarah Taylor, Sophie Jones, Sara Rey, Matthew Bull, Joanne Watkins, Sally Corden, Tom Connor |
| EPI_ISL_560541, EPI_ISL_560542, EPI_ISL_560543, EPI_ISL_560544, EPI_ISL_560546, EPI_ISL_560547, EPI_ISL_560548, EPI_ISL_560549, EPI_ISL_560550, EPI_ISL_560552 | Viollier AG | Department of Biosystems Science and Engineering, ETH Zürich | Christian Beisel, Sarah Nadeau, Ivan Topolsky, Pedro Ferreira, Philipp Jablonski, Susana Posada-Céspedes, Tobias Schär, Ina Nissen, Natascha Santacroce, Elodie Burcklen, Christiane Beckmann, Maurice Redondo, Olivier Kobel, Christoph Noppen, Sophie Seidel, Noemie Santamaria de Souza, Niko Beerenwinkel, Tanja Stadler |
| EPI_ISL_560649, EPI_ISL_560650, EPI_ISL_560651, EPI_ISL_560652, EPI_ISL_560653, EPI_ISL_560654, EPI_ISL_560655, EPI_ISL_560656, EPI_ISL_560657, EPI_ISL_560658, EPI_ISL_560659, EPI_ISL_560660, EPI_ISL_560661, EPI_ISL_560662, EPI_ISL_560663, EPI_ISL_560664, EPI_ISL_560665, EPI_ISL_560666, EPI_ISL_560667, EPI_ISL_560668, EPI_ISL_560669, EPI_ISL_560670, EPI_ISL_560671, EPI_ISL_560672, EPI_ISL_560673, EPI_ISL_560674, EPI_ISL_560675, EPI_ISL_560676, EPI_ISL_560677, EPI_ISL_560678, EPI_ISL_560679, EPI_ISL_560680, EPI_ISL_560681, EPI_ISL_560682, EPI_ISL_560683, EPI_ISL_560684, EPI_ISL_560685, EPI_ISL_560686, EPI_ISL_560687, EPI_ISL_560688, EPI_ISL_560689, EPI_ISL_560690, EPI_ISL_560691, EPI_ISL_560692, EPI_ISL_560693, EPI_ISL_560694, EPI_ISL_560695, EPI_ISL_560696, EPI_ISL_560697, EPI_ISL_560698, EPI_ISL_560699, EPI_ISL_560700, EPI_ISL_560701, EPI_ISL_560702, EPI_ISL_560703, EPI_ISL_560704, EPI_ISL_560705, EPI_ISL_560706, EPI_ISL_560707, EPI_ISL_560708, EPI_ISL_560709, EPI_ISL_560710, EPI_ISL_560711, EPI_ISL_560712, EPI_ISL_560713, EPI_ISL_560714, EPI_ISL_560715, EPI_ISL_560716, EPI_ISL_560717, EPI_ISL_560718, EPI_ISL_560719, EPI_ISL_560720, EPI_ISL_560721, EPI_ISL_560722, EPI_ISL_560723, EPI_ISL_560724, EPI_ISL_560725, EPI_ISL_560726, EPI_ISL_560727, EPI_ISL_560728, EPI_ISL_560729, EPI_ISL_560730, EPI_ISL_560731, EPI_ISL_560732, EPI_ISL_560733, EPI_ISL_560734, EPI_ISL_560735, EPI_ISL_560736, EPI_ISL_560737, EPI_ISL_560738, EPI_ISL_560739, EPI_ISL_560740, EPI_ISL_561363, EPI_ISL_561364, EPI_ISL_561365, EPI_ISL_561366, EPI_ISL_561367, EPI_ISL_561368, EPI_ISL_561369, EPI_ISL_561373, EPI_ISL_561374 | | | |
| see above | Centre for Clinical Infection and Diagnostics Research and Genomics Innovation Unit, Guy's and St. Thomas' NHS Trust | Centre for Clinical Infection and Diagnostics Research and Genomics Innovation Unit, Guy's and St. Thomas' NHS Trust | Chloe Fisher, Luke Snell, Rahul Batra, Jonathan Edgeworth, Ali Raza Awan |
| EPI_ISL_568579 | Virus Molecular Laboratory of the Microbiology and Virology Department | INMI Lazzaro Spallanzani IRCCS | Cesare E.M. Gruber, Martina Rueca, Barbara Bartolini, Francesco Messina, Silvia Meschi, Francesca Colavita, Concetta Castilletti, Elena Percivalle, Irene Cassaniti, Edoardo Vecchio Nepita, Fausto Baldanti, Maria R. Capobianchi, Antonino Di Caro |
| EPI_ISL_572405, EPI_ISL_572407, EPI_ISL_572408, EPI_ISL_572434, EPI_ISL_572446, EPI_ISL_572455, EPI_ISL_572461, EPI_ISL_572465, EPI_ISL_572468, EPI_ISL_572469, EPI_ISL_572474, EPI_ISL_572477, EPI_ISL_572487, EPI_ISL_572494, EPI_ISL_572496, EPI_ISL_572501, EPI_ISL_572512, EPI_ISL_572531, EPI_ISL_572540, EPI_ISL_572545, EPI_ISL_572547, EPI_ISL_572555, EPI_ISL_572558, EPI_ISL_572559, EPI_ISL_572560, EPI_ISL_572561, EPI_ISL_572564, EPI_ISL_572565, EPI_ISL_572566, EPI_ISL_572574, EPI_ISL_572575, EPI_ISL_572576, EPI_ISL_572577, EPI_ISL_572580, EPI_ISL_572586, EPI_ISL_572605, EPI_ISL_572606, EPI_ISL_572611, EPI_ISL_572612, EPI_ISL_572614, EPI_ISL_572643, EPI_ISL_572645, EPI_ISL_572647, EPI_ISL_572652, EPI_ISL_572654, EPI_ISL_572657, EPI_ISL_572659, EPI_ISL_572661, EPI_ISL_572662, EPI_ISL_572667, EPI_ISL_572668, EPI_ISL_572672, EPI_ISL_572674, EPI_ISL_572675, EPI_ISL_572676, EPI_ISL_572680, EPI_ISL_572681, EPI_ISL_572683, EPI_ISL_572690, EPI_ISL_572693, EPI_ISL_572694, EPI_ISL_572697, EPI_ISL_572698, EPI_ISL_572708, EPI_ISL_572731, EPI_ISL_572754, EPI_ISL_572766, EPI_ISL_572772, EPI_ISL_572778, EPI_ISL_572781, EPI_ISL_572782, EPI_ISL_572784, EPI_ISL_572790, EPI_ISL_572805, EPI_ISL_572810, EPI_ISL_572816, EPI_ISL_572818, EPI_ISL_572830, EPI_ISL_572836, EPI_ISL_572853, EPI_ISL_572854, EPI_ISL_572855, EPI_ISL_572856, EPI_ISL_572875, EPI_ISL_572876, EPI_ISL_572877, EPI_ISL_572878, EPI_ISL_572879, EPI_ISL_572880, EPI_ISL_572881, EPI_ISL_572882, EPI_ISL_572883, EPI_ISL_572884, EPI_ISL_572885, EPI_ISL_572886, EPI_ISL_572887, EPI_ISL_572888, EPI_ISL_572889, EPI_ISL_572890, EPI_ISL_572931, EPI_ISL_572932, EPI_ISL_572951, EPI_ISL_573873, EPI_ISL_573874, EPI_ISL_573875, EPI_ISL_573876, EPI_ISL_573877, EPI_ISL_573878, EPI_ISL_573879, EPI_ISL_573880, EPI_ISL_573881, EPI_ISL_573882, EPI_ISL_573883, EPI_ISL_573884, EPI_ISL_573885, EPI_ISL_573886, EPI_ISL_573887, EPI_ISL_573888, EPI_ISL_573889, EPI_ISL_573890, EPI_ISL_573891, EPI_ISL_573892, EPI_ISL_573893, EPI_ISL_573894, EPI_ISL_573895, EPI_ISL_573896, EPI_ISL_573897, EPI_ISL_573898, EPI_ISL_573899, EPI_ISL_573900, EPI_ISL_573901, EPI_ISL_573902, EPI_ISL_573903, EPI_ISL_573904, EPI_ISL_573905, EPI_ISL_573906, EPI_ISL_573907, EPI_ISL_573908, EPI_ISL_573909, EPI_ISL_573910, EPI_ISL_573911, EPI_ISL_573912, EPI_ISL_573913, EPI_ISL_573914, EPI_ISL_573915, EPI_ISL_573916, EPI_ISL_573917, EPI_ISL_573918, EPI_ISL_573919, EPI_ISL_573920, EPI_ISL_573921, EPI_ISL_573922, EPI_ISL_573923, EPI_ISL_573924, EPI_ISL_573925, EPI_ISL_573926, EPI_ISL_573927, EPI_ISL_573928, EPI_ISL_573929, EPI_ISL_573930, EPI_ISL_573931, EPI_ISL_573932, EPI_ISL_573933, EPI_ISL_573934, EPI_ISL_573935, EPI_ISL_573936, EPI_ISL_573937, EPI_ISL_573938, EPI_ISL_573939, EPI_ISL_573940, EPI_ISL_573941, EPI_ISL_573942, EPI_ISL_573943, EPI_ISL_573944, EPI_ISL_573945, EPI_ISL_573946, EPI_ISL_573947, EPI_ISL_573948, EPI_ISL_573949, EPI_ISL_573950, EPI_ISL_573951, EPI_ISL_573952, EPI_ISL_573953, EPI_ISL_573954, EPI_ISL_573955, EPI_ISL_573956, EPI_ISL_573957, EPI_ISL_573958, EPI_ISL_573959, EPI_ISL_573960, EPI_ISL_573961, EPI_ISL_573962, EPI_ISL_573963, EPI_ISL_573964, EPI_ISL_573965, EPI_ISL_573966, EPI_ISL_573967, EPI_ISL_573968, EPI_ISL_573969, EPI_ISL_573970, EPI_ISL_573971, EPI_ISL_573972, EPI_ISL_573973, EPI_ISL_573974, EPI_ISL_573975, EPI_ISL_573976, EPI_ISL_573977, EPI_ISL_573978, EPI_ISL_573979, EPI_ISL_573980, EPI_ISL_573981, EPI_ISL_573982, EPI_ISL_573983, EPI_ISL_573984, EPI_ISL_573985, EPI_ISL_573986, EPI_ISL_573987, EPI_ISL_573988, EPI_ISL_573989, EPI_ISL_573990, EPI_ISL_573991, EPI_ISL_573992, EPI_ISL_573993, EPI_ISL_573994, EPI_ISL_573995, EPI_ISL_573996, EPI_ISL_573997, EPI_ISL_573998, EPI_ISL_573999, EPI_ISL_574000, EPI_ISL_574001, EPI_ISL_574002, EPI_ISL_574003, EPI_ISL_574004, EPI_ISL_574005, EPI_ISL_574006, EPI_ISL_574007, EPI_ISL_574008, EPI_ISL_574009, EPI_ISL_574010, EPI_ISL_574011, EPI_ISL_574012, EPI_ISL_574013, EPI_ISL_574014, EPI_ISL_574015, EPI_ISL_574016, EPI_ISL_574017, EPI_ISL_574018, EPI_ISL_574019, EPI_ISL_574020, EPI_ISL_574021, EPI_ISL_574022, EPI_ISL_574023, EPI_ISL_574024, EPI_ISL_574025, EPI_ISL_574026, EPI_ISL_574027, EPI_ISL_574028, EPI_ISL_574029, EPI_ISL_574030, EPI_ISL_574031, EPI_ISL_574032, EPI_ISL_574033, EPI_ISL_574034, EPI_ISL_574035, EPI_ISL_574036, EPI_ISL_574037, EPI_ISL_574038, EPI_ISL_574039, EPI_ISL_574040, EPI_ISL_574041, EPI_ISL_574042, EPI_ISL_574043, EPI_ISL_574044, EPI_ISL_574045, EPI_ISL_574046, EPI_ISL_574047, EPI_ISL_574048, EPI_ISL_574049, EPI_ISL_574050, EPI_ISL_574051, EPI_ISL_574052, EPI_ISL_574053, EPI_ISL_574054, EPI_ISL_574055, EPI_ISL_574056, EPI_ISL_574057, EPI_ISL_574058, EPI_ISL_574059, EPI_ISL_574060, EPI_ISL_574061, EPI_ISL_574062, EPI_ISL_574063, EPI_ISL_574064, EPI_ISL_574065, EPI_ISL_574066, EPI_ISL_574067, EPI_ISL_574068, EPI_ISL_574069, EPI_ISL_574070, EPI_ISL_574071, EPI_ISL_574072, EPI_ISL_574073, EPI_ISL_574074, EPI_ISL_574075, EPI_ISL_574076, EPI_ISL_574077, EPI_ISL_574078, EPI_ISL_574079, EPI_ISL_574080, EPI_ISL_574081, EPI_ISL_574082, EPI_ISL_574083, EPI_ISL_574084, EPI_ISL_574085, EPI_ISL_574086, EPI_ISL_574087, EPI_ISL_574088, EPI_ISL_574089, EPI_ISL_574090, EPI_ISL_574091, EPI_ISL_574092, EPI_ISL_574093, EPI_ISL_574094, EPI_ISL_574095, EPI_ISL_574096, EPI_ISL_574097, EPI_ISL_574098, EPI_ISL_574099, EPI_ISL_574100, EPI_ISL_574101, EPI_ISL_574102, EPI_ISL_574103, EPI_ISL_574104, EPI_ISL_574105, EPI_ISL_574106, EPI_ISL_574107, EPI_ISL_574108, EPI_ISL_574109, EPI_ISL_574110, EPI_ISL_574111, EPI_ISL_574112, EPI_ISL_574113, EPI_ISL_574114, EPI_ISL_574115, EPI_ISL_574116, EPI_ISL_574117, EPI_ISL_574118, EPI_ISL_574119, EPI_ISL_574120, EPI_ISL_574121, EPI_ISL_574122, EPI_ISL_574123, EPI_ISL_574124, EPI_ISL_574125, EPI_ISL_574126, EPI_ISL_574127, EPI_ISL_574128, EPI_ISL_574129, EPI_ISL_574130, EPI_ISL_574131, EPI_ISL_574132, EPI_ISL_574133, EPI_ISL_574134, EPI_ISL_574135, EPI_ISL_574136, EPI_ISL_574137, EPI_ISL_574138, EPI_ISL_574139, EPI_ISL_574140, EPI_ISL_574141, EPI_ISL_574142, EPI_ISL_574143, EPI_ISL_574144, EPI_ISL_574145, EPI_ISL_574146, EPI_ISL_574147, EPI_ISL_574148, EPI_ISL_574149, EPI_ISL_574150, EPI_ISL_574151, EPI_ISL_574152, EPI_ISL_574153, EPI_ISL_574154, EPI_ISL_574155, EPI_ISL_574156, EPI_ISL_574157, EPI_ISL_574158, EPI_ISL_574159, EPI_ISL_574160, EPI_ISL_574161, EPI_ISL_574162, EPI_ISL_574163, EPI_ISL_574164, EPI_ISL_574165, EPI_ISL_574166, EPI_ISL_574167, EPI_ISL_574168, EPI_ISL_574169, EPI_ISL_574170, EPI_ISL_574171, EPI_ISL_574172, EPI_ISL_574173, EPI_ISL_574174, EPI_ISL_574175, EPI_ISL_574176, EPI_ISL_574177, EPI_ISL_574178, EPI_ISL_574179, EPI_ISL_574180, EPI_ISL_574181, EPI_ISL_574182, EPI_ISL_574183, EPI_ISL_574184, EPI_ISL_574185, EPI_ISL_574186, EPI_ISL_574187, EPI_ISL_574188, EPI_ISL_574189, EPI_ISL_574190, EPI_ISL_574191, EPI_ISL_574192, EPI_ISL_574193, EPI_ISL_574194, EPI_ISL_574195, EPI_ISL_574196, EPI_ISL_574197, EPI_ISL_574198, EPI_ISL_574199, EPI_ISL_574200, EPI_ISL_574201, EPI_ISL_574202, EPI_ISL_574203, EPI_ISL_574204, EPI_ISL_574205, EPI_ISL_574206, EPI_ISL_574207, EPI_ISL_574208, EPI_ISL_574209, EPI_ISL_574210, EPI_ISL_574211, EPI_ISL_574212, EPI_ISL_574213, EPI_ISL_574214, EPI_ISL_574215, EPI_ISL_574216, EPI_ISL_574217, EPI_ISL_574218, EPI_ISL_574219, EPI_ISL_574220, EPI_ISL_574221, EPI_ISL_574222, EPI_ISL_574223, EPI_ISL_574224, EPI_ISL_574225, EPI_ISL_574226, EPI_ISL_574227, EPI_ISL_574228, EPI_ISL_574229, EPI_ISL_574230, EPI_ISL_574231, EPI_ISL_574232, EPI_ISL_574233, EPI_ISL_574234, EPI_ISL_574235, EPI_ISL_574236, EPI_ISL_574237, EPI_ISL_574238, EPI_ISL_574239, EPI_ISL_574240, EPI_ISL_574241, EPI_ISL_574242, EPI_ISL_574243, EPI_ISL_574244, EPI_ISL_574245, EPI_ISL_574246, EPI_ISL_574247, EPI_ISL_574248, EPI_ISL_574249, EPI_ISL_574250, EPI_ISL_574251, EPI_ISL_574252, EPI_ISL_574253, EPI_ISL_574254, EPI_ISL_574255, EPI_ISL_574256 | | | |
| see above | Wales Specialist Virology Centre Sequencing lab: Pathogen Genomics Unit | COVID-19 Genomics UK (COG-UK) Consortium | Catherine Moore, Johnathan Evans, Laura Gifford, Malorie Perry, Simon Cottrell, Angela Marchbank, Alec Birchley, Alexander Adams, Amy Gaskin, Bree Gatica-Wilcox, Jason Coombes, Joel Southgate, Lauren Gilbert, Lee Graham, Nicole Pacchiarini, Sara Kumziene-Summerhayes, Sarah Taylor, Sophie Jones, Sara Rey, Matthew Bull, Joanne Watkins, Sally Corden, Tom Connor |
| EPI_ISL_574876, EPI_ISL_574877, EPI_ISL_574878, EPI_ISL_574879, EPI_ISL_574880, EPI_ISL_574881, EPI_ISL_574882, EPI_ISL_574931, EPI_ISL_574932, EPI_ISL_574933, EPI_ISL_574934, EPI_ISL_574935, EPI_ISL_574936, EPI_ISL_574938, EPI_ISL_574939, EPI_ISL_574940, EPI_ISL_574941, EPI_ISL_574942, EPI_ISL_574943, EPI_ISL_574947 | | | |
| see above | Viollier AG | Department of Biosystems Science and Engineering, ETH Zürich | Christian Beisel, Sarah Nadeau, Ivan Topolsky, Pedro Ferreira, Philipp Jablonski, Susana Posada-Céspedes, Tobias Schär, Ina Nissen, Natascha Santacroce, Elodie Burcklen, Christiane Beckmann, Maurice Redondo, Olivier Kobel, Christoph Noppen, Sophie Seidel, Noemie Santamaria de Souza, Niko Beerenwinkel, Tanja Stadler |
| EPI_ISL_577417, EPI_ISL_577418, EPI_ISL_577419, EPI_ISL_577420, EPI_ISL_577421, EPI_ISL_577422, EPI_ISL_577423, EPI_ISL_577424, EPI_ISL_577425, EPI_ISL_577426, EPI_ISL_577427, EPI_ISL_577428, EPI_ISL_577429, EPI_ISL_577430, EPI_ISL_577431, EPI_ISL_577432, EPI_ISL_577433, EPI_ISL_577434, EPI_ISL_577435, EPI_ISL_577436, EPI_ISL_577437, EPI_ISL_577438, EPI_ISL_577439, EPI_ISL_577440, EPI_ISL_577441, EPI_ISL_577442, EPI_ISL_577443, EPI_ISL_577444, EPI_ISL_577445, EPI_ISL_577446, EPI_ISL_577447, EPI_ISL_577448, EPI_ISL_577449, EPI_ISL_577450, EPI_ISL_577451, EPI_ISL_577452, EPI_ISL_577453, EPI_ISL_577454, EPI_ISL_577455, EPI_ISL_577456, EPI_ISL_577457, EPI_ISL_577458, EPI_ISL_577459, EPI_ISL_577460, EPI_ISL_577461, EPI_ISL_577462, EPI_ISL_577463, EPI_ISL_577464, EPI_ISL_577465, EPI_ISL_577466, EPI_ISL_577467, EPI_ISL_577468, EPI_ISL_577469, EPI_ISL_577470, EPI_ISL_577471, EPI_ISL_577472, EPI_ISL_577473, EPI_ISL_577474, EPI_ISL_577475, EPI_ISL_577476, EPI_ISL_577477, EPI_ISL_577478, EPI_ISL_577479, EPI_ISL_577480, EPI_ISL_577481, EPI_ISL_577482, EPI_ISL_577483, EPI_ISL_577484, EPI_ISL_577485, EPI_ISL_577486, EPI_ISL_577487, EPI_ISL_577488, EPI_ISL_577489, EPI_ISL_577490, EPI_ISL_577491, EPI_ISL_577492, EPI_ISL_577493, EPI_ISL_577494, EPI_ISL_577495, EPI_ISL_577496, EPI_ISL_577497, EPI_ISL_577498, EPI_ISL_577499, EPI_ISL_577500, EPI_ISL_577501, EPI_ISL_577502, EPI_ISL_577503, EPI_ISL_577504, EPI_ISL_577505, EPI_ISL_577506, EPI_ISL_577507, EPI_ISL_577508, EPI_ISL_577509, EPI_ISL_577510, EPI_ISL_577511, EPI_ISL_577512, EPI_ISL_577513, EPI_ISL_577514, EPI_ISL_577515, EPI_ISL_577516, EPI_ISL_577517, EPI_ISL_577518, EPI_ISL_577519, EPI_ISL_577520, EPI_ISL_577521, EPI_ISL_577522, EPI_ISL_577523, EPI_ISL_577524, EPI_ISL_577525, EPI_ISL_577526, EPI_ISL_577527, EPI_ISL_577528, EPI_ISL_577529, EPI_ISL_577530, EPI_ISL_577531, EPI_ISL_577532, EPI_ISL_577533, EPI_ISL_577534, EPI_ISL_577535, EPI_ISL_577536, EPI_ISL_577537, EPI_ISL_577538, EPI_ISL_577539, EPI_ISL_577540, EPI_ISL_577541, EPI_ISL_577542, EPI_ISL_577543 | | | |
| see above | Wales Specialist Virology Centre Sequencing lab: Pathogen Genomics Unit | COVID-19 Genomics UK (COG-UK) Consortium | Catherine Moore, Johnathan Evans, Laura Gifford, Malorie Perry, Simon Cottrell, Angela Marchbank, Alec Birchley, Alexander Adams, Amy Gaskin, Bree Gatica-Wilcox, Jason Coombes, Joel Southgate, Lauren Gilbert, Lee Graham, Nicole Pacchiarini, Sara Kumziene-Summerhayes, Sarah Taylor, Sophie Jones, Sara Rey, Matthew Bull, Joanne Watkins, Sally Corden, Tom Connor |
| EPI_ISL_583914 | DH | Department of Neurovirology, National Institute of Mental Health and Neuroscience (NIMHANS) | Chitra Pattabiraman,Vijayalakshmi Reddy, Harsha PK, Risha Rasheed, Pramada Prasad, Shafeeq S Hameed, Manjunatha Venkataswamy, Anita Desai, Ravi Vasanthapuram |
| EPI_ISL_583915 | Victoria Hospital | Department of Neurovirology, National Institute of Mental Health and Neuroscience (NIMHANS) | Chitra Pattabiraman,Vijayalakshmi Reddy, Harsha PK, Risha Rasheed, Pramada Prasad, Shafeeq S Hameed, Manjunatha Venkataswamy, Anita Desai, Ravi Vasanthapuram |
| EPI_ISL_585630, EPI_ISL_585631, EPI_ISL_585632, EPI_ISL_585633, EPI_ISL_585634, EPI_ISL_585635, EPI_ISL_585636, EPI_ISL_585637, EPI_ISL_585638, EPI_ISL_585639, EPI_ISL_585640, EPI_ISL_585641, EPI_ISL_585642, EPI_ISL_585643, EPI_ISL_585644, EPI_ISL_585645, EPI_ISL_585646, EPI_ISL_585647, EPI_ISL_585648, EPI_ISL_585649, EPI_ISL_585650, EPI_ISL_585651, EPI_ISL_585652, EPI_ISL_585653, EPI_ISL_585654, EPI_ISL_585655, EPI_ISL_585656, EPI_ISL_585657, EPI_ISL_585658, EPI_ISL_585659, EPI_ISL_585660, EPI_ISL_585661, EPI_ISL_585662, EPI_ISL_585663, EPI_ISL_585664, EPI_ISL_585665, EPI_ISL_585666, EPI_ISL_585667, EPI_ISL_585668, EPI_ISL_585669, EPI_ISL_585670, EPI_ISL_585671, EPI_ISL_585672, EPI_ISL_585673, EPI_ISL_585674, EPI_ISL_585675, EPI_ISL_585676, EPI_ISL_585677, EPI_ISL_585678, EPI_ISL_585679, EPI_ISL_585680, EPI_ISL_585681, EPI_ISL_585682, EPI_ISL_585683, | | | |

| | | | |
|---|---|---|---|
| EPI_ISL_585684, EPI_ISL_585685, EPI_ISL_585686, EPI_ISL_585687, EPI_ISL_585688, EPI_ISL_585689, EPI_ISL_585690, EPI_ISL_585691, EPI_ISL_585692, EPI_ISL_585693, EPI_ISL_585694, EPI_ISL_585695, EPI_ISL_585696, EPI_ISL_585697, EPI_ISL_585698, EPI_ISL_585699, EPI_ISL_585700, EPI_ISL_585701, EPI_ISL_585702, EPI_ISL_585703, EPI_ISL_585704, EPI_ISL_585705, EPI_ISL_585706, EPI_ISL_585707, EPI_ISL_585708, EPI_ISL_585709, EPI_ISL_585710, EPI_ISL_585711, EPI_ISL_585712, EPI_ISL_585713, EPI_ISL_585714, EPI_ISL_585715, EPI_ISL_585716, EPI_ISL_585717, EPI_ISL_585718, EPI_ISL_585719, EPI_ISL_585720, EPI_ISL_585721, EPI_ISL_585722, EPI_ISL_585723, EPI_ISL_585724, EPI_ISL_585725, EPI_ISL_585726, EPI_ISL_585727, EPI_ISL_585728, EPI_ISL_585729, EPI_ISL_585730, EPI_ISL_585731, EPI_ISL_585732, EPI_ISL_585733, EPI_ISL_585734, EPI_ISL_585735, EPI_ISL_585736, EPI_ISL_585737, EPI_ISL_585738, EPI_ISL_585739, EPI_ISL_585740, EPI_ISL_585741, EPI_ISL_585742, EPI_ISL_585743, EPI_ISL_585744, EPI_ISL_585745, EPI_ISL_585746, EPI_ISL_585747, EPI_ISL_585748, EPI_ISL_585749, EPI_ISL_585750, EPI_ISL_585751, EPI_ISL_585752, EPI_ISL_585753, EPI_ISL_585754, EPI_ISL_585755, EPI_ISL_585756, EPI_ISL_585757, EPI_ISL_585758, EPI_ISL_585759, EPI_ISL_585760, EPI_ISL_585761, EPI_ISL_585762, EPI_ISL_585763, EPI_ISL_585764, EPI_ISL_585765, EPI_ISL_585766, EPI_ISL_585767, EPI_ISL_585768, EPI_ISL_585769, EPI_ISL_585770, EPI_ISL_585771, EPI_ISL_585772, EPI_ISL_585773, EPI_ISL_585774, EPI_ISL_585775, EPI_ISL_585776, EPI_ISL_585777, EPI_ISL_585778, EPI_ISL_585779, EPI_ISL_585780, EPI_ISL_585781, EPI_ISL_585782, EPI_ISL_585783, EPI_ISL_585784, EPI_ISL_585785, EPI_ISL_585786, EPI_ISL_585787, EPI_ISL_585788, EPI_ISL_585789, EPI_ISL_585790, EPI_ISL_585791, EPI_ISL_585792, EPI_ISL_585793, EPI_ISL_585794, EPI_ISL_585795, EPI_ISL_585796, EPI_ISL_585797, EPI_ISL_585798, EPI_ISL_585799, EPI_ISL_585800, EPI_ISL_585801, EPI_ISL_585802, EPI_ISL_585803, EPI_ISL_585804, EPI_ISL_585805, EPI_ISL_585806, EPI_ISL_585807, EPI_ISL_585808, EPI_ISL_585809, EPI_ISL_585810, EPI_ISL_585811, EPI_ISL_585812, EPI_ISL_585813, EPI_ISL_585814, EPI_ISL_585815, EPI_ISL_585816, EPI_ISL_585817, EPI_ISL_585818, EPI_ISL_585819, EPI_ISL_585820, EPI_ISL_585821, EPI_ISL_585822, EPI_ISL_585823, EPI_ISL_585824, EPI_ISL_585825, EPI_ISL_585826, EPI_ISL_585827, EPI_ISL_585828, EPI_ISL_585829, EPI_ISL_585830, EPI_ISL_585831, EPI_ISL_585832, EPI_ISL_585833, EPI_ISL_585834, EPI_ISL_585835, EPI_ISL_585836, EPI_ISL_585837, EPI_ISL_585838, EPI_ISL_585839, EPI_ISL_585840, EPI_ISL_585841, EPI_ISL_585842, EPI_ISL_585843, EPI_ISL_585844, EPI_ISL_585845, EPI_ISL_585846, EPI_ISL_585847, EPI_ISL_585848, EPI_ISL_585849, EPI_ISL_585850, EPI_ISL_585851, EPI_ISL_585852, EPI_ISL_585853, EPI_ISL_585854, EPI_ISL_585855, EPI_ISL_585856, EPI_ISL_585857, EPI_ISL_585858, EPI_ISL_585859, EPI_ISL_585860, EPI_ISL_585861, EPI_ISL_585862, EPI_ISL_585863, EPI_ISL_585864, EPI_ISL_585865, EPI_ISL_585866, EPI_ISL_585867, EPI_ISL_585868, EPI_ISL_585869, EPI_ISL_585870, EPI_ISL_585871, EPI_ISL_585872, EPI_ISL_585873, EPI_ISL_585874, EPI_ISL_585875, EPI_ISL_585876, EPI_ISL_585877, EPI_ISL_585878, EPI_ISL_585879, EPI_ISL_585880, EPI_ISL_585881, EPI_ISL_585882, EPI_ISL_585883, EPI_ISL_585884, EPI_ISL_585885, EPI_ISL_585886, EPI_ISL_585887, EPI_ISL_585888, EPI_ISL_585889, EPI_ISL_585890, EPI_ISL_585891, EPI_ISL_585892, EPI_ISL_585893, EPI_ISL_585894, EPI_ISL_585895, EPI_ISL_585896, EPI_ISL_585897, EPI_ISL_585898, EPI_ISL_585899, EPI_ISL_585900, EPI_ISL_585901, EPI_ISL_585902, EPI_ISL_585903, EPI_ISL_585904, EPI_ISL_585905, EPI_ISL_585906, EPI_ISL_585907, EPI_ISL_585908, EPI_ISL_585909, EPI_ISL_585910, EPI_ISL_585911, EPI_ISL_585912, EPI_ISL_585913, EPI_ISL_585914, EPI_ISL_585915, EPI_ISL_585916, EPI_ISL_585917, EPI_ISL_585918, EPI_ISL_585919, EPI_ISL_585920, EPI_ISL_585921, EPI_ISL_585922, EPI_ISL_585923, EPI_ISL_585924, EPI_ISL_585925, EPI_ISL_585926, EPI_ISL_585927, EPI_ISL_585928, EPI_ISL_585929, EPI_ISL_585930, EPI_ISL_585931, EPI_ISL_585932, EPI_ISL_585933, EPI_ISL_585934, EPI_ISL_585935, EPI_ISL_585936, EPI_ISL_585937, EPI_ISL_585938, EPI_ISL_585939, EPI_ISL_585940, EPI_ISL_585941, EPI_ISL_585942, EPI_ISL_585943, EPI_ISL_585944, EPI_ISL_585945, EPI_ISL_585946, EPI_ISL_585947, EPI_ISL_585948, EPI_ISL_585949, EPI_ISL_585950, EPI_ISL_585951, EPI_ISL_585952, EPI_ISL_585953, EPI_ISL_585954, EPI_ISL_585955, EPI_ISL_585956, EPI_ISL_585957, EPI_ISL_585958, EPI_ISL_585959, EPI_ISL_585960, EPI_ISL_585961, EPI_ISL_585962, EPI_ISL_585963, EPI_ISL_585964, EPI_ISL_585965, EPI_ISL_585966, EPI_ISL_585967, EPI_ISL_585968, EPI_ISL_585969, EPI_ISL_585970, EPI_ISL_585971, EPI_ISL_585972, EPI_ISL_585973, EPI_ISL_585974, EPI_ISL_585975, EPI_ISL_585976, EPI_ISL_585977, EPI_ISL_585978, EPI_ISL_585979, EPI_ISL_585980, EPI_ISL_585981, EPI_ISL_585982, EPI_ISL_585983, EPI_ISL_585984, EPI_ISL_585985, EPI_ISL_585986, EPI_ISL_585987, EPI_ISL_585988, EPI_ISL_585989, EPI_ISL_585990, EPI_ISL_585991, EPI_ISL_585992, EPI_ISL_585993, EPI_ISL_585994, EPI_ISL_585995, EPI_ISL_585996, EPI_ISL_585997, EPI_ISL_585998, EPI_ISL_585999, EPI_ISL_586000, EPI_ISL_586001, EPI_ISL_586002, EPI_ISL_586003, EPI_ISL_586004, EPI_ISL_586005, EPI_ISL_586006, EPI_ISL_586007, EPI_ISL_586008, EPI_ISL_586009, EPI_ISL_586010, EPI_ISL_586011, EPI_ISL_586012, EPI_ISL_586013, EPI_ISL_586014, EPI_ISL_586015, EPI_ISL_586016, EPI_ISL_586017, EPI_ISL_586018, EPI_ISL_586019, EPI_ISL_586020, EPI_ISL_586021, EPI_ISL_586022, EPI_ISL_586023, EPI_ISL_586024, EPI_ISL_586025, EPI_ISL_586026, EPI_ISL_586027, EPI_ISL_586028, EPI_ISL_586029, EPI_ISL_586030, EPI_ISL_586031, EPI_ISL_586032, EPI_ISL_586033, EPI_ISL_586034, EPI_ISL_586035, EPI_ISL_586036, EPI_ISL_586037, EPI_ISL_586038, EPI_ISL_586039, EPI_ISL_586040, EPI_ISL_586041, EPI_ISL_586042, EPI_ISL_586043, EPI_ISL_586044, EPI_ISL_586045, EPI_ISL_586046, EPI_ISL_586047, EPI_ISL_586048, EPI_ISL_586049, EPI_ISL_586050, EPI_ISL_586051, EPI_ISL_586052, EPI_ISL_586053, EPI_ISL_586054, EPI_ISL_586055, EPI_ISL_586056, EPI_ISL_586057, EPI_ISL_586058, EPI_ISL_586059, EPI_ISL_586060, EPI_ISL_586061, EPI_ISL_586062, EPI_ISL_586063, EPI_ISL_586064, EPI_ISL_586065, EPI_ISL_586066, EPI_ISL_586067, EPI_ISL_586068, EPI_ISL_586069, EPI_ISL_586070, EPI_ISL_586071, EPI_ISL_586072, EPI_ISL_586073, EPI_ISL_586074, EPI_ISL_586075, EPI_ISL_586076, EPI_ISL_586077, EPI_ISL_586078, EPI_ISL_586079, EPI_ISL_586080, EPI_ISL_586081, EPI_ISL_586082, EPI_ISL_586083, EPI_ISL_586084, EPI_ISL_586085, EPI_ISL_586086, EPI_ISL_586087, EPI_ISL_586088, EPI_ISL_586089, EPI_ISL_586090, EPI_ISL_586091, EPI_ISL_586092, EPI_ISL_586093, EPI_ISL_586094, EPI_ISL_586095, EPI_ISL_586096, EPI_ISL_586097, EPI_ISL_586098, EPI_ISL_586099, EPI_ISL_586100, EPI_ISL_586101, EPI_ISL_586102, EPI_ISL_586103, EPI_ISL_586104, EPI_ISL_586105, EPI_ISL_586108, EPI_ISL_586109, EPI_ISL_586110, EPI_ISL_586111, EPI_ISL_586113, EPI_ISL_586114, EPI_ISL_586116, EPI_ISL_586117, EPI_ISL_586118, EPI_ISL_586121, EPI_ISL_586122, EPI_ISL_586123, EPI_ISL_586125, EPI_ISL_586127, EPI_ISL_586128, EPI_ISL_586129, EPI_ISL_586130, EPI_ISL_586132, EPI_ISL_586133, EPI_ISL_586135, EPI_ISL_586136, EPI_ISL_586137, EPI_ISL_586139, EPI_ISL_586142, EPI_ISL_586145, EPI_ISL_586147, EPI_ISL_586149, EPI_ISL_586150, EPI_ISL_586152, EPI_ISL_586153, EPI_ISL_586154, EPI_ISL_586155, EPI_ISL_586156, EPI_ISL_586157, EPI_ISL_586158, EPI_ISL_586159, EPI_ISL_586162, EPI_ISL_586163, EPI_ISL_586164, EPI_ISL_586165, EPI_ISL_586167, EPI_ISL_586168, EPI_ISL_586171, EPI_ISL_586172, EPI_ISL_586176, EPI_ISL_586177, EPI_ISL_586178, EPI_ISL_586179, EPI_ISL_586180, EPI_ISL_586182, EPI_ISL_586184, EPI_ISL_586188, EPI_ISL_586190, EPI_ISL_586191, EPI_ISL_586193, EPI_ISL_586194, EPI_ISL_586198, EPI_ISL_586199, EPI_ISL_586200, EPI_ISL_586202, EPI_ISL_586203, EPI_ISL_586204, EPI_ISL_586205, EPI_ISL_586206, EPI_ISL_586207, EPI_ISL_586210, EPI_ISL_586211, EPI_ISL_586213, EPI_ISL_586214, EPI_ISL_586217, EPI_ISL_586220, EPI_ISL_586221, EPI_ISL_586224, EPI_ISL_586225, EPI_ISL_586227, EPI_ISL_586229, EPI_ISL_586230, EPI_ISL_586231, EPI_ISL_586232, EPI_ISL_586234, EPI_ISL_586235, EPI_ISL_586236, EPI_ISL_586237, EPI_ISL_586238 | | | |
| see above | Wales Specialist Virology Centre Sequencing lab: Pathogen Genomics Unit | COVID-19 Genomics UK (COG-UK) Consortium | Catherine Moore, Johnathan Evans, Laura Gifford, Malorie Perry, Simon Cottrell, Angela Marchbank, Alec Birchley, Alexander Adams, Amy Gaskin, Bree Gatica-Wilcox, Jason Coombes, Joel Southgate, Lauren Gilbert, Lee Graham, Nicole Pacchiarini, Sara Kumziene-Summerhayes, Sarah Taylor, Sophie Jones, Sara Rey, Matthew Bull, Joanne Watkins, Sally Corden, Tom Connor |
| EPI_ISL_590695 | INMI Lazzaro Spallanzani IRCCS | INMI Lazzaro Spallanzani IRCCS | Cesare E.M. Gruber, Francesco Messina, Barbara Bartolini, Martina Rueca, Emanuela Giombini, Beatrice Valli, Eleonora Lalle, Simone Lanini, Francesco Vairo, Antonino Di Caro, Maria R. Capobianchi |
| EPI_ISL_590696 | INMI Lazzaro Spallanzani IRCCS | INMI Lazzaro Spallanzani IRCCS | Cesare E.M. Gruber, Barbara Bartolini, Francesco Messina, Martina Rueca, Emanuela Giombini, Beatrice Valli, Eleonora Lalle, Simone Lanini, Francesco Vairo, Antonino Di Caro, Maria R. Capobianchi |
| EPI_ISL_591540 | St.Johns Hospital | Department of Neurovirology, National Institute of Mental Health and Neuroscience (NIMHANS) | Chitra Pattabiraman,Vijayalakshmi Reddy, Harsha PK, Risha Rasheed, Pramada Prasad, Shafeeq S Hameed, Manjunatha Venkataswamy, Anita Desai, Ravi Vasanthapuram |
| EPI_ISL_595334, EPI_ISL_595335, EPI_ISL_595338, EPI_ISL_595342, EPI_ISL_595347, EPI_ISL_595348, EPI_ISL_595351, EPI_ISL_595353, EPI_ISL_595354, EPI_ISL_595357, EPI_ISL_595360, EPI_ISL_595362, EPI_ISL_595366, EPI_ISL_595367, EPI_ISL_595368, EPI_ISL_595373, EPI_ISL_595379, EPI_ISL_595384, EPI_ISL_595395, EPI_ISL_595398, EPI_ISL_595399, EPI_ISL_595401, EPI_ISL_595411, EPI_ISL_595415, EPI_ISL_595420, EPI_ISL_595421, EPI_ISL_595422, EPI_ISL_595433, EPI_ISL_595436, EPI_ISL_595438, EPI_ISL_595440, EPI_ISL_595442, EPI_ISL_595445, EPI_ISL_595446, EPI_ISL_595454, EPI_ISL_595456, EPI_ISL_595458, EPI_ISL_595461, EPI_ISL_595464, EPI_ISL_595467, EPI_ISL_595468, EPI_ISL_595472, EPI_ISL_595474, EPI_ISL_595477, EPI_ISL_595479, EPI_ISL_595480, EPI_ISL_595482, EPI_ISL_595486, EPI_ISL_595489, EPI_ISL_595499, EPI_ISL_595500, EPI_ISL_595503, EPI_ISL_595507, EPI_ISL_595508, EPI_ISL_595509, EPI_ISL_595516, EPI_ISL_595517, EPI_ISL_595521, EPI_ISL_595522, EPI_ISL_595524, EPI_ISL_595528, EPI_ISL_595531, EPI_ISL_595533, EPI_ISL_595537, EPI_ISL_595539, EPI_ISL_595557, EPI_ISL_595558, EPI_ISL_595560, EPI_ISL_595566, EPI_ISL_595567, EPI_ISL_595568, EPI_ISL_595569, EPI_ISL_595574, EPI_ISL_595575, EPI_ISL_595583, EPI_ISL_595585, EPI_ISL_595591, EPI_ISL_595597, EPI_ISL_595600, EPI_ISL_595602, EPI_ISL_595608, EPI_ISL_595615 | | | |
| see above | Wales Specialist Virology Centre Sequencing lab: Pathogen Genomics Unit | COVID-19 Genomics UK (COG-UK) Consortium | Catherine Moore, Johnathan Evans, Laura Gifford, Malorie Perry, Simon Cottrell, Angela Marchbank, Alec Birchley, Alexander Adams, Amy Gaskin, Bree Gatica-Wilcox, Jason Coombes, Joel Southgate, Lauren Gilbert, Lee Graham, Nicole Pacchiarini, Sara Kumziene-Summerhayes, Sarah Taylor, Sophie Jones, Sara Rey, Matthew Bull, Joanne Watkins, Sally Corden, Tom Connor |
| EPI_ISL_596447, EPI_ISL_596448, EPI_ISL_602208, EPI_ISL_602209, EPI_ISL_602278, EPI_ISL_602279, EPI_ISL_602280, EPI_ISL_602281 | New Mexico Department of Health Scientific Laboratory Division | Los Alamos National Laboratory Bioscience Division | Chien-Chi Lo, Migun Shakya, Cheryl Gleasner, Kim McMurry, Alina Deshpande, Twila Kunde, Joseph Hicks, Michael Edwards, Patrick Chain |
| EPI_ISL_602564 | Centre for Dengue Research, Department of Immunology and Molecular Medicine | Centre for Dengue Research | Chandima Jeewandara, Deshni Jayathilaka, Dinuka Ariyaratne, Diyanath Ranasinghe, Laksiri Gomes, Ananda Wijewickrama, Malika Karunaratne, Gathsaurie Neelika Malavige |
| EPI_ISL_602567, EPI_ISL_602574, EPI_ISL_602575, EPI_ISL_602576 | Centre for Dengue Research, Department of Immunology and Molecular Medicine | Centre for Dengue Research, Department of Immunology and Molecular Medicine | Chandima Jeewandara, Deshni Jayathilaka, Dinuka Ariyaratne, Laksiri Gomes, Diyanath Ranasinghe, Gathsaurie Neelika Malavige |
| EPI_ISL_603137 | INMI Lazzaro Spallanzani IRCCS | INMI Lazzaro Spallanzani IRCCS | Cesare E.M. Gruber, Martina Rueca, Barbara Bartolini, Francesco Messina, Emanuela Giombini, Simone Lanini, Antonino Di Caro, Maria R. Capobianchi |
| EPI_ISL_603144 | INMI Lazzaro Spallanzani IRCCS | INMI Lazzaro Spallanzani IRCCS | Cesare E.M. Gruber, Martina Rueca, Francesco Messina, Barbara Bartolini, Emanuela Giombini, Simone Lanini, Maria R. Capobianchi, Antonino Di Caro |
| EPI_ISL_603146 | INMI Lazzaro Spallanzani IRCCS | INMI Lazzaro Spallanzani IRCCS | Cesare E.M. Gruber, Martina Rueca, Barbara Bartolini, Emanuela Giombini, Francesco Messina, Simone Lanini, Maria R. Capobianchi, Antonino Di Caro |
| EPI_ISL_603147 | INMI Lazzaro Spallanzani IRCCS | INMI Lazzaro Spallanzani IRCCS | Cesare E.M. Gruber, Barbara Bartolini, Francesco Messina, Martina Rueca, Simone Lanini, Emanuela Giombini, Antonino Di Caro, Maria R. Capobianchi |
| EPI_ISL_603148 | INMI Lazzaro Spallanzani IRCCS | INMI Lazzaro Spallanzani IRCCS | Cesare E.M. Gruber, Francesco Messina, Martina Rueca, Simone Lanini, Emanuela Giombini, Barbara Bartolini, Maria R. Capobianchi, Antonino Di Caro |
| EPI_ISL_603161 | INMI Lazzaro Spallanzani IRCCS | INMI Lazzaro Spallanzani IRCCS | Cesare E.M. Gruber, Barbara Bartolini, Francesco Messina, Martina Rueca, Emanuela Giombini, Simone Lanini, Fulvia Pimpinelli, Antonino Di Caro, Maria R. Capobianchi |
| EPI_ISL_603162 | INMI Lazzaro Spallanzani IRCCS | INMI Lazzaro Spallanzani IRCCS | Cesare E.M. Gruber, Francesco Messina, Martina Rueca, Barbara Bartolini, Emanuela Giombini, Simone Lanini, Fulvia Pimpinelli, Antonino Di Caro, Maria R. Capobianchi |
| EPI_ISL_603163 | INMI Lazzaro Spallanzani IRCCS | INMI Lazzaro Spallanzani IRCCS | Cesare E.M. Gruber, Martina Rueca, Emanuela Giombini, Simone Lanini, Francesco Messina, Barbara Bartolini, Fulvia Pimpinelli, Maria R. Capobianchi, Antonino Di Caro |
| EPI_ISL_603164 | INMI Lazzaro Spallanzani IRCCS | INMI Lazzaro Spallanzani IRCCS | Cesare E.M. Gruber, Francesco Messina, Martina Rueca, Barbara Bartolini, Emanuela Giombini, Simone Lanini, Fulvia Pimpinelli, Antonino Di Caro, Maria R. Capobianchi |
| EPI_ISL_603172 | INMI Lazzaro Spallanzani IRCCS | INMI Lazzaro Spallanzani IRCCS | Cesare E.M. Gruber, Martina Rueca, Barbara Bartolini, Simone Lanini, Francesco Messina, Emanuela Giombini, Mariarosa Gaudio, Antonino Di Caro, Maria R. Capobianchi |
| EPI_ISL_603176 | INMI Lazzaro Spallanzani IRCCS | INMI Lazzaro Spallanzani IRCCS | Cesare E.M. Gruber, Barbara Bartolini, Francesco Messina, Martina Rueca, Simone Lanini, Emanuela Giombini, Antonino Di Caro, Maria R. Capobianchi |
| EPI_ISL_603179 | INMI Lazzaro Spallanzani IRCCS | INMI Lazzaro Spallanzani IRCCS | Cesare E.M. Gruber, Barbara Bartolini, Francesco Messina, Martina Rueca, Emanuela Giombini, Simone Lanini, Antonino Di Caro, Maria R. Capobianchi |
| EPI_ISL_603185 | INMI Lazzaro Spallanzani IRCCS | INMI Lazzaro Spallanzani IRCCS | Cesare E.M. Gruber, Martina Rueca, Barbara Bartolini, Francesco Messina, Simone Lanini, Emanuela Giombini, Maria R. Capobianchi, Antonino Di Caro |
| EPI_ISL_603187 | INMI Lazzaro Spallanzani IRCCS | INMI Lazzaro Spallanzani IRCCS | Cesare E.M. Gruber, Martina Rueca, Francesco Messina, Barbara Bartolini, Simone Lanini, Emanuela Giombini, Antonino Di Caro, Maria R. Capobianchi |
| EPI_ISL_603289, EPI_ISL_603344, EPI_ISL_603372, EPI_ISL_603375, EPI_ISL_603384, EPI_ISL_603392, EPI_ISL_603400, EPI_ISL_603401, EPI_ISL_603469, EPI_ISL_603470, EPI_ISL_603475, EPI_ISL_603541, EPI_ISL_603542, EPI_ISL_603543, EPI_ISL_603548, EPI_ISL_603551, EPI_ISL_603595, EPI_ISL_603597, EPI_ISL_603606, EPI_ISL_603622, EPI_ISL_603685, EPI_ISL_603686, EPI_ISL_603687, EPI_ISL_603692, EPI_ISL_603696, EPI_ISL_603718, EPI_ISL_603723, EPI_ISL_603729, EPI_ISL_603752 | | | |

| | | | |
|---|---|---|---|
| see above | Viollier AG | Department of Biosystems Science and Engineering, ETH Zürich | Christian Beisel, Sarah Nadeau, Ivan Topolsky, Pedro Ferreira, Philipp Jablonski, Susana Posada-Céspedes, Tobias Schär, Ina Nissen, Natascha Santacroce, Elodie Burcklen, Christiane Beckmann, Maurice Redondo, Olivier Kobel, Christoph Noppen, Sophie Seidel, Noemie Santamaria de Souza, Niko Beerenwinkel, Tanja Stadler |
| EPI_ISL_611518, EPI_ISL_611519, EPI_ISL_611529, EPI_ISL_611530, EPI_ISL_611534, EPI_ISL_611560, EPI_ISL_611561, EPI_ISL_611562, EPI_ISL_611563, EPI_ISL_611564, EPI_ISL_611566, EPI_ISL_611573, EPI_ISL_611578, EPI_ISL_611581, EPI_ISL_611586, EPI_ISL_611588, EPI_ISL_611601, EPI_ISL_611605, EPI_ISL_611609, EPI_ISL_611624, EPI_ISL_611625, EPI_ISL_611626, EPI_ISL_611631, EPI_ISL_611633, EPI_ISL_611637, EPI_ISL_611640, EPI_ISL_611652, EPI_ISL_611663, EPI_ISL_611666, EPI_ISL_611673, EPI_ISL_611674, EPI_ISL_611684, EPI_ISL_611686, EPI_ISL_611687, EPI_ISL_611688, EPI_ISL_611691, EPI_ISL_611697, EPI_ISL_611704, EPI_ISL_611706, EPI_ISL_611722, EPI_ISL_611725, EPI_ISL_611726, EPI_ISL_611732, EPI_ISL_611735, EPI_ISL_611738, EPI_ISL_611740, EPI_ISL_611742, EPI_ISL_611748, EPI_ISL_611749, EPI_ISL_611809, EPI_ISL_611819, EPI_ISL_611824, EPI_ISL_611828, EPI_ISL_611834, EPI_ISL_611835, EPI_ISL_611849, EPI_ISL_611854, EPI_ISL_611863, EPI_ISL_611865, EPI_ISL_611886, EPI_ISL_611888, EPI_ISL_611896, EPI_ISL_611900, EPI_ISL_611901, EPI_ISL_611902, EPI_ISL_611903, EPI_ISL_611904, EPI_ISL_611906, EPI_ISL_611907, EPI_ISL_611909, EPI_ISL_611912, EPI_ISL_611913, EPI_ISL_611922, EPI_ISL_611923, EPI_ISL_611931, EPI_ISL_611932, EPI_ISL_611968, EPI_ISL_611969, EPI_ISL_611970, EPI_ISL_611973, EPI_ISL_611975, EPI_ISL_611982, EPI_ISL_611986, EPI_ISL_611993, EPI_ISL_611994, EPI_ISL_612002, EPI_ISL_612006, EPI_ISL_612012, EPI_ISL_612013, EPI_ISL_612018, EPI_ISL_612029, EPI_ISL_612031, EPI_ISL_612038, EPI_ISL_612040, EPI_ISL_612045, EPI_ISL_612048, EPI_ISL_612053, EPI_ISL_612076, EPI_ISL_612081, EPI_ISL_612083, EPI_ISL_612087, EPI_ISL_612091, EPI_ISL_612093, EPI_ISL_612098, EPI_ISL_612109, EPI_ISL_612116, EPI_ISL_612120, EPI_ISL_612121, EPI_ISL_612132, EPI_ISL_612135, EPI_ISL_612137, EPI_ISL_612141, EPI_ISL_612143, EPI_ISL_612144, EPI_ISL_612146, EPI_ISL_612148, EPI_ISL_612149, EPI_ISL_612150, EPI_ISL_612152, EPI_ISL_612153, EPI_ISL_612154, EPI_ISL_612155, EPI_ISL_612156, EPI_ISL_612157, EPI_ISL_612165, EPI_ISL_612166, EPI_ISL_612171, EPI_ISL_612173, EPI_ISL_612174, EPI_ISL_612175, EPI_ISL_612176, EPI_ISL_612177, EPI_ISL_612178, EPI_ISL_612179, EPI_ISL_612185, EPI_ISL_612194, EPI_ISL_612195, EPI_ISL_612199, EPI_ISL_612202, EPI_ISL_612203, EPI_ISL_612458, EPI_ISL_612459, EPI_ISL_612460, EPI_ISL_612461, EPI_ISL_612462, EPI_ISL_612463, EPI_ISL_612466, EPI_ISL_612467, EPI_ISL_612468, EPI_ISL_612469, EPI_ISL_612470, EPI_ISL_612471, EPI_ISL_612473, EPI_ISL_612474, EPI_ISL_612476, EPI_ISL_612477, EPI_ISL_612478, EPI_ISL_612479, EPI_ISL_612481, EPI_ISL_612482, EPI_ISL_612483, EPI_ISL_612484, EPI_ISL_612485, EPI_ISL_612486, EPI_ISL_612488, EPI_ISL_612489, EPI_ISL_612490, EPI_ISL_612491, EPI_ISL_612492, EPI_ISL_612493, EPI_ISL_612494, EPI_ISL_612495, EPI_ISL_612496, EPI_ISL_612497, EPI_ISL_612499, EPI_ISL_612500, EPI_ISL_612501, EPI_ISL_612502, EPI_ISL_612503, EPI_ISL_612505, EPI_ISL_612506, EPI_ISL_612507, EPI_ISL_612508, EPI_ISL_612510, EPI_ISL_612512, EPI_ISL_612513, EPI_ISL_612514, EPI_ISL_612515, EPI_ISL_612517, EPI_ISL_612518, EPI_ISL_612519, EPI_ISL_612520, EPI_ISL_612521, EPI_ISL_612522, EPI_ISL_612614, EPI_ISL_612619, EPI_ISL_612620, EPI_ISL_612625, EPI_ISL_612626, EPI_ISL_612627, EPI_ISL_612634, EPI_ISL_612674, EPI_ISL_612675, EPI_ISL_612689, EPI_ISL_612716, EPI_ISL_612718, EPI_ISL_612723, EPI_ISL_612740, EPI_ISL_612765, EPI_ISL_612767, EPI_ISL_612770, EPI_ISL_612775, EPI_ISL_612789, EPI_ISL_612790, EPI_ISL_612807, EPI_ISL_612825, EPI_ISL_612828, EPI_ISL_612840, EPI_ISL_612845, EPI_ISL_612848, EPI_ISL_612849, EPI_ISL_612855, EPI_ISL_612867, EPI_ISL_612868, EPI_ISL_612872, EPI_ISL_612877, EPI_ISL_612879, EPI_ISL_612880, EPI_ISL_612881, EPI_ISL_612882, EPI_ISL_612883, EPI_ISL_612884, EPI_ISL_612885, EPI_ISL_612886, EPI_ISL_612887, EPI_ISL_612888, EPI_ISL_612889, EPI_ISL_612890, EPI_ISL_612891, EPI_ISL_612892, EPI_ISL_612893, EPI_ISL_612894, EPI_ISL_612895, EPI_ISL_612896, EPI_ISL_612897, EPI_ISL_612898, EPI_ISL_612899, EPI_ISL_612900, EPI_ISL_612901, EPI_ISL_612902, EPI_ISL_612909, EPI_ISL_612910, EPI_ISL_612911, EPI_ISL_612912, EPI_ISL_612913, EPI_ISL_612914, EPI_ISL_612915, EPI_ISL_612916, EPI_ISL_612917, EPI_ISL_612918, EPI_ISL_612919, EPI_ISL_612920, EPI_ISL_612921, EPI_ISL_612922, EPI_ISL_612926, EPI_ISL_612933, EPI_ISL_612936, EPI_ISL_612940, EPI_ISL_612957, EPI_ISL_612959, EPI_ISL_612968, EPI_ISL_612969, EPI_ISL_612975, EPI_ISL_612976, EPI_ISL_612981, EPI_ISL_612999, EPI_ISL_613006, EPI_ISL_613010, EPI_ISL_613013, EPI_ISL_613014, EPI_ISL_613015, EPI_ISL_613020, EPI_ISL_613021, EPI_ISL_613022, EPI_ISL_613023, EPI_ISL_613024, EPI_ISL_613025, EPI_ISL_613026, EPI_ISL_613027, EPI_ISL_613028, EPI_ISL_613029, EPI_ISL_613030, EPI_ISL_613031, EPI_ISL_613032, EPI_ISL_613033, EPI_ISL_613034, EPI_ISL_613035, EPI_ISL_613036, EPI_ISL_613037, EPI_ISL_613038, EPI_ISL_613039, EPI_ISL_613040, EPI_ISL_613041, EPI_ISL_613042, EPI_ISL_613043, EPI_ISL_613044, EPI_ISL_613045, EPI_ISL_613046, EPI_ISL_613047, EPI_ISL_613048, EPI_ISL_613049, EPI_ISL_613050, EPI_ISL_613051, EPI_ISL_613052, EPI_ISL_613053, EPI_ISL_613054, EPI_ISL_613055, EPI_ISL_613056, EPI_ISL_613057, EPI_ISL_613058, EPI_ISL_613059, EPI_ISL_613060, EPI_ISL_613061, EPI_ISL_613062, EPI_ISL_613063, EPI_ISL_613064, EPI_ISL_613065, EPI_ISL_613066, EPI_ISL_613067, EPI_ISL_613068, EPI_ISL_613069, EPI_ISL_613070, EPI_ISL_613071, EPI_ISL_613072, EPI_ISL_613073, EPI_ISL_613074, EPI_ISL_613075, EPI_ISL_613076, EPI_ISL_613077, EPI_ISL_613078, EPI_ISL_613079, EPI_ISL_613080, EPI_ISL_613081, EPI_ISL_613082, EPI_ISL_613083, EPI_ISL_613084, EPI_ISL_613085, EPI_ISL_613086, EPI_ISL_613087, EPI_ISL_613088, EPI_ISL_613089, EPI_ISL_613090, EPI_ISL_613091, EPI_ISL_613092, EPI_ISL_613093, EPI_ISL_613094, EPI_ISL_613095, EPI_ISL_613096, EPI_ISL_613097, EPI_ISL_613098, EPI_ISL_613099, EPI_ISL_613100, EPI_ISL_613101, EPI_ISL_613102, EPI_ISL_613103, EPI_ISL_613104, EPI_ISL_613105, EPI_ISL_613106, EPI_ISL_613107, EPI_ISL_613108, EPI_ISL_613110, EPI_ISL_613111, EPI_ISL_613112, EPI_ISL_613113, EPI_ISL_613114, EPI_ISL_613115, EPI_ISL_613116, EPI_ISL_613117, EPI_ISL_613118, EPI_ISL_613119, EPI_ISL_613120, EPI_ISL_613121, EPI_ISL_613122, EPI_ISL_613123, EPI_ISL_613124, EPI_ISL_613125, EPI_ISL_613126, EPI_ISL_613127, EPI_ISL_613128, EPI_ISL_613129, EPI_ISL_613130, EPI_ISL_613131, EPI_ISL_613132, EPI_ISL_613133, EPI_ISL_613134, EPI_ISL_613135, EPI_ISL_613136, EPI_ISL_613137, EPI_ISL_613138, EPI_ISL_613139, EPI_ISL_613140, EPI_ISL_613141, EPI_ISL_613142, EPI_ISL_613143, EPI_ISL_613144, EPI_ISL_613145, EPI_ISL_613146, EPI_ISL_613147, EPI_ISL_613148, EPI_ISL_613149, EPI_ISL_613150, EPI_ISL_613151, EPI_ISL_613152, EPI_ISL_613153, EPI_ISL_613154, EPI_ISL_613155, EPI_ISL_613156, EPI_ISL_613157, EPI_ISL_613158, EPI_ISL_613159, EPI_ISL_613160, EPI_ISL_613161, EPI_ISL_613162, EPI_ISL_613163, EPI_ISL_613164, EPI_ISL_613165, EPI_ISL_613166, EPI_ISL_613167, EPI_ISL_613168, EPI_ISL_613169, EPI_ISL_613170, EPI_ISL_613171, EPI_ISL_613172, EPI_ISL_613173, EPI_ISL_613174, EPI_ISL_613175, EPI_ISL_613176, EPI_ISL_613177, EPI_ISL_613178, EPI_ISL_613179, EPI_ISL_613180, EPI_ISL_613181, EPI_ISL_613182, EPI_ISL_613183, EPI_ISL_613184, EPI_ISL_613185, EPI_ISL_613186, EPI_ISL_613187, EPI_ISL_613188, EPI_ISL_613189, EPI_ISL_613190, EPI_ISL_613191, EPI_ISL_613192, EPI_ISL_613193, EPI_ISL_613194, EPI_ISL_613195, EPI_ISL_613196, EPI_ISL_613197, EPI_ISL_613198, EPI_ISL_613199, EPI_ISL_613200, EPI_ISL_613201, EPI_ISL_613202, EPI_ISL_613203, EPI_ISL_613204, EPI_ISL_613205, EPI_ISL_613206, EPI_ISL_613207, EPI_ISL_613208, EPI_ISL_613209, EPI_ISL_613210, EPI_ISL_613211, EPI_ISL_613212, EPI_ISL_613213, EPI_ISL_613214, EPI_ISL_613215, EPI_ISL_613216, EPI_ISL_613217, EPI_ISL_613218, EPI_ISL_613219, EPI_ISL_613220, EPI_ISL_613221, EPI_ISL_613222, EPI_ISL_613223, EPI_ISL_613224, EPI_ISL_613225, EPI_ISL_613226, EPI_ISL_613227, EPI_ISL_613228, EPI_ISL_613229, EPI_ISL_613230, EPI_ISL_613231, EPI_ISL_613232, EPI_ISL_613233, EPI_ISL_613234, EPI_ISL_613235, EPI_ISL_613236, EPI_ISL_613237, EPI_ISL_613238, EPI_ISL_613239, EPI_ISL_613240, EPI_ISL_613241, EPI_ISL_613242, EPI_ISL_613243, EPI_ISL_613244, EPI_ISL_613245, EPI_ISL_613246, EPI_ISL_613247, EPI_ISL_613248, EPI_ISL_613249, EPI_ISL_613250, EPI_ISL_613251, EPI_ISL_613252, EPI_ISL_613253, EPI_ISL_613254, EPI_ISL_613255, EPI_ISL_613256, EPI_ISL_613257, EPI_ISL_613258, EPI_ISL_613259, EPI_ISL_613260, EPI_ISL_613261, EPI_ISL_613262, EPI_ISL_613263, EPI_ISL_613264, EPI_ISL_613265, EPI_ISL_613266, EPI_ISL_613275 | | | |
| see above | Wales Specialist Virology Centre Sequencing lab: Pathogen Genomics Unit | COVID-19 Genomics UK (COG-UK) Consortium | Catherine Moore, Johnathan Evans, Laura Gifford, Malorie Perry, Simon Cottrell, Angela Marchbank, Alec Birchley, Alexander Adams, Amy Gaskin, Bree Gatica-Wilcox, Jason Coombes, Joel Southgate, Lauren Gilbert, Lee Graham, Nicole Pacchiarini, Sara Kumziene-Summerhayes, Sarah Taylor, Sophie Jones, Sara Rey, Matthew Bull, Joanne Watkins, Sally Corden, Tom Connor |
| EPI_ISL_614347, EPI_ISL_614348, EPI_ISL_614349, EPI_ISL_614350, EPI_ISL_614351, EPI_ISL_614352, EPI_ISL_614353, EPI_ISL_614354, EPI_ISL_614355, EPI_ISL_614356, EPI_ISL_614357, EPI_ISL_614358, EPI_ISL_614359, EPI_ISL_614360, EPI_ISL_614361, EPI_ISL_614362, EPI_ISL_614363, EPI_ISL_614364, EPI_ISL_614365, EPI_ISL_614366, EPI_ISL_614367, EPI_ISL_614368, EPI_ISL_614369, EPI_ISL_614370, EPI_ISL_614371, EPI_ISL_614372, EPI_ISL_614373, EPI_ISL_614374, EPI_ISL_614375, EPI_ISL_614376, EPI_ISL_614377, EPI_ISL_614378, EPI_ISL_614379, EPI_ISL_614380, EPI_ISL_614381, EPI_ISL_614382, EPI_ISL_614383, EPI_ISL_614384, EPI_ISL_614385, EPI_ISL_614386, EPI_ISL_614387, EPI_ISL_614388, EPI_ISL_614389, EPI_ISL_614390, EPI_ISL_614391, EPI_ISL_614392, EPI_ISL_614393, EPI_ISL_614394, EPI_ISL_614395 | | | |
| see above | Molecular diagnostic unit for viral haemorrhagic fevers and emerging viruses, Bouaké CHU Laboratory | Project group Epidemology of Highly Pathogenic Microorganisms, Robert Koch-Institute | Chantal Akoua-Koffi, Diané Bamourou, Etilé Anoh, Essia Belarbi, Safiatou Karidioula, Grit Schubert, Adjaratou Traoré, Soundélé Maïté, Monemo Pacome, Coulibaly Mbegnan, Bamba Fatoumata Touré, Kra Ouffoué, Fabian Leendertz |
| EPI_ISL_614968, EPI_ISL_614969, EPI_ISL_614972, EPI_ISL_614974, EPI_ISL_614975, EPI_ISL_614976, EPI_ISL_614977, EPI_ISL_614978, EPI_ISL_614979 | Viollier AG | Department of Biosystems Science and Engineering, ETH Zürich | Christian Beisel, Sarah Nadeau, Ivan Topolsky, Pedro Ferreira, Philipp Jablonski, Susana Posada-Céspedes, Tobias Schär, Ina Nissen, Natascha Santacroce, Elodie Burcklen, Christiane Beckmann, Maurice Redondo, Olivier Kobel, Christoph Noppen, Sophie Seidel, Noemie Santamaria de Souza, Chaoran Chen, Niko Beerenwinkel, Tanja Stadler |
| EPI_ISL_626658, EPI_ISL_626674, EPI_ISL_626675, EPI_ISL_626696, EPI_ISL_626703, EPI_ISL_626706, EPI_ISL_626707, EPI_ISL_626708, EPI_ISL_626711, EPI_ISL_626733, EPI_ISL_626746, EPI_ISL_626748, EPI_ISL_626761, EPI_ISL_626762, EPI_ISL_626763, EPI_ISL_626767, EPI_ISL_626768, EPI_ISL_626769, EPI_ISL_626780, EPI_ISL_626783, EPI_ISL_626784, EPI_ISL_626787, EPI_ISL_626788, EPI_ISL_626796, EPI_ISL_626820, EPI_ISL_626829, EPI_ISL_626836, EPI_ISL_626850, EPI_ISL_626853, EPI_ISL_626856, EPI_ISL_626857, EPI_ISL_626861, EPI_ISL_626862, EPI_ISL_626863, EPI_ISL_626865, EPI_ISL_626870, EPI_ISL_626871, EPI_ISL_626872, EPI_ISL_626874, EPI_ISL_626876, EPI_ISL_626877, EPI_ISL_626920, EPI_ISL_626921, EPI_ISL_626928, EPI_ISL_626931, EPI_ISL_626960, EPI_ISL_626961, EPI_ISL_626965, EPI_ISL_626993, EPI_ISL_626996, EPI_ISL_626998, EPI_ISL_627002, EPI_ISL_627003, EPI_ISL_627006, EPI_ISL_627007, EPI_ISL_627010, EPI_ISL_627012, EPI_ISL_627013, EPI_ISL_627015, EPI_ISL_627018, EPI_ISL_627021, EPI_ISL_627022, EPI_ISL_627023, EPI_ISL_627025, EPI_ISL_627026, EPI_ISL_627027, EPI_ISL_627028, EPI_ISL_627029, EPI_ISL_627032, EPI_ISL_627033, EPI_ISL_627034, EPI_ISL_627035, EPI_ISL_627037, EPI_ISL_627039, EPI_ISL_627042, EPI_ISL_627043, EPI_ISL_627048, EPI_ISL_627050, EPI_ISL_627051, EPI_ISL_627054, EPI_ISL_627092, EPI_ISL_627100, EPI_ISL_627101, EPI_ISL_627105, EPI_ISL_627113, EPI_ISL_627115, EPI_ISL_627119, EPI_ISL_627120, EPI_ISL_627122, EPI_ISL_627124, EPI_ISL_627126, EPI_ISL_627129, EPI_ISL_627140, EPI_ISL_627146, EPI_ISL_627154, EPI_ISL_627155, EPI_ISL_627157, EPI_ISL_627161, EPI_ISL_627167, EPI_ISL_627168, EPI_ISL_627169, EPI_ISL_627172, EPI_ISL_627192, EPI_ISL_627197, EPI_ISL_627203, EPI_ISL_627204, EPI_ISL_627221, EPI_ISL_627223, EPI_ISL_627224, EPI_ISL_627225, EPI_ISL_627227, EPI_ISL_627228, EPI_ISL_627231, EPI_ISL_627241, EPI_ISL_627242, EPI_ISL_627246, EPI_ISL_627267, EPI_ISL_627270, EPI_ISL_627271, EPI_ISL_627274, EPI_ISL_627293, EPI_ISL_627295, EPI_ISL_627306, EPI_ISL_627308, EPI_ISL_627482, EPI_ISL_627483, EPI_ISL_627484, EPI_ISL_627485, EPI_ISL_627486, EPI_ISL_627489, EPI_ISL_627490, EPI_ISL_627730, EPI_ISL_627731, EPI_ISL_627735, EPI_ISL_627738, EPI_ISL_627739, EPI_ISL_627744, EPI_ISL_627747, EPI_ISL_627748, EPI_ISL_627749, EPI_ISL_627750, EPI_ISL_627751, EPI_ISL_627758, EPI_ISL_627763, EPI_ISL_627781, EPI_ISL_627794, EPI_ISL_627799, EPI_ISL_627811, EPI_ISL_627824, EPI_ISL_627834, EPI_ISL_627841, EPI_ISL_627843, EPI_ISL_627844, EPI_ISL_627846, EPI_ISL_627848, EPI_ISL_627851, EPI_ISL_627859, EPI_ISL_627863, EPI_ISL_627865, EPI_ISL_627868, EPI_ISL_627873, EPI_ISL_627878, EPI_ISL_627880, EPI_ISL_627886, EPI_ISL_627890, EPI_ISL_627912, EPI_ISL_627927, EPI_ISL_627934, EPI_ISL_627936, EPI_ISL_627938, EPI_ISL_627944, EPI_ISL_627946, EPI_ISL_627956, EPI_ISL_627965, EPI_ISL_627966, EPI_ISL_627969, EPI_ISL_627974, EPI_ISL_627979, EPI_ISL_627986, EPI_ISL_627988, EPI_ISL_628001, EPI_ISL_628002, EPI_ISL_628003, EPI_ISL_628008, EPI_ISL_628015, EPI_ISL_628026, EPI_ISL_628027, EPI_ISL_628034, EPI_ISL_628040, EPI_ISL_628061, EPI_ISL_628069, EPI_ISL_628076, EPI_ISL_628088, EPI_ISL_628092, EPI_ISL_628097, EPI_ISL_628102, EPI_ISL_628107, EPI_ISL_628123, EPI_ISL_628128, EPI_ISL_628134, EPI_ISL_628138, EPI_ISL_628141, EPI_ISL_628162, EPI_ISL_628163, EPI_ISL_628183, EPI_ISL_628189, EPI_ISL_628205, EPI_ISL_628211, EPI_ISL_628220, EPI_ISL_628225, EPI_ISL_628236, EPI_ISL_628245, EPI_ISL_628259, EPI_ISL_628263, EPI_ISL_628264, EPI_ISL_628267, EPI_ISL_628272, EPI_ISL_628273, EPI_ISL_628280, EPI_ISL_628292, EPI_ISL_628307, EPI_ISL_628309 | | | |
| see above | Wales Specialist Virology Centre Sequencing lab: Pathogen Genomics Unit | COVID-19 Genomics UK (COG-UK) Consortium | Catherine Moore, Johnathan Evans, Laura Gifford, Malorie Perry, Simon Cottrell, Angela Marchbank, Alec Birchley, Alexander Adams, Amy Gaskin, Bree Gatica-Wilcox, Jason Coombes, Joel Southgate, Lauren Gilbert, Lee Graham, Nicole Pacchiarini, Sara Kumziene-Summerhayes, Sarah Taylor, Sophie Jones, Sara Rey, Matthew Bull, Joanne Watkins, Sally Corden, Tom Connor |

We gratefully acknowledge the following Authors from the Originating laboratories responsible for obtaining the specimens, as well as the Submitting laboratories where the genome data were generated and shared via GISAID, on which this research is based.

All Submitters of data may be contacted directly via www.gisaid.org

Authors are sorted alphabetically.

| Accession ID | Originating Laboratory | Submitting Laboratory | Authors |
|---|---|---|---|
| EPI_ISL_407987 | Singapore General Hospital | Programme in Emerging Infectious Diseases, Duke-NUS Medical School | Danielle E Anderson, Martin Linster, Yan Zhuang, Jayanthi Jayakumar, Kian Sing Chan, Lynette LE Oon, Jenny GH Low, Yvonne CF Su, Linfa Wang, Gavin JD Smith |
| EPI_ISL_407988 | National Centre for Infectious Diseases | Programme in Emerging Infectious Diseases, Duke-NUS Medical School | Danielle E Anderson, Martin Linster, Yan Zhuang, Jayanthi Jayakumar, David CB Lye, Yee Sin Leo, Barnaby E Young, Yvonne CF Su, Linfa Wang, Gavin JD Smith |
| EPI_ISL_409067 | Massachusetts Department of Public Health | Pathogen Discovery, Respiratory Viruses Branch, Division of Viral Diseases, Centers for Dieases Control and Prevention | Clinton R. Paden, Jing Zhang, Krista Queen, Yan Li, Ying Tao, Anna Uehara, Xiaoyan Lu, Brian Lynch, Senthil Kumar K. Sakthivel, Brett L. Whitaker, Shifaq Kamili, Lijuan Wang, Janna' R. Murray, Susan I. Gerber, Stephen Lindstrom, Suxiang Tong |
| EPI_ISL_410535 | National Centre for Infectious Diseases | Programme in Emerging Infectious Diseases, Duke-NUS Medical School | Danielle E Anderson, Martin Linster, Yan Zhuang, Jayanthi Jayakumar, David CB Lye, Yee Sin Leo, Barnaby E Young, Yvonne CF Su, Gavin JD Smith |
| EPI_ISL_410536, EPI_ISL_410537 | Singapore General Hospital, Molecular Laboratory, Division of Pathology | Programme in Emerging Infectious Diseases, Duke-NUS Medical School | Danielle E Anderson, Martin Linster, Yan Zhuang, Jayanthi Jayakumar, Kian Sing Chan, Lynette LE Oon, Shirin Kalimuddin, Jenny GH Low, Yvonne CF Su, Gavin JD Smith |
| EPI_ISL_413456 | Seattle Flu Study, University of Washington Medical Center | Seattle Flu Study, University of Washington Medical Center | Chu et al |
| EPI_ISL_413560 | Seattle Flu Study | Seattle Flu Study | Chu et al |
| EPI_ISL_413618, EPI_ISL_413619, EPI_ISL_413620, EPI_ISL_413621, EPI_ISL_413622, EPI_ISL_413623 | unknown | Pathogen Discovery, Respiratory Viruses Branch, Division of Viral Diseases, Centers for Dieases Control and Prevention | Clinton R. Paden, Ying Tao, Krista Queen, Anna Uehara, Jing Zhang, Yan Li, Haibin Wang, Shifaq Kamili, Xiaoyan Lu, Brian Lynch, Senthil Kumar K. Sakthivel, Brett L. Whitaker, Lijuan Wang, Janna' R. Murray, Jasmine Padilla, Justin Lee, Susan I. Gerber, Stephen Lindstrom, Suxiang Tong |
| EPI_ISL_414014 | Hospital Israelita Albert Einstein | Instituto Adolfo Lutz, Interdiciplinary Procedures Center, Strategic Laboratory | Claudio Tavares Sacchi, Claudia Regina Gonçalves, Katia Correia dos Santos, Carlos Henrique Camargo, Maria do Carmo Sampaito Tavares Timenetsky, Terezinha Maria de Paiva, Ester Cerdeira Sabino |
| EPI_ISL_414015 | Hospital São Joaquim Beneficencia Portuguesa | Instituto Adolfo Lutz, Interdiciplinary Procedures Center, Strategic Laboratory | Claudio Tavares Sacchi, Claudia Regina Gonçalves, SimoneGuadagnucci Morillo, Carlos Henrique Camargo, Maria do Carmo Sampaito Tavares Timenetsky, Fabiana Cristina Pereira dos Santos Terezinha Maria de Paiva, Ester Cerdeira Sabino |
| EPI_ISL_414016 | Hospital São Joaquim Beneficencia Portuguesa | Instituto Adolfo Lutz, Interdiciplinary Procedures Center, Strategic Laboratory | Claudio Tavares Sacchi, Claudia Regina Gonçalves, Audrey Cilli, Carlos Henrique Camargo, Maria do Carmo Sampaito Tavares Timenetsky, Daniela Bernardes Borges da Silva, Terezinha Maria de Paiva, Ester Cerdeira Sabino |
| EPI_ISL_414017 | Hospital São Joaquim Beneficencia Portuguesa | Instituto Adolfo Lutz, Interdiciplinary Procedures Center, Strategic Laboratory | Claudio Tavares Sacchi, Claudia Regina Gonçalves, Fabiana Cristina Pereira dos Santos, Carlos Henrique Camargo, Maria do Carmo Sampaito Tavares Timenetsky, Daniela Bernardes Borges da Silva, Terezinha Maria de Paiva, Ester Cerdeira Sabino |
| EPI_ISL_414378 | National Centre for Infectious Diseases | Programme in Emerging Infectious Diseases, Duke-NUS Medical School | Danielle E Anderson, Martin Linster, Yan Zhuang, Jayanthi Jayakumar, Louisa Sun, David CB Lye, Yee Sin Leo, Barnaby E Young, Yvonne CF Su, Gavin JD Smith |
| EPI_ISL_414379, EPI_ISL_414380 | National Centre for Infectious Diseases | Programme in Emerging Infectious Diseases, Duke-NUS Medical School | Danielle E Anderson, Martin Linster, Yan Zhuang, Jayanthi Jayakumar, David CB Lye, Yee Sin Leo, Barnaby E Young, Yvonne CF Su, Gavin JD Smith |
| EPI_ISL_415152 | Gorgas Memorial Institute for Health Studies | Gorgas Memorial Institute for Health Studies | Danilo Franco, Sandra Lopez-Verges, Elimelec Valdespino, Claudia Gonzalez, Oris Chavarria, Ambar Moreno, Yamilka Diaz, Leyda Abrego, Juan M. Pascale, Alexander A. Martinez. |
| EPI_ISL_416028 | National Influenza Center - Instituto Adolfo Lutz | Instituto Adolfo Lutz, Interdiciplinary Procedures Center, Strategic Laboratory | Claudio Tavares Sacchi, Claudia Regina Gonçalves, Carlos Henrique Camargo, Fabiana Cristina Pereira dos Santos, Daniela Bernardes Borges da Silva, Simone Guadagnucci Morillo, Adriano Abbud, Adriana Bugno, Maria do Carmo Sampaio Tavares Timenetsky, Terezinha Maria de Paiva |
| EPI_ISL_416029 | Laboiratório Fleury | Instituto Adolfo Lutz, Interdiciplinary Procedures Center, Strategic Laboratory | Claudio Tavares Sacchi, Claudia Regina Gonçalves, Carlos Henrique Camargo, Fabiana Cristina Pereira dos Santos, Daniela Bernardes Borges da Silva, Simone Guadagnucci Morillo, Adriano Abbud, Adriana Bugno, Maria do Carmo Sampaio Tavares Timenetsky, Terezinha Maria de Paiva |
| EPI_ISL_416031, EPI_ISL_416032 | National Influenza Center - Instituto Adolfo Lutz | Instituto Adolfo Lutz, Interdiciplinary Procedures Center, Strategic Laboratory | Claudio Tavares Sacchi, Claudia Regina Gonçalves, Carlos Henrique Camargo, Fabiana Cristina Pereira dos Santos, Daniela Bernardes Borges da Silva, Simone Guadagnucci Morillo, Adriano Abbud, Adriana Bugno, Maria do Carmo Sampaio Tavares Timenetsky, Terezinha Maria de Paiva |
| EPI_ISL_416033, EPI_ISL_416034 | Hospital Israelita Albert Einstein | Instituto Adolfo Lutz, Interdiciplinary Procedures Center, Strategic Laboratory | Claudio Tavares Sacchi, Claudia Regina Gonçalves, Carlos Henrique Camargo, Erica Valessa Ramos Gomes, Fabiana Cristina Pereira dos Santos, Daniela Bernardes Borges da Silva, Simone Guadagnucci Morillo, Adriano Abbud, Adriana Bugno, Maria do Carmo Sampaio Tavares Timenetsky, Terezinha Maria de Paiva |
| EPI_ISL_416035, EPI_ISL_416036 | National Influenza Center - Instituto Adolfo Lutz | Instituto Adolfo Lutz, Interdiciplinary Procedures Center, Strategic Laboratory | Claudio Tavares Sacchi, Claudia Regina Gonçalves, Carlos Henrique Camargo, Erica Valessa Ramos Gomes, Fabiana Cristina Pereira dos Santos, Daniela Bernardes Borges da Silva, Simone Guadagnucci Morillo, Adriano Abbud, Adriana Bugno, Maria do Carmo Sampaio Tavares Timenetsky, Terezinha Maria de Paiva |
| EPI_ISL_416460, EPI_ISL_416461, EPI_ISL_416462, EPI_ISL_416465, EPI_ISL_416466 | Seattle Flu Study | Seattle Flu Study | Chu et al |
| EPI_ISL_417065, EPI_ISL_417066, EPI_ISL_417067, EPI_ISL_417068, EPI_ISL_417069, EPI_ISL_417070, EPI_ISL_417071, EPI_ISL_417072, EPI_ISL_417073, EPI_ISL_417074, EPI_ISL_417075, EPI_ISL_417076, EPI_ISL_417077, EPI_ISL_417078, EPI_ISL_417079, EPI_ISL_417080, EPI_ISL_417081, EPI_ISL_417082, EPI_ISL_417083, EPI_ISL_417084, EPI_ISL_417085, EPI_ISL_417086, EPI_ISL_417087, EPI_ISL_417088, EPI_ISL_417089, EPI_ISL_417090, EPI_ISL_417091, EPI_ISL_417092, EPI_ISL_417093, EPI_ISL_417094, EPI_ISL_417095, EPI_ISL_417096, EPI_ISL_417097, EPI_ISL_417098, EPI_ISL_417099, EPI_ISL_417100, EPI_ISL_417101, EPI_ISL_417102, EPI_ISL_417103, EPI_ISL_417104, EPI_ISL_417105, EPI_ISL_417106, EPI_ISL_417107, EPI_ISL_417108, EPI_ISL_417109, EPI_ISL_417110, EPI_ISL_417111, EPI_ISL_417112, EPI_ISL_417113, EPI_ISL_417114, EPI_ISL_417115, EPI_ISL_417116, EPI_ISL_417117, EPI_ISL_417118, EPI_ISL_417119, EPI_ISL_417120, EPI_ISL_417121, EPI_ISL_417122, EPI_ISL_417123, EPI_ISL_417124, EPI_ISL_417125, EPI_ISL_417126, EPI_ISL_417127, EPI_ISL_417128, EPI_ISL_417129, EPI_ISL_417130, EPI_ISL_417131, EPI_ISL_417132, EPI_ISL_417133, EPI_ISL_417134, EPI_ISL_417135, EPI_ISL_417136, EPI_ISL_417137, EPI_ISL_417138, EPI_ISL_417139, EPI_ISL_417140, EPI_ISL_417141, EPI_ISL_417142, EPI_ISL_417143, EPI_ISL_417144, EPI_ISL_417145, EPI_ISL_417146, EPI_ISL_417147, EPI_ISL_417148, EPI_ISL_417149, EPI_ISL_417150, EPI_ISL_417151, EPI_ISL_417152, EPI_ISL_417153, EPI_ISL_417154, EPI_ISL_417155, EPI_ISL_417156, EPI_ISL_417157, EPI_ISL_417158, EPI_ISL_417159, EPI_ISL_417160, EPI_ISL_417161, EPI_ISL_417162 | | | |
| see above | Washington State Department of Health | Seattle Flu Study | Chu etl al |
| EPI_ISL_417163, EPI_ISL_417164, EPI_ISL_417165 | Seattle Flu Study | Seattle Flu Study | Chu etl al |
| EPI_ISL_417166, EPI_ISL_417167, EPI_ISL_417168, EPI_ISL_417169, EPI_ISL_417170, EPI_ISL_417171, EPI_ISL_417172, EPI_ISL_417173, EPI_ISL_417175 | Washington State Department of Health | Seattle Flu Study | Chu etl al |
| EPI_ISL_417481 | deCODE genetics | deCODE genetics | Daniel F Gudbjartsson, Agnar Helgason, Hakon Jonsson, Olafur T Magnusson, Pall Melsted, Gudmundur L Norddahl, Jona Saemundsdottir, Asgeir Sigurdsson, Patrick Sulem, Arna B Agustsdottir, Berglind Eiriksdottir, Run Fridriksdottir, Elisabet E Gardarsdottir, Gudmundur Georgsson, Olafia S Gretarsdottir, Kjartan R Gudmundsson, Thora R Gunnarsdottir, Arnaldur Gylfason, Hilma Holm, Brynjar O Jensson, Aslaug Jonasdottir, Kamilla S |

| | | | |
|---|---|---|---|
| | | | Josefsdottir, Thordur Kristjansson, Droplaug N Magnusdottir, Louise le Roux, Gudrun Sigmundsdottir, Gardar Sveinbjornsson, Kristin E Sveinsdottir, Maney Sveinsdottir, Emil A Thorarensen, Bjarni Thorbjornsson, Gisli Masson, Ingileif Jonsdottir, Alma Moller, Thorolfur Gudnason, Karl G Kristinsson, Unnur Thorsteinsdottir, Kari Stefansson |
| EPI_ISL_417535, EPI_ISL_417536, EPI_ISL_417537, EPI_ISL_417538, EPI_ISL_417539, EPI_ISL_417540, EPI_ISL_417541, EPI_ISL_417542, EPI_ISL_417543, EPI_ISL_417544, EPI_ISL_417545, EPI_ISL_417546, EPI_ISL_417548, EPI_ISL_417549 | | | |
| see above | deCODE genetics | deCODE genetics | Daniel F Gudbjartsson; Agnar Helgason; Hakon Jonsson; Olafur T Magnusson; Pall Melsted; Gudmundur L Norddahl; Jona Saemundsdottir; Asgeir Sigurdsson; Patrick Sulem; Arna B Agustsdottir; Berglind Eiriksdottir; Run Fridriksdottir; Elisabet E Gardarsdottir; Gudmundur Georgsson; Olafia S Gretarsdottir; Kjartan R Gudmundsson; Thora R Gunnarsdottir; Arnaldur Gylfason; Hilma Holm; Brynjar O Jensson; Aslaug Jonasdottir; Kamilla S Josefsdottir; Thordur Kristjansson; Droplaug N Magnusdottir; Louise le Roux; Gudrun Sigmundsdottir; Gardar Sveinbjornsson; Kristin E Sveinsdottir; Maney Sveinsdottir; Emil A Thorarensen; Bjarni Thorbjornsson; Gisli Masson; Ingileif Jonsdottir; Alma Moller; Thorolfur Gudnason; Karl G Kristinsson; Unnur Thorsteinsdottir; Kari Stefansson |
| EPI_ISL_417550, EPI_ISL_417551 | The National University Hospital of Iceland | deCODE genetics | Daniel F Gudbjartsson; Agnar Helgason; Hakon Jonsson; Olafur T Magnusson; Pall Melsted; Gudmundur L Norddahl; Jona Saemundsdottir; Asgeir Sigurdsson; Patrick Sulem; Arna B Agustsdottir; Berglind Eiriksdottir; Run Fridriksdottir; Elisabet E Gardarsdottir; Gudmundur Georgsson; Olafia S Gretarsdottir; Kjartan R Gudmundsson; Thora R Gunnarsdottir; Arnaldur Gylfason; Hilma Holm; Brynjar O Jensson; Aslaug Jonasdottir; Kamilla S Josefsdottir; Thordur Kristjansson; Droplaug N Magnusdottir; Louise le Roux; Gudrun Sigmundsdottir; Gardar Sveinbjornsson; Kristin E Sveinsdottir; Maney Sveinsdottir; Emil A Thorarensen; Bjarni Thorbjornsson; Gisli Masson; Ingileif Jonsdottir; Alma Moller; Thorolfur Gudnason; Karl G Kristinsson; Unnur Thorsteinsdottir; Kari Stefansson |
| EPI_ISL_417552 | deCODE genetics | deCODE genetics | Daniel F Gudbjartsson; Agnar Helgason; Hakon Jonsson; Olafur T Magnusson; Pall Melsted; Gudmundur L Norddahl; Jona Saemundsdottir; Asgeir Sigurdsson; Patrick Sulem; Arna B Agustsdottir; Berglind Eiriksdottir; Run Fridriksdottir; Elisabet E Gardarsdottir; Gudmundur Georgsson; Olafia S Gretarsdottir; Kjartan R Gudmundsson; Thora R Gunnarsdottir; Arnaldur Gylfason; Hilma Holm; Brynjar O Jensson; Aslaug Jonasdottir; Kamilla S Josefsdottir; Thordur Kristjansson; Droplaug N Magnusdottir; Louise le Roux; Gudrun Sigmundsdottir; Gardar Sveinbjornsson; Kristin E Sveinsdottir; Maney Sveinsdottir; Emil A Thorarensen; Bjarni Thorbjornsson; Gisli Masson; Ingileif Jonsdottir; Alma Moller; Thorolfur Gudnason; Karl G Kristinsson; Unnur Thorsteinsdottir; Kari Stefansson |
| EPI_ISL_417553, EPI_ISL_417554, EPI_ISL_417555, EPI_ISL_417556, EPI_ISL_417557, EPI_ISL_417558, EPI_ISL_417559, EPI_ISL_417560, EPI_ISL_417561, EPI_ISL_417562, EPI_ISL_417563, EPI_ISL_417564, EPI_ISL_417565, EPI_ISL_417566, EPI_ISL_417567, EPI_ISL_417568, EPI_ISL_417569, EPI_ISL_417570, EPI_ISL_417571, EPI_ISL_417572, EPI_ISL_417573, EPI_ISL_417574, EPI_ISL_417575, EPI_ISL_417576, EPI_ISL_417577, EPI_ISL_417578, EPI_ISL_417579, EPI_ISL_417580, EPI_ISL_417581, EPI_ISL_417582, EPI_ISL_417583, EPI_ISL_417584, EPI_ISL_417585, EPI_ISL_417586, EPI_ISL_417587, EPI_ISL_417588, EPI_ISL_417589 | | | |
| see above | The National University Hospital of Iceland | deCODE genetics | Daniel F Gudbjartsson; Agnar Helgason; Hakon Jonsson; Olafur T Magnusson; Pall Melsted; Gudmundur L Norddahl; Jona Saemundsdottir; Asgeir Sigurdsson; Patrick Sulem; Arna B Agustsdottir; Berglind Eiriksdottir; Run Fridriksdottir; Elisabet E Gardarsdottir; Gudmundur Georgsson; Olafia S Gretarsdottir; Kjartan R Gudmundsson; Thora R Gunnarsdottir; Arnaldur Gylfason; Hilma Holm; Brynjar O Jensson; Aslaug Jonasdottir; Kamilla S Josefsdottir; Thordur Kristjansson; Droplaug N Magnusdottir; Louise le Roux; Gudrun Sigmundsdottir; Gardar Sveinbjornsson; Kristin E Sveinsdottir; Maney Sveinsdottir; Emil A Thorarensen; Bjarni Thorbjornsson; Gisli Masson; Ingileif Jonsdottir; Alma Moller; Thorolfur Gudnason; Karl G Kristinsson; Unnur Thorsteinsdottir; Kari Stefansson |
| EPI_ISL_417590 | deCODE genetics | deCODE genetics | Daniel F Gudbjartsson; Agnar Helgason; Hakon Jonsson; Olafur T Magnusson; Pall Melsted; Gudmundur L Norddahl; Jona Saemundsdottir; Asgeir Sigurdsson; Patrick Sulem; Arna B Agustsdottir; Berglind Eiriksdottir; Run Fridriksdottir; Elisabet E Gardarsdottir; Gudmundur Georgsson; Olafia S Gretarsdottir; Kjartan R Gudmundsson; Thora R Gunnarsdottir; Arnaldur Gylfason; Hilma Holm; Brynjar O Jensson; Aslaug Jonasdottir; Kamilla S Josefsdottir; Thordur Kristjansson; Droplaug N Magnusdottir; Louise le Roux; Gudrun Sigmundsdottir; Gardar Sveinbjornsson; Kristin E Sveinsdottir; Maney Sveinsdottir; Emil A Thorarensen; Bjarni Thorbjornsson; Gisli Masson; Ingileif Jonsdottir; Alma Moller; Thorolfur Gudnason; Karl G Kristinsson; Unnur Thorsteinsdottir; Kari Stefansson |
| EPI_ISL_417591, EPI_ISL_417592, EPI_ISL_417593, EPI_ISL_417594, EPI_ISL_417595, EPI_ISL_417596, EPI_ISL_417597, EPI_ISL_417598, EPI_ISL_417599, EPI_ISL_417600, EPI_ISL_417601, EPI_ISL_417602, EPI_ISL_417603, EPI_ISL_417604, EPI_ISL_417605, EPI_ISL_417606, EPI_ISL_417607, EPI_ISL_417608, EPI_ISL_417609, EPI_ISL_417610, EPI_ISL_417611, EPI_ISL_417612, EPI_ISL_417613, EPI_ISL_417614, EPI_ISL_417615, EPI_ISL_417616, EPI_ISL_417617 | | | |
| see above | The National University Hospital of Iceland | deCODE genetics | Daniel F Gudbjartsson; Agnar Helgason; Hakon Jonsson; Olafur T Magnusson; Pall Melsted; Gudmundur L Norddahl; Jona Saemundsdottir; Asgeir Sigurdsson; Patrick Sulem; Arna B Agustsdottir; Berglind Eiriksdottir; Run Fridriksdottir; Elisabet E Gardarsdottir; Gudmundur Georgsson; Olafia S Gretarsdottir; Kjartan R Gudmundsson; Thora R Gunnarsdottir; Arnaldur Gylfason; Hilma Holm; Brynjar O Jensson; Aslaug Jonasdottir; Kamilla S Josefsdottir; Thordur Kristjansson; Droplaug N Magnusdottir; Louise le Roux; Gudrun Sigmundsdottir; Gardar Sveinbjornsson; Kristin E Sveinsdottir; Maney Sveinsdottir; Emil A Thorarensen; Bjarni Thorbjornsson; Gisli Masson; Ingileif Jonsdottir; Alma Moller; Thorolfur Gudnason; Karl G Kristinsson; Unnur Thorsteinsdottir; Kari Stefansson |
| EPI_ISL_417618 | deCODE genetics | deCODE genetics | Daniel F Gudbjartsson; Agnar Helgason; Hakon Jonsson; Olafur T Magnusson; Pall Melsted; Gudmundur L Norddahl; Jona Saemundsdottir; Asgeir Sigurdsson; Patrick Sulem; Arna B Agustsdottir; Berglind Eiriksdottir; Run Fridriksdottir; Elisabet E Gardarsdottir; Gudmundur Georgsson; Olafia S Gretarsdottir; Kjartan R Gudmundsson; Thora R Gunnarsdottir; Arnaldur Gylfason; Hilma Holm; Brynjar O Jensson; Aslaug Jonasdottir; Kamilla S Josefsdottir; Thordur Kristjansson; Droplaug N Magnusdottir; Louise le Roux; Gudrun Sigmundsdottir; Gardar Sveinbjornsson; Kristin E Sveinsdottir; Maney Sveinsdottir; Emil A Thorarensen; Bjarni Thorbjornsson; Gisli Masson; Ingileif Jonsdottir; Alma Moller; Thorolfur Gudnason; Karl G Kristinsson; Unnur Thorsteinsdottir; Kari Stefansson |
| EPI_ISL_417619, EPI_ISL_417620, EPI_ISL_417621, EPI_ISL_417622, EPI_ISL_417623, EPI_ISL_417624, EPI_ISL_417625, EPI_ISL_417626, EPI_ISL_417627, EPI_ISL_417628, EPI_ISL_417629, EPI_ISL_417630, EPI_ISL_417631, EPI_ISL_417632, EPI_ISL_417633, EPI_ISL_417634, EPI_ISL_417635, EPI_ISL_417636, EPI_ISL_417637, EPI_ISL_417638, EPI_ISL_417639, EPI_ISL_417640, EPI_ISL_417641, EPI_ISL_417642, EPI_ISL_417643, EPI_ISL_417644, EPI_ISL_417645, EPI_ISL_417646, EPI_ISL_417647, EPI_ISL_417648, EPI_ISL_417649, EPI_ISL_417650, EPI_ISL_417651, EPI_ISL_417652, EPI_ISL_417653, EPI_ISL_417654 | | | |
| see above | The National University Hospital of Iceland | deCODE genetics | Daniel F Gudbjartsson; Agnar Helgason; Hakon Jonsson; Olafur T Magnusson; Pall Melsted; Gudmundur L Norddahl; Jona Saemundsdottir; Asgeir Sigurdsson; Patrick Sulem; Arna B Agustsdottir; Berglind Eiriksdottir; Run Fridriksdottir; Elisabet E Gardarsdottir; Gudmundur Georgsson; Olafia S Gretarsdottir; Kjartan R Gudmundsson; Thora R Gunnarsdottir; Arnaldur Gylfason; Hilma Holm; Brynjar O Jensson; Aslaug Jonasdottir; Kamilla S Josefsdottir; Thordur Kristjansson; Droplaug N Magnusdottir; Louise le Roux; Gudrun Sigmundsdottir; Gardar Sveinbjornsson; Kristin E Sveinsdottir; Maney Sveinsdottir; Emil A Thorarensen; Bjarni Thorbjornsson; Gisli Masson; Ingileif Jonsdottir; Alma Moller; Thorolfur Gudnason; Karl G Kristinsson; Unnur Thorsteinsdottir; Kari Stefansson |
| EPI_ISL_417655, EPI_ISL_417656, EPI_ISL_417657, EPI_ISL_417658, EPI_ISL_417659, EPI_ISL_417660, EPI_ISL_417661, EPI_ISL_417662, EPI_ISL_417663, EPI_ISL_417664, EPI_ISL_417665, EPI_ISL_417666, EPI_ISL_417667, EPI_ISL_417668, EPI_ISL_417669, EPI_ISL_417670, EPI_ISL_417671, EPI_ISL_417672 | | | |
| see above | deCODE genetics | deCODE genetics | Daniel F Gudbjartsson; Agnar Helgason; Hakon Jonsson; Olafur T Magnusson; Pall Melsted; Gudmundur L Norddahl; Jona Saemundsdottir; Asgeir Sigurdsson; Patrick Sulem; Arna B Agustsdottir; Berglind Eiriksdottir; Run Fridriksdottir; Elisabet E Gardarsdottir; Gudmundur Georgsson; Olafia S Gretarsdottir; Kjartan R Gudmundsson; Thora R Gunnarsdottir; Arnaldur Gylfason; Hilma Holm; Brynjar O Jensson; Aslaug Jonasdottir; Kamilla S Josefsdottir; Thordur Kristjansson; Droplaug N Magnusdottir; Louise le Roux; Gudrun Sigmundsdottir; Gardar Sveinbjornsson; Kristin E Sveinsdottir; Maney Sveinsdottir; Emil A Thorarensen; Bjarni Thorbjornsson; Gisli Masson; Ingileif Jonsdottir; Alma Moller; Thorolfur Gudnason; Karl G Kristinsson; Unnur Thorsteinsdottir; Kari Stefansson |
| EPI_ISL_417673, EPI_ISL_417674, EPI_ISL_417675 | The National University Hospital of Iceland | deCODE genetics | Daniel F Gudbjartsson; Agnar Helgason; Hakon Jonsson; Olafur T Magnusson; Pall Melsted; Gudmundur L Norddahl; Jona Saemundsdottir; Asgeir Sigurdsson; Patrick Sulem; Arna B Agustsdottir; Berglind Eiriksdottir; Run Fridriksdottir; Elisabet E Gardarsdottir; Gudmundur Georgsson; Olafia S Gretarsdottir; Kjartan R Gudmundsson; Thora R Gunnarsdottir; Arnaldur Gylfason; Hilma Holm; Brynjar O Jensson; Aslaug Jonasdottir; Kamilla S Josefsdottir; Thordur Kristjansson; Droplaug N Magnusdottir; Louise le Roux; Gudrun Sigmundsdottir; Gardar Sveinbjornsson; Kristin E Sveinsdottir; Maney Sveinsdottir; Emil A Thorarensen; Bjarni Thorbjornsson; Gisli Masson; Ingileif Jonsdottir; Alma Moller; Thorolfur Gudnason; Karl G Kristinsson; Unnur Thorsteinsdottir; Kari Stefansson |
| EPI_ISL_417676 | deCODE genetics | deCODE genetics | Daniel F Gudbjartsson; Agnar Helgason; Hakon Jonsson; Olafur T Magnusson; Pall Melsted; Gudmundur L Norddahl; Jona Saemundsdottir; Asgeir Sigurdsson; Patrick Sulem; Arna B Agustsdottir; Berglind Eiriksdottir; Run Fridriksdottir; Elisabet E Gardarsdottir; Gudmundur Georgsson; Olafia S Gretarsdottir; Kjartan R Gudmundsson; Thora R Gunnarsdottir; Arnaldur Gylfason; Hilma Holm; Brynjar O Jensson; Aslaug Jonasdottir; Kamilla S Josefsdottir; Thordur Kristjansson; Droplaug N Magnusdottir; Louise le Roux; Gudrun Sigmundsdottir; Gardar Sveinbjornsson; Kristin E Sveinsdottir; Maney Sveinsdottir; Emil A Thorarensen; Bjarni Thorbjornsson; Gisli Masson; Ingileif Jonsdottir; Alma Moller; Thorolfur Gudnason; Karl G Kristinsson; Unnur Thorsteinsdottir; Kari Stefansson |
| EPI_ISL_417677, EPI_ISL_417678, EPI_ISL_417679, EPI_ISL_417680, EPI_ISL_417681, EPI_ISL_417682, EPI_ISL_417683, EPI_ISL_417684, EPI_ISL_417685, EPI_ISL_417686, EPI_ISL_417687, EPI_ISL_417688, EPI_ISL_417689, EPI_ISL_417690, EPI_ISL_417691, EPI_ISL_417692, EPI_ISL_417693, EPI_ISL_417694, | | | |

| | | | |
|---|---|---|---|
| EPI_ISL_417695, EPI_ISL_417696, EPI_ISL_417697, EPI_ISL_417698, EPI_ISL_417699, EPI_ISL_417700, EPI_ISL_417701, EPI_ISL_417702, EPI_ISL_417703, EPI_ISL_417704, EPI_ISL_417705, EPI_ISL_417706, EPI_ISL_417707, EPI_ISL_417708, EPI_ISL_417709, EPI_ISL_417710, EPI_ISL_417711, EPI_ISL_417712, EPI_ISL_417713, EPI_ISL_417714, EPI_ISL_417715, EPI_ISL_417716, EPI_ISL_417717, EPI_ISL_417718, EPI_ISL_417719, EPI_ISL_417720, EPI_ISL_417721, EPI_ISL_417722, EPI_ISL_417723, EPI_ISL_417724, EPI_ISL_417725, EPI_ISL_417726, EPI_ISL_417727, EPI_ISL_417728, EPI_ISL_417729, EPI_ISL_417730, EPI_ISL_417731, EPI_ISL_417732, EPI_ISL_417733, EPI_ISL_417734, EPI_ISL_417735, EPI_ISL_417736, EPI_ISL_417737, EPI_ISL_417738, EPI_ISL_417739, EPI_ISL_417740, EPI_ISL_417741, EPI_ISL_417742, EPI_ISL_417743, EPI_ISL_417744, EPI_ISL_417745, EPI_ISL_417746, EPI_ISL_417747, EPI_ISL_417748, EPI_ISL_417749, EPI_ISL_417750, EPI_ISL_417751, EPI_ISL_417752, EPI_ISL_417753, EPI_ISL_417754, EPI_ISL_417755, EPI_ISL_417756, EPI_ISL_417757, EPI_ISL_417758, EPI_ISL_417759, EPI_ISL_417760, EPI_ISL_417761, EPI_ISL_417762, EPI_ISL_417763, EPI_ISL_417764, EPI_ISL_417765, EPI_ISL_417766, EPI_ISL_417767, EPI_ISL_417768, EPI_ISL_417769, EPI_ISL_417770, EPI_ISL_417771, EPI_ISL_417772, EPI_ISL_417773, EPI_ISL_417774, EPI_ISL_417775, EPI_ISL_417776, EPI_ISL_417777, EPI_ISL_417778, EPI_ISL_417779, EPI_ISL_417780, EPI_ISL_417781, EPI_ISL_417782, EPI_ISL_417783, EPI_ISL_417784, EPI_ISL_417785, EPI_ISL_417786, EPI_ISL_417787, EPI_ISL_417788, EPI_ISL_417789, EPI_ISL_417790, EPI_ISL_417791, EPI_ISL_417792, EPI_ISL_417793, EPI_ISL_417794, EPI_ISL_417795, EPI_ISL_417796, EPI_ISL_417797, EPI_ISL_417798, EPI_ISL_417799, EPI_ISL_417800, EPI_ISL_417801, EPI_ISL_417802, EPI_ISL_417803, EPI_ISL_417804, EPI_ISL_417805, EPI_ISL_417806, EPI_ISL_417807, EPI_ISL_417808, EPI_ISL_417809, EPI_ISL_417810, EPI_ISL_417811, EPI_ISL_417812, EPI_ISL_417813, EPI_ISL_417814, EPI_ISL_417815, EPI_ISL_417816, EPI_ISL_417817, EPI_ISL_417818, EPI_ISL_417819, EPI_ISL_417820, EPI_ISL_417821, EPI_ISL_417822, EPI_ISL_417823, EPI_ISL_417824, EPI_ISL_417825, EPI_ISL_417826, EPI_ISL_417827, EPI_ISL_417828, EPI_ISL_417829, EPI_ISL_417830, EPI_ISL_417831, EPI_ISL_417832, EPI_ISL_417833, EPI_ISL_417834, EPI_ISL_417835, EPI_ISL_417836, EPI_ISL_417837, EPI_ISL_417838, EPI_ISL_417839, EPI_ISL_417840, EPI_ISL_417841, EPI_ISL_417842, EPI_ISL_417843, EPI_ISL_417844, EPI_ISL_417845, EPI_ISL_417846, EPI_ISL_417847, EPI_ISL_417848, EPI_ISL_417849, EPI_ISL_417850, EPI_ISL_417851, EPI_ISL_417852, EPI_ISL_417853, EPI_ISL_417854, EPI_ISL_417855, EPI_ISL_417856, EPI_ISL_417857, EPI_ISL_417858, EPI_ISL_417859, EPI_ISL_417860, EPI_ISL_417861, EPI_ISL_417862, EPI_ISL_417863, EPI_ISL_417864, EPI_ISL_417865, EPI_ISL_417866, EPI_ISL_417867, EPI_ISL_417868, EPI_ISL_417869, EPI_ISL_417870, EPI_ISL_417871, EPI_ISL_417872, EPI_ISL_417873, EPI_ISL_417874, EPI_ISL_417875, EPI_ISL_417876 | | | |
| see above | The National University Hospital of Iceland | deCODE genetics | Daniel F Gudbjartsson; Agnar Helgason; Hakon Jonsson; Olafur T Magnusson; Pall Melsted; Gudmundur L Norddahl; Jona Saemundsdottir; Asgeir Sigurdsson; Patrick Sulem; Arna B Agustsdottir; Berglind Eiriksdottir; Run Fridriksdottir; Elisabet E Gardarsdottir; Gudmundur Georgsson; Olafia S Gretarsdottir; Kjartan R Gudmundsson; Thora R Gunnarsdottir; Arnaldur Gylfason; Hilma Holm; Brynjar O Jensson; Aslaug Jonasdottir; Kamilla S Josefsdottir; Thordur Kristjansson; Droplaug N Magnusdottir; Louise le Roux; Gudrun Sigmundsdottir; Gardar Sveinbjornsson; Kristin E Sveinsdottir; Maney Sveinsdottir; Emil A Thorarensen; Bjarni Thorbjornsson; Gisli Masson; Ingileif Jonsdottir; Alma Moller; Thorolfur Gudnason; Karl G Kristinsson; Unnur Thorsteinsdottir; Kari Stefansson |
| EPI_ISL_418184 | Gundersen Molecular Diagnostics Laboratory | Kabara Cancer Research Institute | Craig S. Richmond & Paraic A. Kenny |
| EPI_ISL_418185, EPI_ISL_418186 | Gundersen Molecular Diagnostic Laboratory | Kabara Cancer Research Institute | Craig S. Richmond & Paraic A. Kenny |
| EPI_ISL_418187, EPI_ISL_418188 | Gundersen Molecular Diagnostics Laboratory | Kabara Cancer Research Institute | Craig S. Richmond & Paraic A. Kenny |
| EPI_ISL_418860, EPI_ISL_418861 | Hospital Universitari Vall d'Hebron (HUVH) - Vall d'Hebron Research Institute (VHIR) | Hospital Universitari Vall d'Hebron (HUVH) - Vall d'Hebron Research Institute (VHIR) | Cristina Andrés, Dàmir Garcia-Cehic, Maria Piñana, Mercedes Guerrero-Murillo, Ariadna Rando, Tomàs Pumarola, Maria Gema Codina, Andrés Antón, Josep Quer |
| EPI_ISL_419651, EPI_ISL_419652 | Gundersen Molecular Diagnostics Laboratory | Kabara Cancer Research Institute | Craig S. Richmond & Paraic A. Kenny |
| EPI_ISL_420099, EPI_ISL_420100, EPI_ISL_420101, EPI_ISL_420102, EPI_ISL_420103, EPI_ISL_420104, EPI_ISL_420105, EPI_ISL_420106, EPI_ISL_420107, EPI_ISL_420108, EPI_ISL_420109, EPI_ISL_420110, EPI_ISL_420111 | | | |
| see above | National Centre for Infectious Diseases | Programme in Emerging Infectious Diseases, Duke-NUS Medical School | Danielle E Anderson, Martin Linster, Yan Zhuang, Jayanthi Jayakumar, David CB Lye, Yee Sin Leo, Barnaby E Young, Yvonne CF Su, Gavin JD Smith |
| EPI_ISL_420124 | Servicio de Microbiología. Consorcio Hospital General Universitario de Valencia | Sequencing and Bioinformatics Service and Molecular Epidemiology Research Group. FISABIO-Public Health | Concepcion Gimeno, Maria Alma Bracho, Griselda De Marco, Beatriz Beamud, Lidia Ruiz Roldan, Marta Pla Diaz, Neris Garcia-Gonzalez, Loreto Ferrús Abad, Inma Galán Vendrell, Paula Ruiz-Hueso, Mariana Reyes-Prieto, Vicente Soriano Chirona, Maria Dolores Ocete, Llúcia Martínez-Priego, Giuseppe D'Auria, Fernando Gonzalez-Candelas |
| EPI_ISL_422453, EPI_ISL_422459 | Gundersen Molecular Diagnostics Laboratory | Kabara Cancer Research Institute | Craig S. Richmond & Paraic A. Kenny |
| EPI_ISL_422461, EPI_ISL_422462, EPI_ISL_422463 | Gundersen Molecular Diagnostics Laboratory | Kabara Cancer Research Institute | Craig S. Richmond, Paraic A. Kenny |
| EPI_ISL_422464 | Gundersen Molecular Diagnostics Laboratory | Kabara Cancer Research Institute | Craig S. Richmond; Paraic A. Kenny |
| EPI_ISL_422465 | Gundersen Molecular Diagnostics Laboratory | Kabara Cancer Research Institute | Craig S. Richmond, Paraic A. Kenny |
| EPI_ISL_424342 | INMI Lazzaro Spallanzani IRCCS | Laboratory of Virology, INMI Lazzaro Spallanzani IRCCS | Concetta Castilletti, Barbara Bartolini, Martina Rueca, Cesare Ernesto Maria Gruber, Francesco Messina, Fabrizio Carletti, Eleonora Lalle, Licia Bordi, Giulia Matusali, Francesca Colavita, Maria Rosaria Capobianchi, Francesco Vairo, Giuseppe Ippolito, Antonino Di Caro |
| EPI_ISL_424377, EPI_ISL_424378 | deCODE genetics | deCODE genetics | Daniel F Gudbjartsson; Agnar Helgason; Hakon Jonsson; Olafur T Magnusson; Pall Melsted; Gudmundur L Norddahl; Jona Saemundsdottir; Asgeir Sigurdsson; Patrick Sulem; Arna B Agustsdottir; Berglind Eiriksdottir; Run Fridriksdottir; Elisabet E Gardarsdottir; Gudmundur Georgsson; Olafia S Gretarsdottir; Kjartan R Gudmundsson; Thora R Gunnarsdottir; Arnaldur Gylfason; Hilma Holm; Brynjar O Jensson; Aslaug Jonasdottir; Kamilla S Josefsdottir; Thordur Kristjansson; Droplaug N Magnusdottir; Louise le Roux; Gudrun Sigmundsdottir; Gardar Sveinbjornsson; Kristin E Sveinsdottir; Maney Sveinsdottir; Emil A Thorarensen; Bjarni Thorbjornsson; Gisli Masson; Ingileif Jonsdottir; Alma Moller; Thorolfur Gudnason; Karl G Kristinsson; Unnur Thorsteinsdottir; Kari Stefansson |
| EPI_ISL_424379, EPI_ISL_424380, EPI_ISL_424381, EPI_ISL_424382, EPI_ISL_424383, EPI_ISL_424384, EPI_ISL_424385, EPI_ISL_424386, EPI_ISL_424387, EPI_ISL_424388, EPI_ISL_424389, EPI_ISL_424390, EPI_ISL_424391, EPI_ISL_424392, EPI_ISL_424393, EPI_ISL_424394, EPI_ISL_424395, EPI_ISL_424396, EPI_ISL_424397, EPI_ISL_424398, EPI_ISL_424399, EPI_ISL_424400, EPI_ISL_424401, EPI_ISL_424402, EPI_ISL_424403, EPI_ISL_424404, EPI_ISL_424405, EPI_ISL_424406, EPI_ISL_424407, EPI_ISL_424408, EPI_ISL_424409, EPI_ISL_424410, EPI_ISL_424411, EPI_ISL_424412, EPI_ISL_424413, EPI_ISL_424414, EPI_ISL_424415, EPI_ISL_424416, EPI_ISL_424417, EPI_ISL_424418, EPI_ISL_424419, EPI_ISL_424420, EPI_ISL_424421, EPI_ISL_424422, EPI_ISL_424423, EPI_ISL_424424, EPI_ISL_424425, EPI_ISL_424426, EPI_ISL_424427, EPI_ISL_424428, EPI_ISL_424429, EPI_ISL_424430, EPI_ISL_424431, EPI_ISL_424432, EPI_ISL_424433, EPI_ISL_424434, EPI_ISL_424435, EPI_ISL_424436, EPI_ISL_424437, EPI_ISL_424438, EPI_ISL_424439, EPI_ISL_424440, EPI_ISL_424441, EPI_ISL_424442, EPI_ISL_424443, EPI_ISL_424444, EPI_ISL_424445, EPI_ISL_424446, EPI_ISL_424447, EPI_ISL_424448, EPI_ISL_424449, EPI_ISL_424450, EPI_ISL_424451, EPI_ISL_424452, EPI_ISL_424453, EPI_ISL_424454, EPI_ISL_424455, EPI_ISL_424456, EPI_ISL_424457, EPI_ISL_424458, EPI_ISL_424459, EPI_ISL_424460, EPI_ISL_424461, EPI_ISL_424462, EPI_ISL_424463, EPI_ISL_424464, EPI_ISL_424465, EPI_ISL_424466, EPI_ISL_424467, EPI_ISL_424468, EPI_ISL_424469, EPI_ISL_424470, EPI_ISL_424471, EPI_ISL_424472, EPI_ISL_424473, EPI_ISL_424474, EPI_ISL_424475, EPI_ISL_424476, EPI_ISL_424477, EPI_ISL_424478, EPI_ISL_424479, EPI_ISL_424480, EPI_ISL_424481, EPI_ISL_424482, EPI_ISL_424483, EPI_ISL_424484, EPI_ISL_424485, EPI_ISL_424486, EPI_ISL_424487, EPI_ISL_424488, EPI_ISL_424489, EPI_ISL_424490, EPI_ISL_424491, EPI_ISL_424492, EPI_ISL_424493, EPI_ISL_424494, EPI_ISL_424495, EPI_ISL_424496, EPI_ISL_424497, EPI_ISL_424498, EPI_ISL_424499, EPI_ISL_424500, EPI_ISL_424501, EPI_ISL_424502, EPI_ISL_424503, EPI_ISL_424505, EPI_ISL_424506, EPI_ISL_424507, EPI_ISL_424508, EPI_ISL_424509, EPI_ISL_424510, EPI_ISL_424511, EPI_ISL_424512, EPI_ISL_424513, EPI_ISL_424514, EPI_ISL_424515, EPI_ISL_424516, EPI_ISL_424517, EPI_ISL_424518, EPI_ISL_424519, EPI_ISL_424520, EPI_ISL_424521, EPI_ISL_424522, EPI_ISL_424523, EPI_ISL_424524, EPI_ISL_424525, EPI_ISL_424526, EPI_ISL_424527, EPI_ISL_424528, EPI_ISL_424529, EPI_ISL_424530, EPI_ISL_424531, EPI_ISL_424532 | | | |
| see above | The National University Hospital of Iceland | deCODE genetics | Daniel F Gudbjartsson; Agnar Helgason; Hakon Jonsson; Olafur T Magnusson; Pall Melsted; Gudmundur L Norddahl; Jona Saemundsdottir; Asgeir Sigurdsson; Patrick Sulem; Arna B Agustsdottir; Berglind Eiriksdottir; Run Fridriksdottir; Elisabet E Gardarsdottir; Gudmundur Georgsson; Olafia S Gretarsdottir; Kjartan R Gudmundsson; Thora R Gunnarsdottir; Arnaldur Gylfason; Hilma Holm; Brynjar O Jensson; Aslaug Jonasdottir; Kamilla S Josefsdottir; Thordur Kristjansson; Droplaug N Magnusdottir; Louise le Roux; Gudrun Sigmundsdottir; Gardar Sveinbjornsson; Kristin E Sveinsdottir; Maney Sveinsdottir; Emil A Thorarensen; Bjarni Thorbjornsson; Gisli Masson; Ingileif Jonsdottir; Alma Moller; Thorolfur Gudnason; Karl G Kristinsson; Unnur Thorsteinsdottir; Kari Stefansson |
| EPI_ISL_424533, EPI_ISL_424535, EPI_ISL_424536, EPI_ISL_424538, EPI_ISL_424539, EPI_ISL_424543, EPI_ISL_424544, EPI_ISL_424545, EPI_ISL_424546, EPI_ISL_424551 | deCODE genetics | deCODE genetics | Daniel F Gudbjartsson; Agnar Helgason; Hakon Jonsson; Olafur T Magnusson; Pall Melsted; Gudmundur L Norddahl; Jona Saemundsdottir; Asgeir Sigurdsson; Patrick Sulem; Arna B Agustsdottir; Berglind Eiriksdottir; Run Fridriksdottir; Elisabet E Gardarsdottir; Gudmundur Georgsson; Olafia S Gretarsdottir; Kjartan R Gudmundsson; Thora R Gunnarsdottir; Arnaldur Gylfason; Hilma Holm; Brynjar O Jensson; Aslaug Jonasdottir; Kamilla S Josefsdottir; Thordur Kristjansson; Droplaug N Magnusdottir; Louise le Roux; Gudrun Sigmundsdottir; Gardar Sveinbjornsson; Kristin E Sveinsdottir; Maney Sveinsdottir; Emil A Thorarensen; Bjarni Thorbjornsson; Gisli Masson; Ingileif Jonsdottir; Alma Moller; Thorolfur Gudnason; Karl G Kristinsson; Unnur Thorsteinsdottir; Kari Stefansson |
| EPI_ISL_424552 | The National University Hospital of Iceland | deCODE genetics | Daniel F Gudbjartsson; Agnar Helgason; Hakon Jonsson; Olafur T Magnusson; Pall Melsted; Gudmundur L Norddahl; Jona Saemundsdottir; Asgeir Sigurdsson; Patrick Sulem; Arna B Agustsdottir; Berglind Eiriksdottir; Run Fridriksdottir; Elisabet E Gardarsdottir; Gudmundur Georgsson; Olafia S Gretarsdottir; Kjartan R Gudmundsson; Thora R Gunnarsdottir; Arnaldur Gylfason; Hilma Holm; Brynjar O Jensson; Aslaug Jonasdottir; Kamilla S Josefsdottir; Thordur Kristjansson; Droplaug N Magnusdottir; Louise le Roux; Gudrun Sigmundsdottir; Gardar Sveinbjornsson; Kristin E Sveinsdottir; Maney Sveinsdottir; Emil A Thorarensen; Bjarni Thorbjornsson; Gisli Masson; Ingileif Jonsdottir; Alma Moller; Thorolfur Gudnason; Karl G Kristinsson; Unnur Thorsteinsdottir; Kari Stefansson |
| EPI_ISL_424553 | deCODE genetics | deCODE genetics | Daniel F Gudbjartsson; Agnar Helgason; Hakon Jonsson; Olafur T Magnusson; Pall Melsted; Gudmundur L Norddahl; Jona Saemundsdottir; Asgeir Sigurdsson; Patrick Sulem; Arna B Agustsdottir; Berglind Eiriksdottir; Run Fridriksdottir; Elisabet E Gardarsdottir; Gudmundur Georgsson; Olafia S Gretarsdottir; Kjartan R Gudmundsson; Thora R Gunnarsdottir; Arnaldur Gylfason; Hilma Holm; Brynjar O Jensson; Aslaug Jonasdottir; Kamilla S Josefsdottir; Thordur Kristjansson; Droplaug N Magnusdottir; Louise le Roux; Gudrun Sigmundsdottir; Gardar Sveinbjornsson; Kristin E Sveinsdottir; Maney Sveinsdottir; Emil A Thorarensen; Bjarni Thorbjornsson; Gisli Masson; Ingileif Jonsdottir; Alma Moller; Thorolfur Gudnason; Karl G Kristinsson; Unnur Thorsteinsdottir; Kari Stefansson |
| EPI_ISL_424554, EPI_ISL_424555, EPI_ISL_424556, EPI_ISL_424557, | The National University Hospital of Iceland | deCODE genetics | Daniel F Gudbjartsson; Agnar Helgason; Hakon Jonsson; Olafur T Magnusson; Pall Melsted; Gudmundur L Norddahl; Jona Saemundsdottir; Asgeir Sigurdsson; Patrick Sulem; Arna B Agustsdottir; Berglind Eiriksdottir; Run Fridriksdottir; Elisabet E Gardarsdottir; Gudmundur Georgsson; Olafia S |

| | | | |
|---|---|---|---|
| EPI_ISL_424558, EPI_ISL_424559 | | | Gretarsdottir; Kjartan R Gudmundsson; Thora R Gunnarsdottir; Arnaldur Gylfason; Hilma Holm; Brynjar O Jensson; Aslaug Jonasdottir; Kamilla S Josefsdottir; Thordur Kristjansson; Droplaug N Magnusdottir; Louise le Roux; Gudrun Sigmundsdottir; Gardar Sveinbjornsson; Kristin E Sveinsdottir; Maney Sveinsdottir; Emil A Thorarensen; Bjarni Thorbjornsson; Gisli Masson; Ingileif Jonsdottir; Alma Moller; Thorolfur Gudnason; Karl G Kristinsson; Unnur Thorsteinsdottir; Kari Stefansson |
| EPI_ISL_424560, EPI_ISL_424561, EPI_ISL_424562 | deCODE genetics | deCODE genetics | Daniel F Gudbjartsson; Agnar Helgason; Hakon Jonsson; Olafur T Magnusson; Pall Melsted; Gudmundur L Norddahl; Jona Saemundsdottir; Asgeir Sigurdsson; Patrick Sulem; Arna B Agustsdottir; Berglind Eiriksdottir; Run Fridriksdottir; Elisabet E Gardarsdottir; Gudmundur Georgsson; Olafia S Gretarsdottir; Kjartan R Gudmundsson; Thora R Gunnarsdottir; Arnaldur Gylfason; Hilma Holm; Brynjar O Jensson; Aslaug Jonasdottir; Kamilla S Josefsdottir; Thordur Kristjansson; Droplaug N Magnusdottir; Louise le Roux; Gudrun Sigmundsdottir; Gardar Sveinbjornsson; Kristin E Sveinsdottir; Maney Sveinsdottir; Emil A Thorarensen; Bjarni Thorbjornsson; Gisli Masson; Ingileif Jonsdottir; Alma Moller; Thorolfur Gudnason; Karl G Kristinsson; Unnur Thorsteinsdottir; Kari Stefansson |
| EPI_ISL_424563, EPI_ISL_424564 | The National University Hospital of Iceland | deCODE genetics | Daniel F Gudbjartsson; Agnar Helgason; Hakon Jonsson; Olafur T Magnusson; Pall Melsted; Gudmundur L Norddahl; Jona Saemundsdottir; Asgeir Sigurdsson; Patrick Sulem; Arna B Agustsdottir; Berglind Eiriksdottir; Run Fridriksdottir; Elisabet E Gardarsdottir; Gudmundur Georgsson; Olafia S Gretarsdottir; Kjartan R Gudmundsson; Thora R Gunnarsdottir; Arnaldur Gylfason; Hilma Holm; Brynjar O Jensson; Aslaug Jonasdottir; Kamilla S Josefsdottir; Thordur Kristjansson; Droplaug N Magnusdottir; Louise le Roux; Gudrun Sigmundsdottir; Gardar Sveinbjornsson; Kristin E Sveinsdottir; Maney Sveinsdottir; Emil A Thorarensen; Bjarni Thorbjornsson; Gisli Masson; Ingileif Jonsdottir; Alma Moller; Thorolfur Gudnason; Karl G Kristinsson; Unnur Thorsteinsdottir; Kari Stefansson |
| EPI_ISL_424565, EPI_ISL_424566, EPI_ISL_424567, EPI_ISL_424568, EPI_ISL_424570 | deCODE genetics | deCODE genetics | Daniel F Gudbjartsson; Agnar Helgason; Hakon Jonsson; Olafur T Magnusson; Pall Melsted; Gudmundur L Norddahl; Jona Saemundsdottir; Asgeir Sigurdsson; Patrick Sulem; Arna B Agustsdottir; Berglind Eiriksdottir; Run Fridriksdottir; Elisabet E Gardarsdottir; Gudmundur Georgsson; Olafia S Gretarsdottir; Kjartan R Gudmundsson; Thora R Gunnarsdottir; Arnaldur Gylfason; Hilma Holm; Brynjar O Jensson; Aslaug Jonasdottir; Kamilla S Josefsdottir; Thordur Kristjansson; Droplaug N Magnusdottir; Louise le Roux; Gudrun Sigmundsdottir; Gardar Sveinbjornsson; Kristin E Sveinsdottir; Maney Sveinsdottir; Emil A Thorarensen; Bjarni Thorbjornsson; Gisli Masson; Ingileif Jonsdottir; Alma Moller; Thorolfur Gudnason; Karl G Kristinsson; Unnur Thorsteinsdottir; Kari Stefansson |
| EPI_ISL_424571, EPI_ISL_424572, EPI_ISL_424573, EPI_ISL_424574, EPI_ISL_424575, EPI_ISL_424576, EPI_ISL_424577, EPI_ISL_424578, EPI_ISL_424579, EPI_ISL_424580, EPI_ISL_424581, EPI_ISL_424582, EPI_ISL_424583, EPI_ISL_424584, EPI_ISL_424585, EPI_ISL_424586, EPI_ISL_424587, EPI_ISL_424588, EPI_ISL_424589, EPI_ISL_424590, EPI_ISL_424591, EPI_ISL_424592, EPI_ISL_424593, EPI_ISL_424595, EPI_ISL_424596, EPI_ISL_424597, EPI_ISL_424598, EPI_ISL_424599, EPI_ISL_424600, EPI_ISL_424601, EPI_ISL_424602, EPI_ISL_424603, EPI_ISL_424604, EPI_ISL_424605, EPI_ISL_424606, EPI_ISL_424607 | | | |
| see above | The National University Hospital of Iceland | deCODE genetics | Daniel F Gudbjartsson; Agnar Helgason; Hakon Jonsson; Olafur T Magnusson; Pall Melsted; Gudmundur L Norddahl; Jona Saemundsdottir; Asgeir Sigurdsson; Patrick Sulem; Arna B Agustsdottir; Berglind Eiriksdottir; Run Fridriksdottir; Elisabet E Gardarsdottir; Gudmundur Georgsson; Olafia S Gretarsdottir; Kjartan R Gudmundsson; Thora R Gunnarsdottir; Arnaldur Gylfason; Hilma Holm; Brynjar O Jensson; Aslaug Jonasdottir; Kamilla S Josefsdottir; Thordur Kristjansson; Droplaug N Magnusdottir; Louise le Roux; Gudrun Sigmundsdottir; Gardar Sveinbjornsson; Kristin E Sveinsdottir; Maney Sveinsdottir; Emil A Thorarensen; Bjarni Thorbjornsson; Gisli Masson; Ingileif Jonsdottir; Alma Moller; Thorolfur Gudnason; Karl G Kristinsson; Unnur Thorsteinsdottir; Kari Stefansson |
| EPI_ISL_424608, EPI_ISL_424609 | deCODE genetics | deCODE genetics | Daniel F Gudbjartsson; Agnar Helgason; Hakon Jonsson; Olafur T Magnusson; Pall Melsted; Gudmundur L Norddahl; Jona Saemundsdottir; Asgeir Sigurdsson; Patrick Sulem; Arna B Agustsdottir; Berglind Eiriksdottir; Run Fridriksdottir; Elisabet E Gardarsdottir; Gudmundur Georgsson; Olafia S Gretarsdottir; Kjartan R Gudmundsson; Thora R Gunnarsdottir; Arnaldur Gylfason; Hilma Holm; Brynjar O Jensson; Aslaug Jonasdottir; Kamilla S Josefsdottir; Thordur Kristjansson; Droplaug N Magnusdottir; Louise le Roux; Gudrun Sigmundsdottir; Gardar Sveinbjornsson; Kristin E Sveinsdottir; Maney Sveinsdottir; Emil A Thorarensen; Bjarni Thorbjornsson; Gisli Masson; Ingileif Jonsdottir; Alma Moller; Thorolfur Gudnason; Karl G Kristinsson; Unnur Thorsteinsdottir; Kari Stefansson |
| EPI_ISL_424610, EPI_ISL_424611, EPI_ISL_424612, EPI_ISL_424613, EPI_ISL_424614, EPI_ISL_424615, EPI_ISL_424616, EPI_ISL_424617, EPI_ISL_424618, EPI_ISL_424619, EPI_ISL_424620, EPI_ISL_424621, EPI_ISL_424622, EPI_ISL_424623, EPI_ISL_424624 | | | |
| see above | The National University Hospital of Iceland | deCODE genetics | Daniel F Gudbjartsson; Agnar Helgason; Hakon Jonsson; Olafur T Magnusson; Pall Melsted; Gudmundur L Norddahl; Jona Saemundsdottir; Asgeir Sigurdsson; Patrick Sulem; Arna B Agustsdottir; Berglind Eiriksdottir; Run Fridriksdottir; Elisabet E Gardarsdottir; Gudmundur Georgsson; Olafia S Gretarsdottir; Kjartan R Gudmundsson; Thora R Gunnarsdottir; Arnaldur Gylfason; Hilma Holm; Brynjar O Jensson; Aslaug Jonasdottir; Kamilla S Josefsdottir; Thordur Kristjansson; Droplaug N Magnusdottir; Louise le Roux; Gudrun Sigmundsdottir; Gardar Sveinbjornsson; Kristin E Sveinsdottir; Maney Sveinsdottir; Emil A Thorarensen; Bjarni Thorbjornsson; Gisli Masson; Ingileif Jonsdottir; Alma Moller; Thorolfur Gudnason; Karl G Kristinsson; Unnur Thorsteinsdottir; Kari Stefansson |
| EPI_ISL_426159, EPI_ISL_426160, EPI_ISL_426161 | Gundersen Molecular Diagnostics Laboratory | Kabara Cancer Research Institute | Craig S. Richmond, Paraic A. Kenny |
| EPI_ISL_427526, EPI_ISL_427527, EPI_ISL_427528, EPI_ISL_427529, EPI_ISL_427530, EPI_ISL_427531, EPI_ISL_427532, EPI_ISL_427533, EPI_ISL_427534, EPI_ISL_427535, EPI_ISL_427536, EPI_ISL_427537, EPI_ISL_427538, EPI_ISL_427539, EPI_ISL_427540, EPI_ISL_427541, EPI_ISL_427542, EPI_ISL_427543, EPI_ISL_427544, EPI_ISL_427545, EPI_ISL_427546, EPI_ISL_427547, EPI_ISL_427548, EPI_ISL_427549, EPI_ISL_427550, EPI_ISL_427551, EPI_ISL_427552, EPI_ISL_427553, EPI_ISL_427554, EPI_ISL_427555, EPI_ISL_427556, EPI_ISL_427557, EPI_ISL_427558, EPI_ISL_427559, EPI_ISL_427560, EPI_ISL_427561, EPI_ISL_427562, EPI_ISL_427563, EPI_ISL_427564, EPI_ISL_427565, EPI_ISL_427566, EPI_ISL_427567, EPI_ISL_427568, EPI_ISL_427569, EPI_ISL_427570, EPI_ISL_427571, EPI_ISL_427572, EPI_ISL_427573, EPI_ISL_427574, EPI_ISL_427575, EPI_ISL_427576, EPI_ISL_427577, EPI_ISL_427578, EPI_ISL_427579, EPI_ISL_427580, EPI_ISL_427581, EPI_ISL_427582, EPI_ISL_427583, EPI_ISL_427584, EPI_ISL_427585, EPI_ISL_427586, EPI_ISL_427587, EPI_ISL_427588, EPI_ISL_427589, EPI_ISL_427590, EPI_ISL_427591, EPI_ISL_427592, EPI_ISL_427593, EPI_ISL_427594, EPI_ISL_427595, EPI_ISL_427596, EPI_ISL_427597, EPI_ISL_427598, EPI_ISL_427599, EPI_ISL_427600, EPI_ISL_427601, EPI_ISL_427602, EPI_ISL_427603, EPI_ISL_427604, EPI_ISL_427605, EPI_ISL_427606, EPI_ISL_427607, EPI_ISL_427608, EPI_ISL_427609, EPI_ISL_427610, EPI_ISL_427611, EPI_ISL_427612, EPI_ISL_427613, EPI_ISL_427614, EPI_ISL_427615, EPI_ISL_427616, EPI_ISL_427617, EPI_ISL_427618 | | | |
| see above | NewYork-Presbyterian & Mason Lab | Mason Lab | Daniel J. Butler, Christopher Mozsary, Cem Meydan, David Danko, Jonathan Foox, Joel Rosiene, Alon Shaiber, Matthew MacKay, Ebrahim Afshinnekoo, Fritz J. Sedlazeck, Nikolay A. Ivanov, Maria Sierra, Craig D. Westover, Krista Ryon, Benjamin Young, Chandrima Bhattacharya, Phyllis Ruggiero, Justyna Gawrys, Iman Hajirasouliha, Dmitry Meleshko, Mirella Salvatore, Dong Xu, Jenny Xiang, John Sipley, Lin Cong, Arryn Craney, Priya Velu, Lars F. Westblade, Massimo Loda, Shawn Levy, Melissa Cushing, Marcin Imielinski, Hanna Rennert, Christopher E. Mason |
| EPI_ISL_429843 | Gundersen Molecular Diagnostics Laboratory | Kabara Cancer Research Institute | Craig S. Richmond, Paraic A. Kenny |
| EPI_ISL_429844 | Gundersen Molecular Diagnostics Laboratory | Kabara Cancer Research Institute | Craig S. Richmond; Paraic A. Kenny |
| EPI_ISL_429845, EPI_ISL_429846, EPI_ISL_429847, EPI_ISL_429848 | Gundersen Molecular Diagnostics Laboratory | Kabara Cancer Research Institute | Craig S. Richmond; Paraic A. Kenny |
| EPI_ISL_430018, EPI_ISL_430063 | Molecular Pathology Division, Department of Pathology, Hong Kong Sanatorium & Hospital | Molecular Pathology Division, Department of Pathology, Hong Kong Sanatorium & Hospital | Chun Hang AU, Wai Sing CHAN, Ho Yin LAM, Dona N. HO, Simon Y.M. LAM, Jonpaul S.T. ZEE, Tsun Leung CHAN, Edmond S.K. MA |
| EPI_ISL_430113, EPI_ISL_430114, EPI_ISL_430115, EPI_ISL_430116, EPI_ISL_430117, EPI_ISL_430118, EPI_ISL_430119, EPI_ISL_430121, EPI_ISL_430122, EPI_ISL_430123, EPI_ISL_430124, EPI_ISL_430125, EPI_ISL_430126, EPI_ISL_430127, EPI_ISL_430128, EPI_ISL_430129, EPI_ISL_430130, EPI_ISL_430131, EPI_ISL_430132, EPI_ISL_430133, EPI_ISL_430134, EPI_ISL_430135, EPI_ISL_430136, EPI_ISL_430137, EPI_ISL_430138, EPI_ISL_430139, EPI_ISL_430140, EPI_ISL_430141, EPI_ISL_430142, EPI_ISL_430143, EPI_ISL_430144, EPI_ISL_430145, EPI_ISL_430146, EPI_ISL_430147, EPI_ISL_430148, EPI_ISL_430149, EPI_ISL_430150, EPI_ISL_430151, EPI_ISL_430152, EPI_ISL_430153, EPI_ISL_430154, EPI_ISL_430155, EPI_ISL_430156 | | | |
| see above | Seattle Flu Study | Seattle Flu Study | Chu et al |
| EPI_ISL_430160, EPI_ISL_430161, EPI_ISL_430162, EPI_ISL_430163, EPI_ISL_430164, EPI_ISL_430165, EPI_ISL_430166, EPI_ISL_430167, EPI_ISL_430168, EPI_ISL_430169, EPI_ISL_430170, EPI_ISL_430171, EPI_ISL_430172, EPI_ISL_430173, EPI_ISL_430174, EPI_ISL_430175, EPI_ISL_430176, EPI_ISL_430177, EPI_ISL_430178, EPI_ISL_430179, EPI_ISL_430180, EPI_ISL_430181, EPI_ISL_430182, EPI_ISL_430183, EPI_ISL_430184, EPI_ISL_430185, EPI_ISL_430186, EPI_ISL_430187, EPI_ISL_430188, EPI_ISL_430189, EPI_ISL_430190, EPI_ISL_430191, EPI_ISL_430192, EPI_ISL_430193, EPI_ISL_430194, EPI_ISL_430196, EPI_ISL_430197, EPI_ISL_430198, EPI_ISL_430199, EPI_ISL_430203, EPI_ISL_430204, EPI_ISL_430205, EPI_ISL_430206, EPI_ISL_430207, EPI_ISL_430208, EPI_ISL_430209, EPI_ISL_430210, EPI_ISL_430211, EPI_ISL_430212, EPI_ISL_430213, EPI_ISL_430214, EPI_ISL_430215, EPI_ISL_430216, EPI_ISL_430217, EPI_ISL_430218, EPI_ISL_430219, EPI_ISL_430220, EPI_ISL_430221, EPI_ISL_430222, EPI_ISL_430223, EPI_ISL_430224, EPI_ISL_430225, EPI_ISL_430226, EPI_ISL_430227, EPI_ISL_430228, EPI_ISL_430229, EPI_ISL_430230, EPI_ISL_430231, EPI_ISL_430232, EPI_ISL_430233, EPI_ISL_430234, EPI_ISL_430235, EPI_ISL_430236, EPI_ISL_430237, EPI_ISL_430238, EPI_ISL_430239, EPI_ISL_430240, EPI_ISL_430241, EPI_ISL_430242, EPI_ISL_430243, EPI_ISL_430244, EPI_ISL_430245, EPI_ISL_430246, EPI_ISL_430247, EPI_ISL_430248, EPI_ISL_430249, EPI_ISL_430250, EPI_ISL_430251, EPI_ISL_430252, EPI_ISL_430253, EPI_ISL_430254, EPI_ISL_430255, EPI_ISL_430256, EPI_ISL_430257, EPI_ISL_430258, EPI_ISL_430259, EPI_ISL_430260, EPI_ISL_430261, EPI_ISL_430262, EPI_ISL_430263, EPI_ISL_430264, EPI_ISL_430265, EPI_ISL_430266, EPI_ISL_430267, EPI_ISL_430268, EPI_ISL_430269, EPI_ISL_430270, EPI_ISL_430271, EPI_ISL_430272, EPI_ISL_430273, EPI_ISL_430274, EPI_ISL_430275, EPI_ISL_430276, EPI_ISL_430277, EPI_ISL_430278, EPI_ISL_430279, EPI_ISL_430280, EPI_ISL_430281, EPI_ISL_430282, EPI_ISL_430283, EPI_ISL_430284, EPI_ISL_430285, EPI_ISL_430286, EPI_ISL_430287, EPI_ISL_430288, EPI_ISL_430289, EPI_ISL_430290, EPI_ISL_430291, EPI_ISL_430292, EPI_ISL_430293, EPI_ISL_430294, EPI_ISL_430295, EPI_ISL_430296, EPI_ISL_434063, EPI_ISL_434064, EPI_ISL_434065, EPI_ISL_434066, EPI_ISL_434068, EPI_ISL_434069, EPI_ISL_434070, EPI_ISL_434071, EPI_ISL_434072, EPI_ISL_434073, EPI_ISL_434074, EPI_ISL_434075, EPI_ISL_434076, EPI_ISL_434077, EPI_ISL_434078, EPI_ISL_434079, EPI_ISL_434080, EPI_ISL_434081, EPI_ISL_434082, EPI_ISL_434083, EPI_ISL_434084, EPI_ISL_434085, EPI_ISL_434086, EPI_ISL_434087, EPI_ISL_434088, EPI_ISL_434089, EPI_ISL_434090, EPI_ISL_434091, EPI_ISL_434092, EPI_ISL_434093, EPI_ISL_434094, EPI_ISL_434095, EPI_ISL_434096, EPI_ISL_434097, EPI_ISL_434098, EPI_ISL_434099, EPI_ISL_434100, EPI_ISL_434101, EPI_ISL_434102, EPI_ISL_434103, EPI_ISL_434104, EPI_ISL_434105, EPI_ISL_434106, EPI_ISL_434107, EPI_ISL_434108, EPI_ISL_434109, EPI_ISL_434110, EPI_ISL_434111, EPI_ISL_434112, EPI_ISL_434113, EPI_ISL_434114, EPI_ISL_434115, EPI_ISL_434116, EPI_ISL_434117, EPI_ISL_434118, EPI_ISL_434119, EPI_ISL_434120, EPI_ISL_434121, EPI_ISL_434122, EPI_ISL_434123, EPI_ISL_434124, EPI_ISL_434128, EPI_ISL_434129, EPI_ISL_434130, EPI_ISL_434131, EPI_ISL_434132, EPI_ISL_434133, EPI_ISL_434134, EPI_ISL_434135, EPI_ISL_434136, EPI_ISL_434137, EPI_ISL_434138, EPI_ISL_434139, EPI_ISL_434140, EPI_ISL_434141, EPI_ISL_434142, EPI_ISL_434143, EPI_ISL_434144, EPI_ISL_434145, EPI_ISL_434146, EPI_ISL_434147, EPI_ISL_434148, EPI_ISL_434149, EPI_ISL_434150, EPI_ISL_434151, EPI_ISL_434152, EPI_ISL_434153, EPI_ISL_434154, EPI_ISL_434155, EPI_ISL_434156, EPI_ISL_434157, EPI_ISL_434158, EPI_ISL_434159, EPI_ISL_434160, EPI_ISL_434161, EPI_ISL_434162, EPI_ISL_434163, EPI_ISL_434164, EPI_ISL_434165, EPI_ISL_434166, EPI_ISL_434167, | | | |

| | | | |
|---|---|---|---|
| EPI_ISL_434168, EPI_ISL_434169, EPI_ISL_434170, EPI_ISL_434171, EPI_ISL_434172, EPI_ISL_434173, EPI_ISL_434174, EPI_ISL_434175, EPI_ISL_434176, EPI_ISL_434177, EPI_ISL_434178, EPI_ISL_434179, EPI_ISL_434180, EPI_ISL_434182, EPI_ISL_434183, EPI_ISL_434184, EPI_ISL_434185, EPI_ISL_434186, EPI_ISL_434187, EPI_ISL_434188, EPI_ISL_434189, EPI_ISL_434190, EPI_ISL_434191, EPI_ISL_434192, EPI_ISL_434193, EPI_ISL_434194, EPI_ISL_434196, EPI_ISL_434197, EPI_ISL_434198, EPI_ISL_434199, EPI_ISL_434200, EPI_ISL_434201, EPI_ISL_434202, EPI_ISL_434204, EPI_ISL_434205, EPI_ISL_434206, EPI_ISL_434207, EPI_ISL_434208, EPI_ISL_434209, EPI_ISL_434210, EPI_ISL_434211, EPI_ISL_434212, EPI_ISL_434213, EPI_ISL_434214, EPI_ISL_434215, EPI_ISL_434216, EPI_ISL_434217, EPI_ISL_434218, EPI_ISL_434219, EPI_ISL_434220, EPI_ISL_434221, EPI_ISL_434222, EPI_ISL_434223, EPI_ISL_434224, EPI_ISL_434225, EPI_ISL_434226, EPI_ISL_434227, EPI_ISL_434228, EPI_ISL_434229, EPI_ISL_434230, EPI_ISL_434231, EPI_ISL_434232, EPI_ISL_434233, EPI_ISL_434234, EPI_ISL_434235, EPI_ISL_434236, EPI_ISL_434237, EPI_ISL_434238, EPI_ISL_434239, EPI_ISL_434240, EPI_ISL_434241, EPI_ISL_434242, EPI_ISL_434243, EPI_ISL_434244, EPI_ISL_434245, EPI_ISL_434246, EPI_ISL_434247, EPI_ISL_434248, EPI_ISL_434249, EPI_ISL_434250, EPI_ISL_434251, EPI_ISL_434252, EPI_ISL_434253, EPI_ISL_434254, EPI_ISL_434255, EPI_ISL_434256, EPI_ISL_434257, EPI_ISL_434258, EPI_ISL_434259, EPI_ISL_434260, EPI_ISL_434262, EPI_ISL_434263, EPI_ISL_434264, EPI_ISL_434265, EPI_ISL_434266, EPI_ISL_434268, EPI_ISL_434269, EPI_ISL_434270, EPI_ISL_434271, EPI_ISL_434272, EPI_ISL_434273, EPI_ISL_434274, EPI_ISL_434275, EPI_ISL_434276, EPI_ISL_434277, EPI_ISL_434279, EPI_ISL_434280, EPI_ISL_434281, EPI_ISL_434282, EPI_ISL_434283, EPI_ISL_434284, EPI_ISL_434285, EPI_ISL_434286, EPI_ISL_434287, EPI_ISL_434288, EPI_ISL_434289, EPI_ISL_434290, EPI_ISL_434291, EPI_ISL_434293, EPI_ISL_434294, EPI_ISL_434296, EPI_ISL_434298, EPI_ISL_434299, EPI_ISL_434300, EPI_ISL_434301, EPI_ISL_434302, EPI_ISL_434303, EPI_ISL_434305, EPI_ISL_434306, EPI_ISL_434307, EPI_ISL_434308, EPI_ISL_434309, EPI_ISL_434310, EPI_ISL_434311, EPI_ISL_434313, EPI_ISL_434314, EPI_ISL_434315, EPI_ISL_434316, EPI_ISL_434317, EPI_ISL_434319, EPI_ISL_434320, EPI_ISL_434321, EPI_ISL_434322, EPI_ISL_434323, EPI_ISL_434324, EPI_ISL_434325, EPI_ISL_434327, EPI_ISL_434328, EPI_ISL_434329, EPI_ISL_434330, EPI_ISL_434331, EPI_ISL_434332, EPI_ISL_434333, EPI_ISL_434334, EPI_ISL_434335, EPI_ISL_434336, EPI_ISL_434337, EPI_ISL_434338, EPI_ISL_434339, EPI_ISL_434341, EPI_ISL_434342, EPI_ISL_434343, EPI_ISL_434344, EPI_ISL_434345, EPI_ISL_434346 | | | |
| see above | Washington State Department of Health | Seattle Flu Study | Chu et al |
| EPI_ISL_435144 | Hospital Universitario La Paz | Hospital Universitario 12 de Octubre | Elias Dahdouh, Sara González, Raúl Recio, Fernando Lázaro, Esther Viedma, Natalia Stella, Julio García, Juan Carlos Galán, Rafael Cantón, Mª Dolores Folgueira, Rafael Delgado, Jesús Mingorance |
| EPI_ISL_435394, EPI_ISL_435395, EPI_ISL_435396, EPI_ISL_435397, EPI_ISL_435398, EPI_ISL_435399, EPI_ISL_435400, EPI_ISL_435401, EPI_ISL_435402 | Gundersen Molecular Diagnostics Laboratory | Kabara Cancer Research Institute | Craig S. Richmond, Paraic A. Kenny |
| EPI_ISL_436466, EPI_ISL_436467, EPI_ISL_436468, EPI_ISL_436469, EPI_ISL_436470, EPI_ISL_436471, EPI_ISL_436472, EPI_ISL_436473, EPI_ISL_436474, EPI_ISL_436475, EPI_ISL_436476, EPI_ISL_436477, EPI_ISL_436478, EPI_ISL_436479, EPI_ISL_436480, EPI_ISL_436481, EPI_ISL_436482, EPI_ISL_436483, EPI_ISL_436484, EPI_ISL_436485, EPI_ISL_436486, EPI_ISL_436487, EPI_ISL_436488, EPI_ISL_436489, EPI_ISL_436490, EPI_ISL_436491, EPI_ISL_436492, EPI_ISL_436493, EPI_ISL_436494, EPI_ISL_436495, EPI_ISL_436496, EPI_ISL_436497, EPI_ISL_436498, EPI_ISL_436499, EPI_ISL_436500, EPI_ISL_436501, EPI_ISL_436502, EPI_ISL_436503 | | | |
| see above | UPMC Clinical Laboratory | Microbial Genome Sequencing Center, Microbial Genomic Epidemiological Laboratory | Dan Snyder, Stephanie L Mitchell, Mustapha M Mustapha, Marissa P Griffith, Vatsala R Srinivasa, Kady D Waggle, Chinelo Ezeonwuku, Jane W. Marsh, Lee H. Harrison, Vaughn S. Cooper |
| EPI_ISL_436891, EPI_ISL_436892, EPI_ISL_436893, EPI_ISL_436894, EPI_ISL_436895, EPI_ISL_436896, EPI_ISL_436897, EPI_ISL_436898, EPI_ISL_436899, EPI_ISL_436900 | Gundersen Molecular Diagnostics Laboratory | Kabara Cancer Research Institute | Craig S. Richmond, Paraic A. Kenny |
| EPI_ISL_438140, EPI_ISL_438141, EPI_ISL_438142, EPI_ISL_438143, EPI_ISL_438144, EPI_ISL_438145, EPI_ISL_438146, EPI_ISL_438147, EPI_ISL_438148, EPI_ISL_438149, EPI_ISL_438150, EPI_ISL_438151, EPI_ISL_438152, EPI_ISL_438153, EPI_ISL_438154, EPI_ISL_438155, EPI_ISL_438156, EPI_ISL_438157, EPI_ISL_438158, EPI_ISL_438159, EPI_ISL_438160, EPI_ISL_438161, EPI_ISL_438162, EPI_ISL_438163, EPI_ISL_438164, EPI_ISL_438165, EPI_ISL_438166, EPI_ISL_438167, EPI_ISL_438168, EPI_ISL_438169, EPI_ISL_438170, EPI_ISL_438171, EPI_ISL_438172, EPI_ISL_438173, EPI_ISL_438174, EPI_ISL_438175 | | | |
| see above | Seattle Flu Study | Seattle Flu Study | Chu et al |
| EPI_ISL_438176, EPI_ISL_438177, EPI_ISL_438178, EPI_ISL_438179, EPI_ISL_438180, EPI_ISL_438181, EPI_ISL_438182, EPI_ISL_438183, EPI_ISL_438184, EPI_ISL_438185, EPI_ISL_438186, EPI_ISL_438187, EPI_ISL_438188, EPI_ISL_438189, EPI_ISL_438190, EPI_ISL_438191, EPI_ISL_438192, EPI_ISL_438193, EPI_ISL_438194, EPI_ISL_438195, EPI_ISL_438196, EPI_ISL_438197, EPI_ISL_438198, EPI_ISL_438199, EPI_ISL_438200, EPI_ISL_438201, EPI_ISL_438202, EPI_ISL_438203, EPI_ISL_438204, EPI_ISL_438205, EPI_ISL_438206, EPI_ISL_438207, EPI_ISL_438208, EPI_ISL_438209, EPI_ISL_438210, EPI_ISL_438211, EPI_ISL_438212, EPI_ISL_438213, EPI_ISL_438214, EPI_ISL_438215, EPI_ISL_438216, EPI_ISL_438217, EPI_ISL_438218, EPI_ISL_438219, EPI_ISL_438220, EPI_ISL_438221 | | | |
| see above | Washington State Department of Health | Seattle Flu Study | Chu et al |
| EPI_ISL_441355, EPI_ISL_441356, EPI_ISL_441357, EPI_ISL_441358, EPI_ISL_441359, EPI_ISL_441360, EPI_ISL_441361, EPI_ISL_441362, EPI_ISL_441363, EPI_ISL_441364, EPI_ISL_441365, EPI_ISL_441366, EPI_ISL_441367, EPI_ISL_441368, EPI_ISL_441369, EPI_ISL_441370, EPI_ISL_441371, EPI_ISL_441372, EPI_ISL_441373, EPI_ISL_441374, EPI_ISL_441375, EPI_ISL_441376, EPI_ISL_441377, EPI_ISL_441378, EPI_ISL_441379, EPI_ISL_441380, EPI_ISL_441381, EPI_ISL_441382, EPI_ISL_441383, EPI_ISL_441384, EPI_ISL_441385, EPI_ISL_441386, EPI_ISL_441387, EPI_ISL_441388, EPI_ISL_441389, EPI_ISL_441390, EPI_ISL_441391, EPI_ISL_441392, EPI_ISL_441393, EPI_ISL_441394, EPI_ISL_441395, EPI_ISL_441396, EPI_ISL_441397, EPI_ISL_441398, EPI_ISL_441399, EPI_ISL_441400, EPI_ISL_441401, EPI_ISL_441402, EPI_ISL_441403, EPI_ISL_441404, EPI_ISL_441405, EPI_ISL_441406, EPI_ISL_441407, EPI_ISL_441408, EPI_ISL_441409, EPI_ISL_441410, EPI_ISL_441411, EPI_ISL_441412, EPI_ISL_441413, EPI_ISL_441414, EPI_ISL_441415, EPI_ISL_441416, EPI_ISL_441417, EPI_ISL_441418, EPI_ISL_441419, EPI_ISL_441420, EPI_ISL_441421, EPI_ISL_441422, EPI_ISL_441423, EPI_ISL_441424, EPI_ISL_441425, EPI_ISL_441426, EPI_ISL_441427, EPI_ISL_441428, EPI_ISL_441429, EPI_ISL_441430, EPI_ISL_441431, EPI_ISL_441432, EPI_ISL_441433, EPI_ISL_441434, EPI_ISL_441435, EPI_ISL_441436 | | | |
| see above | Regional Virus Laboratory, Belfast Health and Social Care Trust | COVID-19 Genomics UK (COG-UK) Consortium | Conall McCaughey, James McKenna, Tanya Curran, Susan Feeney, Alison Watt, Ciara Cox, Mairead Connor, Zoltan Molnar, David Simpson, Derek Fairley |
| EPI_ISL_441659, EPI_ISL_441662, EPI_ISL_441663, EPI_ISL_441670, EPI_ISL_441675, EPI_ISL_441676, EPI_ISL_441685, EPI_ISL_441690, EPI_ISL_441691, EPI_ISL_441693, EPI_ISL_441694, EPI_ISL_441696, EPI_ISL_441697, EPI_ISL_441698, EPI_ISL_441699, EPI_ISL_441704, EPI_ISL_441708, EPI_ISL_441712, EPI_ISL_441714, EPI_ISL_441716, EPI_ISL_441717, EPI_ISL_441719, EPI_ISL_441730, EPI_ISL_441733, EPI_ISL_441737, EPI_ISL_441747, EPI_ISL_441751, EPI_ISL_441753, EPI_ISL_441761, EPI_ISL_441765, EPI_ISL_441766, EPI_ISL_441770, EPI_ISL_441771, EPI_ISL_441775, EPI_ISL_441776, EPI_ISL_441781 | | | |
| see above | Regional Virus Laboratory, Belfast Health and Social Care Trust | Wellcome Sanger Institute for the COVID-19 Genomics UK (COG-UK) consortium | Conall McCaughey, James McKenna, Tanya Curran, Susan Feeney, Alison Watt, Ciara Cox, Mairead Connor, Zoltan Molnar, David Simpson, Derek Fairley, Alex Alderton, Roberto Amato, Sonia Goncalves, Ewan Harrison, David K. Jackson, Ian Johnston, Dominic Kwiatkowski, Cordelia Langford, John Sillitoe on behalf of the Wellcome Sanger Institute COVID-19 Surveillance Team (http://www.sanger.ac.uk/covid-team) |
| EPI_ISL_444971 | Hospital Universitari Vall d'Hebron - Vall d'Hebron Institut de Recerca | Hospital Universitari Vall d'Hebron | Cristina Andrés, Maria Piñana, Damir Garcia-Cehic, Mercedes Guerrero-Murillo, Ariadna Rando, Juliana Esperalba, Maria Gema Codina, Tomàs Pumarola, Josep Quer, Andrés Antón |
| EPI_ISL_444972 | Hospital Universitari Vall d'Hebron - Vall d'Hebron Institut de Recerca | Hospital Universitari Vall d'Hebron | Cristina Andrés, Maria Piñana, DAmir Garcia-Cehic, Mercedes Guerrero-Murillo, Ariadna Rando, Juliana Esperalba, Maria Gema Codina, Tomàs Pumarola, Josep Quer, Andrés Antón |
| EPI_ISL_444973 | Hospital Universitari Vall d`Hebron - Vall d'Hebron Institut de Recerca | Hospital Universitari Vall d'Hebron | Cristina Andrés, Maria Piñana, DAmir Garcia-Cehic, Mercedes Guerrero-Murillo, Ariadna Rando, Juliana Esperalba, Maria Gema Codina, Tomàs Pumarola, Josep Quer, Andrés Antón |
| EPI_ISL_444974, EPI_ISL_444975, EPI_ISL_444976, EPI_ISL_444977, EPI_ISL_444978, EPI_ISL_444979 | Hospital Universitari Vall d'Hebron - Vall d'Hebron Institut de Recerca | Hospital Universitari Vall d'Hebron | Cristina Andrés, Maria Piñana, Damir Garcia-Cehic, Mercedes Guerrero-Murillo, Ariadna Rando, Juliana Esperalba, Maria Gema Codina, Tomàs Pumarola, Josep Quer, Andrés Antón |
| EPI_ISL_444980, EPI_ISL_444981, EPI_ISL_444982, EPI_ISL_444983 | Hospital Universitari Vall d'Hebron - Vall d"hebron Institut de Recerca | Hospital Universitari Vall d'Hebron | Cristina Andrés, Maria Piñana, Damir Garcia-Cehic, Mercedes Guerrero-Murillo, Ariadna Rando, Juliana Esperalba, Maria Gema Codina, Tomàs Pumarola, Josep Quer, Andrés Antón |
| EPI_ISL_444984, EPI_ISL_444985, EPI_ISL_444986, EPI_ISL_444987, EPI_ISL_444988, EPI_ISL_444989, EPI_ISL_444990, EPI_ISL_447532, EPI_ISL_447533 | Hospital Universitari Vall d'Hebron - Vall d'Hebron Institut de Recerca | Hospital Universitari Vall d'Hebron | Cristina Andrés, Maria Piñana, Damir Garcia-Cehic, Mercedes Guerrero-Murillo, Ariadna Rando, Juliana Esperalba, Maria Gema Codina, Tomàs Pumarola, Josep Quer, Andrés Antón |
| EPI_ISL_447922, EPI_ISL_447923, EPI_ISL_447924, EPI_ISL_447925, EPI_ISL_447926, EPI_ISL_447927, EPI_ISL_447928, EPI_ISL_447929, EPI_ISL_447930, EPI_ISL_447931, EPI_ISL_447932, EPI_ISL_447933, EPI_ISL_447934, EPI_ISL_447935, EPI_ISL_447936, EPI_ISL_447937, EPI_ISL_447938, EPI_ISL_447939, EPI_ISL_447940, EPI_ISL_447941, EPI_ISL_447942, EPI_ISL_447943, EPI_ISL_447944 | | | |
| see above | University of Birmingham | COVID-19 Genomics UK (COG-UK) Consortium | Claire McMurray, Joanne Stockton, Samuel Nicholls, Radoslaw Poplawski, Will Rowe, Josh Quick, Nicholas Loman, Celina M Whalley, Andrew Bosworth, Charlotte Poxon, Kasun Wanigasooriya, Oliver Pickles, Mike Kidd, Alex Richter, Andrew D Beggs, Husam Osman, Andrew Bosworth |
| EPI_ISL_448918, EPI_ISL_448919, EPI_ISL_448920, EPI_ISL_448921, EPI_ISL_448922, EPI_ISL_448923, EPI_ISL_448924, EPI_ISL_448925, EPI_ISL_448926, EPI_ISL_448927, EPI_ISL_448928, EPI_ISL_448929, EPI_ISL_448930, EPI_ISL_448931, EPI_ISL_448932, EPI_ISL_448933, EPI_ISL_448934, EPI_ISL_448935, EPI_ISL_448936, EPI_ISL_448937, EPI_ISL_448938, EPI_ISL_448939, EPI_ISL_448940, EPI_ISL_448941, EPI_ISL_448942, EPI_ISL_448943, EPI_ISL_448944, EPI_ISL_448945, EPI_ISL_448946, EPI_ISL_448947, EPI_ISL_448948, EPI_ISL_448949, EPI_ISL_448950, EPI_ISL_448951, EPI_ISL_448952, EPI_ISL_448953, EPI_ISL_448954, EPI_ISL_448955, EPI_ISL_448956, EPI_ISL_448957, EPI_ISL_448958, EPI_ISL_448959, EPI_ISL_448960, EPI_ISL_448961, EPI_ISL_448962, EPI_ISL_448963, EPI_ISL_448964, EPI_ISL_448965, EPI_ISL_448966, EPI_ISL_448967, EPI_ISL_448968, EPI_ISL_448969, EPI_ISL_448970, EPI_ISL_448971, EPI_ISL_448972, EPI_ISL_448973, EPI_ISL_448974, EPI_ISL_448975, EPI_ISL_448976, EPI_ISL_448977 | | | |
| see above | Regional Virus Laboratory, Belfast Health and Social Care Trust | COVID-19 Genomics UK (COG-UK) Consortium | Conall McCaughey, James McKenna, Tanya Curran, Susan Feeney, Alison Watt, Ciara Cox, Mairead Connor, Zoltan Molnar, David Simpson, Derek Fairley |
| EPI_ISL_449839, EPI_ISL_449840, EPI_ISL_449841, EPI_ISL_449842, EPI_ISL_449843, EPI_ISL_449844, EPI_ISL_449845, EPI_ISL_449846, EPI_ISL_449847, EPI_ISL_449848, EPI_ISL_449849, EPI_ISL_449850, EPI_ISL_449851, EPI_ISL_449852, EPI_ISL_449853, EPI_ISL_449854, EPI_ISL_449855, EPI_ISL_449856, EPI_ISL_449857, EPI_ISL_449858, EPI_ISL_449859, EPI_ISL_449860, EPI_ISL_449861, EPI_ISL_449862, EPI_ISL_449863, EPI_ISL_449864, EPI_ISL_449865, EPI_ISL_449866, EPI_ISL_449867, EPI_ISL_449868, EPI_ISL_449869, EPI_ISL_449870, EPI_ISL_449871, EPI_ISL_449872, EPI_ISL_449873, EPI_ISL_449874, | | | |

| | | | |
|---|---|---|---|
| EPI_ISL_449875, EPI_ISL_449876, EPI_ISL_449877, EPI_ISL_449878, EPI_ISL_449879, EPI_ISL_449880, EPI_ISL_449881, EPI_ISL_449882, EPI_ISL_449883, EPI_ISL_449884, EPI_ISL_449885, EPI_ISL_449886, EPI_ISL_449887, EPI_ISL_449888, EPI_ISL_449889, EPI_ISL_449890, EPI_ISL_449891, EPI_ISL_449892, EPI_ISL_449893, EPI_ISL_449894, EPI_ISL_449895, EPI_ISL_449896, EPI_ISL_449897, EPI_ISL_449898, EPI_ISL_449899, EPI_ISL_449900, EPI_ISL_449901, EPI_ISL_449902, EPI_ISL_449903, EPI_ISL_449904, EPI_ISL_449905, EPI_ISL_449906, EPI_ISL_449907, EPI_ISL_449908, EPI_ISL_449909, EPI_ISL_449910, EPI_ISL_449911, EPI_ISL_449912, EPI_ISL_449913, EPI_ISL_449914, EPI_ISL_449915, EPI_ISL_449916, EPI_ISL_449917, EPI_ISL_449918, EPI_ISL_449919, EPI_ISL_449920, EPI_ISL_449921, EPI_ISL_449922, EPI_ISL_449923, EPI_ISL_449925, EPI_ISL_449926, EPI_ISL_449927, EPI_ISL_449928, EPI_ISL_449929, EPI_ISL_449930, EPI_ISL_449931, EPI_ISL_449932, EPI_ISL_449933, EPI_ISL_449934, EPI_ISL_449935, EPI_ISL_449936, EPI_ISL_449937, EPI_ISL_449938, EPI_ISL_449939, EPI_ISL_449940, EPI_ISL_449941, EPI_ISL_449942, EPI_ISL_449943, EPI_ISL_449944, EPI_ISL_449945, EPI_ISL_449946, EPI_ISL_449947, EPI_ISL_449948, EPI_ISL_449949, EPI_ISL_449950, EPI_ISL_449951, EPI_ISL_449952, EPI_ISL_449953, EPI_ISL_449954, EPI_ISL_449955, EPI_ISL_449956, EPI_ISL_449957, EPI_ISL_449958, EPI_ISL_449959, EPI_ISL_449960, EPI_ISL_449962, EPI_ISL_449963, EPI_ISL_449964, EPI_ISL_449965, EPI_ISL_449966, EPI_ISL_449967, EPI_ISL_449968, EPI_ISL_449969, EPI_ISL_449970, EPI_ISL_449971, EPI_ISL_449972, EPI_ISL_449973, EPI_ISL_449974, EPI_ISL_449975, EPI_ISL_449976, EPI_ISL_449977, EPI_ISL_449978, EPI_ISL_449979, EPI_ISL_449980, EPI_ISL_449981, EPI_ISL_449982, EPI_ISL_449983, EPI_ISL_449984, EPI_ISL_449985, EPI_ISL_449986, EPI_ISL_449987, EPI_ISL_449988, EPI_ISL_449989, EPI_ISL_449990, EPI_ISL_449991, EPI_ISL_449992, EPI_ISL_449994, EPI_ISL_449995, EPI_ISL_449996, EPI_ISL_449997, EPI_ISL_449998, EPI_ISL_449999, EPI_ISL_450000, EPI_ISL_450001, EPI_ISL_450002, EPI_ISL_450003 | | | |
| see above | Washington State Department of Health | Seattle Flu Study | Chu et al |
| EPI_ISL_450337 | Hospital Universitari Vall d'Hebron - Vall d'Hebron Institut de Recerca | Hospital Universitari Vall d'Hebron | Cristina Andrés, Maria Piñana, Damir Garcia-Cehic, Mercedes Guerrero-Murillo, Ariadna Rando, Juliana Esperalba, Maria Gema Codina, Maria Carmen Martín, Tomàs Pumarola, Josep Quer, Andrés Antón |
| EPI_ISL_451183, EPI_ISL_451184, EPI_ISL_451185, EPI_ISL_451186, EPI_ISL_451187, EPI_ISL_451188, EPI_ISL_451189, EPI_ISL_451190, EPI_ISL_451191, EPI_ISL_451192, EPI_ISL_451193, EPI_ISL_451194, EPI_ISL_451195, EPI_ISL_451196, EPI_ISL_451197, EPI_ISL_451198, EPI_ISL_451199, EPI_ISL_451200, EPI_ISL_451201, EPI_ISL_451202 | | | |
| see above | Uganda Virus Research Institute | MRC/UVRI & LSHTM Uganda Research Unit | Dan Lule Bugembe, John Kayiwa, My V.T Phan, Phionah Tushabe, Stephen Balinandi, Beatrice Dhaala, Deogratius Ssemwanga, Jonas Lexow, Henry Mwebesa, Jane Aceng, Henry Kyobe, Julius Lutwama, Pontiano Kaleebu, Matthew Cotten |
| EPI_ISL_451668, EPI_ISL_451669, EPI_ISL_451670, EPI_ISL_451671, EPI_ISL_451672, EPI_ISL_451673, EPI_ISL_451674, EPI_ISL_451675, EPI_ISL_451676, EPI_ISL_451677, EPI_ISL_451678, EPI_ISL_451679, EPI_ISL_451680, EPI_ISL_451681, EPI_ISL_451682, EPI_ISL_451683, EPI_ISL_451684, EPI_ISL_451685, EPI_ISL_451686, EPI_ISL_451687, EPI_ISL_451688, EPI_ISL_451689, EPI_ISL_451690, EPI_ISL_451691, EPI_ISL_451692, EPI_ISL_451693, EPI_ISL_451695, EPI_ISL_451696, EPI_ISL_451697, EPI_ISL_451698, EPI_ISL_451699, EPI_ISL_451700, EPI_ISL_451701, EPI_ISL_451702, EPI_ISL_451703, EPI_ISL_451704, EPI_ISL_451705, EPI_ISL_451706, EPI_ISL_451707, EPI_ISL_451708, EPI_ISL_451709, EPI_ISL_451710, EPI_ISL_451711, EPI_ISL_451712, EPI_ISL_451713, EPI_ISL_451714, EPI_ISL_451715, EPI_ISL_451716, EPI_ISL_451718, EPI_ISL_451719, EPI_ISL_451720, EPI_ISL_451721, EPI_ISL_451722, EPI_ISL_451723, EPI_ISL_451724, EPI_ISL_451725, EPI_ISL_451726, EPI_ISL_451727, EPI_ISL_451728, EPI_ISL_451729, EPI_ISL_451730, EPI_ISL_451731, EPI_ISL_451732, EPI_ISL_451734, EPI_ISL_451735, EPI_ISL_451736, EPI_ISL_451737, EPI_ISL_451738, EPI_ISL_451739, EPI_ISL_451740, EPI_ISL_451741, EPI_ISL_451742, EPI_ISL_451743, EPI_ISL_451744, EPI_ISL_451745, EPI_ISL_451746, EPI_ISL_451747, EPI_ISL_451748, EPI_ISL_451749, EPI_ISL_451750, EPI_ISL_451751, EPI_ISL_451752, EPI_ISL_451753, EPI_ISL_451754, EPI_ISL_451755, EPI_ISL_451756, EPI_ISL_451757, EPI_ISL_451758, EPI_ISL_451760, EPI_ISL_451761, EPI_ISL_451762, EPI_ISL_451764, EPI_ISL_451765, EPI_ISL_451766, EPI_ISL_451767, EPI_ISL_451769, EPI_ISL_451770, EPI_ISL_451771, EPI_ISL_451772, EPI_ISL_451773, EPI_ISL_451774, EPI_ISL_451775, EPI_ISL_451776, EPI_ISL_451777, EPI_ISL_451778, EPI_ISL_451779, EPI_ISL_451780, EPI_ISL_451781, EPI_ISL_451782, EPI_ISL_451783, EPI_ISL_451784, EPI_ISL_451785, EPI_ISL_451786, EPI_ISL_451787, EPI_ISL_451789, EPI_ISL_451790, EPI_ISL_451792, EPI_ISL_451793, EPI_ISL_451794, EPI_ISL_451795, EPI_ISL_451796, EPI_ISL_451797, EPI_ISL_451798, EPI_ISL_451799, EPI_ISL_451800, EPI_ISL_451801, EPI_ISL_451802, EPI_ISL_451803, EPI_ISL_451804, EPI_ISL_451805, EPI_ISL_451806, EPI_ISL_451807, EPI_ISL_451808, EPI_ISL_451809, EPI_ISL_451811, EPI_ISL_451812, EPI_ISL_451813, EPI_ISL_451814, EPI_ISL_451815, EPI_ISL_451816, EPI_ISL_451817, EPI_ISL_451818, EPI_ISL_451821, EPI_ISL_451822, EPI_ISL_451823, EPI_ISL_451824, EPI_ISL_451825, EPI_ISL_451826, EPI_ISL_451827, EPI_ISL_451828, EPI_ISL_451829, EPI_ISL_451831, EPI_ISL_451833, EPI_ISL_451834, EPI_ISL_451835, EPI_ISL_451837, EPI_ISL_451838, EPI_ISL_451841, EPI_ISL_451842, EPI_ISL_451843, EPI_ISL_451844, EPI_ISL_451845, EPI_ISL_451846, EPI_ISL_451847, EPI_ISL_451848, EPI_ISL_451849, EPI_ISL_451850, EPI_ISL_451851, EPI_ISL_451852, EPI_ISL_451853, EPI_ISL_451854, EPI_ISL_451855, EPI_ISL_451856, EPI_ISL_451857, EPI_ISL_451858, EPI_ISL_451859, EPI_ISL_451860, EPI_ISL_451861, EPI_ISL_451862, EPI_ISL_451863, EPI_ISL_451864, EPI_ISL_451865, EPI_ISL_451866, EPI_ISL_451867, EPI_ISL_451868, EPI_ISL_451869, EPI_ISL_451870, EPI_ISL_451871, EPI_ISL_451872, EPI_ISL_451873, EPI_ISL_451874, EPI_ISL_451875, EPI_ISL_451876, EPI_ISL_451877, EPI_ISL_451878, EPI_ISL_451879, EPI_ISL_451880, EPI_ISL_451881, EPI_ISL_451882, EPI_ISL_451883, EPI_ISL_451884, EPI_ISL_451885, EPI_ISL_451886, EPI_ISL_451887, EPI_ISL_451888, EPI_ISL_451889, EPI_ISL_451890, EPI_ISL_451891, EPI_ISL_451892, EPI_ISL_451893, EPI_ISL_451894, EPI_ISL_451895, EPI_ISL_451896, EPI_ISL_451898, EPI_ISL_451899, EPI_ISL_451900, EPI_ISL_451902, EPI_ISL_451903, EPI_ISL_451904, EPI_ISL_451905, EPI_ISL_451906, EPI_ISL_451907, EPI_ISL_451908, EPI_ISL_451909, EPI_ISL_451910, EPI_ISL_451911, EPI_ISL_451912, EPI_ISL_451913, EPI_ISL_451914, EPI_ISL_451915, EPI_ISL_451916 | | | |
| see above | Viollier AG | Department of Biosystems Science and Engineering, ETH Zürich | Christian Beisel, Sarah Nadeau, Ivan Topolsky, Pedro Ferreira, Philipp Jablonski, Susana Posada-Céspedes, Tobias Schär, Ina Nissen, Natascha Santacroce, Elodie Burcklen, Christiane Beckmann, Maurice Redondo, Olivier Kobel, Christoph Noppen, Sophie Seidel, Noemie Santamaria de Souza, Niko Beerenwinkel, Tanja Stadler |
| EPI_ISL_451957 | Molecular Pathology Division, Department of Pathology, Hong Kong Sanatorium & Hospital | Molecular Pathology Division, Department of Pathology, Hong Kong Sanatorium & Hospital | Chun Hang AU, Wai Sing CHAN, Ho Yin LAM, Dona N. HO, Simon Y.M. LAM, Jonpaul S.T. ZEE, Tsun Leung CHAN, Edmond S.K. MA |
| EPI_ISL_453465, EPI_ISL_453466, EPI_ISL_453467, EPI_ISL_453468, EPI_ISL_453469, EPI_ISL_453470, EPI_ISL_453471, EPI_ISL_453472, EPI_ISL_453473, EPI_ISL_453474, EPI_ISL_453475, EPI_ISL_453476, EPI_ISL_453477, EPI_ISL_453478, EPI_ISL_453479, EPI_ISL_453480, EPI_ISL_453481, EPI_ISL_453482, EPI_ISL_453483, EPI_ISL_453484, EPI_ISL_453485 | | | |
| see above | Regional Virus Laboratory, Belfast Health and Social Care Trust | COVID-19 Genomics UK (COG-UK) Consortium | Conall McCaughey, James McKenna, Tanya Curran, Susan Feeney, Alison Watt, Ciara Cox, Mairead Connor, Zoltan Molnar, David Simpson, Derek Fairley |
| EPI_ISL_454733 | Department of Medical, Biotechnologies University of Siena | Department of Medical, Biotechnologies University of Siena | Cusi,M.G., Pinzauti,D., Gandolfo,C., Anichini,G., Pozzi,G. and Santoro,F. |
| EPI_ISL_455110 | Scania Halsocenter, B288 | The Public Health Agency of Sweden | Christina Lergin, Anna-Malin Linde, Maria Lind Karlberg, Oskar Karlsson Lindsjo, Olov Svartstrom, Anna Risberg, Theresa Enkirch, Mia Brytting, Karin Tegmark-Wisell |
| EPI_ISL_455568, EPI_ISL_455569, EPI_ISL_455570, EPI_ISL_455571, EPI_ISL_455572 | Gundersen Molecular Diagnostics Laboratory | Kabara Cancer Research Institute | Craig S. Richmond, Paraic A. Kenny |
| EPI_ISL_455573, EPI_ISL_455574 | Gundersen Clinical Microbiology Laboratory | Kabara Cancer Research Institute | Craig S. Richmond, Paraic A. Kenny |
| EPI_ISL_455575, EPI_ISL_455576, EPI_ISL_455577, EPI_ISL_455578, EPI_ISL_455579 | Gundersen Molecular Diagnostics Laboratory | Kabara Cancer Research Institute | Craig S. Richmond, Paraic A. Kenny |
| EPI_ISL_455580, EPI_ISL_455581, EPI_ISL_455582 | Gundersen Clinical Microbiology Laboratory | Kabara Cancer Research Institute | Craig S. Richmond, Paraic A. Kenny |
| EPI_ISL_455680 | Institute of pathogenic microbiology, Jiangsu Provincial Center for Disease Control and Prevention | Institute of pathogenic microbiology, Jiangsu Provincial Center for Disease Control and Prevention | Cui,L. |
| EPI_ISL_458130, EPI_ISL_458131 | Hospital Universitari Vall d'Hebron - Vall d'Hebron Institut de Recerca | Hospital Universitari Vall d'Hebron | Cristina Andrés, Maria Piñana, Damir Garcia-Cehic, Mercedes Guerrero-Murillo, Ariadna Rando, Josep Gregori, Juliana Esperalba, Maria Gema Codina, Maria Carmen Martin, Tomàs Pumarola, Josep Quer, Andrés Antón |
| EPI_ISL_458132 | Hospital Universitari Vall d'Hebron - Vall d'Hebron Institut de Rercerca | Hospital Universitari Vall d'Hebron | Cristina Andrés, Maria Piñana, Damir Garcia-Cehic, Mercedes Guerrero-Murillo, Ariadna Rando, Josep Gregori, Juliana Esperalba, Maria Gema Codina, Maria Carmen Martin, Tomàs Pumarola, Josep Quer, Andrés Antón |
| EPI_ISL_459324, EPI_ISL_459325, EPI_ISL_459326, EPI_ISL_459327, EPI_ISL_459328, EPI_ISL_459329, EPI_ISL_459330, EPI_ISL_459331, EPI_ISL_459332, EPI_ISL_459333, EPI_ISL_459334, EPI_ISL_459335, EPI_ISL_459336, EPI_ISL_459337, EPI_ISL_459338, EPI_ISL_459339, EPI_ISL_459340, EPI_ISL_459341, EPI_ISL_459342, EPI_ISL_459343, EPI_ISL_459344, EPI_ISL_459345, EPI_ISL_459346, EPI_ISL_459347, EPI_ISL_459348, EPI_ISL_459349, EPI_ISL_459350, EPI_ISL_459351, EPI_ISL_459352, EPI_ISL_459353, EPI_ISL_459354, EPI_ISL_459355, EPI_ISL_459356, EPI_ISL_459357 | | | |
| see above | Regional Virus Laboratory, Belfast Health and Social Care Trust | Wellcome Sanger Institute for the COVID-19 Genomics UK (COG-UK) consortium | Conall McCaughey, James McKenna, Tanya Curran, Susan Feeney, Alison Watt, Ciara Cox, Mairead Connor, Zoltan Molnar, David Simpson, Derek Fairley; and Alex Alderton, Roberto Amato, Sonia Goncalves, Ewan Harrison, David K. Jackson, Ian Johnston, Dominic Kwiatkowski, Cordelia Langford, John Sillitoe on behalf of the Wellcome Sanger Institute COVID-19 Surveillance Team (http://www.sanger.ac.uk/covid-team) |
| EPI_ISL_459358 | Regional Virus Laboratory, Belfast Health and Social Care Trust | Wellcome Sanger Institute for the COVID-19 Genomics UK (COG-UK) Consortium | Conall McCaughey, James McKenna, Tanya Curran, Susan Feeney, Alison Watt, Ciara Cox, Mairead Connor, Zoltan Molnar, David Simpson, Derek Fairley; and Alex Alderton, Roberto Amato, Sonia Goncalves, Ewan Harrison, David K. Jackson, Ian Johnston, Dominic Kwiatkowski, Cordelia Langford, John Sillitoe on behalf of the Wellcome Sanger Institute COVID-19 Surveillance Team |
| EPI_ISL_459359, EPI_ISL_459360, EPI_ISL_459361, EPI_ISL_459362, EPI_ISL_459363, EPI_ISL_459364, EPI_ISL_459365, EPI_ISL_459366, EPI_ISL_459367, EPI_ISL_459368, EPI_ISL_459369, EPI_ISL_459370, EPI_ISL_459371 | | | |
| see above | Regional Virus Laboratory, Belfast Health and Social Care Trust | Wellcome Sanger Institute for the COVID-19 Genomics UK (COG-UK) consortium | Conall McCaughey, James McKenna, Tanya Curran, Susan Feeney, Alison Watt, Ciara Cox, Mairead Connor, Zoltan Molnar, David Simpson, Derek Fairley; and Alex Alderton, Roberto Amato, Sonia Goncalves, Ewan Harrison, David K. Jackson, Ian Johnston, Dominic Kwiatkowski, Cordelia Langford, John Sillitoe on behalf of the Wellcome Sanger Institute COVID-19 Surveillance Team (http://www.sanger.ac.uk/covid-team) |
| EPI_ISL_459372 | Regional Virus Laboratory, Belfast Health and Social Care Trust | Wellcome Sanger Institute for the COVID-19 Genomics UK (COG-UK) Consortium | Conall McCaughey, James McKenna, Tanya Curran, Susan Feeney, Alison Watt, Ciara Cox, Mairead Connor, Zoltan Molnar, David Simpson, Derek Fairley; and Alex Alderton, Roberto Amato, Sonia Goncalves, Ewan Harrison, David K. Jackson, Ian Johnston, Dominic Kwiatkowski, Cordelia Langford, John Sillitoe on behalf of the Wellcome Sanger Institute COVID-19 Surveillance Team |
| EPI_ISL_459373, EPI_ISL_459374, EPI_ISL_459375, EPI_ISL_459376, EPI_ISL_459377, EPI_ISL_459378, EPI_ISL_459379, EPI_ISL_459380, EPI_ISL_459381, EPI_ISL_459382, EPI_ISL_459383, EPI_ISL_459384, EPI_ISL_459385, EPI_ISL_459386, EPI_ISL_459387, EPI_ISL_459388, EPI_ISL_459389, EPI_ISL_459390, EPI_ISL_459391, EPI_ISL_459392, EPI_ISL_459393, EPI_ISL_459394, EPI_ISL_459395, EPI_ISL_459396, EPI_ISL_459397, EPI_ISL_459398, EPI_ISL_459399, EPI_ISL_459400, EPI_ISL_459401, EPI_ISL_459402, EPI_ISL_459403, EPI_ISL_459404, EPI_ISL_459405, EPI_ISL_459406, EPI_ISL_459407, EPI_ISL_459408, EPI_ISL_459409 | | | |
| see above | Regional Virus Laboratory, Belfast Health and Social Care Trust | Wellcome Sanger Institute for the COVID-19 Genomics UK (COG-UK) consortium | Conall McCaughey, James McKenna, Tanya Curran, Susan Feeney, Alison Watt, Ciara Cox, Mairead Connor, Zoltan Molnar, David Simpson, Derek Fairley; and Alex Alderton, Roberto Amato, Sonia Goncalves, Ewan Harrison, David K. Jackson, Ian Johnston, Dominic Kwiatkowski, Cordelia Langford, |

| | | | |
|---|---|---|---|
| | | | John Sillitoe on behalf of the Wellcome Sanger Institute COVID-19 Surveillance Team (http://www.sanger.ac.uk/covid-team) |
| EPI_ISL_461773, EPI_ISL_461774, EPI_ISL_461775, EPI_ISL_461776, EPI_ISL_461777, EPI_ISL_461778, EPI_ISL_461779, EPI_ISL_461780, EPI_ISL_461781, EPI_ISL_461782, EPI_ISL_461783, EPI_ISL_461784, EPI_ISL_461785, EPI_ISL_461786, EPI_ISL_461787, EPI_ISL_461788, EPI_ISL_461789, EPI_ISL_461790 | | | |
| see above | Regional Virus Laboratory, Belfast Health and Social Care Trust | COVID-19 Genomics UK (COG-UK) Consortium | Conall McCaughey, James McKenna, Tanya Curran, Susan Feeney, Alison Watt, Ciara Cox, Mairead Connor, Zoltan Molnar, David Simpson, Derek Fairley |
| EPI_ISL_463324, EPI_ISL_463325, EPI_ISL_463326, EPI_ISL_463327, EPI_ISL_463328, EPI_ISL_463329, EPI_ISL_463330, EPI_ISL_463331, EPI_ISL_463332, EPI_ISL_463333, EPI_ISL_463334, EPI_ISL_463335, EPI_ISL_463336, EPI_ISL_463337, EPI_ISL_463338, EPI_ISL_463339, EPI_ISL_463340, EPI_ISL_463341, EPI_ISL_463342, EPI_ISL_463343, EPI_ISL_463344, EPI_ISL_463345, EPI_ISL_463346, EPI_ISL_463347, EPI_ISL_463348, EPI_ISL_463349, EPI_ISL_463350, EPI_ISL_463351, EPI_ISL_463352, EPI_ISL_463353, EPI_ISL_463354, EPI_ISL_463355, EPI_ISL_463356, EPI_ISL_463357, EPI_ISL_463358, EPI_ISL_463359, EPI_ISL_463360, EPI_ISL_463361, EPI_ISL_463362, EPI_ISL_463363, EPI_ISL_463364, EPI_ISL_463365, EPI_ISL_463366, EPI_ISL_463367, EPI_ISL_463368, EPI_ISL_463369, EPI_ISL_463370, EPI_ISL_463371, EPI_ISL_463375, EPI_ISL_463378, EPI_ISL_463379, EPI_ISL_463380, EPI_ISL_463381, EPI_ISL_463382, EPI_ISL_463383, EPI_ISL_463384, EPI_ISL_463385, EPI_ISL_463386, EPI_ISL_463387, EPI_ISL_463388, EPI_ISL_463389, EPI_ISL_463390, EPI_ISL_463391, EPI_ISL_463392, EPI_ISL_463393, EPI_ISL_463394, EPI_ISL_463395, EPI_ISL_463396, EPI_ISL_463397, EPI_ISL_463398, EPI_ISL_463399, EPI_ISL_463400, EPI_ISL_463401, EPI_ISL_463402, EPI_ISL_463403, EPI_ISL_463404, EPI_ISL_463405, EPI_ISL_463406, EPI_ISL_463407, EPI_ISL_463408, EPI_ISL_463409, EPI_ISL_463410, EPI_ISL_463411, EPI_ISL_463412, EPI_ISL_463413, EPI_ISL_463414, EPI_ISL_463415, EPI_ISL_463416, EPI_ISL_463417, EPI_ISL_463418, EPI_ISL_463419, EPI_ISL_463420, EPI_ISL_463421, EPI_ISL_463422, EPI_ISL_463423, EPI_ISL_463424, EPI_ISL_463425, EPI_ISL_463426, EPI_ISL_463427, EPI_ISL_463428, EPI_ISL_463429, EPI_ISL_463430, EPI_ISL_463431, EPI_ISL_463432, EPI_ISL_463433, EPI_ISL_463434, EPI_ISL_463435, EPI_ISL_463436, EPI_ISL_463437, EPI_ISL_463438, EPI_ISL_463439, EPI_ISL_463440, EPI_ISL_463441, EPI_ISL_463442, EPI_ISL_463443, EPI_ISL_463444, EPI_ISL_463445, EPI_ISL_463446, EPI_ISL_463449, EPI_ISL_463450, EPI_ISL_463451, EPI_ISL_463452, EPI_ISL_463453, EPI_ISL_463454, EPI_ISL_463455, EPI_ISL_463456, EPI_ISL_463457, EPI_ISL_463458, EPI_ISL_463459, EPI_ISL_463460, EPI_ISL_463461, EPI_ISL_463462, EPI_ISL_463463, EPI_ISL_463464, EPI_ISL_463465, EPI_ISL_463466, EPI_ISL_463467, EPI_ISL_463472, EPI_ISL_463478, EPI_ISL_463479, EPI_ISL_463480, EPI_ISL_463481, EPI_ISL_463482, EPI_ISL_463483, EPI_ISL_463484, EPI_ISL_463485, EPI_ISL_463486, EPI_ISL_463487, EPI_ISL_463488, EPI_ISL_463489, EPI_ISL_463490, EPI_ISL_463491, EPI_ISL_463492, EPI_ISL_463493, EPI_ISL_463494, EPI_ISL_463495, EPI_ISL_463496, EPI_ISL_463497, EPI_ISL_463498, EPI_ISL_463499, EPI_ISL_463500, EPI_ISL_463501, EPI_ISL_463502, EPI_ISL_463503, EPI_ISL_463504, EPI_ISL_463505, EPI_ISL_463506, EPI_ISL_463507, EPI_ISL_463508, EPI_ISL_463509, EPI_ISL_463510, EPI_ISL_463511, EPI_ISL_463512, EPI_ISL_463513, EPI_ISL_463514, EPI_ISL_463515, EPI_ISL_463516, EPI_ISL_463517, EPI_ISL_463518, EPI_ISL_463519, EPI_ISL_463520, EPI_ISL_463521, EPI_ISL_463522, EPI_ISL_463523, EPI_ISL_463524, EPI_ISL_463525, EPI_ISL_463526, EPI_ISL_463527, EPI_ISL_463528, EPI_ISL_463529, EPI_ISL_463530, EPI_ISL_463531, EPI_ISL_463532, EPI_ISL_463533, EPI_ISL_463534, EPI_ISL_463535, EPI_ISL_463536, EPI_ISL_463537, EPI_ISL_463538, EPI_ISL_463539, EPI_ISL_463540, EPI_ISL_463541, EPI_ISL_463542, EPI_ISL_463543, EPI_ISL_463544, EPI_ISL_463545, EPI_ISL_463546, EPI_ISL_463547, EPI_ISL_463548, EPI_ISL_463549, EPI_ISL_463550, EPI_ISL_463551, EPI_ISL_463552, EPI_ISL_463553, EPI_ISL_463554, EPI_ISL_463555, EPI_ISL_463556, EPI_ISL_463557, EPI_ISL_463558, EPI_ISL_463559, EPI_ISL_463560, EPI_ISL_463561, EPI_ISL_463562, EPI_ISL_463563, EPI_ISL_463564, EPI_ISL_463565, EPI_ISL_463566, EPI_ISL_463567, EPI_ISL_463568, EPI_ISL_463569, EPI_ISL_463570, EPI_ISL_463571, EPI_ISL_463572, EPI_ISL_463573, EPI_ISL_463574, EPI_ISL_463575, EPI_ISL_463576, EPI_ISL_463577, EPI_ISL_463578, EPI_ISL_463579, EPI_ISL_463580, EPI_ISL_463581, EPI_ISL_463582, EPI_ISL_463583, EPI_ISL_463584, EPI_ISL_463585, EPI_ISL_463586, EPI_ISL_463587, EPI_ISL_463588, EPI_ISL_463589, EPI_ISL_463590, EPI_ISL_463591, EPI_ISL_463592, EPI_ISL_463593, EPI_ISL_463594, EPI_ISL_463595, EPI_ISL_463596, EPI_ISL_463597, EPI_ISL_463598, EPI_ISL_463599, EPI_ISL_463600, EPI_ISL_463601, EPI_ISL_463602, EPI_ISL_463603, EPI_ISL_463604, EPI_ISL_463605, EPI_ISL_463607, EPI_ISL_463608, EPI_ISL_463609, EPI_ISL_463610, EPI_ISL_463611, EPI_ISL_463612, EPI_ISL_463613, EPI_ISL_463614, EPI_ISL_463615, EPI_ISL_463616, EPI_ISL_463617, EPI_ISL_463618, EPI_ISL_463619, EPI_ISL_463620, EPI_ISL_463621, EPI_ISL_463622, EPI_ISL_463623, EPI_ISL_463624, EPI_ISL_463625, EPI_ISL_463626, EPI_ISL_463627, EPI_ISL_463628, EPI_ISL_463629, EPI_ISL_463630, EPI_ISL_463631, EPI_ISL_463632, EPI_ISL_463633, EPI_ISL_463634, EPI_ISL_463635, EPI_ISL_463636, EPI_ISL_463637, EPI_ISL_463638, EPI_ISL_463639, EPI_ISL_463640, EPI_ISL_463641, EPI_ISL_463642, EPI_ISL_463643, EPI_ISL_463644, EPI_ISL_463645, EPI_ISL_463646, EPI_ISL_463647, EPI_ISL_463648, EPI_ISL_463649, EPI_ISL_463650, EPI_ISL_463651, EPI_ISL_463652, EPI_ISL_463653, EPI_ISL_463654, EPI_ISL_463655, EPI_ISL_463656, EPI_ISL_463657, EPI_ISL_463658, EPI_ISL_463659, EPI_ISL_463660, EPI_ISL_463661, EPI_ISL_463662, EPI_ISL_463663, EPI_ISL_463664, EPI_ISL_463665, EPI_ISL_463666, EPI_ISL_463667, EPI_ISL_463668, EPI_ISL_463669, EPI_ISL_463670, EPI_ISL_463671, EPI_ISL_463672, EPI_ISL_463673, EPI_ISL_463674, EPI_ISL_463675, EPI_ISL_463676, EPI_ISL_463677, EPI_ISL_463678, EPI_ISL_463679, EPI_ISL_463680, EPI_ISL_463681, EPI_ISL_463682, EPI_ISL_463683, EPI_ISL_463684, EPI_ISL_463685, EPI_ISL_463686, EPI_ISL_463687, EPI_ISL_463688, EPI_ISL_463689, EPI_ISL_463690, EPI_ISL_463691, EPI_ISL_463692, EPI_ISL_463693, EPI_ISL_463694, EPI_ISL_463695, EPI_ISL_463696, EPI_ISL_463697, EPI_ISL_463698, EPI_ISL_463699, EPI_ISL_463700, EPI_ISL_463701, EPI_ISL_463702, EPI_ISL_463703, EPI_ISL_463704, EPI_ISL_463705 | | | |
| see above | Washington State Department of Health | Seattle Flu Study | Chu et al |
| EPI_ISL_466927, EPI_ISL_466928, EPI_ISL_466929, EPI_ISL_466930, EPI_ISL_466931, EPI_ISL_466932, EPI_ISL_466933, EPI_ISL_466934, EPI_ISL_466935, EPI_ISL_466936, EPI_ISL_466937, EPI_ISL_466938, EPI_ISL_466940, EPI_ISL_466941, EPI_ISL_466942, EPI_ISL_466943, EPI_ISL_466944, EPI_ISL_466945, EPI_ISL_466946, EPI_ISL_466947, EPI_ISL_466948, EPI_ISL_466949, EPI_ISL_466950, EPI_ISL_466951, EPI_ISL_466952, EPI_ISL_466953, EPI_ISL_466954, EPI_ISL_466955, EPI_ISL_466956, EPI_ISL_466957, EPI_ISL_466958, EPI_ISL_466959, EPI_ISL_466960, EPI_ISL_466961, EPI_ISL_466962, EPI_ISL_466963, EPI_ISL_466964, EPI_ISL_466965, EPI_ISL_466966, EPI_ISL_466967, EPI_ISL_466968, EPI_ISL_466969, EPI_ISL_466970, EPI_ISL_466971, EPI_ISL_466972, EPI_ISL_466973, EPI_ISL_466974, EPI_ISL_466975, EPI_ISL_466976, EPI_ISL_466977, EPI_ISL_466978, EPI_ISL_466979, EPI_ISL_466980, EPI_ISL_466981, EPI_ISL_466982, EPI_ISL_466983, EPI_ISL_466984, EPI_ISL_466985, EPI_ISL_466986, EPI_ISL_466987, EPI_ISL_466988, EPI_ISL_466989, EPI_ISL_466990, EPI_ISL_466991, EPI_ISL_466992, EPI_ISL_466993, EPI_ISL_466994, EPI_ISL_466995, EPI_ISL_466996, EPI_ISL_466997, EPI_ISL_466998, EPI_ISL_466999, EPI_ISL_467000, EPI_ISL_467001, EPI_ISL_467002, EPI_ISL_467003, EPI_ISL_467004, EPI_ISL_467005, EPI_ISL_467006, EPI_ISL_467007, EPI_ISL_467009, EPI_ISL_467010, EPI_ISL_467012, EPI_ISL_467013, EPI_ISL_467014, EPI_ISL_467015, EPI_ISL_467016, EPI_ISL_468202, EPI_ISL_468203, EPI_ISL_468204, EPI_ISL_468205, EPI_ISL_468206, EPI_ISL_468207, EPI_ISL_468209, EPI_ISL_468210, EPI_ISL_468211, EPI_ISL_468212, EPI_ISL_468213, EPI_ISL_468215, EPI_ISL_468216, EPI_ISL_468217, EPI_ISL_468218, EPI_ISL_468219, EPI_ISL_468220, EPI_ISL_468221, EPI_ISL_468222, EPI_ISL_468224, EPI_ISL_468225, EPI_ISL_468226, EPI_ISL_468227, EPI_ISL_468228, EPI_ISL_468229, EPI_ISL_468230, EPI_ISL_468231, EPI_ISL_468232, EPI_ISL_468233, EPI_ISL_468234, EPI_ISL_468235, EPI_ISL_468236, EPI_ISL_468237, EPI_ISL_468238, EPI_ISL_468239, EPI_ISL_468240, EPI_ISL_468241, EPI_ISL_468242, EPI_ISL_468243, EPI_ISL_468244, EPI_ISL_468245, EPI_ISL_468246, EPI_ISL_468248, EPI_ISL_468249, EPI_ISL_468250, EPI_ISL_468251, EPI_ISL_468252, EPI_ISL_468253, EPI_ISL_468254, EPI_ISL_468255, EPI_ISL_468256, EPI_ISL_468258, EPI_ISL_468259, EPI_ISL_468260, EPI_ISL_468261, EPI_ISL_468263, EPI_ISL_468264, EPI_ISL_468265, EPI_ISL_468266, EPI_ISL_468267, EPI_ISL_468268, EPI_ISL_468270, EPI_ISL_468272, EPI_ISL_468273, EPI_ISL_468274, EPI_ISL_468275, EPI_ISL_468276, EPI_ISL_468277, EPI_ISL_468278, EPI_ISL_468279, EPI_ISL_468280, EPI_ISL_468281, EPI_ISL_468282, EPI_ISL_468284, EPI_ISL_468285, EPI_ISL_468287, EPI_ISL_468288, EPI_ISL_468289, EPI_ISL_468290, EPI_ISL_468291, EPI_ISL_468293, EPI_ISL_468294, EPI_ISL_468295, EPI_ISL_468296, EPI_ISL_468298, EPI_ISL_468299, EPI_ISL_468300, EPI_ISL_468301, EPI_ISL_468303, EPI_ISL_468304 | | | |
| see above | Viollier AG | Department of Biosystems Science and Engineering, ETH Zürich | Christian Beisel, Sarah Nadeau, Ivan Topolsky, Pedro Ferreira, Philipp Jablonski, Susana Posada-Céspedes, Tobias Schär, Ina Nissen, Natascha Santacroce, Elodie Burcklen, Christiane Beckmann, Maurice Redondo, Olivier Kobel, Christoph Noppen, Sophie Seidel, Noemie Santamaria de Souza, Niko Beerenwinkel, Tanja Stadler |
| EPI_ISL_468305, EPI_ISL_468307 | Centro de Vigilancia a Saude de Diadema | Instituto Adolfo Lutz, Interdiciplinary Procedures Center, Strategic Laboratory | Claudio Tavares Sacchi, Claudia Regina Gonçalves, Erica Valessa Ramos Gomes |
| EPI_ISL_468308 | Hospital Municipal do Tatuape Carmino Caricchio | Instituto Adolfo Lutz, Interdiciplinary Procedures Center, Strategic Laboratory | Claudio Tavares Sacchi, Claudia Regina Gonçalves, Erica Valessa Ramos Gomes |
| EPI_ISL_468310 | Hospital Sao Paulo de Ensino da UNIFESP | Instituto Adolfo Lutz, Interdiciplinary Procedures Center, Strategic Laboratory | Claudio Tavares Sacchi, Claudia Regina Gonçalves, Erica Valessa Ramos Gomes |
| EPI_ISL_468311, EPI_ISL_468312 | Hospital Municipal Dr Ignacio Proenca de Gouvea | Instituto Adolfo Lutz, Interdiciplinary Procedures Center, Strategic Laboratory | Claudio Tavares Sacchi, Claudia Regina Gonçalves, Erica Valessa Ramos Gomes |
| EPI_ISL_468313 | Vigilancia Epidemiologica de São Bernardo do Campo | Instituto Adolfo Lutz, Interdiciplinary Procedures Center, Strategic Laboratory | Claudio Tavares Sacchi, Claudia Regina Gonçalves, Erica Valessa Ramos Gomes |
| EPI_ISL_468314 | CTA Centro de Testagem e Aconselhamento | Instituto Adolfo Lutz, Interdiciplinary Procedures Center, Strategic Laboratory | Claudio Tavares Sacchi, Claudia Regina Gonçalves, Erica Valessa Ramos Gomes |
| EPI_ISL_468315 | Hospital Municipal do Tatuape Carmino Caricchio | Instituto Adolfo Lutz, Interdiciplinary Procedures Center, Strategic Laboratory | Claudio Tavares Sacchi, Claudia Regina Gonçalves, Erica Valessa Ramos Gomes |
| EPI_ISL_468316 | UPA Vila Assis | Instituto Adolfo Lutz, Interdiciplinary Procedures Center, Strategic Laboratory | Claudio Tavares Sacchi, Claudia Regina Gonçalves, Erica Valessa Ramos Gomes |
| EPI_ISL_468318 | Hospital Universitario da USP | Instituto Adolfo Lutz, Interdiciplinary Procedures Center, Strategic Laboratory | Claudio Tavares Sacchi, Claudia Regina Gonçalves, Erica Valessa Ramos Gomes |
| EPI_ISL_468319 | Vigilancia Epidemiologica de São Bernardo do Campo | Instituto Adolfo Lutz, Interdiciplinary Procedures Center, Strategic Laboratory | Claudio Tavares Sacchi, Claudia Regina Gonçalves, Erica Valessa Ramos Gomes |
| EPI_ISL_468320 | Secretaria Municipal de Saude de Hortolandia | Instituto Adolfo Lutz, Interdiciplinary Procedures Center, Strategic Laboratory | Claudio Tavares Sacchi, Claudia Regina Gonçalves, Erica Valessa Ramos Gomes |
| EPI_ISL_468321 | Hospital Universitario da USP | Instituto Adolfo Lutz, Interdiciplinary Procedures Center, Strategic Laboratory | Claudio Tavares Sacchi, Claudia Regina Gonçalves, Erica Valessa Ramos Gomes |
| EPI_ISL_469844, EPI_ISL_469845, EPI_ISL_469846, EPI_ISL_469847, EPI_ISL_469848, EPI_ISL_469849, EPI_ISL_469850, EPI_ISL_469851, EPI_ISL_469852, EPI_ISL_469853, EPI_ISL_469854, EPI_ISL_469855, EPI_ISL_469856, EPI_ISL_469857, EPI_ISL_469858, EPI_ISL_469859, EPI_ISL_469860, EPI_ISL_469861, EPI_ISL_469862, EPI_ISL_469863, EPI_ISL_469864, EPI_ISL_469865, EPI_ISL_469866, EPI_ISL_469867, EPI_ISL_469868, EPI_ISL_469869, EPI_ISL_469870, EPI_ISL_469871, EPI_ISL_469872, EPI_ISL_469873, EPI_ISL_469874, EPI_ISL_469875, EPI_ISL_469876, EPI_ISL_469877, EPI_ISL_469878, EPI_ISL_469879, EPI_ISL_469880, EPI_ISL_469881, EPI_ISL_469882, EPI_ISL_469883, EPI_ISL_469884, EPI_ISL_469885, EPI_ISL_469886, EPI_ISL_469887, EPI_ISL_469888, EPI_ISL_469889, EPI_ISL_469890, EPI_ISL_470013, EPI_ISL_470014, EPI_ISL_470015, EPI_ISL_470016, EPI_ISL_470017, EPI_ISL_470018, EPI_ISL_470019, EPI_ISL_470020, EPI_ISL_470021, EPI_ISL_470022, EPI_ISL_470023, EPI_ISL_470024, EPI_ISL_470025, EPI_ISL_470026, EPI_ISL_470027, EPI_ISL_470028, EPI_ISL_470029, EPI_ISL_470030, EPI_ISL_470031, EPI_ISL_470032, EPI_ISL_470033, EPI_ISL_470034, EPI_ISL_470035, EPI_ISL_470036, EPI_ISL_470037, EPI_ISL_470038, EPI_ISL_470039, EPI_ISL_470040, EPI_ISL_470041, EPI_ISL_470042, EPI_ISL_470043, EPI_ISL_470044, EPI_ISL_470045, EPI_ISL_470046, EPI_ISL_470047, EPI_ISL_470048, EPI_ISL_470049, EPI_ISL_470050, EPI_ISL_470051, EPI_ISL_470052, EPI_ISL_470053, EPI_ISL_470054, EPI_ISL_470055, EPI_ISL_470056, EPI_ISL_470057, EPI_ISL_470058, EPI_ISL_470059, EPI_ISL_470060, EPI_ISL_470061, EPI_ISL_470062, EPI_ISL_470063, EPI_ISL_470064, EPI_ISL_470065, EPI_ISL_470066, EPI_ISL_470067, EPI_ISL_470068, EPI_ISL_470069, EPI_ISL_470070, EPI_ISL_470071, EPI_ISL_470072, EPI_ISL_470073, EPI_ISL_470074, EPI_ISL_470075, EPI_ISL_470076, EPI_ISL_470077, EPI_ISL_470078, EPI_ISL_470079, EPI_ISL_470080, EPI_ISL_470081, EPI_ISL_470082, EPI_ISL_470083, EPI_ISL_470084, EPI_ISL_470085, EPI_ISL_470086, EPI_ISL_470087, EPI_ISL_470088, EPI_ISL_470089, EPI_ISL_470534 | | | |
| see above | Regional Virus Laboratory, Belfast Health and Social Care Trust | Wellcome Sanger Institute for the COVID-19 Genomics UK (COG-UK) consortium | Conall McCaughey, James McKenna, Tanya Curran, Susan Feeney, Alison Watt, Ciara Cox, Mairead Connor, Zoltan Molnar, David Simpson, Derek Fairley; and Alex Alderton, Roberto Amato, Sonia Goncalves, Ewan Harrison, David K. Jackson, Ian Johnston, Dominic Kwiatkowski, Cordelia Langford, John Sillitoe on behalf of the Wellcome Sanger Institute COVID-19 Surveillance Team (http://www.sanger.ac.uk/covid-team) |
| EPI_ISL_471144, EPI_ISL_471145, EPI_ISL_471146, EPI_ISL_471147, EPI_ISL_471148, EPI_ISL_471149, EPI_ISL_471150, EPI_ISL_471151, EPI_ISL_471152, EPI_ISL_471153, EPI_ISL_471154, EPI_ISL_471155, EPI_ISL_471156 | | | |
| see above | Gundersen Molecular Diagnostics Laboratory | Kabara Cancer Research Institute | Craig S. Richmond, Paraic A. Kenny |
| EPI_ISL_471157 | Gundersen Clinical Microbiology Laboratory | Kabara Cancer Research Institute | Craig S. Richmond, Paraic A. Kenny |

| | | | |
|---|---|---|---|
| EPI_ISL_471539 | Hospital Universitario da USP Sao Paulo | Instituto Adolfo Lutz, Interdiciplinary Procedures Center, Strategic Laboratory | Claudio Tavares Sacchi, Claudia Regina Gonçalves, Erica Valessa Ramos Gomes |
| EPI_ISL_471541 | Hospital Geral Santa Marcelina | Instituto Adolfo Lutz, Interdiciplinary Procedures Center, Strategic Laboratory | Claudio Tavares Sacchi, Claudia Regina Gonçalves, Erica Valessa Ramos Gomes |
| EPI_ISL_471542 | Secretaria de Saude de Mogi das Cruzes | Instituto Adolfo Lutz, Interdiciplinary Procedures Center, Strategic Laboratory | Claudio Tavares Sacchi, Claudia Regina Gonçalves, Erica Valessa Ramos Gomes |
| EPI_ISL_471543 | Centro de Saude I Tacito Leite de Carvalho e Silva | Instituto Adolfo Lutz, Interdiciplinary Procedures Center, Strategic Laboratory | Claudio Tavares Sacchi, Claudia Regina Gonçalves, Erica Valessa Ramos Gomes |
| EPI_ISL_471545 | Hospital Sao Paulo de Ensino da Unifesp | Instituto Adolfo Lutz, Interdiciplinary Procedures Center, Strategic Laboratory | Claudio Tavares Sacchi, Claudia Regina Gonçalves, Erica Valessa Ramos Gomes |
| EPI_ISL_471546 | AMA DR Jose Soares Hungria | Instituto Adolfo Lutz, Interdiciplinary Procedures Center, Strategic Laboratory | Claudio Tavares Sacchi, Claudia Regina Gonçalves, Erica Valessa Ramos Gomes |
| EPI_ISL_471548 | Hospital do Servidor Público Estadual Francisco Morato de Oliveira | Instituto Adolfo Lutz, Interdiciplinary Procedures Center, Strategic Laboratory | Claudio Tavares Sacchi, Claudia Regina Gonçalves, Erica Valessa Ramos Gomes |
| EPI_ISL_471549 | Hospital Municipal Carmen Prudente | Instituto Adolfo Lutz, Interdiciplinary Procedures Center, Strategic Laboratory | Claudio Tavares Sacchi, Claudia Regina Gonçalves, Erica Valessa Ramos Gomes |
| EPI_ISL_471551 | Hospital Sao Paulo de Ensino da Unifesp | Instituto Adolfo Lutz, Interdiciplinary Procedures Center, Strategic Laboratory | Claudio Tavares Sacchi, Claudia Regina Gonçalves, Erica Valessa Ramos Gomes |
| EPI_ISL_471552 | Hospital Sancta Maggiore | Instituto Adolfo Lutz, Interdiciplinary Procedures Center, Strategic Laboratory | Claudio Tavares Sacchi, Claudia Regina Gonçalves, Erica Valessa Ramos Gomes |
| EPI_ISL_471554 | Hospital Bosque da Saúde | Instituto Adolfo Lutz, Interdiciplinary Procedures Center, Strategic Laboratory | Claudio Tavares Sacchi, Claudia Regina Gonçalves, Erica Valessa Ramos Gomes |
| EPI_ISL_471556 | Pronto Socorro Jose Ibrahin | Instituto Adolfo Lutz, Interdiciplinary Procedures Center, Strategic Laboratory | Claudio Tavares Sacchi, Claudia Regina Gonçalves, Erica Valessa Ramos Gomes |
| EPI_ISL_471562, EPI_ISL_471581, EPI_ISL_471582 | Hosp. Municipal Prof. Dr. Alípio Corrêa Netto | Instituto Adolfo Lutz, Interdiciplinary Procedures Center, Strategic Laboratory | Claudio Tavares Sacchi, Claudia Regina Gonçalves, Erica Valessa Ramos Gomes |
| EPI_ISL_471647 | Hospital Municipal de Barueri Dr. Francisco Moran | Instituto Adolfo Lutz, Interdiciplinary Procedures Center, Strategic Laboratory | Claudio Tavares Sacchi, Claudia Regina Gonçalves, Erica Valessa Ramos Gomes |
| EPI_ISL_471648 | UBS e Pronto Socorro Jd. Jacira | Instituto Adolfo Lutz, Interdiciplinary Procedures Center, Strategic Laboratory | Claudio Tavares Sacchi, Claudia Regina Gonçalves, Erica Valessa Ramos Gomes |
| EPI_ISL_472148 | Regional Virus Laboratory, Belfast Health and Social Care Trust | COVID-19 Genomics UK (COG-UK) Consortium | Conall McCaughey, James McKenna, Tanya Curran, Susan Feeney, Alison Watt, Ciara Cox, Mairead Connor, Zoltan Molnar, David Simpson, Derek Fairley |
| EPI_ISL_475717, EPI_ISL_475718, EPI_ISL_475719, EPI_ISL_475720, EPI_ISL_475721 | Microbiology, University Hospital Donostia | Microbiology, University Hospital Donostia | Cilla,G., Montes,M., Pineiro,L., Marimon,J.M. |
| EPI_ISL_476080, EPI_ISL_476081, EPI_ISL_476082, EPI_ISL_476084, EPI_ISL_476085, EPI_ISL_476086, EPI_ISL_476087, EPI_ISL_476088, EPI_ISL_476089, EPI_ISL_476090, EPI_ISL_476091, EPI_ISL_476092, EPI_ISL_476093, EPI_ISL_476094, EPI_ISL_476096, EPI_ISL_476097, EPI_ISL_476098, EPI_ISL_476099, EPI_ISL_476101, EPI_ISL_476102, EPI_ISL_476103, EPI_ISL_476104, EPI_ISL_476105, EPI_ISL_476106, EPI_ISL_476107, EPI_ISL_476108, EPI_ISL_476109, EPI_ISL_476110, EPI_ISL_476112, EPI_ISL_476113, EPI_ISL_476114, EPI_ISL_476115, EPI_ISL_476117, EPI_ISL_476118, EPI_ISL_476119, EPI_ISL_476120, EPI_ISL_476121, EPI_ISL_476122, EPI_ISL_476124, EPI_ISL_476125, EPI_ISL_476126, EPI_ISL_476128, EPI_ISL_476129, EPI_ISL_476130, EPI_ISL_476131, EPI_ISL_476132, EPI_ISL_476133, EPI_ISL_476134 | | | |
| see above | Viollier AG | Department of Biosystems Science and Engineering, ETH Zürich | Christian Beisel, Sarah Nadeau, Ivan Topolsky, Pedro Ferreira, Philipp Jablonski, Susana Posada-Céspedes, Tobias Schär, Ina Nissen, Natascha Santacroce, Elodie Burcklen, Christiane Beckmann, Maurice Redondo, Olivier Kobel, Christoph Noppen, Sophie Seidel, Noemie Santamaria de Souza, Niko Beerenwinkel, Tanja Stadler |
| EPI_ISL_476496 | Hospital Garrahan | Héritas | Dalmacio Pereyra, Roberta Crespo, Mauricio Grisolia, Cristian Rohr, Andrea Mangano, Maria Florencia Fernandez, Fabian Fay, Martin Vazquez |
| EPI_ISL_476563, EPI_ISL_476567 | Hospital de Pediatría "Prof. Dr. Juan P Garrahan" | Héritas | Dalmacio Pereyra, Roberta Crespo, Mauricio Grisolia, Cristian Rohr, Andrea Mangano, Maria Florencia Fernandez, Fabian Fay, Martin Vazquez |
| EPI_ISL_476568 | Hospital de Pediatría "Prof. Dr. Juan P Garrahan" | Héritas | Cristian Rohr, Andrea Mangano, Maria Florencia Fernandez, Dalmacio Pereyra, Roberta Crespo, Mauricio Grisolia, Fabian Fay, Martin Vazquez |
| EPI_ISL_476571, EPI_ISL_476573 | Hospital de Pediatría "Prof. Dr. Juan P Garrahan" | Héritas | Dalmacio Pereyra, Roberta Crespo, Mauricio Grisolia, Cristian Rohr, Andrea Mangano, Maria Florencia Fernandez, Fabian Fay, Martin Vazquez |
| EPI_ISL_478289, EPI_ISL_478290, EPI_ISL_478291, EPI_ISL_478292, EPI_ISL_478293, EPI_ISL_478294, EPI_ISL_478295, EPI_ISL_478296, EPI_ISL_478297, EPI_ISL_478298, EPI_ISL_478299, EPI_ISL_478300, EPI_ISL_478301, EPI_ISL_478302, EPI_ISL_478303, EPI_ISL_478304, EPI_ISL_478305, EPI_ISL_478306, EPI_ISL_478307, EPI_ISL_478308, EPI_ISL_478309, EPI_ISL_478310, EPI_ISL_478311, EPI_ISL_478312, EPI_ISL_478313, EPI_ISL_478314, EPI_ISL_478315, EPI_ISL_478316, EPI_ISL_478317, EPI_ISL_478318, EPI_ISL_478319, EPI_ISL_478320, EPI_ISL_478321, EPI_ISL_478322, EPI_ISL_478323, EPI_ISL_478324, EPI_ISL_478325, EPI_ISL_478326, EPI_ISL_478327, EPI_ISL_478328, EPI_ISL_478329, EPI_ISL_478330, EPI_ISL_478331, EPI_ISL_478332, EPI_ISL_478333, EPI_ISL_478334, EPI_ISL_478335, EPI_ISL_478336, EPI_ISL_478337, EPI_ISL_478338, EPI_ISL_478339, EPI_ISL_478340, EPI_ISL_478341, EPI_ISL_478342, EPI_ISL_478343, EPI_ISL_478344, EPI_ISL_478345, EPI_ISL_478346, EPI_ISL_478347, EPI_ISL_478348, EPI_ISL_478349, EPI_ISL_478350, EPI_ISL_478351, EPI_ISL_478352, EPI_ISL_478353, EPI_ISL_478354, EPI_ISL_478355, EPI_ISL_478356, EPI_ISL_478357, EPI_ISL_478358, EPI_ISL_478359, EPI_ISL_478360, EPI_ISL_478361, EPI_ISL_478362, EPI_ISL_478363, EPI_ISL_478364, EPI_ISL_478365, EPI_ISL_478366, EPI_ISL_478367, EPI_ISL_478368, EPI_ISL_478369, EPI_ISL_478370, EPI_ISL_478371 | | | |
| see above | University Hospitals Of Leicester NHS Trust and DeepSeq Nottingham | COVID-19 Genomics UK (COG-UK) Consortium | Christopher Holmes, Paul Bird, Thomas Helmer, Karlie Fallon, Julian Tang, Jonathan Ball, Patrick McClure, Joeseph Chappell, Nadine Holmes, Matthew Carlisle, Christopher Moore, Fei Sang, Johnny Debebe, Victoria Wright, Matthew Loose |
| EPI_ISL_482032, EPI_ISL_482033, EPI_ISL_482034, EPI_ISL_482035, EPI_ISL_482036, EPI_ISL_482037, EPI_ISL_482038, EPI_ISL_482039, EPI_ISL_482040, EPI_ISL_482041, EPI_ISL_482042, EPI_ISL_482043, EPI_ISL_482044, EPI_ISL_482045, EPI_ISL_482046, EPI_ISL_482047, EPI_ISL_482048, EPI_ISL_482049, EPI_ISL_482050, EPI_ISL_482051, EPI_ISL_482052, EPI_ISL_482053, EPI_ISL_482054, EPI_ISL_482055, EPI_ISL_482056, EPI_ISL_482070, EPI_ISL_482075, EPI_ISL_482077, EPI_ISL_482082, EPI_ISL_482091, EPI_ISL_482096, EPI_ISL_482097, EPI_ISL_482101, EPI_ISL_482102, EPI_ISL_482105, EPI_ISL_482115, EPI_ISL_482132, EPI_ISL_482133, EPI_ISL_482134, EPI_ISL_482135, EPI_ISL_482136, EPI_ISL_482137, EPI_ISL_482139, EPI_ISL_482140, EPI_ISL_482141, EPI_ISL_482142, EPI_ISL_482143, EPI_ISL_482144, EPI_ISL_482145, EPI_ISL_482146, EPI_ISL_482147, EPI_ISL_482148, EPI_ISL_482149, EPI_ISL_482150, EPI_ISL_482151, EPI_ISL_482152, EPI_ISL_482153, EPI_ISL_482154, EPI_ISL_482155, EPI_ISL_482156, EPI_ISL_482157, EPI_ISL_482158, EPI_ISL_482159 | | | |
| see above | Regional Virus Laboratory, Belfast Health and Social Care Trust | Wellcome Sanger Institute for the COVID-19 Genomics UK (COG-UK) consortium | Conall McCaughey, James McKenna, Tanya Curran, Susan Feeney, Alison Watt, Ciara Cox, Mairead Connor, Zoltan Molnar, David Simpson, Derek Fairley; and Alex Alderton, Roberto Amato, Sonia Goncalves, Ewan Harrison, David K. Jackson, Ian Johnston, Dominic Kwiatkowski, Cordelia Langford, John Sillitoe on behalf of the Wellcome Sanger Institute COVID-19 Surveillance Team (http://www.sanger.ac.uk/covid-team) |
| EPI_ISL_483648, EPI_ISL_483649, EPI_ISL_483653, EPI_ISL_483656, EPI_ISL_483657, EPI_ISL_483658, EPI_ISL_483659, EPI_ISL_483661, EPI_ISL_483662, EPI_ISL_483663, EPI_ISL_483664, EPI_ISL_483665, EPI_ISL_483666, EPI_ISL_483667 | | | |
| see above | Viollier AG | Department of Biosystems Science and Engineering, ETH Zürich | Christian Beisel, Sarah Nadeau, Ivan Topolsky, Pedro Ferreira, Philipp Jablonski, Susana Posada-Céspedes, Tobias Schär, Ina Nissen, Natascha Santacroce, Elodie Burcklen, Christiane Beckmann, Maurice Redondo, Olivier Kobel, Christoph Noppen, Sophie Seidel, Noemie Santamaria de Souza, Niko Beerenwinkel, Tanja Stadler |
| EPI_ISL_483668, EPI_ISL_483670, EPI_ISL_483672, EPI_ISL_483674, EPI_ISL_483675, EPI_ISL_483677, EPI_ISL_483682, EPI_ISL_483684, EPI_ISL_483685 | University Hospital Zurich | Department of Biosystems Science and Engineering, ETH Zürich | Christian Beisel, Sarah Nadeau, Ivan Topolsky, Pedro Ferreira, Philipp Jablonski, Susana Posada-Céspedes, Tobias Schär, Ina Nissen, Natascha Santacroce, Elodie Burcklen, Julia Martinez-Gomez, Phil Cheng, Mitch Levesque, Philipp Bosshard, Niko Beerenwinkel, Tanja Stadler |
| EPI_ISL_484221, EPI_ISL_484222, EPI_ISL_484223, EPI_ISL_484224, EPI_ISL_484225, EPI_ISL_484226, EPI_ISL_484227, EPI_ISL_484228, EPI_ISL_484229, EPI_ISL_484230, EPI_ISL_484231, EPI_ISL_484232, EPI_ISL_484233, EPI_ISL_484234, EPI_ISL_484235, EPI_ISL_484236, EPI_ISL_484237, EPI_ISL_484238, EPI_ISL_484239, EPI_ISL_484240, EPI_ISL_484241, EPI_ISL_484242, EPI_ISL_484243, EPI_ISL_484244, EPI_ISL_484245, EPI_ISL_484246, EPI_ISL_484247, EPI_ISL_484248, EPI_ISL_484249, EPI_ISL_484250, EPI_ISL_484251, EPI_ISL_484252, EPI_ISL_484253, EPI_ISL_484376, EPI_ISL_484377, EPI_ISL_484378, EPI_ISL_484379, EPI_ISL_484380, EPI_ISL_484381, EPI_ISL_484382, EPI_ISL_484383, EPI_ISL_484384, EPI_ISL_484385, EPI_ISL_484386, EPI_ISL_484387, EPI_ISL_484388, EPI_ISL_484389 | | | |
| see above | University Hospitals Of Leicester NHS Trust and DeepSeq | COVID-19 Genomics UK (COG-UK) Consortium | Christopher Holmes, Paul Bird, Thomas Helmer, Karlie Fallon, Julian Tang, Jonathan Ball, Patrick McClure, Joeseph Chappell, Nadine Holmes, Matthew |

| | | | |
|---|---|---|---|
| | Nottingham | | Carlisle, Christopher Moore, Fei Sang, Johnny Debebe, Victoria Wright, Matthew Loose |
| EPI_ISL_486442, EPI_ISL_486443, EPI_ISL_486444, EPI_ISL_486445, EPI_ISL_486447, EPI_ISL_486448, EPI_ISL_486449, EPI_ISL_486450, EPI_ISL_486451, EPI_ISL_486452, EPI_ISL_486453, EPI_ISL_486454, EPI_ISL_486455, EPI_ISL_486457, EPI_ISL_486458, EPI_ISL_486459, EPI_ISL_486460, EPI_ISL_486461, EPI_ISL_486462, EPI_ISL_486463, EPI_ISL_486464, EPI_ISL_486465, EPI_ISL_486466, EPI_ISL_486467, EPI_ISL_486468, EPI_ISL_486470, EPI_ISL_486471, EPI_ISL_486472, EPI_ISL_486473, EPI_ISL_486474, EPI_ISL_486475, EPI_ISL_486476, EPI_ISL_486477, EPI_ISL_486478, EPI_ISL_486479, EPI_ISL_486482, EPI_ISL_486483, EPI_ISL_486484, EPI_ISL_486486, EPI_ISL_486487, EPI_ISL_486488, EPI_ISL_486489, EPI_ISL_486490, EPI_ISL_486491, EPI_ISL_486492, EPI_ISL_486493, EPI_ISL_486494, EPI_ISL_486495, EPI_ISL_486496, EPI_ISL_486497, EPI_ISL_486500, EPI_ISL_486501, EPI_ISL_486502, EPI_ISL_486503, EPI_ISL_486504, EPI_ISL_486505, EPI_ISL_486506, EPI_ISL_486507, EPI_ISL_486508, EPI_ISL_486509, EPI_ISL_486511, EPI_ISL_486512, EPI_ISL_486513, EPI_ISL_486514, EPI_ISL_486516, EPI_ISL_486517, EPI_ISL_486519, EPI_ISL_486521, EPI_ISL_486522, EPI_ISL_486523, EPI_ISL_486524, EPI_ISL_486525, EPI_ISL_486526, EPI_ISL_486528, EPI_ISL_486529, EPI_ISL_486530, EPI_ISL_486532, EPI_ISL_486533, EPI_ISL_486534, EPI_ISL_486537, EPI_ISL_486538 | | | |
| see above | Viollier AG | Department of Biosystems Science and Engineering, ETH Zürich | Christian Beisel, Sarah Nadeau, Ivan Topolsky, Pedro Ferreira, Philipp Jablonski, Susana Posada-Céspedes, Tobias Schär, Ina Nissen, Natascha Santacroce, Elodie Burcklen, Christiane Beckmann, Maurice Redondo, Olivier Kobel, Christoph Noppen, Sophie Seidel, Noemie Santamaria de Souza, Niko Beerenwinkel, Tanja Stadler |
| EPI_ISL_488874, EPI_ISL_488875, EPI_ISL_488876, EPI_ISL_489156, EPI_ISL_489157, EPI_ISL_489158, EPI_ISL_489159, EPI_ISL_489160, EPI_ISL_489161, EPI_ISL_489162, EPI_ISL_489163, EPI_ISL_489164, EPI_ISL_489165, EPI_ISL_489166, EPI_ISL_489167, EPI_ISL_489168, EPI_ISL_489169, EPI_ISL_489170, EPI_ISL_489171, EPI_ISL_489172, EPI_ISL_489173, EPI_ISL_489174, EPI_ISL_489175, EPI_ISL_489176, EPI_ISL_489177, EPI_ISL_489178, EPI_ISL_489179, EPI_ISL_489180, EPI_ISL_489181, EPI_ISL_489182, EPI_ISL_489183, EPI_ISL_489184, EPI_ISL_489185, EPI_ISL_489186, EPI_ISL_489187, EPI_ISL_489188, EPI_ISL_489189, EPI_ISL_489190, EPI_ISL_489191, EPI_ISL_489192, EPI_ISL_489193, EPI_ISL_489194, EPI_ISL_489195, EPI_ISL_489196, EPI_ISL_489197, EPI_ISL_489198, EPI_ISL_489199, EPI_ISL_489200, EPI_ISL_489201, EPI_ISL_489202, EPI_ISL_489203, EPI_ISL_489204, EPI_ISL_489205, EPI_ISL_489206, EPI_ISL_489207, EPI_ISL_489208, EPI_ISL_489209, EPI_ISL_489210, EPI_ISL_489211, EPI_ISL_489212, EPI_ISL_489214, EPI_ISL_489215, EPI_ISL_489216, EPI_ISL_489217, EPI_ISL_489218, EPI_ISL_489219, EPI_ISL_489220, EPI_ISL_489221, EPI_ISL_489222, EPI_ISL_489223, EPI_ISL_489224, EPI_ISL_489225, EPI_ISL_489226, EPI_ISL_489227, EPI_ISL_489228, EPI_ISL_489229, EPI_ISL_489230, EPI_ISL_489231, EPI_ISL_489232, EPI_ISL_489233, EPI_ISL_489234, EPI_ISL_489235, EPI_ISL_489236, EPI_ISL_489237, EPI_ISL_489238, EPI_ISL_489239, EPI_ISL_489240, EPI_ISL_489241, EPI_ISL_489242, EPI_ISL_489243, EPI_ISL_489244, EPI_ISL_489245, EPI_ISL_489246, EPI_ISL_489247, EPI_ISL_489248, EPI_ISL_489249, EPI_ISL_489250, EPI_ISL_489251, EPI_ISL_489252, EPI_ISL_489253, EPI_ISL_489254, EPI_ISL_489255, EPI_ISL_489256, EPI_ISL_489257, EPI_ISL_489258, EPI_ISL_489259, EPI_ISL_489260, EPI_ISL_489261, EPI_ISL_489262, EPI_ISL_489263, EPI_ISL_489264, EPI_ISL_489265, EPI_ISL_489266, EPI_ISL_489267, EPI_ISL_489268, EPI_ISL_489269, EPI_ISL_489270, EPI_ISL_489271, EPI_ISL_489272, EPI_ISL_489273, EPI_ISL_489274, EPI_ISL_489275, EPI_ISL_489276, EPI_ISL_489277, EPI_ISL_489278, EPI_ISL_489279, EPI_ISL_489280, EPI_ISL_489281, EPI_ISL_489282, EPI_ISL_489283, EPI_ISL_489284, EPI_ISL_489285, EPI_ISL_489286, EPI_ISL_489287, EPI_ISL_489288, EPI_ISL_489289, EPI_ISL_489290, EPI_ISL_489291, EPI_ISL_489292, EPI_ISL_489293, EPI_ISL_489294, EPI_ISL_489295, EPI_ISL_489296, EPI_ISL_489297, EPI_ISL_489298, EPI_ISL_489299, EPI_ISL_489300, EPI_ISL_489301, EPI_ISL_489302, EPI_ISL_489303, EPI_ISL_489304, EPI_ISL_489305, EPI_ISL_489306, EPI_ISL_489307, EPI_ISL_489308, EPI_ISL_489309, EPI_ISL_489310, EPI_ISL_489311, EPI_ISL_489312, EPI_ISL_489313, EPI_ISL_489314, EPI_ISL_489315, EPI_ISL_489316, EPI_ISL_489317, EPI_ISL_489318, EPI_ISL_489319, EPI_ISL_489320, EPI_ISL_489321, EPI_ISL_489322, EPI_ISL_489323, EPI_ISL_489324, EPI_ISL_489325, EPI_ISL_489326, EPI_ISL_489327, EPI_ISL_489328, EPI_ISL_489329, EPI_ISL_489330, EPI_ISL_489331, EPI_ISL_489332, EPI_ISL_489333, EPI_ISL_489334, EPI_ISL_489335, EPI_ISL_489336, EPI_ISL_489337, EPI_ISL_489338, EPI_ISL_489339, EPI_ISL_489340, EPI_ISL_489341, EPI_ISL_489342, EPI_ISL_489343, EPI_ISL_489344, EPI_ISL_489345, EPI_ISL_489346, EPI_ISL_489347, EPI_ISL_489348, EPI_ISL_489349, EPI_ISL_489350, EPI_ISL_489351, EPI_ISL_489352, EPI_ISL_489353, EPI_ISL_489354, EPI_ISL_489355, EPI_ISL_489356, EPI_ISL_489357, EPI_ISL_489359, EPI_ISL_489360, EPI_ISL_489361, EPI_ISL_489362, EPI_ISL_489363, EPI_ISL_489364, EPI_ISL_489365, EPI_ISL_489366, EPI_ISL_489367, EPI_ISL_489368, EPI_ISL_489369, EPI_ISL_489370, EPI_ISL_489371, EPI_ISL_489372, EPI_ISL_489373, EPI_ISL_489374, EPI_ISL_489375, EPI_ISL_489376, EPI_ISL_489377, EPI_ISL_489378, EPI_ISL_489379 | | | |
| see above | Regional Virus Laboratory, Belfast Health and Social Care Trust | Wellcome Sanger Institute for the COVID-19 Genomics UK (COG-UK) consortium | Conall McCaughey, James McKenna, Tanya Curran, Susan Feeney, Alison Watt, Ciara Cox, Mairead Connor, Zoltan Molnar, David Simpson, Derek Fairley; and Alex Alderton, Roberto Amato, Sonia Goncalves, Ewan Harrison, David K. Jackson, Ian Johnston, Dominic Kwiatkowski, Cordelia Langford, John Sillitoe on behalf of the Wellcome Sanger Institute COVID-19 Surveillance Team (http://www.sanger.ac.uk/covid-team) |
| EPI_ISL_489896, EPI_ISL_489897, EPI_ISL_489898, EPI_ISL_489900, EPI_ISL_489901, EPI_ISL_489902, EPI_ISL_489903 | Gundersen Molecular Diagnostics Laboratory | Kabara Cancer Research Institute | Craig S. Richmond, Paraic A. Kenny |
| EPI_ISL_489904 | Gundersen Clinical Microbiology Laboratory | Kabara Cancer Research Institute | Craig S. Richmond, Paraic A. Kenny |
| EPI_ISL_489905, EPI_ISL_489906, EPI_ISL_489907, EPI_ISL_489908, EPI_ISL_489909, EPI_ISL_489910, EPI_ISL_489911, EPI_ISL_489913, EPI_ISL_489914, EPI_ISL_489915, EPI_ISL_489916, EPI_ISL_489917, EPI_ISL_489918, EPI_ISL_489919, EPI_ISL_489920, EPI_ISL_489921, EPI_ISL_489922, EPI_ISL_489923, EPI_ISL_489925, EPI_ISL_489926, EPI_ISL_489928, EPI_ISL_489929, EPI_ISL_489930, EPI_ISL_489931, EPI_ISL_489932, EPI_ISL_489933, EPI_ISL_489934, EPI_ISL_489935, EPI_ISL_489936, EPI_ISL_489937, EPI_ISL_489938, EPI_ISL_489939, EPI_ISL_489940, EPI_ISL_489941, EPI_ISL_489942, EPI_ISL_489943, EPI_ISL_489944, EPI_ISL_489945, EPI_ISL_489946, EPI_ISL_489947, EPI_ISL_489948, EPI_ISL_489949, EPI_ISL_489950, EPI_ISL_489951, EPI_ISL_489952, EPI_ISL_489953, EPI_ISL_489954, EPI_ISL_489955 | | | |
| see above | Gundersen Molecular Diagnostics Laboratory | Kabara Cancer Research Institute | Craig S. Richmond, Paraic A. Kenny |
| EPI_ISL_489956, EPI_ISL_489957, EPI_ISL_489958 | Gundersen Clinical Microbiology Laboratory | Kabara Cancer Research Institute | Craig S. Richmond, Paraic A. Kenny |
| EPI_ISL_489960, EPI_ISL_489962, EPI_ISL_489963, EPI_ISL_489964, EPI_ISL_489965, EPI_ISL_489966, EPI_ISL_489968, EPI_ISL_489969, EPI_ISL_489970, EPI_ISL_489971, EPI_ISL_489972, EPI_ISL_489973, EPI_ISL_489974, EPI_ISL_489975, EPI_ISL_489976, EPI_ISL_489977, EPI_ISL_489978, EPI_ISL_489979, EPI_ISL_489980, EPI_ISL_489981, EPI_ISL_489982, EPI_ISL_489983, EPI_ISL_489984, EPI_ISL_489985, EPI_ISL_489986 | | | |
| see above | Viollier AG | Department of Biosystems Science and Engineering, ETH Zürich | Christian Beisel, Sarah Nadeau, Ivan Topolsky, Pedro Ferreira, Philipp Jablonski, Susana Posada-Céspedes, Tobias Schär, Ina Nissen, Natascha Santacroce, Elodie Burcklen, Christiane Beckmann, Maurice Redondo, Olivier Kobel, Christoph Noppen, Sophie Seidel, Noemie Santamaria de Souza, Niko Beerenwinkel, Tanja Stadler |
| EPI_ISL_496603, EPI_ISL_496604, EPI_ISL_496605, EPI_ISL_496606, EPI_ISL_496607, EPI_ISL_496608, EPI_ISL_496609, EPI_ISL_496610, EPI_ISL_496611, EPI_ISL_496612, EPI_ISL_496613, EPI_ISL_496614, EPI_ISL_496616, EPI_ISL_496617, EPI_ISL_496618, EPI_ISL_496619, EPI_ISL_496620, EPI_ISL_496621, EPI_ISL_496622, EPI_ISL_496623, EPI_ISL_496624, EPI_ISL_496625, EPI_ISL_496626, EPI_ISL_496627, EPI_ISL_496628, EPI_ISL_496629, EPI_ISL_496630, EPI_ISL_496631, EPI_ISL_496632, EPI_ISL_496633, EPI_ISL_496634, EPI_ISL_496635, EPI_ISL_496636, EPI_ISL_496637, EPI_ISL_496638, EPI_ISL_496639, EPI_ISL_496640, EPI_ISL_496641, EPI_ISL_496642, EPI_ISL_496643, EPI_ISL_496644, EPI_ISL_496645, EPI_ISL_496646, EPI_ISL_496647, EPI_ISL_496648, EPI_ISL_496649, EPI_ISL_496650, EPI_ISL_496651, EPI_ISL_496652, EPI_ISL_496653, EPI_ISL_496654, EPI_ISL_496655, EPI_ISL_496656, EPI_ISL_496657, EPI_ISL_496658, EPI_ISL_496659, EPI_ISL_496660, EPI_ISL_496661, EPI_ISL_496663, EPI_ISL_496664, EPI_ISL_496665, EPI_ISL_496666, EPI_ISL_496668, EPI_ISL_496669, EPI_ISL_496670, EPI_ISL_496671, EPI_ISL_496672, EPI_ISL_496673, EPI_ISL_496675, EPI_ISL_496676, EPI_ISL_496677, EPI_ISL_496678, EPI_ISL_496679, EPI_ISL_496680, EPI_ISL_496681, EPI_ISL_496682, EPI_ISL_496683, EPI_ISL_496685, EPI_ISL_496686, EPI_ISL_496687, EPI_ISL_496688, EPI_ISL_496689, EPI_ISL_496690, EPI_ISL_496691, EPI_ISL_496692, EPI_ISL_496693, EPI_ISL_496694, EPI_ISL_496695, EPI_ISL_496697, EPI_ISL_496698, EPI_ISL_496699, EPI_ISL_496700, EPI_ISL_496701, EPI_ISL_496702, EPI_ISL_496703, EPI_ISL_496704, EPI_ISL_496705, EPI_ISL_496707, EPI_ISL_496708, EPI_ISL_496709, EPI_ISL_496710, EPI_ISL_496711, EPI_ISL_496712, EPI_ISL_496713, EPI_ISL_496714, EPI_ISL_496715, EPI_ISL_496716, EPI_ISL_496717, EPI_ISL_496718, EPI_ISL_496719, EPI_ISL_496720, EPI_ISL_496721, EPI_ISL_496722, EPI_ISL_496723, EPI_ISL_496724, EPI_ISL_496725, EPI_ISL_496726, EPI_ISL_496727, EPI_ISL_496728, EPI_ISL_496729, EPI_ISL_496730, EPI_ISL_496731, EPI_ISL_496732, EPI_ISL_496733, EPI_ISL_496734, EPI_ISL_496735, EPI_ISL_496736, EPI_ISL_496737, EPI_ISL_496738, EPI_ISL_496739, EPI_ISL_496740, EPI_ISL_496741, EPI_ISL_496742, EPI_ISL_496743, EPI_ISL_496744, EPI_ISL_496745, EPI_ISL_496746, EPI_ISL_496747, EPI_ISL_496748, EPI_ISL_496749, EPI_ISL_496750, EPI_ISL_496751, EPI_ISL_496752, EPI_ISL_496753, EPI_ISL_496754, EPI_ISL_496755, EPI_ISL_496756, EPI_ISL_496757, EPI_ISL_496758, EPI_ISL_496759, EPI_ISL_496761, EPI_ISL_496762, EPI_ISL_496763, EPI_ISL_496764, EPI_ISL_496765, EPI_ISL_496766, EPI_ISL_496767, EPI_ISL_496768, EPI_ISL_496769, EPI_ISL_496770, EPI_ISL_496771, EPI_ISL_496772, EPI_ISL_496773, EPI_ISL_496774, EPI_ISL_496775, EPI_ISL_496776, EPI_ISL_496777, EPI_ISL_496778, EPI_ISL_496779, EPI_ISL_496780, EPI_ISL_496781, EPI_ISL_496782, EPI_ISL_496783, EPI_ISL_496784, EPI_ISL_496786, EPI_ISL_496787, EPI_ISL_496788, EPI_ISL_496789, EPI_ISL_496790, EPI_ISL_496791, EPI_ISL_496793, EPI_ISL_496794, EPI_ISL_496795, EPI_ISL_496796, EPI_ISL_496797, EPI_ISL_496798, EPI_ISL_496799, EPI_ISL_496800, EPI_ISL_496801, EPI_ISL_496802, EPI_ISL_496803, EPI_ISL_496804, EPI_ISL_496805, EPI_ISL_496807, EPI_ISL_496809, EPI_ISL_496810, EPI_ISL_496811, EPI_ISL_496812, EPI_ISL_496813, EPI_ISL_496814, EPI_ISL_496815, EPI_ISL_496816, EPI_ISL_496817, EPI_ISL_496818, EPI_ISL_496819, EPI_ISL_496820, EPI_ISL_496821, EPI_ISL_496822, EPI_ISL_496823, EPI_ISL_496824, EPI_ISL_496825, EPI_ISL_496826, EPI_ISL_496827, EPI_ISL_496829, EPI_ISL_496831, EPI_ISL_496833, EPI_ISL_496834, EPI_ISL_496835, EPI_ISL_496836, EPI_ISL_496837, EPI_ISL_496838, EPI_ISL_496839, EPI_ISL_496840, EPI_ISL_496841, EPI_ISL_496842, EPI_ISL_496843, EPI_ISL_496844, EPI_ISL_496845, EPI_ISL_496846, EPI_ISL_496847, EPI_ISL_496848, EPI_ISL_496849, EPI_ISL_496850, EPI_ISL_496851, EPI_ISL_496852, EPI_ISL_496853, EPI_ISL_496854, EPI_ISL_496855, EPI_ISL_496856, EPI_ISL_496857, EPI_ISL_496858, EPI_ISL_496859, EPI_ISL_496860, EPI_ISL_496861, EPI_ISL_496862, EPI_ISL_496863, EPI_ISL_496864, EPI_ISL_496865, EPI_ISL_496866, EPI_ISL_496867, EPI_ISL_496868, EPI_ISL_496869, EPI_ISL_496870, EPI_ISL_496871, EPI_ISL_496872, EPI_ISL_496873, EPI_ISL_496874, EPI_ISL_496875, EPI_ISL_496876, EPI_ISL_496877, EPI_ISL_496878, EPI_ISL_496879, EPI_ISL_496880, EPI_ISL_496881, EPI_ISL_496883, EPI_ISL_496884, EPI_ISL_496885, EPI_ISL_496886, EPI_ISL_496887, EPI_ISL_496888, EPI_ISL_496889, EPI_ISL_496890, EPI_ISL_496891, EPI_ISL_496892, EPI_ISL_496893, EPI_ISL_496894, EPI_ISL_496895, EPI_ISL_496896, EPI_ISL_496897, EPI_ISL_496898, EPI_ISL_496899, EPI_ISL_496900, EPI_ISL_496901, EPI_ISL_496902, EPI_ISL_496903, EPI_ISL_496904, EPI_ISL_496905, EPI_ISL_496906, EPI_ISL_496907, EPI_ISL_496908, EPI_ISL_496909, EPI_ISL_496910, EPI_ISL_496911, EPI_ISL_496912, EPI_ISL_496913, EPI_ISL_496914, EPI_ISL_496915 | | | |
| see above | Gorgas Memorial Laboratory of Health Studies | Gorgas Memorial Laboratory of Health Studies | Danilo Franco, Claudia Gonzalez Sandra Lopez-Verges, Alexander A Martinez |
| EPI_ISL_498193 | Molecular Pathology Division, Department of Pathology, Hong Kong Sanatorium & Hospital | Molecular Pathology Division, Department of Pathology, Hong Kong Sanatorium & Hospital | Chun Hang AU, Wai Sing CHAN, Ho Yin LAM, Dona N. HO, Simon Y.M. LAM, Jonpaul S.T. ZEE, Tsun Leung CHAN, Edmond S.K. MA |
| EPI_ISL_498619, EPI_ISL_498620, EPI_ISL_498621, EPI_ISL_498622, EPI_ISL_498623, EPI_ISL_498624, EPI_ISL_498625, EPI_ISL_498626 | Viollier AG | Department of Biosystems Science and Engineering, ETH Zürich | Christian Beisel, Sarah Nadeau, Ivan Topolsky, Pedro Ferreira, Philipp Jablonski, Susana Posada-Céspedes, Tobias Schär, Ina Nissen, Natascha Santacroce, Elodie Burcklen, Christiane Beckmann, Maurice Redondo, Olivier Kobel, Christoph Noppen, Sophie Seidel, Noemie Santamaria de Souza, Niko Beerenwinkel, Tanja Stadler |
| EPI_ISL_499810, EPI_ISL_499828, EPI_ISL_499829, EPI_ISL_499858, EPI_ISL_499861, EPI_ISL_499869, EPI_ISL_499894, EPI_ISL_499895, EPI_ISL_499897, EPI_ISL_499910, EPI_ISL_499912, EPI_ISL_499913, EPI_ISL_499933, EPI_ISL_499950, EPI_ISL_499974, EPI_ISL_499975, EPI_ISL_499976, EPI_ISL_499977, EPI_ISL_499978, EPI_ISL_499979, EPI_ISL_499980, EPI_ISL_499981, EPI_ISL_499982, EPI_ISL_499983 | | | |
| see above | University Hospitals Of Leicester NHS Trust and DeepSeq Nottingham | COVID-19 Genomics UK (COG-UK) Consortium | Christopher Holmes, Paul Bird, Thomas Helmer, Karlie Fallon, Julian Tang, Jonathan Ball, Patrick McClure, Joeseph Chappell, Nadine Holmes, Matthew Carlisle, Christopher Moore, Fei Sang, Johnny Debebe, Victoria Wright, Matthew Loose |
| EPI_ISL_500877, EPI_ISL_500878, EPI_ISL_500879, EPI_ISL_500880, EPI_ISL_500881, EPI_ISL_500883, EPI_ISL_500884, EPI_ISL_500885, EPI_ISL_500886, EPI_ISL_500887, EPI_ISL_500889, EPI_ISL_500891, EPI_ISL_500892, EPI_ISL_500893, EPI_ISL_500894, EPI_ISL_500895, EPI_ISL_500896, EPI_ISL_500897, EPI_ISL_500898, EPI_ISL_500899, EPI_ISL_500900, EPI_ISL_500901, EPI_ISL_500902, EPI_ISL_500904, EPI_ISL_500906, EPI_ISL_500909, EPI_ISL_500910, EPI_ISL_500911, EPI_ISL_500912, EPI_ISL_500913, EPI_ISL_500914, EPI_ISL_500915, EPI_ISL_500916, EPI_ISL_500917, EPI_ISL_500920, EPI_ISL_500921, EPI_ISL_500922, EPI_ISL_500924, EPI_ISL_500926, EPI_ISL_500927, EPI_ISL_500928, EPI_ISL_500929, EPI_ISL_500930, EPI_ISL_500931, EPI_ISL_500933, EPI_ISL_500934, EPI_ISL_500935, EPI_ISL_500937, EPI_ISL_500939, EPI_ISL_500941, EPI_ISL_500942, EPI_ISL_500943, EPI_ISL_500944, EPI_ISL_500945 | | | |
| see above | Viollier AG | Department of Biosystems Science and Engineering, ETH Zürich | Christian Beisel, Sarah Nadeau, Ivan Topolsky, Pedro Ferreira, Philipp Jablonski, Susana Posada-Céspedes, Tobias Schär, Ina Nissen, Natascha Santacroce, Elodie Burcklen, Christiane Beckmann, Maurice Redondo, Olivier Kobel, Christoph Noppen, Sophie Seidel, Noemie Santamaria de Souza, Niko Beerenwinkel, Tanja Stadler |
| EPI_ISL_500954, EPI_ISL_500955, EPI_ISL_500956, EPI_ISL_500957, EPI_ISL_500958, EPI_ISL_500959, EPI_ISL_500960, EPI_ISL_500961, EPI_ISL_500962, EPI_ISL_500963, EPI_ISL_500964, EPI_ISL_500965, EPI_ISL_500966, EPI_ISL_500967, EPI_ISL_500968, EPI_ISL_500969, EPI_ISL_500970, EPI_ISL_500971, EPI_ISL_500972, EPI_ISL_500973, EPI_ISL_500974, EPI_ISL_500975, EPI_ISL_500976, EPI_ISL_500977, EPI_ISL_500978, EPI_ISL_500979, EPI_ISL_500980, EPI_ISL_500981, EPI_ISL_500982, EPI_ISL_500983, EPI_ISL_500984, EPI_ISL_500985, EPI_ISL_500986, EPI_ISL_500987, EPI_ISL_500988, EPI_ISL_500989, | | | |

| | | | |
|---|---|---|---|
| EPI_ISL_500990, EPI_ISL_500991, EPI_ISL_500992, EPI_ISL_500993, EPI_ISL_500994, EPI_ISL_500995, EPI_ISL_500996, EPI_ISL_500997, EPI_ISL_500998, EPI_ISL_500999, EPI_ISL_501000, EPI_ISL_501001, EPI_ISL_501002, EPI_ISL_501003, EPI_ISL_501004, EPI_ISL_501005, EPI_ISL_501006, EPI_ISL_501007, EPI_ISL_501008, EPI_ISL_501009, EPI_ISL_501010, EPI_ISL_501011, EPI_ISL_501012, EPI_ISL_501013, EPI_ISL_501014, EPI_ISL_501015, EPI_ISL_501016, EPI_ISL_501017, EPI_ISL_501018, EPI_ISL_501019, EPI_ISL_501020, EPI_ISL_501021, EPI_ISL_501022, EPI_ISL_501023, EPI_ISL_501024, EPI_ISL_501025, EPI_ISL_501026, EPI_ISL_501027, EPI_ISL_501028, EPI_ISL_501029, EPI_ISL_501030, EPI_ISL_501031, EPI_ISL_501032, EPI_ISL_501033, EPI_ISL_501034, EPI_ISL_501035, EPI_ISL_501036, EPI_ISL_501037, EPI_ISL_501038, EPI_ISL_501039, EPI_ISL_501040, EPI_ISL_501041, EPI_ISL_501042, EPI_ISL_501043, EPI_ISL_501044, EPI_ISL_501045, EPI_ISL_501046, EPI_ISL_501047, EPI_ISL_501048, EPI_ISL_501049, EPI_ISL_501050, EPI_ISL_501051, EPI_ISL_501052, EPI_ISL_501053, EPI_ISL_501054, EPI_ISL_501055, EPI_ISL_501056, EPI_ISL_501057, EPI_ISL_501058, EPI_ISL_501059, EPI_ISL_501060, EPI_ISL_501061, EPI_ISL_501062, EPI_ISL_501063, EPI_ISL_501064, EPI_ISL_501065, EPI_ISL_501066, EPI_ISL_501067, EPI_ISL_501068, EPI_ISL_501069, EPI_ISL_501070, EPI_ISL_501071 | | | |
| see above | Regional Virus Laboratory, Belfast Health and Social Care Trust | Wellcome Sanger Institute for the COVID-19 Genomics UK (COG-UK) consortium | Conall McCaughey, James McKenna, Tanya Curran, Susan Feeney, Alison Watt, Ciara Cox, Mairead Connor, Zoltan Molnar, David Simpson, Derek Fairley; and Alex Alderton, Roberto Amato, Sonia Goncalves, Ewan Harrison, David K. Jackson, Ian Johnston, Dominic Kwiatkowski, Cordelia Langford, John Sillitoe on behalf of the Wellcome Sanger Institute COVID-19 Surveillance Team (http://www.sanger.ac.uk/covid-team) |
| EPI_ISL_508862, EPI_ISL_508863 | Virology Unit, Institut Pasteur de Madagascar | Virology Unit, Institut Pasteur de Madagascar | Christian Ranaivoson, Cara Brook, Norosoa Razanajatovo, Vida Ahyong, Tsiry Randriambolamanantsoa, Michelle Tan, Vololoniaina Raharinosy, Helisoa Razafimanjato, Cristina M. Tato, Joseph L. DeRisi, Soa Fy Andriamandimby, Jean-Michel Heraud |
| EPI_ISL_510305, EPI_ISL_510306, EPI_ISL_510307, EPI_ISL_510308, EPI_ISL_510309, EPI_ISL_510310, EPI_ISL_510311, EPI_ISL_510312, EPI_ISL_510313, EPI_ISL_510314, EPI_ISL_510315, EPI_ISL_510316, EPI_ISL_510317, EPI_ISL_510318, EPI_ISL_510319, EPI_ISL_510320, EPI_ISL_510321, EPI_ISL_510322, EPI_ISL_510323, EPI_ISL_510324, EPI_ISL_510325, EPI_ISL_510326, EPI_ISL_510327, EPI_ISL_510328, EPI_ISL_510329, EPI_ISL_510330, EPI_ISL_510331, EPI_ISL_510332 | | | |
| see above | Hospital San Pedro de Alcántara (Cáceres) | SeqCOVID-SPAIN consortium/IBV(CSIC) | Cristina Muñoz Cuevas, Guadalupe Rodriguez Rodriguez and SeqCOVID-SPAIN consortium |
| EPI_ISL_510690, EPI_ISL_510691, EPI_ISL_510694, EPI_ISL_510696, EPI_ISL_510697, EPI_ISL_510698, EPI_ISL_510699, EPI_ISL_510701, EPI_ISL_510702, EPI_ISL_510703, EPI_ISL_510704, EPI_ISL_510705, EPI_ISL_510706, EPI_ISL_510707, EPI_ISL_510708, EPI_ISL_510711, EPI_ISL_510712, EPI_ISL_510713, EPI_ISL_510714, EPI_ISL_510715, EPI_ISL_510716, EPI_ISL_510717, EPI_ISL_510718, EPI_ISL_510719, EPI_ISL_510720, EPI_ISL_510722, EPI_ISL_510724, EPI_ISL_510725, EPI_ISL_510726, EPI_ISL_510727, EPI_ISL_510730, EPI_ISL_510731, EPI_ISL_510732, EPI_ISL_510733, EPI_ISL_510734, EPI_ISL_510735, EPI_ISL_510736, EPI_ISL_510737, EPI_ISL_510738, EPI_ISL_510739, EPI_ISL_510740, EPI_ISL_510741, EPI_ISL_510743, EPI_ISL_510744, EPI_ISL_510745, EPI_ISL_510746, EPI_ISL_510747, EPI_ISL_510749, EPI_ISL_510750, EPI_ISL_510751, EPI_ISL_510753, EPI_ISL_510754, EPI_ISL_510755, EPI_ISL_510756, EPI_ISL_510757, EPI_ISL_510758, EPI_ISL_510759, EPI_ISL_510760, EPI_ISL_510761, EPI_ISL_510762, EPI_ISL_510764, EPI_ISL_510765, EPI_ISL_510766, EPI_ISL_510767, EPI_ISL_510768, EPI_ISL_510769, EPI_ISL_510770, EPI_ISL_510771, EPI_ISL_510772, EPI_ISL_510773, EPI_ISL_510774, EPI_ISL_510775, EPI_ISL_510776, EPI_ISL_510777, EPI_ISL_510778, EPI_ISL_510779, EPI_ISL_510780, EPI_ISL_510781, EPI_ISL_510782, EPI_ISL_510783, EPI_ISL_510784, EPI_ISL_510786, EPI_ISL_510787, EPI_ISL_510788, EPI_ISL_510789, EPI_ISL_510790, EPI_ISL_510791, EPI_ISL_510793, EPI_ISL_510794, EPI_ISL_510795, EPI_ISL_510796, EPI_ISL_510797, EPI_ISL_510798, EPI_ISL_510799, EPI_ISL_510800, EPI_ISL_510801, EPI_ISL_510802, EPI_ISL_510803, EPI_ISL_510804, EPI_ISL_510805, EPI_ISL_510806, EPI_ISL_510807, EPI_ISL_510808, EPI_ISL_511985, EPI_ISL_511986, EPI_ISL_511987, EPI_ISL_511988, EPI_ISL_511989, EPI_ISL_511990, EPI_ISL_511992, EPI_ISL_511993, EPI_ISL_511994, EPI_ISL_511995, EPI_ISL_511996, EPI_ISL_511998, EPI_ISL_511999, EPI_ISL_512002, EPI_ISL_512003, EPI_ISL_512004, EPI_ISL_512005, EPI_ISL_512006, EPI_ISL_512008, EPI_ISL_512009, EPI_ISL_512010, EPI_ISL_512011, EPI_ISL_512013, EPI_ISL_512014, EPI_ISL_512015, EPI_ISL_512016, EPI_ISL_512017, EPI_ISL_512018, EPI_ISL_512021, EPI_ISL_512022, EPI_ISL_512023, EPI_ISL_512024, EPI_ISL_512028, EPI_ISL_512029, EPI_ISL_512030, EPI_ISL_512031, EPI_ISL_512032, EPI_ISL_512033, EPI_ISL_512034, EPI_ISL_512035, EPI_ISL_512036, EPI_ISL_512037, EPI_ISL_512042, EPI_ISL_512043, EPI_ISL_512045, EPI_ISL_512046, EPI_ISL_512047, EPI_ISL_512050, EPI_ISL_512051, EPI_ISL_512052, EPI_ISL_512053, EPI_ISL_512054 | | | |
| see above | Viollier AG | Department of Biosystems Science and Engineering, ETH Zürich | Christian Beisel, Sarah Nadeau, Ivan Topolsky, Pedro Ferreira, Philipp Jablonski, Susana Posada-Céspedes, Tobias Schär, Ina Nissen, Natascha Santacroce, Elodie Burcklen, Christiane Beckmann, Maurice Redondo, Olivier Kobel, Christoph Noppen, Sophie Seidel, Noemie Santamaria de Souza, Niko Beerenwinkel, Tanja Stadler |
| EPI_ISL_515520 | Hospital Municipal do Tatuape Carmino Caricchio | Instituto Adolfo Lutz, Interdiciplinary Procedures Center, Strategic Laboratory | Claudio Tavares Sacchi, Claudia Regina Gonçalves, Erica Valessa Ramos Gomes |
| EPI_ISL_515521 | Hospital Municipal Dr Waldemar Tebaldi | Instituto Adolfo Lutz, Interdiciplinary Procedures Center, Strategic Laboratory | Claudio Tavares Sacchi, Claudia Regina Gonçalves, Erica Valessa Ramos Gomes |
| EPI_ISL_515522 | UPA 24HS de Itatiba | Instituto Adolfo Lutz, Interdiciplinary Procedures Center, Strategic Laboratory | Claudio Tavares Sacchi, Claudia Regina Gonçalves, Erica Valessa Ramos Gomes |
| EPI_ISL_515523, EPI_ISL_515524 | PS Municipal Dr Lauro Ribas Braga | Instituto Adolfo Lutz, Interdiciplinary Procedures Center, Strategic Laboratory | Claudio Tavares Sacchi, Claudia Regina Gonçalves, Erica Valessa Ramos Gomes |
| EPI_ISL_515525 | National Influenza Center - Instituto Adolfo Lutz | Instituto Adolfo Lutz, Interdiciplinary Procedures Center, Strategic Laboratory | Claudio Tavares Sacchi, Claudia Regina Gonçalves, Erica Valessa Ramos Gomes |
| EPI_ISL_515526 | Hospital Municipal do Tatuape Carmino Caricchio | Instituto Adolfo Lutz, Interdiciplinary Procedures Center, Strategic Laboratory | Claudio Tavares Sacchi, Claudia Regina Gonçalves, Erica Valessa Ramos Gomes |
| EPI_ISL_515527 | Hospital Santa Clara | Instituto Adolfo Lutz, Interdiciplinary Procedures Center, Strategic Laboratory | Claudio Tavares Sacchi, Claudia Regina Gonçalves, Erica Valessa Ramos Gomes |
| EPI_ISL_515528 | Hospital Sao Paulo de Ensino da Unifesp | Instituto Adolfo Lutz, Interdiciplinary Procedures Center, Strategic Laboratory | Claudio Tavares Sacchi, Claudia Regina Gonçalves, Erica Valessa Ramos Gomes |
| EPI_ISL_515529 | Pronto Socorro Municipal Julio Tupy | Instituto Adolfo Lutz, Interdiciplinary Procedures Center, Strategic Laboratory | Claudio Tavares Sacchi, Claudia Regina Gonçalves, Erica Valessa Ramos Gomes |
| EPI_ISL_515541 | Hospital Montemagno | Instituto Adolfo Lutz, Interdiciplinary Procedures Center, Strategic Laboratory | Claudio Tavares Sacchi, Claudia Regina Gonçalves, Erica Valessa Ramos Gomes |
| EPI_ISL_515542 | Vigilância Epidemiológica de Leme | Instituto Adolfo Lutz, Interdiciplinary Procedures Center, Strategic Laboratory | Claudio Tavares Sacchi, Claudia Regina Gonçalves, Erica Valessa Ramos Gomes |
| EPI_ISL_515543 | Serviço de Vigilância Sanitária e Epidemiológica | Instituto Adolfo Lutz, Interdiciplinary Procedures Center, Strategic Laboratory | Claudio Tavares Sacchi, Claudia Regina Gonçalves, Erica Valessa Ramos Gomes |
| EPI_ISL_515544 | Ama Dr Jose Soares Hungria | Instituto Adolfo Lutz, Interdiciplinary Procedures Center, Strategic Laboratory | Claudio Tavares Sacchi, Claudia Regina Gonçalves, Erica Valessa Ramos Gomes |
| EPI_ISL_515545 | Hospital Sao Paulo de Ensino da Unifesp | Instituto Adolfo Lutz, Interdiciplinary Procedures Center, Strategic Laboratory | Claudio Tavares Sacchi, Claudia Regina Gonçalves, Erica Valessa Ramos Gomes |
| EPI_ISL_515546 | Hospital Municipal do Tatuape Carmino Caricchio | Instituto Adolfo Lutz, Interdiciplinary Procedures Center, Strategic Laboratory | Claudio Tavares Sacchi, Claudia Regina Gonçalves, Erica Valessa Ramos Gomes |
| EPI_ISL_515547 | Centro Medico da Policia Militar do Estado de Sao Paulo | Instituto Adolfo Lutz, Interdiciplinary Procedures Center, Strategic Laboratory | Claudio Tavares Sacchi, Claudia Regina Gonçalves, Erica Valessa Ramos Gomes |
| EPI_ISL_515548 | Hospital Municipal Dr. Jose Soares Hungria | Instituto Adolfo Lutz, Interdiciplinary Procedures Center, Strategic Laboratory | Claudio Tavares Sacchi, Claudia Regina Gonçalves, Erica Valessa Ramos Gomes |
| EPI_ISL_515550 | UPA Vila Santa Catarina | Instituto Adolfo Lutz, Interdiciplinary Procedures Center, Strategic Laboratory | Claudio Tavares Sacchi, Claudia Regina Gonçalves, Erica Valessa Ramos Gomes |
| EPI_ISL_515551, EPI_ISL_515552 | Hospital Municipal do Tatuape Carmino Caricchio | Instituto Adolfo Lutz, Interdiciplinary Procedures Center, Strategic Laboratory | Claudio Tavares Sacchi, Claudia Regina Gonçalves, Erica Valessa Ramos Gomes |
| EPI_ISL_515553 | Hospital Municipal Dr. Ignacio Proença de Gouvea | Instituto Adolfo Lutz, Interdiciplinary Procedures Center, Strategic Laboratory | Claudio Tavares Sacchi, Claudia Regina Gonçalves, Erica Valessa Ramos Gomes |
| EPI_ISL_515554 | Pronto Socorro Municipal de Perus | Instituto Adolfo Lutz, Interdiciplinary Procedures Center, Strategic Laboratory | Claudio Tavares Sacchi, Claudia Regina Gonçalves, Erica Valessa Ramos Gomes |
| EPI_ISL_515555 | Hospital Geral de Vila Nova Cachoeirinha | Instituto Adolfo Lutz, Interdiciplinary Procedures Center, Strategic Laboratory | Claudio Tavares Sacchi, Claudia Regina Gonçalves, Erica Valessa Ramos Gomes |
| EPI_ISL_515557 | Hospital Municipal Dr. Moysés Deutsch | Instituto Adolfo Lutz, Interdiciplinary Procedures Center, Strategic Laboratory | Claudio Tavares Sacchi, Claudia Regina Gonçalves, Erica Valessa Ramos Gomes |
| EPI_ISL_515559, EPI_ISL_515560 | Hospital Sao Paulo de Ensino da Unifesp | Instituto Adolfo Lutz, Interdiciplinary Procedures Center, Strategic Laboratory | Claudio Tavares Sacchi, Claudia Regina Gonçalves, Erica Valessa Ramos Gomes |

| | | | |
|---|---|---|---|
| EPI_ISL_515561 | Hospital Montemagno | Instituto Adolfo Lutz, Interdiciplinary Procedures Center, Strategic Laboratory | Claudio Tavares Sacchi, Claudia Regina Gonçalves, Erica Valessa Ramos Gomes |
| EPI_ISL_515562 | Hospital Municipal Doutor Alexandre Zaio | Instituto Adolfo Lutz, Interdiciplinary Procedures Center, Strategic Laboratory | Claudio Tavares Sacchi, Claudia Regina Gonçalves, Erica Valessa Ramos Gomes |
| EPI_ISL_515563 | Hospital Municipal Dr. Jose Soares Hungria | Instituto Adolfo Lutz, Interdiciplinary Procedures Center, Strategic Laboratory | Claudio Tavares Sacchi, Claudia Regina Gonçalves, Erica Valessa Ramos Gomes |
| EPI_ISL_515564 | Hosp. Municipal Prof. Dr. Alípio Corrêa Netto | Instituto Adolfo Lutz, Interdiciplinary Procedures Center, Strategic Laboratory | Claudio Tavares Sacchi, Claudia Regina Gonçalves, Erica Valessa Ramos Gomes |
| EPI_ISL_515565 | Hospital do Servidor Público Estadual Francisco Morato de Oliveira | Instituto Adolfo Lutz, Interdiciplinary Procedures Center, Strategic Laboratory | Claudio Tavares Sacchi, Claudia Regina Gonçalves, Erica Valessa Ramos Gomes |
| EPI_ISL_515566 | PS Municipal Dr Lauro Ribas Braga | Instituto Adolfo Lutz, Interdiciplinary Procedures Center, Strategic Laboratory | Claudio Tavares Sacchi, Claudia Regina Gonçalves, Erica Valessa Ramos Gomes |
| EPI_ISL_515974 | Department of Virus and Microbiological Special Diagnostics, Statens Serum Institut | Albertsen lab, Department of Chemistry and Bioscience, Aalborg University, Denmark | Danish Corona Danica Consortia |
| EPI_ISL_516553, EPI_ISL_516554, EPI_ISL_516555, EPI_ISL_516556, EPI_ISL_516557, EPI_ISL_516558, EPI_ISL_516559, EPI_ISL_516560, EPI_ISL_516561, EPI_ISL_516562, EPI_ISL_516563, EPI_ISL_516564, EPI_ISL_516565, EPI_ISL_516566, EPI_ISL_516567, EPI_ISL_516568, EPI_ISL_516569, EPI_ISL_516570, EPI_ISL_516571, EPI_ISL_516573, EPI_ISL_516576, EPI_ISL_516577, EPI_ISL_516578, EPI_ISL_516579, EPI_ISL_516580, EPI_ISL_516583, EPI_ISL_516584, EPI_ISL_516585, EPI_ISL_516586, EPI_ISL_516587, EPI_ISL_516588, EPI_ISL_516589, EPI_ISL_516590, EPI_ISL_516591, EPI_ISL_516592, EPI_ISL_516593, EPI_ISL_516594, EPI_ISL_516595, EPI_ISL_516596, EPI_ISL_516597, EPI_ISL_516598, EPI_ISL_516599, EPI_ISL_516601, EPI_ISL_516602, EPI_ISL_516604, EPI_ISL_516605, EPI_ISL_516606 | | | |
| see above | Viollier AG | Department of Biosystems Science and Engineering, ETH Zürich | Christian Beisel, Sarah Nadeau, Ivan Topolsky, Pedro Ferreira, Philipp Jablonski, Susana Posada-Céspedes, Tobias Schär, Ina Nissen, Natascha Santacroce, Elodie Burcklen, Christiane Beckmann, Maurice Redondo, Olivier Kobel, Christoph Noppen, Sophie Seidel, Noemie Santamaria de Souza, Niko Beerenwinkel, Tanja Stadler |
| EPI_ISL_522556, EPI_ISL_522557, EPI_ISL_522558, EPI_ISL_522559, EPI_ISL_522560, EPI_ISL_522561, EPI_ISL_522562, EPI_ISL_522563, EPI_ISL_522564, EPI_ISL_522565, EPI_ISL_522566, EPI_ISL_522567, EPI_ISL_522568, EPI_ISL_522569, EPI_ISL_522570, EPI_ISL_522571, EPI_ISL_522572, EPI_ISL_522573, EPI_ISL_522574, EPI_ISL_522575, EPI_ISL_522576, EPI_ISL_522577, EPI_ISL_522578, EPI_ISL_522579, EPI_ISL_522580, EPI_ISL_522581, EPI_ISL_522582, EPI_ISL_522583, EPI_ISL_522584, EPI_ISL_522585, EPI_ISL_522586, EPI_ISL_522587, EPI_ISL_522588, EPI_ISL_522589, EPI_ISL_522590, EPI_ISL_522591, EPI_ISL_522592, EPI_ISL_522593, EPI_ISL_522594, EPI_ISL_522595, EPI_ISL_522596, EPI_ISL_522597, EPI_ISL_522598, EPI_ISL_522599, EPI_ISL_522600, EPI_ISL_522601, EPI_ISL_522602, EPI_ISL_522603, EPI_ISL_522604, EPI_ISL_522605, EPI_ISL_522606, EPI_ISL_522607, EPI_ISL_522608, EPI_ISL_522609, EPI_ISL_522610, EPI_ISL_522611, EPI_ISL_522612, EPI_ISL_522613, EPI_ISL_522614, EPI_ISL_522615, EPI_ISL_522616, EPI_ISL_522617, EPI_ISL_522618, EPI_ISL_522619, EPI_ISL_522620, EPI_ISL_522621, EPI_ISL_522622, EPI_ISL_522623, EPI_ISL_522624, EPI_ISL_522625, EPI_ISL_522626, EPI_ISL_522628, EPI_ISL_522629, EPI_ISL_522630, EPI_ISL_522631, EPI_ISL_522632, EPI_ISL_522633, EPI_ISL_522635, EPI_ISL_522636, EPI_ISL_522637, EPI_ISL_522638, EPI_ISL_522639, EPI_ISL_522640, EPI_ISL_522641, EPI_ISL_522642, EPI_ISL_522643, EPI_ISL_522644, EPI_ISL_522646, EPI_ISL_522648, EPI_ISL_522649, EPI_ISL_522650, EPI_ISL_522651, EPI_ISL_522652, EPI_ISL_522653, EPI_ISL_522654, EPI_ISL_522655, EPI_ISL_522656, EPI_ISL_522657, EPI_ISL_522658, EPI_ISL_522659, EPI_ISL_522660, EPI_ISL_522661, EPI_ISL_522662, EPI_ISL_522663, EPI_ISL_522664, EPI_ISL_522665, EPI_ISL_522666, EPI_ISL_522667, EPI_ISL_522668, EPI_ISL_522669, EPI_ISL_522670, EPI_ISL_522671, EPI_ISL_522672, EPI_ISL_522673, EPI_ISL_522674, EPI_ISL_522675, EPI_ISL_522676, EPI_ISL_522677, EPI_ISL_522678, EPI_ISL_522679, EPI_ISL_522680, EPI_ISL_522681, EPI_ISL_522683, EPI_ISL_522684, EPI_ISL_522686, EPI_ISL_522687, EPI_ISL_522688, EPI_ISL_522689, EPI_ISL_522690, EPI_ISL_522691, EPI_ISL_522692, EPI_ISL_522693, EPI_ISL_522694, EPI_ISL_522695, EPI_ISL_522696, EPI_ISL_522697, EPI_ISL_522699, EPI_ISL_522700, EPI_ISL_522701, EPI_ISL_522702, EPI_ISL_522703, EPI_ISL_522704, EPI_ISL_522705, EPI_ISL_522706, EPI_ISL_522707, EPI_ISL_522708, EPI_ISL_522709, EPI_ISL_522710, EPI_ISL_522711, EPI_ISL_522712, EPI_ISL_522713, EPI_ISL_522714, EPI_ISL_522715, EPI_ISL_522716, EPI_ISL_522717, EPI_ISL_522718, EPI_ISL_522719, EPI_ISL_522720, EPI_ISL_522721, EPI_ISL_522722, EPI_ISL_522723, EPI_ISL_522724, EPI_ISL_522725, EPI_ISL_522726, EPI_ISL_522727, EPI_ISL_522729, EPI_ISL_522730, EPI_ISL_522731, EPI_ISL_522732, EPI_ISL_522733, EPI_ISL_522734, EPI_ISL_522735, EPI_ISL_522736, EPI_ISL_522737, EPI_ISL_522738, EPI_ISL_522739, EPI_ISL_522740, EPI_ISL_522741, EPI_ISL_522742, EPI_ISL_522743, EPI_ISL_522744, EPI_ISL_522745, EPI_ISL_522746, EPI_ISL_522747, EPI_ISL_522748, EPI_ISL_522749, EPI_ISL_522750, EPI_ISL_522751, EPI_ISL_522752, EPI_ISL_522753, EPI_ISL_522754, EPI_ISL_522755, EPI_ISL_522756, EPI_ISL_522757, EPI_ISL_522758, EPI_ISL_522759, EPI_ISL_522760, EPI_ISL_522761, EPI_ISL_522762, EPI_ISL_522763, EPI_ISL_522764, EPI_ISL_522765, EPI_ISL_522766, EPI_ISL_522767, EPI_ISL_522768, EPI_ISL_522769 | | | |
| see above | Royal Hobart Hospital Microbiology Department | MDU-PHL | Cooley L., van Haeften R., Seemann T., Sait M., Schultz, M.B., Sherry N. |
| EPI_ISL_523856, EPI_ISL_523858, EPI_ISL_523859, EPI_ISL_523860, EPI_ISL_523861, EPI_ISL_523862, EPI_ISL_523863, EPI_ISL_523864, EPI_ISL_523865, EPI_ISL_523866, EPI_ISL_523867, EPI_ISL_523868, EPI_ISL_523869, EPI_ISL_523871, EPI_ISL_523872, EPI_ISL_523873, EPI_ISL_523874, EPI_ISL_523875, EPI_ISL_523876, EPI_ISL_523877, EPI_ISL_523880, EPI_ISL_523881, EPI_ISL_523882, EPI_ISL_523883, EPI_ISL_523884, EPI_ISL_523886, EPI_ISL_523887, EPI_ISL_523888, EPI_ISL_523889, EPI_ISL_523890, EPI_ISL_523891, EPI_ISL_523892, EPI_ISL_523893, EPI_ISL_523894, EPI_ISL_523895, EPI_ISL_523896, EPI_ISL_523897, EPI_ISL_523901, EPI_ISL_523902, EPI_ISL_523905, EPI_ISL_523906, EPI_ISL_523907, EPI_ISL_523910, EPI_ISL_523911, EPI_ISL_523912, EPI_ISL_523913, EPI_ISL_523914, EPI_ISL_523915, EPI_ISL_523916, EPI_ISL_523917, EPI_ISL_523918, EPI_ISL_523919, EPI_ISL_523920, EPI_ISL_523921, EPI_ISL_523922, EPI_ISL_523923, EPI_ISL_523924, EPI_ISL_523925 | | | |
| see above | Viollier AG | Department of Biosystems Science and Engineering, ETH Zürich | Christian Beisel, Sarah Nadeau, Ivan Topolsky, Pedro Ferreira, Philipp Jablonski, Susana Posada-Céspedes, Tobias Schär, Ina Nissen, Natascha Santacroce, Elodie Burcklen, Christiane Beckmann, Maurice Redondo, Olivier Kobel, Christoph Noppen, Sophie Seidel, Noemie Santamaria de Souza, Niko Beerenwinkel, Tanja Stadler |
| EPI_ISL_523955 | Hospital Municipal do Tatuape Carmino Caricchio | Instituto Adolfo Lutz, Interdiciplinary Procedures Center, Strategic Laboratory | Claudio Tavares Sacchi, Claudia Regina Gonçalves, Erica Valessa Ramos Gomes |
| EPI_ISL_523956 | Hospital Regional de Assis | Instituto Adolfo Lutz, Interdiciplinary Procedures Center, Strategic Laboratory | Claudio Tavares Sacchi, Claudia Regina Gonçalves, Erica Valessa Ramos Gomes |
| EPI_ISL_523957 | Hospital Itamaraty | Instituto Adolfo Lutz, Interdiciplinary Procedures Center, Strategic Laboratory | Claudio Tavares Sacchi, Claudia Regina Gonçalves, Erica Valessa Ramos Gomes |
| EPI_ISL_523958, EPI_ISL_523959 | Pronto Socorro Municipal de Perus | Instituto Adolfo Lutz, Interdiciplinary Procedures Center, Strategic Laboratory | Claudio Tavares Sacchi, Claudia Regina Gonçalves, Erica Valessa Ramos Gomes |
| EPI_ISL_523961 | Pronto Socorro Municipal 21 de Junho | Instituto Adolfo Lutz, Interdiciplinary Procedures Center, Strategic Laboratory | Claudio Tavares Sacchi, Claudia Regina Gonçalves, Erica Valessa Ramos Gomes |
| EPI_ISL_523963 | UBS Vila Silvia | Instituto Adolfo Lutz, Interdiciplinary Procedures Center, Strategic Laboratory | Claudio Tavares Sacchi, Claudia Regina Gonçalves, Erica Valessa Ramos Gomes |
| EPI_ISL_523965 | Hospital do Servidor Público Estadual Francisco Morato de Oliveira | Instituto Adolfo Lutz, Interdiciplinary Procedures Center, Strategic Laboratory | Claudio Tavares Sacchi, Claudia Regina Gonçalves, Erica Valessa Ramos Gomes |
| EPI_ISL_523967 | Hospital Sancta Maggiore | Instituto Adolfo Lutz, Interdiciplinary Procedures Center, Strategic Laboratory | Claudio Tavares Sacchi, Claudia Regina Gonçalves, Erica Valessa Ramos Gomes |
| EPI_ISL_523969 | Hospital Sao Paulo de Ensino da Unifesp | Instituto Adolfo Lutz, Interdiciplinary Procedures Center, Strategic Laboratory | Claudio Tavares Sacchi, Claudia Regina Gonçalves, Erica Valessa Ramos Gomes |
| EPI_ISL_523970 | Conjunto Hospitalar do Mandaqui | Instituto Adolfo Lutz, Interdiciplinary Procedures Center, Strategic Laboratory | Claudio Tavares Sacchi, Claudia Regina Gonçalves, Erica Valessa Ramos Gomes |
| EPI_ISL_523971 | Hospital Geral Santa Marcelina | Instituto Adolfo Lutz, Interdiciplinary Procedures Center, Strategic Laboratory | Claudio Tavares Sacchi, Claudia Regina Gonçalves, Erica Valessa Ramos Gomes |
| EPI_ISL_523972 | Hospital do Servidor Público Estadual Francisco Morato de Oliveira | Instituto Adolfo Lutz, Interdiciplinary Procedures Center, Strategic Laboratory | Claudio Tavares Sacchi, Claudia Regina Gonçalves, Erica Valessa Ramos Gomes |
| EPI_ISL_523973 | PS Municipal Dona Maria Antonieta Ferreira de Barros | Instituto Adolfo Lutz, Interdiciplinary Procedures Center, Strategic Laboratory | Claudio Tavares Sacchi, Claudia Regina Gonçalves, Erica Valessa Ramos Gomes |
| EPI_ISL_523974 | Hospital Municipal do Tatuape Carmino Caricchio | Instituto Adolfo Lutz, Interdiciplinary Procedures Center, Strategic Laboratory | Claudio Tavares Sacchi, Claudia Regina Gonçalves, Erica Valessa Ramos Gomes |
| EPI_ISL_523975 | UPA Tito Lopes | Instituto Adolfo Lutz, Interdiciplinary Procedures Center, Strategic Laboratory | Claudio Tavares Sacchi, Claudia Regina Gonçalves, Erica Valessa Ramos Gomes |
| EPI_ISL_523976 | Hospital Municipal do Tatuape Carmino Caricchio | Instituto Adolfo Lutz, Interdiciplinary Procedures Center, Strategic Laboratory | Claudio Tavares Sacchi, Claudia Regina Gonçalves, Erica Valessa Ramos Gomes |
| EPI_ISL_523977 | Hosp. Municipal Prof. Dr. Alípio Corrêa Netto | Instituto Adolfo Lutz, Interdiciplinary Procedures Center, | Claudio Tavares Sacchi, Claudia Regina Gonçalves, Erica Valessa Ramos Gomes |

| | | | |
|---|---|---|---|
| | | Strategic Laboratory | |
| EPI_ISL_523978 | Hospital do Servidor Público Estadual Francisco Morato de Oliveira | Instituto Adolfo Lutz, Interdiciplinary Procedures Center, Strategic Laboratory | Claudio Tavares Sacchi, Claudia Regina Gonçalves, Erica Valessa Ramos Gomes |
| EPI_ISL_523980 | UPA Tito Lopes | Instituto Adolfo Lutz, Interdiciplinary Procedures Center, Strategic Laboratory | Claudio Tavares Sacchi, Claudia Regina Gonçalves, Erica Valessa Ramos Gomes |
| EPI_ISL_523981 | Hospital Sao Paulo de Ensino da Unifesp | Instituto Adolfo Lutz, Interdiciplinary Procedures Center, Strategic Laboratory | Claudio Tavares Sacchi, Claudia Regina Gonçalves, Erica Valessa Ramos Gomes |
| EPI_ISL_523982 | Hospital do Servidor Público Estadual Francisco Morato de Oliveira | Instituto Adolfo Lutz, Interdiciplinary Procedures Center, Strategic Laboratory | Claudio Tavares Sacchi, Claudia Regina Gonçalves, Erica Valessa Ramos Gomes |
| EPI_ISL_523983 | UPA Campo Limpo | Instituto Adolfo Lutz, Interdiciplinary Procedures Center, Strategic Laboratory | Claudio Tavares Sacchi, Claudia Regina Gonçalves, Erica Valessa Ramos Gomes |
| EPI_ISL_523984 | Ama Dr Jose Soares Hungria | Instituto Adolfo Lutz, Interdiciplinary Procedures Center, Strategic Laboratory | Claudio Tavares Sacchi, Claudia Regina Gonçalves, Erica Valessa Ramos Gomes |
| EPI_ISL_523985 | Hospital Municipal Dr. Benedicto Montenegro | Instituto Adolfo Lutz, Interdiciplinary Procedures Center, Strategic Laboratory | Claudio Tavares Sacchi, Claudia Regina Gonçalves, Erica Valessa Ramos Gomes |
| EPI_ISL_523986 | Ama Dr Jose Soares Hungria | Instituto Adolfo Lutz, Interdiciplinary Procedures Center, Strategic Laboratory | Claudio Tavares Sacchi, Claudia Regina Gonçalves, Erica Valessa Ramos Gomes |
| EPI_ISL_523988 | Hospital Sao Paulo de Ensino da Unifesp | Instituto Adolfo Lutz, Interdiciplinary Procedures Center, Strategic Laboratory | Claudio Tavares Sacchi, Claudia Regina Gonçalves, Erica Valessa Ramos Gomes |
| EPI_ISL_523989 | AMA Jardim Joamar | Instituto Adolfo Lutz, Interdiciplinary Procedures Center, Strategic Laboratory | Claudio Tavares Sacchi, Claudia Regina Gonçalves, Erica Valessa Ramos Gomes |
| EPI_ISL_523990 | AMA Jardim Peri | Instituto Adolfo Lutz, Interdiciplinary Procedures Center, Strategic Laboratory | Claudio Tavares Sacchi, Claudia Regina Gonçalves, Erica Valessa Ramos Gomes |
| EPI_ISL_523991, EPI_ISL_523992 | Hospital Municipal Carmen Prudente | Instituto Adolfo Lutz, Interdiciplinary Procedures Center, Strategic Laboratory | Claudio Tavares Sacchi, Claudia Regina Gonçalves, Erica Valessa Ramos Gomes |
| EPI_ISL_523993 | UPA Campo Limpo | Instituto Adolfo Lutz, Interdiciplinary Procedures Center, Strategic Laboratory | Claudio Tavares Sacchi, Claudia Regina Gonçalves, Erica Valessa Ramos Gomes |
| EPI_ISL_524462 | Hospital Metropolitano | Instituto Adolfo Lutz, Interdiciplinary Procedures Center, Strategic Laboratory | Claudio Tavares Sacchi, Claudia Regina Gonçalves, Erica Valessa Ramos Gomes |
| EPI_ISL_524463 | Hospital Regional de Cotia | Instituto Adolfo Lutz, Interdiciplinary Procedures Center, Strategic Laboratory | Claudio Tavares Sacchi, Claudia Regina Gonçalves, Erica Valessa Ramos Gomes |
| EPI_ISL_524464 | Santa Casa de Santa Isabel | Instituto Adolfo Lutz, Interdiciplinary Procedures Center, Strategic Laboratory | Claudio Tavares Sacchi, Claudia Regina Gonçalves, Erica Valessa Ramos Gomes |
| EPI_ISL_524465 | PS Municipal Dr. Caetano Virgílio Neto | Instituto Adolfo Lutz, Interdiciplinary Procedures Center, Strategic Laboratory | Claudio Tavares Sacchi, Claudia Regina Gonçalves, Erica Valessa Ramos Gomes |
| EPI_ISL_524466 | PS Municipal Dr Lauro Ribas Braga | Instituto Adolfo Lutz, Interdiciplinary Procedures Center, Strategic Laboratory | Claudio Tavares Sacchi, Claudia Regina Gonçalves, Erica Valessa Ramos Gomes |
| EPI_ISL_524467 | Hospital Municipal Dr. Moysés Deutsch | Instituto Adolfo Lutz, Interdiciplinary Procedures Center, Strategic Laboratory | Claudio Tavares Sacchi, Claudia Regina Gonçalves, Erica Valessa Ramos Gomes |
| EPI_ISL_524468 | Hospital Municipal Vereador Jose Storopolli | Instituto Adolfo Lutz, Interdiciplinary Procedures Center, Strategic Laboratory | Claudio Tavares Sacchi, Claudia Regina Gonçalves, Erica Valessa Ramos Gomes |
| EPI_ISL_524469 | Santa Casa de Misericordia de Sao Paulo | Instituto Adolfo Lutz, Interdiciplinary Procedures Center, Strategic Laboratory | Claudio Tavares Sacchi, Claudia Regina Gonçalves, Erica Valessa Ramos Gomes |
| EPI_ISL_524470 | Hospital do Servidor Público Estadual Francisco Morato de Oliveira | Instituto Adolfo Lutz, Interdiciplinary Procedures Center, Strategic Laboratory | Claudio Tavares Sacchi, Claudia Regina Gonçalves, Erica Valessa Ramos Gomes |
| EPI_ISL_527856 | Hospital Municipal Prof. Waldomiro de Paula | Instituto Adolfo Lutz, Interdiciplinary Procedures Center, Strategic Laboratory | Claudio Tavares Sacchi, Claudia Regina Gonçalves, Erica Valessa Ramos Gomes |
| EPI_ISL_527857 | Hospital Regional Vale do Ribeira | Instituto Adolfo Lutz, Interdiciplinary Procedures Center, Strategic Laboratory | Claudio Tavares Sacchi, Claudia Regina Gonçalves, Erica Valessa Ramos Gomes |
| EPI_ISL_527858 | Pronto Atendimento Sancta Maggiore Jardim Paulista | Instituto Adolfo Lutz, Interdiciplinary Procedures Center, Strategic Laboratory | Claudio Tavares Sacchi, Claudia Regina Gonçalves, Erica Valessa Ramos Gomes |
| EPI_ISL_527859 | Hospital Municipal Vereador Jose Storopolli | Instituto Adolfo Lutz, Interdiciplinary Procedures Center, Strategic Laboratory | Claudio Tavares Sacchi, Claudia Regina Gonçalves, Erica Valessa Ramos Gomes |
| EPI_ISL_527860 | Hospital Municipal de Parelheiros Josanias Castanha Braga | Instituto Adolfo Lutz, Interdiciplinary Procedures Center, Strategic Laboratory | Claudio Tavares Sacchi, Claudia Regina Gonçalves, Erica Valessa Ramos Gomes |
| EPI_ISL_527862 | Hospital Municipal de Urgência | Instituto Adolfo Lutz, Interdiciplinary Procedures Center, Strategic Laboratory | Claudio Tavares Sacchi, Claudia Regina Gonçalves, Erica Valessa Ramos Gomes |
| EPI_ISL_527863 | Hospital Municipal do Tatuape Carmino Caricchio | Instituto Adolfo Lutz, Interdiciplinary Procedures Center, Strategic Laboratory | Claudio Tavares Sacchi, Claudia Regina Gonçalves, Erica Valessa Ramos Gomes |
| EPI_ISL_527864 | Hospital e Pronto Socorro Comunitário Vila Iolanda | Instituto Adolfo Lutz, Interdiciplinary Procedures Center, Strategic Laboratory | Claudio Tavares Sacchi, Claudia Regina Gonçalves, Erica Valessa Ramos Gomes |
| EPI_ISL_527865 | Hospital e Maternidade São Cristóvão | Instituto Adolfo Lutz, Interdiciplinary Procedures Center, Strategic Laboratory | Claudio Tavares Sacchi, Claudia Regina Gonçalves, Erica Valessa Ramos Gomes |
| EPI_ISL_527867 | Pronto Socorro Municipal - Balneario São José | Instituto Adolfo Lutz, Interdiciplinary Procedures Center, Strategic Laboratory | Claudio Tavares Sacchi, Claudia Regina Gonçalves, Erica Valessa Ramos Gomes |
| EPI_ISL_527868 | Hospital e Maternidade do Braz | Instituto Adolfo Lutz, Interdiciplinary Procedures Center, Strategic Laboratory | Claudio Tavares Sacchi, Claudia Regina Gonçalves, Erica Valessa Ramos Gomes |
| EPI_ISL_527869 | Hospital Municipal Carmen Prudente | Instituto Adolfo Lutz, Interdiciplinary Procedures Center, Strategic Laboratory | Claudio Tavares Sacchi, Claudia Regina Gonçalves, Erica Valessa Ramos Gomes |
| EPI_ISL_527870 | Hospital Municipal Mário Gatti | Instituto Adolfo Lutz, Interdiciplinary Procedures Center, | Claudio Tavares Sacchi, Claudia Regina Gonçalves, Erica Valessa Ramos Gomes |

| | | | |
|---|---|---|---|
| | | Strategic Laboratory | |
| EPI_ISL_534311 | UPA III 26 de Agosto | Instituto Adolfo Lutz, Interdiciplinary Procedures Center, Strategic Laboratory | Claudio Tavares Sacchi, Claudia Regina Gonçalves, Erica Valessa Ramos Gomes |
| EPI_ISL_534312 | Distrito Sanitario Sul | Instituto Adolfo Lutz, Interdiciplinary Procedures Center, Strategic Laboratory | Claudio Tavares Sacchi, Claudia Regina Gonçalves, Erica Valessa Ramos Gomes |
| EPI_ISL_534313 | Hospital da Sta Casa de Sto Amaro | Instituto Adolfo Lutz, Interdiciplinary Procedures Center, Strategic Laboratory | Claudio Tavares Sacchi, Claudia Regina Gonçalves, Erica Valessa Ramos Gomes |
| EPI_ISL_534314 | Hospital Universitario da USP de SP | Instituto Adolfo Lutz, Interdiciplinary Procedures Center, Strategic Laboratory | Claudio Tavares Sacchi, Claudia Regina Gonçalves, Erica Valessa Ramos Gomes |
| EPI_ISL_534315 | Serviço de Verificação de Óbitos SVO Guaruulhos | Instituto Adolfo Lutz, Interdiciplinary Procedures Center, Strategic Laboratory | Claudio Tavares Sacchi, Claudia Regina Gonçalves, Erica Valessa Ramos Gomes |
| EPI_ISL_534316 | OS Mun Santana Lauro Ribas Braga | Instituto Adolfo Lutz, Interdiciplinary Procedures Center, Strategic Laboratory | Claudio Tavares Sacchi, Claudia Regina Gonçalves, Erica Valessa Ramos Gomes |
| EPI_ISL_534317 | Hospital Geral de Itapevi | Instituto Adolfo Lutz, Interdiciplinary Procedures Center, Strategic Laboratory | Claudio Tavares Sacchi, Claudia Regina Gonçalves, Erica Valessa Ramos Gomes |
| EPI_ISL_534318 | Hospital Municipal Antonio Giglio | Instituto Adolfo Lutz, Interdiciplinary Procedures Center, Strategic Laboratory | Claudio Tavares Sacchi, Claudia Regina Gonçalves, Erica Valessa Ramos Gomes |
| EPI_ISL_534319, EPI_ISL_534320 | Hospital do Serv Pub ESTAFCO Morato de Oliveira | Instituto Adolfo Lutz, Interdiciplinary Procedures Center, Strategic Laboratory | Claudio Tavares Sacchi, Claudia Regina Gonçalves, Erica Valessa Ramos Gomes |
| EPI_ISL_534321 | PS e Maternidade Nair Fonseca Leitao Arantes | Instituto Adolfo Lutz, Interdiciplinary Procedures Center, Strategic Laboratory | Claudio Tavares Sacchi, Claudia Regina Gonçalves, Erica Valessa Ramos Gomes |
| EPI_ISL_534322 | PS Mun Julio Tupy | Instituto Adolfo Lutz, Interdiciplinary Procedures Center, Strategic Laboratory | Claudio Tavares Sacchi, Claudia Regina Gonçalves, Erica Valessa Ramos Gomes |
| EPI_ISL_534323 | Hospital e Pronto Socorro Comunitario Vila Yolanda | Instituto Adolfo Lutz, Interdiciplinary Procedures Center, Strategic Laboratory | Claudio Tavares Sacchi, Claudia Regina Gonçalves, Erica Valessa Ramos Gomes |
| EPI_ISL_534324 | Hospital Mun Ver Jose Storopolli | Instituto Adolfo Lutz, Interdiciplinary Procedures Center, Strategic Laboratory | Claudio Tavares Sacchi, Claudia Regina Gonçalves, Erica Valessa Ramos Gomes |
| EPI_ISL_534325 | Unidade de Vigilancia em Saude de Guarulhos | Instituto Adolfo Lutz, Interdiciplinary Procedures Center, Strategic Laboratory | Claudio Tavares Sacchi, Claudia Regina Gonçalves, Erica Valessa Ramos Gomes |
| EPI_ISL_534326 | Notre Dame Intermedica Saude AS | Instituto Adolfo Lutz, Interdiciplinary Procedures Center, Strategic Laboratory | Claudio Tavares Sacchi, Claudia Regina Gonçalves, Erica Valessa Ramos Gomes |
| EPI_ISL_535585, EPI_ISL_535586, EPI_ISL_535587, EPI_ISL_535588, EPI_ISL_535589, EPI_ISL_535590, EPI_ISL_535591, EPI_ISL_535592, EPI_ISL_535593, EPI_ISL_535596, EPI_ISL_535597, EPI_ISL_535598, EPI_ISL_535599, EPI_ISL_535600, EPI_ISL_535601, EPI_ISL_535602, EPI_ISL_535603, EPI_ISL_535604, EPI_ISL_535605, EPI_ISL_535606, EPI_ISL_535607, EPI_ISL_535608, EPI_ISL_535610, EPI_ISL_535611, EPI_ISL_535612, EPI_ISL_535613, EPI_ISL_535614, EPI_ISL_535616, EPI_ISL_535617, EPI_ISL_535618, EPI_ISL_535619, EPI_ISL_535620, EPI_ISL_535621, EPI_ISL_535622, EPI_ISL_535624, EPI_ISL_535625, EPI_ISL_535626, EPI_ISL_535627, EPI_ISL_535628, EPI_ISL_535629, EPI_ISL_535630, EPI_ISL_535631, EPI_ISL_535632, EPI_ISL_535633, EPI_ISL_535634, EPI_ISL_535635, EPI_ISL_535636, EPI_ISL_535637, EPI_ISL_535638, EPI_ISL_535639, EPI_ISL_535640, EPI_ISL_535641, EPI_ISL_535642, EPI_ISL_535644, EPI_ISL_535646, EPI_ISL_535647, EPI_ISL_535648, EPI_ISL_535649, EPI_ISL_539341, EPI_ISL_539342, EPI_ISL_539343, EPI_ISL_539345, EPI_ISL_539347, EPI_ISL_539349, EPI_ISL_539350, EPI_ISL_539351, EPI_ISL_539353, EPI_ISL_539354, EPI_ISL_539356, EPI_ISL_539357, EPI_ISL_539358, EPI_ISL_539359, EPI_ISL_539360, EPI_ISL_539361, EPI_ISL_539362, EPI_ISL_539363, EPI_ISL_539364, EPI_ISL_539365, EPI_ISL_539366, EPI_ISL_539367, EPI_ISL_539368, EPI_ISL_539369, EPI_ISL_539370, EPI_ISL_539371, EPI_ISL_539372, EPI_ISL_539373, EPI_ISL_539374, EPI_ISL_539375, EPI_ISL_539376, EPI_ISL_539377, EPI_ISL_539380, EPI_ISL_539381, EPI_ISL_539383, EPI_ISL_539386, EPI_ISL_539387, EPI_ISL_539388, EPI_ISL_539390, EPI_ISL_539392, EPI_ISL_539393, EPI_ISL_539394, EPI_ISL_539395, EPI_ISL_539396, EPI_ISL_539397, EPI_ISL_539398, EPI_ISL_539399, EPI_ISL_539400, EPI_ISL_539401, EPI_ISL_539402, EPI_ISL_539403, EPI_ISL_539404, EPI_ISL_539405, EPI_ISL_539406, EPI_ISL_539407, EPI_ISL_539408, EPI_ISL_539409, EPI_ISL_539411, EPI_ISL_539412, EPI_ISL_539413, EPI_ISL_539414, EPI_ISL_539415, EPI_ISL_539416, EPI_ISL_539417, EPI_ISL_539419, EPI_ISL_539421, EPI_ISL_539423, EPI_ISL_539424, EPI_ISL_539425, EPI_ISL_539426, EPI_ISL_539427, EPI_ISL_539428, EPI_ISL_539431, EPI_ISL_539432, EPI_ISL_539433, EPI_ISL_539435, EPI_ISL_539436, EPI_ISL_539437, EPI_ISL_539438, EPI_ISL_539440, EPI_ISL_539441, EPI_ISL_539442, EPI_ISL_539443, EPI_ISL_539444, EPI_ISL_539445, EPI_ISL_539447, EPI_ISL_539448, EPI_ISL_539449, EPI_ISL_539450, EPI_ISL_539451, EPI_ISL_539452, EPI_ISL_539453, EPI_ISL_539455, EPI_ISL_539456, EPI_ISL_539457, EPI_ISL_539458, EPI_ISL_539459, EPI_ISL_539460, EPI_ISL_539461, EPI_ISL_539462, EPI_ISL_539463, EPI_ISL_539464, EPI_ISL_539465, EPI_ISL_539467, EPI_ISL_539468, EPI_ISL_539469, EPI_ISL_539470, EPI_ISL_539471, EPI_ISL_539472, EPI_ISL_539475, EPI_ISL_539476, EPI_ISL_539477, EPI_ISL_539478, EPI_ISL_539479, EPI_ISL_539480, EPI_ISL_539481, EPI_ISL_539482, EPI_ISL_541401, EPI_ISL_541402, EPI_ISL_541403, EPI_ISL_541404, EPI_ISL_541405, EPI_ISL_541406, EPI_ISL_541407, EPI_ISL_541408, EPI_ISL_541409, EPI_ISL_541410, EPI_ISL_541411, EPI_ISL_541412, EPI_ISL_541413, EPI_ISL_541414, EPI_ISL_541415, EPI_ISL_541416, EPI_ISL_541417, EPI_ISL_541418, EPI_ISL_541419, EPI_ISL_541420, EPI_ISL_541421, EPI_ISL_541422, EPI_ISL_541423, EPI_ISL_541424, EPI_ISL_541425, EPI_ISL_541426, EPI_ISL_541427, EPI_ISL_541428, EPI_ISL_541429, EPI_ISL_541430, EPI_ISL_541431, EPI_ISL_541432, EPI_ISL_541433, EPI_ISL_541434, EPI_ISL_541435, EPI_ISL_541436, EPI_ISL_541437, EPI_ISL_541438, EPI_ISL_541439, EPI_ISL_541440, EPI_ISL_541441, EPI_ISL_541442, EPI_ISL_541443, EPI_ISL_541444, EPI_ISL_541445, EPI_ISL_541446, EPI_ISL_541447, EPI_ISL_541448, EPI_ISL_541449, EPI_ISL_541450, EPI_ISL_541451, EPI_ISL_541452, EPI_ISL_541453, EPI_ISL_541454, EPI_ISL_541455, EPI_ISL_541456, EPI_ISL_541457, EPI_ISL_541458, EPI_ISL_541459, EPI_ISL_541460, EPI_ISL_541461, EPI_ISL_541462, EPI_ISL_541463, EPI_ISL_541464, EPI_ISL_541465, EPI_ISL_541466, EPI_ISL_541467, EPI_ISL_541468, EPI_ISL_541469, EPI_ISL_541470, EPI_ISL_541471, EPI_ISL_541472, EPI_ISL_541473, EPI_ISL_541474, EPI_ISL_541475, EPI_ISL_541476, EPI_ISL_541477, EPI_ISL_541478, EPI_ISL_541479, EPI_ISL_541480, EPI_ISL_541481, EPI_ISL_541482, EPI_ISL_541483, EPI_ISL_541484, EPI_ISL_541485, EPI_ISL_541486, EPI_ISL_541487, EPI_ISL_541488, EPI_ISL_541489, EPI_ISL_541490, EPI_ISL_541491, EPI_ISL_541492, EPI_ISL_541493, EPI_ISL_541494, EPI_ISL_541495, EPI_ISL_541496, EPI_ISL_541497, EPI_ISL_541498, EPI_ISL_541499, EPI_ISL_541500, EPI_ISL_541501, EPI_ISL_541502, EPI_ISL_541503, EPI_ISL_541504, EPI_ISL_541505, EPI_ISL_541506, EPI_ISL_541507, EPI_ISL_541508, EPI_ISL_541509, EPI_ISL_541511, EPI_ISL_541514, EPI_ISL_541515, EPI_ISL_541518, EPI_ISL_541535, EPI_ISL_541537 | | | |
| see above | Viollier AG | Department of Biosystems Science and Engineering, ETH Zürich | Christian Beisel, Sarah Nadeau, Ivan Topolsky, Pedro Ferreira, Philipp Jablonski, Susana Posada-Céspedes, Tobias Schär, Ina Nissen, Natascha Santacroce, Elodie Burcklen, Christiane Beckmann, Maurice Redondo, Olivier Kobel, Christoph Noppen, Sophie Seidel, Noemie Santamaria de Souza, Niko Beerenwinkel, Tanja Stadler |
| EPI_ISL_542098, EPI_ISL_542099, EPI_ISL_542100, EPI_ISL_542101, EPI_ISL_542102, EPI_ISL_542103, EPI_ISL_542104, EPI_ISL_542105, EPI_ISL_542106, EPI_ISL_542107, EPI_ISL_542108, EPI_ISL_542109, EPI_ISL_542110, EPI_ISL_542111, EPI_ISL_542112, EPI_ISL_542113, EPI_ISL_542114, EPI_ISL_542115, EPI_ISL_542116, EPI_ISL_542117, EPI_ISL_542118, EPI_ISL_542119, EPI_ISL_542120, EPI_ISL_542121, EPI_ISL_542122, EPI_ISL_542123, EPI_ISL_542124, EPI_ISL_542125, EPI_ISL_542126, EPI_ISL_542127, EPI_ISL_542128, EPI_ISL_542129, EPI_ISL_542130, EPI_ISL_542131, EPI_ISL_542132, EPI_ISL_542133, EPI_ISL_542134, EPI_ISL_542135, EPI_ISL_542136, EPI_ISL_542137, EPI_ISL_542138, EPI_ISL_542139, EPI_ISL_542140, EPI_ISL_542141, EPI_ISL_542142, EPI_ISL_542143, EPI_ISL_542144, EPI_ISL_542145, EPI_ISL_542146, EPI_ISL_542147, EPI_ISL_542148, EPI_ISL_542149, EPI_ISL_542150, EPI_ISL_542151, EPI_ISL_542152, EPI_ISL_542153, EPI_ISL_542154, EPI_ISL_542155, EPI_ISL_542156, EPI_ISL_542157, EPI_ISL_542158, EPI_ISL_542159, EPI_ISL_542160, EPI_ISL_542161, EPI_ISL_542162, EPI_ISL_542163, EPI_ISL_542164, EPI_ISL_542165, EPI_ISL_542166, EPI_ISL_542167, EPI_ISL_542168, EPI_ISL_542169, EPI_ISL_542170, EPI_ISL_542171, EPI_ISL_542172, EPI_ISL_542173, EPI_ISL_542174, EPI_ISL_542175, EPI_ISL_542176, EPI_ISL_542177, EPI_ISL_542178, EPI_ISL_542179, EPI_ISL_542180, EPI_ISL_542181, EPI_ISL_542182, EPI_ISL_542183, EPI_ISL_542184, EPI_ISL_542185, EPI_ISL_542186, EPI_ISL_542187, EPI_ISL_542188, EPI_ISL_542189, EPI_ISL_542190, EPI_ISL_542191, EPI_ISL_542192, EPI_ISL_542193, EPI_ISL_542194, EPI_ISL_542195, EPI_ISL_542196, EPI_ISL_542197, EPI_ISL_542198, EPI_ISL_542199, EPI_ISL_542200, EPI_ISL_542201, EPI_ISL_542202, EPI_ISL_542203, EPI_ISL_542204, EPI_ISL_542205, EPI_ISL_542206, EPI_ISL_542207, EPI_ISL_542208, EPI_ISL_542209, EPI_ISL_542210, EPI_ISL_542211, EPI_ISL_542212, EPI_ISL_542213, EPI_ISL_542214, EPI_ISL_542215, EPI_ISL_542216, EPI_ISL_542217, EPI_ISL_542218, EPI_ISL_542219, EPI_ISL_542220, EPI_ISL_542221, EPI_ISL_542222, EPI_ISL_542223, EPI_ISL_542224, EPI_ISL_542225, EPI_ISL_542226, EPI_ISL_542227, EPI_ISL_542228, EPI_ISL_542229, EPI_ISL_542230, EPI_ISL_542231, EPI_ISL_542232, EPI_ISL_542233, EPI_ISL_542234, EPI_ISL_542235, EPI_ISL_542236, EPI_ISL_542237, EPI_ISL_542238, EPI_ISL_542239, EPI_ISL_542240, EPI_ISL_542241, EPI_ISL_542242, EPI_ISL_542243, EPI_ISL_542244, EPI_ISL_542245, EPI_ISL_542246, EPI_ISL_542247, EPI_ISL_542248, EPI_ISL_542249, EPI_ISL_542250, EPI_ISL_542251, EPI_ISL_542252, EPI_ISL_542253, EPI_ISL_542254, EPI_ISL_542255, EPI_ISL_542256, EPI_ISL_542257, EPI_ISL_542258, EPI_ISL_542259, EPI_ISL_542260, EPI_ISL_542261, EPI_ISL_542262, EPI_ISL_542263, EPI_ISL_542264, EPI_ISL_542265, EPI_ISL_542266, EPI_ISL_542267, EPI_ISL_542268, EPI_ISL_542269, EPI_ISL_542270, EPI_ISL_542271, EPI_ISL_542272, EPI_ISL_542273, EPI_ISL_542274, EPI_ISL_542275, EPI_ISL_542276, EPI_ISL_542277 | | | |
| see above | ASST GOM Niguarda | Dep. Of Oncology and Hemato-Oncology University of Milan | Claudia Alteri, Valeria Cento, Antonio Piralla, Valentino Costabile, Monica Tallarita, Luna Colagrossi, Silvia Renica, Federica Giardina, Federica Novazzi, Stefano Gaiarsa, Elisa Matarazzo, Maria Antonello, Chiara Vismara, Roberto Fumagalli, Oscar Massimiliano Epis, Massimo Puoti, Carlo Federico Perno, Fausto Baldanti |
| EPI_ISL_542278, EPI_ISL_542279, EPI_ISL_542280, EPI_ISL_542281, EPI_ISL_542282, EPI_ISL_542283, EPI_ISL_542284, EPI_ISL_542285, EPI_ISL_542286, EPI_ISL_542287, EPI_ISL_542288, EPI_ISL_542289, EPI_ISL_542290, EPI_ISL_542291, EPI_ISL_542292, EPI_ISL_542293, EPI_ISL_542294, EPI_ISL_542295, EPI_ISL_542296, EPI_ISL_542297, EPI_ISL_542298, EPI_ISL_542299, EPI_ISL_542300, EPI_ISL_542301, EPI_ISL_542302, EPI_ISL_542303, EPI_ISL_542304, EPI_ISL_542305, EPI_ISL_542306, EPI_ISL_542307, EPI_ISL_542308, EPI_ISL_542309, EPI_ISL_542310, EPI_ISL_542311, EPI_ISL_542312, EPI_ISL_542313, EPI_ISL_542314, EPI_ISL_542315, EPI_ISL_542316, EPI_ISL_542317, EPI_ISL_542318, EPI_ISL_542319, EPI_ISL_542320, EPI_ISL_542321, EPI_ISL_542322, EPI_ISL_542323, EPI_ISL_542324, EPI_ISL_542325, EPI_ISL_542326, EPI_ISL_542327, EPI_ISL_542328, EPI_ISL_542329, EPI_ISL_542330, EPI_ISL_542331, EPI_ISL_542332, EPI_ISL_542333, EPI_ISL_542334, EPI_ISL_542335, EPI_ISL_542336, EPI_ISL_542337, EPI_ISL_542338, EPI_ISL_542339, EPI_ISL_542340, EPI_ISL_542341, EPI_ISL_542342, EPI_ISL_542343, EPI_ISL_542344, EPI_ISL_542345, EPI_ISL_542346, EPI_ISL_542347, EPI_ISL_542348, EPI_ISL_542349, EPI_ISL_542350, EPI_ISL_542351, EPI_ISL_542352, EPI_ISL_542353, EPI_ISL_542354, EPI_ISL_542355, EPI_ISL_542356, EPI_ISL_542357, EPI_ISL_542358, EPI_ISL_542359, EPI_ISL_542360, EPI_ISL_542361, EPI_ISL_542362, EPI_ISL_542363, EPI_ISL_542364, EPI_ISL_542365, EPI_ISL_542366, EPI_ISL_542367, EPI_ISL_542368, EPI_ISL_542369, EPI_ISL_542370, EPI_ISL_542371, EPI_ISL_542372, EPI_ISL_542373, EPI_ISL_542374, EPI_ISL_542375, EPI_ISL_542376, EPI_ISL_542377, EPI_ISL_542378, EPI_ISL_542379, EPI_ISL_542380, EPI_ISL_542381, EPI_ISL_542382, EPI_ISL_542383, EPI_ISL_542384, EPI_ISL_542385, EPI_ISL_542386, EPI_ISL_542387, EPI_ISL_542388, EPI_ISL_542389, EPI_ISL_542390, EPI_ISL_542391, EPI_ISL_542392, EPI_ISL_542393, EPI_ISL_542394, EPI_ISL_542395, EPI_ISL_542396, EPI_ISL_542397, EPI_ISL_542398, EPI_ISL_542399 | | | |
| see above | San Matteo Hospital Pavia | Dep. Of Oncology and Hemato-Oncology University of Milan | Claudia Alteri, Valeria Cento, Antonio Piralla, Valentino Costabile, Monica Tallarita, Luna Colagrossi, Silvia Renica, Federica Giardina, Federica Novazzi, Stefano Gaiarsa, Elisa Matarazzo, Maria Antonello, Chiara Vismara, Roberto Fumagalli, Oscar Massimiliano Epis, Massimo Puoti, Carlo Federico Perno, Fausto Baldanti |
| EPI_ISL_542400, EPI_ISL_542401, EPI_ISL_542402, EPI_ISL_542403, EPI_ISL_542404, EPI_ISL_542405, EPI_ISL_542406, EPI_ISL_542407, EPI_ISL_542408, EPI_ISL_542409, EPI_ISL_542410, EPI_ISL_542411, EPI_ISL_542412, EPI_ISL_542413, EPI_ISL_542414, EPI_ISL_542415, EPI_ISL_542416, EPI_ISL_542417, EPI_ISL_542418, EPI_ISL_542419, EPI_ISL_542420, EPI_ISL_542421, EPI_ISL_542422, EPI_ISL_542423, EPI_ISL_542424, EPI_ISL_542425, EPI_ISL_542426, EPI_ISL_542427, EPI_ISL_542428, EPI_ISL_542429, EPI_ISL_542430, EPI_ISL_542431, EPI_ISL_542432, EPI_ISL_542433, EPI_ISL_542434, EPI_ISL_542435, | | | |

| | | | |
|---|---|---|---|
| EPI_ISL_542436, EPI_ISL_542437, EPI_ISL_542438, EPI_ISL_542439, EPI_ISL_542440, EPI_ISL_542441, EPI_ISL_542442, EPI_ISL_542443 | | | |
| see above | ASST GOM Niguarda | Dep. Of Oncology and Hemato-Oncology University of Milan | Claudia Alteri, Valeria Cento, Antonio Piralla, Valentino Costabile, Monica Tallarita, Luna Colagrossi, Silvia Renica, Federica Giardina, Federica Novazzi, Stefano Gaiarsa, Elisa Matarazzo, Maria Antonello, Chiara Vismara, Roberto Fumagalli, Oscar Massimiliano Epis, Massimo Puoti, Carlo Federico Perno, Fausto Baldanti |
| EPI_ISL_547570 | PS Municipal Dr Augusto Gomes de Mattos | Instituto Adolfo Lutz, Interdiciplinary Procedures Center, Strategic Laboratory | Claudio Tavares Sacchi, Claudia Regina Gonçalves, Erica Valessa Ramos Gomes, Karoline Rodrigues Campos |
| EPI_ISL_547571 | Hospital Municipal Antônio Giglio | Instituto Adolfo Lutz, Interdiciplinary Procedures Center, Strategic Laboratory | Claudio Tavares Sacchi, Claudia Regina Gonçalves, Erica Valessa Ramos Gomes, Karoline Rodrigues Campos |
| EPI_ISL_547573 | Vigilância em Saúde de Cajamar | Instituto Adolfo Lutz, Interdiciplinary Procedures Center, Strategic Laboratory | Claudio Tavares Sacchi, Claudia Regina Gonçalves, Erica Valessa Ramos Gomes, Karoline Rodrigues Campos |
| EPI_ISL_547574 | Hospital Universitario da USP | Instituto Adolfo Lutz, Interdiciplinary Procedures Center, Strategic Laboratory | Claudio Tavares Sacchi, Claudia Regina Gonçalves, Erica Valessa Ramos Gomes, Karoline Rodrigues Campos |
| EPI_ISL_547575 | SVO Jundiaí | Instituto Adolfo Lutz, Interdiciplinary Procedures Center, Strategic Laboratory | Claudio Tavares Sacchi, Claudia Regina Gonçalves, Erica Valessa Ramos Gomes, Karoline Rodrigues Campos |
| EPI_ISL_547576 | Secretaria Municipal de Saúde | Instituto Adolfo Lutz, Interdiciplinary Procedures Center, Strategic Laboratory | Claudio Tavares Sacchi, Claudia Regina Gonçalves, Erica Valessa Ramos Gomes, Karoline Rodrigues Campos |
| EPI_ISL_547577 | Hospital e Maternidade Nossa Senhora das Graças | Instituto Adolfo Lutz, Interdiciplinary Procedures Center, Strategic Laboratory | Claudio Tavares Sacchi, Claudia Regina Gonçalves, Erica Valessa Ramos Gomes, Karoline Rodrigues Campos |
| EPI_ISL_547578 | Hospital Doutor Domingos Leonardo Cerávolo | Instituto Adolfo Lutz, Interdiciplinary Procedures Center, Strategic Laboratory | Claudio Tavares Sacchi, Claudia Regina Gonçalves, Erica Valessa Ramos Gomes, Karoline Rodrigues Campos |
| EPI_ISL_547579 | Santa Casa de Misericórdia de Araçatuba | Instituto Adolfo Lutz, Interdiciplinary Procedures Center, Strategic Laboratory | Claudio Tavares Sacchi, Claudia Regina Gonçalves, Erica Valessa Ramos Gomes, Karoline Rodrigues Campos |
| EPI_ISL_547580 | Santa Casa da Misericórdia de Presidente Prudente | Instituto Adolfo Lutz, Interdiciplinary Procedures Center, Strategic Laboratory | Claudio Tavares Sacchi, Claudia Regina Gonçalves, Erica Valessa Ramos Gomes, Karoline Rodrigues Campos |
| EPI_ISL_547602, EPI_ISL_547603, EPI_ISL_547604, EPI_ISL_547605, EPI_ISL_547606, EPI_ISL_547607 | Gundersen Molecular Diagnostics Laboratory | Kabara Cancer Research Institute | Craig S. Richmond, Paraic A. Kenny |
| EPI_ISL_547608, EPI_ISL_547609, EPI_ISL_547610 | Gundersen Clinical Microbiology Laboratory | Kabara Cancer Research Institute | Craig S. Richmond, Paraic A. Kenny |
| EPI_ISL_547611, EPI_ISL_547612, EPI_ISL_547613, EPI_ISL_547614, EPI_ISL_547615, EPI_ISL_547616, EPI_ISL_547617, EPI_ISL_547618, EPI_ISL_547619, EPI_ISL_547620, EPI_ISL_547621, EPI_ISL_547622, EPI_ISL_547623, EPI_ISL_547624, EPI_ISL_547625, EPI_ISL_547626, EPI_ISL_547627, EPI_ISL_547628, EPI_ISL_547629, EPI_ISL_547630, EPI_ISL_547631, EPI_ISL_547632, EPI_ISL_547633, EPI_ISL_547634, EPI_ISL_547635, EPI_ISL_547636, EPI_ISL_547637, EPI_ISL_547638, EPI_ISL_547639, EPI_ISL_547640, EPI_ISL_547641, EPI_ISL_547642, EPI_ISL_547643, EPI_ISL_547644, EPI_ISL_547645, EPI_ISL_547646, EPI_ISL_547647, EPI_ISL_547648, EPI_ISL_547649, EPI_ISL_547650, EPI_ISL_547651, EPI_ISL_547652, EPI_ISL_547653, EPI_ISL_547654, EPI_ISL_547655, EPI_ISL_547656, EPI_ISL_547657, EPI_ISL_547658, EPI_ISL_547659, EPI_ISL_547660, EPI_ISL_547661, EPI_ISL_547662, EPI_ISL_547663, EPI_ISL_547664, EPI_ISL_547665 | | | |
| see above | Gundersen Molecular Diagnostics Laboratory | Kabara Cancer Research Institute | Craig S. Richmond, Paraic A. Kenny |
| EPI_ISL_547666, EPI_ISL_547667, EPI_ISL_547668, EPI_ISL_547669, EPI_ISL_547670, EPI_ISL_547671, EPI_ISL_547672, EPI_ISL_547673, EPI_ISL_547674, EPI_ISL_547675, EPI_ISL_547676 | | | |
| see above | Gundersen Clinical Microbiology Laboratory | Kabara Cancer Research Institute | Craig S. Richmond, Paraic A. Kenny |
| EPI_ISL_547677, EPI_ISL_547678, EPI_ISL_547679, EPI_ISL_547680, EPI_ISL_547681, EPI_ISL_547682 | Gundersen Molecular Diagnostics Laboratory | Kabara Cancer Research Institute | Craig S. Richmond, Paraic A. Kenny |
| EPI_ISL_547683, EPI_ISL_547684 | Gundersen Clinical Microbiology Laboratory | Kabara Cancer Research Institute | Craig S. Richmond, Paraic A. Kenny |
| EPI_ISL_547685, EPI_ISL_547686, EPI_ISL_547687, EPI_ISL_547688, EPI_ISL_547689, EPI_ISL_547690, EPI_ISL_547691, EPI_ISL_547692, EPI_ISL_547693, EPI_ISL_547694, EPI_ISL_547695, EPI_ISL_547696, EPI_ISL_547697, EPI_ISL_547698, EPI_ISL_547699, EPI_ISL_547700, EPI_ISL_547701, EPI_ISL_547702, EPI_ISL_547703, EPI_ISL_547704 | | | |
| see above | Gundersen Molecular Diagnostics Laboratory | Kabara Cancer Research Institute | Craig S. Richmond, Paraic A. Kenny |
| EPI_ISL_547705 | Gundersen Clinical Microbiology Laboratory | Kabara Cancer Research Institute | Craig S. Richmond, Paraic A. Kenny |
| EPI_ISL_547706, EPI_ISL_547707, EPI_ISL_547708, EPI_ISL_547709, EPI_ISL_547710, EPI_ISL_547711, EPI_ISL_547712, EPI_ISL_547713, EPI_ISL_547714, EPI_ISL_547715, EPI_ISL_547716, EPI_ISL_547717, EPI_ISL_547718, EPI_ISL_547719, EPI_ISL_547720, EPI_ISL_547721, EPI_ISL_547722, EPI_ISL_547723, EPI_ISL_547724, EPI_ISL_547725, EPI_ISL_547726, EPI_ISL_547727, EPI_ISL_547728, EPI_ISL_547729, EPI_ISL_547730, EPI_ISL_547731, EPI_ISL_547732, EPI_ISL_547733, EPI_ISL_547734, EPI_ISL_547735, EPI_ISL_547736, EPI_ISL_547737, EPI_ISL_547738, EPI_ISL_547739, EPI_ISL_547740, EPI_ISL_547741, EPI_ISL_547742, EPI_ISL_547743, EPI_ISL_547744, EPI_ISL_547745, EPI_ISL_547746, EPI_ISL_547747, EPI_ISL_547748, EPI_ISL_547749, EPI_ISL_547750, EPI_ISL_547751, EPI_ISL_547752, EPI_ISL_547753, EPI_ISL_547754, EPI_ISL_547755, EPI_ISL_547756, EPI_ISL_547757, EPI_ISL_547758, EPI_ISL_547759, EPI_ISL_547760, EPI_ISL_547761, EPI_ISL_547762, EPI_ISL_547763, EPI_ISL_547764, EPI_ISL_547765, EPI_ISL_547766, EPI_ISL_547767, EPI_ISL_547768, EPI_ISL_547769, EPI_ISL_547770, EPI_ISL_547771, EPI_ISL_547772, EPI_ISL_547773, EPI_ISL_547774, EPI_ISL_547775, EPI_ISL_547776, EPI_ISL_547777, EPI_ISL_547778, EPI_ISL_547779, EPI_ISL_547780, EPI_ISL_547781, EPI_ISL_547782, EPI_ISL_547783, EPI_ISL_547784, EPI_ISL_547785, EPI_ISL_547786, EPI_ISL_547787, EPI_ISL_547788, EPI_ISL_547789, EPI_ISL_547790, EPI_ISL_547791, EPI_ISL_547792, EPI_ISL_547793, EPI_ISL_547794, EPI_ISL_547795, EPI_ISL_547796, EPI_ISL_547797, EPI_ISL_547798, EPI_ISL_547799, EPI_ISL_547800, EPI_ISL_547801, EPI_ISL_547802, EPI_ISL_547803, EPI_ISL_547804, EPI_ISL_547805, EPI_ISL_547806, EPI_ISL_547807, EPI_ISL_547808, EPI_ISL_547809, EPI_ISL_547810, EPI_ISL_547811, EPI_ISL_547812, EPI_ISL_547813, EPI_ISL_547814, EPI_ISL_547815, EPI_ISL_547816, EPI_ISL_547817, EPI_ISL_547818, EPI_ISL_547819, EPI_ISL_547820, EPI_ISL_547821, EPI_ISL_547822, EPI_ISL_547823, EPI_ISL_547824, EPI_ISL_547825, EPI_ISL_547826, EPI_ISL_547827, EPI_ISL_547828, EPI_ISL_547829, EPI_ISL_547830, EPI_ISL_547831, EPI_ISL_547832, EPI_ISL_547833, EPI_ISL_547834, EPI_ISL_547835, EPI_ISL_547836, EPI_ISL_547837, EPI_ISL_547838, EPI_ISL_547839, EPI_ISL_547840, EPI_ISL_547841, EPI_ISL_547842, EPI_ISL_547843, EPI_ISL_547844, EPI_ISL_547845, EPI_ISL_547846, EPI_ISL_547847, EPI_ISL_547848, EPI_ISL_547849, EPI_ISL_547850, EPI_ISL_547851, EPI_ISL_547852, EPI_ISL_547853, EPI_ISL_547854, EPI_ISL_547855, EPI_ISL_547856, EPI_ISL_547857, EPI_ISL_547858, EPI_ISL_547859, EPI_ISL_547861, EPI_ISL_547862, EPI_ISL_547863, EPI_ISL_547864, EPI_ISL_547865, EPI_ISL_547866, EPI_ISL_547867, EPI_ISL_547868, EPI_ISL_547869 | | | |
| see above | Gundersen Molecular Diagnostics Laboratory | Kabara Cancer Research Institute | Craig S. Richmond, Paraic A. Kenny |
| EPI_ISL_560416, EPI_ISL_560417, EPI_ISL_560418, EPI_ISL_560419, EPI_ISL_560420, EPI_ISL_560421, EPI_ISL_560422, EPI_ISL_560423, EPI_ISL_560424, EPI_ISL_560425, EPI_ISL_560426, EPI_ISL_560427, EPI_ISL_560428, EPI_ISL_560429, EPI_ISL_560430, EPI_ISL_560431, EPI_ISL_560432, EPI_ISL_560433, EPI_ISL_560434, EPI_ISL_560435, EPI_ISL_560436, EPI_ISL_560437, EPI_ISL_560438, EPI_ISL_560439, EPI_ISL_560440, EPI_ISL_560441, EPI_ISL_560442, EPI_ISL_560443, EPI_ISL_560444, EPI_ISL_560445, EPI_ISL_560446, EPI_ISL_560447, EPI_ISL_560448, EPI_ISL_560449, EPI_ISL_560450, EPI_ISL_560451, EPI_ISL_560452, EPI_ISL_560453, EPI_ISL_560454, EPI_ISL_560455, EPI_ISL_560456, EPI_ISL_560457, EPI_ISL_560458, EPI_ISL_560459, EPI_ISL_560460, EPI_ISL_560461, EPI_ISL_560462, EPI_ISL_560463, EPI_ISL_560464, EPI_ISL_560465, EPI_ISL_560466, EPI_ISL_560467, EPI_ISL_560468, EPI_ISL_560469, EPI_ISL_560470, EPI_ISL_560471, EPI_ISL_560472, EPI_ISL_560473, EPI_ISL_560474, EPI_ISL_560475, EPI_ISL_560476, EPI_ISL_560477, EPI_ISL_560478, EPI_ISL_560479, EPI_ISL_560480, EPI_ISL_560481, EPI_ISL_560482, EPI_ISL_560483, EPI_ISL_560484, EPI_ISL_560485, EPI_ISL_560486, EPI_ISL_560487, EPI_ISL_560488, EPI_ISL_560489, EPI_ISL_560490, EPI_ISL_560491, EPI_ISL_560492, EPI_ISL_560493, EPI_ISL_560494, EPI_ISL_560495, EPI_ISL_560496, EPI_ISL_560497, EPI_ISL_560498, EPI_ISL_560499, EPI_ISL_560500, EPI_ISL_560501, EPI_ISL_560502, EPI_ISL_560503, EPI_ISL_560504, EPI_ISL_560505, EPI_ISL_560506, EPI_ISL_560507, EPI_ISL_560508, EPI_ISL_560509, EPI_ISL_560510, EPI_ISL_560511, EPI_ISL_560512, EPI_ISL_560513, EPI_ISL_560514, EPI_ISL_560515, EPI_ISL_560516, EPI_ISL_560517, EPI_ISL_560518, EPI_ISL_560519, EPI_ISL_560520, EPI_ISL_560521, EPI_ISL_560522, EPI_ISL_560523, EPI_ISL_560524, EPI_ISL_560525, EPI_ISL_560527, EPI_ISL_560528, EPI_ISL_560529, EPI_ISL_560530, EPI_ISL_560531, EPI_ISL_560532, EPI_ISL_560533, EPI_ISL_560534, EPI_ISL_560535, EPI_ISL_560536, EPI_ISL_560537, EPI_ISL_560538, EPI_ISL_560539, EPI_ISL_560540, EPI_ISL_560545, EPI_ISL_560551, EPI_ISL_560553 | | | |
| see above | Viollier AG | Department of Biosystems Science and Engineering, ETH Zürich | Christian Beisel, Sarah Nadeau, Ivan Topolsky, Pedro Ferreira, Philipp Jablonski, Susana Posada-Céspedes, Tobias Schär, Ina Nissen, Natascha Santacroce, Elodie Burcklen, Christiane Beckmann, Maurice Redondo, Olivier Kobel, Christoph Noppen, Sophie Seidel, Noemie Santamaria de Souza, Niko Beerenwinkel, Tanja Stadler |
| EPI_ISL_574577 | Hospital Municipal Dr. Ignacio Proença de Gouvea | Instituto Adolfo Lutz, Interdiciplinary Procedures Center, Strategic Laboratory | Claudio Tavares Sacchi, Claudia Regina Gonçalves, Erica Valessa Ramos Gomes, Karoline Rodrigues Campos |
| EPI_ISL_574578 | Hospital Municipal Mário Gatti | Instituto Adolfo Lutz, Interdiciplinary Procedures Center, Strategic Laboratory | Claudio Tavares Sacchi, Claudia Regina Gonçalves, Erica Valessa Ramos Gomes, Karoline Rodrigues Campos |
| EPI_ISL_574579 | Hospital Municipal Dr. Ignacio Proença de Gouvea | Instituto Adolfo Lutz, Interdiciplinary Procedures Center, Strategic Laboratory | Claudio Tavares Sacchi, Claudia Regina Gonçalves, Erica Valessa Ramos Gomes, Karoline Rodrigues Campos |
| EPI_ISL_574580 | Hospital Cidade Tiradentes Carmen Prudente | Instituto Adolfo Lutz, Interdiciplinary Procedures Center, | Claudio Tavares Sacchi, Claudia Regina Gonçalves, Erica Valessa Ramos Gomes, Karoline Rodrigues Campos |

| | | | |
|---|---|---|---|
| | | Strategic Laboratory | |
| EPI_ISL_574582 | Hospital Municipal Dr. Jose Soares Hungria | Instituto Adolfo Lutz, Interdiciplinary Procedures Center, Strategic Laboratory | Claudio Tavares Sacchi, Claudia Regina Gonçalves, Erica Valessa Ramos Gomes, Karoline Rodrigues Campos |
| EPI_ISL_574583 | Secretaria Municipal de Saude de Jandira | Instituto Adolfo Lutz, Interdiciplinary Procedures Center, Strategic Laboratory | Claudio Tavares Sacchi, Claudia Regina Gonçalves, Erica Valessa Ramos Gomes, Karoline Rodrigues Campos |
| EPI_ISL_574588 | Hospital Estadual Sumare | Instituto Adolfo Lutz, Interdiciplinary Procedures Center, Strategic Laboratory | Claudio Tavares Sacchi, Claudia Regina Gonçalves, Erica Valessa Ramos Gomes, Karoline Rodrigues Campos |
| EPI_ISL_574589 | Hospital Municipal Dr. Jose Soares Hungria | Instituto Adolfo Lutz, Interdiciplinary Procedures Center, Strategic Laboratory | Claudio Tavares Sacchi, Claudia Regina Gonçalves, Erica Valessa Ramos Gomes, Karoline Rodrigues Campos |
| EPI_ISL_574590 | Unidade de Printo Atendimento UPA I Santa Isabel | Instituto Adolfo Lutz, Interdiciplinary Procedures Center, Strategic Laboratory | Claudio Tavares Sacchi, Claudia Regina Gonçalves, Erica Valessa Ramos Gomes, Karoline Rodrigues Campos |
| EPI_ISL_574591, EPI_ISL_574592 | Hospital Domingos Leonardo Ceravolo Presidente Prudente | Instituto Adolfo Lutz, Interdiciplinary Procedures Center, Strategic Laboratory | Claudio Tavares Sacchi, Claudia Regina Gonçalves, Erica Valessa Ramos Gomes, Karoline Rodrigues Campos |
| EPI_ISL_574593 | CS II Dr. Antonio Vicoso Moreira de Rezende Sumare | Instituto Adolfo Lutz, Interdiciplinary Procedures Center, Strategic Laboratory | Claudio Tavares Sacchi, Claudia Regina Gonçalves, Erica Valessa Ramos Gomes, Karoline Rodrigues Campos |
| EPI_ISL_574594 | Hospital Escola da Universidade de Taubate | Instituto Adolfo Lutz, Interdiciplinary Procedures Center, Strategic Laboratory | Claudio Tavares Sacchi, Claudia Regina Gonçalves, Erica Valessa Ramos Gomes, Karoline Rodrigues Campos |
| EPI_ISL_574595 | Hospital Geral de Vila Penteado Dr. Jose Pamgella | Instituto Adolfo Lutz, Interdiciplinary Procedures Center, Strategic Laboratory | Claudio Tavares Sacchi, Claudia Regina Gonçalves, Erica Valessa Ramos Gomes, Karoline Rodrigues Campos |
| EPI_ISL_574596 | CS II Dr. Antonio Vicoso Moreira de Rezende Sumare | Instituto Adolfo Lutz, Interdiciplinary Procedures Center, Strategic Laboratory | Claudio Tavares Sacchi, Claudia Regina Gonçalves, Erica Valessa Ramos Gomes, Karoline Rodrigues Campos |
| EPI_ISL_574597 | Secretaria Municipal de Saude de Jarinu | Instituto Adolfo Lutz, Interdiciplinary Procedures Center, Strategic Laboratory | Claudio Tavares Sacchi, Claudia Regina Gonçalves, Erica Valessa Ramos Gomes, Karoline Rodrigues Campos |
| EPI_ISL_574598 | Servico de Verificacao de Obito SVO | Instituto Adolfo Lutz, Interdiciplinary Procedures Center, Strategic Laboratory | Claudio Tavares Sacchi, Claudia Regina Gonçalves, Erica Valessa Ramos Gomes, Karoline Rodrigues Campos |
| EPI_ISL_574883, EPI_ISL_574884, EPI_ISL_574885, EPI_ISL_574886, EPI_ISL_574887, EPI_ISL_574888, EPI_ISL_574889, EPI_ISL_574890, EPI_ISL_574891, EPI_ISL_574892, EPI_ISL_574893, EPI_ISL_574894, EPI_ISL_574895, EPI_ISL_574896, EPI_ISL_574897, EPI_ISL_574898, EPI_ISL_574899, EPI_ISL_574900, EPI_ISL_574901, EPI_ISL_574902, EPI_ISL_574903, EPI_ISL_574904, EPI_ISL_574905, EPI_ISL_574906, EPI_ISL_574907, EPI_ISL_574908, EPI_ISL_574909, EPI_ISL_574910, EPI_ISL_574911, EPI_ISL_574912, EPI_ISL_574913, EPI_ISL_574914, EPI_ISL_574915, EPI_ISL_574916, EPI_ISL_574917, EPI_ISL_574918, EPI_ISL_574919, EPI_ISL_574920, EPI_ISL_574921, EPI_ISL_574922, EPI_ISL_574923, EPI_ISL_574924, EPI_ISL_574925, EPI_ISL_574926, EPI_ISL_574927, EPI_ISL_574928, EPI_ISL_574929, EPI_ISL_574930, EPI_ISL_574937, EPI_ISL_574944, EPI_ISL_574945, EPI_ISL_574946, EPI_ISL_574948, EPI_ISL_574949, EPI_ISL_574950, EPI_ISL_574951, EPI_ISL_574952, EPI_ISL_574953, EPI_ISL_574954, EPI_ISL_574955, EPI_ISL_574956, EPI_ISL_574957, EPI_ISL_574958, EPI_ISL_574959, EPI_ISL_574960, EPI_ISL_574961, EPI_ISL_574962, EPI_ISL_574963, EPI_ISL_574964, EPI_ISL_574965, EPI_ISL_574966, EPI_ISL_574967, EPI_ISL_574968, EPI_ISL_574969, EPI_ISL_574970, EPI_ISL_574971, EPI_ISL_574972, EPI_ISL_574973, EPI_ISL_574974, EPI_ISL_574975, EPI_ISL_574976, EPI_ISL_574977, EPI_ISL_574978, EPI_ISL_574979, EPI_ISL_574980, EPI_ISL_574981, EPI_ISL_574982, EPI_ISL_574983, EPI_ISL_574984, EPI_ISL_574985, EPI_ISL_574986, EPI_ISL_574987, EPI_ISL_574988, EPI_ISL_574989, EPI_ISL_574990, EPI_ISL_574991, EPI_ISL_574992, EPI_ISL_574993, EPI_ISL_574994, EPI_ISL_574995, EPI_ISL_574996, EPI_ISL_574997, EPI_ISL_574998, EPI_ISL_574999, EPI_ISL_575000, EPI_ISL_575001, EPI_ISL_575002, EPI_ISL_575003, EPI_ISL_575004, EPI_ISL_575005, EPI_ISL_575006, EPI_ISL_575007, EPI_ISL_575008, EPI_ISL_575009, EPI_ISL_575010, EPI_ISL_575011, EPI_ISL_575012, EPI_ISL_575013, EPI_ISL_575014, EPI_ISL_575015, EPI_ISL_575016, EPI_ISL_575017, EPI_ISL_575018, EPI_ISL_575019 | | | |
| see above | Viollier AG | Department of Biosystems Science and Engineering, ETH Zürich | Christian Beisel, Sarah Nadeau, Ivan Topolsky, Pedro Ferreira, Philipp Jablonski, Susana Posada-Céspedes, Tobias Schär, Ina Nissen, Natascha Santacroce, Elodie Burcklen, Christiane Beckmann, Maurice Redondo, Olivier Kobel, Christoph Noppen, Sophie Seidel, Noemie Santamaria de Souza, Niko Beerenwinkel, Tanja Stadler |
| EPI_ISL_577607 | Royal Hobart Hospital | Royal Hobart Hospital | Cooley L., van Haeften R. |
| EPI_ISL_578701, EPI_ISL_578702 | Molecular Pathology Division, Department of Pathology, Hong Kong Sanatorium & Hospital | Molecular Pathology Division, Department of Pathology, Hong Kong Sanatorium & Hospital | Chun Hang AU, Wai Sing CHAN, Ho Yin LAM, Dona N. HO, Simon Y.M. LAM, Jonpaul S.T. ZEE, Tsun Leung CHAN, Edmond S.K. MA |
| EPI_ISL_583490 | Hospital Estadual Sumare | Instituto Adolfo Lutz, Interdiciplinary Procedures Center, Strategic Laboratory | Claudio Tavares Sacchi, Claudia Regina Gonçalves, Erica Valessa Ramos Gomes, Karoline Rodrigues Campos |
| EPI_ISL_583491 | Centro de Saude Esf IV Zona Rual Domingos de SJ Rio Pardo | Instituto Adolfo Lutz, Interdiciplinary Procedures Center, Strategic Laboratory | Claudio Tavares Sacchi, Claudia Regina Gonçalves, Erica Valessa Ramos Gomes, Karoline Rodrigues Campos |
| EPI_ISL_583492 | Santa Casa Anna Cintra | Instituto Adolfo Lutz, Interdiciplinary Procedures Center, Strategic Laboratory | Claudio Tavares Sacchi, Claudia Regina Gonçalves, Erica Valessa Ramos Gomes, Karoline Rodrigues Campos |
| EPI_ISL_583493 | Vigilância em Saúde de Cajamar | Instituto Adolfo Lutz, Interdiciplinary Procedures Center, Strategic Laboratory | Claudio Tavares Sacchi, Claudia Regina Gonçalves, Erica Valessa Ramos Gomes, Karoline Rodrigues Campos |
| EPI_ISL_583494 | CS II Dr. Antonio Vicoso Moreira de Rezende Sumare | Instituto Adolfo Lutz, Interdiciplinary Procedures Center, Strategic Laboratory | Claudio Tavares Sacchi, Claudia Regina Gonçalves, Erica Valessa Ramos Gomes, Karoline Rodrigues Campos |
| EPI_ISL_583495 | Serviço de Verificação de Óbitos SVO Guaruulhos | Instituto Adolfo Lutz, Interdiciplinary Procedures Center, Strategic Laboratory | Claudio Tavares Sacchi, Claudia Regina Gonçalves, Erica Valessa Ramos Gomes, Karoline Rodrigues Campos |
| EPI_ISL_583496 | UPA Jandira | Instituto Adolfo Lutz, Interdiciplinary Procedures Center, Strategic Laboratory | Claudio Tavares Sacchi, Claudia Regina Gonçalves, Erica Valessa Ramos Gomes, Karoline Rodrigues Campos |
| EPI_ISL_583497 | Complexo Hospitalar Ouro Verde de Campinas | Instituto Adolfo Lutz, Interdiciplinary Procedures Center, Strategic Laboratory | Claudio Tavares Sacchi, Claudia Regina Gonçalves, Erica Valessa Ramos Gomes, Karoline Rodrigues Campos |
| EPI_ISL_583498 | Hospital Municipal Dr. Waldemar Tebaldi | Instituto Adolfo Lutz, Interdiciplinary Procedures Center, Strategic Laboratory | Claudio Tavares Sacchi, Claudia Regina Gonçalves, Erica Valessa Ramos Gomes, Karoline Rodrigues Campos |
| EPI_ISL_583499 | Distrito Sanitario Sul Campinas | Instituto Adolfo Lutz, Interdiciplinary Procedures Center, Strategic Laboratory | Claudio Tavares Sacchi, Claudia Regina Gonçalves, Erica Valessa Ramos Gomes, Karoline Rodrigues Campos |
| EPI_ISL_583500 | Centro de Saude I Tacito Leite de Carvalho e Silva | Instituto Adolfo Lutz, Interdiciplinary Procedures Center, Strategic Laboratory | Claudio Tavares Sacchi, Claudia Regina Gonçalves, Erica Valessa Ramos Gomes, Karoline Rodrigues Campos |
| EPI_ISL_583501 | Hospital Estadual de CampanhaCOVID 19 Barradas | Instituto Adolfo Lutz, Interdiciplinary Procedures Center, Strategic Laboratory | Claudio Tavares Sacchi, Claudia Regina Gonçalves, Erica Valessa Ramos Gomes, Karoline Rodrigues Campos |
| EPI_ISL_583502 | Serv de Vig Sanitaria Epidemio e CTRL de Zoonoses Guaruja | Instituto Adolfo Lutz, Interdiciplinary Procedures Center, Strategic Laboratory | Claudio Tavares Sacchi, Claudia Regina Gonçalves, Erica Valessa Ramos Gomes, Karoline Rodrigues Campos |
| EPI_ISL_583503 | CTA Centro de Testagem e Aconselhamento | Instituto Adolfo Lutz, Interdiciplinary Procedures Center, Strategic Laboratory | Claudio Tavares Sacchi, Claudia Regina Gonçalves, Erica Valessa Ramos Gomes, Karoline Rodrigues Campos |
| EPI_ISL_583504, EPI_ISL_583505 | Casa de Saude Stella Maris | Instituto Adolfo Lutz, Interdiciplinary Procedures Center, Strategic Laboratory | Claudio Tavares Sacchi, Claudia Regina Gonçalves, Erica Valessa Ramos Gomes, Karoline Rodrigues Campos |
| EPI_ISL_585092, EPI_ISL_585093, EPI_ISL_585094, EPI_ISL_585095, EPI_ISL_585096, EPI_ISL_585097, EPI_ISL_585098, EPI_ISL_585099, EPI_ISL_585100, EPI_ISL_585101, EPI_ISL_585102, EPI_ISL_585103, EPI_ISL_585104, EPI_ISL_585105, EPI_ISL_585106, EPI_ISL_585107, EPI_ISL_585108, EPI_ISL_585109, EPI_ISL_585110, EPI_ISL_585111, EPI_ISL_585112, EPI_ISL_585113, EPI_ISL_585114, EPI_ISL_585115, EPI_ISL_585116, EPI_ISL_585117, EPI_ISL_585118, EPI_ISL_585119, EPI_ISL_585120, EPI_ISL_585121, EPI_ISL_585122, EPI_ISL_585123, EPI_ISL_585124, EPI_ISL_585125, EPI_ISL_585126, EPI_ISL_585127, EPI_ISL_585128, EPI_ISL_585129, EPI_ISL_585130, EPI_ISL_585131, EPI_ISL_585132, EPI_ISL_585133, EPI_ISL_585134, EPI_ISL_585135, EPI_ISL_585136, EPI_ISL_585137, EPI_ISL_585138, EPI_ISL_585139, EPI_ISL_585140, EPI_ISL_585141, EPI_ISL_585142, EPI_ISL_585143, EPI_ISL_585144, EPI_ISL_585145, EPI_ISL_585146, EPI_ISL_585147, EPI_ISL_585148, EPI_ISL_585149, EPI_ISL_585150, EPI_ISL_585151, EPI_ISL_585152, EPI_ISL_585153, EPI_ISL_585154, EPI_ISL_585155, EPI_ISL_585156, EPI_ISL_585157, EPI_ISL_585158, EPI_ISL_585159, EPI_ISL_585160, EPI_ISL_585161, EPI_ISL_585162, EPI_ISL_585163, | | | |

| | | | |
|---|---|---|---|
| EPI_ISL_585164, EPI_ISL_585165, EPI_ISL_585166, EPI_ISL_585167, EPI_ISL_585168, EPI_ISL_585169, EPI_ISL_585170, EPI_ISL_585171, EPI_ISL_585172, EPI_ISL_585173, EPI_ISL_585174, EPI_ISL_585175, EPI_ISL_585176, EPI_ISL_585177, EPI_ISL_585178, EPI_ISL_585179, EPI_ISL_585180, EPI_ISL_585181, EPI_ISL_585182, EPI_ISL_585183, EPI_ISL_585184, EPI_ISL_585185, EPI_ISL_585186, EPI_ISL_585187, EPI_ISL_585188, EPI_ISL_585189, EPI_ISL_585190, EPI_ISL_585191, EPI_ISL_585192, EPI_ISL_585193, EPI_ISL_585194, EPI_ISL_585195, EPI_ISL_585196, EPI_ISL_585197, EPI_ISL_585199, EPI_ISL_585200, EPI_ISL_585201, EPI_ISL_585202, EPI_ISL_585203, EPI_ISL_585204, EPI_ISL_585205, EPI_ISL_585206, EPI_ISL_585207, EPI_ISL_585208, EPI_ISL_585209, EPI_ISL_585210, EPI_ISL_585211, EPI_ISL_585212, EPI_ISL_585213, EPI_ISL_585214, EPI_ISL_585215, EPI_ISL_585216, EPI_ISL_585217, EPI_ISL_585218, EPI_ISL_585219, EPI_ISL_585220, EPI_ISL_585221, EPI_ISL_585222, EPI_ISL_585223, EPI_ISL_585224, EPI_ISL_585225, EPI_ISL_585226, EPI_ISL_585227, EPI_ISL_585228, EPI_ISL_585229, EPI_ISL_585230, EPI_ISL_585231, EPI_ISL_585232, EPI_ISL_585233, EPI_ISL_585234, EPI_ISL_585235, EPI_ISL_585236, EPI_ISL_585237, EPI_ISL_585238, EPI_ISL_585239, EPI_ISL_585240, EPI_ISL_585241, EPI_ISL_585242, EPI_ISL_585243, EPI_ISL_585244, EPI_ISL_585245, EPI_ISL_585246, EPI_ISL_585247, EPI_ISL_585248, EPI_ISL_585249, EPI_ISL_585250, EPI_ISL_585251, EPI_ISL_585252, EPI_ISL_585253, EPI_ISL_585254, EPI_ISL_585255, EPI_ISL_585256, EPI_ISL_585257, EPI_ISL_585258, EPI_ISL_585259 | | | |
| see above | Regional Virus Laboratory, Belfast Health and Social Care Trust | COVID-19 Genomics UK (COG-UK) Consortium | Conall McCaughey, James McKenna, Tanya Curran, Susan Feeney, Alison Watt, Ciara Cox, Mairead Connor, Zoltan Molnar, David Simpson, Derek Fairley |
| EPI_ISL_591327, EPI_ISL_591328, EPI_ISL_591329, EPI_ISL_591330, EPI_ISL_591331, EPI_ISL_591332, EPI_ISL_591333, EPI_ISL_591334, EPI_ISL_591335, EPI_ISL_591337, EPI_ISL_591338, EPI_ISL_591339, EPI_ISL_591340 | | | |
| see above | Dipartimento di Biotecnologie Mediche, University of Siena | Dipartimento di Biotecnologie Mediche, University of Siena | Cusi,M.G., Pinzauti,D., Gandolfo,C., Anichini,G., Pozzi,G., Santoro,F. |
| EPI_ISL_603021 | Pronto Socorro Dr. Conrado Cesarino Nuvolini | Instituto Adolfo Lutz, Interdiciplinary Procedures Center, Strategic Laboratory | Claudio Tavares Sacchi, Claudia Regina Gonçalves, Erica Valessa Ramos Gomes, Karoline Rodrigues Campos |
| EPI_ISL_603022 | Departamento de Vigilância à Saúde | Instituto Adolfo Lutz, Interdiciplinary Procedures Center, Strategic Laboratory | Claudio Tavares Sacchi, Claudia Regina Gonçalves, Erica Valessa Ramos Gomes, Karoline Rodrigues Campos |
| EPI_ISL_603023 | Vigilância em Saúde Visa Sul | Instituto Adolfo Lutz, Interdiciplinary Procedures Center, Strategic Laboratory | Claudio Tavares Sacchi, Claudia Regina Gonçalves, Erica Valessa Ramos Gomes, Karoline Rodrigues Campos |
| EPI_ISL_603024 | Santa Casa de Misericordia de Araçatuba | Instituto Adolfo Lutz, Interdiciplinary Procedures Center, Strategic Laboratory | Claudio Tavares Sacchi, Claudia Regina Gonçalves, Erica Valessa Ramos Gomes, Karoline Rodrigues Campos |
| EPI_ISL_603025 | UPA Central de Caraguatatuba | Instituto Adolfo Lutz, Interdiciplinary Procedures Center, Strategic Laboratory | Claudio Tavares Sacchi, Claudia Regina Gonçalves, Erica Valessa Ramos Gomes, Karoline Rodrigues Campos |
| EPI_ISL_603026 | Santa Casa da Misericórdia de Presidente Prudente | Instituto Adolfo Lutz, Interdiciplinary Procedures Center, Strategic Laboratory | Claudio Tavares Sacchi, Claudia Regina Gonçalves, Erica Valessa Ramos Gomes, Karoline Rodrigues Campos |
| EPI_ISL_603027 | Santa Casa de Misericordia de Araçatuba | Instituto Adolfo Lutz, Interdiciplinary Procedures Center, Strategic Laboratory | Claudio Tavares Sacchi, Claudia Regina Gonçalves, Erica Valessa Ramos Gomes, Karoline Rodrigues Campos |
| EPI_ISL_603028 | Hospital Municipal Santa Ana | Instituto Adolfo Lutz, Interdiciplinary Procedures Center, Strategic Laboratory | Claudio Tavares Sacchi, Claudia Regina Gonçalves, Erica Valessa Ramos Gomes, Karoline Rodrigues Campos |
| EPI_ISL_603029 | Hospital Municipal Mário Gatti | Instituto Adolfo Lutz, Interdiciplinary Procedures Center, Strategic Laboratory | Claudio Tavares Sacchi, Claudia Regina Gonçalves, Erica Valessa Ramos Gomes, Karoline Rodrigues Campos |
| EPI_ISL_603030 | Hospital Domingos Leonardo Ceravolo Presidente Prudente | Instituto Adolfo Lutz, Interdiciplinary Procedures Center, Strategic Laboratory | Claudio Tavares Sacchi, Claudia Regina Gonçalves, Erica Valessa Ramos Gomes, Karoline Rodrigues Campos |
| EPI_ISL_603031 | Santa Casa de Presidente Epitácio | Instituto Adolfo Lutz, Interdiciplinary Procedures Center, Strategic Laboratory | Claudio Tavares Sacchi, Claudia Regina Gonçalves, Erica Valessa Ramos Gomes, Karoline Rodrigues Campos |
| EPI_ISL_603032 | Santa Casa da Misericórdia de Presidente Prudente | Instituto Adolfo Lutz, Interdiciplinary Procedures Center, Strategic Laboratory | Claudio Tavares Sacchi, Claudia Regina Gonçalves, Erica Valessa Ramos Gomes, Karoline Rodrigues Campos |
| EPI_ISL_603033 | Vigilancia Epidemiologica de São Bernardo do Campo | Instituto Adolfo Lutz, Interdiciplinary Procedures Center, Strategic Laboratory | Claudio Tavares Sacchi, Claudia Regina Gonçalves, Erica Valessa Ramos Gomes, Karoline Rodrigues Campos |
| EPI_ISL_603034 | Departamento de Vigilância à Saúde | Instituto Adolfo Lutz, Interdiciplinary Procedures Center, Strategic Laboratory | Claudio Tavares Sacchi, Claudia Regina Gonçalves, Erica Valessa Ramos Gomes, Karoline Rodrigues Campos |
| EPI_ISL_603035 | Secretaria Municipal de Saúde | Instituto Adolfo Lutz, Interdiciplinary Procedures Center, Strategic Laboratory | Claudio Tavares Sacchi, Claudia Regina Gonçalves, Erica Valessa Ramos Gomes, Karoline Rodrigues Campos |
| EPI_ISL_603036 | Hospital Santa Ana | Instituto Adolfo Lutz, Interdiciplinary Procedures Center, Strategic Laboratory | Claudio Tavares Sacchi, Claudia Regina Gonçalves, Erica Valessa Ramos Gomes, Karoline Rodrigues Campos |
| EPI_ISL_603037 | Hospital Geral de Pedreira | Instituto Adolfo Lutz, Interdiciplinary Procedures Center, Strategic Laboratory | Claudio Tavares Sacchi, Claudia Regina Gonçalves, Erica Valessa Ramos Gomes, Karoline Rodrigues Campos |
| EPI_ISL_603038 | Santa Casa de Misericordia de Araçatuba | Instituto Adolfo Lutz, Interdiciplinary Procedures Center, Strategic Laboratory | Claudio Tavares Sacchi, Claudia Regina Gonçalves, Erica Valessa Ramos Gomes, Karoline Rodrigues Campos |
| EPI_ISL_603039 | Hospital Municipal Mário Gatti | Instituto Adolfo Lutz, Interdiciplinary Procedures Center, Strategic Laboratory | Claudio Tavares Sacchi, Claudia Regina Gonçalves, Erica Valessa Ramos Gomes, Karoline Rodrigues Campos |
| EPI_ISL_603267, EPI_ISL_603268, EPI_ISL_603270, EPI_ISL_603271, EPI_ISL_603272, EPI_ISL_603273, EPI_ISL_603274, EPI_ISL_603275, EPI_ISL_603276, EPI_ISL_603277, EPI_ISL_603278, EPI_ISL_603279, EPI_ISL_603281, EPI_ISL_603282, EPI_ISL_603283, EPI_ISL_603284, EPI_ISL_603285, EPI_ISL_603287, EPI_ISL_603288, EPI_ISL_603290, EPI_ISL_603291, EPI_ISL_603292, EPI_ISL_603293, EPI_ISL_603294, EPI_ISL_603295, EPI_ISL_603296, EPI_ISL_603297, EPI_ISL_603298, EPI_ISL_603299, EPI_ISL_603300, EPI_ISL_603301, EPI_ISL_603302, EPI_ISL_603303, EPI_ISL_603304, EPI_ISL_603305, EPI_ISL_603306, EPI_ISL_603308, EPI_ISL_603309, EPI_ISL_603310, EPI_ISL_603311, EPI_ISL_603312, EPI_ISL_603313, EPI_ISL_603314, EPI_ISL_603315, EPI_ISL_603316, EPI_ISL_603318, EPI_ISL_603319, EPI_ISL_603320, EPI_ISL_603321, EPI_ISL_603322, EPI_ISL_603323, EPI_ISL_603324, EPI_ISL_603325, EPI_ISL_603327, EPI_ISL_603328, EPI_ISL_603330, EPI_ISL_603331, EPI_ISL_603332, EPI_ISL_603333, EPI_ISL_603334, EPI_ISL_603335, EPI_ISL_603336, EPI_ISL_603337, EPI_ISL_603338, EPI_ISL_603339, EPI_ISL_603340, EPI_ISL_603341, EPI_ISL_603342, EPI_ISL_603343, EPI_ISL_603345, EPI_ISL_603346, EPI_ISL_603348, EPI_ISL_603350, EPI_ISL_603351, EPI_ISL_603352, EPI_ISL_603353, EPI_ISL_603354, EPI_ISL_603355, EPI_ISL_603357, EPI_ISL_603358, EPI_ISL_603359, EPI_ISL_603360, EPI_ISL_603361, EPI_ISL_603362, EPI_ISL_603363, EPI_ISL_603364, EPI_ISL_603365, EPI_ISL_603366, EPI_ISL_603367, EPI_ISL_603368, EPI_ISL_603369, EPI_ISL_603370, EPI_ISL_603371, EPI_ISL_603373, EPI_ISL_603374, EPI_ISL_603376, EPI_ISL_603377, EPI_ISL_603378, EPI_ISL_603379, EPI_ISL_603380, EPI_ISL_603381, EPI_ISL_603382, EPI_ISL_603383, EPI_ISL_603385, EPI_ISL_603386, EPI_ISL_603387, EPI_ISL_603388, EPI_ISL_603389, EPI_ISL_603390, EPI_ISL_603391, EPI_ISL_603393, EPI_ISL_603394, EPI_ISL_603395, EPI_ISL_603396, EPI_ISL_603397, EPI_ISL_603398, EPI_ISL_603399, EPI_ISL_603402, EPI_ISL_603403, EPI_ISL_603404, EPI_ISL_603405, EPI_ISL_603406, EPI_ISL_603407, EPI_ISL_603408, EPI_ISL_603409, EPI_ISL_603411, EPI_ISL_603412, EPI_ISL_603413, EPI_ISL_603414, EPI_ISL_603415, EPI_ISL_603416, EPI_ISL_603417, EPI_ISL_603418, EPI_ISL_603419, EPI_ISL_603420, EPI_ISL_603421, EPI_ISL_603422, EPI_ISL_603423, EPI_ISL_603424, EPI_ISL_603426, EPI_ISL_603427, EPI_ISL_603428, EPI_ISL_603429, EPI_ISL_603430, EPI_ISL_603431, EPI_ISL_603432, EPI_ISL_603433, EPI_ISL_603434, EPI_ISL_603435, EPI_ISL_603436, EPI_ISL_603437, EPI_ISL_603438, EPI_ISL_603439, EPI_ISL_603440, EPI_ISL_603441, EPI_ISL_603442, EPI_ISL_603443, EPI_ISL_603444, EPI_ISL_603445, EPI_ISL_603446, EPI_ISL_603447, EPI_ISL_603448, EPI_ISL_603449, EPI_ISL_603450, EPI_ISL_603451, EPI_ISL_603452, EPI_ISL_603453, EPI_ISL_603454, EPI_ISL_603455, EPI_ISL_603456, EPI_ISL_603457, EPI_ISL_603458, EPI_ISL_603459, EPI_ISL_603460, EPI_ISL_603461, EPI_ISL_603462, EPI_ISL_603463, EPI_ISL_603464, EPI_ISL_603465, EPI_ISL_603466, EPI_ISL_603468, EPI_ISL_603471, EPI_ISL_603472, EPI_ISL_603473, EPI_ISL_603474, EPI_ISL_603476, EPI_ISL_603477, EPI_ISL_603478, EPI_ISL_603480, EPI_ISL_603481, EPI_ISL_603482, EPI_ISL_603483, EPI_ISL_603484, EPI_ISL_603485, EPI_ISL_603486, EPI_ISL_603487, EPI_ISL_603488, EPI_ISL_603489, EPI_ISL_603490, EPI_ISL_603491, EPI_ISL_603492, EPI_ISL_603493, EPI_ISL_603494, EPI_ISL_603495, EPI_ISL_603496, EPI_ISL_603497, EPI_ISL_603498, EPI_ISL_603499, EPI_ISL_603500, EPI_ISL_603501, EPI_ISL_603502, EPI_ISL_603503, EPI_ISL_603504, EPI_ISL_603505, EPI_ISL_603506, EPI_ISL_603507, EPI_ISL_603508, EPI_ISL_603509, EPI_ISL_603510, EPI_ISL_603511, EPI_ISL_603512, EPI_ISL_603513, EPI_ISL_603514, EPI_ISL_603515, EPI_ISL_603516, EPI_ISL_603517, EPI_ISL_603518, EPI_ISL_603519, EPI_ISL_603520, EPI_ISL_603521, EPI_ISL_603522, EPI_ISL_603523, EPI_ISL_603524, EPI_ISL_603525, EPI_ISL_603526, EPI_ISL_603527, EPI_ISL_603528, EPI_ISL_603529, EPI_ISL_603530, EPI_ISL_603531, EPI_ISL_603533, EPI_ISL_603534, EPI_ISL_603536, EPI_ISL_603537, EPI_ISL_603538, EPI_ISL_603539, EPI_ISL_603540, EPI_ISL_603544, EPI_ISL_603545, EPI_ISL_603546, EPI_ISL_603547, EPI_ISL_603549, EPI_ISL_603550, EPI_ISL_603552, EPI_ISL_603553, EPI_ISL_603554, EPI_ISL_603555, EPI_ISL_603556, EPI_ISL_603557, EPI_ISL_603558, EPI_ISL_603559, EPI_ISL_603560, EPI_ISL_603561, EPI_ISL_603562, EPI_ISL_603563, EPI_ISL_603564, EPI_ISL_603565, EPI_ISL_603566, EPI_ISL_603567, EPI_ISL_603568, EPI_ISL_603569, EPI_ISL_603570, EPI_ISL_603571, EPI_ISL_603572, EPI_ISL_603573, EPI_ISL_603574, EPI_ISL_603575, EPI_ISL_603576, EPI_ISL_603577, EPI_ISL_603578, EPI_ISL_603579, EPI_ISL_603580, EPI_ISL_603581, EPI_ISL_603582, EPI_ISL_603583, EPI_ISL_603584, EPI_ISL_603585, EPI_ISL_603586, EPI_ISL_603587, EPI_ISL_603588, EPI_ISL_603589, EPI_ISL_603590, EPI_ISL_603591, EPI_ISL_603592, EPI_ISL_603593, EPI_ISL_603594, EPI_ISL_603596, EPI_ISL_603598, EPI_ISL_603599, EPI_ISL_603600, EPI_ISL_603601, EPI_ISL_603602, EPI_ISL_603603, EPI_ISL_603604, EPI_ISL_603605, EPI_ISL_603607, EPI_ISL_603608, EPI_ISL_603609, EPI_ISL_603610, EPI_ISL_603611, EPI_ISL_603612, EPI_ISL_603613, EPI_ISL_603614, EPI_ISL_603615, EPI_ISL_603616, EPI_ISL_603617, EPI_ISL_603618, EPI_ISL_603619, EPI_ISL_603620, EPI_ISL_603621, EPI_ISL_603623, EPI_ISL_603624, EPI_ISL_603625, EPI_ISL_603626, EPI_ISL_603627, EPI_ISL_603628, EPI_ISL_603629, EPI_ISL_603630, EPI_ISL_603631, EPI_ISL_603632, EPI_ISL_603633, EPI_ISL_603634, EPI_ISL_603635, EPI_ISL_603636, EPI_ISL_603637, EPI_ISL_603638, EPI_ISL_603639, EPI_ISL_603640, EPI_ISL_603641, EPI_ISL_603642, EPI_ISL_603643, EPI_ISL_603644, EPI_ISL_603645, EPI_ISL_603646, EPI_ISL_603647, EPI_ISL_603648, EPI_ISL_603649, EPI_ISL_603650, EPI_ISL_603651, EPI_ISL_603652, EPI_ISL_603653, EPI_ISL_603654, EPI_ISL_603655, EPI_ISL_603656, EPI_ISL_603657, EPI_ISL_603658, EPI_ISL_603659, EPI_ISL_603660, EPI_ISL_603661, EPI_ISL_603662, EPI_ISL_603663, EPI_ISL_603664, EPI_ISL_603665, EPI_ISL_603666, EPI_ISL_603667, EPI_ISL_603668, EPI_ISL_603669, EPI_ISL_603670, EPI_ISL_603671, EPI_ISL_603672, EPI_ISL_603673, EPI_ISL_603674, EPI_ISL_603676, EPI_ISL_603677, EPI_ISL_603678, EPI_ISL_603679, EPI_ISL_603680, EPI_ISL_603681, EPI_ISL_603682, EPI_ISL_603683, EPI_ISL_603684, EPI_ISL_603688, EPI_ISL_603689, EPI_ISL_603690, EPI_ISL_603691, EPI_ISL_603693, EPI_ISL_603694, EPI_ISL_603695, EPI_ISL_603697, EPI_ISL_603698, EPI_ISL_603700, EPI_ISL_603702, EPI_ISL_603703, EPI_ISL_603704, EPI_ISL_603705, EPI_ISL_603706, EPI_ISL_603707, EPI_ISL_603708, EPI_ISL_603709, EPI_ISL_603710, EPI_ISL_603711, EPI_ISL_603712, EPI_ISL_603713, EPI_ISL_603714, EPI_ISL_603715, EPI_ISL_603716, EPI_ISL_603717, EPI_ISL_603719, EPI_ISL_603720, EPI_ISL_603721, EPI_ISL_603722, EPI_ISL_603724, EPI_ISL_603725, EPI_ISL_603726, EPI_ISL_603727, EPI_ISL_603728, EPI_ISL_603730, EPI_ISL_603731, EPI_ISL_603732, EPI_ISL_603733, EPI_ISL_603734, EPI_ISL_603735, EPI_ISL_603736, EPI_ISL_603737, EPI_ISL_603738, EPI_ISL_603739, EPI_ISL_603740, EPI_ISL_603741, EPI_ISL_603742, EPI_ISL_603743, EPI_ISL_603744, EPI_ISL_603745, EPI_ISL_603746, EPI_ISL_603747, EPI_ISL_603748, EPI_ISL_603749, EPI_ISL_603750, EPI_ISL_603751, EPI_ISL_603753, EPI_ISL_603754, EPI_ISL_603755, EPI_ISL_603756, EPI_ISL_603758 | | | |
| see above | Viollier AG | Department of Biosystems Science and Engineering, ETH Zürich | Christian Beisel, Sarah Nadeau, Ivan Topolsky, Pedro Ferreira, Philipp Jablonski, Susana Posada-Céspedes, Tobias Schär, Ina Nissen, Natascha Santacroce, Elodie Burcklen, Christiane Beckmann, Maurice Redondo, Olivier Kobel, Christoph Noppen, Sophie Seidel, Noemie Santamaria de Souza, |

| | | | |
|---|---|---|---|
| | | | Niko Beerenwinkel, Tanja Stadler |
| EPI_ISL_611531, EPI_ISL_611628, EPI_ISL_611807, EPI_ISL_611808, EPI_ISL_612546, EPI_ISL_612547, EPI_ISL_612548 | Regional Virus Laboratory, Belfast Health and Social Care Trust | COVID-19 Genomics UK (COG-UK) Consortium | Conall McCaughey, James McKenna, Tanya Curran, Susan Feeney, Alison Watt, Ciara Cox, Mairead Connor, Zoltan Molnar, David Simpson, Derek Fairley |
| EPI_ISL_613410, EPI_ISL_613411 | Hospital San Pedro de Alcántara (Cáceres) | SeqCOVID-SPAIN consortium/IBV(CSIC) | Cristina Muñoz Cuevas, Guadalupe Rodriguez Rodriguez and SeqCOVID-SPAIN consortium |
| EPI_ISL_613541, EPI_ISL_613542, EPI_ISL_613543, EPI_ISL_613544 | Department of Virus and Microbiological Special Diagnostics, Statens Serum Institut, Denmark | Albertsen lab, Department of Chemistry and Bioscience, Aalborg University, Denmark | Danish Corona Genome Consortia |
| EPI_ISL_614404, EPI_ISL_614405, EPI_ISL_614407, EPI_ISL_614409, EPI_ISL_614410, EPI_ISL_614411, EPI_ISL_614414, EPI_ISL_614415, EPI_ISL_614416, EPI_ISL_614422, EPI_ISL_614423, EPI_ISL_614428, EPI_ISL_614429, EPI_ISL_614430, EPI_ISL_614431, EPI_ISL_614435, EPI_ISL_614436, EPI_ISL_614437, EPI_ISL_614438, EPI_ISL_614441, EPI_ISL_614442, EPI_ISL_614445, EPI_ISL_614446, EPI_ISL_614449, EPI_ISL_614461, EPI_ISL_614462, EPI_ISL_614463, EPI_ISL_614464, EPI_ISL_614465, EPI_ISL_614466, EPI_ISL_614467, EPI_ISL_614468, EPI_ISL_614469, EPI_ISL_614470, EPI_ISL_614471, EPI_ISL_614472, EPI_ISL_614515, EPI_ISL_614516, EPI_ISL_614517, EPI_ISL_614518, EPI_ISL_614519, EPI_ISL_614520, EPI_ISL_614521, EPI_ISL_614523, EPI_ISL_614524, EPI_ISL_614525, EPI_ISL_614526, EPI_ISL_614527, EPI_ISL_614528, EPI_ISL_614529, EPI_ISL_614530, EPI_ISL_614531, EPI_ISL_614532, EPI_ISL_614533, EPI_ISL_614534, EPI_ISL_614535, EPI_ISL_614536, EPI_ISL_614537, EPI_ISL_614542, EPI_ISL_614547, EPI_ISL_614548, EPI_ISL_614549, EPI_ISL_614550, EPI_ISL_614551, EPI_ISL_614552, EPI_ISL_614553, EPI_ISL_614554, EPI_ISL_614555, EPI_ISL_614556, EPI_ISL_614557, EPI_ISL_614558, EPI_ISL_614559, EPI_ISL_614560, EPI_ISL_614561, EPI_ISL_614562, EPI_ISL_614563, EPI_ISL_614565, EPI_ISL_614567, EPI_ISL_614568, EPI_ISL_614570, EPI_ISL_614571, EPI_ISL_614572, EPI_ISL_614573, EPI_ISL_614574, EPI_ISL_614575, EPI_ISL_614576, EPI_ISL_614577, EPI_ISL_614578, EPI_ISL_614579, EPI_ISL_614580, EPI_ISL_614581, EPI_ISL_614582, EPI_ISL_614583, EPI_ISL_614584, EPI_ISL_614586, EPI_ISL_614587, EPI_ISL_614590, EPI_ISL_614591, EPI_ISL_614592, EPI_ISL_614595, EPI_ISL_614596, EPI_ISL_614597, EPI_ISL_614598, EPI_ISL_614600, EPI_ISL_614601, EPI_ISL_614603, EPI_ISL_614604, EPI_ISL_614605, EPI_ISL_614606, EPI_ISL_614609, EPI_ISL_614610, EPI_ISL_614611, EPI_ISL_614612, EPI_ISL_614613, EPI_ISL_614614, EPI_ISL_614617, EPI_ISL_614618, EPI_ISL_614619, EPI_ISL_614621, EPI_ISL_614622, EPI_ISL_614623, EPI_ISL_614624, EPI_ISL_614625, EPI_ISL_614626, EPI_ISL_614627, EPI_ISL_614628, EPI_ISL_614629, EPI_ISL_614630, EPI_ISL_614631, EPI_ISL_614632, EPI_ISL_614633, EPI_ISL_614634, EPI_ISL_614635, EPI_ISL_614636, EPI_ISL_614637, EPI_ISL_614638, EPI_ISL_614639, EPI_ISL_614640, EPI_ISL_614641, EPI_ISL_614642, EPI_ISL_614643, EPI_ISL_614644, EPI_ISL_614645, EPI_ISL_614646, EPI_ISL_614647, EPI_ISL_614648, EPI_ISL_614649, EPI_ISL_614650, EPI_ISL_614651, EPI_ISL_614652, EPI_ISL_614653, EPI_ISL_614654, EPI_ISL_614655, EPI_ISL_614656, EPI_ISL_614657, EPI_ISL_614658, EPI_ISL_614659, EPI_ISL_614660, EPI_ISL_614661, EPI_ISL_614662, EPI_ISL_614663, EPI_ISL_614664, EPI_ISL_614665, EPI_ISL_614666, EPI_ISL_614667, EPI_ISL_614668, EPI_ISL_614669, EPI_ISL_614670, EPI_ISL_614695, EPI_ISL_614696, EPI_ISL_614713, EPI_ISL_614733, EPI_ISL_614753, EPI_ISL_614759, EPI_ISL_614760, EPI_ISL_614762, EPI_ISL_614765, EPI_ISL_614766, EPI_ISL_614767, EPI_ISL_614769, EPI_ISL_614771, EPI_ISL_614773, EPI_ISL_614784, EPI_ISL_614796, EPI_ISL_614797, EPI_ISL_614802, EPI_ISL_614803, EPI_ISL_614804, EPI_ISL_614810, EPI_ISL_614821, EPI_ISL_614823, EPI_ISL_614825, EPI_ISL_614830, EPI_ISL_614831, EPI_ISL_614832, EPI_ISL_614833, EPI_ISL_614834, EPI_ISL_614835, EPI_ISL_614836, EPI_ISL_614837, EPI_ISL_614838, EPI_ISL_614839, EPI_ISL_614840, EPI_ISL_614841, EPI_ISL_614842, EPI_ISL_614843, EPI_ISL_614844, EPI_ISL_614845, EPI_ISL_614846, EPI_ISL_614847, EPI_ISL_614848, EPI_ISL_614849, EPI_ISL_614850, EPI_ISL_614851, EPI_ISL_614852, EPI_ISL_614853, EPI_ISL_614854, EPI_ISL_614855, EPI_ISL_614856, EPI_ISL_614858, EPI_ISL_614859, EPI_ISL_614860, EPI_ISL_614861, EPI_ISL_614862, EPI_ISL_614863, EPI_ISL_614864, EPI_ISL_614865, EPI_ISL_614866, EPI_ISL_614867, EPI_ISL_614868, EPI_ISL_614869, EPI_ISL_614870, EPI_ISL_614871, EPI_ISL_614872, EPI_ISL_614873, EPI_ISL_614874, EPI_ISL_614875, EPI_ISL_614884, EPI_ISL_614885, EPI_ISL_614886, EPI_ISL_614887, EPI_ISL_614888 | | | |
| see above | Department of Virus and Microbiological Special Diagnostics, Statens Serum Institut, Denmark | Albertsen lab, Department of Chemistry and Bioscience, Aalborg University, Denmark | Danish Covid-19 Genome Consortia |
| EPI_ISL_614893, EPI_ISL_614894, EPI_ISL_614895, EPI_ISL_614896, EPI_ISL_614897, EPI_ISL_614898, EPI_ISL_614899, EPI_ISL_614900, EPI_ISL_614901, EPI_ISL_614902, EPI_ISL_614903, EPI_ISL_614904, EPI_ISL_614905, EPI_ISL_614906, EPI_ISL_614907, EPI_ISL_614908, EPI_ISL_614909, EPI_ISL_614910, EPI_ISL_614911, EPI_ISL_614912, EPI_ISL_614913, EPI_ISL_614914, EPI_ISL_614915, EPI_ISL_614916, EPI_ISL_614917, EPI_ISL_614918, EPI_ISL_614919, EPI_ISL_614920, EPI_ISL_614921, EPI_ISL_614922, EPI_ISL_614923, EPI_ISL_614924, EPI_ISL_614925, EPI_ISL_614926, EPI_ISL_614927, EPI_ISL_614928, EPI_ISL_614930, EPI_ISL_614931, EPI_ISL_614932, EPI_ISL_614933, EPI_ISL_614934, EPI_ISL_614935, EPI_ISL_614936, EPI_ISL_614937, EPI_ISL_614938, EPI_ISL_614939, EPI_ISL_614940, EPI_ISL_614941, EPI_ISL_614942, EPI_ISL_614943, EPI_ISL_614944, EPI_ISL_614945, EPI_ISL_614946, EPI_ISL_614947, EPI_ISL_614948, EPI_ISL_614949, EPI_ISL_614950, EPI_ISL_614951, EPI_ISL_614952, EPI_ISL_614953, EPI_ISL_614954, EPI_ISL_614955, EPI_ISL_614956, EPI_ISL_614957, EPI_ISL_614958, EPI_ISL_614959, EPI_ISL_614960, EPI_ISL_614961, EPI_ISL_614962, EPI_ISL_614963, EPI_ISL_614964, EPI_ISL_614965, EPI_ISL_614966, EPI_ISL_614967, EPI_ISL_614970, EPI_ISL_614971, EPI_ISL_614973 | | | |
| see above | Viollier AG | Department of Biosystems Science and Engineering, ETH Zürich | Christian Beisel, Sarah Nadeau, Ivan Topolsky, Pedro Ferreira, Philipp Jablonski, Susana Posada-Céspedes, Tobias Schär, Ina Nissen, Natascha Santacroce, Elodie Burcklen, Christiane Beckmann, Maurice Redondo, Olivier Kobel, Christoph Noppen, Sophie Seidel, Noemie Santamaria de Souza, Chaoran Chen, Niko Beerenwinkel, Tanja Stadler |
| EPI_ISL_615121 | Hospital de Pediatría "Prof. Dr. Juan P Garrahan" | Héritas | Cristian Rohr, Bianca Brun, Dalmacio Pereyra, Priscila Aldabe, Andrea Mangano, Maria Florencia Fernandez, Fabian Fay, Martin Vazquez |
| EPI_ISL_615372, EPI_ISL_615373, EPI_ISL_615374, EPI_ISL_615375, EPI_ISL_615404, EPI_ISL_615405, EPI_ISL_615406, EPI_ISL_615407, EPI_ISL_615408, EPI_ISL_615409, EPI_ISL_615410, EPI_ISL_615411, EPI_ISL_615412, EPI_ISL_615413, EPI_ISL_615414, EPI_ISL_615416, EPI_ISL_615417, EPI_ISL_615418, EPI_ISL_615419, EPI_ISL_615420, EPI_ISL_615421, EPI_ISL_615422, EPI_ISL_615423, EPI_ISL_615424, EPI_ISL_615425, EPI_ISL_615426, EPI_ISL_615427, EPI_ISL_615428, EPI_ISL_615429, EPI_ISL_615430, EPI_ISL_615431, EPI_ISL_615432, EPI_ISL_615433, EPI_ISL_615434, EPI_ISL_615435, EPI_ISL_615438, EPI_ISL_615439, EPI_ISL_615440, EPI_ISL_615441, EPI_ISL_615442, EPI_ISL_615443, EPI_ISL_615444, EPI_ISL_615445, EPI_ISL_615446, EPI_ISL_615447, EPI_ISL_615448, EPI_ISL_615449, EPI_ISL_615450, EPI_ISL_615453, EPI_ISL_615454, EPI_ISL_615455, EPI_ISL_615457, EPI_ISL_615458, EPI_ISL_615459, EPI_ISL_615460, EPI_ISL_615461, EPI_ISL_615463, EPI_ISL_615464, EPI_ISL_615465, EPI_ISL_615466, EPI_ISL_615467, EPI_ISL_615468, EPI_ISL_615469, EPI_ISL_615471, EPI_ISL_615472, EPI_ISL_615474, EPI_ISL_615475, EPI_ISL_615477, EPI_ISL_615478, EPI_ISL_615479, EPI_ISL_615480, EPI_ISL_615481, EPI_ISL_615482, EPI_ISL_615483, EPI_ISL_615484, EPI_ISL_615485, EPI_ISL_615486, EPI_ISL_615487, EPI_ISL_615488, EPI_ISL_615489, EPI_ISL_615490, EPI_ISL_615491, EPI_ISL_615492, EPI_ISL_615493, EPI_ISL_615494, EPI_ISL_615495, EPI_ISL_615496, EPI_ISL_615497, EPI_ISL_615498, EPI_ISL_615499, EPI_ISL_615501, EPI_ISL_615502, EPI_ISL_615504, EPI_ISL_615505, EPI_ISL_615506, EPI_ISL_615507, EPI_ISL_615508, EPI_ISL_615509, EPI_ISL_615511, EPI_ISL_615512, EPI_ISL_615513, EPI_ISL_615577, EPI_ISL_615583, EPI_ISL_615584, EPI_ISL_615585, EPI_ISL_615586, EPI_ISL_615587, EPI_ISL_615588, EPI_ISL_615589, EPI_ISL_615590, EPI_ISL_615591, EPI_ISL_615592, EPI_ISL_615593, EPI_ISL_615594, EPI_ISL_615595, EPI_ISL_615596, EPI_ISL_615597, EPI_ISL_615598, EPI_ISL_615599, EPI_ISL_615600, EPI_ISL_615602, EPI_ISL_615603, EPI_ISL_615604, EPI_ISL_615605, EPI_ISL_615606, EPI_ISL_615607, EPI_ISL_615609, EPI_ISL_615617, EPI_ISL_615636, EPI_ISL_615637, EPI_ISL_615906, EPI_ISL_615907, EPI_ISL_615908, EPI_ISL_615909, EPI_ISL_615910, EPI_ISL_615911, EPI_ISL_615912, EPI_ISL_615913, EPI_ISL_615914, EPI_ISL_615915, EPI_ISL_615916, EPI_ISL_615917, EPI_ISL_615918, EPI_ISL_615919, EPI_ISL_615920, EPI_ISL_615921, EPI_ISL_615922, EPI_ISL_615923, EPI_ISL_615924, EPI_ISL_615925, EPI_ISL_615926, EPI_ISL_615927, EPI_ISL_615928, EPI_ISL_615929, EPI_ISL_615930, EPI_ISL_615931, EPI_ISL_615932, EPI_ISL_615933, EPI_ISL_615934, EPI_ISL_615936, EPI_ISL_615937, EPI_ISL_615938, EPI_ISL_615939, EPI_ISL_615940, EPI_ISL_615941, EPI_ISL_615942, EPI_ISL_615943, EPI_ISL_615944, EPI_ISL_615945, EPI_ISL_615946, EPI_ISL_615947, EPI_ISL_615948, EPI_ISL_615949, EPI_ISL_615950, EPI_ISL_615951, EPI_ISL_615952, EPI_ISL_615953, EPI_ISL_615954, EPI_ISL_615955, EPI_ISL_615956, EPI_ISL_615957, EPI_ISL_615958, EPI_ISL_615959, EPI_ISL_615960, EPI_ISL_615961, EPI_ISL_615962, EPI_ISL_615963, EPI_ISL_615964, EPI_ISL_615965, EPI_ISL_615966, EPI_ISL_615967, EPI_ISL_615968, EPI_ISL_615969, EPI_ISL_615970, EPI_ISL_615971, EPI_ISL_615972, EPI_ISL_615973, EPI_ISL_615974, EPI_ISL_615975, EPI_ISL_615976, EPI_ISL_615977, EPI_ISL_615978, EPI_ISL_615979, EPI_ISL_615981, EPI_ISL_615982, EPI_ISL_615983, EPI_ISL_615984, EPI_ISL_615985, EPI_ISL_615986, EPI_ISL_615987, EPI_ISL_615989, EPI_ISL_615990, EPI_ISL_615991, EPI_ISL_615992, EPI_ISL_615993, EPI_ISL_615994, EPI_ISL_615995, EPI_ISL_615997, EPI_ISL_615998, EPI_ISL_615999, EPI_ISL_616000, EPI_ISL_616001, EPI_ISL_616002, EPI_ISL_616003, EPI_ISL_616004, EPI_ISL_616005, EPI_ISL_616006, EPI_ISL_616007, EPI_ISL_616008, EPI_ISL_616009, EPI_ISL_616010, EPI_ISL_616011, EPI_ISL_616012, EPI_ISL_616013, EPI_ISL_616014, EPI_ISL_616015, EPI_ISL_616016, EPI_ISL_616017, EPI_ISL_616018, EPI_ISL_616019, EPI_ISL_616020, EPI_ISL_616021, EPI_ISL_616022, EPI_ISL_616023, EPI_ISL_616024, EPI_ISL_616025, EPI_ISL_616026, EPI_ISL_616027, EPI_ISL_616028, EPI_ISL_616029, EPI_ISL_616030, EPI_ISL_616031, EPI_ISL_616032, EPI_ISL_616033, EPI_ISL_616035, EPI_ISL_616036, EPI_ISL_616037, EPI_ISL_616038, EPI_ISL_616039, EPI_ISL_616040, EPI_ISL_616041, EPI_ISL_616042, EPI_ISL_616043, EPI_ISL_616044, EPI_ISL_616045, EPI_ISL_616046, EPI_ISL_616047, EPI_ISL_616048, EPI_ISL_616050, EPI_ISL_616051, EPI_ISL_616052, EPI_ISL_616053, EPI_ISL_616054, EPI_ISL_616055, EPI_ISL_616056, EPI_ISL_616057, EPI_ISL_616058, EPI_ISL_616059, EPI_ISL_616060, EPI_ISL_616061, EPI_ISL_616062, EPI_ISL_616063, EPI_ISL_616064, EPI_ISL_616065, EPI_ISL_616066, EPI_ISL_616068, EPI_ISL_616069, EPI_ISL_616070, EPI_ISL_616071, EPI_ISL_616072, EPI_ISL_616073, EPI_ISL_616074, EPI_ISL_616075, EPI_ISL_616076, EPI_ISL_616077, EPI_ISL_616078, EPI_ISL_616079, EPI_ISL_616080, EPI_ISL_616081, EPI_ISL_616082, EPI_ISL_616083, EPI_ISL_616084, EPI_ISL_616085, EPI_ISL_616086, EPI_ISL_616087, EPI_ISL_616088, EPI_ISL_616089, EPI_ISL_616090, EPI_ISL_616091, EPI_ISL_616092, EPI_ISL_616093, EPI_ISL_616094, EPI_ISL_616095, EPI_ISL_616096, EPI_ISL_616097, EPI_ISL_616098, EPI_ISL_616099, EPI_ISL_616100, EPI_ISL_616101, EPI_ISL_616102, EPI_ISL_616103, EPI_ISL_616104, EPI_ISL_616105, EPI_ISL_616106, EPI_ISL_616107, EPI_ISL_616108, EPI_ISL_616109, EPI_ISL_616111, EPI_ISL_616112, EPI_ISL_616113, EPI_ISL_616114, EPI_ISL_616116, EPI_ISL_616117, EPI_ISL_616118, EPI_ISL_616119, EPI_ISL_616120, EPI_ISL_616121, EPI_ISL_616122, EPI_ISL_616123, EPI_ISL_616124, EPI_ISL_616125, EPI_ISL_616126, EPI_ISL_616127, EPI_ISL_616129, EPI_ISL_616130, EPI_ISL_616131, EPI_ISL_616132, EPI_ISL_616133, EPI_ISL_616134, EPI_ISL_616135, EPI_ISL_616136, EPI_ISL_616137, EPI_ISL_616138, EPI_ISL_616139, EPI_ISL_616140, EPI_ISL_616141, EPI_ISL_616142, EPI_ISL_616143, EPI_ISL_616144, EPI_ISL_616145, EPI_ISL_616146, EPI_ISL_616147, EPI_ISL_616148, EPI_ISL_616149, EPI_ISL_616150, EPI_ISL_616151, EPI_ISL_616152, EPI_ISL_616153, EPI_ISL_616154, EPI_ISL_616155, EPI_ISL_616156, EPI_ISL_616157, EPI_ISL_616158, EPI_ISL_616159, EPI_ISL_616160, EPI_ISL_616161, EPI_ISL_616162, EPI_ISL_616163, EPI_ISL_616164, EPI_ISL_616165, EPI_ISL_616166, EPI_ISL_616167, EPI_ISL_616168, EPI_ISL_616169, EPI_ISL_616170, EPI_ISL_616171, EPI_ISL_616172, EPI_ISL_616173, EPI_ISL_616174, EPI_ISL_616175, EPI_ISL_616176, EPI_ISL_616177, EPI_ISL_616178, EPI_ISL_616179, EPI_ISL_616180, EPI_ISL_616181, EPI_ISL_616182, EPI_ISL_616183, EPI_ISL_616184, EPI_ISL_616185, EPI_ISL_616186, EPI_ISL_616187, EPI_ISL_616188, EPI_ISL_616189, EPI_ISL_616191, EPI_ISL_616192, EPI_ISL_616193, EPI_ISL_616194, EPI_ISL_616195, EPI_ISL_616196, EPI_ISL_616197, EPI_ISL_616198, EPI_ISL_616199, EPI_ISL_616200, EPI_ISL_616201, EPI_ISL_616202, EPI_ISL_616203, EPI_ISL_616204, EPI_ISL_616205, EPI_ISL_616206, EPI_ISL_616207, EPI_ISL_616208, EPI_ISL_616209, EPI_ISL_616210, EPI_ISL_616211, EPI_ISL_616212, EPI_ISL_616213, EPI_ISL_616214, EPI_ISL_616215, EPI_ISL_616216, EPI_ISL_616217, EPI_ISL_616218, EPI_ISL_616219, EPI_ISL_616220, EPI_ISL_616221, EPI_ISL_616222, EPI_ISL_616223, EPI_ISL_616224, EPI_ISL_616225, EPI_ISL_616227, EPI_ISL_616229, EPI_ISL_616230, EPI_ISL_616231, EPI_ISL_616232, EPI_ISL_616234, EPI_ISL_616235, EPI_ISL_616236, EPI_ISL_616237, EPI_ISL_616238, EPI_ISL_616239, EPI_ISL_616240, EPI_ISL_616241, EPI_ISL_616242, EPI_ISL_616243, EPI_ISL_616244, EPI_ISL_616245, EPI_ISL_616246, EPI_ISL_616247, EPI_ISL_616248, EPI_ISL_616249, EPI_ISL_616250, EPI_ISL_616251, EPI_ISL_616252, EPI_ISL_616253, EPI_ISL_616254, EPI_ISL_616255, EPI_ISL_616256, EPI_ISL_616257, EPI_ISL_616258, EPI_ISL_616259, EPI_ISL_616260, EPI_ISL_616261, EPI_ISL_616262, EPI_ISL_616263, EPI_ISL_616264, EPI_ISL_616265, EPI_ISL_616266, EPI_ISL_616267, EPI_ISL_616268, EPI_ISL_616269, EPI_ISL_616270, EPI_ISL_616271, EPI_ISL_616272, EPI_ISL_616273, EPI_ISL_616274, EPI_ISL_616275, EPI_ISL_616276, EPI_ISL_616277, EPI_ISL_616278, EPI_ISL_616279, EPI_ISL_616280, EPI_ISL_616281, EPI_ISL_616282, EPI_ISL_616283, EPI_ISL_616284, EPI_ISL_616285, EPI_ISL_616286, EPI_ISL_616287, EPI_ISL_616288, EPI_ISL_616289, EPI_ISL_616290, EPI_ISL_616291, EPI_ISL_616292, EPI_ISL_616293, EPI_ISL_616294, EPI_ISL_616295, EPI_ISL_616296, EPI_ISL_616297, EPI_ISL_616298, EPI_ISL_616299, EPI_ISL_616300, EPI_ISL_616301, EPI_ISL_616302, EPI_ISL_616303, EPI_ISL_616304, EPI_ISL_616306, EPI_ISL_616307, EPI_ISL_616308, EPI_ISL_616309, EPI_ISL_616310, EPI_ISL_616311, EPI_ISL_616312, EPI_ISL_616313, EPI_ISL_616314, EPI_ISL_616315, EPI_ISL_616316, EPI_ISL_616317, EPI_ISL_616319, EPI_ISL_616320, EPI_ISL_616321, EPI_ISL_616322, EPI_ISL_616323, EPI_ISL_616324, EPI_ISL_616325, EPI_ISL_616326, EPI_ISL_616327, EPI_ISL_616328, EPI_ISL_616329, EPI_ISL_616330, EPI_ISL_616331, EPI_ISL_616332, EPI_ISL_616333, EPI_ISL_616334, EPI_ISL_616335, EPI_ISL_616337, EPI_ISL_616338, EPI_ISL_616339, EPI_ISL_616340, EPI_ISL_616341, EPI_ISL_616342, EPI_ISL_616343, EPI_ISL_616345, EPI_ISL_616346, EPI_ISL_616347, EPI_ISL_616348, EPI_ISL_616349, EPI_ISL_616350, EPI_ISL_616351, EPI_ISL_616352, EPI_ISL_616353, EPI_ISL_616354, EPI_ISL_616355, EPI_ISL_616356, EPI_ISL_616357, EPI_ISL_616358, EPI_ISL_616359, EPI_ISL_616360, EPI_ISL_616361, EPI_ISL_616362, EPI_ISL_616363, EPI_ISL_616364, EPI_ISL_616365, EPI_ISL_616366, EPI_ISL_616367, EPI_ISL_616368, EPI_ISL_616369, EPI_ISL_616370, EPI_ISL_616371, EPI_ISL_616372, EPI_ISL_616373, EPI_ISL_616374, EPI_ISL_616375, EPI_ISL_616376, EPI_ISL_616378, EPI_ISL_616379, EPI_ISL_616381, EPI_ISL_616382, EPI_ISL_616383, EPI_ISL_616384, EPI_ISL_616385, EPI_ISL_616386, EPI_ISL_616387, EPI_ISL_616388, EPI_ISL_616389, EPI_ISL_616390, EPI_ISL_616391, EPI_ISL_616392, EPI_ISL_616393, EPI_ISL_616394, EPI_ISL_616395, EPI_ISL_616396, EPI_ISL_616397, EPI_ISL_616401, EPI_ISL_616402, EPI_ISL_616403, EPI_ISL_616404, EPI_ISL_616405, EPI_ISL_616406, EPI_ISL_616407, EPI_ISL_616409, EPI_ISL_616415, EPI_ISL_616416, EPI_ISL_616417, EPI_ISL_616418, EPI_ISL_616419, EPI_ISL_616420, EPI_ISL_616421, EPI_ISL_616423, EPI_ISL_616424, EPI_ISL_616425, EPI_ISL_616426, EPI_ISL_616427, EPI_ISL_616428, EPI_ISL_616429, EPI_ISL_616430, EPI_ISL_616431, EPI_ISL_616432, EPI_ISL_616433, EPI_ISL_616434, EPI_ISL_616435, EPI_ISL_616436, EPI_ISL_616438, EPI_ISL_616439, EPI_ISL_616440, EPI_ISL_616441, EPI_ISL_616442, EPI_ISL_616443, EPI_ISL_616444, EPI_ISL_616445, EPI_ISL_616446, EPI_ISL_616447, EPI_ISL_616448, EPI_ISL_616449, EPI_ISL_616462, EPI_ISL_616500, EPI_ISL_616501, EPI_ISL_616502, EPI_ISL_616503, EPI_ISL_616504, EPI_ISL_616505, EPI_ISL_616506, EPI_ISL_616507, EPI_ISL_616508, EPI_ISL_616509, EPI_ISL_616510, EPI_ISL_616512, EPI_ISL_616514, EPI_ISL_616515, EPI_ISL_616516, EPI_ISL_616517, EPI_ISL_616518, EPI_ISL_616519, EPI_ISL_616520, EPI_ISL_616521, EPI_ISL_616522, EPI_ISL_616523, EPI_ISL_616524, EPI_ISL_616525, EPI_ISL_616526, EPI_ISL_616527, EPI_ISL_616528, EPI_ISL_616529, EPI_ISL_616530, EPI_ISL_616531, EPI_ISL_616532, EPI_ISL_616533, EPI_ISL_616569, EPI_ISL_616764, EPI_ISL_616765, EPI_ISL_616766, EPI_ISL_616767, EPI_ISL_616768, EPI_ISL_616769, EPI_ISL_616770, EPI_ISL_616772, EPI_ISL_616773, EPI_ISL_616774, EPI_ISL_616787, EPI_ISL_616789, EPI_ISL_616811, EPI_ISL_616812, EPI_ISL_616813, EPI_ISL_616814, EPI_ISL_616816, EPI_ISL_616817, EPI_ISL_616818, EPI_ISL_616819, EPI_ISL_616820, EPI_ISL_616821, EPI_ISL_616822, EPI_ISL_616823, EPI_ISL_616824, EPI_ISL_616825, EPI_ISL_616827, EPI_ISL_616828, EPI_ISL_616829, EPI_ISL_616830, EPI_ISL_616831, EPI_ISL_616832, EPI_ISL_616833, EPI_ISL_616834, EPI_ISL_616835, EPI_ISL_616836, EPI_ISL_616837, EPI_ISL_616838, EPI_ISL_616839, EPI_ISL_616840, EPI_ISL_616841, EPI_ISL_616842, EPI_ISL_616843, EPI_ISL_616844, EPI_ISL_616845, EPI_ISL_616846, EPI_ISL_616847, EPI_ISL_616848, EPI_ISL_616849, EPI_ISL_616850, EPI_ISL_616851, EPI_ISL_616852, EPI_ISL_616853, EPI_ISL_616854, EPI_ISL_616855, EPI_ISL_616856, EPI_ISL_616858, EPI_ISL_616859, EPI_ISL_616860, EPI_ISL_616861, EPI_ISL_616862, EPI_ISL_616863, EPI_ISL_616864, EPI_ISL_616865, EPI_ISL_616866, EPI_ISL_616867, EPI_ISL_616868, EPI_ISL_616869, EPI_ISL_616870, EPI_ISL_616871, EPI_ISL_616872, EPI_ISL_616873, EPI_ISL_616874, EPI_ISL_616876, EPI_ISL_616877, EPI_ISL_616878, EPI_ISL_616879, EPI_ISL_616880, EPI_ISL_616881, EPI_ISL_616882, EPI_ISL_616883, EPI_ISL_616884, EPI_ISL_616885, EPI_ISL_616886, EPI_ISL_616887, EPI_ISL_616888, EPI_ISL_616889, EPI_ISL_616890, EPI_ISL_616891, EPI_ISL_616892, EPI_ISL_616893, EPI_ISL_616895, EPI_ISL_616896, EPI_ISL_616897, EPI_ISL_616898, EPI_ISL_616899, EPI_ISL_616900, EPI_ISL_616901, EPI_ISL_616902, EPI_ISL_616903, EPI_ISL_616904, EPI_ISL_616905, EPI_ISL_616923, EPI_ISL_616924, EPI_ISL_616975, EPI_ISL_617107, EPI_ISL_617284, EPI_ISL_617382, EPI_ISL_617383, EPI_ISL_617406, EPI_ISL_617407, EPI_ISL_617409, EPI_ISL_617411, EPI_ISL_617412, EPI_ISL_617413, EPI_ISL_617414, EPI_ISL_617416, EPI_ISL_617417, EPI_ISL_617418, EPI_ISL_617419, EPI_ISL_617420, EPI_ISL_617422, EPI_ISL_617423, EPI_ISL_617424, EPI_ISL_617425, EPI_ISL_617427, EPI_ISL_617429, EPI_ISL_617430, EPI_ISL_617431, EPI_ISL_617432, EPI_ISL_617433, EPI_ISL_617434, EPI_ISL_617435, EPI_ISL_617436, EPI_ISL_617437, EPI_ISL_617438, EPI_ISL_617439, EPI_ISL_617440, EPI_ISL_617442, EPI_ISL_617443, EPI_ISL_617444, EPI_ISL_617445, EPI_ISL_617446, EPI_ISL_617447, EPI_ISL_617448, EPI_ISL_617449, EPI_ISL_617450, EPI_ISL_617451, EPI_ISL_617452, EPI_ISL_617453, EPI_ISL_617454, EPI_ISL_617455, EPI_ISL_617456, EPI_ISL_617457, EPI_ISL_617458, EPI_ISL_617459, EPI_ISL_617460, EPI_ISL_617461, EPI_ISL_617462, EPI_ISL_617463, EPI_ISL_617464, EPI_ISL_617465, EPI_ISL_617466, EPI_ISL_617467, EPI_ISL_617468, EPI_ISL_617469, EPI_ISL_617470, EPI_ISL_617472, EPI_ISL_617473, EPI_ISL_617474, EPI_ISL_617475, EPI_ISL_617476, EPI_ISL_617477, EPI_ISL_617478, EPI_ISL_617479, EPI_ISL_617480, EPI_ISL_617481, EPI_ISL_617482, EPI_ISL_617483, EPI_ISL_617484, EPI_ISL_617485, EPI_ISL_617486, EPI_ISL_617487, EPI_ISL_617488, EPI_ISL_617489, EPI_ISL_617490, EPI_ISL_617492, EPI_ISL_617493, EPI_ISL_617494, EPI_ISL_617495, EPI_ISL_617496, EPI_ISL_617497, EPI_ISL_617498, EPI_ISL_617499, EPI_ISL_617500, EPI_ISL_617501, EPI_ISL_617502, EPI_ISL_617503, EPI_ISL_617504, EPI_ISL_617505, EPI_ISL_617506, EPI_ISL_617507, EPI_ISL_617508, EPI_ISL_617509, EPI_ISL_617510, EPI_ISL_617511, EPI_ISL_617512, EPI_ISL_617513, EPI_ISL_617514, EPI_ISL_617515, EPI_ISL_617516, EPI_ISL_617517, EPI_ISL_617518, EPI_ISL_617519, EPI_ISL_617520, EPI_ISL_617521, EPI_ISL_617522, EPI_ISL_617523, EPI_ISL_617525, EPI_ISL_617526, EPI_ISL_617527, EPI_ISL_617528, EPI_ISL_617529, | | | |

EPI_ISL_617530, EPI_ISL_617531, EPI_ISL_617532, EPI_ISL_617533, EPI_ISL_617534, EPI_ISL_617535, EPI_ISL_617536, EPI_ISL_617537, EPI_ISL_617538, EPI_ISL_617539, EPI_ISL_617540, EPI_ISL_617541, EPI_ISL_617542, EPI_ISL_617543, EPI_ISL_617544, EPI_ISL_617545, EPI_ISL_617546, EPI_ISL_617547,
EPI_ISL_617548, EPI_ISL_617549, EPI_ISL_617550, EPI_ISL_617551, EPI_ISL_617552, EPI_ISL_617553, EPI_ISL_617554, EPI_ISL_617555, EPI_ISL_617556, EPI_ISL_617557, EPI_ISL_617558, EPI_ISL_617560, EPI_ISL_617561, EPI_ISL_617562, EPI_ISL_617563, EPI_ISL_617564, EPI_ISL_617565, EPI_ISL_617566,
EPI_ISL_617567, EPI_ISL_617568, EPI_ISL_617569, EPI_ISL_617570, EPI_ISL_617571, EPI_ISL_617572, EPI_ISL_617573, EPI_ISL_617574, EPI_ISL_617575, EPI_ISL_617576, EPI_ISL_617577, EPI_ISL_617578, EPI_ISL_617579, EPI_ISL_617580, EPI_ISL_617581, EPI_ISL_617582, EPI_ISL_617583, EPI_ISL_617584,
EPI_ISL_617585, EPI_ISL_617586, EPI_ISL_617587, EPI_ISL_617588, EPI_ISL_617589, EPI_ISL_617590, EPI_ISL_617591, EPI_ISL_617592, EPI_ISL_617593, EPI_ISL_617594, EPI_ISL_617595, EPI_ISL_617596, EPI_ISL_617597, EPI_ISL_617598, EPI_ISL_617599, EPI_ISL_617600, EPI_ISL_617601, EPI_ISL_617602,
EPI_ISL_617603, EPI_ISL_617604, EPI_ISL_617605, EPI_ISL_617606, EPI_ISL_617607, EPI_ISL_617608, EPI_ISL_617609, EPI_ISL_617610, EPI_ISL_617611, EPI_ISL_617612, EPI_ISL_617613, EPI_ISL_617614, EPI_ISL_617615, EPI_ISL_617616, EPI_ISL_617617, EPI_ISL_617619, EPI_ISL_617620, EPI_ISL_617621,
EPI_ISL_617622, EPI_ISL_617623, EPI_ISL_617624, EPI_ISL_617625, EPI_ISL_617626, EPI_ISL_617627, EPI_ISL_617628, EPI_ISL_617629, EPI_ISL_617630, EPI_ISL_617631, EPI_ISL_617632, EPI_ISL_617633, EPI_ISL_617634, EPI_ISL_617635, EPI_ISL_617636, EPI_ISL_617637, EPI_ISL_617638, EPI_ISL_617639,
EPI_ISL_617640, EPI_ISL_617641, EPI_ISL_617642, EPI_ISL_617643, EPI_ISL_617644, EPI_ISL_617645, EPI_ISL_617646, EPI_ISL_617647, EPI_ISL_617648, EPI_ISL_617650, EPI_ISL_617651, EPI_ISL_617652, EPI_ISL_617653, EPI_ISL_617654, EPI_ISL_617655, EPI_ISL_617656, EPI_ISL_617657, EPI_ISL_617658,
EPI_ISL_617659, EPI_ISL_617660, EPI_ISL_617661, EPI_ISL_617662, EPI_ISL_617663, EPI_ISL_617664, EPI_ISL_617665, EPI_ISL_617666, EPI_ISL_617667, EPI_ISL_617668, EPI_ISL_617669, EPI_ISL_617670, EPI_ISL_617671, EPI_ISL_617672, EPI_ISL_617673, EPI_ISL_617674, EPI_ISL_617675, EPI_ISL_617676,
EPI_ISL_617677, EPI_ISL_617678, EPI_ISL_617679, EPI_ISL_617680, EPI_ISL_617681, EPI_ISL_617682, EPI_ISL_617683, EPI_ISL_617684, EPI_ISL_617685, EPI_ISL_617686, EPI_ISL_617687, EPI_ISL_617688, EPI_ISL_617689, EPI_ISL_617690, EPI_ISL_617691, EPI_ISL_617692, EPI_ISL_617693, EPI_ISL_617694,
EPI_ISL_617695, EPI_ISL_617696, EPI_ISL_617698, EPI_ISL_617699, EPI_ISL_617700, EPI_ISL_617701, EPI_ISL_617702, EPI_ISL_617703, EPI_ISL_617704, EPI_ISL_617705, EPI_ISL_617706, EPI_ISL_617707, EPI_ISL_617708, EPI_ISL_617709, EPI_ISL_617710, EPI_ISL_617711, EPI_ISL_617712, EPI_ISL_617713,
EPI_ISL_617714, EPI_ISL_617716, EPI_ISL_617717, EPI_ISL_617718, EPI_ISL_617719, EPI_ISL_617720, EPI_ISL_617721, EPI_ISL_617722, EPI_ISL_617723, EPI_ISL_617724, EPI_ISL_617725, EPI_ISL_617726, EPI_ISL_617728, EPI_ISL_617729, EPI_ISL_617730, EPI_ISL_617731, EPI_ISL_617732, EPI_ISL_617733,
EPI_ISL_617734, EPI_ISL_617735, EPI_ISL_617736, EPI_ISL_617737, EPI_ISL_617738, EPI_ISL_617739, EPI_ISL_617740, EPI_ISL_617741, EPI_ISL_617742, EPI_ISL_617743, EPI_ISL_617744, EPI_ISL_617745, EPI_ISL_617746, EPI_ISL_617747, EPI_ISL_617748, EPI_ISL_617749, EPI_ISL_617750, EPI_ISL_617751,
EPI_ISL_617752, EPI_ISL_617753, EPI_ISL_617754, EPI_ISL_617755, EPI_ISL_617756, EPI_ISL_617757, EPI_ISL_617758, EPI_ISL_617759, EPI_ISL_617760, EPI_ISL_617761, EPI_ISL_617762, EPI_ISL_617763, EPI_ISL_617764, EPI_ISL_617765, EPI_ISL_617766, EPI_ISL_617767, EPI_ISL_617768, EPI_ISL_617769,
EPI_ISL_617770, EPI_ISL_617771, EPI_ISL_617772, EPI_ISL_617773, EPI_ISL_617774, EPI_ISL_617775, EPI_ISL_617776, EPI_ISL_617777, EPI_ISL_617778, EPI_ISL_617779, EPI_ISL_617780, EPI_ISL_617781, EPI_ISL_617782, EPI_ISL_617783, EPI_ISL_617785, EPI_ISL_617786, EPI_ISL_617787, EPI_ISL_617788,
EPI_ISL_617789, EPI_ISL_617790, EPI_ISL_617791, EPI_ISL_617792, EPI_ISL_617793, EPI_ISL_617794, EPI_ISL_617795, EPI_ISL_617796, EPI_ISL_617797, EPI_ISL_617798, EPI_ISL_617799, EPI_ISL_617800, EPI_ISL_617801, EPI_ISL_617802, EPI_ISL_617803, EPI_ISL_617804, EPI_ISL_617805, EPI_ISL_617806,
EPI_ISL_617807, EPI_ISL_617808, EPI_ISL_617809, EPI_ISL_617810, EPI_ISL_617811, EPI_ISL_617812, EPI_ISL_617813, EPI_ISL_617814, EPI_ISL_617815, EPI_ISL_617816, EPI_ISL_617817, EPI_ISL_617818, EPI_ISL_617819, EPI_ISL_617820, EPI_ISL_617821, EPI_ISL_617822, EPI_ISL_617823, EPI_ISL_617824,
EPI_ISL_617825, EPI_ISL_617826, EPI_ISL_617827, EPI_ISL_617828, EPI_ISL_617829, EPI_ISL_617830, EPI_ISL_617831, EPI_ISL_617832, EPI_ISL_617833, EPI_ISL_617834, EPI_ISL_617835, EPI_ISL_617836, EPI_ISL_617837, EPI_ISL_617838, EPI_ISL_617839, EPI_ISL_617840, EPI_ISL_617841, EPI_ISL_617842,
EPI_ISL_617843, EPI_ISL_617844, EPI_ISL_617845, EPI_ISL_617846, EPI_ISL_617847, EPI_ISL_617848, EPI_ISL_617849, EPI_ISL_617851, EPI_ISL_617852, EPI_ISL_617853, EPI_ISL_617854, EPI_ISL_617855, EPI_ISL_617856, EPI_ISL_617857, EPI_ISL_617858, EPI_ISL_617859, EPI_ISL_617860, EPI_ISL_617861,
EPI_ISL_617862, EPI_ISL_617863, EPI_ISL_617864, EPI_ISL_617865, EPI_ISL_617866, EPI_ISL_617867, EPI_ISL_617868, EPI_ISL_617869, EPI_ISL_617870, EPI_ISL_617871, EPI_ISL_617872, EPI_ISL_617873, EPI_ISL_617874, EPI_ISL_617875, EPI_ISL_617876, EPI_ISL_617877, EPI_ISL_617878, EPI_ISL_617879,
EPI_ISL_617880, EPI_ISL_617881, EPI_ISL_617882, EPI_ISL_617883, EPI_ISL_617884, EPI_ISL_617885, EPI_ISL_617886, EPI_ISL_617887, EPI_ISL_617888, EPI_ISL_617889, EPI_ISL_617890, EPI_ISL_617891, EPI_ISL_617892, EPI_ISL_617893, EPI_ISL_617894, EPI_ISL_617895, EPI_ISL_617896, EPI_ISL_617897,
EPI_ISL_617898, EPI_ISL_617899, EPI_ISL_617900, EPI_ISL_617901, EPI_ISL_617902, EPI_ISL_617903, EPI_ISL_617904, EPI_ISL_617905, EPI_ISL_617939, EPI_ISL_617944, EPI_ISL_617945, EPI_ISL_617947, EPI_ISL_617948, EPI_ISL_617949, EPI_ISL_617950, EPI_ISL_617951, EPI_ISL_617952, EPI_ISL_617953,
EPI_ISL_617954, EPI_ISL_617955, EPI_ISL_617958, EPI_ISL_617961, EPI_ISL_617962, EPI_ISL_617963, EPI_ISL_617964, EPI_ISL_617965, EPI_ISL_617966, EPI_ISL_617967, EPI_ISL_617968, EPI_ISL_617969, EPI_ISL_617970, EPI_ISL_617971, EPI_ISL_617972, EPI_ISL_617973, EPI_ISL_617974, EPI_ISL_617975,
EPI_ISL_617976, EPI_ISL_617977, EPI_ISL_617979, EPI_ISL_617980, EPI_ISL_617981, EPI_ISL_617982, EPI_ISL_617983, EPI_ISL_617984, EPI_ISL_617985, EPI_ISL_617986, EPI_ISL_617987, EPI_ISL_617988, EPI_ISL_617989, EPI_ISL_617990, EPI_ISL_617991, EPI_ISL_617992, EPI_ISL_617993, EPI_ISL_617994,
EPI_ISL_617995, EPI_ISL_617996, EPI_ISL_617997, EPI_ISL_617998, EPI_ISL_617999, EPI_ISL_618000, EPI_ISL_618001, EPI_ISL_618002, EPI_ISL_618003, EPI_ISL_618004, EPI_ISL_618005, EPI_ISL_618006, EPI_ISL_618007, EPI_ISL_618008, EPI_ISL_618009, EPI_ISL_618010, EPI_ISL_618011, EPI_ISL_618012,
EPI_ISL_618013, EPI_ISL_618014, EPI_ISL_618015, EPI_ISL_618016, EPI_ISL_618017, EPI_ISL_618018, EPI_ISL_618019, EPI_ISL_618020, EPI_ISL_618021, EPI_ISL_618022, EPI_ISL_618023, EPI_ISL_618024, EPI_ISL_618025, EPI_ISL_618026, EPI_ISL_618027, EPI_ISL_618028, EPI_ISL_618029, EPI_ISL_618030,
EPI_ISL_618031, EPI_ISL_618039, EPI_ISL_618040, EPI_ISL_618041, EPI_ISL_618043, EPI_ISL_618045, EPI_ISL_618047, EPI_ISL_618048, EPI_ISL_618049, EPI_ISL_618050, EPI_ISL_618051, EPI_ISL_618052, EPI_ISL_618053, EPI_ISL_618054, EPI_ISL_618056, EPI_ISL_618057, EPI_ISL_618058, EPI_ISL_618062,
EPI_ISL_618064, EPI_ISL_618065, EPI_ISL_618066, EPI_ISL_618067, EPI_ISL_618068, EPI_ISL_618069, EPI_ISL_618070, EPI_ISL_618071, EPI_ISL_618072, EPI_ISL_618073, EPI_ISL_618074, EPI_ISL_618076, EPI_ISL_618077, EPI_ISL_618078, EPI_ISL_618079, EPI_ISL_618080, EPI_ISL_618081, EPI_ISL_618082,
EPI_ISL_618083, EPI_ISL_618084, EPI_ISL_618085, EPI_ISL_618086, EPI_ISL_618088, EPI_ISL_618089, EPI_ISL_618090, EPI_ISL_618091, EPI_ISL_618092, EPI_ISL_618093, EPI_ISL_618094, EPI_ISL_618095, EPI_ISL_618096, EPI_ISL_618097, EPI_ISL_618098, EPI_ISL_618099, EPI_ISL_618100, EPI_ISL_618101,
EPI_ISL_618102, EPI_ISL_618103, EPI_ISL_618104, EPI_ISL_618105, EPI_ISL_618106, EPI_ISL_618107, EPI_ISL_618108, EPI_ISL_618109, EPI_ISL_618110, EPI_ISL_618111, EPI_ISL_618112, EPI_ISL_618113, EPI_ISL_618114, EPI_ISL_618115, EPI_ISL_618116, EPI_ISL_618117, EPI_ISL_618118, EPI_ISL_618119,
EPI_ISL_618120, EPI_ISL_618121, EPI_ISL_618122, EPI_ISL_618124, EPI_ISL_618125, EPI_ISL_618127, EPI_ISL_618128, EPI_ISL_618129, EPI_ISL_618130, EPI_ISL_618131, EPI_ISL_618132, EPI_ISL_618133, EPI_ISL_618134, EPI_ISL_618135, EPI_ISL_618136, EPI_ISL_618139, EPI_ISL_618140, EPI_ISL_618141,
EPI_ISL_618143, EPI_ISL_618144, EPI_ISL_618145, EPI_ISL_618146, EPI_ISL_618147, EPI_ISL_618148, EPI_ISL_618149, EPI_ISL_618150, EPI_ISL_618151, EPI_ISL_618152, EPI_ISL_618154, EPI_ISL_618155, EPI_ISL_618156, EPI_ISL_618157, EPI_ISL_618158, EPI_ISL_618159, EPI_ISL_618160, EPI_ISL_618161,
EPI_ISL_618162, EPI_ISL_618163, EPI_ISL_618164, EPI_ISL_618165, EPI_ISL_618166, EPI_ISL_618167, EPI_ISL_618169, EPI_ISL_618170, EPI_ISL_618171, EPI_ISL_618172, EPI_ISL_618173, EPI_ISL_618174, EPI_ISL_618175, EPI_ISL_618176, EPI_ISL_618177, EPI_ISL_618178, EPI_ISL_618179, EPI_ISL_618180,
EPI_ISL_618181, EPI_ISL_618182, EPI_ISL_618183, EPI_ISL_618184, EPI_ISL_618185, EPI_ISL_618186, EPI_ISL_618187, EPI_ISL_618188, EPI_ISL_618189, EPI_ISL_618190, EPI_ISL_618191, EPI_ISL_618192, EPI_ISL_618194, EPI_ISL_618195, EPI_ISL_618196, EPI_ISL_618197, EPI_ISL_618198, EPI_ISL_618199,
EPI_ISL_618200, EPI_ISL_618201, EPI_ISL_618202, EPI_ISL_618203, EPI_ISL_618204, EPI_ISL_618205, EPI_ISL_618206, EPI_ISL_618207, EPI_ISL_618208, EPI_ISL_618209, EPI_ISL_618210, EPI_ISL_618211, EPI_ISL_618212, EPI_ISL_618213, EPI_ISL_618214, EPI_ISL_618215, EPI_ISL_618218, EPI_ISL_618219,
EPI_ISL_618220, EPI_ISL_618221, EPI_ISL_618222, EPI_ISL_618224, EPI_ISL_618225, EPI_ISL_618226, EPI_ISL_618227, EPI_ISL_618228, EPI_ISL_618229, EPI_ISL_618230, EPI_ISL_618231, EPI_ISL_618232, EPI_ISL_618233, EPI_ISL_618234, EPI_ISL_618235, EPI_ISL_618236, EPI_ISL_618237, EPI_ISL_618238,
EPI_ISL_618239, EPI_ISL_618240, EPI_ISL_618241, EPI_ISL_618242, EPI_ISL_618243, EPI_ISL_618244, EPI_ISL_618245, EPI_ISL_618246, EPI_ISL_618247, EPI_ISL_618248, EPI_ISL_618249, EPI_ISL_618250, EPI_ISL_618251, EPI_ISL_618252, EPI_ISL_618253, EPI_ISL_618254, EPI_ISL_618255, EPI_ISL_618256,
EPI_ISL_618257, EPI_ISL_618258, EPI_ISL_618259, EPI_ISL_618260, EPI_ISL_618261, EPI_ISL_618262, EPI_ISL_618263, EPI_ISL_618264, EPI_ISL_618266, EPI_ISL_618267, EPI_ISL_618268, EPI_ISL_618269, EPI_ISL_618270, EPI_ISL_618271, EPI_ISL_618272, EPI_ISL_618273, EPI_ISL_618275, EPI_ISL_618276,
EPI_ISL_618277, EPI_ISL_618278, EPI_ISL_618279, EPI_ISL_618280, EPI_ISL_618281, EPI_ISL_618282, EPI_ISL_618283, EPI_ISL_618284, EPI_ISL_618285, EPI_ISL_618286, EPI_ISL_618287, EPI_ISL_618288, EPI_ISL_618290, EPI_ISL_618291, EPI_ISL_618292, EPI_ISL_618293, EPI_ISL_618294, EPI_ISL_618295,
EPI_ISL_618296, EPI_ISL_618297, EPI_ISL_618298, EPI_ISL_618299, EPI_ISL_618300, EPI_ISL_618301, EPI_ISL_618302, EPI_ISL_618303, EPI_ISL_618304, EPI_ISL_618305, EPI_ISL_618306, EPI_ISL_618307, EPI_ISL_618308, EPI_ISL_618309, EPI_ISL_618310, EPI_ISL_618311, EPI_ISL_618312, EPI_ISL_618313,
EPI_ISL_618314, EPI_ISL_618316, EPI_ISL_618317, EPI_ISL_618318, EPI_ISL_618319, EPI_ISL_618320, EPI_ISL_618321, EPI_ISL_618322, EPI_ISL_618323, EPI_ISL_618324, EPI_ISL_618325, EPI_ISL_618326, EPI_ISL_618327, EPI_ISL_618328, EPI_ISL_618329, EPI_ISL_618330, EPI_ISL_618331, EPI_ISL_618333,
EPI_ISL_618335, EPI_ISL_618336, EPI_ISL_618337, EPI_ISL_618338, EPI_ISL_618339, EPI_ISL_618340, EPI_ISL_618341, EPI_ISL_618342, EPI_ISL_618343, EPI_ISL_618344, EPI_ISL_618345, EPI_ISL_618346, EPI_ISL_618347, EPI_ISL_618348, EPI_ISL_618349, EPI_ISL_618350, EPI_ISL_618351, EPI_ISL_618352,
EPI_ISL_618353, EPI_ISL_618354, EPI_ISL_618355, EPI_ISL_618356, EPI_ISL_618357, EPI_ISL_618358, EPI_ISL_618359, EPI_ISL_618360, EPI_ISL_618361, EPI_ISL_618362, EPI_ISL_618363, EPI_ISL_618364, EPI_ISL_618365, EPI_ISL_618366, EPI_ISL_618367, EPI_ISL_618368, EPI_ISL_618369, EPI_ISL_618370,
EPI_ISL_618371, EPI_ISL_618372, EPI_ISL_618374, EPI_ISL_618375, EPI_ISL_618376, EPI_ISL_618377, EPI_ISL_618378, EPI_ISL_618379, EPI_ISL_618380, EPI_ISL_618381, EPI_ISL_618383, EPI_ISL_618384, EPI_ISL_618385, EPI_ISL_618386, EPI_ISL_618387, EPI_ISL_618388, EPI_ISL_618389, EPI_ISL_618390,
EPI_ISL_618391, EPI_ISL_618392, EPI_ISL_618393, EPI_ISL_618394, EPI_ISL_618395, EPI_ISL_618396, EPI_ISL_618397, EPI_ISL_618398, EPI_ISL_618399, EPI_ISL_618400, EPI_ISL_618402, EPI_ISL_618403, EPI_ISL_618404, EPI_ISL_618405, EPI_ISL_618479, EPI_ISL_618496, EPI_ISL_618511, EPI_ISL_618517,
EPI_ISL_618525, EPI_ISL_618595, EPI_ISL_618609, EPI_ISL_618626, EPI_ISL_618627, EPI_ISL_618628, EPI_ISL_618629, EPI_ISL_618630, EPI_ISL_618631, EPI_ISL_618632, EPI_ISL_618633, EPI_ISL_618634, EPI_ISL_618635, EPI_ISL_618636, EPI_ISL_618637, EPI_ISL_618638, EPI_ISL_618639, EPI_ISL_618640,
EPI_ISL_618641, EPI_ISL_618643, EPI_ISL_618644, EPI_ISL_618645, EPI_ISL_618646, EPI_ISL_618647, EPI_ISL_618648, EPI_ISL_618649, EPI_ISL_618650, EPI_ISL_618651, EPI_ISL_618652, EPI_ISL_618653, EPI_ISL_618654, EPI_ISL_618655, EPI_ISL_618656, EPI_ISL_618657, EPI_ISL_618659, EPI_ISL_618660,
EPI_ISL_618661, EPI_ISL_618720, EPI_ISL_618801, EPI_ISL_618802, EPI_ISL_618803, EPI_ISL_618804, EPI_ISL_618805, EPI_ISL_618806, EPI_ISL_618807, EPI_ISL_618808, EPI_ISL_618969, EPI_ISL_618972, EPI_ISL_618978, EPI_ISL_618979, EPI_ISL_618980, EPI_ISL_619017, EPI_ISL_619018, EPI_ISL_619133,
EPI_ISL_619141, EPI_ISL_619145, EPI_ISL_619151, EPI_ISL_619163, EPI_ISL_619170, EPI_ISL_619176, EPI_ISL_619185, EPI_ISL_619211, EPI_ISL_619226, EPI_ISL_619233, EPI_ISL_619256, EPI_ISL_619260, EPI_ISL_619261, EPI_ISL_619262, EPI_ISL_619263, EPI_ISL_619265, EPI_ISL_619266, EPI_ISL_619267,
EPI_ISL_619268, EPI_ISL_619269, EPI_ISL_619270, EPI_ISL_619271, EPI_ISL_619272, EPI_ISL_619273, EPI_ISL_619292, EPI_ISL_619318, EPI_ISL_619321, EPI_ISL_619328, EPI_ISL_619332, EPI_ISL_619337, EPI_ISL_619344, EPI_ISL_619378, EPI_ISL_619384, EPI_ISL_619389, EPI_ISL_619398, EPI_ISL_619405,
EPI_ISL_619455, EPI_ISL_619492, EPI_ISL_619534, EPI_ISL_619556, EPI_ISL_619751, EPI_ISL_619885, EPI_ISL_619895, EPI_ISL_619906, EPI_ISL_619907, EPI_ISL_619908, EPI_ISL_619909, EPI_ISL_619910, EPI_ISL_619911, EPI_ISL_619912, EPI_ISL_619913, EPI_ISL_619914, EPI_ISL_619915, EPI_ISL_619916,
EPI_ISL_619917, EPI_ISL_619918, EPI_ISL_619920, EPI_ISL_619921, EPI_ISL_619922, EPI_ISL_619923, EPI_ISL_619924, EPI_ISL_619925, EPI_ISL_619926, EPI_ISL_619927, EPI_ISL_619928, EPI_ISL_619929, EPI_ISL_619930, EPI_ISL_619931, EPI_ISL_619932, EPI_ISL_619933, EPI_ISL_619934, EPI_ISL_619935,
EPI_ISL_619936, EPI_ISL_619937, EPI_ISL_619938, EPI_ISL_619939, EPI_ISL_619940, EPI_ISL_619941, EPI_ISL_619942, EPI_ISL_619943, EPI_ISL_619944, EPI_ISL_619945, EPI_ISL_619946, EPI_ISL_619948, EPI_ISL_619949, EPI_ISL_619950, EPI_ISL_619951, EPI_ISL_619952, EPI_ISL_619954, EPI_ISL_619955,
EPI_ISL_619956, EPI_ISL_619957, EPI_ISL_619959, EPI_ISL_619960, EPI_ISL_619962, EPI_ISL_619964, EPI_ISL_619965, EPI_ISL_619966, EPI_ISL_619967, EPI_ISL_619968, EPI_ISL_619969, EPI_ISL_619970, EPI_ISL_619971, EPI_ISL_619973, EPI_ISL_619974, EPI_ISL_619975, EPI_ISL_619977, EPI_ISL_619978,
EPI_ISL_619979, EPI_ISL_619980, EPI_ISL_619981, EPI_ISL_619982, EPI_ISL_619983, EPI_ISL_620056, EPI_ISL_620057, EPI_ISL_620058, EPI_ISL_620059, EPI_ISL_620060, EPI_ISL_620061, EPI_ISL_620062, EPI_ISL_620063, EPI_ISL_620064, EPI_ISL_620065, EPI_ISL_620066, EPI_ISL_620067, EPI_ISL_620068,
EPI_ISL_620069, EPI_ISL_620070, EPI_ISL_620071, EPI_ISL_620072, EPI_ISL_620073, EPI_ISL_620074, EPI_ISL_620075, EPI_ISL_620076, EPI_ISL_620077, EPI_ISL_620078, EPI_ISL_620079, EPI_ISL_620080, EPI_ISL_620081, EPI_ISL_620082, EPI_ISL_620083, EPI_ISL_620084, EPI_ISL_620085, EPI_ISL_620086,
EPI_ISL_620087, EPI_ISL_620088, EPI_ISL_620089, EPI_ISL_620090, EPI_ISL_620092, EPI_ISL_620094, EPI_ISL_620095, EPI_ISL_620096, EPI_ISL_620098, EPI_ISL_620099, EPI_ISL_620100, EPI_ISL_620101, EPI_ISL_620102, EPI_ISL_620103, EPI_ISL_620104, EPI_ISL_620105, EPI_ISL_620106, EPI_ISL_620108,
EPI_ISL_620109, EPI_ISL_620110, EPI_ISL_620111, EPI_ISL_620112, EPI_ISL_620113, EPI_ISL_620114, EPI_ISL_620115, EPI_ISL_620127, EPI_ISL_620128, EPI_ISL_620129, EPI_ISL_620130, EPI_ISL_620131, EPI_ISL_620133, EPI_ISL_620134, EPI_ISL_620135, EPI_ISL_620136, EPI_ISL_620139, EPI_ISL_620141,
EPI_ISL_620142, EPI_ISL_620143, EPI_ISL_620144, EPI_ISL_620145, EPI_ISL_620146, EPI_ISL_620148, EPI_ISL_620149, EPI_ISL_620150, EPI_ISL_620151, EPI_ISL_620153, EPI_ISL_620154, EPI_ISL_620155, EPI_ISL_620156, EPI_ISL_620157, EPI_ISL_620158, EPI_ISL_620159, EPI_ISL_620160, EPI_ISL_620161,
EPI_ISL_620162, EPI_ISL_620163, EPI_ISL_620164, EPI_ISL_620165, EPI_ISL_620166, EPI_ISL_620167, EPI_ISL_620168, EPI_ISL_620169, EPI_ISL_620170, EPI_ISL_620171, EPI_ISL_620172, EPI_ISL_620173, EPI_ISL_620174, EPI_ISL_620175, EPI_ISL_620176, EPI_ISL_620177, EPI_ISL_620178, EPI_ISL_620179,
EPI_ISL_620180, EPI_ISL_620181, EPI_ISL_620182, EPI_ISL_620183, EPI_ISL_620184, EPI_ISL_620185, EPI_ISL_620186, EPI_ISL_620187, EPI_ISL_620188, EPI_ISL_620189, EPI_ISL_620190, EPI_ISL_620192, EPI_ISL_620193, EPI_ISL_620194, EPI_ISL_620195, EPI_ISL_620196, EPI_ISL_620197, EPI_ISL_620198,
EPI_ISL_620199, EPI_ISL_620200, EPI_ISL_620201, EPI_ISL_620202, EPI_ISL_620203, EPI_ISL_620204, EPI_ISL_620205, EPI_ISL_620206, EPI_ISL_620207, EPI_ISL_620208, EPI_ISL_620209, EPI_ISL_620210, EPI_ISL_620211, EPI_ISL_620212, EPI_ISL_620213, EPI_ISL_620214, EPI_ISL_620215, EPI_ISL_620216,
EPI_ISL_620217, EPI_ISL_620218, EPI_ISL_620219, EPI_ISL_620220, EPI_ISL_620222, EPI_ISL_620223, EPI_ISL_620224, EPI_ISL_620225, EPI_ISL_620226, EPI_ISL_620227, EPI_ISL_620228, EPI_ISL_620229, EPI_ISL_620231, EPI_ISL_620232, EPI_ISL_620234, EPI_ISL_620235, EPI_ISL_620236, EPI_ISL_620237,
EPI_ISL_620238, EPI_ISL_620239, EPI_ISL_620240, EPI_ISL_620241, EPI_ISL_620242, EPI_ISL_620243, EPI_ISL_620244, EPI_ISL_620245, EPI_ISL_620246, EPI_ISL_620247, EPI_ISL_620248, EPI_ISL_620249, EPI_ISL_620250, EPI_ISL_620251, EPI_ISL_620252, EPI_ISL_620253, EPI_ISL_620254, EPI_ISL_620255,
EPI_ISL_620256, EPI_ISL_620257, EPI_ISL_620258, EPI_ISL_620259, EPI_ISL_620260, EPI_ISL_620261, EPI_ISL_620262, EPI_ISL_620263, EPI_ISL_620264, EPI_ISL_620266, EPI_ISL_620268, EPI_ISL_620269, EPI_ISL_620270, EPI_ISL_620271, EPI_ISL_620272, EPI_ISL_620273, EPI_ISL_620274, EPI_ISL_620275,
EPI_ISL_620276, EPI_ISL_620277, EPI_ISL_620278, EPI_ISL_620279, EPI_ISL_620280, EPI_ISL_620282, EPI_ISL_620283, EPI_ISL_620284, EPI_ISL_620285, EPI_ISL_620286, EPI_ISL_620287, EPI_ISL_620288, EPI_ISL_620289, EPI_ISL_620292, EPI_ISL_620293, EPI_ISL_620294, EPI_ISL_620295, EPI_ISL_620296,
EPI_ISL_620297, EPI_ISL_620298, EPI_ISL_620299, EPI_ISL_620302, EPI_ISL_620314, EPI_ISL_620808, EPI_ISL_620809, EPI_ISL_620810, EPI_ISL_620811, EPI_ISL_620812, EPI_ISL_620813, EPI_ISL_620815, EPI_ISL_620816, EPI_ISL_620817, EPI_ISL_620818, EPI_ISL_620819, EPI_ISL_620820, EPI_ISL_620821,
EPI_ISL_620822, EPI_ISL_620823, EPI_ISL_620824, EPI_ISL_620825, EPI_ISL_620826, EPI_ISL_620827, EPI_ISL_620828, EPI_ISL_620829, EPI_ISL_620830, EPI_ISL_620831, EPI_ISL_620832, EPI_ISL_620834, EPI_ISL_620835, EPI_ISL_620836, EPI_ISL_620837, EPI_ISL_620838, EPI_ISL_620839, EPI_ISL_620840,
EPI_ISL_620841, EPI_ISL_620842, EPI_ISL_620843, EPI_ISL_620844, EPI_ISL_620845, EPI_ISL_620846, EPI_ISL_620848, EPI_ISL_620850, EPI_ISL_620851, EPI_ISL_620852, EPI_ISL_620853, EPI_ISL_620854, EPI_ISL_620855, EPI_ISL_620856, EPI_ISL_620857, EPI_ISL_620858, EPI_ISL_620859, EPI_ISL_620860,
EPI_ISL_620861, EPI_ISL_620862, EPI_ISL_620863, EPI_ISL_620864, EPI_ISL_620865, EPI_ISL_620866, EPI_ISL_620867, EPI_ISL_620868, EPI_ISL_620869, EPI_ISL_620870, EPI_ISL_620871, EPI_ISL_620872, EPI_ISL_620873, EPI_ISL_620874, EPI_ISL_620875, EPI_ISL_620877, EPI_ISL_620879, EPI_ISL_620880,
EPI_ISL_620881, EPI_ISL_620882, EPI_ISL_620883, EPI_ISL_620884, EPI_ISL_620885, EPI_ISL_620886, EPI_ISL_620887, EPI_ISL_620888, EPI_ISL_620890, EPI_ISL_620891, EPI_ISL_620892, EPI_ISL_620893, EPI_ISL_620894, EPI_ISL_620895, EPI_ISL_620896, EPI_ISL_620897, EPI_ISL_620898, EPI_ISL_620899,
EPI_ISL_620900, EPI_ISL_620901, EPI_ISL_620902, EPI_ISL_620903, EPI_ISL_620904, EPI_ISL_620905, EPI_ISL_620906, EPI_ISL_620907, EPI_ISL_620908, EPI_ISL_620909, EPI_ISL_620910, EPI_ISL_620911, EPI_ISL_620912, EPI_ISL_620914, EPI_ISL_620915, EPI_ISL_620916, EPI_ISL_620918, EPI_ISL_620919,
EPI_ISL_620920, EPI_ISL_620921, EPI_ISL_620922, EPI_ISL_620923, EPI_ISL_620924, EPI_ISL_620925, EPI_ISL_620926, EPI_ISL_620927, EPI_ISL_620929, EPI_ISL_620930, EPI_ISL_620932, EPI_ISL_620933, EPI_ISL_620935, EPI_ISL_620936, EPI_ISL_620937, EPI_ISL_620938, EPI_ISL_620939, EPI_ISL_620940,
EPI_ISL_620941, EPI_ISL_620942, EPI_ISL_620943, EPI_ISL_620944, EPI_ISL_620945, EPI_ISL_620946, EPI_ISL_620947, EPI_ISL_620948, EPI_ISL_620949, EPI_ISL_620950, EPI_ISL_620951, EPI_ISL_620952, EPI_ISL_620953, EPI_ISL_620954, EPI_ISL_620955, EPI_ISL_620956, EPI_ISL_620957, EPI_ISL_620958,
EPI_ISL_620959, EPI_ISL_620960, EPI_ISL_620961, EPI_ISL_620962, EPI_ISL_620963, EPI_ISL_620964, EPI_ISL_620965, EPI_ISL_620966, EPI_ISL_620967, EPI_ISL_620968, EPI_ISL_620969, EPI_ISL_620970, EPI_ISL_620971, EPI_ISL_620972, EPI_ISL_620973, EPI_ISL_620974, EPI_ISL_620975, EPI_ISL_620976,
EPI_ISL_620977, EPI_ISL_620978, EPI_ISL_620979, EPI_ISL_620981, EPI_ISL_620982, EPI_ISL_620983, EPI_ISL_620984, EPI_ISL_620985, EPI_ISL_620986, EPI_ISL_620987, EPI_ISL_620988, EPI_ISL_620989, EPI_ISL_620990, EPI_ISL_620991, EPI_ISL_620992, EPI_ISL_620993, EPI_ISL_620994, EPI_ISL_620995,
EPI_ISL_620996, EPI_ISL_620997, EPI_ISL_620999, EPI_ISL_621000, EPI_ISL_621001, EPI_ISL_621002, EPI_ISL_621003, EPI_ISL_621004, EPI_ISL_621005, EPI_ISL_621006, EPI_ISL_621007, EPI_ISL_621009, EPI_ISL_621010, EPI_ISL_621011, EPI_ISL_621012, EPI_ISL_621013, EPI_ISL_621014, EPI_ISL_621015,
EPI_ISL_621018, EPI_ISL_621019, EPI_ISL_621020, EPI_ISL_621021, EPI_ISL_621022, EPI_ISL_621023, EPI_ISL_621024, EPI_ISL_621025, EPI_ISL_621026, EPI_ISL_621027, EPI_ISL_621028, EPI_ISL_621029, EPI_ISL_621030, EPI_ISL_621031, EPI_ISL_621032, EPI_ISL_621033, EPI_ISL_621034, EPI_ISL_621035,
EPI_ISL_621036, EPI_ISL_621037, EPI_ISL_621038, EPI_ISL_621039, EPI_ISL_621040, EPI_ISL_621041, EPI_ISL_621042, EPI_ISL_621043, EPI_ISL_621044, EPI_ISL_621045, EPI_ISL_621046, EPI_ISL_621047, EPI_ISL_621048, EPI_ISL_621049, EPI_ISL_621050, EPI_ISL_621051, EPI_ISL_621052, EPI_ISL_621053,
EPI_ISL_621054, EPI_ISL_621056, EPI_ISL_621057, EPI_ISL_621058, EPI_ISL_621059, EPI_ISL_621060, EPI_ISL_621061, EPI_ISL_621062, EPI_ISL_621063, EPI_ISL_621064, EPI_ISL_621065, EPI_ISL_621066, EPI_ISL_621067, EPI_ISL_621068, EPI_ISL_621069, EPI_ISL_621070, EPI_ISL_621071, EPI_ISL_621072,
EPI_ISL_621073, EPI_ISL_621074, EPI_ISL_621075, EPI_ISL_621076, EPI_ISL_621077, EPI_ISL_621078, EPI_ISL_621079, EPI_ISL_621080, EPI_ISL_621081, EPI_ISL_621082, EPI_ISL_621083, EPI_ISL_621086, EPI_ISL_621087, EPI_ISL_621088, EPI_ISL_621089, EPI_ISL_621090, EPI_ISL_621091, EPI_ISL_621092,
EPI_ISL_621093, EPI_ISL_621094, EPI_ISL_621095, EPI_ISL_621096, EPI_ISL_621097, EPI_ISL_621098, EPI_ISL_621099, EPI_ISL_621100, EPI_ISL_621101, EPI_ISL_621102, EPI_ISL_621103, EPI_ISL_621104, EPI_ISL_621105, EPI_ISL_621106, EPI_ISL_621107, EPI_ISL_621108, EPI_ISL_621109, EPI_ISL_621110,
EPI_ISL_621111, EPI_ISL_621112, EPI_ISL_621113, EPI_ISL_621114, EPI_ISL_621115, EPI_ISL_621116, EPI_ISL_621117, EPI_ISL_621118, EPI_ISL_621119, EPI_ISL_621120, EPI_ISL_621121, EPI_ISL_621122, EPI_ISL_621124, EPI_ISL_621125, EPI_ISL_621126, EPI_ISL_621127, EPI_ISL_621129, EPI_ISL_621130,
EPI_ISL_621131, EPI_ISL_621132, EPI_ISL_621133, EPI_ISL_621134, EPI_ISL_621135, EPI_ISL_621136, EPI_ISL_621137, EPI_ISL_621138, EPI_ISL_621139, EPI_ISL_621140, EPI_ISL_621141, EPI_ISL_621142, EPI_ISL_621143, EPI_ISL_621144, EPI_ISL_621145, EPI_ISL_621146, EPI_ISL_621149, EPI_ISL_621150,
EPI_ISL_621151, EPI_ISL_621152, EPI_ISL_621153, EPI_ISL_621154, EPI_ISL_621155, EPI_ISL_621156, EPI_ISL_621158, EPI_ISL_621159, EPI_ISL_621160, EPI_ISL_621161, EPI_ISL_621162, EPI_ISL_621163, EPI_ISL_621164, EPI_ISL_621165, EPI_ISL_621166, EPI_ISL_621167, EPI_ISL_621168, EPI_ISL_621169,
EPI_ISL_621170, EPI_ISL_621171, EPI_ISL_621172, EPI_ISL_621173, EPI_ISL_621174, EPI_ISL_621175, EPI_ISL_621176, EPI_ISL_621177, EPI_ISL_621178, EPI_ISL_621179, EPI_ISL_621180, EPI_ISL_621181, EPI_ISL_621182, EPI_ISL_621183, EPI_ISL_621184, EPI_ISL_621185, EPI_ISL_621186, EPI_ISL_621195,
EPI_ISL_621196, EPI_ISL_621197, EPI_ISL_621198, EPI_ISL_621199, EPI_ISL_621200, EPI_ISL_621201, EPI_ISL_621202, EPI_ISL_621203, EPI_ISL_621204, EPI_ISL_621205, EPI_ISL_621206, EPI_ISL_621207, EPI_ISL_621208, EPI_ISL_621209, EPI_ISL_621210, EPI_ISL_621211, EPI_ISL_621212, EPI_ISL_621213,
EPI_ISL_621214, EPI_ISL_621215, EPI_ISL_621216, EPI_ISL_621217, EPI_ISL_621218, EPI_ISL_621219, EPI_ISL_621220, EPI_ISL_621221, EPI_ISL_621222, EPI_ISL_621223, EPI_ISL_621225, EPI_ISL_621226, EPI_ISL_621227, EPI_ISL_621229, EPI_ISL_621230, EPI_ISL_621231, EPI_ISL_621232, EPI_ISL_621233,

| | | | |
|---|---|---|---|
| EPI_ISL_621234, EPI_ISL_621235, EPI_ISL_621236, EPI_ISL_621237, EPI_ISL_621238, EPI_ISL_621239, EPI_ISL_621240, EPI_ISL_621241, EPI_ISL_621242, EPI_ISL_621243, EPI_ISL_621245, EPI_ISL_621246, EPI_ISL_621247, EPI_ISL_621248, EPI_ISL_621249, EPI_ISL_621250, EPI_ISL_621251, EPI_ISL_621252, EPI_ISL_621253, EPI_ISL_621254, EPI_ISL_621255, EPI_ISL_621256, EPI_ISL_621257, EPI_ISL_621259, EPI_ISL_621260, EPI_ISL_621261, EPI_ISL_621273, EPI_ISL_621274, EPI_ISL_621275, EPI_ISL_621276, EPI_ISL_621277, EPI_ISL_621278, EPI_ISL_621279, EPI_ISL_621280, EPI_ISL_621281, EPI_ISL_621282, EPI_ISL_621283, EPI_ISL_621284, EPI_ISL_621285, EPI_ISL_621287, EPI_ISL_621288, EPI_ISL_621291, EPI_ISL_621292, EPI_ISL_621293, EPI_ISL_621295, EPI_ISL_621296, EPI_ISL_621297, EPI_ISL_621299, EPI_ISL_621300, EPI_ISL_621301, EPI_ISL_621302, EPI_ISL_621303, EPI_ISL_621304, EPI_ISL_621305, EPI_ISL_621306, EPI_ISL_621307, EPI_ISL_621308, EPI_ISL_621309, EPI_ISL_621310, EPI_ISL_621311, EPI_ISL_621312, EPI_ISL_621313, EPI_ISL_621314, EPI_ISL_621315, EPI_ISL_621316, EPI_ISL_621318, EPI_ISL_621327, EPI_ISL_621328, EPI_ISL_621329, EPI_ISL_621330, EPI_ISL_621331, EPI_ISL_621332, EPI_ISL_621333, EPI_ISL_621335, EPI_ISL_621336, EPI_ISL_621337, EPI_ISL_621338, EPI_ISL_621339, EPI_ISL_621340, EPI_ISL_621341, EPI_ISL_621342, EPI_ISL_621343, EPI_ISL_621344, EPI_ISL_621345, EPI_ISL_621346, EPI_ISL_621347, EPI_ISL_621348, EPI_ISL_621350, EPI_ISL_621351, EPI_ISL_621352, EPI_ISL_621353, EPI_ISL_621371, EPI_ISL_621535, EPI_ISL_621778, EPI_ISL_621825, EPI_ISL_621874, EPI_ISL_621875, EPI_ISL_621876, EPI_ISL_621877, EPI_ISL_621878, EPI_ISL_621879, EPI_ISL_621880, EPI_ISL_621882, EPI_ISL_621883, EPI_ISL_621884, EPI_ISL_621885, EPI_ISL_621886, EPI_ISL_621887, EPI_ISL_621888, EPI_ISL_621890, EPI_ISL_621891, EPI_ISL_621892, EPI_ISL_621893, EPI_ISL_621894, EPI_ISL_621895, EPI_ISL_621896, EPI_ISL_621897, EPI_ISL_621899, EPI_ISL_621900, EPI_ISL_621901, EPI_ISL_621902, EPI_ISL_621903, EPI_ISL_621904, EPI_ISL_621905, EPI_ISL_621906, EPI_ISL_621907, EPI_ISL_621909, EPI_ISL_621910, EPI_ISL_621911, EPI_ISL_621912, EPI_ISL_621913, EPI_ISL_621914, EPI_ISL_621915, EPI_ISL_621916, EPI_ISL_621917, EPI_ISL_621918, EPI_ISL_621919, EPI_ISL_621920, EPI_ISL_621921, EPI_ISL_621922, EPI_ISL_621923, EPI_ISL_621924, EPI_ISL_621925, EPI_ISL_621926, EPI_ISL_621927, EPI_ISL_621928, EPI_ISL_621929, EPI_ISL_621930, EPI_ISL_621932, EPI_ISL_621933, EPI_ISL_621934, EPI_ISL_621935, EPI_ISL_621936, EPI_ISL_621940, EPI_ISL_622066, EPI_ISL_622201, EPI_ISL_622204, EPI_ISL_622205, EPI_ISL_622206, EPI_ISL_622207, EPI_ISL_622208, EPI_ISL_622209, EPI_ISL_622210, EPI_ISL_622211, EPI_ISL_622212, EPI_ISL_622213, EPI_ISL_622214, EPI_ISL_622215, EPI_ISL_622217, EPI_ISL_622218, EPI_ISL_622220, EPI_ISL_622221, EPI_ISL_622222, EPI_ISL_622223, EPI_ISL_622224, EPI_ISL_622225, EPI_ISL_622226, EPI_ISL_622227, EPI_ISL_622228, EPI_ISL_622229, EPI_ISL_622230, EPI_ISL_622231, EPI_ISL_622232, EPI_ISL_622233, EPI_ISL_622234, EPI_ISL_622235, EPI_ISL_622236, EPI_ISL_622237, EPI_ISL_622238, EPI_ISL_622239, EPI_ISL_622240, EPI_ISL_622241, EPI_ISL_622242, EPI_ISL_622262, EPI_ISL_622263, EPI_ISL_622264, EPI_ISL_622265, EPI_ISL_622266, EPI_ISL_622267, EPI_ISL_622268, EPI_ISL_622269, EPI_ISL_622270, EPI_ISL_622271, EPI_ISL_622272, EPI_ISL_622273, EPI_ISL_622274, EPI_ISL_622275, EPI_ISL_622276, EPI_ISL_622277, EPI_ISL_622403, EPI_ISL_622405, EPI_ISL_622407, EPI_ISL_622408, EPI_ISL_622409, EPI_ISL_622410, EPI_ISL_622411, EPI_ISL_622412, EPI_ISL_622413, EPI_ISL_622414, EPI_ISL_622415, EPI_ISL_622416, EPI_ISL_622417, EPI_ISL_622418, EPI_ISL_622419, EPI_ISL_622420, EPI_ISL_622421, EPI_ISL_622422, EPI_ISL_622423, EPI_ISL_622424, EPI_ISL_622425, EPI_ISL_622426, EPI_ISL_622427, EPI_ISL_622428, EPI_ISL_622429, EPI_ISL_622430, EPI_ISL_622431, EPI_ISL_622432, EPI_ISL_622433, EPI_ISL_622434, EPI_ISL_622435, EPI_ISL_622436, EPI_ISL_622437, EPI_ISL_622438, EPI_ISL_622439, EPI_ISL_622440, EPI_ISL_622441, EPI_ISL_622442, EPI_ISL_622444, EPI_ISL_622445, EPI_ISL_622446, EPI_ISL_622447, EPI_ISL_622448, EPI_ISL_622449, EPI_ISL_622450, EPI_ISL_622451, EPI_ISL_622452, EPI_ISL_622453, EPI_ISL_622454, EPI_ISL_622455, EPI_ISL_622456, EPI_ISL_622457, EPI_ISL_622458, EPI_ISL_622459, EPI_ISL_622460, EPI_ISL_622461, EPI_ISL_622462, EPI_ISL_622463, EPI_ISL_622464, EPI_ISL_622465, EPI_ISL_622466, EPI_ISL_622467, EPI_ISL_622468, EPI_ISL_622469, EPI_ISL_622470, EPI_ISL_622471, EPI_ISL_622472, EPI_ISL_622473, EPI_ISL_622474, EPI_ISL_622475, EPI_ISL_622476, EPI_ISL_622477, EPI_ISL_622478, EPI_ISL_622479, EPI_ISL_622480, EPI_ISL_622481, EPI_ISL_622482, EPI_ISL_622483, EPI_ISL_622484, EPI_ISL_622485, EPI_ISL_622486, EPI_ISL_622487, EPI_ISL_622488, EPI_ISL_622489, EPI_ISL_622490, EPI_ISL_622491, EPI_ISL_622492, EPI_ISL_622493, EPI_ISL_622494, EPI_ISL_622495, EPI_ISL_622496, EPI_ISL_622497, EPI_ISL_622498, EPI_ISL_622499, EPI_ISL_622500, EPI_ISL_622502, EPI_ISL_622503, EPI_ISL_622504, EPI_ISL_622505, EPI_ISL_622506, EPI_ISL_622507, EPI_ISL_622508, EPI_ISL_622509, EPI_ISL_622510, EPI_ISL_622511, EPI_ISL_622512, EPI_ISL_622513, EPI_ISL_622514, EPI_ISL_622516, EPI_ISL_622517, EPI_ISL_622518, EPI_ISL_622519, EPI_ISL_622520, EPI_ISL_622521, EPI_ISL_622522, EPI_ISL_622523, EPI_ISL_622524, EPI_ISL_622525, EPI_ISL_622526, EPI_ISL_622527, EPI_ISL_622528, EPI_ISL_622529, EPI_ISL_622530, EPI_ISL_622531, EPI_ISL_622532, EPI_ISL_622533, EPI_ISL_622534, EPI_ISL_622535, EPI_ISL_622536, EPI_ISL_622537, EPI_ISL_622538, EPI_ISL_622539, EPI_ISL_622540, EPI_ISL_622541, EPI_ISL_622542, EPI_ISL_622543, EPI_ISL_622545, EPI_ISL_622546, EPI_ISL_622547, EPI_ISL_622548, EPI_ISL_622549, EPI_ISL_622550, EPI_ISL_622551, EPI_ISL_622552, EPI_ISL_622553, EPI_ISL_622554, EPI_ISL_622555, EPI_ISL_622556, EPI_ISL_622557, EPI_ISL_622558, EPI_ISL_622559, EPI_ISL_622560, EPI_ISL_622561, EPI_ISL_622562, EPI_ISL_622563, EPI_ISL_622564, EPI_ISL_622565, EPI_ISL_622566, EPI_ISL_622567, EPI_ISL_622568, EPI_ISL_622569, EPI_ISL_622570, EPI_ISL_622571, EPI_ISL_622572, EPI_ISL_622573, EPI_ISL_622574, EPI_ISL_622575, EPI_ISL_622576, EPI_ISL_622577, EPI_ISL_622578, EPI_ISL_622579, EPI_ISL_622580, EPI_ISL_622581, EPI_ISL_622582, EPI_ISL_622583, EPI_ISL_622584, EPI_ISL_622585, EPI_ISL_622586, EPI_ISL_622587, EPI_ISL_622588, EPI_ISL_622589, EPI_ISL_622590, EPI_ISL_622591, EPI_ISL_622592, EPI_ISL_622593, EPI_ISL_622594, EPI_ISL_622595, EPI_ISL_622596, EPI_ISL_622597, EPI_ISL_622598, EPI_ISL_622599, EPI_ISL_622600, EPI_ISL_622601, EPI_ISL_622602, EPI_ISL_622603, EPI_ISL_622604, EPI_ISL_622605, EPI_ISL_622606, EPI_ISL_622607, EPI_ISL_622608, EPI_ISL_622609, EPI_ISL_622610, EPI_ISL_622611, EPI_ISL_622612, EPI_ISL_622613, EPI_ISL_622614, EPI_ISL_622615, EPI_ISL_622616, EPI_ISL_622617, EPI_ISL_622618, EPI_ISL_622619, EPI_ISL_622620, EPI_ISL_622621, EPI_ISL_622622, EPI_ISL_622623, EPI_ISL_622624, EPI_ISL_622625, EPI_ISL_622626, EPI_ISL_622627, EPI_ISL_622628, EPI_ISL_622629, EPI_ISL_622630, EPI_ISL_622631, EPI_ISL_622632, EPI_ISL_622633, EPI_ISL_622634, EPI_ISL_622635, EPI_ISL_622636, EPI_ISL_622637, EPI_ISL_622638, EPI_ISL_622639, EPI_ISL_622640, EPI_ISL_622641, EPI_ISL_622642, EPI_ISL_622644, EPI_ISL_622645, EPI_ISL_622646, EPI_ISL_622647, EPI_ISL_622649, EPI_ISL_622650, EPI_ISL_622652, EPI_ISL_622653, EPI_ISL_622654, EPI_ISL_622655, EPI_ISL_622656, EPI_ISL_622657, EPI_ISL_622658, EPI_ISL_622659, EPI_ISL_622660, EPI_ISL_622661, EPI_ISL_622662, EPI_ISL_622663, EPI_ISL_622664, EPI_ISL_622665, EPI_ISL_622666, EPI_ISL_622667, EPI_ISL_622668, EPI_ISL_622669, EPI_ISL_622670, EPI_ISL_622671, EPI_ISL_622673, EPI_ISL_622674, EPI_ISL_622675, EPI_ISL_622677, EPI_ISL_622678, EPI_ISL_622679, EPI_ISL_622680, EPI_ISL_622681, EPI_ISL_622682, EPI_ISL_622683, EPI_ISL_622684, EPI_ISL_622685, EPI_ISL_622686, EPI_ISL_622687, EPI_ISL_622688, EPI_ISL_622689, EPI_ISL_622690, EPI_ISL_622691, EPI_ISL_622692, EPI_ISL_622693, EPI_ISL_622694, EPI_ISL_622695, EPI_ISL_622696, EPI_ISL_622697, EPI_ISL_622698, EPI_ISL_622699, EPI_ISL_622700, EPI_ISL_622701, EPI_ISL_622702, EPI_ISL_622703, EPI_ISL_622704, EPI_ISL_622705, EPI_ISL_622706, EPI_ISL_622707, EPI_ISL_622708, EPI_ISL_622709, EPI_ISL_622710, EPI_ISL_622711, EPI_ISL_622712, EPI_ISL_622713, EPI_ISL_622714, EPI_ISL_622715, EPI_ISL_622716, EPI_ISL_622717, EPI_ISL_622718, EPI_ISL_622719, EPI_ISL_622720, EPI_ISL_622721, EPI_ISL_622722, EPI_ISL_622723, EPI_ISL_622724, EPI_ISL_622725, EPI_ISL_622726, EPI_ISL_622727, EPI_ISL_622728, EPI_ISL_622729, EPI_ISL_622730, EPI_ISL_622731, EPI_ISL_622732, EPI_ISL_622733, EPI_ISL_622734, EPI_ISL_622735, EPI_ISL_622736, EPI_ISL_622737, EPI_ISL_622738, EPI_ISL_622739, EPI_ISL_622740, EPI_ISL_622741, EPI_ISL_622742, EPI_ISL_622743, EPI_ISL_622744, EPI_ISL_622745, EPI_ISL_622746, EPI_ISL_622747 | | | |
| see above | Department of Virus and Microbiological Special Diagnostics, Statens Serum Institut, Denmark | Albertsen lab, Department of Chemistry and Bioscience, Aalborg University, Denmark | Danish Covid-19 Genome Consortia |
| EPI_ISL_625456 | Virology Unit, Institut Pasteur de Madagascar | Virology Unit, Institut Pasteur de Madagascar | Christian Ranaivoson, Cara Brook, Norosoa Razanajatovo, Vida Ahyong, Tsiry Randriambolamanantsoa, Michelle Tan, Vololoniaina Raharinosy, Helisoa Razafimanjato, Cristina M. Tato, Joseph L. DeRisi, Soa Fy Andriamandimby, Jean-Michel Heraud, Philippe Dussart |
| EPI_ISL_625696, EPI_ISL_625740, EPI_ISL_625756, EPI_ISL_625828, EPI_ISL_625829, EPI_ISL_625830, EPI_ISL_625831, EPI_ISL_625832, EPI_ISL_625833, EPI_ISL_625834, EPI_ISL_625835, EPI_ISL_625837, EPI_ISL_625839, EPI_ISL_625845, EPI_ISL_625848, EPI_ISL_625852, EPI_ISL_625865, EPI_ISL_625871, EPI_ISL_625884, EPI_ISL_626169, EPI_ISL_626170, EPI_ISL_626171, EPI_ISL_626174, EPI_ISL_626175, EPI_ISL_626176, EPI_ISL_626177, EPI_ISL_626179, EPI_ISL_626183, EPI_ISL_626184, EPI_ISL_626185, EPI_ISL_626186, EPI_ISL_626187, EPI_ISL_626190, EPI_ISL_626191, EPI_ISL_626192, EPI_ISL_626193, EPI_ISL_626194, EPI_ISL_626198, EPI_ISL_626202, EPI_ISL_626203, EPI_ISL_626204, EPI_ISL_626205, EPI_ISL_626206, EPI_ISL_626209, EPI_ISL_626210 | | | |
| see above | Department of Virus and Microbiological Special Diagnostics, Statens Serum Institut, Denmark | Albertsen lab, Department of Chemistry and Bioscience, Aalborg University, Denmark | Danish Covid-19 Genome Consortia |
| EPI_ISL_626828, EPI_ISL_626859, EPI_ISL_626953, EPI_ISL_627000, EPI_ISL_627008, EPI_ISL_627020, EPI_ISL_627038, EPI_ISL_627046, EPI_ISL_627070, EPI_ISL_627158, EPI_ISL_627199, EPI_ISL_627518, EPI_ISL_627519, EPI_ISL_627520, EPI_ISL_627521, EPI_ISL_627522, EPI_ISL_627523, EPI_ISL_627524, EPI_ISL_627525, EPI_ISL_627526, EPI_ISL_627527, EPI_ISL_627528, EPI_ISL_627529, EPI_ISL_627530, EPI_ISL_627531, EPI_ISL_627532, EPI_ISL_627533, EPI_ISL_627534, EPI_ISL_627535, EPI_ISL_627536, EPI_ISL_627537, EPI_ISL_627538, EPI_ISL_627539, EPI_ISL_627540, EPI_ISL_627541, EPI_ISL_627542, EPI_ISL_627543, EPI_ISL_627544, EPI_ISL_627545, EPI_ISL_627546, EPI_ISL_627547, EPI_ISL_627548, EPI_ISL_627549, EPI_ISL_627550, EPI_ISL_627551, EPI_ISL_627552, EPI_ISL_627553, EPI_ISL_627554, EPI_ISL_627555, EPI_ISL_627556, EPI_ISL_627557, EPI_ISL_627558, EPI_ISL_627559, EPI_ISL_627560, EPI_ISL_627561, EPI_ISL_627562, EPI_ISL_627563, EPI_ISL_627564, EPI_ISL_627565, EPI_ISL_627566, EPI_ISL_627567, EPI_ISL_627568, EPI_ISL_627569 | | | |
| see above | Regional Virus Laboratory, Belfast Health and Social Care Trust | COVID-19 Genomics UK (COG-UK) Consortium | Conall McCaughey, James McKenna, Tanya Curran, Susan Feeney, Alison Watt, Ciara Cox, Mairead Connor, Zoltan Molnar, David Simpson, Derek Fairley |

We gratefully acknowledge the following Authors from the Originating laboratories responsible for obtaining the specimens, as well as the Submitting laboratories where the genome data were generated and shared via GISAID, on which this research is based.

All Submitters of data may be contacted directly via www.gisaid.org

Authors are sorted alphabetically.

| Accession ID | Originating Laboratory | Submitting Laboratory | Authors |
|---|---|---|---|
| EPI_ISL_413564 | MHC West-Brabant | Erasmus Medical Center | David Nieuwenhuijse, Bas Oude Munnink, Reina Sikkema, Claudia Schapendonk, Irina Chestakova, Anne van der Linden, Mark Pronk, Pascal Lexmond, Corien Swaan, Manon Haverkate, Madelief Mollers, Mart Stein, Sandra Kengne Kamga Mobou, Jeroen van Kampen, Jolanda Voermans, Aura Timen, Corine GeurtsvanKessel, Annemiek van der Eijk, Richard Molenkamp, Marion Koopmans, on behalf of the Dutch national COVID-19 response team. |
| EPI_ISL_413565 | Foundation Pamm | Erasmus Medical Center | David Nieuwenhuijse, Bas Oude Munnink, Reina Sikkema, Claudia Schapendonk, Irina Chestakova, Anne van der Linden, Mark Pronk, Pascal Lexmond, Corien Swaan, Manon Haverkate, Madelief Mollers, Mart Stein, Sandra Kengne Kamga Mobou, Jeroen van Kampen, Jolanda Voermans, Aura Timen, Corine GeurtsvanKessel, Annemiek van der Eijk, Richard Molenkamp, Marion Koopmans, on behalf of the Dutch national COVID-19 response team. |
| EPI_ISL_413566 | MHC Gooi & Vechtstreek | Erasmus Medical Center | David Nieuwenhuijse, Bas Oude Munnink, Reina Sikkema, Claudia Schapendonk, Irina Chestakova, Anne van der Linden, Mark Pronk, Pascal Lexmond, Corien Swaan, Manon Haverkate, Madelief Mollers, Mart Stein, Sandra Kengne Kamga Mobou, Jeroen van Kampen, Jolanda Voermans, Aura Timen, Corine GeurtsvanKessel, Annemiek van der Eijk, Richard Molenkamp, Marion Koopmans, on behalf of the Dutch national COVID-19 response team. |
| EPI_ISL_413568 | MHC Drente | Erasmus Medical Center | David Nieuwenhuijse, Bas Oude Munnink, Reina Sikkema, Claudia Schapendonk, Irina Chestakova, Anne van der Linden, Mark Pronk, Pascal Lexmond, Corien Swaan, Manon Haverkate, Madelief Mollers, Mart Stein, Sandra Kengne Kamga Mobou, Jeroen van Kampen, Jolanda Voermans, Aura Timen, Corine GeurtsvanKessel, Annemiek van der Eijk, Richard Molenkamp, Marion Koopmans, on behalf of the Dutch national COVID-19 response team. |
| EPI_ISL_413569, EPI_ISL_413570 | RIVM | Erasmus Medical Center | David Nieuwenhuijse, Bas Oude Munnink, Reina Sikkema, Claudia Schapendonk, Irina Chestakova, Anne van der Linden, Mark Pronk, Pascal Lexmond, Corien Swaan, Manon Haverkate, Madelief Mollers, Mart Stein, Sandra Kengne Kamga Mobou, Jeroen van Kampen, Jolanda Voermans, Aura Timen, Corine GeurtsvanKessel, Annemiek van der Eijk, Richard Molenkamp, Marion Koopmans, on behalf of the Dutch national COVID-19 response team. |
| EPI_ISL_413571 | MHC Brabant Zuidoost | Erasmus Medical Center | David Nieuwenhuijse, Bas Oude Munnink, Reina Sikkema, Claudia Schapendonk, Irina Chestakova, Anne van der Linden, Mark Pronk, Pascal Lexmond, Corien Swaan, Manon Haverkate, Madelief Mollers, Mart Stein, Sandra Kengne Kamga Mobou, Jeroen van Kampen, Jolanda Voermans, Aura Timen, Corine GeurtsvanKessel, Annemiek van der Eijk, Richard Molenkamp, Marion Koopmans, on behalf of the Dutch national COVID-19 response team. |
| EPI_ISL_413572 | MHC Kennemerland | Erasmus Medical Center | David Nieuwenhuijse, Bas Oude Munnink, Reina Sikkema, Claudia Schapendonk, Irina Chestakova, Anne van der Linden, Mark Pronk, Pascal Lexmond, Corien Swaan, Manon Haverkate, Madelief Mollers, Mart Stein, Sandra Kengne Kamga Mobou, Jeroen van Kampen, Jolanda Voermans, Aura Timen, Corine GeurtsvanKessel, Annemiek van der Eijk, Richard Molenkamp, Marion Koopmans, on behalf of the Dutch national COVID-19 response team. |
| EPI_ISL_413573 | Dienst Gezondheid & Jeugd Zuid-Holland Zuid | Erasmus Medical Center | David Nieuwenhuijse, Bas Oude Munnink, Reina Sikkema, Claudia Schapendonk, Irina Chestakova, Anne van der Linden, Mark Pronk, Pascal Lexmond, Corien Swaan, Manon Haverkate, Madelief Mollers, Mart Stein, Sandra Kengne Kamga Mobou, Jeroen van Kampen, Jolanda Voermans, Aura Timen, Corine GeurtsvanKessel, Annemiek van der Eijk, Richard Molenkamp, Marion Koopmans, on behalf of the Dutch national COVID-19 response team. |
| EPI_ISL_413574 | MHC West-Brabant | Erasmus Medical Center | David Nieuwenhuijse, Bas Oude Munnink, Reina Sikkema, Claudia Schapendonk, Irina Chestakova, Anne van der Linden, Mark Pronk, Pascal Lexmond, Corien Swaan, Manon Haverkate, Madelief Mollers, Mart Stein, Sandra Kengne Kamga Mobou, Jeroen van Kampen, Jolanda Voermans, Aura Timen, Corine GeurtsvanKessel, Annemiek van der Eijk, Richard Molenkamp, Marion Koopmans, on behalf of the Dutch national COVID-19 response team. |
| EPI_ISL_413575, EPI_ISL_413576 | RIVM | Erasmus Medical Center | David Nieuwenhuijse, Bas Oude Munnink, Reina Sikkema, Claudia Schapendonk, Irina Chestakova, Anne van der Linden, Mark Pronk, Pascal Lexmond, Corien Swaan, Manon Haverkate, Madelief Mollers, Mart Stein, Sandra Kengne Kamga Mobou, Jeroen van Kampen, Jolanda Voermans, Aura Timen, Corine GeurtsvanKessel, Annemiek van der Eijk, Richard Molenkamp, Marion Koopmans, on behalf of the Dutch national COVID-19 response team. |
| EPI_ISL_413577 | MHC Gooi & Vechtstreek | Erasmus Medical Center | David Nieuwenhuijse, Bas Oude Munnink, Reina Sikkema, Claudia Schapendonk, Irina Chestakova, Anne van der Linden, Mark Pronk, Pascal Lexmond, Corien Swaan, Manon Haverkate, Madelief Mollers, Mart Stein, Sandra Kengne Kamga Mobou, Jeroen van Kampen, Jolanda Voermans, Aura Timen, Corine GeurtsvanKessel, Annemiek van der Eijk, Richard Molenkamp, Marion Koopmans, on behalf of the Dutch national COVID-19 response team. |
| EPI_ISL_413578 | ErasmusMC | Erasmus Medical Center | David Nieuwenhuijse, Bas Oude Munnink, Reina Sikkema, Claudia Schapendonk, Irina Chestakova, Anne van der Linden, Mark Pronk, Pascal Lexmond, Corien Swaan, Manon Haverkate, Madelief Mollers, Mart Stein, Sandra Kengne Kamga Mobou, Jeroen van Kampen, Jolanda Voermans, Aura Timen, Corine GeurtsvanKessel, Annemiek van der Eijk, Richard Molenkamp, Marion Koopmans, on behalf of the Dutch national COVID-19 response team. |
| EPI_ISL_413579 | MHC Haaglanden | Erasmus Medical Center | David Nieuwenhuijse, Bas Oude Munnink, Reina Sikkema, Claudia Schapendonk, Irina Chestakova, Anne van der Linden, Mark Pronk, Pascal Lexmond, Corien Swaan, Manon Haverkate, Madelief Mollers, Mart Stein, Sandra Kengne Kamga Mobou, Jeroen van Kampen, Jolanda Voermans, Aura Timen, Corine GeurtsvanKessel, Annemiek van der Eijk, Richard Molenkamp, Marion Koopmans, on behalf of the Dutch national COVID-19 response team. |
| EPI_ISL_413580 | MHC Hart voor Brabant | Erasmus Medical Center | David Nieuwenhuijse, Bas Oude Munnink, Reina Sikkema, Claudia Schapendonk, Irina Chestakova, Anne van der Linden, Mark Pronk, Pascal Lexmond, Corien Swaan, Manon Haverkate, Madelief Mollers, Mart Stein, Sandra Kengne Kamga Mobou, Jeroen van Kampen, Jolanda Voermans, Aura Timen, Corine GeurtsvanKessel, Annemiek van der Eijk, Richard Molenkamp, Marion Koopmans, on behalf of the Dutch national COVID-19 response team. |
| EPI_ISL_413581 | RIVM | Erasmus Medical Center | David Nieuwenhuijse, Bas Oude Munnink, Reina Sikkema, Claudia Schapendonk, Irina Chestakova, Anne van der Linden, Mark Pronk, Pascal Lexmond, Corien Swaan, Manon Haverkate, Madelief Mollers, Mart Stein, Sandra Kengne Kamga Mobou, Jeroen van Kampen, Jolanda Voermans, Aura Timen, Corine GeurtsvanKessel, Annemiek van der Eijk, Richard Molenkamp, Marion Koopmans, on behalf of the Dutch national COVID-19 response team. |
| EPI_ISL_413582 | ErasmusMC | Erasmus Medical Center | David Nieuwenhuijse, Bas Oude Munnink, Reina Sikkema, Claudia Schapendonk, Irina Chestakova, Anne van der Linden, Mark Pronk, Pascal Lexmond, Corien Swaan, Manon Haverkate, Madelief Mollers, Mart Stein, Sandra Kengne Kamga Mobou, Jeroen van Kampen, Jolanda Voermans, Aura Timen, Corine GeurtsvanKessel, Annemiek van der Eijk, Richard Molenkamp, Marion Koopmans, on behalf of the Dutch national COVID-19 response team. |
| EPI_ISL_413583 | MHC Rotterdam-Rijnmond | Erasmus Medical Center | David Nieuwenhuijse, Bas Oude Munnink, Reina Sikkema, Claudia Schapendonk, Irina Chestakova, Anne van der Linden, Mark Pronk, Pascal Lexmond, Corien Swaan, Manon Haverkate, Madelief Mollers, Mart Stein, Sandra Kengne Kamga Mobou, Jeroen van Kampen, Jolanda Voermans, Aura Timen, Corine GeurtsvanKessel, Annemiek van der Eijk, Richard Molenkamp, Marion Koopmans, on behalf of the Dutch national COVID-19 response team. |
| EPI_ISL_413584 | unknown | Erasmus Medical Center | David Nieuwenhuijse, Bas Oude Munnink, Reina Sikkema, Claudia Schapendonk, Irina Chestakova, Anne van der Linden, Mark Pronk, Pascal Lexmond, Corien Swaan, Manon Haverkate, Madelief Mollers, Mart Stein, Sandra Kengne Kamga Mobou, Jeroen van Kampen, Jolanda Voermans, Aura Timen, Corine GeurtsvanKessel, Annemiek van der Eijk, Richard Molenkamp, Marion Koopmans, on behalf of the Dutch national COVID-19 response team. |
| EPI_ISL_413586, EPI_ISL_413587 | Foundation Elisabeth-Tweesteden Ziekenhuis | Erasmus Medical Center | David Nieuwenhuijse, Bas Oude Munnink, Reina Sikkema, Claudia Schapendonk, Irina Chestakova, Anne van der Linden, Mark Pronk, Pascal Lexmond, Corien Swaan, Manon Haverkate, Madelief Mollers, Mart Stein, Sandra Kengne Kamga Mobou, Jeroen van Kampen, Jolanda Voermans, Aura Timen, Corine GeurtsvanKessel, Annemiek van der Eijk, Richard Molenkamp, Marion Koopmans, on behalf of the Dutch national COVID-19 response team. |
| EPI_ISL_413588, EPI_ISL_413589, EPI_ISL_413590 | MHC Utrecht | Erasmus Medical Center | David Nieuwenhuijse, Bas Oude Munnink, Reina Sikkema, Claudia Schapendonk, Irina Chestakova, Anne van der Linden, Mark Pronk, Pascal Lexmond, Corien Swaan, Manon Haverkate, Madelief Mollers, Mart Stein, Sandra Kengne Kamga Mobou, Jeroen van Kampen, Jolanda Voermans, Aura Timen, Corine GeurtsvanKessel, Annemiek van der Eijk, Richard Molenkamp, Marion Koopmans, on behalf of the Dutch national COVID-19 response team. |
| EPI_ISL_413591 | MHC Flevoland | Erasmus Medical Center | David Nieuwenhuijse, Bas Oude Munnink, Reina Sikkema, Claudia Schapendonk, Irina Chestakova, Anne van der Linden, Mark Pronk, Pascal Lexmond, Corien Swaan, Manon Haverkate, Madelief Mollers, Mart Stein, Sandra Kengne Kamga Mobou, Jeroen van Kampen, Jolanda Voermans, Aura Timen, Corine GeurtsvanKessel, Annemiek van der Eijk, Richard Molenkamp, Marion Koopmans, on behalf of the Dutch national COVID-19 response team. |
| EPI_ISL_413593 | Laboratoire National de Santé | Erasmus Medical Center | David Nieuwenhuijse, Bas Oude Munnink, Reina Sikkema, Claudia Schapendonk, Irina Chestakova, Anne van der Linden, Mark Pronk, Pascal Lexmond, T. Abdelrahman, G. Fournier, J. Mossong, T. Nguyen, Jeroen van Kampen, Jolanda Voermans, Corine GeurtsvanKessel, Annemiek van der Eijk, Richard |

| | | | |
|---|---|---|---|
| | | | Molenkamp, Marion Koopmans, on behalf of the Dutch national COVID-19 response team. |
| EPI_ISL_414423, EPI_ISL_414424, EPI_ISL_414425, EPI_ISL_414426, EPI_ISL_414427, EPI_ISL_414428, EPI_ISL_414429, EPI_ISL_414430, EPI_ISL_414431, EPI_ISL_414432, EPI_ISL_414433, EPI_ISL_414434, EPI_ISL_414435, EPI_ISL_414436, EPI_ISL_414437, EPI_ISL_414438, EPI_ISL_414439, EPI_ISL_414440, EPI_ISL_414441, EPI_ISL_414442, EPI_ISL_414443, EPI_ISL_414444, EPI_ISL_414445, EPI_ISL_414446, EPI_ISL_414448, EPI_ISL_414449, EPI_ISL_414450, EPI_ISL_414451, EPI_ISL_414452, EPI_ISL_414453, EPI_ISL_414454, EPI_ISL_414455, EPI_ISL_414456, EPI_ISL_414457, EPI_ISL_414458, EPI_ISL_414459, EPI_ISL_414460, EPI_ISL_414461, EPI_ISL_414462, EPI_ISL_414463, EPI_ISL_414464, EPI_ISL_414465, EPI_ISL_414466, EPI_ISL_414467, EPI_ISL_414468, EPI_ISL_414469, EPI_ISL_414470, EPI_ISL_414471 | | | |
| see above | Dutch COVID-19 response team | Erasmus Medical Center | David Nieuwenhuijse, Bas Oude Munnink, Reina Sikkema, Claudia Schapendonk, Irina Chestakova, Anne van der Linden, Mark Pronk, Pascal Lexmond, Corien Swaan, Manon Haverkate, Madelief Mollers, Mart Stein, Sandra Kengne Kamga Mobou, Jeroen van Kampen, Jolanda Voermans, Aura Timen, Corine GeurtsvanKessel, Annemiek van der Eijk, Richard Molenkamp, Marion Koopmans, on behalf of the Dutch national COVID-19 response team. |
| EPI_ISL_414495 | Servicio Microbiología. Hospital Clínico Universitario. Valencia. | Sequencing and Bioinformatics Service. Molecular Epidemiology Laboratory. FISABIO-Public Health | David Navarro, Maria Alma Bracho, Giuseppe D'Auria, Griselda De Marco, Neris Garcia-Gonzalez, Fernando Gonzalez-Candelas |
| EPI_ISL_414496 | Servicio Microbiologia. Hospital Clinico Universitario. Valencia. | Sequencing and Bioinformatics Service. Molecular Epidemiology Laboratory. FISABIO-Public Health | David Navarro, María Alma Bracho, Giuseppe D'Auria, Griselda De Marco, Neris Garcia-Gonzalez, Fernando Gonzalez-Candelas |
| EPI_ISL_414529, EPI_ISL_414530, EPI_ISL_414531, EPI_ISL_414532, EPI_ISL_414533, EPI_ISL_414534, EPI_ISL_414535, EPI_ISL_414536, EPI_ISL_414537, EPI_ISL_414538, EPI_ISL_414539, EPI_ISL_414540, EPI_ISL_414541, EPI_ISL_414544, EPI_ISL_414545, EPI_ISL_414546, EPI_ISL_414547, EPI_ISL_414548, EPI_ISL_414549, EPI_ISL_414550, EPI_ISL_414551, EPI_ISL_414552, EPI_ISL_414553, EPI_ISL_414554, EPI_ISL_414555, EPI_ISL_414556, EPI_ISL_414557, EPI_ISL_414558, EPI_ISL_414559, EPI_ISL_414560, EPI_ISL_414561, EPI_ISL_414562, EPI_ISL_414563, EPI_ISL_414564, EPI_ISL_414565, EPI_ISL_414566 | | | |
| see above | Dutch COVID-19 response team | Erasmus Medical Center | David Nieuwenhuijse, Bas Oude Munnink, Reina Sikkema, Claudia Schapendonk, Irina Chestakova, Anne van der Linden, Mark Pronk, Pascal Lexmond, Corien Swaan, Manon Haverkate, Madelief Mollers, Mart Stein, Sandra Kengne Kamga Mobou, Jeroen van Kampen, Jolanda Voermans, Aura Timen, Corine GeurtsvanKessel, Annemiek van der Eijk, Richard Molenkamp, Marion Koopmans, on behalf of the Dutch national COVID-19 response team. |
| EPI_ISL_414598 | Servicio Microbiologia, Hospital Clinico Universitario, Valencia | Sequencing and Bioinformatics Service and Molecular Epidemiology Research Group. FISABIO-Public Health. | David Navarro, Maria Alma Bracho, Giuseppe D'Auria, Griselda De Marco, Neris Garcia-Gonzalez, Fernando Gonzalez-Candelas |
| EPI_ISL_415460, EPI_ISL_415461, EPI_ISL_415462, EPI_ISL_415463, EPI_ISL_415465, EPI_ISL_415466, EPI_ISL_415467, EPI_ISL_415468, EPI_ISL_415469, EPI_ISL_415470, EPI_ISL_415471, EPI_ISL_415472, EPI_ISL_415473, EPI_ISL_415474, EPI_ISL_415475, EPI_ISL_415476, EPI_ISL_415477, EPI_ISL_415478, EPI_ISL_415479, EPI_ISL_415480, EPI_ISL_415481, EPI_ISL_415482, EPI_ISL_415483, EPI_ISL_415484, EPI_ISL_415485, EPI_ISL_415486, EPI_ISL_415487, EPI_ISL_415488, EPI_ISL_415489, EPI_ISL_415490, EPI_ISL_415491, EPI_ISL_415492, EPI_ISL_415495, EPI_ISL_415496, EPI_ISL_415497, EPI_ISL_415498, EPI_ISL_415500, EPI_ISL_415501, EPI_ISL_415502, EPI_ISL_415503, EPI_ISL_415504, EPI_ISL_415505, EPI_ISL_415506, EPI_ISL_415508, EPI_ISL_415510, EPI_ISL_415511, EPI_ISL_415512, EPI_ISL_415513, EPI_ISL_415514, EPI_ISL_415515, EPI_ISL_415516, EPI_ISL_415524, EPI_ISL_415525, EPI_ISL_415526, EPI_ISL_415527, EPI_ISL_415528, EPI_ISL_415529, EPI_ISL_415530, EPI_ISL_415531, EPI_ISL_415532, EPI_ISL_415533, EPI_ISL_415534, EPI_ISL_415535 | | | |
| see above | Dutch COVID-19 response team | Erasmus Medical Center | David Nieuwenhuijse, Bas Oude Munnink, Reina Sikkema, Claudia Schapendonk, Irina Chestakova, Anne van der Linden, Mark Pronk, Pascal Lexmond, Corien Swaan, Manon Haverkate, Madelief Mollers, Mart Stein, Sandra Kengne Kamga Mobou, Jeroen van Kampen, Jolanda Voermans, Aura Timen, Corine GeurtsvanKessel, Annemiek van der Eijk, Richard Molenkamp, Marion Koopmans, on behalf of the Dutch national COVID-19 response team. |
| EPI_ISL_420125 | Servicio de Microbiologia. Hospital Clinico Universitario de Valencia | Sequencing and Bioinformatics Service and Molecular Epidemiology Research Group. FISABIO-Public Health | David Navarro, Giuseppe D'Auria, Llúcia Martínez-Priego, Maria Alma Bracho, Griselda De Marco, Beatriz Beamud, Lidia Ruiz Roldan, Marta Pla Diaz, Neris Garcia-Gonzalez, Loreto Ferrús Abad, Inma Galán Vendrell, Paula Ruiz-Hueso, Mariana Reyes-Prieto, Vicente Soriano Chirona, Fernando Gonzalez-Candelas |
| EPI_ISL_420130 | Servicio de Microbiologia. Hospital Clinico Universitario de Valencia | Sequencing and Bioinformatics Service and Molecular Epidemiology Research Group. FISABIO-Public Health | David Navarro, Loreto Ferrús Abad, Maria Alma Bracho, Griselda De Marco, Beatriz Beamud, Lidia Ruiz Roldan, Marta Pla Diaz, Neris Garcia-Gonzalez, Inma Galán Vendrell, Paula Ruiz-Hueso, Mariana Reyes-Prieto, Vicente Soriano Chirona, Sandra Carbo, Ivan Ansari, Llúcia Martínez-Priego, Giuseppe D'Auria, Fernando Gonzalez-Candelas |
| EPI_ISL_421272, EPI_ISL_421345, EPI_ISL_421346, EPI_ISL_421347 | Wyoming Public Health Laboratory | Center for Global Health, University of New Mexico Health Sciences Center | Daryl Domman, Kurt Schwalm, Rob Christensen, Wanda Manley, Cari Sloma, Noah Hull, Darrell Dinwiddie |
| EPI_ISL_421519 | Servicio de Microbiologia. Hospital Clinico Universitario de Valencia | Sequencing and Bioinformatics Service and Molecular Epidemiology Research Group. FISABIO-Public Health | David Navarro, Maria Alma Bracho, Griselda De Marco, Beatriz Beamud, Lidia Ruiz Roldan, Marta Pla Diaz, Neris Garcia-Gonzalez, Inma Galán Vendrell, Sandra Carbo, Loreto Ferrús Abad, Paula Ruiz-Hueso, Mariana Reyes-Prieto, Vicente Soriano Chirona, Ivan Ansari, Llúcia Martínez-Priego, Giuseppe D'Auria, Fernando Gonzalez-Candelas |
| EPI_ISL_421543, EPI_ISL_421544, EPI_ISL_421545, EPI_ISL_421546, EPI_ISL_421547, EPI_ISL_421548, EPI_ISL_421549, EPI_ISL_421550, EPI_ISL_421551, EPI_ISL_421552, EPI_ISL_421553, EPI_ISL_421554 | | | |
| see above | Wyoming Public Health Laboratory | Center for Global Health, University of New Mexico Health Sciences Center | Daryl Domman, Kurt Schwalm, Rob Christensen, Wanda Manley, Cari Sloma, Noah Hull, Darrell Dinwiddie |
| EPI_ISL_425178 | Servicio de Microbiología. Consorcio Hospital General Universitario de Valencia | Sequencing and Bioinformatics Service and Molecular Epidemiology Research Group. FISABIO-Public Health | David Navarro, Maria Alma Bracho, Griselda De Marco, Beatriz Beamud, Lidia Ruiz Roldan, Marta Pla Diaz, Neris Garcia-Gonzalez, Inma Galán Vendrell, Sandra Carbo, Loreto Ferrús Abad, Paula Ruiz-Hueso, Mariana Reyes-Prieto, Vicente Soriano Chirona, Ivan Ansari, Llúcia Martínez-Priego, Giuseppe D'Auria, Fernando Gonzalez-Candelas |
| EPI_ISL_425179 | Servicio de Microbiología. Consorcio Hospital General Universitario de Valencia | Sequencing and Bioinformatics Service and Molecular Epidemiology Research Group. FISABIO-Public Health | David Navarro, Maria Alma Bracho, Griselda De Marco, Beatriz Beamud, Lidia Ruiz Roldan, Marta Pla Diaz, Neris Garcia-Gonzalez, Inma Galán Vendrell, Sandra Carbo, Loreto Ferrús Abad, Paula Ruiz-Hueso, Mariana Reyes-Prieto, Vicente Soriano Chirona, Ivan Ansari, David Navarro, Llúcia Martínez-Priego, Giuseppe D'Auria, Fernando Gonzalez-Candelas |
| EPI_ISL_425192, EPI_ISL_425212 | Servicio de Microbiología. Consorcio Hospital General Universitario de Valencia | Sequencing and Bioinformatics Service and Molecular Epidemiology Research Group. FISABIO-Public Health | David Navarro, Maria Alma Bracho, Griselda De Marco, Beatriz Beamud, Lidia Ruiz Roldan, Marta Pla Diaz, Neris Garcia-Gonzalez, Inma Galán Vendrell, Sandra Carbo, Loreto Ferrús Abad, Paula Ruiz-Hueso, Mariana Reyes-Prieto, Vicente Soriano Chirona, Ivan Ansari, Llúcia Martínez-Priego, Giuseppe D'Auria, Fernando Gonzalez-Candelas |
| EPI_ISL_436278 | Servicio de Microbiologia. Hospital Clinico Universitario de Valencia | Sequencing and Bioinformatics Service and Molecular Epidemiology Research Group. FISABIO-Public Health | David Navarro, Maria Alma Bracho, Griselda De Marco, Beatriz Beamud, Lidia Ruiz Roldan, Marta Pla Diaz, Neris Garcia-Gonzalez, Inma Galán Vendrell, Sandra Carbo, Loreto Ferrús Abad, Paula Ruiz-Hueso, Mariana Reyes-Prieto, Vicente Soriano Chirona, Ivan Ansari, David Navarro, Llúcia Martínez-Priego, Giuseppe D'Auria, Fernando Gonzalez-Candelas |
| EPI_ISL_436291 | Servicio de Microbiologia. Hospital Clinico Universitario de Valencia | Sequencing and Bioinformatics Service and Molecular Epidemiology Research Group. FISABIO-Public Health | David Navarro, Maria Alma Bracho, Griselda De Marco, Beatriz Beamud, Lidia Ruiz Roldan, Marta Pla Diaz, Neris Garcia-Gonzalez, Inma Galán Vendrell, Sandra Carbo, Loreto Ferrús Abad, Paula Ruiz-Hueso, Mariana Reyes-Prieto, Vicente Soriano Chirona, Ivan Ansari, Llúcia Martínez-Priego, Giuseppe D'Auria, Fernando Gonzalez-Candelas |
| EPI_ISL_436298 | Servicio de Microbiologia. Hospital Clinico Universitario de Valencia | Sequencing and Bioinformatics Service and Molecular Epidemiology Research Group. FISABIO-Public Health | David Navarro, Maria Alma Bracho, Griselda De Marco, Beatriz Beamud, Lidia Ruiz Roldan, Marta Pla Diaz, Neris Garcia-Gonzalez, Inma Galán Vendrell, Sandra Carbo, Loreto Ferrús Abad, Paula Ruiz-Hueso, Mariana Reyes-Prieto, Vicente Soriano Chirona, Ivan Ansari, David Navarro, Llúcia Martínez-Priego, Giuseppe D'Auria, Fernando Gonzalez-Candelas |
| EPI_ISL_436311 | Servicio de Microbiologia. Hospital Clinico Universitario de Valencia | Sequencing and Bioinformatics Service and Molecular Epidemiology Research Group. FISABIO-Public Health | David Navarro, Maria Alma Bracho, Griselda De Marco, Beatriz Beamud, Lidia Ruiz Roldan, Marta Pla Diaz, Neris Garcia-Gonzalez, Inma Galán Vendrell, Sandra Carbo, Loreto Ferrús Abad, Paula Ruiz-Hueso, Mariana Reyes-Prieto, Vicente Soriano Chirona, Ivan Ansari, Llúcia Martínez-Priego, Giuseppe D'Auria, Fernando Gonzalez-Candelas |
| EPI_ISL_436318 | Servicio de Microbiologia. Hospital Clinico Universitario de Valencia | Sequencing and Bioinformatics Service and Molecular Epidemiology Research Group. FISABIO-Public Health | David Navarro, Maria Alma Bracho, Griselda De Marco, Beatriz Beamud, Lidia Ruiz Roldan, Marta Pla Diaz, Neris Garcia-Gonzalez, Inma Galán Vendrell, Sandra Carbo, Loreto Ferrús Abad, Paula Ruiz-Hueso, Mariana Reyes-Prieto, Vicente Soriano Chirona, Ivan Ansari, David Navarro, Llúcia Martínez-Priego, Giuseppe D'Auria, Fernando Gonzalez-Candelas |
| EPI_ISL_436331 | Servicio de Microbiologia. Hospital Clinico Universitario de Valencia | Sequencing and Bioinformatics Service and Molecular Epidemiology Research Group. FISABIO-Public Health | David Navarro, Maria Alma Bracho, Griselda De Marco, Beatriz Beamud, Lidia Ruiz Roldan, Marta Pla Diaz, Neris Garcia-Gonzalez, Inma Galán Vendrell, Sandra Carbo, Loreto Ferrús Abad, Paula Ruiz-Hueso, Mariana Reyes-Prieto, Vicente Soriano Chirona, Ivan Ansari, Llúcia Martínez-Priego, Giuseppe D'Auria, Fernando Gonzalez-Candelas |
| EPI_ISL_436338 | Servicio de Microbiologia. Hospital Clinico Universitario de Valencia | Sequencing and Bioinformatics Service and Molecular Epidemiology Research Group. FISABIO-Public Health | David Navarro, Maria Alma Bracho, Griselda De Marco, Beatriz Beamud, Lidia Ruiz Roldan, Marta Pla Diaz, Neris Garcia-Gonzalez, Inma Galán Vendrell, Sandra Carbo, Loreto Ferrús Abad, Paula Ruiz-Hueso, Mariana Reyes-Prieto, Vicente Soriano Chirona, Ivan Ansari, David Navarro, Llúcia Martínez-Priego, Giuseppe D'Auria, Fernando Gonzalez-Candelas |
| EPI_ISL_436351 | Servicio de Microbiologia. Hospital Clinico Universitario de Valencia | Sequencing and Bioinformatics Service and Molecular Epidemiology Research Group. FISABIO-Public Health | David Navarro, Maria Alma Bracho, Griselda De Marco, Beatriz Beamud, Lidia Ruiz Roldan, Marta Pla Diaz, Neris Garcia-Gonzalez, Inma Galán Vendrell, Sandra Carbo, Loreto Ferrús Abad, Paula Ruiz-Hueso, Mariana Reyes-Prieto, Vicente Soriano Chirona, Ivan Ansari, Llúcia Martínez-Priego, Giuseppe D'Auria, Fernando Gonzalez-Candelas |
| EPI_ISL_436362 | Servicio de Microbiologia. Hospital Clinico Universitario de Valencia | Sequencing and Bioinformatics Service and Molecular Epidemiology Research Group. FISABIO-Public Health | David Navarro, Maria Alma Bracho, Griselda De Marco, Beatriz Beamud, Lidia Ruiz Roldan, Marta Pla Diaz, Neris Garcia-Gonzalez, Inma Galán Vendrell, Sandra Carbo, Loreto Ferrús Abad, Paula Ruiz-Hueso, Mariana Reyes-Prieto, Vicente Soriano Chirona, Ivan Ansari, David Navarro, Llúcia |

| | | | |
|---|---|---|---|
| | | | Martínez-Priego, Giuseppe D'Auria, Fernando Gonzalez-Candelas |
| EPI_ISL_436402 | Servicio de Microbiologia. Hospital Clinico Universitario de Valencia | Sequencing and Bioinformatics Service and Molecular Epidemiology Research Group. FISABIO-Public Health | David Navarro, Maria Alma Bracho, Griselda De Marco, Beatriz Beamud, Lidia Ruiz Roldan, Marta Pla Diaz, Neris Garcia-Gonzalez, Inma Galán Vendrell, Sandra Carbo, Loreto Ferrús Abad, Paula Ruiz-Hueso, Mariana Reyes-Prieto, Vicente Soriano Chirona, Ivan Ansari, Llúcia Martínez-Priego, Giuseppe D'Auria, Fernando Gonzalez-Candelas |
| EPI_ISL_443183, EPI_ISL_443253, EPI_ISL_443254, EPI_ISL_443255, EPI_ISL_443256, EPI_ISL_443257, EPI_ISL_445077 | M Health Fairview | University of Minnesota Genomics Center | Daryl M. Gohl, John Garbe, Patrick Grady, Jerry Daniel, Ray Watson, Benjamin Auch, Andrew Nelson, Sophia Yohe, and Kenneth B. Beckman |
| EPI_ISL_447470, EPI_ISL_447484, EPI_ISL_447498, EPI_ISL_447512, EPI_ISL_447525 | Servicio de Microbiologia. Hospital Clinico Universitario de Valencia | Sequencing and Bioinformatics Service and Molecular Epidemiology Research Group. FISABIO-Public Health | David Navarro, Eliseo Albert, Maria Alma Bracho, Griselda De Marco, Lidia Ruiz Roldan, Neris Garcia-Gonzalez, Inma Galán Vendrell, Sandra Carbo, Loreto Ferrús Abad, Paula Ruiz-Hueso, Mariana Reyes-Prieto, Vicente Soriano Chirona, Ivan Ansari, Llúcia Martínez-Priego, Giuseppe 'Auria, Fernando Gonzalez-Candelas |
| EPI_ISL_448223, EPI_ISL_448224, EPI_ISL_448225, EPI_ISL_448226, EPI_ISL_448227, EPI_ISL_448229, EPI_ISL_448230, EPI_ISL_448231, EPI_ISL_448232, EPI_ISL_448233, EPI_ISL_448234, EPI_ISL_448235, EPI_ISL_448237, EPI_ISL_448238, EPI_ISL_448239, EPI_ISL_448242, EPI_ISL_448243, EPI_ISL_448244, EPI_ISL_448245, EPI_ISL_448246, EPI_ISL_448248, EPI_ISL_448249, EPI_ISL_448251, EPI_ISL_448252, EPI_ISL_448254, EPI_ISL_448255, EPI_ISL_448256, EPI_ISL_448258, EPI_ISL_448259, EPI_ISL_448262, EPI_ISL_448265, EPI_ISL_448266, EPI_ISL_448267, EPI_ISL_448268, EPI_ISL_448269, EPI_ISL_448270, EPI_ISL_448272, EPI_ISL_448273, EPI_ISL_448274, EPI_ISL_448276, EPI_ISL_448277, EPI_ISL_448278, EPI_ISL_448279, EPI_ISL_448280, EPI_ISL_448281, EPI_ISL_448282, EPI_ISL_448283, EPI_ISL_448284, EPI_ISL_448285, EPI_ISL_448286, EPI_ISL_448287, EPI_ISL_448288, EPI_ISL_448289, EPI_ISL_448290, EPI_ISL_448291, EPI_ISL_448292, EPI_ISL_448293, EPI_ISL_448294, EPI_ISL_448295, EPI_ISL_448296, EPI_ISL_448297, EPI_ISL_448298, EPI_ISL_448299, EPI_ISL_448300, EPI_ISL_448301, EPI_ISL_448304, EPI_ISL_448305, EPI_ISL_448307, EPI_ISL_448309, EPI_ISL_448310, EPI_ISL_448311, EPI_ISL_448312, EPI_ISL_448313, EPI_ISL_448314, EPI_ISL_448315, EPI_ISL_448316, EPI_ISL_448317, EPI_ISL_448318, EPI_ISL_448319, EPI_ISL_448320, EPI_ISL_448321, EPI_ISL_448322, EPI_ISL_448323, EPI_ISL_448324, EPI_ISL_448325, EPI_ISL_448326, EPI_ISL_448327, EPI_ISL_448328, EPI_ISL_448329, EPI_ISL_448330, EPI_ISL_448331, EPI_ISL_448332, EPI_ISL_448333, EPI_ISL_448334, EPI_ISL_448335, EPI_ISL_448336, EPI_ISL_448337, EPI_ISL_448338, EPI_ISL_448339, EPI_ISL_448340, EPI_ISL_448341, EPI_ISL_448342, EPI_ISL_448343, EPI_ISL_448344, EPI_ISL_448345, EPI_ISL_448346, EPI_ISL_448347, EPI_ISL_448348, EPI_ISL_448349, EPI_ISL_448350, EPI_ISL_448351, EPI_ISL_448352, EPI_ISL_448353, EPI_ISL_448354, EPI_ISL_448355, EPI_ISL_448356, EPI_ISL_448357, EPI_ISL_448358, EPI_ISL_448359, EPI_ISL_448360, EPI_ISL_448361, EPI_ISL_448362, EPI_ISL_448363, EPI_ISL_448364, EPI_ISL_448365, EPI_ISL_448366, EPI_ISL_448367, EPI_ISL_448368, EPI_ISL_448369, EPI_ISL_448370, EPI_ISL_448371, EPI_ISL_448372, EPI_ISL_448373, EPI_ISL_448374, EPI_ISL_448375, EPI_ISL_448376, EPI_ISL_448377, EPI_ISL_448378, EPI_ISL_448379, EPI_ISL_448380, EPI_ISL_448381, EPI_ISL_448382, EPI_ISL_448383, EPI_ISL_448384, EPI_ISL_448385, EPI_ISL_448386, EPI_ISL_448387, EPI_ISL_448388, EPI_ISL_448978, EPI_ISL_448979, EPI_ISL_448980, EPI_ISL_448981, EPI_ISL_448982, EPI_ISL_448983, EPI_ISL_448984, EPI_ISL_448985, EPI_ISL_448986, EPI_ISL_448988, EPI_ISL_448989, EPI_ISL_448990, EPI_ISL_448991, EPI_ISL_448992, EPI_ISL_448994, EPI_ISL_448995, EPI_ISL_448996, EPI_ISL_448997, EPI_ISL_448998, EPI_ISL_448999, EPI_ISL_449000, EPI_ISL_449001, EPI_ISL_449002, EPI_ISL_449003, EPI_ISL_449004, EPI_ISL_449005, EPI_ISL_449006, EPI_ISL_449007, EPI_ISL_449008, EPI_ISL_449009, EPI_ISL_449010, EPI_ISL_449011, EPI_ISL_449012, EPI_ISL_449013, EPI_ISL_449014, EPI_ISL_449015, EPI_ISL_449016, EPI_ISL_449017, EPI_ISL_449018, EPI_ISL_449019, EPI_ISL_449020, EPI_ISL_449021, EPI_ISL_449022, EPI_ISL_449023, EPI_ISL_449024, EPI_ISL_449025, EPI_ISL_449026, EPI_ISL_449027, EPI_ISL_449028, EPI_ISL_449029, EPI_ISL_449030, EPI_ISL_449031, EPI_ISL_449032, EPI_ISL_449033, EPI_ISL_449034, EPI_ISL_449035, EPI_ISL_449036, EPI_ISL_449037, EPI_ISL_449038, EPI_ISL_449039, EPI_ISL_449040, EPI_ISL_449041, EPI_ISL_449042, EPI_ISL_449043, EPI_ISL_449044, EPI_ISL_449045, EPI_ISL_449046, EPI_ISL_449047, EPI_ISL_449048, EPI_ISL_449049, EPI_ISL_449050, EPI_ISL_449051, EPI_ISL_449052, EPI_ISL_449053, EPI_ISL_449054, EPI_ISL_449055, EPI_ISL_449056, EPI_ISL_449057, EPI_ISL_449058, EPI_ISL_449059, EPI_ISL_449060, EPI_ISL_449061, EPI_ISL_449062, EPI_ISL_449063, EPI_ISL_449064, EPI_ISL_449065, EPI_ISL_449066, EPI_ISL_449067, EPI_ISL_449068, EPI_ISL_449069, EPI_ISL_449070, EPI_ISL_449071, EPI_ISL_449072, EPI_ISL_449073, EPI_ISL_449074, EPI_ISL_449075, EPI_ISL_449076, EPI_ISL_449077, EPI_ISL_449078, EPI_ISL_449079, EPI_ISL_449080, EPI_ISL_449081, EPI_ISL_449082, EPI_ISL_449083, EPI_ISL_449084, EPI_ISL_449085, EPI_ISL_449086, EPI_ISL_449087, EPI_ISL_449088, EPI_ISL_449089, EPI_ISL_449090, EPI_ISL_449091, EPI_ISL_449092, EPI_ISL_449093, EPI_ISL_449094, EPI_ISL_449095, EPI_ISL_449096, EPI_ISL_449097, EPI_ISL_449098, EPI_ISL_449100, EPI_ISL_449101, EPI_ISL_449102, EPI_ISL_449103, EPI_ISL_449104, EPI_ISL_449105, EPI_ISL_449107, EPI_ISL_449108, EPI_ISL_449109, EPI_ISL_449111, EPI_ISL_449112, EPI_ISL_449113, EPI_ISL_449114, EPI_ISL_449116, EPI_ISL_449117, EPI_ISL_449118, EPI_ISL_449119, EPI_ISL_449120, EPI_ISL_449121, EPI_ISL_449122, EPI_ISL_449123, EPI_ISL_449124, EPI_ISL_449125, EPI_ISL_449127, EPI_ISL_449128, EPI_ISL_449129, EPI_ISL_449130, EPI_ISL_449131, EPI_ISL_449132, EPI_ISL_449133, EPI_ISL_449135, EPI_ISL_449136, EPI_ISL_449139, EPI_ISL_449140, EPI_ISL_449141, EPI_ISL_449143, EPI_ISL_449145, EPI_ISL_449146, EPI_ISL_449147, EPI_ISL_449148, EPI_ISL_449149, EPI_ISL_449150, EPI_ISL_449151, EPI_ISL_449152, EPI_ISL_449153, EPI_ISL_449154, EPI_ISL_449155, EPI_ISL_449156, EPI_ISL_449157, EPI_ISL_449158, EPI_ISL_449159, EPI_ISL_449160, EPI_ISL_449161, EPI_ISL_449162, EPI_ISL_449163, EPI_ISL_449164, EPI_ISL_449165, EPI_ISL_449166, EPI_ISL_449167, EPI_ISL_449168, EPI_ISL_449169, EPI_ISL_449170, EPI_ISL_449171, EPI_ISL_449172, EPI_ISL_449173, EPI_ISL_449174, EPI_ISL_449175 | | | |
| see above | Quadram Institute Bioscience | COVID-19 Genomics UK (COG-UK) Consortium | Dave J. Baker, Gemma L. Kay, Alp Aydin, Thanh Le-Viet, Steven Rudder, Ana P. Tedim, Anastasia Kolyva, Maria Diaz, Leonardo de Oliveira Martins, Nabil-Fareed Alikhan, Lizzie Meadows, Rachael Stanley, Ngozi Elumogo, Muhammed Yasir, Nicholas M. Thomson, Alexander J Trotter, Rachel Gilroy, Samuel Bloomfield, Claire Stuart, Andrew Bell, Reenesh Prakash, Samir Dervisevic, Alison E. Mather, John Wain, Mark Webber, Andrew J. Page, Justin O'Grady |
| EPI_ISL_453486, EPI_ISL_453487, EPI_ISL_453488, EPI_ISL_453489, EPI_ISL_453490, EPI_ISL_453491, EPI_ISL_453492, EPI_ISL_453493, EPI_ISL_453494, EPI_ISL_453495, EPI_ISL_453496, EPI_ISL_453497, EPI_ISL_453498, EPI_ISL_453499, EPI_ISL_453500, EPI_ISL_453501, EPI_ISL_453502, EPI_ISL_453503, EPI_ISL_453504, EPI_ISL_453505, EPI_ISL_453506, EPI_ISL_453507, EPI_ISL_453508, EPI_ISL_453509, EPI_ISL_453510, EPI_ISL_453511, EPI_ISL_453512, EPI_ISL_453513, EPI_ISL_453514, EPI_ISL_453515, EPI_ISL_453516, EPI_ISL_453517, EPI_ISL_453518, EPI_ISL_453519, EPI_ISL_453520, EPI_ISL_453521, EPI_ISL_453522, EPI_ISL_453523, EPI_ISL_453524, EPI_ISL_453525, EPI_ISL_453526, EPI_ISL_453527, EPI_ISL_453528, EPI_ISL_453529, EPI_ISL_453530, EPI_ISL_453531, EPI_ISL_453532, EPI_ISL_453533, EPI_ISL_453534, EPI_ISL_453535, EPI_ISL_453536, EPI_ISL_453537, EPI_ISL_453538, EPI_ISL_453539, EPI_ISL_453540, EPI_ISL_453541, EPI_ISL_453542 | | | |
| see above | Northumbria University / South Tees Hospitals NHS Foundation Trust / North Cumbria Integrated Care NHS Foundation Trust / North Tees and Hartlepool NHS Foundation Trust / Newcastle Hospitals NHS Foundation Trust | COVID-19 Genomics UK (COG-UK) Consortium | Darren L Smith,Andrew Nelson,Matthew Bashton,Greg R Young,Joshua Loh,John Allan,Mohammad A Tariq,Giles S Holt,Gary Black,Wen C Yew,Lynn Dover ,Paul Baker,Steve Liggett,Sarah Essex,Jane Greenaway ,Debra Padgett,Clive Graham,Garren Scott,Edward Barton ,Emma Swindells ,Brendan Payne,Jennifer Collins,Yusri Taha,Gary Eltringham |
| EPI_ISL_453543 | Northumbria University / South Tees Hospitals NHS Foundation Trust / North Cumbria Integrated Care NHS Foundation Trust / North Tees and Hartlepool NHS Foundation Trust / Newcastle Hospitals NHS Foundation Trust | Northumbria University | Darren L Smith,Andrew Nelson,Matthew Bashton,Greg R Young,Joshua Loh,John Allan,Mohammad A Tariq,Giles S Holt,Gary Black,Wen C Yew,Lynn Dover ,Paul Baker,Steve Liggett,Sarah Essex,Jane Greenaway ,Debra Padgett,Clive Graham,Garren Scott,Edward Barton ,Emma Swindells ,Brendan Payne,Jennifer Collins,Yusri Taha,Gary Eltringham |
| EPI_ISL_453544 | Northumbria University / South Tees Hospitals NHS Foundation Trust / North Cumbria Integrated Care NHS Foundation Trust / North Tees and Hartlepool NHS Foundation Trust / Newcastle Hospitals NHS Foundation Trust | COVID-19 Genomics UK (COG-UK) Consortium | Darren L Smith,Andrew Nelson,Matthew Bashton,Greg R Young,Joshua Loh,John Allan,Mohammad A Tariq,Giles S Holt,Gary Black,Wen C Yew,Lynn Dover ,Paul Baker,Steve Liggett,Sarah Essex,Jane Greenaway ,Debra Padgett,Clive Graham,Garren Scott,Edward Barton ,Emma Swindells ,Brendan Payne,Jennifer Collins,Yusri Taha,Gary Eltringham |
| EPI_ISL_453545, EPI_ISL_453546, EPI_ISL_453547, EPI_ISL_453548, EPI_ISL_453549, EPI_ISL_453550, EPI_ISL_453551, EPI_ISL_453552, EPI_ISL_453553, EPI_ISL_453554, EPI_ISL_453555, EPI_ISL_453556, EPI_ISL_453557, EPI_ISL_453558, EPI_ISL_453559, EPI_ISL_453560, EPI_ISL_453561, EPI_ISL_453562, EPI_ISL_453563, EPI_ISL_453564, EPI_ISL_453565, EPI_ISL_453566, EPI_ISL_453567, EPI_ISL_453568, EPI_ISL_453569, EPI_ISL_453570, EPI_ISL_453571, EPI_ISL_453572, EPI_ISL_453573, EPI_ISL_453574, EPI_ISL_453575, EPI_ISL_453576, EPI_ISL_453577, EPI_ISL_453578, EPI_ISL_453579, EPI_ISL_453580, EPI_ISL_453581, EPI_ISL_453582, EPI_ISL_453583, EPI_ISL_453584, EPI_ISL_453585, EPI_ISL_453586, EPI_ISL_453587, EPI_ISL_453588, EPI_ISL_453589, EPI_ISL_453590, EPI_ISL_453591, EPI_ISL_453592, EPI_ISL_453593, EPI_ISL_453594, EPI_ISL_453595, EPI_ISL_453596, EPI_ISL_453597, EPI_ISL_453598, EPI_ISL_453599, EPI_ISL_453600, EPI_ISL_453601, EPI_ISL_453602, EPI_ISL_453603, EPI_ISL_453604, EPI_ISL_453605, EPI_ISL_453606, EPI_ISL_453607, EPI_ISL_453608, EPI_ISL_453609, EPI_ISL_453610, EPI_ISL_453611, EPI_ISL_453612, EPI_ISL_453613 | | | |
| see above | Quadram Institute Bioscience | COVID-19 Genomics UK (COG-UK) Consortium | Dave J. Baker, Gemma L. Kay, Alp Aydin, Thanh Le-Viet, Steven Rudder, Ana P. Tedim, Anastasia Kolyva, Maria Diaz, Leonardo de Oliveira Martins, Nabil-Fareed Alikhan, Lizzie Meadows, Rachael Stanley, Ngozi Elumogo, Muhammed Yasir, Nicholas M. Thomson, Alexander J Trotter, Rachel Gilroy, Samuel Bloomfield, Claire Stuart, Andrew Bell, Reenesh Prakash, Samir Dervisevic, Alison E. Mather, John Wain, Mark Webber, Andrew J. Page, Justin O'Grady |
| EPI_ISL_455727, EPI_ISL_455741 | Servicio de Microbiologia. Hospital Clinico Universitario de Valencia | Sequencing and Bioinformatics Service and Molecular Epidemiology Research Group. FISABIO-Public Health, and SeqCOVID-Spain Consortium | David Navarro, Eliseo Albert, Maria Alma Bracho, Griselda De Marco, Lidia Ruiz Roldan, Neris Garcia-Gonzalez, Inma Galán Vendrell, Sandra Carbo, Loreto Ferrús Abad, Paula Ruiz-Hueso, Mariana Reyes-Prieto, Vicente Soriano Chirona, Ivan Ansari, Llúcia Martínez-Priego, Giuseppe 'Auria, Fernando Gonzalez-Candelas |
| EPI_ISL_457303, EPI_ISL_457304, EPI_ISL_457305, EPI_ISL_457306, EPI_ISL_457307, EPI_ISL_457308, EPI_ISL_457309, EPI_ISL_457310, EPI_ISL_457311, EPI_ISL_457312, EPI_ISL_457313, EPI_ISL_457314, EPI_ISL_457315, EPI_ISL_457316, EPI_ISL_457317, EPI_ISL_457318, EPI_ISL_457319, EPI_ISL_457320, EPI_ISL_457321, EPI_ISL_457322, EPI_ISL_457323, EPI_ISL_457324, EPI_ISL_457325 | | | |
| see above | Northumbria University / South Tees Hospitals NHS Foundation Trust / North Cumbria Integrated Care NHS Foundation Trust / North Tees and Hartlepool NHS Foundation Trust / Newcastle Hospitals NHS Foundation Trust | COVID-19 Genomics UK (COG-UK) Consortium | Darren L Smith,Andrew Nelson,Matthew Bashton,Greg R Young,Joshua Loh,John Allan,Mohammad A Tariq,Giles S Holt,Gary Black,Wen C Yew,Lynn Dover,Paul Baker,Steve Liggett,Sarah Essex,Jane Greenaway,Debra Padgett,Clive Graham,Garren Scott,Edward Barton,Emma Swindells,Brendan Payne,Jennifer Collins,Yusri Taha,Gary Eltringham |
| EPI_ISL_457326, EPI_ISL_457327, EPI_ISL_457328, EPI_ISL_457329, EPI_ISL_457330, EPI_ISL_457331, EPI_ISL_457332, EPI_ISL_457333, EPI_ISL_457334, EPI_ISL_457335, EPI_ISL_457336, EPI_ISL_457337, EPI_ISL_457338, EPI_ISL_457339, EPI_ISL_457340, EPI_ISL_457341, EPI_ISL_457342, EPI_ISL_457343, EPI_ISL_457344, EPI_ISL_457345, EPI_ISL_457346, EPI_ISL_457347, EPI_ISL_457348, EPI_ISL_457349, EPI_ISL_457350, EPI_ISL_457351, EPI_ISL_457352, EPI_ISL_457353, EPI_ISL_457354, EPI_ISL_457355, EPI_ISL_457356, EPI_ISL_457357, EPI_ISL_457358, EPI_ISL_457359, EPI_ISL_457360, EPI_ISL_457361, EPI_ISL_457362, EPI_ISL_457363, EPI_ISL_457364, EPI_ISL_457365, EPI_ISL_457366, EPI_ISL_457367, EPI_ISL_457368, EPI_ISL_457369, EPI_ISL_457370, EPI_ISL_457371, EPI_ISL_457372, EPI_ISL_457373, EPI_ISL_457374, EPI_ISL_457375, EPI_ISL_457376, EPI_ISL_457377, EPI_ISL_457378, EPI_ISL_457379, EPI_ISL_457380, EPI_ISL_457381, EPI_ISL_457382, EPI_ISL_457383, EPI_ISL_457384, EPI_ISL_457385, EPI_ISL_457386, EPI_ISL_457387, EPI_ISL_457388, EPI_ISL_457389, EPI_ISL_457390, EPI_ISL_457391, EPI_ISL_457392, EPI_ISL_457393, EPI_ISL_457394, EPI_ISL_457395, EPI_ISL_457396, EPI_ISL_457397, EPI_ISL_457398, EPI_ISL_457399, EPI_ISL_457400, EPI_ISL_457401, EPI_ISL_457402, EPI_ISL_457403, EPI_ISL_457404, EPI_ISL_457405, EPI_ISL_457406, EPI_ISL_457407, EPI_ISL_457408, EPI_ISL_457409, EPI_ISL_457410, EPI_ISL_457411, EPI_ISL_457412, EPI_ISL_457413, EPI_ISL_457414, EPI_ISL_457415, EPI_ISL_457416, EPI_ISL_457417, EPI_ISL_457418, EPI_ISL_457419, EPI_ISL_457420, EPI_ISL_457421, EPI_ISL_457422, EPI_ISL_457423, EPI_ISL_457424, EPI_ISL_457425, EPI_ISL_457426, EPI_ISL_457427, EPI_ISL_457428, EPI_ISL_457429, EPI_ISL_457430, EPI_ISL_457431, EPI_ISL_457432, EPI_ISL_457433, EPI_ISL_457434, EPI_ISL_457435, EPI_ISL_457436, EPI_ISL_457437, EPI_ISL_457438, EPI_ISL_457439, EPI_ISL_457440, EPI_ISL_457441, EPI_ISL_457442, EPI_ISL_457443, EPI_ISL_457444, EPI_ISL_457445, EPI_ISL_457446, EPI_ISL_457447, EPI_ISL_457448, EPI_ISL_457449, EPI_ISL_457450, EPI_ISL_457451, EPI_ISL_457452, EPI_ISL_457453, EPI_ISL_457454, EPI_ISL_457455, EPI_ISL_457456, EPI_ISL_457457, EPI_ISL_457458, EPI_ISL_457459, EPI_ISL_457460, EPI_ISL_457461, EPI_ISL_457462, EPI_ISL_457463, EPI_ISL_457464, EPI_ISL_457465, EPI_ISL_457466, EPI_ISL_457467, EPI_ISL_457468, EPI_ISL_457469, EPI_ISL_457470, EPI_ISL_457471, EPI_ISL_457472, EPI_ISL_457473, EPI_ISL_457474, EPI_ISL_457475, EPI_ISL_457476, EPI_ISL_457477, EPI_ISL_457478, EPI_ISL_457479, EPI_ISL_457480, EPI_ISL_457481, EPI_ISL_457482, EPI_ISL_457483, EPI_ISL_457484, EPI_ISL_457485, EPI_ISL_457486, EPI_ISL_457487, EPI_ISL_457488, EPI_ISL_457489, EPI_ISL_457490, EPI_ISL_457491, EPI_ISL_457492, EPI_ISL_457493, EPI_ISL_457494, EPI_ISL_457495, EPI_ISL_457496, EPI_ISL_457497, EPI_ISL_457498, EPI_ISL_457499, EPI_ISL_457500, EPI_ISL_457501, EPI_ISL_457502, EPI_ISL_457503, EPI_ISL_457504, EPI_ISL_457505, EPI_ISL_457506, EPI_ISL_457507, EPI_ISL_457508, EPI_ISL_457509, EPI_ISL_457510, EPI_ISL_457511, EPI_ISL_457512, EPI_ISL_457513, EPI_ISL_457514, EPI_ISL_457515, EPI_ISL_457516, EPI_ISL_457517, EPI_ISL_457518, EPI_ISL_457519, EPI_ISL_457520, EPI_ISL_457521, EPI_ISL_457522, EPI_ISL_457523, | | | |

| | | | |
|---|---|---|---|
| EPI_ISL_457524, EPI_ISL_457525, EPI_ISL_457526, EPI_ISL_457527, EPI_ISL_457528, EPI_ISL_457529, EPI_ISL_457530, EPI_ISL_457531, EPI_ISL_457532, EPI_ISL_457533, EPI_ISL_457534, EPI_ISL_457535, EPI_ISL_457536, EPI_ISL_457537, EPI_ISL_457538, EPI_ISL_457539, EPI_ISL_457540, EPI_ISL_457541, EPI_ISL_457542, EPI_ISL_457543, EPI_ISL_457544, EPI_ISL_457545, EPI_ISL_457546, EPI_ISL_457547, EPI_ISL_457548, EPI_ISL_457549, EPI_ISL_457550, EPI_ISL_457551, EPI_ISL_457552, EPI_ISL_457553, EPI_ISL_457555, EPI_ISL_457556, EPI_ISL_457557, EPI_ISL_457558, EPI_ISL_457559, EPI_ISL_457560, EPI_ISL_457561, EPI_ISL_457562, EPI_ISL_457563, EPI_ISL_457564, EPI_ISL_457565, EPI_ISL_457566, EPI_ISL_457567, EPI_ISL_457568, EPI_ISL_457569, EPI_ISL_457570, EPI_ISL_457571, EPI_ISL_457572 | | | |
| see above | Quadram Institute Bioscience | COVID-19 Genomics UK (COG-UK) Consortium | Dave J. Baker, Gemma L. Kay, Alp Aydin, Thanh Le-Viet, Steven Rudder, Ana P. Tedim, Anastasia Kolyva, Maria Diaz, Leonardo de Oliveira Martins, Nabil-Fareed Alikhan, Lizzie Meadows, Rachael Stanley, Ngozi Elumogo, Muhammed Yasir, Nicholas M. Thomson, Alexander J Trotter, Rachel Gilroy, Samuel Bloomfield, Claire Stuart, Andrew Bell, Reenesh Prakash, Samir Dervisevic, Alison E. Mather, John Wain, Mark Webber, Andrew J. Page, Justin O'Grady |
| EPI_ISL_461791, EPI_ISL_461792, EPI_ISL_461793, EPI_ISL_461794, EPI_ISL_461795, EPI_ISL_461796, EPI_ISL_461797, EPI_ISL_461798, EPI_ISL_461799, EPI_ISL_461800, EPI_ISL_461801, EPI_ISL_461802, EPI_ISL_461803, EPI_ISL_461804 | | | |
| see above | Northumbria University / South Tees Hospitals NHS Foundation Trust / North Cumbria Integrated Care NHS Foundation Trust / North Tees and Hartlepool NHS Foundation Trust / Newcastle Hospitals NHS Foundation Trust | COVID-19 Genomics UK (COG-UK) Consortium | Darren L Smith,Andrew Nelson,Matthew Bashton,Greg R Young,Joshua Loh,John Allan,Mohammad A Tariq,Giles S Holt,Gary Black,Wen C Yew,Lynn Dover,Paul Baker,Steve Liggett,Sarah Essex,Jane Greenaway,Debra Padgett,Clive Graham,Garren Scott,Edward Barton,Emma Swindells,Brendan Payne,Jennifer Collins,Yusri Taha,Gary Eltringham |
| EPI_ISL_461805, EPI_ISL_461806, EPI_ISL_461807, EPI_ISL_461808, EPI_ISL_461809, EPI_ISL_461810, EPI_ISL_461811, EPI_ISL_461812, EPI_ISL_461813, EPI_ISL_461814, EPI_ISL_461815, EPI_ISL_461816, EPI_ISL_461817, EPI_ISL_461818, EPI_ISL_461819, EPI_ISL_461820, EPI_ISL_461821, EPI_ISL_461822, EPI_ISL_461823, EPI_ISL_461824, EPI_ISL_461825, EPI_ISL_461826, EPI_ISL_461827, EPI_ISL_461828, EPI_ISL_461829, EPI_ISL_461830, EPI_ISL_461831, EPI_ISL_461832, EPI_ISL_461833, EPI_ISL_461834, EPI_ISL_461835, EPI_ISL_461836, EPI_ISL_461837, EPI_ISL_461838, EPI_ISL_461839, EPI_ISL_461840, EPI_ISL_461841, EPI_ISL_461842, EPI_ISL_461843, EPI_ISL_461844, EPI_ISL_461845, EPI_ISL_461846, EPI_ISL_461847, EPI_ISL_461848, EPI_ISL_461849, EPI_ISL_461850, EPI_ISL_461851, EPI_ISL_461852, EPI_ISL_461853, EPI_ISL_461854, EPI_ISL_461855, EPI_ISL_461856, EPI_ISL_461857, EPI_ISL_461858, EPI_ISL_461859, EPI_ISL_461860, EPI_ISL_461861, EPI_ISL_461862, EPI_ISL_461863, EPI_ISL_461864, EPI_ISL_461865, EPI_ISL_461866, EPI_ISL_461867, EPI_ISL_461868, EPI_ISL_461869, EPI_ISL_461870, EPI_ISL_461871, EPI_ISL_461872, EPI_ISL_461873, EPI_ISL_461874, EPI_ISL_461875, EPI_ISL_461876, EPI_ISL_461877, EPI_ISL_461878, EPI_ISL_461879, EPI_ISL_461880, EPI_ISL_461881, EPI_ISL_461882, EPI_ISL_461883, EPI_ISL_461884, EPI_ISL_461885, EPI_ISL_461886, EPI_ISL_461887, EPI_ISL_461888, EPI_ISL_461889, EPI_ISL_461890, EPI_ISL_461891, EPI_ISL_461892, EPI_ISL_461893 | | | |
| see above | Quadram Institute Bioscience | COVID-19 Genomics UK (COG-UK) Consortium | Dave J. Baker, Gemma L. Kay, Alp Aydin, Thanh Le-Viet, Steven Rudder, Ana P. Tedim, Anastasia Kolyva, Maria Diaz, Leonardo de Oliveira Martins, Nabil-Fareed Alikhan, Lizzie Meadows, Rachael Stanley, Ngozi Elumogo, Muhammed Yasir, Nicholas M. Thomson, Alexander J Trotter, Rachel Gilroy, Samuel Bloomfield, Claire Stuart, Andrew Bell, Reenesh Prakash, Samir Dervisevic, Alison E. Mather, John Wain, Mark Webber, Andrew J. Page, Justin O'Grady |
| EPI_ISL_462912, EPI_ISL_462913, EPI_ISL_462914, EPI_ISL_462915, EPI_ISL_462916, EPI_ISL_462917, EPI_ISL_462918, EPI_ISL_462919, EPI_ISL_462920, EPI_ISL_462921, EPI_ISL_462922, EPI_ISL_462923, EPI_ISL_462924, EPI_ISL_462925, EPI_ISL_462926, EPI_ISL_462927, EPI_ISL_462928, EPI_ISL_462929, EPI_ISL_462930, EPI_ISL_462931, EPI_ISL_462932, EPI_ISL_462933, EPI_ISL_462934, EPI_ISL_462935, EPI_ISL_462936, EPI_ISL_462937, EPI_ISL_462938, EPI_ISL_462939, EPI_ISL_462940, EPI_ISL_462941, EPI_ISL_462942, EPI_ISL_462943, EPI_ISL_462944, EPI_ISL_462945, EPI_ISL_462946, EPI_ISL_462947, EPI_ISL_462948, EPI_ISL_462949, EPI_ISL_462950, EPI_ISL_462951, EPI_ISL_462952, EPI_ISL_462953, EPI_ISL_462954, EPI_ISL_462955, EPI_ISL_462956, EPI_ISL_462957, EPI_ISL_462958, EPI_ISL_462959, EPI_ISL_462960, EPI_ISL_462961, EPI_ISL_462962, EPI_ISL_462963, EPI_ISL_462964, EPI_ISL_462965, EPI_ISL_462966, EPI_ISL_462967, EPI_ISL_462968, EPI_ISL_462969, EPI_ISL_462970, EPI_ISL_462971, EPI_ISL_462972, EPI_ISL_462973, EPI_ISL_462974, EPI_ISL_462975, EPI_ISL_462976, EPI_ISL_462977, EPI_ISL_462978, EPI_ISL_462979, EPI_ISL_462980, EPI_ISL_462981, EPI_ISL_462982, EPI_ISL_462983, EPI_ISL_462984, EPI_ISL_462985, EPI_ISL_462986, EPI_ISL_462987, EPI_ISL_462988, EPI_ISL_462989 | | | |
| see above | Wyoming Public Health Laboratory | Center for Global Health, University of New Mexico Health Sciences Center | Daryl Domman, Kurt Schwalm, Rob Christensen, Wanda Manley, Cari Sloma, Noah Hull, Darrell Dinwiddie |
| EPI_ISL_467302, EPI_ISL_467304 | Washington University in St. Louis | Washington University in St. Louis | David Wang, Carey-Ann Burnham, Scott Handley, Lindsay Droit, Stephen Tahan |
| EPI_ISL_467525, EPI_ISL_467526, EPI_ISL_467527, EPI_ISL_467528, EPI_ISL_467529, EPI_ISL_467530, EPI_ISL_467531, EPI_ISL_467532, EPI_ISL_467533, EPI_ISL_467534, EPI_ISL_467535, EPI_ISL_467536, EPI_ISL_467537, EPI_ISL_467538, EPI_ISL_467539, EPI_ISL_467540, EPI_ISL_467541, EPI_ISL_467542, EPI_ISL_467543, EPI_ISL_467544, EPI_ISL_467545, EPI_ISL_467546, EPI_ISL_467547, EPI_ISL_467548, EPI_ISL_467549, EPI_ISL_467550, EPI_ISL_467551, EPI_ISL_467552, EPI_ISL_467553, EPI_ISL_467554, EPI_ISL_467555, EPI_ISL_467556, EPI_ISL_467557, EPI_ISL_467558, EPI_ISL_467559, EPI_ISL_467560, EPI_ISL_467561, EPI_ISL_467562, EPI_ISL_467563, EPI_ISL_467564, EPI_ISL_467565, EPI_ISL_467566, EPI_ISL_467567, EPI_ISL_467568, EPI_ISL_467569, EPI_ISL_467570, EPI_ISL_467571, EPI_ISL_467572, EPI_ISL_467573, EPI_ISL_467574, EPI_ISL_467575, EPI_ISL_467576, EPI_ISL_467577, EPI_ISL_467578, EPI_ISL_467579, EPI_ISL_467580, EPI_ISL_467581, EPI_ISL_467582, EPI_ISL_467583, EPI_ISL_467584, EPI_ISL_467585, EPI_ISL_467586, EPI_ISL_467587, EPI_ISL_467588, EPI_ISL_467589, EPI_ISL_467590, EPI_ISL_467591, EPI_ISL_467592, EPI_ISL_467593, EPI_ISL_467594, EPI_ISL_467595, EPI_ISL_467596, EPI_ISL_467597, EPI_ISL_467598, EPI_ISL_467599, EPI_ISL_467600, EPI_ISL_467601, EPI_ISL_467602, EPI_ISL_467603, EPI_ISL_467604, EPI_ISL_467605, EPI_ISL_467606, EPI_ISL_467607, EPI_ISL_467608, EPI_ISL_467609, EPI_ISL_467610, EPI_ISL_467611, EPI_ISL_467612, EPI_ISL_467613, EPI_ISL_467614, EPI_ISL_467615, EPI_ISL_467616, EPI_ISL_467617, EPI_ISL_467618, EPI_ISL_467619, EPI_ISL_467620, EPI_ISL_467621, EPI_ISL_467622, EPI_ISL_467623, EPI_ISL_467624, EPI_ISL_467625, EPI_ISL_467626, EPI_ISL_467627, EPI_ISL_467628, EPI_ISL_467629, EPI_ISL_467630, EPI_ISL_467631, EPI_ISL_467632, EPI_ISL_467633, EPI_ISL_467634, EPI_ISL_467635, EPI_ISL_467636, EPI_ISL_467637, EPI_ISL_467638, EPI_ISL_467639, EPI_ISL_467640, EPI_ISL_467641, EPI_ISL_467642, EPI_ISL_467643, EPI_ISL_467644, EPI_ISL_467645, EPI_ISL_467646, EPI_ISL_467647, EPI_ISL_467648, EPI_ISL_467649, EPI_ISL_467650, EPI_ISL_467651, EPI_ISL_467652, EPI_ISL_467653, EPI_ISL_467654, EPI_ISL_467655, EPI_ISL_467656, EPI_ISL_467657, EPI_ISL_467658, EPI_ISL_467659, EPI_ISL_467660, EPI_ISL_467661, EPI_ISL_467662, EPI_ISL_467663, EPI_ISL_467664 | | | |
| see above | New Mexico Department of Health Scientific Laboratory Division | Center for Global Health, University of New Mexico Health Sciences Center | Daryl Domman, Kurt Schwalm, Twila Kunde, Joseph Hicks, Michael Edwards, Darrell Dinwiddie |
| EPI_ISL_472155, EPI_ISL_472156, EPI_ISL_472157, EPI_ISL_472158, EPI_ISL_472159, EPI_ISL_472160, EPI_ISL_472162, EPI_ISL_472163, EPI_ISL_472164, EPI_ISL_472165, EPI_ISL_472166, EPI_ISL_472167, EPI_ISL_472168, EPI_ISL_472169, EPI_ISL_472170, EPI_ISL_472171, EPI_ISL_472172, EPI_ISL_472173, EPI_ISL_472174, EPI_ISL_472175, EPI_ISL_472176, EPI_ISL_472177, EPI_ISL_472178, EPI_ISL_472179, EPI_ISL_472180, EPI_ISL_472181, EPI_ISL_472182, EPI_ISL_472183, EPI_ISL_472184, EPI_ISL_472185, EPI_ISL_472186, EPI_ISL_472187, EPI_ISL_472188, EPI_ISL_472189, EPI_ISL_472190, EPI_ISL_472191, EPI_ISL_472192, EPI_ISL_472193, EPI_ISL_472194, EPI_ISL_472195, EPI_ISL_472196, EPI_ISL_472197, EPI_ISL_472198, EPI_ISL_472199, EPI_ISL_472200, EPI_ISL_472201, EPI_ISL_472202, EPI_ISL_472203, EPI_ISL_472204, EPI_ISL_472205, EPI_ISL_472206, EPI_ISL_472207, EPI_ISL_472208, EPI_ISL_472209, EPI_ISL_472210, EPI_ISL_472211, EPI_ISL_472212, EPI_ISL_472213, EPI_ISL_472214, EPI_ISL_472215, EPI_ISL_472216, EPI_ISL_472217, EPI_ISL_472218, EPI_ISL_472219, EPI_ISL_472220, EPI_ISL_472221, EPI_ISL_472222, EPI_ISL_472223, EPI_ISL_472224, EPI_ISL_472225, EPI_ISL_472226, EPI_ISL_472227, EPI_ISL_472228, EPI_ISL_472229, EPI_ISL_472230, EPI_ISL_472231, EPI_ISL_472232, EPI_ISL_472233, EPI_ISL_472234, EPI_ISL_472235, EPI_ISL_472236, EPI_ISL_472237, EPI_ISL_472238, EPI_ISL_472240, EPI_ISL_472241, EPI_ISL_472242, EPI_ISL_472243, EPI_ISL_472244, EPI_ISL_472245, EPI_ISL_472246, EPI_ISL_472247, EPI_ISL_472248, EPI_ISL_472249, EPI_ISL_472250, EPI_ISL_472251, EPI_ISL_472252, EPI_ISL_472253, EPI_ISL_472254, EPI_ISL_472255, EPI_ISL_472256, EPI_ISL_472257, EPI_ISL_472258, EPI_ISL_472259, EPI_ISL_472260, EPI_ISL_472261, EPI_ISL_472262, EPI_ISL_472263, EPI_ISL_472264, EPI_ISL_472265, EPI_ISL_472266, EPI_ISL_472267, EPI_ISL_472268, EPI_ISL_472269, EPI_ISL_472270, EPI_ISL_472271, EPI_ISL_472272, EPI_ISL_472273, EPI_ISL_472274, EPI_ISL_472275, EPI_ISL_472276, EPI_ISL_472277, EPI_ISL_472278, EPI_ISL_472279, EPI_ISL_472280, EPI_ISL_472281, EPI_ISL_472282, EPI_ISL_472283, EPI_ISL_472284, EPI_ISL_472285, EPI_ISL_472286, EPI_ISL_472287, EPI_ISL_472288, EPI_ISL_472289, EPI_ISL_472290 | | | |
| see above | Northumbria University / South Tees Hospitals NHS Foundation Trust / North Cumbria Integrated Care NHS Foundation Trust / North Tees and Hartlepool NHS Foundation Trust / Newcastle Hospitals NHS Foundation Trust | COVID-19 Genomics UK (COG-UK) Consortium | Darren L Smith,Andrew Nelson,Matthew Bashton,Greg R Young,Joshua Loh,John Allan,Mohammad A Tariq,Giles S Holt,Gary Black,Wen C Yew,Lynn Dover,Paul Baker,Steve Liggett,Sarah Essex,Jane Greenaway,Debra Padgett,Clive Graham,Garren Scott,Edward Barton,Emma Swindells,Brendan Payne,Jennifer Collins,Yusri Taha,Gary Eltringham |
| EPI_ISL_472291, EPI_ISL_472292, EPI_ISL_472293, EPI_ISL_472294, EPI_ISL_472295, EPI_ISL_472296, EPI_ISL_472297, EPI_ISL_472298, EPI_ISL_472299, EPI_ISL_472300, EPI_ISL_472301, EPI_ISL_472302, EPI_ISL_472303, EPI_ISL_472304, EPI_ISL_472305, EPI_ISL_472306, EPI_ISL_472307, EPI_ISL_472308, EPI_ISL_472309, EPI_ISL_472310, EPI_ISL_472311, EPI_ISL_472312, EPI_ISL_472313, EPI_ISL_472314, EPI_ISL_472315, EPI_ISL_472316, EPI_ISL_472317, EPI_ISL_472318, EPI_ISL_472319, EPI_ISL_472320, EPI_ISL_472321, EPI_ISL_472322, EPI_ISL_472323, EPI_ISL_472324, EPI_ISL_472325, EPI_ISL_472326, EPI_ISL_472327, EPI_ISL_472328, EPI_ISL_472329, EPI_ISL_472330, EPI_ISL_472331, EPI_ISL_472332, EPI_ISL_472333, EPI_ISL_472334, EPI_ISL_472335, EPI_ISL_472336, EPI_ISL_472337, EPI_ISL_472338, EPI_ISL_472339, EPI_ISL_472340, EPI_ISL_472341, EPI_ISL_472342, EPI_ISL_472343, EPI_ISL_472344, EPI_ISL_472345, EPI_ISL_472346, EPI_ISL_472347, EPI_ISL_472348, EPI_ISL_472349, EPI_ISL_472350, EPI_ISL_472351, EPI_ISL_472352, EPI_ISL_472353, EPI_ISL_472354, EPI_ISL_472355, EPI_ISL_472356, EPI_ISL_472357, EPI_ISL_472358, EPI_ISL_472359, EPI_ISL_472360, EPI_ISL_472361, EPI_ISL_472362, EPI_ISL_472363, EPI_ISL_472364, EPI_ISL_472365, EPI_ISL_472366, EPI_ISL_472367, EPI_ISL_472368, EPI_ISL_472369, EPI_ISL_472375, EPI_ISL_472377, EPI_ISL_472378, EPI_ISL_472380, EPI_ISL_472382 | | | |
| see above | Quadram Institute Bioscience | COVID-19 Genomics UK (COG-UK) Consortium | Dave J. Baker, Gemma L. Kay, Alp Aydin, Thanh Le-Viet, Steven Rudder, Ana P. Tedim, Anastasia Kolyva, Maria Diaz, Leonardo de Oliveira Martins, Nabil-Fareed Alikhan, Lizzie Meadows, Rachael Stanley, Ngozi Elumogo, Muhammed Yasir, Nicholas M. Thomson, Alexander J Trotter, Rachel Gilroy, Samuel Bloomfield, Claire Stuart, Andrew Bell, Reenesh Prakash, Samir Dervisevic, Alison E. Mather, John Wain, Mark Webber, Andrew J. Page, Justin O'Grady |
| EPI_ISL_476018, EPI_ISL_476019, EPI_ISL_476021 | Washington University in St. Louis | Washington University in St. Louis | David Wang, Carey-Ann Burnham, Scott Handley, Lindsay Droit, Stephen Tahan |
| EPI_ISL_478473, EPI_ISL_478474, EPI_ISL_478475, EPI_ISL_478476, EPI_ISL_478477, EPI_ISL_478478, EPI_ISL_478479, EPI_ISL_478480, EPI_ISL_478481, EPI_ISL_478482, EPI_ISL_478483, EPI_ISL_478484, EPI_ISL_478485, EPI_ISL_478486, EPI_ISL_478487, EPI_ISL_478488, EPI_ISL_478489, EPI_ISL_478490, EPI_ISL_478491, EPI_ISL_478492, EPI_ISL_478493, EPI_ISL_478494, EPI_ISL_478495, EPI_ISL_478496, EPI_ISL_478497, EPI_ISL_478498, EPI_ISL_478500, EPI_ISL_478501, EPI_ISL_478502, EPI_ISL_478503, EPI_ISL_478504, EPI_ISL_478505, EPI_ISL_478506, EPI_ISL_478507, EPI_ISL_478508, EPI_ISL_478509, EPI_ISL_478510, EPI_ISL_478511, EPI_ISL_478513, EPI_ISL_478516, EPI_ISL_478517, EPI_ISL_478518, EPI_ISL_478520, EPI_ISL_478521, EPI_ISL_478522, EPI_ISL_478523, EPI_ISL_478524, EPI_ISL_478525, EPI_ISL_478526, EPI_ISL_478527, EPI_ISL_478528, EPI_ISL_478529, EPI_ISL_478530, EPI_ISL_478531, EPI_ISL_478532, EPI_ISL_478533, EPI_ISL_478534, EPI_ISL_478535, EPI_ISL_478536, EPI_ISL_478537, EPI_ISL_478538, EPI_ISL_478539, EPI_ISL_478540, EPI_ISL_478541, EPI_ISL_478542, EPI_ISL_478543, EPI_ISL_478544, EPI_ISL_478545, EPI_ISL_478546, EPI_ISL_478547, EPI_ISL_478548, EPI_ISL_478549, EPI_ISL_478550, EPI_ISL_478551, EPI_ISL_478552, EPI_ISL_478553, EPI_ISL_478554, EPI_ISL_478555, EPI_ISL_478556, EPI_ISL_478557, EPI_ISL_478558, EPI_ISL_478559, EPI_ISL_478560, EPI_ISL_478561, EPI_ISL_478562, EPI_ISL_478563, EPI_ISL_478564, EPI_ISL_478565, EPI_ISL_478566, EPI_ISL_478567, EPI_ISL_478568, EPI_ISL_478569, EPI_ISL_478570, EPI_ISL_478571, EPI_ISL_478572, EPI_ISL_478573, EPI_ISL_478574, EPI_ISL_478575, EPI_ISL_478576, EPI_ISL_478577, EPI_ISL_478578, EPI_ISL_478579, EPI_ISL_478580, EPI_ISL_478581, EPI_ISL_478582, EPI_ISL_478583, EPI_ISL_478584, EPI_ISL_478585, EPI_ISL_478586, EPI_ISL_478587, EPI_ISL_478588, EPI_ISL_478589, EPI_ISL_478590, EPI_ISL_478591, EPI_ISL_478592, EPI_ISL_478593, EPI_ISL_478594, EPI_ISL_478595, EPI_ISL_478596, EPI_ISL_478597, EPI_ISL_478598, EPI_ISL_478599, EPI_ISL_478600, EPI_ISL_478601, EPI_ISL_478602, EPI_ISL_478603, EPI_ISL_478604, EPI_ISL_478605, EPI_ISL_478606, EPI_ISL_478607, EPI_ISL_478608, EPI_ISL_478609, EPI_ISL_478610, EPI_ISL_478611, EPI_ISL_478612, EPI_ISL_478613, EPI_ISL_478614, EPI_ISL_478615, EPI_ISL_478616, EPI_ISL_478617, EPI_ISL_478618, EPI_ISL_478619, EPI_ISL_478620, EPI_ISL_478621, EPI_ISL_478622, EPI_ISL_478624, EPI_ISL_478625, EPI_ISL_478626, EPI_ISL_478627, EPI_ISL_478628, EPI_ISL_478629, EPI_ISL_478630, EPI_ISL_478631, EPI_ISL_478632, EPI_ISL_478633, EPI_ISL_478634, EPI_ISL_478635, EPI_ISL_478636, EPI_ISL_478637, EPI_ISL_478638, EPI_ISL_478639, EPI_ISL_478640, EPI_ISL_478641, EPI_ISL_478642, EPI_ISL_478643, EPI_ISL_478644, EPI_ISL_478645, EPI_ISL_478646, EPI_ISL_478647, EPI_ISL_478648, EPI_ISL_478649, EPI_ISL_478650, EPI_ISL_478651, EPI_ISL_478652, EPI_ISL_478653, EPI_ISL_478654, EPI_ISL_478655, EPI_ISL_478656, EPI_ISL_478657, EPI_ISL_478658, EPI_ISL_478659, EPI_ISL_478660, EPI_ISL_478661, EPI_ISL_478662, EPI_ISL_478663, EPI_ISL_478664, EPI_ISL_478665, EPI_ISL_478666, EPI_ISL_478667, EPI_ISL_478668 | | | |
| see above | Northumbria University / South Tees Hospitals NHS Foundation Trust / North Cumbria Integrated Care NHS Foundation Trust / North Tees and Hartlepool NHS Foundation Trust / Newcastle Hospitals NHS Foundation Trust | COVID-19 Genomics UK (COG-UK) Consortium | Darren L Smith,Andrew Nelson,Matthew Bashton,Greg R Young,Joshua Loh,John Allan,Mohammad A Tariq,Giles S Holt,Gary Black,Wen C Yew,Lynn Dover,Paul Baker,Steve Liggett,Sarah Essex,Jane Greenaway,Debra Padgett,Clive Graham,Garren Scott,Edward Barton,Emma Swindells,Brendan Payne,Jennifer Collins,Yusri Taha,Gary Eltringham |
| EPI_ISL_478719, EPI_ISL_478720, | Quadram Institute Bioscience | COVID-19 Genomics UK (COG-UK) Consortium | Dave J. Baker, Gemma L. Kay, Alp Aydin, Thanh Le-Viet, Steven Rudder, Ana P. Tedim, Anastasia Kolyva, Maria Diaz, Leonardo de Oliveira Martins, |

| | | | |
|---|---|---|---|
| EPI_ISL_480329, EPI_ISL_480330 | | | Nabil-Fareed Alikhan, Lizzie Meadows, Rachael Stanley, Ngozi Elumogo, Muhammed Yasir, Nicholas M. Thomson, Alexander J Trotter, Rachel Gilroy, Samuel Bloomfield, Claire Stuart, Andrew Bell, Reenesh Prakash, Samir Dervisevic, Alison E. Mather, John Wain, Mark Webber, Andrew J. Page, Justin O'Grady |
| EPI_ISL_484264, EPI_ISL_484265, EPI_ISL_484266, EPI_ISL_484267, EPI_ISL_484268, EPI_ISL_484269, EPI_ISL_484270, EPI_ISL_484271, EPI_ISL_484272, EPI_ISL_484273, EPI_ISL_484274, EPI_ISL_484275, EPI_ISL_484276, EPI_ISL_484277, EPI_ISL_484278, EPI_ISL_484279, EPI_ISL_484280, EPI_ISL_484281, EPI_ISL_484282, EPI_ISL_484283, EPI_ISL_484284, EPI_ISL_484285, EPI_ISL_484286, EPI_ISL_484287, EPI_ISL_484288, EPI_ISL_484289, EPI_ISL_484290, EPI_ISL_484291, EPI_ISL_484292, EPI_ISL_484293, EPI_ISL_484294, EPI_ISL_484295, EPI_ISL_484296, EPI_ISL_484297, EPI_ISL_484298, EPI_ISL_484299, EPI_ISL_484300, EPI_ISL_484301, EPI_ISL_484302, EPI_ISL_484303, EPI_ISL_484304, EPI_ISL_484305, EPI_ISL_484306, EPI_ISL_484307, EPI_ISL_484308, EPI_ISL_484309, EPI_ISL_484310, EPI_ISL_484311, EPI_ISL_484312, EPI_ISL_484313, EPI_ISL_484314, EPI_ISL_484315, EPI_ISL_484316, EPI_ISL_484317, EPI_ISL_484318, EPI_ISL_484319, EPI_ISL_484320, EPI_ISL_484321, EPI_ISL_484322, EPI_ISL_484323, EPI_ISL_484324, EPI_ISL_484325, EPI_ISL_484326, EPI_ISL_484327 | | | |
| see above | Northumbria University / South Tees Hospitals NHS Foundation Trust / North Cumbria Integrated Care NHS Foundation Trust / North Tees and Hartlepool NHS Foundation Trust / Newcastle Hospitals NHS Foundation Trust | COVID-19 Genomics UK (COG-UK) Consortium | Darren L Smith,Andrew Nelson,Matthew Bashton,Greg R Young,Joshua Loh,John Allan,Mohammad A Tariq,Giles S Holt,Gary Black,Wen C Yew,Lynn Dover,Paul Baker,Steve Liggett,Sarah Essex,Jane Greenaway,Debra Padgett,Clive Graham,Garren Scott,Edward Barton,Emma Swindells,Brendan Payne,Jennifer Collins,Yusri Taha,Gary Eltringham |
| EPI_ISL_484328, EPI_ISL_484329, EPI_ISL_484330, EPI_ISL_484331, EPI_ISL_484332, EPI_ISL_484333, EPI_ISL_484334, EPI_ISL_484335, EPI_ISL_484336, EPI_ISL_484337, EPI_ISL_484338 | | | |
| see above | Quadram Institute Bioscience | COVID-19 Genomics UK (COG-UK) Consortium | Dave J. Baker, Gemma L. Kay, Alp Aydin, Thanh Le-Viet, Steven Rudder, Ana P. Tedim, Anastasia Kolyva, Maria Diaz, Leonardo de Oliveira Martins, Nabil-Fareed Alikhan, Lizzie Meadows, Rachael Stanley, Ngozi Elumogo, Muhammed Yasir, Nicholas M. Thomson, Alexander J Trotter, Rachel Gilroy, Samuel Bloomfield, Claire Stuart, Andrew Bell, Reenesh Prakash, Samir Dervisevic, Alison E. Mather, John Wain, Mark Webber, Andrew J. Page, Justin O'Grady |
| EPI_ISL_485872 | New Mexico Department of Health Scientific Laboratory Division | Center for Global Health, University of New Mexico Health Sciences Center | Daryl Domman, Kurt Schwalm, Twila Kunde, Joseph Hicks, Michael Edwards, Darrell Dinwiddie |
| EPI_ISL_490440, EPI_ISL_490441, EPI_ISL_490442, EPI_ISL_490443, EPI_ISL_490444, EPI_ISL_490445, EPI_ISL_490446, EPI_ISL_490447, EPI_ISL_490448, EPI_ISL_490449, EPI_ISL_490450, EPI_ISL_490451, EPI_ISL_490452, EPI_ISL_490453, EPI_ISL_490454, EPI_ISL_490455, EPI_ISL_490456, EPI_ISL_490457, EPI_ISL_490458, EPI_ISL_490459, EPI_ISL_490460, EPI_ISL_490461, EPI_ISL_490462, EPI_ISL_490463, EPI_ISL_490464, EPI_ISL_490465, EPI_ISL_490466, EPI_ISL_490467, EPI_ISL_490468, EPI_ISL_490469, EPI_ISL_490470, EPI_ISL_490471, EPI_ISL_490472, EPI_ISL_490473, EPI_ISL_490474, EPI_ISL_490475, EPI_ISL_490476, EPI_ISL_490477, EPI_ISL_490478, EPI_ISL_490479, EPI_ISL_490480, EPI_ISL_490481, EPI_ISL_490482, EPI_ISL_490483, EPI_ISL_490484, EPI_ISL_490485, EPI_ISL_490486, EPI_ISL_490487, EPI_ISL_490488, EPI_ISL_490489, EPI_ISL_490490, EPI_ISL_490491, EPI_ISL_490492, EPI_ISL_490493, EPI_ISL_490494, EPI_ISL_490495 | | | |
| see above | Northumbria University / South Tees Hospitals NHS Foundation Trust / North Cumbria Integrated Care NHS Foundation Trust / North Tees and Hartlepool NHS Foundation Trust / Newcastle Hospitals NHS Foundation Trust | COVID-19 Genomics UK (COG-UK) Consortium | Darren L Smith,Andrew Nelson,Matthew Bashton,Greg R Young,Joshua Loh,John Allan,Mohammad A Tariq,Giles S Holt,Gary Black,Wen C Yew,Lynn Dover,Paul Baker,Steve Liggett,Sarah Essex,Jane Greenaway,Debra Padgett,Clive Graham,Garren Scott,Edward Barton,Emma Swindells,Brendan Payne,Jennifer Collins,Yusri Taha,Gary Eltringham |
| EPI_ISL_490496, EPI_ISL_490517, EPI_ISL_490518, EPI_ISL_490519, EPI_ISL_490520, EPI_ISL_490521, EPI_ISL_490522, EPI_ISL_490523, EPI_ISL_490525, EPI_ISL_490528, EPI_ISL_490529, EPI_ISL_490532, EPI_ISL_490533, EPI_ISL_490535, EPI_ISL_490539, EPI_ISL_490540, EPI_ISL_490541, EPI_ISL_490542, EPI_ISL_490544, EPI_ISL_490545, EPI_ISL_490547, EPI_ISL_490548, EPI_ISL_490549, EPI_ISL_490550, EPI_ISL_490553, EPI_ISL_490554, EPI_ISL_490555, EPI_ISL_490556, EPI_ISL_490557, EPI_ISL_490558, EPI_ISL_490559 | | | |
| see above | Quadram Institute Bioscience | COVID-19 Genomics UK (COG-UK) Consortium | Dave J. Baker, Gemma L. Kay, Alp Aydin, Thanh Le-Viet, Steven Rudder, Ana P. Tedim, Anastasia Kolyva, Maria Diaz, Leonardo de Oliveira Martins, Nabil-Fareed Alikhan, Lizzie Meadows, Rachael Stanley, Ngozi Elumogo, Muhammed Yasir, Nicholas M. Thomson, Alexander J Trotter, Rachel Gilroy, Samuel Bloomfield, Claire Stuart, Andrew Bell, Reenesh Prakash, Samir Dervisevic, Alison E. Mather, John Wain, Mark Webber, Andrew J. Page, Justin O'Grady |
| EPI_ISL_493069, EPI_ISL_493070, EPI_ISL_493071, EPI_ISL_493072, EPI_ISL_493073, EPI_ISL_493074, EPI_ISL_493075, EPI_ISL_493076, EPI_ISL_493077, EPI_ISL_493079, EPI_ISL_493080, EPI_ISL_493081, EPI_ISL_493082, EPI_ISL_493083, EPI_ISL_493084, EPI_ISL_493085, EPI_ISL_493086 | | | |
| see above | Washington University in St. Louis | Washington University in St. Louis | David Wang, Carey-Ann Burnham, Scott Handley, Lindsay Droit, Stephen Tahan |
| EPI_ISL_493452, EPI_ISL_493453, EPI_ISL_493454, EPI_ISL_493455, EPI_ISL_493456, EPI_ISL_493457, EPI_ISL_493458, EPI_ISL_493459, EPI_ISL_493460, EPI_ISL_493461, EPI_ISL_493462, EPI_ISL_493463, EPI_ISL_493464, EPI_ISL_493465, EPI_ISL_493466, EPI_ISL_493467, EPI_ISL_493468, EPI_ISL_493469, EPI_ISL_493470, EPI_ISL_493471, EPI_ISL_493472, EPI_ISL_493473, EPI_ISL_493474, EPI_ISL_493475, EPI_ISL_493476, EPI_ISL_493477, EPI_ISL_493478, EPI_ISL_493479, EPI_ISL_493480, EPI_ISL_493481, EPI_ISL_493482, EPI_ISL_493483, EPI_ISL_493484, EPI_ISL_493485, EPI_ISL_493486, EPI_ISL_493487, EPI_ISL_493488, EPI_ISL_493489, EPI_ISL_493490, EPI_ISL_493491, EPI_ISL_493492, EPI_ISL_493493, EPI_ISL_493494, EPI_ISL_493495, EPI_ISL_493496, EPI_ISL_493497, EPI_ISL_493498, EPI_ISL_493499, EPI_ISL_493500, EPI_ISL_493501, EPI_ISL_493502, EPI_ISL_493503, EPI_ISL_493504, EPI_ISL_493505, EPI_ISL_493506, EPI_ISL_493507, EPI_ISL_493508, EPI_ISL_493509, EPI_ISL_493510, EPI_ISL_493511, EPI_ISL_493512, EPI_ISL_493513, EPI_ISL_493514, EPI_ISL_493515, EPI_ISL_493516, EPI_ISL_493517, EPI_ISL_493518, EPI_ISL_493519, EPI_ISL_493520, EPI_ISL_493521, EPI_ISL_493522, EPI_ISL_493523, EPI_ISL_493524, EPI_ISL_493525, EPI_ISL_493526, EPI_ISL_493527, EPI_ISL_493528, EPI_ISL_493529, EPI_ISL_493530, EPI_ISL_493531, EPI_ISL_493532, EPI_ISL_493533, EPI_ISL_493534, EPI_ISL_493535, EPI_ISL_493536, EPI_ISL_493537, EPI_ISL_493538, EPI_ISL_493539, EPI_ISL_493540 | | | |
| see above | Northumbria University / South Tees Hospitals NHS Foundation Trust / North Cumbria Integrated Care NHS Foundation Trust / North Tees and Hartlepool NHS Foundation Trust / Newcastle Hospitals NHS Foundation Trust | COVID-19 Genomics UK (COG-UK) Consortium | Darren L Smith,Andrew Nelson,Matthew Bashton,Greg R Young,Joshua Loh,John Allan,Mohammad A Tariq,Giles S Holt,Gary Black,Wen C Yew,Lynn Dover,Paul Baker,Steve Liggett,Sarah Essex,Jane Greenaway,Debra Padgett,Clive Graham,Garren Scott,Edward Barton,Emma Swindells,Brendan Payne,Jennifer Collins,Yusri Taha,Gary Eltringham |
| EPI_ISL_493541, EPI_ISL_493542, EPI_ISL_493543, EPI_ISL_493544, EPI_ISL_493545, EPI_ISL_493546 | Quadram Institute Bioscience | COVID-19 Genomics UK (COG-UK) Consortium | Dave J. Baker, Gemma L. Kay, Alp Aydin, Thanh Le-Viet, Steven Rudder, Ana P. Tedim, Anastasia Kolyva, Maria Diaz, Leonardo de Oliveira Martins, Nabil-Fareed Alikhan, Lizzie Meadows, Rachael Stanley, Ngozi Elumogo, Muhammed Yasir, Nicholas M. Thomson, Alexander J Trotter, Rachel Gilroy, Samuel Bloomfield, Claire Stuart, Andrew Bell, Reenesh Prakash, Samir Dervisevic, Alison E. Mather, John Wain, Mark Webber, Andrew J. Page, Justin O'Grady |
| EPI_ISL_495645 | Washington State Department of Health | Seattle Flu Study | Deborah A. Nickerson, Chris D. Frazar, Jover Lee, Benjamin Pelle, Matthew Richardson, Amanda Adler, Elisabeth Brandstetter, Peter D. Han, Kairsten Fay, Misja Ilcisin, Kirsten Lacombe, Thomas R. Sibley, Melissa Truong, Caitlin R. Wolf, Romesh Gautom, Geoff Melly, Brian Hiatt, Philip Dykema, Scott Lindquist, Michael Boeckh, Janet A. Englund, Michael Famulare, Barry R. Lutz, Mark J. Rieder, Lea M. Starita, Matthew Thompson, Helen Y. Chu, Jay Shendure, Trevor Bedford |
| EPI_ISL_495654, EPI_ISL_495656, EPI_ISL_495657, EPI_ISL_495659 | Seattle Flu Study | Seattle Flu Study | Deborah A. Nickerson, Chris D. Frazar, Jover Lee, Benjamin Pelle, Matthew Richardson, Amanda Adler, Elisabeth Brandstetter, Peter D. Han, Kairsten Fay, Misja Ilcisin, Kirsten Lacombe, Thomas R. Sibley, Melissa Truong, Caitlin R. Wolf, Karen Cowgill, Stephanie Schrag, Jeff Duchin, Michael Boeckh, Janet A. Englund, Michael Famulare, Barry R. Lutz, Mark J. Rieder, Lea M. Starita, Matthew Thompson, Helen Y. Chu, Trevor Bedford, Jay Shendure |
| EPI_ISL_495666, EPI_ISL_495675, EPI_ISL_495679, EPI_ISL_495691, EPI_ISL_495713, EPI_ISL_495724, EPI_ISL_495727, EPI_ISL_495757, EPI_ISL_495767, EPI_ISL_495770, EPI_ISL_495783, EPI_ISL_495807, EPI_ISL_495832, EPI_ISL_495846, EPI_ISL_495850, EPI_ISL_495870, EPI_ISL_495871, EPI_ISL_495879, EPI_ISL_495888, EPI_ISL_495900, EPI_ISL_495910 | | | |
| see above | Washington State Department of Health | Seattle Flu Study | Deborah A. Nickerson, Chris D. Frazar, Jover Lee, Benjamin Pelle, Matthew Richardson, Amanda Adler, Elisabeth Brandstetter, Peter D. Han, Kairsten Fay, Misja Ilcisin, Kirsten Lacombe, Thomas R. Sibley, Melissa Truong, Caitlin R. Wolf, Romesh Gautom, Geoff |
| EPI_ISL_495917, EPI_ISL_495920, EPI_ISL_495932, EPI_ISL_495933, EPI_ISL_495939, EPI_ISL_495965, EPI_ISL_495975, EPI_ISL_495976, EPI_ISL_495977, EPI_ISL_495988, EPI_ISL_495990, EPI_ISL_495993, EPI_ISL_495994, EPI_ISL_496010, EPI_ISL_496014, EPI_ISL_496015, EPI_ISL_496017, EPI_ISL_496020, EPI_ISL_496033, EPI_ISL_496072, EPI_ISL_496075, EPI_ISL_496093, EPI_ISL_496094, EPI_ISL_496095, EPI_ISL_496109, EPI_ISL_496131, EPI_ISL_496149, EPI_ISL_496152, EPI_ISL_496162, EPI_ISL_496170, EPI_ISL_496171, EPI_ISL_496172, EPI_ISL_496177, EPI_ISL_496181, EPI_ISL_496189, EPI_ISL_496198, EPI_ISL_496238, EPI_ISL_496247, EPI_ISL_496251, EPI_ISL_496254, EPI_ISL_496277, EPI_ISL_496278, EPI_ISL_496279, EPI_ISL_496280, EPI_ISL_496284, EPI_ISL_496293, EPI_ISL_496329, EPI_ISL_496931, EPI_ISL_496932, EPI_ISL_496933, EPI_ISL_496934, EPI_ISL_496935, EPI_ISL_496936, EPI_ISL_496937, EPI_ISL_496938, EPI_ISL_496939, EPI_ISL_496940, EPI_ISL_496941, EPI_ISL_496942, EPI_ISL_496943, EPI_ISL_496944, EPI_ISL_496945, EPI_ISL_496946, EPI_ISL_496947, EPI_ISL_496948, EPI_ISL_496949, EPI_ISL_496950, EPI_ISL_496951, EPI_ISL_496952, EPI_ISL_496953, EPI_ISL_496954, EPI_ISL_496955, EPI_ISL_496956, EPI_ISL_496957, EPI_ISL_496958, EPI_ISL_496959, EPI_ISL_496960, EPI_ISL_496961, EPI_ISL_496962, EPI_ISL_496963, EPI_ISL_496964, EPI_ISL_496965, EPI_ISL_496966, EPI_ISL_496967, EPI_ISL_496968, EPI_ISL_496969, EPI_ISL_496970, EPI_ISL_496971, EPI_ISL_496972, EPI_ISL_496973, EPI_ISL_496974, EPI_ISL_496975, EPI_ISL_496976, EPI_ISL_496977, EPI_ISL_496978, EPI_ISL_496979, EPI_ISL_496980, EPI_ISL_496981, EPI_ISL_496982, EPI_ISL_496983, EPI_ISL_496984, EPI_ISL_496985, EPI_ISL_496986, EPI_ISL_496987, EPI_ISL_496988, EPI_ISL_496989, EPI_ISL_496990, EPI_ISL_496991, EPI_ISL_496992, EPI_ISL_496993, EPI_ISL_496994, EPI_ISL_496995, EPI_ISL_496996, EPI_ISL_496997, EPI_ISL_496998, EPI_ISL_496999, EPI_ISL_497000, EPI_ISL_497002, EPI_ISL_497003, EPI_ISL_497004, EPI_ISL_497005, EPI_ISL_497006, EPI_ISL_497007, EPI_ISL_497008, EPI_ISL_497009, EPI_ISL_497010, EPI_ISL_497011, EPI_ISL_497012, EPI_ISL_497013, EPI_ISL_497014, EPI_ISL_497015, EPI_ISL_497016, EPI_ISL_497017, EPI_ISL_497018, EPI_ISL_497019, EPI_ISL_497020, EPI_ISL_497021, EPI_ISL_497022, EPI_ISL_497023, EPI_ISL_497024, EPI_ISL_497025, EPI_ISL_497026, EPI_ISL_497027, EPI_ISL_497028, EPI_ISL_497029, EPI_ISL_497030, EPI_ISL_497031, EPI_ISL_497032, EPI_ISL_497033, EPI_ISL_497034, EPI_ISL_497035, EPI_ISL_497036, EPI_ISL_497037, EPI_ISL_497038, EPI_ISL_497039, EPI_ISL_497040, EPI_ISL_497041, EPI_ISL_497042, EPI_ISL_497043, EPI_ISL_497044, EPI_ISL_497045, EPI_ISL_497046, EPI_ISL_497047, EPI_ISL_497048, EPI_ISL_497049, EPI_ISL_497050, EPI_ISL_497051, EPI_ISL_497052, EPI_ISL_497053, EPI_ISL_497054, EPI_ISL_497055, EPI_ISL_497056, EPI_ISL_497057, EPI_ISL_497058, EPI_ISL_497059, EPI_ISL_497060, EPI_ISL_497061, EPI_ISL_497062, EPI_ISL_497063, EPI_ISL_497064, EPI_ISL_497065, EPI_ISL_497066, EPI_ISL_497067, EPI_ISL_497068, EPI_ISL_497069, EPI_ISL_497070, EPI_ISL_497071, EPI_ISL_497072, EPI_ISL_497073, EPI_ISL_497074, EPI_ISL_497075, EPI_ISL_497076, EPI_ISL_497077, EPI_ISL_497078, EPI_ISL_497079, EPI_ISL_497080, EPI_ISL_497081, EPI_ISL_497082, EPI_ISL_497083, EPI_ISL_497084, EPI_ISL_497085, EPI_ISL_497086, EPI_ISL_497087, EPI_ISL_497088, EPI_ISL_497089, EPI_ISL_497090, EPI_ISL_497091, EPI_ISL_497092, EPI_ISL_497093, EPI_ISL_497094, EPI_ISL_497095, EPI_ISL_497096, EPI_ISL_497097, EPI_ISL_497098, EPI_ISL_497099, EPI_ISL_497100, EPI_ISL_497101, EPI_ISL_497102, EPI_ISL_497103, EPI_ISL_497104, EPI_ISL_497105, EPI_ISL_497106, EPI_ISL_497107, EPI_ISL_497108, EPI_ISL_497109, EPI_ISL_497110, EPI_ISL_497111, EPI_ISL_497112, EPI_ISL_497113, EPI_ISL_497114, EPI_ISL_497115, EPI_ISL_497116, EPI_ISL_497117, EPI_ISL_497118, EPI_ISL_497119, EPI_ISL_497120, EPI_ISL_497121, EPI_ISL_497122, EPI_ISL_497123, EPI_ISL_497124, EPI_ISL_497125, EPI_ISL_497126, EPI_ISL_497127, EPI_ISL_497128, EPI_ISL_497129, EPI_ISL_497130, EPI_ISL_497131, EPI_ISL_497132, EPI_ISL_497133, EPI_ISL_497134, EPI_ISL_497135, EPI_ISL_497136, EPI_ISL_497137, EPI_ISL_497138, EPI_ISL_497139, EPI_ISL_497140, EPI_ISL_497141, EPI_ISL_497142, EPI_ISL_497143, EPI_ISL_497144, EPI_ISL_497145, EPI_ISL_497146, EPI_ISL_497147, EPI_ISL_497148, EPI_ISL_497149, EPI_ISL_497150, EPI_ISL_497151, EPI_ISL_497152, EPI_ISL_497153, EPI_ISL_497154, EPI_ISL_497155, EPI_ISL_497156, EPI_ISL_497157, EPI_ISL_497158, EPI_ISL_497159, EPI_ISL_497160, EPI_ISL_497161, EPI_ISL_497162, EPI_ISL_497163, EPI_ISL_497164, EPI_ISL_497165, EPI_ISL_497166, EPI_ISL_497167, EPI_ISL_497168, EPI_ISL_497169, EPI_ISL_497170, EPI_ISL_497171, EPI_ISL_497172, EPI_ISL_497173, EPI_ISL_497174, EPI_ISL_497175, EPI_ISL_497176, EPI_ISL_497177, EPI_ISL_497178, EPI_ISL_497179, EPI_ISL_497180, EPI_ISL_497181, EPI_ISL_497182, EPI_ISL_497183, EPI_ISL_497184, EPI_ISL_497185, EPI_ISL_497186, EPI_ISL_497187, EPI_ISL_497188, EPI_ISL_497189, EPI_ISL_497190, EPI_ISL_497191, EPI_ISL_497192, EPI_ISL_497193, EPI_ISL_497194, EPI_ISL_497195, EPI_ISL_497196, EPI_ISL_497197, EPI_ISL_497198, EPI_ISL_497199, EPI_ISL_497200, EPI_ISL_497201, EPI_ISL_497202, EPI_ISL_497203, EPI_ISL_497204, EPI_ISL_497205, EPI_ISL_497206, EPI_ISL_497207, EPI_ISL_497208, EPI_ISL_497209, EPI_ISL_497210, EPI_ISL_497211, EPI_ISL_497212, EPI_ISL_497213, EPI_ISL_497214, EPI_ISL_497215, EPI_ISL_497216, EPI_ISL_497217, EPI_ISL_497218, EPI_ISL_497219, EPI_ISL_497220, EPI_ISL_497221, EPI_ISL_497222, EPI_ISL_497223, EPI_ISL_497224, EPI_ISL_497225, EPI_ISL_497226, EPI_ISL_497227, EPI_ISL_497228, EPI_ISL_497229, EPI_ISL_497230, EPI_ISL_497231, EPI_ISL_497232, EPI_ISL_497233, EPI_ISL_497234, EPI_ISL_497235, EPI_ISL_497236, EPI_ISL_497237, EPI_ISL_497238, EPI_ISL_497239, EPI_ISL_497240, EPI_ISL_497241, EPI_ISL_497242, EPI_ISL_497243, EPI_ISL_497244, | | | |

| | | | |
|---|---|---|---|
| EPI_ISL_497245, EPI_ISL_497246, EPI_ISL_497247, EPI_ISL_497248, EPI_ISL_497249, EPI_ISL_497250, EPI_ISL_497251, EPI_ISL_497252, EPI_ISL_497253, EPI_ISL_497254, EPI_ISL_497255, EPI_ISL_497256, EPI_ISL_497257, EPI_ISL_497258, EPI_ISL_497259, EPI_ISL_497260, EPI_ISL_497261, EPI_ISL_497262, EPI_ISL_497263, EPI_ISL_497264, EPI_ISL_497265, EPI_ISL_497266, EPI_ISL_497267, EPI_ISL_497268, EPI_ISL_497269, EPI_ISL_497270, EPI_ISL_497271, EPI_ISL_497272, EPI_ISL_497273, EPI_ISL_497274, EPI_ISL_497275, EPI_ISL_497276, EPI_ISL_497277, EPI_ISL_497278, EPI_ISL_497279, EPI_ISL_497280, EPI_ISL_497281, EPI_ISL_497282, EPI_ISL_497283, EPI_ISL_497284, EPI_ISL_497285, EPI_ISL_497286, EPI_ISL_497287, EPI_ISL_497288, EPI_ISL_497289, EPI_ISL_497290, EPI_ISL_497291, EPI_ISL_497292, EPI_ISL_497293, EPI_ISL_497294, EPI_ISL_497295, EPI_ISL_497296, EPI_ISL_497297, EPI_ISL_497298, EPI_ISL_497299, EPI_ISL_497300, EPI_ISL_497301, EPI_ISL_497302, EPI_ISL_497303, EPI_ISL_497304, EPI_ISL_497305, EPI_ISL_497306, EPI_ISL_497307, EPI_ISL_497308, EPI_ISL_497309, EPI_ISL_497310, EPI_ISL_497311, EPI_ISL_497312, EPI_ISL_497313, EPI_ISL_497314, EPI_ISL_497315, EPI_ISL_497316, EPI_ISL_497317, EPI_ISL_497318, EPI_ISL_497319, EPI_ISL_497320, EPI_ISL_497321, EPI_ISL_497322, EPI_ISL_497323, EPI_ISL_497324, EPI_ISL_497325, EPI_ISL_497326, EPI_ISL_497327, EPI_ISL_497328, EPI_ISL_497329, EPI_ISL_497330, EPI_ISL_497331, EPI_ISL_497332, EPI_ISL_497333, EPI_ISL_497334, EPI_ISL_497335, EPI_ISL_497336, EPI_ISL_497337, EPI_ISL_497338, EPI_ISL_497339, EPI_ISL_497340, EPI_ISL_497341, EPI_ISL_497355, EPI_ISL_497419, EPI_ISL_497439, EPI_ISL_497452, EPI_ISL_497477, EPI_ISL_497566, EPI_ISL_497568, EPI_ISL_497570, EPI_ISL_497571, EPI_ISL_497572, EPI_ISL_497573, EPI_ISL_497574, EPI_ISL_497575, EPI_ISL_497576, EPI_ISL_497577, EPI_ISL_497578, EPI_ISL_497579, EPI_ISL_497580, EPI_ISL_497581, EPI_ISL_497582, EPI_ISL_497583, EPI_ISL_497584, EPI_ISL_497585, EPI_ISL_497586, EPI_ISL_497587, EPI_ISL_497588, EPI_ISL_497589, EPI_ISL_497590, EPI_ISL_497591, EPI_ISL_497592, EPI_ISL_497593, EPI_ISL_497594, EPI_ISL_497595, EPI_ISL_497596, EPI_ISL_497597, EPI_ISL_497598, EPI_ISL_497599, EPI_ISL_497600, EPI_ISL_497601, EPI_ISL_497602, EPI_ISL_497603, EPI_ISL_497604, EPI_ISL_497605, EPI_ISL_497606, EPI_ISL_497607, EPI_ISL_497608, EPI_ISL_497609, EPI_ISL_497610, EPI_ISL_497611, EPI_ISL_497612, EPI_ISL_497613, EPI_ISL_497614, EPI_ISL_497615, EPI_ISL_497616, EPI_ISL_497617, EPI_ISL_497618, EPI_ISL_497619, EPI_ISL_497620, EPI_ISL_497621, EPI_ISL_497622, EPI_ISL_497623, EPI_ISL_497624, EPI_ISL_497625, EPI_ISL_497626, EPI_ISL_497627, EPI_ISL_497628, EPI_ISL_497629, EPI_ISL_497630, EPI_ISL_497631, EPI_ISL_497632, EPI_ISL_497633, EPI_ISL_497634, EPI_ISL_497635, EPI_ISL_497636, EPI_ISL_497637, EPI_ISL_497638, EPI_ISL_497639, EPI_ISL_497640, EPI_ISL_497641, EPI_ISL_497642, EPI_ISL_497643, EPI_ISL_497644, EPI_ISL_497645, EPI_ISL_497646, EPI_ISL_497647, EPI_ISL_497648, EPI_ISL_497649, EPI_ISL_497650, EPI_ISL_497651, EPI_ISL_497652, EPI_ISL_497653, EPI_ISL_497654, EPI_ISL_497655, EPI_ISL_497656, EPI_ISL_497657, EPI_ISL_497658, EPI_ISL_497659, EPI_ISL_497660, EPI_ISL_497661, EPI_ISL_497662, EPI_ISL_497663, EPI_ISL_497664, EPI_ISL_497665, EPI_ISL_497666, EPI_ISL_497667, EPI_ISL_497668, EPI_ISL_497669, EPI_ISL_497670, EPI_ISL_497671, EPI_ISL_497672, EPI_ISL_497673, EPI_ISL_497674, EPI_ISL_497675, EPI_ISL_497676, EPI_ISL_497677, EPI_ISL_497678, EPI_ISL_497679, EPI_ISL_497680, EPI_ISL_497681, EPI_ISL_497682, EPI_ISL_497683, EPI_ISL_497684, EPI_ISL_497685, EPI_ISL_497686, EPI_ISL_497687, EPI_ISL_497688, EPI_ISL_497689, EPI_ISL_497690, EPI_ISL_497691, EPI_ISL_497692, EPI_ISL_497693, EPI_ISL_497694, EPI_ISL_497695, EPI_ISL_497696, EPI_ISL_497697, EPI_ISL_497698, EPI_ISL_497699, EPI_ISL_497700, EPI_ISL_497701, EPI_ISL_497702, EPI_ISL_497703, EPI_ISL_497704, EPI_ISL_497705, EPI_ISL_497706, EPI_ISL_497707, EPI_ISL_497708, EPI_ISL_497709, EPI_ISL_497710, EPI_ISL_497711, EPI_ISL_497712, EPI_ISL_497713, EPI_ISL_497714, EPI_ISL_497715, EPI_ISL_497716, EPI_ISL_497717, EPI_ISL_497718, EPI_ISL_497719, EPI_ISL_497720, EPI_ISL_497721, EPI_ISL_497722, EPI_ISL_497723, EPI_ISL_497724, EPI_ISL_497725, EPI_ISL_497726 | | | |
| see above | Washington State Department of Health | Seattle Flu Study | Deborah A. Nickerson, Chris D. Frazar, Jover Lee, Benjamin Pelle, Matthew Richardson, Amanda Adler, Elisabeth Brandstetter, Peter D. Han, Kairsten Fay, Misja Ilcisin, Kirsten Lacombe, Thomas R. Sibley, Melissa Truong, Caitlin R. Wolf, Romesh Gautom, Geoff Melly, Brian Hiatt, Philip Dykema, Scott Lindquist, Michael Boeckh, Janet A. Englund, Michael Famulare, Barry R. Lutz, Mark J. Rieder, Lea M. Starita, Matthew Thompson, Helen Y. Chu, Jay Shendure, Trevor Bedford |
| EPI_ISL_499472, EPI_ISL_499474, EPI_ISL_499478 | Northumbria University / South Tees Hospitals NHS Foundation Trust / North Cumbria Integrated Care NHS Foundation Trust / North Tees and Hartlepool NHS Foundation Trust / Newcastle Hospitals NHS Foundation Trust | COVID-19 Genomics UK (COG-UK) Consortium | Darren L Smith,Andrew Nelson,Matthew Bashton,Greg R Young,Joshua Loh,John Allan,Mohammad A Tariq,Giles S Holt,Gary Black,Wen C Yew,Lynn Dover,Paul Baker,Steve Liggett,Sarah Essex,Jane Greenaway,Debra Padgett,Clive Graham,Garren Scott,Edward Barton,Emma Swindells,Brendan Payne,Jennifer Collins,Yusri Taha,Gary Eltringham |
| EPI_ISL_499479 | Quadram Institute Bioscience | COVID-19 Genomics UK (COG-UK) Consortium | Dave J. Baker, Gemma L. Kay, Alp Aydin, Thanh Le-Viet, Steven Rudder, Ana P. Tedim, Anastasia Kolyva, Maria Diaz, Leonardo de Oliveira Martins, Nabil-Fareed Alikhan, Lizzie Meadows, Rachael Stanley, Ngozi Elumogo, Muhammed Yasir, Nicholas M. Thomson, Alexander J Trotter, Rachel Gilroy, Samuel Bloomfield, Claire Stuart, Andrew Bell, Reenesh Prakash, Samir Dervisevic, Alison E. Mather, John Wain, Mark Webber, Andrew J. Page, Justin O'Grady |
| EPI_ISL_499482, EPI_ISL_499485 | Northumbria University / South Tees Hospitals NHS Foundation Trust / North Cumbria Integrated Care NHS Foundation Trust / North Tees and Hartlepool NHS Foundation Trust / Newcastle Hospitals NHS Foundation Trust | COVID-19 Genomics UK (COG-UK) Consortium | Darren L Smith,Andrew Nelson,Matthew Bashton,Greg R Young,Joshua Loh,John Allan,Mohammad A Tariq,Giles S Holt,Gary Black,Wen C Yew,Lynn Dover,Paul Baker,Steve Liggett,Sarah Essex,Jane Greenaway,Debra Padgett,Clive Graham,Garren Scott,Edward Barton,Emma Swindells,Brendan Payne,Jennifer Collins,Yusri Taha,Gary Eltringham |
| EPI_ISL_499490 | Quadram Institute Bioscience | COVID-19 Genomics UK (COG-UK) Consortium | Dave J. Baker, Gemma L. Kay, Alp Aydin, Thanh Le-Viet, Steven Rudder, Ana P. Tedim, Anastasia Kolyva, Maria Diaz, Leonardo de Oliveira Martins, Nabil-Fareed Alikhan, Lizzie Meadows, Rachael Stanley, Ngozi Elumogo, Muhammed Yasir, Nicholas M. Thomson, Alexander J Trotter, Rachel Gilroy, Samuel Bloomfield, Claire Stuart, Andrew Bell, Reenesh Prakash, Samir Dervisevic, Alison E. Mather, John Wain, Mark Webber, Andrew J. Page, Justin O'Grady |
| EPI_ISL_499493, EPI_ISL_499494 | Northumbria University / South Tees Hospitals NHS Foundation Trust / North Cumbria Integrated Care NHS Foundation Trust / North Tees and Hartlepool NHS Foundation Trust / Newcastle Hospitals NHS Foundation Trust | COVID-19 Genomics UK (COG-UK) Consortium | Darren L Smith,Andrew Nelson,Matthew Bashton,Greg R Young,Joshua Loh,John Allan,Mohammad A Tariq,Giles S Holt,Gary Black,Wen C Yew,Lynn Dover,Paul Baker,Steve Liggett,Sarah Essex,Jane Greenaway,Debra Padgett,Clive Graham,Garren Scott,Edward Barton,Emma Swindells,Brendan Payne,Jennifer Collins,Yusri Taha,Gary Eltringham |
| EPI_ISL_499495 | Quadram Institute Bioscience | COVID-19 Genomics UK (COG-UK) Consortium | Dave J. Baker, Gemma L. Kay, Alp Aydin, Thanh Le-Viet, Steven Rudder, Ana P. Tedim, Anastasia Kolyva, Maria Diaz, Leonardo de Oliveira Martins, Nabil-Fareed Alikhan, Lizzie Meadows, Rachael Stanley, Ngozi Elumogo, Muhammed Yasir, Nicholas M. Thomson, Alexander J Trotter, Rachel Gilroy, Samuel Bloomfield, Claire Stuart, Andrew Bell, Reenesh Prakash, Samir Dervisevic, Alison E. Mather, John Wain, Mark Webber, Andrew J. Page, Justin O'Grady |
| EPI_ISL_499502, EPI_ISL_499503, EPI_ISL_499504, EPI_ISL_499505, EPI_ISL_499506, EPI_ISL_499507, EPI_ISL_499508, EPI_ISL_499510, EPI_ISL_499515, EPI_ISL_499546, EPI_ISL_499547, EPI_ISL_499548, EPI_ISL_499549, EPI_ISL_499550, EPI_ISL_499551, EPI_ISL_499552, EPI_ISL_499553, EPI_ISL_499554, EPI_ISL_499555, EPI_ISL_499556, EPI_ISL_499558, EPI_ISL_499559, EPI_ISL_499560, EPI_ISL_499561 | | | |
| see above | Northumbria University / South Tees Hospitals NHS Foundation Trust / North Cumbria Integrated Care NHS Foundation Trust / North Tees and Hartlepool NHS Foundation Trust / Newcastle Hospitals NHS Foundation Trust | COVID-19 Genomics UK (COG-UK) Consortium | Darren L Smith,Andrew Nelson,Matthew Bashton,Greg R Young,Joshua Loh,John Allan,Mohammad A Tariq,Giles S Holt,Gary Black,Wen C Yew,Lynn Dover,Paul Baker,Steve Liggett,Sarah Essex,Jane Greenaway,Debra Padgett,Clive Graham,Garren Scott,Edward Barton,Emma Swindells,Brendan Payne,Jennifer Collins,Yusri Taha,Gary Eltringham |
| EPI_ISL_499562 | Quadram Institute Bioscience | COVID-19 Genomics UK (COG-UK) Consortium | Dave J. Baker, Gemma L. Kay, Alp Aydin, Thanh Le-Viet, Steven Rudder, Ana P. Tedim, Anastasia Kolyva, Maria Diaz, Leonardo de Oliveira Martins, Nabil-Fareed Alikhan, Lizzie Meadows, Rachael Stanley, Ngozi Elumogo, Muhammed Yasir, Nicholas M. Thomson, Alexander J Trotter, Rachel Gilroy, Samuel Bloomfield, Claire Stuart, Andrew Bell, Reenesh Prakash, Samir Dervisevic, Alison E. Mather, John Wain, Mark Webber, Andrew J. Page, Justin O'Grady |
| EPI_ISL_499563, EPI_ISL_499568, EPI_ISL_499572, EPI_ISL_499587 | Northumbria University / South Tees Hospitals NHS Foundation Trust / North Cumbria Integrated Care NHS Foundation Trust / North Tees and Hartlepool NHS Foundation Trust / Newcastle Hospitals NHS Foundation Trust | COVID-19 Genomics UK (COG-UK) Consortium | Darren L Smith,Andrew Nelson,Matthew Bashton,Greg R Young,Joshua Loh,John Allan,Mohammad A Tariq,Giles S Holt,Gary Black,Wen C Yew,Lynn Dover,Paul Baker,Steve Liggett,Sarah Essex,Jane Greenaway,Debra Padgett,Clive Graham,Garren Scott,Edward Barton,Emma Swindells,Brendan Payne,Jennifer Collins,Yusri Taha,Gary Eltringham |
| EPI_ISL_499588, EPI_ISL_499591, EPI_ISL_499592 | Quadram Institute Bioscience | COVID-19 Genomics UK (COG-UK) Consortium | Dave J. Baker, Gemma L. Kay, Alp Aydin, Thanh Le-Viet, Steven Rudder, Ana P. Tedim, Anastasia Kolyva, Maria Diaz, Leonardo de Oliveira Martins, Nabil-Fareed Alikhan, Lizzie Meadows, Rachael Stanley, Ngozi Elumogo, Muhammed Yasir, Nicholas M. Thomson, Alexander J Trotter, Rachel Gilroy, Samuel Bloomfield, Claire Stuart, Andrew Bell, Reenesh Prakash, Samir Dervisevic, Alison E. Mather, John Wain, Mark Webber, Andrew J. Page, Justin O'Grady |
| EPI_ISL_499595, EPI_ISL_499596, EPI_ISL_499597, EPI_ISL_499598 | Northumbria University / South Tees Hospitals NHS Foundation Trust / North Cumbria Integrated Care NHS Foundation Trust / North Tees and Hartlepool NHS Foundation Trust / Newcastle Hospitals NHS Foundation Trust | COVID-19 Genomics UK (COG-UK) Consortium | Darren L Smith,Andrew Nelson,Matthew Bashton,Greg R Young,Joshua Loh,John Allan,Mohammad A Tariq,Giles S Holt,Gary Black,Wen C Yew,Lynn Dover,Paul Baker,Steve Liggett,Sarah Essex,Jane Greenaway,Debra Padgett,Clive Graham,Garren Scott,Edward Barton,Emma Swindells,Brendan Payne,Jennifer Collins,Yusri Taha,Gary Eltringham |
| EPI_ISL_499599, EPI_ISL_499601 | Quadram Institute Bioscience | COVID-19 Genomics UK (COG-UK) Consortium | Dave J. Baker, Gemma L. Kay, Alp Aydin, Thanh Le-Viet, Steven Rudder, Ana P. Tedim, Anastasia Kolyva, Maria Diaz, Leonardo de Oliveira Martins, Nabil-Fareed Alikhan, Lizzie Meadows, Rachael Stanley, Ngozi Elumogo, Muhammed Yasir, Nicholas M. Thomson, Alexander J Trotter, Rachel Gilroy, Samuel Bloomfield, Claire Stuart, Andrew Bell, Reenesh Prakash, Samir Dervisevic, Alison E. Mather, John Wain, Mark Webber, Andrew J. Page, Justin O'Grady |
| EPI_ISL_499602 | Northumbria University / South Tees Hospitals NHS Foundation Trust / North Cumbria Integrated Care NHS Foundation Trust / North Tees and Hartlepool NHS Foundation Trust / Newcastle Hospitals NHS Foundation Trust | COVID-19 Genomics UK (COG-UK) Consortium | Darren L Smith,Andrew Nelson,Matthew Bashton,Greg R Young,Joshua Loh,John Allan,Mohammad A Tariq,Giles S Holt,Gary Black,Wen C Yew,Lynn Dover,Paul Baker,Steve Liggett,Sarah Essex,Jane Greenaway,Debra Padgett,Clive Graham,Garren Scott,Edward Barton,Emma Swindells,Brendan Payne,Jennifer Collins,Yusri Taha,Gary Eltringham |
| EPI_ISL_499603 | Quadram Institute Bioscience | COVID-19 Genomics UK (COG-UK) Consortium | Dave J. Baker, Gemma L. Kay, Alp Aydin, Thanh Le-Viet, Steven Rudder, Ana P. Tedim, Anastasia Kolyva, Maria Diaz, Leonardo de Oliveira Martins, Nabil-Fareed Alikhan, Lizzie Meadows, Rachael Stanley, Ngozi Elumogo, Muhammed Yasir, Nicholas M. Thomson, Alexander J Trotter, Rachel Gilroy, Samuel Bloomfield, Claire Stuart, Andrew Bell, Reenesh Prakash, Samir Dervisevic, Alison E. Mather, John Wain, Mark Webber, Andrew J. Page, Justin O'Grady |
| EPI_ISL_499606, EPI_ISL_499609, EPI_ISL_499610 | Northumbria University / South Tees Hospitals NHS Foundation Trust / North Cumbria Integrated Care NHS | COVID-19 Genomics UK (COG-UK) Consortium | Darren L Smith,Andrew Nelson,Matthew Bashton,Greg R Young,Joshua Loh,John Allan,Mohammad A Tariq,Giles S Holt,Gary Black,Wen C Yew,Lynn Dover,Paul Baker,Steve Liggett,Sarah Essex,Jane Greenaway,Debra Padgett,Clive Graham,Garren Scott,Edward Barton,Emma Swindells,Brendan |

| | | | |
|---|---|---|---|
| | Foundation Trust / North Tees and Hartlepool NHS Foundation Trust / Newcastle Hospitals NHS Foundation Trust | | Payne,Jennifer Collins,Yusri Taha,Gary Eltringham |
| EPI_ISL_499623 | Quadram Institute Bioscience | COVID-19 Genomics UK (COG-UK) Consortium | Dave J. Baker, Gemma L. Kay, Alp Aydin, Thanh Le-Viet, Steven Rudder, Ana P. Tedim, Anastasia Kolyva, Maria Diaz, Leonardo de Oliveira Martins, Nabil-Fareed Alikhan, Lizzie Meadows, Rachael Stanley, Ngozi Elumogo, Muhammed Yasir, Nicholas M. Thomson, Alexander J Trotter, Rachel Gilroy, Samuel Bloomfield, Claire Stuart, Andrew Bell, Reenesh Prakash, Samir Dervisevic, Alison E. Mather, John Wain, Mark Webber, Andrew J. Page, Justin O'Grady |
| EPI_ISL_499626 | Northumbria University / South Tees Hospitals NHS Foundation Trust / North Cumbria Integrated Care NHS Foundation Trust / North Tees and Hartlepool NHS Foundation Trust / Newcastle Hospitals NHS Foundation Trust | COVID-19 Genomics UK (COG-UK) Consortium | Darren L Smith,Andrew Nelson,Matthew Bashton,Greg R Young,Joshua Loh,John Allan,Mohammad A Tariq,Giles S Holt,Gary Black,Wen C Yew,Lynn Dover,Paul Baker,Steve Liggett,Sarah Essex,Jane Greenaway,Debra Padgett,Clive Graham,Garren Scott,Edward Barton,Emma Swindells,Brendan Payne,Jennifer Collins,Yusri Taha,Gary Eltringham |
| EPI_ISL_499631 | Quadram Institute Bioscience | COVID-19 Genomics UK (COG-UK) Consortium | Dave J. Baker, Gemma L. Kay, Alp Aydin, Thanh Le-Viet, Steven Rudder, Ana P. Tedim, Anastasia Kolyva, Maria Diaz, Leonardo de Oliveira Martins, Nabil-Fareed Alikhan, Lizzie Meadows, Rachael Stanley, Ngozi Elumogo, Muhammed Yasir, Nicholas M. Thomson, Alexander J Trotter, Rachel Gilroy, Samuel Bloomfield, Claire Stuart, Andrew Bell, Reenesh Prakash, Samir Dervisevic, Alison E. Mather, John Wain, Mark Webber, Andrew J. Page, Justin O'Grady |
| EPI_ISL_499777, EPI_ISL_499778, EPI_ISL_499779, EPI_ISL_499780, EPI_ISL_499781, EPI_ISL_499782, EPI_ISL_499783, EPI_ISL_499784, EPI_ISL_499785, EPI_ISL_499786, EPI_ISL_499787, EPI_ISL_499788, EPI_ISL_499789, EPI_ISL_499790, EPI_ISL_499791, EPI_ISL_499792, EPI_ISL_499793, EPI_ISL_499794, EPI_ISL_499795, EPI_ISL_499796, EPI_ISL_499797, EPI_ISL_499798, EPI_ISL_499799, EPI_ISL_499800, EPI_ISL_499801, EPI_ISL_499802, EPI_ISL_499803, EPI_ISL_499804, EPI_ISL_499805, EPI_ISL_499806, EPI_ISL_499807, EPI_ISL_507110, EPI_ISL_507111, EPI_ISL_507112, EPI_ISL_507113, EPI_ISL_507114, EPI_ISL_507115, EPI_ISL_507116, EPI_ISL_507117, EPI_ISL_507118, EPI_ISL_507119, EPI_ISL_507120, EPI_ISL_507121, EPI_ISL_507122, EPI_ISL_507123, EPI_ISL_507124, EPI_ISL_507125, EPI_ISL_507126, EPI_ISL_507127, EPI_ISL_507128, EPI_ISL_507129 | | | |
| see above | Northumbria University / South Tees Hospitals NHS Foundation Trust / North Cumbria Integrated Care NHS Foundation Trust / North Tees and Hartlepool NHS Foundation Trust / Newcastle Hospitals NHS Foundation Trust | COVID-19 Genomics UK (COG-UK) Consortium | Darren L Smith,Andrew Nelson,Matthew Bashton,Greg R Young,Joshua Loh,John Allan,Mohammad A Tariq,Giles S Holt,Gary Black,Wen C Yew,Lynn Dover,Paul Baker,Steve Liggett,Sarah Essex,Jane Greenaway,Debra Padgett,Clive Graham,Garren Scott,Edward Barton,Emma Swindells,Brendan Payne,Jennifer Collins,Yusri Taha,Gary Eltringham |
| EPI_ISL_507130 | Quadram Institute Bioscience | COVID-19 Genomics UK (COG-UK) Consortium | Dave J. Baker, Gemma L. Kay, Alp Aydin, Thanh Le-Viet, Steven Rudder, Ana P. Tedim, Anastasia Kolyva, Maria Diaz, Leonardo de Oliveira Martins, Nabil-Fareed Alikhan, Lizzie Meadows, Rachael Stanley, Ngozi Elumogo, Muhammed Yasir, Nicholas M. Thomson, Alexander J Trotter, Rachel Gilroy, Samuel Bloomfield, Claire Stuart, Andrew Bell, Reenesh Prakash, Samir Dervisevic, Alison E. Mather, John Wain, Mark Webber, Andrew J. Page, Justin O'Grady |
| EPI_ISL_507984, EPI_ISL_507985, EPI_ISL_507986, EPI_ISL_507987, EPI_ISL_507988, EPI_ISL_507989, EPI_ISL_507990, EPI_ISL_507991, EPI_ISL_507992, EPI_ISL_507993, EPI_ISL_507994, EPI_ISL_507995, EPI_ISL_507996, EPI_ISL_507997, EPI_ISL_507998, EPI_ISL_507999, EPI_ISL_508000, EPI_ISL_508001, EPI_ISL_508002, EPI_ISL_508003, EPI_ISL_508004, EPI_ISL_508005, EPI_ISL_508006, EPI_ISL_508007, EPI_ISL_508008, EPI_ISL_508009, EPI_ISL_508010, EPI_ISL_508011, EPI_ISL_508012, EPI_ISL_508013, EPI_ISL_508014, EPI_ISL_508015, EPI_ISL_508016, EPI_ISL_508017, EPI_ISL_508018, EPI_ISL_508019, EPI_ISL_508020, EPI_ISL_508021, EPI_ISL_508022, EPI_ISL_508023, EPI_ISL_508024, EPI_ISL_508025, EPI_ISL_508026, EPI_ISL_508027, EPI_ISL_508028, EPI_ISL_508029, EPI_ISL_508030, EPI_ISL_508031, EPI_ISL_508032, EPI_ISL_508033, EPI_ISL_508034, EPI_ISL_508035, EPI_ISL_508036, EPI_ISL_508037, EPI_ISL_508038, EPI_ISL_508039, EPI_ISL_508040, EPI_ISL_508041, EPI_ISL_508042, EPI_ISL_508043, EPI_ISL_508044, EPI_ISL_508045, EPI_ISL_508046, EPI_ISL_508047, EPI_ISL_508048, EPI_ISL_508049, EPI_ISL_508050, EPI_ISL_508051, EPI_ISL_508052, EPI_ISL_508053, EPI_ISL_508054, EPI_ISL_508055, EPI_ISL_508056, EPI_ISL_508057, EPI_ISL_508058, EPI_ISL_508059, EPI_ISL_508060, EPI_ISL_508061, EPI_ISL_508062, EPI_ISL_508063, EPI_ISL_508064, EPI_ISL_508065, EPI_ISL_508066, EPI_ISL_508067, EPI_ISL_508068, EPI_ISL_508069, EPI_ISL_508070, EPI_ISL_508071, EPI_ISL_508072, EPI_ISL_508073, EPI_ISL_508074, EPI_ISL_508075, EPI_ISL_508076, EPI_ISL_508077, EPI_ISL_508078, EPI_ISL_508079, EPI_ISL_508080, EPI_ISL_508081, EPI_ISL_508082, EPI_ISL_508083, EPI_ISL_508085, EPI_ISL_508086, EPI_ISL_508087, EPI_ISL_508088, EPI_ISL_508089, EPI_ISL_508090, EPI_ISL_508091, EPI_ISL_508092, EPI_ISL_508093, EPI_ISL_508094, EPI_ISL_508095, EPI_ISL_508096, EPI_ISL_508097, EPI_ISL_508098, EPI_ISL_508099, EPI_ISL_508100, EPI_ISL_508101, EPI_ISL_508102, EPI_ISL_508103, EPI_ISL_508104, EPI_ISL_508105, EPI_ISL_508106, EPI_ISL_508107, EPI_ISL_508108, EPI_ISL_508109, EPI_ISL_508110, EPI_ISL_508111, EPI_ISL_508112, EPI_ISL_508113, EPI_ISL_508114, EPI_ISL_508115, EPI_ISL_508116, EPI_ISL_508117, EPI_ISL_508118, EPI_ISL_508119, EPI_ISL_508120, EPI_ISL_508121 | | | |
| see above | New Mexico Department of Health Scientific Laboratory Division | Center for Global Health, University of New Mexico Health Sciences Center | Daryl Domman, Kurt Schwalm, Twila Kunde, Joseph Hicks, Michael Edwards, Darrell Dinwiddie |
| EPI_ISL_512348, EPI_ISL_512349, EPI_ISL_512350, EPI_ISL_512351, EPI_ISL_512352, EPI_ISL_512353, EPI_ISL_512354, EPI_ISL_512355, EPI_ISL_512356, EPI_ISL_512357, EPI_ISL_512358, EPI_ISL_512359, EPI_ISL_512360, EPI_ISL_512361, EPI_ISL_512362, EPI_ISL_512363, EPI_ISL_512364, EPI_ISL_512365 | | | |
| see above | Northumbria University / South Tees Hospitals NHS Foundation Trust / North Cumbria Integrated Care NHS Foundation Trust / North Tees and Hartlepool NHS Foundation Trust / Newcastle Hospitals NHS Foundation Trust | COVID-19 Genomics UK (COG-UK) Consortium | Darren L Smith,Andrew Nelson,Matthew Bashton,Greg R Young,Joshua Loh,John Allan,Mohammad A Tariq,Giles S Holt,Gary Black,Wen C Yew,Lynn Dover,Paul Baker,Steve Liggett,Sarah Essex,Jane Greenaway,Debra Padgett,Clive Graham,Garren Scott,Edward Barton,Emma Swindells,Brendan Payne,Jennifer Collins,Yusri Taha,Gary Eltringham |
| EPI_ISL_512366, EPI_ISL_512367, EPI_ISL_512368, EPI_ISL_512369, EPI_ISL_512370, EPI_ISL_512371, EPI_ISL_512373, EPI_ISL_512374, EPI_ISL_512375, EPI_ISL_512376, EPI_ISL_512377, EPI_ISL_512378 | | | |
| see above | Quadram Institute Bioscience | COVID-19 Genomics UK (COG-UK) Consortium | Dave J. Baker, Gemma L. Kay, Alp Aydin, Thanh Le-Viet, Steven Rudder, Ana P. Tedim, Anastasia Kolyva, Maria Diaz, Leonardo de Oliveira Martins, Nabil-Fareed Alikhan, Lizzie Meadows, Rachael Stanley, Ngozi Elumogo, Muhammed Yasir, Nicholas M. Thomson, Alexander J Trotter, Rachel Gilroy, Samuel Bloomfield, Claire Stuart, Andrew Bell, Reenesh Prakash, Samir Dervisevic, Alison E. Mather, John Wain, Mark Webber, Andrew J. Page, Justin O'Grady |
| EPI_ISL_522547, EPI_ISL_522548 | Platforme CYROI | UMR PIMIT Université de La Réunion | David Wilkinson, Camille Lebarbenchon, Patrick Mavingui |
| EPI_ISL_522549, EPI_ISL_522550 | Félix Guyon Hospital | UMR PIMIT Université de La Réunion | David Wilkinson, Camille Lebarbenchon, Patrick Mavingui |
| EPI_ISL_525699 | Seattle Flu Study | Seattle Flu Study | Deborah A. Nickerson, Chris D. Frazar, Jover Lee, Benjamin Pelle, Matthew Richardson, Amanda Adler, Elisabeth Brandstetter, Peter D. Han, Kairsten Fay, Misja Ilcisin, Kirsten Lacombe, Thomas R. Sibley, Melissa Truong, Caitlin R. Wolf, Karen Cowgill, Stephanie Schrag, Jeff Duchin, Michael Boeckh, Janet A. Englund, Michael Famulare, Barry R. Lutz, Mark J. Rieder, Lea M. Starita, Matthew Thompson, Helen Y. Chu, Trevor Bedford, Jay Shendure |
| EPI_ISL_525700, EPI_ISL_525701 | Seattle Flu Study | Seattle Flu Study | Deborah A. Nickerson, Chris D. Frazar, Jover Lee, Benjamin Pelle, Matthew Richardson, Amanda Adler, Elisabeth Brandstetter, Peter D. Han, Kairsten Fay, Misja Ilcisin, Kirsten Lacombe, Thomas R. Sibley, Melissa Truong, Caitlin R. Wolf, Michael Boeckh, Janet A. Englund, Michael Famulare, Barry R. Lutz, Mark J. Rieder, Lea M. Starita, Matthew Thompson, Jay Shendure, Trevor Bedford, Helen Y. Chu |
| EPI_ISL_525702, EPI_ISL_525703, EPI_ISL_525704, EPI_ISL_525705 | Seattle Flu Study | Seattle Flu Study | Deborah A. Nickerson, Chris D. Frazar, Jover Lee, Benjamin Pelle, Matthew Richardson, Amanda Adler, Elisabeth Brandstetter, Peter D. Han, Kairsten Fay, Misja Ilcisin, Kirsten Lacombe, Thomas R. Sibley, Melissa Truong, Caitlin R. Wolf, Karen Cowgill, Stephanie Schrag, Jeff Duchin, Michael Boeckh, Janet A. Englund, Michael Famulare, Barry R. Lutz, Mark J. Rieder, Lea M. Starita, Matthew Thompson, Helen Y. Chu, Trevor Bedford, Jay Shendure |
| EPI_ISL_525706 | Seattle Flu Study | Seattle Flu Study | Deborah A. Nickerson, Chris D. Frazar, Jover Lee, Benjamin Pelle, Matthew Richardson, Amanda Adler, Elisabeth Brandstetter, Peter D. Han, Kairsten Fay, Misja Ilcisin, Kirsten Lacombe, Thomas R. Sibley, Melissa Truong, Caitlin R. Wolf, Michael Boeckh, Janet A. Englund, Michael Famulare, Barry R. Lutz, Mark J. Rieder, Lea M. Starita, Matthew Thompson, Jay Shendure, Trevor Bedford, Helen Y. Chu |
| EPI_ISL_525707, EPI_ISL_525708, EPI_ISL_525709, EPI_ISL_525710, EPI_ISL_525711, EPI_ISL_525712, EPI_ISL_525713, EPI_ISL_525714, EPI_ISL_525715, EPI_ISL_525716, EPI_ISL_525717, EPI_ISL_525718, EPI_ISL_525719, EPI_ISL_525720, EPI_ISL_525721, EPI_ISL_525722, EPI_ISL_525723, EPI_ISL_525724, EPI_ISL_525725, EPI_ISL_525726, EPI_ISL_525727 | | | |
| see above | Seattle Flu Study | Seattle Flu Study | Deborah A. Nickerson, Chris D. Frazar, Jover Lee, Benjamin Pelle, Matthew Richardson, Amanda Adler, Elisabeth Brandstetter, Peter D. Han, Kairsten Fay, Misja Ilcisin, Kirsten Lacombe, Thomas R. Sibley, Melissa Truong, Caitlin R. Wolf, Karen Cowgill, Stephanie Schrag, Jeff Duchin, Michael Boeckh, Janet A. Englund, Michael Famulare, Barry R. Lutz, Mark J. Rieder, Lea M. Starita, Matthew Thompson, Helen Y. Chu, Trevor Bedford, Jay Shendure |
| EPI_ISL_525728 | Seattle Flu Study | Seattle Flu Study | Deborah A. Nickerson, Chris D. Frazar, Jover Lee, Benjamin Pelle, Matthew Richardson, Amanda Adler, Elisabeth Brandstetter, Peter D. Han, Kairsten Fay, Misja Ilcisin, Kirsten Lacombe, Thomas R. Sibley, Melissa Truong, Caitlin R. Wolf, Michael Boeckh, Janet A. Englund, Michael Famulare, Barry R. Lutz, Mark J. Rieder, Lea M. Starita, Matthew Thompson, Jay Shendure, Trevor Bedford, Helen Y. Chu |
| EPI_ISL_525729, EPI_ISL_525730, EPI_ISL_525731, EPI_ISL_525732, EPI_ISL_525733, EPI_ISL_525734, EPI_ISL_525735, EPI_ISL_525736, EPI_ISL_525737, EPI_ISL_525738, EPI_ISL_525739, EPI_ISL_525740, EPI_ISL_525741 | | | |
| see above | Seattle Flu Study | Seattle Flu Study | Deborah A. Nickerson, Chris D. Frazar, Jover Lee, Benjamin Pelle, Matthew Richardson, Amanda Adler, Elisabeth Brandstetter, Peter D. Han, Kairsten Fay, Misja Ilcisin, Kirsten Lacombe, Thomas R. Sibley, Melissa Truong, Caitlin R. Wolf, Karen Cowgill, Stephanie Schrag, Jeff Duchin, Michael Boeckh, Janet A. Englund, Michael Famulare, Barry R. Lutz, Mark J. Rieder, Lea M. Starita, Matthew Thompson, Helen Y. Chu, Trevor Bedford, Jay Shendure |
| EPI_ISL_525742 | Seattle Flu Study | Seattle Flu Study | Deborah A. Nickerson, Chris D. Frazar, Jover Lee, Benjamin Pelle, Matthew Richardson, Amanda Adler, Elisabeth Brandstetter, Peter D. Han, Kairsten Fay, Misja Ilcisin, Kirsten Lacombe, Thomas R. Sibley, Melissa Truong, Caitlin R. Wolf, Michael Boeckh, Janet A. Englund, Michael Famulare, Barry R. Lutz, Mark J. Rieder, Lea M. Starita, Matthew Thompson, Jay Shendure, Trevor Bedford, Helen Y. Chu |
| EPI_ISL_525743, EPI_ISL_525744, EPI_ISL_525745, EPI_ISL_525746, EPI_ISL_525747, EPI_ISL_525748, EPI_ISL_525749, EPI_ISL_525750, EPI_ISL_525751, EPI_ISL_525752, EPI_ISL_525753, EPI_ISL_525754, EPI_ISL_525755, EPI_ISL_525756 | | | |

| | | | |
|---|---|---|---|
| see above | Seattle Flu Study | Seattle Flu Study | Deborah A. Nickerson, Chris D. Frazar, Jover Lee, Benjamin Pelle, Matthew Richardson, Amanda Adler, Elisabeth Brandstetter, Peter D. Han, Kairsten Fay, Misja Ilcisin, Kirsten Lacombe, Thomas R. Sibley, Melissa Truong, Caitlin R. Wolf, Karen Cowgill, Stephanie Schrag, Jeff Duchin, Michael Boeckh, Janet A. Englund, Michael Famulare, Barry R. Lutz, Mark J. Rieder, Lea M. Starita, Matthew Thompson, Helen Y. Chu, Trevor Bedford, Jay Shendure |
| EPI_ISL_529267, EPI_ISL_529281, EPI_ISL_529282, EPI_ISL_529378, EPI_ISL_529379, EPI_ISL_529380, EPI_ISL_529381, EPI_ISL_529389, EPI_ISL_529437, EPI_ISL_529451, EPI_ISL_529452, EPI_ISL_529458, EPI_ISL_529460, EPI_ISL_529489, EPI_ISL_529523, EPI_ISL_529557, EPI_ISL_529558, EPI_ISL_529559, EPI_ISL_529560, EPI_ISL_529561, EPI_ISL_529562, EPI_ISL_529563, EPI_ISL_529564, EPI_ISL_529565, EPI_ISL_529566, EPI_ISL_529567, EPI_ISL_529568, EPI_ISL_529569, EPI_ISL_529570, EPI_ISL_529571, EPI_ISL_529572, EPI_ISL_529573, EPI_ISL_529574, EPI_ISL_529575, EPI_ISL_529576, EPI_ISL_529577, EPI_ISL_529578, EPI_ISL_529579, EPI_ISL_529580, EPI_ISL_529581, EPI_ISL_529582, EPI_ISL_529583, EPI_ISL_529584, EPI_ISL_529585, EPI_ISL_529586, EPI_ISL_529587, EPI_ISL_529588, EPI_ISL_529589 | | | |
| see above | Quadram Institute Bioscience | COVID-19 Genomics UK (COG-UK) Consortium | Dave J. Baker, Gemma L. Kay, Alp Aydin, Thanh Le-Viet, Steven Rudder, Ana P. Tedim, Anastasia Kolyva, Maria Diaz, Leonardo de Oliveira Martins, Nabil-Fareed Alikhan, Lizzie Meadows, Rachael Stanley, Ngozi Elumogo, Muhammed Yasir, Nicholas M. Thomson, Alexander J Trotter, Rachel Gilroy, Samuel Bloomfield, Claire Stuart, Andrew Bell, Reenesh Prakash, Samir Dervisevic, Alison E. Mather, John Wain, Mark Webber, Andrew J. Page, Justin O'Grady |
| EPI_ISL_530124, EPI_ISL_530125 | Seattle Flu Study | Seattle Flu Study | Deborah A. Nickerson, Chris D. Frazar, Jover Lee, Benjamin Pelle, Matthew Richardson, Amanda Adler, Elisabeth Brandstetter, Peter D. Han, Kairsten Fay, Misja Ilcisin, Kirsten Lacombe, Thomas R. Sibley, Melissa Truong, Caitlin R. Wolf, Michael Boeckh, Janet A. Englund, Michael Famulare, Barry R. Lutz, Mark J. Rieder, Lea M. Starita, Matthew Thompson, Jay Shendure, Trevor Bedford, Helen Y. Chu |
| EPI_ISL_530126, EPI_ISL_530127, EPI_ISL_530128, EPI_ISL_530129, EPI_ISL_530130, EPI_ISL_530131 | Seattle Flu Study | Seattle Flu Study | Deborah A. Nickerson, Chris D. Frazar, Jover Lee, Benjamin Pelle, Matthew Richardson, Amanda Adler, Elisabeth Brandstetter, Peter D. Han, Kairsten Fay, Misja Ilcisin, Kirsten Lacombe, Thomas R. Sibley, Melissa Truong, Caitlin R. Wolf, Karen Cowgill, Stephanie Schrag, Jeff Duchin, Michael Boeckh, Janet A. Englund, Michael Famulare, Barry R. Lutz, Mark J. Rieder, Lea M. Starita, Matthew Thompson, Helen Y. Chu, Trevor Bedford, Jay Shendure |
| EPI_ISL_530132 | Seattle Flu Study | Seattle Flu Study | Deborah A. Nickerson, Chris D. Frazar, Jover Lee, Benjamin Pelle, Matthew Richardson, Amanda Adler, Elisabeth Brandstetter, Peter D. Han, Kairsten Fay, Misja Ilcisin, Kirsten Lacombe, Thomas R. Sibley, Melissa Truong, Caitlin R. Wolf, Michael Boeckh, Janet A. Englund, Michael Famulare, Barry R. Lutz, Mark J. Rieder, Lea M. Starita, Matthew Thompson, Jay Shendure, Trevor Bedford, Helen Y. Chu |
| EPI_ISL_530133, EPI_ISL_530134, EPI_ISL_530135, EPI_ISL_530136, EPI_ISL_530137, EPI_ISL_530138, EPI_ISL_530139, EPI_ISL_530140, EPI_ISL_530141, EPI_ISL_530142, EPI_ISL_530143, EPI_ISL_530144, EPI_ISL_530145, EPI_ISL_530146, EPI_ISL_530147, EPI_ISL_530148, EPI_ISL_530149, EPI_ISL_530150, EPI_ISL_530151, EPI_ISL_530152, EPI_ISL_530153, EPI_ISL_530154, EPI_ISL_530155, EPI_ISL_530156, EPI_ISL_530157, EPI_ISL_530158, EPI_ISL_530159, EPI_ISL_530160, EPI_ISL_530161, EPI_ISL_530162 | | | |
| see above | Seattle Flu Study | Seattle Flu Study | Deborah A. Nickerson, Chris D. Frazar, Jover Lee, Benjamin Pelle, Matthew Richardson, Amanda Adler, Elisabeth Brandstetter, Peter D. Han, Kairsten Fay, Misja Ilcisin, Kirsten Lacombe, Thomas R. Sibley, Melissa Truong, Caitlin R. Wolf, Karen Cowgill, Stephanie Schrag, Jeff Duchin, Michael Boeckh, Janet A. Englund, Michael Famulare, Barry R. Lutz, Mark J. Rieder, Lea M. Starita, Matthew Thompson, Helen Y. Chu, Trevor Bedford, Jay Shendure |
| EPI_ISL_538233, EPI_ISL_538234, EPI_ISL_538235, EPI_ISL_538236, EPI_ISL_538237, EPI_ISL_538238, EPI_ISL_538239, EPI_ISL_538240, EPI_ISL_538241, EPI_ISL_538242, EPI_ISL_538243, EPI_ISL_538244, EPI_ISL_538245, EPI_ISL_538246, EPI_ISL_538247, EPI_ISL_538248, EPI_ISL_538249, EPI_ISL_538250, EPI_ISL_538251, EPI_ISL_538252, EPI_ISL_538253, EPI_ISL_538254, EPI_ISL_538255, EPI_ISL_538256, EPI_ISL_538257, EPI_ISL_538258, EPI_ISL_538259, EPI_ISL_538260, EPI_ISL_538261, EPI_ISL_538262, EPI_ISL_538263, EPI_ISL_538264 | | | |
| see above | TriCore Reference Laboratories | Center for Global Health, University of New Mexico Health Sciences Center | Daryl Domman, Kurt Schwalm, Twila Kunde, Joseph Hicks, Michael Edwards, Darrell Dinwiddie |
| EPI_ISL_540691, EPI_ISL_540692, EPI_ISL_540693, EPI_ISL_540694, EPI_ISL_540695, EPI_ISL_540696, EPI_ISL_540697, EPI_ISL_540698 | Quadram Institute Bioscience | COVID-19 Genomics UK (COG-UK) Consortium | Dave J. Baker, Gemma L. Kay, Alp Aydin, Thanh Le-Viet, Steven Rudder, Ana P. Tedim, Anastasia Kolyva, Maria Diaz, Leonardo de Oliveira Martins, Nabil-Fareed Alikhan, Lizzie Meadows, Rachael Stanley, Ngozi Elumogo, Muhammed Yasir, Nicholas M. Thomson, Alexander J Trotter, Rachel Gilroy, Samuel Bloomfield, Claire Stuart, Andrew Bell, Reenesh Prakash, Samir Dervisevic, Alison E. Mather, John Wain, Mark Webber, Andrew J. Page, Justin O'Grady |
| EPI_ISL_542935, EPI_ISL_542936, EPI_ISL_542937, EPI_ISL_542938, EPI_ISL_542939, EPI_ISL_542940, EPI_ISL_542941, EPI_ISL_542942, EPI_ISL_542943, EPI_ISL_542944, EPI_ISL_542945, EPI_ISL_542946, EPI_ISL_542947, EPI_ISL_542948, EPI_ISL_542949, EPI_ISL_542950, EPI_ISL_542951, EPI_ISL_542952, EPI_ISL_542953, EPI_ISL_542954, EPI_ISL_542955, EPI_ISL_542956, EPI_ISL_542957, EPI_ISL_542958, EPI_ISL_542959, EPI_ISL_542960, EPI_ISL_542961, EPI_ISL_542962, EPI_ISL_542963, EPI_ISL_542964, EPI_ISL_542965, EPI_ISL_542966, EPI_ISL_542967, EPI_ISL_542968, EPI_ISL_542969, EPI_ISL_542970, EPI_ISL_542971, EPI_ISL_542972, EPI_ISL_542973, EPI_ISL_542974, EPI_ISL_542975, EPI_ISL_542976, EPI_ISL_542977, EPI_ISL_542978, EPI_ISL_542979, EPI_ISL_542980, EPI_ISL_542981, EPI_ISL_542982, EPI_ISL_542983, EPI_ISL_542984, EPI_ISL_542985, EPI_ISL_542986, EPI_ISL_542987, EPI_ISL_542988, EPI_ISL_542989, EPI_ISL_542990, EPI_ISL_542991, EPI_ISL_542992, EPI_ISL_542993, EPI_ISL_542994, EPI_ISL_542995, EPI_ISL_542996, EPI_ISL_542997, EPI_ISL_542998, EPI_ISL_542999, EPI_ISL_543000, EPI_ISL_543001, EPI_ISL_543002, EPI_ISL_543003, EPI_ISL_543004, EPI_ISL_543005, EPI_ISL_543006 | | | |
| see above | TriCore Reference Laboratories | Center for Global Health, University of New Mexico Health Sciences Center | Daryl Domman, Kurt Schwalm, Twila Kunde, Joseph Hicks, Michael Edwards, Darrell Dinwiddie |
| EPI_ISL_549344, EPI_ISL_549350, EPI_ISL_549351, EPI_ISL_549355, EPI_ISL_549367, EPI_ISL_549382, EPI_ISL_549479, EPI_ISL_549480, EPI_ISL_549481, EPI_ISL_549488 | Quadram Institute Bioscience | COVID-19 Genomics UK (COG-UK) Consortium | Dave J. Baker, Gemma L. Kay, Alp Aydin, Thanh Le-Viet, Steven Rudder, Ana P. Tedim, Anastasia Kolyva, Maria Diaz, Leonardo de Oliveira Martins, Nabil-Fareed Alikhan, Lizzie Meadows, Rachael Stanley, Ngozi Elumogo, Muhammed Yasir, Nicholas M. Thomson, Alexander J Trotter, Rachel Gilroy, Samuel Bloomfield, Claire Stuart, Andrew Bell, Reenesh Prakash, Samir Dervisevic, Alison E. Mather, John Wain, Mark Webber, Andrew J. Page, Justin O'Grady |
| EPI_ISL_560325, EPI_ISL_560326, EPI_ISL_560327, EPI_ISL_560328, EPI_ISL_560329, EPI_ISL_560330, EPI_ISL_560331, EPI_ISL_560332, EPI_ISL_560333, EPI_ISL_560334, EPI_ISL_560335, EPI_ISL_560336, EPI_ISL_560337, EPI_ISL_560338, EPI_ISL_560339, EPI_ISL_560340, EPI_ISL_560341, EPI_ISL_560342, EPI_ISL_560343, EPI_ISL_560344, EPI_ISL_560345, EPI_ISL_560346, EPI_ISL_560347, EPI_ISL_560348, EPI_ISL_560349, EPI_ISL_560350, EPI_ISL_560351, EPI_ISL_560352, EPI_ISL_560353, EPI_ISL_560354, EPI_ISL_560356, EPI_ISL_560357, EPI_ISL_560358, EPI_ISL_560359, EPI_ISL_560361, EPI_ISL_560362 | | | |
| see above | TriCore Reference Laboratories | Center for Global Health, University of New Mexico Health Sciences Center | Daryl Domman, Kurt Schwalm, Twila Kunde, Joseph Hicks, Michael Edwards, Darrell Dinwiddie |
| EPI_ISL_572399, EPI_ISL_572400, EPI_ISL_572401, EPI_ISL_572403, EPI_ISL_572406 | Quadram Institute Bioscience | COVID-19 Genomics UK (COG-UK) Consortium | Dave J. Baker, Gemma L. Kay, Alp Aydin, Thanh Le-Viet, Steven Rudder, Ana P. Tedim, Anastasia Kolyva, Maria Diaz, Leonardo de Oliveira Martins, Nabil-Fareed Alikhan, Lizzie Meadows, Rachael Stanley, Ngozi Elumogo, Muhammed Yasir, Nicholas M. Thomson, Alexander J Trotter, Rachel Gilroy, Samuel Bloomfield, Claire Stuart, Andrew Bell, Reenesh Prakash, Samir Dervisevic, Alison E. Mather, John Wain, Mark Webber, Andrew J. Page, Justin O'Grady |
| EPI_ISL_572409, EPI_ISL_572410, EPI_ISL_572412 | Northumbria University / South Tees Hospitals NHS Foundation Trust / North Cumbria Integrated Care NHS Foundation Trust / North Tees and Hartlepool NHS Foundation Trust / Newcastle Hospitals NHS Foundation Trust | COVID-19 Genomics UK (COG-UK) Consortium | Darren L Smith,Andrew Nelson,Matthew Bashton,Greg R Young,Joshua Loh,John Allan,Mohammad A Tariq,Giles S Holt,Gary Black,Wen C Yew,Lynn Dover,Paul Baker,Steve Liggett,Sarah Essex,Jane Greenaway,Debra Padgett,Clive Graham,Garren Scott,Edward Barton,Emma Swindells,Brendan Payne,Jennifer Collins,Yusri Taha,Gary Eltringham |
| EPI_ISL_572419, EPI_ISL_572423 | Quadram Institute Bioscience | COVID-19 Genomics UK (COG-UK) Consortium | Dave J. Baker, Gemma L. Kay, Alp Aydin, Thanh Le-Viet, Steven Rudder, Ana P. Tedim, Anastasia Kolyva, Maria Diaz, Leonardo de Oliveira Martins, Nabil-Fareed Alikhan, Lizzie Meadows, Rachael Stanley, Ngozi Elumogo, Muhammed Yasir, Nicholas M. Thomson, Alexander J Trotter, Rachel Gilroy, Samuel Bloomfield, Claire Stuart, Andrew Bell, Reenesh Prakash, Samir Dervisevic, Alison E. Mather, John Wain, Mark Webber, Andrew J. Page, Justin O'Grady |
| EPI_ISL_572424 | Northumbria University / South Tees Hospitals NHS Foundation Trust / North Cumbria Integrated Care NHS Foundation Trust / North Tees and Hartlepool NHS Foundation Trust / Newcastle Hospitals NHS Foundation Trust | COVID-19 Genomics UK (COG-UK) Consortium | Darren L Smith,Andrew Nelson,Matthew Bashton,Greg R Young,Joshua Loh,John Allan,Mohammad A Tariq,Giles S Holt,Gary Black,Wen C Yew,Lynn Dover,Paul Baker,Steve Liggett,Sarah Essex,Jane Greenaway,Debra Padgett,Clive Graham,Garren Scott,Edward Barton,Emma Swindells,Brendan Payne,Jennifer Collins,Yusri Taha,Gary Eltringham |
| EPI_ISL_572430, EPI_ISL_572437 | Quadram Institute Bioscience | COVID-19 Genomics UK (COG-UK) Consortium | Dave J. Baker, Gemma L. Kay, Alp Aydin, Thanh Le-Viet, Steven Rudder, Ana P. Tedim, Anastasia Kolyva, Maria Diaz, Leonardo de Oliveira Martins, Nabil-Fareed Alikhan, Lizzie Meadows, Rachael Stanley, Ngozi Elumogo, Muhammed Yasir, Nicholas M. Thomson, Alexander J Trotter, Rachel Gilroy, Samuel Bloomfield, Claire Stuart, Andrew Bell, Reenesh Prakash, Samir Dervisevic, Alison E. Mather, John Wain, Mark Webber, Andrew J. Page, Justin O'Grady |
| EPI_ISL_572438, EPI_ISL_572439 | Northumbria University / South Tees Hospitals NHS Foundation Trust / North Cumbria Integrated Care NHS Foundation Trust / North Tees and Hartlepool NHS Foundation Trust / Newcastle Hospitals NHS Foundation Trust | COVID-19 Genomics UK (COG-UK) Consortium | Darren L Smith,Andrew Nelson,Matthew Bashton,Greg R Young,Joshua Loh,John Allan,Mohammad A Tariq,Giles S Holt,Gary Black,Wen C Yew,Lynn Dover,Paul Baker,Steve Liggett,Sarah Essex,Jane Greenaway,Debra Padgett,Clive Graham,Garren Scott,Edward Barton,Emma Swindells,Brendan Payne,Jennifer Collins,Yusri Taha,Gary Eltringham |
| EPI_ISL_572440, EPI_ISL_572441, EPI_ISL_572447, EPI_ISL_572448, EPI_ISL_572449 | Quadram Institute Bioscience | COVID-19 Genomics UK (COG-UK) Consortium | Dave J. Baker, Gemma L. Kay, Alp Aydin, Thanh Le-Viet, Steven Rudder, Ana P. Tedim, Anastasia Kolyva, Maria Diaz, Leonardo de Oliveira Martins, Nabil-Fareed Alikhan, Lizzie Meadows, Rachael Stanley, Ngozi Elumogo, Muhammed Yasir, Nicholas M. Thomson, Alexander J Trotter, Rachel Gilroy, Samuel Bloomfield, Claire Stuart, Andrew Bell, Reenesh Prakash, Samir Dervisevic, Alison E. Mather, John Wain, Mark Webber, Andrew J. Page, Justin O'Grady |
| EPI_ISL_572450, EPI_ISL_572452 | Northumbria University / South Tees Hospitals NHS | COVID-19 Genomics UK (COG-UK) Consortium | Darren L Smith,Andrew Nelson,Matthew Bashton,Greg R Young,Joshua Loh,John Allan,Mohammad A Tariq,Giles S Holt,Gary Black,Wen C Yew,Lynn |

| | | | |
|---|---|---|---|
| | Foundation Trust / North Cumbria Integrated Care NHS Foundation Trust / North Tees and Hartlepool NHS Foundation Trust / Newcastle Hospitals NHS Foundation Trust | | Dover,Paul Baker,Steve Liggett,Sarah Essex,Jane Greenaway,Debra Padgett,Clive Graham,Garren Scott,Edward Barton,Emma Swindells,Brendan Payne,Jennifer Collins,Yusri Taha,Gary Eltringham |
| EPI_ISL_572453, EPI_ISL_572454, EPI_ISL_572456, EPI_ISL_572459, EPI_ISL_572462, EPI_ISL_572463 | Quadram Institute Bioscience | COVID-19 Genomics UK (COG-UK) Consortium | Dave J. Baker, Gemma L. Kay, Alp Aydin, Thanh Le-Viet, Steven Rudder, Ana P. Tedim, Anastasia Kolyva, Maria Diaz, Leonardo de Oliveira Martins, Nabil-Fareed Alikhan, Lizzie Meadows, Rachael Stanley, Ngozi Elumogo, Muhammed Yasir, Nicholas M. Thomson, Alexander J Trotter, Rachel Gilroy, Samuel Bloomfield, Claire Stuart, Andrew Bell, Reenesh Prakash, Samir Dervisevic, Alison E. Mather, John Wain, Mark Webber, Andrew J. Page, Justin O'Grady |
| EPI_ISL_572464, EPI_ISL_572480 | Northumbria University / South Tees Hospitals NHS Foundation Trust / North Cumbria Integrated Care NHS Foundation Trust / North Tees and Hartlepool NHS Foundation Trust / Newcastle Hospitals NHS Foundation Trust | COVID-19 Genomics UK (COG-UK) Consortium | Darren L Smith,Andrew Nelson,Matthew Bashton,Greg R Young,Joshua Loh,John Allan,Mohammad A Tariq,Giles S Holt,Gary Black,Wen C Yew,Lynn Dover,Paul Baker,Steve Liggett,Sarah Essex,Jane Greenaway,Debra Padgett,Clive Graham,Garren Scott,Edward Barton,Emma Swindells,Brendan Payne,Jennifer Collins,Yusri Taha,Gary Eltringham |
| EPI_ISL_572485, EPI_ISL_572486 | Quadram Institute Bioscience | COVID-19 Genomics UK (COG-UK) Consortium | Dave J. Baker, Gemma L. Kay, Alp Aydin, Thanh Le-Viet, Steven Rudder, Ana P. Tedim, Anastasia Kolyva, Maria Diaz, Leonardo de Oliveira Martins, Nabil-Fareed Alikhan, Lizzie Meadows, Rachael Stanley, Ngozi Elumogo, Muhammed Yasir, Nicholas M. Thomson, Alexander J Trotter, Rachel Gilroy, Samuel Bloomfield, Claire Stuart, Andrew Bell, Reenesh Prakash, Samir Dervisevic, Alison E. Mather, John Wain, Mark Webber, Andrew J. Page, Justin O'Grady |
| EPI_ISL_572488, EPI_ISL_572489, EPI_ISL_572491 | Northumbria University / South Tees Hospitals NHS Foundation Trust / North Cumbria Integrated Care NHS Foundation Trust / North Tees and Hartlepool NHS Foundation Trust / Newcastle Hospitals NHS Foundation Trust | COVID-19 Genomics UK (COG-UK) Consortium | Darren L Smith,Andrew Nelson,Matthew Bashton,Greg R Young,Joshua Loh,John Allan,Mohammad A Tariq,Giles S Holt,Gary Black,Wen C Yew,Lynn Dover,Paul Baker,Steve Liggett,Sarah Essex,Jane Greenaway,Debra Padgett,Clive Graham,Garren Scott,Edward Barton,Emma Swindells,Brendan Payne,Jennifer Collins,Yusri Taha,Gary Eltringham |
| EPI_ISL_572492, EPI_ISL_572493, EPI_ISL_572495, EPI_ISL_572497, EPI_ISL_572498, EPI_ISL_572499, EPI_ISL_572503, EPI_ISL_572504, EPI_ISL_572505 | Quadram Institute Bioscience | COVID-19 Genomics UK (COG-UK) Consortium | Dave J. Baker, Gemma L. Kay, Alp Aydin, Thanh Le-Viet, Steven Rudder, Ana P. Tedim, Anastasia Kolyva, Maria Diaz, Leonardo de Oliveira Martins, Nabil-Fareed Alikhan, Lizzie Meadows, Rachael Stanley, Ngozi Elumogo, Muhammed Yasir, Nicholas M. Thomson, Alexander J Trotter, Rachel Gilroy, Samuel Bloomfield, Claire Stuart, Andrew Bell, Reenesh Prakash, Samir Dervisevic, Alison E. Mather, John Wain, Mark Webber, Andrew J. Page, Justin O'Grady |
| EPI_ISL_572513, EPI_ISL_572515, EPI_ISL_572517 | Northumbria University / South Tees Hospitals NHS Foundation Trust / North Cumbria Integrated Care NHS Foundation Trust / North Tees and Hartlepool NHS Foundation Trust / Newcastle Hospitals NHS Foundation Trust | COVID-19 Genomics UK (COG-UK) Consortium | Darren L Smith,Andrew Nelson,Matthew Bashton,Greg R Young,Joshua Loh,John Allan,Mohammad A Tariq,Giles S Holt,Gary Black,Wen C Yew,Lynn Dover,Paul Baker,Steve Liggett,Sarah Essex,Jane Greenaway,Debra Padgett,Clive Graham,Garren Scott,Edward Barton,Emma Swindells,Brendan Payne,Jennifer Collins,Yusri Taha,Gary Eltringham |
| EPI_ISL_572520 | Quadram Institute Bioscience | COVID-19 Genomics UK (COG-UK) Consortium | Dave J. Baker, Gemma L. Kay, Alp Aydin, Thanh Le-Viet, Steven Rudder, Ana P. Tedim, Anastasia Kolyva, Maria Diaz, Leonardo de Oliveira Martins, Nabil-Fareed Alikhan, Lizzie Meadows, Rachael Stanley, Ngozi Elumogo, Muhammed Yasir, Nicholas M. Thomson, Alexander J Trotter, Rachel Gilroy, Samuel Bloomfield, Claire Stuart, Andrew Bell, Reenesh Prakash, Samir Dervisevic, Alison E. Mather, John Wain, Mark Webber, Andrew J. Page, Justin O'Grady |
| EPI_ISL_572525 | Northumbria University / South Tees Hospitals NHS Foundation Trust / North Cumbria Integrated Care NHS Foundation Trust / North Tees and Hartlepool NHS Foundation Trust / Newcastle Hospitals NHS Foundation Trust | COVID-19 Genomics UK (COG-UK) Consortium | Darren L Smith,Andrew Nelson,Matthew Bashton,Greg R Young,Joshua Loh,John Allan,Mohammad A Tariq,Giles S Holt,Gary Black,Wen C Yew,Lynn Dover,Paul Baker,Steve Liggett,Sarah Essex,Jane Greenaway,Debra Padgett,Clive Graham,Garren Scott,Edward Barton,Emma Swindells,Brendan Payne,Jennifer Collins,Yusri Taha,Gary Eltringham |
| EPI_ISL_572526 | Quadram Institute Bioscience | COVID-19 Genomics UK (COG-UK) Consortium | Dave J. Baker, Gemma L. Kay, Alp Aydin, Thanh Le-Viet, Steven Rudder, Ana P. Tedim, Anastasia Kolyva, Maria Diaz, Leonardo de Oliveira Martins, Nabil-Fareed Alikhan, Lizzie Meadows, Rachael Stanley, Ngozi Elumogo, Muhammed Yasir, Nicholas M. Thomson, Alexander J Trotter, Rachel Gilroy, Samuel Bloomfield, Claire Stuart, Andrew Bell, Reenesh Prakash, Samir Dervisevic, Alison E. Mather, John Wain, Mark Webber, Andrew J. Page, Justin O'Grady |
| EPI_ISL_572530, EPI_ISL_572532 | Northumbria University / South Tees Hospitals NHS Foundation Trust / North Cumbria Integrated Care NHS Foundation Trust / North Tees and Hartlepool NHS Foundation Trust / Newcastle Hospitals NHS Foundation Trust | COVID-19 Genomics UK (COG-UK) Consortium | Darren L Smith,Andrew Nelson,Matthew Bashton,Greg R Young,Joshua Loh,John Allan,Mohammad A Tariq,Giles S Holt,Gary Black,Wen C Yew,Lynn Dover,Paul Baker,Steve Liggett,Sarah Essex,Jane Greenaway,Debra Padgett,Clive Graham,Garren Scott,Edward Barton,Emma Swindells,Brendan Payne,Jennifer Collins,Yusri Taha,Gary Eltringham |
| EPI_ISL_572533, EPI_ISL_572546, EPI_ISL_572548, EPI_ISL_572549, EPI_ISL_572550, EPI_ISL_572551, EPI_ISL_572553, EPI_ISL_572556, EPI_ISL_572563 | Quadram Institute Bioscience | COVID-19 Genomics UK (COG-UK) Consortium | Dave J. Baker, Gemma L. Kay, Alp Aydin, Thanh Le-Viet, Steven Rudder, Ana P. Tedim, Anastasia Kolyva, Maria Diaz, Leonardo de Oliveira Martins, Nabil-Fareed Alikhan, Lizzie Meadows, Rachael Stanley, Ngozi Elumogo, Muhammed Yasir, Nicholas M. Thomson, Alexander J Trotter, Rachel Gilroy, Samuel Bloomfield, Claire Stuart, Andrew Bell, Reenesh Prakash, Samir Dervisevic, Alison E. Mather, John Wain, Mark Webber, Andrew J. Page, Justin O'Grady |
| EPI_ISL_572568 | Northumbria University / South Tees Hospitals NHS Foundation Trust / North Cumbria Integrated Care NHS Foundation Trust / North Tees and Hartlepool NHS Foundation Trust / Newcastle Hospitals NHS Foundation Trust | COVID-19 Genomics UK (COG-UK) Consortium | Darren L Smith,Andrew Nelson,Matthew Bashton,Greg R Young,Joshua Loh,John Allan,Mohammad A Tariq,Giles S Holt,Gary Black,Wen C Yew,Lynn Dover,Paul Baker,Steve Liggett,Sarah Essex,Jane Greenaway,Debra Padgett,Clive Graham,Garren Scott,Edward Barton,Emma Swindells,Brendan Payne,Jennifer Collins,Yusri Taha,Gary Eltringham |
| EPI_ISL_572569 | Quadram Institute Bioscience | COVID-19 Genomics UK (COG-UK) Consortium | Dave J. Baker, Gemma L. Kay, Alp Aydin, Thanh Le-Viet, Steven Rudder, Ana P. Tedim, Anastasia Kolyva, Maria Diaz, Leonardo de Oliveira Martins, Nabil-Fareed Alikhan, Lizzie Meadows, Rachael Stanley, Ngozi Elumogo, Muhammed Yasir, Nicholas M. Thomson, Alexander J Trotter, Rachel Gilroy, Samuel Bloomfield, Claire Stuart, Andrew Bell, Reenesh Prakash, Samir Dervisevic, Alison E. Mather, John Wain, Mark Webber, Andrew J. Page, Justin O'Grady |
| EPI_ISL_572570, EPI_ISL_572571, EPI_ISL_572572, EPI_ISL_572579, EPI_ISL_572581 | Northumbria University / South Tees Hospitals NHS Foundation Trust / North Cumbria Integrated Care NHS Foundation Trust / North Tees and Hartlepool NHS Foundation Trust / Newcastle Hospitals NHS Foundation Trust | COVID-19 Genomics UK (COG-UK) Consortium | Darren L Smith,Andrew Nelson,Matthew Bashton,Greg R Young,Joshua Loh,John Allan,Mohammad A Tariq,Giles S Holt,Gary Black,Wen C Yew,Lynn Dover,Paul Baker,Steve Liggett,Sarah Essex,Jane Greenaway,Debra Padgett,Clive Graham,Garren Scott,Edward Barton,Emma Swindells,Brendan Payne,Jennifer Collins,Yusri Taha,Gary Eltringham |
| EPI_ISL_572583, EPI_ISL_572591 | Quadram Institute Bioscience | COVID-19 Genomics UK (COG-UK) Consortium | Dave J. Baker, Gemma L. Kay, Alp Aydin, Thanh Le-Viet, Steven Rudder, Ana P. Tedim, Anastasia Kolyva, Maria Diaz, Leonardo de Oliveira Martins, Nabil-Fareed Alikhan, Lizzie Meadows, Rachael Stanley, Ngozi Elumogo, Muhammed Yasir, Nicholas M. Thomson, Alexander J Trotter, Rachel Gilroy, Samuel Bloomfield, Claire Stuart, Andrew Bell, Reenesh Prakash, Samir Dervisevic, Alison E. Mather, John Wain, Mark Webber, Andrew J. Page, Justin O'Grady |
| EPI_ISL_572594, EPI_ISL_572595, EPI_ISL_572596, EPI_ISL_572597 | Northumbria University / South Tees Hospitals NHS Foundation Trust / North Cumbria Integrated Care NHS Foundation Trust / North Tees and Hartlepool NHS Foundation Trust / Newcastle Hospitals NHS Foundation Trust | COVID-19 Genomics UK (COG-UK) Consortium | Darren L Smith,Andrew Nelson,Matthew Bashton,Greg R Young,Joshua Loh,John Allan,Mohammad A Tariq,Giles S Holt,Gary Black,Wen C Yew,Lynn Dover,Paul Baker,Steve Liggett,Sarah Essex,Jane Greenaway,Debra Padgett,Clive Graham,Garren Scott,Edward Barton,Emma Swindells,Brendan Payne,Jennifer Collins,Yusri Taha,Gary Eltringham |
| EPI_ISL_572600, EPI_ISL_572602, EPI_ISL_572607, EPI_ISL_572615 | Quadram Institute Bioscience | COVID-19 Genomics UK (COG-UK) Consortium | Dave J. Baker, Gemma L. Kay, Alp Aydin, Thanh Le-Viet, Steven Rudder, Ana P. Tedim, Anastasia Kolyva, Maria Diaz, Leonardo de Oliveira Martins, Nabil-Fareed Alikhan, Lizzie Meadows, Rachael Stanley, Ngozi Elumogo, Muhammed Yasir, Nicholas M. Thomson, Alexander J Trotter, Rachel Gilroy, Samuel Bloomfield, Claire Stuart, Andrew Bell, Reenesh Prakash, Samir Dervisevic, Alison E. Mather, John Wain, Mark Webber, Andrew J. Page, Justin O'Grady |
| EPI_ISL_572644 | Northumbria University / South Tees Hospitals NHS Foundation Trust / North Cumbria Integrated Care NHS Foundation Trust / North Tees and Hartlepool NHS Foundation Trust / Newcastle Hospitals NHS Foundation Trust | COVID-19 Genomics UK (COG-UK) Consortium | Darren L Smith,Andrew Nelson,Matthew Bashton,Greg R Young,Joshua Loh,John Allan,Mohammad A Tariq,Giles S Holt,Gary Black,Wen C Yew,Lynn Dover,Paul Baker,Steve Liggett,Sarah Essex,Jane Greenaway,Debra Padgett,Clive Graham,Garren Scott,Edward Barton,Emma Swindells,Brendan Payne,Jennifer Collins,Yusri Taha,Gary Eltringham |
| EPI_ISL_572646 | Quadram Institute Bioscience | COVID-19 Genomics UK (COG-UK) Consortium | Dave J. Baker, Gemma L. Kay, Alp Aydin, Thanh Le-Viet, Steven Rudder, Ana P. Tedim, Anastasia Kolyva, Maria Diaz, Leonardo de Oliveira Martins, |

| | | | |
|---|---|---|---|
| | | | Nabil-Fareed Alikhan, Lizzie Meadows, Rachael Stanley, Ngozi Elumogo, Muhammed Yasir, Nicholas M. Thomson, Alexander J Trotter, Rachel Gilroy, Samuel Bloomfield, Claire Stuart, Andrew Bell, Reenesh Prakash, Samir Dervisevic, Alison E. Mather, John Wain, Mark Webber, Andrew J. Page, Justin O'Grady |
| EPI_ISL_572648 | Northumbria University / South Tees Hospitals NHS Foundation Trust / North Cumbria Integrated Care NHS Foundation Trust / North Tees and Hartlepool NHS Foundation Trust / Newcastle Hospitals NHS Foundation Trust | COVID-19 Genomics UK (COG-UK) Consortium | Darren L Smith,Andrew Nelson,Matthew Bashton,Greg R Young,Joshua Loh,John Allan,Mohammad A Tariq,Giles S Holt,Gary Black,Wen C Yew,Lynn Dover,Paul Baker,Steve Liggett,Sarah Essex,Jane Greenaway,Debra Padgett,Clive Graham,Garren Scott,Edward Barton,Emma Swindells,Brendan Payne,Jennifer Collins,Yusri Taha,Gary Eltringham |
| EPI_ISL_572649, EPI_ISL_572651, EPI_ISL_572653, EPI_ISL_572655, EPI_ISL_572656, EPI_ISL_572658, EPI_ISL_572660 | Quadram Institute Bioscience | COVID-19 Genomics UK (COG-UK) Consortium | Dave J. Baker, Gemma L. Kay, Alp Aydin, Thanh Le-Viet, Steven Rudder, Ana P. Tedim, Anastasia Kolyva, Maria Diaz, Leonardo de Oliveira Martins, Nabil-Fareed Alikhan, Lizzie Meadows, Rachael Stanley, Ngozi Elumogo, Muhammed Yasir, Nicholas M. Thomson, Alexander J Trotter, Rachel Gilroy, Samuel Bloomfield, Claire Stuart, Andrew Bell, Reenesh Prakash, Samir Dervisevic, Alison E. Mather, John Wain, Mark Webber, Andrew J. Page, Justin O'Grady |
| EPI_ISL_572663, EPI_ISL_572664 | Northumbria University / South Tees Hospitals NHS Foundation Trust / North Cumbria Integrated Care NHS Foundation Trust / North Tees and Hartlepool NHS Foundation Trust / Newcastle Hospitals NHS Foundation Trust | COVID-19 Genomics UK (COG-UK) Consortium | Darren L Smith,Andrew Nelson,Matthew Bashton,Greg R Young,Joshua Loh,John Allan,Mohammad A Tariq,Giles S Holt,Gary Black,Wen C Yew,Lynn Dover,Paul Baker,Steve Liggett,Sarah Essex,Jane Greenaway,Debra Padgett,Clive Graham,Garren Scott,Edward Barton,Emma Swindells,Brendan Payne,Jennifer Collins,Yusri Taha,Gary Eltringham |
| EPI_ISL_572665, EPI_ISL_572666, EPI_ISL_572670, EPI_ISL_572671, EPI_ISL_572673 | Quadram Institute Bioscience | COVID-19 Genomics UK (COG-UK) Consortium | Dave J. Baker, Gemma L. Kay, Alp Aydin, Thanh Le-Viet, Steven Rudder, Ana P. Tedim, Anastasia Kolyva, Maria Diaz, Leonardo de Oliveira Martins, Nabil-Fareed Alikhan, Lizzie Meadows, Rachael Stanley, Ngozi Elumogo, Muhammed Yasir, Nicholas M. Thomson, Alexander J Trotter, Rachel Gilroy, Samuel Bloomfield, Claire Stuart, Andrew Bell, Reenesh Prakash, Samir Dervisevic, Alison E. Mather, John Wain, Mark Webber, Andrew J. Page, Justin O'Grady |
| EPI_ISL_572700, EPI_ISL_572701, EPI_ISL_572702, EPI_ISL_572703, EPI_ISL_572704 | Northumbria University / South Tees Hospitals NHS Foundation Trust / North Cumbria Integrated Care NHS Foundation Trust / North Tees and Hartlepool NHS Foundation Trust / Newcastle Hospitals NHS Foundation Trust | COVID-19 Genomics UK (COG-UK) Consortium | Darren L Smith,Andrew Nelson,Matthew Bashton,Greg R Young,Joshua Loh,John Allan,Mohammad A Tariq,Giles S Holt,Gary Black,Wen C Yew,Lynn Dover,Paul Baker,Steve Liggett,Sarah Essex,Jane Greenaway,Debra Padgett,Clive Graham,Garren Scott,Edward Barton,Emma Swindells,Brendan Payne,Jennifer Collins,Yusri Taha,Gary Eltringham |
| EPI_ISL_572711 | Quadram Institute Bioscience | COVID-19 Genomics UK (COG-UK) Consortium | Dave J. Baker, Gemma L. Kay, Alp Aydin, Thanh Le-Viet, Steven Rudder, Ana P. Tedim, Anastasia Kolyva, Maria Diaz, Leonardo de Oliveira Martins, Nabil-Fareed Alikhan, Lizzie Meadows, Rachael Stanley, Ngozi Elumogo, Muhammed Yasir, Nicholas M. Thomson, Alexander J Trotter, Rachel Gilroy, Samuel Bloomfield, Claire Stuart, Andrew Bell, Reenesh Prakash, Samir Dervisevic, Alison E. Mather, John Wain, Mark Webber, Andrew J. Page, Justin O'Grady |
| EPI_ISL_572712, EPI_ISL_572713 | Northumbria University / South Tees Hospitals NHS Foundation Trust / North Cumbria Integrated Care NHS Foundation Trust / North Tees and Hartlepool NHS Foundation Trust / Newcastle Hospitals NHS Foundation Trust | COVID-19 Genomics UK (COG-UK) Consortium | Darren L Smith,Andrew Nelson,Matthew Bashton,Greg R Young,Joshua Loh,John Allan,Mohammad A Tariq,Giles S Holt,Gary Black,Wen C Yew,Lynn Dover,Paul Baker,Steve Liggett,Sarah Essex,Jane Greenaway,Debra Padgett,Clive Graham,Garren Scott,Edward Barton,Emma Swindells,Brendan Payne,Jennifer Collins,Yusri Taha,Gary Eltringham |
| EPI_ISL_572717, EPI_ISL_572718, EPI_ISL_572719, EPI_ISL_572720 | Quadram Institute Bioscience | COVID-19 Genomics UK (COG-UK) Consortium | Dave J. Baker, Gemma L. Kay, Alp Aydin, Thanh Le-Viet, Steven Rudder, Ana P. Tedim, Anastasia Kolyva, Maria Diaz, Leonardo de Oliveira Martins, Nabil-Fareed Alikhan, Lizzie Meadows, Rachael Stanley, Ngozi Elumogo, Muhammed Yasir, Nicholas M. Thomson, Alexander J Trotter, Rachel Gilroy, Samuel Bloomfield, Claire Stuart, Andrew Bell, Reenesh Prakash, Samir Dervisevic, Alison E. Mather, John Wain, Mark Webber, Andrew J. Page, Justin O'Grady |
| EPI_ISL_572729 | Northumbria University / South Tees Hospitals NHS Foundation Trust / North Cumbria Integrated Care NHS Foundation Trust / North Tees and Hartlepool NHS Foundation Trust / Newcastle Hospitals NHS Foundation Trust | COVID-19 Genomics UK (COG-UK) Consortium | Darren L Smith,Andrew Nelson,Matthew Bashton,Greg R Young,Joshua Loh,John Allan,Mohammad A Tariq,Giles S Holt,Gary Black,Wen C Yew,Lynn Dover,Paul Baker,Steve Liggett,Sarah Essex,Jane Greenaway,Debra Padgett,Clive Graham,Garren Scott,Edward Barton,Emma Swindells,Brendan Payne,Jennifer Collins,Yusri Taha,Gary Eltringham |
| EPI_ISL_572730, EPI_ISL_572732, EPI_ISL_572735, EPI_ISL_572742 | Quadram Institute Bioscience | COVID-19 Genomics UK (COG-UK) Consortium | Dave J. Baker, Gemma L. Kay, Alp Aydin, Thanh Le-Viet, Steven Rudder, Ana P. Tedim, Anastasia Kolyva, Maria Diaz, Leonardo de Oliveira Martins, Nabil-Fareed Alikhan, Lizzie Meadows, Rachael Stanley, Ngozi Elumogo, Muhammed Yasir, Nicholas M. Thomson, Alexander J Trotter, Rachel Gilroy, Samuel Bloomfield, Claire Stuart, Andrew Bell, Reenesh Prakash, Samir Dervisevic, Alison E. Mather, John Wain, Mark Webber, Andrew J. Page, Justin O'Grady |
| EPI_ISL_572744 | Northumbria University / South Tees Hospitals NHS Foundation Trust / North Cumbria Integrated Care NHS Foundation Trust / North Tees and Hartlepool NHS Foundation Trust / Newcastle Hospitals NHS Foundation Trust | COVID-19 Genomics UK (COG-UK) Consortium | Darren L Smith,Andrew Nelson,Matthew Bashton,Greg R Young,Joshua Loh,John Allan,Mohammad A Tariq,Giles S Holt,Gary Black,Wen C Yew,Lynn Dover,Paul Baker,Steve Liggett,Sarah Essex,Jane Greenaway,Debra Padgett,Clive Graham,Garren Scott,Edward Barton,Emma Swindells,Brendan Payne,Jennifer Collins,Yusri Taha,Gary Eltringham |
| EPI_ISL_572745 | Quadram Institute Bioscience | COVID-19 Genomics UK (COG-UK) Consortium | Dave J. Baker, Gemma L. Kay, Alp Aydin, Thanh Le-Viet, Steven Rudder, Ana P. Tedim, Anastasia Kolyva, Maria Diaz, Leonardo de Oliveira Martins, Nabil-Fareed Alikhan, Lizzie Meadows, Rachael Stanley, Ngozi Elumogo, Muhammed Yasir, Nicholas M. Thomson, Alexander J Trotter, Rachel Gilroy, Samuel Bloomfield, Claire Stuart, Andrew Bell, Reenesh Prakash, Samir Dervisevic, Alison E. Mather, John Wain, Mark Webber, Andrew J. Page, Justin O'Grady |
| EPI_ISL_572747 | Northumbria University / South Tees Hospitals NHS Foundation Trust / North Cumbria Integrated Care NHS Foundation Trust / North Tees and Hartlepool NHS Foundation Trust / Newcastle Hospitals NHS Foundation Trust | COVID-19 Genomics UK (COG-UK) Consortium | Darren L Smith,Andrew Nelson,Matthew Bashton,Greg R Young,Joshua Loh,John Allan,Mohammad A Tariq,Giles S Holt,Gary Black,Wen C Yew,Lynn Dover,Paul Baker,Steve Liggett,Sarah Essex,Jane Greenaway,Debra Padgett,Clive Graham,Garren Scott,Edward Barton,Emma Swindells,Brendan Payne,Jennifer Collins,Yusri Taha,Gary Eltringham |
| EPI_ISL_572749 | Quadram Institute Bioscience | COVID-19 Genomics UK (COG-UK) Consortium | Dave J. Baker, Gemma L. Kay, Alp Aydin, Thanh Le-Viet, Steven Rudder, Ana P. Tedim, Anastasia Kolyva, Maria Diaz, Leonardo de Oliveira Martins, Nabil-Fareed Alikhan, Lizzie Meadows, Rachael Stanley, Ngozi Elumogo, Muhammed Yasir, Nicholas M. Thomson, Alexander J Trotter, Rachel Gilroy, Samuel Bloomfield, Claire Stuart, Andrew Bell, Reenesh Prakash, Samir Dervisevic, Alison E. Mather, John Wain, Mark Webber, Andrew J. Page, Justin O'Grady |
| EPI_ISL_572751, EPI_ISL_572753 | Northumbria University / South Tees Hospitals NHS Foundation Trust / North Cumbria Integrated Care NHS Foundation Trust / North Tees and Hartlepool NHS Foundation Trust / Newcastle Hospitals NHS Foundation Trust | COVID-19 Genomics UK (COG-UK) Consortium | Darren L Smith,Andrew Nelson,Matthew Bashton,Greg R Young,Joshua Loh,John Allan,Mohammad A Tariq,Giles S Holt,Gary Black,Wen C Yew,Lynn Dover,Paul Baker,Steve Liggett,Sarah Essex,Jane Greenaway,Debra Padgett,Clive Graham,Garren Scott,Edward Barton,Emma Swindells,Brendan Payne,Jennifer Collins,Yusri Taha,Gary Eltringham |
| EPI_ISL_572755, EPI_ISL_572758, EPI_ISL_572759, EPI_ISL_572764, EPI_ISL_572768, EPI_ISL_572775, EPI_ISL_572780 | Quadram Institute Bioscience | COVID-19 Genomics UK (COG-UK) Consortium | Dave J. Baker, Gemma L. Kay, Alp Aydin, Thanh Le-Viet, Steven Rudder, Ana P. Tedim, Anastasia Kolyva, Maria Diaz, Leonardo de Oliveira Martins, Nabil-Fareed Alikhan, Lizzie Meadows, Rachael Stanley, Ngozi Elumogo, Muhammed Yasir, Nicholas M. Thomson, Alexander J Trotter, Rachel Gilroy, Samuel Bloomfield, Claire Stuart, Andrew Bell, Reenesh Prakash, Samir Dervisevic, Alison E. Mather, John Wain, Mark Webber, Andrew J. Page, Justin O'Grady |
| EPI_ISL_572783 | Northumbria University / South Tees Hospitals NHS Foundation Trust / North Cumbria Integrated Care NHS Foundation Trust / North Tees and Hartlepool NHS Foundation Trust / Newcastle Hospitals NHS Foundation Trust | COVID-19 Genomics UK (COG-UK) Consortium | Darren L Smith,Andrew Nelson,Matthew Bashton,Greg R Young,Joshua Loh,John Allan,Mohammad A Tariq,Giles S Holt,Gary Black,Wen C Yew,Lynn Dover,Paul Baker,Steve Liggett,Sarah Essex,Jane Greenaway,Debra Padgett,Clive Graham,Garren Scott,Edward Barton,Emma Swindells,Brendan Payne,Jennifer Collins,Yusri Taha,Gary Eltringham |
| EPI_ISL_572785, EPI_ISL_572786, EPI_ISL_572787, EPI_ISL_572789, EPI_ISL_572791, EPI_ISL_572793 | Quadram Institute Bioscience | COVID-19 Genomics UK (COG-UK) Consortium | Dave J. Baker, Gemma L. Kay, Alp Aydin, Thanh Le-Viet, Steven Rudder, Ana P. Tedim, Anastasia Kolyva, Maria Diaz, Leonardo de Oliveira Martins, Nabil-Fareed Alikhan, Lizzie Meadows, Rachael Stanley, Ngozi Elumogo, Muhammed Yasir, Nicholas M. Thomson, Alexander J Trotter, Rachel Gilroy, Samuel Bloomfield, Claire Stuart, Andrew Bell, Reenesh Prakash, Samir Dervisevic, Alison E. Mather, John Wain, Mark Webber, Andrew J. Page, Justin O'Grady |
| EPI_ISL_572795 | Northumbria University / South Tees Hospitals NHS Foundation Trust / North Cumbria Integrated Care NHS Foundation Trust / North Tees and Hartlepool NHS | COVID-19 Genomics UK (COG-UK) Consortium | Darren L Smith,Andrew Nelson,Matthew Bashton,Greg R Young,Joshua Loh,John Allan,Mohammad A Tariq,Giles S Holt,Gary Black,Wen C Yew,Lynn Dover,Paul Baker,Steve Liggett,Sarah Essex,Jane Greenaway,Debra Padgett,Clive Graham,Garren Scott,Edward Barton,Emma Swindells,Brendan Payne,Jennifer Collins,Yusri Taha,Gary Eltringham |

| | | | |
|---|---|---|---|
| | Foundation Trust / Newcastle Hospitals NHS Foundation Trust | | |
| EPI_ISL_572797 | Quadram Institute Bioscience | COVID-19 Genomics UK (COG-UK) Consortium | Dave J. Baker, Gemma L. Kay, Alp Aydin, Thanh Le-Viet, Steven Rudder, Ana P. Tedim, Anastasia Kolyva, Maria Diaz, Leonardo de Oliveira Martins, Nabil-Fareed Alikhan, Lizzie Meadows, Rachael Stanley, Ngozi Elumogo, Muhammed Yasir, Nicholas M. Thomson, Alexander J Trotter, Rachel Gilroy, Samuel Bloomfield, Claire Stuart, Andrew Bell, Reenesh Prakash, Samir Dervisevic, Alison E. Mather, John Wain, Mark Webber, Andrew J. Page, Justin O'Grady |
| EPI_ISL_572798, EPI_ISL_572802 | Northumbria University / South Tees Hospitals NHS Foundation Trust / North Cumbria Integrated Care NHS Foundation Trust / North Tees and Hartlepool NHS Foundation Trust / Newcastle Hospitals NHS Foundation Trust | COVID-19 Genomics UK (COG-UK) Consortium | Darren L Smith,Andrew Nelson,Matthew Bashton,Greg R Young,Joshua Loh,John Allan,Mohammad A Tariq,Giles S Holt,Gary Black,Wen C Yew,Lynn Dover,Paul Baker,Steve Liggett,Sarah Essex,Jane Greenaway,Debra Padgett,Clive Graham,Garren Scott,Edward Barton,Emma Swindells,Brendan Payne,Jennifer Collins,Yusri Taha,Gary Eltringham |
| EPI_ISL_572804, EPI_ISL_572806, EPI_ISL_572811, EPI_ISL_572812, EPI_ISL_572814, EPI_ISL_572819, EPI_ISL_572823, EPI_ISL_572825, EPI_ISL_572826, EPI_ISL_572831, EPI_ISL_572835 | | | |
| see above | Quadram Institute Bioscience | COVID-19 Genomics UK (COG-UK) Consortium | Dave J. Baker, Gemma L. Kay, Alp Aydin, Thanh Le-Viet, Steven Rudder, Ana P. Tedim, Anastasia Kolyva, Maria Diaz, Leonardo de Oliveira Martins, Nabil-Fareed Alikhan, Lizzie Meadows, Rachael Stanley, Ngozi Elumogo, Muhammed Yasir, Nicholas M. Thomson, Alexander J Trotter, Rachel Gilroy, Samuel Bloomfield, Claire Stuart, Andrew Bell, Reenesh Prakash, Samir Dervisevic, Alison E. Mather, John Wain, Mark Webber, Andrew J. Page, Justin O'Grady |
| EPI_ISL_572840 | Northumbria University / South Tees Hospitals NHS Foundation Trust / North Cumbria Integrated Care NHS Foundation Trust / North Tees and Hartlepool NHS Foundation Trust / Newcastle Hospitals NHS Foundation Trust | COVID-19 Genomics UK (COG-UK) Consortium | Darren L Smith,Andrew Nelson,Matthew Bashton,Greg R Young,Joshua Loh,John Allan,Mohammad A Tariq,Giles S Holt,Gary Black,Wen C Yew,Lynn Dover,Paul Baker,Steve Liggett,Sarah Essex,Jane Greenaway,Debra Padgett,Clive Graham,Garren Scott,Edward Barton,Emma Swindells,Brendan Payne,Jennifer Collins,Yusri Taha,Gary Eltringham |
| EPI_ISL_572842, EPI_ISL_572844 | Quadram Institute Bioscience | COVID-19 Genomics UK (COG-UK) Consortium | Dave J. Baker, Gemma L. Kay, Alp Aydin, Thanh Le-Viet, Steven Rudder, Ana P. Tedim, Anastasia Kolyva, Maria Diaz, Leonardo de Oliveira Martins, Nabil-Fareed Alikhan, Lizzie Meadows, Rachael Stanley, Ngozi Elumogo, Muhammed Yasir, Nicholas M. Thomson, Alexander J Trotter, Rachel Gilroy, Samuel Bloomfield, Claire Stuart, Andrew Bell, Reenesh Prakash, Samir Dervisevic, Alison E. Mather, John Wain, Mark Webber, Andrew J. Page, Justin O'Grady |
| EPI_ISL_572845 | Northumbria University / South Tees Hospitals NHS Foundation Trust / North Cumbria Integrated Care NHS Foundation Trust / North Tees and Hartlepool NHS Foundation Trust / Newcastle Hospitals NHS Foundation Trust | COVID-19 Genomics UK (COG-UK) Consortium | Darren L Smith,Andrew Nelson,Matthew Bashton,Greg R Young,Joshua Loh,John Allan,Mohammad A Tariq,Giles S Holt,Gary Black,Wen C Yew,Lynn Dover,Paul Baker,Steve Liggett,Sarah Essex,Jane Greenaway,Debra Padgett,Clive Graham,Garren Scott,Edward Barton,Emma Swindells,Brendan Payne,Jennifer Collins,Yusri Taha,Gary Eltringham |
| EPI_ISL_572847, EPI_ISL_572898, EPI_ISL_572899, EPI_ISL_572900, EPI_ISL_572901 | Quadram Institute Bioscience | COVID-19 Genomics UK (COG-UK) Consortium | Dave J. Baker, Gemma L. Kay, Alp Aydin, Thanh Le-Viet, Steven Rudder, Ana P. Tedim, Anastasia Kolyva, Maria Diaz, Leonardo de Oliveira Martins, Nabil-Fareed Alikhan, Lizzie Meadows, Rachael Stanley, Ngozi Elumogo, Muhammed Yasir, Nicholas M. Thomson, Alexander J Trotter, Rachel Gilroy, Samuel Bloomfield, Claire Stuart, Andrew Bell, Reenesh Prakash, Samir Dervisevic, Alison E. Mather, John Wain, Mark Webber, Andrew J. Page, Justin O'Grady |
| EPI_ISL_572909, EPI_ISL_572910, EPI_ISL_572911 | Northumbria University / South Tees Hospitals NHS Foundation Trust / North Cumbria Integrated Care NHS Foundation Trust / North Tees and Hartlepool NHS Foundation Trust / Newcastle Hospitals NHS Foundation Trust | COVID-19 Genomics UK (COG-UK) Consortium | Darren L Smith,Andrew Nelson,Matthew Bashton,Greg R Young,Joshua Loh,John Allan,Mohammad A Tariq,Giles S Holt,Gary Black,Wen C Yew,Lynn Dover,Paul Baker,Steve Liggett,Sarah Essex,Jane Greenaway,Debra Padgett,Clive Graham,Garren Scott,Edward Barton,Emma Swindells,Brendan Payne,Jennifer Collins,Yusri Taha,Gary Eltringham |
| EPI_ISL_572939, EPI_ISL_572940, EPI_ISL_572941 | Quadram Institute Bioscience | COVID-19 Genomics UK (COG-UK) Consortium | Dave J. Baker, Gemma L. Kay, Alp Aydin, Thanh Le-Viet, Steven Rudder, Ana P. Tedim, Anastasia Kolyva, Maria Diaz, Leonardo de Oliveira Martins, Nabil-Fareed Alikhan, Lizzie Meadows, Rachael Stanley, Ngozi Elumogo, Muhammed Yasir, Nicholas M. Thomson, Alexander J Trotter, Rachel Gilroy, Samuel Bloomfield, Claire Stuart, Andrew Bell, Reenesh Prakash, Samir Dervisevic, Alison E. Mather, John Wain, Mark Webber, Andrew J. Page, Justin O'Grady |
| EPI_ISL_572942, EPI_ISL_572950 | Northumbria University / South Tees Hospitals NHS Foundation Trust / North Cumbria Integrated Care NHS Foundation Trust / North Tees and Hartlepool NHS Foundation Trust / Newcastle Hospitals NHS Foundation Trust | COVID-19 Genomics UK (COG-UK) Consortium | Darren L Smith,Andrew Nelson,Matthew Bashton,Greg R Young,Joshua Loh,John Allan,Mohammad A Tariq,Giles S Holt,Gary Black,Wen C Yew,Lynn Dover,Paul Baker,Steve Liggett,Sarah Essex,Jane Greenaway,Debra Padgett,Clive Graham,Garren Scott,Edward Barton,Emma Swindells,Brendan Payne,Jennifer Collins,Yusri Taha,Gary Eltringham |
| EPI_ISL_572954, EPI_ISL_573085, EPI_ISL_573086, EPI_ISL_573087, EPI_ISL_573088, EPI_ISL_573089, EPI_ISL_573090, EPI_ISL_573091, EPI_ISL_573092, EPI_ISL_573093, EPI_ISL_573094, EPI_ISL_573095, EPI_ISL_573096, EPI_ISL_573097, EPI_ISL_573098, EPI_ISL_573099, EPI_ISL_573100, EPI_ISL_573101, EPI_ISL_573102, EPI_ISL_573103, EPI_ISL_573104, EPI_ISL_573105, EPI_ISL_573106, EPI_ISL_573107, EPI_ISL_573108, EPI_ISL_573109, EPI_ISL_573110, EPI_ISL_573111, EPI_ISL_573112, EPI_ISL_573113, EPI_ISL_573114, EPI_ISL_573115, EPI_ISL_573116, EPI_ISL_573117, EPI_ISL_573118, EPI_ISL_573119, EPI_ISL_573120, EPI_ISL_573121, EPI_ISL_573122, EPI_ISL_573123, EPI_ISL_573124, EPI_ISL_573125, EPI_ISL_573126, EPI_ISL_573127, EPI_ISL_573128, EPI_ISL_573129, EPI_ISL_573130, EPI_ISL_573131 | | | |
| see above | Quadram Institute Bioscience | COVID-19 Genomics UK (COG-UK) Consortium | Dave J. Baker, Gemma L. Kay, Alp Aydin, Thanh Le-Viet, Steven Rudder, Ana P. Tedim, Anastasia Kolyva, Maria Diaz, Leonardo de Oliveira Martins, Nabil-Fareed Alikhan, Lizzie Meadows, Rachael Stanley, Ngozi Elumogo, Muhammed Yasir, Nicholas M. Thomson, Alexander J Trotter, Rachel Gilroy, Samuel Bloomfield, Claire Stuart, Andrew Bell, Reenesh Prakash, Samir Dervisevic, Alison E. Mather, John Wain, Mark Webber, Andrew J. Page, Justin O'Grady |
| EPI_ISL_573266, EPI_ISL_573267, EPI_ISL_573268, EPI_ISL_573269, EPI_ISL_573270, EPI_ISL_573271, EPI_ISL_573272, EPI_ISL_573273, EPI_ISL_573274, EPI_ISL_573275, EPI_ISL_573276, EPI_ISL_573277, EPI_ISL_573278, EPI_ISL_573279, EPI_ISL_573280, EPI_ISL_573281, EPI_ISL_573282, EPI_ISL_573283, EPI_ISL_573284, EPI_ISL_573285, EPI_ISL_573286, EPI_ISL_573287, EPI_ISL_573288, EPI_ISL_573289, EPI_ISL_573290, EPI_ISL_573291, EPI_ISL_573292, EPI_ISL_573293, EPI_ISL_573294, EPI_ISL_573295, EPI_ISL_573296, EPI_ISL_573297, EPI_ISL_573298, EPI_ISL_573299, EPI_ISL_573300, EPI_ISL_573301, EPI_ISL_573302, EPI_ISL_573303, EPI_ISL_573304, EPI_ISL_573305, EPI_ISL_573306, EPI_ISL_573307, EPI_ISL_573308, EPI_ISL_573309, EPI_ISL_573310, EPI_ISL_573311, EPI_ISL_573312, EPI_ISL_573313, EPI_ISL_573314, EPI_ISL_573315, EPI_ISL_573316, EPI_ISL_573317, EPI_ISL_573318, EPI_ISL_573319, EPI_ISL_573320, EPI_ISL_573321, EPI_ISL_573322, EPI_ISL_573323, EPI_ISL_573324, EPI_ISL_573325, EPI_ISL_573326, EPI_ISL_573327, EPI_ISL_573328, EPI_ISL_573329, EPI_ISL_573330, EPI_ISL_573331, EPI_ISL_573332, EPI_ISL_573333, EPI_ISL_573334, EPI_ISL_573335, EPI_ISL_573336, EPI_ISL_573337, EPI_ISL_573338, EPI_ISL_573339, EPI_ISL_573340, EPI_ISL_573341, EPI_ISL_573342, EPI_ISL_573343, EPI_ISL_573344, EPI_ISL_573345, EPI_ISL_573346, EPI_ISL_573347, EPI_ISL_573348, EPI_ISL_573349, EPI_ISL_573350, EPI_ISL_573351, EPI_ISL_573352, EPI_ISL_573353, EPI_ISL_573354, EPI_ISL_573355, EPI_ISL_573356, EPI_ISL_573357, EPI_ISL_573358, EPI_ISL_573359, EPI_ISL_573360, EPI_ISL_573361, EPI_ISL_573362, EPI_ISL_573363, EPI_ISL_573364, EPI_ISL_573365, EPI_ISL_573366, EPI_ISL_573367, EPI_ISL_573368, EPI_ISL_573369, EPI_ISL_573370, EPI_ISL_573371, EPI_ISL_573372, EPI_ISL_573373, EPI_ISL_573374, EPI_ISL_573375, EPI_ISL_573376, EPI_ISL_573377, EPI_ISL_573378, EPI_ISL_573379, EPI_ISL_573380, EPI_ISL_573381, EPI_ISL_573382, EPI_ISL_573383, EPI_ISL_573384, EPI_ISL_573385, EPI_ISL_573386, EPI_ISL_573387, EPI_ISL_573388, EPI_ISL_573389, EPI_ISL_573390, EPI_ISL_573391, EPI_ISL_573392, EPI_ISL_573393, EPI_ISL_573394, EPI_ISL_573395, EPI_ISL_573396, EPI_ISL_573397, EPI_ISL_573398, EPI_ISL_573399, EPI_ISL_573400, EPI_ISL_573401, EPI_ISL_573402, EPI_ISL_573403, EPI_ISL_573404, EPI_ISL_573405, EPI_ISL_573406, EPI_ISL_573407, EPI_ISL_573408, EPI_ISL_573409, EPI_ISL_573410, EPI_ISL_573411, EPI_ISL_573412, EPI_ISL_573413 | | | |
| see above | Northumbria University / South Tees Hospitals NHS Foundation Trust / North Cumbria Integrated Care NHS Foundation Trust / North Tees and Hartlepool NHS Foundation Trust / Newcastle Hospitals NHS Foundation Trust | COVID-19 Genomics UK (COG-UK) Consortium | Darren L Smith,Andrew Nelson,Matthew Bashton,Greg R Young,Joshua Loh,John Allan,Mohammad A Tariq,Giles S Holt,Gary Black,Wen C Yew,Lynn Dover,Paul Baker,Steve Liggett,Sarah Essex,Jane Greenaway,Debra Padgett,Clive Graham,Garren Scott,Edward Barton,Emma Swindells,Brendan Payne,Jennifer Collins,Yusri Taha,Gary Eltringham |
| EPI_ISL_573414, EPI_ISL_573415, EPI_ISL_573416, EPI_ISL_573417 | Quadram Institute Bioscience | COVID-19 Genomics UK (COG-UK) Consortium | Dave J. Baker, Gemma L. Kay, Alp Aydin, Thanh Le-Viet, Steven Rudder, Ana P. Tedim, Anastasia Kolyva, Maria Diaz, Leonardo de Oliveira Martins, Nabil-Fareed Alikhan, Lizzie Meadows, Rachael Stanley, Ngozi Elumogo, Muhammed Yasir, Nicholas M. Thomson, Alexander J Trotter, Rachel Gilroy, Samuel Bloomfield, Claire Stuart, Andrew Bell, Reenesh Prakash, Samir Dervisevic, Alison E. Mather, John Wain, Mark Webber, Andrew J. Page, Justin O'Grady |
| EPI_ISL_573760, EPI_ISL_573761, EPI_ISL_573799, EPI_ISL_573871, EPI_ISL_573872 | Northumbria University / South Tees Hospitals NHS Foundation Trust / North Cumbria Integrated Care NHS Foundation Trust / North Tees and Hartlepool NHS Foundation Trust / Newcastle Hospitals NHS Foundation Trust | COVID-19 Genomics UK (COG-UK) Consortium | Darren L Smith,Andrew Nelson,Matthew Bashton,Greg R Young,Joshua Loh,John Allan,Mohammad A Tariq,Giles S Holt,Gary Black,Wen C Yew,Lynn Dover,Paul Baker,Steve Liggett,Sarah Essex,Jane Greenaway,Debra Padgett,Clive Graham,Garren Scott,Edward Barton,Emma Swindells,Brendan Payne,Jennifer Collins,Yusri Taha,Gary Eltringham |
| EPI_ISL_574648 | Seattle Flu Study | Seattle Flu Study | Deborah A. Nickerson, Chris D. Frazar, Jover Lee, Benjamin Pelle, Matthew Richardson, Amanda Adler, Elisabeth Brandstetter, Peter D. Han, Kairsten Fay, Misja Ilcisin, Kirsten Lacombe, Thomas R. Sibley, Melissa Truong, Caitlin R. Wolf, Michael Boeckh, Janet A. Englund, Michael Famulare, Barry R. Lutz, Mark J. Rieder, Lea M. Starita, Matthew Thompson, Jay Shendure, Trevor Bedford, Helen Y. Chu |
| EPI_ISL_574649, EPI_ISL_574650, EPI_ISL_574651, EPI_ISL_574652, EPI_ISL_574653, EPI_ISL_574654, EPI_ISL_574655, EPI_ISL_574656, EPI_ISL_574657, EPI_ISL_574658, EPI_ISL_574659, EPI_ISL_574660, EPI_ISL_574661, EPI_ISL_574662 | | | |
| see above | Seattle Flu Study | Seattle Flu Study | Deborah A. Nickerson, Chris D. Frazar, Jover Lee, Benjamin Pelle, Matthew Richardson, Amanda Adler, Elisabeth Brandstetter, Peter D. Han, Kairsten Fay, Misja Ilcisin, Kirsten Lacombe, Thomas R. Sibley, Melissa Truong, Caitlin R. Wolf, Karen Cowgill, Stephanie Schrag, Jeff Duchin, Michael Boeckh, Janet A. Englund, Michael Famulare, Barry R. Lutz, Mark J. Rieder, Lea M. Starita, Matthew Thompson, Helen Y. Chu, Trevor Bedford, Jay Shendure |

| | | | |
|---|---|---|---|
| EPI_ISL_574663, EPI_ISL_574664, EPI_ISL_574665, EPI_ISL_574666, EPI_ISL_574667 | Seattle Flu Study | Seattle Flu Study | Deborah A. Nickerson, Chris D. Frazar, Jover Lee, Benjamin Pelle, Matthew Richardson, Amanda Adler, Elisabeth Brandstetter, Peter D. Han, Kairsten Fay, Misja Ilcisin, Kirsten Lacombe, Thomas R. Sibley, Melissa Truong, Caitlin R. Wolf, Michael Boeckh, Janet A. Englund, Michael Famulare, Barry R. Lutz, Mark J. Rieder, Lea M. Starita, Matthew Thompson, Jay Shendure, Trevor Bedford, Helen Y. Chu |
| EPI_ISL_574668, EPI_ISL_574669, EPI_ISL_574670, EPI_ISL_574671, EPI_ISL_574672, EPI_ISL_574673, EPI_ISL_574674, EPI_ISL_574675, EPI_ISL_574676, EPI_ISL_574677, EPI_ISL_574678 | | | |
| see above | Seattle Flu Study | Seattle Flu Study | Deborah A. Nickerson, Chris D. Frazar, Jover Lee, Benjamin Pelle, Matthew Richardson, Amanda Adler, Elisabeth Brandstetter, Peter D. Han, Kairsten Fay, Misja Ilcisin, Kirsten Lacombe, Thomas R. Sibley, Melissa Truong, Caitlin R. Wolf, Karen Cowgill, Stephanie Schrag, Jeff Duchin, Michael Boeckh, Janet A. Englund, Michael Famulare, Barry R. Lutz, Mark J. Rieder, Lea M. Starita, Matthew Thompson, Helen Y. Chu, Trevor Bedford, Jay Shendure |
| EPI_ISL_574679 | Seattle Flu Study | Seattle Flu Study | Deborah A. Nickerson, Chris D. Frazar, Jover Lee, Benjamin Pelle, Matthew Richardson, Amanda Adler, Elisabeth Brandstetter, Peter D. Han, Kairsten Fay, Misja Ilcisin, Kirsten Lacombe, Thomas R. Sibley, Melissa Truong, Caitlin R. Wolf, Michael Boeckh, Janet A. Englund, Michael Famulare, Barry R. Lutz, Mark J. Rieder, Lea M. Starita, Matthew Thompson, Jay Shendure, Trevor Bedford, Helen Y. Chu |
| EPI_ISL_574680, EPI_ISL_574681 | Seattle Flu Study | Seattle Flu Study | Deborah A. Nickerson, Chris D. Frazar, Jover Lee, Benjamin Pelle, Matthew Richardson, Amanda Adler, Elisabeth Brandstetter, Peter D. Han, Kairsten Fay, Misja Ilcisin, Kirsten Lacombe, Thomas R. Sibley, Melissa Truong, Caitlin R. Wolf, Karen Cowgill, Stephanie Schrag, Jeff Duchin, Michael Boeckh, Janet A. Englund, Michael Famulare, Barry R. Lutz, Mark J. Rieder, Lea M. Starita, Matthew Thompson, Helen Y. Chu, Trevor Bedford, Jay Shendure |
| EPI_ISL_574682 | Seattle Flu Study | Seattle Flu Study | Deborah A. Nickerson, Chris D. Frazar, Jover Lee, Benjamin Pelle, Matthew Richardson, Amanda Adler, Elisabeth Brandstetter, Peter D. Han, Kairsten Fay, Misja Ilcisin, Kirsten Lacombe, Thomas R. Sibley, Melissa Truong, Caitlin R. Wolf, Michael Boeckh, Janet A. Englund, Michael Famulare, Barry R. Lutz, Mark J. Rieder, Lea M. Starita, Matthew Thompson, Jay Shendure, Trevor Bedford, Helen Y. Chu |
| EPI_ISL_574683 | Seattle Flu Study | Seattle Flu Study | Deborah A. Nickerson, Chris D. Frazar, Jover Lee, Benjamin Pelle, Matthew Richardson, Amanda Adler, Elisabeth Brandstetter, Peter D. Han, Kairsten Fay, Misja Ilcisin, Kirsten Lacombe, Thomas R. Sibley, Melissa Truong, Caitlin R. Wolf, Karen Cowgill, Stephanie Schrag, Jeff Duchin, Michael Boeckh, Janet A. Englund, Michael Famulare, Barry R. Lutz, Mark J. Rieder, Lea M. Starita, Matthew Thompson, Helen Y. Chu, Trevor Bedford, Jay Shendure |
| EPI_ISL_574684 | Seattle Flu Study | Seattle Flu Study | Deborah A. Nickerson, Chris D. Frazar, Jover Lee, Benjamin Pelle, Matthew Richardson, Amanda Adler, Elisabeth Brandstetter, Peter D. Han, Kairsten Fay, Misja Ilcisin, Kirsten Lacombe, Thomas R. Sibley, Melissa Truong, Caitlin R. Wolf, Michael Boeckh, Janet A. Englund, Michael Famulare, Barry R. Lutz, Mark J. Rieder, Lea M. Starita, Matthew Thompson, Jay Shendure, Trevor Bedford, Helen Y. Chu |
| EPI_ISL_574685, EPI_ISL_574686, EPI_ISL_574687, EPI_ISL_574688, EPI_ISL_574689, EPI_ISL_574690, EPI_ISL_574691, EPI_ISL_575028, EPI_ISL_575029, EPI_ISL_575030, EPI_ISL_575031, EPI_ISL_575032 | | | |
| see above | Seattle Flu Study | Seattle Flu Study | Deborah A. Nickerson, Chris D. Frazar, Jover Lee, Benjamin Pelle, Matthew Richardson, Amanda Adler, Elisabeth Brandstetter, Peter D. Han, Kairsten Fay, Misja Ilcisin, Kirsten Lacombe, Thomas R. Sibley, Melissa Truong, Caitlin R. Wolf, Karen Cowgill, Stephanie Schrag, Jeff Duchin, Michael Boeckh, Janet A. Englund, Michael Famulare, Barry R. Lutz, Mark J. Rieder, Lea M. Starita, Matthew Thompson, Helen Y. Chu, Trevor Bedford, Jay Shendure |
| EPI_ISL_575033 | Seattle Flu Study | Seattle Flu Study | Deborah A. Nickerson, Chris D. Frazar, Jover Lee, Benjamin Pelle, Matthew Richardson, Amanda Adler, Elisabeth Brandstetter, Peter D. Han, Kairsten Fay, Misja Ilcisin, Kirsten Lacombe, Thomas R. Sibley, Melissa Truong, Caitlin R. Wolf, Romesh Gautom, Geoff Melly, Brian Hiatt, Philip Dykema, Scott Lindquist, Michael Boeckh, Janet A. Englund, Michael Famulare, Barry R. Lutz, Mark J. Rieder, Lea M. Starita, Matthew Thompson, Helen Y. Chu, Jay Shendure, Trevor Bedford |
| EPI_ISL_576986, EPI_ISL_576987, EPI_ISL_576988, EPI_ISL_576989, EPI_ISL_576990, EPI_ISL_576991, EPI_ISL_576992, EPI_ISL_576993, EPI_ISL_576994, EPI_ISL_576995, EPI_ISL_576996, EPI_ISL_576997, EPI_ISL_576998, EPI_ISL_576999, EPI_ISL_577000, EPI_ISL_577001, EPI_ISL_577002, EPI_ISL_577003, EPI_ISL_577004, EPI_ISL_577005, EPI_ISL_577006, EPI_ISL_577007, EPI_ISL_577008, EPI_ISL_577009, EPI_ISL_577010, EPI_ISL_577011, EPI_ISL_577012, EPI_ISL_577013, EPI_ISL_577014, EPI_ISL_577015, EPI_ISL_577016, EPI_ISL_577017, EPI_ISL_577018, EPI_ISL_577019, EPI_ISL_577020, EPI_ISL_577021, EPI_ISL_577022, EPI_ISL_577023, EPI_ISL_577024, EPI_ISL_577025, EPI_ISL_577026, EPI_ISL_577027, EPI_ISL_577028, EPI_ISL_577029, EPI_ISL_577030, EPI_ISL_577031, EPI_ISL_577032, EPI_ISL_577033, EPI_ISL_577034, EPI_ISL_577035, EPI_ISL_577036, EPI_ISL_577037, EPI_ISL_577038, EPI_ISL_577039 | | | |
| see above | Northumbria University / South Tees Hospitals NHS Foundation Trust / North Cumbria Integrated Care NHS Foundation Trust / North Tees and Hartlepool NHS Foundation Trust / Newcastle Hospitals NHS Foundation Trust | COVID-19 Genomics UK (COG-UK) Consortium | Darren L Smith,Andrew Nelson,Matthew Bashton,Greg R Young,Joshua Loh,John Allan,Mohammad A Tariq,Giles S Holt,Gary Black,Wen C Yew,Lynn Dover,Paul Baker,Steve Liggett,Sarah Essex,Jane Greenaway,Debra Padgett,Clive Graham,Garren Scott,Edward Barton,Emma Swindells,Brendan Payne,Jennifer Collins,Yusri Taha,Gary Eltringham |
| EPI_ISL_577096, EPI_ISL_577097, EPI_ISL_577098, EPI_ISL_577099, EPI_ISL_577100, EPI_ISL_577101, EPI_ISL_577102, EPI_ISL_577103, EPI_ISL_577104, EPI_ISL_577105, EPI_ISL_577106, EPI_ISL_577107, EPI_ISL_577108, EPI_ISL_577109 | | | |
| see above | Quadram Institute Bioscience | COVID-19 Genomics UK (COG-UK) Consortium | Dave J. Baker, Gemma L. Kay, Alp Aydin, Thanh Le-Viet, Steven Rudder, Ana P. Tedim, Anastasia Kolyva, Maria Diaz, Leonardo de Oliveira Martins, Nabil-Fareed Alikhan, Lizzie Meadows, Rachael Stanley, Ngozi Elumogo, Muhammed Yasir, Nicholas M. Thomson, Alexander J Trotter, Rachel Gilroy, Samuel Bloomfield, Claire Stuart, Andrew Bell, Reenesh Prakash, Samir Dervisevic, Alison E. Mather, John Wain, Mark Webber, Andrew J. Page, Justin O'Grady |
| EPI_ISL_582148, EPI_ISL_582149, EPI_ISL_582150, EPI_ISL_582151, EPI_ISL_582152, EPI_ISL_582153, EPI_ISL_582154, EPI_ISL_582155, EPI_ISL_582156, EPI_ISL_582157, EPI_ISL_582158, EPI_ISL_582159, EPI_ISL_582160, EPI_ISL_582161, EPI_ISL_582162, EPI_ISL_582163, EPI_ISL_582164, EPI_ISL_582165, EPI_ISL_582166, EPI_ISL_582167, EPI_ISL_582168, EPI_ISL_582169, EPI_ISL_582170, EPI_ISL_582171, EPI_ISL_582172, EPI_ISL_582173, EPI_ISL_582174, EPI_ISL_582175, EPI_ISL_582176, EPI_ISL_582177, EPI_ISL_582178, EPI_ISL_582179, EPI_ISL_582180, EPI_ISL_582181, EPI_ISL_582182, EPI_ISL_582183, EPI_ISL_582184, EPI_ISL_582185, EPI_ISL_582186, EPI_ISL_582187, EPI_ISL_582188, EPI_ISL_582189, EPI_ISL_582190, EPI_ISL_582191, EPI_ISL_582192, EPI_ISL_582193, EPI_ISL_582194, EPI_ISL_582195, EPI_ISL_582196, EPI_ISL_582197, EPI_ISL_582198, EPI_ISL_582199, EPI_ISL_582200, EPI_ISL_582201, EPI_ISL_582202, EPI_ISL_582203, EPI_ISL_582205, EPI_ISL_582206, EPI_ISL_582207, EPI_ISL_582208, EPI_ISL_582209, EPI_ISL_582210, EPI_ISL_582211, EPI_ISL_582212, EPI_ISL_582213, EPI_ISL_582214, EPI_ISL_582215, EPI_ISL_582216, EPI_ISL_582217, EPI_ISL_582218, EPI_ISL_582219 | | | |
| see above | TriCore Reference Laboratories | Center for Global Health, University of New Mexico Health Sciences Center | Daryl Domman, Kurt Schwalm, Twila Kunde, Joseph Hicks, Michael Edwards, Darrell Dinwiddie |
| EPI_ISL_582220, EPI_ISL_582221, EPI_ISL_582222, EPI_ISL_582223, EPI_ISL_582224, EPI_ISL_582225, EPI_ISL_582226, EPI_ISL_582227, EPI_ISL_582228, EPI_ISL_582229, EPI_ISL_582230, EPI_ISL_582231, EPI_ISL_582232, EPI_ISL_582233, EPI_ISL_582234, EPI_ISL_582235, EPI_ISL_582236, EPI_ISL_582237, EPI_ISL_582238, EPI_ISL_582239, EPI_ISL_582240 | | | |
| see above | Wyoming Public Health Laboratory | Center for Global Health, University of New Mexico Health Sciences Center | Daryl Domman, Kurt Schwalm, Rob Christensen, Wanda Manley, Cari Sloma, Noah Hull, Darrell Dinwiddie |
| EPI_ISL_582812, EPI_ISL_582813, EPI_ISL_582814, EPI_ISL_582815, EPI_ISL_582816, EPI_ISL_582817, EPI_ISL_582818, EPI_ISL_582819, EPI_ISL_582820, EPI_ISL_582821, EPI_ISL_582822, EPI_ISL_582823, EPI_ISL_582824, EPI_ISL_582825, EPI_ISL_582826, EPI_ISL_582827, EPI_ISL_582828, EPI_ISL_582829, EPI_ISL_582830, EPI_ISL_582831, EPI_ISL_582832, EPI_ISL_582843, EPI_ISL_582844 | | | |
| see above | Hospital General Universitario Gregorio Marañón | SeqCOVID-SPAIN consortium/IBV(CSIC) | Darío García de Viedma, Laura Pérez-Lago, Marta Herranz, Jon Sicilia, Julia Suárez, Pilar Catalán, Patricia Muñoz and SeqCOVID-SPAIN consortium |
| EPI_ISL_584165, EPI_ISL_584166 | Quadram Institute Bioscience | COVID-19 Genomics UK (COG-UK) Consortium | Dave J. Baker, Gemma L. Kay, Alp Aydin, Thanh Le-Viet, Steven Rudder, Ana P. Tedim, Anastasia Kolyva, Maria Diaz, Leonardo de Oliveira Martins, Nabil-Fareed Alikhan, Lizzie Meadows, Rachael Stanley, Ngozi Elumogo, Muhammed Yasir, Nicholas M. Thomson, Alexander J Trotter, Rachel Gilroy, Samuel Bloomfield, Claire Stuart, Andrew Bell, Reenesh Prakash, Samir Dervisevic, Alison E. Mather, John Wain, Mark Webber, Andrew J. Page, Justin O'Grady |
| EPI_ISL_584681, EPI_ISL_584682, EPI_ISL_584683, EPI_ISL_584684, EPI_ISL_584685, EPI_ISL_584686, EPI_ISL_584687, EPI_ISL_584688, EPI_ISL_584689, EPI_ISL_584690, EPI_ISL_584691, EPI_ISL_584692, EPI_ISL_584693, EPI_ISL_584694, EPI_ISL_584695, EPI_ISL_584696, EPI_ISL_584697, EPI_ISL_584698, EPI_ISL_584699, EPI_ISL_584700, EPI_ISL_584701, EPI_ISL_584702, EPI_ISL_584703, EPI_ISL_584704, EPI_ISL_584705, EPI_ISL_584706, EPI_ISL_584707, EPI_ISL_584708, EPI_ISL_584709, EPI_ISL_584710, EPI_ISL_584711, EPI_ISL_584712, EPI_ISL_584713, EPI_ISL_584714, EPI_ISL_584715 | | | |
| see above | Northumbria University / South Tees Hospitals NHS Foundation Trust / North Cumbria Integrated Care NHS Foundation Trust / North Tees and Hartlepool NHS Foundation Trust / Newcastle Hospitals NHS Foundation Trust | COVID-19 Genomics UK (COG-UK) Consortium | Darren L Smith,Andrew Nelson,Matthew Bashton,Greg R Young,Joshua Loh,John Allan,Mohammad A Tariq,Giles S Holt,Gary Black,Wen C Yew,Lynn Dover,Paul Baker,Steve Liggett,Sarah Essex,Jane Greenaway,Debra Padgett,Clive Graham,Garren Scott,Edward Barton,Emma Swindells,Brendan Payne,Jennifer Collins,Yusri Taha,Gary Eltringham |
| EPI_ISL_584716, EPI_ISL_584717, EPI_ISL_584718, EPI_ISL_584719, EPI_ISL_584720, EPI_ISL_584721, EPI_ISL_584722, EPI_ISL_584723, EPI_ISL_584724, EPI_ISL_584725, EPI_ISL_584726, EPI_ISL_584727, EPI_ISL_584728, EPI_ISL_584729, EPI_ISL_584730, EPI_ISL_584731, EPI_ISL_584732, EPI_ISL_584733, EPI_ISL_584734, EPI_ISL_584735, EPI_ISL_584736, EPI_ISL_584737, EPI_ISL_584738, EPI_ISL_584739, EPI_ISL_584740, EPI_ISL_584741, EPI_ISL_584742, EPI_ISL_584743, EPI_ISL_584744, EPI_ISL_584745, EPI_ISL_584746, EPI_ISL_584747, EPI_ISL_584748, EPI_ISL_584749 | | | |
| see above | Quadram Institute Bioscience | COVID-19 Genomics UK (COG-UK) Consortium | Dave J. Baker, Gemma L. Kay, Alp Aydin, Thanh Le-Viet, Steven Rudder, Ana P. Tedim, Anastasia Kolyva, Maria Diaz, Leonardo de Oliveira Martins, Nabil-Fareed Alikhan, Lizzie Meadows, Rachael Stanley, Ngozi Elumogo, Muhammed Yasir, Nicholas M. Thomson, Alexander J Trotter, Rachel Gilroy, Samuel Bloomfield, Claire Stuart, Andrew Bell, Reenesh Prakash, Samir Dervisevic, Alison E. Mather, John Wain, Mark Webber, Andrew J. Page, Justin O'Grady |
| EPI_ISL_584847, EPI_ISL_584848, EPI_ISL_584849, EPI_ISL_584850, EPI_ISL_584851, EPI_ISL_584852, EPI_ISL_584853, EPI_ISL_584854, EPI_ISL_584855, EPI_ISL_584856, EPI_ISL_584857, EPI_ISL_584858, EPI_ISL_584859, EPI_ISL_584860, EPI_ISL_584861, EPI_ISL_584862, EPI_ISL_584863, EPI_ISL_584864, EPI_ISL_584865, EPI_ISL_584866, EPI_ISL_584867, EPI_ISL_584868, EPI_ISL_584869, EPI_ISL_584870, EPI_ISL_584871, EPI_ISL_584872, EPI_ISL_584873, EPI_ISL_584874, EPI_ISL_584875, EPI_ISL_584876, EPI_ISL_584877, EPI_ISL_584878, EPI_ISL_584879, EPI_ISL_584880, EPI_ISL_584881, EPI_ISL_584882 | | | |
| see above | Northumbria University / South Tees Hospitals NHS | COVID-19 Genomics UK (COG-UK) Consortium | Darren L Smith,Andrew Nelson,Matthew Bashton,Greg R Young,Joshua Loh,John Allan,Mohammad A Tariq,Giles S Holt,Gary Black,Wen C Yew,Lynn |

| | | | |
|---|---|---|---|
| | Foundation Trust / North Cumbria Integrated Care NHS Foundation Trust / North Tees and Hartlepool NHS Foundation Trust / Newcastle Hospitals NHS Foundation Trust | | Dover,Paul Baker,Steve Liggett,Sarah Essex,Jane Greenaway,Debra Padgett,Clive Graham,Garren Scott,Edward Barton,Emma Swindells,Brendan Payne,Jennifer Collins,Yusri Taha,Gary Eltringham |
| EPI_ISL_584906, EPI_ISL_584907, EPI_ISL_584908, EPI_ISL_584909, EPI_ISL_584910, EPI_ISL_584911, EPI_ISL_584912, EPI_ISL_584913, EPI_ISL_584914, EPI_ISL_584915, EPI_ISL_584916, EPI_ISL_584917, EPI_ISL_584918, EPI_ISL_584919, EPI_ISL_584920, EPI_ISL_584921, EPI_ISL_584922, EPI_ISL_584923, EPI_ISL_584924, EPI_ISL_584925, EPI_ISL_584926, EPI_ISL_584927, EPI_ISL_584928, EPI_ISL_584929, EPI_ISL_584930, EPI_ISL_584931, EPI_ISL_584932, EPI_ISL_584933, EPI_ISL_584934, EPI_ISL_584935, EPI_ISL_584936, EPI_ISL_584937, EPI_ISL_584938, EPI_ISL_584939, EPI_ISL_584940, EPI_ISL_584941, EPI_ISL_584942, EPI_ISL_584943, EPI_ISL_584944, EPI_ISL_584945, EPI_ISL_584946, EPI_ISL_584947, EPI_ISL_584948, EPI_ISL_584949, EPI_ISL_584950, EPI_ISL_584951, EPI_ISL_584952, EPI_ISL_584953, EPI_ISL_584954, EPI_ISL_584955, EPI_ISL_584956, EPI_ISL_584957, EPI_ISL_584958, EPI_ISL_584959, EPI_ISL_584960, EPI_ISL_584961, EPI_ISL_584962, EPI_ISL_584963, EPI_ISL_584964, EPI_ISL_584965, EPI_ISL_584966, EPI_ISL_584967, EPI_ISL_584968, EPI_ISL_584969, EPI_ISL_584970, EPI_ISL_584971, EPI_ISL_584972, EPI_ISL_584973, EPI_ISL_584974, EPI_ISL_584975, EPI_ISL_584976, EPI_ISL_584977, EPI_ISL_584978, EPI_ISL_584979, EPI_ISL_584980, EPI_ISL_584981, EPI_ISL_584982, EPI_ISL_584983, EPI_ISL_584984, EPI_ISL_584985, EPI_ISL_584986, EPI_ISL_584987, EPI_ISL_584988, EPI_ISL_584989, EPI_ISL_584990, EPI_ISL_584991, EPI_ISL_584992, EPI_ISL_584993, EPI_ISL_584994, EPI_ISL_584995, EPI_ISL_584996, EPI_ISL_584997, EPI_ISL_584998, EPI_ISL_584999, EPI_ISL_585000, EPI_ISL_585001, EPI_ISL_585002, EPI_ISL_585003, EPI_ISL_585004, EPI_ISL_585005, EPI_ISL_585006, EPI_ISL_585007, EPI_ISL_585008, EPI_ISL_585260 | | | |
| see above | Quadram Institute Bioscience | COVID-19 Genomics UK (COG-UK) Consortium | Dave J. Baker, Gemma L. Kay, Alp Aydin, Thanh Le-Viet, Steven Rudder, Ana P. Tedim, Anastasia Kolyva, Maria Diaz, Leonardo de Oliveira Martins, Nabil-Fareed Alikhan, Lizzie Meadows, Rachael Stanley, Ngozi Elumogo, Muhammed Yasir, Nicholas M. Thomson, Alexander J Trotter, Rachel Gilroy, Samuel Bloomfield, Claire Stuart, Andrew Bell, Reenesh Prakash, Samir Dervisevic, Alison E. Mather, John Wain, Mark Webber, Andrew J. Page, Justin O'Grady |
| EPI_ISL_585504, EPI_ISL_585505, EPI_ISL_585506, EPI_ISL_585507, EPI_ISL_585508, EPI_ISL_585509, EPI_ISL_585510, EPI_ISL_585511, EPI_ISL_585512, EPI_ISL_585513, EPI_ISL_585514, EPI_ISL_585515, EPI_ISL_585516, EPI_ISL_585517, EPI_ISL_585518, EPI_ISL_585519, EPI_ISL_585520, EPI_ISL_585521, EPI_ISL_585522, EPI_ISL_585523, EPI_ISL_585524, EPI_ISL_585525, EPI_ISL_585526, EPI_ISL_585527, EPI_ISL_585528, EPI_ISL_585529, EPI_ISL_585530, EPI_ISL_585531, EPI_ISL_585532 | | | |
| see above | Northumbria University / South Tees Hospitals NHS Foundation Trust / North Cumbria Integrated Care NHS Foundation Trust / North Tees and Hartlepool NHS Foundation Trust / Newcastle Hospitals NHS Foundation Trust | COVID-19 Genomics UK (COG-UK) Consortium | Darren L Smith,Andrew Nelson,Matthew Bashton,Greg R Young,Joshua Loh,John Allan,Mohammad A Tariq,Giles S Holt,Gary Black,Wen C Yew,Lynn Dover,Paul Baker,Steve Liggett,Sarah Essex,Jane Greenaway,Debra Padgett,Clive Graham,Garren Scott,Edward Barton,Emma Swindells,Brendan Payne,Jennifer Collins,Yusri Taha,Gary Eltringham |
| EPI_ISL_585583, EPI_ISL_585584, EPI_ISL_585585, EPI_ISL_585586, EPI_ISL_585587, EPI_ISL_585588, EPI_ISL_585589, EPI_ISL_585590, EPI_ISL_585591, EPI_ISL_585592, EPI_ISL_585593, EPI_ISL_585594, EPI_ISL_585595, EPI_ISL_585596, EPI_ISL_585597, EPI_ISL_585598, EPI_ISL_585599, EPI_ISL_585600, EPI_ISL_585601, EPI_ISL_585602, EPI_ISL_585603, EPI_ISL_585604, EPI_ISL_585605, EPI_ISL_585606, EPI_ISL_585607, EPI_ISL_585608, EPI_ISL_585609, EPI_ISL_585610, EPI_ISL_585611, EPI_ISL_585612, EPI_ISL_585613, EPI_ISL_585614, EPI_ISL_585615, EPI_ISL_585616, EPI_ISL_585617, EPI_ISL_585618, EPI_ISL_585619, EPI_ISL_586240, EPI_ISL_586241, EPI_ISL_586242 | | | |
| see above | Quadram Institute Bioscience | COVID-19 Genomics UK (COG-UK) Consortium | Dave J. Baker, Gemma L. Kay, Alp Aydin, Thanh Le-Viet, Steven Rudder, Ana P. Tedim, Anastasia Kolyva, Maria Diaz, Leonardo de Oliveira Martins, Nabil-Fareed Alikhan, Lizzie Meadows, Rachael Stanley, Ngozi Elumogo, Muhammed Yasir, Nicholas M. Thomson, Alexander J Trotter, Rachel Gilroy, Samuel Bloomfield, Claire Stuart, Andrew Bell, Reenesh Prakash, Samir Dervisevic, Alison E. Mather, John Wain, Mark Webber, Andrew J. Page, Justin O'Grady |
| EPI_ISL_594987, EPI_ISL_594988, EPI_ISL_594989, EPI_ISL_594990, EPI_ISL_594991, EPI_ISL_594992, EPI_ISL_594993, EPI_ISL_594994, EPI_ISL_594995, EPI_ISL_594996, EPI_ISL_594997, EPI_ISL_594998, EPI_ISL_594999, EPI_ISL_595000, EPI_ISL_595001, EPI_ISL_595002, EPI_ISL_595003, EPI_ISL_595004, EPI_ISL_595005, EPI_ISL_595006, EPI_ISL_595007, EPI_ISL_595008, EPI_ISL_595009, EPI_ISL_595010, EPI_ISL_595011, EPI_ISL_595012, EPI_ISL_595013, EPI_ISL_595014, EPI_ISL_595015, EPI_ISL_595016, EPI_ISL_595017, EPI_ISL_595018, EPI_ISL_595019, EPI_ISL_595020, EPI_ISL_595021, EPI_ISL_595022, EPI_ISL_595023, EPI_ISL_595024, EPI_ISL_595025, EPI_ISL_595026, EPI_ISL_595027, EPI_ISL_595028, EPI_ISL_595029, EPI_ISL_595030, EPI_ISL_595031, EPI_ISL_595032, EPI_ISL_595033, EPI_ISL_595034, EPI_ISL_595035, EPI_ISL_595036, EPI_ISL_595037, EPI_ISL_595038, EPI_ISL_595039, EPI_ISL_595040, EPI_ISL_595041, EPI_ISL_595042, EPI_ISL_595043, EPI_ISL_595044, EPI_ISL_595045 | | | |
| see above | Northumbria University / South Tees Hospitals NHS Foundation Trust / North Cumbria Integrated Care NHS Foundation Trust / North Tees and Hartlepool NHS Foundation Trust / Newcastle Hospitals NHS Foundation Trust | COVID-19 Genomics UK (COG-UK) Consortium | Darren L Smith,Andrew Nelson,Matthew Bashton,Greg R Young,Joshua Loh,John Allan,Mohammad A Tariq,Giles S Holt,Gary Black,Wen C Yew,Lynn Dover,Paul Baker,Steve Liggett,Sarah Essex,Jane Greenaway,Debra Padgett,Clive Graham,Garren Scott,Edward Barton,Emma Swindells,Brendan Payne,Jennifer Collins,Yusri Taha,Gary Eltringham |
| EPI_ISL_595115, EPI_ISL_595116, EPI_ISL_595117, EPI_ISL_595118, EPI_ISL_595119, EPI_ISL_595120, EPI_ISL_595121, EPI_ISL_595122, EPI_ISL_595123, EPI_ISL_595124, EPI_ISL_595125, EPI_ISL_595126, EPI_ISL_595127, EPI_ISL_595128, EPI_ISL_595129, EPI_ISL_595130, EPI_ISL_595131, EPI_ISL_595132, EPI_ISL_595133, EPI_ISL_595134, EPI_ISL_595135, EPI_ISL_595136, EPI_ISL_595137, EPI_ISL_595138, EPI_ISL_595139, EPI_ISL_595140, EPI_ISL_595141, EPI_ISL_595142, EPI_ISL_595143, EPI_ISL_595144, EPI_ISL_595145, EPI_ISL_595146, EPI_ISL_595147, EPI_ISL_595148, EPI_ISL_595149, EPI_ISL_595150, EPI_ISL_595151, EPI_ISL_595152, EPI_ISL_595153, EPI_ISL_595154, EPI_ISL_595155, EPI_ISL_595156, EPI_ISL_595157, EPI_ISL_595158, EPI_ISL_595159, EPI_ISL_595160, EPI_ISL_595161, EPI_ISL_595162, EPI_ISL_595163, EPI_ISL_595164, EPI_ISL_595165, EPI_ISL_595166, EPI_ISL_595167, EPI_ISL_595168, EPI_ISL_595169, EPI_ISL_595170, EPI_ISL_595171, EPI_ISL_595172, EPI_ISL_595173, EPI_ISL_595174, EPI_ISL_595175, EPI_ISL_595176, EPI_ISL_595177, EPI_ISL_595178, EPI_ISL_595179, EPI_ISL_595180, EPI_ISL_595181, EPI_ISL_595182, EPI_ISL_595183, EPI_ISL_595184, EPI_ISL_595185, EPI_ISL_595186, EPI_ISL_595187, EPI_ISL_595188, EPI_ISL_595189, EPI_ISL_595190, EPI_ISL_595191, EPI_ISL_595192, EPI_ISL_595193, EPI_ISL_595194, EPI_ISL_595195, EPI_ISL_595196, EPI_ISL_595197, EPI_ISL_595198, EPI_ISL_595199, EPI_ISL_595200, EPI_ISL_595201, EPI_ISL_595202, EPI_ISL_595203, EPI_ISL_595204, EPI_ISL_595205, EPI_ISL_595206, EPI_ISL_595207, EPI_ISL_595208, EPI_ISL_595221, EPI_ISL_595223, EPI_ISL_595225, EPI_ISL_595226, EPI_ISL_595229, EPI_ISL_595837, EPI_ISL_595838, EPI_ISL_595839, EPI_ISL_595840, EPI_ISL_595841, EPI_ISL_595842, EPI_ISL_595843, EPI_ISL_595844, EPI_ISL_595845, EPI_ISL_595846, EPI_ISL_595847, EPI_ISL_595848, EPI_ISL_595849, EPI_ISL_595850, EPI_ISL_595851, EPI_ISL_595852, EPI_ISL_595853, EPI_ISL_595854, EPI_ISL_595855, EPI_ISL_595856, EPI_ISL_595857, EPI_ISL_595858, EPI_ISL_595859, EPI_ISL_595860, EPI_ISL_595861, EPI_ISL_595862, EPI_ISL_595863, EPI_ISL_595864, EPI_ISL_595865, EPI_ISL_595866, EPI_ISL_595867, EPI_ISL_595868, EPI_ISL_595869, EPI_ISL_595870, EPI_ISL_595871, EPI_ISL_595872, EPI_ISL_595873, EPI_ISL_595874, EPI_ISL_595875, EPI_ISL_595876, EPI_ISL_595877, EPI_ISL_595878, EPI_ISL_595879, EPI_ISL_595880, EPI_ISL_595881, EPI_ISL_595882, EPI_ISL_595883, EPI_ISL_595884, EPI_ISL_595885, EPI_ISL_595886, EPI_ISL_595887, EPI_ISL_595888, EPI_ISL_595889, EPI_ISL_595890, EPI_ISL_595891, EPI_ISL_595892, EPI_ISL_595893, EPI_ISL_595894, EPI_ISL_595895, EPI_ISL_595896, EPI_ISL_595897, EPI_ISL_595898, EPI_ISL_595899, EPI_ISL_595900, EPI_ISL_595901, EPI_ISL_595902, EPI_ISL_595903, EPI_ISL_595904, EPI_ISL_595905, EPI_ISL_595906, EPI_ISL_595907, EPI_ISL_595908, EPI_ISL_595909, EPI_ISL_595910, EPI_ISL_595911, EPI_ISL_595912, EPI_ISL_595913, EPI_ISL_595914, EPI_ISL_595915, EPI_ISL_595916, EPI_ISL_595917, EPI_ISL_595918, EPI_ISL_595919, EPI_ISL_595920, EPI_ISL_595921, EPI_ISL_595922, EPI_ISL_595923, EPI_ISL_595924, EPI_ISL_595925, EPI_ISL_595926, EPI_ISL_595927, EPI_ISL_595928, EPI_ISL_595929, EPI_ISL_595930, EPI_ISL_595931, EPI_ISL_595932, EPI_ISL_595933, EPI_ISL_595934, EPI_ISL_595935, EPI_ISL_595936, EPI_ISL_595937, EPI_ISL_595938, EPI_ISL_595939, EPI_ISL_595940, EPI_ISL_595941, EPI_ISL_595942, EPI_ISL_595943, EPI_ISL_595944, EPI_ISL_595945, EPI_ISL_595946, EPI_ISL_595947, EPI_ISL_595948, EPI_ISL_595949, EPI_ISL_595950, EPI_ISL_595951, EPI_ISL_595952, EPI_ISL_595953, EPI_ISL_595954, EPI_ISL_595955, EPI_ISL_595956, EPI_ISL_595957, EPI_ISL_595958, EPI_ISL_595959, EPI_ISL_595960, EPI_ISL_595961, EPI_ISL_595962, EPI_ISL_595963, EPI_ISL_595964, EPI_ISL_595965, EPI_ISL_595966, EPI_ISL_595967, EPI_ISL_595968, EPI_ISL_595969, EPI_ISL_595970, EPI_ISL_595971, EPI_ISL_595972, EPI_ISL_595973, EPI_ISL_595974, EPI_ISL_595975, EPI_ISL_595976, EPI_ISL_595977, EPI_ISL_595978, EPI_ISL_595979, EPI_ISL_595980, EPI_ISL_595981, EPI_ISL_595982, EPI_ISL_595983, EPI_ISL_595984, EPI_ISL_595985, EPI_ISL_595986, EPI_ISL_595987, EPI_ISL_595988, EPI_ISL_595989, EPI_ISL_595990, EPI_ISL_595991, EPI_ISL_595992, EPI_ISL_595993, EPI_ISL_595994, EPI_ISL_595995, EPI_ISL_595996, EPI_ISL_595997, EPI_ISL_595998, EPI_ISL_595999, EPI_ISL_596000, EPI_ISL_596001, EPI_ISL_596002, EPI_ISL_596003, EPI_ISL_596004, EPI_ISL_596005, EPI_ISL_596006, EPI_ISL_596007, EPI_ISL_596008, EPI_ISL_596009, EPI_ISL_596010, EPI_ISL_596011, EPI_ISL_596012, EPI_ISL_596013, EPI_ISL_596014, EPI_ISL_596015, EPI_ISL_596016, EPI_ISL_596017, EPI_ISL_596018, EPI_ISL_596019, EPI_ISL_596020, EPI_ISL_596021, EPI_ISL_596022, EPI_ISL_596023, EPI_ISL_596024, EPI_ISL_596025, EPI_ISL_596026, EPI_ISL_596027, EPI_ISL_596028, EPI_ISL_596029, EPI_ISL_596030, EPI_ISL_596031, EPI_ISL_596032, EPI_ISL_596033, EPI_ISL_596034, EPI_ISL_596035, EPI_ISL_596036, EPI_ISL_596037, EPI_ISL_596038, EPI_ISL_596039, EPI_ISL_596040, EPI_ISL_596041, EPI_ISL_596042, EPI_ISL_596043, EPI_ISL_596044, EPI_ISL_596045, EPI_ISL_596046, EPI_ISL_596047, EPI_ISL_596048, EPI_ISL_596049, EPI_ISL_596050, EPI_ISL_596051, EPI_ISL_596052, EPI_ISL_596053, EPI_ISL_596054, EPI_ISL_596055, EPI_ISL_596056, EPI_ISL_596057, EPI_ISL_596058, EPI_ISL_596059, EPI_ISL_596060, EPI_ISL_596061, EPI_ISL_596062, EPI_ISL_596063, EPI_ISL_596064, EPI_ISL_596065, EPI_ISL_596066, EPI_ISL_596067, EPI_ISL_596068, EPI_ISL_596069, EPI_ISL_596070, EPI_ISL_596071, EPI_ISL_596072, EPI_ISL_596073, EPI_ISL_596074, EPI_ISL_596075, EPI_ISL_596076, EPI_ISL_596077, EPI_ISL_596078, EPI_ISL_596079, EPI_ISL_596080, EPI_ISL_596081, EPI_ISL_596082, EPI_ISL_596083, EPI_ISL_596084, EPI_ISL_596085, EPI_ISL_596086, EPI_ISL_596087, EPI_ISL_596088, EPI_ISL_596089, EPI_ISL_596090, EPI_ISL_596091, EPI_ISL_596092, EPI_ISL_596093, EPI_ISL_596094, EPI_ISL_596095, EPI_ISL_596096, EPI_ISL_596097, EPI_ISL_596098, EPI_ISL_596099, EPI_ISL_596100, EPI_ISL_596101, EPI_ISL_596102, EPI_ISL_596103, EPI_ISL_596104, EPI_ISL_596105, EPI_ISL_596106, EPI_ISL_596107, EPI_ISL_596108, EPI_ISL_596109, EPI_ISL_596110, EPI_ISL_596111, EPI_ISL_596112, EPI_ISL_596113, EPI_ISL_596114, EPI_ISL_596115, EPI_ISL_596116, EPI_ISL_596117, EPI_ISL_596118, EPI_ISL_596119, EPI_ISL_596120, EPI_ISL_596121, EPI_ISL_596122, EPI_ISL_596123, EPI_ISL_596124, EPI_ISL_596125, EPI_ISL_596126, EPI_ISL_596127, EPI_ISL_596128, EPI_ISL_596129, EPI_ISL_596130, EPI_ISL_596131, EPI_ISL_596132, EPI_ISL_596133, EPI_ISL_596134, EPI_ISL_596135, EPI_ISL_596136, EPI_ISL_596137, EPI_ISL_596138, EPI_ISL_596139, EPI_ISL_596140, EPI_ISL_596141, EPI_ISL_596142, EPI_ISL_596143, EPI_ISL_596144, EPI_ISL_596145, EPI_ISL_596146, EPI_ISL_596147, EPI_ISL_596148, EPI_ISL_596149, EPI_ISL_596150, EPI_ISL_596151, EPI_ISL_596152, EPI_ISL_596153, EPI_ISL_596154, EPI_ISL_596155, EPI_ISL_596156, EPI_ISL_596157, EPI_ISL_596158, EPI_ISL_596159, EPI_ISL_596160, EPI_ISL_596161, EPI_ISL_596162, EPI_ISL_596163, EPI_ISL_596164, EPI_ISL_596165, EPI_ISL_596166, EPI_ISL_596167, EPI_ISL_596168, EPI_ISL_596169, EPI_ISL_596170, EPI_ISL_596171, EPI_ISL_596172, EPI_ISL_596173, EPI_ISL_596174, EPI_ISL_596175, EPI_ISL_596176, EPI_ISL_596177, EPI_ISL_596178, EPI_ISL_596179, EPI_ISL_596180, EPI_ISL_596181, EPI_ISL_596182, EPI_ISL_596183, EPI_ISL_596184, EPI_ISL_596185, EPI_ISL_596186, EPI_ISL_596187, EPI_ISL_596188, EPI_ISL_596189, EPI_ISL_596190 | | | |
| see above | Quadram Institute Bioscience | COVID-19 Genomics UK (COG-UK) Consortium | Dave J. Baker, Gemma L. Kay, Alp Aydin, Thanh Le-Viet, Steven Rudder, Ana P. Tedim, Anastasia Kolyva, Maria Diaz, Leonardo de Oliveira Martins, Nabil-Fareed Alikhan, Lizzie Meadows, Rachael Stanley, Ngozi Elumogo, Muhammed Yasir, Nicholas M. Thomson, Alexander J Trotter, Rachel Gilroy, Samuel Bloomfield, Claire Stuart, Andrew Bell, Reenesh Prakash, Samir Dervisevic, Alison E. Mather, John Wain, Mark Webber, Andrew J. Page, Justin O'Grady |
| EPI_ISL_596387 | Seattle Flu Study | Seattle Flu Study | Deborah A. Nickerson, Chris D. Frazar, Jover Lee, Benjamin Pelle, Matthew Richardson, Amanda Adler, Elisabeth Brandstetter, Peter D. Han, Kairsten Fay, Misja Ilcisin, Kirsten Lacombe, Thomas R. Sibley, Melissa Truong, Caitlin R. Wolf, Michael Boeckh, Janet A. Englund, Michael Famulare, Barry R. Lutz, Mark J. Rieder, Lea M. Starita, Matthew Thompson, Jay Shendure, Trevor Bedford, Helen Y. Chu |
| EPI_ISL_596388, EPI_ISL_596389, EPI_ISL_596390, EPI_ISL_596391, EPI_ISL_596392 | Seattle Flu Study | Seattle Flu Study | Deborah A. Nickerson, Chris D. Frazar, Jover Lee, Benjamin Pelle, Matthew Richardson, Amanda Adler, Elisabeth Brandstetter, Peter D. Han, Kairsten Fay, Misja Ilcisin, Kirsten Lacombe, Thomas R. Sibley, Melissa Truong, Caitlin R. Wolf, Karen Cowgill, Stephanie Schrag, Jeff Duchin, Michael Boeckh, Janet A. Englund, Michael Famulare, Barry R. Lutz, Mark J. Rieder, Lea M. Starita, Matthew Thompson, Helen Y. Chu, Trevor Bedford, Jay Shendure |
| EPI_ISL_596393 | Seattle Flu Study | Seattle Flu Study | Deborah A. Nickerson, Chris D. Frazar, Jover Lee, Benjamin Pelle, Matthew Richardson, Amanda Adler, Elisabeth Brandstetter, Peter D. Han, Kairsten Fay, Misja Ilcisin, Kirsten Lacombe, Thomas R. Sibley, Melissa Truong, Caitlin R. Wolf, Michael Boeckh, Janet A. Englund, Michael Famulare, Barry R. Lutz, Mark J. Rieder, Lea M. Starita, Matthew Thompson, Jay Shendure, Trevor Bedford, Helen Y. Chu |
| EPI_ISL_596394, EPI_ISL_596395, EPI_ISL_596396, EPI_ISL_596397, EPI_ISL_596398, EPI_ISL_596399, EPI_ISL_596400, EPI_ISL_596401, EPI_ISL_596402, EPI_ISL_596403, EPI_ISL_596404, EPI_ISL_596405, EPI_ISL_596406, EPI_ISL_596407, EPI_ISL_596408, EPI_ISL_596409 | | | |
| see above | Seattle Flu Study | Seattle Flu Study | Deborah A. Nickerson, Chris D. Frazar, Jover Lee, Benjamin Pelle, Matthew Richardson, Amanda Adler, Elisabeth Brandstetter, Peter D. Han, Kairsten Fay, Misja Ilcisin, Kirsten Lacombe, Thomas R. Sibley, Melissa Truong, Caitlin R. Wolf, Karen Cowgill, Stephanie Schrag, Jeff Duchin, Michael Boeckh, Janet A. Englund, Michael Famulare, Barry R. Lutz, Mark J. Rieder, Lea M. Starita, Matthew Thompson, Helen Y. Chu, Trevor Bedford, Jay Shendure |
| EPI_ISL_596410 | Seattle Flu Study | Seattle Flu Study | Deborah A. Nickerson, Chris D. Frazar, Jover Lee, Benjamin Pelle, Matthew Richardson, Amanda Adler, Elisabeth Brandstetter, Peter D. Han, Kairsten Fay, Misja Ilcisin, Kirsten Lacombe, Thomas R. Sibley, Melissa Truong, Caitlin R. Wolf, Michael Boeckh, Janet A. Englund, Michael Famulare, Barry R. |

| | | | |
|---|---|---|---|
| | | | Lutz, Mark J. Rieder, Lea M. Starita, Matthew Thompson, Jay Shendure, Trevor Bedford, Helen Y. Chu |
| EPI_ISL_596411 | Seattle Flu Study | Seattle Flu Study | Deborah A. Nickerson, Chris D. Frazar, Jover Lee, Benjamin Pelle, Matthew Richardson, Amanda Adler, Elisabeth Brandstetter, Peter D. Han, Kairsten Fay, Misja Ilcisin, Kirsten Lacombe, Thomas R. Sibley, Melissa Truong, Caitlin R. Wolf, Karen Cowgill, Stephanie Schrag, Jeff Duchin, Michael Boeckh, Janet A. Englund, Michael Famulare, Barry R. Lutz, Mark J. Rieder, Lea M. Starita, Matthew Thompson, Helen Y. Chu, Trevor Bedford, Jay Shendure |
| EPI_ISL_596412, EPI_ISL_596413, EPI_ISL_596414 | Seattle Flu Study | Seattle Flu Study | Deborah A. Nickerson, Chris D. Frazar, Jover Lee, Benjamin Pelle, Matthew Richardson, Amanda Adler, Elisabeth Brandstetter, Peter D. Han, Kairsten Fay, Misja Ilcisin, Kirsten Lacombe, Thomas R. Sibley, Melissa Truong, Caitlin R. Wolf, Michael Boeckh, Janet A. Englund, Michael Famulare, Barry R. Lutz, Mark J. Rieder, Lea M. Starita, Matthew Thompson, Jay Shendure, Trevor Bedford, Helen Y. Chu |
| EPI_ISL_596415, EPI_ISL_596416 | Seattle Flu Study | Seattle Flu Study | Deborah A. Nickerson, Chris D. Frazar, Jover Lee, Benjamin Pelle, Matthew Richardson, Amanda Adler, Elisabeth Brandstetter, Peter D. Han, Kairsten Fay, Misja Ilcisin, Kirsten Lacombe, Thomas R. Sibley, Melissa Truong, Caitlin R. Wolf, Karen Cowgill, Stephanie Schrag, Jeff Duchin, Michael Boeckh, Janet A. Englund, Michael Famulare, Barry R. Lutz, Mark J. Rieder, Lea M. Starita, Matthew Thompson, Helen Y. Chu, Trevor Bedford, Jay Shendure |
| EPI_ISL_596417, EPI_ISL_596418, EPI_ISL_596419, EPI_ISL_596420 | Seattle Flu Study | Seattle Flu Study | Deborah A. Nickerson, Chris D. Frazar, Jover Lee, Benjamin Pelle, Matthew Richardson, Amanda Adler, Elisabeth Brandstetter, Peter D. Han, Kairsten Fay, Misja Ilcisin, Kirsten Lacombe, Thomas R. Sibley, Melissa Truong, Caitlin R. Wolf, Michael Boeckh, Janet A. Englund, Michael Famulare, Barry R. Lutz, Mark J. Rieder, Lea M. Starita, Matthew Thompson, Jay Shendure, Trevor Bedford, Helen Y. Chu |
| EPI_ISL_596421, EPI_ISL_596422, EPI_ISL_596423, EPI_ISL_596424, EPI_ISL_596425 | Seattle Flu Study | Seattle Flu Study | Deborah A. Nickerson, Chris D. Frazar, Jover Lee, Benjamin Pelle, Matthew Richardson, Amanda Adler, Elisabeth Brandstetter, Peter D. Han, Kairsten Fay, Misja Ilcisin, Kirsten Lacombe, Thomas R. Sibley, Melissa Truong, Caitlin R. Wolf, Karen Cowgill, Stephanie Schrag, Jeff Duchin, Michael Boeckh, Janet A. Englund, Michael Famulare, Barry R. Lutz, Mark J. Rieder, Lea M. Starita, Matthew Thompson, Helen Y. Chu, Trevor Bedford, Jay Shendure |
| EPI_ISL_596426, EPI_ISL_596427, EPI_ISL_596428, EPI_ISL_596429, EPI_ISL_596430, EPI_ISL_596431, EPI_ISL_596432, EPI_ISL_596433, EPI_ISL_596434, EPI_ISL_596435, EPI_ISL_596436 | | | |
| see above | Seattle Flu Study | Seattle Flu Study | Deborah A. Nickerson, Chris D. Frazar, Jover Lee, Benjamin Pelle, Matthew Richardson, Amanda Adler, Elisabeth Brandstetter, Peter D. Han, Kairsten Fay, Misja Ilcisin, Kirsten Lacombe, Thomas R. Sibley, Melissa Truong, Caitlin R. Wolf, Michael Boeckh, Janet A. Englund, Michael Famulare, Barry R. Lutz, Mark J. Rieder, Lea M. Starita, Matthew Thompson, Jay Shendure, Trevor Bedford, Helen Y. Chu |
| EPI_ISL_596437, EPI_ISL_596438, EPI_ISL_596439 | Seattle Flu Study | Seattle Flu Study | Deborah A. Nickerson, Chris D. Frazar, Jover Lee, Benjamin Pelle, Matthew Richardson, Amanda Adler, Elisabeth Brandstetter, Peter D. Han, Kairsten Fay, Misja Ilcisin, Kirsten Lacombe, Thomas R. Sibley, Melissa Truong, Caitlin R. Wolf, Karen Cowgill, Stephanie Schrag, Jeff Duchin, Michael Boeckh, Janet A. Englund, Michael Famulare, Barry R. Lutz, Mark J. Rieder, Lea M. Starita, Matthew Thompson, Helen Y. Chu, Trevor Bedford, Jay Shendure |
| EPI_ISL_596440, EPI_ISL_596441, EPI_ISL_596442, EPI_ISL_596443, EPI_ISL_596444, EPI_ISL_596445 | Seattle Flu Study | Seattle Flu Study | Deborah A. Nickerson, Chris D. Frazar, Jover Lee, Benjamin Pelle, Matthew Richardson, Amanda Adler, Elisabeth Brandstetter, Peter D. Han, Kairsten Fay, Misja Ilcisin, Kirsten Lacombe, Thomas R. Sibley, Melissa Truong, Caitlin R. Wolf, Michael Boeckh, Janet A. Englund, Michael Famulare, Barry R. Lutz, Mark J. Rieder, Lea M. Starita, Matthew Thompson, Jay Shendure, Trevor Bedford, Helen Y. Chu |
| EPI_ISL_596446 | Seattle Flu Study | Seattle Flu Study | Deborah A. Nickerson, Chris D. Frazar, Jover Lee, Benjamin Pelle, Matthew Richardson, Amanda Adler, Elisabeth Brandstetter, Peter D. Han, Kairsten Fay, Misja Ilcisin, Kirsten Lacombe, Thomas R. Sibley, Melissa Truong, Caitlin R. Wolf, Karen Cowgill, Stephanie Schrag, Jeff Duchin, Michael Boeckh, Janet A. Englund, Michael Famulare, Barry R. Lutz, Mark J. Rieder, Lea M. Starita, Matthew Thompson, Helen Y. Chu, Trevor Bedford, Jay Shendure |
| EPI_ISL_602282, EPI_ISL_602283, EPI_ISL_602284, EPI_ISL_602285, EPI_ISL_602286, EPI_ISL_602287, EPI_ISL_602288, EPI_ISL_602289, EPI_ISL_602290, EPI_ISL_602291, EPI_ISL_602292, EPI_ISL_602293, EPI_ISL_602294, EPI_ISL_602295, EPI_ISL_602296, EPI_ISL_602297, EPI_ISL_602298, EPI_ISL_602299, EPI_ISL_602300, EPI_ISL_602301, EPI_ISL_602302, EPI_ISL_602303 | | | |
| see above | Evangelisches Klinikum Bethel, Institut für Laboratoriumsmedizin, Mikrobiologie und Hygiene | Bielefeld University | David Brandt, Tobias Busche, Markus Haak, Jörn Kalinowski, Levin-Joe Klages, Christiane Scherer, Alexander Sczyrba, Marina Simunovic, Svenja Vinke |
| EPI_ISL_602559, EPI_ISL_602560, EPI_ISL_602561 | Alaska State Virology Laboratory | Alaska State Virology Laboratory | DeRonde,S., Deuling,H., Chen,J. |
| EPI_ISL_610157, EPI_ISL_610159, EPI_ISL_610160 | Washington University in St. Louis | Washington University in St. Louis | David Wang, Carey-Ann Burnham, Bijal Parikh, Scott Handley, Lindsay Droit, Stephen Tahan |
| EPI_ISL_610250 | New Mexico Department of Health Scientific Laboratory Division | Center for Global Health, University of New Mexico Health Sciences Center | Daryl Domman, Kurt Schwalm, Twila Kunde, Joseph Hicks, Michael Edwards, Darrell Dinwiddie |
| EPI_ISL_610251, EPI_ISL_610252 | TriCore Reference Laboratories | Center for Global Health, University of New Mexico Health Sciences Center | Daryl Domman, Kurt Schwalm, Twila Kunde, Joseph Hicks, Michael Edwards, Darrell Dinwiddie |
| EPI_ISL_610253 | New Mexico Department of Health Scientific Laboratory Division | Center for Global Health, University of New Mexico Health Sciences Center | Daryl Domman, Kurt Schwalm, Twila Kunde, Joseph Hicks, Michael Edwards, Darrell Dinwiddie |
| EPI_ISL_610254, EPI_ISL_610255, EPI_ISL_610256, EPI_ISL_610257, EPI_ISL_610258, EPI_ISL_610259, EPI_ISL_610260, EPI_ISL_610261, EPI_ISL_610262, EPI_ISL_610263, EPI_ISL_610264, EPI_ISL_610265, EPI_ISL_610266, EPI_ISL_610267 | | | |
| see above | TriCore Reference Laboratories | Center for Global Health, University of New Mexico Health Sciences Center | Daryl Domman, Kurt Schwalm, Twila Kunde, Joseph Hicks, Michael Edwards, Darrell Dinwiddie |
| EPI_ISL_610268, EPI_ISL_610269, EPI_ISL_610270, EPI_ISL_610271, EPI_ISL_610272, EPI_ISL_610273, EPI_ISL_610274, EPI_ISL_610275, EPI_ISL_610276, EPI_ISL_610277, EPI_ISL_610278, EPI_ISL_610279, EPI_ISL_610280, EPI_ISL_610281 | | | |
| see above | New Mexico Department of Health Scientific Laboratory Division | Center for Global Health, University of New Mexico Health Sciences Center | Daryl Domman, Kurt Schwalm, Twila Kunde, Joseph Hicks, Michael Edwards, Darrell Dinwiddie |
| EPI_ISL_610282 | TriCore Reference Laboratories | Center for Global Health, University of New Mexico Health Sciences Center | Daryl Domman, Kurt Schwalm, Twila Kunde, Joseph Hicks, Michael Edwards, Darrell Dinwiddie |
| EPI_ISL_611524, EPI_ISL_611572, EPI_ISL_611574, EPI_ISL_611576, EPI_ISL_611577, EPI_ISL_611591, EPI_ISL_611643, EPI_ISL_611644, EPI_ISL_611645, EPI_ISL_611646, EPI_ISL_611647, EPI_ISL_611648, EPI_ISL_611649, EPI_ISL_611657, EPI_ISL_611668, EPI_ISL_611746, EPI_ISL_611747, EPI_ISL_611752, EPI_ISL_611756, EPI_ISL_611757, EPI_ISL_611758, EPI_ISL_611759, EPI_ISL_611760, EPI_ISL_611766, EPI_ISL_611767, EPI_ISL_611768, EPI_ISL_611769, EPI_ISL_611770, EPI_ISL_611839, EPI_ISL_611846, EPI_ISL_611864, EPI_ISL_611868, EPI_ISL_611917, EPI_ISL_611930, EPI_ISL_611941, EPI_ISL_611942, EPI_ISL_611943, EPI_ISL_611944, EPI_ISL_611945, EPI_ISL_611946, EPI_ISL_611947, EPI_ISL_611948, EPI_ISL_611949, EPI_ISL_611950, EPI_ISL_611951, EPI_ISL_611952, EPI_ISL_611953, EPI_ISL_611980, EPI_ISL_612008, EPI_ISL_612026, EPI_ISL_612027, EPI_ISL_612030, EPI_ISL_612054, EPI_ISL_612085, EPI_ISL_612090, EPI_ISL_612097, EPI_ISL_612100, EPI_ISL_612104, EPI_ISL_612110, EPI_ISL_612134, EPI_ISL_612523, EPI_ISL_612524, EPI_ISL_612525, EPI_ISL_612526, EPI_ISL_612527, EPI_ISL_612528, EPI_ISL_612529, EPI_ISL_612530, EPI_ISL_612531, EPI_ISL_612532, EPI_ISL_612533, EPI_ISL_612534, EPI_ISL_612535, EPI_ISL_612536, EPI_ISL_612537, EPI_ISL_612538, EPI_ISL_612539, EPI_ISL_612540, EPI_ISL_612541, EPI_ISL_612542, EPI_ISL_612543, EPI_ISL_612544, EPI_ISL_612545, EPI_ISL_612549, EPI_ISL_612550, EPI_ISL_612551, EPI_ISL_612552, EPI_ISL_612553, EPI_ISL_612554, EPI_ISL_612555, EPI_ISL_612556, EPI_ISL_612557, EPI_ISL_612558, EPI_ISL_612559, EPI_ISL_612560, EPI_ISL_612561, EPI_ISL_612562, EPI_ISL_612563, EPI_ISL_612564, EPI_ISL_612565, EPI_ISL_612566, EPI_ISL_612567, EPI_ISL_612568, EPI_ISL_612569, EPI_ISL_612570 | | | |
| see above | Northumbria University / South Tees Hospitals NHS Foundation Trust / North Cumbria Integrated Care NHS Foundation Trust / North Tees and Hartlepool NHS Foundation Trust / Newcastle Hospitals NHS Foundation Trust | COVID-19 Genomics UK (COG-UK) Consortium | Darren L Smith,Andrew Nelson,Matthew Bashton,Greg R Young,Joshua Loh,John Allan,Mohammad A Tariq,Giles S Holt,Gary Black,Wen C Yew,Lynn Dover,Paul Baker,Steve Liggett,Sarah Essex,Jane Greenaway,Debra Padgett,Clive Graham,Garren Scott,Edward Barton,Emma Swindells,Brendan Payne,Jennifer Collins,Yusri Taha,Gary Eltringham |
| EPI_ISL_614248, EPI_ISL_614249, EPI_ISL_614250, EPI_ISL_614251, EPI_ISL_614252, EPI_ISL_614253, EPI_ISL_614254, EPI_ISL_614255, EPI_ISL_614256, EPI_ISL_614257, EPI_ISL_614258, EPI_ISL_614259, EPI_ISL_614260, EPI_ISL_614261, EPI_ISL_614262, EPI_ISL_614263, EPI_ISL_614264 | | | |
| see above | Wyoming Public Health Laboratory | Center for Global Health, University of New Mexico Health Sciences Center | Daryl Domman, Kurt Schwalm, Rob Christensen, Wanda Manley, Cari Sloma, Noah Hull, Darrell Dinwiddie |
| EPI_ISL_614403, EPI_ISL_614406, EPI_ISL_614408, EPI_ISL_614412, EPI_ISL_614413, EPI_ISL_614417, EPI_ISL_614418, EPI_ISL_614419, EPI_ISL_614420, EPI_ISL_614421, EPI_ISL_614424, EPI_ISL_614425, EPI_ISL_614426, EPI_ISL_614427, EPI_ISL_614432, EPI_ISL_614433, EPI_ISL_614434, EPI_ISL_614439, EPI_ISL_614440, EPI_ISL_614443, EPI_ISL_614444, EPI_ISL_614447, EPI_ISL_614448, EPI_ISL_614450, EPI_ISL_614451, EPI_ISL_614452, EPI_ISL_614453, EPI_ISL_614454, EPI_ISL_614455, EPI_ISL_614456, EPI_ISL_614457, EPI_ISL_614458, EPI_ISL_614459, EPI_ISL_614460, EPI_ISL_614473, EPI_ISL_614474, EPI_ISL_614475, EPI_ISL_614476, EPI_ISL_614477, EPI_ISL_614478, EPI_ISL_614479, EPI_ISL_614480, EPI_ISL_614481, EPI_ISL_614482, EPI_ISL_614483, EPI_ISL_614484, EPI_ISL_614485, EPI_ISL_614486, EPI_ISL_614487, EPI_ISL_614488, EPI_ISL_614489, EPI_ISL_614490, EPI_ISL_614491, EPI_ISL_614492, EPI_ISL_614493, EPI_ISL_614494, EPI_ISL_614495, EPI_ISL_614496, EPI_ISL_614497, EPI_ISL_614498, EPI_ISL_614499, EPI_ISL_614500, EPI_ISL_614501, EPI_ISL_614502, EPI_ISL_614503, EPI_ISL_614504, EPI_ISL_614505, EPI_ISL_614506, EPI_ISL_614507, EPI_ISL_614508, EPI_ISL_614509, EPI_ISL_614510, EPI_ISL_614511, EPI_ISL_614512, EPI_ISL_614513, EPI_ISL_614514, EPI_ISL_614522, EPI_ISL_614538, EPI_ISL_614539, EPI_ISL_614540, EPI_ISL_614541, EPI_ISL_614543, EPI_ISL_614544, EPI_ISL_614545, EPI_ISL_614546, EPI_ISL_614564, EPI_ISL_614566, EPI_ISL_614569, EPI_ISL_614585, EPI_ISL_614588, EPI_ISL_614589, EPI_ISL_614593, EPI_ISL_614594, EPI_ISL_614599, EPI_ISL_614602, EPI_ISL_614607, EPI_ISL_614608, EPI_ISL_614615, EPI_ISL_614616, EPI_ISL_614620, EPI_ISL_614671, EPI_ISL_614672, EPI_ISL_614673, EPI_ISL_614674, EPI_ISL_614675, EPI_ISL_614676, EPI_ISL_614677, EPI_ISL_614678, EPI_ISL_614679, EPI_ISL_614680, EPI_ISL_614681, EPI_ISL_614682, EPI_ISL_614683, EPI_ISL_614684, EPI_ISL_614685, EPI_ISL_614686, EPI_ISL_614687, EPI_ISL_614688, EPI_ISL_614689, EPI_ISL_614690, EPI_ISL_614691, EPI_ISL_614692, EPI_ISL_614693, EPI_ISL_614694, EPI_ISL_614697, EPI_ISL_614698, EPI_ISL_614699, EPI_ISL_614700, EPI_ISL_614701, EPI_ISL_614702, EPI_ISL_614703, EPI_ISL_614704, EPI_ISL_614705, EPI_ISL_614706, EPI_ISL_614707, EPI_ISL_614708, EPI_ISL_614709, EPI_ISL_614710, EPI_ISL_614711, EPI_ISL_614712, EPI_ISL_614714, EPI_ISL_614715, EPI_ISL_614716, EPI_ISL_614717, EPI_ISL_614718, EPI_ISL_614719, EPI_ISL_614720, EPI_ISL_614721, EPI_ISL_614722, EPI_ISL_614723, EPI_ISL_614724, EPI_ISL_614725, EPI_ISL_614726, EPI_ISL_614727, EPI_ISL_614728, EPI_ISL_614729, EPI_ISL_614730, EPI_ISL_614731, EPI_ISL_614732, EPI_ISL_614734, EPI_ISL_614735, EPI_ISL_614736, EPI_ISL_614737, EPI_ISL_614738, EPI_ISL_614739, EPI_ISL_614740, EPI_ISL_614741, EPI_ISL_614742, EPI_ISL_614743, EPI_ISL_614744, EPI_ISL_614745, EPI_ISL_614746, EPI_ISL_614747, EPI_ISL_614748, EPI_ISL_614749, EPI_ISL_614750, EPI_ISL_614751, EPI_ISL_614752, EPI_ISL_614754, EPI_ISL_614755, | | | |

EPI_ISL_614756, EPI_ISL_614757, EPI_ISL_614758, EPI_ISL_614761, EPI_ISL_614764, EPI_ISL_614768, EPI_ISL_614770, EPI_ISL_614772, EPI_ISL_614774, EPI_ISL_614775, EPI_ISL_614776, EPI_ISL_614777, EPI_ISL_614778, EPI_ISL_614779, EPI_ISL_614780, EPI_ISL_614781, EPI_ISL_614782, EPI_ISL_614783,
EPI_ISL_614785, EPI_ISL_614786, EPI_ISL_614787, EPI_ISL_614788, EPI_ISL_614789, EPI_ISL_614790, EPI_ISL_614791, EPI_ISL_614792, EPI_ISL_614793, EPI_ISL_614794, EPI_ISL_614795, EPI_ISL_614798, EPI_ISL_614799, EPI_ISL_614800, EPI_ISL_614801, EPI_ISL_614805, EPI_ISL_614806, EPI_ISL_614807,
EPI_ISL_614808, EPI_ISL_614809, EPI_ISL_614811, EPI_ISL_614812, EPI_ISL_614813, EPI_ISL_614814, EPI_ISL_614815, EPI_ISL_614816, EPI_ISL_614817, EPI_ISL_614818, EPI_ISL_614819, EPI_ISL_614820, EPI_ISL_614822, EPI_ISL_614824, EPI_ISL_614826, EPI_ISL_614827, EPI_ISL_614828, EPI_ISL_614829,
EPI_ISL_614857, EPI_ISL_614876, EPI_ISL_614877, EPI_ISL_614878, EPI_ISL_614879, EPI_ISL_614880, EPI_ISL_614881, EPI_ISL_614882, EPI_ISL_614883, EPI_ISL_615166, EPI_ISL_615167, EPI_ISL_615168, EPI_ISL_615169, EPI_ISL_615170, EPI_ISL_615171, EPI_ISL_615172, EPI_ISL_615173, EPI_ISL_615174,
EPI_ISL_615175, EPI_ISL_615176, EPI_ISL_615177, EPI_ISL_615178, EPI_ISL_615179, EPI_ISL_615180, EPI_ISL_615181, EPI_ISL_615182, EPI_ISL_615183, EPI_ISL_615184, EPI_ISL_615185, EPI_ISL_615186, EPI_ISL_615187, EPI_ISL_615188, EPI_ISL_615189, EPI_ISL_615190, EPI_ISL_615191, EPI_ISL_615192,
EPI_ISL_615193, EPI_ISL_615194, EPI_ISL_615195, EPI_ISL_615196, EPI_ISL_615197, EPI_ISL_615198, EPI_ISL_615199, EPI_ISL_615200, EPI_ISL_615201, EPI_ISL_615202, EPI_ISL_615203, EPI_ISL_615204, EPI_ISL_615205, EPI_ISL_615206, EPI_ISL_615207, EPI_ISL_615208, EPI_ISL_615209, EPI_ISL_615210,
EPI_ISL_615211, EPI_ISL_615212, EPI_ISL_615213, EPI_ISL_615214, EPI_ISL_615215, EPI_ISL_615216, EPI_ISL_615217, EPI_ISL_615218, EPI_ISL_615219, EPI_ISL_615220, EPI_ISL_615221, EPI_ISL_615222, EPI_ISL_615223, EPI_ISL_615224, EPI_ISL_615225, EPI_ISL_615226, EPI_ISL_615227, EPI_ISL_615228,
EPI_ISL_615229, EPI_ISL_615230, EPI_ISL_615231, EPI_ISL_615232, EPI_ISL_615233, EPI_ISL_615234, EPI_ISL_615235, EPI_ISL_615236, EPI_ISL_615237, EPI_ISL_615238, EPI_ISL_615239, EPI_ISL_615240, EPI_ISL_615241, EPI_ISL_615242, EPI_ISL_615243, EPI_ISL_615244, EPI_ISL_615245, EPI_ISL_615246,
EPI_ISL_615247, EPI_ISL_615248, EPI_ISL_615249, EPI_ISL_615250, EPI_ISL_615251, EPI_ISL_615252, EPI_ISL_615253, EPI_ISL_615254, EPI_ISL_615255, EPI_ISL_615256, EPI_ISL_615257, EPI_ISL_615258, EPI_ISL_615259, EPI_ISL_615260, EPI_ISL_615261, EPI_ISL_615262, EPI_ISL_615263, EPI_ISL_615264,
EPI_ISL_615265, EPI_ISL_615266, EPI_ISL_615267, EPI_ISL_615268, EPI_ISL_615269, EPI_ISL_615270, EPI_ISL_615271, EPI_ISL_615272, EPI_ISL_615273, EPI_ISL_615274, EPI_ISL_615275, EPI_ISL_615276, EPI_ISL_615277, EPI_ISL_615278, EPI_ISL_615279, EPI_ISL_615280, EPI_ISL_615281, EPI_ISL_615282,
EPI_ISL_615283, EPI_ISL_615284, EPI_ISL_615285, EPI_ISL_615286, EPI_ISL_615287, EPI_ISL_615288, EPI_ISL_615289, EPI_ISL_615290, EPI_ISL_615291, EPI_ISL_615292, EPI_ISL_615293, EPI_ISL_615294, EPI_ISL_615295, EPI_ISL_615296, EPI_ISL_615297, EPI_ISL_615298, EPI_ISL_615299, EPI_ISL_615300,
EPI_ISL_615301, EPI_ISL_615302, EPI_ISL_615303, EPI_ISL_615304, EPI_ISL_615305, EPI_ISL_615306, EPI_ISL_615307, EPI_ISL_615308, EPI_ISL_615309, EPI_ISL_615310, EPI_ISL_615311, EPI_ISL_615312, EPI_ISL_615313, EPI_ISL_615314, EPI_ISL_615315, EPI_ISL_615316, EPI_ISL_615317, EPI_ISL_615318,
EPI_ISL_615319, EPI_ISL_615320, EPI_ISL_615321, EPI_ISL_615322, EPI_ISL_615323, EPI_ISL_615324, EPI_ISL_615325, EPI_ISL_615326, EPI_ISL_615327, EPI_ISL_615328, EPI_ISL_615329, EPI_ISL_615330, EPI_ISL_615331, EPI_ISL_615332, EPI_ISL_615333, EPI_ISL_615334, EPI_ISL_615335, EPI_ISL_615336,
EPI_ISL_615337, EPI_ISL_615338, EPI_ISL_615339, EPI_ISL_615340, EPI_ISL_615341, EPI_ISL_615342, EPI_ISL_615343, EPI_ISL_615344, EPI_ISL_615345, EPI_ISL_615346, EPI_ISL_615347, EPI_ISL_615348, EPI_ISL_615349, EPI_ISL_615350, EPI_ISL_615351, EPI_ISL_615352, EPI_ISL_615353, EPI_ISL_615354,
EPI_ISL_615355, EPI_ISL_615356, EPI_ISL_615357, EPI_ISL_615358, EPI_ISL_615359, EPI_ISL_615360, EPI_ISL_615361, EPI_ISL_615362, EPI_ISL_615363, EPI_ISL_615364, EPI_ISL_615365, EPI_ISL_615366, EPI_ISL_615367, EPI_ISL_615368, EPI_ISL_615369, EPI_ISL_615370, EPI_ISL_615371, EPI_ISL_615376,
EPI_ISL_615377, EPI_ISL_615378, EPI_ISL_615379, EPI_ISL_615380, EPI_ISL_615381, EPI_ISL_615382, EPI_ISL_615383, EPI_ISL_615384, EPI_ISL_615385, EPI_ISL_615386, EPI_ISL_615387, EPI_ISL_615388, EPI_ISL_615389, EPI_ISL_615390, EPI_ISL_615391, EPI_ISL_615392, EPI_ISL_615393, EPI_ISL_615394,
EPI_ISL_615395, EPI_ISL_615396, EPI_ISL_615397, EPI_ISL_615398, EPI_ISL_615399, EPI_ISL_615400, EPI_ISL_615401, EPI_ISL_615402, EPI_ISL_615403, EPI_ISL_615415, EPI_ISL_615436, EPI_ISL_615437, EPI_ISL_615451, EPI_ISL_615452, EPI_ISL_615456, EPI_ISL_615462, EPI_ISL_615470, EPI_ISL_615473,
EPI_ISL_615476, EPI_ISL_615500, EPI_ISL_615503, EPI_ISL_615510, EPI_ISL_615514, EPI_ISL_615515, EPI_ISL_615516, EPI_ISL_615517, EPI_ISL_615518, EPI_ISL_615519, EPI_ISL_615520, EPI_ISL_615521, EPI_ISL_615522, EPI_ISL_615523, EPI_ISL_615524, EPI_ISL_615525, EPI_ISL_615526, EPI_ISL_615527,
EPI_ISL_615528, EPI_ISL_615529, EPI_ISL_615530, EPI_ISL_615531, EPI_ISL_615532, EPI_ISL_615533, EPI_ISL_615534, EPI_ISL_615535, EPI_ISL_615536, EPI_ISL_615537, EPI_ISL_615538, EPI_ISL_615539, EPI_ISL_615540, EPI_ISL_615541, EPI_ISL_615542, EPI_ISL_615543, EPI_ISL_615544, EPI_ISL_615545,
EPI_ISL_615546, EPI_ISL_615547, EPI_ISL_615548, EPI_ISL_615549, EPI_ISL_615550, EPI_ISL_615551, EPI_ISL_615552, EPI_ISL_615553, EPI_ISL_615554, EPI_ISL_615555, EPI_ISL_615556, EPI_ISL_615557, EPI_ISL_615558, EPI_ISL_615559, EPI_ISL_615560, EPI_ISL_615561, EPI_ISL_615562, EPI_ISL_615563,
EPI_ISL_615564, EPI_ISL_615565, EPI_ISL_615566, EPI_ISL_615567, EPI_ISL_615568, EPI_ISL_615569, EPI_ISL_615570, EPI_ISL_615571, EPI_ISL_615572, EPI_ISL_615573, EPI_ISL_615574, EPI_ISL_615575, EPI_ISL_615576, EPI_ISL_615578, EPI_ISL_615579, EPI_ISL_615580, EPI_ISL_615581, EPI_ISL_615582,
EPI_ISL_615601, EPI_ISL_615608, EPI_ISL_615610, EPI_ISL_615611, EPI_ISL_615612, EPI_ISL_615613, EPI_ISL_615614, EPI_ISL_615615, EPI_ISL_615616, EPI_ISL_615618, EPI_ISL_615619, EPI_ISL_615620, EPI_ISL_615621, EPI_ISL_615622, EPI_ISL_615623, EPI_ISL_615624, EPI_ISL_615625, EPI_ISL_615626,
EPI_ISL_615627, EPI_ISL_615628, EPI_ISL_615629, EPI_ISL_615630, EPI_ISL_615631, EPI_ISL_615632, EPI_ISL_615633, EPI_ISL_615634, EPI_ISL_615635, EPI_ISL_615638, EPI_ISL_615639, EPI_ISL_615640, EPI_ISL_615641, EPI_ISL_615642, EPI_ISL_615643, EPI_ISL_615644, EPI_ISL_615645, EPI_ISL_615646,
EPI_ISL_615647, EPI_ISL_615648, EPI_ISL_615649, EPI_ISL_615650, EPI_ISL_615651, EPI_ISL_615652, EPI_ISL_615653, EPI_ISL_615654, EPI_ISL_615655, EPI_ISL_615656, EPI_ISL_615657, EPI_ISL_615658, EPI_ISL_615659, EPI_ISL_615660, EPI_ISL_615661, EPI_ISL_615662, EPI_ISL_615663, EPI_ISL_615664,
EPI_ISL_615665, EPI_ISL_615666, EPI_ISL_615667, EPI_ISL_615668, EPI_ISL_615669, EPI_ISL_615670, EPI_ISL_615671, EPI_ISL_615672, EPI_ISL_615673, EPI_ISL_615674, EPI_ISL_615675, EPI_ISL_615676, EPI_ISL_615677, EPI_ISL_615678, EPI_ISL_615679, EPI_ISL_615680, EPI_ISL_615681, EPI_ISL_615682,
EPI_ISL_615683, EPI_ISL_615684, EPI_ISL_615685, EPI_ISL_615686, EPI_ISL_615687, EPI_ISL_615688, EPI_ISL_615689, EPI_ISL_615690, EPI_ISL_615691, EPI_ISL_615692, EPI_ISL_615693, EPI_ISL_615694, EPI_ISL_615695, EPI_ISL_615696, EPI_ISL_615697, EPI_ISL_615698, EPI_ISL_615699, EPI_ISL_615700,
EPI_ISL_615701, EPI_ISL_615702, EPI_ISL_615703, EPI_ISL_615704, EPI_ISL_615705, EPI_ISL_615706, EPI_ISL_615707, EPI_ISL_615708, EPI_ISL_615709, EPI_ISL_615710, EPI_ISL_615711, EPI_ISL_615712, EPI_ISL_615713, EPI_ISL_615714, EPI_ISL_615715, EPI_ISL_615716, EPI_ISL_615717, EPI_ISL_615718,
EPI_ISL_615719, EPI_ISL_615720, EPI_ISL_615721, EPI_ISL_615722, EPI_ISL_615723, EPI_ISL_615724, EPI_ISL_615725, EPI_ISL_615726, EPI_ISL_615727, EPI_ISL_615728, EPI_ISL_615729, EPI_ISL_615730, EPI_ISL_615731, EPI_ISL_615732, EPI_ISL_615733, EPI_ISL_615734, EPI_ISL_615735, EPI_ISL_615736,
EPI_ISL_615737, EPI_ISL_615738, EPI_ISL_615739, EPI_ISL_615740, EPI_ISL_615741, EPI_ISL_615742, EPI_ISL_615743, EPI_ISL_615744, EPI_ISL_615745, EPI_ISL_615746, EPI_ISL_615747, EPI_ISL_615748, EPI_ISL_615749, EPI_ISL_615750, EPI_ISL_615751, EPI_ISL_615752, EPI_ISL_615753, EPI_ISL_615754,
EPI_ISL_615755, EPI_ISL_615756, EPI_ISL_615757, EPI_ISL_615758, EPI_ISL_615759, EPI_ISL_615760, EPI_ISL_615761, EPI_ISL_615762, EPI_ISL_615763, EPI_ISL_615764, EPI_ISL_615765, EPI_ISL_615766, EPI_ISL_615767, EPI_ISL_615768, EPI_ISL_615769, EPI_ISL_615770, EPI_ISL_615771, EPI_ISL_615772,
EPI_ISL_615773, EPI_ISL_615774, EPI_ISL_615775, EPI_ISL_615776, EPI_ISL_615777, EPI_ISL_615778, EPI_ISL_615779, EPI_ISL_615780, EPI_ISL_615781, EPI_ISL_615782, EPI_ISL_615783, EPI_ISL_615784, EPI_ISL_615785, EPI_ISL_615786, EPI_ISL_615787, EPI_ISL_615788, EPI_ISL_615789, EPI_ISL_615790,
EPI_ISL_615791, EPI_ISL_615792, EPI_ISL_615793, EPI_ISL_615794, EPI_ISL_615795, EPI_ISL_615796, EPI_ISL_615797, EPI_ISL_615798, EPI_ISL_615799, EPI_ISL_615800, EPI_ISL_615801, EPI_ISL_615802, EPI_ISL_615803, EPI_ISL_615804, EPI_ISL_615805, EPI_ISL_615806, EPI_ISL_615807, EPI_ISL_615808,
EPI_ISL_615809, EPI_ISL_615810, EPI_ISL_615811, EPI_ISL_615812, EPI_ISL_615813, EPI_ISL_615814, EPI_ISL_615815, EPI_ISL_615816, EPI_ISL_615817, EPI_ISL_615818, EPI_ISL_615819, EPI_ISL_615820, EPI_ISL_615821, EPI_ISL_615822, EPI_ISL_615823, EPI_ISL_615824, EPI_ISL_615825, EPI_ISL_615826,
EPI_ISL_615827, EPI_ISL_615828, EPI_ISL_615829, EPI_ISL_615830, EPI_ISL_615831, EPI_ISL_615832, EPI_ISL_615833, EPI_ISL_615834, EPI_ISL_615835, EPI_ISL_615836, EPI_ISL_615837, EPI_ISL_615838, EPI_ISL_615839, EPI_ISL_615840, EPI_ISL_615841, EPI_ISL_615842, EPI_ISL_615843, EPI_ISL_615844,
EPI_ISL_615845, EPI_ISL_615846, EPI_ISL_615847, EPI_ISL_615848, EPI_ISL_615849, EPI_ISL_615850, EPI_ISL_615851, EPI_ISL_615852, EPI_ISL_615853, EPI_ISL_615854, EPI_ISL_615855, EPI_ISL_615856, EPI_ISL_615857, EPI_ISL_615858, EPI_ISL_615859, EPI_ISL_615860, EPI_ISL_615861, EPI_ISL_615862,
EPI_ISL_615863, EPI_ISL_615864, EPI_ISL_615865, EPI_ISL_615866, EPI_ISL_615867, EPI_ISL_615868, EPI_ISL_615869, EPI_ISL_615870, EPI_ISL_615871, EPI_ISL_615872, EPI_ISL_615873, EPI_ISL_615874, EPI_ISL_615875, EPI_ISL_615876, EPI_ISL_615877, EPI_ISL_615878, EPI_ISL_615879, EPI_ISL_615880,
EPI_ISL_615881, EPI_ISL_615882, EPI_ISL_615883, EPI_ISL_615884, EPI_ISL_615885, EPI_ISL_615886, EPI_ISL_615887, EPI_ISL_615888, EPI_ISL_615889, EPI_ISL_615890, EPI_ISL_615891, EPI_ISL_615892, EPI_ISL_615893, EPI_ISL_615894, EPI_ISL_615895, EPI_ISL_615896, EPI_ISL_615897, EPI_ISL_615898,
EPI_ISL_615899, EPI_ISL_615900, EPI_ISL_615901, EPI_ISL_615902, EPI_ISL_615903, EPI_ISL_615904, EPI_ISL_615905, EPI_ISL_615935, EPI_ISL_615980, EPI_ISL_615988, EPI_ISL_615996, EPI_ISL_616034, EPI_ISL_616067, EPI_ISL_616110, EPI_ISL_616115, EPI_ISL_616128, EPI_ISL_616190, EPI_ISL_616226,
EPI_ISL_616228, EPI_ISL_616233, EPI_ISL_616305, EPI_ISL_616318, EPI_ISL_616336, EPI_ISL_616344, EPI_ISL_616377, EPI_ISL_616380, EPI_ISL_616398, EPI_ISL_616399, EPI_ISL_616400, EPI_ISL_616408, EPI_ISL_616410, EPI_ISL_616411, EPI_ISL_616412, EPI_ISL_616413, EPI_ISL_616414, EPI_ISL_616422,
EPI_ISL_616437, EPI_ISL_616450, EPI_ISL_616451, EPI_ISL_616452, EPI_ISL_616453, EPI_ISL_616454, EPI_ISL_616455, EPI_ISL_616456, EPI_ISL_616457, EPI_ISL_616458, EPI_ISL_616459, EPI_ISL_616460, EPI_ISL_616461, EPI_ISL_616463, EPI_ISL_616464, EPI_ISL_616465, EPI_ISL_616466, EPI_ISL_616467,
EPI_ISL_616468, EPI_ISL_616469, EPI_ISL_616470, EPI_ISL_616471, EPI_ISL_616472, EPI_ISL_616473, EPI_ISL_616474, EPI_ISL_616475, EPI_ISL_616476, EPI_ISL_616477, EPI_ISL_616478, EPI_ISL_616479, EPI_ISL_616480, EPI_ISL_616481, EPI_ISL_616482, EPI_ISL_616483, EPI_ISL_616484, EPI_ISL_616485,
EPI_ISL_616486, EPI_ISL_616487, EPI_ISL_616488, EPI_ISL_616489, EPI_ISL_616490, EPI_ISL_616491, EPI_ISL_616492, EPI_ISL_616493, EPI_ISL_616494, EPI_ISL_616495, EPI_ISL_616496, EPI_ISL_616497, EPI_ISL_616498, EPI_ISL_616499, EPI_ISL_616511, EPI_ISL_616513, EPI_ISL_616534, EPI_ISL_616535,
EPI_ISL_616536, EPI_ISL_616537, EPI_ISL_616538, EPI_ISL_616539, EPI_ISL_616540, EPI_ISL_616541, EPI_ISL_616542, EPI_ISL_616543, EPI_ISL_616544, EPI_ISL_616545, EPI_ISL_616546, EPI_ISL_616547, EPI_ISL_616548, EPI_ISL_616549, EPI_ISL_616550, EPI_ISL_616551, EPI_ISL_616552, EPI_ISL_616553,
EPI_ISL_616554, EPI_ISL_616555, EPI_ISL_616556, EPI_ISL_616557, EPI_ISL_616558, EPI_ISL_616559, EPI_ISL_616560, EPI_ISL_616561, EPI_ISL_616562, EPI_ISL_616563, EPI_ISL_616564, EPI_ISL_616565, EPI_ISL_616566, EPI_ISL_616567, EPI_ISL_616568, EPI_ISL_616570, EPI_ISL_616571, EPI_ISL_616572,
EPI_ISL_616573, EPI_ISL_616574, EPI_ISL_616575, EPI_ISL_616576, EPI_ISL_616577, EPI_ISL_616578, EPI_ISL_616579, EPI_ISL_616580, EPI_ISL_616581, EPI_ISL_616582, EPI_ISL_616583, EPI_ISL_616584, EPI_ISL_616585, EPI_ISL_616586, EPI_ISL_616587, EPI_ISL_616588, EPI_ISL_616589, EPI_ISL_616590,
EPI_ISL_616591, EPI_ISL_616592, EPI_ISL_616593, EPI_ISL_616594, EPI_ISL_616595, EPI_ISL_616596, EPI_ISL_616597, EPI_ISL_616598, EPI_ISL_616599, EPI_ISL_616600, EPI_ISL_616601, EPI_ISL_616602, EPI_ISL_616603, EPI_ISL_616604, EPI_ISL_616605, EPI_ISL_616606, EPI_ISL_616607, EPI_ISL_616608,
EPI_ISL_616609, EPI_ISL_616610, EPI_ISL_616611, EPI_ISL_616612, EPI_ISL_616613, EPI_ISL_616614, EPI_ISL_616615, EPI_ISL_616616, EPI_ISL_616617, EPI_ISL_616618, EPI_ISL_616619, EPI_ISL_616620, EPI_ISL_616621, EPI_ISL_616622, EPI_ISL_616623, EPI_ISL_616624, EPI_ISL_616625, EPI_ISL_616626,
EPI_ISL_616627, EPI_ISL_616628, EPI_ISL_616629, EPI_ISL_616630, EPI_ISL_616631, EPI_ISL_616632, EPI_ISL_616633, EPI_ISL_616634, EPI_ISL_616635, EPI_ISL_616636, EPI_ISL_616637, EPI_ISL_616638, EPI_ISL_616639, EPI_ISL_616640, EPI_ISL_616641, EPI_ISL_616642, EPI_ISL_616643, EPI_ISL_616644,
EPI_ISL_616645, EPI_ISL_616646, EPI_ISL_616647, EPI_ISL_616648, EPI_ISL_616649, EPI_ISL_616650, EPI_ISL_616651, EPI_ISL_616652, EPI_ISL_616653, EPI_ISL_616654, EPI_ISL_616655, EPI_ISL_616656, EPI_ISL_616657, EPI_ISL_616658, EPI_ISL_616659, EPI_ISL_616660, EPI_ISL_616661, EPI_ISL_616662,
EPI_ISL_616663, EPI_ISL_616664, EPI_ISL_616665, EPI_ISL_616666, EPI_ISL_616667, EPI_ISL_616668, EPI_ISL_616669, EPI_ISL_616670, EPI_ISL_616671, EPI_ISL_616672, EPI_ISL_616673, EPI_ISL_616674, EPI_ISL_616675, EPI_ISL_616676, EPI_ISL_616677, EPI_ISL_616678, EPI_ISL_616679, EPI_ISL_616680,
EPI_ISL_616681, EPI_ISL_616682, EPI_ISL_616683, EPI_ISL_616684, EPI_ISL_616685, EPI_ISL_616686, EPI_ISL_616687, EPI_ISL_616688, EPI_ISL_616689, EPI_ISL_616690, EPI_ISL_616691, EPI_ISL_616692, EPI_ISL_616693, EPI_ISL_616694, EPI_ISL_616695, EPI_ISL_616696, EPI_ISL_616697, EPI_ISL_616698,
EPI_ISL_616699, EPI_ISL_616700, EPI_ISL_616701, EPI_ISL_616702, EPI_ISL_616703, EPI_ISL_616704, EPI_ISL_616705, EPI_ISL_616706, EPI_ISL_616707, EPI_ISL_616708, EPI_ISL_616709, EPI_ISL_616710, EPI_ISL_616711, EPI_ISL_616712, EPI_ISL_616713, EPI_ISL_616714, EPI_ISL_616715, EPI_ISL_616716,
EPI_ISL_616717, EPI_ISL_616718, EPI_ISL_616719, EPI_ISL_616720, EPI_ISL_616721, EPI_ISL_616722, EPI_ISL_616723, EPI_ISL_616724, EPI_ISL_616725, EPI_ISL_616726, EPI_ISL_616727, EPI_ISL_616728, EPI_ISL_616729, EPI_ISL_616730, EPI_ISL_616731, EPI_ISL_616732, EPI_ISL_616733, EPI_ISL_616734,
EPI_ISL_616735, EPI_ISL_616736, EPI_ISL_616737, EPI_ISL_616738, EPI_ISL_616739, EPI_ISL_616740, EPI_ISL_616741, EPI_ISL_616742, EPI_ISL_616743, EPI_ISL_616744, EPI_ISL_616745, EPI_ISL_616746, EPI_ISL_616747, EPI_ISL_616748, EPI_ISL_616749, EPI_ISL_616750, EPI_ISL_616751, EPI_ISL_616752,
EPI_ISL_616753, EPI_ISL_616754, EPI_ISL_616755, EPI_ISL_616756, EPI_ISL_616757, EPI_ISL_616758, EPI_ISL_616759, EPI_ISL_616760, EPI_ISL_616761, EPI_ISL_616762, EPI_ISL_616763, EPI_ISL_616771, EPI_ISL_616775, EPI_ISL_616776, EPI_ISL_616777, EPI_ISL_616778, EPI_ISL_616779, EPI_ISL_616780,
EPI_ISL_616781, EPI_ISL_616782, EPI_ISL_616783, EPI_ISL_616784, EPI_ISL_616785, EPI_ISL_616786, EPI_ISL_616788, EPI_ISL_616790, EPI_ISL_616791, EPI_ISL_616792, EPI_ISL_616793, EPI_ISL_616794, EPI_ISL_616795, EPI_ISL_616796, EPI_ISL_616797, EPI_ISL_616798, EPI_ISL_616799, EPI_ISL_616800,
EPI_ISL_616801, EPI_ISL_616802, EPI_ISL_616803, EPI_ISL_616804, EPI_ISL_616805, EPI_ISL_616806, EPI_ISL_616807, EPI_ISL_616808, EPI_ISL_616809, EPI_ISL_616810, EPI_ISL_616815, EPI_ISL_616826, EPI_ISL_616857, EPI_ISL_616875, EPI_ISL_616894, EPI_ISL_616906, EPI_ISL_616907, EPI_ISL_616908,
EPI_ISL_616909, EPI_ISL_616910, EPI_ISL_616911, EPI_ISL_616912, EPI_ISL_616913, EPI_ISL_616914, EPI_ISL_616915, EPI_ISL_616916, EPI_ISL_616917, EPI_ISL_616918, EPI_ISL_616919, EPI_ISL_616920, EPI_ISL_616921, EPI_ISL_616922, EPI_ISL_616925, EPI_ISL_616926, EPI_ISL_616927, EPI_ISL_616928,
EPI_ISL_616929, EPI_ISL_616930, EPI_ISL_616931, EPI_ISL_616932, EPI_ISL_616933, EPI_ISL_616934, EPI_ISL_616935, EPI_ISL_616936, EPI_ISL_616937, EPI_ISL_616938, EPI_ISL_616939, EPI_ISL_616940, EPI_ISL_616941, EPI_ISL_616942, EPI_ISL_616943, EPI_ISL_616944, EPI_ISL_616945, EPI_ISL_616946,
EPI_ISL_616947, EPI_ISL_616948, EPI_ISL_616949, EPI_ISL_616950, EPI_ISL_616951, EPI_ISL_616952, EPI_ISL_616953, EPI_ISL_616954, EPI_ISL_616955, EPI_ISL_616956, EPI_ISL_616957, EPI_ISL_616958, EPI_ISL_616959, EPI_ISL_616960, EPI_ISL_616961, EPI_ISL_616962, EPI_ISL_616963, EPI_ISL_616964,
EPI_ISL_616965, EPI_ISL_616966, EPI_ISL_616967, EPI_ISL_616968, EPI_ISL_616969, EPI_ISL_616970, EPI_ISL_616971, EPI_ISL_616972, EPI_ISL_616973, EPI_ISL_616974, EPI_ISL_616976, EPI_ISL_616977, EPI_ISL_616978, EPI_ISL_616979, EPI_ISL_616980, EPI_ISL_616981, EPI_ISL_616982, EPI_ISL_616983,
EPI_ISL_616984, EPI_ISL_616985, EPI_ISL_616986, EPI_ISL_616987, EPI_ISL_616988, EPI_ISL_616989, EPI_ISL_616990, EPI_ISL_616991, EPI_ISL_616992, EPI_ISL_616993, EPI_ISL_616994, EPI_ISL_616995, EPI_ISL_616996, EPI_ISL_616997, EPI_ISL_616998, EPI_ISL_616999, EPI_ISL_617000, EPI_ISL_617001,
EPI_ISL_617002, EPI_ISL_617003, EPI_ISL_617004, EPI_ISL_617005, EPI_ISL_617006, EPI_ISL_617007, EPI_ISL_617008, EPI_ISL_617009, EPI_ISL_617010, EPI_ISL_617011, EPI_ISL_617012, EPI_ISL_617013, EPI_ISL_617014, EPI_ISL_617015, EPI_ISL_617016, EPI_ISL_617017, EPI_ISL_617018, EPI_ISL_617019,
EPI_ISL_617020, EPI_ISL_617021, EPI_ISL_617022, EPI_ISL_617023, EPI_ISL_617024, EPI_ISL_617025, EPI_ISL_617026, EPI_ISL_617027, EPI_ISL_617028, EPI_ISL_617029, EPI_ISL_617030, EPI_ISL_617031, EPI_ISL_617032, EPI_ISL_617033, EPI_ISL_617034, EPI_ISL_617035, EPI_ISL_617036, EPI_ISL_617037,
EPI_ISL_617038, EPI_ISL_617039, EPI_ISL_617040, EPI_ISL_617041, EPI_ISL_617042, EPI_ISL_617043, EPI_ISL_617044, EPI_ISL_617045, EPI_ISL_617046, EPI_ISL_617047, EPI_ISL_617048, EPI_ISL_617049, EPI_ISL_617050, EPI_ISL_617051, EPI_ISL_617052, EPI_ISL_617053, EPI_ISL_617054, EPI_ISL_617055,
EPI_ISL_617056, EPI_ISL_617057, EPI_ISL_617058, EPI_ISL_617059, EPI_ISL_617060, EPI_ISL_617061, EPI_ISL_617062, EPI_ISL_617063, EPI_ISL_617064, EPI_ISL_617065, EPI_ISL_617066, EPI_ISL_617067, EPI_ISL_617068, EPI_ISL_617069, EPI_ISL_617070, EPI_ISL_617071, EPI_ISL_617072, EPI_ISL_617073,
EPI_ISL_617074, EPI_ISL_617075, EPI_ISL_617076, EPI_ISL_617077, EPI_ISL_617078, EPI_ISL_617079, EPI_ISL_617080, EPI_ISL_617081, EPI_ISL_617082, EPI_ISL_617083, EPI_ISL_617084, EPI_ISL_617085, EPI_ISL_617086, EPI_ISL_617087, EPI_ISL_617088, EPI_ISL_617089, EPI_ISL_617090, EPI_ISL_617091,
EPI_ISL_617092, EPI_ISL_617093, EPI_ISL_617094, EPI_ISL_617095, EPI_ISL_617096, EPI_ISL_617097, EPI_ISL_617098, EPI_ISL_617099, EPI_ISL_617100, EPI_ISL_617101, EPI_ISL_617102, EPI_ISL_617103, EPI_ISL_617104, EPI_ISL_617105, EPI_ISL_617106, EPI_ISL_617108, EPI_ISL_617109, EPI_ISL_617110,
EPI_ISL_617111, EPI_ISL_617112, EPI_ISL_617113, EPI_ISL_617114, EPI_ISL_617115, EPI_ISL_617116, EPI_ISL_617117, EPI_ISL_617118, EPI_ISL_617119, EPI_ISL_617120, EPI_ISL_617121, EPI_ISL_617122, EPI_ISL_617123, EPI_ISL_617124, EPI_ISL_617125, EPI_ISL_617126, EPI_ISL_617127, EPI_ISL_617128,
EPI_ISL_617129, EPI_ISL_617130, EPI_ISL_617131, EPI_ISL_617132, EPI_ISL_617133, EPI_ISL_617134, EPI_ISL_617135, EPI_ISL_617136, EPI_ISL_617137, EPI_ISL_617138, EPI_ISL_617139, EPI_ISL_617140, EPI_ISL_617141, EPI_ISL_617142, EPI_ISL_617143, EPI_ISL_617144, EPI_ISL_617145, EPI_ISL_617146,
EPI_ISL_617147, EPI_ISL_617148, EPI_ISL_617149, EPI_ISL_617150, EPI_ISL_617151, EPI_ISL_617152, EPI_ISL_617153, EPI_ISL_617154, EPI_ISL_617155, EPI_ISL_617156, EPI_ISL_617157, EPI_ISL_617158, EPI_ISL_617159, EPI_ISL_617160, EPI_ISL_617161, EPI_ISL_617162, EPI_ISL_617163, EPI_ISL_617164,
EPI_ISL_617165, EPI_ISL_617166, EPI_ISL_617167, EPI_ISL_617168, EPI_ISL_617169, EPI_ISL_617170, EPI_ISL_617171, EPI_ISL_617172, EPI_ISL_617173, EPI_ISL_617174, EPI_ISL_617175, EPI_ISL_617176, EPI_ISL_617177, EPI_ISL_617178, EPI_ISL_617179, EPI_ISL_617180, EPI_ISL_617181, EPI_ISL_617182,
EPI_ISL_617183, EPI_ISL_617184, EPI_ISL_617185, EPI_ISL_617186, EPI_ISL_617187, EPI_ISL_617188, EPI_ISL_617189, EPI_ISL_617190, EPI_ISL_617191, EPI_ISL_617192, EPI_ISL_617193, EPI_ISL_617194, EPI_ISL_617195, EPI_ISL_617196, EPI_ISL_617197, EPI_ISL_617198, EPI_ISL_617199, EPI_ISL_617200,
EPI_ISL_617201, EPI_ISL_617202, EPI_ISL_617203, EPI_ISL_617204, EPI_ISL_617205, EPI_ISL_617206, EPI_ISL_617207, EPI_ISL_617208, EPI_ISL_617209, EPI_ISL_617210, EPI_ISL_617211, EPI_ISL_617212, EPI_ISL_617213, EPI_ISL_617214, EPI_ISL_617215, EPI_ISL_617216, EPI_ISL_617217, EPI_ISL_617218,
EPI_ISL_617219, EPI_ISL_617220, EPI_ISL_617221, EPI_ISL_617222, EPI_ISL_617223, EPI_ISL_617224, EPI_ISL_617225, EPI_ISL_617226, EPI_ISL_617227, EPI_ISL_617228, EPI_ISL_617229, EPI_ISL_617230, EPI_ISL_617231, EPI_ISL_617232, EPI_ISL_617233, EPI_ISL_617234, EPI_ISL_617235, EPI_ISL_617236,
EPI_ISL_617237, EPI_ISL_617238, EPI_ISL_617239, EPI_ISL_617240, EPI_ISL_617241, EPI_ISL_617242, EPI_ISL_617243, EPI_ISL_617244, EPI_ISL_617245, EPI_ISL_617246, EPI_ISL_617247, EPI_ISL_617248, EPI_ISL_617249, EPI_ISL_617250, EPI_ISL_617251, EPI_ISL_617252, EPI_ISL_617253, EPI_ISL_617254,
EPI_ISL_617255, EPI_ISL_617256, EPI_ISL_617257, EPI_ISL_617258, EPI_ISL_617259, EPI_ISL_617260, EPI_ISL_617261, EPI_ISL_617262, EPI_ISL_617263, EPI_ISL_617264, EPI_ISL_617265, EPI_ISL_617266, EPI_ISL_617267, EPI_ISL_617268, EPI_ISL_617269, EPI_ISL_617270, EPI_ISL_617271, EPI_ISL_617272,
EPI_ISL_617273, EPI_ISL_617274, EPI_ISL_617275, EPI_ISL_617276, EPI_ISL_617277, EPI_ISL_617278, EPI_ISL_617279, EPI_ISL_617280, EPI_ISL_617281, EPI_ISL_617282, EPI_ISL_617283, EPI_ISL_617285, EPI_ISL_617286, EPI_ISL_617287, EPI_ISL_617288, EPI_ISL_617289, EPI_ISL_617290, EPI_ISL_617291,
EPI_ISL_617292, EPI_ISL_617293, EPI_ISL_617294, EPI_ISL_617295, EPI_ISL_617296, EPI_ISL_617297, EPI_ISL_617298, EPI_ISL_617299, EPI_ISL_617300, EPI_ISL_617301, EPI_ISL_617302, EPI_ISL_617303, EPI_ISL_617304, EPI_ISL_617305, EPI_ISL_617306, EPI_ISL_617307, EPI_ISL_617308, EPI_ISL_617309,
EPI_ISL_617310, EPI_ISL_617311, EPI_ISL_617312, EPI_ISL_617313, EPI_ISL_617314, EPI_ISL_617315, EPI_ISL_617316, EPI_ISL_617317, EPI_ISL_617318, EPI_ISL_617319, EPI_ISL_617320, EPI_ISL_617321, EPI_ISL_617322, EPI_ISL_617323, EPI_ISL_617324, EPI_ISL_617325, EPI_ISL_617326, EPI_ISL_617327,
EPI_ISL_617328, EPI_ISL_617329, EPI_ISL_617330, EPI_ISL_617331, EPI_ISL_617332, EPI_ISL_617333, EPI_ISL_617334, EPI_ISL_617335, EPI_ISL_617336, EPI_ISL_617337, EPI_ISL_617338, EPI_ISL_617339, EPI_ISL_617340, EPI_ISL_617341, EPI_ISL_617342, EPI_ISL_617343, EPI_ISL_617344, EPI_ISL_617345,
EPI_ISL_617346, EPI_ISL_617347, EPI_ISL_617348, EPI_ISL_617349, EPI_ISL_617350, EPI_ISL_617351, EPI_ISL_617352, EPI_ISL_617353, EPI_ISL_617354, EPI_ISL_617355, EPI_ISL_617356, EPI_ISL_617357, EPI_ISL_617358, EPI_ISL_617359, EPI_ISL_617360, EPI_ISL_617361, EPI_ISL_617362, EPI_ISL_617363,
EPI_ISL_617364, EPI_ISL_617365, EPI_ISL_617366, EPI_ISL_617367, EPI_ISL_617368, EPI_ISL_617369, EPI_ISL_617370, EPI_ISL_617371, EPI_ISL_617372, EPI_ISL_617373, EPI_ISL_617374, EPI_ISL_617375, EPI_ISL_617376, EPI_ISL_617377, EPI_ISL_617378, EPI_ISL_617379, EPI_ISL_617380, EPI_ISL_617381,
EPI_ISL_617384, EPI_ISL_617385, EPI_ISL_617386, EPI_ISL_617387, EPI_ISL_617388, EPI_ISL_617389, EPI_ISL_617390, EPI_ISL_617391, EPI_ISL_617392, EPI_ISL_617393, EPI_ISL_617394, EPI_ISL_617395, EPI_ISL_617396, EPI_ISL_617397, EPI_ISL_617398, EPI_ISL_617399, EPI_ISL_617400, EPI_ISL_617401,
EPI_ISL_617402, EPI_ISL_617403, EPI_ISL_617404, EPI_ISL_617405, EPI_ISL_617408, EPI_ISL_617410, EPI_ISL_617415, EPI_ISL_617421, EPI_ISL_617426, EPI_ISL_617428, EPI_ISL_617441, EPI_ISL_617471, EPI_ISL_617491, EPI_ISL_617524, EPI_ISL_617559, EPI_ISL_617618, EPI_ISL_617649, EPI_ISL_617697,
EPI_ISL_617715, EPI_ISL_617727, EPI_ISL_617784, EPI_ISL_617850, EPI_ISL_617906, EPI_ISL_617907, EPI_ISL_617908, EPI_ISL_617909, EPI_ISL_617910, EPI_ISL_617911, EPI_ISL_617912, EPI_ISL_617913, EPI_ISL_617914, EPI_ISL_617915, EPI_ISL_617916, EPI_ISL_617917, EPI_ISL_617918, EPI_ISL_617919,
EPI_ISL_617920, EPI_ISL_617921, EPI_ISL_617922, EPI_ISL_617923, EPI_ISL_617924, EPI_ISL_617925, EPI_ISL_617926, EPI_ISL_617927, EPI_ISL_617928, EPI_ISL_617929, EPI_ISL_617930, EPI_ISL_617931, EPI_ISL_617932, EPI_ISL_617933, EPI_ISL_617934, EPI_ISL_617935, EPI_ISL_617936, EPI_ISL_617937,

EPI_ISL_617938, EPI_ISL_617940, EPI_ISL_617941, EPI_ISL_617942, EPI_ISL_617943, EPI_ISL_617946, EPI_ISL_617956, EPI_ISL_617957, EPI_ISL_617959, EPI_ISL_617960, EPI_ISL_617978, EPI_ISL_618032, EPI_ISL_618033, EPI_ISL_618034, EPI_ISL_618035, EPI_ISL_618036, EPI_ISL_618037, EPI_ISL_618038,
EPI_ISL_618042, EPI_ISL_618044, EPI_ISL_618046, EPI_ISL_618055, EPI_ISL_618059, EPI_ISL_618060, EPI_ISL_618061, EPI_ISL_618063, EPI_ISL_618075, EPI_ISL_618087, EPI_ISL_618123, EPI_ISL_618126, EPI_ISL_618137, EPI_ISL_618138, EPI_ISL_618142, EPI_ISL_618153, EPI_ISL_618168, EPI_ISL_618193,
EPI_ISL_618216, EPI_ISL_618217, EPI_ISL_618223, EPI_ISL_618265, EPI_ISL_618274, EPI_ISL_618289, EPI_ISL_618315, EPI_ISL_618332, EPI_ISL_618334, EPI_ISL_618373, EPI_ISL_618382, EPI_ISL_618401, EPI_ISL_618406, EPI_ISL_618407, EPI_ISL_618408, EPI_ISL_618409, EPI_ISL_618410, EPI_ISL_618411,
EPI_ISL_618412, EPI_ISL_618413, EPI_ISL_618414, EPI_ISL_618415, EPI_ISL_618416, EPI_ISL_618417, EPI_ISL_618418, EPI_ISL_618419, EPI_ISL_618420, EPI_ISL_618421, EPI_ISL_618422, EPI_ISL_618423, EPI_ISL_618424, EPI_ISL_618425, EPI_ISL_618426, EPI_ISL_618427, EPI_ISL_618428, EPI_ISL_618429,
EPI_ISL_618430, EPI_ISL_618431, EPI_ISL_618432, EPI_ISL_618433, EPI_ISL_618434, EPI_ISL_618435, EPI_ISL_618436, EPI_ISL_618437, EPI_ISL_618438, EPI_ISL_618439, EPI_ISL_618440, EPI_ISL_618441, EPI_ISL_618442, EPI_ISL_618443, EPI_ISL_618444, EPI_ISL_618445, EPI_ISL_618446, EPI_ISL_618447,
EPI_ISL_618448, EPI_ISL_618449, EPI_ISL_618450, EPI_ISL_618451, EPI_ISL_618452, EPI_ISL_618453, EPI_ISL_618454, EPI_ISL_618455, EPI_ISL_618456, EPI_ISL_618457, EPI_ISL_618458, EPI_ISL_618459, EPI_ISL_618460, EPI_ISL_618461, EPI_ISL_618462, EPI_ISL_618463, EPI_ISL_618464, EPI_ISL_618465,
EPI_ISL_618466, EPI_ISL_618467, EPI_ISL_618468, EPI_ISL_618469, EPI_ISL_618470, EPI_ISL_618471, EPI_ISL_618472, EPI_ISL_618473, EPI_ISL_618474, EPI_ISL_618475, EPI_ISL_618476, EPI_ISL_618477, EPI_ISL_618478, EPI_ISL_618480, EPI_ISL_618481, EPI_ISL_618482, EPI_ISL_618483, EPI_ISL_618484,
EPI_ISL_618485, EPI_ISL_618486, EPI_ISL_618487, EPI_ISL_618488, EPI_ISL_618489, EPI_ISL_618490, EPI_ISL_618491, EPI_ISL_618492, EPI_ISL_618493, EPI_ISL_618494, EPI_ISL_618495, EPI_ISL_618497, EPI_ISL_618498, EPI_ISL_618499, EPI_ISL_618500, EPI_ISL_618501, EPI_ISL_618502, EPI_ISL_618503,
EPI_ISL_618504, EPI_ISL_618505, EPI_ISL_618506, EPI_ISL_618507, EPI_ISL_618508, EPI_ISL_618509, EPI_ISL_618510, EPI_ISL_618512, EPI_ISL_618513, EPI_ISL_618514, EPI_ISL_618515, EPI_ISL_618516, EPI_ISL_618518, EPI_ISL_618519, EPI_ISL_618520, EPI_ISL_618521, EPI_ISL_618522, EPI_ISL_618523,
EPI_ISL_618524, EPI_ISL_618526, EPI_ISL_618527, EPI_ISL_618528, EPI_ISL_618529, EPI_ISL_618530, EPI_ISL_618531, EPI_ISL_618532, EPI_ISL_618533, EPI_ISL_618534, EPI_ISL_618535, EPI_ISL_618536, EPI_ISL_618537, EPI_ISL_618538, EPI_ISL_618539, EPI_ISL_618540, EPI_ISL_618541, EPI_ISL_618542,
EPI_ISL_618543, EPI_ISL_618544, EPI_ISL_618545, EPI_ISL_618546, EPI_ISL_618547, EPI_ISL_618548, EPI_ISL_618549, EPI_ISL_618550, EPI_ISL_618551, EPI_ISL_618552, EPI_ISL_618553, EPI_ISL_618554, EPI_ISL_618555, EPI_ISL_618556, EPI_ISL_618557, EPI_ISL_618558, EPI_ISL_618559, EPI_ISL_618560,
EPI_ISL_618561, EPI_ISL_618562, EPI_ISL_618563, EPI_ISL_618564, EPI_ISL_618565, EPI_ISL_618566, EPI_ISL_618567, EPI_ISL_618568, EPI_ISL_618569, EPI_ISL_618570, EPI_ISL_618571, EPI_ISL_618572, EPI_ISL_618573, EPI_ISL_618574, EPI_ISL_618575, EPI_ISL_618576, EPI_ISL_618577, EPI_ISL_618578,
EPI_ISL_618579, EPI_ISL_618580, EPI_ISL_618581, EPI_ISL_618582, EPI_ISL_618583, EPI_ISL_618584, EPI_ISL_618585, EPI_ISL_618586, EPI_ISL_618587, EPI_ISL_618588, EPI_ISL_618589, EPI_ISL_618590, EPI_ISL_618591, EPI_ISL_618592, EPI_ISL_618593, EPI_ISL_618594, EPI_ISL_618596, EPI_ISL_618597,
EPI_ISL_618598, EPI_ISL_618599, EPI_ISL_618600, EPI_ISL_618601, EPI_ISL_618602, EPI_ISL_618603, EPI_ISL_618604, EPI_ISL_618605, EPI_ISL_618606, EPI_ISL_618607, EPI_ISL_618608, EPI_ISL_618610, EPI_ISL_618611, EPI_ISL_618612, EPI_ISL_618613, EPI_ISL_618614, EPI_ISL_618615, EPI_ISL_618616,
EPI_ISL_618617, EPI_ISL_618618, EPI_ISL_618619, EPI_ISL_618620, EPI_ISL_618621, EPI_ISL_618622, EPI_ISL_618623, EPI_ISL_618624, EPI_ISL_618625, EPI_ISL_618642, EPI_ISL_618658, EPI_ISL_618662, EPI_ISL_618663, EPI_ISL_618664, EPI_ISL_618665, EPI_ISL_618666, EPI_ISL_618667, EPI_ISL_618668,
EPI_ISL_618669, EPI_ISL_618670, EPI_ISL_618671, EPI_ISL_618672, EPI_ISL_618673, EPI_ISL_618674, EPI_ISL_618675, EPI_ISL_618676, EPI_ISL_618677, EPI_ISL_618678, EPI_ISL_618679, EPI_ISL_618680, EPI_ISL_618681, EPI_ISL_618682, EPI_ISL_618683, EPI_ISL_618684, EPI_ISL_618685, EPI_ISL_618686,
EPI_ISL_618687, EPI_ISL_618688, EPI_ISL_618689, EPI_ISL_618690, EPI_ISL_618691, EPI_ISL_618692, EPI_ISL_618693, EPI_ISL_618694, EPI_ISL_618695, EPI_ISL_618696, EPI_ISL_618697, EPI_ISL_618698, EPI_ISL_618699, EPI_ISL_618700, EPI_ISL_618701, EPI_ISL_618702, EPI_ISL_618703, EPI_ISL_618704,
EPI_ISL_618705, EPI_ISL_618706, EPI_ISL_618707, EPI_ISL_618708, EPI_ISL_618709, EPI_ISL_618710, EPI_ISL_618711, EPI_ISL_618712, EPI_ISL_618713, EPI_ISL_618714, EPI_ISL_618715, EPI_ISL_618716, EPI_ISL_618717, EPI_ISL_618718, EPI_ISL_618719, EPI_ISL_618721, EPI_ISL_618722, EPI_ISL_618723,
EPI_ISL_618724, EPI_ISL_618725, EPI_ISL_618726, EPI_ISL_618727, EPI_ISL_618728, EPI_ISL_618729, EPI_ISL_618730, EPI_ISL_618731, EPI_ISL_618732, EPI_ISL_618733, EPI_ISL_618734, EPI_ISL_618735, EPI_ISL_618736, EPI_ISL_618737, EPI_ISL_618738, EPI_ISL_618739, EPI_ISL_618740, EPI_ISL_618741,
EPI_ISL_618742, EPI_ISL_618743, EPI_ISL_618744, EPI_ISL_618745, EPI_ISL_618746, EPI_ISL_618747, EPI_ISL_618748, EPI_ISL_618749, EPI_ISL_618750, EPI_ISL_618751, EPI_ISL_618752, EPI_ISL_618753, EPI_ISL_618754, EPI_ISL_618755, EPI_ISL_618756, EPI_ISL_618757, EPI_ISL_618758, EPI_ISL_618759,
EPI_ISL_618760, EPI_ISL_618761, EPI_ISL_618762, EPI_ISL_618763, EPI_ISL_618764, EPI_ISL_618765, EPI_ISL_618766, EPI_ISL_618767, EPI_ISL_618768, EPI_ISL_618769, EPI_ISL_618770, EPI_ISL_618771, EPI_ISL_618772, EPI_ISL_618773, EPI_ISL_618774, EPI_ISL_618775, EPI_ISL_618776, EPI_ISL_618777,
EPI_ISL_618778, EPI_ISL_618779, EPI_ISL_618780, EPI_ISL_618781, EPI_ISL_618782, EPI_ISL_618783, EPI_ISL_618784, EPI_ISL_618785, EPI_ISL_618786, EPI_ISL_618787, EPI_ISL_618788, EPI_ISL_618789, EPI_ISL_618790, EPI_ISL_618791, EPI_ISL_618792, EPI_ISL_618793, EPI_ISL_618794, EPI_ISL_618795,
EPI_ISL_618796, EPI_ISL_618797, EPI_ISL_618798, EPI_ISL_618799, EPI_ISL_618800, EPI_ISL_618809, EPI_ISL_618810, EPI_ISL_618811, EPI_ISL_618812, EPI_ISL_618813, EPI_ISL_618814, EPI_ISL_618815, EPI_ISL_618816, EPI_ISL_618817, EPI_ISL_618818, EPI_ISL_618819, EPI_ISL_618820, EPI_ISL_618821,
EPI_ISL_618822, EPI_ISL_618823, EPI_ISL_618824, EPI_ISL_618825, EPI_ISL_618826, EPI_ISL_618827, EPI_ISL_618828, EPI_ISL_618829, EPI_ISL_618830, EPI_ISL_618831, EPI_ISL_618832, EPI_ISL_618833, EPI_ISL_618834, EPI_ISL_618835, EPI_ISL_618836, EPI_ISL_618837, EPI_ISL_618838, EPI_ISL_618839,
EPI_ISL_618840, EPI_ISL_618841, EPI_ISL_618842, EPI_ISL_618843, EPI_ISL_618844, EPI_ISL_618845, EPI_ISL_618846, EPI_ISL_618847, EPI_ISL_618848, EPI_ISL_618849, EPI_ISL_618850, EPI_ISL_618851, EPI_ISL_618852, EPI_ISL_618853, EPI_ISL_618854, EPI_ISL_618855, EPI_ISL_618856, EPI_ISL_618857,
EPI_ISL_618858, EPI_ISL_618859, EPI_ISL_618860, EPI_ISL_618861, EPI_ISL_618862, EPI_ISL_618863, EPI_ISL_618864, EPI_ISL_618865, EPI_ISL_618866, EPI_ISL_618867, EPI_ISL_618868, EPI_ISL_618869, EPI_ISL_618870, EPI_ISL_618871, EPI_ISL_618872, EPI_ISL_618873, EPI_ISL_618874, EPI_ISL_618875,
EPI_ISL_618876, EPI_ISL_618877, EPI_ISL_618878, EPI_ISL_618879, EPI_ISL_618880, EPI_ISL_618881, EPI_ISL_618882, EPI_ISL_618883, EPI_ISL_618884, EPI_ISL_618885, EPI_ISL_618886, EPI_ISL_618887, EPI_ISL_618888, EPI_ISL_618889, EPI_ISL_618890, EPI_ISL_618891, EPI_ISL_618892, EPI_ISL_618893,
EPI_ISL_618894, EPI_ISL_618895, EPI_ISL_618896, EPI_ISL_618897, EPI_ISL_618898, EPI_ISL_618899, EPI_ISL_618900, EPI_ISL_618901, EPI_ISL_618902, EPI_ISL_618903, EPI_ISL_618904, EPI_ISL_618905, EPI_ISL_618906, EPI_ISL_618907, EPI_ISL_618908, EPI_ISL_618909, EPI_ISL_618910, EPI_ISL_618911,
EPI_ISL_618912, EPI_ISL_618913, EPI_ISL_618914, EPI_ISL_618915, EPI_ISL_618916, EPI_ISL_618917, EPI_ISL_618918, EPI_ISL_618919, EPI_ISL_618920, EPI_ISL_618921, EPI_ISL_618922, EPI_ISL_618923, EPI_ISL_618924, EPI_ISL_618925, EPI_ISL_618926, EPI_ISL_618927, EPI_ISL_618928, EPI_ISL_618929,
EPI_ISL_618930, EPI_ISL_618931, EPI_ISL_618932, EPI_ISL_618933, EPI_ISL_618934, EPI_ISL_618935, EPI_ISL_618936, EPI_ISL_618937, EPI_ISL_618938, EPI_ISL_618939, EPI_ISL_618940, EPI_ISL_618941, EPI_ISL_618942, EPI_ISL_618943, EPI_ISL_618944, EPI_ISL_618945, EPI_ISL_618946, EPI_ISL_618947,
EPI_ISL_618948, EPI_ISL_618949, EPI_ISL_618950, EPI_ISL_618951, EPI_ISL_618952, EPI_ISL_618953, EPI_ISL_618954, EPI_ISL_618955, EPI_ISL_618956, EPI_ISL_618957, EPI_ISL_618958, EPI_ISL_618959, EPI_ISL_618960, EPI_ISL_618961, EPI_ISL_618962, EPI_ISL_618963, EPI_ISL_618964, EPI_ISL_618965,
EPI_ISL_618966, EPI_ISL_618967, EPI_ISL_618968, EPI_ISL_618970, EPI_ISL_618971, EPI_ISL_618973, EPI_ISL_618974, EPI_ISL_618975, EPI_ISL_618976, EPI_ISL_618977, EPI_ISL_618981, EPI_ISL_618982, EPI_ISL_618983, EPI_ISL_618984, EPI_ISL_618985, EPI_ISL_618986, EPI_ISL_618987, EPI_ISL_618988,
EPI_ISL_618989, EPI_ISL_618990, EPI_ISL_618991, EPI_ISL_618992, EPI_ISL_618993, EPI_ISL_618994, EPI_ISL_618995, EPI_ISL_618996, EPI_ISL_618997, EPI_ISL_618998, EPI_ISL_618999, EPI_ISL_619000, EPI_ISL_619001, EPI_ISL_619002, EPI_ISL_619003, EPI_ISL_619004, EPI_ISL_619005, EPI_ISL_619006,
EPI_ISL_619007, EPI_ISL_619008, EPI_ISL_619009, EPI_ISL_619010, EPI_ISL_619011, EPI_ISL_619012, EPI_ISL_619013, EPI_ISL_619014, EPI_ISL_619015, EPI_ISL_619016, EPI_ISL_619019, EPI_ISL_619020, EPI_ISL_619021, EPI_ISL_619022, EPI_ISL_619023, EPI_ISL_619024, EPI_ISL_619025, EPI_ISL_619026,
EPI_ISL_619027, EPI_ISL_619028, EPI_ISL_619029, EPI_ISL_619030, EPI_ISL_619031, EPI_ISL_619032, EPI_ISL_619033, EPI_ISL_619034, EPI_ISL_619035, EPI_ISL_619036, EPI_ISL_619037, EPI_ISL_619038, EPI_ISL_619039, EPI_ISL_619040, EPI_ISL_619041, EPI_ISL_619042, EPI_ISL_619043, EPI_ISL_619044,
EPI_ISL_619045, EPI_ISL_619046, EPI_ISL_619047, EPI_ISL_619048, EPI_ISL_619049, EPI_ISL_619050, EPI_ISL_619051, EPI_ISL_619052, EPI_ISL_619053, EPI_ISL_619054, EPI_ISL_619055, EPI_ISL_619056, EPI_ISL_619057, EPI_ISL_619058, EPI_ISL_619059, EPI_ISL_619060, EPI_ISL_619061, EPI_ISL_619062,
EPI_ISL_619063, EPI_ISL_619064, EPI_ISL_619065, EPI_ISL_619066, EPI_ISL_619067, EPI_ISL_619068, EPI_ISL_619069, EPI_ISL_619070, EPI_ISL_619071, EPI_ISL_619072, EPI_ISL_619073, EPI_ISL_619074, EPI_ISL_619075, EPI_ISL_619076, EPI_ISL_619077, EPI_ISL_619078, EPI_ISL_619079, EPI_ISL_619080,
EPI_ISL_619081, EPI_ISL_619082, EPI_ISL_619083, EPI_ISL_619084, EPI_ISL_619085, EPI_ISL_619086, EPI_ISL_619087, EPI_ISL_619088, EPI_ISL_619089, EPI_ISL_619090, EPI_ISL_619091, EPI_ISL_619092, EPI_ISL_619093, EPI_ISL_619094, EPI_ISL_619095, EPI_ISL_619096, EPI_ISL_619097, EPI_ISL_619098,
EPI_ISL_619099, EPI_ISL_619100, EPI_ISL_619101, EPI_ISL_619102, EPI_ISL_619103, EPI_ISL_619104, EPI_ISL_619105, EPI_ISL_619106, EPI_ISL_619107, EPI_ISL_619108, EPI_ISL_619109, EPI_ISL_619110, EPI_ISL_619111, EPI_ISL_619112, EPI_ISL_619113, EPI_ISL_619114, EPI_ISL_619115, EPI_ISL_619116,
EPI_ISL_619117, EPI_ISL_619118, EPI_ISL_619119, EPI_ISL_619120, EPI_ISL_619121, EPI_ISL_619122, EPI_ISL_619123, EPI_ISL_619124, EPI_ISL_619125, EPI_ISL_619126, EPI_ISL_619127, EPI_ISL_619128, EPI_ISL_619129, EPI_ISL_619130, EPI_ISL_619131, EPI_ISL_619132, EPI_ISL_619134, EPI_ISL_619135,
EPI_ISL_619136, EPI_ISL_619137, EPI_ISL_619138, EPI_ISL_619139, EPI_ISL_619140, EPI_ISL_619142, EPI_ISL_619143, EPI_ISL_619144, EPI_ISL_619146, EPI_ISL_619147, EPI_ISL_619148, EPI_ISL_619149, EPI_ISL_619150, EPI_ISL_619152, EPI_ISL_619153, EPI_ISL_619154, EPI_ISL_619155, EPI_ISL_619156,
EPI_ISL_619157, EPI_ISL_619158, EPI_ISL_619159, EPI_ISL_619160, EPI_ISL_619161, EPI_ISL_619162, EPI_ISL_619164, EPI_ISL_619165, EPI_ISL_619166, EPI_ISL_619167, EPI_ISL_619168, EPI_ISL_619169, EPI_ISL_619171, EPI_ISL_619172, EPI_ISL_619173, EPI_ISL_619174, EPI_ISL_619175, EPI_ISL_619177,
EPI_ISL_619178, EPI_ISL_619179, EPI_ISL_619180, EPI_ISL_619181, EPI_ISL_619182, EPI_ISL_619183, EPI_ISL_619184, EPI_ISL_619186, EPI_ISL_619187, EPI_ISL_619188, EPI_ISL_619189, EPI_ISL_619190, EPI_ISL_619191, EPI_ISL_619192, EPI_ISL_619193, EPI_ISL_619194, EPI_ISL_619195, EPI_ISL_619196,
EPI_ISL_619197, EPI_ISL_619198, EPI_ISL_619199, EPI_ISL_619200, EPI_ISL_619201, EPI_ISL_619202, EPI_ISL_619203, EPI_ISL_619204, EPI_ISL_619205, EPI_ISL_619206, EPI_ISL_619207, EPI_ISL_619208, EPI_ISL_619209, EPI_ISL_619210, EPI_ISL_619212, EPI_ISL_619213, EPI_ISL_619214, EPI_ISL_619215,
EPI_ISL_619216, EPI_ISL_619217, EPI_ISL_619218, EPI_ISL_619219, EPI_ISL_619220, EPI_ISL_619221, EPI_ISL_619222, EPI_ISL_619223, EPI_ISL_619224, EPI_ISL_619225, EPI_ISL_619227, EPI_ISL_619228, EPI_ISL_619229, EPI_ISL_619230, EPI_ISL_619231, EPI_ISL_619232, EPI_ISL_619234, EPI_ISL_619235,
EPI_ISL_619236, EPI_ISL_619237, EPI_ISL_619238, EPI_ISL_619239, EPI_ISL_619240, EPI_ISL_619241, EPI_ISL_619242, EPI_ISL_619243, EPI_ISL_619244, EPI_ISL_619245, EPI_ISL_619246, EPI_ISL_619247, EPI_ISL_619248, EPI_ISL_619249, EPI_ISL_619250, EPI_ISL_619251, EPI_ISL_619252, EPI_ISL_619253,
EPI_ISL_619254, EPI_ISL_619255, EPI_ISL_619257, EPI_ISL_619258, EPI_ISL_619259, EPI_ISL_619264, EPI_ISL_619274, EPI_ISL_619275, EPI_ISL_619276, EPI_ISL_619277, EPI_ISL_619278, EPI_ISL_619279, EPI_ISL_619280, EPI_ISL_619281, EPI_ISL_619282, EPI_ISL_619283, EPI_ISL_619284, EPI_ISL_619285,
EPI_ISL_619286, EPI_ISL_619287, EPI_ISL_619288, EPI_ISL_619289, EPI_ISL_619290, EPI_ISL_619291, EPI_ISL_619293, EPI_ISL_619294, EPI_ISL_619295, EPI_ISL_619296, EPI_ISL_619297, EPI_ISL_619298, EPI_ISL_619299, EPI_ISL_619300, EPI_ISL_619301, EPI_ISL_619302, EPI_ISL_619303, EPI_ISL_619304,
EPI_ISL_619305, EPI_ISL_619306, EPI_ISL_619307, EPI_ISL_619308, EPI_ISL_619309, EPI_ISL_619310, EPI_ISL_619311, EPI_ISL_619312, EPI_ISL_619313, EPI_ISL_619314, EPI_ISL_619315, EPI_ISL_619316, EPI_ISL_619317, EPI_ISL_619319, EPI_ISL_619320, EPI_ISL_619322, EPI_ISL_619323, EPI_ISL_619324,
EPI_ISL_619325, EPI_ISL_619326, EPI_ISL_619327, EPI_ISL_619329, EPI_ISL_619330, EPI_ISL_619331, EPI_ISL_619333, EPI_ISL_619334, EPI_ISL_619335, EPI_ISL_619336, EPI_ISL_619338, EPI_ISL_619339, EPI_ISL_619340, EPI_ISL_619341, EPI_ISL_619342, EPI_ISL_619343, EPI_ISL_619345, EPI_ISL_619346,
EPI_ISL_619347, EPI_ISL_619348, EPI_ISL_619349, EPI_ISL_619350, EPI_ISL_619351, EPI_ISL_619352, EPI_ISL_619353, EPI_ISL_619354, EPI_ISL_619355, EPI_ISL_619356, EPI_ISL_619357, EPI_ISL_619358, EPI_ISL_619359, EPI_ISL_619360, EPI_ISL_619361, EPI_ISL_619362, EPI_ISL_619363, EPI_ISL_619364,
EPI_ISL_619365, EPI_ISL_619366, EPI_ISL_619367, EPI_ISL_619368, EPI_ISL_619369, EPI_ISL_619370, EPI_ISL_619371, EPI_ISL_619372, EPI_ISL_619373, EPI_ISL_619374, EPI_ISL_619375, EPI_ISL_619376, EPI_ISL_619377, EPI_ISL_619379, EPI_ISL_619380, EPI_ISL_619381, EPI_ISL_619382, EPI_ISL_619383,
EPI_ISL_619385, EPI_ISL_619386, EPI_ISL_619387, EPI_ISL_619388, EPI_ISL_619390, EPI_ISL_619391, EPI_ISL_619392, EPI_ISL_619393, EPI_ISL_619394, EPI_ISL_619395, EPI_ISL_619396, EPI_ISL_619397, EPI_ISL_619399, EPI_ISL_619400, EPI_ISL_619401, EPI_ISL_619402, EPI_ISL_619403, EPI_ISL_619404,
EPI_ISL_619406, EPI_ISL_619407, EPI_ISL_619408, EPI_ISL_619409, EPI_ISL_619410, EPI_ISL_619411, EPI_ISL_619412, EPI_ISL_619413, EPI_ISL_619414, EPI_ISL_619415, EPI_ISL_619416, EPI_ISL_619417, EPI_ISL_619418, EPI_ISL_619419, EPI_ISL_619420, EPI_ISL_619421, EPI_ISL_619422, EPI_ISL_619423,
EPI_ISL_619424, EPI_ISL_619425, EPI_ISL_619426, EPI_ISL_619427, EPI_ISL_619428, EPI_ISL_619429, EPI_ISL_619430, EPI_ISL_619431, EPI_ISL_619432, EPI_ISL_619433, EPI_ISL_619434, EPI_ISL_619435, EPI_ISL_619436, EPI_ISL_619437, EPI_ISL_619438, EPI_ISL_619439, EPI_ISL_619440, EPI_ISL_619441,
EPI_ISL_619442, EPI_ISL_619443, EPI_ISL_619444, EPI_ISL_619445, EPI_ISL_619446, EPI_ISL_619447, EPI_ISL_619448, EPI_ISL_619449, EPI_ISL_619450, EPI_ISL_619451, EPI_ISL_619452, EPI_ISL_619453, EPI_ISL_619454, EPI_ISL_619456, EPI_ISL_619457, EPI_ISL_619458, EPI_ISL_619459, EPI_ISL_619460,
EPI_ISL_619461, EPI_ISL_619462, EPI_ISL_619463, EPI_ISL_619464, EPI_ISL_619465, EPI_ISL_619466, EPI_ISL_619467, EPI_ISL_619468, EPI_ISL_619469, EPI_ISL_619470, EPI_ISL_619471, EPI_ISL_619472, EPI_ISL_619473, EPI_ISL_619474, EPI_ISL_619475, EPI_ISL_619476, EPI_ISL_619477, EPI_ISL_619478,
EPI_ISL_619479, EPI_ISL_619480, EPI_ISL_619481, EPI_ISL_619482, EPI_ISL_619483, EPI_ISL_619484, EPI_ISL_619485, EPI_ISL_619486, EPI_ISL_619487, EPI_ISL_619488, EPI_ISL_619489, EPI_ISL_619490, EPI_ISL_619491, EPI_ISL_619493, EPI_ISL_619494, EPI_ISL_619495, EPI_ISL_619496, EPI_ISL_619497,
EPI_ISL_619498, EPI_ISL_619499, EPI_ISL_619500, EPI_ISL_619501, EPI_ISL_619502, EPI_ISL_619503, EPI_ISL_619504, EPI_ISL_619505, EPI_ISL_619506, EPI_ISL_619507, EPI_ISL_619508, EPI_ISL_619509, EPI_ISL_619510, EPI_ISL_619511, EPI_ISL_619512, EPI_ISL_619513, EPI_ISL_619514, EPI_ISL_619515,
EPI_ISL_619516, EPI_ISL_619517, EPI_ISL_619518, EPI_ISL_619519, EPI_ISL_619520, EPI_ISL_619521, EPI_ISL_619522, EPI_ISL_619523, EPI_ISL_619524, EPI_ISL_619525, EPI_ISL_619526, EPI_ISL_619527, EPI_ISL_619528, EPI_ISL_619529, EPI_ISL_619530, EPI_ISL_619531, EPI_ISL_619532, EPI_ISL_619533,
EPI_ISL_619535, EPI_ISL_619536, EPI_ISL_619537, EPI_ISL_619538, EPI_ISL_619539, EPI_ISL_619540, EPI_ISL_619541, EPI_ISL_619542, EPI_ISL_619543, EPI_ISL_619544, EPI_ISL_619545, EPI_ISL_619546, EPI_ISL_619547, EPI_ISL_619548, EPI_ISL_619549, EPI_ISL_619550, EPI_ISL_619551, EPI_ISL_619552,
EPI_ISL_619553, EPI_ISL_619554, EPI_ISL_619555, EPI_ISL_619557, EPI_ISL_619558, EPI_ISL_619559, EPI_ISL_619560, EPI_ISL_619561, EPI_ISL_619562, EPI_ISL_619563, EPI_ISL_619564, EPI_ISL_619565, EPI_ISL_619566, EPI_ISL_619567, EPI_ISL_619568, EPI_ISL_619569, EPI_ISL_619570, EPI_ISL_619571,
EPI_ISL_619572, EPI_ISL_619573, EPI_ISL_619574, EPI_ISL_619575, EPI_ISL_619576, EPI_ISL_619577, EPI_ISL_619578, EPI_ISL_619579, EPI_ISL_619580, EPI_ISL_619581, EPI_ISL_619582, EPI_ISL_619583, EPI_ISL_619584, EPI_ISL_619585, EPI_ISL_619586, EPI_ISL_619587, EPI_ISL_619588, EPI_ISL_619589,
EPI_ISL_619590, EPI_ISL_619591, EPI_ISL_619592, EPI_ISL_619593, EPI_ISL_619594, EPI_ISL_619595, EPI_ISL_619596, EPI_ISL_619597, EPI_ISL_619598, EPI_ISL_619599, EPI_ISL_619600, EPI_ISL_619601, EPI_ISL_619602, EPI_ISL_619603, EPI_ISL_619604, EPI_ISL_619605, EPI_ISL_619606, EPI_ISL_619607,
EPI_ISL_619608, EPI_ISL_619609, EPI_ISL_619610, EPI_ISL_619611, EPI_ISL_619612, EPI_ISL_619613, EPI_ISL_619614, EPI_ISL_619615, EPI_ISL_619616, EPI_ISL_619617, EPI_ISL_619618, EPI_ISL_619619, EPI_ISL_619620, EPI_ISL_619621, EPI_ISL_619622, EPI_ISL_619623, EPI_ISL_619624, EPI_ISL_619625,
EPI_ISL_619626, EPI_ISL_619627, EPI_ISL_619628, EPI_ISL_619629, EPI_ISL_619630, EPI_ISL_619631, EPI_ISL_619632, EPI_ISL_619633, EPI_ISL_619634, EPI_ISL_619635, EPI_ISL_619636, EPI_ISL_619637, EPI_ISL_619638, EPI_ISL_619639, EPI_ISL_619640, EPI_ISL_619641, EPI_ISL_619642, EPI_ISL_619643,
EPI_ISL_619644, EPI_ISL_619645, EPI_ISL_619646, EPI_ISL_619647, EPI_ISL_619648, EPI_ISL_619649, EPI_ISL_619650, EPI_ISL_619651, EPI_ISL_619652, EPI_ISL_619653, EPI_ISL_619654, EPI_ISL_619655, EPI_ISL_619656, EPI_ISL_619657, EPI_ISL_619658, EPI_ISL_619659, EPI_ISL_619660, EPI_ISL_619661,
EPI_ISL_619662, EPI_ISL_619663, EPI_ISL_619664, EPI_ISL_619665, EPI_ISL_619666, EPI_ISL_619667, EPI_ISL_619668, EPI_ISL_619669, EPI_ISL_619670, EPI_ISL_619671, EPI_ISL_619672, EPI_ISL_619673, EPI_ISL_619674, EPI_ISL_619675, EPI_ISL_619676, EPI_ISL_619677, EPI_ISL_619678, EPI_ISL_619679,
EPI_ISL_619680, EPI_ISL_619681, EPI_ISL_619682, EPI_ISL_619683, EPI_ISL_619684, EPI_ISL_619685, EPI_ISL_619686, EPI_ISL_619687, EPI_ISL_619688, EPI_ISL_619689, EPI_ISL_619690, EPI_ISL_619691, EPI_ISL_619692, EPI_ISL_619693, EPI_ISL_619694, EPI_ISL_619695, EPI_ISL_619696, EPI_ISL_619697,
EPI_ISL_619698, EPI_ISL_619699, EPI_ISL_619700, EPI_ISL_619701, EPI_ISL_619702, EPI_ISL_619703, EPI_ISL_619704, EPI_ISL_619705, EPI_ISL_619706, EPI_ISL_619707, EPI_ISL_619708, EPI_ISL_619709, EPI_ISL_619710, EPI_ISL_619711, EPI_ISL_619712, EPI_ISL_619713, EPI_ISL_619714, EPI_ISL_619715,
EPI_ISL_619716, EPI_ISL_619717, EPI_ISL_619718, EPI_ISL_619719, EPI_ISL_619720, EPI_ISL_619721, EPI_ISL_619722, EPI_ISL_619723, EPI_ISL_619724, EPI_ISL_619725, EPI_ISL_619726, EPI_ISL_619727, EPI_ISL_619728, EPI_ISL_619729, EPI_ISL_619730, EPI_ISL_619731, EPI_ISL_619732, EPI_ISL_619733,
EPI_ISL_619734, EPI_ISL_619735, EPI_ISL_619736, EPI_ISL_619737, EPI_ISL_619738, EPI_ISL_619739, EPI_ISL_619740, EPI_ISL_619741, EPI_ISL_619742, EPI_ISL_619743, EPI_ISL_619744, EPI_ISL_619745, EPI_ISL_619746, EPI_ISL_619747, EPI_ISL_619748, EPI_ISL_619749, EPI_ISL_619750, EPI_ISL_619752,
EPI_ISL_619753, EPI_ISL_619754, EPI_ISL_619755, EPI_ISL_619756, EPI_ISL_619757, EPI_ISL_619758, EPI_ISL_619759, EPI_ISL_619760, EPI_ISL_619761, EPI_ISL_619762, EPI_ISL_619763, EPI_ISL_619764, EPI_ISL_619765, EPI_ISL_619766, EPI_ISL_619767, EPI_ISL_619768, EPI_ISL_619769, EPI_ISL_619770,
EPI_ISL_619771, EPI_ISL_619772, EPI_ISL_619773, EPI_ISL_619774, EPI_ISL_619775, EPI_ISL_619776, EPI_ISL_619777, EPI_ISL_619778, EPI_ISL_619779, EPI_ISL_619780, EPI_ISL_619781, EPI_ISL_619782, EPI_ISL_619783, EPI_ISL_619784, EPI_ISL_619785, EPI_ISL_619786, EPI_ISL_619787, EPI_ISL_619788,
EPI_ISL_619789, EPI_ISL_619790, EPI_ISL_619791, EPI_ISL_619792, EPI_ISL_619793, EPI_ISL_619794, EPI_ISL_619795, EPI_ISL_619796, EPI_ISL_619797, EPI_ISL_619798, EPI_ISL_619799, EPI_ISL_619800, EPI_ISL_619801, EPI_ISL_619802, EPI_ISL_619803, EPI_ISL_619804, EPI_ISL_619805, EPI_ISL_619806,
EPI_ISL_619807, EPI_ISL_619808, EPI_ISL_619809, EPI_ISL_619810, EPI_ISL_619811, EPI_ISL_619812, EPI_ISL_619813, EPI_ISL_619814, EPI_ISL_619815, EPI_ISL_619816, EPI_ISL_619817, EPI_ISL_619818, EPI_ISL_619819, EPI_ISL_619820, EPI_ISL_619821, EPI_ISL_619822, EPI_ISL_619823, EPI_ISL_619824,
EPI_ISL_619825, EPI_ISL_619826, EPI_ISL_619827, EPI_ISL_619828, EPI_ISL_619829, EPI_ISL_619830, EPI_ISL_619831, EPI_ISL_619832, EPI_ISL_619833, EPI_ISL_619834, EPI_ISL_619835, EPI_ISL_619836, EPI_ISL_619837, EPI_ISL_619838, EPI_ISL_619839, EPI_ISL_619840, EPI_ISL_619841, EPI_ISL_619842,
EPI_ISL_619843, EPI_ISL_619844, EPI_ISL_619845, EPI_ISL_619846, EPI_ISL_619847, EPI_ISL_619848, EPI_ISL_619849, EPI_ISL_619850, EPI_ISL_619851, EPI_ISL_619852, EPI_ISL_619853, EPI_ISL_619854, EPI_ISL_619855, EPI_ISL_619856, EPI_ISL_619857, EPI_ISL_619858, EPI_ISL_619859, EPI_ISL_619860,
EPI_ISL_619861, EPI_ISL_619862, EPI_ISL_619863, EPI_ISL_619864, EPI_ISL_619865, EPI_ISL_619866, EPI_ISL_619867, EPI_ISL_619868, EPI_ISL_619869, EPI_ISL_619870, EPI_ISL_619871, EPI_ISL_619872, EPI_ISL_619873, EPI_ISL_619874, EPI_ISL_619875, EPI_ISL_619876, EPI_ISL_619877, EPI_ISL_619878,
EPI_ISL_619879, EPI_ISL_619880, EPI_ISL_619881, EPI_ISL_619882, EPI_ISL_619883, EPI_ISL_619884, EPI_ISL_619886, EPI_ISL_619887, EPI_ISL_619888, EPI_ISL_619889, EPI_ISL_619890, EPI_ISL_619891, EPI_ISL_619892, EPI_ISL_619893, EPI_ISL_619894, EPI_ISL_619896, EPI_ISL_619897, EPI_ISL_619898,
EPI_ISL_619899, EPI_ISL_619900, EPI_ISL_619901, EPI_ISL_619902, EPI_ISL_619903, EPI_ISL_619904, EPI_ISL_619905, EPI_ISL_619919, EPI_ISL_619947, EPI_ISL_619953, EPI_ISL_619958, EPI_ISL_619961, EPI_ISL_619963, EPI_ISL_619972, EPI_ISL_619976, EPI_ISL_619984, EPI_ISL_619985, EPI_ISL_619986,
EPI_ISL_619987, EPI_ISL_619988, EPI_ISL_619989, EPI_ISL_619990, EPI_ISL_619991, EPI_ISL_619992, EPI_ISL_619993, EPI_ISL_619994, EPI_ISL_619995, EPI_ISL_619996, EPI_ISL_619997, EPI_ISL_619998, EPI_ISL_619999, EPI_ISL_620000, EPI_ISL_620001, EPI_ISL_620002, EPI_ISL_620003, EPI_ISL_620004,
EPI_ISL_620005, EPI_ISL_620006, EPI_ISL_620007, EPI_ISL_620008, EPI_ISL_620009, EPI_ISL_620010, EPI_ISL_620011, EPI_ISL_620012, EPI_ISL_620013, EPI_ISL_620014, EPI_ISL_620015, EPI_ISL_620016, EPI_ISL_620017, EPI_ISL_620018, EPI_ISL_620019, EPI_ISL_620020, EPI_ISL_620021, EPI_ISL_620022,
EPI_ISL_620023, EPI_ISL_620024, EPI_ISL_620025, EPI_ISL_620026, EPI_ISL_620027, EPI_ISL_620028, EPI_ISL_620029, EPI_ISL_620030, EPI_ISL_620031, EPI_ISL_620032, EPI_ISL_620033, EPI_ISL_620034, EPI_ISL_620035, EPI_ISL_620036, EPI_ISL_620037, EPI_ISL_620038, EPI_ISL_620039, EPI_ISL_620040,
EPI_ISL_620041, EPI_ISL_620042, EPI_ISL_620043, EPI_ISL_620044, EPI_ISL_620045, EPI_ISL_620046, EPI_ISL_620047, EPI_ISL_620048, EPI_ISL_620049, EPI_ISL_620050, EPI_ISL_620051, EPI_ISL_620052, EPI_ISL_620053, EPI_ISL_620054, EPI_ISL_620055, EPI_ISL_620091, EPI_ISL_620093, EPI_ISL_620097,
EPI_ISL_620107, EPI_ISL_620116, EPI_ISL_620117, EPI_ISL_620118, EPI_ISL_620119, EPI_ISL_620120, EPI_ISL_620121, EPI_ISL_620122, EPI_ISL_620123, EPI_ISL_620124, EPI_ISL_620125, EPI_ISL_620126, EPI_ISL_620132, EPI_ISL_620137, EPI_ISL_620138, EPI_ISL_620140, EPI_ISL_620147, EPI_ISL_620152,
EPI_ISL_620191, EPI_ISL_620221, EPI_ISL_620230, EPI_ISL_620233, EPI_ISL_620265, EPI_ISL_620267, EPI_ISL_620281, EPI_ISL_620290, EPI_ISL_620291, EPI_ISL_620300, EPI_ISL_620301, EPI_ISL_620303, EPI_ISL_620304, EPI_ISL_620305, EPI_ISL_620306, EPI_ISL_620307, EPI_ISL_620308, EPI_ISL_620309,

EPI_ISL_620310, EPI_ISL_620311, EPI_ISL_620312, EPI_ISL_620313, EPI_ISL_620315, EPI_ISL_620316, EPI_ISL_620317, EPI_ISL_620318, EPI_ISL_620319, EPI_ISL_620320, EPI_ISL_620321, EPI_ISL_620322, EPI_ISL_620323, EPI_ISL_620324, EPI_ISL_620325, EPI_ISL_620326, EPI_ISL_620327, EPI_ISL_620328,
EPI_ISL_620329, EPI_ISL_620330, EPI_ISL_620331, EPI_ISL_620332, EPI_ISL_620333, EPI_ISL_620334, EPI_ISL_620335, EPI_ISL_620336, EPI_ISL_620337, EPI_ISL_620338, EPI_ISL_620339, EPI_ISL_620340, EPI_ISL_620341, EPI_ISL_620342, EPI_ISL_620343, EPI_ISL_620344, EPI_ISL_620345, EPI_ISL_620346,
EPI_ISL_620347, EPI_ISL_620348, EPI_ISL_620349, EPI_ISL_620350, EPI_ISL_620351, EPI_ISL_620352, EPI_ISL_620353, EPI_ISL_620354, EPI_ISL_620355, EPI_ISL_620356, EPI_ISL_620357, EPI_ISL_620358, EPI_ISL_620359, EPI_ISL_620360, EPI_ISL_620361, EPI_ISL_620362, EPI_ISL_620363, EPI_ISL_620364,
EPI_ISL_620365, EPI_ISL_620366, EPI_ISL_620367, EPI_ISL_620368, EPI_ISL_620369, EPI_ISL_620370, EPI_ISL_620371, EPI_ISL_620372, EPI_ISL_620373, EPI_ISL_620374, EPI_ISL_620375, EPI_ISL_620376, EPI_ISL_620377, EPI_ISL_620378, EPI_ISL_620379, EPI_ISL_620380, EPI_ISL_620381, EPI_ISL_620382,
EPI_ISL_620383, EPI_ISL_620384, EPI_ISL_620385, EPI_ISL_620386, EPI_ISL_620387, EPI_ISL_620388, EPI_ISL_620389, EPI_ISL_620390, EPI_ISL_620391, EPI_ISL_620392, EPI_ISL_620393, EPI_ISL_620394, EPI_ISL_620395, EPI_ISL_620396, EPI_ISL_620397, EPI_ISL_620398, EPI_ISL_620399, EPI_ISL_620400,
EPI_ISL_620401, EPI_ISL_620402, EPI_ISL_620403, EPI_ISL_620404, EPI_ISL_620405, EPI_ISL_620762, EPI_ISL_620763, EPI_ISL_620764, EPI_ISL_620765, EPI_ISL_620766, EPI_ISL_620767, EPI_ISL_620768, EPI_ISL_620769, EPI_ISL_620770, EPI_ISL_620771, EPI_ISL_620772, EPI_ISL_620773, EPI_ISL_620774,
EPI_ISL_620775, EPI_ISL_620776, EPI_ISL_620777, EPI_ISL_620778, EPI_ISL_620779, EPI_ISL_620780, EPI_ISL_620781, EPI_ISL_620782, EPI_ISL_620783, EPI_ISL_620784, EPI_ISL_620785, EPI_ISL_620786, EPI_ISL_620787, EPI_ISL_620788, EPI_ISL_620789, EPI_ISL_620790, EPI_ISL_620791, EPI_ISL_620792,
EPI_ISL_620793, EPI_ISL_620794, EPI_ISL_620795, EPI_ISL_620796, EPI_ISL_620797, EPI_ISL_620798, EPI_ISL_620799, EPI_ISL_620800, EPI_ISL_620801, EPI_ISL_620802, EPI_ISL_620803, EPI_ISL_620804, EPI_ISL_620805, EPI_ISL_620806, EPI_ISL_620807, EPI_ISL_620814, EPI_ISL_620833, EPI_ISL_620847,
EPI_ISL_620849, EPI_ISL_620876, EPI_ISL_620878, EPI_ISL_620889, EPI_ISL_620913, EPI_ISL_620917, EPI_ISL_620928, EPI_ISL_620931, EPI_ISL_620934, EPI_ISL_620980, EPI_ISL_620998, EPI_ISL_621008, EPI_ISL_621016, EPI_ISL_621017, EPI_ISL_621055, EPI_ISL_621084, EPI_ISL_621085, EPI_ISL_621123,
EPI_ISL_621128, EPI_ISL_621147, EPI_ISL_621148, EPI_ISL_621157, EPI_ISL_621187, EPI_ISL_621188, EPI_ISL_621189, EPI_ISL_621190, EPI_ISL_621191, EPI_ISL_621192, EPI_ISL_621193, EPI_ISL_621194, EPI_ISL_621224, EPI_ISL_621228, EPI_ISL_621244, EPI_ISL_621258, EPI_ISL_621262, EPI_ISL_621263,
EPI_ISL_621264, EPI_ISL_621265, EPI_ISL_621266, EPI_ISL_621267, EPI_ISL_621268, EPI_ISL_621269, EPI_ISL_621270, EPI_ISL_621271, EPI_ISL_621272, EPI_ISL_621286, EPI_ISL_621289, EPI_ISL_621290, EPI_ISL_621294, EPI_ISL_621298, EPI_ISL_621317, EPI_ISL_621319, EPI_ISL_621320, EPI_ISL_621321,
EPI_ISL_621322, EPI_ISL_621323, EPI_ISL_621324, EPI_ISL_621325, EPI_ISL_621326, EPI_ISL_621334, EPI_ISL_621349, EPI_ISL_621354, EPI_ISL_621355, EPI_ISL_621356, EPI_ISL_621357, EPI_ISL_621358, EPI_ISL_621359, EPI_ISL_621360, EPI_ISL_621361, EPI_ISL_621362, EPI_ISL_621363, EPI_ISL_621364,
EPI_ISL_621365, EPI_ISL_621366, EPI_ISL_621367, EPI_ISL_621368, EPI_ISL_621369, EPI_ISL_621370, EPI_ISL_621372, EPI_ISL_621373, EPI_ISL_621374, EPI_ISL_621375, EPI_ISL_621376, EPI_ISL_621377, EPI_ISL_621378, EPI_ISL_621379, EPI_ISL_621380, EPI_ISL_621381, EPI_ISL_621382, EPI_ISL_621383,
EPI_ISL_621384, EPI_ISL_621385, EPI_ISL_621386, EPI_ISL_621387, EPI_ISL_621388, EPI_ISL_621389, EPI_ISL_621390, EPI_ISL_621391, EPI_ISL_621392, EPI_ISL_621393, EPI_ISL_621394, EPI_ISL_621395, EPI_ISL_621396, EPI_ISL_621397, EPI_ISL_621398, EPI_ISL_621399, EPI_ISL_621400, EPI_ISL_621401,
EPI_ISL_621402, EPI_ISL_621403, EPI_ISL_621404, EPI_ISL_621405, EPI_ISL_621406, EPI_ISL_621407, EPI_ISL_621408, EPI_ISL_621409, EPI_ISL_621410, EPI_ISL_621411, EPI_ISL_621412, EPI_ISL_621413, EPI_ISL_621414, EPI_ISL_621415, EPI_ISL_621416, EPI_ISL_621417, EPI_ISL_621418, EPI_ISL_621419,
EPI_ISL_621420, EPI_ISL_621421, EPI_ISL_621422, EPI_ISL_621423, EPI_ISL_621424, EPI_ISL_621425, EPI_ISL_621426, EPI_ISL_621427, EPI_ISL_621428, EPI_ISL_621429, EPI_ISL_621430, EPI_ISL_621431, EPI_ISL_621432, EPI_ISL_621433, EPI_ISL_621434, EPI_ISL_621435, EPI_ISL_621436, EPI_ISL_621437,
EPI_ISL_621438, EPI_ISL_621439, EPI_ISL_621440, EPI_ISL_621441, EPI_ISL_621442, EPI_ISL_621443, EPI_ISL_621444, EPI_ISL_621445, EPI_ISL_621446, EPI_ISL_621447, EPI_ISL_621448, EPI_ISL_621449, EPI_ISL_621450, EPI_ISL_621451, EPI_ISL_621452, EPI_ISL_621453, EPI_ISL_621454, EPI_ISL_621455,
EPI_ISL_621456, EPI_ISL_621457, EPI_ISL_621458, EPI_ISL_621459, EPI_ISL_621460, EPI_ISL_621461, EPI_ISL_621462, EPI_ISL_621463, EPI_ISL_621464, EPI_ISL_621465, EPI_ISL_621466, EPI_ISL_621467, EPI_ISL_621468, EPI_ISL_621469, EPI_ISL_621470, EPI_ISL_621471, EPI_ISL_621472, EPI_ISL_621473,
EPI_ISL_621474, EPI_ISL_621475, EPI_ISL_621476, EPI_ISL_621477, EPI_ISL_621478, EPI_ISL_621479, EPI_ISL_621480, EPI_ISL_621481, EPI_ISL_621482, EPI_ISL_621483, EPI_ISL_621484, EPI_ISL_621485, EPI_ISL_621486, EPI_ISL_621487, EPI_ISL_621488, EPI_ISL_621489, EPI_ISL_621490, EPI_ISL_621491,
EPI_ISL_621492, EPI_ISL_621493, EPI_ISL_621494, EPI_ISL_621495, EPI_ISL_621496, EPI_ISL_621497, EPI_ISL_621498, EPI_ISL_621499, EPI_ISL_621500, EPI_ISL_621501, EPI_ISL_621502, EPI_ISL_621503, EPI_ISL_621504, EPI_ISL_621505, EPI_ISL_621506, EPI_ISL_621507, EPI_ISL_621508, EPI_ISL_621509,
EPI_ISL_621510, EPI_ISL_621511, EPI_ISL_621512, EPI_ISL_621513, EPI_ISL_621514, EPI_ISL_621515, EPI_ISL_621516, EPI_ISL_621517, EPI_ISL_621518, EPI_ISL_621519, EPI_ISL_621520, EPI_ISL_621521, EPI_ISL_621522, EPI_ISL_621523, EPI_ISL_621524, EPI_ISL_621525, EPI_ISL_621526, EPI_ISL_621527,
EPI_ISL_621528, EPI_ISL_621529, EPI_ISL_621530, EPI_ISL_621531, EPI_ISL_621532, EPI_ISL_621533, EPI_ISL_621534, EPI_ISL_621536, EPI_ISL_621537, EPI_ISL_621538, EPI_ISL_621539, EPI_ISL_621540, EPI_ISL_621541, EPI_ISL_621542, EPI_ISL_621543, EPI_ISL_621544, EPI_ISL_621545, EPI_ISL_621546,
EPI_ISL_621547, EPI_ISL_621548, EPI_ISL_621549, EPI_ISL_621550, EPI_ISL_621551, EPI_ISL_621552, EPI_ISL_621553, EPI_ISL_621554, EPI_ISL_621555, EPI_ISL_621556, EPI_ISL_621557, EPI_ISL_621558, EPI_ISL_621559, EPI_ISL_621560, EPI_ISL_621561, EPI_ISL_621562, EPI_ISL_621563, EPI_ISL_621564,
EPI_ISL_621565, EPI_ISL_621566, EPI_ISL_621567, EPI_ISL_621568, EPI_ISL_621569, EPI_ISL_621570, EPI_ISL_621571, EPI_ISL_621572, EPI_ISL_621573, EPI_ISL_621574, EPI_ISL_621575, EPI_ISL_621576, EPI_ISL_621577, EPI_ISL_621578, EPI_ISL_621579, EPI_ISL_621580, EPI_ISL_621581, EPI_ISL_621582,
EPI_ISL_621583, EPI_ISL_621584, EPI_ISL_621585, EPI_ISL_621586, EPI_ISL_621587, EPI_ISL_621588, EPI_ISL_621589, EPI_ISL_621590, EPI_ISL_621591, EPI_ISL_621592, EPI_ISL_621593, EPI_ISL_621594, EPI_ISL_621595, EPI_ISL_621596, EPI_ISL_621597, EPI_ISL_621598, EPI_ISL_621599, EPI_ISL_621600,
EPI_ISL_621601, EPI_ISL_621602, EPI_ISL_621603, EPI_ISL_621604, EPI_ISL_621605, EPI_ISL_621606, EPI_ISL_621607, EPI_ISL_621608, EPI_ISL_621609, EPI_ISL_621610, EPI_ISL_621611, EPI_ISL_621612, EPI_ISL_621613, EPI_ISL_621614, EPI_ISL_621615, EPI_ISL_621616, EPI_ISL_621617, EPI_ISL_621618,
EPI_ISL_621619, EPI_ISL_621620, EPI_ISL_621621, EPI_ISL_621622, EPI_ISL_621623, EPI_ISL_621624, EPI_ISL_621625, EPI_ISL_621626, EPI_ISL_621627, EPI_ISL_621628, EPI_ISL_621629, EPI_ISL_621630, EPI_ISL_621631, EPI_ISL_621632, EPI_ISL_621633, EPI_ISL_621634, EPI_ISL_621635, EPI_ISL_621636,
EPI_ISL_621637, EPI_ISL_621638, EPI_ISL_621639, EPI_ISL_621640, EPI_ISL_621641, EPI_ISL_621642, EPI_ISL_621643, EPI_ISL_621644, EPI_ISL_621645, EPI_ISL_621646, EPI_ISL_621647, EPI_ISL_621648, EPI_ISL_621649, EPI_ISL_621650, EPI_ISL_621651, EPI_ISL_621652, EPI_ISL_621653, EPI_ISL_621654,
EPI_ISL_621655, EPI_ISL_621656, EPI_ISL_621657, EPI_ISL_621658, EPI_ISL_621659, EPI_ISL_621660, EPI_ISL_621661, EPI_ISL_621662, EPI_ISL_621663, EPI_ISL_621664, EPI_ISL_621665, EPI_ISL_621666, EPI_ISL_621667, EPI_ISL_621668, EPI_ISL_621669, EPI_ISL_621670, EPI_ISL_621671, EPI_ISL_621672,
EPI_ISL_621673, EPI_ISL_621674, EPI_ISL_621675, EPI_ISL_621676, EPI_ISL_621677, EPI_ISL_621678, EPI_ISL_621679, EPI_ISL_621680, EPI_ISL_621681, EPI_ISL_621682, EPI_ISL_621683, EPI_ISL_621684, EPI_ISL_621685, EPI_ISL_621686, EPI_ISL_621687, EPI_ISL_621688, EPI_ISL_621689, EPI_ISL_621690,
EPI_ISL_621691, EPI_ISL_621692, EPI_ISL_621693, EPI_ISL_621694, EPI_ISL_621695, EPI_ISL_621696, EPI_ISL_621697, EPI_ISL_621698, EPI_ISL_621699, EPI_ISL_621700, EPI_ISL_621701, EPI_ISL_621702, EPI_ISL_621703, EPI_ISL_621704, EPI_ISL_621705, EPI_ISL_621706, EPI_ISL_621707, EPI_ISL_621708,
EPI_ISL_621709, EPI_ISL_621710, EPI_ISL_621711, EPI_ISL_621712, EPI_ISL_621713, EPI_ISL_621714, EPI_ISL_621715, EPI_ISL_621716, EPI_ISL_621717, EPI_ISL_621718, EPI_ISL_621719, EPI_ISL_621720, EPI_ISL_621721, EPI_ISL_621722, EPI_ISL_621723, EPI_ISL_621724, EPI_ISL_621725, EPI_ISL_621726,
EPI_ISL_621727, EPI_ISL_621728, EPI_ISL_621729, EPI_ISL_621730, EPI_ISL_621731, EPI_ISL_621732, EPI_ISL_621733, EPI_ISL_621734, EPI_ISL_621735, EPI_ISL_621736, EPI_ISL_621737, EPI_ISL_621738, EPI_ISL_621739, EPI_ISL_621740, EPI_ISL_621741, EPI_ISL_621742, EPI_ISL_621743, EPI_ISL_621744,
EPI_ISL_621745, EPI_ISL_621746, EPI_ISL_621747, EPI_ISL_621748, EPI_ISL_621749, EPI_ISL_621750, EPI_ISL_621751, EPI_ISL_621752, EPI_ISL_621753, EPI_ISL_621754, EPI_ISL_621755, EPI_ISL_621756, EPI_ISL_621757, EPI_ISL_621758, EPI_ISL_621759, EPI_ISL_621760, EPI_ISL_621761, EPI_ISL_621762,
EPI_ISL_621763, EPI_ISL_621764, EPI_ISL_621765, EPI_ISL_621766, EPI_ISL_621767, EPI_ISL_621768, EPI_ISL_621769, EPI_ISL_621770, EPI_ISL_621771, EPI_ISL_621772, EPI_ISL_621773, EPI_ISL_621774, EPI_ISL_621775, EPI_ISL_621776, EPI_ISL_621777, EPI_ISL_621779, EPI_ISL_621780, EPI_ISL_621781,
EPI_ISL_621782, EPI_ISL_621783, EPI_ISL_621784, EPI_ISL_621785, EPI_ISL_621786, EPI_ISL_621787, EPI_ISL_621788, EPI_ISL_621789, EPI_ISL_621790, EPI_ISL_621791, EPI_ISL_621792, EPI_ISL_621793, EPI_ISL_621794, EPI_ISL_621795, EPI_ISL_621796, EPI_ISL_621797, EPI_ISL_621798, EPI_ISL_621799,
EPI_ISL_621800, EPI_ISL_621801, EPI_ISL_621802, EPI_ISL_621803, EPI_ISL_621804, EPI_ISL_621805, EPI_ISL_621806, EPI_ISL_621807, EPI_ISL_621808, EPI_ISL_621809, EPI_ISL_621810, EPI_ISL_621811, EPI_ISL_621812, EPI_ISL_621813, EPI_ISL_621814, EPI_ISL_621815, EPI_ISL_621816, EPI_ISL_621817,
EPI_ISL_621818, EPI_ISL_621819, EPI_ISL_621820, EPI_ISL_621821, EPI_ISL_621822, EPI_ISL_621823, EPI_ISL_621824, EPI_ISL_621826, EPI_ISL_621827, EPI_ISL_621828, EPI_ISL_621829, EPI_ISL_621830, EPI_ISL_621831, EPI_ISL_621832, EPI_ISL_621833, EPI_ISL_621834, EPI_ISL_621835, EPI_ISL_621836,
EPI_ISL_621837, EPI_ISL_621838, EPI_ISL_621839, EPI_ISL_621840, EPI_ISL_621841, EPI_ISL_621842, EPI_ISL_621843, EPI_ISL_621844, EPI_ISL_621845, EPI_ISL_621846, EPI_ISL_621847, EPI_ISL_621848, EPI_ISL_621849, EPI_ISL_621850, EPI_ISL_621851, EPI_ISL_621852, EPI_ISL_621853, EPI_ISL_621854,
EPI_ISL_621855, EPI_ISL_621856, EPI_ISL_621857, EPI_ISL_621858, EPI_ISL_621859, EPI_ISL_621860, EPI_ISL_621861, EPI_ISL_621862, EPI_ISL_621863, EPI_ISL_621864, EPI_ISL_621865, EPI_ISL_621866, EPI_ISL_621867, EPI_ISL_621868, EPI_ISL_621869, EPI_ISL_621870, EPI_ISL_621871, EPI_ISL_621872,
EPI_ISL_621873, EPI_ISL_621881, EPI_ISL_621889, EPI_ISL_621898, EPI_ISL_621908, EPI_ISL_621931, EPI_ISL_621937, EPI_ISL_621938, EPI_ISL_621939, EPI_ISL_621941, EPI_ISL_621942, EPI_ISL_621943, EPI_ISL_621944, EPI_ISL_621945, EPI_ISL_621946, EPI_ISL_621947, EPI_ISL_621948, EPI_ISL_621949,
EPI_ISL_621950, EPI_ISL_621951, EPI_ISL_621952, EPI_ISL_621953, EPI_ISL_621954, EPI_ISL_621955, EPI_ISL_621956, EPI_ISL_621957, EPI_ISL_621958, EPI_ISL_621959, EPI_ISL_621960, EPI_ISL_621961, EPI_ISL_621962, EPI_ISL_621963, EPI_ISL_621964, EPI_ISL_621965, EPI_ISL_621966, EPI_ISL_621967,
EPI_ISL_621968, EPI_ISL_621969, EPI_ISL_621970, EPI_ISL_621971, EPI_ISL_621972, EPI_ISL_621973, EPI_ISL_621974, EPI_ISL_621975, EPI_ISL_621976, EPI_ISL_621977, EPI_ISL_621978, EPI_ISL_621979, EPI_ISL_621980, EPI_ISL_621981, EPI_ISL_621982, EPI_ISL_621983, EPI_ISL_621984, EPI_ISL_621985,
EPI_ISL_621986, EPI_ISL_621987, EPI_ISL_621988, EPI_ISL_621989, EPI_ISL_621990, EPI_ISL_621991, EPI_ISL_621992, EPI_ISL_621993, EPI_ISL_621994, EPI_ISL_621995, EPI_ISL_621996, EPI_ISL_621997, EPI_ISL_621998, EPI_ISL_621999, EPI_ISL_622000, EPI_ISL_622001, EPI_ISL_622002, EPI_ISL_622003,
EPI_ISL_622004, EPI_ISL_622005, EPI_ISL_622006, EPI_ISL_622007, EPI_ISL_622008, EPI_ISL_622009, EPI_ISL_622010, EPI_ISL_622011, EPI_ISL_622012, EPI_ISL_622013, EPI_ISL_622014, EPI_ISL_622015, EPI_ISL_622016, EPI_ISL_622017, EPI_ISL_622018, EPI_ISL_622019, EPI_ISL_622020, EPI_ISL_622021,
EPI_ISL_622022, EPI_ISL_622023, EPI_ISL_622024, EPI_ISL_622025, EPI_ISL_622026, EPI_ISL_622027, EPI_ISL_622028, EPI_ISL_622029, EPI_ISL_622030, EPI_ISL_622031, EPI_ISL_622032, EPI_ISL_622033, EPI_ISL_622034, EPI_ISL_622035, EPI_ISL_622036, EPI_ISL_622037, EPI_ISL_622038, EPI_ISL_622039,
EPI_ISL_622040, EPI_ISL_622041, EPI_ISL_622042, EPI_ISL_622043, EPI_ISL_622044, EPI_ISL_622045, EPI_ISL_622046, EPI_ISL_622047, EPI_ISL_622048, EPI_ISL_622049, EPI_ISL_622050, EPI_ISL_622051, EPI_ISL_622052, EPI_ISL_622053, EPI_ISL_622054, EPI_ISL_622055, EPI_ISL_622056, EPI_ISL_622057,
EPI_ISL_622058, EPI_ISL_622059, EPI_ISL_622060, EPI_ISL_622061, EPI_ISL_622062, EPI_ISL_622063, EPI_ISL_622064, EPI_ISL_622065, EPI_ISL_622067, EPI_ISL_622068, EPI_ISL_622069, EPI_ISL_622070, EPI_ISL_622071, EPI_ISL_622072, EPI_ISL_622073, EPI_ISL_622074, EPI_ISL_622075, EPI_ISL_622076,
EPI_ISL_622077, EPI_ISL_622078, EPI_ISL_622079, EPI_ISL_622080, EPI_ISL_622081, EPI_ISL_622082, EPI_ISL_622083, EPI_ISL_622084, EPI_ISL_622085, EPI_ISL_622086, EPI_ISL_622087, EPI_ISL_622088, EPI_ISL_622089, EPI_ISL_622090, EPI_ISL_622091, EPI_ISL_622092, EPI_ISL_622093, EPI_ISL_622094,
EPI_ISL_622095, EPI_ISL_622096, EPI_ISL_622097, EPI_ISL_622098, EPI_ISL_622099, EPI_ISL_622100, EPI_ISL_622101, EPI_ISL_622102, EPI_ISL_622103, EPI_ISL_622104, EPI_ISL_622105, EPI_ISL_622106, EPI_ISL_622107, EPI_ISL_622108, EPI_ISL_622109, EPI_ISL_622110, EPI_ISL_622111, EPI_ISL_622112,
EPI_ISL_622113, EPI_ISL_622114, EPI_ISL_622115, EPI_ISL_622116, EPI_ISL_622117, EPI_ISL_622118, EPI_ISL_622119, EPI_ISL_622120, EPI_ISL_622121, EPI_ISL_622122, EPI_ISL_622123, EPI_ISL_622124, EPI_ISL_622125, EPI_ISL_622126, EPI_ISL_622127, EPI_ISL_622128, EPI_ISL_622129, EPI_ISL_622130,
EPI_ISL_622131, EPI_ISL_622132, EPI_ISL_622133, EPI_ISL_622134, EPI_ISL_622135, EPI_ISL_622136, EPI_ISL_622137, EPI_ISL_622138, EPI_ISL_622139, EPI_ISL_622140, EPI_ISL_622141, EPI_ISL_622142, EPI_ISL_622143, EPI_ISL_622144, EPI_ISL_622145, EPI_ISL_622146, EPI_ISL_622147, EPI_ISL_622148,
EPI_ISL_622149, EPI_ISL_622150, EPI_ISL_622151, EPI_ISL_622152, EPI_ISL_622153, EPI_ISL_622154, EPI_ISL_622155, EPI_ISL_622156, EPI_ISL_622157, EPI_ISL_622158, EPI_ISL_622159, EPI_ISL_622160, EPI_ISL_622161, EPI_ISL_622162, EPI_ISL_622163, EPI_ISL_622164, EPI_ISL_622165, EPI_ISL_622166,
EPI_ISL_622167, EPI_ISL_622168, EPI_ISL_622169, EPI_ISL_622170, EPI_ISL_622171, EPI_ISL_622172, EPI_ISL_622173, EPI_ISL_622174, EPI_ISL_622175, EPI_ISL_622176, EPI_ISL_622177, EPI_ISL_622178, EPI_ISL_622179, EPI_ISL_622180, EPI_ISL_622181, EPI_ISL_622182, EPI_ISL_622183, EPI_ISL_622184,
EPI_ISL_622185, EPI_ISL_622186, EPI_ISL_622187, EPI_ISL_622188, EPI_ISL_622189, EPI_ISL_622190, EPI_ISL_622191, EPI_ISL_622192, EPI_ISL_622193, EPI_ISL_622194, EPI_ISL_622195, EPI_ISL_622196, EPI_ISL_622197, EPI_ISL_622198, EPI_ISL_622199, EPI_ISL_622200, EPI_ISL_622202, EPI_ISL_622203,
EPI_ISL_622216, EPI_ISL_622219, EPI_ISL_622243, EPI_ISL_622244, EPI_ISL_622245, EPI_ISL_622246, EPI_ISL_622247, EPI_ISL_622248, EPI_ISL_622249, EPI_ISL_622250, EPI_ISL_622251, EPI_ISL_622252, EPI_ISL_622253, EPI_ISL_622254, EPI_ISL_622255, EPI_ISL_622256, EPI_ISL_622257, EPI_ISL_622258,
EPI_ISL_622259, EPI_ISL_622260, EPI_ISL_622261, EPI_ISL_622278, EPI_ISL_622279, EPI_ISL_622280, EPI_ISL_622281, EPI_ISL_622282, EPI_ISL_622283, EPI_ISL_622284, EPI_ISL_622285, EPI_ISL_622286, EPI_ISL_622287, EPI_ISL_622288, EPI_ISL_622289, EPI_ISL_622290, EPI_ISL_622291, EPI_ISL_622292,
EPI_ISL_622293, EPI_ISL_622294, EPI_ISL_622295, EPI_ISL_622296, EPI_ISL_622297, EPI_ISL_622298, EPI_ISL_622299, EPI_ISL_622300, EPI_ISL_622301, EPI_ISL_622302, EPI_ISL_622303, EPI_ISL_622304, EPI_ISL_622305, EPI_ISL_622306, EPI_ISL_622307, EPI_ISL_622308, EPI_ISL_622309, EPI_ISL_622310,
EPI_ISL_622311, EPI_ISL_622312, EPI_ISL_622313, EPI_ISL_622314, EPI_ISL_622315, EPI_ISL_622316, EPI_ISL_622317, EPI_ISL_622318, EPI_ISL_622319, EPI_ISL_622320, EPI_ISL_622321, EPI_ISL_622322, EPI_ISL_622323, EPI_ISL_622324, EPI_ISL_622325, EPI_ISL_622326, EPI_ISL_622327, EPI_ISL_622328,
EPI_ISL_622329, EPI_ISL_622330, EPI_ISL_622331, EPI_ISL_622332, EPI_ISL_622333, EPI_ISL_622334, EPI_ISL_622335, EPI_ISL_622336, EPI_ISL_622337, EPI_ISL_622338, EPI_ISL_622339, EPI_ISL_622340, EPI_ISL_622341, EPI_ISL_622342, EPI_ISL_622343, EPI_ISL_622344, EPI_ISL_622345, EPI_ISL_622346,
EPI_ISL_622347, EPI_ISL_622348, EPI_ISL_622349, EPI_ISL_622350, EPI_ISL_622351, EPI_ISL_622352, EPI_ISL_622353, EPI_ISL_622354, EPI_ISL_622355, EPI_ISL_622356, EPI_ISL_622357, EPI_ISL_622358, EPI_ISL_622359, EPI_ISL_622360, EPI_ISL_622361, EPI_ISL_622362, EPI_ISL_622363, EPI_ISL_622364,
EPI_ISL_622365, EPI_ISL_622366, EPI_ISL_622367, EPI_ISL_622368, EPI_ISL_622369, EPI_ISL_622370, EPI_ISL_622371, EPI_ISL_622372, EPI_ISL_622373, EPI_ISL_622374, EPI_ISL_622375, EPI_ISL_622376, EPI_ISL_622377, EPI_ISL_622378, EPI_ISL_622379, EPI_ISL_622380, EPI_ISL_622381, EPI_ISL_622382,
EPI_ISL_622383, EPI_ISL_622384, EPI_ISL_622385, EPI_ISL_622386, EPI_ISL_622387, EPI_ISL_622388, EPI_ISL_622389, EPI_ISL_622390, EPI_ISL_622391, EPI_ISL_622392, EPI_ISL_622393, EPI_ISL_622394, EPI_ISL_622395, EPI_ISL_622396, EPI_ISL_622397, EPI_ISL_622398, EPI_ISL_622399, EPI_ISL_622400,
EPI_ISL_622401, EPI_ISL_622402, EPI_ISL_622404, EPI_ISL_622406, EPI_ISL_622443, EPI_ISL_622501, EPI_ISL_622515, EPI_ISL_622544, EPI_ISL_622643, EPI_ISL_622648, EPI_ISL_622651, EPI_ISL_622672, EPI_ISL_622676, EPI_ISL_622748, EPI_ISL_622749, EPI_ISL_622750, EPI_ISL_622751, EPI_ISL_622752,
EPI_ISL_622753, EPI_ISL_622754, EPI_ISL_622755, EPI_ISL_622756, EPI_ISL_622757, EPI_ISL_622758, EPI_ISL_622759, EPI_ISL_622760, EPI_ISL_625684, EPI_ISL_625685, EPI_ISL_625686, EPI_ISL_625687, EPI_ISL_625688, EPI_ISL_625689, EPI_ISL_625690, EPI_ISL_625691, EPI_ISL_625692, EPI_ISL_625693,
EPI_ISL_625694, EPI_ISL_625695, EPI_ISL_625697, EPI_ISL_625698, EPI_ISL_625699, EPI_ISL_625700, EPI_ISL_625701, EPI_ISL_625702, EPI_ISL_625703, EPI_ISL_625704, EPI_ISL_625705, EPI_ISL_625706, EPI_ISL_625707, EPI_ISL_625708, EPI_ISL_625709, EPI_ISL_625710, EPI_ISL_625711, EPI_ISL_625712,
EPI_ISL_625713, EPI_ISL_625714, EPI_ISL_625715, EPI_ISL_625716, EPI_ISL_625717, EPI_ISL_625718, EPI_ISL_625719, EPI_ISL_625720, EPI_ISL_625721, EPI_ISL_625722, EPI_ISL_625723, EPI_ISL_625724, EPI_ISL_625725, EPI_ISL_625726, EPI_ISL_625727, EPI_ISL_625728, EPI_ISL_625729, EPI_ISL_625730,
EPI_ISL_625731, EPI_ISL_625732, EPI_ISL_625733, EPI_ISL_625734, EPI_ISL_625735, EPI_ISL_625736, EPI_ISL_625737, EPI_ISL_625738, EPI_ISL_625739, EPI_ISL_625741, EPI_ISL_625742, EPI_ISL_625743, EPI_ISL_625744, EPI_ISL_625745, EPI_ISL_625746, EPI_ISL_625747, EPI_ISL_625748, EPI_ISL_625749,
EPI_ISL_625750, EPI_ISL_625751, EPI_ISL_625752, EPI_ISL_625753, EPI_ISL_625754, EPI_ISL_625755, EPI_ISL_625757, EPI_ISL_625758, EPI_ISL_625759, EPI_ISL_625760, EPI_ISL_625761, EPI_ISL_625762, EPI_ISL_625763, EPI_ISL_625764, EPI_ISL_625765, EPI_ISL_625766, EPI_ISL_625767, EPI_ISL_625768,
EPI_ISL_625769, EPI_ISL_625770, EPI_ISL_625771, EPI_ISL_625772, EPI_ISL_625773, EPI_ISL_625774, EPI_ISL_625775, EPI_ISL_625776, EPI_ISL_625777, EPI_ISL_625778, EPI_ISL_625779, EPI_ISL_625780, EPI_ISL_625781, EPI_ISL_625782, EPI_ISL_625783, EPI_ISL_625784, EPI_ISL_625785, EPI_ISL_625786,
EPI_ISL_625787, EPI_ISL_625788, EPI_ISL_625789, EPI_ISL_625790, EPI_ISL_625791, EPI_ISL_625792, EPI_ISL_625793, EPI_ISL_625794, EPI_ISL_625795, EPI_ISL_625796, EPI_ISL_625797, EPI_ISL_625798, EPI_ISL_625799, EPI_ISL_625800, EPI_ISL_625801, EPI_ISL_625802, EPI_ISL_625803, EPI_ISL_625804,
EPI_ISL_625805, EPI_ISL_625806, EPI_ISL_625807, EPI_ISL_625808, EPI_ISL_625809, EPI_ISL_625810, EPI_ISL_625811, EPI_ISL_625812, EPI_ISL_625813, EPI_ISL_625814, EPI_ISL_625815, EPI_ISL_625816, EPI_ISL_625817, EPI_ISL_625818, EPI_ISL_625819, EPI_ISL_625820, EPI_ISL_625821, EPI_ISL_625822,
EPI_ISL_625823, EPI_ISL_625824, EPI_ISL_625825, EPI_ISL_625826, EPI_ISL_625827, EPI_ISL_625836, EPI_ISL_625838, EPI_ISL_625840, EPI_ISL_625841, EPI_ISL_625842, EPI_ISL_625843, EPI_ISL_625844, EPI_ISL_625846, EPI_ISL_625847, EPI_ISL_625849, EPI_ISL_625850, EPI_ISL_625851, EPI_ISL_625853,
EPI_ISL_625854, EPI_ISL_625855, EPI_ISL_625856, EPI_ISL_625857, EPI_ISL_625858, EPI_ISL_625859, EPI_ISL_625860, EPI_ISL_625861, EPI_ISL_625862, EPI_ISL_625863, EPI_ISL_625864, EPI_ISL_625866, EPI_ISL_625867, EPI_ISL_625868, EPI_ISL_625869, EPI_ISL_625870, EPI_ISL_625872, EPI_ISL_625873,
EPI_ISL_625874, EPI_ISL_625875, EPI_ISL_625876, EPI_ISL_625877, EPI_ISL_625878, EPI_ISL_625879, EPI_ISL_625880, EPI_ISL_625881, EPI_ISL_625882, EPI_ISL_625883, EPI_ISL_625885, EPI_ISL_625886, EPI_ISL_625887, EPI_ISL_625888, EPI_ISL_625889, EPI_ISL_625890, EPI_ISL_625891, EPI_ISL_625892,
EPI_ISL_625893, EPI_ISL_625894, EPI_ISL_625895, EPI_ISL_625896, EPI_ISL_625897, EPI_ISL_625898, EPI_ISL_625899, EPI_ISL_625900, EPI_ISL_625901, EPI_ISL_625902, EPI_ISL_625903, EPI_ISL_625904, EPI_ISL_625905, EPI_ISL_625906, EPI_ISL_625907, EPI_ISL_625908, EPI_ISL_625909, EPI_ISL_625910,
EPI_ISL_625911, EPI_ISL_625912, EPI_ISL_625913, EPI_ISL_625914, EPI_ISL_625915, EPI_ISL_625916, EPI_ISL_625917, EPI_ISL_625918, EPI_ISL_625919, EPI_ISL_625920, EPI_ISL_625921, EPI_ISL_625922, EPI_ISL_625923, EPI_ISL_625924, EPI_ISL_625925, EPI_ISL_625926, EPI_ISL_625927, EPI_ISL_625928,
EPI_ISL_625929, EPI_ISL_625930, EPI_ISL_625931, EPI_ISL_625932, EPI_ISL_625934, EPI_ISL_625935, EPI_ISL_625936, EPI_ISL_625937, EPI_ISL_625938, EPI_ISL_625939, EPI_ISL_625940, EPI_ISL_625941, EPI_ISL_625942, EPI_ISL_625943, EPI_ISL_625944, EPI_ISL_625945, EPI_ISL_625946, EPI_ISL_625947,
EPI_ISL_625948, EPI_ISL_625949, EPI_ISL_625950, EPI_ISL_625951, EPI_ISL_625952, EPI_ISL_625953, EPI_ISL_625954, EPI_ISL_625955, EPI_ISL_625956, EPI_ISL_625957, EPI_ISL_625958, EPI_ISL_625959, EPI_ISL_625961, EPI_ISL_625962, EPI_ISL_625963, EPI_ISL_625964, EPI_ISL_625965, EPI_ISL_625966,
EPI_ISL_625967, EPI_ISL_625968, EPI_ISL_625969, EPI_ISL_625970, EPI_ISL_625971, EPI_ISL_625974, EPI_ISL_625975, EPI_ISL_625976, EPI_ISL_625977, EPI_ISL_625978, EPI_ISL_625979, EPI_ISL_625980, EPI_ISL_625981, EPI_ISL_625982, EPI_ISL_625983, EPI_ISL_625984, EPI_ISL_625985, EPI_ISL_625986,
EPI_ISL_625987, EPI_ISL_625988, EPI_ISL_625989, EPI_ISL_625990, EPI_ISL_625991, EPI_ISL_625992, EPI_ISL_625993, EPI_ISL_625994, EPI_ISL_625995, EPI_ISL_625996, EPI_ISL_625997, EPI_ISL_625999, EPI_ISL_626000, EPI_ISL_626001, EPI_ISL_626002, EPI_ISL_626003, EPI_ISL_626004, EPI_ISL_626005,
EPI_ISL_626006, EPI_ISL_626007, EPI_ISL_626008, EPI_ISL_626009, EPI_ISL_626011, EPI_ISL_626012, EPI_ISL_626013, EPI_ISL_626014, EPI_ISL_626015, EPI_ISL_626016, EPI_ISL_626017, EPI_ISL_626018, EPI_ISL_626019, EPI_ISL_626020, EPI_ISL_626021, EPI_ISL_626022, EPI_ISL_626023, EPI_ISL_626024,
EPI_ISL_626025, EPI_ISL_626026, EPI_ISL_626027, EPI_ISL_626028, EPI_ISL_626029, EPI_ISL_626030, EPI_ISL_626031, EPI_ISL_626032, EPI_ISL_626033, EPI_ISL_626034, EPI_ISL_626035, EPI_ISL_626036, EPI_ISL_626037, EPI_ISL_626038, EPI_ISL_626039, EPI_ISL_626040, EPI_ISL_626041, EPI_ISL_626042,
EPI_ISL_626043, EPI_ISL_626044, EPI_ISL_626045, EPI_ISL_626046, EPI_ISL_626047, EPI_ISL_626048, EPI_ISL_626049, EPI_ISL_626050, EPI_ISL_626051, EPI_ISL_626052, EPI_ISL_626053, EPI_ISL_626054, EPI_ISL_626055, EPI_ISL_626056, EPI_ISL_626057, EPI_ISL_626058, EPI_ISL_626059, EPI_ISL_626060,
EPI_ISL_626061, EPI_ISL_626062, EPI_ISL_626063, EPI_ISL_626064, EPI_ISL_626065, EPI_ISL_626066, EPI_ISL_626067, EPI_ISL_626068, EPI_ISL_626069, EPI_ISL_626070, EPI_ISL_626071, EPI_ISL_626072, EPI_ISL_626073, EPI_ISL_626074, EPI_ISL_626075, EPI_ISL_626076, EPI_ISL_626077, EPI_ISL_626078,
EPI_ISL_626079, EPI_ISL_626080, EPI_ISL_626081, EPI_ISL_626082, EPI_ISL_626083, EPI_ISL_626084, EPI_ISL_626085, EPI_ISL_626086, EPI_ISL_626087, EPI_ISL_626088, EPI_ISL_626089, EPI_ISL_626090, EPI_ISL_626091, EPI_ISL_626092, EPI_ISL_626093, EPI_ISL_626094, EPI_ISL_626095, EPI_ISL_626096,
EPI_ISL_626097, EPI_ISL_626098, EPI_ISL_626099, EPI_ISL_626100, EPI_ISL_626101, EPI_ISL_626102, EPI_ISL_626103, EPI_ISL_626104, EPI_ISL_626105, EPI_ISL_626106, EPI_ISL_626107, EPI_ISL_626108, EPI_ISL_626109, EPI_ISL_626110, EPI_ISL_626111, EPI_ISL_626112, EPI_ISL_626113, EPI_ISL_626114,

| | | | |
|---|---|---|---|
| EPI_ISL_626115, EPI_ISL_626116, EPI_ISL_626117, EPI_ISL_626118, EPI_ISL_626119, EPI_ISL_626120, EPI_ISL_626121, EPI_ISL_626122, EPI_ISL_626123, EPI_ISL_626124, EPI_ISL_626125, EPI_ISL_626126, EPI_ISL_626127, EPI_ISL_626128, EPI_ISL_626129, EPI_ISL_626130, EPI_ISL_626131, EPI_ISL_626132, EPI_ISL_626133, EPI_ISL_626134, EPI_ISL_626135, EPI_ISL_626136, EPI_ISL_626137, EPI_ISL_626138, EPI_ISL_626139, EPI_ISL_626140, EPI_ISL_626141, EPI_ISL_626142, EPI_ISL_626143, EPI_ISL_626144, EPI_ISL_626145, EPI_ISL_626146, EPI_ISL_626147, EPI_ISL_626148, EPI_ISL_626149, EPI_ISL_626150, EPI_ISL_626151, EPI_ISL_626152, EPI_ISL_626153, EPI_ISL_626154, EPI_ISL_626155, EPI_ISL_626156, EPI_ISL_626157, EPI_ISL_626158, EPI_ISL_626159, EPI_ISL_626160, EPI_ISL_626161, EPI_ISL_626162, EPI_ISL_626163, EPI_ISL_626164, EPI_ISL_626165, EPI_ISL_626166, EPI_ISL_626167, EPI_ISL_626168, EPI_ISL_626172, EPI_ISL_626173, EPI_ISL_626178, EPI_ISL_626180, EPI_ISL_626181, EPI_ISL_626182, EPI_ISL_626188, EPI_ISL_626189, EPI_ISL_626195, EPI_ISL_626196, EPI_ISL_626197, EPI_ISL_626199, EPI_ISL_626200, EPI_ISL_626201, EPI_ISL_626207, EPI_ISL_626208 | | | |
| see above | Department of Virus and Microbiological Special Diagnostics, Statens Serum Institut, Denmark | Albertsen lab, Department of Chemistry and Bioscience, Aalborg University, Denmark | Danish Covid-19 Genome Consortia |
| EPI_ISL_626656, EPI_ISL_626657, EPI_ISL_626660, EPI_ISL_626663, EPI_ISL_626667, EPI_ISL_626681, EPI_ISL_626693, EPI_ISL_626698, EPI_ISL_626700, EPI_ISL_626701, EPI_ISL_626702, EPI_ISL_626704, EPI_ISL_626705, EPI_ISL_626737, EPI_ISL_626741, EPI_ISL_626744, EPI_ISL_626745, EPI_ISL_626751, EPI_ISL_626753, EPI_ISL_626756, EPI_ISL_626759, EPI_ISL_626765, EPI_ISL_626779, EPI_ISL_626790, EPI_ISL_626795, EPI_ISL_626808, EPI_ISL_626814, EPI_ISL_626819, EPI_ISL_626860, EPI_ISL_626864, EPI_ISL_626868, EPI_ISL_626893, EPI_ISL_626894, EPI_ISL_626914, EPI_ISL_626954, EPI_ISL_626955, EPI_ISL_626956, EPI_ISL_626964, EPI_ISL_627009, EPI_ISL_627024, EPI_ISL_627041, EPI_ISL_627080, EPI_ISL_627081, EPI_ISL_627098, EPI_ISL_627099, EPI_ISL_627148, EPI_ISL_627170, EPI_ISL_627180, EPI_ISL_627187, EPI_ISL_627193, EPI_ISL_627198, EPI_ISL_627213, EPI_ISL_627214, EPI_ISL_627300, EPI_ISL_627570, EPI_ISL_627571, EPI_ISL_627572, EPI_ISL_627573, EPI_ISL_627574, EPI_ISL_627575, EPI_ISL_627576, EPI_ISL_627577, EPI_ISL_627578, EPI_ISL_627579, EPI_ISL_627580, EPI_ISL_627581, EPI_ISL_627582, EPI_ISL_627583, EPI_ISL_627586, EPI_ISL_627587, EPI_ISL_627588, EPI_ISL_627589, EPI_ISL_627590, EPI_ISL_627591, EPI_ISL_627592, EPI_ISL_627593, EPI_ISL_627594, EPI_ISL_627595, EPI_ISL_627596, EPI_ISL_627597, EPI_ISL_627598, EPI_ISL_627599, EPI_ISL_627600, EPI_ISL_627601, EPI_ISL_627602, EPI_ISL_627603, EPI_ISL_627604, EPI_ISL_627605, EPI_ISL_627606, EPI_ISL_627607, EPI_ISL_627608, EPI_ISL_627609, EPI_ISL_627610, EPI_ISL_627611, EPI_ISL_627612, EPI_ISL_627613, EPI_ISL_627614, EPI_ISL_627615, EPI_ISL_627616, EPI_ISL_627617, EPI_ISL_627618, EPI_ISL_627619, EPI_ISL_627620, EPI_ISL_627621, EPI_ISL_627622, EPI_ISL_627623, EPI_ISL_627624, EPI_ISL_627625, EPI_ISL_627626, EPI_ISL_627627, EPI_ISL_627628, EPI_ISL_627629, EPI_ISL_627630, EPI_ISL_627631, EPI_ISL_627632, EPI_ISL_627633, EPI_ISL_627634, EPI_ISL_627635, EPI_ISL_627636, EPI_ISL_627637, EPI_ISL_627638, EPI_ISL_627639, EPI_ISL_627640, EPI_ISL_627641, EPI_ISL_627642, EPI_ISL_627643, EPI_ISL_627644, EPI_ISL_627645, EPI_ISL_627646, EPI_ISL_627647, EPI_ISL_627648, EPI_ISL_627649, EPI_ISL_627650, EPI_ISL_627651, EPI_ISL_627652, EPI_ISL_627653, EPI_ISL_627654, EPI_ISL_627655, EPI_ISL_627656, EPI_ISL_627657, EPI_ISL_627658 | | | |
| see above | Quadram Institute Bioscience | COVID-19 Genomics UK (COG-UK) Consortium | Dave J. Baker, Gemma L. Kay, Alp Aydin, Thanh Le-Viet, Steven Rudder, Ana P. Tedim, Anastasia Kolyva, Maria Diaz, Leonardo de Oliveira Martins, Nabil-Fareed Alikhan, Lizzie Meadows, Rachael Stanley, Ngozi Elumogo, Muhammed Yasir, Nicholas M. Thomson, Alexander J Trotter, Rachel Gilroy, Samuel Bloomfield, Claire Stuart, Andrew Bell, Reenesh Prakash, Samir Dervisevic, Alison E. Mather, John Wain, Mark Webber, Andrew J. Page, Justin O'Grady |
| EPI_ISL_631277, EPI_ISL_631278, EPI_ISL_631279, EPI_ISL_631280, EPI_ISL_631281, EPI_ISL_631282, EPI_ISL_631283, EPI_ISL_631284, EPI_ISL_631285, EPI_ISL_631286, EPI_ISL_631287, EPI_ISL_631288, EPI_ISL_631289, EPI_ISL_631290, EPI_ISL_631291, EPI_ISL_631292, EPI_ISL_631293, EPI_ISL_631294, EPI_ISL_631295, EPI_ISL_631296, EPI_ISL_631297, EPI_ISL_631298, EPI_ISL_631299, EPI_ISL_631300, EPI_ISL_631301, EPI_ISL_631302, EPI_ISL_631303, EPI_ISL_632311 | | | |
| see above | MVZ DIAMEDIS Diagnostische Medizin Sennestadt GmbH | Bielefeld University | David Brandt, Tobias Busche, Markus Haak, Jörn Kalinowski, Levin-Joe Klages, Christiane Scherer, Alexander Sczyrba, Marina Simunovic, Svenja Vinke |
| EPI_ISL_632847, EPI_ISL_632848, EPI_ISL_632849, EPI_ISL_632850, EPI_ISL_632851, EPI_ISL_632852, EPI_ISL_632853, EPI_ISL_632854, EPI_ISL_632855, EPI_ISL_632856, EPI_ISL_632857, EPI_ISL_632858, EPI_ISL_632859, EPI_ISL_632860, EPI_ISL_632861, EPI_ISL_632862, EPI_ISL_632863, EPI_ISL_632864, EPI_ISL_632865, EPI_ISL_632866, EPI_ISL_632867, EPI_ISL_632868, EPI_ISL_632869, EPI_ISL_632870, EPI_ISL_632871, EPI_ISL_632872, EPI_ISL_632873, EPI_ISL_632874, EPI_ISL_632875, EPI_ISL_632876, EPI_ISL_632877, EPI_ISL_632878, EPI_ISL_632879, EPI_ISL_632880, EPI_ISL_632881, EPI_ISL_632882, EPI_ISL_632883, EPI_ISL_632884, EPI_ISL_632885, EPI_ISL_632886, EPI_ISL_632887, EPI_ISL_632888, EPI_ISL_632889 | | | |
| see above | Idaho Bureau of Laboratories | Center for Global Health, University of New Mexico Health Sciences Center | Daryl Domman, Kurt Schwalm, Matthew Burns, Robert Voermans, Christopher Ball, Darrell Dinwiddie |

We gratefully acknowledge the following Authors from the Originating laboratories responsible for obtaining the specimens, as well as the Submitting laboratories where the genome data were generated and shared via GISAID, on which this research is based.

All Submitters of data may be contacted directly via www.gisaid.org

Authors are sorted alphabetically.

| Accession ID | Originating Laboratory | Submitting Laboratory | Authors |
|---|---|---|---|
| EPI_ISL_407893 | Centre for Infectious Diseases and Microbiology Laboratory Services | NSW Health Pathology - Institute of Clinical Pathology and Medical Research; Westmead Hospital; University of Sydney | Eden J-S, Carter I, Rahman H, Holmes EC, Rockett R, O'Sullivan MV, Sintchenko V, Chen SC, Maddocks S, Kok J and Dwyer DE for the 2019-nCoV Study Group |
| EPI_ISL_408977 | Serology, Virology and OTDS Laboratories (SAViD), NSW Health Pathology Randwick | NSW Health Pathology - Institute of Clinical Pathology and Medical Research; Centre for Infectious Diseases and Microbiology Laboratory Services; Westmead Hospital; University of Sydney | Eden J-S, Carter I, Rahman H, Rawlinson W, Holmes EC, Rockett R, O'Sullivan MV, Sintchenko V, Chen SC, Maddocks S, Kok J and Dwyer DE for the 2019-nCoV Study Group* |
| EPI_ISL_411902 | Virology Unit, Institut Pasteur du Cambodge. | Virology Unit, Institut Pasteur du Cambodge (Sequencing done by: Jessica E Manning/Jennifer A Bohl at Malaria and Vector Research Research Laboratory, National Institute of Allergy and Infectious Diseases and Vida Ahyong from Chan-Zuckerberg Biohub) | Erik A Karlsson, Jennifer A Bohl, Vida Ahyong, Veasna Duong, Philippe Dussart, Jessica E Manning. |
| EPI_ISL_412028 | Hong Kong Department of Health | School of Public Health, The University of Hon g Kong | Dominic N.C. Tsang, Daniel K.W. Chu, Leo L.M. Poon, Malik Peiris |
| EPI_ISL_412029 | Hong Kong Department of Health | The University of Hong Kong | Dominic N.C. Tsang, Daniel K.W. Chu, Leo L.M. Poon, Malik Peiris |
| EPI_ISL_412030 | Hong Kong Department of Health | School of Public Health, The University of Hon g Kong | Dominic N.C. Tsang, Daniel K.W. Chu, Leo L.M. Poon, Malik Peiris |
| EPI_ISL_412975 | Centre for Infectious Diseases and Microbiology Laboratory Services | NSW Health Pathology - Institute of Clinical Pathology and Medical Research; Westmead Hospital; University of Sydney | Eden J-S, Carter I, Rahman H, Holmes EC, Rockett R, O'Sullivan MV, Sintchenko V, Chen SC, Maddocks S, Kok J and Dwyer DE for the 2019-nCoV Study Group |
| EPI_ISL_413213, EPI_ISL_413214 | Centre for Infectious Diseases and Microbiology Laboratory Services | NSW Health Pathology - Institute of Clinical Pathology and Medical Research; Westmead Hospital; University of Sydney | Eden J-S, Carter I, Rahman H, Holmes EC, Rockett R, O'Sullivan MV, Sintchenko V, Chen SC, Maddocks S, Kok J and Dwyer DE for the 2019-nCoV Study Group* |
| EPI_ISL_413221 | West of Scotland Specialist Virology Centre, NHSGGC | MRC-University of Glasgow Centre for Virus Research | Emma Thomson, Antonia Ho; James Shephard, Shirin Ashraf; Kathy Smollett, Daniel Mair, Stephen Carmichael, Ana da Silva Filipe; Richard Orton, Josh Singer, David L Robertson; Andrew Rambaut; Alasdair MacLean, Rory Gunson. |
| EPI_ISL_413600 | Centre for Infectious Diseases and Microbiology - Public Health | NSW Health Pathology - Institute of Clinical Pathology and Medical Research; Westmead Hospital; University of Sydney | Gall, M, Eden J-S, Lam C, Gray K, Timms, V, Rockett R, Carter I, Rahman H, Holmes EC, O'Sullivan MV, Sintchenko V, Chen SC, Maddocks S, Kok J and Dwyer DE for the 2019-nCoV Study Group* |
| EPI_ISL_414024, EPI_ISL_414025, EPI_ISL_414026, EPI_ISL_414027 | West of Scotland Specialist Virology Centre, NHSGGC | MRC-University of Glasgow Centre for Virus Research | Emma Thomson, Antonia Ho; Kathy Smollett, Daniel Mair, Stephen Carmichael, Ana da Silva Filipe; Richard Orton, David L Robertson; Alasdair MacLean, Rory Gunson. |
| EPI_ISL_414517, EPI_ISL_414519, EPI_ISL_414527, EPI_ISL_414528, EPI_ISL_414567, EPI_ISL_414569, EPI_ISL_414571 | Hong Kong Department of Health | School of Public Health, The University of Hong Kong | Dominic N.C. Tsang, Daniel K.W. Chu, Leo L.M. Poon, Malik Peiris |
| EPI_ISL_415539, EPI_ISL_415541, EPI_ISL_415542, EPI_ISL_415543, EPI_ISL_415544 | Utah Public Health Laboratory | Utah Public Health Laboratory | Erin Young, Kelly Oakeson |
| EPI_ISL_416458 | Virology laboratory Ministry of Health Kuwait sequenced at Dasman Diabetes Institute | Dasman Diabetes Institute | Fahd Al-Mulla, Sumi John, Sara Alqabandi, Rasheeba iqbal, Motasem Melhem, Ebaa alOzairi, Qais Al-Duwairi |
| EPI_ISL_416489, EPI_ISL_416491, EPI_ISL_416492, EPI_ISL_416523 | University of Wisconsin-Madison AIDS Vaccine Research Laboratories | University of Wisconsin-Madison AIDS Vaccine Research Laboratories | Gage Moreno, Katarina Braun, et al. AIDS Vaccine Research Laboratories |
| EPI_ISL_416541 | Dasman Diabetes Institute and Virology Laboratory Ministry of Health | Dasman Diabetes Institute | Fahd Al-Mulla, Sumi John, Rasheeba Iqbal, Motasem Melhem, Ebaa AlOzairi, Sara Al-Qabandi, Qais Al-Duwairi |
| EPI_ISL_416542 | Dasman Diabetes Institute | Dasman Diabetes Institute | Fahd Al-Mulla, Sumi John, Rasheeba Iqbal, Motasem Melhem, Ebaa AlOzairi, Sara Al-Qabandi, Qais Al-Duwairi |
| EPI_ISL_416543 | Dasman Diabetes Institute | Dasman Diabetes Institute | Fahd Al-Mulla, Rasheeba Iqbal, Sumi John, Motasem Melhem, Ebaa AlOzairi, Sara Al-Qabandi, Qais Al-Duwairi |
| EPI_ISL_416997, EPI_ISL_417004, EPI_ISL_417005, EPI_ISL_417006, EPI_ISL_417008, EPI_ISL_417009, EPI_ISL_417011, EPI_ISL_417012, EPI_ISL_417013, EPI_ISL_417014, EPI_ISL_417015, EPI_ISL_417016, EPI_ISL_417017, EPI_ISL_417018, EPI_ISL_417019, EPI_ISL_417020, EPI_ISL_417021, EPI_ISL_417022, EPI_ISL_417023, EPI_ISL_417025 | | | |
| see above | Department of Clinical Microbiology | GIGA Medical Genomics | Durkin Keith, Artesi Maria, Bontems Sébastien, Boreux Raphaël, Meex Cécile, Melin Pierrette, Hayette Marie-Pierre, Bours Vincent. |
| EPI_ISL_417026, EPI_ISL_417027, EPI_ISL_417028 | Utah Public Health Laboratory | Utah Public Health Laboratory | Erin Young, Kelly Oakeson |
| EPI_ISL_417030 | Centre for Infectious Diseases and Microbiology Laboratory Services | NSW Health Pathology - Institute of Clinical Pathology and Medical Research; Westmead Hospital; University of Sydney | Eden J-S, Rockett R, Carter I, Rahman H, Holmes EC, O'Sullivan MV, Sintchenko V, Chen SC, Maddocks S, Kok J and Dwyer DE for the 2019-nCoV Study Group* |
| EPI_ISL_417200, EPI_ISL_417201, EPI_ISL_417202, EPI_ISL_417203, EPI_ISL_417204 | University of Wisconsin-Madison AIDS Vaccine Research Laboratories | University of Wisconsin-Madison AIDS Vaccine Research Laboratories | Gage Moreno, Katarina Braun, et al. AIDS Vaccine Research Laboratories |
| EPI_ISL_417384 | Centre for Infectious Diseases and Microbiology Public Health | NSW Health Pathology - Institute of Clinical Pathology and Medical Research; Westmead Hospital; University of Sydney | Eden J-S, Lam C, Gray K, Timms V, Gall M, Arnott A, Sadsad R, Carter I, Rahman H, Holmes EC, O'Sullivan MV, Sintchenko V, Chen SC, Maddocks S, Kok J, Dwyer DE and Rockett R for the 2019-nCoV Study Group |
| EPI_ISL_417388 | Centre for Infectious Diseases and Microbiology Public Health | NSW Health Pathology - Institute of Clinical Pathology and Medical Research; Westmead Hospital; University of Sydney | Gall M, Arnott A, Sadsad R, Carter I, Rahman H, Holmes EC, O'Sullivan MV, Sintchenko V, Chen SC, Maddocks S, Kok J, Dwyer DE, Rockett R, Eden J-S, Lam C, Gray K and Timms V for the 2019-nCoV Study Group |
| EPI_ISL_417399 | Centre for Infectious Diseases and Microbiology Public Health | NSW Health Pathology - Institute of Clinical Pathology and Medical Research; Westmead Hospital; University of Sydney | Dwyer DE, Rockett R, Eden J-S, Lam C, Gray K, Timms V, Gall M, Arnott A, Sadsad R, Carter I, Rahman H, Holmes EC, O'Sullivan MV, Sintchenko V, Chen SC, Maddocks S and Kok J for the 2019-nCoV Study Group |
| EPI_ISL_417401 | Centre for Infectious Diseases and Microbiology Public Health | NSW Health Pathology - Institute of Clinical Pathology and Medical Research; Westmead Hospital; University of Sydney | Eden J-S, Lam C, Gray K, Timms V, Gall M, Arnott A, Sadsad R, Carter I, Rahman H, Holmes EC, O'Sullivan MV, Sintchenko V, Chen SC, Maddocks S, Kok J, Dwyer DE and Rockett R for the 2019-nCoV Study Group |
| EPI_ISL_417405 | Centre for Infectious Diseases and Microbiology Public Health | NSW Health Pathology - Institute of Clinical Pathology and Medical Research; Westmead Hospital; University of Sydney | Gall M, Arnott A, Sadsad R, Carter I, Rahman H, Holmes EC, O'Sullivan MV, Sintchenko V, Chen SC, Maddocks S, Kok J, Dwyer DE, Rockett R, Eden J-S, Lam C, Gray K and Timms V for the 2019-nCoV Study Group |
| EPI_ISL_417413 | Ministry of Health Turkey | Ministry of Health Turkey | Fatma Bayrakdar,Aye Baak Alta,Yasemin Cogun,Gülay Korukluolu,Selçuk Klç |
| EPI_ISL_417504, EPI_ISL_417505, EPI_ISL_417506, EPI_ISL_417507, EPI_ISL_417508, EPI_ISL_417509, EPI_ISL_417510, EPI_ISL_417512, EPI_ISL_417513, EPI_ISL_417514, EPI_ISL_417515, EPI_ISL_417516, EPI_ISL_417517 | | | |

| | | | |
|---|---|---|---|
| see above | University of Wisconsin-Madison AIDS Vaccine Research Laboratories | University of Wisconsin-Madison AIDS Vaccine Research Laboratories | Gage Moreno, Katarina Braun, et al. AIDS Vaccine Research Laboratories |
| EPI_ISL_417923 | INMI Lazzaro Spallanzani IRCCS | Laboratory of Virology, INMI Lazzaro Spallanzani IRCCS | Francesco Messina, Barbara Bartolini, Martina Rueca, Cesare E. M. Gruber, Emanuela Giombini, Maria R. Capobianchi, Fabrizio Carletti, Francesca Colavita, Concetta Castilletti, Eleonora Lalle, Daniele Lapa, Giuseppe Ippolito. |
| EPI_ISL_417925 | Laboratório Simili | Bioinformatics Laboratory / LNCC | Filipe Romero, Ana Paula Guimarães, Mariane Talon, Luiz Gonzaga Paula de Almeida, Ronaldo da Silva, Francisco Junior, Diana Mariani, Lídia Boullosa, Alexandra Gerber, Jaqueline Goes de Jesus, Ingra Morales Claro, Ester Cerdeira Sabino, Nuno Rodrigues Faria, Terezinha Marta Pereira, Pinto Castiñeiras, Isabela de Carvalho Leitão, Rafael de Mello Galliez, Cássia Cristina Alves Gonçalves, Érica Ramos dos Santos Nascimento, Richard Araújo Maia, Mauro Teixeira,Cristiano Xavier Lima, Orlando Ferreira Jr., Rodrigo Brindeiro, Luciana Jesus Costa e André Felipe Santos, Laboratorio Hermes Pardini, Laboratorio Simile, Amilcar Tanuri, Renato Santana Aguiar e Ana Tereza Vasconcelos |
| EPI_ISL_417926 | Laboratório Simili | Bioinformatics Laboratory / LNCC | Filipe Romero, Ana Paula Guimarães, Mariane Talon, Luiz Gonzaga Paula de Almeida, Ronaldo da Silva Francisco Junior, Diana Mariani, Lídia Boullosa, Alexandra Gerber, Jaqueline Goes de Jesus, Ingra Morales Claro, Ester Cerdeira Sabino, Nuno Rodrigues Faria, Terezinha Marta Pereira, Pinto Castiñeiras, Isabela de Carvalho Leitão, Rafael de Mello Galliez, Cássia Cristina Alves Gonçalves, Érica Ramos dos Santos Nascimento, Richard Araújo Maia, Mauro Teixeira,Cristiano Xavier Lima, Orlando Ferreira Jr., Rodrigo Brindeiro, Luciana Jesus Costa e André Felipe Santos, Laboratorio Hermes Pardini, Laboratorio Simile, Amilcar Tanuri, Renato Santana Aguiar e Ana Tereza Vasconcelos |
| EPI_ISL_417928, EPI_ISL_417929, EPI_ISL_417930, EPI_ISL_417934, EPI_ISL_417936 | Laboratório Hermes Pardini | Bioinformatics Laboratory - LNCC | Filipe Romero, Ana Paula Guimarães, Mariane Talon, Luiz Gonzaga Paula de Almeida, Ronaldo da Silva Francisco Junior, Diana Mariani, Lídia Boullosa, Alexandra Gerber, Jaqueline Goes de Jesus, Ingra Morales Claro, Ester Cerdeira Sabino, Nuno Rodrigues Faria, Terezinha Marta Pereira, Pinto Castiñeiras, Isabela de Carvalho Leitão, Rafael de Mello Galliez, Cássia Cristina Alves Gonçalves, Érica Ramos dos Santos Nascimento, Richard Araújo Maia, Mauro Teixeira,Cristiano Xavier Lima, Orlando Ferreira Jr., Rodrigo Brindeiro, Luciana Jesus Costa e André Felipe Santos, Laboratorio Hermes Pardini, Laboratorio Simile, Amilcar Tanuri, Renato Santana Aguiar e Ana Tereza Vasconcelos |
| EPI_ISL_417940 | Laboratório Hermes Pardini | Bioinformatics Laboratory - LNCC | Filipe Romero, Ana Paula Guimarães, Mariane Talon, Luiz Gonzaga Paula de Almeida, Ronaldo da Silva Francisco Junior, Diana Mariani, Lídia Boullosa,Alexandra Gerber, Jaqueline Goes de Jesus, Ingra Morales Claro, Ester Cerdeira Sabino, Nuno Rodrigues Faria, Terezinha Marta Pereira, Pinto Castiñeiras, Isabela de Carvalho Leitão, Rafael de Mello Galliez, Cássia Alves Gonçalves, Érica Ramos dos Santos Nascimento, Richard Araújo Maia, Mauro Teixeira,Cristiano Xavier Lima, Orlando Ferreira Jr., Rodrigo Brindeiro, Luciana Jesus Costa e André Felipe Santos, Laboratorio Hermes Pardini, Laboratorio Simile, Amilcar Tanuri, Renato Santana Aguiar e Ana Tereza Vasconcelos |
| EPI_ISL_417943 | Laboratório Hermes Pardini | Bioinformatics Laboratory | Filipe Romero, Ana Paula Guimarães, Mariane Talon, Luiz Gonzaga Paula de Almeida, Ronaldo da Silva Francisco Junior, Diana Mariani, Lídia Boullosa,Alexandra Gerber, Jaqueline Goes de Jesus, Ingra Morales Claro, Ester Cerdeira Sabino, Nuno Rodrigues Faria, Terezinha Marta Pereira, Pinto Castiñeiras, Isabela de Carvalho Leitão, Rafael de Mello Galliez, Cássia Alves Gonçalves, Érica Ramos dos Santos Nascimento, Richard Araújo Maia, Mauro Teixeira,Cristiano Xavier Lima, Orlando Ferreira Jr., Rodrigo Brindeiro, Luciana Jesus Costa e André Felipe Santos, Laboratorio Hermes Pardini, Laboratorio Simile, Amilcar Tanuri, Renato Santana Aguiar e Ana Tereza Vasconcelos |
| EPI_ISL_417945, EPI_ISL_417949 | Laboratório Hermes Pardini | Bioinformatics Laboratory - LNCC | Filipe Romero, Ana Paula Guimarães, Mariane Talon, Luiz Gonzaga Paula de Almeida, Ronaldo da Silva Francisco Junior, Diana Mariani, Lídia Boullosa,Alexandra Gerber, Jaqueline Goes de Jesus, Ingra Morales Claro, Ester Cerdeira Sabino, Nuno Rodrigues Faria, Terezinha Marta Pereira, Pinto Castiñeiras, Isabela de Carvalho Leitão, Rafael de Mello Galliez, Cássia Alves Gonçalves, Érica Ramos dos Santos Nascimento, Richard Araújo Maia, Mauro Teixeira,Cristiano Xavier Lima, Orlando Ferreira Jr., Rodrigo Brindeiro, Luciana Jesus Costa e André Felipe Santos, Laboratorio Hermes Pardini, Laboratorio Simile, Amilcar Tanuri, Renato Santana Aguiar e Ana Tereza Vasconcelos |
| EPI_ISL_417951 | Universidade Federal do Rio de Janeiro | Bioinformatics Laboratory - LNCC | Filipe Romero, Ana Paula Guimarães, Mariane Talon, Luiz Gonzaga Paula de Almeida, Ronaldo da Silva Francisco Junior, Diana Mariani, Lídia Boullosa,Alexandra Gerber, Jaqueline Goes de Jesus, Ingra Morales Claro, Ester Cerdeira Sabino, Nuno Rodrigues Faria, Terezinha Marta Pereira, Pinto Castiñeiras, Isabela de Carvalho Leitão, Rafael de Mello Galliez, Cássia Alves Gonçalves, Érica Ramos dos Santos Nascimento, Richard Araújo Maia, Mauro Teixeira,Cristiano Xavier Lima, Orlando Ferreira Jr., Rodrigo Brindeiro, Luciana Jesus Costa e André Felipe Santos, Laboratorio Hermes Pardini, Laboratorio Simile, Amilcar Tanuri, Renato Santana Aguiar e Ana Tereza Vasconcelos |
| EPI_ISL_417952 | Hospital Universitario 12 de Octubre | Hospital Universitario La Paz | Elias Dahdouh, Sara González, Fernando Lázaro, Esther Viedma, Natalia Stella, Julio García, Juan Carlos Galán, Rafael Cantón, Mª Dolores Folgueira, Rafael Delgado, Jesús Mingorance |
| EPI_ISL_417953 | Universidade Federal do Rio de Janeiro | Bioinformatics Laboratory - LNCC | Filipe Romero, Ana Paula Guimarães, Mariane Talon, Luiz Gonzaga Paula de Almeida, Ronaldo da Silva Francisco Junior, Diana Mariani, Lídia Boullosa,Alexandra Gerber, Jaqueline Goes de Jesus, Ingra Morales Claro, Ester Cerdeira Sabino, Nuno Rodrigues Faria, Terezinha Marta Pereira, Pinto Castiñeiras, Isabela de Carvalho Leitão, Rafael de Mello Galliez, Cássia Alves Gonçalves, Érica Ramos dos Santos Nascimento, Richard Araújo Maia, Mauro Teixeira,Cristiano Xavier Lima, Orlando Ferreira Jr., Rodrigo Brindeiro, Luciana Jesus Costa e André Felipe Santos, Laboratorio Hermes Pardini, Laboratorio Simile, Amilcar Tanuri, Renato Santana Aguiar e Ana Tereza Vasconcelos |
| EPI_ISL_417954, EPI_ISL_417956, EPI_ISL_417957 | Hospital Universitario 12 de Octubre | Hospital Universitario La Paz | Elias Dahdouh, Sara González, Fernando Lázaro, Esther Viedma, Natalia Stella, Julio García, Juan Carlos Galán, Rafael Cantón, Mª Dolores Folgueira, Rafael Delgado, Jesús Mingorance |
| EPI_ISL_417958, EPI_ISL_417959, EPI_ISL_417960 | Utah Public Health Laboratory | Utah Public Health Laboratory | Erin Young, Kelly Oakeson |
| EPI_ISL_417961, EPI_ISL_417963 | Hospital Universitario 12 de Octubre | Hospital Universitario La Paz | Elias Dahdouh, Sara González, Fernando Lázaro, Esther Viedma, Natalia Stella, Julio García, Juan Carlos Galán, Rafael Cantón, Mª Dolores Folgueira, Rafael Delgado, Jesús Mingorance |
| EPI_ISL_417964, EPI_ISL_417966 | Utah Public Health Laboratory | Utah Public Health Laboratory | Erin Young, Kelly Oakeson |
| EPI_ISL_417967 | Hospital Universitario 12 de Octubre | Hospital Universitario La Paz | Elias Dahdouh, Sara González, Fernando Lázaro, Esther Viedma, Natalia Stella, Julio García, Juan Carlos Galán, Rafael Cantón, Mª Dolores Folgueira, Rafael Delgado, Jesús Mingorance |
| EPI_ISL_417969 | Hospital Universitario La Paz | Hospital Universitario La Paz | Elias Dahdouh, Sara González, Fernando Lázaro, Esther Viedma, Natalia Stella, Julio García, Juan Carlos Galán, Rafael Cantón, Mª Dolores Folgueira, Rafael Delgado, Jesús Mingorance |
| EPI_ISL_417970, EPI_ISL_417971 | Utah Public Health Laboratory | Utah Public Health Laboratory | Erin Young, Kelly Oakeson |
| EPI_ISL_417972 | Hospital Universitario La Paz | Hospital Universitario La Paz | Elias Dahdouh, Sara González, Fernando Lázaro, Esther Viedma, Natalia Stella, Julio García, Juan Carlos Galán, Rafael Cantón, Mª Dolores Folgueira, Rafael Delgado, Jesús Mingorance |
| EPI_ISL_417973, EPI_ISL_417974 | Utah Public Health Laboratory | Utah Public Health Laboratory | Erin Young, Kelly Oakeson |
| EPI_ISL_417975 | Hospital Universitario La Paz | Hospital Universitario La Paz | Elias Dahdouh, Sara González, Fernando Lázaro, Esther Viedma, Natalia Stella, Julio García, Juan Carlos Galán, Rafael Cantón, Mª Dolores Folgueira, Rafael Delgado, Jesús Mingorance |
| EPI_ISL_417976, EPI_ISL_417977 | Utah Public Health Laboratory | Utah Public Health Laboratory | Erin Young, Kelly Oakeson |
| EPI_ISL_417978 | Hospital Universitario La Paz | Hospital Universitario La Paz | Elias Dahdouh, Sara González, Fernando Lázaro, Esther Viedma, Natalia Stella, Julio García, Juan Carlos Galán, Rafael Cantón, Mª Dolores Folgueira, Rafael Delgado, Jesús Mingorance |
| EPI_ISL_417979, EPI_ISL_417980, EPI_ISL_417981 | Hospital Universitario Ramón y Cajal | Hospital Universitario La Paz | Elias Dahdouh, Sara González, Fernando Lázaro, Esther Viedma, Natalia Stella, Julio García, Juan Carlos Galán, Rafael Cantón, Mª Dolores Folgueira, Rafael Delgado, Jesús Mingorance |
| EPI_ISL_417982, EPI_ISL_417983, EPI_ISL_417984, EPI_ISL_417985 | Universidade Federal do Rio de Janeiro | Bioinformatics Laboratory - LNCC | Filipe Romero, Ana Paula Guimarães, Mariane Talon, Luiz Gonzaga Paula de Almeida, Ronaldo da Silva Francisco Junior, Diana Mariani, Lídia Boullosa,Alexandra Gerber, Jaqueline Goes de Jesus, Ingra Morales Claro, Ester Cerdeira Sabino, Nuno Rodrigues Faria, Terezinha Marta Pereira, Pinto Castiñeiras, Isabela de Carvalho Leitão, Rafael de Mello Galliez, Cássia Alves Gonçalves, Érica Ramos dos Santos Nascimento, Richard Araújo Maia, Mauro Teixeira,Cristiano Xavier Lima, Orlando Ferreira Jr., Rodrigo Brindeiro, Luciana Jesus Costa e André Felipe Santos, Laboratorio Hermes Pardini, Laboratorio Simile, Amilcar Tanuri, Renato Santana Aguiar e Ana Tereza Vasconcelos |
| EPI_ISL_418182 | Hospital Universitario 12 de Octubre | Hospital Universitario La Paz | Elias Dahdouh, Sara González, Fernando Lázaro, Esther Viedma, Natalia Stella, Julio García, Juan Carlos Galán, Rafael Cantón, Mª Dolores Folgueira, Rafael Delgado, Jesús Mingorance |

| | | | |
|---|---|---|---|
| EPI_ISL_418959 | Universidade Federal do Rio de Janeiro - UFRJ | Bioinformatics Laboratory - LNCC | Filipe Romero, Ana Paula Guimarães, Mariane Talon, Luiz Gonzaga Paula de Almeida, Ronaldo da Silva Francisco Junior, Diana Mariani, Lídia Boullosa, Alexandra Gerber, Jaqueline Goes de Jesus, Ingra Morales Claro, Ester Cerdeira Sabino, Nuno Rodrigues Faria, Terezinha Marta Pereira, Pinto Castiñeiras, Isabela de Carvalho Leitão, Rafael de Mello Galliez, Cássia Cristina Alves Gonçalves, Érica Ramos dos Santos Nascimento, Richard Araújo Maia, Mauro Teixeira,Cristiano Xavier Lima, Orlando Ferreira Jr., Rodrigo Brindeiro, Luciana Jesus Costa e André Felipe Santos, Laboratorio Hermes Pardini, Laboratorio Simile, Amilcar Tanuri, Renato Santana Aguiar, e Ana Tereza Vasconcelos |
| EPI_ISL_418962, EPI_ISL_418963, EPI_ISL_418964, EPI_ISL_418965, EPI_ISL_418967, EPI_ISL_420803, EPI_ISL_420804, EPI_ISL_420805, EPI_ISL_420806, EPI_ISL_420807, EPI_ISL_420808, EPI_ISL_420809, EPI_ISL_420810, EPI_ISL_420811, EPI_ISL_420812, EPI_ISL_420813, EPI_ISL_420814, EPI_ISL_420815, EPI_ISL_420816, EPI_ISL_420817, EPI_ISL_420818, EPI_ISL_420819, EPI_ISL_420820, EPI_ISL_420821, EPI_ISL_420822, EPI_ISL_420823, EPI_ISL_420824, EPI_ISL_420825 | | | |
| see above | Utah Public Health Laboratory | Utah Public Health Laboratory | Erin Young, Kelly Oakeson |
| EPI_ISL_421171, EPI_ISL_421172, EPI_ISL_421173, EPI_ISL_421174, EPI_ISL_421175, EPI_ISL_421176, EPI_ISL_421177, EPI_ISL_421178, EPI_ISL_421179, EPI_ISL_421180 | Hospital Universitario 12 de Octubre | Hospital Universitario 12 de Octubre | Esther Viedma, Sara González, Elias Dahdouh, Raúl Recio, Fernando Lázaro, Julio García, Mª Dolores Folgueira, Jesús Mingorance, Rafael Delgado |
| EPI_ISL_421283, EPI_ISL_421284, EPI_ISL_421285, EPI_ISL_421286, EPI_ISL_421287, EPI_ISL_421288, EPI_ISL_421289, EPI_ISL_421290, EPI_ISL_421291, EPI_ISL_421292, EPI_ISL_421293, EPI_ISL_421294, EPI_ISL_421295, EPI_ISL_421296, EPI_ISL_421297, EPI_ISL_421298, EPI_ISL_421299, EPI_ISL_421300, EPI_ISL_421301, EPI_ISL_421302, EPI_ISL_421303, EPI_ISL_421304, EPI_ISL_421305, EPI_ISL_421306, EPI_ISL_421307, EPI_ISL_421308, EPI_ISL_421309, EPI_ISL_421310, EPI_ISL_421311, EPI_ISL_421312, EPI_ISL_421313, EPI_ISL_421314, EPI_ISL_421315, EPI_ISL_421316, EPI_ISL_421317, EPI_ISL_421318, EPI_ISL_421319, EPI_ISL_421320, EPI_ISL_421321, EPI_ISL_421322, EPI_ISL_421323, EPI_ISL_421324, EPI_ISL_421325, EPI_ISL_421326, EPI_ISL_421327, EPI_ISL_421328, EPI_ISL_421329, EPI_ISL_421330, EPI_ISL_421331, EPI_ISL_421332, EPI_ISL_421333, EPI_ISL_421334, EPI_ISL_421335, EPI_ISL_421336, EPI_ISL_421338, EPI_ISL_421339, EPI_ISL_421340, EPI_ISL_421341, EPI_ISL_421342, EPI_ISL_421343 | | | |
| see above | University of Wisconsin-Madison AIDS Vaccine Research Laboratories | University of Wisconsin-Madison AIDS Vaccine Research Laboratories | Gage Moreno, Katarina Braun, et al. AIDS Vaccine Research Laboratories |
| EPI_ISL_421560, EPI_ISL_421561, EPI_ISL_421562, EPI_ISL_421563 | Utah Public Health Laboratory | Utah Public Health Laboratory | Erin Young, Kelly Oakeson |
| EPI_ISL_421572 | Molecular Diagnostic Services and FLowpath | KRISP, KZN Research Innovation and Sequencing Platform | Giandhari J, Pillay S, Ngcapu S, Samsunder N, Lessells R, Chimukangara B, Deforche K, Tegally H, Wilkinson E, de Oliveira T |
| EPI_ISL_421573, EPI_ISL_421574, EPI_ISL_421575 | Molecular Diagnostic Services | KRISP, KZN Research Innovation and Sequencing Platform | Giandhari J, Pillay S, Ngcapu S, Samsunder N, Lessells R, Chimukangara B, Deforche K, Tegally H, Wilkinson E, de Oliveira T |
| EPI_ISL_421652 | Dasman Diabetes Institute | Dasman Diabetes Institute | Fahd Al-Mulla, Rasheeba Iqbal, Sumi John, Ebaa Al-Ozairi, Qais Al-Duwairi |
| EPI_ISL_422424 | Jaber Al Ahmad Al Sabah Hospital | Dasman diabetes Institute | Fahd Al-Mulla, Rasheeba Iqbal, Sumi John, Ebaa Al-Ozairi, Qais Al-Duwairi |
| EPI_ISL_422426, EPI_ISL_422427 | JABER AL AHMAD AL SABAH HOSPITAL - KUWAIT CITY | Dasman Diabetes Institute | Fahd Al-Mulla, Rasheeba Iqbal, Sumi John, Ebaa Al-Ozairi, Qais Al-Duwairi |
| EPI_ISL_423039, EPI_ISL_423040, EPI_ISL_423041, EPI_ISL_423042, EPI_ISL_423043 | Ramathibodi Hospital | COVID-19 Network Investigations (CONI) Alliance | Elizabeth Batty, Wasun Chantratita, Thanat Chookajorn, Stefan Fernandez, Angkana Huang, Anthony R. Jones, Khajohn Joonsalak, Chonticha Klungtong, Theerarat Kochakarn, Namfon Kotanan, Krittikorn Kumpornsin, Wudtichai Manasatienkij, Bhakbhoom Panthan, Ekawat Pasomsub, Insee Sensorn, Arporn Wangwiwatsin |
| EPI_ISL_424343 | INMI Lazzaro Spallanzani IRCCS | Laboratory of Virology, INMI Lazzaro Spallanzani IRCCS | Fabrizio Carletti, Barbara Bartolini, Martina Rueca, Cesare Ernesto Maria Gruber, Francesco Messina, Eleonora Lalle, Licia Bordi, Giulia Matusali, Francesca Colavita, Maria Rosaria Capobianchi, Concetta Castilletti, Francesco Vairo, Giuseppe Ippolito, Antonino Di Caro |
| EPI_ISL_424344 | INMI Lazzaro Spallanzani IRCCS | Laboratory of Virology, INMI Lazzaro Spallanzani IRCCS | Eleonora Lalle, Barbara Bartolini, Martina Rueca, Cesare Ernesto Maria Gruber, Francesco Messina, Fabrizio Carletti, Licia Bordi, Giulia Matusali, Francesca Colavita, Maria Rosaria Capobianchi, Concetta Castilletti, Francesco Vairo, Giuseppe Ippolito, Antonino Di Caro |
| EPI_ISL_424352 | Clinical Laboratory, Fuyang City Center for Disease Control and Prevention | Clinical Laboratory, Fuyang City Center for Disease Control and Prevention | Ge,B. |
| EPI_ISL_424355, EPI_ISL_424356, EPI_ISL_424357, EPI_ISL_424358, EPI_ISL_424359, EPI_ISL_424360 | Beijing Institute of Microbiology and Epidemiology | Beijing Institute of Microbiology and Epidemiology | Fan,H., Qin,E., Wu,Y., Guo,Y., Zhang,X., Yong,Y., Hou,J., Xu,Z., Mu,J., Teng,Y., Mi,Z., Yang,R., Song,Y., Li,B. and Cui,Y. |
| EPI_ISL_424672 | Laboratorio Estatal de Salud Publica del Estado de Puebla | Instituto de Diagnostico y Referencia Epidemiologicos | Fabiola Garcés Ayala, Gisela Barrera Badillo, Irma López Martínez, Lucia Hernández Rivas, Edgar Mendieta Condado, Adnan Araiza Rodríguez, Celia Boukadida, Santiago Avila Ríos, Mario Mújica Sánchez, José Arturo Martínez Orozco,Eduardo Becerril Vargas, Joel Armando Vázquez Pérez, Victor Hugo Borja Aburto, Concepción Grajales Muñiz, Cesar Raúl González Bonilla, Carolina González Torres, Francisco Javier Gaytán Cervantes, José Esteban Muñoz Medina, Guillermo M. Ruiz-Palacios, Pilar Ramos Cervantes, Violeta Ibarra Gonzalez, Fernando Ledesma Barrientos, Luis Alberto García Andrade, Alfredo Ponce de León Garduño, Blanca Taboada, Alejandro Sánchez, Pavel Isa, Ricardo Grande, Gloria Vázquez, Francisco Pulido, Carlos F. Arias,José Ernesto Ramírez González |
| EPI_ISL_424703 | Klinisk mikrobiologi, Region Västerbotten | Unit for Biological Agents, Department for CBRN Defence and Security, Swedish Defence Research Agency | FOI Bioinformatics team |
| EPI_ISL_425142, EPI_ISL_425143, EPI_ISL_425144, EPI_ISL_425145, EPI_ISL_425146, EPI_ISL_425147, EPI_ISL_425148, EPI_ISL_425149, EPI_ISL_425150, EPI_ISL_425151, EPI_ISL_425152, EPI_ISL_425153, EPI_ISL_425154, EPI_ISL_425155, EPI_ISL_425156, EPI_ISL_425157, EPI_ISL_425158, EPI_ISL_425159, EPI_ISL_425160, EPI_ISL_425161, EPI_ISL_425162, EPI_ISL_425163, EPI_ISL_425164, EPI_ISL_425165, EPI_ISL_425166, EPI_ISL_425167, EPI_ISL_425168, EPI_ISL_425169, EPI_ISL_425170, EPI_ISL_425171, EPI_ISL_425172, EPI_ISL_425173, EPI_ISL_425174, EPI_ISL_425175 | | | |
| see above | University of Wisconsin-Madison AIDS Vaccine Research Laboratories | University of Wisconsin-Madison AIDS Vaccine Research Laboratories | Gage Moreno, Katarina Braun, et al. AIDS Vaccine Research Laboratories |
| EPI_ISL_425462, EPI_ISL_425463, EPI_ISL_425464, EPI_ISL_425465, EPI_ISL_425466, EPI_ISL_425467, EPI_ISL_425468, EPI_ISL_425469, EPI_ISL_425470, EPI_ISL_425471, EPI_ISL_425472, EPI_ISL_425473, EPI_ISL_425474, EPI_ISL_425475, EPI_ISL_425476, EPI_ISL_425477, EPI_ISL_425478, EPI_ISL_425479, EPI_ISL_425480, EPI_ISL_425481, EPI_ISL_425482, EPI_ISL_425483, EPI_ISL_425484, EPI_ISL_425485, EPI_ISL_425486, EPI_ISL_425487, EPI_ISL_425488, EPI_ISL_425489, EPI_ISL_425490, EPI_ISL_425491, EPI_ISL_425492, EPI_ISL_425493, EPI_ISL_425494, EPI_ISL_425495, EPI_ISL_425496, EPI_ISL_425497, EPI_ISL_425498, EPI_ISL_425499, EPI_ISL_425500, EPI_ISL_425501, EPI_ISL_425502, EPI_ISL_425503, EPI_ISL_425504, EPI_ISL_425505, EPI_ISL_425506, EPI_ISL_425507, EPI_ISL_425508, EPI_ISL_425509, EPI_ISL_425510, EPI_ISL_425511, EPI_ISL_425512, EPI_ISL_425513, EPI_ISL_425514, EPI_ISL_425515, EPI_ISL_425516, EPI_ISL_425517, EPI_ISL_425518, EPI_ISL_425519, EPI_ISL_425520, EPI_ISL_425521, EPI_ISL_425522, EPI_ISL_425523, EPI_ISL_425524, EPI_ISL_425525, EPI_ISL_425526, EPI_ISL_425527, EPI_ISL_425528, EPI_ISL_425529, EPI_ISL_425530, EPI_ISL_425531, EPI_ISL_425532, EPI_ISL_425533, EPI_ISL_425534, EPI_ISL_425535, EPI_ISL_425536, EPI_ISL_425537, EPI_ISL_425538, EPI_ISL_425539, EPI_ISL_425540, EPI_ISL_425541, EPI_ISL_425542, EPI_ISL_425543, EPI_ISL_425544, EPI_ISL_425545, EPI_ISL_425546, EPI_ISL_425547, EPI_ISL_425548, EPI_ISL_425549, EPI_ISL_425550, EPI_ISL_425551, EPI_ISL_425552, EPI_ISL_425553, EPI_ISL_425554, EPI_ISL_425555, EPI_ISL_425556, EPI_ISL_425557, EPI_ISL_425558, EPI_ISL_425559, EPI_ISL_425560, EPI_ISL_425561, EPI_ISL_425562, EPI_ISL_425563, EPI_ISL_425564, EPI_ISL_425565, EPI_ISL_425566, EPI_ISL_425567, EPI_ISL_425568, EPI_ISL_425569, EPI_ISL_425570, EPI_ISL_425571, EPI_ISL_425572, EPI_ISL_425573, EPI_ISL_425574, EPI_ISL_425575, EPI_ISL_425576, EPI_ISL_425577, EPI_ISL_425579, EPI_ISL_425580, EPI_ISL_425581, EPI_ISL_425582, EPI_ISL_425583, EPI_ISL_425584, EPI_ISL_425585, EPI_ISL_425586, EPI_ISL_425587, EPI_ISL_425588, EPI_ISL_425589, EPI_ISL_425590, EPI_ISL_425591, EPI_ISL_425592, EPI_ISL_425593, EPI_ISL_425594, EPI_ISL_425595, EPI_ISL_425596, EPI_ISL_425597, EPI_ISL_425598, EPI_ISL_425599, EPI_ISL_425600, EPI_ISL_425601, EPI_ISL_425602, EPI_ISL_425603, EPI_ISL_425604, EPI_ISL_425605, EPI_ISL_425606, EPI_ISL_425607, EPI_ISL_425608, EPI_ISL_425609, EPI_ISL_425610, EPI_ISL_425611, EPI_ISL_425612, EPI_ISL_425613, EPI_ISL_425614, EPI_ISL_425615, EPI_ISL_425616, EPI_ISL_425617, EPI_ISL_425618, EPI_ISL_425619, EPI_ISL_425620, EPI_ISL_425621, EPI_ISL_425622, EPI_ISL_425623, EPI_ISL_425624, EPI_ISL_425625, EPI_ISL_425626, EPI_ISL_425627, EPI_ISL_425628, EPI_ISL_425629, EPI_ISL_425630, EPI_ISL_425631, EPI_ISL_425632, EPI_ISL_425633, EPI_ISL_425634, EPI_ISL_425635, EPI_ISL_425636, EPI_ISL_425637, EPI_ISL_425638, EPI_ISL_425639, EPI_ISL_425640, EPI_ISL_425641, EPI_ISL_425642, EPI_ISL_425643, EPI_ISL_425644, EPI_ISL_425645, EPI_ISL_425646 | | | |
| see above | Queens Medical Centre, Clinical Microbiology Department / DeepSeq Nottingham | COVID-19 Genomics UK (COG-UK) Consortium | Gemma Clark, Wendy Smith, Manjinder Khakh, Hannah Howson-Wells, Jonathan Ball, Patrick McClure, Joseph Chappell, Theocharis Tsoleridis, Nadine Holmes, Matthew Carlisle, Christopher Moore, Fei Sang, Johnny Debebe, Victoria Wright, Matthew Loose |
| EPI_ISL_426580 | Instituto Sabin | Laboratory of Virology | Fernando L Melo,Gustavo Barra, Ticiane H Santa-Rita, Pedro G Mesquita, Ikaro A Andrade, Tatsuya Nagata, Bergmann M Ribeiro |
| EPI_ISL_427427, EPI_ISL_427428, EPI_ISL_427429, EPI_ISL_427430, EPI_ISL_427431, EPI_ISL_427432, EPI_ISL_427433, EPI_ISL_427434, EPI_ISL_427435, EPI_ISL_427436, EPI_ISL_427437, EPI_ISL_427438, EPI_ISL_427439, EPI_ISL_427440, EPI_ISL_427441, EPI_ISL_427442, EPI_ISL_427443, EPI_ISL_427444, EPI_ISL_427445, EPI_ISL_427446, EPI_ISL_427447, EPI_ISL_427448, EPI_ISL_427449, EPI_ISL_427450, EPI_ISL_427451, EPI_ISL_427452, EPI_ISL_427453, EPI_ISL_427454, EPI_ISL_427455, EPI_ISL_427456, EPI_ISL_427457, EPI_ISL_427458, EPI_ISL_427459, EPI_ISL_427460, EPI_ISL_427461, EPI_ISL_427462 | | | |
| see above | University of Wisconsin-Madison AIDS Vaccine Research Laboratories | University of Wisconsin-Madison AIDS Vaccine Research Laboratories | Gage Moreno, Katarina Braun, et al. AIDS Vaccine Research Laboratories |
| EPI_ISL_427650 | Centre for Infectious Diseases and Microbiology Public Health | NSW Health Pathology - Institute of Clinical Pathology and Medical Research; Westmead Hospital; University of Sydney | Gall M, Arnott A, Sadsad R, Draper J, Sim E, Bachmann N, Rockett R, Lam C, Gray K, Timms V, Carter I, Holmes EC, O'Sullivan MV, Byun R, Sintchenko V, Chen SC, Eden JS, Maddocks S, Kok J, Propenko M, Sorrell T, Chang S, Basile K, Dwyer DE for the 2019-nCoV Study Group |
| EPI_ISL_427655 | Centre for Infectious Diseases and Microbiology Public Health | NSW Health Pathology - Institute of Clinical Pathology and Medical Research; Westmead Hospital; University of Sydney | Draper J, Sim E, Bachmann N, Rockett R, Lam C, Gray K, Timms V, Gall M, Arnott A, Sadsad R, Carter I, Holmes EC, O'Sullivan MV, Byun R, Sintchenko V, Chen SC, Eden JS, Maddocks S, Kok J, Propenko M, Sorrell T, Chang S, Basile K, Dwyer DE for the 2019-nCoV Study Group |

| | | | |
|---|---|---|---|
| EPI_ISL_427667, EPI_ISL_427674 | Centre for Infectious Diseases and Microbiology Public Health | NSW Health Pathology - Institute of Clinical Pathology and Medical Research; Westmead Hospital; University of Sydney | Gall M, Arnott A, Sadsad R, Draper J, Sim E, Bachmann N, Rockett R, Lam C, Gray K, Timms V, Carter I, Holmes EC, O'Sullivan MV, Byun R, Sintchenko V, Chen SC, Eden JS, Maddocks S, Kok J, Propenko M, Sorrell T, Chang S, Basile K, Dwyer DE for the 2019-nCoV Study Group |
| EPI_ISL_427676, EPI_ISL_427679, EPI_ISL_427680, EPI_ISL_427684, EPI_ISL_427690 | Centre for Infectious Diseases and Microbiology Public Health | NSW Health Pathology - Institute of Clinical Pathology and Medical Research; Westmead Hospital; University of Sydney | Draper J, Sim E, Bachmann N, Rockett R, Lam C, Gray K, Timms V, Gall M, Arnott A, Sadsad R, Carter I, Holmes EC, O'Sullivan MV, Byun R, Sintchenko V, Chen SC, Eden JS, Maddocks S, Kok J, Propenko M, Sorrell T, Chang S, Basile K, Dwyer DE for the 2019-nCoV Study Group |
| EPI_ISL_427691, EPI_ISL_427697, EPI_ISL_427700 | Centre for Infectious Diseases and Microbiology Public Health | NSW Health Pathology - Institute of Clinical Pathology and Medical Research; Westmead Hospital; University of Sydney | Gall M, Arnott A, Sadsad R, Draper J, Sim E, Bachmann N, Rockett R, Lam C, Gray K, Timms V, Carter I, Holmes EC, O'Sullivan MV, Byun R, Sintchenko V, Chen SC, Eden JS, Maddocks S, Kok J, Propenko M, Sorrell T, Chang S, Basile K, Dwyer DE for the 2019-nCoV Study Group |
| EPI_ISL_427701, EPI_ISL_427722 | Centre for Infectious Diseases and Microbiology Public Health | NSW Health Pathology - Institute of Clinical Pathology and Medical Research; Westmead Hospital; University of Sydney | Draper J, Sim E, Bachmann N, Rockett R, Lam C, Gray K, Timms V, Gall M, Arnott A, Sadsad R, Carter I, Holmes EC, O'Sullivan MV, Byun R, Sintchenko V, Chen SC, Eden JS, Maddocks S, Kok J, Propenko M, Sorrell T, Chang S, Basile K, Dwyer DE for the 2019-nCoV Study Group |
| EPI_ISL_427729, EPI_ISL_427733 | Centre for Infectious Diseases and Microbiology Public Health | NSW Health Pathology - Institute of Clinical Pathology and Medical Research; Westmead Hospital; University of Sydney | Gall M, Arnott A, Sadsad R, Draper J, Sim E, Bachmann N, Rockett R, Lam C, Gray K, Timms V, Carter I, Holmes EC, O'Sullivan MV, Byun R, Sintchenko V, Chen SC, Eden JS, Maddocks S, Kok J, Propenko M, Sorrell T, Chang S, Basile K, Dwyer DE for the 2019-nCoV Study Group |
| EPI_ISL_427736, EPI_ISL_427741 | Centre for Infectious Diseases and Microbiology Public Health | NSW Health Pathology - Institute of Clinical Pathology and Medical Research; Westmead Hospital; University of Sydney | Draper J, Sim E, Bachmann N, Rockett R, Lam C, Gray K, Timms V, Gall M, Arnott A, Sadsad R, Carter I, Holmes EC, O'Sullivan MV, Byun R, Sintchenko V, Chen SC, Eden JS, Maddocks S, Kok J, Propenko M, Sorrell T, Chang S, Basile K, Dwyer DE for the 2019-nCoV Study Group |
| EPI_ISL_427746, EPI_ISL_427756 | Centre for Infectious Diseases and Microbiology Public Health | NSW Health Pathology - Institute of Clinical Pathology and Medical Research; Westmead Hospital; University of Sydney | Gall M, Arnott A, Sadsad R, Draper J, Sim E, Bachmann N, Rockett R, Lam C, Gray K, Timms V, Carter I, Holmes EC, O'Sullivan MV, Byun R, Sintchenko V, Chen SC, Eden JS, Maddocks S, Kok J, Propenko M, Sorrell T, Chang S, Basile K, Dwyer DE for the 2019-nCoV Study Group |
| EPI_ISL_427757 | Centre for Infectious Diseases and Microbiology Public Health | NSW Health Pathology - Institute of Clinical Pathology and Medical Research; Westmead Hospital; University of Sydney | Draper J, Sim E, Bachmann N, Rockett R, Lam C, Gray K, Timms V, Gall M, Arnott A, Sadsad R, Carter I, Holmes EC, O'Sullivan MV, Byun R, Sintchenko V, Chen SC, Eden JS, Maddocks S, Kok J, Propenko M, Sorrell T, Chang S, Basile K, Dwyer DE for the 2019-nCoV Study Group |
| EPI_ISL_427764 | Centre for Infectious Diseases and Microbiology Public Health | NSW Health Pathology - Institute of Clinical Pathology and Medical Research; Westmead Hospital; University of Sydney | Gall M, Arnott A, Sadsad R, Draper J, Sim E, Bachmann N, Rockett R, Lam C, Gray K, Timms V, Carter I, Holmes EC, O'Sullivan MV, Byun R, Sintchenko V, Chen SC, Eden JS, Maddocks S, Kok J, Propenko M, Sorrell T, Chang S, Basile K, Dwyer DE for the 2019-nCoV Study Group |
| EPI_ISL_427766, EPI_ISL_427768 | Centre for Infectious Diseases and Microbiology Public Health | NSW Health Pathology - Institute of Clinical Pathology and Medical Research; Westmead Hospital; University of Sydney | Draper J, Sim E, Bachmann N, Rockett R, Lam C, Gray K, Timms V, Gall M, Arnott A, Sadsad R, Carter I, Holmes EC, O'Sullivan MV, Byun R, Sintchenko V, Chen SC, Eden JS, Maddocks S, Kok J, Propenko M, Sorrell T, Chang S, Basile K, Dwyer DE for the 2019-nCoV Study Group |
| EPI_ISL_427769, EPI_ISL_427772 | Centre for Infectious Diseases and Microbiology Public Health | NSW Health Pathology - Institute of Clinical Pathology and Medical Research; Westmead Hospital; University of Sydney | Gall M, Arnott A, Sadsad R, Draper J, Sim E, Bachmann N, Rockett R, Lam C, Gray K, Timms V, Carter I, Holmes EC, O'Sullivan MV, Byun R, Sintchenko V, Chen SC, Eden JS, Maddocks S, Kok J, Propenko M, Sorrell T, Chang S, Basile K, Dwyer DE for the 2019-nCoV Study Group |
| EPI_ISL_427784, EPI_ISL_427796, EPI_ISL_427799 | Centre for Infectious Diseases and Microbiology Public Health | NSW Health Pathology - Institute of Clinical Pathology and Medical Research; Westmead Hospital; University of Sydney | Draper J, Sim E, Bachmann N, Rockett R, Lam C, Gray K, Timms V, Gall M, Arnott A, Sadsad R, Carter I, Holmes EC, O'Sullivan MV, Byun R, Sintchenko V, Chen SC, Eden JS, Maddocks S, Kok J, Propenko M, Sorrell T, Chang S, Basile K, Dwyer DE for the 2019-nCoV Study Group |
| EPI_ISL_427800, EPI_ISL_427807, EPI_ISL_427808 | Centre for Infectious Diseases and Microbiology Public Health | NSW Health Pathology - Institute of Clinical Pathology and Medical Research; Westmead Hospital; University of Sydney | Gall M, Arnott A, Sadsad R, Draper J, Sim E, Bachmann N, Rockett R, Lam C, Gray K, Timms V, Carter I, Holmes EC, O'Sullivan MV, Byun R, Sintchenko V, Chen SC, Eden JS, Maddocks S, Kok J, Propenko M, Sorrell T, Chang S, Basile K, Dwyer DE for the 2019-nCoV Study Group |
| EPI_ISL_428148 | Klinisk mikrobiologi, Region Västerbotten | Unit for Biological Agents, Department for CBRN Defence and Security, Swedish Defence Research Agency | FOI bioinformatics team |
| EPI_ISL_428201 | Klinisk mikrobiologi, Region Västerbotten | Unit for Biological Agents, Department for CBRN Defence and Security, Swedish Defence Research Agency | FOI Bioinformatics team |
| EPI_ISL_428252, EPI_ISL_428253, EPI_ISL_428254, EPI_ISL_428255, EPI_ISL_428256, EPI_ISL_428257, EPI_ISL_428258, EPI_ISL_428259, EPI_ISL_428260, EPI_ISL_428261, EPI_ISL_428262, EPI_ISL_428263, EPI_ISL_428264, EPI_ISL_428265, EPI_ISL_428266, EPI_ISL_428267, EPI_ISL_428268, EPI_ISL_428269, EPI_ISL_428270, EPI_ISL_428271, EPI_ISL_428272, EPI_ISL_428273, EPI_ISL_428274, EPI_ISL_428275, EPI_ISL_428276, EPI_ISL_428277, EPI_ISL_428278, EPI_ISL_428279, EPI_ISL_428280, EPI_ISL_428281, EPI_ISL_428282, EPI_ISL_428283, EPI_ISL_428284, EPI_ISL_428285, EPI_ISL_428286, EPI_ISL_428287, EPI_ISL_428288, EPI_ISL_428289, EPI_ISL_428290, EPI_ISL_428291, EPI_ISL_428292, EPI_ISL_428293, EPI_ISL_428294, EPI_ISL_428295, EPI_ISL_428296, EPI_ISL_428297, EPI_ISL_428298, EPI_ISL_428299, EPI_ISL_428300, EPI_ISL_428301, EPI_ISL_428302, EPI_ISL_428303, EPI_ISL_428304, EPI_ISL_428305, EPI_ISL_428306, EPI_ISL_428307, EPI_ISL_428308, EPI_ISL_428309, EPI_ISL_428310, EPI_ISL_428311, EPI_ISL_428312, EPI_ISL_428313, EPI_ISL_428314, EPI_ISL_428315, EPI_ISL_428316, EPI_ISL_428317, EPI_ISL_428318, EPI_ISL_428319, EPI_ISL_428320, EPI_ISL_428321, EPI_ISL_428322, EPI_ISL_428323, EPI_ISL_428324, EPI_ISL_428325, EPI_ISL_428326, EPI_ISL_428327, EPI_ISL_428328, EPI_ISL_428329, EPI_ISL_428330, EPI_ISL_428331, EPI_ISL_428332, EPI_ISL_428333, EPI_ISL_428334, EPI_ISL_428335, EPI_ISL_428336, EPI_ISL_428337, EPI_ISL_428338, EPI_ISL_428339, EPI_ISL_428340, EPI_ISL_428341, EPI_ISL_428342, EPI_ISL_428343, EPI_ISL_428344, EPI_ISL_428345 | | | |
| see above | University of Wisconsin-Madison AIDS Vaccine Research Laboratories | University of Wisconsin-Madison AIDS Vaccine Research Laboratories | Gage Moreno, Katarina Braun, et al. AIDS Vaccine Research Laboratories |
| EPI_ISL_428674, EPI_ISL_428675, EPI_ISL_428676, EPI_ISL_428677, EPI_ISL_428678, EPI_ISL_428679, EPI_ISL_428680, EPI_ISL_428681, EPI_ISL_428682 | Hospital Universitario La Paz | Hospital Universitario 12 de Octubre | Elias Dahdouh, Sara González, Raúl Recio, Fernando Lázaro, Esther Viedma, Natalia Stella, Julio García, Juan Carlos Galán, Rafael Cantón, Mª Dolores Folgueira, Rafael Delgado, Jesús Mingorance |
| EPI_ISL_428700, EPI_ISL_428701, EPI_ISL_428702, EPI_ISL_428703, EPI_ISL_428704, EPI_ISL_428705, EPI_ISL_428706, EPI_ISL_428707, EPI_ISL_428708, EPI_ISL_428709, EPI_ISL_428710, EPI_ISL_428711 | | | |
| see above | Hospital Universitario 12 de Octubre | Hospital Universitario 12 de Octubre | Esther Viedma, Sara González, Raúl Recio, Elias Dahdouh, Fernando Lázaro, Julio García, Mª Dolores Folgueira, Jesús Mingorance, Rafael Delgado |
| EPI_ISL_428712, EPI_ISL_428713, EPI_ISL_428714, EPI_ISL_428715, EPI_ISL_428716, EPI_ISL_428717, EPI_ISL_428718, EPI_ISL_428719, EPI_ISL_428720, EPI_ISL_428721, EPI_ISL_428722, EPI_ISL_428723 | | | |
| see above | Ministry of Health Turkey | Ministry of Health Turkey | Fatma Bayrakdar,Aye Baak Alta,Yasemin Cogun,Gülay Korukluolu,Selçuk Klç |
| EPI_ISL_428724, EPI_ISL_428725, EPI_ISL_428726, EPI_ISL_428727, EPI_ISL_428728, EPI_ISL_428729, EPI_ISL_428730, EPI_ISL_428731, EPI_ISL_428732, EPI_ISL_428935, EPI_ISL_428936 | | | |
| see above | University of Wisconsin-Madison AIDS Vaccine Research Laboratories | University of Wisconsin-Madison AIDS Vaccine Research Laboratories | Gage Moreno, Katarina Braun, et al. AIDS Vaccine Research Laboratories |
| EPI_ISL_429164, EPI_ISL_429165, EPI_ISL_429166, EPI_ISL_429167, EPI_ISL_429168, EPI_ISL_429169, EPI_ISL_429170, EPI_ISL_429171, EPI_ISL_429172, EPI_ISL_429173, EPI_ISL_429174, EPI_ISL_429175, EPI_ISL_429176, EPI_ISL_429177, EPI_ISL_429178, EPI_ISL_429179, EPI_ISL_429180, EPI_ISL_429181, EPI_ISL_429182, EPI_ISL_429183, EPI_ISL_429184 | | | |
| see above | Ramathibodi Hospital | COVID-19 Network Investigations (CONI) Alliance | Elizabeth Batty, Wasun Chantratita, Thanat Chookajorn, Stefan Fernandez, Angkana Huang, Poramate Jiaranai, Anthony R. Jones, Khajohn Joonsalak, Chonticha Klungtong, Theerarat Kochakarn, Namfon Kotanan, Krittikorn Kumpornsin, Wudtichai Manasatienkij, Bhakbhoom Panthan, Ekawat Pasomsub, Kingkan Rakmanee, Insee Sensorn, Janjira Thaipadungpanit, Arporn Wangwiwatsin,Treewat Watthanachockchai |
| EPI_ISL_429861, EPI_ISL_429862, EPI_ISL_429863, EPI_ISL_429864, EPI_ISL_429865, EPI_ISL_429866, EPI_ISL_429867, EPI_ISL_429868, EPI_ISL_429869, EPI_ISL_429870, EPI_ISL_429871, EPI_ISL_429872, EPI_ISL_429873 | | | |
| see above | Ministry of Health Turkey | Ministry of Health Turkey | Fatma Bayrakdar,Aye Baak Alta,Yasemin Cogun,Gülay Korukluolu,Selçuk Klç |
| EPI_ISL_430019, EPI_ISL_430020, EPI_ISL_430021, EPI_ISL_430022, EPI_ISL_430023, EPI_ISL_430024, EPI_ISL_430025, EPI_ISL_430026, EPI_ISL_430027, EPI_ISL_430028, EPI_ISL_430029, EPI_ISL_430030, EPI_ISL_430031, EPI_ISL_430032, EPI_ISL_430033, EPI_ISL_430034, EPI_ISL_430035, EPI_ISL_430036, EPI_ISL_430037, EPI_ISL_430038, EPI_ISL_430039, EPI_ISL_430040, EPI_ISL_430041, EPI_ISL_430042, EPI_ISL_430043, EPI_ISL_430044, EPI_ISL_430045, EPI_ISL_430046, EPI_ISL_430047, EPI_ISL_430048, EPI_ISL_430049, EPI_ISL_430050, EPI_ISL_430051, EPI_ISL_430052, EPI_ISL_430053, EPI_ISL_430054, EPI_ISL_430055, EPI_ISL_430056, EPI_ISL_430057, EPI_ISL_430058, EPI_ISL_430059, EPI_ISL_430060, EPI_ISL_430061, EPI_ISL_430062 | | | |
| see above | Utah Public Health Laboratory | Utah Public Health Laboratory | Erin Young, Kelly Oakeson |
| EPI_ISL_430719, EPI_ISL_430720, EPI_ISL_430721 | Hospital Universitario La Paz | Hospital Universitario 12 de Octubre | Elias Dahdouh, Sara González, Raúl Recio, Fernando Lázaro, Esther Viedma, Natalia Stella, Julio García, Juan Carlos Galán, Rafael Cantón, Mª Dolores Folgueira, Rafael Delgado, Jesús Mingorance |
| EPI_ISL_432668, EPI_ISL_432669, EPI_ISL_432670, EPI_ISL_432671, EPI_ISL_432672, EPI_ISL_432673, EPI_ISL_432674, EPI_ISL_432675, EPI_ISL_432676, EPI_ISL_432677, EPI_ISL_432678, EPI_ISL_432679, EPI_ISL_432680, EPI_ISL_432681, EPI_ISL_432682, EPI_ISL_432683, EPI_ISL_432684, EPI_ISL_432685, EPI_ISL_432686, EPI_ISL_432687, EPI_ISL_432688, EPI_ISL_432689, EPI_ISL_432690, EPI_ISL_432691, EPI_ISL_432692, EPI_ISL_432693, EPI_ISL_432694, EPI_ISL_432695, EPI_ISL_432696, EPI_ISL_432697, EPI_ISL_432698, EPI_ISL_432699, EPI_ISL_432700, EPI_ISL_432701, EPI_ISL_432702, EPI_ISL_432703, EPI_ISL_432704, EPI_ISL_432705, EPI_ISL_432706, EPI_ISL_432707, EPI_ISL_432708, EPI_ISL_432709, EPI_ISL_432900, EPI_ISL_432901, EPI_ISL_432902, EPI_ISL_432903, EPI_ISL_432904, EPI_ISL_432905, EPI_ISL_432906, EPI_ISL_432907, EPI_ISL_432908, EPI_ISL_432909, EPI_ISL_432910, EPI_ISL_432911, EPI_ISL_432912, EPI_ISL_432913, EPI_ISL_432914, EPI_ISL_432915, EPI_ISL_432916, EPI_ISL_432917, EPI_ISL_432918, EPI_ISL_432919, EPI_ISL_432920, EPI_ISL_432921, EPI_ISL_432922, EPI_ISL_432923, EPI_ISL_432924, EPI_ISL_432925, EPI_ISL_432926, EPI_ISL_432927, EPI_ISL_432928, EPI_ISL_432929, EPI_ISL_432930, EPI_ISL_432931, EPI_ISL_432932, EPI_ISL_432933, EPI_ISL_432934, EPI_ISL_432935, EPI_ISL_432936, EPI_ISL_432937, EPI_ISL_432938, EPI_ISL_432939, EPI_ISL_432940, EPI_ISL_432941, EPI_ISL_432942, EPI_ISL_432943, EPI_ISL_432944, EPI_ISL_432945, EPI_ISL_432946, EPI_ISL_432947, EPI_ISL_432948, EPI_ISL_432949, EPI_ISL_432950, EPI_ISL_432951, EPI_ISL_432952, EPI_ISL_432953, EPI_ISL_432954, EPI_ISL_432955, EPI_ISL_432956, EPI_ISL_432957, EPI_ISL_432958, EPI_ISL_432959, EPI_ISL_432960, EPI_ISL_432961, EPI_ISL_432962, EPI_ISL_432963, EPI_ISL_432964, EPI_ISL_432965, EPI_ISL_432966, EPI_ISL_432967, EPI_ISL_432968, EPI_ISL_432969, EPI_ISL_432970, EPI_ISL_432971, EPI_ISL_432972, EPI_ISL_432973, EPI_ISL_432974, EPI_ISL_432975, EPI_ISL_432976, EPI_ISL_432977, EPI_ISL_432978, EPI_ISL_432979, EPI_ISL_432980, EPI_ISL_432981, EPI_ISL_432982, EPI_ISL_432983, EPI_ISL_432984, EPI_ISL_432985, EPI_ISL_432986, EPI_ISL_432987, EPI_ISL_432988, EPI_ISL_432989, EPI_ISL_432990, EPI_ISL_432991, EPI_ISL_432992, EPI_ISL_432993, EPI_ISL_432994, EPI_ISL_432995, EPI_ISL_432996, EPI_ISL_432997, EPI_ISL_432998, EPI_ISL_432999, EPI_ISL_433000, EPI_ISL_433001, | | | |

| | | | |
|---|---|---|---|
| EPI_ISL_433002, EPI_ISL_433003, EPI_ISL_433004, EPI_ISL_433005, EPI_ISL_433006, EPI_ISL_433007, EPI_ISL_433008, EPI_ISL_433009, EPI_ISL_433010, EPI_ISL_433011, EPI_ISL_433012, EPI_ISL_433013, EPI_ISL_433014, EPI_ISL_433015, EPI_ISL_433016, EPI_ISL_433017, EPI_ISL_433018, EPI_ISL_433019, EPI_ISL_433020, EPI_ISL_433021, EPI_ISL_433022, EPI_ISL_433023, EPI_ISL_433024, EPI_ISL_433025, EPI_ISL_433026, EPI_ISL_433027, EPI_ISL_433028, EPI_ISL_433029, EPI_ISL_433030, EPI_ISL_433031, EPI_ISL_433032, EPI_ISL_433033, EPI_ISL_433034, EPI_ISL_433035, EPI_ISL_433036, EPI_ISL_433037, EPI_ISL_433038, EPI_ISL_433039, EPI_ISL_433040, EPI_ISL_433041, EPI_ISL_433042, EPI_ISL_433043, EPI_ISL_433044, EPI_ISL_433045, EPI_ISL_433046, EPI_ISL_433047, EPI_ISL_433048, EPI_ISL_433049, EPI_ISL_433050, EPI_ISL_433051, EPI_ISL_433052, EPI_ISL_433053, EPI_ISL_433054, EPI_ISL_433055, EPI_ISL_433056, EPI_ISL_433057, EPI_ISL_433058, EPI_ISL_433059, EPI_ISL_433060, EPI_ISL_433061, EPI_ISL_433062, EPI_ISL_433063, EPI_ISL_433064, EPI_ISL_433065, EPI_ISL_433066, EPI_ISL_433067 | | | |
| see above | Queens Medical Centre, Clinical Microbiology Department / DeepSeq Nottingham | COVID-19 Genomics UK (COG-UK) Consortium | Gemma Clark, Wendy Smith, Manjinder Khakh, Hannah Howson-Wells, Jonathan Ball, Patrick McClure, Joseph Chappell, Theocharis Tsoleridis, Nadine Holmes, Matthew Carlisle, Christopher Moore, Fei Sang, Johnny Debebe, Victoria Wright, Matthew Loose |
| EPI_ISL_434533, EPI_ISL_434535 | Area de Salud Alajuela Sur | Inciensa, Instituto Costarricense de Investigación y Enseñanza en Nutrición y Salud | Francisco Duarte, Hebleen Porras, Claudio Soto-Garita, Estela Cordero, Adriana Godinez & Melany Calderon |
| EPI_ISL_434536 | Hospital San Vicente de Paul | Inciensa, Instituto Costarricense de Investigación y Enseñanza en Nutrición y Salud | Francisco Duarte, Hebleen Porras, Claudio Soto-Garita, Estela Cordero, Adriana Godinez & Melany Calderon |
| EPI_ISL_434538 | COOPESAIN | Inciensa, Instituto Costarricense de Investigación y Enseñanza en Nutrición y Salud | Francisco Duarte, Hebleen Porras, Claudio Soto-Garita, Estela Cordero, Adriana Godinez & Melany Calderon |
| EPI_ISL_434539 | Area de Salud Orotina | Inciensa, Instituto Costarricense de Investigación y Enseñanza en Nutrición y Salud | Francisco Duarte, Hebleen Porras, Claudio Soto-Garita, Estela Cordero, Adriana Godinez & Melany Calderon |
| EPI_ISL_434540 | EBAIS Concepción Norte | Inciensa, Instituto Costarricense de Investigación y Enseñanza en Nutrición y Salud | Francisco Duarte, Hebleen Porras, Claudio Soto-Garita, Estela Cordero, Adriana Godinez & Melany Calderon |
| EPI_ISL_434607, EPI_ISL_434608, EPI_ISL_434609, EPI_ISL_434610, EPI_ISL_434611, EPI_ISL_434612, EPI_ISL_434613, EPI_ISL_434614, EPI_ISL_434615 | University of Wisconsin-Madison AIDS Vaccine Research Laboratories | University of Wisconsin-Madison AIDS Vaccine Research Laboratories | Gage Moreno, Katarina Braun, et al. AIDS Vaccine Research Laboratories |
| EPI_ISL_434636 | Lednicky Laboratory, Emerging Pathogens Institute, University of Florida | Lednicky Laboratory at Emerging Pathogens Institute, University of Florida | Elbadry,M.A., Subramaniam,K., Waltzek,T.B., Stephenson,C.J.,Gibson,J.C., Alam,M., Morris,J.G. Jr. and Lednicky,J.A. |
| EPI_ISL_434637 | Lednicky Laboratory, Emerging Pathogens Institute, University of Florida. | Lednicky Laboratory, Emerging Pathogens Institute, University of Florida. | Elbadry,M.A.; Subramaniam,K.; Waltzek,T.B.; Gibson,J.C.; Stephenson,C.J.; Morris,J.G. Jr. and Lednicky,J.A. |
| EPI_ISL_434652 | Ektorps Vardcentral | The Public Health Agency of Sweden | Eva Espmark, Oskar Karlsson Lindsjo, Maria Lind Karlberg, Anna-Malin Linde, Olov Svartstrom, Anna Risberg, Theresa Enkirch, Mia Brytting, Karin Tegmark-Wisell |
| EPI_ISL_434663, EPI_ISL_434672, EPI_ISL_434675, EPI_ISL_434676 | Surbrunns VC | The Public Health Agency of Sweden | Erik Embring, Oskar Karlsson Lindsjo, Maria Lind Karlberg, Anna-Malin Linde, Olov Svartstrom, Anna Risberg, Theresa Enkirch, Mia Brytting, Karin Tegmark-Wisell |
| EPI_ISL_435281 | Medistra Hospital Jakarta | Eijkman Institute for Molecular Biology, Ministry of Research and Technology/National Agency for Research and Innovation | Edison Johar, Frilasita A Yudhaputri, Hidayat Trimarsanto, David H Muljono, Safarina G Malik, Khin Saw Myint, Amin Soebandrio |
| EPI_ISL_435282, EPI_ISL_435283 | RS Pondok Indah Hospital - Pondok Indah | Eijkman Institute for Molecular Biology, Ministry of Research and Technology/National Agency for Research and Innovation | Edison Johar, Frilasita A Yudhaputri, Hidayat Trimarsanto, David H Muljono, Safarina G Malik, Khin Saw Myint, Amin Soebandrio |
| EPI_ISL_435348, EPI_ISL_435349, EPI_ISL_435350, EPI_ISL_435351, EPI_ISL_435352, EPI_ISL_435353, EPI_ISL_435354, EPI_ISL_435355, EPI_ISL_435356, EPI_ISL_435357, EPI_ISL_435358, EPI_ISL_435359, EPI_ISL_435360, EPI_ISL_435361, EPI_ISL_435362, EPI_ISL_435363, EPI_ISL_435364, EPI_ISL_435365, EPI_ISL_435366, EPI_ISL_435367, EPI_ISL_435368, EPI_ISL_435369, EPI_ISL_435370, EPI_ISL_435371, EPI_ISL_435372, EPI_ISL_435373, EPI_ISL_435374, EPI_ISL_435375, EPI_ISL_435376, EPI_ISL_435377, EPI_ISL_435378, EPI_ISL_435379, EPI_ISL_435380, EPI_ISL_435381, EPI_ISL_435382, EPI_ISL_435383, EPI_ISL_435384, EPI_ISL_435385, EPI_ISL_435386, EPI_ISL_435387, EPI_ISL_435388, EPI_ISL_435389, EPI_ISL_435390, EPI_ISL_435391, EPI_ISL_435392, EPI_ISL_435393 | | | |
| see above | Utah Public Health Laboratory | Utah Public Health Laboratory | Erin Young, Kelly Oakeson |
| EPI_ISL_436564, EPI_ISL_436565, EPI_ISL_436566, EPI_ISL_436567, EPI_ISL_436568, EPI_ISL_436569, EPI_ISL_436570, EPI_ISL_436571, EPI_ISL_436572, EPI_ISL_436573, EPI_ISL_436574, EPI_ISL_436575, EPI_ISL_436576, EPI_ISL_436577, EPI_ISL_436578, EPI_ISL_436579, EPI_ISL_436580, EPI_ISL_436581, EPI_ISL_436582, EPI_ISL_436583, EPI_ISL_436584, EPI_ISL_436585, EPI_ISL_436586, EPI_ISL_436587, EPI_ISL_436588, EPI_ISL_436589, EPI_ISL_436590, EPI_ISL_436591, EPI_ISL_436592, EPI_ISL_436593, EPI_ISL_436594, EPI_ISL_436595, EPI_ISL_436596, EPI_ISL_436597, EPI_ISL_436598, EPI_ISL_436599, EPI_ISL_436600, EPI_ISL_436601, EPI_ISL_436602, EPI_ISL_436603, EPI_ISL_436604, EPI_ISL_436605, EPI_ISL_436606, EPI_ISL_436607, EPI_ISL_436608, EPI_ISL_436609, EPI_ISL_436610, EPI_ISL_436611, EPI_ISL_436612, EPI_ISL_436613, EPI_ISL_436614, EPI_ISL_436615, EPI_ISL_436616, EPI_ISL_436617, EPI_ISL_436618, EPI_ISL_436619, EPI_ISL_436620, EPI_ISL_436621, EPI_ISL_436622, EPI_ISL_436623, EPI_ISL_436624, EPI_ISL_436625, EPI_ISL_436626, EPI_ISL_436627, EPI_ISL_436628, EPI_ISL_436629, EPI_ISL_436630, EPI_ISL_436631, EPI_ISL_436632, EPI_ISL_436633, EPI_ISL_436634, EPI_ISL_436635, EPI_ISL_436636, EPI_ISL_436637, EPI_ISL_436638, EPI_ISL_436639, EPI_ISL_436640 | | | |
| see above | University of Wisconsin-Madison AIDS Vaccine Research Laboratories | University of Wisconsin-Madison AIDS Vaccine Research Laboratories | Gage Moreno, Katarina Braun, et al. AIDS Vaccine Research Laboratories |
| EPI_ISL_436684, EPI_ISL_436685, EPI_ISL_436686, EPI_ISL_436687 | KRISP, KZN Research Innovation and Sequencing Platform | KRISP, KZN Research Innovation and Sequencing Platform | Giandhari J, Pillay S, Lessells R, Chimukangara B, Deforche K, Tegally H, Wilkinson E, de Oliveira T |
| EPI_ISL_436901, EPI_ISL_436902, EPI_ISL_436903, EPI_ISL_436904, EPI_ISL_436905, EPI_ISL_436906, EPI_ISL_436907, EPI_ISL_436908, EPI_ISL_436909, EPI_ISL_436910, EPI_ISL_436911, EPI_ISL_436912, EPI_ISL_436913, EPI_ISL_436914, EPI_ISL_436915, EPI_ISL_436916, EPI_ISL_436917, EPI_ISL_436918, EPI_ISL_436919, EPI_ISL_436920, EPI_ISL_436921, EPI_ISL_436922, EPI_ISL_436923, EPI_ISL_436924, EPI_ISL_436925 | | | |
| see above | Utah Public Health Laboratory | Utah Public Health Laboratory | Erin Young, Kelly Oakeson |
| EPI_ISL_436926 | $x^2$ | Utah Public Health Laboratory | Erin Young, Kelly Oakeson |
| EPI_ISL_436927, EPI_ISL_436928, EPI_ISL_436929, EPI_ISL_436930, EPI_ISL_436931, EPI_ISL_436932, EPI_ISL_436933, EPI_ISL_436934, EPI_ISL_436935, EPI_ISL_436936, EPI_ISL_436937, EPI_ISL_436938 | | | |
| see above | Utah Public Health Laboratory | Utah Public Health Laboratory | Erin Young, Kelly Oakeson |
| EPI_ISL_437189 | Pusat Pertamina Hospital | Eijkman Institute for Molecular Biology, Ministry of Research and Technology/National Agency for Research and Innovation | Edison Johar, Frilasita A Yudhaputri, Hidayat Trimarsanto, David H Muljono, Safarina G Malik, Khin Saw Myint, Amin Soebandrio |
| EPI_ISL_437190, EPI_ISL_437191 | RS Pondok Indah Hospital - Pondok Indah | Eijkman Institute for Molecular Biology, Ministry of Research and Technology/National Agency for Research and Innovation | Edison Johar, Frilasita A Yudhaputri, Hidayat Trimarsanto, David H Muljono, Safarina G Malik, Khin Saw Myint, Amin Soebandrio |
| EPI_ISL_437192 | Mitra Keluarga Kelapa Gading Hospital | Eijkman Institute for Molecular Biology, Ministry of Research and Technology/National Agency for Research and Innovation | Edison Johar, Frilasita A Yudhaputri, Hidayat Trimarsanto, David H Muljono, Safarina G Malik, Khin Saw Myint, Amin Soebandrio |
| EPI_ISL_437304, EPI_ISL_437305, EPI_ISL_437306, EPI_ISL_437307, EPI_ISL_437308, EPI_ISL_437309, EPI_ISL_437310, EPI_ISL_437311, EPI_ISL_437312, EPI_ISL_437313, EPI_ISL_437314, EPI_ISL_437315, EPI_ISL_437316, EPI_ISL_437317, EPI_ISL_437318 | | | |
| see above | Ministry of Health Turkey | Ministry of Health Turkey | Fatma Bayrakdar,Tülin Demir,Süleyman Yalçn, Selçuk Klç |
| EPI_ISL_437319, EPI_ISL_437320, EPI_ISL_437321 | Ministry of Health Turkey | Ministry of Health Turkey | Fatma Bayrakdar,Aye Baak Alta,Yasemin Cogun,Süleyman Yalçn, Gülay Korukluolu,Selçuk Klç |
| EPI_ISL_437322 | Ministry of Health Turkey | Ministry of Health Turkey | Fatma Bayrakdar,Tülin Demir,Süleyman Yalçn, Selçuk Klç |
| EPI_ISL_437323, EPI_ISL_437324, EPI_ISL_437325, EPI_ISL_437326, EPI_ISL_437327, EPI_ISL_437328, EPI_ISL_437329, EPI_ISL_437330 | Ministry of Health Turkey | Ministry of Health Turkey | Fatma Bayrakdar,Aye Baak Alta,Yasemin Cogun,Süleyman Yalçn, Gülay Korukluolu,Selçuk Klç |
| EPI_ISL_437331 | Ministry of Health Turkey | Ministry of Health Turkey | Fatma Bayrakdar,Tülin Demir,Süleyman Yalçn, Selçuk Klç |
| EPI_ISL_437332, EPI_ISL_437333, EPI_ISL_437334, EPI_ISL_437335 | Ministry of Health Turkey | Ministry of Health Turkey | Fatma Bayrakdar,Aye Baak Alta,Yasemin Cogun,Süleyman Yalçn, Gülay Korukluolu,Selçuk Klç |
| EPI_ISL_437435, EPI_ISL_437436 | Veterinary Specialized Institue Kraljevo | Veterinary Specialized Institue Kraljevo | Dejan Vidanovic. Bojana Tesovic, Milanko Sekler, Marko Dmitric, Kazimir Matovic, Zoran Debeljak, Nikola Vaskovic, Tamas Petrovic, Jeremy Volkening, |

| | | | |
|---|---|---|---|
| | | | Claudio L Afonso |
| EPI_ISL_437442 | Department of MicroBiology, Government Medical College, Surat | Gujarat Biotechnology Research Centre | Dinesh Kumar, Zuber Saiyed, Dipa Kinariwala, Disha Patel, Binita Aring, Neeta Khandelwal, Geeta Vaghela, Sonia Barve, Bhavesh Modi, Kairavi Joshi, Gaurishankar Shrimali, Nidhi Sood, Pranay Shah, R D Dixit, Snehal Bagatharia, Kamlesh J Upadhyay, Ramesh Pandit, Tejas Shah, Ankit Hinsu, Pritesh Sabara, Apurvasinh Puvar, Janvi Raval, Monika Gandhi, Pinal Trivedi, Maharshi Pandya, Amit Kanani, Akanksha Verma, Nitin Savaliya, Raghawendra Kumar, Dipeshwari Shewale, Chaitanya Joshi, Madhvi Joshi |
| EPI_ISL_437444 | Department of MicroBiology, Government Medical College, Surat | Gujarat Biotechnology Research Centre | Dipa Kinariwala, Disha Patel, Binita Aring, Neeta Khandelwal, Geeta Vaghela, Sonia Barve, Bhavesh Modi, Kairavi Joshi, Gaurishankar Shrimali, Nidhi Sood, Pranay Shah, R D Dixit, Snehal Bagatharia, Kamlesh J Upadhyay, Ramesh Pandit, Tejas Shah, Ankit Hinsu, Pritesh Sabara, Apurvasinh Puvar, Janvi Raval, Monika Gandhi, Pinal Trivedi, Maharshi Pandya, Amit Kanani, Akanksha Verma, Nitin Savaliya, Raghawendra Kumar, Dinesh Kumar, Zuber Saiyed, Pooja P Doshi, Chaitanya Joshi, Madhvi Joshi |
| EPI_ISL_437445 | B.J. Medical College and Civil hospital | Gujarat Biotechnology Research Centre | Disha Patel, Binita Aring, Neeta Khandelwal, Geeta Vaghela, Sonia Barve, Bhavesh Modi, Kairavi Joshi, Gaurishankar Shrimali, Nidhi Sood, Pranay Shah, R D Dixit, Snehal Bagatharia, Kamlesh J Upadhyay, Ramesh Pandit, Tejas Shah, Ankit Hinsu, Pritesh Sabara, Apurvasinh Puvar, Janvi Raval, Monika Gandhi, Pinal Trivedi, Maharshi Pandya, Amit Kanani, Akanksha Verma, Nitin Savaliya, Raghawendra Kumar, Dinesh Kumar, Zuber Saiyed, Dipa Kinariwala, Nidhi Patel, Chaitanya Joshi, Madhvi Joshi |
| EPI_ISL_437448 | B.J. Medical College and Civil hospital | Gujarat Biotechnology Research Centre | Geeta Vaghela, Sonia Barve, Bhavesh Modi, Kairavi Joshi, Gaurishankar Shrimali, Nidhi Sood, Pranay Shah, R D Dixit, Snehal Bagatharia, Kamlesh J Upadhyay, Ramesh Pandit, Tejas Shah, Ankit Hinsu, Pritesh Sabara, Apurvasinh Puvar, Janvi Raval, Monika Gandhi, Pinal Trivedi, Maharshi Pandya, Amit Kanani, Akanksha Verma, Nitin Savaliya, Raghawendra Kumar, Dinesh Kumar, Zuber Saiyed, Dipa Kinariwala, Disha Patel, Binita Aring, Neeta Khandelwal, Afzal Ansari, Chaitanya Joshi, Madhvi Joshi |
| EPI_ISL_437452 | B.J. Medical College and Civil hospital | Gujarat Biotechnology Research Centre | Gaurishankar Shrimali, Nidhi Sood, Pranay Shah, R D Dixit, Snehal Bagatharia, Kamlesh J Upadhyay, Ramesh Pandit, Tejas Shah, Ankit Hinsu, Pritesh Sabara, Apurvasinh Puvar, Janvi Raval, Monika Gandhi, Pinal Trivedi, Maharshi Pandya, Amit Kanani, Akanksha Verma, Nitin Savaliya, Raghawendra Kumar, Dinesh Kumar, Zuber Saiyed, Dipa Kinariwala, Disha Patel, Binita Aring, Neeta Khandelwal, Geeta Vaghela, Sonia Barve, Bhavesh Modi, Kairavi Joshi, Chaitanya Joshi, Anjali Rajwar, Madhvi Joshi |
| EPI_ISL_437625 | Laboratory of Genomics & Bioinformatics, Institute of Immunology and Experimental Therapy, Polish Academy of Sciences Oddzial Mikrobiologii Wojewodzkiej Stacji Sanitarno-Epidemiologicznej. | Laboratory of Genomics & Bioinformatics, Institute of Immunology and Experimental Therapy, Polish Academy of Sciences | Dorota Kujawa, Aleksandra Herud, Dariusz Martynowski, Krzysztof Jakub Pawlik, Joanna Sikorska, Paulina Zebrowska, Grazyna Zalewska, Oskar Karpinski and Lukasz Laczmanski |
| EPI_ISL_441437, EPI_ISL_441438, EPI_ISL_441439, EPI_ISL_441440, EPI_ISL_441441, EPI_ISL_441442, EPI_ISL_441443, EPI_ISL_441444, EPI_ISL_441445, EPI_ISL_441446, EPI_ISL_441447, EPI_ISL_441448, EPI_ISL_441449, EPI_ISL_441450, EPI_ISL_441451, EPI_ISL_441452, EPI_ISL_441453, EPI_ISL_441454, EPI_ISL_441455, EPI_ISL_441456, EPI_ISL_441457, EPI_ISL_441458, EPI_ISL_441459, EPI_ISL_441460, EPI_ISL_441461, EPI_ISL_441462, EPI_ISL_441463, EPI_ISL_441464, EPI_ISL_441465, EPI_ISL_441466, EPI_ISL_441467, EPI_ISL_441468, EPI_ISL_441469, EPI_ISL_441470, EPI_ISL_441471, EPI_ISL_441472, EPI_ISL_441473, EPI_ISL_441474, EPI_ISL_441475, EPI_ISL_441476, EPI_ISL_441477, EPI_ISL_441478, EPI_ISL_441479, EPI_ISL_441480, EPI_ISL_441481, EPI_ISL_441482, EPI_ISL_441483, EPI_ISL_441484, EPI_ISL_441485, EPI_ISL_441486, EPI_ISL_441487, EPI_ISL_441488, EPI_ISL_441489, EPI_ISL_441490, EPI_ISL_441491, EPI_ISL_441492, EPI_ISL_441493, EPI_ISL_441494, EPI_ISL_441495, EPI_ISL_441496, EPI_ISL_441497, EPI_ISL_441498, EPI_ISL_441499, EPI_ISL_441500, EPI_ISL_441501, EPI_ISL_441502, EPI_ISL_441503, EPI_ISL_441504, EPI_ISL_441505, EPI_ISL_441506, EPI_ISL_441507, EPI_ISL_441508, EPI_ISL_441509, EPI_ISL_441510, EPI_ISL_441511, EPI_ISL_441512, EPI_ISL_441513, EPI_ISL_441514, EPI_ISL_441515, EPI_ISL_441516, EPI_ISL_441517, EPI_ISL_441518, EPI_ISL_441519, EPI_ISL_441520, EPI_ISL_441521, EPI_ISL_441522, EPI_ISL_441523, EPI_ISL_441524, EPI_ISL_441525, EPI_ISL_441526, EPI_ISL_441527, EPI_ISL_441528, EPI_ISL_441529, EPI_ISL_441530, EPI_ISL_441531, EPI_ISL_441532, EPI_ISL_441533, EPI_ISL_441534, EPI_ISL_441535, EPI_ISL_441536, EPI_ISL_441537, EPI_ISL_441538, EPI_ISL_441539, EPI_ISL_441540, EPI_ISL_441541, EPI_ISL_441542, EPI_ISL_441543, EPI_ISL_441544, EPI_ISL_441545, EPI_ISL_441546, EPI_ISL_441847, EPI_ISL_441848, EPI_ISL_441849, EPI_ISL_441850, EPI_ISL_441851, EPI_ISL_441852, EPI_ISL_441853, EPI_ISL_441854, EPI_ISL_441855, EPI_ISL_441856, EPI_ISL_441857, EPI_ISL_441858, EPI_ISL_441859, EPI_ISL_441860, EPI_ISL_441861, EPI_ISL_441862, EPI_ISL_441863, EPI_ISL_441864, EPI_ISL_441865, EPI_ISL_441866, EPI_ISL_441867, EPI_ISL_441868, EPI_ISL_441869, EPI_ISL_441870, EPI_ISL_441871, EPI_ISL_441872, EPI_ISL_441873, EPI_ISL_441874, EPI_ISL_441875, EPI_ISL_441876, EPI_ISL_441877, EPI_ISL_441878, EPI_ISL_441879, EPI_ISL_441880, EPI_ISL_441881, EPI_ISL_441882, EPI_ISL_441883, EPI_ISL_441884, EPI_ISL_441885, EPI_ISL_441886, EPI_ISL_441887, EPI_ISL_441888, EPI_ISL_441889, EPI_ISL_441890, EPI_ISL_441891, EPI_ISL_441892, EPI_ISL_441893, EPI_ISL_441894, EPI_ISL_441895, EPI_ISL_441896, EPI_ISL_441897 | | | |
| see above | Queens Medical Centre, Clinical Microbiology Department / DeepSeq Nottingham | COVID-19 Genomics UK (COG-UK) Consortium | Gemma Clark, Wendy Smith, Manjinder Khakh, Hannah Howson-Wells, Jonathan Ball, Patrick McClure, Joseph Chappell, Theocharis Tsoleridis, Nadine Holmes, Matthew Carlisle, Christopher Moore, Fei Sang, Johnny Debebe, Victoria Wright, Matthew Loose |
| EPI_ISL_444472 | B.J. Medical College and Civil hospital | Gujarat Biotechnology Research Centre | Dinesh Kumar, Zuber Saiyed, Dipa Kinariwala, Disha Patel, Binita Aring, Neeta Khandelwal, Geeta Vaghela, Sonia Barve, Bhavesh Modi, Kairavi Joshi, Gaurishankar Shrimali, Nidhi Sood, Pranay Shah, R D Dixit, Snehal Bagatharia, Kamlesh J Upadhyay, Ramesh Pandit, Tejas Shah, Ankit Hinsu, Pritesh Sabara, Apurvasinh Puvar, Janvi Raval, Monika Gandhi, Pinal Trivedi, Maharshi Pandya, Amit Kanani, Akanksha Verma, Nitin Savaliya, Raghawendra Kumar, Neelam Nathani, Chaitanya Joshi, Madhvi Joshi |
| EPI_ISL_444474 | B.J. Medical College and Civil hospital | Gujarat Biotechnology Research Centre | Dipa Kinariwala, Disha Patel, Binita Aring, Neeta Khandelwal, Geeta Vaghela, Sonia Barve, Bhavesh Modi, Kairavi Joshi, Gaurishankar Shrimali, Nidhi Sood, Pranay Shah, R D Dixit, Snehal Bagatharia, Kamlesh J Upadhyay, Ramesh Pandit, Tejas Shah, Ankit Hinsu, Pritesh Sabara, Apurvasinh Puvar, Janvi Raval, Monika Gandhi, Pinal Trivedi, Maharshi Pandya, Amit Kanani, Akanksha Verma, Nitin Savaliya, Raghawendra Kumar, Dinesh Kumar, Zuber Saiyed, Bhavya Jindal, Chaitanya Joshi, Madhvi Joshi |
| EPI_ISL_444475 | B.J. Medical College and Civil hospital | Gujarat Biotechnology Research Centre | Disha Patel, Binita Aring, Neeta Khandelwal, Geeta Vaghela, Sonia Barve, Bhavesh Modi, Kairavi Joshi, Gaurishankar Shrimali, Nidhi Sood, Pranay Shah, R D Dixit, Snehal Bagatharia, Kamlesh J Upadhyay, Ramesh Pandit, Tejas Shah, Ankit Hinsu, Pritesh Sabara, Apurvasinh Puvar, Janvi Raval, Monika Gandhi, Pinal Trivedi, Maharshi Pandya, Amit Kanani, Akanksha Verma, Nitin Savaliya, Raghawendra Kumar, Dinesh Kumar, Zuber Saiyed, Dipa Kinariwala, Dipeshwari Shewale, Chaitanya Joshi, Madhvi Joshi |
| EPI_ISL_444478 | B.J. Medical College and Civil hospital | Gujarat Biotechnology Research Centre | Geeta Vaghela, Sonia Barve, Bhavesh Modi, Kairavi Joshi, Gaurishankar Shrimali, Nidhi Sood, Pranay Shah, R D Dixit, Snehal Bagatharia, Kamlesh J Upadhyay, Ramesh Pandit, Tejas Shah, Ankit Hinsu, Pritesh Sabara, Apurvasinh Puvar, Janvi Raval, Monika Gandhi, Pinal Trivedi, Maharshi Pandya, Amit Kanani, Akanksha Verma, Nitin Savaliya, Raghawendra Kumar, Dinesh Kumar, Zuber Saiyed, Dipa Kinariwala, Disha Patel, Binita Aring, Neeta Khandelwal, Pooja P Doshi, Chaitanya Joshi, Madhvi Joshi |
| EPI_ISL_444482 | Gujarat Biotechnology Research Centre | Gujarat Biotechnology Research Centre | Gaurishankar Shrimali, Nidhi Sood, Pranay Shah, R D Dixit, Snehal Bagatharia, Kamlesh J Upadhyay, Ramesh Pandit, Tejas Shah, Ankit Hinsu, Pritesh Sabara, Apurvasinh Puvar, Janvi Raval, Monika Gandhi, Pinal Trivedi, Maharshi Pandya, Amit Kanani, Akanksha Verma, Nitin Savaliya, Raghawendra Kumar, Dinesh Kumar, Zuber Saiyed, Dipa Kinariwala, Disha Patel, Binita Aring, Neeta Khandelwal, Geeta Vaghela, Sonia Barve, Bhavesh Modi, Kairavi Joshi, Afzal Ansari, Chaitanya Joshi, Madhvi Joshi |
| EPI_ISL_445221 | Wasterlakarna | The Public Health Agency of Sweden | Frida Ahlfors, Oskar Karlsson Lindsjo, Maria Lind Karlberg, Anna-Malin Linde, Olov Svartstrom, Anna Risberg, Theresa Enkirch, Mia Brytting, Karin Tegmark-Wisell |
| EPI_ISL_445225 | Surbrunns VC | The Public Health Agency of Sweden | Erik Embring, Oskar Karlsson Lindsjo, Maria Lind Karlberg, Anna-Malin Linde, Olov Svartstrom, Anna Risberg, Theresa Enkirch, Mia Brytting, Karin Tegmark-Wisell |
| EPI_ISL_445234, EPI_ISL_445235 | Wasterlakarna | The Public Health Agency of Sweden | Frida Ahlfors, Oskar Karlsson Lindsjo, Maria Lind Karlberg, Anna-Malin Linde, Olov Svartstrom, Anna Risberg, Theresa Enkirch, Mia Brytting, Karin Tegmark-Wisell |
| EPI_ISL_445238 | Ã-resundslakarna | The Public Health Agency of Sweden | Del Akrawi, Oskar Karlsson Lindsjo, Maria Lind Karlberg, Anna-Malin Linde, Olov Svartstrom, Anna Risberg, Theresa Enkirch, Mia Brytting, Karin Tegmark-Wisell |
| EPI_ISL_445380 | Ramathibodi Hospital | COVID-19 Network Investigations (CONI) Alliance | Elizabeth Batty, Wasun Chantratita, Thanat Chookajorn, Stefan Fernandez, Angkana Huang, Anthony R. Jones, Khajohn Joonsalak, Chonticha Klungtong, Theerarat Kochakarn, Namfon Kotanan, Krittikorn Kumpornsin, Wudtichai Manasatienkij, Bhakbhoom Panthan, Ekawat Pasomsub, Kingkan Rakmanee, Insee Sensorn, Janjira Thaipadungpanit, Arporn Wangwiwatsin, Treewat Watthanachockchai |
| EPI_ISL_446996, EPI_ISL_446997, EPI_ISL_446998, EPI_ISL_446999, EPI_ISL_447000, EPI_ISL_447001, EPI_ISL_447002, EPI_ISL_447003, EPI_ISL_447004, EPI_ISL_447005, EPI_ISL_447006, EPI_ISL_447007, EPI_ISL_447008, EPI_ISL_447009, EPI_ISL_447010, EPI_ISL_447011, EPI_ISL_447012, EPI_ISL_447013, EPI_ISL_447014, EPI_ISL_447015, EPI_ISL_447016, EPI_ISL_447017, EPI_ISL_447018, EPI_ISL_447019, EPI_ISL_447020, EPI_ISL_447021, EPI_ISL_447022, EPI_ISL_447023, EPI_ISL_447024, EPI_ISL_447025, EPI_ISL_447026, EPI_ISL_447027, EPI_ISL_447028, EPI_ISL_447029 | | | |
| see above | Ramathibodi Hospital | COVID-19 Network Investigations (CONI) Alliance | Elizabeth Batty, Wasun Chantratita, Thanat Chookajorn, Stefan Fernandez, Angkana Huang, Anthony R. Jones, Khajohn Joonsalak, Chonticha Klungtong, Theerarat Kochakarn, Namfon Kotanan, Krittikorn Kumpornsin, Wudtichai Manasatienkij, Bhakbhoom Panthan, Ekawat Pasomsub, Kingkan Rakmanee, Insee Sensorn, Janjira Thaipadungpanit, Arporn Wangwiwatsin,Treewat Watthanachockchai |
| EPI_ISL_447044 | B.J. Medical College and Civil hospital | Gujarat Biotechnology Research Centre | Dinesh Kumar, Zuber Saiyed, Dipa Kinariwala, Disha Patel, Binita Aring, Neeta Khandelwal, Geeta Vaghela, Sonia Barve, Bhavesh Modi, Kairavi Joshi, Gaurishankar Shrimali, Nidhi Sood, Pranay Shah, R D Dixit, Snehal Bagatharia, Kamlesh J Upadhyay, Ramesh Pandit, Tejas Shah, Ankit Hinsu, Pritesh Sabara, Apurvasinh Puvar, Janvi Raval, Monika Gandhi, Pinal Trivedi, Maharshi Pandya, Amit Kanani, Akanksha Verma, Nitin Savaliya, Raghawendra Kumar, Priti Pandita, Chaitanya Joshi, Madhvi Joshi |

| | | | |
|---|---|---|---|
| EPI_ISL_447046 | B.J. Medical College and Civil hospital | Gujarat Biotechnology Research Centre | Dipa Kinariwala, Disha Patel, Binita Aring, Neeta Khandelwal, Geeta Vaghela, Sonia Barve, Bhavesh Modi, Kairavi Joshi, Gaurishankar Shrimali, Nidhi Sood, Pranay Shah, R D Dixit, Snehal Bagatharia, Kamlesh J Upadhyay, Ramesh Pandit, Tejas Shah, Ankit Hinsu, Pritesh Sabara, Apurvasinh Puvar, Janvi Raval, Monika Gandhi, Pinal Trivedi, Maharshi Pandya, Amit Kanani, Akanksha Verma, Nitin Savaliya, Raghawendra Kumar, Dinesh Kumar, Zuber Saiyed, Afzal Ansari, Chaitanya Joshi, Madhvi Joshi |
| EPI_ISL_447047 | GMERS Medical College and Hospital, Gandhinagar | Gujarat Biotechnology Research Centre | Disha Patel, Binita Aring, Neeta Khandelwal, Geeta Vaghela, Sonia Barve, Bhavesh Modi, Kairavi Joshi, Gaurishankar Shrimali, Nidhi Sood, Pranay Shah, R D Dixit, Snehal Bagatharia, Kamlesh J Upadhyay, Ramesh Pandit, Tejas Shah, Ankit Hinsu, Pritesh Sabara, Apurvasinh Puvar, Janvi Raval, Monika Gandhi, Pinal Trivedi, Maharshi Pandya, Amit Kanani, Akanksha Verma, Nitin Savaliya, Raghawendra Kumar, Dinesh Kumar, Zuber Saiyed, Dipa Kinariwala, Neelam Nathani, Chaitanya Joshi, Madhvi Joshi |
| EPI_ISL_447050 | GMERS Medical College and Hospital, Gandhinagar | Gujarat Biotechnology Research Centre | Geeta Vaghela, Sonia Barve, Bhavesh Modi, Kairavi Joshi, Gaurishankar Shrimali, Nidhi Sood, Pranay Shah, R D Dixit, Snehal Bagatharia, Kamlesh J Upadhyay, Ramesh Pandit, Tejas Shah, Ankit Hinsu, Pritesh Sabara, Apurvasinh Puvar, Janvi Raval, Monika Gandhi, Pinal Trivedi, Maharshi Pandya, Amit Kanani, Akanksha Verma, Nitin Savaliya, Raghawendra Kumar, Dinesh Kumar, Zuber Saiyed, Dipa Kinariwala, Disha Patel, Binita Aring, Neeta Khandelwal, Dipeshwari Shewale, Chaitanya Joshi, Madhvi Joshi |
| EPI_ISL_447471, EPI_ISL_447485, EPI_ISL_447499, EPI_ISL_447513, EPI_ISL_447526 | Servicio de Microbiologia. Hospital Clinico Universitario de Valencia | Sequencing and Bioinformatics Service and Molecular Epidemiology Research Group. FISABIO-Public Health | Eliseo Albert, Maria Alma Bracho, Griselda De Marco, Lidia Ruiz Roldan, Neris Garcia-Gonzalez, Inma Galán Vendrell, Sandra Carbo, Loreto Ferrús Abad, Paula Ruiz-Hueso, Mariana Reyes-Prieto, Vicente Soriano Chirona, Ivan Ansari, Llúcia Martínez-Priego, Giuseppe 'Auria, David Navarro, Fernando Gonzalez-Candelas |
| EPI_ISL_447534 | Gujarat Biotechnology Research Centre | Gujarat Biotechnology Research Centre | Gaurishankar Shrimali, Nidhi Sood, Pranay Shah, R D Dixit, Snehal Bagatharia, Kamlesh J Upadhyay, Ramesh Pandit, Tejas Shah, Ankit Hinsu, Pritesh Sabara, Apurvasinh Puvar, Janvi Raval, Monika Gandhi, Pinal Trivedi, Maharshi Pandya, Amit Kanani, Akanksha Verma, Nitin Savaliya, Raghawendra Kumar, Dinesh Kumar, Zuber Saiyed, Dipa Kinariwala, Disha Patel, Binita Aring, Neeta Khandelwal, Geeta Vaghela, Sonia Barve, Bhavesh Modi, Kairavi Joshi, Nidhi Patel, Chaitanya Joshi, Madhvi Joshi |
| EPI_ISL_447553 | GMERS Medical College and Hospital, Gandhinagar | Gujarat Biotechnology Research Centre | Dinesh Kumar, Zuber Saiyed, Dipa Kinariwala, Disha Patel, Binita Aring, Neeta Khandelwal, Geeta Vaghela, Sonia Barve, Bhavesh Modi, Kairavi Joshi, Gaurishankar Shrimali, Nidhi Sood, Pranay Shah, R D Dixit, Snehal Bagatharia, Kamlesh J Upadhyay, Ramesh Pandit, Tejas Shah, Ankit Hinsu, Pritesh Sabara, Apurvasinh Puvar, Janvi Raval, Monika Gandhi, Pinal Trivedi, Maharshi Pandya, Amit Kanani, Akanksha Verma, Nitin Savaliya, Raghawendra Kumar, Dipeshwari Shewale, Chaitanya Joshi, Madhvi Joshi |
| EPI_ISL_447555 | GMERS Medical College and Hospital, Gandhinagar | Gujarat Biotechnology Research Centre | Dipa Kinariwala, Disha Patel, Binita Aring, Neeta Khandelwal, Geeta Vaghela, Sonia Barve, Bhavesh Modi, Kairavi Joshi, Gaurishankar Shrimali, Nidhi Sood, Pranay Shah, R D Dixit, Snehal Bagatharia, Kamlesh J Upadhyay, Ramesh Pandit, Tejas Shah, Ankit Hinsu, Pritesh Sabara, Apurvasinh Puvar, Janvi Raval, Monika Gandhi, Pinal Trivedi, Maharshi Pandya, Amit Kanani, Akanksha Verma, Nitin Savaliya, Raghawendra Kumar, Dinesh Kumar, Zuber Saiyed, Pooja P Doshi, Chaitanya Joshi, Madhvi Joshi |
| EPI_ISL_447588 | Lednicky Lab | Lednicky lab | Elbadry,M.A., Subramaniam,K., Waltzek,T.B., Gibson,J.C., Stephenson,C.J., Alam,M.M., Morris,J.G. Jr. and Lednicky,J.A. |
| EPI_ISL_447589 | University of Florida, Lednicky Lab | University of Florida, Lednicky Lab | Elbadry,M.A., Subramaniam,K., Waltzek,T.B., Gibson,J.C., Stephenson,C.J., Alam,M.M., Morris,J.G. Jr. and Lednicky,J.A. |
| EPI_ISL_447903 | University of Florida | University of Florida | Elbadry,M.A., Subramaniam,K., Waltzek,T.B., Lauzardo,M., Gibson,J.C., Stephenson,C.J., Alam,M.M., Morris,J.G. Jr. and Lednicky,J.A. |
| EPI_ISL_447905 | University of Florida | University of Florida | Elbadry,M.A., Subramaniam,K., Waltzek,T.B., Gibson,J.C., Stephenson,C.J., Alam,M.M., Morris,J.G. Jr. and Lednicky,J.A. |
| EPI_ISL_448389, EPI_ISL_448390, EPI_ISL_448391, EPI_ISL_448392, EPI_ISL_448393, EPI_ISL_448394, EPI_ISL_448395, EPI_ISL_448396, EPI_ISL_448397, EPI_ISL_448398, EPI_ISL_448399, EPI_ISL_448400, EPI_ISL_448401, EPI_ISL_448402, EPI_ISL_448403, EPI_ISL_448404, EPI_ISL_448405, EPI_ISL_448406, EPI_ISL_448407, EPI_ISL_448408, EPI_ISL_448409, EPI_ISL_448410, EPI_ISL_448411, EPI_ISL_448412, EPI_ISL_448413, EPI_ISL_448414, EPI_ISL_448415, EPI_ISL_448416, EPI_ISL_448417, EPI_ISL_448418, EPI_ISL_448419, EPI_ISL_448420, EPI_ISL_448421, EPI_ISL_448422, EPI_ISL_448423, EPI_ISL_448424, EPI_ISL_448425, EPI_ISL_448426, EPI_ISL_448427, EPI_ISL_448428, EPI_ISL_448429, EPI_ISL_448430, EPI_ISL_448431, EPI_ISL_448432, EPI_ISL_448433, EPI_ISL_448434, EPI_ISL_448435, EPI_ISL_448436, EPI_ISL_448437, EPI_ISL_448438, EPI_ISL_448439, EPI_ISL_448440, EPI_ISL_448441, EPI_ISL_448442, EPI_ISL_448443, EPI_ISL_448444, EPI_ISL_448445, EPI_ISL_448446, EPI_ISL_448447, EPI_ISL_448448 | | | |
| see above | Queens Medical Centre, Clinical Microbiology Department / DeepSeq Nottingham | COVID-19 Genomics UK (COG-UK) Consortium | Gemma Clark, Wendy Smith, Manjinder Khakh, Hannah Howson-Wells, Jonathan Ball, Patrick McClure, Joseph Chappell, Theocharis Tsoleridis, Nadine Holmes, Matthew Carlisle, Christopher Moore, Fei Sang, Johnny Debebe, Victoria Wright, Matthew Loose |
| EPI_ISL_449801, EPI_ISL_449802, EPI_ISL_449803, EPI_ISL_449804, EPI_ISL_449805, EPI_ISL_449806, EPI_ISL_449807, EPI_ISL_449808, EPI_ISL_449809, EPI_ISL_449810, EPI_ISL_449811, EPI_ISL_449812, EPI_ISL_449813, EPI_ISL_449815, EPI_ISL_449816, EPI_ISL_449817, EPI_ISL_449818, EPI_ISL_449819, EPI_ISL_449820, EPI_ISL_449821, EPI_ISL_449822, EPI_ISL_449823, EPI_ISL_449824, EPI_ISL_449825, EPI_ISL_449826, EPI_ISL_449827, EPI_ISL_449828, EPI_ISL_449829, EPI_ISL_449830, EPI_ISL_449831, EPI_ISL_449832, EPI_ISL_449833, EPI_ISL_449834, EPI_ISL_449835, EPI_ISL_449836, EPI_ISL_449837, EPI_ISL_449838 | | | |
| see above | Utah Public Health Laboratory | Utah Public Health Laboratory | Erin Young, Kelly Oakeson |
| EPI_ISL_450321 | NIV Pune | CSIR-Centre for Cellular and Molecular Biology | Dr V A Potdar, Dr ML Choudhary,Dr Priya Abraham,V. Vipat, S. Jadhav, U. Saha, H. Kengle, A. Awhale, A. Jagtap, A. Gondhalikar, V Malik, N Srivastava, S. Digraskar, P. Malsane, S. Hundekar, K. Patel, Yogesh Balakartik, M. Kakade, S. Jadhav, R. Gunjikar, V. Awtade, S. Bhorekar, P Shinde, S. Salve, B. Minhas S. Bharadwaj, H Kaushal Y. Gurav, S. Tomar,Payel Mukherjee, Sofia Banu, Priya Singh, Dhiviya Vedagiri, Divya Gupta, Vishal Sah, Santosh Kumar Kuncha, Krishnan Harinivas Harshan, Archana Bharadwaj Siva, Karthik Bharadwaj Tallapaka, Shagufta Khan, Lamuk Zaveri, Namami Gaur, Sakshi Shambhavi, Tulasi Nagabandi, Purushotham Vodnala,G. Aditya Kumar, Koushick Sivakumar, Pooja Ramesh Gupta, Rajan Kumar Jha, Shraddha Vijay Lahoti, Deepak Kumar, Devi Prasad Vijayashankara, Disha Nanda, Divya Das, Jotin Gogoi, Manish |
| EPI_ISL_450322 | NIV Pune | CSIR-Centre for Cellular and Molecular Biology | Dr V A Potdar, Dr ML Choudhary,Dr Priya Abraham,V. Vipat, S. Jadhav, U. Saha, H. Kengle, A. Awhale, A. Jagtap, A. Gondhalikar, V Malik, N Srivastava, S. Digraskar, P. Malsane, S. Hundekar, K. Patel, Yogesh Balakartik, M. Kakade, S. Jadhav, R. Gunjikar, V. Awtade, S. Bhorekar, P Shinde, S. Salve, B. Minhas S. Bharadwaj, H Kaushal Y. Gurav, S. Tomar,Sofia Banu, Payel Mukherjee, Priya Singh, Dhiviya Vedagiri, Divya Gupta, Vishal Sah, Santosh Kumar Kuncha, Krishnan Harinivas Harshan, Archana Bharadwaj Siva, Karthik Bharadwaj Tallapaka, Shagufta Khan, Lamuk Zaveri, Namami Gaur, Sakshi Shambhavi, Tulasi Nagabandi, Purushotham Vodnala, Disha Nanda, Divya Das, Jotin Gogoi, Manish Bhattacharjee, Ravi Prasad Mukku, Renu Sudhakar, Somesh Gorde, Gangumala Srinivas Reddy, Sujoy Deb, Swati Bayyana, Zeba Rizvi, Rakesh K Mishra |
| EPI_ISL_450323 | NIV Pune | CSIR-Centre for Cellular and Molecular Biology | Dr V A Potdar, Dr ML Choudhary,Dr Priya Abraham,V. Vipat, S. Jadhav, U. Saha, H. Kengle, A. Awhale, A. Jagtap, A. Gondhalikar, V Malik, N Srivastava, S. Digraskar, P. Malsane, S. Hundekar, K. Patel, Yogesh Balakartik, M. Kakade, S. Jadhav, R. Gunjikar, V. Awtade, S. Bhorekar, P Shinde, S. Salve, B. Minhas S. Bharadwaj, H Kaushal Y. Gurav, S. Tomar,Payel Mukherjee, Sofia Banu, Priya Singh, Dhiviya Vedagiri, Divya Gupta, Vishal Sah, Santosh Kumar Kuncha, Krishnan Harinivas Harshan, Archana Bharadwaj Siva, Karthik Bharadwaj Tallapaka, Shagufta Khan, Lamuk Zaveri, Namami Gaur, Sakshi Shambhavi, Tulasi Nagabandi, Purushotham Vodnala,G. Aditya Kumar, Koushick Sivakumar, Pooja Ramesh Gupta, Rajan Kumar Jha, Shraddha Vijay Lahoti, Deepak Kumar, Devi Prasad Vijayashankara, Disha Nanda, Divya Das, Jotin Gogoi, Manish |
| EPI_ISL_450324 | NIV Pune | CSIR-Centre for Cellular and Molecular Biology | Dr V A Potdar, Dr ML Choudhary,Dr Priya Abraham,V. Vipat, S. Jadhav, U. Saha, H. Kengle, A. Awhale, A. Jagtap, A. Gondhalikar, V Malik, N Srivastava, S. Digraskar, P. Malsane, S. Hundekar, K. Patel, Yogesh Balakartik, M. Kakade, S. Jadhav, R. Gunjikar, V. Awtade, S. Bhorekar, P Shinde, S. Salve, B. Minhas S. Bharadwaj, H Kaushal Y. Gurav, S. Tomar,Sofia Banu, Payel Mukherjee, Priya Singh, Dhiviya Vedagiri, Divya Gupta, Vishal Sah, Santosh Kumar Kuncha, Krishnan Harinivas Harshan, Archana Bharadwaj Siva, Karthik Bharadwaj Tallapaka, Shagufta Khan, Lamuk Zaveri, Namami Gaur, Sakshi Shambhavi, Tulasi Nagabandi, Purushotham Vodnala, Disha Nanda, Divya Das, Jotin Gogoi, Manish Bhattacharjee, Ravi Prasad Mukku, Renu Sudhakar, Somesh Gorde, Gangumala Srinivas Reddy, Sujoy Deb, Swati Bayyana, Zeba Rizvi, Rakesh K Mishra |
| EPI_ISL_450325 | NIV Pune | CSIR-Centre for Cellular and Molecular Biology | Dr V A Potdar, Dr ML Choudhary,Dr Priya Abraham,V. Vipat, S. Jadhav, U. Saha, H. Kengle, A. Awhale, A. Jagtap, A. Gondhalikar, V Malik, N Srivastava, S. Digraskar, P. Malsane, S. Hundekar, K. Patel, Yogesh Balakartik, M. Kakade, S. Jadhav, R. Gunjikar, V. Awtade, S. Bhorekar, P Shinde, S. Salve, B. Minhas S. Bharadwaj, H Kaushal Y. Gurav, S. Tomar,Payel Mukherjee, Sofia Banu, Priya Singh, Dhiviya Vedagiri, Divya Gupta, Vishal Sah, Santosh Kumar Kuncha, Krishnan Harinivas Harshan, Archana Bharadwaj Siva, Karthik Bharadwaj Tallapaka, Shagufta Khan, Lamuk Zaveri, Namami Gaur, Sakshi Shambhavi, Tulasi Nagabandi, Purushotham Vodnala,G. Aditya Kumar, Koushick Sivakumar, Pooja Ramesh Gupta, Rajan Kumar Jha, Shraddha Vijay Lahoti, Deepak Kumar, Devi Prasad Vijayashankara, Disha Nanda, Divya Das, Jotin Gogoi, Manish |
| EPI_ISL_450538, EPI_ISL_450539, EPI_ISL_450540, EPI_ISL_450541, EPI_ISL_450542, EPI_ISL_450543, EPI_ISL_450544, EPI_ISL_450545, EPI_ISL_450546, EPI_ISL_450547, EPI_ISL_450548, EPI_ISL_450549, EPI_ISL_450550, EPI_ISL_450551, EPI_ISL_450552, EPI_ISL_450553, EPI_ISL_450554, EPI_ISL_450555, EPI_ISL_450557, EPI_ISL_450559, EPI_ISL_450560, EPI_ISL_450561, EPI_ISL_450562, EPI_ISL_450563, EPI_ISL_450564, EPI_ISL_450565, EPI_ISL_450566, EPI_ISL_450567, EPI_ISL_450568 | | | |
| see above | Utah Public Health Laboratory | Utah Public Health Laboratory | Erin Young, Kelly Oakeson |
| EPI_ISL_450700, EPI_ISL_450701, EPI_ISL_450702, EPI_ISL_450703, EPI_ISL_450704, EPI_ISL_450705, EPI_ISL_450706, EPI_ISL_450707, EPI_ISL_450708, EPI_ISL_450709, EPI_ISL_450710, EPI_ISL_450711, EPI_ISL_450712, EPI_ISL_450713, EPI_ISL_450714, EPI_ISL_450715, EPI_ISL_450716, EPI_ISL_450717, EPI_ISL_450718, EPI_ISL_450719, EPI_ISL_450720, EPI_ISL_450721, EPI_ISL_450722 | | | |

| | | | |
|---|---|---|---|
| see above | University of Wisconsin-Madison AIDS Vaccine Research Laboratories | University of Wisconsin-Madison AIDS Vaccine Research Laboratories | Gage Moreno, Katarina Braun, et al. AIDS Vaccine Research Laboratories |
| EPI_ISL_450723 | Ramathibodi Hospital | COVID-19 Network Investigations (CONI) Alliance | Elizabeth Batty, Wasun Chantratita, Thanat Chookajorn, Stefan Fernandez, Angkana Huang, Anthony R. Jones, Khajohn Joonsalak, Chonticha Klungtong, Theerarat Kochakarn, Namfon Kotanan, Krittikorn Kumpornsin, Wudtichai Manasatienkij, Bhakbhoom Panthan, Ekawat Pasomsub, Insee Sensorn, Arporn Wangwiwatsin |
| EPI_ISL_450838 | Utah Public Health Laboratory | Utah Public Health Laboratory | Erin Young, Kelly Oakeson |
| EPI_ISL_451160 | Government Medical College, Vadodara | Gujarat Biotechnology Research Centre | Dinesh Kumar, Zuber Saiyed, Komal Patel, Labdhi Pandya, Snehal Bagatharia, Ramesh Pandit, Tejas Shah, Ankit Hinsu, Pritesh Sabara, Apurvasinh Puvar, Janvi Raval, Zarna Patel, Monika Gandhi, Pinal Trivedi, Maharshi Pandya, Manish Pattani, Tanuja Javadekar , Amit Kanani, Nidhi Patel, Nitin Savaliya, Raghawendra Kumar, Bhavesh Modi, Gaurishankar Shrimali, R D Dixit, A M Kadri, Pooja P Doshi, Chaitanya Joshi, Madhvi Joshi |
| EPI_ISL_451307 | Molecular Virology Unit, Fondazione IRCCS Policlinico San Matteo , Pavia | Laboratory of Virology, INMI Lazzaro Spallanzani IRCCS | Fausto Baldanti, Antonio Piralla, Antonino Di Caro, Cesare E.M. Gruber, Martina Rueca, Barbara Bartolini, Maria R. Capobianchi |
| EPI_ISL_451309 | Molecular Virology Unit, Fondazione IRCCS Policlinico San Matteo , Pavia | Laboratory of Virology, INMI Lazzaro Spallanzani IRCCS | Fausto Baldanti, Antonio Piralla, Cesare E.M. Gruber, Maria R. Capobianchi, Antonino Di Caro, Martina Rueca, Barbara Bartolini |
| EPI_ISL_451963, EPI_ISL_451964, EPI_ISL_451965, EPI_ISL_451966, EPI_ISL_451967, EPI_ISL_451968, EPI_ISL_451969, EPI_ISL_451970 | Federal Budget Institution of Science, State Research Center for Applied Microbiology & Biotechnology | Federal Budget Institution of Science, State Research Center for Applied Microbiology & Biotechnology | Dyatlov I, Shemyakin I, Khramov M, Bogun A, Kislichkina A, Frolov V, Shishkina L, Sizova A, Chekan L, Blagodatskikh S, Podkopaev Y, Kosilova I, Koroleva-Ushakova A, Tyurin E, Galkina E, Slukina N, Shaikhutdinova R, Kalmantayev T, Kalmantayeva O, Fursova N, Silkina M, Gorbatov A, Titareva G, Firstova V, Makarova M, Gapelchenkova T, Solovieva A, Slukin P, Dentovskaya S, Detushev K, Vagayskaya A, Kartsev N, Detusheva E, Zeninskaya N, Ivanov S, Kartseva A, Platonov M, Hlyntseva A, Khomyakov A, Chernysh S, Krasilnikova E, Ryabko A, Solomentsev V, Teymurazov M, Bakhteeva I, Borzilov A, Skryabin Y, Kanashenko M, Abaimova A, Kolchanova A, Novikova T, Goncharova J, Timofeev V, Kuzina E, Fursov M, Zhumakaev R, Marin M, Denisenko E, Trunyakova A, Kuzin V |
| EPI_ISL_452153, EPI_ISL_452154, EPI_ISL_452155, EPI_ISL_452156, EPI_ISL_452157, EPI_ISL_452158, EPI_ISL_452159, EPI_ISL_452160, EPI_ISL_452161, EPI_ISL_452162, EPI_ISL_452163, EPI_ISL_452164, EPI_ISL_452165, EPI_ISL_452166, EPI_ISL_452167, EPI_ISL_452168, EPI_ISL_452169, EPI_ISL_452170, EPI_ISL_452171, EPI_ISL_452172, EPI_ISL_452173, EPI_ISL_452174, EPI_ISL_452175, EPI_ISL_452176, EPI_ISL_452177 | | | |
| see above | Utah Public Health Laboratory | Utah Public Health Laboratory | Erin Young, Kelly Oakeson |
| EPI_ISL_452242 | Wernstedt Medical AB | The Public Health Agency of Sweden | Eva Sandberg, Anna-Malin Linde, Maria Lind Karlberg, Oskar Karlsson Lindsjo, Olov Svartstrom, Anna Risberg, Theresa Enkirch, Mia Brytting, Karin Tegmark-Wisell |
| EPI_ISL_453614, EPI_ISL_453615, EPI_ISL_453616, EPI_ISL_453617, EPI_ISL_453618, EPI_ISL_453619, EPI_ISL_453620, EPI_ISL_453621, EPI_ISL_453622, EPI_ISL_453623, EPI_ISL_453624, EPI_ISL_453625, EPI_ISL_453626, EPI_ISL_453627, EPI_ISL_453628, EPI_ISL_453629, EPI_ISL_453630, EPI_ISL_453631, EPI_ISL_453632, EPI_ISL_453633, EPI_ISL_453634, EPI_ISL_453635, EPI_ISL_453636, EPI_ISL_453637, EPI_ISL_453638, EPI_ISL_453639, EPI_ISL_453640, EPI_ISL_453641, EPI_ISL_453642, EPI_ISL_453643, EPI_ISL_453644, EPI_ISL_453645, EPI_ISL_453646, EPI_ISL_453647, EPI_ISL_453648, EPI_ISL_453649, EPI_ISL_453650, EPI_ISL_453651, EPI_ISL_453652, EPI_ISL_453653, EPI_ISL_453654, EPI_ISL_453655, EPI_ISL_453656, EPI_ISL_453657, EPI_ISL_453658, EPI_ISL_453659, EPI_ISL_453660, EPI_ISL_453661, EPI_ISL_453662, EPI_ISL_453663, EPI_ISL_453664 | | | |
| see above | Queens Medical Centre, Clinical Microbiology Department / DeepSeq Nottingham | COVID-19 Genomics UK (COG-UK) Consortium | Gemma Clark, Wendy Smith, Manjinder Khakh, Hannah Howson-Wells, Jonathan Ball, Patrick McClure, Joseph Chappell, Theocharis Tsoleridis, Nadine Holmes, Matthew Carlisle, Christopher Moore, Fei Sang, Johnny Debebe, Victoria Wright, Matthew Loose |
| EPI_ISL_454574 | nstitute for Public Health | Laboratory for advanced genomics | Filip Roki, Lovro Trgovec-Greif, Neven Sui, Tomislav Rukavina, Igor Jurak, Oliver Vugrek |
| EPI_ISL_454605, EPI_ISL_454606 | Institute for Public Health | Laboratory for advanced genomics | Filip Roki, Lovro Trgovec-Greif, Neven Sui, Tomislav Rukavina, Igor Jurak, Oliver Vugrek |
| EPI_ISL_454707, EPI_ISL_454708, EPI_ISL_454709, EPI_ISL_454710, EPI_ISL_454711, EPI_ISL_454712, EPI_ISL_454713, EPI_ISL_454714, EPI_ISL_454715, EPI_ISL_454716, EPI_ISL_454717, EPI_ISL_454718, EPI_ISL_454719, EPI_ISL_454720, EPI_ISL_454721, EPI_ISL_454722, EPI_ISL_454723, EPI_ISL_454724, EPI_ISL_454725, EPI_ISL_454726, EPI_ISL_454727, EPI_ISL_454728, EPI_ISL_454729, EPI_ISL_454730, EPI_ISL_454731 | | | |
| see above | Utah Public Health Laboratory | Utah Public Health Laboratory | Erin Young, Kelly Oakeson |
| EPI_ISL_454732 | Russian State Collection of Viruses | Pathogenic Microorganisms Variability Laboratory | Denis Protsenko, Alexey Shchetinin, Maria Nikiforova, Elena Shidlovskaya, Nadezhda Kuznetsova, Vladimir Gushchin, Inna Dolzhikova, Daria Grousova, Andrey Botikov, Denis Logunov, Alexander Gintsburg, Alexey Mazus |
| EPI_ISL_455106 | Surbrunns VC | The Public Health Agency of Sweden | Erik Embring, Anna-Malin Linde, Maria Lind Karlberg, Oskar Karlsson Lindsjo, Olov Svartstrom, Anna Risberg, Theresa Enkirch, Mia Brytting, Karin Tegmark-Wisell |
| EPI_ISL_455435 | Instituto de Diagnostico y Referencia Epidemiologicos (INDRE) | Instituto de Diagnostico y Referencia Epidemiologicos (INDRE) | Garces-Ayala Fabiola. Taboada Ramírez Blanca. Ramirez-Gonzalez Ernesto, Araiza-Rodriguez Adnan , Mendieta-Condado Edgar, Rodriguez-Maldonado Abril, Wong-Arambula Claudia, Barrera-Badillo Gisela, Hernandez-Rivas Lucia, Lopez-Martinez Irma |
| EPI_ISL_455436 | Instituto de Diagnostico y Referencia Epidemiologicos (INDRE) | Instituto de Diagnostico y Referencia Epidemiologicos (INDRE) | Garces-Ayala Fabiola. Taboada Ramírez Blanca. Ramirez-Gonzalez Ernesto, Araiza-Rodriguez Adnan, Mendieta-Condado Edgar, Rodriguez-Maldonado Abril, Wong-Arambula Claudia, Barrera-Badillo Gisela, Hernandez-Rivas Lucia, Lopez-Martinez Irma. |
| EPI_ISL_455566, EPI_ISL_455567 | Institute for Public Health | Laboratory for advanced genomics | Filip Roki, Lovro Trgovec-Greif, Neven Sui, Tomislav Rukavina, Igor Jurak, Oliver Vugrek |
| EPI_ISL_455629, EPI_ISL_455630, EPI_ISL_455631, EPI_ISL_455632, EPI_ISL_455633, EPI_ISL_455634, EPI_ISL_455635, EPI_ISL_455636, EPI_ISL_455637, EPI_ISL_455638, EPI_ISL_455639 | | | |
| see above | KRISP, KZN Research Innovation and Sequencing Platform | KRISP, KZN Research Innovation and Sequencing Platform | Giandhari J, Pillay S, Lessells R, Chimukangara B, Deforche K, Tegally H, Wilkinson E, de Oliveira T |
| EPI_ISL_455683, EPI_ISL_455684, EPI_ISL_455685, EPI_ISL_455686, EPI_ISL_455687, EPI_ISL_455688, EPI_ISL_455689, EPI_ISL_455690, EPI_ISL_455691, EPI_ISL_455692, EPI_ISL_455693 | | | |
| see above | unknown | Department of Microbiology | Gao,Q., Bao,L., Mao,H., Wang,L., Xu,K., Yang,M., Li,Y., Zhu,L., Wang,N., Lv,Z., Gao,H., Ge,X., Kan,B., Hu,Y., Liu,J., Cai,F., Jiang,D., Yin,Y., Qin,C., Li,J., Gong,X., Lou,X., Shi,W., Wu,D., Zhang,H., Deng,W., Lu,J., Li,C., Wang,X., Yin,W., Zhang,Y., Sun,Y. |
| EPI_ISL_455725, EPI_ISL_455743 | Servicio de Microbiologia. Hospital Clinico Universitario de Valencia | Sequencing and Bioinformatics Service and Molecular Epidemiology Research Group. FISABIO-Public Health, and SeqCOVID-Spain Consortium | Eliseo Albert, Maria Alma Bracho, Griselda De Marco, Lidia Ruiz Roldan, Neris Garcia-Gonzalez, Inma Galán Vendrell, Sandra Carbo, Loreto Ferrús Abad, Paula Ruiz-Hueso, Mariana Reyes-Prieto, Vicente Soriano Chirona, Ivan Ansari, Llúcia Martínez-Priego, Giuseppe 'Auria, David Navarro, Fernando Gonzalez-Candelas |
| EPI_ISL_455908, EPI_ISL_455909, EPI_ISL_455910, EPI_ISL_455911, EPI_ISL_455912, EPI_ISL_455913, EPI_ISL_455914, EPI_ISL_455915, EPI_ISL_455916, EPI_ISL_455917, EPI_ISL_455918, EPI_ISL_455919, EPI_ISL_455920, EPI_ISL_455921, EPI_ISL_455922, EPI_ISL_455923, EPI_ISL_455924, EPI_ISL_455925, EPI_ISL_455926, EPI_ISL_455927, EPI_ISL_455928, EPI_ISL_455929, EPI_ISL_455930, EPI_ISL_455931, EPI_ISL_455932, EPI_ISL_455933, EPI_ISL_455934, EPI_ISL_455935, EPI_ISL_455936, EPI_ISL_455937, EPI_ISL_455938, EPI_ISL_455939, EPI_ISL_455940, EPI_ISL_455941, EPI_ISL_455942, EPI_ISL_455943, EPI_ISL_455944, EPI_ISL_455945, EPI_ISL_455946, EPI_ISL_455947, EPI_ISL_455948 | | | |
| see above | Ramathibodi Hospital | COVID-19 Network Investigations (CONI) Alliance | Elizabeth Batty, Wasun Chantratita, Thanat Chookajorn, Stefan Fernandez, Angkana Huang, Anthony R. Jones, Khajohn Joonsalak, Chonticha Klungtong, Theerarat Kochakarn, Namfon Kotanan, Krittikorn Kumpornsin, Wudtichai Manasatienkij, Bhakbhoom Panthan, Ekawat Pasomsub, Kingkan Rakmanee, Insee Sensorn, Janjira Thaipadungpanit, Arporn Wangwiwatsin,Treewat Watthanachockchai |
| EPI_ISL_457573, EPI_ISL_457574, EPI_ISL_457575, EPI_ISL_457576, EPI_ISL_457577, EPI_ISL_457578, EPI_ISL_457579, EPI_ISL_457580 | Queens Medical Centre, Clinical Microbiology Department / DeepSeq Nottingham | COVID-19 Genomics UK (COG-UK) Consortium | Gemma Clark, Wendy Smith, Manjinder Khakh, Hannah Howson-Wells, Jonathan Ball, Patrick McClure, Joseph Chappell, Theocharis Tsoleridis, Nadine Holmes, Matthew Carlisle, Christopher Moore, Fei Sang, Johnny Debebe, Victoria Wright, Matthew Loose |
| EPI_ISL_457702 | Oman-NIC | Microbiology laboratory- Sultan Qaboos University Hospital | Fahad Zadjali, Samira Al-Maruqi, Amina Al Jardani, Khulood Al-Mammary, Hanan Al-kindi, Fatma BaAlawi, Hamida AL Barwani, Zeyana AL-Dahmani, Intisar Al-Shukri, Aisha Al-Busaidi, Aisha Al-Amri, Ahlam Al-Amri, Mohammed Al-Tobi, Samiha Al Kharusi, Abdulla Balkhair |
| EPI_ISL_457703 | Oman-NIC | Department of Microbiology and Immunology- SQUH | Fahad Zadjali, Samira Al-Maruqi, Amina Al Jardani, Khulood Al-Mammary, Hanan Al-kindi, Fatma BaAlawi, Hamida AL Barwani, Zeyana AL-Dahmani, Intisar Al-Shukri, Aisha Al-Busaidi, Aisha Al-Amri, Ahlam Al-Amri, Mohammed Al-Tobi, Samiha Al Kharusi, Abdulla Balkhair |
| EPI_ISL_457705 | OMAN-NIC | Department of Microbiology and Immunology- SQUH | Fahad Zadjali, Samira Al-Maruqi, Amina Al Jardani, Khulood Al-Mammary, Hanan Al-kindi, Fatma BaAlawi, Hamida AL Barwani, Zeyana AL-Dahmani, Intisar Al-Shukri, Aisha Al-Busaidi, Aisha Al-Amri, Ahlam Al-Amri, Mohammed Al-Tobi, Samiha Al Kharusi, Abdulla Balkhair |
| EPI_ISL_457707 | Oman-NIC | Department of Microbiology and Immunology- SQUH | Fahad Zadjali, Samira Al-Maruqi, Amina Al Jardani, Khulood Al-Mammary, Hanan Al-kindi, Fatma BaAlawi, Hamida AL Barwani, Zeyana AL-Dahmani, Intisar Al-Shukri, Aisha Al-Busaidi, Aisha Al-Amri, Ahlam Al-Amri, Mohammed Al-Tobi, Samiha Al Kharusi, Abdulla Balkhair |
| EPI_ISL_457981, EPI_ISL_457982, EPI_ISL_457983, EPI_ISL_457984 | Oman-NIC | Department of Microbiology and Immunology-SQUH | Fahad Zadjali, Samira Al-Maruqi, Amina Al Jardani, Khulood Al-Mammary, Hanan Al-kindi, Fatma BaAlawi, Hamida AL Barwani, Zeyana AL-Dahmani, Intisar Al-Shukri, Aisha Al-Busaidi, Aisha Al-Amri, Ahlam Al-Amri, Mohammed Al-Tobi, Samiha Al Kharusi, Abdulla Balkhair |

| | | | |
|---|---|---|---|
| EPI_ISL_458023, EPI_ISL_458024, EPI_ISL_458025, EPI_ISL_458026, EPI_ISL_458027, EPI_ISL_458028 | Hospital for Tropical Diseases | COVID-19 Network Investigations (CONI) Alliance | Elizabeth Batty, Nantarat Chantawat, Wasun Chantratita, Thanat Chookajorn, Stefan Fernandez, Angkana Huang, Weena Janwitthayanan, Akanitt Jittmittraphap, Anthony R. Jones, Khajohn Joonsalak, Chonticha Klungtong, Theerarat Kochakarn, Namfon Kotanan, Krittikorn Kumpornsin, Pornsawan Leaungwutiwong, Wudtichai Manasatienkij, Bhakbhoom Panthan, Ekawat Pasomsub, Kingkan Rakmanee, Insee Sensorn, Janjira Thaipadungpanit, Arporn Wangwiwatsin,Treewat Watthanachockchai |
| EPI_ISL_458072, EPI_ISL_458073, EPI_ISL_458074, EPI_ISL_458075, EPI_ISL_458076, EPI_ISL_458077 | CSIR-Centre for Cellular and Molecular Biology | CSIR-Centre for Cellular and Molecular Biology | Dhiviya Vedagiri, Divya Gupta, Vishal Sah, Payel Mukherjee, Sofia Banu, Priya Singh, Santosh Kumar Kuncha, Archana Bharadwaj Siva, Karthik Bharadwaj Tallapaka, Shagufta Khan, Lamuk Zaveri, Namami Gaur, Sakshi Shambhavi, Tulasi Nagabandi, Purushotham Vodnala, Rakesh K Mishra, Divya Tej Sowpati, Krishnan Harinivas Harshan |
| EPI_ISL_458086 | B.J. Medical College and Civil hospital | Gujarat Biotechnology Research Centre | Dhaval Vaghela, Ramesh Patel, Pranay Shah, Kamlesh J Upadhyay, Ramesh Pandit, Tejas Shah, Ankit Hinsu, Pritesh Sabara, Apurvasinh Puvar, Janvi Raval, Zarna Patel, Monika Gandhi, Pinal Trivedi, Maharshi Pandya, Amit Kanani, Nidhi Patel, Nitin Savaliya, Raghawendra Kumar, Dinesh Kumar, Zuber Saiyed, Komal Patel, Labdhi Pandya, Snehal Bagatharia, Neha Rajpara, Bhavesh Modi, Gaurishankar Shrimali, R D Dixit, A M Kadri, Umang Mishra, Chaitanya Joshi, Madhvi Joshi |
| EPI_ISL_458096 | B.J. Medical College and Civil hospital | Gujarat Biotechnology Research Centre | Dinesh Kumar, Zuber Saiyed, Komal Patel, Labdhi Pandya, Snehal Bagatharia, Dhaval Vaghela, Ramesh Patel, Pranay Shah, Kamlesh J Upadhyay, Ramesh Pandit, Tejas Shah, Ankit Hinsu, Pritesh Sabara, Apurvasinh Puvar, Janvi Raval, Zarna Patel, Monika Gandhi, Pinal Trivedi, Maharshi Pandya, Amit Kanani, Nidhi Patel, Nitin Savaliya, Raghawendra Kumar, Akanksha Verma, Bhavesh Modi, Gaurishankar Shrimali, R D Dixit, A M Kadri, Umang Mishra, Chaitanya Joshi, Madhvi Joshi |
| EPI_ISL_458101 | B.J. Medical College and Civil hospital | Gujarat Biotechnology Research Centre | Dhaval Vaghela, Ramesh Patel, Pranay Shah, Kamlesh J Upadhyay, Ramesh Pandit, Tejas Shah, Ankit Hinsu, Pritesh Sabara, Apurvasinh Puvar, Janvi Raval, Zarna Patel, Monika Gandhi, Pinal Trivedi, Maharshi Pandya, Amit Kanani, Nidhi Patel, Nitin Savaliya, Raghawendra Kumar, Dinesh Kumar, Zuber Saiyed, Komal Patel, Labdhi Pandya, Snehal Bagatharia, Fenil Patel, Bhavesh Modi, Gaurishankar Shrimali, R D Dixit, A M Kadri, Umang Mishra, Chaitanya Joshi, Madhvi Joshi |
| EPI_ISL_458116, EPI_ISL_458117, EPI_ISL_458118, EPI_ISL_458119, EPI_ISL_458120, EPI_ISL_458121, EPI_ISL_458122, EPI_ISL_458123, EPI_ISL_458124, EPI_ISL_458125, EPI_ISL_458126, EPI_ISL_458127, EPI_ISL_458128 | | | |
| see above | Oman National Influenza Centre | Department of Microbiology and Immunology-SQUH | Fahad Zadjali, Samira Al-Maruqi, Amina Al Jardani, Khulood Al-Mammary, Hanan Al-kindi, Fatma BaAlawi, Hamida AL Barwani, Zeyana AL-Dahmani, Intisar Al-Shukri, Aisha Al-Busaidi, Aisha Al-Amri, Ahlam Al-Amri, Mohammed Al-Tobi, Samiha Al Kharusi, Abdulla Balkhair |
| EPI_ISL_458137 | Oman National Influenza Centre | Department of Microbiology and Immunology | Fahad Zadjali, Samira Al-Maruqi, Amina Al Jardani, Khulood Al-Mammary, Hanan Al-kindi, Fatma BaAlawi, Hamida AL Barwani, Zeyana AL-Dahmani, Intisar Al-Shukri, Aisha Al-Busaidi, Aisha Al-Amri, Ahlam Al-Amri, Mohammed Al-Tobi, Samiha Al Kharusi, Abdulla Balkhair |
| EPI_ISL_459856, EPI_ISL_459857, EPI_ISL_459858, EPI_ISL_459859, EPI_ISL_459860, EPI_ISL_459861, EPI_ISL_459862, EPI_ISL_459863, EPI_ISL_459864 | Center for Genome Regulation (CRG) | Center for Mathematical Modeling and Center for Genome Regulation. Santiago, Chile | Gaete A, Travisany D, Palma R, Urra C, Varas M, Allende ML, Maass A, González M. |
| EPI_ISL_460081 | Molecular Virology Unit, Fondazione IRCCS Policlinico San Matteo , Pavia | Laboratory of Virology, INMI Lazzaro Spallanzani IRCCS | Fausto Baldanti, Martina Rueca, Antonio Piralla, Antonino Di Caro, Maria R. Capobianchi, Cesare E.M. Gruber, Barbara Bartolini |
| EPI_ISL_460084 | Molecular Virology Unit, Fondazione IRCCS Policlinico San Matteo , Pavia | Laboratory of Virology, INMI Lazzaro Spallanzani IRCCS | Fausto Baldanti, Antonio Piralla, Martina Rueca, Barbara Bartolini, Maria R. Capobianchi, Cesare E.M. Gruber, Antonino Di Caro |
| EPI_ISL_461478 | Government Medical College, Vadodara | Gujarat Biotechnology Research Centre | Fenil Patel, Nidhi Patel, Nitin Savaliya, Raghawendra Kumar, Dinesh Kumar, Zuber Saiyed, Komal Patel, Labdhi Pandya, Snehal Bagatharia, Tanuja Javadekar , R N Daveshwar, Tejas Shah, Ankit Hinsu, Pritesh Sabara, Apurvasinh Puvar, Janvi Raval, Zarna Patel, Monika Gandhi, Pinal Trivedi, Maharshi Pandya, R D Dixit, A M Kadri, Harsh Bakshi, Chaitanya Joshi, Madhvi Joshi, |
| EPI_ISL_461483 | B.J. Medical College and Civil hospital | Gujarat Biotechnology Research Centre | Dipeshwari Shewale, Komal Patel, Labdhi Pandya, Snehal Bagatharia, Pranay Shah, Kamlesh J Upadhyay, Tejas Shah, Ankit Hinsu, Pritesh Sabara, Apurvasinh Puvar, Janvi Raval, Zarna Patel, Monika Gandhi, Pinal Trivedi, Maharshi Pandya, Nidhi Patel, Nitin Savaliya, Raghawendra Kumar, Dinesh Kumar, Zuber Saiyed, R D Dixit, A M Kadri, Harsh Bakshi, Chaitanya Joshi, Madhvi Joshi, |
| EPI_ISL_461491 | B.J. Medical College and Civil hospital | Gujarat Biotechnology Research Centre | Fenil Patel, Apurvasinh Puvar, Janvi Raval, Zarna Patel, Monika Gandhi, Pinal Trivedi, Maharshi Pandya, Nidhi Patel, Nitin Savaliya, Raghawendra Kumar, Dinesh Kumar, Zuber Saiyed, Komal Patel, Labdhi Pandya, Snehal Bagatharia, Pranay Shah, Kamlesh J Upadhyay, Tejas Shah, Ankit Hinsu, Pritesh Sabara, R D Dixit, A M Kadri, Harsh Bakshi, Chaitanya Joshi, Madhvi Joshi, |
| EPI_ISL_461496 | B.J. Medical College and Civil hospital | Gujarat Biotechnology Research Centre | Dipeshwari Shewale, Maharshi Pandya, Nidhi Patel, Nitin Savaliya, Raghawendra Kumar, Dinesh Kumar, Zuber Saiyed, Komal Patel, Labdhi Pandya, Snehal Bagatharia, Pranay Shah, Kamlesh J Upadhyay, Tejas Shah, Ankit Hinsu, Pritesh Sabara, Apurvasinh Puvar, Janvi Raval, Zarna Patel, Monika Gandhi, Pinal Trivedi, R D Dixit, A M Kadri, Harsh Bakshi, Chaitanya Joshi, Madhvi Joshi, |
| EPI_ISL_461894, EPI_ISL_461895, EPI_ISL_461896, EPI_ISL_461897, EPI_ISL_461898, EPI_ISL_461899, EPI_ISL_461900, EPI_ISL_461901, EPI_ISL_461902, EPI_ISL_461903, EPI_ISL_461904, EPI_ISL_461905, EPI_ISL_461906, EPI_ISL_461907, EPI_ISL_461908, EPI_ISL_461909, EPI_ISL_461910, EPI_ISL_461911, EPI_ISL_461912, EPI_ISL_461913, EPI_ISL_461914, EPI_ISL_461915, EPI_ISL_461916, EPI_ISL_461917, EPI_ISL_461918, EPI_ISL_461919, EPI_ISL_461920, EPI_ISL_461921, EPI_ISL_461922, EPI_ISL_461923, EPI_ISL_461924, EPI_ISL_461925, EPI_ISL_461926, EPI_ISL_461927, EPI_ISL_461928, EPI_ISL_461929, EPI_ISL_461930, EPI_ISL_461931, EPI_ISL_461932, EPI_ISL_461933, EPI_ISL_461934, EPI_ISL_461935, EPI_ISL_461936, EPI_ISL_461937, EPI_ISL_461938, EPI_ISL_461939, EPI_ISL_461940, EPI_ISL_461941, EPI_ISL_461942, EPI_ISL_461943, EPI_ISL_461944, EPI_ISL_461945, EPI_ISL_461946, EPI_ISL_461947, EPI_ISL_461948, EPI_ISL_461949, EPI_ISL_461950, EPI_ISL_461951, EPI_ISL_461952, EPI_ISL_461953, EPI_ISL_461954, EPI_ISL_461955, EPI_ISL_461956, EPI_ISL_461957, EPI_ISL_461958, EPI_ISL_461959, EPI_ISL_461960, EPI_ISL_461961, EPI_ISL_461962, EPI_ISL_461963, EPI_ISL_461964, EPI_ISL_461965, EPI_ISL_461966, EPI_ISL_461967, EPI_ISL_461968, EPI_ISL_461969 | | | |
| see above | Queens Medical Centre, Clinical Microbiology Department / DeepSeq Nottingham | COVID-19 Genomics UK (COG-UK) Consortium | Gemma Clark, Wendy Smith, Manjinder Khakh, Hannah Howson-Wells, Jonathan Ball, Patrick McClure, Joseph Chappell, Theocharis Tsoleridis, Nadine Holmes, Matthew Carlisle, Christopher Moore, Fei Sang, Johnny Debebe, Victoria Wright, Matthew Loose |
| EPI_ISL_462991 | Microbiology Division | Microbiology Division | Flores,H. |
| EPI_ISL_463001, EPI_ISL_463002, EPI_ISL_463003, EPI_ISL_463004, EPI_ISL_463005, EPI_ISL_463006 | unknown | Clinical virology | Fares,W., Triki,H. |
| EPI_ISL_467030 | GMERS Medical College and Hospital, Gandhinagar | Gujarat Biotechnology Research Centre | Gaurishankar Shrimali, Bhavesh Modi, Bharti Rajani, Tejas Shah, Ankit Hinsu, Pritesh Sabara, Apurvasinh Puvar, Janvi Raval, Zarna Patel, Monika Gandhi, Pinal Trivedi, Maharshi Pandya, Nidhi Patel, Nitin Savaliya, Raghawendra Kumar, Dinesh Kumar, Zuber Saiyed, Komal Patel, Labdhi Pandya, Snehal Bagatharia, Seema Bhatt, Priyanka P Vatsa, R D Dixit, A M Kadri, Harsh Bakshi, Chaitanya Joshi, Madhvi Joshi |
| EPI_ISL_467047 | B.J. Medical College and Civil hospital | Gujarat Biotechnology Research Centre | Dinesh Kumar, Zuber Saiyed, Komal Patel, Labdhi Pandya, Snehal Bagatharia, Pranay Shah, Kamlesh J Upadhyay, Nirav Mungalpara, Tejas Shah, Ankit Hinsu, Pritesh Sabara, Apurvasinh Puvar, Janvi Raval, Zarna Patel, Monika Gandhi, Pinal Trivedi, Maharshi Pandya, Nidhi Patel, Nitin Savaliya, Raghawendra Kumar, Afzal Ansari, R D Dixit, A M Kadri, Harsh Bakshi, Chaitanya Joshi, Madhvi Joshi, |
| EPI_ISL_467374 | Dinkes Samarinda | Eijkman Institute for Molecular Biology, Ministry of Research and Technology/National Agency for Research and Innovation | Edison Johar, Frilasita A Yudhaputri, Hidayat Trimarsanto, David H Muljono, Safarina G Malik, Khin Saw Myint, Amin Soebandrio |
| EPI_ISL_467375 | RSUP Prof. Dr. R. Kandou Manado | Eijkman Institute for Molecular Biology, Ministry of Research and Technology/National Agency for Research and Innovation | Edison Johar, Frilasita A Yudhaputri, Hidayat Trimarsanto, David H Muljono, Safarina G Malik, Khin Saw Myint, Amin Soebandrio |
| EPI_ISL_467376 | RSUP Fatmawati | Eijkman Institute for Molecular Biology, Ministry of Research and Technology/National Agency for Research and Innovation | Edison Johar, Frilasita A Yudhaputri, Hidayat Trimarsanto, David H Muljono, Safarina G Malik, Khin Saw Myint, Amin Soebandrio |
| EPI_ISL_467436, EPI_ISL_467438, EPI_ISL_467439 | NHLS-IALCH | KRISP, KZN Research Innovation and Sequencing Platform | Giandhari J, Pillay S, Lessells R, Chimukangara B, Mdlalose K, York D, Khan S, Tegally H, Wilkinson E, de Oliveira T |
| EPI_ISL_467447 | Molecular Diagnostics Services (MDS) | KRISP, KZN Research Innovation and Sequencing Platform | Giandhari J, Pillay S, Lessells R, Chimukangara B, Mdlalose K, York D, Khan S, Tegally H, Wilkinson E, de Oliveira T |
| EPI_ISL_467451, EPI_ISL_467452, EPI_ISL_467453, EPI_ISL_467454, EPI_ISL_467455, EPI_ISL_467456, EPI_ISL_467457, EPI_ISL_467458, EPI_ISL_467459, EPI_ISL_467460, EPI_ISL_467461, EPI_ISL_467462, EPI_ISL_467463 | | | |
| see above | AMPATH-DBN | KRISP, KZN Research Innovation and Sequencing Platform | Giandhari J, Pillay S, Lessells R, Chimukangara B, Mdlalose K, York D, Khan S, Tegally H, Wilkinson E, de Oliveira T |

| | | | |
|---|---|---|---|
| EPI_ISL_467492, EPI_ISL_467510, EPI_ISL_467517, EPI_ISL_467518, EPI_ISL_467519, EPI_ISL_467520, EPI_ISL_467521, EPI_ISL_467522, EPI_ISL_467523, EPI_ISL_467524 | NHLS-IALCH | KRISP, KZN Research Innovation and Sequencing Platform | Giandhari J, Pillay S, Lessells R, Chimukangara B, Mdlalose K, York D, Khan S, Tegally H, Wilkinson E, de Oliveira T |
| EPI_ISL_468591, EPI_ISL_468656 | Institute for Public Health | Laboratory for advanced genomics | Filip Roki, Lovro Trgovec-Greif, Neven Sui, Tomislav Rukavina, Igor Jurak, Oliver Vugrek |
| EPI_ISL_468719 | University of Florida | University of Florida | Elbadry,M.A., Subramaniam,K., Waltzek,T.B., Stephenson,C.J., Gibson,J.C., Alam,M.M., Lauzardo,M., Morris,J.G., Lednicky,J.A. |
| EPI_ISL_468722 | University of Florida | University of Florida | Elbadry,M.A., Subramaniam,K., Waltzek,T.B., Lauzardo,M., Morris,J.G., Lednicky,J.A. |
| EPI_ISL_468747, EPI_ISL_468748, EPI_ISL_468749, EPI_ISL_468750, EPI_ISL_468751 | Facultad de Medicina UC | Center for Mathematical Modeling and Center for Genome Regulation. Santiago, Chile | Gaete A, Travisany D, Palma R, Urra C, Varas M, Allende ML, Maass A, González M, Ferres M. |
| EPI_ISL_468752 | Center for Genome Regulation (CRG) | Center for Mathematical Modeling and Center for Genome Regulation. Santiago, Chile | Gaete A, Travisany D, Palma R, Urra C, Varas M, Allende ML, Maass A, González M. |
| EPI_ISL_468753, EPI_ISL_468754, EPI_ISL_468755, EPI_ISL_468756, EPI_ISL_468757, EPI_ISL_468758, EPI_ISL_468759 | Laboratorio de Biologia Molecular, Facultad de Medicina, Universidad de Atacama | Center for Mathematical Modeling and Center for Genome Regulation. Santiago, Chile | Gaete A, Travisany D, Palma R, Urra C, Varas M, Allende ML, Maass A, González M, C Echeverria |
| EPI_ISL_468760 | Center for Genome Regulation (CRG) | Center for Mathematical Modeling and Center for Genome Regulation. Santiago, Chile | Gaete A, Travisany D, Palma R, Urra C, Varas M, Allende ML, Maass A, González M. |
| EPI_ISL_469036 | Government Medical College, Vadodara | Gujarat Biotechnology Research Centre | Dinesh Kumar, Zuber Saiyed, Komal Patel, Labdhi Pandya, Snehal Bagatharia, Meenakshi Shah, Neena Doshi, Varsha Godbole, Tejas Shah, Ankit Hinsu, Pritesh Sabara, Apurvasinh Puvar, Janvi Raval, Zarna Patel, Monika Gandhi, Pinal Trivedi, Maharshi Pandya, Nidhi Patel, Nitin Savaliya, Raghawendra Kumar, Priti Pandita, R D Dixit, A M Kadri, Harsh Bakshi, Chaitanya Joshi, Madhvi Joshi |
| EPI_ISL_470541, EPI_ISL_470542, EPI_ISL_470543, EPI_ISL_470544, EPI_ISL_470545, EPI_ISL_470546, EPI_ISL_470547, EPI_ISL_470548, EPI_ISL_470549, EPI_ISL_470550, EPI_ISL_470551, EPI_ISL_470552, EPI_ISL_470553, EPI_ISL_470554, EPI_ISL_470555, EPI_ISL_470556, EPI_ISL_470557, EPI_ISL_470558, EPI_ISL_470559, EPI_ISL_470560, EPI_ISL_470561, EPI_ISL_470562, EPI_ISL_470563, EPI_ISL_470564, EPI_ISL_470565, EPI_ISL_470566, EPI_ISL_470567, EPI_ISL_470656, EPI_ISL_470657, EPI_ISL_470658, EPI_ISL_470659, EPI_ISL_470660, EPI_ISL_470661, EPI_ISL_470662, EPI_ISL_470663, EPI_ISL_470664, EPI_ISL_470665, EPI_ISL_470666, EPI_ISL_470667, EPI_ISL_470668, EPI_ISL_470669, EPI_ISL_470670, EPI_ISL_470671, EPI_ISL_470672, EPI_ISL_470673, EPI_ISL_470674, EPI_ISL_470675, EPI_ISL_470676, EPI_ISL_470677, EPI_ISL_470678, EPI_ISL_470679, EPI_ISL_470680, EPI_ISL_470681, EPI_ISL_470682, EPI_ISL_470683, EPI_ISL_470684, EPI_ISL_470685, EPI_ISL_470686, EPI_ISL_470687, EPI_ISL_470688, EPI_ISL_470689, EPI_ISL_470690, EPI_ISL_470691, EPI_ISL_470692, EPI_ISL_470693, EPI_ISL_470694, EPI_ISL_470695, EPI_ISL_470696, EPI_ISL_470697, EPI_ISL_470698, EPI_ISL_470699, EPI_ISL_470700, EPI_ISL_470701, EPI_ISL_470702, EPI_ISL_470703, EPI_ISL_470704, EPI_ISL_470705, EPI_ISL_470706, EPI_ISL_470707, EPI_ISL_470708, EPI_ISL_470709, EPI_ISL_470710, EPI_ISL_470711, EPI_ISL_470712, EPI_ISL_470713, EPI_ISL_470714, EPI_ISL_470715, EPI_ISL_470716, EPI_ISL_470717, EPI_ISL_470718 | | | |
| see above | Utah Public Health Laboratory | Utah Public Health Laboratory | Erin Young, Kelly Oakeson |
| EPI_ISL_471585, EPI_ISL_471587, EPI_ISL_471641, EPI_ISL_471642, EPI_ISL_471645, EPI_ISL_471646 | CSIR-Centre for Cellular and Molecular Biology | CSIR-Centre for Cellular and Molecular Biology | Dhiviya Vedagiri, Divya Gupta, Vishal Sah, Payel Mukherjee, Sofia Banu, Priya Singh, Santosh Kumar Kuncha, Archana Bharadwaj Siva, Karthik Bharadwaj Tallapaka, Shagufta Khan, Lamuk Zaveri, Namami Gaur, Sakshi Shambhavi, Tulasi Nagabandi, Purushotham Vodnala, Rakesh K Mishra, Divya Tej Sowpati, Krishnan Harinivas Harshan |
| EPI_ISL_472384, EPI_ISL_472385, EPI_ISL_472386, EPI_ISL_472387, EPI_ISL_472388, EPI_ISL_472389, EPI_ISL_472390, EPI_ISL_472391, EPI_ISL_472392, EPI_ISL_472393, EPI_ISL_472394, EPI_ISL_472395, EPI_ISL_472396, EPI_ISL_472397, EPI_ISL_472398, EPI_ISL_472399, EPI_ISL_472400, EPI_ISL_472401, EPI_ISL_472402, EPI_ISL_472403, EPI_ISL_472404, EPI_ISL_472405, EPI_ISL_472406, EPI_ISL_472407, EPI_ISL_472408, EPI_ISL_472409, EPI_ISL_472410, EPI_ISL_472411, EPI_ISL_472412, EPI_ISL_472413, EPI_ISL_472414, EPI_ISL_472415, EPI_ISL_472416, EPI_ISL_472417, EPI_ISL_472418, EPI_ISL_472419, EPI_ISL_472420, EPI_ISL_472421, EPI_ISL_472422, EPI_ISL_472423, EPI_ISL_472424, EPI_ISL_472425, EPI_ISL_472426, EPI_ISL_472427, EPI_ISL_472428, EPI_ISL_472429 | | | |
| see above | Queens Medical Centre, Clinical Microbiology Department / DeepSeq Nottingham | COVID-19 Genomics UK (COG-UK) Consortium | Gemma Clark, Wendy Smith, Manjinder Khakh, Vicki M Fleming, Michelle M Lister, Hannah Howson-Wells, Jonathan Ball, Patrick McClure, Joseph Chappell, Theocharis Tsoleridis, Nadine Holmes, Matthew Carlisle, Christopher Moore, Fei Sang, Johnny Debebe, Victoria Wright, Matthew Loose |
| EPI_ISL_474798, EPI_ISL_474799, EPI_ISL_474800, EPI_ISL_474801, EPI_ISL_474802, EPI_ISL_474803, EPI_ISL_474804, EPI_ISL_474805, EPI_ISL_474806, EPI_ISL_474807, EPI_ISL_474808, EPI_ISL_474809, EPI_ISL_474810, EPI_ISL_474811, EPI_ISL_474812, EPI_ISL_474813, EPI_ISL_474814, EPI_ISL_474815, EPI_ISL_474816, EPI_ISL_474817, EPI_ISL_474818, EPI_ISL_474819, EPI_ISL_474820, EPI_ISL_474821, EPI_ISL_474822, EPI_ISL_474823, EPI_ISL_474824, EPI_ISL_474825, EPI_ISL_474826, EPI_ISL_474827, EPI_ISL_474828, EPI_ISL_474829, EPI_ISL_474830, EPI_ISL_474831, EPI_ISL_474837, EPI_ISL_474838, EPI_ISL_474839, EPI_ISL_474840, EPI_ISL_474847, EPI_ISL_474848, EPI_ISL_474849, EPI_ISL_474853, EPI_ISL_474901, EPI_ISL_474902, EPI_ISL_474903, EPI_ISL_474904, EPI_ISL_474905, EPI_ISL_474906, EPI_ISL_474919, EPI_ISL_474921, EPI_ISL_474933, EPI_ISL_474940, EPI_ISL_474941, EPI_ISL_474945, EPI_ISL_474946, EPI_ISL_474947, EPI_ISL_474951, EPI_ISL_474952, EPI_ISL_474953, EPI_ISL_474954, EPI_ISL_474955, EPI_ISL_474956 | | | |
| see above | Complejo Hospitalario Universitario de Albacete | SeqCOVID-SPAIN consortium/IBV(CSIC) | Encarnacion Simarro Córdoba, Julia Lozano Serra, Lorena Robles Fonseca , Monica Parra Grandes, Caridad Sainz de Baranda Camino and SeqCOVID-SPAIN consortium |
| EPI_ISL_475038 | Department of MicroBiology, Government Medical College, Surat | Gujarat Biotechnology Research Centre | Dinesh Kumar, Zuber Saiyed, Komal Patel, Labdhi Pandya, Snehal Bagatharia, Naresh Chauhan, Summaiya Mullan, Amit gamit, Pritesh Sabara, Apurvasinh Puvar, Janvi Raval, Zarna Patel, Monika Gandhi, Pinal Trivedi, Maharshi Pandya, Nidhi Patel, Nitin Savaliya, Raghawendra Kumar, Afzal Ansari, R D Dixit, A M Kadri, Harsh Bakshi, Chaitanya Joshi, Madhvi Joshi |
| EPI_ISL_475056 | Dr. N. D. Desai Medical College & Hospital | Gujarat Biotechnology Research Centre | Dinesh Kumar, Zuber Saiyed, Komal Patel, Labdhi Pandya, Snehal Bagatharia, J G Buch, Jigar Gusani, Supreet Prabhu, Pritesh Sabara, Apurvasinh Puvar, Janvi Raval, Zarna Patel, Monika Gandhi, Pinal Trivedi, Maharshi Pandya, Nidhi Patel, Nitin Savaliya, Raghawendra Kumar, Akanksha Verma, R D Dixit, A M Kadri, Harsh Bakshi, Chaitanya Joshi, Madhvi Joshi |
| EPI_ISL_475725, EPI_ISL_475726, EPI_ISL_475727, EPI_ISL_475728, EPI_ISL_475729, EPI_ISL_475730, EPI_ISL_475731, EPI_ISL_475732, EPI_ISL_475733, EPI_ISL_475735, EPI_ISL_475736, EPI_ISL_475737, EPI_ISL_475738, EPI_ISL_475739, EPI_ISL_475740, EPI_ISL_475741, EPI_ISL_475742, EPI_ISL_475743, EPI_ISL_475744 | | | |
| see above | Utah Public Health Laboratory | Utah Public Health Laboratory | Erin Young, Kelly Oakeson |
| EPI_ISL_476068, EPI_ISL_476069, EPI_ISL_476070, EPI_ISL_476071, EPI_ISL_476072, EPI_ISL_476073, EPI_ISL_476074, EPI_ISL_476075, EPI_ISL_476076, EPI_ISL_476077 | University of Debrecen, Department of Medical Microbiology | National Laboratory of Virology, Szentágothai Research Centre | Endre Gábor Tóth, Balázs Somogyi, Brigitta Zana, Eszter Csoma, Ferenc Jakab, Gábor Kemenesi |
| EPI_ISL_476078 | University of Szeged, Institute of Clinical Microbiology | National Laboratory of Virology, Szentágothai Research Centre | Endre Gábor Tóth, Balázs Somogyi, Brigitta Zana, Terhes Gabriella, Ferenc Jakab, Gábor Kemenesi |
| EPI_ISL_476801, EPI_ISL_476802, EPI_ISL_476803, EPI_ISL_476804 | Hong Kong Department of Health | School of Public Health, The University of Hong Kong | Dominic N.C. Tsang, Daniel K.W. Chu, Leo L.M. Poon, Malik Peiris |
| EPI_ISL_476864 | GMERS Medical College and Hospital, Gandhinagar | Gujarat Biotechnology Research Centre | Dinesh Kumar, Zuber Saiyed, Komal Patel, Labdhi Pandya, Afzal Ansari, Nikha Trivedi, Seema Bhatt, Gaurishankar Shrimali, Bhavesh Modi, Bharti Rajani, Apurvasinh Puvar, Janvi Raval, Zarna Patel, Monika Gandhi, Pinal Trivedi, Maharshi Pandya, Nidhi Patel, Nitin Savaliya, Raghawendra Kumar, R D Dixit, A M Kadri, Harsh Bakshi, Chaitanya Joshi, Madhvi Joshi |
| EPI_ISL_476881 | Department of MicroBiology, Government Medical College, Surat | Gujarat Biotechnology Research Centre | Dinesh Kumar, Zuber Saiyed, Komal Patel, Labdhi Pandya, Afzal Ansari, Nikha Trivedi, Naresh Chauhan, Summaiya Mullan, Amit gamit, Apurvasinh Puvar, Janvi Raval, Zarna Patel, Monika Gandhi, Pinal Trivedi, Maharshi Pandya, Nidhi Patel, Nitin Savaliya, Raghawendra Kumar, R D Dixit, A M Kadri, Harsh Bakshi, Chaitanya Joshi, Madhvi Joshi |
| EPI_ISL_477008, EPI_ISL_477009, EPI_ISL_477010, EPI_ISL_477011, EPI_ISL_477012, EPI_ISL_477013 | University of Debrecen, Department of Medical Microbiology | National Laboratory of Virology, Szentágothai Research Centre | Endre Gábor Tóth, Balázs Somogyi, Brigitta Zana, Eszter Csoma, Ferenc Jakab, Gábor Kemenesi |
| EPI_ISL_477205, EPI_ISL_477207, EPI_ISL_477208, EPI_ISL_477210, EPI_ISL_477211, EPI_ISL_477233, EPI_ISL_477237, EPI_ISL_477238, EPI_ISL_477239, EPI_ISL_477241, EPI_ISL_477242, EPI_ISL_477246, EPI_ISL_477249, EPI_ISL_477250, EPI_ISL_477255, EPI_ISL_477256, EPI_ISL_477258, EPI_ISL_477261 | | | |
| see above | Institute for Stem Cell Science and Regenerative Medicine | National Centre for Biological Sciences | Farhan Ali, Vanessa Molin Paynter, Srikar Krishna, Mohak Sharda, Shah-e-Jahan Gulzar, Awadhesh Pandit, Varadha Sundarmurthy, Uma Ramakrishnan, Dasaradhi Palakodeti, Aswin Seshasayee |
| EPI_ISL_477615, EPI_ISL_477616, EPI_ISL_477617, EPI_ISL_477618, EPI_ISL_477619, EPI_ISL_477620, EPI_ISL_477621, EPI_ISL_477622, EPI_ISL_477623, EPI_ISL_477624, EPI_ISL_477625 | | | |

| | | | |
|---|---|---|---|
| see above | University of Szeged, Institute of Clinical Microbiology | National Laboratory of Virology, Szentágothai Research Centre | Endre Gábor Tóth, Balázs Somogyi, Brigitta Zana, Terhes Gabriella, Ferenc Jakab, Gábor Kemenesi |
| EPI_ISL_478721, EPI_ISL_478722, EPI_ISL_478723, EPI_ISL_478724, EPI_ISL_478725, EPI_ISL_478726, EPI_ISL_478727, EPI_ISL_478728 | Queens Medical Centre, Clinical Microbiology Department / DeepSeq Nottingham | COVID-19 Genomics UK (COG-UK) Consortium | Gemma Clark, Wendy Smith, Manjinder Khakh, Vicki M Fleming, Michelle M Lister, Hannah Howson-Wells, Jonathan Ball, Patrick McClure, Joseph Chappell, Theocharis Tsoleridis, Nadine Holmes, Matthew Carlisle, Christopher Moore, Fei Sang, Johnny Debebe, Victoria Wright, Matthew Loose |
| EPI_ISL_479759, EPI_ISL_479760, EPI_ISL_479761, EPI_ISL_479762, EPI_ISL_479763, EPI_ISL_479764, EPI_ISL_479765, EPI_ISL_479766, EPI_ISL_479767, EPI_ISL_479768, EPI_ISL_479769, EPI_ISL_479770, EPI_ISL_479771, EPI_ISL_479772, EPI_ISL_479773, EPI_ISL_479774, EPI_ISL_479775 | | | |
| see above | University of Miami Immunology and Histocompatibility Laboratory | University of Miami Immunology and Histocompatibility Laboratory | Emilio Margolles-Clark, PhD and Phillip Ruiz, MD, PhD |
| EPI_ISL_480351, EPI_ISL_480352, EPI_ISL_480353, EPI_ISL_480354, EPI_ISL_480355, EPI_ISL_480356, EPI_ISL_480357, EPI_ISL_480358, EPI_ISL_480359, EPI_ISL_480360, EPI_ISL_480361, EPI_ISL_480362, EPI_ISL_480363, EPI_ISL_480364, EPI_ISL_480365, EPI_ISL_480366, EPI_ISL_480367, EPI_ISL_480368, EPI_ISL_480369, EPI_ISL_480370, EPI_ISL_480371, EPI_ISL_480372, EPI_ISL_480373, EPI_ISL_480374, EPI_ISL_480375, EPI_ISL_480376, EPI_ISL_480377, EPI_ISL_480378, EPI_ISL_480379, EPI_ISL_480380, EPI_ISL_480381, EPI_ISL_480382, EPI_ISL_480383, EPI_ISL_480384, EPI_ISL_480385, EPI_ISL_480386, EPI_ISL_480387, EPI_ISL_480388, EPI_ISL_480389, EPI_ISL_480390, EPI_ISL_480391, EPI_ISL_480392, EPI_ISL_480393, EPI_ISL_480394, EPI_ISL_480395, EPI_ISL_480396, EPI_ISL_480397, EPI_ISL_480398, EPI_ISL_480399, EPI_ISL_480400, EPI_ISL_480401, EPI_ISL_480402, EPI_ISL_480403, EPI_ISL_480404, EPI_ISL_480405, EPI_ISL_480406, EPI_ISL_480407, EPI_ISL_480408, EPI_ISL_480409, EPI_ISL_480410, EPI_ISL_480411, EPI_ISL_480412, EPI_ISL_480413 | | | |
| see above | University of Wisconsin-Madison AIDS Vaccine Research Laboratories | University of Wisconsin-Madison AIDS Vaccine Research Laboratories | Gage Moreno, Katarina Braun, et al. AIDS Vaccine Research Laboratories |
| EPI_ISL_482702, EPI_ISL_482703, EPI_ISL_482704, EPI_ISL_482705, EPI_ISL_482708, EPI_ISL_482709 | Molecular Diagnostics Services (MDS) | KRISP, KZN Research Innovation and Sequencing Platform | Giandhari J, Pillay S, Lessells R, Chimukangara B, Mdlalose K, York D, Khan S, Tegally H, Wilkinson E, de Oliveira T |
| EPI_ISL_482710, EPI_ISL_482711, EPI_ISL_482712, EPI_ISL_482713 | NHLS-IALCH | KRISP, KZN Research Innovation and Sequencing Platform | Giandhari J, Pillay S, Lessells R, Chimukangara B, Mdlalose K, York D, Khan S, Tegally H, Wilkinson E, de Oliveira T |
| EPI_ISL_482714, EPI_ISL_482715, EPI_ISL_482716, EPI_ISL_482717, EPI_ISL_482718, EPI_ISL_482719, EPI_ISL_482720, EPI_ISL_482721, EPI_ISL_482722, EPI_ISL_482723 | Molecular Diagnostics Services (MDS) | KRISP, KZN Research Innovation and Sequencing Platform | Giandhari J, Pillay S, Lessells R, Chimukangara B, Mdlalose K, York D, Khan S, Tegally H, Wilkinson E, de Oliveira T |
| EPI_ISL_482724, EPI_ISL_482725, EPI_ISL_482726, EPI_ISL_482727, EPI_ISL_482728, EPI_ISL_482729, EPI_ISL_482730, EPI_ISL_482731, EPI_ISL_482848, EPI_ISL_482849, EPI_ISL_482850 | | | |
| see above | NHLS-IALCH | KRISP, KZN Research Innovation and Sequencing Platform | Giandhari J, Pillay S, Lessells R, Chimukangara B, Mdlalose K, York D, Khan S, Tegally H, Wilkinson E, de Oliveira T |
| EPI_ISL_482851, EPI_ISL_482852, EPI_ISL_482853, EPI_ISL_482854, EPI_ISL_482855, EPI_ISL_482856, EPI_ISL_482857, EPI_ISL_482858, EPI_ISL_482859, EPI_ISL_482860, EPI_ISL_482861, EPI_ISL_482862, EPI_ISL_482863, EPI_ISL_482864, EPI_ISL_482865, EPI_ISL_482866, EPI_ISL_482867, EPI_ISL_482868, EPI_ISL_482869, EPI_ISL_482870, EPI_ISL_482871, EPI_ISL_482872 | | | |
| see above | Molecular Diagnostics Services (MDS) | KRISP, KZN Research Innovation and Sequencing Platform | Giandhari J, Pillay S, Lessells R, Chimukangara B, Mdlalose K, York D, Khan S, Tegally H, Wilkinson E, de Oliveira T |
| EPI_ISL_483060 | unknown | Microbiology, Canterbury Health Laboratories | Dilcher,M., Anderson,T. |
| EPI_ISL_483637 | National Laboratory of Virology, Szentágothai Research Centre | National Laboratory of Virology, Szentágothai Research Centre | Endre Gábor Tóth, Balázs Somogyi, Ferenc Jakab, Gábor Kemenesi |
| EPI_ISL_483835 | Pandit Deendayal Upadhyay Government Medical College, Rajkot | Gujarat Biotechnology Research Centre | Dinesh Kumar, Zuber Saiyed, Komal Patel, Labdhi Pandya, Afzal Ansari, Nikha Trivedi, Gauravi Dhruv, Arti Trivedi, Apurvasinh Puvar, Janvi Raval, Zarna Patel, Monika Gandhi, Pinal Trivedi, Maharshi Pandya, Nidhi Patel, Nitin Savaliya, Raghawendra Kumar, R D Dixit, A M Kadri, Harsh Bakshi, Chaitanya Joshi, Madhvi Joshi |
| EPI_ISL_483841 | Pandit Deendayal Upadhyay Government Medical College, Rajkot | Gujarat Biotechnology Research Centre | Gauravi Dhruv, Arti Trivedi, Apurvasinh Puvar, Janvi Raval, Zarna Patel, Monika Gandhi, Pinal Trivedi, Maharshi Pandya, Nidhi Patel, Nitin Savaliya, Raghawendra Kumar, Dinesh Kumar, Zuber Saiyed, Komal Patel, Labdhi Pandya, Afzal Ansari, Nikha Trivedi, R D Dixit, A M Kadri, Harsh Bakshi, Chaitanya Joshi, Madhvi Joshi |
| EPI_ISL_483852, EPI_ISL_483870 | Department of Microbiology, Government Medical College, Surat | Gujarat Biotechnology Research Centre | Dinesh Kumar, Zuber Saiyed, Komal Patel, Labdhi Pandya, Afzal Ansari, Nikha Trivedi, Naresh Chauhan, Summaiya Mullan, Amit gamit, Apurvasinh Puvar, Janvi Raval, Zarna Patel, Monika Gandhi, Pinal Trivedi, Maharshi Pandya, Nidhi Patel, Nitin Savaliya, Raghawendra Kumar, R D Dixit, A M Kadri, Harsh Bakshi, Chaitanya Joshi, Madhvi Joshi |
| EPI_ISL_484339, EPI_ISL_484340, EPI_ISL_484341, EPI_ISL_484342, EPI_ISL_484343, EPI_ISL_484344, EPI_ISL_484345, EPI_ISL_484346, EPI_ISL_484347, EPI_ISL_484348, EPI_ISL_484349, EPI_ISL_484350, EPI_ISL_484351, EPI_ISL_484352, EPI_ISL_484353, EPI_ISL_484374, EPI_ISL_484375, EPI_ISL_484390, EPI_ISL_484391 | | | |
| see above | Queens Medical Centre, Clinical Microbiology Department / DeepSeq Nottingham | COVID-19 Genomics UK (COG-UK) Consortium | Gemma Clark, Wendy Smith, Manjinder Khakh, Vicki M Fleming, Michelle M Lister, Hannah Howson-Wells, Jonathan Ball, Patrick McClure, Joseph Chappell, Theocharis Tsoleridis, Nadine Holmes, Matthew Carlisle, Christopher Moore, Fei Sang, Johnny Debebe, Victoria Wright, Matthew Loose |
| EPI_ISL_484807, EPI_ISL_484808, EPI_ISL_484809, EPI_ISL_484810, EPI_ISL_484811, EPI_ISL_484812, EPI_ISL_484813, EPI_ISL_484814, EPI_ISL_484815, EPI_ISL_484816, EPI_ISL_484817, EPI_ISL_484818, EPI_ISL_484819, EPI_ISL_484820, EPI_ISL_484821, EPI_ISL_484822, EPI_ISL_484823, EPI_ISL_484824, EPI_ISL_484825, EPI_ISL_484826, EPI_ISL_484827, EPI_ISL_484828, EPI_ISL_484829, EPI_ISL_484830, EPI_ISL_484831, EPI_ISL_484832, EPI_ISL_484833, EPI_ISL_484834, EPI_ISL_484835, EPI_ISL_484836, EPI_ISL_484837, EPI_ISL_484838, EPI_ISL_484839, EPI_ISL_484840, EPI_ISL_484841, EPI_ISL_484842, EPI_ISL_484843, EPI_ISL_484844, EPI_ISL_484845, EPI_ISL_484846, EPI_ISL_484847, EPI_ISL_484848, EPI_ISL_484849, EPI_ISL_484850, EPI_ISL_484851, EPI_ISL_484852, EPI_ISL_484853, EPI_ISL_484854, EPI_ISL_484855, EPI_ISL_484856, EPI_ISL_484857, EPI_ISL_484858, EPI_ISL_484859, EPI_ISL_484860, EPI_ISL_484861, EPI_ISL_484862, EPI_ISL_484863, EPI_ISL_484864, EPI_ISL_484865, EPI_ISL_484866, EPI_ISL_484867, EPI_ISL_484868, EPI_ISL_484869, EPI_ISL_484870, EPI_ISL_484871, EPI_ISL_484872, EPI_ISL_484873, EPI_ISL_484874, EPI_ISL_484875, EPI_ISL_484876, EPI_ISL_484877, EPI_ISL_484878, EPI_ISL_484879, EPI_ISL_484880, EPI_ISL_484881, EPI_ISL_484882, EPI_ISL_484883, EPI_ISL_484884, EPI_ISL_484885, EPI_ISL_484886, EPI_ISL_484887, EPI_ISL_484888, EPI_ISL_484889, EPI_ISL_484890, EPI_ISL_484891, EPI_ISL_484892, EPI_ISL_484893, EPI_ISL_484894, EPI_ISL_484895, EPI_ISL_484896, EPI_ISL_484897, EPI_ISL_484898, EPI_ISL_484899, EPI_ISL_484900, EPI_ISL_484901, EPI_ISL_484902, EPI_ISL_484903, EPI_ISL_484904, EPI_ISL_484905, EPI_ISL_484906, EPI_ISL_484907, EPI_ISL_484908, EPI_ISL_484909, EPI_ISL_484910, EPI_ISL_484911, EPI_ISL_484912, EPI_ISL_484913, EPI_ISL_484914, EPI_ISL_484915, EPI_ISL_484916, EPI_ISL_484917, EPI_ISL_484918, EPI_ISL_484919, EPI_ISL_484920, EPI_ISL_484921, EPI_ISL_484922, EPI_ISL_484923, EPI_ISL_484924, EPI_ISL_484925, EPI_ISL_484926, EPI_ISL_484927, EPI_ISL_484928, EPI_ISL_484929, EPI_ISL_484930, EPI_ISL_484931, EPI_ISL_484932, EPI_ISL_484933, EPI_ISL_484934, EPI_ISL_484935, EPI_ISL_484936, EPI_ISL_484937, EPI_ISL_484938, EPI_ISL_484939, EPI_ISL_484940, EPI_ISL_484941, EPI_ISL_484942, EPI_ISL_484943, EPI_ISL_484944, EPI_ISL_484945, EPI_ISL_484946, EPI_ISL_484947, EPI_ISL_484948, EPI_ISL_484949, EPI_ISL_484950, EPI_ISL_484951, EPI_ISL_484952, EPI_ISL_484953, EPI_ISL_484954, EPI_ISL_484955, EPI_ISL_484956, EPI_ISL_484957, EPI_ISL_484958, EPI_ISL_484959, EPI_ISL_484960, EPI_ISL_484961, EPI_ISL_484962, EPI_ISL_484963, EPI_ISL_484964, EPI_ISL_484965, EPI_ISL_484966, EPI_ISL_484967, EPI_ISL_484968, EPI_ISL_484969, EPI_ISL_484970, EPI_ISL_484971, EPI_ISL_484972, EPI_ISL_484973, EPI_ISL_484974, EPI_ISL_484975, EPI_ISL_484976, EPI_ISL_484977, EPI_ISL_484978, EPI_ISL_484979, EPI_ISL_484980, EPI_ISL_484981, EPI_ISL_484982, EPI_ISL_484983, EPI_ISL_484984, EPI_ISL_484985, EPI_ISL_484986, EPI_ISL_484987, EPI_ISL_484988, EPI_ISL_484989, EPI_ISL_484990, EPI_ISL_484991, EPI_ISL_484992, EPI_ISL_484993, EPI_ISL_484994, EPI_ISL_484995, EPI_ISL_484996, EPI_ISL_484997, EPI_ISL_484998, EPI_ISL_484999, EPI_ISL_485000 | | | |
| see above | University of Wisconsin-Madison AIDS Vaccine Research Laboratories | University of Wisconsin-Madison AIDS Vaccine Research Laboratories | Gage Moreno, Katarina Braun, et al. AIDS Vaccine Research Laboratories |
| EPI_ISL_487276 | Department of Food Safety, Nutrition and Veterinary public health, Istituto Superiore di Sanita' | Department of Biomedical, Surgical and Dental Sciences and Department of Biomedical Sciences for Health | Delbue,S., Ferrante,P., Basilico,N., Parapini,S., Binda,S., D'Alessandro,S., Galli,C., Signorini,L., Primache,V., Anselmi,G., Pariani,E. |
| EPI_ISL_487277, EPI_ISL_487278, EPI_ISL_487279, EPI_ISL_487280, EPI_ISL_487281, EPI_ISL_487282, EPI_ISL_487283, EPI_ISL_487284, EPI_ISL_487285, EPI_ISL_487286, EPI_ISL_487287, EPI_ISL_487288, EPI_ISL_487289, EPI_ISL_487290, EPI_ISL_487291, EPI_ISL_487292, EPI_ISL_487293, EPI_ISL_487294, EPI_ISL_487295, EPI_ISL_487296, EPI_ISL_487297, EPI_ISL_487298, EPI_ISL_487299, EPI_ISL_487300, EPI_ISL_487301, EPI_ISL_487302, EPI_ISL_487303, EPI_ISL_487304, EPI_ISL_487305, EPI_ISL_487306, EPI_ISL_487307, EPI_ISL_487308, EPI_ISL_487309, EPI_ISL_487310, EPI_ISL_487311, EPI_ISL_487312, EPI_ISL_487313, EPI_ISL_487314, EPI_ISL_487315, EPI_ISL_487316, EPI_ISL_487317, EPI_ISL_487318, EPI_ISL_487319, EPI_ISL_487320, EPI_ISL_487321, EPI_ISL_487322, EPI_ISL_487323, EPI_ISL_487324, EPI_ISL_487325, EPI_ISL_487326, EPI_ISL_487327, EPI_ISL_487328 | | | |
| see above | NHLS-IALCH | KRISP, KZN Research Innovation and Sequencing Platform | Giandhari J, Pillay S, Lessells R, Chimukangara B, Mdlalose K, York D, Khan S, Tegally H, Wilkinson E, de Oliveira T |
| EPI_ISL_487329, EPI_ISL_487330, EPI_ISL_487331, EPI_ISL_487332, EPI_ISL_487333, EPI_ISL_487334, EPI_ISL_487335, EPI_ISL_487336, EPI_ISL_487337, EPI_ISL_487338, EPI_ISL_487339, EPI_ISL_487340, EPI_ISL_487341 | | | |
| see above | Molecular Diagnostics Services (MDS) | KRISP, KZN Research Innovation and Sequencing Platform | Giandhari J, Pillay S, Lessells R, Chimukangara B, Mdlalose K, York D, Khan S, Tegally H, Wilkinson E, de Oliveira T |
| EPI_ISL_487348 | NHLS-IALCH | KRISP, KZN Research Innovation and Sequencing Platform | Giandhari J, Pillay S, Lessells R, Chimukangara B, Mdlalose K, York D, Khan S, Tegally H, Wilkinson E, de Oliveira T |
| EPI_ISL_490205, EPI_ISL_490206, EPI_ISL_490207, EPI_ISL_490208 | München Klinik Schwabing | MGZ Medical Genetics Center | Dieter A. Wolf, Elke Holinski-Feder |
| EPI_ISL_490560, EPI_ISL_490561, EPI_ISL_490562 | Queens Medical Centre, Clinical Microbiology Department / DeepSeq Nottingham | COVID-19 Genomics UK (COG-UK) Consortium | Gemma Clark, Wendy Smith, Manjinder Khakh, Vicki M Fleming, Michelle M Lister, Hannah Howson-Wells, Jonathan Ball, Patrick McClure, Joseph Chappell, Theocharis Tsoleridis, Nadine Holmes, Matthew Carlisle, Christopher Moore, Fei Sang, Johnny Debebe, Victoria Wright, Matthew Loose |
| EPI_ISL_491097, EPI_ISL_491098, EPI_ISL_491099, EPI_ISL_491100, EPI_ISL_491101, EPI_ISL_491102, EPI_ISL_491103, EPI_ISL_491104, EPI_ISL_491105, EPI_ISL_491106, EPI_ISL_491107, EPI_ISL_491108, EPI_ISL_491109, EPI_ISL_491110 | | | |

| | | | |
|---|---|---|---|
| see above | SC Department of Health and Environmental Control | SC Department of Health and Environmental Control | Flores,H. |
| EPI_ISL_491299, EPI_ISL_491300, EPI_ISL_491301, EPI_ISL_491302, EPI_ISL_491303, EPI_ISL_491304, EPI_ISL_491305, EPI_ISL_491306, EPI_ISL_491307, EPI_ISL_491308, EPI_ISL_491309, EPI_ISL_491310, EPI_ISL_491311, EPI_ISL_491312, EPI_ISL_491313, EPI_ISL_491314, EPI_ISL_491315, EPI_ISL_491316, EPI_ISL_491317, EPI_ISL_491318, EPI_ISL_491319, EPI_ISL_491320, EPI_ISL_491321, EPI_ISL_491322, EPI_ISL_491323, EPI_ISL_491324, EPI_ISL_491325, EPI_ISL_491326, EPI_ISL_491327, EPI_ISL_491328, EPI_ISL_491329, EPI_ISL_491330, EPI_ISL_491331, EPI_ISL_491332, EPI_ISL_491333, EPI_ISL_491334, EPI_ISL_491335, EPI_ISL_491336, EPI_ISL_491337, EPI_ISL_491338, EPI_ISL_491339, EPI_ISL_491340, EPI_ISL_491341, EPI_ISL_491342, EPI_ISL_491343, EPI_ISL_491344, EPI_ISL_491345, EPI_ISL_491346, EPI_ISL_491347, EPI_ISL_491348, EPI_ISL_491349, EPI_ISL_491350, EPI_ISL_491351, EPI_ISL_491352, EPI_ISL_491353, EPI_ISL_491354, EPI_ISL_491355, EPI_ISL_491356, EPI_ISL_491357, EPI_ISL_491358, EPI_ISL_491359, EPI_ISL_491360, EPI_ISL_491361, EPI_ISL_491362, EPI_ISL_491363, EPI_ISL_491364, EPI_ISL_491365, EPI_ISL_491366, EPI_ISL_491367, EPI_ISL_491368, EPI_ISL_491369, EPI_ISL_491370, EPI_ISL_491371, EPI_ISL_491372, EPI_ISL_491373, EPI_ISL_491374, EPI_ISL_491375, EPI_ISL_491376, EPI_ISL_491377, EPI_ISL_491378, EPI_ISL_491379, EPI_ISL_491380, EPI_ISL_491381, EPI_ISL_491382, EPI_ISL_491383, EPI_ISL_491384, EPI_ISL_491385, EPI_ISL_491386, EPI_ISL_491387, EPI_ISL_491388, EPI_ISL_491389, EPI_ISL_491390, EPI_ISL_491391, EPI_ISL_491392, EPI_ISL_491393, EPI_ISL_491394, EPI_ISL_491395, EPI_ISL_491396, EPI_ISL_491397, EPI_ISL_491398, EPI_ISL_491399, EPI_ISL_491400, EPI_ISL_491401, EPI_ISL_491402, EPI_ISL_491403, EPI_ISL_491404, EPI_ISL_491405, EPI_ISL_491406, EPI_ISL_491407, EPI_ISL_491408, EPI_ISL_491409, EPI_ISL_491410, EPI_ISL_491411, EPI_ISL_491412, EPI_ISL_491413, EPI_ISL_491414, EPI_ISL_491415, EPI_ISL_491416, EPI_ISL_491417, EPI_ISL_491418, EPI_ISL_491419, EPI_ISL_491420, EPI_ISL_491421, EPI_ISL_491422, EPI_ISL_491423, EPI_ISL_491424, EPI_ISL_491425, EPI_ISL_491426 | | | |
| see above | University of Wisconsin-Madison AIDS Vaccine Research Laboratories | University of Wisconsin-Madison AIDS Vaccine Research Laboratories | Gage Moreno, Katarina Braun, et al. AIDS Vaccine Research Laboratories |
| EPI_ISL_491437 | Area de Salud Escazu (Coopesana) | Inciensa, Instituto Costarricense de Investigación y Enseñanza en Nutrición y Salud | Francisco Duarte, Hebleen Brenes, Claudio Soto-Garita, Estela Cordero, Adriana Godinez & Melany Calderon |
| EPI_ISL_491438 | Hospital San Rafael de Alajuela | Inciensa, Instituto Costarricense de Investigación y Enseñanza en Nutrición y Salud | Francisco Duarte, Hebleen Brenes, Claudio Soto-Garita, Estela Cordero, Adriana Godinez & Melany Calderon |
| EPI_ISL_491439 | Hospital Calderon Guardia | Inciensa, Instituto Costarricense de Investigación y Enseñanza en Nutrición y Salud | Francisco Duarte, Hebleen Brenes, Claudio Soto-Garita, Estela Cordero, Adriana Godinez & Melany Calderon |
| EPI_ISL_491440 | Hospital San Rafael de Alajuela | Inciensa, Instituto Costarricense de Investigación y Enseñanza en Nutrición y Salud | Francisco Duarte, Hebleen Brenes, Claudio Soto-Garita, Estela Cordero, Adriana Godinez & Melany Calderon |
| EPI_ISL_491441, EPI_ISL_491442 | Hospital Clinica Biblica | Inciensa, Instituto Costarricense de Investigación y Enseñanza en Nutrición y Salud | Francisco Duarte, Hebleen Brenes, Claudio Soto-Garita, Estela Cordero, Adriana Godinez & Melany Calderon |
| EPI_ISL_491443 | Hospital Fernando Escalante Pradilla | Inciensa, Instituto Costarricense de Investigación y Enseñanza en Nutrición y Salud | Francisco Duarte, Hebleen Brenes, Claudio Soto-Garita, Estela Cordero, Adriana Godinez & Melany Calderon |
| EPI_ISL_491444 | Area de Salud Escazu (Coopesana) | Inciensa, Instituto Costarricense de Investigación y Enseñanza en Nutrición y Salud | Francisco Duarte, Hebleen Brenes, Claudio Soto-Garita, Estela Cordero, Adriana Godinez & Melany Calderon |
| EPI_ISL_491445 | Area de Salud Mata Redonda | Inciensa, Instituto Costarricense de Investigación y Enseñanza en Nutrición y Salud | Francisco Duarte, Hebleen Brenes, Claudio Soto-Garita, Estela Cordero, Adriana Godinez & Melany Calderon |
| EPI_ISL_491446 | Area de Salud Alajuela Central | Inciensa, Instituto Costarricense de Investigación y Enseñanza en Nutrición y Salud | Francisco Duarte, Hebleen Brenes, Claudio Soto-Garita, Estela Cordero, Adriana Godinez & Melany Calderon |
| EPI_ISL_491447 | Hospital Fernando Escalante Pradilla | Inciensa, Instituto Costarricense de Investigación y Enseñanza en Nutrición y Salud | Francisco Duarte, Hebleen Brenes, Claudio Soto-Garita, Estela Cordero, Adriana Godinez & Melany Calderon |
| EPI_ISL_491448 | Hospital San Rafael de Alajuela | Inciensa, Instituto Costarricense de Investigación y Enseñanza en Nutrición y Salud | Francisco Duarte, Hebleen Brenes, Claudio Soto-Garita, Estela Cordero, Adriana Godinez & Melany Calderon |
| EPI_ISL_491449 | Area de Salud Alajuela Sur | Inciensa, Instituto Costarricense de Investigación y Enseñanza en Nutrición y Salud | Francisco Duarte, Hebleen Brenes, Claudio Soto-Garita, Estela Cordero, Adriana Godinez & Melany Calderon |
| EPI_ISL_491450 | Hospital San Juan de Dios | Inciensa, Instituto Costarricense de Investigación y Enseñanza en Nutrición y Salud | Francisco Duarte, Hebleen Brenes, Claudio Soto-Garita, Estela Cordero, Adriana Godinez & Melany Calderon |
| EPI_ISL_491451 | Hospital México | Inciensa, Instituto Costarricense de Investigación y Enseñanza en Nutrición y Salud | Francisco Duarte, Hebleen Brenes, Claudio Soto-Garita, Estela Cordero, Adriana Godinez & Melany Calderon |
| EPI_ISL_491452 | Hospital San Rafael de Alajuela | Inciensa, Instituto Costarricense de Investigación y Enseñanza en Nutrición y Salud | Francisco Duarte, Hebleen Brenes, Claudio Soto-Garita, Estela Cordero, Adriana Godinez & Melany Calderon |
| EPI_ISL_491453 | Hospital México | Inciensa, Instituto Costarricense de Investigación y Enseñanza en Nutrición y Salud | Francisco Duarte, Hebleen Brenes, Claudio Soto-Garita, Estela Cordero, Adriana Godinez & Melany Calderon |
| EPI_ISL_491454 | Hospital San Juan de Dios | Inciensa, Instituto Costarricense de Investigación y Enseñanza en Nutrición y Salud | Francisco Duarte, Hebleen Brenes, Claudio Soto-Garita, Estela Cordero, Adriana Godinez & Melany Calderon |
| EPI_ISL_491455 | Hospital Clinica Biblica | Inciensa, Instituto Costarricense de Investigación y Enseñanza en Nutrición y Salud | Francisco Duarte, Hebleen Brenes, Claudio Soto-Garita, Estela Cordero, Adriana Godinez & Melany Calderon |
| EPI_ISL_491456 | Hospital San Juan de Dios | Inciensa, Instituto Costarricense de Investigación y Enseñanza en Nutrición y Salud | Francisco Duarte, Hebleen Brenes, Claudio Soto-Garita, Estela Cordero, Adriana Godinez & Melany Calderon |
| EPI_ISL_491457 | Area de Salud Los Santos | Inciensa, Instituto Costarricense de Investigación y Enseñanza en Nutrición y Salud | Francisco Duarte, Hebleen Brenes, Claudio Soto-Garita, Estela Cordero, Adriana Godinez & Melany Calderon |
| EPI_ISL_491933, EPI_ISL_491934, EPI_ISL_491935, EPI_ISL_491941 | Centro de Investigaciones, Universidad de Especialidades Espíritu Santo | Institute of Microbiology, Universidad San Francisco de Quito | Derly Andrade, Juan Carlos Fernandez, Belén Prado-Vivar, Sully Márquez, Juan José Guadalupe, Monica Becerra-Wong, Bernardo Gutiérrez, Gabriel Morey, Ruben Armas, Jose Pedro Barberan, Fernando Espinoza, Edith Lopez, Verónica Barragán, Patricio Rojas-Silva, Gabriel Trueba, Michelle Grunauer, Paúl Cárdenas |
| EPI_ISL_491968, EPI_ISL_491969, EPI_ISL_491970, EPI_ISL_491971, EPI_ISL_491972, EPI_ISL_491973, EPI_ISL_491974, EPI_ISL_491975, EPI_ISL_491976, EPI_ISL_491977, EPI_ISL_491978, EPI_ISL_491979, EPI_ISL_491980, EPI_ISL_491981, EPI_ISL_491982, EPI_ISL_491983, EPI_ISL_491984, EPI_ISL_491985, EPI_ISL_491986, EPI_ISL_491988, EPI_ISL_491989, EPI_ISL_491990, EPI_ISL_491991, EPI_ISL_491992, EPI_ISL_491993, EPI_ISL_491994, EPI_ISL_491995, EPI_ISL_491996, EPI_ISL_491997, EPI_ISL_491998, EPI_ISL_491999, EPI_ISL_492000, EPI_ISL_492001, EPI_ISL_492002, EPI_ISL_492003, EPI_ISL_492004, EPI_ISL_492005, EPI_ISL_492006, EPI_ISL_492007, EPI_ISL_492008, EPI_ISL_492009, EPI_ISL_492010, EPI_ISL_492011, EPI_ISL_492012, EPI_ISL_492013, EPI_ISL_492014, EPI_ISL_492015, EPI_ISL_492016, EPI_ISL_492017, EPI_ISL_492018, EPI_ISL_492019, EPI_ISL_492020, EPI_ISL_492021, EPI_ISL_492022, EPI_ISL_492023, EPI_ISL_492024, EPI_ISL_492025, EPI_ISL_492026 | | | |
| see above | Oman-NIC | Department of Microbiology and Immunology-SQUH | Fahad Zadjali, Samira Al-Maruqi, Amina Al Jardani, Khulood Al-Mammary, Hanan Al-kindi, Fatma BaAlawi, Hamida AL Barwani, Zeyana AL-Dahmani, Intisar Al-Shukri, Aisha Al-Busaidi, Aisha Al-Amri, Ahlam Al-Amri, Mohammed Al-Tobi, Samiha Al Kharusi, Abdulla Balkhair |
| EPI_ISL_493342, EPI_ISL_493343 | Instituto de Diagnostico y Referencia Epidemiologicos (INDRE) | Instituto de Diagnostico y Referencia Epidemiologicos (INDRE) | Ernesto Ramirez-Gonzalez, Abril Rodriguez-Maldonado, Claudia Wong-Arambula , Natividad Cruz-Ortiz, Tatiana Nunez-Garcia, Dayanira Arellano-Suarez, Adnan Araiza-Rodriguez, Edgar Mendieta-Condado, Lucia Hernandez-Rivas, Irma Lopez-Martinez, Gisela Barrera-Badillo. |
| EPI_ISL_493344, EPI_ISL_493345, EPI_ISL_493346, EPI_ISL_493347, EPI_ISL_493348, EPI_ISL_493349 | Instituto de Diagnostico y Referencia Epidemiologicos (INDRE) | Instituto de Diagnostico y Referencia Epidemiologicos (INDRE) | Ernesto Ramirez-Gonzalez, Abril Rodriguez-Maldonado, Claudia Wong-Arambula , Natividad Cruz-Ortiz, Tatiana Nunez-Garcia, Dayanira Arellano-Suarez, Adnan Araiza-Rodriguez, Fabiola Garces-Ayala, Lucia Hernandez-Rivas, Irma Lopez-Martinez, Gisela Barrera-Badillo. |
| EPI_ISL_493547, EPI_ISL_493548, EPI_ISL_493601, EPI_ISL_493602, EPI_ISL_493606, EPI_ISL_493607, EPI_ISL_493608, EPI_ISL_493609 | Queens Medical Centre, Clinical Microbiology Department / DeepSeq Nottingham | COVID-19 Genomics UK (COG-UK) Consortium | Gemma Clark, Wendy Smith, Manjinder Khakh, Vicki M Fleming, Michelle M Lister, Hannah Howson-Wells, Jonathan Ball, Patrick McClure, Joseph Chappell, Theocharis Tsoleridis, Nadine Holmes, Matthew Carlisle, Christopher Moore, Fei Sang, Johnny Debebe, Victoria Wright, Matthew Loose |
| EPI_ISL_495025 | Government Medical College, Bhavnagar | Gujarat Biotechnology Research Centre | Dinesh Kumar, Zuber Saiyed, Komal Patel, Labdhi Pandya, Afzal Ansari, Nikha Trivedi, Kairavi Desai, Saklin Malek, Shirish Patel, Apurvasinh Puvar, Janvi Raval, Zarna Patel, Monika Gandhi, Pinal Trivedi, Maharshi Pandya, Nidhi Patel, Nitin Savaliya, Raghawendra Kumar, R D Dixit, A M Kadri, Harsh Bakshi, Chaitanya Joshi, Madhvi Joshi |

| | | | |
|---|---|---|---|
| EPI_ISL_495042, EPI_ISL_495060 | GMERS Medical College & Hospital, Gotri, Vadodara | Gujarat Biotechnology Research Centre | Dinesh Kumar, Zuber Saiyed, Komal Patel, Labdhi Pandya, Afzal Ansari, Nikha Trivedi, Meenakshi Shah, Neena Doshi, Varsha Godbole, Apurvasinh Puvar, Janvi Raval, Zarna Patel, Monika Gandhi, Pinal Trivedi, Maharshi Pandya, Nidhi Patel, Nitin Savaliya, Raghawendra Kumar, R D Dixit, A M Kadri, Harsh Bakshi, Chaitanya Joshi, Madhvi Joshi |
| EPI_ISL_495078 | Department of MicroBiology, Government Medical College, Surat | Gujarat Biotechnology Research Centre | Dinesh Kumar, Zuber Saiyed, Komal Patel, Labdhi Pandya, Afzal Ansari, Nikha Trivedi, Naresh Chauhan, Summaiya Mullan, Amit gamit, Apurvasinh Puvar, Janvi Raval, Zarna Patel, Monika Gandhi, Pinal Trivedi, Maharshi Pandya, Nidhi Patel, Nitin Savaliya, Raghawendra Kumar, R D Dixit, A M Kadri, Harsh Bakshi, Chaitanya Joshi, Madhvi Joshi |
| EPI_ISL_495460, EPI_ISL_495461, EPI_ISL_495462, EPI_ISL_495463, EPI_ISL_495464, EPI_ISL_495465, EPI_ISL_495466, EPI_ISL_495467, EPI_ISL_495468, EPI_ISL_495469, EPI_ISL_495470, EPI_ISL_495471, EPI_ISL_495472, EPI_ISL_495473, EPI_ISL_495474, EPI_ISL_495475, EPI_ISL_495476, EPI_ISL_495477, EPI_ISL_495478, EPI_ISL_495479, EPI_ISL_495480, EPI_ISL_495481, EPI_ISL_495482, EPI_ISL_495483, EPI_ISL_495484, EPI_ISL_495485, EPI_ISL_495486, EPI_ISL_495487, EPI_ISL_495488, EPI_ISL_495489, EPI_ISL_495490, EPI_ISL_495491, EPI_ISL_495492, EPI_ISL_495493, EPI_ISL_495494, EPI_ISL_495495, EPI_ISL_495496, EPI_ISL_495497, EPI_ISL_495498, EPI_ISL_495499, EPI_ISL_495500, EPI_ISL_495501, EPI_ISL_495502, EPI_ISL_495503, EPI_ISL_495504, EPI_ISL_495505, EPI_ISL_495506, EPI_ISL_495507, EPI_ISL_495508, EPI_ISL_495509 | | | |
| see above | University of Wisconsin-Madison AIDS Vaccine Research Laboratories | University of Wisconsin-Madison AIDS Vaccine Research Laboratories | Gage Moreno, Katarina Braun, et al. AIDS Vaccine Research Laboratories |
| EPI_ISL_495516, EPI_ISL_495517, EPI_ISL_495518, EPI_ISL_495519, EPI_ISL_495520, EPI_ISL_495521, EPI_ISL_495522, EPI_ISL_495523, EPI_ISL_495524, EPI_ISL_495525, EPI_ISL_495526, EPI_ISL_495527, EPI_ISL_495528, EPI_ISL_495529, EPI_ISL_495530, EPI_ISL_495531, EPI_ISL_495532, EPI_ISL_495533, EPI_ISL_495534 | | | |
| see above | NHLS-IALCH | KRISP, KZN Research Innovation and Sequencing Platform | Giandhari J, Pillay S, Lessells R, Chimukangara B, Mdlalose K, York D, Khan S, Tegally H, Wilkinson E, de Oliveira T |
| EPI_ISL_495535, EPI_ISL_495536, EPI_ISL_495537, EPI_ISL_495538, EPI_ISL_495539, EPI_ISL_495540, EPI_ISL_495541, EPI_ISL_495542 | Medical Disagnostics Services (MDS) | KRISP, KZN Research Innovation and Sequencing Platform | Giandhari J, Pillay S, Lessells R, Chimukangara B, Mdlalose K, York D, Khan S, Tegally H, Wilkinson E, de Oliveira T |
| EPI_ISL_495543, EPI_ISL_495544, EPI_ISL_495545, EPI_ISL_495546, EPI_ISL_495547, EPI_ISL_495548, EPI_ISL_495549, EPI_ISL_495550, EPI_ISL_495551, EPI_ISL_495552, EPI_ISL_495553, EPI_ISL_495555, EPI_ISL_495556, EPI_ISL_495557, EPI_ISL_495558, EPI_ISL_495559, EPI_ISL_495560, EPI_ISL_495561, EPI_ISL_495562 | | | |
| see above | NHLS-IALCH | KRISP, KZN Research Innovation and Sequencing Platform | Giandhari J, Pillay S, Lessells R, Chimukangara B, Mdlalose K, York D, Khan S, Tegally H, Wilkinson E, de Oliveira T |
| EPI_ISL_495642, EPI_ISL_495643, EPI_ISL_495644, EPI_ISL_495646, EPI_ISL_495647, EPI_ISL_495648, EPI_ISL_495649, EPI_ISL_495650, EPI_ISL_495651, EPI_ISL_495652, EPI_ISL_495653 | | | |
| see above | Washington State Department of Health | Seattle Flu Study | Deborah A. Nickerson, Chris D. Frazar, Jover Lee, Benjamin Pelle, Matthew Richardson, Amanda Adler, Elisabeth Brandstetter, Peter D. Han, Kairsten Fay, Misja Ilcisin, Kirsten Lacombe, Thomas R. Sibley, Melissa Truong, Caitlin R. Wolf, Romesh Gautom, Geoff Melly, Brian Hiatt, Philip Dykema, Scott Lindquist, Michael Boeckh, Janet A. Englund, Michael Famulare, Barry R. Lutz, Mark J. Rieder, Lea M. Starita, Matthew Thompson, Helen Y. Chu, Jay Shendure, Trevor Bedford |
| EPI_ISL_495655, EPI_ISL_495658, EPI_ISL_495660, EPI_ISL_495661, EPI_ISL_495662 | Seattle Flu Study | Seattle Flu Study | Deborah A. Nickerson, Chris D. Frazar, Jover Lee, Benjamin Pelle, Matthew Richardson, Amanda Adler, Elisabeth Brandstetter, Peter D. Han, Kairsten Fay, Misja Ilcisin, Kirsten Lacombe, Thomas R. Sibley, Melissa Truong, Caitlin R. Wolf, Karen Cowgill, Stephanie Schrag, Jeff Duchin, Michael Boeckh, Janet A. Englund, Michael Famulare, Barry R. Lutz, Mark J. Rieder, Lea M. Starita, Matthew Thompson, Helen Y. Chu, Trevor Bedford, Jay Shendure |
| EPI_ISL_495663 | Seattle Flu Study | Seattle Flu Study | Deborah A. Nickerson, Chris D. Frazar, Jover Lee, Benjamin Pelle, Matthew Richardson, Amanda Adler, Elisabeth Brandstetter, Peter D. Han, Kairsten Fay, Misja Ilcisin, Kirsten Lacombe, Thomas R. Sibley, Melissa Truong, Caitlin R. Wolf, Michael Boeckh, Janet A. Englund, Michael Famulare, Barry R. Lutz, Mark J. Rieder, Lea M. Starita, Matthew Thompson, Jay Shendure, Trevor Bedford, Helen Y. Chu |
| EPI_ISL_495664, EPI_ISL_495665 | Seattle Flu Study | Seattle Flu Study | Deborah A. Nickerson, Chris D. Frazar, Jover Lee, Benjamin Pelle, Matthew Richardson, Amanda Adler, Elisabeth Brandstetter, Peter D. Han, Kairsten Fay, Misja Ilcisin, Kirsten Lacombe, Thomas R. Sibley, Melissa Truong, Caitlin R. Wolf, Karen Cowgill, Stephanie Schrag, Jeff Duchin, Michael Boeckh, Janet A. Englund, Michael Famulare, Barry R. Lutz, Mark J. Rieder, Lea M. Starita, Matthew Thompson, Helen Y. Chu, Trevor Bedford, Jay Shendure |
| EPI_ISL_495667, EPI_ISL_495668, EPI_ISL_495669, EPI_ISL_495670, EPI_ISL_495671, EPI_ISL_495672, EPI_ISL_495673, EPI_ISL_495674, EPI_ISL_495676, EPI_ISL_495677, EPI_ISL_495680, EPI_ISL_495681, EPI_ISL_495682, EPI_ISL_495683, EPI_ISL_495684, EPI_ISL_495685, EPI_ISL_495686, EPI_ISL_495687, EPI_ISL_495688, EPI_ISL_495689, EPI_ISL_495690, EPI_ISL_495692, EPI_ISL_495693, EPI_ISL_495694, EPI_ISL_495695, EPI_ISL_495696, EPI_ISL_495697, EPI_ISL_495698, EPI_ISL_495699, EPI_ISL_495700, EPI_ISL_495701, EPI_ISL_495702, EPI_ISL_495703, EPI_ISL_495704, EPI_ISL_495705, EPI_ISL_495706, EPI_ISL_495707, EPI_ISL_495708, EPI_ISL_495709, EPI_ISL_495710, EPI_ISL_495711, EPI_ISL_495712, EPI_ISL_495714, EPI_ISL_495715, EPI_ISL_495716, EPI_ISL_495717, EPI_ISL_495718, EPI_ISL_495719, EPI_ISL_495720, EPI_ISL_495721, EPI_ISL_495722, EPI_ISL_495723, EPI_ISL_495725, EPI_ISL_495726, EPI_ISL_495728, EPI_ISL_495729, EPI_ISL_495730, EPI_ISL_495731, EPI_ISL_495732, EPI_ISL_495733, EPI_ISL_495734, EPI_ISL_495735, EPI_ISL_495736, EPI_ISL_495737, EPI_ISL_495738, EPI_ISL_495739, EPI_ISL_495740, EPI_ISL_495741, EPI_ISL_495742, EPI_ISL_495743, EPI_ISL_495744, EPI_ISL_495745, EPI_ISL_495746, EPI_ISL_495747, EPI_ISL_495748, EPI_ISL_495749, EPI_ISL_495750, EPI_ISL_495751, EPI_ISL_495752, EPI_ISL_495753, EPI_ISL_495754, EPI_ISL_495755, EPI_ISL_495756, EPI_ISL_495758, EPI_ISL_495759, EPI_ISL_495760, EPI_ISL_495761, EPI_ISL_495762, EPI_ISL_495763, EPI_ISL_495764, EPI_ISL_495765, EPI_ISL_495766, EPI_ISL_495768, EPI_ISL_495769, EPI_ISL_495771, EPI_ISL_495772, EPI_ISL_495773, EPI_ISL_495774, EPI_ISL_495775, EPI_ISL_495776, EPI_ISL_495777, EPI_ISL_495778, EPI_ISL_495779, EPI_ISL_495780, EPI_ISL_495781, EPI_ISL_495782, EPI_ISL_495784, EPI_ISL_495785, EPI_ISL_495786, EPI_ISL_495787, EPI_ISL_495788, EPI_ISL_495789, EPI_ISL_495790, EPI_ISL_495791, EPI_ISL_495792, EPI_ISL_495793, EPI_ISL_495794, EPI_ISL_495795, EPI_ISL_495796, EPI_ISL_495797, EPI_ISL_495798, EPI_ISL_495799, EPI_ISL_495800, EPI_ISL_495801, EPI_ISL_495802, EPI_ISL_495803, EPI_ISL_495804, EPI_ISL_495805, EPI_ISL_495806, EPI_ISL_495808, EPI_ISL_495809, EPI_ISL_495810, EPI_ISL_495811, EPI_ISL_495812, EPI_ISL_495813, EPI_ISL_495814, EPI_ISL_495815, EPI_ISL_495816, EPI_ISL_495817, EPI_ISL_495818, EPI_ISL_495819, EPI_ISL_495820, EPI_ISL_495821, EPI_ISL_495822, EPI_ISL_495823, EPI_ISL_495824, EPI_ISL_495825, EPI_ISL_495826, EPI_ISL_495827, EPI_ISL_495828, EPI_ISL_495829, EPI_ISL_495830, EPI_ISL_495831, EPI_ISL_495833, EPI_ISL_495834, EPI_ISL_495835, EPI_ISL_495836, EPI_ISL_495837, EPI_ISL_495838, EPI_ISL_495839, EPI_ISL_495840, EPI_ISL_495841, EPI_ISL_495842, EPI_ISL_495843, EPI_ISL_495844, EPI_ISL_495845, EPI_ISL_495847, EPI_ISL_495848, EPI_ISL_495849, EPI_ISL_495851, EPI_ISL_495852, EPI_ISL_495853, EPI_ISL_495854, EPI_ISL_495855, EPI_ISL_495856, EPI_ISL_495857, EPI_ISL_495858, EPI_ISL_495859, EPI_ISL_495860, EPI_ISL_495861, EPI_ISL_495862, EPI_ISL_495863, EPI_ISL_495864, EPI_ISL_495865, EPI_ISL_495866, EPI_ISL_495867, EPI_ISL_495868, EPI_ISL_495869, EPI_ISL_495874, EPI_ISL_495875, EPI_ISL_495876, EPI_ISL_495877, EPI_ISL_495878, EPI_ISL_495880, EPI_ISL_495881, EPI_ISL_495882, EPI_ISL_495883, EPI_ISL_495884, EPI_ISL_495889, EPI_ISL_495890, EPI_ISL_495891, EPI_ISL_495892, EPI_ISL_495893, EPI_ISL_495894, EPI_ISL_495895, EPI_ISL_495896, EPI_ISL_495897, EPI_ISL_495898, EPI_ISL_495899, EPI_ISL_495901, EPI_ISL_495902, EPI_ISL_495903, EPI_ISL_495904, EPI_ISL_495905, EPI_ISL_495906, EPI_ISL_495907, EPI_ISL_495908, EPI_ISL_495909, EPI_ISL_495911, EPI_ISL_495912, EPI_ISL_495913, EPI_ISL_495914, EPI_ISL_495915, EPI_ISL_495916 | | | |
| see above | Washington State Department of Health | Seattle Flu Study | Deborah A. Nickerson, Chris D. Frazar, Jover Lee, Benjamin Pelle, Matthew Richardson, Amanda Adler, Elisabeth Brandstetter, Peter D. Han, Kairsten Fay, Misja Ilcisin, Kirsten Lacombe, Thomas R. Sibley, Melissa Truong, Caitlin R. Wolf, Romesh Gautom, Geoff |
| EPI_ISL_495918, EPI_ISL_495919, EPI_ISL_495921, EPI_ISL_495922, EPI_ISL_495923, EPI_ISL_495924, EPI_ISL_495925, EPI_ISL_495926, EPI_ISL_495927, EPI_ISL_495928, EPI_ISL_495929, EPI_ISL_495930, EPI_ISL_495931, EPI_ISL_495934, EPI_ISL_495935, EPI_ISL_495936, EPI_ISL_495937, EPI_ISL_495938, EPI_ISL_495940, EPI_ISL_495941, EPI_ISL_495942, EPI_ISL_495943, EPI_ISL_495944, EPI_ISL_495945, EPI_ISL_495946, EPI_ISL_495947, EPI_ISL_495948, EPI_ISL_495949, EPI_ISL_495950, EPI_ISL_495951, EPI_ISL_495952, EPI_ISL_495953, EPI_ISL_495954, EPI_ISL_495955, EPI_ISL_495956, EPI_ISL_495957, EPI_ISL_495958, EPI_ISL_495959, EPI_ISL_495960, EPI_ISL_495961, EPI_ISL_495962, EPI_ISL_495963, EPI_ISL_495964, EPI_ISL_495966, EPI_ISL_495967, EPI_ISL_495968, EPI_ISL_495969, EPI_ISL_495970, EPI_ISL_495971, EPI_ISL_495972, EPI_ISL_495973, EPI_ISL_495974, EPI_ISL_495978, EPI_ISL_495979, EPI_ISL_495980, EPI_ISL_495981, EPI_ISL_495982, EPI_ISL_495983, EPI_ISL_495984, EPI_ISL_495985, EPI_ISL_495986, EPI_ISL_495987, EPI_ISL_495989, EPI_ISL_495991, EPI_ISL_495992, EPI_ISL_495995, EPI_ISL_495996, EPI_ISL_495997, EPI_ISL_495998, EPI_ISL_495999, EPI_ISL_496000, EPI_ISL_496001, EPI_ISL_496002, EPI_ISL_496003, EPI_ISL_496004, EPI_ISL_496005, EPI_ISL_496006, EPI_ISL_496007, EPI_ISL_496008, EPI_ISL_496009, EPI_ISL_496011, EPI_ISL_496012, EPI_ISL_496013, EPI_ISL_496016, EPI_ISL_496018, EPI_ISL_496019, EPI_ISL_496021, EPI_ISL_496022, EPI_ISL_496023, EPI_ISL_496024, EPI_ISL_496025, EPI_ISL_496026, EPI_ISL_496027, EPI_ISL_496028, EPI_ISL_496029, EPI_ISL_496030, EPI_ISL_496031, EPI_ISL_496032, EPI_ISL_496034, EPI_ISL_496035, EPI_ISL_496036, EPI_ISL_496037, EPI_ISL_496038, EPI_ISL_496039, EPI_ISL_496040, EPI_ISL_496041, EPI_ISL_496042, EPI_ISL_496043, EPI_ISL_496044, EPI_ISL_496045, EPI_ISL_496046, EPI_ISL_496047, EPI_ISL_496048, EPI_ISL_496049, EPI_ISL_496050, EPI_ISL_496051, EPI_ISL_496052, EPI_ISL_496053, EPI_ISL_496054, EPI_ISL_496055, EPI_ISL_496056, EPI_ISL_496057, EPI_ISL_496058, EPI_ISL_496059, EPI_ISL_496060, EPI_ISL_496061, EPI_ISL_496062, EPI_ISL_496063, EPI_ISL_496064, EPI_ISL_496065, EPI_ISL_496066, EPI_ISL_496067, EPI_ISL_496068, EPI_ISL_496069, EPI_ISL_496070, EPI_ISL_496071, EPI_ISL_496073, EPI_ISL_496074, EPI_ISL_496076, EPI_ISL_496077, EPI_ISL_496078, EPI_ISL_496079, EPI_ISL_496080, EPI_ISL_496081, EPI_ISL_496082, EPI_ISL_496083, EPI_ISL_496084, EPI_ISL_496085, EPI_ISL_496086, EPI_ISL_496087, EPI_ISL_496088, EPI_ISL_496089, EPI_ISL_496090, EPI_ISL_496091, EPI_ISL_496092, EPI_ISL_496096, EPI_ISL_496097, EPI_ISL_496098, EPI_ISL_496099, EPI_ISL_496100, EPI_ISL_496101, EPI_ISL_496102, EPI_ISL_496103, EPI_ISL_496104, EPI_ISL_496105, EPI_ISL_496106, EPI_ISL_496107, EPI_ISL_496108, EPI_ISL_496110, EPI_ISL_496111, EPI_ISL_496112, EPI_ISL_496113, EPI_ISL_496114, EPI_ISL_496115, EPI_ISL_496116, EPI_ISL_496117, EPI_ISL_496118, EPI_ISL_496119, EPI_ISL_496120, EPI_ISL_496121, EPI_ISL_496122, EPI_ISL_496123, EPI_ISL_496124, EPI_ISL_496125, EPI_ISL_496126, EPI_ISL_496127, EPI_ISL_496128, EPI_ISL_496129, EPI_ISL_496130, EPI_ISL_496132, EPI_ISL_496133, EPI_ISL_496134, EPI_ISL_496135, EPI_ISL_496136, EPI_ISL_496137, EPI_ISL_496138, EPI_ISL_496139, EPI_ISL_496140, EPI_ISL_496141, EPI_ISL_496142, EPI_ISL_496143, EPI_ISL_496144, EPI_ISL_496145, EPI_ISL_496146, EPI_ISL_496147, EPI_ISL_496148, EPI_ISL_496150, EPI_ISL_496151, EPI_ISL_496153, EPI_ISL_496154, EPI_ISL_496155, EPI_ISL_496156, EPI_ISL_496157, EPI_ISL_496158, EPI_ISL_496159, EPI_ISL_496160, EPI_ISL_496161, EPI_ISL_496163, EPI_ISL_496164, EPI_ISL_496165, EPI_ISL_496166, EPI_ISL_496167, EPI_ISL_496168, EPI_ISL_496169, EPI_ISL_496173, EPI_ISL_496174, EPI_ISL_496175, EPI_ISL_496176, EPI_ISL_496178, EPI_ISL_496179, EPI_ISL_496180, EPI_ISL_496182, EPI_ISL_496183, EPI_ISL_496184, EPI_ISL_496185, EPI_ISL_496186, EPI_ISL_496187, EPI_ISL_496188, EPI_ISL_496190, EPI_ISL_496191, EPI_ISL_496192, EPI_ISL_496193, EPI_ISL_496194, EPI_ISL_496195, EPI_ISL_496196, EPI_ISL_496197, EPI_ISL_496199, EPI_ISL_496200, EPI_ISL_496201, EPI_ISL_496202, EPI_ISL_496203, EPI_ISL_496204, EPI_ISL_496205, EPI_ISL_496206, EPI_ISL_496207, EPI_ISL_496208, EPI_ISL_496209, EPI_ISL_496210, EPI_ISL_496211, EPI_ISL_496212, EPI_ISL_496213, EPI_ISL_496214, EPI_ISL_496215, EPI_ISL_496216, EPI_ISL_496217, EPI_ISL_496218, EPI_ISL_496219, EPI_ISL_496220, EPI_ISL_496221, EPI_ISL_496222, EPI_ISL_496223, EPI_ISL_496224, EPI_ISL_496225, EPI_ISL_496226, EPI_ISL_496227, EPI_ISL_496228, EPI_ISL_496229, EPI_ISL_496230, EPI_ISL_496231, EPI_ISL_496232, EPI_ISL_496233, EPI_ISL_496234, EPI_ISL_496235, EPI_ISL_496236, EPI_ISL_496237, EPI_ISL_496239, EPI_ISL_496240, EPI_ISL_496241, EPI_ISL_496242, EPI_ISL_496243, EPI_ISL_496244, EPI_ISL_496245, EPI_ISL_496246, EPI_ISL_496248, EPI_ISL_496249, EPI_ISL_496250, EPI_ISL_496252, EPI_ISL_496253, EPI_ISL_496255, EPI_ISL_496256, EPI_ISL_496257, EPI_ISL_496258, EPI_ISL_496259, EPI_ISL_496260, EPI_ISL_496261, EPI_ISL_496262, EPI_ISL_496263, EPI_ISL_496264, EPI_ISL_496265, EPI_ISL_496266, EPI_ISL_496267, EPI_ISL_496268, EPI_ISL_496269, EPI_ISL_496270, EPI_ISL_496271, EPI_ISL_496272, EPI_ISL_496273, EPI_ISL_496274, EPI_ISL_496275, EPI_ISL_496276, EPI_ISL_496281, EPI_ISL_496282, EPI_ISL_496283, EPI_ISL_496285, EPI_ISL_496286, EPI_ISL_496287, EPI_ISL_496288, EPI_ISL_496289, EPI_ISL_496290, EPI_ISL_496291, EPI_ISL_496292, EPI_ISL_496294, EPI_ISL_496295, EPI_ISL_496296, EPI_ISL_496297, EPI_ISL_496298, EPI_ISL_496299, EPI_ISL_496300, EPI_ISL_496301, EPI_ISL_496302, EPI_ISL_496303, EPI_ISL_496304, EPI_ISL_496305, EPI_ISL_496306, EPI_ISL_496307, EPI_ISL_496308, EPI_ISL_496309, EPI_ISL_496310, EPI_ISL_496311, EPI_ISL_496312, EPI_ISL_496313, EPI_ISL_496314, EPI_ISL_496315, EPI_ISL_496316, EPI_ISL_496317, EPI_ISL_496318, EPI_ISL_496319, EPI_ISL_496320, EPI_ISL_496321, EPI_ISL_496322, EPI_ISL_496323, EPI_ISL_496324, EPI_ISL_496325, EPI_ISL_496326, EPI_ISL_496327, EPI_ISL_496328, EPI_ISL_496330, EPI_ISL_496331, EPI_ISL_496332, EPI_ISL_496333, EPI_ISL_496334, EPI_ISL_496335, EPI_ISL_496336, EPI_ISL_496337, EPI_ISL_496338 | | | |
| see above | Washington State Department of Health | Seattle Flu Study | Deborah A. Nickerson, Chris D. Frazar, Jover Lee, Benjamin Pelle, Matthew Richardson, Amanda Adler, Elisabeth Brandstetter, Peter D. Han, Kairsten Fay, Misja Ilcisin, Kirsten Lacombe, Thomas R. Sibley, Melissa Truong, Caitlin R. Wolf, Romesh Gautom, Geoff Melly, Brian Hiatt, Philip Dykema, Scott Lindquist, Michael Boeckh, Janet A. Englund, Michael Famulare, Barry R. Lutz, Mark J. Rieder, Lea M. Starita, Matthew Thompson, Helen Y. Chu, Jay Shendure, Trevor Bedford |
| EPI_ISL_496518, EPI_ISL_496519, EPI_ISL_496520 | Armed Forces Medical College | National Centre For Cell Science | Dhiraj Paul, Kunal Jani, Radha Chauhan, Janesh Kumar, Vasudevan Seshadri, Girdhari Lal, Rajesh Karyakarte, Suvarna Joshi, Murlidhar Tambe, Sourav Sen, Santosh Karade, Kavita Bala Anand, Shelinder Pal Singh Shergill, Rajiv Mohan Gupta, Manoj Kumar Bhat, Arvind Sahu, Maharashtra COVID-19 Study Group, DBT's PAN-INDIA 1000 SARS-CoV2 RNA genome sequencing consortium, Yogesh S Shouche |

| | | | |
|---|---|---|---|
| EPI_ISL_496521, EPI_ISL_496522, EPI_ISL_496523 | B.J. Govt. Medical College | National Centre For Cell Science | Dhiraj Paul, Kunal Jani, Radha Chauhan, Janesh Kumar, Vasudevan Seshadri, Girdhari Lal, Rajesh Karyakarte, Suvarna Joshi, Murlidhar Tambe, Sourav Sen, Santosh Karade, Kavita Bala Anand, Shelinder Pal Singh Shergill, Rajiv Mohan Gupta, Manoj Kumar Bhat, Arvind Sahu, Maharashtra COVID-19 Study Group, DBT's PAN-INDIA 1000 SARS-CoV2 RNA genome sequencing consortium, Yogesh S Shouche |
| EPI_ISL_496524, EPI_ISL_496525, EPI_ISL_496526 | National Centre For Cell Science | National Centre For Cell Science | Dhiraj Paul, Kunal Jani, Radha Chauhan, Janesh Kumar, Vasudevan Seshadri, Girdhari Lal, Rajesh Karyakarte, Suvarna Joshi, Murlidhar Tambe, Sourav Sen, Santosh Karade, Kavita Bala Anand, Shelinder Pal Singh Shergill, Rajiv Mohan Gupta, Manoj Kumar Bhat, Arvind Sahu, Maharashtra COVID-19 Study Group, DBT's PAN-INDIA 1000 SARS-CoV2 RNA genome sequencing consortium, Yogesh S Shouche |
| EPI_ISL_496527, EPI_ISL_496528 | B.J. Govt. Medical College | National Centre For Cell Science | Dhiraj Paul, Kunal Jani, Radha Chauhan, Janesh Kumar, Vasudevan Seshadri, Girdhari Lal, Rajesh Karyakarte, Suvarna Joshi, Murlidhar Tambe, Sourav Sen, Santosh Karade, Kavita Bala Anand, Shelinder Pal Singh Shergill, Rajiv Mohan Gupta, Manoj Kumar Bhat, Arvind Sahu, Maharashtra COVID-19 Study Group, DBT's PAN-INDIA 1000 SARS-CoV2 RNA genome sequencing consortium, Yogesh S Shouche |
| EPI_ISL_496529 | Armed Forces Medical College | National Centre For Cell Science | Dhiraj Paul, Kunal Jani, Radha Chauhan, Janesh Kumar, Vasudevan Seshadri, Girdhari Lal, Rajesh Karyakarte, Suvarna Joshi, Murlidhar Tambe, Sourav Sen, Santosh Karade, Kavita Bala Anand, Shelinder Pal Singh Shergill, Rajiv Mohan Gupta, Manoj Kumar Bhat, Arvind Sahu, Maharashtra COVID-19 Study Group, DBT's PAN-INDIA 1000 SARS-CoV2 RNA genome sequencing consortium, Yogesh S Shouche |
| EPI_ISL_496530 | B.J. Govt. Medical College | National Centre For Cell Science | Dhiraj Paul, Kunal Jani, Radha Chauhan, Janesh Kumar, Vasudevan Seshadri, Girdhari Lal, Rajesh Karyakarte, Suvarna Joshi, Murlidhar Tambe, Sourav Sen, Santosh Karade, Kavita Bala Anand, Shelinder Pal Singh Shergill, Rajiv Mohan Gupta, Manoj Kumar Bhat, Arvind Sahu, Maharashtra COVID-19 Study Group, DBT's PAN-INDIA 1000 SARS-CoV2 RNA genome sequencing consortium, Yogesh S Shouche |
| EPI_ISL_496531, EPI_ISL_496532 | National Centre For Cell Science | National Centre For Cell Science | Dhiraj Paul, Kunal Jani, Radha Chauhan, Janesh Kumar, Vasudevan Seshadri, Girdhari Lal, Rajesh Karyakarte, Suvarna Joshi, Murlidhar Tambe, Sourav Sen, Santosh Karade, Kavita Bala Anand, Shelinder Pal Singh Shergill, Rajiv Mohan Gupta, Manoj Kumar Bhat, Arvind Sahu, Maharashtra COVID-19 Study Group, DBT's PAN-INDIA 1000 SARS-CoV2 RNA genome sequencing consortium, Yogesh S Shouche |
| EPI_ISL_496533 | Armed Forces Medical College | National Centre For Cell Science | Dhiraj Paul, Kunal Jani, Radha Chauhan, Janesh Kumar, Vasudevan Seshadri, Girdhari Lal, Rajesh Karyakarte, Suvarna Joshi, Murlidhar Tambe, Sourav Sen, Santosh Karade, Kavita Bala Anand, Shelinder Pal Singh Shergill, Rajiv Mohan Gupta, Manoj Kumar Bhat, Arvind Sahu, Maharashtra COVID-19 Study Group, DBT's PAN-INDIA 1000 SARS-CoV2 RNA genome sequencing consortium, Yogesh S Shouche |
| EPI_ISL_496534 | B.J. Govt. Medical College | National Centre For Cell Science | Dhiraj Paul, Kunal Jani, Radha Chauhan, Janesh Kumar, Vasudevan Seshadri, Girdhari Lal, Rajesh Karyakarte, Suvarna Joshi, Murlidhar Tambe, Sourav Sen, Santosh Karade, Kavita Bala Anand, Shelinder Pal Singh Shergill, Rajiv Mohan Gupta, Manoj Kumar Bhat, Arvind Sahu, Maharashtra COVID-19 Study Group, DBT's PAN-INDIA 1000 SARS-CoV2 RNA genome sequencing consortium, Yogesh S Shouche |
| EPI_ISL_496535, EPI_ISL_496536 | National Centre For Cell Science | National Centre For Cell Science | Dhiraj Paul, Kunal Jani, Radha Chauhan, Janesh Kumar, Vasudevan Seshadri, Girdhari Lal, Rajesh Karyakarte, Suvarna Joshi, Murlidhar Tambe, Sourav Sen, Santosh Karade, Kavita Bala Anand, Shelinder Pal Singh Shergill, Rajiv Mohan Gupta, Manoj Kumar Bhat, Arvind Sahu, Maharashtra COVID-19 Study Group, DBT's PAN-INDIA 1000 SARS-CoV2 RNA genome sequencing consortium, Yogesh S Shouche |
| EPI_ISL_496537, EPI_ISL_496538, EPI_ISL_496539, EPI_ISL_496540, EPI_ISL_496541, EPI_ISL_496542, EPI_ISL_496543, EPI_ISL_496544, EPI_ISL_496545 | Armed Forces Medical College | National Centre For Cell Science | Dhiraj Paul, Kunal Jani, Radha Chauhan, Janesh Kumar, Vasudevan Seshadri, Girdhari Lal, Rajesh Karyakarte, Suvarna Joshi, Murlidhar Tambe, Sourav Sen, Santosh Karade, Kavita Bala Anand, Shelinder Pal Singh Shergill, Rajiv Mohan Gupta, Manoj Kumar Bhat, Arvind Sahu, Maharashtra COVID-19 Study Group, DBT's PAN-INDIA 1000 SARS-CoV2 RNA genome sequencing consortium, Yogesh S Shouche |
| EPI_ISL_496546, EPI_ISL_496547, EPI_ISL_496548, EPI_ISL_496549, EPI_ISL_496550, EPI_ISL_496551, EPI_ISL_496552, EPI_ISL_496553, EPI_ISL_496554 | B.J. Govt. Medical College | National Centre For Cell Science | Dhiraj Paul, Kunal Jani, Radha Chauhan, Janesh Kumar, Vasudevan Seshadri, Girdhari Lal, Rajesh Karyakarte, Suvarna Joshi, Murlidhar Tambe, Sourav Sen, Santosh Karade, Kavita Bala Anand, Shelinder Pal Singh Shergill, Rajiv Mohan Gupta, Manoj Kumar Bhat, Arvind Sahu, Maharashtra COVID-19 Study Group, DBT's PAN-INDIA 1000 SARS-CoV2 RNA genome sequencing consortium, Yogesh S Shouche |
| EPI_ISL_496555, EPI_ISL_496556, EPI_ISL_496557, EPI_ISL_496558, EPI_ISL_496559, EPI_ISL_496560, EPI_ISL_496561, EPI_ISL_496562, EPI_ISL_496563, EPI_ISL_496564, EPI_ISL_496565, EPI_ISL_496566, EPI_ISL_496567, EPI_ISL_496568, EPI_ISL_496569, EPI_ISL_496570, EPI_ISL_496571, EPI_ISL_496572, EPI_ISL_496573, EPI_ISL_496574, EPI_ISL_496575, EPI_ISL_496576, EPI_ISL_496577, EPI_ISL_496578, EPI_ISL_496579, EPI_ISL_496580, EPI_ISL_496581, EPI_ISL_496582, EPI_ISL_496583, EPI_ISL_496584, EPI_ISL_496585, EPI_ISL_496586, EPI_ISL_496587 | | | |
| see above | National Centre For Cell Science | National Centre For Cell Science | Dhiraj Paul, Kunal Jani, Radha Chauhan, Janesh Kumar, Vasudevan Seshadri, Girdhari Lal, Rajesh Karyakarte, Suvarna Joshi, Murlidhar Tambe, Sourav Sen, Santosh Karade, Kavita Bala Anand, Shelinder Pal Singh Shergill, Rajiv Mohan Gupta, Manoj Kumar Bhat, Arvind Sahu, Maharashtra COVID-19 Study Group, DBT's PAN-INDIA 1000 SARS-CoV2 RNA genome sequencing consortium, Yogesh S Shouche |
| EPI_ISL_496602 | Armed Forces Medical College | National Centre For Cell Science | Dhiraj Paul, Kunal Jani, Radha Chauhan, Janesh Kumar, Vasudevan Seshadri, Girdhari Lal, Rajesh Karyakarte, Suvarna Joshi, Murlidhar Tambe, Sourav Sen, Santosh Karade, Kavita Bala Anand, Shelinder Pal Singh Shergill, Rajiv Mohan Gupta, Manoj Kumar Bhat, Arvind Sahu, Maharashtra COVID-19 Study Group, DBT's PAN-INDIA 1000 SARS-CoV2 RNA genome sequencing consortium, Yogesh S Shouche |
| EPI_ISL_497342, EPI_ISL_497343, EPI_ISL_497344, EPI_ISL_497345, EPI_ISL_497346, EPI_ISL_497347, EPI_ISL_497348, EPI_ISL_497349, EPI_ISL_497350, EPI_ISL_497351, EPI_ISL_497352, EPI_ISL_497353, EPI_ISL_497354, EPI_ISL_497356, EPI_ISL_497357, EPI_ISL_497358, EPI_ISL_497359, EPI_ISL_497360, EPI_ISL_497361, EPI_ISL_497362, EPI_ISL_497363, EPI_ISL_497364, EPI_ISL_497365, EPI_ISL_497366, EPI_ISL_497367, EPI_ISL_497368, EPI_ISL_497369, EPI_ISL_497370, EPI_ISL_497371, EPI_ISL_497372, EPI_ISL_497373, EPI_ISL_497374, EPI_ISL_497375, EPI_ISL_497376, EPI_ISL_497377, EPI_ISL_497378, EPI_ISL_497379, EPI_ISL_497380, EPI_ISL_497381, EPI_ISL_497382, EPI_ISL_497383, EPI_ISL_497384, EPI_ISL_497385, EPI_ISL_497386, EPI_ISL_497387, EPI_ISL_497388, EPI_ISL_497389, EPI_ISL_497390, EPI_ISL_497391, EPI_ISL_497392, EPI_ISL_497393, EPI_ISL_497394, EPI_ISL_497395, EPI_ISL_497396, EPI_ISL_497397, EPI_ISL_497398, EPI_ISL_497399, EPI_ISL_497400, EPI_ISL_497401, EPI_ISL_497402, EPI_ISL_497403, EPI_ISL_497404, EPI_ISL_497405, EPI_ISL_497406, EPI_ISL_497407, EPI_ISL_497408, EPI_ISL_497409, EPI_ISL_497410, EPI_ISL_497411, EPI_ISL_497412, EPI_ISL_497413, EPI_ISL_497414, EPI_ISL_497415, EPI_ISL_497416, EPI_ISL_497417, EPI_ISL_497418, EPI_ISL_497420, EPI_ISL_497421, EPI_ISL_497422, EPI_ISL_497423, EPI_ISL_497424, EPI_ISL_497425, EPI_ISL_497426, EPI_ISL_497427, EPI_ISL_497428, EPI_ISL_497429, EPI_ISL_497430, EPI_ISL_497431, EPI_ISL_497432, EPI_ISL_497433, EPI_ISL_497434, EPI_ISL_497435, EPI_ISL_497436, EPI_ISL_497437, EPI_ISL_497438, EPI_ISL_497440, EPI_ISL_497441, EPI_ISL_497442, EPI_ISL_497443, EPI_ISL_497444, EPI_ISL_497445, EPI_ISL_497446, EPI_ISL_497447, EPI_ISL_497448, EPI_ISL_497449, EPI_ISL_497450, EPI_ISL_497451, EPI_ISL_497453, EPI_ISL_497454, EPI_ISL_497455, EPI_ISL_497456, EPI_ISL_497457, EPI_ISL_497458, EPI_ISL_497459, EPI_ISL_497460, EPI_ISL_497461, EPI_ISL_497462, EPI_ISL_497463, EPI_ISL_497464, EPI_ISL_497465, EPI_ISL_497466, EPI_ISL_497467, EPI_ISL_497468, EPI_ISL_497469, EPI_ISL_497470, EPI_ISL_497471, EPI_ISL_497472, EPI_ISL_497473, EPI_ISL_497474, EPI_ISL_497475, EPI_ISL_497476, EPI_ISL_497478, EPI_ISL_497479, EPI_ISL_497480, EPI_ISL_497481, EPI_ISL_497482, EPI_ISL_497483, EPI_ISL_497485, EPI_ISL_497486, EPI_ISL_497487, EPI_ISL_497488, EPI_ISL_497489, EPI_ISL_497490, EPI_ISL_497491, EPI_ISL_497492, EPI_ISL_497493, EPI_ISL_497494, EPI_ISL_497495, EPI_ISL_497496, EPI_ISL_497497, EPI_ISL_497498, EPI_ISL_497499, EPI_ISL_497500, EPI_ISL_497501, EPI_ISL_497502, EPI_ISL_497503, EPI_ISL_497504, EPI_ISL_497505, EPI_ISL_497506, EPI_ISL_497507, EPI_ISL_497508, EPI_ISL_497509, EPI_ISL_497510, EPI_ISL_497511, EPI_ISL_497512, EPI_ISL_497513, EPI_ISL_497514, EPI_ISL_497515, EPI_ISL_497516, EPI_ISL_497517, EPI_ISL_497518, EPI_ISL_497519, EPI_ISL_497520, EPI_ISL_497521, EPI_ISL_497522, EPI_ISL_497523, EPI_ISL_497524, EPI_ISL_497525, EPI_ISL_497526, EPI_ISL_497527, EPI_ISL_497528, EPI_ISL_497529, EPI_ISL_497530, EPI_ISL_497531, EPI_ISL_497532, EPI_ISL_497533, EPI_ISL_497534, EPI_ISL_497535, EPI_ISL_497536, EPI_ISL_497537, EPI_ISL_497538, EPI_ISL_497539, EPI_ISL_497540, EPI_ISL_497541, EPI_ISL_497542, EPI_ISL_497543, EPI_ISL_497544, EPI_ISL_497545, EPI_ISL_497546, EPI_ISL_497547, EPI_ISL_497548, EPI_ISL_497549, EPI_ISL_497550, EPI_ISL_497551, EPI_ISL_497552, EPI_ISL_497553, EPI_ISL_497554, EPI_ISL_497555, EPI_ISL_497556, EPI_ISL_497557, EPI_ISL_497558, EPI_ISL_497559, EPI_ISL_497560, EPI_ISL_497561, EPI_ISL_497562, EPI_ISL_497563, EPI_ISL_497564, EPI_ISL_497565, EPI_ISL_497567, EPI_ISL_497569 | | | |
| see above | Washington State Department of Health | Seattle Flu Study | Deborah A. Nickerson, Chris D. Frazar, Jover Lee, Benjamin Pelle, Matthew Richardson, Amanda Adler, Elisabeth Brandstetter, Peter D. Han, Kairsten Fay, Misja Ilcisin, Kirsten Lacombe, Thomas R. Sibley, Melissa Truong, Caitlin R. Wolf, Romesh Gautom, Geoff Melly, Brian Hiatt, Philip Dykema, Scott Lindquist, Michael Boeckh, Janet A. Englund, Michael Famulare, Barry R. Lutz, Mark J. Rieder, Lea M. Starita, Matthew Thompson, Helen Y. Chu, Jay Shendure, Trevor Bedford |
| EPI_ISL_497873, EPI_ISL_497874, EPI_ISL_497875, EPI_ISL_497876, EPI_ISL_497877, EPI_ISL_497878, EPI_ISL_497879 | National Centre For Cell Science | National Centre For Cell Science | Dhiraj Paul, Kunal Jani, Radha Chauhan, Janesh Kumar, Vasudevan Seshadri, Girdhari Lal, Rajesh Karyakarte, Suvarna Joshi, Murlidhar Tambe, Sourav Sen, Santosh Karade, Kavita Bala Anand, Shelinder Pal Singh Shergill, Rajiv Mohan Gupta, Manoj Kumar Bhat, Arvind Sahu, Maharashtra COVID-19 Study Group, DBT's PAN-INDIA 1000 SARS-CoV2 RNA genome sequencing consortium, Yogesh S Shouche |
| EPI_ISL_497880, EPI_ISL_497881, EPI_ISL_497882, EPI_ISL_497883, EPI_ISL_497884, EPI_ISL_497885, EPI_ISL_497886, EPI_ISL_497887 | Armed Forces Medical College | National Centre For Cell Science | Dhiraj Paul, Kunal Jani, Radha Chauhan, Janesh Kumar, Vasudevan Seshadri, Girdhari Lal, Rajesh Karyakarte, Suvarna Joshi, Murlidhar Tambe, Sourav Sen, Santosh Karade, Kavita Bala Anand, Shelinder Pal Singh Shergill, Rajiv Mohan Gupta, Manoj Kumar Bhat, Arvind Sahu, Maharashtra COVID-19 Study Group, DBT's PAN-INDIA 1000 SARS-CoV2 RNA genome sequencing consortium, Yogesh S Shouche |
| EPI_ISL_497888, EPI_ISL_497889, EPI_ISL_497890, EPI_ISL_497891 | B.J. Govt. Medical College | National Centre For Cell Science | Dhiraj Paul, Kunal Jani, Radha Chauhan, Janesh Kumar, Vasudevan Seshadri, Girdhari Lal, Rajesh Karyakarte, Suvarna Joshi, Murlidhar Tambe, Sourav Sen, Santosh Karade, Kavita Bala Anand, Shelinder Pal Singh Shergill, Rajiv Mohan Gupta, Manoj Kumar Bhat, Arvind Sahu, Maharashtra COVID-19 Study Group, DBT's PAN-INDIA 1000 SARS-CoV2 RNA genome sequencing consortium, Yogesh S Shouche |
| EPI_ISL_498054, EPI_ISL_498055, EPI_ISL_498056, EPI_ISL_498057, EPI_ISL_498058, EPI_ISL_498059, EPI_ISL_498060, EPI_ISL_498061, EPI_ISL_498062, EPI_ISL_498063, EPI_ISL_498064, EPI_ISL_498065, EPI_ISL_498066, EPI_ISL_498067, EPI_ISL_498068, EPI_ISL_498069, EPI_ISL_498070, EPI_ISL_498071, EPI_ISL_498072, EPI_ISL_498073, EPI_ISL_498074, EPI_ISL_498075, EPI_ISL_498077, EPI_ISL_498078, EPI_ISL_498079, EPI_ISL_498080, EPI_ISL_498081, EPI_ISL_498082, EPI_ISL_498083, EPI_ISL_498084, EPI_ISL_498085, EPI_ISL_498086, EPI_ISL_498087, EPI_ISL_498088, EPI_ISL_498089, EPI_ISL_498090, EPI_ISL_498091, EPI_ISL_498092, EPI_ISL_498093, EPI_ISL_498094, EPI_ISL_498095, EPI_ISL_498096, EPI_ISL_498097, EPI_ISL_498098, EPI_ISL_498099, EPI_ISL_498100, EPI_ISL_498101, EPI_ISL_498102, EPI_ISL_498103, EPI_ISL_498104, EPI_ISL_498105, EPI_ISL_498106, EPI_ISL_498107, EPI_ISL_498108, EPI_ISL_498109, EPI_ISL_498110, EPI_ISL_498111, EPI_ISL_498112, EPI_ISL_498113, EPI_ISL_498114, EPI_ISL_498115, EPI_ISL_498116, EPI_ISL_498117, EPI_ISL_498118, EPI_ISL_498119, EPI_ISL_498120, EPI_ISL_498121, EPI_ISL_498122, EPI_ISL_498123, EPI_ISL_498124, EPI_ISL_498125, EPI_ISL_498126 | | | |

| | | | |
|---|---|---|---|
| see above | NHLS-IALCH | KRISP, KZN Research Innovation and Sequencing Platform | Giandhari J, Pillay S, Lessells R, Chimukangara B, Mdlalose K, York D, Khan S, Tegally H, Wilkinson E, de Oliveira T |
| EPI_ISL_498255, EPI_ISL_498256, EPI_ISL_498257, EPI_ISL_498258, EPI_ISL_498259, EPI_ISL_498260, EPI_ISL_498261 | Hospital for Tropical Diseases | COVID-19 Network Investigations (CONI) Alliance | Elizabeth Batty, Nantarat Chantawat, Wasun Chantratita, Thanat Chookajorn, Stefan Fernandez, Angkana Huang, Weena Janwitthayanan, Akanitt Jittmittraphap, Anthony R. Jones, Khajohn Joonsalak, Chonticha Klungthong, Theerarat Kochakarn, Namfon Kotanan, Krittikorn Kumpornsin, Pornsawan Leaungwutiwong, Wudtichai Manasatienkij, Bhakbhoom Panthan, Ekawat Pasomsub, Kingkan Rakmanee, Insee Sensorn, Janjira Thaipadungpanit, Arporn Wangwiwatsin,Treewat Watthanachockchai |
| EPI_ISL_498262, EPI_ISL_498263, EPI_ISL_498264, EPI_ISL_498265, EPI_ISL_498266 | Ramathibodi Hospital | COVID-19 Network Investigations (CONI) Alliance | Elizabeth Batty, Wasun Chantratita, Thanat Chookajorn, Stefan Fernandez, Angkana Huang, Anthony R. Jones, Khajohn Joonsalak, Chonticha Klungtong, Theerarat Kochakarn, Namfon Kotanan, Krittikorn Kumpornsin, Wudtichai Manasatienkij, Bhakbhoom Panthan, Ekawat Pasomsub, Kingkan Rakmanee, Insee Sensorn, Janjira Thaipadungpanit, Arporn Wangwiwatsin,Treewat Watthanachockchai |
| EPI_ISL_499266, EPI_ISL_499267, EPI_ISL_499268, EPI_ISL_499269, EPI_ISL_499808, EPI_ISL_499809 | Queens Medical Centre, Clinical Microbiology Department / DeepSeq Nottingham | COVID-19 Genomics UK (COG-UK) Consortium | Gemma Clark, Wendy Smith, Manjinder Khakh, Vicki M Fleming, Michelle M Lister, Hannah Howson-Wells, Jonathan Ball, Patrick McClure, Joseph Chappell, Theocharis Tsoleridis, Nadine Holmes, Matthew Carlisle, Christopher Moore, Fei Sang, Johnny Debebe, Victoria Wright, Matthew Loose |
| EPI_ISL_500157, EPI_ISL_500158, EPI_ISL_500159, EPI_ISL_500166, EPI_ISL_500207, EPI_ISL_500218 | Complejo Hospitalario Universitario de Albacete | SeqCOVID-SPAIN consortium/IBV(CSIC) | Encarnacion Simarro Córdoba, Julia Lozano Serra, Lorena Robles Fonseca , Monica Parra Grandes, Caridad Sainz de Baranda Camino and SeqCOVID-SPAIN consortium |
| EPI_ISL_504185 | Discovery DNA | Discovery DNA | Dustin Hittel, Marina Kerr, Leo Dimnik, Desmond Koo, Alice Li, Aneal Khan |
| EPI_ISL_505003 | Biology Dpt | Microbiology and Infections Diseases | Emmanuelle Billon-Denis, Audrey Ferrier-Rembert, Annabelle Garnier, Laurence Cheutin, Clarisse Vigne, Emilie Tessier, Jessica Denis, Olivier Gorgé, Flora Nolent, Isabelle Drouet, Noémie Verguet, Olivier Ferraris, Jean-Nicolas Tournier |
| EPI_ISL_506041 | Biology Dpt, HIA Percy | Microbiology and Infectious Diseases Dpt | Emmanuelle Billon-Denis, Audrey Ferrier-Rembert, Annabelle Garnier, Laurence Cheutin, Clarisse Vigne, Emilie Tessier, Jessica Denis, Olivier Gorgé, Flora Nolent, Isabelle Drouet, Noémie Verguet, Olivier Ferraris, Jean-Nicolas Tournier |
| EPI_ISL_509223, EPI_ISL_509224, EPI_ISL_509225, EPI_ISL_509226, EPI_ISL_509227, EPI_ISL_509228, EPI_ISL_509229, EPI_ISL_509230, EPI_ISL_509231, EPI_ISL_509232, EPI_ISL_509233, EPI_ISL_509234, EPI_ISL_509235, EPI_ISL_509236, EPI_ISL_509237, EPI_ISL_509238, EPI_ISL_509239, EPI_ISL_509240, EPI_ISL_509241, EPI_ISL_509242, EPI_ISL_509243, EPI_ISL_509244, EPI_ISL_509245, EPI_ISL_509246, EPI_ISL_509247, EPI_ISL_509248, EPI_ISL_509249, EPI_ISL_509250, EPI_ISL_509251, EPI_ISL_509252, EPI_ISL_509253, EPI_ISL_509254, EPI_ISL_509255, EPI_ISL_509256, EPI_ISL_509257, EPI_ISL_509258, EPI_ISL_509259, EPI_ISL_509260, EPI_ISL_509261, EPI_ISL_509262, EPI_ISL_509263, EPI_ISL_509264, EPI_ISL_509265, EPI_ISL_509266, EPI_ISL_509267, EPI_ISL_509268, EPI_ISL_509269, EPI_ISL_509270, EPI_ISL_509271, EPI_ISL_509272, EPI_ISL_509273, EPI_ISL_509274, EPI_ISL_509275, EPI_ISL_509276, EPI_ISL_509277, EPI_ISL_509278, EPI_ISL_509279, EPI_ISL_509280, EPI_ISL_509281, EPI_ISL_509282, EPI_ISL_509283, EPI_ISL_509284, EPI_ISL_509285, EPI_ISL_509286, EPI_ISL_509287, EPI_ISL_509288, EPI_ISL_509289, EPI_ISL_509290, EPI_ISL_509291, EPI_ISL_509292, EPI_ISL_509293, EPI_ISL_509294, EPI_ISL_509295, EPI_ISL_509296, EPI_ISL_509297, EPI_ISL_509298, EPI_ISL_509299, EPI_ISL_509300, EPI_ISL_509301, EPI_ISL_509302, EPI_ISL_509303, EPI_ISL_509304, EPI_ISL_509305, EPI_ISL_509306, EPI_ISL_509307, EPI_ISL_509308, EPI_ISL_509309, EPI_ISL_509310, EPI_ISL_509311, EPI_ISL_509312, EPI_ISL_509313, EPI_ISL_509314, EPI_ISL_509315, EPI_ISL_509316, EPI_ISL_509317, EPI_ISL_509318, EPI_ISL_509319, EPI_ISL_509320, EPI_ISL_509321, EPI_ISL_509322, EPI_ISL_509323, EPI_ISL_509324, EPI_ISL_509325, EPI_ISL_509326, EPI_ISL_509327, EPI_ISL_509328, EPI_ISL_509329, EPI_ISL_509330, EPI_ISL_509331, EPI_ISL_509332, EPI_ISL_509333, EPI_ISL_509334, EPI_ISL_509335, EPI_ISL_509336, EPI_ISL_509337, EPI_ISL_509338, EPI_ISL_509339, EPI_ISL_509340, EPI_ISL_509341, EPI_ISL_509342, EPI_ISL_509343, EPI_ISL_509344, EPI_ISL_509345, EPI_ISL_509346, EPI_ISL_509347, EPI_ISL_509348, EPI_ISL_509349, EPI_ISL_509350, EPI_ISL_509351, EPI_ISL_509352, EPI_ISL_509353, EPI_ISL_509354, EPI_ISL_509355, EPI_ISL_509356, EPI_ISL_509357, EPI_ISL_509358, EPI_ISL_509359, EPI_ISL_509360, EPI_ISL_509361, EPI_ISL_509362, EPI_ISL_509363, EPI_ISL_509364, EPI_ISL_509365, EPI_ISL_509366, EPI_ISL_509367, EPI_ISL_509368, EPI_ISL_509369, EPI_ISL_509370, EPI_ISL_509371 | | | |
| see above | NHLS-IALCH | KRISP, KZN Research Innovation and Sequencing Platform | Giandhari J, Pillay S, Lessells R, Mdlalose K, York D, Tegally H, Wilkinson E, de Oliveira T |
| EPI_ISL_509799, EPI_ISL_509800, EPI_ISL_509801, EPI_ISL_509802, EPI_ISL_509803, EPI_ISL_509804, EPI_ISL_509805, EPI_ISL_509806, EPI_ISL_509807, EPI_ISL_509808, EPI_ISL_509809, EPI_ISL_509810, EPI_ISL_509811, EPI_ISL_509812, EPI_ISL_509813, EPI_ISL_509814, EPI_ISL_509815, EPI_ISL_509816, EPI_ISL_509817, EPI_ISL_509818, EPI_ISL_509819, EPI_ISL_509820, EPI_ISL_509821, EPI_ISL_509822, EPI_ISL_509823, EPI_ISL_509824, EPI_ISL_509825, EPI_ISL_509826, EPI_ISL_509827, EPI_ISL_509828, EPI_ISL_509829, EPI_ISL_509830, EPI_ISL_509831, EPI_ISL_509832, EPI_ISL_509833, EPI_ISL_509834, EPI_ISL_509835, EPI_ISL_509836, EPI_ISL_509837, EPI_ISL_509838, EPI_ISL_509839, EPI_ISL_509840, EPI_ISL_509841, EPI_ISL_509842, EPI_ISL_509843, EPI_ISL_509844, EPI_ISL_509845, EPI_ISL_509846, EPI_ISL_509847, EPI_ISL_509848, EPI_ISL_509849, EPI_ISL_509850, EPI_ISL_509851, EPI_ISL_509852, EPI_ISL_509853, EPI_ISL_509854, EPI_ISL_509855, EPI_ISL_509856, EPI_ISL_509857, EPI_ISL_509858, EPI_ISL_509859, EPI_ISL_509860, EPI_ISL_509861, EPI_ISL_509862, EPI_ISL_509863, EPI_ISL_509864, EPI_ISL_509865, EPI_ISL_509866, EPI_ISL_509867, EPI_ISL_509868, EPI_ISL_509869, EPI_ISL_509870, EPI_ISL_509871, EPI_ISL_509872, EPI_ISL_509873, EPI_ISL_509874, EPI_ISL_509875, EPI_ISL_509876, EPI_ISL_509877, EPI_ISL_509878, EPI_ISL_509879, EPI_ISL_509880, EPI_ISL_509881, EPI_ISL_509882, EPI_ISL_509883, EPI_ISL_509884, EPI_ISL_509885, EPI_ISL_509886, EPI_ISL_509887, EPI_ISL_509888, EPI_ISL_509889, EPI_ISL_509890, EPI_ISL_509891, EPI_ISL_509892, EPI_ISL_509893, EPI_ISL_509894, EPI_ISL_509895, EPI_ISL_509896, EPI_ISL_509897, EPI_ISL_509898, EPI_ISL_509899, EPI_ISL_509900, EPI_ISL_509901, EPI_ISL_509902, EPI_ISL_509903, EPI_ISL_509904, EPI_ISL_509905, EPI_ISL_509906, EPI_ISL_509907, EPI_ISL_509908, EPI_ISL_509909, EPI_ISL_509910, EPI_ISL_509911, EPI_ISL_509912, EPI_ISL_509914, EPI_ISL_509915, EPI_ISL_509916, EPI_ISL_509917, EPI_ISL_509918, EPI_ISL_509919, EPI_ISL_509920, EPI_ISL_509921, EPI_ISL_509922, EPI_ISL_509923, EPI_ISL_509924, EPI_ISL_509925, EPI_ISL_509926, EPI_ISL_509927, EPI_ISL_509928, EPI_ISL_509929, EPI_ISL_509930, EPI_ISL_509931, EPI_ISL_509932, EPI_ISL_509933, EPI_ISL_509934, EPI_ISL_509935, EPI_ISL_509936, EPI_ISL_509937, EPI_ISL_509938, EPI_ISL_509939, EPI_ISL_509940, EPI_ISL_509941, EPI_ISL_509942, EPI_ISL_509943, EPI_ISL_509944, EPI_ISL_509945, EPI_ISL_509946, EPI_ISL_509947, EPI_ISL_509948, EPI_ISL_509949, EPI_ISL_509950, EPI_ISL_509951, EPI_ISL_509952, EPI_ISL_509953, EPI_ISL_509954, EPI_ISL_509955, EPI_ISL_509956, EPI_ISL_509957, EPI_ISL_509958, EPI_ISL_509959, EPI_ISL_509960, EPI_ISL_509961, EPI_ISL_509962, EPI_ISL_509963, EPI_ISL_509964, EPI_ISL_509965, EPI_ISL_509966, EPI_ISL_509967, EPI_ISL_509968, EPI_ISL_509969, EPI_ISL_509970, EPI_ISL_509971, EPI_ISL_509972, EPI_ISL_509973, EPI_ISL_509974, EPI_ISL_509975, EPI_ISL_509976, EPI_ISL_509977, EPI_ISL_509978, EPI_ISL_509979, EPI_ISL_509980, EPI_ISL_509981, EPI_ISL_509982, EPI_ISL_509983, EPI_ISL_509984, EPI_ISL_509985, EPI_ISL_509986, EPI_ISL_509987, EPI_ISL_509988, EPI_ISL_509989, EPI_ISL_509990, EPI_ISL_509991, EPI_ISL_509992, EPI_ISL_509993, EPI_ISL_509994, EPI_ISL_509995, EPI_ISL_509996, EPI_ISL_509997, EPI_ISL_509998 | | | |
| see above | University of Wisconsin-Madison AIDS Vaccine Research Laboratories | University of Wisconsin-Madison AIDS Vaccine Research Laboratories | Gage Moreno, Katarina Braun, et al. AIDS Vaccine Research Laboratories |
| EPI_ISL_510246, EPI_ISL_510247, EPI_ISL_510248, EPI_ISL_510249, EPI_ISL_510250, EPI_ISL_510251, EPI_ISL_510252, EPI_ISL_510253, EPI_ISL_510254, EPI_ISL_510255, EPI_ISL_510256, EPI_ISL_510257, EPI_ISL_510258, EPI_ISL_510259, EPI_ISL_510260, EPI_ISL_510261, EPI_ISL_510262, EPI_ISL_510263, EPI_ISL_510264, EPI_ISL_510265, EPI_ISL_510266, EPI_ISL_510267 | | | |
| see above | Hospital de la Santa Creu i Sant Pau. Servicio de Microbiología | SeqCOVID-SPAIN consortium/IBV(CSIC) | Ferran Navarro, Núria Rabella, Elisenda Miró and SeqCOVID-SPAIN consortium |
| EPI_ISL_510510, EPI_ISL_510511, EPI_ISL_510512, EPI_ISL_510513, EPI_ISL_510514, EPI_ISL_510515, EPI_ISL_510516, EPI_ISL_510517, EPI_ISL_510518, EPI_ISL_510519, EPI_ISL_510520, EPI_ISL_510521, EPI_ISL_510522, EPI_ISL_510523 | | | |
| see above | Servicio de Microbiología, Laboratori Clínic Metropolitana Nord. Hospital Universitari Germans Trias i Pujol. Institut d'Investigació en Ciències de la Salut Germans Trias i Pujol (IGTP) | SeqCOVID-SPAIN consortium/IBV(CSIC) | Elisa Martró, Antoni E. Bordoy, Anna Not, Adrián Antuori, Anabel Fernández, Nona Romaní and SeqCOVID-SPAIN consortium |
| EPI_ISL_510524, EPI_ISL_510525 | University of Wisconsin-Madison AIDS Vaccine Research Laboratories | University of Wisconsin-Madison AIDS Vaccine Research Laboratories | Gage Moreno, Katarina Braun, et al. AIDS Vaccine Research Laboratories |
| EPI_ISL_510537 | Microbiology Division, SC DHEC | Microbiology Division, SC DHEC | Flores,H. |
| EPI_ISL_510878, EPI_ISL_510879, EPI_ISL_510880, EPI_ISL_510881, EPI_ISL_510882, EPI_ISL_510883, EPI_ISL_510884, EPI_ISL_510885, EPI_ISL_510886, EPI_ISL_510887, EPI_ISL_510888 | | | |
| see above | New Mexico Department of Health Scientific Laboratory | New Mexico Department of Health Scientific Laboratory | Ellie Johnson, Anastacia Griego-Fisher |
| EPI_ISL_511878 | Santosa Hospital Bandung Kopo | Laboratorium Kesehatan Provinsi Jawa Barat; Molecular Genetics Laboratory-Faculty of Medicine-Universitas Padjadjaran; School of Life Sciences and Technology & School of Pharmacy-Institut Teknologi Bandung | Ema Rahmawati, Marselina Irasonia Tan, Yunia Sribudiani, Catur Riani, Azzania Fibriani, Husna Nugrahapraja, Tarwadi, Hesti Lina Wiraswati, Lia Faridah, Savira Ekawardhani, Ryan Bayusantika Ristandi, Rifky Waluyajati Rachman, Cut Nur Cinthia Alamanda, Hammam Riza, Soni Solistia Wirawan, Agung Eru Wibowo |
| EPI_ISL_512064 | B.J. Medical College and Civil hospital, Ahmedabad | Gujarat Biotechnology Research Centre | Dinesh Kumar, Zuber Saiyed, Komal Patel, Labdhi Pandya, Afzal Ansari, Nikha Trivedi, Pranay Shah, Kamlesh J Upadhyay, Sanjay Kapadia, Apurvasinh Puvar, Janvi Raval, Zarna Patel, Monika Gandhi, Pinal Trivedi, Maharshi Pandya, Nidhi Patel, Nitin Savaliya, Raghawendra Kumar, R D Dixit, A M Kadri, Harsh Bakshi, Chaitanya Joshi, Madhvi Joshi |
| EPI_ISL_512379, EPI_ISL_512380, EPI_ISL_512381, EPI_ISL_512382, EPI_ISL_512383 | Queens Medical Centre, Clinical Microbiology Department / DeepSeq Nottingham | COVID-19 Genomics UK (COG-UK) Consortium | Gemma Clark, Wendy Smith, Manjinder Khakh, Vicki M Fleming, Michelle M Lister, Hannah Howson-Wells, Jonathan Ball, Patrick McClure, Joseph Chappell, Theocharis Tsoleridis, Nadine Holmes, Matthew Carlisle, Christopher Moore, Fei Sang, Johnny Debebe, Victoria Wright, Matthew Loose |
| EPI_ISL_512653 | Area De Salud Desamparados 1 - Clinica Dr. Marcial Fallas [Grifo Alto/Desampara | Inciensa, Instituto Costarricense de Investigación y Enseñanza en Nutrición y Salud | Francisco Duarte, Hebleen Porras, Claudio Soto-Garita, Estela Cordero, Adriana Godinez & Melany Calderon |
| EPI_ISL_512654 | Hospital Dr. Rafael A. Calderon Guardia [San Jose/San Jose] | Inciensa, Instituto Costarricense de Investigación y Enseñanza en Nutrición y Salud | Francisco Duarte, Hebleen Porras, Claudio Soto-Garita, Estela Cordero, Adriana Godinez & Melany Calderon |
| EPI_ISL_512655 | Hospital De Niños Dr. Carlos Saenz Herrera [San Jose/San Jose] | Inciensa, Instituto Costarricense de Investigación y Enseñanza en Nutrición y Salud | Francisco Duarte, Hebleen Porras, Claudio Soto-Garita, Estela Cordero, Adriana Godinez & Melany Calderon |
| EPI_ISL_512656 | Area De Salud Pavas (Coopesalud) [Pavas/San Jose] | Inciensa, Instituto Costarricense de Investigación y | Francisco Duarte, Hebleen Porras, Claudio Soto-Garita, Estela Cordero, Adriana Godinez & Melany Calderon |

| | | | |
|---|---|---|---|
| | | Enseñanza en Nutrición y Salud | |
| EPI_ISL_512657 | Area De Salud Tibas-Uruca-Merced - Clinica Dr. Clorito Picado [Tibas/San Jose] | Inciensa, Instituto Costarricense de Investigación y Enseñanza en Nutrición y Salud | Francisco Duarte, Hebleen Porras, Claudio Soto-Garita, Estela Cordero, Adriana Godinez & Melany Calderon |
| EPI_ISL_512658 | Area De Salud Orotina-San Mateo [Orotina/Alajuela] | Inciensa, Instituto Costarricense de Investigación y Enseñanza en Nutrición y Salud | Francisco Duarte, Hebleen Porras, Claudio Soto-Garita, Estela Cordero, Adriana Godinez & Melany Calderon |
| EPI_ISL_512659 | Area De Salud Fortuna | Inciensa, Instituto Costarricense de Investigación y Enseñanza en Nutrición y Salud | Francisco Duarte, Hebleen Porras, Claudio Soto-Garita, Estela Cordero, Adriana Godinez & Melany Calderon |
| EPI_ISL_512660, EPI_ISL_512661 | Area De Salud Los Chiles | Inciensa, Instituto Costarricense de Investigación y Enseñanza en Nutrición y Salud | Francisco Duarte, Hebleen Porras, Claudio Soto-Garita, Estela Cordero, Adriana Godinez & Melany Calderon |
| EPI_ISL_512662 | Area De Salud La Cruz | Inciensa, Instituto Costarricense de Investigación y Enseñanza en Nutrición y Salud | Francisco Duarte, Hebleen Porras, Claudio Soto-Garita, Estela Cordero, Adriana Godinez & Melany Calderon |
| EPI_ISL_512663, EPI_ISL_512664 | Area De Salud Alajuela Norte - Clinica Dr. Marcial Rodriguez | Inciensa, Instituto Costarricense de Investigación y Enseñanza en Nutrición y Salud | Francisco Duarte, Hebleen Porras, Claudio Soto-Garita, Estela Cordero, Adriana Godinez & Melany Calderon |
| EPI_ISL_512665 | Hospital De Las Mujeres Dr. Adolfo Carit | Inciensa, Instituto Costarricense de Investigación y Enseñanza en Nutrición y Salud | Francisco Duarte, Hebleen Porras, Claudio Soto-Garita, Estela Cordero, Adriana Godinez & Melany Calderon |
| EPI_ISL_512666, EPI_ISL_512667 | Area De Salud La Cruz | Inciensa, Instituto Costarricense de Investigación y Enseñanza en Nutrición y Salud | Francisco Duarte, Hebleen Porras, Claudio Soto-Garita, Estela Cordero, Adriana Godinez & Melany Calderon |
| EPI_ISL_512668 | Area De Salud Corredores | Inciensa, Instituto Costarricense de Investigación y Enseñanza en Nutrición y Salud | Francisco Duarte, Hebleen Porras, Claudio Soto-Garita, Estela Cordero, Adriana Godinez & Melany Calderon |
| EPI_ISL_512669 | Area De Salud Alajuela Norte - Clinica Dr. Marcial Rodriguez | Inciensa, Instituto Costarricense de Investigación y Enseñanza en Nutrición y Salud | Francisco Duarte, Hebleen Porras, Claudio Soto-Garita, Estela Cordero, Adriana Godinez & Melany Calderon |
| EPI_ISL_512670 | Centro Nacional De Rehabilitacion Humberto Araya Rojas (Cenare) | Inciensa, Instituto Costarricense de Investigación y Enseñanza en Nutrición y Salud | Francisco Duarte, Hebleen Porras, Claudio Soto-Garita, Estela Cordero, Adriana Godinez & Melany Calderon |
| EPI_ISL_512671 | Area De Salud La Cruz | Inciensa, Instituto Costarricense de Investigación y Enseñanza en Nutrición y Salud | Francisco Duarte, Hebleen Porras, Claudio Soto-Garita, Estela Cordero, Adriana Godinez & Melany Calderon |
| EPI_ISL_512672, EPI_ISL_512673, EPI_ISL_512674, EPI_ISL_512675 | Hospital De Niños Dr. Carlos Saenz Herrera [San Jose/San Jose] | Inciensa, Instituto Costarricense de Investigación y Enseñanza en Nutrición y Salud | Francisco Duarte, Hebleen Porras, Claudio Soto-Garita, Estela Cordero, Adriana Godinez & Melany Calderon |
| EPI_ISL_512785, EPI_ISL_512786, EPI_ISL_512787, EPI_ISL_512788, EPI_ISL_512789, EPI_ISL_512790, EPI_ISL_512791, EPI_ISL_512792, EPI_ISL_512793, EPI_ISL_512794, EPI_ISL_512795, EPI_ISL_512796, EPI_ISL_512797, EPI_ISL_512798, EPI_ISL_512799, EPI_ISL_512800, EPI_ISL_512801, EPI_ISL_512802, EPI_ISL_512803, EPI_ISL_512804, EPI_ISL_512805, EPI_ISL_512806, EPI_ISL_512807, EPI_ISL_512808, EPI_ISL_512809 | | | |
| see above | Public Health, United States Air Force School of Aerospace Medicine | Public Health, United States Air Force School of Aerospace Medicine | Fries,A.C., Purves,S.M., Meyer,J.R., Javorina,A.K., Connors,B.C., Macias,E.A., Lambert,A.W., Chapleau,R.R., Starr,C.R. |
| EPI_ISL_512846 | O.I.J. MORGUE JUDICIAL | Inciensa, Instituto Costarricense de Investigación y Enseñanza en Nutrición y Salud | Francisco Duarte, Hebleen Porras, Claudio Soto-Garita, Estela Cordero, Adriana Godinez & Melany Calderon |
| EPI_ISL_512847, EPI_ISL_512848, EPI_ISL_512849, EPI_ISL_512850, EPI_ISL_512851, EPI_ISL_512852, EPI_ISL_512853, EPI_ISL_512854, EPI_ISL_512855, EPI_ISL_512856, EPI_ISL_512857, EPI_ISL_512858, EPI_ISL_512859, EPI_ISL_512860, EPI_ISL_512861, EPI_ISL_512862, EPI_ISL_512863, EPI_ISL_512864, EPI_ISL_512865, EPI_ISL_512866, EPI_ISL_512867, EPI_ISL_512868, EPI_ISL_512869, EPI_ISL_512870, EPI_ISL_512871, EPI_ISL_512872 | | | |
| see above | Ramathibodi Hospital | COVID-19 Network Investigations (CONI) Alliance | Elizabeth Batty, Wasun Chantratita, Thanat Chookajorn, Stefan Fernandez, Angkana Huang, Anthony R. Jones, Khajohn Joonsalak, Chonticha Klungtong, Theerarat Kochakarn, Namfon Kotanan, Krittikorn Kumpornsin, Wudtichai Manasatienkij, Bhakbhoom Panthan, Ekawat Pasomsub, Kingkan Rakmanee, Insee Sensorn, Janjira Thaipadungpanit, Arporn Wangwiwatsin,Treewat Watthanachockchai |
| EPI_ISL_512904, EPI_ISL_512905, EPI_ISL_512906, EPI_ISL_512907 | Pathogen Genomics Lab King Abdullah University of Science and Technology(KAUST) | Pathogen Genomics Lab King Abdullah University of Science and Technology(KAUST) | Fathia Ben Rached, Raeece Naeem, Sharif Hala, Fadwa Alofi, Rahul P Salunke, Sara Mfarrej, Amit Kumar Subudhi, Afrah Alsomali, Asim Khogeer, Ahmad Bakur Mahmoud, Anwar Hashem, Naif Almontashiri, Arnab Pain |
| EPI_ISL_512918, EPI_ISL_512919, EPI_ISL_512920, EPI_ISL_512921, EPI_ISL_512922, EPI_ISL_512923, EPI_ISL_512924, EPI_ISL_512925, EPI_ISL_512926, EPI_ISL_512927, EPI_ISL_512928, EPI_ISL_512929, EPI_ISL_512930, EPI_ISL_512931, EPI_ISL_512932, EPI_ISL_512933, EPI_ISL_512934, EPI_ISL_512935, EPI_ISL_512936, EPI_ISL_512937, EPI_ISL_512938, EPI_ISL_512939, EPI_ISL_512940, EPI_ISL_512941, EPI_ISL_512942, EPI_ISL_512943, EPI_ISL_512944, EPI_ISL_512945 | | | |
| see above | Pathogen Genomics Lab King Abdullah University of Science and Technology(KAUST) | Pathogen Genomics Lab King Abdullah University of Science and Technology(KAUST) | Fadwa Alofi, Sharif Hala, Rahul P Salunke, Sara Mfarrej, Amit Kumar Subudhi, Fathia Ben Rached, Amanda, Luke, Afrah Alsomali, Asim Khogeer, Jumana Taha, Abdulaziz Alahmadi, Kahled Algithami, Raeece Naeem, Anwar Hashem, Naif Almontashiri, Arnab Pain |
| EPI_ISL_513077, EPI_ISL_513078, EPI_ISL_513079, EPI_ISL_513080, EPI_ISL_513081, EPI_ISL_513082, EPI_ISL_513083, EPI_ISL_513084, EPI_ISL_513085, EPI_ISL_513086, EPI_ISL_513087, EPI_ISL_513088, EPI_ISL_513089, EPI_ISL_513090, EPI_ISL_513091, EPI_ISL_513092, EPI_ISL_513093, EPI_ISL_513094, EPI_ISL_513095, EPI_ISL_513096, EPI_ISL_513097, EPI_ISL_513098, EPI_ISL_513099, EPI_ISL_513100, EPI_ISL_513101, EPI_ISL_513102, EPI_ISL_513103, EPI_ISL_513104, EPI_ISL_513105, EPI_ISL_513106, EPI_ISL_513107, EPI_ISL_513108, EPI_ISL_513109, EPI_ISL_513110, EPI_ISL_513111, EPI_ISL_513112, EPI_ISL_513113, EPI_ISL_513114, EPI_ISL_513115, EPI_ISL_513116, EPI_ISL_513117, EPI_ISL_513118, EPI_ISL_513119, EPI_ISL_513120 | | | |
| see above | Pathogen Genomics Lab King Abdullah University of Science and Technology(KAUST) | Pathogen Genomics Lab King Abdullah University of Science and Technology(KAUST) | Fathia Ben Rached, Raeece Naeem, Sharif Hala, Fadwa Alofi, Rahul P Salunke, Sara Mfarrej, Amit Kumar Subudhi, Afrah Alsomali, Asim Khogeer, Ahmad Bakur Mahmoud, Anwar Hashem, Naif Almontashiri, Arnab Pain |
| EPI_ISL_513147, EPI_ISL_513148, EPI_ISL_513149, EPI_ISL_513150, EPI_ISL_513151, EPI_ISL_513152, EPI_ISL_513153, EPI_ISL_513154, EPI_ISL_513155, EPI_ISL_513156, EPI_ISL_513157, EPI_ISL_513158, EPI_ISL_513159, EPI_ISL_513160, EPI_ISL_513161, EPI_ISL_513162, EPI_ISL_513163, EPI_ISL_513164, EPI_ISL_513165, EPI_ISL_513166, EPI_ISL_513167 | | | |
| see above | Pathogen Genomics Lab King Abdullah University of Science and Technology(KAUST) | Pathogen Genomics Lab King Abdullah University of Science and Technology(KAUST) | Fadwa Alofi, Sharif Hala, Rahul P Salunke, Sara Mfarrej, Amit Kumar Subudhi, Fathia Ben Rached, Amanda, Luke, Afrah Alsomali, Asim Khogeer, Jumana Taha, Abdulaziz Alahmadi, Kahled Algithami, Raeece Naeem, Anwar Hashem, Naif Almontashiri, Arnab Pain |
| EPI_ISL_513265, EPI_ISL_513266, EPI_ISL_513267, EPI_ISL_513268, EPI_ISL_513269, EPI_ISL_513270, EPI_ISL_513271, EPI_ISL_513272, EPI_ISL_513273, EPI_ISL_513274, EPI_ISL_513275, EPI_ISL_513276, EPI_ISL_513277, EPI_ISL_513278, EPI_ISL_513279, EPI_ISL_513280, EPI_ISL_513281, EPI_ISL_513282, EPI_ISL_513283, EPI_ISL_513284, EPI_ISL_513285, EPI_ISL_513286, EPI_ISL_513287, EPI_ISL_513288, EPI_ISL_513289, EPI_ISL_513290, EPI_ISL_513291, EPI_ISL_513292, EPI_ISL_513293 | | | |
| see above | University of Miami Immunology and Histocompatibility Laboratory | University of Miami Immunology and Histocompatibility Laboratory | Emilio Margolles-Clark, PhD and Phillip Ruiz, MD, PhD |
| EPI_ISL_513298, EPI_ISL_513299, EPI_ISL_513300, EPI_ISL_513301, EPI_ISL_513302, EPI_ISL_513303, EPI_ISL_513304, EPI_ISL_513305, EPI_ISL_513306, EPI_ISL_513307 | Department of Infection Prevention and Infectious Diseases, University Hospital Regensburg | University Hospital Regensburg | Fritsch,J., Holzmann,T., Schneider-Brachert,W. |
| EPI_ISL_513310 | Public Health, United States Air Force School of Aerospace Medicine | Public Health, United States Air Force School of Aerospace Medicine | Fries,A.C., Purves,S.M., Meyer,J.R., Javorina,A.K., Connors,B.C., Macias,E.A., Lambert,A.W., Chapleau,R.R. and Starr,C.R. |
| EPI_ISL_513311, EPI_ISL_513312, EPI_ISL_513313 | Public Health, United States Air Force School of Aerospace Medicine | Public Health, United States Air Force School of Aerospace Medicine | Fries,A.C., Purves,S.M., Meyer,J.R., Javorina,A.K., Connors,B.C., Macias,E.A., Lambert,A.W., Chapleau,R.R. and Starr,C.R. |
| EPI_ISL_514434 | NSTU COVID-19 Diagnostic Center, | NSU Genome Research Institute (NGRI), North South University | Dr. Muhammad Maqsud Hossain, Aura Rahman, Prof. Firoz Ahmed, Tahrima Huq, Abdus Sadique, Jahidul Alam, Md Aminul Islam, Prof. Md. Didar-Ul-Alam, Prof. Kazi Nadim Hasan, Prof. Abdul Khaleque, Prof, Hasan Mahmud Reza |
| EPI_ISL_514440 | NSTU COVID-19 Diagnostic Center | NSU Genome Research Institute (NGRI), North South University | Dr. Muhammad Maqsud Hossain, Aura Rahman, Prof. Firoz Ahmed, Tahrima Huq, Abdus Sadique, Tamanna Afroze, Jahidul Alam, Md Aminul Islam, Prof. Md. Didar-Ul-Alam, Prof. Kazi Nadim Hasan, Prof. Abdul Khaleque, Prof, Hasan Mahmud Reza |
| EPI_ISL_514441 | NSTU COVID-19 Diagnostic Center | NSU Genome Research Institute (NGRI), North South | Dr. Muhammad Maqsud Hossain, Aura Rahman, Prof. Firoz Ahmed, Tahrima Huq, Abdus Sadique, Jahidul Alam, Md Aminul Islam, Prof. Md. |

| | | | |
|---|---|---|---|
| | | University | Didar-Ul-Alam, Prof. Kazi Nadim Hasan, Prof. Abdul Khaleque, Prof, Hasan Mahmud Reza |
| EPI_ISL_514442 | NSTU COVID-19 Diagnostic Center | NSU Genome Research Institute (NGRI), North South University | Dr. Muhammad Maqsud Hossain, Aura Rahman, Prof. Firoz Ahmed, Tahrima Huq, Abdus Sadique, Jahidul Alam, Tamanna Afroze, Md Aminul Islam, Prof. Md. Didar-Ul-Alam, Prof. Kazi Nadim Hasan, Prof. Abdul Khaleque, Prof, Hasan Mahmud Reza |
| EPI_ISL_514452, EPI_ISL_514453, EPI_ISL_514454 | Queens Medical Centre, Clinical Microbiology Department / DeepSeq Nottingham | COVID-19 Genomics UK (COG-UK) Consortium | Gemma Clark, Wendy Smith, Manjinder Khakh, Vicki M Fleming, Michelle M Lister, Hannah Howson-Wells, Jonathan Ball, Patrick McClure, Joseph Chappell, Theocharis Tsoleridis, Nadine Holmes, Matthew Carlisle, Christopher Moore, Fei Sang, Johnny Debebe, Victoria Wright, Matthew Loose |
| EPI_ISL_514580 | NSTU COVID-19 Diagnostic Center | NSU Genome Research Institute (NGRI), North South University | Dr. Muhammad Maqsud Hossain, Aura Rahman, Prof. Firoz Ahmed, Tahrima Huq, Abdus Sadique, Jahidul Alam, Tamanna Afroze, Md Aminul Islam, Prof. Md. Didar-Ul-Alam, Prof. Kazi Nadim Hasan, Prof. Abdul Khaleque, Prof, Hasan Mahmud Reza |
| EPI_ISL_514583, EPI_ISL_514601 | B.J. Medical College and Civil hospital, Ahmedabad | Gujarat Biotechnology Research Centre | Dinesh Kumar, Zuber Saiyed, Komal Patel, Labdhi Pandya, Afzal Ansari, Nikha Trivedi, Pranay Shah, Kamlesh J Upadhyay, Sanjay Kapadia, Apurvasinh Puvar, Janvi Raval, Zarna Patel, Monika Gandhi, Pinal Trivedi, Maharshi Pandya, Nidhi Patel, Nitin Savaliya, Raghawendra Kumar, R D Dixit, A M Kadri, Harsh Bakshi, Chaitanya Joshi, Madhvi Joshi |
| EPI_ISL_514612 | GMERS Medical College and Hospital, Gandhinagar | Gujarat Biotechnology Research Centre | Gaurishankar Shrimali, Bhavesh Modi, Bharti Rajani, Apurvasinh Puvar, Janvi Raval, Zarna Patel, Monika Gandhi, Pinal Trivedi, Maharshi Pandya, Nidhi Patel, Nitin Savaliya, Raghawendra Kumar, Dinesh Kumar, Zuber Saiyed, Komal Patel, Labdhi Pandya, Afzal Ansari, Nikha Trivedi, Pranay Shah, Kamlesh J Upadhyay, Sanjay Kapadia, Seema Bhatt, R D Dixit, A M Kadri, Harsh Bakshi, Chaitanya Joshi, Madhvi Joshi |
| EPI_ISL_514613, EPI_ISL_514614, EPI_ISL_514615 | NSTU COVID-19 Diagnostic Center | NSU Genome Research Institute (NGRI), North South University | Dr. Muhammad Maqsud Hossain, Aura Rahman, Prof. Firoz Ahmed, Tahrima Huq, Abdus Sadique, Jahidul Alam, Tamanna Afroze, Md Aminul Islam, Prof. Md. Didar-Ul-Alam, Prof. Kazi Nadim Hasan, Prof. Abdul Khaleque, Prof, Hasan Mahmud Reza |
| EPI_ISL_515466 | Discovery DNA | Discovery DNA | Dustin Hittel, Marina Kerr, Leo Dimnik, Desmond Koo, Alice Li, Aneal Khan |
| EPI_ISL_515572, EPI_ISL_515577, EPI_ISL_515578, EPI_ISL_515583, EPI_ISL_515587, EPI_ISL_515588, EPI_ISL_515590, EPI_ISL_515591, EPI_ISL_515593, EPI_ISL_515594, EPI_ISL_515595, EPI_ISL_515596, EPI_ISL_515597, EPI_ISL_515598, EPI_ISL_515599, EPI_ISL_515600, EPI_ISL_515601, EPI_ISL_515602, EPI_ISL_515603, EPI_ISL_515604, EPI_ISL_515605, EPI_ISL_515606, EPI_ISL_515607, EPI_ISL_515608, EPI_ISL_515609, EPI_ISL_515610, EPI_ISL_515611, EPI_ISL_515612, EPI_ISL_515613, EPI_ISL_515614, EPI_ISL_515615, EPI_ISL_515616, EPI_ISL_515617, EPI_ISL_515618, EPI_ISL_515619, EPI_ISL_515620, EPI_ISL_515621, EPI_ISL_515622, EPI_ISL_515623, EPI_ISL_515624, EPI_ISL_515625, EPI_ISL_515626, EPI_ISL_515627, EPI_ISL_515628, EPI_ISL_515629, EPI_ISL_515630, EPI_ISL_515631, EPI_ISL_515632, EPI_ISL_515633, EPI_ISL_515634, EPI_ISL_515635, EPI_ISL_515636, EPI_ISL_515637, EPI_ISL_515638, EPI_ISL_515639, EPI_ISL_515640, EPI_ISL_515641, EPI_ISL_515642, EPI_ISL_515643, EPI_ISL_515644, EPI_ISL_515645, EPI_ISL_515646, EPI_ISL_515647, EPI_ISL_515648, EPI_ISL_515649, EPI_ISL_515650, EPI_ISL_515651, EPI_ISL_515652, EPI_ISL_515653, EPI_ISL_515654, EPI_ISL_515655, EPI_ISL_515656, EPI_ISL_515657, EPI_ISL_515658, EPI_ISL_515659, EPI_ISL_515660, EPI_ISL_515661, EPI_ISL_515662, EPI_ISL_515663, EPI_ISL_515664, EPI_ISL_515665, EPI_ISL_515666, EPI_ISL_515667, EPI_ISL_515668, EPI_ISL_515669, EPI_ISL_515670, EPI_ISL_515671, EPI_ISL_515672, EPI_ISL_515673, EPI_ISL_515674, EPI_ISL_515675, EPI_ISL_515676, EPI_ISL_515677, EPI_ISL_515678, EPI_ISL_515679, EPI_ISL_515680, EPI_ISL_515681, EPI_ISL_515682, EPI_ISL_515683, EPI_ISL_515684, EPI_ISL_515685, EPI_ISL_515686, EPI_ISL_515687, EPI_ISL_515688, EPI_ISL_515689, EPI_ISL_515690, EPI_ISL_515691, EPI_ISL_515692, EPI_ISL_515693, EPI_ISL_515694, EPI_ISL_515696, EPI_ISL_515697, EPI_ISL_515698, EPI_ISL_515699, EPI_ISL_515700, EPI_ISL_515701, EPI_ISL_515702, EPI_ISL_515703, EPI_ISL_515704, EPI_ISL_515705, EPI_ISL_515706, EPI_ISL_515707, EPI_ISL_515708, EPI_ISL_515709, EPI_ISL_515710, EPI_ISL_515711, EPI_ISL_515712, EPI_ISL_515714, EPI_ISL_515715, EPI_ISL_515716, EPI_ISL_515718, EPI_ISL_515719, EPI_ISL_515722, EPI_ISL_515723, EPI_ISL_515724, EPI_ISL_515725, EPI_ISL_515728, EPI_ISL_515729, EPI_ISL_515730, EPI_ISL_515731, EPI_ISL_515732, EPI_ISL_515733, EPI_ISL_515734, EPI_ISL_515735, EPI_ISL_515736, EPI_ISL_515737, EPI_ISL_515738, EPI_ISL_515739, EPI_ISL_515740, EPI_ISL_515741, EPI_ISL_515742, EPI_ISL_515743, EPI_ISL_515744, EPI_ISL_515745, EPI_ISL_515748, EPI_ISL_515749, EPI_ISL_515750, EPI_ISL_515751, EPI_ISL_515752, EPI_ISL_515753, EPI_ISL_515754, EPI_ISL_515755, EPI_ISL_515757, EPI_ISL_515758, EPI_ISL_515759, EPI_ISL_515760, EPI_ISL_515761, EPI_ISL_515762, EPI_ISL_515763, EPI_ISL_515764, EPI_ISL_515767, EPI_ISL_515768, EPI_ISL_515769, EPI_ISL_515770, EPI_ISL_515771, EPI_ISL_515773, EPI_ISL_515775, EPI_ISL_515776, EPI_ISL_515777, EPI_ISL_515778, EPI_ISL_515779, EPI_ISL_515780, EPI_ISL_515781, EPI_ISL_515782, EPI_ISL_515783, EPI_ISL_515784, EPI_ISL_515785, EPI_ISL_515786, EPI_ISL_515787, EPI_ISL_515789, EPI_ISL_515790, EPI_ISL_515791, EPI_ISL_515792, EPI_ISL_515794, EPI_ISL_515795, EPI_ISL_515796, EPI_ISL_515797 | | | |
| see above | NHLS-IALCH | KRISP, KZN Research Innovation and Sequencing Platform | Giandhari J, Pillay S, Lessells R, Mdlalose K, York D, Khan S, Tegally H, Wilkinson E, de Oliveira T |
| EPI_ISL_515800, EPI_ISL_515801, EPI_ISL_515802, EPI_ISL_515803, EPI_ISL_515804, EPI_ISL_515805, EPI_ISL_515806, EPI_ISL_515807, EPI_ISL_515808, EPI_ISL_515809, EPI_ISL_515810, EPI_ISL_515811, EPI_ISL_515812, EPI_ISL_515813, EPI_ISL_515814, EPI_ISL_515815, EPI_ISL_515816, EPI_ISL_515817, EPI_ISL_515818, EPI_ISL_515819, EPI_ISL_515820, EPI_ISL_515821, EPI_ISL_515822, EPI_ISL_515823, EPI_ISL_515824, EPI_ISL_515825, EPI_ISL_515826, EPI_ISL_515827, EPI_ISL_515828, EPI_ISL_515829, EPI_ISL_515859, EPI_ISL_515879, EPI_ISL_515888 | | | |
| see above | Medical Disagnostics Services (MDS) | KRISP, KZN Research Innovation and Sequencing Platform | Giandhari J, Pillay S, Lessells R, ChimukangaraB, Mdlalose K, York D, Khan S, Tegally H, Wilkinson E, de Oliveira T |
| EPI_ISL_516432, EPI_ISL_516433, EPI_ISL_516434, EPI_ISL_516435, EPI_ISL_516436, EPI_ISL_516437, EPI_ISL_516438, EPI_ISL_516439, EPI_ISL_516440, EPI_ISL_516441, EPI_ISL_516442, EPI_ISL_516443, EPI_ISL_516444, EPI_ISL_516445, EPI_ISL_516446, EPI_ISL_516447, EPI_ISL_516448, EPI_ISL_516449, EPI_ISL_516450, EPI_ISL_516451, EPI_ISL_516452, EPI_ISL_516453, EPI_ISL_516454, EPI_ISL_516455, EPI_ISL_516456, EPI_ISL_516457, EPI_ISL_516458, EPI_ISL_516459, EPI_ISL_516460, EPI_ISL_516461, EPI_ISL_516462, EPI_ISL_516463, EPI_ISL_516464, EPI_ISL_516465, EPI_ISL_516466, EPI_ISL_516467, EPI_ISL_516468, EPI_ISL_516469, EPI_ISL_516470, EPI_ISL_516471, EPI_ISL_516472, EPI_ISL_516473, EPI_ISL_516474, EPI_ISL_516475, EPI_ISL_516476, EPI_ISL_516477, EPI_ISL_516478, EPI_ISL_516479, EPI_ISL_516480, EPI_ISL_516481, EPI_ISL_516482, EPI_ISL_516483, EPI_ISL_516484, EPI_ISL_516485, EPI_ISL_516486, EPI_ISL_516487, EPI_ISL_516488, EPI_ISL_516489, EPI_ISL_516490, EPI_ISL_516491, EPI_ISL_516492, EPI_ISL_516493, EPI_ISL_516494, EPI_ISL_516495, EPI_ISL_516496, EPI_ISL_516497, EPI_ISL_516498, EPI_ISL_516499, EPI_ISL_516500, EPI_ISL_516501, EPI_ISL_516502, EPI_ISL_516503, EPI_ISL_516504, EPI_ISL_516505, EPI_ISL_516506, EPI_ISL_516507, EPI_ISL_516508, EPI_ISL_516509, EPI_ISL_516510, EPI_ISL_516511, EPI_ISL_516512, EPI_ISL_516513, EPI_ISL_516514, EPI_ISL_516515, EPI_ISL_516516, EPI_ISL_516517, EPI_ISL_516518, EPI_ISL_516519, EPI_ISL_516520, EPI_ISL_516521, EPI_ISL_516522, EPI_ISL_516523, EPI_ISL_516524, EPI_ISL_516525, EPI_ISL_516526, EPI_ISL_516527, EPI_ISL_516528, EPI_ISL_516529, EPI_ISL_516530, EPI_ISL_516531, EPI_ISL_516532, EPI_ISL_516533, EPI_ISL_516534, EPI_ISL_516535, EPI_ISL_516536, EPI_ISL_516537, EPI_ISL_516538, EPI_ISL_516539, EPI_ISL_516540, EPI_ISL_516541, EPI_ISL_516542, EPI_ISL_516543, EPI_ISL_516544, EPI_ISL_516545, EPI_ISL_516546, EPI_ISL_516547, EPI_ISL_516548, EPI_ISL_516549 | | | |
| see above | University of Wisconsin-Madison AIDS Vaccine Research Laboratories | University of Wisconsin-Madison AIDS Vaccine Research Laboratories | Gage Moreno, Katarina Braun, et al. AIDS Vaccine Research Laboratories |
| EPI_ISL_516608 | Instituto de Diagnostico y Referencia Epidemiologicos (INDRE) | Instituto de Diagnostico y Referencia Epidemiologicos (INDRE) | Ernesto Ramirez-Gonzalez, Abril Rodriguez-Maldonado, Claudia Wong-Arambula , Natividad Cruz-Ortiz, Tatiana Nunez-Garcia, Dayanira Arellano-Suarez, Adnan Araiza-Rodriguez, Edgar Mendieta-Condado, Lucia Hernandez-Rivas, Irma Lopez-Martinez, Gisela Barrera-Badillo. |
| EPI_ISL_516609, EPI_ISL_516610, EPI_ISL_516612 | Instituto de Diagnostico y Referencia Epidemiologicos (INDRE) | Instituto de Diagnostico y Referencia Epidemiologicos (INDRE) | Ernesto Ramirez-Gonzalez, Abril Rodriguez-Maldonado, Claudia Wong-Arambula , Natividad Cruz-Ortiz, Tatiana Nunez-Garcia, Dayanira Arellano-Suarez, Adnan Araiza-Rodriguez, Fabiola Garces-Ayala, Lucia Hernandez-Rivas, Irma Lopez-Martinez, Gisela Barrera-Badillo. |
| EPI_ISL_516613 | Instituto de Diagnostico y Referencia Epidemiologicos (INDRE) | Instituto de Diagnostico y Referencia Epidemiologicos (INDRE) | Ernesto Ramirez-Gonzalez, Abril Rodriguez-Maldonado, Claudia Wong-Arambula , Natividad Cruz-Ortiz, Tatiana Nunez-Garcia, Dayanira Arellano-Suarez, Adnan Araiza-Rodriguez, Edgar Mendieta-Condado, Lucia Hernandez-Rivas, Irma Lopez-Martinez, Gisela Barrera-Badillo. |
| EPI_ISL_516614, EPI_ISL_516615, EPI_ISL_516616, EPI_ISL_516617 | Instituto de Diagnostico y Referencia Epidemiologicos (INDRE) | Instituto de Diagnostico y Referencia Epidemiologicos (INDRE) | Ernesto Ramirez-Gonzalez, Abril Rodriguez-Maldonado, Claudia Wong-Arambula , Natividad Cruz-Ortiz, Tatiana Nunez-Garcia, Dayanira Arellano-Suarez, Adnan Araiza-Rodriguez, Fabiola Garces-Ayala, Lucia Hernandez-Rivas, Irma Lopez-Martinez, Gisela Barrera-Badillo. |
| EPI_ISL_518033, EPI_ISL_518034, EPI_ISL_518035, EPI_ISL_518037, EPI_ISL_518038, EPI_ISL_518039, EPI_ISL_518040, EPI_ISL_518041, EPI_ISL_518043, EPI_ISL_518044, EPI_ISL_518045, EPI_ISL_518046, EPI_ISL_518047, EPI_ISL_518048, EPI_ISL_518049, EPI_ISL_518050, EPI_ISL_518051, EPI_ISL_518052 | | | |
| see above | NHLS-IALCH | KRISP, KZN Research Innovation and Sequencing Platform | Giandhari J, Pillay S, Lessells R, Mdlalose K, York D, Khan S, Tegally H, Wilkinson E, de Oliveira T |
| EPI_ISL_522254, EPI_ISL_522255, EPI_ISL_522256, EPI_ISL_522257, EPI_ISL_522258, EPI_ISL_522259, EPI_ISL_522260, EPI_ISL_522261, EPI_ISL_522262, EPI_ISL_522263, EPI_ISL_522264, EPI_ISL_522265, EPI_ISL_522266, EPI_ISL_522267, EPI_ISL_522268, EPI_ISL_522269, EPI_ISL_522270, EPI_ISL_522271, EPI_ISL_522272, EPI_ISL_522273, EPI_ISL_522274, EPI_ISL_522275, EPI_ISL_522276, EPI_ISL_522277, EPI_ISL_522278, EPI_ISL_522279, EPI_ISL_522280, EPI_ISL_522281, EPI_ISL_522282, EPI_ISL_522283, EPI_ISL_522284, EPI_ISL_522285, EPI_ISL_522286, EPI_ISL_522287, EPI_ISL_522288, EPI_ISL_522289, EPI_ISL_522290, EPI_ISL_522291, EPI_ISL_522292, EPI_ISL_522293, EPI_ISL_522294, EPI_ISL_522295, EPI_ISL_522296, EPI_ISL_522297, EPI_ISL_522298, EPI_ISL_522299, EPI_ISL_522300, EPI_ISL_522301, EPI_ISL_522302, EPI_ISL_522303, EPI_ISL_522304, EPI_ISL_522305, EPI_ISL_522306, EPI_ISL_522307, EPI_ISL_522308, EPI_ISL_522309, EPI_ISL_522310, EPI_ISL_522311, EPI_ISL_522312, EPI_ISL_522313, EPI_ISL_522314, EPI_ISL_522315, EPI_ISL_522316, EPI_ISL_522317, EPI_ISL_522318, EPI_ISL_522319, EPI_ISL_522320, EPI_ISL_522321, EPI_ISL_522322, EPI_ISL_522323, EPI_ISL_522324, EPI_ISL_522325, EPI_ISL_522326, EPI_ISL_522327, EPI_ISL_522328, EPI_ISL_522329, EPI_ISL_522330, EPI_ISL_522331, EPI_ISL_522332, EPI_ISL_522333, EPI_ISL_522334, EPI_ISL_522335, EPI_ISL_522336, EPI_ISL_522337, EPI_ISL_522338, EPI_ISL_522339, EPI_ISL_522340, EPI_ISL_522341, EPI_ISL_522342, EPI_ISL_522343, EPI_ISL_522344, EPI_ISL_522345, EPI_ISL_522346, EPI_ISL_522347, EPI_ISL_522348 | | | |
| see above | Utah Public Health Laboratory | Utah Public Health Laboratory | Erin Young, Kelly Oakeson |
| EPI_ISL_522407, EPI_ISL_522408 | Department of Infection Prevention and Infectious Diseases, University Hospital Regensburg | Department of Infection Prevention and Infectious Diseases, University Hospital Regensburg | Fritsch,J., Holzmann,T., Schneider-Brachert,W. |
| EPI_ISL_522881, EPI_ISL_522882, EPI_ISL_522883, EPI_ISL_522884, EPI_ISL_522885, EPI_ISL_522886, EPI_ISL_522887, EPI_ISL_522888, EPI_ISL_522889, EPI_ISL_522890, EPI_ISL_522891, EPI_ISL_522892, EPI_ISL_522893, EPI_ISL_522894, EPI_ISL_522895, EPI_ISL_522896, EPI_ISL_522897, EPI_ISL_522898, EPI_ISL_522899, EPI_ISL_522900, EPI_ISL_522901, EPI_ISL_522902, EPI_ISL_522903, EPI_ISL_522904, EPI_ISL_522905, EPI_ISL_522906, EPI_ISL_522907, EPI_ISL_522908, EPI_ISL_522909, EPI_ISL_522910, EPI_ISL_522911, EPI_ISL_522912, EPI_ISL_522913, EPI_ISL_522914, EPI_ISL_522915, EPI_ISL_522916, EPI_ISL_522917, EPI_ISL_522918, EPI_ISL_522919, EPI_ISL_522920, EPI_ISL_522921, EPI_ISL_522922, EPI_ISL_522923, EPI_ISL_522924, EPI_ISL_522925, EPI_ISL_522926, EPI_ISL_522927, EPI_ISL_522928, EPI_ISL_522929, EPI_ISL_522930, EPI_ISL_522931, EPI_ISL_522932, EPI_ISL_522933, EPI_ISL_522934, EPI_ISL_522935, EPI_ISL_522936, EPI_ISL_522937, EPI_ISL_522938, EPI_ISL_522939 | | | |
| see above | University of Wisconsin-Madison AIDS Vaccine Research Laboratories | University of Wisconsin-Madison AIDS Vaccine Research Laboratories | Gage Moreno, Katarina Braun, et al. AIDS Vaccine Research Laboratories |
| EPI_ISL_524074, EPI_ISL_524075, EPI_ISL_524076, EPI_ISL_524077, EPI_ISL_524078, EPI_ISL_524079, EPI_ISL_524080, EPI_ISL_524081, EPI_ISL_524082, EPI_ISL_524083, EPI_ISL_524084, EPI_ISL_524085, EPI_ISL_524086, EPI_ISL_524087, EPI_ISL_524088, EPI_ISL_524089, EPI_ISL_524090, EPI_ISL_524091, EPI_ISL_524092, EPI_ISL_524093, EPI_ISL_524094, EPI_ISL_524095, EPI_ISL_524096, EPI_ISL_524097, EPI_ISL_524098, EPI_ISL_524099, EPI_ISL_524100, EPI_ISL_524101, EPI_ISL_524102, EPI_ISL_524103, EPI_ISL_524104, EPI_ISL_524105, EPI_ISL_524106, EPI_ISL_524107, EPI_ISL_524108, EPI_ISL_524109, EPI_ISL_524110, EPI_ISL_524111, EPI_ISL_524112, EPI_ISL_524113, EPI_ISL_524114, EPI_ISL_524115, EPI_ISL_524116, EPI_ISL_524117, EPI_ISL_524118, EPI_ISL_524119, EPI_ISL_524120, EPI_ISL_524121, EPI_ISL_524122, EPI_ISL_524123, EPI_ISL_524124, EPI_ISL_524125, EPI_ISL_524126, EPI_ISL_524127, EPI_ISL_524128, EPI_ISL_524129, EPI_ISL_524130, EPI_ISL_524131, EPI_ISL_524132, EPI_ISL_524133, EPI_ISL_524134, EPI_ISL_524135, EPI_ISL_524136, EPI_ISL_524137, EPI_ISL_524138, EPI_ISL_524139, EPI_ISL_524140, EPI_ISL_524141, EPI_ISL_524142, EPI_ISL_524143, EPI_ISL_524144, EPI_ISL_524145, EPI_ISL_524146, EPI_ISL_524147, EPI_ISL_524148, EPI_ISL_524149, EPI_ISL_524150, EPI_ISL_524151, EPI_ISL_524152, EPI_ISL_524153, EPI_ISL_524154, EPI_ISL_524155, EPI_ISL_524156, EPI_ISL_524157, EPI_ISL_524158, EPI_ISL_524159, EPI_ISL_524160, EPI_ISL_524161, EPI_ISL_524162, EPI_ISL_524163, EPI_ISL_524164, EPI_ISL_524165, EPI_ISL_524166, EPI_ISL_524167, EPI_ISL_524168, EPI_ISL_524169, EPI_ISL_524170, EPI_ISL_524171, EPI_ISL_524172, EPI_ISL_524173, EPI_ISL_524174, EPI_ISL_524175, EPI_ISL_524176, EPI_ISL_524177, EPI_ISL_524178, EPI_ISL_524179, EPI_ISL_524180, EPI_ISL_524181, EPI_ISL_524182, EPI_ISL_524183, EPI_ISL_524184, EPI_ISL_524185, EPI_ISL_524186, EPI_ISL_524187, EPI_ISL_524188, EPI_ISL_524189, EPI_ISL_524190, EPI_ISL_524191, EPI_ISL_524192, EPI_ISL_524193, EPI_ISL_524194, EPI_ISL_524195, EPI_ISL_524196, EPI_ISL_524197, EPI_ISL_524198, EPI_ISL_524199, EPI_ISL_524200, EPI_ISL_524201, EPI_ISL_524202, EPI_ISL_524203, EPI_ISL_524204, EPI_ISL_524205, EPI_ISL_524206, EPI_ISL_524207, EPI_ISL_524208, EPI_ISL_524209, EPI_ISL_524210, EPI_ISL_524211, EPI_ISL_524212, EPI_ISL_524213, EPI_ISL_524214, EPI_ISL_524215, EPI_ISL_524216, EPI_ISL_524217, EPI_ISL_524218, EPI_ISL_524219, EPI_ISL_524220, EPI_ISL_524221, EPI_ISL_524222, EPI_ISL_524223, EPI_ISL_524224, EPI_ISL_524225, EPI_ISL_524226, EPI_ISL_524227, EPI_ISL_524228, EPI_ISL_524229, EPI_ISL_524230, EPI_ISL_524231, EPI_ISL_524232, EPI_ISL_524233, EPI_ISL_524234, EPI_ISL_524235, EPI_ISL_524236, EPI_ISL_524237, EPI_ISL_524238, EPI_ISL_524239, EPI_ISL_524240, EPI_ISL_524241, EPI_ISL_524242, EPI_ISL_524243, EPI_ISL_524244, EPI_ISL_524245, EPI_ISL_524246, EPI_ISL_524247, EPI_ISL_524248, EPI_ISL_524249, EPI_ISL_524250, EPI_ISL_524251, EPI_ISL_524252, EPI_ISL_524253, | | | |

| | | | |
|---|---|---|---|
| EPI_ISL_524254, EPI_ISL_524255, EPI_ISL_524256, EPI_ISL_524257, EPI_ISL_524258, EPI_ISL_524259, EPI_ISL_524260, EPI_ISL_524261, EPI_ISL_524262, EPI_ISL_524263, EPI_ISL_524264, EPI_ISL_524265, EPI_ISL_524266, EPI_ISL_524267, EPI_ISL_524268, EPI_ISL_524269, EPI_ISL_524270, EPI_ISL_524271, EPI_ISL_524272, EPI_ISL_524273, EPI_ISL_524274, EPI_ISL_524275, EPI_ISL_524276, EPI_ISL_524277, EPI_ISL_524278, EPI_ISL_524279, EPI_ISL_524280, EPI_ISL_524281, EPI_ISL_524282, EPI_ISL_524283, EPI_ISL_524284, EPI_ISL_524285, EPI_ISL_524286, EPI_ISL_524287, EPI_ISL_524288, EPI_ISL_524289, EPI_ISL_524290, EPI_ISL_524291, EPI_ISL_524292, EPI_ISL_524293, EPI_ISL_524294, EPI_ISL_524295, EPI_ISL_524296, EPI_ISL_524297, EPI_ISL_524298, EPI_ISL_524299, EPI_ISL_524300, EPI_ISL_524301, EPI_ISL_524302, EPI_ISL_524303, EPI_ISL_524304, EPI_ISL_524305, EPI_ISL_524306, EPI_ISL_524307, EPI_ISL_524308, EPI_ISL_524309, EPI_ISL_524310, EPI_ISL_524311, EPI_ISL_524312, EPI_ISL_524313, EPI_ISL_524314, EPI_ISL_524315, EPI_ISL_524316, EPI_ISL_524317, EPI_ISL_524318, EPI_ISL_524319, EPI_ISL_524320, EPI_ISL_524321, EPI_ISL_524322, EPI_ISL_524323, EPI_ISL_524324, EPI_ISL_524325, EPI_ISL_524326, EPI_ISL_524327, EPI_ISL_524328, EPI_ISL_524329, EPI_ISL_524330, EPI_ISL_524331, EPI_ISL_524332, EPI_ISL_524333, EPI_ISL_524334, EPI_ISL_524335, EPI_ISL_524336, EPI_ISL_524337, EPI_ISL_524338, EPI_ISL_524339, EPI_ISL_524340, EPI_ISL_524341, EPI_ISL_524342, EPI_ISL_524343, EPI_ISL_524344, EPI_ISL_524345, EPI_ISL_524346, EPI_ISL_524347, EPI_ISL_524348, EPI_ISL_524349, EPI_ISL_524350, EPI_ISL_524351, EPI_ISL_524352, EPI_ISL_524353, EPI_ISL_524354, EPI_ISL_524355, EPI_ISL_524356, EPI_ISL_524357, EPI_ISL_524358, EPI_ISL_524359, EPI_ISL_524360, EPI_ISL_524361, EPI_ISL_524362, EPI_ISL_524363, EPI_ISL_524364, EPI_ISL_524365, EPI_ISL_524366, EPI_ISL_524367, EPI_ISL_524368, EPI_ISL_524369, EPI_ISL_524370, EPI_ISL_524371, EPI_ISL_524372, EPI_ISL_524373, EPI_ISL_524374, EPI_ISL_524375, EPI_ISL_524376, EPI_ISL_524377, EPI_ISL_524378, EPI_ISL_524379, EPI_ISL_524380, EPI_ISL_524381, EPI_ISL_524382, EPI_ISL_524383, EPI_ISL_524384, EPI_ISL_524385, EPI_ISL_524386, EPI_ISL_524387, EPI_ISL_524388, EPI_ISL_524389, EPI_ISL_524390, EPI_ISL_524391, EPI_ISL_524392, EPI_ISL_524393, EPI_ISL_524394, EPI_ISL_524395, EPI_ISL_524396, EPI_ISL_524397, EPI_ISL_524398, EPI_ISL_524399, EPI_ISL_524400, EPI_ISL_524401, EPI_ISL_524402, EPI_ISL_524403, EPI_ISL_524404, EPI_ISL_524405, EPI_ISL_524406, EPI_ISL_524407, EPI_ISL_524408, EPI_ISL_524409, EPI_ISL_524410, EPI_ISL_524411, EPI_ISL_524412, EPI_ISL_524413, EPI_ISL_524414, EPI_ISL_524415, EPI_ISL_524416, EPI_ISL_524417, EPI_ISL_524418, EPI_ISL_524419, EPI_ISL_524420, EPI_ISL_524421, EPI_ISL_524422, EPI_ISL_524423, EPI_ISL_524424, EPI_ISL_524425 | | | |
| see above | Utah Public Health Laboratory | Utah Public Health Laboratory | Erin L. Young, Kelly Oakeson, Tara Gallagher, Michael T. Pyne, E. Susan Slechta, Melanie A. Mallory, Jeffrey B. Stevenson, Salika M. Shakir, David R. Hillyard |
| EPI_ISL_524433, EPI_ISL_524434 | Environmental and Global Health, University of Florida - Gainesville | University of Florida | Elbadry,M.A., Subramaniam,K., Waltzek,T.B., Gibson,J.C., Stephenson,C.J., Alam,M.M., Morris,J.G. Jr., Lednicky,J.A. |
| EPI_ISL_524722 | B.J. Medical College and Civil hospital, Ahmedabad | Gujarat Biotechnology Research Centre | Dinesh Kumar, Zuber Saiyed, Komal Patel, Labdhi Pandya, Afzal Ansari, Nikha Trivedi, Pranay Shah, Kamlesh J Upadhyay, Sanjay Kapadia, Apurvasinh Puvar, Janvi Raval, Zarna Patel, Monika Gandhi, Pinal Trivedi, Maharshi Pandya, Nidhi Patel, Nitin Savaliya, Raghawendra Kumar, R D Dixit, A M Kadri, Harsh Bakshi, Chaitanya Joshi, Madhvi Joshi |
| EPI_ISL_524742 | GMERS Medical College and Hospital, Gandhinagar | Gujarat Biotechnology Research Centre | Dinesh Kumar, Zuber Saiyed, Komal Patel, Labdhi Pandya, Afzal Ansari, Nikha Trivedi, Seema Bhatt, Gaurishankar Shrimali, Bhavesh Modi, Bharti Rajani, Apurvasinh Puvar, Janvi Raval, Zarna Patel, Monika Gandhi, Pinal Trivedi, Maharshi Pandya, Nidhi Patel, Nitin Savaliya, Raghawendra Kumar, R D Dixit, A M Kadri, Harsh Bakshi, Chaitanya Joshi, Madhvi Joshi |
| EPI_ISL_524758 | GMERS Medical College and Hospital, Dharpur, Patan | Gujarat Biotechnology Research Centre | Dinesh Kumar, Zuber Saiyed, Komal Patel, Labdhi Pandya, Afzal Ansari, Nikha Trivedi, A N Parmar, Apurvasinh Puvar, Janvi Raval, Zarna Patel, Monika Gandhi, Pinal Trivedi, Maharshi Pandya, Nidhi Patel, Nitin Savaliya, Raghawendra Kumar, R D Dixit, A M Kadri, Harsh Bakshi, Chaitanya Joshi, Madhvi Joshi |
| EPI_ISL_524767, EPI_ISL_524768, EPI_ISL_524769, EPI_ISL_524770, EPI_ISL_524771, EPI_ISL_524772, EPI_ISL_524773, EPI_ISL_524774, EPI_ISL_524775, EPI_ISL_524776, EPI_ISL_524777, EPI_ISL_524778, EPI_ISL_524779, EPI_ISL_524780, EPI_ISL_524781, EPI_ISL_524802, EPI_ISL_524803, EPI_ISL_524804, EPI_ISL_524805, EPI_ISL_524806, EPI_ISL_524807, EPI_ISL_524808, EPI_ISL_524809, EPI_ISL_524810, EPI_ISL_524811, EPI_ISL_524812, EPI_ISL_524813, EPI_ISL_524814, EPI_ISL_524815, EPI_ISL_524816, EPI_ISL_524817, EPI_ISL_524818, EPI_ISL_524819, EPI_ISL_524820, EPI_ISL_524821, EPI_ISL_524822, EPI_ISL_524823, EPI_ISL_524824, EPI_ISL_524825, EPI_ISL_524826, EPI_ISL_524827, EPI_ISL_524828, EPI_ISL_524829, EPI_ISL_524830, EPI_ISL_524831, EPI_ISL_524832, EPI_ISL_524833, EPI_ISL_524834, EPI_ISL_524835, EPI_ISL_524836, EPI_ISL_524837, EPI_ISL_524838, EPI_ISL_524839, EPI_ISL_524840, EPI_ISL_524841, EPI_ISL_524842, EPI_ISL_524843, EPI_ISL_524844, EPI_ISL_524845, EPI_ISL_524846, EPI_ISL_524847, EPI_ISL_524848, EPI_ISL_524849, EPI_ISL_524850, EPI_ISL_524851, EPI_ISL_524852, EPI_ISL_524853, EPI_ISL_524854, EPI_ISL_524855, EPI_ISL_524856, EPI_ISL_524857, EPI_ISL_524858, EPI_ISL_524859, EPI_ISL_524860, EPI_ISL_524861, EPI_ISL_524862, EPI_ISL_524863, EPI_ISL_524864, EPI_ISL_524865, EPI_ISL_524866, EPI_ISL_524867, EPI_ISL_524868, EPI_ISL_524869, EPI_ISL_524870, EPI_ISL_524871, EPI_ISL_524872, EPI_ISL_524904, EPI_ISL_524905, EPI_ISL_524906, EPI_ISL_524907, EPI_ISL_524908, EPI_ISL_524909, EPI_ISL_524910, EPI_ISL_524911, EPI_ISL_524912, EPI_ISL_524913, EPI_ISL_524914, EPI_ISL_524915, EPI_ISL_524916, EPI_ISL_524917, EPI_ISL_524918, EPI_ISL_524919, EPI_ISL_524920, EPI_ISL_524921, EPI_ISL_524922, EPI_ISL_524923, EPI_ISL_524924, EPI_ISL_524925, EPI_ISL_524926, EPI_ISL_524927, EPI_ISL_524928, EPI_ISL_524929, EPI_ISL_524930, EPI_ISL_524931, EPI_ISL_524932, EPI_ISL_524933, EPI_ISL_524934, EPI_ISL_524935, EPI_ISL_524936, EPI_ISL_524937, EPI_ISL_524938, EPI_ISL_524939, EPI_ISL_524940, EPI_ISL_524941, EPI_ISL_524942, EPI_ISL_524943, EPI_ISL_524944, EPI_ISL_524945, EPI_ISL_524946, EPI_ISL_524947, EPI_ISL_524948, EPI_ISL_524949, EPI_ISL_524950, EPI_ISL_524951, EPI_ISL_524952, EPI_ISL_524953, EPI_ISL_524954, EPI_ISL_524955, EPI_ISL_524956, EPI_ISL_524957, EPI_ISL_524958, EPI_ISL_524959, EPI_ISL_524960, EPI_ISL_524961, EPI_ISL_524962, EPI_ISL_524963, EPI_ISL_524964, EPI_ISL_524965, EPI_ISL_524966, EPI_ISL_524967, EPI_ISL_524968, EPI_ISL_524969, EPI_ISL_524970, EPI_ISL_524971, EPI_ISL_524972, EPI_ISL_524973, EPI_ISL_524974, EPI_ISL_524975, EPI_ISL_524976, EPI_ISL_524977, EPI_ISL_524978, EPI_ISL_524979, EPI_ISL_524980, EPI_ISL_524981, EPI_ISL_524982, EPI_ISL_524983, EPI_ISL_524984, EPI_ISL_524985, EPI_ISL_524986, EPI_ISL_524987, EPI_ISL_524988, EPI_ISL_524989, EPI_ISL_524990, EPI_ISL_524991, EPI_ISL_524992, EPI_ISL_524993, EPI_ISL_524994, EPI_ISL_524995, EPI_ISL_524996, EPI_ISL_524997, EPI_ISL_524998, EPI_ISL_524999, EPI_ISL_525000, EPI_ISL_525001, EPI_ISL_525002, EPI_ISL_525003, EPI_ISL_525004, EPI_ISL_525005, EPI_ISL_525006, EPI_ISL_525007, EPI_ISL_525008, EPI_ISL_525009, EPI_ISL_525010, EPI_ISL_525011, EPI_ISL_525012, EPI_ISL_525013, EPI_ISL_525014, EPI_ISL_525015, EPI_ISL_525016, EPI_ISL_525017, EPI_ISL_525018, EPI_ISL_525019, EPI_ISL_525020, EPI_ISL_525021, EPI_ISL_525022, EPI_ISL_525023, EPI_ISL_525024, EPI_ISL_525025, EPI_ISL_525026, EPI_ISL_525027, EPI_ISL_525028, EPI_ISL_525029, EPI_ISL_525030, EPI_ISL_525031, EPI_ISL_525032, EPI_ISL_525033, EPI_ISL_525034, EPI_ISL_525035, EPI_ISL_525036, EPI_ISL_525037, EPI_ISL_525038, EPI_ISL_525039, EPI_ISL_525040, EPI_ISL_525041, EPI_ISL_525042, EPI_ISL_525043, EPI_ISL_525044, EPI_ISL_525045, EPI_ISL_525046, EPI_ISL_525047, EPI_ISL_525048, EPI_ISL_525049, EPI_ISL_525050, EPI_ISL_525051, EPI_ISL_525052, EPI_ISL_525053, EPI_ISL_525054, EPI_ISL_525055, EPI_ISL_525056, EPI_ISL_525057, EPI_ISL_525058, EPI_ISL_525059, EPI_ISL_525060, EPI_ISL_525061, EPI_ISL_525062, EPI_ISL_525063, EPI_ISL_525064, EPI_ISL_525065, EPI_ISL_525066, EPI_ISL_525067, EPI_ISL_525068, EPI_ISL_525069, EPI_ISL_525070, EPI_ISL_525071, EPI_ISL_525072, EPI_ISL_525073, EPI_ISL_525074, EPI_ISL_525075, EPI_ISL_525076, EPI_ISL_525077, EPI_ISL_525078, EPI_ISL_525079, EPI_ISL_525080, EPI_ISL_525081, EPI_ISL_525082, EPI_ISL_525083, EPI_ISL_525084, EPI_ISL_525085, EPI_ISL_525086, EPI_ISL_525087, EPI_ISL_525088, EPI_ISL_525089, EPI_ISL_525090, EPI_ISL_525091, EPI_ISL_525092, EPI_ISL_525093, EPI_ISL_525094, EPI_ISL_525095, EPI_ISL_525096, EPI_ISL_525097, EPI_ISL_525098, EPI_ISL_525099, EPI_ISL_525100, EPI_ISL_525101, EPI_ISL_525102, EPI_ISL_525103, EPI_ISL_525104, EPI_ISL_525105, EPI_ISL_525106, EPI_ISL_525107, EPI_ISL_525108, EPI_ISL_525109, EPI_ISL_525110, EPI_ISL_525111, EPI_ISL_525112, EPI_ISL_525113, EPI_ISL_525114, EPI_ISL_525115, EPI_ISL_525116, EPI_ISL_525117, EPI_ISL_525119, EPI_ISL_525120, EPI_ISL_525121, EPI_ISL_525122, EPI_ISL_525123, EPI_ISL_525124, EPI_ISL_525125, EPI_ISL_525126, EPI_ISL_525127, EPI_ISL_525128, EPI_ISL_525129, EPI_ISL_525130, EPI_ISL_525131, EPI_ISL_525132, EPI_ISL_525133, EPI_ISL_525134, EPI_ISL_525135, EPI_ISL_525136, EPI_ISL_525137, EPI_ISL_525138, EPI_ISL_525139, EPI_ISL_525140, EPI_ISL_525141, EPI_ISL_525142, EPI_ISL_525143, EPI_ISL_525144, EPI_ISL_525145, EPI_ISL_525146, EPI_ISL_525147, EPI_ISL_525148, EPI_ISL_525149, EPI_ISL_525150, EPI_ISL_525151, EPI_ISL_525152, EPI_ISL_525153, EPI_ISL_525154, EPI_ISL_525155, EPI_ISL_525156, EPI_ISL_525157, EPI_ISL_525158, EPI_ISL_525159, EPI_ISL_525160, EPI_ISL_525161, EPI_ISL_525162, EPI_ISL_525163, EPI_ISL_525164, EPI_ISL_525165, EPI_ISL_525166, EPI_ISL_525167, EPI_ISL_525168, EPI_ISL_525169, EPI_ISL_525170, EPI_ISL_525171, EPI_ISL_525172, EPI_ISL_525173, EPI_ISL_525174, EPI_ISL_525175, EPI_ISL_525176, EPI_ISL_525177, EPI_ISL_525178, EPI_ISL_525179, EPI_ISL_525180, EPI_ISL_525181, EPI_ISL_525182, EPI_ISL_525183, EPI_ISL_525184, EPI_ISL_525185, EPI_ISL_525186, EPI_ISL_525187, EPI_ISL_525188, EPI_ISL_525207, EPI_ISL_525208, EPI_ISL_525209, EPI_ISL_525210, EPI_ISL_525211, EPI_ISL_525212, EPI_ISL_525213, EPI_ISL_525214, EPI_ISL_525215, EPI_ISL_525216, EPI_ISL_525217, EPI_ISL_525218, EPI_ISL_525219, EPI_ISL_525220, EPI_ISL_525221, EPI_ISL_525222, EPI_ISL_525223, EPI_ISL_525224, EPI_ISL_525225, EPI_ISL_525226, EPI_ISL_525227, EPI_ISL_525228, EPI_ISL_525229, EPI_ISL_525230, EPI_ISL_525231, EPI_ISL_525232, EPI_ISL_525233, EPI_ISL_525234, EPI_ISL_525235, EPI_ISL_525236, EPI_ISL_525237, EPI_ISL_525238, EPI_ISL_525239, EPI_ISL_525240, EPI_ISL_525241, EPI_ISL_525242, EPI_ISL_525243, EPI_ISL_525244, EPI_ISL_525245, EPI_ISL_525246, EPI_ISL_525247, EPI_ISL_525248, EPI_ISL_525249, EPI_ISL_525250, EPI_ISL_525251, EPI_ISL_525252, EPI_ISL_525253, EPI_ISL_525254, EPI_ISL_525255, EPI_ISL_525256, EPI_ISL_525257, EPI_ISL_525258, EPI_ISL_525259, EPI_ISL_525260, EPI_ISL_525261, EPI_ISL_525262, EPI_ISL_525263, EPI_ISL_525264, EPI_ISL_525265, EPI_ISL_525266, EPI_ISL_525267, EPI_ISL_525268, EPI_ISL_525269, EPI_ISL_525270, EPI_ISL_525271, EPI_ISL_525272, EPI_ISL_525273, EPI_ISL_525274, EPI_ISL_525275, EPI_ISL_525276, EPI_ISL_525277, EPI_ISL_525278, EPI_ISL_525279, EPI_ISL_525280, EPI_ISL_525281, EPI_ISL_525282, EPI_ISL_525283, EPI_ISL_525284, EPI_ISL_525285, EPI_ISL_525286, EPI_ISL_525287, EPI_ISL_525288, EPI_ISL_525289, EPI_ISL_525290, EPI_ISL_525291, EPI_ISL_525292, EPI_ISL_525293, EPI_ISL_525294, EPI_ISL_525295, EPI_ISL_525296, EPI_ISL_525297, EPI_ISL_525298, EPI_ISL_525299, EPI_ISL_525300, EPI_ISL_525301, EPI_ISL_525302, EPI_ISL_525303, EPI_ISL_525304, EPI_ISL_525305, EPI_ISL_525306, EPI_ISL_525307, EPI_ISL_525308, EPI_ISL_525309, EPI_ISL_525310, EPI_ISL_525311, EPI_ISL_525312, EPI_ISL_525313, EPI_ISL_525314, EPI_ISL_525315, EPI_ISL_525316, EPI_ISL_525317, EPI_ISL_525318, EPI_ISL_525319, EPI_ISL_525320, EPI_ISL_525321, EPI_ISL_525322, EPI_ISL_525323, EPI_ISL_525324, EPI_ISL_525325, EPI_ISL_525326, EPI_ISL_525327, EPI_ISL_525328, EPI_ISL_525329, EPI_ISL_525330, EPI_ISL_525331, EPI_ISL_525332, EPI_ISL_525333, EPI_ISL_525334, EPI_ISL_525335, EPI_ISL_525336 | | | |
| see above | Utah Public Health Laboratory | Utah Public Health Laboratory | Erin L. Young, Kelly Oakeson, Tara Gallagher, Michael T. Pyne, E. Susan Slechta, Melanie A. Mallory, Jeffrey B. Stevenson, Salika M. Shakir, David R. Hillyard |
| EPI_ISL_525337, EPI_ISL_525338, EPI_ISL_525339, EPI_ISL_525340, EPI_ISL_525341, EPI_ISL_525342, EPI_ISL_525343, EPI_ISL_525344, EPI_ISL_525345, EPI_ISL_525346, EPI_ISL_525347, EPI_ISL_525348, EPI_ISL_525349, EPI_ISL_525350, EPI_ISL_525351 | | | |
| see above | Utah Public Health Laboratory | Utah Public Health Laboratory | Erin Young, Kelly Oakeson |
| EPI_ISL_525438 | Institute of Microbiology, Universidad San Francisco de Quito | Institute of Microbiology, Universidad San Francisco de Quito | Diego Egas, Juan José Guadalupe, Monica Becerra-Wong, Belén Prado-Vivar, Sully Márquez, Bernardo Gutiérrez, Stalin Castillo, Eddy Chavez, Francisco Rodriguez, Verónica Barragán, Patricio Rojas-Silva, Gabriel Trueba, Michelle Grunauer, Paúl Cárdenas |
| EPI_ISL_526215, EPI_ISL_526216, EPI_ISL_526217, EPI_ISL_526218, EPI_ISL_526219, EPI_ISL_526220, EPI_ISL_526221, EPI_ISL_526222, EPI_ISL_526223, EPI_ISL_526224, EPI_ISL_526225, EPI_ISL_526226, EPI_ISL_526227, EPI_ISL_526228, EPI_ISL_526229, EPI_ISL_526230, EPI_ISL_526231, EPI_ISL_526232, EPI_ISL_526233, EPI_ISL_526234, EPI_ISL_526235, EPI_ISL_526236, EPI_ISL_526237, EPI_ISL_526238 | | | |
| see above | Hungarian Defence Forces Military Medical Centre | National Laboratory of Virology, Szentágothai Research Centre | Endre Gábor Tóth, Balázs Somogyi, Bálint Eszenyi, Ferenc Jakab, Gábor Kemenesi |
| EPI_ISL_526337, EPI_ISL_526338, EPI_ISL_526339, EPI_ISL_526340, EPI_ISL_526341, EPI_ISL_526342, EPI_ISL_526343, EPI_ISL_526344, EPI_ISL_526345, EPI_ISL_526346, EPI_ISL_526347, EPI_ISL_526348, EPI_ISL_526349, EPI_ISL_526350, EPI_ISL_526351, EPI_ISL_526352, EPI_ISL_526353, EPI_ISL_526354, EPI_ISL_526355, EPI_ISL_526356, EPI_ISL_526357, EPI_ISL_526358, EPI_ISL_526359, EPI_ISL_526360, EPI_ISL_526361, EPI_ISL_526362, EPI_ISL_526363, EPI_ISL_526364, EPI_ISL_526365, EPI_ISL_526366, EPI_ISL_526395, EPI_ISL_526396, EPI_ISL_526397, EPI_ISL_526398, EPI_ISL_526399, EPI_ISL_526400, EPI_ISL_526401, EPI_ISL_526402, EPI_ISL_526403, EPI_ISL_526404, EPI_ISL_526405, EPI_ISL_526406, EPI_ISL_526407, EPI_ISL_526408, EPI_ISL_526409, EPI_ISL_526410, EPI_ISL_526411, EPI_ISL_526412, EPI_ISL_526413, EPI_ISL_526414, EPI_ISL_526415, EPI_ISL_526416, EPI_ISL_526417, EPI_ISL_526418, EPI_ISL_526419, EPI_ISL_526420, EPI_ISL_526421, EPI_ISL_526422, EPI_ISL_526423, EPI_ISL_526424, EPI_ISL_526425, EPI_ISL_526426, EPI_ISL_526427, EPI_ISL_526428, EPI_ISL_526429, EPI_ISL_526430 | | | |
| see above | Queens Medical Centre, Clinical Microbiology Department / DeepSeq Nottingham | COVID-19 Genomics UK (COG-UK) Consortium | Gemma Clark, Wendy Smith, Manjinder Khakh, Vicki M Fleming, Michelle M Lister, Hannah Howson-Wells, Jonathan Ball, Patrick McClure, Joseph Chappell, Theocharis Tsoleridis, Nadine Holmes, Matthew Carlisle, Christopher Moore, Fei Sang, Johnny Debebe, Victoria Wright, Matthew Loose |
| EPI_ISL_527381, EPI_ISL_527382, EPI_ISL_527383, EPI_ISL_527384, EPI_ISL_527385, EPI_ISL_527386, EPI_ISL_527387, EPI_ISL_527388, EPI_ISL_527389, EPI_ISL_527390, EPI_ISL_527391, EPI_ISL_527392, EPI_ISL_527393, EPI_ISL_527394, EPI_ISL_527395, EPI_ISL_527396, EPI_ISL_527397, EPI_ISL_527398, EPI_ISL_527399 | | | |
| see above | University of Miami Immunology and Histocompatibility Laboratory | University of Miami Immunology and Histocompatibility Laboratory | Emilio Margolles-Clark, PhD and Phillip Ruiz, MD, PhD |
| EPI_ISL_527739 | Hospital Mexico [San Jose/San Jose] | Inciensa, Instituto Costarricense de Investigación y Enseñanza en Nutrición y Salud | Francisco Duarte, Hebleen Porras, Claudio Soto-Garita, Estela Cordero, Adriana Godinez & Melany Calderon |
| EPI_ISL_527740 | Area De Salud Alajuela Norte - Clinica Dr. Marcial Rodriguez | Inciensa, Instituto Costarricense de Investigación y Enseñanza en Nutrición y Salud | Francisco Duarte, Hebleen Porras, Claudio Soto-Garita, Estela Cordero, Adriana Godinez & Melany Calderon |
| EPI_ISL_527741 | Hospital Metropolitano | Inciensa, Instituto Costarricense de Investigación y Enseñanza en Nutrición y Salud | Francisco Duarte, Hebleen Porras, Claudio Soto-Garita, Estela Cordero, Adriana Godinez & Melany Calderon |

| | | | |
|---|---|---|---|
| EPI_ISL_527742 | Centro Nacional De Rehabilitacion Humberto Araya Rojas (Cenare) | Inciensa, Instituto Costarricense de Investigación y Enseñanza en Nutrición y Salud | Francisco Duarte, Hebleen Porras, Claudio Soto-Garita, Estela Cordero, Adriana Godinez & Melany Calderon |
| EPI_ISL_527743, EPI_ISL_527744 | Hospital Cima | Inciensa, Instituto Costarricense de Investigación y Enseñanza en Nutrición y Salud | Francisco Duarte, Hebleen Porras, Claudio Soto-Garita, Estela Cordero, Adriana Godinez & Melany Calderon |
| EPI_ISL_527745 | Area De Salud Corredores | Inciensa, Instituto Costarricense de Investigación y Enseñanza en Nutrición y Salud | Francisco Duarte, Hebleen Porras, Claudio Soto-Garita, Estela Cordero, Adriana Godinez & Melany Calderon |
| EPI_ISL_527746, EPI_ISL_527747 | Area De Salud La Cruz | Inciensa, Instituto Costarricense de Investigación y Enseñanza en Nutrición y Salud | Francisco Duarte, Hebleen Porras, Claudio Soto-Garita, Estela Cordero, Adriana Godinez & Melany Calderon |
| EPI_ISL_527748 | Area De Salud Corredores | Inciensa, Instituto Costarricense de Investigación y Enseñanza en Nutrición y Salud | Francisco Duarte, Hebleen Porras, Claudio Soto-Garita, Estela Cordero, Adriana Godinez & Melany Calderon |
| EPI_ISL_527749 | Area De Salud Alajuela Norte - Clinica Dr. Marcial Rodriguez | Inciensa, Instituto Costarricense de Investigación y Enseñanza en Nutrición y Salud | Francisco Duarte, Hebleen Porras, Claudio Soto-Garita, Estela Cordero, Adriana Godinez & Melany Calderon |
| EPI_ISL_527750 | Hospital De Niños Dr. Carlos Saenz Herrera [San Jose/San Jose] | Inciensa, Instituto Costarricense de Investigación y Enseñanza en Nutrición y Salud | Francisco Duarte, Hebleen Porras, Claudio Soto-Garita, Estela Cordero, Adriana Godinez & Melany Calderon |
| EPI_ISL_527751 | Area De Salud Corredores | Inciensa, Instituto Costarricense de Investigación y Enseñanza en Nutrición y Salud | Francisco Duarte, Hebleen Porras, Claudio Soto-Garita, Estela Cordero, Adriana Godinez & Melany Calderon |
| EPI_ISL_527752 | Hospital Dr. Rafael A. Calderon Guardia | Inciensa, Instituto Costarricense de Investigación y Enseñanza en Nutrición y Salud | Francisco Duarte, Hebleen Porras, Claudio Soto-Garita, Estela Cordero, Adriana Godinez & Melany Calderon |
| EPI_ISL_527753, EPI_ISL_527754 | Hospital San Vicente De Paul | Inciensa, Instituto Costarricense de Investigación y Enseñanza en Nutrición y Salud | Francisco Duarte, Hebleen Porras, Claudio Soto-Garita, Estela Cordero, Adriana Godinez & Melany Calderon |
| EPI_ISL_527755 | Centro Nacional De Rehabilitacion Humberto Araya Rojas (Cenare) | Inciensa, Instituto Costarricense de Investigación y Enseñanza en Nutrición y Salud | Francisco Duarte, Hebleen Porras, Claudio Soto-Garita, Estela Cordero, Adriana Godinez & Melany Calderon |
| EPI_ISL_527756 | Area De Salud Aserri | Inciensa, Instituto Costarricense de Investigación y Enseñanza en Nutrición y Salud | Francisco Duarte, Hebleen Porras, Claudio Soto-Garita, Estela Cordero, Adriana Godinez & Melany Calderon |
| EPI_ISL_527757 | Area De Salud Goicoechea 1 | Inciensa, Instituto Costarricense de Investigación y Enseñanza en Nutrición y Salud | Francisco Duarte, Hebleen Porras, Claudio Soto-Garita, Estela Cordero, Adriana Godinez & Melany Calderon |
| EPI_ISL_527794, EPI_ISL_527795, EPI_ISL_527796, EPI_ISL_527797, EPI_ISL_527798, EPI_ISL_527799, EPI_ISL_527800, EPI_ISL_527801, EPI_ISL_527802, EPI_ISL_527803, EPI_ISL_527804, EPI_ISL_527805, EPI_ISL_527806, EPI_ISL_527807, EPI_ISL_527808 | | | |
| see above | University of Wisconsin-Madison AIDS Vaccine Research Laboratories | University of Wisconsin-Madison AIDS Vaccine Research Laboratories | Gage Moreno, Katarina Braun, et al. AIDS Vaccine Research Laboratories |
| EPI_ISL_527818, EPI_ISL_527819 | Centro de Investigaciones, Universidad de Especialidades Espíritu Santo | Institute of Microbiology, Universidad San Francisco de Quito | Derly Andrade, Juan Carlos Fernandez, Belén Prado-Vivar, Sully Márquez, Juan José Guadalupe, Monica Becerra-Wong, Bernardo Gutiérrez, Gabriel Morey, Ruben Armas, Jose Pedro Barberan, Fernando Espinoza, Edith Lopez, Verónica Barragán, Patricio Rojas-Silva, Gabriel Trueba, Michelle Grunauer, Paúl Cárdenas |
| EPI_ISL_528540, EPI_ISL_528541 | National Genomics Core-Center for DNA Fingerprinting and Diagnostics | National Genomics Core- Center for DNA Fingerprinting and Diagnostics (NGC-CDFD)- DBT's PAN-INDIA-1000 Genome consortium | G Shashikanth, Heena Shah, Bala Pratyusha, Vinay Donipadi, K.Manohar, Madhumohan Rao, Shubhra Ganguli, Suchitra Upreti, Swathi Chodisetty, Vani Singh, R Harinarayanan, Rashna Bhandari, Murali Dharan Bashyam, Debashish Mitra, Divya Vashisht, Ashwin Dalal |
| EPI_ISL_528542, EPI_ISL_528543, EPI_ISL_528544, EPI_ISL_528545, EPI_ISL_528546, EPI_ISL_528547, EPI_ISL_528548, EPI_ISL_528549 | National Genomics Core-Center for DNA Fingerprinting and Diagnostics | National Genomics Core- Center for DNA Fingerprinting and Diagnostics (NGC-CDFD)- DBT's PAN-INDIA-1000 Genome consortium | G Shashikanth, Heena Shah, Bala Pratyusha, Vinay Donipadi, S Vasantha Rani, M Sri Lalitha, R. Angalena, Usha Rani Dutta, Nimmala Naresh, Nanci Rani K, Ch Venkateshwar Goud, Devinder Singh Negi, R Harinarayanan, Rashna Bhandari, Murali Dharan Bashyam, Debashish Mitra, Divya Vashisht, Ashwin Dalal |
| EPI_ISL_528565, EPI_ISL_528566, EPI_ISL_528567, EPI_ISL_528568, EPI_ISL_528569, EPI_ISL_528570, EPI_ISL_528571, EPI_ISL_528572 | National Genomics Core-Center for DNA Fingerprinting and Diagnostics | National Genomics Core- Center for DNA Fingerprinting and Diagnostics (NGC-CDFD)- DBT's PAN-INDIA-1000 Genome consortium | G Shashikanth, Heena Shah, Bala Pratyusha, Vinay Donipadi, Rajitha Ponnala, Seyed Khaja Ali, B. Krishna Murthy, Akruti Shah, Jayashree Ladke, Shivangi Wagh, Asodu Sandeep Sarma, Sunu Joseph, R Harinarayanan, Rashna Bhandari, Murali Dharan Bashyam, Debashish Mitra, Divya Vashisht, Ashwin Dalal |
| EPI_ISL_528581, EPI_ISL_528582, EPI_ISL_528583, EPI_ISL_528584, EPI_ISL_528585, EPI_ISL_528586, EPI_ISL_528587 | National Genomics Core-Center for DNA Fingerprinting and Diagnostics | National Genomics Core- Center for DNA Fingerprinting and Diagnostics (NGC-CDFD)- DBT's PAN-INDIA-1000 Genome consortium | G Shashikanth, Heena Shah, Bala Pratyusha, Vinay Donipadi, Bathula Siddardha, Vineesha Oddi, Lavanya Banda, Surya Chodisetty, Abhijeeth Singh Thakur, Mohammad Mudassir, Nalini Raghunathan, Rajeshree Sanyal, R Harinarayanan, Rashna Bhandari, Murali Dharan Bashyam, Debashish Mitra, Divya Vashisht, Ashwin Dalal |
| EPI_ISL_528609, EPI_ISL_528610, EPI_ISL_528611, EPI_ISL_528612, EPI_ISL_528613, EPI_ISL_528614, EPI_ISL_528615, EPI_ISL_528616 | National Genomics Core-Center for DNA Fingerprinting and Diagnostics | National Genomics Core- Center for DNA Fingerprinting and Diagnostics (NGC-CDFD)- DBT's PAN-INDIA-1000 Genome consortium | Divya Vashisht, Bala Pratyusha, Heena Shah, G Shashikanth, Vinay Donipadi, K.Manohar, Madhumohan Rao, SPR Prasad, Yogesh Patidar, Arjita Jaiswal, Arpita Singh, Devanshi Gupta, Romila Moirangthem, Sanjana Sarkar, Shivani Yadav, R Harinarayanan, Rashna Bhandari, Murali Dharan Bashyam, Debashish Mitra, Ashwin Dalal |
| EPI_ISL_528625, EPI_ISL_528626, EPI_ISL_528627, EPI_ISL_528628, EPI_ISL_528629, EPI_ISL_528630 | National Genomics Core-Center for DNA Fingerprinting and Diagnostics | National Genomics Core- Center for DNA Fingerprinting and Diagnostics (NGC-CDFD)- DBT's PAN-INDIA-1000 Genome consortium | G Shashikanth, Heena Shah, Bala Pratyusha, Vinay Donipadi, K.Manohar, Madhumohan Rao, Shubhra Ganguli, Suchitra Upreti, Swathi Chodisetty, Vani Singh, R Harinarayanan, Rashna Bhandari, Murali Dharan Bashyam, Debashish Mitra, Divya Vashisht, Ashwin Dalal |
| EPI_ISL_528880 | National Genomics Core-Center for DNA Fingerprinting and Diagnostics | National Genomics Core- Center for DNA Fingerprinting and Diagnostics (NGC-CDFD)- DBT's PAN-INDIA-1000 Genome consortium | G Shashikanth, Heena Shah, Bala Pratyusha, Vinay Donipadi, Bathula Siddardha, Vineesha Oddi, Lavanya Banda, Surya Chodisetty, Abhijeeth Singh Thakur, Mohammad Mudassir, Nalini Raghunathan, Rajeshree Sanyal, R Harinarayanan, Rashna Bhandari, Murali Dharan Bashyam, Debashish Mitra, Divya Vashisht, Ashwin Dalal |
| EPI_ISL_529080, EPI_ISL_529081, EPI_ISL_529082, EPI_ISL_529083, EPI_ISL_529084, EPI_ISL_529085, EPI_ISL_529086, EPI_ISL_529087, EPI_ISL_529088, EPI_ISL_529089, EPI_ISL_529090, EPI_ISL_529091, EPI_ISL_529092, EPI_ISL_529093, EPI_ISL_529094, EPI_ISL_529095, EPI_ISL_529096, EPI_ISL_529097, EPI_ISL_529098, EPI_ISL_529099, EPI_ISL_529100, EPI_ISL_529101, EPI_ISL_529102, EPI_ISL_529103, EPI_ISL_529104, EPI_ISL_529105, EPI_ISL_529106, EPI_ISL_529107, EPI_ISL_529108 | | | |
| see above | Microbiology Division, SC DHEC | Microbiology Division, SC DHEC | Flores,H. |
| EPI_ISL_529167, EPI_ISL_529168 | Maryland Department of Health Laboratory | Maryland Department of Health Laboratory | Eric Keller, Maryland Department of Health Laboratories Administration |
| EPI_ISL_529202, EPI_ISL_529203 | Utah Public Health Laboratory | Utah Public Health Laboratory | Erin Young, Kelly Oakeson |
| EPI_ISL_529213, EPI_ISL_529214, EPI_ISL_529215, EPI_ISL_529216, EPI_ISL_529217 | Beijing Institute of Microbiology and Epidemiology | Beijing Institute of Microbiology and Epidemiology | Fan, Hang; Qin, E.; Wu, Y.; Guo, Y.; Zhang, X.; Yong, Y.; Hou, J.; Xu, Z.; Mu, J.; Teng, Yue; Mi, Z.; Yang, R.; Song, Yajun.; Li, B.; Cui, Y. |
| EPI_ISL_529264, EPI_ISL_529266, EPI_ISL_529276, EPI_ISL_529279, EPI_ISL_529288, EPI_ISL_529306, EPI_ISL_529307, EPI_ISL_529308, EPI_ISL_529309, EPI_ISL_529310, EPI_ISL_529311, EPI_ISL_529383, EPI_ISL_529385, EPI_ISL_529387, EPI_ISL_529410, EPI_ISL_529411, EPI_ISL_529412, EPI_ISL_529414, EPI_ISL_529423, EPI_ISL_529436, EPI_ISL_529450, EPI_ISL_529454, EPI_ISL_529455, EPI_ISL_529456, EPI_ISL_529463, EPI_ISL_529464, EPI_ISL_529467, EPI_ISL_529479, EPI_ISL_529483, EPI_ISL_529486, EPI_ISL_529504, EPI_ISL_529519, EPI_ISL_529528, EPI_ISL_529529, EPI_ISL_529530, EPI_ISL_529531, EPI_ISL_529532, EPI_ISL_529533, EPI_ISL_529534, EPI_ISL_529535, EPI_ISL_529536, EPI_ISL_529537, EPI_ISL_529538, EPI_ISL_529539, EPI_ISL_529540, EPI_ISL_529541, EPI_ISL_529542, EPI_ISL_529543, EPI_ISL_529544, EPI_ISL_529545, EPI_ISL_529546, EPI_ISL_529547, EPI_ISL_529548, EPI_ISL_529549, EPI_ISL_529550, EPI_ISL_529551, EPI_ISL_529552, EPI_ISL_529553, EPI_ISL_529554, EPI_ISL_529555, EPI_ISL_529556, EPI_ISL_529667 | | | |
| see above | Queens Medical Centre, Clinical Microbiology Department / DeepSeq Nottingham | COVID-19 Genomics UK (COG-UK) Consortium | Gemma Clark, Wendy Smith, Manjinder Khakh, Vicki M Fleming, Michelle M Lister, Hannah Howson-Wells, Jonathan Ball, Patrick McClure, Joseph Chappell, Theocharis Tsoleridis, Nadine Holmes, Matthew Carlisle, Christopher Moore, Fei Sang, Johnny Debebe, Victoria Wright, Matthew Loose |
| EPI_ISL_529720, EPI_ISL_529721, EPI_ISL_529722, EPI_ISL_529723, EPI_ISL_529724, EPI_ISL_529726, EPI_ISL_529728, EPI_ISL_529729, EPI_ISL_529730, EPI_ISL_529731, EPI_ISL_529733, EPI_ISL_529734, EPI_ISL_529735, EPI_ISL_529736, EPI_ISL_529737, EPI_ISL_529739, EPI_ISL_529740, EPI_ISL_529741, EPI_ISL_529742, EPI_ISL_529743, EPI_ISL_529744, EPI_ISL_529747, EPI_ISL_529748, EPI_ISL_529749, EPI_ISL_529750, EPI_ISL_529751, EPI_ISL_529752, EPI_ISL_529753, EPI_ISL_529754, EPI_ISL_529755, EPI_ISL_529756, EPI_ISL_529757, EPI_ISL_529758, EPI_ISL_529759, EPI_ISL_529760, EPI_ISL_529761, | | | |

| | | | |
|---|---|---|---|
| EPI_ISL_529762, EPI_ISL_529764, EPI_ISL_529765, EPI_ISL_529766, EPI_ISL_529767, EPI_ISL_529769, EPI_ISL_529770, EPI_ISL_529771, EPI_ISL_529772, EPI_ISL_529773, EPI_ISL_529774, EPI_ISL_529775, EPI_ISL_529776, EPI_ISL_529777, EPI_ISL_529778, EPI_ISL_529779, EPI_ISL_529780, EPI_ISL_529782, EPI_ISL_529783, EPI_ISL_529784, EPI_ISL_529785, EPI_ISL_529787, EPI_ISL_529788, EPI_ISL_529789, EPI_ISL_529790, EPI_ISL_529791, EPI_ISL_529792, EPI_ISL_529794, EPI_ISL_529796, EPI_ISL_529798, EPI_ISL_529799, EPI_ISL_529800, EPI_ISL_529801, EPI_ISL_529802 | | | |
| see above | NHLS-IALCH | KRISP, KZN Research Innovation and Sequencing Platform | Giandhari J, Pillay S, Lessells R, Mdlalose K, York D, Khan S, Tegally H, Wilkinson E, de Oliveira T |
| EPI_ISL_530007, EPI_ISL_530008, EPI_ISL_530009, EPI_ISL_530010, EPI_ISL_530011, EPI_ISL_530012, EPI_ISL_530013, EPI_ISL_530014, EPI_ISL_530015, EPI_ISL_530016, EPI_ISL_530017, EPI_ISL_530018, EPI_ISL_530019, EPI_ISL_530020, EPI_ISL_530021, EPI_ISL_530022, EPI_ISL_530023, EPI_ISL_530024 | | | |
| see above | Hospital Universitario 12 de Octubre | Hospital Universitario 12 de Octubre | Esther Viedma, Raúl Recio, Sara González, Elias Dahdouh, Fernando Lázaro, Natalia Stella, Julio García, Juan Carlos Galán, Rafael Cantón, Mª Dolores Folgueira, Rafael Delgado, Jesús Mingorance |
| EPI_ISL_530044, EPI_ISL_530045, EPI_ISL_530046, EPI_ISL_530047, EPI_ISL_530048, EPI_ISL_530049, EPI_ISL_530050, EPI_ISL_530051, EPI_ISL_530052, EPI_ISL_530053, EPI_ISL_530054, EPI_ISL_530055, EPI_ISL_530056, EPI_ISL_530057, EPI_ISL_530058, EPI_ISL_530059, EPI_ISL_530060, EPI_ISL_530061, EPI_ISL_530062, EPI_ISL_530063, EPI_ISL_530064, EPI_ISL_530065, EPI_ISL_530066, EPI_ISL_530067, EPI_ISL_530068, EPI_ISL_530069 | | | |
| see above | Hospital Universitario La Paz | Hospital Universitario La Paz | Elias Dahdouh, Sara González, Raúl Recio, Fernando Lázaro, Esther Viedma, Natalia Stella, Julio García, Juan Carlos Galán, Rafael Cantón, Mª Dolores Folgueira, Rafael Delgado, Jesús Mingorance |
| EPI_ISL_534329 | Hospital Universitario 12 de Octubre | Hospital Universitario 12 de Octubre | Esther Viedma, Raúl Recio, Sara González, Elias Dahdouh, Fernando Lázaro, Natalia Stella, Julio García, Juan Carlos Galán, Rafael Cantón, Ma Dolores Folgueira, Rafael Delgado, Jesús Mingorance |
| EPI_ISL_534693, EPI_ISL_534694, EPI_ISL_534695, EPI_ISL_534696, EPI_ISL_534697, EPI_ISL_534698 | University of Miami Immunology and Histocompatibility Laboratory | University of Miami Immunology and Histocompatibility Laboratory | Emilio Margolles-Clark, PhD and Phillip Ruiz, MD, PhD |
| EPI_ISL_534756, EPI_ISL_534757, EPI_ISL_534758, EPI_ISL_534759, EPI_ISL_534760, EPI_ISL_534761, EPI_ISL_534762, EPI_ISL_535183 | Queens Medical Centre, Clinical Microbiology Department / DeepSeq Nottingham | COVID-19 Genomics UK (COG-UK) Consortium | Gemma Clark, Wendy Smith, Manjinder Khakh, Vicki M Fleming, Michelle M Lister, Hannah Howson-Wells, Jonathan Ball, Patrick McClure, Joseph Chappell, Theocharis Tsoleridis, Nadine Holmes, Matthew Carlisle, Christopher Moore, Fei Sang, Johnny Debebe, Victoria Wright, Matthew Loose |
| EPI_ISL_535263, EPI_ISL_535264, EPI_ISL_535265, EPI_ISL_535266, EPI_ISL_535267, EPI_ISL_535268, EPI_ISL_535269, EPI_ISL_535270, EPI_ISL_535271, EPI_ISL_535272, EPI_ISL_535273, EPI_ISL_535274, EPI_ISL_535275, EPI_ISL_535276, EPI_ISL_535277, EPI_ISL_535278, EPI_ISL_535279, EPI_ISL_535280, EPI_ISL_535281, EPI_ISL_535282, EPI_ISL_535283, EPI_ISL_535284, EPI_ISL_535285, EPI_ISL_535286, EPI_ISL_535287, EPI_ISL_535288, EPI_ISL_535289, EPI_ISL_535290, EPI_ISL_535291, EPI_ISL_535292, EPI_ISL_535293, EPI_ISL_535294, EPI_ISL_535295, EPI_ISL_535296, EPI_ISL_535297, EPI_ISL_535298, EPI_ISL_535299, EPI_ISL_535300, EPI_ISL_535301, EPI_ISL_535302, EPI_ISL_535303 | | | |
| see above | New Mexico Department of Health Scientific Laboratory | New Mexico Department of Health Scientific Laboratory | Ellie Johnson, Anastacia Griego-Fisher, D'Eldra Malone |
| EPI_ISL_535390, EPI_ISL_535392, EPI_ISL_535393, EPI_ISL_535394, EPI_ISL_535396, EPI_ISL_535397, EPI_ISL_535398, EPI_ISL_535399, EPI_ISL_535400, EPI_ISL_535402, EPI_ISL_535404, EPI_ISL_535405, EPI_ISL_535406, EPI_ISL_535408, EPI_ISL_535409, EPI_ISL_535411, EPI_ISL_535412, EPI_ISL_535413, EPI_ISL_535414, EPI_ISL_535415, EPI_ISL_535416, EPI_ISL_535417, EPI_ISL_535418, EPI_ISL_535420, EPI_ISL_535421, EPI_ISL_535422, EPI_ISL_535424, EPI_ISL_535425, EPI_ISL_535426, EPI_ISL_535427, EPI_ISL_535428, EPI_ISL_535430, EPI_ISL_535431, EPI_ISL_535432, EPI_ISL_535433, EPI_ISL_535434, EPI_ISL_535435, EPI_ISL_535436, EPI_ISL_535438, EPI_ISL_535439, EPI_ISL_535440, EPI_ISL_535441, EPI_ISL_535442, EPI_ISL_535443, EPI_ISL_535444, EPI_ISL_535445, EPI_ISL_535446, EPI_ISL_535447, EPI_ISL_535448, EPI_ISL_535449, EPI_ISL_535450, EPI_ISL_535451, EPI_ISL_535452, EPI_ISL_535454, EPI_ISL_535455, EPI_ISL_535456, EPI_ISL_535457, EPI_ISL_535458, EPI_ISL_535459, EPI_ISL_535460, EPI_ISL_535461, EPI_ISL_535462, EPI_ISL_535463, EPI_ISL_535464, EPI_ISL_535465, EPI_ISL_535466, EPI_ISL_535468, EPI_ISL_535470, EPI_ISL_535471, EPI_ISL_535472, EPI_ISL_535473, EPI_ISL_535475, EPI_ISL_535476, EPI_ISL_535477, EPI_ISL_535479, EPI_ISL_535480, EPI_ISL_535481, EPI_ISL_535483, EPI_ISL_535484, EPI_ISL_535485, EPI_ISL_535486, EPI_ISL_535487, EPI_ISL_535488, EPI_ISL_535496, EPI_ISL_535497, EPI_ISL_535498, EPI_ISL_535499, EPI_ISL_535500, EPI_ISL_535501, EPI_ISL_535502, EPI_ISL_535503, EPI_ISL_535504, EPI_ISL_535505, EPI_ISL_535506, EPI_ISL_535507, EPI_ISL_535508, EPI_ISL_535509, EPI_ISL_535510, EPI_ISL_535511, EPI_ISL_535512, EPI_ISL_535513, EPI_ISL_535514, EPI_ISL_535515, EPI_ISL_535516, EPI_ISL_535518, EPI_ISL_535519, EPI_ISL_535520, EPI_ISL_535522, EPI_ISL_535523, EPI_ISL_535526, EPI_ISL_535527, EPI_ISL_535529, EPI_ISL_535530, EPI_ISL_535531, EPI_ISL_535532, EPI_ISL_535533, EPI_ISL_535534, EPI_ISL_535536, EPI_ISL_535537, EPI_ISL_535539, EPI_ISL_535541, EPI_ISL_535542, EPI_ISL_535543, EPI_ISL_535544, EPI_ISL_535545, EPI_ISL_535546, EPI_ISL_535547, EPI_ISL_535550, EPI_ISL_535552, EPI_ISL_535553, EPI_ISL_535554, EPI_ISL_535555, EPI_ISL_535556, EPI_ISL_535557, EPI_ISL_535558, EPI_ISL_535559, EPI_ISL_535560, EPI_ISL_535561, EPI_ISL_535562, EPI_ISL_535563, EPI_ISL_535564, EPI_ISL_535565, EPI_ISL_535566, EPI_ISL_535567, EPI_ISL_535568, EPI_ISL_535569, EPI_ISL_535570, EPI_ISL_535571, EPI_ISL_535572 | | | |
| see above | NHLS-IALCH | KRISP, KZN Research Innovation and Sequencing Platform | Giandhari J, Pillay S, Lessells R, Mdlalose K, York D, Khan S, Tegally H, Wilkinson E, de Oliveira T |
| EPI_ISL_536411 | Medtimes Molecular Laboratory | Medtimes Molecular Laboratory | Eric Chan, Winsome Wong, Jacqueline Tam, Isaac Chow |
| EPI_ISL_536477, EPI_ISL_536478, EPI_ISL_536479, EPI_ISL_536480, EPI_ISL_536481, EPI_ISL_536482, EPI_ISL_536483, EPI_ISL_536484, EPI_ISL_536485, EPI_ISL_536486, EPI_ISL_536487, EPI_ISL_536488, EPI_ISL_536489, EPI_ISL_536490, EPI_ISL_536491, EPI_ISL_536492, EPI_ISL_536493, EPI_ISL_536494, EPI_ISL_536495, EPI_ISL_536496, EPI_ISL_536497, EPI_ISL_536498, EPI_ISL_536499, EPI_ISL_536500, EPI_ISL_536501, EPI_ISL_536502, EPI_ISL_536503, EPI_ISL_536504, EPI_ISL_536505, EPI_ISL_536506, EPI_ISL_536507, EPI_ISL_536508, EPI_ISL_536509, EPI_ISL_536510, EPI_ISL_536511, EPI_ISL_536512, EPI_ISL_536513, EPI_ISL_536514, EPI_ISL_536515, EPI_ISL_536516, EPI_ISL_536517, EPI_ISL_536518, EPI_ISL_536519, EPI_ISL_536520, EPI_ISL_536521, EPI_ISL_536522, EPI_ISL_536523, EPI_ISL_536524, EPI_ISL_536525, EPI_ISL_536526, EPI_ISL_536527, EPI_ISL_536528, EPI_ISL_536529, EPI_ISL_536530, EPI_ISL_536531, EPI_ISL_536532, EPI_ISL_536533, EPI_ISL_536534, EPI_ISL_536535, EPI_ISL_536536, EPI_ISL_536537, EPI_ISL_536538, EPI_ISL_536539, EPI_ISL_536540, EPI_ISL_536541, EPI_ISL_536542, EPI_ISL_536543, EPI_ISL_536544, EPI_ISL_536545, EPI_ISL_536546, EPI_ISL_536547, EPI_ISL_536548, EPI_ISL_536549, EPI_ISL_536550, EPI_ISL_536551, EPI_ISL_536552, EPI_ISL_536553, EPI_ISL_536554, EPI_ISL_536555, EPI_ISL_536556, EPI_ISL_536557, EPI_ISL_536558, EPI_ISL_536559, EPI_ISL_536560, EPI_ISL_536561, EPI_ISL_536562, EPI_ISL_536563, EPI_ISL_536564, EPI_ISL_536565 | | | |
| see above | Instituto Nacional de Salud | Laboratorio de Infecciones Respiratorias Agudas | Eduardo Juscamayta Lopez, David Tarazona, Faviola Valdivia Guerrero, Nancy Rojas Serrano, Dennis Carhuaricra, Lenin Maturrano Hernandez, Ronnie Gavilan Chavez |
| EPI_ISL_536572, EPI_ISL_536573, EPI_ISL_536574, EPI_ISL_536575, EPI_ISL_536576, EPI_ISL_536577, EPI_ISL_536578, EPI_ISL_536579, EPI_ISL_536580, EPI_ISL_536581, EPI_ISL_536582, EPI_ISL_536583, EPI_ISL_536584, EPI_ISL_536585, EPI_ISL_536586, EPI_ISL_536587, EPI_ISL_536588, EPI_ISL_536589, EPI_ISL_536590, EPI_ISL_536591, EPI_ISL_536592, EPI_ISL_536593, EPI_ISL_536594, EPI_ISL_536595, EPI_ISL_536596, EPI_ISL_536597, EPI_ISL_536598, EPI_ISL_536599, EPI_ISL_536600, EPI_ISL_536601, EPI_ISL_536602, EPI_ISL_536603, EPI_ISL_536604, EPI_ISL_536605, EPI_ISL_536606, EPI_ISL_536607, EPI_ISL_536608, EPI_ISL_536609, EPI_ISL_536610, EPI_ISL_536611, EPI_ISL_536612, EPI_ISL_536613, EPI_ISL_536614, EPI_ISL_536615, EPI_ISL_536616, EPI_ISL_536617, EPI_ISL_536618, EPI_ISL_536619, EPI_ISL_536620, EPI_ISL_536621, EPI_ISL_536622, EPI_ISL_536623, EPI_ISL_536624, EPI_ISL_536625, EPI_ISL_536626, EPI_ISL_536627, EPI_ISL_536628, EPI_ISL_536629, EPI_ISL_536630, EPI_ISL_536631, EPI_ISL_536632, EPI_ISL_536633, EPI_ISL_536634, EPI_ISL_536635, EPI_ISL_536636, EPI_ISL_536637, EPI_ISL_536638, EPI_ISL_536639, EPI_ISL_536640, EPI_ISL_536641, EPI_ISL_536642, EPI_ISL_536643, EPI_ISL_536644, EPI_ISL_536645, EPI_ISL_536646, EPI_ISL_536647, EPI_ISL_536648, EPI_ISL_536649, EPI_ISL_536650, EPI_ISL_536651, EPI_ISL_536652, EPI_ISL_536653, EPI_ISL_536654, EPI_ISL_536655 | | | |
| see above | University of Wisconsin-Madison AIDS Vaccine Research Laboratories | University of Wisconsin-Madison AIDS Vaccine Research Laboratories | Gage Moreno, Katarina Braun, et al. AIDS Vaccine Research Laboratories |
| EPI_ISL_536656, EPI_ISL_536657 | University of Wisconsin-Madison Campus AIDS Vaccine Research Laboratories | University of Wisconsin-Madison AIDS Vaccine Research Laboratories | Gage Moreno, Katarina Braun, et al. AIDS Vaccine Research Laboratories |
| EPI_ISL_536658, EPI_ISL_536659, EPI_ISL_536660, EPI_ISL_536661, EPI_ISL_536662, EPI_ISL_536663, EPI_ISL_536664, EPI_ISL_536665, EPI_ISL_536666, EPI_ISL_536667, EPI_ISL_536668, EPI_ISL_536669, EPI_ISL_536670, EPI_ISL_536671, EPI_ISL_536672, EPI_ISL_536673, EPI_ISL_536674, EPI_ISL_536675, EPI_ISL_536676, EPI_ISL_536677, EPI_ISL_536678, EPI_ISL_536679, EPI_ISL_536680, EPI_ISL_536681, EPI_ISL_536682, EPI_ISL_536683, EPI_ISL_536684, EPI_ISL_536685, EPI_ISL_536686, EPI_ISL_536687, EPI_ISL_536688, EPI_ISL_536689, EPI_ISL_536690, EPI_ISL_536691, EPI_ISL_536692, EPI_ISL_536693, EPI_ISL_536694, EPI_ISL_536695, EPI_ISL_536696, EPI_ISL_536697, EPI_ISL_536698, EPI_ISL_536699, EPI_ISL_536700, EPI_ISL_536701, EPI_ISL_536702, EPI_ISL_536703, EPI_ISL_536704, EPI_ISL_536705, EPI_ISL_536706, EPI_ISL_536707, EPI_ISL_536708, EPI_ISL_536709, EPI_ISL_536710, EPI_ISL_536711, EPI_ISL_536712, EPI_ISL_536713, EPI_ISL_536714, EPI_ISL_536715, EPI_ISL_536716, EPI_ISL_536717, EPI_ISL_536718, EPI_ISL_536719, EPI_ISL_536720, EPI_ISL_536721, EPI_ISL_536722, EPI_ISL_536723, EPI_ISL_536724, EPI_ISL_536725, EPI_ISL_536726, EPI_ISL_536727, EPI_ISL_536728, EPI_ISL_536729, EPI_ISL_536730, EPI_ISL_536731, EPI_ISL_536732, EPI_ISL_536733, EPI_ISL_536734, EPI_ISL_536735, EPI_ISL_536736, EPI_ISL_536737, EPI_ISL_536738, EPI_ISL_536739, EPI_ISL_536740, EPI_ISL_536741, EPI_ISL_536742, EPI_ISL_536743, EPI_ISL_536744, EPI_ISL_536745, EPI_ISL_536746, EPI_ISL_536747, EPI_ISL_536748, EPI_ISL_536749, EPI_ISL_536750, EPI_ISL_536751, EPI_ISL_536752, EPI_ISL_536753, EPI_ISL_536754, EPI_ISL_536755, EPI_ISL_536756, EPI_ISL_536757, EPI_ISL_536758, EPI_ISL_536759, EPI_ISL_536760, EPI_ISL_536761, EPI_ISL_536762, EPI_ISL_536763, EPI_ISL_536764, EPI_ISL_536765, EPI_ISL_536766, EPI_ISL_536767, EPI_ISL_536768, EPI_ISL_536769, EPI_ISL_536770, EPI_ISL_536771, EPI_ISL_536772, EPI_ISL_536773, EPI_ISL_536774, EPI_ISL_536775, EPI_ISL_536776, EPI_ISL_536777, EPI_ISL_536778, EPI_ISL_536779, EPI_ISL_536780, EPI_ISL_536781, EPI_ISL_536782, EPI_ISL_536783, EPI_ISL_536784 | | | |
| see above | University of Wisconsin-Madison AIDS Vaccine Research Laboratories | University of Wisconsin-Madison AIDS Vaccine Research Laboratories | Gage Moreno, Katarina Braun, et al. AIDS Vaccine Research Laboratories |
| EPI_ISL_536785 | University of Wisconsin-Madison Campus AIDS Vaccine Research Laboratories | University of Wisconsin-Madison AIDS Vaccine Research Laboratories | Gage Moreno, Katarina Braun, et al. AIDS Vaccine Research Laboratories |
| EPI_ISL_536786, EPI_ISL_536787 | University of Wisconsin-Madison AIDS Vaccine Research Laboratories | University of Wisconsin-Madison AIDS Vaccine Research Laboratories | Gage Moreno, Katarina Braun, et al. AIDS Vaccine Research Laboratories |
| EPI_ISL_536793 | Medtimes Molecular Laboratory | Medtimes Molecular Laboratory | Eric Chan, Winsome Wong, Jacqueline Tam, Isaac Chow |
| EPI_ISL_538390, EPI_ISL_538391, EPI_ISL_538392, EPI_ISL_538393, EPI_ISL_538394, EPI_ISL_538395, EPI_ISL_538396, EPI_ISL_538397, EPI_ISL_538398, EPI_ISL_538399, EPI_ISL_538400, EPI_ISL_538401, EPI_ISL_538402, EPI_ISL_538403, EPI_ISL_538404, EPI_ISL_538405, EPI_ISL_538406, EPI_ISL_538407, EPI_ISL_538408, EPI_ISL_538409, EPI_ISL_538410, EPI_ISL_538411, EPI_ISL_538412, EPI_ISL_538413, EPI_ISL_538414, EPI_ISL_538415, EPI_ISL_538416, EPI_ISL_538417, EPI_ISL_538418, EPI_ISL_538419, EPI_ISL_538420, EPI_ISL_538421, EPI_ISL_538422, EPI_ISL_538423, EPI_ISL_538424, EPI_ISL_538425, EPI_ISL_538426, EPI_ISL_538427, EPI_ISL_538428, EPI_ISL_538429, EPI_ISL_538430, EPI_ISL_538431, EPI_ISL_538432, EPI_ISL_538433, EPI_ISL_538434 | | | |
| see above | Microbiology Division, South Carolina Department of Health and Environmental Control | Microbiology Division, South Carolina Department of Health and Environmental Control | Flores,H. |
| EPI_ISL_539783, EPI_ISL_539784 | Universidad Regional Amazonica IKIAM | Institute of Microbiology, Universidad San Francisco de Quito | Fabian Aguilar, Katherine Apunte, Andrea Carrera, Nina Espinoza de los Monteros, Giovanna Moran, Marcelo Ortiz, Yeimy Rojas, Sonia Sislema, Carolina Proaño-Bolaños, Belén Prado-Vivar, Sully Márquez, Juan José Guadalupe, Monica Becerra-Wong, Bernardo Gutiérrez, Verónica Barragán, Patricio Rojas-Silva, Gabriel Trueba, Michelle Grunauer, Paúl Cárdenas |
| EPI_ISL_539879 | Microbiology Division, South Carolina Department of Health | Microbiology Division, South Carolina Department of Health | Flores,H. |

| | | | |
|---|---|---|---|
| | and Environmental Control | and Environmental Control | |
| EPI_ISL_540640, EPI_ISL_540641, EPI_ISL_540642, EPI_ISL_540643, EPI_ISL_540644, EPI_ISL_540645, EPI_ISL_540646, EPI_ISL_540647, EPI_ISL_540648, EPI_ISL_540649, EPI_ISL_540650, EPI_ISL_540651, EPI_ISL_540652, EPI_ISL_540653, EPI_ISL_540654, EPI_ISL_540655, EPI_ISL_540656, EPI_ISL_540657, EPI_ISL_540658, EPI_ISL_540659, EPI_ISL_540660, EPI_ISL_540661, EPI_ISL_540662, EPI_ISL_540663, EPI_ISL_540664, EPI_ISL_540665, EPI_ISL_540666, EPI_ISL_540667, EPI_ISL_540668, EPI_ISL_540669, EPI_ISL_540670, EPI_ISL_540671, EPI_ISL_540672, EPI_ISL_540673, EPI_ISL_540674, EPI_ISL_540675, EPI_ISL_540676, EPI_ISL_540677, EPI_ISL_540678, EPI_ISL_540679, EPI_ISL_540680, EPI_ISL_540681, EPI_ISL_540682, EPI_ISL_540683, EPI_ISL_540684, EPI_ISL_540685, EPI_ISL_540686, EPI_ISL_540687, EPI_ISL_540688, EPI_ISL_540689, EPI_ISL_540690, EPI_ISL_540699, EPI_ISL_540700, EPI_ISL_540701, EPI_ISL_540702, EPI_ISL_540703, EPI_ISL_540704, EPI_ISL_540705, EPI_ISL_540706, EPI_ISL_540707, EPI_ISL_540708, EPI_ISL_540709, EPI_ISL_540710, EPI_ISL_540711, EPI_ISL_540712, EPI_ISL_540713, EPI_ISL_540714, EPI_ISL_540715, EPI_ISL_540716, EPI_ISL_540717, EPI_ISL_540718, EPI_ISL_540894, EPI_ISL_540895, EPI_ISL_540896 | | | |
| see above | Queens Medical Centre, Clinical Microbiology Department / DeepSeq Nottingham | COVID-19 Genomics UK (COG-UK) Consortium | Gemma Clark, Wendy Smith, Manjinder Khakh, Vicki M Fleming, Michelle M Lister, Hannah Howson-Wells, Jonathan Ball, Patrick McClure, Joseph Chappell, Theocharis Tsoleridis, Nadine Holmes, Matthew Carlisle, Christopher Moore, Fei Sang, Johnny Debebe, Victoria Wright, Matthew Loose |
| EPI_ISL_541067, EPI_ISL_541068, EPI_ISL_541069, EPI_ISL_541070, EPI_ISL_541071, EPI_ISL_541072, EPI_ISL_541073, EPI_ISL_541074, EPI_ISL_541075, EPI_ISL_541076, EPI_ISL_541077, EPI_ISL_541078, EPI_ISL_541079, EPI_ISL_541080, EPI_ISL_541081 | | | |
| see above | Hospital de la Santa Creu i Sant Pau. Servicio de Microbiología | SeqCOVID-SPAIN consortium/Institute of Biomedicine of Valencia, IBV-CSIC | Ferran Navarro, Núria Rabella, Elisenda Miró and SeqCOVID-SPAIN consortium |
| EPI_ISL_541540, EPI_ISL_541541, EPI_ISL_541542, EPI_ISL_541543, EPI_ISL_541544, EPI_ISL_541545, EPI_ISL_541546, EPI_ISL_541547, EPI_ISL_541548, EPI_ISL_541549, EPI_ISL_541550 | | | |
| see above | University of Wisconsin-Madison AIDS Vaccine Research Laboratories | University of Wisconsin-Madison AIDS Vaccine Research Laboratories | Gage Moreno, Katarina Braun, et al. AIDS Vaccine Research Laboratories |
| EPI_ISL_541551, EPI_ISL_541552, EPI_ISL_541553, EPI_ISL_541554, EPI_ISL_541555, EPI_ISL_541556, EPI_ISL_541557, EPI_ISL_541558, EPI_ISL_541559, EPI_ISL_541560, EPI_ISL_541561, EPI_ISL_541562, EPI_ISL_541563, EPI_ISL_541564, EPI_ISL_541565 | | | |
| see above | University of Wisconsin-Madison Campus AIDS Vaccine Research Laboratories | University of Wisconsin-Madison AIDS Vaccine Research Laboratories | Gage Moreno, Katarina Braun, et al. AIDS Vaccine Research Laboratories |
| EPI_ISL_541566, EPI_ISL_541567 | University of Wisconsin-Madison AIDS Vaccine Research Laboratories | University of Wisconsin-Madison AIDS Vaccine Research Laboratories | Gage Moreno, Katarina Braun, et al. AIDS Vaccine Research Laboratories |
| EPI_ISL_541568 | University of Wisconsin-Madison Campus AIDS Vaccine Research Laboratories | University of Wisconsin-Madison AIDS Vaccine Research Laboratories | Gage Moreno, Katarina Braun, et al. AIDS Vaccine Research Laboratories |
| EPI_ISL_541569, EPI_ISL_541570, EPI_ISL_541571, EPI_ISL_541572, EPI_ISL_541573, EPI_ISL_541574, EPI_ISL_541575, EPI_ISL_541576, EPI_ISL_541577, EPI_ISL_541578, EPI_ISL_541579, EPI_ISL_541580, EPI_ISL_541581, EPI_ISL_541582, EPI_ISL_541583, EPI_ISL_541584, EPI_ISL_541585, EPI_ISL_541586, EPI_ISL_541587, EPI_ISL_541588, EPI_ISL_541589, EPI_ISL_541590, EPI_ISL_541591, EPI_ISL_541592, EPI_ISL_541593, EPI_ISL_541594, EPI_ISL_541595, EPI_ISL_541596, EPI_ISL_541597, EPI_ISL_541598, EPI_ISL_541599, EPI_ISL_541600, EPI_ISL_541601, EPI_ISL_541602, EPI_ISL_541603, EPI_ISL_541604, EPI_ISL_541605, EPI_ISL_541606, EPI_ISL_541607, EPI_ISL_541608, EPI_ISL_541609, EPI_ISL_541610, EPI_ISL_541611, EPI_ISL_541612, EPI_ISL_541613, EPI_ISL_541614, EPI_ISL_541615, EPI_ISL_541616, EPI_ISL_541617, EPI_ISL_541618, EPI_ISL_541619, EPI_ISL_541620, EPI_ISL_541621, EPI_ISL_541622, EPI_ISL_541623, EPI_ISL_541624, EPI_ISL_541625, EPI_ISL_541626, EPI_ISL_541627, EPI_ISL_541628, EPI_ISL_541629, EPI_ISL_541630, EPI_ISL_541631, EPI_ISL_541632, EPI_ISL_541633, EPI_ISL_541634, EPI_ISL_541635, EPI_ISL_541636, EPI_ISL_541637, EPI_ISL_541638, EPI_ISL_541639, EPI_ISL_541640, EPI_ISL_541641, EPI_ISL_541642, EPI_ISL_541643, EPI_ISL_541644, EPI_ISL_541645, EPI_ISL_541646, EPI_ISL_541647, EPI_ISL_541648 | | | |
| see above | University of Wisconsin-Madison AIDS Vaccine Research Laboratories | University of Wisconsin-Madison AIDS Vaccine Research Laboratories | Gage Moreno, Katarina Braun, et al. AIDS Vaccine Research Laboratories |
| EPI_ISL_541663, EPI_ISL_541664, EPI_ISL_541665, EPI_ISL_541666, EPI_ISL_541667, EPI_ISL_541668, EPI_ISL_541669, EPI_ISL_541670, EPI_ISL_541671, EPI_ISL_541672, EPI_ISL_541673, EPI_ISL_541674, EPI_ISL_541675, EPI_ISL_541676, EPI_ISL_541677, EPI_ISL_541678, EPI_ISL_541679 | | | |
| see above | Microbiology Division, South Carolina Department of Health and Environmental Control | Microbiology Division, South Carolina Department of Health and Environmental Control | Flores,H. |
| EPI_ISL_541970 | Influenza Centre, University of Bergen | Norwegian Institute of Public Health, Department of Virology | Fan Zhou, Rebecca J Cox, Karl A Brokstad, Bjørn Blomberg, Kathrine Stene-Johansen, Kamilla Heddeland Instefjord, Hilde Elshaug, Rasmus Riis Kopperud, Hilde Synnøve Vollan, Karoline Bragstad, Olav Hungnes |
| EPI_ISL_542012, EPI_ISL_542013, EPI_ISL_542014, EPI_ISL_542015, EPI_ISL_542016, EPI_ISL_542017, EPI_ISL_542018, EPI_ISL_542019, EPI_ISL_542020, EPI_ISL_542021, EPI_ISL_542022, EPI_ISL_542023, EPI_ISL_542024, EPI_ISL_542025, EPI_ISL_542026, EPI_ISL_542027, EPI_ISL_542028, EPI_ISL_542029, EPI_ISL_542030, EPI_ISL_542031, EPI_ISL_542032, EPI_ISL_542033, EPI_ISL_542034, EPI_ISL_542035, EPI_ISL_542036, EPI_ISL_542037, EPI_ISL_542038, EPI_ISL_542039, EPI_ISL_542040, EPI_ISL_542041, EPI_ISL_542042, EPI_ISL_542043, EPI_ISL_542044, EPI_ISL_542045, EPI_ISL_542046, EPI_ISL_542047, EPI_ISL_542048, EPI_ISL_542049, EPI_ISL_542050, EPI_ISL_542051, EPI_ISL_542052, EPI_ISL_542053, EPI_ISL_542054, EPI_ISL_542055, EPI_ISL_542056, EPI_ISL_542057, EPI_ISL_542058, EPI_ISL_542059, EPI_ISL_542060, EPI_ISL_542061, EPI_ISL_542062, EPI_ISL_542063, EPI_ISL_542064, EPI_ISL_542065, EPI_ISL_542066, EPI_ISL_542067, EPI_ISL_542068, EPI_ISL_542069, EPI_ISL_542070, EPI_ISL_542071, EPI_ISL_542072, EPI_ISL_542073, EPI_ISL_542074, EPI_ISL_542075, EPI_ISL_542076, EPI_ISL_542077, EPI_ISL_542078, EPI_ISL_542079, EPI_ISL_542080, EPI_ISL_542081, EPI_ISL_542082, EPI_ISL_542083, EPI_ISL_542084, EPI_ISL_542085, EPI_ISL_542086, EPI_ISL_542087, EPI_ISL_542088, EPI_ISL_542089, EPI_ISL_542090, EPI_ISL_542091, EPI_ISL_542092, EPI_ISL_542093, EPI_ISL_542094, EPI_ISL_542095, EPI_ISL_542096, EPI_ISL_542097 | | | |
| see above | New Mexico Department of Health Scientific Laboratory | New Mexico Department of Health Scientific Laboratory | Ellie Johnson, Anastacia Griego-Fisher, D'Eldra Malone |
| EPI_ISL_547581, EPI_ISL_547582, EPI_ISL_547583 | University of Wisconsin-Madison AIDS Vaccine Research Laboratories | University of Wisconsin-Madison AIDS Vaccine Research Laboratories | Gage Moreno, Katarina Braun, et al. AIDS Vaccine Research Laboratories |
| EPI_ISL_548962, EPI_ISL_548963, EPI_ISL_549024, EPI_ISL_549026, EPI_ISL_549176 | Klinisk mikrobiologi, Region Västerbotten | Unit for Biological Agents, Department for CBRN Defence and Security, Swedish Defence Research Agency | FOI Bioinformatics team |
| EPI_ISL_549334, EPI_ISL_549357, EPI_ISL_549361, EPI_ISL_549376, EPI_ISL_549380, EPI_ISL_549385, EPI_ISL_549491, EPI_ISL_549492, EPI_ISL_549493, EPI_ISL_549494, EPI_ISL_549495, EPI_ISL_549496, EPI_ISL_549497, EPI_ISL_549498, EPI_ISL_549499, EPI_ISL_549500, EPI_ISL_549501, EPI_ISL_549502, EPI_ISL_549503, EPI_ISL_549504, EPI_ISL_549505, EPI_ISL_549506, EPI_ISL_549507, EPI_ISL_549508, EPI_ISL_549509, EPI_ISL_549510, EPI_ISL_549511, EPI_ISL_549512, EPI_ISL_549513, EPI_ISL_549514, EPI_ISL_549515, EPI_ISL_549516 | | | |
| see above | Queens Medical Centre, Clinical Microbiology Department / DeepSeq Nottingham | COVID-19 Genomics UK (COG-UK) Consortium | Gemma Clark, Wendy Smith, Manjinder Khakh, Vicki M Fleming, Michelle M Lister, Hannah Howson-Wells, Jonathan Ball, Patrick McClure, Joseph Chappell, Theocharis Tsoleridis, Nadine Holmes, Matthew Carlisle, Christopher Moore, Fei Sang, Johnny Debebe, Victoria Wright, Matthew Loose |
| EPI_ISL_560833, EPI_ISL_560834, EPI_ISL_560835, EPI_ISL_560836, EPI_ISL_560837, EPI_ISL_560838, EPI_ISL_560839, EPI_ISL_560840, EPI_ISL_560841, EPI_ISL_560842, EPI_ISL_560843, EPI_ISL_560844, EPI_ISL_560845, EPI_ISL_560846, EPI_ISL_560847, EPI_ISL_560848, EPI_ISL_560849, EPI_ISL_560850, EPI_ISL_560851, EPI_ISL_560852, EPI_ISL_560853, EPI_ISL_560854, EPI_ISL_560855, EPI_ISL_560856, EPI_ISL_560857, EPI_ISL_560858, EPI_ISL_560859, EPI_ISL_560860, EPI_ISL_560861, EPI_ISL_560862, EPI_ISL_560863, EPI_ISL_560864, EPI_ISL_560865, EPI_ISL_560866, EPI_ISL_560867, EPI_ISL_560868, EPI_ISL_560869, EPI_ISL_560870, EPI_ISL_560871, EPI_ISL_560872, EPI_ISL_560873, EPI_ISL_560874, EPI_ISL_560875, EPI_ISL_560876, EPI_ISL_560877, EPI_ISL_560878, EPI_ISL_560879, EPI_ISL_560880, EPI_ISL_560881, EPI_ISL_560882, EPI_ISL_560883, EPI_ISL_560884, EPI_ISL_560885, EPI_ISL_560886, EPI_ISL_560887, EPI_ISL_560888, EPI_ISL_560889, EPI_ISL_560890, EPI_ISL_560891, EPI_ISL_560892, EPI_ISL_560893, EPI_ISL_560894, EPI_ISL_560895, EPI_ISL_560896, EPI_ISL_560897, EPI_ISL_560898, EPI_ISL_560899, EPI_ISL_560900, EPI_ISL_560901, EPI_ISL_560902, EPI_ISL_560903, EPI_ISL_560904, EPI_ISL_560905, EPI_ISL_560906, EPI_ISL_560907, EPI_ISL_560908, EPI_ISL_560909, EPI_ISL_560910, EPI_ISL_560911, EPI_ISL_560912, EPI_ISL_560913, EPI_ISL_560914, EPI_ISL_560915, EPI_ISL_560916, EPI_ISL_560917, EPI_ISL_560918, EPI_ISL_560919, EPI_ISL_560920, EPI_ISL_560921, EPI_ISL_560922, EPI_ISL_560923, EPI_ISL_560924, EPI_ISL_560925 | | | |
| see above | Utah Public Health Laboratory | Utah Public Health Laboratory | Erin Young, Kelly Oakeson |
| EPI_ISL_568580, EPI_ISL_568581, EPI_ISL_568582, EPI_ISL_568583, EPI_ISL_568584, EPI_ISL_568585, EPI_ISL_568586 | Ramathibodi Hospital | COVID-19 Network Investigations (CONI) Alliance | Elizabeth Batty, Wasun Chantratita, Thanat Chookajorn, Stefan Fernandez, Angkana Huang, Anthony R. Jones, Khajohn Joonsalak, Chonticha Klungtong, Theerarat Kochakarn, Namfon Kotanan, Krittikorn Kumpornsin, Wudtichai Manasatienkij, Bhakbhoom Panthan, Ekawat Pasomsub, Kingkan Rakmanee, Insee Sensorn, Janjira Thaipadungpanit, Arporn Wangwiwatsin,Treewat Watthanachockchai |
| EPI_ISL_568686 | RSUD Prof. DR. Margono Soekarjo | Eijkman Institute for Molecular Biology, Ministry of Research and Technology/National Agency for Research and Innovation | Frilasita A Yudhaputri, Edison Johar, Hidayat Trimarsanto, Iskandar A Adnan, Willy Agustine, David H Muljono, Safarina G Malik, Herawati Sudoyo, Khin Saw Myint, Amin Soebandrio |
| EPI_ISL_568687 | RSUP Fatmawati | Eijkman Institute for Molecular Biology, Ministry of Research and Technology/National Agency for Research and Innovation | Frilasita A Yudhaputri, Edison Johar, Hidayat Trimarsanto, Iskandar A Adnan, Willy Agustine, David H Muljono, Safarina G Malik, Herawati Sudoyo, Khin Saw Myint, Amin Soebandrio |
| EPI_ISL_568688 | RS Pelni | Eijkman Institute for Molecular Biology, Ministry of Research and Technology/National Agency for Research and Innovation | Frilasita A Yudhaputri, Edison Johar, Hidayat Trimarsanto, Iskandar A Adnan, Willy Agustine, David H Muljono, Safarina G Malik, Herawati Sudoyo, Khin Saw Myint, Amin Soebandrio |
| EPI_ISL_568689 | RS Pondok Indah Puri Indah | Eijkman Institute for Molecular Biology, Ministry of Research and Technology/National Agency for Research and Innovation | Frilasita A Yudhaputri, Edison Johar, Hidayat Trimarsanto, Iskandar A Adnan, Willy Agustine, David H Muljono, Safarina G Malik, Herawati Sudoyo, Khin Saw Myint, Amin Soebandrio |
| EPI_ISL_568690, EPI_ISL_568691 | RSUP Fatmawati | Eijkman Institute for Molecular Biology, Ministry of Research and Technology/National Agency for Research and Innovation | Frilasita A Yudhaputri, Edison Johar, Hidayat Trimarsanto, Iskandar A Adnan, Willy Agustine, David H Muljono, Safarina G Malik, Herawati Sudoyo, Khin Saw Myint, Amin Soebandrio |
| EPI_ISL_568692 | RS Kramat 128 | Eijkman Institute for Molecular Biology, Ministry of Research and Technology/National Agency for Research and Innovation | Frilasita A Yudhaputri, Edison Johar, Hidayat Trimarsanto, Iskandar A Adnan, Willy Agustine, David H Muljono, Safarina G Malik, Herawati Sudoyo, Khin Saw Myint, Amin Soebandrio |
| EPI_ISL_568693 | RS Freeport Tembagapura | Eijkman Institute for Molecular Biology, Ministry of Research | Frilasita A Yudhaputri, Edison Johar, Hidayat Trimarsanto, Iskandar A Adnan, Willy Agustine, David H Muljono, Safarina G Malik, Herawati Sudoyo, Khin |

| | | | |
|---|---|---|---|
| | | and Technology/National Agency for Research and Innovation | Saw Myint, Amin Soebandrio |
| EPI_ISL_568694 | RSUP Prof. Dr. R. Kandou Manado | Eijkman Institute for Molecular Biology, Ministry of Research and Technology/National Agency for Research and Innovation | Frilasita A Yudhaputri, Edison Johar, Hidayat Trimarsanto, Iskandar A Adnan, Willy Agustine, David H Muljono, Safarina G Malik, Herawati Sudoyo, Khin Saw Myint, Amin Soebandrio |
| EPI_ISL_569625, EPI_ISL_569626, EPI_ISL_569627, EPI_ISL_569628, EPI_ISL_569629, EPI_ISL_569630, EPI_ISL_569631, EPI_ISL_569632, EPI_ISL_569633, EPI_ISL_569634, EPI_ISL_569635, EPI_ISL_569636, EPI_ISL_569637, EPI_ISL_569638, EPI_ISL_569639, EPI_ISL_569640, EPI_ISL_569641, EPI_ISL_569642, EPI_ISL_569643, EPI_ISL_569644, EPI_ISL_569645, EPI_ISL_569646, EPI_ISL_569647, EPI_ISL_569648, EPI_ISL_569649, EPI_ISL_569650, EPI_ISL_569651, EPI_ISL_569652, EPI_ISL_569653, EPI_ISL_569654, EPI_ISL_569655, EPI_ISL_569656, EPI_ISL_569657, EPI_ISL_569658, EPI_ISL_569659, EPI_ISL_569660, EPI_ISL_569661, EPI_ISL_569662, EPI_ISL_569663 | | | |
| see above | New Mexico Department of Health Scientific Laboratory | New Mexico Department of Health Scientific Laboratory | Ellie Johnson, Anastacia Griego-Fisher, D'Eldra Malone |
| EPI_ISL_572329 | Public Health Authority of the Slovak Republic, Bratislava | Faculty of Natural Sciences, Comenius University in Bratislava | Dominika Friová, Viktória Hodorová, Kristína Boršová, Broa Brejová, Viktória abanová, Sabina Fumaová Havlíková, Juraj Kopáek, Martina Liková, ubomíra Lukáiková, Martina Neboháová, Monika Sláviková, Edita Staroová, Elena Tichá, Tomáš Vina, Boris Klempa, Jozef Nosek |
| EPI_ISL_572444, EPI_ISL_572460, EPI_ISL_572475, EPI_ISL_572476, EPI_ISL_572509, EPI_ISL_572582, EPI_ISL_572598, EPI_ISL_572603, EPI_ISL_572706, EPI_ISL_572799, EPI_ISL_572833, EPI_ISL_572872, EPI_ISL_573418, EPI_ISL_573419, EPI_ISL_573420, EPI_ISL_573421, EPI_ISL_573422, EPI_ISL_573423, EPI_ISL_573424, EPI_ISL_573425, EPI_ISL_573426, EPI_ISL_573427, EPI_ISL_573428, EPI_ISL_573429, EPI_ISL_573430, EPI_ISL_573431, EPI_ISL_573432, EPI_ISL_573433, EPI_ISL_573434, EPI_ISL_573435, EPI_ISL_573436, EPI_ISL_573450, EPI_ISL_573451, EPI_ISL_573452, EPI_ISL_573453, EPI_ISL_573454, EPI_ISL_573455, EPI_ISL_573456, EPI_ISL_573457, EPI_ISL_573458, EPI_ISL_573459, EPI_ISL_573460, EPI_ISL_573461, EPI_ISL_573462, EPI_ISL_573463, EPI_ISL_573464, EPI_ISL_573465, EPI_ISL_573466, EPI_ISL_573467, EPI_ISL_573468, EPI_ISL_573469, EPI_ISL_573470, EPI_ISL_573471, EPI_ISL_573472, EPI_ISL_573473, EPI_ISL_573474, EPI_ISL_573475, EPI_ISL_573476, EPI_ISL_573477 | | | |
| see above | Queens Medical Centre, Clinical Microbiology Department / DeepSeq Nottingham | COVID-19 Genomics UK (COG-UK) Consortium | Gemma Clark, Wendy Smith, Manjinder Khakh, Vicki M Fleming, Michelle M Lister, Hannah Howson-Wells, Jonathan Ball, Patrick McClure, Joseph Chappell, Theocharis Tsoleridis, Nadine Holmes, Matthew Carlisle, Christopher Moore, Fei Sang, Johnny Debebe, Victoria Wright, Matthew Loose |
| EPI_ISL_574260, EPI_ISL_574261, EPI_ISL_574262, EPI_ISL_574263, EPI_ISL_574264, EPI_ISL_574265, EPI_ISL_574266, EPI_ISL_574267, EPI_ISL_574268, EPI_ISL_574269, EPI_ISL_574270, EPI_ISL_574271, EPI_ISL_574272, EPI_ISL_574273, EPI_ISL_574274, EPI_ISL_574275, EPI_ISL_574276, EPI_ISL_574277, EPI_ISL_574278, EPI_ISL_574279, EPI_ISL_574280, EPI_ISL_574281, EPI_ISL_574282, EPI_ISL_574283, EPI_ISL_574284, EPI_ISL_574285, EPI_ISL_574286, EPI_ISL_574287, EPI_ISL_574288, EPI_ISL_574289, EPI_ISL_574290, EPI_ISL_574291 | | | |
| see above | New Mexico Department of Health Scientific Laboratory | New Mexico Department of Health Scientific Laboratory | Ellie Johnson, Anastacia Griego-Fisher, D'Eldra Malone |
| EPI_ISL_574543, EPI_ISL_574544, EPI_ISL_574545, EPI_ISL_574546, EPI_ISL_574547, EPI_ISL_574548, EPI_ISL_574549, EPI_ISL_574550, EPI_ISL_574551, EPI_ISL_574552, EPI_ISL_574553, EPI_ISL_574554, EPI_ISL_574555, EPI_ISL_574556, EPI_ISL_574557, EPI_ISL_574558, EPI_ISL_574559, EPI_ISL_574560, EPI_ISL_574561, EPI_ISL_574562, EPI_ISL_574563, EPI_ISL_574564, EPI_ISL_574565, EPI_ISL_574566, EPI_ISL_574567, EPI_ISL_574568, EPI_ISL_574569, EPI_ISL_574570, EPI_ISL_574571, EPI_ISL_574572, EPI_ISL_574573, EPI_ISL_574574, EPI_ISL_574575 | | | |
| see above | Microbiology Division, South Carolina Department of Health and Environmental Control | Microbiology Division, South Carolina Department of Health and Environmental Control | Flores,H. |
| EPI_ISL_574603 | RS Freeport Tembagapura | Eijkman Institute for Molecular Biology, Ministry of Research and Technology/National Agency for Research and Innovation | Frilasita A Yudhaputri, Edison Johar, Hidayat Trimarsanto, Iskandar A Adnan, Willy Agustine, David H Muljono, Safarina G Malik, Herawati Sudoyo, Khin Saw Myint, Amin Soebandrio |
| EPI_ISL_574604 | RSUP Prof. Dr. R. Kandou Manado | Eijkman Institute for Molecular Biology, Ministry of Research and Technology/National Agency for Research and Innovation | Frilasita A Yudhaputri, Edison Johar, Hidayat Trimarsanto, Iskandar A Adnan, Willy Agustine, David H Muljono, Safarina G Malik, Herawati Sudoyo, Khin Saw Myint, Amin Soebandrio |
| EPI_ISL_574605 | RSUP dr. Sardjito | Eijkman Institute for Molecular Biology, Ministry of Research and Technology/National Agency for Research and Innovation | Frilasita A Yudhaputri, Edison Johar, Hidayat Trimarsanto, Iskandar A Adnan, Willy Agustine, David H Muljono, Safarina G Malik, Herawati Sudoyo, Khin Saw Myint, Amin Soebandrio |
| EPI_ISL_574606 | RS Siloam Purwakarta | Eijkman Institute for Molecular Biology, Ministry of Research and Technology/National Agency for Research and Innovation | Frilasita A Yudhaputri, Edison Johar, Hidayat Trimarsanto, Iskandar A Adnan, Willy Agustine, David H Muljono, Safarina G Malik, Herawati Sudoyo, Khin Saw Myint, Amin Soebandrio |
| EPI_ISL_574607 | RS Husada | Eijkman Institute for Molecular Biology, Ministry of Research and Technology/National Agency for Research and Innovation | Frilasita A Yudhaputri, Edison Johar, Hidayat Trimarsanto, Iskandar A Adnan, Willy Agustine, David H Muljono, Safarina G Malik, Herawati Sudoyo, Khin Saw Myint, Amin Soebandrio |
| EPI_ISL_574608 | RS Mitra Keluarga Kelapa Gading | Eijkman Institute for Molecular Biology, Ministry of Research and Technology/National Agency for Research and Innovation | Frilasita A Yudhaputri, Edison Johar, Hidayat Trimarsanto, Iskandar A Adnan, Willy Agustine, David H Muljono, Safarina G Malik, Herawati Sudoyo, Khin Saw Myint, Amin Soebandrio |
| EPI_ISL_574609 | RSJPD Harapan Kita | Eijkman Institute for Molecular Biology, Ministry of Research and Technology/National Agency for Research and Innovation | Frilasita A Yudhaputri, Edison Johar, Hidayat Trimarsanto, Iskandar A Adnan, Willy Agustine, David H Muljono, Safarina G Malik, Herawati Sudoyo, Khin Saw Myint, Amin Soebandrio |
| EPI_ISL_574610 | Puskesmas Tambora | Eijkman Institute for Molecular Biology, Ministry of Research and Technology/National Agency for Research and Innovation | Frilasita A Yudhaputri, Edison Johar, Hidayat Trimarsanto, Iskandar A Adnan, Willy Agustine, David H Muljono, Safarina G Malik, Herawati Sudoyo, Khin Saw Myint, Amin Soebandrio |
| EPI_ISL_574611 | RS Premier Bintaro | Eijkman Institute for Molecular Biology, Ministry of Research and Technology/National Agency for Research and Innovation | Frilasita A Yudhaputri, Edison Johar, Hidayat Trimarsanto, Iskandar A Adnan, Willy Agustine, David H Muljono, Safarina G Malik, Herawati Sudoyo, Khin Saw Myint, Amin Soebandrio |
| EPI_ISL_574612 | RS Hermina Mekarsari | Eijkman Institute for Molecular Biology, Ministry of Research and Technology/National Agency for Research and Innovation | Frilasita A Yudhaputri, Edison Johar, Hidayat Trimarsanto, Iskandar A Adnan, Willy Agustine, David H Muljono, Safarina G Malik, Herawati Sudoyo, Khin Saw Myint, Amin Soebandrio |
| EPI_ISL_574613 | RS Harapan Bunda | Eijkman Institute for Molecular Biology, Ministry of Research and Technology/National Agency for Research and Innovation | Frilasita A Yudhaputri, Edison Johar, Hidayat Trimarsanto, Iskandar A Adnan, Willy Agustine, David H Muljono, Safarina G Malik, Herawati Sudoyo, Khin Saw Myint, Amin Soebandrio |
| EPI_ISL_574614 | RS Kramat 128 | Eijkman Institute for Molecular Biology, Ministry of Research and Technology/National Agency for Research and Innovation | Frilasita A Yudhaputri, Edison Johar, Hidayat Trimarsanto, Iskandar A Adnan, Willy Agustine, David H Muljono, Safarina G Malik, Herawati Sudoyo, Khin Saw Myint, Amin Soebandrio |
| EPI_ISL_574615 | RS Harapan Bunda | Eijkman Institute for Molecular Biology, Ministry of Research and Technology/National Agency for Research and Innovation | Frilasita A Yudhaputri, Edison Johar, Hidayat Trimarsanto, Iskandar A Adnan, Willy Agustine, David H Muljono, Safarina G Malik, Herawati Sudoyo, Khin Saw Myint, Amin Soebandrio |
| EPI_ISL_574616, EPI_ISL_574617 | RS Kramat 128 | Eijkman Institute for Molecular Biology, Ministry of Research and Technology/National Agency for Research and Innovation | Frilasita A Yudhaputri, Edison Johar, Hidayat Trimarsanto, Iskandar A Adnan, Willy Agustine, David H Muljono, Safarina G Malik, Herawati Sudoyo, Khin Saw Myint, Amin Soebandrio |
| EPI_ISL_575034, EPI_ISL_575035, EPI_ISL_575036, EPI_ISL_575037, EPI_ISL_575038, EPI_ISL_575039, EPI_ISL_575040, EPI_ISL_575041, EPI_ISL_575042, EPI_ISL_575043, EPI_ISL_575044, EPI_ISL_575045, EPI_ISL_575046, EPI_ISL_575047, EPI_ISL_575048, EPI_ISL_575049, EPI_ISL_575050, EPI_ISL_575051, EPI_ISL_575052, EPI_ISL_575053, EPI_ISL_575054, EPI_ISL_575055, EPI_ISL_575056, EPI_ISL_575057, EPI_ISL_575058, EPI_ISL_575059, EPI_ISL_575060, EPI_ISL_575061, EPI_ISL_575062, EPI_ISL_575063, EPI_ISL_575064, EPI_ISL_575065, EPI_ISL_575066, EPI_ISL_575067, EPI_ISL_575068, EPI_ISL_575069, EPI_ISL_575070, EPI_ISL_575071, EPI_ISL_575072, EPI_ISL_575073, EPI_ISL_575074, EPI_ISL_575075, EPI_ISL_575076, EPI_ISL_575077, EPI_ISL_575078, EPI_ISL_575079, EPI_ISL_575080, EPI_ISL_575081, EPI_ISL_575082, EPI_ISL_575083, EPI_ISL_575084, EPI_ISL_575085, EPI_ISL_575086, EPI_ISL_575087, EPI_ISL_575088, EPI_ISL_575089, EPI_ISL_575090, EPI_ISL_575091, EPI_ISL_575092, EPI_ISL_575093, EPI_ISL_575094, EPI_ISL_575095, EPI_ISL_575096, EPI_ISL_575097, EPI_ISL_575098, EPI_ISL_575099, EPI_ISL_575100, EPI_ISL_575101, EPI_ISL_575102, EPI_ISL_575103, EPI_ISL_575104, EPI_ISL_575105, EPI_ISL_575106, EPI_ISL_575107, EPI_ISL_575108, EPI_ISL_575109, EPI_ISL_575110, EPI_ISL_575111, EPI_ISL_575112, EPI_ISL_575113, EPI_ISL_575114, EPI_ISL_575115, EPI_ISL_575116, EPI_ISL_575117, EPI_ISL_575118, EPI_ISL_575119, EPI_ISL_575120, EPI_ISL_575121, EPI_ISL_575122, EPI_ISL_575123, EPI_ISL_575124, EPI_ISL_575125, EPI_ISL_575126, EPI_ISL_575127, EPI_ISL_575128, EPI_ISL_575129, EPI_ISL_575130, EPI_ISL_575131, EPI_ISL_575132, EPI_ISL_575133, EPI_ISL_575134, EPI_ISL_575135, EPI_ISL_575136, EPI_ISL_575137, EPI_ISL_575138, EPI_ISL_575139, EPI_ISL_575140, EPI_ISL_575141, EPI_ISL_575142, EPI_ISL_575143, EPI_ISL_575144, EPI_ISL_575145, EPI_ISL_575146, EPI_ISL_575147, EPI_ISL_575148, EPI_ISL_575149, EPI_ISL_575150, EPI_ISL_575151, EPI_ISL_575152, EPI_ISL_575153, EPI_ISL_575154, EPI_ISL_575155, EPI_ISL_575156, EPI_ISL_575157, EPI_ISL_575158, EPI_ISL_575159, EPI_ISL_575160, EPI_ISL_575161, EPI_ISL_575162, EPI_ISL_575163, EPI_ISL_575164, EPI_ISL_575165, EPI_ISL_575166, EPI_ISL_575167, EPI_ISL_575168, EPI_ISL_575169, EPI_ISL_575170, EPI_ISL_575171, EPI_ISL_575172, EPI_ISL_575173, EPI_ISL_575174, EPI_ISL_575175, EPI_ISL_575176, EPI_ISL_575177, EPI_ISL_575178, EPI_ISL_575179, EPI_ISL_575180, EPI_ISL_575181, EPI_ISL_575182, EPI_ISL_575183, EPI_ISL_575184, EPI_ISL_575185, EPI_ISL_575186, EPI_ISL_575187, EPI_ISL_575188, EPI_ISL_575189, EPI_ISL_575190, EPI_ISL_575191, EPI_ISL_575192, EPI_ISL_575193, EPI_ISL_575194, EPI_ISL_575195, EPI_ISL_575196, EPI_ISL_575197, EPI_ISL_575198, EPI_ISL_575199, EPI_ISL_575200, EPI_ISL_575201, EPI_ISL_575202, EPI_ISL_575203, EPI_ISL_575204, EPI_ISL_575205, EPI_ISL_575206, EPI_ISL_575207, EPI_ISL_575208, EPI_ISL_575209, EPI_ISL_575210, EPI_ISL_575211, EPI_ISL_575212, EPI_ISL_575213, EPI_ISL_575214, EPI_ISL_575215, EPI_ISL_575216, EPI_ISL_575217, EPI_ISL_575218, EPI_ISL_575219, EPI_ISL_575220, EPI_ISL_575221, EPI_ISL_575222, EPI_ISL_575223, EPI_ISL_575224, EPI_ISL_575225, EPI_ISL_575226, EPI_ISL_575227, EPI_ISL_575228, EPI_ISL_575229, EPI_ISL_575230, EPI_ISL_575231, EPI_ISL_575232, EPI_ISL_575233, EPI_ISL_575234, EPI_ISL_575235, EPI_ISL_575236, EPI_ISL_575237, EPI_ISL_575238, EPI_ISL_575239, EPI_ISL_575240, EPI_ISL_575241, EPI_ISL_575242, EPI_ISL_575243, EPI_ISL_575244, EPI_ISL_575245, EPI_ISL_575246, EPI_ISL_575247, EPI_ISL_575248, EPI_ISL_575249, EPI_ISL_575250, EPI_ISL_575251, EPI_ISL_575252, EPI_ISL_575253, EPI_ISL_575254, EPI_ISL_575255, EPI_ISL_575256, EPI_ISL_575257, EPI_ISL_575258, EPI_ISL_575259, EPI_ISL_575260, EPI_ISL_575261, EPI_ISL_575262, EPI_ISL_575263, EPI_ISL_575264, EPI_ISL_575265, EPI_ISL_575266, EPI_ISL_575267, EPI_ISL_575268, EPI_ISL_575269, EPI_ISL_575270, EPI_ISL_575271, EPI_ISL_575272, EPI_ISL_575273, EPI_ISL_575274, EPI_ISL_575275, EPI_ISL_575276, EPI_ISL_575277, EPI_ISL_575278, EPI_ISL_575279, EPI_ISL_575280, EPI_ISL_575281, EPI_ISL_575282, EPI_ISL_575283, EPI_ISL_575284, EPI_ISL_575285, EPI_ISL_575286, EPI_ISL_575287, EPI_ISL_575288 | | | |
| see above | Utah Public Health Laboratory | Utah Public Health Laboratory | Erin Young, Kelly Oakeson |
| EPI_ISL_576264, EPI_ISL_576265 | Instituto de Diagnostico y Referencia Epidemiologicos (INDRE) | Instituto de Diagnostico y Referencia Epidemiologicos (INDRE) | Ernesto Ramirez-Gonzalez, Abril Rodriguez-Maldonado, Claudia Wong-Arambula , Natividad Cruz-Ortiz, Tatiana Nunez-Garcia, Dayanira Arellano-Suarez, Adnan Araiza-Rodriguez, Edgar Mendieta-Condado, Lucia Hernandez-Rivas, Irma Lopez-Martinez, Gisela Barrera-Badillo. |
| EPI_ISL_576266, EPI_ISL_576267, EPI_ISL_576268, EPI_ISL_576269, EPI_ISL_576270, EPI_ISL_576271, EPI_ISL_576273 | Instituto de Diagnostico y Referencia Epidemiologicos (INDRE) | Instituto de Diagnostico y Referencia Epidemiologicos (INDRE) | Ernesto Ramirez-Gonzalez, Abril Rodriguez-Maldonado, Claudia Wong-Arambula , Natividad Cruz-Ortiz, Tatiana Nunez-Garcia, Dayanira Arellano-Suarez, Adnan Araiza-Rodriguez, Fabiola Garces-Ayala, Lucia Hernandez-Rivas, Irma Lopez-Martinez, Gisela Barrera-Badillo. |
| EPI_ISL_576277, EPI_ISL_576278 | Instituto de Diagnostico y Referencia Epidemiologicos | Instituto de Diagnostico y Referencia Epidemiologicos | Ernesto Ramirez-Gonzalez, Abril Rodriguez-Maldonado, Claudia Wong-Arambula , Natividad Cruz-Ortiz, Tatiana Nunez-Garcia, Dayanira Arellano-Suarez, |

| | | | |
|---|---|---|---|
| | (INDRE) | (INDRE) | Adnan Araiza-Rodriguez, Edgar Mendieta-Condado, Lucia Hernandez-Rivas, Irma Lopez-Martinez, Gisela Barrera-Badillo. |
| EPI_ISL_576279 | Instituto de Diagnostico y Referencia Epidemiologicos (INDRE) | Instituto de Diagnostico y Referencia Epidemiologicos (INDRE) | Ernesto Ramirez-Gonzalez, Abril Rodriguez-Maldonado, Claudia Wong-Arambula , Natividad Cruz-Ortiz, Tatiana Nunez-Garcia, Dayanira Arellano-Suarez, Adnan Araiza-Rodriguez, Fabiola Garces-Ayala, Lucia Hernandez-Rivas, Irma Lopez-Martinez, Gisela Barrera-Badillo. |
| EPI_ISL_577110, EPI_ISL_577111, EPI_ISL_577112, EPI_ISL_577113, EPI_ISL_577114, EPI_ISL_577115, EPI_ISL_577116, EPI_ISL_577117, EPI_ISL_577118, EPI_ISL_577119, EPI_ISL_577120, EPI_ISL_577121, EPI_ISL_577122, EPI_ISL_577123, EPI_ISL_577124, EPI_ISL_577125, EPI_ISL_577126, EPI_ISL_577127, EPI_ISL_577128, EPI_ISL_577129, EPI_ISL_577130, EPI_ISL_577131, EPI_ISL_577132, EPI_ISL_577133, EPI_ISL_577134, EPI_ISL_577135, EPI_ISL_577136, EPI_ISL_577137, EPI_ISL_577138, EPI_ISL_577139, EPI_ISL_577140, EPI_ISL_577141, EPI_ISL_577142, EPI_ISL_577143, EPI_ISL_577144, EPI_ISL_577145, EPI_ISL_577146, EPI_ISL_577147, EPI_ISL_577148, EPI_ISL_577149, EPI_ISL_577150, EPI_ISL_577151, EPI_ISL_577152, EPI_ISL_577153, EPI_ISL_577154, EPI_ISL_577155, EPI_ISL_577156, EPI_ISL_577157, EPI_ISL_577158, EPI_ISL_577159, EPI_ISL_577160, EPI_ISL_577161, EPI_ISL_577162, EPI_ISL_577163, EPI_ISL_577164, EPI_ISL_577165, EPI_ISL_577166, EPI_ISL_577167, EPI_ISL_577168, EPI_ISL_577169, EPI_ISL_577170, EPI_ISL_577171, EPI_ISL_577172, EPI_ISL_577173, EPI_ISL_577174, EPI_ISL_577175, EPI_ISL_577176, EPI_ISL_577177, EPI_ISL_577178, EPI_ISL_577179, EPI_ISL_577180, EPI_ISL_577181, EPI_ISL_577182, EPI_ISL_577183, EPI_ISL_577184, EPI_ISL_577185, EPI_ISL_577186, EPI_ISL_577187, EPI_ISL_577188, EPI_ISL_577189, EPI_ISL_577190, EPI_ISL_577191, EPI_ISL_577192 | | | |
| see above | Queens Medical Centre, Clinical Microbiology Department / DeepSeq Nottingham | COVID-19 Genomics UK (COG-UK) Consortium | Gemma Clark, Wendy Smith, Manjinder Khakh, Vicki M Fleming, Michelle M Lister, Hannah Howson-Wells, Jonathan Ball, Patrick McClure, Joseph Chappell, Theocharis Tsoleridis, Nadine Holmes, Matthew Carlisle, Christopher Moore, Fei Sang, Johnny Debebe, Victoria Wright, Matthew Loose |
| EPI_ISL_578697 | Clinical Virology Laboratory, Institute of Liver and Biliary Sciences | ILBS - IGIB | Ekta Gupta, Sheetalnath Rooge, Abhishek Padhi, Reshu Agarwal, Jaswinder Singh Maras, Shridhar Sivasubbu, Vinod Scaria, Shvetank Sharma |
| EPI_ISL_584048 | Laboratory of Molecular Virology, Department of Biomedical, Surgical and Dental Sciences University of Milano | Laboratory of Molecular Virology, Department of Biomedical, Surgical and Dental Sciences University of Milano | Delbue,S., Modenese,A., Bianchi,M., Fattori,M., D'Alessandro,S., Pariani,E., Basilico,N., Galli,C. and Ferrante,P. |
| EPI_ISL_584049 | Laboratory of Molecular Virology, Department of Biomedical, Surgical and Dental Sciences University of Milano | Laboratory of Molecular Virology, Department of Biomedical, Surgical and Dental Sciences University of Milano | Delbue,S., Modenese,A., Bianchi,M., Fattori,M., D'Alessandro,S.,Pariani,E., Basilico,N., Galli,C. and Ferrante,P. |
| EPI_ISL_584051 | Laboratory of Molecular Virology, Department of Biomedical, Surgical and Dental Sciences University of Milano | Laboratory of Molecular Virology, Department of Biomedical, Surgical and Dental Sciences University of Milano | Delbue,S., Ferrante,P., Basilico,N., Parapini,S., Binda,S., D'Alessandro,S., Galli,C., Signorini,L., Primache,V., Anselmi,G. and Pariani,E |
| EPI_ISL_584052 | Laboratory of Molecular Virology, Department of Biomedical, Surgical and Dental Sciences University of Milano | Laboratory of Molecular Virology, Department of Biomedical, Surgical and Dental Sciences University of Milano | Delbue,S., D'Alessandro,S., Modenese,A., Signorini,L., Parapini,S.,Dolci,M., Binda,S., Primache,V., Taramelli,D., Incorvaia,B. and Ferrante,P. |
| EPI_ISL_584381, EPI_ISL_584382, EPI_ISL_584383, EPI_ISL_584384, EPI_ISL_584385, EPI_ISL_584386, EPI_ISL_584387, EPI_ISL_584388, EPI_ISL_584389, EPI_ISL_584390, EPI_ISL_584391, EPI_ISL_584392, EPI_ISL_584393, EPI_ISL_584394, EPI_ISL_584395, EPI_ISL_584396, EPI_ISL_584397, EPI_ISL_584398, EPI_ISL_584399, EPI_ISL_584400, EPI_ISL_584401, EPI_ISL_584402, EPI_ISL_584403, EPI_ISL_584404, EPI_ISL_584405, EPI_ISL_584406, EPI_ISL_584407, EPI_ISL_584408, EPI_ISL_584409, EPI_ISL_584410, EPI_ISL_584411, EPI_ISL_584412, EPI_ISL_584413, EPI_ISL_584414, EPI_ISL_584415, EPI_ISL_584416, EPI_ISL_584417, EPI_ISL_584418, EPI_ISL_584419, EPI_ISL_584420, EPI_ISL_584421, EPI_ISL_584422, EPI_ISL_584423, EPI_ISL_584424, EPI_ISL_584425, EPI_ISL_584426, EPI_ISL_584427, EPI_ISL_584428, EPI_ISL_584429, EPI_ISL_584430, EPI_ISL_584431, EPI_ISL_584432, EPI_ISL_584433, EPI_ISL_584434, EPI_ISL_584435, EPI_ISL_584436, EPI_ISL_584437, EPI_ISL_584438, EPI_ISL_584439, EPI_ISL_584750, EPI_ISL_584751, EPI_ISL_584752, EPI_ISL_584753, EPI_ISL_584754, EPI_ISL_584755, EPI_ISL_584756, EPI_ISL_584757, EPI_ISL_584758, EPI_ISL_584759, EPI_ISL_584760, EPI_ISL_584761, EPI_ISL_584762, EPI_ISL_584763, EPI_ISL_584764, EPI_ISL_584765, EPI_ISL_584766, EPI_ISL_584767, EPI_ISL_584768, EPI_ISL_584769, EPI_ISL_584770, EPI_ISL_584771, EPI_ISL_584772, EPI_ISL_584773, EPI_ISL_584774, EPI_ISL_584775, EPI_ISL_584776, EPI_ISL_584777, EPI_ISL_584778, EPI_ISL_584779, EPI_ISL_584780, EPI_ISL_584781, EPI_ISL_584782, EPI_ISL_584783, EPI_ISL_584784, EPI_ISL_584785, EPI_ISL_584786, EPI_ISL_584787, EPI_ISL_584788, EPI_ISL_584789, EPI_ISL_584790, EPI_ISL_584791, EPI_ISL_584792, EPI_ISL_584793, EPI_ISL_584794, EPI_ISL_584795, EPI_ISL_584796, EPI_ISL_584797, EPI_ISL_584798, EPI_ISL_584799, EPI_ISL_584800, EPI_ISL_584801, EPI_ISL_584802, EPI_ISL_584803, EPI_ISL_584804, EPI_ISL_584805, EPI_ISL_584806, EPI_ISL_584807, EPI_ISL_584808, EPI_ISL_585627, EPI_ISL_585628, EPI_ISL_585629 | | | |
| see above | Queens Medical Centre, Clinical Microbiology Department / DeepSeq Nottingham | COVID-19 Genomics UK (COG-UK) Consortium | Gemma Clark, Wendy Smith, Manjinder Khakh, Vicki M Fleming, Michelle M Lister, Hannah Howson-Wells, Jonathan Ball, Patrick McClure, Joseph Chappell, Theocharis Tsoleridis, Nadine Holmes, Matthew Carlisle, Christopher Moore, Fei Sang, Johnny Debebe, Victoria Wright, Matthew Loose |
| EPI_ISL_586518 | B.J. Medical College and Civil hospital, Ahmedabad | Gujarat Biotechnology Research Centre | Dinesh Kumar, Zuber Saiyed, Komal Patel, Labdhi Pandya, Afzal Ansari, Nikha Trivedi, Pranay Shah, Kamlesh J Upadhyay, Sanjay Kapadia, Apurvasinh Puvar, Janvi Raval, Zarna Patel, Monika Gandhi, Pinal Trivedi, Maharshi Pandya, Nidhi Patel, Nitin Savaliya, Raghawendra Kumar, R D Dixit, A M Kadri, Harsh Bakshi, Chaitanya Joshi, Madhvi Joshi |
| EPI_ISL_586533 | General Hospital,Chhotaudepur | Gujarat Biotechnology Research Centre | Dharak Pandya, Maharshi Pandya, Nidhi Patel, Nitin Savaliya, Raghawendra Kumar, Dinesh Kumar, Zuber Saiyed, Komal Patel, Labdhi Pandya, Afzal Ansari, Nikha Trivedi, Apurvasinh Puvar, Janvi Raval, Zarna Patel, Monika Gandhi, Pinal Trivedi, Yogesh Parmar, Arti Thakur, R D Dixit, A M Kadri, Harsh Bakshi, Chaitanya Joshi, Madhvi Joshi |
| EPI_ISL_586540 | Zydus Medical Hospital,Dahod | Gujarat Biotechnology Research Centre | Dinesh Kumar, Zuber Saiyed, Komal Patel, Labdhi Pandya, Afzal Ansari, Nikha Trivedi, Apurvasinh Puvar, Janvi Raval, Zarna Patel, Monika Gandhi, Pinal Trivedi, Maharshi Pandya, Nidhi Patel, Kalpesh Labana, Mohit Desai, Nitin Savaliya, Raghawendra Kumar, R D Dixit, A M Kadri, Harsh Bakshi, Chaitanya Joshi, Madhvi Joshi |
| EPI_ISL_586557 | CCC,Veraval | Gujarat Biotechnology Research Centre | Dinesh Kumar, Zuber Saiyed, Komal Patel, Labdhi Pandya, Afzal Ansari, Nikha Trivedi, Jignesh Parmar, Jitendra Bamrotia, Apurvasinh Puvar, Janvi Raval, Zarna Patel, Monika Gandhi, Pinal Trivedi, Maharshi Pandya, Nidhi Patel, Nitin Savaliya, Raghawendra Kumar, R D Dixit, A M Kadri, Harsh Bakshi, Chaitanya Joshi, Madhvi Joshi |
| EPI_ISL_591281, EPI_ISL_591282, EPI_ISL_591283, EPI_ISL_591284, EPI_ISL_591285, EPI_ISL_591286, EPI_ISL_591287, EPI_ISL_591288, EPI_ISL_591289, EPI_ISL_591290, EPI_ISL_591291, EPI_ISL_591292, EPI_ISL_591293, EPI_ISL_591294, EPI_ISL_591295, EPI_ISL_591296, EPI_ISL_591297, EPI_ISL_591298, EPI_ISL_591299, EPI_ISL_591300, EPI_ISL_591301, EPI_ISL_591302, EPI_ISL_591303, EPI_ISL_591304, EPI_ISL_591305, EPI_ISL_591306, EPI_ISL_591307, EPI_ISL_591308, EPI_ISL_591309, EPI_ISL_591310, EPI_ISL_591311, EPI_ISL_591312, EPI_ISL_591313, EPI_ISL_591314, EPI_ISL_591315 | | | |
| see above | Utah Public Health Laboratory | Utah Public Health Laboratory | Erin Young, Kelly Oakeson |
| EPI_ISL_591519, EPI_ISL_591520, EPI_ISL_591521 | Programa de Virología, Facultad de Medicina UC | Center for Mathematical Modeling and Center for Genome Regulation. Santiago, Chile | Gaggero A, Valiente F, Gaete A, Travisany D, Palma R, Urra C, Varas M, Allende ML, Maass A, González M, Ferres M. |
| EPI_ISL_591522, EPI_ISL_591523, EPI_ISL_591524, EPI_ISL_591525, EPI_ISL_591526, EPI_ISL_591527, EPI_ISL_591528, EPI_ISL_591529, EPI_ISL_591530 | Medicina Norte U Chile - Servicio Medico Legal | Center for Mathematical Modeling and Center for Genome Regulation. Santiago, Chile | Gaggero A, Valiente F, Gaete A, Travisany D, Palma R, Urra C, Varas M, Allende ML, Maass A, González M, Ferres M. |
| EPI_ISL_593772, EPI_ISL_593773, EPI_ISL_593774, EPI_ISL_593775, EPI_ISL_593776 | HOSPITAL REGIONAL LAMBAYEQUE | GENOMA MAYOR | Franklin R. Aguilar-Gamboa, Luis M. López-Serquén, Heber Silva-Díaz, Percy O. Tullume-Vergara, Luis Salcedo-Mejía, Juan J. Bonifacio-Briceño, Ramsés Salas-Asencios, Marco E. Mechán-Llontop and Juan P. Cárdenas. |
| EPI_ISL_593999, EPI_ISL_594000, EPI_ISL_594001, EPI_ISL_594002, EPI_ISL_594003, EPI_ISL_594004, EPI_ISL_594005, EPI_ISL_594006, EPI_ISL_594007, EPI_ISL_594008, EPI_ISL_594009, EPI_ISL_594010, EPI_ISL_594011, EPI_ISL_594012, EPI_ISL_594013, EPI_ISL_594014, EPI_ISL_594015, EPI_ISL_594016, EPI_ISL_594017, EPI_ISL_594018, EPI_ISL_594019, EPI_ISL_594020, EPI_ISL_594021, EPI_ISL_594022, EPI_ISL_594023, EPI_ISL_594024, EPI_ISL_594025, EPI_ISL_594026, EPI_ISL_594027, EPI_ISL_594028, EPI_ISL_594029, EPI_ISL_594030, EPI_ISL_594031, EPI_ISL_594032, EPI_ISL_594033, EPI_ISL_594034, EPI_ISL_594035, EPI_ISL_594036, EPI_ISL_594037, EPI_ISL_594038, EPI_ISL_594039, EPI_ISL_594040, EPI_ISL_594041, EPI_ISL_594042, EPI_ISL_594043, EPI_ISL_594044, EPI_ISL_594045, EPI_ISL_594046, EPI_ISL_594047, EPI_ISL_594048, EPI_ISL_594049, EPI_ISL_594050, EPI_ISL_594051, EPI_ISL_594052, EPI_ISL_594053, EPI_ISL_594054, EPI_ISL_594055, EPI_ISL_594056, EPI_ISL_594057, EPI_ISL_594058, EPI_ISL_594059, EPI_ISL_594060, EPI_ISL_594061, EPI_ISL_594062, EPI_ISL_594063, EPI_ISL_594064, EPI_ISL_594065, EPI_ISL_594066, EPI_ISL_594067, EPI_ISL_594068, EPI_ISL_594069, EPI_ISL_594070, EPI_ISL_594071, EPI_ISL_594072, EPI_ISL_594073, EPI_ISL_594074, EPI_ISL_594075, EPI_ISL_594076, EPI_ISL_594077, EPI_ISL_594078, EPI_ISL_594079, EPI_ISL_594080, EPI_ISL_594081, EPI_ISL_594082, EPI_ISL_594083, EPI_ISL_594084, EPI_ISL_594085, EPI_ISL_594086, EPI_ISL_594087, EPI_ISL_594088, EPI_ISL_594089, EPI_ISL_594090, EPI_ISL_594091, EPI_ISL_594092, EPI_ISL_594093, EPI_ISL_594094, EPI_ISL_594095, EPI_ISL_594096, EPI_ISL_594097, EPI_ISL_594098, EPI_ISL_594099, EPI_ISL_594100, EPI_ISL_594101, EPI_ISL_594102, EPI_ISL_594103, EPI_ISL_594104, EPI_ISL_594105, EPI_ISL_594106, EPI_ISL_594107, EPI_ISL_594108, EPI_ISL_594109, EPI_ISL_594110, EPI_ISL_594111, EPI_ISL_594112, EPI_ISL_594113, EPI_ISL_594114, EPI_ISL_594115, EPI_ISL_594116, EPI_ISL_594117 | | | |
| see above | Utah Public Health Laboratory | Utah Public Health Laboratory | Erin Young, Kelly Oakeson |
| EPI_ISL_594118 | Centro de Investigaciones, Universidad de Especialidades Espíritu Santo | Institute of Microbiology, Universidad San Francisco de Quito | Derly Andrade, Juan Carlos Fernandez, Belén Prado-Vivar, Sully Márquez, Juan José Guadalupe, Monica Becerra-Wong, Bernardo Gutiérrez, Gabriel Morey, Ruben Armas, Jose Pedro Barberan, Fernando Espinoza, Edith Lopez, Verónica Barragán, Patricio Rojas-Silva, Gabriel Trueba, Michelle Grunauer, Paúl Cárdenas |
| EPI_ISL_594157, EPI_ISL_594158, EPI_ISL_594159, EPI_ISL_594160, EPI_ISL_594162 | Israel Institute for Biological Research | Israel Institute for Biological Research | Galia Zaide, Inbar Cohen-Gihon, Ofir Israeli, Dana Stein, Shay Weiss, Orly Laskar, Yoav Gal, Libby Weiss, Emanuelle Mamroud, Adi Beth-Din and Anat Zvi |
| EPI_ISL_594248, EPI_ISL_594249, EPI_ISL_594250, EPI_ISL_594251, EPI_ISL_594252, EPI_ISL_594253, EPI_ISL_594254, EPI_ISL_594255, EPI_ISL_594256, EPI_ISL_594257, EPI_ISL_594258, EPI_ISL_594259, EPI_ISL_594260, EPI_ISL_594261, EPI_ISL_594262, EPI_ISL_594263, EPI_ISL_594264, EPI_ISL_594265, EPI_ISL_594266, EPI_ISL_594267, EPI_ISL_594268, EPI_ISL_594269, EPI_ISL_594270, EPI_ISL_594271, EPI_ISL_594272, EPI_ISL_594273, EPI_ISL_594274, EPI_ISL_594275, EPI_ISL_594276, EPI_ISL_594394, EPI_ISL_594395, EPI_ISL_594396, EPI_ISL_594397, EPI_ISL_594398, EPI_ISL_594399, EPI_ISL_594400, EPI_ISL_594401, EPI_ISL_594402, EPI_ISL_594403, EPI_ISL_594404, EPI_ISL_594405, EPI_ISL_594406, EPI_ISL_594407, EPI_ISL_594408, EPI_ISL_594409, EPI_ISL_594410, EPI_ISL_594411, EPI_ISL_594412, EPI_ISL_594413, EPI_ISL_594414, EPI_ISL_594415, EPI_ISL_594416, EPI_ISL_594417, EPI_ISL_594418, EPI_ISL_594419, EPI_ISL_594420, EPI_ISL_594421, EPI_ISL_594422, EPI_ISL_594423, EPI_ISL_594424, EPI_ISL_594425, EPI_ISL_594426, EPI_ISL_594427, EPI_ISL_594428, EPI_ISL_594429, EPI_ISL_594430, EPI_ISL_594431, EPI_ISL_594432, EPI_ISL_594433, EPI_ISL_594434, EPI_ISL_594435, EPI_ISL_594436, EPI_ISL_594437, EPI_ISL_594438, EPI_ISL_594439, EPI_ISL_594440, EPI_ISL_594441, EPI_ISL_594442, EPI_ISL_594443, EPI_ISL_594444, EPI_ISL_594445, EPI_ISL_594446 | | | |
| see above | Utah Public Health Laboratory | Utah Public Health Laboratory | Erin Young, Kelly Oakeson |

| | | | |
|---|---|---|---|
| EPI_ISL_594923, EPI_ISL_594924, EPI_ISL_594925, EPI_ISL_594926, EPI_ISL_594927, EPI_ISL_594928, EPI_ISL_594929, EPI_ISL_594930, EPI_ISL_594931, EPI_ISL_594932, EPI_ISL_594933, EPI_ISL_594934, EPI_ISL_594935, EPI_ISL_594936, EPI_ISL_594937, EPI_ISL_594938, EPI_ISL_594939, EPI_ISL_594940, EPI_ISL_594941, EPI_ISL_594942, EPI_ISL_594943, EPI_ISL_594944, EPI_ISL_594945, EPI_ISL_594946, EPI_ISL_594947, EPI_ISL_594948, EPI_ISL_594949, EPI_ISL_594950, EPI_ISL_594951, EPI_ISL_594952, EPI_ISL_594953, EPI_ISL_594954, EPI_ISL_594955, EPI_ISL_594956, EPI_ISL_594957, EPI_ISL_594958, EPI_ISL_594959, EPI_ISL_594960, EPI_ISL_594961, EPI_ISL_594962, EPI_ISL_594963, EPI_ISL_594964, EPI_ISL_594965, EPI_ISL_594966, EPI_ISL_594967, EPI_ISL_594968, EPI_ISL_594969, EPI_ISL_594970, EPI_ISL_594971, EPI_ISL_594972, EPI_ISL_594973, EPI_ISL_594974, EPI_ISL_594975, EPI_ISL_594976, EPI_ISL_594977, EPI_ISL_594978, EPI_ISL_594979, EPI_ISL_594980, EPI_ISL_594981, EPI_ISL_594982, EPI_ISL_594983, EPI_ISL_594984, EPI_ISL_594985, EPI_ISL_594986, EPI_ISL_595046, EPI_ISL_595047, EPI_ISL_595048, EPI_ISL_595049, EPI_ISL_595050, EPI_ISL_595051, EPI_ISL_595052, EPI_ISL_595053, EPI_ISL_595054, EPI_ISL_595055, EPI_ISL_595056, EPI_ISL_595057, EPI_ISL_595058, EPI_ISL_595059, EPI_ISL_595060, EPI_ISL_595061, EPI_ISL_595062, EPI_ISL_595063, EPI_ISL_595064, EPI_ISL_595065, EPI_ISL_595066, EPI_ISL_595067, EPI_ISL_595068, EPI_ISL_595069, EPI_ISL_595070, EPI_ISL_595071, EPI_ISL_595072, EPI_ISL_595073, EPI_ISL_595074, EPI_ISL_595075, EPI_ISL_595076, EPI_ISL_595077, EPI_ISL_595078, EPI_ISL_595079, EPI_ISL_595080, EPI_ISL_595081, EPI_ISL_595082, EPI_ISL_595083, EPI_ISL_595084, EPI_ISL_595085, EPI_ISL_595086, EPI_ISL_595087, EPI_ISL_595088, EPI_ISL_595089, EPI_ISL_595090, EPI_ISL_595091, EPI_ISL_595092, EPI_ISL_595093, EPI_ISL_595094, EPI_ISL_595095, EPI_ISL_595096, EPI_ISL_595097, EPI_ISL_595098, EPI_ISL_595099, EPI_ISL_595232, EPI_ISL_595233, EPI_ISL_595234, EPI_ISL_595235, EPI_ISL_595236, EPI_ISL_595237, EPI_ISL_595238, EPI_ISL_595239, EPI_ISL_595240, EPI_ISL_595241, EPI_ISL_595242, EPI_ISL_595243, EPI_ISL_595244, EPI_ISL_595245, EPI_ISL_595246, EPI_ISL_595247, EPI_ISL_595248, EPI_ISL_595249, EPI_ISL_595250, EPI_ISL_595251, EPI_ISL_595252, EPI_ISL_595253, EPI_ISL_595254, EPI_ISL_595255, EPI_ISL_595256, EPI_ISL_595257, EPI_ISL_595258, EPI_ISL_595259, EPI_ISL_595260, EPI_ISL_595261, EPI_ISL_595262, EPI_ISL_595263, EPI_ISL_595264, EPI_ISL_595265, EPI_ISL_595266, EPI_ISL_595267, EPI_ISL_595268, EPI_ISL_595269, EPI_ISL_595270, EPI_ISL_595271, EPI_ISL_595272, EPI_ISL_595273, EPI_ISL_595274, EPI_ISL_595275, EPI_ISL_595276, EPI_ISL_595277, EPI_ISL_595278, EPI_ISL_595279, EPI_ISL_595280, EPI_ISL_595281, EPI_ISL_595282, EPI_ISL_595283, EPI_ISL_595284, EPI_ISL_595285, EPI_ISL_595286, EPI_ISL_595287, EPI_ISL_595288, EPI_ISL_595289, EPI_ISL_595290, EPI_ISL_595291, EPI_ISL_595292, EPI_ISL_595293, EPI_ISL_595294, EPI_ISL_595295, EPI_ISL_595296, EPI_ISL_595297, EPI_ISL_595298, EPI_ISL_595299, EPI_ISL_595300, EPI_ISL_595301, EPI_ISL_595302, EPI_ISL_595303, EPI_ISL_595304, EPI_ISL_595305, EPI_ISL_595306, EPI_ISL_595307, EPI_ISL_595308, EPI_ISL_595309, EPI_ISL_595310, EPI_ISL_595311, EPI_ISL_595312, EPI_ISL_595313, EPI_ISL_595314, EPI_ISL_595315, EPI_ISL_595316, EPI_ISL_595317, EPI_ISL_595318, EPI_ISL_595319, EPI_ISL_595320, EPI_ISL_595321, EPI_ISL_595322, EPI_ISL_595323, EPI_ISL_595324, EPI_ISL_595325 | | | |
| see above | Queens Medical Centre, Clinical Microbiology Department / DeepSeq Nottingham | COVID-19 Genomics UK (COG-UK) Consortium | Gemma Clark, Wendy Smith, Manjinder Khakh, Vicki M Fleming, Michelle M Lister, Hannah Howson-Wells, Jonathan Ball, Patrick McClure, Joseph Chappell, Theocharis Tsoleridis, Nadine Holmes, Matthew Carlisle, Christopher Moore, Fei Sang, Johnny Debebe, Victoria Wright, Matthew Loose |
| EPI_ISL_602162, EPI_ISL_602163, EPI_ISL_602164, EPI_ISL_602165, EPI_ISL_602166, EPI_ISL_602167, EPI_ISL_602168, EPI_ISL_602169, EPI_ISL_602170, EPI_ISL_602171, EPI_ISL_602172, EPI_ISL_602173, EPI_ISL_602174, EPI_ISL_602175, EPI_ISL_602176, EPI_ISL_602177, EPI_ISL_602178, EPI_ISL_602179, EPI_ISL_602180, EPI_ISL_602181, EPI_ISL_602182, EPI_ISL_602183, EPI_ISL_602184, EPI_ISL_602185, EPI_ISL_602186, EPI_ISL_602187, EPI_ISL_602188, EPI_ISL_602189, EPI_ISL_602190, EPI_ISL_602191, EPI_ISL_602192, EPI_ISL_602193, EPI_ISL_602194, EPI_ISL_602195, EPI_ISL_602196, EPI_ISL_602197, EPI_ISL_602198, EPI_ISL_602199, EPI_ISL_602200, EPI_ISL_602201, EPI_ISL_602202, EPI_ISL_602203, EPI_ISL_602204, EPI_ISL_602205, EPI_ISL_602206, EPI_ISL_602207 | | | |
| see above | New Mexico Department of Health Scientific Laboratory | New Mexico Department of Health Scientific Laboratory | Ellie Johnson, Anastacia Griego-Fisher, D'Eldra Malone |
| EPI_ISL_602306, EPI_ISL_602307, EPI_ISL_602308, EPI_ISL_602309, EPI_ISL_602310, EPI_ISL_602311, EPI_ISL_602312, EPI_ISL_602313, EPI_ISL_602314, EPI_ISL_602315, EPI_ISL_602316, EPI_ISL_602317, EPI_ISL_602318, EPI_ISL_602319, EPI_ISL_602320, EPI_ISL_602321, EPI_ISL_602322, EPI_ISL_602323, EPI_ISL_602324, EPI_ISL_602325, EPI_ISL_602326, EPI_ISL_602327, EPI_ISL_602328 | | | |
| see above | University of Miami Immunology and Histocompatibility Laboratory | University of Miami Immunology and Histocompatibility Laboratory | Emilio Margolles-Clark, PhD and Phillip Ruiz, MD, PhD |
| EPI_ISL_602622 | AHRI-Sigal | KRISP, KZN Research Innovation and Sequencing Platform | Gazy I, Sigla, Karim F, Cele S, Giandhari J, Pillay S, Tegally H, Wilkinson E, de Oliveira T |
| EPI_ISL_602623, EPI_ISL_602624, EPI_ISL_602625, EPI_ISL_602626, EPI_ISL_602627, EPI_ISL_602628, EPI_ISL_602629, EPI_ISL_602630, EPI_ISL_602631 | AHRI-Sigal | KRISP, KZN Research Innovation and Sequencing Platform | Gazy I, Sigl A, Karim F, Cele S, Giandhari J, Pillay S, Tegally H, Wilkinson E, de Oliveira T |
| EPI_ISL_602632, EPI_ISL_602633, EPI_ISL_602634, EPI_ISL_602635, EPI_ISL_602637, EPI_ISL_602638, EPI_ISL_602639, EPI_ISL_602640, EPI_ISL_602641, EPI_ISL_602642, EPI_ISL_602644, EPI_ISL_602646, EPI_ISL_602647, EPI_ISL_602648, EPI_ISL_602649, EPI_ISL_602650, EPI_ISL_602651, EPI_ISL_602652, EPI_ISL_602653, EPI_ISL_602656, EPI_ISL_602657, EPI_ISL_602658, EPI_ISL_602659, EPI_ISL_602660, EPI_ISL_602661, EPI_ISL_602662, EPI_ISL_602663, EPI_ISL_602664, EPI_ISL_602665, EPI_ISL_602666, EPI_ISL_602667, EPI_ISL_602668, EPI_ISL_602669, EPI_ISL_602670, EPI_ISL_602671, EPI_ISL_602672, EPI_ISL_602673, EPI_ISL_602674, EPI_ISL_602675, EPI_ISL_602676, EPI_ISL_602677, EPI_ISL_602678, EPI_ISL_602679, EPI_ISL_602680, EPI_ISL_602681, EPI_ISL_602683, EPI_ISL_602684, EPI_ISL_602686, EPI_ISL_602687, EPI_ISL_602688, EPI_ISL_602689, EPI_ISL_602690, EPI_ISL_602691, EPI_ISL_602692, EPI_ISL_602693, EPI_ISL_602694, EPI_ISL_602695, EPI_ISL_602696, EPI_ISL_602697, EPI_ISL_602698, EPI_ISL_602700, EPI_ISL_602701, EPI_ISL_602703, EPI_ISL_602704, EPI_ISL_602705, EPI_ISL_602707, EPI_ISL_602708, EPI_ISL_602709, EPI_ISL_602710, EPI_ISL_602711, EPI_ISL_602712, EPI_ISL_602713, EPI_ISL_602714, EPI_ISL_602715, EPI_ISL_602716, EPI_ISL_602717, EPI_ISL_602718, EPI_ISL_602719, EPI_ISL_602720, EPI_ISL_602721, EPI_ISL_602725, EPI_ISL_602727, EPI_ISL_602729, EPI_ISL_602730, EPI_ISL_602731, EPI_ISL_602732, EPI_ISL_602733, EPI_ISL_602734, EPI_ISL_602735, EPI_ISL_602736, EPI_ISL_602737, EPI_ISL_602738, EPI_ISL_602739, EPI_ISL_602740, EPI_ISL_602741, EPI_ISL_602742, EPI_ISL_602743, EPI_ISL_602744, EPI_ISL_602747, EPI_ISL_602748, EPI_ISL_602749, EPI_ISL_602750, EPI_ISL_602751, EPI_ISL_602752, EPI_ISL_602753, EPI_ISL_602754, EPI_ISL_602755, EPI_ISL_602756, EPI_ISL_602757, EPI_ISL_602758, EPI_ISL_602759, EPI_ISL_602760, EPI_ISL_602761, EPI_ISL_602762, EPI_ISL_602763, EPI_ISL_602764, EPI_ISL_602767, EPI_ISL_602768, EPI_ISL_602769, EPI_ISL_602770, EPI_ISL_602771, EPI_ISL_602772, EPI_ISL_602773, EPI_ISL_602774, EPI_ISL_602775, EPI_ISL_602776, EPI_ISL_602778, EPI_ISL_602779, EPI_ISL_602780, EPI_ISL_602782, EPI_ISL_602783, EPI_ISL_602784, EPI_ISL_602785, EPI_ISL_602786, EPI_ISL_602787, EPI_ISL_602788, EPI_ISL_602789, EPI_ISL_602791, EPI_ISL_602795, EPI_ISL_602796, EPI_ISL_602797, EPI_ISL_602798, EPI_ISL_602799, EPI_ISL_602800, EPI_ISL_602801, EPI_ISL_602803, EPI_ISL_602804, EPI_ISL_602806, EPI_ISL_602807, EPI_ISL_602808, EPI_ISL_602809, EPI_ISL_602810, EPI_ISL_602811, EPI_ISL_602812, EPI_ISL_602813, EPI_ISL_602814, EPI_ISL_602816, EPI_ISL_602817, EPI_ISL_602818, EPI_ISL_602820, EPI_ISL_602821, EPI_ISL_602822, EPI_ISL_602824, EPI_ISL_602826, EPI_ISL_602831, EPI_ISL_602832, EPI_ISL_602833, EPI_ISL_602834, EPI_ISL_602835, EPI_ISL_602836, EPI_ISL_602837, EPI_ISL_602838, EPI_ISL_602839, EPI_ISL_602841, EPI_ISL_602842, EPI_ISL_602844, EPI_ISL_602846, EPI_ISL_602848, EPI_ISL_602849, EPI_ISL_602850, EPI_ISL_602851, EPI_ISL_602853, EPI_ISL_602854, EPI_ISL_602855, EPI_ISL_602858, EPI_ISL_602859, EPI_ISL_602860, EPI_ISL_602861, EPI_ISL_602862, EPI_ISL_602863, EPI_ISL_602865, EPI_ISL_602867, EPI_ISL_602870, EPI_ISL_602871, EPI_ISL_602872, EPI_ISL_602873, EPI_ISL_602878, EPI_ISL_602881, EPI_ISL_602883, EPI_ISL_602884, EPI_ISL_602890, EPI_ISL_602892, EPI_ISL_602896, EPI_ISL_602899, EPI_ISL_602900, EPI_ISL_602902, EPI_ISL_602907, EPI_ISL_602908, EPI_ISL_602916, EPI_ISL_602917, EPI_ISL_602919, EPI_ISL_602920, EPI_ISL_602923, EPI_ISL_602926, EPI_ISL_602928, EPI_ISL_602929 | | | |
| see above | NHLS-IALCH | KRISP, KZN Research Innovation and Sequencing Platform | Giandhari J, Pillay S, Lessells R, Mdlalose K, York D, Khan S, Tegally H, Wilkinson E, de Oliveira T |
| EPI_ISL_602931, EPI_ISL_602932, EPI_ISL_602933, EPI_ISL_602934, EPI_ISL_602935, EPI_ISL_602936, EPI_ISL_602937, EPI_ISL_602938, EPI_ISL_602939, EPI_ISL_602940, EPI_ISL_602941, EPI_ISL_602942, EPI_ISL_602943, EPI_ISL_602944, EPI_ISL_602945, EPI_ISL_602946, EPI_ISL_602947, EPI_ISL_602948, EPI_ISL_602949, EPI_ISL_602950, EPI_ISL_602951, EPI_ISL_602952, EPI_ISL_602953, EPI_ISL_602954, EPI_ISL_602955 | | | |
| see above | Utah Public Health Laboratory | Utah Public Health Laboratory | Erin Young, Kelly Oakeson |
| EPI_ISL_603141 | INMI Lazzaro Spallanzani IRCCS | INMI Lazzaro Spallanzani IRCCS | Francesco Messina, Cesare E.M. Gruber, Martina Rueca, Barbara Bartolini, Emanuela Giombini, Simone Lanini, Maria R. Capobianchi, Antonino Di Caro |
| EPI_ISL_603145 | INMI Lazzaro Spallanzani IRCCS | INMI Lazzaro Spallanzani IRCCS | Francesco Messina, Cesare E.M. Gruber, Martina Rueca, Barbara Bartolini, Emanuela Giombini, Simone Lanini, Antonino Di Caro, Maria R. Capobianchi |
| EPI_ISL_603149 | INMI Lazzaro Spallanzani IRCCS | INMI Lazzaro Spallanzani IRCCS | Francesco Messina, Barbara Bartolini, Cesare E.M. Gruber, Martina Rueca, Simone Lanini, Emanuela Giombini, Patrizia Massarelli, Antonino Di Caro, Maria R. Capobianchi |
| EPI_ISL_603151 | INMI Lazzaro Spallanzani IRCCS | INMI Lazzaro Spallanzani IRCCS | Francesco Messina, Barbara Bartolini, Cesare E.M. Gruber, Martina Rueca, Emanuela Giombini, Simone Lanini, Patrizia Massarelli, Maria R. Capobianchi, Antonino Di Caro |
| EPI_ISL_603153 | INMI Lazzaro Spallanzani IRCCS | INMI Lazzaro Spallanzani IRCCS | Francesco Messina, Cesare E.M. Gruber, Barbara Bartolini, Simone Lanini, Emanuela Giombini, Martina Rueca, Marcello Meledandri, Maria Letizia Schiavone, Annunziata Tamburro, Antonino Di Caro, Maria R. Capobianchi |
| EPI_ISL_603157 | INMI Lazzaro Spallanzani IRCCS | INMI Lazzaro Spallanzani IRCCS | Francesco Messina, Martina Rueca, Barbara Bartolini, Cesare E.M. Gruber, Simone Lanini, Emanuela Giombini, Fulvia Pimpinelli, Antonino Di Caro, Maria R. Capobianchi |
| EPI_ISL_603169 | INMI Lazzaro Spallanzani IRCCS | INMI Lazzaro Spallanzani IRCCS | Francesco Messina, Barbara Bartolini, Cesare E.M. Gruber, Martina Rueca, Simone Lanini, Emanuela Giombini, Fulvia Pimpinelli, Maria R. Capobianchi, Antonino Di Caro |
| EPI_ISL_603174 | INMI Lazzaro Spallanzani IRCCS | INMI Lazzaro Spallanzani IRCCS | Francesco Messina, Barbara Bartolini, Martina Rueca, Emanuela Giombini, Simone Lanini, Cesare E.M. Gruber, Mariarosa Gaudio, Maria R. Capobianchi, Antonino Di Caro |
| EPI_ISL_603181 | INMI Lazzaro Spallanzani IRCCS | INMI Lazzaro Spallanzani IRCCS | Francesco Messina, Cesare E.M. Gruber, Barbara Bartolini, Martina Rueca, Simone Lanini, Emanuela Giombini, Maria R. Capobianchi, Antonino Di Caro |
| EPI_ISL_603186 | INMI Lazzaro Spallanzani IRCCS | INMI Lazzaro Spallanzani IRCCS | Francesco Messina, Martina Rueca, Barbara Bartolini, Cesare E.M. Gruber, Emanuela Giombini, Simone Lanini, Maria R. Capobianchi, Antonino Di Caro |
| EPI_ISL_605147 | University of Miami Immunology and Histocompatibility Laboratory | University of Miami Immunology and Histocompatibility Laboratory | Emilio Margolles-Clark, PhD and Phillip Ruiz, MD, PhD |
| EPI_ISL_605148, EPI_ISL_605149, EPI_ISL_605150, EPI_ISL_605151, EPI_ISL_605153, EPI_ISL_605154, EPI_ISL_605155, EPI_ISL_605156, EPI_ISL_605157, EPI_ISL_605158, EPI_ISL_605159, EPI_ISL_605160, EPI_ISL_605161, EPI_ISL_605162, EPI_ISL_605163, EPI_ISL_605164, EPI_ISL_605165, EPI_ISL_605166, EPI_ISL_605167, EPI_ISL_605168, EPI_ISL_605169, EPI_ISL_605170, EPI_ISL_605171, EPI_ISL_605172, EPI_ISL_605173, EPI_ISL_605174, EPI_ISL_605175, EPI_ISL_605176, EPI_ISL_605177, EPI_ISL_605178, EPI_ISL_605179, EPI_ISL_605180, EPI_ISL_605181, EPI_ISL_605182, EPI_ISL_605183, EPI_ISL_605184, EPI_ISL_605185, EPI_ISL_605186, EPI_ISL_605187, EPI_ISL_605188, EPI_ISL_605189, EPI_ISL_605190, EPI_ISL_605191, EPI_ISL_605192, EPI_ISL_605193, EPI_ISL_605194, EPI_ISL_605195, EPI_ISL_605196, EPI_ISL_605197, EPI_ISL_605198, EPI_ISL_605199, EPI_ISL_605200, EPI_ISL_605201, EPI_ISL_605202, EPI_ISL_605203, EPI_ISL_605204, EPI_ISL_605205, EPI_ISL_605206, EPI_ISL_605207, EPI_ISL_605208, EPI_ISL_605209, EPI_ISL_605210, EPI_ISL_605211, EPI_ISL_605212, EPI_ISL_605213, EPI_ISL_605214, EPI_ISL_605215, EPI_ISL_605216, EPI_ISL_605217, EPI_ISL_605218, EPI_ISL_605219, EPI_ISL_605220, EPI_ISL_605221, EPI_ISL_605222, EPI_ISL_605223, EPI_ISL_605224, EPI_ISL_605225, EPI_ISL_605226, EPI_ISL_605227, EPI_ISL_605228, EPI_ISL_605229, EPI_ISL_605230, EPI_ISL_605231, EPI_ISL_605232, EPI_ISL_605233, EPI_ISL_605234, EPI_ISL_605235, EPI_ISL_605236, EPI_ISL_605237, EPI_ISL_605238, EPI_ISL_605239, EPI_ISL_605240, EPI_ISL_605241, EPI_ISL_605242, EPI_ISL_605243, EPI_ISL_605244, EPI_ISL_605245, EPI_ISL_605246, EPI_ISL_605247, EPI_ISL_605248, EPI_ISL_605249, EPI_ISL_605250, EPI_ISL_605251, EPI_ISL_605252, EPI_ISL_605253, EPI_ISL_605254, EPI_ISL_605255, EPI_ISL_605256, EPI_ISL_605257, EPI_ISL_605258, EPI_ISL_605259, EPI_ISL_605260, EPI_ISL_605261, EPI_ISL_605262, EPI_ISL_605263, EPI_ISL_605264, EPI_ISL_605265, EPI_ISL_605266, EPI_ISL_605267, EPI_ISL_605268, EPI_ISL_605269, EPI_ISL_605270, EPI_ISL_605271, EPI_ISL_605272, EPI_ISL_605273, EPI_ISL_605274, EPI_ISL_605275, EPI_ISL_605276, EPI_ISL_605277, EPI_ISL_605278, EPI_ISL_605279, EPI_ISL_605280, EPI_ISL_605281, EPI_ISL_605282, EPI_ISL_605283, EPI_ISL_605284, EPI_ISL_605285, EPI_ISL_605286, EPI_ISL_605287, EPI_ISL_605288, EPI_ISL_605289, EPI_ISL_605290, EPI_ISL_605291, EPI_ISL_605292, EPI_ISL_605293, EPI_ISL_605294, EPI_ISL_605295, EPI_ISL_605296, EPI_ISL_605297, EPI_ISL_605298, EPI_ISL_605299, EPI_ISL_605300, EPI_ISL_605301, EPI_ISL_605302, EPI_ISL_605303, EPI_ISL_605304, EPI_ISL_605305, EPI_ISL_605306, EPI_ISL_605307, EPI_ISL_605308, EPI_ISL_605309, EPI_ISL_605310, EPI_ISL_605311, EPI_ISL_605312, EPI_ISL_605313, EPI_ISL_605314, EPI_ISL_605315, EPI_ISL_605316, EPI_ISL_605317, EPI_ISL_605318, EPI_ISL_605319, EPI_ISL_605320, EPI_ISL_605321, EPI_ISL_605322, EPI_ISL_605323, EPI_ISL_605324, EPI_ISL_605325, EPI_ISL_605326, EPI_ISL_605327, EPI_ISL_605328, EPI_ISL_605329, EPI_ISL_605330, EPI_ISL_605331, EPI_ISL_605332, EPI_ISL_605333, EPI_ISL_605334, EPI_ISL_605335, EPI_ISL_605336, EPI_ISL_605337, EPI_ISL_605338, EPI_ISL_605339, EPI_ISL_605340, EPI_ISL_605341, EPI_ISL_605342, EPI_ISL_605343, EPI_ISL_605344, EPI_ISL_605345, EPI_ISL_605346, EPI_ISL_605347, EPI_ISL_605348, EPI_ISL_605349, EPI_ISL_605350, EPI_ISL_605351, EPI_ISL_605352, EPI_ISL_605353, EPI_ISL_605354, EPI_ISL_605355, EPI_ISL_605356, EPI_ISL_605357, EPI_ISL_605358, EPI_ISL_605359, EPI_ISL_605360, EPI_ISL_605361, EPI_ISL_605362, EPI_ISL_605363, EPI_ISL_605364, | | | |

| | | | |
|---|---|---|---|
| EPI_ISL_605365, EPI_ISL_605366, EPI_ISL_605367, EPI_ISL_605368, EPI_ISL_605369, EPI_ISL_605370, EPI_ISL_605371, EPI_ISL_605372, EPI_ISL_605373, EPI_ISL_605374, EPI_ISL_605375, EPI_ISL_605376, EPI_ISL_605377, EPI_ISL_605378, EPI_ISL_605379, EPI_ISL_605380, EPI_ISL_605381, EPI_ISL_605382, EPI_ISL_605383, EPI_ISL_605384, EPI_ISL_605385, EPI_ISL_605386, EPI_ISL_605387, EPI_ISL_605388, EPI_ISL_605389, EPI_ISL_605390, EPI_ISL_605391, EPI_ISL_605392, EPI_ISL_605393, EPI_ISL_605394, EPI_ISL_605395, EPI_ISL_605396, EPI_ISL_605397, EPI_ISL_605398, EPI_ISL_605399, EPI_ISL_605400, EPI_ISL_605401, EPI_ISL_605402, EPI_ISL_605403, EPI_ISL_605404 | | | |
| see above | Utah Public Health Laboratory | Utah Public Health Laboratory | Erin L. Young, Kelly Oakeson, Tara Gallagher, Michael T. Pyne, E. Susan Slechta, Melanie A. Mallory, Jeffrey B. Stevenson, Salika M. Shakir, David R. Hillyard |
| EPI_ISL_605405, EPI_ISL_605406, EPI_ISL_605407, EPI_ISL_605408, EPI_ISL_605409, EPI_ISL_605410, EPI_ISL_605411, EPI_ISL_605412, EPI_ISL_605413, EPI_ISL_605414, EPI_ISL_605415, EPI_ISL_605416, EPI_ISL_605417, EPI_ISL_605418, EPI_ISL_605419, EPI_ISL_605420, EPI_ISL_605421, EPI_ISL_605422, EPI_ISL_605423, EPI_ISL_605424, EPI_ISL_605425, EPI_ISL_605426, EPI_ISL_605427, EPI_ISL_605428, EPI_ISL_605429, EPI_ISL_605430, EPI_ISL_605431, EPI_ISL_605432, EPI_ISL_605433, EPI_ISL_605434, EPI_ISL_605435, EPI_ISL_605436, EPI_ISL_605437, EPI_ISL_605438, EPI_ISL_605439, EPI_ISL_605440, EPI_ISL_605441, EPI_ISL_605442, EPI_ISL_605443, EPI_ISL_605444, EPI_ISL_605445, EPI_ISL_605446, EPI_ISL_605447, EPI_ISL_605448, EPI_ISL_605449, EPI_ISL_605450, EPI_ISL_605451, EPI_ISL_605452, EPI_ISL_605453, EPI_ISL_605454, EPI_ISL_605455, EPI_ISL_605456, EPI_ISL_605457, EPI_ISL_605458, EPI_ISL_605459, EPI_ISL_605460, EPI_ISL_605461, EPI_ISL_605462, EPI_ISL_605463, EPI_ISL_605464, EPI_ISL_605465, EPI_ISL_605466, EPI_ISL_605467, EPI_ISL_605468, EPI_ISL_605469, EPI_ISL_605470, EPI_ISL_605471, EPI_ISL_605472, EPI_ISL_605473, EPI_ISL_605474, EPI_ISL_605475, EPI_ISL_605476, EPI_ISL_605477, EPI_ISL_605478, EPI_ISL_605479, EPI_ISL_605480, EPI_ISL_605481, EPI_ISL_605482, EPI_ISL_605483, EPI_ISL_605484, EPI_ISL_605485, EPI_ISL_605486, EPI_ISL_605487, EPI_ISL_605488, EPI_ISL_605489, EPI_ISL_605490, EPI_ISL_605491, EPI_ISL_605492, EPI_ISL_605493, EPI_ISL_605494, EPI_ISL_605495, EPI_ISL_605496, EPI_ISL_605497, EPI_ISL_605498, EPI_ISL_605499, EPI_ISL_605500, EPI_ISL_605501, EPI_ISL_605504, EPI_ISL_605505, EPI_ISL_605506, EPI_ISL_605507, EPI_ISL_605508, EPI_ISL_605509, EPI_ISL_605510, EPI_ISL_605511, EPI_ISL_605512, EPI_ISL_605513, EPI_ISL_605514, EPI_ISL_605515, EPI_ISL_605516, EPI_ISL_605517, EPI_ISL_605518, EPI_ISL_605519, EPI_ISL_605520, EPI_ISL_605521, EPI_ISL_605522, EPI_ISL_605523, EPI_ISL_605524, EPI_ISL_605525, EPI_ISL_605526, EPI_ISL_605527, EPI_ISL_605528, EPI_ISL_605529, EPI_ISL_605530, EPI_ISL_605531, EPI_ISL_605532, EPI_ISL_605533, EPI_ISL_605534, EPI_ISL_605535, EPI_ISL_605536, EPI_ISL_605537, EPI_ISL_605538, EPI_ISL_605539, EPI_ISL_605540, EPI_ISL_605541, EPI_ISL_605542, EPI_ISL_605543, EPI_ISL_605544, EPI_ISL_605545, EPI_ISL_605546, EPI_ISL_605547, EPI_ISL_605548, EPI_ISL_605549, EPI_ISL_605550, EPI_ISL_605551, EPI_ISL_605552, EPI_ISL_605553, EPI_ISL_605554, EPI_ISL_605555, EPI_ISL_605556, EPI_ISL_605557, EPI_ISL_605558, EPI_ISL_605559, EPI_ISL_605560, EPI_ISL_605561, EPI_ISL_605562, EPI_ISL_605563, EPI_ISL_605564, EPI_ISL_605565, EPI_ISL_605566, EPI_ISL_605567, EPI_ISL_605568, EPI_ISL_605569, EPI_ISL_605570, EPI_ISL_605571, EPI_ISL_605572, EPI_ISL_605573, EPI_ISL_605574, EPI_ISL_605575, EPI_ISL_605576, EPI_ISL_605577, EPI_ISL_605578, EPI_ISL_605579, EPI_ISL_605580, EPI_ISL_605581, EPI_ISL_605582, EPI_ISL_605583, EPI_ISL_605584, EPI_ISL_605585, EPI_ISL_605586, EPI_ISL_605587, EPI_ISL_605588, EPI_ISL_605589, EPI_ISL_605590, EPI_ISL_605591, EPI_ISL_605592, EPI_ISL_605593, EPI_ISL_605594, EPI_ISL_605595, EPI_ISL_605596, EPI_ISL_605597, EPI_ISL_605598, EPI_ISL_605599, EPI_ISL_605600, EPI_ISL_605601, EPI_ISL_605602, EPI_ISL_605603, EPI_ISL_605604, EPI_ISL_605605, EPI_ISL_605606, EPI_ISL_605607, EPI_ISL_605608, EPI_ISL_605609, EPI_ISL_605610, EPI_ISL_605611, EPI_ISL_605612, EPI_ISL_605613, EPI_ISL_605614, EPI_ISL_605615, EPI_ISL_605616, EPI_ISL_605617, EPI_ISL_605618, EPI_ISL_605619, EPI_ISL_605620, EPI_ISL_605621, EPI_ISL_605622, EPI_ISL_605623, EPI_ISL_605624, EPI_ISL_605625, EPI_ISL_605626, EPI_ISL_605627, EPI_ISL_605628, EPI_ISL_605629, EPI_ISL_605630, EPI_ISL_605631, EPI_ISL_605632, EPI_ISL_605633, EPI_ISL_605634, EPI_ISL_605635, EPI_ISL_605636, EPI_ISL_605637, EPI_ISL_605638, EPI_ISL_605639, EPI_ISL_605640, EPI_ISL_605641, EPI_ISL_605642, EPI_ISL_605643, EPI_ISL_605644, EPI_ISL_605645, EPI_ISL_605646, EPI_ISL_605647, EPI_ISL_605648, EPI_ISL_605649, EPI_ISL_605650, EPI_ISL_605651, EPI_ISL_605652, EPI_ISL_605653, EPI_ISL_605654, EPI_ISL_605655, EPI_ISL_605656, EPI_ISL_605657, EPI_ISL_605658, EPI_ISL_605659, EPI_ISL_605660, EPI_ISL_605661, EPI_ISL_605662, EPI_ISL_605663, EPI_ISL_605664, EPI_ISL_605665, EPI_ISL_605666, EPI_ISL_605667, EPI_ISL_605668, EPI_ISL_605669, EPI_ISL_605670, EPI_ISL_605671, EPI_ISL_605672, EPI_ISL_605673, EPI_ISL_605674, EPI_ISL_605675, EPI_ISL_605676, EPI_ISL_605677, EPI_ISL_605678, EPI_ISL_605679, EPI_ISL_605680, EPI_ISL_605681, EPI_ISL_605682, EPI_ISL_605683, EPI_ISL_605684, EPI_ISL_605685, EPI_ISL_605686, EPI_ISL_605687, EPI_ISL_605688, EPI_ISL_605689, EPI_ISL_605690, EPI_ISL_605691, EPI_ISL_605692, EPI_ISL_605693, EPI_ISL_605694, EPI_ISL_605695, EPI_ISL_605696, EPI_ISL_605697, EPI_ISL_605698, EPI_ISL_605699, EPI_ISL_605700, EPI_ISL_605701, EPI_ISL_605702, EPI_ISL_605703, EPI_ISL_605704, EPI_ISL_605705, EPI_ISL_605706, EPI_ISL_605707, EPI_ISL_605708, EPI_ISL_605709, EPI_ISL_605710, EPI_ISL_605711, EPI_ISL_605712, EPI_ISL_605713, EPI_ISL_605714, EPI_ISL_605715, EPI_ISL_605716, EPI_ISL_605717, EPI_ISL_605718, EPI_ISL_605719, EPI_ISL_605720, EPI_ISL_605721, EPI_ISL_605722, EPI_ISL_605723, EPI_ISL_605724, EPI_ISL_605725, EPI_ISL_605726, EPI_ISL_605727, EPI_ISL_605728, EPI_ISL_605729, EPI_ISL_605730, EPI_ISL_605731, EPI_ISL_605732, EPI_ISL_605733, EPI_ISL_605734, EPI_ISL_605735, EPI_ISL_605736, EPI_ISL_605737, EPI_ISL_605738, EPI_ISL_605739, EPI_ISL_605740, EPI_ISL_605741, EPI_ISL_605742, EPI_ISL_605743, EPI_ISL_605744, EPI_ISL_605745, EPI_ISL_605746, EPI_ISL_605747, EPI_ISL_605748, EPI_ISL_605749, EPI_ISL_605750, EPI_ISL_605751, EPI_ISL_605752, EPI_ISL_605753, EPI_ISL_605754, EPI_ISL_605755, EPI_ISL_605756, EPI_ISL_605757, EPI_ISL_605758, EPI_ISL_605759, EPI_ISL_605760, EPI_ISL_605761, EPI_ISL_605762, EPI_ISL_605763, EPI_ISL_605764, EPI_ISL_605765, EPI_ISL_605766, EPI_ISL_605767, EPI_ISL_605768, EPI_ISL_605769, EPI_ISL_605770, EPI_ISL_605771, EPI_ISL_605772, EPI_ISL_605773, EPI_ISL_605774, EPI_ISL_605775, EPI_ISL_605776, EPI_ISL_605777, EPI_ISL_605778, EPI_ISL_605779 | | | |
| see above | University of Wisconsin-Madison AIDS Vaccine Research Laboratories | University of Wisconsin-Madison AIDS Vaccine Research Laboratories | Gage Moreno, Katarina Braun, et al. AIDS Vaccine Research Laboratories |
| EPI_ISL_605784 | AHRI-Sigal | KRISP, KZN Research Innovation and Sequencing Platform | Gazy I, Sigal A, Karim F, Cele S, Giandhari J, Pillay S, Tegally H, Wilkinson E, de Oliveira T |
| EPI_ISL_605785, EPI_ISL_605786, EPI_ISL_605787, EPI_ISL_605788, EPI_ISL_605790 | NHLS-IALCH | KRISP, KZN Research Innovation and Sequencing Platform | Giandhari J, Pillay S, Lessells R, Mdlalose K, York D, Khan S, Tegally H, Wilkinson E, de Oliveira T |
| EPI_ISL_605799, EPI_ISL_605800, EPI_ISL_605801, EPI_ISL_605802, EPI_ISL_605803, EPI_ISL_605804, EPI_ISL_605805, EPI_ISL_605806, EPI_ISL_605807, EPI_ISL_605808, EPI_ISL_605809, EPI_ISL_605810, EPI_ISL_605811, EPI_ISL_605812, EPI_ISL_605813, EPI_ISL_605814, EPI_ISL_605815, EPI_ISL_605816 | | | |
| see above | Clinical Virology Laboratory, Institute of Liver and Biliary Sciences | ILBS - IGIB | Ekta Gupta, Sheetalnath Rooge, Abhishek Padhi, Reshu Agarwal, Jaswinder Singh Maras, Shridhar Sivasubbu, Vinod Scaria, Shvetank Sharma |
| EPI_ISL_609992 | INMI Lazzaro Spallanzani IRCCS | INMI Lazzaro Spallanzani IRCCS | F Messina, E Giombini, M Rueca, B Bartolini, C.E.M Gruber, MR Capobianchi, A Di Caro |
| EPI_ISL_609993 | INMI Lazzaro Spallanzani IRCCS | INMI Lazzaro Spallanzani IRCCS | E Giombini, M Rueca, B Bartolini, C.E.M Gruber, F Messina, A Di Caro, MR Capobianchi |
| EPI_ISL_609995 | INMI Lazzaro Spallanzani IRCCS | INMI Lazzaro Spallanzani IRCCS | E Giombini, C.E.M Gruber, M Rueca, B Bartolini, F Messina, A Di Caro, MR Capobianchi |
| EPI_ISL_609996 | INMI Lazzaro Spallanzani IRCCS | INMI Lazzaro Spallanzani IRCCS | F Messina, M Rueca, B Bartolini, C.E.M Gruber, E Giombini, MR Capobianchi, A Di Caro |
| EPI_ISL_609998 | INMI Lazzaro Spallanzani IRCCS | INMI Lazzaro Spallanzani IRCCS | F Messina, B Bartolini, M Rueca, C.E.M Gruber, E Giombini, A Di Caro, MR Capobianchi |
| EPI_ISL_611485, EPI_ISL_611486, EPI_ISL_611487, EPI_ISL_611488, EPI_ISL_611489, EPI_ISL_611490, EPI_ISL_611491, EPI_ISL_611492, EPI_ISL_611493, EPI_ISL_611494, EPI_ISL_611495, EPI_ISL_611496, EPI_ISL_611497, EPI_ISL_611498, EPI_ISL_611499, EPI_ISL_611500, EPI_ISL_611501, EPI_ISL_611502, EPI_ISL_611503, EPI_ISL_611504, EPI_ISL_611505, EPI_ISL_611506, EPI_ISL_611507, EPI_ISL_611508, EPI_ISL_611509, EPI_ISL_611510, EPI_ISL_611511, EPI_ISL_611512, EPI_ISL_611513 | | | |
| see above | Utah Public Health Laboratory | Utah Public Health Laboratory | Erin Young, Kelly Oakeson |
| EPI_ISL_611546, EPI_ISL_611582, EPI_ISL_611604, EPI_ISL_611659, EPI_ISL_611662, EPI_ISL_611670, EPI_ISL_611676, EPI_ISL_611677, EPI_ISL_611680, EPI_ISL_611753, EPI_ISL_611791, EPI_ISL_611810, EPI_ISL_611812, EPI_ISL_611813, EPI_ISL_611831, EPI_ISL_611833, EPI_ISL_611840, EPI_ISL_611919, EPI_ISL_611954, EPI_ISL_611955, EPI_ISL_611956, EPI_ISL_611963, EPI_ISL_612032, EPI_ISL_612077, EPI_ISL_612107, EPI_ISL_612158, EPI_ISL_612159, EPI_ISL_612181, EPI_ISL_612182, EPI_ISL_612183, EPI_ISL_612192, EPI_ISL_612197, EPI_ISL_612571, EPI_ISL_612572, EPI_ISL_612573, EPI_ISL_612574, EPI_ISL_612575, EPI_ISL_612576, EPI_ISL_612577, EPI_ISL_612578, EPI_ISL_612579, EPI_ISL_612580, EPI_ISL_612581, EPI_ISL_612582, EPI_ISL_612583, EPI_ISL_612584, EPI_ISL_612585, EPI_ISL_612586, EPI_ISL_612587, EPI_ISL_612588, EPI_ISL_612589, EPI_ISL_612590, EPI_ISL_612591, EPI_ISL_612592, EPI_ISL_612593, EPI_ISL_612594, EPI_ISL_612595, EPI_ISL_612596, EPI_ISL_612597, EPI_ISL_612598, EPI_ISL_612599, EPI_ISL_612600, EPI_ISL_612601, EPI_ISL_612602, EPI_ISL_612603, EPI_ISL_612604, EPI_ISL_612605, EPI_ISL_612606, EPI_ISL_613307, EPI_ISL_613308, EPI_ISL_613309, EPI_ISL_613310, EPI_ISL_613311, EPI_ISL_613312, EPI_ISL_613313, EPI_ISL_613314, EPI_ISL_613315, EPI_ISL_613316, EPI_ISL_613317, EPI_ISL_613318, EPI_ISL_613319, EPI_ISL_613320, EPI_ISL_613321, EPI_ISL_613322, EPI_ISL_613323, EPI_ISL_613324, EPI_ISL_613325, EPI_ISL_613326, EPI_ISL_613327, EPI_ISL_613328, EPI_ISL_613329, EPI_ISL_613330, EPI_ISL_613331, EPI_ISL_613332, EPI_ISL_613333, EPI_ISL_613334, EPI_ISL_613335, EPI_ISL_613336, EPI_ISL_613337, EPI_ISL_613338, EPI_ISL_613339, EPI_ISL_613340, EPI_ISL_613341, EPI_ISL_613342, EPI_ISL_613343, EPI_ISL_613344, EPI_ISL_613345, EPI_ISL_613346, EPI_ISL_613347, EPI_ISL_613348, EPI_ISL_613349, EPI_ISL_613350, EPI_ISL_613351, EPI_ISL_613352, EPI_ISL_613353, EPI_ISL_613354, EPI_ISL_613355 | | | |
| see above | Queens Medical Centre, Clinical Microbiology Department / DeepSeq Nottingham | COVID-19 Genomics UK (COG-UK) Consortium | Gemma Clark, Wendy Smith, Manjinder Khakh, Vicki M Fleming, Michelle M Lister, Hannah Howson-Wells, Jonathan Ball, Patrick McClure, Joseph Chappell, Theocharis Tsoleridis, Nadine Holmes, Matthew Carlisle, Christopher Moore, Fei Sang, Johnny Debebe, Victoria Wright, Matthew Loose |
| EPI_ISL_623170, EPI_ISL_623171, EPI_ISL_623172, EPI_ISL_623173, EPI_ISL_623174, EPI_ISL_623175, EPI_ISL_623176, EPI_ISL_623177, EPI_ISL_623178, EPI_ISL_623179, EPI_ISL_623180, EPI_ISL_623181, EPI_ISL_623182, EPI_ISL_623183, EPI_ISL_623184, EPI_ISL_623185, EPI_ISL_623186, EPI_ISL_623187, EPI_ISL_623188, EPI_ISL_623189, EPI_ISL_623190, EPI_ISL_623191, EPI_ISL_623192, EPI_ISL_623193, EPI_ISL_623194, EPI_ISL_623195, EPI_ISL_623196, EPI_ISL_623197, EPI_ISL_623198, EPI_ISL_623199, EPI_ISL_623200, EPI_ISL_623201, EPI_ISL_623202, EPI_ISL_623203, EPI_ISL_623204, EPI_ISL_623205, EPI_ISL_623206, EPI_ISL_623207, EPI_ISL_623208, EPI_ISL_623209, EPI_ISL_623210 | | | |
| see above | Utah Public Health Laboratory | Utah Public Health Laboratory | Erin Young, Kelly Oakeson |
| EPI_ISL_626549, EPI_ISL_626550, EPI_ISL_626551, EPI_ISL_626552, EPI_ISL_626553, EPI_ISL_626554, EPI_ISL_626555, EPI_ISL_626556, EPI_ISL_626557, EPI_ISL_626558, EPI_ISL_626559, EPI_ISL_626560, EPI_ISL_626561, EPI_ISL_626562, EPI_ISL_626563, EPI_ISL_626564, EPI_ISL_626565 | | | |
| see above | Laboratorio de Biología Molecular, Facultad de Medicina, Universidad de Atacama, Copiapo, Chile/ FONDAP CRG, Universidad Andrés Bello, Santiago, Chile | Center for Mathematical Modeling and Center for Genome Regulation. Santiago, Chile | Echeverría C, Manríquez R, Bastias M, Sanhueza D, Travisany D, Allende ML, Maass A, González M, Montecino, M, Orellana A, Castro E, Meneses C. |
| EPI_ISL_626659, EPI_ISL_626670, EPI_ISL_626688, EPI_ISL_626695, EPI_ISL_626740, EPI_ISL_626750, EPI_ISL_626770, EPI_ISL_626811, EPI_ISL_626816, EPI_ISL_626834, EPI_ISL_626881, EPI_ISL_626919, EPI_ISL_627053, EPI_ISL_627068, EPI_ISL_627147, EPI_ISL_627215, EPI_ISL_627216, EPI_ISL_627304, EPI_ISL_627309, EPI_ISL_627310, EPI_ISL_627666, EPI_ISL_627667, EPI_ISL_627668, EPI_ISL_627669, EPI_ISL_627670, EPI_ISL_627671, EPI_ISL_627672, EPI_ISL_627673, EPI_ISL_627674, EPI_ISL_627675, EPI_ISL_627676, EPI_ISL_627677, EPI_ISL_627678, EPI_ISL_627679, EPI_ISL_627680, EPI_ISL_627681, EPI_ISL_627682, EPI_ISL_627683, EPI_ISL_627684, EPI_ISL_627685, EPI_ISL_627686, EPI_ISL_627687, EPI_ISL_627688, EPI_ISL_627689, EPI_ISL_627690, EPI_ISL_627691, EPI_ISL_627692, EPI_ISL_627693, EPI_ISL_627694, EPI_ISL_627695, EPI_ISL_627696, EPI_ISL_627697, EPI_ISL_627698, EPI_ISL_627699, EPI_ISL_627700, EPI_ISL_627701, EPI_ISL_627702, EPI_ISL_627703, EPI_ISL_627704, EPI_ISL_627705, EPI_ISL_627706, EPI_ISL_627707, EPI_ISL_627708, EPI_ISL_627709, EPI_ISL_627710, EPI_ISL_627711, EPI_ISL_627712, EPI_ISL_627713, EPI_ISL_627714, EPI_ISL_627715, EPI_ISL_627716, EPI_ISL_627717, EPI_ISL_627718, EPI_ISL_627719, EPI_ISL_627720, EPI_ISL_627721, EPI_ISL_627722, EPI_ISL_627723, EPI_ISL_627724, EPI_ISL_627725, EPI_ISL_627726, EPI_ISL_627727, EPI_ISL_627728, EPI_ISL_627729 | | | |
| see above | Queens Medical Centre, Clinical Microbiology Department / DeepSeq Nottingham | COVID-19 Genomics UK (COG-UK) Consortium | Gemma Clark, Wendy Smith, Manjinder Khakh, Vicki M Fleming, Michelle M Lister, Hannah Howson-Wells, Jonathan Ball, Patrick McClure, Joseph Chappell, Theocharis Tsoleridis, Nadine Holmes, Matthew Carlisle, Christopher Moore, Fei Sang, Johnny Debebe, Victoria Wright, Matthew Loose |
| EPI_ISL_628932, EPI_ISL_628933, EPI_ISL_628934, EPI_ISL_628935, EPI_ISL_628936, EPI_ISL_628937, EPI_ISL_628938, EPI_ISL_628939, EPI_ISL_628940, EPI_ISL_628941, EPI_ISL_628942, EPI_ISL_628943, EPI_ISL_628944, EPI_ISL_628945, EPI_ISL_628946, EPI_ISL_628947, EPI_ISL_628948, EPI_ISL_628949, EPI_ISL_628950, EPI_ISL_628951, EPI_ISL_628952, EPI_ISL_628953, EPI_ISL_628954, EPI_ISL_628955, EPI_ISL_628956, EPI_ISL_628957, EPI_ISL_628958, EPI_ISL_628959, EPI_ISL_628960, EPI_ISL_628961, EPI_ISL_628962, EPI_ISL_628963, EPI_ISL_628964, EPI_ISL_628965, EPI_ISL_628966, EPI_ISL_628967, EPI_ISL_628968, EPI_ISL_628969, EPI_ISL_628970, EPI_ISL_628971, EPI_ISL_628972, EPI_ISL_628973, EPI_ISL_628974, EPI_ISL_628975, EPI_ISL_628976, EPI_ISL_628977, EPI_ISL_628978, EPI_ISL_628979, EPI_ISL_628980, EPI_ISL_628981, EPI_ISL_628982, EPI_ISL_628983, EPI_ISL_628984, EPI_ISL_628985, EPI_ISL_628986, EPI_ISL_628987, EPI_ISL_628988, EPI_ISL_628989, EPI_ISL_628990, EPI_ISL_628991, EPI_ISL_628992, EPI_ISL_628993, EPI_ISL_628994, EPI_ISL_628995, EPI_ISL_628996, EPI_ISL_628997, EPI_ISL_628998, EPI_ISL_628999 | | | |
| see above | Utah Public Health Laboratory | Utah Public Health Laboratory | Erin Young, Kelly Oakeson |
| EPI_ISL_632823, EPI_ISL_632824, EPI_ISL_632825, EPI_ISL_632826, EPI_ISL_632827, EPI_ISL_632828, EPI_ISL_632829, EPI_ISL_632830, EPI_ISL_632831, EPI_ISL_632832, EPI_ISL_632833, EPI_ISL_632834, EPI_ISL_632835, EPI_ISL_632836, EPI_ISL_632837, EPI_ISL_632838, EPI_ISL_632839, EPI_ISL_632840, EPI_ISL_632841, EPI_ISL_632842, EPI_ISL_632843, EPI_ISL_632844, EPI_ISL_632845 | | | |

| | | | |
|---|---|---|---|
| see above | New Mexico Department of Health Scientific Laboratory | New Mexico Department of Health Scientific Laboratory | Ellie Johnson, Anastacia Griego-Fisher, D'Eldra Malone |
| EPI_ISL_632890, EPI_ISL_632891, EPI_ISL_632892, EPI_ISL_632893, EPI_ISL_632894, EPI_ISL_632895, EPI_ISL_632896, EPI_ISL_632897, EPI_ISL_632898 | University of Wisconsin-Madison AIDS Vaccine Research Laboratories | University of Wisconsin-Madison AIDS Vaccine Research Laboratories | Gage Moreno, Katarina Braun, et al. AIDS Vaccine Research Laboratories |

We gratefully acknowledge the following Authors from the Originating laboratories responsible for obtaining the specimens, as well as the Submitting laboratories where the genome data were generated and shared via GISAID, on which this research is based.

All Submitters of data may be contacted directly via www.gisaid.org

Authors are sorted alphabetically.

| Accession ID | Originating Laboratory | Submitting Laboratory | Authors |
|---|---|---|---|
| EPI_ISL_413598 | Centre for Infectious Diseases and Microbiology - Public Health | NSW Health Pathology - Institute of Clinical Pathology and Medical Research; Westmead Hospital; University of Sydney | Gray K, Eden J-S, Lam C, Rockett R, Timms, V, Gall, M, Carter I, Rahman H, Holmes EC, O'Sullivan MV, Sintchenko V, Chen SC, Maddocks S, Kok J and Dwyer DE for the 2019-nCoV Study Group* |
| EPI_ISL_414476 | MSHS Clinical Microbiology Laboratories | MSHS Pathogen Surveillance Program | Gopi Patel, Emilia Sordillo, Melissa Gitman, Alberto Paniz-mondolfi, Matthew Hernandez, Shelcie Fabre, Jose Polanco, Ana Sylvia Gonzalez-Reiche, Zenab Khan, Nancy Francoeur, Melissa Smith, Robert Sebra, Lisa Miorin, Wen-chun Liu, Randy Albrecht, Judith Aberg, Florian Krammer, Adolfo Garcia-Sarstre, Viviana Simon, Harm van Bakel |
| EPI_ISL_415151 | MSHS Clinical Microbiology Laboratories | MSHS Pathogen Surveillance Program | Gopi Patel, Emilia Sordillo, Melissa Gitman, Alberto Paniz-mondolfi, Matthew Hernandez, Shelcie Fabre, Jose Polanco, Ana Silvia Gonzalez-Reiche, Zenab Khan, Nancy Francoeur, Melissa Smith, Robert Sebra, Lisa Miorin, Wen-chun Liu, Randy Albrecht, Judith Aberg, Florian Krammer, Adolfo Garcia-Sarstre, Viviana Simon, Harm van Bakel |
| EPI_ISL_415709, EPI_ISL_415711, EPI_ISL_416042 | State Key Laboratory for Diagnosis and Treatment of Infectious Diseases, National Clinical Research Center for Infectious Diseases, First Affiliated Hospital, Zhejiang University School of Medicine, Hangzhou, China. 310003 | State Key Laboratory for Diagnosis and Treatment of Infectious Diseases, National Clinical Research Center for Infectious Diseases, First Affiliated Hospital, Zhejiang University School of Medicine, Hangzhou, China. 310003 | Hangping Yao, Nanping Wu, Chao Jiang, Xiangyun Lu, Linfang Cheng, Fumin Liu, Zhigang Wu, Haibo Wu, Changzhong Jin, Min Zheng, Lanjuan Li |
| EPI_ISL_416044, EPI_ISL_416046, EPI_ISL_416047, EPI_ISL_416425, EPI_ISL_416473, EPI_ISL_416474 | State Key Laboratory for Diagnosis and Treatment of Infectious Diseases, National Clinical Research Center for Infectious Diseases, First Affiliated Hospital, Zhejiang University School of Medicine, Hangzhou, China 310003 | State Key Laboratory for Diagnosis and Treatment of Infectious Diseases, National Clinical Research Center for Infectious Diseases, First Affiliated Hospital, Zhejiang University School of Medicine, Hangzhou, China 310003 | Hangping Yao, Nanping Wu, Chao Jiang, Xiangyun Lu, Linfang Cheng, Fumin Liu, Zhigang Wu, Haibo Wu, Changzhong Jin, Min Zheng, Lanjuan Li |
| EPI_ISL_416481 | R. G. Lugar Center for Public Health Research, National Center for Disease Control and Public Health (NCDC) of Georgia. | R. G. Lugar Center for Public Health Research, National Center for Disease Control and Public Health (NCDC) of Georgia. | Gvantsa Chanturia, Marine Murtskhvaladze, Nato Kotaria, Ann Machablishvili, Lela Sabadze, Mari Gavashelidze, Ana Papkiauri, Meri Pantsulaia, Gvantsa Brachveli, Tata Imnadze, Tamar Jashiashvili, Tea Tevdoradze, Ketevan Sidamonidze, Ekaterine Khmaladze, Ekaterine Zhghenti, Roena Sukhiashvili, Mariam Zakalashvili, Lela Urushadze, Magda Dgebuadze, Giorgi Tomashvili, Davit Tsaguria, Ekaterine Zangaladze, Nino Berishvili, Adam Kotorashvili, Maia Alkhazashvili, Irma Burjanadze, Anna Kasradze, Khatuna Zakhashvili, Paata Imnadze, Amiran Gamkrelidze. |
| EPI_ISL_416485 | Servicio de Microbiologia. Consorcio Hospital General Universitario de Valencia | Sequencing and Bioinformatics Service and Molecular Epidemiology Research Group. FISABIO-Public Health | Griselda De Marco, Neris Garcia-Gonzalez, Maria Alma Bracho, Maria Dolores Ocete, Concepcion Gimeno, Giuseppe D'Auria, Fernando Gonzalez-Candelas |
| EPI_ISL_416487 | Servicio de Microbiología. Consorcio Hospital General Universitario de Valencia | Sequencing and Bioinformatics Service and Molecular Epidemiology Research Group. FISABIO-Public Health | Giuseppe D'Auria, Griselda De Marco, Neris Garcia-Gonzalez, Maria Alma Bracho, Maria Dolores Ocete, Concepcion Gimeno, Fernando Gonzalez-Candelas |
| EPI_ISL_417386, EPI_ISL_417403 | Centre for Infectious Diseases and Microbiology Public Health | NSW Health Pathology - Institute of Clinical Pathology and Medical Research; Westmead Hospital; University of Sydney | Gray K, Timms V, Gall M, Arnott A, Sadsad R, Carter I, Rahman H, Holmes EC, O'Sullivan MV, Sintchenko V, Chen SC, Maddocks S, Kok J, Dwyer DE, Rockett R, Eden J-S and Lam C for the 2019-nCoV Study Group |
| EPI_ISL_417445, EPI_ISL_417446, EPI_ISL_417447 | Laboratory of Infectious Diseases, Department of Biomedical and Clinical Sciences L. Sacco, University of Milan | Laboratory of Infectious Diseases, Department of Biomedical and Clinical Sciences L. Sacco, University of Milan | Gianguglielmo Zehender, Alessia Lai, Annalisa Bergna, Luca Meroni, Agostino Riva, Claudia Balotta, Maciej Tarkowski, Arianna Gabrieli, Dario Bernacchia, Stefano Rusconi, Giuliano Rizzardini, Spinello Antinori, Massimo Galli |
| EPI_ISL_417986, EPI_ISL_417987 | Centro Hospitalar e Universitario de Sao Joao, Porto | Instituto Nacional de Saude (INSA) | Guiomar et al |
| EPI_ISL_417988 | CHULC - H Curry Cabral | Instituto Nacional de Saude (INSA) | Guiomar et al |
| EPI_ISL_417989 | Centro Hospitalar e Universitario de Sao Joao, Porto | Instituto Nacional de Saude (INSA) | Guiomar et al |
| EPI_ISL_417990, EPI_ISL_417991 | CHULC - H Curry Cabral | Instituto Nacional de Saude (INSA) | Guiomar et al |
| EPI_ISL_417992, EPI_ISL_417993 | CHULC - H D Estefania | Instituto Nacional de Saude (INSA) | Guiomar et al |
| EPI_ISL_417994, EPI_ISL_417995, EPI_ISL_417996 | CHULC - H Curry Cabral | Instituto Nacional de Saude (INSA) | Guiomar et al |
| EPI_ISL_417997, EPI_ISL_417998, EPI_ISL_417999 | Centro Hospital do Porto, E.P.E. - H. Geral de Santo Antonio | Instituto Nacional de Saude (INSA) | Guiomar et al |
| EPI_ISL_418000, EPI_ISL_418001 | ARS Algarve - Laboratório Laura Ayres | Instituto Nacional de Saude (INSA) | Guiomar et al |
| EPI_ISL_418002 | CHU Coimbra | Instituto Nacional de Saude (INSA) | Guiomar et al |
| EPI_ISL_418003 | H Braga | Instituto Nacional de Saude (INSA) | Guiomar et al |
| EPI_ISL_418004 | ARS Algarve - Laboratório Laura Ayres | Instituto Nacional de Saude (INSA) | Guiomar et al |
| EPI_ISL_418005 | CHU Coimbra - Pediátrico | Instituto Nacional de Saude (INSA) | Guiomar et al |
| EPI_ISL_418006 | CHBarreiro Montijo | Instituto Nacional de Saude (INSA) | Guiomar et al |
| EPI_ISL_418007, EPI_ISL_418008 | H Braga | Instituto Nacional de Saude (INSA) | Guiomar et al |
| EPI_ISL_418009 | HSE Ilha Terceira - Angra do Heroismo | Instituto Nacional de Saude (INSA) | Guiomar et al |
| EPI_ISL_418010, EPI_ISL_418011, EPI_ISL_418012, EPI_ISL_418013, EPI_ISL_418014, EPI_ISL_418015, EPI_ISL_418016 | CHULC - H Curry Cabral | Instituto Nacional de Saude (INSA) | Guiomar et al |
| EPI_ISL_418017 | CHMT | Instituto Nacional de Saude (INSA) | Guiomar et al |
| EPI_ISL_418018 | H Garcia de Orta | Instituto Nacional de Saude (INSA) | Guiomar et al |
| EPI_ISL_418019, EPI_ISL_418020, EPI_ISL_418021, EPI_ISL_418022 | H Braga | Instituto Nacional de Saude (INSA) | Guiomar et al |
| EPI_ISL_418023 | H Evora | Instituto Nacional de Saude (INSA) | Guiomar et al |
| EPI_ISL_418024 | CHUA - Faro | Instituto Nacional de Saude (INSA) | Guiomar et al |
| EPI_ISL_418025 | H Santarem | Instituto Nacional de Saude (INSA) | Guiomar et al |
| EPI_ISL_418026 | H Dr. Nelio Mendonca - Funchal | Instituto Nacional de Saude (INSA) | Guiomar et al |
| EPI_ISL_418027 | CHTMAD | Instituto Nacional de Saude (INSA) | Guiomar et al |

| | | | |
|---|---|---|---|
| EPI_ISL_418990, EPI_ISL_418991 | State Key Laboratory for Diagnosis and Treatment of Infectious Diseases, National Clinical Research Center for Infectious Diseases, First Affiliated Hospital, Zhejiang University School of Medicine, Hangzhou, China 310003 | State Key Laboratory for Diagnosis and Treatment of Infectious Diseases, National Clinical Research Center for Infectious Diseases, First Affiliated Hospital, Zhejiang University School of Medicine, Hangzhou, China 310003 | Hangping Yao, Nanping Wu, Chao Jiang, Xiangyun Lu, Linfang Cheng, Fumin Liu, Zhigang Wu, Haibo Wu, Changzhong Jin, Min Zheng, Lanjuan Li |
| EPI_ISL_419677 | Servicio de Microbiología. Consorcio Hospital General Universitario de Valencia | Sequencing and Bioinformatics Service and Molecular Epidemiology Research Group. FISABIO-Public Health | Giuseppe D'Auria, Griselda De Marco, Neris Garcia-Gonzalez, Maria Alma Bracho, Maria Dolores Ocete, Concepcion Gimeno, Fernando Gonzalez-Candelas |
| EPI_ISL_419678 | Servicio de Microbiología. Consorcio Hospital General Universitario de Valencia | Sequencing and Bioinformatics Service and Molecular Epidemiology Research Group. FISABIO-Public Health | Griselda De Marco, Neris Garcia-Gonzalez, Maria Alma Bracho, Maria Dolores Ocete, Giuseppe D'Auria, Concepcion Gimeno, Fernando Gonzalez-Candelas |
| EPI_ISL_419682 | Servicio de Microbiología. Consorcio Hospital General Universitario de Valencia | Sequencing and Bioinformatics Service and Molecular Epidemiology Research Group. FISABIO-Public Health | Giuseppe D'Auria, Griselda De Marco, Neris Garcia-Gonzalez, Maria Alma Bracho, Maria Dolores Ocete, Concepcion Gimeno, Fernando Gonzalez-Candelas |
| EPI_ISL_419683 | Servicio de Microbiología. Consorcio Hospital General Universitario de Valencia | Sequencing and Bioinformatics Service and Molecular Epidemiology Research Group. FISABIO-Public Health | Griselda De Marco, Neris Garcia-Gonzalez, Maria Alma Bracho, Maria Dolores Ocete, Giuseppe D'Auria, Concepcion Gimeno, Fernando Gonzalez-Candelas |
| EPI_ISL_419687 | Servicio de Microbiología. Consorcio Hospital General Universitario de Valencia | Sequencing and Bioinformatics Service and Molecular Epidemiology Research Group. FISABIO-Public Health | Giuseppe D'Auria, Griselda De Marco, Neris Garcia-Gonzalez, Maria Alma Bracho, Maria Dolores Ocete, Concepcion Gimeno, Fernando Gonzalez-Candelas |
| EPI_ISL_419688 | Servicio de Microbiología. Consorcio Hospital General Universitario de Valencia | Sequencing and Bioinformatics Service and Molecular Epidemiology Research Group. FISABIO-Public Health | Griselda De Marco, Neris Garcia-Gonzalez, Maria Alma Bracho, Maria Dolores Ocete, Giuseppe D'Auria, Concepcion Gimeno, Fernando Gonzalez-Candelas |
| EPI_ISL_420114 | Servicio de Microbiología. Consorcio Hospital General Universitario de Valencia | Sequencing and Bioinformatics Service and Molecular Epidemiology Research Group. FISABIO-Public Health | Griselda De Marco, Beatriz Beamud, Lidia Ruiz Roldan, Marta Pla Diaz, Neris Garcia-Gonzalez, Loreto Ferrús Abad, Inma Galán Vendrell, Paula Ruiz-Hueso, Mariana Reyes-Prieto, Vicente Soriano Chirona, Maria Alma Bracho, Maria Dolores Ocete, Llúcia Martínez-Priego, Concepcion Gimeno, Giuseppe D'Auria, Fernando Gonzalez-Candelas |
| EPI_ISL_420128 | Servicio de Microbiologia. Hospital Clinico Universitario de Valencia | Sequencing and Bioinformatics Service and Molecular Epidemiology Research Group. FISABIO-Public Health | Griselda De Marco, Beatriz Beamud, Lidia Ruiz Roldan, Marta Pla Diaz, Neris Garcia-Gonzalez, Loreto Ferrús Abad, Inma Galán Vendrell, Paula Ruiz-Hueso, Mariana Reyes-Prieto, Vicente Soriano Chirona, David Navarro, Maria Alma Bracho, Llúcia Martínez-Priego, Giuseppe D'Auria, Fernando Gonzalez-Candelas |
| EPI_ISL_420132 | Servicio de Microbiologia. Hospital Clinico Universitario de Valencia | Sequencing and Bioinformatics Service and Molecular Epidemiology Research Group. FISABIO-Public Health | Giuseppe D'Auria,Sandra Carbo, Loreto Ferrús Abad, Maria Alma Bracho, Griselda De Marco, Beatriz Beamud, Lidia Ruiz Roldan, Marta Pla Diaz, Neris Garcia-Gonzalez, Inma Galán Vendrell, Paula Ruiz-Hueso, Mariana Reyes-Prieto, Vicente Soriano Chirona, Ivan Ansari, David Navarro, Llúcia Martínez-Priego, Fernando Gonzalez-Candelas |
| EPI_ISL_420144 | Department for Virology, Molecular Biology and Genome Research, R. G. Lugar Center for Public Health Research, National Center for Disease Control and Public Health (NCDC) of Georgia. | Department for Virology, Molecular Biology and Genome Research, R. G. Lugar Center for Public Health Research, National Center for Disease Control and Public Health (NCDC) of Georgia. | Gvantsa Chanturia, Ann Machablishvili, Nato Kotaria, Marine Murtskhvaladze, Lela Sabadze, Mari Gavashelidze, Ana Papkiauri, Meri Pantsulaia, Gvantsa Brachveli, Tata Imnadze, Tamar Jashiashvili, Tea Tevdoradze, Ketevan Sidamonidze, Ekaterine Khmaladze, Ekaterine Zhghenti, Roena Sukhiashvili, Mariam Zakalashvili, Lela Urushadze, Magda Dgebuadze, Giorgi Tomashvili, Davit Tsaguria, Ekaterine Zangaladze, Nino Berishvili, Adam Kotorashvili, Maia Alkhazashvili, Irma Burjanadze, Anna Kasradze, Khatuna Zakhashvili, Paata Imnadze, Amiran Gamkrelidze. |
| EPI_ISL_421446, EPI_ISL_421447, EPI_ISL_421448 | H Guimaraes | Instituto Nacional de Saude (INSA) | Guiomar et al |
| EPI_ISL_421449 | H Dr. Nelio Mendonca - Funchal | Instituto Nacional de Saude (INSA) | Guiomar et al |
| EPI_ISL_421450, EPI_ISL_421451, EPI_ISL_421452 | Instituto Nacional de Saude (INSA) | Instituto Nacional de Saude (INSA) | Guiomar et al |
| EPI_ISL_421453 | CHTMAD | Instituto Nacional de Saude (INSA) | Guiomar et al |
| EPI_ISL_421454 | H Beatriz Angelo | Instituto Nacional de Saude (INSA) | Guiomar et al |
| EPI_ISL_421455 | CH Barreiro Montijo | Instituto Nacional de Saude (INSA) | Guiomar et al |
| EPI_ISL_421456 | Instituto Nacional de Saude (INSA) | Instituto Nacional de Saude (INSA) | Guiomar et al |
| EPI_ISL_421457 | H Dr Nelio Mendonca - Funchal | Instituto Nacional de Saude (INSA) | Guiomar et al |
| EPI_ISL_421458 | H Beatriz Angelo | Instituto Nacional de Saude (INSA) | Guiomar et al |
| EPI_ISL_421459, EPI_ISL_421460, EPI_ISL_421461 | H Dr. Nelio Mendonca - Funchal | Instituto Nacional de Saude (INSA) | Guiomar et al |
| EPI_ISL_421462 | H Santarem | Instituto Nacional de Saude (INSA) | Guiomar et al |
| EPI_ISL_421463 | HSE Ilha Terceira - Angra do Heroismo | Instituto Nacional de Saude (INSA) | Guiomar et al |
| EPI_ISL_421464, EPI_ISL_421465 | CHTMAD | Instituto Nacional de Saude (INSA) | Guiomar et al |
| EPI_ISL_421466, EPI_ISL_421467 | H Evora | Instituto Nacional de Saude (INSA) | Guiomar et al |
| EPI_ISL_421468, EPI_ISL_421469, EPI_ISL_421470, EPI_ISL_421471 | H Santarem | Instituto Nacional de Saude (INSA) | Guiomar et al |
| EPI_ISL_421472, EPI_ISL_421473, EPI_ISL_421474, EPI_ISL_421475, EPI_ISL_421476, EPI_ISL_421477, EPI_ISL_421478 | Instituto Nacional de Saude (INSA) | Instituto Nacional de Saude (INSA) | Guiomar et al |
| EPI_ISL_421479, EPI_ISL_421480 | CH Barreiro Montijo | Instituto Nacional de Saude (INSA) | Guiomar et al |
| EPI_ISL_421481 | H Beatriz Angelo | Instituto Nacional de Saude (INSA) | Guiomar et al |
| EPI_ISL_421482, EPI_ISL_421483, EPI_ISL_421484 | CH VN Gaia - Espinho | Instituto Nacional de Saude (INSA) | Guiomar et al |
| EPI_ISL_421485, EPI_ISL_421486 | CH Barreiro Montijo | Instituto Nacional de Saude (INSA) | Guiomar et al |
| EPI_ISL_421487 | H Beatriz Angelo | Instituto Nacional de Saude (INSA) | Guiomar et al |
| EPI_ISL_421488 | H Santarem | Instituto Nacional de Saude (INSA) | Guiomar et al |
| EPI_ISL_421489 | HSE Ilha Terceira - Angra do Heroismo | Instituto Nacional de Saude (INSA) | Guiomar et al |
| EPI_ISL_421490 | H Santarem | Instituto Nacional de Saude (INSA) | Guiomar et al |
| EPI_ISL_421491 | H Beatriz Angelo | Instituto Nacional de Saude (INSA) | Guiomar et al |
| EPI_ISL_421492 | H Santarem | Instituto Nacional de Saude (INSA) | Guiomar et al |
| EPI_ISL_421493, EPI_ISL_421494 | HSE Ilha Terceira - Angra do Heroismo | Instituto Nacional de Saude (INSA) | Guiomar et al |
| EPI_ISL_421495 | H Santarem | Instituto Nacional de Saude (INSA) | Guiomar et al |
| EPI_ISL_421496, EPI_ISL_421497, EPI_ISL_421498, EPI_ISL_421499 | Instituto Nacional de Saude (INSA) | Instituto Nacional de Saude (INSA) | Guiomar et al |
| EPI_ISL_421515 | Servicio de Microbiologia. Hospital Clinico Universitario de | Sequencing and Bioinformatics Service and Molecular | Giuseppe D'Auria, Llúcia Martínez-Priego, Maria Alma Bracho, Griselda De Marco, Beatriz Beamud, Lidia Ruiz Roldan, Marta Pla Diaz, Neris |

| | | | |
|---|---|---|---|
| | Valencia | Epidemiology Research Group. FISABIO-Public Health | Garcia-Gonzalez, Loreto Ferrús Abad, Inma Galán Vendrell, Paula Ruiz-Hueso, Mariana Reyes-Prieto, Vicente Soriano Chirona, David Navarro, Fernando Gonzalez-Candelas |
| EPI_ISL_421576 | Molecular Diagnostic Services | KRISP, KZN Research Innovation and Sequencing Platform | Giandhari J, Pillay S, Ngcapu S, Samsunder N, Lessells R, Chimukangara B, Deforche K, Tegally H, Wilkinson E, de Oliveira T |
| EPI_ISL_424670 | Laboratorio Estatal de Salud Publica del Estado de Queretaro | Instituto de Diagnóstico y Referencia Epidemiologicos | Gisela Barrera Badillo, Irma López Martínez, Lucia Hernández Rivas, Edgar Mendieta Condado, Fabiola Garcés Ayala, Adnan Araiza Rodríguez, Celia Boukadida, Santiago Avila Ríos, Mario Mújica Sánchez, José Arturo Martínez Orozco,Eduardo Becerril Vargas, Joel Armando Vázquez Pérez, Victor Hugo Borja Aburto, Concepción Grajales Muñiz, Cesar Raúl González Bonilla, Carolina González Torres, Francisco Javier Gaytán Cervantes, José Esteban Muñoz Medina, Guillermo M. Ruiz-Palacios, Pilar Ramos Cervantes, Violeta Ibarra Gonzalez, Fernando Ledesma Barrientos, Luis Alberto García Andrade, Alfredo Ponce de León Garduño, Blanca Taboada, Alejandro Sánchez, Pavel Isa, Ricardo Grande, Gloria Vázquez, Francisco Pulido, Carlos F. Arias,José Ernesto Ramírez González |
| EPI_ISL_425180, EPI_ISL_425194 | Servicio de Microbiología. Consorcio Hospital General Universitario de Valencia | Sequencing and Bioinformatics Service and Molecular Epidemiology Research Group. FISABIO-Public Health | Griselda De Marco, Beatriz Beamud, Lidia Ruiz Roldan, Marta Pla Diaz, Neris Garcia-Gonzalez, Inma Galán Vendrell, Sandra Carbo, Loreto Ferrús Abad, Paula Ruiz-Hueso, Mariana Reyes-Prieto, Vicente Soriano Chirona, Ivan Ansari, David Navarro, Maria Alma Bracho, Llúcia Martínez-Priego, Giuseppe D'Auria, Fernando Gonzalez-Candelas |
| EPI_ISL_425205 | Servicio de Microbiologia. Hospital Clinico Universitario de Valencia | Sequencing and Bioinformatics Service and Molecular Epidemiology Research Group. FISABIO-Public Health | Griselda De Marco, Beatriz Beamud, Lidia Ruiz Roldan, Marta Pla Diaz,Neris Garcia-Gonzalez, Loreto Ferrús Abad, Maria Dolores Ocete, Inma Galán Vendrell, Paula Ruiz-Hueso, Mariana Reyes-Prieto, Vicente Soriano Chirona, Maria Alma Bracho, Llúcia Martínez-Priego, Concepcion Gimeno, Giuseppe D'Auria, Fernando Gonzalez-Candelas |
| EPI_ISL_425214 | Servicio de Microbiología. Consorcio Hospital General Universitario de Valencia | Sequencing and Bioinformatics Service and Molecular Epidemiology Research Group. FISABIO-Public Health | Griselda De Marco, Beatriz Beamud, Lidia Ruiz Roldan, Marta Pla Diaz, Neris Garcia-Gonzalez, Inma Galán Vendrell, Sandra Carbo, Loreto Ferrús Abad, Paula Ruiz-Hueso, Mariana Reyes-Prieto, Vicente Soriano Chirona, Ivan Ansari, David Navarro, Maria Alma Bracho, Llúcia Martínez-Priego, Giuseppe D'Auria, Fernando Gonzalez-Candelas |
| EPI_ISL_426361, EPI_ISL_426362, EPI_ISL_426363 | Instituto Nacional de Ciencias Medicas y Nutricion Salvador Zubiran | Instituto Nacional de Ciencias Medicas y Nutricion Salvador Zubiran | Guillermo M. Ruiz-Palacios, Pilar Ramos Cervantes, Violeta Ibarra Gonzalez, Fernando Ledesma Barrientos, Luis Alberto García Andrade, Alfredo Ponce de León Garduño, Irma López Martínez, Lucia Hernández Rivas, Gisela Barrera Badillo, Edgar Mendieta Condado, Fabiola Garcés Ayala, Adnan Araiza Rodríguez, José Ernesto Ramírez González, Celia Boukadida, Santiago Avila Ríos, Mario Mújica Sánchez, José Arturo Martínez Orozco, Eduardo Becerril Vargas, Joel Armando Vázquez Pérez, Victor Hugo Borja Aburto, Concepción Grajales Muñiz, Cesar Raúl González Bonilla, Carolina González Torres, Francisco Javier Gaytán Cervantes, José Esteban Muñoz Medina, Blanca Taboada, Alejandro Sánchez, Pavel Isa, Ricardo Grande, Gloria Vázquez, Francisco Pulido, Carlos F. Arias |
| EPI_ISL_426364 | Instituto Nacional de Ciencias Medicas y Nutricion Salvador Zubiran | Instituto Nacional de Ciencias Medicas y Nutricion | Guillermo M. Ruiz-Palacios, Pilar Ramos Cervantes, Violeta Ibarra Gonzalez, Fernando Ledesma Barrientos, Luis Alberto García Andrade, Alfredo Ponce de León Garduño, Irma López Martínez, Lucia Hernández Rivas, Gisela Barrera Badillo, Edgar Mendieta Condado, Fabiola Garcés Ayala, Adnan Araiza Rodríguez, José Ernesto Ramírez González, Celia Boukadida, Santiago Avila Ríos, Mario Mújica Sánchez, José Arturo Martínez Orozco, Eduardo Becerril Vargas, Joel Armando Vázquez Pérez, Victor Hugo Borja Aburto, Concepción Grajales Muñiz, Cesar Raúl González Bonilla, Carolina González Torres, Francisco Javier Gaytán Cervantes, José Esteban Muñoz Medina, Blanca Taboada, Alejandro Sánchez, Pavel Isa, Ricardo Grande, Gloria Vázquez, Francisco Pulido, Carlos F. Arias |
| EPI_ISL_426365 | Instituto Nacional de Ciencias Medicas y Nutricion Salvador Zubiran | Instituto Nacional de Ciencias Medicas y Nutricion Salvador Zubiran | Guillermo M. Ruiz-Palacios, Pilar Ramos Cervantes, Violeta Ibarra Gonzalez, Fernando Ledesma Barrientos, Luis Alberto García Andrade, Alfredo Ponce de León Garduño, Irma López Martínez, Lucia Hernández Rivas, Gisela Barrera Badillo, Edgar Mendieta Condado, Fabiola Garcés Ayala, Adnan Araiza Rodríguez, José Ernesto Ramírez González, Celia Boukadida, Santiago Avila Ríos, Mario Mújica Sánchez, José Arturo Martínez Orozco, Eduardo Becerril Vargas, Joel Armando Vázquez Pérez, Victor Hugo Borja Aburto, Concepción Grajales Muñiz, Cesar Raúl González Bonilla, Carolina González Torres, Francisco Javier Gaytán Cervantes, José Esteban Muñoz Medina, Blanca Taboada, Alejandro Sánchez, Pavel Isa, Ricardo Grande, Gloria Vázquez, Francisco Pulido, Carlos F. Arias |
| EPI_ISL_427645, EPI_ISL_427647, EPI_ISL_427672, EPI_ISL_427673, EPI_ISL_427686, EPI_ISL_427687, EPI_ISL_427696, EPI_ISL_427703, EPI_ISL_427708 | Centre for Infectious Diseases and Microbiology Public Health | NSW Health Pathology - Institute of Clinical Pathology and Medical Research; Westmead Hospital; University of Sydney | Gray K, Timms V, Gall M, Arnott A, Sadsad R, Draper J, Sim E, Bachmann N, Rockett R, Lam C, Carter I, Holmes EC, O'Sullivan MV, Byun R, Sintchenko V, Chen SC, Eden JS, Maddocks S, Kok J, Propenko M, Sorrell T, Chang S, Basile K, Dwyer DE for the 2019-nCoV Study Group |
| EPI_ISL_427718 | ACT Pathology, The Canberra Hospital | NSW Health Pathology - Institute of Clinical Pathology and Medical Research; Westmead Hospital; University of Sydney | Gray K, Timms V, Gall M, Arnott A, Sadsad R, Draper J, Sim E, Bachmann N, Rockett R, Lam C, Carter I, Holmes EC, O'Sullivan MV, Byun R, Sintchenko V, Chen SC, Eden JS, Maddocks S, Kok J, Propenko M, Sorrell T, Chang S, Basile K, Dwyer DE for the 2019-nCoV Study Group |
| EPI_ISL_427730, EPI_ISL_427731, EPI_ISL_427732, EPI_ISL_427745, EPI_ISL_427767, EPI_ISL_427785, EPI_ISL_427793 | Centre for Infectious Diseases and Microbiology Public Health | NSW Health Pathology - Institute of Clinical Pathology and Medical Research; Westmead Hospital; University of Sydney | Gray K, Timms V, Gall M, Arnott A, Sadsad R, Draper J, Sim E, Bachmann N, Rockett R, Lam C, Carter I, Holmes EC, O'Sullivan MV, Byun R, Sintchenko V, Chen SC, Eden JS, Maddocks S, Kok J, Propenko M, Sorrell T, Chang S, Basile K, Dwyer DE for the 2019-nCoV Study Group |
| EPI_ISL_434541, EPI_ISL_434542, EPI_ISL_434543, EPI_ISL_434544, EPI_ISL_434545, EPI_ISL_434546, EPI_ISL_434547, EPI_ISL_434548, EPI_ISL_434549, EPI_ISL_434550, EPI_ISL_434551, EPI_ISL_434552, EPI_ISL_434553 | | | |
| see above | Puerto Rico Department of Health | Centers for Disease Control and Prevention, Dengue Branch | Gilberto A. Santiago, Glenda Gonzalez, Betzabel Flores, Keyla Charriez, Fabiola Cruz, Chaney Kalinich, Joseph Fauver, Jessica I. Falcon, Nathan Grubaugh, Jorge L. Munoz-Jordan |
| EPI_ISL_434626, EPI_ISL_434627, EPI_ISL_434628, EPI_ISL_434629, EPI_ISL_434630, EPI_ISL_434631, EPI_ISL_434632, EPI_ISL_434633, EPI_ISL_434634, EPI_ISL_434635 | CHU Purpan - Laboratoire de Virologie - Institut Fédératif de Biologie | Laboratoire de virologie - École Nationale Vétérinaire de Toulouse | Guillaume Croville, Jean-Luc Guérin, Jacques Izopet |
| EPI_ISL_436196, EPI_ISL_436204, EPI_ISL_436212, EPI_ISL_436220, EPI_ISL_436227, EPI_ISL_436271 | Servicio de Microbiología. Consorcio Hospital General Universitario de Valencia | Sequencing and Bioinformatics Service and Molecular Epidemiology Research Group. FISABIO-Public Health | Griselda De Marco, Beatriz Beamud, Lidia Ruiz Roldan, Marta Pla Diaz,Neris Garcia-Gonzalez, Loreto Ferrús Abad, Maria Dolores Ocete, Inma Galán Vendrell, Paula Ruiz-Hueso, Mariana Reyes-Prieto, Vicente Soriano Chirona, Maria Alma Bracho, Llúcia Martínez-Priego, Concepcion Gimeno, Giuseppe D'Auria, Fernando Gonzalez-Candelas |
| EPI_ISL_436279 | Servicio de Microbiologia. Hospital Clinico Universitario de Valencia | Sequencing and Bioinformatics Service and Molecular Epidemiology Research Group. FISABIO-Public Health | Griselda De Marco, Beatriz Beamud, Lidia Ruiz Roldan, Marta Pla Diaz, Neris Garcia-Gonzalez, Inma Galán Vendrell, Sandra Carbo, Loreto Ferrús Abad, Paula Ruiz-Hueso, Mariana Reyes-Prieto, Vicente Soriano Chirona, Ivan Ansari, David Navarro, Maria Alma Bracho, Llúcia Martínez-Priego, Giuseppe D'Auria, Fernando Gonzalez-Candelas |
| EPI_ISL_436293 | Servicio de Microbiologia. Hospital Clinico Universitario de Valencia | Sequencing and Bioinformatics Service and Molecular Epidemiology Research Group. FISABIO-Public Health | Griselda De Marco, Beatriz Beamud, Lidia Ruiz Roldan, Marta Pla Diaz, Neris Garcia-Gonzalez, Inma Galán Vendrell, Sandra Carbo, Loreto Ferrús Abad, Paula Ruiz-Hueso, Mariana Reyes-Prieto, Vicente Soriano Chirona, Ivan Ansari, David Navarro, Maria Alma Bracho, Llúcia Martínez-Priego, Giuseppe D'Auria, Fernando Gonzalez-Candelas |
| EPI_ISL_436299 | Servicio de Microbiologia. Hospital Clinico Universitario de Valencia | Sequencing and Bioinformatics Service and Molecular Epidemiology Research Group. FISABIO-Public Health | Griselda De Marco, Beatriz Beamud, Lidia Ruiz Roldan, Marta Pla Diaz, Neris Garcia-Gonzalez, Inma Galán Vendrell, Sandra Carbo, Loreto Ferrús Abad, Paula Ruiz-Hueso, Mariana Reyes-Prieto, Vicente Soriano Chirona, Ivan Ansari, David Navarro, Maria Alma Bracho, Llúcia Martínez-Priego, Giuseppe D'Auria, Fernando Gonzalez-Candelas |
| EPI_ISL_436313 | Servicio de Microbiologia. Hospital Clinico Universitario de Valencia | Sequencing and Bioinformatics Service and Molecular Epidemiology Research Group. FISABIO-Public Health | Griselda De Marco, Beatriz Beamud, Lidia Ruiz Roldan, Marta Pla Diaz, Neris Garcia-Gonzalez, Inma Galán Vendrell, Sandra Carbo, Loreto Ferrús Abad, Paula Ruiz-Hueso, Mariana Reyes-Prieto, Vicente Soriano Chirona, Ivan Ansari, David Navarro, Maria Alma Bracho, Llúcia Martínez-Priego, Giuseppe D'Auria, Fernando Gonzalez-Candelas |
| EPI_ISL_436319 | Servicio de Microbiologia. Hospital Clinico Universitario de Valencia | Sequencing and Bioinformatics Service and Molecular Epidemiology Research Group. FISABIO-Public Health | Griselda De Marco, Beatriz Beamud, Lidia Ruiz Roldan, Marta Pla Diaz, Neris Garcia-Gonzalez, Inma Galán Vendrell, Sandra Carbo, Loreto Ferrús Abad, Paula Ruiz-Hueso, Mariana Reyes-Prieto, Vicente Soriano Chirona, Ivan Ansari, David Navarro, Maria Alma Bracho, Llúcia Martínez-Priego, Giuseppe D'Auria, Fernando Gonzalez-Candelas |
| EPI_ISL_436333 | Servicio de Microbiologia. Hospital Clinico Universitario de Valencia | Sequencing and Bioinformatics Service and Molecular Epidemiology Research Group. FISABIO-Public Health | Griselda De Marco, Beatriz Beamud, Lidia Ruiz Roldan, Marta Pla Diaz, Neris Garcia-Gonzalez, Inma Galán Vendrell, Sandra Carbo, Loreto Ferrús Abad, Paula Ruiz-Hueso, Mariana Reyes-Prieto, Vicente Soriano Chirona, Ivan Ansari, David Navarro, Maria Alma Bracho, Llúcia Martínez-Priego, Giuseppe D'Auria, Fernando Gonzalez-Candelas |

| | | | |
|---|---|---|---|
| EPI_ISL_436339 | Servicio de Microbiologia. Hospital Clinico Universitario de Valencia | Sequencing and Bioinformatics Service and Molecular Epidemiology Research Group. FISABIO-Public Health | Griselda De Marco, Beatriz Beamud, Lidia Ruiz Roldan, Marta Pla Diaz, Neris Garcia-Gonzalez, Inma Galán Vendrell, Sandra Carbo, Loreto Ferrús Abad, Paula Ruiz-Hueso, Mariana Reyes-Prieto, Vicente Soriano Chirona, Ivan Ansari, David Navarro, Maria Alma Bracho, Llúcia Martínez-Priego, Giuseppe D'Auria, Fernando Gonzalez-Candelas |
| EPI_ISL_436353 | Servicio de Microbiologia. Hospital Clinico Universitario de Valencia | Sequencing and Bioinformatics Service and Molecular Epidemiology Research Group. FISABIO-Public Health | Griselda De Marco, Beatriz Beamud, Lidia Ruiz Roldan, Marta Pla Diaz, Neris Garcia-Gonzalez, Inma Galán Vendrell, Sandra Carbo, Loreto Ferrús Abad, Paula Ruiz-Hueso, Mariana Reyes-Prieto, Vicente Soriano Chirona, Ivan Ansari, David Navarro, Maria Alma Bracho, Llúcia Martínez-Priego, Giuseppe D'Auria, Fernando Gonzalez-Candelas |
| EPI_ISL_436363 | Servicio de Microbiologia. Hospital Clinico Universitario de Valencia | Sequencing and Bioinformatics Service and Molecular Epidemiology Research Group. FISABIO-Public Health | Griselda De Marco, Beatriz Beamud, Lidia Ruiz Roldan, Marta Pla Diaz, Neris Garcia-Gonzalez, Inma Galán Vendrell, Sandra Carbo, Loreto Ferrús Abad, Paula Ruiz-Hueso, Mariana Reyes-Prieto, Vicente Soriano Chirona, Ivan Ansari, David Navarro, Maria Alma Bracho, Llúcia Martínez-Priego, Giuseppe D'Auria, Fernando Gonzalez-Candelas |
| EPI_ISL_436384, EPI_ISL_436391 | Servicio de Microbiología. Consorcio Hospital General Universitario de Valencia | Sequencing and Bioinformatics Service and Molecular Epidemiology Research Group. FISABIO-Public Health | Griselda De Marco, Beatriz Beamud, Lidia Ruiz Roldan, Marta Pla Diaz,Neris Garcia-Gonzalez, Loreto Ferrús Abad, Maria Dolores Ocete, Inma Galán Vendrell, Paula Ruiz-Hueso, Mariana Reyes-Prieto, Vicente Soriano Chirona, Maria Alma Bracho, Llúcia Martínez-Priego, Concepcion Gimeno, Giuseppe D'Auria, Fernando Gonzalez-Candelas |
| EPI_ISL_436404 | Servicio de Microbiologia. Hospital Clinico Universitario de Valencia | Sequencing and Bioinformatics Service and Molecular Epidemiology Research Group. FISABIO-Public Health | Griselda De Marco, Beatriz Beamud, Lidia Ruiz Roldan, Marta Pla Diaz, Neris Garcia-Gonzalez, Inma Galán Vendrell, Sandra Carbo, Loreto Ferrús Abad, Paula Ruiz-Hueso, Mariana Reyes-Prieto, Vicente Soriano Chirona, Ivan Ansari, David Navarro, Maria Alma Bracho, Llúcia Martínez-Priego, Giuseppe D'Auria, Fernando Gonzalez-Candelas |
| EPI_ISL_447473 | Servicio de Microbiologia. Hospital Clinico Universitario de Valencia | Sequencing and Bioinformatics Service and Molecular Epidemiology Research Group. FISABIO-Public Health | Griselda De Marco, Lidia Ruiz Roldan, Neris Garcia-Gonzalez, Inma Galán Vendrell, Sandra Carbo, Loreto Ferrús Abad, Paula Ruiz-Hueso, Mariana Reyes-Prieto, Vicente Soriano Chirona, Ivan Ansari, Llúcia Martínez-Priego, Giuseppe 'Auria, David Navarro, Eliseo Albert, Maria Alma Bracho, Fernando Gonzalez-Candelas |
| EPI_ISL_447482 | Servicio de Microbiologia. Hospital Clinico Universitario de Valencia | Sequencing and Bioinformatics Service and Molecular Epidemiology Research Group. FISABIO-Public Health | Giuseppe 'Auria, David Navarro, Eliseo Albert, Maria Alma Bracho, Lidia Ruiz Roldan, Neris Garcia-Gonzalez, Inma Galán Vendrell, Sandra Carbo, Loreto Ferrús Abad, Paula Ruiz-Hueso, Mariana Reyes-Prieto, Vicente Soriano Chirona, Ivan Ansari, Llúcia Martínez-Priego, Fernando Gonzalez-Candelas |
| EPI_ISL_447487 | Servicio de Microbiologia. Hospital Clinico Universitario de Valencia | Sequencing and Bioinformatics Service and Molecular Epidemiology Research Group. FISABIO-Public Health | Griselda De Marco, Lidia Ruiz Roldan, Neris Garcia-Gonzalez, Inma Galán Vendrell, Sandra Carbo, Loreto Ferrús Abad, Paula Ruiz-Hueso, Mariana Reyes-Prieto, Vicente Soriano Chirona, Ivan Ansari, Llúcia Martínez-Priego, Giuseppe 'Auria, David Navarro, Eliseo Albert, Maria Alma Bracho, Fernando Gonzalez-Candelas |
| EPI_ISL_447496 | Servicio de Microbiologia. Hospital Clinico Universitario de Valencia | Sequencing and Bioinformatics Service and Molecular Epidemiology Research Group. FISABIO-Public Health | Giuseppe 'Auria, David Navarro, Eliseo Albert, Maria Alma Bracho, Lidia Ruiz Roldan, Neris Garcia-Gonzalez, Inma Galán Vendrell, Sandra Carbo, Loreto Ferrús Abad, Paula Ruiz-Hueso, Mariana Reyes-Prieto, Vicente Soriano Chirona, Ivan Ansari, Llúcia Martínez-Priego, Fernando Gonzalez-Candelas |
| EPI_ISL_447501 | Servicio de Microbiologia. Hospital Clinico Universitario de Valencia | Sequencing and Bioinformatics Service and Molecular Epidemiology Research Group. FISABIO-Public Health | Griselda De Marco, Lidia Ruiz Roldan, Neris Garcia-Gonzalez, Inma Galán Vendrell, Sandra Carbo, Loreto Ferrús Abad, Paula Ruiz-Hueso, Mariana Reyes-Prieto, Vicente Soriano Chirona, Ivan Ansari, Llúcia Martínez-Priego, Giuseppe 'Auria, David Navarro, Eliseo Albert, Maria Alma Bracho, Fernando Gonzalez-Candelas |
| EPI_ISL_447510 | Servicio de Microbiologia. Hospital Clinico Universitario de Valencia | Sequencing and Bioinformatics Service and Molecular Epidemiology Research Group. FISABIO-Public Health | Giuseppe 'Auria, David Navarro, Eliseo Albert, Maria Alma Bracho, Lidia Ruiz Roldan, Neris Garcia-Gonzalez, Inma Galán Vendrell, Sandra Carbo, Loreto Ferrús Abad, Paula Ruiz-Hueso, Mariana Reyes-Prieto, Vicente Soriano Chirona, Ivan Ansari, Llúcia Martínez-Priego, Fernando Gonzalez-Candelas |
| EPI_ISL_447515 | Servicio de Microbiologia. Hospital Clinico Universitario de Valencia | Sequencing and Bioinformatics Service and Molecular Epidemiology Research Group. FISABIO-Public Health | Griselda De Marco, Lidia Ruiz Roldan, Neris Garcia-Gonzalez, Inma Galán Vendrell, Sandra Carbo, Loreto Ferrús Abad, Paula Ruiz-Hueso, Mariana Reyes-Prieto, Vicente Soriano Chirona, Ivan Ansari, Llúcia Martínez-Priego, Giuseppe 'Auria, David Navarro, Eliseo Albert, Maria Alma Bracho, Fernando Gonzalez-Candelas |
| EPI_ISL_447523 | Servicio de Microbiologia. Hospital Clinico Universitario de Valencia | Sequencing and Bioinformatics Service and Molecular Epidemiology Research Group. FISABIO-Public Health | Giuseppe 'Auria, David Navarro, Eliseo Albert, Maria Alma Bracho, Lidia Ruiz Roldan, Neris Garcia-Gonzalez, Inma Galán Vendrell, Sandra Carbo, Loreto Ferrús Abad, Paula Ruiz-Hueso, Mariana Reyes-Prieto, Vicente Soriano Chirona, Ivan Ansari, Llúcia Martínez-Priego, Fernando Gonzalez-Candelas |
| EPI_ISL_447528 | Servicio de Microbiologia. Hospital Clinico Universitario de Valencia | Sequencing and Bioinformatics Service and Molecular Epidemiology Research Group. FISABIO-Public Health | Griselda De Marco, Lidia Ruiz Roldan, Neris Garcia-Gonzalez, Inma Galán Vendrell, Sandra Carbo, Loreto Ferrús Abad, Paula Ruiz-Hueso, Mariana Reyes-Prieto, Vicente Soriano Chirona, Ivan Ansari, Llúcia Martínez-Priego, Giuseppe 'Auria, David Navarro, Eliseo Albert, Maria Alma Bracho, Fernando Gonzalez-Candelas |
| EPI_ISL_450821, EPI_ISL_450822 | Jarpens HC | The Public Health Agency of Sweden | Gunilla Johansson, Anna-Malin Linde, Maria Lind Karlberg, Oskar Karlsson Lindsjo, Olov Svartstrom, Anna Risberg, Theresa Enkirch, Mia Brytting, Karin Tegmark-Wisell |
| EPI_ISL_452544, EPI_ISL_452545, EPI_ISL_452546, EPI_ISL_452547, EPI_ISL_452548, EPI_ISL_452549, EPI_ISL_452550, EPI_ISL_452551, EPI_ISL_452552, EPI_ISL_452553, EPI_ISL_452554, EPI_ISL_452555, EPI_ISL_452556, EPI_ISL_452557, EPI_ISL_452558, EPI_ISL_452559, EPI_ISL_452560, EPI_ISL_452561, EPI_ISL_452562, EPI_ISL_452563, EPI_ISL_452564, EPI_ISL_452565, EPI_ISL_452566, EPI_ISL_452567, EPI_ISL_452568, EPI_ISL_452569, EPI_ISL_452570, EPI_ISL_452571, EPI_ISL_452572, EPI_ISL_452573, EPI_ISL_452574, EPI_ISL_452575, EPI_ISL_452576, EPI_ISL_452577, EPI_ISL_452578, EPI_ISL_452579, EPI_ISL_452580, EPI_ISL_452581, EPI_ISL_452582, EPI_ISL_452583, EPI_ISL_452584, EPI_ISL_452585, EPI_ISL_452586, EPI_ISL_452587, EPI_ISL_452588, EPI_ISL_452589, EPI_ISL_452590, EPI_ISL_452592, EPI_ISL_452593, EPI_ISL_452594, EPI_ISL_452595, EPI_ISL_452596, EPI_ISL_452597, EPI_ISL_452598, EPI_ISL_452599, EPI_ISL_452600, EPI_ISL_452601, EPI_ISL_452602, EPI_ISL_452603, EPI_ISL_452604, EPI_ISL_452605, EPI_ISL_452606, EPI_ISL_452607, EPI_ISL_452608, EPI_ISL_452609, EPI_ISL_452610, EPI_ISL_452611, EPI_ISL_452612, EPI_ISL_452613, EPI_ISL_452614, EPI_ISL_452615, EPI_ISL_452616 | | | |
| see above | Servicio de Microbiología y Parasitología clínica. UCEIMP. Hospital Universitario Virgen del Rocío/IBIS/CSIC/US. | SeqCOVID-SPAIN consortium/IBV(CSIC) | Guillermo Martí-n Gutiérrez, Ángel Rodrí-guez Villodres, Lidia Gálvez Benitez, Verónica González Galán, Javier Aznar Martí-n and SeqCOVID-SPAIN consortium |
| EPI_ISL_452617, EPI_ISL_452618, EPI_ISL_452620, EPI_ISL_452621, EPI_ISL_452622, EPI_ISL_452623, EPI_ISL_452624, EPI_ISL_452625, EPI_ISL_452626, EPI_ISL_452627, EPI_ISL_452628, EPI_ISL_452629, EPI_ISL_452630, EPI_ISL_452631, EPI_ISL_452632, EPI_ISL_452633, EPI_ISL_452634, EPI_ISL_452635, EPI_ISL_452636, EPI_ISL_452637, EPI_ISL_452638, EPI_ISL_452639, EPI_ISL_452640, EPI_ISL_452641, EPI_ISL_452642, EPI_ISL_452643, EPI_ISL_452644, EPI_ISL_452645, EPI_ISL_452646, EPI_ISL_452647, EPI_ISL_452648, EPI_ISL_452649, EPI_ISL_452650, EPI_ISL_452651, EPI_ISL_452652, EPI_ISL_452653, EPI_ISL_452654, EPI_ISL_452655, EPI_ISL_452656, EPI_ISL_452657, EPI_ISL_452658, EPI_ISL_452659, EPI_ISL_452660, EPI_ISL_452661, EPI_ISL_452662, EPI_ISL_452663, EPI_ISL_452664, EPI_ISL_452665, EPI_ISL_452666, EPI_ISL_452667, EPI_ISL_452668, EPI_ISL_452669, EPI_ISL_452670, EPI_ISL_452671, EPI_ISL_452672, EPI_ISL_452673, EPI_ISL_452674, EPI_ISL_452675, EPI_ISL_452676, EPI_ISL_452677, EPI_ISL_452678, EPI_ISL_452679, EPI_ISL_452680, EPI_ISL_452681, EPI_ISL_452682, EPI_ISL_452683, EPI_ISL_452684, EPI_ISL_452685, EPI_ISL_452686, EPI_ISL_452687, EPI_ISL_452688, EPI_ISL_452689, EPI_ISL_452690, EPI_ISL_452691 | | | |
| see above | Servicio de Microbiología. Hospital Universitario Donostia. OSI Donostialdea. Área de Enfermedades Infecciosas, Grupo de Infección Respiratoria y Resistencia Antimicrobiana. Instituto de Investigación Sanitaria Biodonostia. | SeqCOVID-SPAIN consortium/IBV(CSIC) | Gustavo Cilla, Milagrosa Montes, Luis Piñeiro, Jose Maria Marimón and SeqCOVID-SPAIN consortium |
| EPI_ISL_454417, EPI_ISL_454418 | Research and Experiment Center, Meizhou People Hospital | Research and Experiment Center, Meizhou People Hospital | Guo,X., Zeng,L. and Yu,Z. |
| EPI_ISL_455313 | Microbiology Unit, Department of Pathology & Laboratory Medicine, IIUM Medical Centre | SEA Microbiome Unit, Faculty of Industrial Sciences & Technology, Universiti Malaysia Pahang | Hajar Fauzan Ahmad, Norhidayah Kamarudin, Ahmad Hafiz Zulkifly, IIUM Medical Centre Covid19 Taskforce, UMP Covid19 Team |
| EPI_ISL_455722 | Servicio de Microbiologia. Hospital Clinico Universitario de Valencia | Sequencing and Bioinformatics Service and Molecular Epidemiology Research Group. FISABIO-Public Health, and SeqCOVID-Spain Consortium | Giuseppe 'Auria, David Navarro, Eliseo Albert, Maria Alma Bracho, Lidia Ruiz Roldan, Neris Garcia-Gonzalez, Inma Galán Vendrell, Sandra Carbo, Loreto Ferrús Abad, Paula Ruiz-Hueso, Mariana Reyes-Prieto, Vicente Soriano Chirona, Ivan Ansari, Llúcia Martínez-Priego, Fernando Gonzalez-Candelas |
| EPI_ISL_455734, EPI_ISL_455737 | Servicio de Microbiologia. Hospital Clinico Universitario de Valencia | Sequencing and Bioinformatics Service and Molecular Epidemiology Research Group. FISABIO-Public Health, and SeqCOVID-Spain Consortium | Griselda De Marco, Lidia Ruiz Roldan, Neris Garcia-Gonzalez, Inma Galán Vendrell, Sandra Carbo, Loreto Ferrús Abad, Paula Ruiz-Hueso, Mariana Reyes-Prieto, Vicente Soriano Chirona, Ivan Ansari, Llúcia Martínez-Priego, Giuseppe 'Auria, David Navarro, Eliseo Albert, Maria Alma Bracho, Fernando Gonzalez-Candelas |
| EPI_ISL_457833, EPI_ISL_457834, EPI_ISL_457835, EPI_ISL_457836, EPI_ISL_457837, EPI_ISL_457838, EPI_ISL_457842, EPI_ISL_457843, EPI_ISL_457844 | National Public Health Laboratory | KEMRI-Wellcome Trust Research Programme/KEMRI-CGMR-C Kilifi | Githinji G. et al 2020 |
| EPI_ISL_457845, EPI_ISL_457846, EPI_ISL_457847, EPI_ISL_457848, EPI_ISL_457849, EPI_ISL_457850, EPI_ISL_457851, EPI_ISL_457852, EPI_ISL_457853, EPI_ISL_457854, EPI_ISL_457855, EPI_ISL_457856, EPI_ISL_457857, EPI_ISL_457858, EPI_ISL_457859, EPI_ISL_457860, EPI_ISL_457861, EPI_ISL_457862, EPI_ISL_457863, EPI_ISL_457864, EPI_ISL_457865, EPI_ISL_457866, EPI_ISL_457867, EPI_ISL_457868, EPI_ISL_457869, EPI_ISL_457870, EPI_ISL_457871, EPI_ISL_457872, EPI_ISL_457873, EPI_ISL_457874, EPI_ISL_457875, EPI_ISL_457876, EPI_ISL_457877, EPI_ISL_457878, EPI_ISL_457879, EPI_ISL_457880, EPI_ISL_457881, EPI_ISL_457882, EPI_ISL_457883, EPI_ISL_457884, EPI_ISL_457885, EPI_ISL_457886, EPI_ISL_457887, EPI_ISL_457888, EPI_ISL_457889, EPI_ISL_457890, EPI_ISL_457891, EPI_ISL_457892, EPI_ISL_457893, EPI_ISL_457894, EPI_ISL_457895, EPI_ISL_457896, EPI_ISL_457897, EPI_ISL_457898, EPI_ISL_457899, EPI_ISL_457900, EPI_ISL_457901, EPI_ISL_457902, EPI_ISL_457903, EPI_ISL_457904, EPI_ISL_457905, EPI_ISL_457906, EPI_ISL_457907, EPI_ISL_457908, EPI_ISL_457909, EPI_ISL_457910, EPI_ISL_457911, EPI_ISL_457912, EPI_ISL_457913, EPI_ISL_457914, EPI_ISL_457915, EPI_ISL_457916, EPI_ISL_457917, EPI_ISL_457918, EPI_ISL_457919, EPI_ISL_457920, EPI_ISL_457921, EPI_ISL_457922, EPI_ISL_457923, EPI_ISL_457924, EPI_ISL_457925, EPI_ISL_457926, EPI_ISL_457927, EPI_ISL_457928, EPI_ISL_457929, EPI_ISL_457930, EPI_ISL_457931 | | | |
| see above | KEMRI-CGMR-C | KEMRI-Wellcome Trust Research | Githinji G. et al 2020 |

| | | | |
|---|---|---|---|
| | | Programme/KEMRI-CGMR-C Kilifi | |
| EPI_ISL_457932, EPI_ISL_457933, EPI_ISL_457934, EPI_ISL_457935, EPI_ISL_457936 | KEMRI-Centre for Virus Research | KEMRI-Wellcome Trust Research Programme/KEMRI-CGMR-C Kilifi | Githinji G. et al 2020 |
| EPI_ISL_458285, EPI_ISL_458286 | unknown | Bundeswehr Institute of Microbiology | Handrick,S., Bestehorn-Willmann,M.S., Eckstein,S., Walter,M.C., Antwerpen,M.H., Rehn,A., Naija,H., Stoecker,K., Woelfel,R. and Ben Moussa,M. |
| EPI_ISL_462753 | University Clinical Hospital of Mostar | University of Sarajevo Veterinary Faculty | Goletic, T., Softic, A., Goletic, S., Ostojic, M., Hukic, M., Eterovic, T., Seho-Alic, A. |
| EPI_ISL_462995, EPI_ISL_462996, EPI_ISL_462997, EPI_ISL_462998, EPI_ISL_462999, EPI_ISL_463000 | Molecular Genetics | Molecular Genetics | Gomez,J., Coto,E. |
| EPI_ISL_467431 | Molecular Diagnostics Services (MDS) | KRISP, KZN Research Innovation and Sequencing Platform | Giandhari J, Pillay S, Lessells R, Chimukangara B, Mdlalose K, York D, Khan S, Tegally H, Wilkinson E, de Oliveira T |
| EPI_ISL_467432, EPI_ISL_467433, EPI_ISL_467434, EPI_ISL_467435 | AMPATH-DBN | KRISP, KZN Research Innovation and Sequencing Platform | Giandhari J, Pillay S, Lessells R, Chimukangara B, Mdlalose K, York D, Khan S, Tegally H, Wilkinson E, de Oliveira T |
| EPI_ISL_467437, EPI_ISL_467440, EPI_ISL_467441, EPI_ISL_467442, EPI_ISL_467443 | NHLS-IALCH | KRISP, KZN Research Innovation and Sequencing Platform | Giandhari J, Pillay S, Lessells R, Chimukangara B, Mdlalose K, York D, Khan S, Tegally H, Wilkinson E, de Oliveira T |
| EPI_ISL_467444, EPI_ISL_467445, EPI_ISL_467446, EPI_ISL_467448 | Molecular Diagnostics Services (MDS) | KRISP, KZN Research Innovation and Sequencing Platform | Giandhari J, Pillay S, Lessells R, Chimukangara B, Mdlalose K, York D, Khan S, Tegally H, Wilkinson E, de Oliveira T |
| EPI_ISL_467449, EPI_ISL_467450, EPI_ISL_467465, EPI_ISL_467466, EPI_ISL_467467, EPI_ISL_467468, EPI_ISL_467469, EPI_ISL_467470, EPI_ISL_467471, EPI_ISL_467472, EPI_ISL_467473, EPI_ISL_467474 | | | |
| see above | AMPATH-DBN | KRISP, KZN Research Innovation and Sequencing Platform | Giandhari J, Pillay S, Lessells R, Chimukangara B, Mdlalose K, York D, Khan S, Tegally H, Wilkinson E, de Oliveira T |
| EPI_ISL_467475, EPI_ISL_467476, EPI_ISL_467477, EPI_ISL_467478, EPI_ISL_467479, EPI_ISL_467480, EPI_ISL_467481, EPI_ISL_467482, EPI_ISL_467483, EPI_ISL_467484, EPI_ISL_467485, EPI_ISL_467486, EPI_ISL_467487, EPI_ISL_467488, EPI_ISL_467489, EPI_ISL_467490, EPI_ISL_467491 | | | |
| see above | Molecular Diagnostics Services (MDS) | KRISP, KZN Research Innovation and Sequencing Platform | Giandhari J, Pillay S, Lessells R, Chimukangara B, Mdlalose K, York D, Khan S, Tegally H, Wilkinson E, de Oliveira T |
| EPI_ISL_467493 | NHLS-IALCH | KRISP, KZN Research Innovation and Sequencing Platform | Giandhari J, Pillay S, Lessells R, Chimukangara B, Mdlalose K, York D, Khan S, Tegally H, Wilkinson E, de Oliveira T |
| EPI_ISL_467494, EPI_ISL_467495, EPI_ISL_467496, EPI_ISL_467497, EPI_ISL_467498, EPI_ISL_467499, EPI_ISL_467500, EPI_ISL_467501, EPI_ISL_467502, EPI_ISL_467503, EPI_ISL_467504, EPI_ISL_467505, EPI_ISL_467506 | | | |
| see above | Molecular Diagnostics Services (MDS) | KRISP, KZN Research Innovation and Sequencing Platform | Giandhari J, Pillay S, Lessells R, Chimukangara B, Mdlalose K, York D, Khan S, Tegally H, Wilkinson E, de Oliveira T |
| EPI_ISL_467507, EPI_ISL_467508, EPI_ISL_467509, EPI_ISL_467511, EPI_ISL_467512, EPI_ISL_467513, EPI_ISL_467514, EPI_ISL_467515 | NHLS-IALCH | KRISP, KZN Research Innovation and Sequencing Platform | Giandhari J, Pillay S, Lessells R, Chimukangara B, Mdlalose K, York D, Khan S, Tegally H, Wilkinson E, de Oliveira T |
| EPI_ISL_467516 | CAPRISA | KRISP, KZN Research Innovation and Sequencing Platform | Giandhari J, Pillay S, Lessells R, Chimukangara B, Mdlalose K, York D, Khan S, Tegally H, Wilkinson E, de Oliveira T |
| EPI_ISL_468860, EPI_ISL_468861, EPI_ISL_468862, EPI_ISL_468863, EPI_ISL_468864, EPI_ISL_468865, EPI_ISL_468866, EPI_ISL_468867, EPI_ISL_468868, EPI_ISL_468869, EPI_ISL_468870, EPI_ISL_468871, EPI_ISL_468872, EPI_ISL_468873, EPI_ISL_468874, EPI_ISL_468875, EPI_ISL_468876, EPI_ISL_468877, EPI_ISL_468878, EPI_ISL_468879, EPI_ISL_468880, EPI_ISL_468881, EPI_ISL_468882, EPI_ISL_468883, EPI_ISL_468884, EPI_ISL_468885, EPI_ISL_468886, EPI_ISL_468887, EPI_ISL_468888, EPI_ISL_468889, EPI_ISL_468890, EPI_ISL_468891, EPI_ISL_468892, EPI_ISL_468893, EPI_ISL_468894, EPI_ISL_468895, EPI_ISL_468896, EPI_ISL_468897, EPI_ISL_468898, EPI_ISL_468899, EPI_ISL_468900, EPI_ISL_468901, EPI_ISL_468902, EPI_ISL_468903, EPI_ISL_468904, EPI_ISL_468905, EPI_ISL_468906, EPI_ISL_468907, EPI_ISL_468908, EPI_ISL_468909, EPI_ISL_468910, EPI_ISL_468911, EPI_ISL_468912, EPI_ISL_468913, EPI_ISL_468915, EPI_ISL_468916, EPI_ISL_468917, EPI_ISL_468918, EPI_ISL_468919, EPI_ISL_468920, EPI_ISL_468921, EPI_ISL_468922, EPI_ISL_468923, EPI_ISL_468924, EPI_ISL_468925, EPI_ISL_468926, EPI_ISL_468927, EPI_ISL_468928, EPI_ISL_468929, EPI_ISL_468930, EPI_ISL_468931, EPI_ISL_468932, EPI_ISL_468933, EPI_ISL_468934, EPI_ISL_468935, EPI_ISL_468936, EPI_ISL_468937, EPI_ISL_468938, EPI_ISL_468939, EPI_ISL_468940, EPI_ISL_468941, EPI_ISL_468942, EPI_ISL_468943, EPI_ISL_468944, EPI_ISL_468945, EPI_ISL_468946, EPI_ISL_468947, EPI_ISL_468948, EPI_ISL_468949, EPI_ISL_468950, EPI_ISL_468951 | | | |
| see above | Servicio de Microbiología. Hospital Universitario Donostia. OSI Donostialdea. Área de Enfermedades Infecciosas, Grupo de Infección Respiratoria y Resistencia Antimicrobiana. Instituto de Investigación Sanitaria Biodonostia. | SeqCOVID-SPAIN consortium/IBV(CSIC) | Gustavo Cilla, Milagrosa Montes, Luis Piñeiro, Jose Maria Marimón and SeqCOVID-SPAIN consortium |
| EPI_ISL_470876 | Department for Virology, Molecular Biology and Genome Research, R. G. Lugar Center for Public Health Research, National Center for Disease Control and Public Health (NCDC) of Georgia. | Department for Virology, Molecular Biology and Genome Research, R. G. Lugar Center for Public Health Research, National Center for Disease Control and Public Health (NCDC) of Georgia. | Giorgi Tomashvili, Meri Pantsulaia, Gvantsa Brachveli, Gvantsa Chanturia, Ann Machablishvili, Nato Kotaria, Marine Murtskhvaladze, Lela Sabadze, Mari Gavashelidze, Ana Papkiauri, Tata Imnadze, Tamar Jashiashvili, Tea Tevdoradze, Ketevan Sidamonidze, Ekaterine Khmaladze, Ekaterine Zhghenti, Roena Sukhiashvili, Mariam Zakalashvili, Lela Urushadze, Magda Dgebuadze, Davit Tsaguria, Ekaterine Zangaladze, Nino Berishvili, Adam Kotorashvili, Maia Alkhazashvili, Irma Burjanadze, Anna Kasradze, Khatuna Zakhashvili, Paata Imnadze, Amiran Gamkrelidze. |
| EPI_ISL_470877 | Department for Virology, Molecular Biology and Genome Research, R. G. Lugar Center for Public Health Research, National Center for Disease Control and Public Health (NCDC) of Georgia. | Department for Virology, Molecular Biology and Genome Research, R. G. Lugar Center for Public Health Research, National Center for Disease Control and Public Health (NCDC) of Georgia. | Gvantsa Brachveli, Meri Pantsulaia, Giorgi Tomashvili, Gvantsa Chanturia, Ann Machablishvili, Nato Kotaria, Marine Murtskhvaladze, Lela Sabadze, Mari Gavashelidze, Ana Papkiauri, Tata Imnadze, Tamar Jashiashvili, Tea Tevdoradze, Ketevan Sidamonidze, Ekaterine Khmaladze, Ekaterine Zhghenti, Roena Sukhiashvili, Mariam Zakalashvili, Lela Urushadze, Magda Dgebuadze, Davit Tsaguria, Ekaterine Zangaladze, Nino Berishvili, Adam Kotorashvili, Maia Alkhazashvili, Irma Burjanadze, Anna Kasradze, Khatuna Zakhashvili, Paata Imnadze, Amiran Gamkrelidze. |
| EPI_ISL_476559 | unknown | Laboratoire Sciences et Technologies de la Santé (STS) Institut Supérieur des Sciences de la Santé Université Hassan 1er, Settat, Morocco | Hajar Lemriss, Sanaâ Lemriss, Amal Souiri, Narjis Amar, Mustapha Mouallif, Touria Essayagh, Jawad Bouzid, Saâd EL Kabbaj, Abderraouf Hilali |
| EPI_ISL_480952, EPI_ISL_480953, EPI_ISL_480954, EPI_ISL_480955, EPI_ISL_480956, EPI_ISL_480957, EPI_ISL_480958, EPI_ISL_480959, EPI_ISL_480960, EPI_ISL_480962, EPI_ISL_480963, EPI_ISL_480964, EPI_ISL_480965, EPI_ISL_480966, EPI_ISL_480967, EPI_ISL_480968, EPI_ISL_480969, EPI_ISL_480970, EPI_ISL_480971, EPI_ISL_480972, EPI_ISL_480973, EPI_ISL_480974, EPI_ISL_480976, EPI_ISL_480977, EPI_ISL_480978, EPI_ISL_480979, EPI_ISL_480980, EPI_ISL_480982, EPI_ISL_480983, EPI_ISL_480984, EPI_ISL_480985, EPI_ISL_480986, EPI_ISL_480987, EPI_ISL_480988, EPI_ISL_480990, EPI_ISL_480991, EPI_ISL_480992, EPI_ISL_480993, EPI_ISL_480994, EPI_ISL_480996, EPI_ISL_480997, EPI_ISL_480998, EPI_ISL_480999, EPI_ISL_481000, EPI_ISL_481001, EPI_ISL_481002, EPI_ISL_481004, EPI_ISL_481005, EPI_ISL_481006, EPI_ISL_481007, EPI_ISL_481008, EPI_ISL_481009, EPI_ISL_481010, EPI_ISL_481011, EPI_ISL_481012, EPI_ISL_481013, EPI_ISL_481014, EPI_ISL_481015, EPI_ISL_481016, EPI_ISL_481018, EPI_ISL_481019, EPI_ISL_481020, EPI_ISL_481021, EPI_ISL_481022, EPI_ISL_481023, EPI_ISL_481024, EPI_ISL_481026, EPI_ISL_481027, EPI_ISL_481028, EPI_ISL_481030, EPI_ISL_481031, EPI_ISL_481032, EPI_ISL_481033, EPI_ISL_481036, EPI_ISL_481037, EPI_ISL_481038, EPI_ISL_481039, EPI_ISL_481040 | | | |
| see above | Servicio de Microbiología. Hospital Universitario Donostia. OSI Donostialdea. Área de Enfermedades Infecciosas, Grupo de Infección Respiratoria y Resistencia Antimicrobiana. Instituto de Investigación Sanitaria Biodonostia | SeqCOVID-SPAIN consortium/IBV(CSIC) | Gustavo Cilla, Milagrosa Montes, Luis Piñeiro, Jose Maria Marimón and SeqCOVID-SPAIN consortium |
| EPI_ISL_482879, EPI_ISL_482880, EPI_ISL_482881, EPI_ISL_482882, EPI_ISL_482883, EPI_ISL_482884, EPI_ISL_482885, EPI_ISL_482886, EPI_ISL_482887, EPI_ISL_482888, EPI_ISL_482889 | | | |
| see above | CHU Purpan - Laboratoire de Virologie - Institut Fédératif de Biologie | Laboratoire de virologie - École Nationale Vétérinaire de Toulouse | Guillaume Croville, Jean-Luc Guérin, Jacques Izopet |
| EPI_ISL_491481 | Functional Genomics Core University of South Carolina / Prisma Health-Midlands | Functional Genomics Core, Center For Targeted Therapeutics, | Hao Ji, Diego Altomare, B.Celia Cui, Mengqian Chen, Alyssa Clay-Glimour, Michael Wyatt, Phillip Buckhaults, Helmut Albrecht, Michael Shtutman |
| EPI_ISL_492074 | Functional Genomics Core University of South Carolina / Prisma Health-Midlands | Functional Genomics Core, University of South Carolina, | Hao Ji, Diego Altomare, B.Celia Cui, Mengqian Chen, Alyssa Clay-Glimour, Michael Wyatt, Phillip Buckhaults, Helmut Albrecht, Michael Shtutman |
| EPI_ISL_493129, EPI_ISL_493130 | Functional Genomics Core University of South Carolina / Prisma Health-Midlands | Functional Genomics Core, University of South Carolina, | Hao Ji, Diego Altomare, B.Celia Cui, Mengqian Chen, Alyssa Clay-Glimour, Michael Wyatt, Phillip Buckhaults, Helmut Albrecht, Michael Shtutman |
| EPI_ISL_493209, EPI_ISL_493210, EPI_ISL_493211, EPI_ISL_493212 | Functional Genomics Core University of South Carolina / Prisma Health-Midlands | Functional Genomics Core, University of South Carolina | Hao Ji, Diego Altomare, B.Celia Cui, Mengqian Chen, Alyssa Clay-Glimour, Michael Wyatt, Phillip Buckhaults, Helmut Albrecht, Michael Shtutman |
| EPI_ISL_493334, EPI_ISL_493335, EPI_ISL_493336, EPI_ISL_493337, | Instituto de Diagnostico y Referencia Epidemiologicos (INDRE) | Instituto de Diagnostico y Referencia Epidemiologicos (INDRE) | Gisela Barrera-Badillo , Abril Rodriguez-Maldonado, Claudia Wong-Arambula , Natividad Cruz-Ortiz, Tatiana Nunez-Garcia, Dayanira Arellano-Suarez, Fabiola Garces-Ayala, Edgar Mendieta-Condado, Lucia Hernandez-Rivas, Irma Lopez-Martinez, Ernesto Ramirez-Gonzalez. |

| | | | |
|---|---|---|---|
| EPI_ISL_493338 | | | |
| EPI_ISL_493339, EPI_ISL_493340, EPI_ISL_493341 | Instituto de Diagnostico y Referencia Epidemiologicos (INDRE) | Instituto de Diagnostico y Referencia Epidemiologicos (INDRE) | Gisela Barrera-Badillo , Abril Rodriguez-Maldonado, Claudia Wong-Arambula , Natividad Cruz-Ortiz, Tatiana Nunez-Garcia, Dayanira Arellano-Suarez, Adnan Araiza-Rodriguez, Edgar Mendieta-Condado, Lucia Hernandez-Rivas, Irma Lopez-Martinez, Ernesto Ramirez-Gonzalez. |
| EPI_ISL_493426 | Functional Genomics Core University of South Carolina / Prisma Health-Midlands | Functional Genomics Core, University of South Carolina, | Hao Ji, Diego Altomare, B.Celia Cui, Mengqian Chen, Alyssa Clay-Glimour, Michael Wyatt, Phillip Buckhaults, Helmut Albrecht, Michael Shtutman |
| EPI_ISL_493427, EPI_ISL_494553, EPI_ISL_494554, EPI_ISL_494555, EPI_ISL_494558 | Functional Genomics Core University of South Carolina / Prisma Health-Midlands | Functional Genomics Core, University of South Carolina | Hao Ji, Diego Altomare, B.Celia Cui, Mengqian Chen, Alyssa Clay-Glimour, Michael Wyatt, Phillip Buckhaults, Helmut Albrecht, Michael Shtutman |
| EPI_ISL_500292, EPI_ISL_500293, EPI_ISL_500294, EPI_ISL_500295, EPI_ISL_500296, EPI_ISL_500297, EPI_ISL_500298, EPI_ISL_500299, EPI_ISL_500300, EPI_ISL_500301, EPI_ISL_500302, EPI_ISL_500303, EPI_ISL_500304, EPI_ISL_500305, EPI_ISL_500306, EPI_ISL_500307, EPI_ISL_500308, EPI_ISL_500309, EPI_ISL_500310, EPI_ISL_500311, EPI_ISL_500312, EPI_ISL_500313, EPI_ISL_500314, EPI_ISL_500315, EPI_ISL_500316, EPI_ISL_500317, EPI_ISL_500318, EPI_ISL_500319, EPI_ISL_500320, EPI_ISL_500321, EPI_ISL_500322, EPI_ISL_500323, EPI_ISL_500324, EPI_ISL_500325, EPI_ISL_500326, EPI_ISL_500327, EPI_ISL_500328, EPI_ISL_500329, EPI_ISL_500330, EPI_ISL_500331, EPI_ISL_500332, EPI_ISL_500333, EPI_ISL_500334, EPI_ISL_500335, EPI_ISL_509601, EPI_ISL_509610, EPI_ISL_509612, EPI_ISL_509613, EPI_ISL_509614, EPI_ISL_509633, EPI_ISL_509634, EPI_ISL_509635, EPI_ISL_509636, EPI_ISL_509637, EPI_ISL_509638, EPI_ISL_509639, EPI_ISL_509640, EPI_ISL_509641, EPI_ISL_509642, EPI_ISL_509643, EPI_ISL_509644, EPI_ISL_509645, EPI_ISL_509646, EPI_ISL_509647, EPI_ISL_509648, EPI_ISL_509649, EPI_ISL_509650, EPI_ISL_509651, EPI_ISL_509652, EPI_ISL_509653, EPI_ISL_509654, EPI_ISL_510391, EPI_ISL_510392, EPI_ISL_510393, EPI_ISL_510394, EPI_ISL_510395, EPI_ISL_510396, EPI_ISL_510397, EPI_ISL_510398, EPI_ISL_510399, EPI_ISL_510400, EPI_ISL_510401, EPI_ISL_510402, EPI_ISL_510403, EPI_ISL_510404, EPI_ISL_510405, EPI_ISL_510406, EPI_ISL_510407, EPI_ISL_510408, EPI_ISL_510409, EPI_ISL_510410, EPI_ISL_510411, EPI_ISL_510412, EPI_ISL_510413, EPI_ISL_510414, EPI_ISL_510415, EPI_ISL_510416, EPI_ISL_510417, EPI_ISL_510418, EPI_ISL_510419, EPI_ISL_510420, EPI_ISL_510421, EPI_ISL_510422, EPI_ISL_510423, EPI_ISL_510467, EPI_ISL_510468, EPI_ISL_510469, EPI_ISL_510470, EPI_ISL_510471, EPI_ISL_510472, EPI_ISL_510473, EPI_ISL_510474, EPI_ISL_510475, EPI_ISL_510476, EPI_ISL_510477, EPI_ISL_510478, EPI_ISL_510479, EPI_ISL_510480, EPI_ISL_510481, EPI_ISL_510482, EPI_ISL_510483, EPI_ISL_510484, EPI_ISL_510485, EPI_ISL_510486, EPI_ISL_510487, EPI_ISL_510488, EPI_ISL_510489, EPI_ISL_510490, EPI_ISL_510491, EPI_ISL_510492, EPI_ISL_510493, EPI_ISL_510494, EPI_ISL_510495, EPI_ISL_510496, EPI_ISL_510497, EPI_ISL_510498, EPI_ISL_510499, EPI_ISL_510500, EPI_ISL_510501, EPI_ISL_510502, EPI_ISL_510503, EPI_ISL_510504, EPI_ISL_510505, EPI_ISL_510506, EPI_ISL_510507, EPI_ISL_510508, EPI_ISL_510509 | | | |
| see above | Servicio de Microbiología. Hospital Universitario Donostia. OSI Donostialdea. Área de Enfermedades Infecciosas, Grupo de Infección Respiratoria y Resistencia Antimicrobiana. Instituto de Investigación Sanitaria Biodonostia | SeqCOVID-SPAIN consortium/IBV(CSIC) | Gustavo Cilla, Milagrosa Montes, Luis Piñeiro, Jose Maria Marimón and SeqCOVID-SPAIN consortium |
| EPI_ISL_512811, EPI_ISL_512812, EPI_ISL_512813, EPI_ISL_512814, EPI_ISL_512815, EPI_ISL_512816, EPI_ISL_512817, EPI_ISL_512818, EPI_ISL_512819, EPI_ISL_512820, EPI_ISL_512821 | | | |
| see above | Kenema Government Hospital, Ministry of Health and Sanitation | Kenema Government Hospital, Ministry of Health and Sanitation | Goba,A., Momoh,M., Sandi,J., Tomkins-Tinch,C., Siddle,K., Mehta,S., Oluniyi,P., Jalloh,S., Park,D., Andersen,K., Garry,R., Happi,C., Grant,D., Olawoye,I. |
| EPI_ISL_515568, EPI_ISL_515569, EPI_ISL_515570, EPI_ISL_515571, EPI_ISL_515573, EPI_ISL_515574, EPI_ISL_515575, EPI_ISL_515576, EPI_ISL_515579, EPI_ISL_515580, EPI_ISL_515581, EPI_ISL_515582, EPI_ISL_515584, EPI_ISL_515585, EPI_ISL_515586, EPI_ISL_515589, EPI_ISL_515592, EPI_ISL_515695, EPI_ISL_515713, EPI_ISL_515717, EPI_ISL_515720, EPI_ISL_515721, EPI_ISL_515726, EPI_ISL_515727, EPI_ISL_515746, EPI_ISL_515747, EPI_ISL_515756, EPI_ISL_515765, EPI_ISL_515766, EPI_ISL_515772, EPI_ISL_515774, EPI_ISL_515788, EPI_ISL_515793, EPI_ISL_515798, EPI_ISL_515799 | | | |
| see above | NHLS-IALCH | KRISP, KZN Research Innovation and Sequencing Platform | Giandhari J, Pillay S, Lessells R, Mdlalose K, York D, Khan S, Tegally H, Wilkinson E, de Oliveira T |
| EPI_ISL_515830, EPI_ISL_515831, EPI_ISL_515832, EPI_ISL_515833, EPI_ISL_515834, EPI_ISL_515835, EPI_ISL_515836, EPI_ISL_515837, EPI_ISL_515838, EPI_ISL_515839, EPI_ISL_515840, EPI_ISL_515841, EPI_ISL_515842, EPI_ISL_515843, EPI_ISL_515844, EPI_ISL_515845, EPI_ISL_515846, EPI_ISL_515847, EPI_ISL_515848, EPI_ISL_515849, EPI_ISL_515850, EPI_ISL_515851, EPI_ISL_515852, EPI_ISL_515853, EPI_ISL_515854, EPI_ISL_515855, EPI_ISL_515856, EPI_ISL_515857, EPI_ISL_515858, EPI_ISL_515860, EPI_ISL_515861, EPI_ISL_515862, EPI_ISL_515863, EPI_ISL_515864, EPI_ISL_515865, EPI_ISL_515866, EPI_ISL_515867, EPI_ISL_515868, EPI_ISL_515869, EPI_ISL_515870, EPI_ISL_515871, EPI_ISL_515872, EPI_ISL_515873, EPI_ISL_515874, EPI_ISL_515875, EPI_ISL_515876, EPI_ISL_515877, EPI_ISL_515878, EPI_ISL_515880, EPI_ISL_515881, EPI_ISL_515882, EPI_ISL_515883, EPI_ISL_515884, EPI_ISL_515885, EPI_ISL_515886, EPI_ISL_515887, EPI_ISL_515889, EPI_ISL_515890, EPI_ISL_515891, EPI_ISL_515892, EPI_ISL_515893 | | | |
| see above | Medical Disagnostics Services (MDS) | KRISP, KZN Research Innovation and Sequencing Platform | Giandhari J, Pillay S, Lessells R, ChimukangaraB, Mdlalose K, York D, Khan S, Tegally H, Wilkinson E, de Oliveira T |
| EPI_ISL_516611, EPI_ISL_516618, EPI_ISL_516619 | Instituto de Diagnostico y Referencia Epidemiologicos (INDRE) | Instituto de Diagnostico y Referencia Epidemiologicos (INDRE) | Gisela Barrera-Badillo , Abril Rodriguez-Maldonado, Claudia Wong-Arambula , Natividad Cruz-Ortiz, Tatiana Nunez-Garcia, Dayanira Arellano-Suarez, Fabiola Garces-Ayala, Edgar Mendieta-Condado, Lucia Hernandez-Rivas, Irma Lopez-Martinez, Ernesto Ramirez-Gonzalez. |
| EPI_ISL_516620, EPI_ISL_516621 | Instituto de Diagnostico y Referencia Epidemiologicos (INDRE) | Instituto de Diagnostico y Referencia Epidemiologicos (INDRE) | Gisela Barrera-Badillo , Abril Rodriguez-Maldonado, Claudia Wong-Arambula , Natividad Cruz-Ortiz, Tatiana Nunez-Garcia, Dayanira Arellano-Suarez, Adnan Araiza-Rodriguez, Edgar Mendieta-Condado, Lucia Hernandez-Rivas, Irma Lopez-Martinez, Ernesto Ramirez-Gonzalez. |
| EPI_ISL_516622 | Instituto de Diagnostico y Referencia Epidemiologicos (INDRE) | Instituto de Diagnostico y Referencia Epidemiologicos (INDRE) | Gisela Barrera-Badillo , Abril Rodriguez-Maldonado, Claudia Wong-Arambula , Natividad Cruz-Ortiz, Tatiana Nunez-Garcia, Dayanira Arellano-Suarez, Fabiola Garces-Ayala, Edgar Mendieta-Condado, Lucia Hernandez-Rivas, Irma Lopez-Martinez, Ernesto Ramirez-Gonzalez. |
| EPI_ISL_516623 | Instituto de Diagnostico y Referencia Epidemiologicos (INDRE) | Instituto de Diagnostico y Referencia Epidemiologicos (INDRE) | Gisela Barrera-Badillo , Abril Rodriguez-Maldonado, Claudia Wong-Arambula , Natividad Cruz-Ortiz, Tatiana Nunez-Garcia, Dayanira Arellano-Suarez, Adnan Araiza-Rodriguez, Edgar Mendieta-Condado, Lucia Hernandez-Rivas, Irma Lopez-Martinez, Ernesto Ramirez-Gonzalez. |
| EPI_ISL_516624, EPI_ISL_516625 | Instituto de Diagnostico y Referencia Epidemiologicos (INDRE) | Instituto de Diagnostico y Referencia Epidemiologicos (INDRE) | Gisela Barrera-Badillo , Abril Rodriguez-Maldonado, Claudia Wong-Arambula , Natividad Cruz-Ortiz, Tatiana Nunez-Garcia, Dayanira Arellano-Suarez, Fabiola Garces-Ayala, Edgar Mendieta-Condado, Lucia Hernandez-Rivas, Irma Lopez-Martinez, Ernesto Ramirez-Gonzalez. |
| EPI_ISL_516800 | Rumah Sakit Akademik Universitas Gadjah Mada | Genetics Working Group (Pokja Genetik) Faculty of Medicine, Public Health and Nursing Universitas Gadjah Mada (FK-KMK UGM); Disease Investigation Center Wates Ministry of Agriculture Indonesia; Department of Microbiology FK-KMK UGM; Laboratorium Diagnostik Yayasan Tahija World Mosquito Program (WMP) Yogyakarta Center for Tropical Medicine FK-KMK UGM; Integrated Research center FK-KMK UGM; Department of Computer Science and Electronics FMIPA UGM | Gunadi, Hendra Wibawa, . Marcellus, Mohamad S. Hakim, Edwin W. Daniwijaya, Ludhang P. Rizki, Endah Supriyati, Eggi Arguni, Titik Nuryastuti, Tri Wibawa, Dwi AA Nugrahaningsih, Afiahayati, Siswanto, Alvin S. Kalim, Desyifa Mursalin |
| EPI_ISL_516806 | Rumah Sakit PKU Gamping | Genetics Working Group (Pokja Genetik) Faculty of Medicine, Public Health and Nursing Universitas Gadjah Mada (FK-KMK UGM); Disease Investigation Center Wates Ministry of Agriculture Indonesia; Department of Microbiology FK-KMK UGM; Laboratorium Diagnostik Yayasan Tahija World Mosquito Program (WMP) Yogyakarta Center for Tropical Medicine FK-KMK UGM; Integrated Research center FK-KMK UGM; Department of Computer Science and Electronics FMIPA UGM | Gunadi, Hendra Wibawa, . Marcellus, Mohamad S. Hakim, Edwin W. Daniwijaya, Ludhang P. Rizki, Endah Supriyati, Eggi Arguni, Titik Nuryastuti, Tri Wibawa, Dwi AA Nugrahaningsih, . Afiahayati , . Siswanto, Ardorisye Saptaty Fornia, Kemala Athollah |
| EPI_ISL_516829 | RSUD Nyi Ageng Serang | Genetics Working Group (Pokja Genetik) Faculty of Medicine, Public Health and Nursing Universitas Gadjah Mada (FK-KMK UGM), Disease Investigation Center Wates Ministry of Agriculture Indonesia, Department of Microbiology FK-KMK UGM, Laboratorium Diagnostik Yayasan Tahija World Mosquito Program (WMP) Yogyakarta Center for Tropical Medicine FK-KMK UGM, Integrated Research Center FK-KMK UGM, Department of Computer Science and Electronics FMIPA UGM | Gunadi, Hendra Wibawa, . Marcellus, Mohamad S. Hakim, Edwin W. Daniwijaya, Ludhang P. Rizki, Endah Supriyati, Eggi Arguni, Titik Nuryastuti, Tri Wibawa, Dwi AA Nugrahaningsih, Afiahayati , . Siswanto, Beby Dewi Sartika, Dyah Ayu Puspitarani |
| EPI_ISL_518036, EPI_ISL_518042, EPI_ISL_518053 | NHLS-IALCH | KRISP, KZN Research Innovation and Sequencing Platform | Giandhari J, Pillay S, Lessells R, Mdlalose K, York D, Khan S, Tegally H, Wilkinson E, de Oliveira T |
| EPI_ISL_524735 | Government General Hospital, Jam Khambhaliya, Devbhoomi Dwarka | Gujarat Biotechnology Research Centre | Harish Matani, Janvi Raval, Zarna Patel, Monika Gandhi, Pinal Trivedi, Maharshi Pandya, Nidhi Patel, Nitin Savaliya, Raghawendra Kumar, Dinesh Kumar, Zuber Saiyed, Komal Patel, Labdhi Pandya, Afzal Ansari, Nikha Trivedi, Apurvasinh Puvar, R D Dixit, A M Kadri, Harsh Bakshi, Chaitanya Joshi, Madhvi Joshi |
| EPI_ISL_525492 | RSUP dr. SOERADJI TIRTONEGORO | Genetics Working Group (Pokja Genetik) Faculty of Medicine, Public Health and Nursing Universitas Gadjah Mada (FK-KMK | Gunadi, Hendra Wibawa, . Marcellus, Mohamad S. Hakim, Edwin W. Daniwijaya, Ludhang P. Rizki, Endah Supriyati, Eggi Arguni, Titik Nuryastuti, Tri Wibawa, Dwi AA Nugrahaningsih, Afiahayati , . Siswanto, Kurniyanto, Indah Juliana, Alvin S. Kalim, Dwiki Afandy |

| | | | |
|---|---|---|---|
| | | UGM), Disease Investigation Center Wates Ministry of Agriculture Indonesia, Department of Microbiology FK-KMK UGM, Laboratorium Diagnostik Yayasan Tahija World Mosquito Program (WMP) Yogyakarta Center for Tropical Medicine FK-KMK UGM, Integrated Research Center FK-KMK UGM, Department of Computer Science and Electronics FMIPA UGM | |
| EPI_ISL_526686, EPI_ISL_526687, EPI_ISL_526688 | Faith Laboratory, Immunology Institute, Icahn School of Medicine at Mount Sinai | van Bakel Laboratory, Genetics and Genomics Sciences, Icahn School of Medicine at Mount Sinai | Graham J. Britton, Alice Chen-Liaw, Francesca Cossarini, Alexandra Livanos, Matthew P. Spindler, Tamar Plitt, Joseph Eggers, Ilaria Mogno, Ana S. Gonzalez-Reiche, Sophia Sui, Michael Tankelevich, Lauren Tal Grinspan, Rebekah E. Dixon, Divya Jha, Gustavo Martinez-Delgado, Fatima Amanat, Daisy Hoagland, Benjamin R. tenOever, Marla C. Dubinsky, Miriam Merad, Harm Van Bakel, Florian Krammer, Gerold Bongers, Saurabh Mehandru and Jeremiah J. Faith |
| EPI_ISL_527401, EPI_ISL_527402, EPI_ISL_527403, EPI_ISL_527404, EPI_ISL_527405, EPI_ISL_527406, EPI_ISL_527407, EPI_ISL_527408, EPI_ISL_527409, EPI_ISL_527410, EPI_ISL_527411, EPI_ISL_527412, EPI_ISL_527413, EPI_ISL_527414, EPI_ISL_527415, EPI_ISL_527416, EPI_ISL_527417, EPI_ISL_527418, EPI_ISL_527419, EPI_ISL_527420, EPI_ISL_527421, EPI_ISL_527422, EPI_ISL_527423, EPI_ISL_527424, EPI_ISL_527425, EPI_ISL_527426, EPI_ISL_527427, EPI_ISL_527428, EPI_ISL_527429, EPI_ISL_527430, EPI_ISL_527431, EPI_ISL_527432, EPI_ISL_527433, EPI_ISL_527434, EPI_ISL_527435, EPI_ISL_527436, EPI_ISL_527437, EPI_ISL_527438, EPI_ISL_527439, EPI_ISL_527440, EPI_ISL_527441, EPI_ISL_527442, EPI_ISL_527443, EPI_ISL_527444, EPI_ISL_527445, EPI_ISL_527446, EPI_ISL_527447, EPI_ISL_527448, EPI_ISL_527449, EPI_ISL_527450, EPI_ISL_527451, EPI_ISL_527452, EPI_ISL_527453, EPI_ISL_527454, EPI_ISL_527455, EPI_ISL_527456, EPI_ISL_527457, EPI_ISL_527458, EPI_ISL_527459, EPI_ISL_527460, EPI_ISL_527461, EPI_ISL_527462, EPI_ISL_527463, EPI_ISL_527464, EPI_ISL_527465, EPI_ISL_527466, EPI_ISL_527467, EPI_ISL_527468, EPI_ISL_527469, EPI_ISL_527470, EPI_ISL_527471, EPI_ISL_527472, EPI_ISL_527473, EPI_ISL_527474, EPI_ISL_527475, EPI_ISL_527476, EPI_ISL_527477, EPI_ISL_527478, EPI_ISL_527479, EPI_ISL_527480, EPI_ISL_527481, EPI_ISL_527482, EPI_ISL_527483, EPI_ISL_527484, EPI_ISL_527485, EPI_ISL_527486, EPI_ISL_527487, EPI_ISL_527488 | | | |
| see above | Colorado State University - Ebel Lab | Colorado State University - Ebel Lab | Greg Ebel et al. |
| EPI_ISL_528539 | LVM/UFRJ | LNCC | Gustavo M. Romário M. de Souza; Bruno B. Bezerra; Lucio A. Caldas; Fabio Limonte; Elena Cobos; Sharton V. A. Coelho; Luiz Almeida; Luiza Higga; Isadora A. Correa; Diana Marianni; Luciana B. Arruda; Marcelo Bozza; Orlando Ferreira; Wanderley de Souza; Ana Teresa R. Vasconcelos; Terezinha M. Castineiras; Amilcar Tanuri; Luciana J. Costa. |
| EPI_ISL_528637, EPI_ISL_528638 | LVM/UFRJ | Bioinformatics Laboratory / LNCC | Gustavo D. P. Silva; M. Romário M. de Souza; Bruno B. Bezerra; Lucio A. Caldas; Fabio Limonte; Elena Cobos; Sharton V. A. Coelho; Luiz Almeida; Luiza Higga; Isadora A. Correa; Diana Marianni; Luciana B. Arruda; Marcelo Bozza; Orlando Ferreira; Wanderley de Souza; Ana Teresa R. Vasconcelos; Terezinha M. Castineiras; Amilcar Tanuri; Luciana J. Costa |
| EPI_ISL_529171, EPI_ISL_529172, EPI_ISL_529173, EPI_ISL_529174, EPI_ISL_529175, EPI_ISL_529176, EPI_ISL_529177, EPI_ISL_529178, EPI_ISL_529180, EPI_ISL_529181, EPI_ISL_529182, EPI_ISL_529183, EPI_ISL_529184, EPI_ISL_529185, EPI_ISL_529186, EPI_ISL_529187, EPI_ISL_529188, EPI_ISL_529189, EPI_ISL_529190, EPI_ISL_529191, EPI_ISL_529192, EPI_ISL_529193, EPI_ISL_529194, EPI_ISL_529195, EPI_ISL_529196, EPI_ISL_529197, EPI_ISL_529198, EPI_ISL_529199 | | | |
| see above | South Carolina Department of Health and Environmental Control | South Carolina Department of Health and Environmental Control | Haley V. Flores |
| EPI_ISL_529725, EPI_ISL_529727, EPI_ISL_529732, EPI_ISL_529738, EPI_ISL_529745, EPI_ISL_529746, EPI_ISL_529763, EPI_ISL_529768, EPI_ISL_529781, EPI_ISL_529786, EPI_ISL_529793, EPI_ISL_529795, EPI_ISL_529797, EPI_ISL_529803 | | | |
| see above | NHLS-IALCH | KRISP, KZN Research Innovation and Sequencing Platform | Giandhari J, Pillay S, Lessells R, Mdlalose K, York D, Khan S, Tegally H, Wilkinson E, de Oliveira T |
| EPI_ISL_530351, EPI_ISL_530352, EPI_ISL_530353, EPI_ISL_530354, EPI_ISL_530355, EPI_ISL_530356, EPI_ISL_530357, EPI_ISL_530358, EPI_ISL_530359, EPI_ISL_530360, EPI_ISL_530362, EPI_ISL_530363, EPI_ISL_530364, EPI_ISL_530365, EPI_ISL_530367, EPI_ISL_530368, EPI_ISL_530369, EPI_ISL_530371, EPI_ISL_530372, EPI_ISL_530373, EPI_ISL_530374, EPI_ISL_530375, EPI_ISL_530376, EPI_ISL_530378, EPI_ISL_530379, EPI_ISL_530380, EPI_ISL_530381, EPI_ISL_530382, EPI_ISL_530383, EPI_ISL_530384, EPI_ISL_530385, EPI_ISL_530387, EPI_ISL_530389, EPI_ISL_530390, EPI_ISL_530392 | | | |
| see above | Lighthouse Lab in Glasgow | Wellcome Sanger Institute for the COVID-19 Genomics UK (COG-UK) consortium | Harper VanSteenhouse, Yumi Kasai, David Gray, Carol Clugston, Anna Dominiczak and Alex Alderton, Roberto Amato, Sonia Goncalves, Ewan Harrison, David K. Jackson, Ian Johnston, Dominic Kwiatkowski, Cordelia Langford, John Sillitoe |
| EPI_ISL_530393 | Lighthouse Lab in Glasgow | Wellcome Sanger Institute for the COVID-19 Genomics UK (COG-UK) consortium | Harper VanSteenhouse, Yumi Kasai, David Gray, Carol Clugston, Anna Dominiczak and Alex Alderton, Roberto Amato, Sonia Goncalves, Ewan Harrison, David K. Jackson, Ian Johnston, Dominic Kwiatkowski, Cordelia Langford, John Sillitoe on behalf of the Wellcome Sanger Institute COVID-19 Surveillance Team (http://www.sanger.ac.uk/covid-team) |
| EPI_ISL_530394, EPI_ISL_530395, EPI_ISL_530396, EPI_ISL_530397, EPI_ISL_530399, EPI_ISL_530400, EPI_ISL_530401, EPI_ISL_530402, EPI_ISL_530403, EPI_ISL_530404, EPI_ISL_530405, EPI_ISL_530406, EPI_ISL_530407, EPI_ISL_530408, EPI_ISL_530409, EPI_ISL_530411, EPI_ISL_530412, EPI_ISL_530413, EPI_ISL_530414, EPI_ISL_530415, EPI_ISL_530416, EPI_ISL_530417, EPI_ISL_530418, EPI_ISL_530419, EPI_ISL_530420, EPI_ISL_530421, EPI_ISL_530422, EPI_ISL_530423, EPI_ISL_530424, EPI_ISL_530425, EPI_ISL_530426, EPI_ISL_530427, EPI_ISL_530428, EPI_ISL_530430, EPI_ISL_530431, EPI_ISL_530432, EPI_ISL_530433, EPI_ISL_530434, EPI_ISL_530436, EPI_ISL_530437, EPI_ISL_530438, EPI_ISL_530439, EPI_ISL_530441, EPI_ISL_530442, EPI_ISL_530443, EPI_ISL_530444, EPI_ISL_530445, EPI_ISL_530446, EPI_ISL_530447, EPI_ISL_530448, EPI_ISL_530449, EPI_ISL_530450, EPI_ISL_530451, EPI_ISL_530452, EPI_ISL_530453, EPI_ISL_530454, EPI_ISL_530455, EPI_ISL_530456, EPI_ISL_530457, EPI_ISL_530458, EPI_ISL_530459, EPI_ISL_530460, EPI_ISL_530461, EPI_ISL_530462, EPI_ISL_530463, EPI_ISL_530464, EPI_ISL_530465, EPI_ISL_530466, EPI_ISL_530467 | | | |
| see above | Lighthouse Lab in Glasgow | Wellcome Sanger Institute for the COVID-19 Genomics UK (COG-UK) consortium | Harper VanSteenhouse, Yumi Kasai, David Gray, Carol Clugston, Anna Dominiczak and Alex Alderton, Roberto Amato, Sonia Goncalves, Ewan Harrison, David K. Jackson, Ian Johnston, Dominic Kwiatkowski, Cordelia Langford, John Sillitoe |
| EPI_ISL_530469 | Lighthouse Lab in Glasgow | Wellcome Sanger Institute for the COVID-19 Genomics UK (COG-UK) consortium | Harper VanSteenhouse, Yumi Kasai, David Gray, Carol Clugston, Anna Dominiczak and Alex Alderton, Roberto Amato, Sonia Goncalves, Ewan Harrison, David K. Jackson, Ian Johnston, Dominic Kwiatkowski, Cordelia Langford, John Sillitoe on behalf of the Wellcome Sanger Institute COVID-19 Surveillance Team (http://www.sanger.ac.uk/covid-team) |
| EPI_ISL_530470, EPI_ISL_530471, EPI_ISL_530472, EPI_ISL_530473, EPI_ISL_530474, EPI_ISL_530475, EPI_ISL_530476, EPI_ISL_530477, EPI_ISL_530478, EPI_ISL_530479, EPI_ISL_530481, EPI_ISL_530483, EPI_ISL_530484, EPI_ISL_530485, EPI_ISL_530486, EPI_ISL_530487, EPI_ISL_530490, EPI_ISL_530491, EPI_ISL_530492, EPI_ISL_530493, EPI_ISL_530494, EPI_ISL_530495, EPI_ISL_530496 | | | |
| see above | Lighthouse Lab in Glasgow | Wellcome Sanger Institute for the COVID-19 Genomics UK (COG-UK) consortium | Harper VanSteenhouse, Yumi Kasai, David Gray, Carol Clugston, Anna Dominiczak and Alex Alderton, Roberto Amato, Sonia Goncalves, Ewan Harrison, David K. Jackson, Ian Johnston, Dominic Kwiatkowski, Cordelia Langford, John Sillitoe |
| EPI_ISL_530497 | Lighthouse Lab in Glasgow | Wellcome Sanger Institute for the COVID-19 Genomics UK (COG-UK) consortium | Harper VanSteenhouse, Yumi Kasai, David Gray, Carol Clugston, Anna Dominiczak and Alex Alderton, Roberto Amato, Sonia Goncalves, Ewan Harrison, David K. Jackson, Ian Johnston, Dominic Kwiatkowski, Cordelia Langford, John Sillitoe on behalf of the Wellcome Sanger Institute COVID-19 Surveillance Team (http://www.sanger.ac.uk/covid-team) |
| EPI_ISL_530498, EPI_ISL_530499, EPI_ISL_530500, EPI_ISL_530501, EPI_ISL_530502, EPI_ISL_530503, EPI_ISL_530504, EPI_ISL_530505, EPI_ISL_530506, EPI_ISL_530507, EPI_ISL_530508, EPI_ISL_530509, EPI_ISL_530510, EPI_ISL_530511, EPI_ISL_530512, EPI_ISL_530513, EPI_ISL_530514, EPI_ISL_530515, EPI_ISL_530516, EPI_ISL_530517, EPI_ISL_530518, EPI_ISL_530519, EPI_ISL_530520, EPI_ISL_530521, EPI_ISL_530522, EPI_ISL_530524, EPI_ISL_530525 | | | |
| see above | Lighthouse Lab in Glasgow | Wellcome Sanger Institute for the COVID-19 Genomics UK (COG-UK) consortium | Harper VanSteenhouse, Yumi Kasai, David Gray, Carol Clugston, Anna Dominiczak and Alex Alderton, Roberto Amato, Sonia Goncalves, Ewan Harrison, David K. Jackson, Ian Johnston, Dominic Kwiatkowski, Cordelia Langford, John Sillitoe |
| EPI_ISL_530526, EPI_ISL_530527 | Lighthouse Lab in Glasgow | Wellcome Sanger Institute for the COVID-19 Genomics UK (COG-UK) consortium | Harper VanSteenhouse, Yumi Kasai, David Gray, Carol Clugston, Anna Dominiczak and Alex Alderton, Roberto Amato, Sonia Goncalves, Ewan Harrison, David K. Jackson, Ian Johnston, Dominic Kwiatkowski, Cordelia Langford, John Sillitoe on behalf of the Wellcome Sanger Institute COVID-19 Surveillance Team (http://www.sanger.ac.uk/covid-team) |
| EPI_ISL_530529, EPI_ISL_530530, EPI_ISL_530531, EPI_ISL_530532, EPI_ISL_530534, EPI_ISL_530536, EPI_ISL_530537, EPI_ISL_530538, EPI_ISL_530539, EPI_ISL_530540, EPI_ISL_530541, EPI_ISL_530543, EPI_ISL_530544, EPI_ISL_530545, EPI_ISL_530546, EPI_ISL_530547, EPI_ISL_530548, EPI_ISL_530549, EPI_ISL_530550, EPI_ISL_530551, EPI_ISL_530552, EPI_ISL_530554, EPI_ISL_530555, EPI_ISL_530556, EPI_ISL_530557, EPI_ISL_530558, EPI_ISL_530559, EPI_ISL_530560, EPI_ISL_530561, EPI_ISL_530563, EPI_ISL_530564, EPI_ISL_530565, EPI_ISL_530566, EPI_ISL_530567, EPI_ISL_530568, EPI_ISL_530569, EPI_ISL_530570, EPI_ISL_530571, EPI_ISL_530572, EPI_ISL_530573, EPI_ISL_530574, EPI_ISL_530575, EPI_ISL_530576 | | | |
| see above | Lighthouse Lab in Glasgow | Wellcome Sanger Institute for the COVID-19 Genomics UK (COG-UK) consortium | Harper VanSteenhouse, Yumi Kasai, David Gray, Carol Clugston, Anna Dominiczak and Alex Alderton, Roberto Amato, Sonia Goncalves, Ewan Harrison, David K. Jackson, Ian Johnston, Dominic Kwiatkowski, Cordelia Langford, John Sillitoe |
| EPI_ISL_530577 | Lighthouse Lab in Glasgow | Wellcome Sanger Institute for the COVID-19 Genomics UK (COG-UK) consortium | Harper VanSteenhouse, Yumi Kasai, David Gray, Carol Clugston, Anna Dominiczak and Alex Alderton, Roberto Amato, Sonia Goncalves, Ewan Harrison, David K. Jackson, Ian Johnston, Dominic Kwiatkowski, Cordelia Langford, John Sillitoe on behalf of the Wellcome Sanger Institute COVID-19 Surveillance Team (http://www.sanger.ac.uk/covid-team) |
| EPI_ISL_530578, EPI_ISL_530579, EPI_ISL_530580, EPI_ISL_530581, EPI_ISL_530582, EPI_ISL_530583, EPI_ISL_530584, EPI_ISL_530585, EPI_ISL_530586, EPI_ISL_530587, EPI_ISL_530588, EPI_ISL_530589, EPI_ISL_530590, EPI_ISL_530591, EPI_ISL_530592, EPI_ISL_530593, EPI_ISL_530594, EPI_ISL_530595, EPI_ISL_530597, EPI_ISL_530598, EPI_ISL_530599, EPI_ISL_530600, EPI_ISL_530601, EPI_ISL_530602, EPI_ISL_530603, EPI_ISL_530604, EPI_ISL_530605, EPI_ISL_530606, EPI_ISL_530607, EPI_ISL_530608, EPI_ISL_530609, EPI_ISL_530610, EPI_ISL_530611, EPI_ISL_530612, EPI_ISL_530613, EPI_ISL_530614, EPI_ISL_530615, EPI_ISL_530616, EPI_ISL_530617, EPI_ISL_530618, EPI_ISL_530619, EPI_ISL_530620, EPI_ISL_530621, EPI_ISL_530622, EPI_ISL_530623, EPI_ISL_530624, EPI_ISL_530625, EPI_ISL_530626, EPI_ISL_530627, EPI_ISL_530628, EPI_ISL_530629, EPI_ISL_530630, EPI_ISL_530631, EPI_ISL_530632, EPI_ISL_530633, EPI_ISL_530634, EPI_ISL_530635, EPI_ISL_530636, EPI_ISL_530637, EPI_ISL_530638, EPI_ISL_530639, EPI_ISL_530640, EPI_ISL_530641, EPI_ISL_530643, EPI_ISL_530644, EPI_ISL_530645, EPI_ISL_530646, EPI_ISL_530647, EPI_ISL_530648, EPI_ISL_530649, EPI_ISL_530650, EPI_ISL_530651, EPI_ISL_530652, EPI_ISL_530653, EPI_ISL_530654, EPI_ISL_530655, EPI_ISL_530656, EPI_ISL_530657, EPI_ISL_530658, EPI_ISL_530660, EPI_ISL_530661, EPI_ISL_530662, EPI_ISL_530663, EPI_ISL_530664, EPI_ISL_530665, EPI_ISL_530666, EPI_ISL_530667, EPI_ISL_530668, EPI_ISL_530669, EPI_ISL_530670, EPI_ISL_530671, EPI_ISL_530672, EPI_ISL_530674, EPI_ISL_530675, EPI_ISL_530676, EPI_ISL_530677, EPI_ISL_530678, EPI_ISL_530679, EPI_ISL_530680, EPI_ISL_530681, EPI_ISL_530683, EPI_ISL_530684, EPI_ISL_530685 | | | |
| see above | Lighthouse Lab in Glasgow | Wellcome Sanger Institute for the COVID-19 Genomics UK | Harper VanSteenhouse, Yumi Kasai, David Gray, Carol Clugston, Anna Dominiczak and Alex Alderton, Roberto Amato, Sonia Goncalves, Ewan Harrison, |

| | | | |
|---|---|---|---|
| | | (COG-UK) consortium | David K. Jackson, Ian Johnston, Dominic Kwiatkowski, Cordelia Langford, John Sillitoe |
| EPI_ISL_530686 | Lighthouse Lab in Glasgow | Wellcome Sanger Institute for the COVID-19 Genomics UK (COG-UK) consortium | Harper VanSteenhouse, Yumi Kasai, David Gray, Carol Clugston, Anna Dominiczak and Alex Alderton, Roberto Amato, Sonia Goncalves, Ewan Harrison, David K. Jackson, Ian Johnston, Dominic Kwiatkowski, Cordelia Langford, John Sillitoe on behalf of the Wellcome Sanger Institute COVID-19 Surveillance Team (http://www.sanger.ac.uk/covid-team) |
| EPI_ISL_530687, EPI_ISL_530688, EPI_ISL_530690, EPI_ISL_530691, EPI_ISL_530692, EPI_ISL_530693, EPI_ISL_530694, EPI_ISL_530695, EPI_ISL_530696, EPI_ISL_530697, EPI_ISL_530698, EPI_ISL_530699, EPI_ISL_530700, EPI_ISL_530701, EPI_ISL_530702, EPI_ISL_530703, EPI_ISL_530704, EPI_ISL_530705, EPI_ISL_530706, EPI_ISL_530707, EPI_ISL_530709, EPI_ISL_530710, EPI_ISL_530711, EPI_ISL_530712, EPI_ISL_530713, EPI_ISL_530714, EPI_ISL_530715, EPI_ISL_530716, EPI_ISL_530717, EPI_ISL_530718, EPI_ISL_530719, EPI_ISL_530720, EPI_ISL_530721, EPI_ISL_530723, EPI_ISL_530724, EPI_ISL_530725, EPI_ISL_530726, EPI_ISL_530727, EPI_ISL_530728, EPI_ISL_530729, EPI_ISL_530730, EPI_ISL_530731, EPI_ISL_530732, EPI_ISL_530733, EPI_ISL_530734, EPI_ISL_530735, EPI_ISL_530736, EPI_ISL_530737, EPI_ISL_530738, EPI_ISL_530739, EPI_ISL_530741, EPI_ISL_530742, EPI_ISL_530744, EPI_ISL_530745, EPI_ISL_530747, EPI_ISL_530748, EPI_ISL_530749, EPI_ISL_530750, EPI_ISL_530751, EPI_ISL_530752, EPI_ISL_530753, EPI_ISL_530755, EPI_ISL_530756, EPI_ISL_530757, EPI_ISL_530758, EPI_ISL_530759, EPI_ISL_530760, EPI_ISL_530761, EPI_ISL_530762, EPI_ISL_530763, EPI_ISL_530765, EPI_ISL_530769, EPI_ISL_530770, EPI_ISL_530771, EPI_ISL_530772, EPI_ISL_530773, EPI_ISL_530774, EPI_ISL_530775, EPI_ISL_530776, EPI_ISL_530777, EPI_ISL_530778, EPI_ISL_530779, EPI_ISL_530780, EPI_ISL_530781, EPI_ISL_530782, EPI_ISL_530783, EPI_ISL_530784, EPI_ISL_530785, EPI_ISL_530786, EPI_ISL_530787, EPI_ISL_530788, EPI_ISL_530789, EPI_ISL_530790, EPI_ISL_530791, EPI_ISL_530792, EPI_ISL_530793, EPI_ISL_530794 | | | |
| see above | Lighthouse Lab in Glasgow | Wellcome Sanger Institute for the COVID-19 Genomics UK (COG-UK) consortium | Harper VanSteenhouse, Yumi Kasai, David Gray, Carol Clugston, Anna Dominiczak and Alex Alderton, Roberto Amato, Sonia Goncalves, Ewan Harrison, David K. Jackson, Ian Johnston, Dominic Kwiatkowski, Cordelia Langford, John Sillitoe |
| EPI_ISL_530795 | Lighthouse Lab in Glasgow | Wellcome Sanger Institute for the COVID-19 Genomics UK (COG-UK) consortium | Harper VanSteenhouse, Yumi Kasai, David Gray, Carol Clugston, Anna Dominiczak and Alex Alderton, Roberto Amato, Sonia Goncalves, Ewan Harrison, David K. Jackson, Ian Johnston, Dominic Kwiatkowski, Cordelia Langford, John Sillitoe on behalf of the Wellcome Sanger Institute COVID-19 Surveillance Team (http://www.sanger.ac.uk/covid-team) |
| EPI_ISL_530796, EPI_ISL_530797, EPI_ISL_530798, EPI_ISL_530799 | Lighthouse Lab in Glasgow | Wellcome Sanger Institute for the COVID-19 Genomics UK (COG-UK) consortium | Harper VanSteenhouse, Yumi Kasai, David Gray, Carol Clugston, Anna Dominiczak and Alex Alderton, Roberto Amato, Sonia Goncalves, Ewan Harrison, David K. Jackson, Ian Johnston, Dominic Kwiatkowski, Cordelia Langford, John Sillitoe |
| EPI_ISL_530800 | Lighthouse Lab in Glasgow | Wellcome Sanger Institute for the COVID-19 Genomics UK (COG-UK) consortium | Harper VanSteenhouse, Yumi Kasai, David Gray, Carol Clugston, Anna Dominiczak and Alex Alderton, Roberto Amato, Sonia Goncalves, Ewan Harrison, David K. Jackson, Ian Johnston, Dominic Kwiatkowski, Cordelia Langford, John Sillitoe on behalf of the Wellcome Sanger Institute COVID-19 Surveillance Team (http://www.sanger.ac.uk/covid-team) |
| EPI_ISL_530801, EPI_ISL_530802, EPI_ISL_530803, EPI_ISL_530804, EPI_ISL_530805, EPI_ISL_530806, EPI_ISL_530807, EPI_ISL_530808, EPI_ISL_530809, EPI_ISL_530810, EPI_ISL_530811, EPI_ISL_530812, EPI_ISL_530813, EPI_ISL_530814, EPI_ISL_530815, EPI_ISL_530816, EPI_ISL_530817, EPI_ISL_530818, EPI_ISL_530819, EPI_ISL_530820, EPI_ISL_530821, EPI_ISL_530822, EPI_ISL_530823, EPI_ISL_530824, EPI_ISL_530825, EPI_ISL_530827, EPI_ISL_530828, EPI_ISL_530829, EPI_ISL_530830, EPI_ISL_530831, EPI_ISL_530832, EPI_ISL_530833, EPI_ISL_530834, EPI_ISL_530835, EPI_ISL_530837, EPI_ISL_530838, EPI_ISL_530839, EPI_ISL_530840, EPI_ISL_530841, EPI_ISL_530842, EPI_ISL_530843, EPI_ISL_530844, EPI_ISL_530846, EPI_ISL_530847, EPI_ISL_530848, EPI_ISL_530849, EPI_ISL_530850, EPI_ISL_530851, EPI_ISL_530852, EPI_ISL_530853, EPI_ISL_530854, EPI_ISL_530855, EPI_ISL_530856, EPI_ISL_530857, EPI_ISL_530858, EPI_ISL_530859, EPI_ISL_530860, EPI_ISL_530861, EPI_ISL_530863, EPI_ISL_530864, EPI_ISL_530865, EPI_ISL_530866, EPI_ISL_530867 | | | |
| see above | Lighthouse Lab in Glasgow | Wellcome Sanger Institute for the COVID-19 Genomics UK (COG-UK) consortium | Harper VanSteenhouse, Yumi Kasai, David Gray, Carol Clugston, Anna Dominiczak and Alex Alderton, Roberto Amato, Sonia Goncalves, Ewan Harrison, David K. Jackson, Ian Johnston, Dominic Kwiatkowski, Cordelia Langford, John Sillitoe |
| EPI_ISL_530868 | Lighthouse Lab in Glasgow | Wellcome Sanger Institute for the COVID-19 Genomics UK (COG-UK) consortium | Harper VanSteenhouse, Yumi Kasai, David Gray, Carol Clugston, Anna Dominiczak and Alex Alderton, Roberto Amato, Sonia Goncalves, Ewan Harrison, David K. Jackson, Ian Johnston, Dominic Kwiatkowski, Cordelia Langford, John Sillitoe on behalf of the Wellcome Sanger Institute COVID-19 Surveillance Team (http://www.sanger.ac.uk/covid-team) |
| EPI_ISL_530869, EPI_ISL_530870, EPI_ISL_530872 | Lighthouse Lab in Glasgow | Wellcome Sanger Institute for the COVID-19 Genomics UK (COG-UK) consortium | Harper VanSteenhouse, Yumi Kasai, David Gray, Carol Clugston, Anna Dominiczak and Alex Alderton, Roberto Amato, Sonia Goncalves, Ewan Harrison, David K. Jackson, Ian Johnston, Dominic Kwiatkowski, Cordelia Langford, John Sillitoe |
| EPI_ISL_530874 | Lighthouse Lab in Glasgow | Wellcome Sanger Institute for the COVID-19 Genomics UK (COG-UK) consortium | Harper VanSteenhouse, Yumi Kasai, David Gray, Carol Clugston, Anna Dominiczak and Alex Alderton, Roberto Amato, Sonia Goncalves, Ewan Harrison, David K. Jackson, Ian Johnston, Dominic Kwiatkowski, Cordelia Langford, John Sillitoe on behalf of the Wellcome Sanger Institute COVID-19 Surveillance Team (http://www.sanger.ac.uk/covid-team) |
| EPI_ISL_530875, EPI_ISL_530876, EPI_ISL_530877, EPI_ISL_530878, EPI_ISL_530879, EPI_ISL_530880, EPI_ISL_530881, EPI_ISL_530882, EPI_ISL_530883, EPI_ISL_530884, EPI_ISL_530885, EPI_ISL_530886, EPI_ISL_530887, EPI_ISL_530888, EPI_ISL_530889, EPI_ISL_530890, EPI_ISL_530891, EPI_ISL_530892, EPI_ISL_530893, EPI_ISL_530894, EPI_ISL_530895, EPI_ISL_530896, EPI_ISL_530897 | | | |
| see above | Lighthouse Lab in Glasgow | Wellcome Sanger Institute for the COVID-19 Genomics UK (COG-UK) consortium | Harper VanSteenhouse, Yumi Kasai, David Gray, Carol Clugston, Anna Dominiczak and Alex Alderton, Roberto Amato, Sonia Goncalves, Ewan Harrison, David K. Jackson, Ian Johnston, Dominic Kwiatkowski, Cordelia Langford, John Sillitoe |
| EPI_ISL_530898 | Lighthouse Lab in Glasgow | Wellcome Sanger Institute for the COVID-19 Genomics UK (COG-UK) consortium | Harper VanSteenhouse, Yumi Kasai, David Gray, Carol Clugston, Anna Dominiczak and Alex Alderton, Roberto Amato, Sonia Goncalves, Ewan Harrison, David K. Jackson, Ian Johnston, Dominic Kwiatkowski, Cordelia Langford, John Sillitoe on behalf of the Wellcome Sanger Institute COVID-19 Surveillance Team (http://www.sanger.ac.uk/covid-team) |
| EPI_ISL_530900, EPI_ISL_530901, EPI_ISL_530902, EPI_ISL_530903, EPI_ISL_530905, EPI_ISL_530907, EPI_ISL_530908, EPI_ISL_530909, EPI_ISL_530910, EPI_ISL_530911, EPI_ISL_530912, EPI_ISL_530913, EPI_ISL_530914, EPI_ISL_530915, EPI_ISL_530916, EPI_ISL_530917, EPI_ISL_530918, EPI_ISL_530919, EPI_ISL_530920, EPI_ISL_530921, EPI_ISL_530922, EPI_ISL_530924, EPI_ISL_530925, EPI_ISL_530926, EPI_ISL_530927, EPI_ISL_530928, EPI_ISL_530930, EPI_ISL_530931, EPI_ISL_530932, EPI_ISL_530933, EPI_ISL_530934, EPI_ISL_530935, EPI_ISL_530936, EPI_ISL_530937, EPI_ISL_530938, EPI_ISL_530939, EPI_ISL_530940, EPI_ISL_530941, EPI_ISL_530942, EPI_ISL_530943, EPI_ISL_530944, EPI_ISL_530945, EPI_ISL_530946, EPI_ISL_530947, EPI_ISL_530948, EPI_ISL_530949, EPI_ISL_530950, EPI_ISL_530951, EPI_ISL_530952, EPI_ISL_530954, EPI_ISL_530955, EPI_ISL_530956, EPI_ISL_530957, EPI_ISL_530958, EPI_ISL_530960, EPI_ISL_530964, EPI_ISL_530965, EPI_ISL_530966, EPI_ISL_530967, EPI_ISL_530968, EPI_ISL_530969, EPI_ISL_530970, EPI_ISL_530971, EPI_ISL_530972, EPI_ISL_530974, EPI_ISL_530975, EPI_ISL_530977, EPI_ISL_530978, EPI_ISL_530980, EPI_ISL_530981, EPI_ISL_530982, EPI_ISL_530984, EPI_ISL_530987, EPI_ISL_530988, EPI_ISL_530989, EPI_ISL_530991, EPI_ISL_530992, EPI_ISL_530993, EPI_ISL_530994, EPI_ISL_530995, EPI_ISL_530998, EPI_ISL_530999, EPI_ISL_531000, EPI_ISL_531001, EPI_ISL_531003, EPI_ISL_531005, EPI_ISL_531007, EPI_ISL_531008, EPI_ISL_531009, EPI_ISL_531010, EPI_ISL_531011, EPI_ISL_531012, EPI_ISL_531013, EPI_ISL_531014, EPI_ISL_531015, EPI_ISL_531016, EPI_ISL_531018, EPI_ISL_531019, EPI_ISL_531021, EPI_ISL_531022, EPI_ISL_531023, EPI_ISL_531024 | | | |
| see above | Lighthouse Lab in Glasgow | Wellcome Sanger Institute for the COVID-19 Genomics UK (COG-UK) consortium | Harper VanSteenhouse, Yumi Kasai, David Gray, Carol Clugston, Anna Dominiczak and Alex Alderton, Roberto Amato, Sonia Goncalves, Ewan Harrison, David K. Jackson, Ian Johnston, Dominic Kwiatkowski, Cordelia Langford, John Sillitoe |
| EPI_ISL_531025 | Lighthouse Lab in Glasgow | Wellcome Sanger Institute for the COVID-19 Genomics UK (COG-UK) consortium | Harper VanSteenhouse, Yumi Kasai, David Gray, Carol Clugston, Anna Dominiczak and Alex Alderton, Roberto Amato, Sonia Goncalves, Ewan Harrison, David K. Jackson, Ian Johnston, Dominic Kwiatkowski, Cordelia Langford, John Sillitoe on behalf of the Wellcome Sanger Institute COVID-19 Surveillance Team (http://www.sanger.ac.uk/covid-team) |
| EPI_ISL_531026, EPI_ISL_531027, EPI_ISL_531028, EPI_ISL_531029, EPI_ISL_531030, EPI_ISL_531031, EPI_ISL_531034, EPI_ISL_531035, EPI_ISL_531036, EPI_ISL_531037, EPI_ISL_531039, EPI_ISL_531041, EPI_ISL_531042, EPI_ISL_531043, EPI_ISL_531044, EPI_ISL_531046, EPI_ISL_531047, EPI_ISL_531050 | | | |
| see above | Lighthouse Lab in Glasgow | Wellcome Sanger Institute for the COVID-19 Genomics UK (COG-UK) consortium | Harper VanSteenhouse, Yumi Kasai, David Gray, Carol Clugston, Anna Dominiczak and Alex Alderton, Roberto Amato, Sonia Goncalves, Ewan Harrison, David K. Jackson, Ian Johnston, Dominic Kwiatkowski, Cordelia Langford, John Sillitoe |
| EPI_ISL_531051 | Lighthouse Lab in Glasgow | Wellcome Sanger Institute for the COVID-19 Genomics UK (COG-UK) consortium | Harper VanSteenhouse, Yumi Kasai, David Gray, Carol Clugston, Anna Dominiczak and Alex Alderton, Roberto Amato, Sonia Goncalves, Ewan Harrison, David K. Jackson, Ian Johnston, Dominic Kwiatkowski, Cordelia Langford, John Sillitoe on behalf of the Wellcome Sanger Institute COVID-19 Surveillance Team (http://www.sanger.ac.uk/covid-team) |
| EPI_ISL_531053, EPI_ISL_531055, EPI_ISL_531058, EPI_ISL_531059, EPI_ISL_531060, EPI_ISL_531061, EPI_ISL_531062, EPI_ISL_531063, EPI_ISL_531064, EPI_ISL_531066, EPI_ISL_531067, EPI_ISL_531068, EPI_ISL_531069, EPI_ISL_531070, EPI_ISL_531071, EPI_ISL_531072, EPI_ISL_531073, EPI_ISL_531074, EPI_ISL_531075, EPI_ISL_531076, EPI_ISL_531077, EPI_ISL_531078, EPI_ISL_531080, EPI_ISL_531081, EPI_ISL_531082, EPI_ISL_531084, EPI_ISL_531085, EPI_ISL_531086, EPI_ISL_531088, EPI_ISL_531089, EPI_ISL_531090, EPI_ISL_531092, EPI_ISL_531093, EPI_ISL_531094, EPI_ISL_531095, EPI_ISL_531097, EPI_ISL_531098, EPI_ISL_531101, EPI_ISL_531102, EPI_ISL_531105, EPI_ISL_531106, EPI_ISL_531107, EPI_ISL_531108, EPI_ISL_531109, EPI_ISL_531110, EPI_ISL_531111, EPI_ISL_531112, EPI_ISL_531113, EPI_ISL_531115, EPI_ISL_531117, EPI_ISL_531119, EPI_ISL_531120, EPI_ISL_531121, EPI_ISL_531122, EPI_ISL_531124, EPI_ISL_531125, EPI_ISL_531127, EPI_ISL_531128, EPI_ISL_531129, EPI_ISL_531132, EPI_ISL_531133, EPI_ISL_531134, EPI_ISL_531135, EPI_ISL_531136, EPI_ISL_531137, EPI_ISL_531138, EPI_ISL_531139, EPI_ISL_531140, EPI_ISL_531141, EPI_ISL_531143, EPI_ISL_531144, EPI_ISL_531145, EPI_ISL_531146, EPI_ISL_531148, EPI_ISL_531149, EPI_ISL_531150, EPI_ISL_531151, EPI_ISL_531152, EPI_ISL_531153, EPI_ISL_531158, EPI_ISL_531160, EPI_ISL_531161, EPI_ISL_531162, EPI_ISL_531163, EPI_ISL_531164, EPI_ISL_531165, EPI_ISL_531167, EPI_ISL_531168, EPI_ISL_531169, EPI_ISL_531170, EPI_ISL_531171, EPI_ISL_531172, EPI_ISL_531173, EPI_ISL_531176, EPI_ISL_531177, EPI_ISL_531178, EPI_ISL_531179, EPI_ISL_531180, EPI_ISL_531181, EPI_ISL_531182, EPI_ISL_531183, EPI_ISL_531184, EPI_ISL_531185, EPI_ISL_531186, EPI_ISL_531187, EPI_ISL_531190, EPI_ISL_531191, EPI_ISL_531192, EPI_ISL_531193, EPI_ISL_531194, EPI_ISL_531195, EPI_ISL_531196, EPI_ISL_531197, EPI_ISL_531199, EPI_ISL_531200, EPI_ISL_531201, EPI_ISL_531202, EPI_ISL_531203, EPI_ISL_531204, EPI_ISL_531205, EPI_ISL_531206, EPI_ISL_531208, EPI_ISL_531209, EPI_ISL_531210, EPI_ISL_531213, EPI_ISL_531215, EPI_ISL_531216, EPI_ISL_531217, EPI_ISL_531218, EPI_ISL_531219, EPI_ISL_531220, EPI_ISL_531221, EPI_ISL_531222, EPI_ISL_531224 | | | |
| see above | Lighthouse Lab in Glasgow | Wellcome Sanger Institute for the COVID-19 Genomics UK (COG-UK) consortium | Harper VanSteenhouse, Yumi Kasai, David Gray, Carol Clugston, Anna Dominiczak and Alex Alderton, Roberto Amato, Sonia Goncalves, Ewan Harrison, David K. Jackson, Ian Johnston, Dominic Kwiatkowski, Cordelia Langford, John Sillitoe |
| EPI_ISL_531225 | Lighthouse Lab in Glasgow | Wellcome Sanger Institute for the COVID-19 Genomics UK (COG-UK) consortium | Harper VanSteenhouse, Yumi Kasai, David Gray, Carol Clugston, Anna Dominiczak and Alex Alderton, Roberto Amato, Sonia Goncalves, Ewan Harrison, David K. Jackson, Ian Johnston, Dominic Kwiatkowski, Cordelia Langford, John Sillitoe on behalf of the Wellcome Sanger Institute COVID-19 Surveillance Team (http://www.sanger.ac.uk/covid-team) |
| EPI_ISL_531226, EPI_ISL_531227, EPI_ISL_531228, EPI_ISL_531229, EPI_ISL_531230 | Lighthouse Lab in Glasgow | Wellcome Sanger Institute for the COVID-19 Genomics UK (COG-UK) consortium | Harper VanSteenhouse, Yumi Kasai, David Gray, Carol Clugston, Anna Dominiczak and Alex Alderton, Roberto Amato, Sonia Goncalves, Ewan Harrison, David K. Jackson, Ian Johnston, Dominic Kwiatkowski, Cordelia Langford, John Sillitoe |
| EPI_ISL_531231 | Lighthouse Lab in Glasgow | Wellcome Sanger Institute for the COVID-19 Genomics UK | Harper VanSteenhouse, Yumi Kasai, David Gray, Carol Clugston, Anna Dominiczak and Alex Alderton, Roberto Amato, Sonia Goncalves, Ewan Harrison, |

| | | | |
|---|---|---|---|
| | | (COG-UK) consortium | David K. Jackson, Ian Johnston, Dominic Kwiatkowski, Cordelia Langford, John Sillitoe on behalf of the Wellcome Sanger Institute COVID-19 Surveillance Team (http://www.sanger.ac.uk/covid-team) |
| EPI_ISL_531232, EPI_ISL_531233, EPI_ISL_531234, EPI_ISL_531235, EPI_ISL_531236, EPI_ISL_531237, EPI_ISL_531238, EPI_ISL_531239, EPI_ISL_531240, EPI_ISL_531241, EPI_ISL_531242, EPI_ISL_531243, EPI_ISL_531244, EPI_ISL_531245, EPI_ISL_531246, EPI_ISL_531247, EPI_ISL_531248, EPI_ISL_531250, EPI_ISL_531251, EPI_ISL_531252, EPI_ISL_531253, EPI_ISL_531254, EPI_ISL_531255, EPI_ISL_531256, EPI_ISL_531257, EPI_ISL_531258, EPI_ISL_531259, EPI_ISL_531260, EPI_ISL_531261, EPI_ISL_531262, EPI_ISL_531263, EPI_ISL_531264, EPI_ISL_531265, EPI_ISL_531267, EPI_ISL_531268, EPI_ISL_531269, EPI_ISL_531270, EPI_ISL_531271, EPI_ISL_531272, EPI_ISL_531273, EPI_ISL_531274, EPI_ISL_531275, EPI_ISL_531276, EPI_ISL_531277, EPI_ISL_531278, EPI_ISL_531279, EPI_ISL_531280, EPI_ISL_531281, EPI_ISL_531282, EPI_ISL_531283, EPI_ISL_531284, EPI_ISL_531285, EPI_ISL_531286, EPI_ISL_531287, EPI_ISL_531288, EPI_ISL_531289, EPI_ISL_531290, EPI_ISL_531291, EPI_ISL_531292, EPI_ISL_531293, EPI_ISL_531294, EPI_ISL_531295, EPI_ISL_531296, EPI_ISL_531297, EPI_ISL_531298, EPI_ISL_531299, EPI_ISL_531300, EPI_ISL_531301, EPI_ISL_531302, EPI_ISL_531303, EPI_ISL_531304, EPI_ISL_531305, EPI_ISL_531306, EPI_ISL_531307, EPI_ISL_531308, EPI_ISL_531309, EPI_ISL_531310, EPI_ISL_531311, EPI_ISL_531312, EPI_ISL_531313, EPI_ISL_531314, EPI_ISL_531315, EPI_ISL_531316, EPI_ISL_531317, EPI_ISL_531318, EPI_ISL_531319, EPI_ISL_531320, EPI_ISL_531322, EPI_ISL_531323, EPI_ISL_531324, EPI_ISL_531325, EPI_ISL_531326, EPI_ISL_531327, EPI_ISL_531328, EPI_ISL_531329, EPI_ISL_531331, EPI_ISL_531332, EPI_ISL_531333, EPI_ISL_531334, EPI_ISL_531335, EPI_ISL_531336, EPI_ISL_531337, EPI_ISL_531338, EPI_ISL_531339, EPI_ISL_531340, EPI_ISL_531341, EPI_ISL_531342, EPI_ISL_531343, EPI_ISL_531344, EPI_ISL_531345, EPI_ISL_531346, EPI_ISL_531347, EPI_ISL_531348, EPI_ISL_531349, EPI_ISL_531350, EPI_ISL_531351, EPI_ISL_531352, EPI_ISL_531353, EPI_ISL_531354, EPI_ISL_531355, EPI_ISL_531356, EPI_ISL_531357, EPI_ISL_531358, EPI_ISL_531359, EPI_ISL_531360, EPI_ISL_531361, EPI_ISL_531362, EPI_ISL_531363, EPI_ISL_531364, EPI_ISL_531366, EPI_ISL_531367, EPI_ISL_531368, EPI_ISL_531369, EPI_ISL_531370, EPI_ISL_531371, EPI_ISL_531372, EPI_ISL_531373, EPI_ISL_531375, EPI_ISL_531376, EPI_ISL_531377, EPI_ISL_531378, EPI_ISL_531379, EPI_ISL_531380 | | | |
| see above | Lighthouse Lab in Glasgow | Wellcome Sanger Institute for the COVID-19 Genomics UK (COG-UK) consortium | Harper VanSteenhouse, Yumi Kasai, David Gray, Carol Clugston, Anna Dominiczak and Alex Alderton, Roberto Amato, Sonia Goncalves, Ewan Harrison, David K. Jackson, Ian Johnston, Dominic Kwiatkowski, Cordelia Langford, John Sillitoe |
| EPI_ISL_531381 | Lighthouse Lab in Glasgow | Wellcome Sanger Institute for the COVID-19 Genomics UK (COG-UK) consortium | Harper VanSteenhouse, Yumi Kasai, David Gray, Carol Clugston, Anna Dominiczak and Alex Alderton, Roberto Amato, Sonia Goncalves, Ewan Harrison, David K. Jackson, Ian Johnston, Dominic Kwiatkowski, Cordelia Langford, John Sillitoe on behalf of the Wellcome Sanger Institute COVID-19 Surveillance Team (http://www.sanger.ac.uk/covid-team) |
| EPI_ISL_531383, EPI_ISL_531384, EPI_ISL_531385, EPI_ISL_531386, EPI_ISL_531387, EPI_ISL_531388, EPI_ISL_531389, EPI_ISL_531390, EPI_ISL_531391, EPI_ISL_531392, EPI_ISL_531393, EPI_ISL_531394, EPI_ISL_531395, EPI_ISL_531396, EPI_ISL_531397, EPI_ISL_531398, EPI_ISL_531399, EPI_ISL_531400, EPI_ISL_531401, EPI_ISL_531402, EPI_ISL_531403, EPI_ISL_531404, EPI_ISL_531405, EPI_ISL_531406, EPI_ISL_531407, EPI_ISL_531408, EPI_ISL_531409, EPI_ISL_531410, EPI_ISL_531411, EPI_ISL_531413, EPI_ISL_531414, EPI_ISL_531415, EPI_ISL_531417, EPI_ISL_531418, EPI_ISL_531419, EPI_ISL_531421, EPI_ISL_531423, EPI_ISL_531424, EPI_ISL_531425, EPI_ISL_531426, EPI_ISL_531427, EPI_ISL_531428, EPI_ISL_531429, EPI_ISL_531430, EPI_ISL_531431, EPI_ISL_531433, EPI_ISL_531434, EPI_ISL_531435, EPI_ISL_531436, EPI_ISL_531437, EPI_ISL_531438, EPI_ISL_531439, EPI_ISL_531440, EPI_ISL_531441, EPI_ISL_531443, EPI_ISL_531444, EPI_ISL_531445, EPI_ISL_531446, EPI_ISL_531447, EPI_ISL_531448, EPI_ISL_531449, EPI_ISL_531450, EPI_ISL_531451, EPI_ISL_531452, EPI_ISL_531453, EPI_ISL_531454, EPI_ISL_531455, EPI_ISL_531456, EPI_ISL_531457, EPI_ISL_531458, EPI_ISL_531459, EPI_ISL_531460, EPI_ISL_531461, EPI_ISL_531462, EPI_ISL_531463, EPI_ISL_531465, EPI_ISL_531466, EPI_ISL_531467, EPI_ISL_531468, EPI_ISL_531469, EPI_ISL_531471, EPI_ISL_531472, EPI_ISL_531474, EPI_ISL_531475, EPI_ISL_531478, EPI_ISL_531479, EPI_ISL_531481, EPI_ISL_531482, EPI_ISL_531486, EPI_ISL_531487, EPI_ISL_531488, EPI_ISL_531489, EPI_ISL_531490, EPI_ISL_531491, EPI_ISL_531493, EPI_ISL_531494, EPI_ISL_531496, EPI_ISL_531497, EPI_ISL_531498, EPI_ISL_531499, EPI_ISL_531501, EPI_ISL_531502, EPI_ISL_531503, EPI_ISL_531505, EPI_ISL_531506, EPI_ISL_531507, EPI_ISL_531508, EPI_ISL_531509, EPI_ISL_531510, EPI_ISL_531511, EPI_ISL_531512, EPI_ISL_531513, EPI_ISL_531514, EPI_ISL_531515, EPI_ISL_531516, EPI_ISL_531517, EPI_ISL_531518, EPI_ISL_531519, EPI_ISL_531520, EPI_ISL_531521, EPI_ISL_531522, EPI_ISL_531523, EPI_ISL_531525, EPI_ISL_531526, EPI_ISL_531527, EPI_ISL_531528, EPI_ISL_531529, EPI_ISL_531530, EPI_ISL_531531, EPI_ISL_531532, EPI_ISL_531533, EPI_ISL_531534, EPI_ISL_531535, EPI_ISL_531536, EPI_ISL_531537, EPI_ISL_531538, EPI_ISL_531539, EPI_ISL_531540, EPI_ISL_531541, EPI_ISL_531542, EPI_ISL_531543, EPI_ISL_531544, EPI_ISL_531545, EPI_ISL_531547, EPI_ISL_531549, EPI_ISL_531550, EPI_ISL_531551, EPI_ISL_531552, EPI_ISL_531553, EPI_ISL_531554, EPI_ISL_531555, EPI_ISL_531556, EPI_ISL_531558, EPI_ISL_531559, EPI_ISL_531560, EPI_ISL_531561, EPI_ISL_531562, EPI_ISL_531566, EPI_ISL_531567, EPI_ISL_531568, EPI_ISL_531571, EPI_ISL_531572, EPI_ISL_531573, EPI_ISL_531574, EPI_ISL_531576, EPI_ISL_531578, EPI_ISL_531579, EPI_ISL_531580, EPI_ISL_531581, EPI_ISL_531582, EPI_ISL_531584 | | | |
| see above | Lighthouse Lab in Glasgow | Wellcome Sanger Institute for the COVID-19 Genomics UK (COG-UK) consortium | Harper VanSteenhouse, Yumi Kasai, David Gray, Carol Clugston, Anna Dominiczak and Alex Alderton, Roberto Amato, Sonia Goncalves, Ewan Harrison, David K. Jackson, Ian Johnston, Dominic Kwiatkowski, Cordelia Langford, John Sillitoe |
| EPI_ISL_531585 | Lighthouse Lab in Glasgow | Wellcome Sanger Institute for the COVID-19 Genomics UK (COG-UK) consortium | Harper VanSteenhouse, Yumi Kasai, David Gray, Carol Clugston, Anna Dominiczak and Alex Alderton, Roberto Amato, Sonia Goncalves, Ewan Harrison, David K. Jackson, Ian Johnston, Dominic Kwiatkowski, Cordelia Langford, John Sillitoe on behalf of the Wellcome Sanger Institute COVID-19 Surveillance Team (http://www.sanger.ac.uk/covid-team) |
| EPI_ISL_531586, EPI_ISL_531587, EPI_ISL_531588, EPI_ISL_531589, EPI_ISL_531591, EPI_ISL_531592, EPI_ISL_531593, EPI_ISL_531594, EPI_ISL_531596, EPI_ISL_531597, EPI_ISL_531600, EPI_ISL_531601, EPI_ISL_531602, EPI_ISL_531603, EPI_ISL_531604, EPI_ISL_531605, EPI_ISL_531606, EPI_ISL_531608, EPI_ISL_531610, EPI_ISL_531611, EPI_ISL_531612, EPI_ISL_531613, EPI_ISL_531614, EPI_ISL_531615, EPI_ISL_531616, EPI_ISL_531619, EPI_ISL_531620, EPI_ISL_531621, EPI_ISL_531622, EPI_ISL_531623, EPI_ISL_531624, EPI_ISL_531626, EPI_ISL_531627, EPI_ISL_531628, EPI_ISL_531629, EPI_ISL_531630, EPI_ISL_531632, EPI_ISL_531633, EPI_ISL_531635, EPI_ISL_531636, EPI_ISL_531637, EPI_ISL_531640, EPI_ISL_531641, EPI_ISL_531642, EPI_ISL_531643, EPI_ISL_531644, EPI_ISL_531645, EPI_ISL_531646, EPI_ISL_531647, EPI_ISL_531648, EPI_ISL_531650, EPI_ISL_531651, EPI_ISL_531652, EPI_ISL_531653, EPI_ISL_531654, EPI_ISL_531655, EPI_ISL_531656, EPI_ISL_531657, EPI_ISL_531658, EPI_ISL_531659, EPI_ISL_531661, EPI_ISL_531662, EPI_ISL_531663, EPI_ISL_531664, EPI_ISL_531665, EPI_ISL_531666, EPI_ISL_531667, EPI_ISL_531669, EPI_ISL_531670, EPI_ISL_531671, EPI_ISL_531673, EPI_ISL_531674, EPI_ISL_531675, EPI_ISL_531676, EPI_ISL_531679, EPI_ISL_531680, EPI_ISL_531681, EPI_ISL_531682, EPI_ISL_531683, EPI_ISL_531684, EPI_ISL_531685, EPI_ISL_531686, EPI_ISL_531692, EPI_ISL_531693, EPI_ISL_531694, EPI_ISL_531695, EPI_ISL_531696, EPI_ISL_531697, EPI_ISL_531698, EPI_ISL_531699, EPI_ISL_531700, EPI_ISL_531701, EPI_ISL_531704, EPI_ISL_531705, EPI_ISL_531706, EPI_ISL_531707, EPI_ISL_531708, EPI_ISL_531709, EPI_ISL_531710, EPI_ISL_531711, EPI_ISL_531712, EPI_ISL_531713, EPI_ISL_531715, EPI_ISL_531716, EPI_ISL_531717, EPI_ISL_531718, EPI_ISL_531719, EPI_ISL_531720, EPI_ISL_531721, EPI_ISL_531722, EPI_ISL_531723, EPI_ISL_531724, EPI_ISL_531725 | | | |
| see above | Lighthouse Lab in Glasgow | Wellcome Sanger Institute for the COVID-19 Genomics UK (COG-UK) consortium | Harper VanSteenhouse, Yumi Kasai, David Gray, Carol Clugston, Anna Dominiczak and Alex Alderton, Roberto Amato, Sonia Goncalves, Ewan Harrison, David K. Jackson, Ian Johnston, Dominic Kwiatkowski, Cordelia Langford, John Sillitoe |
| EPI_ISL_531726 | Lighthouse Lab in Glasgow | Wellcome Sanger Institute for the COVID-19 Genomics UK (COG-UK) consortium | Harper VanSteenhouse, Yumi Kasai, David Gray, Carol Clugston, Anna Dominiczak and Alex Alderton, Roberto Amato, Sonia Goncalves, Ewan Harrison, David K. Jackson, Ian Johnston, Dominic Kwiatkowski, Cordelia Langford, John Sillitoe on behalf of the Wellcome Sanger Institute COVID-19 Surveillance Team (http://www.sanger.ac.uk/covid-team) |
| EPI_ISL_531728, EPI_ISL_531729, EPI_ISL_531731, EPI_ISL_531733 | Lighthouse Lab in Glasgow | Wellcome Sanger Institute for the COVID-19 Genomics UK (COG-UK) consortium | Harper VanSteenhouse, Yumi Kasai, David Gray, Carol Clugston, Anna Dominiczak and Alex Alderton, Roberto Amato, Sonia Goncalves, Ewan Harrison, David K. Jackson, Ian Johnston, Dominic Kwiatkowski, Cordelia Langford, John Sillitoe |
| EPI_ISL_531734 | Lighthouse Lab in Glasgow | Wellcome Sanger Institute for the COVID-19 Genomics UK (COG-UK) consortium | Harper VanSteenhouse, Yumi Kasai, David Gray, Carol Clugston, Anna Dominiczak and Alex Alderton, Roberto Amato, Sonia Goncalves, Ewan Harrison, David K. Jackson, Ian Johnston, Dominic Kwiatkowski, Cordelia Langford, John Sillitoe on behalf of the Wellcome Sanger Institute COVID-19 Surveillance Team (http://www.sanger.ac.uk/covid-team) |
| EPI_ISL_531735, EPI_ISL_531736, EPI_ISL_531737, EPI_ISL_531738, EPI_ISL_531739, EPI_ISL_531740, EPI_ISL_531741, EPI_ISL_531742, EPI_ISL_531744, EPI_ISL_531745, EPI_ISL_531746, EPI_ISL_531747, EPI_ISL_531748, EPI_ISL_531749, EPI_ISL_531750, EPI_ISL_531751, EPI_ISL_531752, EPI_ISL_531756, EPI_ISL_531757, EPI_ISL_531759, EPI_ISL_531760, EPI_ISL_531761, EPI_ISL_531762, EPI_ISL_531763, EPI_ISL_531764, EPI_ISL_531765, EPI_ISL_531766, EPI_ISL_531767, EPI_ISL_531769, EPI_ISL_531770, EPI_ISL_531771, EPI_ISL_531772, EPI_ISL_531773, EPI_ISL_531774, EPI_ISL_531775, EPI_ISL_531776, EPI_ISL_531777, EPI_ISL_531778, EPI_ISL_531780, EPI_ISL_531781, EPI_ISL_531782, EPI_ISL_531783, EPI_ISL_531784, EPI_ISL_531785, EPI_ISL_531786, EPI_ISL_531787, EPI_ISL_531789, EPI_ISL_531790, EPI_ISL_531791, EPI_ISL_531793, EPI_ISL_531794, EPI_ISL_531795, EPI_ISL_531796, EPI_ISL_531797, EPI_ISL_531798, EPI_ISL_531799, EPI_ISL_531800, EPI_ISL_531803, EPI_ISL_531804, EPI_ISL_531806, EPI_ISL_531807, EPI_ISL_531808, EPI_ISL_531809, EPI_ISL_531810, EPI_ISL_531811, EPI_ISL_531812, EPI_ISL_531813, EPI_ISL_531816, EPI_ISL_531817, EPI_ISL_531818, EPI_ISL_531819, EPI_ISL_531820, EPI_ISL_531822, EPI_ISL_531823, EPI_ISL_531825 | | | |
| see above | Lighthouse Lab in Glasgow | Wellcome Sanger Institute for the COVID-19 Genomics UK (COG-UK) consortium | Harper VanSteenhouse, Yumi Kasai, David Gray, Carol Clugston, Anna Dominiczak and Alex Alderton, Roberto Amato, Sonia Goncalves, Ewan Harrison, David K. Jackson, Ian Johnston, Dominic Kwiatkowski, Cordelia Langford, John Sillitoe |
| EPI_ISL_531826 | Lighthouse Lab in Glasgow | Wellcome Sanger Institute for the COVID-19 Genomics UK (COG-UK) consortium | Harper VanSteenhouse, Yumi Kasai, David Gray, Carol Clugston, Anna Dominiczak and Alex Alderton, Roberto Amato, Sonia Goncalves, Ewan Harrison, David K. Jackson, Ian Johnston, Dominic Kwiatkowski, Cordelia Langford, John Sillitoe on behalf of the Wellcome Sanger Institute COVID-19 Surveillance Team (http://www.sanger.ac.uk/covid-team) |
| EPI_ISL_531827, EPI_ISL_531828, EPI_ISL_531829, EPI_ISL_531830, EPI_ISL_531831, EPI_ISL_531833, EPI_ISL_531834, EPI_ISL_531835, EPI_ISL_531836, EPI_ISL_531837, EPI_ISL_531838, EPI_ISL_531839, EPI_ISL_531840, EPI_ISL_531841, EPI_ISL_531843, EPI_ISL_531844, EPI_ISL_531845, EPI_ISL_531846, EPI_ISL_531847, EPI_ISL_531848, EPI_ISL_531849, EPI_ISL_531850, EPI_ISL_531852, EPI_ISL_531853, EPI_ISL_531854, EPI_ISL_531855, EPI_ISL_531859, EPI_ISL_531860, EPI_ISL_531861, EPI_ISL_531866, EPI_ISL_531870, EPI_ISL_531878, EPI_ISL_531895, EPI_ISL_531896, EPI_ISL_531897, EPI_ISL_531903, EPI_ISL_531911, EPI_ISL_531921, EPI_ISL_531924, EPI_ISL_531939, EPI_ISL_531940, EPI_ISL_531941, EPI_ISL_531946, EPI_ISL_531947, EPI_ISL_531948, EPI_ISL_531955, EPI_ISL_531958, EPI_ISL_531960, EPI_ISL_531964, EPI_ISL_531966, EPI_ISL_531967, EPI_ISL_531970, EPI_ISL_531977, EPI_ISL_531983, EPI_ISL_531985, EPI_ISL_531986, EPI_ISL_531992, EPI_ISL_532000, EPI_ISL_532009, EPI_ISL_532016, EPI_ISL_532017, EPI_ISL_532019, EPI_ISL_532020, EPI_ISL_532025, EPI_ISL_532028, EPI_ISL_532030, EPI_ISL_532031, EPI_ISL_532035, EPI_ISL_532036, EPI_ISL_532039, EPI_ISL_532042, EPI_ISL_532046, EPI_ISL_532051, EPI_ISL_532058, EPI_ISL_532060, EPI_ISL_532064, EPI_ISL_532069, EPI_ISL_532082, EPI_ISL_532087, EPI_ISL_532089, EPI_ISL_532090, EPI_ISL_532096, EPI_ISL_532100, EPI_ISL_532106, EPI_ISL_532107, EPI_ISL_532110, EPI_ISL_532113, EPI_ISL_532128, EPI_ISL_532135, EPI_ISL_532142, EPI_ISL_532145, EPI_ISL_532154, EPI_ISL_532161, EPI_ISL_532163, EPI_ISL_532169, EPI_ISL_532174, EPI_ISL_532177, EPI_ISL_532179, EPI_ISL_532187, EPI_ISL_532189, EPI_ISL_532207, EPI_ISL_532208, EPI_ISL_532211, EPI_ISL_532218, EPI_ISL_532229, EPI_ISL_532240, EPI_ISL_532284, EPI_ISL_532285, EPI_ISL_532286, EPI_ISL_532287, EPI_ISL_532289, EPI_ISL_532290, EPI_ISL_532291, EPI_ISL_532296, EPI_ISL_532297, EPI_ISL_532298, EPI_ISL_532299, EPI_ISL_532300, EPI_ISL_532303, EPI_ISL_532304, EPI_ISL_532305, EPI_ISL_532306, EPI_ISL_532307, EPI_ISL_532308, EPI_ISL_532310, EPI_ISL_532315, EPI_ISL_532318, EPI_ISL_532319, EPI_ISL_532320, EPI_ISL_532322, EPI_ISL_532324, EPI_ISL_532326, EPI_ISL_532330, EPI_ISL_532331, EPI_ISL_532332, EPI_ISL_532337, EPI_ISL_532341, EPI_ISL_532344, EPI_ISL_532345, EPI_ISL_532346, EPI_ISL_532348, EPI_ISL_532350, EPI_ISL_532354, EPI_ISL_532357, EPI_ISL_532358, EPI_ISL_532359, EPI_ISL_532360, EPI_ISL_532362 | | | |
| see above | Lighthouse Lab in Glasgow | Wellcome Sanger Institute for the COVID-19 Genomics UK (COG-UK) consortium | Harper VanSteenhouse, Yumi Kasai, David Gray, Carol Clugston, Anna Dominiczak and Alex Alderton, Roberto Amato, Sonia Goncalves, Ewan Harrison, David K. Jackson, Ian Johnston, Dominic Kwiatkowski, Cordelia Langford, John Sillitoe |
| EPI_ISL_532365 | Lighthouse Lab in Glasgow | Wellcome Sanger Institute for the COVID-19 Genomics UK (COG-UK) consortium | Harper VanSteenhouse, Yumi Kasai, David Gray, Carol Clugston, Anna Dominiczak and Alex Alderton, Roberto Amato, Sonia Goncalves, Ewan Harrison, David K. Jackson, Ian Johnston, Dominic Kwiatkowski, Cordelia Langford, John Sillitoe on behalf of the Wellcome Sanger Institute COVID-19 Surveillance Team (http://www.sanger.ac.uk/covid-team) |
| EPI_ISL_532367, EPI_ISL_532368, EPI_ISL_532370, EPI_ISL_532371, EPI_ISL_532372, EPI_ISL_532373, EPI_ISL_532375, EPI_ISL_532376, EPI_ISL_532377, EPI_ISL_532378, EPI_ISL_532380, EPI_ISL_532381, EPI_ISL_532382, EPI_ISL_532383, EPI_ISL_532385, EPI_ISL_532389, EPI_ISL_532391, EPI_ISL_532392, EPI_ISL_532393, EPI_ISL_532397, EPI_ISL_532398, EPI_ISL_532401, EPI_ISL_532405, EPI_ISL_532406, EPI_ISL_532407, EPI_ISL_532408, EPI_ISL_532409, EPI_ISL_532410, EPI_ISL_532411, EPI_ISL_532413, EPI_ISL_532416, EPI_ISL_532417, EPI_ISL_532418, EPI_ISL_532419, EPI_ISL_532420, EPI_ISL_532422, EPI_ISL_532427, EPI_ISL_532429, EPI_ISL_532434, EPI_ISL_532435, EPI_ISL_532436, EPI_ISL_532441, EPI_ISL_532443, EPI_ISL_532445, EPI_ISL_532447, EPI_ISL_532448, EPI_ISL_532450, EPI_ISL_532451, EPI_ISL_532453, EPI_ISL_532457, EPI_ISL_532458, EPI_ISL_532459, EPI_ISL_532460, EPI_ISL_532461, EPI_ISL_532462, EPI_ISL_532465, EPI_ISL_532466, EPI_ISL_532468, EPI_ISL_532470, EPI_ISL_532471, EPI_ISL_532474, EPI_ISL_532476, EPI_ISL_532478, EPI_ISL_532481, EPI_ISL_532482, EPI_ISL_532486, EPI_ISL_532488, EPI_ISL_532489, EPI_ISL_532495, EPI_ISL_532496, EPI_ISL_532498, EPI_ISL_532499, EPI_ISL_532500, EPI_ISL_532501, EPI_ISL_532502, EPI_ISL_532503, EPI_ISL_532504, EPI_ISL_532507, EPI_ISL_532508, EPI_ISL_532513, EPI_ISL_532516, EPI_ISL_532517, EPI_ISL_532519, EPI_ISL_532520, EPI_ISL_532521, EPI_ISL_532522, EPI_ISL_532524, EPI_ISL_532525, EPI_ISL_532526, EPI_ISL_532528, EPI_ISL_532529, EPI_ISL_532530, EPI_ISL_532532, EPI_ISL_532534, EPI_ISL_532535, EPI_ISL_532537, EPI_ISL_532539, EPI_ISL_532541, EPI_ISL_532542, EPI_ISL_532543, EPI_ISL_532544, EPI_ISL_532545, EPI_ISL_532546, EPI_ISL_532547, EPI_ISL_532548, EPI_ISL_532549, EPI_ISL_532550, EPI_ISL_532551, EPI_ISL_532552, EPI_ISL_532553, EPI_ISL_532554, EPI_ISL_532555, EPI_ISL_532556, EPI_ISL_532557, EPI_ISL_532558, EPI_ISL_532559, EPI_ISL_532560, EPI_ISL_532561, EPI_ISL_532562, EPI_ISL_532565, EPI_ISL_532566, EPI_ISL_532567, EPI_ISL_532568, EPI_ISL_532569, EPI_ISL_532570, EPI_ISL_532571, | | | |

| | | | |
|---|---|---|---|
| EPI_ISL_532573, EPI_ISL_532574, EPI_ISL_532575, EPI_ISL_532576, EPI_ISL_532577, EPI_ISL_532578, EPI_ISL_532579, EPI_ISL_532580, EPI_ISL_532581, EPI_ISL_532584, EPI_ISL_532585, EPI_ISL_532586, EPI_ISL_532588, EPI_ISL_532589, EPI_ISL_532590, EPI_ISL_532591, EPI_ISL_532592, EPI_ISL_532593, EPI_ISL_532595, EPI_ISL_532596, EPI_ISL_532597, EPI_ISL_532599, EPI_ISL_532600, EPI_ISL_532601, EPI_ISL_532602, EPI_ISL_532603, EPI_ISL_532604, EPI_ISL_532605, EPI_ISL_532606, EPI_ISL_532607, EPI_ISL_532608, EPI_ISL_532609, EPI_ISL_532610, EPI_ISL_532611, EPI_ISL_532612, EPI_ISL_532613, EPI_ISL_532614, EPI_ISL_532615, EPI_ISL_532616, EPI_ISL_532617, EPI_ISL_532618, EPI_ISL_532619, EPI_ISL_532622, EPI_ISL_532623, EPI_ISL_532624, EPI_ISL_532627, EPI_ISL_532628, EPI_ISL_532629, EPI_ISL_532630, EPI_ISL_532631, EPI_ISL_532632, EPI_ISL_532634, EPI_ISL_532635, EPI_ISL_532636, EPI_ISL_532637, EPI_ISL_532638, EPI_ISL_532640, EPI_ISL_532641, EPI_ISL_532642, EPI_ISL_532643, EPI_ISL_532644, EPI_ISL_532645, EPI_ISL_532646, EPI_ISL_532647, EPI_ISL_532649, EPI_ISL_532650, EPI_ISL_532651, EPI_ISL_532652, EPI_ISL_532653, EPI_ISL_532654, EPI_ISL_532655, EPI_ISL_532656, EPI_ISL_532657, EPI_ISL_532658, EPI_ISL_532660, EPI_ISL_532661, EPI_ISL_532662, EPI_ISL_532663, EPI_ISL_532664, EPI_ISL_532665, EPI_ISL_532666, EPI_ISL_532667, EPI_ISL_532668, EPI_ISL_532669, EPI_ISL_532671, EPI_ISL_532672, EPI_ISL_532673, EPI_ISL_532674, EPI_ISL_532675, EPI_ISL_532676, EPI_ISL_532677, EPI_ISL_532678, EPI_ISL_532679, EPI_ISL_532680, EPI_ISL_532682, EPI_ISL_532684, EPI_ISL_532685, EPI_ISL_532686, EPI_ISL_532687, EPI_ISL_532688, EPI_ISL_532690, EPI_ISL_532691, EPI_ISL_532692, EPI_ISL_532693, EPI_ISL_532694, EPI_ISL_532695, EPI_ISL_532696, EPI_ISL_532698, EPI_ISL_532699, EPI_ISL_532700, EPI_ISL_532701, EPI_ISL_532702, EPI_ISL_532703, EPI_ISL_532704, EPI_ISL_532706, EPI_ISL_532707, EPI_ISL_532708, EPI_ISL_532709, EPI_ISL_532710, EPI_ISL_532711, EPI_ISL_532712, EPI_ISL_532713, EPI_ISL_532714, EPI_ISL_532716, EPI_ISL_532717, EPI_ISL_532718, EPI_ISL_532719, EPI_ISL_532720, EPI_ISL_532721, EPI_ISL_532722, EPI_ISL_532723, EPI_ISL_532724, EPI_ISL_532725, EPI_ISL_532726, EPI_ISL_532727, EPI_ISL_532728, EPI_ISL_532729, EPI_ISL_532730, EPI_ISL_532731, EPI_ISL_532734, EPI_ISL_532735, EPI_ISL_532736, EPI_ISL_532737, EPI_ISL_532739, EPI_ISL_532741, EPI_ISL_532742, EPI_ISL_532743, EPI_ISL_532745, EPI_ISL_532747, EPI_ISL_532748, EPI_ISL_532749, EPI_ISL_532751, EPI_ISL_532752, EPI_ISL_532754, EPI_ISL_532756, EPI_ISL_532757, EPI_ISL_532758, EPI_ISL_532759, EPI_ISL_532761, EPI_ISL_532762, EPI_ISL_532763, EPI_ISL_532764, EPI_ISL_532766, EPI_ISL_532768, EPI_ISL_532769, EPI_ISL_532770, EPI_ISL_532771, EPI_ISL_532772, EPI_ISL_532773, EPI_ISL_532774, EPI_ISL_532775, EPI_ISL_532776, EPI_ISL_532777, EPI_ISL_532778, EPI_ISL_532779, EPI_ISL_532780, EPI_ISL_532781, EPI_ISL_532782, EPI_ISL_532783, EPI_ISL_532785, EPI_ISL_532787, EPI_ISL_532788, EPI_ISL_532789, EPI_ISL_532790, EPI_ISL_532792, EPI_ISL_532794, EPI_ISL_532795, EPI_ISL_532796, EPI_ISL_532797, EPI_ISL_532798, EPI_ISL_532799, EPI_ISL_532800, EPI_ISL_532801, EPI_ISL_532802, EPI_ISL_532803, EPI_ISL_532804, EPI_ISL_532805, EPI_ISL_532806, EPI_ISL_532807, EPI_ISL_532808, EPI_ISL_532809, EPI_ISL_532810, EPI_ISL_532812, EPI_ISL_532813, EPI_ISL_532814, EPI_ISL_532815, EPI_ISL_532816, EPI_ISL_532817, EPI_ISL_532818, EPI_ISL_532819, EPI_ISL_532820, EPI_ISL_532821, EPI_ISL_532822, EPI_ISL_532823, EPI_ISL_532824, EPI_ISL_532825, EPI_ISL_532826, EPI_ISL_532827, EPI_ISL_532829, EPI_ISL_532830, EPI_ISL_532831, EPI_ISL_532832, EPI_ISL_532833, EPI_ISL_532834, EPI_ISL_532836, EPI_ISL_532839, EPI_ISL_532842, EPI_ISL_532843, EPI_ISL_532850, EPI_ISL_532852, EPI_ISL_532854, EPI_ISL_532857, EPI_ISL_532862, EPI_ISL_532872, EPI_ISL_532873, EPI_ISL_532880, EPI_ISL_532881, EPI_ISL_532897, EPI_ISL_532899, EPI_ISL_532902, EPI_ISL_532903, EPI_ISL_532907, EPI_ISL_532909, EPI_ISL_532910, EPI_ISL_532913, EPI_ISL_532916, EPI_ISL_532919, EPI_ISL_532920, EPI_ISL_532921, EPI_ISL_532922, EPI_ISL_532924, EPI_ISL_532925, EPI_ISL_532929, EPI_ISL_532937, EPI_ISL_532940, EPI_ISL_532943, EPI_ISL_532944, EPI_ISL_532946, EPI_ISL_532949, EPI_ISL_532950, EPI_ISL_532951, EPI_ISL_532976, EPI_ISL_532979, EPI_ISL_532982, EPI_ISL_532984, EPI_ISL_532985, EPI_ISL_532997, EPI_ISL_533001, EPI_ISL_533007, EPI_ISL_533013, EPI_ISL_533015, EPI_ISL_533017, EPI_ISL_533021, EPI_ISL_533025, EPI_ISL_533026, EPI_ISL_533029, EPI_ISL_533030, EPI_ISL_533038, EPI_ISL_533046, EPI_ISL_533049, EPI_ISL_533052, EPI_ISL_533054, EPI_ISL_533065, EPI_ISL_533066, EPI_ISL_533067, EPI_ISL_533070, EPI_ISL_533074, EPI_ISL_533077, EPI_ISL_533079, EPI_ISL_533082, EPI_ISL_533089, EPI_ISL_533091, EPI_ISL_533097, EPI_ISL_533101, EPI_ISL_533107, EPI_ISL_533117, EPI_ISL_533130, EPI_ISL_533132, EPI_ISL_533133, EPI_ISL_533138, EPI_ISL_533139, EPI_ISL_533141, EPI_ISL_533144, EPI_ISL_533151, EPI_ISL_533161, EPI_ISL_533166, EPI_ISL_533169, EPI_ISL_533177, EPI_ISL_533188, EPI_ISL_533190, EPI_ISL_533201, EPI_ISL_533202, EPI_ISL_533204, EPI_ISL_533207, EPI_ISL_533222, EPI_ISL_533234, EPI_ISL_533240, EPI_ISL_533244, EPI_ISL_533250, EPI_ISL_533251, EPI_ISL_533266, EPI_ISL_533267, EPI_ISL_533272, EPI_ISL_533281, EPI_ISL_533291, EPI_ISL_533305, EPI_ISL_533307, EPI_ISL_533312, EPI_ISL_533316, EPI_ISL_533317, EPI_ISL_533328, EPI_ISL_533331, EPI_ISL_533357, EPI_ISL_533363, EPI_ISL_533380, EPI_ISL_533411, EPI_ISL_533413, EPI_ISL_533417, EPI_ISL_533418, EPI_ISL_533427 | | | |
| see above | Lighthouse Lab in Glasgow | Wellcome Sanger Institute for the COVID-19 Genomics UK (COG-UK) consortium | Harper VanSteenhouse, Yumi Kasai, David Gray, Carol Clugston, Anna Dominiczak and Alex Alderton, Roberto Amato, Sonia Goncalves, Ewan Harrison, David K. Jackson, Ian Johnston, Dominic Kwiatkowski, Cordelia Langford, John Sillitoe |
| EPI_ISL_535391, EPI_ISL_535395, EPI_ISL_535401, EPI_ISL_535403, EPI_ISL_535407, EPI_ISL_535410, EPI_ISL_535419, EPI_ISL_535423, EPI_ISL_535429, EPI_ISL_535437, EPI_ISL_535453, EPI_ISL_535467, EPI_ISL_535469, EPI_ISL_535474, EPI_ISL_535478, EPI_ISL_535482, EPI_ISL_535517, EPI_ISL_535521, EPI_ISL_535524, EPI_ISL_535525, EPI_ISL_535528, EPI_ISL_535535, EPI_ISL_535538, EPI_ISL_535540, EPI_ISL_535548, EPI_ISL_535549, EPI_ISL_535551 | | | |
| see above | NHLS-IALCH | KRISP, KZN Research Innovation and Sequencing Platform | Giandhari J, Pillay S, Lessells R, Mdlalose K, York D, Khan S, Tegally H, Wilkinson E, de Oliveira T |
| EPI_ISL_536819, EPI_ISL_536820, EPI_ISL_536821, EPI_ISL_536822, EPI_ISL_536823, EPI_ISL_536824, EPI_ISL_536825, EPI_ISL_536826, EPI_ISL_536827, EPI_ISL_536828, EPI_ISL_536829, EPI_ISL_536830, EPI_ISL_536831, EPI_ISL_536832, EPI_ISL_536833, EPI_ISL_536834, EPI_ISL_536835, EPI_ISL_536836, EPI_ISL_536837, EPI_ISL_536838, EPI_ISL_536839, EPI_ISL_536840, EPI_ISL_536841, EPI_ISL_536842, EPI_ISL_536843, EPI_ISL_536844, EPI_ISL_536845, EPI_ISL_536846, EPI_ISL_536847, EPI_ISL_536848, EPI_ISL_536849, EPI_ISL_536850, EPI_ISL_536851, EPI_ISL_536852, EPI_ISL_536853, EPI_ISL_536854, EPI_ISL_536855, EPI_ISL_536856, EPI_ISL_536857, EPI_ISL_536858, EPI_ISL_536859, EPI_ISL_536860, EPI_ISL_536861, EPI_ISL_536862, EPI_ISL_536863, EPI_ISL_536864, EPI_ISL_536865, EPI_ISL_536866, EPI_ISL_536867, EPI_ISL_536868, EPI_ISL_536869, EPI_ISL_536870, EPI_ISL_536871, EPI_ISL_536872, EPI_ISL_536873, EPI_ISL_536874, EPI_ISL_536875, EPI_ISL_536876, EPI_ISL_536877, EPI_ISL_536878, EPI_ISL_536879, EPI_ISL_536880, EPI_ISL_536881, EPI_ISL_536882, EPI_ISL_536883, EPI_ISL_536884, EPI_ISL_536885, EPI_ISL_536886, EPI_ISL_536887, EPI_ISL_536888, EPI_ISL_536890, EPI_ISL_536891, EPI_ISL_536892, EPI_ISL_536893, EPI_ISL_536894, EPI_ISL_536895, EPI_ISL_536896, EPI_ISL_536897, EPI_ISL_536898, EPI_ISL_536899, EPI_ISL_536900, EPI_ISL_536901, EPI_ISL_536902, EPI_ISL_536903, EPI_ISL_536904, EPI_ISL_536905, EPI_ISL_536906, EPI_ISL_536907, EPI_ISL_536908, EPI_ISL_536909, EPI_ISL_536910, EPI_ISL_536911, EPI_ISL_536912, EPI_ISL_536913, EPI_ISL_536914, EPI_ISL_536915, EPI_ISL_536916, EPI_ISL_536917, EPI_ISL_536918, EPI_ISL_536919, EPI_ISL_536920, EPI_ISL_536921, EPI_ISL_536922, EPI_ISL_536923, EPI_ISL_536924, EPI_ISL_536925, EPI_ISL_536926, EPI_ISL_536927, EPI_ISL_536928, EPI_ISL_536929, EPI_ISL_536930, EPI_ISL_536931, EPI_ISL_536932, EPI_ISL_536933, EPI_ISL_536934, EPI_ISL_536935, EPI_ISL_536936, EPI_ISL_536937, EPI_ISL_536938, EPI_ISL_536939, EPI_ISL_536940, EPI_ISL_536941, EPI_ISL_536942, EPI_ISL_536943, EPI_ISL_536944, EPI_ISL_536945, EPI_ISL_536946, EPI_ISL_536947, EPI_ISL_536948, EPI_ISL_536949, EPI_ISL_536950, EPI_ISL_536951, EPI_ISL_536952, EPI_ISL_536953, EPI_ISL_536954, EPI_ISL_536955, EPI_ISL_536956, EPI_ISL_536957, EPI_ISL_536958, EPI_ISL_536959, EPI_ISL_536960, EPI_ISL_536961, EPI_ISL_536962, EPI_ISL_536963, EPI_ISL_536964, EPI_ISL_536965, EPI_ISL_536966, EPI_ISL_536967, EPI_ISL_536968, EPI_ISL_536969, EPI_ISL_536970, EPI_ISL_536971, EPI_ISL_536972, EPI_ISL_536973, EPI_ISL_536974, EPI_ISL_536975, EPI_ISL_536976, EPI_ISL_536977, EPI_ISL_536978, EPI_ISL_536979, EPI_ISL_536980, EPI_ISL_536981, EPI_ISL_536982, EPI_ISL_536983, EPI_ISL_536984, EPI_ISL_536985, EPI_ISL_536986, EPI_ISL_536987, EPI_ISL_536988, EPI_ISL_536989, EPI_ISL_536990, EPI_ISL_536991, EPI_ISL_536992, EPI_ISL_536993, EPI_ISL_536994, EPI_ISL_536995, EPI_ISL_536996, EPI_ISL_536997, EPI_ISL_536998, EPI_ISL_536999, EPI_ISL_537000, EPI_ISL_537001, EPI_ISL_537002, EPI_ISL_537003, EPI_ISL_537004, EPI_ISL_537005, EPI_ISL_537006, EPI_ISL_537007, EPI_ISL_537008, EPI_ISL_537009, EPI_ISL_537010, EPI_ISL_537011, EPI_ISL_537012, EPI_ISL_537013, EPI_ISL_537014, EPI_ISL_537015, EPI_ISL_537016, EPI_ISL_537018, EPI_ISL_537019, EPI_ISL_537020, EPI_ISL_537021, EPI_ISL_537022, EPI_ISL_537023, EPI_ISL_537024, EPI_ISL_537025, EPI_ISL_537026, EPI_ISL_537027, EPI_ISL_537028, EPI_ISL_537029, EPI_ISL_537030, EPI_ISL_537031, EPI_ISL_537032, EPI_ISL_537033, EPI_ISL_537034, EPI_ISL_537035, EPI_ISL_537036, EPI_ISL_537037, EPI_ISL_537038, EPI_ISL_537039, EPI_ISL_537040, EPI_ISL_537041, EPI_ISL_537042, EPI_ISL_537044, EPI_ISL_537045, EPI_ISL_537046, EPI_ISL_537047, EPI_ISL_537048, EPI_ISL_537049, EPI_ISL_537050, EPI_ISL_537051, EPI_ISL_537052, EPI_ISL_537053, EPI_ISL_537054, EPI_ISL_537055, EPI_ISL_537056, EPI_ISL_537057, EPI_ISL_537058, EPI_ISL_537059, EPI_ISL_537060, EPI_ISL_537061, EPI_ISL_537062, EPI_ISL_537063, EPI_ISL_537064, EPI_ISL_537065, EPI_ISL_537066, EPI_ISL_537067, EPI_ISL_537068, EPI_ISL_537069, EPI_ISL_537070, EPI_ISL_537071, EPI_ISL_537072, EPI_ISL_537073, EPI_ISL_537074, EPI_ISL_537075, EPI_ISL_537076, EPI_ISL_537077, EPI_ISL_537078, EPI_ISL_537079, EPI_ISL_537080, EPI_ISL_537081, EPI_ISL_537082, EPI_ISL_537083, EPI_ISL_537084, EPI_ISL_537085, EPI_ISL_537086, EPI_ISL_537087, EPI_ISL_537088, EPI_ISL_537089, EPI_ISL_537090, EPI_ISL_537091, EPI_ISL_537092, EPI_ISL_537093, EPI_ISL_537094, EPI_ISL_537095, EPI_ISL_537096, EPI_ISL_537097, EPI_ISL_537098, EPI_ISL_537099, EPI_ISL_537100, EPI_ISL_537101, EPI_ISL_537102, EPI_ISL_537103, EPI_ISL_537104, EPI_ISL_537105, EPI_ISL_537106, EPI_ISL_537107, EPI_ISL_537108, EPI_ISL_537109, EPI_ISL_537110, EPI_ISL_537111, EPI_ISL_537112, EPI_ISL_537113, EPI_ISL_537114, EPI_ISL_537115, EPI_ISL_537116, EPI_ISL_537117, EPI_ISL_537118, EPI_ISL_537119, EPI_ISL_537120, EPI_ISL_537121, EPI_ISL_537122, EPI_ISL_537123, EPI_ISL_537124, EPI_ISL_537125, EPI_ISL_537126, EPI_ISL_537127, EPI_ISL_537128, EPI_ISL_537129, EPI_ISL_537130, EPI_ISL_537131, EPI_ISL_537132, EPI_ISL_537133, EPI_ISL_537134, EPI_ISL_537135, EPI_ISL_537136, EPI_ISL_537137, EPI_ISL_537138, EPI_ISL_537139, EPI_ISL_537140, EPI_ISL_537141, EPI_ISL_537142, EPI_ISL_537143, EPI_ISL_537144, EPI_ISL_537145, EPI_ISL_537146, EPI_ISL_537147, EPI_ISL_537148, EPI_ISL_537149, EPI_ISL_537150, EPI_ISL_537151, EPI_ISL_537152, EPI_ISL_537153, EPI_ISL_537154, EPI_ISL_537155, EPI_ISL_537156, EPI_ISL_537157, EPI_ISL_537158, EPI_ISL_537159, EPI_ISL_537160, EPI_ISL_537161, EPI_ISL_537162, EPI_ISL_537163, EPI_ISL_537164, EPI_ISL_537165, EPI_ISL_537166, EPI_ISL_537167, EPI_ISL_537168, EPI_ISL_537169, EPI_ISL_537170, EPI_ISL_537171, EPI_ISL_537172, EPI_ISL_537173, EPI_ISL_537174, EPI_ISL_537175, EPI_ISL_537176, EPI_ISL_537177, EPI_ISL_537178, EPI_ISL_537179, EPI_ISL_537180, EPI_ISL_537181, EPI_ISL_537182, EPI_ISL_537183, EPI_ISL_537184, EPI_ISL_537185, EPI_ISL_537186, EPI_ISL_537187, EPI_ISL_537188, EPI_ISL_537189, EPI_ISL_537190, EPI_ISL_537191, EPI_ISL_537192, EPI_ISL_537193, EPI_ISL_537194, EPI_ISL_537195, EPI_ISL_537196, EPI_ISL_537197, EPI_ISL_537198, EPI_ISL_537199, EPI_ISL_537200, EPI_ISL_537201, EPI_ISL_537202, EPI_ISL_537203, EPI_ISL_537204, EPI_ISL_537205 | | | |
| see above | Lighthouse Lab in Glasgow | Wellcome Sanger Institute for the COVID-19 Genomics UK (COG-UK) consortium | Harper VanSteenhouse, Yumi Kasai, David Gray, Carol Clugston, Anna Dominiczak and Alex Alderton, Roberto Amato, Sonia Goncalves, Ewan Harrison, David K. Jackson, Ian Johnston, Dominic Kwiatkowski, Cordelia Langford, John Sillitoe on behalf of the Wellcome Sanger Institute COVID-19 Surveillance Team |
| EPI_ISL_537467, EPI_ISL_537468, EPI_ISL_537469, EPI_ISL_537470, EPI_ISL_537471, EPI_ISL_537472, EPI_ISL_537473, EPI_ISL_537474, EPI_ISL_537475, EPI_ISL_537476, EPI_ISL_537477, EPI_ISL_537478, EPI_ISL_537479, EPI_ISL_537480, EPI_ISL_537481, EPI_ISL_537482, EPI_ISL_537483, EPI_ISL_537484, EPI_ISL_537485, EPI_ISL_537486, EPI_ISL_537487, EPI_ISL_537488, EPI_ISL_537489, EPI_ISL_537490, EPI_ISL_537491, EPI_ISL_537492, EPI_ISL_537493, EPI_ISL_537494, EPI_ISL_537495, EPI_ISL_537496, EPI_ISL_537497, EPI_ISL_537498, EPI_ISL_537499, EPI_ISL_537500, EPI_ISL_537501, EPI_ISL_537502, EPI_ISL_537503, EPI_ISL_537504, EPI_ISL_537505, EPI_ISL_537506, EPI_ISL_537507, EPI_ISL_537508, EPI_ISL_537509, EPI_ISL_537510, EPI_ISL_537512, EPI_ISL_537513, EPI_ISL_537514, EPI_ISL_537515, EPI_ISL_537516, EPI_ISL_537517, EPI_ISL_537518, EPI_ISL_537519, EPI_ISL_537520, EPI_ISL_537521, EPI_ISL_537522, EPI_ISL_537523, EPI_ISL_537524, EPI_ISL_537525, EPI_ISL_537526, EPI_ISL_537527, EPI_ISL_537528, EPI_ISL_537529, EPI_ISL_537530, EPI_ISL_537531, EPI_ISL_537532, EPI_ISL_537533, EPI_ISL_537534, EPI_ISL_537535, EPI_ISL_537536, EPI_ISL_537537, EPI_ISL_537538, EPI_ISL_537539, EPI_ISL_537540, EPI_ISL_537541, EPI_ISL_537542, EPI_ISL_537543, EPI_ISL_537544, EPI_ISL_537545, EPI_ISL_537546, EPI_ISL_537547, EPI_ISL_537548, EPI_ISL_537549, EPI_ISL_537550, EPI_ISL_537551, EPI_ISL_537552, EPI_ISL_537553, EPI_ISL_537554, EPI_ISL_537555, EPI_ISL_537556, EPI_ISL_537557, EPI_ISL_537558, EPI_ISL_537559, EPI_ISL_537560, EPI_ISL_537561, EPI_ISL_537562, EPI_ISL_537563, EPI_ISL_537564, EPI_ISL_537565, EPI_ISL_537566, EPI_ISL_537567, EPI_ISL_537568, EPI_ISL_537569, EPI_ISL_537570, EPI_ISL_537571, EPI_ISL_537572, EPI_ISL_537573, EPI_ISL_537575, EPI_ISL_537576, EPI_ISL_537577, EPI_ISL_537578, EPI_ISL_537579, EPI_ISL_537582, EPI_ISL_537583, EPI_ISL_537584, EPI_ISL_537585, EPI_ISL_537586, EPI_ISL_537587, EPI_ISL_537588, EPI_ISL_537589, EPI_ISL_537590, EPI_ISL_537591, EPI_ISL_537592, EPI_ISL_537593, EPI_ISL_537594, EPI_ISL_537595, EPI_ISL_537596, EPI_ISL_537597, EPI_ISL_537598, EPI_ISL_537599, EPI_ISL_537601, EPI_ISL_537602, EPI_ISL_537603, EPI_ISL_537604, EPI_ISL_537605, EPI_ISL_537606, EPI_ISL_537607 | | | |
| see above | UCLA Pathology Clinical Microbiology Lab | Kruglyak Lab | Guo et al. |
| EPI_ISL_538055, EPI_ISL_538056, EPI_ISL_538057, EPI_ISL_538058, EPI_ISL_538059, EPI_ISL_538060, EPI_ISL_538061 | Servicio de Microbiología y Parasitología clínica. UCEIMP. Hospital Universitario Virgen del Rocío/IBIS/CSIC/US | SeqCOVID-SPAIN consortium/IBV(CSIC) | Guillermo Martín Gutiérrez, Ángel Rodríguez Villodres, Lidia Gálvez Benitez, Verónica González Galán, Javier Aznar Martín and SeqCOVID-SPAIN consortium |
| EPI_ISL_538062, EPI_ISL_538063, EPI_ISL_538064, EPI_ISL_538065, EPI_ISL_538066, EPI_ISL_538067, EPI_ISL_538068, EPI_ISL_538069, EPI_ISL_538070, EPI_ISL_538071, EPI_ISL_538072, EPI_ISL_538073, EPI_ISL_538074, EPI_ISL_538075, EPI_ISL_538076, EPI_ISL_538077, EPI_ISL_538078, EPI_ISL_538079, EPI_ISL_538080, EPI_ISL_538081 | | | |
| see above | Servicio de Microbiología. Hospital Universitario Donostia. OSI Donostialdea. Área de Enfermedades Infecciosas, Grupo de Infección Respiratoria y Resistencia Antimicrobiana. Instituto de Investigación Sanitaria Biodonostia | SeqCOVID-SPAIN consortium/IBV(CSIC) | Gustavo Cilla, Milagrosa Montes, Luis Piñeiro, Jose Maria Marimón and SeqCOVID-SPAIN consortium |
| EPI_ISL_538119, EPI_ISL_538120, EPI_ISL_538121, EPI_ISL_538122, EPI_ISL_538123, EPI_ISL_538124, EPI_ISL_538125, EPI_ISL_538126, EPI_ISL_538127, EPI_ISL_538128, EPI_ISL_538129, EPI_ISL_538130, EPI_ISL_538131, EPI_ISL_538132, EPI_ISL_538133, EPI_ISL_538134, EPI_ISL_538135, EPI_ISL_538136, EPI_ISL_538137, EPI_ISL_538138, EPI_ISL_538139, EPI_ISL_538140, EPI_ISL_538141, EPI_ISL_538142, EPI_ISL_538143, EPI_ISL_538144, EPI_ISL_538145, EPI_ISL_538146, EPI_ISL_538147, EPI_ISL_538148, EPI_ISL_538149, EPI_ISL_538150, EPI_ISL_538151, EPI_ISL_538152, EPI_ISL_538153, EPI_ISL_538154, EPI_ISL_538155, EPI_ISL_538156, EPI_ISL_538157, EPI_ISL_538158, EPI_ISL_538159, EPI_ISL_538160, EPI_ISL_538161, EPI_ISL_538162, EPI_ISL_538163, EPI_ISL_538164, EPI_ISL_538165, EPI_ISL_538166, EPI_ISL_538167, EPI_ISL_538168, EPI_ISL_538169, EPI_ISL_538170, EPI_ISL_538171, EPI_ISL_538172, EPI_ISL_538173, EPI_ISL_538174, EPI_ISL_538175, EPI_ISL_538176, EPI_ISL_538177, EPI_ISL_538178, EPI_ISL_538179, EPI_ISL_538180, EPI_ISL_538181, EPI_ISL_538182, EPI_ISL_538183, EPI_ISL_538184, EPI_ISL_538185, EPI_ISL_538186, EPI_ISL_538187, EPI_ISL_538188, EPI_ISL_538189, EPI_ISL_538190, EPI_ISL_538191, EPI_ISL_538192, EPI_ISL_538193, EPI_ISL_538194, EPI_ISL_538195, EPI_ISL_538196, EPI_ISL_538197, EPI_ISL_538198, EPI_ISL_538199, EPI_ISL_538200, EPI_ISL_538201, EPI_ISL_538202, EPI_ISL_538203, EPI_ISL_538204, EPI_ISL_538205, EPI_ISL_538206, EPI_ISL_538207, EPI_ISL_538208, EPI_ISL_538209, EPI_ISL_538210, EPI_ISL_538211, EPI_ISL_538212, EPI_ISL_538213, EPI_ISL_538214, EPI_ISL_538215, EPI_ISL_538216, EPI_ISL_538217, EPI_ISL_538218, EPI_ISL_538219, EPI_ISL_538220, EPI_ISL_538221, EPI_ISL_538222, EPI_ISL_538223, EPI_ISL_538224, EPI_ISL_538225, EPI_ISL_538226, EPI_ISL_538227, EPI_ISL_538228, EPI_ISL_538229, EPI_ISL_538230, EPI_ISL_538568, EPI_ISL_538569, EPI_ISL_538570, EPI_ISL_538571, EPI_ISL_538572, EPI_ISL_538573, EPI_ISL_538574, EPI_ISL_538575, EPI_ISL_538576, EPI_ISL_538577, EPI_ISL_538578, EPI_ISL_538579, EPI_ISL_538580, EPI_ISL_538581, EPI_ISL_538582, EPI_ISL_538583, EPI_ISL_538584, EPI_ISL_538585, EPI_ISL_538586, EPI_ISL_538587, EPI_ISL_538588, EPI_ISL_538589, EPI_ISL_538590, EPI_ISL_538591, EPI_ISL_538592, EPI_ISL_538593, EPI_ISL_538594, EPI_ISL_538595, EPI_ISL_538596, EPI_ISL_538597, EPI_ISL_538598, EPI_ISL_538599, EPI_ISL_538600, EPI_ISL_538601, EPI_ISL_538602, EPI_ISL_538603, EPI_ISL_538604, EPI_ISL_538605, EPI_ISL_538606, EPI_ISL_538607 | | | |
| see above | Servicio de Microbiología y Parasitología clínica. UCEIMP. Hospital Universitario Virgen del Rocío/IBIS/CSIC/US | SeqCOVID-SPAIN consortium/IBV(CSIC) | Guillermo Martín Gutiérrez, Ángel Rodríguez Villodres, Lidia Gálvez Benitez, Verónica González Galán, Javier Aznar Martín and SeqCOVID-SPAIN consortium |
| EPI_ISL_539495 | Centers for Disease Control and Prevention, Dengue Branch | Centers for Disease Control and Prevention, Dengue Branch | Gilberto A. Santiago, Glenda Gonzalez, Betzabel Flores, Keyla Charriez, Jorge L. Munoz-Jordan, Gabriela Paz-Bailey, Janice Perez, Vanessa Rivera-Amill, Diego Sainz de la Peña, Jorge Bertran |

| | | | |
|---|---|---|---|
| EPI_ISL_539897, EPI_ISL_539898, EPI_ISL_539899, EPI_ISL_539900, EPI_ISL_539901, EPI_ISL_539902, EPI_ISL_539903, EPI_ISL_539904, EPI_ISL_539905, EPI_ISL_539907, EPI_ISL_539908, EPI_ISL_539909, EPI_ISL_539910, EPI_ISL_539911, EPI_ISL_539912, EPI_ISL_539913, EPI_ISL_539914, EPI_ISL_539915, EPI_ISL_539916, EPI_ISL_539917, EPI_ISL_539918, EPI_ISL_539919, EPI_ISL_539920, EPI_ISL_539921, EPI_ISL_539922, EPI_ISL_539923, EPI_ISL_539924, EPI_ISL_539925, EPI_ISL_539926, EPI_ISL_539927, EPI_ISL_539928, EPI_ISL_539929, EPI_ISL_539930, EPI_ISL_539931, EPI_ISL_539932, EPI_ISL_539933, EPI_ISL_539934, EPI_ISL_539935, EPI_ISL_539936, EPI_ISL_539937, EPI_ISL_539938, EPI_ISL_539939, EPI_ISL_539940, EPI_ISL_539941, EPI_ISL_539942, EPI_ISL_539943, EPI_ISL_539944, EPI_ISL_539945, EPI_ISL_539946, EPI_ISL_539947, EPI_ISL_539948, EPI_ISL_539949, EPI_ISL_539950, EPI_ISL_539951, EPI_ISL_539952, EPI_ISL_539953, EPI_ISL_539954, EPI_ISL_539955, EPI_ISL_539956, EPI_ISL_539957, EPI_ISL_539958, EPI_ISL_539959, EPI_ISL_539960, EPI_ISL_539961, EPI_ISL_539962, EPI_ISL_539963, EPI_ISL_539964, EPI_ISL_539965, EPI_ISL_539966, EPI_ISL_539967, EPI_ISL_539968, EPI_ISL_539969, EPI_ISL_539970, EPI_ISL_539971, EPI_ISL_539972, EPI_ISL_539973, EPI_ISL_539974, EPI_ISL_539975, EPI_ISL_539976, EPI_ISL_539977, EPI_ISL_539978, EPI_ISL_539979, EPI_ISL_539980, EPI_ISL_539981, EPI_ISL_539983, EPI_ISL_539984, EPI_ISL_539985, EPI_ISL_539986, EPI_ISL_539987, EPI_ISL_539988, EPI_ISL_539989, EPI_ISL_539990, EPI_ISL_539991, EPI_ISL_539992, EPI_ISL_539993, EPI_ISL_539994, EPI_ISL_539995, EPI_ISL_539996, EPI_ISL_539997, EPI_ISL_539998, EPI_ISL_539999, EPI_ISL_540000, EPI_ISL_540001, EPI_ISL_540002, EPI_ISL_540003, EPI_ISL_540004, EPI_ISL_540005, EPI_ISL_540006, EPI_ISL_540007, EPI_ISL_540008, EPI_ISL_540009, EPI_ISL_540010, EPI_ISL_540011, EPI_ISL_540012, EPI_ISL_540013, EPI_ISL_540014, EPI_ISL_540015, EPI_ISL_540016, EPI_ISL_540017, EPI_ISL_540018, EPI_ISL_540019, EPI_ISL_540020, EPI_ISL_540021, EPI_ISL_540022, EPI_ISL_540023, EPI_ISL_540024, EPI_ISL_540025, EPI_ISL_540026, EPI_ISL_540027, EPI_ISL_540028, EPI_ISL_540029, EPI_ISL_540031, EPI_ISL_540032, EPI_ISL_540033, EPI_ISL_540034, EPI_ISL_540035, EPI_ISL_540036, EPI_ISL_540037, EPI_ISL_540038, EPI_ISL_540039, EPI_ISL_540040, EPI_ISL_540041, EPI_ISL_540042, EPI_ISL_540043, EPI_ISL_540044, EPI_ISL_540045, EPI_ISL_540046, EPI_ISL_540047, EPI_ISL_540048, EPI_ISL_540049, EPI_ISL_540050, EPI_ISL_540051, EPI_ISL_540052, EPI_ISL_540053, EPI_ISL_540054, EPI_ISL_540055, EPI_ISL_540056, EPI_ISL_540057, EPI_ISL_540058, EPI_ISL_540059, EPI_ISL_540060, EPI_ISL_540061, EPI_ISL_540062, EPI_ISL_540063, EPI_ISL_540064, EPI_ISL_540065, EPI_ISL_540066, EPI_ISL_540067, EPI_ISL_540068, EPI_ISL_540069, EPI_ISL_540070, EPI_ISL_540071, EPI_ISL_540072, EPI_ISL_540073, EPI_ISL_540074, EPI_ISL_540075, EPI_ISL_540076, EPI_ISL_540077, EPI_ISL_540078, EPI_ISL_540079, EPI_ISL_540080, EPI_ISL_540081, EPI_ISL_540082, EPI_ISL_540083, EPI_ISL_540084, EPI_ISL_540085, EPI_ISL_540086, EPI_ISL_540087, EPI_ISL_540088, EPI_ISL_540089, EPI_ISL_540090, EPI_ISL_540091, EPI_ISL_540092, EPI_ISL_540093, EPI_ISL_540094, EPI_ISL_540095, EPI_ISL_540096, EPI_ISL_540097, EPI_ISL_540098, EPI_ISL_540099, EPI_ISL_540100, EPI_ISL_540101, EPI_ISL_540102, EPI_ISL_540103, EPI_ISL_540104, EPI_ISL_540105, EPI_ISL_540106, EPI_ISL_540107, EPI_ISL_540108, EPI_ISL_540109, EPI_ISL_540110, EPI_ISL_540111, EPI_ISL_540112, EPI_ISL_540113, EPI_ISL_540114, EPI_ISL_540115, EPI_ISL_540116, EPI_ISL_540117, EPI_ISL_540118, EPI_ISL_540119, EPI_ISL_540120, EPI_ISL_540121, EPI_ISL_540122, EPI_ISL_540123, EPI_ISL_540124, EPI_ISL_540125, EPI_ISL_540126, EPI_ISL_540127, EPI_ISL_540128, EPI_ISL_540129, EPI_ISL_540130, EPI_ISL_540131, EPI_ISL_540132, EPI_ISL_540133, EPI_ISL_540134, EPI_ISL_540135, EPI_ISL_540136, EPI_ISL_540137, EPI_ISL_540138, EPI_ISL_540139, EPI_ISL_540140, EPI_ISL_540141, EPI_ISL_540142, EPI_ISL_540143, EPI_ISL_540144, EPI_ISL_540145, EPI_ISL_540146, EPI_ISL_540147, EPI_ISL_540148, EPI_ISL_540149, EPI_ISL_540150, EPI_ISL_540151, EPI_ISL_540152, EPI_ISL_540153, EPI_ISL_540154, EPI_ISL_540155, EPI_ISL_540156, EPI_ISL_540157, EPI_ISL_540158, EPI_ISL_540159, EPI_ISL_540160, EPI_ISL_540161, EPI_ISL_540162, EPI_ISL_540163, EPI_ISL_540164, EPI_ISL_540165, EPI_ISL_540166, EPI_ISL_540167, EPI_ISL_540168, EPI_ISL_540169, EPI_ISL_540170, EPI_ISL_540171, EPI_ISL_540172, EPI_ISL_540173, EPI_ISL_540174, EPI_ISL_540175, EPI_ISL_540176, EPI_ISL_540177, EPI_ISL_540178, EPI_ISL_540179, EPI_ISL_540180, EPI_ISL_540181, EPI_ISL_540182, EPI_ISL_540183, EPI_ISL_540184, EPI_ISL_540185, EPI_ISL_540186, EPI_ISL_540187, EPI_ISL_540188, EPI_ISL_540189, EPI_ISL_540190, EPI_ISL_540191, EPI_ISL_540192, EPI_ISL_540193, EPI_ISL_540194, EPI_ISL_540195, EPI_ISL_540196, EPI_ISL_540197, EPI_ISL_540198, EPI_ISL_540199, EPI_ISL_540200, EPI_ISL_540201, EPI_ISL_540202, EPI_ISL_540203, EPI_ISL_540204, EPI_ISL_540205, EPI_ISL_540206, EPI_ISL_540207, EPI_ISL_540208, EPI_ISL_540209, EPI_ISL_540210, EPI_ISL_540211, EPI_ISL_540212, EPI_ISL_540213, EPI_ISL_540214, EPI_ISL_540215, EPI_ISL_540216, EPI_ISL_540217, EPI_ISL_540218, EPI_ISL_540219, EPI_ISL_540220, EPI_ISL_540221, EPI_ISL_540222, EPI_ISL_540223, EPI_ISL_540224, EPI_ISL_540225, EPI_ISL_540226, EPI_ISL_540227, EPI_ISL_540228, EPI_ISL_540229, EPI_ISL_540230, EPI_ISL_540231, EPI_ISL_540232, EPI_ISL_540233, EPI_ISL_540234, EPI_ISL_540235, EPI_ISL_540236, EPI_ISL_540237, EPI_ISL_540238, EPI_ISL_540239, EPI_ISL_540240, EPI_ISL_540241, EPI_ISL_540242, EPI_ISL_540243, EPI_ISL_540244, EPI_ISL_540245, EPI_ISL_540246, EPI_ISL_540247, EPI_ISL_540248, EPI_ISL_540249, EPI_ISL_540250, EPI_ISL_540251, EPI_ISL_540252, EPI_ISL_540253, EPI_ISL_540254, EPI_ISL_540255, EPI_ISL_540256, EPI_ISL_540257, EPI_ISL_540258, EPI_ISL_540259, EPI_ISL_540260, EPI_ISL_540261, EPI_ISL_540262, EPI_ISL_540263, EPI_ISL_540264, EPI_ISL_540265, EPI_ISL_540266, EPI_ISL_540267, EPI_ISL_540268, EPI_ISL_540269, EPI_ISL_540270, EPI_ISL_540271, EPI_ISL_540272, EPI_ISL_540273, EPI_ISL_540274, EPI_ISL_540275, EPI_ISL_540276, EPI_ISL_540277, EPI_ISL_540279, EPI_ISL_540280, EPI_ISL_540282, EPI_ISL_540283, EPI_ISL_540284, EPI_ISL_540285, EPI_ISL_540286, EPI_ISL_540287, EPI_ISL_540288, EPI_ISL_540289, EPI_ISL_540290, EPI_ISL_540291, EPI_ISL_540292, EPI_ISL_540293, EPI_ISL_540294, EPI_ISL_540295, EPI_ISL_540296, EPI_ISL_540297, EPI_ISL_540298, EPI_ISL_540299, EPI_ISL_540300, EPI_ISL_540301, EPI_ISL_540302, EPI_ISL_540303, EPI_ISL_540304, EPI_ISL_540305, EPI_ISL_540306, EPI_ISL_540307, EPI_ISL_540308, EPI_ISL_540309, EPI_ISL_540310, EPI_ISL_540311, EPI_ISL_540312, EPI_ISL_540313, EPI_ISL_540314, EPI_ISL_540315, EPI_ISL_540316, EPI_ISL_540317, EPI_ISL_540318, EPI_ISL_540319, EPI_ISL_540320, EPI_ISL_540321, EPI_ISL_540322, EPI_ISL_540323, EPI_ISL_540324, EPI_ISL_540325, EPI_ISL_540326, EPI_ISL_540327, EPI_ISL_540328, EPI_ISL_540329, EPI_ISL_540330, EPI_ISL_540331, EPI_ISL_540332, EPI_ISL_540333, EPI_ISL_540334, EPI_ISL_540335, EPI_ISL_540336, EPI_ISL_540337, EPI_ISL_540338, EPI_ISL_540339, EPI_ISL_540340, EPI_ISL_540341, EPI_ISL_540342, EPI_ISL_540343, EPI_ISL_540344, EPI_ISL_540345, EPI_ISL_540346, EPI_ISL_540347, EPI_ISL_540348, EPI_ISL_540349, EPI_ISL_540350, EPI_ISL_540352, EPI_ISL_540353, EPI_ISL_540354, EPI_ISL_540355, EPI_ISL_540356, EPI_ISL_540357, EPI_ISL_540358, EPI_ISL_540359, EPI_ISL_540360, EPI_ISL_540361, EPI_ISL_540362, EPI_ISL_540363, EPI_ISL_540364, EPI_ISL_540365, EPI_ISL_540366, EPI_ISL_540367, EPI_ISL_540368, EPI_ISL_540369, EPI_ISL_540370, EPI_ISL_540371, EPI_ISL_540372, EPI_ISL_540373, EPI_ISL_540374, EPI_ISL_540375, EPI_ISL_540376, EPI_ISL_540377, EPI_ISL_540378, EPI_ISL_540379, EPI_ISL_540380, EPI_ISL_540381, EPI_ISL_540382, EPI_ISL_540383, EPI_ISL_540384, EPI_ISL_540385, EPI_ISL_540386, EPI_ISL_540387, EPI_ISL_540388, EPI_ISL_540389, EPI_ISL_540390, EPI_ISL_540391, EPI_ISL_540392, EPI_ISL_540393, EPI_ISL_540394, EPI_ISL_540395, EPI_ISL_540396, EPI_ISL_540397, EPI_ISL_540398, EPI_ISL_540399, EPI_ISL_540400, EPI_ISL_540401, EPI_ISL_540402, EPI_ISL_540403, EPI_ISL_540404, EPI_ISL_540405, EPI_ISL_540406, EPI_ISL_540407, EPI_ISL_540408, EPI_ISL_540409, EPI_ISL_540410, EPI_ISL_540411, EPI_ISL_540412, EPI_ISL_540413, EPI_ISL_540414, EPI_ISL_540415, EPI_ISL_540416, EPI_ISL_540417, EPI_ISL_540418, EPI_ISL_540419, EPI_ISL_540420 | | | |
| see above | Lighthouse Lab in Glasgow | Wellcome Sanger Institute for the COVID-19 Genomics UK (COG-UK) consortium | Harper VanSteenhouse, Yumi Kasai, David Gray, Carol Clugston, Anna Dominiczak and Alex Alderton, Roberto Amato, Sonia Goncalves, Ewan Harrison, David K. Jackson, Ian Johnston, Dominic Kwiatkowski, Cordelia Langford, John Sillitoe on behalf of the Wellcome Sanger Institute COVID-19 Surveillance Team |
| EPI_ISL_541083, EPI_ISL_541084, EPI_ISL_541085, EPI_ISL_541086, EPI_ISL_541087, EPI_ISL_541088, EPI_ISL_541089, EPI_ISL_541090, EPI_ISL_541091, EPI_ISL_541092, EPI_ISL_541093, EPI_ISL_541094, EPI_ISL_541095, EPI_ISL_541096, EPI_ISL_541097, EPI_ISL_541098, EPI_ISL_541099, EPI_ISL_541100, EPI_ISL_541101, EPI_ISL_541102, EPI_ISL_541103, EPI_ISL_541104, EPI_ISL_541105, EPI_ISL_541106, EPI_ISL_541107, EPI_ISL_541108, EPI_ISL_541109, EPI_ISL_541110, EPI_ISL_541111, EPI_ISL_541112, EPI_ISL_541113, EPI_ISL_541114, EPI_ISL_541115, EPI_ISL_541116, EPI_ISL_541117, EPI_ISL_541118, EPI_ISL_541119, EPI_ISL_541120, EPI_ISL_541121, EPI_ISL_541122, EPI_ISL_541123, EPI_ISL_541124, EPI_ISL_541125, EPI_ISL_541126, EPI_ISL_541127, EPI_ISL_541128, EPI_ISL_541129, EPI_ISL_541130, EPI_ISL_541131, EPI_ISL_541132, EPI_ISL_541133, EPI_ISL_541134, EPI_ISL_541135, EPI_ISL_541136, EPI_ISL_541137 | | | |
| see above | Servicio de Microbiología. Hospital Universitario Donostia. OSI Donostialdea. Área de Enfermedades Infecciosas, Grupo de Infección Respiratoria y Resistencia Antimicrobiana. Instituto de Investigación Sanitaria Biodonostia | SeqCOVID-SPAIN consortium/Institute of Biomedicine of Valencia, IBV-CSIC | Gustavo Cilla, Milagrosa Montes, Luis Piñeiro, Jose Maria Marimón and SeqCOVID-SPAIN consortium |
| EPI_ISL_541784, EPI_ISL_541785, EPI_ISL_541786, EPI_ISL_541787, EPI_ISL_541788, EPI_ISL_541789, EPI_ISL_541790, EPI_ISL_541791, EPI_ISL_541792, EPI_ISL_541793, EPI_ISL_541794, EPI_ISL_541795, EPI_ISL_541796, EPI_ISL_541797, EPI_ISL_541798, EPI_ISL_541799, EPI_ISL_541800, EPI_ISL_541801, EPI_ISL_541802, EPI_ISL_541803, EPI_ISL_541804, EPI_ISL_541805, EPI_ISL_541806, EPI_ISL_541807, EPI_ISL_541808, EPI_ISL_541809, EPI_ISL_541810, EPI_ISL_541811, EPI_ISL_541812, EPI_ISL_541813, EPI_ISL_541814, EPI_ISL_541815, EPI_ISL_541816, EPI_ISL_541817, EPI_ISL_541818, EPI_ISL_541819, EPI_ISL_541820, EPI_ISL_541821, EPI_ISL_541822, EPI_ISL_541823, EPI_ISL_541824, EPI_ISL_541825, EPI_ISL_541826, EPI_ISL_541827, EPI_ISL_541828, EPI_ISL_541829, EPI_ISL_541830, EPI_ISL_541831, EPI_ISL_541832, EPI_ISL_541833, EPI_ISL_541834, EPI_ISL_541835, EPI_ISL_541836, EPI_ISL_541837, EPI_ISL_541838, EPI_ISL_541839, EPI_ISL_541840, EPI_ISL_541841, EPI_ISL_541842, EPI_ISL_541843, EPI_ISL_541844, EPI_ISL_541845, EPI_ISL_541846 | | | |
| see above | Lighthouse Lab in Glasgow | Wellcome Sanger Institute for the COVID-19 Genomics UK (COG-UK) consortium | Harper VanSteenhouse, Yumi Kasai, David Gray, Carol Clugston, Anna Dominiczak and Alex Alderton, Roberto Amato, Sonia Goncalves, Ewan Harrison, David K. Jackson, Ian Johnston, Dominic Kwiatkowski, Cordelia Langford, John Sillitoe on behalf of the Wellcome Sanger Institute COVID-19 Surveillance Team |
| EPI_ISL_548243, EPI_ISL_548244 | Faith Laboratory, Immunology Institute, Icahn School of Medicine at Mount Sinai | van Bakel Laboratory, Genetics and Genomics Sciences, Icahn School of Medicine at Mount Sinai | Graham J. Britton, Alice Chen-Liaw, Francesca Cossarini, Alexandra Livanos, Matthew P. Spindler, Tamar Plitt, Joseph Eggers, Ilaria Mogno, Ana S. Gonzalez-Reiche, Sophia Sui, Michael Tankelevich, Lauren Tal Grinspan, Rebekah E. Dixon, Divya Jha, Gustavo Martinez-Delgado, Fatima Amanat, Daisy Hoagland, Benjamin R. tenOever, Marla C. Dubinsky, Miriam Merad, Harm Van Bakel, Florian Krammer, Gerold Bongers, Saurabh Mehandru and Jeremiah J. Faith |
| EPI_ISL_549547, EPI_ISL_549548, EPI_ISL_549549, EPI_ISL_549550, EPI_ISL_549551, EPI_ISL_549552, EPI_ISL_549553, EPI_ISL_549554, EPI_ISL_549555, EPI_ISL_549556, EPI_ISL_549557, EPI_ISL_549558, EPI_ISL_549559, EPI_ISL_549560, EPI_ISL_549561, EPI_ISL_549562, EPI_ISL_549563, EPI_ISL_549564, EPI_ISL_549565, EPI_ISL_549566, EPI_ISL_549567, EPI_ISL_549568, EPI_ISL_549569, EPI_ISL_549570, EPI_ISL_549571, EPI_ISL_549572, EPI_ISL_549573, EPI_ISL_549574, EPI_ISL_549575, EPI_ISL_549576, EPI_ISL_549577, EPI_ISL_549578, EPI_ISL_549579, EPI_ISL_549580, EPI_ISL_549581, EPI_ISL_549582, EPI_ISL_549583, EPI_ISL_549584, EPI_ISL_549585, EPI_ISL_549586, EPI_ISL_549587, EPI_ISL_549588, EPI_ISL_549589, EPI_ISL_549590, EPI_ISL_549591, EPI_ISL_549592, EPI_ISL_549593, EPI_ISL_549594, EPI_ISL_549595, EPI_ISL_549596, EPI_ISL_549597, EPI_ISL_549598, EPI_ISL_549599, EPI_ISL_549600, EPI_ISL_549601, EPI_ISL_549602, EPI_ISL_549603, EPI_ISL_549604, EPI_ISL_549605, EPI_ISL_549606, EPI_ISL_549607, EPI_ISL_549608, EPI_ISL_549609, EPI_ISL_549610, EPI_ISL_549611, EPI_ISL_549612, EPI_ISL_549613, EPI_ISL_549614, EPI_ISL_549615, EPI_ISL_549616, EPI_ISL_549617, EPI_ISL_549618, EPI_ISL_549619, EPI_ISL_549620, EPI_ISL_549621, EPI_ISL_549622, EPI_ISL_549623, EPI_ISL_549624, EPI_ISL_549625, EPI_ISL_549626, EPI_ISL_549627, EPI_ISL_549628, EPI_ISL_549629, EPI_ISL_549630, EPI_ISL_549631, EPI_ISL_549632, EPI_ISL_549633, EPI_ISL_549634, EPI_ISL_549635, EPI_ISL_549636, EPI_ISL_549637, EPI_ISL_549638, EPI_ISL_549639, EPI_ISL_549640, EPI_ISL_549641, EPI_ISL_549642, EPI_ISL_549643, EPI_ISL_549644, EPI_ISL_549645, EPI_ISL_549646, EPI_ISL_549647, EPI_ISL_549648, EPI_ISL_549649, EPI_ISL_549650, EPI_ISL_549651, EPI_ISL_549652, EPI_ISL_549653, EPI_ISL_549654, EPI_ISL_549655, EPI_ISL_549656, EPI_ISL_549657, EPI_ISL_549658, EPI_ISL_549659, EPI_ISL_549660, EPI_ISL_549661, EPI_ISL_549662, EPI_ISL_549663, EPI_ISL_549664, EPI_ISL_549665, EPI_ISL_549666, EPI_ISL_549667, EPI_ISL_549668, EPI_ISL_549669, EPI_ISL_549670, EPI_ISL_549671, EPI_ISL_549672, EPI_ISL_549673, EPI_ISL_549674, EPI_ISL_549675, EPI_ISL_549676, EPI_ISL_549677, EPI_ISL_549678, EPI_ISL_549679, EPI_ISL_549680, EPI_ISL_549681, EPI_ISL_549682, EPI_ISL_549683, EPI_ISL_549684, EPI_ISL_549685, EPI_ISL_549686, EPI_ISL_549687, EPI_ISL_549688, EPI_ISL_549689, EPI_ISL_549690, EPI_ISL_549691, EPI_ISL_549692, EPI_ISL_549693, EPI_ISL_549694, EPI_ISL_549695, EPI_ISL_549696, EPI_ISL_549697, EPI_ISL_549698, EPI_ISL_549699, EPI_ISL_549700, EPI_ISL_549701, EPI_ISL_549702, EPI_ISL_549703, EPI_ISL_549704, EPI_ISL_549705, EPI_ISL_549706, EPI_ISL_549707, EPI_ISL_549708, EPI_ISL_549709, EPI_ISL_549710, EPI_ISL_549711, EPI_ISL_549712, EPI_ISL_549713, EPI_ISL_549714, EPI_ISL_549715, EPI_ISL_549716, EPI_ISL_549717, EPI_ISL_549718, EPI_ISL_549719, EPI_ISL_549720, EPI_ISL_549721, EPI_ISL_549722, EPI_ISL_549723, EPI_ISL_549724, EPI_ISL_549725, EPI_ISL_549726, EPI_ISL_549727, EPI_ISL_549728, EPI_ISL_549729, EPI_ISL_549730, EPI_ISL_549731, EPI_ISL_549732, EPI_ISL_549733, EPI_ISL_549734, EPI_ISL_549735, EPI_ISL_549736, EPI_ISL_549737, EPI_ISL_549738, EPI_ISL_549739, EPI_ISL_549740, EPI_ISL_549741, EPI_ISL_549742, EPI_ISL_549743, EPI_ISL_549744, EPI_ISL_549745, EPI_ISL_549746, EPI_ISL_549747, EPI_ISL_549748, EPI_ISL_549749, EPI_ISL_549750, EPI_ISL_549751, EPI_ISL_549752, EPI_ISL_549753, EPI_ISL_549754, EPI_ISL_549755, EPI_ISL_549756, EPI_ISL_549757, EPI_ISL_549758, EPI_ISL_549759, EPI_ISL_549760, EPI_ISL_549761, EPI_ISL_549762, EPI_ISL_549763, EPI_ISL_549764, EPI_ISL_549765, EPI_ISL_549766, EPI_ISL_549767, EPI_ISL_549768, EPI_ISL_549769, EPI_ISL_549770, EPI_ISL_549771, EPI_ISL_549773, EPI_ISL_549774, EPI_ISL_549775, EPI_ISL_549776, EPI_ISL_549777, EPI_ISL_549778, EPI_ISL_549779, EPI_ISL_549780, EPI_ISL_549781, EPI_ISL_549782, EPI_ISL_549783, EPI_ISL_549785, EPI_ISL_549786, EPI_ISL_549787, EPI_ISL_549788, EPI_ISL_549789, EPI_ISL_549790, EPI_ISL_549791, EPI_ISL_549792, EPI_ISL_549793, EPI_ISL_549794, EPI_ISL_549795, EPI_ISL_549796, EPI_ISL_549797, EPI_ISL_549798, EPI_ISL_549799, EPI_ISL_549800, EPI_ISL_549801, EPI_ISL_549802, EPI_ISL_549803, EPI_ISL_549804, EPI_ISL_549805, EPI_ISL_549806, EPI_ISL_549807, EPI_ISL_553732 | | | |
| see above | Lighthouse Lab in Glasgow | Wellcome Sanger Institute for the COVID-19 Genomics UK (COG-UK) consortium | Harper VanSteenhouse, Yumi Kasai, David Gray, Carol Clugston, Anna Dominiczak and Alex Alderton, Roberto Amato, Sonia Goncalves, Ewan Harrison, David K. Jackson, Ian Johnston, Dominic Kwiatkowski, Cordelia Langford, John Sillitoe on behalf of the Wellcome Sanger Institute COVID-19 Surveillance Team |
| EPI_ISL_560307, EPI_ISL_560406, EPI_ISL_560567, EPI_ISL_560647, EPI_ISL_560648, EPI_ISL_560741, EPI_ISL_560742, EPI_ISL_561335, EPI_ISL_561336, EPI_ISL_561345, EPI_ISL_561346, EPI_ISL_561347, EPI_ISL_561349, EPI_ISL_561350, EPI_ISL_561351, EPI_ISL_561352, EPI_ISL_561353, EPI_ISL_561356, EPI_ISL_561357, EPI_ISL_561358, EPI_ISL_561359, EPI_ISL_561360, EPI_ISL_561362, EPI_ISL_565830, EPI_ISL_565831, EPI_ISL_565832 | | | |
| see above | Delaware Public Health Lab | Delaware Public Health Lab | Gregory Hovan |
| EPI_ISL_565920, EPI_ISL_565921, EPI_ISL_565922, EPI_ISL_565923, EPI_ISL_565924, EPI_ISL_565925 | Servicio de Microbiología. Hospital Universitario Donostia. OSI Donostialdea. Área de Enfermedades Infecciosas, Grupo de Infección Respiratoria y Resistencia Antimicrobiana. Instituto de Investigación Sanitaria Biodonostia | SeqCOVID-SPAIN consortium/IBV(CSIC) | Gustavo Cilla, Milagrosa Montes, Luis Piñeiro, Jose Maria Marimón and SeqCOVID-SPAIN consortium |
| EPI_ISL_566084, EPI_ISL_566108, EPI_ISL_566109 | Delaware Public Health Lab | Delaware Public Health Lab | Gregory Hovan |
| EPI_ISL_566998, EPI_ISL_566999, EPI_ISL_567000, EPI_ISL_567001, EPI_ISL_567002, EPI_ISL_567003, EPI_ISL_567004, EPI_ISL_567005, EPI_ISL_567006, EPI_ISL_567007, EPI_ISL_567008, EPI_ISL_567009, EPI_ISL_567010, EPI_ISL_567011, EPI_ISL_567012, EPI_ISL_567013, EPI_ISL_567014, EPI_ISL_567015, EPI_ISL_567016, EPI_ISL_567017, EPI_ISL_567018, EPI_ISL_567019, EPI_ISL_567020, EPI_ISL_567021, EPI_ISL_567022, EPI_ISL_567023, EPI_ISL_567025, EPI_ISL_567026, EPI_ISL_567027, EPI_ISL_567028, EPI_ISL_567029, EPI_ISL_567030, EPI_ISL_567031, EPI_ISL_567032, EPI_ISL_567033, EPI_ISL_567034, EPI_ISL_567035, EPI_ISL_567036, EPI_ISL_567037, EPI_ISL_567038, EPI_ISL_567039, EPI_ISL_567040, EPI_ISL_567041, EPI_ISL_567042, EPI_ISL_567043, EPI_ISL_567044, EPI_ISL_567045, EPI_ISL_567046, EPI_ISL_567047, EPI_ISL_567048, EPI_ISL_567049, EPI_ISL_567050, EPI_ISL_567051, EPI_ISL_567052, | | | |

| | | | |
|---|---|---|---|
| EPI_ISL_567053, EPI_ISL_567054, EPI_ISL_567055, EPI_ISL_567056, EPI_ISL_567057, EPI_ISL_567058, EPI_ISL_567060, EPI_ISL_567061, EPI_ISL_567062, EPI_ISL_567063, EPI_ISL_567064, EPI_ISL_567065, EPI_ISL_567066, EPI_ISL_567067, EPI_ISL_567068, EPI_ISL_567069, EPI_ISL_567070, EPI_ISL_567071, EPI_ISL_567072, EPI_ISL_567073, EPI_ISL_567074, EPI_ISL_567075, EPI_ISL_567076, EPI_ISL_567077, EPI_ISL_567078, EPI_ISL_567079, EPI_ISL_567080, EPI_ISL_567081, EPI_ISL_567082, EPI_ISL_567083, EPI_ISL_567084, EPI_ISL_567085, EPI_ISL_567086, EPI_ISL_567087, EPI_ISL_567088, EPI_ISL_567089, EPI_ISL_567090, EPI_ISL_567091, EPI_ISL_567092, EPI_ISL_567093, EPI_ISL_567094, EPI_ISL_567095, EPI_ISL_567096, EPI_ISL_567097, EPI_ISL_567098, EPI_ISL_567099, EPI_ISL_567100, EPI_ISL_567101, EPI_ISL_567102, EPI_ISL_567103, EPI_ISL_567104, EPI_ISL_567105, EPI_ISL_567106, EPI_ISL_567107, EPI_ISL_567108, EPI_ISL_567109, EPI_ISL_567110, EPI_ISL_567111, EPI_ISL_567112, EPI_ISL_567113, EPI_ISL_567114, EPI_ISL_567115, EPI_ISL_567116, EPI_ISL_567117, EPI_ISL_567118, EPI_ISL_567119, EPI_ISL_567120, EPI_ISL_567121, EPI_ISL_567122, EPI_ISL_567123, EPI_ISL_567124, EPI_ISL_567125, EPI_ISL_567126, EPI_ISL_567127, EPI_ISL_567128, EPI_ISL_567129, EPI_ISL_567130, EPI_ISL_567131, EPI_ISL_567132, EPI_ISL_567133, EPI_ISL_567134, EPI_ISL_567135, EPI_ISL_567136, EPI_ISL_567137, EPI_ISL_567138, EPI_ISL_567139, EPI_ISL_567140, EPI_ISL_567141, EPI_ISL_567142, EPI_ISL_567143, EPI_ISL_567144, EPI_ISL_567145, EPI_ISL_567146, EPI_ISL_567147, EPI_ISL_567148, EPI_ISL_567149, EPI_ISL_567150, EPI_ISL_567151, EPI_ISL_567152, EPI_ISL_567153, EPI_ISL_567154, EPI_ISL_567155, EPI_ISL_567156, EPI_ISL_567157, EPI_ISL_567158, EPI_ISL_567159, EPI_ISL_567160, EPI_ISL_567161, EPI_ISL_567162, EPI_ISL_567163, EPI_ISL_567164, EPI_ISL_567165, EPI_ISL_567166, EPI_ISL_567167, EPI_ISL_567168, EPI_ISL_567169, EPI_ISL_567170, EPI_ISL_567171, EPI_ISL_567172, EPI_ISL_567173, EPI_ISL_567174, EPI_ISL_567175, EPI_ISL_567176, EPI_ISL_567177, EPI_ISL_567178, EPI_ISL_567179, EPI_ISL_567180, EPI_ISL_567181, EPI_ISL_567182, EPI_ISL_567183, EPI_ISL_567185, EPI_ISL_567186, EPI_ISL_567187, EPI_ISL_567188, EPI_ISL_567189, EPI_ISL_567190, EPI_ISL_567191, EPI_ISL_567192, EPI_ISL_567193, EPI_ISL_567194, EPI_ISL_567195, EPI_ISL_567196, EPI_ISL_567197, EPI_ISL_567198, EPI_ISL_567199, EPI_ISL_567200, EPI_ISL_567201, EPI_ISL_567203, EPI_ISL_567204, EPI_ISL_567205, EPI_ISL_567206, EPI_ISL_567207, EPI_ISL_567208, EPI_ISL_567209, EPI_ISL_567210, EPI_ISL_567211, EPI_ISL_567212, EPI_ISL_567213, EPI_ISL_567214, EPI_ISL_567215, EPI_ISL_567216, EPI_ISL_567217, EPI_ISL_567218, EPI_ISL_567219, EPI_ISL_567220, EPI_ISL_567221, EPI_ISL_567222, EPI_ISL_567223, EPI_ISL_567224, EPI_ISL_567225, EPI_ISL_567226, EPI_ISL_567227, EPI_ISL_567228, EPI_ISL_567229, EPI_ISL_567230, EPI_ISL_567232, EPI_ISL_567233, EPI_ISL_567234, EPI_ISL_567235, EPI_ISL_567236, EPI_ISL_567237, EPI_ISL_567238, EPI_ISL_567239, EPI_ISL_567240, EPI_ISL_567241, EPI_ISL_567242, EPI_ISL_567243, EPI_ISL_567244, EPI_ISL_567245, EPI_ISL_567246, EPI_ISL_567247, EPI_ISL_567248, EPI_ISL_567249, EPI_ISL_567250, EPI_ISL_567251, EPI_ISL_567252, EPI_ISL_567253, EPI_ISL_567254, EPI_ISL_567255, EPI_ISL_567256, EPI_ISL_567257, EPI_ISL_567258, EPI_ISL_567259, EPI_ISL_567260, EPI_ISL_567261, EPI_ISL_567262, EPI_ISL_567263, EPI_ISL_567264, EPI_ISL_567265, EPI_ISL_567266, EPI_ISL_567267, EPI_ISL_567268, EPI_ISL_567269, EPI_ISL_567270, EPI_ISL_567271, EPI_ISL_567272, EPI_ISL_567273, EPI_ISL_567274, EPI_ISL_567275, EPI_ISL_567276, EPI_ISL_567277, EPI_ISL_567278, EPI_ISL_567279, EPI_ISL_567280, EPI_ISL_567281, EPI_ISL_567282, EPI_ISL_567283, EPI_ISL_567284, EPI_ISL_567285, EPI_ISL_567286, EPI_ISL_567287, EPI_ISL_567288, EPI_ISL_567289, EPI_ISL_567290, EPI_ISL_567291, EPI_ISL_567292, EPI_ISL_567293, EPI_ISL_567294, EPI_ISL_567295, EPI_ISL_567296, EPI_ISL_567297, EPI_ISL_567298, EPI_ISL_567299, EPI_ISL_567300, EPI_ISL_567301, EPI_ISL_567302, EPI_ISL_567303, EPI_ISL_567304, EPI_ISL_567305, EPI_ISL_567306, EPI_ISL_567308, EPI_ISL_567309, EPI_ISL_567310, EPI_ISL_567311, EPI_ISL_567312, EPI_ISL_567313, EPI_ISL_567314, EPI_ISL_567315, EPI_ISL_567316, EPI_ISL_567317, EPI_ISL_567318, EPI_ISL_567319, EPI_ISL_567320, EPI_ISL_567321, EPI_ISL_567322, EPI_ISL_567323, EPI_ISL_567324, EPI_ISL_567325, EPI_ISL_567326, EPI_ISL_567327, EPI_ISL_567328, EPI_ISL_567329, EPI_ISL_567330, EPI_ISL_567331, EPI_ISL_567332, EPI_ISL_567333, EPI_ISL_567334, EPI_ISL_567335, EPI_ISL_567336, EPI_ISL_567337, EPI_ISL_567338, EPI_ISL_567339, EPI_ISL_567340, EPI_ISL_567341, EPI_ISL_567342, EPI_ISL_567343, EPI_ISL_567344, EPI_ISL_567345, EPI_ISL_567346, EPI_ISL_567347, EPI_ISL_567348, EPI_ISL_567349, EPI_ISL_567350, EPI_ISL_567351, EPI_ISL_567352, EPI_ISL_567353, EPI_ISL_567354, EPI_ISL_567355, EPI_ISL_567356, EPI_ISL_567357, EPI_ISL_567358, EPI_ISL_567359, EPI_ISL_567360, EPI_ISL_567361, EPI_ISL_567362, EPI_ISL_567363, EPI_ISL_567364, EPI_ISL_567365, EPI_ISL_567366, EPI_ISL_567367, EPI_ISL_567368, EPI_ISL_567369, EPI_ISL_567370, EPI_ISL_567371, EPI_ISL_567372, EPI_ISL_567373, EPI_ISL_567374, EPI_ISL_567375, EPI_ISL_567376, EPI_ISL_567377, EPI_ISL_567378, EPI_ISL_567379, EPI_ISL_567380, EPI_ISL_567381, EPI_ISL_567382, EPI_ISL_567383, EPI_ISL_567384, EPI_ISL_567385, EPI_ISL_567386, EPI_ISL_567387, EPI_ISL_567388, EPI_ISL_567389, EPI_ISL_567390, EPI_ISL_567391, EPI_ISL_567392, EPI_ISL_567393, EPI_ISL_567394, EPI_ISL_567395, EPI_ISL_567396, EPI_ISL_567397, EPI_ISL_567398, EPI_ISL_567399, EPI_ISL_567400, EPI_ISL_567401, EPI_ISL_567402, EPI_ISL_567403, EPI_ISL_567404, EPI_ISL_567405, EPI_ISL_567406, EPI_ISL_567407, EPI_ISL_567408, EPI_ISL_567409, EPI_ISL_567410, EPI_ISL_567411, EPI_ISL_567412, EPI_ISL_567413, EPI_ISL_567414, EPI_ISL_567415, EPI_ISL_567416, EPI_ISL_567417, EPI_ISL_567418, EPI_ISL_567419, EPI_ISL_567420, EPI_ISL_567421, EPI_ISL_567422, EPI_ISL_567423, EPI_ISL_567424, EPI_ISL_567425, EPI_ISL_567426, EPI_ISL_567427, EPI_ISL_567428, EPI_ISL_567429, EPI_ISL_567430, EPI_ISL_567431, EPI_ISL_567432, EPI_ISL_567433, EPI_ISL_567434, EPI_ISL_567435, EPI_ISL_567436, EPI_ISL_567437, EPI_ISL_567438, EPI_ISL_567439, EPI_ISL_567440, EPI_ISL_567441, EPI_ISL_567442, EPI_ISL_567443, EPI_ISL_567444, EPI_ISL_567445, EPI_ISL_567446, EPI_ISL_567447, EPI_ISL_567448, EPI_ISL_567449, EPI_ISL_567450, EPI_ISL_567451, EPI_ISL_567452, EPI_ISL_567453, EPI_ISL_567454, EPI_ISL_567455, EPI_ISL_567456, EPI_ISL_567457, EPI_ISL_567458, EPI_ISL_567459, EPI_ISL_567460, EPI_ISL_567461, EPI_ISL_567462, EPI_ISL_567463, EPI_ISL_567464, EPI_ISL_567465, EPI_ISL_567466, EPI_ISL_567467, EPI_ISL_567468, EPI_ISL_567469, EPI_ISL_567470, EPI_ISL_567471, EPI_ISL_567472, EPI_ISL_567473, EPI_ISL_567474, EPI_ISL_567475, EPI_ISL_567476, EPI_ISL_567477, EPI_ISL_567478, EPI_ISL_567479, EPI_ISL_567480, EPI_ISL_567481, EPI_ISL_567482, EPI_ISL_567483, EPI_ISL_567484, EPI_ISL_567485, EPI_ISL_567486, EPI_ISL_567487, EPI_ISL_567488, EPI_ISL_567489, EPI_ISL_567490, EPI_ISL_567491, EPI_ISL_567492, EPI_ISL_567493, EPI_ISL_567494, EPI_ISL_567495, EPI_ISL_567496, EPI_ISL_567497, EPI_ISL_567498, EPI_ISL_567499, EPI_ISL_567500, EPI_ISL_567501, EPI_ISL_567502, EPI_ISL_567503, EPI_ISL_567504, EPI_ISL_567505, EPI_ISL_567506, EPI_ISL_567507, EPI_ISL_567508, EPI_ISL_567509, EPI_ISL_567510, EPI_ISL_567511, EPI_ISL_567512, EPI_ISL_567513, EPI_ISL_567514, EPI_ISL_567515, EPI_ISL_567516, EPI_ISL_567517, EPI_ISL_567518, EPI_ISL_567519, EPI_ISL_567520, EPI_ISL_567521, EPI_ISL_567522, EPI_ISL_567523, EPI_ISL_567524, EPI_ISL_567525, EPI_ISL_567526, EPI_ISL_567527, EPI_ISL_567528, EPI_ISL_567529, EPI_ISL_567530, EPI_ISL_567531, EPI_ISL_567532, EPI_ISL_568223, EPI_ISL_568224, EPI_ISL_568225, EPI_ISL_568226, EPI_ISL_568227, EPI_ISL_568228, EPI_ISL_568229, EPI_ISL_568230, EPI_ISL_568231, EPI_ISL_568232, EPI_ISL_568233, EPI_ISL_568234, EPI_ISL_568235, EPI_ISL_568236, EPI_ISL_568237, EPI_ISL_568238, EPI_ISL_568239, EPI_ISL_568240, EPI_ISL_568241, EPI_ISL_568242, EPI_ISL_568243, EPI_ISL_568244, EPI_ISL_568245, EPI_ISL_568246, EPI_ISL_568247, EPI_ISL_568248, EPI_ISL_568249, EPI_ISL_568250, EPI_ISL_568251, EPI_ISL_568252, EPI_ISL_568253, EPI_ISL_568254, EPI_ISL_568255, EPI_ISL_568256, EPI_ISL_568257, EPI_ISL_568258, EPI_ISL_568259, EPI_ISL_568260, EPI_ISL_568261, EPI_ISL_568262, EPI_ISL_568263, EPI_ISL_568264, EPI_ISL_568265, EPI_ISL_568266, EPI_ISL_568267, EPI_ISL_568268, EPI_ISL_568269, EPI_ISL_568270, EPI_ISL_568271, EPI_ISL_568272, EPI_ISL_568273, EPI_ISL_568274, EPI_ISL_568275, EPI_ISL_568276, EPI_ISL_568277, EPI_ISL_568278, EPI_ISL_568279, EPI_ISL_568280, EPI_ISL_568281, EPI_ISL_568282, EPI_ISL_568283, EPI_ISL_568284, EPI_ISL_568285, EPI_ISL_568286, EPI_ISL_568287, EPI_ISL_568288, EPI_ISL_568291, EPI_ISL_568292, EPI_ISL_568293, EPI_ISL_568294, EPI_ISL_568295, EPI_ISL_568296, EPI_ISL_568297, EPI_ISL_568298, EPI_ISL_568299, EPI_ISL_568300, EPI_ISL_568302, EPI_ISL_568303, EPI_ISL_568304, EPI_ISL_568305, EPI_ISL_568306, EPI_ISL_568307, EPI_ISL_568308, EPI_ISL_568309, EPI_ISL_568310, EPI_ISL_568311, EPI_ISL_568312, EPI_ISL_568313, EPI_ISL_568314, EPI_ISL_568316, EPI_ISL_568318, EPI_ISL_568319, EPI_ISL_568320, EPI_ISL_568321, EPI_ISL_568322, EPI_ISL_568323, EPI_ISL_568324, EPI_ISL_568325, EPI_ISL_568326, EPI_ISL_568327, EPI_ISL_568328, EPI_ISL_568329, EPI_ISL_568330, EPI_ISL_568331, EPI_ISL_568332, EPI_ISL_568333, EPI_ISL_568334, EPI_ISL_568335, EPI_ISL_568336, EPI_ISL_568337, EPI_ISL_568338, EPI_ISL_568339, EPI_ISL_568340, EPI_ISL_568341, EPI_ISL_568342, EPI_ISL_568343, EPI_ISL_568345, EPI_ISL_568346, EPI_ISL_568347, EPI_ISL_568348, EPI_ISL_568349, EPI_ISL_568350, EPI_ISL_568351, EPI_ISL_568352, EPI_ISL_568353, EPI_ISL_568354, EPI_ISL_568355, EPI_ISL_568356, EPI_ISL_568357, EPI_ISL_568358, EPI_ISL_568359, EPI_ISL_568360, EPI_ISL_568361, EPI_ISL_568362, EPI_ISL_568363, EPI_ISL_568366, EPI_ISL_568367, EPI_ISL_568368, EPI_ISL_568369, EPI_ISL_568370, EPI_ISL_568371, EPI_ISL_568372, EPI_ISL_568373, EPI_ISL_568374, EPI_ISL_568375, EPI_ISL_568376, EPI_ISL_568377, EPI_ISL_568378, EPI_ISL_568379, EPI_ISL_568380, EPI_ISL_568381, EPI_ISL_568382, EPI_ISL_568383, EPI_ISL_568384, EPI_ISL_568388, EPI_ISL_568389, EPI_ISL_568390, EPI_ISL_568391, EPI_ISL_568393, EPI_ISL_568394, EPI_ISL_568396, EPI_ISL_568397, EPI_ISL_568399, EPI_ISL_568400, EPI_ISL_568401, EPI_ISL_568402, EPI_ISL_568403, EPI_ISL_568404, EPI_ISL_568405, EPI_ISL_568406, EPI_ISL_568407, EPI_ISL_568408, EPI_ISL_568409, EPI_ISL_568410, EPI_ISL_568411, EPI_ISL_568412, EPI_ISL_568413, EPI_ISL_568414, EPI_ISL_568415, EPI_ISL_568416, EPI_ISL_568417, EPI_ISL_568418, EPI_ISL_568419, EPI_ISL_568420, EPI_ISL_568421, EPI_ISL_568422, EPI_ISL_568423, EPI_ISL_568424, EPI_ISL_568425, EPI_ISL_568426, EPI_ISL_568427, EPI_ISL_568428, EPI_ISL_568429, EPI_ISL_568430, EPI_ISL_568431, EPI_ISL_568432, EPI_ISL_568433, EPI_ISL_568434, EPI_ISL_568435, EPI_ISL_568437, EPI_ISL_568438, EPI_ISL_568439, EPI_ISL_568440, EPI_ISL_568441, EPI_ISL_568442, EPI_ISL_568443, EPI_ISL_568444, EPI_ISL_568445, EPI_ISL_568446, EPI_ISL_568447, EPI_ISL_568448, EPI_ISL_568449, EPI_ISL_568450, EPI_ISL_568451, EPI_ISL_568452, EPI_ISL_568455, EPI_ISL_568456, EPI_ISL_568457, EPI_ISL_568459, EPI_ISL_568460, EPI_ISL_568462, EPI_ISL_568463, EPI_ISL_568464, EPI_ISL_568465 | | | |
| see above | Lighthouse Lab in Glasgow | Wellcome Sanger Institute for the COVID-19 Genomics UK (COG-UK) consortium | Harper VanSteenhouse, Yumi Kasai, David Gray, Carol Clugston, Anna Dominiczak and Alex Alderton, Roberto Amato, Sonia Goncalves, Ewan Harrison, David K. Jackson, Ian Johnston, Dominic Kwiatkowski, Cordelia Langford, John Sillitoe on behalf of the Wellcome Sanger Institute COVID-19 Surveillance Team |
| EPI_ISL_568695, EPI_ISL_568696, EPI_ISL_568697, EPI_ISL_568698, EPI_ISL_568699, EPI_ISL_568700, EPI_ISL_568701, EPI_ISL_568702, EPI_ISL_568703, EPI_ISL_568704, EPI_ISL_568705, EPI_ISL_568706, EPI_ISL_568707, EPI_ISL_568708, EPI_ISL_568709, EPI_ISL_568710, EPI_ISL_568711, EPI_ISL_568712, EPI_ISL_568713, EPI_ISL_568714, EPI_ISL_568715, EPI_ISL_568716, EPI_ISL_568717, EPI_ISL_568718, EPI_ISL_568719, EPI_ISL_568720, EPI_ISL_568721, EPI_ISL_568722, EPI_ISL_568723, EPI_ISL_568724, EPI_ISL_568725, EPI_ISL_568726, EPI_ISL_568727, EPI_ISL_568728, EPI_ISL_568729, EPI_ISL_568730, EPI_ISL_568731, EPI_ISL_568732, EPI_ISL_568733, EPI_ISL_568734, EPI_ISL_568735, EPI_ISL_568736, EPI_ISL_568737, EPI_ISL_568738, EPI_ISL_568739, EPI_ISL_568740, EPI_ISL_568741, EPI_ISL_568742, EPI_ISL_568743, EPI_ISL_568744, EPI_ISL_568745, EPI_ISL_568746, EPI_ISL_568747, EPI_ISL_568748, EPI_ISL_568749, EPI_ISL_568750, EPI_ISL_568751, EPI_ISL_568752, EPI_ISL_568753, EPI_ISL_568754, EPI_ISL_568755, EPI_ISL_568756, EPI_ISL_568757, EPI_ISL_568758, EPI_ISL_568759, EPI_ISL_568760, EPI_ISL_568761, EPI_ISL_568762, EPI_ISL_568763, EPI_ISL_568764, EPI_ISL_568765, EPI_ISL_568766, EPI_ISL_568767, EPI_ISL_568768, EPI_ISL_568769, EPI_ISL_568770, EPI_ISL_568771, EPI_ISL_568772, EPI_ISL_568773, EPI_ISL_568774, EPI_ISL_568775, EPI_ISL_568776, EPI_ISL_568777, EPI_ISL_568778, EPI_ISL_568779, EPI_ISL_568780, EPI_ISL_568781, EPI_ISL_568782, EPI_ISL_568783, EPI_ISL_568784, EPI_ISL_568785, EPI_ISL_568786, EPI_ISL_568787, EPI_ISL_568788, EPI_ISL_568789, EPI_ISL_568790, EPI_ISL_568791, EPI_ISL_568792, EPI_ISL_568793, EPI_ISL_568794, EPI_ISL_568795, EPI_ISL_568796, EPI_ISL_568797, EPI_ISL_568798, EPI_ISL_568799, EPI_ISL_568800, EPI_ISL_568801, EPI_ISL_568802, EPI_ISL_568803, EPI_ISL_568804, EPI_ISL_568805, EPI_ISL_568806, EPI_ISL_568807, EPI_ISL_568808, EPI_ISL_568809, EPI_ISL_568810, EPI_ISL_568811, EPI_ISL_568812, EPI_ISL_568813, EPI_ISL_568814, EPI_ISL_568815, EPI_ISL_568816, EPI_ISL_568817, EPI_ISL_568818, EPI_ISL_568819, EPI_ISL_568820, EPI_ISL_568821, EPI_ISL_568822, EPI_ISL_568823, EPI_ISL_568824, EPI_ISL_568825, EPI_ISL_568826, EPI_ISL_568827, EPI_ISL_568828, EPI_ISL_568829, EPI_ISL_568830, EPI_ISL_568831, EPI_ISL_568832, EPI_ISL_568833, EPI_ISL_568834, EPI_ISL_568835, EPI_ISL_568836, EPI_ISL_568837, EPI_ISL_568838, EPI_ISL_568839, EPI_ISL_568840, EPI_ISL_568841, EPI_ISL_568842, EPI_ISL_568843, EPI_ISL_568844, EPI_ISL_568845, EPI_ISL_568846, EPI_ISL_568847, EPI_ISL_568848, EPI_ISL_568849, EPI_ISL_568850, EPI_ISL_568851, EPI_ISL_568852, EPI_ISL_568853, EPI_ISL_568854, EPI_ISL_568855, EPI_ISL_568856, EPI_ISL_568857, EPI_ISL_568858, EPI_ISL_568859, EPI_ISL_568860, EPI_ISL_568861, EPI_ISL_568862, EPI_ISL_568863, EPI_ISL_568864, EPI_ISL_568865, EPI_ISL_568866, EPI_ISL_568867, EPI_ISL_568868, EPI_ISL_568869, EPI_ISL_568870, EPI_ISL_568871, EPI_ISL_568872 | | | |
| see above | KEMRI-Wellcome Trust Research Programme/KEMRI-CGMR-C Kilifi | KEMRI-Wellcome Trust Research Programme/KEMRI-CGMR-C Kilifi | Githinji et al 2020 |
| EPI_ISL_574581, EPI_ISL_574584, EPI_ISL_574585, EPI_ISL_574586, EPI_ISL_574599 | Delaware Public Health Lab | Delaware Public Health Lab | Gregory Hovan |
| EPI_ISL_575331 | RSUD Wates | Genetics Working Group (Pokja Genetik) Faculty of Medicine, Public Health and Nursing Universitas Gadjah Mada (FK-KMK UGM); Disease Investigation Center Wates Ministry of Agriculture Indonesia; Department of Microbiology FK-KMK UGM; Laboratorium Diagnostik Yayasan Tahija World Mosquito Program (WMP) Yogyakarta Center for Tropical Medicine FK-KMK UGM; Integrated Research Center FK-KMK UGM; Department of Computer Science and Electronics FMIPA UGM | Gunadi, Hendra Wibawa, . Marcellus, Mohamad S. Hakim, Edwin W. Daniwijaya, Ludhang P. Rizki, Endah Supriyati, Eggi Arguni, Titik Nuryastuti, Tri Wibawa, Dwi AA Nugrahaningsih, Afiahayati, Siswanto, Kristy Iskandar, Nungki Anggorowati, Bastianto Kusumajaya, Zumrati Ahmad, Alvin S. Kalim, Susan Simanjaya |
| EPI_ISL_575335 | Delaware Public Health Lab | Delaware Public Health Lab | Gregory Hovan |
| EPI_ISL_575339, EPI_ISL_575348, EPI_ISL_575364, EPI_ISL_575378, EPI_ISL_575383, EPI_ISL_575389 | Lighthouse Lab in Glasgow | Wellcome Sanger Institute for the COVID-19 Genomics UK (COG-UK) consortium | Harper VanSteenhouse, Yumi Kasai, David Gray, Carol Clugston, Anna Dominiczak and Alex Alderton, Roberto Amato, Sonia Goncalves, Ewan Harrison, David K. Jackson, Ian Johnston, Dominic Kwiatkowski, Cordelia Langford, John Sillitoe on behalf of the Wellcome Sanger Institute COVID-19 Surveillance Team |
| EPI_ISL_575995, EPI_ISL_575996, EPI_ISL_575997, EPI_ISL_575998, EPI_ISL_575999, EPI_ISL_576000, EPI_ISL_576001, EPI_ISL_576002, EPI_ISL_576003, EPI_ISL_576004, EPI_ISL_576005, EPI_ISL_576006, EPI_ISL_576007, EPI_ISL_576008, EPI_ISL_576009, EPI_ISL_576010, EPI_ISL_576011, EPI_ISL_576012, EPI_ISL_576013, EPI_ISL_576014, EPI_ISL_576015, EPI_ISL_576016, EPI_ISL_576017, EPI_ISL_576018, EPI_ISL_576019, EPI_ISL_576020, EPI_ISL_576021, EPI_ISL_576022, EPI_ISL_576023, EPI_ISL_576024, EPI_ISL_576025, EPI_ISL_576026, EPI_ISL_576027, EPI_ISL_576028, EPI_ISL_576029, EPI_ISL_576030, EPI_ISL_576031, EPI_ISL_576032, EPI_ISL_576033, EPI_ISL_576034, EPI_ISL_576035, EPI_ISL_576036, EPI_ISL_576037, EPI_ISL_576038, EPI_ISL_576039, EPI_ISL_576040, EPI_ISL_576041, EPI_ISL_576042, EPI_ISL_576043, EPI_ISL_576044, EPI_ISL_576045, EPI_ISL_576046, EPI_ISL_576047, EPI_ISL_576048, EPI_ISL_576049, EPI_ISL_576051, EPI_ISL_576052, EPI_ISL_576053, EPI_ISL_576054, EPI_ISL_576055, EPI_ISL_576056, EPI_ISL_576057, EPI_ISL_576058, EPI_ISL_576059, EPI_ISL_576060, EPI_ISL_576061, EPI_ISL_576062, EPI_ISL_576063, EPI_ISL_576065, EPI_ISL_576066, EPI_ISL_576067, EPI_ISL_576068, EPI_ISL_576069, EPI_ISL_576071, EPI_ISL_576072, EPI_ISL_576073, EPI_ISL_576074, EPI_ISL_576075, EPI_ISL_576076, EPI_ISL_576077, EPI_ISL_576078, EPI_ISL_576079, EPI_ISL_576080, EPI_ISL_576081, EPI_ISL_576082, EPI_ISL_576083, EPI_ISL_576084, EPI_ISL_576085, EPI_ISL_576086, EPI_ISL_576087, EPI_ISL_576089, EPI_ISL_576090, EPI_ISL_576091, EPI_ISL_576092, EPI_ISL_576093 | | | |
| see above | Lighthouse Lab in Glasgow | Wellcome Sanger Institute for the COVID-19 Genomics UK (COG-UK) consortium | Harper VanSteenhouse, Yumi Kasai, David Gray, Carol Clugston, Anna Dominiczak and Alex Alderton, Roberto Amato, Sonia Goncalves, Ewan Harrison, David K. Jackson, Ian Johnston, Dominic Kwiatkowski, Cordelia Langford, John Sillitoe on behalf of the Wellcome Sanger Institute COVID-19 Surveillance Team (http://www.sanger.ac.uk/covid-team) |
| EPI_ISL_576113 | RSUP Dr. Sardjito | Genetics Working Group (Pokja Genetik) Faculty of Medicine, Public Health and Nursing Universitas Gadjah Mada (FK-KMK UGM); Disease Investigation Center Wates Ministry of | Gunadi, Hendra Wibawa, . Marcellus, Mohamad S. Hakim, Edwin W. Daniwijaya, Ludhang P. Rizki, Endah Supriyati, Eggi Arguni, Titik Nuryastuti, Tri Wibawa, Dwi AA Nugrahaningsih, Afiahayati, Siswanto, Kristy Iskandar, Nungki Anggorowati, Ika Trisnawati, Riat El Khair, William Widitjiarso, Fadil Fahri |

| | | | |
|---|---|---|---|
| | | Agriculture Indonesia; Department of Microbiology FK-KMK UGM; Laboratorium Diagnostik Yayasan Tahija World Mosquito Program (WMP) Yogyakarta Center for Tropical Medicine FK-KMK UGM; Integrated Research Center FK-KMK UGM; Department of Computer Science and Electronics FMIPA UGM | |
| EPI_ISL_576114 | RSUP Dr. Sardjito | Genetics Working Group (Pokja Genetik) Faculty of Medicine, Public Health and Nursing Universitas Gadjah Mada (FK-KMK UGM); Disease Investigation Center Wates Ministry of Agriculture Indonesia; Department of Microbiology FK-KMK UGM; Laboratorium Diagnostik Yayasan Tahija World Mosquito Program (WMP) Yogyakarta Center for Tropical Medicine FK-KMK UGM; Integrated Research Center FK-KMK UGM; Department of Computer Science and Electronics FMIPA UGM | Gunadi, Hendra Wibawa, . Marcellus, Mohamad S. Hakim, Edwin W. Daniwijaya, Ludhang P. Rizki, Endah Supriyati, Eggi Arguni, Titik Nuryastuti, Tri Wibawa, Dwi AA Nugrahaningsih, Afiahayati, Siswanto, Kristy Iskandar, Nungki Anggorowati, Ika Trisnawati, Riat El Khair, Audric Kenny Tedja, Aditya Rifqi Fauzi |
| EPI_ISL_576115 | RSUP Dr Sardjito | Genetics Working Group (Pokja Genetik) Faculty of Medicine, Public Health and Nursing Universitas Gadjah Mada (FK-KMK UGM); Disease Investigation Center Wates Ministry of Agriculture Indonesia; Department of Microbiology FK-KMK UGM; Laboratorium Diagnostik Yayasan Tahija World Mosquito Program (WMP) Yogyakarta Center for Tropical Medicine FK-KMK UGM; Integrated Research Center FK-KMK UGM; Department of Computer Science and Electronics FMIPA UGM | Gunadi, Hendra Wibawa, . Marcellus, Mohamad S. Hakim, Edwin W. Daniwijaya, Ludhang P. Rizki, Endah Supriyati, Eggi Arguni, Titik Nuryastuti, Tri Wibawa, Dwi AA Nugrahaningsih, Afiahayati, Siswanto, Kristy Iskandar, Nungki Anggorowati, Ika Trisnawati, Riat El Khair, Yunika Puspadewi, Osman Sianipar, Umi Solekhah Intansari, Elizabeth Henny Herningtiyas, Ira Puspitawati, Nur Imma Fatimah Harahap, Untung Wirawan, Maria Patricia Inggriani |
| EPI_ISL_576116 | RSUP Dr. Sardjito | Genetics Working Group (Pokja Genetik) Faculty of Medicine, Public Health and Nursing Universitas Gadjah Mada (FK-KMK UGM); Disease Investigation Center Wates Ministry of Agriculture Indonesia; Department of Microbiology FK-KMK UGM; Laboratorium Diagnostik Yayasan Tahija World Mosquito Program (WMP) Yogyakarta Center for Tropical Medicine FK-KMK UGM; Integrated Research Center FK-KMK UGM; Department of Computer Science and Electronics FMIPA UGM | Gunadi, Hendra Wibawa, . Marcellus, Mohamad S. Hakim, Edwin W. Daniwijaya, Ludhang P. Rizki, Endah Supriyati, Eggi Arguni, Titik Nuryastuti, Tri Wibawa, Dwi AA Nugrahaningsih, Afiahayati, Siswanto, Kristy Iskandar, Nungki Anggorowati, Ika Trisnawati, Sumardi, Eko Budiono, Bambang Sigit Riyanto, Heni Retnowulan, Munawar Gani, Satria Maulana, Nur Rahmi Ananda, Riat El Khair, Yunika Puspadewi, Osman Sianipar, Umi Solekhah Intansari, Elizabeth Henny Herningtiyas, Ira Puspitawati, Nur Imma Fatimah Harahap, Ririn Enggy Yuliyanti, Alvin Santoso Kalim |
| EPI_ISL_576128 | RSUP Dr. Sardjito | Genetics Working Group (Pokja Genetik) Faculty of Medicine, Public Health and Nursing Universitas Gadjah Mada (FK-KMK UGM); Disease Investigation Center Wates Ministry of Agriculture Indonesia; Department of Microbiology FK-KMK UGM; Laboratorium Diagnostik Yayasan Tahija World Mosquito Program (WMP) Yogyakarta Center for Tropical Medicine FK-KMK UGM; Integrated Research Center FK-KMK UGM; Department of Computer Science and Electronics FMIPA UGM | Gunadi, Hendra Wibawa, Marcellus, Mohamad S. Hakim, Edwin W. Daniwijaya, Ludhang P. Rizki, Endah Supriyati, Eggi Arguni, Titik Nuryastuti, Tri Wibawa, Dwi AA Nugrahaningsih, Afiahayati, Siswanto, Kristy Iskandar, Nungki Anggorowati, Ika Trisnawati, Sumardi, Eko Budiono, Bambang Sigit Riyanto, Heni Retnowulan, Munawar Gani, Satria Maulana, Nur Rahmi Ananda, Riat El Khair, Yunika Puspadewi, Osman Sianipar, Umi Solekhah Intansari, Elizabeth Henny Herningtiyas, Ira Puspitawati, Nur Imma Fatimah Harahap, Dyah Ayu Puspitarani, Kemala Athollah |
| EPI_ISL_576130 | RSUP Dr. Sardjito | Genetics Working Group (Pokja Genetik) Faculty of Medicine, Public Health and Nursing Universitas Gadjah Mada (FK-KMK UGM); Disease Investigation Center Wates Ministry of Agriculture Indonesia; Department of Microbiology FK-KMK UGM; Laboratorium Diagnostik Yayasan Tahija World Mosquito Program (WMP) Yogyakarta Center for Tropical Medicine FK-KMK UGM; Integrated Research Center FK-KMK UGM; Department of Computer Science and Electronics FMIPA UGM | Gunadi, Hendra Wibawa, Marcellus, Mohamad S. Hakim, Edwin W. Daniwijaya, Ludhang P. Rizki, Endah Supriyati, Eggi Arguni, Titik Nuryastuti, Tri Wibawa, Dwi AA Nugrahaningsih, Afiahayati, Siswanto, Kristy Iskandar, Nungki Anggorowati, Rina Triasih, Amalia Setyati, Dwikisworo Setyowireni, Ida Safitri Laksanawati, Cahya Dewi Satria, Titis Widowati, Elisabeth Siti Herini, Riat El Khair, Yunika Puspadewi, Osman Sianipar, Umi Solekhah Intansari, Elizabeth Henny Herningtiyas, Ira Puspitawati, Nur Imma Fatimah Harahap, Alvin S. Kalim, Susan Simanjaya, Desyifa Mursalin |
| EPI_ISL_576145 | RSA Universitas Gadjah Mada | Genetics Working Group (Pokja Genetik) Faculty of Medicine, Public Health and Nursing Universitas Gadjah Mada (FK-KMK UGM); Disease Investigation Center Wates Ministry of Agriculture Indonesia; Department of Microbiology FK-KMK UGM; Laboratorium Diagnostik Yayasan Tahija World Mosquito Program (WMP) Yogyakarta Center for Tropical Medicine FK-KMK UGM; Integrated Research Center FK-KMK UGM; Department of Computer Science and Electronics FMIPA UGM | Gunadi, Hendra Wibawa, Marcellus, Mohamad S. Hakim, Edwin W. Daniwijaya, Ludhang P. Rizki, Endah Supriyati, Eggi Arguni, Titik Nuryastuti, Tri Wibawa, Dwi AA Nugrahaningsih, Afiahayati, Siswanto, Kristy Iskandar, Nungki Anggorowati, William Widitjiarso, Fadil Fahri |
| EPI_ISL_576150, EPI_ISL_576151, EPI_ISL_576152, EPI_ISL_576153, EPI_ISL_576154, EPI_ISL_576155, EPI_ISL_576156, EPI_ISL_576157, EPI_ISL_576158, EPI_ISL_576159, EPI_ISL_576160, EPI_ISL_576161, EPI_ISL_576162, EPI_ISL_576163, EPI_ISL_576164, EPI_ISL_576165, EPI_ISL_576166, EPI_ISL_576167, EPI_ISL_576168, EPI_ISL_576169, EPI_ISL_576222, EPI_ISL_576223, EPI_ISL_576224, EPI_ISL_576225, EPI_ISL_576226, EPI_ISL_576227 | | | |
| see above | Delaware Public Health Lab | Delaware Public Health Lab | Gregory Hovan |
| EPI_ISL_576257, EPI_ISL_576258, EPI_ISL_576259, EPI_ISL_576260 | Instituto de Diagnostico y Referencia Epidemiologicos (INDRE) | Instituto de Diagnostico y Referencia Epidemiologicos (INDRE) | Gisela Barrera-Badillo , Abril Rodriguez-Maldonado, Claudia Wong-Arambula , Natividad Cruz-Ortiz, Tatiana Nunez-Garcia, Dayanira Arellano-Suarez, Fabiola Garces-Ayala, Edgar Mendieta-Condado, Lucia Hernandez-Rivas, Irma Lopez-Martinez, Ernesto Ramirez-Gonzalez. |
| EPI_ISL_576261, EPI_ISL_576262, EPI_ISL_576263 | Instituto de Diagnostico y Referencia Epidemiologicos (INDRE) | Instituto de Diagnostico y Referencia Epidemiologicos (INDRE) | Gisela Barrera-Badillo , Abril Rodriguez-Maldonado, Claudia Wong-Arambula , Natividad Cruz-Ortiz, Tatiana Nunez-Garcia, Dayanira Arellano-Suarez, Adnan Araiza-Rodriguez, Edgar Mendieta-Condado, Lucia Hernandez-Rivas, Irma Lopez-Martinez, Ernesto Ramirez-Gonzalez. |
| EPI_ISL_576272 | Instituto de Diagnostico y Referencia Epidemiologicos (INDRE) | Instituto de Diagnostico y Referencia Epidemiologicos (INDRE) | Gisela Barrera-Badillo , Abril Rodriguez-Maldonado, Claudia Wong-Arambula , Natividad Cruz-Ortiz, Tatiana Nunez-Garcia, Dayanira Arellano-Suarez, Fabiola Garces-Ayala, Edgar Mendieta-Condado, Lucia Hernandez-Rivas, Irma Lopez-Martinez, Ernesto Ramirez-Gonzalez. |
| EPI_ISL_576274, EPI_ISL_576275, EPI_ISL_576276 | Instituto de Diagnostico y Referencia Epidemiologicos (INDRE) | Instituto de Diagnostico y Referencia Epidemiologicos (INDRE) | Gisela Barrera-Badillo , Abril Rodriguez-Maldonado, Claudia Wong-Arambula , Natividad Cruz-Ortiz, Tatiana Nunez-Garcia, Dayanira Arellano-Suarez, Adnan Araiza-Rodriguez, Edgar Mendieta-Condado, Lucia Hernandez-Rivas, Irma Lopez-Martinez, Ernesto Ramirez-Gonzalez. |
| EPI_ISL_576383 | RSUD Budi Rahayu Kota Magelang | Genetics Working Group (Pokja Genetik) Faculty of Medicine, Public Health and Nursing Universitas Gadjah Mada (FK-KMK UGM); Disease Investigation Center Wates Ministry of Agriculture Indonesia; Department of Microbiology FK-KMK UGM; Laboratorium Diagnostik Yayasan Tahija World Mosquito Program (WMP) Yogyakarta Center for Tropical Medicine FK-KMK UGM; Integrated Research Center FK-KMK UGM; Department of Computer Science and Electronics FMIPA UGM; Balai Besar Teknik Kesehatan Lingkungan dan Pengendalian Penyakit (BBTKLPP) Yogyakarta | Gunadi, Hendra Wibawa, Marcellus, Mohamad S. Hakim, Edwin W. Daniwijaya, Ludhang P. Rizki, Endah Supriyati, Eggi Arguni, Titik Nuryastuti, Tri Wibawa, Dwi AA Nugrahaningsih, Afiahayati, Siswanto, Kristy Iskandar, Nungki Anggorowati, Irene, Indaryati, Havid Setyawan, Ari Meliyanti, Merliana Sari Situmeang, Audric Kenny Tedja, Aditya Rifqi Fauzi |

EPI_ISL_579759, EPI_ISL_579760, EPI_ISL_579761, EPI_ISL_579762, EPI_ISL_579763, EPI_ISL_579764, EPI_ISL_579765, EPI_ISL_579766, EPI_ISL_579767, EPI_ISL_579768, EPI_ISL_579769, EPI_ISL_579770, EPI_ISL_579771, EPI_ISL_579772, EPI_ISL_579773, EPI_ISL_579774, EPI_ISL_579775, EPI_ISL_579776, EPI_ISL_579777, EPI_ISL_579778, EPI_ISL_579779, EPI_ISL_579780, EPI_ISL_579781, EPI_ISL_579782, EPI_ISL_579783, EPI_ISL_579784, EPI_ISL_579785, EPI_ISL_579786, EPI_ISL_579787, EPI_ISL_579788, EPI_ISL_579789, EPI_ISL_579790, EPI_ISL_579791, EPI_ISL_579792, EPI_ISL_579793, EPI_ISL_579795, EPI_ISL_579796, EPI_ISL_579797, EPI_ISL_579798, EPI_ISL_579799, EPI_ISL_579800, EPI_ISL_579801, EPI_ISL_579802, EPI_ISL_579803, EPI_ISL_579804, EPI_ISL_579805, EPI_ISL_579806, EPI_ISL_579807, EPI_ISL_579808, EPI_ISL_579809, EPI_ISL_579810, EPI_ISL_579811, EPI_ISL_579812, EPI_ISL_579813, EPI_ISL_579814, EPI_ISL_579815, EPI_ISL_579816, EPI_ISL_579817, EPI_ISL_579818, EPI_ISL_579819, EPI_ISL_579820, EPI_ISL_579821, EPI_ISL_579822, EPI_ISL_579823, EPI_ISL_579824, EPI_ISL_579825, EPI_ISL_579826, EPI_ISL_579827, EPI_ISL_579828, EPI_ISL_579829, EPI_ISL_579830, EPI_ISL_579831, EPI_ISL_579832, EPI_ISL_579833, EPI_ISL_579834, EPI_ISL_579836, EPI_ISL_579837, EPI_ISL_579838, EPI_ISL_579839, EPI_ISL_579840, EPI_ISL_579841, EPI_ISL_579842, EPI_ISL_579844, EPI_ISL_579845, EPI_ISL_579846, EPI_ISL_579847, EPI_ISL_579848, EPI_ISL_579849, EPI_ISL_579850, EPI_ISL_579851, EPI_ISL_579852, EPI_ISL_579853, EPI_ISL_579854, EPI_ISL_579855, EPI_ISL_579856, EPI_ISL_579857, EPI_ISL_579858, EPI_ISL_579859, EPI_ISL_579860, EPI_ISL_579861, EPI_ISL_579862, EPI_ISL_579863, EPI_ISL_579864, EPI_ISL_579865, EPI_ISL_579866, EPI_ISL_579867, EPI_ISL_579868, EPI_ISL_579869, EPI_ISL_579870, EPI_ISL_579871, EPI_ISL_579872, EPI_ISL_579873, EPI_ISL_579874, EPI_ISL_579875, EPI_ISL_579876, EPI_ISL_579877, EPI_ISL_579878, EPI_ISL_579879, EPI_ISL_579880, EPI_ISL_579881, EPI_ISL_579882, EPI_ISL_579883, EPI_ISL_579884, EPI_ISL_579885, EPI_ISL_579886, EPI_ISL_579887, EPI_ISL_579888, EPI_ISL_579889, EPI_ISL_579890, EPI_ISL_579891, EPI_ISL_579892, EPI_ISL_579893, EPI_ISL_579894, EPI_ISL_579895, EPI_ISL_579896, EPI_ISL_579897, EPI_ISL_579898, EPI_ISL_579899, EPI_ISL_579900, EPI_ISL_579901, EPI_ISL_579902, EPI_ISL_579903, EPI_ISL_579904, EPI_ISL_579905, EPI_ISL_579906, EPI_ISL_579907, EPI_ISL_579908, EPI_ISL_579909, EPI_ISL_579910, EPI_ISL_579911, EPI_ISL_579912, EPI_ISL_579913, EPI_ISL_579914, EPI_ISL_579915, EPI_ISL_579916, EPI_ISL_579917, EPI_ISL_579918, EPI_ISL_580858, EPI_ISL_580859, EPI_ISL_580860, EPI_ISL_580862, EPI_ISL_580863, EPI_ISL_580865, EPI_ISL_580866, EPI_ISL_580867, EPI_ISL_580868, EPI_ISL_580869, EPI_ISL_580870, EPI_ISL_580871, EPI_ISL_580872, EPI_ISL_580874, EPI_ISL_580876, EPI_ISL_580878, EPI_ISL_580879, EPI_ISL_580880, EPI_ISL_580881, EPI_ISL_580883, EPI_ISL_580884, EPI_ISL_580886, EPI_ISL_580887, EPI_ISL_580888, EPI_ISL_580889, EPI_ISL_580890, EPI_ISL_580891, EPI_ISL_580893, EPI_ISL_580894, EPI_ISL_580895, EPI_ISL_580899, EPI_ISL_580901, EPI_ISL_580905, EPI_ISL_580906, EPI_ISL_580907, EPI_ISL_580908, EPI_ISL_580911, EPI_ISL_580912, EPI_ISL_580914, EPI_ISL_580916, EPI_ISL_580919, EPI_ISL_580920, EPI_ISL_580922, EPI_ISL_580923, EPI_ISL_580924, EPI_ISL_580925, EPI_ISL_580926, EPI_ISL_580927, EPI_ISL_580928, EPI_ISL_580930, EPI_ISL_580931, EPI_ISL_580932, EPI_ISL_580933, EPI_ISL_580934, EPI_ISL_580936, EPI_ISL_580937, EPI_ISL_580938, EPI_ISL_580939, EPI_ISL_580941, EPI_ISL_580942, EPI_ISL_580943, EPI_ISL_580944, EPI_ISL_580945, EPI_ISL_580946, EPI_ISL_580947, EPI_ISL_580949, EPI_ISL_580952, EPI_ISL_580953, EPI_ISL_580954, EPI_ISL_580955, EPI_ISL_580956, EPI_ISL_580957, EPI_ISL_580958, EPI_ISL_580960, EPI_ISL_580962, EPI_ISL_580963, EPI_ISL_580964, EPI_ISL_580965, EPI_ISL_580966, EPI_ISL_580967, EPI_ISL_580968, EPI_ISL_580969, EPI_ISL_580970, EPI_ISL_580971, EPI_ISL_580972, EPI_ISL_580973, EPI_ISL_580974, EPI_ISL_580975, EPI_ISL_580977, EPI_ISL_580978, EPI_ISL_580981, EPI_ISL_580982, EPI_ISL_580983, EPI_ISL_580984, EPI_ISL_580985, EPI_ISL_580986, EPI_ISL_580987, EPI_ISL_580988, EPI_ISL_580989, EPI_ISL_580992, EPI_ISL_580994, EPI_ISL_580997, EPI_ISL_580999, EPI_ISL_581001, EPI_ISL_581002, EPI_ISL_581003, EPI_ISL_581005, EPI_ISL_581006, EPI_ISL_581007, EPI_ISL_581010, EPI_ISL_581011, EPI_ISL_581012, EPI_ISL_581013, EPI_ISL_581014, EPI_ISL_581015, EPI_ISL_581016, EPI_ISL_581017, EPI_ISL_581019, EPI_ISL_581020, EPI_ISL_581021, EPI_ISL_581022, EPI_ISL_581024, EPI_ISL_581027, EPI_ISL_581030, EPI_ISL_581031, EPI_ISL_581033, EPI_ISL_581034, EPI_ISL_581036, EPI_ISL_581037, EPI_ISL_581039, EPI_ISL_581042, EPI_ISL_581043, EPI_ISL_581044, EPI_ISL_581047, EPI_ISL_581048, EPI_ISL_581049, EPI_ISL_581050, EPI_ISL_581051, EPI_ISL_581052, EPI_ISL_581053, EPI_ISL_581055, EPI_ISL_581057, EPI_ISL_581059, EPI_ISL_581060, EPI_ISL_581061, EPI_ISL_581062, EPI_ISL_581063, EPI_ISL_581066, EPI_ISL_581068, EPI_ISL_581070, EPI_ISL_581071, EPI_ISL_581072, EPI_ISL_581073, EPI_ISL_581075, EPI_ISL_581077, EPI_ISL_581079, EPI_ISL_581080, EPI_ISL_581081, EPI_ISL_581082, EPI_ISL_581083, EPI_ISL_581085, EPI_ISL_581087, EPI_ISL_581088, EPI_ISL_581089, EPI_ISL_581090, EPI_ISL_581091, EPI_ISL_581092, EPI_ISL_581094, EPI_ISL_581095, EPI_ISL_581096, EPI_ISL_581098, EPI_ISL_581100, EPI_ISL_581101, EPI_ISL_581102, EPI_ISL_581104, EPI_ISL_581105, EPI_ISL_581108, EPI_ISL_581109, EPI_ISL_581110, EPI_ISL_581112, EPI_ISL_581127, EPI_ISL_581132, EPI_ISL_581133, EPI_ISL_581141, EPI_ISL_581146, EPI_ISL_581154, EPI_ISL_581166, EPI_ISL_581174, EPI_ISL_581182, EPI_ISL_581183, EPI_ISL_581189, EPI_ISL_581193, EPI_ISL_581194, EPI_ISL_581196, EPI_ISL_581214, EPI_ISL_581217, EPI_ISL_581223, EPI_ISL_581226, EPI_ISL_581229, EPI_ISL_581230, EPI_ISL_581245, EPI_ISL_581247, EPI_ISL_581261, EPI_ISL_581262, EPI_ISL_581267, EPI_ISL_581285, EPI_ISL_581291, EPI_ISL_581310, EPI_ISL_581323, EPI_ISL_581324, EPI_ISL_581329, EPI_ISL_581334, EPI_ISL_581337, EPI_ISL_581341, EPI_ISL_581344, EPI_ISL_581346, EPI_ISL_581350, EPI_ISL_581354, EPI_ISL_581358, EPI_ISL_581377, EPI_ISL_581381, EPI_ISL_581384, EPI_ISL_581387, EPI_ISL_581392, EPI_ISL_581395, EPI_ISL_581402, EPI_ISL_581403, EPI_ISL_581405, EPI_ISL_581406, EPI_ISL_581407, EPI_ISL_581408, EPI_ISL_581410

| | | | |
|---|---|---|---|
| see above | Lighthouse Lab in Glasgow | Wellcome Sanger Institute for the COVID-19 Genomics UK (COG-UK) consortium | Harper VanSteenhouse, Yumi Kasai, David Gray, Carol Clugston, Anna Dominiczak and Alex Alderton, Roberto Amato, Sonia Goncalves, Ewan Harrison, David K. Jackson, Ian Johnston, Dominic Kwiatkowski, Cordelia Langford, John Sillitoe on behalf of the Wellcome Sanger Institute COVID-19 Surveillance Team |

EPI_ISL_582034, EPI_ISL_582035, EPI_ISL_582036, EPI_ISL_582037, EPI_ISL_582038, EPI_ISL_582039, EPI_ISL_582040, EPI_ISL_582041, EPI_ISL_582042, EPI_ISL_582043, EPI_ISL_582044, EPI_ISL_582045, EPI_ISL_582046, EPI_ISL_582047, EPI_ISL_582048, EPI_ISL_582049, EPI_ISL_582050, EPI_ISL_582051, EPI_ISL_582052, EPI_ISL_582053, EPI_ISL_582054, EPI_ISL_582055, EPI_ISL_582056, EPI_ISL_582057, EPI_ISL_582058, EPI_ISL_582059, EPI_ISL_582060, EPI_ISL_582061, EPI_ISL_582062, EPI_ISL_582063, EPI_ISL_582064, EPI_ISL_582065, EPI_ISL_582066, EPI_ISL_582067, EPI_ISL_582068, EPI_ISL_582069, EPI_ISL_582070, EPI_ISL_582071, EPI_ISL_582072, EPI_ISL_582073, EPI_ISL_582074, EPI_ISL_582075, EPI_ISL_582076, EPI_ISL_582077, EPI_ISL_582078, EPI_ISL_582079, EPI_ISL_582080, EPI_ISL_582081, EPI_ISL_582082, EPI_ISL_582083, EPI_ISL_582084, EPI_ISL_582085, EPI_ISL_582086, EPI_ISL_582087, EPI_ISL_582088, EPI_ISL_582089, EPI_ISL_582090

| | | | |
|---|---|---|---|
| see above | Servicio de Microbiología. Hospital Universitario Donostia. OSI Donostialdea. Área de Enfermedades Infecciosas, Grupo de Infección Respiratoria y Resistencia Antimicrobiana. Instituto de Investigación Sanitaria Biodonostia | SeqCOVID-SPAIN consortium/IBV(CSIC) | Gustavo Cilla, Milagrosa Montes, Luis Piñeiro, Jose Maria Marimón and SeqCOVID-SPAIN consortium |
| EPI_ISL_582133, EPI_ISL_583246, EPI_ISL_583247, EPI_ISL_583248, EPI_ISL_583994, EPI_ISL_583995, EPI_ISL_584013, EPI_ISL_584014 | Delaware Public Health Lab | Delaware Public Health Lab | Gregory Hovan |

EPI_ISL_587304, EPI_ISL_587305, EPI_ISL_587306, EPI_ISL_587307, EPI_ISL_587308, EPI_ISL_587309, EPI_ISL_587310, EPI_ISL_587311, EPI_ISL_587312, EPI_ISL_587314, EPI_ISL_587315, EPI_ISL_587316, EPI_ISL_587317, EPI_ISL_587318, EPI_ISL_587319, EPI_ISL_587320, EPI_ISL_587321, EPI_ISL_587322, EPI_ISL_587323, EPI_ISL_587324, EPI_ISL_587325, EPI_ISL_587326, EPI_ISL_587327, EPI_ISL_587328, EPI_ISL_587329, EPI_ISL_587330, EPI_ISL_587331, EPI_ISL_587332, EPI_ISL_587333, EPI_ISL_587334, EPI_ISL_587335, EPI_ISL_587336, EPI_ISL_587337, EPI_ISL_587338, EPI_ISL_587339, EPI_ISL_587342, EPI_ISL_587343, EPI_ISL_587344, EPI_ISL_587346, EPI_ISL_587347, EPI_ISL_587348, EPI_ISL_587349, EPI_ISL_587350, EPI_ISL_587351, EPI_ISL_587352, EPI_ISL_587353, EPI_ISL_587354, EPI_ISL_587355, EPI_ISL_587356, EPI_ISL_587357, EPI_ISL_587358, EPI_ISL_587359, EPI_ISL_587360, EPI_ISL_587361, EPI_ISL_587362, EPI_ISL_587363, EPI_ISL_587364, EPI_ISL_587365, EPI_ISL_587367, EPI_ISL_587368, EPI_ISL_587369, EPI_ISL_587370, EPI_ISL_587371, EPI_ISL_587372, EPI_ISL_587373, EPI_ISL_587374, EPI_ISL_587375, EPI_ISL_587376, EPI_ISL_587377, EPI_ISL_587378, EPI_ISL_587379, EPI_ISL_587380, EPI_ISL_587381, EPI_ISL_587382, EPI_ISL_587383, EPI_ISL_587384, EPI_ISL_587385, EPI_ISL_587387, EPI_ISL_587388, EPI_ISL_587389, EPI_ISL_587390, EPI_ISL_587392, EPI_ISL_587393, EPI_ISL_587394, EPI_ISL_587395, EPI_ISL_587396, EPI_ISL_587397, EPI_ISL_587398, EPI_ISL_587399, EPI_ISL_587400, EPI_ISL_587401, EPI_ISL_587405, EPI_ISL_587406, EPI_ISL_587407, EPI_ISL_587408, EPI_ISL_587409, EPI_ISL_587410, EPI_ISL_587412, EPI_ISL_587413, EPI_ISL_587414, EPI_ISL_587415, EPI_ISL_587417, EPI_ISL_587418, EPI_ISL_587419, EPI_ISL_587420, EPI_ISL_587421, EPI_ISL_587422, EPI_ISL_587423, EPI_ISL_587424, EPI_ISL_587426, EPI_ISL_587429, EPI_ISL_587430, EPI_ISL_587431, EPI_ISL_587432, EPI_ISL_587433, EPI_ISL_587434, EPI_ISL_587435, EPI_ISL_587436, EPI_ISL_587437, EPI_ISL_587438, EPI_ISL_587439, EPI_ISL_587440, EPI_ISL_587441, EPI_ISL_587442, EPI_ISL_587443, EPI_ISL_587444, EPI_ISL_587445, EPI_ISL_587446, EPI_ISL_587447, EPI_ISL_587448, EPI_ISL_587449, EPI_ISL_587450, EPI_ISL_587452, EPI_ISL_587453, EPI_ISL_587454, EPI_ISL_587455, EPI_ISL_587456, EPI_ISL_587457, EPI_ISL_587458, EPI_ISL_587460, EPI_ISL_587461, EPI_ISL_587462, EPI_ISL_587463, EPI_ISL_587464, EPI_ISL_587465, EPI_ISL_587466, EPI_ISL_587467, EPI_ISL_587468, EPI_ISL_587469, EPI_ISL_587470, EPI_ISL_587471, EPI_ISL_587472, EPI_ISL_587473, EPI_ISL_587474, EPI_ISL_587475, EPI_ISL_587476, EPI_ISL_587478, EPI_ISL_587479, EPI_ISL_587480, EPI_ISL_587481, EPI_ISL_587482, EPI_ISL_587483, EPI_ISL_587484, EPI_ISL_587485, EPI_ISL_587486, EPI_ISL_587488, EPI_ISL_587489, EPI_ISL_587490, EPI_ISL_587491, EPI_ISL_587492, EPI_ISL_587493, EPI_ISL_587494, EPI_ISL_587495, EPI_ISL_587496, EPI_ISL_587497, EPI_ISL_587499, EPI_ISL_587501, EPI_ISL_587502, EPI_ISL_587503, EPI_ISL_587504, EPI_ISL_587505, EPI_ISL_587506, EPI_ISL_587507, EPI_ISL_587508, EPI_ISL_587509, EPI_ISL_587510, EPI_ISL_587511, EPI_ISL_587512, EPI_ISL_587513, EPI_ISL_587514, EPI_ISL_587515, EPI_ISL_587516, EPI_ISL_587517, EPI_ISL_587518, EPI_ISL_587520, EPI_ISL_587521, EPI_ISL_587522, EPI_ISL_587523, EPI_ISL_587524, EPI_ISL_587526, EPI_ISL_587527, EPI_ISL_587528, EPI_ISL_587529, EPI_ISL_587530, EPI_ISL_587531, EPI_ISL_587532, EPI_ISL_587533, EPI_ISL_587535, EPI_ISL_587537, EPI_ISL_587538, EPI_ISL_587539, EPI_ISL_587541, EPI_ISL_587542, EPI_ISL_587543, EPI_ISL_587544, EPI_ISL_587545, EPI_ISL_587546, EPI_ISL_587547, EPI_ISL_587548, EPI_ISL_587549, EPI_ISL_587550, EPI_ISL_587551, EPI_ISL_587552, EPI_ISL_587553, EPI_ISL_587554, EPI_ISL_587555, EPI_ISL_587556, EPI_ISL_587557, EPI_ISL_587558, EPI_ISL_587559, EPI_ISL_587561, EPI_ISL_587562, EPI_ISL_587563, EPI_ISL_587564, EPI_ISL_587565, EPI_ISL_587566, EPI_ISL_587567, EPI_ISL_587568, EPI_ISL_587569, EPI_ISL_587570, EPI_ISL_587571, EPI_ISL_587572, EPI_ISL_587573, EPI_ISL_587574, EPI_ISL_587576, EPI_ISL_587578, EPI_ISL_587579, EPI_ISL_587580, EPI_ISL_587581, EPI_ISL_587582, EPI_ISL_587583, EPI_ISL_587584, EPI_ISL_587586, EPI_ISL_587589, EPI_ISL_587590, EPI_ISL_587591, EPI_ISL_587592, EPI_ISL_587593, EPI_ISL_587595, EPI_ISL_587596, EPI_ISL_587597, EPI_ISL_587599, EPI_ISL_587600, EPI_ISL_587602, EPI_ISL_587603, EPI_ISL_587604, EPI_ISL_587607, EPI_ISL_587608, EPI_ISL_587610, EPI_ISL_587611, EPI_ISL_587612, EPI_ISL_587613, EPI_ISL_587614, EPI_ISL_587615, EPI_ISL_587616, EPI_ISL_587618, EPI_ISL_587620, EPI_ISL_587621, EPI_ISL_587624, EPI_ISL_587625, EPI_ISL_587626, EPI_ISL_587627, EPI_ISL_587628, EPI_ISL_587629, EPI_ISL_587631, EPI_ISL_587632, EPI_ISL_587633, EPI_ISL_587634, EPI_ISL_587635, EPI_ISL_587636, EPI_ISL_587637, EPI_ISL_587638, EPI_ISL_587639, EPI_ISL_587641, EPI_ISL_587642, EPI_ISL_587644, EPI_ISL_587645, EPI_ISL_587646, EPI_ISL_587647, EPI_ISL_587648, EPI_ISL_587649, EPI_ISL_587650, EPI_ISL_587651, EPI_ISL_587652, EPI_ISL_587653, EPI_ISL_587654, EPI_ISL_587655, EPI_ISL_587656, EPI_ISL_587657, EPI_ISL_587659, EPI_ISL_587660, EPI_ISL_587661, EPI_ISL_587662, EPI_ISL_587663, EPI_ISL_587664, EPI_ISL_587666, EPI_ISL_587667, EPI_ISL_587668, EPI_ISL_587669, EPI_ISL_587671, EPI_ISL_587672, EPI_ISL_587673, EPI_ISL_587675, EPI_ISL_587676, EPI_ISL_587677, EPI_ISL_587679, EPI_ISL_587680, EPI_ISL_587681, EPI_ISL_587682, EPI_ISL_587683, EPI_ISL_587684, EPI_ISL_587686, EPI_ISL_587687, EPI_ISL_587688, EPI_ISL_587689, EPI_ISL_587690, EPI_ISL_587691, EPI_ISL_587692, EPI_ISL_587693, EPI_ISL_587694, EPI_ISL_587695, EPI_ISL_587696, EPI_ISL_587697, EPI_ISL_587698, EPI_ISL_587699, EPI_ISL_587700, EPI_ISL_587701, EPI_ISL_587703, EPI_ISL_587704, EPI_ISL_587706, EPI_ISL_587707, EPI_ISL_587708, EPI_ISL_587709, EPI_ISL_587710, EPI_ISL_588069, EPI_ISL_588079, EPI_ISL_588081, EPI_ISL_588095, EPI_ISL_588111, EPI_ISL_588137, EPI_ISL_588141, EPI_ISL_588144, EPI_ISL_588181, EPI_ISL_588189, EPI_ISL_588199, EPI_ISL_588211, EPI_ISL_588218, EPI_ISL_588220, EPI_ISL_588228, EPI_ISL_588233, EPI_ISL_588236, EPI_ISL_588244, EPI_ISL_588280, EPI_ISL_588282, EPI_ISL_588284, EPI_ISL_588289, EPI_ISL_588313, EPI_ISL_588327, EPI_ISL_588349, EPI_ISL_588360, EPI_ISL_588364, EPI_ISL_588373, EPI_ISL_588374, EPI_ISL_588380, EPI_ISL_588396, EPI_ISL_588409, EPI_ISL_588443, EPI_ISL_588445, EPI_ISL_588455, EPI_ISL_588469, EPI_ISL_588476, EPI_ISL_588477, EPI_ISL_588478, EPI_ISL_588479, EPI_ISL_588480, EPI_ISL_588481, EPI_ISL_588482, EPI_ISL_588483, EPI_ISL_588484, EPI_ISL_588485, EPI_ISL_588486, EPI_ISL_588487, EPI_ISL_588488, EPI_ISL_588489, EPI_ISL_588490, EPI_ISL_588491, EPI_ISL_588492, EPI_ISL_588493, EPI_ISL_588494, EPI_ISL_588495, EPI_ISL_588497, EPI_ISL_588498, EPI_ISL_588499, EPI_ISL_588500, EPI_ISL_588501, EPI_ISL_588502, EPI_ISL_588503, EPI_ISL_588504, EPI_ISL_588505, EPI_ISL_588506, EPI_ISL_588507, EPI_ISL_588508, EPI_ISL_588509, EPI_ISL_588510, EPI_ISL_588511, EPI_ISL_588512, EPI_ISL_588513, EPI_ISL_588514, EPI_ISL_588515, EPI_ISL_588516, EPI_ISL_588517, EPI_ISL_588518, EPI_ISL_588520, EPI_ISL_588521, EPI_ISL_588522, EPI_ISL_588523, EPI_ISL_588524, EPI_ISL_588525, EPI_ISL_588526, EPI_ISL_588527, EPI_ISL_588528, EPI_ISL_588529, EPI_ISL_588530, EPI_ISL_588531, EPI_ISL_588532, EPI_ISL_588533, EPI_ISL_588534, EPI_ISL_588535, EPI_ISL_588536, EPI_ISL_588537, EPI_ISL_588538, EPI_ISL_588539, EPI_ISL_588540, EPI_ISL_588543, EPI_ISL_588544, EPI_ISL_588545, EPI_ISL_588546, EPI_ISL_588547, EPI_ISL_588548, EPI_ISL_588549, EPI_ISL_588550, EPI_ISL_588551, EPI_ISL_588552, EPI_ISL_588553, EPI_ISL_588554, EPI_ISL_588555, EPI_ISL_588556, EPI_ISL_588557, EPI_ISL_588558, EPI_ISL_588559, EPI_ISL_588560, EPI_ISL_588561, EPI_ISL_588562, EPI_ISL_588563, EPI_ISL_588564, EPI_ISL_588565, EPI_ISL_588566, EPI_ISL_588567, EPI_ISL_588568, EPI_ISL_588569, EPI_ISL_588570, EPI_ISL_588571, EPI_ISL_588573, EPI_ISL_588574, EPI_ISL_588575, EPI_ISL_588576, EPI_ISL_588577, EPI_ISL_588578, EPI_ISL_588579, EPI_ISL_588580, EPI_ISL_588582, EPI_ISL_588583, EPI_ISL_588584, EPI_ISL_588585, EPI_ISL_588586, EPI_ISL_588587, EPI_ISL_588588, EPI_ISL_588590, EPI_ISL_588591, EPI_ISL_588592, EPI_ISL_588593, EPI_ISL_588594, EPI_ISL_588595, EPI_ISL_588596, EPI_ISL_588597, EPI_ISL_588599, EPI_ISL_588600, EPI_ISL_588601, EPI_ISL_588602, EPI_ISL_588603, EPI_ISL_588604, EPI_ISL_588605, EPI_ISL_588606, EPI_ISL_588607, EPI_ISL_588608, EPI_ISL_588609, EPI_ISL_588610, EPI_ISL_588611, EPI_ISL_588612, EPI_ISL_588613, EPI_ISL_588614, EPI_ISL_588615, EPI_ISL_588616, EPI_ISL_588617, EPI_ISL_588618, EPI_ISL_588620, EPI_ISL_588621, EPI_ISL_588622, EPI_ISL_588623, EPI_ISL_588624, EPI_ISL_588625, EPI_ISL_588627, EPI_ISL_588629, EPI_ISL_588630, EPI_ISL_588631, EPI_ISL_588632, EPI_ISL_588633, EPI_ISL_588634, EPI_ISL_588635, EPI_ISL_588636, EPI_ISL_588637, EPI_ISL_588638, EPI_ISL_588639, EPI_ISL_588640, EPI_ISL_588641, EPI_ISL_588642, EPI_ISL_588644, EPI_ISL_588645, EPI_ISL_588646, EPI_ISL_588647, EPI_ISL_588648, EPI_ISL_588650, EPI_ISL_588651, EPI_ISL_588652, EPI_ISL_588653, EPI_ISL_588654, EPI_ISL_588655, EPI_ISL_588656, EPI_ISL_588657, EPI_ISL_588658, EPI_ISL_588659, EPI_ISL_588660, EPI_ISL_588661, EPI_ISL_588662, EPI_ISL_588663, EPI_ISL_588664, EPI_ISL_588665, EPI_ISL_588666, EPI_ISL_588667, EPI_ISL_588668, EPI_ISL_588669, EPI_ISL_588670, EPI_ISL_588671, EPI_ISL_588673, EPI_ISL_588676, EPI_ISL_588677, EPI_ISL_588679, EPI_ISL_588680, EPI_ISL_588681, EPI_ISL_588682, EPI_ISL_588683, EPI_ISL_588684, EPI_ISL_588685, EPI_ISL_588686, EPI_ISL_588687, EPI_ISL_588688, EPI_ISL_588689, EPI_ISL_588690, EPI_ISL_588692, EPI_ISL_588693, EPI_ISL_588694, EPI_ISL_588695, EPI_ISL_588696, EPI_ISL_588697, EPI_ISL_588698, EPI_ISL_588700, EPI_ISL_588701, EPI_ISL_588702, EPI_ISL_588703, EPI_ISL_588704, EPI_ISL_588705, EPI_ISL_588706, EPI_ISL_588707, EPI_ISL_588708, EPI_ISL_588709, EPI_ISL_588710, EPI_ISL_588711, EPI_ISL_588712, EPI_ISL_588713, EPI_ISL_588714, EPI_ISL_588715, EPI_ISL_588716, EPI_ISL_588718, EPI_ISL_588719, EPI_ISL_588720, EPI_ISL_588721, EPI_ISL_588722, EPI_ISL_588723, EPI_ISL_588724, EPI_ISL_588728, EPI_ISL_588745, EPI_ISL_588770, EPI_ISL_588772, EPI_ISL_588792, EPI_ISL_588827, EPI_ISL_588855, EPI_ISL_588867, EPI_ISL_588901, EPI_ISL_588905, EPI_ISL_588909, EPI_ISL_588912, EPI_ISL_588918, EPI_ISL_588932, EPI_ISL_588944, EPI_ISL_588947, EPI_ISL_588966, EPI_ISL_588975, EPI_ISL_588986, EPI_ISL_588988, EPI_ISL_589012, EPI_ISL_589023, EPI_ISL_589037, EPI_ISL_589102, EPI_ISL_589119, EPI_ISL_589149, EPI_ISL_589193, EPI_ISL_589208

| | | | |
|---|---|---|---|
| see above | Lighthouse Lab in Glasgow | Wellcome Sanger Institute for the COVID-19 Genomics UK (COG-UK) consortium | Harper VanSteenhouse, Yumi Kasai, David Gray, Carol Clugston, Anna Dominiczak and Alex Alderton, Roberto Amato, Sonia Goncalves, Ewan Harrison, David K. Jackson, Ian Johnston, Dominic Kwiatkowski, Cordelia Langford, John Sillitoe on behalf of the Wellcome Sanger Institute COVID-19 Surveillance Team |

EPI_ISL_589774, EPI_ISL_589775, EPI_ISL_589776, EPI_ISL_589777, EPI_ISL_589788, EPI_ISL_589790, EPI_ISL_589793, EPI_ISL_589803, EPI_ISL_589804, EPI_ISL_589805, EPI_ISL_589806, EPI_ISL_589807, EPI_ISL_589808, EPI_ISL_589809, EPI_ISL_589810, EPI_ISL_589811, EPI_ISL_589812, EPI_ISL_589813, EPI_ISL_589814, EPI_ISL_589815, EPI_ISL_589816, EPI_ISL_589817, EPI_ISL_589818, EPI_ISL_589819, EPI_ISL_589820, EPI_ISL_589821, EPI_ISL_589822, EPI_ISL_589823, EPI_ISL_589824, EPI_ISL_589825, EPI_ISL_589826, EPI_ISL_589827, EPI_ISL_589828, EPI_ISL_589829, EPI_ISL_589830, EPI_ISL_589831, EPI_ISL_589832, EPI_ISL_589833, EPI_ISL_589834, EPI_ISL_589835, EPI_ISL_589836, EPI_ISL_589837, EPI_ISL_589838, EPI_ISL_589839, EPI_ISL_589840, EPI_ISL_589841, EPI_ISL_589842, EPI_ISL_589843, EPI_ISL_589844, EPI_ISL_589845, EPI_ISL_589846, EPI_ISL_589847, EPI_ISL_589848, EPI_ISL_589849, EPI_ISL_589850, EPI_ISL_589851, EPI_ISL_589852, EPI_ISL_589853, EPI_ISL_589854, EPI_ISL_589855, EPI_ISL_589856, EPI_ISL_589857, EPI_ISL_589858, EPI_ISL_589859, EPI_ISL_589860, EPI_ISL_589861, EPI_ISL_589862, EPI_ISL_589863, EPI_ISL_589864, EPI_ISL_589865, EPI_ISL_589866, EPI_ISL_589867, EPI_ISL_589868, EPI_ISL_589869, EPI_ISL_589870, EPI_ISL_589871, EPI_ISL_589872, EPI_ISL_589873, EPI_ISL_589874, EPI_ISL_589875, EPI_ISL_589876, EPI_ISL_589877, EPI_ISL_589878, EPI_ISL_589879, EPI_ISL_589880, EPI_ISL_589881, EPI_ISL_589882, EPI_ISL_589883, EPI_ISL_589884, EPI_ISL_589885, EPI_ISL_589886, EPI_ISL_589887, EPI_ISL_589888, EPI_ISL_589889, EPI_ISL_589890, EPI_ISL_589891, EPI_ISL_589892, EPI_ISL_589893, EPI_ISL_589894, EPI_ISL_589895, EPI_ISL_589896, EPI_ISL_589897, EPI_ISL_589898, EPI_ISL_589899, EPI_ISL_589900, EPI_ISL_589901, EPI_ISL_589902, EPI_ISL_589903, EPI_ISL_589904, EPI_ISL_589905, EPI_ISL_589906, EPI_ISL_589907, EPI_ISL_589908, EPI_ISL_589909, EPI_ISL_589910, EPI_ISL_589911, EPI_ISL_589912, EPI_ISL_589913, EPI_ISL_589914, EPI_ISL_589915, EPI_ISL_589916, EPI_ISL_589917, EPI_ISL_589918, EPI_ISL_589919, EPI_ISL_589920, EPI_ISL_589921, EPI_ISL_589922, EPI_ISL_589923, EPI_ISL_589924, EPI_ISL_589925, EPI_ISL_589926, EPI_ISL_589927, EPI_ISL_589928, EPI_ISL_589929, EPI_ISL_589930, EPI_ISL_589931, EPI_ISL_589932, EPI_ISL_589933, EPI_ISL_589934, EPI_ISL_589935, EPI_ISL_589937, EPI_ISL_589938, EPI_ISL_589939, EPI_ISL_589940,

EPI_ISL_589941, EPI_ISL_589942, EPI_ISL_589943, EPI_ISL_589944, EPI_ISL_589945, EPI_ISL_589946, EPI_ISL_589947, EPI_ISL_589948, EPI_ISL_589949, EPI_ISL_589950, EPI_ISL_589951, EPI_ISL_589952, EPI_ISL_589953, EPI_ISL_589954, EPI_ISL_589955, EPI_ISL_589956, EPI_ISL_589957, EPI_ISL_589958, EPI_ISL_589959, EPI_ISL_589960, EPI_ISL_589961, EPI_ISL_589962, EPI_ISL_589963, EPI_ISL_589964, EPI_ISL_589965, EPI_ISL_589966, EPI_ISL_589967, EPI_ISL_589968, EPI_ISL_589969, EPI_ISL_589970, EPI_ISL_589971, EPI_ISL_589972, EPI_ISL_589973, EPI_ISL_589974, EPI_ISL_589975, EPI_ISL_589976, EPI_ISL_589977, EPI_ISL_589978, EPI_ISL_589979, EPI_ISL_589980, EPI_ISL_589981, EPI_ISL_589982, EPI_ISL_589983, EPI_ISL_589984, EPI_ISL_589985, EPI_ISL_589986, EPI_ISL_589987, EPI_ISL_589988, EPI_ISL_589989, EPI_ISL_589990, EPI_ISL_589991, EPI_ISL_589992, EPI_ISL_589993, EPI_ISL_589994, EPI_ISL_589995, EPI_ISL_589996, EPI_ISL_589997, EPI_ISL_589998, EPI_ISL_589999, EPI_ISL_590000, EPI_ISL_590001, EPI_ISL_590002, EPI_ISL_590003, EPI_ISL_590004, EPI_ISL_590005, EPI_ISL_590006, EPI_ISL_590007, EPI_ISL_590008, EPI_ISL_590009, EPI_ISL_590010, EPI_ISL_590011, EPI_ISL_590012, EPI_ISL_590013, EPI_ISL_590014, EPI_ISL_590015, EPI_ISL_590016, EPI_ISL_590017, EPI_ISL_590018, EPI_ISL_590019, EPI_ISL_590020, EPI_ISL_590021, EPI_ISL_590022, EPI_ISL_590023, EPI_ISL_590024, EPI_ISL_590025, EPI_ISL_590026, EPI_ISL_590027, EPI_ISL_590028, EPI_ISL_590029, EPI_ISL_590030, EPI_ISL_590031, EPI_ISL_590032, EPI_ISL_590033, EPI_ISL_590034, EPI_ISL_590035, EPI_ISL_590036, EPI_ISL_590037, EPI_ISL_590038, EPI_ISL_590039, EPI_ISL_590040, EPI_ISL_590041, EPI_ISL_590042, EPI_ISL_590043, EPI_ISL_590044, EPI_ISL_590045, EPI_ISL_590047, EPI_ISL_590048, EPI_ISL_590049, EPI_ISL_590050, EPI_ISL_590051, EPI_ISL_590052, EPI_ISL_590053, EPI_ISL_590054, EPI_ISL_590055, EPI_ISL_590056, EPI_ISL_590057, EPI_ISL_590058, EPI_ISL_590059, EPI_ISL_590060, EPI_ISL_590061, EPI_ISL_590062, EPI_ISL_590063, EPI_ISL_590064, EPI_ISL_590065, EPI_ISL_590066, EPI_ISL_590067, EPI_ISL_590068, EPI_ISL_590069, EPI_ISL_590070, EPI_ISL_590071, EPI_ISL_590072, EPI_ISL_590073, EPI_ISL_590074, EPI_ISL_590075, EPI_ISL_590076, EPI_ISL_590077, EPI_ISL_590078, EPI_ISL_590079, EPI_ISL_590080, EPI_ISL_590081, EPI_ISL_590082, EPI_ISL_590083, EPI_ISL_590084, EPI_ISL_590085, EPI_ISL_590086, EPI_ISL_590087, EPI_ISL_590088, EPI_ISL_590089, EPI_ISL_590090, EPI_ISL_590091, EPI_ISL_590092, EPI_ISL_590093, EPI_ISL_590094, EPI_ISL_590095, EPI_ISL_590096, EPI_ISL_590097, EPI_ISL_590098, EPI_ISL_590099, EPI_ISL_590100, EPI_ISL_590101, EPI_ISL_590102, EPI_ISL_590103, EPI_ISL_590104, EPI_ISL_590105, EPI_ISL_590106, EPI_ISL_590107, EPI_ISL_590108, EPI_ISL_590109, EPI_ISL_590110, EPI_ISL_590111, EPI_ISL_590112, EPI_ISL_590113, EPI_ISL_590114, EPI_ISL_590115, EPI_ISL_590116, EPI_ISL_590117, EPI_ISL_590118, EPI_ISL_590119, EPI_ISL_590120, EPI_ISL_590121, EPI_ISL_590122, EPI_ISL_590123, EPI_ISL_590124, EPI_ISL_590125, EPI_ISL_590126, EPI_ISL_590127, EPI_ISL_590128, EPI_ISL_590129, EPI_ISL_590130, EPI_ISL_590131, EPI_ISL_590132, EPI_ISL_590133, EPI_ISL_590134, EPI_ISL_590135, EPI_ISL_590136, EPI_ISL_590137, EPI_ISL_590138, EPI_ISL_590139, EPI_ISL_590140, EPI_ISL_590141, EPI_ISL_590142, EPI_ISL_590143, EPI_ISL_590144, EPI_ISL_590145, EPI_ISL_590146, EPI_ISL_590147, EPI_ISL_590148, EPI_ISL_590149, EPI_ISL_590150, EPI_ISL_590151, EPI_ISL_590152, EPI_ISL_590153, EPI_ISL_590154, EPI_ISL_590155, EPI_ISL_590156, EPI_ISL_590157, EPI_ISL_590158, EPI_ISL_590159, EPI_ISL_590160, EPI_ISL_590161, EPI_ISL_590162, EPI_ISL_590163, EPI_ISL_590164, EPI_ISL_590165, EPI_ISL_590166, EPI_ISL_590167, EPI_ISL_590168, EPI_ISL_590169, EPI_ISL_590170, EPI_ISL_590171, EPI_ISL_590176, EPI_ISL_590177, EPI_ISL_590178, EPI_ISL_590179, EPI_ISL_590180, EPI_ISL_590182, EPI_ISL_590183, EPI_ISL_590184, EPI_ISL_590185, EPI_ISL_590187, EPI_ISL_590188, EPI_ISL_590189, EPI_ISL_590190, EPI_ISL_590191, EPI_ISL_590192, EPI_ISL_590195, EPI_ISL_590196, EPI_ISL_590197, EPI_ISL_590199, EPI_ISL_590201, EPI_ISL_590202, EPI_ISL_590203, EPI_ISL_590204, EPI_ISL_590205, EPI_ISL_590207, EPI_ISL_590208, EPI_ISL_590209, EPI_ISL_590211, EPI_ISL_590212, EPI_ISL_590216, EPI_ISL_590217, EPI_ISL_590218, EPI_ISL_590219, EPI_ISL_590221, EPI_ISL_590222, EPI_ISL_590223, EPI_ISL_590227, EPI_ISL_590229, EPI_ISL_590230, EPI_ISL_590231, EPI_ISL_590232, EPI_ISL_590233, EPI_ISL_590236, EPI_ISL_590237, EPI_ISL_590238, EPI_ISL_590240, EPI_ISL_590241, EPI_ISL_590242, EPI_ISL_590243, EPI_ISL_590244, EPI_ISL_590245, EPI_ISL_590247, EPI_ISL_590248, EPI_ISL_590249, EPI_ISL_590251, EPI_ISL_590252, EPI_ISL_590253, EPI_ISL_590254, EPI_ISL_590256, EPI_ISL_590258, EPI_ISL_590262, EPI_ISL_590264, EPI_ISL_590266, EPI_ISL_590267, EPI_ISL_590268, EPI_ISL_590269, EPI_ISL_590270, EPI_ISL_590271, EPI_ISL_590273, EPI_ISL_590278, EPI_ISL_590279, EPI_ISL_590280, EPI_ISL_590281, EPI_ISL_590285, EPI_ISL_590287, EPI_ISL_590289, EPI_ISL_590290, EPI_ISL_590291, EPI_ISL_590292, EPI_ISL_590293, EPI_ISL_590294, EPI_ISL_590297, EPI_ISL_590298, EPI_ISL_590299, EPI_ISL_590301, EPI_ISL_590302, EPI_ISL_590303, EPI_ISL_590304, EPI_ISL_590305, EPI_ISL_590308, EPI_ISL_590310, EPI_ISL_590311, EPI_ISL_590312, EPI_ISL_590313, EPI_ISL_590315, EPI_ISL_590316, EPI_ISL_590317, EPI_ISL_590318, EPI_ISL_590320, EPI_ISL_590322, EPI_ISL_590323, EPI_ISL_590324, EPI_ISL_590325, EPI_ISL_590326, EPI_ISL_590330, EPI_ISL_590331, EPI_ISL_590332, EPI_ISL_590333, EPI_ISL_590334, EPI_ISL_590336, EPI_ISL_590338, EPI_ISL_590339, EPI_ISL_590340, EPI_ISL_590341, EPI_ISL_590342, EPI_ISL_590343, EPI_ISL_590345, EPI_ISL_590348, EPI_ISL_590349, EPI_ISL_590350, EPI_ISL_590353, EPI_ISL_590354, EPI_ISL_590356, EPI_ISL_590358, EPI_ISL_590360, EPI_ISL_590362, EPI_ISL_590363, EPI_ISL_590365, EPI_ISL_590368, EPI_ISL_590370, EPI_ISL_590371, EPI_ISL_590372, EPI_ISL_590374, EPI_ISL_590375, EPI_ISL_590376, EPI_ISL_590377, EPI_ISL_590378, EPI_ISL_590381, EPI_ISL_590382, EPI_ISL_590384, EPI_ISL_590385, EPI_ISL_590387, EPI_ISL_590389, EPI_ISL_590391, EPI_ISL_590393, EPI_ISL_590394, EPI_ISL_590395, EPI_ISL_590396, EPI_ISL_590400, EPI_ISL_590401, EPI_ISL_590402, EPI_ISL_590404, EPI_ISL_590406, EPI_ISL_590407, EPI_ISL_590408, EPI_ISL_590409, EPI_ISL_590410, EPI_ISL_590411, EPI_ISL_590412, EPI_ISL_590413, EPI_ISL_590414, EPI_ISL_590415, EPI_ISL_590416, EPI_ISL_590417, EPI_ISL_590418, EPI_ISL_590420, EPI_ISL_590421, EPI_ISL_590424, EPI_ISL_590425, EPI_ISL_590426, EPI_ISL_590427, EPI_ISL_590428, EPI_ISL_590429, EPI_ISL_590430, EPI_ISL_590431, EPI_ISL_590432, EPI_ISL_590433, EPI_ISL_590434, EPI_ISL_590435, EPI_ISL_590436, EPI_ISL_590437, EPI_ISL_590439, EPI_ISL_590440, EPI_ISL_590443, EPI_ISL_590444, EPI_ISL_590445, EPI_ISL_590447, EPI_ISL_590449, EPI_ISL_590450, EPI_ISL_590451, EPI_ISL_590454, EPI_ISL_590455, EPI_ISL_590456, EPI_ISL_590457, EPI_ISL_590458, EPI_ISL_590459, EPI_ISL_590460, EPI_ISL_590461, EPI_ISL_590463, EPI_ISL_590464, EPI_ISL_590465, EPI_ISL_590466, EPI_ISL_590467, EPI_ISL_590468, EPI_ISL_590469, EPI_ISL_590470, EPI_ISL_590471, EPI_ISL_590472, EPI_ISL_590473, EPI_ISL_590474, EPI_ISL_590475, EPI_ISL_590476, EPI_ISL_590477, EPI_ISL_590478, EPI_ISL_590479, EPI_ISL_590480, EPI_ISL_590481, EPI_ISL_590482, EPI_ISL_590483, EPI_ISL_590484, EPI_ISL_590485, EPI_ISL_590486, EPI_ISL_590487, EPI_ISL_590488, EPI_ISL_590489, EPI_ISL_590490, EPI_ISL_590491, EPI_ISL_590492, EPI_ISL_590493, EPI_ISL_590494, EPI_ISL_590495, EPI_ISL_590496, EPI_ISL_590498, EPI_ISL_590499, EPI_ISL_590500, EPI_ISL_590501, EPI_ISL_590502, EPI_ISL_590503, EPI_ISL_590504, EPI_ISL_590505, EPI_ISL_590506, EPI_ISL_590507, EPI_ISL_590508, EPI_ISL_590509, EPI_ISL_590510, EPI_ISL_590511, EPI_ISL_590512, EPI_ISL_590513, EPI_ISL_590514, EPI_ISL_590515, EPI_ISL_590516, EPI_ISL_590517, EPI_ISL_590518, EPI_ISL_590519, EPI_ISL_590520, EPI_ISL_590521, EPI_ISL_590522, EPI_ISL_590523, EPI_ISL_590524, EPI_ISL_590525, EPI_ISL_590526, EPI_ISL_590527, EPI_ISL_590528, EPI_ISL_590529, EPI_ISL_590530, EPI_ISL_590531, EPI_ISL_590532, EPI_ISL_590533, EPI_ISL_590534, EPI_ISL_590535, EPI_ISL_590536, EPI_ISL_590537, EPI_ISL_590538, EPI_ISL_590539, EPI_ISL_590540, EPI_ISL_590541, EPI_ISL_590542, EPI_ISL_590543, EPI_ISL_590544, EPI_ISL_590545, EPI_ISL_590546, EPI_ISL_590547, EPI_ISL_590548, EPI_ISL_590549, EPI_ISL_590550, EPI_ISL_590551, EPI_ISL_590552, EPI_ISL_590553, EPI_ISL_590554, EPI_ISL_590555, EPI_ISL_590556, EPI_ISL_590557, EPI_ISL_590558, EPI_ISL_590559, EPI_ISL_590560, EPI_ISL_590561, EPI_ISL_590562, EPI_ISL_590563, EPI_ISL_590564, EPI_ISL_590565, EPI_ISL_590566, EPI_ISL_590567, EPI_ISL_590568, EPI_ISL_590569, EPI_ISL_590570, EPI_ISL_590571, EPI_ISL_590572, EPI_ISL_590573, EPI_ISL_590574, EPI_ISL_590575, EPI_ISL_590576, EPI_ISL_590577, EPI_ISL_590578, EPI_ISL_590579, EPI_ISL_590580, EPI_ISL_590581, EPI_ISL_590582, EPI_ISL_590583, EPI_ISL_590584, EPI_ISL_590585, EPI_ISL_590586, EPI_ISL_590587, EPI_ISL_590588, EPI_ISL_590589, EPI_ISL_590590, EPI_ISL_590591, EPI_ISL_590592, EPI_ISL_590593, EPI_ISL_590594, EPI_ISL_590595, EPI_ISL_590596, EPI_ISL_590597, EPI_ISL_590598, EPI_ISL_590599, EPI_ISL_590600, EPI_ISL_590601, EPI_ISL_590602, EPI_ISL_590603, EPI_ISL_590604, EPI_ISL_590605, EPI_ISL_590606, EPI_ISL_590607, EPI_ISL_590608, EPI_ISL_590609, EPI_ISL_590610, EPI_ISL_590611, EPI_ISL_590612, EPI_ISL_590613, EPI_ISL_590614, EPI_ISL_590615, EPI_ISL_590616, EPI_ISL_590617, EPI_ISL_590618, EPI_ISL_590619, EPI_ISL_590620, EPI_ISL_590621, EPI_ISL_590622, EPI_ISL_590623, EPI_ISL_590624, EPI_ISL_590625, EPI_ISL_590626, EPI_ISL_590627, EPI_ISL_590628, EPI_ISL_590629, EPI_ISL_590630, EPI_ISL_590631, EPI_ISL_590632, EPI_ISL_590633, EPI_ISL_590634, EPI_ISL_590635, EPI_ISL_590636, EPI_ISL_590637, EPI_ISL_590638, EPI_ISL_590639, EPI_ISL_590640, EPI_ISL_590641, EPI_ISL_590642, EPI_ISL_590643, EPI_ISL_590644, EPI_ISL_590645, EPI_ISL_590646, EPI_ISL_590647, EPI_ISL_590648, EPI_ISL_590649, EPI_ISL_590650, EPI_ISL_590651, EPI_ISL_590652, EPI_ISL_590653, EPI_ISL_590654, EPI_ISL_590655, EPI_ISL_590656, EPI_ISL_590657, EPI_ISL_590658, EPI_ISL_590659, EPI_ISL_590660, EPI_ISL_590661, EPI_ISL_590662, EPI_ISL_590663, EPI_ISL_590664, EPI_ISL_590665, EPI_ISL_590666, EPI_ISL_590667, EPI_ISL_590668, EPI_ISL_590669, EPI_ISL_590670, EPI_ISL_590671, EPI_ISL_590672, EPI_ISL_590673, EPI_ISL_590674, EPI_ISL_590675, EPI_ISL_590676, EPI_ISL_590677, EPI_ISL_590678, EPI_ISL_590679, EPI_ISL_590680, EPI_ISL_590681, EPI_ISL_590682, EPI_ISL_590683, EPI_ISL_590684, EPI_ISL_590685, EPI_ISL_590686, EPI_ISL_590687, EPI_ISL_590688

| | | | |
|---|---|---|---|
| see above | Lighthouse Lab in Glasgow | Wellcome Sanger Institute for the COVID-19 Genomics UK (COG-UK) consortium | Harper VanSteenhouse, Yumi Kasai, David Gray, Carol Clugston, Anna Dominiczak and Alex Alderton, Roberto Amato, Sonia Goncalves, Ewan Harrison, David K. Jackson, Ian Johnston, Dominic Kwiatkowski, Cordelia Langford, John Sillitoe on behalf of the Wellcome Sanger Institute COVID-19 Surveillance Team (http://www.sanger.ac.uk/covid-team) |

EPI_ISL_593937, EPI_ISL_593938, EPI_ISL_593939, EPI_ISL_593940, EPI_ISL_593941, EPI_ISL_593942, EPI_ISL_593943, EPI_ISL_593944, EPI_ISL_593945, EPI_ISL_593946, EPI_ISL_593947, EPI_ISL_593948, EPI_ISL_593949, EPI_ISL_593950, EPI_ISL_593951, EPI_ISL_593952, EPI_ISL_593953, EPI_ISL_593954, EPI_ISL_593955, EPI_ISL_593956, EPI_ISL_593957, EPI_ISL_593958, EPI_ISL_593959, EPI_ISL_593960, EPI_ISL_593961, EPI_ISL_593962, EPI_ISL_593963, EPI_ISL_593964, EPI_ISL_593965, EPI_ISL_593966, EPI_ISL_593967, EPI_ISL_593968, EPI_ISL_593969, EPI_ISL_593970, EPI_ISL_593971, EPI_ISL_593972, EPI_ISL_593973, EPI_ISL_593974, EPI_ISL_593975, EPI_ISL_593976, EPI_ISL_593977, EPI_ISL_593978, EPI_ISL_593979, EPI_ISL_593980, EPI_ISL_593981

| | | | |
|---|---|---|---|
| see above | Delaware Public Health Lab | Delaware Public Health Lab | Gregory Hovan |

EPI_ISL_597234, EPI_ISL_597235, EPI_ISL_597236, EPI_ISL_597240, EPI_ISL_597242, EPI_ISL_597244, EPI_ISL_597245, EPI_ISL_597249, EPI_ISL_597252, EPI_ISL_597260, EPI_ISL_597261, EPI_ISL_597263, EPI_ISL_597267, EPI_ISL_597268, EPI_ISL_597272, EPI_ISL_597275, EPI_ISL_597276, EPI_ISL_597277, EPI_ISL_597278, EPI_ISL_597280, EPI_ISL_597281, EPI_ISL_597288, EPI_ISL_597290, EPI_ISL_597291, EPI_ISL_597292, EPI_ISL_597296, EPI_ISL_597298, EPI_ISL_597299, EPI_ISL_597302, EPI_ISL_597303, EPI_ISL_597311, EPI_ISL_597312, EPI_ISL_597313, EPI_ISL_597314, EPI_ISL_597315, EPI_ISL_597318, EPI_ISL_597322, EPI_ISL_597326, EPI_ISL_597328, EPI_ISL_597329, EPI_ISL_597330, EPI_ISL_597332, EPI_ISL_597333, EPI_ISL_597334, EPI_ISL_597336, EPI_ISL_597338, EPI_ISL_597339, EPI_ISL_597347, EPI_ISL_597348, EPI_ISL_597351, EPI_ISL_597352, EPI_ISL_597353, EPI_ISL_597355, EPI_ISL_597356, EPI_ISL_597357, EPI_ISL_597359, EPI_ISL_597360, EPI_ISL_597362, EPI_ISL_597363, EPI_ISL_597364, EPI_ISL_597369, EPI_ISL_597370, EPI_ISL_597376, EPI_ISL_597379, EPI_ISL_597382, EPI_ISL_597383, EPI_ISL_597384, EPI_ISL_597385, EPI_ISL_597386, EPI_ISL_597387, EPI_ISL_597389, EPI_ISL_597391, EPI_ISL_597393, EPI_ISL_597394, EPI_ISL_597395, EPI_ISL_597400, EPI_ISL_597404, EPI_ISL_597406, EPI_ISL_597407, EPI_ISL_597408, EPI_ISL_597409, EPI_ISL_597410, EPI_ISL_597411, EPI_ISL_597413, EPI_ISL_597415, EPI_ISL_597416, EPI_ISL_597417, EPI_ISL_597419, EPI_ISL_597420, EPI_ISL_597421, EPI_ISL_597422, EPI_ISL_597423, EPI_ISL_597429, EPI_ISL_597430, EPI_ISL_597431, EPI_ISL_597433, EPI_ISL_597434, EPI_ISL_597436, EPI_ISL_597439, EPI_ISL_597440, EPI_ISL_597442, EPI_ISL_597443, EPI_ISL_597444, EPI_ISL_597447, EPI_ISL_597448, EPI_ISL_597452, EPI_ISL_597454, EPI_ISL_597455, EPI_ISL_597456, EPI_ISL_597457, EPI_ISL_597463, EPI_ISL_597465, EPI_ISL_597471, EPI_ISL_597474, EPI_ISL_597475, EPI_ISL_597481, EPI_ISL_597482, EPI_ISL_597484, EPI_ISL_597485, EPI_ISL_597486, EPI_ISL_597491, EPI_ISL_597492, EPI_ISL_597493, EPI_ISL_597496, EPI_ISL_597498, EPI_ISL_597499, EPI_ISL_597500, EPI_ISL_597501, EPI_ISL_597503, EPI_ISL_597506, EPI_ISL_597509, EPI_ISL_597511, EPI_ISL_597513, EPI_ISL_597514, EPI_ISL_597515, EPI_ISL_597516, EPI_ISL_597518, EPI_ISL_597521, EPI_ISL_597522, EPI_ISL_598831, EPI_ISL_598832, EPI_ISL_598833, EPI_ISL_598834, EPI_ISL_598835, EPI_ISL_598836, EPI_ISL_598837, EPI_ISL_598838, EPI_ISL_598839, EPI_ISL_598840, EPI_ISL_598841, EPI_ISL_598842, EPI_ISL_598843, EPI_ISL_598844, EPI_ISL_598845, EPI_ISL_598847, EPI_ISL_598848, EPI_ISL_598849, EPI_ISL_598850, EPI_ISL_598851, EPI_ISL_598852, EPI_ISL_598853, EPI_ISL_598854, EPI_ISL_598855, EPI_ISL_598856, EPI_ISL_598857, EPI_ISL_598858, EPI_ISL_598859, EPI_ISL_598860, EPI_ISL_598861, EPI_ISL_598862, EPI_ISL_598863, EPI_ISL_598864, EPI_ISL_598865, EPI_ISL_598866, EPI_ISL_598867, EPI_ISL_598868, EPI_ISL_598869, EPI_ISL_598870, EPI_ISL_598871, EPI_ISL_598872, EPI_ISL_598873, EPI_ISL_598874, EPI_ISL_598875, EPI_ISL_598876, EPI_ISL_598877, EPI_ISL_598878, EPI_ISL_598879, EPI_ISL_598880, EPI_ISL_598881, EPI_ISL_598882, EPI_ISL_598883, EPI_ISL_598884, EPI_ISL_598885, EPI_ISL_598886, EPI_ISL_598887, EPI_ISL_598888, EPI_ISL_598889, EPI_ISL_598890, EPI_ISL_598891, EPI_ISL_598892, EPI_ISL_598893, EPI_ISL_598894, EPI_ISL_598895, EPI_ISL_598896, EPI_ISL_598897, EPI_ISL_598898, EPI_ISL_598899, EPI_ISL_598900, EPI_ISL_598901, EPI_ISL_598902, EPI_ISL_598903, EPI_ISL_598904, EPI_ISL_598905, EPI_ISL_598906, EPI_ISL_598907, EPI_ISL_598908, EPI_ISL_598909, EPI_ISL_598910, EPI_ISL_598911, EPI_ISL_598912, EPI_ISL_598913, EPI_ISL_598914, EPI_ISL_598915, EPI_ISL_598916, EPI_ISL_598917, EPI_ISL_598918, EPI_ISL_598919, EPI_ISL_598920, EPI_ISL_598921, EPI_ISL_598922, EPI_ISL_598923, EPI_ISL_598924, EPI_ISL_598925, EPI_ISL_598926, EPI_ISL_598927, EPI_ISL_598928, EPI_ISL_598929, EPI_ISL_598930, EPI_ISL_598931, EPI_ISL_598932, EPI_ISL_598933, EPI_ISL_598934, EPI_ISL_598935, EPI_ISL_598936, EPI_ISL_598937, EPI_ISL_598938, EPI_ISL_598939, EPI_ISL_598940, EPI_ISL_598941, EPI_ISL_598942, EPI_ISL_598943, EPI_ISL_598944, EPI_ISL_598945, EPI_ISL_598946, EPI_ISL_598947, EPI_ISL_598948, EPI_ISL_598949, EPI_ISL_598950, EPI_ISL_598951, EPI_ISL_598952, EPI_ISL_598953, EPI_ISL_598954, EPI_ISL_598955, EPI_ISL_598956, EPI_ISL_598957, EPI_ISL_598958, EPI_ISL_598959, EPI_ISL_598960, EPI_ISL_598961, EPI_ISL_598962, EPI_ISL_598963, EPI_ISL_598964, EPI_ISL_598965, EPI_ISL_598966, EPI_ISL_598967, EPI_ISL_598968, EPI_ISL_598969, EPI_ISL_598971, EPI_ISL_598972, EPI_ISL_598973, EPI_ISL_598974, EPI_ISL_598975, EPI_ISL_598976, EPI_ISL_598977, EPI_ISL_598978, EPI_ISL_598979, EPI_ISL_598980, EPI_ISL_598981, EPI_ISL_598982, EPI_ISL_598983, EPI_ISL_598984, EPI_ISL_598985, EPI_ISL_598986, EPI_ISL_598987, EPI_ISL_598988, EPI_ISL_598989, EPI_ISL_598990, EPI_ISL_598991, EPI_ISL_598992, EPI_ISL_598993, EPI_ISL_598994, EPI_ISL_598995, EPI_ISL_598996, EPI_ISL_598997, EPI_ISL_598998, EPI_ISL_598999, EPI_ISL_599000, EPI_ISL_599001, EPI_ISL_599002, EPI_ISL_599003, EPI_ISL_599004, EPI_ISL_599005, EPI_ISL_599006, EPI_ISL_599007, EPI_ISL_599008, EPI_ISL_599009, EPI_ISL_599011, EPI_ISL_599012, EPI_ISL_599013, EPI_ISL_599014, EPI_ISL_599015, EPI_ISL_599016, EPI_ISL_599017, EPI_ISL_599018, EPI_ISL_599019, EPI_ISL_599020, EPI_ISL_599021, EPI_ISL_599022, EPI_ISL_599023, EPI_ISL_599024, EPI_ISL_599025, EPI_ISL_599026, EPI_ISL_599027, EPI_ISL_599028, EPI_ISL_599029, EPI_ISL_599030, EPI_ISL_599031, EPI_ISL_599032, EPI_ISL_599033, EPI_ISL_599034, EPI_ISL_599035, EPI_ISL_599036, EPI_ISL_599037, EPI_ISL_599038, EPI_ISL_599039, EPI_ISL_599040, EPI_ISL_599041, EPI_ISL_599042, EPI_ISL_599043, EPI_ISL_599044, EPI_ISL_599045, EPI_ISL_599046, EPI_ISL_599047, EPI_ISL_599048, EPI_ISL_599049, EPI_ISL_599050, EPI_ISL_599051, EPI_ISL_599052, EPI_ISL_599053, EPI_ISL_599054, EPI_ISL_599055, EPI_ISL_599056, EPI_ISL_599057, EPI_ISL_599058, EPI_ISL_599059, EPI_ISL_599060, EPI_ISL_599061, EPI_ISL_599062, EPI_ISL_599063, EPI_ISL_599064, EPI_ISL_599065, EPI_ISL_599066, EPI_ISL_599067, EPI_ISL_599068, EPI_ISL_599069, EPI_ISL_599070, EPI_ISL_599071, EPI_ISL_599072, EPI_ISL_599073, EPI_ISL_599074, EPI_ISL_599075, EPI_ISL_599076, EPI_ISL_599078, EPI_ISL_599079, EPI_ISL_599080, EPI_ISL_599081, EPI_ISL_599082, EPI_ISL_599083, EPI_ISL_599084, EPI_ISL_599085, EPI_ISL_599086, EPI_ISL_599087, EPI_ISL_599088, EPI_ISL_599089, EPI_ISL_599090, EPI_ISL_599091, EPI_ISL_599092, EPI_ISL_599093, EPI_ISL_599094, EPI_ISL_599095, EPI_ISL_599096, EPI_ISL_599097, EPI_ISL_599098, EPI_ISL_599099, EPI_ISL_599100, EPI_ISL_599101, EPI_ISL_599102, EPI_ISL_599103, EPI_ISL_599104, EPI_ISL_599105, EPI_ISL_599106, EPI_ISL_599107, EPI_ISL_599108, EPI_ISL_599109, EPI_ISL_599110, EPI_ISL_599111, EPI_ISL_599112, EPI_ISL_599113, EPI_ISL_599114, EPI_ISL_599115, EPI_ISL_599116, EPI_ISL_599117, EPI_ISL_599118, EPI_ISL_599119, EPI_ISL_599120, EPI_ISL_599121, EPI_ISL_599122, EPI_ISL_599123, EPI_ISL_599124, EPI_ISL_599125, EPI_ISL_599126, EPI_ISL_599127, EPI_ISL_599128, EPI_ISL_599129, EPI_ISL_599130, EPI_ISL_599131, EPI_ISL_599132, EPI_ISL_599133, EPI_ISL_599134, EPI_ISL_599135, EPI_ISL_599136, EPI_ISL_599137, EPI_ISL_599145, EPI_ISL_599154, EPI_ISL_599160, EPI_ISL_599161, EPI_ISL_599164, EPI_ISL_599168, EPI_ISL_599169, EPI_ISL_599171, EPI_ISL_599172, EPI_ISL_599173, EPI_ISL_599176, EPI_ISL_599177, EPI_ISL_599179, EPI_ISL_599180, EPI_ISL_599189, EPI_ISL_599191, EPI_ISL_599197, EPI_ISL_599206, EPI_ISL_599209, EPI_ISL_599211, EPI_ISL_599212, EPI_ISL_599214, EPI_ISL_599215, EPI_ISL_599216, EPI_ISL_599219, EPI_ISL_599220, EPI_ISL_599221, EPI_ISL_599223, EPI_ISL_599225, EPI_ISL_599227, EPI_ISL_599228, EPI_ISL_599231, EPI_ISL_599233, EPI_ISL_599235, EPI_ISL_599236, EPI_ISL_599237, EPI_ISL_599238, EPI_ISL_599239, EPI_ISL_599241, EPI_ISL_599243, EPI_ISL_599247, EPI_ISL_599248, EPI_ISL_599249, EPI_ISL_599250, EPI_ISL_599253, EPI_ISL_599255, EPI_ISL_599256, EPI_ISL_599257, EPI_ISL_599261, EPI_ISL_599262, EPI_ISL_599267, EPI_ISL_599268, EPI_ISL_599271, EPI_ISL_599272, EPI_ISL_599276, EPI_ISL_599279, EPI_ISL_599286, EPI_ISL_599291, EPI_ISL_599293, EPI_ISL_599294, EPI_ISL_599295, EPI_ISL_599298, EPI_ISL_599299, EPI_ISL_599300, EPI_ISL_599306, EPI_ISL_599310, EPI_ISL_599311, EPI_ISL_599316, EPI_ISL_599317, EPI_ISL_599323, EPI_ISL_599324, EPI_ISL_599325, EPI_ISL_599326, EPI_ISL_599329, EPI_ISL_599331, EPI_ISL_599332, EPI_ISL_599333, EPI_ISL_599341, EPI_ISL_599342, EPI_ISL_599343, EPI_ISL_599344, EPI_ISL_599345, EPI_ISL_599353, EPI_ISL_599354, EPI_ISL_599357, EPI_ISL_599361, EPI_ISL_599362, EPI_ISL_599365, EPI_ISL_599366, EPI_ISL_599367, EPI_ISL_599369, EPI_ISL_599372, EPI_ISL_599376, EPI_ISL_599377, EPI_ISL_599378, EPI_ISL_599379, EPI_ISL_599383, EPI_ISL_599384, EPI_ISL_599385, EPI_ISL_599386, EPI_ISL_599390, EPI_ISL_599391, EPI_ISL_599393, EPI_ISL_599394, EPI_ISL_599395, EPI_ISL_599398, EPI_ISL_599399, EPI_ISL_599402, EPI_ISL_599403, EPI_ISL_599404, EPI_ISL_599408, EPI_ISL_599409, EPI_ISL_599411, EPI_ISL_599413, EPI_ISL_599414, EPI_ISL_599415, EPI_ISL_599416, EPI_ISL_599419, EPI_ISL_599425, EPI_ISL_599426, EPI_ISL_599429, EPI_ISL_599431, EPI_ISL_599435, EPI_ISL_599436, EPI_ISL_599437, EPI_ISL_599444, EPI_ISL_599445, EPI_ISL_599449, EPI_ISL_599450, EPI_ISL_599452, EPI_ISL_599457, EPI_ISL_599463, EPI_ISL_599466, EPI_ISL_599468, EPI_ISL_599472, EPI_ISL_599474, EPI_ISL_599475, EPI_ISL_599477, EPI_ISL_599480, EPI_ISL_599482, EPI_ISL_599484, EPI_ISL_599485, EPI_ISL_599486, EPI_ISL_599487, EPI_ISL_599488, EPI_ISL_599490, EPI_ISL_599493, EPI_ISL_599494, EPI_ISL_599495, EPI_ISL_599496, EPI_ISL_599497, EPI_ISL_599498, EPI_ISL_599499, EPI_ISL_599500, EPI_ISL_599501, EPI_ISL_599502, EPI_ISL_599503, EPI_ISL_599504, EPI_ISL_599505, EPI_ISL_599506, EPI_ISL_599507, EPI_ISL_599509, EPI_ISL_599510, EPI_ISL_599512, EPI_ISL_599513, EPI_ISL_599514, EPI_ISL_599515, EPI_ISL_599516, EPI_ISL_599517, EPI_ISL_599518, EPI_ISL_599519, EPI_ISL_599520, EPI_ISL_599522, EPI_ISL_599523, EPI_ISL_599524, EPI_ISL_599526, EPI_ISL_599527, EPI_ISL_599528, EPI_ISL_599529, EPI_ISL_599530, EPI_ISL_599531, EPI_ISL_599532, EPI_ISL_599533, EPI_ISL_599534, EPI_ISL_599535, EPI_ISL_599536, EPI_ISL_599538, EPI_ISL_599541, EPI_ISL_599542, EPI_ISL_599543, EPI_ISL_599544, EPI_ISL_599545, EPI_ISL_599547, EPI_ISL_599548, EPI_ISL_599549, EPI_ISL_599550, EPI_ISL_599551, EPI_ISL_599553, EPI_ISL_599554, EPI_ISL_599555, EPI_ISL_599556, EPI_ISL_599558, EPI_ISL_599559, EPI_ISL_599560, EPI_ISL_599561, EPI_ISL_599562, EPI_ISL_599563, EPI_ISL_599564, EPI_ISL_599565, EPI_ISL_599566, EPI_ISL_599567, EPI_ISL_599568, EPI_ISL_599569, EPI_ISL_599570, EPI_ISL_599571, EPI_ISL_599573, EPI_ISL_599574, EPI_ISL_599575, EPI_ISL_599576, EPI_ISL_599577, EPI_ISL_599579, EPI_ISL_599580, EPI_ISL_599581, EPI_ISL_599582, EPI_ISL_599583, EPI_ISL_599584, EPI_ISL_599585, EPI_ISL_599586, EPI_ISL_599587, EPI_ISL_599589, EPI_ISL_599591, EPI_ISL_599592, EPI_ISL_599594, EPI_ISL_599595, EPI_ISL_599598, EPI_ISL_599599, EPI_ISL_599600, EPI_ISL_599601, EPI_ISL_599602, EPI_ISL_599603, EPI_ISL_599604, EPI_ISL_599605, EPI_ISL_599606, EPI_ISL_599607, EPI_ISL_599608, EPI_ISL_599609, EPI_ISL_599610, EPI_ISL_599611, EPI_ISL_599612, EPI_ISL_599613, EPI_ISL_599614, EPI_ISL_599615, EPI_ISL_599616, EPI_ISL_599617, EPI_ISL_599618, EPI_ISL_599619, EPI_ISL_599620, EPI_ISL_599621, EPI_ISL_599622, EPI_ISL_599623, EPI_ISL_599624, EPI_ISL_599625, EPI_ISL_599626, EPI_ISL_599628, EPI_ISL_599629, EPI_ISL_599630, EPI_ISL_599631, EPI_ISL_599632, EPI_ISL_599633, EPI_ISL_599634, EPI_ISL_599636, EPI_ISL_599637, EPI_ISL_599638, EPI_ISL_599639, EPI_ISL_599640, EPI_ISL_599641, EPI_ISL_599642, EPI_ISL_599643, EPI_ISL_599644, EPI_ISL_599645, EPI_ISL_599648, EPI_ISL_599649, EPI_ISL_599650, EPI_ISL_599651, EPI_ISL_599652, EPI_ISL_599653, EPI_ISL_599654, EPI_ISL_599655, EPI_ISL_599656, EPI_ISL_599657, EPI_ISL_599659, EPI_ISL_599660, EPI_ISL_599661, EPI_ISL_599662, EPI_ISL_599663,

EPI_ISL_599664, EPI_ISL_599665, EPI_ISL_599666, EPI_ISL_599667, EPI_ISL_599668, EPI_ISL_599669, EPI_ISL_599670, EPI_ISL_599672, EPI_ISL_599673, EPI_ISL_599674, EPI_ISL_599675, EPI_ISL_599678, EPI_ISL_599679, EPI_ISL_599680, EPI_ISL_599681, EPI_ISL_599682, EPI_ISL_599684, EPI_ISL_599686, EPI_ISL_599687, EPI_ISL_599688, EPI_ISL_599689, EPI_ISL_599690, EPI_ISL_599692, EPI_ISL_599693, EPI_ISL_599694, EPI_ISL_599695, EPI_ISL_599696, EPI_ISL_599699, EPI_ISL_599701, EPI_ISL_599703, EPI_ISL_599705, EPI_ISL_599706, EPI_ISL_599707, EPI_ISL_599708, EPI_ISL_599709, EPI_ISL_599710, EPI_ISL_599711, EPI_ISL_599712, EPI_ISL_599713, EPI_ISL_599714, EPI_ISL_599715, EPI_ISL_599716, EPI_ISL_599717, EPI_ISL_599719, EPI_ISL_599720, EPI_ISL_599724, EPI_ISL_599727, EPI_ISL_599728, EPI_ISL_599729, EPI_ISL_599730, EPI_ISL_599731, EPI_ISL_599732, EPI_ISL_599733, EPI_ISL_599734, EPI_ISL_599736, EPI_ISL_599738, EPI_ISL_599739, EPI_ISL_599740, EPI_ISL_599741, EPI_ISL_599743, EPI_ISL_599744, EPI_ISL_599747, EPI_ISL_599748, EPI_ISL_599749, EPI_ISL_599750, EPI_ISL_599751, EPI_ISL_599752, EPI_ISL_599753, EPI_ISL_599754, EPI_ISL_599755, EPI_ISL_599756, EPI_ISL_599757, EPI_ISL_599758, EPI_ISL_599759, EPI_ISL_599760, EPI_ISL_599761, EPI_ISL_599762, EPI_ISL_599763, EPI_ISL_599764, EPI_ISL_599765, EPI_ISL_599766, EPI_ISL_599767, EPI_ISL_599768, EPI_ISL_599769, EPI_ISL_599771, EPI_ISL_599772, EPI_ISL_599773, EPI_ISL_599774, EPI_ISL_599775, EPI_ISL_599777, EPI_ISL_599778, EPI_ISL_599780, EPI_ISL_599781, EPI_ISL_599782, EPI_ISL_599783, EPI_ISL_599784, EPI_ISL_599785, EPI_ISL_599787, EPI_ISL_599788, EPI_ISL_599789, EPI_ISL_599790, EPI_ISL_599791, EPI_ISL_599792, EPI_ISL_599794, EPI_ISL_599795, EPI_ISL_599796, EPI_ISL_599797, EPI_ISL_599798, EPI_ISL_599799, EPI_ISL_599800, EPI_ISL_599801, EPI_ISL_599802, EPI_ISL_599803, EPI_ISL_599804, EPI_ISL_599805, EPI_ISL_599806, EPI_ISL_599808, EPI_ISL_599809, EPI_ISL_599810, EPI_ISL_599811, EPI_ISL_599812, EPI_ISL_599813, EPI_ISL_599814, EPI_ISL_599815, EPI_ISL_599817, EPI_ISL_600129, EPI_ISL_600132, EPI_ISL_600141, EPI_ISL_600142, EPI_ISL_600147, EPI_ISL_600152, EPI_ISL_600153, EPI_ISL_600155, EPI_ISL_600169, EPI_ISL_600171, EPI_ISL_600172, EPI_ISL_600174, EPI_ISL_600175, EPI_ISL_600181, EPI_ISL_600184, EPI_ISL_600186, EPI_ISL_600190, EPI_ISL_600191, EPI_ISL_600203, EPI_ISL_600205, EPI_ISL_600207, EPI_ISL_600213, EPI_ISL_600214, EPI_ISL_600228, EPI_ISL_600233, EPI_ISL_600235, EPI_ISL_600237, EPI_ISL_600238, EPI_ISL_600239, EPI_ISL_600244, EPI_ISL_600245, EPI_ISL_600246, EPI_ISL_600253, EPI_ISL_600258, EPI_ISL_600268, EPI_ISL_600269, EPI_ISL_600273, EPI_ISL_600289, EPI_ISL_600296, EPI_ISL_600297, EPI_ISL_600300, EPI_ISL_600316, EPI_ISL_600318, EPI_ISL_600320, EPI_ISL_600329, EPI_ISL_600333, EPI_ISL_600335, EPI_ISL_600339, EPI_ISL_600341, EPI_ISL_600351, EPI_ISL_600353, EPI_ISL_600362, EPI_ISL_600365, EPI_ISL_600369, EPI_ISL_600370, EPI_ISL_600376, EPI_ISL_600384, EPI_ISL_600394, EPI_ISL_600400, EPI_ISL_600401, EPI_ISL_600405, EPI_ISL_600408, EPI_ISL_600411, EPI_ISL_600414, EPI_ISL_600426, EPI_ISL_600450, EPI_ISL_600453, EPI_ISL_600457, EPI_ISL_600462, EPI_ISL_600465, EPI_ISL_600467, EPI_ISL_600469, EPI_ISL_600470, EPI_ISL_600479, EPI_ISL_600481, EPI_ISL_600488, EPI_ISL_600494, EPI_ISL_600511, EPI_ISL_600516, EPI_ISL_600539, EPI_ISL_600540, EPI_ISL_600542, EPI_ISL_600544, EPI_ISL_600545, EPI_ISL_600547, EPI_ISL_600548, EPI_ISL_600550, EPI_ISL_600551, EPI_ISL_600553, EPI_ISL_600555, EPI_ISL_600556, EPI_ISL_600558, EPI_ISL_600560, EPI_ISL_600561, EPI_ISL_600563, EPI_ISL_600565, EPI_ISL_600566, EPI_ISL_600568, EPI_ISL_600569, EPI_ISL_600571, EPI_ISL_600572, EPI_ISL_600573, EPI_ISL_600574, EPI_ISL_600575, EPI_ISL_600576, EPI_ISL_600577, EPI_ISL_600578, EPI_ISL_600579, EPI_ISL_600580, EPI_ISL_600581, EPI_ISL_600582, EPI_ISL_600583, EPI_ISL_600584, EPI_ISL_600585, EPI_ISL_600586, EPI_ISL_600587, EPI_ISL_600588, EPI_ISL_600589, EPI_ISL_600590, EPI_ISL_600591, EPI_ISL_600592, EPI_ISL_600593, EPI_ISL_600594, EPI_ISL_600595, EPI_ISL_600596, EPI_ISL_600597, EPI_ISL_600598, EPI_ISL_600599, EPI_ISL_600600, EPI_ISL_600601, EPI_ISL_600602, EPI_ISL_600603, EPI_ISL_600604, EPI_ISL_600605, EPI_ISL_600606, EPI_ISL_600607, EPI_ISL_600608, EPI_ISL_600609, EPI_ISL_600610, EPI_ISL_600611, EPI_ISL_600612, EPI_ISL_600613, EPI_ISL_600614, EPI_ISL_600615, EPI_ISL_600616, EPI_ISL_600617, EPI_ISL_600618, EPI_ISL_600619, EPI_ISL_600620, EPI_ISL_600621, EPI_ISL_600622, EPI_ISL_600623, EPI_ISL_600624, EPI_ISL_600625, EPI_ISL_600626, EPI_ISL_600627, EPI_ISL_600628, EPI_ISL_600629, EPI_ISL_600630, EPI_ISL_600631, EPI_ISL_600632, EPI_ISL_600633, EPI_ISL_600634, EPI_ISL_600635, EPI_ISL_600636, EPI_ISL_600637, EPI_ISL_600638, EPI_ISL_600639, EPI_ISL_600640, EPI_ISL_600641, EPI_ISL_600642, EPI_ISL_600644, EPI_ISL_600645, EPI_ISL_600646, EPI_ISL_600647, EPI_ISL_600648, EPI_ISL_600649, EPI_ISL_600650, EPI_ISL_600651, EPI_ISL_600652, EPI_ISL_600653, EPI_ISL_600654, EPI_ISL_600655, EPI_ISL_600656, EPI_ISL_600657, EPI_ISL_600658, EPI_ISL_600659, EPI_ISL_600660, EPI_ISL_600661, EPI_ISL_600662, EPI_ISL_600663, EPI_ISL_600664, EPI_ISL_600665, EPI_ISL_600666, EPI_ISL_600667, EPI_ISL_600668, EPI_ISL_600669, EPI_ISL_600670, EPI_ISL_600671, EPI_ISL_600672, EPI_ISL_600673, EPI_ISL_600674, EPI_ISL_600675, EPI_ISL_600676, EPI_ISL_600677, EPI_ISL_600678, EPI_ISL_600679, EPI_ISL_600680, EPI_ISL_600681, EPI_ISL_600682, EPI_ISL_600683, EPI_ISL_600684, EPI_ISL_600685, EPI_ISL_600686, EPI_ISL_600687, EPI_ISL_600688, EPI_ISL_600689, EPI_ISL_600690, EPI_ISL_600691, EPI_ISL_600692, EPI_ISL_600693, EPI_ISL_600694, EPI_ISL_600695, EPI_ISL_600696, EPI_ISL_600697, EPI_ISL_600698, EPI_ISL_600699, EPI_ISL_600700, EPI_ISL_600701, EPI_ISL_600702, EPI_ISL_600703, EPI_ISL_600704, EPI_ISL_600705, EPI_ISL_600706, EPI_ISL_600707, EPI_ISL_600709, EPI_ISL_600710, EPI_ISL_600711, EPI_ISL_600712, EPI_ISL_600713, EPI_ISL_600714, EPI_ISL_600715, EPI_ISL_600716, EPI_ISL_600717, EPI_ISL_600718, EPI_ISL_600719, EPI_ISL_600720, EPI_ISL_600721, EPI_ISL_600722, EPI_ISL_600723, EPI_ISL_600724, EPI_ISL_600725, EPI_ISL_600726, EPI_ISL_600727, EPI_ISL_600728, EPI_ISL_600729, EPI_ISL_600730, EPI_ISL_600731, EPI_ISL_600732, EPI_ISL_600733, EPI_ISL_600734, EPI_ISL_600735, EPI_ISL_600736, EPI_ISL_600737, EPI_ISL_600738, EPI_ISL_600740, EPI_ISL_600741, EPI_ISL_600742, EPI_ISL_600743, EPI_ISL_600744, EPI_ISL_600745, EPI_ISL_600746, EPI_ISL_600747, EPI_ISL_600748, EPI_ISL_600749, EPI_ISL_600750, EPI_ISL_600751, EPI_ISL_600752, EPI_ISL_600753, EPI_ISL_600754, EPI_ISL_600755, EPI_ISL_600756, EPI_ISL_600757, EPI_ISL_600758, EPI_ISL_600759, EPI_ISL_600760, EPI_ISL_600761, EPI_ISL_600762, EPI_ISL_600763, EPI_ISL_600764, EPI_ISL_600765, EPI_ISL_600766, EPI_ISL_600767, EPI_ISL_600768, EPI_ISL_600769, EPI_ISL_600770, EPI_ISL_600771, EPI_ISL_600772, EPI_ISL_600773, EPI_ISL_600774, EPI_ISL_600775, EPI_ISL_600776, EPI_ISL_600777, EPI_ISL_600778, EPI_ISL_600779, EPI_ISL_600780, EPI_ISL_600781, EPI_ISL_600782, EPI_ISL_600783, EPI_ISL_600784, EPI_ISL_600785, EPI_ISL_600786, EPI_ISL_600787, EPI_ISL_600788, EPI_ISL_600789, EPI_ISL_600790, EPI_ISL_600791, EPI_ISL_600792, EPI_ISL_600793, EPI_ISL_600794, EPI_ISL_600795, EPI_ISL_600796, EPI_ISL_600797, EPI_ISL_600798, EPI_ISL_600799, EPI_ISL_600800, EPI_ISL_600801, EPI_ISL_600802, EPI_ISL_600803, EPI_ISL_600804, EPI_ISL_600805, EPI_ISL_600806, EPI_ISL_600807, EPI_ISL_600808, EPI_ISL_600809, EPI_ISL_600810, EPI_ISL_600811, EPI_ISL_600812, EPI_ISL_600813, EPI_ISL_600814, EPI_ISL_600815, EPI_ISL_600816, EPI_ISL_600817, EPI_ISL_600818, EPI_ISL_600819, EPI_ISL_600820, EPI_ISL_600821, EPI_ISL_600822, EPI_ISL_600823, EPI_ISL_600825, EPI_ISL_600826, EPI_ISL_600827, EPI_ISL_600828, EPI_ISL_600829, EPI_ISL_600830, EPI_ISL_600831, EPI_ISL_600832, EPI_ISL_600833, EPI_ISL_600834, EPI_ISL_600835, EPI_ISL_600836, EPI_ISL_600837, EPI_ISL_600838, EPI_ISL_600839, EPI_ISL_600840, EPI_ISL_600841, EPI_ISL_600842, EPI_ISL_600843, EPI_ISL_600844, EPI_ISL_600845, EPI_ISL_600846, EPI_ISL_600847, EPI_ISL_600848, EPI_ISL_600849, EPI_ISL_600850, EPI_ISL_600851, EPI_ISL_600852, EPI_ISL_600853, EPI_ISL_600854, EPI_ISL_600855, EPI_ISL_600856, EPI_ISL_600857, EPI_ISL_600858, EPI_ISL_600859, EPI_ISL_600860, EPI_ISL_600861, EPI_ISL_600862, EPI_ISL_600863, EPI_ISL_600864, EPI_ISL_600865, EPI_ISL_600866, EPI_ISL_600867, EPI_ISL_600868, EPI_ISL_600869, EPI_ISL_600870, EPI_ISL_600871, EPI_ISL_600872, EPI_ISL_600873, EPI_ISL_600874, EPI_ISL_600875, EPI_ISL_600876, EPI_ISL_600877, EPI_ISL_600878, EPI_ISL_600879, EPI_ISL_600880, EPI_ISL_600881, EPI_ISL_600882, EPI_ISL_600883, EPI_ISL_600884, EPI_ISL_600885, EPI_ISL_600886, EPI_ISL_600887, EPI_ISL_600888, EPI_ISL_600889, EPI_ISL_600890, EPI_ISL_600891, EPI_ISL_600893, EPI_ISL_600894, EPI_ISL_600895, EPI_ISL_600896, EPI_ISL_600897, EPI_ISL_600898, EPI_ISL_600899, EPI_ISL_600900, EPI_ISL_600902, EPI_ISL_600903, EPI_ISL_600905, EPI_ISL_600906, EPI_ISL_600907, EPI_ISL_600908, EPI_ISL_600909, EPI_ISL_600910, EPI_ISL_600911, EPI_ISL_600914, EPI_ISL_600916, EPI_ISL_600917, EPI_ISL_600918, EPI_ISL_600919, EPI_ISL_600920, EPI_ISL_600921, EPI_ISL_600922, EPI_ISL_600924, EPI_ISL_600932, EPI_ISL_600970, EPI_ISL_600974, EPI_ISL_600980, EPI_ISL_600983, EPI_ISL_600999, EPI_ISL_601002, EPI_ISL_601023, EPI_ISL_601025, EPI_ISL_601032, EPI_ISL_601047, EPI_ISL_601061, EPI_ISL_601071, EPI_ISL_601099, EPI_ISL_601110, EPI_ISL_601118, EPI_ISL_601136, EPI_ISL_601139, EPI_ISL_601172, EPI_ISL_601203, EPI_ISL_601236, EPI_ISL_601237, EPI_ISL_601238, EPI_ISL_601241, EPI_ISL_601244, EPI_ISL_601245, EPI_ISL_601246, EPI_ISL_601250, EPI_ISL_601251, EPI_ISL_601252, EPI_ISL_601253, EPI_ISL_601254, EPI_ISL_601255, EPI_ISL_601257, EPI_ISL_601259, EPI_ISL_601260, EPI_ISL_601261, EPI_ISL_601265, EPI_ISL_601266, EPI_ISL_601269, EPI_ISL_601271, EPI_ISL_601274, EPI_ISL_601276, EPI_ISL_601279, EPI_ISL_601284, EPI_ISL_601285, EPI_ISL_601286, EPI_ISL_601288, EPI_ISL_601295, EPI_ISL_601296, EPI_ISL_601300, EPI_ISL_601308, EPI_ISL_601309, EPI_ISL_601315, EPI_ISL_601318, EPI_ISL_601319, EPI_ISL_601323, EPI_ISL_601328, EPI_ISL_601331, EPI_ISL_601332, EPI_ISL_601333, EPI_ISL_601334, EPI_ISL_601335, EPI_ISL_601336, EPI_ISL_601339, EPI_ISL_601341, EPI_ISL_601342, EPI_ISL_601344, EPI_ISL_601347, EPI_ISL_601348, EPI_ISL_601351, EPI_ISL_601352, EPI_ISL_601353, EPI_ISL_601355, EPI_ISL_601356, EPI_ISL_601357, EPI_ISL_601358, EPI_ISL_601361, EPI_ISL_601362, EPI_ISL_601364, EPI_ISL_601372, EPI_ISL_601374, EPI_ISL_601377, EPI_ISL_601380, EPI_ISL_601384, EPI_ISL_601386, EPI_ISL_601387, EPI_ISL_601389, EPI_ISL_601391, EPI_ISL_601393, EPI_ISL_601400, EPI_ISL_601401, EPI_ISL_601402, EPI_ISL_601403, EPI_ISL_601405, EPI_ISL_601407, EPI_ISL_601408, EPI_ISL_601411, EPI_ISL_601413, EPI_ISL_601414, EPI_ISL_601416, EPI_ISL_601421, EPI_ISL_601423, EPI_ISL_601424, EPI_ISL_601425, EPI_ISL_601426, EPI_ISL_601429, EPI_ISL_601430, EPI_ISL_601432, EPI_ISL_601433, EPI_ISL_601435, EPI_ISL_601436, EPI_ISL_601445, EPI_ISL_601446, EPI_ISL_601453, EPI_ISL_601454, EPI_ISL_601459, EPI_ISL_601460, EPI_ISL_601461, EPI_ISL_601462, EPI_ISL_601466, EPI_ISL_601467, EPI_ISL_601469, EPI_ISL_601471, EPI_ISL_601477, EPI_ISL_601479, EPI_ISL_601481, EPI_ISL_601482, EPI_ISL_601483, EPI_ISL_601486, EPI_ISL_601488, EPI_ISL_601489, EPI_ISL_601789, EPI_ISL_601790, EPI_ISL_601791, EPI_ISL_601796, EPI_ISL_601799, EPI_ISL_601801, EPI_ISL_601803, EPI_ISL_601804, EPI_ISL_601808, EPI_ISL_601810, EPI_ISL_601811, EPI_ISL_601813, EPI_ISL_601818, EPI_ISL_601819, EPI_ISL_601820, EPI_ISL_601821, EPI_ISL_601824, EPI_ISL_601826, EPI_ISL_601830, EPI_ISL_601833, EPI_ISL_601838, EPI_ISL_601841, EPI_ISL_601844, EPI_ISL_601845, EPI_ISL_601847, EPI_ISL_601848, EPI_ISL_601850, EPI_ISL_601851, EPI_ISL_601854, EPI_ISL_601856, EPI_ISL_601857, EPI_ISL_601864, EPI_ISL_601866, EPI_ISL_601868, EPI_ISL_601869, EPI_ISL_601870, EPI_ISL_601871, EPI_ISL_601872, EPI_ISL_601873, EPI_ISL_601874, EPI_ISL_601875, EPI_ISL_601876, EPI_ISL_601877, EPI_ISL_601878, EPI_ISL_601879, EPI_ISL_601880, EPI_ISL_601881, EPI_ISL_601882, EPI_ISL_601883, EPI_ISL_601884, EPI_ISL_601885, EPI_ISL_601886, EPI_ISL_601887, EPI_ISL_601888, EPI_ISL_601889, EPI_ISL_601890, EPI_ISL_601891, EPI_ISL_601892, EPI_ISL_601893, EPI_ISL_601894, EPI_ISL_601895, EPI_ISL_601896, EPI_ISL_601897, EPI_ISL_601898, EPI_ISL_601899, EPI_ISL_601900, EPI_ISL_601901, EPI_ISL_601902, EPI_ISL_601903, EPI_ISL_601904, EPI_ISL_601905, EPI_ISL_601906, EPI_ISL_601907, EPI_ISL_601908, EPI_ISL_601909, EPI_ISL_601910, EPI_ISL_601911, EPI_ISL_601912, EPI_ISL_601913, EPI_ISL_601914, EPI_ISL_601915, EPI_ISL_601916, EPI_ISL_601917, EPI_ISL_601918, EPI_ISL_601919, EPI_ISL_601920, EPI_ISL_601921, EPI_ISL_601922, EPI_ISL_601923, EPI_ISL_601924, EPI_ISL_601925, EPI_ISL_601926, EPI_ISL_601927, EPI_ISL_601928, EPI_ISL_601929, EPI_ISL_601930, EPI_ISL_601931, EPI_ISL_601932, EPI_ISL_601933, EPI_ISL_601934, EPI_ISL_601935, EPI_ISL_601936, EPI_ISL_601937, EPI_ISL_601938, EPI_ISL_601939, EPI_ISL_601940, EPI_ISL_601941, EPI_ISL_601942, EPI_ISL_601943, EPI_ISL_601944, EPI_ISL_601945, EPI_ISL_601946, EPI_ISL_601947, EPI_ISL_601948, EPI_ISL_601949, EPI_ISL_601951, EPI_ISL_601952, EPI_ISL_601953, EPI_ISL_601954, EPI_ISL_601955, EPI_ISL_601956, EPI_ISL_601957, EPI_ISL_601958, EPI_ISL_601959, EPI_ISL_601960, EPI_ISL_601961, EPI_ISL_601962, EPI_ISL_601963, EPI_ISL_601964, EPI_ISL_601965, EPI_ISL_601966, EPI_ISL_601967, EPI_ISL_601968, EPI_ISL_601969, EPI_ISL_601970, EPI_ISL_601971, EPI_ISL_601972, EPI_ISL_601973, EPI_ISL_601974, EPI_ISL_601975, EPI_ISL_601976, EPI_ISL_601977, EPI_ISL_601978, EPI_ISL_601979, EPI_ISL_601980, EPI_ISL_601981, EPI_ISL_601982, EPI_ISL_601983, EPI_ISL_601984, EPI_ISL_601985, EPI_ISL_601986, EPI_ISL_601987, EPI_ISL_601988, EPI_ISL_601989, EPI_ISL_601990, EPI_ISL_601991, EPI_ISL_601992, EPI_ISL_601993, EPI_ISL_601994, EPI_ISL_601995, EPI_ISL_601996, EPI_ISL_601997, EPI_ISL_601998, EPI_ISL_601999, EPI_ISL_602000, EPI_ISL_602001, EPI_ISL_602002, EPI_ISL_602003, EPI_ISL_602004, EPI_ISL_602005, EPI_ISL_602007, EPI_ISL_602008, EPI_ISL_602009, EPI_ISL_602010, EPI_ISL_602011, EPI_ISL_602012, EPI_ISL_602013, EPI_ISL_602014, EPI_ISL_602016, EPI_ISL_602017, EPI_ISL_602018, EPI_ISL_602019, EPI_ISL_602020, EPI_ISL_602021, EPI_ISL_602022, EPI_ISL_602023, EPI_ISL_602024, EPI_ISL_602025, EPI_ISL_602026, EPI_ISL_602030, EPI_ISL_602031, EPI_ISL_602032, EPI_ISL_602033, EPI_ISL_602034, EPI_ISL_602035, EPI_ISL_602036, EPI_ISL_602037, EPI_ISL_602038, EPI_ISL_602039, EPI_ISL_602040, EPI_ISL_602041, EPI_ISL_602042, EPI_ISL_602043, EPI_ISL_602044, EPI_ISL_602045, EPI_ISL_602046, EPI_ISL_602047, EPI_ISL_602048, EPI_ISL_602049, EPI_ISL_602050, EPI_ISL_602051, EPI_ISL_602052, EPI_ISL_602053, EPI_ISL_602054, EPI_ISL_602055, EPI_ISL_602056, EPI_ISL_602057, EPI_ISL_602058, EPI_ISL_602059, EPI_ISL_602060, EPI_ISL_602061, EPI_ISL_602062, EPI_ISL_602063, EPI_ISL_602064, EPI_ISL_602065, EPI_ISL_602066, EPI_ISL_602067, EPI_ISL_602068, EPI_ISL_602069, EPI_ISL_602070, EPI_ISL_602071, EPI_ISL_602072, EPI_ISL_602073, EPI_ISL_602074, EPI_ISL_602075, EPI_ISL_602076, EPI_ISL_602078, EPI_ISL_602079, EPI_ISL_602080, EPI_ISL_602081, EPI_ISL_602082, EPI_ISL_602084, EPI_ISL_602085, EPI_ISL_602086, EPI_ISL_602087, EPI_ISL_602088, EPI_ISL_602089, EPI_ISL_602092, EPI_ISL_602093, EPI_ISL_602094, EPI_ISL_602095, EPI_ISL_602096, EPI_ISL_602097, EPI_ISL_602098, EPI_ISL_602099, EPI_ISL_602100, EPI_ISL_602101, EPI_ISL_602103, EPI_ISL_602104, EPI_ISL_602105, EPI_ISL_602107, EPI_ISL_602108, EPI_ISL_602110, EPI_ISL_602111, EPI_ISL_602112, EPI_ISL_602113, EPI_ISL_602114, EPI_ISL_602115, EPI_ISL_602116, EPI_ISL_602117, EPI_ISL_602118, EPI_ISL_602119, EPI_ISL_602120, EPI_ISL_602121, EPI_ISL_602123, EPI_ISL_602124, EPI_ISL_602125, EPI_ISL_602126, EPI_ISL_602127, EPI_ISL_602128, EPI_ISL_602129, EPI_ISL_602130, EPI_ISL_602131, EPI_ISL_602132, EPI_ISL_602133, EPI_ISL_602134, EPI_ISL_602135, EPI_ISL_602136, EPI_ISL_602137, EPI_ISL_602138, EPI_ISL_602139, EPI_ISL_602140, EPI_ISL_602141, EPI_ISL_602142, EPI_ISL_602143, EPI_ISL_602144, EPI_ISL_602156, EPI_ISL_602157

| | | | |
|---|---|---|---|
| see above | Lighthouse Lab in Glasgow | Wellcome Sanger Institute for the COVID-19 Genomics UK (COG-UK) consortium | Harper VanSteenhouse, Yumi Kasai, David Gray, Carol Clugston, Anna Dominiczak and Alex Alderton, Roberto Amato, Sonia Goncalves, Ewan Harrison, David K. Jackson, Ian Johnston, Dominic Kwiatkowski, Cordelia Langford, John Sillitoe on behalf of the Wellcome Sanger Institute COVID-19 Surveillance Team (http://www.sanger.ac.uk/covid-team) |

EPI_ISL_602636, EPI_ISL_602643, EPI_ISL_602645, EPI_ISL_602654, EPI_ISL_602655, EPI_ISL_602682, EPI_ISL_602685, EPI_ISL_602699, EPI_ISL_602702, EPI_ISL_602706, EPI_ISL_602722, EPI_ISL_602723, EPI_ISL_602724, EPI_ISL_602726, EPI_ISL_602728, EPI_ISL_602745, EPI_ISL_602746, EPI_ISL_602765, EPI_ISL_602766, EPI_ISL_602777, EPI_ISL_602781, EPI_ISL_602790, EPI_ISL_602792, EPI_ISL_602793, EPI_ISL_602794, EPI_ISL_602802, EPI_ISL_602805, EPI_ISL_602815, EPI_ISL_602819, EPI_ISL_602823, EPI_ISL_602825, EPI_ISL_602827, EPI_ISL_602828, EPI_ISL_602829, EPI_ISL_602830, EPI_ISL_602840, EPI_ISL_602843, EPI_ISL_602845, EPI_ISL_602847, EPI_ISL_602852, EPI_ISL_602856, EPI_ISL_602857, EPI_ISL_602864, EPI_ISL_602866, EPI_ISL_602868, EPI_ISL_602869, EPI_ISL_602874, EPI_ISL_602875, EPI_ISL_602876, EPI_ISL_602877, EPI_ISL_602879, EPI_ISL_602880, EPI_ISL_602882, EPI_ISL_602885, EPI_ISL_602886, EPI_ISL_602887, EPI_ISL_602888, EPI_ISL_602889, EPI_ISL_602891, EPI_ISL_602893, EPI_ISL_602894, EPI_ISL_602895, EPI_ISL_602897, EPI_ISL_602898, EPI_ISL_602901, EPI_ISL_602903, EPI_ISL_602904, EPI_ISL_602905, EPI_ISL_602906, EPI_ISL_602909, EPI_ISL_602910, EPI_ISL_602911, EPI_ISL_602912, EPI_ISL_602913, EPI_ISL_602914, EPI_ISL_602915, EPI_ISL_602918, EPI_ISL_602921, EPI_ISL_602922, EPI_ISL_602924, EPI_ISL_602925, EPI_ISL_602927, EPI_ISL_602930, EPI_ISL_605789

| | | | |
|---|---|---|---|
| see above | NHLS-IALCH | KRISP, KZN Research Innovation and Sequencing Platform | Giandhari J, Pillay S, Lessells R, Mdlalose K, York D, Khan S, Tegally H, Wilkinson E, de Oliveira T |

EPI_ISL_609178, EPI_ISL_609179, EPI_ISL_609182, EPI_ISL_609184, EPI_ISL_609185, EPI_ISL_609186, EPI_ISL_609189, EPI_ISL_609190, EPI_ISL_609191, EPI_ISL_609194, EPI_ISL_609197, EPI_ISL_609198, EPI_ISL_609199, EPI_ISL_609202, EPI_ISL_609206

| | | | |
|---|---|---|---|
| see above | Lighthouse Lab in Glasgow | Wellcome Sanger Institute for the COVID-19 Genomics UK (COG-UK) consortium | Harper VanSteenhouse, Yumi Kasai, David Gray, Carol Clugston, Anna Dominiczak and Alex Alderton, Roberto Amato, Sonia Goncalves, Ewan Harrison, David K. Jackson, Ian Johnston, Dominic Kwiatkowski, Cordelia Langford, John Sillitoe on behalf of the Wellcome Sanger Institute COVID-19 Surveillance Team |
| EPI_ISL_610155 | Puskesmas Cangkrep | Genetics Working Group (Pokja Genetik) Faculty of Medicine, Public Health and Nursing Universitas Gadjah Mada (FK-KMK UGM); Disease Investigation Center Wates Ministry of Agriculture Indonesia; Department of Microbiology FK-KMK UGM; Laboratorium Diagnostik Yayasan Tahija World Mosquito Program (WMP) Yogyakarta Center for Tropical Medicine FK-KMK UGM; Integrated Research Center FK-KMK UGM; Department of Computer Science and Electronics FMIPA UGM; Balai Besar Teknik Kesehatan Lingkungan dan Pengendalian Penyakit (BBTKLPP) Yogyakarta | Gunadi, Hendra Wibawa, Marcellus, Mohamad S. Hakim, Edwin W. Daniwijaya, Ludhang P. Rizki, Endah Supriyati, Eggi Arguni, Titik Nuryastuti, Tri Wibawa, Dwi AA Nugrahaningsih, Afiahayati, Siswanto, Kristy Iskandar, Nungki Anggorowati, Irene, Indaryati, Havid Setyawan, Nawang Sukestiningsih, Fadil Fahri, Ririn Enggy Yuliyanti |
| EPI_ISL_610161 | RSUD Dr. Tjitrowardojo | Genetics Working Group (Pokja Genetik) Faculty of Medicine, Public Health and Nursing Universitas Gadjah Mada (FK-KMK UGM); Disease Investigation Center Wates Ministry of Agriculture Indonesia; Department of Microbiology FK-KMK UGM; Laboratorium Diagnostik Yayasan Tahija World | Gunadi, Hendra Wibawa, Marcellus, Mohamad S. Hakim, Edwin W. Daniwijaya, Ludhang P. Rizki, Endah Supriyati, Eggi Arguni, Titik Nuryastuti, Tri Wibawa, Dwi AA Nugrahaningsih, Afiahayati, Siswanto, Kristy Iskandar, Nungki Anggorowati, Irene, Indaryati, Havid Setyawan, Wuryanto, Kemala Athollah, Desyifa Mursalin |

| | | | |
|---|---|---|---|
| | | Mosquito Program (WMP) Yogyakarta Center for Tropical Medicine FK-KMK UGM; Integrated Research Center FK-KMK UGM; Department of Computer Science and Electronics FMIPA UGM; Balai Besar Teknik Kesehatan Lingkungan dan Pengendalian Penyakit (BBTKLPP) Yogyakarta | |
| EPI_ISL_610162 | RSUD Dr. Tjitrowardojo | Genetics Working Group (Pokja Genetik) Faculty of Medicine, Public Health and Nursing Universitas Gadjah Mada (FK-KMK UGM); Disease Investigation Center Wates Ministry of Agriculture Indonesia; Department of Microbiology FK-KMK UGM; Laboratorium Diagnostik Yayasan Tahija World Mosquito Program (WMP) Yogyakarta Center for Tropical Medicine FK-KMK UGM; Integrated Research Center FK-KMK UGM; Department of Computer Science and Electronics FMIPA UGM; Balai Besar Teknik Kesehatan Lingkungan dan Pengendalian Penyakit (BBTKLPP) Yogyakarta | Gunadi, Hendra Wibawa, Marcellus, Mohamad S. Hakim, Edwin W. Daniwijaya, Ludhang P. Rizki, Endah Supriyati, Eggi Arguni, Titik Nuryastuti, Tri Wibawa, Dwi AA Nugrahaningsih, Afiahayati, Siswanto, Kristy Iskandar, Nungki Anggorowati, Irene, Indaryati, Havid Setyawan, Wuryanto, Susan Simanjaya, Alvin Santoso Kalim |
| EPI_ISL_611287, EPI_ISL_611288, EPI_ISL_611289, EPI_ISL_611290, EPI_ISL_611291, EPI_ISL_611292, EPI_ISL_611293, EPI_ISL_611294, EPI_ISL_611295, EPI_ISL_611296, EPI_ISL_611297, EPI_ISL_611298, EPI_ISL_611299, EPI_ISL_611300, EPI_ISL_611301, EPI_ISL_611302, EPI_ISL_611303, EPI_ISL_611305, EPI_ISL_611306, EPI_ISL_611307, EPI_ISL_611308, EPI_ISL_611309, EPI_ISL_611310, EPI_ISL_611311, EPI_ISL_611312, EPI_ISL_611313, EPI_ISL_611314, EPI_ISL_611315, EPI_ISL_611316, EPI_ISL_611317, EPI_ISL_611318, EPI_ISL_611319, EPI_ISL_611320, EPI_ISL_611321, EPI_ISL_611322, EPI_ISL_611323, EPI_ISL_611324, EPI_ISL_611325, EPI_ISL_611326, EPI_ISL_611327, EPI_ISL_611328, EPI_ISL_611329, EPI_ISL_611330, EPI_ISL_611331, EPI_ISL_611332, EPI_ISL_611333, EPI_ISL_611334, EPI_ISL_611335, EPI_ISL_611336, EPI_ISL_611337, EPI_ISL_611338, EPI_ISL_611339, EPI_ISL_611340, EPI_ISL_611341, EPI_ISL_611342, EPI_ISL_611343, EPI_ISL_611344, EPI_ISL_611345, EPI_ISL_611346, EPI_ISL_611347, EPI_ISL_611348, EPI_ISL_611349, EPI_ISL_611350, EPI_ISL_611351, EPI_ISL_611352, EPI_ISL_611353, EPI_ISL_611354, EPI_ISL_611355, EPI_ISL_611356, EPI_ISL_611357, EPI_ISL_611358, EPI_ISL_611359, EPI_ISL_611360, EPI_ISL_611361, EPI_ISL_611362, EPI_ISL_611363, EPI_ISL_611364, EPI_ISL_611365, EPI_ISL_611366, EPI_ISL_611367, EPI_ISL_611368, EPI_ISL_611369, EPI_ISL_611370, EPI_ISL_611371, EPI_ISL_611372, EPI_ISL_611373, EPI_ISL_611374, EPI_ISL_611375, EPI_ISL_611376, EPI_ISL_611377, EPI_ISL_611379, EPI_ISL_611380, EPI_ISL_611381, EPI_ISL_611382, EPI_ISL_611383, EPI_ISL_611384, EPI_ISL_611385, EPI_ISL_611386, EPI_ISL_611387, EPI_ISL_611388, EPI_ISL_611389, EPI_ISL_611390, EPI_ISL_611391, EPI_ISL_611392, EPI_ISL_611393, EPI_ISL_611394, EPI_ISL_611395, EPI_ISL_611396, EPI_ISL_611397, EPI_ISL_611398, EPI_ISL_611399, EPI_ISL_611400, EPI_ISL_611401, EPI_ISL_611402, EPI_ISL_611403, EPI_ISL_611404, EPI_ISL_611405, EPI_ISL_611406, EPI_ISL_611407, EPI_ISL_611408, EPI_ISL_611409, EPI_ISL_611410, EPI_ISL_611411, EPI_ISL_611412, EPI_ISL_611413, EPI_ISL_611414, EPI_ISL_611415, EPI_ISL_611416, EPI_ISL_611417, EPI_ISL_611418, EPI_ISL_611419, EPI_ISL_611420, EPI_ISL_611421, EPI_ISL_611422, EPI_ISL_611423, EPI_ISL_611424, EPI_ISL_611425, EPI_ISL_611426, EPI_ISL_611427, EPI_ISL_611428, EPI_ISL_611429, EPI_ISL_611430, EPI_ISL_611431, EPI_ISL_611432, EPI_ISL_611433, EPI_ISL_611434, EPI_ISL_611435, EPI_ISL_611436, EPI_ISL_611437, EPI_ISL_611438, EPI_ISL_611439, EPI_ISL_611440, EPI_ISL_611441, EPI_ISL_611442, EPI_ISL_611443, EPI_ISL_611444, EPI_ISL_611445, EPI_ISL_611446, EPI_ISL_611447, EPI_ISL_611448, EPI_ISL_611449, EPI_ISL_611450, EPI_ISL_611451, EPI_ISL_611452, EPI_ISL_611453, EPI_ISL_611454, EPI_ISL_611455, EPI_ISL_611456, EPI_ISL_611457, EPI_ISL_611458, EPI_ISL_611459, EPI_ISL_611460, EPI_ISL_611461, EPI_ISL_611462, EPI_ISL_611463, EPI_ISL_611464, EPI_ISL_611465, EPI_ISL_611466, EPI_ISL_611467, EPI_ISL_611468, EPI_ISL_611469, EPI_ISL_611470, EPI_ISL_611471, EPI_ISL_611472, EPI_ISL_611473, EPI_ISL_611474, EPI_ISL_611475, EPI_ISL_611476, EPI_ISL_611477, EPI_ISL_611478, EPI_ISL_611479, EPI_ISL_611480, EPI_ISL_611481, EPI_ISL_611482, EPI_ISL_611483, EPI_ISL_611484, EPI_ISL_623863, EPI_ISL_623865, EPI_ISL_623868, EPI_ISL_623872, EPI_ISL_623873, EPI_ISL_623874, EPI_ISL_623876, EPI_ISL_623883, EPI_ISL_623884, EPI_ISL_623886, EPI_ISL_623893, EPI_ISL_623895, EPI_ISL_623903, EPI_ISL_623907, EPI_ISL_623908, EPI_ISL_623911, EPI_ISL_623916, EPI_ISL_623917, EPI_ISL_623921, EPI_ISL_623924, EPI_ISL_623930, EPI_ISL_623933, EPI_ISL_623934, EPI_ISL_623942, EPI_ISL_623944, EPI_ISL_623949, EPI_ISL_623954, EPI_ISL_623956, EPI_ISL_623959, EPI_ISL_623963, EPI_ISL_623975, EPI_ISL_623976, EPI_ISL_623977, EPI_ISL_623986, EPI_ISL_623990, EPI_ISL_623991, EPI_ISL_623994, EPI_ISL_623995, EPI_ISL_624010, EPI_ISL_624012, EPI_ISL_624018, EPI_ISL_624019, EPI_ISL_624022, EPI_ISL_624027, EPI_ISL_624028, EPI_ISL_624029, EPI_ISL_624032, EPI_ISL_624034, EPI_ISL_624043, EPI_ISL_624044, EPI_ISL_624049, EPI_ISL_624067, EPI_ISL_624073, EPI_ISL_624077, EPI_ISL_624079, EPI_ISL_624092, EPI_ISL_624105, EPI_ISL_624112, EPI_ISL_624114, EPI_ISL_624120, EPI_ISL_624122, EPI_ISL_624128, EPI_ISL_624132, EPI_ISL_624140, EPI_ISL_624145, EPI_ISL_624147, EPI_ISL_624148, EPI_ISL_624158, EPI_ISL_624163, EPI_ISL_624164, EPI_ISL_624165, EPI_ISL_624176, EPI_ISL_624181, EPI_ISL_624188, EPI_ISL_624189, EPI_ISL_625315, EPI_ISL_625316, EPI_ISL_625317, EPI_ISL_625318, EPI_ISL_625319, EPI_ISL_625320, EPI_ISL_625321, EPI_ISL_625322, EPI_ISL_625323, EPI_ISL_625324, EPI_ISL_625325, EPI_ISL_625326, EPI_ISL_625327, EPI_ISL_625328, EPI_ISL_625329, EPI_ISL_625330, EPI_ISL_625331, EPI_ISL_625332, EPI_ISL_625333, EPI_ISL_625334, EPI_ISL_625335, EPI_ISL_625336, EPI_ISL_625337, EPI_ISL_625338, EPI_ISL_625339, EPI_ISL_625340, EPI_ISL_625341, EPI_ISL_625342, EPI_ISL_625343, EPI_ISL_625344, EPI_ISL_625345, EPI_ISL_625346, EPI_ISL_625347, EPI_ISL_625348, EPI_ISL_625349, EPI_ISL_625350, EPI_ISL_625351, EPI_ISL_625352, EPI_ISL_625353, EPI_ISL_625354, EPI_ISL_625355, EPI_ISL_625356, EPI_ISL_625357, EPI_ISL_625358, EPI_ISL_625359, EPI_ISL_625360, EPI_ISL_625361, EPI_ISL_625362, EPI_ISL_625363, EPI_ISL_625364, EPI_ISL_625365, EPI_ISL_625366, EPI_ISL_625367, EPI_ISL_625368, EPI_ISL_625369, EPI_ISL_625370, EPI_ISL_625371, EPI_ISL_625372, EPI_ISL_625373, EPI_ISL_625374, EPI_ISL_625375, EPI_ISL_625376, EPI_ISL_625377, EPI_ISL_625378, EPI_ISL_625379, EPI_ISL_625380, EPI_ISL_625381, EPI_ISL_625382, EPI_ISL_625383, EPI_ISL_625384, EPI_ISL_625385, EPI_ISL_625386, EPI_ISL_625387, EPI_ISL_625388, EPI_ISL_625389, EPI_ISL_625390, EPI_ISL_625391, EPI_ISL_625392, EPI_ISL_625393, EPI_ISL_625394, EPI_ISL_625395 | | | |
| see above | Lighthouse Lab in Glasgow | Wellcome Sanger Institute for the COVID-19 Genomics UK (COG-UK) consortium | Harper VanSteenhouse, Yumi Kasai, David Gray, Carol Clugston, Anna Dominiczak and Alex Alderton, Roberto Amato, Sonia Goncalves, Ewan Harrison, David K. Jackson, Ian Johnston, Dominic Kwiatkowski, Cordelia Langford, John Sillitoe on behalf of the Wellcome Sanger Institute COVID-19 Surveillance Team (http://www.sanger.ac.uk/covid-team) |
| EPI_ISL_629755, EPI_ISL_629758, EPI_ISL_629764, EPI_ISL_629771, EPI_ISL_629774, EPI_ISL_629776, EPI_ISL_629777, EPI_ISL_629795, EPI_ISL_629811, EPI_ISL_629821, EPI_ISL_629822, EPI_ISL_629836, EPI_ISL_629846, EPI_ISL_629851, EPI_ISL_629877, EPI_ISL_629913, EPI_ISL_629926, EPI_ISL_629932, EPI_ISL_629943, EPI_ISL_629950, EPI_ISL_629951, EPI_ISL_629952, EPI_ISL_629969, EPI_ISL_629970, EPI_ISL_629977, EPI_ISL_629983, EPI_ISL_629986, EPI_ISL_630702, EPI_ISL_630703, EPI_ISL_630704, EPI_ISL_630705, EPI_ISL_630712, EPI_ISL_630718, EPI_ISL_630719, EPI_ISL_630725, EPI_ISL_630731, EPI_ISL_630733, EPI_ISL_630741, EPI_ISL_630745, EPI_ISL_630749, EPI_ISL_630761, EPI_ISL_630764, EPI_ISL_630775, EPI_ISL_630778, EPI_ISL_630782, EPI_ISL_630787, EPI_ISL_630804, EPI_ISL_630813, EPI_ISL_630815, EPI_ISL_630824, EPI_ISL_630827, EPI_ISL_630833, EPI_ISL_630838, EPI_ISL_630850, EPI_ISL_630851, EPI_ISL_630853, EPI_ISL_630859, EPI_ISL_630862, EPI_ISL_630864, EPI_ISL_630870, EPI_ISL_630871, EPI_ISL_630872, EPI_ISL_630873, EPI_ISL_630875, EPI_ISL_630876, EPI_ISL_630883, EPI_ISL_630888, EPI_ISL_630900, EPI_ISL_630915, EPI_ISL_630919, EPI_ISL_630923, EPI_ISL_630926, EPI_ISL_630931, EPI_ISL_630933, EPI_ISL_630941, EPI_ISL_630948, EPI_ISL_630949, EPI_ISL_630951, EPI_ISL_630955, EPI_ISL_630957, EPI_ISL_630959, EPI_ISL_630965 | | | |
| see above | Lighthouse Lab in Glasgow | Wellcome Sanger Institute for the COVID-19 Genomics UK (COG-UK) consortium | Harper VanSteenhouse, Yumi Kasai, David Gray, Carol Clugston, Anna Dominiczak and Alex Alderton, Roberto Amato, Sonia Goncalves, Ewan Harrison, David K. Jackson, Ian Johnston, Dominic Kwiatkowski, Cordelia Langford, John Sillitoe on behalf of the Wellcome Sanger Institute COVID-19 Surveillance Team |

We gratefully acknowledge the following Authors from the Originating laboratories responsible for obtaining the specimens, as well as the Submitting laboratories where the genome data were generated and shared via GISAID, on which this research is based.

All Submitters of data may be contacted directly via www.gisaid.org

Authors are sorted alphabetically.

| Accession ID | Originating Laboratory | Submitting Laboratory | Authors |
|---|---|---|---|
| EPI_ISL_412965 | BCCDC Public Health Laboratory | BCCDC Public Health Laboratory | Harrigan, Prystajecky, Krajden, Lee, Kamelian, Lapointe, Choi, Hoang, Sekirov, Levett, Tyson, Loman, Quick, Li, Gilmour |
| EPI_ISL_412968, EPI_ISL_412969 | Takayuki Hishiki Kanagawa Prefectural Institute of Public Health | Takayuki Hishiki Kanagawa Prefectural Institute of Public Health | Hishiki,T., Suzuki,R., Sakuragi,J., Usui,K., Tanaka,Y., Kawai,J., Kogo,Y., Matsuki,Y., An,T., Hayashizaki,Y. and Takasaki,T. |
| EPI_ISL_412970 | Washington State Department of Health | Seattle Flu Study | Helen Chu, Michael Boeckh, Janet Englund, Michael Famulare, Barry Lutz, Deborah Nickerson, Mark Rieder, Lea Starita, Matthew Thompson, Jay Shendure, and Trevor Bedford |
| EPI_ISL_415577, EPI_ISL_415578, EPI_ISL_415579, EPI_ISL_415580, EPI_ISL_415581, EPI_ISL_415582, EPI_ISL_415583, EPI_ISL_415584, EPI_ISL_415585, EPI_ISL_415586, EPI_ISL_415587, EPI_ISL_415588, EPI_ISL_415589, EPI_ISL_415590 | | | |
| see above | BCCDC Public Health Laboratory | BCCDC Public Health Laboratory | Harrigan, Prystajecky, Krajden, Lee, Kamelian, Lapointe, Choi, Hoang, Sekirov, Levett, Tyson, Snutch, Loman, Quick, Li, Gilmour |
| EPI_ISL_416994 | COMPLEJO ASISTENCIAL UNIVERSITARIO DE BURGOS | Instituto de Salud Carlos III | Iglesias-Caballero, M. Molinero Calamita, M. González-Esguevillas, M. Camarero S. Pozo F. Casas I. Jiménez P. Jiménez M. Zaballos A. Monzón, S. Varona, S. Juliá M. Cuesta I. Megias Lobón, G. Hospital: ------ |
| EPI_ISL_417007 | HOSPITAL SANTA MARIA NAI | Instituto de Salud Carlos III | Iglesias-Caballero, M. Molinero Calamita, M. González-Esguevillas, M. Camarero S. Pozo F. Casas I. Jiménez, P. Jiménez, M. Zaballos, A. Monzón, S. Varona, S. Juliá, M. Cuesta, I. García Costa, J. |
| EPI_ISL_417010 | FUNDACION JIMENEZ DIAZ | Instituto de Salud Carlos III | Iglesias-Caballero, M. Molinero Calamita, M. González-Esguevillas, M. Camarero, S. Pozo, F. Casas, I. Jiménez, P. Jiménez, M. Zaballos, A. Monzón, S. Varona, S. Juliá, M. Cuesta, I. Fernández Roblas, R. |
| EPI_ISL_417393, EPI_ISL_417410 | Centre for Infectious Diseases and Microbiology Public Health | NSW Health Pathology - Institute of Clinical Pathology and Medical Research; Westmead Hospital; University of Sydney | Holmes EC, O'Sullivan MV, Sintchenko V, Chen SC, Maddocks S, Kok J, Dwyer DE, Rockett R, Eden J-S, Lam C, Gray K, Timms V, Gall M, Arnott A, Sadsad R, Carter I and Rahman H for the 2019-nCoV Study Group |
| EPI_ISL_418243, EPI_ISL_418244 | HOSPITAL UNIVERSITARIO VIRGEN DE LAS NIEVES | Instituto de Salud Carlos III | Iglesias-Caballero, M. Molinero Calamita, M. González-Esguevillas, M. Camarero, S. Pozo, F. Casas, I. Jiménez, P. Jiménez, M. Zaballos, A. Monzón, S. Varona, S. Juliá, M. Cuesta, I. Sanbonmatsu S. |
| EPI_ISL_418245, EPI_ISL_418246 | Hospital General y Universitario de Guadalajara | Instituto de Salud Carlos III | Iglesias-Caballero, M. Molinero Calamita, M. González-Esguevillas, M. Camarero, S. Pozo, F. Casas, I. Jiménez, P. Jiménez, M. Zaballos, A. Monzón, S. Varona, S. Juliá, M. Cuesta, I. Gonzalez-Praetorius A. |
| EPI_ISL_418247 | HOSPITAL GENERAL DE SEGOVIA | Instituto de Salud Carlos III | Iglesias-Caballero, M. Molinero Calamita, M. González-Esguevillas, M. Camarero, S. Pozo, F. Casas, I. Jiménez, P. Jiménez, M. Zaballos, A. Monzón, S. Varona, S. Juliá, M. Cuesta, I. Hernando-Real S. |
| EPI_ISL_418248, EPI_ISL_418249 | COMPLEJO ASISTENCIAL UNIVERSITARIO DE BURGOS | Instituto de Salud Carlos III | Iglesias-Caballero, M. Molinero Calamita, M. González-Esguevillas, M. Camarero, S. Pozo, F. Casas, I. Jiménez, P. Jiménez, M. Zaballos, A. Monzón, S. Varona, S. Juliá, M. Cuesta, I. Megias-Lobon G. |
| EPI_ISL_418251 | HOSPITAL UNIVERSITARIO LA PAZ | Instituto de Salud Carlos III | Iglesias-Caballero, M. Molinero Calamita, M. González-Esguevillas, M. Camarero, S. Pozo, F. Casas, I. Jiménez, P. Jiménez, M. Zaballos, A. Monzón, S. Varona, S. Juliá, M. Cuesta, I. Romero P. |
| EPI_ISL_418252 | FUNDACION JIMENEZ DIAZ | Instituto de Salud Carlos III | Iglesias-Caballero, M. Molinero Calamita, M. González-Esguevillas, M. Camarero, S. Pozo, F. Casas, I. Jiménez, P. Jiménez, M. Zaballos, A. Monzón, S. Varona, S. Juliá, M. Cuesta, I. Fernández Roblas, R. |
| EPI_ISL_418253 | HOSPITAL TXAGORRITXU | Instituto de Salud Carlos III | Iglesias-Caballero, M. Molinero Calamita, M. González-Esguevillas, M. Camarero, S. Pozo, F. Casas, I. Jiménez, P. Jiménez, M. Zaballos, A. Monzón, S. Varona, S. Juliá, M. Cuesta, I. Gomez-Gonzalez C. |
| EPI_ISL_418271, EPI_ISL_418273 | University Hospital Basel, Clinical Virology | University Hospital Basel, Labormedizin | Hirsch, H., Leuzinger, K., Seth-Smith, H., Mari, A., Roloff, T., Egli, A. |
| EPI_ISL_418275, EPI_ISL_418277, EPI_ISL_418278, EPI_ISL_418279, EPI_ISL_418280, EPI_ISL_418281, EPI_ISL_418282, EPI_ISL_418283, EPI_ISL_418284, EPI_ISL_418433, EPI_ISL_418434, EPI_ISL_418435 | | | |
| see above | University Hospital Basel, Clinical Virology | University Hospital Basel, Clinical Bacteriology | Hirsch, H., Leuzinger, K., Seth-Smith, H., Mari, A., Roloff, T., Egli, A. |
| EPI_ISL_418436 | University Hospital Basel, Clinical Virology | University Hospital Basel, Clinical Virology | Hirsch, H., Leuzinger, K., Seth-Smith, H., Mari, A., Roloff, T., Egli, A. |
| EPI_ISL_418438, EPI_ISL_418439, EPI_ISL_418440 | University Hospital Basel, Clinical Virology | University Hospital Basel, Clinical Bacteriology | Hirsch, H., Leuzinger, K., Seth-Smith, H., Mari, A., Roloff, T., Egli, A. |
| EPI_ISL_418816, EPI_ISL_418817, EPI_ISL_418818, EPI_ISL_418819, EPI_ISL_418820, EPI_ISL_418821, EPI_ISL_418822, EPI_ISL_418823, EPI_ISL_418824, EPI_ISL_418825, EPI_ISL_418826, EPI_ISL_418827, EPI_ISL_418828, EPI_ISL_418829, EPI_ISL_418830, EPI_ISL_418831, EPI_ISL_418832, EPI_ISL_418833, EPI_ISL_418834, EPI_ISL_418835, EPI_ISL_418836, EPI_ISL_418837, EPI_ISL_418838, EPI_ISL_418839, EPI_ISL_418840, EPI_ISL_418841, EPI_ISL_418842, EPI_ISL_418843, EPI_ISL_418844, EPI_ISL_418845, EPI_ISL_418846, EPI_ISL_418847, EPI_ISL_418848, EPI_ISL_418849, EPI_ISL_418850, EPI_ISL_418851, EPI_ISL_418852, EPI_ISL_418853, EPI_ISL_418854, EPI_ISL_418855, EPI_ISL_418856, EPI_ISL_418857, EPI_ISL_418858, EPI_ISL_418859 | | | |
| see above | BCCDC Public Health Laboratory | BCCDC Public Health Laboratory | Harrigan, Prystajecky, Krajden, Lee, Kamelian, Lapointe, Choi, Hoang, Sekirov, Levett, Tyson, Snutch, Loman, Quick, Li, Gilmour |
| EPI_ISL_419210 | The Chaim Sheba Medical Center | Israel Institute for Biological Research | Inbar Cohen-Gihon, Ofir Israeli, Ohad Shifman, Dana Stein, Sharon Melamed, Nir Paran, Tomer Israely, Hagit Achdout, Yfat Yahalom Ronen, Hadas Tamir, Boaz Politi, Lilach Cherry, Einat Vitner, Orly Laskar, Shay Weiss, Michal Mandelboim, Oran Erster, Gili Regev-Yochay, Gadi Segal, Shmuel Yitzhaki, Shmuel C. Shapira, Adi Beth-Din, Anat Zvi |
| EPI_ISL_419211 | Central Virology Laboratory | Israel Institute for Biological Research | Inbar Cohen-Gihon, Ofir Israeli, Ohad Shifman, Dana Stein, Sharon Melamed, Nir Paran, Tomer Israely, Hagit Achdout, Yfat Yahalom Ronen, Hadas Tamir, Boaz Politi, Lilach Cherry, Einat Vitner, Orly Laskar, Shay Weiss, Michal Mandelboim, Oran Erster, Gili Regev-Yochay, Gadi Segal, Shmuel Yitzhaki, Shmuel C. Shapira, Adi Beth-Din, Anat Zvi |
| EPI_ISL_419230 | Hospital Universitario Virgen de las Nieves | Instituto de Salud Carlos III | Iglesias-Caballero, M.; Molinero Calamita, M.; González-Esguevillas, M.; Camarero, S.; Pozo, F.; Casas, I.; Jiménez, P.; Jiménez, M.; Zaballos, A.; Monzón, S.; Varona, S.; Juliá, M.; Cuesta, I.; Sanbonmatsu, S. |
| EPI_ISL_419233 | Hospital Universitario de Canarias | Instituto de Salud Carlos III | Iglesias-Caballero, M.; Molinero Calamita, M.; González-Esguevillas, M.; Camarero, S.; Pozo, F.; Casas, I.; Jiménez, P.; Jiménez, M.; Zaballos, A.; Monzón, S.; Varona, S.; Juliá, M.; Cuesta, I.; Castro, B. |
| EPI_ISL_419234 | Hospital San Pedro | Instituto de Salud Carlos III | Iglesias-Caballero, M.; Molinero Calamita, M.; González-Esguevillas, M.; Camarero, S.; Pozo, F.; Casas, I.; Jiménez, P.; Jiménez, M.; Zaballos, A.; Monzón, S.; Varona, S.; Juliá, M.; Cuesta, I.; Alonso, C. |
| EPI_ISL_419235, EPI_ISL_419236, EPI_ISL_419237 | Fundacion Jimenez Diaz | Instituto de Salud Carlos III | Iglesias-Caballero, M.; Molinero Calamita, M.; González-Esguevillas, M.; Camarero, S.; Pozo, F.; Casas, I.; Jiménez, P.; Jiménez, M.; Zaballos, A.; Monzón, S.; Varona, S.; Juliá, M.; Cuesta, I.; Fernández, R. |
| EPI_ISL_419238 | HOSPITAL DE CRUCES. | Instituto de Salud Carlos III | Iglesias-Caballero, M. Molinero Calamita, M. González-Esguevillas, M. Camarero, S. Pozo, F. Casas, I. Jiménez, P. Jiménez, M. Zaballos, A. Monzón, S. Varona, S. Juliá, M. Cuesta, I. Aranzamendi, M. |
| EPI_ISL_419240 | HOSPITAL TXAGORRITXU | Instituto de Salud Carlos III | Iglesias-Caballero, M. Molinero Calamita, M. González-Esguevillas, M. Camarero, S. Pozo, F. Casas, I. Jiménez, P. Jiménez, M. Zaballos, A. Monzón, S. Varona, S. Juliá, M. Cuesta, I. Gómez, C |
| EPI_ISL_419709 | HOSPITAL TXAGORRITXU | Instituto de Salud Carlos III | Iglesias-Caballero, M. Molinero Calamita, M. González-Esguevillas, M. Camarero S. Pozo F. Casas I. Jiménez, P. Jiménez, M. Zaballos, A. Monzón, S. Varona, S. Juliá, M. Cuesta, I. Gómez, C. |

| | | | |
|---|---|---|---|
| EPI_ISL_420118 | Servicio de Microbiología. Consorcio Hospital General Universitario de Valencia | Sequencing and Bioinformatics Service and Molecular Epidemiology Research Group. FISABIO-Public Health | Inma Galán Vendrell, Paula Ruiz-Hueso, Mariana Reyes-Prieto, Vicente Soriano Chirona, Maria Alma Bracho, Griselda De Marco, Beatriz Beamud, Lidia Ruiz Roldan, Marta Pla Diaz,Neris Garcia-Gonzalez, Loreto Ferrús Abad, Maria Dolores Ocete, Llúcia Martínez-Priego, Concepcion Gimeno, Giuseppe D'Auria, Fernando Gonzalez-Candelas |
| EPI_ISL_420129 | Servicio de Microbiologia. Hospital Clinico Universitario de Valencia | Sequencing and Bioinformatics Service and Molecular Epidemiology Research Group. FISABIO-Public Health | Inma Galán Vendrell, Loreto Ferrús Abad, Maria Alma Bracho, Griselda De Marco, Beatriz Beamud, Lidia Ruiz Roldan, Marta Pla Diaz, Neris Garcia-Gonzalez, Paula Ruiz-Hueso, Mariana Reyes-Prieto, Vicente Soriano Chirona, David Navarro, Llúcia Martínez-Priego, Giuseppe D'Auria, Fernando Gonzalez-Candelas |
| EPI_ISL_420889, EPI_ISL_420890 | Takayuki Hishiki Kanagawa Prefectural Institute of Public Health | Takayuki Hishiki Kanagawa Prefectural Institute of Public Health | Hishiki,T., Suzuki,R., Sakuragi,J., Usui,K., Tanaka,Y., Kawai,J., Kogo,Y., Matsuki,Y., An,T., Hayashizaki,Y. and Takasaki,T. |
| EPI_ISL_421518 | Servicio de Microbiologia. Hospital Clinico Universitario de Valencia | Sequencing and Bioinformatics Service and Molecular Epidemiology Research Group. FISABIO-Public Health | Inma Galán Vendrell, Paula Ruiz-Hueso, Sandra Carbo, Loreto Ferrús Abad, Maria Alma Bracho, Griselda De Marco, Beatriz Beamud, Lidia Ruiz Roldan, Marta Pla Diaz, Neris Garcia-Gonzalez, Mariana Reyes-Prieto, Vicente Soriano Chirona, Ivan Ansari, David Navarro, Llúcia Martínez-Priego, Giuseppe D'Auria, Fernando Gonzalez-Candelas |
| EPI_ISL_421653, EPI_ISL_421654, EPI_ISL_421655, EPI_ISL_421656 | E. Gulbja Laboratorija | Latvian Biomedical Research and Study Centre | Ivars Silamielis, Kaspars Megnis, Monta Ustinova, ikita Zrelovs, Vita Rovte, Mikus Gavars, Dmitrijs Perminovs, Uga Dumpis, Jnis Kloviš |
| EPI_ISL_421660 | The Ohio State University | The Ohio State University-James Molecular Lab at Polaris | Huolin Tu, Matthew Avenarius, Preeti Panchioli, Sean Caruthers, Joan-Miquel Balada-Llasat, Jason Garee, Matt Hunt, Xiaokang Pan, Dan Jones |
| EPI_ISL_421675 | The Ohio State University Wexner Medical Center | The Ohio State University James Molecular lab | Huolin Tu, Sean Caruthers, Matthew Avenarius, Joan-Miquel Balada-Llasat, Matthew Hunt, Preeti Panchioli, Xiaokang Pen, Jason Garee, Pam Snyder, Dan Jones |
| EPI_ISL_424667 | Laboratorio Estatal de Salud Publica del Estado de México | Instituto de Diagnóstico y Referencia Epidemiológicos | Irma López Martínez, José Ernesto Ramírez González, Lucia Hernández Rivas, Gisela Barrera Badillo, Edgar Mendieta Condado, Fabiola Garcés Ayala, Adnan Araiza Rodríguez, Celia Boukadida, Santiago Avila Ríos, Mario Mújica Sánchez, José Arturo Martínez Orozco,Eduardo Becerril Vargas, Joel Armando Vázquez Pérez, Victor Hugo Borja Aburto, Concepción Grajales Muñiz, Cesar Raúl González Bonilla, Carolina González Torres, Francisco Javier Gaytán Cervantes, José Esteban Muñoz Medina, Guillermo M. Ruiz-Palacios, Pilar Ramos Cervantes, Violeta Ibarra Gonzalez, Fernando Ledesma Barrientos, Luis Alberto García Andrade, Alfredo Ponce de León Garduño, Blanca Taboada, Alejandro Sánchez, Pavel Isa, Ricardo Grande, Gloria Vázquez, Francisco Pulido, Carlos F. Arias. |
| EPI_ISL_425185 | Servicio de Microbiología. Consorcio Hospital General Universitario de Valencia | Sequencing and Bioinformatics Service and Molecular Epidemiology Research Group. FISABIO-Public Health | Inma Galán Vendrell, Sandra Carbo, Loreto Ferrús Abad, Paula Ruiz-Hueso, Mariana Reyes-Prieto, Vicente Soriano Chirona, Ivan Ansari, David Navarro, Maria Alma Bracho, Griselda De Marco, Beatriz Beamud, Lidia Ruiz Roldan, Marta Pla Diaz, Neris Garcia-Gonzalez, Llúcia Martínez-Priego, Giuseppe D'Auria, Fernando Gonzalez-Candelas |
| EPI_ISL_425191 | Servicio de Microbiología. Consorcio Hospital General Universitario de Valencia | Sequencing and Bioinformatics Service and Molecular Epidemiology Research Group. FISABIO-Public Health | Ivan Ansari, David Navarro, Maria Alma Bracho, Griselda De Marco, Beatriz Beamud, Lidia Ruiz Roldan, Marta Pla Diaz, Neris Garcia-Gonzalez, Inma Galán Vendrell, Sandra Carbo, Loreto Ferrús Abad, Paula Ruiz-Hueso, Mariana Reyes-Prieto, Vicente Soriano Chirona, Llúcia Martínez-Priego, Giuseppe D'Auria, Fernando Gonzalez-Candelas |
| EPI_ISL_425199 | Servicio de Microbiologia. Hospital Clinico Universitario de Valencia | Sequencing and Bioinformatics Service and Molecular Epidemiology Research Group. FISABIO-Public Health | Inma Galán Vendrell, Paula Ruiz-Hueso, Mariana Reyes-Prieto, Vicente Soriano Chirona, Maria Alma Bracho, Griselda De Marco, Beatriz Beamud, Lidia Ruiz Roldan, Marta Pla Diaz,Neris Garcia-Gonzalez, Loreto Ferrús Abad, Maria Dolores Ocete, Llúcia Martínez-Priego, Concepcion Gimeno, Giuseppe D'Auria, Fernando Gonzalez-Candelas |
| EPI_ISL_425220 | Servicio de Microbiología. Consorcio Hospital General Universitario de Valencia | Sequencing and Bioinformatics Service and Molecular Epidemiology Research Group. FISABIO-Public Health | Inma Galán Vendrell, Sandra Carbo, Loreto Ferrús Abad, Paula Ruiz-Hueso, Mariana Reyes-Prieto, Vicente Soriano Chirona, Ivan Ansari, David Navarro, Maria Alma Bracho, Griselda De Marco, Beatriz Beamud, Lidia Ruiz Roldan, Marta Pla Diaz, Neris Garcia-Gonzalez, Llúcia Martínez-Priego, Giuseppe D'Auria, Fernando Gonzalez-Candelas |
| EPI_ISL_426285, EPI_ISL_426286, EPI_ISL_426287, EPI_ISL_426288, EPI_ISL_426289 | E. Gulbja Laboratorija | Latvian Biomedical Research and Study Centre | Ivars Silamielis, Kaspars Megnis, Monta Ustinova, ikita Zrelovs, Vita Rovte, Mikus Gavars, Dmitrijs Perminovs, Uga Dumpis, Jnis Kloviš |
| EPI_ISL_427288 | The Ohio State University | The Ohio State University-James Molecular Lab at Polaris | Huolin Tu, Preeti Panchioli, Jason Garee, Matthew Hunt, Joan-Miquel Balada-Llasat, Erica Vincent, Weiqiang Zhao, Dan Jones |
| EPI_ISL_427289 | The Ohio State University | The Ohio State University-James Molecular Lab at Polaris | Huolin Tu, Joan-Miquel Balada-Llasat, Jason Garee, Matthew Hunt, Preeti Pancholi, Erica Vincent, Xiaokang Zhao, Dan Jones |
| EPI_ISL_427290 | The Ohio State University | The Ohio State University-James Molecular Lab at Polaris | Huolin Tu, Jason Garee, Matthew Hunt, Joan-Miquel Balada-Llasat, Preeti Pancholi, Erica Vincent, Rongqin Ren, Dan Jones |
| EPI_ISL_427291 | The Ohio State University | The Ohio State University-James Molecular Lab at Polaris | Huolin Tu, Matthew Hunt, Preeti Pancholi, Jason Garee, Joan-Miquel Balada-Llasat, Erica Vincent, Weiqiang Zhao, Dan Jones |
| EPI_ISL_427391 | Genomic Laboratory (GLAB) (Conjoint lab of Health Directorate of Istanbul and Istanbul Technical University) | Genomic Laboratory (GLAB), Istanbul Technical University | Ilker Karacan, Tugba Kizilboga Akgun, Bugra Agaoglu, Gizem Alkurt, Jale Yildiz, Betsi Köse, Elifnaz Çelik, Mehtap Aydn, Levent Doganay, Gizem Dinler Doganay |
| EPI_ISL_428346, EPI_ISL_428368 | Genomic Laboratory (GLAB) (Conjoint lab of Health Directorate of Istanbul and Istanbul Technical University) | Genomic Laboratory (GLAB), Istanbul Technical University | Ilker Karacan, Tugba Kizilboga Akgun, Bugra Agaoglu, Gizem Alkurt, Jale Yildiz, Betsi Köse, Elifnaz Çelik, Arzu Irvem, Yasemin Kendir Demirkol, Ozlem Akgun Dogan, Mehtap Aydn, Levent Doganay, Gizem Dinler Doganay |
| EPI_ISL_429992, EPI_ISL_429993, EPI_ISL_429994, EPI_ISL_429995, EPI_ISL_429996, EPI_ISL_429997, EPI_ISL_429998, EPI_ISL_429999, EPI_ISL_430000, EPI_ISL_430001, EPI_ISL_430002, EPI_ISL_430003, EPI_ISL_430004, EPI_ISL_430005, EPI_ISL_430006, EPI_ISL_430007, EPI_ISL_430008, EPI_ISL_430009 | | | |
| see above | Biolab Diagnostic Laboratories | Andersen lab at Scripps Research | Issa Abu-Dayyeh, Ahmad Tibi, Lama Hussein, Lina Mohammad, Zein Naber, Amid Abdelnour with SEARCH Alliance San Diego |
| EPI_ISL_430010 | OSU Wexner Medical Center | James Molecular Lab - OSUWMC | Huolin Tu, Preeti Pancholi, Matt Avenarius, Erica Vincent, Matt Hunt, Dan Jones |
| EPI_ISL_430011, EPI_ISL_430012, EPI_ISL_430013, EPI_ISL_430014, EPI_ISL_430015 | Biolab Diagnostic Laboratories | Andersen lab at Scripps Research | Issa Abu-Dayyeh, Ahmad Tibi, Lama Hussein, Lina Mohammad, Zein Naber, Amid Abdelnour with SEARCH Alliance San Diego |
| EPI_ISL_431013 | Alaska State Virology Laboratory | Alaska State Virology Laboratory | Jack Chen |
| EPI_ISL_431014, EPI_ISL_431015, EPI_ISL_431016, EPI_ISL_431017, EPI_ISL_431018, EPI_ISL_431019 | Alaska State Virology Laboratory | Alaska State Virology Laboratory | Jack Chen, Ph.D. |
| EPI_ISL_434516 | Biolab Diagnostic Laboratories | Andersen lab at Scripps Research | Issa Abu-Dayyeh, Ahmad Tibi, Lama Hussein, Lina Mohammad, Zein Naber, Amid Abdelnour with SEARCH Alliance San Diego |
| EPI_ISL_434646, EPI_ISL_434667 | Ulltuna Vardcentral | The Public Health Agency of Sweden | Heidi Lindback, Oskar Karlsson Lindsjo, Maria Lind Karlberg, Anna-Malin Linde, Olov Svartstrom, Anna Risberg, Theresa Enkirch, Mia Brytting, Karin Tegmark-Wisell |
| EPI_ISL_435441, EPI_ISL_435442, EPI_ISL_435443, EPI_ISL_435444 | Alaska State Virology Laboratory | Alaska State Virology Laboratory | Jack Chen, Ph.D. |
| EPI_ISL_436284 | Servicio de Microbiologia. Hospital Clinico Universitario de Valencia | Sequencing and Bioinformatics Service and Molecular Epidemiology Research Group. FISABIO-Public Health | Inma Galán Vendrell, Sandra Carbo, Loreto Ferrús Abad, Paula Ruiz-Hueso, Mariana Reyes-Prieto, Vicente Soriano Chirona, Ivan Ansari, David Navarro, Maria Alma Bracho, Griselda De Marco, Beatriz Beamud, Lidia Ruiz Roldan, Marta Pla Diaz, Neris Garcia-Gonzalez, Llúcia Martínez-Priego, Giuseppe D'Auria, Fernando Gonzalez-Candelas |
| EPI_ISL_436290 | Servicio de Microbiologia. Hospital Clinico Universitario de Valencia | Sequencing and Bioinformatics Service and Molecular Epidemiology Research Group. FISABIO-Public Health | Ivan Ansari, David Navarro, Maria Alma Bracho, Griselda De Marco, Beatriz Beamud, Lidia Ruiz Roldan, Marta Pla Diaz, Neris Garcia-Gonzalez, Inma Galán Vendrell, Sandra Carbo, Loreto Ferrús Abad, Paula Ruiz-Hueso, Mariana Reyes-Prieto, Vicente Soriano Chirona, Llúcia Martínez-Priego, Giuseppe D'Auria, Fernando Gonzalez-Candelas |
| EPI_ISL_436304 | Servicio de Microbiologia. Hospital Clinico Universitario de Valencia | Sequencing and Bioinformatics Service and Molecular Epidemiology Research Group. FISABIO-Public Health | Inma Galán Vendrell, Sandra Carbo, Loreto Ferrús Abad, Paula Ruiz-Hueso, Mariana Reyes-Prieto, Vicente Soriano Chirona, Ivan Ansari, David Navarro, Maria Alma Bracho, Griselda De Marco, Beatriz Beamud, Lidia Ruiz Roldan, Marta Pla Diaz, Neris Garcia-Gonzalez, Llúcia Martínez-Priego, Giuseppe D'Auria, Fernando Gonzalez-Candelas |
| EPI_ISL_436310 | Servicio de Microbiologia. Hospital Clinico Universitario de Valencia | Sequencing and Bioinformatics Service and Molecular Epidemiology Research Group. FISABIO-Public Health | Ivan Ansari, David Navarro, Maria Alma Bracho, Griselda De Marco, Beatriz Beamud, Lidia Ruiz Roldan, Marta Pla Diaz, Neris Garcia-Gonzalez, Inma Galán Vendrell, Sandra Carbo, Loreto Ferrús Abad, Paula Ruiz-Hueso, Mariana Reyes-Prieto, Vicente Soriano Chirona, Llúcia Martínez-Priego, Giuseppe D'Auria, Fernando Gonzalez-Candelas |

| | | | |
|---|---|---|---|
| EPI_ISL_436324 | Servicio de Microbiologia. Hospital Clinico Universitario de Valencia | Sequencing and Bioinformatics Service and Molecular Epidemiology Research Group. FISABIO-Public Health | Inma Galán Vendrell, Sandra Carbo, Loreto Ferrús Abad, Paula Ruiz-Hueso, Mariana Reyes-Prieto, Vicente Soriano Chirona, Ivan Ansari, David Navarro, Maria Alma Bracho, Griselda De Marco, Beatriz Beamud, Lidia Ruiz Roldan, Marta Pla Diaz, Neris Garcia-Gonzalez, Llúcia Martínez-Priego, Giuseppe D'Auria, Fernando Gonzalez-Candelas |
| EPI_ISL_436330 | Servicio de Microbiologia. Hospital Clinico Universitario de Valencia | Sequencing and Bioinformatics Service and Molecular Epidemiology Research Group. FISABIO-Public Health | Ivan Ansari, David Navarro, Maria Alma Bracho, Griselda De Marco, Beatriz Beamud, Lidia Ruiz Roldan, Marta Pla Diaz, Neris Garcia-Gonzalez, Inma Galán Vendrell, Sandra Carbo, Loreto Ferrús Abad, Paula Ruiz-Hueso, Mariana Reyes-Prieto, Vicente Soriano Chirona, Llúcia Martínez-Priego, Giuseppe D'Auria, Fernando Gonzalez-Candelas |
| EPI_ISL_436344 | Servicio de Microbiologia. Hospital Clinico Universitario de Valencia | Sequencing and Bioinformatics Service and Molecular Epidemiology Research Group. FISABIO-Public Health | Inma Galán Vendrell, Sandra Carbo, Loreto Ferrús Abad, Paula Ruiz-Hueso, Mariana Reyes-Prieto, Vicente Soriano Chirona, Ivan Ansari, David Navarro, Maria Alma Bracho, Griselda De Marco, Beatriz Beamud, Lidia Ruiz Roldan, Marta Pla Diaz, Neris Garcia-Gonzalez, Llúcia Martínez-Priego, Giuseppe D'Auria, Fernando Gonzalez-Candelas |
| EPI_ISL_436350 | Servicio de Microbiologia. Hospital Clinico Universitario de Valencia | Sequencing and Bioinformatics Service and Molecular Epidemiology Research Group. FISABIO-Public Health | Ivan Ansari, David Navarro, Maria Alma Bracho, Griselda De Marco, Beatriz Beamud, Lidia Ruiz Roldan, Marta Pla Diaz, Neris Garcia-Gonzalez, Inma Galán Vendrell, Sandra Carbo, Loreto Ferrús Abad, Paula Ruiz-Hueso, Mariana Reyes-Prieto, Vicente Soriano Chirona, Llúcia Martínez-Priego, Giuseppe D'Auria, Fernando Gonzalez-Candelas |
| EPI_ISL_436368 | Servicio de Microbiologia. Hospital Clinico Universitario de Valencia | Sequencing and Bioinformatics Service and Molecular Epidemiology Research Group. FISABIO-Public Health | Inma Galán Vendrell, Sandra Carbo, Loreto Ferrús Abad, Paula Ruiz-Hueso, Mariana Reyes-Prieto, Vicente Soriano Chirona, Ivan Ansari, David Navarro, Maria Alma Bracho, Griselda De Marco, Beatriz Beamud, Lidia Ruiz Roldan, Marta Pla Diaz, Neris Garcia-Gonzalez, Llúcia Martínez-Priego, Giuseppe D'Auria, Fernando Gonzalez-Candelas |
| EPI_ISL_436401 | Servicio de Microbiologia. Hospital Clinico Universitario de Valencia | Sequencing and Bioinformatics Service and Molecular Epidemiology Research Group. FISABIO-Public Health | Ivan Ansari, David Navarro, Maria Alma Bracho, Griselda De Marco, Beatriz Beamud, Lidia Ruiz Roldan, Marta Pla Diaz, Neris Garcia-Gonzalez, Inma Galán Vendrell, Sandra Carbo, Loreto Ferrús Abad, Paula Ruiz-Hueso, Mariana Reyes-Prieto, Vicente Soriano Chirona, Llúcia Martínez-Priego, Giuseppe D'Auria, Fernando Gonzalez-Candelas |
| EPI_ISL_436464 | Alaska State Virology Laboratory | Alaska State Virology Laboratory | Jack Chen |
| EPI_ISL_437089, EPI_ISL_437090, EPI_ISL_437091, EPI_ISL_437092, EPI_ISL_437093, EPI_ISL_437094, EPI_ISL_437095, EPI_ISL_437096 | Latvijas Infektoloijas centrs | Latvian Biomedical Research and Study Centre | Ivars Silamielis, Kaspars Megnis, Monta Ustinova, ikita Zrelovs, Vita Rovte, Jeena Storoženko, Tatjana Kolupajeva, Oksana Savicka, Uga Dumpis, Jnis Kloviš |
| EPI_ISL_437437, EPI_ISL_437513, EPI_ISL_437514, EPI_ISL_437515, EPI_ISL_437516, EPI_ISL_437517, EPI_ISL_437518, EPI_ISL_437873 | Alaska State Virology Laboratory | Alaska State Virology Laboratory | Jack Chen, Ph.D. |
| EPI_ISL_445183 | Takayuki Hishiki Kanagawa Prefectural Institute of Public Health | Takayuki Hishiki Kanagawa Prefectural Institute of Public Health | Hishiki,T., Suzuki,R., Sakuragi,J., Usui,K., Tanaka,Y., Kawai,J., Kogo,Y., Matsuki,Y., An,T., Hayashizaki,Y. and Takasaki,T. |
| EPI_ISL_445228, EPI_ISL_445240 | Ulltuna Vardcentral | The Public Health Agency of Sweden | Heidi Lindback, Oskar Karlsson Lindsjo, Maria Lind Karlberg, Anna-Malin Linde, Olov Svartstrom, Anna Risberg, Theresa Enkirch, Mia Brytting, Karin Tegmark-Wisell |
| EPI_ISL_447476, EPI_ISL_447490, EPI_ISL_447504, EPI_ISL_447518, EPI_ISL_447531 | Servicio de Microbiologia. Hospital Clinico Universitario de Valencia | Sequencing and Bioinformatics Service and Molecular Epidemiology Research Group. FISABIO-Public Health | Inma Galán Vendrell, Sandra Carbo, Loreto Ferrús Abad, Paula Ruiz-Hueso, Mariana Reyes-Prieto, Vicente Soriano Chirona, Ivan Ansari, Llúcia Martínez-Priego, Giuseppe 'Auria, David Navarro, Eliseo Albert, Maria Alma Bracho, Lidia Ruiz Roldan, Neris Garcia-Gonzalez, Fernando Gonzalez-Candelas |
| EPI_ISL_450186, EPI_ISL_450187, EPI_ISL_450188, EPI_ISL_450189 | Biolab Diagnostic Laboratories | Andersen lab at Scripps Research | Issa Abu-Dayyeh, Ahmad Tibi, Lama Hussein, Lina Mohammad, Zein Naber, Amid Abdelnour with SEARCH Alliance San Diego |
| EPI_ISL_450413 | Center for Diagnostics, Institute of Medical Microbiology, Virology and Hygiene | University Medical Center Hamburg-Eppendorf | Huang,J., Pfefferle,S. and Fischer,N. |
| EPI_ISL_450518, EPI_ISL_450519 | E. Gulbja Laboratorija | Latvian Biomedical Research and Study Centre | Ivars Silamielis, Kaspars Megnis, Monta Ustinova, ikita Zrelovs, Vita Rovte, Mikus Gavars, Dmitrijs Perminovs, Uga Dumpis, Jnis Kloviš |
| EPI_ISL_450520, EPI_ISL_450521, EPI_ISL_450522, EPI_ISL_450523, EPI_ISL_450524 | Centrl Laboratorija | Latvian Biomedical Research and Study Centre | Ivars Silamielis, Kaspars Megnis, Monta Ustinova, ikita Zrelovs, Vita Rovte, Stella Lapia, Jana Oste, Marta Priedte, Uga Dumpis, Jnis Kloviš |
| EPI_ISL_450824 | Ulltuna Vardcentral | The Public Health Agency of Sweden | Heidi Lindback, Anna-Malin Linde, Maria Lind Karlberg, Oskar Karlsson Lindsjo, Olov Svartstrom, Anna Risberg, Theresa Enkirch, Mia Brytting, Karin Tegmark-Wisell |
| EPI_ISL_452140 | CUB Hopital Erasme Laboratoire d'Anatomie Pathologique | CUB Hopital Erasme Laboratoire d'Anatomie Pathologique | Isabelle Salmon, Nicky D'Haene |
| EPI_ISL_452142 | CUB Hopital Erasme Laboratoire d'Anatomie Pathologique | CUB Hopital Erasme Laboratoire d'Anatomie Pathologique | Isabelle Salmon, Nikcy D'Haene |
| EPI_ISL_452148, EPI_ISL_452149 | CUB Hopital Erasme Laboratoire d'Anatomie Pathologique | CUB Hopital Erasme Laboratoire d'Anatomie Pathologique | Isabelle Salmon, Nicky D'Haene |
| EPI_ISL_452234 | Din Klinik | The Public Health Agency of Sweden | Helene Warnborg, Anna-Malin Linde, Maria Lind Karlberg, Oskar Karlsson Lindsjo, Olov Svartstrom, Anna Risberg, Theresa Enkirch, Mia Brytting, Karin Tegmark-Wisell |
| EPI_ISL_452365, EPI_ISL_452366, EPI_ISL_452367, EPI_ISL_452368, EPI_ISL_452369, EPI_ISL_452370, EPI_ISL_452371, EPI_ISL_452372, EPI_ISL_452373, EPI_ISL_452374, EPI_ISL_452375, EPI_ISL_452376, EPI_ISL_452377, EPI_ISL_452378, EPI_ISL_452379, EPI_ISL_452380, EPI_ISL_452381, EPI_ISL_452382, EPI_ISL_452383, EPI_ISL_452384, EPI_ISL_452385, EPI_ISL_452386, EPI_ISL_452387, EPI_ISL_452388, EPI_ISL_452389, EPI_ISL_452390, EPI_ISL_452391, EPI_ISL_452392, EPI_ISL_452393, EPI_ISL_452394, EPI_ISL_452395, EPI_ISL_452396, EPI_ISL_452397, EPI_ISL_452398, EPI_ISL_452399, EPI_ISL_452400, EPI_ISL_452401, EPI_ISL_452402, EPI_ISL_452403, EPI_ISL_452404, EPI_ISL_452405, EPI_ISL_452406, EPI_ISL_452407, EPI_ISL_452408, EPI_ISL_452409, EPI_ISL_452410, EPI_ISL_452411, EPI_ISL_452412, EPI_ISL_452413, EPI_ISL_452414, EPI_ISL_452415, EPI_ISL_452416, EPI_ISL_452417, EPI_ISL_452418, EPI_ISL_452419, EPI_ISL_452420, EPI_ISL_452421, EPI_ISL_452422, EPI_ISL_452423, EPI_ISL_452424, EPI_ISL_452425, EPI_ISL_452426, EPI_ISL_452427, EPI_ISL_452428, EPI_ISL_452429, EPI_ISL_452430, EPI_ISL_452431, EPI_ISL_452432, EPI_ISL_452433, EPI_ISL_452434, EPI_ISL_452435, EPI_ISL_452436, EPI_ISL_452437, EPI_ISL_452438, EPI_ISL_452439, EPI_ISL_452440, EPI_ISL_452441, EPI_ISL_452442, EPI_ISL_452443, EPI_ISL_452444, EPI_ISL_452445, EPI_ISL_452446, EPI_ISL_452447, EPI_ISL_452448, EPI_ISL_452449, EPI_ISL_452450, EPI_ISL_452451, EPI_ISL_452452 | | | |
| see above | Servicio de Microbiología. HRU de Málaga. Servicio Andaluz de Salud | SeqCOVID-SPAIN consortium/IBV(CSIC) | Inmaculada de Toro Peinado, Maria Concepción Mediavilla Gradolph, Begoña Palop Borrás and SeqCOVID-SPAIN consortium |
| EPI_ISL_454578, EPI_ISL_454581, EPI_ISL_454583, EPI_ISL_454588, EPI_ISL_454592, EPI_ISL_454595 | University Hospital for Infectious Diseases "Dr. Fran Mihaljevi", Research Unit | University of Zagreb, Centre for research and knowledge transfer in biotechnology | Ivan-Christian Kurolt, Jelena Ivancic Jelecki, Anamarija Slovic |
| EPI_ISL_454602 | Croatian Institute of Public Health | University of Zagreb, Centre for research and knowledge transfer in biotechnology | Irena Tabain, Tatjana Vilibic-Cavlek, Jelena Ivancic Jelecki, Anamarija Slovic |
| EPI_ISL_455104, EPI_ISL_455109 | Ulltuna Vardcentral | The Public Health Agency of Sweden | Heidi Lindback, Anna-Malin Linde, Maria Lind Karlberg, Oskar Karlsson Lindsjo, Olov Svartstrom, Anna Risberg, Theresa Enkirch, Mia Brytting, Karin Tegmark-Wisell |
| EPI_ISL_455314 | Hospital Virgen del Rocio | Instituto de Salud Carlos III | Iglesias-Caballero, M. Molinero Calamita, M. González-Esguevillas, M. Camarero, S. Pozo, F. Casas, I. Jiménez, P. Jiménez, M. Zaballos, A. Monzón, S. Varona, S. Juliá, M. Cuesta, I, J. Lepe |
| EPI_ISL_455315, EPI_ISL_455316, EPI_ISL_455317, EPI_ISL_455318, EPI_ISL_455319, EPI_ISL_455320, EPI_ISL_455321, EPI_ISL_455322 | Hospital Virgen de las Nieves | Instituto de Salud Carlos III | Iglesias-Caballero, M. Molinero Calamita, M. González-Esguevillas, M. Camarero, S. Pozo, F. Casas, I. Jiménez, P. Jiménez, M. Zaballos, A. Monzón, S. Varona, S. Juliá, M. Cuesta, I, S. Sanbonmatsu |
| EPI_ISL_455323 | Hospital Virgen del Rocio | Instituto de Salud Carlos III | Iglesias-Caballero, M. Molinero Calamita, M. González-Esguevillas, M. Camarero, S. Pozo, F. Casas, I. Jiménez, P. Jiménez, M. Zaballos, A. Monzón, S. Varona, S. Juliá, M. Cuesta, I, J. Lepe |
| EPI_ISL_455324 | Hospital Virgen de las Nieves | Instituto de Salud Carlos III | Iglesias-Caballero, M. Molinero Calamita, M. González-Esguevillas, M. Camarero, S. Pozo, F. Casas, I. Jiménez, P. Jiménez, M. Zaballos, A. Monzón, S. Varona, S. Juliá, M. Cuesta, I, S. Sanbonmatsu |

| | | | |
|---|---|---|---|
| EPI_ISL_455325 | Hospital Universitario de Canarias | Instituto de Salud Carlos III | Iglesias-Caballero, M. Molinero Calamita, M. González-Esguevillas, M. Camarero, S. Pozo, F. Casas, I. Jiménez, P. Jiménez, M. Zaballos, A. Monzón, S. Varona, S. Juliá, M. Cuesta, I, B. Castro |
| EPI_ISL_455326 | Hospital Universitario Insular de Gran Canaria | Instituto de Salud Carlos III | Iglesias-Caballero, M. Molinero Calamita, M. González-Esguevillas, M. Camarero, S. Pozo, F. Casas, I. Jiménez, P. Jiménez, M. Zaballos, A. Monzón, S. Varona, S. Juliá, M. Cuesta, I, A. Hernández |
| EPI_ISL_455327 | Consejeria de Sanidad y Asuntos Sociales | Instituto de Salud Carlos III | Iglesias-Caballero, M. Molinero Calamita, M. González-Esguevillas, M. Camarero, S. Pozo, F. Casas, I. Jiménez, P. Jiménez, M. Zaballos, A. Monzón, S. Varona, S. Juliá, M. Cuesta, I, G. Gutiérrez |
| EPI_ISL_455328, EPI_ISL_455329, EPI_ISL_455330, EPI_ISL_455331 | Complejo Hospitalario Universitario La Coruna | Instituto de Salud Carlos III | Iglesias-Caballero, M. Molinero Calamita, M. González-Esguevillas, M. Camarero, S. Pozo, F. Casas, I. Jiménez, P. Jiménez, M. Zaballos, A. Monzón, S. Varona, S. Juliá, M. Cuesta, I, M.A Canizares |
| EPI_ISL_455332 | Xerencia de Xestión Integrada de Pontevedra e o Salnés | Instituto de Salud Carlos III | Iglesias-Caballero, M. Molinero Calamita, M. González-Esguevillas, M. Camarero, S. Pozo, F. Casas, I. Jiménez, P. Jiménez, M. Zaballos, A. Monzón, S. Varona, S. Juliá, M. Cuesta, I, M. Garcia |
| EPI_ISL_455333 | Complejo Hospitalario Universitario de Santiago | Instituto de Salud Carlos III | Iglesias-Caballero, M. Molinero Calamita, M. González-Esguevillas, M. Camarero, S. Pozo, F. Casas, I. Jiménez, P. Jiménez, M. Zaballos, A. Monzón, S. Varona, S. Juliá, M. Cuesta, I, J. Llovo |
| EPI_ISL_455334, EPI_ISL_455335 | Complejo Hospitalario de Orense | Instituto de Salud Carlos III | Iglesias-Caballero, M. Molinero Calamita, M. González-Esguevillas, M. Camarero, S. Pozo, F. Casas, I. Jiménez, P. Jiménez, M. Zaballos, A. Monzón, S. Varona, S. Juliá, M. Cuesta, I, M. Paz |
| EPI_ISL_455336, EPI_ISL_455337, EPI_ISL_455338, EPI_ISL_455339, EPI_ISL_455340, EPI_ISL_455341, EPI_ISL_455342, EPI_ISL_455343 | Hospital San Pedro | Instituto de Salud Carlos III | Iglesias-Caballero, M. Molinero Calamita, M. González-Esguevillas, M. Camarero, S. Pozo, F. Casas, I. Jiménez, P. Jiménez, M. Zaballos, A. Monzón, S. Varona, S. Juliá, M. Cuesta, I, C. Alonso |
| EPI_ISL_455344, EPI_ISL_455345, EPI_ISL_455346, EPI_ISL_455347, EPI_ISL_455348, EPI_ISL_455349 | Hospital Comarcal de Melilla | Instituto de Salud Carlos III | Iglesias-Caballero, M. Molinero Calamita, M. González-Esguevillas, M. Camarero, S. Pozo, F. Casas, I. Jiménez, P. Jiménez, M. Zaballos, A. Monzón, S. Varona, S. Juliá, M. Cuesta, I, I. Pérez |
| EPI_ISL_455350, EPI_ISL_455351 | Hospital Txagorritxu | Instituto de Salud Carlos III | Iglesias-Caballero, M. Molinero Calamita, M. González-Esguevillas, M. Camarero, S. Pozo, F. Casas, I. Jiménez, P. Jiménez, M. Zaballos, A. Monzón, S. Varona, S. Juliá, M. Cuesta, I, C. Gómez |
| EPI_ISL_455352, EPI_ISL_455353, EPI_ISL_455354 | Hospital de Cruces | Instituto de Salud Carlos III | Iglesias-Caballero, M. Molinero Calamita, M. González-Esguevillas, M. Camarero, S. Pozo, F. Casas, I. Jiménez, P. Jiménez, M. Zaballos, A. Monzón, S. Varona, S. Juliá, M. Cuesta, I, M. Aranzamendi |
| EPI_ISL_455726 | Servicio de Microbiologia. Hospital Clinico Universitario de Valencia | Sequencing and Bioinformatics Service and Molecular Epidemiology Research Group. FISABIO-Public Health, and SeqCOVID-Spain Consortium | Inma Galán Vendrell, Sandra Carbo, Loreto Ferrús Abad, Paula Ruiz-Hueso, Mariana Reyes-Prieto, Vicente Soriano Chirona, Ivan Ansari, Llúcia Martínez-Priego, Giuseppe 'Auria, David Navarro, Eliseo Albert, Maria Alma Bracho, Lidia Ruiz Roldan, Neris Garcia-Gonzalez, Fernando Gonzalez-Candelas |
| EPI_ISL_462091 | National Institute of Laboratory Medicine and Referral Center | Genomic Research Lab, BCSIR | Iffat Jahan, Abu Sayeed Mohammad Mahmud, Mohammad Samir Uzzaman, Eshrar Osman, Md. Ahasan Habib, Shahina Akter, Tanjina Akhter Banu, Barna Goswami, Md. Saddam Hossain, Tasnim Nafisa, Md. Maruf Ahmed Molla, Mahmuda Yeasmin, Asish Kumar Ghos, Bayzid Bin Monir, Arifa Akram, Sheikh Md. Selim Al Din, Salek Ahmed Sajib, Utpal Chandra Ray, Md. Salim Khan |
| EPI_ISL_462477 | Hospital Costa del Sol | Instituto de Salud Carlos III | Iglesias-Caballero, M. Molinero Calamita, M. González-Esguevillas, M. Camarero, S. Pozo, F. Casas, I. Jiménez, P. Jiménez, M. Zaballos, A. Monzón, S. Varona, S. Juliá, M. Cuesta, I, F. Fernández |
| EPI_ISL_462478 | Fundación Jiménez Díaz | Instituto de Salud Carlos III | Iglesias-Caballero, M. Molinero Calamita, M. González-Esguevillas, M. Camarero, S. Pozo, F. Casas, I. Jiménez, P. Jiménez, M. Zaballos, A. Monzón, S. Varona, S. Juliá, M. Cuesta, I, R. Fernández |
| EPI_ISL_462479 | Hospital Clinic | Instituto de Salud Carlos III | Iglesias-Caballero, M. Molinero Calamita, M. González-Esguevillas, M. Camarero, S. Pozo, F. Casas, I. Jiménez, P. Jiménez, M. Zaballos, A. Monzón, S. Varona, S. Juliá, M. Cuesta, I, M.A Marcos |
| EPI_ISL_465163, EPI_ISL_465164 | National Institute of Laboratory Medicine and Referral Center | Genomic Research Lab, BCSIR | Iffat Jahan, Abu Sayeed Mohammad Mahmud, Mohammad Samir Uzzaman, Eshrar Osman, Md. Ahasan Habib, Shahina Akter, Tanjina Akhter Banu, Md. Murshed Hasan Sarker, Barna Goswami, Md. Saddam Hossain, Tasnim Nafisa, Md. Maruf Ahmed Molla, Mahmuda Yeasmin, Asish Kumar Ghosh, Arifa Akram, A. K. M. Shamsuzzaman, Sheikh Md. Selim Al Din, Utpal Chandra Ray, Salek Ahmed Sajib, Md. Salim Khan |
| EPI_ISL_467945 | Montefiore Medical Center, Dept. of Pathology, Clinical Virology | Albert Einstein College of Medicine, Dept. of Microbiology & Immunology, Chandran lab | J. Maximilian Fels, Saad Khan, Ryan Forster, Karin A. Skalina, Ariel S. Wirchnianski, Denise Haslwanter, Catalina Florez, Robert H. Bortz III, M. Eugenia Dieterle, Ethan Laudermilch, Rohit K. Jangra, Amanda Mengotto, Duncan Kimmel, Shahina B. Maqbool, John M. Greally, Wendy A. Szymczak, Amy S. Fox, Michael B. Prystowsky, D. Yitzchak Goldstein, Johanna P. Daily, Libusha Kelly, Kartik Chandran |
| EPI_ISL_469043 | Dr. N. D. Desai Medical College & Hospital | Gujarat Biotechnology Research Centre | J G Buch, Jigar Gusani, Supreet Prabhu, Tejas Shah, Ankit Hinsu, Pritesh Sabara, Apurvasinh Puvar, Janvi Raval, Zarna Patel, Monika Gandhi, Pinal Trivedi, Maharshi Pandya, Nidhi Patel, Nitin Savaliya, Raghawendra Kumar, Dinesh Kumar, Zuber Saiyed, Komal Patel, Labdhi Pandya, Snehal Bagatharia, Bhavya Jindal, R D Dixit, A M Kadri, Harsh Bakshi, Chaitanya Joshi, Madhvi Joshi |
| EPI_ISL_469299 | National Institute of Laboratory Medicine and Referral Center | Genomic Research Lab, BCSIR | Iffat Jahan, Abu Sayeed Mohammad Mahmud, Mohammad Samir Uzzaman, Eshrar Osman, Md. Ahasan Habib, Shahina Akter, Tanjina Akhter Banu, Md. Murshed Hasan Sarkar, Barna Goswami, Md. Saddam Hossain, Tasnim Nafisa, Md. Maruf Ahmed Molla, Mahmuda Yeasmin, Asish Kumar Ghosh, Bayzid Bin Monir, A. K. M. Shamsuzzaman, Sheikh Md. Selim Al Din, Utpal Chandra Ray, Salek Ahmed Sajib, Md. Salim Khan |
| EPI_ISL_470719, EPI_ISL_470720, EPI_ISL_470721, EPI_ISL_470722, EPI_ISL_470723, EPI_ISL_470724, EPI_ISL_470725, EPI_ISL_470726, EPI_ISL_470727, EPI_ISL_470728, EPI_ISL_470729, EPI_ISL_470730, EPI_ISL_470731, EPI_ISL_470732, EPI_ISL_470733, EPI_ISL_470734, EPI_ISL_470735, EPI_ISL_470736, EPI_ISL_470737, EPI_ISL_470738, EPI_ISL_470740, EPI_ISL_470741, EPI_ISL_470742, EPI_ISL_470743, EPI_ISL_470744, EPI_ISL_470745, EPI_ISL_470746 | | | |
| see above | Utah Public Health Laboratory | Utah Public Health Laboratory | Heidi Butz, Erin Young, Kelly Oakeson |
| EPI_ISL_470801 | Virology | Virology | Hossain,M.E., Hasan,R., Miah,M., Hasan,M.M., Sumaiya,M.K., Rahman,M.M., Alam,M.S., Clemens,J.D., Ahmed,T., Rahman,M.Z. and Rahman,M. |
| EPI_ISL_471456, EPI_ISL_471457, EPI_ISL_471458, EPI_ISL_471459, EPI_ISL_471460 | Centre de Virologie des Maladies Tropicales | Functional Genomic Platform/Service Analyses Biologique/UATRS/ Centre National Pour la Recherche Scientifique Et Technique (CNRST) | Hicham EL ANNAZ, Elmostafa EL FAHIME, Marouane MELLOUL, Youssef AKHOUAD, Mly Abdelaziz ELALAOUI, Ahmed REGGAD, Sanaa ALAOUI-Amine, Rachid ABI, Rida TAGAJDID, Zohour KASMY, Safae ELKOCHRI, Nadia TOUIL, Farida HILALI, Abdelkader LAATIRIS, Abdelillah LARAQUI, Tahar BAJJOU , Yassine SEKHSOKH , Idriss-Amine LAHLOU, Mostafa ELOUENNASS, Khalid ENNIBI |
| EPI_ISL_473306, EPI_ISL_473307, EPI_ISL_473308, EPI_ISL_473309, EPI_ISL_473310, EPI_ISL_473311, EPI_ISL_473312, EPI_ISL_473313, EPI_ISL_473314, EPI_ISL_473315, EPI_ISL_473316, EPI_ISL_473317, EPI_ISL_473318, EPI_ISL_473319, EPI_ISL_473320, EPI_ISL_473321, EPI_ISL_473322, EPI_ISL_473323, EPI_ISL_473324, EPI_ISL_473325, EPI_ISL_473326, EPI_ISL_473327, EPI_ISL_473328, EPI_ISL_473329, EPI_ISL_473330, EPI_ISL_473331, EPI_ISL_473332, EPI_ISL_473333, EPI_ISL_473334, EPI_ISL_473335, EPI_ISL_473336, EPI_ISL_473337, EPI_ISL_473338, EPI_ISL_473339, EPI_ISL_473340, EPI_ISL_473341, EPI_ISL_473342, EPI_ISL_473343, EPI_ISL_473344, EPI_ISL_473345, EPI_ISL_473346, EPI_ISL_473347, EPI_ISL_473348, EPI_ISL_473349, EPI_ISL_473350, EPI_ISL_473351, EPI_ISL_473352, EPI_ISL_473353, EPI_ISL_473354, EPI_ISL_473355, EPI_ISL_473356, EPI_ISL_473357, EPI_ISL_473358, EPI_ISL_473359, EPI_ISL_473360, EPI_ISL_473361, EPI_ISL_473362, EPI_ISL_473363, EPI_ISL_473364, EPI_ISL_473365, EPI_ISL_473366, EPI_ISL_473367, EPI_ISL_473368, EPI_ISL_473369, EPI_ISL_473370, EPI_ISL_473371, EPI_ISL_473372, EPI_ISL_473373, EPI_ISL_473374, EPI_ISL_473375, EPI_ISL_473376, EPI_ISL_473377, EPI_ISL_473378, EPI_ISL_473379, EPI_ISL_473380, EPI_ISL_473381, EPI_ISL_473382, EPI_ISL_473383, EPI_ISL_473384, EPI_ISL_473385, EPI_ISL_473386, EPI_ISL_473387, EPI_ISL_473388, EPI_ISL_473389, EPI_ISL_473390, EPI_ISL_473391, EPI_ISL_473392, EPI_ISL_473393, EPI_ISL_473394, EPI_ISL_473395, EPI_ISL_473396, EPI_ISL_473397, EPI_ISL_473398, EPI_ISL_473399, EPI_ISL_473400, EPI_ISL_473401, EPI_ISL_473402, EPI_ISL_473403, EPI_ISL_473404, EPI_ISL_473405, EPI_ISL_473406, EPI_ISL_473407, EPI_ISL_473408, EPI_ISL_473409, EPI_ISL_473410, EPI_ISL_473411, EPI_ISL_473412, EPI_ISL_473413, EPI_ISL_473414, EPI_ISL_473415, EPI_ISL_473416, EPI_ISL_473417, EPI_ISL_473418, EPI_ISL_473419, EPI_ISL_473420, EPI_ISL_473421, EPI_ISL_473422, EPI_ISL_473423, EPI_ISL_473424, EPI_ISL_473425, EPI_ISL_473426, EPI_ISL_473427, EPI_ISL_473428, EPI_ISL_473429, EPI_ISL_473430, EPI_ISL_473431, EPI_ISL_473432, EPI_ISL_473433, EPI_ISL_473434, EPI_ISL_473435, EPI_ISL_473436, EPI_ISL_473437, EPI_ISL_473438, EPI_ISL_473439, EPI_ISL_473440, EPI_ISL_473441, EPI_ISL_473442, EPI_ISL_473443, EPI_ISL_473444, EPI_ISL_473445, EPI_ISL_473446, EPI_ISL_473447, EPI_ISL_473448, EPI_ISL_473449, EPI_ISL_473450 | | | |
| see above | University of Birmingham | COVID-19 Genomics UK (COG-UK) Consortium | Institute of Microbiology, University of Birmingham: Claire McMurray, Joanne Stockton, Samuel Nicholls, Radoslaw Poplawski, Will Rowe, Josh Quick, Nicholas Loman, University of Birmingham Testing Laboratory: Celina M Whalley, Andrew Bosworth, Charlotte Poxon, Kasun Wanigasooriya, Oliver Pickles, Mike Kidd, Alex Richter, Andrew D Beggs PHE Heartlands Lab: Husam Osman, Andrew Bosworth. Queen Elizabeth Hospital: Anna Casey |
| EPI_ISL_475059 | GAIMS & G K General Hospital | Gujarat Biotechnology Research Centre | Hitesh Assudani, Babulal Babhoria, Labdhi Pandya, Snehal Bagatharia, Pritesh Sabara, Apurvasinh Puvar, Janvi Raval, Zarna Patel, Monika Gandhi, Pinal Trivedi, Maharshi Pandya, Nidhi Patel, Nitin Savaliya, Raghawendra Kumar, Dinesh Kumar, Zuber Saiyed, Komal Patel, Neha Rajpara, R D Dixit, A M Kadri, Harsh Bakshi, Chaitanya Joshi, Madhvi Joshi |
| EPI_ISL_475168 | National Institute of Laboratory Medicine and Referral Center | Genomic Research Lab, BCSIR | Iffat Jahan, Abu Sayeed Mohammad Mahmud, Mohammad Samir Uzzaman, Eshrar Osman, Md. Ahasan Habib, Shahina Akter, Tanjina Akhter Banu, Md. Murshed Hasan Sarkar, Barna Goswami, Md. Saddam Hossain, Tasnim Nafisa, Md. Maruf Ahmed Molla, Mahmuda Yeasmin, Asish Kumar Ghosh, Bayzid Bin Monir, A. K. M. Shamsuzzaman, Sheikh Md. Selim Al Din, Utpal Chandra Ray, Salek Ahmed Sajib, Md. Salim Khan |

| | | | |
|---|---|---|---|
| EPI_ISL_475571 | Genome Center | Genome Center | Hassan M. Al-Emran, Md. Shazid Hasan, Ovinu Kibria Islam, A. S. M. Rubayet- Ul- Alam, Pravas Chandra Roy, Selina Akter, Shireen Nigar, Shovon Lal Sarkar, Md. Tanvir Islam, Mithun Talukder Md. Tawyabur, Md. Taijul Islam, Provakar Mondol, Md. Muzahidul Islam, Md. Iqbal Kabir Jahid Md. Anwar Hossain |
| EPI_ISL_475758 | National Institute of Laboratory Medicine and Referral Center | Genomic Research Lab, BCSIR | Iffat Jahan, Abu Sayeed Mohammad Mahmud, Mohammad Samir Uzzaman, Eshrar Osman, Md. Ahasan Habib, Shahina Akter, Tanjina Akhter Banu, Md. Murshed Hasan Sarkar, Barna Goswami, Md. Saddam Hossain, Tasnim Nafisa, Md. Maruf Ahmed Molla, Mahmuda Yeasmin, Asish Kumar Ghosh, Arifa Akram, A. K. M. Shamsuzzaman, Sheikh Md. Selim Al Din, Utpal Chandra Ray, Salek Ahmed Sajib, Md. Salim Khan |
| EPI_ISL_476898, EPI_ISL_476899 | Alaska State Virology Laboratory | Alaska State Virology Laboratory | Jack Chen, Ph.D. |
| EPI_ISL_477727, EPI_ISL_477728, EPI_ISL_477729, EPI_ISL_477730, EPI_ISL_477731, EPI_ISL_477732, EPI_ISL_477733, EPI_ISL_477734, EPI_ISL_477735, EPI_ISL_477736, EPI_ISL_477737, EPI_ISL_477738, EPI_ISL_477739, EPI_ISL_477740, EPI_ISL_477741, EPI_ISL_477742, EPI_ISL_477743, EPI_ISL_477744, EPI_ISL_477745, EPI_ISL_477746, EPI_ISL_477747, EPI_ISL_477748, EPI_ISL_477749, EPI_ISL_477750, EPI_ISL_477751, EPI_ISL_477752, EPI_ISL_477753, EPI_ISL_477754, EPI_ISL_477755, EPI_ISL_477756, EPI_ISL_477757, EPI_ISL_477758, EPI_ISL_477759, EPI_ISL_477760, EPI_ISL_477761, EPI_ISL_477762, EPI_ISL_477763, EPI_ISL_477764, EPI_ISL_477765, EPI_ISL_477766, EPI_ISL_477767, EPI_ISL_477768, EPI_ISL_477769, EPI_ISL_477770, EPI_ISL_477771, EPI_ISL_477772, EPI_ISL_477773, EPI_ISL_477774, EPI_ISL_477775, EPI_ISL_477776, EPI_ISL_477777, EPI_ISL_477778, EPI_ISL_477779, EPI_ISL_477780, EPI_ISL_477781, EPI_ISL_477782 | | | |
| see above | University of Birmingham | COVID-19 Genomics UK (COG-UK) Consortium | Institute of Microbiology, University of Birmingham: Claire McMurray, Joanne Stockton, Samuel Nicholls, Radoslaw Poplawski, Will Rowe, Josh Quick, Nicholas Loman. University of Birmingham Testing Laboratory: Celina M Whalley, Andrew Bosworth, Charlotte Poxon, Kasun Wanigasooriya, Oliver Pickles, Mike Kidd, Alex Richter, Andrew D Beggs PHE Heartlands Lab: Husam Osman, Andrew Bosworth. Queen Elizabeth Hospital: Anna Casey |
| EPI_ISL_480224 | National Reference Laboratory "Influenza and acute respiratory diseases" | NRL-HIV | Ivan Ivanov, Ivailo Alexiev, Ivva Philipova |
| EPI_ISL_480228, EPI_ISL_480229, EPI_ISL_480230, EPI_ISL_480231, EPI_ISL_480232, EPI_ISL_480233, EPI_ISL_480234, EPI_ISL_480235, EPI_ISL_480236, EPI_ISL_480237, EPI_ISL_480238, EPI_ISL_480239, EPI_ISL_480240, EPI_ISL_480241, EPI_ISL_480242, EPI_ISL_480243, EPI_ISL_480244, EPI_ISL_480245, EPI_ISL_480246, EPI_ISL_480247, EPI_ISL_480248, EPI_ISL_480249, EPI_ISL_480250, EPI_ISL_480251, EPI_ISL_480252, EPI_ISL_480253, EPI_ISL_480254, EPI_ISL_480255, EPI_ISL_480256, EPI_ISL_480257, EPI_ISL_480258, EPI_ISL_480259, EPI_ISL_480260, EPI_ISL_480261, EPI_ISL_480262, EPI_ISL_480263, EPI_ISL_480264, EPI_ISL_480265, EPI_ISL_480266, EPI_ISL_480267, EPI_ISL_480268, EPI_ISL_480269, EPI_ISL_480270, EPI_ISL_480271, EPI_ISL_480272, EPI_ISL_480273, EPI_ISL_480274, EPI_ISL_480275, EPI_ISL_480276, EPI_ISL_480277, EPI_ISL_480278, EPI_ISL_480279, EPI_ISL_480280, EPI_ISL_480281, EPI_ISL_480282, EPI_ISL_480283, EPI_ISL_480284, EPI_ISL_480285, EPI_ISL_480286, EPI_ISL_480287, EPI_ISL_480288, EPI_ISL_480289, EPI_ISL_480290, EPI_ISL_480291, EPI_ISL_480292 | | | |
| see above | Genomic Laboratory (GLAB) (Conjoint lab of Health Directorate of Istanbul and Istanbul Technical University) | Genomic Laboratory (GLAB), Istanbul Technical University | Ilker Karacan, Tugba Kizilboga Akgun, Bugra Agaoglu, Gizem Alkurt, Jale Yildiz, Betsi Köse, Elifnaz Çelik, Arzu Irvem, Yasemin Kendir Demirkol, Ozlem Akgun Dogan, Mehtap Aydn, Levent Doganay, Gizem Dinler Doganay |
| EPI_ISL_480297, EPI_ISL_480298, EPI_ISL_480299, EPI_ISL_480300, EPI_ISL_480301, EPI_ISL_480302, EPI_ISL_480303, EPI_ISL_480304, EPI_ISL_480305, EPI_ISL_480306, EPI_ISL_480307, EPI_ISL_480308, EPI_ISL_480309, EPI_ISL_480310 | | | |
| see above | National Reference Laboratory "Influenza and acute respiratory diseases" | NRL-HIV | Ivan Ivanov, Ivailo Alexiev, Ivva Philipova |
| EPI_ISL_480443, EPI_ISL_480444 | National Institute of Laboratory Medicine and Referral Center | Bangladesh Council of Scientific and Industrial Research | Iffat Jahan, Abu Sayeed Mohammad Mahmud, Mohammad Samir Uzzaman, Eshrar Osman, Md. Ahasan Habib, Shahina Akter, Tanjina Akhter Banu, Md. Murshed Hasan Sarkar, Barna Goswami, Md. Saddam Hossain, Tasnim Nafisa, Md. Maruf Ahmed Molla, Mahmuda Yeasmin, Asish Kumar Ghosh, Shahjahan Siddike, A. K. M. Shamsuzzaman, Sheikh Md. Selim Al Din, Utpal Chandra Ray, Salek Ahmed Sajib, Md. Salim Khan |
| EPI_ISL_482759, EPI_ISL_482760, EPI_ISL_482761, EPI_ISL_482762, EPI_ISL_482763, EPI_ISL_482764, EPI_ISL_482765, EPI_ISL_482766, EPI_ISL_482767, EPI_ISL_482768, EPI_ISL_482769, EPI_ISL_482770, EPI_ISL_482771, EPI_ISL_482772, EPI_ISL_482773, EPI_ISL_482774, EPI_ISL_482775 | | | |
| see above | Medical Ain Shams Research Institute (MASRI), Ain Shams University | Medical Ain Shams Research Institute (MASRI), Ain Shams University | Hesham Elghazaly, Sara Hassan Agwa, Ahmad Moustafa, Hala Hafez, Sara Elnakeep, Shaimaa Moustafa, Aya Mohamed, Reham Mamdouh, Ghada Ismael, Ashraf Omar, Osama Mansour, Mahmoud Elmeitini |
| EPI_ISL_483018, EPI_ISL_483019, EPI_ISL_483020, EPI_ISL_483021, EPI_ISL_483022, EPI_ISL_483023, EPI_ISL_483024, EPI_ISL_483025, EPI_ISL_483026, EPI_ISL_483027, EPI_ISL_483028, EPI_ISL_483029, EPI_ISL_483030, EPI_ISL_483031, EPI_ISL_483032, EPI_ISL_483033 | | | |
| see above | Utah Public Health Laboratory | Utah Public Health Laboratory | Heidi Butz, Erin Young, Kelly Oakeson |
| EPI_ISL_483035, EPI_ISL_483036, EPI_ISL_483037, EPI_ISL_483038 | Medical Ain Shams Research Institute (MASRI), Ain Shams University | Medical Ain Shams Research Institute (MASRI), Ain Shams University | Hesham Elghazaly, Sara Hassan Agwa, Ahmad Moustafa, Hala Hafez, Sara Elnakeep, Shaimaa Moustafa, Aya Mohamed, Reham Mamdouh, Ghada Ismael, Ashraf Omar, Osama Mansour, Mahmoud Elmeitini |
| EPI_ISL_483059 | Hospital Universitari Germans Trias i Pujol | IrsiCaixa AIDS Research Lab | J. Segalés, M. Puig, J. Rodon, C. Avila-Nieto, J. Carrillo, G. Cantero, M.T. Terrón, S. Cruz, M. Parera ,M. Noguera-Julián, N. Izquierdo-Useros, V. Guallar, E. Vidal, A. Valencia, I. Blanco, J. Blanco, B. Clotet, J. Vergara-Alert |
| EPI_ISL_483643, EPI_ISL_483644 | National Institute of Laboratory Medicine and Referral Center | Genomic Research Lab, BCSIR | Iffat Jahan, Abu Sayeed Mohammad Mahmud, Mohammad Samir Uzzaman, Eshrar Osman, Md. Ahasan Habib, Shahina Akter, Tanjina Akhter Banu, Md. Murshed Hasan Sarkar, Barna Goswami, Md. Saddam Hossain, Tasnim Nafisa, Md. Maruf Ahmed Molla, Mahmuda Yeasmin, Asish Kumar Ghosh, A. K. M. Shamsuzzaman, Sheikh Md. Selim Al Din, Utpal Chandra Ray, Salek Ahmed Sajib, Md. Salim Khan |
| EPI_ISL_483824 | GMERS Medical College Himmatnagar | Gujarat Biotechnology Research Centre | Himanshu Khatri, Mayur Gandhi, Apurvasinh Puvar, Janvi Raval, Zarna Patel, Monika Gandhi, Pinal Trivedi, Maharshi Pandya, Nidhi Patel, Nitin Savaliya, Raghawendra Kumar, Dinesh Kumar, Zuber Saiyed, Komal Patel, Labdhi Pandya, Afzal Ansari, Nikha Trivedi, R D Dixit, A M Kadri, Harsh Bakshi, Chaitanya Joshi, Madhvi Joshi |
| EPI_ISL_483880, EPI_ISL_483881, EPI_ISL_483882, EPI_ISL_483883, EPI_ISL_483884, EPI_ISL_483885, EPI_ISL_483886, EPI_ISL_483887, EPI_ISL_483888, EPI_ISL_483889, EPI_ISL_483890, EPI_ISL_483891, EPI_ISL_483892, EPI_ISL_483893, EPI_ISL_483894, EPI_ISL_483895, EPI_ISL_483896, EPI_ISL_483897, EPI_ISL_483898, EPI_ISL_483899, EPI_ISL_483900, EPI_ISL_483901, EPI_ISL_483902, EPI_ISL_483903, EPI_ISL_483904, EPI_ISL_483905, EPI_ISL_483906, EPI_ISL_483907, EPI_ISL_483908, EPI_ISL_483909, EPI_ISL_483910, EPI_ISL_483911, EPI_ISL_483912, EPI_ISL_484218, EPI_ISL_484219, EPI_ISL_484220 | | | |
| see above | University of Birmingham | COVID-19 Genomics UK (COG-UK) Consortium | Institute of Microbiology, University of Birmingham: Claire McMurray, Joanne Stockton, Samuel Nicholls, Radoslaw Poplawski, Will Rowe, Josh Quick, Nicholas Loman. University of Birmingham Testing Laboratory: Celina M Whalley, Andrew Bosworth, Charlotte Poxon, Kasun Wanigasooriya, Oliver Pickles, Mike Kidd, Alex Richter, Andrew D Beggs PHE Heartlands Lab: Husam Osman, Andrew Bosworth. Queen Elizabeth Hospital: Anna Casey |
| EPI_ISL_486390, EPI_ISL_486391 | Centrl laboratorija | Latvian Biomedical Research and Study Centre | Ivars Silamielis, Kaspars Megnis, Monta Ustinova, ikita Zrelovs, Vita Rovte, Stella Lapia, Jana Oste, Marta Priedte, Uga Dumpis, Jnis Kloviš |
| EPI_ISL_486410 | Centrala laboratorija | Latvian Biomedical Research and Study Centre | Ivars Silamielis, Kaspars Megnis, Monta Ustinova, ikita Zrelovs, Vita Rovte, Stella Lapia, Jana Oste, Marta Priedte, Uga Dumpis, Jnis Kloviš |
| EPI_ISL_486411, EPI_ISL_486412, EPI_ISL_486413, EPI_ISL_486414, EPI_ISL_486415, EPI_ISL_486416 | Centrl laboratorija | Latvian Biomedical Research and Study Centre | Ivars Silamielis, Kaspars Megnis, Monta Ustinova, ikita Zrelovs, Vita Rovte, Stella Lapia, Jana Oste, Marta Priedte, Uga Dumpis, Jnis Kloviš |
| EPI_ISL_486417 | Centrala laboratorija | Latvian Biomedical Research and Study Centre | Ivars Silamielis, Kaspars Megnis, Monta Ustinova, ikita Zrelovs, Vita Rovte, Stella Lapia, Jana Oste, Marta Priedte, Uga Dumpis, Jnis Kloviš |
| EPI_ISL_486418, EPI_ISL_486419, EPI_ISL_486420, EPI_ISL_486421 | Centrl laboratorija | Latvian Biomedical Research and Study Centre | Ivars Silamielis, Kaspars Megnis, Monta Ustinova, ikita Zrelovs, Vita Rovte, Stella Lapia, Jana Oste, Marta Priedte, Uga Dumpis, Jnis Kloviš |
| EPI_ISL_486422, EPI_ISL_486423, EPI_ISL_486424, EPI_ISL_486425, EPI_ISL_486426, EPI_ISL_486428, EPI_ISL_486430, EPI_ISL_486431, EPI_ISL_486432, EPI_ISL_486433, EPI_ISL_486434, EPI_ISL_486435, EPI_ISL_486436 | | | |
| see above | Latvijas Infektoloijas centrs | Latvian Biomedical Research and Study Centre | Ivars Silamielis, Kaspars Megnis, Monta Ustinova, ikita Zrelovs, Vita Rovte, Jeena Storoženko, Tatjana Kolupajeva, Oksana Savicka, Uga Dumpis, Jnis Kloviš |
| EPI_ISL_486437 | Centrl laboratorija | Latvian Biomedical Research and Study Centre | Ivars Silamielis, Kaspars Megnis, Monta Ustinova, ikita Zrelovs, Vita Rovte, Stella Lapia, Jana Oste, Marta Priedte, Uga Dumpis, Jnis Kloviš |
| EPI_ISL_486438 | E. Gulbja laboratorija | Latvian Biomedical Research and Study Centre | Ivars Silamielis, Kaspars Megnis, Monta Ustinova, ikita Zrelovs, Vita Rovte, Mikus Gavars, Dmitrijs Perminovs, Uga Dumpis, Jnis Kloviš |
| EPI_ISL_487193, EPI_ISL_487194, EPI_ISL_487195, EPI_ISL_487196, EPI_ISL_487197, EPI_ISL_487198, EPI_ISL_487199, EPI_ISL_487200, EPI_ISL_487201, EPI_ISL_487202, EPI_ISL_487203, EPI_ISL_487204, EPI_ISL_487205, EPI_ISL_487206, EPI_ISL_487207, EPI_ISL_487208, EPI_ISL_487209, EPI_ISL_487210, EPI_ISL_487211, EPI_ISL_487212, EPI_ISL_487213, EPI_ISL_487214, EPI_ISL_487215, EPI_ISL_487216, EPI_ISL_487217, EPI_ISL_487218, EPI_ISL_487219, EPI_ISL_487220, EPI_ISL_487221, EPI_ISL_487222, EPI_ISL_487223, EPI_ISL_487224, EPI_ISL_487225, EPI_ISL_487226, EPI_ISL_487227, EPI_ISL_487228, EPI_ISL_487244, EPI_ISL_487245, EPI_ISL_487246, EPI_ISL_487247, EPI_ISL_487248, EPI_ISL_487249, EPI_ISL_487250, EPI_ISL_487251, EPI_ISL_487252, EPI_ISL_487253, EPI_ISL_487254, EPI_ISL_487255, EPI_ISL_487256, EPI_ISL_487257, EPI_ISL_487258, EPI_ISL_487259, EPI_ISL_487260, EPI_ISL_487261, EPI_ISL_487262, EPI_ISL_487263, EPI_ISL_487264, EPI_ISL_487265, EPI_ISL_487266, EPI_ISL_487267, EPI_ISL_487268 | | | |
| see above | Utah Public Health Laboratory | Utah Public Health Laboratory | Heidi Butz, Erin Young, Kelly Oakeson |
| EPI_ISL_487372, EPI_ISL_487373, EPI_ISL_490164 | National Institute of Laboratory Medicine and Referral Center | Genomic Research Lab, BCSIR | Iffat Jahan, Abu Sayeed Mohammad Mahmud, Mohammad Samir Uzzaman, Eshrar Osman, Md. Ahasan Habib, Shahina Akter, Tanjina Akhter Banu, Md. Murshed Hasan Sarkar, Barna Goswami, Md. Saddam Hossain, Tasnim Nafisa, Md. Maruf Ahmed Molla, Mahmuda Yeasmin, Asish Kumar Ghosh, A. K. M. Shamsuzzaman, Sheikh Md. Selim Al Din, Utpal Chandra Ray, Salek Ahmed Sajib, Md. Salim Khan |
| EPI_ISL_492988, EPI_ISL_492989, EPI_ISL_492990, EPI_ISL_492991, | Centrl laboratorija | Latvian Biomedical Research and Study Centre | Ivars Silamielis, Kaspars Megnis, Monta Ustinova, ikita Zrelovs, Vita Rovte, Stella Lapia, Jana Oste, Marta Priedte, Uga Dumpis, Jnis Kloviš |

| | | | |
|---|---|---|---|
| EPI_ISL_492992 | | | |
| EPI_ISL_492993, EPI_ISL_492994, EPI_ISL_492995, EPI_ISL_492996, EPI_ISL_492997, EPI_ISL_492998, EPI_ISL_492999, EPI_ISL_493000 | E. Gulbja Laboratorija | Latvian Biomedical Research and Study Centre | Ivars Silamielis, Kaspars Megnis, Monta Ustinova, ikita Zrelovs, Vita Rovte, Mikus Gavars, Dmitrijs Perminovs, Uga Dumpis, Jnis Kloviš |
| EPI_ISL_493004, EPI_ISL_493005, EPI_ISL_493006, EPI_ISL_493007, EPI_ISL_493008, EPI_ISL_493009, EPI_ISL_493010, EPI_ISL_493011, EPI_ISL_493012, EPI_ISL_493013, EPI_ISL_493014, EPI_ISL_493015, EPI_ISL_493016, EPI_ISL_493017, EPI_ISL_493018, EPI_ISL_493019, EPI_ISL_493020, EPI_ISL_493021, EPI_ISL_493022, EPI_ISL_493023, EPI_ISL_493024, EPI_ISL_493025, EPI_ISL_493026, EPI_ISL_493027, EPI_ISL_493028, EPI_ISL_493029, EPI_ISL_493030, EPI_ISL_493031, EPI_ISL_493032, EPI_ISL_493033, EPI_ISL_493034, EPI_ISL_493035, EPI_ISL_493036, EPI_ISL_493037, EPI_ISL_493038, EPI_ISL_493039, EPI_ISL_493040, EPI_ISL_493041, EPI_ISL_493042, EPI_ISL_493043, EPI_ISL_493044, EPI_ISL_493045, EPI_ISL_493046, EPI_ISL_493047, EPI_ISL_493048, EPI_ISL_493049, EPI_ISL_493050, EPI_ISL_493051, EPI_ISL_493052, EPI_ISL_493054, EPI_ISL_493055, EPI_ISL_493056, EPI_ISL_493057, EPI_ISL_493058, EPI_ISL_493059, EPI_ISL_493060, EPI_ISL_493087, EPI_ISL_493088, EPI_ISL_493089, EPI_ISL_493090, EPI_ISL_493091, EPI_ISL_493092, EPI_ISL_493093, EPI_ISL_493094, EPI_ISL_493095, EPI_ISL_493096, EPI_ISL_493097, EPI_ISL_493098, EPI_ISL_493099, EPI_ISL_493100, EPI_ISL_493101, EPI_ISL_493102, EPI_ISL_493103, EPI_ISL_493104, EPI_ISL_493105, EPI_ISL_493106, EPI_ISL_493107, EPI_ISL_493108, EPI_ISL_493109, EPI_ISL_493110, EPI_ISL_493111, EPI_ISL_493112, EPI_ISL_493113, EPI_ISL_493114, EPI_ISL_493115, EPI_ISL_493116, EPI_ISL_493117, EPI_ISL_493118, EPI_ISL_493119, EPI_ISL_493120, EPI_ISL_493121, EPI_ISL_493122, EPI_ISL_493123, EPI_ISL_493124 | | | |
| see above | Utah Public Health Laboratory | Utah Public Health Laboratory | Heidi Butz, Erin Young, Kelly Oakeson |
| EPI_ISL_493198, EPI_ISL_493199, EPI_ISL_493200, EPI_ISL_493202, EPI_ISL_493205, EPI_ISL_493206, EPI_ISL_493207 | Virology Lab,Department of Pathology, National Cheng Kung University Hospital | Virology Lab,Department of Pathology, National Cheng Kung University Hospital | Huey-Pin Tsai, et al |
| EPI_ISL_493208 | Virology Lab,Department of Pathology, National Cheng Kung University Hospital | Virology Lab,Department of Pathology, National Cheng Kung University Hospital | Huey-Pin Tsai et al |
| EPI_ISL_493441, EPI_ISL_493442, EPI_ISL_493443, EPI_ISL_493444, EPI_ISL_493445, EPI_ISL_493446, EPI_ISL_493447, EPI_ISL_493448, EPI_ISL_493449, EPI_ISL_493450, EPI_ISL_493451 | | | |
| see above | University of Birmingham | COVID-19 Genomics UK (COG-UK) Consortium | Institute of Microbiology, University of Birmingham: Claire McMurray, Joanne Stockton, Samuel Nicholls, Radoslaw Poplawski, Will Rowe, Josh Quick, Nicholas Loman. University of Birmingham Testing Laboratory: Celina M Whalley, Andrew Bosworth, Charlotte Poxon, Kasun Wanigasooriya, Oliver Pickles, Mike Kidd, Alex Richter, Andrew D Beggs PHE Heartlands Lab: Husam Osman, Andrew Bosworth. Queen Elizabeth Hospital: Anna Casey |
| EPI_ISL_494747 | Virology Lab,Department of Pathology, National Cheng Kung University Hospital | Virology Lab,Department of Pathology, National Cheng Kung University Hospital | Huey-Pin Tsai, et al |
| EPI_ISL_495067 | Dr. N. D. Desai Medical College & Hospital | Gujarat Biotechnology Research Centre | J G Buch, Jigar Gusani, Supreet Prabhu, Apurvasinh Puvar, Janvi Raval, Zarna Patel, Monika Gandhi, Pinal Trivedi, Maharshi Pandya, Nidhi Patel, Nitin Savaliya, Raghawendra Kumar, Dinesh Kumar, Zuber Saiyed, Komal Patel, Labdhi Pandya, Afzal Ansari, Nikha Trivedi, R D Dixit, A M Kadri, Harsh Bakshi, Chaitanya Joshi, Madhvi Joshi |
| EPI_ISL_498272, EPI_ISL_498273 | National Institute of Laboratory Medicine and Referral Center | Genomic Research Lab, BCSIR | Iffat Jahan, Abu Sayeed Mohammad Mahmud, Mohammad Samir Uzzaman, Eshrar Osman, Md. Ahasan Habib, Shahina Akter, Tanjina Akhter Banu, Md. Murshed Hasan Sarkar, Barna Goswami, Md. Saddam Hossain, Tasnim Nafisa, Md. Maruf Ahmed Molla, Mahmuda Yeasmin, Asish Kumar Ghosh, A. K. M. Shamsuzzaman, Sheikh Md. Selim Al Din, Utpal Chandra Ray, Salek Ahmed Sajib, Md. Salim Khan |
| EPI_ISL_498630, EPI_ISL_498631, EPI_ISL_498632, EPI_ISL_498633, EPI_ISL_498634, EPI_ISL_498635, EPI_ISL_498637, EPI_ISL_498638, EPI_ISL_498639, EPI_ISL_498640, EPI_ISL_498641, EPI_ISL_498642, EPI_ISL_498643, EPI_ISL_498644, EPI_ISL_498645, EPI_ISL_498646, EPI_ISL_498647, EPI_ISL_498648, EPI_ISL_498649, EPI_ISL_498650, EPI_ISL_498651, EPI_ISL_498652, EPI_ISL_498653, EPI_ISL_498654, EPI_ISL_498655, EPI_ISL_498656, EPI_ISL_498657, EPI_ISL_498658, EPI_ISL_498659, EPI_ISL_498660, EPI_ISL_498661, EPI_ISL_498662, EPI_ISL_498663, EPI_ISL_498664, EPI_ISL_498665, EPI_ISL_498666, EPI_ISL_498667, EPI_ISL_498668, EPI_ISL_498669, EPI_ISL_498670, EPI_ISL_498671, EPI_ISL_498672, EPI_ISL_498673, EPI_ISL_498674, EPI_ISL_498675, EPI_ISL_498676, EPI_ISL_498677, EPI_ISL_498678, EPI_ISL_498679, EPI_ISL_498680, EPI_ISL_498681, EPI_ISL_498682, EPI_ISL_498683, EPI_ISL_498684, EPI_ISL_498685, EPI_ISL_498686, EPI_ISL_498687, EPI_ISL_498688, EPI_ISL_498689, EPI_ISL_498690 | | | |
| see above | Utah Public Health Laboratory | Utah Public Health Laboratory | Heidi Butz, Erin Young, Kelly Oakeson |
| EPI_ISL_501275, EPI_ISL_501284, EPI_ISL_501285 | E. Gulbja Laboratorija | Latvian Biomedical Research and Study Centre | Ivars Silamielis, Kaspars Megnis, Monta Ustinova, ikita Zrelovs, Vita Rovte, Mikus Gavars, Dmitrijs Perminovs, Uga Dumpis, Jnis Kloviš |
| EPI_ISL_501286, EPI_ISL_501287, EPI_ISL_501288, EPI_ISL_501289, EPI_ISL_501808, EPI_ISL_501817 | Centrl laboratorija | Latvian Biomedical Research and Study Centre | Ivars Silamielis, Kaspars Megnis, Monta Ustinova, ikita Zrelovs, Vita Rovte, Stella Lapia, Jana Oste, Marta Priedte, Uga Dumpis, Jnis Kloviš |
| EPI_ISL_501823, EPI_ISL_501829, EPI_ISL_501833, EPI_ISL_501839, EPI_ISL_501849, EPI_ISL_501894 | E. Gulbja Laboratorija | Latvian Biomedical Research and Study Centre | Ivars Silamielis, Kaspars Megnis, Monta Ustinova, ikita Zrelovs, Vita Rovte, Mikus Gavars, Dmitrijs Perminovs, Uga Dumpis, Jnis Kloviš |
| EPI_ISL_501895, EPI_ISL_501896, EPI_ISL_501915, EPI_ISL_501922 | Centrl laboratorija | Latvian Biomedical Research and Study Centre | Ivars Silamielis, Kaspars Megnis, Monta Ustinova, ikita Zrelovs, Vita Rovte, Stella Lapia, Jana Oste, Marta Priedte, Uga Dumpis, Jnis Kloviš |
| EPI_ISL_501929, EPI_ISL_501936 | E. Gulbja Laboratorija | Latvian Biomedical Research and Study Centre | Ivars Silamielis, Kaspars Megnis, Monta Ustinova, ikita Zrelovs, Vita Rovte, Mikus Gavars, Dmitrijs Perminovs, Uga Dumpis, Jnis Kloviš |
| EPI_ISL_509527, EPI_ISL_509528, EPI_ISL_509529, EPI_ISL_509530, EPI_ISL_509531, EPI_ISL_509532, EPI_ISL_509533, EPI_ISL_509534, EPI_ISL_509535, EPI_ISL_509536, EPI_ISL_509537, EPI_ISL_509538, EPI_ISL_509539, EPI_ISL_509540, EPI_ISL_509541, EPI_ISL_509542, EPI_ISL_509543, EPI_ISL_509544, EPI_ISL_509545, EPI_ISL_509546, EPI_ISL_509547, EPI_ISL_509548, EPI_ISL_509549, EPI_ISL_509550, EPI_ISL_509551, EPI_ISL_509552, EPI_ISL_509553, EPI_ISL_509554, EPI_ISL_509555, EPI_ISL_509556, EPI_ISL_509557, EPI_ISL_509558, EPI_ISL_509559, EPI_ISL_509560, EPI_ISL_509561, EPI_ISL_509562, EPI_ISL_509563, EPI_ISL_509567, EPI_ISL_509568, EPI_ISL_509569, EPI_ISL_509570, EPI_ISL_509571, EPI_ISL_509572, EPI_ISL_509573, EPI_ISL_509574, EPI_ISL_509575, EPI_ISL_509576, EPI_ISL_509577, EPI_ISL_509578, EPI_ISL_509579, EPI_ISL_509580, EPI_ISL_509581, EPI_ISL_509582, EPI_ISL_509583, EPI_ISL_509584, EPI_ISL_509585, EPI_ISL_509586, EPI_ISL_509587, EPI_ISL_509588, EPI_ISL_509589, EPI_ISL_509590, EPI_ISL_509591, EPI_ISL_509592, EPI_ISL_509593, EPI_ISL_509594, EPI_ISL_509595, EPI_ISL_509596, EPI_ISL_509597, EPI_ISL_509598, EPI_ISL_509599, EPI_ISL_509600 | | | |
| see above | Utah Public Health Laboratory | Utah Public Health Laboratory | Heidi Butz, Erin Young, Kelly Oakeson |
| EPI_ISL_509602, EPI_ISL_509603, EPI_ISL_509604, EPI_ISL_509605, EPI_ISL_509606, EPI_ISL_509607, EPI_ISL_509608, EPI_ISL_509609, EPI_ISL_509611, EPI_ISL_509619, EPI_ISL_509620, EPI_ISL_509621, EPI_ISL_509622, EPI_ISL_509623, EPI_ISL_509624, EPI_ISL_509625, EPI_ISL_509626, EPI_ISL_509627, EPI_ISL_509628, EPI_ISL_509629, EPI_ISL_509630, EPI_ISL_509631, EPI_ISL_509632, EPI_ISL_510044, EPI_ISL_510045, EPI_ISL_510046, EPI_ISL_510047, EPI_ISL_510048, EPI_ISL_510049, EPI_ISL_510050, EPI_ISL_510051, EPI_ISL_510052, EPI_ISL_510053, EPI_ISL_510054, EPI_ISL_510055, EPI_ISL_510056, EPI_ISL_510057, EPI_ISL_510058, EPI_ISL_510059, EPI_ISL_510060, EPI_ISL_510061, EPI_ISL_510062, EPI_ISL_510063 | | | |
| see above | Servicio de Microbiología. HRU de Málaga. Servicio Andaluz de Salud | SeqCOVID-SPAIN consortium/IBV(CSIC) | Inmaculada de Toro Peinado. MªConcepción Mediavilla Gradolph. Begoña Palop Borrás and SeqCOVID-SPAIN consortium |
| EPI_ISL_512072 | Saikrishna Hospital, Mehsana | Gujarat Biotechnology Research Centre | Harshadbhai Parmar, Apurvasinh Puvar, Janvi Raval, Zarna Patel, Monika Gandhi, Pinal Trivedi, Maharshi Pandya, Nidhi Patel, Nitin Savaliya, Raghawendra Kumar, Dinesh Kumar, Zuber Saiyed, Komal Patel, Labdhi Pandya, Afzal Ansari, Nikha Trivedi, R D Dixit, A M Kadri, Harsh Bakshi, Chaitanya Joshi, Madhvi Joshi |
| EPI_ISL_512313, EPI_ISL_512314, EPI_ISL_512645, EPI_ISL_512646 | E. Gulbja Laboratorija | Latvian Biomedical Research and Study Centre | Ivars Silamielis, Kaspars Megnis, Monta Ustinova, ikita Zrelovs, Vita Rovte, Mikus Gavars, Dmitrijs Perminovs, Uga Dumpis, Jnis Kloviš |
| EPI_ISL_512647, EPI_ISL_512648, EPI_ISL_512649, EPI_ISL_512650, EPI_ISL_512651, EPI_ISL_512652 | Latvijas Infektoloijas centrs | Latvian Biomedical Research and Study Centre | Ivars Silamielis, Kaspars Megnis, Monta Ustinova, ikita Zrelovs, Vita Rovte, Jeena Storoženko, Tatjana Kolupajeva, Oksana Savicka, Uga Dumpis, Jnis Kloviš |
| EPI_ISL_513640, EPI_ISL_513641, EPI_ISL_513642, EPI_ISL_513643, EPI_ISL_513644, EPI_ISL_513645, EPI_ISL_513646, EPI_ISL_513647, EPI_ISL_513648, EPI_ISL_513649, EPI_ISL_513650, EPI_ISL_513651, EPI_ISL_513652, EPI_ISL_513653, EPI_ISL_513654, EPI_ISL_513655, EPI_ISL_513656, EPI_ISL_513657, EPI_ISL_513658, EPI_ISL_513659, EPI_ISL_513660, EPI_ISL_513661, EPI_ISL_513662, EPI_ISL_513663, EPI_ISL_513664, EPI_ISL_513665, EPI_ISL_513666, EPI_ISL_513667, EPI_ISL_513668, EPI_ISL_513669, EPI_ISL_513670, EPI_ISL_513671, EPI_ISL_513672, EPI_ISL_513673, EPI_ISL_513674, EPI_ISL_513675, EPI_ISL_513676, EPI_ISL_513677, EPI_ISL_513678, EPI_ISL_513679, EPI_ISL_513680, EPI_ISL_513681, EPI_ISL_513682, EPI_ISL_513683, EPI_ISL_513684, EPI_ISL_513685, EPI_ISL_513686, EPI_ISL_513687, EPI_ISL_513688, EPI_ISL_513689, EPI_ISL_513690, EPI_ISL_513691, EPI_ISL_513692, EPI_ISL_513693, EPI_ISL_513694, EPI_ISL_513695, EPI_ISL_513696, EPI_ISL_513697, EPI_ISL_513698, EPI_ISL_513699, EPI_ISL_513700, EPI_ISL_513701, EPI_ISL_513702, EPI_ISL_513703, EPI_ISL_513704, EPI_ISL_513705, EPI_ISL_513706 | | | |
| see above | Utah Public Health Laboratory | Utah Public Health Laboratory | Heidi Butz, Erin Young, Kelly Oakeson |
| EPI_ISL_514245, EPI_ISL_514246, EPI_ISL_514247 | National Institute of Laboratory Medicine and Referral Center | Genomic Research Lab, BCSIR | Iffat Jahan, Abu Sayeed Mohammad Mahmud, Mohammad Samir Uzzaman, Eshrar Osman, Md. Ahashan Habib, Shahina Akter, Tanjina Akhter Banu, Md. Murshed Hasan Sarkar, Barna Goswami, Md. Saddam Hossain, Tasnim Nafisa, Md. Maruf Ahmed Molla, Mahmuda Yeasmin, Asish Kumar Ghosh, A. K. M. Shamsuzzaman, Sheikh Md. Selim Al Din, Utpal Chandra Ray, Salek Ahmed Sajib, Md. Salim Khan |
| EPI_ISL_514435 | Dr. RSS Hospital, Modasa | Gujarat Biotechnology Research Centre | Harsh Chaudhari, Zarna Patel, Monika Gandhi, Pinal Trivedi, Maharshi Pandya, Nidhi Patel, Nitin Savaliya, Raghawendra Kumar, Dinesh Kumar, Zuber |

| | | | |
|---|---|---|---|
| | | | Saiyed, Komal Patel, Labdhi Pandya, Afzal Ansari, Nikha Trivedi, Apurvasinh Puvar, Janvi Raval, R D Dixit, A M Kadri, Harsh Bakshi, Chaitanya Joshi, Madhvi Joshi |
| EPI_ISL_515185, EPI_ISL_515186 | Latvijas Infektoloijas centrs | Latvian Biomedical Research and Study Centre | Ivars Silamielis, Kaspars Megnis, Monta Ustinova, ikita Zrelovs, Vita Rovte, Jeena Storoženko, Tatjana Kolupajeva, Oksana Savicka, Uga Dumpis, Jnis Kloviš |
| EPI_ISL_515187, EPI_ISL_515188, EPI_ISL_515189, EPI_ISL_515190, EPI_ISL_515191, EPI_ISL_515192, EPI_ISL_515193, EPI_ISL_515194, EPI_ISL_515195 | E. Gulbja Laboratorija | Latvian Biomedical Research and Study Centre | Ivars Silamielis, Kaspars Megnis, Monta Ustinova, ikita Zrelovs, Vita Rovte, Mikus Gavars, Dmitrijs Perminovs, Uga Dumpis, Jnis Kloviš |
| EPI_ISL_515196 | Centrl laboratorija | Latvian Biomedical Research and Study Centre | Ivars Silamielis, Kaspars Megnis, Monta Ustinova, ikita Zrelovs, Vita Rovte, Stella Lapia, Jana Oste, Marta Priedte, Uga Dumpis, Jnis Kloviš |
| EPI_ISL_522396, EPI_ISL_522397, EPI_ISL_522398, EPI_ISL_522399, EPI_ISL_522400, EPI_ISL_522401, EPI_ISL_522402, EPI_ISL_522403, EPI_ISL_522404, EPI_ISL_522405 | Alaska State Virology Laboratory | Alaska State Virology Laboratory | Jack Chen, Ph.D. |
| EPI_ISL_522872 | Instituto Nacional de Medicina Genómica | Instituto Nacional de Medicina Genómica | Hidalgo-Miranda A, Mendoza-Vargas A, Reyes-Grajeda JP, Cisneros-Villanueva M, Cedro-Tanda A, Hurtado-Cordova E, Peñaloza-Figueroa F, Herrera-Montalvo LA |
| EPI_ISL_522873, EPI_ISL_522874, EPI_ISL_522875, EPI_ISL_522876, EPI_ISL_522877 | Instituto Nacional de Medicina Genomica | Instituto Nacional de Medicina Genomica | Hidalgo-Miranda A, Mendoza-Vargas A, Reyes-Grajeda JP, Cisneros-Villanueva M, Cedro-Tanda A, Hurtado-Cordova E, Peñaloza-Figueroa F, Herrera-Montalvo LA |
| EPI_ISL_522878 | Instituto Nacional de Medicina Genomica | Instituto Nacional de Medicina Genomica | Hidalgo-Miranda A, Mendoza-Vargas A, Reyes-Grajeda JP, Cisneros-Villanueva M, Hurtado-Cordova E, Cedro-Tanda A, Peñaloza-Figueroa F, Herrera-Montalvo LA |
| EPI_ISL_522879, EPI_ISL_522880, EPI_ISL_522940, EPI_ISL_522941, EPI_ISL_522942, EPI_ISL_522978, EPI_ISL_522979, EPI_ISL_522980, EPI_ISL_522981, EPI_ISL_522982, EPI_ISL_522983, EPI_ISL_522984, EPI_ISL_522985, EPI_ISL_522986, EPI_ISL_523500 | | | |
| see above | Instituto Nacional de Medicina Genómica | Instituto Nacional de Medicina Genómica | Hidalgo-Miranda A, Mendoza-Vargas A, Reyes-Grajeda JP, Cisneros-Villanueva M, Cedro-Tanda A, Hurtado-Cordova E, Peñaloza-Figueroa F, Herrera-Montalvo LA |
| EPI_ISL_524732 | Dr. RSS Hospital, Modasa | Gujarat Biotechnology Research Centre | Harsh Chaudhari, Apurvasinh Puvar, Janvi Raval, Zarna Patel, Monika Gandhi, Pinal Trivedi, Maharshi Pandya, Nidhi Patel, Nitin Savaliya, Raghawendra Kumar, Dinesh Kumar, Zuber Saiyed, Komal Patel, Labdhi Pandya, Afzal Ansari, Nikha Trivedi, R D Dixit, A M Kadri, Harsh Bakshi, Chaitanya Joshi, Madhvi Joshi, Neena Doshi, Varsha Godbole |
| EPI_ISL_525472, EPI_ISL_525473 | Institute of Clinical Microbiology and Hygiene, University Hospital Regensburg | Institute of Clinical Microbiology and Hygiene, University Hospital Regensburg | Hiergeist, A. |
| EPI_ISL_525758, EPI_ISL_525759, EPI_ISL_525760 | Alaska State Virology Laboratory | Alaska State Virology Laboratory | Jack Chen, Ph.D. |
| EPI_ISL_526287, EPI_ISL_526288, EPI_ISL_526289, EPI_ISL_526290, EPI_ISL_526291, EPI_ISL_526292, EPI_ISL_526293, EPI_ISL_526294, EPI_ISL_526295, EPI_ISL_526296, EPI_ISL_526297, EPI_ISL_526298, EPI_ISL_526299, EPI_ISL_526300, EPI_ISL_526301, EPI_ISL_526302, EPI_ISL_526303, EPI_ISL_526304, EPI_ISL_526305, EPI_ISL_526306, EPI_ISL_526307, EPI_ISL_526308, EPI_ISL_526309, EPI_ISL_526310, EPI_ISL_526311, EPI_ISL_526312, EPI_ISL_526313, EPI_ISL_526314, EPI_ISL_526315, EPI_ISL_526316, EPI_ISL_526317, EPI_ISL_526318, EPI_ISL_526319, EPI_ISL_526320, EPI_ISL_526321, EPI_ISL_526322, EPI_ISL_526323, EPI_ISL_526324, EPI_ISL_526325, EPI_ISL_526326, EPI_ISL_526327, EPI_ISL_526328, EPI_ISL_526329, EPI_ISL_526330, EPI_ISL_526331, EPI_ISL_526332, EPI_ISL_526333, EPI_ISL_526334, EPI_ISL_526335, EPI_ISL_526336 | | | |
| see above | University of Birmingham | COVID-19 Genomics UK (COG-UK) Consortium | Institute of Microbiology, University of Birmingham: Claire McMurray, Joanne Stockton, Samuel Nicholls, Radoslaw Poplawski, Will Rowe, Josh Quick, Nicholas Loman. University of Birmingham Testing Laboratory: Celina M Whalley, Andrew Bosworth, Charlotte Poxon, Kasun Wanigasooriya, Oliver Pickles, Mike Kidd, Alex Richter, Andrew D Beggs PHE Heartlands Lab: Husam Osman, Andrew Bosworth. Queen Elizabeth Hospital: Anna Casey |
| EPI_ISL_528550, EPI_ISL_528551, EPI_ISL_528552, EPI_ISL_528553, EPI_ISL_528554, EPI_ISL_528555, EPI_ISL_528556 | National Genomics Core-Center for DNA Fingerprinting and Diagnostics | National Genomics Core- Center for DNA Fingerprinting and Diagnostics (NGC-CDFD)- DBT's PAN-INDIA-1000 Genome consortium | Heena Shah, G Shashikanth, Bala Pratyusha, Vinay Donipadi, Shruti Dasgupta, Kandali Sreethi Sreenivasulu Reddy, Chandra Shekhar Singh, Sunke Vijayakumar, R Lakshmi Vaishna, Jenige Aravindh Kumar, Muthulakshmi, V Naga Sailaja, R Harinarayanan, Rashna Bhandari, Murali Dharan Bashyam, Debashish Mitra, Divya Vashisht, Ashwin Dalal |
| EPI_ISL_528557, EPI_ISL_528558, EPI_ISL_528559, EPI_ISL_528560, EPI_ISL_528561, EPI_ISL_528562, EPI_ISL_528563, EPI_ISL_528564 | National Genomics Core-Center for DNA Fingerprinting and Diagnostics | National Genomics Core- Center for DNA Fingerprinting and Diagnostics (NGC-CDFD)- DBT's PAN-INDIA-1000 Genome consortium | Heena Shah, G Shashikanth, Bala Pratyusha, Vinay Donipadi, Binod Bihari Pradhan, Jamal Md Nurul Jain, Srinivas G, C Bala Maddeleti, R. Manorama, T. Navaneetha, Surya Vamshi, Chendra Shekar P, R Harinarayanan, Rashna Bhandari, Murali Dharan Bashyam, Debashish Mitra, Divya Vashisht, Ashwin Dalal |
| EPI_ISL_528588, EPI_ISL_528589, EPI_ISL_528590, EPI_ISL_528591, EPI_ISL_528592, EPI_ISL_528593, EPI_ISL_528594, EPI_ISL_528595 | National Genomics Core-Center for DNA Fingerprinting and Diagnostics | National Genomics Core- Center for DNA Fingerprinting and Diagnostics (NGC-CDFD)- DBT's PAN-INDIA-1000 Genome consortium | Heena Shah, G Shashikanth, Bala Pratyusha, Vinay Donipadi, Raju Kumar, Ajay Kumar Chaudhary, Akash Chinchole, Brahmaji Sontyana, C. Arun Kumar, Chandra Shekhar V, Chilakala Gangi Reddy, Chinthakindi KrishnaPrasad, R Harinarayanan, Rashna Bhandari, Murali Dharan Bashyam, Debashish Mitra, Divya Vashisht, Ashwin Dalal |
| EPI_ISL_528617, EPI_ISL_528618, EPI_ISL_528619, EPI_ISL_528620, EPI_ISL_528621, EPI_ISL_528622, EPI_ISL_528623, EPI_ISL_528624 | National Genomics Core-Center for DNA Fingerprinting and Diagnostics | National Genomics Core- Center for DNA Fingerprinting and Diagnostics (NGC-CDFD)- DBT's PAN-INDIA-1000 Genome consortium | Heena Shah, G Shashikanth, Bala Pratyusha, Vinay Donipadi, K.Manohar, Madhumohan Rao, Shubhra Ganguli, Suchitra Upreti, Swathi Chodisetty, Vani Singh, R Harinarayanan, Rashna Bhandari, Murali Dharan Bashyam, Debashish Mitra, Divya Vashisht, Ashwin Dalal |
| EPI_ISL_528631, EPI_ISL_528632, EPI_ISL_528633, EPI_ISL_528634, EPI_ISL_528635, EPI_ISL_528636 | National Genomics Core-Center for DNA Fingerprinting and Diagnostics | National Genomics Core- Center for DNA Fingerprinting and Diagnostics (NGC-CDFD)- DBT's PAN-INDIA-1000 Genome consortium | Heena Shah, G Shashikanth, Bala Pratyusha, Vinay Donipadi, K.Manohar, Madhumohan Rao, Shruti Dasgupta, Kandali Sreethi Sreenivasulu Reddy, Chandra Shekhar Singh, Sunke Vijayakumar, R Lakshmi Vaishna, Jenige Aravindh Kumar, Muthulakshmi, V Naga Sailaja, R Harinarayanan, Rashna Bhandari, Murali Dharan Bashyam, Debashish Mitra, Divya Vashisht, Ashwin Dalal |
| EPI_ISL_528749 | Santosa Hospital Bandung Central | School of Life Sciences and Technology & School of Pharmacy-Institut Teknologi Bandung; Molecular Genetics Laboratory-Faculty of Medicine-Universitas Padjadjaran; Laboratorium Kesehatan Provinsi Jawa Barat | Husna Nugrahapraja, Azzania Fibriani, Catur Riani, Marselina Irasonia Tan, Yunia Sribudiani, Tarwadi, Ema Rahmawati, Savira Ekawardhani, Hesti Lina Wiraswati, Ryan Bayusantika Ristandi, Rifky Waluyajati Rachman, Cut Nur Cinthia Alamanda, Lia Faridah, Davin H. E. Setiamarga, Rizki Mardian, Hammam Riza, Sony Solistia Wirawan, Agung Eru Wibowo, Irvan Faizal |
| EPI_ISL_528759 | Santo Borromeus Hospital | School of Life Sciences and Technology & School of Pharmacy-Institut Teknologi Bandung; Molecular Genetics Laboratory-Faculty of Medicine-Universitas Padjadjaran; Laboratorium Kesehatan Provinsi Jawa Barat | Husna Nugrahapraja, Azzania Fibriani, Catur Riani, Marselina Irasonia Tan, Yunia Sribudiani, Tarwadi, Ema Rahmawati, Savira Ekawardhani, Hesti Lina Wiraswati, Ryan Bayusantika Ristandi, Rifky Waluyajati Rachman, Cut Nur Cinthia Alamanda, Lia Faridah, Tri Hanggono Achmad, Mas Rizky A.A. Syamsunarno, Fensi Amalina, Hammam Riza, Sony Solistia Wirawan, Agung Eru Wibowo, Irvan Faizal |
| EPI_ISL_529149, EPI_ISL_529150 | Technology Centre, Guangzhou Customs | Technology Centre, Guangzhou Customs | Huang, J., Shi, Y., Sun, J., Zheng, K., Zhu, ., Sun, F., Zhuang, Z., Dai, J., Zhang, Z., Huang, S., Wang, Y., Li, X. |
| EPI_ISL_529218, EPI_ISL_529219, EPI_ISL_529220, EPI_ISL_529221, EPI_ISL_529222, EPI_ISL_529223, EPI_ISL_529234, EPI_ISL_529235, EPI_ISL_529237, EPI_ISL_529239, EPI_ISL_529251, EPI_ISL_529252, EPI_ISL_529255, EPI_ISL_529256, EPI_ISL_529262, EPI_ISL_529268, EPI_ISL_529275, EPI_ISL_529277, EPI_ISL_529278, EPI_ISL_529289, EPI_ISL_529291, EPI_ISL_529298, EPI_ISL_529300, EPI_ISL_529312, EPI_ISL_529313, EPI_ISL_529314, EPI_ISL_529320, EPI_ISL_529321, EPI_ISL_529322, EPI_ISL_529323, EPI_ISL_529324, EPI_ISL_529325, EPI_ISL_529326, EPI_ISL_529327, EPI_ISL_529328, EPI_ISL_529329, EPI_ISL_529330, EPI_ISL_529331, EPI_ISL_529332, EPI_ISL_529333, EPI_ISL_529334, EPI_ISL_529335, EPI_ISL_529336, EPI_ISL_529337, EPI_ISL_529338, EPI_ISL_529339, EPI_ISL_529340, EPI_ISL_529357, EPI_ISL_529358, EPI_ISL_529359, EPI_ISL_529360, EPI_ISL_529361, EPI_ISL_529362, EPI_ISL_529363, EPI_ISL_529364, EPI_ISL_529365, EPI_ISL_529366, EPI_ISL_529367, EPI_ISL_529368, EPI_ISL_529369, EPI_ISL_529370, EPI_ISL_529371, EPI_ISL_529372, EPI_ISL_529384, EPI_ISL_529386, EPI_ISL_529390, EPI_ISL_529391, EPI_ISL_529392, EPI_ISL_529393, EPI_ISL_529394, EPI_ISL_529395, EPI_ISL_529396, EPI_ISL_529397, EPI_ISL_529398, EPI_ISL_529399, EPI_ISL_529400, EPI_ISL_529401, EPI_ISL_529402, EPI_ISL_529403, EPI_ISL_529404, EPI_ISL_529405, EPI_ISL_529406, EPI_ISL_529408, EPI_ISL_529420, EPI_ISL_529421, EPI_ISL_529424, EPI_ISL_529425, EPI_ISL_529426, EPI_ISL_529427, EPI_ISL_529428, EPI_ISL_529429, EPI_ISL_529430, EPI_ISL_529434, EPI_ISL_529435, EPI_ISL_529438, EPI_ISL_529439, EPI_ISL_529440, EPI_ISL_529441, EPI_ISL_529442, EPI_ISL_529443, EPI_ISL_529444, EPI_ISL_529448, EPI_ISL_529453, EPI_ISL_529461, EPI_ISL_529462, EPI_ISL_529468, EPI_ISL_529470, EPI_ISL_529471, EPI_ISL_529472, EPI_ISL_529473, EPI_ISL_529474, EPI_ISL_529475, EPI_ISL_529480, EPI_ISL_529482, EPI_ISL_529485, EPI_ISL_529493, EPI_ISL_529494, EPI_ISL_529495, EPI_ISL_529496, EPI_ISL_529497, EPI_ISL_529499, EPI_ISL_529508, EPI_ISL_529512, EPI_ISL_529514, EPI_ISL_529515, EPI_ISL_529517, EPI_ISL_529518, EPI_ISL_529520, EPI_ISL_529522, EPI_ISL_529590, EPI_ISL_529591, EPI_ISL_529592, EPI_ISL_529593, EPI_ISL_529594, EPI_ISL_529595, EPI_ISL_529596, EPI_ISL_529597, EPI_ISL_529598, EPI_ISL_529599, EPI_ISL_529600, EPI_ISL_529601, EPI_ISL_529602, EPI_ISL_529603, EPI_ISL_529604, EPI_ISL_529605, EPI_ISL_529606, EPI_ISL_529607, EPI_ISL_529608, EPI_ISL_529609, EPI_ISL_529610, EPI_ISL_529611, EPI_ISL_529612, EPI_ISL_529613, EPI_ISL_529614, EPI_ISL_529615, EPI_ISL_529616, EPI_ISL_529617, EPI_ISL_529618, EPI_ISL_529619, EPI_ISL_529620, EPI_ISL_529621, EPI_ISL_529622, EPI_ISL_529623, EPI_ISL_529624, EPI_ISL_529625, EPI_ISL_529626, EPI_ISL_529627, EPI_ISL_529628, EPI_ISL_529629, EPI_ISL_529630, EPI_ISL_529631, EPI_ISL_529632, EPI_ISL_529633, EPI_ISL_529634, EPI_ISL_529635, EPI_ISL_529636, EPI_ISL_529637, EPI_ISL_529638, EPI_ISL_529639, EPI_ISL_529640, | | | |

EPI_ISL_529641, EPI_ISL_529642, EPI_ISL_529643, EPI_ISL_529644, EPI_ISL_529645, EPI_ISL_529646, EPI_ISL_529647, EPI_ISL_529648, EPI_ISL_529649, EPI_ISL_529650, EPI_ISL_529651, EPI_ISL_529652, EPI_ISL_529653, EPI_ISL_529654, EPI_ISL_529655, EPI_ISL_529661, EPI_ISL_529662, EPI_ISL_529663, EPI_ISL_529664, EPI_ISL_529665, EPI_ISL_529666

| | | | |
|---|---|---|---|
| see above | University of Birmingham | COVID-19 Genomics UK (COG-UK) Consortium | Institute of Microbiology, University of Birmingham: Claire McMurray, Joanne Stockton, Samuel Nicholls, Radoslaw Poplawski, Will Rowe, Josh Quick, Nicholas Loman. University of Birmingham Testing Laboratory: Celina M Whalley, Andrew Bosworth, Charlotte Poxon, Kasun Wanigasooriya, Oliver Pickles, Mike Kidd, Alex Richter, Andrew D Beggs PHE Heartlands Lab: Husam Osman, Andrew Bosworth. Queen Elizabeth Hospital: Anna Casey |

EPI_ISL_531864, EPI_ISL_531867, EPI_ISL_531868, EPI_ISL_531869, EPI_ISL_531871, EPI_ISL_531874, EPI_ISL_531875, EPI_ISL_531876, EPI_ISL_531879, EPI_ISL_531882, EPI_ISL_531883, EPI_ISL_531885, EPI_ISL_531886, EPI_ISL_531888, EPI_ISL_531891, EPI_ISL_531893, EPI_ISL_531898, EPI_ISL_531899, EPI_ISL_531900, EPI_ISL_531901, EPI_ISL_531902, EPI_ISL_531904, EPI_ISL_531906, EPI_ISL_531907, EPI_ISL_531908, EPI_ISL_531909, EPI_ISL_531910, EPI_ISL_531914, EPI_ISL_531915, EPI_ISL_531916, EPI_ISL_531917, EPI_ISL_531920, EPI_ISL_531922, EPI_ISL_531923, EPI_ISL_531926, EPI_ISL_531927, EPI_ISL_531928, EPI_ISL_531930, EPI_ISL_531931, EPI_ISL_531934, EPI_ISL_531937, EPI_ISL_531938, EPI_ISL_531942, EPI_ISL_531944, EPI_ISL_531949, EPI_ISL_531950, EPI_ISL_531952, EPI_ISL_531953, EPI_ISL_531956, EPI_ISL_531957, EPI_ISL_531959, EPI_ISL_531961, EPI_ISL_531962, EPI_ISL_531965, EPI_ISL_531968, EPI_ISL_531969, EPI_ISL_531971, EPI_ISL_531972, EPI_ISL_531974, EPI_ISL_531975, EPI_ISL_531976, EPI_ISL_531980, EPI_ISL_531981, EPI_ISL_531988, EPI_ISL_531989, EPI_ISL_531991, EPI_ISL_531993, EPI_ISL_531994, EPI_ISL_531996, EPI_ISL_531997, EPI_ISL_531998, EPI_ISL_531999, EPI_ISL_532001, EPI_ISL_532002, EPI_ISL_532006, EPI_ISL_532008, EPI_ISL_532010, EPI_ISL_532011, EPI_ISL_532013, EPI_ISL_532014, EPI_ISL_532015, EPI_ISL_532021, EPI_ISL_532022, EPI_ISL_532023, EPI_ISL_532026, EPI_ISL_532027, EPI_ISL_532029, EPI_ISL_532033, EPI_ISL_532034, EPI_ISL_532037, EPI_ISL_532040, EPI_ISL_532041, EPI_ISL_532043, EPI_ISL_532044, EPI_ISL_532045, EPI_ISL_532047, EPI_ISL_532048, EPI_ISL_532050, EPI_ISL_532052, EPI_ISL_532054, EPI_ISL_532056, EPI_ISL_532059, EPI_ISL_532061, EPI_ISL_532063, EPI_ISL_532065, EPI_ISL_532066, EPI_ISL_532067, EPI_ISL_532068, EPI_ISL_532072, EPI_ISL_532073, EPI_ISL_532074, EPI_ISL_532075, EPI_ISL_532079, EPI_ISL_532083, EPI_ISL_532084, EPI_ISL_532086, EPI_ISL_532088, EPI_ISL_532091, EPI_ISL_532092, EPI_ISL_532095, EPI_ISL_532097, EPI_ISL_532099, EPI_ISL_532101, EPI_ISL_532103, EPI_ISL_532104, EPI_ISL_532108, EPI_ISL_532109, EPI_ISL_532111, EPI_ISL_532112, EPI_ISL_532114, EPI_ISL_532115, EPI_ISL_532116, EPI_ISL_532117, EPI_ISL_532119, EPI_ISL_532120, EPI_ISL_532121, EPI_ISL_532122, EPI_ISL_532124, EPI_ISL_532126, EPI_ISL_532127, EPI_ISL_532132, EPI_ISL_532133, EPI_ISL_532134, EPI_ISL_532136, EPI_ISL_532137, EPI_ISL_532139, EPI_ISL_532140, EPI_ISL_532141, EPI_ISL_532143, EPI_ISL_532144, EPI_ISL_532146, EPI_ISL_532147, EPI_ISL_532148, EPI_ISL_532149, EPI_ISL_532150, EPI_ISL_532151, EPI_ISL_532152, EPI_ISL_532155, EPI_ISL_532156, EPI_ISL_532157, EPI_ISL_532158, EPI_ISL_532159, EPI_ISL_532160, EPI_ISL_532162, EPI_ISL_532164, EPI_ISL_532165, EPI_ISL_532166, EPI_ISL_532167, EPI_ISL_532170, EPI_ISL_532171, EPI_ISL_532172, EPI_ISL_532173, EPI_ISL_532175, EPI_ISL_532176, EPI_ISL_532180, EPI_ISL_532181, EPI_ISL_532182, EPI_ISL_532183, EPI_ISL_532184, EPI_ISL_532185, EPI_ISL_532188, EPI_ISL_532190, EPI_ISL_532191, EPI_ISL_532193, EPI_ISL_532198, EPI_ISL_532199, EPI_ISL_532201, EPI_ISL_532204, EPI_ISL_532205, EPI_ISL_532206, EPI_ISL_532209, EPI_ISL_532212, EPI_ISL_532213, EPI_ISL_532215, EPI_ISL_532216, EPI_ISL_532217, EPI_ISL_532220, EPI_ISL_532222, EPI_ISL_532223, EPI_ISL_532224, EPI_ISL_532225, EPI_ISL_532226, EPI_ISL_532227, EPI_ISL_532228, EPI_ISL_532230, EPI_ISL_532231, EPI_ISL_532232, EPI_ISL_532233, EPI_ISL_532234, EPI_ISL_532236, EPI_ISL_532237, EPI_ISL_532238, EPI_ISL_532239, EPI_ISL_532242, EPI_ISL_532243, EPI_ISL_532245, EPI_ISL_532246, EPI_ISL_532247, EPI_ISL_532248, EPI_ISL_532250, EPI_ISL_532251, EPI_ISL_532252, EPI_ISL_532253, EPI_ISL_532254, EPI_ISL_532255, EPI_ISL_532256, EPI_ISL_532257, EPI_ISL_532259, EPI_ISL_532260, EPI_ISL_532261, EPI_ISL_532262, EPI_ISL_532264, EPI_ISL_532265, EPI_ISL_532266, EPI_ISL_532267, EPI_ISL_532268, EPI_ISL_532269, EPI_ISL_532270, EPI_ISL_532273, EPI_ISL_532274, EPI_ISL_532277, EPI_ISL_532279, EPI_ISL_532280, EPI_ISL_532281, EPI_ISL_532282, EPI_ISL_532292, EPI_ISL_532295, EPI_ISL_532302, EPI_ISL_532309, EPI_ISL_532312, EPI_ISL_532313, EPI_ISL_532314, EPI_ISL_532323, EPI_ISL_532325, EPI_ISL_532329, EPI_ISL_532333, EPI_ISL_532336, EPI_ISL_532339, EPI_ISL_532343, EPI_ISL_532352, EPI_ISL_532361, EPI_ISL_532366, EPI_ISL_532369, EPI_ISL_532374, EPI_ISL_532386, EPI_ISL_532387, EPI_ISL_532395, EPI_ISL_532396, EPI_ISL_532400, EPI_ISL_532403, EPI_ISL_532412, EPI_ISL_532415, EPI_ISL_532421, EPI_ISL_532423, EPI_ISL_532424, EPI_ISL_532425, EPI_ISL_532426, EPI_ISL_532428, EPI_ISL_532433, EPI_ISL_532437, EPI_ISL_532439, EPI_ISL_532442, EPI_ISL_532449, EPI_ISL_532455, EPI_ISL_532467, EPI_ISL_532475, EPI_ISL_532477, EPI_ISL_532479, EPI_ISL_532483, EPI_ISL_532485, EPI_ISL_532487, EPI_ISL_532491, EPI_ISL_532494, EPI_ISL_532497, EPI_ISL_532505, EPI_ISL_532509, EPI_ISL_532527, EPI_ISL_532533, EPI_ISL_532536, EPI_ISL_532681, EPI_ISL_532732, EPI_ISL_532740, EPI_ISL_532760, EPI_ISL_532793, EPI_ISL_532828, EPI_ISL_532837, EPI_ISL_532840, EPI_ISL_532841, EPI_ISL_532845, EPI_ISL_532846, EPI_ISL_532847, EPI_ISL_532848, EPI_ISL_532849, EPI_ISL_532853, EPI_ISL_532855, EPI_ISL_532856, EPI_ISL_532859, EPI_ISL_532860, EPI_ISL_532861, EPI_ISL_532863, EPI_ISL_532866, EPI_ISL_532867, EPI_ISL_532868, EPI_ISL_532869, EPI_ISL_532870, EPI_ISL_532871, EPI_ISL_532877, EPI_ISL_532879, EPI_ISL_532882, EPI_ISL_532883, EPI_ISL_532884, EPI_ISL_532887, EPI_ISL_532888, EPI_ISL_532889, EPI_ISL_532890, EPI_ISL_532892, EPI_ISL_532893, EPI_ISL_532895, EPI_ISL_532898, EPI_ISL_532900, EPI_ISL_532901, EPI_ISL_532904, EPI_ISL_532908, EPI_ISL_532911, EPI_ISL_532912, EPI_ISL_532914, EPI_ISL_532917, EPI_ISL_532918, EPI_ISL_532923, EPI_ISL_532927, EPI_ISL_532928, EPI_ISL_532930, EPI_ISL_532931, EPI_ISL_532934, EPI_ISL_532935, EPI_ISL_532936, EPI_ISL_532938, EPI_ISL_532939, EPI_ISL_532941, EPI_ISL_532942, EPI_ISL_532945, EPI_ISL_532947, EPI_ISL_532952, EPI_ISL_532953, EPI_ISL_532954, EPI_ISL_532955, EPI_ISL_532956, EPI_ISL_532957, EPI_ISL_532958, EPI_ISL_532959, EPI_ISL_532960, EPI_ISL_532961, EPI_ISL_532963, EPI_ISL_532964, EPI_ISL_532965, EPI_ISL_532966, EPI_ISL_532967, EPI_ISL_532968, EPI_ISL_532969, EPI_ISL_532970, EPI_ISL_532971, EPI_ISL_532972, EPI_ISL_532973, EPI_ISL_532974, EPI_ISL_532975, EPI_ISL_532977, EPI_ISL_532980, EPI_ISL_532981, EPI_ISL_532986, EPI_ISL_532987, EPI_ISL_532989, EPI_ISL_532991, EPI_ISL_532993, EPI_ISL_532995, EPI_ISL_532996, EPI_ISL_532998, EPI_ISL_532999, EPI_ISL_533002, EPI_ISL_533003, EPI_ISL_533004, EPI_ISL_533005, EPI_ISL_533006, EPI_ISL_533008, EPI_ISL_533009, EPI_ISL_533010, EPI_ISL_533011, EPI_ISL_533012, EPI_ISL_533014, EPI_ISL_533018, EPI_ISL_533019, EPI_ISL_533020, EPI_ISL_533022, EPI_ISL_533023, EPI_ISL_533024, EPI_ISL_533028, EPI_ISL_533031, EPI_ISL_533032, EPI_ISL_533033, EPI_ISL_533034, EPI_ISL_533035, EPI_ISL_533036, EPI_ISL_533037, EPI_ISL_533039, EPI_ISL_533040, EPI_ISL_533041, EPI_ISL_533042, EPI_ISL_533043, EPI_ISL_533044, EPI_ISL_533045, EPI_ISL_533047, EPI_ISL_533048, EPI_ISL_533050, EPI_ISL_533051, EPI_ISL_533053, EPI_ISL_533055, EPI_ISL_533056, EPI_ISL_533057, EPI_ISL_533058, EPI_ISL_533059, EPI_ISL_533060, EPI_ISL_533061, EPI_ISL_533062, EPI_ISL_533063, EPI_ISL_533064, EPI_ISL_533068, EPI_ISL_533069, EPI_ISL_533071, EPI_ISL_533073, EPI_ISL_533075, EPI_ISL_533076, EPI_ISL_533078, EPI_ISL_533080, EPI_ISL_533081, EPI_ISL_533083, EPI_ISL_533084, EPI_ISL_533085, EPI_ISL_533087, EPI_ISL_533088, EPI_ISL_533090, EPI_ISL_533092, EPI_ISL_533093, EPI_ISL_533094, EPI_ISL_533095, EPI_ISL_533096, EPI_ISL_533098, EPI_ISL_533099, EPI_ISL_533100, EPI_ISL_533102, EPI_ISL_533103, EPI_ISL_533104, EPI_ISL_533105, EPI_ISL_533106, EPI_ISL_533108, EPI_ISL_533109, EPI_ISL_533110, EPI_ISL_533111, EPI_ISL_533112, EPI_ISL_533113, EPI_ISL_533114, EPI_ISL_533115, EPI_ISL_533116, EPI_ISL_533118, EPI_ISL_533119, EPI_ISL_533120, EPI_ISL_533121, EPI_ISL_533122, EPI_ISL_533123, EPI_ISL_533124, EPI_ISL_533125, EPI_ISL_533126, EPI_ISL_533127, EPI_ISL_533128, EPI_ISL_533129, EPI_ISL_533131, EPI_ISL_533134, EPI_ISL_533135, EPI_ISL_533136, EPI_ISL_533137, EPI_ISL_533140, EPI_ISL_533142, EPI_ISL_533143, EPI_ISL_533145, EPI_ISL_533146, EPI_ISL_533147, EPI_ISL_533148, EPI_ISL_533149, EPI_ISL_533150, EPI_ISL_533152, EPI_ISL_533153, EPI_ISL_533154, EPI_ISL_533155, EPI_ISL_533156, EPI_ISL_533157, EPI_ISL_533158, EPI_ISL_533159, EPI_ISL_533160, EPI_ISL_533162, EPI_ISL_533163, EPI_ISL_533164, EPI_ISL_533165, EPI_ISL_533167, EPI_ISL_533168, EPI_ISL_533170, EPI_ISL_533171, EPI_ISL_533172, EPI_ISL_533173, EPI_ISL_533174, EPI_ISL_533175, EPI_ISL_533176, EPI_ISL_533178, EPI_ISL_533179, EPI_ISL_533180, EPI_ISL_533181, EPI_ISL_533182, EPI_ISL_533183, EPI_ISL_533184, EPI_ISL_533185, EPI_ISL_533186, EPI_ISL_533187, EPI_ISL_533189, EPI_ISL_533191, EPI_ISL_533192, EPI_ISL_533193, EPI_ISL_533194, EPI_ISL_533195, EPI_ISL_533196, EPI_ISL_533197, EPI_ISL_533198, EPI_ISL_533199, EPI_ISL_533200, EPI_ISL_533203, EPI_ISL_533205, EPI_ISL_533206, EPI_ISL_533208, EPI_ISL_533209, EPI_ISL_533210, EPI_ISL_533211, EPI_ISL_533212, EPI_ISL_533213, EPI_ISL_533214, EPI_ISL_533215, EPI_ISL_533216, EPI_ISL_533217, EPI_ISL_533218, EPI_ISL_533219, EPI_ISL_533220, EPI_ISL_533221, EPI_ISL_533223, EPI_ISL_533224, EPI_ISL_533225, EPI_ISL_533226, EPI_ISL_533227, EPI_ISL_533229, EPI_ISL_533233, EPI_ISL_533235, EPI_ISL_533238, EPI_ISL_533239, EPI_ISL_533242, EPI_ISL_533243, EPI_ISL_533245, EPI_ISL_533246, EPI_ISL_533247, EPI_ISL_533248, EPI_ISL_533249, EPI_ISL_533253, EPI_ISL_533254, EPI_ISL_533255, EPI_ISL_533257, EPI_ISL_533258, EPI_ISL_533259, EPI_ISL_533260, EPI_ISL_533261, EPI_ISL_533263, EPI_ISL_533264, EPI_ISL_533265, EPI_ISL_533268, EPI_ISL_533269, EPI_ISL_533273, EPI_ISL_533274, EPI_ISL_533275, EPI_ISL_533276, EPI_ISL_533277, EPI_ISL_533278, EPI_ISL_533279, EPI_ISL_533280, EPI_ISL_533282, EPI_ISL_533284, EPI_ISL_533285, EPI_ISL_533286, EPI_ISL_533287, EPI_ISL_533288, EPI_ISL_533290, EPI_ISL_533292, EPI_ISL_533293, EPI_ISL_533294, EPI_ISL_533295, EPI_ISL_533296, EPI_ISL_533297, EPI_ISL_533299, EPI_ISL_533300, EPI_ISL_533301, EPI_ISL_533302, EPI_ISL_533303, EPI_ISL_533304, EPI_ISL_533306, EPI_ISL_533308, EPI_ISL_533309, EPI_ISL_533310, EPI_ISL_533313, EPI_ISL_533314, EPI_ISL_533315, EPI_ISL_533318, EPI_ISL_533319, EPI_ISL_533320, EPI_ISL_533321, EPI_ISL_533322, EPI_ISL_533323, EPI_ISL_533324, EPI_ISL_533325, EPI_ISL_533326, EPI_ISL_533327, EPI_ISL_533329, EPI_ISL_533330, EPI_ISL_533332, EPI_ISL_533333, EPI_ISL_533334, EPI_ISL_533335, EPI_ISL_533336, EPI_ISL_533337, EPI_ISL_533338, EPI_ISL_533339, EPI_ISL_533340, EPI_ISL_533341, EPI_ISL_533342, EPI_ISL_533344, EPI_ISL_533345, EPI_ISL_533347, EPI_ISL_533348, EPI_ISL_533349, EPI_ISL_533352, EPI_ISL_533353, EPI_ISL_533354, EPI_ISL_533356, EPI_ISL_533359, EPI_ISL_533360, EPI_ISL_533361, EPI_ISL_533362, EPI_ISL_533364, EPI_ISL_533366, EPI_ISL_533367, EPI_ISL_533368, EPI_ISL_533369, EPI_ISL_533370, EPI_ISL_533371, EPI_ISL_533372, EPI_ISL_533373, EPI_ISL_533374, EPI_ISL_533375, EPI_ISL_533377, EPI_ISL_533378, EPI_ISL_533379, EPI_ISL_533381, EPI_ISL_533382, EPI_ISL_533383, EPI_ISL_533384, EPI_ISL_533385, EPI_ISL_533386, EPI_ISL_533387, EPI_ISL_533388, EPI_ISL_533389, EPI_ISL_533390, EPI_ISL_533391, EPI_ISL_533392, EPI_ISL_533393, EPI_ISL_533395, EPI_ISL_533397, EPI_ISL_533398, EPI_ISL_533399

| | | | |
|---|---|---|---|
| see above | Lighthouse Lab in Glasgow | Wellcome Sanger Institute for the COVID-19 Genomics UK (COG-UK) consortium | Harper VanSteenhouse, Yumi Kasai, David Gray, Carol Clugston, Anna Dominiczak and Alex Alderton, Roberto Amato, Sonia Goncalves, Ewan Harrison, David K. Jackson, Ian Johnston, Dominic Kwiatkowski, Cordelia Langford, John Sillitoe |
| EPI_ISL_533400 | Lighthouse Lab in Glasgow | Wellcome Sanger Institute for the COVID-19 Genomics UK (COG-UK) consortium | Harper VanSteenhouse, Yumi Kasai, David Gray, Carol Clugston, Anna Dominiczak and Alex Alderton, Roberto Amato, Sonia Goncalves, Ewan Harrison, David K. Jackson, Ian Johnston, Dominic Kwiatkowski, Cordelia Langford, John Sillitoe on behalf of the Wellcome Sanger Institute COVID-19 Surveillance Team |

EPI_ISL_533401, EPI_ISL_533402, EPI_ISL_533404, EPI_ISL_533405, EPI_ISL_533407, EPI_ISL_533408, EPI_ISL_533409, EPI_ISL_533410, EPI_ISL_533412, EPI_ISL_533414, EPI_ISL_533415, EPI_ISL_533416, EPI_ISL_533420, EPI_ISL_533422, EPI_ISL_533423, EPI_ISL_533424, EPI_ISL_533425, EPI_ISL_533426, EPI_ISL_533428, EPI_ISL_533429, EPI_ISL_533430, EPI_ISL_533431

| | | | |
|---|---|---|---|
| see above | Lighthouse Lab in Glasgow | Wellcome Sanger Institute for the COVID-19 Genomics UK (COG-UK) consortium | Harper VanSteenhouse, Yumi Kasai, David Gray, Carol Clugston, Anna Dominiczak and Alex Alderton, Roberto Amato, Sonia Goncalves, Ewan Harrison, David K. Jackson, Ian Johnston, Dominic Kwiatkowski, Cordelia Langford, John Sillitoe |
| EPI_ISL_533432, EPI_ISL_533433, EPI_ISL_533434 | Lighthouse Lab in Glasgow | Wellcome Sanger Institute for the COVID-19 Genomics UK (COG-UK) consortium | Harper VanSteenhouse, Yumi Kasai, David Gray, Carol Clugston, Anna Dominiczak and Alex Alderton, Roberto Amato, Sonia Goncalves, Ewan Harrison, David K. Jackson, Ian Johnston, Dominic Kwiatkowski, Cordelia Langford, John Sillitoe on behalf of the Wellcome Sanger Institute COVID-19 Surveillance Team |
| EPI_ISL_534199 | E. Gulbja Laboratorija | Latvian Biomedical Research and Study Centre | Ivars Silamielis, Jnis Pjalkovskis, Kaspars Megnis, Monta Ustinova, ikita Zrelovs, Vita Rovte, Mikus Gavars, Dmitrijs Perminovs, Uga Dumpis, Jnis Kloviš |
| EPI_ISL_534200, EPI_ISL_534201, EPI_ISL_534202 | Centrl laboratorija | Latvian Biomedical Research and Study Centre | Ivars Silamielis, Jnis Pjalkovskis, Kaspars Megnis, Monta Ustinova, ikita Zrelovs, Vita Rovte, Stella Lapia, Jana Oste, Marta Priedte, Uga Dumpis, Jnis Kloviš |
| EPI_ISL_534203 | E. Gulbja Laboratorija | Latvian Biomedical Research and Study Centre | Ivars Silamielis, Jnis Pjalkovskis, Kaspars Megnis, Monta Ustinova, ikita Zrelovs, Vita Rovte, Mikus Gavars, Dmitrijs Perminovs, Uga Dumpis, Jnis Kloviš |
| EPI_ISL_534204, EPI_ISL_534205 | Latvijas Infektoloijas centrs | Latvian Biomedical Research and Study Centre | Ivars Silamielis, Jnis Pjalkovskis, Kaspars Megnis, Monta Ustinova, ikita Zrelovs, Vita Rovte, Jeena Storoženko, Tatjana Kolupajeva, Oksana Savicka, Uga Dumpis, Jnis Kloviš |
| EPI_ISL_534206, EPI_ISL_534207, EPI_ISL_534208 | Centrl laboratorija | Latvian Biomedical Research and Study Centre | Ivars Silamielis, Jnis Pjalkovskis, Kaspars Megnis, Monta Ustinova, ikita Zrelovs, Vita Rovte, Stella Lapia, Jana Oste, Marta Priedte, Uga Dumpis, Jnis Kloviš |
| EPI_ISL_534209 | Latvijas Infektoloijas centrs | Latvian Biomedical Research and Study Centre | Ivars Silamielis, Jnis Pjalkovskis, Kaspars Megnis, Monta Ustinova, ikita Zrelovs, Vita Rovte, Jeena Storoženko, Tatjana Kolupajeva, Oksana Savicka, Uga Dumpis, Jnis Kloviš |
| EPI_ISL_534210, EPI_ISL_534211 | Centrl laboratorija | Latvian Biomedical Research and Study Centre | Ivars Silamielis, Jnis Pjalkovskis, Kaspars Megnis, Monta Ustinova, ikita Zrelovs, Vita Rovte, Stella Lapia, Jana Oste, Marta Priedte, Uga Dumpis, Jnis Kloviš |
| EPI_ISL_534212, EPI_ISL_534213, EPI_ISL_534214, EPI_ISL_534215, EPI_ISL_534216, EPI_ISL_534218, EPI_ISL_534219 | E. Gulbja Laboratorija | Latvian Biomedical Research and Study Centre | Ivars Silamielis, Jnis Pjalkovskis, Kaspars Megnis, Monta Ustinova, ikita Zrelovs, Vita Rovte, Mikus Gavars, Dmitrijs Perminovs, Uga Dumpis, Jnis Kloviš |
| EPI_ISL_534220 | Centrl laboratorija | Latvian Biomedical Research and Study Centre | Ivars Silamielis, Jnis Pjalkovskis, Kaspars Megnis, Monta Ustinova, ikita Zrelovs, Vita Rovte, Stella Lapia, Jana Oste, Marta Priedte, Uga Dumpis, Jnis Kloviš |
| EPI_ISL_534221 | E. Gulbja Laboratorija | Latvian Biomedical Research and Study Centre | Ivars Silamielis, Jnis Pjalkovskis, Kaspars Megnis, Monta Ustinova, ikita Zrelovs, Vita Rovte, Mikus Gavars, Dmitrijs Perminovs, Uga Dumpis, Jnis Kloviš |
| EPI_ISL_534222, EPI_ISL_534223 | Centrl laboratorija | Latvian Biomedical Research and Study Centre | Ivars Silamielis, Jnis Pjalkovskis, Kaspars Megnis, Monta Ustinova, ikita Zrelovs, Vita Rovte, Stella Lapia, Jana Oste, Marta Priedte, Uga Dumpis, Jnis Kloviš |
| EPI_ISL_537043 | Lighthouse Lab in Glasgow | Wellcome Sanger Institute for the COVID-19 Genomics UK | Harper VanSteenhouse, Yumi Kasai, David Gray, Carol Clugston, Anna Dominiczak and Alex Alderton, Roberto Amato, Sonia Goncalves, Ewan Harrison, |

| | | | |
|---|---|---|---|
| | | (COG-UK) consortium | David K. Jackson, Ian Johnston, Dominic Kwiatkowski, Cordelia Langford, John Sillitoe on behalf of the Wellcome Sanger Institute COVID-19 Surveillance Team |
| EPI_ISL_539496 | Hospital Nostra Senyora de Meritxell | Instituto de Salud Carlos III | Iglesias-Caballero, M. Molinero Calamita, M. González-Esguevillas, M. Camarero, S. Pozo, F. Casas, I. Jiménez, P. Jiménez, M. Zaballos, A. Monzón, S. Varona, S. Juliá, M. Cuesta, I, F. Fernández |
| EPI_ISL_539497, EPI_ISL_539498 | Hospital Virgen de las Nieves | Instituto de Salud Carlos III | Iglesias-Caballero, M. Molinero Calamita, M. González-Esguevillas, M. Camarero, S. Pozo, F. Casas, I. Jiménez, P. Jiménez, M. Zaballos, A. Monzón, S. Varona, S. Juliá, M. Cuesta, I, J. Lepe |
| EPI_ISL_539499 | Hospital Costa del Sol | Instituto de Salud Carlos III | Iglesias-Caballero, M. Molinero Calamita, M. González-Esguevillas, M. Camarero, S. Pozo, F. Casas, I. Jiménez, P. Jiménez, M. Zaballos, A. Monzón, S. Varona, S. Juliá, M. Cuesta, I, F. Fernández |
| EPI_ISL_539500, EPI_ISL_539501, EPI_ISL_539502 | Hospital Clínico Universitario Lozano Blesa | Instituto de Salud Carlos III | Iglesias-Caballero, M. Molinero Calamita, M. González-Esguevillas, M. Camarero, S. Pozo, F. Casas, I. Jiménez, P. Jiménez, M. Zaballos, A. Monzón, S. Varona, S. Juliá, M. Cuesta, I, R. Benito |
| EPI_ISL_539503 | Hospital Universitario Miguel Servet | Instituto de Salud Carlos III | Iglesias-Caballero, M. Molinero Calamita, M. González-Esguevillas, M. Camarero, S. Pozo, F. Casas, I. Jiménez, P. Jiménez, M. Zaballos, A. Monzón, S. Varona, S. Juliá, M. Cuesta, I, A. Rezusta |
| EPI_ISL_539504 | Hospital Universitario Miguel Servet | Instituto de Salud Carlos III | Iglesias-Caballero, M. Molinero Calamita, M. González-Esguevillas, M. Camarero, S. Pozo, F. Casas, I. Jiménez, P. Jiménez, M. Zaballos, A. Monzón, S. Varona, S. Juliá, M. Cuesta, I, R. Benito |
| EPI_ISL_539505, EPI_ISL_539506, EPI_ISL_539507, EPI_ISL_539508 | Hospital Universitario Miguel Servet | Instituto de Salud Carlos III | Iglesias-Caballero, M. Molinero Calamita, M. González-Esguevillas, M. Camarero, S. Pozo, F. Casas, I. Jiménez, P. Jiménez, M. Zaballos, A. Monzón, S. Varona, S. Juliá, M. Cuesta, I, A. Rezusta |
| EPI_ISL_539509, EPI_ISL_539510, EPI_ISL_539511, EPI_ISL_539512 | Hospital Clínico Universitario Lozano Blesa | Instituto de Salud Carlos III | Iglesias-Caballero, M. Molinero Calamita, M. González-Esguevillas, M. Camarero, S. Pozo, F. Casas, I. Jiménez, P. Jiménez, M. Zaballos, A. Monzón, S. Varona, S. Juliá, M. Cuesta, I, R. Benito |
| EPI_ISL_539513, EPI_ISL_539514, EPI_ISL_539515, EPI_ISL_539516, EPI_ISL_539517, EPI_ISL_539518, EPI_ISL_539519 | Hospital Universitario Miguel Servet | Instituto de Salud Carlos III | Iglesias-Caballero, M. Molinero Calamita, M. González-Esguevillas, M. Camarero, S. Pozo, F. Casas, I. Jiménez, P. Jiménez, M. Zaballos, A. Monzón, S. Varona, S. Juliá, M. Cuesta, I, A. Rezusta |
| EPI_ISL_539520, EPI_ISL_539521 | Hospital Universitario de Ceuta | Instituto de Salud Carlos III | Iglesias-Caballero, M. Molinero Calamita, M. González-Esguevillas, M. Camarero, S. Pozo, F. Casas, I. Jiménez, P. Jiménez, M. Zaballos, A. Monzón, S. Varona, S. Juliá, M. Cuesta, I, J. López |
| EPI_ISL_539522 | Hospital Universitario de Ceuta | Instituto de Salud Carlos III | Iglesias-Caballero, M. Molinero Calamita, M. González-Esguevillas, M. Camarero, S. Pozo, F. Casas, I. Jiménez, P. Jiménez, M. Zaballos, A. Monzón, S. Varona, S. Juliá, M. Cuesta, I, G. Sánchez |
| EPI_ISL_539523 | Hospital General de Segovia | Instituto de Salud Carlos III | Iglesias-Caballero, M. Molinero Calamita, M. González-Esguevillas, M. Camarero, S. Pozo, F. Casas, I. Jiménez, P. Jiménez, M. Zaballos, A. Monzón, S. Varona, S. Juliá, M. Cuesta, I, S. Hernando |
| EPI_ISL_539524 | Gerencia de Asistencia Sanitaria de Soria | Instituto de Salud Carlos III | Iglesias-Caballero, M. Molinero Calamita, M. González-Esguevillas, M. Camarero, S. Pozo, F. Casas, I. Jiménez, P. Jiménez, M. Zaballos, A. Monzón, S. Varona, S. Juliá, M. Cuesta, I, C. Aldea |
| EPI_ISL_539525 | Hospital Nuestra Señora de Sonsoles | Instituto de Salud Carlos III | Iglesias-Caballero, M. Molinero Calamita, M. González-Esguevillas, M. Camarero, S. Pozo, F. Casas, I. Jiménez, P. Jiménez, M. Zaballos, A. Monzón, S. Varona, S. Juliá, M. Cuesta, I, A. San Pedro |
| EPI_ISL_539526, EPI_ISL_539527, EPI_ISL_539528, EPI_ISL_539529, EPI_ISL_539530 | Consejeria de Sanidad y Asuntos Sociales | Instituto de Salud Carlos III | Iglesias-Caballero, M. Molinero Calamita, M. González-Esguevillas, M. Camarero, S. Pozo, F. Casas, I. Jiménez, P. Jiménez, M. Zaballos, A. Monzón, S. Varona, S. Juliá, M. Cuesta, I, G. Gutiérrez |
| EPI_ISL_539531 | C.H.U Nuestra Señora de Candelaria | Instituto de Salud Carlos III | Iglesias-Caballero, M. Molinero Calamita, M. González-Esguevillas, M. Camarero, S. Pozo, F. Casas, I. Jiménez, P. Jiménez, M. Zaballos, A. Monzón, S. Varona, S. Juliá, M. Cuesta, I, O. Díez |
| EPI_ISL_539532 | Hospital Universitario de Canarias | Instituto de Salud Carlos III | Iglesias-Caballero, M. Molinero Calamita, M. González-Esguevillas, M. Camarero, S. Pozo, F. Casas, I. Jiménez, P. Jiménez, M. Zaballos, A. Monzón, S. Varona, S. Juliá, M. Cuesta, I, B. Castro |
| EPI_ISL_539533, EPI_ISL_539534, EPI_ISL_539535, EPI_ISL_539536, EPI_ISL_539537, EPI_ISL_539538, EPI_ISL_539539, EPI_ISL_539540, EPI_ISL_539541, EPI_ISL_539542, EPI_ISL_539543, EPI_ISL_539544, EPI_ISL_539545, EPI_ISL_539546, EPI_ISL_539547, EPI_ISL_539548, EPI_ISL_539549, EPI_ISL_539550, EPI_ISL_539551, EPI_ISL_539552, EPI_ISL_539553, EPI_ISL_539554, EPI_ISL_539555, EPI_ISL_539556 | | | |
| see above | Hospital Clínic | Instituto de Salud Carlos III | Iglesias-Caballero, M. Molinero Calamita, M. González-Esguevillas, M. Camarero, S. Pozo, F. Casas, I. Jiménez, P. Jiménez, M. Zaballos, A. Monzón, S. Varona, S. Juliá, M. Cuesta, I, M.A Marcos |
| EPI_ISL_539557 | Gerencia del área de salud de Badajoz, Llerena y Zafra | Instituto de Salud Carlos III | Iglesias-Caballero, M. Molinero Calamita, M. González-Esguevillas, M. Camarero, S. Pozo, F. Casas, I. Jiménez, P. Jiménez, M. Zaballos, A. Monzón, S. Varona, S. Juliá, M. Cuesta, I, C. Pazos |
| EPI_ISL_539558 | Hospital San Pedro de Alcántara | Instituto de Salud Carlos III | Iglesias-Caballero, M. Molinero Calamita, M. González-Esguevillas, M. Camarero, S. Pozo, F. Casas, I. Jiménez, P. Jiménez, M. Zaballos, A. Monzón, S. Varona, S. Juliá, M. Cuesta, I, E. Cerro |
| EPI_ISL_539559 | Hospital Campo Arañuelo | Instituto de Salud Carlos III | Iglesias-Caballero, M. Molinero Calamita, M. González-Esguevillas, M. Camarero, S. Pozo, F. Casas, I. Jiménez, P. Jiménez, M. Zaballos, A. Monzón, S. Varona, S. Juliá, M. Cuesta, I, J. López |
| EPI_ISL_539560 | Hospital Campo Arañuelo | Instituto de Salud Carlos III | Iglesias-Caballero, M. Molinero Calamita, M. González-Esguevillas, M. Camarero, S. Pozo, F. Casas, I. Jiménez, P. Jiménez, M. Zaballos, A. Monzón, S. Varona, S. Juliá, M. Cuesta, I, G. Rodríguez |
| EPI_ISL_539561 | Hospital San Pedro de Alcántara | Instituto de Salud Carlos III | Iglesias-Caballero, M. Molinero Calamita, M. González-Esguevillas, M. Camarero, S. Pozo, F. Casas, I. Jiménez, P. Jiménez, M. Zaballos, A. Monzón, S. Varona, S. Juliá, M. Cuesta, I, J. López |
| EPI_ISL_539562 | Hospital San Pedro de Alcántara | Instituto de Salud Carlos III | Iglesias-Caballero, M. Molinero Calamita, M. González-Esguevillas, M. Camarero, S. Pozo, F. Casas, I. Jiménez, P. Jiménez, M. Zaballos, A. Monzón, S. Varona, S. Juliá, M. Cuesta, I, M.A Cañizares |
| EPI_ISL_539563 | Complejo Hospitalario Universitario La Coruna | Instituto de Salud Carlos III | Iglesias-Caballero, M. Molinero Calamita, M. González-Esguevillas, M. Camarero, S. Pozo, F. Casas, I. Jiménez, P. Jiménez, M. Zaballos, A. Monzón, S. Varona, S. Juliá, M. Cuesta, I, J. López |
| EPI_ISL_539564 | Complejo Hospitalario Universitario La Coruna | Instituto de Salud Carlos III | Iglesias-Caballero, M. Molinero Calamita, M. González-Esguevillas, M. Camarero, S. Pozo, F. Casas, I. Jiménez, P. Jiménez, M. Zaballos, A. Monzón, S. Varona, S. Juliá, M. Cuesta, I, M.I Paz |
| EPI_ISL_539565 | Xerencia de Xestión Integrada de Pontevedra e o Salnés | Instituto de Salud Carlos III | Iglesias-Caballero, M. Molinero Calamita, M. González-Esguevillas, M. Camarero, S. Pozo, F. Casas, I. Jiménez, P. Jiménez, M. Zaballos, A. Monzón, S. Varona, S. Juliá, M. Cuesta, I, J. López |
| EPI_ISL_539566 | Xerencia de Xestión Integrada de Pontevedra e o Salnés | Instituto de Salud Carlos III | Iglesias-Caballero, M. Molinero Calamita, M. González-Esguevillas, M. Camarero, S. Pozo, F. Casas, I. Jiménez, P. Jiménez, M. Zaballos, A. Monzón, S. Varona, S. Juliá, M. Cuesta, I, I. Pérez |
| EPI_ISL_539567 | Hospital Comarcal de Melilla | Instituto de Salud Carlos III | Iglesias-Caballero, M. Molinero Calamita, M. González-Esguevillas, M. Camarero, S. Pozo, F. Casas, I. Jiménez, P. Jiménez, M. Zaballos, A. Monzón, S. Varona, S. Juliá, M. Cuesta, I, J. López |
| EPI_ISL_539568 | Hospital Comarcal de Melilla | Instituto de Salud Carlos III | Iglesias-Caballero, M. Molinero Calamita, M. González-Esguevillas, M. Camarero, S. Pozo, F. Casas, I. Jiménez, P. Jiménez, M. Zaballos, A. Monzón, S. Varona, S. Juliá, M. Cuesta, I, C. Ezpeleta |
| EPI_ISL_539569, EPI_ISL_539570, EPI_ISL_539571, EPI_ISL_539572 | Complejo Hospitalario de Navarra | Instituto de Salud Carlos III | Iglesias-Caballero, M. Molinero Calamita, M. González-Esguevillas, M. Camarero, S. Pozo, F. Casas, I. Jiménez, P. Jiménez, M. Zaballos, A. Monzón, S. Varona, S. Juliá, M. Cuesta, I, J. López |
| EPI_ISL_539573, EPI_ISL_539574, EPI_ISL_539575, EPI_ISL_539576 | Centre de Recherches Medicales de Lambarene (CERMEL) | Department of Emerging Infectious Diseases, Institute of Tropical Medicine, Nagasaki University | Haruka Abe, Yuri Ushijima, Rodrigue Bikangui, Akim A. Adegnika, Bertrand Lell, Jiro Yasuda |

| | | | |
|---|---|---|---|
| EPI_ISL_560554, EPI_ISL_560555, EPI_ISL_560556, EPI_ISL_560557, EPI_ISL_560558, EPI_ISL_560559, EPI_ISL_560560, EPI_ISL_560561, EPI_ISL_560562, EPI_ISL_560563, EPI_ISL_560564, EPI_ISL_560565, EPI_ISL_560566 | | | |
| see above | Alaska State Virology Laboratory | Alaska State Virology Laboratory | Jack Chen, Ph.D. |
| EPI_ISL_565909, EPI_ISL_565910, EPI_ISL_565911, EPI_ISL_565912, EPI_ISL_565913, EPI_ISL_565914, EPI_ISL_565915, EPI_ISL_565916, EPI_ISL_565917, EPI_ISL_565918, EPI_ISL_565919 | | | |
| see above | Servicio de Microbiología. HRU de Málaga. Servicio Andaluz de Salud | SeqCOVID-SPAIN consortium/IBV(CSIC) | Inmaculada de Toro Peinado, Ma Concepción Mediavilla Gradolph, Begoña Palop Borrás and SeqCOVID-SPAIN consortium |
| EPI_ISL_566110, EPI_ISL_566111, EPI_ISL_566112, EPI_ISL_566113, EPI_ISL_566115, EPI_ISL_566116, EPI_ISL_566117, EPI_ISL_566118, EPI_ISL_566120, EPI_ISL_566121, EPI_ISL_566122, EPI_ISL_566123, EPI_ISL_566124, EPI_ISL_566125, EPI_ISL_566126, EPI_ISL_566127, EPI_ISL_566128, EPI_ISL_566130, EPI_ISL_566131, EPI_ISL_566132, EPI_ISL_566133, EPI_ISL_566134, EPI_ISL_566135, EPI_ISL_566136, EPI_ISL_566138, EPI_ISL_566139, EPI_ISL_566140, EPI_ISL_566141, EPI_ISL_566143, EPI_ISL_566144, EPI_ISL_566147, EPI_ISL_566148, EPI_ISL_566149, EPI_ISL_566150, EPI_ISL_566151, EPI_ISL_566152, EPI_ISL_566153, EPI_ISL_566154, EPI_ISL_566155, EPI_ISL_566156, EPI_ISL_566157, EPI_ISL_566158, EPI_ISL_566159, EPI_ISL_566160, EPI_ISL_566161, EPI_ISL_566162, EPI_ISL_566163, EPI_ISL_566164, EPI_ISL_566165, EPI_ISL_566166, EPI_ISL_566167, EPI_ISL_566168, EPI_ISL_566170, EPI_ISL_566171, EPI_ISL_566172, EPI_ISL_566173, EPI_ISL_566174, EPI_ISL_566175, EPI_ISL_566176, EPI_ISL_566177, EPI_ISL_566178, EPI_ISL_566179, EPI_ISL_566180, EPI_ISL_566181, EPI_ISL_566183, EPI_ISL_566184, EPI_ISL_566185, EPI_ISL_566186, EPI_ISL_566187, EPI_ISL_566188, EPI_ISL_566189, EPI_ISL_566190, EPI_ISL_566191, EPI_ISL_566192, EPI_ISL_566193, EPI_ISL_566195, EPI_ISL_566196, EPI_ISL_566197, EPI_ISL_566198, EPI_ISL_566199, EPI_ISL_566200, EPI_ISL_566201, EPI_ISL_566202, EPI_ISL_566203, EPI_ISL_566204, EPI_ISL_566205, EPI_ISL_566206, EPI_ISL_566208, EPI_ISL_566209, EPI_ISL_566211, EPI_ISL_566212, EPI_ISL_566213, EPI_ISL_566214, EPI_ISL_566215, EPI_ISL_566216, EPI_ISL_566217, EPI_ISL_566218, EPI_ISL_566219, EPI_ISL_566220, EPI_ISL_566221, EPI_ISL_566222, EPI_ISL_566223, EPI_ISL_566224, EPI_ISL_566225, EPI_ISL_566227, EPI_ISL_566228, EPI_ISL_566229, EPI_ISL_566230, EPI_ISL_566231, EPI_ISL_566233, EPI_ISL_566234, EPI_ISL_566235, EPI_ISL_566236, EPI_ISL_566237, EPI_ISL_566239, EPI_ISL_566240, EPI_ISL_566241, EPI_ISL_566243, EPI_ISL_566244, EPI_ISL_566245, EPI_ISL_566246, EPI_ISL_566247, EPI_ISL_566248, EPI_ISL_566250, EPI_ISL_566251, EPI_ISL_566252, EPI_ISL_566254, EPI_ISL_566255, EPI_ISL_566256, EPI_ISL_566257, EPI_ISL_566258, EPI_ISL_566259, EPI_ISL_566261, EPI_ISL_566262, EPI_ISL_566263, EPI_ISL_566264, EPI_ISL_566265, EPI_ISL_566266, EPI_ISL_566267, EPI_ISL_566268, EPI_ISL_566269, EPI_ISL_566270, EPI_ISL_566271, EPI_ISL_566272, EPI_ISL_566273, EPI_ISL_566275, EPI_ISL_566276, EPI_ISL_566277, EPI_ISL_566278, EPI_ISL_566279, EPI_ISL_566280, EPI_ISL_566281, EPI_ISL_566282, EPI_ISL_566283, EPI_ISL_566284, EPI_ISL_566286, EPI_ISL_566287, EPI_ISL_566288, EPI_ISL_566289, EPI_ISL_566290, EPI_ISL_566291, EPI_ISL_566293, EPI_ISL_566294, EPI_ISL_566295, EPI_ISL_566296, EPI_ISL_566297, EPI_ISL_566298, EPI_ISL_566300, EPI_ISL_566301, EPI_ISL_566302, EPI_ISL_566303, EPI_ISL_566304, EPI_ISL_566305, EPI_ISL_566306, EPI_ISL_566307, EPI_ISL_566308, EPI_ISL_566309, EPI_ISL_566310, EPI_ISL_566311, EPI_ISL_566312, EPI_ISL_566313, EPI_ISL_566314, EPI_ISL_566315, EPI_ISL_566316, EPI_ISL_566318, EPI_ISL_566320, EPI_ISL_566321, EPI_ISL_566323, EPI_ISL_566324, EPI_ISL_566325, EPI_ISL_566326, EPI_ISL_566327, EPI_ISL_566328, EPI_ISL_566329, EPI_ISL_566330, EPI_ISL_566331, EPI_ISL_566332, EPI_ISL_566333, EPI_ISL_566334, EPI_ISL_566335, EPI_ISL_566336, EPI_ISL_566337, EPI_ISL_566338, EPI_ISL_566339, EPI_ISL_566340, EPI_ISL_566341, EPI_ISL_566342, EPI_ISL_566343, EPI_ISL_566344, EPI_ISL_566345, EPI_ISL_566346, EPI_ISL_566347, EPI_ISL_566349, EPI_ISL_566350, EPI_ISL_566351, EPI_ISL_566352, EPI_ISL_566353, EPI_ISL_566354, EPI_ISL_566355, EPI_ISL_566356, EPI_ISL_566357, EPI_ISL_566358, EPI_ISL_566359, EPI_ISL_566360, EPI_ISL_566361, EPI_ISL_566363, EPI_ISL_566364, EPI_ISL_566365, EPI_ISL_566366, EPI_ISL_566367, EPI_ISL_566368, EPI_ISL_566369, EPI_ISL_566370, EPI_ISL_566371, EPI_ISL_566372, EPI_ISL_566373, EPI_ISL_566374, EPI_ISL_566375, EPI_ISL_566376, EPI_ISL_566377, EPI_ISL_566378, EPI_ISL_566379, EPI_ISL_566381, EPI_ISL_566382, EPI_ISL_566383, EPI_ISL_566384, EPI_ISL_566386, EPI_ISL_566387, EPI_ISL_566388, EPI_ISL_566389, EPI_ISL_566390, EPI_ISL_566391, EPI_ISL_566392, EPI_ISL_566393, EPI_ISL_566394, EPI_ISL_566398, EPI_ISL_566399, EPI_ISL_566400, EPI_ISL_566401, EPI_ISL_566402, EPI_ISL_566403, EPI_ISL_566404, EPI_ISL_566406, EPI_ISL_566407, EPI_ISL_566408, EPI_ISL_566409, EPI_ISL_566410, EPI_ISL_566411, EPI_ISL_566412, EPI_ISL_566413, EPI_ISL_566415, EPI_ISL_566416, EPI_ISL_566417, EPI_ISL_566418, EPI_ISL_566420 | | | |
| see above | Lighthouse Lab in Alderley Park | Wellcome Sanger Institute for the COVID-19 Genomics UK (COG-UK) consortium | Jacquelyn Wynn, Mairead Hyland, The Lighthouse Lab in Alderley Park and Alex Alderton, Roberto Amato, Sonia Goncalves, Ewan Harrison, David K. Jackson, Ian Johnston, Dominic Kwiatkowski, Cordelia Langford, John Sillitoe on behalf of the Wellcome Sanger Institute COVID-19 Surveillance Team |
| EPI_ISL_567307 | Lighthouse Lab in Glasgow | Wellcome Sanger Institute for the COVID-19 Genomics UK (COG-UK) consortium | Harper VanSteenhouse, Yumi Kasai, David Gray, Carol Clugston, Anna Dominiczak and Alex Alderton, Roberto Amato, Sonia Goncalves, Ewan Harrison, David K. Jackson, Ian Johnston, Dominic Kwiatkowski, Cordelia Langford, John Sillitoe on behalf of the Wellcome Sanger Institute COVID-19 Surveillance Team |
| EPI_ISL_567547, EPI_ISL_567570, EPI_ISL_567596, EPI_ISL_567609, EPI_ISL_567667, EPI_ISL_567688, EPI_ISL_567720, EPI_ISL_567722, EPI_ISL_567833, EPI_ISL_567840, EPI_ISL_567879, EPI_ISL_567887, EPI_ISL_567919, EPI_ISL_567931, EPI_ISL_567968, EPI_ISL_568079, EPI_ISL_568134, EPI_ISL_568151, EPI_ISL_568158, EPI_ISL_568180, EPI_ISL_568467, EPI_ISL_568468, EPI_ISL_568474, EPI_ISL_568475, EPI_ISL_568477 | | | |
| see above | Lighthouse Lab in Alderley Park | Wellcome Sanger Institute for the COVID-19 Genomics UK (COG-UK) consortium | Jacquelyn Wynn, Mairead Hyland, The Lighthouse Lab in Alderley Park and Alex Alderton, Roberto Amato, Sonia Goncalves, Ewan Harrison, David K. Jackson, Ian Johnston, Dominic Kwiatkowski, Cordelia Langford, John Sillitoe on behalf of the Wellcome Sanger Institute COVID-19 Surveillance Team |
| EPI_ISL_574618 | RS Kramat 128 | Eijkman Institute for Molecular Biology, Ministry of Research and Technology/National Agency for Research and Innovation | Hidayat Trimarsanto, Frilasita A Yudhaputri, Edison Johar, Iskandar A Adnan, Willy Agustine, David H Muljono, Safarina G Malik, Herawati Sudoyo, Khin Saw Myint, Amin Soebandrio |
| EPI_ISL_574619, EPI_ISL_574620, EPI_ISL_574621, EPI_ISL_574622, EPI_ISL_574623 | BTKLPP Kelas I Manado | Eijkman Institute for Molecular Biology, Ministry of Research and Technology/National Agency for Research and Innovation | Hidayat Trimarsanto, Frilasita A Yudhaputri, Edison Johar, Iskandar A Adnan, Willy Agustine, David H Muljono, Safarina G Malik, Herawati Sudoyo, Khin Saw Myint, Amin Soebandrio |
| EPI_ISL_575338, EPI_ISL_575340, EPI_ISL_575341, EPI_ISL_575342, EPI_ISL_575343, EPI_ISL_575344, EPI_ISL_575345, EPI_ISL_575346, EPI_ISL_575347, EPI_ISL_575349, EPI_ISL_575350, EPI_ISL_575351, EPI_ISL_575352, EPI_ISL_575353, EPI_ISL_575354, EPI_ISL_575355, EPI_ISL_575356, EPI_ISL_575357, EPI_ISL_575358, EPI_ISL_575359, EPI_ISL_575360, EPI_ISL_575361, EPI_ISL_575362, EPI_ISL_575363, EPI_ISL_575365, EPI_ISL_575366, EPI_ISL_575368, EPI_ISL_575369, EPI_ISL_575370, EPI_ISL_575371, EPI_ISL_575372, EPI_ISL_575373, EPI_ISL_575374, EPI_ISL_575375, EPI_ISL_575376, EPI_ISL_575377, EPI_ISL_575379, EPI_ISL_575380, EPI_ISL_575381, EPI_ISL_575382, EPI_ISL_575384, EPI_ISL_575385, EPI_ISL_575386, EPI_ISL_575387, EPI_ISL_575388, EPI_ISL_575390, EPI_ISL_575391, EPI_ISL_575392, EPI_ISL_575393, EPI_ISL_575394, EPI_ISL_575395, EPI_ISL_575396, EPI_ISL_575397, EPI_ISL_575398 | | | |
| see above | Lighthouse Lab in Glasgow | Wellcome Sanger Institute for the COVID-19 Genomics UK (COG-UK) consortium | Harper VanSteenhouse, Yumi Kasai, David Gray, Carol Clugston, Anna Dominiczak and Alex Alderton, Roberto Amato, Sonia Goncalves, Ewan Harrison, David K. Jackson, Ian Johnston, Dominic Kwiatkowski, Cordelia Langford, John Sillitoe on behalf of the Wellcome Sanger Institute COVID-19 Surveillance Team |
| EPI_ISL_575404, EPI_ISL_575417, EPI_ISL_575440, EPI_ISL_575458, EPI_ISL_575483, EPI_ISL_575489, EPI_ISL_575552, EPI_ISL_575556, EPI_ISL_575611, EPI_ISL_575618, EPI_ISL_575645, EPI_ISL_575659, EPI_ISL_575670, EPI_ISL_575699, EPI_ISL_575710, EPI_ISL_575759 | | | |
| see above | Lighthouse Lab in Alderley Park | Wellcome Sanger Institute for the COVID-19 Genomics UK (COG-UK) consortium | Jacquelyn Wynn, Mairead Hyland, The Lighthouse Lab in Alderley Park and Alex Alderton, Roberto Amato, Sonia Goncalves, Ewan Harrison, David K. Jackson, Ian Johnston, Dominic Kwiatkowski, Cordelia Langford, John Sillitoe on behalf of the Wellcome Sanger Institute COVID-19 Surveillance Team |
| EPI_ISL_575982, EPI_ISL_575983 | Lighthouse Lab in Alderley Park | Wellcome Sanger Institute for the COVID-19 Genomics UK (COG-UK) consortium | Jacquelyn Wynn, Mairead Hyland, The Lighthouse Lab in Alderley Park and Alex Alderton, Roberto Amato, Sonia Goncalves, Ewan Harrison, David K. Jackson, Ian Johnston, Dominic Kwiatkowski, Cordelia Langford, John Sillitoe on behalf of the Wellcome Sanger Institute COVID-19 Surveillance Team (http://www.sanger.ac.uk/covid-team) |
| EPI_ISL_576111, EPI_ISL_576112 | Alaska State Virology Laboratory | Alaska State Virology Laboratory | Jack Chen, Ph.D. |
| EPI_ISL_576877, EPI_ISL_576878, EPI_ISL_576879, EPI_ISL_576880, EPI_ISL_576881, EPI_ISL_576882, EPI_ISL_576883, EPI_ISL_576884, EPI_ISL_576885, EPI_ISL_576886, EPI_ISL_576887, EPI_ISL_576888, EPI_ISL_576889, EPI_ISL_576890, EPI_ISL_576891, EPI_ISL_576892, EPI_ISL_576893, EPI_ISL_576894, EPI_ISL_576895, EPI_ISL_576896 | | | |
| see above | University of Birmingham | COVID-19 Genomics UK (COG-UK) Consortium | Institute of Microbiology, University of Birmingham: Claire McMurray, Joanne Stockton, Samuel Nicholls, Radoslaw Poplawski, Will Rowe, Josh Quick, Nicholas Loman. University of Birmingham Testing Laboratory: Celina M Whalley, Andrew Bosworth, Charlotte Poxon, Kasun Wanigasooriya, Oliver Pickles, Mike Kidd, Alex Richter, Andrew D Beggs PHE Heartlands Lab: Husam Osman, Andrew Bosworth. Queen Elizabeth Hospital: Anna Casey |
| EPI_ISL_578186 | Hospital General Juan Ramón Jiménez | Instituto de Salud Carlos III | Iglesias-Caballero, M. Molinero Calamita, M. González-Esguevillas, M. Camarero, S. Pozo, F. Casas, I. Jiménez, P. Jiménez, M. Zaballos, A. Monzón, S. Varona, S. Juliá, M. Cuesta, I, J. Saavedra |
| EPI_ISL_578187, EPI_ISL_578188 | Hospital Virgen de las Nieves | Instituto de Salud Carlos III | Iglesias-Caballero, M. Molinero Calamita, M. González-Esguevillas, M. Camarero, S. Pozo, F. Casas, I. Jiménez, P. Jiménez, M. Zaballos, A. Monzón, S. Varona, S. Juliá, M. Cuesta, I, J.M Navarro |
| EPI_ISL_578189 | Hospital Universitario Miguel Servet | Instituto de Salud Carlos III | Iglesias-Caballero, M. Molinero Calamita, M. González-Esguevillas, M. Camarero, S. Pozo, F. Casas, I. Jiménez, P. Jiménez, M. Zaballos, A. Monzón, S. Varona, S. Juliá, M. Cuesta, I, A. Rezusta |
| EPI_ISL_578190, EPI_ISL_578191 | Hospital Clínico Universitario Lozano Blesa | Instituto de Salud Carlos III | Iglesias-Caballero, M. Molinero Calamita, M. González-Esguevillas, M. Camarero, S. Pozo, F. Casas, I. Jiménez, P. Jiménez, M. Zaballos, A. Monzón, S. Varona, S. Juliá, M. Cuesta, I, R. Benito |
| EPI_ISL_578192, EPI_ISL_578193 | Hospital Universitario Miguel Servet | Instituto de Salud Carlos III | Iglesias-Caballero, M. Molinero Calamita, M. González-Esguevillas, M. Camarero, S. Pozo, F. Casas, I. Jiménez, P. Jiménez, M. Zaballos, A. Monzón, S. Varona, S. Juliá, M. Cuesta, I, A. Rezusta |
| EPI_ISL_578194 | Complejo Hospitalario Universitario La Coruna | Instituto de Salud Carlos III | Iglesias-Caballero, M. Molinero Calamita, M. González-Esguevillas, M. Camarero, S. Pozo, F. Casas, I. Jiménez, P. Jiménez, M. Zaballos, A. Monzón, S. Varona, S. Juliá, M. Cuesta, I, J. López |
| EPI_ISL_578195 | Complejo Hospitalario de Orense | Instituto de Salud Carlos III | Iglesias-Caballero, M. Molinero Calamita, M. González-Esguevillas, M. Camarero, S. Pozo, F. Casas, I. Jiménez, P. Jiménez, M. Zaballos, A. Monzón, S. Varona, S. Juliá, M. Cuesta, I, M. García |
| EPI_ISL_578196 | Hospital San Pedro | Instituto de Salud Carlos III | Iglesias-Caballero, M. Molinero Calamita, M. González-Esguevillas, M. Camarero, S. Pozo, F. Casas, I. Jiménez, P. Jiménez, M. Zaballos, A. Monzón, S. Varona, S. Juliá, M. Cuesta, I, J.M Azcona |
| EPI_ISL_578197 | Hospital San Pedro | Instituto de Salud Carlos III | Iglesias-Caballero, M. Molinero Calamita, M. González-Esguevillas, M. Camarero, S. Pozo, F. Casas, I. Jiménez, P. Jiménez, M. Zaballos, A. Monzón, S. Varona, S. Juliá, M. Cuesta, I, C. Alonso |
| EPI_ISL_578198, EPI_ISL_578199 | Hospital San Pedro | Instituto de Salud Carlos III | Iglesias-Caballero, M. Molinero Calamita, M. González-Esguevillas, M. Camarero, S. Pozo, F. Casas, I. Jiménez, P. Jiménez, M. Zaballos, A. Monzón, S. Varona, S. Juliá, M. Cuesta, I, M. Blasco |

| | | | |
|---|---|---|---|
| EPI_ISL_578200, EPI_ISL_578201 | Hospital San Pedro | Instituto de Salud Carlos III | Iglesias-Caballero, M. Molinero Calamita, M. González-Esguevillas, M. Camarero, S. Pozo, F. Casas, I. Jiménez, P. Jiménez, M. Zaballos, A. Monzón, S. Varona, S. Juliá, M. Cuesta, I, C. Alonso |
| EPI_ISL_579919, EPI_ISL_579920, EPI_ISL_579921, EPI_ISL_579922, EPI_ISL_579923, EPI_ISL_579924, EPI_ISL_579925, EPI_ISL_579927, EPI_ISL_579928, EPI_ISL_579929, EPI_ISL_579930, EPI_ISL_579931, EPI_ISL_579932, EPI_ISL_579933, EPI_ISL_579934, EPI_ISL_579935, EPI_ISL_579936, EPI_ISL_579937, EPI_ISL_579938, EPI_ISL_579939, EPI_ISL_579940, EPI_ISL_579941, EPI_ISL_579942, EPI_ISL_579943, EPI_ISL_579944, EPI_ISL_579945, EPI_ISL_579946, EPI_ISL_579947, EPI_ISL_579948, EPI_ISL_579949, EPI_ISL_579950, EPI_ISL_579952, EPI_ISL_579953, EPI_ISL_579954, EPI_ISL_579955, EPI_ISL_579956, EPI_ISL_579957, EPI_ISL_579958, EPI_ISL_579959, EPI_ISL_579960, EPI_ISL_579961, EPI_ISL_579962, EPI_ISL_579963, EPI_ISL_579964, EPI_ISL_579965, EPI_ISL_579966, EPI_ISL_579967, EPI_ISL_579968, EPI_ISL_579969, EPI_ISL_579970, EPI_ISL_579972, EPI_ISL_579973, EPI_ISL_579974, EPI_ISL_579975, EPI_ISL_579976, EPI_ISL_579977, EPI_ISL_579978, EPI_ISL_579979, EPI_ISL_579980, EPI_ISL_579981, EPI_ISL_579982, EPI_ISL_579983, EPI_ISL_579984, EPI_ISL_579985, EPI_ISL_579986, EPI_ISL_579987, EPI_ISL_579988, EPI_ISL_579989, EPI_ISL_579990, EPI_ISL_579991, EPI_ISL_579992, EPI_ISL_579993, EPI_ISL_579994, EPI_ISL_579995, EPI_ISL_579996, EPI_ISL_579997, EPI_ISL_579998, EPI_ISL_579999, EPI_ISL_580000, EPI_ISL_580001, EPI_ISL_580002, EPI_ISL_580003, EPI_ISL_580004, EPI_ISL_580005, EPI_ISL_580006, EPI_ISL_580007, EPI_ISL_580008, EPI_ISL_580009, EPI_ISL_580010, EPI_ISL_580011, EPI_ISL_580012, EPI_ISL_580013, EPI_ISL_580014, EPI_ISL_580015, EPI_ISL_580016, EPI_ISL_580017, EPI_ISL_580018, EPI_ISL_580019, EPI_ISL_580020, EPI_ISL_580021, EPI_ISL_580022, EPI_ISL_580023, EPI_ISL_580024, EPI_ISL_580025, EPI_ISL_580026, EPI_ISL_580027, EPI_ISL_580028, EPI_ISL_580029, EPI_ISL_580030, EPI_ISL_580031, EPI_ISL_580032, EPI_ISL_580033, EPI_ISL_580034, EPI_ISL_580035, EPI_ISL_580036, EPI_ISL_580037, EPI_ISL_580038, EPI_ISL_580039, EPI_ISL_580040, EPI_ISL_580041, EPI_ISL_580042, EPI_ISL_580043, EPI_ISL_580044, EPI_ISL_580045, EPI_ISL_580046, EPI_ISL_580047, EPI_ISL_580048, EPI_ISL_580049, EPI_ISL_580050, EPI_ISL_580051, EPI_ISL_580052, EPI_ISL_580053, EPI_ISL_580054, EPI_ISL_580055, EPI_ISL_580056, EPI_ISL_580057, EPI_ISL_580058, EPI_ISL_580059, EPI_ISL_580060, EPI_ISL_580061, EPI_ISL_580063, EPI_ISL_580064, EPI_ISL_580065, EPI_ISL_580066, EPI_ISL_580067, EPI_ISL_580068, EPI_ISL_580069, EPI_ISL_580070, EPI_ISL_580071, EPI_ISL_580072, EPI_ISL_580073, EPI_ISL_580074, EPI_ISL_580075, EPI_ISL_580076, EPI_ISL_580077, EPI_ISL_580078, EPI_ISL_580079, EPI_ISL_580080, EPI_ISL_580081, EPI_ISL_580082, EPI_ISL_580083, EPI_ISL_580084, EPI_ISL_580085, EPI_ISL_580086, EPI_ISL_580087, EPI_ISL_580088, EPI_ISL_580089, EPI_ISL_580090, EPI_ISL_580091, EPI_ISL_580092, EPI_ISL_580093, EPI_ISL_580094, EPI_ISL_580095, EPI_ISL_580096, EPI_ISL_580097, EPI_ISL_580098, EPI_ISL_580099, EPI_ISL_580100, EPI_ISL_580101, EPI_ISL_580102, EPI_ISL_580103, EPI_ISL_580104, EPI_ISL_580105, EPI_ISL_580106, EPI_ISL_580107, EPI_ISL_580108, EPI_ISL_580109, EPI_ISL_580110, EPI_ISL_580111, EPI_ISL_580112, EPI_ISL_580113, EPI_ISL_580114, EPI_ISL_580115, EPI_ISL_580116, EPI_ISL_580117, EPI_ISL_580118, EPI_ISL_580119, EPI_ISL_580120, EPI_ISL_580121, EPI_ISL_580122, EPI_ISL_580123, EPI_ISL_580124, EPI_ISL_580125, EPI_ISL_580126, EPI_ISL_580127, EPI_ISL_580128, EPI_ISL_580129, EPI_ISL_580130, EPI_ISL_580131, EPI_ISL_580132, EPI_ISL_580133, EPI_ISL_580134, EPI_ISL_580135, EPI_ISL_580136, EPI_ISL_580137, EPI_ISL_580139, EPI_ISL_580140, EPI_ISL_580141, EPI_ISL_580142, EPI_ISL_580143, EPI_ISL_580144, EPI_ISL_580145, EPI_ISL_580146, EPI_ISL_580147, EPI_ISL_580148, EPI_ISL_580149, EPI_ISL_580150, EPI_ISL_580151, EPI_ISL_580152, EPI_ISL_580153, EPI_ISL_580154, EPI_ISL_580155, EPI_ISL_580156, EPI_ISL_580157, EPI_ISL_580158, EPI_ISL_580159, EPI_ISL_580160, EPI_ISL_580161, EPI_ISL_580162, EPI_ISL_580163, EPI_ISL_580164, EPI_ISL_580165, EPI_ISL_580166, EPI_ISL_580167, EPI_ISL_580168, EPI_ISL_580169, EPI_ISL_580170, EPI_ISL_580171, EPI_ISL_580172, EPI_ISL_580173, EPI_ISL_580174, EPI_ISL_580175, EPI_ISL_580176, EPI_ISL_580177, EPI_ISL_580178, EPI_ISL_580179, EPI_ISL_580180, EPI_ISL_580181, EPI_ISL_580182, EPI_ISL_580183, EPI_ISL_580184, EPI_ISL_580185, EPI_ISL_580186, EPI_ISL_580188, EPI_ISL_580189, EPI_ISL_580190, EPI_ISL_580191, EPI_ISL_580192, EPI_ISL_580193, EPI_ISL_580194, EPI_ISL_580195, EPI_ISL_580196, EPI_ISL_580197, EPI_ISL_580198, EPI_ISL_580199, EPI_ISL_580200, EPI_ISL_580201, EPI_ISL_580202, EPI_ISL_580203, EPI_ISL_580204, EPI_ISL_580205, EPI_ISL_580206, EPI_ISL_580207, EPI_ISL_580208, EPI_ISL_580209, EPI_ISL_580211, EPI_ISL_580212, EPI_ISL_580213, EPI_ISL_580214, EPI_ISL_580215, EPI_ISL_580216, EPI_ISL_580217, EPI_ISL_580218, EPI_ISL_580219, EPI_ISL_580220, EPI_ISL_580221, EPI_ISL_580222, EPI_ISL_580223, EPI_ISL_580224, EPI_ISL_580225, EPI_ISL_580226, EPI_ISL_580227, EPI_ISL_580228, EPI_ISL_580229, EPI_ISL_580230, EPI_ISL_580231, EPI_ISL_580232, EPI_ISL_580233, EPI_ISL_580234, EPI_ISL_580235, EPI_ISL_580236, EPI_ISL_580237, EPI_ISL_580238, EPI_ISL_580239, EPI_ISL_580240, EPI_ISL_580241, EPI_ISL_580242, EPI_ISL_580243, EPI_ISL_580244, EPI_ISL_580245, EPI_ISL_580246, EPI_ISL_580247, EPI_ISL_580248, EPI_ISL_580249, EPI_ISL_580250, EPI_ISL_580251, EPI_ISL_580252, EPI_ISL_580253, EPI_ISL_580254, EPI_ISL_580255, EPI_ISL_580256, EPI_ISL_580257, EPI_ISL_580258, EPI_ISL_580259, EPI_ISL_580261, EPI_ISL_580262, EPI_ISL_580263, EPI_ISL_580264, EPI_ISL_580265, EPI_ISL_580266, EPI_ISL_580267, EPI_ISL_580268, EPI_ISL_580269, EPI_ISL_580270, EPI_ISL_580271, EPI_ISL_580272, EPI_ISL_580273, EPI_ISL_580274, EPI_ISL_580275, EPI_ISL_580276, EPI_ISL_580277, EPI_ISL_580278, EPI_ISL_580279, EPI_ISL_580280, EPI_ISL_580281, EPI_ISL_580284, EPI_ISL_580288, EPI_ISL_580289, EPI_ISL_580290, EPI_ISL_580291, EPI_ISL_580292, EPI_ISL_580293, EPI_ISL_580294, EPI_ISL_580295, EPI_ISL_580296, EPI_ISL_580297, EPI_ISL_580298, EPI_ISL_580302, EPI_ISL_580303, EPI_ISL_580304, EPI_ISL_580306, EPI_ISL_580307, EPI_ISL_580308, EPI_ISL_580309, EPI_ISL_580310, EPI_ISL_580311, EPI_ISL_580312, EPI_ISL_580314, EPI_ISL_580315, EPI_ISL_580316, EPI_ISL_580317, EPI_ISL_580318, EPI_ISL_580319, EPI_ISL_580320, EPI_ISL_580321, EPI_ISL_580323, EPI_ISL_580324, EPI_ISL_580326, EPI_ISL_580328, EPI_ISL_580329, EPI_ISL_580330, EPI_ISL_580331, EPI_ISL_580334, EPI_ISL_580335, EPI_ISL_580336, EPI_ISL_580337, EPI_ISL_580339, EPI_ISL_580342, EPI_ISL_580343, EPI_ISL_580344, EPI_ISL_580345, EPI_ISL_580346, EPI_ISL_580347, EPI_ISL_580348, EPI_ISL_580349, EPI_ISL_580350, EPI_ISL_580351, EPI_ISL_580352, EPI_ISL_580355, EPI_ISL_580356, EPI_ISL_580357, EPI_ISL_580358, EPI_ISL_580359, EPI_ISL_580360, EPI_ISL_580361, EPI_ISL_580363, EPI_ISL_580365, EPI_ISL_580366, EPI_ISL_580370, EPI_ISL_580372, EPI_ISL_580374, EPI_ISL_580375, EPI_ISL_580376, EPI_ISL_580377, EPI_ISL_580378, EPI_ISL_580379, EPI_ISL_580380, EPI_ISL_580381, EPI_ISL_580382, EPI_ISL_580383, EPI_ISL_580384, EPI_ISL_580386, EPI_ISL_580387, EPI_ISL_580388, EPI_ISL_580389, EPI_ISL_580390, EPI_ISL_580392, EPI_ISL_580395, EPI_ISL_580397, EPI_ISL_580398, EPI_ISL_580399, EPI_ISL_580401, EPI_ISL_580402, EPI_ISL_580403, EPI_ISL_580404, EPI_ISL_580405, EPI_ISL_580406, EPI_ISL_580407, EPI_ISL_580408, EPI_ISL_580409, EPI_ISL_580410, EPI_ISL_580411, EPI_ISL_580412, EPI_ISL_580413, EPI_ISL_580415, EPI_ISL_580416, EPI_ISL_580417, EPI_ISL_580419, EPI_ISL_580420, EPI_ISL_580422, EPI_ISL_580423, EPI_ISL_580424, EPI_ISL_580425, EPI_ISL_580426, EPI_ISL_580427, EPI_ISL_580428, EPI_ISL_580429, EPI_ISL_580430, EPI_ISL_580431, EPI_ISL_580432, EPI_ISL_580433, EPI_ISL_580434, EPI_ISL_580436, EPI_ISL_580437, EPI_ISL_580438, EPI_ISL_580439, EPI_ISL_580440, EPI_ISL_580441, EPI_ISL_580442, EPI_ISL_580443, EPI_ISL_580445, EPI_ISL_580446, EPI_ISL_580447, EPI_ISL_580448, EPI_ISL_580450, EPI_ISL_580451, EPI_ISL_580452, EPI_ISL_580454, EPI_ISL_580455, EPI_ISL_580456, EPI_ISL_580457, EPI_ISL_580458, EPI_ISL_580460, EPI_ISL_580461, EPI_ISL_580462, EPI_ISL_580463, EPI_ISL_580464, EPI_ISL_580466, EPI_ISL_580468, EPI_ISL_580469, EPI_ISL_580470, EPI_ISL_580472, EPI_ISL_580473, EPI_ISL_580474, EPI_ISL_580475, EPI_ISL_580476, EPI_ISL_580477, EPI_ISL_580482, EPI_ISL_580483, EPI_ISL_580487, EPI_ISL_580488, EPI_ISL_580489, EPI_ISL_580490, EPI_ISL_580491, EPI_ISL_580492, EPI_ISL_580493, EPI_ISL_580494, EPI_ISL_580496, EPI_ISL_580497, EPI_ISL_580498, EPI_ISL_580500, EPI_ISL_580501, EPI_ISL_580503, EPI_ISL_580504, EPI_ISL_580505, EPI_ISL_580506, EPI_ISL_580507, EPI_ISL_580510, EPI_ISL_580512, EPI_ISL_580513, EPI_ISL_580514, EPI_ISL_580516, EPI_ISL_580517, EPI_ISL_580518, EPI_ISL_580519, EPI_ISL_580520, EPI_ISL_580521, EPI_ISL_580522, EPI_ISL_580523, EPI_ISL_580524, EPI_ISL_580525, EPI_ISL_580526, EPI_ISL_580527, EPI_ISL_580528, EPI_ISL_580529, EPI_ISL_580530, EPI_ISL_580531, EPI_ISL_580534, EPI_ISL_580535, EPI_ISL_580537, EPI_ISL_580538, EPI_ISL_580539, EPI_ISL_580540, EPI_ISL_580541, EPI_ISL_580542, EPI_ISL_580543, EPI_ISL_580544, EPI_ISL_580545, EPI_ISL_580546, EPI_ISL_580547, EPI_ISL_580548, EPI_ISL_580549, EPI_ISL_580551, EPI_ISL_580552, EPI_ISL_580553, EPI_ISL_580554, EPI_ISL_580555, EPI_ISL_580556, EPI_ISL_580558, EPI_ISL_580560, EPI_ISL_580561, EPI_ISL_580562, EPI_ISL_580566, EPI_ISL_580568, EPI_ISL_580570, EPI_ISL_580571, EPI_ISL_580572, EPI_ISL_580574, EPI_ISL_580575, EPI_ISL_580578, EPI_ISL_580580, EPI_ISL_580581, EPI_ISL_580583, EPI_ISL_580584, EPI_ISL_580585, EPI_ISL_580588, EPI_ISL_580589, EPI_ISL_580590, EPI_ISL_580591, EPI_ISL_580592, EPI_ISL_580595, EPI_ISL_580596, EPI_ISL_580597, EPI_ISL_580599, EPI_ISL_580600, EPI_ISL_580601, EPI_ISL_580602, EPI_ISL_580604, EPI_ISL_580605, EPI_ISL_580606, EPI_ISL_580608, EPI_ISL_580609, EPI_ISL_580611, EPI_ISL_580612, EPI_ISL_580613, EPI_ISL_580614, EPI_ISL_580615, EPI_ISL_580616, EPI_ISL_580617, EPI_ISL_580619, EPI_ISL_580620, EPI_ISL_580622, EPI_ISL_580623, EPI_ISL_580624, EPI_ISL_580627, EPI_ISL_580628, EPI_ISL_580629, EPI_ISL_580630, EPI_ISL_580633, EPI_ISL_580635, EPI_ISL_580636, EPI_ISL_580640, EPI_ISL_580645, EPI_ISL_580648, EPI_ISL_580650, EPI_ISL_580651, EPI_ISL_580655, EPI_ISL_580660, EPI_ISL_580663, EPI_ISL_580674, EPI_ISL_580675, EPI_ISL_580677, EPI_ISL_580678, EPI_ISL_580679, EPI_ISL_580680, EPI_ISL_580681, EPI_ISL_580682, EPI_ISL_580689, EPI_ISL_580690, EPI_ISL_580691, EPI_ISL_580692, EPI_ISL_580695, EPI_ISL_580696, EPI_ISL_580697, EPI_ISL_580699, EPI_ISL_580706, EPI_ISL_580710, EPI_ISL_580720, EPI_ISL_580722, EPI_ISL_580723, EPI_ISL_580730, EPI_ISL_580734, EPI_ISL_580737, EPI_ISL_580738, EPI_ISL_580740, EPI_ISL_580754, EPI_ISL_580763, EPI_ISL_580764, EPI_ISL_580769, EPI_ISL_580771, EPI_ISL_580784, EPI_ISL_580787, EPI_ISL_580788, EPI_ISL_580789, EPI_ISL_580791, EPI_ISL_580821, EPI_ISL_580822, EPI_ISL_580823, EPI_ISL_580825, EPI_ISL_580828, EPI_ISL_580833, EPI_ISL_580835, EPI_ISL_580849, EPI_ISL_580851, EPI_ISL_580853, EPI_ISL_581379, EPI_ISL_581391 | | | |
| see above | Lighthouse Lab in Alderley Park | Wellcome Sanger Institute for the COVID-19 Genomics UK (COG-UK) consortium | Jacquelyn Wynn, Mairead Hyland, The Lighthouse Lab in Alderley Park and Alex Alderton, Roberto Amato, Sonia Goncalves, Ewan Harrison, David K. Jackson, Ian Johnston, Dominic Kwiatkowski, Cordelia Langford, John Sillitoe on behalf of the Wellcome Sanger Institute COVID-19 Surveillance Team |
| EPI_ISL_581418, EPI_ISL_581419, EPI_ISL_581420 | Lighthouse Lab in Glasgow | Wellcome Sanger Institute for the COVID-19 Genomics UK (COG-UK) consortium | Harper VanSteenhouse, Yumi Kasai, David Gray, Carol Clugston, Anna Dominiczak and Alex Alderton, Roberto Amato, Sonia Goncalves, Ewan Harrison, David K. Jackson, Ian Johnston, Dominic Kwiatkowski, Cordelia Langford, John Sillitoe on behalf of the Wellcome Sanger Institute COVID-19 Surveillance Team |
| EPI_ISL_581425, EPI_ISL_581433 | Lighthouse Lab in Alderley Park | Wellcome Sanger Institute for the COVID-19 Genomics UK (COG-UK) consortium | Jacquelyn Wynn, Mairead Hyland, The Lighthouse Lab in Alderley Park and Alex Alderton, Roberto Amato, Sonia Goncalves, Ewan Harrison, David K. Jackson, Ian Johnston, Dominic Kwiatkowski, Cordelia Langford, John Sillitoe on behalf of the Wellcome Sanger Institute COVID-19 Surveillance Team |
| EPI_ISL_581437, EPI_ISL_581439, EPI_ISL_581441, EPI_ISL_581443 | Lighthouse Lab in Glasgow | Wellcome Sanger Institute for the COVID-19 Genomics UK (COG-UK) consortium | Harper VanSteenhouse, Yumi Kasai, David Gray, Carol Clugston, Anna Dominiczak and Alex Alderton, Roberto Amato, Sonia Goncalves, Ewan Harrison, David K. Jackson, Ian Johnston, Dominic Kwiatkowski, Cordelia Langford, John Sillitoe on behalf of the Wellcome Sanger Institute COVID-19 Surveillance Team |
| EPI_ISL_581445 | Lighthouse Lab in Alderley Park | Wellcome Sanger Institute for the COVID-19 Genomics UK (COG-UK) consortium | Jacquelyn Wynn, Mairead Hyland, The Lighthouse Lab in Alderley Park and Alex Alderton, Roberto Amato, Sonia Goncalves, Ewan Harrison, David K. Jackson, Ian Johnston, Dominic Kwiatkowski, Cordelia Langford, John Sillitoe on behalf of the Wellcome Sanger Institute COVID-19 Surveillance Team |
| EPI_ISL_586243, EPI_ISL_586244, EPI_ISL_586245, EPI_ISL_586246, EPI_ISL_586247, EPI_ISL_586248, EPI_ISL_586249, EPI_ISL_586250, EPI_ISL_586251, EPI_ISL_586252, EPI_ISL_586253, EPI_ISL_586254, EPI_ISL_586255, EPI_ISL_586256, EPI_ISL_586257, EPI_ISL_586258, EPI_ISL_586259, EPI_ISL_586260, EPI_ISL_586261, EPI_ISL_586262, EPI_ISL_586263, EPI_ISL_586264, EPI_ISL_586265, EPI_ISL_586266 | | | |
| see above | Alaska State Virology Laboratory | Alaska State Virology Laboratory | Jack Chen, Ph.D. |
| EPI_ISL_586570 | GCRI,Ahmedabad | Gujarat Biotechnology Research Centre | Harsha Panchal, Apurvasinh Puvar, Janvi Raval, Zarna Patel, Monika Gandhi, Pinal Trivedi, Maharshi Pandya, Nidhi Patel, Nitin Savaliya, Raghawendra Kumar, Dinesh Kumar, Zuber Saiyed, Komal Patel, Labdhi Pandya, Afzal Ansari, Nikha Trivedi, Shashank Pandya, R D Dixit, A M Kadri, Harsh Bakshi, Chaitanya Joshi, Madhvi Joshi |
| EPI_ISL_586572, EPI_ISL_586573, EPI_ISL_586574, EPI_ISL_586575, EPI_ISL_586576, EPI_ISL_586577, EPI_ISL_586578, EPI_ISL_586579, EPI_ISL_586580, EPI_ISL_586581, EPI_ISL_586582, EPI_ISL_586583, EPI_ISL_586584, EPI_ISL_586585, EPI_ISL_586586, EPI_ISL_586587, EPI_ISL_586588, EPI_ISL_586589, EPI_ISL_586590, EPI_ISL_586591, EPI_ISL_586592, EPI_ISL_586593, EPI_ISL_586594, EPI_ISL_586595, EPI_ISL_586596, EPI_ISL_586597, EPI_ISL_586598, EPI_ISL_586599, EPI_ISL_586600, EPI_ISL_586601, EPI_ISL_586602, EPI_ISL_586603, EPI_ISL_586604, EPI_ISL_586605, EPI_ISL_586606, EPI_ISL_586607, EPI_ISL_586608, EPI_ISL_586609, EPI_ISL_586610, EPI_ISL_586611, EPI_ISL_586613, EPI_ISL_586614, EPI_ISL_586615, EPI_ISL_586616, EPI_ISL_586618, EPI_ISL_586619, EPI_ISL_586620, EPI_ISL_586622, EPI_ISL_586623, EPI_ISL_586624, EPI_ISL_586625, EPI_ISL_586626, EPI_ISL_586627, EPI_ISL_586628, EPI_ISL_586629, EPI_ISL_586630, EPI_ISL_586631, EPI_ISL_586632, EPI_ISL_586633, EPI_ISL_586634, EPI_ISL_586635, EPI_ISL_586636, EPI_ISL_586637, EPI_ISL_586638, EPI_ISL_586639, EPI_ISL_586640, EPI_ISL_586641, EPI_ISL_586642, EPI_ISL_586643, EPI_ISL_586644, EPI_ISL_586645, EPI_ISL_586646, EPI_ISL_586647, EPI_ISL_586648, EPI_ISL_586649, EPI_ISL_586650, EPI_ISL_586651, EPI_ISL_586652, EPI_ISL_586653, EPI_ISL_586654, EPI_ISL_586655, EPI_ISL_586656, EPI_ISL_586657, EPI_ISL_586658, EPI_ISL_586659, EPI_ISL_586661, EPI_ISL_586662, EPI_ISL_586663, EPI_ISL_586664, EPI_ISL_586665, EPI_ISL_586666, EPI_ISL_586667, EPI_ISL_586668, EPI_ISL_586669, EPI_ISL_586670, EPI_ISL_586671, EPI_ISL_586672, EPI_ISL_586673, EPI_ISL_586674, EPI_ISL_586675, EPI_ISL_586676, EPI_ISL_586677, EPI_ISL_586678, EPI_ISL_586679, EPI_ISL_586680, EPI_ISL_586681, EPI_ISL_586682, EPI_ISL_586683, EPI_ISL_586684, EPI_ISL_586685, EPI_ISL_586686, EPI_ISL_586687, EPI_ISL_586688, EPI_ISL_586689, EPI_ISL_586690, EPI_ISL_586691, EPI_ISL_586692, EPI_ISL_586693, EPI_ISL_586694, EPI_ISL_586695, EPI_ISL_586696, EPI_ISL_586697, EPI_ISL_586698, EPI_ISL_586699, EPI_ISL_586700, EPI_ISL_586701, EPI_ISL_586702, EPI_ISL_586703, EPI_ISL_586704, EPI_ISL_586705, EPI_ISL_586706, EPI_ISL_586707, EPI_ISL_586708, EPI_ISL_586710, EPI_ISL_586711, EPI_ISL_586712, EPI_ISL_586713, EPI_ISL_586714, EPI_ISL_586715, EPI_ISL_586716, EPI_ISL_586717, EPI_ISL_586718, EPI_ISL_586719, EPI_ISL_586720, EPI_ISL_586721, EPI_ISL_586722, EPI_ISL_586723, EPI_ISL_586724, EPI_ISL_586725, EPI_ISL_586726, EPI_ISL_586727, EPI_ISL_586728, EPI_ISL_586729, EPI_ISL_586730, EPI_ISL_586731, EPI_ISL_586732, EPI_ISL_586733, EPI_ISL_586734, EPI_ISL_586735, EPI_ISL_586736, EPI_ISL_586737, EPI_ISL_586738, EPI_ISL_586739, EPI_ISL_586740, EPI_ISL_586741, EPI_ISL_586742, EPI_ISL_586743, EPI_ISL_586744, EPI_ISL_586745, EPI_ISL_586746, EPI_ISL_586747, EPI_ISL_586748, EPI_ISL_586749, EPI_ISL_586750, EPI_ISL_586751, EPI_ISL_586752, EPI_ISL_586753, EPI_ISL_586754, EPI_ISL_586755, EPI_ISL_586756, EPI_ISL_586757, EPI_ISL_586758, EPI_ISL_586759, EPI_ISL_586760, EPI_ISL_586761, EPI_ISL_586762, EPI_ISL_586763, EPI_ISL_586764, EPI_ISL_586765, EPI_ISL_586766, EPI_ISL_586767, EPI_ISL_586768, EPI_ISL_586769, EPI_ISL_586770, EPI_ISL_586771, EPI_ISL_586772, EPI_ISL_586773, EPI_ISL_586774, EPI_ISL_586775, EPI_ISL_586776, EPI_ISL_586777, EPI_ISL_586778, EPI_ISL_586779, EPI_ISL_586780, EPI_ISL_586781, EPI_ISL_586782, EPI_ISL_586783, EPI_ISL_586784, EPI_ISL_586785, EPI_ISL_586786, EPI_ISL_586787, EPI_ISL_586788, EPI_ISL_586789, EPI_ISL_586790, EPI_ISL_586791, EPI_ISL_586792, EPI_ISL_586793, EPI_ISL_586794, EPI_ISL_586795, EPI_ISL_586796, EPI_ISL_586797, EPI_ISL_586798, EPI_ISL_586799, EPI_ISL_586800, EPI_ISL_586801, EPI_ISL_586802, EPI_ISL_586803, EPI_ISL_586804, EPI_ISL_586805, EPI_ISL_586806, EPI_ISL_586807, EPI_ISL_586808, EPI_ISL_586809, EPI_ISL_586810, EPI_ISL_586811, EPI_ISL_586812, EPI_ISL_586813, EPI_ISL_586814, EPI_ISL_586815, EPI_ISL_586816, EPI_ISL_586817, EPI_ISL_586818, EPI_ISL_586819, EPI_ISL_586820, EPI_ISL_586821, EPI_ISL_586822, EPI_ISL_586823, EPI_ISL_586824, EPI_ISL_586825, EPI_ISL_586826, EPI_ISL_586827, EPI_ISL_586828, EPI_ISL_586829, EPI_ISL_586830, EPI_ISL_586831, EPI_ISL_586832, EPI_ISL_586833, EPI_ISL_586834, EPI_ISL_586835, EPI_ISL_586836, EPI_ISL_586837, EPI_ISL_586838, EPI_ISL_586839, EPI_ISL_586840, EPI_ISL_586841, EPI_ISL_586842, EPI_ISL_586843, EPI_ISL_586844, EPI_ISL_586845, EPI_ISL_586846, EPI_ISL_586847, EPI_ISL_586848, EPI_ISL_586849, EPI_ISL_586850, EPI_ISL_586851, EPI_ISL_586852, EPI_ISL_586853, EPI_ISL_586854, EPI_ISL_586855, EPI_ISL_586856, EPI_ISL_586857, EPI_ISL_586858, EPI_ISL_586859, EPI_ISL_586860, EPI_ISL_586861, EPI_ISL_586862, EPI_ISL_586863, EPI_ISL_586864, EPI_ISL_586865, EPI_ISL_586866, EPI_ISL_586867, EPI_ISL_586868, EPI_ISL_586869, EPI_ISL_586870, EPI_ISL_586871, EPI_ISL_586872, EPI_ISL_586873, EPI_ISL_586874, EPI_ISL_586875, EPI_ISL_586876, EPI_ISL_586877, EPI_ISL_586878, EPI_ISL_586879, EPI_ISL_586880, EPI_ISL_586881, EPI_ISL_586882, EPI_ISL_586883, EPI_ISL_586884, EPI_ISL_586885, EPI_ISL_586886, EPI_ISL_586887, EPI_ISL_586888, EPI_ISL_586889, EPI_ISL_586890, EPI_ISL_586891, EPI_ISL_586892, EPI_ISL_586893, EPI_ISL_586894, EPI_ISL_586895, EPI_ISL_586896, EPI_ISL_586897, EPI_ISL_586898, EPI_ISL_586899, EPI_ISL_586900, EPI_ISL_586901, EPI_ISL_586902, EPI_ISL_586903, EPI_ISL_586904, EPI_ISL_586905, EPI_ISL_586906, EPI_ISL_586908, EPI_ISL_586909, EPI_ISL_586910, EPI_ISL_586911, EPI_ISL_586912, EPI_ISL_586913, EPI_ISL_586914, EPI_ISL_586915, EPI_ISL_586916, EPI_ISL_586917, EPI_ISL_586918, EPI_ISL_586919, EPI_ISL_586920, EPI_ISL_586921, EPI_ISL_586922, EPI_ISL_586923, EPI_ISL_586924, EPI_ISL_586925, EPI_ISL_586927, EPI_ISL_586928, EPI_ISL_586929, EPI_ISL_586930, EPI_ISL_586931, EPI_ISL_586932, EPI_ISL_586933, EPI_ISL_586934, EPI_ISL_586935, EPI_ISL_586936, EPI_ISL_586937, EPI_ISL_586938, EPI_ISL_586939, EPI_ISL_586940, EPI_ISL_586941, EPI_ISL_586942, EPI_ISL_586943, EPI_ISL_586944, EPI_ISL_586945, EPI_ISL_586946, EPI_ISL_586947, EPI_ISL_586948, EPI_ISL_586949, EPI_ISL_586950, EPI_ISL_586951, EPI_ISL_586952, EPI_ISL_586953, EPI_ISL_586954, EPI_ISL_586955, EPI_ISL_586956, EPI_ISL_586957, EPI_ISL_586958, EPI_ISL_586959, EPI_ISL_586960, EPI_ISL_586961, EPI_ISL_586962, EPI_ISL_586963, EPI_ISL_586964, EPI_ISL_586965, EPI_ISL_586966, EPI_ISL_586967, EPI_ISL_586968, EPI_ISL_586969, EPI_ISL_586970, EPI_ISL_586971, EPI_ISL_586972, EPI_ISL_586973, EPI_ISL_586974, EPI_ISL_586975, EPI_ISL_586976, EPI_ISL_586977, EPI_ISL_586978, EPI_ISL_586979, EPI_ISL_586980, EPI_ISL_586981, EPI_ISL_586982, EPI_ISL_586983, EPI_ISL_586984, EPI_ISL_586985, EPI_ISL_586986, EPI_ISL_586987, EPI_ISL_586988, EPI_ISL_586989, EPI_ISL_586990, EPI_ISL_586991, EPI_ISL_586992, EPI_ISL_586993, EPI_ISL_586994, EPI_ISL_586995, EPI_ISL_586996, EPI_ISL_586997, EPI_ISL_586998, EPI_ISL_586999, EPI_ISL_587000, EPI_ISL_587001, EPI_ISL_587002, EPI_ISL_587003, EPI_ISL_587004, EPI_ISL_587005, EPI_ISL_587006, EPI_ISL_587007, EPI_ISL_587008, EPI_ISL_587009, EPI_ISL_587010, | | | |

EPI_ISL_587011, EPI_ISL_587012, EPI_ISL_587013, EPI_ISL_587014, EPI_ISL_587015, EPI_ISL_587016, EPI_ISL_587017, EPI_ISL_587018, EPI_ISL_587019, EPI_ISL_587020, EPI_ISL_587021, EPI_ISL_587022, EPI_ISL_587023, EPI_ISL_587024, EPI_ISL_587025, EPI_ISL_587026, EPI_ISL_587027, EPI_ISL_587028, EPI_ISL_587029, EPI_ISL_587030, EPI_ISL_587031, EPI_ISL_587032, EPI_ISL_587033, EPI_ISL_587034, EPI_ISL_587035, EPI_ISL_587036, EPI_ISL_587037, EPI_ISL_587038, EPI_ISL_587039, EPI_ISL_587040, EPI_ISL_587041, EPI_ISL_587042, EPI_ISL_587043, EPI_ISL_587044, EPI_ISL_587045, EPI_ISL_587046, EPI_ISL_587047, EPI_ISL_587048, EPI_ISL_587049, EPI_ISL_587050, EPI_ISL_587051, EPI_ISL_587052, EPI_ISL_587053, EPI_ISL_587054, EPI_ISL_587055, EPI_ISL_587056, EPI_ISL_587057, EPI_ISL_587058, EPI_ISL_587059, EPI_ISL_587060, EPI_ISL_587061, EPI_ISL_587062, EPI_ISL_587063, EPI_ISL_587064, EPI_ISL_587065, EPI_ISL_587066, EPI_ISL_587067, EPI_ISL_587068, EPI_ISL_587069, EPI_ISL_587070, EPI_ISL_587071

| | | | |
|---|---|---|---|
| see above | Lighthouse Lab in Alderley Park | Wellcome Sanger Institute for the COVID-19 Genomics UK (COG-UK) consortium | Jacquelyn Wynn, Mairead Hyland, The Lighthouse Lab in Alderley Park and Alex Alderton, Roberto Amato, Sonia Goncalves, Ewan Harrison, David K. Jackson, Ian Johnston, Dominic Kwiatkowski, Cordelia Langford, John Sillitoe on behalf of the Wellcome Sanger Institute COVID-19 Surveillance Team (http://www.sanger.ac.uk/covid-team) |

EPI_ISL_587072, EPI_ISL_587073, EPI_ISL_587074, EPI_ISL_587075, EPI_ISL_587076, EPI_ISL_587077, EPI_ISL_587078, EPI_ISL_587079, EPI_ISL_587080, EPI_ISL_587081, EPI_ISL_587082, EPI_ISL_587083, EPI_ISL_587084, EPI_ISL_587086, EPI_ISL_587087, EPI_ISL_587088, EPI_ISL_587089, EPI_ISL_587090, EPI_ISL_587091, EPI_ISL_587092, EPI_ISL_587093, EPI_ISL_587094, EPI_ISL_587095, EPI_ISL_587096, EPI_ISL_587097, EPI_ISL_587098, EPI_ISL_587099, EPI_ISL_587100, EPI_ISL_587101, EPI_ISL_587102, EPI_ISL_587103, EPI_ISL_587104, EPI_ISL_587105, EPI_ISL_587106, EPI_ISL_587107, EPI_ISL_587108, EPI_ISL_587109, EPI_ISL_587110, EPI_ISL_587111, EPI_ISL_587112, EPI_ISL_587113, EPI_ISL_587114, EPI_ISL_587115, EPI_ISL_587116, EPI_ISL_587117, EPI_ISL_587118, EPI_ISL_587119, EPI_ISL_587120, EPI_ISL_587121, EPI_ISL_587122, EPI_ISL_587123, EPI_ISL_587124, EPI_ISL_587125, EPI_ISL_587126, EPI_ISL_587127, EPI_ISL_587128, EPI_ISL_587129, EPI_ISL_587130, EPI_ISL_587131, EPI_ISL_587132, EPI_ISL_587133, EPI_ISL_587134, EPI_ISL_587135, EPI_ISL_587136, EPI_ISL_587137, EPI_ISL_587138, EPI_ISL_587139, EPI_ISL_587140, EPI_ISL_587141, EPI_ISL_587142, EPI_ISL_587143, EPI_ISL_587144, EPI_ISL_587145, EPI_ISL_587146, EPI_ISL_587147, EPI_ISL_587148, EPI_ISL_587149, EPI_ISL_587150, EPI_ISL_587152, EPI_ISL_587153, EPI_ISL_587154, EPI_ISL_587155, EPI_ISL_587156, EPI_ISL_587157, EPI_ISL_587158, EPI_ISL_587159, EPI_ISL_587160, EPI_ISL_587161, EPI_ISL_587162, EPI_ISL_587163, EPI_ISL_587164, EPI_ISL_587165, EPI_ISL_587166, EPI_ISL_587167, EPI_ISL_587168, EPI_ISL_587169, EPI_ISL_587170, EPI_ISL_587171, EPI_ISL_587172, EPI_ISL_587173, EPI_ISL_587174, EPI_ISL_587175, EPI_ISL_587176, EPI_ISL_587177, EPI_ISL_587178, EPI_ISL_587179, EPI_ISL_587180, EPI_ISL_587181, EPI_ISL_587182, EPI_ISL_587183, EPI_ISL_587184, EPI_ISL_587185, EPI_ISL_587186, EPI_ISL_587187, EPI_ISL_587188, EPI_ISL_587189, EPI_ISL_587190, EPI_ISL_587191, EPI_ISL_587192, EPI_ISL_587193, EPI_ISL_587194, EPI_ISL_587195, EPI_ISL_587196, EPI_ISL_587197, EPI_ISL_587198, EPI_ISL_587199, EPI_ISL_587200, EPI_ISL_587201, EPI_ISL_587202, EPI_ISL_587203, EPI_ISL_587204, EPI_ISL_587205, EPI_ISL_587206, EPI_ISL_587207, EPI_ISL_587208, EPI_ISL_587209, EPI_ISL_587210, EPI_ISL_587211, EPI_ISL_587212, EPI_ISL_587213, EPI_ISL_587214, EPI_ISL_587215, EPI_ISL_587216, EPI_ISL_587217, EPI_ISL_587218, EPI_ISL_587219, EPI_ISL_587220, EPI_ISL_587221, EPI_ISL_587222, EPI_ISL_587223, EPI_ISL_587224, EPI_ISL_587225, EPI_ISL_587226, EPI_ISL_587227, EPI_ISL_587228, EPI_ISL_587229, EPI_ISL_587230, EPI_ISL_587231, EPI_ISL_587232, EPI_ISL_587233, EPI_ISL_587234, EPI_ISL_587235, EPI_ISL_587236, EPI_ISL_587237, EPI_ISL_587238, EPI_ISL_587239, EPI_ISL_587240, EPI_ISL_587241, EPI_ISL_587242, EPI_ISL_587243, EPI_ISL_587244, EPI_ISL_587245, EPI_ISL_587246, EPI_ISL_587247, EPI_ISL_587248, EPI_ISL_587249, EPI_ISL_587250, EPI_ISL_587251, EPI_ISL_587252, EPI_ISL_587253, EPI_ISL_587254, EPI_ISL_587255, EPI_ISL_587256, EPI_ISL_587257, EPI_ISL_587258, EPI_ISL_587259, EPI_ISL_587260, EPI_ISL_587261, EPI_ISL_587262, EPI_ISL_587263, EPI_ISL_587264, EPI_ISL_587266, EPI_ISL_587267, EPI_ISL_587268, EPI_ISL_587269, EPI_ISL_587270, EPI_ISL_587271, EPI_ISL_587272, EPI_ISL_587273, EPI_ISL_587274, EPI_ISL_587275, EPI_ISL_587276, EPI_ISL_587277, EPI_ISL_587278, EPI_ISL_587279, EPI_ISL_587280, EPI_ISL_587281, EPI_ISL_587282, EPI_ISL_587283, EPI_ISL_587284, EPI_ISL_587285, EPI_ISL_587286, EPI_ISL_587287, EPI_ISL_587288, EPI_ISL_587289, EPI_ISL_587290, EPI_ISL_587291, EPI_ISL_587292, EPI_ISL_587293, EPI_ISL_587294, EPI_ISL_587295, EPI_ISL_587296, EPI_ISL_587297, EPI_ISL_587298, EPI_ISL_587299, EPI_ISL_587300, EPI_ISL_587301, EPI_ISL_587302

| | | | |
|---|---|---|---|
| see above | Lighthouse Lab in Alderley Park | Wellcome Sanger Institute for the COVID-19 Genomics UK (COG-UK) consortium | Jacquelyn Wynn, Mairead Hyland, The Lighthouse Lab in Alderley Park and Alex Alderton, Roberto Amato, Sonia Goncalves, Ewan Harrison, David K. Jackson, Ian Johnston, Dominic Kwiatkowski, Cordelia Langford, John Sillitoe on behalf of the Wellcome Sanger Institute COVID-19 Surveillance Team |

EPI_ISL_587313, EPI_ISL_587340, EPI_ISL_587341, EPI_ISL_587345, EPI_ISL_587366, EPI_ISL_587391, EPI_ISL_587402, EPI_ISL_587403, EPI_ISL_587404, EPI_ISL_587411, EPI_ISL_587416, EPI_ISL_587425, EPI_ISL_587427, EPI_ISL_587428, EPI_ISL_587451, EPI_ISL_587459, EPI_ISL_587487, EPI_ISL_587498, EPI_ISL_587500, EPI_ISL_587519, EPI_ISL_587525, EPI_ISL_587534, EPI_ISL_587540, EPI_ISL_587560, EPI_ISL_587575, EPI_ISL_587577, EPI_ISL_587585, EPI_ISL_587587, EPI_ISL_587588, EPI_ISL_587594, EPI_ISL_587598, EPI_ISL_587601, EPI_ISL_587605, EPI_ISL_587606, EPI_ISL_587609, EPI_ISL_587617, EPI_ISL_587619, EPI_ISL_587622, EPI_ISL_587623, EPI_ISL_587630, EPI_ISL_587640, EPI_ISL_587643, EPI_ISL_587658, EPI_ISL_587665, EPI_ISL_587670, EPI_ISL_587674, EPI_ISL_587678, EPI_ISL_587685, EPI_ISL_587702, EPI_ISL_587705

| | | | |
|---|---|---|---|
| see above | Lighthouse Lab in Glasgow | Wellcome Sanger Institute for the COVID-19 Genomics UK (COG-UK) consortium | Harper VanSteenhouse, Yumi Kasai, David Gray, Carol Clugston, Anna Dominiczak and Alex Alderton, Roberto Amato, Sonia Goncalves, Ewan Harrison, David K. Jackson, Ian Johnston, Dominic Kwiatkowski, Cordelia Langford, John Sillitoe on behalf of the Wellcome Sanger Institute COVID-19 Surveillance Team |

EPI_ISL_587711, EPI_ISL_587712, EPI_ISL_587713, EPI_ISL_587714, EPI_ISL_587715, EPI_ISL_587716, EPI_ISL_587717, EPI_ISL_587718, EPI_ISL_587719, EPI_ISL_587720, EPI_ISL_587721, EPI_ISL_587722, EPI_ISL_587723, EPI_ISL_587724, EPI_ISL_587725, EPI_ISL_587726, EPI_ISL_587727, EPI_ISL_587728, EPI_ISL_587729, EPI_ISL_587730, EPI_ISL_587731, EPI_ISL_587732, EPI_ISL_587733, EPI_ISL_587734, EPI_ISL_587735, EPI_ISL_587736, EPI_ISL_587737, EPI_ISL_587738, EPI_ISL_587739, EPI_ISL_587740, EPI_ISL_587741, EPI_ISL_587742, EPI_ISL_587743, EPI_ISL_587744, EPI_ISL_587745, EPI_ISL_587746, EPI_ISL_587747, EPI_ISL_587748, EPI_ISL_587749, EPI_ISL_587750, EPI_ISL_587751, EPI_ISL_587752, EPI_ISL_587753, EPI_ISL_587754, EPI_ISL_587755, EPI_ISL_587756, EPI_ISL_587757, EPI_ISL_587758, EPI_ISL_587759, EPI_ISL_587760, EPI_ISL_587761, EPI_ISL_587762, EPI_ISL_587763, EPI_ISL_587764, EPI_ISL_587765, EPI_ISL_587766, EPI_ISL_587767, EPI_ISL_587768, EPI_ISL_587769, EPI_ISL_587770, EPI_ISL_587771, EPI_ISL_587772, EPI_ISL_587773, EPI_ISL_587774, EPI_ISL_587775, EPI_ISL_587776, EPI_ISL_587777, EPI_ISL_587778, EPI_ISL_587779, EPI_ISL_587780, EPI_ISL_587781, EPI_ISL_587782, EPI_ISL_587783, EPI_ISL_587784, EPI_ISL_587785, EPI_ISL_587786, EPI_ISL_587787, EPI_ISL_587788, EPI_ISL_587789, EPI_ISL_587790, EPI_ISL_587791, EPI_ISL_587792, EPI_ISL_587793, EPI_ISL_587794, EPI_ISL_587795, EPI_ISL_587796, EPI_ISL_587797, EPI_ISL_587798, EPI_ISL_587799, EPI_ISL_587800, EPI_ISL_587801, EPI_ISL_587802, EPI_ISL_587803, EPI_ISL_587804, EPI_ISL_587805, EPI_ISL_587806, EPI_ISL_587807, EPI_ISL_587808, EPI_ISL_587809, EPI_ISL_587810, EPI_ISL_587811, EPI_ISL_587812, EPI_ISL_587813, EPI_ISL_587814, EPI_ISL_587815, EPI_ISL_587816, EPI_ISL_587817, EPI_ISL_587818, EPI_ISL_587819, EPI_ISL_587820, EPI_ISL_587821, EPI_ISL_587822, EPI_ISL_587823, EPI_ISL_587824, EPI_ISL_587825, EPI_ISL_587826, EPI_ISL_587827, EPI_ISL_587828, EPI_ISL_587829, EPI_ISL_587830, EPI_ISL_587831, EPI_ISL_587832, EPI_ISL_587833, EPI_ISL_587834, EPI_ISL_587835, EPI_ISL_587836, EPI_ISL_587837, EPI_ISL_587838, EPI_ISL_587839, EPI_ISL_587840, EPI_ISL_587841, EPI_ISL_587842, EPI_ISL_587843, EPI_ISL_587844, EPI_ISL_587845, EPI_ISL_587846, EPI_ISL_587847, EPI_ISL_587848, EPI_ISL_587849, EPI_ISL_587850, EPI_ISL_587851, EPI_ISL_587852, EPI_ISL_587853, EPI_ISL_587854, EPI_ISL_587855, EPI_ISL_587856, EPI_ISL_587857, EPI_ISL_587858, EPI_ISL_587859, EPI_ISL_587860, EPI_ISL_587861, EPI_ISL_587862, EPI_ISL_587863, EPI_ISL_587864, EPI_ISL_587865, EPI_ISL_587866, EPI_ISL_587867, EPI_ISL_587868, EPI_ISL_587869, EPI_ISL_587870, EPI_ISL_587871, EPI_ISL_587872, EPI_ISL_587873, EPI_ISL_587874, EPI_ISL_587875, EPI_ISL_587876, EPI_ISL_587877, EPI_ISL_587878, EPI_ISL_587879, EPI_ISL_587880, EPI_ISL_587881, EPI_ISL_587882, EPI_ISL_587883, EPI_ISL_587884, EPI_ISL_587885, EPI_ISL_587886, EPI_ISL_587887, EPI_ISL_587888, EPI_ISL_587889, EPI_ISL_587890, EPI_ISL_587891, EPI_ISL_587892, EPI_ISL_587893, EPI_ISL_587894, EPI_ISL_587895, EPI_ISL_587896, EPI_ISL_587897, EPI_ISL_587898, EPI_ISL_587899, EPI_ISL_587900, EPI_ISL_587901, EPI_ISL_587902, EPI_ISL_587903, EPI_ISL_587904, EPI_ISL_587905, EPI_ISL_587906, EPI_ISL_587907, EPI_ISL_587908, EPI_ISL_587909, EPI_ISL_587910, EPI_ISL_587911, EPI_ISL_587912, EPI_ISL_587913, EPI_ISL_587914, EPI_ISL_587915, EPI_ISL_587916, EPI_ISL_587917, EPI_ISL_587918, EPI_ISL_587919, EPI_ISL_587920, EPI_ISL_587921, EPI_ISL_587922, EPI_ISL_587923, EPI_ISL_587924, EPI_ISL_587925, EPI_ISL_587926, EPI_ISL_587927, EPI_ISL_587928, EPI_ISL_587929, EPI_ISL_587930, EPI_ISL_587931, EPI_ISL_587932, EPI_ISL_587933, EPI_ISL_587934, EPI_ISL_587935, EPI_ISL_587936, EPI_ISL_587937, EPI_ISL_587938, EPI_ISL_587939, EPI_ISL_587940, EPI_ISL_587941, EPI_ISL_587942, EPI_ISL_587943, EPI_ISL_587944, EPI_ISL_587945, EPI_ISL_587946, EPI_ISL_587947, EPI_ISL_587948, EPI_ISL_587949, EPI_ISL_587950, EPI_ISL_587951, EPI_ISL_587952, EPI_ISL_587953, EPI_ISL_587954, EPI_ISL_587955, EPI_ISL_587956, EPI_ISL_587957, EPI_ISL_587958, EPI_ISL_587959, EPI_ISL_587960, EPI_ISL_587961, EPI_ISL_587962, EPI_ISL_587963, EPI_ISL_587964, EPI_ISL_587965, EPI_ISL_587966, EPI_ISL_587967, EPI_ISL_587968, EPI_ISL_587969, EPI_ISL_587970, EPI_ISL_587971, EPI_ISL_587972, EPI_ISL_587973, EPI_ISL_587974, EPI_ISL_587975, EPI_ISL_587976, EPI_ISL_587978, EPI_ISL_587979, EPI_ISL_587980, EPI_ISL_587981, EPI_ISL_587982, EPI_ISL_587983, EPI_ISL_587984, EPI_ISL_587985, EPI_ISL_587986, EPI_ISL_587987, EPI_ISL_587988, EPI_ISL_587989, EPI_ISL_587990, EPI_ISL_587991, EPI_ISL_587992, EPI_ISL_587993, EPI_ISL_587994, EPI_ISL_587995, EPI_ISL_587996, EPI_ISL_587997, EPI_ISL_587998, EPI_ISL_587999, EPI_ISL_588000, EPI_ISL_588001, EPI_ISL_588002, EPI_ISL_588003, EPI_ISL_588004, EPI_ISL_588005, EPI_ISL_588006, EPI_ISL_588008, EPI_ISL_588009, EPI_ISL_588010, EPI_ISL_588011, EPI_ISL_588012, EPI_ISL_588013, EPI_ISL_588014, EPI_ISL_588015, EPI_ISL_588016, EPI_ISL_588017, EPI_ISL_588018, EPI_ISL_588019, EPI_ISL_588020, EPI_ISL_588021, EPI_ISL_588022, EPI_ISL_588023, EPI_ISL_588024, EPI_ISL_588025, EPI_ISL_588027, EPI_ISL_588028, EPI_ISL_588029, EPI_ISL_588030, EPI_ISL_588031, EPI_ISL_588032, EPI_ISL_588033, EPI_ISL_588034, EPI_ISL_588035, EPI_ISL_588036, EPI_ISL_588037, EPI_ISL_588038, EPI_ISL_588039, EPI_ISL_588040, EPI_ISL_588041, EPI_ISL_588042, EPI_ISL_588043, EPI_ISL_588044, EPI_ISL_588045, EPI_ISL_588046, EPI_ISL_588047, EPI_ISL_588048, EPI_ISL_588049, EPI_ISL_588050, EPI_ISL_588051, EPI_ISL_588052, EPI_ISL_588053, EPI_ISL_588054, EPI_ISL_588055, EPI_ISL_588056, EPI_ISL_588057, EPI_ISL_588058, EPI_ISL_588059

| | | | |
|---|---|---|---|
| see above | Lighthouse Lab in Alderley Park | Wellcome Sanger Institute for the COVID-19 Genomics UK (COG-UK) consortium | Jacquelyn Wynn, Mairead Hyland, The Lighthouse Lab in Alderley Park and Alex Alderton, Roberto Amato, Sonia Goncalves, Ewan Harrison, David K. Jackson, Ian Johnston, Dominic Kwiatkowski, Cordelia Langford, John Sillitoe on behalf of the Wellcome Sanger Institute COVID-19 Surveillance Team |

EPI_ISL_588060, EPI_ISL_588061, EPI_ISL_588062, EPI_ISL_588063, EPI_ISL_588064, EPI_ISL_588065, EPI_ISL_588066, EPI_ISL_588067, EPI_ISL_588068, EPI_ISL_588070, EPI_ISL_588071, EPI_ISL_588072, EPI_ISL_588073, EPI_ISL_588074, EPI_ISL_588076, EPI_ISL_588077, EPI_ISL_588078, EPI_ISL_588080, EPI_ISL_588082, EPI_ISL_588083, EPI_ISL_588084, EPI_ISL_588085, EPI_ISL_588086, EPI_ISL_588087, EPI_ISL_588088, EPI_ISL_588089, EPI_ISL_588090, EPI_ISL_588091, EPI_ISL_588092, EPI_ISL_588093, EPI_ISL_588094, EPI_ISL_588096, EPI_ISL_588097, EPI_ISL_588098, EPI_ISL_588099, EPI_ISL_588100, EPI_ISL_588101, EPI_ISL_588102, EPI_ISL_588103, EPI_ISL_588104, EPI_ISL_588105, EPI_ISL_588106, EPI_ISL_588107, EPI_ISL_588108, EPI_ISL_588109, EPI_ISL_588110, EPI_ISL_588112, EPI_ISL_588113, EPI_ISL_588114, EPI_ISL_588115, EPI_ISL_588116, EPI_ISL_588117, EPI_ISL_588118, EPI_ISL_588119, EPI_ISL_588120, EPI_ISL_588121, EPI_ISL_588122, EPI_ISL_588123, EPI_ISL_588124, EPI_ISL_588126, EPI_ISL_588127, EPI_ISL_588129, EPI_ISL_588130, EPI_ISL_588131, EPI_ISL_588132, EPI_ISL_588133, EPI_ISL_588134, EPI_ISL_588135, EPI_ISL_588136, EPI_ISL_588138, EPI_ISL_588139, EPI_ISL_588140, EPI_ISL_588142, EPI_ISL_588143, EPI_ISL_588145, EPI_ISL_588146, EPI_ISL_588147, EPI_ISL_588148, EPI_ISL_588149, EPI_ISL_588150, EPI_ISL_588151, EPI_ISL_588152, EPI_ISL_588153, EPI_ISL_588154, EPI_ISL_588155, EPI_ISL_588156, EPI_ISL_588157, EPI_ISL_588158, EPI_ISL_588159, EPI_ISL_588160, EPI_ISL_588161, EPI_ISL_588162, EPI_ISL_588163, EPI_ISL_588164, EPI_ISL_588165, EPI_ISL_588166, EPI_ISL_588167, EPI_ISL_588168, EPI_ISL_588169, EPI_ISL_588170, EPI_ISL_588171, EPI_ISL_588172, EPI_ISL_588173, EPI_ISL_588174, EPI_ISL_588176, EPI_ISL_588177, EPI_ISL_588178, EPI_ISL_588179, EPI_ISL_588180, EPI_ISL_588182, EPI_ISL_588183, EPI_ISL_588184, EPI_ISL_588185, EPI_ISL_588186, EPI_ISL_588187, EPI_ISL_588188, EPI_ISL_588190, EPI_ISL_588191, EPI_ISL_588192, EPI_ISL_588193, EPI_ISL_588194, EPI_ISL_588195, EPI_ISL_588196, EPI_ISL_588197, EPI_ISL_588198, EPI_ISL_588200, EPI_ISL_588201, EPI_ISL_588202, EPI_ISL_588203, EPI_ISL_588204, EPI_ISL_588205, EPI_ISL_588206, EPI_ISL_588207, EPI_ISL_588208, EPI_ISL_588209, EPI_ISL_588210, EPI_ISL_588212, EPI_ISL_588213, EPI_ISL_588214, EPI_ISL_588215, EPI_ISL_588216, EPI_ISL_588217, EPI_ISL_588219, EPI_ISL_588221, EPI_ISL_588222, EPI_ISL_588223, EPI_ISL_588224, EPI_ISL_588225, EPI_ISL_588226, EPI_ISL_588227, EPI_ISL_588229, EPI_ISL_588230, EPI_ISL_588231, EPI_ISL_588232, EPI_ISL_588234, EPI_ISL_588235, EPI_ISL_588237, EPI_ISL_588238, EPI_ISL_588239, EPI_ISL_588240, EPI_ISL_588241, EPI_ISL_588242, EPI_ISL_588243, EPI_ISL_588245, EPI_ISL_588246, EPI_ISL_588247, EPI_ISL_588248, EPI_ISL_588249, EPI_ISL_588250, EPI_ISL_588251, EPI_ISL_588252, EPI_ISL_588253, EPI_ISL_588254, EPI_ISL_588255, EPI_ISL_588256, EPI_ISL_588257, EPI_ISL_588258, EPI_ISL_588259, EPI_ISL_588260, EPI_ISL_588261, EPI_ISL_588262, EPI_ISL_588263, EPI_ISL_588264, EPI_ISL_588265, EPI_ISL_588266, EPI_ISL_588267, EPI_ISL_588268, EPI_ISL_588269, EPI_ISL_588270, EPI_ISL_588271, EPI_ISL_588272, EPI_ISL_588273, EPI_ISL_588274, EPI_ISL_588275, EPI_ISL_588276, EPI_ISL_588277, EPI_ISL_588278, EPI_ISL_588279, EPI_ISL_588281, EPI_ISL_588283, EPI_ISL_588285, EPI_ISL_588286, EPI_ISL_588287, EPI_ISL_588288, EPI_ISL_588290, EPI_ISL_588291, EPI_ISL_588292, EPI_ISL_588293, EPI_ISL_588294, EPI_ISL_588295, EPI_ISL_588296, EPI_ISL_588297, EPI_ISL_588298, EPI_ISL_588299, EPI_ISL_588300, EPI_ISL_588301, EPI_ISL_588302, EPI_ISL_588303, EPI_ISL_588304, EPI_ISL_588305, EPI_ISL_588306, EPI_ISL_588307, EPI_ISL_588308, EPI_ISL_588309, EPI_ISL_588310, EPI_ISL_588311, EPI_ISL_588312, EPI_ISL_588314, EPI_ISL_588315, EPI_ISL_588316, EPI_ISL_588317, EPI_ISL_588318, EPI_ISL_588319, EPI_ISL_588320, EPI_ISL_588321, EPI_ISL_588322, EPI_ISL_588323, EPI_ISL_588324, EPI_ISL_588325, EPI_ISL_588326, EPI_ISL_588328, EPI_ISL_588329, EPI_ISL_588330, EPI_ISL_588331, EPI_ISL_588332, EPI_ISL_588333, EPI_ISL_588334, EPI_ISL_588335, EPI_ISL_588336, EPI_ISL_588337, EPI_ISL_588338, EPI_ISL_588339, EPI_ISL_588340, EPI_ISL_588341, EPI_ISL_588342, EPI_ISL_588343, EPI_ISL_588344, EPI_ISL_588345, EPI_ISL_588346, EPI_ISL_588347, EPI_ISL_588348, EPI_ISL_588350, EPI_ISL_588351, EPI_ISL_588352, EPI_ISL_588353, EPI_ISL_588354, EPI_ISL_588355, EPI_ISL_588356, EPI_ISL_588357, EPI_ISL_588358, EPI_ISL_588359, EPI_ISL_588361, EPI_ISL_588362, EPI_ISL_588363, EPI_ISL_588365, EPI_ISL_588366, EPI_ISL_588367, EPI_ISL_588368, EPI_ISL_588369, EPI_ISL_588370, EPI_ISL_588371, EPI_ISL_588372, EPI_ISL_588375, EPI_ISL_588376, EPI_ISL_588377, EPI_ISL_588378, EPI_ISL_588379, EPI_ISL_588381, EPI_ISL_588382, EPI_ISL_588383, EPI_ISL_588384, EPI_ISL_588385, EPI_ISL_588386, EPI_ISL_588387, EPI_ISL_588388, EPI_ISL_588389, EPI_ISL_588390, EPI_ISL_588391, EPI_ISL_588392, EPI_ISL_588393, EPI_ISL_588394, EPI_ISL_588395, EPI_ISL_588397, EPI_ISL_588398, EPI_ISL_588399, EPI_ISL_588400, EPI_ISL_588401, EPI_ISL_588402, EPI_ISL_588403, EPI_ISL_588404, EPI_ISL_588405, EPI_ISL_588406, EPI_ISL_588407, EPI_ISL_588408, EPI_ISL_588410, EPI_ISL_588411, EPI_ISL_588412, EPI_ISL_588413, EPI_ISL_588414, EPI_ISL_588415, EPI_ISL_588416, EPI_ISL_588417, EPI_ISL_588418, EPI_ISL_588419, EPI_ISL_588420, EPI_ISL_588421, EPI_ISL_588422, EPI_ISL_588423, EPI_ISL_588424, EPI_ISL_588425, EPI_ISL_588426, EPI_ISL_588427, EPI_ISL_588428, EPI_ISL_588429, EPI_ISL_588430, EPI_ISL_588431, EPI_ISL_588432, EPI_ISL_588433, EPI_ISL_588434, EPI_ISL_588435, EPI_ISL_588436, EPI_ISL_588437, EPI_ISL_588438, EPI_ISL_588439, EPI_ISL_588440, EPI_ISL_588441, EPI_ISL_588442, EPI_ISL_588444, EPI_ISL_588446, EPI_ISL_588447, EPI_ISL_588448, EPI_ISL_588449, EPI_ISL_588450, EPI_ISL_588451, EPI_ISL_588452, EPI_ISL_588453, EPI_ISL_588454, EPI_ISL_588456, EPI_ISL_588457, EPI_ISL_588458, EPI_ISL_588459, EPI_ISL_588460, EPI_ISL_588461, EPI_ISL_588462, EPI_ISL_588463, EPI_ISL_588464, EPI_ISL_588465, EPI_ISL_588466, EPI_ISL_588467, EPI_ISL_588468, EPI_ISL_588470, EPI_ISL_588471, EPI_ISL_588472, EPI_ISL_588473, EPI_ISL_588474, EPI_ISL_588475, EPI_ISL_588519, EPI_ISL_588542, EPI_ISL_588581, EPI_ISL_588589, EPI_ISL_588598, EPI_ISL_588619, EPI_ISL_588626, EPI_ISL_588628, EPI_ISL_588643, EPI_ISL_588649, EPI_ISL_588672, EPI_ISL_588675, EPI_ISL_588678, EPI_ISL_588691, EPI_ISL_588699, EPI_ISL_588725, EPI_ISL_588726, EPI_ISL_588727, EPI_ISL_588729, EPI_ISL_588730, EPI_ISL_588731, EPI_ISL_588732, EPI_ISL_588733, EPI_ISL_588734, EPI_ISL_588735, EPI_ISL_588736, EPI_ISL_588737, EPI_ISL_588738, EPI_ISL_588739, EPI_ISL_588740, EPI_ISL_588741, EPI_ISL_588742, EPI_ISL_588743, EPI_ISL_588744, EPI_ISL_588746, EPI_ISL_588747, EPI_ISL_588748, EPI_ISL_588749, EPI_ISL_588750, EPI_ISL_588751, EPI_ISL_588752, EPI_ISL_588753, EPI_ISL_588754, EPI_ISL_588755, EPI_ISL_588756, EPI_ISL_588757, EPI_ISL_588758, EPI_ISL_588759, EPI_ISL_588760, EPI_ISL_588761, EPI_ISL_588762, EPI_ISL_588763, EPI_ISL_588764, EPI_ISL_588765, EPI_ISL_588766, EPI_ISL_588767, EPI_ISL_588768, EPI_ISL_588769, EPI_ISL_588771, EPI_ISL_588773, EPI_ISL_588774, EPI_ISL_588775, EPI_ISL_588776, EPI_ISL_588777, EPI_ISL_588778, EPI_ISL_588779, EPI_ISL_588780, EPI_ISL_588781, EPI_ISL_588782, EPI_ISL_588783, EPI_ISL_588784, EPI_ISL_588785, EPI_ISL_588786, EPI_ISL_588787, EPI_ISL_588788, EPI_ISL_588789, EPI_ISL_588790, EPI_ISL_588791, EPI_ISL_588793, EPI_ISL_588794, EPI_ISL_588795, EPI_ISL_588796, EPI_ISL_588797, EPI_ISL_588798, EPI_ISL_588799, EPI_ISL_588800, EPI_ISL_588801, EPI_ISL_588802, EPI_ISL_588803, EPI_ISL_588804, EPI_ISL_588805, EPI_ISL_588806, EPI_ISL_588807, EPI_ISL_588808, EPI_ISL_588809, EPI_ISL_588810, EPI_ISL_588811, EPI_ISL_588812, EPI_ISL_588813, EPI_ISL_588814, EPI_ISL_588815, EPI_ISL_588816, EPI_ISL_588817, EPI_ISL_588818, EPI_ISL_588819, EPI_ISL_588820, EPI_ISL_588821, EPI_ISL_588822, EPI_ISL_588823, EPI_ISL_588824, EPI_ISL_588825, EPI_ISL_588826, EPI_ISL_588828, EPI_ISL_588829, EPI_ISL_588830, EPI_ISL_588831, EPI_ISL_588832, EPI_ISL_588833, EPI_ISL_588834, EPI_ISL_588835, EPI_ISL_588836, EPI_ISL_588837, EPI_ISL_588838, EPI_ISL_588839, EPI_ISL_588840, EPI_ISL_588841, EPI_ISL_588842, EPI_ISL_588843, EPI_ISL_588846, EPI_ISL_588847, EPI_ISL_588848, EPI_ISL_588849, EPI_ISL_588850, EPI_ISL_588851, EPI_ISL_588852, EPI_ISL_588853, EPI_ISL_588854, EPI_ISL_588856, EPI_ISL_588857, EPI_ISL_588858, EPI_ISL_588859, EPI_ISL_588860, EPI_ISL_588861, EPI_ISL_588862, EPI_ISL_588863, EPI_ISL_588864, EPI_ISL_588865, EPI_ISL_588866, EPI_ISL_588868, EPI_ISL_588869, EPI_ISL_588870, EPI_ISL_588871, EPI_ISL_588872, EPI_ISL_588873, EPI_ISL_588874, EPI_ISL_588875, EPI_ISL_588876, EPI_ISL_588877, EPI_ISL_588878, EPI_ISL_588879, EPI_ISL_588880, EPI_ISL_588881, EPI_ISL_588882, EPI_ISL_588883, EPI_ISL_588884, EPI_ISL_588885, EPI_ISL_588886, EPI_ISL_588887, EPI_ISL_588888, EPI_ISL_588889, EPI_ISL_588890, EPI_ISL_588891, EPI_ISL_588892, EPI_ISL_588893, EPI_ISL_588894, EPI_ISL_588895, EPI_ISL_588896, EPI_ISL_588897, EPI_ISL_588898, EPI_ISL_588899, EPI_ISL_588900, EPI_ISL_588902, EPI_ISL_588903, EPI_ISL_588904, EPI_ISL_588906, EPI_ISL_588907, EPI_ISL_588908, EPI_ISL_588910, EPI_ISL_588911, EPI_ISL_588913, EPI_ISL_588914, EPI_ISL_588915, EPI_ISL_588916, EPI_ISL_588917, EPI_ISL_588919, EPI_ISL_588920, EPI_ISL_588921, EPI_ISL_588922, EPI_ISL_588923, EPI_ISL_588924, EPI_ISL_588925, EPI_ISL_588926, EPI_ISL_588928, EPI_ISL_588929, EPI_ISL_588930, EPI_ISL_588931, EPI_ISL_588933, EPI_ISL_588934, EPI_ISL_588935, EPI_ISL_588936, EPI_ISL_588937, EPI_ISL_588938, EPI_ISL_588939, EPI_ISL_588941, EPI_ISL_588942, EPI_ISL_588943, EPI_ISL_588945, EPI_ISL_588946,

EPI_ISL_588948, EPI_ISL_588949, EPI_ISL_588950, EPI_ISL_588951, EPI_ISL_588952, EPI_ISL_588953, EPI_ISL_588954, EPI_ISL_588955, EPI_ISL_588956, EPI_ISL_588957, EPI_ISL_588958, EPI_ISL_588959, EPI_ISL_588960, EPI_ISL_588961, EPI_ISL_588963, EPI_ISL_588964, EPI_ISL_588965, EPI_ISL_588967, EPI_ISL_588968, EPI_ISL_588969, EPI_ISL_588970, EPI_ISL_588971, EPI_ISL_588972, EPI_ISL_588973, EPI_ISL_588974, EPI_ISL_588976, EPI_ISL_588977, EPI_ISL_588978, EPI_ISL_588979, EPI_ISL_588980, EPI_ISL_588981, EPI_ISL_588982, EPI_ISL_588983, EPI_ISL_588984, EPI_ISL_588985, EPI_ISL_588987, EPI_ISL_588989, EPI_ISL_588990, EPI_ISL_588991, EPI_ISL_588992, EPI_ISL_588993, EPI_ISL_588994, EPI_ISL_588995, EPI_ISL_588996, EPI_ISL_588997, EPI_ISL_588998, EPI_ISL_588999, EPI_ISL_589000, EPI_ISL_589001, EPI_ISL_589002, EPI_ISL_589003, EPI_ISL_589004, EPI_ISL_589005, EPI_ISL_589006, EPI_ISL_589007, EPI_ISL_589008, EPI_ISL_589009, EPI_ISL_589010, EPI_ISL_589011, EPI_ISL_589013, EPI_ISL_589014, EPI_ISL_589016, EPI_ISL_589020, EPI_ISL_589022, EPI_ISL_589025, EPI_ISL_589027, EPI_ISL_589031, EPI_ISL_589032, EPI_ISL_589033, EPI_ISL_589035, EPI_ISL_589036, EPI_ISL_589038, EPI_ISL_589042, EPI_ISL_589044, EPI_ISL_589045, EPI_ISL_589046, EPI_ISL_589047, EPI_ISL_589048, EPI_ISL_589049, EPI_ISL_589052, EPI_ISL_589053, EPI_ISL_589054, EPI_ISL_589055, EPI_ISL_589056, EPI_ISL_589058, EPI_ISL_589059, EPI_ISL_589060, EPI_ISL_589061, EPI_ISL_589062, EPI_ISL_589064, EPI_ISL_589065, EPI_ISL_589066, EPI_ISL_589068, EPI_ISL_589069, EPI_ISL_589071, EPI_ISL_589072, EPI_ISL_589073, EPI_ISL_589076, EPI_ISL_589078, EPI_ISL_589082, EPI_ISL_589084, EPI_ISL_589085, EPI_ISL_589087, EPI_ISL_589088, EPI_ISL_589089, EPI_ISL_589092, EPI_ISL_589093, EPI_ISL_589094, EPI_ISL_589096, EPI_ISL_589100, EPI_ISL_589105, EPI_ISL_589106, EPI_ISL_589107, EPI_ISL_589112, EPI_ISL_589113, EPI_ISL_589116, EPI_ISL_589120, EPI_ISL_589124, EPI_ISL_589125, EPI_ISL_589126, EPI_ISL_589128, EPI_ISL_589129, EPI_ISL_589131, EPI_ISL_589132, EPI_ISL_589134, EPI_ISL_589136, EPI_ISL_589138, EPI_ISL_589139, EPI_ISL_589140, EPI_ISL_589141, EPI_ISL_589142, EPI_ISL_589143, EPI_ISL_589144, EPI_ISL_589145, EPI_ISL_589146, EPI_ISL_589148, EPI_ISL_589150, EPI_ISL_589151, EPI_ISL_589152, EPI_ISL_589154, EPI_ISL_589155, EPI_ISL_589156, EPI_ISL_589158, EPI_ISL_589159, EPI_ISL_589162, EPI_ISL_589163, EPI_ISL_589164, EPI_ISL_589165, EPI_ISL_589175, EPI_ISL_589177, EPI_ISL_589178, EPI_ISL_589180, EPI_ISL_589181, EPI_ISL_589186, EPI_ISL_589188, EPI_ISL_589189, EPI_ISL_589190, EPI_ISL_589191, EPI_ISL_589192, EPI_ISL_589194, EPI_ISL_589197, EPI_ISL_589201, EPI_ISL_589202, EPI_ISL_589203, EPI_ISL_589204, EPI_ISL_589206, EPI_ISL_589207, EPI_ISL_589209, EPI_ISL_589210, EPI_ISL_589213, EPI_ISL_589214, EPI_ISL_589215, EPI_ISL_589216, EPI_ISL_589217, EPI_ISL_589218, EPI_ISL_589219, EPI_ISL_589220, EPI_ISL_589222, EPI_ISL_589223, EPI_ISL_589228, EPI_ISL_589230, EPI_ISL_589232, EPI_ISL_589234, EPI_ISL_589236, EPI_ISL_589238, EPI_ISL_589239, EPI_ISL_589240, EPI_ISL_589241, EPI_ISL_589244, EPI_ISL_589245, EPI_ISL_589246, EPI_ISL_589249, EPI_ISL_589250, EPI_ISL_589252, EPI_ISL_589254, EPI_ISL_589255, EPI_ISL_589257, EPI_ISL_589258, EPI_ISL_589263

| | | | |
|---|---|---|---|
| see above | Lighthouse Lab in Glasgow | Wellcome Sanger Institute for the COVID-19 Genomics UK (COG-UK) consortium | Harper VanSteenhouse, Yumi Kasai, David Gray, Carol Clugston, Anna Dominiczak and Alex Alderton, Roberto Amato, Sonia Goncalves, Ewan Harrison, David K. Jackson, Ian Johnston, Dominic Kwiatkowski, Cordelia Langford, John Sillitoe on behalf of the Wellcome Sanger Institute COVID-19 Surveillance Team |

EPI_ISL_589267, EPI_ISL_589268, EPI_ISL_589269, EPI_ISL_589270, EPI_ISL_589271, EPI_ISL_589272, EPI_ISL_589273, EPI_ISL_589274, EPI_ISL_589275, EPI_ISL_589276, EPI_ISL_589277, EPI_ISL_589278, EPI_ISL_589279, EPI_ISL_589280, EPI_ISL_589281, EPI_ISL_589282, EPI_ISL_589283, EPI_ISL_589284, EPI_ISL_589285, EPI_ISL_589286, EPI_ISL_589287, EPI_ISL_589288, EPI_ISL_589290, EPI_ISL_589291, EPI_ISL_589292, EPI_ISL_589293, EPI_ISL_589294, EPI_ISL_589295, EPI_ISL_589296, EPI_ISL_589298, EPI_ISL_589299, EPI_ISL_589300, EPI_ISL_589301, EPI_ISL_589303, EPI_ISL_589305, EPI_ISL_589306, EPI_ISL_589307, EPI_ISL_589308, EPI_ISL_589309, EPI_ISL_589310, EPI_ISL_589311, EPI_ISL_589312, EPI_ISL_589313, EPI_ISL_589314, EPI_ISL_589315, EPI_ISL_589316, EPI_ISL_589317, EPI_ISL_589318, EPI_ISL_589319, EPI_ISL_589320, EPI_ISL_589321, EPI_ISL_589322, EPI_ISL_589323, EPI_ISL_589324, EPI_ISL_589325, EPI_ISL_589326, EPI_ISL_589327, EPI_ISL_589328, EPI_ISL_589330, EPI_ISL_589331, EPI_ISL_589332, EPI_ISL_589333, EPI_ISL_589336, EPI_ISL_589337, EPI_ISL_589338, EPI_ISL_589339, EPI_ISL_589340, EPI_ISL_589341, EPI_ISL_589343, EPI_ISL_589345, EPI_ISL_589346, EPI_ISL_589347, EPI_ISL_589348, EPI_ISL_589349, EPI_ISL_589350, EPI_ISL_589351, EPI_ISL_589352, EPI_ISL_589353, EPI_ISL_589354, EPI_ISL_589356, EPI_ISL_589357, EPI_ISL_589358, EPI_ISL_589359, EPI_ISL_589360, EPI_ISL_589361, EPI_ISL_589362, EPI_ISL_589363, EPI_ISL_589364, EPI_ISL_589365, EPI_ISL_589366, EPI_ISL_589367, EPI_ISL_589368, EPI_ISL_589369, EPI_ISL_589370, EPI_ISL_589371, EPI_ISL_589372, EPI_ISL_589373, EPI_ISL_589374, EPI_ISL_589375, EPI_ISL_589378, EPI_ISL_589379, EPI_ISL_589380, EPI_ISL_589381, EPI_ISL_589382, EPI_ISL_589383, EPI_ISL_589384, EPI_ISL_589385, EPI_ISL_589386, EPI_ISL_589387, EPI_ISL_589388, EPI_ISL_589389, EPI_ISL_589390, EPI_ISL_589391, EPI_ISL_589392, EPI_ISL_589393, EPI_ISL_589394, EPI_ISL_589395, EPI_ISL_589396, EPI_ISL_589397, EPI_ISL_589398, EPI_ISL_589399, EPI_ISL_589400, EPI_ISL_589401, EPI_ISL_589402, EPI_ISL_589403, EPI_ISL_589404, EPI_ISL_589405, EPI_ISL_589406, EPI_ISL_589408, EPI_ISL_589409, EPI_ISL_589410, EPI_ISL_589411, EPI_ISL_589412, EPI_ISL_589413, EPI_ISL_589414, EPI_ISL_589415, EPI_ISL_589416, EPI_ISL_589418, EPI_ISL_589419, EPI_ISL_589420, EPI_ISL_589421, EPI_ISL_589422, EPI_ISL_589424, EPI_ISL_589425, EPI_ISL_589426, EPI_ISL_589427, EPI_ISL_589428, EPI_ISL_589429, EPI_ISL_589431, EPI_ISL_589432, EPI_ISL_589433, EPI_ISL_589434, EPI_ISL_589435, EPI_ISL_589436, EPI_ISL_589437, EPI_ISL_589438, EPI_ISL_589439, EPI_ISL_589441, EPI_ISL_589442, EPI_ISL_589443, EPI_ISL_589444, EPI_ISL_589445, EPI_ISL_589446, EPI_ISL_589447, EPI_ISL_589448, EPI_ISL_589449, EPI_ISL_589450, EPI_ISL_589451, EPI_ISL_589452, EPI_ISL_589453, EPI_ISL_589454, EPI_ISL_589455, EPI_ISL_589456, EPI_ISL_589457, EPI_ISL_589458, EPI_ISL_589459, EPI_ISL_589460, EPI_ISL_589461, EPI_ISL_589462, EPI_ISL_589463, EPI_ISL_589464, EPI_ISL_589465, EPI_ISL_589466, EPI_ISL_589467, EPI_ISL_589468, EPI_ISL_589469, EPI_ISL_589470, EPI_ISL_589471, EPI_ISL_589472, EPI_ISL_589473, EPI_ISL_589474, EPI_ISL_589475, EPI_ISL_589476, EPI_ISL_589477, EPI_ISL_589478, EPI_ISL_589479, EPI_ISL_589480, EPI_ISL_589481, EPI_ISL_589482, EPI_ISL_589483, EPI_ISL_589484, EPI_ISL_589485, EPI_ISL_589486, EPI_ISL_589487, EPI_ISL_589488, EPI_ISL_589489, EPI_ISL_589490, EPI_ISL_589491, EPI_ISL_589492, EPI_ISL_589494, EPI_ISL_589495, EPI_ISL_589496, EPI_ISL_589497, EPI_ISL_589498, EPI_ISL_589499, EPI_ISL_589500, EPI_ISL_589501, EPI_ISL_589502, EPI_ISL_589503, EPI_ISL_589504, EPI_ISL_589505, EPI_ISL_589506, EPI_ISL_589507, EPI_ISL_589508, EPI_ISL_589509, EPI_ISL_589510, EPI_ISL_589511, EPI_ISL_589512, EPI_ISL_589513, EPI_ISL_589514, EPI_ISL_589515, EPI_ISL_589516, EPI_ISL_589517, EPI_ISL_589518, EPI_ISL_589522, EPI_ISL_589523, EPI_ISL_589524, EPI_ISL_589525, EPI_ISL_589527, EPI_ISL_589528, EPI_ISL_589529, EPI_ISL_589530, EPI_ISL_589531, EPI_ISL_589532, EPI_ISL_589533, EPI_ISL_589534, EPI_ISL_589535, EPI_ISL_589536, EPI_ISL_589537, EPI_ISL_589538, EPI_ISL_589539, EPI_ISL_589540, EPI_ISL_589541, EPI_ISL_589542, EPI_ISL_589543, EPI_ISL_589544, EPI_ISL_589545, EPI_ISL_589546, EPI_ISL_589547, EPI_ISL_589548, EPI_ISL_589549, EPI_ISL_589550, EPI_ISL_589551, EPI_ISL_589552, EPI_ISL_589553, EPI_ISL_589554, EPI_ISL_589555, EPI_ISL_589556, EPI_ISL_589557, EPI_ISL_589558, EPI_ISL_589559, EPI_ISL_589560, EPI_ISL_589561, EPI_ISL_589562, EPI_ISL_589563, EPI_ISL_589564, EPI_ISL_589565, EPI_ISL_589566, EPI_ISL_589568, EPI_ISL_589569, EPI_ISL_589570, EPI_ISL_589571

| | | | |
|---|---|---|---|
| see above | Lighthouse Lab in Alderley Park | Wellcome Sanger Institute for the COVID-19 Genomics UK (COG-UK) consortium | Jacquelyn Wynn, Mairead Hyland, The Lighthouse Lab in Alderley Park and Alex Alderton, Roberto Amato, Sonia Goncalves, Ewan Harrison, David K. Jackson, Ian Johnston, Dominic Kwiatkowski, Cordelia Langford, John Sillitoe on behalf of the Wellcome Sanger Institute COVID-19 Surveillance Team |

EPI_ISL_589762, EPI_ISL_589765, EPI_ISL_589778, EPI_ISL_589779, EPI_ISL_589782, EPI_ISL_589783, EPI_ISL_589786, EPI_ISL_590198, EPI_ISL_590213, EPI_ISL_590234, EPI_ISL_590307, EPI_ISL_590364, EPI_ISL_590419

| | | | |
|---|---|---|---|
| see above | Lighthouse Lab in Alderley Park | Wellcome Sanger Institute for the COVID-19 Genomics UK (COG-UK) consortium | Jacquelyn Wynn, Mairead Hyland, The Lighthouse Lab in Alderley Park and Alex Alderton, Roberto Amato, Sonia Goncalves, Ewan Harrison, David K. Jackson, Ian Johnston, Dominic Kwiatkowski, Cordelia Langford, John Sillitoe on behalf of the Wellcome Sanger Institute COVID-19 Surveillance Team (http://www.sanger.ac.uk/covid-team) |
| EPI_ISL_591270, EPI_ISL_591271 | National Institute for Viral Disease Control and Prevention, China CDC | National Institute for Viral Disease Control and Prevention, China CDC | Huilai Ma, Zhaoguo Wang, Xiang Zhao, Jun Han, Yong Zhang, Hong Wang, Cao Chen, Ji Wang, Jingdong Song, Yao Meng, Yuchao Wu, Zhixiao Chen, Dayan Wang, Ruqin Gao, George F.Gao, Wenbo Xu |

EPI_ISL_594484, EPI_ISL_594485, EPI_ISL_594486, EPI_ISL_594487, EPI_ISL_594488, EPI_ISL_594489, EPI_ISL_594490, EPI_ISL_594491, EPI_ISL_594492, EPI_ISL_594493, EPI_ISL_594494, EPI_ISL_594495, EPI_ISL_594496, EPI_ISL_594497, EPI_ISL_594498, EPI_ISL_594499, EPI_ISL_594500, EPI_ISL_594501, EPI_ISL_594502, EPI_ISL_594503, EPI_ISL_594504, EPI_ISL_594505, EPI_ISL_594506, EPI_ISL_594507, EPI_ISL_594508, EPI_ISL_594509, EPI_ISL_594510, EPI_ISL_594511, EPI_ISL_594512, EPI_ISL_594513, EPI_ISL_594514, EPI_ISL_594515, EPI_ISL_594516, EPI_ISL_594517, EPI_ISL_594518, EPI_ISL_594519, EPI_ISL_594520, EPI_ISL_594521, EPI_ISL_594522, EPI_ISL_594523, EPI_ISL_594524, EPI_ISL_594525, EPI_ISL_594526, EPI_ISL_594527, EPI_ISL_594528, EPI_ISL_594529, EPI_ISL_594530, EPI_ISL_594531, EPI_ISL_594532, EPI_ISL_594533, EPI_ISL_594534, EPI_ISL_594535, EPI_ISL_594536, EPI_ISL_594537, EPI_ISL_594538, EPI_ISL_594539, EPI_ISL_594540, EPI_ISL_594541, EPI_ISL_594542, EPI_ISL_594543, EPI_ISL_594544, EPI_ISL_594545, EPI_ISL_594546, EPI_ISL_594547, EPI_ISL_594548, EPI_ISL_594549, EPI_ISL_594550, EPI_ISL_594551, EPI_ISL_594552, EPI_ISL_594553, EPI_ISL_594554, EPI_ISL_594555, EPI_ISL_594556, EPI_ISL_594557, EPI_ISL_594558, EPI_ISL_594559, EPI_ISL_594560, EPI_ISL_594561, EPI_ISL_594562, EPI_ISL_594563, EPI_ISL_594564, EPI_ISL_594565, EPI_ISL_594566, EPI_ISL_594567, EPI_ISL_594568, EPI_ISL_594569, EPI_ISL_594570, EPI_ISL_594571, EPI_ISL_594572, EPI_ISL_594573, EPI_ISL_594574, EPI_ISL_594575, EPI_ISL_594576, EPI_ISL_594577, EPI_ISL_594578, EPI_ISL_594579, EPI_ISL_594580, EPI_ISL_594581, EPI_ISL_594582, EPI_ISL_594583, EPI_ISL_594584, EPI_ISL_594585, EPI_ISL_594586, EPI_ISL_594587, EPI_ISL_594588, EPI_ISL_594589, EPI_ISL_594590, EPI_ISL_594591, EPI_ISL_594592, EPI_ISL_594593, EPI_ISL_594594, EPI_ISL_594595, EPI_ISL_594596, EPI_ISL_594597, EPI_ISL_594598, EPI_ISL_594599, EPI_ISL_594600, EPI_ISL_594601, EPI_ISL_594602, EPI_ISL_594603, EPI_ISL_594604, EPI_ISL_594605, EPI_ISL_594606, EPI_ISL_594607, EPI_ISL_594608, EPI_ISL_594609, EPI_ISL_594610, EPI_ISL_594611, EPI_ISL_594612, EPI_ISL_594613, EPI_ISL_594614, EPI_ISL_594615, EPI_ISL_594616

| | | | |
|---|---|---|---|
| see above | University of Birmingham | COVID-19 Genomics UK (COG-UK) Consortium | Institute of Microbiology, University of Birmingham: Claire McMurray, Joanne Stockton, Samuel Nicholls, Radoslaw Poplawski, Will Rowe, Josh Quick, Nicholas Loman. University of Birmingham Testing Laboratory: Celina M Whalley, Andrew Bosworth, Charlotte Poxon, Kasun Wanigasooriya, Oliver Pickles, Mike Kidd, Alex Richter, Andrew D Beggs PHE Heartlands Lab: Husam Osman, Andrew Bosworth. Queen Elizabeth Hospital: Anna Casey |

EPI_ISL_597525, EPI_ISL_597526, EPI_ISL_597527, EPI_ISL_597528, EPI_ISL_597529, EPI_ISL_597530, EPI_ISL_597531, EPI_ISL_597532, EPI_ISL_597533, EPI_ISL_597535, EPI_ISL_597536, EPI_ISL_597537, EPI_ISL_597538, EPI_ISL_597540, EPI_ISL_597542, EPI_ISL_597543, EPI_ISL_597544, EPI_ISL_597545, EPI_ISL_597546, EPI_ISL_597548, EPI_ISL_597549, EPI_ISL_597550, EPI_ISL_597552, EPI_ISL_597553, EPI_ISL_597554, EPI_ISL_597555, EPI_ISL_597556, EPI_ISL_597557, EPI_ISL_597561, EPI_ISL_597562, EPI_ISL_597563, EPI_ISL_597564, EPI_ISL_597565, EPI_ISL_597566, EPI_ISL_597568, EPI_ISL_597569, EPI_ISL_597571, EPI_ISL_597572, EPI_ISL_597573, EPI_ISL_597574, EPI_ISL_597575, EPI_ISL_597576, EPI_ISL_597577, EPI_ISL_597579, EPI_ISL_597582, EPI_ISL_597583, EPI_ISL_597585, EPI_ISL_597587, EPI_ISL_597588, EPI_ISL_597589, EPI_ISL_597590, EPI_ISL_597591, EPI_ISL_597592, EPI_ISL_597593, EPI_ISL_597594, EPI_ISL_597595, EPI_ISL_597598, EPI_ISL_597599, EPI_ISL_597601, EPI_ISL_597602, EPI_ISL_597603, EPI_ISL_597605, EPI_ISL_597606, EPI_ISL_597607, EPI_ISL_597609, EPI_ISL_597610, EPI_ISL_597611, EPI_ISL_597612, EPI_ISL_597613, EPI_ISL_597614, EPI_ISL_597615, EPI_ISL_597617, EPI_ISL_597618, EPI_ISL_597619, EPI_ISL_597622, EPI_ISL_597623, EPI_ISL_597624, EPI_ISL_597625, EPI_ISL_597627, EPI_ISL_597628, EPI_ISL_597630, EPI_ISL_597631, EPI_ISL_597632, EPI_ISL_597635, EPI_ISL_597636, EPI_ISL_597637, EPI_ISL_597638, EPI_ISL_597640, EPI_ISL_597641, EPI_ISL_597642, EPI_ISL_597643, EPI_ISL_597644, EPI_ISL_597646, EPI_ISL_597648, EPI_ISL_597649, EPI_ISL_597651, EPI_ISL_597653, EPI_ISL_597657, EPI_ISL_597658, EPI_ISL_597659, EPI_ISL_597661, EPI_ISL_597662, EPI_ISL_597664, EPI_ISL_597665, EPI_ISL_597667, EPI_ISL_597668, EPI_ISL_597669, EPI_ISL_597670, EPI_ISL_597671, EPI_ISL_597672, EPI_ISL_597673, EPI_ISL_597674, EPI_ISL_597675, EPI_ISL_597676, EPI_ISL_597677, EPI_ISL_597678, EPI_ISL_597679, EPI_ISL_597680, EPI_ISL_597681, EPI_ISL_597682, EPI_ISL_597683, EPI_ISL_597684, EPI_ISL_597685, EPI_ISL_597686, EPI_ISL_597687, EPI_ISL_597688, EPI_ISL_597690, EPI_ISL_597691, EPI_ISL_597693, EPI_ISL_597696, EPI_ISL_597697, EPI_ISL_597698, EPI_ISL_597699, EPI_ISL_597701, EPI_ISL_597702, EPI_ISL_597703, EPI_ISL_597704, EPI_ISL_597706, EPI_ISL_597707, EPI_ISL_597708, EPI_ISL_597709, EPI_ISL_597710, EPI_ISL_597714, EPI_ISL_597716, EPI_ISL_597717, EPI_ISL_597718, EPI_ISL_597719, EPI_ISL_597720, EPI_ISL_597724, EPI_ISL_597726, EPI_ISL_597727, EPI_ISL_597728, EPI_ISL_597729, EPI_ISL_597730, EPI_ISL_597734, EPI_ISL_597735, EPI_ISL_597736, EPI_ISL_597737, EPI_ISL_597739, EPI_ISL_597740, EPI_ISL_597741, EPI_ISL_597742, EPI_ISL_597743, EPI_ISL_597744, EPI_ISL_597745, EPI_ISL_597746, EPI_ISL_597748, EPI_ISL_597749, EPI_ISL_597750, EPI_ISL_597751, EPI_ISL_597752, EPI_ISL_597753, EPI_ISL_597758, EPI_ISL_597759, EPI_ISL_597761, EPI_ISL_597762, EPI_ISL_597764, EPI_ISL_597765, EPI_ISL_597766, EPI_ISL_597767, EPI_ISL_597768, EPI_ISL_597769, EPI_ISL_597770, EPI_ISL_597772, EPI_ISL_597774, EPI_ISL_597775, EPI_ISL_597776, EPI_ISL_597777, EPI_ISL_597779, EPI_ISL_597780, EPI_ISL_597782, EPI_ISL_597783, EPI_ISL_597784, EPI_ISL_597786, EPI_ISL_597788, EPI_ISL_597789, EPI_ISL_597790, EPI_ISL_597791, EPI_ISL_597792, EPI_ISL_597793, EPI_ISL_597795, EPI_ISL_597797, EPI_ISL_597799, EPI_ISL_597800, EPI_ISL_597801, EPI_ISL_597802, EPI_ISL_597803, EPI_ISL_597804, EPI_ISL_597805, EPI_ISL_597806, EPI_ISL_597807, EPI_ISL_597809, EPI_ISL_597810, EPI_ISL_597811, EPI_ISL_597813, EPI_ISL_597814, EPI_ISL_597815, EPI_ISL_597816, EPI_ISL_597817, EPI_ISL_597818, EPI_ISL_597819, EPI_ISL_597820, EPI_ISL_597821, EPI_ISL_597822, EPI_ISL_597823, EPI_ISL_597824, EPI_ISL_597825, EPI_ISL_597828, EPI_ISL_597831, EPI_ISL_597832, EPI_ISL_597833, EPI_ISL_597835, EPI_ISL_597837, EPI_ISL_597841, EPI_ISL_597842, EPI_ISL_597843, EPI_ISL_597844, EPI_ISL_597845, EPI_ISL_597847, EPI_ISL_597848, EPI_ISL_597849, EPI_ISL_597850, EPI_ISL_597851, EPI_ISL_597852, EPI_ISL_597854, EPI_ISL_597857, EPI_ISL_597859, EPI_ISL_597860, EPI_ISL_597861, EPI_ISL_597862, EPI_ISL_597864, EPI_ISL_597865, EPI_ISL_597866, EPI_ISL_597869, EPI_ISL_597870, EPI_ISL_597872, EPI_ISL_597873, EPI_ISL_597874, EPI_ISL_597875, EPI_ISL_597877, EPI_ISL_597880, EPI_ISL_597883, EPI_ISL_597884, EPI_ISL_597885, EPI_ISL_597886, EPI_ISL_597887, EPI_ISL_597888, EPI_ISL_597889, EPI_ISL_597890, EPI_ISL_597891, EPI_ISL_597892, EPI_ISL_597893, EPI_ISL_597894, EPI_ISL_597895, EPI_ISL_597896, EPI_ISL_597897, EPI_ISL_597898, EPI_ISL_597899, EPI_ISL_597900, EPI_ISL_597901, EPI_ISL_597902, EPI_ISL_597903, EPI_ISL_597904, EPI_ISL_597905, EPI_ISL_597906, EPI_ISL_597907, EPI_ISL_597908, EPI_ISL_597909, EPI_ISL_597910, EPI_ISL_597911, EPI_ISL_597912, EPI_ISL_597913, EPI_ISL_597914, EPI_ISL_597915, EPI_ISL_597916, EPI_ISL_597917, EPI_ISL_597918, EPI_ISL_597919, EPI_ISL_597920, EPI_ISL_597921, EPI_ISL_597922, EPI_ISL_597923, EPI_ISL_597924, EPI_ISL_597925, EPI_ISL_597926, EPI_ISL_597927, EPI_ISL_597928, EPI_ISL_597929, EPI_ISL_597930, EPI_ISL_597931, EPI_ISL_597932, EPI_ISL_597933, EPI_ISL_597934, EPI_ISL_597935, EPI_ISL_597936, EPI_ISL_597937, EPI_ISL_597938, EPI_ISL_597939, EPI_ISL_597940, EPI_ISL_597941, EPI_ISL_597942, EPI_ISL_597943, EPI_ISL_597944, EPI_ISL_597945, EPI_ISL_597946, EPI_ISL_597947, EPI_ISL_597948, EPI_ISL_597949, EPI_ISL_597950, EPI_ISL_597951, EPI_ISL_597952, EPI_ISL_597953, EPI_ISL_597954, EPI_ISL_597955, EPI_ISL_597956, EPI_ISL_597957, EPI_ISL_597958, EPI_ISL_597959, EPI_ISL_597960, EPI_ISL_597961, EPI_ISL_597962, EPI_ISL_597963, EPI_ISL_597964, EPI_ISL_597965, EPI_ISL_597966, EPI_ISL_597967, EPI_ISL_597968, EPI_ISL_597969, EPI_ISL_597970, EPI_ISL_597971, EPI_ISL_597972, EPI_ISL_597973, EPI_ISL_597974, EPI_ISL_597975, EPI_ISL_597976, EPI_ISL_597977, EPI_ISL_597978, EPI_ISL_597979, EPI_ISL_597980, EPI_ISL_597981, EPI_ISL_597982, EPI_ISL_597983, EPI_ISL_597985, EPI_ISL_597986, EPI_ISL_597987, EPI_ISL_597988, EPI_ISL_597989, EPI_ISL_597990, EPI_ISL_597991, EPI_ISL_597992, EPI_ISL_597993, EPI_ISL_597994, EPI_ISL_597995, EPI_ISL_597996, EPI_ISL_597997, EPI_ISL_597998, EPI_ISL_597999, EPI_ISL_598000, EPI_ISL_598001, EPI_ISL_598002, EPI_ISL_598003, EPI_ISL_598004, EPI_ISL_598005, EPI_ISL_598006, EPI_ISL_598007, EPI_ISL_598008, EPI_ISL_598009, EPI_ISL_598010, EPI_ISL_598011, EPI_ISL_598012, EPI_ISL_598013, EPI_ISL_598014, EPI_ISL_598015, EPI_ISL_598016, EPI_ISL_598017, EPI_ISL_598018, EPI_ISL_598019, EPI_ISL_598020, EPI_ISL_598021, EPI_ISL_598022, EPI_ISL_598023, EPI_ISL_598024, EPI_ISL_598025, EPI_ISL_598026, EPI_ISL_598027, EPI_ISL_598028, EPI_ISL_598029, EPI_ISL_598030, EPI_ISL_598031, EPI_ISL_598032, EPI_ISL_598033, EPI_ISL_598034, EPI_ISL_598035, EPI_ISL_598036, EPI_ISL_598037, EPI_ISL_598038, EPI_ISL_598039, EPI_ISL_598040, EPI_ISL_598041, EPI_ISL_598042, EPI_ISL_598043, EPI_ISL_598044, EPI_ISL_598045, EPI_ISL_598046, EPI_ISL_598047, EPI_ISL_598048, EPI_ISL_598049, EPI_ISL_598050, EPI_ISL_598051, EPI_ISL_598052, EPI_ISL_598053, EPI_ISL_598054, EPI_ISL_598055, EPI_ISL_598056, EPI_ISL_598057, EPI_ISL_598058, EPI_ISL_598059, EPI_ISL_598060, EPI_ISL_598061, EPI_ISL_598062, EPI_ISL_598063, EPI_ISL_598064, EPI_ISL_598065, EPI_ISL_598066, EPI_ISL_598067, EPI_ISL_598068, EPI_ISL_598069, EPI_ISL_598070, EPI_ISL_598071, EPI_ISL_598072, EPI_ISL_598073, EPI_ISL_598074, EPI_ISL_598075, EPI_ISL_598076, EPI_ISL_598077, EPI_ISL_598078, EPI_ISL_598079, EPI_ISL_598080, EPI_ISL_598081, EPI_ISL_598082, EPI_ISL_598083, EPI_ISL_598084, EPI_ISL_598085, EPI_ISL_598086, EPI_ISL_598087, EPI_ISL_598088, EPI_ISL_598089, EPI_ISL_598090, EPI_ISL_598091, EPI_ISL_598092, EPI_ISL_598093, EPI_ISL_598094, EPI_ISL_598095, EPI_ISL_598096, EPI_ISL_598097, EPI_ISL_598098, EPI_ISL_598099, EPI_ISL_598100, EPI_ISL_598101, EPI_ISL_598102, EPI_ISL_598103, EPI_ISL_598104, EPI_ISL_598105, EPI_ISL_598106, EPI_ISL_598107, EPI_ISL_598108, EPI_ISL_598109, EPI_ISL_598110, EPI_ISL_598111, EPI_ISL_598112, EPI_ISL_598113, EPI_ISL_598114, EPI_ISL_598115, EPI_ISL_598116, EPI_ISL_598117, EPI_ISL_598118, EPI_ISL_598119, EPI_ISL_598120, EPI_ISL_598121, EPI_ISL_598122, EPI_ISL_598123, EPI_ISL_598124, EPI_ISL_598125, EPI_ISL_598126, EPI_ISL_598127, EPI_ISL_598128, EPI_ISL_598129, EPI_ISL_598130, EPI_ISL_598131, EPI_ISL_598132, EPI_ISL_598133, EPI_ISL_598134, EPI_ISL_598135, EPI_ISL_598136, EPI_ISL_598137, EPI_ISL_598138, EPI_ISL_598139, EPI_ISL_598140, EPI_ISL_598141, EPI_ISL_598142, EPI_ISL_598143, EPI_ISL_598144, EPI_ISL_598145, EPI_ISL_598146, EPI_ISL_598147, EPI_ISL_598148, EPI_ISL_598149, EPI_ISL_598150, EPI_ISL_598151, EPI_ISL_598152, EPI_ISL_598153, EPI_ISL_598154, EPI_ISL_598155, EPI_ISL_598156, EPI_ISL_598157, EPI_ISL_598158, EPI_ISL_598159, EPI_ISL_598160, EPI_ISL_598161, EPI_ISL_598162, EPI_ISL_598164, EPI_ISL_598165, EPI_ISL_598166, EPI_ISL_598167, EPI_ISL_598168, EPI_ISL_598169, EPI_ISL_598170, EPI_ISL_598171, EPI_ISL_598172, EPI_ISL_598173, EPI_ISL_598174, EPI_ISL_598175, EPI_ISL_598176, EPI_ISL_598177, EPI_ISL_598178, EPI_ISL_598179, EPI_ISL_598180, EPI_ISL_598181, EPI_ISL_598182, EPI_ISL_598183, EPI_ISL_598184, EPI_ISL_598185, EPI_ISL_598186, EPI_ISL_598187, EPI_ISL_598188, EPI_ISL_598189, EPI_ISL_598190, EPI_ISL_598191, EPI_ISL_598192, EPI_ISL_598193, EPI_ISL_598194, EPI_ISL_598195, EPI_ISL_598196, EPI_ISL_598197, EPI_ISL_598198, EPI_ISL_598199, EPI_ISL_598200, EPI_ISL_598201, EPI_ISL_598202, EPI_ISL_598203, EPI_ISL_598205, EPI_ISL_598206, EPI_ISL_598207, EPI_ISL_598208, EPI_ISL_598209, EPI_ISL_598210, EPI_ISL_598211, EPI_ISL_598212, EPI_ISL_598213, EPI_ISL_598214, EPI_ISL_598215, EPI_ISL_598216, EPI_ISL_598217,

| | | | |
|---|---|---|---|
| EPI_ISL_598218, EPI_ISL_598219, EPI_ISL_598220, EPI_ISL_598221, EPI_ISL_598222, EPI_ISL_598223, EPI_ISL_598224, EPI_ISL_598225, EPI_ISL_598226, EPI_ISL_598227, EPI_ISL_598228, EPI_ISL_598229, EPI_ISL_598230, EPI_ISL_598231, EPI_ISL_598232, EPI_ISL_598233, EPI_ISL_598234, EPI_ISL_598235, EPI_ISL_598236, EPI_ISL_598237, EPI_ISL_598238, EPI_ISL_598239, EPI_ISL_598240, EPI_ISL_598241, EPI_ISL_598243, EPI_ISL_598244, EPI_ISL_598245, EPI_ISL_598246, EPI_ISL_598247, EPI_ISL_598248, EPI_ISL_598828 | | | |
| see above | Lighthouse Lab in Alderley Park | Wellcome Sanger Institute for the COVID-19 Genomics UK (COG-UK) consortium | Jacquelyn Wynn, Mairead Hyland, The Lighthouse Lab in Alderley Park and Alex Alderton, Roberto Amato, Sonia Goncalves, Ewan Harrison, David K. Jackson, Ian Johnston, Dominic Kwiatkowski, Cordelia Langford, John Sillitoe on behalf of the Wellcome Sanger Institute COVID-19 Surveillance Team (http://www.sanger.ac.uk/covid-team) |
| EPI_ISL_599138, EPI_ISL_599139, EPI_ISL_599140, EPI_ISL_599141, EPI_ISL_599142, EPI_ISL_599143, EPI_ISL_599144, EPI_ISL_599146, EPI_ISL_599147, EPI_ISL_599148, EPI_ISL_599149, EPI_ISL_599150, EPI_ISL_599151, EPI_ISL_599152, EPI_ISL_599153, EPI_ISL_599155, EPI_ISL_599156, EPI_ISL_599157, EPI_ISL_599158, EPI_ISL_599159, EPI_ISL_599162, EPI_ISL_599163, EPI_ISL_599165, EPI_ISL_599167, EPI_ISL_599170, EPI_ISL_599174, EPI_ISL_599175, EPI_ISL_599178, EPI_ISL_599181, EPI_ISL_599182, EPI_ISL_599184, EPI_ISL_599185, EPI_ISL_599186, EPI_ISL_599187, EPI_ISL_599188, EPI_ISL_599190, EPI_ISL_599192, EPI_ISL_599193, EPI_ISL_599195, EPI_ISL_599196, EPI_ISL_599198, EPI_ISL_599199, EPI_ISL_599200, EPI_ISL_599202, EPI_ISL_599203, EPI_ISL_599204, EPI_ISL_599205, EPI_ISL_599207, EPI_ISL_599208, EPI_ISL_599210, EPI_ISL_599213, EPI_ISL_599217, EPI_ISL_599218, EPI_ISL_599222, EPI_ISL_599224, EPI_ISL_599226, EPI_ISL_599229, EPI_ISL_599230, EPI_ISL_599232, EPI_ISL_599234, EPI_ISL_599240, EPI_ISL_599242, EPI_ISL_599244, EPI_ISL_599245, EPI_ISL_599246, EPI_ISL_599251, EPI_ISL_599252, EPI_ISL_599254, EPI_ISL_599258, EPI_ISL_599259, EPI_ISL_599260, EPI_ISL_599263, EPI_ISL_599264, EPI_ISL_599265, EPI_ISL_599266, EPI_ISL_599269, EPI_ISL_599273, EPI_ISL_599274, EPI_ISL_599275, EPI_ISL_599277, EPI_ISL_599278, EPI_ISL_599280, EPI_ISL_599281, EPI_ISL_599282, EPI_ISL_599283, EPI_ISL_599284, EPI_ISL_599285, EPI_ISL_599287, EPI_ISL_599288, EPI_ISL_599289, EPI_ISL_599290, EPI_ISL_599292, EPI_ISL_599296, EPI_ISL_599297, EPI_ISL_599301, EPI_ISL_599302, EPI_ISL_599303, EPI_ISL_599304, EPI_ISL_599305, EPI_ISL_599307, EPI_ISL_599308, EPI_ISL_599309, EPI_ISL_599312, EPI_ISL_599313, EPI_ISL_599315, EPI_ISL_599318, EPI_ISL_599319, EPI_ISL_599320, EPI_ISL_599321, EPI_ISL_599322, EPI_ISL_599327, EPI_ISL_599328, EPI_ISL_599330, EPI_ISL_599334, EPI_ISL_599335, EPI_ISL_599336, EPI_ISL_599337, EPI_ISL_599338, EPI_ISL_599339, EPI_ISL_599340, EPI_ISL_599346, EPI_ISL_599347, EPI_ISL_599348, EPI_ISL_599349, EPI_ISL_599350, EPI_ISL_599351, EPI_ISL_599352, EPI_ISL_599355, EPI_ISL_599356, EPI_ISL_599358, EPI_ISL_599359, EPI_ISL_599360, EPI_ISL_599363, EPI_ISL_599364, EPI_ISL_599368, EPI_ISL_599370, EPI_ISL_599371, EPI_ISL_599373, EPI_ISL_599374, EPI_ISL_599375, EPI_ISL_599380, EPI_ISL_599381, EPI_ISL_599382, EPI_ISL_599387, EPI_ISL_599388, EPI_ISL_599389, EPI_ISL_599392, EPI_ISL_599396, EPI_ISL_599397, EPI_ISL_599400, EPI_ISL_599401, EPI_ISL_599405, EPI_ISL_599406, EPI_ISL_599407, EPI_ISL_599410, EPI_ISL_599412, EPI_ISL_599417, EPI_ISL_599418, EPI_ISL_599420, EPI_ISL_599421, EPI_ISL_599422, EPI_ISL_599423, EPI_ISL_599424, EPI_ISL_599427, EPI_ISL_599428, EPI_ISL_599430, EPI_ISL_599432, EPI_ISL_599433, EPI_ISL_599434, EPI_ISL_599438, EPI_ISL_599439, EPI_ISL_599440, EPI_ISL_599441, EPI_ISL_599442, EPI_ISL_599443, EPI_ISL_599446, EPI_ISL_599447, EPI_ISL_599448, EPI_ISL_599451, EPI_ISL_599453, EPI_ISL_599454, EPI_ISL_599455, EPI_ISL_599456, EPI_ISL_599458, EPI_ISL_599459, EPI_ISL_599460, EPI_ISL_599461, EPI_ISL_599462, EPI_ISL_599464, EPI_ISL_599465, EPI_ISL_599467, EPI_ISL_599469, EPI_ISL_599470, EPI_ISL_599471, EPI_ISL_599473, EPI_ISL_599476, EPI_ISL_599478, EPI_ISL_599479, EPI_ISL_599481, EPI_ISL_599483, EPI_ISL_599489, EPI_ISL_599491, EPI_ISL_599492, EPI_ISL_599508, EPI_ISL_599511, EPI_ISL_599521, EPI_ISL_599525, EPI_ISL_599537, EPI_ISL_599539, EPI_ISL_599540, EPI_ISL_599546, EPI_ISL_599552, EPI_ISL_599557, EPI_ISL_599572, EPI_ISL_599578, EPI_ISL_599588, EPI_ISL_599593, EPI_ISL_599596, EPI_ISL_599597, EPI_ISL_599635, EPI_ISL_599646, EPI_ISL_599647, EPI_ISL_599658, EPI_ISL_599671, EPI_ISL_599676, EPI_ISL_599677, EPI_ISL_599683, EPI_ISL_599685, EPI_ISL_599691, EPI_ISL_599697, EPI_ISL_599698, EPI_ISL_599700, EPI_ISL_599702, EPI_ISL_599704, EPI_ISL_599718, EPI_ISL_599721, EPI_ISL_599722, EPI_ISL_599723, EPI_ISL_599725, EPI_ISL_599726, EPI_ISL_599735, EPI_ISL_599737, EPI_ISL_599742, EPI_ISL_599745, EPI_ISL_599746, EPI_ISL_599770, EPI_ISL_599776, EPI_ISL_599779, EPI_ISL_599786, EPI_ISL_599793, EPI_ISL_599807, EPI_ISL_600128, EPI_ISL_600130, EPI_ISL_600131, EPI_ISL_600133, EPI_ISL_600134, EPI_ISL_600135, EPI_ISL_600136, EPI_ISL_600137, EPI_ISL_600138, EPI_ISL_600140, EPI_ISL_600143, EPI_ISL_600144, EPI_ISL_600145, EPI_ISL_600146, EPI_ISL_600148, EPI_ISL_600149, EPI_ISL_600150, EPI_ISL_600151, EPI_ISL_600154, EPI_ISL_600156, EPI_ISL_600157, EPI_ISL_600158, EPI_ISL_600159, EPI_ISL_600160, EPI_ISL_600161, EPI_ISL_600162, EPI_ISL_600163, EPI_ISL_600164, EPI_ISL_600165, EPI_ISL_600166, EPI_ISL_600167, EPI_ISL_600168, EPI_ISL_600170, EPI_ISL_600173, EPI_ISL_600176, EPI_ISL_600177, EPI_ISL_600178, EPI_ISL_600179, EPI_ISL_600180, EPI_ISL_600182, EPI_ISL_600183, EPI_ISL_600185, EPI_ISL_600187, EPI_ISL_600188, EPI_ISL_600189, EPI_ISL_600192, EPI_ISL_600193, EPI_ISL_600194, EPI_ISL_600195, EPI_ISL_600196, EPI_ISL_600197, EPI_ISL_600198, EPI_ISL_600199, EPI_ISL_600200, EPI_ISL_600201, EPI_ISL_600202, EPI_ISL_600204, EPI_ISL_600206, EPI_ISL_600208, EPI_ISL_600209, EPI_ISL_600210, EPI_ISL_600211, EPI_ISL_600212, EPI_ISL_600215, EPI_ISL_600216, EPI_ISL_600217, EPI_ISL_600218, EPI_ISL_600219, EPI_ISL_600220, EPI_ISL_600222, EPI_ISL_600223, EPI_ISL_600224, EPI_ISL_600225, EPI_ISL_600226, EPI_ISL_600227, EPI_ISL_600229, EPI_ISL_600230, EPI_ISL_600231, EPI_ISL_600232, EPI_ISL_600234, EPI_ISL_600236, EPI_ISL_600240, EPI_ISL_600241, EPI_ISL_600242, EPI_ISL_600243, EPI_ISL_600247, EPI_ISL_600248, EPI_ISL_600249, EPI_ISL_600250, EPI_ISL_600251, EPI_ISL_600252, EPI_ISL_600254, EPI_ISL_600255, EPI_ISL_600256, EPI_ISL_600257, EPI_ISL_600259, EPI_ISL_600260, EPI_ISL_600261, EPI_ISL_600262, EPI_ISL_600263, EPI_ISL_600264, EPI_ISL_600265, EPI_ISL_600266, EPI_ISL_600267, EPI_ISL_600270, EPI_ISL_600271, EPI_ISL_600272, EPI_ISL_600274, EPI_ISL_600275, EPI_ISL_600276, EPI_ISL_600277, EPI_ISL_600278, EPI_ISL_600279, EPI_ISL_600280, EPI_ISL_600281, EPI_ISL_600282, EPI_ISL_600283, EPI_ISL_600284, EPI_ISL_600285, EPI_ISL_600286, EPI_ISL_600287, EPI_ISL_600288, EPI_ISL_600290, EPI_ISL_600291, EPI_ISL_600292, EPI_ISL_600293, EPI_ISL_600294, EPI_ISL_600295, EPI_ISL_600298, EPI_ISL_600299, EPI_ISL_600301, EPI_ISL_600302, EPI_ISL_600303, EPI_ISL_600304, EPI_ISL_600305, EPI_ISL_600306, EPI_ISL_600307, EPI_ISL_600308, EPI_ISL_600309, EPI_ISL_600310, EPI_ISL_600311, EPI_ISL_600312, EPI_ISL_600313, EPI_ISL_600314, EPI_ISL_600315, EPI_ISL_600317, EPI_ISL_600319, EPI_ISL_600321, EPI_ISL_600322, EPI_ISL_600323, EPI_ISL_600324, EPI_ISL_600325, EPI_ISL_600326, EPI_ISL_600327, EPI_ISL_600328, EPI_ISL_600330, EPI_ISL_600331, EPI_ISL_600332, EPI_ISL_600334, EPI_ISL_600336, EPI_ISL_600337, EPI_ISL_600338, EPI_ISL_600340, EPI_ISL_600342, EPI_ISL_600343, EPI_ISL_600344, EPI_ISL_600345, EPI_ISL_600346, EPI_ISL_600347, EPI_ISL_600348, EPI_ISL_600349, EPI_ISL_600350, EPI_ISL_600352, EPI_ISL_600354, EPI_ISL_600355, EPI_ISL_600356, EPI_ISL_600357, EPI_ISL_600358, EPI_ISL_600359, EPI_ISL_600360, EPI_ISL_600361, EPI_ISL_600363, EPI_ISL_600364, EPI_ISL_600366, EPI_ISL_600367, EPI_ISL_600368, EPI_ISL_600371, EPI_ISL_600372, EPI_ISL_600373, EPI_ISL_600374, EPI_ISL_600375, EPI_ISL_600377, EPI_ISL_600378, EPI_ISL_600379, EPI_ISL_600380, EPI_ISL_600381, EPI_ISL_600382, EPI_ISL_600383, EPI_ISL_600385, EPI_ISL_600386, EPI_ISL_600387, EPI_ISL_600388, EPI_ISL_600389, EPI_ISL_600390, EPI_ISL_600391, EPI_ISL_600392, EPI_ISL_600393, EPI_ISL_600395, EPI_ISL_600396, EPI_ISL_600397, EPI_ISL_600398, EPI_ISL_600399, EPI_ISL_600402, EPI_ISL_600403, EPI_ISL_600404, EPI_ISL_600406, EPI_ISL_600407, EPI_ISL_600409, EPI_ISL_600410, EPI_ISL_600412, EPI_ISL_600413, EPI_ISL_600415, EPI_ISL_600416, EPI_ISL_600417, EPI_ISL_600418, EPI_ISL_600419, EPI_ISL_600420, EPI_ISL_600421, EPI_ISL_600422, EPI_ISL_600423, EPI_ISL_600424, EPI_ISL_600425, EPI_ISL_600427, EPI_ISL_600447, EPI_ISL_600448, EPI_ISL_600452, EPI_ISL_600455, EPI_ISL_600458, EPI_ISL_600460, EPI_ISL_600464, EPI_ISL_600472, EPI_ISL_600474, EPI_ISL_600475, EPI_ISL_600477, EPI_ISL_600482, EPI_ISL_600483, EPI_ISL_600485, EPI_ISL_600486, EPI_ISL_600490, EPI_ISL_600491, EPI_ISL_600493, EPI_ISL_600496, EPI_ISL_600497, EPI_ISL_600499, EPI_ISL_600501, EPI_ISL_600503, EPI_ISL_600504, EPI_ISL_600506, EPI_ISL_600508, EPI_ISL_600509, EPI_ISL_600513, EPI_ISL_600514, EPI_ISL_600518, EPI_ISL_600520 | | | |
| see above | Lighthouse Lab in Glasgow | Wellcome Sanger Institute for the COVID-19 Genomics UK (COG-UK) consortium | Harper VanSteenhouse, Yumi Kasai, David Gray, Carol Clugston, Anna Dominiczak and Alex Alderton, Roberto Amato, Sonia Goncalves, Ewan Harrison, David K. Jackson, Ian Johnston, Dominic Kwiatkowski, Cordelia Langford, John Sillitoe on behalf of the Wellcome Sanger Institute COVID-19 Surveillance Team (http://www.sanger.ac.uk/covid-team) |
| EPI_ISL_600523, EPI_ISL_600536 | Lighthouse Lab in Alderley Park | Wellcome Sanger Institute for the COVID-19 Genomics UK (COG-UK) consortium | Jacquelyn Wynn, Mairead Hyland, The Lighthouse Lab in Alderley Park and Alex Alderton, Roberto Amato, Sonia Goncalves, Ewan Harrison, David K. Jackson, Ian Johnston, Dominic Kwiatkowski, Cordelia Langford, John Sillitoe on behalf of the Wellcome Sanger Institute COVID-19 Surveillance Team (http://www.sanger.ac.uk/covid-team) |
| EPI_ISL_600708, EPI_ISL_600739, EPI_ISL_600933, EPI_ISL_600934, EPI_ISL_600935, EPI_ISL_600936, EPI_ISL_600937, EPI_ISL_600938, EPI_ISL_600939, EPI_ISL_600940, EPI_ISL_600941, EPI_ISL_600942, EPI_ISL_600943, EPI_ISL_600944, EPI_ISL_600945, EPI_ISL_600946, EPI_ISL_600947, EPI_ISL_600948, EPI_ISL_600949, EPI_ISL_600950, EPI_ISL_600951, EPI_ISL_600952, EPI_ISL_600953, EPI_ISL_600954, EPI_ISL_600955, EPI_ISL_600956, EPI_ISL_600957, EPI_ISL_600958, EPI_ISL_600959, EPI_ISL_600960, EPI_ISL_600961, EPI_ISL_600962, EPI_ISL_600963, EPI_ISL_600964, EPI_ISL_600965, EPI_ISL_600966, EPI_ISL_600967, EPI_ISL_600968, EPI_ISL_600969, EPI_ISL_600971, EPI_ISL_600972, EPI_ISL_600973, EPI_ISL_600975, EPI_ISL_600976, EPI_ISL_600977, EPI_ISL_600978, EPI_ISL_600979, EPI_ISL_600981, EPI_ISL_600982, EPI_ISL_600984, EPI_ISL_600985, EPI_ISL_600986, EPI_ISL_600987, EPI_ISL_600988, EPI_ISL_600989, EPI_ISL_600990, EPI_ISL_600991, EPI_ISL_600992, EPI_ISL_600993, EPI_ISL_600994, EPI_ISL_600995, EPI_ISL_600996, EPI_ISL_600997, EPI_ISL_600998, EPI_ISL_601000, EPI_ISL_601001, EPI_ISL_601003, EPI_ISL_601004, EPI_ISL_601005, EPI_ISL_601006, EPI_ISL_601007, EPI_ISL_601008, EPI_ISL_601009, EPI_ISL_601010, EPI_ISL_601011, EPI_ISL_601012, EPI_ISL_601013, EPI_ISL_601014, EPI_ISL_601015, EPI_ISL_601016, EPI_ISL_601017, EPI_ISL_601018, EPI_ISL_601019, EPI_ISL_601020, EPI_ISL_601021, EPI_ISL_601024, EPI_ISL_601026, EPI_ISL_601027, EPI_ISL_601028, EPI_ISL_601029, EPI_ISL_601030, EPI_ISL_601031, EPI_ISL_601033, EPI_ISL_601034, EPI_ISL_601035, EPI_ISL_601036, EPI_ISL_601037, EPI_ISL_601038, EPI_ISL_601039, EPI_ISL_601040, EPI_ISL_601041, EPI_ISL_601042, EPI_ISL_601043, EPI_ISL_601044, EPI_ISL_601045, EPI_ISL_601046, EPI_ISL_601048, EPI_ISL_601049, EPI_ISL_601050, EPI_ISL_601051, EPI_ISL_601052, EPI_ISL_601053, EPI_ISL_601054, EPI_ISL_601055, EPI_ISL_601056, EPI_ISL_601057, EPI_ISL_601058, EPI_ISL_601059, EPI_ISL_601060, EPI_ISL_601062, EPI_ISL_601063, EPI_ISL_601064, EPI_ISL_601065, EPI_ISL_601066, EPI_ISL_601067, EPI_ISL_601068, EPI_ISL_601069, EPI_ISL_601070, EPI_ISL_601072, EPI_ISL_601073, EPI_ISL_601074, EPI_ISL_601075, EPI_ISL_601076, EPI_ISL_601077, EPI_ISL_601078, EPI_ISL_601079, EPI_ISL_601080, EPI_ISL_601081, EPI_ISL_601082, EPI_ISL_601083, EPI_ISL_601084, EPI_ISL_601085, EPI_ISL_601086, EPI_ISL_601087, EPI_ISL_601088, EPI_ISL_601089, EPI_ISL_601090, EPI_ISL_601091, EPI_ISL_601092, EPI_ISL_601093, EPI_ISL_601094, EPI_ISL_601095, EPI_ISL_601096, EPI_ISL_601097, EPI_ISL_601098, EPI_ISL_601100, EPI_ISL_601101, EPI_ISL_601102, EPI_ISL_601103, EPI_ISL_601104, EPI_ISL_601105, EPI_ISL_601106, EPI_ISL_601107, EPI_ISL_601108, EPI_ISL_601109, EPI_ISL_601111, EPI_ISL_601112, EPI_ISL_601113, EPI_ISL_601114, EPI_ISL_601115, EPI_ISL_601116, EPI_ISL_601117, EPI_ISL_601119, EPI_ISL_601120, EPI_ISL_601121, EPI_ISL_601122, EPI_ISL_601123, EPI_ISL_601124, EPI_ISL_601125, EPI_ISL_601126, EPI_ISL_601127, EPI_ISL_601128, EPI_ISL_601129, EPI_ISL_601130, EPI_ISL_601131, EPI_ISL_601132, EPI_ISL_601133, EPI_ISL_601134, EPI_ISL_601135, EPI_ISL_601137, EPI_ISL_601138, EPI_ISL_601140, EPI_ISL_601141, EPI_ISL_601142, EPI_ISL_601143, EPI_ISL_601144, EPI_ISL_601145, EPI_ISL_601146, EPI_ISL_601147, EPI_ISL_601148, EPI_ISL_601149, EPI_ISL_601151, EPI_ISL_601152, EPI_ISL_601153, EPI_ISL_601154, EPI_ISL_601155, EPI_ISL_601156, EPI_ISL_601157, EPI_ISL_601158, EPI_ISL_601159, EPI_ISL_601160, EPI_ISL_601161, EPI_ISL_601162, EPI_ISL_601163, EPI_ISL_601164, EPI_ISL_601165, EPI_ISL_601166, EPI_ISL_601167, EPI_ISL_601168, EPI_ISL_601169, EPI_ISL_601170, EPI_ISL_601171, EPI_ISL_601173, EPI_ISL_601174, EPI_ISL_601175, EPI_ISL_601176, EPI_ISL_601177, EPI_ISL_601178, EPI_ISL_601179, EPI_ISL_601180, EPI_ISL_601181, EPI_ISL_601182, EPI_ISL_601183, EPI_ISL_601184, EPI_ISL_601185, EPI_ISL_601186, EPI_ISL_601187, EPI_ISL_601188, EPI_ISL_601189, EPI_ISL_601190, EPI_ISL_601191, EPI_ISL_601193, EPI_ISL_601194, EPI_ISL_601195, EPI_ISL_601196, EPI_ISL_601197, EPI_ISL_601198, EPI_ISL_601199, EPI_ISL_601200, EPI_ISL_601201, EPI_ISL_601202, EPI_ISL_601204, EPI_ISL_601205, EPI_ISL_601206, EPI_ISL_601207, EPI_ISL_601208, EPI_ISL_601209, EPI_ISL_601210, EPI_ISL_601211, EPI_ISL_601212, EPI_ISL_601213, EPI_ISL_601215, EPI_ISL_601216, EPI_ISL_601217, EPI_ISL_601218, EPI_ISL_601219, EPI_ISL_601220, EPI_ISL_601221, EPI_ISL_601222, EPI_ISL_601223, EPI_ISL_601225, EPI_ISL_601226, EPI_ISL_601227, EPI_ISL_601228, EPI_ISL_601229, EPI_ISL_601230, EPI_ISL_601231, EPI_ISL_601232, EPI_ISL_601233, EPI_ISL_601787, EPI_ISL_601788, EPI_ISL_601792, EPI_ISL_601793, EPI_ISL_601794, EPI_ISL_601795, EPI_ISL_601798, EPI_ISL_601800, EPI_ISL_601802, EPI_ISL_601805, EPI_ISL_601806, EPI_ISL_601807, EPI_ISL_601809, EPI_ISL_601812, EPI_ISL_601814, EPI_ISL_601815, EPI_ISL_601816, EPI_ISL_601817, EPI_ISL_601822, EPI_ISL_601823, EPI_ISL_601825, EPI_ISL_601827, EPI_ISL_601828, EPI_ISL_601829, EPI_ISL_601831, EPI_ISL_601832, EPI_ISL_601835, EPI_ISL_601836, EPI_ISL_601837, EPI_ISL_601839, EPI_ISL_601840, EPI_ISL_601842, EPI_ISL_601843, EPI_ISL_601846, EPI_ISL_601849, EPI_ISL_601852, EPI_ISL_601853, EPI_ISL_601855, EPI_ISL_601858, EPI_ISL_601859, EPI_ISL_601860, EPI_ISL_601861, EPI_ISL_601862, EPI_ISL_601863, EPI_ISL_601865, EPI_ISL_601867 | | | |
| see above | Lighthouse Lab in Glasgow | Wellcome Sanger Institute for the COVID-19 Genomics UK (COG-UK) consortium | Harper VanSteenhouse, Yumi Kasai, David Gray, Carol Clugston, Anna Dominiczak and Alex Alderton, Roberto Amato, Sonia Goncalves, Ewan Harrison, David K. Jackson, Ian Johnston, Dominic Kwiatkowski, Cordelia Langford, John Sillitoe on behalf of the Wellcome Sanger Institute COVID-19 Surveillance Team (http://www.sanger.ac.uk/covid-team) |
| EPI_ISL_602159, EPI_ISL_602160 | Lighthouse Lab in Alderley Park | Wellcome Sanger Institute for the COVID-19 Genomics UK (COG-UK) consortium | Jacquelyn Wynn, Mairead Hyland, The Lighthouse Lab in Alderley Park and Alex Alderton, Roberto Amato, Sonia Goncalves, Ewan Harrison, David K. Jackson, Ian Johnston, Dominic Kwiatkowski, Cordelia Langford, John Sillitoe on behalf of the Wellcome Sanger Institute COVID-19 Surveillance Team (http://www.sanger.ac.uk/covid-team) |
| EPI_ISL_603244, EPI_ISL_603245 | National Institute of Laboratory Medicine and Referral Center | Genomic Research Lab, BCSIR | Iffat Jahan, Abu Sayeed Mohammad Mahmud, Mohammad Samir Uzzaman, Eshrar Osman, Md. Ahashan Habib, Shahina Akter, Tanjina Akhter Banu, Md. Murshed Hasan Sarkar, Barna Goswami, Md. Saddam Hossain, Tasnim Nafisa, Md. Maruf Ahmed Molla, Mahmuda Yeasmin, Asish Kumar Ghosh, A. K. M. Shamsuzzaman, Monira Parveen, Md. Masum Hossain Arif, Md. Salim Khan |
| EPI_ISL_605932, EPI_ISL_605933, EPI_ISL_605934, EPI_ISL_605935, EPI_ISL_605936, EPI_ISL_605937, EPI_ISL_605938, EPI_ISL_605939, EPI_ISL_605940, EPI_ISL_605941, EPI_ISL_605942, EPI_ISL_605943, EPI_ISL_605944, EPI_ISL_605945, EPI_ISL_605946, EPI_ISL_605947, EPI_ISL_605948, EPI_ISL_605949, EPI_ISL_605950, EPI_ISL_605951, EPI_ISL_605952, EPI_ISL_605953, EPI_ISL_605954, EPI_ISL_605955, EPI_ISL_605956, EPI_ISL_605957, EPI_ISL_605958, EPI_ISL_605959, EPI_ISL_605960, EPI_ISL_605961, EPI_ISL_605962, EPI_ISL_605963, EPI_ISL_605964, EPI_ISL_605965, EPI_ISL_605966, EPI_ISL_605967, EPI_ISL_605968, EPI_ISL_605969, EPI_ISL_605970, EPI_ISL_605971, EPI_ISL_605972, EPI_ISL_605973, EPI_ISL_605974, EPI_ISL_605975, EPI_ISL_605976, EPI_ISL_605977, EPI_ISL_605978, EPI_ISL_605979, EPI_ISL_605980, EPI_ISL_605981, EPI_ISL_605982, EPI_ISL_605983, EPI_ISL_605984, EPI_ISL_605985, EPI_ISL_605986, EPI_ISL_605987, EPI_ISL_605988, EPI_ISL_605989, EPI_ISL_605990, EPI_ISL_605991, EPI_ISL_605992, EPI_ISL_605993, EPI_ISL_605994, EPI_ISL_605995, EPI_ISL_605996, EPI_ISL_605997, EPI_ISL_605998, EPI_ISL_605999, EPI_ISL_606000, EPI_ISL_606001, EPI_ISL_606002, EPI_ISL_606003, EPI_ISL_606004, EPI_ISL_606005, EPI_ISL_606006, EPI_ISL_606007, EPI_ISL_606008, EPI_ISL_606009, EPI_ISL_606010, EPI_ISL_606011, EPI_ISL_606012, EPI_ISL_606013, EPI_ISL_606014, EPI_ISL_606015, EPI_ISL_606016, EPI_ISL_606017, EPI_ISL_606018, EPI_ISL_606019, EPI_ISL_606020, EPI_ISL_606021, EPI_ISL_606022, EPI_ISL_606023, EPI_ISL_606024, EPI_ISL_606025, EPI_ISL_606026, EPI_ISL_606027, EPI_ISL_606028, EPI_ISL_606029, EPI_ISL_606030, EPI_ISL_606031, EPI_ISL_606032, EPI_ISL_606033, EPI_ISL_606034, EPI_ISL_606035, EPI_ISL_606036, EPI_ISL_606037, EPI_ISL_606038, EPI_ISL_606039, EPI_ISL_606040, EPI_ISL_606041, EPI_ISL_606042, EPI_ISL_606043, EPI_ISL_606044, EPI_ISL_606045, EPI_ISL_606046, EPI_ISL_606047, EPI_ISL_606048, EPI_ISL_606049, EPI_ISL_606050, EPI_ISL_606051, EPI_ISL_606052, EPI_ISL_606053, EPI_ISL_606054, EPI_ISL_606055, EPI_ISL_606056, EPI_ISL_606057, EPI_ISL_606058, EPI_ISL_606059, EPI_ISL_606060, EPI_ISL_606061, EPI_ISL_606062, EPI_ISL_606063, EPI_ISL_606064, EPI_ISL_606065, EPI_ISL_606066, EPI_ISL_606067, EPI_ISL_606068, EPI_ISL_606069, EPI_ISL_606070, EPI_ISL_606071, EPI_ISL_606072, EPI_ISL_606073, EPI_ISL_606074, EPI_ISL_606075, EPI_ISL_606076, EPI_ISL_606077, EPI_ISL_606078, EPI_ISL_606079, EPI_ISL_606080, EPI_ISL_606081, EPI_ISL_606082, EPI_ISL_606083, EPI_ISL_606084, EPI_ISL_606085, EPI_ISL_606086, EPI_ISL_606087, EPI_ISL_606088, EPI_ISL_606089, EPI_ISL_606090, EPI_ISL_606091, EPI_ISL_606092, EPI_ISL_606093, EPI_ISL_606094, EPI_ISL_606095, EPI_ISL_606096, EPI_ISL_606097, EPI_ISL_606098, EPI_ISL_606099, EPI_ISL_606100, EPI_ISL_606101, EPI_ISL_606102, EPI_ISL_606103, EPI_ISL_606104, EPI_ISL_606105, EPI_ISL_606106, EPI_ISL_606107, EPI_ISL_606108, EPI_ISL_606109, EPI_ISL_606110, EPI_ISL_606111, EPI_ISL_606112, EPI_ISL_606113, EPI_ISL_606114, EPI_ISL_606115, EPI_ISL_606117, EPI_ISL_606118, EPI_ISL_606119, EPI_ISL_606120, EPI_ISL_606121, EPI_ISL_606122, EPI_ISL_606123, EPI_ISL_606124, EPI_ISL_606125, EPI_ISL_606126, EPI_ISL_606127, EPI_ISL_606128, EPI_ISL_606129, EPI_ISL_606130, EPI_ISL_606131, EPI_ISL_606132, EPI_ISL_606133, EPI_ISL_606134, EPI_ISL_606135, EPI_ISL_606136, EPI_ISL_606137, EPI_ISL_606138, EPI_ISL_606139, EPI_ISL_606140, EPI_ISL_606141, EPI_ISL_606142, EPI_ISL_606143, EPI_ISL_606144, EPI_ISL_606145, EPI_ISL_606146, EPI_ISL_606147, EPI_ISL_606148, EPI_ISL_606149, EPI_ISL_606150, EPI_ISL_606151, EPI_ISL_606152, EPI_ISL_606153, EPI_ISL_606154, EPI_ISL_606155, EPI_ISL_606156, EPI_ISL_606157, EPI_ISL_606158, EPI_ISL_606159, EPI_ISL_606160, EPI_ISL_606161, EPI_ISL_606162, EPI_ISL_606163, EPI_ISL_606164, EPI_ISL_606165, EPI_ISL_606166, EPI_ISL_606167, EPI_ISL_606168, EPI_ISL_606169, EPI_ISL_606170, EPI_ISL_606171, EPI_ISL_606172, EPI_ISL_606173, EPI_ISL_606174, EPI_ISL_606175, EPI_ISL_606176, EPI_ISL_606177, EPI_ISL_606178, EPI_ISL_606179, EPI_ISL_606180, EPI_ISL_606181, EPI_ISL_606182, EPI_ISL_606183, EPI_ISL_606184, EPI_ISL_606185, EPI_ISL_606186, EPI_ISL_606187, EPI_ISL_606188, EPI_ISL_606190, EPI_ISL_606191, EPI_ISL_606192, EPI_ISL_606193, EPI_ISL_606194, EPI_ISL_606195, EPI_ISL_606196, EPI_ISL_606197, EPI_ISL_606198, EPI_ISL_606199, EPI_ISL_606200, EPI_ISL_606201, EPI_ISL_606202, EPI_ISL_606203, | | | |

EPI_ISL_606204, EPI_ISL_606205, EPI_ISL_606206, EPI_ISL_606207, EPI_ISL_606208, EPI_ISL_606209, EPI_ISL_606210, EPI_ISL_606211, EPI_ISL_606212, EPI_ISL_606213, EPI_ISL_606214, EPI_ISL_606215, EPI_ISL_606216, EPI_ISL_606217, EPI_ISL_606218, EPI_ISL_606219, EPI_ISL_606220, EPI_ISL_606221,
EPI_ISL_606222, EPI_ISL_606223, EPI_ISL_606224, EPI_ISL_606225, EPI_ISL_606226, EPI_ISL_606227, EPI_ISL_606229, EPI_ISL_606230, EPI_ISL_606231, EPI_ISL_606232, EPI_ISL_606233, EPI_ISL_606234, EPI_ISL_606235, EPI_ISL_606236, EPI_ISL_606237, EPI_ISL_606239, EPI_ISL_606240, EPI_ISL_606241,
EPI_ISL_606242, EPI_ISL_606243, EPI_ISL_606244, EPI_ISL_606245, EPI_ISL_606246, EPI_ISL_606247, EPI_ISL_606248, EPI_ISL_606249, EPI_ISL_606250, EPI_ISL_606251, EPI_ISL_606252, EPI_ISL_606253, EPI_ISL_606254, EPI_ISL_606255, EPI_ISL_606256, EPI_ISL_606257, EPI_ISL_606258, EPI_ISL_606259,
EPI_ISL_606260, EPI_ISL_606261, EPI_ISL_606262, EPI_ISL_606263, EPI_ISL_606264, EPI_ISL_606265, EPI_ISL_606266, EPI_ISL_606267, EPI_ISL_606268, EPI_ISL_606269, EPI_ISL_606270, EPI_ISL_606271, EPI_ISL_606272, EPI_ISL_606273, EPI_ISL_606274, EPI_ISL_606275, EPI_ISL_606276, EPI_ISL_606277,
EPI_ISL_606278, EPI_ISL_606279, EPI_ISL_606280, EPI_ISL_606281, EPI_ISL_606282, EPI_ISL_606283, EPI_ISL_606284, EPI_ISL_606285, EPI_ISL_606286, EPI_ISL_606287, EPI_ISL_606288, EPI_ISL_606289, EPI_ISL_606290, EPI_ISL_606291, EPI_ISL_606292, EPI_ISL_606293, EPI_ISL_606294, EPI_ISL_606295,
EPI_ISL_606926, EPI_ISL_606930, EPI_ISL_606953, EPI_ISL_606964, EPI_ISL_606979, EPI_ISL_607017, EPI_ISL_607020, EPI_ISL_607028, EPI_ISL_607080, EPI_ISL_607086, EPI_ISL_607113, EPI_ISL_607148, EPI_ISL_607193, EPI_ISL_607196, EPI_ISL_607209, EPI_ISL_607256, EPI_ISL_607259, EPI_ISL_607260,
EPI_ISL_607625, EPI_ISL_607639, EPI_ISL_607643, EPI_ISL_607657, EPI_ISL_607662, EPI_ISL_607671, EPI_ISL_607675, EPI_ISL_607698, EPI_ISL_607748, EPI_ISL_607758, EPI_ISL_607780, EPI_ISL_607813, EPI_ISL_607826, EPI_ISL_607829, EPI_ISL_607860, EPI_ISL_607887, EPI_ISL_607905, EPI_ISL_607917,
EPI_ISL_607948, EPI_ISL_607950, EPI_ISL_607990, EPI_ISL_607991, EPI_ISL_607993, EPI_ISL_607998, EPI_ISL_608008, EPI_ISL_608015, EPI_ISL_608020, EPI_ISL_608023, EPI_ISL_608061, EPI_ISL_608074, EPI_ISL_608091, EPI_ISL_608117, EPI_ISL_608132, EPI_ISL_608174, EPI_ISL_608175, EPI_ISL_608215,
EPI_ISL_608239, EPI_ISL_608297, EPI_ISL_608308, EPI_ISL_608358, EPI_ISL_608366, EPI_ISL_608383, EPI_ISL_608414, EPI_ISL_608421, EPI_ISL_608425, EPI_ISL_608444, EPI_ISL_608453, EPI_ISL_608469, EPI_ISL_608474, EPI_ISL_608483, EPI_ISL_608520, EPI_ISL_608533, EPI_ISL_608556, EPI_ISL_608563,
EPI_ISL_608564, EPI_ISL_608574, EPI_ISL_608582, EPI_ISL_608591, EPI_ISL_608593, EPI_ISL_608603, EPI_ISL_608614, EPI_ISL_608621, EPI_ISL_608635, EPI_ISL_608652, EPI_ISL_608664, EPI_ISL_608677, EPI_ISL_608687, EPI_ISL_608691, EPI_ISL_608864, EPI_ISL_608872, EPI_ISL_608889, EPI_ISL_608891,
EPI_ISL_608896, EPI_ISL_608903, EPI_ISL_608904, EPI_ISL_608909, EPI_ISL_608915, EPI_ISL_608920, EPI_ISL_608932, EPI_ISL_608955, EPI_ISL_608973, EPI_ISL_608979, EPI_ISL_608980, EPI_ISL_608981, EPI_ISL_608985, EPI_ISL_608989, EPI_ISL_608994, EPI_ISL_609002, EPI_ISL_609019, EPI_ISL_609020,
EPI_ISL_609055, EPI_ISL_609073, EPI_ISL_609074, EPI_ISL_609107, EPI_ISL_609112, EPI_ISL_609141, EPI_ISL_609148, EPI_ISL_609150, EPI_ISL_609214, EPI_ISL_610286, EPI_ISL_610287, EPI_ISL_610288, EPI_ISL_610290, EPI_ISL_610292, EPI_ISL_610293, EPI_ISL_610294, EPI_ISL_610295, EPI_ISL_610297,
EPI_ISL_610298, EPI_ISL_610300, EPI_ISL_610303, EPI_ISL_610304, EPI_ISL_610305, EPI_ISL_610306, EPI_ISL_610308, EPI_ISL_610309, EPI_ISL_610310, EPI_ISL_610311, EPI_ISL_610314, EPI_ISL_610315, EPI_ISL_610316, EPI_ISL_610317, EPI_ISL_610319, EPI_ISL_610320, EPI_ISL_610321, EPI_ISL_610322,
EPI_ISL_610323, EPI_ISL_610324, EPI_ISL_610327, EPI_ISL_610329, EPI_ISL_610331, EPI_ISL_610334, EPI_ISL_610336, EPI_ISL_610337, EPI_ISL_610339, EPI_ISL_610341, EPI_ISL_610343, EPI_ISL_610344, EPI_ISL_610346, EPI_ISL_610348, EPI_ISL_610349, EPI_ISL_610350, EPI_ISL_610353, EPI_ISL_610354,
EPI_ISL_610355, EPI_ISL_610356, EPI_ISL_610359, EPI_ISL_610361, EPI_ISL_610363, EPI_ISL_610364, EPI_ISL_610367, EPI_ISL_610368, EPI_ISL_610369, EPI_ISL_610370, EPI_ISL_610372, EPI_ISL_610373, EPI_ISL_610374, EPI_ISL_610375, EPI_ISL_610378, EPI_ISL_610381, EPI_ISL_610382, EPI_ISL_610383,
EPI_ISL_610384, EPI_ISL_610385, EPI_ISL_610387, EPI_ISL_610389, EPI_ISL_610392, EPI_ISL_610393, EPI_ISL_610395, EPI_ISL_610397, EPI_ISL_610398, EPI_ISL_610399, EPI_ISL_610400, EPI_ISL_610401, EPI_ISL_610402, EPI_ISL_610403, EPI_ISL_610405, EPI_ISL_610407, EPI_ISL_610408, EPI_ISL_610409,
EPI_ISL_610410, EPI_ISL_610411, EPI_ISL_610415, EPI_ISL_610416, EPI_ISL_610417, EPI_ISL_610419, EPI_ISL_610421, EPI_ISL_610423, EPI_ISL_610426, EPI_ISL_610427, EPI_ISL_610428, EPI_ISL_610429, EPI_ISL_610430, EPI_ISL_610431, EPI_ISL_610434, EPI_ISL_610435, EPI_ISL_610437, EPI_ISL_610443,
EPI_ISL_610444, EPI_ISL_610445, EPI_ISL_610446, EPI_ISL_610447, EPI_ISL_610448, EPI_ISL_610449, EPI_ISL_610451, EPI_ISL_610452, EPI_ISL_610453, EPI_ISL_610454, EPI_ISL_610456, EPI_ISL_610459, EPI_ISL_610462, EPI_ISL_610463, EPI_ISL_610464, EPI_ISL_610465, EPI_ISL_610467, EPI_ISL_610468,
EPI_ISL_610469, EPI_ISL_610470, EPI_ISL_610472, EPI_ISL_610473, EPI_ISL_610474, EPI_ISL_610475, EPI_ISL_610476, EPI_ISL_610478, EPI_ISL_610483, EPI_ISL_610486, EPI_ISL_610487, EPI_ISL_610488, EPI_ISL_610489, EPI_ISL_610492, EPI_ISL_610493, EPI_ISL_610494, EPI_ISL_610495, EPI_ISL_610497,
EPI_ISL_610500, EPI_ISL_610501, EPI_ISL_610504, EPI_ISL_610507, EPI_ISL_610508, EPI_ISL_610509, EPI_ISL_610510, EPI_ISL_610511, EPI_ISL_610512, EPI_ISL_610513, EPI_ISL_610516, EPI_ISL_610519, EPI_ISL_610522, EPI_ISL_610526, EPI_ISL_610527, EPI_ISL_610530, EPI_ISL_610531, EPI_ISL_610532,
EPI_ISL_610534, EPI_ISL_610535, EPI_ISL_610536, EPI_ISL_610537, EPI_ISL_610538, EPI_ISL_610540, EPI_ISL_610543, EPI_ISL_610545, EPI_ISL_610546, EPI_ISL_610547, EPI_ISL_610548, EPI_ISL_610549, EPI_ISL_610550, EPI_ISL_610555, EPI_ISL_610556, EPI_ISL_610557, EPI_ISL_610558, EPI_ISL_610559,
EPI_ISL_610560, EPI_ISL_610561, EPI_ISL_610563, EPI_ISL_610564, EPI_ISL_610566, EPI_ISL_610567, EPI_ISL_610568, EPI_ISL_610570, EPI_ISL_610571, EPI_ISL_610572, EPI_ISL_610574, EPI_ISL_610576, EPI_ISL_610577, EPI_ISL_610581, EPI_ISL_610583, EPI_ISL_610584, EPI_ISL_610585, EPI_ISL_610587,
EPI_ISL_610588, EPI_ISL_610590, EPI_ISL_610591, EPI_ISL_610592, EPI_ISL_610593, EPI_ISL_610595, EPI_ISL_610596, EPI_ISL_610597, EPI_ISL_610598, EPI_ISL_610601, EPI_ISL_610603, EPI_ISL_610605, EPI_ISL_610606, EPI_ISL_610608, EPI_ISL_610610, EPI_ISL_610612, EPI_ISL_610613, EPI_ISL_610614,
EPI_ISL_610616, EPI_ISL_610617, EPI_ISL_610619, EPI_ISL_610621, EPI_ISL_610624, EPI_ISL_610627, EPI_ISL_610628

| | | | |
|---|---|---|---|
| see above | Lighthouse Lab in Alderley Park | Wellcome Sanger Institute for the COVID-19 Genomics UK (COG-UK) consortium | Jacquelyn Wynn, Mairead Hyland, The Lighthouse Lab in Alderley Park and Alex Alderton, Roberto Amato, Sonia Goncalves, Ewan Harrison, David K. Jackson, Ian Johnston, Dominic Kwiatkowski, Cordelia Langford, John Sillitoe on behalf of the Wellcome Sanger Institute COVID-19 Surveillance Team |

EPI_ISL_610922, EPI_ISL_610923, EPI_ISL_610924, EPI_ISL_610925, EPI_ISL_610926, EPI_ISL_610927, EPI_ISL_610928, EPI_ISL_610929, EPI_ISL_610930, EPI_ISL_610931, EPI_ISL_610932, EPI_ISL_610933, EPI_ISL_610934, EPI_ISL_610935, EPI_ISL_610936, EPI_ISL_610937, EPI_ISL_610938, EPI_ISL_610939,
EPI_ISL_610940, EPI_ISL_610941, EPI_ISL_610943, EPI_ISL_610944, EPI_ISL_610945, EPI_ISL_610946, EPI_ISL_610947, EPI_ISL_610948, EPI_ISL_610949, EPI_ISL_610950, EPI_ISL_610951, EPI_ISL_610952, EPI_ISL_610953, EPI_ISL_610954, EPI_ISL_610955, EPI_ISL_610956, EPI_ISL_610957, EPI_ISL_610958,
EPI_ISL_610959, EPI_ISL_610960, EPI_ISL_610962, EPI_ISL_610963, EPI_ISL_610965, EPI_ISL_610966, EPI_ISL_610967, EPI_ISL_610969, EPI_ISL_610970, EPI_ISL_610971, EPI_ISL_610972, EPI_ISL_610973, EPI_ISL_610974, EPI_ISL_610975, EPI_ISL_610976, EPI_ISL_610977, EPI_ISL_610978,
EPI_ISL_610979, EPI_ISL_610980, EPI_ISL_610981, EPI_ISL_610982, EPI_ISL_610983, EPI_ISL_610984, EPI_ISL_610985, EPI_ISL_610986, EPI_ISL_610987, EPI_ISL_610988, EPI_ISL_610989, EPI_ISL_610990, EPI_ISL_610991, EPI_ISL_610992, EPI_ISL_610993, EPI_ISL_610994, EPI_ISL_610995, EPI_ISL_610997,
EPI_ISL_610999, EPI_ISL_611000, EPI_ISL_611001, EPI_ISL_611002, EPI_ISL_611003, EPI_ISL_611004, EPI_ISL_611005, EPI_ISL_611006, EPI_ISL_611007, EPI_ISL_611008, EPI_ISL_611009, EPI_ISL_611010, EPI_ISL_611011, EPI_ISL_611012, EPI_ISL_611013, EPI_ISL_611014, EPI_ISL_611015, EPI_ISL_611016,
EPI_ISL_611017, EPI_ISL_611018, EPI_ISL_611019, EPI_ISL_611020, EPI_ISL_611021, EPI_ISL_611022, EPI_ISL_611023, EPI_ISL_611025, EPI_ISL_611026, EPI_ISL_611027, EPI_ISL_611028, EPI_ISL_611029, EPI_ISL_611031, EPI_ISL_611032, EPI_ISL_611033, EPI_ISL_611034, EPI_ISL_611035, EPI_ISL_611036,
EPI_ISL_611037, EPI_ISL_611039, EPI_ISL_611040, EPI_ISL_611041, EPI_ISL_611042, EPI_ISL_611043, EPI_ISL_611044, EPI_ISL_611045, EPI_ISL_611046, EPI_ISL_611047, EPI_ISL_611048, EPI_ISL_611049, EPI_ISL_611050, EPI_ISL_611051, EPI_ISL_611052, EPI_ISL_611053, EPI_ISL_611054, EPI_ISL_611055,
EPI_ISL_611056, EPI_ISL_611057, EPI_ISL_611058, EPI_ISL_611059, EPI_ISL_611062, EPI_ISL_611063, EPI_ISL_611066, EPI_ISL_611067, EPI_ISL_611068, EPI_ISL_611069, EPI_ISL_611070, EPI_ISL_611071, EPI_ISL_611072, EPI_ISL_611074, EPI_ISL_611075, EPI_ISL_611076, EPI_ISL_611077, EPI_ISL_611078,
EPI_ISL_611079, EPI_ISL_611080, EPI_ISL_611081, EPI_ISL_611082, EPI_ISL_611083, EPI_ISL_611084, EPI_ISL_611085, EPI_ISL_611086, EPI_ISL_611087, EPI_ISL_611088, EPI_ISL_611089, EPI_ISL_611090, EPI_ISL_611091, EPI_ISL_611092, EPI_ISL_611094, EPI_ISL_611095, EPI_ISL_611096, EPI_ISL_611097,
EPI_ISL_611098, EPI_ISL_611099, EPI_ISL_611100, EPI_ISL_611101, EPI_ISL_611102, EPI_ISL_611103, EPI_ISL_611104, EPI_ISL_611105, EPI_ISL_611106, EPI_ISL_611107, EPI_ISL_611108, EPI_ISL_611112, EPI_ISL_611113, EPI_ISL_611114, EPI_ISL_611115, EPI_ISL_611116, EPI_ISL_611117, EPI_ISL_611118,
EPI_ISL_611119, EPI_ISL_611120, EPI_ISL_611121, EPI_ISL_611122, EPI_ISL_611123, EPI_ISL_611124, EPI_ISL_611125, EPI_ISL_611126, EPI_ISL_611127, EPI_ISL_611128, EPI_ISL_611129, EPI_ISL_611131, EPI_ISL_611132, EPI_ISL_611133, EPI_ISL_611134, EPI_ISL_611135, EPI_ISL_611136, EPI_ISL_611137,
EPI_ISL_611138, EPI_ISL_611139, EPI_ISL_611140, EPI_ISL_611141, EPI_ISL_611142, EPI_ISL_611143, EPI_ISL_611144, EPI_ISL_611145, EPI_ISL_611146, EPI_ISL_611147, EPI_ISL_611148, EPI_ISL_611149, EPI_ISL_611150, EPI_ISL_611151, EPI_ISL_611152, EPI_ISL_611153, EPI_ISL_611154, EPI_ISL_611155,
EPI_ISL_611156, EPI_ISL_611157, EPI_ISL_611158, EPI_ISL_611159, EPI_ISL_611160, EPI_ISL_611161, EPI_ISL_611162, EPI_ISL_611163, EPI_ISL_611164, EPI_ISL_611165, EPI_ISL_611166, EPI_ISL_611167, EPI_ISL_611168, EPI_ISL_611169, EPI_ISL_611170, EPI_ISL_611171, EPI_ISL_611172, EPI_ISL_611174,
EPI_ISL_611175, EPI_ISL_611176, EPI_ISL_611177, EPI_ISL_611178, EPI_ISL_611179, EPI_ISL_611181, EPI_ISL_611182, EPI_ISL_611183, EPI_ISL_611184, EPI_ISL_611185, EPI_ISL_611187, EPI_ISL_611188, EPI_ISL_611189, EPI_ISL_611190, EPI_ISL_611191, EPI_ISL_611192, EPI_ISL_611193, EPI_ISL_611194,
EPI_ISL_611195, EPI_ISL_611196, EPI_ISL_611197, EPI_ISL_611198, EPI_ISL_611199, EPI_ISL_611200, EPI_ISL_611201, EPI_ISL_611203, EPI_ISL_611204, EPI_ISL_611205, EPI_ISL_611206, EPI_ISL_611207, EPI_ISL_611208, EPI_ISL_611209, EPI_ISL_611210, EPI_ISL_611212, EPI_ISL_611213, EPI_ISL_611214,
EPI_ISL_611215, EPI_ISL_611217, EPI_ISL_611218, EPI_ISL_611219, EPI_ISL_611220, EPI_ISL_611221, EPI_ISL_611222, EPI_ISL_611223, EPI_ISL_611224, EPI_ISL_611225, EPI_ISL_611226, EPI_ISL_611227, EPI_ISL_611228, EPI_ISL_611229, EPI_ISL_611230, EPI_ISL_611232, EPI_ISL_611233, EPI_ISL_611234,
EPI_ISL_611235, EPI_ISL_611236, EPI_ISL_611237, EPI_ISL_611238, EPI_ISL_611239, EPI_ISL_611240, EPI_ISL_611241, EPI_ISL_611242, EPI_ISL_611243, EPI_ISL_611244, EPI_ISL_611245, EPI_ISL_611246, EPI_ISL_611247, EPI_ISL_611248, EPI_ISL_611249, EPI_ISL_611250, EPI_ISL_611251, EPI_ISL_611252,
EPI_ISL_611253, EPI_ISL_611254, EPI_ISL_611255, EPI_ISL_611256, EPI_ISL_611257, EPI_ISL_611258, EPI_ISL_611259, EPI_ISL_611260, EPI_ISL_611261, EPI_ISL_611262, EPI_ISL_611263, EPI_ISL_611264, EPI_ISL_611265, EPI_ISL_611266, EPI_ISL_611267, EPI_ISL_611268, EPI_ISL_611269, EPI_ISL_611270,
EPI_ISL_611271, EPI_ISL_611272, EPI_ISL_611273, EPI_ISL_611274, EPI_ISL_611275, EPI_ISL_611276, EPI_ISL_611277, EPI_ISL_611278, EPI_ISL_611279, EPI_ISL_611280, EPI_ISL_611281, EPI_ISL_611282, EPI_ISL_611283, EPI_ISL_611284, EPI_ISL_611285, EPI_ISL_611286

| | | | |
|---|---|---|---|
| see above | Lighthouse Lab in Alderley Park | Wellcome Sanger Institute for the COVID-19 Genomics UK (COG-UK) consortium | Jacquelyn Wynn, Mairead Hyland, The Lighthouse Lab in Alderley Park and Alex Alderton, Roberto Amato, Sonia Goncalves, Ewan Harrison, David K. Jackson, Ian Johnston, Dominic Kwiatkowski, Cordelia Langford, John Sillitoe on behalf of the Wellcome Sanger Institute COVID-19 Surveillance Team (http://www.sanger.ac.uk/covid-team) |

EPI_ISL_611550, EPI_ISL_611554, EPI_ISL_611556, EPI_ISL_611608, EPI_ISL_611612, EPI_ISL_611671, EPI_ISL_611692, EPI_ISL_611696, EPI_ISL_611700, EPI_ISL_611705, EPI_ISL_611776, EPI_ISL_611814, EPI_ISL_611825, EPI_ISL_611847, EPI_ISL_611855, EPI_ISL_611856, EPI_ISL_611858, EPI_ISL_611859,
EPI_ISL_611860, EPI_ISL_612011, EPI_ISL_612015, EPI_ISL_612047, EPI_ISL_612073, EPI_ISL_612207, EPI_ISL_612208, EPI_ISL_612209, EPI_ISL_612210, EPI_ISL_612211, EPI_ISL_612212, EPI_ISL_612213, EPI_ISL_612214, EPI_ISL_612215, EPI_ISL_612216, EPI_ISL_612217, EPI_ISL_612218, EPI_ISL_612219,
EPI_ISL_612220, EPI_ISL_612221, EPI_ISL_612222, EPI_ISL_612223, EPI_ISL_612224, EPI_ISL_612225, EPI_ISL_612226, EPI_ISL_612227, EPI_ISL_612228, EPI_ISL_612229, EPI_ISL_612230, EPI_ISL_612231, EPI_ISL_612232, EPI_ISL_612233, EPI_ISL_612234, EPI_ISL_612235, EPI_ISL_612236, EPI_ISL_612237,
EPI_ISL_612238, EPI_ISL_612239, EPI_ISL_612240, EPI_ISL_612241, EPI_ISL_612242, EPI_ISL_612243, EPI_ISL_612244, EPI_ISL_612245, EPI_ISL_612246, EPI_ISL_612247, EPI_ISL_612248, EPI_ISL_612249, EPI_ISL_612250, EPI_ISL_612251, EPI_ISL_612252, EPI_ISL_612253, EPI_ISL_612254, EPI_ISL_612255,
EPI_ISL_612256, EPI_ISL_612257, EPI_ISL_612258, EPI_ISL_612259, EPI_ISL_612260

| | | | |
|---|---|---|---|
| see above | University of Birmingham | COVID-19 Genomics UK (COG-UK) Consortium | Institute of Microbiology, University of Birmingham: Claire McMurray, Joanne Stockton, Samuel Nicholls, Radoslaw Poplawski, Will Rowe, Josh Quick, Nicholas Loman. University of Birmingham Testing Laboratory: Celina M Whalley, Andrew Bosworth, Charlotte Poxon, Kasun Wanigasooriya, Oliver Pickles, Mike Kidd, Alex Richter, Andrew D Beggs PHE Heartlands Lab: Husam Osman, Andrew Bosworth. Queen Elizabeth Hospital: Anna Casey |

EPI_ISL_623858, EPI_ISL_623859, EPI_ISL_623860, EPI_ISL_623861, EPI_ISL_623862, EPI_ISL_623866, EPI_ISL_623867, EPI_ISL_623869, EPI_ISL_623870, EPI_ISL_623875, EPI_ISL_623878, EPI_ISL_623880, EPI_ISL_623881, EPI_ISL_623882, EPI_ISL_623887, EPI_ISL_623888, EPI_ISL_623889, EPI_ISL_623890,
EPI_ISL_623891, EPI_ISL_623892, EPI_ISL_623894, EPI_ISL_623897, EPI_ISL_623898, EPI_ISL_623899, EPI_ISL_623900, EPI_ISL_623901, EPI_ISL_623902, EPI_ISL_623906, EPI_ISL_623909, EPI_ISL_623910, EPI_ISL_623912, EPI_ISL_623913, EPI_ISL_623914, EPI_ISL_623915, EPI_ISL_623918, EPI_ISL_623920,
EPI_ISL_623922, EPI_ISL_623923, EPI_ISL_623926, EPI_ISL_623927, EPI_ISL_623928, EPI_ISL_623929, EPI_ISL_623931, EPI_ISL_623932, EPI_ISL_623935, EPI_ISL_623936, EPI_ISL_623937, EPI_ISL_623938, EPI_ISL_623939, EPI_ISL_623940, EPI_ISL_623941, EPI_ISL_623943, EPI_ISL_623945, EPI_ISL_623946,
EPI_ISL_623947, EPI_ISL_623950, EPI_ISL_623952, EPI_ISL_623955, EPI_ISL_623957, EPI_ISL_623958, EPI_ISL_623960, EPI_ISL_623961, EPI_ISL_623962, EPI_ISL_623965, EPI_ISL_623966, EPI_ISL_623967, EPI_ISL_623968, EPI_ISL_623969, EPI_ISL_623971, EPI_ISL_623972, EPI_ISL_623978, EPI_ISL_623979,
EPI_ISL_623980, EPI_ISL_623981, EPI_ISL_623982, EPI_ISL_623984, EPI_ISL_623985, EPI_ISL_623987, EPI_ISL_623988, EPI_ISL_623992, EPI_ISL_623993, EPI_ISL_623996, EPI_ISL_623997, EPI_ISL_623999, EPI_ISL_624000, EPI_ISL_624001, EPI_ISL_624002, EPI_ISL_624003, EPI_ISL_624004, EPI_ISL_624005,
EPI_ISL_624007, EPI_ISL_624008, EPI_ISL_624011, EPI_ISL_624013, EPI_ISL_624014, EPI_ISL_624015, EPI_ISL_624016, EPI_ISL_624020, EPI_ISL_624023, EPI_ISL_624024, EPI_ISL_624025, EPI_ISL_624026, EPI_ISL_624030, EPI_ISL_624031, EPI_ISL_624033, EPI_ISL_624035, EPI_ISL_624036, EPI_ISL_624038,
EPI_ISL_624040, EPI_ISL_624041, EPI_ISL_624042, EPI_ISL_624045, EPI_ISL_624046, EPI_ISL_624047, EPI_ISL_624050, EPI_ISL_624051, EPI_ISL_624052, EPI_ISL_624054, EPI_ISL_624055, EPI_ISL_624057, EPI_ISL_624058, EPI_ISL_624060, EPI_ISL_624061, EPI_ISL_624063, EPI_ISL_624064, EPI_ISL_624065,
EPI_ISL_624066, EPI_ISL_624068, EPI_ISL_624069, EPI_ISL_624070, EPI_ISL_624072, EPI_ISL_624074, EPI_ISL_624075, EPI_ISL_624076, EPI_ISL_624078, EPI_ISL_624080, EPI_ISL_624081, EPI_ISL_624082, EPI_ISL_624083, EPI_ISL_624084, EPI_ISL_624085, EPI_ISL_624086, EPI_ISL_624087, EPI_ISL_624088,
EPI_ISL_624089, EPI_ISL_624090, EPI_ISL_624091, EPI_ISL_624095, EPI_ISL_624096, EPI_ISL_624098, EPI_ISL_624099, EPI_ISL_624101, EPI_ISL_624102, EPI_ISL_624103, EPI_ISL_624104, EPI_ISL_624106, EPI_ISL_624108, EPI_ISL_624109, EPI_ISL_624110, EPI_ISL_624113, EPI_ISL_624115, EPI_ISL_624116,
EPI_ISL_624117, EPI_ISL_624118, EPI_ISL_624121, EPI_ISL_624123, EPI_ISL_624124, EPI_ISL_624126, EPI_ISL_624127, EPI_ISL_624129, EPI_ISL_624130, EPI_ISL_624131, EPI_ISL_624133, EPI_ISL_624136, EPI_ISL_624137, EPI_ISL_624138, EPI_ISL_624139, EPI_ISL_624142, EPI_ISL_624144, EPI_ISL_624149,
EPI_ISL_624150, EPI_ISL_624151, EPI_ISL_624152, EPI_ISL_624155, EPI_ISL_624156, EPI_ISL_624159, EPI_ISL_624160, EPI_ISL_624161, EPI_ISL_624162, EPI_ISL_624166, EPI_ISL_624167, EPI_ISL_624168, EPI_ISL_624169, EPI_ISL_624170, EPI_ISL_624171, EPI_ISL_624172, EPI_ISL_624173, EPI_ISL_624174,
EPI_ISL_624175, EPI_ISL_624177, EPI_ISL_624179, EPI_ISL_624180, EPI_ISL_624182, EPI_ISL_624183, EPI_ISL_624184, EPI_ISL_624185, EPI_ISL_624186, EPI_ISL_624187, EPI_ISL_624190, EPI_ISL_624191, EPI_ISL_624194, EPI_ISL_624195, EPI_ISL_624196, EPI_ISL_624198, EPI_ISL_624200, EPI_ISL_624201,
EPI_ISL_624202, EPI_ISL_624203, EPI_ISL_624204, EPI_ISL_624207, EPI_ISL_624209, EPI_ISL_624210, EPI_ISL_624211, EPI_ISL_624212, EPI_ISL_624213, EPI_ISL_624214, EPI_ISL_624215, EPI_ISL_624216, EPI_ISL_624219, EPI_ISL_624222, EPI_ISL_624223, EPI_ISL_624224, EPI_ISL_624225, EPI_ISL_624229,
EPI_ISL_624233, EPI_ISL_624237, EPI_ISL_624240, EPI_ISL_624242, EPI_ISL_624243, EPI_ISL_624245, EPI_ISL_624246, EPI_ISL_624247, EPI_ISL_624248, EPI_ISL_624249, EPI_ISL_624250, EPI_ISL_624252, EPI_ISL_624253, EPI_ISL_624254, EPI_ISL_624259, EPI_ISL_624260, EPI_ISL_624261, EPI_ISL_624262,
EPI_ISL_624263, EPI_ISL_624265, EPI_ISL_624267, EPI_ISL_624269, EPI_ISL_624270, EPI_ISL_624271, EPI_ISL_624273, EPI_ISL_624274, EPI_ISL_624275, EPI_ISL_624276, EPI_ISL_624278, EPI_ISL_624279, EPI_ISL_624282, EPI_ISL_624283, EPI_ISL_624285, EPI_ISL_624290, EPI_ISL_624297, EPI_ISL_624298,
EPI_ISL_624300, EPI_ISL_624302, EPI_ISL_624303, EPI_ISL_624304, EPI_ISL_624305, EPI_ISL_624306, EPI_ISL_624307, EPI_ISL_624308, EPI_ISL_624312, EPI_ISL_624314, EPI_ISL_624318, EPI_ISL_624321, EPI_ISL_624322, EPI_ISL_624324, EPI_ISL_624325, EPI_ISL_624328, EPI_ISL_624330, EPI_ISL_624332,
EPI_ISL_624335, EPI_ISL_624336, EPI_ISL_624339, EPI_ISL_624342, EPI_ISL_624345, EPI_ISL_624348, EPI_ISL_624351, EPI_ISL_624353, EPI_ISL_624355, EPI_ISL_624356, EPI_ISL_624360, EPI_ISL_624361, EPI_ISL_624363, EPI_ISL_624366, EPI_ISL_624367, EPI_ISL_624368, EPI_ISL_624369, EPI_ISL_624371,
EPI_ISL_624373, EPI_ISL_624374, EPI_ISL_624376, EPI_ISL_624378, EPI_ISL_624379, EPI_ISL_624382, EPI_ISL_624383, EPI_ISL_624384, EPI_ISL_624388, EPI_ISL_624389, EPI_ISL_624390, EPI_ISL_624391, EPI_ISL_624392, EPI_ISL_624393, EPI_ISL_624395, EPI_ISL_624396, EPI_ISL_624398, EPI_ISL_624399,
EPI_ISL_624401, EPI_ISL_624404, EPI_ISL_624406, EPI_ISL_624408, EPI_ISL_624412, EPI_ISL_624414, EPI_ISL_624415, EPI_ISL_624417, EPI_ISL_624418, EPI_ISL_624421, EPI_ISL_624422, EPI_ISL_624428, EPI_ISL_624429, EPI_ISL_624430, EPI_ISL_624433, EPI_ISL_624435, EPI_ISL_624436, EPI_ISL_624437,
EPI_ISL_624448, EPI_ISL_624449, EPI_ISL_624452, EPI_ISL_624454, EPI_ISL_624455, EPI_ISL_624456, EPI_ISL_624457, EPI_ISL_624458, EPI_ISL_624459, EPI_ISL_624461, EPI_ISL_624464, EPI_ISL_624466, EPI_ISL_624467, EPI_ISL_624468, EPI_ISL_624469, EPI_ISL_624470, EPI_ISL_624475, EPI_ISL_624477,
EPI_ISL_624480, EPI_ISL_624481, EPI_ISL_624482, EPI_ISL_624485, EPI_ISL_624486, EPI_ISL_624487, EPI_ISL_624491, EPI_ISL_624494, EPI_ISL_624495, EPI_ISL_624496, EPI_ISL_624499, EPI_ISL_624503, EPI_ISL_624507, EPI_ISL_624508, EPI_ISL_624511, EPI_ISL_624512, EPI_ISL_624513, EPI_ISL_624514,
EPI_ISL_624516, EPI_ISL_624520, EPI_ISL_624525, EPI_ISL_624527, EPI_ISL_624530, EPI_ISL_624532, EPI_ISL_624536, EPI_ISL_624537, EPI_ISL_624538, EPI_ISL_624541, EPI_ISL_624542, EPI_ISL_624545, EPI_ISL_624546, EPI_ISL_624551, EPI_ISL_624552, EPI_ISL_624553, EPI_ISL_624555, EPI_ISL_624557,
EPI_ISL_624560, EPI_ISL_624561, EPI_ISL_624562, EPI_ISL_624563, EPI_ISL_624564, EPI_ISL_624566, EPI_ISL_624570, EPI_ISL_624571, EPI_ISL_624572, EPI_ISL_624575, EPI_ISL_624577, EPI_ISL_624579, EPI_ISL_624580, EPI_ISL_624583, EPI_ISL_624585, EPI_ISL_624586, EPI_ISL_624587, EPI_ISL_624589,
EPI_ISL_624590, EPI_ISL_624592, EPI_ISL_624593, EPI_ISL_624595, EPI_ISL_624597, EPI_ISL_624598, EPI_ISL_624599, EPI_ISL_624600, EPI_ISL_624601, EPI_ISL_624605, EPI_ISL_624608, EPI_ISL_624609, EPI_ISL_624610, EPI_ISL_624612, EPI_ISL_624614, EPI_ISL_624615, EPI_ISL_624616, EPI_ISL_624619,
EPI_ISL_624621, EPI_ISL_624623, EPI_ISL_624624, EPI_ISL_624625, EPI_ISL_624628, EPI_ISL_624630, EPI_ISL_624631, EPI_ISL_624632, EPI_ISL_624633, EPI_ISL_624634, EPI_ISL_624637, EPI_ISL_624638, EPI_ISL_624640, EPI_ISL_624643, EPI_ISL_624644, EPI_ISL_624646, EPI_ISL_624648, EPI_ISL_624654,
EPI_ISL_624655, EPI_ISL_624656, EPI_ISL_624657, EPI_ISL_624658, EPI_ISL_624659, EPI_ISL_624660, EPI_ISL_624661, EPI_ISL_624663, EPI_ISL_624665, EPI_ISL_624666, EPI_ISL_624667, EPI_ISL_624668, EPI_ISL_624669, EPI_ISL_624670, EPI_ISL_624671, EPI_ISL_624672, EPI_ISL_624673, EPI_ISL_624677,
EPI_ISL_624678, EPI_ISL_624679, EPI_ISL_624680, EPI_ISL_624682, EPI_ISL_624683, EPI_ISL_624685, EPI_ISL_624688, EPI_ISL_624689, EPI_ISL_624690, EPI_ISL_624698, EPI_ISL_624699, EPI_ISL_624700, EPI_ISL_624701, EPI_ISL_624703, EPI_ISL_624709, EPI_ISL_624710, EPI_ISL_624711, EPI_ISL_624712,
EPI_ISL_624714, EPI_ISL_624716, EPI_ISL_624717, EPI_ISL_624719, EPI_ISL_624720, EPI_ISL_624721, EPI_ISL_624722, EPI_ISL_624723, EPI_ISL_624724, EPI_ISL_624725, EPI_ISL_624729, EPI_ISL_624730, EPI_ISL_624733, EPI_ISL_624736, EPI_ISL_624740, EPI_ISL_624741, EPI_ISL_624742, EPI_ISL_624743,

EPI_ISL_624744, EPI_ISL_624746, EPI_ISL_624747, EPI_ISL_624748, EPI_ISL_624749, EPI_ISL_624750, EPI_ISL_624751, EPI_ISL_624752, EPI_ISL_624755, EPI_ISL_624756, EPI_ISL_624758, EPI_ISL_624759, EPI_ISL_624760, EPI_ISL_624761, EPI_ISL_624764, EPI_ISL_624765, EPI_ISL_624766, EPI_ISL_624767, EPI_ISL_624768, EPI_ISL_624769, EPI_ISL_624771, EPI_ISL_624774, EPI_ISL_624775, EPI_ISL_624779, EPI_ISL_624782, EPI_ISL_624783, EPI_ISL_624785, EPI_ISL_624786, EPI_ISL_624787, EPI_ISL_624788, EPI_ISL_624789, EPI_ISL_624792, EPI_ISL_624793, EPI_ISL_624797, EPI_ISL_624798, EPI_ISL_624799, EPI_ISL_624801, EPI_ISL_624802, EPI_ISL_624803, EPI_ISL_624806, EPI_ISL_624807, EPI_ISL_624808, EPI_ISL_624809, EPI_ISL_624810, EPI_ISL_624811, EPI_ISL_624813, EPI_ISL_624814, EPI_ISL_624815, EPI_ISL_624816, EPI_ISL_624817, EPI_ISL_624819, EPI_ISL_624820, EPI_ISL_624821, EPI_ISL_624822, EPI_ISL_624823, EPI_ISL_624825, EPI_ISL_624827, EPI_ISL_624828, EPI_ISL_624830, EPI_ISL_624832, EPI_ISL_624833, EPI_ISL_624834, EPI_ISL_624837, EPI_ISL_624838, EPI_ISL_624839, EPI_ISL_624842, EPI_ISL_624843, EPI_ISL_624845, EPI_ISL_624846, EPI_ISL_624850, EPI_ISL_624851, EPI_ISL_624852, EPI_ISL_624853, EPI_ISL_624854, EPI_ISL_624855, EPI_ISL_624856, EPI_ISL_624857, EPI_ISL_624860, EPI_ISL_624862, EPI_ISL_624863, EPI_ISL_624864, EPI_ISL_624865, EPI_ISL_624866, EPI_ISL_624868, EPI_ISL_624869, EPI_ISL_624870, EPI_ISL_624872, EPI_ISL_624874, EPI_ISL_624875, EPI_ISL_624876, EPI_ISL_624877, EPI_ISL_624878, EPI_ISL_624880, EPI_ISL_624884, EPI_ISL_624885, EPI_ISL_624886, EPI_ISL_624888, EPI_ISL_624891, EPI_ISL_624892, EPI_ISL_624893, EPI_ISL_624894, EPI_ISL_624897, EPI_ISL_624898, EPI_ISL_624899, EPI_ISL_624900, EPI_ISL_624902, EPI_ISL_624905, EPI_ISL_624907, EPI_ISL_624908, EPI_ISL_624910, EPI_ISL_624911, EPI_ISL_624912, EPI_ISL_624914, EPI_ISL_624915, EPI_ISL_624917, EPI_ISL_624919, EPI_ISL_624920, EPI_ISL_624921, EPI_ISL_624924, EPI_ISL_624926, EPI_ISL_624927, EPI_ISL_624928, EPI_ISL_624932, EPI_ISL_624933, EPI_ISL_624934, EPI_ISL_624935, EPI_ISL_624936, EPI_ISL_624940, EPI_ISL_624942, EPI_ISL_624943, EPI_ISL_624945, EPI_ISL_624946, EPI_ISL_624947, EPI_ISL_624949, EPI_ISL_624950, EPI_ISL_624951, EPI_ISL_624952, EPI_ISL_624953, EPI_ISL_624955, EPI_ISL_624956, EPI_ISL_624957, EPI_ISL_624958, EPI_ISL_624959, EPI_ISL_624960, EPI_ISL_624961, EPI_ISL_624962, EPI_ISL_624964, EPI_ISL_624966, EPI_ISL_624967, EPI_ISL_624968, EPI_ISL_624970, EPI_ISL_624971, EPI_ISL_624973, EPI_ISL_624974, EPI_ISL_624975, EPI_ISL_624976, EPI_ISL_624977, EPI_ISL_624979, EPI_ISL_624980, EPI_ISL_624982, EPI_ISL_624983, EPI_ISL_624985, EPI_ISL_624986, EPI_ISL_624987, EPI_ISL_624988, EPI_ISL_624990, EPI_ISL_624991, EPI_ISL_624992, EPI_ISL_624993, EPI_ISL_624994, EPI_ISL_624995, EPI_ISL_624997, EPI_ISL_624999, EPI_ISL_625001, EPI_ISL_625003, EPI_ISL_625004, EPI_ISL_625005, EPI_ISL_625007, EPI_ISL_625008, EPI_ISL_625009, EPI_ISL_625010, EPI_ISL_625011, EPI_ISL_625014, EPI_ISL_625016, EPI_ISL_625017, EPI_ISL_625018, EPI_ISL_625020, EPI_ISL_625021, EPI_ISL_625022, EPI_ISL_625027, EPI_ISL_625028, EPI_ISL_625029, EPI_ISL_625030, EPI_ISL_625034, EPI_ISL_625035, EPI_ISL_625036, EPI_ISL_625037, EPI_ISL_625039, EPI_ISL_625040, EPI_ISL_625043, EPI_ISL_625045, EPI_ISL_625046, EPI_ISL_625047, EPI_ISL_625048, EPI_ISL_625049, EPI_ISL_625050, EPI_ISL_625051, EPI_ISL_625052, EPI_ISL_625053, EPI_ISL_625054, EPI_ISL_625055, EPI_ISL_625056, EPI_ISL_625059, EPI_ISL_625061, EPI_ISL_625063, EPI_ISL_625067, EPI_ISL_625069, EPI_ISL_625070, EPI_ISL_625071, EPI_ISL_625072, EPI_ISL_625073, EPI_ISL_625074, EPI_ISL_625075, EPI_ISL_625078, EPI_ISL_625082, EPI_ISL_625084, EPI_ISL_625086, EPI_ISL_625087, EPI_ISL_625088, EPI_ISL_625089, EPI_ISL_625090, EPI_ISL_625091, EPI_ISL_625092, EPI_ISL_625095, EPI_ISL_625096, EPI_ISL_625098, EPI_ISL_625099, EPI_ISL_625100, EPI_ISL_625101, EPI_ISL_625103, EPI_ISL_625104, EPI_ISL_625107, EPI_ISL_625108, EPI_ISL_625109, EPI_ISL_625110, EPI_ISL_625111, EPI_ISL_625112, EPI_ISL_625115, EPI_ISL_625118, EPI_ISL_625120, EPI_ISL_625121, EPI_ISL_625123, EPI_ISL_625128, EPI_ISL_625129, EPI_ISL_625130, EPI_ISL_625131, EPI_ISL_625132, EPI_ISL_625134, EPI_ISL_625135, EPI_ISL_625136, EPI_ISL_625137, EPI_ISL_625139, EPI_ISL_625140, EPI_ISL_625141, EPI_ISL_625142, EPI_ISL_625144, EPI_ISL_625147, EPI_ISL_625148, EPI_ISL_625151, EPI_ISL_625152, EPI_ISL_625154, EPI_ISL_625156, EPI_ISL_625158, EPI_ISL_625159, EPI_ISL_625161, EPI_ISL_625162, EPI_ISL_625165, EPI_ISL_625168, EPI_ISL_625169, EPI_ISL_625170, EPI_ISL_625171, EPI_ISL_625174, EPI_ISL_625175, EPI_ISL_625179, EPI_ISL_625180, EPI_ISL_625181, EPI_ISL_625187, EPI_ISL_625188, EPI_ISL_625191, EPI_ISL_625193, EPI_ISL_625195, EPI_ISL_625196, EPI_ISL_625197, EPI_ISL_625198, EPI_ISL_625200, EPI_ISL_625202, EPI_ISL_625204, EPI_ISL_625205, EPI_ISL_625206, EPI_ISL_625208, EPI_ISL_625209, EPI_ISL_625211, EPI_ISL_625214, EPI_ISL_625216, EPI_ISL_625218, EPI_ISL_625219, EPI_ISL_625220, EPI_ISL_625222, EPI_ISL_625223, EPI_ISL_625224, EPI_ISL_625225, EPI_ISL_625226, EPI_ISL_625228, EPI_ISL_625229, EPI_ISL_625231, EPI_ISL_625232, EPI_ISL_625235, EPI_ISL_625236, EPI_ISL_625238, EPI_ISL_625239, EPI_ISL_625241, EPI_ISL_625245, EPI_ISL_625246, EPI_ISL_625247, EPI_ISL_625250, EPI_ISL_625251, EPI_ISL_625253, EPI_ISL_625254, EPI_ISL_625256, EPI_ISL_625258, EPI_ISL_625261, EPI_ISL_625262, EPI_ISL_625263, EPI_ISL_625264, EPI_ISL_625401, EPI_ISL_625402, EPI_ISL_625423

see above | Lighthouse Lab in Alderley Park | Wellcome Sanger Institute for the COVID-19 Genomics UK (COG-UK) consortium | Jacquelyn Wynn, Mairead Hyland, The Lighthouse Lab in Alderley Park and Alex Alderton, Roberto Amato, Sonia Goncalves, Ewan Harrison, David K. Jackson, Ian Johnston, Dominic Kwiatkowski, Cordelia Langford, John Sillitoe on behalf of the Wellcome Sanger Institute COVID-19 Surveillance Team (http://www.sanger.ac.uk/covid-team)

EPI_ISL_629754, EPI_ISL_629756, EPI_ISL_629757, EPI_ISL_629759, EPI_ISL_629760, EPI_ISL_629761, EPI_ISL_629762, EPI_ISL_629763, EPI_ISL_629765, EPI_ISL_629766, EPI_ISL_629767, EPI_ISL_629768, EPI_ISL_629769, EPI_ISL_629770, EPI_ISL_629772, EPI_ISL_629773, EPI_ISL_629775, EPI_ISL_629778, EPI_ISL_629779, EPI_ISL_629780, EPI_ISL_629781, EPI_ISL_629782, EPI_ISL_629783, EPI_ISL_629784, EPI_ISL_629785, EPI_ISL_629786, EPI_ISL_629787, EPI_ISL_629788, EPI_ISL_629789, EPI_ISL_629790, EPI_ISL_629791, EPI_ISL_629792, EPI_ISL_629793, EPI_ISL_629794, EPI_ISL_629796, EPI_ISL_629797, EPI_ISL_629798, EPI_ISL_629799, EPI_ISL_629800, EPI_ISL_629801, EPI_ISL_629802, EPI_ISL_629803, EPI_ISL_629804, EPI_ISL_629805, EPI_ISL_629806, EPI_ISL_629807, EPI_ISL_629808, EPI_ISL_629809, EPI_ISL_629810, EPI_ISL_629812, EPI_ISL_629813, EPI_ISL_629814, EPI_ISL_629815, EPI_ISL_629816, EPI_ISL_629817, EPI_ISL_629818, EPI_ISL_629819, EPI_ISL_629820, EPI_ISL_629823, EPI_ISL_629824, EPI_ISL_629825, EPI_ISL_629826, EPI_ISL_629827, EPI_ISL_629828, EPI_ISL_629829, EPI_ISL_629830, EPI_ISL_629831, EPI_ISL_629832, EPI_ISL_629833, EPI_ISL_629834, EPI_ISL_629835, EPI_ISL_629837, EPI_ISL_629838, EPI_ISL_629839, EPI_ISL_629840, EPI_ISL_629841, EPI_ISL_629842, EPI_ISL_629843, EPI_ISL_629844, EPI_ISL_629845, EPI_ISL_629847, EPI_ISL_629848, EPI_ISL_629849, EPI_ISL_629850, EPI_ISL_629852, EPI_ISL_629853, EPI_ISL_629854, EPI_ISL_629855, EPI_ISL_629856, EPI_ISL_629857, EPI_ISL_629858, EPI_ISL_629859, EPI_ISL_629860, EPI_ISL_629861, EPI_ISL_629862, EPI_ISL_629863, EPI_ISL_629864, EPI_ISL_629865, EPI_ISL_629866, EPI_ISL_629867, EPI_ISL_629868, EPI_ISL_629869, EPI_ISL_629870, EPI_ISL_629871, EPI_ISL_629872, EPI_ISL_629873, EPI_ISL_629874, EPI_ISL_629875, EPI_ISL_629876, EPI_ISL_629878, EPI_ISL_629879, EPI_ISL_629880, EPI_ISL_629881, EPI_ISL_629882, EPI_ISL_629883, EPI_ISL_629884, EPI_ISL_629885, EPI_ISL_629886, EPI_ISL_629887, EPI_ISL_629888, EPI_ISL_629889, EPI_ISL_629890, EPI_ISL_629891, EPI_ISL_629892, EPI_ISL_629893, EPI_ISL_629894, EPI_ISL_629895, EPI_ISL_629896, EPI_ISL_629897, EPI_ISL_629898, EPI_ISL_629899, EPI_ISL_629900, EPI_ISL_629901, EPI_ISL_629902, EPI_ISL_629903, EPI_ISL_629904, EPI_ISL_629905, EPI_ISL_629906, EPI_ISL_629907, EPI_ISL_629908, EPI_ISL_629909, EPI_ISL_629910, EPI_ISL_629911, EPI_ISL_629912, EPI_ISL_629914, EPI_ISL_629915, EPI_ISL_629916, EPI_ISL_629917, EPI_ISL_629918, EPI_ISL_629919, EPI_ISL_629920, EPI_ISL_629921, EPI_ISL_629922, EPI_ISL_629923, EPI_ISL_629924, EPI_ISL_629925, EPI_ISL_629927, EPI_ISL_629928, EPI_ISL_629929, EPI_ISL_629930, EPI_ISL_629931, EPI_ISL_629933, EPI_ISL_629934, EPI_ISL_629935, EPI_ISL_629936, EPI_ISL_629937, EPI_ISL_629938, EPI_ISL_629939, EPI_ISL_629941, EPI_ISL_629942, EPI_ISL_629944, EPI_ISL_629945, EPI_ISL_629946, EPI_ISL_629947, EPI_ISL_629948, EPI_ISL_629949, EPI_ISL_629953, EPI_ISL_629954, EPI_ISL_629955, EPI_ISL_629956, EPI_ISL_629957, EPI_ISL_629958, EPI_ISL_629959, EPI_ISL_629960, EPI_ISL_629961, EPI_ISL_629962, EPI_ISL_629963, EPI_ISL_629964, EPI_ISL_629965, EPI_ISL_629966, EPI_ISL_629967, EPI_ISL_629968, EPI_ISL_629971, EPI_ISL_629972, EPI_ISL_629973, EPI_ISL_629974, EPI_ISL_629975, EPI_ISL_629976, EPI_ISL_629978, EPI_ISL_629979, EPI_ISL_629980, EPI_ISL_629981, EPI_ISL_629982, EPI_ISL_629984, EPI_ISL_629985, EPI_ISL_629987, EPI_ISL_629988, EPI_ISL_629989, EPI_ISL_629990, EPI_ISL_629991, EPI_ISL_629992, EPI_ISL_629993, EPI_ISL_629994, EPI_ISL_629995, EPI_ISL_629996, EPI_ISL_629997, EPI_ISL_629998, EPI_ISL_629999, EPI_ISL_630000, EPI_ISL_630001, EPI_ISL_630002, EPI_ISL_630003, EPI_ISL_630004, EPI_ISL_630005, EPI_ISL_630006

see above | Lighthouse Lab in Glasgow | Wellcome Sanger Institute for the COVID-19 Genomics UK (COG-UK) consortium | Harper VanSteenhouse, Yumi Kasai, David Gray, Carol Clugston, Anna Dominiczak and Alex Alderton, Roberto Amato, Sonia Goncalves, Ewan Harrison, David K. Jackson, Ian Johnston, Dominic Kwiatkowski, Cordelia Langford, John Sillitoe on behalf of the Wellcome Sanger Institute COVID-19 Surveillance Team

EPI_ISL_630348, EPI_ISL_630349, EPI_ISL_630350, EPI_ISL_630351, EPI_ISL_630353, EPI_ISL_630354, EPI_ISL_630355, EPI_ISL_630356, EPI_ISL_630357, EPI_ISL_630358, EPI_ISL_630359, EPI_ISL_630360, EPI_ISL_630361, EPI_ISL_630362, EPI_ISL_630363, EPI_ISL_630364, EPI_ISL_630365, EPI_ISL_630366, EPI_ISL_630367, EPI_ISL_630368, EPI_ISL_630369, EPI_ISL_630370, EPI_ISL_630371, EPI_ISL_630373, EPI_ISL_630374, EPI_ISL_630375, EPI_ISL_630376, EPI_ISL_630377, EPI_ISL_630378, EPI_ISL_630379, EPI_ISL_630383, EPI_ISL_630384, EPI_ISL_630386, EPI_ISL_630387, EPI_ISL_630388, EPI_ISL_630389, EPI_ISL_630390, EPI_ISL_630391, EPI_ISL_630393, EPI_ISL_630394, EPI_ISL_630395, EPI_ISL_630396, EPI_ISL_630397, EPI_ISL_630398, EPI_ISL_630399, EPI_ISL_630400, EPI_ISL_630401, EPI_ISL_630403, EPI_ISL_630404, EPI_ISL_630405, EPI_ISL_630407, EPI_ISL_630408, EPI_ISL_630410, EPI_ISL_630411, EPI_ISL_630412, EPI_ISL_630413, EPI_ISL_630414, EPI_ISL_630415, EPI_ISL_630417, EPI_ISL_630418, EPI_ISL_630419, EPI_ISL_630420, EPI_ISL_630421, EPI_ISL_630422, EPI_ISL_630423, EPI_ISL_630426, EPI_ISL_630427, EPI_ISL_630428, EPI_ISL_630429, EPI_ISL_630430, EPI_ISL_630431, EPI_ISL_630432, EPI_ISL_630433, EPI_ISL_630434, EPI_ISL_630435, EPI_ISL_630437, EPI_ISL_630438, EPI_ISL_630439, EPI_ISL_630440, EPI_ISL_630441, EPI_ISL_630442, EPI_ISL_630443, EPI_ISL_630444, EPI_ISL_630445, EPI_ISL_630446, EPI_ISL_630447, EPI_ISL_630448, EPI_ISL_630450, EPI_ISL_630451, EPI_ISL_630452, EPI_ISL_630453, EPI_ISL_630454, EPI_ISL_630456, EPI_ISL_630457, EPI_ISL_630458, EPI_ISL_630459, EPI_ISL_630460, EPI_ISL_630461, EPI_ISL_630462, EPI_ISL_630463, EPI_ISL_630464, EPI_ISL_630465, EPI_ISL_630466, EPI_ISL_630467, EPI_ISL_630468, EPI_ISL_630470, EPI_ISL_630471, EPI_ISL_630472, EPI_ISL_630473, EPI_ISL_630475, EPI_ISL_630477, EPI_ISL_630478, EPI_ISL_630479, EPI_ISL_630480, EPI_ISL_630481, EPI_ISL_630482, EPI_ISL_630484, EPI_ISL_630485, EPI_ISL_630486, EPI_ISL_630488, EPI_ISL_630489, EPI_ISL_630490, EPI_ISL_630491, EPI_ISL_630493, EPI_ISL_630494, EPI_ISL_630496, EPI_ISL_630497, EPI_ISL_630498, EPI_ISL_630499, EPI_ISL_630500, EPI_ISL_630501, EPI_ISL_630502, EPI_ISL_630503, EPI_ISL_630504, EPI_ISL_630505, EPI_ISL_630507, EPI_ISL_630508, EPI_ISL_630509, EPI_ISL_630510, EPI_ISL_630511, EPI_ISL_630512, EPI_ISL_630513, EPI_ISL_630514, EPI_ISL_630515, EPI_ISL_630516, EPI_ISL_630517, EPI_ISL_630518, EPI_ISL_630519, EPI_ISL_630520, EPI_ISL_630522, EPI_ISL_630524, EPI_ISL_630525, EPI_ISL_630526, EPI_ISL_630527, EPI_ISL_630529, EPI_ISL_630530, EPI_ISL_630532, EPI_ISL_630534, EPI_ISL_630535, EPI_ISL_630536, EPI_ISL_630537, EPI_ISL_630538, EPI_ISL_630539, EPI_ISL_630540, EPI_ISL_630541, EPI_ISL_630542, EPI_ISL_630543, EPI_ISL_630544, EPI_ISL_630546, EPI_ISL_630548, EPI_ISL_630549, EPI_ISL_630550, EPI_ISL_630552, EPI_ISL_630553, EPI_ISL_630554, EPI_ISL_630555, EPI_ISL_630557, EPI_ISL_630558, EPI_ISL_630559, EPI_ISL_630560, EPI_ISL_630561, EPI_ISL_630562, EPI_ISL_630563, EPI_ISL_630564, EPI_ISL_630566, EPI_ISL_630567, EPI_ISL_630568, EPI_ISL_630569, EPI_ISL_630570, EPI_ISL_630571, EPI_ISL_630572, EPI_ISL_630573, EPI_ISL_630574, EPI_ISL_630575, EPI_ISL_630577, EPI_ISL_630578, EPI_ISL_630580, EPI_ISL_630581, EPI_ISL_630582, EPI_ISL_630583, EPI_ISL_630584, EPI_ISL_630585, EPI_ISL_630586, EPI_ISL_630587, EPI_ISL_630588, EPI_ISL_630589, EPI_ISL_630591, EPI_ISL_630592, EPI_ISL_630593, EPI_ISL_630594, EPI_ISL_630595, EPI_ISL_630596, EPI_ISL_630597, EPI_ISL_630599, EPI_ISL_630600, EPI_ISL_630601, EPI_ISL_630602, EPI_ISL_630603, EPI_ISL_630604, EPI_ISL_630605, EPI_ISL_630606, EPI_ISL_630607, EPI_ISL_630608, EPI_ISL_630609, EPI_ISL_630610, EPI_ISL_630611, EPI_ISL_630612, EPI_ISL_630614, EPI_ISL_630615, EPI_ISL_630616, EPI_ISL_630618, EPI_ISL_630619, EPI_ISL_630620, EPI_ISL_630621, EPI_ISL_630622, EPI_ISL_630623, EPI_ISL_630624, EPI_ISL_630625, EPI_ISL_630626, EPI_ISL_630627, EPI_ISL_630628, EPI_ISL_630629, EPI_ISL_630630, EPI_ISL_630631, EPI_ISL_630632, EPI_ISL_630634, EPI_ISL_630635, EPI_ISL_630637, EPI_ISL_630638, EPI_ISL_630639, EPI_ISL_630641, EPI_ISL_630642, EPI_ISL_630643, EPI_ISL_630644, EPI_ISL_630645, EPI_ISL_630646, EPI_ISL_630647, EPI_ISL_630648, EPI_ISL_630649, EPI_ISL_630650, EPI_ISL_630651, EPI_ISL_630652, EPI_ISL_630653, EPI_ISL_630655, EPI_ISL_630656, EPI_ISL_630658, EPI_ISL_630659, EPI_ISL_630660, EPI_ISL_630661, EPI_ISL_630662, EPI_ISL_630663, EPI_ISL_630664, EPI_ISL_630665, EPI_ISL_630666, EPI_ISL_630667, EPI_ISL_630668, EPI_ISL_630669, EPI_ISL_630670, EPI_ISL_630671, EPI_ISL_630672, EPI_ISL_630673, EPI_ISL_630674, EPI_ISL_630675, EPI_ISL_630676, EPI_ISL_630677, EPI_ISL_630678, EPI_ISL_630679, EPI_ISL_630680, EPI_ISL_630681, EPI_ISL_630682, EPI_ISL_630683, EPI_ISL_630685, EPI_ISL_630686, EPI_ISL_630687, EPI_ISL_630688, EPI_ISL_630689, EPI_ISL_630690, EPI_ISL_630691, EPI_ISL_630692, EPI_ISL_630693, EPI_ISL_630694, EPI_ISL_630695, EPI_ISL_630696, EPI_ISL_630697, EPI_ISL_630698, EPI_ISL_630699, EPI_ISL_630700, EPI_ISL_630701, EPI_ISL_630709, EPI_ISL_630710, EPI_ISL_630711, EPI_ISL_630713, EPI_ISL_630715, EPI_ISL_630716, EPI_ISL_630717, EPI_ISL_630720, EPI_ISL_630721, EPI_ISL_630722, EPI_ISL_630727, EPI_ISL_630728, EPI_ISL_630730, EPI_ISL_630732, EPI_ISL_630734, EPI_ISL_630738, EPI_ISL_630739, EPI_ISL_630740, EPI_ISL_630743, EPI_ISL_630746, EPI_ISL_630750, EPI_ISL_630751, EPI_ISL_630752, EPI_ISL_630756, EPI_ISL_630757, EPI_ISL_630758, EPI_ISL_630760, EPI_ISL_630763, EPI_ISL_630765, EPI_ISL_630768, EPI_ISL_630769, EPI_ISL_630770, EPI_ISL_630771, EPI_ISL_630772, EPI_ISL_630773, EPI_ISL_630774, EPI_ISL_630776, EPI_ISL_630777, EPI_ISL_630783, EPI_ISL_630786, EPI_ISL_630789, EPI_ISL_630791, EPI_ISL_630792, EPI_ISL_630794, EPI_ISL_630795, EPI_ISL_630797, EPI_ISL_630799, EPI_ISL_630801, EPI_ISL_630802, EPI_ISL_630805, EPI_ISL_630806, EPI_ISL_630811, EPI_ISL_630814, EPI_ISL_630817, EPI_ISL_630818, EPI_ISL_630819, EPI_ISL_630821, EPI_ISL_630823, EPI_ISL_630826, EPI_ISL_630831, EPI_ISL_630834, EPI_ISL_630835, EPI_ISL_630836, EPI_ISL_630837, EPI_ISL_630841, EPI_ISL_630842, EPI_ISL_630843, EPI_ISL_630846, EPI_ISL_630852, EPI_ISL_630854, EPI_ISL_630860, EPI_ISL_630865, EPI_ISL_630868, EPI_ISL_630874, EPI_ISL_630878, EPI_ISL_630881, EPI_ISL_630884, EPI_ISL_630885, EPI_ISL_630886, EPI_ISL_630887, EPI_ISL_630889, EPI_ISL_630890, EPI_ISL_630892, EPI_ISL_630895, EPI_ISL_630897, EPI_ISL_630898, EPI_ISL_630903, EPI_ISL_630906, EPI_ISL_630907, EPI_ISL_630909, EPI_ISL_630910, EPI_ISL_630916, EPI_ISL_630920, EPI_ISL_630922, EPI_ISL_630924, EPI_ISL_630928, EPI_ISL_630930, EPI_ISL_630935, EPI_ISL_630936, EPI_ISL_630937, EPI_ISL_630938, EPI_ISL_630947, EPI_ISL_630953, EPI_ISL_630961, EPI_ISL_630962, EPI_ISL_630964, EPI_ISL_630968, EPI_ISL_630970, EPI_ISL_631163, EPI_ISL_631175

see above | Lighthouse Lab in Alderley Park | Wellcome Sanger Institute for the COVID-19 Genomics UK (COG-UK) consortium | Jacquelyn Wynn, Mairead Hyland, The Lighthouse Lab in Alderley Park and Alex Alderton, Roberto Amato, Sonia Goncalves, Ewan Harrison, David K. Jackson, Ian Johnston, Dominic Kwiatkowski, Cordelia Langford, John Sillitoe on behalf of the Wellcome Sanger Institute COVID-19 Surveillance Team

We gratefully acknowledge the following Authors from the Originating laboratories responsible for obtaining the specimens, as well as the Submitting laboratories where the genome data were generated and shared via GISAID, on which this research is based.

All Submitters of data may be contacted directly via www.gisaid.org

Authors are sorted alphabetically.

| Accession ID | Originating Laboratory | Submitting Laboratory | Authors |
|---|---|---|---|
| EPI_ISL_406031 | Centers for Disease Control, R.O.C. (Taiwan) | Centers for Disease Control, R.O.C. (Taiwan) | Ji-Rong Yang, Yu-Chi Lin, Jung-Jung Mu, Ming-Tsan Liu, Shu-Ying Li |
| EPI_ISL_407193 | Korea Centers for Disease Control & Prevention (KCDC) Center for Laboratory Control of Infectious Diseases Division of Viral Diseases | Korea Centers for Disease Control & Prevention (KCDC) Center for Laboratory Control of Infectious Diseases Division of Viral Diseases | Jeong-Min Kim, Yoon-Seok Chung, Namjoo Lee, Mi-Seon Kim, SangHee Woo, Hye-Joon Jo, Sehee Park, Heui Man Kim, Myung Guk Han |
| EPI_ISL_407214, EPI_ISL_407215 | Washington State Department of Health | Pathogen Discovery, Respiratory Viruses Branch, Division of Viral Diseases, Centers for Dieases Control and Prevention | Krista Queen, Azaibi Tamin, Jennifer Harcourt, Ying Tao, Clinton R. Paden, Jing Zhang, Yan Li, Anna Uehara, Xiaoyan Lu, Shifaq Kamili, Rashi Gautam, Haibin Wang, Janna' R. Murray, Susan I. Gerber, Stephen Lindstrom, Natalie Thornburg, Suxiang Tong |
| EPI_ISL_407313 | Hangzhou Center for Disease Control and Prevention | Hangzhou Center for Disease Control and Prevention | Jun Li, Haoqiu Wang, Hua Yu, Lingfeng Mao, Xinfen Yu, Zhou Sun, Qingxin Kong, Xin Qian, Shuchang Chen, Xuchu Wang |
| EPI_ISL_408008 | California Department of Health | Pathogen Discovery, Respiratory Viruses Branch, Division of Viral Diseases, Centers for Disease Control and Prevention | Krista Queen, Jing Zhang, Yan Li, Ying Tao, Anna Uehara, Clinton Paden, Xiaoyan Lu, Brian Lynch, Senthil Kumar K. Sakthivel, Brett L. Whitaker, Shifaq Kamili, Lijuan Wang, Janna' R. Murray, Susan I. Gerber, Stephen Lindstrom, Suxiang Tong |
| EPI_ISL_408009 | California Department of Health | Pathogen Discovery, Respiratory Viruses Branch, Division of Viral Diseases, Centers for Dieases Control and Prevention | Krista Queen, Jing Zhang, Yan Li, Ying Tao, Anna Uehara, Clinton Paden, Xiaoyan Lu, Brian Lynch, Senthil Kumar K. Sakthivel, Brett L. Whitaker, Shifaq Kamili, Lijuan Wang, Janna' R. Murray, Susan I. Gerber, Stephen Lindstrom, Suxiang Tong |
| EPI_ISL_408670 | Wisconsin Department of Health Services | Pathogen Discovery, Respiratory Viruses Branch, Division of Viral Diseases, Centers for Dieases Control and Prevention | Jing Zhang, Anna Uehara, Krista Queen, Yan Li, Ying Tao, Clinton R. Paden, Xiaoyan Lu, Brian Lynch, Senthil Kumar K. Sakthivel, Brett L. Whitaker, Shifaq Kamili, Lijuan Wang, Janna' R. Murray, Susan I. Gerber, Stephen Lindstrom, Suxiang Tong |
| EPI_ISL_410044 | California Department of Public Health | Pathogen Discovery, Respiratory Viruses Branch, Division of Viral Diseases, Centers for Dieases Control and Prevention | Jing Zhang, Krista Queen, Yan Li, Ying Tao, Anna Uehara, Clinton R. Paden, Xiaoyan Lu, Brian Lynch, Senthil Kumar K. Sakthivel, Brett L. Whitaker, Shifaq Kamili, Lijuan Wang, Janna' R. Murray, Susan I. Gerber, Stephen Lindstrom, Suxiang Tong |
| EPI_ISL_411915 | Laboratory Medicine | Department of Laboratory Medicine, Lin-Kou Chang Gung Memorial Hospital, Taoyuan, Taiwan. | Kuo-Chien Tsao, Yu-Nong Gong, Shu-Li Yang, Yi-Chun Li, Chung-Guei Huang, Yhu-Chering Huang, Shin-Ru Shih |
| EPI_ISL_411926, EPI_ISL_411927 | Taiwan Centers for Disease Control | Taiwan Centers for Disease Control | Ji-Rong Yang, Yu-Chi-Lin, Jung-Jung Mu, Ming-Tsan-Liu |
| EPI_ISL_411952, EPI_ISL_411953 | NHC Key laboratory of Enteric Pathogenic Microbiology, Institute of Pathogenic Microbiology | Jiangsu Provincial Center for Disease Control & Prevention | Kangchen Zhao, Xiaojuan Zhu, Lunbiao Cui, Tao Wu, Yiyue Ge, Bin Wu, Yin Chen, Fengcai Zhu, Baoli Zhu, Ming Wu |
| EPI_ISL_411954, EPI_ISL_411955 | California Department of Public Health | Pathogen Discovery, Respiratory Viruses Branch, Division of Viral Diseases, Centers for Dieases Control and Prevention | Krista Queen, Anna Uehara, Jing Zhang, Yan Li, Ying Tao, Clinton R. Paden, Haibin Wang, Shifaq Kamili, Xiaoyan Lu, Brian Lynch, Senthil Kumar K. Sakthivel, Brett L. Whitaker, Lijuan Wang, Janna' R. Murray, Susan I. Gerber, Stephen Lindstrom, Suxiang Tong |
| EPI_ISL_411956 | Texas Department of State Health Services | Pathogen Discovery, Respiratory Viruses Branch, Division of Viral Diseases, Centers for Dieases Control and Prevention | Krista Queen, Anna Uehara, Jing Zhang, Yan Li, Ying Tao, Clinton R. Paden, Haibin Wang, Shifaq Kamili, Xiaoyan Lu, Brian Lynch, Senthil Kumar K. Sakthivel, Brett L. Whitaker, Lijuan Wang, Janna' R. Murray, Susan I. Gerber, Stephen Lindstrom, Suxiang Tong |
| EPI_ISL_412862 | California Department of Public Health | Pathogen Discovery, Respiratory Viruses Branch, Division of Viral Diseases, Centers for Disease Control and Prevention | Krista Queen, Anna Uehara, Jing Zhang, Yan Li, Ying Tao, Clinton R. Paden, Haibin Wang, Shifaq Kamili, Xiaoyan Lu, Brian Lynch, Senthil Kumar K. Sakthivel, Brett L. Whitaker, Lijuan Wang, Janna' R. Murray, Jasmine Padilla, Justin Lee, Susan I. Gerber, Stephen Lindstrom, Suxiang Tong |
| EPI_ISL_412869, EPI_ISL_412870, EPI_ISL_412871, EPI_ISL_412872, EPI_ISL_412873 | Division of Viral Diseases, Center for Laboratory Control of Infectious Diseases, Korea Centers for Diseases Control and Prevention | Division of Viral Diseases, Center for Laboratory Control of Infectious Diseases, Korea Centers for Diseases Control and Prevention | Jeong-Min Kim, Yoon-Seok Chung, Namjoo Lee, Mi-Seon Kim, Sang Hee Woo, Hye-Jun Jo, Sehee Park, Heui Man Kim, Myung Guk Han |
| EPI_ISL_412964 | Hospital Israelita Albert Einstein | Instituto Adolfo Lutz Interdisciplinary Procedures Center Strategic Laboratory | Jaqueline Goes de Jesus, Claudio Tavares Sacchi, Daniela Bernardes Borges da Silva, Ingra Morales Claro, Flávia Cristina da Silva Sales, Claudia Regina Gonçalves, Joshua Quick, Maria do Carmo, Sampaio Tavares Timenetsky, Nicholas James Loman, Andrew Rambaut, Ester Cerdeira Sabino, Nuno Rodrigues Faria |
| EPI_ISL_413016 | Hospital Israelita Albert Einstein | Instituto Adolfo Lutz, Interdisciplinary Procedures Center, Strategic Laboratory | Jaqueline Goes de Jesus, Claudio Tavares Sacchi, Fabiana Cristina Pereira dos Santos, Ingra Morales Claro, Flávia Cristina da Silva Sales, Claudia Regina Gonçalves, Joshua Quick, Maria do Carmo Sampaio Tavares Timenetsky, Nicholas James Loman, Andrew Rambaut, Ester Cerdeira Sabino, Nuno Rodrigues Faria |
| EPI_ISL_413597 | Centre for Infectious Diseases and Microbiology- Public Health | NSW Health Pathology - Institute of Clinical Pathology and Medical Research; Westmead Hospital; University of Sydney | Lam C, Eden J-S, Rockett R, Gray K, Timms, V, Gall, M, Carter I, Rahman H, Holmes EC, O'Sullivan MV, Sintchenko V, Chen SC, Maddocks S, Kok J and Dwyer DE for the 2019-nCoV Study Group* |
| EPI_ISL_413850, EPI_ISL_413851, EPI_ISL_413852, EPI_ISL_413853, EPI_ISL_413854, EPI_ISL_413855, EPI_ISL_413856, EPI_ISL_413857, EPI_ISL_413858, EPI_ISL_413859, EPI_ISL_413860, EPI_ISL_413861, EPI_ISL_413862, EPI_ISL_413863, EPI_ISL_413864, EPI_ISL_413866, EPI_ISL_413867, EPI_ISL_413868, EPI_ISL_413870, EPI_ISL_413871, EPI_ISL_413872, EPI_ISL_413873, EPI_ISL_413874, EPI_ISL_413875, EPI_ISL_413876, EPI_ISL_413877, EPI_ISL_413878, EPI_ISL_413879, EPI_ISL_413880, EPI_ISL_413881, EPI_ISL_413882, EPI_ISL_413883, EPI_ISL_413884, EPI_ISL_413885, EPI_ISL_413886, EPI_ISL_413887, EPI_ISL_413888, EPI_ISL_413889, EPI_ISL_413890, EPI_ISL_413891, EPI_ISL_413892, EPI_ISL_413893, EPI_ISL_413894, EPI_ISL_413895, EPI_ISL_413896, EPI_ISL_413897, EPI_ISL_413898, EPI_ISL_413899, EPI_ISL_413900, EPI_ISL_413902 | | | |
| see above | Guangdong Provincial Institution of Public Health, Guangdong Provinical Center for Disease Control and Prevention | Guangdong Provincial Institution of Public Health | Jing Lu, Louis du Plessis, Liu Zhe, Jiufeng Sun, Sarah François, Huifang Lin, Moritz Kraemer, Jingju Peng, Qianlin Xiong, Runyu Yuan, Lilian Zeng, Pingping Zhou, Chuming Liang, Tao Liu, Wei Li, Juan Su, Huanying Zheng, Kang Min, Song Tie, Bo Peng, Shisong Fang, Wenzhe Su, Kuibiao Li, Ruilin Sun, Ru bai, Xi Tang, Minfeng Liang, Nuno Faria, Josh Quick, Andrew Rambaut, Verity Hill, Wenjun Ma, Nick Loman, Oliver Pybus, Changwen Ke |
| EPI_ISL_413996, EPI_ISL_413997, EPI_ISL_413999, EPI_ISL_414019, EPI_ISL_414020, EPI_ISL_414021, EPI_ISL_414022, EPI_ISL_414023 | Laboratoire de Virologie, HUG | Swiss National Reference Centre for Influenza | LAUBSCHER Florian et al. |
| EPI_ISL_414482, EPI_ISL_414483, EPI_ISL_414484, EPI_ISL_414485 | unknown | Pathogen Discovery, Respiratory Viruses Branch, Division of Viral Diseases, Centers for Disease Control and Prevention | Krista Queen, Anna Uehara, Ying Tao, Clinton R. Paden, Jing Zhang, Yan Li, Haibin Wang, Shifaq Kamili, Xiaoyan Lu, Brian Lynch, Senthil Kumar K. Sakthivel, Brett L. Whitaker, Lijuan Wang, Janna' R. Murray, Jasmine Padilla, Justin Lee, Susan I. Gerber, Stephen Lindstrom, Suxiang Tong |
| EPI_ISL_414639 | NYU Langone Health | Departments of Pathology and Medicine, New York University School of Medicine | John Chen, Dacia Dimartino, Xiaojun Feng, Adriana Heguy, Megan Hogan, Emily Huang, George Jour, Christian Marier, Matthew T. Maurano, Mark J. Mulligan, Peter Meyn, Marie Samanovic-Golden, Amy Rapkiewicz, Guomiao Shen, Matija Snuderl, Gael Westby, Paul Zappile |
| EPI_ISL_415628, EPI_ISL_415630, EPI_ISL_415631 | West of Scotland Specialist Virology Centre, NHSGGC | MRC-University of Glasgow Centre for Virus Research | Kathy Smollett, Daniel Mair, Stephen Carmichael, Ana da Silva Filipe; Richard Orton, David L Robertson; Alasdair MacLean, Rory Gunson; Natasha Jesudason, Kathy Li, Antonia Ho; Emma Thomson. |
| EPI_ISL_415741, EPI_ISL_415742, EPI_ISL_415743 | Laboratory Medicine | Department of Laboratory Medicine, Lin-Kou Chang Gung Memorial Hospital, Taoyuan, Taiwan | Kuo-Chien Tsao, Yu-Nong Gong, Shu-Li Yang, Yi-Chun Liu, Chung-Guei Huang, Po-Wei Huang, Mei-Jen Hsiao, Cheng-Ta Yang, Cheng-Hsun Chiu, Chi-Hsien Huang, Kuang-Tso Le, Shu-Min Lin, Peng-Nien Huang, Kuo-Ming Lee, Guang-Wu Chen, Shin-Ru Shih |
| EPI_ISL_416416, EPI_ISL_416417, EPI_ISL_416418, EPI_ISL_416419 | Connecticut State Department of Public Health | Grubaugh Lab - Yale School of Public Health | Joseph Fauver, Chantal Vogels, Anderson Brito, Tara Alpert, Nagarjuna Cheemarla, Ellen Foxman, Anthony Muyombwe, Jafar Razeq, Richard Martinello, Albert Ko, Marie-Louise Landry, Nathan Grubaugh |
| EPI_ISL_416420, EPI_ISL_416421, EPI_ISL_416422, EPI_ISL_416423, EPI_ISL_416424 | Yale Clinical Virology Laboratory | Grubaugh Lab - Yale School of Public Health | Joseph Fauver, Chantal Vogels, Anderson Brito, Tara Alpert, Nagarjuna Cheemarla, Ellen Foxman, Anthony Muyombwe, Jafar Razeq, Richard Martinello, Albert Ko, Marie-Louise Landry, Nathan Grubaugh |
| EPI_ISL_416524 | Saitama Medical University Hospital | Saitama Medical University | Kazuo Imai |

| | | | |
|---|---|---|---|
| EPI_ISL_416525 | Saitama Medical University | Saitama Medical University | Kazuo Imai |
| EPI_ISL_416830, EPI_ISL_416831, EPI_ISL_416832 | NYU Langone Health | Department of Pathology and Medicine, New York University School of Medicine | John Chen, Dacia Dimartino, Xiaojun Feng, Adriana Heguy, Megan Hogan, Emily Huang, George Jour, Christian Marier, Matt Maurano, Mark Mulligan, Peter Meyn, Marie Samanovic-Golden, Amy Rapkiewicz, Guomiao Shen, Matija Snuderl, Gael Westby, Paul Zappile |
| EPI_ISL_417176, EPI_ISL_417177, EPI_ISL_417178, EPI_ISL_417179 | Department of Pathology, Princess Margaret Hospital | Department of Health Technology and Informatics, Faculty of Health and Social Science, The Hong Kong Polytechnic University | Kenneth Siu-Sing LEUNG, Timothy Ting-Leung NG, Alan Ka-Lun WU, Miranda Chong-Yee YAU, Hiu-Yin LAO, Ming-Pan CHOI, Kingsley King-Gee TAM, Lam-Kwong LEE, Barry Kin-Chung WONG, Alex Yat-Man HO, Kam-Tong Yip, Kwok-Cheung LUNG, Raymond Wai-To LIU, Eugene Yuk-Keung TSO, Wai-Shing LEUNG, Man-Chun CHAN, Yuk-Yung NG, Kit-Man SIN, Kitty Sau-Chun FUNG, Sandy Ka-Yee CHAU, Wing-Kin TO, Tak-Lun Que, David Ho-Keung SHUM, Shea Ping YIP, Wing Cheong YAM, Gilman Kit-Hang SIU |
| EPI_ISL_417180, EPI_ISL_417181, EPI_ISL_417182 | Department of Pathology, United Christian Hospital | Department of Health Technology and Informatics, Faculty of Health and Social Science, The Hong Kong Polytechnic University | Kenneth Siu-Sing LEUNG, Timothy Ting-Leung NG, Alan Ka-Lun WU, Miranda Chong-Yee YAU, Hiu-Yin LAO, Ming-Pan CHOI, Kingsley King-Gee TAM, Lam-Kwong LEE, Barry Kin-Chung WONG, Alex Yat-Man HO, Kam-Tong Yip, Kwok-Cheung LUNG, Raymond Wai-To LIU, Eugene Yuk-Keung TSO, Wai-Shing LEUNG, Man-Chun CHAN, Yuk-Yung NG, Kit-Man SIN, Kitty Sau-Chun FUNG, Sandy Ka-Yee CHAU, Wing-Kin TO, Tak-Lun Que, David Ho-Keung SHUM, Shea Ping YIP, Wing Cheong YAM, Gilman Kit-Hang SIU |
| EPI_ISL_417183, EPI_ISL_417184 | Department of Clinical Pathology, Pamela Youde Nethersole Eastern Hospital | Department of Health Technology and Informatics, Faculty of Health and Social Science, The Hong Kong Polytechnic University | Kenneth Siu-Sing LEUNG, Timothy Ting-Leung NG, Alan Ka-Lun WU, Miranda Chong-Yee YAU, Hiu-Yin LAO, Ming-Pan CHOI, Kingsley King-Gee TAM, Lam-Kwong LEE, Barry Kin-Chung WONG, Alex Yat-Man HO, Kam-Tong Yip, Kwok-Cheung LUNG, Raymond Wai-To LIU, Eugene Yuk-Keung TSO, Wai-Shing LEUNG, Man-Chun CHAN, Yuk-Yung NG, Kit-Man SIN, Kitty Sau-Chun FUNG, Sandy Ka-Yee CHAU, Wing-Kin TO, Tak-Lun Que, David Ho-Keung SHUM, Shea Ping YIP, Wing Cheong YAM, Gilman Kit-Hang SIU |
| EPI_ISL_417185 | Department of Pathology, United Christian Hospital | Department of Health Technology and Informatics, Faculty of Health and Social Science, The Hong Kong Polytechnic University | Kenneth Siu-Sing LEUNG, Timothy Ting-Leung NG, Alan Ka-Lun WU, Miranda Chong-Yee YAU, Hiu-Yin LAO, Ming-Pan CHOI, Kingsley King-Gee TAM, Lam-Kwong LEE, Barry Kin-Chung WONG, Alex Yat-Man HO, Kam-Tong Yip, Kwok-Cheung LUNG, Raymond Wai-To LIU, Eugene Yuk-Keung TSO, Wai-Shing LEUNG, Man-Chun CHAN, Yuk-Yung NG, Kit-Man SIN, Kitty Sau-Chun FUNG, Sandy Ka-Yee CHAU, Wing-Kin TO, Tak-Lun Que, David Ho-Keung SHUM, Shea Ping YIP, Wing Cheong YAM, Gilman Kit-Hang SIU |
| EPI_ISL_417187, EPI_ISL_417188, EPI_ISL_417190, EPI_ISL_417193, EPI_ISL_417195, EPI_ISL_417197 | Department of Clinical Pathology, Pamela Youde Nethersole Eastern Hospital | Department of Health Technology and Informatics, Faculty of Health and Social Science, The Hong Kong Polytechnic University | Kenneth Siu-Sing LEUNG, Timothy Ting-Leung NG, Alan Ka-Lun WU, Miranda Chong-Yee YAU, Hiu-Yin LAO, Ming-Pan CHOI, Kingsley King-Gee TAM, Lam-Kwong LEE, Barry Kin-Chung WONG, Alex Yat-Man HO, Kam-Tong Yip, Kwok-Cheung LUNG, Raymond Wai-To LIU, Eugene Yuk-Keung TSO, Wai-Shing LEUNG, Man-Chun CHAN, Yuk-Yung NG, Kit-Man SIN, Kitty Sau-Chun FUNG, Sandy Ka-Yee CHAU, Wing-Kin TO, Tak-Lun Que, David Ho-Keung SHUM, Shea Ping YIP, Wing Cheong YAM, Gilman Kit-Hang SIU |
| EPI_ISL_417385 | Centre for Infectious Diseases and Microbiology Public Health | NSW Health Pathology - Institute of Clinical Pathology and Medical Research; Westmead Hospital; University of Sydney | Lam C, Gray K, Timms V, Gall M, Arnott A, Sadsad R, Carter I, Rahman H, Holmes EC, O'Sullivan MV, Sintchenko V, Chen SC, Maddocks S, Kok J, Dwyer DE, Rockett R and Eden J-S for the 2019-nCoV Study Group |
| EPI_ISL_417398 | Centre for Infectious Diseases and Microbiology Public Health | NSW Health Pathology - Institute of Clinical Pathology and Medical Research; Westmead Hospital; University of Sydney | Kok J, Dwyer DE, Rockett R, Eden J-S, Lam C, Gray K, Timms V, Gall M, Arnott A, Sadsad R, Carter I, Rahman H, Holmes EC, O'Sullivan MV, Sintchenko V, Chen SC and Maddocks S for the 2019-nCoV Study Group |
| EPI_ISL_417402 | Centre for Infectious Diseases and Microbiology Public Health | NSW Health Pathology - Institute of Clinical Pathology and Medical Research; Westmead Hospital; University of Sydney | Lam C, Gray K, Timms V, Gall M, Arnott A, Sadsad R, Carter I, Rahman H, Holmes EC, O'Sullivan MV, Sintchenko V, Chen SC, Maddocks S, Kok J, Dwyer DE, Rockett R and Eden J-S for the 2019-nCoV Study Group |
| EPI_ISL_417422, EPI_ISL_417424, EPI_ISL_417425, EPI_ISL_417426, EPI_ISL_417427, EPI_ISL_417428, EPI_ISL_417429, EPI_ISL_417430 | KU Leuven, Clinical and Epidemiological Virology | KU Leuven, Clinical and Epidemiological Virology | Joan Marti-Carerras, Tony Wawina, Bert Vanmechelen, Piet Maes |
| EPI_ISL_417444 | Department of Healthcare Biotechnology, National University of Sciences and Technology (NUST) | Department of Healthcare Biotechnology, National University of Sciences and Technology (NUST) | Javed,A., Niazi,S.K., Ghani,E., Saqib,M., Janjua,H.A., Corman,V.M. and Zohaib,A. |
| EPI_ISL_417483 | Oslo University Hospital, Department of Medical Microbiology | Norwegian Institute of Public Health | Kathrine Stene-Johansen, Kamilla Heddeland Instefjord, Hilde Elshaug, Karoline Bragstad, Olav Hungnes |
| EPI_ISL_417484 | Oslo University Hospital, Department of Medical Microbiology | Norwegian Institute of Public Health, Department of Virology | Kathrine Stene-Johansen, Kamilla Heddeland Instefjord, Hilde Elshaug, Karoline Bragstad, Olav Hungnes |
| EPI_ISL_417485 | University Hospital of Northern Norway, Department for Microbiology and Infectious Disease Control | Norwegian Institute of Public Health, Department of Virology | Kathrine Stene-Johansen, Kamilla Heddeland Instefjord, Hilde Elshaug, Karoline Bragstad, Olav Hungnes |
| EPI_ISL_417486, EPI_ISL_417487 | Hospital of Southern Norway - Kristiansand, Department of Medical Microbiology | Norwegian Institute of Public Health, Department of Virology | Kathrine Stene-Johansen, Kamilla Heddeland Instefjord, Hilde Elshaug, Karoline Bragstad, Olav Hungnes |
| EPI_ISL_417488, EPI_ISL_417489, EPI_ISL_417490 | Oslo University Hospital, Department of Medical Microbiology | Norwegian Institute of Public Health, Department of Virology | Kathrine Stene-Johansen, Kamilla Heddeland Instefjord, Hilde Elshaug, Karoline Bragstad, Olav Hungnes |
| EPI_ISL_417518, EPI_ISL_417519, EPI_ISL_417520, EPI_ISL_417521, EPI_ISL_417522, EPI_ISL_417523, EPI_ISL_417524, EPI_ISL_417525 | Laboratory Medicine | Department of Laboratory Medicine, Lin-Kou Chang Gung Memorial Hospital, Taoyuan, Taiwan | Kuo-Chien Tsao, Yu-Nong Gong, Shu-Li Yang, Yi-Chun Liu, Chung-Guei Huang, Po-Wei Huang, Mei-Jen Hsiao, Cheng-Ta Yang, Cheng-Hsun Chiu, Peng-Nien Huang, Kuo-Ming Lee, Guang-Wu Chen , Shin-Ru Shih |
| EPI_ISL_418623, EPI_ISL_418624, EPI_ISL_418625, EPI_ISL_418626, EPI_ISL_418627, EPI_ISL_418628, EPI_ISL_418629, EPI_ISL_418630, EPI_ISL_418631, EPI_ISL_418632, EPI_ISL_418633, EPI_ISL_418634, EPI_ISL_418635, EPI_ISL_418636, EPI_ISL_418637, EPI_ISL_418638, EPI_ISL_418639, EPI_ISL_418640, EPI_ISL_418641, EPI_ISL_418642, EPI_ISL_418643, EPI_ISL_418644, EPI_ISL_418645, EPI_ISL_418646, EPI_ISL_418647, EPI_ISL_418648, EPI_ISL_418649, EPI_ISL_418650, EPI_ISL_418651, EPI_ISL_418652, EPI_ISL_418653, EPI_ISL_418654, EPI_ISL_418655, EPI_ISL_418656, EPI_ISL_418657, EPI_ISL_418658, EPI_ISL_418659, EPI_ISL_418660, EPI_ISL_418661, EPI_ISL_418662, EPI_ISL_418663, EPI_ISL_418664, EPI_ISL_418665, EPI_ISL_418666 | | | |
| see above | Department of Clinical Microbiology | GIGA Medical Genomics | Keith Durkin, Maria Artesi, Sébastien Bontems, Raphaël Boreux, Cécile Meex, Pierrette Melin, Marie-Pierre Hayette, Vincent Bours. |
| EPI_ISL_418771, EPI_ISL_418772, EPI_ISL_418773, EPI_ISL_418774, EPI_ISL_418775, EPI_ISL_418776, EPI_ISL_418777 | WA State Department of Health | Pathogen Discovery, Respiratory Viruses Branch, Division of Viral Diseases, Centers for Disease Control and Prevention | Jing Zhang, Ying Tao, Clinton R. Paden, Krista Queen, Anna Uehara, Yan Li, Haibin Wang, Jesica Jacobs, Denny Russell, Brian Hiatt, Jessica Gant, Suxiang Tong |
| EPI_ISL_418809 | University of Wisconsin - Madison: Influenza Research Institute | University of Wisconsin Madison, AIDS Vaccine Research Laboratories | Katarina Braun, Gage Moreno, Peter Halfmann, et al. |
| EPI_ISL_418815, EPI_ISL_419213, EPI_ISL_419214, EPI_ISL_419215, EPI_ISL_419216 | Department of Clinical Pathology, Pamela Youde Nethersole Eastern Hospital | Department of Health Technology and Informatics, Faculty of Health and Social Science, The Hong Kong Polytechnic University | Kenneth Siu-Sing LEUNG, Timothy Ting-Leung NG, Alan Ka-Lun WU, Miranda Chong-Yee YAU, Hiu-Yin LAO, Ming-Pan CHOI, Kingsley King-Gee TAM, Lam-Kwong LEE, Barry Kin-Chung WONG, Alex Yat-Man HO, Kam-Tong YIP, Kwok-Cheung LUNG, Raymond Wai-To LIU, Eugene Yuk-Keung TSO, Wai-Shing LEUNG, Man-Chun CHAN, Yuk-Yung NG, Kit-Man SIN, Kitty Sau-Chun FUNG, Sandy Ka-Yee CHAU, Wing-Kin TO, Tak-Lun QUE, David Ho-Keung SHUM, Shea Ping YIP, Wing Cheong YAM, Gilman Kit-Hang SIU |
| EPI_ISL_419217 | Department of Pathology, Princess Margaret Hospital | Department of Health Technology and Informatics, Faculty of Health and Social Science, The Hong Kong Polytechnic University | Kenneth Siu-Sing LEUNG, Timothy Ting-Leung NG, Alan Ka-Lun WU, Miranda Chong-Yee YAU, Hiu-Yin LAO, Ming-Pan CHOI, Kingsley King-Gee TAM, Lam-Kwong LEE, Barry Kin-Chung WONG, Alex Yat-Man HO, Kam-Tong YIP, Kwok-Cheung LUNG, Raymond Wai-To LIU, Eugene Yuk-Keung TSO, Wai-Shing LEUNG, Man-Chun CHAN, Yuk-Yung NG, Kit-Man SIN, Kitty Sau-Chun FUNG, Sandy Ka-Yee CHAU, Wing-Kin TO, Tak-Lun QUE, David Ho-Keung SHUM, Shea Ping YIP, Wing Cheong YAM, Gilman Kit-Hang SIU |
| EPI_ISL_419218, EPI_ISL_419219 | Department of Clinical Pathology, Pamela Youde Nethersole Eastern Hospital | Department of Health Technology and Informatics, Faculty of Health and Social Science, The Hong Kong Polytechnic University | Kenneth Siu-Sing LEUNG, Timothy Ting-Leung NG, Alan Ka-Lun WU, Miranda Chong-Yee YAU, Hiu-Yin LAO, Ming-Pan CHOI, Kingsley King-Gee TAM, Lam-Kwong LEE, Barry Kin-Chung WONG, Alex Yat-Man HO, Kam-Tong YIP, Kwok-Cheung LUNG, Raymond Wai-To LIU, Eugene Yuk-Keung TSO, Wai-Shing LEUNG, Man-Chun CHAN, Yuk-Yung NG, Kit-Man SIN, Kitty Sau-Chun FUNG, Sandy Ka-Yee CHAU, Wing-Kin TO, Tak-Lun QUE, David Ho-Keung SHUM, Shea Ping YIP, Wing Cheong YAM, Gilman Kit-Hang SIU |
| EPI_ISL_419221 | Department of Pathology, United Christian Hospital | Department of Health Technology and Informatics, Faculty of Health and Social Science, The Hong Kong Polytechnic University | Kenneth Siu-Sing LEUNG, Timothy Ting-Leung NG, Alan Ka-Lun WU, Miranda Chong-Yee YAU, Hiu-Yin LAO, Ming-Pan CHOI, Kingsley King-Gee TAM, Lam-Kwong LEE, Barry Kin-Chung WONG, Alex Yat-Man HO, Kam-Tong YIP, Kwok-Cheung LUNG, Raymond Wai-To LIU, Eugene Yuk-Keung TSO, Wai-Shing LEUNG, Man-Chun CHAN, Yuk-Yung NG, Kit-Man SIN, Kitty Sau-Chun FUNG, Sandy Ka-Yee CHAU, Wing-Kin TO, Tak-Lun QUE, David Ho-Keung SHUM, Shea Ping YIP, Wing Cheong YAM, Gilman Kit-Hang SIU |
| EPI_ISL_419222, EPI_ISL_419223 | Department of Pathology, Princess Margaret Hospital | Department of Health Technology and Informatics, Faculty of | Kenneth Siu-Sing LEUNG, Timothy Ting-Leung NG, Alan Ka-Lun WU, Miranda Chong-Yee YAU, Hiu-Yin LAO, Ming-Pan CHOI, Kingsley King-Gee TAM, |

| | | | |
|---|---|---|---|
| | | Health and Social Science, The Hong Kong Polytechnic University | Lam-Kwong LEE, Barry Kin-Chung WONG, Alex Yat-Man HO, Kam-Tong YIP, Kwok-Cheung LUNG, Raymond Wai-To LIU, Eugene Yuk-Keung TSO, Wai-Shing LEUNG, Man-Chun CHAN, Yuk-Yung NG, Kit-Man SIN, Kitty Sau-Chun FUNG, Sandy Ka-Yee CHAU, Wing-Kin TO, Tak-Lun QUE, David Ho-Keung SHUM, Shea Ping YIP, Wing Cheong YAM, Gilman Kit-Hang SIU |
| EPI_ISL_419224, EPI_ISL_419225, EPI_ISL_419226, EPI_ISL_419227, EPI_ISL_419228, EPI_ISL_419229 | Department of Clinical Pathology, Pamela Youde Nethersole Eastern Hospital | Department of Health Technology and Informatics, Faculty of Health and Social Science, The Hong Kong Polytechnic University | Kenneth Siu-Sing LEUNG, Timothy Ting-Leung NG, Alan Ka-Lun WU, Miranda Chong-Yee YAU, Hiu-Yin LAO, Ming-Pan CHOI, Kingsley King-Gee TAM, Lam-Kwong LEE, Barry Kin-Chung WONG, Alex Yat-Man HO, Kam-Tong YIP, Kwok-Cheung LUNG, Raymond Wai-To LIU, Eugene Yuk-Keung TSO, Wai-Shing LEUNG, Man-Chun CHAN, Yuk-Yung NG, Kit-Man SIN, Kitty Sau-Chun FUNG, Sandy Ka-Yee CHAU, Wing-Kin TO, Tak-Lun QUE, David Ho-Keung SHUM, Shea Ping YIP, Wing Cheong YAM, Gilman Kit-Hang SIU |
| EPI_ISL_419231 | Department of Clinical Pathology, Tuen Mun Hospital, 23 Tsing Chung Koon Road, Tuen Mun, N.T. | Department of Health Technology and Informatics, Faculty of Health and Social Science, The Hong Kong Polytechnic University | Kenneth Siu-Sing LEUNG, Timothy Ting-Leung NG, Alan Ka-Lun WU, Miranda Chong-Yee YAU, Hiu-Yin LAO, Ming-Pan CHOI, Kingsley King-Gee TAM, Lam-Kwong LEE, Barry Kin-Chung WONG, Alex Yat-Man HO, Kam-Tong YIP, Kwok-Cheung LUNG, Raymond Wai-To LIU, Eugene Yuk-Keung TSO, Wai-Shing LEUNG, Man-Chun CHAN, Yuk-Yung NG, Kit-Man SIN, Kitty Sau-Chun FUNG, Sandy Ka-Yee CHAU, Wing-Kin TO, Tak-Lun QUE, David Ho-Keung SHUM, Shea Ping YIP, Wing Cheong YAM, Gilman Kit-Hang SIU |
| EPI_ISL_419232, EPI_ISL_419241 | Department of Clinical Pathology, Pamela Youde Nethersole Eastern Hospital | Department of Health Technology and Informatics, Faculty of Health and Social Science, The Hong Kong Polytechnic University | Kenneth Siu-Sing LEUNG, Timothy Ting-Leung NG, Alan Ka-Lun WU, Miranda Chong-Yee YAU, Hiu-Yin LAO, Ming-Pan CHOI, Kingsley King-Gee TAM, Lam-Kwong LEE, Barry Kin-Chung WONG, Alex Yat-Man HO, Kam-Tong YIP, Kwok-Cheung LUNG, Raymond Wai-To LIU, Eugene Yuk-Keung TSO, Wai-Shing LEUNG, Man-Chun CHAN, Yuk-Yung NG, Kit-Man SIN, Kitty Sau-Chun FUNG, Sandy Ka-Yee CHAU, Wing-Kin TO, Tak-Lun QUE, David Ho-Keung SHUM, Shea Ping YIP, Wing Cheong YAM, Gilman Kit-Hang SIU |
| EPI_ISL_419242 | Department of Clinical Pathology, Tuen Mun Hospital | Department of Health Technology and Informatics, Faculty of Health and Social Science, The Hong Kong Polytechnic University | Kenneth Siu-Sing LEUNG, Timothy Ting-Leung NG, Alan Ka-Lun WU, Miranda Chong-Yee YAU, Hiu-Yin LAO, Ming-Pan CHOI, Kingsley King-Gee TAM, Lam-Kwong LEE, Barry Kin-Chung WONG, Alex Yat-Man HO, Kam-Tong YIP, Kwok-Cheung LUNG, Raymond Wai-To LIU, Eugene Yuk-Keung TSO, Wai-Shing LEUNG, Man-Chun CHAN, Yuk-Yung NG, Kit-Man SIN, Kitty Sau-Chun FUNG, Sandy Ka-Yee CHAU, Wing-Kin TO, Tak-Lun QUE, David Ho-Keung SHUM, Shea Ping YIP, Wing Cheong YAM, Gilman Kit-Hang SIU |
| EPI_ISL_419243, EPI_ISL_419244, EPI_ISL_419245, EPI_ISL_419246, EPI_ISL_419247, EPI_ISL_419248, EPI_ISL_419249, EPI_ISL_419250, EPI_ISL_419251, EPI_ISL_419252 | Department of Clinical Pathology, Pamela Youde Nethersole Eastern Hospital | Department of Health Technology and Informatics, Faculty of Health and Social Science, The Hong Kong Polytechnic University | Kenneth Siu-Sing LEUNG, Timothy Ting-Leung NG, Alan Ka-Lun WU, Miranda Chong-Yee YAU, Hiu-Yin LAO, Ming-Pan CHOI, Kingsley King-Gee TAM, Lam-Kwong LEE, Barry Kin-Chung WONG, Alex Yat-Man HO, Kam-Tong YIP, Kwok-Cheung LUNG, Raymond Wai-To LIU, Eugene Yuk-Keung TSO, Wai-Shing LEUNG, Man-Chun CHAN, Yuk-Yung NG, Kit-Man SIN, Kitty Sau-Chun FUNG, Sandy Ka-Yee CHAU, Wing-Kin TO, Tak-Lun QUE, David Ho-Keung SHUM, Shea Ping YIP, Wing Cheong YAM, Gilman Kit-Hang SIU |
| EPI_ISL_419253 | Department of Pathology, United Christian Hospital | Department of Health Technology and Informatics, Faculty of Health and Social Science, The Hong Kong Polytechnic University | Kenneth Siu-Sing LEUNG, Timothy Ting-Leung NG, Alan Ka-Lun WU, Miranda Chong-Yee YAU, Hiu-Yin LAO, Ming-Pan CHOI, Kingsley King-Gee TAM, Lam-Kwong LEE, Barry Kin-Chung WONG, Alex Yat-Man HO, Kam-Tong YIP, Kwok-Cheung LUNG, Raymond Wai-To LIU, Eugene Yuk-Keung TSO, Wai-Shing LEUNG, Man-Chun CHAN, Yuk-Yung NG, Kit-Man SIN, Kitty Sau-Chun FUNG, Sandy Ka-Yee CHAU, Wing-Kin TO, Tak-Lun QUE, David Ho-Keung SHUM, Shea Ping YIP, Wing Cheong YAM, Gilman Kit-Hang SIU |
| EPI_ISL_419386, EPI_ISL_419387 | Hospital Prof. Doutor Fernando Fonseca, EPE | Instituto Gulbenkian de Ciência | João Costa, Cathy Paulino, Joao Sobral, Susana Ladeiro, Ricardo Leite |
| EPI_ISL_419512, EPI_ISL_419513, EPI_ISL_419514, EPI_ISL_419515, EPI_ISL_419516 | Yale COVID-19 Biorepository | Grubaugh Lab - Yale School of Public Health | Joseph Fauver, Tara Alpert, Anderson Brito, Anne Wyllie, Chantal Vogels, Mary Petrone, Chaney Kalinich, Isabel Ott, Arnau Casanovas, Catherine Muenker, Adam Moore, Alice Lu, Maria Tokuyama, Patrick Wong, Peiwen Lu, Saad Omer, Richard Martinello, Allison Nelson, Shelli Farhadian, Akiko Iwasaki, Charlese Dela Cruz, Albert Ko, Nathan Grubaugh |
| EPI_ISL_419517, EPI_ISL_419518, EPI_ISL_419519, EPI_ISL_419520, EPI_ISL_419521, EPI_ISL_419522, EPI_ISL_419523, EPI_ISL_419524, EPI_ISL_419525, EPI_ISL_419526, EPI_ISL_419527, EPI_ISL_419528 | | | |
| see above | Yale Clinical Virology Laboratory | Grubaugh Lab - Yale School of Public Health | Joseph Fauver, Anderson Brito, Tara Alpert, Chantal Vogels, Ellen Foxman, Albert Ko, Marie Landry, Nathan Grubaugh |
| EPI_ISL_420082 | Centers for Disease Control, R.O.C. (Taiwan) | Centers for Disease Control, R.O.C. (Taiwan) | Ji-Rong Yang, Yu-Chi-Lin, Jung-Jung Mu, Ming-Tsan Liu |
| EPI_ISL_420083, EPI_ISL_420084, EPI_ISL_420085 | Centers for Disease Control, R.O.C. (Taiwan) | Centers for Disease Control, R.O.C. (Taiwan) | Ji-Rong Yang, Yu-Chi Lin, Jung-Jung Mu, Ming-Tsan Liu |
| EPI_ISL_420086, EPI_ISL_420087, EPI_ISL_420088, EPI_ISL_420089, EPI_ISL_420090, EPI_ISL_420091, EPI_ISL_420092, EPI_ISL_420093, EPI_ISL_420094, EPI_ISL_420095, EPI_ISL_420096, EPI_ISL_420097, EPI_ISL_420098 | | | |
| see above | Yale Clinical Virology Laboratory | Grubaugh Lab - Yale School of Public Health | Joseph Fauver, Anderson Brito, Tara Alpert, Chantal Vogels, Ellen Foxman, Albert Ko, Marie Landry, Nathan Grubaugh |
| EPI_ISL_420134 | Akershus University Hospital, Department for Microbiology and Infectious Disease Control | Norwegian Institute of Public Health, Department of Virology | Kathrine Stene-Johansen, Kamilla Heddeland Instefjord, Hilde Elshaug, Karoline Bragstad, Olav Hungnes |
| EPI_ISL_420135 | Oslo University Hospital, Department of Medical Microbiology | Norwegian Institute of Public Health, Department of Virology | Kathrine Stene-Johansen, Kamilla Heddeland Instefjord, Hilde Elshaug, Karoline Bragstad, Olav Hungnes |
| EPI_ISL_420136 | Akershus University Hospital, Department for Microbiology and Infectious Disease Control | Norwegian Institute of Public Health, Department of Virology | Kathrine Stene-Johansen, Kamilla Heddeland Instefjord, Hilde Elshaug, Karoline Bragstad, Olav Hungnes |
| EPI_ISL_420137 | Vestfold Hospital, Tonsberg Department of Microbiology | Norwegian Institute of Public Health, Department of Virology | Kathrine Stene-Johansen, Kamilla Heddeland Instefjord, Hilde Elshaug, Karoline Bragstad, Olav Hungnes |
| EPI_ISL_420138 | University Hospital of Northern Norway, Department for Microbiology and Infectious Disease Control | Norwegian Institute of Public Health, Department of Virology | Kathrine Stene-Johansen, Kamilla Heddeland Instefjord, Hilde Elshaug, Karoline Bragstad, Olav Hungnes |
| EPI_ISL_420139 | Akershus University Hospital, Department for Microbiology and Infectious Disease Control | Norwegian Institute of Public Health, Department of Virology | Kathrine Stene-Johansen, Kamilla Heddeland Instefjord, Hilde Elshaug, Karoline Bragstad, Olav Hungnes |
| EPI_ISL_420141 | Furst Medical Laboratory | Norwegian Institute of Public Health, Department of Virology | Kathrine Stene-Johansen, Kamilla Heddeland Instefjord, Hilde Elshaug, Karoline Bragstad, Olav Hungnes |
| EPI_ISL_420143 | Unilabs Laboratory Medicine | Norwegian Institute of Public Health, Department of Virology | Kathrine Stene-Johansen, Kamilla Heddeland Instefjord, Hilde Elshaug, Karoline Bragstad, Olav Hungnes |
| EPI_ISL_420145 | Forde Hospital Department of Microbiology | Norwegian Institute of Public Health, Department of Virology | Kathrine Stene-Johansen, Kamilla Heddeland Instefjord, Hilde Elshaug, Karoline Bragstad, Olav Hungnes |
| EPI_ISL_420146 | Furst Medical Laboratory | Norwegian Institute of Public Health, Department of Virology | Kathrine Stene-Johansen, Kamilla Heddeland Instefjord, Hilde Elshaug, Karoline Bragstad, Olav Hungnes |
| EPI_ISL_420147 | Hospital of Southern Norway - Kristiansand, Department of Medical Microbiology | Norwegian Institute of Public Health, Department of Virology | Kathrine Stene-Johansen, Kamilla Heddeland Instefjord, Hilde Elshaug, Karoline Bragstad, Olav Hungnes |
| EPI_ISL_420148 | Ostfold Hospital Trust -Kalnes Centre for Laboratory Medicine Section for gene technology and infection serology | Norwegian Institute of Public Health, Department of Virology | Kathrine Stene-Johansen, Kamilla Heddeland Instefjord, Hilde Elshaug, Karoline Bragstad, Olav Hungnes |
| EPI_ISL_420149 | Stavanger University Hospital, Department of Medical Microbiology | Norwegian Institute of Public Health, Department of Virology | Kathrine Stene-Johansen, Kamilla Heddeland Instefjord, Hilde Elshaug, Karoline Bragstad, Olav Hungnes |
| EPI_ISL_420150 | Oslo University Hospital, Department of Medical Microbiology | Norwegian Institute of Public Health, Department of Virology | Kathrine Stene-Johansen, Kamilla Heddeland Instefjord, Hilde Elshaug, Karoline Bragstad, Olav Hungnes |
| EPI_ISL_420151 | Nordland Hospital - Bodo, Laboratory Department, Molecular Biology Unit | Norwegian Institute of Public Health, Department of Virology | Kathrine Stene-Johansen, Kamilla Heddeland Instefjord, Hilde Elshaug, Karoline Bragstad, Olav Hungnes |
| EPI_ISL_420152, EPI_ISL_420153, EPI_ISL_420310 | University Hospital of Northern Norway, Department for Microbiology and Infectious Disease Control | Norwegian Institute of Public Health, Department of Virology | Kathrine Stene-Johansen, Kamilla Heddeland Instefjord, Hilde Elshaug, Karoline Bragstad, Olav Hungnes |
| EPI_ISL_420311, EPI_ISL_420312 | Akershus University Hospital, Department for Microbiology and Infectious Disease Control | Norwegian Institute of Public Health, Department of Virology | Kathrine Stene-Johansen, Kamilla Heddeland Instefjord, Hilde Elshaug, Karoline Bragstad, Olav Hungnes |
| EPI_ISL_420313 | Furst Medical Laboratory | Norwegian Institute of Public Health, Department of Virology | Kathrine Stene-Johansen, Kamilla Heddeland Instefjord, Hilde Elshaug, Karoline Bragstad, Olav Hungnes |
| EPI_ISL_420314, EPI_ISL_420315, EPI_ISL_420316, EPI_ISL_420317, EPI_ISL_420318, EPI_ISL_420319, EPI_ISL_420320, EPI_ISL_420321, EPI_ISL_420322, EPI_ISL_420323, EPI_ISL_420324, EPI_ISL_420325, EPI_ISL_420326, EPI_ISL_420327, EPI_ISL_420328, EPI_ISL_420329, EPI_ISL_420330, EPI_ISL_420331, EPI_ISL_420332, EPI_ISL_420333, EPI_ISL_420334, EPI_ISL_420335, EPI_ISL_420336, EPI_ISL_420337, EPI_ISL_420338, EPI_ISL_420339, EPI_ISL_420340, EPI_ISL_420341, EPI_ISL_420342, EPI_ISL_420343, EPI_ISL_420344, EPI_ISL_420345, EPI_ISL_420346, EPI_ISL_420347, EPI_ISL_420348, EPI_ISL_420349, EPI_ISL_420350, EPI_ISL_420351, EPI_ISL_420352, EPI_ISL_420353, EPI_ISL_420354, EPI_ISL_420355, EPI_ISL_420356, EPI_ISL_420357, EPI_ISL_420358, EPI_ISL_420359, EPI_ISL_420360, EPI_ISL_420361, EPI_ISL_420362, EPI_ISL_420363, EPI_ISL_420364, EPI_ISL_420365, EPI_ISL_420366, EPI_ISL_420367, EPI_ISL_420368, EPI_ISL_420369, EPI_ISL_420370, EPI_ISL_420371, EPI_ISL_420372, EPI_ISL_420373, EPI_ISL_420374, EPI_ISL_420375, EPI_ISL_420376, EPI_ISL_420377, EPI_ISL_420378, EPI_ISL_420379, EPI_ISL_420380, EPI_ISL_420381, EPI_ISL_420382, EPI_ISL_420383, EPI_ISL_420384, EPI_ISL_420385, | | | |

| | | | |
|---|---|---|---|
| EPI_ISL_420386, EPI_ISL_420387, EPI_ISL_420388, EPI_ISL_420389, EPI_ISL_420390, EPI_ISL_420391, EPI_ISL_420392, EPI_ISL_420393, EPI_ISL_420394, EPI_ISL_420395, EPI_ISL_420396, EPI_ISL_420397, EPI_ISL_420398, EPI_ISL_420399, EPI_ISL_420400, EPI_ISL_420401, EPI_ISL_420402, EPI_ISL_420403, EPI_ISL_420404, EPI_ISL_420405, EPI_ISL_420406, EPI_ISL_420407, EPI_ISL_420408, EPI_ISL_420409, EPI_ISL_420410, EPI_ISL_420411, EPI_ISL_420412, EPI_ISL_420413, EPI_ISL_420414, EPI_ISL_420415, EPI_ISL_420416, EPI_ISL_420417, EPI_ISL_420418, EPI_ISL_420419, EPI_ISL_420420, EPI_ISL_420421, EPI_ISL_420422, EPI_ISL_420423, EPI_ISL_420424, EPI_ISL_420425, EPI_ISL_420426, EPI_ISL_420427, EPI_ISL_420428, EPI_ISL_420429, EPI_ISL_420430, EPI_ISL_420431, EPI_ISL_420432, EPI_ISL_420433, EPI_ISL_420434, EPI_ISL_420435, EPI_ISL_420436, EPI_ISL_420437, EPI_ISL_420438, EPI_ISL_420439, EPI_ISL_420440, EPI_ISL_420441, EPI_ISL_420442, EPI_ISL_420443, EPI_ISL_420444, EPI_ISL_420445, EPI_ISL_420446, EPI_ISL_420447, EPI_ISL_420448, EPI_ISL_420449, EPI_ISL_420450, EPI_ISL_420451, EPI_ISL_420452, EPI_ISL_420453, EPI_ISL_420454 | | | |
| see above | KU Leuven, Clinical and Epidemiological Virology | KU Leuven, Clinical and Epidemiological Virology | Joan Marti-Carreras, Bert Vanmechelen, Tony Wawina, Piet Maes |
| EPI_ISL_420455 | Department of Clinical Pathology, Pamela Youde Nethersole Eastern Hospital | Department of Health Technology and Informatics, Faculty of Health and Social Science, The Hong Kong Polytechnic University | Kenneth Siu-Sing LEUNG, Timothy Ting-Leung NG, Alan Ka-Lun WU, Miranda Chong-Yee YAU, Hiu-Yin LAO, Ming-Pan CHOI, Kingsley King-Gee TAM, Lam-Kwong LEE, Barry Kin-Chung WONG, Alex Yat-Man HO, Kam-Tong YIP, Kwok-Cheung LUNG, Raymond Wai-To LIU, Eugene Yuk-Keung TSO, Wai-Shing LEUNG, Man-Chun CHAN, Yuk-Yung NG, Kit-Man SIN, Kitty Sau-Chun FUNG, Sandy Ka-Yee CHAU, Wing-Kin TO, Tak-Lun QUE, David Ho-Keung SHUM, Shea Ping YIP, Wing Cheong YAM, Gilman Kit-Hang SIU |
| EPI_ISL_420784 | AZ Department of Health Services | Pathogen Discovery, Respiratory Viruses Branch, Division of Viral Diseases, Centers for Disease Control and Prevention | Krista Queen, Yan Li, Ying Tao, Jing Zhang, Anne Uehara, Clinton R. Paden, Haibin Wang, Rachel Marine, Mary S. Keckler, Alison S. Laufer Halpin, Jasmine Padilla, Justin Lee, Christopher A. Elkins, Suxiang Tong |
| EPI_ISL_420785 | FL Bureau of Health Laboratories Tampa | Pathogen Discovery, Respiratory Viruses Branch, Division of Viral Diseases, Centers for Disease Control and Prevention | Krista Queen, Yan Li, Ying Tao, Jing Zhang, Anne Uehara, Clinton R. Paden, Haibin Wang, Rachel Marine, Mary S. Keckler, Alison S. Laufer Halpin, Jasmine Padilla, Justin Lee, Christopher A. Elkins, Suxiang Tong |
| EPI_ISL_420786, EPI_ISL_420787, EPI_ISL_420788 | GA Department of Public Health | Pathogen Discovery, Respiratory Viruses Branch, Division of Viral Diseases, Centers for Disease Control and Prevention | Krista Queen, Yan Li, Ying Tao, Jing Zhang, Anne Uehara, Clinton R. Paden, Haibin Wang, Rachel Marine, Mary S. Keckler, Alison S. Laufer Halpin, Jasmine Padilla, Justin Lee, Christopher A. Elkins, Suxiang Tong |
| EPI_ISL_420789, EPI_ISL_420790 | Illinois Department of Public Health Chicago Laboratory | Pathogen Discovery, Respiratory Viruses Branch, Division of Viral Diseases, Centers for Disease Control and Prevention | Krista Queen, Yan Li, Ying Tao, Jing Zhang, Anne Uehara, Clinton R. Paden, Haibin Wang, Rachel Marine, Mary S. Keckler, Alison S. Laufer Halpin, Jasmine Padilla, Justin Lee, Christopher A. Elkins, Suxiang Tong |
| EPI_ISL_420791, EPI_ISL_420792 | NH Department of Health and Human Services Public Health Labs | Pathogen Discovery, Respiratory Viruses Branch, Division of Viral Diseases, Centers for Disease Control and Prevention | Krista Queen, Yan Li, Ying Tao, Jing Zhang, Anne Uehara, Clinton R. Paden, Haibin Wang, Rachel Marine, Mary S. Keckler, Alison S. Laufer Halpin, Jasmine Padilla, Justin Lee, Christopher A. Elkins, Suxiang Tong |
| EPI_ISL_420793 | NYC Department of Health and Mental Hygiene | Pathogen Discovery, Respiratory Viruses Branch, Division of Viral Diseases, Centers for Disease Control and Prevention | Krista Queen, Yan Li, Ying Tao, Jing Zhang, Anne Uehara, Clinton R. Paden, Haibin Wang, Rachel Marine, Mary S. Keckler, Alison S. Laufer Halpin, Jasmine Padilla, Justin Lee, Christopher A. Elkins, Suxiang Tong |
| EPI_ISL_420794 | Oregon State Public Health- Virology section | Pathogen Discovery, Respiratory Viruses Branch, Division of Viral Diseases, Centers for Disease Control and Prevention | Krista Queen, Yan Li, Ying Tao, Jing Zhang, Anne Uehara, Clinton R. Paden, Haibin Wang, Rachel Marine, Mary S. Keckler, Alison S. Laufer Halpin, Jasmine Padilla, Justin Lee, Christopher A. Elkins, Suxiang Tong |
| EPI_ISL_420795 | RI State Health Laboratory | Pathogen Discovery, Respiratory Viruses Branch, Division of Viral Diseases, Centers for Disease Control and Prevention | Krista Queen, Yan Li, Ying Tao, Jing Zhang, Anne Uehara, Clinton R. Paden, Haibin Wang, Rachel Marine, Mary S. Keckler, Alison S. Laufer Halpin, Jasmine Padilla, Justin Lee, Christopher A. Elkins, Suxiang Tong |
| EPI_ISL_420796, EPI_ISL_420797, EPI_ISL_420798 | Texas DSHS Lab Services | Pathogen Discovery, Respiratory Viruses Branch, Division of Viral Diseases, Centers for Disease Control and Prevention | Krista Queen, Yan Li, Ying Tao, Jing Zhang, Anne Uehara, Clinton R. Paden, Haibin Wang, Rachel Marine, Mary S. Keckler, Alison S. Laufer Halpin, Jasmine Padilla, Justin Lee, Christopher A. Elkins, Suxiang Tong |
| EPI_ISL_420799, EPI_ISL_420800, EPI_ISL_420801 | Brian D. Allgood Army Community Hospital | Pathogen Discovery, Respiratory Viruses Branch, Division of Viral Diseases, Centers for Disease Control and Prevention | Krista Queen, Yan Li, Ying Tao, Jing Zhang, Anne Uehara, Clinton R. Paden, Haibin Wang, Rachel Marine, Mary S. Keckler, Alison S. Laufer Halpin, Jasmine Padilla, Justin Lee, Christopher A. Elkins, Suxiang Tong |
| EPI_ISL_421182, EPI_ISL_421183, EPI_ISL_421184, EPI_ISL_421185, EPI_ISL_421186, EPI_ISL_421187, EPI_ISL_421188, EPI_ISL_421189, EPI_ISL_421190, EPI_ISL_421191, EPI_ISL_421192, EPI_ISL_421193, EPI_ISL_421194, EPI_ISL_421195, EPI_ISL_421196, EPI_ISL_421197, EPI_ISL_421198, EPI_ISL_421199, EPI_ISL_421200, EPI_ISL_421201, EPI_ISL_421202, EPI_ISL_421203, EPI_ISL_421204, EPI_ISL_421205, EPI_ISL_421206, EPI_ISL_421207, EPI_ISL_421208, EPI_ISL_421209, EPI_ISL_421210, EPI_ISL_421211, EPI_ISL_421212, EPI_ISL_421213, EPI_ISL_421214 | | | |
| see above | Department of Clinical Microbiology | GIGA Medical Genomics | Keith Durkin, Maria Artesi, Sébastien Bontems, Raphaël Boreux, Cécile Meex, Pierrette Melin, Marie-Pierre Hayette, Vincent Bours. |
| EPI_ISL_421221, EPI_ISL_421222, EPI_ISL_421223, EPI_ISL_421224, EPI_ISL_421225, EPI_ISL_421226, EPI_ISL_421227, EPI_ISL_421228, EPI_ISL_421229, EPI_ISL_421230, EPI_ISL_421231, EPI_ISL_421232, EPI_ISL_421233, EPI_ISL_421234, EPI_ISL_421235, EPI_ISL_421236 | | | |
| see above | Hangzhou Center for Diseases Control and Prevention | Hangzhou Center for Diseases Control and Prevention | Jun Li, Haoqiu Wang, Lingfeng Mao, Hua Yu, Xinfen Yu, Zhou Sun, Xin Qian, Shuchang Chen, Junfang Chen, Xuchu Wang |
| EPI_ISL_421237, EPI_ISL_421238, EPI_ISL_421239, EPI_ISL_421240, EPI_ISL_421241, EPI_ISL_421242, EPI_ISL_421243, EPI_ISL_421244, EPI_ISL_421245, EPI_ISL_421246, EPI_ISL_421247, EPI_ISL_421248, EPI_ISL_421249, EPI_ISL_421250, EPI_ISL_421251, EPI_ISL_421252, EPI_ISL_421253, EPI_ISL_421254, EPI_ISL_421256, EPI_ISL_421257, EPI_ISL_421258, EPI_ISL_421259, EPI_ISL_421260, EPI_ISL_421261, EPI_ISL_421262 | | | |
| see above | Jiangxi Province Center for Disease Control and Prevention | Jiangxi Province Center for Disease Control and Prevention | JianXiong Li,Ying Xiong,Tian Gong,Yong Shi,Jun Zhou,Fang Xiao,ShiWen Liu,XiaoQing Liu,Gang Xu,DaJin Xiao,Xin Ran,YanNi Zhang |
| EPI_ISL_421641, EPI_ISL_421651 | Centers for Disease Control, R.O.C. (Taiwan) | Centers for Disease Control, R.O.C. (Taiwan) | Ji-Rong Yang, Yu-Chi Lin, Jung-Jung Mu, Ming-Tsan Liu |
| EPI_ISL_422382 | NMIMR, Department of Virology | WACCBIP, University of Ghana | Joyce M. Ngoi, Bright Adu, Collins M. Misita, Selassie Kumordjie, Miriam Eshun, Linda Boatemaa, Vanessa Magnussen, Erasmus Kotey, Fred Tei-Maya, Dominic S. Y. Amuzu, Peter Quashie, Augustina Arjaquah, Ivy Asante, Evelyn Bonney, George B. Kyei, Kofi Bonney, Gordon A. Awandare, William Ampofo |
| EPI_ISL_422384, EPI_ISL_422387, EPI_ISL_422390, EPI_ISL_422394, EPI_ISL_422397, EPI_ISL_422398, EPI_ISL_422399, EPI_ISL_422400, EPI_ISL_422401, EPI_ISL_422402, EPI_ISL_422403, EPI_ISL_422404, EPI_ISL_422405, EPI_ISL_422406 | | | |
| see above | NMIMR, Department of Virology | WACCBIP, University of Ghana | Joyce M. Ngoi, Bright Adu, Collins M. Morang'a, Selassie Kumordjie, Miriam Eshun, Linda Boatemaa, Vanessa Magnussen, Erasmus Kotey, Fred Tei-Maya, Dominic S. Y. Amuzu, Peter Quashie, Augustina Arjarquah, Ivy Asante, Evelyn Bonney, George B. Kyei, Kofi Bonney, Abraham Kwabena Anang, Gordon A. Awandare, William Ampofo |
| EPI_ISL_422564 | Institute of Microbiology Universidad San Francisco de Quito | Institute of Microbiology Universidad San Francisco de Quito | Juan Jose Guadalupe, Belen Prado-Vivar, Sully Marquez, Bernardo Gutierrez, Francisco Mora, Juan Gaviria, Alejandra Ramones, Franklin Espinoza, Edison Ligña, Jorge Reyes, Patricio Rojas-Silva, Veronica Barragan, Gabriel Trueba, Michelle Grunauer, Paul Cardenas |
| EPI_ISL_424345, EPI_ISL_424348, EPI_ISL_424625, EPI_ISL_424626, EPI_ISL_424627 | Instituto Nacional de Enfermedades Respiratorias | Instituto Nacional de Enfermedades Respiratorias | Joel Armando Vázquez Pérez, Celia Boukadida, Santiago Avila Ríos, Mario Mújica Sánchez, José Arturo Martínez Orozco, Eduardo Becerril Vargas, Jorge Salas Hernández, Irma López Martínez, Lucia Hernández Rivas, Gisela Barrera Badillo, Edgar Mendieta Condado, Fabiola Garcés Ayala, Adnan Araiza Rodríguez, José Ernesto Ramírez González, Victor Hugo Borja Aburto, Concepción Grajales Muñiz, Cesar Raúl González Bonilla, Carolina González Torres, Francisco Javier Gaytán Cervantes, José Esteban Muñoz Medina, Guillermo M. Ruiz-Palacios, Pilar Ramos Cervantes, Violeta Ibarra Gonzalez, Fernando Ledesma Barrientos, Luis Alberto García Andrade, Alfredo Ponce de León Garduño, Blanca Taboada, Alejandro Sánchez, Pavel Isa, Ricardo Grande, Gloria Vázquez, Francisco Pulido, Carlos F. Arias. |
| EPI_ISL_424628, EPI_ISL_424629, EPI_ISL_424630, EPI_ISL_424631, EPI_ISL_424632, EPI_ISL_424633, EPI_ISL_424634, EPI_ISL_424635, EPI_ISL_424636, EPI_ISL_424637, EPI_ISL_424638, EPI_ISL_424639, EPI_ISL_424640, EPI_ISL_424641, EPI_ISL_424642, EPI_ISL_424643, EPI_ISL_424644, EPI_ISL_424645, EPI_ISL_424646, EPI_ISL_424647, EPI_ISL_424648, EPI_ISL_424649, EPI_ISL_424650, EPI_ISL_424651, EPI_ISL_424652, EPI_ISL_424653, EPI_ISL_424654, EPI_ISL_424655, EPI_ISL_424656, EPI_ISL_424657, EPI_ISL_424658, EPI_ISL_424659, EPI_ISL_424660, EPI_ISL_424661, EPI_ISL_424662, EPI_ISL_424663, EPI_ISL_424664, EPI_ISL_424665 | | | |
| see above | Department of Clinical Microbiology | GIGA Medical Genomics | Keith Durkin, Maria Artesi, Sébastien Bontems, Raphaël Boreux, Cécile Meex, Pierrette Melin, Marie-Pierre Hayette, Vincent Bours. |
| EPI_ISL_424666 | Instituto de Diagnóstico y Referencia Epidemiológicos | Instituto de Diagnóstico y Referencia Epidemiológicos | José Ernesto Ramírez González, Irma López Martínez, Lucia Hernández Rivas, Gisela Barrera Badillo, Edgar Mendieta Condado, Fabiola Garcés Ayala, Adnan Araiza Rodríguez, Celia Boukadida, Santiago Avila Ríos, Mario Mújica Sánchez, José Arturo Martínez Orozco,Eduardo Becerril Vargas, Joel Armando Vázquez Pérez, Victor Hugo Borja Aburto, Concepción Grajales Muñiz, Cesar Raúl González Bonilla, Carolina González Torres, Francisco Javier Gaytán Cervantes, José Esteban Muñoz Medina, Guillermo M. Ruiz-Palacios, Pilar Ramos Cervantes, Violeta Ibarra Gonzalez, Fernando Ledesma Barrientos, Luis Alberto García Andrade, Alfredo Ponce de León Garduño, Blanca Taboada, Alejandro Sánchez, Pavel Isa, Ricardo Grande, Gloria Vázquez, Francisco Pulido, Carlos F. Arias. |
| EPI_ISL_424969, EPI_ISL_424970, EPI_ISL_424971, EPI_ISL_424972, EPI_ISL_424973, EPI_ISL_424974, EPI_ISL_424975, EPI_ISL_424978 | Laboratory Medicine | Department of Laboratory Medicine, Lin-Kou Chang Gung Memorial Hospital, Taoyuan, Taiwan | Kuo-Chien Tsao, Yu-Nong Gong, Shu-Li Yang, Yi-Chun Liu, Chung-Guei Huang, Mei-Jen Hsiao, Po-Wei Huang, Cheng-Ta Yang, Cheng-Hsun Chiu, Peng-Nien Huang, Kuo-Ming Lee, Guang-Wu Chen, Shin-Ru Shih |
| EPI_ISL_425117, EPI_ISL_425118 | Division of Viral Diseases, Center for Laboratory Control of Infectious Diseases, Korea Centers for Diseases Control and Prevention | Division of Viral Diseases, Center for Laboratory Control of Infectious Diseases, Korea Centers for Diseases Control and Prevention | Jeong-Min Kim, Yoon-Seok Chung, Namjoo Lee, Mi-Seon Kim, Sang Hee Woo, Hye-Jun Jo, Sehee Park, Heui Man Kim, Jun-Sub Kim, Junhyeong Jang, Dong Hyun Song, Daesang Lee, Seong Tae Jeong, Myung Guk Han |
| EPI_ISL_425224 | Wadsworth Center, New York State Department of Health | Wadsworth Center, New York State Department of Health | Kirsten St. George, Daryl M. Lamson, Sara Griesemer, Jonathan Plitnick, Navjot Singh, Matthew D. Shudt, Erica Lasek-Nesselquist |
| EPI_ISL_426025, EPI_ISL_426026, EPI_ISL_426027, EPI_ISL_426028, EPI_ISL_426029, EPI_ISL_426030, EPI_ISL_426031, EPI_ISL_426032, EPI_ISL_426033, EPI_ISL_426034, EPI_ISL_426035, EPI_ISL_426036, EPI_ISL_426037, EPI_ISL_426038, EPI_ISL_426039, EPI_ISL_426040, EPI_ISL_426041, EPI_ISL_426042, EPI_ISL_426043, EPI_ISL_426044, EPI_ISL_426045, EPI_ISL_426046, EPI_ISL_426047, EPI_ISL_426048, EPI_ISL_426049, EPI_ISL_426050 | | | |

| | | | |
|---|---|---|---|
| see above | Wadsworth Center, New York State Department.of Health | Wadsworth Center, New York State Department.of Health | Kirsten St. George, Daryl M. Lamson, Sara Griesemer, Jonathan Plitnick, Navjot Singh, Matthew D. Shudt, Erica Lasek-Nesselquist |
| EPI_ISL_426163 | Division of Viral Diseases, Center for Laboratory Control of Infectious Diseases, Korea Centers for Diseases Control and Prevention | Division of Viral Diseases, Center for Laboratory Control of Infectious Diseases, Korea Centers for Diseases Control and Prevention | Jeong-Min Kim, Yoon-Seok Chung, Namjoo Lee, Mi-Seon Kim, Sang Hee Woo, Hye-Jun Jo, Sehee Park, Heui Man Kim, Jun-Sub Kim, Junhyeong Jang, Myung Guk Han |
| EPI_ISL_426164 | Division of Viral Diseases, Center for Laboratory Control of Infectious Diseases, Korea Centers for Diseases Control and Prevention | Division of Viral Diseases, Center for Laboratory Control of Infectious Diseases, Korea Centers for Diseases Control and Prevention | Jeong-Min Kim, Yoon-Seok Chung, Namjoo Lee, Mi-Seon Kim, Sang Hee Woo, Hye-Jun Jo, Sehee Park, Heui Man Kim, Jun-Sub Kim, Junhyeong Jang, Dong Hyun Song, Daesang Lee, Seong Tae Jeong, Myung Guk Han |
| EPI_ISL_426166, EPI_ISL_426168 | Division of Viral Diseases, Center for Laboratory Control of Infectious Diseases, Korea Centers for Diseases Control and Prevention | Division of Viral Diseases, Center for Laboratory Control of Infectious Diseases, Korea Centers for Diseases Control and Prevention | Jeong-Min Kim, Yoon-Seok Chung, Namjoo Lee, Mi-Seon Kim, Sang Hee Woo, Hye-Jun Jo, Sehee Park, Heui Man Kim, Jun-Sub Kim, Junhyeong Jang, Myung Guk Han |
| EPI_ISL_426169, EPI_ISL_426171 | Division of Viral Diseases, Center for Laboratory Control of Infectious Diseases, Korea Centers for Diseases Control and Prevention | Division of Viral Diseases, Center for Laboratory Control of Infectious Diseases, Korea Centers for Diseases Control and Prevention | Jeong-Min Kim, Yoon-Seok Chung, Namjoo Lee, Mi-Seon Kim, Sang Hee Woo, Hye-Jun Jo, Sehee Park, Heui Man Kim, Jun-Sub Kim, Junhyeong Jang, Dong Hyun Song, Daesang Lee, Seong Tae Jeong, Myung Guk Han |
| EPI_ISL_426173, EPI_ISL_426180, EPI_ISL_426181, EPI_ISL_426182, EPI_ISL_426183, EPI_ISL_426187 | Division of Viral Diseases, Center for Laboratory Control of Infectious Diseases, Korea Centers for Diseases Control and Prevention | Division of Viral Diseases, Center for Laboratory Control of Infectious Diseases, Korea Centers for Diseases Control and Prevention | Jeong-Min Kim, Yoon-Seok Chung, Namjoo Lee, Mi-Seon Kim, Sang Hee Woo, Hye-Jun Jo, Sehee Park, Heui Man Kim, Jun-Sub Kim, Junhyeong Jang, Myung Guk Han |
| EPI_ISL_426290 | Wadsworth Center, New York State Department of Health | Wadsworth Center, New York State Department.of Health | Kirsten St. George, Daryl M. Lamson, Sara Griesemer, Jonathan Plitnick, Navjot Singh, Matthew D. Shudt, Erica Lasek-Nesselquist |
| EPI_ISL_426291, EPI_ISL_426292, EPI_ISL_426293, EPI_ISL_426294, EPI_ISL_426295, EPI_ISL_426296, EPI_ISL_426297, EPI_ISL_426298, EPI_ISL_426299, EPI_ISL_426300, EPI_ISL_426301, EPI_ISL_426302, EPI_ISL_426303, EPI_ISL_426304, EPI_ISL_426305, EPI_ISL_426306, EPI_ISL_426307, EPI_ISL_426308, EPI_ISL_426309, EPI_ISL_426310, EPI_ISL_426311, EPI_ISL_426312, EPI_ISL_426313, EPI_ISL_426314, EPI_ISL_426315, EPI_ISL_426316, EPI_ISL_426317, EPI_ISL_426318, EPI_ISL_426319, EPI_ISL_426320, EPI_ISL_426321, EPI_ISL_426322, EPI_ISL_426323, EPI_ISL_426324, EPI_ISL_426325, EPI_ISL_426326, EPI_ISL_426327, EPI_ISL_426328 | | | |
| see above | Wadsworth Center, New York State Department.of Health | Wadsworth Center, New York State Department.of Health | Kirsten St. George, Daryl M. Lamson, Sara Griesemer, Jonathan Plitnick, Navjot Singh, Matthew D. Shudt, Erica Lasek-Nesselquist |
| EPI_ISL_426422, EPI_ISL_426423, EPI_ISL_426424 | IN State Department of Health Laboratory Services | Pathogen Discovery, Respiratory Viruses Branch, Division of Viral Diseases, Centers for Disease Control and Prevention | Krista Queen, Yan Li, Anna Uehara, Clinton R. Paden, Rachel Marine, Ying Tao, Jing Zhang, Haibin Wang, Mary S. Keckler, Alison S. Laufer Halpin, Christopher A. Elkins, Suxiang Tong |
| EPI_ISL_426425 | MD DOH Laboratories Administration | Pathogen Discovery, Respiratory Viruses Branch, Division of Viral Diseases, Centers for Disease Control and Prevention | Krista Queen, Yan Li, Anna Uehara, Clinton R. Paden, Rachel Marine, Ying Tao, Jing Zhang, Haibin Wang, Mary S. Keckler, Alison S. Laufer Halpin, Christopher A. Elkins, Suxiang Tong |
| EPI_ISL_426426, EPI_ISL_426427 | MN PHL Division, Minnesota Department of Health | Pathogen Discovery, Respiratory Viruses Branch, Division of Viral Diseases, Centers for Disease Control and Prevention | Krista Queen, Yan Li, Anna Uehara, Clinton R. Paden, Rachel Marine, Ying Tao, Jing Zhang, Haibin Wang, Mary S. Keckler, Alison S. Laufer Halpin, Christopher A. Elkins, Suxiang Tong |
| EPI_ISL_426428 | NC State Laboratory of Public Health | Pathogen Discovery, Respiratory Viruses Branch, Division of Viral Diseases, Centers for Disease Control and Prevention | Krista Queen, Yan Li, Anna Uehara, Clinton R. Paden, Rachel Marine, Ying Tao, Jing Zhang, Haibin Wang, Mary S. Keckler, Alison S. Laufer Halpin, Christopher A. Elkins, Suxiang Tong |
| EPI_ISL_426429 | NV State Public Health Laboratory | Pathogen Discovery, Respiratory Viruses Branch, Division of Viral Diseases, Centers for Disease Control and Prevention | Krista Queen, Yan Li, Anna Uehara, Clinton R. Paden, Rachel Marine, Ying Tao, Jing Zhang, Haibin Wang, Mary S. Keckler, Alison S. Laufer Halpin, Christopher A. Elkins, Suxiang Tong |
| EPI_ISL_426430, EPI_ISL_426431 | OH Department of Health Laboratory | Pathogen Discovery, Respiratory Viruses Branch, Division of Viral Diseases, Centers for Disease Control and Prevention | Krista Queen, Yan Li, Anna Uehara, Clinton R. Paden, Rachel Marine, Ying Tao, Jing Zhang, Haibin Wang, Mary S. Keckler, Alison S. Laufer Halpin, Christopher A. Elkins, Suxiang Tong |
| EPI_ISL_426432, EPI_ISL_426433, EPI_ISL_426434 | PA Department of Health, Bureau of Laboratories | Pathogen Discovery, Respiratory Viruses Branch, Division of Viral Diseases, Centers for Disease Control and Prevention | Krista Queen, Yan Li, Anna Uehara, Clinton R. Paden, Rachel Marine, Ying Tao, Jing Zhang, Haibin Wang, Mary S. Keckler, Alison S. Laufer Halpin, Christopher A. Elkins, Suxiang Tong |
| EPI_ISL_426435 | RI State Health Laboratories | Pathogen Discovery, Respiratory Viruses Branch, Division of Viral Diseases, Centers for Disease Control and Prevention | Krista Queen, Yan Li, Anna Uehara, Clinton R. Paden, Rachel Marine, Ying Tao, Jing Zhang, Haibin Wang, Mary S. Keckler, Alison S. Laufer Halpin, Christopher A. Elkins, Suxiang Tong |
| EPI_ISL_426436, EPI_ISL_426438, EPI_ISL_426439, EPI_ISL_426442, EPI_ISL_426445, EPI_ISL_426450 | WA State Department of Health | Pathogen Discovery, Respiratory Viruses Branch, Division of Viral Diseases, Centers for Disease Control and Prevention | Jing Zhang, Ying Tao, Clinton R. Paden, Krista Queen, Anna Uehara, Yan Li, Haibin Wang, Jesica Jacobs, Denny Russell, Brian Hiatt, Jessica Gant, Suxiang Tong |
| EPI_ISL_426483, EPI_ISL_426485 | AZ SPHL, Arizona Department of Health Services | TGen North | Jolene Bowers, Megan Folkerts, Darrin Lemmer, Dave Engelthaler |
| EPI_ISL_426499, EPI_ISL_426500, EPI_ISL_426501, EPI_ISL_426502, EPI_ISL_426503, EPI_ISL_426504, EPI_ISL_426505, EPI_ISL_426506, EPI_ISL_426507, EPI_ISL_426509, EPI_ISL_426510, EPI_ISL_426511 | | | |
| see above | TGen North | TGen North | Jolene Bowers, Megan Folkerts, Darrin Lemmer, Dave Engelthaler |
| EPI_ISL_426512, EPI_ISL_426513, EPI_ISL_426514, EPI_ISL_426515, EPI_ISL_426516, EPI_ISL_426517, EPI_ISL_426518, EPI_ISL_426519 | AZ SPHL, Arizona Department of Health Services | TGen North | Jolene Bowers, Megan Folkerts, Darrin Lemmer, Dave Engelthaler |
| EPI_ISL_426520, EPI_ISL_426521, EPI_ISL_426522, EPI_ISL_426523, EPI_ISL_426524, EPI_ISL_426525, EPI_ISL_426526 | TGen North | TGen North | Jolene Bowers, Megan Folkerts, Darrin Lemmer, Dave Engelthaler |
| EPI_ISL_426527, EPI_ISL_426528, EPI_ISL_426529, EPI_ISL_426530, EPI_ISL_426531 | AZ SPHL, Arizona Department of Health Services | TGen North | Jolene Bowers, Megan Folkerts, Darrin Lemmer, Dave Engelthaler |
| EPI_ISL_426532, EPI_ISL_426533, EPI_ISL_426534, EPI_ISL_426535, EPI_ISL_426536 | TGen North | TGen North | Jolene Bowers, Megan Folkerts, Darrin Lemmer, Dave Engelthaler |
| EPI_ISL_426537, EPI_ISL_426538, EPI_ISL_426539, EPI_ISL_426540, EPI_ISL_426541, EPI_ISL_426542, EPI_ISL_426543, EPI_ISL_426544, EPI_ISL_426545, EPI_ISL_426546, EPI_ISL_426547, EPI_ISL_426548, EPI_ISL_426549, EPI_ISL_426550, EPI_ISL_426551, EPI_ISL_426552, EPI_ISL_426553, EPI_ISL_426554, EPI_ISL_426555 | | | |
| see above | AZ SPHL, Arizona Department of Health Services | TGen North | Jolene Bowers, Megan Folkerts, Darrin Lemmer, Dave Engelthaler |
| EPI_ISL_426556, EPI_ISL_426557 | TGen North | TGen North | Jolene Bowers, Megan Folkerts, Darrin Lemmer, Dave Engelthaler |
| EPI_ISL_426558, EPI_ISL_426559, EPI_ISL_426560, EPI_ISL_426561, EPI_ISL_426562, EPI_ISL_426563, EPI_ISL_426564, EPI_ISL_426565, EPI_ISL_426566, EPI_ISL_426567, EPI_ISL_426568, EPI_ISL_426569, EPI_ISL_427271, EPI_ISL_427272 | | | |
| see above | AZ SPHL, Arizona Department of Health Services | TGen North | Jolene Bowers, Megan Folkerts, Darrin Lemmer, Dave Engelthaler |
| EPI_ISL_427340, EPI_ISL_427341, EPI_ISL_427342, EPI_ISL_427343, EPI_ISL_427344, EPI_ISL_427345, EPI_ISL_427346, EPI_ISL_427347, EPI_ISL_427348, EPI_ISL_427349, EPI_ISL_427350, EPI_ISL_427351, EPI_ISL_427352, EPI_ISL_427353, EPI_ISL_427354, EPI_ISL_427355, EPI_ISL_427356, EPI_ISL_427357, EPI_ISL_427358, EPI_ISL_427359, EPI_ISL_427360, EPI_ISL_427361, EPI_ISL_427362, EPI_ISL_427363, EPI_ISL_427364, EPI_ISL_427365, EPI_ISL_427366, EPI_ISL_427367, EPI_ISL_427368, EPI_ISL_427369, EPI_ISL_427370, EPI_ISL_427371, EPI_ISL_427372, EPI_ISL_427373, EPI_ISL_427374, EPI_ISL_427375, EPI_ISL_427376, EPI_ISL_427377, EPI_ISL_427378, EPI_ISL_427379, EPI_ISL_427380, EPI_ISL_427381, EPI_ISL_427382, EPI_ISL_427383, EPI_ISL_427384, EPI_ISL_427385, EPI_ISL_427386, EPI_ISL_427387, EPI_ISL_427388, EPI_ISL_427389, EPI_ISL_427390 | | | |
| see above | Department of Clinical Microbiology | GIGA Medical Genomics | Keith Durkin, Maria Artesi, Sébastien Bontems, Raphaël Boreux, Cécile Meex, Pierrette Melin, Marie-Pierre Hayette, Vincent Bours. |
| EPI_ISL_427652, EPI_ISL_427662, EPI_ISL_427668, EPI_ISL_427677, EPI_ISL_427706, EPI_ISL_427715, EPI_ISL_427716, EPI_ISL_427723, EPI_ISL_427724, EPI_ISL_427727, EPI_ISL_427738, EPI_ISL_427742, EPI_ISL_427752, EPI_ISL_427763, EPI_ISL_427773, EPI_ISL_427782, EPI_ISL_427792 | | | |
| see above | Centre for Infectious Diseases and Microbiology Public Health | NSW Health Pathology - Institute of Clinical Pathology and Medical Research; Westmead Hospital; University of Sydney | Lam C, Gray K, Timms, V, Gall M, Arnott A, Sadsad R, Draper J, Sim E, Bachmann N, Rockett R, Carter I, Holmes EC, O'Sullivan MV, Byun R, Sintchenko V, Chen SC, Eden JS, Maddocks S, Kok J, Propenko M, Sorrell T, Chang S, Basile K, Dwyer DE for the 2019-nCoV Study Group |
| EPI_ISL_427809 | Division of Viral Diseases, Center for Laboratory Control of | Division of Viral Diseases, Center for Laboratory Control of | Jeong-Min Kim, Yoon-Seok Chung, Namjoo Lee, Mi-Seon Kim, Sang Hee Woo, Hye-Jun Jo, Sehee Park, Heui Man Kim, Jun-Sub Kim, Junhyeong Jang, |

| | | | |
|---|---|---|---|
| | Infectious Diseases, Korea Centers for Diseases Control and Prevention | Infectious Diseases, Korea Centers for Diseases Control and Prevention | Myung Guk Han |
| EPI_ISL_427810, EPI_ISL_427811, EPI_ISL_427812, EPI_ISL_427813 | Division of Viral Diseases, Center for Laboratory Control of Infectious Diseases, Korea Centers for Diseases Control and Prevention | Division of Viral Diseases, Center for Laboratory Control of Infectious Diseases, Korea Centers for Diseases Control and Prevention | Jeong-Min Kim, Yoon-Seok Chung, Namjoo Lee, Mi-Seon Kim, Sang Hee Woo, Hye-Jun Jo, Sehee Park, Heui Man Kim, Jun-Sub Kim, Junhyeong Jang, Dong Hyun Song, Daesang Lee, Seong Tae Jeong, Myung Guk Han |
| EPI_ISL_428250, EPI_ISL_428370, EPI_ISL_428371, EPI_ISL_428372 | Yale Clinical Virology Laboratory | Grubaugh Lab - Yale School of Public Health | Joseph Fauver, Anderson Brito, Tara Alpert, Chantal Vogels, Ellen Foxman, Albert Ko, Marie Landry, Nathan Grubaugh |
| EPI_ISL_428373, EPI_ISL_428374, EPI_ISL_428375, EPI_ISL_428376, EPI_ISL_428377, EPI_ISL_428378, EPI_ISL_428379, EPI_ISL_428380 | Yale COVID-19 Biorepository | Grubaugh Lab - Yale School of Public Health | Joseph Fauver, Tara Alpert, Anderson Brito, Anne Wyllie, Chantal Vogels, Mary Petrone, Chaney Kalinich, Isabel Ott, Arnau Casanovas, Catherine Muenker, Adam Moore, Alice Lu, Maria Tokuyama, Patrick Wong, Peiwen Lu, Saad Omer, Richard Martinello, Allison Nelson, Shelli Farhadian, Akiko Iwasaki, Charlese Dela Cruz, Albert Ko, Nathan Grubaugh |
| EPI_ISL_428381, EPI_ISL_428382 | Yale Clinical Virology Laboratory | Grubaugh Lab - Yale School of Public Health | Joseph Fauver, Anderson Brito, Tara Alpert, Chantal Vogels, Ellen Foxman, Albert Ko, Marie Landry, Nathan Grubaugh |
| EPI_ISL_428384, EPI_ISL_428385, EPI_ISL_428386, EPI_ISL_428387, EPI_ISL_428388, EPI_ISL_428389, EPI_ISL_428390, EPI_ISL_428391, EPI_ISL_428392, EPI_ISL_428393, EPI_ISL_428394, EPI_ISL_428395, EPI_ISL_428396, EPI_ISL_428397, EPI_ISL_428398 | | | |
| see above | Yale COVID-19 Biorepository | Grubaugh Lab - Yale School of Public Health | Joseph Fauver, Tara Alpert, Anderson Brito, Anne Wyllie, Chantal Vogels, Mary Petrone, Chaney Kalinich, Isabel Ott, Arnau Casanovas, Catherine Muenker, Adam Moore, Alice Lu, Maria Tokuyama, Patrick Wong, Peiwen Lu, Saad Omer, Richard Martinello, Allison Nelson, Shelli Farhadian, Akiko Iwasaki, Charlese Dela Cruz, Albert Ko, Nathan Grubaugh |
| EPI_ISL_428399, EPI_ISL_428400, EPI_ISL_428401, EPI_ISL_428402, EPI_ISL_428403, EPI_ISL_428404, EPI_ISL_428405 | Yale COVID-19 Biorepository | Grubaugh Lab - Yale School of Public Health | Joseph Fauver, Tara Alpert, Anderson Brito, Anne Wyllie, Chantal Vogels, Mary Petrone, Cole Jensen, Chaney Kalinich, Isabel Ott, Arnau Casanovas, Catherine Muenker, Adam Moore, Alice Lu, Maria Tokuyama, Patrick Wong, Peiwen Lu, Saad Omer, Richard Martinello, Allison Nelson, Shelli Farhadian, Akiko Iwasaki, Charlese Dela Cruz, Albert Ko, Nathan Grubaugh |
| EPI_ISL_428488, EPI_ISL_428489, EPI_ISL_428490, EPI_ISL_428491 | Centers for Disease Control, R.O.C. (Taiwan) | Centers for Disease Control, R.O.C. (Taiwan) | Ji-Rong Yang, Yu-Chi Lin, Jung-Jung Mu, Ming-Tsan Liu |
| EPI_ISL_428740 | Yale Clinical Virology Laboratory | Grubaugh Lab - Yale School of Public Health | Joseph Fauver, Anderson Brito, Tara Alpert, Chantal Vogels, Ellen Foxman, Albert Ko, Marie Landry, Nathan Grubaugh |
| EPI_ISL_428745, EPI_ISL_428746, EPI_ISL_428747, EPI_ISL_428748, EPI_ISL_428749, EPI_ISL_428750, EPI_ISL_428751, EPI_ISL_428752 | Yale COVID-19 Biorepository | Grubaugh Lab - Yale School of Public Health | Joseph Fauver, Tara Alpert, Anderson Brito, Anne Wyllie, Chantal Vogels, Mary Petrone, Cole Jensen, Chaney Kalinich, Isabel Ott, Arnau Casanovas, Catherine Muenker, Adam Moore, Alice Lu, Maria Tokuyama, Patrick Wong, Peiwen Lu, Saad Omer, Richard Martinello, Allison Nelson, Shelli Farhadian, Akiko Iwasaki, Charlese Dela Cruz, Albert Ko, Nathan Grubaugh |
| EPI_ISL_429256 | Health Sciences Technology Park, Avicena, 8, 18016 Granada. Spain | Sequencing and Bioinformatics Service FISABIO-Public Health | Joaquín Mendoza, Almudena Rojas, Pablo Mendoza |
| EPI_ISL_429882 | Centers for Disease Control, R.O.C. (Taiwan) | Centers for Disease Control, R.O.C. (Taiwan) | Ji-Rong Yang, Yu-Chi Lin, Jung-Jung Mu, Ming-Tsan Liu |
| EPI_ISL_429883 | Centers for Disease Control, R.O.C. (Taiwan) | Centers for Disease Control, R.O.C. (Taiwan) | Ji-Rong Yang, Yu-Chi Lin, Jung-Jung Mu, Ming-Tsan-Liu |
| EPI_ISL_429884 | Centers for Disease Control, R.O.C. (Taiwan) | Centers for Disease Control, R.O.C. (Taiwan) | Ji-Rong Yang, Yu-Chi Lin, Jung-Jung Mu, Ming-Tsan Liu |
| EPI_ISL_431080, EPI_ISL_431081, EPI_ISL_431082, EPI_ISL_431083, EPI_ISL_431084, EPI_ISL_431085, EPI_ISL_431086, EPI_ISL_431087, EPI_ISL_431088, EPI_ISL_431089, EPI_ISL_431090, EPI_ISL_431091, EPI_ISL_431092, EPI_ISL_431093, EPI_ISL_431094, EPI_ISL_431095, EPI_ISL_431096 | | | |
| see above | Yale COVID-19 Biorepository | Grubaugh Lab - Yale School of Public Health | Joseph Fauver, Tara Alpert, Anderson Brito, Anne Wyllie, Chantal Vogels, Mary Petrone, Cole Jensen, Chaney Kalinich, Isabel Ott, Arnau Casanovas, Catherine Muenker, Adam Moore, Alice Lu, Maria Tokuyama, Patrick Wong, Peiwen Lu, Saad Omer, Richard Martinello, Allison Nelson, Shelli Farhadian, Akiko Iwasaki, Charlese Dela Cruz, Albert Ko, Nathan Grubaugh |
| EPI_ISL_434455, EPI_ISL_434456, EPI_ISL_434457, EPI_ISL_434458, EPI_ISL_434459, EPI_ISL_434460, EPI_ISL_434461, EPI_ISL_434462, EPI_ISL_434463, EPI_ISL_434464, EPI_ISL_434465, EPI_ISL_434466, EPI_ISL_434467, EPI_ISL_434468, EPI_ISL_434469, EPI_ISL_434470, EPI_ISL_434471, EPI_ISL_434472, EPI_ISL_434473, EPI_ISL_434474, EPI_ISL_434475, EPI_ISL_434476, EPI_ISL_434477, EPI_ISL_434478, EPI_ISL_434479, EPI_ISL_434480, EPI_ISL_434481, EPI_ISL_434482, EPI_ISL_434483, EPI_ISL_434484, EPI_ISL_434485, EPI_ISL_434486 | | | |
| see above | Laboratory of Microbiology, Medical School, National and Kapodistrian University of Athens | Laboratory of Biology, Department of Medicine, Democritus University of Thrace | Kassela K., Bampali,M., Dovrolis,N., Gatzidou,E., Froukala,E., Stavropoulou,A., Veletza,S., Tsakris,A., Spanakis,N. and Karakasiliotis,I. |
| EPI_ISL_434650 | Follinge Halsocentral | The Public Health Agency of Sweden | Kerstin Persson Moberg, Oskar Karlsson Lindsjo, Maria Lind Karlberg, Anna-Malin Linde, Olov Svartstrom, Anna Risberg, Theresa Enkirch, Mia Brytting, Karin Tegmark-Wisell |
| EPI_ISL_434654 | Saroledens Familjelakare | The Public Health Agency of Sweden | Katarina Jarbur, Oskar Karlsson Lindsjo, Maria Lind Karlberg, Anna-Malin Linde, Olov Svartstrom, Anna Risberg, Theresa Enkirch, Mia Brytting, Karin Tegmark-Wisell |
| EPI_ISL_434659 | Knivsta VC | The Public Health Agency of Sweden | Johanna Carlson, Oskar Karlsson Lindsjo, Maria Lind Karlberg, Anna-Malin Linde, Olov Svartstrom, Anna Risberg, Theresa Enkirch, Mia Brytting, Karin Tegmark-Wisell |
| EPI_ISL_434668 | Kungsors VC | The Public Health Agency of Sweden | Jessica Karlsson, Oskar Karlsson Lindsjo, Maria Lind Karlberg, Anna-Malin Linde, Olov Svartstrom, Anna Risberg, Theresa Enkirch, Mia Brytting, Karin Tegmark-Wisell |
| EPI_ISL_435034 | LSUHS Emerging Viral Threat Laboratory | Microbial Genome Sequencing Center | Jeremy P. Kamil, John A. Vanchiere, Rona S. Scott, Camille F. Abshire, Abida Siddiqa, Byeong-Jae Lee, Chan-ki Min, Md Maksudul Alam, Monica Gestal-Cartele, Edna Ondari, Adam Greer, Malgorzata Bienkowska-Haba, Katarzyna Zwolinska, Michelle M. Arnold, Jason M. Bodily, Andrew D. Yurochko, Paul M. Weinberger, Christopher G. Kevil, Martin J. Sapp, Daniel J. Snyder, Vaughn S. Cooper |
| EPI_ISL_435035, EPI_ISL_435036, EPI_ISL_435037, EPI_ISL_435038, EPI_ISL_435039, EPI_ISL_435040, EPI_ISL_435041, EPI_ISL_435042, EPI_ISL_435043, EPI_ISL_435044 | LSUHS Emerging Viral Threat Laboratory | Microbial Genome Sequencing Center | Jeremy P. Kamil, John A. Vanchiere, Rona S. Scott, Camille F. Abshire, Abida Siddiqa, Byeong-Jae Lee, Chan-ki Min, Md Maksudul Alam, Monica Gestal-Cartele, Edna Ondari, Adam Greer, Malgorzata Bienkowska-Haba, Katarzyna Zwolinska, Jason M. Bodily, Andrew D. Yurochko, Paul M. Weinberger, Christopher G. Kevil, Martin J. Sapp, Daniel J. Snyder, Vaughn S. Cooper |
| EPI_ISL_435053 | B.J. Medical College and Civil hospital | Gujarat Biotechnology Research Centre | Janvi Raval, Monika Gandhi, Pinal Trivedi, Maharshi Pandya, Amit Kanani, Akanksha Verma, Nitin Savaliya, Raghawendra Kumar, Dinesh Kumar, Zuber Saiyed, Dipa Kinariwala, Disha Patel, Binita Aring, Geeta Vaghela, Sonia Barve, Bhavesh Modi, Kairavi Joshi, Gaurishankar Shrimali, Nidhi Sood, Pranay Shah, R D Dixit, Snehal Bagatharia, Kamlesh J Upadhyay, Ramesh Pandit, Tejas Shah, Ankit Hinsu, Pritesh Sabara, Apurvasinh Puvar, Nidhi Patel, Chaitanya Joshi, Madhvi Joshi |
| EPI_ISL_435555, EPI_ISL_435556, EPI_ISL_435557, EPI_ISL_435558, EPI_ISL_435559, EPI_ISL_435560, EPI_ISL_435561, EPI_ISL_435562, EPI_ISL_435563, EPI_ISL_435564, EPI_ISL_435565, EPI_ISL_435566, EPI_ISL_435567, EPI_ISL_435568 | | | |
| see above | LSUHS Emerging Viral Threat Laboratory | Microbial Genome Sequencing Center | John A. Vanchiere, Jeremy P. Kamil, Rona S. Scott, Camille F. Abshire, Abida Siddiqa, Byeong-Jae Lee, Chan-ki Min, Md Maksudul Alam, Monica Gestal-Cartele, Edna Ondari, Adam Greer, Malgorzata Bienkowska-Haba, Katarzyna Zwolinska, Jason M. Bodily, Andrew D. Yurochko, Paul M. Weinberger, Christopher G. Kevil, Martin J. Sapp, Daniel J. Snyder, Vaughn S. Cooper |
| EPI_ISL_435569, EPI_ISL_435570, EPI_ISL_435571, EPI_ISL_435572, EPI_ISL_435573, EPI_ISL_435574, EPI_ISL_435575, EPI_ISL_435576, EPI_ISL_435577, EPI_ISL_435578, EPI_ISL_435579 | | | |
| see above | LSUHS Emerging Viral Threat Laboratory | Microbial Genome Sequencing Center | Jeremy P. Kamil, John A. Vanchiere, Rona S. Scott, Camille F. Abshire, Abida Siddiqa, Byeong-Jae Lee, Chan-ki Min, Md Maksudul Alam, Monica Gestal-Cartele, Edna Ondari, Adam Greer, Malgorzata Bienkowska-Haba, Katarzyna Zwolinska, Jason M. Bodily, Andrew D. Yurochko, Paul M. Weinberger, Christopher G. Kevil, Martin J. Sapp, Daniel J. Snyder, Vaughn S. Cooper |
| EPI_ISL_435702, EPI_ISL_435703, EPI_ISL_435704, EPI_ISL_435705, EPI_ISL_435706, EPI_ISL_435707, EPI_ISL_435708, EPI_ISL_435709 | Yale COVID-19 Biorepository | Grubaugh Lab - Yale School of Public Health | Joseph Fauver, Tara Alpert, Anderson Brito, Anne Wyllie, Chantal Vogels, Mary Petrone, Cole Jensen, Chaney Kalinich, Isabel Ott, Arnau Casanovas, Catherine Muenker, Adam Moore, Alice Lu, Maria Tokuyama, Patrick Wong, Peiwen Lu, Saad Omer, Richard Martinello, Allison Nelson, Shelli Farhadian, Akiko Iwasaki, Charlese Dela Cruz, Albert Ko, Nathan Grubaugh |

| | | | |
|---|---|---|---|
| EPI_ISL_435710, EPI_ISL_435711, EPI_ISL_435712, EPI_ISL_435713, EPI_ISL_435714, EPI_ISL_435715, EPI_ISL_435716, EPI_ISL_435717, EPI_ISL_435718, EPI_ISL_435719 | Connecticut State Department of Public Health | Grubaugh Lab - Yale School of Public Health | Joseph Fauver, Tara Alpert, Anderson Brito, Anne Wyllie, Chantal Vogels, Mary Petrone, Cole Jensen, Chaney Kalinich, Isabel Ott, Arnau Casanovas, Catherine Muenker, Adam Moore, Alice Lu, Maria Tokuyama, Patrick Wong, Peiwen Lu, Saad Omer, Richard Martinello, Allison Nelson, Shelli Farhadian, Akiko Iwasaki, Charlese Dela Cruz, Albert Ko, Nathan Grubaugh |
| EPI_ISL_435720 | Yale Clinical Virology Laboratory | Grubaugh Lab - Yale School of Public Health | Joseph Fauver, Tara Alpert, Anderson Brito, Anne Wyllie, Chantal Vogels, Mary Petrone, Cole Jensen, Chaney Kalinich, Isabel Ott, Arnau Casanovas, Catherine Muenker, Adam Moore, Alice Lu, Maria Tokuyama, Patrick Wong, Peiwen Lu, Saad Omer, Richard Martinello, Allison Nelson, Shelli Farhadian, Akiko Iwasaki, Charlese Dela Cruz, Albert Ko, Nathan Grubaugh |
| EPI_ISL_436235, EPI_ISL_436236, EPI_ISL_436237, EPI_ISL_436238, EPI_ISL_436239, EPI_ISL_436240, EPI_ISL_436241, EPI_ISL_436242, EPI_ISL_436243, EPI_ISL_436244, EPI_ISL_436245, EPI_ISL_436246, EPI_ISL_436247, EPI_ISL_436248, EPI_ISL_436249, EPI_ISL_436250, EPI_ISL_436251, EPI_ISL_436252, EPI_ISL_436253, EPI_ISL_436254, EPI_ISL_436255, EPI_ISL_436256, EPI_ISL_436257, EPI_ISL_436258, EPI_ISL_436259, EPI_ISL_436260, EPI_ISL_436261, EPI_ISL_436262, EPI_ISL_436263, EPI_ISL_436264, EPI_ISL_436265, EPI_ISL_436266, EPI_ISL_436267, EPI_ISL_436268, EPI_ISL_436269, EPI_ISL_436270, EPI_ISL_436372, EPI_ISL_436373, EPI_ISL_436374, EPI_ISL_436375, EPI_ISL_436376, EPI_ISL_436377, EPI_ISL_436378, EPI_ISL_436379, EPI_ISL_436380, EPI_ISL_436381 | | | |
| see above | Servicio de Microbiologia. Hospital Universitario Doctor Peset | Sequencing and Bioinformatics Service and Molecular Epidemiology Research Group. FISABIO-Public Health | Juan Alberola Enguídanos, Juan Jose Camarena Miñana, Rosa González Pellicer, Neris Garcia-Gonzalez, Inma Galán Vendrell, Sandra Carbo, Loreto Ferrús Abad, Paula Ruiz-Hueso, Mariana Reyes-Prieto, Vicente Soriano Chirona, Ivan Ansari, Maria Alma Bracho, Griselda De Marco, Beatriz Beamud, Lidia Ruiz Roldan, Marta Pla Diaz, Llúcia Martínez-Priego, Giuseppe D'Auria, Jose Miguel Nogueira Coito, Fernando Gonzalez-Candelas |
| EPI_ISL_437187 | Siloam Hospitals | Institute of Tropical Disease, Universitas Airlangga | Kazufumi Shimizu, Krisnoadi Rahardjo, Aldise M Nastri, Jezzy R Dewantari, Rima R Prasetya, Maria M Padmidewi, Gatot Soegiarto, Laksmi Wulandari, Retno A Setyoningrum, Resti Y Meliana, Yohko K Shimizu, Mitsuhiro Nishimura, Yasuko Mori, Soetjipto, Maria I Lusida |
| EPI_ISL_437188 | RSUD Dr. Soetomo | Institute of Tropical Disease, Universitas Airlangga | Krisnoadi Rahardjo, Aldise M Nastri, Jezzy R Dewantari, Rima R Prasetya, Joni Wahyuhadi, Gatot Soegiarto, Laksmi Wulandari, Retno A Setyoningrum, Resti Y Meliana, Yohko K Shimizu, Mitsuhiro Nishimura, Yasuko Mori, Soetjipto, Kazufumi Shimizu, Maria I Lusida |
| EPI_ISL_437197, EPI_ISL_437198, EPI_ISL_437199, EPI_ISL_437200, EPI_ISL_437201, EPI_ISL_437202, EPI_ISL_437203, EPI_ISL_437298, EPI_ISL_437299, EPI_ISL_437300, EPI_ISL_437301, EPI_ISL_437302, EPI_ISL_437303 | | | |
| see above | Diagnostic- and Research Institute of Pathology, Medical University of Graz | Diagnostic- and Research Institute of Pathology, Medical University of Graz | Karl Kashofer, Peter Regitnig, Martin Zacharias, Gregor Gorkiewicz |
| EPI_ISL_437451 | B.J. Medical College and Civil hospital | Gujarat Biotechnology Research Centre | Kairavi Joshi, Gaurishankar Shrimali, Nidhi Sood, Pranay Shah, R D Dixit, Snehal Bagatharia, Kamlesh J Upadhyay, Ramesh Pandit, Tejas Shah, Ankit Hinsu, Pritesh Sabara, Apurvasinh Puvar, Janvi Raval, Monika Gandhi, Pinal Trivedi, Maharshi Pandya, Amit Kanani, Akanksha Verma, Nitin Savaliya, Raghawendra Kumar, Dinesh Kumar, Zuber Saiyed, Dipa Kinariwala, Disha Patel, Binita Aring, Neeta Khandelwal, Geeta Vaghela, Sonia Barve, Bhavesh Modi, Bhavya Jindal, Chaitanya Joshi, Madhvi Joshi |
| EPI_ISL_437512 | Human Genetic Research Center, Kawsar Biotech Company | Human Genetic Research Center, Kawsar Biotech Company | Khosravi,M.A., Abbasalipour,M., Zeinali,S., Sabeghi,S., Kehsvar,Y., Hosseini,F. and Haghdoost,Y. |
| EPI_ISL_437601 | Keio University School of Medicine | Keio University School of Medicine | Kenjiro Kosaki, Yuka Iwasaki, Toshiki Takenouchi, Haruhiko Siomi, |
| EPI_ISL_437626 | Department of Microbiology,Gandhi Medical College and Hospital | Department of Veterinary Biotechnalogy, College of Veterinary Science, Rajendranagar, PV Narsimha Rao Telengana Veterinary University | Kalyani Putty, Muttineni Radhakrishna, Nagamani K, Thrilok Chander B, Raja Rao M, Ravikumar P, Sunitha P, Pankaj Singh D, Anand Kumar K, Amit A. Upadhyay, Steven Bosinger, Rama Amara |
| EPI_ISL_437874, EPI_ISL_437875, EPI_ISL_437876, EPI_ISL_437877, EPI_ISL_437878, EPI_ISL_437879, EPI_ISL_437880, EPI_ISL_437881, EPI_ISL_437882, EPI_ISL_437883, EPI_ISL_437884, EPI_ISL_437885, EPI_ISL_437886, EPI_ISL_437887, EPI_ISL_437888, EPI_ISL_437889, EPI_ISL_437890, EPI_ISL_437891, EPI_ISL_437892, EPI_ISL_437893, EPI_ISL_437894, EPI_ISL_437895, EPI_ISL_437896, EPI_ISL_437897, EPI_ISL_437898, EPI_ISL_437899, EPI_ISL_437900, EPI_ISL_437901, EPI_ISL_437902, EPI_ISL_437903, EPI_ISL_437904, EPI_ISL_437905, EPI_ISL_437906, EPI_ISL_437907, EPI_ISL_437908, EPI_ISL_437909, EPI_ISL_437910, EPI_ISL_437911 | | | |
| see above | Laboratory of Microbiology, Medical School, National and Kapodistrian University of Athens | Laboratory of Biology, Department of Medicine, Democritus University of Thrace | Kassela K., Dovrolis,N., Bampali,M., Gatzidou,E., Froukala,E., Stavropoulou,A., Veletza,S., Tsakris,A., Spanakis,N. and Karakasiliotis,I. |
| EPI_ISL_438947, EPI_ISL_438948, EPI_ISL_438949, EPI_ISL_438951, EPI_ISL_438952, EPI_ISL_438953, EPI_ISL_438954, EPI_ISL_438955, EPI_ISL_438956, EPI_ISL_438958, EPI_ISL_438959, EPI_ISL_438961, EPI_ISL_438962, EPI_ISL_438963, EPI_ISL_438964, EPI_ISL_438965, EPI_ISL_438966, EPI_ISL_438967, EPI_ISL_438968, EPI_ISL_438969, EPI_ISL_438970, EPI_ISL_438971, EPI_ISL_438972 | | | |
| see above | Keio University School of Medicine | Keio University School of Medicine | Kenjiro Kosaki |
| EPI_ISL_444274, EPI_ISL_444275, EPI_ISL_444276, EPI_ISL_444277, EPI_ISL_444278 | Laboratory Medicine | Department of Laboratory Medicine, Lin-Kou Chang Gung Memorial Hospital, Taoyuan, Taiwan | Kuo-Chien Tsao, Yu-Nong Gong, Shu-Li Yang, Yi-Chun Liu, Chung-Guei Huang, Mei-Jen Hsiao, Po-Wei Huang, Cheng-Ta Yang, Cheng-Hsun Chiu, Peng-Nien Huang, Kuo-Ming Lee, Guang-Wu Chen, Shin-Ru Shih |
| EPI_ISL_444458 | B.J. Medical College and Civil hospital | Gujarat Biotechnology Research Centre | Kamlesh J Upadhyay, Ramesh Pandit, Tejas Shah, Ankit Hinsu, Pritesh Sabara, Apurvasinh Puvar, Janvi Raval, Monika Gandhi, Pinal Trivedi, Maharshi Pandya, Amit Kanani, Akanksha Verma, Nitin Savaliya, Raghawendra Kumar, Dinesh Kumar, Zuber Saiyed, Dipa Kinariwala, Disha Patel, Binita Aring, Neeta Khandelwal, Geeta Vaghela, Sonia Barve, Bhavesh Modi, Kairavi Joshi, Gaurishankar Shrimali, Nidhi Sood, Pranay Shah, R D Dixit, Snehal Bagatharia, Priti Pandita, Chaitanya Joshi, Madhvi Joshi |
| EPI_ISL_444464 | B.J. Medical College and Civil hospital | Gujarat Biotechnology Research Centre | Janvi Raval, Monika Gandhi, Pinal Trivedi, Maharshi Pandya, Amit Kanani, Akanksha Verma, Nitin Savaliya, Raghawendra Kumar, Dinesh Kumar, Zuber Saiyed, Dipa Kinariwala, Disha Patel, Binita Aring, Neeta Khandelwal, Geeta Vaghela, Sonia Barve, Bhavesh Modi, Kairavi Joshi, Gaurishankar Shrimali, Nidhi Sood, Pranay Shah, R D Dixit, Snehal Bagatharia, Kamlesh J Upadhyay, Ramesh Pandit, Tejas Shah, Ankit Hinsu, Pritesh Sabara, Apurvasinh Puvar, Dipeshwari Shewale, Chaitanya Joshi, Madhvi Joshi |
| EPI_ISL_444481 | B.J. Medical College and Civil hospital | Gujarat Biotechnology Research Centre | Kairavi Joshi, Gaurishankar Shrimali, Nidhi Sood, Pranay Shah, R D Dixit, Snehal Bagatharia, Kamlesh J Upadhyay, Ramesh Pandit, Tejas Shah, Ankit Hinsu, Pritesh Sabara, Apurvasinh Puvar, Janvi Raval, Monika Gandhi, Pinal Trivedi, Maharshi Pandya, Amit Kanani, Akanksha Verma, Nitin Savaliya, Raghawendra Kumar, Dinesh Kumar, Zuber Saiyed, Dipa Kinariwala, Disha Patel, Binita Aring, Neeta Khandelwal, Geeta Vaghela, Sonia Barve, Bhavesh Modi, Neha Rajpara, Chaitanya Joshi, Madhvi Joshi |
| EPI_ISL_444969 | Guangzhou Eighth People's Hospital (Jiahe Sector) | Institute of Human Virology, Zhongshan School of Medicine, Sun Yat-sen University | Junsong Zhang, Fei Yu, Jun Liu, Huimin Fan, Ruosu Ying, Feng Huang, Ting Pan, Bingfeng Liu,Yiwen Zhang, Xu Zhang, Mang Shi, Fengyu Hu, Fang Li, Kai Deng, Hui Zhang |
| EPI_ISL_445222, EPI_ISL_445226 | Saroledens Familjelakare | The Public Health Agency of Sweden | Katarina Jarbur, Oskar Karlsson Lindsjo, Maria Lind Karlberg, Anna-Malin Linde, Olov Svartstrom, Anna Risberg, Theresa Enkirch, Mia Brytting, Karin Tegmark-Wisell |
| EPI_ISL_445232 | Kungsors VC | The Public Health Agency of Sweden | Jessica Karlsson, Oskar Karlsson Lindsjo, Maria Lind Karlberg, Anna-Malin Linde, Olov Svartstrom, Anna Risberg, Theresa Enkirch, Mia Brytting, Karin Tegmark-Wisell |
| EPI_ISL_447030 | B.J. Medical College and Civil hospital | Gujarat Biotechnology Research Centre | Kamlesh J Upadhyay, Ramesh Pandit, Tejas Shah, Ankit Hinsu, Pritesh Sabara, Apurvasinh Puvar, Janvi Raval, Monika Gandhi, Pinal Trivedi, Maharshi Pandya, Amit Kanani, Akanksha Verma, Nitin Savaliya, Raghawendra Kumar, Dinesh Kumar, Zuber Saiyed, Dipa Kinariwala, Disha Patel, Binita Aring, Neeta Khandelwal, Geeta Vaghela, Sonia Barve, Bhavesh Modi, Kairavi Joshi, Gaurishankar Shrimali, Nidhi Sood, Pranay Shah, R D Dixit, Snehal Bagatharia, Anjali Rajwar, Chaitanya Joshi, Madhvi Joshi |
| EPI_ISL_447036 | B.J. Medical College and Civil hospital | Gujarat Biotechnology Research Centre | Janvi Raval, Monika Gandhi, Pinal Trivedi, Maharshi Pandya, Amit Kanani, Akanksha Verma, Nitin Savaliya, Raghawendra Kumar, Dinesh Kumar, Zuber Saiyed, Dipa Kinariwala, Disha Patel, Binita Aring, Neeta Khandelwal, Geeta Vaghela, Sonia Barve, Bhavesh Modi, Kairavi Joshi, Gaurishankar Shrimali, Nidhi Sood, Pranay Shah, R D Dixit, Snehal Bagatharia, Kamlesh J Upadhyay, Ramesh Pandit, Tejas Shah, Ankit Hinsu, Pritesh Sabara, Apurvasinh Puvar, Afzal Ansari, Chaitanya Joshi, Madhvi Joshi |
| EPI_ISL_447053 | GMERS Medical College and Hospital, Gandhinagar | Gujarat Biotechnology Research Centre | Kairavi Joshi, Gaurishankar Shrimali, Nidhi Sood, Pranay Shah, R D Dixit, Snehal Bagatharia, Kamlesh J Upadhyay, Ramesh Pandit, Tejas Shah, Ankit Hinsu, Pritesh Sabara, Apurvasinh Puvar, Janvi Raval, Monika Gandhi, Pinal Trivedi, Maharshi Pandya, Amit Kanani, Akanksha Verma, Nitin Savaliya, Raghawendra Kumar, Dinesh Kumar, Zuber Saiyed, Dipa Kinariwala, Disha Patel, Binita Aring, Neeta Khandelwal, Geeta Vaghela, Sonia Barve, Bhavesh Modi, Pooja P Doshi, Chaitanya Joshi, Madhvi Joshi |
| EPI_ISL_447120, EPI_ISL_447121, EPI_ISL_447122, EPI_ISL_447123, EPI_ISL_447124, EPI_ISL_447125, EPI_ISL_447126, EPI_ISL_447127, EPI_ISL_447128, EPI_ISL_447129, EPI_ISL_447130, EPI_ISL_447131, EPI_ISL_447132, EPI_ISL_447133, EPI_ISL_447134, EPI_ISL_447135, EPI_ISL_447136, EPI_ISL_447137, EPI_ISL_447138, EPI_ISL_447139, EPI_ISL_447140, EPI_ISL_447141, EPI_ISL_447142, EPI_ISL_447143, EPI_ISL_447144, EPI_ISL_447145, EPI_ISL_447146, EPI_ISL_447147, EPI_ISL_447148, EPI_ISL_447149, EPI_ISL_447150, EPI_ISL_447151, EPI_ISL_447152, EPI_ISL_447153, EPI_ISL_447154, EPI_ISL_447155, EPI_ISL_447156, EPI_ISL_447157, EPI_ISL_447158, EPI_ISL_447159, EPI_ISL_447160, EPI_ISL_447161, EPI_ISL_447162 | | | |

| | | | |
|---|---|---|---|
| see above | Department of Clinical Microbiology | GIGA Medical Genomics | Keith Durkin, Maria Artesi, Sébastien Bontems, Raphaël Boreux, Cécile Meex, Pierrette Melin, Marie-Pierre Hayette, Vincent Bours. |
| EPI_ISL_447539 | Gujarat Biotechnology Research Centre | Gujarat Biotechnology Research Centre | Kamlesh J Upadhyay, Ramesh Pandit, Tejas Shah, Ankit Hinsu, Pritesh Sabara, Apurvasinh Puvar, Janvi Raval, Monika Gandhi, Pinal Trivedi, Maharshi Pandya, Amit Kanani, Akanksha Verma, Nitin Savaliya, Raghawendra Kumar, Dinesh Kumar, Zuber Saiyed, Dipa Kinariwala, Disha Patel, Binita Aring, Neeta Khandelwal, Geeta Vaghela, Sonia Barve, Bhavesh Modi, Kairavi Joshi, Gaurishankar Shrimali, Nidhi Sood, Pranay Shah, R D Dixit, Snehal Bagatharia, Armi Chaudhari, Chaitanya Joshi, Madhvi Joshi |
| EPI_ISL_447545 | Gujarat Biotechnology Research Centre | Gujarat Biotechnology Research Centre | Janvi Raval, Monika Gandhi, Pinal Trivedi, Maharshi Pandya, Amit Kanani, Akanksha Verma, Nitin Savaliya, Raghawendra Kumar, Dinesh Kumar, Zuber Saiyed, Dipa Kinariwala, Disha Patel, Binita Aring, Neeta Khandelwal, Geeta Vaghela, Sonia Barve, Bhavesh Modi, Kairavi Joshi, Gaurishankar Shrimali, Nidhi Sood, Pranay Shah, R D Dixit, Snehal Bagatharia, Kamlesh J Upadhyay, Ramesh Pandit, Tejas Shah, Ankit Hinsu, Pritesh Sabara, Apurvasinh Puvar, Nidhi Patel, Chaitanya Joshi, Madhvi Joshi |
| EPI_ISL_447570, EPI_ISL_447571, EPI_ISL_447572 | CSIR-Centre for Cellular and Molecular Biology | CSIR-Centre for Cellular and Molecular Biology | Lamuk Zaveri, Shagufta Khan, Namami Gaur, Sakshi Shambhavi, Tulasi Nagabandi, Purushotham Vodnala, Payel Mukherjee, Sofia Banu, Priya Singh, Dhiviya Vedagiri, Divya Gupta, Vishal Sah, Santosh Kumar Kuncha, Krishnan Harinivas Harshan, Archana Bharadwaj Siva, Karthik Bharadwaj Tallapaka, Rakesh K Mishra, Divya Tej Sowpati |
| EPI_ISL_447584, EPI_ISL_447585, EPI_ISL_447586, EPI_ISL_447587 | Tamil Nadu Veterinary and Animal Sciences University | CSIR-Centre for Cellular and Molecular Biology | K Kaveri, S Sivasubramanian, S Vennila, P Padmapriya, R Kiruba, S Magesh, G Dhinakar Raj, G Ravi Kumar, Payel Mukherjee, Tulasi Nagabandi, Namami Gaur, Sakshi Shambhavi, Lamuk Zaveri, Shagufta Khan, Purushotham Vodnala, Sofia Banu, Priya Singh, Dhiviya Vedagiri, Divya Gupta, Vishal Sah, Santosh Kumar Kuncha, Krishnan Harinivas Harshan, Archana Bharadwaj Siva, Karthik Bharadwaj Tallapaka, Kumarasamy Thangaraj, Rakesh K Mishra, Divya Tej Sowpati |
| EPI_ISL_447635, EPI_ISL_447636, EPI_ISL_447637, EPI_ISL_447638, EPI_ISL_447639, EPI_ISL_447640, EPI_ISL_447641, EPI_ISL_447642, EPI_ISL_447643, EPI_ISL_447644, EPI_ISL_447645, EPI_ISL_447646, EPI_ISL_447647, EPI_ISL_447648, EPI_ISL_447649, EPI_ISL_447650, EPI_ISL_447651, EPI_ISL_447652, EPI_ISL_447653 | | | |
| see above | unknown | Department of Medicine | Kassela,K., Dovrolis,N., Bampali,M., Gatzidou,E., Froukala,E., Stavropoulou,A., Veletza,S., Tsakris,A., Spanakis,N. and Karakasiliotis,I. |
| EPI_ISL_447734, EPI_ISL_447735, EPI_ISL_447736, EPI_ISL_447737, EPI_ISL_447738, EPI_ISL_447739, EPI_ISL_447740, EPI_ISL_447741, EPI_ISL_447742, EPI_ISL_447743, EPI_ISL_447744, EPI_ISL_447745, EPI_ISL_447746, EPI_ISL_447747, EPI_ISL_447748, EPI_ISL_447749, EPI_ISL_447750, EPI_ISL_447751, EPI_ISL_447752, EPI_ISL_447753, EPI_ISL_447754 | | | |
| see above | Grupo de Investigaciones Microbiológicas-UR (GIMUR), Departamento de Biología, Facultad de Ciencias Naturales, Universidad del Rosario, Bogotá, Colombia | Grupo de Investigaciones Microbiológicas-UR (GIMUR), Departamento de Biología, Facultad de Ciencias Naturales, Universidad del Rosario, Bogotá, Colombia Instituto Nacional de Salud, Bogotá, Colombia Icahn School of Medicine at Mount Sinai, New York, USA | Juan David Ramírez, Carolina Florez, Marina Muñoz, Carolina Hernandez, Adriana Castillo, Sergio Castañeda, Nathalia Ballesteros, David Martínez, Laura Vega, Jesús E. Jaimes, Sergio Gomez, Angelica Rico, Lisseth Pardo, Esther C. Barros, Martha L. Ospina, Aníbal A. Teherán, Ana S. Gonzalez-Reiche, Matthew M. Hernandez, Emilia Mia Sordillo, Viviana Simon, Harm van Bakel, Alberto Paniz-Mondolfi |
| EPI_ISL_447755, EPI_ISL_447756, EPI_ISL_447757, EPI_ISL_447758, EPI_ISL_447759, EPI_ISL_447760, EPI_ISL_447761, EPI_ISL_447762, EPI_ISL_447763, EPI_ISL_447764, EPI_ISL_447765, EPI_ISL_447766, EPI_ISL_447767, EPI_ISL_447768, EPI_ISL_447769, EPI_ISL_447771, EPI_ISL_447772, EPI_ISL_447774, EPI_ISL_447775, EPI_ISL_447776, EPI_ISL_447777, EPI_ISL_447778, EPI_ISL_447779, EPI_ISL_447780, EPI_ISL_447781, EPI_ISL_447782, EPI_ISL_447783, EPI_ISL_447784, EPI_ISL_447785, EPI_ISL_447786, EPI_ISL_447787, EPI_ISL_447789, EPI_ISL_447790, EPI_ISL_447791, EPI_ISL_447792, EPI_ISL_447793, EPI_ISL_447794, EPI_ISL_447795, EPI_ISL_447796, EPI_ISL_447797, EPI_ISL_447798, EPI_ISL_447799, EPI_ISL_447800, EPI_ISL_447801, EPI_ISL_447802, EPI_ISL_447803, EPI_ISL_447804, EPI_ISL_447805, EPI_ISL_447806, EPI_ISL_447807, EPI_ISL_447808, EPI_ISL_447809, EPI_ISL_447810, EPI_ISL_447811, EPI_ISL_447812, EPI_ISL_447813, EPI_ISL_447814, EPI_ISL_447815, EPI_ISL_447816, EPI_ISL_447817 | | | |
| see above | Instituto Nacional de Salud, Bogotá, Colombia | Grupo de Investigaciones Microbiológicas-UR (GIMUR), Departamento de Biología, Facultad de Ciencias Naturales, Universidad del Rosario, Bogotá, Colombia Instituto Nacional de Salud, Bogotá, Colombia Icahn School of Medicine at Mount Sinai, New York, USA | Juan David Ramírez, Carolina Florez, Marina Muñoz, Carolina Hernandez, Adriana Castillo, Sergio Castañeda, Nathalia Ballesteros, David Martínez, Laura Vega, Jesús E. Jaimes, Sergio Gomez, Angelica Rico, Lisseth Pardo, Esther C. Barros, Martha L. Ospina, Aníbal A. Teherán, Ana S. Gonzalez-Reiche, Matthew M. Hernandez, Emilia Mia Sordillo, Viviana Simon, Harm van Bakel, Alberto Paniz-Mondolfi |
| EPI_ISL_447832, EPI_ISL_447833, EPI_ISL_447834, EPI_ISL_447835, EPI_ISL_447836 | unknown | Department of Medicine | Kassela,K., Dovrolis,N., Bampali,M., Gatzidou,E., Froukala,E., Stavropoulou,A., Veletza,S., Tsakris,A., Spanakis,N. and Karakasiliotis,I. |
| EPI_ISL_447837 | Dept. of Medical Microbiology, Stavanger University Hospital, Helse Stavanger HF, | Norwegian Institute of Public Health, Department of Virology | Kathrine Stene-Johansen, Kamilla Heddeland Instefjord, Hilde Elshaug, Rasmus Riis Kopperud, Karoline Bragstad, Olav Hungnes |
| EPI_ISL_447838, EPI_ISL_447839 | Medical Microbiology Unit, Department for Laboratory Medicine, Drammen Hospital, Vestre Viken Health Trust, | Norwegian Institute of Public Health, Department of Virology | Kathrine Stene-Johansen, Kamilla Heddeland Instefjord, Hilde Elshaug, Rasmus Riis Kopperud, Karoline Bragstad, Olav Hungnes |
| EPI_ISL_447840 | DC Public Health Lab/ Dept. of Forensic Sciences | Pathogen Discovery, Respiratory Viruses Branch, Division of Viral Diseases, Centers for Disease Control and Prevention | Krista Queen, Yan Li, Anna Uehara, Jing Zhang, Ying Tao, Clinton R. Paden, Haibin Wang, Jasmine Padilla, Mary S. Keckler, Alison S. Laufer Halpin, Justin Lee, Christopher A. Elkins, Suxiang Tong |
| EPI_ISL_447841 | FL Bureau of Public Health Laboratories-Tampa | Pathogen Discovery, Respiratory Viruses Branch, Division of Viral Diseases, Centers for Disease Control and Prevention | Krista Queen, Yan Li, Anna Uehara, Jing Zhang, Ying Tao, Clinton R. Paden, Haibin Wang, Jasmine Padilla, Mary S. Keckler, Alison S. Laufer Halpin, Justin Lee, Christopher A. Elkins, Suxiang Tong |
| EPI_ISL_447842 | IA State Hygienic Laboratory | Pathogen Discovery, Respiratory Viruses Branch, Division of Viral Diseases, Centers for Disease Control and Prevention | Krista Queen, Yan Li, Anna Uehara, Jing Zhang, Ying Tao, Clinton R. Paden, Haibin Wang, Jasmine Padilla, Mary S. Keckler, Alison S. Laufer Halpin, Justin Lee, Christopher A. Elkins, Suxiang Tong |
| EPI_ISL_447843 | MD DOH Laboratories Administration | Pathogen Discovery, Respiratory Viruses Branch, Division of Viral Diseases, Centers for Disease Control and Prevention | Krista Queen, Yan Li, Anna Uehara, Jing Zhang, Ying Tao, Clinton R. Paden, Haibin Wang, Jasmine Padilla, Mary S. Keckler, Alison S. Laufer Halpin, Justin Lee, Christopher A. Elkins, Suxiang Tong |
| EPI_ISL_447844 | PA Department of Health, Bureau of Laboratories | Pathogen Discovery, Respiratory Viruses Branch, Division of Viral Diseases, Centers for Disease Control and Prevention | Krista Queen, Yan Li, Anna Uehara, Jing Zhang, Ying Tao, Clinton R. Paden, Haibin Wang, Jasmine Padilla, Mary S. Keckler, Alison S. Laufer Halpin, Justin Lee, Christopher A. Elkins, Suxiang Tong |
| EPI_ISL_447845 | PR - Biological and Chemical Emergencies Lab Office of Public Health Preparedness and Response | Pathogen Discovery, Respiratory Viruses Branch, Division of Viral Diseases, Centers for Disease Control and Prevention | Krista Queen, Yan Li, Anna Uehara, Jing Zhang, Ying Tao, Clinton R. Paden, Haibin Wang, Jasmine Padilla, Mary S. Keckler, Alison S. Laufer Halpin, Justin Lee, Christopher A. Elkins, Suxiang Tong |
| EPI_ISL_447846 | VT Dept. of Health Laboratory | Pathogen Discovery, Respiratory Viruses Branch, Division of Viral Diseases, Centers for Disease Control and Prevention | Krista Queen, Yan Li, Anna Uehara, Jing Zhang, Ying Tao, Clinton R. Paden, Haibin Wang, Jasmine Padilla, Mary S. Keckler, Alison S. Laufer Halpin, Justin Lee, Christopher A. Elkins, Suxiang Tong |
| EPI_ISL_447851, EPI_ISL_447852, EPI_ISL_447855 | CSIR-Centre for Cellular and Molecular Biology | CSIR-Centre for Cellular and Molecular Biology | Lamuk Zaveri, Shagufta Khan, Namami Gaur, Sakshi Shambhavi, Tulasi Nagabandi, Purushotham Vodnala, Payel Mukherjee, Sofia Banu, Priya Singh, Dhiviya Vedagiri, Divya Gupta, Vishal Sah, Santosh Kumar Kuncha, Krishnan Harinivas Harshan, Archana Bharadwaj Siva, Karthik Bharadwaj Tallapaka, Rakesh K Mishra, Divya Tej Sowpati |
| EPI_ISL_449786 | Ostfold Hospital Trust - Kalnes, Centre for Laboratory Medicine, Section for gene technology and infection serology | Norwegian Institute of Public Health, Department of Virology | Kathrine Stene-Johansen, Kamilla Heddeland Instefjord, Hilde Elshaug, Rasmus Riis Kopperud, Karoline Bragstad, Olav Hungnes |
| EPI_ISL_449787, EPI_ISL_449788 | Furst Medical Laboratory | Norwegian Institute of Public Health, Department of Virology | Kathrine Stene-Johansen, Kamilla Heddeland Instefjord, Hilde Elshaug, Rasmus Riis Kopperud, Karoline Bragstad, Olav Hungnes |
| EPI_ISL_449789, EPI_ISL_449790, EPI_ISL_449791, EPI_ISL_449792, EPI_ISL_449793, EPI_ISL_449794 | Dept. of Medical Microbiology, Stavanger University Hospital, Helse Stavanger HF | Norwegian Institute of Public Health, Department of Virology | Kathrine Stene-Johansen, Kamilla Heddeland Instefjord, Hilde Elshaug, Rasmus Riis Kopperud, Karoline Bragstad, Olav Hungnes |
| EPI_ISL_450171, EPI_ISL_450172 | National Influenza centre, Sri Lanka | The University of Hong Kong | Jude Jayamaha, Daniel KW Chu, Malik Peiris |
| EPI_ISL_450747 | Sunnybrook Health Sciences Centre | Department of Laboratory Medicine and Molecular Diagnostics, Sunnybrook Health Sciences Centre | Jalees A. Nasir, Robert A. Kozak, Patryk Aftanas, Amogelang R. Raphenya, Kendrick M. Smith, Finlay Maguire, Hassaan Maan, Muhannad Alruwaili, Arinjay Banerjee, Hamza Mbareche, Brian P. Alcock, Natalie C. Knox, Karen Mossman, Bo Wang, Julian A. Hiscox, Andrew G. McArthur, Samira Mubareka |
| EPI_ISL_450781 | Government Medical College-Bhavnagar | Gujarat Biotechnology Research Centre | Kairavi Desai, Saklain Malek, Shirish Patel, Ramesh Pandit, Tejas Shah, Ankit Hinsu, Pritesh Sabara, Apurvasinh Puvar, Janvi Raval, Zarna Patel, Monika Gandhi, Pinal Trivedi, Maharshi Pandya, Amit Kanani, Nidhi Patel, Nitin Savaliya, Raghawendra Kumar, Dinesh Kumar, Zuber Saiyed, Komal Patel, Labdhi Pandya, Snehal Bagatharia, Bhavesh Modi, Gaurishankar Shrimali, R D Dixit, A M Kadri, Akanksha Verma, Chaitanya Joshi, Madhvi Joshi |
| EPI_ISL_450792, EPI_ISL_450798 | Jamaica Ministry of Health and Wellness | Pathogen Discovery, Respiratory Viruses Branch, Division of Viral Diseases, Centers for Disease Control and Prevention | Krista Queen, Yan Li, Anna Montmayeur, Ying Tao, Jing Zhang, Anna Uehara, Clinton R. Paden, Rachel Marine, Haibin Wang, Jasmine Padilla, Justin Lee, Zachary Weiner, Bettina Bankamp, Suxiang Tong |

| | | | |
|---|---|---|---|
| EPI_ISL_450805 | VI-US Virgin Islands Department of Health | Pathogen Discovery, Respiratory Viruses Branch, Division of Viral Diseases, Centers for Disease Control and Prevention | Krista Queen, Yan Li, Anna Montmayeur, Ying Tao, Jing Zhang, Anna Uehara, Clinton R. Paden, Rachel Marine, Haibin Wang, Jasmine Padilla, Justin Lee, Zachary Weiner, Bettina Bankamp, Suxiang Tong |
| EPI_ISL_450808, EPI_ISL_450810 | Saroledens Familjelakare | The Public Health Agency of Sweden | Katarina Jarbur, Anna-Malin Linde, Maria Lind Karlberg, Oskar Karlsson Lindsjo, Olov Svartstrom, Anna Risberg, Theresa Enkirch, Mia Brytting, Karin Tegmark-Wisell |
| EPI_ISL_450811 | Knivsta VC | The Public Health Agency of Sweden | Johanna Carlson, Anna-Malin Linde, Maria Lind Karlberg, Oskar Karlsson Lindsjo, Olov Svartstrom, Anna Risberg, Theresa Enkirch, Mia Brytting, Karin Tegmark-Wisell |
| EPI_ISL_450814 | Huslakarna Varmbadhuset Varberg | The Public Health Agency of Sweden | Johanna Hilmersson, Anna-Malin Linde, Maria Lind Karlberg, Oskar Karlsson Lindsjo, Olov Svartstrom, Anna Risberg, Theresa Enkirch, Mia Brytting, Karin Tegmark-Wisell |
| EPI_ISL_450818 | Aneby VC | The Public Health Agency of Sweden | Ken Granath, Anna-Malin Linde, Maria Lind Karlberg, Oskar Karlsson Lindsjo, Olov Svartstrom, Anna Risberg, Theresa Enkirch, Mia Brytting, Karin Tegmark-Wisell |
| EPI_ISL_451149 | M.P Shah Government Medocal college Jamnagar | Gujarat Biotechnology Research Centre | Janvi Raval, Zarna Patel, Monika Gandhi, Pinal Trivedi, Maharshi Pandya, Amit Kanani, Nidhi Patel, Nitin Savaliya, Raghawendra Kumar, Dinesh Kumar, Zuber Saiyed, Komal Patel, Labdhi Pandya, Snehal Bagatharia, Ramesh Pandit, Tejas Shah, Ankit Hinsu, Pritesh Sabara, Apurvasinh Puvar, Binita Aring, Bhavesh Modi, Gaurishankar Shrimali, R D Dixit, A M Kadri, Priti Pandita, Chaitanya Joshi, Madhvi Joshi, |
| EPI_ISL_451162 | Government Medical College, Vadodara | Gujarat Biotechnology Research Centre | Komal Patel, Labdhi Pandya, Snehal Bagatharia, Ramesh Pandit, Tejas Shah, Ankit Hinsu, Pritesh Sabara, Apurvasinh Puvar, Janvi Raval, Zarna Patel, Monika Gandhi, Pinal Trivedi, Maharshi Pandya, Manish Pattani, Tanuja Javadekar , Amit Kanani, Nidhi Patel, Nitin Savaliya, Raghawendra Kumar, Dinesh Kumar, Zuber Saiyed, Bhavesh Modi, Gaurishankar Shrimali, R D Dixit, A M Kadri, Priti Pandita, Chaitanya Joshi, Madhvi Joshi |
| EPI_ISL_451163 | Government Medical College, Vadodara | Gujarat Biotechnology Research Centre | Labdhi Pandya, Snehal Bagatharia, Ramesh Pandit, Tejas Shah, Ankit Hinsu, Pritesh Sabara, Apurvasinh Puvar, Janvi Raval, Zarna Patel, Monika Gandhi, Pinal Trivedi, Maharshi Pandya, Manish Pattani, Tanuja Javadekar , Amit Kanani, Nidhi Patel, Nitin Savaliya, Raghawendra Kumar, Dinesh Kumar, Zuber Saiyed, Komal Patel, Bhavesh Modi, Gaurishankar Shrimali, R D Dixit, A M Kadri, Pragya Sharma, Chaitanya Joshi, Madhvi Joshi |
| EPI_ISL_451203, EPI_ISL_451204, EPI_ISL_451205, EPI_ISL_451206, EPI_ISL_451207, EPI_ISL_451208, EPI_ISL_451209, EPI_ISL_451210, EPI_ISL_451211, EPI_ISL_451212, EPI_ISL_451213, EPI_ISL_451214, EPI_ISL_451215, EPI_ISL_451216, EPI_ISL_451217, EPI_ISL_451218 | | | |
| see above | LSUHS Emerging Viral Threat Laboratory | Microbial Genome Sequencing Center | Jeremy P. Kamil, John A. Vanchiere, Rona S. Scott, Camille F. Abshire, Abida Siddiqa, Byeong-Jae Lee, Chan-ki Min, Md Maksudul Alam, Monica Gestal-Cartele, Edna Ondari, Adam Greer, Malgorzata Bienkowska-Haba, Katarzyna Zwolinska, Jason M. Bodily, Andrew D. Yurochko, Paul M. Weinberger, Christopher G. Kevil, Martin J. Sapp, Daniel J. Snyder, Vaughn S. Cooper |
| EPI_ISL_451230, EPI_ISL_451231, EPI_ISL_451232, EPI_ISL_451233, EPI_ISL_451234, EPI_ISL_451235, EPI_ISL_451236, EPI_ISL_451237, EPI_ISL_451238, EPI_ISL_451239, EPI_ISL_451240, EPI_ISL_451241, EPI_ISL_451242, EPI_ISL_451243, EPI_ISL_451244, EPI_ISL_451245, EPI_ISL_451246, EPI_ISL_451247 | | | |
| see above | LSUHS Emerging Viral Threat Laboratory | Microbial Genome Sequencing Center | John A. Vanchiere, Jeremy P. Kamil, Rona S. Scott, Camille F. Abshire, Abida Siddiqa, Byeong-Jae Lee, Chan-ki Min, Md Maksudul Alam, Monica Gestal-Cartele, Edna Ondari, Adam Greer, Malgorzata Bienkowska-Haba, Katarzyna Zwolinska, Jason M. Bodily, Andrew D. Yurochko, Paul M. Weinberger, Christopher G. Kevil, Martin J. Sapp, Daniel J. Snyder, Vaughn S. Cooper |
| EPI_ISL_451248, EPI_ISL_451249, EPI_ISL_451250, EPI_ISL_451251, EPI_ISL_451252, EPI_ISL_451253, EPI_ISL_451254, EPI_ISL_451255, EPI_ISL_451256, EPI_ISL_451257, EPI_ISL_451258, EPI_ISL_451259, EPI_ISL_451260, EPI_ISL_451261, EPI_ISL_451262, EPI_ISL_451263, EPI_ISL_451264, EPI_ISL_451265, EPI_ISL_451266, EPI_ISL_451267, EPI_ISL_451268, EPI_ISL_451269, EPI_ISL_451270, EPI_ISL_451271, EPI_ISL_451272, EPI_ISL_451273, EPI_ISL_451274, EPI_ISL_451275, EPI_ISL_451276, EPI_ISL_451277, EPI_ISL_451278, EPI_ISL_451279, EPI_ISL_451280, EPI_ISL_451281, EPI_ISL_451282, EPI_ISL_451283, EPI_ISL_451284, EPI_ISL_451285, EPI_ISL_451286, EPI_ISL_451287, EPI_ISL_451288, EPI_ISL_451289, EPI_ISL_451290, EPI_ISL_451291, EPI_ISL_451292, EPI_ISL_451293, EPI_ISL_451294, EPI_ISL_451295, EPI_ISL_451296, EPI_ISL_451297 | | | |
| see above | LSUHS Emerging Viral Threat Laboratory | Microbial Genome Sequencing Center | Jeremy P. Kamil, John A. Vanchiere, Rona S. Scott, Camille F. Abshire, Abida Siddiqa, Byeong-Jae Lee, Chan-ki Min, Md Maksudul Alam, Monica Gestal-Cartele, Edna Ondari, Adam Greer, Malgorzata Bienkowska-Haba, Katarzyna Zwolinska, Jason M. Bodily, Andrew D. Yurochko, Paul M. Weinberger, Christopher G. Kevil, Martin J. Sapp, Daniel J. Snyder, Vaughn S. Cooper |
| EPI_ISL_452120 | Georgia Department of Health | Pathogen Discovery, Respiratory Viruses Branch, Division of Viral Diseases, Centers for Disease Control and Prevention | Jing Zhang, Anna Montmayeur, Yan Li, Ying Tao, Krista Queen, Anna Uehara, Clinton R. Paden, Rachel Marine, Mary S. Keckler, Alison S. Laufer Halpin, Haibin Wang, Christopher A. Elkins, Zachary Weiner, Suxiang Tong |
| EPI_ISL_452121, EPI_ISL_452122, EPI_ISL_452123 | VI-US Virgin Islands Department of Health | Pathogen Discovery, Respiratory Viruses Branch, Division of Viral Diseases, Centers for Disease Control and Prevention | Jing Zhang, Anna Montmayeur, Yan Li, Ying Tao, Krista Queen, Anna Uehara, Clinton R. Paden, Rachel Marine, Mary S. Keckler, Alison S. Laufer Halpin, Haibin Wang, Christopher A. Elkins, Zachary Weiner, Suxiang Tong |
| EPI_ISL_452124 | NC State Laboratory of Public Health | Pathogen Discovery, Respiratory Viruses Branch, Division of Viral Diseases, Centers for Disease Control and Prevention | Jing Zhang, Anna Montmayeur, Yan Li, Ying Tao, Krista Queen, Anna Uehara, Clinton R. Paden, Rachel Marine, Mary S. Keckler, Alison S. Laufer Halpin, Haibin Wang, Christopher A. Elkins, Zachary Weiner, Suxiang Tong |
| EPI_ISL_452125 | Georgia Department of Health | Pathogen Discovery, Respiratory Viruses Branch, Division of Viral Diseases, Centers for Disease Control and Prevention | Jing Zhang, Anna Montmayeur, Yan Li, Ying Tao, Krista Queen, Anna Uehara, Clinton R. Paden, Rachel Marine, Mary S. Keckler, Alison S. Laufer Halpin, Haibin Wang, Christopher A. Elkins, Zachary Weiner, Suxiang Tong |
| EPI_ISL_452126 | NC State Laboratory of Public Health | Pathogen Discovery, Respiratory Viruses Branch, Division of Viral Diseases, Centers for Disease Control and Prevention | Jing Zhang, Anna Montmayeur, Yan Li, Ying Tao, Krista Queen, Anna Uehara, Clinton R. Paden, Rachel Marine, Mary S. Keckler, Alison S. Laufer Halpin, Haibin Wang, Christopher A. Elkins, Zachary Weiner, Suxiang Tong |
| EPI_ISL_452127, EPI_ISL_452128, EPI_ISL_452129, EPI_ISL_452130 | CO Department of Public Health and Environment | Pathogen Discovery, Respiratory Viruses Branch, Division of Viral Diseases, Centers for Disease Control and Prevention | Jing Zhang, Anna Montmayeur, Yan Li, Ying Tao, Krista Queen, Anna Uehara, Clinton R. Paden, Rachel Marine, Mary S. Keckler, Alison S. Laufer Halpin, Haibin Wang, Christopher A. Elkins, Zachary Weiner, Suxiang Tong |
| EPI_ISL_452131 | IN State Department of Health Laboratory Services | Pathogen Discovery, Respiratory Viruses Branch, Division of Viral Diseases, Centers for Disease Control and Prevention | Jing Zhang, Anna Montmayeur, Yan Li, Ying Tao, Krista Queen, Anna Uehara, Clinton R. Paden, Rachel Marine, Mary S. Keckler, Alison S. Laufer Halpin, Haibin Wang, Christopher A. Elkins, Zachary Weiner, Suxiang Tong |
| EPI_ISL_452132 | FL Bureau of Public Health Laboratories | Pathogen Discovery, Respiratory Viruses Branch, Division of Viral Diseases, Centers for Disease Control and Prevention | Krista Queen, Yan Li, Anna Montmayeur, Ying Tao, Jing Zhang, Anna Uehara, Clinton R. Paden, Rachel Marine, Mary S. Keckler, Alison S. Laufer Halpin, Haibin Wang, Christopher A. Elkins, Zachary Weiner, Suxiang Tong |
| EPI_ISL_452133 | MN Department of Health | Pathogen Discovery, Respiratory Viruses Branch, Division of Viral Diseases, Centers for Disease Control and Prevention | Krista Queen, Yan Li, Anna Montmayeur, Ying Tao, Jing Zhang, Anna Uehara, Clinton R. Paden, Rachel Marine, Mary S. Keckler, Alison S. Laufer Halpin, Haibin Wang, Christopher A. Elkins, Zachary Weiner, Suxiang Tong |
| EPI_ISL_452134 | IL Department of Public Health Chicago Laboratory | Pathogen Discovery, Respiratory Viruses Branch, Division of Viral Diseases, Centers for Disease Control and Prevention | Krista Queen, Yan Li, Anna Montmayeur, Ying Tao, Jing Zhang, Anna Uehara, Clinton R. Paden, Rachel Marine, Mary S. Keckler, Alison S. Laufer Halpin, Haibin Wang, Christopher A. Elkins, Zachary Weiner, Suxiang Tong |
| EPI_ISL_452135, EPI_ISL_452136 | IN State Department of Health Laboratory Services | Pathogen Discovery, Respiratory Viruses Branch, Division of Viral Diseases, Centers for Disease Control and Prevention | Krista Queen, Yan Li, Anna Montmayeur, Ying Tao, Jing Zhang, Anna Uehara, Clinton R. Paden, Rachel Marine, Mary S. Keckler, Alison S. Laufer Halpin, Haibin Wang, Christopher A. Elkins, Zachary Weiner, Suxiang Tong |
| EPI_ISL_452143, EPI_ISL_452144, EPI_ISL_452145, EPI_ISL_452146, EPI_ISL_452147 | Yale COVID-19 Biorepository | Grubaugh Lab - Yale School of Public Health | Joseph Fauver, Tara Alpert, Anderson Brito, Anne Wyllie, Chantal Vogels, Mary Petrone, Cole Jensen, Chaney Kalinich, Isabel Ott, Arnau Casanovas, Catherine Muenker, Adam Moore, Alice Lu, Maria Tokuyama, Patrick Wong, Peiwen Lu, Saad Omer, Richard Martinello, Allison Nelson, Shelli Farhadian, Akiko Iwasaki, Charlese Dela Cruz, Albert Ko, Nathan Grubaugh |
| EPI_ISL_452178, EPI_ISL_452179 | Laboratory Medicine | Department of Laboratory Medicine, Lin-Kou Chang Gung Memorial Hospital, Taoyuan, Taiwan | Kuo-Chien Tsao, Yu-Nong Gong, Shu-Li Yang, Yi-Chun Liu, Chung-Guei Huang, Mei-Jen Hsiao, Po-Wei Huang, Cheng-Ta Yang, Cheng-Hsun Chiu, Peng-Nien Huang, Kuo-Ming Lee, Guang-Wu Chen, Shin-Ru Shih |
| EPI_ISL_452231, EPI_ISL_452232 | Saroledens Familjelakare | The Public Health Agency of Sweden | Katarina Jarbur, Anna-Malin Linde, Maria Lind Karlberg, Oskar Karlsson Lindsjo, Olov Svartstrom, Anna Risberg, Theresa Enkirch, Mia Brytting, Karin Tegmark-Wisell |
| EPI_ISL_452241 | Huslakarna Varmbadhuset Varberg | The Public Health Agency of Sweden | Johanna Hilmersson, Anna-Malin Linde, Maria Lind Karlberg, Oskar Karlsson Lindsjo, Olov Svartstrom, Anna Risberg, Theresa Enkirch, Mia Brytting, Karin Tegmark-Wisell |
| EPI_ISL_454642 | CT-Dr. Katherine A. Kelley State Public Health Lab | Pathogen Discovery, Respiratory Viruses Branch, Division of Viral Diseases, Centers for Disease Control and Prevention | Jing Zhang, Ying Tao, Clinton R. Paden, Anna Uehara, Krista Queen, Yan Li, Haibin Wang, Zachary Weiner, Bettina Bankamp, Suxiang Tong |
| EPI_ISL_454643, EPI_ISL_454644 | VI-US Virgin Islands Department of Health | Pathogen Discovery, Respiratory Viruses Branch, Division of Viral Diseases, Centers for Disease Control and Prevention | Jing Zhang, Ying Tao, Clinton R. Paden, Anna Uehara, Krista Queen, Yan Li, Haibin Wang, Zachary Weiner, Bettina Bankamp, Suxiang Tong |
| EPI_ISL_454645 | CT-Dr. Katherine A. Kelley State Public Health Lab | Pathogen Discovery, Respiratory Viruses Branch, Division of Viral Diseases, Centers for Disease Control and Prevention | Jing Zhang, Ying Tao, Clinton R. Paden, Anna Uehara, Krista Queen, Yan Li, Haibin Wang, Zachary Weiner, Bettina Bankamp, Suxiang Tong |
| EPI_ISL_455024 | Government Medical College, Vadodara | Gujarat Biotechnology Research Centre | Janvi Raval, Zarna Patel, Monika Gandhi, Pinal Trivedi, Maharshi Pandya, Amit Kanani, Nidhi Patel, Nitin Savaliya, Raghawendra Kumar, Dinesh Kumar, Zuber Saiyed, Komal Patel, Labdhi Pandya, Snehal Bagatharia, Tanuja Javadekar , R N Daveshwar, Ramesh Pandit, Tejas Shah, Ankit Hinsu, Pritesh |

| | | | |
|---|---|---|---|
| | | | Sabara, Apurvasinh Puvar, Pooja P Doshi, Bhavesh Modi, Gaurishankar Shrimali, R D Dixit, A M Kadri, Umang Mishra, Chaitanya Joshi, Madhvi Joshi, |
| EPI_ISL_455102 | Kungsors VC | The Public Health Agency of Sweden | Jessica Karlsson, Anna-Malin Linde, Maria Lind Karlberg, Oskar Karlsson Lindsjo, Olov Svartstrom, Anna Risberg, Theresa Enkirch, Mia Brytting, Karin Tegmark-Wisell |
| EPI_ISL_455103 | Huslakarna Varmbadhuset Varberg | The Public Health Agency of Sweden | Johanna Hilmersson, Anna-Malin Linde, Maria Lind Karlberg, Oskar Karlsson Lindsjo, Olov Svartstrom, Anna Risberg, Theresa Enkirch, Mia Brytting, Karin Tegmark-Wisell |
| EPI_ISL_455440, EPI_ISL_455441, EPI_ISL_455442, EPI_ISL_455443, EPI_ISL_455444, EPI_ISL_455445, EPI_ISL_455446, EPI_ISL_455447, EPI_ISL_455448, EPI_ISL_455449, EPI_ISL_455450, EPI_ISL_455451, EPI_ISL_455452, EPI_ISL_455453 | | | |
| see above | 1. ViroGenetics - BSL3 Laboratory of Virology, Maopolska Centre of Biotechnology, Jagiellonian University; 2. II Department of Internal Medicine, Faculty of Medicine, Jagiellonian University Medical College; 3. Narodowy Instytut Zdrowia Publicznego - Pastwowy Zakad Higieny (NIZP-PZH) | 1. ViroGenetics - BSL3 Laboratory of Virology, Maopolska Centre of Biotechnology, Jagiellonian University; 2. II Department of Internal Medicine, Faculty of Medicine, Jagiellonian University Medical College; 3. Narodowy Instytut Zdrowia Publicznego - Pastwowy Zakad Higieny (NIZP-PZH). | Katarzyna Pancer, Marek Sanak, Aleksandra A. Zasada, Magdalena Rzeczkowska, Tomasz Wokowicz, Katarzyna Zacharczuk, Agnieszka Koakowska-Kulesza, Katarzyna Owczarek, Aleksandra Milewska, Natalia Wolaniuk, Ewelina Hallman-Szeliska, Pawe P abaj, Wojciech Branicki, Krzysztof Pyr |
| EPI_ISL_455460, EPI_ISL_455461, EPI_ISL_455462, EPI_ISL_455463, EPI_ISL_455464, EPI_ISL_455465, EPI_ISL_455466, EPI_ISL_455467 | Jiangxi Province Center for Disease Control and Prevention | Jiangxi Province Center for Disease Control and Prevention | JianXiong Li,Ying Xiong,Tian Gong,Yong Shi,Jun Zhou,Fang Xiao,ShiWen Liu,XiaoQing Liu,Gang Xu,DaJin Xiao,Xin Ran,YanNi Zhang |
| EPI_ISL_455958, EPI_ISL_455959, EPI_ISL_455960, EPI_ISL_455961, EPI_ISL_455962, EPI_ISL_455963, EPI_ISL_455964, EPI_ISL_455965, EPI_ISL_455966, EPI_ISL_455967, EPI_ISL_455968, EPI_ISL_455969, EPI_ISL_455970, EPI_ISL_455971, EPI_ISL_455972, EPI_ISL_455973, EPI_ISL_455974, EPI_ISL_455975, EPI_ISL_455976, EPI_ISL_455977, EPI_ISL_455978, EPI_ISL_455979 | | | |
| see above | Department of Clinical Microbiology | GIGA Medical Genomics | Keith Durkin, Maria Artesi, Sébastien Bontems, Raphaël Boreux, Cécile Meex, Pierrette Melin, Marie-Pierre Hayette, Vincent Bours. |
| EPI_ISL_455980, EPI_ISL_455981, EPI_ISL_455982, EPI_ISL_455983, EPI_ISL_455984, EPI_ISL_455985, EPI_ISL_455986, EPI_ISL_455987, EPI_ISL_455988, EPI_ISL_455989 | LSUHS Emerging Viral Threat Laboratory | Microbial Genome Sequencing Center | Jeremy P. Kamil, John A. Vanchiere, Rona S. Scott, Camille F. Abshire, Abida Siddiqa, Byeong-Jae Lee, Chan-ki Min, Md Maksudul Alam, Monica Gestal-Cartele, Edna Ondari, Adam Greer, Malgorzata Bienkowska-Haba, Katarzyna Zwolinska, Michelle M. Arnold, Jason M. Bodily, Andrew D. Yurochko, Paul M. Weinberger, Christopher G. Kevil, Martin J. Sapp, Daniel J. Snyder, Vaughn S. Cooper |
| EPI_ISL_455990, EPI_ISL_455991, EPI_ISL_455992, EPI_ISL_455993, EPI_ISL_455994, EPI_ISL_455995, EPI_ISL_455996, EPI_ISL_455997, EPI_ISL_455998, EPI_ISL_455999 | LSUHS Emerging Viral Threat Laboratory | Microbial Genome Sequencing Center | John A. Vanchiere, Jeremy P. Kamil, Rona S. Scott, Camille F. Abshire, Abida Siddiqa, Byeong-Jae Lee, Chan-ki Min, Md Maksudul Alam, Monica Gestal-Cartele, Edna Ondari, Adam Greer, Malgorzata Bienkowska-Haba, Katarzyna Zwolinska, Michelle M. Arnold, Jason M. Bodily, Andrew D. Yurochko, Paul M. Weinberger, Christopher G. Kevil, Martin J. Sapp, Daniel J. Snyder, Vaughn S. Cooper |
| EPI_ISL_456116, EPI_ISL_456117, EPI_ISL_456118, EPI_ISL_456119, EPI_ISL_456120, EPI_ISL_456121, EPI_ISL_456122, EPI_ISL_456123, EPI_ISL_456124, EPI_ISL_456125, EPI_ISL_456126, EPI_ISL_456127, EPI_ISL_456128, EPI_ISL_456129, EPI_ISL_456130, EPI_ISL_456131, EPI_ISL_456132, EPI_ISL_456133, EPI_ISL_456134, EPI_ISL_456135, EPI_ISL_456136, EPI_ISL_456137, EPI_ISL_456138, EPI_ISL_456139, EPI_ISL_456140, EPI_ISL_456141, EPI_ISL_456142, EPI_ISL_456143, EPI_ISL_456144, EPI_ISL_456145, EPI_ISL_456146, EPI_ISL_456147, EPI_ISL_456148, EPI_ISL_456149, EPI_ISL_456150, EPI_ISL_456151, EPI_ISL_456152, EPI_ISL_456153, EPI_ISL_456154, EPI_ISL_456155, EPI_ISL_456156 | | | |
| see above | Instituto Nacional de Salud - Unidad de Secuenciación y Análisis Genómico | Instituto Nacional de Salud, Universidad Cooperativa de Colombia, Instituto Alexander von Humboldt, Imperial College-London, London School of Hygiene & Tropical Medicine | Katherine Laiton-Donato, Diego A. Álvarez-Díaz, Carlos Franco-Muñoz, Jose A. Usme-Ciro, Gloria Puerto, Nicolas D. Franco-Sierra, Mailyn A.Gonzalez, Zulma M. Cucunubá, Christian Julian VillabonaArenas, Liz Villabona-Arenas, Sussy Echeverria, Astrid C. Flórez, Sergio Gomez-Rangel, Luz Dary Rodriguez, Juliana Barbosa, Erika Ospitia, Diana Marcela Walteros-Acero, Martha Lucia Ospina Martinez, Marcela Mercado-Reyes. |
| EPI_ISL_458030 | King Institute of Preventive Medicine & Research | CSIR-Centre for Cellular and Molecular Biology | K.Kaveri,S.Sivasubramanian,S.Vennila,P.Padmapriya,R.Kiruba,S.Magesh,G. Dhinakar Raj, G. Ravikumar, P. Azhahianambi,K Thangaraj,Payel Mukherjee, Sofia Banu, Priya Singh, Dhiviya Vedagiri, Divya Gupta, Vishal Sah, Santosh Kumar Kuncha, Krishnan Harinivas Harshan, Archana Bharadwaj Siva, Karthik Bharadwaj Tallapaka, Shagufta Khan, Lamuk Zaveri, Namami Gaur, Sakshi Shambhavi, Tulasi Nagabandi, Purushotham Vodnala, Rakesh K Mishra, Divya Tej Sowpati |
| EPI_ISL_458031 | King Institute of Preventive Medicine & Research | CSIR-Centre for Cellular and Molecular Biology | K.Kaveri,S.Sivasubramanian,S.Vennila,P.Padmapriya,R.Kiruba,S.Magesh,G. Dhinakar Raj, G. Ravikumar, P. Azhahianambi, K Thangaraj,Sofia Banu, Payel Mukherjee, Priya Singh, Dhiviya Vedagiri, Divya Gupta, Vishal Sah, Santosh Kumar Kuncha, Krishnan Harinivas Harshan, Archana Bharadwaj Siva, Karthik Bharadwaj Tallapaka, Shagufta Khan, Lamuk Zaveri, Namami Gaur, Sakshi Shambhavi, Tulasi Nagabandi, Purushotham Vodnala, Rakesh K Mishra, Divya Tej Sowpati |
| EPI_ISL_458032 | King Institute of Preventive Medicine & Research | CSIR-Centre for Cellular and Molecular Biology | K.Kaveri,S.Sivasubramanian,S.Vennila,P.Padmapriya,R.Kiruba,S.Magesh,G. Dhinakar Raj, G. Ravikumar, P. Azhahianambi, K Thangaraj,Shagufta Khan, Lamuk Zaveri, Namami Gaur, Sakshi Shambhavi, Tulasi Nagabandi, Purushotham Vodnala, Payel Mukherjee, Sofia Banu, Priya Singh, Dhiviya Vedagiri, Divya Gupta, Vishal Sah, Santosh Kumar Kuncha, Krishnan Harinivas Harshan, Archana Bharadwaj Siva, Karthik Bharadwaj Tallapaka, Rakesh K Mishra, Divya Tej Sowpati |
| EPI_ISL_458033 | King Institute of Preventive Medicine & Research | CSIR-Centre for Cellular and Molecular Biology | K.Kaveri,S.Sivasubramanian,S.Vennila,P.Padmapriya,R.Kiruba,S.Magesh,G. Dhinakar Raj, G. Ravikumar, P. Azhahianambi, K Thangaraj,Lamuk Zaveri, Shagufta Khan, Namami Gaur, Sakshi Shambhavi, Tulasi Nagabandi, Purushotham Vodnala, Payel Mukherjee, Sofia Banu, Priya Singh, Dhiviya Vedagiri, Divya Gupta, Vishal Sah, Santosh Kumar Kuncha, Krishnan Harinivas Harshan, Archana Bharadwaj Siva, Karthik Bharadwaj Tallapaka, Rakesh K Mishra, Divya Tej Sowpati |
| EPI_ISL_458034 | King Institute of Preventive Medicine & Research | CSIR-Centre for Cellular and Molecular Biology | K.Kaveri,S.Sivasubramanian,S.Vennila,P.Padmapriya,R.Kiruba,S.Magesh,G. Dhinakar Raj, G. Ravikumar, P. Azhahianambi, K Thangaraj, Namami Gaur, Sakshi Shambhavi, Lamuk Zaveri, Shagufta Khan, Tulasi Nagabandi, Purushotham Vodnala, Payel Mukherjee, Sofia Banu, Priya Singh, Dhiviya Vedagiri, Divya Gupta, Vishal Sah, Santosh Kumar Kuncha, Krishnan Harinivas Harshan, Archana Bharadwaj Siva, Karthik Bharadwaj Tallapaka, Rakesh K Mishra, Divya Tej Sowpati |
| EPI_ISL_458035 | King Institute of Preventive Medicine & Research | CSIR-Centre for Cellular and Molecular Biology | K.Kaveri,S.Sivasubramanian,S.Vennila,P.Padmapriya,R.Kiruba,S.Magesh,G. Dhinakar Raj, G. Ravikumar, P. Azhahianambi, K Thangaraj, Tulasi Nagabandi, Namami Gaur, Sakshi Shambhavi, Lamuk Zaveri, Shagufta Khan, Purushotham Vodnala, Payel Mukherjee, Sofia Banu, Priya Singh, Dhiviya Vedagiri, Divya Gupta, Vishal Sah, Santosh Kumar Kuncha, Krishnan Harinivas Harshan, Archana Bharadwaj Siva, Karthik Bharadwaj Tallapaka, Rakesh K Mishra, Divya Tej Sowpati |
| EPI_ISL_458036 | King Institute of Preventive Medicine & Research | CSIR-Centre for Cellular and Molecular Biology | K.Kaveri,S.Sivasubramanian,S.Vennila,P.Padmapriya,R.Kiruba,S.Magesh,G. Dhinakar Raj, G. Ravikumar, R. P. Aravindh Babu, K Thangaraj, Payel Mukherjee, Sofia Banu, Priya Singh, Dhiviya Vedagiri, Divya Gupta, Vishal Sah, Santosh Kumar Kuncha, Krishnan Harinivas Harshan, Archana Bharadwaj Siva, Karthik Bharadwaj Tallapaka, Shagufta Khan, Lamuk Zaveri, Namami Gaur, Sakshi Shambhavi, Tulasi Nagabandi, Purushotham Vodnala, Rakesh K Mishra, Divya Tej Sowpati |
| EPI_ISL_458037 | King Institute of Preventive Medicine & Research | CSIR-Centre for Cellular and Molecular Biology | K.Kaveri,S.Sivasubramanian,S.Vennila,P.Padmapriya,R.Kiruba,S.Magesh,G. Dhinakar Raj, G. Ravikumar, R. P. Aravindh Babu, K Thangaraj, Sofia Banu, Payel Mukherjee, Priya Singh, Dhiviya Vedagiri, Divya Gupta, Vishal Sah, Santosh Kumar Kuncha, Krishnan Harinivas Harshan, Archana Bharadwaj Siva, Karthik Bharadwaj Tallapaka, Shagufta Khan, Lamuk Zaveri, Namami Gaur, Sakshi Shambhavi, Tulasi Nagabandi, Purushotham Vodnala, Rakesh K Mishra, Divya Tej Sowpati |
| EPI_ISL_458038 | King Institute of Preventive Medicine & Research | CSIR-Centre for Cellular and Molecular Biology | K.Kaveri,S.Sivasubramanian,S.Vennila,P.Padmapriya,R.Kiruba,S.Magesh,G. Dhinakar Raj, G. Ravikumar, R. P. Aravindh Babu, K Thangaraj, Shagufta Khan, Lamuk Zaveri, Namami Gaur, Sakshi Shambhavi, Tulasi Nagabandi, Purushotham Vodnala, Payel Mukherjee, Sofia Banu, Priya Singh, Dhiviya Vedagiri, Divya Gupta, Vishal Sah, Santosh Kumar Kuncha, Krishnan Harinivas Harshan, Archana Bharadwaj Siva, Karthik Bharadwaj Tallapaka, Rakesh K Mishra, Divya Tej Sowpati |
| EPI_ISL_458039 | King Institute of Preventive Medicine & Research | CSIR-Centre for Cellular and Molecular Biology | K.Kaveri,S.Sivasubramanian,S.Vennila,P.Padmapriya,R.Kiruba,S.Magesh,G. Dhinakar Raj, G. Ravikumar, R. P. Aravindh Babu, K Thangaraj, Lamuk Zaveri, Shagufta Khan, Namami Gaur, Sakshi Shambhavi, Tulasi Nagabandi, Purushotham Vodnala, Payel Mukherjee, Sofia Banu, Priya Singh, Dhiviya Vedagiri, Divya Gupta, Vishal Sah, Santosh Kumar Kuncha, Krishnan Harinivas Harshan, Archana Bharadwaj Siva, Karthik Bharadwaj Tallapaka, Rakesh K Mishra, Divya Tej Sowpati |
| EPI_ISL_458040 | King Institute of Preventive Medicine & Research | CSIR-Centre for Cellular and Molecular Biology | K.Kaveri,S.Sivasubramanian,S.Vennila,P.Padmapriya,R.Kiruba,S.Magesh,G. Dhinakar Raj, G. Ravikumar, R. P. Aravindh Babu, K Thangaraj, Namami |

| | | | |
|---|---|---|---|
| | | | Gaur, Sakshi Shambhavi, Lamuk Zaveri, Shagufta Khan, Tulasi Nagabandi, Purushotham Vodnala, Payel Mukherjee, Sofia Banu, Priya Singh, Dhiviya Vedagiri, Divya Gupta, Vishal Sah, Santosh Kumar Kuncha, Krishnan Harinivas Harshan, Archana Bharadwaj Siva, Karthik Bharadwaj Tallapaka, Rakesh K Mishra, Divya Tej Sowpati |
| EPI_ISL_458041 | King Institute of Preventive Medicine & Research | CSIR-Centre for Cellular and Molecular Biology | K.Kaveri,S.Sivasubramanian,S.Vennila,P.Padmapriya,R.Kiruba,S.Magesh,G. Dhinakar Raj, G. Ravikumar, R. P. Aravindh Babu, K Thangaraj, Tulasi Nagabandi, Namami Gaur, Sakshi Shambhavi, Lamuk Zaveri, Shagufta Khan, Purushotham Vodnala, Payel Mukherjee, Sofia Banu, Priya Singh, Dhiviya Vedagiri, Divya Gupta, Vishal Sah, Santosh Kumar Kuncha, Krishnan Harinivas Harshan, Archana Bharadwaj Siva, Karthik Bharadwaj Tallapaka, Rakesh K Mishra, Divya Tej Sowpati |
| EPI_ISL_458042 | King Institute of Preventive Medicine & Research | CSIR-Centre for Cellular and Molecular Biology | K.Kaveri,S.Sivasubramanian,S.Vennila,P.Padmapriya,R.Kiruba,S.Magesh,G. Dhinakar Raj, G. Ravikumar, M. Sekar, K Thangaraj, Payel Mukherjee, Sofia Banu, Priya Singh, Dhiviya Vedagiri, Divya Gupta, Vishal Sah, Santosh Kumar Kuncha, Krishnan Harinivas Harshan, Archana Bharadwaj Siva, Karthik Bharadwaj Tallapaka, Shagufta Khan, Lamuk Zaveri, Namami Gaur, Sakshi Shambhavi, Tulasi Nagabandi, Purushotham Vodnala, Rakesh K Mishra, Divya Tej Sowpati |
| EPI_ISL_458043 | King Institute of Preventive Medicine & Research | CSIR-Centre for Cellular and Molecular Biology | K.Kaveri,S.Sivasubramanian,S.Vennila,P.Padmapriya,R.Kiruba,S.Magesh,G. Dhinakar Raj, G. Ravikumar, M. Sekar, K Thangaraj,Sofia Banu, Payel Mukherjee, Priya Singh, Dhiviya Vedagiri, Divya Gupta, Vishal Sah, Santosh Kumar Kuncha, Krishnan Harinivas Harshan, Archana Bharadwaj Siva, Karthik Bharadwaj Tallapaka, Shagufta Khan, Lamuk Zaveri, Namami Gaur, Sakshi Shambhavi, Tulasi Nagabandi, Purushotham Vodnala, Rakesh K Mishra, Divya Tej Sowpati |
| EPI_ISL_458044 | King Institute of Preventive Medicine & Research | CSIR-Centre for Cellular and Molecular Biology | K.Kaveri,S.Sivasubramanian,S.Vennila,P.Padmapriya,R.Kiruba,S.Magesh,G. Dhinakar Raj, G. Ravikumar, M. Sekar, K Thangaraj,Shagufta Khan, Lamuk Zaveri, Namami Gaur, Sakshi Shambhavi, Tulasi Nagabandi, Purushotham Vodnala, Payel Mukherjee, Sofia Banu, Priya Singh, Dhiviya Vedagiri, Divya Gupta, Vishal Sah, Santosh Kumar Kuncha, Krishnan Harinivas Harshan, Archana Bharadwaj Siva, Karthik Bharadwaj Tallapaka, Rakesh K Mishra, Divya Tej Sowpati |
| EPI_ISL_458048 | CSIR-Centre for Cellular and Molecular Biology | CSIR-Centre for Cellular and Molecular Biology | Lamuk Zaveri, Shagufta Khan, Namami Gaur, Sakshi Shambhavi, Tulasi Nagabandi, Purushotham Vodnala, Payel Mukherjee, Sofia Banu, Priya Singh, Dhiviya Vedagiri, Divya Gupta, Vishal Sah, Santosh Kumar Kuncha, Krishnan Harinivas Harshan, Archana Bharadwaj Siva, Karthik Bharadwaj Tallapaka, Renu Sudhakar, Somesh Gorde, Gangumala Srinivas Reddy, Sujoy Deb, Swati Bayyana, Rakesh K Mishra, Divya Tej Sowpati |
| EPI_ISL_458054 | CSIR-Centre for Cellular and Molecular Biology | CSIR-Centre for Cellular and Molecular Biology | Lamuk Zaveri, Shagufta Khan, Namami Gaur, Sakshi Shambhavi, Tulasi Nagabandi, Purushotham Vodnala, Payel Mukherjee, Sofia Banu, Priya Singh, Dhiviya Vedagiri, Divya Gupta, Vishal Sah, Santosh Kumar Kuncha, Krishnan Harinivas Harshan, Archana Bharadwaj Siva, Karthik Bharadwaj Tallapaka,Umesh Kumar, Unis Ahmad Bhat, Ajay Sarawagi, Priyanka Pant, Rajkanwar Nathawat, Rakesh K Mishra, Divya Tej Sowpati |
| EPI_ISL_458060 | CSIR-Centre for Cellular and Molecular Biology | CSIR-Centre for Cellular and Molecular Biology | Lamuk Zaveri, Shagufta Khan, Namami Gaur, Sakshi Shambhavi, Tulasi Nagabandi, Purushotham Vodnala, Payel Mukherjee, Sofia Banu, Priya Singh, Dhiviya Vedagiri, Divya Gupta, Vishal Sah, Santosh Kumar Kuncha, Krishnan Harinivas Harshan, Archana Bharadwaj Siva, Karthik Bharadwaj Tallapaka,Zeba Rizvi, Zuberwasim Sayyad, Kakade Aishwarya Arun, Amrutha H C, Ananga Ghosh, Rakesh K Mishra, Divya Tej Sowpati |
| EPI_ISL_458065 | CSIR-Centre for Cellular and Molecular Biology | CSIR-Centre for Cellular and Molecular Biology | Lamuk Zaveri, Shagufta Khan, Namami Gaur, Sakshi Shambhavi, Tulasi Nagabandi, Purushotham Vodnala, Payel Mukherjee, Sofia Banu, Priya Singh, Dhiviya Vedagiri, Divya Gupta, Vishal Sah, Santosh Kumar Kuncha, Krishnan Harinivas Harshan, Archana Bharadwaj Siva, Karthik Bharadwaj Tallapaka,Umesh Kumar, Unis Ahmad Bhat, Ajay Sarawagi, Priyanka Pant, Rajkanwar Nathawat, Rakesh K Mishra, Divya Tej Sowpati |
| EPI_ISL_458081 | RSUD Bangil Pasuruan | Institute of Tropical Disease, Universitas Airlangga | Jezzy R Dewantari, Rima R Prasetya, Krisnoadi Rahardjo, Aldise M Nastri, Arma Roosalina, Gatot Soegiarto, Laksmi Wulandari, Retno A Setyoningrum, Resti Yudhawati, Yohko K Shimizu, Mitsuhiro Nishimura, Yasuko Mori, Soetjipto, Kazufumi Shimizu, Maria I Lusida |
| EPI_ISL_458082 | Universitas Airlangga Hospital | Institute of Tropical Disease, Universitas Airlangga | Kazufumi Shimizu, Krisnoadi Rahardjo, Aldise M Nastri, Jezzy R Dewantari, Rima R Prasetya, Nasronudin, Gatot Soegiarto, Laksmi Wulandari, Retno A Setyoningrum, Resti Yudhawati, Yohko K Shimizu, Mitsuhiro Nishimura, Yasuko Mori, Soetjipto, Maria I Lusida |
| EPI_ISL_458090 | B.J. Medical College and Civil hospital | Gujarat Biotechnology Research Centre | Kamlesh J Upadhyay, Ramesh Pandit, Tejas Shah, Ankit Hinsu, Pritesh Sabara, Apurvasinh Puvar, Janvi Raval, Zarna Patel, Monika Gandhi, Pinal Trivedi, Maharshi Pandya, Amit Kanani, Nidhi Patel, Nitin Savaliya, Raghawendra Kumar, Dinesh Kumar, Zuber Saiyed, Komal Patel, Labdhi Pandya, Snehal Bagatharia, Dhaval Vaghela, Ramesh Patel, Pranay Shah, Armi Chaudhari, Bhavesh Modi, Gaurishankar Shrimali, R D Dixit, A M Kadri, Umang Mishra, Chaitanya Joshi, Madhvi Joshi |
| EPI_ISL_458098 | B.J. Medical College and Civil hospital | Gujarat Biotechnology Research Centre | Komal Patel, Labdhi Pandya, Snehal Bagatharia, Dhaval Vaghela, Ramesh Patel, Pranay Shah, Kamlesh J Upadhyay, Ramesh Pandit, Tejas Shah, Ankit Hinsu, Pritesh Sabara, Apurvasinh Puvar, Janvi Raval, Zarna Patel, Monika Gandhi, Pinal Trivedi, Maharshi Pandya, Amit Kanani, Nidhi Patel, Nitin Savaliya, Raghawendra Kumar, Dinesh Kumar, Zuber Saiyed, Pragya Sharma, Bhavesh Modi, Gaurishankar Shrimali, R D Dixit, A M Kadri, Umang Mishra, Chaitanya Joshi, Madhvi Joshi |
| EPI_ISL_458099 | B.J. Medical College and Civil hospital | Gujarat Biotechnology Research Centre | Labdhi Pandya, Snehal Bagatharia, Dhaval Vaghela, Ramesh Patel, Pranay Shah, Kamlesh J Upadhyay, Ramesh Pandit, Tejas Shah, Ankit Hinsu, Pritesh Sabara, Apurvasinh Puvar, Janvi Raval, Zarna Patel, Monika Gandhi, Pinal Trivedi, Maharshi Pandya, Amit Kanani, Nidhi Patel, Nitin Savaliya, Raghawendra Kumar, Dinesh Kumar, Zuber Saiyed, Komal Patel, Neha Rajpara, Bhavesh Modi, Gaurishankar Shrimali, R D Dixit, A M Kadri, Umang Mishra, Chaitanya Joshi, Madhvi Joshi |
| EPI_ISL_458108 | Gujarat Biotechnology Research Centre | Gujarat Biotechnology Research Centre | Janvi Raval, Zarna Patel, Monika Gandhi, Pinal Trivedi, Maharshi Pandya, Amit Kanani, Nidhi Patel, Nitin Savaliya, Raghawendra Kumar, Dinesh Kumar, Zuber Saiyed, Komal Patel, Labdhi Pandya, Snehal Bagatharia, Ramesh Pandit, Tejas Shah, Ankit Hinsu, Pritesh Sabara, Apurvasinh Puvar, Pooja P Doshi, Bhavesh Modi, Gaurishankar Shrimali, R D Dixit, A M Kadri, Umang Mishra, Chaitanya Joshi, Madhvi Joshi, , , , , |
| EPI_ISL_458150 | ANOUAL | ANOUAL | Jouali Farah, El Ansari Fatima Zahra, Marchoudi Nabila, Kasmi Yassine, Chenaoui Mohamed, El Aliani Aissam, Benhida Rachid, Azami Nawfel, Kitane Driss Lahlou, Loukman Salma, Fekkak Jamal |
| EPI_ISL_461506 | B.J. Medical College and Civil hospital | Gujarat Biotechnology Research Centre | Kamlesh J Upadhyay, Tejas Shah, Ankit Hinsu, Pritesh Sabara, Apurvasinh Puvar, Janvi Raval, Zarna Patel, Monika Gandhi, Pinal Trivedi, Maharshi Pandya, Nidhi Patel, Nitin Savaliya, Raghawendra Kumar, Dinesh Kumar, Zuber Saiyed, Komal Patel, Labdhi Pandya, Snehal Bagatharia, Pranay Shah, Armi Chaudhari, R D Dixit, A M Kadri, Harsh Bakshi, Chaitanya Joshi, Madhvi Joshi, |
| EPI_ISL_463138, EPI_ISL_463139, EPI_ISL_463140, EPI_ISL_463141, EPI_ISL_463142, EPI_ISL_463143, EPI_ISL_463144, EPI_ISL_463145, EPI_ISL_463146, EPI_ISL_463147, EPI_ISL_463148, EPI_ISL_463149, EPI_ISL_463150, EPI_ISL_463151, EPI_ISL_463152, EPI_ISL_463153, EPI_ISL_463154, EPI_ISL_463155, EPI_ISL_463156, EPI_ISL_463157, EPI_ISL_463158, EPI_ISL_463159, EPI_ISL_463160, EPI_ISL_463161, EPI_ISL_463162, EPI_ISL_463163, EPI_ISL_463164, EPI_ISL_463165, EPI_ISL_463166, EPI_ISL_463167, EPI_ISL_463168, EPI_ISL_463169, EPI_ISL_463170, EPI_ISL_463171, EPI_ISL_463172, EPI_ISL_463173, EPI_ISL_463174 | | | |
| see above | Yale Clinical Virology Laboratory | Grubaugh Lab - Yale School of Public Health | Joseph Fauver, Tara Alpert, Anderson Brito, Anne Wyllie, Chantal Vogels, Mary Petrone, Cole Jensen, Chaney Kalinich, Isabel Ott, Arnau Casanovas, Catherine Muenker, Adam Moore, Alice Lu, Maria Tokuyama, Patrick Wong, Peiwen Lu, Saad Omer, Richard Martinello, Allison Nelson, Shelli Farhadian, Akiko Iwasaki, Charlese Dela Cruz, Albert Ko, Nathan Grubaugh |
| EPI_ISL_463741, EPI_ISL_463742, EPI_ISL_463743, EPI_ISL_463744, EPI_ISL_463745, EPI_ISL_463746, EPI_ISL_463747, EPI_ISL_463748 | Department of Molecular Virology, Cyprus Institute of Neurology and Genetics | Department of Molecular Virology, Cyprus Institute of Neurology and Genetics | Jan Richter, George Krashias, Christina Tryfonos, Stavros Bashiardes, Dana Koptides, Christina Christodoulou |
| EPI_ISL_463889 | Shaoxing Center for Disease Control and Prevention | Department of Pathology and Laboratory Medicine, University of California Los Angeles | Jinkun Chen, Evann E. Hilt, Huan Wu, Zhuojing Jiang, QinChao Zhang, JiLing Wang, Yifang Wang, Fan Li, Ziqin Li, Jialiang Tang, Shangxin Yang |
| EPI_ISL_463893 | University Clinical Center Tuzla | Alea Genetiki Centar | Konjhodži,R;Salihefendi,L;Goleti,T;Pear,D;Tihi,N;Marjanovi,D;Huki,M. |
| EPI_ISL_463894, EPI_ISL_463895, EPI_ISL_463896, EPI_ISL_463897, EPI_ISL_463899, EPI_ISL_463902 | Shaoxing Center for Disease Control and Prevention | Department of Pathology and Laboratory Medicine, University of California Los Angeles | Jinkun Chen, Evann E. Hilt, Huan Wu, Zhuojing Jiang, QinChao Zhang, JiLing Wang, Yifang Wang, Fan Li, Ziqin Li, Jialiang Tang, Shangxin Yang |
| EPI_ISL_464092, EPI_ISL_464093, EPI_ISL_464094 | Laboratory Medicine | Department of Laboratory Medicine, Lin-Kou Chang Gung Memorial Hospital, Taoyuan, Taiwan | Kuo-Chien Tsao, Yu-Nong Gong, Shu-Li Yang, Yi-Chun Liu, Chung-Guei Huang, Mei-Jen Hsiao, Po-Wei Huang, Cheng-Ta Yang, Cheng-Hsun Chiu, Peng-Nien Huang, Kuo-Ming Lee, Guang-Wu Chen, Shin-Ru Shih |
| EPI_ISL_464167 | VI-US Virgin Islands Department of Health | Pathogen Discovery, Respiratory Viruses Branch, Division of Viral Diseases, Centers for Disease Control and Prevention | Krista Queen, Ying Tao, Jing Zhang, Yan Li, Anna Uehara, Clinton R. Paden, Mary S. Keckler, Alison S. Laufer Halpin, Haibin Wang, Jasmine Padilla, Justin Lee, Christopher A. Elkins, Suxiang Tong |

| | | | |
|---|---|---|---|
| EPI_ISL_467037 | GMERS Medical College and Hospital, Gandhinagar | Gujarat Biotechnology Research Centre | Janvi Raval, Zarna Patel, Monika Gandhi, Pinal Trivedi, Maharshi Pandya, Nidhi Patel, Nitin Savaliya, Raghawendra Kumar, Dinesh Kumar, Zuber Saiyed, Komal Patel, Labdhi Pandya, Snehal Bagatharia, Seema Bhatt, Gaurishankar Shrimali, Bhavesh Modi, Bharti Rajani, Tejas Shah, Ankit Hinsu, Pritesh Sabara, Apurvasinh Puvar, Fenil Patel, R D Dixit, A M Kadri, Harsh Bakshi, Chaitanya Joshi, Madhvi Joshi |
| EPI_ISL_467049 | B.J. Medical College and Civil hospital | Gujarat Biotechnology Research Centre | Komal Patel, Labdhi Pandya, Snehal Bagatharia, Pranay Shah, Kamlesh J Upadhyay, Nirav Mungalpara, Tejas Shah, Ankit Hinsu, Pritesh Sabara, Apurvasinh Puvar, Janvi Raval, Zarna Patel, Monika Gandhi, Pinal Trivedi, Maharshi Pandya, Nidhi Patel, Nitin Savaliya, Raghawendra Kumar, Dinesh Kumar, Zuber Saiyed, Neelam Nathani, R D Dixit, A M Kadri, Harsh Bakshi, Chaitanya Joshi, Madhvi Joshi, |
| EPI_ISL_467050 | B.J. Medical College and Civil hospital | Gujarat Biotechnology Research Centre | Labdhi Pandya, Snehal Bagatharia, Pranay Shah, Kamlesh J Upadhyay, Nirav Mungalpara, Tejas Shah, Ankit Hinsu, Pritesh Sabara, Apurvasinh Puvar, Janvi Raval, Zarna Patel, Monika Gandhi, Pinal Trivedi, Maharshi Pandya, Nidhi Patel, Nitin Savaliya, Raghawendra Kumar, Dinesh Kumar, Zuber Saiyed, Komal Patel, Armi Chaudhari, R D Dixit, A M Kadri, Harsh Bakshi, Chaitanya Joshi, Madhvi Joshi, |
| EPI_ISL_467053 | B.J. Medical College and Civil hospital | Gujarat Biotechnology Research Centre | Kamlesh J Upadhyay, Nirav Mungalpara, Tejas Shah, Ankit Hinsu, Pritesh Sabara, Apurvasinh Puvar, Janvi Raval, Zarna Patel, Monika Gandhi, Pinal Trivedi, Maharshi Pandya, Nidhi Patel, Nitin Savaliya, Raghawendra Kumar, Dinesh Kumar, Zuber Saiyed, Komal Patel, Labdhi Pandya, Snehal Bagatharia, Pranay Shah, Pooja P Doshi, R D Dixit, A M Kadri, Harsh Bakshi, Chaitanya Joshi, Madhvi Joshi, |
| EPI_ISL_467262, EPI_ISL_467263, EPI_ISL_467264, EPI_ISL_467265, EPI_ISL_467266, EPI_ISL_467267, EPI_ISL_467268, EPI_ISL_467269, EPI_ISL_467270, EPI_ISL_467271, EPI_ISL_467272, EPI_ISL_467273, EPI_ISL_467274, EPI_ISL_467275, EPI_ISL_467276, EPI_ISL_467277, EPI_ISL_467278, EPI_ISL_467279, EPI_ISL_467280, EPI_ISL_467281, EPI_ISL_467282, EPI_ISL_467283, EPI_ISL_467284, EPI_ISL_467285, EPI_ISL_467286, EPI_ISL_467287, EPI_ISL_467288, EPI_ISL_467289, EPI_ISL_467290, EPI_ISL_467291, EPI_ISL_467292, EPI_ISL_467293, EPI_ISL_467294, EPI_ISL_467295, EPI_ISL_467296, EPI_ISL_467297 | | | |
| see above | Hospital Clínico Universitario de Santiago de Compostela | SeqCOVID-SPAIN consortium/IBV(CSIC) | José Javier Costa Alcalde, Antonio Aguilera Guirao, Mª Luisa Pérez del Molino Bernal, Amparo Coira Nieto, Gema Barbeito Castiñeiras, Rocio Trastoy Pena and SeqCOVID-SPAIN consortium |
| EPI_ISL_467430 | Zoonotic and Exotic infection Diseases Division | Zoontic and Exotic infection Diseases Division | Jinlang Wang, Zhigao Bu |
| EPI_ISL_468331, EPI_ISL_468332, EPI_ISL_468333, EPI_ISL_468334, EPI_ISL_468335, EPI_ISL_468336, EPI_ISL_468337, EPI_ISL_468338, EPI_ISL_468339, EPI_ISL_468340, EPI_ISL_468341, EPI_ISL_468342, EPI_ISL_468343, EPI_ISL_468344 | | | |
| see above | Microbiology Service, University Hospital of A Coruna-Biomedical Research Institute | Genomes & Disease, Center for Research in Molecular Medicine and Chronic Diseases, University of Santiago de Compostela | Kelly Conde, Jorge Arca, Soraya Rumbo, Juan A. Vallejo, M Poza, G Bou, Ana Pequeno-Valtierra, Jorge Rodriguez-Castro, Javier Temes, Daniel Garcia-Souto, Martin Santamarina, Cristina Gomez, Jose M. C. Tubio |
| EPI_ISL_469028 | B.J. Medical College and Civil hospital | Gujarat Biotechnology Research Centre | Janvi Raval, Zarna Patel, Monika Gandhi, Pinal Trivedi, Maharshi Pandya, Nidhi Patel, Nitin Savaliya, Raghawendra Kumar, Dinesh Kumar, Zuber Saiyed, Komal Patel, Labdhi Pandya, Snehal Bagatharia, Pranay Shah, Kamlesh J Upadhyay, Nirav Mungalpara, Tejas Shah, Ankit Hinsu, Pritesh Sabara, Apurvasinh Puvar, Fenil Patel, R D Dixit, A M Kadri, Harsh Bakshi, Chaitanya Joshi, Madhvi Joshi |
| EPI_ISL_469038 | GMERS Medical College & Hospital | Gujarat Biotechnology Research Centre | Komal Patel, Labdhi Pandya, Snehal Bagatharia, Meenakshi Shah, Neena Doshi, Varsha Godbole, Tejas Shah, Ankit Hinsu, Pritesh Sabara, Apurvasinh Puvar, Janvi Raval, Zarna Patel, Monika Gandhi, Pinal Trivedi, Maharshi Pandya, Nidhi Patel, Nitin Savaliya, Raghawendra Kumar, Dinesh Kumar, Zuber Saiyed, Neha Rajpara, R D Dixit, A M Kadri, Harsh Bakshi, Chaitanya Joshi, Madhvi Joshi |
| EPI_ISL_469039 | GMERS Medical College & Hospital | Gujarat Biotechnology Research Centre | Labdhi Pandya, Snehal Bagatharia, Meenakshi Shah, Neena Doshi, Varsha Godbole, Tejas Shah, Ankit Hinsu, Pritesh Sabara, Apurvasinh Puvar, Janvi Raval, Zarna Patel, Monika Gandhi, Pinal Trivedi, Maharshi Pandya, Nidhi Patel, Nitin Savaliya, Raghawendra Kumar, Dinesh Kumar, Zuber Saiyed, Komal Patel, Afzal Ansari, R D Dixit, A M Kadri, Harsh Bakshi, Chaitanya Joshi, Madhvi Joshi |
| EPI_ISL_469044 | Dr. N. D. Desai Medical College & Hospital | Gujarat Biotechnology Research Centre | Jigar Gusani, Supreet Prabhu, Tejas Shah, Ankit Hinsu, Pritesh Sabara, Apurvasinh Puvar, Janvi Raval, Zarna Patel, Monika Gandhi, Pinal Trivedi, Maharshi Pandya, Nidhi Patel, Nitin Savaliya, Raghawendra Kumar, Dinesh Kumar, Zuber Saiyed, Komal Patel, Labdhi Pandya, Snehal Bagatharia, J G Buch, Neha Rajpara, R D Dixit, A M Kadri, Harsh Bakshi, Chaitanya Joshi, Madhvi Joshi |
| EPI_ISL_469171, EPI_ISL_469172, EPI_ISL_469173, EPI_ISL_469174, EPI_ISL_469175, EPI_ISL_469176, EPI_ISL_469177, EPI_ISL_469178, EPI_ISL_469179, EPI_ISL_469180, EPI_ISL_469181, EPI_ISL_469182, EPI_ISL_469183, EPI_ISL_469184, EPI_ISL_469185, EPI_ISL_469186, EPI_ISL_469187, EPI_ISL_469188, EPI_ISL_469189, EPI_ISL_469190, EPI_ISL_469191, EPI_ISL_469192, EPI_ISL_469193, EPI_ISL_469194, EPI_ISL_469195, EPI_ISL_469196, EPI_ISL_469197, EPI_ISL_469198, EPI_ISL_469199, EPI_ISL_469200, EPI_ISL_469201, EPI_ISL_469202, EPI_ISL_469203, EPI_ISL_469204, EPI_ISL_469205, EPI_ISL_469206, EPI_ISL_469207, EPI_ISL_469208 | | | |
| see above | Yale Clinical Virology Laboratory | Grubaugh Lab - Yale School of Public Health | Joseph Fauver, Tara Alpert, Anderson Brito, Anne Wyllie, Chantal Vogels, Mary Petrone, Cole Jensen, Chaney Kalinich, Isabel Ott, Arnau Casanovas, Catherine Muenker, Adam Moore, Alice Lu, Maria Tokuyama, Patrick Wong, Peiwen Lu, Saad Omer, Richard Martinello, Allison Nelson, Shelli Farhadian, Akiko Iwasaki, Charlese Dela Cruz, Albert Ko, Nathan Grubaugh |
| EPI_ISL_469287, EPI_ISL_469288, EPI_ISL_469289, EPI_ISL_469290, EPI_ISL_469291, EPI_ISL_469292, EPI_ISL_469293, EPI_ISL_469294, EPI_ISL_469295, EPI_ISL_469296 | Keio University Hospital | Keio University Hospital | Kenjiro Kosaki |
| EPI_ISL_470540 | Wisconsin State Laboratory of Hygiene Communicable Disease Division | Wisconsin State Laboratory of Hygiene Communicable Disease Division | Kelsey R Florek |
| EPI_ISL_470882 | Foerde Hospital, Department of Microbiology | Norwegian Institute of Public Health, Department of Virology | Kathrine Stene-Johansen, Kamilla Heddeland Instefjord, Hilde Elshaug, Rasmus Riis Kopperud, Karoline Bragstad, Olav Hungnes |
| EPI_ISL_471172 | Unilabs Laboratory Medicine | Norwegian Institute of Public Health, Department of Virology | Kathrine Stene-Johansen, Kamilla Heddeland Instefjord, Hilde Elshaug, Rasmus Riis Kopperud, Karoline Bragstad, Olav Hungnes |
| EPI_ISL_471173 | Hospital of Southern Norway - Kristiansand, Department of Medical Microbiology | Norwegian Institute of Public Health, Department of Virology | Kathrine Stene-Johansen, Kamilla Heddeland Instefjord, Hilde Elshaug, Rasmus Riis Kopperud, Karoline Bragstad, Olav Hungnes |
| EPI_ISL_471174 | Ostfold Hospital Trust - Kalnes, Centre for Laboratory Medicine, Section for gene technology and infection serology | Norwegian Institute of Public Health, Department of Virology | Kathrine Stene-Johansen, Kamilla Heddeland Instefjord, Hilde Elshaug, Rasmus Riis Kopperud, Karoline Bragstad, Olav Hungnes |
| EPI_ISL_471175 | Oslo University Hospital, Department of Medical Microbiology | Norwegian Institute of Public Health, Department of Virology | Kathrine Stene-Johansen, Kamilla Heddeland Instefjord, Hilde Elshaug, Rasmus Riis Kopperud, Karoline Bragstad, Olav Hungnes |
| EPI_ISL_471176 | Hospital of Southern Norway - Kristiansand, Department of Medical Microbiology | Norwegian Institute of Public Health, Department of Virology | Kathrine Stene-Johansen, Kamilla Heddeland Instefjord, Hilde Elshaug, Rasmus Riis Kopperud, Karoline Bragstad, Olav Hungnes |
| EPI_ISL_471177 | Oslo University Hospital, Department of Medical Microbiology | Norwegian Institute of Public Health, Department of Virology | Kathrine Stene-Johansen, Kamilla Heddeland Instefjord, Hilde Elshaug, Rasmus Riis Kopperud, Karoline Bragstad, Olav Hungnes |
| EPI_ISL_471178, EPI_ISL_471179, EPI_ISL_471180, EPI_ISL_471181, EPI_ISL_471182, EPI_ISL_471183, EPI_ISL_471184, EPI_ISL_471185, EPI_ISL_471186, EPI_ISL_471187, EPI_ISL_471188, EPI_ISL_471189, EPI_ISL_471190, EPI_ISL_471191, EPI_ISL_471192, EPI_ISL_471193, EPI_ISL_471194, EPI_ISL_471195, EPI_ISL_471196, EPI_ISL_471197, EPI_ISL_471198, EPI_ISL_471199, EPI_ISL_471200, EPI_ISL_471201, EPI_ISL_471202, EPI_ISL_471203, EPI_ISL_471204, EPI_ISL_471205, EPI_ISL_471206, EPI_ISL_471207, EPI_ISL_471208, EPI_ISL_471209, EPI_ISL_471210, EPI_ISL_471211, EPI_ISL_471212, EPI_ISL_471213, EPI_ISL_471214, EPI_ISL_471215, EPI_ISL_471216, EPI_ISL_471217, EPI_ISL_471218, EPI_ISL_471219, EPI_ISL_471220, EPI_ISL_471221, EPI_ISL_471222, EPI_ISL_471223, EPI_ISL_471224, EPI_ISL_471225, EPI_ISL_471226, EPI_ISL_471227, EPI_ISL_471228, EPI_ISL_471229, EPI_ISL_471230, EPI_ISL_471231, EPI_ISL_471232, EPI_ISL_471233, EPI_ISL_471234, EPI_ISL_471235, EPI_ISL_471236, EPI_ISL_471237, EPI_ISL_471238, EPI_ISL_471239, EPI_ISL_471240, EPI_ISL_471241, EPI_ISL_471242, EPI_ISL_471243, EPI_ISL_471244, EPI_ISL_471245, EPI_ISL_471246, EPI_ISL_471247, EPI_ISL_471248, EPI_ISL_471249, EPI_ISL_471250, EPI_ISL_471251, EPI_ISL_471252, EPI_ISL_471253, EPI_ISL_471254, EPI_ISL_471255, EPI_ISL_471256, EPI_ISL_471257, EPI_ISL_471258, EPI_ISL_471259, EPI_ISL_471260, EPI_ISL_471261, EPI_ISL_471262, EPI_ISL_471263, EPI_ISL_471264, EPI_ISL_471265, EPI_ISL_471266 | | | |
| see above | Wisconsin State Laboratory of Hygiene Communicable Disease Division | Wisconsin State Laboratory of Hygiene Communicable Disease Division | Kelsey R. Florek, Abigail C. Shockey |
| EPI_ISL_471425, EPI_ISL_471426 | Division of Viral Diseases, Center for Laboratory Control of Infectious Diseases, Korea Centers for Diseases Control and Prevention | Division of Viral Diseases, Center for Laboratory Control of Infectious Diseases, Korea Centers for Diseases Control and Prevention | Jeong-Min Kim, Yoon-Seok Chung, Namjoo Lee, Mi-Seon Kim, Sang Hee Woo, Hye-Jun Jo, Sehee Park, Heui Man Kim, Jun-Sub Kim, Junhyeong Jang, Dong Hyun Song, Daesang Lee, Seong Tae Jeong, Myung Guk Han |
| EPI_ISL_471427, EPI_ISL_471428, EPI_ISL_471429, EPI_ISL_471430, EPI_ISL_471431, EPI_ISL_471432, EPI_ISL_471433, EPI_ISL_471434, EPI_ISL_471435, EPI_ISL_471436, EPI_ISL_471437 | | | |
| see above | Department of Clinical Microbiology | GIGA Medical Genomics | Keith Durkin, Maria Artesi, Sébastien Bontems, Raphaël Boreux, Cécile Meex, Axelle Chaslain, Céline Fombellida-Lopez, Pierrette Melin, Marie-Pierre Hayette, Vincent Bours. |
| EPI_ISL_471438, EPI_ISL_471439, EPI_ISL_471440, EPI_ISL_471441, EPI_ISL_471442, EPI_ISL_471443, EPI_ISL_471444 | Division of Viral Diseases, Center for Laboratory Control of Infectious Diseases, Korea Centers for Diseases Control and Prevention | Division of Viral Diseases, Center for Laboratory Control of Infectious Diseases, Korea Centers for Diseases Control and Prevention | Jeong-Min Kim, Yoon-Seok Chung, Namjoo Lee, Sang Hee Woo, Hye-Jun Jo, Heui Man Kim, Jun-Sub Kim, Dong Hyun Song, Daesang Lee, Seong Tae Jeong, Myung Guk Han |

| | | | |
|---|---|---|---|
| EPI_ISL_471445 | Division of Viral Diseases, Center for Laboratory Control of Infectious Diseases, Korea Centers for Diseases Control and Prevention | Division of Viral Diseases, Center for Laboratory Control of Infectious Diseases, Korea Centers for Diseases Control and Prevention | Jeong-Min Kim, Yoon-Seok Chung, Namjoo Lee, Sang Hee Woo, Hye-Jun Jo, Heui Man Kim, Jun-Sub Kim, Myung Guk Han |
| EPI_ISL_471446, EPI_ISL_471447, EPI_ISL_471448, EPI_ISL_471449, EPI_ISL_471450, EPI_ISL_471451, EPI_ISL_471452 | Division of Viral Diseases, Center for Laboratory Control of Infectious Diseases, Korea Centers for Diseases Control and Prevention | Division of Viral Diseases, Center for Laboratory Control of Infectious Diseases, Korea Centers for Diseases Control and Prevention | Jeong-Min Kim, Yoon-Seok Chung, Namjoo Lee, Sang Hee Woo, Hye-Jun Jo, Heui Man Kim, Jun-Sub Kim, Dong Hyun Song, Daesang Lee, Seong Tae Jeong, Myung Guk Han |
| EPI_ISL_471453 | Division of Viral Diseases, Center for Laboratory Control of Infectious Diseases, Korea Centers for Diseases Control and Prevention | Division of Viral Diseases, Center for Laboratory Control of Infectious Diseases, Korea Centers for Diseases Control and Prevention | Jeong-Min Kim, Yoon-Seok Chung, Namjoo Lee, Sang Hee Woo, Hye-Jun Jo, Heui Man Kim, Jun-Sub Kim, Myung Guk Han |
| EPI_ISL_471454, EPI_ISL_471455 | Division of Viral Diseases, Center for Laboratory Control of Infectious Diseases, Korea Centers for Diseases Control and Prevention | Division of Viral Diseases, Center for Laboratory Control of Infectious Diseases, Korea Centers for Diseases Control and Prevention | Jeong-Min Kim, Yoon-Seok Chung, Namjoo Lee, Sang Hee Woo, Hye-Jun Jo, Heui Man Kim, Jun-Sub Kim, Dong Hyun Song, Daesang Lee, Seong Tae Jeong, Myung Guk Han |
| EPI_ISL_471583, EPI_ISL_471584 | King Institute of Preventive Medicine & Research | CSIR-Centre for Cellular and Molecular Biology | K.Kaveri,S.Sivasubramanian,S.Vennila,P.Padmapriya,R.Kiruba,S.Magesh,G. Dhinakar Raj, G. Ravikumar, P. Azhahianambi,K Thangaraj,Payel Mukherjee, Sofia Banu, Priya Singh, Dhiviya Vedagiri, Divya Gupta, Vishal Sah, Santosh Kumar Kuncha, Krishnan Harinivas Harshan, Archana Bharadwaj Siva, Karthik Bharadwaj Tallapaka, Shagufta Khan, Lamuk Zaveri, Namami Gaur, Sakshi Shambhavi, Tulasi Nagabandi, Purushotham Vodnala, Rakesh K Mishra, Divya Tej Sowpati |
| EPI_ISL_471586 | CSIR-Centre for Cellular and Molecular Biology | CSIR-Centre for Cellular and Molecular Biology | Lamuk Zaveri, Shagufta Khan, Namami Gaur, Sakshi Shambhavi, Tulasi Nagabandi, Purushotham Vodnala, Payel Mukherjee, Sofia Banu, Priya Singh, Dhiviya Vedagiri, Divya Gupta, Vishal Sah, Santosh Kumar Kuncha, Krishnan Harinivas Harshan, Archana Bharadwaj Siva, Karthik Bharadwaj Tallapaka,Zeba Rizvi, Zuberwasim Sayyad, Kakade Aishwarya Arun, Amrutha H C, Ananga Ghosh, Rakesh K Mishra, Divya Tej Sowpati |
| EPI_ISL_471588 | CSIR-Centre for Cellular and Molecular Biology | CSIR-Centre for Cellular and Molecular Biology | Lamuk Zaveri, Shagufta Khan, Namami Gaur, Sakshi Shambhavi, Tulasi Nagabandi, Purushotham Vodnala, Payel Mukherjee, Sofia Banu, Priya Singh, Dhiviya Vedagiri, Divya Gupta, Vishal Sah, Santosh Kumar Kuncha, Krishnan Harinivas Harshan, Archana Bharadwaj Siva, Karthik Bharadwaj Tallapaka, Renu Sudhakar, Somesh Gorde, Gangumala Srinivas Reddy, Sujoy Deb, Swati Bayyana, Rakesh K Mishra, Divya Tej Sowpati |
| EPI_ISL_471589 | CSIR-Centre for Cellular and Molecular Biology | CSIR-Centre for Cellular and Molecular Biology | Lamuk Zaveri, Shagufta Khan, Namami Gaur, Sakshi Shambhavi, Tulasi Nagabandi, Purushotham Vodnala, Payel Mukherjee, Sofia Banu, Priya Singh, Dhiviya Vedagiri, Divya Gupta, Vishal Sah, Santosh Kumar Kuncha, Krishnan Harinivas Harshan, Archana Bharadwaj Siva, Karthik Bharadwaj Tallapaka,Umesh Kumar, Unis Ahmad Bhat, Ajay Sarawagi, Priyanka Pant, Rajkanwar Nathawat, Rakesh K Mishra, Divya Tej Sowpati |
| EPI_ISL_471590, EPI_ISL_471615 | CSIR-Centre for Cellular and Molecular Biology | CSIR-Centre for Cellular and Molecular Biology | Lamuk Zaveri, Shagufta Khan, Namami Gaur, Sakshi Shambhavi, Tulasi Nagabandi, Purushotham Vodnala, Payel Mukherjee, Sofia Banu, Priya Singh, Dhiviya Vedagiri, Divya Gupta, Vishal Sah, Santosh Kumar Kuncha, Krishnan Harinivas Harshan, Archana Bharadwaj Siva, Karthik Bharadwaj Tallapaka,Zeba Rizvi, Zuberwasim Sayyad, Kakade Aishwarya Arun, Amrutha H C, Ananga Ghosh, Rakesh K Mishra, Divya Tej Sowpati |
| EPI_ISL_471616 | CSIR-Centre for Cellular and Molecular Biology | CSIR-Centre for Cellular and Molecular Biology | Lamuk Zaveri, Shagufta Khan, Namami Gaur, Sakshi Shambhavi, Tulasi Nagabandi, Purushotham Vodnala, Payel Mukherjee, Sofia Banu, Priya Singh, Dhiviya Vedagiri, Divya Gupta, Vishal Sah, Santosh Kumar Kuncha, Krishnan Harinivas Harshan, Archana Bharadwaj Siva, Karthik Bharadwaj Tallapaka, Renu Sudhakar, Somesh Gorde, Gangumala Srinivas Reddy, Sujoy Deb, Swati Bayyana, Rakesh K Mishra, Divya Tej Sowpati |
| EPI_ISL_471617 | CSIR-Centre for Cellular and Molecular Biology | CSIR-Centre for Cellular and Molecular Biology | Lamuk Zaveri, Shagufta Khan, Namami Gaur, Sakshi Shambhavi, Tulasi Nagabandi, Purushotham Vodnala, Payel Mukherjee, Sofia Banu, Priya Singh, Dhiviya Vedagiri, Divya Gupta, Vishal Sah, Santosh Kumar Kuncha, Krishnan Harinivas Harshan, Archana Bharadwaj Siva, Karthik Bharadwaj Tallapaka,Umesh Kumar, Unis Ahmad Bhat, Ajay Sarawagi, Priyanka Pant, Rajkanwar Nathawat, Rakesh K Mishra, Divya Tej Sowpati |
| EPI_ISL_471618 | CSIR-Centre for Cellular and Molecular Biology | CSIR-Centre for Cellular and Molecular Biology | Lamuk Zaveri, Shagufta Khan, Namami Gaur, Sakshi Shambhavi, Tulasi Nagabandi, Purushotham Vodnala, Payel Mukherjee, Sofia Banu, Priya Singh, Dhiviya Vedagiri, Divya Gupta, Vishal Sah, Santosh Kumar Kuncha, Krishnan Harinivas Harshan, Archana Bharadwaj Siva, Karthik Bharadwaj Tallapaka,Zeba Rizvi, Zuberwasim Sayyad, Kakade Aishwarya Arun, Amrutha H C, Ananga Ghosh, Rakesh K Mishra, Divya Tej Sowpati |
| EPI_ISL_475030 | Department of MicroBiology, Government Medical College, Surat | Gujarat Biotechnology Research Centre | Janvi Raval, Zarna Patel, Monika Gandhi, Pinal Trivedi, Maharshi Pandya, Nidhi Patel, Nitin Savaliya, Raghawendra Kumar, Dinesh Kumar, Zuber Saiyed, Komal Patel, Labdhi Pandya, Snehal Bagatharia, Naresh Chauhan, Summaiya Mullan, Amit gamit, Pritesh Sabara, Apurvasinh Puvar, Armi Chaudhari, R D Dixit, A M Kadri, Harsh Bakshi, Chaitanya Joshi, Madhvi Joshi |
| EPI_ISL_475040 | Department of MicroBiology, Government Medical College, Surat | Gujarat Biotechnology Research Centre | Komal Patel, Labdhi Pandya, Snehal Bagatharia, Naresh Chauhan, Summaiya Mullan, Amit gamit, Pritesh Sabara, Apurvasinh Puvar, Janvi Raval, Zarna Patel, Monika Gandhi, Pinal Trivedi, Maharshi Pandya, Nidhi Patel, Nitin Savaliya, Raghawendra Kumar, Dinesh Kumar, Zuber Saiyed, Neelam Nathani, R D Dixit, A M Kadri, Harsh Bakshi, Chaitanya Joshi, Madhvi Joshi |
| EPI_ISL_475041 | Department of MicroBiology, Government Medical College, Surat | Gujarat Biotechnology Research Centre | Labdhi Pandya, Snehal Bagatharia, Naresh Chauhan, Summaiya Mullan, Amit gamit, Pritesh Sabara, Apurvasinh Puvar, Janvi Raval, Zarna Patel, Monika Gandhi, Pinal Trivedi, Maharshi Pandya, Nidhi Patel, Nitin Savaliya, Raghawendra Kumar, Dinesh Kumar, Zuber Saiyed, Komal Patel, Armi Chaudhari, R D Dixit, A M Kadri, Harsh Bakshi, Chaitanya Joshi, Madhvi Joshi |
| EPI_ISL_475048 | GMERS Medical College & Hospital | Gujarat Biotechnology Research Centre | Janvi Raval, Zarna Patel, Monika Gandhi, Pinal Trivedi, Maharshi Pandya, Nidhi Patel, Nitin Savaliya, Raghawendra Kumar, Dinesh Kumar, Zuber Saiyed, Komal Patel, Labdhi Pandya, Snehal Bagatharia, Meenakshi Shah, Neena Doshi, Varsha Godbole, Pritesh Sabara, Apurvasinh Puvar, Neha Rajpara, R D Dixit, A M Kadri, Harsh Bakshi, Chaitanya Joshi, Madhvi Joshi |
| EPI_ISL_475572 | Imperial College London | Imperial College London | Jie Zhou, Wendy Barclay |
| EPI_ISL_476152, EPI_ISL_476153, EPI_ISL_476154, EPI_ISL_476155, EPI_ISL_476156, EPI_ISL_476157, EPI_ISL_476158, EPI_ISL_476159, EPI_ISL_476160, EPI_ISL_476161, EPI_ISL_476162, EPI_ISL_476163, EPI_ISL_476164, EPI_ISL_476165, EPI_ISL_476166, EPI_ISL_476167, EPI_ISL_476168, EPI_ISL_476169, EPI_ISL_476170, EPI_ISL_476337, EPI_ISL_476338, EPI_ISL_476339, EPI_ISL_476340, EPI_ISL_476341, EPI_ISL_476342, EPI_ISL_476343, EPI_ISL_476344, EPI_ISL_476345, EPI_ISL_476346, EPI_ISL_476347, EPI_ISL_476348, EPI_ISL_476349, EPI_ISL_476387, EPI_ISL_476388, EPI_ISL_476389, EPI_ISL_476390, EPI_ISL_476391, EPI_ISL_476392, EPI_ISL_476393, EPI_ISL_476394, EPI_ISL_476395, EPI_ISL_476396, EPI_ISL_476397, EPI_ISL_476398, EPI_ISL_476399, EPI_ISL_476400, EPI_ISL_476401, EPI_ISL_476402, EPI_ISL_476408, EPI_ISL_476409, EPI_ISL_476410, EPI_ISL_476411, EPI_ISL_476412, EPI_ISL_476413, EPI_ISL_476414, EPI_ISL_476415, EPI_ISL_476416, EPI_ISL_476417, EPI_ISL_476418, EPI_ISL_476419, EPI_ISL_476420, EPI_ISL_476421, EPI_ISL_476422, EPI_ISL_476423, EPI_ISL_476424, EPI_ISL_476425 | | | |
| see above | Laboratório de Patologia Clínica - UNICAMP | Laboratório de Estudos de Vírus Emergentes - UNICAMP | José Luiz Proença-Modena, Magnun Nueldo Nunes dos Santos, Angelica Schreiber, Julia Forato,Camila Simeoni, Marcilio Jorge Fumagalli, Mariene Ribeiro Amorim, Darlan da Silva Candido, Nuno Rodrigues Faria, Julien Theze, Luiz Gonzaga,Jaqueline Goes Jesus e William Marciel de Souza |
| EPI_ISL_476517, EPI_ISL_476518, EPI_ISL_476519, EPI_ISL_476520, EPI_ISL_476521, EPI_ISL_476522, EPI_ISL_476523, EPI_ISL_476524, EPI_ISL_476525, EPI_ISL_476526, EPI_ISL_476527, EPI_ISL_476528, EPI_ISL_476529, EPI_ISL_476530, EPI_ISL_476531, EPI_ISL_476532, EPI_ISL_476533, EPI_ISL_476534, EPI_ISL_476535, EPI_ISL_476536, EPI_ISL_476537, EPI_ISL_476538, EPI_ISL_476539, EPI_ISL_476540, EPI_ISL_476541, EPI_ISL_476542, EPI_ISL_476543, EPI_ISL_476544, EPI_ISL_476545, EPI_ISL_476546, EPI_ISL_476547, EPI_ISL_476548, EPI_ISL_476549, EPI_ISL_476550, EPI_ISL_476551, EPI_ISL_476552, EPI_ISL_476553, EPI_ISL_476554, EPI_ISL_476555, EPI_ISL_476556, EPI_ISL_476557 | | | |
| see above | Yale Clinical Virology Laboratory | Grubaugh Lab - Yale School of Public Health | Joseph Fauver, Tara Alpert, Anderson Brito, Anne Wyllie, Chantal Vogels, Mary Petrone, Cole Jensen, Chaney Kalinich, Isabel Ott, Arnau Casanovas, Catherine Muenker, Adam Moore, Alice Lu, Maria Tokuyama, Patrick Wong, Peiwen Lu, Saad Omer, Richard Martinello, Allison Nelson, Shelli Farhadian, Akiko Iwasaki, Charlese Dela Cruz, Albert Ko, Nathan Grubaugh |
| EPI_ISL_476856 | GMERS Medical College & Hospital, Gotri, Vadodara | Gujarat Biotechnology Research Centre | Janvi Raval, Zarna Patel, Monika Gandhi, Pinal Trivedi, Maharshi Pandya, Nidhi Patel, Nitin Savaliya, Raghawendra Kumar, Dinesh Kumar, Zuber Saiyed, Komal Patel, Labdhi Pandya, Afzal Ansari, Nikha Trivedi, Meenakshi Shah, Neena Doshi, Varsha Godbole, Apurvasinh Puvar, R D Dixit, A M Kadri, Harsh Bakshi, Chaitanya Joshi, Madhvi Joshi |
| EPI_ISL_476866 | GMERS Medical College and Hospital, Gandhinagar | Gujarat Biotechnology Research Centre | Komal Patel, Labdhi Pandya, Afzal Ansari, Nikha Trivedi, Seema Bhatt, Gaurishankar Shrimali, Bhavesh Modi, Bharti Rajani, Apurvasinh Puvar, Janvi Raval, Zarna Patel, Monika Gandhi, Pinal Trivedi, Maharshi Pandya, Nidhi Patel, Nitin Savaliya, Raghawendra Kumar, Dinesh Kumar, Zuber Saiyed, R D Dixit, A M Kadri, Harsh Bakshi, Chaitanya Joshi, Madhvi Joshi |
| EPI_ISL_476867 | Banas Medical College and Research Institute | Gujarat Biotechnology Research Centre | Labdhi Pandya, Afzal Ansari, Nikha Trivedi, Radhika Khara, Sunil R Joshi, Viren s Doshi, Apurvasinh Puvar, Janvi Raval, Zarna Patel, Monika Gandhi, Pinal Trivedi, Maharshi Pandya, Nidhi Patel, Nitin Savaliya, Raghawendra Kumar, Dinesh Kumar, Zuber Saiyed, Komal Patel, R D Dixit, A M Kadri, Harsh Bakshi, Chaitanya Joshi, Madhvi Joshi |
| EPI_ISL_476874 | Department of MicroBiology, Government Medical College, Surat | Gujarat Biotechnology Research Centre | Janvi Raval, Zarna Patel, Monika Gandhi, Pinal Trivedi, Maharshi Pandya, Nidhi Patel, Nitin Savaliya, Raghawendra Kumar, Dinesh Kumar, Zuber Saiyed, Komal Patel, Labdhi Pandya, Afzal Ansari, Nikha Trivedi, Naresh Chauhan, Summaiya Mullan, Amit gamit, Apurvasinh Puvar, R D Dixit, A M Kadri, Harsh Bakshi, Chaitanya Joshi, Madhvi Joshi |

| | | | |
|---|---|---|---|
| EPI_ISL_477016 | Institute of Microbiology, Universidad San Francisco de Quito | Institute of Microbiology, Universidad San Francisco de Quito | Juan José Guadalupe, Sully Márquez, Belén Prado-Vivar, Monica Becerra-Wong, Carla Torres, Bernardo Gutiérrez, Jorge Luis Velez, Verónica Barragán, Patricio Rojas-Silva, Gabriel Trueba, Michelle Grunauer, Paúl Cárdenas |
| EPI_ISL_477163 | Laboratory of Dr. John Lednicky | University of Florida | John A. Lednicky, Maha A. Elbadry, Kuttichantran Subramaniam, Thomas B. Waltzek, John Glenn Morris, Jr. |
| EPI_ISL_480784, EPI_ISL_480785 | NYC Department of Health and Mental Hygiene | Pathogen Discovery, Respiratory Viruses Branch, Division of Viral Diseases, Centers for Disease Control and Prevention | Krista Queen, Christine Mahl, Jennifer Rakeman, Anna Uehara, Ying Tao, Jing Zhang, Yan Li, Clinton R. Paden, Haibin Wang, Jasmine Padilla, Justin Lee, Sally Slavinski, Suxiang Tong |
| EPI_ISL_481207 | Ostfold Hospital Trust - Kalnes, Centre for Laboratory Medicine, Section for gene technology and infection serology | Norwegian Institute of Public Health, Department of Virology | Kathrine Stene-Johansen, Kamilla Heddeland Instefjord, Hilde Elshaug, Rasmus Riis Kopperud, Karoline Bragstad, Olav Hungnes |
| EPI_ISL_481208 | Furst Medical Laboratory | Norwegian Institute of Public Health, Department of Virology | Kathrine Stene-Johansen, Kamilla Heddeland Instefjord, Hilde Elshaug, Rasmus Riis Kopperud, Karoline Bragstad, Olav Hungnes |
| EPI_ISL_481209, EPI_ISL_481210, EPI_ISL_481211, EPI_ISL_481212, EPI_ISL_481213 | Ostfold Hospital Trust - Kalnes, Centre for Laboratory Medicine, Section for gene technology and infection serology | Norwegian Institute of Public Health, Department of Virology | Kathrine Stene-Johansen, Kamilla Heddeland Instefjord, Hilde Elshaug, Rasmus Riis Kopperud, Karoline Bragstad, Olav Hungnes |
| EPI_ISL_481215 | Oslo University Hospital, Department of Medical Microbiology | Norwegian Institute of Public Health, Department of Virology | Kathrine Stene-Johansen, Kamilla Heddeland Instefjord, Hilde Elshaug, Rasmus Riis Kopperud, Karoline Bragstad, Olav Hungnes |
| EPI_ISL_481216 | Medical Microbiology Unit, Department for Laboratory Medicine, Drammen Hospital, Vestre Viken Health Trust, | Norwegian Institute of Public Health, Department of Virology | Kathrine Stene-Johansen, Kamilla Heddeland Instefjord, Hilde Elshaug, Rasmus Riis Kopperud, Karoline Bragstad, Olav Hungnes |
| EPI_ISL_481217, EPI_ISL_481218 | Oslo University Hospital, Department of Medical Microbiology | Norwegian Institute of Public Health, Department of Virology | Kathrine Stene-Johansen, Kamilla Heddeland Instefjord, Hilde Elshaug, Rasmus Riis Kopperud, Karoline Bragstad, Olav Hungnes |
| EPI_ISL_481251, EPI_ISL_481252, EPI_ISL_481254, EPI_ISL_481255, EPI_ISL_481257, EPI_ISL_481258, EPI_ISL_481259, EPI_ISL_481260, EPI_ISL_481261, EPI_ISL_481263 | Department of Emerging Infectious Diseases, Institute of Tropical Medicine, Nagasaki University | Department of Emerging Infectious Diseases, Institute of Tropical Medicine, Nagasaki University | Jiro Yasuda, Rokusuke Yoshikawa, Yuichiro Furusato, Haruka Abe |
| EPI_ISL_481265, EPI_ISL_481266, EPI_ISL_481267, EPI_ISL_481268, EPI_ISL_481269, EPI_ISL_481270, EPI_ISL_481271, EPI_ISL_481272, EPI_ISL_481273, EPI_ISL_481274, EPI_ISL_481275, EPI_ISL_481276, EPI_ISL_481277, EPI_ISL_481278, EPI_ISL_481279, EPI_ISL_481280, EPI_ISL_481281, EPI_ISL_481282 | | | |
| see above | Maryland Department of Health | Maryland Department of Health | Keller,E. |
| EPI_ISL_481370 | Division of Viral Diseases, Center for Laboratory Control of Infectious Diseases, Korea Centers for Diseases Control and Prevention | Division of Viral Diseases, Center for Laboratory Control of Infectious Diseases, Korea Centers for Diseases Control and Prevention | Jeong-Min Kim, Yoon-Seok Chung, Namjoo Lee, Sang Hee Woo, Hye-Jun Jo, Heui Man Kim, Jun-Sub Kim, Myung Guk Han |
| EPI_ISL_481371, EPI_ISL_481372, EPI_ISL_481373, EPI_ISL_481374, EPI_ISL_481375, EPI_ISL_481376, EPI_ISL_481377, EPI_ISL_481378, EPI_ISL_481379 | Division of Viral Diseases, Center for Laboratory Control of Infectious Diseases, Korea Centers for Diseases Control and Prevention | Division of Viral Diseases, Center for Laboratory Control of Infectious Diseases, Korea Centers for Diseases Control and Prevention | Jeong-Min Kim, Yoon-Seok Chung, Namjoo Lee, Sang Hee Woo, Hye-Jun Jo, Heui Man Kim, Jun-Sub Kim, Dong Hyun Song, Daesang Lee, Seong Tae Jeong, Myung Guk Han |
| EPI_ISL_482575, EPI_ISL_482576, EPI_ISL_482577, EPI_ISL_482578, EPI_ISL_482579, EPI_ISL_482580, EPI_ISL_482581, EPI_ISL_482582, EPI_ISL_482583, EPI_ISL_482584, EPI_ISL_482585, EPI_ISL_482586 | | | |
| see above | Hangzhou Center for Diseases Control and Prevention | Hangzhou Center for Diseases Control and Prevention | Jun Li, Haoqiu Wang, Lingfeng Mao, Hua Yu, Xinfen Yu, Zhou Sun, Xin Qian, Shuchang Chen, Junfang Chen, Xuchu Wang |
| EPI_ISL_483139, EPI_ISL_483140, EPI_ISL_483141, EPI_ISL_483142, EPI_ISL_483143, EPI_ISL_483144, EPI_ISL_483145, EPI_ISL_483146, EPI_ISL_483147, EPI_ISL_483148, EPI_ISL_483149, EPI_ISL_483150, EPI_ISL_483151, EPI_ISL_483152, EPI_ISL_483153, EPI_ISL_483154, EPI_ISL_483155, EPI_ISL_483156, EPI_ISL_483157 | | | |
| see above | Robert Koch Institute, ZBS1 Highly Pathogenic Viruses, Berlin, Germany | Robert Koch Institute, Bioinformatics MF1, Berlin, Germany | Janine Michel, Andrea Thuermer, Oliver Drechsel, Rene Kmiecinski, Stephan Fuchs, Max v. Kleist, Andreas Nitsche |
| EPI_ISL_483820 | GMERS Medical College and Hospital, Gandhinagar | Gujarat Biotechnology Research Centre | Komal Patel, Labdhi Pandya, Afzal Ansari, Nikha Trivedi, Seema Bhatt, Gaurishankar Shrimali, Bhavesh Modi, Bharti Rajani, Apurvasinh Puvar, Janvi Raval, Zarna Patel, Monika Gandhi, Pinal Trivedi, Maharshi Pandya, Nidhi Patel, Nitin Savaliya, Raghawendra Kumar, Dinesh Kumar, Zuber Saiyed, R D Dixit, A M Kadri, Harsh Bakshi, Chaitanya Joshi, Madhvi Joshi |
| EPI_ISL_483821 | Government Medical College, Vadodara | Gujarat Biotechnology Research Centre | Labdhi Pandya, Afzal Ansari, Nikha Trivedi, Meenakshi Shah, Neena Doshi, Varsha Godbole, Apurvasinh Puvar, Janvi Raval, Zarna Patel, Monika Gandhi, Pinal Trivedi, Maharshi Pandya, Nidhi Patel, Nitin Savaliya, Raghawendra Kumar, Dinesh Kumar, Zuber Saiyed, Komal Patel, R D Dixit, A M Kadri, Harsh Bakshi, Chaitanya Joshi, Madhvi Joshi |
| EPI_ISL_483827 | GMERS Medical College Himmatnagar | Gujarat Biotechnology Research Centre | Janvi Raval, Zarna Patel, Monika Gandhi, Pinal Trivedi, Maharshi Pandya, Nidhi Patel, Nitin Savaliya, Raghawendra Kumar, Dinesh Kumar, Zuber Saiyed, Komal Patel, Labdhi Pandya, Afzal Ansari, Nikha Trivedi, Himanshu Khatri, Mayur Gandhi, Apurvasinh Puvar, R D Dixit, A M Kadri, Harsh Bakshi, Chaitanya Joshi, Madhvi Joshi |
| EPI_ISL_483837 | Pandit Deendayal Upadhyay Government Medical College, Rajkot | Gujarat Biotechnology Research Centre | Komal Patel, Labdhi Pandya, Afzal Ansari, Nikha Trivedi, Gauravi Dhruv, Arti Trivedi, Apurvasinh Puvar, Janvi Raval, Zarna Patel, Monika Gandhi, Pinal Trivedi, Maharshi Pandya, Nidhi Patel, Nitin Savaliya, Raghawendra Kumar, Dinesh Kumar, Zuber Saiyed, R D Dixit, A M Kadri, Harsh Bakshi, Chaitanya Joshi, Madhvi Joshi |
| EPI_ISL_483838 | Pandit Deendayal Upadhyay Government Medical College, Rajkot | Gujarat Biotechnology Research Centre | Labdhi Pandya, Afzal Ansari, Nikha Trivedi, Gauravi Dhruv, Arti Trivedi, Apurvasinh Puvar, Janvi Raval, Zarna Patel, Monika Gandhi, Pinal Trivedi, Maharshi Pandya, Nidhi Patel, Nitin Savaliya, Raghawendra Kumar, Dinesh Kumar, Zuber Saiyed, Komal Patel, R D Dixit, A M Kadri, Harsh Bakshi, Chaitanya Joshi, Madhvi Joshi |
| EPI_ISL_483844 | Pandit Deendayal Upadhyay Government Medical College, Rajkot | Gujarat Biotechnology Research Centre | Janvi Raval, Zarna Patel, Monika Gandhi, Pinal Trivedi, Maharshi Pandya, Nidhi Patel, Nitin Savaliya, Raghawendra Kumar, Dinesh Kumar, Zuber Saiyed, Komal Patel, Labdhi Pandya, Afzal Ansari, Nikha Trivedi, Gauravi Dhruv, Arti Trivedi, Apurvasinh Puvar, R D Dixit, A M Kadri, Harsh Bakshi, Chaitanya Joshi, Madhvi Joshi |
| EPI_ISL_483854 | Department of Microbiology, Government Medical College, Surat | Gujarat Biotechnology Research Centre | Komal Patel, Labdhi Pandya, Afzal Ansari, Nikha Trivedi, Naresh Chauhan, Summaiya Mullan, Amit gamit, Apurvasinh Puvar, Janvi Raval, Zarna Patel, Monika Gandhi, Pinal Trivedi, Maharshi Pandya, Nidhi Patel, Nitin Savaliya, Raghawendra Kumar, Dinesh Kumar, Zuber Saiyed, R D Dixit, A M Kadri, Harsh Bakshi, Chaitanya Joshi, Madhvi Joshi |
| EPI_ISL_483855 | Department of Microbiology, Government Medical College, Surat | Gujarat Biotechnology Research Centre | Labdhi Pandya, Afzal Ansari, Nikha Trivedi, Naresh Chauhan, Summaiya Mullan, Amit gamit, Apurvasinh Puvar, Janvi Raval, Zarna Patel, Monika Gandhi, Pinal Trivedi, Maharshi Pandya, Nidhi Patel, Nitin Savaliya, Raghawendra Kumar, Dinesh Kumar, Zuber Saiyed, Komal Patel, R D Dixit, A M Kadri, Harsh Bakshi, Chaitanya Joshi, Madhvi Joshi |
| EPI_ISL_483862 | Department of Microbiology, Government Medical College, Surat | Gujarat Biotechnology Research Centre | Janvi Raval, Zarna Patel, Monika Gandhi, Pinal Trivedi, Maharshi Pandya, Nidhi Patel, Nitin Savaliya, Raghawendra Kumar, Dinesh Kumar, Zuber Saiyed, Komal Patel, Labdhi Pandya, Afzal Ansari, Nikha Trivedi, Naresh Chauhan, Summaiya Mullan, Amit gamit, Apurvasinh Puvar, R D Dixit, A M Kadri, Harsh Bakshi, Chaitanya Joshi, Madhvi Joshi |
| EPI_ISL_483872 | Department of Microbiology, Government Medical College, Surat | Gujarat Biotechnology Research Centre | Komal Patel, Labdhi Pandya, Afzal Ansari, Nikha Trivedi, Naresh Chauhan, Summaiya Mullan, Amit gamit, Apurvasinh Puvar, Janvi Raval, Zarna Patel, Monika Gandhi, Pinal Trivedi, Maharshi Pandya, Nidhi Patel, Nitin Savaliya, Raghawendra Kumar, Dinesh Kumar, Zuber Saiyed, R D Dixit, A M Kadri, Harsh Bakshi, Chaitanya Joshi, Madhvi Joshi |
| EPI_ISL_483873 | Department of Microbiology, Government Medical College, Surat | Gujarat Biotechnology Research Centre | Labdhi Pandya, Afzal Ansari, Nikha Trivedi, Naresh Chauhan, Summaiya Mullan, Amit gamit, Apurvasinh Puvar, Janvi Raval, Zarna Patel, Monika Gandhi, Pinal Trivedi, Maharshi Pandya, Nidhi Patel, Nitin Savaliya, Raghawendra Kumar, Dinesh Kumar, Zuber Saiyed, Komal Patel, R D Dixit, A M Kadri, Harsh Bakshi, Chaitanya Joshi, Madhvi Joshi |
| EPI_ISL_484693, EPI_ISL_484694, EPI_ISL_484695, EPI_ISL_484696, EPI_ISL_484697, EPI_ISL_484698, EPI_ISL_484699, EPI_ISL_484700, EPI_ISL_484701, EPI_ISL_484702, EPI_ISL_484703, EPI_ISL_484704, EPI_ISL_484705, EPI_ISL_484706, EPI_ISL_484707, EPI_ISL_484708 | | | |
| see above | Department of Clinical Microbiology | GIGA Medical Genomics | Keith Durkin, Maria Artesi, Sébastien Bontems, Raphaël Boreux, Cécile Meex, Axelle Chaslain, Céline Fombellida-Lopez, Pierrette Melin, Marie-Pierre Hayette, Vincent Bours. |

| | | | |
|---|---|---|---|
| EPI_ISL_485001, EPI_ISL_485002, EPI_ISL_485388, EPI_ISL_485389, EPI_ISL_485390, EPI_ISL_485391, EPI_ISL_485392, EPI_ISL_485393, EPI_ISL_485394, EPI_ISL_485395, EPI_ISL_485396 | | | |
| see above | University of Ulsan College of Medicine and Asan Medical Center | University of Ulsan College of Medicine and Asan Medical Center | Kuenyoul Park, Jaewoong Lee, Kihyun Lee, Jiwon Jung, Sung-Han Kim, Jina Lee, Mauricio Chalita, Seok-Hwan Yoon, Jongsik Chun, Kyu-Hwa Hur, Heungsup Sung, Mi-Na Kim, and Hae Kyung Lee |
| EPI_ISL_485398 | Department of Internal Medicine, College of Medicine, Chosun University | Department of Internal Medicine, College of Medicine, Chosun University | Kim,D.-M. |
| EPI_ISL_485603 | Division of Infectious Disease | Steininger Lab | Jakob Thannesberger, Ingeborg Klymiuk, Nicolas Rascovan, Lorenz Schubert, Oliver Robak, Christoph Steininger |
| EPI_ISL_486912, EPI_ISL_486913, EPI_ISL_486914, EPI_ISL_486915, EPI_ISL_486916, EPI_ISL_486917 | Maryland Department of Health | Maryland Department of Health | Keller,E. |
| EPI_ISL_487446, EPI_ISL_487447, EPI_ISL_487448, EPI_ISL_487449, EPI_ISL_487450, EPI_ISL_487451, EPI_ISL_487452, EPI_ISL_487453, EPI_ISL_487454, EPI_ISL_487455, EPI_ISL_487456, EPI_ISL_487457, EPI_ISL_487458, EPI_ISL_487459, EPI_ISL_487460, EPI_ISL_487461, EPI_ISL_487462, EPI_ISL_487463, EPI_ISL_487464, EPI_ISL_487465, EPI_ISL_487466 | | | |
| see above | CICM-Mali | Bundeswehr Institut of Microbiology | Kouriba, Dürr, Sangaré, Rehn, Traoré, Bestehorn-Willmann, Walter, Quedraogo, Zimmermann, Maiga, Heitzer, Sogodogo, Antwerpen, Wölfel |
| EPI_ISL_491174, EPI_ISL_491175, EPI_ISL_491176, EPI_ISL_491177, EPI_ISL_491178, EPI_ISL_491179, EPI_ISL_491180, EPI_ISL_491181, EPI_ISL_491182, EPI_ISL_491183, EPI_ISL_491184, EPI_ISL_491185, EPI_ISL_491186, EPI_ISL_491187, EPI_ISL_491188, EPI_ISL_491189, EPI_ISL_491190, EPI_ISL_491191, EPI_ISL_491192, EPI_ISL_491193, EPI_ISL_491194, EPI_ISL_491195, EPI_ISL_491196, EPI_ISL_491197, EPI_ISL_491198, EPI_ISL_491199, EPI_ISL_491200, EPI_ISL_491201 | | | |
| see above | Instituto Gulbenkian de Ciência | Instituto Gulbenkian de Ciência | João Costa, Cathy Paulino, Joao Sobral, Susana Ladeiro, Ricardo Leite |
| EPI_ISL_491231, EPI_ISL_491232, EPI_ISL_491233, EPI_ISL_491234, EPI_ISL_491235, EPI_ISL_491236, EPI_ISL_491237, EPI_ISL_491238, EPI_ISL_491239, EPI_ISL_491240, EPI_ISL_491241, EPI_ISL_491242, EPI_ISL_491243, EPI_ISL_491244, EPI_ISL_491245, EPI_ISL_491246, EPI_ISL_491247, EPI_ISL_491248, EPI_ISL_491249, EPI_ISL_491250, EPI_ISL_491251, EPI_ISL_491252, EPI_ISL_491253, EPI_ISL_491254, EPI_ISL_491255, EPI_ISL_491256, EPI_ISL_491257, EPI_ISL_491258, EPI_ISL_491259, EPI_ISL_491260, EPI_ISL_491261, EPI_ISL_491262, EPI_ISL_491263, EPI_ISL_491264 | | | |
| see above | Instituto Gulbenkian de Ciência | Instituto Gulbenkian de Ciência | Joao Sobral, Susana Ladeiro, João Costa, Cathy Paulino, Ricardo Leite |
| EPI_ISL_492066, EPI_ISL_492067, EPI_ISL_492068, EPI_ISL_492069, EPI_ISL_492070, EPI_ISL_492071, EPI_ISL_492072, EPI_ISL_492073 | 1. ViroGenetics - BSL3 Laboratory of Virology, Maopolska Centre of Biotechnology, Jagiellonian University; 2. II Department of Internal Medicine, Faculty of Medicine, Jagiellonian University Medical College; 3. Narodowy Instytut Zdrowia Publicznego - Pastwowy Zakad Higieny (NIZP-PZH) | 1. ViroGenetics - BSL3 Laboratory of Virology, Maopolska Centre of Biotechnology, Jagiellonian University; 2. II Department of Internal Medicine, Faculty of Medicine, Jagiellonian University Medical College; 3. Narodowy Instytut Zdrowia Publicznego - Pastwowy Zakad Higieny (NIZP-PZH). | Katarzyna Pancer, Marek Sanak, Aleksandra A. Zasada, Magdalena Rzeczkowska, Tomasz Wokowicz, Katarzyna Zacharczuk, Agnieszka Koakowska-Kulesza, Katarzyna Owczarek, Aleksandra Milewska, Natalia Wolaniuk, Ewelina Hallman-Szeliska, Pawe P abaj, Wojciech Branicki, Krzysztof Pyr |
| EPI_ISL_493137, EPI_ISL_493139 | Center for Research and Innovation, Faculty of Medical Technology, Mahidol University | Center for Research and Innovation, Faculty of Medical Technology, Mahidol University | Kantima Sangsiriwut; Hatairat Lerdsamran; Jarunee Prasertsopon; Tipsuda Chanmanee; Anek Mungaomklang; Kamolthip Atsawawaranunt; Prabda Praphasiri; Somrak Sirikhetkon; Nattakan Thinpan; Pilaipan Puthavathana |
| EPI_ISL_493149, EPI_ISL_493150, EPI_ISL_493151, EPI_ISL_493152, EPI_ISL_493153, EPI_ISL_493154, EPI_ISL_493155, EPI_ISL_493156, EPI_ISL_493157, EPI_ISL_493158, EPI_ISL_493159, EPI_ISL_493160, EPI_ISL_493161, EPI_ISL_493162, EPI_ISL_493163, EPI_ISL_493164, EPI_ISL_493165, EPI_ISL_493166, EPI_ISL_493167, EPI_ISL_493168, EPI_ISL_493169, EPI_ISL_493170, EPI_ISL_493171, EPI_ISL_493172, EPI_ISL_493173, EPI_ISL_493174, EPI_ISL_493175, EPI_ISL_493176, EPI_ISL_493177, EPI_ISL_493178, EPI_ISL_493179, EPI_ISL_493180, EPI_ISL_493181, EPI_ISL_493182, EPI_ISL_493183, EPI_ISL_493184, EPI_ISL_493185, EPI_ISL_493186, EPI_ISL_493188, EPI_ISL_493189, EPI_ISL_493190 | | | |
| see above | National Virus Resource Center, Chinese Academy of Sciences, Wuhan 430071, China | Computational Virology Group, Center for Bacteria and Viruses Resources and Bioinformation, Wuhan Institute of Virology, Chinese Academy of SciencesWuhan 430071, China | Jianjun Chen, Yi Yan, Yi Huang, Jin Xiong, Hongping Wei, Di Liu |
| EPI_ISL_493350 | Ostfold Hospital Trust - Kalnes, Centre for Laboratory Medicine, Section for gene technology and infection serology | Norwegian Institute of Public Health, Department of Virology | Kathrine Stene-Johansen, Kamilla Heddeland Instefjord, Hilde Elshaug, Rasmus Riis Kopperud, Karoline Bragstad, Olav Hungnes |
| EPI_ISL_493351, EPI_ISL_493352, EPI_ISL_493353, EPI_ISL_493354, EPI_ISL_493355 | Oslo University Hospital, Department of Medical Microbiology | Norwegian Institute of Public Health, Department of Virology | Kathrine Stene-Johansen, Kamilla Heddeland Instefjord, Hilde Elshaug, Rasmus Riis Kopperud, Karoline Bragstad, Olav Hungnes |
| EPI_ISL_493356, EPI_ISL_493357, EPI_ISL_493358, EPI_ISL_493359, EPI_ISL_493360, EPI_ISL_493361, EPI_ISL_493362, EPI_ISL_493363, EPI_ISL_493364, EPI_ISL_493365, EPI_ISL_493366, EPI_ISL_493367, EPI_ISL_493368, EPI_ISL_493369, EPI_ISL_493370, EPI_ISL_493371, EPI_ISL_493372, EPI_ISL_493373, EPI_ISL_493374, EPI_ISL_493375, EPI_ISL_493376, EPI_ISL_493377, EPI_ISL_493378, EPI_ISL_493379, EPI_ISL_493380 | | | |
| see above | Furst Medical Laboratory | Norwegian Institute of Public Health, Department of Virology | Kathrine Stene-Johansen, Kamilla Heddeland Instefjord, Hilde Elshaug, Rasmus Riis Kopperud, Karoline Bragstad, Olav Hungnes |
| EPI_ISL_493381, EPI_ISL_493382, EPI_ISL_493383 | Medical Microbiology Unit, Department for Laboratory Medicine, Drammen Hospital, Vestre Viken Health Trust, | Norwegian Institute of Public Health, Department of Virology | Kathrine Stene-Johansen, Kamilla Heddeland Instefjord, Hilde Elshaug, Rasmus Riis Kopperud, Karoline Bragstad, Olav Hungnes |
| EPI_ISL_493384 | Hospital of Southern Norway - Kristiansand, Department of Medical Microbiology | Norwegian Institute of Public Health, Department of Virology | Kathrine Stene-Johansen, Kamilla Heddeland Instefjord, Hilde Elshaug, Rasmus Riis Kopperud, Karoline Bragstad, Olav Hungnes |
| EPI_ISL_493385, EPI_ISL_493386, EPI_ISL_493387, EPI_ISL_493388, EPI_ISL_493389 | Oslo University Hospital, Department of Medical Microbiology | Norwegian Institute of Public Health, Department of Virology | Kathrine Stene-Johansen, Kamilla Heddeland Instefjord, Hilde Elshaug, Rasmus Riis Kopperud, Karoline Bragstad, Olav Hungnes |
| EPI_ISL_494970 | Dr. Tony Mazzulli Microbiologist-in-Chief | Dr. Jeff Wrana, Senior Investigator | Jeff Wrana, Jess Shen, Seda Barutcu, Kin Chan, Dan Trcka, Marie-Ming Aynaud, Javier Hernandez, Jessica Bourke, Christine Bruce, Bryn Hazlett, Laurence Pelletier, Sue Poutanen, Tony Mazzulli |
| EPI_ISL_495012 | Dr. Tony Mazzulli Microbiologist-in-Chief | Dr. Jeff Wrana, Senior Investigator | Jeff Wrana, Jess Shen, Seda Barutcu, Kin Chan, Dan Trcka, Marie-Ming Aynaud, Javier Hernandez, Jessica Bourke, Christine Bruce, Bryn Hazlett, Laurence Pelletier, Sue Poutanen, Tony Mazzulli |
| EPI_ISL_495013 | Dr. Tony Mazzulli Microbiologist-in-Chief | Dr. Jeff Wrana, Senior Investigator | Jeff Wrana, Jess Shen, Seda Barutcu, Kin Chan, Dan Trcka, Marie-Ming Aynaud, Javier Hernandez, Jessica Bourke, Christine Bruce, Bryn Hazlett, Laurence Pelletier, Sue Poutanen, Tony Mazzulli |
| EPI_ISL_495014 | B.J. Medical College and Civil hospital | Gujarat Biotechnology Research Centre | Janvi Raval, Zarna Patel, Monika Gandhi, Pinal Trivedi, Maharshi Pandya, Nidhi Patel, Nitin Savaliya, Raghawendra Kumar, Dinesh Kumar, Zuber Saiyed, Komal Patel, Labdhi Pandya, Afzal Ansari, Nikha Trivedi, Pranay Shah, Kamlesh J Upadhyay, Sanjay Kapadia, Apurvasinh Puvar, R D Dixit, A M Kadri, Harsh Bakshi, Chaitanya Joshi, Madhvi Joshi |
| EPI_ISL_495020 | Government Medical College, Bhavnagar | Gujarat Biotechnology Research Centre | Kairavi Desai, Saklin Malek, Shirish Patel, Nitin Savaliya, Raghawendra Kumar, Dinesh Kumar, Zuber Saiyed, Komal Patel, Labdhi Pandya, Afzal Ansari, Nikha Trivedi, Apurvasinh Puvar, Janvi Raval, Zarna Patel, Monika Gandhi, Pinal Trivedi, Maharshi Pandya, Nidhi Patel, R D Dixit, A M Kadri, Harsh Bakshi, Chaitanya Joshi, Madhvi Joshi |
| EPI_ISL_495026 | GMERS Medical College & Hospital, Gotri, Vadodara | Gujarat Biotechnology Research Centre | Komal Patel, Labdhi Pandya, Afzal Ansari, Nikha Trivedi, Meenakshi Shah, Neena Doshi, Varsha Godbole, Apurvasinh Puvar, Janvi Raval, Zarna Patel, Monika Gandhi, Pinal Trivedi, Maharshi Pandya, Nidhi Patel, Nitin Savaliya, Raghawendra Kumar, Dinesh Kumar, Zuber Saiyed, R D Dixit, A M Kadri, Harsh Bakshi, Chaitanya Joshi, Madhvi Joshi |
| EPI_ISL_495027 | GMERS Medical College & Hospital, Gotri, Vadodara | Gujarat Biotechnology Research Centre | Labdhi Pandya, Afzal Ansari, Nikha Trivedi, Meenakshi Shah, Neena Doshi, Varsha Godbole, Apurvasinh Puvar, Janvi Raval, Zarna Patel, Monika Gandhi, Pinal Trivedi, Maharshi Pandya, Nidhi Patel, Nitin Savaliya, Raghawendra Kumar, Dinesh Kumar, Zuber Saiyed, Komal Patel, R D Dixit, A M Kadri, Harsh Bakshi, Chaitanya Joshi, Madhvi Joshi |
| EPI_ISL_495034 | GMERS Medical College & Hospital, Gotri, Vadodara | Gujarat Biotechnology Research Centre | Janvi Raval, Zarna Patel, Monika Gandhi, Pinal Trivedi, Maharshi Pandya, Nidhi Patel, Nitin Savaliya, Raghawendra Kumar, Dinesh Kumar, Zuber Saiyed, Komal Patel, Labdhi Pandya, Afzal Ansari, Nikha Trivedi, Meenakshi Shah, Neena Doshi, Varsha Godbole, Apurvasinh Puvar, R D Dixit, A M Kadri, Harsh Bakshi, Chaitanya Joshi, Madhvi Joshi |
| EPI_ISL_495044 | GMERS Medical College & Hospital, Gotri, Vadodara | Gujarat Biotechnology Research Centre | Komal Patel, Labdhi Pandya, Afzal Ansari, Nikha Trivedi, Meenakshi Shah, Neena Doshi, Varsha Godbole, Apurvasinh Puvar, Janvi Raval, Zarna Patel, Monika Gandhi, Pinal Trivedi, Maharshi Pandya, Nidhi Patel, Nitin Savaliya, Raghawendra Kumar, Dinesh Kumar, Zuber Saiyed, R D Dixit, A M Kadri, Harsh Bakshi, Chaitanya Joshi, Madhvi Joshi |
| EPI_ISL_495045 | GMERS Medical College & Hospital, Gotri, Vadodara | Gujarat Biotechnology Research Centre | Labdhi Pandya, Afzal Ansari, Nikha Trivedi, Meenakshi Shah, Neena Doshi, Varsha Godbole, Apurvasinh Puvar, Janvi Raval, Zarna Patel, Monika Gandhi, |

| | | | |
|---|---|---|---|
| | | | Pinal Trivedi, Maharshi Pandya, Nidhi Patel, Nitin Savaliya, Raghawendra Kumar, Dinesh Kumar, Zuber Saiyed, Komal Patel, R D Dixit, A M Kadri, Harsh Bakshi, Chaitanya Joshi, Madhvi Joshi |
| EPI_ISL_495052 | GMERS Medical College & Hospital, Gotri, Vadodara | Gujarat Biotechnology Research Centre | Janvi Raval, Zarna Patel, Monika Gandhi, Pinal Trivedi, Maharshi Pandya, Nidhi Patel, Nitin Savaliya, Raghawendra Kumar, Dinesh Kumar, Zuber Saiyed, Komal Patel, Labdhi Pandya, Afzal Ansari, Nikha Trivedi, Meenakshi Shah, Neena Doshi, Varsha Godbole, Apurvasinh Puvar, R D Dixit, A M Kadri, Harsh Bakshi, Chaitanya Joshi, Madhvi Joshi |
| EPI_ISL_495062 | GMERS Medical College & Hospital, Gotri, Vadodara | Gujarat Biotechnology Research Centre | Komal Patel, Labdhi Pandya, Afzal Ansari, Nikha Trivedi, Meenakshi Shah, Neena Doshi, Varsha Godbole, Apurvasinh Puvar, Janvi Raval, Zarna Patel, Monika Gandhi, Pinal Trivedi, Maharshi Pandya, Nidhi Patel, Nitin Savaliya, Raghawendra Kumar, Dinesh Kumar, Zuber Saiyed, R D Dixit, A M Kadri, Harsh Bakshi, Chaitanya Joshi, Madhvi Joshi |
| EPI_ISL_495063 | GMERS Medical College & Hospital, Himmatnagar | Gujarat Biotechnology Research Centre | Labdhi Pandya, Afzal Ansari, Nikha Trivedi, Himanshu Khatri, Mayur Gandhi, Apurvasinh Puvar, Janvi Raval, Zarna Patel, Monika Gandhi, Pinal Trivedi, Maharshi Pandya, Nidhi Patel, Nitin Savaliya, Raghawendra Kumar, Dinesh Kumar, Zuber Saiyed, Komal Patel, R D Dixit, A M Kadri, Harsh Bakshi, Chaitanya Joshi, Madhvi Joshi, |
| EPI_ISL_495068 | Dr. N. D. Desai Medical College & Hospital | Gujarat Biotechnology Research Centre | Jigar Gusani, Supreet Prabhu, Apurvasinh Puvar, Janvi Raval, Zarna Patel, Monika Gandhi, Pinal Trivedi, Maharshi Pandya, Nidhi Patel, Nitin Savaliya, Raghawendra Kumar, Dinesh Kumar, Zuber Saiyed, Komal Patel, Labdhi Pandya, Afzal Ansari, Nikha Trivedi, J G Buch, R D Dixit, A M Kadri, Harsh Bakshi, Chaitanya Joshi, Madhvi Joshi |
| EPI_ISL_495071 | Department of MicroBiology, Government Medical College, Surat | Gujarat Biotechnology Research Centre | Janvi Raval, Zarna Patel, Monika Gandhi, Pinal Trivedi, Maharshi Pandya, Nidhi Patel, Nitin Savaliya, Raghawendra Kumar, Dinesh Kumar, Zuber Saiyed, Komal Patel, Labdhi Pandya, Afzal Ansari, Nikha Trivedi, Naresh Chauhan, Summaiya Mullan, Amit gamit, Apurvasinh Puvar, R D Dixit, A M Kadri, Harsh Bakshi, Chaitanya Joshi, Madhvi Joshi |
| EPI_ISL_495200 | CSIR-Centre for Cellular and Molecular Biology | CSIR-Centre for Cellular and Molecular Biology | Lamuk Zaveri, Shagufta Khan, Namami Gaur, Sakshi Shambhavi, Nikhil Hajirnis, M Soujanya Reddy, Pratheusa Maccha, Tulasi Nagabandi, Purushotham Vodnala, Payel Mukherjee, Sofia Banu, Priya Singh, Onkar Kulkarni, Dhiviya Vedagiri, Divya Gupta, Vishal Sah, Santosh Kumar Kuncha, Krishnan Harinivas Harshan, Archana Bharadwaj Siva, Karthik Bharadwaj Tallapaka, Renu Sudhakar, Somesh Gorde, Gangumala Srinivas Reddy, Sujoy Deb, Swati Bayyana, Rakesh K Mishra, Divya Tej Sowpati |
| EPI_ISL_495215, EPI_ISL_495233 | CSIR-Centre for Cellular and Molecular Biology | CSIR-Centre for Cellular and Molecular Biology | Lamuk Zaveri, Shagufta Khan,Nikhil Hajirnis, M Soujanya Reddy, Pratheusa Maccha, Namami Gaur, Sakshi Shambhavi, Tulasi Nagabandi, Purushotham Vodnala, Payel Mukherjee, Sofia Banu, Priya Singh, Onkar Kulkarni, Dhiviya Vedagiri, Divya Gupta, Vishal Sah, Santosh Kumar Kuncha, Krishnan Harinivas Harshan, Archana Bharadwaj Siva, Karthik Bharadwaj Tallapaka,Umesh Kumar, Unis Ahmad Bhat, Ajay Sarawagi, Priyanka Pant, Rajkanwar Nathawat, Rakesh K Mishra, Divya Tej Sowpati |
| EPI_ISL_495234, EPI_ISL_495243 | CSIR-Centre for Cellular and Molecular Biology | CSIR-Centre for Cellular and Molecular Biology | Lamuk Zaveri, Shagufta Khan,Nikhil Hajirnis, M Soujanya Reddy, Pratheusa Maccha, Namami Gaur, Sakshi Shambhavi, Tulasi Nagabandi, Purushotham Vodnala, Payel Mukherjee, Sofia Banu, Priya Singh, Onkar Kulkarni, Dhiviya Vedagiri, Divya Gupta, Vishal Sah, Santosh Kumar Kuncha, Krishnan Harinivas Harshan, Archana Bharadwaj Siva, Karthik Bharadwaj Tallapaka,Zeba Rizvi, Zuberwasim Sayyad, Kakade Aishwarya Arun, Amrutha H C, Ananga Ghosh, Rakesh K Mishra, Divya Tej Sowpati |
| EPI_ISL_495246 | CSIR-Centre for Cellular and Molecular Biology | CSIR-Centre for Cellular and Molecular Biology | Lamuk Zaveri, Shagufta Khan, Namami Gaur, Sakshi Shambhavi, Nikhil Hajirnis, M Soujanya Reddy, Pratheusa Maccha, Tulasi Nagabandi, Purushotham Vodnala, Payel Mukherjee, Sofia Banu, Priya Singh, Onkar Kulkarni, Dhiviya Vedagiri, Divya Gupta, Vishal Sah, Santosh Kumar Kuncha, Krishnan Harinivas Harshan, Archana Bharadwaj Siva, Karthik Bharadwaj Tallapaka, Renu Sudhakar, Somesh Gorde, Gangumala Srinivas Reddy, Sujoy Deb, Swati Bayyana, Rakesh K Mishra, Divya Tej Sowpati |
| EPI_ISL_495247 | CSIR-Centre for Cellular and Molecular Biology | CSIR-Centre for Cellular and Molecular Biology | Lamuk Zaveri, Shagufta Khan,Nikhil Hajirnis, M Soujanya Reddy, Pratheusa Maccha, Namami Gaur, Sakshi Shambhavi, Tulasi Nagabandi, Purushotham Vodnala, Payel Mukherjee, Sofia Banu, Priya Singh, Onkar Kulkarni, Dhiviya Vedagiri, Divya Gupta, Vishal Sah, Santosh Kumar Kuncha, Krishnan Harinivas Harshan, Archana Bharadwaj Siva, Karthik Bharadwaj Tallapaka,Zeba Rizvi, Zuberwasim Sayyad, Kakade Aishwarya Arun, Amrutha H C, Ananga Ghosh, Rakesh K Mishra, Divya Tej Sowpati |
| EPI_ISL_495250 | CSIR-Centre for Cellular and Molecular Biology | CSIR-Centre for Cellular and Molecular Biology | Lamuk Zaveri, Shagufta Khan,Nikhil Hajirnis, M Soujanya Reddy, Pratheusa Maccha, Namami Gaur, Sakshi Shambhavi, Tulasi Nagabandi, Purushotham Vodnala, Payel Mukherjee, Sofia Banu, Priya Singh, Onkar Kulkarni, Dhiviya Vedagiri, Divya Gupta, Vishal Sah, Santosh Kumar Kuncha, Krishnan Harinivas Harshan, Archana Bharadwaj Siva, Karthik Bharadwaj Tallapaka,Umesh Kumar, Unis Ahmad Bhat, Ajay Sarawagi, Priyanka Pant, Rajkanwar Nathawat, Rakesh K Mishra, Divya Tej Sowpati |
| EPI_ISL_495258, EPI_ISL_495265 | CSIR-Centre for Cellular and Molecular Biology | CSIR-Centre for Cellular and Molecular Biology | Lamuk Zaveri, Shagufta Khan, Namami Gaur, Sakshi Shambhavi, Nikhil Hajirnis, M Soujanya Reddy, Pratheusa Maccha, Tulasi Nagabandi, Purushotham Vodnala, Payel Mukherjee, Sofia Banu, Priya Singh, Onkar Kulkarni, Dhiviya Vedagiri, Divya Gupta, Vishal Sah, Santosh Kumar Kuncha, Krishnan Harinivas Harshan, Archana Bharadwaj Siva, Karthik Bharadwaj Tallapaka, Renu Sudhakar, Somesh Gorde, Gangumala Srinivas Reddy, Sujoy Deb, Swati Bayyana, Rakesh K Mishra, Divya Tej Sowpati |
| EPI_ISL_497736, EPI_ISL_497738, EPI_ISL_497744, EPI_ISL_497745 | Instituto Nacional de Salud, Bogotá, Colombia | Instituto Nacional de Salud, Bogotá, Colombia | Katherine Laiton-Donato, Diego A. Álvarez-Díaz, Carlos Franco-Muñoz, Jonathan Reales, Diego Andrés Prada, Jose A. Usme-Ciro, Nicolas D. Franco-Sierra, Zulma M. Cucunubá, Christian Julian VillabonaArenas, Liz Villabona-Arenas, Sussy Echeverria, Astrid C. Flórez, Carolina Ferro, Diana Marcela Walteros-Acero, Franklin Prieto, Carlos Andrés Durán, Martha Lucia Ospina Martinez, Marcela Mercado-Reyes |
| EPI_ISL_497768, EPI_ISL_497769, EPI_ISL_497770, EPI_ISL_497771, EPI_ISL_497772, EPI_ISL_497773, EPI_ISL_497774, EPI_ISL_497775, EPI_ISL_497776, EPI_ISL_497777, EPI_ISL_497778, EPI_ISL_497779, EPI_ISL_497780, EPI_ISL_497781, EPI_ISL_497782, EPI_ISL_497783, EPI_ISL_497784, EPI_ISL_497785, EPI_ISL_497786, EPI_ISL_497787, EPI_ISL_497788, EPI_ISL_497789, EPI_ISL_497790, EPI_ISL_497791, EPI_ISL_497792, EPI_ISL_497793, EPI_ISL_497794, EPI_ISL_497795, EPI_ISL_497796, EPI_ISL_497797, EPI_ISL_497798, EPI_ISL_497799, EPI_ISL_497800, EPI_ISL_497801, EPI_ISL_497802, EPI_ISL_497803, EPI_ISL_497804, EPI_ISL_497805, EPI_ISL_497806, EPI_ISL_497807, EPI_ISL_497808, EPI_ISL_497809, EPI_ISL_497810, EPI_ISL_497811, EPI_ISL_497812, EPI_ISL_497813, EPI_ISL_497814, EPI_ISL_497815, EPI_ISL_497816, EPI_ISL_497817, EPI_ISL_497818, EPI_ISL_497819, EPI_ISL_497820, EPI_ISL_497821, EPI_ISL_497822, EPI_ISL_497823, EPI_ISL_497824, EPI_ISL_497825, EPI_ISL_497826, EPI_ISL_497827, EPI_ISL_497828, EPI_ISL_497829, EPI_ISL_497830, EPI_ISL_497831, EPI_ISL_497832, EPI_ISL_497833, EPI_ISL_497834, EPI_ISL_497835, EPI_ISL_497836, EPI_ISL_497837, EPI_ISL_497838, EPI_ISL_497839, EPI_ISL_497840, EPI_ISL_497841, EPI_ISL_497842, EPI_ISL_497843, EPI_ISL_497844, EPI_ISL_497845, EPI_ISL_497846, EPI_ISL_497847, EPI_ISL_497848, EPI_ISL_497849, EPI_ISL_497850, EPI_ISL_497851, EPI_ISL_497852, EPI_ISL_497853, EPI_ISL_497854, EPI_ISL_497855, EPI_ISL_497856, EPI_ISL_497857, EPI_ISL_497858, EPI_ISL_497859, EPI_ISL_497860, EPI_ISL_497861, EPI_ISL_497862, EPI_ISL_497863, EPI_ISL_497864, EPI_ISL_497865, EPI_ISL_497866, EPI_ISL_497867, EPI_ISL_497868, EPI_ISL_497869, EPI_ISL_497870 | | | |
| see above | Department of Microbiology, The University of Hong Kong | Department of Microbiology, The University of Hong Kong | Kelvin K.W. To, Kwok-Yung Yuen |
| EPI_ISL_497951 | Division of Viral Diseases, Center for Laboratory Control of Infectious Diseases, Korea Centers for Diseases Control and Prevention | Division of Viral Diseases, Center for Laboratory Control of Infectious Diseases, Korea Centers for Diseases Control and Prevention | Jeong-Min Kim, Yoon-Seok Chung, Namjoo Lee, Sang Hee Woo, Hye-Jun Jo, Heui Man Kim, Jun-Sub Kim, Myung Guk Han |
| EPI_ISL_497953, EPI_ISL_497954, EPI_ISL_497955, EPI_ISL_497956, EPI_ISL_497957, EPI_ISL_497958, EPI_ISL_497959, EPI_ISL_497960 | Division of Viral Diseases, Center for Laboratory Control of Infectious Diseases, Korea Centers for Diseases Control and Prevention | Division of Viral Diseases, Center for Laboratory Control of Infectious Diseases, Korea Centers for Diseases Control and Prevention | Jeong-Min Kim, Yoon-Seok Chung, Namjoo Lee, Sang Hee Woo, Hye-Jun Jo, Heui Man Kim, Jun-Sub Kim, Dong Hyun Song, Daesang Lee, Seong Tae Jeong, Myung Guk Han |
| EPI_ISL_497961, EPI_ISL_497962, EPI_ISL_497963, EPI_ISL_497964, EPI_ISL_497965, EPI_ISL_497966, EPI_ISL_497967 | Division of Viral Diseases, Center for Laboratory Control of Infectious Diseases, Korea Centers for Diseases Control and Prevention | Division of Viral Diseases, Center for Laboratory Control of Infectious Diseases, Korea Centers for Diseases Control and Prevention | Jeong-Min Kim, Yoon-Seok Chung, Namjoo Lee, Sang Hee Woo, Hye-Jun Jo, Heui Man Kim, Jun-Sub Kim, Myung Guk Han |
| EPI_ISL_497968, EPI_ISL_497969, EPI_ISL_497970, EPI_ISL_497971 | Division of Viral Diseases, Center for Laboratory Control of Infectious Diseases, Korea Centers for Diseases Control and Prevention | Division of Viral Diseases, Center for Laboratory Control of Infectious Diseases, Korea Centers for Diseases Control and Prevention | Jeong-Min Kim, Yoon-Seok Chung, Namjoo Lee, Sang Hee Woo, Hye-Jun Jo, Heui Man Kim, Jun-Sub Kim, Dong Hyun Song, Daesang Lee, Seong Tae Jeong, Myung Guk Han |
| EPI_ISL_497972, EPI_ISL_497973 | Division of Viral Diseases, Center for Laboratory Control of Infectious Diseases, Korea Centers for Diseases Control and Prevention | Division of Viral Diseases, Center for Laboratory Control of Infectious Diseases, Korea Centers for Diseases Control and Prevention | Jeong-Min Kim, Yoon-Seok Chung, Namjoo Lee, Sang Hee Woo, Hye-Jun Jo, Heui Man Kim, Jun-Sub Kim, Myung Guk Han |
| EPI_ISL_497974 | Division of Viral Diseases, Center for Laboratory Control of Infectious Diseases, Korea Centers for Diseases Control and Prevention | Division of Viral Diseases, Center for Laboratory Control of Infectious Diseases, Korea Centers for Diseases Control and Prevention | Jeong-Min Kim, Yoon-Seok Chung, Namjoo Lee, Sang Hee Woo, Hye-Jun Jo, Heui Man Kim, Jun-Sub Kim, Dong Hyun Song, Daesang Lee, Seong Tae Jeong, Myung Guk Han |
| EPI_ISL_497975, EPI_ISL_497976, | Division of Viral Diseases, Center for Laboratory Control of | Division of Viral Diseases, Center for Laboratory Control of | Jeong-Min Kim, Yoon-Seok Chung, Namjoo Lee, Sang Hee Woo, Hye-Jun Jo, Heui Man Kim, Jun-Sub Kim, Myung Guk Han |

| | | | |
|---|---|---|---|
| EPI_ISL_497977, EPI_ISL_497978, EPI_ISL_497979, EPI_ISL_497980, EPI_ISL_497981, EPI_ISL_497982, EPI_ISL_497983, EPI_ISL_497984 | Infectious Diseases, Korea Centers for Diseases Control and Prevention | Infectious Diseases, Korea Centers for Diseases Control and Prevention | |
| EPI_ISL_497985 | Division of Viral Diseases, Center for Laboratory Control of Infectious Diseases, Korea Centers for Diseases Control and Prevention | Division of Viral Diseases, Center for Laboratory Control of Infectious Diseases, Korea Centers for Diseases Control and Prevention | Jeong-Min Kim, Yoon-Seok Chung, Namjoo Lee, Sang Hee Woo, Hye-Jun Jo, Heui Man Kim, Jun-Sub Kim, Dong Hyun Song, Daesang Lee, Seong Tae Jeong, Myung Guk Han |
| EPI_ISL_497986 | Division of Viral Diseases, Center for Laboratory Control of Infectious Diseases, Korea Centers for Diseases Control and Prevention | Division of Viral Diseases, Center for Laboratory Control of Infectious Diseases, Korea Centers for Diseases Control and Prevention | Jeong-Min Kim, Yoon-Seok Chung, Namjoo Lee, Sang Hee Woo, Hye-Jun Jo, Heui Man Kim, Jun-Sub Kim, Myung Guk Han |
| EPI_ISL_497987, EPI_ISL_497988, EPI_ISL_497989, EPI_ISL_497990 | Division of Viral Diseases, Center for Laboratory Control of Infectious Diseases, Korea Centers for Diseases Control and Prevention | Division of Viral Diseases, Center for Laboratory Control of Infectious Diseases, Korea Centers for Diseases Control and Prevention | Jeong-Min Kim, Yoon-Seok Chung, Namjoo Lee, Sang Hee Woo, Hye-Jun Jo, Heui Man Kim, Jun-Sub Kim, Dong Hyun Song, Daesang Lee, Seong Tae Jeong, Myung Guk Han |
| EPI_ISL_497991, EPI_ISL_497992, EPI_ISL_497993 | Division of Viral Diseases, Center for Laboratory Control of Infectious Diseases, Korea Centers for Diseases Control and Prevention | Division of Viral Diseases, Center for Laboratory Control of Infectious Diseases, Korea Centers for Diseases Control and Prevention | Jeong-Min Kim, Yoon-Seok Chung, Namjoo Lee, Sang Hee Woo, Hye-Jun Jo, Heui Man Kim, Jun-Sub Kim, Myung Guk Han |
| EPI_ISL_497994, EPI_ISL_497995 | Division of Viral Diseases, Center for Laboratory Control of Infectious Diseases, Korea Centers for Diseases Control and Prevention | Division of Viral Diseases, Center for Laboratory Control of Infectious Diseases, Korea Centers for Diseases Control and Prevention | Jeong-Min Kim, Yoon-Seok Chung, Namjoo Lee, Sang Hee Woo, Hye-Jun Jo, Heui Man Kim, Jun-Sub Kim, Dong Hyun Song, Daesang Lee, Seong Tae Jeong, Myung Guk Han |
| EPI_ISL_497996 | Division of Viral Diseases, Center for Laboratory Control of Infectious Diseases, Korea Centers for Diseases Control and Prevention | Division of Viral Diseases, Center for Laboratory Control of Infectious Diseases, Korea Centers for Diseases Control and Prevention | Jeong-Min Kim, Yoon-Seok Chung, Namjoo Lee, Sang Hee Woo, Hye-Jun Jo, Heui Man Kim, Jun-Sub Kim, Myung Guk Han |
| EPI_ISL_497997 | Division of Viral Diseases, Center for Laboratory Control of Infectious Diseases, Korea Centers for Diseases Control and Prevention | Division of Viral Diseases, Center for Laboratory Control of Infectious Diseases, Korea Centers for Diseases Control and Prevention | Jeong-Min Kim, Yoon-Seok Chung, Namjoo Lee, Sang Hee Woo, Hye-Jun Jo, Heui Man Kim, Jun-Sub Kim, Dong Hyun Song, Daesang Lee, Seong Tae Jeong, Myung Guk Han |
| EPI_ISL_497998, EPI_ISL_497999, EPI_ISL_498000, EPI_ISL_498001, EPI_ISL_498002, EPI_ISL_498003 | Division of Viral Diseases, Center for Laboratory Control of Infectious Diseases, Korea Centers for Diseases Control and Prevention | Division of Viral Diseases, Center for Laboratory Control of Infectious Diseases, Korea Centers for Diseases Control and Prevention | Jeong-Min Kim, Yoon-Seok Chung, Namjoo Lee, Sang Hee Woo, Hye-Jun Jo, Heui Man Kim, Jun-Sub Kim, Myung Guk Han |
| EPI_ISL_498004 | Division of Viral Diseases, Center for Laboratory Control of Infectious Diseases, Korea Centers for Diseases Control and Prevention | Division of Viral Diseases, Center for Laboratory Control of Infectious Diseases, Korea Centers for Diseases Control and Prevention | Jeong-Min Kim, Yoon-Seok Chung, Namjoo Lee, Sang Hee Woo, Hye-Jun Jo, Heui Man Kim, Jun-Sub Kim, Dong Hyun Song, Daesang Lee, Seong Tae Jeong, Myung Guk Han |
| EPI_ISL_498005 | Division of Viral Diseases, Center for Laboratory Control of Infectious Diseases, Korea Centers for Diseases Control and Prevention | Division of Viral Diseases, Center for Laboratory Control of Infectious Diseases, Korea Centers for Diseases Control and Prevention | Jeong-Min Kim, Yoon-Seok Chung, Namjoo Lee, Sang Hee Woo, Hye-Jun Jo, Heui Man Kim, Jun-Sub Kim, Myung Guk Han |
| EPI_ISL_498006, EPI_ISL_498007, EPI_ISL_498008, EPI_ISL_498009, EPI_ISL_498010, EPI_ISL_498011, EPI_ISL_498012, EPI_ISL_498013, EPI_ISL_498014, EPI_ISL_498015, EPI_ISL_498016 | | | |
| see above | Division of Viral Diseases, Center for Laboratory Control of Infectious Diseases, Korea Centers for Diseases Control and Prevention | Division of Viral Diseases, Center for Laboratory Control of Infectious Diseases, Korea Centers for Diseases Control and Prevention | Jeong-Min Kim, Yoon-Seok Chung, Namjoo Lee, Sang Hee Woo, Hye-Jun Jo, Heui Man Kim, Jun-Sub Kim, Dong Hyun Song, Daesang Lee, Seong Tae Jeong, Myung Guk Han |
| EPI_ISL_498017 | Division of Viral Diseases, Center for Laboratory Control of Infectious Diseases, Korea Centers for Diseases Control and Prevention | Division of Viral Diseases, Center for Laboratory Control of Infectious Diseases, Korea Centers for Diseases Control and Prevention | Jeong-Min Kim, Yoon-Seok Chung, Namjoo Lee, Sang Hee Woo, Hye-Jun Jo, Heui Man Kim, Jun-Sub Kim, Myung Guk Han |
| EPI_ISL_498018, EPI_ISL_498019, EPI_ISL_498020, EPI_ISL_498021, EPI_ISL_498022, EPI_ISL_498023, EPI_ISL_498024, EPI_ISL_498025, EPI_ISL_498026, EPI_ISL_498027, EPI_ISL_498028, EPI_ISL_498029, EPI_ISL_498030, EPI_ISL_498031, EPI_ISL_498032, EPI_ISL_498033, EPI_ISL_498034, EPI_ISL_498035 | | | |
| see above | Division of Viral Diseases, Center for Laboratory Control of Infectious Diseases, Korea Centers for Diseases Control and Prevention | Division of Viral Diseases, Center for Laboratory Control of Infectious Diseases, Korea Centers for Diseases Control and Prevention | Jeong-Min Kim, Yoon-Seok Chung, Namjoo Lee, Sang Hee Woo, Hye-Jun Jo, Heui Man Kim, Jun-Sub Kim, Dong Hyun Song, Daesang Lee, Seong Tae Jeong, Myung Guk Han |
| EPI_ISL_498036 | Division of Viral Diseases, Center for Laboratory Control of Infectious Diseases, Korea Centers for Diseases Control and Prevention | Division of Viral Diseases, Center for Laboratory Control of Infectious Diseases, Korea Centers for Diseases Control and Prevention | Jeong-Min Kim, Yoon-Seok Chung, Namjoo Lee, Sang Hee Woo, Hye-Jun Jo, Heui Man Kim, Jun-Sub Kim, Myung Guk Han |
| EPI_ISL_498037, EPI_ISL_498038, EPI_ISL_498039, EPI_ISL_498040, EPI_ISL_498041, EPI_ISL_498042, EPI_ISL_498043, EPI_ISL_498044, EPI_ISL_498045, EPI_ISL_498046, EPI_ISL_498047, EPI_ISL_498048, EPI_ISL_498049, EPI_ISL_498050, EPI_ISL_498051, EPI_ISL_498052, EPI_ISL_498053 | | | |
| see above | Division of Viral Diseases, Center for Laboratory Control of Infectious Diseases, Korea Centers for Diseases Control and Prevention | Division of Viral Diseases, Center for Laboratory Control of Infectious Diseases, Korea Centers for Diseases Control and Prevention | Jeong-Min Kim, Yoon-Seok Chung, Namjoo Lee, Sang Hee Woo, Hye-Jun Jo, Heui Man Kim, Jun-Sub Kim, Dong Hyun Song, Daesang Lee, Seong Tae Jeong, Myung Guk Han |
| EPI_ISL_498127, EPI_ISL_498128, EPI_ISL_498129, EPI_ISL_498130, EPI_ISL_498131, EPI_ISL_498132, EPI_ISL_498133, EPI_ISL_498134, EPI_ISL_498135, EPI_ISL_498136, EPI_ISL_498137, EPI_ISL_498138, EPI_ISL_498139, EPI_ISL_498140, EPI_ISL_498141, EPI_ISL_498142, EPI_ISL_498143, EPI_ISL_498144, EPI_ISL_498145, EPI_ISL_498146, EPI_ISL_498147, EPI_ISL_498148, EPI_ISL_498149, EPI_ISL_498150, EPI_ISL_498151 | | | |
| see above | Department of Clinical Microbiology | GIGA Medical Genomics | Keith Durkin, Maria Artesi, Sébastien Bontems, Raphaël Boreux, Cécile Meex, Axelle Chaslain, Céline Fombellida-Lopez, Pierrette Melin, Marie-Pierre Hayette, Vincent Bours. |
| EPI_ISL_498152, EPI_ISL_498153, EPI_ISL_498154, EPI_ISL_498155, EPI_ISL_498156, EPI_ISL_498157, EPI_ISL_498158, EPI_ISL_498159, EPI_ISL_498160, EPI_ISL_498161, EPI_ISL_498162, EPI_ISL_498163, EPI_ISL_498164, EPI_ISL_498165, EPI_ISL_498166, EPI_ISL_498167, EPI_ISL_498168, EPI_ISL_498169, EPI_ISL_498170 | | | |
| see above | Instituto Nacional de Salud, Bogotá, Colombia | Instituto Nacional de Salud, Bogotá, Colombia | Katherine Laiton-Donato, Diego A. Álvarez-Díaz, Carlos Franco-Muñoz, Jonathan Reales, Diego Andrés Prada, Jose A. Usme-Ciro, Nicolas D. Franco-Sierra, Zulma M. Cucunubá, Christian Julian VillabonaArenas, Liz Villabona-Arenas, Sussy Echeverria, Astrid C. Flórez, Carolina Ferro, Diana Marcela Walteros-Acero, Franklin Prieto, Carlos Andrés Durán, Martha Lucia Ospina Martinez, Marcela Mercado-Reyes |
| EPI_ISL_498226 | LIC | LIC | LIC |
| EPI_ISL_498269, EPI_ISL_498270, EPI_ISL_498271 | Department of Microbiology, The University of Hong Kong | Department of Microbiology, The University of Hong Kong | Kelvin K.W. To, Kwok-Yung Yuen |
| EPI_ISL_498628, EPI_ISL_498629 | Department of Clinical Microbiology | GIGA Medical Genomics | Keith Durkin, Maria Artesi, Sébastien Bontems, Raphaël Boreux, Cécile Meex, Axelle Chaslain, Céline Fombellida-Lopez, Pierrette Melin, Marie-Pierre Hayette, Vincent Bours. |
| EPI_ISL_500162, EPI_ISL_500167, EPI_ISL_500209, EPI_ISL_500219, EPI_ISL_500221 | Hospital Clínico Universitario de Santiago de Compostela | SeqCOVID-SPAIN consortium/IBV(CSIC) | José Javier Costa Alcalde, Antonio Aguilera Guirao, Mª Luisa Pérez del Molino Bernal, Amparo Coira Nieto, Gema Barbeito Castiñeiras, Rocio Trastoy Pena and SeqCOVID-SPAIN consortium |
| EPI_ISL_500768, EPI_ISL_500769, EPI_ISL_500770, EPI_ISL_500771, EPI_ISL_500772, EPI_ISL_500773, EPI_ISL_500774 | Furst Medical Laboratory | Norwegian Institute of Public Health, Department of Virology | Kathrine Stene-Johansen, Kamilla Heddeland Instefjord, Hilde Elshaug, Rasmus Riis Kopperud, Karoline Bragstad, Olav Hungnes |

| | | | |
|---|---|---|---|
| EPI_ISL_500775 | Hospital of Southern Norway - Kristiansand, Department of Medical Microbiology | Norwegian Institute of Public Health, Department of Virology | Kathrine Stene-Johansen, Kamilla Heddeland Instefjord, Hilde Elshaug, Rasmus Riis Kopperud, Karoline Bragstad, Olav Hungnes |
| EPI_ISL_500776, EPI_ISL_500777, EPI_ISL_500778 | Foerde Hospital, Department of Microbiology | Norwegian Institute of Public Health, Department of Virology | Kathrine Stene-Johansen, Kamilla Heddeland Instefjord, Hilde Elshaug, Rasmus Riis Kopperud, Karoline Bragstad, Olav Hungnes |
| EPI_ISL_500779, EPI_ISL_500780, EPI_ISL_500781, EPI_ISL_500782, EPI_ISL_500783 | Akershus University Hospital, Department for Microbiology and Infectious Disease Control | Norwegian Institute of Public Health, Department of Virology | Kathrine Stene-Johansen, Kamilla Heddeland Instefjord, Hilde Elshaug, Rasmus Riis Kopperud, Karoline Bragstad, Olav Hungnes |
| EPI_ISL_500784, EPI_ISL_500785, EPI_ISL_500786, EPI_ISL_500787, EPI_ISL_500788, EPI_ISL_500789, EPI_ISL_500790 | Furst Medical Laboratory | Norwegian Institute of Public Health, Department of Virology | Kathrine Stene-Johansen, Kamilla Heddeland Instefjord, Hilde Elshaug, Rasmus Riis Kopperud, Karoline Bragstad, Olav Hungnes |
| EPI_ISL_500791, EPI_ISL_500792 | Hospital of Southern Norway - Kristiansand, Department of Medical Microbiology | Norwegian Institute of Public Health, Department of Virology | Kathrine Stene-Johansen, Kamilla Heddeland Instefjord, Hilde Elshaug, Rasmus Riis Kopperud, Karoline Bragstad, Olav Hungnes |
| EPI_ISL_500793, EPI_ISL_500794, EPI_ISL_500795, EPI_ISL_500796 | Akershus University Hospital, Department for Microbiology and Infectious Disease Control | Norwegian Institute of Public Health, Department of Virology | Kathrine Stene-Johansen, Kamilla Heddeland Instefjord, Hilde Elshaug, Rasmus Riis Kopperud, Karoline Bragstad, Olav Hungnes |
| EPI_ISL_500797, EPI_ISL_500798, EPI_ISL_500799 | Hospital of Southern Norway - Kristiansand, Department of Medical Microbiology | Norwegian Institute of Public Health, Department of Virology | Kathrine Stene-Johansen, Kamilla Heddeland Instefjord, Hilde Elshaug, Rasmus Riis Kopperud, Karoline Bragstad, Olav Hungnes |
| EPI_ISL_500800 | Oslo University Hospital, Department of Medical Microbiology | Norwegian Institute of Public Health, Department of Virology | Kathrine Stene-Johansen, Kamilla Heddeland Instefjord, Hilde Elshaug, Rasmus Riis Kopperud, Karoline Bragstad, Olav Hungnes |
| EPI_ISL_500949 | Department of MicroBiology, Government Medical College, Surat | Gujarat Biotechnology Research Centre | Komal Patel, Labdhi Pandya, Afzal Ansari, Nikha Trivedi, Naresh Chauhan, Summaiya Mullan, Amit gamit, Apurvasinh Puvar, Janvi Raval, Zarna Patel, Monika Gandhi, Pinal Trivedi, Maharshi Pandya, Nidhi Patel, Nitin Savaliya, Raghawendra Kumar, Dinesh Kumar, Zuber Saiyed, R D Dixit, A M Kadri, Harsh Bakshi, Chaitanya Joshi, Madhvi Joshi |
| EPI_ISL_500950 | Department of MicroBiology, Government Medical College, Surat | Gujarat Biotechnology Research Centre | Labdhi Pandya, Afzal Ansari, Nikha Trivedi, Naresh Chauhan, Summaiya Mullan, Amit gamit, Apurvasinh Puvar, Janvi Raval, Zarna Patel, Monika Gandhi, Pinal Trivedi, Maharshi Pandya, Nidhi Patel, Nitin Savaliya, Raghawendra Kumar, Dinesh Kumar, Zuber Saiyed, Komal Patel, R D Dixit, A M Kadri, Harsh Bakshi, Chaitanya Joshi, Madhvi Joshi |
| EPI_ISL_506947, EPI_ISL_506948, EPI_ISL_506949, EPI_ISL_506950, EPI_ISL_506951, EPI_ISL_506952, EPI_ISL_506953, EPI_ISL_506954, EPI_ISL_506955, EPI_ISL_506956 | Division of Viral Diseases, Center for Laboratory Control of Infectious Diseases, Korea Centers for Diseases Control and Prevention | Division of Viral Diseases, Center for Laboratory Control of Infectious Diseases, Korea Centers for Diseases Control and Prevention | Jeong-Min Kim, Yoon-Seok Chung, Namjoo Lee, Sang Hee Woo, Hye-Jun Jo, Heui Man Kim, Jun-Sub Kim, Dong Hyun Song, Daesang Lee, Seong Tae Jeong, Myung Guk Han |
| EPI_ISL_506957, EPI_ISL_506958, EPI_ISL_506959, EPI_ISL_506960, EPI_ISL_506961, EPI_ISL_506962, EPI_ISL_506963, EPI_ISL_506964 | Division of Viral Diseases, Center for Laboratory Control of Infectious Diseases, Korea Centers for Diseases Control and Prevention | Division of Viral Diseases, Center for Laboratory Control of Infectious Diseases, Korea Centers for Diseases Control and Prevention | Jeong-Min Kim, Yoon-Seok Chung, Namjoo Lee, Sang Hee Woo, Hye-Jun Jo, Heui Man Kim, Jun-Sub Kim, Myung Guk Han |
| EPI_ISL_506965, EPI_ISL_506966, EPI_ISL_506967, EPI_ISL_506968, EPI_ISL_506969, EPI_ISL_506970, EPI_ISL_506971, EPI_ISL_506972, EPI_ISL_506973, EPI_ISL_506974 | Division of Viral Diseases, Center for Laboratory Control of Infectious Diseases, Korea Centers for Diseases Control and Prevention | Division of Viral Diseases, Center for Laboratory Control of Infectious Diseases, Korea Centers for Diseases Control and Prevention | Jeong-Min Kim, Yoon-Seok Chung, Namjoo Lee, Sang Hee Woo, Hye-Jun Jo, Heui Man Kim, Jun-Sub Kim, Dong Hyun Song, Daesang Lee, Seong Tae Jeong, Myung Guk Han |
| EPI_ISL_506975 | Division of Viral Diseases, Center for Laboratory Control of Infectious Diseases, Korea Centers for Diseases Control and Prevention | Division of Viral Diseases, Center for Laboratory Control of Infectious Diseases, Korea Centers for Diseases Control and Prevention | Jeong-Min Kim, Yoon-Seok Chung, Namjoo Lee, Sang Hee Woo, Hye-Jun Jo, Heui Man Kim, Jun-Sub Kim, Myung Guk Han |
| EPI_ISL_506976, EPI_ISL_506977, EPI_ISL_506978, EPI_ISL_506979, EPI_ISL_506980, EPI_ISL_506981, EPI_ISL_506982, EPI_ISL_506983, EPI_ISL_506984, EPI_ISL_506985, EPI_ISL_506986, EPI_ISL_506987, EPI_ISL_506988, EPI_ISL_506989, EPI_ISL_506990, EPI_ISL_506991, EPI_ISL_506992, EPI_ISL_506993 | | | |
| see above | Division of Viral Diseases, Center for Laboratory Control of Infectious Diseases, Korea Centers for Diseases Control and Prevention | Division of Viral Diseases, Center for Laboratory Control of Infectious Diseases, Korea Centers for Diseases Control and Prevention | Jeong-Min Kim, Yoon-Seok Chung, Namjoo Lee, Sang Hee Woo, Hye-Jun Jo, Heui Man Kim, Jun-Sub Kim, Dong Hyun Song, Daesang Lee, Seong Tae Jeong, Myung Guk Han |
| EPI_ISL_506994, EPI_ISL_506995 | Division of Viral Diseases, Center for Laboratory Control of Infectious Diseases, Korea Centers for Diseases Control and Prevention | Division of Viral Diseases, Center for Laboratory Control of Infectious Diseases, Korea Centers for Diseases Control and Prevention | Jeong-Min Kim, Yoon-Seok Chung, Namjoo Lee, Sang Hee Woo, Hye-Jun Jo, Heui Man Kim, Jun-Sub Kim, Myung Guk Han |
| EPI_ISL_507040, EPI_ISL_507041, EPI_ISL_507042, EPI_ISL_507043, EPI_ISL_507044, EPI_ISL_507045, EPI_ISL_507046, EPI_ISL_507047, EPI_ISL_507048, EPI_ISL_507049, EPI_ISL_507050, EPI_ISL_507051, EPI_ISL_507052, EPI_ISL_507053, EPI_ISL_507054, EPI_ISL_507055, EPI_ISL_507056, EPI_ISL_507057, EPI_ISL_507058, EPI_ISL_507059, EPI_ISL_507060, EPI_ISL_507061, EPI_ISL_507062, EPI_ISL_507063, EPI_ISL_507064, EPI_ISL_507065, EPI_ISL_507066, EPI_ISL_507067, EPI_ISL_507068, EPI_ISL_507069, EPI_ISL_507070, EPI_ISL_507071, EPI_ISL_507072, EPI_ISL_507073, EPI_ISL_507074, EPI_ISL_507075, EPI_ISL_507076, EPI_ISL_507077, EPI_ISL_507078, EPI_ISL_507079, EPI_ISL_507080, EPI_ISL_507081, EPI_ISL_507082, EPI_ISL_507083, EPI_ISL_507084, EPI_ISL_507085, EPI_ISL_507086, EPI_ISL_507087, EPI_ISL_507088, EPI_ISL_507089, EPI_ISL_507090, EPI_ISL_507091, EPI_ISL_507092, EPI_ISL_507093, EPI_ISL_507094, EPI_ISL_507095, EPI_ISL_507096, EPI_ISL_507097, EPI_ISL_507098, EPI_ISL_507099, EPI_ISL_507100, EPI_ISL_507101, EPI_ISL_507102, EPI_ISL_507103, EPI_ISL_507104, EPI_ISL_507105, EPI_ISL_507106, EPI_ISL_507107, EPI_ISL_507108, EPI_ISL_507109 | | | |
| see above | University College London Hospital | COVID-19 Genomics UK (COG-UK) Consortium | Judith Heaney, Matthew Byott, Catherine Houlihan, Dan Frampton, Stuart Kirk, Moira Spyer and Eleni Nastouli |
| EPI_ISL_509444, EPI_ISL_509445, EPI_ISL_509446, EPI_ISL_509447, EPI_ISL_509448, EPI_ISL_509449, EPI_ISL_509450, EPI_ISL_509451, EPI_ISL_509452, EPI_ISL_509453, EPI_ISL_509454, EPI_ISL_509455, EPI_ISL_509456, EPI_ISL_509457, EPI_ISL_509458, EPI_ISL_509459, EPI_ISL_509460, EPI_ISL_509461, EPI_ISL_509462, EPI_ISL_509463, EPI_ISL_509464, EPI_ISL_509465, EPI_ISL_509466, EPI_ISL_509467, EPI_ISL_509468, EPI_ISL_509469, EPI_ISL_509470, EPI_ISL_509471, EPI_ISL_509472, EPI_ISL_509473, EPI_ISL_509474, EPI_ISL_509475, EPI_ISL_509476, EPI_ISL_509477, EPI_ISL_509478, EPI_ISL_509479, EPI_ISL_509480, EPI_ISL_509481, EPI_ISL_509482, EPI_ISL_509483, EPI_ISL_509484, EPI_ISL_509485, EPI_ISL_509486, EPI_ISL_509487, EPI_ISL_509488, EPI_ISL_509489, EPI_ISL_509490, EPI_ISL_509491 | | | |
| see above | Maryland Department of Health | Maryland Department of Health | Keller,E. |
| EPI_ISL_509683, EPI_ISL_509684, EPI_ISL_509685 | AR Dept. of Health-Public Health Lab | Pathogen Discovery, Respiratory Viruses Branch, Division of Viral Diseases, Centers for Disease Control and Prevention | Jing Zhang, Yan Li, Anna Montmayeur, Krista Queen, Anna Uehara, Ying Tao, Rachel Marine, Clinton R. Paden, Haibin Wang, Suxiang Tong |
| EPI_ISL_509707, EPI_ISL_509708, EPI_ISL_509709 | Utah Public Health Laboratory | Pathogen Discovery, Respiratory Viruses Branch, Division of Viral Diseases, Centers for Disease Control and Prevention | Jing Zhang, Ying Tao, Krista Queen, Anna Uehara, Yan Li, Clinton Paden, Haibin Wang, Suxiang Tong |
| EPI_ISL_509710 | Guatemala Ministry of Public Health | Pathogen Discovery, Respiratory Viruses Branch, Division of Viral Diseases, Centers for Disease Control and Prevention | Jing Zhang, Ying Tao, Krista Queen, Anna Uehara, Yan Li, Clinton Paden, Haibin Wang, Suxiang Tong |
| EPI_ISL_509711, EPI_ISL_509712, EPI_ISL_509713, EPI_ISL_509714 | Belize Ministry of Health | Pathogen Discovery, Respiratory Viruses Branch, Division of Viral Diseases, Centers for Disease Control and Prevention | Jing Zhang, Ying Tao, Krista Queen, Anna Uehara, Yan Li, Clinton Paden, Haibin Wang, Suxiang Tong |
| EPI_ISL_509715 | Wisconsin Department of Health Services | Pathogen Discovery, Respiratory Viruses Branch, Division of Viral Diseases, Centers for Disease Control and Prevention | Jing Zhang, Ying Tao, Krista Queen, Anna Uehara, Yan Li, Clinton Paden, Haibin Wang, Suxiang Tong |
| EPI_ISL_510268, EPI_ISL_510269, EPI_ISL_510270, EPI_ISL_510271, EPI_ISL_510272, EPI_ISL_510273, EPI_ISL_510274, EPI_ISL_510275, EPI_ISL_510276, EPI_ISL_510277, EPI_ISL_510278, EPI_ISL_510279, EPI_ISL_510280, EPI_ISL_510281, EPI_ISL_510282, EPI_ISL_510283, EPI_ISL_510284, EPI_ISL_510285, EPI_ISL_510286, EPI_ISL_510287, EPI_ISL_510288, EPI_ISL_510289, EPI_ISL_510290, EPI_ISL_510291, EPI_ISL_510292, EPI_ISL_510293, EPI_ISL_510294, EPI_ISL_510295, EPI_ISL_510296, EPI_ISL_510297, EPI_ISL_510298, EPI_ISL_510299, EPI_ISL_510300, EPI_ISL_510301, EPI_ISL_510302, EPI_ISL_510303, EPI_ISL_510304 | | | |
| see above | Hospital Clínico Universitario de Santiago de Compostela | SeqCOVID-SPAIN consortium/IBV(CSIC) | José Javier Costa Alcalde, Antonio Aguilera Guirao, Mª Luisa Pérez del Molino Bernal, Amparo Coira Nieto, Gema Barbeito Castiñeiras, Rocio Trastoy Pena and SeqCOVID-SPAIN consortium |
| EPI_ISL_510538, EPI_ISL_510539, | Department of Microbiology, The University of Hong Kong | Department of Microbiology, The University of Hong Kong | Kelvin K.W. To, Kwok-Yung Yuen |

| | | | |
|---|---|---|---|
| EPI_ISL_510540 | | | |
| EPI_ISL_510547, EPI_ISL_510548, EPI_ISL_510549, EPI_ISL_510550, EPI_ISL_510551, EPI_ISL_510552, EPI_ISL_510553, EPI_ISL_510554, EPI_ISL_510555, EPI_ISL_510556, EPI_ISL_510557, EPI_ISL_510558, EPI_ISL_510559, EPI_ISL_510560, EPI_ISL_510561, EPI_ISL_510562, EPI_ISL_510563, EPI_ISL_510564, EPI_ISL_510565, EPI_ISL_510566, EPI_ISL_510567, EPI_ISL_510568, EPI_ISL_510569, EPI_ISL_510570, EPI_ISL_510571, EPI_ISL_510572, EPI_ISL_510573, EPI_ISL_510574, EPI_ISL_510575, EPI_ISL_510576, EPI_ISL_510577, EPI_ISL_510578, EPI_ISL_510579, EPI_ISL_510580, EPI_ISL_510581, EPI_ISL_510582, EPI_ISL_510583, EPI_ISL_510584, EPI_ISL_510585, EPI_ISL_510586, EPI_ISL_510587, EPI_ISL_510588, EPI_ISL_510589, EPI_ISL_510590, EPI_ISL_510591, EPI_ISL_510592, EPI_ISL_510593, EPI_ISL_510594, EPI_ISL_510595, EPI_ISL_510596, EPI_ISL_510597, EPI_ISL_510598, EPI_ISL_510599, EPI_ISL_510600, EPI_ISL_510601, EPI_ISL_510602, EPI_ISL_510603, EPI_ISL_510604, EPI_ISL_510605, EPI_ISL_510606, EPI_ISL_510607, EPI_ISL_510608, EPI_ISL_510609, EPI_ISL_510610, EPI_ISL_510611, EPI_ISL_510612, EPI_ISL_510613, EPI_ISL_510614, EPI_ISL_510615, EPI_ISL_510616, EPI_ISL_510617, EPI_ISL_510618, EPI_ISL_510619, EPI_ISL_510620, EPI_ISL_510621, EPI_ISL_510622, EPI_ISL_510623, EPI_ISL_510624, EPI_ISL_510625, EPI_ISL_510626, EPI_ISL_510627, EPI_ISL_510628, EPI_ISL_510629, EPI_ISL_510630, EPI_ISL_510631, EPI_ISL_510632, EPI_ISL_510633, EPI_ISL_510634, EPI_ISL_510635, EPI_ISL_510636, EPI_ISL_510637, EPI_ISL_510638, EPI_ISL_510639, EPI_ISL_510640, EPI_ISL_510641, EPI_ISL_510642, EPI_ISL_510643, EPI_ISL_510644, EPI_ISL_510645, EPI_ISL_510646, EPI_ISL_510647, EPI_ISL_510648, EPI_ISL_510649, EPI_ISL_510650, EPI_ISL_510651, EPI_ISL_510652, EPI_ISL_510653, EPI_ISL_510654, EPI_ISL_510655, EPI_ISL_510656, EPI_ISL_510657, EPI_ISL_510658, EPI_ISL_510659, EPI_ISL_510660, EPI_ISL_510661, EPI_ISL_510662, EPI_ISL_510663, EPI_ISL_510664, EPI_ISL_510665, EPI_ISL_510666, EPI_ISL_510667, EPI_ISL_510668, EPI_ISL_510669, EPI_ISL_510670, EPI_ISL_510671, EPI_ISL_510672, EPI_ISL_510673, EPI_ISL_510674, EPI_ISL_510675, EPI_ISL_510676, EPI_ISL_510677, EPI_ISL_510678, EPI_ISL_510679, EPI_ISL_510680, EPI_ISL_510681, EPI_ISL_510682, EPI_ISL_510683, EPI_ISL_510684, EPI_ISL_510685, EPI_ISL_510686, EPI_ISL_510687, EPI_ISL_510688 | | | |
| see above | Division of Viral Diseases, Center for Laboratory Control of Infectious Diseases, Korea Centers for Diseases Control and Prevention | Division of Viral Diseases, Center for Laboratory Control of Infectious Diseases, Korea Centers for Diseases Control and Prevention | Jeong-Min Kim, Yoon-Seok Chung, Namjoo Lee, Sang Hee Woo, Hye-Jun Jo, Heui Man Kim, Jun-Sub Kim, Myung Guk Han |
| EPI_ISL_512066 | Sardar Vallabhbhai Patel Institute of Medical Sciences & Research | Gujarat Biotechnology Research Centre | Komal Patel, Labdhi Pandya, Afzal Ansari, Nikha Trivedi, Pranay Shah, Kamlesh J Upadhyay, Sanjay Kapadia, Apurvasinh Puvar, Janvi Raval, Zarna Patel, Monika Gandhi, Pinal Trivedi, Maharshi Pandya, Nidhi Patel, Nitin Savaliya, Raghawendra Kumar, Dinesh Kumar, Zuber Saiyed, R D Dixit, A M Kadri, Harsh Bakshi, Chaitanya Joshi, Madhvi Joshi |
| EPI_ISL_512067 | Sardar Vallabhbhai Patel Institute of Medical Sciences & Research | Gujarat Biotechnology Research Centre | Labdhi Pandya, Afzal Ansari, Nikha Trivedi, Pranay Shah, Kamlesh J Upadhyay, Sanjay Kapadia, Apurvasinh Puvar, Janvi Raval, Zarna Patel, Monika Gandhi, Pinal Trivedi, Maharshi Pandya, Nidhi Patel, Nitin Savaliya, Raghawendra Kumar, Dinesh Kumar, Zuber Saiyed, Komal Patel, R D Dixit, A M Kadri, Harsh Bakshi, Chaitanya Joshi, Madhvi Joshi |
| EPI_ISL_512074 | Saikrishna Hospital, Mehsana | Gujarat Biotechnology Research Centre | Janvi Raval, Zarna Patel, Monika Gandhi, Pinal Trivedi, Maharshi Pandya, Nidhi Patel, Nitin Savaliya, Raghawendra Kumar, Dinesh Kumar, Zuber Saiyed, Komal Patel, Labdhi Pandya, Afzal Ansari, Nikha Trivedi, Harshadbhai Parmar, Apurvasinh Puvar, R D Dixit, A M Kadri, Harsh Bakshi, Chaitanya Joshi, Madhvi Joshi |
| EPI_ISL_513513, EPI_ISL_513514, EPI_ISL_513515, EPI_ISL_513516, EPI_ISL_513517, EPI_ISL_513518, EPI_ISL_513519, EPI_ISL_513520, EPI_ISL_513521, EPI_ISL_513522, EPI_ISL_513523, EPI_ISL_513524, EPI_ISL_513525, EPI_ISL_513526, EPI_ISL_513527, EPI_ISL_513528, EPI_ISL_513529, EPI_ISL_513530, EPI_ISL_513531, EPI_ISL_513532, EPI_ISL_513533, EPI_ISL_513534, EPI_ISL_513535, EPI_ISL_513536, EPI_ISL_513537, EPI_ISL_513538, EPI_ISL_513539, EPI_ISL_513540, EPI_ISL_513541, EPI_ISL_513542, EPI_ISL_513543, EPI_ISL_513544, EPI_ISL_513545, EPI_ISL_513546, EPI_ISL_513547, EPI_ISL_513548, EPI_ISL_513549, EPI_ISL_513550, EPI_ISL_513551, EPI_ISL_513552, EPI_ISL_513553, EPI_ISL_513554, EPI_ISL_513555, EPI_ISL_513556, EPI_ISL_513557, EPI_ISL_513558, EPI_ISL_513559, EPI_ISL_513560, EPI_ISL_513561, EPI_ISL_513562, EPI_ISL_513563, EPI_ISL_513564, EPI_ISL_513565, EPI_ISL_513566, EPI_ISL_513567, EPI_ISL_513568, EPI_ISL_513569, EPI_ISL_513570, EPI_ISL_513571, EPI_ISL_513572, EPI_ISL_513573, EPI_ISL_513574, EPI_ISL_513575, EPI_ISL_513576, EPI_ISL_513577, EPI_ISL_513578, EPI_ISL_513579, EPI_ISL_513580, EPI_ISL_513581, EPI_ISL_513582, EPI_ISL_513583 | | | |
| see above | Programa de Oncovirologia, Instituto Nacional de Câncer | Programa de Oncovirologia, Instituto Nacional de Câncer | Juliana D. Siqueira, Livia R. Goes, Brunna M. Alves, Claudia Cicala,James Arthos, João P.B. Viola, Andreia C. de Melo, Marcelo A. Soares |
| EPI_ISL_513925 | Microbiology & Bioinformatics and Biostatistics, Kohat University of Science and Technology (Pakistan) & Shanghai Jiao Tong University (China) | Microbiology & Bioinformatics and Biostatistics, Kohat University of Science and Technology (Pakistan) & Shanghai Jiao Tong University (China) | Khan,M.T., Khan,T.A., Ali,S., Khan,A.S., Muhammad,N. and Wei,D.Q. |
| EPI_ISL_514065, EPI_ISL_514067, EPI_ISL_514068, EPI_ISL_514071, EPI_ISL_514072, EPI_ISL_514075, EPI_ISL_514076, EPI_ISL_514080, EPI_ISL_514081, EPI_ISL_514082, EPI_ISL_514083, EPI_ISL_514085, EPI_ISL_514086, EPI_ISL_514087, EPI_ISL_514089 | | | |
| see above | Navy and Marine Corps Public Health Center | Pathogen Discovery, Respiratory Viruses Branch, Division of Viral Diseases, Centers for Disease Control and Prevention | Krista Queen, Yan Li, Anna Montmayeur, Jing Zhang, Ying Tao, Anna Uehara, Rachel Marine, Clinton R. Paden, Haibin Wang, Suxiang Tong |
| EPI_ISL_514128 | Navy and Marine Corps Public Health Center | Centers for Disease Control and Prevention | Krista Queen, Yan Li, Anna Montmayeur, Jing Zhang, Ying Tao, Anna Uehara, Rachel Marine, Clinton R. Paden, Haibin Wang, Suxiang Tong |
| EPI_ISL_514585 | B.J. Medical College and Civil hospital, Ahmedabad | Gujarat Biotechnology Research Centre | Komal Patel, Labdhi Pandya, Afzal Ansari, Nikha Trivedi, Pranay Shah, Kamlesh J Upadhyay, Sanjay Kapadia, Apurvasinh Puvar, Janvi Raval, Zarna Patel, Monika Gandhi, Pinal Trivedi, Maharshi Pandya, Nidhi Patel, Nitin Savaliya, Raghawendra Kumar, Dinesh Kumar, Zuber Saiyed, R D Dixit, A M Kadri, Harsh Bakshi, Chaitanya Joshi, Madhvi Joshi |
| EPI_ISL_514586 | B.J. Medical College and Civil hospital, Ahmedabad | Gujarat Biotechnology Research Centre | Labdhi Pandya, Afzal Ansari, Nikha Trivedi, Pranay Shah, Kamlesh J Upadhyay, Sanjay Kapadia, Apurvasinh Puvar, Janvi Raval, Zarna Patel, Monika Gandhi, Pinal Trivedi, Maharshi Pandya, Nidhi Patel, Nitin Savaliya, Raghawendra Kumar, Dinesh Kumar, Zuber Saiyed, Komal Patel, R D Dixit, A M Kadri, Harsh Bakshi, Chaitanya Joshi, Madhvi Joshi |
| EPI_ISL_514590 | B.J. Medical College and Civil hospital, Ahmedabad | Gujarat Biotechnology Research Centre | Kamlesh J Upadhyay, Sanjay Kapadia, Apurvasinh Puvar, Janvi Raval, Zarna Patel, Monika Gandhi, Pinal Trivedi, Maharshi Pandya, Nidhi Patel, Nitin Savaliya, Raghawendra Kumar, Dinesh Kumar, Zuber Saiyed, Komal Patel, Labdhi Pandya, Afzal Ansari, Nikha Trivedi, Pranay Shah, R D Dixit, A M Kadri, Harsh Bakshi, Chaitanya Joshi, Madhvi Joshi |
| EPI_ISL_514593 | B.J. Medical College and Civil hospital, Ahmedabad | Gujarat Biotechnology Research Centre | Janvi Raval, Zarna Patel, Monika Gandhi, Pinal Trivedi, Maharshi Pandya, Nidhi Patel, Nitin Savaliya, Raghawendra Kumar, Dinesh Kumar, Zuber Saiyed, Komal Patel, Labdhi Pandya, Afzal Ansari, Nikha Trivedi, Pranay Shah, Kamlesh J Upadhyay, Sanjay Kapadia, Apurvasinh Puvar, R D Dixit, A M Kadri, Harsh Bakshi, Chaitanya Joshi, Madhvi Joshi |
| EPI_ISL_514603 | B.J. Medical College and Civil hospital, Ahmedabad | Gujarat Biotechnology Research Centre | Komal Patel, Labdhi Pandya, Afzal Ansari, Nikha Trivedi, Pranay Shah, Kamlesh J Upadhyay, Sanjay Kapadia, Apurvasinh Puvar, Janvi Raval, Zarna Patel, Monika Gandhi, Pinal Trivedi, Maharshi Pandya, Nidhi Patel, Nitin Savaliya, Raghawendra Kumar, Dinesh Kumar, Zuber Saiyed, R D Dixit, A M Kadri, Harsh Bakshi, Chaitanya Joshi, Madhvi Joshi |
| EPI_ISL_514604 | B.J. Medical College and Civil hospital, Ahmedabad | Gujarat Biotechnology Research Centre | Labdhi Pandya, Afzal Ansari, Nikha Trivedi, Pranay Shah, Kamlesh J Upadhyay, Sanjay Kapadia, Apurvasinh Puvar, Janvi Raval, Zarna Patel, Monika Gandhi, Pinal Trivedi, Maharshi Pandya, Nidhi Patel, Nitin Savaliya, Raghawendra Kumar, Dinesh Kumar, Zuber Saiyed, Komal Patel, R D Dixit, A M Kadri, Harsh Bakshi, Chaitanya Joshi, Madhvi Joshi |
| EPI_ISL_514608 | B.J. Medical College and Civil hospital, Ahmedabad | Gujarat Biotechnology Research Centre | Kamlesh J Upadhyay, Sanjay Kapadia, Apurvasinh Puvar, Janvi Raval, Zarna Patel, Monika Gandhi, Pinal Trivedi, Maharshi Pandya, Nidhi Patel, Nitin Savaliya, Raghawendra Kumar, Dinesh Kumar, Zuber Saiyed, Komal Patel, Labdhi Pandya, Afzal Ansari, Nikha Trivedi, Pranay Shah, R D Dixit, A M Kadri, Harsh Bakshi, Chaitanya Joshi, Madhvi Joshi |
| EPI_ISL_514756, EPI_ISL_514757, EPI_ISL_514758, EPI_ISL_514759, EPI_ISL_514760, EPI_ISL_514761, EPI_ISL_514762, EPI_ISL_514763, EPI_ISL_514764, EPI_ISL_514765, EPI_ISL_514766, EPI_ISL_514767, EPI_ISL_514768, EPI_ISL_514769, EPI_ISL_514770, EPI_ISL_514771, EPI_ISL_514772, EPI_ISL_514773, EPI_ISL_514774, EPI_ISL_514775, EPI_ISL_514776, EPI_ISL_514777, EPI_ISL_514778, EPI_ISL_514779, EPI_ISL_514780, EPI_ISL_514781, EPI_ISL_514782, EPI_ISL_514783, EPI_ISL_514784, EPI_ISL_514785, EPI_ISL_514786, EPI_ISL_514787, EPI_ISL_514788, EPI_ISL_514789, EPI_ISL_514790, EPI_ISL_514791, EPI_ISL_514792, EPI_ISL_514793, EPI_ISL_514794, EPI_ISL_514795, EPI_ISL_514796, EPI_ISL_514797, EPI_ISL_514798, EPI_ISL_514799, EPI_ISL_514800, EPI_ISL_514801, EPI_ISL_514802, EPI_ISL_514803, EPI_ISL_514804, EPI_ISL_514805, EPI_ISL_514806, EPI_ISL_514807, EPI_ISL_514808, EPI_ISL_514809, EPI_ISL_514810, EPI_ISL_514811, EPI_ISL_514812, EPI_ISL_514813, EPI_ISL_514814, EPI_ISL_514815, EPI_ISL_514816, EPI_ISL_514817, EPI_ISL_514818, EPI_ISL_514819, EPI_ISL_514820, EPI_ISL_514821, EPI_ISL_514822, EPI_ISL_514823, EPI_ISL_514824, EPI_ISL_514825, EPI_ISL_514826, EPI_ISL_514827, EPI_ISL_514828, EPI_ISL_514829, EPI_ISL_514830, EPI_ISL_514831, EPI_ISL_514832, EPI_ISL_514833, EPI_ISL_514834, EPI_ISL_514835, EPI_ISL_514836, EPI_ISL_514837, EPI_ISL_514838, EPI_ISL_514839, EPI_ISL_514840, EPI_ISL_514841, EPI_ISL_514842, EPI_ISL_514843, EPI_ISL_514844, EPI_ISL_514845, EPI_ISL_514846, EPI_ISL_514847, EPI_ISL_514848, EPI_ISL_514849, EPI_ISL_514850, EPI_ISL_514851, EPI_ISL_514852, EPI_ISL_514853, EPI_ISL_514854, EPI_ISL_514855, EPI_ISL_514856, EPI_ISL_514857, EPI_ISL_514858, EPI_ISL_514859, EPI_ISL_514860, EPI_ISL_514861, EPI_ISL_514862, EPI_ISL_514863, EPI_ISL_514864, EPI_ISL_514865, EPI_ISL_514866, EPI_ISL_514867, EPI_ISL_514868, EPI_ISL_514869, EPI_ISL_514870, EPI_ISL_514871, EPI_ISL_514872, EPI_ISL_514873, EPI_ISL_514874, EPI_ISL_514875, EPI_ISL_514876, EPI_ISL_514877, EPI_ISL_514878, EPI_ISL_514879, EPI_ISL_514880, EPI_ISL_514881, EPI_ISL_514882, EPI_ISL_514883, EPI_ISL_514884, EPI_ISL_514885, EPI_ISL_514886, EPI_ISL_514887, EPI_ISL_514888, EPI_ISL_514889, EPI_ISL_514890, EPI_ISL_514891, EPI_ISL_514892, EPI_ISL_514893, EPI_ISL_514894, EPI_ISL_514895, EPI_ISL_514896, EPI_ISL_514897, EPI_ISL_514898, EPI_ISL_514899, EPI_ISL_514900, EPI_ISL_514901, EPI_ISL_514902, EPI_ISL_514903, EPI_ISL_514904, EPI_ISL_514905, EPI_ISL_514906, EPI_ISL_514907, EPI_ISL_514908, EPI_ISL_514909, EPI_ISL_514910, EPI_ISL_514911, EPI_ISL_514912, EPI_ISL_514913, EPI_ISL_514914, EPI_ISL_514915, EPI_ISL_514916, EPI_ISL_514917, EPI_ISL_514918, EPI_ISL_514919, EPI_ISL_514920, EPI_ISL_514921, EPI_ISL_514922, EPI_ISL_514923, EPI_ISL_514924, EPI_ISL_514925, EPI_ISL_514926, EPI_ISL_514927, EPI_ISL_514928, EPI_ISL_514929, EPI_ISL_514930, EPI_ISL_514931, EPI_ISL_514932, EPI_ISL_514933, EPI_ISL_514934, EPI_ISL_514935, EPI_ISL_514936, EPI_ISL_514937, EPI_ISL_514938, EPI_ISL_514939, EPI_ISL_514940, EPI_ISL_514941, EPI_ISL_514942, EPI_ISL_514943, EPI_ISL_514944, EPI_ISL_514945, EPI_ISL_514946, EPI_ISL_514947, EPI_ISL_514948, EPI_ISL_514949, EPI_ISL_514950, EPI_ISL_514951, EPI_ISL_514952, EPI_ISL_514953, EPI_ISL_514954, EPI_ISL_514955, EPI_ISL_514956, EPI_ISL_514957, EPI_ISL_514958, EPI_ISL_514959, EPI_ISL_514960, EPI_ISL_514961, EPI_ISL_514962, EPI_ISL_514963, EPI_ISL_514964, EPI_ISL_514965, EPI_ISL_514966, EPI_ISL_514967, EPI_ISL_514968, EPI_ISL_514969, EPI_ISL_514970, EPI_ISL_514971, EPI_ISL_514972, EPI_ISL_514973, EPI_ISL_514974, EPI_ISL_514975, EPI_ISL_514976, EPI_ISL_514977, EPI_ISL_514978, EPI_ISL_514979, EPI_ISL_514980, EPI_ISL_514981, EPI_ISL_514982, EPI_ISL_514983, EPI_ISL_514984, EPI_ISL_514985, EPI_ISL_514986, EPI_ISL_514987, EPI_ISL_514988, EPI_ISL_514989, EPI_ISL_514990, EPI_ISL_514991, EPI_ISL_514992, EPI_ISL_514993, EPI_ISL_514994, EPI_ISL_514995, EPI_ISL_514996, EPI_ISL_514997, EPI_ISL_514998, EPI_ISL_514999, EPI_ISL_515000, EPI_ISL_515001, EPI_ISL_515002, EPI_ISL_515003, EPI_ISL_515004, EPI_ISL_515005, EPI_ISL_515006, EPI_ISL_515007, EPI_ISL_515008, EPI_ISL_515009, EPI_ISL_515010, EPI_ISL_515011, EPI_ISL_515012, EPI_ISL_515013, EPI_ISL_515014, EPI_ISL_515015, EPI_ISL_515016, EPI_ISL_515017, EPI_ISL_515018, EPI_ISL_515019, EPI_ISL_515020, EPI_ISL_515021, EPI_ISL_515022, EPI_ISL_515023, EPI_ISL_515024, EPI_ISL_515025, EPI_ISL_515026, EPI_ISL_515027, EPI_ISL_515028, EPI_ISL_515029, EPI_ISL_515030, EPI_ISL_515031, EPI_ISL_515032, EPI_ISL_515033, EPI_ISL_515034, EPI_ISL_515035, EPI_ISL_515036, EPI_ISL_515037, EPI_ISL_515038, EPI_ISL_515039, EPI_ISL_515040, EPI_ISL_515041, EPI_ISL_515042, EPI_ISL_515043, EPI_ISL_515044, EPI_ISL_515045, EPI_ISL_515046, EPI_ISL_515047, EPI_ISL_515048, EPI_ISL_515049, EPI_ISL_515050, EPI_ISL_515051, EPI_ISL_515052, EPI_ISL_515053, EPI_ISL_515054 | | | |
| see above | Division of Viral Diseases, Center for Laboratory Control of Infectious Diseases, Korea Centers for Diseases Control and Prevention | Division of Viral Diseases, Center for Laboratory Control of Infectious Diseases, Korea Centers for Diseases Control and Prevention | Jeong-Min Kim, Yoon-Seok Chung, Namjoo Lee, Sang Hee Woo, Hye-Jun Jo, Heui Man Kim, Jun-Sub Kim, Myung Guk Han |
| EPI_ISL_515055, EPI_ISL_515056, EPI_ISL_515057, EPI_ISL_515058, EPI_ISL_515059, EPI_ISL_515060, EPI_ISL_515061, EPI_ISL_515062, EPI_ISL_515063, EPI_ISL_515064, EPI_ISL_515065, EPI_ISL_515066, EPI_ISL_515067, EPI_ISL_515068, EPI_ISL_515069, EPI_ISL_515070, EPI_ISL_515071, EPI_ISL_515072, EPI_ISL_515073, EPI_ISL_515074, EPI_ISL_515075, EPI_ISL_515076, EPI_ISL_515077, EPI_ISL_515078, EPI_ISL_515079, EPI_ISL_515080, EPI_ISL_515081 | | | |

| | | | |
|---|---|---|---|
| see above | Department of Clinical Microbiology | GIGA Medical Genomics | Keith Durkin, Maria Artesi, Sebastien Bontems, Raphael Boreux, Cecile Meex, Axelle Chaslain, Celine Fombellida-Lopez, Pierrette Melin, Marie-Pierre Hayette, Vincent Bours. |
| EPI_ISL_516648 | Institute of Microbiology, Universidad San Francisco de Quito | Institute of Microbiology, Universidad San Francisco de Quito | Juan José Guadalupe, Monica Becerra-Wong, Prado-Vivar, Sully Márquez, Bernardo Gutiérrez, Nabih Dahik, Carlos Mena, Ligia Briceño, Verónica Barragán, Patricio Rojas-Silva, Gabriel Trueba, Michelle Grunauer, Paúl Cárdenas |
| EPI_ISL_516649 | Institute of Microbiology, Universidad San Francisco de Quito | Institute of Microbiology, Universidad San Francisco de Quito | Juan José Guadalupe, Monica Becerra-Wong, Bernardo Gutiérrez, Nabih Dahik, Carlos Mena, Ligia Briceño, Verónica Barragán, Patricio Rojas-Silva, Gabriel Trueba, Michelle Grunauer, Paúl Cárdenas |
| EPI_ISL_516783, EPI_ISL_516784, EPI_ISL_516785 | Center for Laboratory Control of Infectious Diseases, Korea Centers for Diseases Control and Prevention | Center for Laboratory Control of Infectious Diseases, Korea Centers for Diseases Control and Prevention | Junyoung Kim, Ae Kyung Park, Eunkyung Shin, Jin Sun No, Jeong-Min Kim, Yoon-Seok Chung, Heui Man Kim, Myung Guk Han |
| EPI_ISL_516798, EPI_ISL_516799 | Department of Microbiology, The University of Hong Kong | Department of Microbiology, The University of Hong Kong | Kelvin K.W. To, Kwok-Yung Yuen |
| EPI_ISL_522439, EPI_ISL_522440, EPI_ISL_522441, EPI_ISL_522442, EPI_ISL_522443, EPI_ISL_522444, EPI_ISL_522445, EPI_ISL_522446, EPI_ISL_522447, EPI_ISL_522448, EPI_ISL_522449, EPI_ISL_522450, EPI_ISL_522451, EPI_ISL_522452, EPI_ISL_522453 | | | |
| see above | Center for Laboratory Control of Infectious Diseases, Korea Centers for Diseases Control and Prevention | Center for Laboratory Control of Infectious Diseases, Korea Centers for Diseases Control and Prevention | Junyoung Kim, Ae Kyung Park, Eunkyung Shin, Jin Sun No, Jeong-Min Kim, Yoon-Seok Chung, Heui Man Kim, Myung Guk Han |
| EPI_ISL_522454 | Division of Viral Diseases, Center for Laboratory Control of Infectious Diseases, Korea Centers for Diseases Control and Prevention | Division of Viral Diseases, Center for Laboratory Control of Infectious Diseases, Korea Centers for Diseases Control and Prevention | Jeong-Min Kim, Yoon-Seok Chung, Namjoo Lee, Sang Hee Woo, Hye-Jun Jo, Heui Man Kim, Jun-Sub Kim, Myung Guk Han |
| EPI_ISL_522455, EPI_ISL_522456, EPI_ISL_522457, EPI_ISL_522458, EPI_ISL_522459, EPI_ISL_522460, EPI_ISL_522461, EPI_ISL_522462, EPI_ISL_522463, EPI_ISL_522464 | Center for Laboratory Control of Infectious Diseases, Korea Centers for Diseases Control and Prevention | Center for Laboratory Control of Infectious Diseases, Korea Centers for Diseases Control and Prevention | Junyoung Kim, Ae Kyung Park, Eunkyung Shin, Jin Sun No, Jeong-Min Kim, Yoon-Seok Chung, Heui Man Kim, Myung Guk Han |
| EPI_ISL_522465, EPI_ISL_522466 | Division of Viral Diseases, Center for Laboratory Control of Infectious Diseases, Korea Centers for Diseases Control and Prevention | Division of Viral Diseases, Center for Laboratory Control of Infectious Diseases, Korea Centers for Diseases Control and Prevention | Jeong-Min Kim, Yoon-Seok Chung, Namjoo Lee, Sang Hee Woo, Hye-Jun Jo, Heui Man Kim, Jun-Sub Kim, Myung Guk Han |
| EPI_ISL_522467, EPI_ISL_522468, EPI_ISL_522469, EPI_ISL_522470, EPI_ISL_522471, EPI_ISL_522472, EPI_ISL_522473 | Center for Laboratory Control of Infectious Diseases, Korea Centers for Diseases Control and Prevention | Center for Laboratory Control of Infectious Diseases, Korea Centers for Diseases Control and Prevention | Junyoung Kim, Ae Kyung Park, Eunkyung Shin, Jin Sun No, Jeong-Min Kim, Yoon-Seok Chung, Heui Man Kim, Myung Guk Han |
| EPI_ISL_522474, EPI_ISL_522475 | Division of Viral Diseases, Center for Laboratory Control of Infectious Diseases, Korea Centers for Diseases Control and Prevention | Division of Viral Diseases, Center for Laboratory Control of Infectious Diseases, Korea Centers for Diseases Control and Prevention | Jeong-Min Kim, Yoon-Seok Chung, Namjoo Lee, Sang Hee Woo, Hye-Jun Jo, Heui Man Kim, Jun-Sub Kim, Myung Guk Han |
| EPI_ISL_522476 | Center for Laboratory Control of Infectious Diseases, Korea Centers for Diseases Control and Prevention | Center for Laboratory Control of Infectious Diseases, Korea Centers for Diseases Control and Prevention | Junyoung Kim, Ae Kyung Park, Eunkyung Shin, Jin Sun No, Jeong-Min Kim, Yoon-Seok Chung, Heui Man Kim, Myung Guk Han |
| EPI_ISL_522477, EPI_ISL_522478 | Division of Viral Diseases, Center for Laboratory Control of Infectious Diseases, Korea Centers for Diseases Control and Prevention | Division of Viral Diseases, Center for Laboratory Control of Infectious Diseases, Korea Centers for Diseases Control and Prevention | Jeong-Min Kim, Yoon-Seok Chung, Namjoo Lee, Sang Hee Woo, Hye-Jun Jo, Heui Man Kim, Jun-Sub Kim, Myung Guk Han |
| EPI_ISL_522479, EPI_ISL_522480, EPI_ISL_522481, EPI_ISL_522482, EPI_ISL_522483 | Center for Laboratory Control of Infectious Diseases, Korea Centers for Diseases Control and Prevention | Center for Laboratory Control of Infectious Diseases, Korea Centers for Diseases Control and Prevention | Junyoung Kim, Ae Kyung Park, Eunkyung Shin, Jin Sun No, Jeong-Min Kim, Yoon-Seok Chung, Heui Man Kim, Myung Guk Han |
| EPI_ISL_522484 | Division of Viral Diseases, Center for Laboratory Control of Infectious Diseases, Korea Centers for Diseases Control and Prevention | Division of Viral Diseases, Center for Laboratory Control of Infectious Diseases, Korea Centers for Diseases Control and Prevention | Jeong-Min Kim, Yoon-Seok Chung, Namjoo Lee, Sang Hee Woo, Hye-Jun Jo, Heui Man Kim, Jun-Sub Kim, Myung Guk Han |
| EPI_ISL_522485 | Center for Laboratory Control of Infectious Diseases, Korea Centers for Diseases Control and Prevention | Center for Laboratory Control of Infectious Diseases, Korea Centers for Diseases Control and Prevention | Junyoung Kim, Ae Kyung Park, Eunkyung Shin, Jin Sun No, Jeong-Min Kim, Yoon-Seok Chung, Heui Man Kim, Myung Guk Han |
| EPI_ISL_522486, EPI_ISL_522487, EPI_ISL_522488, EPI_ISL_522489, EPI_ISL_522490 | Division of Viral Diseases, Center for Laboratory Control of Infectious Diseases, Korea Centers for Diseases Control and Prevention | Division of Viral Diseases, Center for Laboratory Control of Infectious Diseases, Korea Centers for Diseases Control and Prevention | Jeong-Min Kim, Yoon-Seok Chung, Namjoo Lee, Sang Hee Woo, Hye-Jun Jo, Heui Man Kim, Jun-Sub Kim, Myung Guk Han |
| EPI_ISL_522491, EPI_ISL_522492, EPI_ISL_522493, EPI_ISL_522494, EPI_ISL_522495, EPI_ISL_522496, EPI_ISL_522497 | Center for Laboratory Control of Infectious Diseases, Korea Centers for Diseases Control and Prevention | Center for Laboratory Control of Infectious Diseases, Korea Centers for Diseases Control and Prevention | Junyoung Kim, Ae Kyung Park, Eunkyung Shin, Jin Sun No, Jeong-Min Kim, Yoon-Seok Chung, Heui Man Kim, Myung Guk Han |
| EPI_ISL_522498, EPI_ISL_522499, EPI_ISL_522500, EPI_ISL_522501 | Division of Viral Diseases, Center for Laboratory Control of Infectious Diseases, Korea Centers for Diseases Control and Prevention | Division of Viral Diseases, Center for Laboratory Control of Infectious Diseases, Korea Centers for Diseases Control and Prevention | Jeong-Min Kim, Yoon-Seok Chung, Namjoo Lee, Sang Hee Woo, Hye-Jun Jo, Heui Man Kim, Jun-Sub Kim, Myung Guk Han |
| EPI_ISL_522502 | Center for Laboratory Control of Infectious Diseases, Korea Centers for Diseases Control and Prevention | Center for Laboratory Control of Infectious Diseases, Korea Centers for Diseases Control and Prevention | Junyoung Kim, Ae Kyung Park, Eunkyung Shin, Jin Sun No, Jeong-Min Kim, Yoon-Seok Chung, Heui Man Kim, Myung Guk Han |
| EPI_ISL_522503, EPI_ISL_522504 | Division of Viral Diseases, Center for Laboratory Control of Infectious Diseases, Korea Centers for Diseases Control and Prevention | Division of Viral Diseases, Center for Laboratory Control of Infectious Diseases, Korea Centers for Diseases Control and Prevention | Jeong-Min Kim, Yoon-Seok Chung, Namjoo Lee, Sang Hee Woo, Hye-Jun Jo, Heui Man Kim, Jun-Sub Kim, Myung Guk Han |
| EPI_ISL_522505 | Center for Laboratory Control of Infectious Diseases, Korea Centers for Diseases Control and Prevention | Center for Laboratory Control of Infectious Diseases, Korea Centers for Diseases Control and Prevention | Junyoung Kim, Ae Kyung Park, Eunkyung Shin, Jin Sun No, Jeong-Min Kim, Yoon-Seok Chung, Heui Man Kim, Myung Guk Han |
| EPI_ISL_522506, EPI_ISL_522507 | Division of Viral Diseases, Center for Laboratory Control of Infectious Diseases, Korea Centers for Diseases Control and Prevention | Division of Viral Diseases, Center for Laboratory Control of Infectious Diseases, Korea Centers for Diseases Control and Prevention | Jeong-Min Kim, Yoon-Seok Chung, Namjoo Lee, Sang Hee Woo, Hye-Jun Jo, Heui Man Kim, Jun-Sub Kim, Myung Guk Han |
| EPI_ISL_522508 | Center for Laboratory Control of Infectious Diseases, Korea Centers for Diseases Control and Prevention | Center for Laboratory Control of Infectious Diseases, Korea Centers for Diseases Control and Prevention | Junyoung Kim, Ae Kyung Park, Eunkyung Shin, Jin Sun No, Jeong-Min Kim, Yoon-Seok Chung, Heui Man Kim, Myung Guk Han |
| EPI_ISL_522509 | Division of Viral Diseases, Center for Laboratory Control of Infectious Diseases, Korea Centers for Diseases Control and Prevention | Division of Viral Diseases, Center for Laboratory Control of Infectious Diseases, Korea Centers for Diseases Control and Prevention | Jeong-Min Kim, Yoon-Seok Chung, Namjoo Lee, Sang Hee Woo, Hye-Jun Jo, Heui Man Kim, Jun-Sub Kim, Myung Guk Han |
| EPI_ISL_522510 | Center for Laboratory Control of Infectious Diseases, Korea Centers for Diseases Control and Prevention | Center for Laboratory Control of Infectious Diseases, Korea Centers for Diseases Control and Prevention | Junyoung Kim, Ae Kyung Park, Eunkyung Shin, Jin Sun No, Jeong-Min Kim, Yoon-Seok Chung, Heui Man Kim, Myung Guk Han |
| EPI_ISL_522511, EPI_ISL_522512, EPI_ISL_522513, EPI_ISL_522514 | Division of Viral Diseases, Center for Laboratory Control of Infectious Diseases, Korea Centers for Diseases Control and Prevention | Division of Viral Diseases, Center for Laboratory Control of Infectious Diseases, Korea Centers for Diseases Control and Prevention | Jeong-Min Kim, Yoon-Seok Chung, Namjoo Lee, Sang Hee Woo, Hye-Jun Jo, Heui Man Kim, Jun-Sub Kim, Myung Guk Han |

| | | | |
|---|---|---|---|
| EPI_ISL_522515, EPI_ISL_522516 | Center for Laboratory Control of Infectious Diseases, Korea Centers for Diseases Control and Prevention | Center for Laboratory Control of Infectious Diseases, Korea Centers for Diseases Control and Prevention | Junyoung Kim, Ae Kyung Park, Eunkyung Shin, Jin Sun No, Jeong-Min Kim, Yoon-Seok Chung, Heui Man Kim, Myung Guk Han |
| EPI_ISL_522517 | Division of Viral Diseases, Center for Laboratory Control of Infectious Diseases, Korea Centers for Diseases Control and Prevention | Division of Viral Diseases, Center for Laboratory Control of Infectious Diseases, Korea Centers for Diseases Control and Prevention | Jeong-Min Kim, Yoon-Seok Chung, Namjoo Lee, Sang Hee Woo, Hye-Jun Jo, Heui Man Kim, Jun-Sub Kim, Myung Guk Han |
| EPI_ISL_522518 | Center for Laboratory Control of Infectious Diseases, Korea Centers for Diseases Control and Prevention | Center for Laboratory Control of Infectious Diseases, Korea Centers for Diseases Control and Prevention | Junyoung Kim, Ae Kyung Park, Eunkyung Shin, Jin Sun No, Jeong-Min Kim, Yoon-Seok Chung, Heui Man Kim, Myung Guk Han |
| EPI_ISL_522519, EPI_ISL_522520, EPI_ISL_522521, EPI_ISL_522522, EPI_ISL_522523, EPI_ISL_522524, EPI_ISL_522525, EPI_ISL_522526, EPI_ISL_522527, EPI_ISL_522528, EPI_ISL_522529, EPI_ISL_522530, EPI_ISL_522531, EPI_ISL_522532, EPI_ISL_522533, EPI_ISL_522534, EPI_ISL_522535, EPI_ISL_522536, EPI_ISL_522537, EPI_ISL_522538, EPI_ISL_522539, EPI_ISL_522540, EPI_ISL_522541, EPI_ISL_522542, EPI_ISL_522543, EPI_ISL_522544 | | | |
| see above | Division of Viral Diseases, Center for Laboratory Control of Infectious Diseases, Korea Centers for Diseases Control and Prevention | Division of Viral Diseases, Center for Laboratory Control of Infectious Diseases, Korea Centers for Diseases Control and Prevention | Jeong-Min Kim, Yoon-Seok Chung, Namjoo Lee, Sang Hee Woo, Hye-Jun Jo, Heui Man Kim, Jun-Sub Kim, Myung Guk Han |
| EPI_ISL_524714 | B.J. Medical College and Civil hospital, Ahmedabad | Gujarat Biotechnology Research Centre | Janvi Raval, Zarna Patel, Monika Gandhi, Pinal Trivedi, Maharshi Pandya, Nidhi Patel, Nitin Savaliya, Raghawendra Kumar, Dinesh Kumar, Zuber Saiyed, Komal Patel, Labdhi Pandya, Afzal Ansari, Nikha Trivedi, Pranay Shah, Kamlesh J Upadhyay, Sanjay Kapadia, Apurvasinh Puvar, R D Dixit, A M Kadri, Harsh Bakshi, Chaitanya Joshi, Madhvi Joshi |
| EPI_ISL_524724 | B.J. Medical College and Civil hospital, Ahmedabad | Gujarat Biotechnology Research Centre | Komal Patel, Labdhi Pandya, Afzal Ansari, Nikha Trivedi, Pranay Shah, Kamlesh J Upadhyay, Sanjay Kapadia, Apurvasinh Puvar, Janvi Raval, Zarna Patel, Monika Gandhi, Pinal Trivedi, Maharshi Pandya, Nidhi Patel, Nitin Savaliya, Raghawendra Kumar, Dinesh Kumar, Zuber Saiyed, R D Dixit, A M Kadri, Harsh Bakshi, Chaitanya Joshi, Madhvi Joshi |
| EPI_ISL_524725 | B.J. Medical College and Civil hospital, Ahmedabad | Gujarat Biotechnology Research Centre | Labdhi Pandya, Afzal Ansari, Nikha Trivedi, Pranay Shah, Kamlesh J Upadhyay, Sanjay Kapadia, Apurvasinh Puvar, Janvi Raval, Zarna Patel, Monika Gandhi, Pinal Trivedi, Maharshi Pandya, Nidhi Patel, Nitin Savaliya, Raghawendra Kumar, Dinesh Kumar, Zuber Saiyed, Komal Patel, R D Dixit, A M Kadri, Harsh Bakshi, Chaitanya Joshi, Madhvi Joshi |
| EPI_ISL_524729 | B.J. Medical College and Civil hospital, Ahmedabad | Gujarat Biotechnology Research Centre | Kamlesh J Upadhyay, Sanjay Kapadia, Apurvasinh Puvar, Janvi Raval, Zarna Patel, Monika Gandhi, Pinal Trivedi, Maharshi Pandya, Nidhi Patel, Nitin Savaliya, Raghawendra Kumar, Dinesh Kumar, Zuber Saiyed, Komal Patel, Labdhi Pandya, Afzal Ansari, Nikha Trivedi, Pranay Shah, R D Dixit, A M Kadri, Harsh Bakshi, Chaitanya Joshi, Madhvi Joshi |
| EPI_ISL_524736 | Government General Hospital, Jam Khambhaliya, Devbhoomi Dwarka | Gujarat Biotechnology Research Centre | Janvi Raval, Zarna Patel, Monika Gandhi, Pinal Trivedi, Maharshi Pandya, Nidhi Patel, Nitin Savaliya, Raghawendra Kumar, Dinesh Kumar, Zuber Saiyed, Komal Patel, Labdhi Pandya, Afzal Ansari, Nikha Trivedi, Harish Matani, Apurvasinh Puvar, R D Dixit, A M Kadri, Harsh Bakshi, Chaitanya Joshi, Madhvi Joshi |
| EPI_ISL_524744 | GMERS Medical College and Hospital, Gandhinagar | Gujarat Biotechnology Research Centre | Komal Patel, Labdhi Pandya, Afzal Ansari, Nikha Trivedi, Seema Bhatt, Gaurishankar Shrimali, Bhavesh Modi, Bharti Rajani, Apurvasinh Puvar, Janvi Raval, Zarna Patel, Monika Gandhi, Pinal Trivedi, Maharshi Pandya, Nidhi Patel, Nitin Savaliya, Raghawendra Kumar, Dinesh Kumar, Zuber Saiyed, R D Dixit, A M Kadri, Harsh Bakshi, Chaitanya Joshi, Madhvi Joshi |
| EPI_ISL_524745 | GMERS Medical College and Hospital, Gandhinagar | Gujarat Biotechnology Research Centre | Labdhi Pandya, Afzal Ansari, Nikha Trivedi, Seema Bhatt, Gaurishankar Shrimali, Bhavesh Modi, Bharti Rajani, Apurvasinh Puvar, Janvi Raval, Zarna Patel, Monika Gandhi, Pinal Trivedi, Maharshi Pandya, Nidhi Patel, Nitin Savaliya, Raghawendra Kumar, Dinesh Kumar, Zuber Saiyed, Komal Patel, R D Dixit, A M Kadri, Harsh Bakshi, Chaitanya Joshi, Madhvi Joshi |
| EPI_ISL_524750 | Sir M P Shah Government Medical College, Jamnagar | Gujarat Biotechnology Research Centre | Janvi Raval, Zarna Patel, Monika Gandhi, Pinal Trivedi, Maharshi Pandya, Nidhi Patel, Nitin Savaliya, Raghawendra Kumar, Dinesh Kumar, Zuber Saiyed, Komal Patel, Labdhi Pandya, Afzal Ansari, Nikha Trivedi, Binita Aring, Apurvasinh Puvar, R D Dixit, A M Kadri, Harsh Bakshi, Chaitanya Joshi, Madhvi Joshi |
| EPI_ISL_524760 | GMERS Medical College and Hospital, Dharpur, Patan | Gujarat Biotechnology Research Centre | Komal Patel, Labdhi Pandya, Afzal Ansari, Nikha Trivedi, A N Parmar, Apurvasinh Puvar, Janvi Raval, Zarna Patel, Monika Gandhi, Pinal Trivedi, Maharshi Pandya, Nidhi Patel, Nitin Savaliya, Raghawendra Kumar, Dinesh Kumar, Zuber Saiyed, R D Dixit, A M Kadri, Harsh Bakshi, Chaitanya Joshi, Madhvi Joshi |
| EPI_ISL_524761 | GMERS Medical College and Hospital, Dharpur, Patan | Gujarat Biotechnology Research Centre | Labdhi Pandya, Afzal Ansari, Nikha Trivedi, A N Parmar, Apurvasinh Puvar, Janvi Raval, Zarna Patel, Monika Gandhi, Pinal Trivedi, Maharshi Pandya, Nidhi Patel, Nitin Savaliya, Raghawendra Kumar, Dinesh Kumar, Zuber Saiyed, Komal Patel, R D Dixit, A M Kadri, Harsh Bakshi, Chaitanya Joshi, Madhvi Joshi |
| EPI_ISL_524766 | GMERS Medical College and Hospital, Dharpur, Patan | Gujarat Biotechnology Research Centre | Janvi Raval, Zarna Patel, Monika Gandhi, Pinal Trivedi, Maharshi Pandya, Nidhi Patel, Nitin Savaliya, Raghawendra Kumar, Dinesh Kumar, Zuber Saiyed, Komal Patel, Labdhi Pandya, Afzal Ansari, Nikha Trivedi, Apurvasinh Puvar, A N Parmar, R D Dixit, A M Kadri, Harsh Bakshi, Chaitanya Joshi, Madhvi Joshi |
| EPI_ISL_525430 | Institute of Microbiology, Universidad San Francisco de Quito | Institute of Microbiology, Universidad San Francisco de Quito | Juan José Guadalupe, Monica Becerra-Wong, Belén Prado-Vivar, Sully Márquez, Bernardo Gutiérrez, Verónica Barragán, Patricio Rojas-Silva, Gabriel Trueba, Michelle Grunauer, Paúl Cárdenas |
| EPI_ISL_525431, EPI_ISL_525432 | Institute of Microbiology, Universidad San Francisco de Quito | Institute of Microbiology, Universidad San Francisco de Quito | Juan José Guadalupe, Monica Becerra-Wong, Belén Prado-Vivar, Sully Márquez, Ligia Briceño, Carlos Mena, Nabih Dahik, Bernardo Gutiérrez, Verónica Barragán, Patricio Rojas-Silva, Gabriel Trueba, Michelle Grunauer, Paúl Cárdenas |
| EPI_ISL_525433, EPI_ISL_525434 | Institute of Microbiology, Universidad San Francisco de Quito | Institute of Microbiology, Universidad San Francisco de Quito | Juan José Guadalupe, Monica Becerra-Wong, Belén Prado-Vivar, Sully Márquez, Bernardo Gutiérrez, Eulalia Pazmiño, Carolina Pacheco, Damaris Zandoya, Carlos Mena, Nabih Dahik, Verónica Barragán, Patricio Rojas-Silva, Gabriel Trueba, Michelle Grunauer, Paúl Cárdenas |
| EPI_ISL_525435 | Institute of Microbiology, Universidad San Francisco de Quito | Institute of Microbiology, Universidad San Francisco de Quito | Juan José Guadalupe, Monica Becerra-Wong, Belén Prado-Vivar, Sully Márquez, Bernardo Gutiérrez, Yomara Napa, Edy Quizhpe, Carlos Mena, Nabih Dahik, Verónica Barragán, Patricio Rojas-Silva, Gabriel Trueba, Michelle Grunauer, Paúl Cárdenas |
| EPI_ISL_525436, EPI_ISL_525437 | Institute of Microbiology, Universidad San Francisco de Quito | Institute of Microbiology, Universidad San Francisco de Quito | Juan José Guadalupe, Monica Becerra-Wong, Belén Prado-Vivar, Sully Márquez, Bernardo Gutiérrez, Dayron Brossard, Carlos Mena, Nabih Dahik, Verónica Barragán, Patricio Rojas-Silva, Gabriel Trueba, Michelle Grunauer, Paúl Cárdenas |
| EPI_ISL_525575, EPI_ISL_525576, EPI_ISL_525577, EPI_ISL_525578, EPI_ISL_525579, EPI_ISL_525580, EPI_ISL_525581, EPI_ISL_525582, EPI_ISL_525583, EPI_ISL_525584, EPI_ISL_525585, EPI_ISL_525586, EPI_ISL_525587, EPI_ISL_525588, EPI_ISL_525589, EPI_ISL_525590, EPI_ISL_525591, EPI_ISL_525592, EPI_ISL_525593, EPI_ISL_525594, EPI_ISL_525595, EPI_ISL_525596, EPI_ISL_525597, EPI_ISL_525598, EPI_ISL_525599, EPI_ISL_525600, EPI_ISL_525601, EPI_ISL_525602, EPI_ISL_525603, EPI_ISL_525604, EPI_ISL_525605, EPI_ISL_525606, EPI_ISL_525607, EPI_ISL_525608, EPI_ISL_525609, EPI_ISL_525610, EPI_ISL_525611, EPI_ISL_525612, EPI_ISL_525613, EPI_ISL_525614, EPI_ISL_525615, EPI_ISL_525616, EPI_ISL_525617, EPI_ISL_525618, EPI_ISL_525619, EPI_ISL_525620, EPI_ISL_525621, EPI_ISL_525622, EPI_ISL_525623, EPI_ISL_525624, EPI_ISL_525625, EPI_ISL_525626, EPI_ISL_525627, EPI_ISL_525628, EPI_ISL_525629, EPI_ISL_525630, EPI_ISL_525631, EPI_ISL_525632, EPI_ISL_525633, EPI_ISL_525634, EPI_ISL_525635, EPI_ISL_525636, EPI_ISL_525637, EPI_ISL_525638, EPI_ISL_525639, EPI_ISL_525640, EPI_ISL_525641, EPI_ISL_525642, EPI_ISL_525643, EPI_ISL_525644, EPI_ISL_525645, EPI_ISL_525646, EPI_ISL_525647, EPI_ISL_525648, EPI_ISL_525649, EPI_ISL_525650, EPI_ISL_525651, EPI_ISL_525652, EPI_ISL_525653, EPI_ISL_525654, EPI_ISL_525655, EPI_ISL_525656, EPI_ISL_525657, EPI_ISL_525658, EPI_ISL_525659, EPI_ISL_525660, EPI_ISL_525661, EPI_ISL_525662, EPI_ISL_525663, EPI_ISL_525664, EPI_ISL_525665, EPI_ISL_525666, EPI_ISL_525667, EPI_ISL_525668, EPI_ISL_525669, EPI_ISL_525670, EPI_ISL_525671, EPI_ISL_525672, EPI_ISL_525673, EPI_ISL_525674, EPI_ISL_525675, EPI_ISL_525676, EPI_ISL_525677, EPI_ISL_525678, EPI_ISL_525679, EPI_ISL_525680, EPI_ISL_525681, EPI_ISL_525682, EPI_ISL_525683, EPI_ISL_525684, EPI_ISL_525685, EPI_ISL_525686 | | | |
| see above | Wadsworth Center, New York State Department of Health | Wadsworth Center, New York State Department of Health | Kirsten St. George, Daryl M. Lamson, Sara Griesemer, Jonathan Plitnick, Navjot Singh, Matthew D. Shudt, Erica Lasek-Nesselquist |
| EPI_ISL_525763, EPI_ISL_525764, EPI_ISL_525765, EPI_ISL_525766, EPI_ISL_525767, EPI_ISL_525768, EPI_ISL_525769, EPI_ISL_525770, EPI_ISL_525771, EPI_ISL_525772, EPI_ISL_525773, EPI_ISL_525774, EPI_ISL_525775, EPI_ISL_525776, EPI_ISL_525777, EPI_ISL_525778, EPI_ISL_525779, EPI_ISL_525780, EPI_ISL_525781, EPI_ISL_525782, EPI_ISL_525783, EPI_ISL_525784, EPI_ISL_525785, EPI_ISL_525786, EPI_ISL_525787, EPI_ISL_525788, EPI_ISL_525789 | | | |
| see above | Texas Department of State Health Services | Texas Department of State Health Services | Jenny Zhang, Rashmi Tuladhar, Bonnie Oh, Maliha Rahman, Anita Pokharel, Myong Koag, Chun Wang, Rachel Lee, Grace Kubin |
| EPI_ISL_526700, EPI_ISL_526701, EPI_ISL_526702, EPI_ISL_526703, EPI_ISL_526704, EPI_ISL_526705, EPI_ISL_526706, EPI_ISL_526707, EPI_ISL_526708, EPI_ISL_526709, EPI_ISL_526710, EPI_ISL_526711, EPI_ISL_526712, EPI_ISL_526713, EPI_ISL_526714, EPI_ISL_526715, EPI_ISL_526716, EPI_ISL_526717, EPI_ISL_526718, EPI_ISL_526719, EPI_ISL_526720, EPI_ISL_526721, EPI_ISL_526722, EPI_ISL_526723, EPI_ISL_526724, EPI_ISL_526725, EPI_ISL_526726, EPI_ISL_526727, EPI_ISL_526728, EPI_ISL_526729 | | | |
| see above | Division of Viral Diseases, Center for Laboratory Control of Infectious Diseases, Korea Centers for Diseases Control and Prevention | Division of Viral Diseases, Center for Laboratory Control of Infectious Diseases, Korea Centers for Diseases Control and Prevention | Jeong-Min Kim, Yoon-Seok Chung, Namjoo Lee, Sang Hee Woo, Hye-Jun Jo, Heui Man Kim, Jun-Sub Kim, Myung Guk Han |
| EPI_ISL_526730, EPI_ISL_526731 | Center for Laboratory Control of Infectious Diseases, Korea Centers for Diseases Control and Prevention | Center for Laboratory Control of Infectious Diseases, Korea Centers for Diseases Control and Prevention | Junyoung Kim, Ae Kyung Park, Eunkyung Shin, Jin Sun No, Jeong-Min Kim, Yoon-Seok Chung, Heui Man Kim, Myung Guk Han |
| EPI_ISL_526732, EPI_ISL_526733 | Division of Viral Diseases, Center for Laboratory Control of Infectious Diseases, Korea Centers for Diseases Control and Prevention | Division of Viral Diseases, Center for Laboratory Control of Infectious Diseases, Korea Centers for Diseases Control and Prevention | Jeong-Min Kim, Yoon-Seok Chung, Namjoo Lee, Sang Hee Woo, Hye-Jun Jo, Heui Man Kim, Jun-Sub Kim, Myung Guk Han |
| EPI_ISL_526734, EPI_ISL_526735, EPI_ISL_526736, EPI_ISL_526737, EPI_ISL_526738, EPI_ISL_526739, EPI_ISL_526740, EPI_ISL_526741, EPI_ISL_526742, EPI_ISL_526743, EPI_ISL_526744, EPI_ISL_526745, EPI_ISL_526746 | | | |

| | | | |
|---|---|---|---|
| see above | Center for Laboratory Control of Infectious Diseases, Korea Centers for Diseases Control and Prevention | Center for Laboratory Control of Infectious Diseases, Korea Centers for Diseases Control and Prevention | Junyoung Kim, Ae Kyung Park, Eunkyung Shin, Jin Sun No, Jeong-Min Kim, Yoon-Seok Chung, Heui Man Kim, Myung Guk Han |
| EPI_ISL_526932, EPI_ISL_526933, EPI_ISL_526934, EPI_ISL_526949, EPI_ISL_526950, EPI_ISL_526951, EPI_ISL_526953, EPI_ISL_526954, EPI_ISL_526955, EPI_ISL_526956, EPI_ISL_526957, EPI_ISL_526958, EPI_ISL_526960 | | | |
| see above | Instituto Nacional de Salud, Bogotá, Colombia | Instituto Nacional de Salud, Bogotá, Colombia | Katherine Laiton-Donato, Diego A. Álvarez-Díaz, Carlos Franco-Muñoz, Mauricio Pacheco-Montealegre, Jonathan Reales, Diego Andrés Prada, Jose A. Usme-Ciro, Zulma M. Cucunubá, Christian Julian VillabonaArenas, Liz Villabona-Arenas, Sussy Echeverria, Astrid C. Flórez, Carolina Ferro, Diana Marcela Walteros-Acero, Franklin Prieto, Carlos Andrés Durán, Martha Lucia Ospina Martinez, Marcela Mercado-Reyes |
| EPI_ISL_526962, EPI_ISL_526963, EPI_ISL_526964, EPI_ISL_526965 | Instituto Nacional de Salud, Bogotá, Colombia | Instituto Nacional de Salud, Bogotá, Colombia | Katherine Laiton-Donato, Diego A. Álvarez-Díaz, Carlos Franco-Muñoz, Jonathan Reales, Diego Andrés Prada, Jeadran Malagón-Rojas, Felix Betzler, Wendy K. Jo, Edmilson F. de Oliveira-Filho, Carolina Ferro, Diana Marcela Walteros-Acero, Franklin Prieto, Carlos Andrés Durán, Martha Lucia Ospina Martinez, Marcela Mercado-Reyes |
| EPI_ISL_526967, EPI_ISL_526968, EPI_ISL_526969, EPI_ISL_526970, EPI_ISL_526971, EPI_ISL_526973, EPI_ISL_526974 | Instituto Nacional de Salud, Bogotá, Colombia | Instituto Nacional de Salud, Bogotá, Colombia | Katherine Laiton-Donato, Diego A. Álvarez-Díaz, Carlos Franco-Muñoz, Mauricio Pacheco-Montealegre, Jonathan Reales, Diego Andrés Prada, Jose A. Usme-Ciro, Zulma M. Cucunubá, Christian Julian VillabonaArenas, Liz Villabona-Arenas, Sussy Echeverria, Astrid C. Flórez, Carolina Ferro, Diana Marcela Walteros-Acero, Franklin Prieto, Carlos Andrés Durán, Martha Lucia Ospina Martinez, Marcela Mercado-Reyes |
| EPI_ISL_527666 | IN State Department of Health Laboratory Services | Pathogen Discovery, Respiratory Viruses Branch, Division of Viral Diseases, Centers for Disease Control and Prevention | Krista Queen, Brian Lynch, Yan Li, Anna Montmayeur, Jing Zhang, Ying Tao, Anna Uehara, Rachel Marine, Clinton R. Paden, Haibin Wang, Suxiang Tong |
| EPI_ISL_527692, EPI_ISL_527699, EPI_ISL_527701, EPI_ISL_527714, EPI_ISL_527716, EPI_ISL_527717, EPI_ISL_527720, EPI_ISL_527724, EPI_ISL_527731, EPI_ISL_527734, EPI_ISL_527735 | | | |
| see above | MN PHL Division, Minnesota Department of Health | Pathogen Discovery, Respiratory Viruses Branch, Division of Viral Diseases, Centers for Disease Control and Prevention | Krista Queen, Brian Lynch, Yan Li, Anna Montmayeur, Jing Zhang, Ying Tao, Anna Uehara, Rachel Marine, Clinton R. Paden, Haibin Wang, Suxiang Tong |
| EPI_ISL_527738 | Connecticut State Department of Public Health | Grubaugh Lab - Yale School of Public Health | Joseph Fauver, Mary Petrone, Anderson Brito, Tara Alpert, Nathan Grubaugh |
| EPI_ISL_527761, EPI_ISL_527762, EPI_ISL_527763, EPI_ISL_527764, EPI_ISL_527765, EPI_ISL_527766, EPI_ISL_527767, EPI_ISL_527768, EPI_ISL_527769, EPI_ISL_527770, EPI_ISL_527771, EPI_ISL_527772, EPI_ISL_527773, EPI_ISL_527774, EPI_ISL_527775, EPI_ISL_527776, EPI_ISL_527777, EPI_ISL_527778, EPI_ISL_527779, EPI_ISL_527780, EPI_ISL_527781, EPI_ISL_527782, EPI_ISL_527783, EPI_ISL_527784, EPI_ISL_527785 | | | |
| see above | Connecticut State Department of Public Health | Grubaugh Lab - Yale School of Public Health | Joseph Fauver, Mary Petrone, Chantal Vogels, Anderson Brito, Tara Alpert, Anthony Muyombwe, Jafar Razeq, Albert Ko, Nathan Grubaugh |
| EPI_ISL_528831, EPI_ISL_528833, EPI_ISL_528850, EPI_ISL_528862 | CSIR-Centre for Cellular and Molecular Biology | CSIR-Centre for Cellular and Molecular Biology | Lamuk Zaveri, Shagufta Khan, Namami Gaur, Sakshi Shambhavi, Nikhil Hajirnis, M Soujanya Reddy, Pratheusa Maccha, Tulasi Nagabandi, Purushotham Vodnala, Payel Mukherjee, Sofia Banu, Priya Singh, Onkar Kulkarni, Dhiviya Vedagiri, Divya Gupta, Vishal Sah, Santosh Kumar Kuncha, Krishnan Harinivas Harshan, Archana Bharadwaj Siva, Karthik Bharadwaj Tallapaka, Renu Sudhakar, Somesh Gorde, Gangumala Srinivas Reddy, Sujoy Deb, Swati Bayyana, Rakesh K Mishra, Divya Tej Sowpati |
| EPI_ISL_529033, EPI_ISL_529034, EPI_ISL_529035, EPI_ISL_529036 | Wadsworth Center, New York State Department of Health | Wadsworth Center, New York State Department of Health | Kirsten St. George, Daryl M. Lamson, Sara Griesemer, Jonathan Plitnick, Navjot Singh, Matthew D. Shudt, Erica Lasek-Nesselquist |
| EPI_ISL_529138 | RSAL Dr. Ramelan Surabaya | Institute of Tropical Disease, Universitas Airlangga | Jezzy R Dewantari, Rima R Prasetya, Krisnoadi Rahardjo, Aldise M Nastri, Radito Soesanto, Gatot Soegiarto, Laksmi Wulandari, Retno A Setyoningrum, Resti Yudhawati, Yohko K Shimizu, Mitsuhiro Nishimura, Yasuko Mori, Soetjipto, Kazufumi Shimizu, Maria I Lusida |
| EPI_ISL_529147, EPI_ISL_529148 | Democritus University of Thrace, Department of Medicine | Democritus University of Thrace, Department of Medicine | Kassela,K., Dovrolis,N., Bampali,M., Gatzidou,E., Froukala,E., Stavropoulou,A., Veletza,S., Tsakris,A., Spanakis,N., Karakasiliotis,I. |
| EPI_ISL_529961 | Husada Utama Hospital | Institute of Tropical Disease, Universitas Airlangga | Krisnoadi Rahardjo, Aldise M Nastri, Jezzy R Dewantari, Rima R Prasetya, Didi Dewanto, Gatot Soegiarto, Laksmi Wulandari, Retno A Setyoningrum, Resti Yudhawati, Yohko K Shimizu, Mitsuhiro Nishimura, Yasuko Mori, Soetjipto, Kazufumi Shimizu, Maria I Lusida |
| EPI_ISL_529962 | Universitas Airlangga Hospital | Institute of Tropical Disease, Universitas Airlangga | Jezzy R Dewantari, Rima R Prasetya, Krisnoadi Rahardjo, Aldise M Nastri, Nasronudin, Gatot Soegiarto, Laksmi Wulandari, Retno A Setyoningrum, Resti Yudhawati, Yohko K Shimizu, Mitsuhiro Nishimura, Yasuko Mori, Soetjipto, Kazufumi Shimizu, Maria I Lusida |
| EPI_ISL_529965 | RSUD Dr. Soetomo | Institute of Tropical Disease, Universitas Airlangga | Krisnoadi Rahardjo, Aldise M Nastri, Jezzy R Dewantari, Rima R Prasetya, Joni Wahyuhadi, Gatot Soegiarto, Laksmi Wulandari, Retno A Setyoningrum, Resti Yudhawati, Yohko K Shimizu, Mitsuhiro Nishimura, Yasuko Mori, Soetjipto, Kazufumi Shimizu, Maria I Lusida |
| EPI_ISL_529966 | RSUD Sidoarjo | Institute of Tropical Disease, Universitas Airlangga | Kazufumi Shimizu, Krisnoadi Rahardjo, Aldise M Nastri, Jezzy R Dewantari, Rima R Prasetya, Atok Irawan, Gatot Soegiarto, Laksmi Wulandari, Retno A Setyoningrum, Resti Yudhawati, Yohko K Shimizu, Mitsuhiro Nishimura, Yasuko Mori, Soetjipto, Maria I Lusida |
| EPI_ISL_534731 | Wadsworth Center, New York State Department of Health | Wadsworth Center, New York State Department of Health | Kirsten St. George, Daryl M. Lamson, Sara Griesemer, Jonathan Plitnick, Navjot Singh, Matthew D. Shudt, Erica Lasek-Nesselquist |
| EPI_ISL_535341, EPI_ISL_535342, EPI_ISL_535344, EPI_ISL_535345, EPI_ISL_535346, EPI_ISL_535348, EPI_ISL_535351, EPI_ISL_535357 | LA Office of Public Health Laboratories | Pathogen Discovery, Respiratory Viruses Branch, Division of Viral Diseases, Centers for Disease Control and Prevention | Jing Zhang, Ying Tao, Yan Li, Krista Queen, Anna Uehara, Clinton Paden, Haibin Wang, Suxiang Tong |
| EPI_ISL_535359, EPI_ISL_535360 | RI State Health Laboratories | Pathogen Discovery, Respiratory Viruses Branch, Division of Viral Diseases, Centers for Disease Control and Prevention | Jing Zhang, Ying Tao, Yan Li, Krista Queen, Anna Uehara, Clinton Paden, Haibin Wang, Suxiang Tong |
| EPI_ISL_539483, EPI_ISL_539484, EPI_ISL_539485, EPI_ISL_539486, EPI_ISL_539487, EPI_ISL_539488, EPI_ISL_539489, EPI_ISL_539490 | Civil Hospital, Rupnagar | CSIR-Institute of Microbial Technology | Kanika Bansal, Sanjeet Kumar, Anu Singh, Debarghya Ghose, Amandeep Kaur, Rajesh Kumar Mishra, Poushali Chakraborty, Harsh Goar, Navin Baid, Ashwani Kumar, Dipak Dutta, Sanjeev Khosla, Prabhu B. Patil |
| EPI_ISL_539491 | IDSP unit, Dehradun | CSIR-Institute of Microbial Technology | Kanika Bansal, Sanjeet Kumar, Anu Singh, Debarghya Ghose, Amandeep Kaur, Rajesh Kumar Mishra, Poushali Chakraborty, Harsh Goar, Navin Baid, Ashwani Kumar, Dipak Dutta, Sanjeev Khosla, Prabhu B. Patil |
| EPI_ISL_539492 | CMS, Roorkee | CSIR-Institute of Microbial Technology | Kanika Bansal, Sanjeet Kumar, Anu Singh, Debarghya Ghose, Amandeep Kaur, Rajesh Kumar Mishra, Poushali Chakraborty, Harsh Goar, Navin Baid, Ashwani Kumar, Dipak Dutta, Sanjeev Khosla, Prabhu B. Patil |
| EPI_ISL_539616 | CSIR-Centre for Cellular and Molecular Biology | CSIR-Centre for Cellular and Molecular Biology | Lamuk Zaveri, Shagufta Khan, Namami Gaur, Sakshi Shambhavi, Nikhil Hajirnis, M Soujanya Reddy, Pratheusa Maccha, Tulasi Nagabandi, Purushotham Vodnala, Payel Mukherjee, Sofia Banu, Priya Singh, Onkar Kulkarni, Dhiviya Vedagiri, Divya Gupta, Vishal Sah, Santosh Kumar Kuncha, Krishnan Harinivas Harshan, Archana Bharadwaj Siva, Karthik Bharadwaj Tallapaka, Renu Sudhakar, Somesh Gorde, Gangumala Srinivas Reddy, Sujoy Deb, Swati Bayyana, Rakesh K Mishra, Divya Tej Sowpati |
| EPI_ISL_539617 | CSIR-Centre for Cellular and Molecular Biology | CSIR-Centre for Cellular and Molecular Biology | Lamuk Zaveri, Shagufta Khan,Nikhil Hajirnis, M Soujanya Reddy, Pratheusa Maccha, Namami Gaur, Sakshi Shambhavi, Tulasi Nagabandi, Purushotham Vodnala, Payel Mukherjee, Sofia Banu, Priya Singh, Onkar Kulkarni, Dhiviya Vedagiri, Divya Gupta, Vishal Sah, Santosh Kumar Kuncha, Krishnan Harinivas Harshan, Archana Bharadwaj Siva, Karthik Bharadwaj Tallapaka,Umesh Kumar, Unis Ahmad Bhat, Ajay Sarawagi, Priyanka Pant, Rajkanwar Nathawat, Rakesh K Mishra, Divya Tej Sowpati |
| EPI_ISL_539618 | CSIR-Centre for Cellular and Molecular Biology | CSIR-Centre for Cellular and Molecular Biology | Lamuk Zaveri, Shagufta Khan,Nikhil Hajirnis, M Soujanya Reddy, Pratheusa Maccha, Namami Gaur, Sakshi Shambhavi, Tulasi Nagabandi, Purushotham Vodnala, Payel Mukherjee, Sofia Banu, Priya Singh, Onkar Kulkarni, Dhiviya Vedagiri, Divya Gupta, Vishal Sah, Santosh Kumar Kuncha, Krishnan Harinivas Harshan, Archana Bharadwaj Siva, Karthik Bharadwaj Tallapaka,Zeba Rizvi, Zuberwasim Sayyad, Kakade Aishwarya Arun, Amrutha H C, Ananga Ghosh, Rakesh K Mishra, Divya Tej Sowpati |
| EPI_ISL_539646 | CSIR-Centre for Cellular and Molecular Biology | CSIR-Centre for Cellular and Molecular Biology | Lamuk Zaveri, Shagufta Khan, Namami Gaur, Sakshi Shambhavi, Nikhil Hajirnis, M Soujanya Reddy, Pratheusa Maccha, Tulasi Nagabandi, Purushotham Vodnala, Payel Mukherjee, Sofia Banu, Priya Singh, Onkar Kulkarni, Dhiviya Vedagiri, Divya Gupta, Vishal Sah, Santosh Kumar Kuncha, Krishnan Harinivas Harshan, Archana Bharadwaj Siva, Karthik Bharadwaj Tallapaka, Renu Sudhakar, Somesh Gorde, Gangumala Srinivas Reddy, Sujoy Deb, Swati Bayyana, Rakesh K Mishra, Divya Tej Sowpati |
| EPI_ISL_539647 | CSIR-Centre for Cellular and Molecular Biology | CSIR-Centre for Cellular and Molecular Biology | Lamuk Zaveri, Shagufta Khan,Nikhil Hajirnis, M Soujanya Reddy, Pratheusa Maccha, Namami Gaur, Sakshi Shambhavi, Tulasi Nagabandi, Purushotham Vodnala, Payel Mukherjee, Sofia Banu, Priya Singh, Onkar Kulkarni, Dhiviya Vedagiri, Divya Gupta, Vishal Sah, Santosh Kumar Kuncha, Krishnan Harinivas Harshan, Archana Bharadwaj Siva, Karthik Bharadwaj Tallapaka,Umesh Kumar, Unis Ahmad Bhat, Ajay Sarawagi, Priyanka Pant, Rajkanwar Nathawat, |

| | | | |
|---|---|---|---|
| | | | Rakesh K Mishra, Divya Tej Sowpati |
| EPI_ISL_539648 | CSIR-Centre for Cellular and Molecular Biology | CSIR-Centre for Cellular and Molecular Biology | Lamuk Zaveri, Shagufta Khan,Nikhil Hajirnis, M Soujanya Reddy, Pratheusa Maccha, Namami Gaur, Sakshi Shambhavi, Tulasi Nagabandi, Purushotham Vodnala, Payel Mukherjee, Sofia Banu, Priya Singh, Onkar Kulkarni, Dhiviya Vedagiri, Divya Gupta, Vishal Sah, Santosh Kumar Kuncha, Krishnan Harinivas Harshan, Archana Bharadwaj Siva, Karthik Bharadwaj Tallapaka,Zeba Rizvi, Zuberwasim Sayyad, Kakade Aishwarya Arun, Amrutha H C, Ananga Ghosh, Rakesh K Mishra, Divya Tej Sowpati |
| EPI_ISL_539676 | CSIR-Centre for Cellular and Molecular Biology | CSIR-Centre for Cellular and Molecular Biology | Lamuk Zaveri, Shagufta Khan, Namami Gaur, Sakshi Shambhavi, Nikhil Hajirnis, M Soujanya Reddy, Pratheusa Maccha, Tulasi Nagabandi, Purushotham Vodnala, Payel Mukherjee, Sofia Banu, Priya Singh, Onkar Kulkarni, Dhiviya Vedagiri, Divya Gupta, Vishal Sah, Santosh Kumar Kuncha, Krishnan Harinivas Harshan, Archana Bharadwaj Siva, Karthik Bharadwaj Tallapaka, Renu Sudhakar, Somesh Gorde, Gangumala Srinivas Reddy, Sujoy Deb, Swati Bayyana, Rakesh K Mishra, Divya Tej Sowpati |
| EPI_ISL_539677 | CSIR-Centre for Cellular and Molecular Biology | CSIR-Centre for Cellular and Molecular Biology | Lamuk Zaveri, Shagufta Khan,Nikhil Hajirnis, M Soujanya Reddy, Pratheusa Maccha, Namami Gaur, Sakshi Shambhavi, Tulasi Nagabandi, Purushotham Vodnala, Payel Mukherjee, Sofia Banu, Priya Singh, Onkar Kulkarni, Dhiviya Vedagiri, Divya Gupta, Vishal Sah, Santosh Kumar Kuncha, Krishnan Harinivas Harshan, Archana Bharadwaj Siva, Karthik Bharadwaj Tallapaka,Umesh Kumar, Unis Ahmad Bhat, Ajay Sarawagi, Priyanka Pant, Rajkanwar Nathawat, Rakesh K Mishra, Divya Tej Sowpati |
| EPI_ISL_539678 | CSIR-Centre for Cellular and Molecular Biology | CSIR-Centre for Cellular and Molecular Biology | Lamuk Zaveri, Shagufta Khan,Nikhil Hajirnis, M Soujanya Reddy, Pratheusa Maccha, Namami Gaur, Sakshi Shambhavi, Tulasi Nagabandi, Purushotham Vodnala, Payel Mukherjee, Sofia Banu, Priya Singh, Onkar Kulkarni, Dhiviya Vedagiri, Divya Gupta, Vishal Sah, Santosh Kumar Kuncha, Krishnan Harinivas Harshan, Archana Bharadwaj Siva, Karthik Bharadwaj Tallapaka,Zeba Rizvi, Zuberwasim Sayyad, Kakade Aishwarya Arun, Amrutha H C, Ananga Ghosh, Rakesh K Mishra, Divya Tej Sowpati |
| EPI_ISL_539706 | CSIR-Centre for Cellular and Molecular Biology | CSIR-Centre for Cellular and Molecular Biology | Lamuk Zaveri, Shagufta Khan, Namami Gaur, Sakshi Shambhavi, Nikhil Hajirnis, M Soujanya Reddy, Pratheusa Maccha, Tulasi Nagabandi, Purushotham Vodnala, Payel Mukherjee, Sofia Banu, Priya Singh, Onkar Kulkarni, Dhiviya Vedagiri, Divya Gupta, Vishal Sah, Santosh Kumar Kuncha, Krishnan Harinivas Harshan, Archana Bharadwaj Siva, Karthik Bharadwaj Tallapaka, Renu Sudhakar, Somesh Gorde, Gangumala Srinivas Reddy, Sujoy Deb, Swati Bayyana, Rakesh K Mishra, Divya Tej Sowpati |
| EPI_ISL_539707 | CSIR-Centre for Cellular and Molecular Biology | CSIR-Centre for Cellular and Molecular Biology | Lamuk Zaveri, Shagufta Khan,Nikhil Hajirnis, M Soujanya Reddy, Pratheusa Maccha, Namami Gaur, Sakshi Shambhavi, Tulasi Nagabandi, Purushotham Vodnala, Payel Mukherjee, Sofia Banu, Priya Singh, Onkar Kulkarni, Dhiviya Vedagiri, Divya Gupta, Vishal Sah, Santosh Kumar Kuncha, Krishnan Harinivas Harshan, Archana Bharadwaj Siva, Karthik Bharadwaj Tallapaka,Umesh Kumar, Unis Ahmad Bhat, Ajay Sarawagi, Priyanka Pant, Rajkanwar Nathawat, Rakesh K Mishra, Divya Tej Sowpati |
| EPI_ISL_539708 | CSIR-Centre for Cellular and Molecular Biology | CSIR-Centre for Cellular and Molecular Biology | Lamuk Zaveri, Shagufta Khan,Nikhil Hajirnis, M Soujanya Reddy, Pratheusa Maccha, Namami Gaur, Sakshi Shambhavi, Tulasi Nagabandi, Purushotham Vodnala, Payel Mukherjee, Sofia Banu, Priya Singh, Onkar Kulkarni, Dhiviya Vedagiri, Divya Gupta, Vishal Sah, Santosh Kumar Kuncha, Krishnan Harinivas Harshan, Archana Bharadwaj Siva, Karthik Bharadwaj Tallapaka,Zeba Rizvi, Zuberwasim Sayyad, Kakade Aishwarya Arun, Amrutha H C, Ananga Ghosh, Rakesh K Mishra, Divya Tej Sowpati |
| EPI_ISL_539736 | CSIR-Centre for Cellular and Molecular Biology | CSIR-Centre for Cellular and Molecular Biology | Lamuk Zaveri, Shagufta Khan, Namami Gaur, Sakshi Shambhavi, Nikhil Hajirnis, M Soujanya Reddy, Pratheusa Maccha, Tulasi Nagabandi, Purushotham Vodnala, Payel Mukherjee, Sofia Banu, Priya Singh, Onkar Kulkarni, Dhiviya Vedagiri, Divya Gupta, Vishal Sah, Santosh Kumar Kuncha, Krishnan Harinivas Harshan, Archana Bharadwaj Siva, Karthik Bharadwaj Tallapaka, Renu Sudhakar, Somesh Gorde, Gangumala Srinivas Reddy, Sujoy Deb, Swati Bayyana, Rakesh K Mishra, Divya Tej Sowpati |
| EPI_ISL_539737 | CSIR-Centre for Cellular and Molecular Biology | CSIR-Centre for Cellular and Molecular Biology | Lamuk Zaveri, Shagufta Khan,Nikhil Hajirnis, M Soujanya Reddy, Pratheusa Maccha, Namami Gaur, Sakshi Shambhavi, Tulasi Nagabandi, Purushotham Vodnala, Payel Mukherjee, Sofia Banu, Priya Singh, Onkar Kulkarni, Dhiviya Vedagiri, Divya Gupta, Vishal Sah, Santosh Kumar Kuncha, Krishnan Harinivas Harshan, Archana Bharadwaj Siva, Karthik Bharadwaj Tallapaka,Umesh Kumar, Unis Ahmad Bhat, Ajay Sarawagi, Priyanka Pant, Rajkanwar Nathawat, Rakesh K Mishra, Divya Tej Sowpati |
| EPI_ISL_539738 | CSIR-Centre for Cellular and Molecular Biology | CSIR-Centre for Cellular and Molecular Biology | Lamuk Zaveri, Shagufta Khan,Nikhil Hajirnis, M Soujanya Reddy, Pratheusa Maccha, Namami Gaur, Sakshi Shambhavi, Tulasi Nagabandi, Purushotham Vodnala, Payel Mukherjee, Sofia Banu, Priya Singh, Onkar Kulkarni, Dhiviya Vedagiri, Divya Gupta, Vishal Sah, Santosh Kumar Kuncha, Krishnan Harinivas Harshan, Archana Bharadwaj Siva, Karthik Bharadwaj Tallapaka,Zeba Rizvi, Zuberwasim Sayyad, Kakade Aishwarya Arun, Amrutha H C, Ananga Ghosh, Rakesh K Mishra, Divya Tej Sowpati |
| EPI_ISL_539769 | CSIR-Centre for Cellular and Molecular Biology | CSIR-Centre for Cellular and Molecular Biology | Lamuk Zaveri, Shagufta Khan,Nikhil Hajirnis, M Soujanya Reddy, Pratheusa Maccha, Namami Gaur, Sakshi Shambhavi, Tulasi Nagabandi, Purushotham Vodnala, Payel Mukherjee, Sofia Banu, Priya Singh, Onkar Kulkarni, Dhiviya Vedagiri, Divya Gupta, Vishal Sah, Santosh Kumar Kuncha, Krishnan Harinivas Harshan, Archana Bharadwaj Siva, Karthik Bharadwaj Tallapaka,Umesh Kumar, Unis Ahmad Bhat, Ajay Sarawagi, Priyanka Pant, Rajkanwar Nathawat, Rakesh K Mishra, Divya Tej Sowpati |
| EPI_ISL_539770 | CSIR-Centre for Cellular and Molecular Biology | CSIR-Centre for Cellular and Molecular Biology | Lamuk Zaveri, Shagufta Khan,Nikhil Hajirnis, M Soujanya Reddy, Pratheusa Maccha, Namami Gaur, Sakshi Shambhavi, Tulasi Nagabandi, Purushotham Vodnala, Payel Mukherjee, Sofia Banu, Priya Singh, Onkar Kulkarni, Dhiviya Vedagiri, Divya Gupta, Vishal Sah, Santosh Kumar Kuncha, Krishnan Harinivas Harshan, Archana Bharadwaj Siva, Karthik Bharadwaj Tallapaka,Zeba Rizvi, Zuberwasim Sayyad, Kakade Aishwarya Arun, Amrutha H C, Ananga Ghosh, Rakesh K Mishra, Divya Tej Sowpati |
| EPI_ISL_540435 | VA-Division of Consolidated Laboratory Services | Pathogen Discovery, Respiratory Viruses Branch, Division of Viral Diseases, Centers for Disease Control and Prevention | Krista Queen, Yan Li, Jing Zhang, Anna Montmayeur, Krista Queen,Ying Tao, Anna Uehara, Clinton R. Paden, Rachel Marine, Haibin Wang, Suxiang Tong |
| EPI_ISL_540442, EPI_ISL_540443, EPI_ISL_540444, EPI_ISL_540445, EPI_ISL_540446, EPI_ISL_540447, EPI_ISL_540448, EPI_ISL_540449, EPI_ISL_540450, EPI_ISL_540451, EPI_ISL_540452, EPI_ISL_540453, EPI_ISL_540454, EPI_ISL_540455, EPI_ISL_540456, EPI_ISL_540457, EPI_ISL_540458, EPI_ISL_540459, EPI_ISL_540460, EPI_ISL_540461, EPI_ISL_540462, EPI_ISL_540463, EPI_ISL_540464, EPI_ISL_540465, EPI_ISL_540466, EPI_ISL_540467, EPI_ISL_540468 | | | |
| see above | University of Liège COVID-19 testing center | GIGA Medical Genomics | Keith Durkin, Maria Artesi, Emmanuel André, Marc Van Ranst, Fabrice Bureau, Laurent Gillet, Wouter Coppieters, Vincent Bours |
| EPI_ISL_540469, EPI_ISL_540470, EPI_ISL_540471, EPI_ISL_540472, EPI_ISL_540473, EPI_ISL_540474, EPI_ISL_540475, EPI_ISL_540476, EPI_ISL_540477, EPI_ISL_540478, EPI_ISL_540479, EPI_ISL_540480, EPI_ISL_540481, EPI_ISL_540482, EPI_ISL_540483, EPI_ISL_540484, EPI_ISL_540485, EPI_ISL_540486, EPI_ISL_540487, EPI_ISL_540488, EPI_ISL_540489, EPI_ISL_540490, EPI_ISL_540491, EPI_ISL_540492, EPI_ISL_540493, EPI_ISL_540494, EPI_ISL_540495, EPI_ISL_540496, EPI_ISL_540497, EPI_ISL_540498, EPI_ISL_540499, EPI_ISL_540500, EPI_ISL_540501, EPI_ISL_540502, EPI_ISL_540503, EPI_ISL_540504, EPI_ISL_540505, EPI_ISL_540506, EPI_ISL_540507, EPI_ISL_540508, EPI_ISL_540509, EPI_ISL_540510, EPI_ISL_540511, EPI_ISL_540512, EPI_ISL_540513, EPI_ISL_540514, EPI_ISL_540515, EPI_ISL_540516, EPI_ISL_540517, EPI_ISL_540518, EPI_ISL_540519, EPI_ISL_540520, EPI_ISL_540521, EPI_ISL_540522, EPI_ISL_540523, EPI_ISL_540524, EPI_ISL_540525, EPI_ISL_540526, EPI_ISL_540527, EPI_ISL_540528, EPI_ISL_540529, EPI_ISL_540530, EPI_ISL_540531, EPI_ISL_540532, EPI_ISL_540533, EPI_ISL_540534, EPI_ISL_540535, EPI_ISL_540536, EPI_ISL_540537, EPI_ISL_540538, EPI_ISL_540539, EPI_ISL_540540, EPI_ISL_540541, EPI_ISL_540542, EPI_ISL_540543, EPI_ISL_540544, EPI_ISL_540545, EPI_ISL_540546, EPI_ISL_540547, EPI_ISL_540548, EPI_ISL_540549, EPI_ISL_540550, EPI_ISL_540551, EPI_ISL_540552, EPI_ISL_540553, EPI_ISL_540554, EPI_ISL_540555, EPI_ISL_540556, EPI_ISL_540557, EPI_ISL_540558, EPI_ISL_540559, EPI_ISL_540560, EPI_ISL_540561, EPI_ISL_540562, EPI_ISL_540563, EPI_ISL_540564, EPI_ISL_540565, EPI_ISL_540566, EPI_ISL_540567, EPI_ISL_540568, EPI_ISL_540569, EPI_ISL_540570, EPI_ISL_540571, EPI_ISL_540572, EPI_ISL_540573, EPI_ISL_540574, EPI_ISL_540575, EPI_ISL_540576, EPI_ISL_540577, EPI_ISL_540578 | | | |
| see above | Department of Clinical Microbiology | GIGA Medical Genomics | Keith Durkin, Maria Artesi, Sébastien Bontems, Raphaël Boreux, Bouchra Boujemla, Cécile Meex, Axelle Chaslain, Céline Fombellida-Lopez, Pierrette Melin, Marie-Pierre Hayette, Vincent Bours |
| EPI_ISL_541037, EPI_ISL_541038, EPI_ISL_541039, EPI_ISL_541040, EPI_ISL_541041, EPI_ISL_541042, EPI_ISL_541043, EPI_ISL_541044, EPI_ISL_541045, EPI_ISL_541046, EPI_ISL_541047, EPI_ISL_541048, EPI_ISL_541049, EPI_ISL_541050, EPI_ISL_541051, EPI_ISL_541052, EPI_ISL_541053, EPI_ISL_541054, EPI_ISL_541055, EPI_ISL_541056, EPI_ISL_541057, EPI_ISL_541058, EPI_ISL_541059, EPI_ISL_541060, EPI_ISL_541061, EPI_ISL_541062, EPI_ISL_541063, EPI_ISL_541064, EPI_ISL_541065, EPI_ISL_541066 | | | |
| see above | Hospital Clínico Universitario de Santiago de Compostela | SeqCOVID-SPAIN consortium/Institute of Biomedicine of Valencia, IBV-CSIC | José Javier Costa Alcalde, Antonio Aguilera Guirao, Mª Luisa Pérez del Molino Bernal, Amparo Coira Nieto, Gema Barbeito Castiñeiras, Rocio Trastoy Pena and SeqCOVID-SPAIN consortium |
| EPI_ISL_545955 | Laboratorio de Infecciones Respiratorias Agudas. Centro Nacional de Salud Publica,Instituto Nacional de Salud | Laboratorio de Infecciones Respiratorias Agudas. Centro Nacional de Salud Publica,Instituto Nacional de Salud | Juscamayta,E. |
| EPI_ISL_547584 | CMS, Roorkee | CSIR-Institute of Microbial Technology | Kanika Bansal, Sanjeet Kumar, Anu Singh, Debarghya Ghose, Rajesh Kumar Mishra, Dipak Dutta, Sanjeev Khosla, Prabhu B. Patil |
| EPI_ISL_547585, EPI_ISL_547586, EPI_ISL_547587, EPI_ISL_547588, EPI_ISL_547589, EPI_ISL_547590, EPI_ISL_547591, EPI_ISL_547592, EPI_ISL_547593 | Civil Hospital, Panchkula | CSIR-Institute of Microbial Technology | Kanika Bansal, Sanjeet Kumar, Anu Singh, Debarghya Ghose, Rajesh Kumar Mishra, Dipak Dutta, Sanjeev Khosla, Prabhu B. Patil |
| EPI_ISL_548147, EPI_ISL_548148, EPI_ISL_548149, EPI_ISL_548150, EPI_ISL_548151, EPI_ISL_548152, EPI_ISL_548153, EPI_ISL_548154, EPI_ISL_548155, EPI_ISL_548156, EPI_ISL_548157, EPI_ISL_548158, EPI_ISL_548159, EPI_ISL_548160, EPI_ISL_548161, EPI_ISL_548162, EPI_ISL_548163, EPI_ISL_548164, | | | |

| | | | |
|---|---|---|---|
| EPI_ISL_548165, EPI_ISL_548166, EPI_ISL_548167, EPI_ISL_548168, EPI_ISL_548169, EPI_ISL_548170, EPI_ISL_548171, EPI_ISL_548172, EPI_ISL_548173, EPI_ISL_548174, EPI_ISL_548175, EPI_ISL_548176, EPI_ISL_548177, EPI_ISL_548178, EPI_ISL_548179, EPI_ISL_548180, EPI_ISL_548181, EPI_ISL_548182, EPI_ISL_548183, EPI_ISL_548184, EPI_ISL_548185, EPI_ISL_548186, EPI_ISL_548187, EPI_ISL_548188, EPI_ISL_548189, EPI_ISL_548190, EPI_ISL_548191, EPI_ISL_548192, EPI_ISL_548193, EPI_ISL_548194, EPI_ISL_548195, EPI_ISL_548196, EPI_ISL_548197, EPI_ISL_548198, EPI_ISL_548199, EPI_ISL_548200, EPI_ISL_548201, EPI_ISL_548202, EPI_ISL_548203, EPI_ISL_548204, EPI_ISL_548205, EPI_ISL_548206, EPI_ISL_548207, EPI_ISL_548208, EPI_ISL_548209, EPI_ISL_548210, EPI_ISL_548211, EPI_ISL_548212, EPI_ISL_548213, EPI_ISL_548214, EPI_ISL_548215, EPI_ISL_548216, EPI_ISL_548217, EPI_ISL_548218, EPI_ISL_548219, EPI_ISL_548220, EPI_ISL_548221, EPI_ISL_548222, EPI_ISL_548223, EPI_ISL_548224, EPI_ISL_548225, EPI_ISL_548226, EPI_ISL_548227, EPI_ISL_548228, EPI_ISL_548229, EPI_ISL_548230, EPI_ISL_548231, EPI_ISL_548232, EPI_ISL_548233, EPI_ISL_548234, EPI_ISL_548235, EPI_ISL_548236, EPI_ISL_548237, EPI_ISL_548238, EPI_ISL_548239, EPI_ISL_548240, EPI_ISL_548241, EPI_ISL_548242 | | | |
| see above | Laboratoire de Virologie, HUG | Swiss National Reference Centre for Influenza | LAUBSCHER F. |
| EPI_ISL_549027, EPI_ISL_549028 | Furst Medical Laboratory | Norwegian Institute of Public Health, Department of Virology | Kathrine Stene-Johansen, Kamilla Heddeland Instefjord, Hilde Elshaug, Rasmus Riis Kopperud, Hilde Synnøve Vollan, Karoline Bragstad, Olav Hungnes |
| EPI_ISL_549029, EPI_ISL_549030, EPI_ISL_549031, EPI_ISL_549032, EPI_ISL_549033, EPI_ISL_549034, EPI_ISL_549035 | Oslo University Hospital, Department of Medical Microbiology | Norwegian Institute of Public Health, Department of Virology | Kathrine Stene-Johansen, Kamilla Heddeland Instefjord, Hilde Elshaug, Rasmus Riis Kopperud, Hilde Synnøve Vollan, Karoline Bragstad, Olav Hungnes |
| EPI_ISL_549036, EPI_ISL_549037 | Hospital of Southern Norway - Kristiansand, Department of Medical Microbiology | Norwegian Institute of Public Health, Department of Virology | Kathrine Stene-Johansen, Kamilla Heddeland Instefjord, Hilde Elshaug, Rasmus Riis Kopperud, Hilde Synnøve Vollan, Karoline Bragstad, Olav Hungnes |
| EPI_ISL_549038 | Furst Medical Laboratory | Norwegian Institute of Public Health, Department of Virology | Kathrine Stene-Johansen, Kamilla Heddeland Instefjord, Hilde Elshaug, Rasmus Riis Kopperud, Hilde Synnøve Vollan, Karoline Bragstad, Olav Hungnes |
| EPI_ISL_549039, EPI_ISL_549040, EPI_ISL_549041 | Medical Microbiology Unit, Department for Laboratory Medicine, Drammen Hospital, Vestre Viken Health Trust, | Norwegian Institute of Public Health, Department of Virology | Kathrine Stene-Johansen, Kamilla Heddeland Instefjord, Hilde Elshaug, Rasmus Riis Kopperud, Hilde Synnøve Vollan, Karoline Bragstad, Olav Hungnes |
| EPI_ISL_549042, EPI_ISL_549043, EPI_ISL_549044, EPI_ISL_549045, EPI_ISL_549046, EPI_ISL_549047 | Ostfold Hospital Trust - Kalnes, Centre for Laboratory Medicine, Section for gene technology and infection serology | Norwegian Institute of Public Health, Department of Virology | Kathrine Stene-Johansen, Kamilla Heddeland Instefjord, Hilde Elshaug, Rasmus Riis Kopperud, Hilde Synnøve Vollan, Karoline Bragstad, Olav Hungnes |
| EPI_ISL_549048, EPI_ISL_549049 | Furst Medical Laboratory | Norwegian Institute of Public Health, Department of Virology | Kathrine Stene-Johansen, Kamilla Heddeland Instefjord, Hilde Elshaug, Rasmus Riis Kopperud, Hilde Synnøve Vollan, Karoline Bragstad, Olav Hungnes |
| EPI_ISL_549050 | Unilabs Laboratory Medicine | Norwegian Institute of Public Health, Department of Virology | Kathrine Stene-Johansen, Kamilla Heddeland Instefjord, Hilde Elshaug, Rasmus Riis Kopperud, Hilde Synnøve Vollan, Karoline Bragstad, Olav Hungnes |
| EPI_ISL_549051 | Vestfold Hospital, Toensberg Department of Microbiology | Norwegian Institute of Public Health, Department of Virology | Kathrine Stene-Johansen, Kamilla Heddeland Instefjord, Hilde Elshaug, Rasmus Riis Kopperud, Hilde Synnøve Vollan, Karoline Bragstad, Olav Hungnes |
| EPI_ISL_549052 | Hospital of Southern Norway - Kristiansand, Department of Medical Microbiology | Norwegian Institute of Public Health, Department of Virology | Kathrine Stene-Johansen, Kamilla Heddeland Instefjord, Hilde Elshaug, Rasmus Riis Kopperud, Hilde Synnøve Vollan, Karoline Bragstad, Olav Hungnes |
| EPI_ISL_549053, EPI_ISL_549054, EPI_ISL_549055, EPI_ISL_549056, EPI_ISL_549057, EPI_ISL_549058 | Furst Medical Laboratory | Norwegian Institute of Public Health, Department of Virology | Kathrine Stene-Johansen, Kamilla Heddeland Instefjord, Hilde Elshaug, Rasmus Riis Kopperud, Hilde Synnøve Vollan, Karoline Bragstad, Olav Hungnes |
| EPI_ISL_549059 | Medical Microbiology Unit, Department for Laboratory Medicine, Drammen Hospital, Vestre Viken Health Trust, | Norwegian Institute of Public Health, Department of Virology | Kathrine Stene-Johansen, Kamilla Heddeland Instefjord, Hilde Elshaug, Rasmus Riis Kopperud, Hilde Synnøve Vollan, Karoline Bragstad, Olav Hungnes |
| EPI_ISL_549060, EPI_ISL_549061, EPI_ISL_549062, EPI_ISL_549063, EPI_ISL_549064, EPI_ISL_549065, EPI_ISL_549066, EPI_ISL_549067, EPI_ISL_549068, EPI_ISL_549069 | Furst Medical Laboratory | Norwegian Institute of Public Health, Department of Virology | Kathrine Stene-Johansen, Kamilla Heddeland Instefjord, Hilde Elshaug, Rasmus Riis Kopperud, Hilde Synnøve Vollan, Karoline Bragstad, Olav Hungnes |
| EPI_ISL_549070 | Medical Microbiology Unit, Department for Laboratory Medicine, Drammen Hospital, Vestre Viken Health Trust, | Norwegian Institute of Public Health, Department of Virology | Kathrine Stene-Johansen, Kamilla Heddeland Instefjord, Hilde Elshaug, Rasmus Riis Kopperud, Hilde Synnøve Vollan, Karoline Bragstad, Olav Hungnes |
| EPI_ISL_549071, EPI_ISL_549072, EPI_ISL_549073, EPI_ISL_549074, EPI_ISL_549075, EPI_ISL_549076, EPI_ISL_549077, EPI_ISL_549078, EPI_ISL_549079, EPI_ISL_549080 | Furst Medical Laboratory | Norwegian Institute of Public Health, Department of Virology | Kathrine Stene-Johansen, Kamilla Heddeland Instefjord, Hilde Elshaug, Rasmus Riis Kopperud, Hilde Synnøve Vollan, Karoline Bragstad, Olav Hungnes |
| EPI_ISL_549081 | Medical Microbiology Unit, Department for Laboratory Medicine, Drammen Hospital, Vestre Viken Health Trust, | Norwegian Institute of Public Health, Department of Virology | Kathrine Stene-Johansen, Kamilla Heddeland Instefjord, Hilde Elshaug, Rasmus Riis Kopperud, Hilde Synnøve Vollan, Karoline Bragstad, Olav Hungnes |
| EPI_ISL_549082 | Furst Medical Laboratory | Norwegian Institute of Public Health, Department of Virology | Kathrine Stene-Johansen, Kamilla Heddeland Instefjord, Hilde Elshaug, Rasmus Riis Kopperud, Hilde Synnøve Vollan, Karoline Bragstad, Olav Hungnes |
| EPI_ISL_549083, EPI_ISL_549084 | Akershus University Hospital, Department for Microbiology and Infectious Disease Control | Norwegian Institute of Public Health, Department of Virology | Kathrine Stene-Johansen, Kamilla Heddeland Instefjord, Hilde Elshaug, Rasmus Riis Kopperud, Hilde Synnøve Vollan, Karoline Bragstad, Olav Hungnes |
| EPI_ISL_549085, EPI_ISL_549086 | Medical Microbiology Unit, Department for Laboratory Medicine, Drammen Hospital, Vestre Viken Health Trust, | Norwegian Institute of Public Health, Department of Virology | Kathrine Stene-Johansen, Kamilla Heddeland Instefjord, Hilde Elshaug, Rasmus Riis Kopperud, Hilde Synnøve Vollan, Karoline Bragstad, Olav Hungnes |
| EPI_ISL_549087 | Vestfold Hospital, Toensberg Department of Microbiology | Norwegian Institute of Public Health, Department of Virology | Kathrine Stene-Johansen, Kamilla Heddeland Instefjord, Hilde Elshaug, Rasmus Riis Kopperud, Hilde Synnøve Vollan, Karoline Bragstad, Olav Hungnes |
| EPI_ISL_549088 | Furst Medical Laboratory | Norwegian Institute of Public Health, Department of Virology | Kathrine Stene-Johansen, Kamilla Heddeland Instefjord, Hilde Elshaug, Rasmus Riis Kopperud, Hilde Synnøve Vollan, Karoline Bragstad, Olav Hungnes |
| EPI_ISL_549089, EPI_ISL_549090, EPI_ISL_549091 | Akershus University Hospital, Department for Microbiology and Infectious Disease Control | Norwegian Institute of Public Health, Department of Virology | Kathrine Stene-Johansen, Kamilla Heddeland Instefjord, Hilde Elshaug, Rasmus Riis Kopperud, Hilde Synnøve Vollan, Karoline Bragstad, Olav Hungnes |
| EPI_ISL_549092, EPI_ISL_549093, EPI_ISL_549094, EPI_ISL_549095, EPI_ISL_549096, EPI_ISL_549097, EPI_ISL_549098, EPI_ISL_549099, EPI_ISL_549100, EPI_ISL_549101, EPI_ISL_549102, EPI_ISL_549103, EPI_ISL_549104, EPI_ISL_549105, EPI_ISL_549106, EPI_ISL_549107, EPI_ISL_549108, EPI_ISL_549109, EPI_ISL_549110, EPI_ISL_549111 | | | |
| see above | Ostfold Hospital Trust - Kalnes, Centre for Laboratory Medicine, Section for gene technology and infection serology | Norwegian Institute of Public Health, Department of Virology | Kathrine Stene-Johansen, Kamilla Heddeland Instefjord, Hilde Elshaug, Rasmus Riis Kopperud, Hilde Synnøve Vollan, Karoline Bragstad, Olav Hungnes |
| EPI_ISL_549112 | Furst Medical Laboratory | Norwegian Institute of Public Health, Department of Virology | Kathrine Stene-Johansen, Kamilla Heddeland Instefjord, Hilde Elshaug, Rasmus Riis Kopperud, Hilde Synnøve Vollan, Karoline Bragstad, Olav Hungnes |
| EPI_ISL_549113, EPI_ISL_549114, EPI_ISL_549115, EPI_ISL_549116, EPI_ISL_549117, EPI_ISL_549118 | Ostfold Hospital Trust - Kalnes, Centre for Laboratory Medicine, Section for gene technology and infection serology | Norwegian Institute of Public Health, Department of Virology | Kathrine Stene-Johansen, Kamilla Heddeland Instefjord, Hilde Elshaug, Rasmus Riis Kopperud, Hilde Synnøve Vollan, Karoline Bragstad, Olav Hungnes |
| EPI_ISL_549119, EPI_ISL_549120, EPI_ISL_549121, EPI_ISL_549122 | Furst Medical Laboratory | Norwegian Institute of Public Health, Department of Virology | Kathrine Stene-Johansen, Kamilla Heddeland Instefjord, Hilde Elshaug, Rasmus Riis Kopperud, Hilde Synnøve Vollan, Karoline Bragstad, Olav Hungnes |
| EPI_ISL_549123 | Unilabs Laboratory Medicine | Norwegian Institute of Public Health, Department of Virology | Kathrine Stene-Johansen, Kamilla Heddeland Instefjord, Hilde Elshaug, Rasmus Riis Kopperud, Hilde Synnøve Vollan, Karoline Bragstad, Olav Hungnes |
| EPI_ISL_549124 | Furst Medical Laboratory | Norwegian Institute of Public Health, Department of Virology | Kathrine Stene-Johansen, Kamilla Heddeland Instefjord, Hilde Elshaug, Rasmus Riis Kopperud, Hilde Synnøve Vollan, Karoline Bragstad, Olav Hungnes |
| EPI_ISL_549125 | Unilabs Laboratory Medicine | Norwegian Institute of Public Health, Department of Virology | Kathrine Stene-Johansen, Kamilla Heddeland Instefjord, Hilde Elshaug, Rasmus Riis Kopperud, Hilde Synnøve Vollan, Karoline Bragstad, Olav Hungnes |
| EPI_ISL_549126, EPI_ISL_549127, EPI_ISL_549128, EPI_ISL_549129, EPI_ISL_549130, EPI_ISL_549131 | Ostfold Hospital Trust - Kalnes, Centre for Laboratory Medicine, Section for gene technology and infection serology | Norwegian Institute of Public Health, Department of Virology | Kathrine Stene-Johansen, Kamilla Heddeland Instefjord, Hilde Elshaug, Rasmus Riis Kopperud, Hilde Synnøve Vollan, Karoline Bragstad, Olav Hungnes |
| EPI_ISL_549132 | Furst Medical Laboratory | Norwegian Institute of Public Health, Department of Virology | Kathrine Stene-Johansen, Kamilla Heddeland Instefjord, Hilde Elshaug, Rasmus Riis Kopperud, Hilde Synnøve Vollan, Karoline Bragstad, Olav Hungnes |
| EPI_ISL_549133, EPI_ISL_549134, EPI_ISL_549135, EPI_ISL_549136, | Ostfold Hospital Trust - Kalnes, Centre for Laboratory Medicine, Section for gene technology and infection serology | Norwegian Institute of Public Health, Department of Virology | Kathrine Stene-Johansen, Kamilla Heddeland Instefjord, Hilde Elshaug, Rasmus Riis Kopperud, Hilde Synnøve Vollan, Karoline Bragstad, Olav Hungnes |

| | | | |
|---|---|---|---|
| EPI_ISL_549137, EPI_ISL_549138, EPI_ISL_549139, EPI_ISL_549140, EPI_ISL_549141, EPI_ISL_549142 | | | |
| EPI_ISL_549143 | Furst Medical Laboratory | Norwegian Institute of Public Health, Department of Virology | Kathrine Stene-Johansen, Kamilla Heddeland Instefjord, Hilde Elshaug, Rasmus Riis Kopperud, Hilde Synnøve Vollan, Karoline Bragstad, Olav Hungnes |
| EPI_ISL_549144, EPI_ISL_549145, EPI_ISL_549146, EPI_ISL_549147, EPI_ISL_549148, EPI_ISL_549149, EPI_ISL_549150, EPI_ISL_549151, EPI_ISL_549152, EPI_ISL_549153 | Ostfold Hospital Trust - Kalnes, Centre for Laboratory Medicine, Section for gene technology and infection serology | Norwegian Institute of Public Health, Department of Virology | Kathrine Stene-Johansen, Kamilla Heddeland Instefjord, Hilde Elshaug, Rasmus Riis Kopperud, Hilde Synnøve Vollan, Karoline Bragstad, Olav Hungnes |
| EPI_ISL_549154 | Furst Medical Laboratory | Norwegian Institute of Public Health, Department of Virology | Kathrine Stene-Johansen, Kamilla Heddeland Instefjord, Hilde Elshaug, Rasmus Riis Kopperud, Hilde Synnøve Vollan, Karoline Bragstad, Olav Hungnes |
| EPI_ISL_549155, EPI_ISL_549156, EPI_ISL_549157, EPI_ISL_549158, EPI_ISL_549159, EPI_ISL_549160, EPI_ISL_549161, EPI_ISL_549162, EPI_ISL_549163 | Ostfold Hospital Trust - Kalnes, Centre for Laboratory Medicine, Section for gene technology and infection serology | Norwegian Institute of Public Health, Department of Virology | Kathrine Stene-Johansen, Kamilla Heddeland Instefjord, Hilde Elshaug, Rasmus Riis Kopperud, Hilde Synnøve Vollan, Karoline Bragstad, Olav Hungnes |
| EPI_ISL_549164 | Medical Microbiology Unit, Department for Laboratory Medicine, Drammen Hospital, Vestre Viken Health Trust, | Norwegian Institute of Public Health, Department of Virology | Kathrine Stene-Johansen, Kamilla Heddeland Instefjord, Hilde Elshaug, Rasmus Riis Kopperud, Hilde Synnøve Vollan, Karoline Bragstad, Olav Hungnes |
| EPI_ISL_549165 | Furst Medical Laboratory | Norwegian Institute of Public Health, Department of Virology | Kathrine Stene-Johansen, Kamilla Heddeland Instefjord, Hilde Elshaug, Rasmus Riis Kopperud, Hilde Synnøve Vollan, Karoline Bragstad, Olav Hungnes |
| EPI_ISL_549166 | Medical Microbiology Unit, Department for Laboratory Medicine, Drammen Hospital, Vestre Viken Health Trust, | Norwegian Institute of Public Health, Department of Virology | Kathrine Stene-Johansen, Kamilla Heddeland Instefjord, Hilde Elshaug, Rasmus Riis Kopperud, Hilde Synnøve Vollan, Karoline Bragstad, Olav Hungnes |
| EPI_ISL_549167 | Hospital of Southern Norway - Kristiansand, Department of Medical Microbiology | Norwegian Institute of Public Health, Department of Virology | Kathrine Stene-Johansen, Kamilla Heddeland Instefjord, Hilde Elshaug, Rasmus Riis Kopperud, Hilde Synnøve Vollan, Karoline Bragstad, Olav Hungnes |
| EPI_ISL_549168 | Furst Medical Laboratory | Norwegian Institute of Public Health, Department of Virology | Kathrine Stene-Johansen, Kamilla Heddeland Instefjord, Hilde Elshaug, Rasmus Riis Kopperud, Hilde Synnøve Vollan, Karoline Bragstad, Olav Hungnes |
| EPI_ISL_549169, EPI_ISL_549170, EPI_ISL_549171 | Akershus University Hospital, Department for Microbiology and Infectious Disease Control | Norwegian Institute of Public Health, Department of Virology | Kathrine Stene-Johansen, Kamilla Heddeland Instefjord, Hilde Elshaug, Rasmus Riis Kopperud, Hilde Synnøve Vollan, Karoline Bragstad, Olav Hungnes |
| EPI_ISL_549172 | Unilabs Laboratory Medicine | Norwegian Institute of Public Health, Department of Virology | Kathrine Stene-Johansen, Kamilla Heddeland Instefjord, Hilde Elshaug, Rasmus Riis Kopperud, Hilde Synnøve Vollan, Karoline Bragstad, Olav Hungnes |
| EPI_ISL_549173, EPI_ISL_549174 | Vestfold Hospital, Toensberg Department of Microbiology | Norwegian Institute of Public Health, Department of Virology | Kathrine Stene-Johansen, Kamilla Heddeland Instefjord, Hilde Elshaug, Rasmus Riis Kopperud, Hilde Synnøve Vollan, Karoline Bragstad, Olav Hungnes |
| EPI_ISL_549175 | Unilabs Laboratory Medicine | Norwegian Institute of Public Health, Department of Virology | Kathrine Stene-Johansen, Kamilla Heddeland Instefjord, Hilde Elshaug, Rasmus Riis Kopperud, Hilde Synnøve Vollan, Karoline Bragstad, Olav Hungnes |
| EPI_ISL_560318 | Civil Hospital, Panchkula | CSIR-Institute of Microbial Technology | Kanika Bansal, Sanjeet Kumar, Anu Singh, Debarghya Ghose, Amandeep Kaur, Rajesh Kumar Mishra, Poushali Chakraborty, Harsh Goar, Navin Baid, Ashwani Kumar, Dipak Dutta, Sanjeev Khosla, Prabhu B. Patil |
| EPI_ISL_560319 | IDSP unit, Dehradun | CSIR-Institute of Microbial Technology | Kanika Bansal, Sanjeet Kumar, Anu Singh, Debarghya Ghose, Amandeep Kaur, Rajesh Kumar Mishra, Poushali Chakraborty, Harsh Goar, Navin Baid, Ashwani Kumar, Dipak Dutta, Sanjeev Khosla, Prabhu B. Patil |
| EPI_ISL_560320, EPI_ISL_560321 | CMS, Roorkee | CSIR-Institute of Microbial Technology | Kanika Bansal, Sanjeet Kumar, Anu Singh, Debarghya Ghose, Amandeep Kaur, Rajesh Kumar Mishra, Poushali Chakraborty, Harsh Goar, Navin Baid, Ashwani Kumar, Dipak Dutta, Sanjeev Khosla, Prabhu B. Patil |
| EPI_ISL_560322, EPI_ISL_560323, EPI_ISL_560324 | Civil Hospital, Panchkula | CSIR-Institute of Microbial Technology | Kanika Bansal, Sanjeet Kumar, Anu Singh, Debarghya Ghose, Rajesh Kumar Mishra, Dipak Dutta, Sanjeev Khosla, Prabhu B. Patil |
| EPI_ISL_560385 | National Health Laboratory | Botswana Institute for Technology Research and Innovation | Kefentse Arnold Tumedi, Madisa Mine, Dineo Emang Tshiamo, Gape Nyepetsi, Thongbotho Mphoyakgosi, Maitshwarelo Ignatius Matsheka |
| EPI_ISL_560386 | National Health Laboratory | Botswana Institute for Technology Research and innovation | Kefentse Arnold Tumedi, Madisa Mine, Dineo Emang Tshiamo. Gape Nyepetsi, Thongbotho Mphoyakgosi, Maitshwarelo Ignatius Matsheka |
| EPI_ISL_560387, EPI_ISL_560388, EPI_ISL_560389, EPI_ISL_560390 | National Health Laboratory | Botswana Institute for Technology Research and Innovation | Kefentse Arnold Tumedi, Madisa Mine, Dineo Emang Tshiamo. Gape Nyepetsi, Thongbotho Mphoyakgosi, Maitshwarelo Ignatius Matsheka |
| EPI_ISL_560391, EPI_ISL_560392, EPI_ISL_560393, EPI_ISL_560394, EPI_ISL_560395, EPI_ISL_560396, EPI_ISL_560397, EPI_ISL_560398, EPI_ISL_560399, EPI_ISL_560400, EPI_ISL_560401, EPI_ISL_560402, EPI_ISL_560403, EPI_ISL_560404, EPI_ISL_560405 | | | |
| see above | Vilnius University Hospital Santaros Klinikos, Vilnius University | Institute of Biotechnology, Life Sciences Center, Vilnius University and Thermo Fisher Scientific | Justinas Slikas, Albertas Timinskas, Alma Gedvilaite, Aurelija Zvirbliene, Daniel Naumovas, Laimonas Griskevicius, Ligita Janciorienе, Mindaugas Paulauskas |
| EPI_ISL_561337, EPI_ISL_561338, EPI_ISL_561339, EPI_ISL_561340, EPI_ISL_561341, EPI_ISL_561342, EPI_ISL_561343 | Civil Hospital, Panchkula | CSIR-Institute of Microbial Technology | Kanika Bansal, Sanjeet Kumar, Anu Singh, Debarghya Ghose, Rajesh Kumar Mishra, Dipak Dutta, Sanjeev Khosla, Prabhu B. Patil |
| EPI_ISL_561344 | Civil Hospital, Rupnagar | CSIR-Institute of Microbial Technology | Kanika Bansal, Sanjeet Kumar, Anu Singh, Debarghya Ghose, Amandeep Kaur, Rajesh Kumar Mishra, Poushali Chakraborty, Harsh Goar, Navin Baid, Ashwani Kumar, Dipak Dutta, Sanjeev Khosla, Prabhu B. Patil |
| EPI_ISL_566085, EPI_ISL_566086, EPI_ISL_566087, EPI_ISL_566088, EPI_ISL_566089, EPI_ISL_566090, EPI_ISL_566091, EPI_ISL_566092, EPI_ISL_566093, EPI_ISL_566094, EPI_ISL_566095, EPI_ISL_566096, EPI_ISL_566097, EPI_ISL_566098, EPI_ISL_566099, EPI_ISL_566100, EPI_ISL_566101, EPI_ISL_566102, EPI_ISL_566103, EPI_ISL_566104, EPI_ISL_566105, EPI_ISL_566106, EPI_ISL_566107 | | | |
| see above | GA Department of Public Health Laboratory | Pathogen Discovery, Respiratory Viruses Branch, Division of Viral Diseases, Centers for Disease Control and Prevention | Jing Zhang, Brian Lynch, Yan Li, Anna Montmayeur, Krista Queen, Ying Tao, Anna Uehara, Clinton R. Paden, Rachel Marine, Haibin Wang, Suxiang Tong |
| EPI_ISL_566114, EPI_ISL_566119, EPI_ISL_566129, EPI_ISL_566137, EPI_ISL_566142, EPI_ISL_566145, EPI_ISL_566146, EPI_ISL_566169, EPI_ISL_566182, EPI_ISL_566194, EPI_ISL_566207, EPI_ISL_566210, EPI_ISL_566226, EPI_ISL_566232, EPI_ISL_566238, EPI_ISL_566242, EPI_ISL_566249, EPI_ISL_566253, EPI_ISL_566260, EPI_ISL_566274, EPI_ISL_566285, EPI_ISL_566292, EPI_ISL_566299, EPI_ISL_566317, EPI_ISL_566319, EPI_ISL_566322, EPI_ISL_566348, EPI_ISL_566362, EPI_ISL_566380, EPI_ISL_566385, EPI_ISL_566395, EPI_ISL_566396, EPI_ISL_566397, EPI_ISL_566405, EPI_ISL_566414, EPI_ISL_566419, EPI_ISL_567533, EPI_ISL_567534, EPI_ISL_567535, EPI_ISL_567536, EPI_ISL_567537, EPI_ISL_567538, EPI_ISL_567539, EPI_ISL_567540, EPI_ISL_567541, EPI_ISL_567542, EPI_ISL_567543, EPI_ISL_567544, EPI_ISL_567545, EPI_ISL_567546, EPI_ISL_567548, EPI_ISL_567549, EPI_ISL_567550, EPI_ISL_567551, EPI_ISL_567552, EPI_ISL_567553, EPI_ISL_567554, EPI_ISL_567555, EPI_ISL_567556, EPI_ISL_567557, EPI_ISL_567558, EPI_ISL_567559, EPI_ISL_567560, EPI_ISL_567561, EPI_ISL_567562, EPI_ISL_567563, EPI_ISL_567564, EPI_ISL_567565, EPI_ISL_567566, EPI_ISL_567567, EPI_ISL_567568, EPI_ISL_567569, EPI_ISL_567571, EPI_ISL_567572, EPI_ISL_567573, EPI_ISL_567574, EPI_ISL_567575, EPI_ISL_567576, EPI_ISL_567577, EPI_ISL_567578, EPI_ISL_567579, EPI_ISL_567580, EPI_ISL_567581, EPI_ISL_567582, EPI_ISL_567583, EPI_ISL_567584, EPI_ISL_567585, EPI_ISL_567586, EPI_ISL_567587, EPI_ISL_567588, EPI_ISL_567589, EPI_ISL_567590, EPI_ISL_567591, EPI_ISL_567592, EPI_ISL_567593, EPI_ISL_567594, EPI_ISL_567595, EPI_ISL_567597, EPI_ISL_567598, EPI_ISL_567599, EPI_ISL_567600, EPI_ISL_567601, EPI_ISL_567602, EPI_ISL_567603, EPI_ISL_567604, EPI_ISL_567605, EPI_ISL_567606, EPI_ISL_567607, EPI_ISL_567608, EPI_ISL_567610, EPI_ISL_567611, EPI_ISL_567612, EPI_ISL_567613, EPI_ISL_567614, EPI_ISL_567615, EPI_ISL_567616, EPI_ISL_567617, EPI_ISL_567618, EPI_ISL_567619, EPI_ISL_567620, EPI_ISL_567621, EPI_ISL_567622, EPI_ISL_567623, EPI_ISL_567624, EPI_ISL_567625, EPI_ISL_567626, EPI_ISL_567627, EPI_ISL_567628, EPI_ISL_567629, EPI_ISL_567630, EPI_ISL_567631, EPI_ISL_567632, EPI_ISL_567633, EPI_ISL_567634, EPI_ISL_567635, EPI_ISL_567636, EPI_ISL_567637, EPI_ISL_567638, EPI_ISL_567639, EPI_ISL_567640, EPI_ISL_567641, EPI_ISL_567642, EPI_ISL_567643, EPI_ISL_567644, EPI_ISL_567645, EPI_ISL_567646, EPI_ISL_567647, EPI_ISL_567648, EPI_ISL_567649, EPI_ISL_567650, EPI_ISL_567651, EPI_ISL_567652, EPI_ISL_567653, EPI_ISL_567654, EPI_ISL_567655, EPI_ISL_567656, EPI_ISL_567657, EPI_ISL_567658, EPI_ISL_567659, EPI_ISL_567660, EPI_ISL_567661, EPI_ISL_567662, EPI_ISL_567663, EPI_ISL_567664, EPI_ISL_567665, EPI_ISL_567666, EPI_ISL_567668, EPI_ISL_567669, EPI_ISL_567670, EPI_ISL_567671, EPI_ISL_567672, EPI_ISL_567673, EPI_ISL_567674, EPI_ISL_567675, EPI_ISL_567676, EPI_ISL_567677, EPI_ISL_567678, EPI_ISL_567679, EPI_ISL_567680, EPI_ISL_567681, EPI_ISL_567682, EPI_ISL_567683, EPI_ISL_567684, EPI_ISL_567685, EPI_ISL_567686, EPI_ISL_567687, EPI_ISL_567689, EPI_ISL_567690, EPI_ISL_567691, EPI_ISL_567692, EPI_ISL_567693, EPI_ISL_567694, EPI_ISL_567695, EPI_ISL_567696, EPI_ISL_567697, EPI_ISL_567698, EPI_ISL_567699, EPI_ISL_567700, EPI_ISL_567701, EPI_ISL_567702, EPI_ISL_567703, EPI_ISL_567704, EPI_ISL_567705, EPI_ISL_567706, EPI_ISL_567707, EPI_ISL_567708, EPI_ISL_567709, EPI_ISL_567710, EPI_ISL_567711, EPI_ISL_567712, EPI_ISL_567713, EPI_ISL_567714, EPI_ISL_567715, EPI_ISL_567716, EPI_ISL_567717, EPI_ISL_567718, EPI_ISL_567719, EPI_ISL_567721, EPI_ISL_567723, EPI_ISL_567724, EPI_ISL_567725, EPI_ISL_567726, EPI_ISL_567727, EPI_ISL_567728, EPI_ISL_567729, EPI_ISL_567730, EPI_ISL_567731, EPI_ISL_567732, EPI_ISL_567733, EPI_ISL_567734, EPI_ISL_567735, EPI_ISL_567736, EPI_ISL_567737, EPI_ISL_567738, EPI_ISL_567739, EPI_ISL_567740, EPI_ISL_567741, EPI_ISL_567742, EPI_ISL_567743, EPI_ISL_567744, EPI_ISL_567745, EPI_ISL_567746, EPI_ISL_567747, EPI_ISL_567748, EPI_ISL_567749, EPI_ISL_567750, EPI_ISL_567751, EPI_ISL_567752, EPI_ISL_567753, EPI_ISL_567754, EPI_ISL_567755, EPI_ISL_567756, EPI_ISL_567757, EPI_ISL_567758, EPI_ISL_567759, EPI_ISL_567760, EPI_ISL_567761, EPI_ISL_567762, EPI_ISL_567763, EPI_ISL_567764, EPI_ISL_567765, EPI_ISL_567766, EPI_ISL_567767, EPI_ISL_567768, EPI_ISL_567769, EPI_ISL_567770, EPI_ISL_567771, EPI_ISL_567772, EPI_ISL_567773, EPI_ISL_567774, EPI_ISL_567775, EPI_ISL_567776, EPI_ISL_567777, EPI_ISL_567778, EPI_ISL_567779, EPI_ISL_567780, EPI_ISL_567781, EPI_ISL_567782, EPI_ISL_567783, EPI_ISL_567784, EPI_ISL_567785, EPI_ISL_567786, EPI_ISL_567787, EPI_ISL_567788, EPI_ISL_567789, EPI_ISL_567790, EPI_ISL_567791, EPI_ISL_567792, EPI_ISL_567793, EPI_ISL_567794, EPI_ISL_567795, EPI_ISL_567796, EPI_ISL_567797, EPI_ISL_567798, EPI_ISL_567799, EPI_ISL_567800, EPI_ISL_567801, EPI_ISL_567802, EPI_ISL_567804, EPI_ISL_567805, EPI_ISL_567806, EPI_ISL_567807, EPI_ISL_567808, EPI_ISL_567809, EPI_ISL_567810, EPI_ISL_567812, EPI_ISL_567813, EPI_ISL_567814, EPI_ISL_567815, EPI_ISL_567816, EPI_ISL_567817, EPI_ISL_567818, EPI_ISL_567819, EPI_ISL_567820, EPI_ISL_567821, EPI_ISL_567822, EPI_ISL_567823, EPI_ISL_567824, EPI_ISL_567825, EPI_ISL_567826, EPI_ISL_567827, EPI_ISL_567828, EPI_ISL_567829, EPI_ISL_567830, EPI_ISL_567831, EPI_ISL_567832, EPI_ISL_567834, EPI_ISL_567835, EPI_ISL_567836, EPI_ISL_567837, EPI_ISL_567838, EPI_ISL_567839, EPI_ISL_567841, EPI_ISL_567842, EPI_ISL_567843, EPI_ISL_567844, EPI_ISL_567845, EPI_ISL_567846, EPI_ISL_567847, EPI_ISL_567848, EPI_ISL_567849, EPI_ISL_567850, EPI_ISL_567851, EPI_ISL_567852, EPI_ISL_567853, EPI_ISL_567854, EPI_ISL_567855, EPI_ISL_567856, EPI_ISL_567857, EPI_ISL_567858, EPI_ISL_567859, EPI_ISL_567860, EPI_ISL_567861, EPI_ISL_567862, EPI_ISL_567863, EPI_ISL_567864, EPI_ISL_567865, EPI_ISL_567866, EPI_ISL_567867, EPI_ISL_567868, EPI_ISL_567869, EPI_ISL_567870, EPI_ISL_567871, EPI_ISL_567872, EPI_ISL_567873, EPI_ISL_567874, EPI_ISL_567875, EPI_ISL_567876, EPI_ISL_567877, EPI_ISL_567878, EPI_ISL_567880, EPI_ISL_567881, EPI_ISL_567882, EPI_ISL_567883, EPI_ISL_567884, EPI_ISL_567885, EPI_ISL_567886, EPI_ISL_567888, | | | |

EPI_ISL_567889, EPI_ISL_567890, EPI_ISL_567891, EPI_ISL_567892, EPI_ISL_567893, EPI_ISL_567894, EPI_ISL_567895, EPI_ISL_567896, EPI_ISL_567897, EPI_ISL_567898, EPI_ISL_567899, EPI_ISL_567900, EPI_ISL_567901, EPI_ISL_567902, EPI_ISL_567903, EPI_ISL_567904, EPI_ISL_567905, EPI_ISL_567906, EPI_ISL_567907, EPI_ISL_567908, EPI_ISL_567909, EPI_ISL_567910, EPI_ISL_567911, EPI_ISL_567912, EPI_ISL_567913, EPI_ISL_567914, EPI_ISL_567915, EPI_ISL_567916, EPI_ISL_567917, EPI_ISL_567918, EPI_ISL_567920, EPI_ISL_567921, EPI_ISL_567922, EPI_ISL_567923, EPI_ISL_567924, EPI_ISL_567925, EPI_ISL_567926, EPI_ISL_567927, EPI_ISL_567928, EPI_ISL_567929, EPI_ISL_567930, EPI_ISL_567932, EPI_ISL_567933, EPI_ISL_567934, EPI_ISL_567935, EPI_ISL_567936, EPI_ISL_567937, EPI_ISL_567938, EPI_ISL_567939, EPI_ISL_567940, EPI_ISL_567941, EPI_ISL_567942, EPI_ISL_567943, EPI_ISL_567944, EPI_ISL_567945, EPI_ISL_567946, EPI_ISL_567947, EPI_ISL_567948, EPI_ISL_567949, EPI_ISL_567950, EPI_ISL_567951, EPI_ISL_567952, EPI_ISL_567953, EPI_ISL_567954, EPI_ISL_567955, EPI_ISL_567956, EPI_ISL_567957, EPI_ISL_567958, EPI_ISL_567959, EPI_ISL_567960, EPI_ISL_567961, EPI_ISL_567962, EPI_ISL_567963, EPI_ISL_567964, EPI_ISL_567965, EPI_ISL_567966, EPI_ISL_567967, EPI_ISL_567969, EPI_ISL_567970, EPI_ISL_567971, EPI_ISL_567972, EPI_ISL_567973, EPI_ISL_567974, EPI_ISL_567975, EPI_ISL_567976, EPI_ISL_567977, EPI_ISL_567978, EPI_ISL_567979, EPI_ISL_567980, EPI_ISL_567981, EPI_ISL_567982, EPI_ISL_567983, EPI_ISL_567984, EPI_ISL_567985, EPI_ISL_567986, EPI_ISL_567987, EPI_ISL_567988, EPI_ISL_567989, EPI_ISL_567990, EPI_ISL_567991, EPI_ISL_567992, EPI_ISL_567993, EPI_ISL_567994, EPI_ISL_567995, EPI_ISL_567996, EPI_ISL_567997, EPI_ISL_567998, EPI_ISL_567999, EPI_ISL_568000, EPI_ISL_568001, EPI_ISL_568002, EPI_ISL_568003, EPI_ISL_568004, EPI_ISL_568005, EPI_ISL_568006, EPI_ISL_568007, EPI_ISL_568008, EPI_ISL_568009, EPI_ISL_568010, EPI_ISL_568011, EPI_ISL_568012, EPI_ISL_568013, EPI_ISL_568014, EPI_ISL_568015, EPI_ISL_568016, EPI_ISL_568017, EPI_ISL_568018, EPI_ISL_568019, EPI_ISL_568020, EPI_ISL_568021, EPI_ISL_568022, EPI_ISL_568023, EPI_ISL_568024, EPI_ISL_568025, EPI_ISL_568026, EPI_ISL_568027, EPI_ISL_568028, EPI_ISL_568029, EPI_ISL_568030, EPI_ISL_568031, EPI_ISL_568032, EPI_ISL_568033, EPI_ISL_568034, EPI_ISL_568035, EPI_ISL_568036, EPI_ISL_568037, EPI_ISL_568038, EPI_ISL_568039, EPI_ISL_568040, EPI_ISL_568041, EPI_ISL_568042, EPI_ISL_568043, EPI_ISL_568044, EPI_ISL_568045, EPI_ISL_568046, EPI_ISL_568047, EPI_ISL_568048, EPI_ISL_568049, EPI_ISL_568050, EPI_ISL_568051, EPI_ISL_568052, EPI_ISL_568053, EPI_ISL_568054, EPI_ISL_568055, EPI_ISL_568056, EPI_ISL_568057, EPI_ISL_568058, EPI_ISL_568059, EPI_ISL_568060, EPI_ISL_568061, EPI_ISL_568062, EPI_ISL_568063, EPI_ISL_568064, EPI_ISL_568065, EPI_ISL_568066, EPI_ISL_568067, EPI_ISL_568068, EPI_ISL_568069, EPI_ISL_568070, EPI_ISL_568071, EPI_ISL_568072, EPI_ISL_568073, EPI_ISL_568074, EPI_ISL_568075, EPI_ISL_568076, EPI_ISL_568077, EPI_ISL_568078, EPI_ISL_568080, EPI_ISL_568081, EPI_ISL_568082, EPI_ISL_568083, EPI_ISL_568084, EPI_ISL_568085, EPI_ISL_568086, EPI_ISL_568087, EPI_ISL_568088, EPI_ISL_568089, EPI_ISL_568090, EPI_ISL_568091, EPI_ISL_568092, EPI_ISL_568093, EPI_ISL_568094, EPI_ISL_568095, EPI_ISL_568096, EPI_ISL_568097, EPI_ISL_568098, EPI_ISL_568099, EPI_ISL_568100, EPI_ISL_568101, EPI_ISL_568102, EPI_ISL_568103, EPI_ISL_568104, EPI_ISL_568105, EPI_ISL_568106, EPI_ISL_568107, EPI_ISL_568108, EPI_ISL_568109, EPI_ISL_568110, EPI_ISL_568111, EPI_ISL_568112, EPI_ISL_568113, EPI_ISL_568114, EPI_ISL_568115, EPI_ISL_568116, EPI_ISL_568117, EPI_ISL_568118, EPI_ISL_568119, EPI_ISL_568120, EPI_ISL_568121, EPI_ISL_568122, EPI_ISL_568123, EPI_ISL_568124, EPI_ISL_568125, EPI_ISL_568126, EPI_ISL_568127, EPI_ISL_568128, EPI_ISL_568129, EPI_ISL_568130, EPI_ISL_568131, EPI_ISL_568132, EPI_ISL_568133, EPI_ISL_568135, EPI_ISL_568136, EPI_ISL_568137, EPI_ISL_568138, EPI_ISL_568139, EPI_ISL_568140, EPI_ISL_568141, EPI_ISL_568142, EPI_ISL_568143, EPI_ISL_568144, EPI_ISL_568145, EPI_ISL_568146, EPI_ISL_568147, EPI_ISL_568148, EPI_ISL_568149, EPI_ISL_568150, EPI_ISL_568152, EPI_ISL_568153, EPI_ISL_568154, EPI_ISL_568155, EPI_ISL_568156, EPI_ISL_568157, EPI_ISL_568159, EPI_ISL_568160, EPI_ISL_568161, EPI_ISL_568162, EPI_ISL_568163, EPI_ISL_568164, EPI_ISL_568165, EPI_ISL_568166, EPI_ISL_568167, EPI_ISL_568168, EPI_ISL_568169, EPI_ISL_568170, EPI_ISL_568171, EPI_ISL_568172, EPI_ISL_568173, EPI_ISL_568174, EPI_ISL_568175, EPI_ISL_568176, EPI_ISL_568177, EPI_ISL_568178, EPI_ISL_568179, EPI_ISL_568181, EPI_ISL_568182, EPI_ISL_568183, EPI_ISL_568184, EPI_ISL_568185, EPI_ISL_568186, EPI_ISL_568187, EPI_ISL_568188, EPI_ISL_568189, EPI_ISL_568190, EPI_ISL_568191, EPI_ISL_568192, EPI_ISL_568193, EPI_ISL_568194, EPI_ISL_568195, EPI_ISL_568196, EPI_ISL_568197, EPI_ISL_568198, EPI_ISL_568199, EPI_ISL_568200, EPI_ISL_568201, EPI_ISL_568202, EPI_ISL_568203, EPI_ISL_568204, EPI_ISL_568205, EPI_ISL_568206, EPI_ISL_568207, EPI_ISL_568208, EPI_ISL_568209, EPI_ISL_568210, EPI_ISL_568211, EPI_ISL_568212, EPI_ISL_568213, EPI_ISL_568214, EPI_ISL_568215, EPI_ISL_568216, EPI_ISL_568217, EPI_ISL_568218, EPI_ISL_568219, EPI_ISL_568220, EPI_ISL_568221, EPI_ISL_568222

| | | | |
|---|---|---|---|
| see above | Lighthouse Lab in Alderley Park | Wellcome Sanger Institute for the COVID-19 Genomics UK (COG-UK) consortium | Jacquelyn Wynn, Mairead Hyland, The Lighthouse Lab in Alderley Park and Alex Alderton, Roberto Amato, Sonia Goncalves, Ewan Harrison, David K. Jackson, Ian Johnston, Dominic Kwiatkowski, Cordelia Langford, John Sillitoe on behalf of the Wellcome Sanger Institute COVID-19 Surveillance Team |

EPI_ISL_574295, EPI_ISL_574296, EPI_ISL_574297, EPI_ISL_574298, EPI_ISL_574299, EPI_ISL_574300, EPI_ISL_574320, EPI_ISL_574321, EPI_ISL_574322, EPI_ISL_574323, EPI_ISL_574324, EPI_ISL_574325, EPI_ISL_574326, EPI_ISL_574327, EPI_ISL_574328, EPI_ISL_574329, EPI_ISL_574330, EPI_ISL_574331, EPI_ISL_574332, EPI_ISL_574333, EPI_ISL_574334, EPI_ISL_574335, EPI_ISL_574336, EPI_ISL_574337, EPI_ISL_574338, EPI_ISL_574339, EPI_ISL_574340, EPI_ISL_574341, EPI_ISL_574342, EPI_ISL_574343, EPI_ISL_574344, EPI_ISL_574345, EPI_ISL_574346, EPI_ISL_574347, EPI_ISL_574348, EPI_ISL_574349, EPI_ISL_574350, EPI_ISL_574351, EPI_ISL_574352, EPI_ISL_574353, EPI_ISL_574354, EPI_ISL_574355, EPI_ISL_574356, EPI_ISL_574357, EPI_ISL_574358, EPI_ISL_574359, EPI_ISL_574360, EPI_ISL_574361, EPI_ISL_574362, EPI_ISL_574363, EPI_ISL_574364, EPI_ISL_574365, EPI_ISL_574366, EPI_ISL_574367, EPI_ISL_574368, EPI_ISL_574369, EPI_ISL_574370, EPI_ISL_574371, EPI_ISL_574372, EPI_ISL_574373, EPI_ISL_574374, EPI_ISL_574375, EPI_ISL_574376, EPI_ISL_574377, EPI_ISL_574378, EPI_ISL_574379, EPI_ISL_574380, EPI_ISL_574381, EPI_ISL_574382, EPI_ISL_574383, EPI_ISL_574384, EPI_ISL_574385, EPI_ISL_574386, EPI_ISL_574387, EPI_ISL_574388, EPI_ISL_574389, EPI_ISL_574390, EPI_ISL_574391, EPI_ISL_574392, EPI_ISL_574393, EPI_ISL_574394, EPI_ISL_574395, EPI_ISL_574396, EPI_ISL_574397, EPI_ISL_574398, EPI_ISL_574399, EPI_ISL_574400, EPI_ISL_574401, EPI_ISL_574402, EPI_ISL_574403, EPI_ISL_574404, EPI_ISL_574405, EPI_ISL_574406, EPI_ISL_574407, EPI_ISL_574408, EPI_ISL_574409, EPI_ISL_574410, EPI_ISL_574411, EPI_ISL_574412, EPI_ISL_574413, EPI_ISL_574414, EPI_ISL_574415, EPI_ISL_574416, EPI_ISL_574417, EPI_ISL_574418, EPI_ISL_574419, EPI_ISL_574420, EPI_ISL_574421, EPI_ISL_574422, EPI_ISL_574423, EPI_ISL_574424, EPI_ISL_574425, EPI_ISL_574426, EPI_ISL_574427, EPI_ISL_574428, EPI_ISL_574429, EPI_ISL_574430

| | | | |
|---|---|---|---|
| see above | LSUHS Emerging Viral Threat Laboratory | Microbial Genome Sequencing Center | Jeremy P. Kamil, Rona S. Scott, Maarten Van Diest, Malgorzata Bienkowska-Haba, Katarzyna Zwolinska, Andrew D. Yurochko, Christopher G. Kevil, Martin J. Sapp, Daniel J. Snyder, Vaughn S. Cooper, John A. Vanchiere |

EPI_ISL_574576, EPI_ISL_575289, EPI_ISL_575290, EPI_ISL_575291, EPI_ISL_575292, EPI_ISL_575293, EPI_ISL_575294, EPI_ISL_575295, EPI_ISL_575296, EPI_ISL_575297, EPI_ISL_575298, EPI_ISL_575299, EPI_ISL_575300, EPI_ISL_575301, EPI_ISL_575302, EPI_ISL_575303, EPI_ISL_575304, EPI_ISL_575305, EPI_ISL_575306, EPI_ISL_575307, EPI_ISL_575308, EPI_ISL_575309, EPI_ISL_575310, EPI_ISL_575311, EPI_ISL_575314, EPI_ISL_575322

| | | | |
|---|---|---|---|
| see above | Molecular Biology, New Mexico Department of Health Scientific Laboratory | Molecular Biology, New Mexico Department of Health Scientific Laboratory | Johnson,E.J., Griego-Fisher,A.M., Malone,D. |

EPI_ISL_575399, EPI_ISL_575400, EPI_ISL_575401, EPI_ISL_575402, EPI_ISL_575403, EPI_ISL_575405, EPI_ISL_575406, EPI_ISL_575407, EPI_ISL_575408, EPI_ISL_575409, EPI_ISL_575410, EPI_ISL_575411, EPI_ISL_575412, EPI_ISL_575413, EPI_ISL_575414, EPI_ISL_575415, EPI_ISL_575416, EPI_ISL_575418, EPI_ISL_575419, EPI_ISL_575420, EPI_ISL_575421, EPI_ISL_575422, EPI_ISL_575423, EPI_ISL_575424, EPI_ISL_575425, EPI_ISL_575426, EPI_ISL_575427, EPI_ISL_575428, EPI_ISL_575429, EPI_ISL_575430, EPI_ISL_575431, EPI_ISL_575432, EPI_ISL_575433, EPI_ISL_575434, EPI_ISL_575435, EPI_ISL_575436, EPI_ISL_575437, EPI_ISL_575438, EPI_ISL_575439, EPI_ISL_575441, EPI_ISL_575442, EPI_ISL_575443, EPI_ISL_575444, EPI_ISL_575445, EPI_ISL_575446, EPI_ISL_575447, EPI_ISL_575448, EPI_ISL_575449, EPI_ISL_575450, EPI_ISL_575451, EPI_ISL_575452, EPI_ISL_575453, EPI_ISL_575454, EPI_ISL_575455, EPI_ISL_575456, EPI_ISL_575457, EPI_ISL_575459, EPI_ISL_575460, EPI_ISL_575461, EPI_ISL_575462, EPI_ISL_575463, EPI_ISL_575464, EPI_ISL_575465, EPI_ISL_575466, EPI_ISL_575467, EPI_ISL_575468, EPI_ISL_575469, EPI_ISL_575470, EPI_ISL_575471, EPI_ISL_575472, EPI_ISL_575473, EPI_ISL_575474, EPI_ISL_575475, EPI_ISL_575476, EPI_ISL_575477, EPI_ISL_575478, EPI_ISL_575479, EPI_ISL_575480, EPI_ISL_575481, EPI_ISL_575482, EPI_ISL_575484, EPI_ISL_575485, EPI_ISL_575486, EPI_ISL_575487, EPI_ISL_575488, EPI_ISL_575490, EPI_ISL_575491, EPI_ISL_575492, EPI_ISL_575493, EPI_ISL_575494, EPI_ISL_575495, EPI_ISL_575496, EPI_ISL_575497, EPI_ISL_575498, EPI_ISL_575499, EPI_ISL_575500, EPI_ISL_575501, EPI_ISL_575502, EPI_ISL_575503, EPI_ISL_575504, EPI_ISL_575505, EPI_ISL_575506, EPI_ISL_575507, EPI_ISL_575508, EPI_ISL_575509, EPI_ISL_575510, EPI_ISL_575511, EPI_ISL_575512, EPI_ISL_575513, EPI_ISL_575514, EPI_ISL_575515, EPI_ISL_575516, EPI_ISL_575517, EPI_ISL_575518, EPI_ISL_575519, EPI_ISL_575520, EPI_ISL_575521, EPI_ISL_575522, EPI_ISL_575523, EPI_ISL_575524, EPI_ISL_575525, EPI_ISL_575526, EPI_ISL_575527, EPI_ISL_575528, EPI_ISL_575529, EPI_ISL_575530, EPI_ISL_575531, EPI_ISL_575532, EPI_ISL_575533, EPI_ISL_575534, EPI_ISL_575535, EPI_ISL_575537, EPI_ISL_575538, EPI_ISL_575539, EPI_ISL_575540, EPI_ISL_575541, EPI_ISL_575542, EPI_ISL_575543, EPI_ISL_575544, EPI_ISL_575545, EPI_ISL_575546, EPI_ISL_575547, EPI_ISL_575548, EPI_ISL_575549, EPI_ISL_575550, EPI_ISL_575551, EPI_ISL_575553, EPI_ISL_575554, EPI_ISL_575555, EPI_ISL_575557, EPI_ISL_575558, EPI_ISL_575559, EPI_ISL_575560, EPI_ISL_575561, EPI_ISL_575562, EPI_ISL_575563, EPI_ISL_575564, EPI_ISL_575565, EPI_ISL_575566, EPI_ISL_575567, EPI_ISL_575568, EPI_ISL_575569, EPI_ISL_575570, EPI_ISL_575571, EPI_ISL_575572, EPI_ISL_575573, EPI_ISL_575574, EPI_ISL_575575, EPI_ISL_575576, EPI_ISL_575577, EPI_ISL_575578, EPI_ISL_575579, EPI_ISL_575580, EPI_ISL_575581, EPI_ISL_575582, EPI_ISL_575583, EPI_ISL_575584, EPI_ISL_575585, EPI_ISL_575586, EPI_ISL_575587, EPI_ISL_575588, EPI_ISL_575589, EPI_ISL_575590, EPI_ISL_575591, EPI_ISL_575592, EPI_ISL_575593, EPI_ISL_575594, EPI_ISL_575595, EPI_ISL_575596, EPI_ISL_575597, EPI_ISL_575598, EPI_ISL_575599, EPI_ISL_575600, EPI_ISL_575601, EPI_ISL_575602, EPI_ISL_575603, EPI_ISL_575604, EPI_ISL_575605, EPI_ISL_575606, EPI_ISL_575607, EPI_ISL_575608, EPI_ISL_575609, EPI_ISL_575610, EPI_ISL_575612, EPI_ISL_575613, EPI_ISL_575614, EPI_ISL_575615, EPI_ISL_575616, EPI_ISL_575617, EPI_ISL_575619, EPI_ISL_575620, EPI_ISL_575621, EPI_ISL_575622, EPI_ISL_575623, EPI_ISL_575624, EPI_ISL_575625, EPI_ISL_575626, EPI_ISL_575627, EPI_ISL_575628, EPI_ISL_575629, EPI_ISL_575630, EPI_ISL_575631, EPI_ISL_575632, EPI_ISL_575633, EPI_ISL_575634, EPI_ISL_575635, EPI_ISL_575636, EPI_ISL_575637, EPI_ISL_575638, EPI_ISL_575639, EPI_ISL_575640, EPI_ISL_575641, EPI_ISL_575642, EPI_ISL_575643, EPI_ISL_575644, EPI_ISL_575646, EPI_ISL_575647, EPI_ISL_575648, EPI_ISL_575649, EPI_ISL_575650, EPI_ISL_575651, EPI_ISL_575652, EPI_ISL_575653, EPI_ISL_575654, EPI_ISL_575655, EPI_ISL_575656, EPI_ISL_575657, EPI_ISL_575658, EPI_ISL_575660, EPI_ISL_575661, EPI_ISL_575662, EPI_ISL_575663, EPI_ISL_575664, EPI_ISL_575665, EPI_ISL_575666, EPI_ISL_575667, EPI_ISL_575668, EPI_ISL_575669, EPI_ISL_575671, EPI_ISL_575672, EPI_ISL_575673, EPI_ISL_575674, EPI_ISL_575675, EPI_ISL_575676, EPI_ISL_575677, EPI_ISL_575678, EPI_ISL_575679, EPI_ISL_575680, EPI_ISL_575681, EPI_ISL_575682, EPI_ISL_575683, EPI_ISL_575684, EPI_ISL_575685, EPI_ISL_575686, EPI_ISL_575687, EPI_ISL_575688, EPI_ISL_575689, EPI_ISL_575690, EPI_ISL_575691, EPI_ISL_575692, EPI_ISL_575693, EPI_ISL_575694, EPI_ISL_575695, EPI_ISL_575696, EPI_ISL_575697, EPI_ISL_575698, EPI_ISL_575700, EPI_ISL_575701, EPI_ISL_575702, EPI_ISL_575703, EPI_ISL_575704, EPI_ISL_575705, EPI_ISL_575706, EPI_ISL_575707, EPI_ISL_575708, EPI_ISL_575709, EPI_ISL_575711, EPI_ISL_575712, EPI_ISL_575713, EPI_ISL_575714, EPI_ISL_575715, EPI_ISL_575716, EPI_ISL_575717, EPI_ISL_575718, EPI_ISL_575719, EPI_ISL_575720, EPI_ISL_575721, EPI_ISL_575722, EPI_ISL_575723, EPI_ISL_575724, EPI_ISL_575725, EPI_ISL_575726, EPI_ISL_575727, EPI_ISL_575728, EPI_ISL_575729, EPI_ISL_575730, EPI_ISL_575731, EPI_ISL_575732, EPI_ISL_575733, EPI_ISL_575734, EPI_ISL_575735, EPI_ISL_575736, EPI_ISL_575737, EPI_ISL_575738, EPI_ISL_575739, EPI_ISL_575740, EPI_ISL_575741, EPI_ISL_575742, EPI_ISL_575743, EPI_ISL_575744, EPI_ISL_575745, EPI_ISL_575746, EPI_ISL_575747, EPI_ISL_575748, EPI_ISL_575749, EPI_ISL_575750, EPI_ISL_575751, EPI_ISL_575752, EPI_ISL_575753, EPI_ISL_575754, EPI_ISL_575755, EPI_ISL_575756, EPI_ISL_575757, EPI_ISL_575758, EPI_ISL_575760

| | | | |
|---|---|---|---|
| see above | Lighthouse Lab in Alderley Park | Wellcome Sanger Institute for the COVID-19 Genomics UK (COG-UK) consortium | Jacquelyn Wynn, Mairead Hyland, The Lighthouse Lab in Alderley Park and Alex Alderton, Roberto Amato, Sonia Goncalves, Ewan Harrison, David K. Jackson, Ian Johnston, Dominic Kwiatkowski, Cordelia Langford, John Sillitoe on behalf of the Wellcome Sanger Institute COVID-19 Surveillance Team |
| EPI_ISL_577735, EPI_ISL_577737, EPI_ISL_577739 | Institute of Virology, Biomedical Research Center of the Slovak Academy of Sciences, Bratislava | Faculty of Natural Sciences, Comenius University, Bratislava | Kristína Boršová, Viktória Hodorová, Broa Brejová, Viktória abanová, Dominika Friová, Sabina Fumaová Havlíková, Juraj Kopáek, Martina Liková, ubomíra Lukáiková, Martina Neboháová, Monika Sláviková, Edita Staroová, Elena Tichá, Tomáš Vina, Boris Klempa, Jozef Nosek |

EPI_ISL_578335, EPI_ISL_578336, EPI_ISL_578337, EPI_ISL_578338, EPI_ISL_578339, EPI_ISL_578340, EPI_ISL_578341, EPI_ISL_578342, EPI_ISL_578343, EPI_ISL_578344, EPI_ISL_578345, EPI_ISL_578346, EPI_ISL_578347, EPI_ISL_578348, EPI_ISL_578349, EPI_ISL_578350, EPI_ISL_578351, EPI_ISL_578352, EPI_ISL_578353, EPI_ISL_578354, EPI_ISL_578355, EPI_ISL_578356, EPI_ISL_578357, EPI_ISL_578358, EPI_ISL_578359, EPI_ISL_578360, EPI_ISL_578361, EPI_ISL_578362, EPI_ISL_578363, EPI_ISL_578364, EPI_ISL_578365, EPI_ISL_578366, EPI_ISL_578367, EPI_ISL_578368, EPI_ISL_578369, EPI_ISL_578370, EPI_ISL_578371, EPI_ISL_578372, EPI_ISL_578373, EPI_ISL_578374, EPI_ISL_578375, EPI_ISL_578376, EPI_ISL_578377, EPI_ISL_578378, EPI_ISL_578379, EPI_ISL_578380, EPI_ISL_578381, EPI_ISL_578382, EPI_ISL_578383, EPI_ISL_578384, EPI_ISL_578385, EPI_ISL_578386, EPI_ISL_578387, EPI_ISL_578388, EPI_ISL_578389, EPI_ISL_578390, EPI_ISL_578391, EPI_ISL_578392, EPI_ISL_578393, EPI_ISL_578394, EPI_ISL_578395, EPI_ISL_578396, EPI_ISL_578397, EPI_ISL_578398, EPI_ISL_578399, EPI_ISL_578400, EPI_ISL_578401, EPI_ISL_578402, EPI_ISL_578403, EPI_ISL_578404, EPI_ISL_578405, EPI_ISL_578406, EPI_ISL_578407, EPI_ISL_578408, EPI_ISL_578409, EPI_ISL_578410, EPI_ISL_578411, EPI_ISL_578412, EPI_ISL_578413, EPI_ISL_578414, EPI_ISL_578415, EPI_ISL_578416, EPI_ISL_578417, EPI_ISL_578418, EPI_ISL_578419, EPI_ISL_578420, EPI_ISL_578421, EPI_ISL_578422, EPI_ISL_578423, EPI_ISL_578424, EPI_ISL_578425, EPI_ISL_578426, EPI_ISL_578427, EPI_ISL_578428, EPI_ISL_578429, EPI_ISL_578430, EPI_ISL_578431, EPI_ISL_578432, EPI_ISL_578433, EPI_ISL_578434, EPI_ISL_578435, EPI_ISL_578436, EPI_ISL_578437, EPI_ISL_578438, EPI_ISL_578439, EPI_ISL_578440, EPI_ISL_578441, EPI_ISL_578442, EPI_ISL_578443, EPI_ISL_578444, EPI_ISL_578445, EPI_ISL_578446, EPI_ISL_578447, EPI_ISL_578448, EPI_ISL_578449, EPI_ISL_578450, EPI_ISL_578451, EPI_ISL_578452, EPI_ISL_578453, EPI_ISL_578454, EPI_ISL_578455, EPI_ISL_578456, EPI_ISL_578457, EPI_ISL_578458, EPI_ISL_578459, EPI_ISL_578460, EPI_ISL_578461, EPI_ISL_578462, EPI_ISL_578463, EPI_ISL_578464, EPI_ISL_578465, EPI_ISL_578466, EPI_ISL_578467, EPI_ISL_578468, EPI_ISL_578469, EPI_ISL_578470, EPI_ISL_578471, EPI_ISL_578472, EPI_ISL_578473, EPI_ISL_578474, EPI_ISL_578475, EPI_ISL_578476, EPI_ISL_578477, EPI_ISL_578478, EPI_ISL_578479, EPI_ISL_578480, EPI_ISL_578481, EPI_ISL_578482, EPI_ISL_578483, EPI_ISL_578484, EPI_ISL_578485, EPI_ISL_578486, EPI_ISL_578487, EPI_ISL_578488, EPI_ISL_578489, EPI_ISL_578490, EPI_ISL_578491, EPI_ISL_578492, EPI_ISL_578493, EPI_ISL_578494, EPI_ISL_578495, EPI_ISL_578496, EPI_ISL_578497, EPI_ISL_578498, EPI_ISL_578499, EPI_ISL_578500, EPI_ISL_578501, EPI_ISL_578502, EPI_ISL_578503, EPI_ISL_578504, EPI_ISL_578505, EPI_ISL_578506, EPI_ISL_578507, EPI_ISL_578508, EPI_ISL_578509, EPI_ISL_578510, EPI_ISL_578511, EPI_ISL_578512, EPI_ISL_578513, EPI_ISL_578514, EPI_ISL_578515, EPI_ISL_578516, EPI_ISL_578517, EPI_ISL_578518, EPI_ISL_578519, EPI_ISL_578520, EPI_ISL_578521, EPI_ISL_578522, EPI_ISL_578523, EPI_ISL_578524, EPI_ISL_578525, EPI_ISL_578526, EPI_ISL_578527, EPI_ISL_578528, EPI_ISL_578529, EPI_ISL_578530, EPI_ISL_578531, EPI_ISL_578532, EPI_ISL_578533, EPI_ISL_578534, EPI_ISL_578535, EPI_ISL_578536, EPI_ISL_578537, EPI_ISL_578538, EPI_ISL_578539, EPI_ISL_578540, EPI_ISL_578541, EPI_ISL_578542, EPI_ISL_578543, EPI_ISL_578544, EPI_ISL_578545, EPI_ISL_578546, EPI_ISL_578547, EPI_ISL_578548, EPI_ISL_578549, EPI_ISL_578550, EPI_ISL_578551, EPI_ISL_578552, EPI_ISL_578553, EPI_ISL_578554, EPI_ISL_578555, EPI_ISL_578556, EPI_ISL_578557, EPI_ISL_578558, EPI_ISL_578559, EPI_ISL_578560, EPI_ISL_578561, EPI_ISL_578562, EPI_ISL_578563, EPI_ISL_578564, EPI_ISL_578565, EPI_ISL_578566, EPI_ISL_578567, EPI_ISL_578568, EPI_ISL_578569, EPI_ISL_578570, EPI_ISL_578571, EPI_ISL_578572, EPI_ISL_578573, EPI_ISL_578574, EPI_ISL_578575, EPI_ISL_578576, EPI_ISL_578577, EPI_ISL_578578, EPI_ISL_578579, EPI_ISL_578580, EPI_ISL_578581, EPI_ISL_578582, EPI_ISL_578583, EPI_ISL_578584, EPI_ISL_578585, EPI_ISL_578586, EPI_ISL_578587, EPI_ISL_578588, EPI_ISL_578589, EPI_ISL_578590, EPI_ISL_578591, EPI_ISL_578592, EPI_ISL_578593, EPI_ISL_578594, EPI_ISL_578595, EPI_ISL_578596, EPI_ISL_578597, EPI_ISL_578598, EPI_ISL_578599, EPI_ISL_578600, EPI_ISL_578601, EPI_ISL_578602, EPI_ISL_578603, EPI_ISL_578604, EPI_ISL_578605, EPI_ISL_578606, EPI_ISL_578607, EPI_ISL_578608, EPI_ISL_578609, EPI_ISL_578610, EPI_ISL_578611, EPI_ISL_578612, EPI_ISL_578613, EPI_ISL_578614, EPI_ISL_578615, EPI_ISL_578616, EPI_ISL_578617, EPI_ISL_578618, EPI_ISL_578619, EPI_ISL_578620, EPI_ISL_578621, EPI_ISL_578622, EPI_ISL_578623, EPI_ISL_578624, EPI_ISL_578625, EPI_ISL_578626, EPI_ISL_578627, EPI_ISL_578628, EPI_ISL_578629, EPI_ISL_578630, EPI_ISL_578631, EPI_ISL_578632, EPI_ISL_578633, EPI_ISL_578634, EPI_ISL_578635, EPI_ISL_578636, EPI_ISL_578637, EPI_ISL_578638, EPI_ISL_578639, EPI_ISL_578640, EPI_ISL_578641, EPI_ISL_578642, EPI_ISL_578643, EPI_ISL_578644, EPI_ISL_578645, EPI_ISL_578646, EPI_ISL_578647, EPI_ISL_578648, EPI_ISL_578649, EPI_ISL_578650, EPI_ISL_578651, EPI_ISL_578652, EPI_ISL_578653, EPI_ISL_578654, EPI_ISL_578655, EPI_ISL_578656, EPI_ISL_578657, EPI_ISL_578658, EPI_ISL_578659, EPI_ISL_578660, EPI_ISL_578661, EPI_ISL_578662, EPI_ISL_578663, EPI_ISL_578664, EPI_ISL_578665, EPI_ISL_578666, EPI_ISL_578667, EPI_ISL_578668, EPI_ISL_578669, EPI_ISL_578670, EPI_ISL_578671, EPI_ISL_578672, EPI_ISL_578673, EPI_ISL_578674, EPI_ISL_578675, EPI_ISL_578676, EPI_ISL_578677, EPI_ISL_578678, EPI_ISL_578679, EPI_ISL_578680, EPI_ISL_578681, EPI_ISL_578682, EPI_ISL_578683, EPI_ISL_578684, EPI_ISL_578685, EPI_ISL_578686, EPI_ISL_578687, EPI_ISL_578688, EPI_ISL_578689, EPI_ISL_578690, EPI_ISL_578691, EPI_ISL_578692, EPI_ISL_578693, EPI_ISL_578694, EPI_ISL_578695

| | | | |
|---|---|---|---|
| see above | Wisconsin State Laboratory of Hygiene Communicable Disease Division | Wisconsin State Laboratory of Hygiene Communicable Disease Division | Kelsey R. Florek, Abigail C. Shockey |

EPI_ISL_578699, EPI_ISL_578700, EPI_ISL_578753, EPI_ISL_578754, EPI_ISL_578755, EPI_ISL_578756, EPI_ISL_578757, EPI_ISL_578758, EPI_ISL_578759, EPI_ISL_578760, EPI_ISL_578761, EPI_ISL_578762, EPI_ISL_578763, EPI_ISL_578764, EPI_ISL_578765, EPI_ISL_578766, EPI_ISL_578767, EPI_ISL_578768,

| | | | |
|---|---|---|---|
| EPI_ISL_578769, EPI_ISL_578770, EPI_ISL_578771, EPI_ISL_578772, EPI_ISL_578773, EPI_ISL_578774, EPI_ISL_578775, EPI_ISL_578776, EPI_ISL_578777, EPI_ISL_578778, EPI_ISL_578779, EPI_ISL_578780, EPI_ISL_578781, EPI_ISL_578782, EPI_ISL_578783, EPI_ISL_578784, EPI_ISL_578785, EPI_ISL_578786, EPI_ISL_578787, EPI_ISL_578788, EPI_ISL_578789, EPI_ISL_578790, EPI_ISL_578791, EPI_ISL_578792, EPI_ISL_578793, EPI_ISL_578794, EPI_ISL_578795, EPI_ISL_578796, EPI_ISL_578797, EPI_ISL_578798, EPI_ISL_578799, EPI_ISL_578800, EPI_ISL_578801, EPI_ISL_578802, EPI_ISL_578803, EPI_ISL_578804, EPI_ISL_578805, EPI_ISL_578806, EPI_ISL_578807, EPI_ISL_578808, EPI_ISL_578809, EPI_ISL_578810, EPI_ISL_578811, EPI_ISL_578812, EPI_ISL_578813, EPI_ISL_578814, EPI_ISL_578815, EPI_ISL_578816, EPI_ISL_578817, EPI_ISL_578818, EPI_ISL_578819, EPI_ISL_578820, EPI_ISL_578821, EPI_ISL_578822, EPI_ISL_578823, EPI_ISL_578824, EPI_ISL_578825, EPI_ISL_578826, EPI_ISL_578827, EPI_ISL_578828, EPI_ISL_578829, EPI_ISL_578830, EPI_ISL_578831, EPI_ISL_578832, EPI_ISL_578833, EPI_ISL_578834, EPI_ISL_578835, EPI_ISL_578836, EPI_ISL_578837, EPI_ISL_578838, EPI_ISL_578839, EPI_ISL_578840, EPI_ISL_578841, EPI_ISL_578842, EPI_ISL_578843, EPI_ISL_578844, EPI_ISL_578845, EPI_ISL_578846, EPI_ISL_578847, EPI_ISL_578848, EPI_ISL_578849, EPI_ISL_578850, EPI_ISL_578851, EPI_ISL_578852, EPI_ISL_578853, EPI_ISL_578854, EPI_ISL_578855, EPI_ISL_578856, EPI_ISL_578857, EPI_ISL_578858, EPI_ISL_578859, EPI_ISL_578860, EPI_ISL_578861, EPI_ISL_578862, EPI_ISL_578863, EPI_ISL_578864, EPI_ISL_578865, EPI_ISL_578866, EPI_ISL_578867, EPI_ISL_578868, EPI_ISL_578869, EPI_ISL_578870, EPI_ISL_578871, EPI_ISL_578872, EPI_ISL_578873, EPI_ISL_578874, EPI_ISL_578875, EPI_ISL_578876, EPI_ISL_578877, EPI_ISL_578878, EPI_ISL_578879, EPI_ISL_578880, EPI_ISL_578881, EPI_ISL_578882, EPI_ISL_578883, EPI_ISL_578884, EPI_ISL_578885, EPI_ISL_578886, EPI_ISL_578887, EPI_ISL_578888, EPI_ISL_578889, EPI_ISL_578890, EPI_ISL_578891, EPI_ISL_578892, EPI_ISL_578893, EPI_ISL_578894, EPI_ISL_578895, EPI_ISL_578896, EPI_ISL_578897, EPI_ISL_578898, EPI_ISL_578899, EPI_ISL_578900, EPI_ISL_578901, EPI_ISL_578902, EPI_ISL_578903, EPI_ISL_578904, EPI_ISL_578905, EPI_ISL_578906, EPI_ISL_578907, EPI_ISL_578908, EPI_ISL_578909, EPI_ISL_578910, EPI_ISL_578911, EPI_ISL_578912, EPI_ISL_578913, EPI_ISL_578914, EPI_ISL_578915, EPI_ISL_578916, EPI_ISL_578917, EPI_ISL_578918, EPI_ISL_578919, EPI_ISL_578920, EPI_ISL_578921, EPI_ISL_578922, EPI_ISL_578923, EPI_ISL_578924, EPI_ISL_578925, EPI_ISL_578926, EPI_ISL_578927, EPI_ISL_578928, EPI_ISL_578929, EPI_ISL_578930, EPI_ISL_578931, EPI_ISL_578932, EPI_ISL_578933, EPI_ISL_578934, EPI_ISL_578935, EPI_ISL_578936, EPI_ISL_578937, EPI_ISL_578938, EPI_ISL_578939, EPI_ISL_578940, EPI_ISL_578941, EPI_ISL_578942, EPI_ISL_578943, EPI_ISL_578944, EPI_ISL_578945, EPI_ISL_578946, EPI_ISL_578947, EPI_ISL_578948, EPI_ISL_578949, EPI_ISL_578950, EPI_ISL_578951, EPI_ISL_578952, EPI_ISL_578953, EPI_ISL_578954, EPI_ISL_578955, EPI_ISL_578956, EPI_ISL_578957, EPI_ISL_578958, EPI_ISL_578959, EPI_ISL_578960, EPI_ISL_578961, EPI_ISL_578962, EPI_ISL_578963, EPI_ISL_578964, EPI_ISL_578965, EPI_ISL_578966, EPI_ISL_578967, EPI_ISL_578968, EPI_ISL_578969, EPI_ISL_578970, EPI_ISL_578971, EPI_ISL_578972, EPI_ISL_578973, EPI_ISL_578974, EPI_ISL_578975, EPI_ISL_578976, EPI_ISL_578977, EPI_ISL_578978, EPI_ISL_578979, EPI_ISL_578980, EPI_ISL_578981, EPI_ISL_578982, EPI_ISL_578983, EPI_ISL_578984, EPI_ISL_578985, EPI_ISL_578986, EPI_ISL_578987, EPI_ISL_578988, EPI_ISL_578989, EPI_ISL_578990, EPI_ISL_578991, EPI_ISL_578992, EPI_ISL_578993, EPI_ISL_578994, EPI_ISL_578995, EPI_ISL_578996, EPI_ISL_578997, EPI_ISL_578998, EPI_ISL_578999, EPI_ISL_579000, EPI_ISL_579001, EPI_ISL_579002, EPI_ISL_579003, EPI_ISL_579004, EPI_ISL_579005, EPI_ISL_579006, EPI_ISL_579007, EPI_ISL_579008, EPI_ISL_579009, EPI_ISL_579010, EPI_ISL_579011, EPI_ISL_579012, EPI_ISL_579013, EPI_ISL_579014, EPI_ISL_579015, EPI_ISL_579016, EPI_ISL_579017, EPI_ISL_579018, EPI_ISL_579019, EPI_ISL_579020, EPI_ISL_579021, EPI_ISL_579022, EPI_ISL_579023, EPI_ISL_579024, EPI_ISL_579025, EPI_ISL_579026, EPI_ISL_579027, EPI_ISL_579028, EPI_ISL_579029, EPI_ISL_579030, EPI_ISL_579031, EPI_ISL_579032, EPI_ISL_579033, EPI_ISL_579034, EPI_ISL_579035, EPI_ISL_579036, EPI_ISL_579037, EPI_ISL_579038, EPI_ISL_579039, EPI_ISL_579040, EPI_ISL_579041, EPI_ISL_579042, EPI_ISL_579043, EPI_ISL_579044, EPI_ISL_579045, EPI_ISL_579046, EPI_ISL_579047, EPI_ISL_579048, EPI_ISL_579049, EPI_ISL_579050, EPI_ISL_579051, EPI_ISL_579052, EPI_ISL_579053, EPI_ISL_579054, EPI_ISL_579055, EPI_ISL_579056, EPI_ISL_579057 | | | |
| see above | LSUHS Emerging Viral Threat Laboratory | Microbial Genome Sequencing Center | Jeremy P. Kamil, Rona S. Scott, Maarten Van Diest, Malgorzata Bienkowska-Haba, Katarzyna Zwolinska, Andrew D. Yurochko, Christopher G. Kevil, Martin J. Sapp, Daniel J. Snyder, Vaughn S. Cooper, John A. Vanchiere |
| EPI_ISL_579926, EPI_ISL_580260, EPI_ISL_580484, EPI_ISL_580598, EPI_ISL_581367, EPI_ISL_581378, EPI_ISL_581380, EPI_ISL_581404, EPI_ISL_581409, EPI_ISL_581411, EPI_ISL_581412, EPI_ISL_581413, EPI_ISL_581414, EPI_ISL_581416, EPI_ISL_581417, EPI_ISL_581424, EPI_ISL_581426, EPI_ISL_581427, EPI_ISL_581429, EPI_ISL_581430, EPI_ISL_581431, EPI_ISL_581432, EPI_ISL_581435, EPI_ISL_581436, EPI_ISL_581442, EPI_ISL_581444, EPI_ISL_581446, EPI_ISL_581448 | | | |
| see above | Lighthouse Lab in Alderley Park | Wellcome Sanger Institute for the COVID-19 Genomics UK (COG-UK) consortium | Jacquelyn Wynn, Mairead Hyland, The Lighthouse Lab in Alderley Park and Alex Alderton, Roberto Amato, Sonia Goncalves, Ewan Harrison, David K. Jackson, Ian Johnston, Dominic Kwiatkowski, Cordelia Langford, John Sillitoe on behalf of the Wellcome Sanger Institute COVID-19 Surveillance Team |
| EPI_ISL_581575, EPI_ISL_581576, EPI_ISL_581577, EPI_ISL_581578, EPI_ISL_581579, EPI_ISL_581580, EPI_ISL_581581, EPI_ISL_581582, EPI_ISL_581583, EPI_ISL_581584, EPI_ISL_581585, EPI_ISL_581586, EPI_ISL_581587, EPI_ISL_581588, EPI_ISL_581589, EPI_ISL_581590, EPI_ISL_581591, EPI_ISL_581592, EPI_ISL_581593, EPI_ISL_581594, EPI_ISL_581595, EPI_ISL_581596, EPI_ISL_581597, EPI_ISL_581598, EPI_ISL_581599, EPI_ISL_581600, EPI_ISL_581601, EPI_ISL_581602, EPI_ISL_581603, EPI_ISL_581604, EPI_ISL_581605, EPI_ISL_581606, EPI_ISL_581607, EPI_ISL_581608, EPI_ISL_581609, EPI_ISL_581610, EPI_ISL_581611, EPI_ISL_581612, EPI_ISL_581613, EPI_ISL_581614, EPI_ISL_581615, EPI_ISL_581616, EPI_ISL_581617, EPI_ISL_581618, EPI_ISL_581619, EPI_ISL_581620, EPI_ISL_581621, EPI_ISL_581622, EPI_ISL_581623, EPI_ISL_581624, EPI_ISL_581625, EPI_ISL_581626, EPI_ISL_581627, EPI_ISL_581628, EPI_ISL_581629, EPI_ISL_581630, EPI_ISL_581631, EPI_ISL_581632, EPI_ISL_581633, EPI_ISL_581634, EPI_ISL_581635, EPI_ISL_581636, EPI_ISL_581637, EPI_ISL_581638, EPI_ISL_581639, EPI_ISL_581640, EPI_ISL_581641, EPI_ISL_581642, EPI_ISL_581643, EPI_ISL_581644, EPI_ISL_581645, EPI_ISL_581646, EPI_ISL_581647, EPI_ISL_581648, EPI_ISL_581649, EPI_ISL_581650, EPI_ISL_581651, EPI_ISL_581652, EPI_ISL_581653, EPI_ISL_581654, EPI_ISL_581655, EPI_ISL_581656, EPI_ISL_581657, EPI_ISL_581658, EPI_ISL_581659, EPI_ISL_581660, EPI_ISL_581661, EPI_ISL_581662, EPI_ISL_581663, EPI_ISL_581664, EPI_ISL_581665, EPI_ISL_581666, EPI_ISL_581667, EPI_ISL_582027 | | | |
| see above | Department of Clinical Microbiology | GIGA Medical Genomics | Keith Durkin, Maria Artesi, Sébastien Bontems, Raphaël Boreux, Bouchra Boujemla, Cécile Meex, Pierrette Melin, Marie-Pierre Hayette, Vincent Bours |
| EPI_ISL_583482, EPI_ISL_583484, EPI_ISL_583486 | Institute of Virology, Biomedical Research Center of the Slovak Academy of Sciences, Bratislava | Faculty of Natural Sciences, Comenius University, Bratislava | Kristína Boršová, Viktória Hodorová, Broa Brejová, Viktória abanová, Dominika Friová, Sabina Fumaová Havlíková, Juraj Kopáek, Martina Liková, ubomíra Lukáiková, Martina Neboháová, Monika Sláviková, Edita Staroová, Elena Tichá, Tomáš Vina, Boris Klempa, Jozef Nosek |
| EPI_ISL_586520 | B.J. Medical College and Civil hospital, Ahmedabad | Gujarat Biotechnology Research Centre | Komal Patel, Labdhi Pandya, Afzal Ansari, Nikha Trivedi, Pranay Shah, Kamlesh J Upadhyay, Sanjay Kapadia, Apurvasinh Puvar, Janvi Raval, Zarna Patel, Monika Gandhi, Pinal Trivedi, Maharshi Pandya, Nidhi Patel, Nitin Savaliya, Raghawendra Kumar, Dinesh Kumar, Zuber Saiyed, R D Dixit, A M Kadri, Harsh Bakshi, Chaitanya Joshi, Madhvi Joshi |
| EPI_ISL_586521 | B.J. Medical College and Civil hospital, Ahmedabad | Gujarat Biotechnology Research Centre | Labdhi Pandya, Afzal Ansari, Nikha Trivedi, Pranay Shah, Kamlesh J Upadhyay, Sanjay Kapadia, Apurvasinh Puvar, Janvi Raval, Zarna Patel, Monika Gandhi, Pinal Trivedi, Maharshi Pandya, Nidhi Patel, Nitin Savaliya, Raghawendra Kumar, Dinesh Kumar, Zuber Saiyed, Komal Patel, R D Dixit, A M Kadri, Harsh Bakshi, Chaitanya Joshi, Madhvi Joshi |
| EPI_ISL_586525 | B.J. Medical College and Civil hospital, Ahmedabad | Gujarat Biotechnology Research Centre | Kamlesh J Upadhyay, Sanjay Kapadia, Apurvasinh Puvar, Janvi Raval, Zarna Patel, Monika Gandhi, Pinal Trivedi, Maharshi Pandya, Nidhi Patel, Nitin Savaliya, Raghawendra Kumar, Dinesh Kumar, Zuber Saiyed, Komal Patel, Labdhi Pandya, Afzal Ansari, Nikha Trivedi, Pranay Shah, R D Dixit, A M Kadri, Harsh Bakshi, Chaitanya Joshi, Madhvi Joshi |
| EPI_ISL_586536 | Zydus Medical Hospital,Dahod | Gujarat Biotechnology Research Centre | Kalpesh Labana, Mohit Desai, Nitin Savaliya, Raghawendra Kumar, Dinesh Kumar, Zuber Saiyed, Komal Patel, Labdhi Pandya, Afzal Ansari, Nikha Trivedi, Apurvasinh Puvar, Janvi Raval, Zarna Patel, Monika Gandhi, Pinal Trivedi, Maharshi Pandya, Nidhi Patel, R D Dixit, A M Kadri, Harsh Bakshi, Chaitanya Joshi, Madhvi Joshi |
| EPI_ISL_586542 | Government General Hospital, Jam Khambhaliya, Devbhoomi Dwarka | Gujarat Biotechnology Research Centre | Komal Patel, Labdhi Pandya, Afzal Ansari, Nikha Trivedi, Harish Matani, Apurvasinh Puvar, Janvi Raval, Zarna Patel, Monika Gandhi, Pinal Trivedi, Maharshi Pandya, Nidhi Patel, Nitin Savaliya, Raghawendra Kumar, Dinesh Kumar, Zuber Saiyed, R D Dixit, A M Kadri, Harsh Bakshi, Chaitanya Joshi, Madhvi Joshi |
| EPI_ISL_586543 | Government General Hospital, Jam Khambhaliya, Devbhoomi Dwarka | Gujarat Biotechnology Research Centre | Labdhi Pandya, Afzal Ansari, Nikha Trivedi, Harish Matani, Apurvasinh Puvar, Janvi Raval, Zarna Patel, Monika Gandhi, Pinal Trivedi, Maharshi Pandya, Nidhi Patel, Nitin Savaliya, Raghawendra Kumar, Dinesh Kumar, Zuber Saiyed, Komal Patel, R D Dixit, A M Kadri, Harsh Bakshi, Chaitanya Joshi, Madhvi Joshi |
| EPI_ISL_586549 | CCC,Veraval | Gujarat Biotechnology Research Centre | Jignesh Parmar, Jitendra Bamrotia, Apurvasinh Puvar, Janvi Raval, Zarna Patel, Monika Gandhi, Pinal Trivedi, Maharshi Pandya, Nidhi Patel, Nitin Savaliya, Raghawendra Kumar, Dinesh Kumar, Zuber Saiyed, Komal Patel, Labdhi Pandya, Afzal Ansari, Nikha Trivedi, R D Dixit, A M Kadri, Harsh Bakshi, Chaitanya Joshi, Madhvi Joshi |
| EPI_ISL_586550 | CCC,Veraval | Gujarat Biotechnology Research Centre | Jitendra Bamrotia, Apurvasinh Puvar, Janvi Raval, Zarna Patel, Monika Gandhi, Pinal Trivedi, Maharshi Pandya, Nidhi Patel, Nitin Savaliya, Raghawendra Kumar, Dinesh Kumar, Zuber Saiyed, Komal Patel, Labdhi Pandya, Afzal Ansari, Nikha Trivedi, Jignesh Parmar, R D Dixit, A M Kadri, Harsh Bakshi, Chaitanya Joshi, Madhvi Joshi |
| EPI_ISL_586559 | CCC,Veraval | Gujarat Biotechnology Research Centre | Komal Patel, Labdhi Pandya, Afzal Ansari, Nikha Trivedi, Jignesh Parmar, Jitendra Bamrotia, Apurvasinh Puvar, Janvi Raval, Zarna Patel, Monika Gandhi, Pinal Trivedi, Maharshi Pandya, Nidhi Patel, Nitin Savaliya, Raghawendra Kumar, Dinesh Kumar, Zuber Saiyed, R D Dixit, A M Kadri, Harsh Bakshi, Chaitanya Joshi, Madhvi Joshi |
| EPI_ISL_586560 | CCC,Veraval | Gujarat Biotechnology Research Centre | Labdhi Pandya, Afzal Ansari, Nikha Trivedi, Jignesh Parmar, Jitendra Bamrotia, Apurvasinh Puvar, Janvi Raval, Zarna Patel, Monika Gandhi, Pinal Trivedi, Maharshi Pandya, Nidhi Patel, Nitin Savaliya, Raghawendra Kumar, Dinesh Kumar, Zuber Saiyed, Komal Patel, R D Dixit, A M Kadri, Harsh Bakshi, Chaitanya Joshi, Madhvi Joshi |
| EPI_ISL_586566 | GMERS GMERS Medical College and Hospital, Junagadh | Gujarat Biotechnology Research Centre | Janvi Raval, Zarna Patel, Monika Gandhi, Pinal Trivedi, Maharshi Pandya, Nidhi Patel, Nitin Savaliya, Raghawendra Kumar, Dinesh Kumar, Zuber Saiyed, Komal Patel, Labdhi Pandya, Afzal Ansari, Nikha Trivedi, Bhavesh Bagda , Ravi Dedaniya, Apurvasinh Puvar, R D Dixit, A M Kadri, Harsh Bakshi, Chaitanya Joshi, Madhvi Joshi |
| EPI_ISL_586660, EPI_ISL_586926 | Lighthouse Lab in Alderley Park | Wellcome Sanger Institute for the COVID-19 Genomics UK (COG-UK) consortium | Jacquelyn Wynn, Mairead Hyland, The Lighthouse Lab in Alderley Park and Alex Alderton, Roberto Amato, Sonia Goncalves, Ewan Harrison, David K. Jackson, Ian Johnston, Dominic Kwiatkowski, Cordelia Langford, John Sillitoe on behalf of the Wellcome Sanger Institute COVID-19 Surveillance Team (http://www.sanger.ac.uk/covid-team) |
| EPI_ISL_587151 | Lighthouse Lab in Alderley Park | Wellcome Sanger Institute for the COVID-19 Genomics UK (COG-UK) consortium | Jacquelyn Wynn, Mairead Hyland, The Lighthouse Lab in Alderley Park and Alex Alderton, Roberto Amato, Sonia Goncalves, Ewan Harrison, David K. Jackson, Ian Johnston, Dominic Kwiatkowski, Cordelia Langford, John Sillitoe on behalf of the Wellcome Sanger Institute COVID-19 Surveillance Team |
| EPI_ISL_589780, EPI_ISL_589781, EPI_ISL_589784, EPI_ISL_589785, EPI_ISL_589802, EPI_ISL_590181, EPI_ISL_590186, EPI_ISL_590193, EPI_ISL_590194, EPI_ISL_590200, EPI_ISL_590206, EPI_ISL_590210, EPI_ISL_590214, EPI_ISL_590215, EPI_ISL_590220, EPI_ISL_590224, EPI_ISL_590225, EPI_ISL_590226, EPI_ISL_590228, EPI_ISL_590235, EPI_ISL_590239, EPI_ISL_590246, EPI_ISL_590250, EPI_ISL_590255, EPI_ISL_590257, EPI_ISL_590259, EPI_ISL_590260, EPI_ISL_590261, EPI_ISL_590263, EPI_ISL_590265, EPI_ISL_590272, EPI_ISL_590274, EPI_ISL_590275, EPI_ISL_590276, EPI_ISL_590277, EPI_ISL_590282, EPI_ISL_590283, EPI_ISL_590284, EPI_ISL_590286, EPI_ISL_590288, EPI_ISL_590295, EPI_ISL_590296, EPI_ISL_590300, EPI_ISL_590306, EPI_ISL_590309, EPI_ISL_590314, EPI_ISL_590319, EPI_ISL_590321, EPI_ISL_590327, EPI_ISL_590328, EPI_ISL_590329, EPI_ISL_590335, EPI_ISL_590337, EPI_ISL_590346, EPI_ISL_590347, EPI_ISL_590351, EPI_ISL_590352, EPI_ISL_590355, EPI_ISL_590357, EPI_ISL_590359, EPI_ISL_590361, EPI_ISL_590366, EPI_ISL_590367, EPI_ISL_590369, EPI_ISL_590373, EPI_ISL_590379, EPI_ISL_590380, EPI_ISL_590383, EPI_ISL_590386, EPI_ISL_590388, EPI_ISL_590390, EPI_ISL_590392, EPI_ISL_590397, EPI_ISL_590398, EPI_ISL_590399, EPI_ISL_590403, EPI_ISL_590405, EPI_ISL_590422, EPI_ISL_590423, EPI_ISL_590438, EPI_ISL_590441, EPI_ISL_590442, EPI_ISL_590446, EPI_ISL_590448, EPI_ISL_590452, EPI_ISL_590453, EPI_ISL_590462 | | | |
| see above | Lighthouse Lab in Alderley Park | Wellcome Sanger Institute for the COVID-19 Genomics UK (COG-UK) consortium | Jacquelyn Wynn, Mairead Hyland, The Lighthouse Lab in Alderley Park and Alex Alderton, Roberto Amato, Sonia Goncalves, Ewan Harrison, David K. Jackson, Ian Johnston, Dominic Kwiatkowski, Cordelia Langford, John Sillitoe on behalf of the Wellcome Sanger Institute COVID-19 Surveillance Team |

| | | | |
|---|---|---|---|
| | | | (http://www.sanger.ac.uk/covid-team) |
| EPI_ISL_590689 | B.J. Medical College and Civil hospital, Ahmedabad | Gujarat Biotechnology Research Centre | Janvi Raval, Zarna Patel, Monika Gandhi, Pinal Trivedi, Maharshi Pandya, Nidhi Patel, Nitin Savaliya, Raghawendra Kumar, Dinesh Kumar, Zuber Saiyed, Komal Patel, Labdhi Pandya, Afzal Ansari, Nikha Trivedi, Pranay Shah, Kamlesh J Upadhyay, Sanjay Kapadia, Apurvasinh Puvar, R D Dixit, A M Kadri, Harsh Bakshi, Chaitanya Joshi, Madhvi Joshi |
| EPI_ISL_590691 | Government Hospital,Veraval | Gujarat Biotechnology Research Centre | Janvi Raval, Zarna Patel, Monika Gandhi, Pinal Trivedi, Maharshi Pandya, Nidhi Patel, Nitin Savaliya, Raghawendra Kumar, Dinesh Kumar, Zuber Saiyed, Komal Patel, Labdhi Pandya, Afzal Ansari, Nikha Trivedi, Jignesh Parmar, Jitendra Bamrotia, Apurvasinh Puvar, R D Dixit, A M Kadri, Harsh Bakshi, Chaitanya Joshi, Madhvi Joshi |
| EPI_ISL_590877 | Dept. of Medical Microbiology, Stavanger University Hospital, Helse Stavanger HF | Norwegian Institute of Public Health, Department of Virology | Kathrine Stene-Johansen, Kamilla Heddeland Instefjord, Hilde Elshaug, Marie Paulsen Madsen, Rasmus Riis Kopperud, Hilde Vollan, Karoline Bragstad, Olav Hungnes |
| EPI_ISL_590878 | Oslo University Hospital, Department of Medical Microbiology | Norwegian Institute of Public Health, Department of Virology | Kathrine Stene-Johansen, Kamilla Heddeland Instefjord, Hilde Elshaug, Rasmus Riis Kopperud, Hilde Vollan, Karoline Bragstad, Olav Hungnes |
| EPI_ISL_590879, EPI_ISL_590880 | Medical Microbiology Unit, Department for Laboratory Medicine, Drammen Hospital, Vestre Viken Health Trust, | Norwegian Institute of Public Health, Department of Virology | Kathrine Stene-Johansen, Kamilla Heddeland Instefjord, Hilde Elshaug, Rasmus Riis Kopperud, Hilde Vollan, Karoline Bragstad, Olav Hungnes |
| EPI_ISL_590881 | Furst Medical Laboratory | Norwegian Institute of Public Health, Department of Virology | Kathrine Stene-Johansen, Kamilla Heddeland Instefjord, Hilde Elshaug, Rasmus Riis Kopperud, Hilde Vollan, Karoline Bragstad, Olav Hungnes |
| EPI_ISL_590882 | Unilabs Laboratory Medicine | Norwegian Institute of Public Health, Department of Virology | Kathrine Stene-Johansen, Kamilla Heddeland Instefjord, Hilde Elshaug, Rasmus Riis Kopperud, Hilde Vollan, Karoline Bragstad, Olav Hungnes |
| EPI_ISL_590883, EPI_ISL_590884 | Foerde Hospital, Department of Microbiology | Norwegian Institute of Public Health, Department of Virology | Kathrine Stene-Johansen, Kamilla Heddeland Instefjord, Hilde Elshaug, Rasmus Riis Kopperud, Hilde Vollan, Karoline Bragstad, Olav Hungnes |
| EPI_ISL_590885, EPI_ISL_590886, EPI_ISL_590887, EPI_ISL_590888 | Vestfold Hospital, Toensberg Department of Microbiology | Norwegian Institute of Public Health, Department of Virology | Kathrine Stene-Johansen, Kamilla Heddeland Instefjord, Hilde Elshaug, Rasmus Riis Kopperud, Hilde Vollan, Karoline Bragstad, Olav Hungnes |
| EPI_ISL_590889, EPI_ISL_590890, EPI_ISL_590891, EPI_ISL_590892 | Akershus University Hospital, Department for Microbiology and Infectious Disease Control | Norwegian Institute of Public Health, Department of Virology | Kathrine Stene-Johansen, Kamilla Heddeland Instefjord, Hilde Elshaug, Rasmus Riis Kopperud, Hilde Vollan, Karoline Bragstad, Olav Hungnes |
| EPI_ISL_590893 | Medical Microbiology Unit, Department for Laboratory Medicine, Drammen Hospital, Vestre Viken Health Trust, | Norwegian Institute of Public Health, Department of Virology | Kathrine Stene-Johansen, Kamilla Heddeland Instefjord, Hilde Elshaug, Rasmus Riis Kopperud, Hilde Vollan, Karoline Bragstad, Olav Hungnes |
| EPI_ISL_590894, EPI_ISL_590895, EPI_ISL_590896, EPI_ISL_590897 | Ostfold Hospital Trust - Kalnes, Centre for Laboratory Medicine, Section for gene technology and infection serology | Norwegian Institute of Public Health, Department of Virology | Kathrine Stene-Johansen, Kamilla Heddeland Instefjord, Hilde Elshaug, Rasmus Riis Kopperud, Hilde Vollan, Karoline Bragstad, Olav Hungnes |
| EPI_ISL_590898 | Furst Medical Laboratory | Norwegian Institute of Public Health, Department of Virology | Kathrine Stene-Johansen, Kamilla Heddeland Instefjord, Hilde Elshaug, Rasmus Riis Kopperud, Hilde Vollan, Karoline Bragstad, Olav Hungnes |
| EPI_ISL_590899 | Foerde Hospital, Department of Microbiology | Norwegian Institute of Public Health, Department of Virology | Kathrine Stene-Johansen, Kamilla Heddeland Instefjord, Hilde Elshaug, Rasmus Riis Kopperud, Hilde Vollan, Karoline Bragstad, Olav Hungnes |
| EPI_ISL_590900 | Oslo University Hospital, Department of Medical Microbiology | Norwegian Institute of Public Health, Department of Virology | Kathrine Stene-Johansen, Kamilla Heddeland Instefjord, Hilde Elshaug, Rasmus Riis Kopperud, Hilde Vollan, Karoline Bragstad, Olav Hungnes |
| EPI_ISL_590901, EPI_ISL_590902, EPI_ISL_590903 | Furst Medical Laboratory | Norwegian Institute of Public Health, Department of Virology | Kathrine Stene-Johansen, Kamilla Heddeland Instefjord, Hilde Elshaug, Rasmus Riis Kopperud, Hilde Vollan, Karoline Bragstad, Olav Hungnes |
| EPI_ISL_590904, EPI_ISL_590905, EPI_ISL_590906, EPI_ISL_590907 | Hospital of Southern Norway - Kristiansand, Department of Medical Microbiology | Norwegian Institute of Public Health, Department of Virology | Kathrine Stene-Johansen, Kamilla Heddeland Instefjord, Hilde Elshaug, Rasmus Riis Kopperud, Hilde Vollan, Karoline Bragstad, Olav Hungnes |
| EPI_ISL_590908, EPI_ISL_590909, EPI_ISL_590910 | Oslo University Hospital, Department of Medical Microbiology | Norwegian Institute of Public Health, Department of Virology | Kathrine Stene-Johansen, Kamilla Heddeland Instefjord, Hilde Elshaug, Rasmus Riis Kopperud, Hilde Vollan, Karoline Bragstad, Olav Hungnes |
| EPI_ISL_590911 | Unilabs Laboratory Medicine | Norwegian Institute of Public Health, Department of Virology | Kathrine Stene-Johansen, Kamilla Heddeland Instefjord, Hilde Elshaug, Rasmus Riis Kopperud, Hilde Vollan, Karoline Bragstad, Olav Hungnes |
| EPI_ISL_590912 | Furst Medical Laboratory | Norwegian Institute of Public Health, Department of Virology | Kathrine Stene-Johansen, Kamilla Heddeland Instefjord, Hilde Elshaug, Rasmus Riis Kopperud, Hilde Vollan, Karoline Bragstad, Olav Hungnes |
| EPI_ISL_590913, EPI_ISL_590914 | Oslo University Hospital, Department of Medical Microbiology | Norwegian Institute of Public Health, Department of Virology | Kathrine Stene-Johansen, Kamilla Heddeland Instefjord, Hilde Elshaug, Rasmus Riis Kopperud, Hilde Vollan, Karoline Bragstad, Olav Hungnes |
| EPI_ISL_590915 | Hospital of Southern Norway - Kristiansand, Department of Medical Microbiology | Norwegian Institute of Public Health, Department of Virology | Kathrine Stene-Johansen, Kamilla Heddeland Instefjord, Hilde Elshaug, Rasmus Riis Kopperud, Hilde Vollan, Karoline Bragstad, Olav Hungnes |
| EPI_ISL_590916 | Oslo University Hospital, Department of Medical Microbiology | Norwegian Institute of Public Health, Department of Virology | Kathrine Stene-Johansen, Kamilla Heddeland Instefjord, Hilde Elshaug, Rasmus Riis Kopperud, Hilde Vollan, Karoline Bragstad, Olav Hungnes |
| EPI_ISL_590917 | Unilabs Laboratory Medicine | Norwegian Institute of Public Health, Department of Virology | Kathrine Stene-Johansen, Kamilla Heddeland Instefjord, Hilde Elshaug, Rasmus Riis Kopperud, Hilde Vollan, Karoline Bragstad, Olav Hungnes |
| EPI_ISL_590918 | Oslo University Hospital, Department of Medical Microbiology | Norwegian Institute of Public Health, Department of Virology | Kathrine Stene-Johansen, Kamilla Heddeland Instefjord, Hilde Elshaug, Rasmus Riis Kopperud, Hilde Vollan, Karoline Bragstad, Olav Hungnes |
| EPI_ISL_590919, EPI_ISL_590920 | Vestfold Hospital, Toensberg Department of Microbiology | Norwegian Institute of Public Health, Department of Virology | Kathrine Stene-Johansen, Kamilla Heddeland Instefjord, Hilde Elshaug, Rasmus Riis Kopperud, Hilde Vollan, Karoline Bragstad, Olav Hungnes |
| EPI_ISL_590921, EPI_ISL_590922 | Ostfold Hospital Trust - Kalnes, Centre for Laboratory Medicine, Section for gene technology and infection serology | Norwegian Institute of Public Health, Department of Virology | Kathrine Stene-Johansen, Kamilla Heddeland Instefjord, Hilde Elshaug, Rasmus Riis Kopperud, Hilde Vollan, Karoline Bragstad, Olav Hungnes |
| EPI_ISL_590923 | Oslo University Hospital, Department of Medical Microbiology | Norwegian Institute of Public Health, Department of Virology | Kathrine Stene-Johansen, Kamilla Heddeland Instefjord, Hilde Elshaug, Rasmus Riis Kopperud, Hilde Vollan, Karoline Bragstad, Olav Hungnes |
| EPI_ISL_590924, EPI_ISL_590925, EPI_ISL_590926 | University Hospital of Northern Norway, Department for Microbiology and Infectious Disease Control | Norwegian Institute of Public Health, Department of Virology | Kathrine Stene-Johansen, Kamilla Heddeland Instefjord, Hilde Elshaug, Rasmus Riis Kopperud, Hilde Vollan, Karoline Bragstad, Olav Hungnes |
| EPI_ISL_590927, EPI_ISL_590928 | Oslo University Hospital, Department of Medical Microbiology | Norwegian Institute of Public Health, Department of Virology | Kathrine Stene-Johansen, Kamilla Heddeland Instefjord, Hilde Elshaug, Rasmus Riis Kopperud, Hilde Vollan, Karoline Bragstad, Olav Hungnes |
| EPI_ISL_590929 | Medical Microbiology Unit, Department for Laboratory Medicine, Drammen Hospital, Vestre Viken Health Trust, | Norwegian Institute of Public Health, Department of Virology | Kathrine Stene-Johansen, Kamilla Heddeland Instefjord, Hilde Elshaug, Rasmus Riis Kopperud, Hilde Vollan, Karoline Bragstad, Olav Hungnes |
| EPI_ISL_590930 | Oslo University Hospital, Department of Medical Microbiology | Norwegian Institute of Public Health, Department of Virology | Kathrine Stene-Johansen, Kamilla Heddeland Instefjord, Hilde Elshaug, Rasmus Riis Kopperud, Hilde Vollan, Karoline Bragstad, Olav Hungnes |
| EPI_ISL_590931 | Ostfold Hospital Trust - Kalnes, Centre for Laboratory Medicine, Section for gene technology and infection serology | Norwegian Institute of Public Health, Department of Virology | Kathrine Stene-Johansen, Kamilla Heddeland Instefjord, Hilde Elshaug, Rasmus Riis Kopperud, Hilde Vollan, Karoline Bragstad, Olav Hungnes |
| EPI_ISL_590932, EPI_ISL_590933 | Medical Microbiology Unit, Department for Laboratory Medicine, Drammen Hospital, Vestre Viken Health Trust, | Norwegian Institute of Public Health, Department of Virology | Kathrine Stene-Johansen, Kamilla Heddeland Instefjord, Hilde Elshaug, Rasmus Riis Kopperud, Hilde Vollan, Karoline Bragstad, Olav Hungnes |
| EPI_ISL_590934, EPI_ISL_590935 | Oslo University Hospital, Department of Medical Microbiology | Norwegian Institute of Public Health, Department of Virology | Kathrine Stene-Johansen, Kamilla Heddeland Instefjord, Hilde Elshaug, Rasmus Riis Kopperud, Hilde Vollan, Karoline Bragstad, Olav Hungnes |
| EPI_ISL_590936, EPI_ISL_590937 | Hospital of Southern Norway - Kristiansand, Department of Medical Microbiology | Norwegian Institute of Public Health, Department of Virology | Kathrine Stene-Johansen, Kamilla Heddeland Instefjord, Hilde Elshaug, Rasmus Riis Kopperud, Hilde Vollan, Karoline Bragstad, Olav Hungnes |
| EPI_ISL_590938, EPI_ISL_590939, EPI_ISL_590940, EPI_ISL_590941, EPI_ISL_590942, EPI_ISL_590943, EPI_ISL_590944 | University Hospital of Northern Norway, Department for Microbiology and Infectious Disease Control | Norwegian Institute of Public Health, Department of Virology | Kathrine Stene-Johansen, Kamilla Heddeland Instefjord, Hilde Elshaug, Rasmus Riis Kopperud, Hilde Vollan, Karoline Bragstad, Olav Hungnes |
| EPI_ISL_590945 | Medical Microbiology Unit, Department for Laboratory Medicine, Drammen Hospital, Vestre Viken Health Trust, | Norwegian Institute of Public Health, Department of Virology | Kathrine Stene-Johansen, Kamilla Heddeland Instefjord, Hilde Elshaug, Rasmus Riis Kopperud, Hilde Vollan, Karoline Bragstad, Olav Hungnes |
| EPI_ISL_590946, EPI_ISL_590947 | University Hospital of Northern Norway, Department for Microbiology and Infectious Disease Control | Norwegian Institute of Public Health, Department of Virology | Kathrine Stene-Johansen, Kamilla Heddeland Instefjord, Hilde Elshaug, Rasmus Riis Kopperud, Hilde Vollan, Karoline Bragstad, Olav Hungnes |
| EPI_ISL_590948 | Ostfold Hospital Trust - Kalnes, Centre for Laboratory Medicine, Section for gene technology and infection serology | Norwegian Institute of Public Health, Department of Virology | Kathrine Stene-Johansen, Kamilla Heddeland Instefjord, Hilde Elshaug, Rasmus Riis Kopperud, Hilde Vollan, Karoline Bragstad, Olav Hungnes |

| | | | |
|---|---|---|---|
| EPI_ISL_590949, EPI_ISL_590950 | University Hospital of Northern Norway, Department for Microbiology and Infectious Disease Control | Norwegian Institute of Public Health, Department of Virology | Kathrine Stene-Johansen, Kamilla Heddeland Instefjord, Hilde Elshaug, Rasmus Riis Kopperud, Hilde Vollan, Karoline Bragstad, Olav Hungnes |
| EPI_ISL_590951, EPI_ISL_590952 | Akershus University Hospital, Department for Microbiology and Infectious Disease Control | Norwegian Institute of Public Health, Department of Virology | Kathrine Stene-Johansen, Kamilla Heddeland Instefjord, Hilde Elshaug, Rasmus Riis Kopperud, Hilde Vollan, Karoline Bragstad, Olav Hungnes |
| EPI_ISL_590953, EPI_ISL_590954, EPI_ISL_590955, EPI_ISL_590956, EPI_ISL_590957, EPI_ISL_590958, EPI_ISL_590959, EPI_ISL_590960, EPI_ISL_590961, EPI_ISL_590962, EPI_ISL_590963, EPI_ISL_590964, EPI_ISL_590965, EPI_ISL_590966, EPI_ISL_590967, EPI_ISL_590968, EPI_ISL_590969, EPI_ISL_590970, EPI_ISL_590971, EPI_ISL_590972, EPI_ISL_590973, EPI_ISL_590974, EPI_ISL_590975 | | | |
| see above | Dept. of Medical Microbiology, Stavanger University Hospital, Helse Stavanger HF | Norwegian Institute of Public Health, Department of Virology | Kathrine Stene-Johansen, Iren Löhr, Kamilla Heddeland Instefjord, Hilde Elshaug, Rasmus Riis Kopperud, Hilde Vollan, Karoline Bragstad, Olav Hungnes |
| EPI_ISL_590976, EPI_ISL_590977 | Dept. of Medical Microbiology, Stavanger University Hospital, Helse Stavanger HF | Norwegian Institute of Public Health, Department of Virology | Kathrine Stene-Johansen, Kamilla Heddeland Instefjord, Hilde Elshaug, Rasmus Riis Kopperud, Hilde Vollan, Karoline Bragstad, Olav Hungnes |
| EPI_ISL_590978, EPI_ISL_590979, EPI_ISL_590980 | Oslo University Hospital, Department of Medical Microbiology | Norwegian Institute of Public Health, Department of Virology | Kathrine Stene-Johansen, Kamilla Heddeland Instefjord, Hilde Elshaug, Rasmus Riis Kopperud, Hilde Vollan, Karoline Bragstad, Olav Hungnes |
| EPI_ISL_590981 | Foerde Hospital, Department of Microbiology | Norwegian Institute of Public Health, Department of Virology | Kathrine Stene-Johansen, Kamilla Heddeland Instefjord, Hilde Elshaug, Rasmus Riis Kopperud, Hilde Vollan, Karoline Bragstad, Olav Hungnes |
| EPI_ISL_590982, EPI_ISL_590983 | Vestfold Hospital, Toensberg Department of Microbiology | Norwegian Institute of Public Health, Department of Virology | Kathrine Stene-Johansen, Kamilla Heddeland Instefjord, Hilde Elshaug, Rasmus Riis Kopperud, Hilde Vollan, Karoline Bragstad, Olav Hungnes |
| EPI_ISL_590984 | Hospital of Southern Norway - Kristiansand, Department of Medical Microbiology | Norwegian Institute of Public Health, Department of Virology | Kathrine Stene-Johansen, Kamilla Heddeland Instefjord, Hilde Elshaug, Rasmus Riis Kopperud, Hilde Vollan, Karoline Bragstad, Olav Hungnes |
| EPI_ISL_590985 | Foerde Hospital, Department of Microbiology | Norwegian Institute of Public Health, Department of Virology | Kathrine Stene-Johansen, Kamilla Heddeland Instefjord, Hilde Elshaug, Rasmus Riis Kopperud, Hilde Vollan, Karoline Bragstad, Olav Hungnes |
| EPI_ISL_590986, EPI_ISL_590987, EPI_ISL_590988, EPI_ISL_590989, EPI_ISL_590990, EPI_ISL_590991, EPI_ISL_590992, EPI_ISL_590993, EPI_ISL_590994, EPI_ISL_590995, EPI_ISL_590996, EPI_ISL_590997, EPI_ISL_590998, EPI_ISL_590999, EPI_ISL_591000, EPI_ISL_591001, EPI_ISL_591002, EPI_ISL_591003, EPI_ISL_591004, EPI_ISL_591005, EPI_ISL_591006 | | | |
| see above | Ostfold Hospital Trust - Kalnes, Centre for Laboratory Medicine, Section for gene technology and infection serology | Norwegian Institute of Public Health, Department of Virology | Kathrine Stene-Johansen, Kamilla Heddeland Instefjord, Hilde Elshaug, Rasmus Riis Kopperud, Hilde Vollan, Karoline Bragstad, Olav Hungnes |
| EPI_ISL_591007 | Department of Medical Microbiology, St. Olavs hospital | Norwegian Institute of Public Health, Department of Virology | Kathrine Stene-Johansen, Kamilla Heddeland Instefjord, Hilde Elshaug, Rasmus Riis Kopperud, Hilde Vollan, Karoline Bragstad, Olav Hungnes |
| EPI_ISL_591008 | Foerde Hospital, Department of Microbiology | Norwegian Institute of Public Health, Department of Virology | Kathrine Stene-Johansen, Kamilla Heddeland Instefjord, Hilde Elshaug, Rasmus Riis Kopperud, Hilde Vollan, Karoline Bragstad, Olav Hungnes |
| EPI_ISL_591009 | Ostfold Hospital Trust - Kalnes, Centre for Laboratory Medicine, Section for gene technology and infection serology | Norwegian Institute of Public Health, Department of Virology | Kathrine Stene-Johansen, Kamilla Heddeland Instefjord, Hilde Elshaug, Rasmus Riis Kopperud, Hilde Vollan, Karoline Bragstad, Olav Hungnes |
| EPI_ISL_591010 | Unilabs Laboratory Medicine | Norwegian Institute of Public Health, Department of Virology | Kathrine Stene-Johansen, Kamilla Heddeland Instefjord, Hilde Elshaug, Rasmus Riis Kopperud, Hilde Vollan, Karoline Bragstad, Olav Hungnes |
| EPI_ISL_591011 | Oslo University Hospital, Department of Medical Microbiology | Norwegian Institute of Public Health, Department of Virology | Kathrine Stene-Johansen, Kamilla Heddeland Instefjord, Hilde Elshaug, Rasmus Riis Kopperud, Hilde Vollan, Karoline Bragstad, Olav Hungnes |
| EPI_ISL_591012 | Department of Medical Microbiology - section Molde, Molde Hospital | Norwegian Institute of Public Health, Department of Virology | Kathrine Stene-Johansen, Kamilla Heddeland Instefjord, Hilde Elshaug, Rasmus Riis Kopperud, Hilde Vollan, Karoline Bragstad, Olav Hungnes |
| EPI_ISL_591013 | Unilabs Laboratory Medicine | Norwegian Institute of Public Health, Department of Virology | Kathrine Stene-Johansen, Kamilla Heddeland Instefjord, Hilde Elshaug, Rasmus Riis Kopperud, Hilde Vollan, Karoline Bragstad, Olav Hungnes |
| EPI_ISL_591014 | Department of Medical Microbiology, St. Olavs hospital | Norwegian Institute of Public Health, Department of Virology | Kathrine Stene-Johansen, Kamilla Heddeland Instefjord, Hilde Elshaug, Rasmus Riis Kopperud, Hilde Vollan, Karoline Bragstad, Olav Hungnes |
| EPI_ISL_591015, EPI_ISL_591016, EPI_ISL_591017 | Ostfold Hospital Trust - Kalnes, Centre for Laboratory Medicine, Section for gene technology and infection serology | Norwegian Institute of Public Health, Department of Virology | Kathrine Stene-Johansen, Kamilla Heddeland Instefjord, Hilde Elshaug, Rasmus Riis Kopperud, Hilde Vollan, Karoline Bragstad, Olav Hungnes |
| EPI_ISL_591018, EPI_ISL_591019 | Department of Medical Microbiology - section Molde, Molde Hospital | Norwegian Institute of Public Health, Department of Virology | Kathrine Stene-Johansen, Kamilla Heddeland Instefjord, Hilde Elshaug, Rasmus Riis Kopperud, Hilde Vollan, Karoline Bragstad, Olav Hungnes |
| EPI_ISL_591020 | Oslo University Hospital, Department of Medical Microbiology | Norwegian Institute of Public Health, Department of Virology | Kathrine Stene-Johansen, Kamilla Heddeland Instefjord, Hilde Elshaug, Rasmus Riis Kopperud, Hilde Vollan, Karoline Bragstad, Olav Hungnes |
| EPI_ISL_591021 | Dept. of Medical Microbiology, Stavanger University Hospital, Helse Stavanger HF | Norwegian Institute of Public Health, Department of Virology | Kathrine Stene-Johansen, Kamilla Heddeland Instefjord, Hilde Elshaug, Rasmus Riis Kopperud, Hilde Vollan, Karoline Bragstad, Olav Hungnes |
| EPI_ISL_591099, EPI_ISL_591541, EPI_ISL_593856, EPI_ISL_593875, EPI_ISL_593876, EPI_ISL_593877, EPI_ISL_593878, EPI_ISL_593879, EPI_ISL_593880, EPI_ISL_593881, EPI_ISL_593882, EPI_ISL_593883, EPI_ISL_593886, EPI_ISL_593887, EPI_ISL_593888, EPI_ISL_593889 | | | |
| see above | CHU Purpan - Laboratoire de Virologie - Institut Fédératif de Biologie | CHU Purpan - Laboratoire de Virologie - Institut Fédératif de Biologie | Latour J., Ranger N., Dubois M., Carcenac R., Harter A., Boyer P., Tremeaux P., Izopet J. |
| EPI_ISL_594119, EPI_ISL_594120, EPI_ISL_594121, EPI_ISL_594122, EPI_ISL_594123, EPI_ISL_594124, EPI_ISL_594125, EPI_ISL_594126, EPI_ISL_594127, EPI_ISL_594128, EPI_ISL_594129, EPI_ISL_594130, EPI_ISL_594131, EPI_ISL_594132, EPI_ISL_594133 | | | |
| see above | Yale COVID-19 Biorepository | Grubaugh Lab - Yale School of Public Health | Joseph Fauver, Tara Alpert, Anderson Brito, Anne Wyllie, Chantal Vogels, Mary Petrone, Chaney Kalinich, Isabel Ott, Arnau Casanovas, Catherine Muenker, Adam Moore, Alice Lu, Maria Tokuyama, Patrick Wong, Peiwen Lu, Saad Omer, Richard Martinello, Allison Nelson, Shelli Farhadian, Akiko Iwasaki, Charlese Dela Cruz, Albert Ko, Nathan Grubaugh |
| EPI_ISL_596386 | National Institute for Allergy and Infectious Diseases Integrated Research Facility - Frederick (NIAID IRF-Frederick), National Institutes of Health (NIH) | National Institute for Allergy and Infectious Diseases Integrated Research Facility - Frederick (NIAID IRF-Frederick), National Institutes of Health (NIH) | Kocher,G., Kugelman,J.R., Beitzel,B. and Palacios,G. |
| EPI_ISL_597523, EPI_ISL_597524, EPI_ISL_597534, EPI_ISL_597539, EPI_ISL_597541, EPI_ISL_597547, EPI_ISL_597551, EPI_ISL_597558, EPI_ISL_597559, EPI_ISL_597560, EPI_ISL_597567, EPI_ISL_597570, EPI_ISL_597578, EPI_ISL_597580, EPI_ISL_597581, EPI_ISL_597584, EPI_ISL_597586, EPI_ISL_597596, EPI_ISL_597597, EPI_ISL_597600, EPI_ISL_597604, EPI_ISL_597608, EPI_ISL_597616, EPI_ISL_597620, EPI_ISL_597621, EPI_ISL_597626, EPI_ISL_597629, EPI_ISL_597634, EPI_ISL_597639, EPI_ISL_597645, EPI_ISL_597647, EPI_ISL_597650, EPI_ISL_597652, EPI_ISL_597654, EPI_ISL_597655, EPI_ISL_597656, EPI_ISL_597660, EPI_ISL_597663, EPI_ISL_597666, EPI_ISL_597689, EPI_ISL_597692, EPI_ISL_597694, EPI_ISL_597695, EPI_ISL_597700, EPI_ISL_597705, EPI_ISL_597711, EPI_ISL_597712, EPI_ISL_597713, EPI_ISL_597715, EPI_ISL_597721, EPI_ISL_597722, EPI_ISL_597723, EPI_ISL_597725, EPI_ISL_597731, EPI_ISL_597732, EPI_ISL_597733, EPI_ISL_597738, EPI_ISL_597747, EPI_ISL_597754, EPI_ISL_597755, EPI_ISL_597756, EPI_ISL_597757, EPI_ISL_597760, EPI_ISL_597763, EPI_ISL_597771, EPI_ISL_597773, EPI_ISL_597778, EPI_ISL_597781, EPI_ISL_597785, EPI_ISL_597787, EPI_ISL_597794, EPI_ISL_597796, EPI_ISL_597798, EPI_ISL_597808, EPI_ISL_597812, EPI_ISL_597826, EPI_ISL_597827, EPI_ISL_597829, EPI_ISL_597830, EPI_ISL_597834, EPI_ISL_597836, EPI_ISL_597838, EPI_ISL_597839, EPI_ISL_597840, EPI_ISL_597846, EPI_ISL_597853, EPI_ISL_597855, EPI_ISL_597858, EPI_ISL_597863, EPI_ISL_597867, EPI_ISL_597868, EPI_ISL_597871, EPI_ISL_597876, EPI_ISL_597878, EPI_ISL_597879, EPI_ISL_597881, EPI_ISL_597882, EPI_ISL_598204, EPI_ISL_598242, EPI_ISL_598824, EPI_ISL_598825, EPI_ISL_598826, EPI_ISL_598827, EPI_ISL_598829, EPI_ISL_598830, EPI_ISL_600521, EPI_ISL_600525, EPI_ISL_600526, EPI_ISL_600528, EPI_ISL_600529, EPI_ISL_600531, EPI_ISL_600532, EPI_ISL_600534, EPI_ISL_600537, EPI_ISL_602155, EPI_ISL_602158, EPI_ISL_602161 | | | |
| see above | Lighthouse Lab in Alderley Park | Wellcome Sanger Institute for the COVID-19 Genomics UK (COG-UK) consortium | Jacquelyn Wynn, Mairead Hyland, The Lighthouse Lab in Alderley Park and Alex Alderton, Roberto Amato, Sonia Goncalves, Ewan Harrison, David K. Jackson, Ian Johnston, Dominic Kwiatkowski, Cordelia Langford, John Sillitoe on behalf of the Wellcome Sanger Institute COVID-19 Surveillance Team (http://www.sanger.ac.uk/covid-team) |
| EPI_ISL_603216 | CHU Purpan - Laboratoire de Virologie - Institut Fédératif de Biologie | CHU Purpan - Laboratoire de Virologie - Institut Fédératif de Biologie | Latour J., Ranger N., Dubois M., Carcenac R., Harter A., Boyer P., Tremeaux P., Izopet J. |
| EPI_ISL_605882, EPI_ISL_605883, EPI_ISL_605884, EPI_ISL_605885, EPI_ISL_605886, EPI_ISL_605887, EPI_ISL_605888, EPI_ISL_605889, EPI_ISL_605890, EPI_ISL_605891, EPI_ISL_605892, EPI_ISL_605893, EPI_ISL_605894, EPI_ISL_605895, EPI_ISL_605896, EPI_ISL_605897, EPI_ISL_605898, EPI_ISL_605899, EPI_ISL_605900, EPI_ISL_605901, EPI_ISL_605902, EPI_ISL_605903, EPI_ISL_605904, EPI_ISL_605905, EPI_ISL_605906, EPI_ISL_605907, EPI_ISL_605908 | | | |
| see above | NGS Lab, DNA SOLUTION LTD. | NGS Lab, DNA SOLUTION LTD. | Khan,M.I., Hasan,K.N., Sufian,A., Polol,M.N.I., Khaleque,A., Rahman,M., Chowdhury,M., Haider,H.U., Razu,M.H., Khan,M., Rabbi,M.F.A. |
| EPI_ISL_605909, EPI_ISL_605910, EPI_ISL_605911, EPI_ISL_605912, EPI_ISL_605913 | NGS Lab, DNA SOLUTION LTD. | NGS Lab, DNA SOLUTION LTD. | Khan,M.I., Hasan,K.N., Sufian,A., Hosen,M.B., Polol,M.N.I., Khaleque,A., Rahman,M., Chowdhury,M., Haider,H.U., Razu,M.H., Khan,M., Rabbi,M.F.A. |
| EPI_ISL_605914, EPI_ISL_605915, EPI_ISL_605916, EPI_ISL_605917, | NGS Lab, DNA SOLUTION LTD. | NGS Lab, DNA SOLUTION LTD. | Khan,M.I., Hasan,K.N., Sufian,A., Hosen,M.B., Khaleque,A., Rahman,M., Chowdhury,M., Haider,H.U., Razu,M.H., Khan,M., Rabbi,M.F.A. |

EPI_ISL_605918, EPI_ISL_605919,
EPI_ISL_605920, EPI_ISL_605921,
EPI_ISL_605922, EPI_ISL_605923

EPI_ISL_606116, EPI_ISL_606189, EPI_ISL_606228, EPI_ISL_606238, EPI_ISL_606924, EPI_ISL_606925, EPI_ISL_606927, EPI_ISL_606928, EPI_ISL_606929, EPI_ISL_606931, EPI_ISL_606932, EPI_ISL_606933, EPI_ISL_606934, EPI_ISL_606935, EPI_ISL_606936, EPI_ISL_606937, EPI_ISL_606938, EPI_ISL_606939,
EPI_ISL_606940, EPI_ISL_606941, EPI_ISL_606942, EPI_ISL_606943, EPI_ISL_606944, EPI_ISL_606945, EPI_ISL_606946, EPI_ISL_606947, EPI_ISL_606949, EPI_ISL_606950, EPI_ISL_606951, EPI_ISL_606952, EPI_ISL_606954, EPI_ISL_606955, EPI_ISL_606956, EPI_ISL_606957, EPI_ISL_606958, EPI_ISL_606959,
EPI_ISL_606961, EPI_ISL_606962, EPI_ISL_606963, EPI_ISL_606965, EPI_ISL_606966, EPI_ISL_606967, EPI_ISL_606968, EPI_ISL_606969, EPI_ISL_606970, EPI_ISL_606971, EPI_ISL_606972, EPI_ISL_606973, EPI_ISL_606974, EPI_ISL_606975, EPI_ISL_606976, EPI_ISL_606977, EPI_ISL_606978, EPI_ISL_606980,
EPI_ISL_606981, EPI_ISL_606982, EPI_ISL_606983, EPI_ISL_606984, EPI_ISL_606985, EPI_ISL_606986, EPI_ISL_606987, EPI_ISL_606988, EPI_ISL_606989, EPI_ISL_606990, EPI_ISL_606991, EPI_ISL_606992, EPI_ISL_606993, EPI_ISL_606994, EPI_ISL_606995, EPI_ISL_606996, EPI_ISL_606997, EPI_ISL_606998,
EPI_ISL_606999, EPI_ISL_607000, EPI_ISL_607001, EPI_ISL_607002, EPI_ISL_607003, EPI_ISL_607004, EPI_ISL_607005, EPI_ISL_607006, EPI_ISL_607007, EPI_ISL_607008, EPI_ISL_607009, EPI_ISL_607010, EPI_ISL_607011, EPI_ISL_607012, EPI_ISL_607013, EPI_ISL_607014, EPI_ISL_607015, EPI_ISL_607016,
EPI_ISL_607018, EPI_ISL_607019, EPI_ISL_607021, EPI_ISL_607022, EPI_ISL_607023, EPI_ISL_607024, EPI_ISL_607025, EPI_ISL_607026, EPI_ISL_607027, EPI_ISL_607029, EPI_ISL_607030, EPI_ISL_607031, EPI_ISL_607032, EPI_ISL_607033, EPI_ISL_607034, EPI_ISL_607035, EPI_ISL_607036, EPI_ISL_607037,
EPI_ISL_607038, EPI_ISL_607039, EPI_ISL_607040, EPI_ISL_607041, EPI_ISL_607042, EPI_ISL_607043, EPI_ISL_607044, EPI_ISL_607045, EPI_ISL_607046, EPI_ISL_607047, EPI_ISL_607048, EPI_ISL_607049, EPI_ISL_607050, EPI_ISL_607051, EPI_ISL_607052, EPI_ISL_607053, EPI_ISL_607054, EPI_ISL_607055,
EPI_ISL_607056, EPI_ISL_607057, EPI_ISL_607058, EPI_ISL_607059, EPI_ISL_607060, EPI_ISL_607061, EPI_ISL_607062, EPI_ISL_607063, EPI_ISL_607064, EPI_ISL_607065, EPI_ISL_607066, EPI_ISL_607067, EPI_ISL_607068, EPI_ISL_607069, EPI_ISL_607070, EPI_ISL_607071, EPI_ISL_607072, EPI_ISL_607073,
EPI_ISL_607074, EPI_ISL_607075, EPI_ISL_607076, EPI_ISL_607077, EPI_ISL_607078, EPI_ISL_607079, EPI_ISL_607081, EPI_ISL_607082, EPI_ISL_607083, EPI_ISL_607084, EPI_ISL_607085, EPI_ISL_607087, EPI_ISL_607088, EPI_ISL_607089, EPI_ISL_607090, EPI_ISL_607091, EPI_ISL_607092, EPI_ISL_607093,
EPI_ISL_607094, EPI_ISL_607095, EPI_ISL_607096, EPI_ISL_607097, EPI_ISL_607098, EPI_ISL_607099, EPI_ISL_607100, EPI_ISL_607101, EPI_ISL_607102, EPI_ISL_607103, EPI_ISL_607104, EPI_ISL_607105, EPI_ISL_607106, EPI_ISL_607107, EPI_ISL_607108, EPI_ISL_607109, EPI_ISL_607110, EPI_ISL_607111,
EPI_ISL_607112, EPI_ISL_607114, EPI_ISL_607115, EPI_ISL_607116, EPI_ISL_607117, EPI_ISL_607118, EPI_ISL_607119, EPI_ISL_607120, EPI_ISL_607121, EPI_ISL_607122, EPI_ISL_607123, EPI_ISL_607124, EPI_ISL_607125, EPI_ISL_607126, EPI_ISL_607127, EPI_ISL_607128, EPI_ISL_607129, EPI_ISL_607130,
EPI_ISL_607131, EPI_ISL_607132, EPI_ISL_607133, EPI_ISL_607134, EPI_ISL_607135, EPI_ISL_607136, EPI_ISL_607137, EPI_ISL_607138, EPI_ISL_607139, EPI_ISL_607140, EPI_ISL_607141, EPI_ISL_607142, EPI_ISL_607143, EPI_ISL_607144, EPI_ISL_607145, EPI_ISL_607146, EPI_ISL_607147, EPI_ISL_607149,
EPI_ISL_607150, EPI_ISL_607151, EPI_ISL_607152, EPI_ISL_607153, EPI_ISL_607154, EPI_ISL_607155, EPI_ISL_607156, EPI_ISL_607157, EPI_ISL_607158, EPI_ISL_607159, EPI_ISL_607160, EPI_ISL_607161, EPI_ISL_607162, EPI_ISL_607163, EPI_ISL_607164, EPI_ISL_607165, EPI_ISL_607166, EPI_ISL_607167,
EPI_ISL_607168, EPI_ISL_607169, EPI_ISL_607170, EPI_ISL_607171, EPI_ISL_607172, EPI_ISL_607173, EPI_ISL_607174, EPI_ISL_607175, EPI_ISL_607176, EPI_ISL_607177, EPI_ISL_607178, EPI_ISL_607179, EPI_ISL_607180, EPI_ISL_607181, EPI_ISL_607182, EPI_ISL_607183, EPI_ISL_607184, EPI_ISL_607185,
EPI_ISL_607186, EPI_ISL_607187, EPI_ISL_607188, EPI_ISL_607190, EPI_ISL_607191, EPI_ISL_607192, EPI_ISL_607194, EPI_ISL_607195, EPI_ISL_607197, EPI_ISL_607198, EPI_ISL_607199, EPI_ISL_607201, EPI_ISL_607202, EPI_ISL_607203, EPI_ISL_607204, EPI_ISL_607205, EPI_ISL_607206, EPI_ISL_607207,
EPI_ISL_607208, EPI_ISL_607210, EPI_ISL_607211, EPI_ISL_607212, EPI_ISL_607213, EPI_ISL_607214, EPI_ISL_607215, EPI_ISL_607216, EPI_ISL_607217, EPI_ISL_607218, EPI_ISL_607219, EPI_ISL_607220, EPI_ISL_607221, EPI_ISL_607222, EPI_ISL_607223, EPI_ISL_607224, EPI_ISL_607225, EPI_ISL_607226,
EPI_ISL_607227, EPI_ISL_607228, EPI_ISL_607229, EPI_ISL_607230, EPI_ISL_607231, EPI_ISL_607232, EPI_ISL_607233, EPI_ISL_607234, EPI_ISL_607235, EPI_ISL_607236, EPI_ISL_607237, EPI_ISL_607238, EPI_ISL_607239, EPI_ISL_607240, EPI_ISL_607241, EPI_ISL_607242, EPI_ISL_607243, EPI_ISL_607244,
EPI_ISL_607245, EPI_ISL_607246, EPI_ISL_607247, EPI_ISL_607248, EPI_ISL_607249, EPI_ISL_607250, EPI_ISL_607251, EPI_ISL_607252, EPI_ISL_607253, EPI_ISL_607254, EPI_ISL_607255, EPI_ISL_607257, EPI_ISL_607258, EPI_ISL_607261, EPI_ISL_607262, EPI_ISL_607263, EPI_ISL_607264, EPI_ISL_607265,
EPI_ISL_607266, EPI_ISL_607267, EPI_ISL_607268, EPI_ISL_607269, EPI_ISL_607270, EPI_ISL_607271, EPI_ISL_607272, EPI_ISL_607273, EPI_ISL_607274, EPI_ISL_607275, EPI_ISL_607276, EPI_ISL_607277, EPI_ISL_607278, EPI_ISL_607279, EPI_ISL_607280, EPI_ISL_607281, EPI_ISL_607282, EPI_ISL_607283,
EPI_ISL_607284, EPI_ISL_607285, EPI_ISL_607286, EPI_ISL_607287, EPI_ISL_607288, EPI_ISL_607289, EPI_ISL_607290, EPI_ISL_607623, EPI_ISL_607624, EPI_ISL_607626, EPI_ISL_607627, EPI_ISL_607628, EPI_ISL_607629, EPI_ISL_607630, EPI_ISL_607631, EPI_ISL_607632, EPI_ISL_607633, EPI_ISL_607634,
EPI_ISL_607635, EPI_ISL_607636, EPI_ISL_607637, EPI_ISL_607638, EPI_ISL_607640, EPI_ISL_607641, EPI_ISL_607642, EPI_ISL_607644, EPI_ISL_607645, EPI_ISL_607646, EPI_ISL_607647, EPI_ISL_607648, EPI_ISL_607649, EPI_ISL_607650, EPI_ISL_607651, EPI_ISL_607652, EPI_ISL_607653, EPI_ISL_607654,
EPI_ISL_607655, EPI_ISL_607656, EPI_ISL_607658, EPI_ISL_607659, EPI_ISL_607660, EPI_ISL_607661, EPI_ISL_607663, EPI_ISL_607664, EPI_ISL_607665, EPI_ISL_607666, EPI_ISL_607667, EPI_ISL_607668, EPI_ISL_607669, EPI_ISL_607670, EPI_ISL_607672, EPI_ISL_607673, EPI_ISL_607674, EPI_ISL_607676,
EPI_ISL_607677, EPI_ISL_607678, EPI_ISL_607679, EPI_ISL_607680, EPI_ISL_607681, EPI_ISL_607682, EPI_ISL_607683, EPI_ISL_607684, EPI_ISL_607685, EPI_ISL_607686, EPI_ISL_607687, EPI_ISL_607688, EPI_ISL_607689, EPI_ISL_607690, EPI_ISL_607691, EPI_ISL_607692, EPI_ISL_607693, EPI_ISL_607694,
EPI_ISL_607695, EPI_ISL_607696, EPI_ISL_607697, EPI_ISL_607699, EPI_ISL_607700, EPI_ISL_607701, EPI_ISL_607702, EPI_ISL_607703, EPI_ISL_607704, EPI_ISL_607705, EPI_ISL_607706, EPI_ISL_607707, EPI_ISL_607708, EPI_ISL_607709, EPI_ISL_607710, EPI_ISL_607711, EPI_ISL_607712, EPI_ISL_607713,
EPI_ISL_607714, EPI_ISL_607715, EPI_ISL_607716, EPI_ISL_607717, EPI_ISL_607718, EPI_ISL_607719, EPI_ISL_607720, EPI_ISL_607721, EPI_ISL_607722, EPI_ISL_607723, EPI_ISL_607724, EPI_ISL_607725, EPI_ISL_607726, EPI_ISL_607727, EPI_ISL_607728, EPI_ISL_607729, EPI_ISL_607730, EPI_ISL_607731,
EPI_ISL_607732, EPI_ISL_607733, EPI_ISL_607734, EPI_ISL_607735, EPI_ISL_607736, EPI_ISL_607737, EPI_ISL_607738, EPI_ISL_607739, EPI_ISL_607740, EPI_ISL_607741, EPI_ISL_607742, EPI_ISL_607743, EPI_ISL_607744, EPI_ISL_607745, EPI_ISL_607746, EPI_ISL_607747, EPI_ISL_607749, EPI_ISL_607750,
EPI_ISL_607751, EPI_ISL_607752, EPI_ISL_607753, EPI_ISL_607754, EPI_ISL_607755, EPI_ISL_607756, EPI_ISL_607757, EPI_ISL_607759, EPI_ISL_607760, EPI_ISL_607761, EPI_ISL_607762, EPI_ISL_607763, EPI_ISL_607764, EPI_ISL_607765, EPI_ISL_607766, EPI_ISL_607767, EPI_ISL_607768, EPI_ISL_607769,
EPI_ISL_607770, EPI_ISL_607771, EPI_ISL_607772, EPI_ISL_607773, EPI_ISL_607774, EPI_ISL_607775, EPI_ISL_607776, EPI_ISL_607777, EPI_ISL_607778, EPI_ISL_607779, EPI_ISL_607781, EPI_ISL_607782, EPI_ISL_607783, EPI_ISL_607784, EPI_ISL_607785, EPI_ISL_607786, EPI_ISL_607787, EPI_ISL_607788,
EPI_ISL_607789, EPI_ISL_607790, EPI_ISL_607791, EPI_ISL_607792, EPI_ISL_607793, EPI_ISL_607794, EPI_ISL_607795, EPI_ISL_607796, EPI_ISL_607797, EPI_ISL_607798, EPI_ISL_607799, EPI_ISL_607800, EPI_ISL_607801, EPI_ISL_607802, EPI_ISL_607803, EPI_ISL_607804, EPI_ISL_607805, EPI_ISL_607806,
EPI_ISL_607807, EPI_ISL_607808, EPI_ISL_607809, EPI_ISL_607810, EPI_ISL_607811, EPI_ISL_607812, EPI_ISL_607814, EPI_ISL_607815, EPI_ISL_607816, EPI_ISL_607817, EPI_ISL_607818, EPI_ISL_607819, EPI_ISL_607820, EPI_ISL_607821, EPI_ISL_607822, EPI_ISL_607823, EPI_ISL_607824, EPI_ISL_607825,
EPI_ISL_607827, EPI_ISL_607828, EPI_ISL_607830, EPI_ISL_607831, EPI_ISL_607832, EPI_ISL_607833, EPI_ISL_607834, EPI_ISL_607835, EPI_ISL_607836, EPI_ISL_607837, EPI_ISL_607838, EPI_ISL_607839, EPI_ISL_607840, EPI_ISL_607841, EPI_ISL_607842, EPI_ISL_607843, EPI_ISL_607844, EPI_ISL_607845,
EPI_ISL_607846, EPI_ISL_607847, EPI_ISL_607848, EPI_ISL_607849, EPI_ISL_607850, EPI_ISL_607851, EPI_ISL_607852, EPI_ISL_607853, EPI_ISL_607854, EPI_ISL_607855, EPI_ISL_607856, EPI_ISL_607857, EPI_ISL_607858, EPI_ISL_607859, EPI_ISL_607861, EPI_ISL_607862, EPI_ISL_607863, EPI_ISL_607864,
EPI_ISL_607865, EPI_ISL_607866, EPI_ISL_607867, EPI_ISL_607868, EPI_ISL_607869, EPI_ISL_607870, EPI_ISL_607871, EPI_ISL_607872, EPI_ISL_607873, EPI_ISL_607874, EPI_ISL_607875, EPI_ISL_607876, EPI_ISL_607877, EPI_ISL_607878, EPI_ISL_607879, EPI_ISL_607880, EPI_ISL_607881, EPI_ISL_607882,
EPI_ISL_607883, EPI_ISL_607884, EPI_ISL_607885, EPI_ISL_607886, EPI_ISL_607888, EPI_ISL_607889, EPI_ISL_607890, EPI_ISL_607891, EPI_ISL_607892, EPI_ISL_607893, EPI_ISL_607895, EPI_ISL_607896, EPI_ISL_607897, EPI_ISL_607898, EPI_ISL_607899, EPI_ISL_607900, EPI_ISL_607901, EPI_ISL_607902,
EPI_ISL_607903, EPI_ISL_607904, EPI_ISL_607906, EPI_ISL_607907, EPI_ISL_607908, EPI_ISL_607909, EPI_ISL_607910, EPI_ISL_607911, EPI_ISL_607912, EPI_ISL_607913, EPI_ISL_607914, EPI_ISL_607915, EPI_ISL_607916, EPI_ISL_607918, EPI_ISL_607919, EPI_ISL_607920, EPI_ISL_607921, EPI_ISL_607922,
EPI_ISL_607923, EPI_ISL_607924, EPI_ISL_607925, EPI_ISL_607926, EPI_ISL_607927, EPI_ISL_607928, EPI_ISL_607929, EPI_ISL_607930, EPI_ISL_607931, EPI_ISL_607932, EPI_ISL_607933, EPI_ISL_607934, EPI_ISL_607935, EPI_ISL_607936, EPI_ISL_607937, EPI_ISL_607938, EPI_ISL_607939, EPI_ISL_607940,
EPI_ISL_607941, EPI_ISL_607942, EPI_ISL_607943, EPI_ISL_607944, EPI_ISL_607945, EPI_ISL_607946, EPI_ISL_607947, EPI_ISL_607949, EPI_ISL_607951, EPI_ISL_607952, EPI_ISL_607953, EPI_ISL_607954, EPI_ISL_607955, EPI_ISL_607956, EPI_ISL_607957, EPI_ISL_607958, EPI_ISL_607959, EPI_ISL_607960,
EPI_ISL_607961, EPI_ISL_607962, EPI_ISL_607963, EPI_ISL_607964, EPI_ISL_607965, EPI_ISL_607966, EPI_ISL_607967, EPI_ISL_607968, EPI_ISL_607969, EPI_ISL_607970, EPI_ISL_607971, EPI_ISL_607972, EPI_ISL_607973, EPI_ISL_607974, EPI_ISL_607975, EPI_ISL_607976, EPI_ISL_607977, EPI_ISL_607978,
EPI_ISL_607979, EPI_ISL_607980, EPI_ISL_607981, EPI_ISL_607982, EPI_ISL_607983, EPI_ISL_607984, EPI_ISL_607985, EPI_ISL_607986, EPI_ISL_607987, EPI_ISL_607988, EPI_ISL_607989, EPI_ISL_607992, EPI_ISL_607994, EPI_ISL_607995, EPI_ISL_607996, EPI_ISL_607997, EPI_ISL_607999, EPI_ISL_608000,
EPI_ISL_608001, EPI_ISL_608002, EPI_ISL_608003, EPI_ISL_608004, EPI_ISL_608005, EPI_ISL_608006, EPI_ISL_608007, EPI_ISL_608009, EPI_ISL_608010, EPI_ISL_608011, EPI_ISL_608012, EPI_ISL_608013, EPI_ISL_608014, EPI_ISL_608016, EPI_ISL_608017, EPI_ISL_608018, EPI_ISL_608019, EPI_ISL_608021,
EPI_ISL_608022, EPI_ISL_608024, EPI_ISL_608025, EPI_ISL_608026, EPI_ISL_608027, EPI_ISL_608028, EPI_ISL_608029, EPI_ISL_608030, EPI_ISL_608031, EPI_ISL_608032, EPI_ISL_608033, EPI_ISL_608034, EPI_ISL_608035, EPI_ISL_608036, EPI_ISL_608037, EPI_ISL_608038, EPI_ISL_608039, EPI_ISL_608040,
EPI_ISL_608041, EPI_ISL_608042, EPI_ISL_608043, EPI_ISL_608044, EPI_ISL_608045, EPI_ISL_608046, EPI_ISL_608047, EPI_ISL_608048, EPI_ISL_608049, EPI_ISL_608050, EPI_ISL_608051, EPI_ISL_608052, EPI_ISL_608053, EPI_ISL_608054, EPI_ISL_608055, EPI_ISL_608056, EPI_ISL_608057, EPI_ISL_608058,
EPI_ISL_608059, EPI_ISL_608060, EPI_ISL_608062, EPI_ISL_608063, EPI_ISL_608064, EPI_ISL_608065, EPI_ISL_608066, EPI_ISL_608067, EPI_ISL_608068, EPI_ISL_608069, EPI_ISL_608070, EPI_ISL_608071, EPI_ISL_608072, EPI_ISL_608073, EPI_ISL_608075, EPI_ISL_608076, EPI_ISL_608077, EPI_ISL_608078,
EPI_ISL_608079, EPI_ISL_608080, EPI_ISL_608081, EPI_ISL_608082, EPI_ISL_608083, EPI_ISL_608084, EPI_ISL_608085, EPI_ISL_608086, EPI_ISL_608087, EPI_ISL_608088, EPI_ISL_608089, EPI_ISL_608090, EPI_ISL_608092, EPI_ISL_608093, EPI_ISL_608094, EPI_ISL_608095, EPI_ISL_608096, EPI_ISL_608097,
EPI_ISL_608098, EPI_ISL_608099, EPI_ISL_608100, EPI_ISL_608101, EPI_ISL_608102, EPI_ISL_608103, EPI_ISL_608104, EPI_ISL_608105, EPI_ISL_608106, EPI_ISL_608107, EPI_ISL_608108, EPI_ISL_608109, EPI_ISL_608110, EPI_ISL_608111, EPI_ISL_608112, EPI_ISL_608113, EPI_ISL_608114, EPI_ISL_608115,
EPI_ISL_608116, EPI_ISL_608118, EPI_ISL_608119, EPI_ISL_608120, EPI_ISL_608121, EPI_ISL_608122, EPI_ISL_608123, EPI_ISL_608124, EPI_ISL_608125, EPI_ISL_608126, EPI_ISL_608127, EPI_ISL_608128, EPI_ISL_608129, EPI_ISL_608130, EPI_ISL_608131, EPI_ISL_608133, EPI_ISL_608134, EPI_ISL_608135,
EPI_ISL_608136, EPI_ISL_608137, EPI_ISL_608138, EPI_ISL_608139, EPI_ISL_608140, EPI_ISL_608141, EPI_ISL_608142, EPI_ISL_608143, EPI_ISL_608144, EPI_ISL_608145, EPI_ISL_608146, EPI_ISL_608148, EPI_ISL_608149, EPI_ISL_608150, EPI_ISL_608151, EPI_ISL_608152, EPI_ISL_608153, EPI_ISL_608154,
EPI_ISL_608155, EPI_ISL_608156, EPI_ISL_608157, EPI_ISL_608158, EPI_ISL_608159, EPI_ISL_608160, EPI_ISL_608161, EPI_ISL_608162, EPI_ISL_608163, EPI_ISL_608164, EPI_ISL_608165, EPI_ISL_608166, EPI_ISL_608167, EPI_ISL_608168, EPI_ISL_608169, EPI_ISL_608170, EPI_ISL_608171, EPI_ISL_608172,
EPI_ISL_608173, EPI_ISL_608176, EPI_ISL_608177, EPI_ISL_608178, EPI_ISL_608179, EPI_ISL_608180, EPI_ISL_608181, EPI_ISL_608182, EPI_ISL_608183, EPI_ISL_608184, EPI_ISL_608185, EPI_ISL_608186, EPI_ISL_608187, EPI_ISL_608188, EPI_ISL_608189, EPI_ISL_608190, EPI_ISL_608191, EPI_ISL_608192,
EPI_ISL_608193, EPI_ISL_608194, EPI_ISL_608195, EPI_ISL_608196, EPI_ISL_608197, EPI_ISL_608198, EPI_ISL_608199, EPI_ISL_608200, EPI_ISL_608201, EPI_ISL_608202, EPI_ISL_608203, EPI_ISL_608204, EPI_ISL_608205, EPI_ISL_608206, EPI_ISL_608207, EPI_ISL_608208, EPI_ISL_608209, EPI_ISL_608210,
EPI_ISL_608211, EPI_ISL_608212, EPI_ISL_608213, EPI_ISL_608214, EPI_ISL_608217, EPI_ISL_608218, EPI_ISL_608219, EPI_ISL_608220, EPI_ISL_608221, EPI_ISL_608222, EPI_ISL_608223, EPI_ISL_608224, EPI_ISL_608225, EPI_ISL_608226, EPI_ISL_608227, EPI_ISL_608228, EPI_ISL_608229, EPI_ISL_608230,
EPI_ISL_608231, EPI_ISL_608232, EPI_ISL_608233, EPI_ISL_608234, EPI_ISL_608235, EPI_ISL_608236, EPI_ISL_608237, EPI_ISL_608238, EPI_ISL_608240, EPI_ISL_608241, EPI_ISL_608242, EPI_ISL_608243, EPI_ISL_608244, EPI_ISL_608245, EPI_ISL_608246, EPI_ISL_608247, EPI_ISL_608248, EPI_ISL_608249,
EPI_ISL_608250, EPI_ISL_608251, EPI_ISL_608252, EPI_ISL_608253, EPI_ISL_608254, EPI_ISL_608255, EPI_ISL_608256, EPI_ISL_608257, EPI_ISL_608258, EPI_ISL_608259, EPI_ISL_608260, EPI_ISL_608261, EPI_ISL_608262, EPI_ISL_608263, EPI_ISL_608264, EPI_ISL_608265, EPI_ISL_608266, EPI_ISL_608267,
EPI_ISL_608268, EPI_ISL_608269, EPI_ISL_608270, EPI_ISL_608271, EPI_ISL_608272, EPI_ISL_608273, EPI_ISL_608274, EPI_ISL_608275, EPI_ISL_608276, EPI_ISL_608277, EPI_ISL_608278, EPI_ISL_608279, EPI_ISL_608280, EPI_ISL_608281, EPI_ISL_608282, EPI_ISL_608283, EPI_ISL_608284, EPI_ISL_608285,
EPI_ISL_608286, EPI_ISL_608287, EPI_ISL_608288, EPI_ISL_608289, EPI_ISL_608290, EPI_ISL_608291, EPI_ISL_608292, EPI_ISL_608293, EPI_ISL_608294, EPI_ISL_608295, EPI_ISL_608296, EPI_ISL_608298, EPI_ISL_608299, EPI_ISL_608300, EPI_ISL_608301, EPI_ISL_608302, EPI_ISL_608303, EPI_ISL_608304,
EPI_ISL_608305, EPI_ISL_608306, EPI_ISL_608307, EPI_ISL_608309, EPI_ISL_608311, EPI_ISL_608312, EPI_ISL_608313, EPI_ISL_608314, EPI_ISL_608315, EPI_ISL_608316, EPI_ISL_608317, EPI_ISL_608318, EPI_ISL_608319, EPI_ISL_608320, EPI_ISL_608321, EPI_ISL_608322, EPI_ISL_608323, EPI_ISL_608324,
EPI_ISL_608325, EPI_ISL_608326, EPI_ISL_608327, EPI_ISL_608328, EPI_ISL_608329, EPI_ISL_608330, EPI_ISL_608331, EPI_ISL_608332, EPI_ISL_608333, EPI_ISL_608334, EPI_ISL_608335, EPI_ISL_608336, EPI_ISL_608337, EPI_ISL_608338, EPI_ISL_608339, EPI_ISL_608340, EPI_ISL_608341, EPI_ISL_608342,
EPI_ISL_608343, EPI_ISL_608344, EPI_ISL_608345, EPI_ISL_608346, EPI_ISL_608350, EPI_ISL_608351, EPI_ISL_608352, EPI_ISL_608355, EPI_ISL_608356, EPI_ISL_608361, EPI_ISL_608362, EPI_ISL_608364, EPI_ISL_608365, EPI_ISL_608367, EPI_ISL_608369, EPI_ISL_608371, EPI_ISL_608372, EPI_ISL_608374,
EPI_ISL_608375, EPI_ISL_608376, EPI_ISL_608377, EPI_ISL_608378, EPI_ISL_608379, EPI_ISL_608380, EPI_ISL_608384, EPI_ISL_608385, EPI_ISL_608386, EPI_ISL_608387, EPI_ISL_608390, EPI_ISL_608399, EPI_ISL_608401, EPI_ISL_608403, EPI_ISL_608405, EPI_ISL_608406, EPI_ISL_608409, EPI_ISL_608413,
EPI_ISL_608415, EPI_ISL_608418, EPI_ISL_608419, EPI_ISL_608420, EPI_ISL_608422, EPI_ISL_608431, EPI_ISL_608434, EPI_ISL_608435, EPI_ISL_608438, EPI_ISL_608440, EPI_ISL_608441, EPI_ISL_608443, EPI_ISL_608445, EPI_ISL_608448, EPI_ISL_608452, EPI_ISL_608454, EPI_ISL_608457, EPI_ISL_608460,
EPI_ISL_608461, EPI_ISL_608462, EPI_ISL_608465, EPI_ISL_608468, EPI_ISL_608470, EPI_ISL_608473, EPI_ISL_608479, EPI_ISL_608481, EPI_ISL_608487, EPI_ISL_608488, EPI_ISL_608489, EPI_ISL_608491, EPI_ISL_608496, EPI_ISL_608499, EPI_ISL_608500, EPI_ISL_608502, EPI_ISL_608503, EPI_ISL_608505,
EPI_ISL_608511, EPI_ISL_608512, EPI_ISL_608513, EPI_ISL_608516, EPI_ISL_608517, EPI_ISL_608518, EPI_ISL_608519, EPI_ISL_608526, EPI_ISL_608528, EPI_ISL_608534, EPI_ISL_608539, EPI_ISL_608541, EPI_ISL_608542, EPI_ISL_608543, EPI_ISL_608548, EPI_ISL_608549, EPI_ISL_608551, EPI_ISL_608553,
EPI_ISL_608554, EPI_ISL_608555, EPI_ISL_608562, EPI_ISL_608567, EPI_ISL_608569, EPI_ISL_608575, EPI_ISL_608579, EPI_ISL_608580, EPI_ISL_608581, EPI_ISL_608583, EPI_ISL_608584, EPI_ISL_608586, EPI_ISL_608587, EPI_ISL_608589, EPI_ISL_608590, EPI_ISL_608592, EPI_ISL_608594, EPI_ISL_608595,
EPI_ISL_608599, EPI_ISL_608600, EPI_ISL_608601, EPI_ISL_608605, EPI_ISL_608606, EPI_ISL_608609, EPI_ISL_608610, EPI_ISL_608617, EPI_ISL_608622, EPI_ISL_608623, EPI_ISL_608625, EPI_ISL_608630, EPI_ISL_608634, EPI_ISL_608639, EPI_ISL_608640, EPI_ISL_608646, EPI_ISL_608647, EPI_ISL_608648,
EPI_ISL_608649, EPI_ISL_608655, EPI_ISL_608657, EPI_ISL_608658, EPI_ISL_608659, EPI_ISL_608661, EPI_ISL_608662, EPI_ISL_608663, EPI_ISL_608665, EPI_ISL_608667, EPI_ISL_608668, EPI_ISL_608669, EPI_ISL_608670, EPI_ISL_608672, EPI_ISL_608675, EPI_ISL_608676, EPI_ISL_608680, EPI_ISL_608684,
EPI_ISL_608685, EPI_ISL_608686, EPI_ISL_608690, EPI_ISL_608692, EPI_ISL_608694, EPI_ISL_608847, EPI_ISL_608849, EPI_ISL_608851, EPI_ISL_608853, EPI_ISL_608855, EPI_ISL_608856, EPI_ISL_608858, EPI_ISL_608860, EPI_ISL_608862, EPI_ISL_608863, EPI_ISL_608865, EPI_ISL_608866, EPI_ISL_608869,
EPI_ISL_608874, EPI_ISL_608877, EPI_ISL_608878, EPI_ISL_608880, EPI_ISL_608883, EPI_ISL_608884, EPI_ISL_608885, EPI_ISL_608886, EPI_ISL_608887, EPI_ISL_608888, EPI_ISL_608890, EPI_ISL_608895, EPI_ISL_608897, EPI_ISL_608898, EPI_ISL_608899, EPI_ISL_608900, EPI_ISL_608911, EPI_ISL_608913,
EPI_ISL_608914, EPI_ISL_608916, EPI_ISL_608917, EPI_ISL_608919, EPI_ISL_608921, EPI_ISL_608922, EPI_ISL_608925, EPI_ISL_608926, EPI_ISL_608928, EPI_ISL_608929, EPI_ISL_608930, EPI_ISL_608931, EPI_ISL_608933, EPI_ISL_608934, EPI_ISL_608938, EPI_ISL_608941, EPI_ISL_608942, EPI_ISL_608944,
EPI_ISL_608945, EPI_ISL_608946, EPI_ISL_608948, EPI_ISL_608950, EPI_ISL_608952, EPI_ISL_608956, EPI_ISL_608957, EPI_ISL_608961, EPI_ISL_608962, EPI_ISL_608963, EPI_ISL_608964, EPI_ISL_608966, EPI_ISL_608970, EPI_ISL_608971, EPI_ISL_608972, EPI_ISL_608976, EPI_ISL_608983, EPI_ISL_608984,
EPI_ISL_608988, EPI_ISL_608991, EPI_ISL_608992, EPI_ISL_608993, EPI_ISL_608995, EPI_ISL_608998, EPI_ISL_608999, EPI_ISL_609000, EPI_ISL_609004, EPI_ISL_609007, EPI_ISL_609009, EPI_ISL_609010, EPI_ISL_609011, EPI_ISL_609012, EPI_ISL_609013, EPI_ISL_609016, EPI_ISL_609022, EPI_ISL_609028,
EPI_ISL_609031, EPI_ISL_609032, EPI_ISL_609034, EPI_ISL_609035, EPI_ISL_609039, EPI_ISL_609040, EPI_ISL_609041, EPI_ISL_609043, EPI_ISL_609047, EPI_ISL_609051, EPI_ISL_609052, EPI_ISL_609053, EPI_ISL_609056, EPI_ISL_609057, EPI_ISL_609060, EPI_ISL_609061, EPI_ISL_609064, EPI_ISL_609065,
EPI_ISL_609068, EPI_ISL_609069, EPI_ISL_609070, EPI_ISL_609075, EPI_ISL_609078, EPI_ISL_609079, EPI_ISL_609080, EPI_ISL_609083, EPI_ISL_609085, EPI_ISL_609087, EPI_ISL_609089, EPI_ISL_609090, EPI_ISL_609092, EPI_ISL_609093, EPI_ISL_609094, EPI_ISL_609095, EPI_ISL_609096, EPI_ISL_609098,
EPI_ISL_609102, EPI_ISL_609103, EPI_ISL_609104, EPI_ISL_609105, EPI_ISL_609106, EPI_ISL_609109, EPI_ISL_609120, EPI_ISL_609124, EPI_ISL_609125, EPI_ISL_609127, EPI_ISL_609130, EPI_ISL_609132, EPI_ISL_609133, EPI_ISL_609134, EPI_ISL_609135, EPI_ISL_609144, EPI_ISL_609146, EPI_ISL_609147,
EPI_ISL_609152, EPI_ISL_609153, EPI_ISL_609154, EPI_ISL_609157, EPI_ISL_609207, EPI_ISL_609208, EPI_ISL_609210, EPI_ISL_609211, EPI_ISL_609212, EPI_ISL_609213, EPI_ISL_609215, EPI_ISL_609216, EPI_ISL_609217, EPI_ISL_609218, EPI_ISL_610289, EPI_ISL_610291, EPI_ISL_610296, EPI_ISL_610299,
EPI_ISL_610301, EPI_ISL_610302, EPI_ISL_610307, EPI_ISL_610312, EPI_ISL_610313, EPI_ISL_610318, EPI_ISL_610325, EPI_ISL_610326, EPI_ISL_610328, EPI_ISL_610330, EPI_ISL_610333, EPI_ISL_610335, EPI_ISL_610338, EPI_ISL_610340, EPI_ISL_610342, EPI_ISL_610345, EPI_ISL_610347, EPI_ISL_610351,
EPI_ISL_610352, EPI_ISL_610357, EPI_ISL_610358, EPI_ISL_610360, EPI_ISL_610362, EPI_ISL_610365, EPI_ISL_610366, EPI_ISL_610371, EPI_ISL_610376, EPI_ISL_610377, EPI_ISL_610379, EPI_ISL_610380, EPI_ISL_610386, EPI_ISL_610388, EPI_ISL_610390, EPI_ISL_610391, EPI_ISL_610394, EPI_ISL_610396,
EPI_ISL_610404, EPI_ISL_610406, EPI_ISL_610412, EPI_ISL_610413, EPI_ISL_610414, EPI_ISL_610418, EPI_ISL_610420, EPI_ISL_610422, EPI_ISL_610424, EPI_ISL_610425, EPI_ISL_610432, EPI_ISL_610433, EPI_ISL_610436, EPI_ISL_610438, EPI_ISL_610439, EPI_ISL_610440, EPI_ISL_610441, EPI_ISL_610442,
EPI_ISL_610450, EPI_ISL_610455, EPI_ISL_610457, EPI_ISL_610458, EPI_ISL_610460, EPI_ISL_610461, EPI_ISL_610466, EPI_ISL_610471, EPI_ISL_610477, EPI_ISL_610479, EPI_ISL_610480, EPI_ISL_610481, EPI_ISL_610482, EPI_ISL_610484, EPI_ISL_610485, EPI_ISL_610490, EPI_ISL_610491, EPI_ISL_610496,
EPI_ISL_610498, EPI_ISL_610499, EPI_ISL_610505, EPI_ISL_610506, EPI_ISL_610514, EPI_ISL_610515, EPI_ISL_610517, EPI_ISL_610518, EPI_ISL_610520, EPI_ISL_610521, EPI_ISL_610523, EPI_ISL_610524, EPI_ISL_610525, EPI_ISL_610528, EPI_ISL_610529, EPI_ISL_610533, EPI_ISL_610539, EPI_ISL_610541,
EPI_ISL_610542, EPI_ISL_610544, EPI_ISL_610551, EPI_ISL_610552, EPI_ISL_610553, EPI_ISL_610554, EPI_ISL_610562, EPI_ISL_610565, EPI_ISL_610569, EPI_ISL_610573, EPI_ISL_610575, EPI_ISL_610578, EPI_ISL_610579, EPI_ISL_610580, EPI_ISL_610582, EPI_ISL_610586, EPI_ISL_610589, EPI_ISL_610594,
EPI_ISL_610599, EPI_ISL_610600, EPI_ISL_610602, EPI_ISL_610604, EPI_ISL_610607, EPI_ISL_610609, EPI_ISL_610611, EPI_ISL_610615, EPI_ISL_610618, EPI_ISL_610620, EPI_ISL_610622, EPI_ISL_610623, EPI_ISL_610625, EPI_ISL_610626

| | | | |
|---|---|---|---|
| see above | Lighthouse Lab in Alderley Park | Wellcome Sanger Institute for the COVID-19 Genomics UK (COG-UK) consortium | Jacquelyn Wynn, Mairead Hyland, The Lighthouse Lab in Alderley Park and Alex Alderton, Roberto Amato, Sonia Goncalves, Ewan Harrison, David K. Jackson, Ian Johnston, Dominic Kwiatkowski, Cordelia Langford, John Sillitoe on behalf of the Wellcome Sanger Institute COVID-19 Surveillance Team |

| | | | |
|---|---|---|---|
| EPI_ISL_610942, EPI_ISL_610961, EPI_ISL_610968, EPI_ISL_610996, EPI_ISL_610998, EPI_ISL_611024, EPI_ISL_611030, EPI_ISL_611038, EPI_ISL_611060, EPI_ISL_611061, EPI_ISL_611064, EPI_ISL_611065, EPI_ISL_611073, EPI_ISL_611109, EPI_ISL_611110, EPI_ISL_611111, EPI_ISL_611130, EPI_ISL_611173, EPI_ISL_611180, EPI_ISL_611186, EPI_ISL_611202, EPI_ISL_611211, EPI_ISL_611216, EPI_ISL_611231 | | | |
| see above | Lighthouse Lab in Alderley Park | Wellcome Sanger Institute for the COVID-19 Genomics UK (COG-UK) consortium | Jacquelyn Wynn, Mairead Hyland, The Lighthouse Lab in Alderley Park and Alex Alderton, Roberto Amato, Sonia Goncalves, Ewan Harrison, David K. Jackson, Ian Johnston, Dominic Kwiatkowski, Cordelia Langford, John Sillitoe on behalf of the Wellcome Sanger Institute COVID-19 Surveillance Team (http://www.sanger.ac.uk/covid-team) |
| EPI_ISL_614294 | Maimonides Medical Center | New York City Public Health Laboratory | Jade Wang, et al. |
| EPI_ISL_623871, EPI_ISL_623877, EPI_ISL_623879, EPI_ISL_623885, EPI_ISL_623896, EPI_ISL_623905, EPI_ISL_623919, EPI_ISL_623925, EPI_ISL_623948, EPI_ISL_623951, EPI_ISL_623953, EPI_ISL_623964, EPI_ISL_623970, EPI_ISL_623973, EPI_ISL_623989, EPI_ISL_623998, EPI_ISL_624006, EPI_ISL_624009, EPI_ISL_624017, EPI_ISL_624021, EPI_ISL_624037, EPI_ISL_624039, EPI_ISL_624048, EPI_ISL_624053, EPI_ISL_624056, EPI_ISL_624059, EPI_ISL_624062, EPI_ISL_624071, EPI_ISL_624093, EPI_ISL_624097, EPI_ISL_624100, EPI_ISL_624107, EPI_ISL_624111, EPI_ISL_624119, EPI_ISL_624125, EPI_ISL_624134, EPI_ISL_624135, EPI_ISL_624141, EPI_ISL_624143, EPI_ISL_624146, EPI_ISL_624153, EPI_ISL_624154, EPI_ISL_624157, EPI_ISL_624178, EPI_ISL_624192, EPI_ISL_624193, EPI_ISL_624199, EPI_ISL_624205, EPI_ISL_624206, EPI_ISL_624208, EPI_ISL_624217, EPI_ISL_624218, EPI_ISL_624220, EPI_ISL_624221, EPI_ISL_624226, EPI_ISL_624227, EPI_ISL_624228, EPI_ISL_624230, EPI_ISL_624231, EPI_ISL_624232, EPI_ISL_624234, EPI_ISL_624235, EPI_ISL_624236, EPI_ISL_624238, EPI_ISL_624239, EPI_ISL_624241, EPI_ISL_624251, EPI_ISL_624255, EPI_ISL_624256, EPI_ISL_624257, EPI_ISL_624258, EPI_ISL_624264, EPI_ISL_624266, EPI_ISL_624268, EPI_ISL_624272, EPI_ISL_624277, EPI_ISL_624280, EPI_ISL_624281, EPI_ISL_624284, EPI_ISL_624286, EPI_ISL_624287, EPI_ISL_624288, EPI_ISL_624289, EPI_ISL_624292, EPI_ISL_624293, EPI_ISL_624294, EPI_ISL_624295, EPI_ISL_624296, EPI_ISL_624299, EPI_ISL_624301, EPI_ISL_624309, EPI_ISL_624310, EPI_ISL_624311, EPI_ISL_624313, EPI_ISL_624315, EPI_ISL_624316, EPI_ISL_624317, EPI_ISL_624319, EPI_ISL_624320, EPI_ISL_624323, EPI_ISL_624326, EPI_ISL_624327, EPI_ISL_624329, EPI_ISL_624331, EPI_ISL_624333, EPI_ISL_624334, EPI_ISL_624337, EPI_ISL_624338, EPI_ISL_624340, EPI_ISL_624341, EPI_ISL_624343, EPI_ISL_624344, EPI_ISL_624346, EPI_ISL_624347, EPI_ISL_624349, EPI_ISL_624352, EPI_ISL_624357, EPI_ISL_624358, EPI_ISL_624359, EPI_ISL_624362, EPI_ISL_624364, EPI_ISL_624365, EPI_ISL_624370, EPI_ISL_624372, EPI_ISL_624375, EPI_ISL_624377, EPI_ISL_624380, EPI_ISL_624381, EPI_ISL_624385, EPI_ISL_624386, EPI_ISL_624387, EPI_ISL_624394, EPI_ISL_624397, EPI_ISL_624400, EPI_ISL_624402, EPI_ISL_624403, EPI_ISL_624405, EPI_ISL_624407, EPI_ISL_624409, EPI_ISL_624410, EPI_ISL_624411, EPI_ISL_624413, EPI_ISL_624416, EPI_ISL_624419, EPI_ISL_624420, EPI_ISL_624423, EPI_ISL_624424, EPI_ISL_624425, EPI_ISL_624426, EPI_ISL_624427, EPI_ISL_624431, EPI_ISL_624432, EPI_ISL_624434, EPI_ISL_624438, EPI_ISL_624439, EPI_ISL_624440, EPI_ISL_624441, EPI_ISL_624442, EPI_ISL_624443, EPI_ISL_624444, EPI_ISL_624445, EPI_ISL_624446, EPI_ISL_624447, EPI_ISL_624450, EPI_ISL_624451, EPI_ISL_624453, EPI_ISL_624460, EPI_ISL_624462, EPI_ISL_624463, EPI_ISL_624465, EPI_ISL_624471, EPI_ISL_624472, EPI_ISL_624473, EPI_ISL_624474, EPI_ISL_624476, EPI_ISL_624478, EPI_ISL_624479, EPI_ISL_624483, EPI_ISL_624484, EPI_ISL_624488, EPI_ISL_624489, EPI_ISL_624490, EPI_ISL_624492, EPI_ISL_624493, EPI_ISL_624497, EPI_ISL_624498, EPI_ISL_624500, EPI_ISL_624501, EPI_ISL_624502, EPI_ISL_624504, EPI_ISL_624505, EPI_ISL_624506, EPI_ISL_624509, EPI_ISL_624510, EPI_ISL_624515, EPI_ISL_624517, EPI_ISL_624518, EPI_ISL_624519, EPI_ISL_624521, EPI_ISL_624522, EPI_ISL_624523, EPI_ISL_624524, EPI_ISL_624526, EPI_ISL_624528, EPI_ISL_624529, EPI_ISL_624531, EPI_ISL_624533, EPI_ISL_624534, EPI_ISL_624535, EPI_ISL_624539, EPI_ISL_624540, EPI_ISL_624543, EPI_ISL_624544, EPI_ISL_624547, EPI_ISL_624548, EPI_ISL_624549, EPI_ISL_624550, EPI_ISL_624554, EPI_ISL_624556, EPI_ISL_624558, EPI_ISL_624559, EPI_ISL_624565, EPI_ISL_624567, EPI_ISL_624568, EPI_ISL_624569, EPI_ISL_624573, EPI_ISL_624574, EPI_ISL_624576, EPI_ISL_624578, EPI_ISL_624581, EPI_ISL_624582, EPI_ISL_624584, EPI_ISL_624588, EPI_ISL_624591, EPI_ISL_624594, EPI_ISL_624596, EPI_ISL_624602, EPI_ISL_624603, EPI_ISL_624604, EPI_ISL_624606, EPI_ISL_624607, EPI_ISL_624611, EPI_ISL_624613, EPI_ISL_624617, EPI_ISL_624618, EPI_ISL_624620, EPI_ISL_624622, EPI_ISL_624626, EPI_ISL_624627, EPI_ISL_624629, EPI_ISL_624635, EPI_ISL_624636, EPI_ISL_624639, EPI_ISL_624641, EPI_ISL_624642, EPI_ISL_624645, EPI_ISL_624647, EPI_ISL_624649, EPI_ISL_624650, EPI_ISL_624651, EPI_ISL_624652, EPI_ISL_624653, EPI_ISL_624662, EPI_ISL_624664, EPI_ISL_624674, EPI_ISL_624675, EPI_ISL_624676, EPI_ISL_624681, EPI_ISL_624684, EPI_ISL_624686, EPI_ISL_624687, EPI_ISL_624691, EPI_ISL_624692, EPI_ISL_624693, EPI_ISL_624694, EPI_ISL_624695, EPI_ISL_624696, EPI_ISL_624697, EPI_ISL_624702, EPI_ISL_624704, EPI_ISL_624705, EPI_ISL_624706, EPI_ISL_624707, EPI_ISL_624708, EPI_ISL_624713, EPI_ISL_624715, EPI_ISL_624718, EPI_ISL_624726, EPI_ISL_624727, EPI_ISL_624728, EPI_ISL_624731, EPI_ISL_624732, EPI_ISL_624734, EPI_ISL_624735, EPI_ISL_624737, EPI_ISL_624738, EPI_ISL_624739, EPI_ISL_624745, EPI_ISL_624753, EPI_ISL_624754, EPI_ISL_624757, EPI_ISL_624762, EPI_ISL_624763, EPI_ISL_624770, EPI_ISL_624772, EPI_ISL_624773, EPI_ISL_624776, EPI_ISL_624777, EPI_ISL_624778, EPI_ISL_624780, EPI_ISL_624781, EPI_ISL_624784, EPI_ISL_624790, EPI_ISL_624791, EPI_ISL_624794, EPI_ISL_624795, EPI_ISL_624796, EPI_ISL_624800, EPI_ISL_624804, EPI_ISL_624805, EPI_ISL_624812, EPI_ISL_624818, EPI_ISL_624824, EPI_ISL_624826, EPI_ISL_624829, EPI_ISL_624831, EPI_ISL_624835, EPI_ISL_624836, EPI_ISL_624840, EPI_ISL_624841, EPI_ISL_624844, EPI_ISL_624847, EPI_ISL_624848, EPI_ISL_624849, EPI_ISL_624858, EPI_ISL_624859, EPI_ISL_624861, EPI_ISL_624867, EPI_ISL_624871, EPI_ISL_624873, EPI_ISL_624879, EPI_ISL_624881, EPI_ISL_624882, EPI_ISL_624883, EPI_ISL_624887, EPI_ISL_624889, EPI_ISL_624890, EPI_ISL_624895, EPI_ISL_624896, EPI_ISL_624901, EPI_ISL_624903, EPI_ISL_624904, EPI_ISL_624906, EPI_ISL_624913, EPI_ISL_624916, EPI_ISL_624918, EPI_ISL_624922, EPI_ISL_624923, EPI_ISL_624925, EPI_ISL_624929, EPI_ISL_624930, EPI_ISL_624931, EPI_ISL_624937, EPI_ISL_624938, EPI_ISL_624939, EPI_ISL_624941, EPI_ISL_624944, EPI_ISL_624948, EPI_ISL_624954, EPI_ISL_624963, EPI_ISL_624965, EPI_ISL_624969, EPI_ISL_624972, EPI_ISL_624978, EPI_ISL_624981, EPI_ISL_624984, EPI_ISL_624989, EPI_ISL_624996, EPI_ISL_624998, EPI_ISL_625000, EPI_ISL_625002, EPI_ISL_625006, EPI_ISL_625012, EPI_ISL_625013, EPI_ISL_625015, EPI_ISL_625019, EPI_ISL_625023, EPI_ISL_625024, EPI_ISL_625025, EPI_ISL_625026, EPI_ISL_625031, EPI_ISL_625032, EPI_ISL_625033, EPI_ISL_625038, EPI_ISL_625041, EPI_ISL_625042, EPI_ISL_625044, EPI_ISL_625057, EPI_ISL_625058, EPI_ISL_625060, EPI_ISL_625062, EPI_ISL_625064, EPI_ISL_625065, EPI_ISL_625066, EPI_ISL_625068, EPI_ISL_625076, EPI_ISL_625077, EPI_ISL_625079, EPI_ISL_625080, EPI_ISL_625081, EPI_ISL_625083, EPI_ISL_625085, EPI_ISL_625093, EPI_ISL_625094, EPI_ISL_625097, EPI_ISL_625102, EPI_ISL_625105, EPI_ISL_625106, EPI_ISL_625113, EPI_ISL_625114, EPI_ISL_625116, EPI_ISL_625117, EPI_ISL_625119, EPI_ISL_625122, EPI_ISL_625124, EPI_ISL_625125, EPI_ISL_625126, EPI_ISL_625127, EPI_ISL_625133, EPI_ISL_625138, EPI_ISL_625143, EPI_ISL_625145, EPI_ISL_625146, EPI_ISL_625149, EPI_ISL_625150, EPI_ISL_625153, EPI_ISL_625155, EPI_ISL_625157, EPI_ISL_625160, EPI_ISL_625163, EPI_ISL_625164, EPI_ISL_625166, EPI_ISL_625167, EPI_ISL_625172, EPI_ISL_625173, EPI_ISL_625176, EPI_ISL_625177, EPI_ISL_625178, EPI_ISL_625182, EPI_ISL_625183, EPI_ISL_625184, EPI_ISL_625185, EPI_ISL_625186, EPI_ISL_625189, EPI_ISL_625190, EPI_ISL_625192, EPI_ISL_625194, EPI_ISL_625199, EPI_ISL_625201, EPI_ISL_625203, EPI_ISL_625207, EPI_ISL_625210, EPI_ISL_625212, EPI_ISL_625215, EPI_ISL_625217, EPI_ISL_625221, EPI_ISL_625227, EPI_ISL_625230, EPI_ISL_625233, EPI_ISL_625234, EPI_ISL_625237, EPI_ISL_625242, EPI_ISL_625243, EPI_ISL_625244, EPI_ISL_625248, EPI_ISL_625249, EPI_ISL_625252, EPI_ISL_625255, EPI_ISL_625257, EPI_ISL_625259, EPI_ISL_625260, EPI_ISL_625265, EPI_ISL_625266, EPI_ISL_625267, EPI_ISL_625268, EPI_ISL_625269, EPI_ISL_625270, EPI_ISL_625271, EPI_ISL_625272, EPI_ISL_625273, EPI_ISL_625274, EPI_ISL_625275, EPI_ISL_625276, EPI_ISL_625277, EPI_ISL_625278, EPI_ISL_625279, EPI_ISL_625280, EPI_ISL_625396, EPI_ISL_625397, EPI_ISL_625398, EPI_ISL_625399, EPI_ISL_625400, EPI_ISL_625403, EPI_ISL_625404, EPI_ISL_625422, EPI_ISL_625425, EPI_ISL_625427, EPI_ISL_625429, EPI_ISL_625431, EPI_ISL_625433, EPI_ISL_625434, EPI_ISL_625435, EPI_ISL_625436 | | | |
| see above | Lighthouse Lab in Alderley Park | Wellcome Sanger Institute for the COVID-19 Genomics UK (COG-UK) consortium | Jacquelyn Wynn, Mairead Hyland, The Lighthouse Lab in Alderley Park and Alex Alderton, Roberto Amato, Sonia Goncalves, Ewan Harrison, David K. Jackson, Ian Johnston, Dominic Kwiatkowski, Cordelia Langford, John Sillitoe on behalf of the Wellcome Sanger Institute COVID-19 Surveillance Team (http://www.sanger.ac.uk/covid-team) |
| EPI_ISL_625673, EPI_ISL_625674, EPI_ISL_625675, EPI_ISL_625676, EPI_ISL_625677, EPI_ISL_625678, EPI_ISL_625679, EPI_ISL_625680, EPI_ISL_625681, EPI_ISL_625682 | Laboratory of Molecular Medicine, University of Magallanes | Centro Asistencial Docente y de Investigacion, Universidad de Magallanes | Jorge Gonzalez, Jacqueline Aldridge, Diego Alvarez, Marcelo Navarrete |
| EPI_ISL_626231, EPI_ISL_626232, EPI_ISL_626233, EPI_ISL_626234, EPI_ISL_626235, EPI_ISL_626236, EPI_ISL_626237, EPI_ISL_626238, EPI_ISL_626239, EPI_ISL_626240, EPI_ISL_626241, EPI_ISL_626242, EPI_ISL_626243, EPI_ISL_626244, EPI_ISL_626245, EPI_ISL_626246, EPI_ISL_626247, EPI_ISL_626248, EPI_ISL_626249, EPI_ISL_626250, EPI_ISL_626251, EPI_ISL_626252, EPI_ISL_626253, EPI_ISL_626254, EPI_ISL_626255, EPI_ISL_626256, EPI_ISL_626257, EPI_ISL_626258, EPI_ISL_626259, EPI_ISL_626260, EPI_ISL_626261, EPI_ISL_626262, EPI_ISL_626263, EPI_ISL_626264, EPI_ISL_626265, EPI_ISL_626266, EPI_ISL_626267, EPI_ISL_626268, EPI_ISL_626269, EPI_ISL_626270, EPI_ISL_626271, EPI_ISL_626272, EPI_ISL_626273, EPI_ISL_626274, EPI_ISL_626275, EPI_ISL_626276, EPI_ISL_626277, EPI_ISL_626278, EPI_ISL_626279, EPI_ISL_626280, EPI_ISL_626281, EPI_ISL_626282, EPI_ISL_626283, EPI_ISL_626284, EPI_ISL_626285, EPI_ISL_626286, EPI_ISL_626287, EPI_ISL_626288, EPI_ISL_626289, EPI_ISL_626290, EPI_ISL_626291, EPI_ISL_626292, EPI_ISL_626293, EPI_ISL_626294, EPI_ISL_626295, EPI_ISL_626296, EPI_ISL_626297, EPI_ISL_626298, EPI_ISL_626299, EPI_ISL_626300, EPI_ISL_626301, EPI_ISL_626302, EPI_ISL_626303, EPI_ISL_626304, EPI_ISL_626305, EPI_ISL_626306, EPI_ISL_626307, EPI_ISL_626308, EPI_ISL_626309, EPI_ISL_626310, EPI_ISL_626311, EPI_ISL_626312, EPI_ISL_626313, EPI_ISL_626314, EPI_ISL_626315, EPI_ISL_626316, EPI_ISL_626317, EPI_ISL_626318, EPI_ISL_626319, EPI_ISL_626320, EPI_ISL_626321, EPI_ISL_626322, EPI_ISL_626323, EPI_ISL_626324, EPI_ISL_626325, EPI_ISL_626326, EPI_ISL_626327, EPI_ISL_626328, EPI_ISL_626329, EPI_ISL_626330, EPI_ISL_626331, EPI_ISL_626332, EPI_ISL_626333, EPI_ISL_626334, EPI_ISL_626335, EPI_ISL_626336, EPI_ISL_626337, EPI_ISL_626338, EPI_ISL_626339 | | | |
| see above | Department of Clinical Microbiology | GIGA Medical Genomics | Keith Durkin, Maria Artesi, Sébastien Bontems, Raphaël Boreux, Bouchra Boujemla, Cécile Meex, Pierrette Melin, Marie-Pierre Hayette, Vincent Bours |
| EPI_ISL_628750, EPI_ISL_628751, EPI_ISL_628752, EPI_ISL_628754, EPI_ISL_628755, EPI_ISL_628756, EPI_ISL_628758, EPI_ISL_628760, EPI_ISL_628761 | UHAS COVID-19 Lab | UHAS COVID-19 Lab | Kwabena O. Duedu, Jones Gyamfi, Reuben Ayivor-Djanie, John O. Gyapong and the UHAS COVID-19 Lab Team |
| EPI_ISL_630352, EPI_ISL_630372, EPI_ISL_630380, EPI_ISL_630381, EPI_ISL_630382, EPI_ISL_630385, EPI_ISL_630392, EPI_ISL_630402, EPI_ISL_630406, EPI_ISL_630409, EPI_ISL_630416, EPI_ISL_630424, EPI_ISL_630425, EPI_ISL_630436, EPI_ISL_630449, EPI_ISL_630455, EPI_ISL_630469, EPI_ISL_630476, EPI_ISL_630483, EPI_ISL_630487, EPI_ISL_630492, EPI_ISL_630495, EPI_ISL_630506, EPI_ISL_630521, EPI_ISL_630523, EPI_ISL_630528, EPI_ISL_630531, EPI_ISL_630545, EPI_ISL_630551, EPI_ISL_630556, EPI_ISL_630565, EPI_ISL_630576, EPI_ISL_630579, EPI_ISL_630590, EPI_ISL_630598, EPI_ISL_630613, EPI_ISL_630617, EPI_ISL_630633, EPI_ISL_630636, EPI_ISL_630640, EPI_ISL_630654, EPI_ISL_630657, EPI_ISL_630684, EPI_ISL_630706, EPI_ISL_630707, EPI_ISL_630708, EPI_ISL_630714, EPI_ISL_630723, EPI_ISL_630724, EPI_ISL_630726, EPI_ISL_630729, EPI_ISL_630735, EPI_ISL_630736, EPI_ISL_630737, EPI_ISL_630742, EPI_ISL_630744, EPI_ISL_630747, EPI_ISL_630748, EPI_ISL_630753, EPI_ISL_630754, EPI_ISL_630755, EPI_ISL_630759, EPI_ISL_630762, EPI_ISL_630766, EPI_ISL_630767, EPI_ISL_630779, EPI_ISL_630781, EPI_ISL_630784, EPI_ISL_630785, EPI_ISL_630788, EPI_ISL_630790, EPI_ISL_630793, EPI_ISL_630796, EPI_ISL_630798, EPI_ISL_630800, EPI_ISL_630803, EPI_ISL_630807, EPI_ISL_630808, EPI_ISL_630809, EPI_ISL_630810, EPI_ISL_630812, EPI_ISL_630816, EPI_ISL_630820, EPI_ISL_630822, EPI_ISL_630825, EPI_ISL_630828, EPI_ISL_630829, EPI_ISL_630830, EPI_ISL_630832, EPI_ISL_630839, EPI_ISL_630840, EPI_ISL_630844, EPI_ISL_630845, EPI_ISL_630847, EPI_ISL_630848, EPI_ISL_630849, EPI_ISL_630855, EPI_ISL_630856, EPI_ISL_630857, EPI_ISL_630858, EPI_ISL_630861, EPI_ISL_630863, EPI_ISL_630866, EPI_ISL_630867, EPI_ISL_630869, EPI_ISL_630877, EPI_ISL_630879, EPI_ISL_630880, EPI_ISL_630882, EPI_ISL_630893, EPI_ISL_630894, EPI_ISL_630896, EPI_ISL_630899, EPI_ISL_630901, EPI_ISL_630902, EPI_ISL_630904, EPI_ISL_630905, EPI_ISL_630908, EPI_ISL_630911, EPI_ISL_630912, EPI_ISL_630913, EPI_ISL_630914, EPI_ISL_630917, EPI_ISL_630918, EPI_ISL_630921, EPI_ISL_630925, EPI_ISL_630927, EPI_ISL_630929, EPI_ISL_630932, EPI_ISL_630934, EPI_ISL_630939, EPI_ISL_630940, EPI_ISL_630942, EPI_ISL_630943, EPI_ISL_630944, EPI_ISL_630945, EPI_ISL_630946, EPI_ISL_630950, EPI_ISL_630952, EPI_ISL_630954, EPI_ISL_630956, EPI_ISL_630958, EPI_ISL_630960, EPI_ISL_630963, EPI_ISL_630966, EPI_ISL_630967, EPI_ISL_630969, EPI_ISL_630971, EPI_ISL_630975, EPI_ISL_630976, EPI_ISL_630982, EPI_ISL_630983, EPI_ISL_630985, EPI_ISL_630992, EPI_ISL_630994, EPI_ISL_630999, EPI_ISL_631011, EPI_ISL_631012, EPI_ISL_631014, EPI_ISL_631018, EPI_ISL_631023, EPI_ISL_631027, EPI_ISL_631031, EPI_ISL_631033, EPI_ISL_631035, EPI_ISL_631041, EPI_ISL_631044, EPI_ISL_631046, EPI_ISL_631050, EPI_ISL_631056, EPI_ISL_631057, EPI_ISL_631059, EPI_ISL_631060, EPI_ISL_631074, EPI_ISL_631081, EPI_ISL_631084, EPI_ISL_631089, EPI_ISL_631093, EPI_ISL_631096, EPI_ISL_631099, EPI_ISL_631107, EPI_ISL_631111, EPI_ISL_631113, EPI_ISL_631116, EPI_ISL_631127, EPI_ISL_631128, EPI_ISL_631131, EPI_ISL_631133, EPI_ISL_631150, EPI_ISL_631165, EPI_ISL_631173, EPI_ISL_631184, EPI_ISL_631189, EPI_ISL_631190, EPI_ISL_631202, EPI_ISL_631212, EPI_ISL_631215, EPI_ISL_631217, EPI_ISL_631222, EPI_ISL_631228, EPI_ISL_631231, EPI_ISL_631241, EPI_ISL_631250, EPI_ISL_631265, EPI_ISL_631269 | | | |
| see above | Lighthouse Lab in Alderley Park | Wellcome Sanger Institute for the COVID-19 Genomics UK (COG-UK) consortium | Jacquelyn Wynn, Mairead Hyland, The Lighthouse Lab in Alderley Park and Alex Alderton, Roberto Amato, Sonia Goncalves, Ewan Harrison, David K. Jackson, Ian Johnston, Dominic Kwiatkowski, Cordelia Langford, John Sillitoe on behalf of the Wellcome Sanger Institute COVID-19 Surveillance Team |
| EPI_ISL_631389, EPI_ISL_631390, EPI_ISL_631391, EPI_ISL_631392, EPI_ISL_631393, EPI_ISL_631394, EPI_ISL_631395, EPI_ISL_631396, EPI_ISL_631397, EPI_ISL_631398, EPI_ISL_631399, EPI_ISL_631400, EPI_ISL_631401, EPI_ISL_631402, EPI_ISL_631403, EPI_ISL_631404, EPI_ISL_631405, EPI_ISL_631406, EPI_ISL_631407, EPI_ISL_631408, EPI_ISL_631409, EPI_ISL_631410, EPI_ISL_631411, EPI_ISL_631412, EPI_ISL_631413, EPI_ISL_631414, EPI_ISL_631415, EPI_ISL_631416, EPI_ISL_631417, EPI_ISL_631418, EPI_ISL_631419, EPI_ISL_631420, EPI_ISL_631421, EPI_ISL_631422, EPI_ISL_631423, EPI_ISL_631424, EPI_ISL_631425, EPI_ISL_631426, EPI_ISL_631427, EPI_ISL_631428, EPI_ISL_631429, EPI_ISL_631430, EPI_ISL_631431, EPI_ISL_631432, EPI_ISL_631433, EPI_ISL_631434, EPI_ISL_631435, EPI_ISL_631436, EPI_ISL_631437, EPI_ISL_631438, EPI_ISL_631439, EPI_ISL_631440, EPI_ISL_631441, EPI_ISL_631442, EPI_ISL_631443, EPI_ISL_631444, EPI_ISL_631445, EPI_ISL_631446, EPI_ISL_631447, EPI_ISL_631448, EPI_ISL_631449, EPI_ISL_631450, EPI_ISL_631451, EPI_ISL_631452, EPI_ISL_631453, EPI_ISL_631454, EPI_ISL_631455, EPI_ISL_631456, EPI_ISL_631457, EPI_ISL_631458, EPI_ISL_631459, EPI_ISL_631460, EPI_ISL_631461, EPI_ISL_631462, EPI_ISL_631463, EPI_ISL_631464, EPI_ISL_631465, EPI_ISL_631466, EPI_ISL_631467, EPI_ISL_631468, EPI_ISL_631469, EPI_ISL_631470, EPI_ISL_631471, EPI_ISL_631472, EPI_ISL_631473, EPI_ISL_631474, EPI_ISL_631475, EPI_ISL_631476, EPI_ISL_631477, EPI_ISL_631478, EPI_ISL_631479, EPI_ISL_631480, EPI_ISL_631481, EPI_ISL_631482, EPI_ISL_631483, EPI_ISL_631484, EPI_ISL_631485, EPI_ISL_631486, EPI_ISL_631487, EPI_ISL_631488, EPI_ISL_631489, EPI_ISL_631490, EPI_ISL_631491, EPI_ISL_631492, EPI_ISL_631493, EPI_ISL_631494, EPI_ISL_631495, EPI_ISL_631496, EPI_ISL_631497, EPI_ISL_631498, EPI_ISL_631499, EPI_ISL_631500, EPI_ISL_631501 | | | |
| see above | Wisconsin State Laboratory of Hygiene Communicable Disease Division | Wisconsin State Laboratory of Hygiene Communicable Disease Division | Kelsey R. Florek, Abigail C. Shockey |
| EPI_ISL_631502, EPI_ISL_631503 | New York-Presbyterian-Columbia University Medical Center | New York City Public Health Laboratory | Jade Wang, et al. |

| | | | |
|---|---|---|---|
| EPI_ISL_631504 | Wyckoff Heights Medical Center | New York City Public Health Laboratory | Jade Wang, et al. |
| EPI_ISL_631505 | Northwell Health-GoHealth Urgent Care 10025 | New York City Public Health Laboratory | Jade Wang, et al. |
| EPI_ISL_631506 | NYC Department Of Health And Mental Hygiene | New York City Public Health Laboratory | Jade Wang, et al. |
| EPI_ISL_631507 | Mount Sinai West | New York City Public Health Laboratory | Jade Wang, et al. |
| EPI_ISL_631508 | Brooklyn Hospital Center | New York City Public Health Laboratory | Jade Wang, et al. |
| EPI_ISL_631509 | New York Presbyterian Lower Manhattan | New York City Public Health Laboratory | Jade Wang, et al. |
| EPI_ISL_631510, EPI_ISL_631511 | New York Presbyterian/ Weill Cornell Medical Center | New York City Public Health Laboratory | Jade Wang, et al. |
| EPI_ISL_631512 | Mount Sinai Hospital | New York City Public Health Laboratory | Jade Wang, et al. |
| EPI_ISL_631513 | Montefiore Medical Center | New York City Public Health Laboratory | Jade Wang, et al. |
| EPI_ISL_631514 | New York Community Hospital | New York City Public Health Laboratory | Jade Wang, et al. |
| EPI_ISL_631515 | NYU Langone Health | New York City Public Health Laboratory | Jade Wang, et al. |
| EPI_ISL_631516 | Jamaica Hospital Medical Center | New York City Public Health Laboratory | Jade Wang, et al. |
| EPI_ISL_631517, EPI_ISL_631518, EPI_ISL_631519 | New York Presbyterian-Brooklyn Methodist Hospital | New York City Public Health Laboratory | Jade Wang, et al. |
| EPI_ISL_631520 | NYC HH Lincoln Medical And Mental Health Center | New York City Public Health Laboratory | Jade Wang, et al. |
| EPI_ISL_631521 | Mount Sinai Hospital | New York City Public Health Laboratory | Jade Wang, et al. |
| EPI_ISL_631522 | Robert Kutnick | New York City Public Health Laboratory | Jade Wang, et al. |
| EPI_ISL_631523 | NYU Langone Health | New York City Public Health Laboratory | Jade Wang, et al. |
| EPI_ISL_631524 | Jamaica Hospital Medical Center | New York City Public Health Laboratory | Jade Wang, et al. |
| EPI_ISL_631525 | Mount Sinai Hospital | New York City Public Health Laboratory | Jade Wang, et al. |
| EPI_ISL_631526, EPI_ISL_631527 | NYU Langone Health | New York City Public Health Laboratory | Jade Wang, et al. |
| EPI_ISL_631528 | Maimonides Medical Center | New York City Public Health Laboratory | Jade Wang, et al. |
| EPI_ISL_631529 | St. Johns Episcopal Hospital | New York City Public Health Laboratory | Jade Wang, et al. |
| EPI_ISL_631530 | New York Presbyterian Queens | New York City Public Health Laboratory | Jade Wang, et al. |
| EPI_ISL_631531 | NYU Langone Health | New York City Public Health Laboratory | Jade Wang, et al. |
| EPI_ISL_631532 | Mount Sinai Hospital | New York City Public Health Laboratory | Jade Wang, et al. |
| EPI_ISL_631533 | New York Presbyterian Lower Manhattan | New York City Public Health Laboratory | Jade Wang, et al. |
| EPI_ISL_631534 | Mount Sinai Hospital | New York City Public Health Laboratory | Jade Wang, et al. |
| EPI_ISL_631535 | NYC Department Of Health And Mental Hygiene | New York City Public Health Laboratory | Jade Wang, et al. |
| EPI_ISL_631536 | New York Presbyterian Lower Manhattan | New York City Public Health Laboratory | Jade Wang, et al. |
| EPI_ISL_631537 | Mount Sinai Hospital | New York City Public Health Laboratory | Jade Wang, et al. |
| EPI_ISL_631538 | Maimonides Medical Center | New York City Public Health Laboratory | Jade Wang, et al. |
| EPI_ISL_631539, EPI_ISL_631540 | New York Presbyterian Lower Manhattan | New York City Public Health Laboratory | Jade Wang, et al. |
| EPI_ISL_631541 | NYC Department Of Health And Mental Hygiene | New York City Public Health Laboratory | Jade Wang, et al. |
| EPI_ISL_631542 | New York Presbyterian Lower Manhattan | New York City Public Health Laboratory | Jade Wang, et al. |
| EPI_ISL_631543 | Robert Kutnick | New York City Public Health Laboratory | Jade Wang, et al. |
| EPI_ISL_631544, EPI_ISL_631545 | New York Presbyterian-Brooklyn Methodist Hospital | New York City Public Health Laboratory | Jade Wang, et al. |
| EPI_ISL_631546 | Maimonides Medical Center | New York City Public Health Laboratory | Jade Wang, et al. |
| EPI_ISL_631547 | New York Presbyterian Queens | New York City Public Health Laboratory | Jade Wang, et al. |
| EPI_ISL_631548 | New York Presbyterian-Brooklyn Methodist Hospital | New York City Public Health Laboratory | Jade Wang, et al. |
| EPI_ISL_631549 | Mount Sinai Hospital | New York City Public Health Laboratory | Jade Wang, et al. |
| EPI_ISL_631550, EPI_ISL_631551 | Jamaica Hospital Medical Center | New York City Public Health Laboratory | Jade Wang, et al. |
| EPI_ISL_631552, EPI_ISL_631553 | Brookdale University Hospital Medical Center | New York City Public Health Laboratory | Jade Wang, et al. |
| EPI_ISL_631554 | Jamaica Hospital Medical Center | New York City Public Health Laboratory | Jade Wang, et al. |
| EPI_ISL_631555 | Richmond University Medical Center | New York City Public Health Laboratory | Jade Wang, et al. |
| EPI_ISL_631556 | NYC HH Jacobi Medical Center | New York City Public Health Laboratory | Jade Wang, et al. |
| EPI_ISL_631557 | NYC HH Lincoln Medical And Mental Health Center | New York City Public Health Laboratory | Jade Wang, et al. |
| EPI_ISL_631558, EPI_ISL_631559 | NYU Langone Health | New York City Public Health Laboratory | Jade Wang, et al. |
| EPI_ISL_631560, EPI_ISL_631561 | Montefiore Medical Center | New York City Public Health Laboratory | Jade Wang, et al. |
| EPI_ISL_631562 | Brookdale University Hospital Medical Center | New York City Public Health Laboratory | Jade Wang, et al. |
| EPI_ISL_631563 | Flushing Hospital Medical Center | New York City Public Health Laboratory | Jade Wang, et al. |
| EPI_ISL_631564 | NYC HH Elmhurst Hospital Medical Center | New York City Public Health Laboratory | Jade Wang, et al. |
| EPI_ISL_631565 | Brookdale University Hospital Medical Center | New York City Public Health Laboratory | Jade Wang, et al. |
| EPI_ISL_631566, EPI_ISL_631567 | New York Presbyterian-Brooklyn Methodist Hospital | New York City Public Health Laboratory | Jade Wang, et al. |
| EPI_ISL_631568, EPI_ISL_631569 | New York Presbyterian Queens | New York City Public Health Laboratory | Jade Wang, et al. |
| EPI_ISL_631570 | OCME Office Of Chief Medical Examiner | New York City Public Health Laboratory | Jade Wang, et al. |
| EPI_ISL_631571 | Jamaica Hospital Medical Center | New York City Public Health Laboratory | Jade Wang, et al. |
| EPI_ISL_631572 | NYC Department Of Health And Mental Hygiene | New York City Public Health Laboratory | Jade Wang, et al. |
| EPI_ISL_631573 | Brookdale University Hospital Medical Center | New York City Public Health Laboratory | Jade Wang, et al. |

| | | | |
|---|---|---|---|
| EPI_ISL_631574, EPI_ISL_631575 | Jamaica Hospital Medical Center | New York City Public Health Laboratory | Jade Wang, et al. |
| EPI_ISL_631576 | Richmond University Medical Center | New York City Public Health Laboratory | Jade Wang, et al. |
| EPI_ISL_631577 | NYC HH Lincoln Medical And Mental Health Center | New York City Public Health Laboratory | Jade Wang, et al. |
| EPI_ISL_631578 | Jamaica Hospital Medical Center | New York City Public Health Laboratory | Jade Wang, et al. |
| EPI_ISL_631579 | Flushing Hospital Medical Center | New York City Public Health Laboratory | Jade Wang, et al. |
| EPI_ISL_631580 | Bellevue Hospital Center | New York City Public Health Laboratory | Jade Wang, et al. |
| EPI_ISL_631581 | SUNY Downstate Medical Center | New York City Public Health Laboratory | Jade Wang, et al. |
| EPI_ISL_631582 | Jamaica Hospital Medical Center | New York City Public Health Laboratory | Jade Wang, et al. |
| EPI_ISL_631583 | Richmond University Medical Center | New York City Public Health Laboratory | Jade Wang, et al. |
| EPI_ISL_631584 | NYC HH Lincoln Medical And Mental Health Center | New York City Public Health Laboratory | Jade Wang, et al. |
| EPI_ISL_631585 | Jamaica Hospital Medical Center | New York City Public Health Laboratory | Jade Wang, et al. |
| EPI_ISL_631586 | Flushing Hospital Medical Center | New York City Public Health Laboratory | Jade Wang, et al. |
| EPI_ISL_631587 | NYC HH Lincoln Medical And Mental Health Center | New York City Public Health Laboratory | Jade Wang, et al. |
| EPI_ISL_631588 | SUNY Downstate Medical Center | New York City Public Health Laboratory | Jade Wang, et al. |
| EPI_ISL_631589 | Flushing Hospital Medical Center | New York City Public Health Laboratory | Jade Wang, et al. |
| EPI_ISL_631590 | Jamaica Hospital Medical Center | New York City Public Health Laboratory | Jade Wang, et al. |
| EPI_ISL_631591 | OCME Office Of Chief Medical Examiner | New York City Public Health Laboratory | Jade Wang, et al. |
| EPI_ISL_631592, EPI_ISL_631593 | Jamaica Hospital Medical Center | New York City Public Health Laboratory | Jade Wang, et al. |
| EPI_ISL_631594 | OCME Office Of Chief Medical Examiner | New York City Public Health Laboratory | Jade Wang, et al. |
| EPI_ISL_631595 | Jamaica Hospital Medical Center | New York City Public Health Laboratory | Jade Wang, et al. |
| EPI_ISL_631596 | James J. Peters VA Medical Center | New York City Public Health Laboratory | Jade Wang, et al. |
| EPI_ISL_631597 | Jamaica Hospital Medical Center | New York City Public Health Laboratory | Jade Wang, et al. |
| EPI_ISL_631598 | OCME Office Of Chief Medical Examiner | New York City Public Health Laboratory | Jade Wang, et al. |
| EPI_ISL_631599 | Jamaica Hospital Medical Center | New York City Public Health Laboratory | Jade Wang, et al. |
| EPI_ISL_631600 | James J. Peters VA Medical Center | New York City Public Health Laboratory | Jade Wang, et al. |
| EPI_ISL_631601 | Jamaica Hospital Medical Center | New York City Public Health Laboratory | Jade Wang, et al. |
| EPI_ISL_631602 | Flushing Hospital Medical Center | New York City Public Health Laboratory | Jade Wang, et al. |
| EPI_ISL_631603 | NYC HH Lincoln Medical And Mental Health Center | New York City Public Health Laboratory | Jade Wang, et al. |
| EPI_ISL_631604 | Jamaica Hospital Medical Center | New York City Public Health Laboratory | Jade Wang, et al. |
| EPI_ISL_631605 | OCME Office Of Chief Medical Examiner | New York City Public Health Laboratory | Jade Wang, et al. |
| EPI_ISL_631606 | Flushing Hospital Medical Center | New York City Public Health Laboratory | Jade Wang, et al. |
| EPI_ISL_631607 | NYC HH Lincoln Medical And Mental Health Center | New York City Public Health Laboratory | Jade Wang, et al. |
| EPI_ISL_631608, EPI_ISL_631609, EPI_ISL_631610 | Jamaica Hospital Medical Center | New York City Public Health Laboratory | Jade Wang, et al. |
| EPI_ISL_631611 | OCME Office Of Chief Medical Examiner | New York City Public Health Laboratory | Jade Wang, et al. |
| EPI_ISL_631612, EPI_ISL_631613 | Jamaica Hospital Medical Center | New York City Public Health Laboratory | Jade Wang, et al. |
| EPI_ISL_631614 | Flushing Hospital Medical Center | New York City Public Health Laboratory | Jade Wang, et al. |
| EPI_ISL_631615 | Jamaica Hospital Medical Center | New York City Public Health Laboratory | Jade Wang, et al. |
| EPI_ISL_631616 | Flushing Hospital Medical Center | New York City Public Health Laboratory | Jade Wang, et al. |
| EPI_ISL_631617, EPI_ISL_631618, EPI_ISL_631619 | OCME Office Of Chief Medical Examiner | New York City Public Health Laboratory | Jade Wang, et al. |
| EPI_ISL_631620 | Flushing Hospital Medical Center | New York City Public Health Laboratory | Jade Wang, et al. |
| EPI_ISL_631621 | Jamaica Hospital Medical Center | New York City Public Health Laboratory | Jade Wang, et al. |
| EPI_ISL_631622, EPI_ISL_631623, EPI_ISL_631624, EPI_ISL_631625, EPI_ISL_631626, EPI_ISL_631627, EPI_ISL_631628, EPI_ISL_631629, EPI_ISL_631630, EPI_ISL_631631, EPI_ISL_631632, EPI_ISL_631633 | | | |
| see above | Richmond University Medical Center | New York City Public Health Laboratory | Jade Wang, et al. |
| EPI_ISL_631634 | Jamaica Hospital Medical Center | New York City Public Health Laboratory | Jade Wang, et al. |
| EPI_ISL_631635 | OCME Office Of Chief Medical Examiner | New York City Public Health Laboratory | Jade Wang, et al. |
| EPI_ISL_631636, EPI_ISL_631637 | Jamaica Hospital Medical Center | New York City Public Health Laboratory | Jade Wang, et al. |
| EPI_ISL_631638 | Flushing Hospital Medical Center | New York City Public Health Laboratory | Jade Wang, et al. |
| EPI_ISL_631639, EPI_ISL_631685 | OCME Office Of Chief Medical Examiner | New York City Public Health Laboratory | Jade Wang, et al. |
| EPI_ISL_631686 | NYC HH Lincoln Medical And Mental Health Center | New York City Public Health Laboratory | Jade Wang, et al. |
| EPI_ISL_631687 | OCME Office Of Chief Medical Examiner | New York City Public Health Laboratory | Jade Wang, et al. |
| EPI_ISL_631688 | Wildlife Conservation Society - Bronx Zoo | New York City Public Health Laboratory | Jade Wang, et al. |
| EPI_ISL_631689, EPI_ISL_631690 | Lincoln Medical Center | New York City Public Health Laboratory | Jade Wang, et al. |
| EPI_ISL_631691 | NYC HH Lincoln Medical And Mental Health Center | New York City Public Health Laboratory | Jade Wang, et al. |
| EPI_ISL_631692 | Flushing Hospital Medical Center | New York City Public Health Laboratory | Jade Wang, et al. |
| EPI_ISL_631693 | NYC HH Lincoln Medical And Mental Health Center | New York City Public Health Laboratory | Jade Wang, et al. |
| EPI_ISL_631694, EPI_ISL_631695 | OCME Office Of Chief Medical Examiner | New York City Public Health Laboratory | Jade Wang, et al. |
| EPI_ISL_631696 | Flushing Hospital Medical Center | New York City Public Health Laboratory | Jade Wang, et al. |

| | | | |
|---|---|---|---|
| EPI_ISL_631697, EPI_ISL_631698 | Jamaica Hospital Medical Center | New York City Public Health Laboratory | Jade Wang, et al. |
| EPI_ISL_631699, EPI_ISL_631700, EPI_ISL_631701 | Flushing Hospital Medical Center | New York City Public Health Laboratory | Jade Wang, et al. |
| EPI_ISL_631702, EPI_ISL_631703, EPI_ISL_631704, EPI_ISL_631705, EPI_ISL_631706 | Jamaica Hospital Medical Center | New York City Public Health Laboratory | Jade Wang, et al. |
| EPI_ISL_631707, EPI_ISL_631708 | Flushing Hospital Medical Center | New York City Public Health Laboratory | Jade Wang, et al. |
| EPI_ISL_631709 | Jamaica Hospital Medical Center | New York City Public Health Laboratory | Jade Wang, et al. |
| EPI_ISL_631710 | Flushing Hospital Medical Center | New York City Public Health Laboratory | Jade Wang, et al. |
| EPI_ISL_631711, EPI_ISL_631712, EPI_ISL_631713 | Jamaica Hospital Medical Center | New York City Public Health Laboratory | Jade Wang, et al. |
| EPI_ISL_631714, EPI_ISL_631715 | Flushing Hospital Medical Center | New York City Public Health Laboratory | Jade Wang, et al. |
| EPI_ISL_631716, EPI_ISL_631717, EPI_ISL_631718, EPI_ISL_631719, EPI_ISL_631720, EPI_ISL_631721, EPI_ISL_631722, EPI_ISL_631723 | Jamaica Hospital Medical Center | New York City Public Health Laboratory | Jade Wang, et al. |
| EPI_ISL_631724, EPI_ISL_631725 | Flushing Hospital Medical Center | New York City Public Health Laboratory | Jade Wang, et al. |
| EPI_ISL_631726 | Jamaica Hospital Medical Center | New York City Public Health Laboratory | Jade Wang, et al. |
| EPI_ISL_631727 | Flushing Hospital Medical Center | New York City Public Health Laboratory | Jade Wang, et al. |
| EPI_ISL_631728, EPI_ISL_631729, EPI_ISL_631730, EPI_ISL_631731 | Jamaica Hospital Medical Center | New York City Public Health Laboratory | Jade Wang, et al. |
| EPI_ISL_631732 | Flushing Hospital Medical Center | New York City Public Health Laboratory | Jade Wang, et al. |
| EPI_ISL_631733, EPI_ISL_631734, EPI_ISL_631735, EPI_ISL_631736, EPI_ISL_631737, EPI_ISL_631738, EPI_ISL_631739, EPI_ISL_631740, EPI_ISL_631741, EPI_ISL_631742, EPI_ISL_631743, EPI_ISL_631744, EPI_ISL_631745, EPI_ISL_631746, EPI_ISL_631747 | | | |
| see above | Jamaica Hospital Medical Center | New York City Public Health Laboratory | Jade Wang, et al. |
| EPI_ISL_631748, EPI_ISL_631749, EPI_ISL_631750, EPI_ISL_631751 | St Barnabas Hospital | New York City Public Health Laboratory | Jade Wang, et al. |
| EPI_ISL_631752, EPI_ISL_631753, EPI_ISL_631754 | NYC HH Lincoln Medical And Mental Health Center | New York City Public Health Laboratory | Jade Wang, et al. |
| EPI_ISL_631755 | Richmond University Medical Center | New York City Public Health Laboratory | Jade Wang, et al. |
| EPI_ISL_631756 | Flushing Hospital Medical Center | New York City Public Health Laboratory | Jade Wang, et al. |
| EPI_ISL_631757 | Richmond University Medical Center | New York City Public Health Laboratory | Jade Wang, et al. |
| EPI_ISL_631758, EPI_ISL_631759, EPI_ISL_631760, EPI_ISL_631761, EPI_ISL_631762, EPI_ISL_631763 | OCME Office Of Chief Medical Examiner | New York City Public Health Laboratory | Jade Wang, et al. |
| EPI_ISL_631764 | Mount Sinai Hospital | New York City Public Health Laboratory | Jade Wang, et al. |
| EPI_ISL_631765 | Bellevue Hospital Center | New York City Public Health Laboratory | Jade Wang, et al. |
| EPI_ISL_631766 | Flushing Hospital Medical Center | New York City Public Health Laboratory | Jade Wang, et al. |
| EPI_ISL_631767, EPI_ISL_631768 | Jamaica Hospital Medical Center | New York City Public Health Laboratory | Jade Wang, et al. |
| EPI_ISL_631769, EPI_ISL_631770, EPI_ISL_631771 | Flushing Hospital Medical Center | New York City Public Health Laboratory | Jade Wang, et al. |
| EPI_ISL_631772 | Jamaica Hospital Medical Center | New York City Public Health Laboratory | Jade Wang, et al. |
| EPI_ISL_631773, EPI_ISL_631774 | Flushing Hospital Medical Center | New York City Public Health Laboratory | Jade Wang, et al. |
| EPI_ISL_631775, EPI_ISL_631776, EPI_ISL_631777 | Jamaica Hospital Medical Center | New York City Public Health Laboratory | Jade Wang, et al. |
| EPI_ISL_631778, EPI_ISL_631779, EPI_ISL_631780, EPI_ISL_631781, EPI_ISL_631782, EPI_ISL_631783, EPI_ISL_631784 | St Barnabas Hospital | New York City Public Health Laboratory | Jade Wang, et al. |
| EPI_ISL_631785, EPI_ISL_631786 | Mount Sinai Hospital | New York City Public Health Laboratory | Jade Wang, et al. |
| EPI_ISL_631787 | Flushing Hospital Medical Center | New York City Public Health Laboratory | Jade Wang, et al. |
| EPI_ISL_631788, EPI_ISL_631789, EPI_ISL_631790, EPI_ISL_631791 | Jamaica Hospital Medical Center | New York City Public Health Laboratory | Jade Wang, et al. |
| EPI_ISL_631792 | Richmond University Medical Center | New York City Public Health Laboratory | Jade Wang, et al. |
| EPI_ISL_631793, EPI_ISL_631794 | SUNY Downstate Medical Center | New York City Public Health Laboratory | Jade Wang, et al. |
| EPI_ISL_631795 | Jamaica Hospital Medical Center | New York City Public Health Laboratory | Jade Wang, et al. |
| EPI_ISL_631796 | Maimonides Medical Center | New York City Public Health Laboratory | Jade Wang, et al. |
| EPI_ISL_631797 | OCME Office Of Chief Medical Examiner | New York City Public Health Laboratory | Jade Wang, et al. |
| EPI_ISL_631798 | Jamaica Hospital Medical Center | New York City Public Health Laboratory | Jade Wang, et al. |
| EPI_ISL_631799 | NYC HH Lincoln Medical And Mental Health Center | New York City Public Health Laboratory | Jade Wang, et al. |
| EPI_ISL_631800 | Brookdale University Hospital Medical Center | New York City Public Health Laboratory | Jade Wang, et al. |
| EPI_ISL_631801 | Richmond University Medical Center | New York City Public Health Laboratory | Jade Wang, et al. |
| EPI_ISL_631802 | SUNY Downstate Medical Center | New York City Public Health Laboratory | Jade Wang, et al. |
| EPI_ISL_631803 | Brookdale University Hospital Medical Center | New York City Public Health Laboratory | Jade Wang, et al. |
| EPI_ISL_631804 | Bellevue Hospital Center | New York City Public Health Laboratory | Jade Wang, et al. |
| EPI_ISL_631805, EPI_ISL_631806, | Brookdale University Hospital Medical Center | New York City Public Health Laboratory | Jade Wang, et al. |

| | | | |
|---|---|---|---|
| EPI_ISL_631807 | | | |
| EPI_ISL_631808 | Wyckoff Heights Medical Center | New York City Public Health Laboratory | Jade Wang, et al. |
| EPI_ISL_631809, EPI_ISL_631810 | Jamaica Hospital Medical Center | New York City Public Health Laboratory | Jade Wang, et al. |
| EPI_ISL_631811 | Wyckoff Heights Medical Center | New York City Public Health Laboratory | Jade Wang, et al. |
| EPI_ISL_631812, EPI_ISL_631813 | Jamaica Hospital Medical Center | New York City Public Health Laboratory | Jade Wang, et al. |
| EPI_ISL_631814, EPI_ISL_631815, EPI_ISL_631816, EPI_ISL_631817, EPI_ISL_631818 | Richmond University Medical Center | New York City Public Health Laboratory | Jade Wang, et al. |
| EPI_ISL_631819 | Brookdale University Hospital Medical Center | New York City Public Health Laboratory | Jade Wang, et al. |
| EPI_ISL_631820 | Jamaica Hospital Medical Center | New York City Public Health Laboratory | Jade Wang, et al. |
| EPI_ISL_631821 | Richmond University Medical Center | New York City Public Health Laboratory | Jade Wang, et al. |
| EPI_ISL_631822, EPI_ISL_631823, EPI_ISL_631825 | Jamaica Hospital Medical Center | New York City Public Health Laboratory | Jade Wang, et al. |
| EPI_ISL_631826 | Flushing Hospital Medical Center | New York City Public Health Laboratory | Jade Wang, et al. |
| EPI_ISL_631827 | NYC HH Lincoln Medical And Mental Health Center | New York City Public Health Laboratory | Jade Wang, et al. |
| EPI_ISL_631828 | Richmond University Medical Center | New York City Public Health Laboratory | Jade Wang, et al. |
| EPI_ISL_631829 | Brookdale University Hospital Medical Center | New York City Public Health Laboratory | Jade Wang, et al. |
| EPI_ISL_631830 | Mount Sinai Hospital | New York City Public Health Laboratory | Jade Wang, et al. |
| EPI_ISL_631831 | VA NY Harbor Healthcare System | New York City Public Health Laboratory | Jade Wang, et al. |
| EPI_ISL_631832 | Metro Urgicare | New York City Public Health Laboratory | Jade Wang, et al. |
| EPI_ISL_631833 | Jamaica Hospital Medical Center | New York City Public Health Laboratory | Jade Wang, et al. |
| EPI_ISL_631834 | Mount Sinai Hospital | New York City Public Health Laboratory | Jade Wang, et al. |
| EPI_ISL_631835, EPI_ISL_631836 | Jamaica Hospital Medical Center | New York City Public Health Laboratory | Jade Wang, et al. |
| EPI_ISL_631837 | Richmond University Medical Center | New York City Public Health Laboratory | Jade Wang, et al. |
| EPI_ISL_631838 | Bellevue Hospital Center | New York City Public Health Laboratory | Jade Wang, et al. |
| EPI_ISL_631839 | Mount Sinai Hospital | New York City Public Health Laboratory | Jade Wang, et al. |
| EPI_ISL_631840, EPI_ISL_631841, EPI_ISL_631842, EPI_ISL_631843, EPI_ISL_631844, EPI_ISL_631845, EPI_ISL_631846 | OCME Office Of Chief Medical Examiner | New York City Public Health Laboratory | Jade Wang, et al. |
| EPI_ISL_631847 | Jamaica Hospital Medical Center | New York City Public Health Laboratory | Jade Wang, et al. |
| EPI_ISL_631848 | Flushing Hospital Medical Center | New York City Public Health Laboratory | Jade Wang, et al. |
| EPI_ISL_631849, EPI_ISL_631850 | Jamaica Hospital Medical Center | New York City Public Health Laboratory | Jade Wang, et al. |
| EPI_ISL_631851 | Flushing Hospital Medical Center | New York City Public Health Laboratory | Jade Wang, et al. |
| EPI_ISL_631852, EPI_ISL_631853, EPI_ISL_631854 | Jamaica Hospital Medical Center | New York City Public Health Laboratory | Jade Wang, et al. |
| EPI_ISL_631855 | Flushing Hospital Medical Center | New York City Public Health Laboratory | Jade Wang, et al. |
| EPI_ISL_631856, EPI_ISL_631857, EPI_ISL_631858, EPI_ISL_631859 | St Barnabas Hospital | New York City Public Health Laboratory | Jade Wang, et al. |
| EPI_ISL_631860 | New York Presbyterian/ Weill Cornell Medical Center | New York City Public Health Laboratory | Jade Wang, et al. |
| EPI_ISL_631861 | New York Presbyterian Queens | New York City Public Health Laboratory | Jade Wang, et al. |
| EPI_ISL_631862 | NYC Department Of Health And Mental Hygiene | New York City Public Health Laboratory | Jade Wang, et al. |
| EPI_ISL_631863 | Maimonides Medical Center | New York City Public Health Laboratory | Jade Wang, et al. |
| EPI_ISL_631864 | Brookdale University Hospital Medical Center | New York City Public Health Laboratory | Jade Wang, et al. |
| EPI_ISL_631865 | NYC HH Elmhurst Hospital Medical Center | New York City Public Health Laboratory | Jade Wang, et al. |
| EPI_ISL_631866, EPI_ISL_631867 | Jamaica Hospital Medical Center | New York City Public Health Laboratory | Jade Wang, et al. |
| EPI_ISL_631868, EPI_ISL_631869 | Richmond University Medical Center | New York City Public Health Laboratory | Jade Wang, et al. |
| EPI_ISL_631870, EPI_ISL_631871, EPI_ISL_631872 | Jamaica Hospital Medical Center | New York City Public Health Laboratory | Jade Wang, et al. |
| EPI_ISL_631873 | Richmond University Medical Center | New York City Public Health Laboratory | Jade Wang, et al. |
| EPI_ISL_631874 | James J. Peters VA Medical Center | New York City Public Health Laboratory | Jade Wang, et al. |
| EPI_ISL_631875 | Richmond University Medical Center | New York City Public Health Laboratory | Jade Wang, et al. |
| EPI_ISL_631876, EPI_ISL_631877, EPI_ISL_631878, EPI_ISL_631879 | Jamaica Hospital Medical Center | New York City Public Health Laboratory | Jade Wang, et al. |
| EPI_ISL_631880 | Mount Sinai West | New York City Public Health Laboratory | Jade Wang, et al. |
| EPI_ISL_631881 | Maimonides Medical Center | New York City Public Health Laboratory | Jade Wang, et al. |
| EPI_ISL_631882, EPI_ISL_631883 | Jamaica Hospital Medical Center | New York City Public Health Laboratory | Jade Wang, et al. |
| EPI_ISL_631884 | Brookdale University Hospital Medical Center | New York City Public Health Laboratory | Jade Wang, et al. |
| EPI_ISL_631885 | Flushing Hospital Medical Center | New York City Public Health Laboratory | Jade Wang, et al. |
| EPI_ISL_631886 | NYC HH Lincoln Medical And Mental Health Center | New York City Public Health Laboratory | Jade Wang, et al. |
| EPI_ISL_631887 | Jamaica Hospital Medical Center | New York City Public Health Laboratory | Jade Wang, et al. |
| EPI_ISL_631888 | St Barnabas Hospital | New York City Public Health Laboratory | Jade Wang, et al. |

| | | | |
|---|---|---|---|
| EPI_ISL_631889, EPI_ISL_631890 | Jamaica Hospital Medical Center | New York City Public Health Laboratory | Jade Wang, et al. |
| EPI_ISL_631891 | Flushing Hospital Medical Center | New York City Public Health Laboratory | Jade Wang, et al. |
| EPI_ISL_631892 | Richmond University Medical Center | New York City Public Health Laboratory | Jade Wang, et al. |
| EPI_ISL_631893, EPI_ISL_631894 | Flushing Hospital Medical Center | New York City Public Health Laboratory | Jade Wang, et al. |
| EPI_ISL_631895, EPI_ISL_631896 | Jamaica Hospital Medical Center | New York City Public Health Laboratory | Jade Wang, et al. |
| EPI_ISL_631897 | Richmond University Medical Center | New York City Public Health Laboratory | Jade Wang, et al. |
| EPI_ISL_631898 | OCME Office Of Chief Medical Examiner | New York City Public Health Laboratory | Jade Wang, et al. |
| EPI_ISL_631899 | Richmond University Medical Center | New York City Public Health Laboratory | Jade Wang, et al. |
| EPI_ISL_631900 | St Barnabas Hospital | New York City Public Health Laboratory | Jade Wang, et al. |
| EPI_ISL_631901, EPI_ISL_631902 | Jamaica Hospital Medical Center | New York City Public Health Laboratory | Jade Wang, et al. |
| EPI_ISL_631903 | Richmond University Medical Center | New York City Public Health Laboratory | Jade Wang, et al. |
| EPI_ISL_631904 | Jamaica Hospital Medical Center | New York City Public Health Laboratory | Jade Wang, et al. |
| EPI_ISL_631905 | Richmond University Medical Center | New York City Public Health Laboratory | Jade Wang, et al. |
| EPI_ISL_631906, EPI_ISL_631907, EPI_ISL_631908 | Flushing Hospital Medical Center | New York City Public Health Laboratory | Jade Wang, et al. |
| EPI_ISL_631909, EPI_ISL_631910 | Jamaica Hospital Medical Center | New York City Public Health Laboratory | Jade Wang, et al. |
| EPI_ISL_631911, EPI_ISL_631912 | St Barnabas Hospital | New York City Public Health Laboratory | Jade Wang, et al. |
| EPI_ISL_631913, EPI_ISL_631914 | Mount Sinai Hospital | New York City Public Health Laboratory | Jade Wang, et al. |
| EPI_ISL_631915 | Maimonides Medical Center | New York City Public Health Laboratory | Jade Wang, et al. |
| EPI_ISL_631916 | Flushing Hospital Medical Center | New York City Public Health Laboratory | Jade Wang, et al. |
| EPI_ISL_631917 | St Barnabas Hospital | New York City Public Health Laboratory | Jade Wang, et al. |
| EPI_ISL_631918 | Flushing Hospital Medical Center | New York City Public Health Laboratory | Jade Wang, et al. |
| EPI_ISL_631919, EPI_ISL_631920 | Richmond University Medical Center | New York City Public Health Laboratory | Jade Wang, et al. |
| EPI_ISL_631921, EPI_ISL_631922 | Jamaica Hospital Medical Center | New York City Public Health Laboratory | Jade Wang, et al. |
| EPI_ISL_631923 | Flushing Hospital Medical Center | New York City Public Health Laboratory | Jade Wang, et al. |
| EPI_ISL_631924 | St Barnabas Hospital | New York City Public Health Laboratory | Jade Wang, et al. |
| EPI_ISL_631925, EPI_ISL_631926, EPI_ISL_631927, EPI_ISL_631928 | Jamaica Hospital Medical Center | New York City Public Health Laboratory | Jade Wang, et al. |
| EPI_ISL_631929 | Flushing Hospital Medical Center | New York City Public Health Laboratory | Jade Wang, et al. |
| EPI_ISL_631930 | Jamaica Hospital Medical Center | New York City Public Health Laboratory | Jade Wang, et al. |
| EPI_ISL_631931 | Flushing Hospital Medical Center | New York City Public Health Laboratory | Jade Wang, et al. |
| EPI_ISL_631932 | OCME Office Of Chief Medical Examiner | New York City Public Health Laboratory | Jade Wang, et al. |
| EPI_ISL_631933 | St Barnabas Hospital | New York City Public Health Laboratory | Jade Wang, et al. |
| EPI_ISL_631934 | OCME Office Of Chief Medical Examiner | New York City Public Health Laboratory | Jade Wang, et al. |
| EPI_ISL_631935 | St Barnabas Hospital | New York City Public Health Laboratory | Jade Wang, et al. |
| EPI_ISL_631936, EPI_ISL_631937, EPI_ISL_631938 | OCME Office Of Chief Medical Examiner | New York City Public Health Laboratory | Jade Wang, et al. |
| EPI_ISL_631939, EPI_ISL_631940, EPI_ISL_631941, EPI_ISL_631942, EPI_ISL_631943, EPI_ISL_631944, EPI_ISL_631945, EPI_ISL_631946, EPI_ISL_631947, EPI_ISL_631948, EPI_ISL_631949 | | | |
| see above | St Barnabas Hospital | New York City Public Health Laboratory | Jade Wang, et al. |
| EPI_ISL_631950, EPI_ISL_631951 | Jamaica Hospital Medical Center | New York City Public Health Laboratory | Jade Wang, et al. |
| EPI_ISL_631952 | St Barnabas Hospital | New York City Public Health Laboratory | Jade Wang, et al. |
| EPI_ISL_631953 | NYC HH Lincoln Medical And Mental Health Center | New York City Public Health Laboratory | Jade Wang, et al. |
| EPI_ISL_631954 | Richmond University Medical Center | New York City Public Health Laboratory | Jade Wang, et al. |
| EPI_ISL_631955 | Flushing Hospital Medical Center | New York City Public Health Laboratory | Jade Wang, et al. |
| EPI_ISL_631956 | OCME Office Of Chief Medical Examiner | New York City Public Health Laboratory | Jade Wang, et al. |
| EPI_ISL_631957, EPI_ISL_631958 | St Barnabas Hospital | New York City Public Health Laboratory | Jade Wang, et al. |
| EPI_ISL_631959, EPI_ISL_631960, EPI_ISL_631961 | Flushing Hospital Medical Center | New York City Public Health Laboratory | Jade Wang, et al. |
| EPI_ISL_631962 | Richmond University Medical Center | New York City Public Health Laboratory | Jade Wang, et al. |
| EPI_ISL_631963 | Jamaica Hospital Medical Center | New York City Public Health Laboratory | Jade Wang, et al. |
| EPI_ISL_631964 | Flushing Hospital Medical Center | New York City Public Health Laboratory | Jade Wang, et al. |
| EPI_ISL_631965 | Richmond University Medical Center | New York City Public Health Laboratory | Jade Wang, et al. |
| EPI_ISL_631966 | Jamaica Hospital Medical Center | New York City Public Health Laboratory | Jade Wang, et al. |
| EPI_ISL_631967 | Richmond University Medical Center | New York City Public Health Laboratory | Jade Wang, et al. |
| EPI_ISL_631968 | Flushing Hospital Medical Center | New York City Public Health Laboratory | Jade Wang, et al. |
| EPI_ISL_631969, EPI_ISL_631970 | Jamaica Hospital Medical Center | New York City Public Health Laboratory | Jade Wang, et al. |
| EPI_ISL_631971 | St Barnabas Hospital | New York City Public Health Laboratory | Jade Wang, et al. |
| EPI_ISL_631972 | Richmond University Medical Center | New York City Public Health Laboratory | Jade Wang, et al. |
| EPI_ISL_631973 | Jamaica Hospital Medical Center | New York City Public Health Laboratory | Jade Wang, et al. |
| EPI_ISL_631974 | NYC HH Coney Island Hospital | New York City Public Health Laboratory | Jade Wang, et al. |

| | | | |
|---|---|---|---|
| EPI_ISL_631975 | Richmond University Medical Center | New York City Public Health Laboratory | Jade Wang, et al. |
| EPI_ISL_631976 | Brookdale University Hospital Medical Center | New York City Public Health Laboratory | Jade Wang, et al. |
| EPI_ISL_631977 | SUNY Downstate Medical Center | New York City Public Health Laboratory | Jade Wang, et al. |
| EPI_ISL_631978 | Brookdale University Hospital Medical Center | New York City Public Health Laboratory | Jade Wang, et al. |
| EPI_ISL_631979 | Jamaica Hospital Medical Center | New York City Public Health Laboratory | Jade Wang, et al. |
| EPI_ISL_631980 | St Barnabas Hospital | New York City Public Health Laboratory | Jade Wang, et al. |
| EPI_ISL_631981 | Brookdale University Hospital Medical Center | New York City Public Health Laboratory | Jade Wang, et al. |
| EPI_ISL_631982 | Montefiore Medical Center | New York City Public Health Laboratory | Jade Wang, et al. |
| EPI_ISL_631983 | Mount Sinai Hospital | New York City Public Health Laboratory | Jade Wang, et al. |
| EPI_ISL_631984 | Brookdale University Hospital Medical Center | New York City Public Health Laboratory | Jade Wang, et al. |
| EPI_ISL_631985 | St Barnabas Hospital | New York City Public Health Laboratory | Jade Wang, et al. |
| EPI_ISL_631986 | Flushing Hospital Medical Center | New York City Public Health Laboratory | Jade Wang, et al. |
| EPI_ISL_631987, EPI_ISL_631988 | St Barnabas Hospital | New York City Public Health Laboratory | Jade Wang, et al. |
| EPI_ISL_631989 | Flushing Hospital Medical Center | New York City Public Health Laboratory | Jade Wang, et al. |
| EPI_ISL_631990 | St Barnabas Hospital | New York City Public Health Laboratory | Jade Wang, et al. |
| EPI_ISL_631991 | Flushing Hospital Medical Center | New York City Public Health Laboratory | Jade Wang, et al. |
| EPI_ISL_631992 | Richmond University Medical Center | New York City Public Health Laboratory | Jade Wang, et al. |
| EPI_ISL_631993 | Maimonides Medical Center | New York City Public Health Laboratory | Jade Wang, et al. |
| EPI_ISL_631994 | Jamaica Hospital Medical Center | New York City Public Health Laboratory | Jade Wang, et al. |
| EPI_ISL_631995 | Wyckoff Heights Medical Center | New York City Public Health Laboratory | Jade Wang, et al. |
| EPI_ISL_631996, EPI_ISL_631997 | Jamaica Hospital Medical Center | New York City Public Health Laboratory | Jade Wang, et al. |
| EPI_ISL_631998, EPI_ISL_631999, EPI_ISL_632000, EPI_ISL_632001, EPI_ISL_632002 | Richmond University Medical Center | New York City Public Health Laboratory | Jade Wang, et al. |
| EPI_ISL_632003, EPI_ISL_632004 | Flushing Hospital Medical Center | New York City Public Health Laboratory | Jade Wang, et al. |
| EPI_ISL_632005 | Jamaica Hospital Medical Center | New York City Public Health Laboratory | Jade Wang, et al. |
| EPI_ISL_632006 | OCME Office Of Chief Medical Examiner | New York City Public Health Laboratory | Jade Wang, et al. |
| EPI_ISL_632007, EPI_ISL_632008, EPI_ISL_632009 | Jamaica Hospital Medical Center | New York City Public Health Laboratory | Jade Wang, et al. |
| EPI_ISL_632010 | Flushing Hospital Medical Center | New York City Public Health Laboratory | Jade Wang, et al. |
| EPI_ISL_632011, EPI_ISL_632012 | Jamaica Hospital Medical Center | New York City Public Health Laboratory | Jade Wang, et al. |
| EPI_ISL_632013, EPI_ISL_632014, EPI_ISL_632015, EPI_ISL_632016 | Richmond University Medical Center | New York City Public Health Laboratory | Jade Wang, et al. |
| EPI_ISL_632017 | Jamaica Hospital Medical Center | New York City Public Health Laboratory | Jade Wang, et al. |
| EPI_ISL_632018, EPI_ISL_632019, EPI_ISL_632020 | Flushing Hospital Medical Center | New York City Public Health Laboratory | Jade Wang, et al. |
| EPI_ISL_632021 | Jamaica Hospital Medical Center | New York City Public Health Laboratory | Jade Wang, et al. |
| EPI_ISL_632022 | Flushing Hospital Medical Center | New York City Public Health Laboratory | Jade Wang, et al. |
| EPI_ISL_632023 | Jamaica Hospital Medical Center | New York City Public Health Laboratory | Jade Wang, et al. |
| EPI_ISL_632024 | Flushing Hospital Medical Center | New York City Public Health Laboratory | Jade Wang, et al. |
| EPI_ISL_632025 | Jamaica Hospital Medical Center | New York City Public Health Laboratory | Jade Wang, et al. |
| EPI_ISL_632026 | NYC HH Lincoln Medical And Mental Health Center | New York City Public Health Laboratory | Jade Wang, et al. |
| EPI_ISL_632027 | Flushing Hospital Medical Center | New York City Public Health Laboratory | Jade Wang, et al. |
| EPI_ISL_632028, EPI_ISL_632029 | Jamaica Hospital Medical Center | New York City Public Health Laboratory | Jade Wang, et al. |
| EPI_ISL_632030, EPI_ISL_632031, EPI_ISL_632032 | Richmond University Medical Center | New York City Public Health Laboratory | Jade Wang, et al. |
| EPI_ISL_632034 | Flushing Hospital Medical Center | New York City Public Health Laboratory | Jade Wang, et al. |
| EPI_ISL_632035 | Jamaica Hospital Medical Center | New York City Public Health Laboratory | Jade Wang, et al. |
| EPI_ISL_632036 | Richmond University Medical Center | New York City Public Health Laboratory | Jade Wang, et al. |
| EPI_ISL_632037 | Jamaica Hospital Medical Center | New York City Public Health Laboratory | Jade Wang, et al. |
| EPI_ISL_632038 | New York Presbyterian Lower Manhattan | New York City Public Health Laboratory | Jade Wang, et al. |
| EPI_ISL_632039 | NYC Department Of Health And Mental Hygiene | New York City Public Health Laboratory | Jade Wang, et al. |
| EPI_ISL_632040 | New York Presbyterian Queens | New York City Public Health Laboratory | Jade Wang, et al. |
| EPI_ISL_632041, EPI_ISL_632042 | New York Presbyterian Lower Manhattan | New York City Public Health Laboratory | Jade Wang, et al. |
| EPI_ISL_632043, EPI_ISL_632044 | NYU Langone Health | New York City Public Health Laboratory | Jade Wang, et al. |
| EPI_ISL_632045 | New York Presbyterian Lower Manhattan | New York City Public Health Laboratory | Jade Wang, et al. |
| EPI_ISL_632046 | New York Presbyterian-Brooklyn Methodist Hospital | New York City Public Health Laboratory | Jade Wang, et al. |
| EPI_ISL_632047 | New York Presbyterian Lower Manhattan | New York City Public Health Laboratory | Jade Wang, et al. |
| EPI_ISL_632048 | New York Presbyterian-Brooklyn Methodist Hospital | New York City Public Health Laboratory | Jade Wang, et al. |
| EPI_ISL_632049 | New York Presbyterian Queens | New York City Public Health Laboratory | Jade Wang, et al. |
| EPI_ISL_632050 | Long Island Jewish Medical Center | New York City Public Health Laboratory | Jade Wang, et al. |

| | | | |
|---|---|---|---|
| EPI_ISL_632051 | Jamaica Hospital Medical Center | New York City Public Health Laboratory | Jade Wang, et al. |
| EPI_ISL_632052, EPI_ISL_632053 | New York Presbyterian Queens | New York City Public Health Laboratory | Jade Wang, et al. |
| EPI_ISL_632054 | New York Presbyterian-Brooklyn Methodist Hospital | New York City Public Health Laboratory | Jade Wang, et al. |
| EPI_ISL_632055, EPI_ISL_632056 | New York Community Hospital | New York City Public Health Laboratory | Jade Wang, et al. |
| EPI_ISL_632057 | New York-Presbyterian-Columbia University Medical Center | New York City Public Health Laboratory | Jade Wang, et al. |
| EPI_ISL_632058, EPI_ISL_632059 | Proximedical Urgent Care | New York City Public Health Laboratory | Jade Wang, et al. |
| EPI_ISL_632060 | NYC HH Queens Hospital Center | New York City Public Health Laboratory | Jade Wang, et al. |
| EPI_ISL_632061 | Jamaica Hospital Medical Center | New York City Public Health Laboratory | Jade Wang, et al. |
| EPI_ISL_632062 | NYC Department Of Health And Mental Hygiene | New York City Public Health Laboratory | Jade Wang, et al. |
| EPI_ISL_632063, EPI_ISL_632064 | New York Presbyterian Queens | New York City Public Health Laboratory | Jade Wang, et al. |
| EPI_ISL_632065, EPI_ISL_632066 | Jamaica Hospital Medical Center | New York City Public Health Laboratory | Jade Wang, et al. |
| EPI_ISL_632067 | NYU Langone Health | New York City Public Health Laboratory | Jade Wang, et al. |
| EPI_ISL_632068, EPI_ISL_632069 | Bellevue Hospital Center | New York City Public Health Laboratory | Jade Wang, et al. |
| EPI_ISL_632070, EPI_ISL_632071 | New York Presbyterian Queens | New York City Public Health Laboratory | Jade Wang, et al. |
| EPI_ISL_632072 | NYC HH Lincoln Medical And Mental Health Center | New York City Public Health Laboratory | Jade Wang, et al. |
| EPI_ISL_632073 | OCME Office Of Chief Medical Examiner | New York City Public Health Laboratory | Jade Wang, et al. |
| EPI_ISL_632074 | NYC Department Of Health And Mental Hygiene | New York City Public Health Laboratory | Jade Wang, et al. |
| EPI_ISL_632075 | Jamaica Hospital Medical Center | New York City Public Health Laboratory | Jade Wang, et al. |
| EPI_ISL_632076 | NYU Langone Health | New York City Public Health Laboratory | Jade Wang, et al. |
| EPI_ISL_632077, EPI_ISL_632078 | Brookdale University Hospital Medical Center | New York City Public Health Laboratory | Jade Wang, et al. |
| EPI_ISL_632079 | NYU Langone Health | New York City Public Health Laboratory | Jade Wang, et al. |
| EPI_ISL_632080 | SUNY Downstate Medical Center | New York City Public Health Laboratory | Jade Wang, et al. |
| EPI_ISL_632081 | New York Presbyterian-Brooklyn Methodist Hospital | New York City Public Health Laboratory | Jade Wang, et al. |
| EPI_ISL_632082 | New York Presbyterian/ Weill Cornell Medical Center | New York City Public Health Laboratory | Jade Wang, et al. |
| EPI_ISL_632083, EPI_ISL_632084 | NYC Department Of Health And Mental Hygiene | New York City Public Health Laboratory | Jade Wang, et al. |
| EPI_ISL_632085 | Richmond University Medical Center | New York City Public Health Laboratory | Jade Wang, et al. |
| EPI_ISL_632086 | Jamaica Hospital Medical Center | New York City Public Health Laboratory | Jade Wang, et al. |
| EPI_ISL_632087 | Flushing Hospital Medical Center | New York City Public Health Laboratory | Jade Wang, et al. |
| EPI_ISL_632088 | Jamaica Hospital Medical Center | New York City Public Health Laboratory | Jade Wang, et al. |
| EPI_ISL_632089 | Brookdale University Hospital Medical Center | New York City Public Health Laboratory | Jade Wang, et al. |
| EPI_ISL_632090 | Jamaica Hospital Medical Center | New York City Public Health Laboratory | Jade Wang, et al. |
| EPI_ISL_632091, EPI_ISL_632092 | SUNY Downstate Medical Center | New York City Public Health Laboratory | Jade Wang, et al. |
| EPI_ISL_632093, EPI_ISL_632094, EPI_ISL_632095, EPI_ISL_632096 | Jamaica Hospital Medical Center | New York City Public Health Laboratory | Jade Wang, et al. |
| EPI_ISL_632097 | Flushing Hospital Medical Center | New York City Public Health Laboratory | Jade Wang, et al. |
| EPI_ISL_632098 | Bellevue Hospital Center | New York City Public Health Laboratory | Jade Wang, et al. |
| EPI_ISL_632099 | VA NY Harbor Healthcare System | New York City Public Health Laboratory | Jade Wang, et al. |
| EPI_ISL_632100 | Brookdale University Hospital Medical Center | New York City Public Health Laboratory | Jade Wang, et al. |
| EPI_ISL_632101, EPI_ISL_632102 | Mount Sinai Queens | New York City Public Health Laboratory | Jade Wang, et al. |
| EPI_ISL_632103, EPI_ISL_632104, EPI_ISL_632105 | Flushing Hospital Medical Center | New York City Public Health Laboratory | Jade Wang, et al. |
| EPI_ISL_632106, EPI_ISL_632107, EPI_ISL_632108, EPI_ISL_632109, EPI_ISL_632110 | Jamaica Hospital Medical Center | New York City Public Health Laboratory | Jade Wang, et al. |
| EPI_ISL_632111 | Montefiore Medical Center | New York City Public Health Laboratory | Jade Wang, et al. |
| EPI_ISL_632112, EPI_ISL_632113 | Richmond University Medical Center | New York City Public Health Laboratory | Jade Wang, et al. |
| EPI_ISL_632114 | Brookdale University Hospital Medical Center | New York City Public Health Laboratory | Jade Wang, et al. |
| EPI_ISL_632115 | Richmond University Medical Center | New York City Public Health Laboratory | Jade Wang, et al. |
| EPI_ISL_632116 | NYC HH Lincoln Medical And Mental Health Center | New York City Public Health Laboratory | Jade Wang, et al. |
| EPI_ISL_632117, EPI_ISL_632118 | Richmond University Medical Center | New York City Public Health Laboratory | Jade Wang, et al. |
| EPI_ISL_632119, EPI_ISL_632120 | Flushing Hospital Medical Center | New York City Public Health Laboratory | Jade Wang, et al. |
| EPI_ISL_632121 | Bellevue Hospital Center | New York City Public Health Laboratory | Jade Wang, et al. |
| EPI_ISL_632122 | Flushing Hospital Medical Center | New York City Public Health Laboratory | Jade Wang, et al. |
| EPI_ISL_632123 | Brookdale University Hospital Medical Center | New York City Public Health Laboratory | Jade Wang, et al. |
| EPI_ISL_632124, EPI_ISL_632125, EPI_ISL_632126, EPI_ISL_632127 | Flushing Hospital Medical Center | New York City Public Health Laboratory | Jade Wang, et al. |
| EPI_ISL_632128 | Jamaica Hospital Medical Center | New York City Public Health Laboratory | Jade Wang, et al. |
| EPI_ISL_632129 | Wyckoff Heights Medical Center | New York City Public Health Laboratory | Jade Wang, et al. |
| EPI_ISL_632130 | Jamaica Hospital Medical Center | New York City Public Health Laboratory | Jade Wang, et al. |
| EPI_ISL_632131 | NYC HH Lincoln Medical And Mental Health Center | New York City Public Health Laboratory | Jade Wang, et al. |

| | | | |
|---|---|---|---|
| EPI_ISL_632132, EPI_ISL_632133, EPI_ISL_632134 | Flushing Hospital Medical Center | New York City Public Health Laboratory | Jade Wang, et al. |
| EPI_ISL_632135 | SUNY Downstate Medical Center | New York City Public Health Laboratory | Jade Wang, et al. |
| EPI_ISL_632136 | Bellevue Hospital Center | New York City Public Health Laboratory | Jade Wang, et al. |
| EPI_ISL_632137, EPI_ISL_632138, EPI_ISL_632139, EPI_ISL_632140 | Flushing Hospital Medical Center | New York City Public Health Laboratory | Jade Wang, et al. |
| EPI_ISL_632141 | Wyckoff Heights Medical Center | New York City Public Health Laboratory | Jade Wang, et al. |
| EPI_ISL_632142 | Brookdale University Hospital Medical Center | New York City Public Health Laboratory | Jade Wang, et al. |
| EPI_ISL_632143 | Jamaica Hospital Medical Center | New York City Public Health Laboratory | Jade Wang, et al. |
| EPI_ISL_632144 | Flushing Hospital Medical Center | New York City Public Health Laboratory | Jade Wang, et al. |
| EPI_ISL_632145 | Jamaica Hospital Medical Center | New York City Public Health Laboratory | Jade Wang, et al. |
| EPI_ISL_632146, EPI_ISL_632147 | NYC HH Lincoln Medical And Mental Health Center | New York City Public Health Laboratory | Jade Wang, et al. |
| EPI_ISL_632148 | SUNY Downstate Medical Center | New York City Public Health Laboratory | Jade Wang, et al. |
| EPI_ISL_632149, EPI_ISL_632150, EPI_ISL_632151, EPI_ISL_632152 | Jamaica Hospital Medical Center | New York City Public Health Laboratory | Jade Wang, et al. |
| EPI_ISL_632153 | SUNY Downstate Medical Center | New York City Public Health Laboratory | Jade Wang, et al. |
| EPI_ISL_632154 | Richmond University Medical Center | New York City Public Health Laboratory | Jade Wang, et al. |
| EPI_ISL_632155, EPI_ISL_632156 | Brookdale University Hospital Medical Center | New York City Public Health Laboratory | Jade Wang, et al. |
| EPI_ISL_632157 | Wyckoff Heights Medical Center | New York City Public Health Laboratory | Jade Wang, et al. |
| EPI_ISL_632158 | SUNY Downstate Medical Center | New York City Public Health Laboratory | Jade Wang, et al. |
| EPI_ISL_632159 | Wyckoff Heights Medical Center | New York City Public Health Laboratory | Jade Wang, et al. |
| EPI_ISL_632160, EPI_ISL_632161 | SUNY Downstate Medical Center | New York City Public Health Laboratory | Jade Wang, et al. |
| EPI_ISL_632162 | Jamaica Hospital Medical Center | New York City Public Health Laboratory | Jade Wang, et al. |
| EPI_ISL_632163, EPI_ISL_632164, EPI_ISL_632165 | Richmond University Medical Center | New York City Public Health Laboratory | Jade Wang, et al. |
| EPI_ISL_632166, EPI_ISL_632167, EPI_ISL_632168, EPI_ISL_632169, EPI_ISL_632170, EPI_ISL_632171, EPI_ISL_632172 | Jamaica Hospital Medical Center | New York City Public Health Laboratory | Jade Wang, et al. |
| EPI_ISL_632173 | Richmond University Medical Center | New York City Public Health Laboratory | Jade Wang, et al. |
| EPI_ISL_632174 | Flushing Hospital Medical Center | New York City Public Health Laboratory | Jade Wang, et al. |
| EPI_ISL_632175, EPI_ISL_632176 | OCME Office Of Chief Medical Examiner | New York City Public Health Laboratory | Jade Wang, et al. |
| EPI_ISL_632177 | NYC HH Lincoln Medical And Mental Health Center | New York City Public Health Laboratory | Jade Wang, et al. |
| EPI_ISL_632178 | OCME Office Of Chief Medical Examiner | New York City Public Health Laboratory | Jade Wang, et al. |
| EPI_ISL_632179 | James J. Peters VA Medical Center | New York City Public Health Laboratory | Jade Wang, et al. |
| EPI_ISL_632180 | Jamaica Hospital Medical Center | New York City Public Health Laboratory | Jade Wang, et al. |
| EPI_ISL_632181 | Flushing Hospital Medical Center | New York City Public Health Laboratory | Jade Wang, et al. |
| EPI_ISL_632182 | Jamaica Hospital Medical Center | New York City Public Health Laboratory | Jade Wang, et al. |
| EPI_ISL_632183 | OCME Office Of Chief Medical Examiner | New York City Public Health Laboratory | Jade Wang, et al. |
| EPI_ISL_632184 | Jamaica Hospital Medical Center | New York City Public Health Laboratory | Jade Wang, et al. |
| EPI_ISL_632185 | Flushing Hospital Medical Center | New York City Public Health Laboratory | Jade Wang, et al. |
| EPI_ISL_632186, EPI_ISL_632187 | NYC HH Lincoln Medical And Mental Health Center | New York City Public Health Laboratory | Jade Wang, et al. |
| EPI_ISL_632188, EPI_ISL_632189 | Jamaica Hospital Medical Center | New York City Public Health Laboratory | Jade Wang, et al. |
| EPI_ISL_632190 | Flushing Hospital Medical Center | New York City Public Health Laboratory | Jade Wang, et al. |
| EPI_ISL_632191 | Jamaica Hospital Medical Center | New York City Public Health Laboratory | Jade Wang, et al. |
| EPI_ISL_632192, EPI_ISL_632193 | NYC HH Lincoln Medical And Mental Health Center | New York City Public Health Laboratory | Jade Wang, et al. |
| EPI_ISL_632194 | Flushing Hospital Medical Center | New York City Public Health Laboratory | Jade Wang, et al. |
| EPI_ISL_632195 | Jamaica Hospital Medical Center | New York City Public Health Laboratory | Jade Wang, et al. |
| EPI_ISL_632196, EPI_ISL_632197 | Flushing Hospital Medical Center | New York City Public Health Laboratory | Jade Wang, et al. |
| EPI_ISL_632198 | Jamaica Hospital Medical Center | New York City Public Health Laboratory | Jade Wang, et al. |
| EPI_ISL_632199 | Richmond University Medical Center | New York City Public Health Laboratory | Jade Wang, et al. |
| EPI_ISL_632200 | James J. Peters VA Medical Center | New York City Public Health Laboratory | Jade Wang, et al. |
| EPI_ISL_632201 | Richmond University Medical Center | New York City Public Health Laboratory | Jade Wang, et al. |
| EPI_ISL_632202 | OCME Office Of Chief Medical Examiner | New York City Public Health Laboratory | Jade Wang, et al. |
| EPI_ISL_632203 | Richmond University Medical Center | New York City Public Health Laboratory | Jade Wang, et al. |
| EPI_ISL_632204 | Jamaica Hospital Medical Center | New York City Public Health Laboratory | Jade Wang, et al. |
| EPI_ISL_632205, EPI_ISL_632206, EPI_ISL_632207, EPI_ISL_632208 | Richmond University Medical Center | New York City Public Health Laboratory | Jade Wang, et al. |
| EPI_ISL_632209, EPI_ISL_632210 | Jamaica Hospital Medical Center | New York City Public Health Laboratory | Jade Wang, et al. |
| EPI_ISL_632211, EPI_ISL_632212, EPI_ISL_632213 | Flushing Hospital Medical Center | New York City Public Health Laboratory | Jade Wang, et al. |
| EPI_ISL_632214 | Jamaica Hospital Medical Center | New York City Public Health Laboratory | Jade Wang, et al. |

| | | | |
|---|---|---|---|
| EPI_ISL_632215, EPI_ISL_632216, EPI_ISL_632217, EPI_ISL_632218 | Richmond University Medical Center | New York City Public Health Laboratory | Jade Wang, et al. |
| EPI_ISL_632219 | Jamaica Hospital Medical Center | New York City Public Health Laboratory | Jade Wang, et al. |
| EPI_ISL_632220 | Flushing Hospital Medical Center | New York City Public Health Laboratory | Jade Wang, et al. |
| EPI_ISL_632221 | Jamaica Hospital Medical Center | New York City Public Health Laboratory | Jade Wang, et al. |
| EPI_ISL_632222, EPI_ISL_632223, EPI_ISL_632224, EPI_ISL_632225, EPI_ISL_632226 | Flushing Hospital Medical Center | New York City Public Health Laboratory | Jade Wang, et al. |
| EPI_ISL_632227, EPI_ISL_632228, EPI_ISL_632229 | Jamaica Hospital Medical Center | New York City Public Health Laboratory | Jade Wang, et al. |
| EPI_ISL_632230, EPI_ISL_632231, EPI_ISL_632232, EPI_ISL_632233, EPI_ISL_632234 | Richmond University Medical Center | New York City Public Health Laboratory | Jade Wang, et al. |
| EPI_ISL_632235 | NYC HH Lincoln Medical And Mental Health Center | New York City Public Health Laboratory | Jade Wang, et al. |
| EPI_ISL_632236, EPI_ISL_632237 | Flushing Hospital Medical Center | New York City Public Health Laboratory | Jade Wang, et al. |
| EPI_ISL_632238 | OCME Office Of Chief Medical Examiner | New York City Public Health Laboratory | Jade Wang, et al. |
| EPI_ISL_632239, EPI_ISL_632240, EPI_ISL_632241 | Jamaica Hospital Medical Center | New York City Public Health Laboratory | Jade Wang, et al. |
| EPI_ISL_632312 | NU-sjukvården | Clinical microbiology, Sahlgrenska University Hospital | Johan Ringlander, Josefin Olausson, Hedvig Engström Jakobsson, Magnus Lindh |
| EPI_ISL_632313 | NU-sjukvården | Clinical Microbiology, Sahlgrenska University Hospital, | Johan Ringlander, Josefin Olausson, Hedvig Engström Jakobsson, Magnus Lindh |

We gratefully acknowledge the following Authors from the Originating laboratories responsible for obtaining the specimens, as well as the Submitting laboratories where the genome data were generated and shared via GISAID, on which this research is based.

All Submitters of data may be contacted directly via www.gisaid.org

Authors are sorted alphabetically.

| Accession ID | Originating Laboratory | Submitting Laboratory | Authors |
|---|---|---|---|
| EPI_ISL_402123, EPI_ISL_403928, EPI_ISL_403929, EPI_ISL_403930, EPI_ISL_403931 | Institute of Pathogen Biology, Chinese Academy of Medical Sciences & Peking Union Medical College | Institute of Pathogen Biology, Chinese Academy of Medical Sciences & Peking Union Medical College | Lili Ren, Jianwei Wang, Qi Jin, Zichun Xiang, Zhiqiang Wu, Chao Wu, Yiwei Liu |
| EPI_ISL_406973 | Singapore General Hospital | National Public Health Laboratory | Mak, TM; Octavia S; Chavatte JM; Zhou, ZY; Cui, L; Lin, RTP |
| EPI_ISL_411950 | NHC Key laboratory of Enteric Pathogenic Microbiology, Institute of Pathogenic Microbiology | Jiangsu Provincial Center for Disease Control & Prevention | Lunbiao Cui,Kangchen Zhao,Xiaojuan Zhu,Yiyue Ge,Tao Wu,Bin Wu,Yin Chen,Fengcai Zhu,Baoli Zhu,Ming Wu |
| EPI_ISL_411957 | Key Laboratory of Human Diseases, Comparative Medicine, Institute of Laboratory Animal Science | Key Laboratory of Human Diseases, Comparative Medicine, Institute of Laboratory Animal Science | Linlin,B., Lili,R., Shuran,G., Jiangning,L., Feifei,Q., Qi,L., Fengdi,L., Jing,X., Wei,D., Pin,Y., Yanfeng,X., Yajin,Q., Hong,G., Qiang,W., Mingya,L., Guanpeng,W., Shunyi,W., Zhiqi,S., Li,G., Lan,C., Conghui,W., Ying,W., Xinming,W., Yan,X., Qi,J. and Chuan,Q. |
| EPI_ISL_413518, EPI_ISL_413519, EPI_ISL_413520, EPI_ISL_413521 | Infectious Disease Control Center, Center for Disease Control and Prevention of PLA | Infectious Disease Control Center, Center for Disease Control and Prevention of PLA | Li,J., Li,L., Li,Z., Qiu,S., Song,H., Li,P. and Li,P. |
| EPI_ISL_415454, EPI_ISL_415455, EPI_ISL_415456, EPI_ISL_415457, EPI_ISL_415458, EPI_ISL_415459, EPI_ISL_415698, EPI_ISL_415699, EPI_ISL_415700, EPI_ISL_415701, EPI_ISL_415702, EPI_ISL_415703, EPI_ISL_415704, EPI_ISL_415705, EPI_ISL_415706, EPI_ISL_415707, EPI_ISL_415708 | | | |
| see above | University Hospitals of Geneva Laboratory of Virology | University Hospitals of Geneva Laboratory of Virology | Laubscher F. |
| EPI_ISL_416427, EPI_ISL_416428, EPI_ISL_416429, EPI_ISL_416430, EPI_ISL_416431 | National Influenza Center, National Institute of Hygiene and Epidemiology (NIHE) | National Influenza Center, National Institute of Hygiene and Epidemiology (NIHE) | Le Quynh Mai, Taichiro Takemura, Meng Ling Moi, Takeshi Nabeshima, Nguyen Le Khanh Hang, Hoang Vu Mai Phuong, Ung Thi Hong Trang, Le Thi Thanh, Nguyen Vu Son, Vuong Duc Cuong, Pham Thi Hien, Tran Thu Huong, Nguyen Phuong Anh, Pham Hong Quynh Anh, Kouichi Morita, Futoshi Hasebe, Dang Duc Anh |
| EPI_ISL_416432 | Clinical Microbiology Lab | Infectious Disease Research Department, King Abdullah International Medical Research Center (KAIMRC) | Majed Alghoribi, Sadeem Alhayli, Abdulrahman Alswaji, Liliane Okdah, Sameera Al Johani, Michel Doumith |
| EPI_ISL_417211, EPI_ISL_417212 | Dunedin Hospital | University of Otago | M.E. Quiñones-Mateu, B. Lawley, J. Grant, R. Harfoot, J. Ussher |
| EPI_ISL_417397 | Centre for Infectious Diseases and Microbiology Public Health | NSW Health Pathology - Institute of Clinical Pathology and Medical Research; Westmead Hospital; University of Sydney | Maddocks S, Kok J, Dwyer DE, Rockett R, Eden J-S, Lam C, Gray K, Timms V, Gall M, Arnott A, Sadsad R, Carter I, Rahman H, Holmes EC, O'Sullivan MV, Sintchenko V and Chen SC for the 2019-nCoV Study Group |
| EPI_ISL_417418 | Laboratory of Molecular Virology International Center fro Genetic Engineering and Biotechnology (ICGEB) | ARGO Open Lab Platform for Genome sequencing | Licastro D, Rajasekharan S, Dal Monego S, Segat L, D'Agaro P, Marcello A |
| EPI_ISL_417419 | Laboratory of Molecular Virology International Center for Genetic Engineering and Biotechnology (ICGEB) | ARGO Open Lab Platform for Genome sequencing | Licastro D, Rajasekharan S, Dal Monego S, Segat L, D'Agaro P, Marcello A |
| EPI_ISL_417420 | Jiangxi province Center for Disease Control and Prevention | Jiangxi province Center for Disease Control and Prevention | Li jian Xiong |
| EPI_ISL_417421 | Laboratory of Molecular Virology International Center for Genetic Engineering and Biotechnology (ICGEB) | ARGO Open Lab Platform for Genome sequencing | Licastro D, Rajasekharan S, Dal Monego S, Segat L, D'Agaro P, Marcello A |
| EPI_ISL_417423 | Laboratory of Molecular Virology International Center for Genetic Engineering and Biotechnology (ICGEB) | ARGO Open Lab Platform for Genome sequencing | Licastro D, Rajasekharan, Dal Monego S, Segat L, D'Agaro P, Marcello A |
| EPI_ISL_417924 | Secretaría de Salud Medellín | Instituto Nacional de Salud, Universidad Cooperativa de Colombia, Instituto Alexander von Humboldt, Imperial College-London, London School of Hygiene & Tropical Medicine | Marcela Mercado-Reyes, Katherine Laiton-Donato, Diego A. Álvarez-Díaz, Carlos Franco-Muñoz, Jose A. Usme-Ciro, Gloria Puerto, Nicolás D. Franco-Sierra, Mailyn A. Gonzalez, Zulma M. Cucunubá, Christian Julian VillabonaArenas, Liz Villabona-Arenas, Sussy Echeverria-Londoño, Astrid C. Flórez, Sergio Gomez Rangel, Luz Dary Rodriguez, Juliana Barbosa, Erika Ospitia, Diana Marcela Walteros-Acero, Martha Lucia Ospina Martinez |
| EPI_ISL_418190, EPI_ISL_418192, EPI_ISL_418193, EPI_ISL_418194, EPI_ISL_418196, EPI_ISL_418197, EPI_ISL_418198, EPI_ISL_418199, EPI_ISL_418200, EPI_ISL_418201, EPI_ISL_418202, EPI_ISL_418203, EPI_ISL_418204, EPI_ISL_418205, EPI_ISL_418254 | | | |
| see above | NYU Langone Health | Department of Pathology and Medicine, New York University School of Medicine | Margaret Black, John Cadley, Paolo Cotzia, John Chen, Dacia Dimartino, Xiaojun Feng, Adriana Heguy, Megan Hogan, Emily Huang, George Jour, Christian Marier, Matthew T. Maurano, Mark J. Mulligan, Peter Meyn, Jared Pinnell, Amy Rapkiewicz, Marie Samanovic-Golden, Antonio Serrano, Guomiao Shen, Matija Snuderl, Nick Vulpescu, Gael Westby, Paul Zappile |
| EPI_ISL_418255 | Presidio Ospedaliero "S. Spirito" - PESCARA | Istituto Zooprofilattico Sperimentale dell'Abruzzo e Molise "G. Caporale" | Lorusso A, Marcacci M, Cammà C, Monaco F, Puglia I, Di Pasquale A, Rinaldi A, Mangone I, Savini G |
| EPI_ISL_418256 | Ospedale "San Liberatore" di Atri | Istituto Zooprofilattico Sperimentale dell'Abruzzo e Molise "G. Caporale" | Lorusso A, Marcacci M, Di Domenico M, Puglia I, Curini V, Ancora M, Di Pasquale A, Rinaldi A, Mangone I, Cammà C, Savini G. |
| EPI_ISL_418257 | Ospedale Civile Giuseppe Mazzini, Teramo | Istituto Zooprofilattico Sperimentale dell'Abruzzo e Molise "G.Caporale" | Lorusso A, Marcacci M, Di Domenico M, Puglia I, Curini V, Ancora M, Di Pasquale A, Rinaldi A, Mangone I, Cammà C, Savini G. |
| EPI_ISL_418258, EPI_ISL_418259 | Presidio ospedaliero "Santo Spirito" | Istituto Zooprofilattico Sperimentale dell'Abruzzo e Molise "G. Caporale" | Lorusso A, Marcacci M, Di Domenico M, Puglia I, Curini V, Ancora M, Di Pasquale A, Rinaldi A, Mangone I, Cammà C, Savini G. |
| EPI_ISL_418260, EPI_ISL_418261 | Ospedale Civile Giuseppe Mazzini | Istituto Zooprofilattico Sperimentale dell'Abruzzo e Molise "G. Caporale" | Lorusso A, Marcacci M, Di Domenico M, Puglia I, Curini V, Ancora M, Di Pasquale A, Rinaldi A, Mangone I, Cammà C, Savini G. |
| EPI_ISL_418262 | Instituto Nacional de Salud | Instituto Nacional de Salud Universidad Cooperativa de Colombia Instituto Alexander von Humboldt Imperial College-London London School of Hygiene & Tropical Medicine | Marcela Mercado-Reyes, Katherine Laiton-Donato, Diego A. Álvarez-Díaz, Carlos Franco-Muñoz, Jose A. Usme-Ciro, Gloria Puerto, Nicolas D. Franco-Sierra, Mailyn A.Gonzalez, Zulma M. Cucunubá, Christian Julian VillabonaArenas, Liz Villabona-Arenas, Sussy Echeverria, Astrid C. Flórez, Sergio Gomez Rangel, Luz Dary Rodriguez, Juliana Barbosa, Erika Ospitia, Diana Marcela Walteros-Acero, Nuno Rodrigues Faria, Martha Lucia Ospina Martinez |
| EPI_ISL_418992, EPI_ISL_418993, EPI_ISL_418994, EPI_ISL_418995, EPI_ISL_418996, EPI_ISL_418997, EPI_ISL_418998, EPI_ISL_418999, EPI_ISL_419000, EPI_ISL_419001 | National Public Health Laboratory, National Centre for Infectious Diseases | National Public Health Laboratory, National Centre for Infectious Diseases | Mak TM, Octavia S, Cui L, Lin RTP |
| EPI_ISL_419259, EPI_ISL_419265 | Lab voor klinische biologie | Onderzoeksgroep Virologie | Laurens Lambrechts, Nick Vereecke, Marthe Pauwels, Basiel Cole, Bruno Verhasselt, Linos Vandekerckhove, Hans Nauwynck, Sebastiaan Theuns |
| EPI_ISL_419703 | NYU Langone Health | Departments of Pathology and Medicine, New York University School of Medicine | Maria Aguero-Rosenfeld, Margaret Black, John Cadley, Paolo Cotzia, John Chen, Dacia Dimartino, Xiaojun Feng, Adriana Heguy, Megan Hogan, Emily Huang, George Jour, Christian Marier, Matthew T. Maurano, Mark J. Mulligan, Peter Meyn, Jared Pinnell, Sitharam Ramaswami, Amy Rapkiewicz, Marie Samanovic-Golden, Antonio Serrano, Guomiao Shen, Matija Snuderl, Nick Vulpescu, Gael Westby, Paul Zappile, Yutong Zhang |
| EPI_ISL_420112 | Servicio de Microbiología. Consorcio Hospital General | Sequencing and Bioinformatics Service and Molecular | Lidia Ruiz Roldan, Marta Pla Diaz, Neris Garcia-Gonzalez, Loreto Ferrús Abad, Inma Galán Vendrell, Paula Ruiz-Hueso, Mariana Reyes-Prieto, Vicente |

| | | | |
|---|---|---|---|
| | Universitario de Valencia | Epidemiology Research Group. FISABIO-Public Health | Soriano Chirona, Maria Alma Bracho, Griselda De Marco, Beatriz Beamud, Maria Dolores Ocete, Llúcia Martínez-Priego, Concepcion Gimeno, Giuseppe D'Auria, Fernando Gonzalez-Candelas |
| EPI_ISL_420117 | Servicio de Microbiología. Consorcio Hospital General Universitario de Valencia | Sequencing and Bioinformatics Service and Molecular Epidemiology Research Group. FISABIO-Public Health | Loreto Ferrús Abad, Inma Galán Vendrell, Paula Ruiz-Hueso, Mariana Reyes-Prieto, Vicente Soriano Chirona, Maria Alma Bracho, Griselda De Marco, Beatriz Beamud, Lidia Ruiz Roldan, Marta Pla Diaz,Neris Garcia-Gonzalez, Maria Dolores Ocete, Llúcia Martínez-Priego, Concepcion Gimeno, Giuseppe D'Auria, Fernando Gonzalez-Candelas |
| EPI_ISL_420126 | Servicio de Microbiologia. Hospital Clinico Universitario de Valencia | Sequencing and Bioinformatics Service and Molecular Epidemiology Research Group. FISABIO-Public Health | Llúcia Martínez-Priego, Maria Alma Bracho, Griselda De Marco, Beatriz Beamud, Lidia Ruiz Roldan, Marta Pla Diaz, Neris Garcia-Gonzalez, Loreto Ferrús Abad, Inma Galán Vendrell, Paula Ruiz-Hueso, Mariana Reyes-Prieto, Vicente Soriano Chirona, David Navarro, Giuseppe D'Auria, Fernando Gonzalez-Candelas |
| EPI_ISL_420563 | Ospedale Civile Giuseppe Mazzini | Istituto Zooprofilattico Sperimentale dell'Abruzzo e Molise "G. Caporale" | Lorusso A, Marcacci M, Di Domenico M, Ancora M, Curini V, Mangone I, Rinaldi A, Di Pasquale A, Cammà C, Puglia I, Savini G |
| EPI_ISL_420564 | Ospedale Civile Castel Di Sangro | Istituto Zooprofilattico Sperimentale dell'Abruzzo e Molise "G.Caporale" | Lorusso A, Marcacci M, Di Domenico M, Ancora M, Curini V, Mangone I, Rinaldi A, Di Pasquale A, Cammà C, Puglia I, Savini G |
| EPI_ISL_420565 | Ospedale Civile Giuseppe Mazzini | Istituto Zooprofilattico Sperimentale dell'Abruzzo e Molise "G.Caporale" | Lorusso A, Marcacci M, Di Domenico M, Ancora M, Curini V, Mangone I, Rinaldi A, Di Pasquale A, Cammà C, Puglia I, Savini G |
| EPI_ISL_420566, EPI_ISL_420567 | Ospedale Regionale San Salvatore | Istituto Zooprofilattico Sperimentale dell'Abruzzo e Molise "G.Caporale" | Lorusso A, Marcacci M, Di Domenico M, Ancora M, Curini V, Mangone I, Rinaldi A, Di Pasquale A, Cammà C, Puglia I, Savini G |
| EPI_ISL_420568, EPI_ISL_420569, EPI_ISL_420583, EPI_ISL_420592 | Ospedale Civile Giuseppe Mazzini | Istituto Zooprofilattico Sperimentale dell'Abruzzo e Molise "G.Caporale" | Lorusso A, Marcacci M, Di Domenico M, Ancora M, Curini V, Mangone I, Rinaldi A, Di Pasquale A, Cammà C, Puglia I, Savini G |
| EPI_ISL_421516 | Servicio de Microbiologia. Hospital Clinico Universitario de Valencia | Sequencing and Bioinformatics Service and Molecular Epidemiology Research Group. FISABIO-Public Health | Loreto Ferrús Abad, Maria Alma Bracho, Griselda De Marco, Beatriz Beamud, Lidia Ruiz Roldan, Marta Pla Diaz, Neris Garcia-Gonzalez, Inma Galán Vendrell, Paula Ruiz-Hueso, Mariana Reyes-Prieto, Vicente Soriano Chirona, David Navarro, Llúcia Martínez-Priego, Giuseppe D'Auria, Fernando Gonzalez-Candelas |
| EPI_ISL_421520 | Servicio de Microbiologia. Hospital Clinico Universitario de Valencia | Sequencing and Bioinformatics Service and Molecular Epidemiology Research Group. FISABIO-Public Health | Loreto Ferrús Abad, Maria Alma Bracho, Griselda De Marco, Sandra Carbo, Beatriz Beamud, Lidia Ruiz Roldan, Marta Pla Diaz, Neris Garcia-Gonzalez, Inma Galán Vendrell, Paula Ruiz-Hueso, Mariana Reyes-Prieto, Vicente Soriano Chirona, Ivan Ansari, David Navarro, Llúcia Martínez-Priego, Giuseppe D'Auria, Fernando Gonzalez-Candelas |
| EPI_ISL_422428, EPI_ISL_422429, EPI_ISL_422430, EPI_ISL_422431, EPI_ISL_422432, EPI_ISL_422433, EPI_ISL_422434, EPI_ISL_422435 | National Public Health Laboratory, National Centre for Infectious Diseases | National Public Health Laboratory, National Centre for Infectious Diseases | Mak TM, Octavia S, Cui L, Lin RTP |
| EPI_ISL_425048, EPI_ISL_425050, EPI_ISL_425056, EPI_ISL_425057, EPI_ISL_425058 | Lab voor klinische biologie | Onderzoeksgroep Virologie | Laurens Lambrechts, Nick Vereecke, Marthe Pauwels, Basiel Cole, Bruno Verhasselt, Linos Vandekerckhove, Hans Nauwynck, Sebastiaan Theuns |
| EPI_ISL_425062, EPI_ISL_425063, EPI_ISL_425064 | Lab voor klinische biologie | Onderzoeksgroep Virologie | Laurens Lambrechts, Nick Vereecke, Marthe Pauwels, Jozefien De Clercq, Bruno Verhasselt, Linos Vandekerckhove, Hans Nauwynck, Sebastiaan Theuns |
| EPI_ISL_425182 | Servicio de Microbiología. Consorcio Hospital General Universitario de Valencia | Sequencing and Bioinformatics Service and Molecular Epidemiology Research Group. FISABIO-Public Health | Lidia Ruiz Roldan, Marta Pla Diaz, Neris Garcia-Gonzalez, Inma Galán Vendrell, Sandra Carbo, Loreto Ferrús Abad, Paula Ruiz-Hueso, Mariana Reyes-Prieto, Vicente Soriano Chirona, Ivan Ansari, David Navarro, Maria Alma Bracho, Griselda De Marco, Beatriz Beamud, Llúcia Martínez-Priego, Giuseppe D'Auria, Fernando Gonzalez-Candelas |
| EPI_ISL_425187 | Servicio de Microbiología. Consorcio Hospital General Universitario de Valencia | Sequencing and Bioinformatics Service and Molecular Epidemiology Research Group. FISABIO-Public Health | Loreto Ferrús Abad, Paula Ruiz-Hueso, Mariana Reyes-Prieto, Vicente Soriano Chirona, Ivan Ansari, David Navarro, Maria Alma Bracho, Griselda De Marco, Beatriz Beamud, Lidia Ruiz Roldan, Marta Pla Diaz, Neris Garcia-Gonzalez, Inma Galán Vendrell, Sandra Carbo, Llúcia Martínez-Priego, Giuseppe D'Auria, Fernando Gonzalez-Candelas |
| EPI_ISL_425196 | Servicio de Microbiología. Consorcio Hospital General Universitario de Valencia | Sequencing and Bioinformatics Service and Molecular Epidemiology Research Group. FISABIO-Public Health | Lidia Ruiz Roldan, Marta Pla Diaz, Neris Garcia-Gonzalez, Inma Galán Vendrell, Sandra Carbo, Loreto Ferrús Abad, Paula Ruiz-Hueso, Mariana Reyes-Prieto, Vicente Soriano Chirona, Ivan Ansari, David Navarro, Maria Alma Bracho, Griselda De Marco, Beatriz Beamud, Llúcia Martínez-Priego, Giuseppe D'Auria, Fernando Gonzalez-Candelas |
| EPI_ISL_425207 | Servicio de Microbiologia. Hospital Clinico Universitario de Valencia | Sequencing and Bioinformatics Service and Molecular Epidemiology Research Group. FISABIO-Public Health | Lidia Ruiz Roldan, Marta Pla Diaz,Neris Garcia-Gonzalez, Loreto Ferrús Abad, Maria Dolores Ocete, Inma Galán Vendrell, Paula Ruiz-Hueso, Mariana Reyes-Prieto, Vicente Soriano Chirona, Maria Alma Bracho, Griselda De Marco, Beatriz Beamud, Llúcia Martínez-Priego, Concepcion Gimeno, Giuseppe D'Auria, Fernando Gonzalez-Candelas |
| EPI_ISL_425210 | Servicio de Microbiologia. Hospital Clinico Universitario de Valencia | Sequencing and Bioinformatics Service and Molecular Epidemiology Research Group. FISABIO-Public Health | Loreto Ferrús Abad, Maria Dolores Ocete, Inma Galán Vendrell, Paula Ruiz-Hueso, Mariana Reyes-Prieto, Vicente Soriano Chirona, Maria Alma Bracho, Griselda De Marco, Beatriz Beamud, Lidia Ruiz Roldan, Marta Pla Diaz,Neris Garcia-Gonzalez, Llúcia Martínez-Priego, Concepcion Gimeno, Giuseppe D'Auria, Fernando Gonzalez-Candelas |
| EPI_ISL_425211 | Servicio de Microbiologia. Hospital Clinico Universitario de Valencia | Sequencing and Bioinformatics Service and Molecular Epidemiology Research Group. FISABIO-Public Health | Loreto Ferrús Abad, Maria Dolores Ocete,Inma Galán Vendrell, Paula Ruiz-Hueso, Mariana Reyes-Prieto, Vicente Soriano Chirona, Maria Alma Bracho, Griselda De Marco, Beatriz Beamud, Lidia Ruiz Roldan, Marta Pla Diaz,Neris Garcia-Gonzalez, Llúcia Martínez-Priego, Concepcion Gimeno, Giuseppe D'Auria, Fernando Gonzalez-Candelas |
| EPI_ISL_425216 | Servicio de Microbiología. Consorcio Hospital General Universitario de Valencia | Sequencing and Bioinformatics Service and Molecular Epidemiology Research Group. FISABIO-Public Health | Lidia Ruiz Roldan, Marta Pla Diaz, Neris Garcia-Gonzalez, Inma Galán Vendrell, Sandra Carbo, Loreto Ferrús Abad, Paula Ruiz-Hueso, Mariana Reyes-Prieto, Vicente Soriano Chirona, Ivan Ansari, David Navarro, Maria Alma Bracho, Griselda De Marco, Beatriz Beamud, Llúcia Martínez-Priego, Giuseppe D'Auria, Fernando Gonzalez-Candelas |
| EPI_ISL_425222 | Servicio de Microbiología. Consorcio Hospital General Universitario de Valencia | Sequencing and Bioinformatics Service and Molecular Epidemiology Research Group. FISABIO-Public Health | Loreto Ferrús Abad, Paula Ruiz-Hueso, Mariana Reyes-Prieto, Vicente Soriano Chirona, Ivan Ansari, David Navarro, Maria Alma Bracho, Griselda De Marco, Beatriz Beamud, Lidia Ruiz Roldan, Marta Pla Diaz, Neris Garcia-Gonzalez, Inma Galán Vendrell, Sandra Carbo, Llúcia Martínez-Priego, Giuseppe D'Auria, Fernando Gonzalez-Candelas |
| EPI_ISL_425228, EPI_ISL_425229 | Laboratory of Molecular Genetics, 2nd Faculty of Medicine, Charles University in Prague, Prague, Czech Republic | Laboratory of Molecular Genetics, 2nd Faculty of Medicine, Charles University in Prague, Prague, Czech Republic | Lenka Kramná, Kateina Poláková, Ondej Cinek |
| EPI_ISL_425231, EPI_ISL_425232, EPI_ISL_425233, EPI_ISL_425234, EPI_ISL_425235, EPI_ISL_425236, EPI_ISL_425237, EPI_ISL_425238, EPI_ISL_425239, EPI_ISL_425240, EPI_ISL_425241, EPI_ISL_425242, EPI_ISL_425243, EPI_ISL_425244, EPI_ISL_425245, EPI_ISL_425246, EPI_ISL_425247, EPI_ISL_425248, EPI_ISL_425249, EPI_ISL_425250, EPI_ISL_425251, EPI_ISL_425252, EPI_ISL_425253, EPI_ISL_425254, EPI_ISL_425255, EPI_ISL_425256, EPI_ISL_425257, EPI_ISL_425260, EPI_ISL_425262, EPI_ISL_425263, EPI_ISL_425264, EPI_ISL_425265, EPI_ISL_425266, EPI_ISL_425267, EPI_ISL_425268, EPI_ISL_425269, EPI_ISL_425270, EPI_ISL_425271, EPI_ISL_425272, EPI_ISL_425273, EPI_ISL_425275, EPI_ISL_425276, EPI_ISL_425277, EPI_ISL_425278, EPI_ISL_425279, EPI_ISL_425281, EPI_ISL_425282, EPI_ISL_425283, EPI_ISL_425284, EPI_ISL_425285, EPI_ISL_425286, EPI_ISL_425287, EPI_ISL_425288, EPI_ISL_425289, EPI_ISL_425290, EPI_ISL_425291, EPI_ISL_425292, EPI_ISL_425293, EPI_ISL_425294, EPI_ISL_425295, EPI_ISL_425296, EPI_ISL_425297, EPI_ISL_425298, EPI_ISL_425299, EPI_ISL_425300, EPI_ISL_425301, EPI_ISL_425302, EPI_ISL_425303, EPI_ISL_425304, EPI_ISL_425305, EPI_ISL_425306, EPI_ISL_425307, EPI_ISL_425308, EPI_ISL_425309, EPI_ISL_425310, EPI_ISL_425311, EPI_ISL_425312, EPI_ISL_425313, EPI_ISL_425314, EPI_ISL_425315, EPI_ISL_425316, EPI_ISL_425317, EPI_ISL_425318, EPI_ISL_425319, EPI_ISL_425320, EPI_ISL_425321, EPI_ISL_425322, EPI_ISL_425323, EPI_ISL_425324, EPI_ISL_425325, EPI_ISL_425326, EPI_ISL_425327, EPI_ISL_425328, EPI_ISL_425329, EPI_ISL_425330, EPI_ISL_425331, EPI_ISL_425332, EPI_ISL_425333, EPI_ISL_425334, EPI_ISL_425335, EPI_ISL_425336, EPI_ISL_425337, EPI_ISL_425338, EPI_ISL_425339, EPI_ISL_425340, EPI_ISL_425341, EPI_ISL_425342, EPI_ISL_425343, EPI_ISL_425344, EPI_ISL_425345, EPI_ISL_425347, EPI_ISL_425348, EPI_ISL_425349, EPI_ISL_425350, EPI_ISL_425351, EPI_ISL_425352, EPI_ISL_425353, EPI_ISL_425354, EPI_ISL_425355, EPI_ISL_425356, EPI_ISL_425357, EPI_ISL_425358, EPI_ISL_425359, EPI_ISL_425360, EPI_ISL_425361, EPI_ISL_425362, EPI_ISL_425363, EPI_ISL_425364, EPI_ISL_425365, EPI_ISL_425366, EPI_ISL_425367, EPI_ISL_425368, EPI_ISL_425369, EPI_ISL_425370, EPI_ISL_425371, EPI_ISL_425373, EPI_ISL_425374, EPI_ISL_425375, EPI_ISL_425376, EPI_ISL_425377, EPI_ISL_425378, EPI_ISL_425379, EPI_ISL_425380, EPI_ISL_425381, EPI_ISL_425382, EPI_ISL_425383, EPI_ISL_425384, EPI_ISL_425385, EPI_ISL_425387, EPI_ISL_425388, EPI_ISL_425389, EPI_ISL_425390, EPI_ISL_425391, EPI_ISL_425392, EPI_ISL_425393, EPI_ISL_425394, EPI_ISL_425395, EPI_ISL_425396, EPI_ISL_425397, EPI_ISL_425398, EPI_ISL_425399, EPI_ISL_425400, EPI_ISL_425401, EPI_ISL_425402, EPI_ISL_425403, EPI_ISL_425404, EPI_ISL_425406, EPI_ISL_425407, EPI_ISL_425408, EPI_ISL_425409, EPI_ISL_425411, EPI_ISL_425412, EPI_ISL_425413, EPI_ISL_425414, EPI_ISL_425415, EPI_ISL_425416, EPI_ISL_425417, EPI_ISL_425418, EPI_ISL_425420, EPI_ISL_425421, EPI_ISL_425423, EPI_ISL_425424, EPI_ISL_425425, EPI_ISL_425426, EPI_ISL_425427, EPI_ISL_425428, EPI_ISL_425429, EPI_ISL_425430, EPI_ISL_425431, EPI_ISL_425432, EPI_ISL_425433, EPI_ISL_425434, EPI_ISL_425435, EPI_ISL_425436, EPI_ISL_425437, EPI_ISL_425438, EPI_ISL_425441, EPI_ISL_425442, EPI_ISL_425443, EPI_ISL_425444, EPI_ISL_425445, EPI_ISL_425446, EPI_ISL_425447, EPI_ISL_425448, EPI_ISL_425449, EPI_ISL_425450, EPI_ISL_425451, EPI_ISL_425452, EPI_ISL_425453, EPI_ISL_425454, EPI_ISL_425455, EPI_ISL_425456, EPI_ISL_425457, EPI_ISL_425458, EPI_ISL_425459, EPI_ISL_425460, EPI_ISL_425461 | | | |
| see above | Department of Pathology, University of Cambridge | COVID-19 Genomics UK (COG-UK) Consortium | Luke W Meredith, M. Estee Torok , Myra Hosmillo, William L. Hamilton, Martin D. Curran, Theresa Feltwell, Anna Yakovleva, Charlotte J. Houldcroft, Aminu S. Jahun, Sarah L. Caddy, Ian Goodfellow |
| EPI_ISL_426051 | Laboratory of Molecular Genetics, 2nd Faculty of Medicine, | Laboratory of Molecular Genetics, 2nd Faculty of Medicine, | Lenka Kramná, Kateina Poláková, Ondej Cinek |

| | | | |
|---|---|---|---|
| | Charles University in Prague, Prague, Czech Republic | Charles University in Prague, Prague, Czech Republic | |
| EPI_ISL_426356, EPI_ISL_426357, EPI_ISL_426358, EPI_ISL_426359, EPI_ISL_426360 | Laboratory of Molecular Genetics, 2nd Faculty of Medicine, Charles University in Prague, Prague, Czech Republic | Laboratory of Molecular Genetics, 2nd Faculty of Medicine, Charles University in Prague, Prague, Czech Republic | Lenka Kramna, Katerina Polackova, Ondrej Cinek |
| EPI_ISL_426379 | The National Laboratory of Health, Environment and Food, Maribor, Slovenia | The National Laboratory of Health, Environment and Food, Maribor, Slovenia | Mahnic A., Hedzet S., Janezic S., Duh D., Zavrsnik J., Blazun Vosner H., Rupnik M. |
| EPI_ISL_428209 | Laboratory of Molecular Biology, Diagnostyka sp. z o.o. | Laboratory of Recombinant Vaccines | Lukasz Rabalski, Anna Piotrowska-Mietelska, Boguslaw Szewczyk, Krystyna Bienkowska-Szewczyk |
| EPI_ISL_428822, EPI_ISL_428823, EPI_ISL_428824, EPI_ISL_428825, EPI_ISL_428826, EPI_ISL_428827, EPI_ISL_428828, EPI_ISL_428829, EPI_ISL_428830, EPI_ISL_428831, EPI_ISL_428832, EPI_ISL_428833, EPI_ISL_428834, EPI_ISL_428835, EPI_ISL_428836, EPI_ISL_428837, EPI_ISL_428838, EPI_ISL_428839, EPI_ISL_428840, EPI_ISL_428841, EPI_ISL_428842, EPI_ISL_428843, EPI_ISL_428844, EPI_ISL_428845, EPI_ISL_428846, EPI_ISL_428847, EPI_ISL_428848, EPI_ISL_428849, EPI_ISL_428850 | | | |
| see above | National Public Health Laboratory, National Centre for Infectious Diseases | National Public Health Laboratory, National Centre for Infectious Diseases | Mak TM, Octavia S, Chavatte JM, Cui L, Lin RTP |
| EPI_ISL_428851, EPI_ISL_428852 | FSBSI "Chumakov Federal Scientific Center for Research and Development of Immune-and-Biological Products of Russian Academy of Sciences" | FSBSI "Chumakov Federal Scientific Center for Research and Development of Immune-and-Biological Products of Russian Academy of Sciences" & NRC "Kurchatov institute" | Liubov Kozlovskaya, Anastasia Piniaeva, Georgy Ignatyev, Anna Shishova, Aydar Ishmukhametov, Mikhail Rychev, Egor Prokhorchuk, Denis Protsenko, Anastasia Berestovskaya |
| EPI_ISL_428853 | Laboratory of Molecular Virology International Center for Genetic Engineering and Biotechnology (ICGEB) | ARGO Open Lab Platform for Genome Sequencing | Licastro D, Rajasekharan S, Dal Monego S, Segat L, D'Agaro P, Marcello A |
| EPI_ISL_428854 | Laboratory of Molecular Virology International Center for Genetic Engineering and Biotechnology (ICGEB) | ARGO Open Lab Platform for Genome sequencing | Licastro D, Rajasekharan S, Dal Monego S, Segat L, D'Agaro P, Marcello A |
| EPI_ISL_429196, EPI_ISL_429197, EPI_ISL_429199, EPI_ISL_429200, EPI_ISL_429201, EPI_ISL_429203, EPI_ISL_429204, EPI_ISL_429206, EPI_ISL_429207, EPI_ISL_429208, EPI_ISL_429209, EPI_ISL_429210, EPI_ISL_429211, EPI_ISL_429212, EPI_ISL_429213, EPI_ISL_429214, EPI_ISL_429215, EPI_ISL_429217, EPI_ISL_429218, EPI_ISL_429219, EPI_ISL_429220, EPI_ISL_429221, EPI_ISL_429222, EPI_ISL_429223 | | | |
| see above | University Hospitals of Geneva Laboratory of Virology | University Hospitals of Geneva Laboratory of Virology | Laubscher F. |
| EPI_ISL_429226, EPI_ISL_429227 | Presidio Ospedaliero Santo Spirito | Istituto Zooprofilattico Sperimentale dell'Abruzzo e Molise "G. Caporale" | Lorusso A, Marcacci M, Di Domenico M, Ancora M, Curini V, Mangone I, Rinaldi A, Di Pasquale A, Camma C, Puglia I, Savini G |
| EPI_ISL_429228 | Ospedale Civile Giuseppe Mazzini | Istituto Zooprofilattico Sperimentale dell'Abruzzo e Molise "G. Caporale" | Lorusso A, Marcacci M, Di Domenico M, Ancora M, Curini V, Mangone I, Rinaldi A, Di Pasquale A, Camma C, Puglia I, Savini G |
| EPI_ISL_429229 | Ospedale Regionale San Salvatore | Istituto Zooprofilattico Sperimentale dell'Abruzzo e Molise "G. Caporale" | Lorusso A, Marcacci M, Di Domenico M, Ancora M, Curini V, Mangone I, Rinaldi A, Di Pasquale A, Camma C, Puglia I, Savini G |
| EPI_ISL_429230, EPI_ISL_429231, EPI_ISL_429232, EPI_ISL_429233, EPI_ISL_429234, EPI_ISL_429235 | Ospedale Civile Giuseppe Mazzini | Istituto Zooprofilattico Sperimentale dell'Abruzzo e Molise "G. Caporale" | Lorusso A, Marcacci M, Di Domenico M, Ancora M, Curini V, Mangone I, Rinaldi A, Di Pasquale A, Camma C, Puglia I, Savini G |
| EPI_ISL_429236 | Ospedale Civile S. Liberatore di Atri | Istituto Zooprofilattico Sperimentale dell'Abruzzo e Molise "G. Caporale" | Lorusso A, Marcacci M, Di Domenico M, Ancora M, Curini V, Mangone I, Rinaldi A, Di Pasquale A, Camma C, Puglia I, Savini G |
| EPI_ISL_430419, EPI_ISL_430420, EPI_ISL_430421, EPI_ISL_430422, EPI_ISL_430423, EPI_ISL_430424, EPI_ISL_430425, EPI_ISL_430426, EPI_ISL_430427, EPI_ISL_430429 | NYU Langone Health | Departments of Pathology and Medicine, New York University School of Medicine | Maria Aguero-Rosenfeld, Brendan Belovarac, Margaret Black, Ludovic Boytard, John Cadley, Paolo Cotzia, John Chen, Dacia Dimartino, Xiaojun Feng, Tatyana Gindin, Emily Guzman, Adriana Heguy, Megan Hogan, Emily Huang, George Jour, Lawrence H. Lin, Raven Luther, Andrew Lytle, Christian Marier, Matthew T. Maurano, Mark J. Mulligan, Peter Meyn, Raquel Ordonez Ciriza, Iman Osman, Jared Pinnell, Vanessa Raabe, Sitharam Ramaswami, Amy Rapkiewicz, Andre M. Ribeiro-dos-Santos, Marie Samanovic-Golden, Antonio Serrano, Guomiao Shen, Matija Snuderl, Theodore Vougiouklakis, Nick Vulpescu, Gael Westby, Paul Zappile, Yutong Zhang |
| EPI_ISL_431118, EPI_ISL_431180, EPI_ISL_431240, EPI_ISL_431292, EPI_ISL_431779, EPI_ISL_431780, EPI_ISL_431781, EPI_ISL_431782, EPI_ISL_431783, EPI_ISL_431784, EPI_ISL_431785 | | | |
| see above | Fujian Center for Disease Control and Prevention | Fujian Center for Disease Control and Prevention | Lin Qi, Huang Zhimiao, Zhang Yanhua, Weng Yuwei |
| EPI_ISL_433466, EPI_ISL_433467, EPI_ISL_433468, EPI_ISL_433469, EPI_ISL_433470, EPI_ISL_433471, EPI_ISL_433472, EPI_ISL_433473, EPI_ISL_433474, EPI_ISL_433475, EPI_ISL_433476, EPI_ISL_433477, EPI_ISL_433478, EPI_ISL_433479, EPI_ISL_433480, EPI_ISL_433481, EPI_ISL_433482, EPI_ISL_433483, EPI_ISL_433484, EPI_ISL_433485, EPI_ISL_433486, EPI_ISL_433487, EPI_ISL_433489, EPI_ISL_433490, EPI_ISL_433491, EPI_ISL_433492, EPI_ISL_433493, EPI_ISL_433494, EPI_ISL_433495, EPI_ISL_433496, EPI_ISL_433497, EPI_ISL_433666, EPI_ISL_433667, EPI_ISL_433668, EPI_ISL_433669, EPI_ISL_433670, EPI_ISL_433671, EPI_ISL_433672, EPI_ISL_433673, EPI_ISL_433674, EPI_ISL_433675, EPI_ISL_433676, EPI_ISL_433677, EPI_ISL_433678, EPI_ISL_433679, EPI_ISL_433680, EPI_ISL_433681, EPI_ISL_433682, EPI_ISL_433683, EPI_ISL_433684, EPI_ISL_433685, EPI_ISL_433686, EPI_ISL_433688, EPI_ISL_433689, EPI_ISL_433690, EPI_ISL_433691, EPI_ISL_433692, EPI_ISL_433693, EPI_ISL_433694, EPI_ISL_433695, EPI_ISL_433696, EPI_ISL_433697, EPI_ISL_433698, EPI_ISL_433699, EPI_ISL_433700, EPI_ISL_433701, EPI_ISL_433702, EPI_ISL_433703, EPI_ISL_433704, EPI_ISL_433705, EPI_ISL_433706, EPI_ISL_433707, EPI_ISL_433709, EPI_ISL_433710, EPI_ISL_433711, EPI_ISL_433712, EPI_ISL_433713, EPI_ISL_433714, EPI_ISL_433715, EPI_ISL_433717, EPI_ISL_433718, EPI_ISL_433719, EPI_ISL_433720, EPI_ISL_433721, EPI_ISL_433722, EPI_ISL_433723, EPI_ISL_433724, EPI_ISL_433725, EPI_ISL_433726, EPI_ISL_433727, EPI_ISL_433728, EPI_ISL_433729, EPI_ISL_433730, EPI_ISL_433731, EPI_ISL_433732, EPI_ISL_433733, EPI_ISL_433734, EPI_ISL_433735, EPI_ISL_433736, EPI_ISL_433737, EPI_ISL_433738, EPI_ISL_433739, EPI_ISL_433741, EPI_ISL_433742, EPI_ISL_433743, EPI_ISL_433744, EPI_ISL_433745, EPI_ISL_433746, EPI_ISL_433748, EPI_ISL_433749, EPI_ISL_433750, EPI_ISL_433751, EPI_ISL_433752, EPI_ISL_433753, EPI_ISL_433754, EPI_ISL_433755, EPI_ISL_433756, EPI_ISL_433757, EPI_ISL_433758, EPI_ISL_433759, EPI_ISL_433760, EPI_ISL_433761, EPI_ISL_433762, EPI_ISL_433763, EPI_ISL_433764, EPI_ISL_433765, EPI_ISL_433767, EPI_ISL_433768, EPI_ISL_433769, EPI_ISL_433770, EPI_ISL_433771, EPI_ISL_433772, EPI_ISL_433773, EPI_ISL_433774, EPI_ISL_433775, EPI_ISL_433776, EPI_ISL_433777, EPI_ISL_433778, EPI_ISL_433779, EPI_ISL_433780, EPI_ISL_433781, EPI_ISL_433782, EPI_ISL_433783, EPI_ISL_433784, EPI_ISL_433785, EPI_ISL_433786, EPI_ISL_433787, EPI_ISL_433788, EPI_ISL_433789, EPI_ISL_433790, EPI_ISL_433791, EPI_ISL_433792, EPI_ISL_433793, EPI_ISL_433794, EPI_ISL_433795, EPI_ISL_433796, EPI_ISL_433797, EPI_ISL_433798, EPI_ISL_433799, EPI_ISL_433800, EPI_ISL_433801, EPI_ISL_433802, EPI_ISL_433803, EPI_ISL_433804, EPI_ISL_433805, EPI_ISL_433806, EPI_ISL_433808, EPI_ISL_433809, EPI_ISL_433810, EPI_ISL_433811, EPI_ISL_433812, EPI_ISL_433813, EPI_ISL_433814, EPI_ISL_433815, EPI_ISL_433816, EPI_ISL_433817, EPI_ISL_433818, EPI_ISL_433819, EPI_ISL_433820, EPI_ISL_433821, EPI_ISL_433822, EPI_ISL_433823, EPI_ISL_433824, EPI_ISL_433825, EPI_ISL_433826, EPI_ISL_433827, EPI_ISL_433828, EPI_ISL_433829, EPI_ISL_433830, EPI_ISL_433831, EPI_ISL_433832, EPI_ISL_433833, EPI_ISL_433834, EPI_ISL_433835, EPI_ISL_433836, EPI_ISL_433837, EPI_ISL_433839, EPI_ISL_433840, EPI_ISL_433841, EPI_ISL_433842, EPI_ISL_433843, EPI_ISL_433844, EPI_ISL_433845, EPI_ISL_433846, EPI_ISL_433847, EPI_ISL_433848, EPI_ISL_433849, EPI_ISL_433850, EPI_ISL_433851, EPI_ISL_433852, EPI_ISL_433853, EPI_ISL_433854, EPI_ISL_433855, EPI_ISL_433856, EPI_ISL_433857, EPI_ISL_433858, EPI_ISL_433859, EPI_ISL_433860, EPI_ISL_433861, EPI_ISL_433862, EPI_ISL_433863, EPI_ISL_433864, EPI_ISL_433865, EPI_ISL_433866, EPI_ISL_433867, EPI_ISL_433868, EPI_ISL_433869, EPI_ISL_433870, EPI_ISL_433871, EPI_ISL_433872, EPI_ISL_433873, EPI_ISL_433874, EPI_ISL_433875, EPI_ISL_433876, EPI_ISL_433877, EPI_ISL_433878, EPI_ISL_433879, EPI_ISL_433880, EPI_ISL_433881, EPI_ISL_433882, EPI_ISL_433883, EPI_ISL_433884, EPI_ISL_433885, EPI_ISL_433886, EPI_ISL_433887, EPI_ISL_433888, EPI_ISL_433889, EPI_ISL_433890, EPI_ISL_433891, EPI_ISL_433892, EPI_ISL_433893, EPI_ISL_433894, EPI_ISL_433895, EPI_ISL_433896, EPI_ISL_433897, EPI_ISL_433898, EPI_ISL_433899, EPI_ISL_433900, EPI_ISL_433901, EPI_ISL_433902, EPI_ISL_433903, EPI_ISL_433904, EPI_ISL_433905, EPI_ISL_433906, EPI_ISL_433907, EPI_ISL_433908, EPI_ISL_433909, EPI_ISL_433910, EPI_ISL_433911, EPI_ISL_433912, EPI_ISL_433913, EPI_ISL_433914, EPI_ISL_433915, EPI_ISL_433916, EPI_ISL_433917, EPI_ISL_433918, EPI_ISL_433919, EPI_ISL_433920, EPI_ISL_433921, EPI_ISL_433922, EPI_ISL_433923, EPI_ISL_433924, EPI_ISL_433925, EPI_ISL_433926, EPI_ISL_433927, EPI_ISL_433928, EPI_ISL_433929, EPI_ISL_433930, EPI_ISL_433931, EPI_ISL_433932, EPI_ISL_433933, EPI_ISL_433934, EPI_ISL_433935, EPI_ISL_433936, EPI_ISL_433937, EPI_ISL_433938, EPI_ISL_433939, EPI_ISL_433940, EPI_ISL_433941, EPI_ISL_433942, EPI_ISL_433943, EPI_ISL_433944, EPI_ISL_433945, EPI_ISL_433946, EPI_ISL_433947, EPI_ISL_433948, EPI_ISL_433949, EPI_ISL_433950, EPI_ISL_433951, EPI_ISL_433952, EPI_ISL_433953, EPI_ISL_433954, EPI_ISL_433955, EPI_ISL_433956, EPI_ISL_433957, EPI_ISL_433958, EPI_ISL_433959, EPI_ISL_433960, EPI_ISL_433961, EPI_ISL_433962, EPI_ISL_433963, EPI_ISL_433964, EPI_ISL_433965, EPI_ISL_433966, EPI_ISL_433967, EPI_ISL_433968, EPI_ISL_433969, EPI_ISL_433970, EPI_ISL_433971, EPI_ISL_433974, EPI_ISL_433975, EPI_ISL_433976, EPI_ISL_433977, EPI_ISL_433978, EPI_ISL_433979, EPI_ISL_433980, EPI_ISL_433981, EPI_ISL_433982, EPI_ISL_433983, EPI_ISL_433986, EPI_ISL_433987, EPI_ISL_433988, EPI_ISL_433989, EPI_ISL_433990, EPI_ISL_433991, EPI_ISL_433992, EPI_ISL_433993, EPI_ISL_433994, EPI_ISL_433995, EPI_ISL_433996, EPI_ISL_433997, EPI_ISL_433999, EPI_ISL_434000, EPI_ISL_434001, EPI_ISL_434002, EPI_ISL_434003, EPI_ISL_434004, EPI_ISL_434005, EPI_ISL_434006, EPI_ISL_434008, EPI_ISL_434009, EPI_ISL_434010, EPI_ISL_434011, EPI_ISL_434012, EPI_ISL_434013, EPI_ISL_434014, EPI_ISL_434015, EPI_ISL_434016, EPI_ISL_434017, EPI_ISL_434018, EPI_ISL_434019, EPI_ISL_434020, EPI_ISL_434021, EPI_ISL_434022, EPI_ISL_434023, EPI_ISL_434024, EPI_ISL_434025, EPI_ISL_434026, EPI_ISL_434027, EPI_ISL_434028, EPI_ISL_434029, EPI_ISL_434030, EPI_ISL_434031, EPI_ISL_434032, EPI_ISL_434033, EPI_ISL_434034, EPI_ISL_434035, EPI_ISL_434036, EPI_ISL_434037, EPI_ISL_434038, EPI_ISL_434039, EPI_ISL_434040, EPI_ISL_434041, EPI_ISL_434042, EPI_ISL_434043, EPI_ISL_434044, EPI_ISL_434045, EPI_ISL_434046, EPI_ISL_434047, EPI_ISL_434048, EPI_ISL_434049, EPI_ISL_434050, EPI_ISL_434051, EPI_ISL_434052, EPI_ISL_434053, EPI_ISL_434054, EPI_ISL_434055, EPI_ISL_434056, EPI_ISL_434057, EPI_ISL_434058, EPI_ISL_434059, EPI_ISL_434060, EPI_ISL_434061, EPI_ISL_434062 | | | |
| see above | Department of Pathology, University of Cambridge | COVID-19 Genomics UK (COG-UK) Consortium | Luke W Meredith, M. Estee Torok , Myra Hosmillo, William L. Hamilton, Martin D. Curran, Theresa Feltwell, Grant Hall, Anna Yakovleva, Fahad A Khokhar, Charlotte J. Houldcroft, Laura G Caller, Aminu S. Jahun, Sarah L. Caddy, Ian Goodfellow |
| EPI_ISL_434347, EPI_ISL_434348, EPI_ISL_434349, EPI_ISL_434350, EPI_ISL_434351, EPI_ISL_434352, EPI_ISL_434353, EPI_ISL_434354, EPI_ISL_434355 | Lab voor klinische biologie | Onderzoeksgroep Virologie | Laurens Lambrechts, Nick Vereecke, Marthe Pauwels, Jozefien De Clercq, Bruno Verhasselt, Linos Vandekerckhove, Hans Nauwynck, Sebastiaan Theuns |
| EPI_ISL_434644 | Kungsholmsdoktorn | The Public Health Agency of Sweden | Linus Hammar, Oskar Karlsson Lindsjo, Maria Lind Karlberg, Anna-Malin Linde, Olov Svartstrom, Anna Risberg, Theresa Enkirch, Mia Brytting, Karin Tegmark-Wisell |
| EPI_ISL_434671 | Narhalsan Sjobo vardcentral | The Public Health Agency of Sweden | Lovisa Hjerten, Oskar Karlsson Lindsjo, Maria Lind Karlberg, Anna-Malin Linde, Olov Svartstrom, Anna Risberg, Theresa Enkirch, Mia Brytting, Karin Tegmark-Wisell |
| EPI_ISL_435056 | Gujarat Biotechnology Research Centre | Gujarat Biotechnology Research Centre | Maharshi Pandya, Amit Kanani, Akanksha Verma, Nitin Savaliya, Raghawendra Kumar, Dinesh Kumar, Zuber Saiyed, Dipa Kinariwala, Disha Patel, Binita Aring, Geeta Vaghela, Sonia Barve, Bhavesh Modi, Kairavi Joshi, Gaurishankar Shrimali, Nidhi Sood, Pranay Shah, R D Dixit, Snehal Bagatharia, Kamlesh |

| | | | |
|---|---|---|---|
| | | | J Upadhyay, Ramesh Pandit, Tejas Shah, Ankit Hinsu, Pritesh Sabara, Apurvasinh Puvar, Janvi Raval, Monika Gandhi, Pinal Trivedi, Afzal Ansari, Chaitanya Joshi, Madhvi Joshi |
| EPI_ISL_435145 | Ospedale Civile Giuseppe Mazzini | Istituto Zooprofilattico Sperimentale dell'Abruzzo e Molise "G.Caporale" | Lorusso A, Marcacci M, Di Domenico M, Ancora M, Curini V, Mangone I, Rinaldi A, Di Pasquale A, Cammà C, Puglia I, Savini G |
| EPI_ISL_435146, EPI_ISL_435147 | Villa Serena del Dr. Leonardo Petruzzi | Istituto Zooprofilattico Sperimentale dell'Abruzzo e Molise "G.Caporale" | Lorusso A, Marcacci M, Di Domenico M, Ancora M, Curini V, Mangone I, Rinaldi A, Di Pasquale A, Cammà C, Puglia I, Savini G |
| EPI_ISL_435148 | Ospedale SS Annunziata | Istituto Zooprofilattico Sperimentale dell'Abruzzo e Molise "G.Caporale" | Lorusso A, Marcacci M, Di Domenico M, Ancora M, Curini V, Mangone I, Rinaldi A, Di Pasquale A, Cammà C, Puglia I, Savini G |
| EPI_ISL_435149 | SERVIZIO DI IGIENE E SANITÀ PUBBLICA ASL Teramo | Istituto Zooprofilattico Sperimentale dell'Abruzzo e Molise "G.Caporale" | Lorusso A, Marcacci M, Di Domenico M, Ancora M, Curini V, Mangone I, Rinaldi A, Di Pasquale A, Cammà C, Puglia I, Savini G |
| EPI_ISL_435150, EPI_ISL_435151 | Ospedale SS Annunziata | Istituto Zooprofilattico Sperimentale dell'Abruzzo e Molise "G.Caporale" | Lorusso A, Marcacci M, Di Domenico M, Ancora M, Curini V, Mangone I, Rinaldi A, Di Pasquale A, Cammà C, Puglia I, Savini G |
| EPI_ISL_435152 | Servizio di Igiene, Epidemiologia e Sanità Pubblica (SIESP) Avezzano | Istituto Zooprofilattico Sperimentale dell'Abruzzo e Molise "G.Caporale" | Lorusso A, Marcacci M, Di Domenico M, Ancora M, Curini V, Mangone I, Rinaldi A, Di Pasquale A, Cammà C, Puglia I, Savini G |
| EPI_ISL_435153, EPI_ISL_435154, EPI_ISL_435155 | SERVIZIO DI IGIENE E SANITÀ PUBBLICA ASL Teramo | Istituto Zooprofilattico Sperimentale dell'Abruzzo e Molise "G.Caporale" | Lorusso A, Marcacci M, Di Domenico M, Ancora M, Curini V, Mangone I, Rinaldi A, Di Pasquale A, Cammà C, Puglia I, Savini G |
| EPI_ISL_435481, EPI_ISL_435482, EPI_ISL_435484, EPI_ISL_435485, EPI_ISL_435486, EPI_ISL_435487, EPI_ISL_435489, EPI_ISL_435490, EPI_ISL_435491, EPI_ISL_435492 | NYU Langone Health | Departments of Pathology and Medicine, New York University School of Medicine | Maria Aguero-Rosenfeld, Brendan Belovarac, Margaret Black, Ludovic Boytard, John Cadley, Paolo Cotzia, John Chen, Dacia Dimartino, Xiaojun Feng, Tatyana Gindin, Emily Guzman, Adriana Heguy, Megan Hogan, Emily Huang, George Jour, Lawrence H. Lin, Raven Luther, Andrew Lytle, Christian Marier, Matthew T. Maurano, Mark J. Mulligan, Peter Meyn, Raquel Ordonez Ciriza, Iman Osman, Jared Pinnell, Vanessa Raabe, Sitharam Ramaswami, Amy Rapkiewicz, Andre M. Ribeiro-dos-Santos, Marie Samanovic-Golden, Antonio Serrano, Guomiao Shen, Matija Snuderl, Theodore Vougiouklakis, Nick Vulpescu, Gael Westby, Paul Zappile, Yutong Zhang |
| EPI_ISL_435674, EPI_ISL_435675, EPI_ISL_435676, EPI_ISL_435677 | National Virology Reference Laboratory | National Public Health Laboratory, National Centre for Infectious Diseases | Mak Tze Minn, Octavia Sophie, Chavatte Jean-Marc, Zaini Zainun, Taib Surita, Cui Lin, Lin Raymond Tzer Pin |
| EPI_ISL_435678, EPI_ISL_435679, EPI_ISL_435680, EPI_ISL_435681, EPI_ISL_435682, EPI_ISL_435683, EPI_ISL_435684, EPI_ISL_435685, EPI_ISL_435686, EPI_ISL_435687, EPI_ISL_435688, EPI_ISL_435689, EPI_ISL_435690, EPI_ISL_435691, EPI_ISL_435692, EPI_ISL_435693, EPI_ISL_435694, EPI_ISL_435695, EPI_ISL_435696, EPI_ISL_435697, EPI_ISL_435698, EPI_ISL_435699, EPI_ISL_435700 | | | |
| see above | National Public Health Laboratory, National Centre for Infectious Diseases | National Public Health Laboratory, National Centre for Infectious Diseases | Mak Tze Minn, Octavia Sophie, Chavatte Jean-Marc, Cui Lin, Lin Raymond Tzer Pin |
| EPI_ISL_436198 | Servicio de Microbiología. Consorcio Hospital General Universitario de Valencia | Sequencing and Bioinformatics Service and Molecular Epidemiology Research Group. FISABIO-Public Health | Lidia Ruiz Roldan, Marta Pla Diaz,Neris Garcia-Gonzalez, Loreto Ferrús Abad, Maria Dolores Ocete, Inma Galán Vendrell, Paula Ruiz-Hueso, Mariana Reyes-Prieto, Vicente Soriano Chirona, Maria Alma Bracho, Griselda De Marco, Beatriz Beamud, Llúcia Martínez-Priego, Concepcion Gimeno, Giuseppe D'Auria, Fernando Gonzalez-Candelas |
| EPI_ISL_436201 | Servicio de Microbiología. Consorcio Hospital General Universitario de Valencia | Sequencing and Bioinformatics Service and Molecular Epidemiology Research Group. FISABIO-Public Health | Loreto Ferrús Abad, Maria Dolores Ocete, Inma Galán Vendrell, Paula Ruiz-Hueso, Mariana Reyes-Prieto, Vicente Soriano Chirona, Maria Alma Bracho, Griselda De Marco, Beatriz Beamud, Lidia Ruiz Roldan, Marta Pla Diaz,Neris Garcia-Gonzalez, Llúcia Martínez-Priego, Concepcion Gimeno, Giuseppe D'Auria, Fernando Gonzalez-Candelas |
| EPI_ISL_436202 | Servicio de Microbiología. Consorcio Hospital General Universitario de Valencia | Sequencing and Bioinformatics Service and Molecular Epidemiology Research Group. FISABIO-Public Health | Loreto Ferrús Abad, Maria Dolores Ocete,Inma Galán Vendrell, Paula Ruiz-Hueso, Mariana Reyes-Prieto, Vicente Soriano Chirona, Maria Alma Bracho, Griselda De Marco, Beatriz Beamud, Lidia Ruiz Roldan, Marta Pla Diaz,Neris Garcia-Gonzalez, Llúcia Martínez-Priego, Concepcion Gimeno, Giuseppe D'Auria, Fernando Gonzalez-Candelas |
| EPI_ISL_436206 | Servicio de Microbiología. Consorcio Hospital General Universitario de Valencia | Sequencing and Bioinformatics Service and Molecular Epidemiology Research Group. FISABIO-Public Health | Lidia Ruiz Roldan, Marta Pla Diaz,Neris Garcia-Gonzalez, Loreto Ferrús Abad, Maria Dolores Ocete, Inma Galán Vendrell, Paula Ruiz-Hueso, Mariana Reyes-Prieto, Vicente Soriano Chirona, Maria Alma Bracho, Griselda De Marco, Beatriz Beamud, Llúcia Martínez-Priego, Concepcion Gimeno, Giuseppe D'Auria, Fernando Gonzalez-Candelas |
| EPI_ISL_436209 | Servicio de Microbiología. Consorcio Hospital General Universitario de Valencia | Sequencing and Bioinformatics Service and Molecular Epidemiology Research Group. FISABIO-Public Health | Loreto Ferrús Abad, Maria Dolores Ocete, Inma Galán Vendrell, Paula Ruiz-Hueso, Mariana Reyes-Prieto, Vicente Soriano Chirona, Maria Alma Bracho, Griselda De Marco, Beatriz Beamud, Lidia Ruiz Roldan, Marta Pla Diaz,Neris Garcia-Gonzalez, Llúcia Martínez-Priego, Concepcion Gimeno, Giuseppe D'Auria, Fernando Gonzalez-Candelas |
| EPI_ISL_436210 | Servicio de Microbiología. Consorcio Hospital General Universitario de Valencia | Sequencing and Bioinformatics Service and Molecular Epidemiology Research Group. FISABIO-Public Health | Loreto Ferrús Abad, Maria Dolores Ocete,Inma Galán Vendrell, Paula Ruiz-Hueso, Mariana Reyes-Prieto, Vicente Soriano Chirona, Maria Alma Bracho, Griselda De Marco, Beatriz Beamud, Lidia Ruiz Roldan, Marta Pla Diaz,Neris Garcia-Gonzalez, Llúcia Martínez-Priego, Concepcion Gimeno, Giuseppe D'Auria, Fernando Gonzalez-Candelas |
| EPI_ISL_436214 | Servicio de Microbiología. Consorcio Hospital General Universitario de Valencia | Sequencing and Bioinformatics Service and Molecular Epidemiology Research Group. FISABIO-Public Health | Lidia Ruiz Roldan, Marta Pla Diaz,Neris Garcia-Gonzalez, Loreto Ferrús Abad, Maria Dolores Ocete, Inma Galán Vendrell, Paula Ruiz-Hueso, Mariana Reyes-Prieto, Vicente Soriano Chirona, Maria Alma Bracho, Griselda De Marco, Beatriz Beamud, Llúcia Martínez-Priego, Concepcion Gimeno, Giuseppe D'Auria, Fernando Gonzalez-Candelas |
| EPI_ISL_436217 | Servicio de Microbiología. Consorcio Hospital General Universitario de Valencia | Sequencing and Bioinformatics Service and Molecular Epidemiology Research Group. FISABIO-Public Health | Loreto Ferrús Abad, Maria Dolores Ocete, Inma Galán Vendrell, Paula Ruiz-Hueso, Mariana Reyes-Prieto, Vicente Soriano Chirona, Maria Alma Bracho, Griselda De Marco, Beatriz Beamud, Lidia Ruiz Roldan, Marta Pla Diaz,Neris Garcia-Gonzalez, Llúcia Martínez-Priego, Concepcion Gimeno, Giuseppe D'Auria, Fernando Gonzalez-Candelas |
| EPI_ISL_436218 | Servicio de Microbiología. Consorcio Hospital General Universitario de Valencia | Sequencing and Bioinformatics Service and Molecular Epidemiology Research Group. FISABIO-Public Health | Loreto Ferrús Abad, Maria Dolores Ocete,Inma Galán Vendrell, Paula Ruiz-Hueso, Mariana Reyes-Prieto, Vicente Soriano Chirona, Maria Alma Bracho, Griselda De Marco, Beatriz Beamud, Lidia Ruiz Roldan, Marta Pla Diaz,Neris Garcia-Gonzalez, Llúcia Martínez-Priego, Concepcion Gimeno, Giuseppe D'Auria, Fernando Gonzalez-Candelas |
| EPI_ISL_436222 | Servicio de Microbiología. Consorcio Hospital General Universitario de Valencia | Sequencing and Bioinformatics Service and Molecular Epidemiology Research Group. FISABIO-Public Health | Lidia Ruiz Roldan, Marta Pla Diaz,Neris Garcia-Gonzalez, Loreto Ferrús Abad, Maria Dolores Ocete, Inma Galán Vendrell, Paula Ruiz-Hueso, Mariana Reyes-Prieto, Vicente Soriano Chirona, Maria Alma Bracho, Griselda De Marco, Beatriz Beamud, Llúcia Martínez-Priego, Concepcion Gimeno, Giuseppe D'Auria, Fernando Gonzalez-Candelas |
| EPI_ISL_436225 | Servicio de Microbiología. Consorcio Hospital General Universitario de Valencia | Sequencing and Bioinformatics Service and Molecular Epidemiology Research Group. FISABIO-Public Health | Loreto Ferrús Abad, Maria Dolores Ocete, Inma Galán Vendrell, Paula Ruiz-Hueso, Mariana Reyes-Prieto, Vicente Soriano Chirona, Maria Alma Bracho, Griselda De Marco, Beatriz Beamud, Lidia Ruiz Roldan, Marta Pla Diaz,Neris Garcia-Gonzalez, Llúcia Martínez-Priego, Concepcion Gimeno, Giuseppe D'Auria, Fernando Gonzalez-Candelas |
| EPI_ISL_436229 | Servicio de Microbiología. Consorcio Hospital General Universitario de Valencia | Sequencing and Bioinformatics Service and Molecular Epidemiology Research Group. FISABIO-Public Health | Lidia Ruiz Roldan, Marta Pla Diaz,Neris Garcia-Gonzalez, Loreto Ferrús Abad, Maria Dolores Ocete, Inma Galán Vendrell, Paula Ruiz-Hueso, Mariana Reyes-Prieto, Vicente Soriano Chirona, Maria Alma Bracho, Griselda De Marco, Beatriz Beamud, Llúcia Martínez-Priego, Concepcion Gimeno, Giuseppe D'Auria, Fernando Gonzalez-Candelas |
| EPI_ISL_436232 | Servicio de Microbiología. Consorcio Hospital General Universitario de Valencia | Sequencing and Bioinformatics Service and Molecular Epidemiology Research Group. FISABIO-Public Health | Loreto Ferrús Abad, Maria Dolores Ocete, Inma Galán Vendrell, Paula Ruiz-Hueso, Mariana Reyes-Prieto, Vicente Soriano Chirona, Maria Alma Bracho, Griselda De Marco, Beatriz Beamud, Lidia Ruiz Roldan, Marta Pla Diaz,Neris Garcia-Gonzalez, Llúcia Martínez-Priego, Concepcion Gimeno, Giuseppe D'Auria, Fernando Gonzalez-Candelas |
| EPI_ISL_436233 | Servicio de Microbiología. Consorcio Hospital General Universitario de Valencia | Sequencing and Bioinformatics Service and Molecular Epidemiology Research Group. FISABIO-Public Health | Loreto Ferrús Abad, Maria Dolores Ocete,Inma Galán Vendrell, Paula Ruiz-Hueso, Mariana Reyes-Prieto, Vicente Soriano Chirona, Maria Alma Bracho, Griselda De Marco, Beatriz Beamud, Lidia Ruiz Roldan, Marta Pla Diaz,Neris Garcia-Gonzalez, Llúcia Martínez-Priego, Concepcion Gimeno, Giuseppe D'Auria, Fernando Gonzalez-Candelas |
| EPI_ISL_436273 | Servicio de Microbiología. Consorcio Hospital General Universitario de Valencia | Sequencing and Bioinformatics Service and Molecular Epidemiology Research Group. FISABIO-Public Health | Lidia Ruiz Roldan, Marta Pla Diaz,Neris Garcia-Gonzalez, Loreto Ferrús Abad, Maria Dolores Ocete, Inma Galán Vendrell, Paula Ruiz-Hueso, Mariana Reyes-Prieto, Vicente Soriano Chirona, Maria Alma Bracho, Griselda De Marco, Beatriz Beamud, Llúcia Martínez-Priego, Concepcion Gimeno, Giuseppe D'Auria, Fernando Gonzalez-Candelas |
| EPI_ISL_436276 | Servicio de Microbiología. Consorcio Hospital General Universitario de Valencia | Sequencing and Bioinformatics Service and Molecular Epidemiology Research Group. FISABIO-Public Health | Loreto Ferrús Abad, Maria Dolores Ocete, Inma Galán Vendrell, Paula Ruiz-Hueso, Mariana Reyes-Prieto, Vicente Soriano Chirona, Maria Alma Bracho, Griselda De Marco, Beatriz Beamud, Lidia Ruiz Roldan, Marta Pla Diaz,Neris Garcia-Gonzalez, Llúcia Martínez-Priego, Concepcion Gimeno, Giuseppe |

| | | | |
|---|---|---|---|
| | | | D'Auria, Fernando Gonzalez-Candelas |
| EPI_ISL_436281 | Servicio de Microbiologia. Hospital Clinico Universitario de Valencia | Sequencing and Bioinformatics Service and Molecular Epidemiology Research Group. FISABIO-Public Health | Lidia Ruiz Roldan, Marta Pla Diaz, Neris Garcia-Gonzalez, Inma Galán Vendrell, Sandra Carbo, Loreto Ferrús Abad, Paula Ruiz-Hueso, Mariana Reyes-Prieto, Vicente Soriano Chirona, Ivan Ansari, David Navarro, Maria Alma Bracho, Griselda De Marco, Beatriz Beamud, Llúcia Martínez-Priego, Giuseppe D'Auria, Fernando Gonzalez-Candelas |
| EPI_ISL_436286 | Servicio de Microbiologia. Hospital Clinico Universitario de Valencia | Sequencing and Bioinformatics Service and Molecular Epidemiology Research Group. FISABIO-Public Health | Loreto Ferrús Abad, Paula Ruiz-Hueso, Mariana Reyes-Prieto, Vicente Soriano Chirona, Ivan Ansari, David Navarro, Maria Alma Bracho, Griselda De Marco, Beatriz Beamud, Lidia Ruiz Roldan, Marta Pla Diaz, Neris Garcia-Gonzalez, Inma Galán Vendrell, Sandra Carbo, Llúcia Martínez-Priego, Giuseppe D'Auria, Fernando Gonzalez-Candelas |
| EPI_ISL_436295, EPI_ISL_436301 | Servicio de Microbiologia. Hospital Clinico Universitario de Valencia | Sequencing and Bioinformatics Service and Molecular Epidemiology Research Group. FISABIO-Public Health | Lidia Ruiz Roldan, Marta Pla Diaz, Neris Garcia-Gonzalez, Inma Galán Vendrell, Sandra Carbo, Loreto Ferrús Abad, Paula Ruiz-Hueso, Mariana Reyes-Prieto, Vicente Soriano Chirona, Ivan Ansari, David Navarro, Maria Alma Bracho, Griselda De Marco, Beatriz Beamud, Llúcia Martínez-Priego, Giuseppe D'Auria, Fernando Gonzalez-Candelas |
| EPI_ISL_436306 | Servicio de Microbiologia. Hospital Clinico Universitario de Valencia | Sequencing and Bioinformatics Service and Molecular Epidemiology Research Group. FISABIO-Public Health | Loreto Ferrús Abad, Paula Ruiz-Hueso, Mariana Reyes-Prieto, Vicente Soriano Chirona, Ivan Ansari, David Navarro, Maria Alma Bracho, Griselda De Marco, Beatriz Beamud, Lidia Ruiz Roldan, Marta Pla Diaz, Neris Garcia-Gonzalez, Inma Galán Vendrell, Sandra Carbo, Llúcia Martínez-Priego, Giuseppe D'Auria, Fernando Gonzalez-Candelas |
| EPI_ISL_436315, EPI_ISL_436321 | Servicio de Microbiologia. Hospital Clinico Universitario de Valencia | Sequencing and Bioinformatics Service and Molecular Epidemiology Research Group. FISABIO-Public Health | Lidia Ruiz Roldan, Marta Pla Diaz, Neris Garcia-Gonzalez, Inma Galán Vendrell, Sandra Carbo, Loreto Ferrús Abad, Paula Ruiz-Hueso, Mariana Reyes-Prieto, Vicente Soriano Chirona, Ivan Ansari, David Navarro, Maria Alma Bracho, Griselda De Marco, Beatriz Beamud, Llúcia Martínez-Priego, Giuseppe D'Auria, Fernando Gonzalez-Candelas |
| EPI_ISL_436326 | Servicio de Microbiologia. Hospital Clinico Universitario de Valencia | Sequencing and Bioinformatics Service and Molecular Epidemiology Research Group. FISABIO-Public Health | Loreto Ferrús Abad, Paula Ruiz-Hueso, Mariana Reyes-Prieto, Vicente Soriano Chirona, Ivan Ansari, David Navarro, Maria Alma Bracho, Griselda De Marco, Beatriz Beamud, Lidia Ruiz Roldan, Marta Pla Diaz, Neris Garcia-Gonzalez, Inma Galán Vendrell, Sandra Carbo, Llúcia Martínez-Priego, Giuseppe D'Auria, Fernando Gonzalez-Candelas |
| EPI_ISL_436335, EPI_ISL_436341 | Servicio de Microbiologia. Hospital Clinico Universitario de Valencia | Sequencing and Bioinformatics Service and Molecular Epidemiology Research Group. FISABIO-Public Health | Lidia Ruiz Roldan, Marta Pla Diaz, Neris Garcia-Gonzalez, Inma Galán Vendrell, Sandra Carbo, Loreto Ferrús Abad, Paula Ruiz-Hueso, Mariana Reyes-Prieto, Vicente Soriano Chirona, Ivan Ansari, David Navarro, Maria Alma Bracho, Griselda De Marco, Beatriz Beamud, Llúcia Martínez-Priego, Giuseppe D'Auria, Fernando Gonzalez-Candelas |
| EPI_ISL_436346 | Servicio de Microbiologia. Hospital Clinico Universitario de Valencia | Sequencing and Bioinformatics Service and Molecular Epidemiology Research Group. FISABIO-Public Health | Loreto Ferrús Abad, Paula Ruiz-Hueso, Mariana Reyes-Prieto, Vicente Soriano Chirona, Ivan Ansari, David Navarro, Maria Alma Bracho, Griselda De Marco, Beatriz Beamud, Lidia Ruiz Roldan, Marta Pla Diaz, Neris Garcia-Gonzalez, Inma Galán Vendrell, Sandra Carbo, Llúcia Martínez-Priego, Giuseppe D'Auria, Fernando Gonzalez-Candelas |
| EPI_ISL_436355, EPI_ISL_436358, EPI_ISL_436365 | Servicio de Microbiologia. Hospital Clinico Universitario de Valencia | Sequencing and Bioinformatics Service and Molecular Epidemiology Research Group. FISABIO-Public Health | Lidia Ruiz Roldan, Marta Pla Diaz, Neris Garcia-Gonzalez, Inma Galán Vendrell, Sandra Carbo, Loreto Ferrús Abad, Paula Ruiz-Hueso, Mariana Reyes-Prieto, Vicente Soriano Chirona, Ivan Ansari, David Navarro, Maria Alma Bracho, Griselda De Marco, Beatriz Beamud, Llúcia Martínez-Priego, Giuseppe D'Auria, Fernando Gonzalez-Candelas |
| EPI_ISL_436370 | Servicio de Microbiologia. Hospital Clinico Universitario de Valencia | Sequencing and Bioinformatics Service and Molecular Epidemiology Research Group. FISABIO-Public Health | Loreto Ferrús Abad, Paula Ruiz-Hueso, Mariana Reyes-Prieto, Vicente Soriano Chirona, Ivan Ansari, David Navarro, Maria Alma Bracho, Griselda De Marco, Beatriz Beamud, Lidia Ruiz Roldan, Marta Pla Diaz, Neris Garcia-Gonzalez, Inma Galán Vendrell, Sandra Carbo, Llúcia Martínez-Priego, Giuseppe D'Auria, Fernando Gonzalez-Candelas |
| EPI_ISL_436382 | Servicio de Microbiología. Consorcio Hospital General Universitario de Valencia | Sequencing and Bioinformatics Service and Molecular Epidemiology Research Group. FISABIO-Public Health | Loreto Ferrús Abad, Maria Dolores Ocete,Inma Galán Vendrell, Paula Ruiz-Hueso, Mariana Reyes-Prieto, Vicente Soriano Chirona, Maria Alma Bracho, Griselda De Marco, Beatriz Beamud, Lidia Ruiz Roldan, Marta Pla Diaz,Neris Garcia-Gonzalez, Llúcia Martínez-Priego, Concepcion Gimeno, Giuseppe D'Auria, Fernando Gonzalez-Candelas |
| EPI_ISL_436386 | Servicio de Microbiología. Consorcio Hospital General Universitario de Valencia | Sequencing and Bioinformatics Service and Molecular Epidemiology Research Group. FISABIO-Public Health | Lidia Ruiz Roldan, Marta Pla Diaz,Neris Garcia-Gonzalez, Loreto Ferrús Abad, Maria Dolores Ocete, Inma Galán Vendrell, Paula Ruiz-Hueso, Mariana Reyes-Prieto, Vicente Soriano Chirona, Maria Alma Bracho, Griselda De Marco, Beatriz Beamud, Llúcia Martínez-Priego, Concepcion Gimeno, Giuseppe D'Auria, Fernando Gonzalez-Candelas |
| EPI_ISL_436389 | Servicio de Microbiología. Consorcio Hospital General Universitario de Valencia | Sequencing and Bioinformatics Service and Molecular Epidemiology Research Group. FISABIO-Public Health | Loreto Ferrús Abad, Maria Dolores Ocete, Inma Galán Vendrell, Paula Ruiz-Hueso, Mariana Reyes-Prieto, Vicente Soriano Chirona, Maria Alma Bracho, Griselda De Marco, Beatriz Beamud, Lidia Ruiz Roldan, Marta Pla Diaz,Neris Garcia-Gonzalez, Llúcia Martínez-Priego, Concepcion Gimeno, Giuseppe D'Auria, Fernando Gonzalez-Candelas |
| EPI_ISL_436393 | Servicio de Microbiología. Consorcio Hospital General Universitario de Valencia | Sequencing and Bioinformatics Service and Molecular Epidemiology Research Group. FISABIO-Public Health | Lidia Ruiz Roldan, Marta Pla Diaz,Neris Garcia-Gonzalez, Loreto Ferrús Abad, Maria Dolores Ocete, Inma Galán Vendrell, Paula Ruiz-Hueso, Mariana Reyes-Prieto, Vicente Soriano Chirona, Maria Alma Bracho, Griselda De Marco, Beatriz Beamud, Llúcia Martínez-Priego, Concepcion Gimeno, Giuseppe D'Auria, Fernando Gonzalez-Candelas |
| EPI_ISL_436397 | Servicio de Microbiologia. Hospital Clinico Universitario de Valencia | Sequencing and Bioinformatics Service and Molecular Epidemiology Research Group. FISABIO-Public Health | Loreto Ferrús Abad, Paula Ruiz-Hueso, Mariana Reyes-Prieto, Vicente Soriano Chirona, Ivan Ansari, David Navarro, Maria Alma Bracho, Griselda De Marco, Beatriz Beamud, Lidia Ruiz Roldan, Marta Pla Diaz, Neris Garcia-Gonzalez, Inma Galán Vendrell, Sandra Carbo, Llúcia Martínez-Priego, Giuseppe D'Auria, Fernando Gonzalez-Candelas |
| EPI_ISL_436406, EPI_ISL_436409 | Servicio de Microbiologia. Hospital Clinico Universitario de Valencia | Sequencing and Bioinformatics Service and Molecular Epidemiology Research Group. FISABIO-Public Health | Lidia Ruiz Roldan, Marta Pla Diaz, Neris Garcia-Gonzalez, Inma Galán Vendrell, Sandra Carbo, Loreto Ferrús Abad, Paula Ruiz-Hueso, Mariana Reyes-Prieto, Vicente Soriano Chirona, Ivan Ansari, David Navarro, Maria Alma Bracho, Griselda De Marco, Beatriz Beamud, Llúcia Martínez-Priego, Giuseppe D'Auria, Fernando Gonzalez-Candelas |
| EPI_ISL_436718 | Ospedale Regionale San Salvatore | Istituto Zooprofilattico Sperimentale dell'Abruzzo e Molise "G.Caporale" | Lorusso A, Marcacci M, Di Domenico M, Ancora M, Curini V, Mangone I, Rinaldi A, Di Pasquale A, Cammà C, Puglia I, Savini G |
| EPI_ISL_436719, EPI_ISL_436720, EPI_ISL_436721, EPI_ISL_436722 | Ospedale Civile S. Liberatore di Atri | Istituto Zooprofilattico Sperimentale dell'Abruzzo e Molise "G.Caporale" | Lorusso A, Marcacci M, Di Domenico M, Ancora M, Curini V, Mangone I, Rinaldi A, Di Pasquale A, Cammà C, Puglia I, Savini G |
| EPI_ISL_436723 | Ospedale Civile Giuseppe Mazzini | Istituto Zooprofilattico Sperimentale dell'Abruzzo e Molise "G.Caporale" | Lorusso A, Marcacci M, Di Domenico M, Ancora M, Curini V, Mangone I, Rinaldi A, Di Pasquale A, Cammà C, Puglia I, Savini G |
| EPI_ISL_436724 | Ospedale Civile S. Liberatore di Atri | Istituto Zooprofilattico Sperimentale dell'Abruzzo e Molise "G.Caporale" | Lorusso A, Marcacci M, Di Domenico M, Ancora M, Curini V, Mangone I, Rinaldi A, Di Pasquale A, Cammà C, Puglia I, Savini G |
| EPI_ISL_436725 | RSA/RP Villa San Giovanni - Gruppo Edos | Istituto Zooprofilattico Sperimentale dell'Abruzzo e Molise "G.Caporale" | Lorusso A, Marcacci M, Di Domenico M, Ancora M, Curini V, Mangone I, Rinaldi A, Di Pasquale A, Cammà C, Puglia I, Savini G |
| EPI_ISL_436726, EPI_ISL_436727, EPI_ISL_436728, EPI_ISL_436729 | SERVIZIO DI IGIENE E SANITÀ PUBBLICA ASL Teramo | Istituto Zooprofilattico Sperimentale dell'Abruzzo e Molise "G.Caporale" | Lorusso A, Marcacci M, Di Domenico M, Ancora M, Curini V, Mangone I, Rinaldi A, Di Pasquale A, Cammà C, Puglia I, Savini G |
| EPI_ISL_436730 | Servizio di igiene epidemiologia e sanità pubblica (Siesp) Chieti | Istituto Zooprofilattico Sperimentale dell'Abruzzo e Molise "G.Caporale" | Lorusso A, Marcacci M, Di Domenico M, Ancora M, Curini V, Mangone I, Rinaldi A, Di Pasquale A, Cammà C, Puglia I, Savini G |
| EPI_ISL_436731, EPI_ISL_436732 | Ospedale Civile S. Liberatore di Atri | Istituto Zooprofilattico Sperimentale dell'Abruzzo e Molise "G.Caporale" | Lorusso A, Marcacci M, Di Domenico M, Ancora M, Curini V, Mangone I, Rinaldi A, Di Pasquale A, Cammà C, Puglia I, Savini G |

EPI_ISL_438248, EPI_ISL_438249, EPI_ISL_438250, EPI_ISL_438251, EPI_ISL_438252, EPI_ISL_438253, EPI_ISL_438254, EPI_ISL_438255, EPI_ISL_438256, EPI_ISL_438257, EPI_ISL_438258, EPI_ISL_438259, EPI_ISL_438260, EPI_ISL_438261, EPI_ISL_438262, EPI_ISL_438263, EPI_ISL_438264, EPI_ISL_438265, EPI_ISL_438266, EPI_ISL_438267, EPI_ISL_438268, EPI_ISL_438269, EPI_ISL_438270, EPI_ISL_438271, EPI_ISL_438272, EPI_ISL_438273, EPI_ISL_438274, EPI_ISL_438275, EPI_ISL_438276, EPI_ISL_438277, EPI_ISL_438278, EPI_ISL_438279, EPI_ISL_438280, EPI_ISL_438281, EPI_ISL_438282, EPI_ISL_438283, EPI_ISL_438284, EPI_ISL_438285, EPI_ISL_438286, EPI_ISL_438287, EPI_ISL_438288, EPI_ISL_438289, EPI_ISL_438290, EPI_ISL_438291, EPI_ISL_438292, EPI_ISL_438293, EPI_ISL_438294, EPI_ISL_438295, EPI_ISL_438296, EPI_ISL_438297, EPI_ISL_438298, EPI_ISL_438299, EPI_ISL_438300, EPI_ISL_438301, EPI_ISL_438302, EPI_ISL_438303, EPI_ISL_438304, EPI_ISL_438305, EPI_ISL_438306, EPI_ISL_438307, EPI_ISL_438308, EPI_ISL_438309, EPI_ISL_438310, EPI_ISL_438311, EPI_ISL_438312, EPI_ISL_438313, EPI_ISL_438314, EPI_ISL_438315, EPI_ISL_438316, EPI_ISL_438317, EPI_ISL_438318, EPI_ISL_438319, EPI_ISL_438320, EPI_ISL_438321, EPI_ISL_438322, EPI_ISL_438323, EPI_ISL_438324, EPI_ISL_438325, EPI_ISL_438326, EPI_ISL_438327, EPI_ISL_438328, EPI_ISL_438329, EPI_ISL_438330, EPI_ISL_438331, EPI_ISL_438332, EPI_ISL_438333, EPI_ISL_438334, EPI_ISL_438335, EPI_ISL_438336, EPI_ISL_438337, EPI_ISL_438338, EPI_ISL_438339, EPI_ISL_438340, EPI_ISL_438341, EPI_ISL_438342, EPI_ISL_438343, EPI_ISL_438344, EPI_ISL_438345, EPI_ISL_438346, EPI_ISL_438347, EPI_ISL_438348, EPI_ISL_438349, EPI_ISL_438350, EPI_ISL_438351, EPI_ISL_438352, EPI_ISL_438353, EPI_ISL_438354, EPI_ISL_438355, EPI_ISL_438356, EPI_ISL_438357, EPI_ISL_438358, EPI_ISL_438359, EPI_ISL_438360, EPI_ISL_438361, EPI_ISL_438362, EPI_ISL_438363, EPI_ISL_438364, EPI_ISL_438365, EPI_ISL_438366, EPI_ISL_438367, EPI_ISL_438368, EPI_ISL_438369, EPI_ISL_438370, EPI_ISL_438371, EPI_ISL_438372, EPI_ISL_438373, EPI_ISL_438374, EPI_ISL_438375, EPI_ISL_438376, EPI_ISL_438377, EPI_ISL_438378, EPI_ISL_438379, EPI_ISL_438380, EPI_ISL_438381, EPI_ISL_438382, EPI_ISL_438383, EPI_ISL_438384, EPI_ISL_438385, EPI_ISL_438386, EPI_ISL_438387, EPI_ISL_438388, EPI_ISL_438389, EPI_ISL_438390, EPI_ISL_438391, EPI_ISL_438392, EPI_ISL_438393, EPI_ISL_438394, EPI_ISL_438395, EPI_ISL_438396, EPI_ISL_438397, EPI_ISL_438398, EPI_ISL_438399, EPI_ISL_438400, EPI_ISL_438401, EPI_ISL_438402, EPI_ISL_438403, EPI_ISL_438404, EPI_ISL_438405, EPI_ISL_438406, EPI_ISL_438407, EPI_ISL_438408, EPI_ISL_438409, EPI_ISL_438410, EPI_ISL_438411, EPI_ISL_438412, EPI_ISL_438413, EPI_ISL_438414, EPI_ISL_438415, EPI_ISL_438416, EPI_ISL_438417, EPI_ISL_438418, EPI_ISL_438419, EPI_ISL_438420, EPI_ISL_438421, EPI_ISL_438422, EPI_ISL_438423, EPI_ISL_438424, EPI_ISL_438425, EPI_ISL_438426, EPI_ISL_438427,

EPI_ISL_438428, EPI_ISL_438429, EPI_ISL_438430, EPI_ISL_438431, EPI_ISL_438432, EPI_ISL_438433, EPI_ISL_438434, EPI_ISL_438435, EPI_ISL_438436, EPI_ISL_438437, EPI_ISL_438438, EPI_ISL_438439, EPI_ISL_438440, EPI_ISL_438441, EPI_ISL_438442, EPI_ISL_438443, EPI_ISL_438444, EPI_ISL_438445,
EPI_ISL_438446, EPI_ISL_438447, EPI_ISL_438448, EPI_ISL_438449, EPI_ISL_438450, EPI_ISL_438451, EPI_ISL_438452, EPI_ISL_438453, EPI_ISL_438454, EPI_ISL_438455, EPI_ISL_438456, EPI_ISL_438457, EPI_ISL_438458, EPI_ISL_438459, EPI_ISL_438460, EPI_ISL_438461, EPI_ISL_438462, EPI_ISL_438463,
EPI_ISL_438464, EPI_ISL_438465, EPI_ISL_438466, EPI_ISL_438467, EPI_ISL_438468, EPI_ISL_438469, EPI_ISL_438470, EPI_ISL_438471, EPI_ISL_438472, EPI_ISL_438473, EPI_ISL_438474, EPI_ISL_438475, EPI_ISL_438476, EPI_ISL_438477, EPI_ISL_438478, EPI_ISL_438479, EPI_ISL_438480, EPI_ISL_438481,
EPI_ISL_438482, EPI_ISL_438483, EPI_ISL_438484, EPI_ISL_438485, EPI_ISL_438486, EPI_ISL_438487, EPI_ISL_438488, EPI_ISL_438489, EPI_ISL_438490, EPI_ISL_438491, EPI_ISL_438492, EPI_ISL_438493, EPI_ISL_438494, EPI_ISL_438495, EPI_ISL_438496, EPI_ISL_438497, EPI_ISL_438498, EPI_ISL_438499,
EPI_ISL_438500, EPI_ISL_438501, EPI_ISL_438502, EPI_ISL_438503, EPI_ISL_438504, EPI_ISL_438505, EPI_ISL_438506, EPI_ISL_438507, EPI_ISL_438508, EPI_ISL_438509, EPI_ISL_438510, EPI_ISL_438511, EPI_ISL_438512, EPI_ISL_438513, EPI_ISL_438514, EPI_ISL_438515, EPI_ISL_438516, EPI_ISL_438517,
EPI_ISL_438518, EPI_ISL_438519, EPI_ISL_438520, EPI_ISL_438521, EPI_ISL_438522, EPI_ISL_438523, EPI_ISL_438524, EPI_ISL_438525, EPI_ISL_438526, EPI_ISL_438527, EPI_ISL_438528, EPI_ISL_438529, EPI_ISL_438530, EPI_ISL_438531, EPI_ISL_438532, EPI_ISL_438533, EPI_ISL_438534, EPI_ISL_438535,
EPI_ISL_438536, EPI_ISL_438537, EPI_ISL_438538, EPI_ISL_438539, EPI_ISL_438540, EPI_ISL_438541, EPI_ISL_438542, EPI_ISL_438543, EPI_ISL_438544, EPI_ISL_438545

| | | | |
|---|---|---|---|
| see above | Department of Pathology, University of Cambridge | Wellcome Sanger Institute for the COVID-19 Genomics UK (COG-UK) consortium | Luke W Meredith, M. Estée Török , Myra Hosmillo, William L. Hamilton, Martin D. Curran, Theresa Feltwell, Grant Hall, Anna Yakovleva, Fahad A Khokhar, Charlotte J. Houldcroft, Laura G Caller, Aminu S. Jahun, Sarah L. Caddy, Ian Goodfellow, Alex Alderton, Roberto Amato, Sonia Goncalves, Ewan Harrison, David K. Jackson, Ian Johnston, Dominic Kwiatkowski, Cordelia Langford, John Sillitoe on behalf of the Wellcome Sanger Institute COVID-19 Surveillance Team (http://www.sanger.ac.uk/covid-team) |

EPI_ISL_438550, EPI_ISL_438551, EPI_ISL_438552, EPI_ISL_438553, EPI_ISL_438554, EPI_ISL_438555, EPI_ISL_438556, EPI_ISL_438557, EPI_ISL_438558, EPI_ISL_438559, EPI_ISL_438560, EPI_ISL_438561, EPI_ISL_438562, EPI_ISL_438563, EPI_ISL_438564, EPI_ISL_438565, EPI_ISL_438566, EPI_ISL_438567,
EPI_ISL_438568, EPI_ISL_438569, EPI_ISL_438570, EPI_ISL_438571, EPI_ISL_438573, EPI_ISL_438574, EPI_ISL_438575, EPI_ISL_438576, EPI_ISL_438577, EPI_ISL_438578, EPI_ISL_438579, EPI_ISL_438580, EPI_ISL_438581, EPI_ISL_438582, EPI_ISL_438583, EPI_ISL_438584, EPI_ISL_438585, EPI_ISL_438586,
EPI_ISL_438587, EPI_ISL_438588, EPI_ISL_438589, EPI_ISL_438590, EPI_ISL_438591, EPI_ISL_438592, EPI_ISL_438593, EPI_ISL_438594, EPI_ISL_438595, EPI_ISL_438596, EPI_ISL_438597, EPI_ISL_438598, EPI_ISL_438599, EPI_ISL_438600, EPI_ISL_438601, EPI_ISL_438602, EPI_ISL_438603, EPI_ISL_438604,
EPI_ISL_438605, EPI_ISL_438606, EPI_ISL_438607, EPI_ISL_438608, EPI_ISL_438609, EPI_ISL_438610, EPI_ISL_438612, EPI_ISL_438613, EPI_ISL_438614, EPI_ISL_438615, EPI_ISL_438616, EPI_ISL_438617, EPI_ISL_438618, EPI_ISL_438619, EPI_ISL_438620, EPI_ISL_438621, EPI_ISL_438622, EPI_ISL_438623,
EPI_ISL_438624, EPI_ISL_438625, EPI_ISL_438626, EPI_ISL_438627, EPI_ISL_438628, EPI_ISL_438629, EPI_ISL_438630, EPI_ISL_438631, EPI_ISL_438632, EPI_ISL_438633, EPI_ISL_438634, EPI_ISL_438635, EPI_ISL_438636, EPI_ISL_438637, EPI_ISL_438638, EPI_ISL_438639, EPI_ISL_438640, EPI_ISL_438641,
EPI_ISL_438642, EPI_ISL_438643, EPI_ISL_438644, EPI_ISL_438645, EPI_ISL_438646, EPI_ISL_438647, EPI_ISL_438648, EPI_ISL_438649, EPI_ISL_438650, EPI_ISL_438651, EPI_ISL_438652, EPI_ISL_438653, EPI_ISL_438654, EPI_ISL_438656, EPI_ISL_438657, EPI_ISL_438658, EPI_ISL_438659, EPI_ISL_438660,
EPI_ISL_438661, EPI_ISL_438662, EPI_ISL_438663, EPI_ISL_438664, EPI_ISL_438665, EPI_ISL_438666, EPI_ISL_438667, EPI_ISL_438668, EPI_ISL_438669, EPI_ISL_438670, EPI_ISL_438671, EPI_ISL_438672, EPI_ISL_438673, EPI_ISL_438674, EPI_ISL_438675, EPI_ISL_438676, EPI_ISL_438677, EPI_ISL_438678,
EPI_ISL_438679, EPI_ISL_438680, EPI_ISL_438681, EPI_ISL_438682, EPI_ISL_438683, EPI_ISL_438684, EPI_ISL_438685, EPI_ISL_438686, EPI_ISL_438687, EPI_ISL_438689, EPI_ISL_438690, EPI_ISL_438691, EPI_ISL_438692, EPI_ISL_438693, EPI_ISL_438694, EPI_ISL_438695, EPI_ISL_438696, EPI_ISL_438697,
EPI_ISL_438698, EPI_ISL_438699, EPI_ISL_438700, EPI_ISL_438701, EPI_ISL_438702, EPI_ISL_438703, EPI_ISL_438704, EPI_ISL_438705, EPI_ISL_438706, EPI_ISL_438707, EPI_ISL_438708, EPI_ISL_438709, EPI_ISL_438710, EPI_ISL_438711, EPI_ISL_438712, EPI_ISL_438713, EPI_ISL_438714, EPI_ISL_438715,
EPI_ISL_438716, EPI_ISL_438717, EPI_ISL_438718, EPI_ISL_438719, EPI_ISL_438720, EPI_ISL_438721, EPI_ISL_438722, EPI_ISL_438723, EPI_ISL_438724, EPI_ISL_438725, EPI_ISL_438726, EPI_ISL_438727, EPI_ISL_438728, EPI_ISL_438729, EPI_ISL_438730, EPI_ISL_438731, EPI_ISL_438732, EPI_ISL_438733,
EPI_ISL_438734, EPI_ISL_438735, EPI_ISL_438736, EPI_ISL_438737, EPI_ISL_438738, EPI_ISL_438739, EPI_ISL_438740, EPI_ISL_438741, EPI_ISL_438742, EPI_ISL_438743, EPI_ISL_438744, EPI_ISL_438745, EPI_ISL_438746, EPI_ISL_438747, EPI_ISL_438748

| | | | |
|---|---|---|---|
| see above | Department of Pathology, University of Cambridge | COVID-19 Genomics UK (COG-UK) Consortium | Luke W Meredith, M. Est©e Trk , Myra Hosmillo, William L. Hamilton, Martin D. Curran, Theresa Feltwell, Grant Hall, Anna Yakovleva, Fahad A Khokhar, Charlotte J. Houldcroft, Laura G Caller, Aminu S. Jahun, Sarah L. Caddy, Ian Goodfellow |

EPI_ISL_439368, EPI_ISL_439369, EPI_ISL_439370, EPI_ISL_439371, EPI_ISL_439372, EPI_ISL_439373, EPI_ISL_439374, EPI_ISL_439375, EPI_ISL_439376, EPI_ISL_439377, EPI_ISL_439378, EPI_ISL_439379, EPI_ISL_439380, EPI_ISL_439381, EPI_ISL_439382, EPI_ISL_439383, EPI_ISL_439384, EPI_ISL_439385,
EPI_ISL_439386, EPI_ISL_439387, EPI_ISL_439388, EPI_ISL_439389, EPI_ISL_439390, EPI_ISL_439391, EPI_ISL_439392, EPI_ISL_439393, EPI_ISL_439394, EPI_ISL_439395, EPI_ISL_439396, EPI_ISL_439397, EPI_ISL_439398, EPI_ISL_439399, EPI_ISL_439400, EPI_ISL_439401, EPI_ISL_439402, EPI_ISL_439403,
EPI_ISL_439404, EPI_ISL_439405, EPI_ISL_439406, EPI_ISL_439407, EPI_ISL_439408, EPI_ISL_439409, EPI_ISL_439410, EPI_ISL_439411, EPI_ISL_439412, EPI_ISL_439413, EPI_ISL_439414, EPI_ISL_439415, EPI_ISL_439416, EPI_ISL_439417, EPI_ISL_439418, EPI_ISL_439419, EPI_ISL_439420, EPI_ISL_439421,
EPI_ISL_439422, EPI_ISL_439423, EPI_ISL_439424, EPI_ISL_439425, EPI_ISL_439426, EPI_ISL_439427, EPI_ISL_439428, EPI_ISL_439429, EPI_ISL_439430, EPI_ISL_439431, EPI_ISL_439432, EPI_ISL_439433, EPI_ISL_439434, EPI_ISL_439435, EPI_ISL_439436, EPI_ISL_439437, EPI_ISL_439438, EPI_ISL_439439,
EPI_ISL_439440, EPI_ISL_439441, EPI_ISL_439442, EPI_ISL_439443, EPI_ISL_439444, EPI_ISL_439445, EPI_ISL_439446, EPI_ISL_439447, EPI_ISL_439448, EPI_ISL_439449, EPI_ISL_439450, EPI_ISL_439451, EPI_ISL_439452, EPI_ISL_439453, EPI_ISL_439454, EPI_ISL_439455, EPI_ISL_439456, EPI_ISL_439457,
EPI_ISL_439458, EPI_ISL_439459, EPI_ISL_439460, EPI_ISL_439461, EPI_ISL_439462, EPI_ISL_439463, EPI_ISL_439464, EPI_ISL_439465, EPI_ISL_439466, EPI_ISL_439467, EPI_ISL_439468, EPI_ISL_439469, EPI_ISL_439470, EPI_ISL_439471, EPI_ISL_439472, EPI_ISL_439473, EPI_ISL_439474, EPI_ISL_439475,
EPI_ISL_439476, EPI_ISL_439477, EPI_ISL_439478, EPI_ISL_439479, EPI_ISL_439480, EPI_ISL_439481, EPI_ISL_439482, EPI_ISL_439483, EPI_ISL_439484, EPI_ISL_439485, EPI_ISL_439486, EPI_ISL_439487, EPI_ISL_439488, EPI_ISL_439489, EPI_ISL_439490, EPI_ISL_439491, EPI_ISL_439492, EPI_ISL_439493,
EPI_ISL_439494, EPI_ISL_439495, EPI_ISL_439496, EPI_ISL_439497, EPI_ISL_439498, EPI_ISL_439499, EPI_ISL_439500, EPI_ISL_439501, EPI_ISL_439502, EPI_ISL_439503, EPI_ISL_439504, EPI_ISL_439505, EPI_ISL_439506, EPI_ISL_439507, EPI_ISL_439508, EPI_ISL_439509, EPI_ISL_439510, EPI_ISL_439511,
EPI_ISL_439512, EPI_ISL_439513, EPI_ISL_439514, EPI_ISL_439515, EPI_ISL_439516, EPI_ISL_439517, EPI_ISL_439518, EPI_ISL_439519, EPI_ISL_439520, EPI_ISL_439521, EPI_ISL_439522, EPI_ISL_439523, EPI_ISL_439524, EPI_ISL_439525, EPI_ISL_439526, EPI_ISL_439527, EPI_ISL_439528, EPI_ISL_439529,
EPI_ISL_439530, EPI_ISL_439531, EPI_ISL_439532, EPI_ISL_439533, EPI_ISL_439534, EPI_ISL_439535, EPI_ISL_439536, EPI_ISL_439537, EPI_ISL_439538, EPI_ISL_439539, EPI_ISL_439540, EPI_ISL_439541, EPI_ISL_439542, EPI_ISL_439543, EPI_ISL_439544, EPI_ISL_439545, EPI_ISL_439546, EPI_ISL_439547,
EPI_ISL_439548, EPI_ISL_439549, EPI_ISL_439550, EPI_ISL_439551, EPI_ISL_439552, EPI_ISL_439553, EPI_ISL_439554, EPI_ISL_439555, EPI_ISL_439556, EPI_ISL_439557, EPI_ISL_439558, EPI_ISL_439559, EPI_ISL_439560, EPI_ISL_439561, EPI_ISL_439562, EPI_ISL_439563, EPI_ISL_439564, EPI_ISL_439565,
EPI_ISL_439566, EPI_ISL_439567, EPI_ISL_439568, EPI_ISL_439569, EPI_ISL_439570, EPI_ISL_439571, EPI_ISL_439572, EPI_ISL_439573, EPI_ISL_439574, EPI_ISL_439575, EPI_ISL_439576, EPI_ISL_439577, EPI_ISL_439578, EPI_ISL_439579, EPI_ISL_439580, EPI_ISL_439581, EPI_ISL_439582, EPI_ISL_439583,
EPI_ISL_439584, EPI_ISL_439585, EPI_ISL_439586, EPI_ISL_439587, EPI_ISL_439588, EPI_ISL_439589, EPI_ISL_439590, EPI_ISL_439591, EPI_ISL_439592, EPI_ISL_439593, EPI_ISL_439594, EPI_ISL_439595, EPI_ISL_439596, EPI_ISL_439597, EPI_ISL_439598, EPI_ISL_439599, EPI_ISL_439600, EPI_ISL_439601,
EPI_ISL_439602, EPI_ISL_439603, EPI_ISL_439604, EPI_ISL_439605, EPI_ISL_439606, EPI_ISL_439607, EPI_ISL_439608, EPI_ISL_439609, EPI_ISL_439610, EPI_ISL_439611, EPI_ISL_439612, EPI_ISL_439613, EPI_ISL_439614, EPI_ISL_439615, EPI_ISL_439616, EPI_ISL_439617, EPI_ISL_439618, EPI_ISL_439619,
EPI_ISL_439620, EPI_ISL_439621, EPI_ISL_439622, EPI_ISL_439623, EPI_ISL_439624, EPI_ISL_439625, EPI_ISL_439626, EPI_ISL_439627, EPI_ISL_439628, EPI_ISL_439629, EPI_ISL_439630, EPI_ISL_439631, EPI_ISL_439632, EPI_ISL_439633, EPI_ISL_439634, EPI_ISL_439635, EPI_ISL_439636, EPI_ISL_439637,
EPI_ISL_439638, EPI_ISL_439639, EPI_ISL_439640, EPI_ISL_439641, EPI_ISL_439642, EPI_ISL_439643, EPI_ISL_439644, EPI_ISL_439645, EPI_ISL_439646, EPI_ISL_439647, EPI_ISL_439648, EPI_ISL_439649, EPI_ISL_439650, EPI_ISL_439651, EPI_ISL_439652, EPI_ISL_439653, EPI_ISL_439654, EPI_ISL_439655,
EPI_ISL_439656, EPI_ISL_439657, EPI_ISL_439658, EPI_ISL_439659, EPI_ISL_439660, EPI_ISL_439661, EPI_ISL_439662, EPI_ISL_439663, EPI_ISL_439664, EPI_ISL_439863, EPI_ISL_439864, EPI_ISL_439865, EPI_ISL_439866, EPI_ISL_439867, EPI_ISL_439868, EPI_ISL_439869, EPI_ISL_439870, EPI_ISL_439871,
EPI_ISL_439872, EPI_ISL_439873, EPI_ISL_439874, EPI_ISL_439875, EPI_ISL_439876, EPI_ISL_439877, EPI_ISL_439878, EPI_ISL_439879, EPI_ISL_439880, EPI_ISL_439881, EPI_ISL_439882, EPI_ISL_439883, EPI_ISL_439884, EPI_ISL_439885, EPI_ISL_439886, EPI_ISL_439887, EPI_ISL_439888, EPI_ISL_439889,
EPI_ISL_439890, EPI_ISL_439891, EPI_ISL_439892, EPI_ISL_439893, EPI_ISL_439894, EPI_ISL_439895, EPI_ISL_439896, EPI_ISL_439897, EPI_ISL_439898, EPI_ISL_439899, EPI_ISL_439900, EPI_ISL_439901, EPI_ISL_439902, EPI_ISL_439903, EPI_ISL_439904, EPI_ISL_439905, EPI_ISL_439906, EPI_ISL_439907,
EPI_ISL_439908, EPI_ISL_439909, EPI_ISL_439910, EPI_ISL_439911, EPI_ISL_439912, EPI_ISL_439913, EPI_ISL_439914, EPI_ISL_439915, EPI_ISL_439916, EPI_ISL_439917, EPI_ISL_439918, EPI_ISL_439919, EPI_ISL_439920, EPI_ISL_439921, EPI_ISL_439922, EPI_ISL_439923, EPI_ISL_439924, EPI_ISL_439925,
EPI_ISL_439926, EPI_ISL_439927, EPI_ISL_439928, EPI_ISL_439929, EPI_ISL_439930, EPI_ISL_439931, EPI_ISL_439932, EPI_ISL_439933, EPI_ISL_439934, EPI_ISL_439935, EPI_ISL_439936, EPI_ISL_439937, EPI_ISL_439938, EPI_ISL_439939, EPI_ISL_439940, EPI_ISL_439941, EPI_ISL_439942, EPI_ISL_439943,
EPI_ISL_439944, EPI_ISL_439945, EPI_ISL_439946, EPI_ISL_439947, EPI_ISL_439948, EPI_ISL_439949, EPI_ISL_439950, EPI_ISL_439951, EPI_ISL_439952, EPI_ISL_439954, EPI_ISL_439955, EPI_ISL_439956, EPI_ISL_439958, EPI_ISL_439961, EPI_ISL_439963, EPI_ISL_439968, EPI_ISL_439970, EPI_ISL_439975,
EPI_ISL_439977, EPI_ISL_439980, EPI_ISL_439987, EPI_ISL_439989, EPI_ISL_439990, EPI_ISL_439991, EPI_ISL_439993, EPI_ISL_439994, EPI_ISL_439995, EPI_ISL_439996, EPI_ISL_439997, EPI_ISL_439999, EPI_ISL_440003, EPI_ISL_440004, EPI_ISL_440007, EPI_ISL_440009, EPI_ISL_440010, EPI_ISL_440011,
EPI_ISL_440012, EPI_ISL_440014, EPI_ISL_440015, EPI_ISL_440017, EPI_ISL_440019, EPI_ISL_440024, EPI_ISL_440025, EPI_ISL_440030, EPI_ISL_440034, EPI_ISL_440040, EPI_ISL_440041, EPI_ISL_440044, EPI_ISL_440045, EPI_ISL_440046, EPI_ISL_440047, EPI_ISL_440048, EPI_ISL_440049, EPI_ISL_440053,
EPI_ISL_440055, EPI_ISL_440057, EPI_ISL_440059, EPI_ISL_440060, EPI_ISL_440061, EPI_ISL_440062, EPI_ISL_440064, EPI_ISL_440066, EPI_ISL_440067, EPI_ISL_440069, EPI_ISL_440072, EPI_ISL_440073, EPI_ISL_440074, EPI_ISL_440075, EPI_ISL_440076, EPI_ISL_440077, EPI_ISL_440078, EPI_ISL_440079,
EPI_ISL_440080, EPI_ISL_440082, EPI_ISL_440083, EPI_ISL_440085, EPI_ISL_440086, EPI_ISL_440088, EPI_ISL_440089, EPI_ISL_440090, EPI_ISL_440096, EPI_ISL_440097, EPI_ISL_440099, EPI_ISL_440104, EPI_ISL_440107, EPI_ISL_440108, EPI_ISL_440109, EPI_ISL_440110, EPI_ISL_440111, EPI_ISL_440112,
EPI_ISL_440117, EPI_ISL_440120, EPI_ISL_440121, EPI_ISL_440122, EPI_ISL_440125, EPI_ISL_440129, EPI_ISL_440132, EPI_ISL_440133, EPI_ISL_440138, EPI_ISL_440143, EPI_ISL_440144, EPI_ISL_440146, EPI_ISL_440147, EPI_ISL_440150, EPI_ISL_440152, EPI_ISL_440155, EPI_ISL_440159, EPI_ISL_440161,
EPI_ISL_440163, EPI_ISL_440164, EPI_ISL_440166, EPI_ISL_440169, EPI_ISL_440170, EPI_ISL_440171, EPI_ISL_440173, EPI_ISL_440176, EPI_ISL_440177, EPI_ISL_440179, EPI_ISL_440180, EPI_ISL_440181, EPI_ISL_440182, EPI_ISL_440183, EPI_ISL_440184, EPI_ISL_440185, EPI_ISL_440186, EPI_ISL_440188,
EPI_ISL_440193, EPI_ISL_440195, EPI_ISL_440197, EPI_ISL_440202, EPI_ISL_440203, EPI_ISL_440207, EPI_ISL_440208, EPI_ISL_440209, EPI_ISL_440211, EPI_ISL_440213, EPI_ISL_440217, EPI_ISL_440223, EPI_ISL_440226, EPI_ISL_440231, EPI_ISL_440234, EPI_ISL_440237, EPI_ISL_440239, EPI_ISL_440240,
EPI_ISL_440244, EPI_ISL_440246, EPI_ISL_440247, EPI_ISL_440254, EPI_ISL_440257, EPI_ISL_440258, EPI_ISL_440259, EPI_ISL_440260, EPI_ISL_440261, EPI_ISL_440263, EPI_ISL_440264, EPI_ISL_440265, EPI_ISL_440267, EPI_ISL_440268, EPI_ISL_440270, EPI_ISL_440271, EPI_ISL_440278, EPI_ISL_440279,
EPI_ISL_440280, EPI_ISL_440281, EPI_ISL_440282, EPI_ISL_440283, EPI_ISL_440284, EPI_ISL_440285, EPI_ISL_440287, EPI_ISL_440288, EPI_ISL_440289, EPI_ISL_440290, EPI_ISL_440293, EPI_ISL_440295, EPI_ISL_440296, EPI_ISL_440297, EPI_ISL_440300, EPI_ISL_440301, EPI_ISL_440302, EPI_ISL_440303,
EPI_ISL_440304, EPI_ISL_440305, EPI_ISL_440306, EPI_ISL_440307, EPI_ISL_440308, EPI_ISL_440309, EPI_ISL_440310, EPI_ISL_440311, EPI_ISL_440312, EPI_ISL_440313, EPI_ISL_440314, EPI_ISL_440315, EPI_ISL_440316, EPI_ISL_440317, EPI_ISL_440318, EPI_ISL_440319, EPI_ISL_440320, EPI_ISL_440321,
EPI_ISL_440322, EPI_ISL_440323, EPI_ISL_440324, EPI_ISL_440325, EPI_ISL_440326, EPI_ISL_440327, EPI_ISL_440328, EPI_ISL_440329, EPI_ISL_440330, EPI_ISL_440331, EPI_ISL_440332, EPI_ISL_440333, EPI_ISL_440334, EPI_ISL_440335, EPI_ISL_440336, EPI_ISL_440337, EPI_ISL_440338, EPI_ISL_440339,
EPI_ISL_440340, EPI_ISL_440341, EPI_ISL_440342, EPI_ISL_440343, EPI_ISL_440344, EPI_ISL_440345, EPI_ISL_440346, EPI_ISL_440347, EPI_ISL_440348, EPI_ISL_440349, EPI_ISL_440350, EPI_ISL_440351, EPI_ISL_440352, EPI_ISL_440353, EPI_ISL_440354, EPI_ISL_440355, EPI_ISL_440356, EPI_ISL_440357,
EPI_ISL_440358, EPI_ISL_440359, EPI_ISL_440360, EPI_ISL_440361, EPI_ISL_440362, EPI_ISL_440363, EPI_ISL_440364, EPI_ISL_440365, EPI_ISL_440366, EPI_ISL_440367, EPI_ISL_440368, EPI_ISL_440369, EPI_ISL_440370, EPI_ISL_440371, EPI_ISL_440372, EPI_ISL_440373, EPI_ISL_440374, EPI_ISL_440375,
EPI_ISL_440376, EPI_ISL_440377, EPI_ISL_440378, EPI_ISL_440379, EPI_ISL_440380, EPI_ISL_440381, EPI_ISL_440382, EPI_ISL_440383, EPI_ISL_440384, EPI_ISL_440385, EPI_ISL_440386, EPI_ISL_440387, EPI_ISL_440388, EPI_ISL_440389, EPI_ISL_440390, EPI_ISL_440391, EPI_ISL_440392, EPI_ISL_440393,
EPI_ISL_440394, EPI_ISL_440395, EPI_ISL_440396, EPI_ISL_440397, EPI_ISL_440398, EPI_ISL_440399, EPI_ISL_440400, EPI_ISL_440401, EPI_ISL_440402, EPI_ISL_440403, EPI_ISL_440404, EPI_ISL_440405, EPI_ISL_440406, EPI_ISL_440407, EPI_ISL_440408, EPI_ISL_440409, EPI_ISL_440410, EPI_ISL_440411,
EPI_ISL_440412, EPI_ISL_440413, EPI_ISL_440414, EPI_ISL_440415, EPI_ISL_440416, EPI_ISL_440417, EPI_ISL_440418, EPI_ISL_440419, EPI_ISL_440420, EPI_ISL_440421, EPI_ISL_440422, EPI_ISL_440423, EPI_ISL_440424, EPI_ISL_440425, EPI_ISL_440426, EPI_ISL_440427, EPI_ISL_440428, EPI_ISL_440429,
EPI_ISL_440430, EPI_ISL_440431, EPI_ISL_440432, EPI_ISL_440433, EPI_ISL_440434, EPI_ISL_440435, EPI_ISL_440436, EPI_ISL_440437, EPI_ISL_440438, EPI_ISL_440439, EPI_ISL_440440, EPI_ISL_440441, EPI_ISL_440442, EPI_ISL_440443, EPI_ISL_440444, EPI_ISL_440445, EPI_ISL_440446, EPI_ISL_440447,
EPI_ISL_440448, EPI_ISL_440449, EPI_ISL_440450, EPI_ISL_440451, EPI_ISL_440452, EPI_ISL_440453, EPI_ISL_440454, EPI_ISL_440455, EPI_ISL_440456, EPI_ISL_440457, EPI_ISL_440458, EPI_ISL_440459, EPI_ISL_440460, EPI_ISL_440461, EPI_ISL_440462, EPI_ISL_440463, EPI_ISL_440464, EPI_ISL_440465,
EPI_ISL_440466, EPI_ISL_440467, EPI_ISL_440468, EPI_ISL_440469, EPI_ISL_440470, EPI_ISL_440471, EPI_ISL_440472, EPI_ISL_440473, EPI_ISL_440474, EPI_ISL_440475, EPI_ISL_440476, EPI_ISL_440477, EPI_ISL_440478, EPI_ISL_440479, EPI_ISL_440480, EPI_ISL_440481, EPI_ISL_440482, EPI_ISL_440483,
EPI_ISL_440484, EPI_ISL_440485, EPI_ISL_440486, EPI_ISL_440487, EPI_ISL_440488, EPI_ISL_440489, EPI_ISL_440490, EPI_ISL_440491, EPI_ISL_440492, EPI_ISL_440493, EPI_ISL_440494, EPI_ISL_440495, EPI_ISL_440496, EPI_ISL_440497, EPI_ISL_440498, EPI_ISL_440499, EPI_ISL_440500, EPI_ISL_440501,
EPI_ISL_440502, EPI_ISL_440503, EPI_ISL_440504, EPI_ISL_440505, EPI_ISL_440506, EPI_ISL_440507, EPI_ISL_440508, EPI_ISL_440509, EPI_ISL_440510, EPI_ISL_440511, EPI_ISL_440512, EPI_ISL_440513, EPI_ISL_440514, EPI_ISL_440515, EPI_ISL_440516, EPI_ISL_440517, EPI_ISL_440518, EPI_ISL_440519,
EPI_ISL_440520, EPI_ISL_440521, EPI_ISL_440522, EPI_ISL_440523, EPI_ISL_440524, EPI_ISL_440525, EPI_ISL_440526, EPI_ISL_440527, EPI_ISL_440528, EPI_ISL_440529, EPI_ISL_440530, EPI_ISL_440531, EPI_ISL_440532, EPI_ISL_440533, EPI_ISL_440534, EPI_ISL_440535, EPI_ISL_440536, EPI_ISL_440537,
EPI_ISL_440538, EPI_ISL_440539, EPI_ISL_440540, EPI_ISL_440541, EPI_ISL_440542, EPI_ISL_440543, EPI_ISL_440544, EPI_ISL_440545, EPI_ISL_440546, EPI_ISL_440547, EPI_ISL_440548, EPI_ISL_440549, EPI_ISL_440550, EPI_ISL_440551, EPI_ISL_440552, EPI_ISL_440553, EPI_ISL_440554, EPI_ISL_440555,
EPI_ISL_440556, EPI_ISL_440557, EPI_ISL_440558, EPI_ISL_440559, EPI_ISL_440560, EPI_ISL_440561, EPI_ISL_440562, EPI_ISL_440563, EPI_ISL_440564, EPI_ISL_440565, EPI_ISL_440566, EPI_ISL_440567, EPI_ISL_440568, EPI_ISL_440569, EPI_ISL_440570, EPI_ISL_440571, EPI_ISL_440572, EPI_ISL_440573,
EPI_ISL_440574, EPI_ISL_440575, EPI_ISL_440576, EPI_ISL_440577, EPI_ISL_440578, EPI_ISL_440579, EPI_ISL_440580, EPI_ISL_440581, EPI_ISL_440582, EPI_ISL_440583, EPI_ISL_440584, EPI_ISL_440585, EPI_ISL_440586, EPI_ISL_440587, EPI_ISL_440588, EPI_ISL_440589, EPI_ISL_440590, EPI_ISL_440591,
EPI_ISL_440592, EPI_ISL_440593, EPI_ISL_440594, EPI_ISL_440595, EPI_ISL_440596, EPI_ISL_440597, EPI_ISL_440598, EPI_ISL_440599, EPI_ISL_440600, EPI_ISL_440601, EPI_ISL_440602, EPI_ISL_440603, EPI_ISL_440604, EPI_ISL_440605, EPI_ISL_440606, EPI_ISL_440607, EPI_ISL_440608, EPI_ISL_440609,
EPI_ISL_440610, EPI_ISL_440611, EPI_ISL_440612, EPI_ISL_440613, EPI_ISL_440614, EPI_ISL_440615, EPI_ISL_440616, EPI_ISL_440617, EPI_ISL_440618, EPI_ISL_440619, EPI_ISL_440620, EPI_ISL_440621, EPI_ISL_440622, EPI_ISL_440810, EPI_ISL_440811, EPI_ISL_440812, EPI_ISL_440813, EPI_ISL_440814,
EPI_ISL_440815, EPI_ISL_440816, EPI_ISL_440817, EPI_ISL_440818, EPI_ISL_440819, EPI_ISL_440820, EPI_ISL_440821, EPI_ISL_440822, EPI_ISL_440823, EPI_ISL_440824, EPI_ISL_440825, EPI_ISL_440826, EPI_ISL_440827, EPI_ISL_440828, EPI_ISL_440829, EPI_ISL_440830, EPI_ISL_440831, EPI_ISL_440832,
EPI_ISL_440833, EPI_ISL_440834, EPI_ISL_440835, EPI_ISL_440836, EPI_ISL_440837, EPI_ISL_440838, EPI_ISL_440839, EPI_ISL_440840, EPI_ISL_440841, EPI_ISL_440842, EPI_ISL_440843, EPI_ISL_440844, EPI_ISL_440845, EPI_ISL_440846, EPI_ISL_440847, EPI_ISL_440848, EPI_ISL_440849, EPI_ISL_440850,
EPI_ISL_440851, EPI_ISL_441052, EPI_ISL_441053, EPI_ISL_441054, EPI_ISL_441055, EPI_ISL_441056, EPI_ISL_441057, EPI_ISL_441058, EPI_ISL_441059, EPI_ISL_441060, EPI_ISL_441061, EPI_ISL_441062, EPI_ISL_441063, EPI_ISL_441064, EPI_ISL_441065, EPI_ISL_441066, EPI_ISL_441067, EPI_ISL_441068,
EPI_ISL_441069, EPI_ISL_441070, EPI_ISL_441071, EPI_ISL_441072, EPI_ISL_441073, EPI_ISL_441074, EPI_ISL_441075, EPI_ISL_441076, EPI_ISL_441077, EPI_ISL_441078, EPI_ISL_441079, EPI_ISL_441080, EPI_ISL_441081, EPI_ISL_441082, EPI_ISL_441083, EPI_ISL_441084, EPI_ISL_441085, EPI_ISL_441086,
EPI_ISL_441087, EPI_ISL_441088, EPI_ISL_441089, EPI_ISL_441090, EPI_ISL_441091, EPI_ISL_441092, EPI_ISL_441093, EPI_ISL_441094, EPI_ISL_441095, EPI_ISL_441096, EPI_ISL_441097, EPI_ISL_441098, EPI_ISL_441099, EPI_ISL_441100, EPI_ISL_441101, EPI_ISL_441102, EPI_ISL_441103, EPI_ISL_441104,
EPI_ISL_441105, EPI_ISL_441106, EPI_ISL_441107, EPI_ISL_441108, EPI_ISL_441109, EPI_ISL_441110, EPI_ISL_441111, EPI_ISL_441112, EPI_ISL_441113, EPI_ISL_441114, EPI_ISL_441115, EPI_ISL_441116, EPI_ISL_441117, EPI_ISL_441118, EPI_ISL_441119, EPI_ISL_441120, EPI_ISL_441121, EPI_ISL_441122,
EPI_ISL_441123, EPI_ISL_441124, EPI_ISL_441125, EPI_ISL_441126, EPI_ISL_441127, EPI_ISL_441128, EPI_ISL_441129, EPI_ISL_441130, EPI_ISL_441131, EPI_ISL_441132, EPI_ISL_441133, EPI_ISL_441134, EPI_ISL_441135, EPI_ISL_441136, EPI_ISL_441137, EPI_ISL_441138, EPI_ISL_441139, EPI_ISL_441140,
EPI_ISL_441141, EPI_ISL_441142, EPI_ISL_441143, EPI_ISL_441144, EPI_ISL_441145, EPI_ISL_441146, EPI_ISL_441147, EPI_ISL_441148, EPI_ISL_441149, EPI_ISL_441150, EPI_ISL_441151, EPI_ISL_441152, EPI_ISL_441153, EPI_ISL_441154, EPI_ISL_441155, EPI_ISL_441156, EPI_ISL_441157, EPI_ISL_441158,
EPI_ISL_441159, EPI_ISL_441160, EPI_ISL_441161, EPI_ISL_441162, EPI_ISL_441163, EPI_ISL_441164, EPI_ISL_441165, EPI_ISL_441166, EPI_ISL_441167, EPI_ISL_441168, EPI_ISL_441169, EPI_ISL_441170, EPI_ISL_441171, EPI_ISL_441172, EPI_ISL_441173, EPI_ISL_441174, EPI_ISL_441175, EPI_ISL_441176,
EPI_ISL_441177, EPI_ISL_441178, EPI_ISL_441179, EPI_ISL_441180, EPI_ISL_441181, EPI_ISL_441182, EPI_ISL_441183, EPI_ISL_441184, EPI_ISL_441185, EPI_ISL_441186, EPI_ISL_441187, EPI_ISL_441188, EPI_ISL_441189, EPI_ISL_441190, EPI_ISL_441191, EPI_ISL_441192, EPI_ISL_441193, EPI_ISL_441194,
EPI_ISL_441195, EPI_ISL_441196, EPI_ISL_441197, EPI_ISL_441198, EPI_ISL_441199, EPI_ISL_441200, EPI_ISL_441201, EPI_ISL_441202, EPI_ISL_441203, EPI_ISL_441204, EPI_ISL_441205, EPI_ISL_441206, EPI_ISL_441207, EPI_ISL_441208, EPI_ISL_441209, EPI_ISL_441210, EPI_ISL_441211, EPI_ISL_441212,
EPI_ISL_441213, EPI_ISL_441214, EPI_ISL_441215, EPI_ISL_441216, EPI_ISL_441217, EPI_ISL_441218, EPI_ISL_441219, EPI_ISL_441220, EPI_ISL_441221, EPI_ISL_441222, EPI_ISL_441223, EPI_ISL_441224, EPI_ISL_441225, EPI_ISL_441226, EPI_ISL_441227, EPI_ISL_441228, EPI_ISL_441229, EPI_ISL_441230,

| | | | |
|---|---|---|---|
| EPI_ISL_441231, EPI_ISL_441232, EPI_ISL_441233, EPI_ISL_441234, EPI_ISL_441235, EPI_ISL_441236, EPI_ISL_441237, EPI_ISL_441238, EPI_ISL_441239, EPI_ISL_441240, EPI_ISL_441241, EPI_ISL_441242, EPI_ISL_441243, EPI_ISL_441244, EPI_ISL_441245, EPI_ISL_441246, EPI_ISL_441247, EPI_ISL_441248, EPI_ISL_441249, EPI_ISL_441250, EPI_ISL_441251, EPI_ISL_441252, EPI_ISL_441253, EPI_ISL_441254, EPI_ISL_441255, EPI_ISL_441256, EPI_ISL_441257, EPI_ISL_441258, EPI_ISL_441259, EPI_ISL_441260, EPI_ISL_441261, EPI_ISL_441262, EPI_ISL_441263, EPI_ISL_441264, EPI_ISL_441265, EPI_ISL_441266, EPI_ISL_441267, EPI_ISL_441268, EPI_ISL_441269, EPI_ISL_441270, EPI_ISL_441271, EPI_ISL_441272, EPI_ISL_441273, EPI_ISL_441274, EPI_ISL_441275, EPI_ISL_441276, EPI_ISL_441277, EPI_ISL_441278, EPI_ISL_441279, EPI_ISL_441280, EPI_ISL_441281, EPI_ISL_441282, EPI_ISL_441283, EPI_ISL_441284, EPI_ISL_441285, EPI_ISL_441286, EPI_ISL_441287, EPI_ISL_441288, EPI_ISL_441289, EPI_ISL_441290, EPI_ISL_441291, EPI_ISL_441292, EPI_ISL_441293, EPI_ISL_441294, EPI_ISL_441295, EPI_ISL_441296, EPI_ISL_441297, EPI_ISL_441298, EPI_ISL_441299, EPI_ISL_441300, EPI_ISL_441301, EPI_ISL_441302, EPI_ISL_441303, EPI_ISL_441304, EPI_ISL_441305, EPI_ISL_441306, EPI_ISL_441307, EPI_ISL_441308, EPI_ISL_441309, EPI_ISL_441310, EPI_ISL_441311, EPI_ISL_441312, EPI_ISL_441313, EPI_ISL_441314, EPI_ISL_441315, EPI_ISL_441316, EPI_ISL_441317, EPI_ISL_441318, EPI_ISL_441319, EPI_ISL_441320, EPI_ISL_441321, EPI_ISL_441322, EPI_ISL_441323, EPI_ISL_441324, EPI_ISL_441325, EPI_ISL_441326, EPI_ISL_441327, EPI_ISL_441328, EPI_ISL_441329, EPI_ISL_441330, EPI_ISL_441331, EPI_ISL_441332, EPI_ISL_441333, EPI_ISL_441334, EPI_ISL_441335, EPI_ISL_441336, EPI_ISL_441337, EPI_ISL_441338, EPI_ISL_441339, EPI_ISL_441340, EPI_ISL_441341, EPI_ISL_441342, EPI_ISL_441343, EPI_ISL_441344, EPI_ISL_441345, EPI_ISL_441346, EPI_ISL_441347, EPI_ISL_441348, EPI_ISL_441349, EPI_ISL_441547, EPI_ISL_441548, EPI_ISL_441549, EPI_ISL_441550, EPI_ISL_441551, EPI_ISL_441552, EPI_ISL_441553, EPI_ISL_441554, EPI_ISL_441555, EPI_ISL_441556, EPI_ISL_441557, EPI_ISL_441558, EPI_ISL_441559, EPI_ISL_441560, EPI_ISL_441561, EPI_ISL_441562, EPI_ISL_441563, EPI_ISL_441564, EPI_ISL_441565, EPI_ISL_441566, EPI_ISL_441567, EPI_ISL_441568, EPI_ISL_441569, EPI_ISL_441570, EPI_ISL_441571, EPI_ISL_441572, EPI_ISL_441573, EPI_ISL_441574, EPI_ISL_441575, EPI_ISL_441576, EPI_ISL_441577, EPI_ISL_441578, EPI_ISL_441579, EPI_ISL_441580, EPI_ISL_441581, EPI_ISL_441582, EPI_ISL_441583, EPI_ISL_441584, EPI_ISL_441585, EPI_ISL_441586, EPI_ISL_441587, EPI_ISL_441588, EPI_ISL_441589, EPI_ISL_441590, EPI_ISL_441591, EPI_ISL_441592, EPI_ISL_441593, EPI_ISL_441594, EPI_ISL_441595, EPI_ISL_441596, EPI_ISL_441597, EPI_ISL_441598, EPI_ISL_441599, EPI_ISL_441600, EPI_ISL_441601, EPI_ISL_441602, EPI_ISL_441603, EPI_ISL_441604, EPI_ISL_441605, EPI_ISL_441606, EPI_ISL_441607, EPI_ISL_441608, EPI_ISL_441609, EPI_ISL_441610, EPI_ISL_441611, EPI_ISL_441612, EPI_ISL_441613, EPI_ISL_441614, EPI_ISL_441615, EPI_ISL_441616, EPI_ISL_441617, EPI_ISL_441618, EPI_ISL_441619, EPI_ISL_441620, EPI_ISL_441621, EPI_ISL_441622, EPI_ISL_441623, EPI_ISL_441624, EPI_ISL_441625, EPI_ISL_441626, EPI_ISL_441627, EPI_ISL_441628, EPI_ISL_441629, EPI_ISL_441630, EPI_ISL_441631, EPI_ISL_441632, EPI_ISL_441633, EPI_ISL_441634, EPI_ISL_441635, EPI_ISL_441636, EPI_ISL_441637, EPI_ISL_441638, EPI_ISL_441639, EPI_ISL_441640, EPI_ISL_441641, EPI_ISL_441642, EPI_ISL_441643, EPI_ISL_441644, EPI_ISL_441645, EPI_ISL_441646, EPI_ISL_441647, EPI_ISL_441648, EPI_ISL_441649, EPI_ISL_441650, EPI_ISL_441651, EPI_ISL_441652, EPI_ISL_441653, EPI_ISL_441654, EPI_ISL_441655, EPI_ISL_441656, EPI_ISL_441657, EPI_ISL_441658, EPI_ISL_441660, EPI_ISL_441661, EPI_ISL_441664, EPI_ISL_441665, EPI_ISL_441666, EPI_ISL_441667, EPI_ISL_441668, EPI_ISL_441669, EPI_ISL_441671, EPI_ISL_441672, EPI_ISL_441673, EPI_ISL_441674, EPI_ISL_441677, EPI_ISL_441678, EPI_ISL_441679, EPI_ISL_441680, EPI_ISL_441681, EPI_ISL_441682, EPI_ISL_441683, EPI_ISL_441684, EPI_ISL_441686, EPI_ISL_441687, EPI_ISL_441688, EPI_ISL_441689, EPI_ISL_441692, EPI_ISL_441695, EPI_ISL_441700, EPI_ISL_441701, EPI_ISL_441702, EPI_ISL_441703, EPI_ISL_441705, EPI_ISL_441706, EPI_ISL_441707, EPI_ISL_441709, EPI_ISL_441710, EPI_ISL_441711, EPI_ISL_441713, EPI_ISL_441715, EPI_ISL_441718, EPI_ISL_441720, EPI_ISL_441721, EPI_ISL_441722, EPI_ISL_441723, EPI_ISL_441724, EPI_ISL_441725, EPI_ISL_441726, EPI_ISL_441727, EPI_ISL_441728, EPI_ISL_441729, EPI_ISL_441731, EPI_ISL_441732, EPI_ISL_441734, EPI_ISL_441735, EPI_ISL_441736, EPI_ISL_441738, EPI_ISL_441739, EPI_ISL_441740, EPI_ISL_441741, EPI_ISL_441742, EPI_ISL_441743, EPI_ISL_441744, EPI_ISL_441745, EPI_ISL_441746, EPI_ISL_441748, EPI_ISL_441749, EPI_ISL_441750, EPI_ISL_441752, EPI_ISL_441754, EPI_ISL_441755, EPI_ISL_441756, EPI_ISL_441757, EPI_ISL_441758, EPI_ISL_441759, EPI_ISL_441760, EPI_ISL_441762, EPI_ISL_441763, EPI_ISL_441764, EPI_ISL_441767, EPI_ISL_441768, EPI_ISL_441769, EPI_ISL_441772, EPI_ISL_441773, EPI_ISL_441774, EPI_ISL_441777, EPI_ISL_441778, EPI_ISL_441779, EPI_ISL_441780, EPI_ISL_441782, EPI_ISL_441783, EPI_ISL_441784, EPI_ISL_441785, EPI_ISL_441786, EPI_ISL_441787, EPI_ISL_441788, EPI_ISL_441789, EPI_ISL_441790, EPI_ISL_441791, EPI_ISL_441792, EPI_ISL_441793, EPI_ISL_441794, EPI_ISL_441795, EPI_ISL_441796, EPI_ISL_441797, EPI_ISL_441798, EPI_ISL_441799, EPI_ISL_441800, EPI_ISL_441801, EPI_ISL_441802, EPI_ISL_441803, EPI_ISL_441804, EPI_ISL_441805, EPI_ISL_441806, EPI_ISL_441807, EPI_ISL_441808, EPI_ISL_441809, EPI_ISL_441810, EPI_ISL_441811, EPI_ISL_441812, EPI_ISL_441813, EPI_ISL_441814, EPI_ISL_441815, EPI_ISL_441816, EPI_ISL_441817, EPI_ISL_441818, EPI_ISL_441819, EPI_ISL_441820, EPI_ISL_441821, EPI_ISL_441822, EPI_ISL_441823, EPI_ISL_441824, EPI_ISL_441825, EPI_ISL_441826, EPI_ISL_441827, EPI_ISL_441828, EPI_ISL_441829, EPI_ISL_441830, EPI_ISL_441831, EPI_ISL_441832, EPI_ISL_441833, EPI_ISL_441834, EPI_ISL_441835, EPI_ISL_441836, EPI_ISL_441837, EPI_ISL_441838, EPI_ISL_441839, EPI_ISL_441840, EPI_ISL_441841, EPI_ISL_441842, EPI_ISL_441843, EPI_ISL_441844, EPI_ISL_441845, EPI_ISL_441846, EPI_ISL_442045, EPI_ISL_442046, EPI_ISL_442047, EPI_ISL_442048, EPI_ISL_442049, EPI_ISL_442050, EPI_ISL_442051, EPI_ISL_442052, EPI_ISL_442053, EPI_ISL_442054, EPI_ISL_442055, EPI_ISL_442056, EPI_ISL_442057, EPI_ISL_442058, EPI_ISL_442059, EPI_ISL_442060, EPI_ISL_442061, EPI_ISL_442062, EPI_ISL_442063, EPI_ISL_442064, EPI_ISL_442065, EPI_ISL_442066, EPI_ISL_442067, EPI_ISL_442068, EPI_ISL_442069, EPI_ISL_442070, EPI_ISL_442071, EPI_ISL_442072, EPI_ISL_442073, EPI_ISL_442074, EPI_ISL_442075, EPI_ISL_442076, EPI_ISL_442077, EPI_ISL_442078, EPI_ISL_442079, EPI_ISL_442080, EPI_ISL_442081, EPI_ISL_442082, EPI_ISL_442083, EPI_ISL_442084, EPI_ISL_442085, EPI_ISL_442086, EPI_ISL_442087, EPI_ISL_442088, EPI_ISL_442089, EPI_ISL_442090, EPI_ISL_442091, EPI_ISL_442092, EPI_ISL_442093, EPI_ISL_442094, EPI_ISL_442095, EPI_ISL_442096, EPI_ISL_442097, EPI_ISL_442098, EPI_ISL_442099, EPI_ISL_442100, EPI_ISL_442101, EPI_ISL_442102, EPI_ISL_442103, EPI_ISL_442104, EPI_ISL_442105, EPI_ISL_442106, EPI_ISL_442107, EPI_ISL_442108, EPI_ISL_442109, EPI_ISL_442110, EPI_ISL_442111, EPI_ISL_442112, EPI_ISL_442113, EPI_ISL_442114, EPI_ISL_442115, EPI_ISL_442116, EPI_ISL_442117, EPI_ISL_442118, EPI_ISL_442119, EPI_ISL_442120, EPI_ISL_442121, EPI_ISL_442122, EPI_ISL_442123, EPI_ISL_442124, EPI_ISL_442125, EPI_ISL_442126, EPI_ISL_442127, EPI_ISL_442128, EPI_ISL_442129, EPI_ISL_442130, EPI_ISL_442131, EPI_ISL_442132, EPI_ISL_442133, EPI_ISL_442134, EPI_ISL_442135, EPI_ISL_442136, EPI_ISL_442137, EPI_ISL_442138, EPI_ISL_442139, EPI_ISL_442140, EPI_ISL_442141, EPI_ISL_442142, EPI_ISL_442143, EPI_ISL_442144, EPI_ISL_442145, EPI_ISL_442146, EPI_ISL_442147, EPI_ISL_442148, EPI_ISL_442149, EPI_ISL_442150, EPI_ISL_442151, EPI_ISL_442152, EPI_ISL_442153, EPI_ISL_442154, EPI_ISL_442155, EPI_ISL_442156, EPI_ISL_442157, EPI_ISL_442158, EPI_ISL_442159, EPI_ISL_442160, EPI_ISL_442161, EPI_ISL_442162, EPI_ISL_442163, EPI_ISL_442164, EPI_ISL_442165, EPI_ISL_442166, EPI_ISL_442167, EPI_ISL_442168, EPI_ISL_442169, EPI_ISL_442170, EPI_ISL_442171, EPI_ISL_442172, EPI_ISL_442173, EPI_ISL_442174, EPI_ISL_442175, EPI_ISL_442176, EPI_ISL_442177, EPI_ISL_442178, EPI_ISL_442179, EPI_ISL_442180, EPI_ISL_442181, EPI_ISL_442182, EPI_ISL_442183, EPI_ISL_442184, EPI_ISL_442185, EPI_ISL_442186, EPI_ISL_442187, EPI_ISL_442188, EPI_ISL_442189, EPI_ISL_442190, EPI_ISL_442191, EPI_ISL_442192, EPI_ISL_442193, EPI_ISL_442194, EPI_ISL_442195, EPI_ISL_442196, EPI_ISL_442197, EPI_ISL_442198, EPI_ISL_442199, EPI_ISL_442200, EPI_ISL_442201, EPI_ISL_442202, EPI_ISL_442203, EPI_ISL_442204, EPI_ISL_442205, EPI_ISL_442206, EPI_ISL_442207, EPI_ISL_442208, EPI_ISL_442209, EPI_ISL_442210, EPI_ISL_442211, EPI_ISL_442212, EPI_ISL_442213, EPI_ISL_442214, EPI_ISL_442215, EPI_ISL_442216, EPI_ISL_442217, EPI_ISL_442218, EPI_ISL_442219, EPI_ISL_442220, EPI_ISL_442221, EPI_ISL_442222, EPI_ISL_442223, EPI_ISL_442224, EPI_ISL_442225, EPI_ISL_442226, EPI_ISL_442227, EPI_ISL_442228, EPI_ISL_442229, EPI_ISL_442230, EPI_ISL_442231, EPI_ISL_442232, EPI_ISL_442233, EPI_ISL_442234, EPI_ISL_442235, EPI_ISL_442236, EPI_ISL_442237, EPI_ISL_442238, EPI_ISL_442239, EPI_ISL_442240, EPI_ISL_442241, EPI_ISL_442242, EPI_ISL_442243, EPI_ISL_442244, EPI_ISL_442245, EPI_ISL_442246, EPI_ISL_442247, EPI_ISL_442248, EPI_ISL_442249, EPI_ISL_442250, EPI_ISL_442251, EPI_ISL_442252, EPI_ISL_442253, EPI_ISL_442254, EPI_ISL_442255, EPI_ISL_442256, EPI_ISL_442257, EPI_ISL_442258, EPI_ISL_442259, EPI_ISL_442260, EPI_ISL_442261, EPI_ISL_442262, EPI_ISL_442263, EPI_ISL_442264, EPI_ISL_442265, EPI_ISL_442266, EPI_ISL_442267, EPI_ISL_442268, EPI_ISL_442269, EPI_ISL_442270, EPI_ISL_442271, EPI_ISL_442272, EPI_ISL_442273, EPI_ISL_442274, EPI_ISL_442275, EPI_ISL_442276, EPI_ISL_442277, EPI_ISL_442278, EPI_ISL_442279, EPI_ISL_442280, EPI_ISL_442281, EPI_ISL_442282, EPI_ISL_442283, EPI_ISL_442284, EPI_ISL_442285, EPI_ISL_442286, EPI_ISL_442287, EPI_ISL_442288, EPI_ISL_442289, EPI_ISL_442290, EPI_ISL_442291, EPI_ISL_442292, EPI_ISL_442293, EPI_ISL_442294, EPI_ISL_442295, EPI_ISL_442296, EPI_ISL_442297, EPI_ISL_442298, EPI_ISL_442299, EPI_ISL_442300, EPI_ISL_442301, EPI_ISL_442302, EPI_ISL_442303, EPI_ISL_442304, EPI_ISL_442305, EPI_ISL_442306, EPI_ISL_442307, EPI_ISL_442308, EPI_ISL_442309, EPI_ISL_442310, EPI_ISL_442311, EPI_ISL_442312, EPI_ISL_442313, EPI_ISL_442314, EPI_ISL_442315, EPI_ISL_442316, EPI_ISL_442317, EPI_ISL_442318, EPI_ISL_442319, EPI_ISL_442320, EPI_ISL_442321, EPI_ISL_442322, EPI_ISL_442323, EPI_ISL_442324, EPI_ISL_442325, EPI_ISL_442326, EPI_ISL_442327, EPI_ISL_442328, EPI_ISL_442329, EPI_ISL_442330, EPI_ISL_442331, EPI_ISL_442332, EPI_ISL_442333, EPI_ISL_442334, EPI_ISL_442335, EPI_ISL_442336, EPI_ISL_442337, EPI_ISL_442338, EPI_ISL_442339, EPI_ISL_442340, EPI_ISL_442341, EPI_ISL_442342, EPI_ISL_442343, EPI_ISL_442524, EPI_ISL_442525, EPI_ISL_442526, EPI_ISL_442527, EPI_ISL_442528, EPI_ISL_442529, EPI_ISL_442530, EPI_ISL_442531, EPI_ISL_442532, EPI_ISL_442533, EPI_ISL_442534, EPI_ISL_442535, EPI_ISL_442536, EPI_ISL_442537, EPI_ISL_442538, EPI_ISL_442539, EPI_ISL_442540, EPI_ISL_442541, EPI_ISL_442542, EPI_ISL_442543, EPI_ISL_442544, EPI_ISL_442545, EPI_ISL_442546, EPI_ISL_442547, EPI_ISL_442548, EPI_ISL_442549, EPI_ISL_442550, EPI_ISL_442551, EPI_ISL_442552, EPI_ISL_442553, EPI_ISL_442554, EPI_ISL_442555, EPI_ISL_442556, EPI_ISL_442557, EPI_ISL_442558, EPI_ISL_442559, EPI_ISL_442560, EPI_ISL_442561, EPI_ISL_442562, EPI_ISL_442563, EPI_ISL_442564, EPI_ISL_442565, EPI_ISL_442566, EPI_ISL_442567, EPI_ISL_442568, EPI_ISL_442569, EPI_ISL_442570, EPI_ISL_442571, EPI_ISL_442572, EPI_ISL_442573, EPI_ISL_442574, EPI_ISL_442575, EPI_ISL_442576, EPI_ISL_442577, EPI_ISL_442578, EPI_ISL_442579, EPI_ISL_442580, EPI_ISL_442581, EPI_ISL_442582, EPI_ISL_442583, EPI_ISL_442584, EPI_ISL_442585, EPI_ISL_442586, EPI_ISL_442587, EPI_ISL_442588, EPI_ISL_442589, EPI_ISL_442590, EPI_ISL_442591, EPI_ISL_442592, EPI_ISL_442593, EPI_ISL_442594, EPI_ISL_442595, EPI_ISL_442596, EPI_ISL_442597, EPI_ISL_442598, EPI_ISL_442599, EPI_ISL_442600, EPI_ISL_442601, EPI_ISL_442602, EPI_ISL_442603, EPI_ISL_442604, EPI_ISL_442605, EPI_ISL_442606, EPI_ISL_442607, EPI_ISL_442608, EPI_ISL_442609, EPI_ISL_442610, EPI_ISL_442611, EPI_ISL_442612, EPI_ISL_442613, EPI_ISL_442614, EPI_ISL_442615, EPI_ISL_442616, EPI_ISL_442617, EPI_ISL_442618, EPI_ISL_442619, EPI_ISL_442620, EPI_ISL_442621, EPI_ISL_442622, EPI_ISL_442623, EPI_ISL_442811, EPI_ISL_442812, EPI_ISL_442813, EPI_ISL_442814, EPI_ISL_442815, EPI_ISL_442816, EPI_ISL_442817, EPI_ISL_442818, EPI_ISL_442819, EPI_ISL_442820, EPI_ISL_442821, EPI_ISL_442822, EPI_ISL_442823, EPI_ISL_442824, EPI_ISL_442825, EPI_ISL_442826, EPI_ISL_442827, EPI_ISL_442828, EPI_ISL_442829, EPI_ISL_442830, EPI_ISL_442831, EPI_ISL_442832, EPI_ISL_442833, EPI_ISL_442834, EPI_ISL_442835, EPI_ISL_442836, EPI_ISL_442837, EPI_ISL_442838, EPI_ISL_442839, EPI_ISL_442840, EPI_ISL_442841, EPI_ISL_442842, EPI_ISL_442843, EPI_ISL_442844, EPI_ISL_442845, EPI_ISL_442846, EPI_ISL_442847, EPI_ISL_442848, EPI_ISL_442849, EPI_ISL_442850, EPI_ISL_442851, EPI_ISL_442852, EPI_ISL_442853, EPI_ISL_442854, EPI_ISL_442855, EPI_ISL_442856, EPI_ISL_442857, EPI_ISL_442858, EPI_ISL_442859, EPI_ISL_442860, EPI_ISL_442861, EPI_ISL_442862, EPI_ISL_442863, EPI_ISL_442864, EPI_ISL_442865, EPI_ISL_442866, EPI_ISL_442867, EPI_ISL_442868, EPI_ISL_442869, EPI_ISL_442870, EPI_ISL_442871, EPI_ISL_442872, EPI_ISL_442873, EPI_ISL_442874, EPI_ISL_442875, EPI_ISL_442876, EPI_ISL_442877, EPI_ISL_442878, EPI_ISL_442879, EPI_ISL_442880, EPI_ISL_442881, EPI_ISL_442882, EPI_ISL_442883, EPI_ISL_442884, EPI_ISL_442885, EPI_ISL_442886, EPI_ISL_442887, EPI_ISL_442888, EPI_ISL_442889, EPI_ISL_442890, EPI_ISL_442891, EPI_ISL_442892, EPI_ISL_442893, EPI_ISL_442894, EPI_ISL_442895, EPI_ISL_442896, EPI_ISL_442897, EPI_ISL_442898, EPI_ISL_442899, EPI_ISL_442900, EPI_ISL_442901, EPI_ISL_442902, EPI_ISL_442903, EPI_ISL_442904, EPI_ISL_442905, EPI_ISL_442906, EPI_ISL_442907, EPI_ISL_442908, EPI_ISL_442909, EPI_ISL_442910, EPI_ISL_442911, EPI_ISL_442912, EPI_ISL_442913, EPI_ISL_442914, EPI_ISL_442915, EPI_ISL_442916, EPI_ISL_442917, EPI_ISL_442918, EPI_ISL_442919, EPI_ISL_442920, EPI_ISL_442921, EPI_ISL_442922, EPI_ISL_442923, EPI_ISL_442924, EPI_ISL_442925, EPI_ISL_442926, EPI_ISL_442927, EPI_ISL_442928, EPI_ISL_442929, EPI_ISL_442930, EPI_ISL_442931, EPI_ISL_442932, EPI_ISL_442934, EPI_ISL_442935, EPI_ISL_442936, EPI_ISL_442937, EPI_ISL_442938, EPI_ISL_442939, EPI_ISL_442940, EPI_ISL_442941, EPI_ISL_442942, EPI_ISL_442943, EPI_ISL_442944, EPI_ISL_442945, EPI_ISL_442946, EPI_ISL_442947, EPI_ISL_442948, EPI_ISL_442949, EPI_ISL_442950, EPI_ISL_442951, EPI_ISL_442952, EPI_ISL_442953, EPI_ISL_442954, EPI_ISL_442955, EPI_ISL_442956, EPI_ISL_442957, EPI_ISL_442958, EPI_ISL_442959, EPI_ISL_442960, EPI_ISL_442961, EPI_ISL_442962, EPI_ISL_442963, EPI_ISL_442964, EPI_ISL_442965, EPI_ISL_442966, EPI_ISL_442967, EPI_ISL_442968, EPI_ISL_442969, EPI_ISL_442970, EPI_ISL_442971, EPI_ISL_442972, EPI_ISL_442973, EPI_ISL_442974, EPI_ISL_442975, EPI_ISL_442976, EPI_ISL_442977, EPI_ISL_442978, EPI_ISL_442979, EPI_ISL_442980, EPI_ISL_442981, EPI_ISL_442982, EPI_ISL_442983, EPI_ISL_442984, EPI_ISL_442985, EPI_ISL_442986, EPI_ISL_442987, EPI_ISL_442988, EPI_ISL_442989, EPI_ISL_442990, EPI_ISL_442991, EPI_ISL_442992, EPI_ISL_442993, EPI_ISL_442994, EPI_ISL_442995, EPI_ISL_442996, EPI_ISL_442997, EPI_ISL_442998, EPI_ISL_442999, EPI_ISL_443000, EPI_ISL_443001, EPI_ISL_443002, EPI_ISL_443003, EPI_ISL_443004, EPI_ISL_443005, EPI_ISL_443006, EPI_ISL_443007, EPI_ISL_443008, EPI_ISL_443009, EPI_ISL_443010, EPI_ISL_443011, EPI_ISL_443012, EPI_ISL_443013, EPI_ISL_443014, EPI_ISL_443015, EPI_ISL_443016, EPI_ISL_443017, EPI_ISL_443018, EPI_ISL_443019, EPI_ISL_443020, EPI_ISL_443021, EPI_ISL_443022, EPI_ISL_443023, EPI_ISL_443024, EPI_ISL_443025, EPI_ISL_443026, EPI_ISL_443027, EPI_ISL_443028, EPI_ISL_443029, EPI_ISL_443030, EPI_ISL_443031, EPI_ISL_443032, EPI_ISL_443033, EPI_ISL_443034, EPI_ISL_443035, EPI_ISL_443036, EPI_ISL_443037, EPI_ISL_443038, EPI_ISL_443039, EPI_ISL_443040, EPI_ISL_443041, EPI_ISL_443042, EPI_ISL_443043, EPI_ISL_443044, EPI_ISL_443045, EPI_ISL_443046, EPI_ISL_443047, EPI_ISL_443048, EPI_ISL_443049, EPI_ISL_443050, EPI_ISL_443051, EPI_ISL_443052, EPI_ISL_443053, EPI_ISL_443054, EPI_ISL_443055, EPI_ISL_443056, EPI_ISL_443057, EPI_ISL_443058, EPI_ISL_443059, EPI_ISL_443060, EPI_ISL_443061, EPI_ISL_443062, EPI_ISL_443063, EPI_ISL_443064, EPI_ISL_443065, EPI_ISL_443066, EPI_ISL_443067, EPI_ISL_443068, EPI_ISL_443069, EPI_ISL_443070, EPI_ISL_443071, EPI_ISL_443072, EPI_ISL_443073, EPI_ISL_443074, EPI_ISL_443075, EPI_ISL_443076, EPI_ISL_443077, EPI_ISL_443078, EPI_ISL_443079, EPI_ISL_443080, EPI_ISL_443081, EPI_ISL_443082, EPI_ISL_443083, EPI_ISL_443084, EPI_ISL_443085, EPI_ISL_443086, EPI_ISL_443087, EPI_ISL_443088, EPI_ISL_443089, EPI_ISL_443090, EPI_ISL_443091, EPI_ISL_443092, EPI_ISL_443093, EPI_ISL_443094, EPI_ISL_443095, EPI_ISL_443096, EPI_ISL_443097, EPI_ISL_443098, EPI_ISL_443099, EPI_ISL_443100, EPI_ISL_443101, EPI_ISL_443102, EPI_ISL_443103, EPI_ISL_443104, EPI_ISL_443105, EPI_ISL_443106, EPI_ISL_443107, EPI_ISL_443108, EPI_ISL_443109, EPI_ISL_443110, EPI_ISL_443111, EPI_ISL_443112, EPI_ISL_443113, EPI_ISL_443114, EPI_ISL_443115, EPI_ISL_443116, EPI_ISL_443117, EPI_ISL_443118 | | | |
| see above | Department of Pathology, University of Cambridge | Wellcome Sanger Institute for the COVID-19 Genomics UK (COG-UK) consortium | Luke W Meredith, M. Estée Török , Myra Hosmillo, William L. Hamilton, Martin D. Curran, Theresa Feltwell, Grant Hall, Anna Yakovleva, Fahad A Khokhar, Charlotte J. Houldcroft, Laura G Caller, Aminu S. Jahun, Sarah L. Caddy, Ian Goodfellow, Alex Alderton, Roberto Amato, Sonia Goncalves, Ewan Harrison, David K. Jackson, Ian Johnston, Dominic Kwiatkowski, Cordelia Langford, John Sillitoe on behalf of the Wellcome Sanger Institute COVID-19 Surveillance Team (http://www.sanger.ac.uk/covid-team) |
| EPI_ISL_443187 | National Virology Reference Laboratory | National Public Health Laboratory, National Centre for Infectious Diseases | Mak Tze Minn, Octavia Sophie, Chavatte Jean-Marc, Zaini Zainun, Taib Surita, Cui Lin, Lin Raymond Tzer Pin |
| EPI_ISL_443188, EPI_ISL_443189, EPI_ISL_443190, EPI_ISL_443191, EPI_ISL_443192, EPI_ISL_443193, EPI_ISL_443194, EPI_ISL_443195, EPI_ISL_443196, EPI_ISL_443197, EPI_ISL_443198, EPI_ISL_443199, EPI_ISL_443200, EPI_ISL_443201, EPI_ISL_443202, EPI_ISL_443203, EPI_ISL_443204, EPI_ISL_443205, EPI_ISL_443206, EPI_ISL_443207, EPI_ISL_443208, EPI_ISL_443209, EPI_ISL_443210, EPI_ISL_443211, EPI_ISL_443212, EPI_ISL_443213, EPI_ISL_443214, EPI_ISL_443215, EPI_ISL_443216, EPI_ISL_443217, EPI_ISL_443218, EPI_ISL_443219, EPI_ISL_443220, EPI_ISL_443221, EPI_ISL_443222, EPI_ISL_443223, EPI_ISL_443224, EPI_ISL_443225, EPI_ISL_443226, EPI_ISL_443227, EPI_ISL_443228, EPI_ISL_443229, EPI_ISL_443230, EPI_ISL_443231, EPI_ISL_443232, EPI_ISL_443233, EPI_ISL_443234, EPI_ISL_443235, EPI_ISL_443236, EPI_ISL_443237, EPI_ISL_443238, EPI_ISL_443239, EPI_ISL_443240, EPI_ISL_443241, EPI_ISL_443242, EPI_ISL_443243, EPI_ISL_443244, EPI_ISL_443245, EPI_ISL_443246, EPI_ISL_443247, EPI_ISL_443248, EPI_ISL_443249 | | | |
| see above | National Public Health Laboratory, National Centre for Infectious Diseases | National Public Health Laboratory, National Centre for Infectious Diseases | Mak Tze Minn, Octavia Sophie, Chavatte Jean-Marc, Cui Lin, Lin Raymond Tzer Pin |
| EPI_ISL_443318, EPI_ISL_443319, EPI_ISL_443320, EPI_ISL_443321, EPI_ISL_443322, EPI_ISL_443323, EPI_ISL_443324, EPI_ISL_443325, EPI_ISL_443326, EPI_ISL_443327, EPI_ISL_443328, EPI_ISL_443329, EPI_ISL_443330, EPI_ISL_443331, EPI_ISL_443332, EPI_ISL_443333, EPI_ISL_443334, EPI_ISL_443335, EPI_ISL_443336, EPI_ISL_443337, EPI_ISL_443338, EPI_ISL_443339, EPI_ISL_443340, EPI_ISL_443341, EPI_ISL_443342, EPI_ISL_443343, EPI_ISL_443344, EPI_ISL_443345, EPI_ISL_443346, EPI_ISL_443347, EPI_ISL_443348, EPI_ISL_443349, EPI_ISL_443350, EPI_ISL_443351, EPI_ISL_443352, EPI_ISL_443353, EPI_ISL_443354, EPI_ISL_443355, EPI_ISL_443356, EPI_ISL_443357, EPI_ISL_443358, EPI_ISL_443359, EPI_ISL_443360, EPI_ISL_443361, EPI_ISL_443362, EPI_ISL_443363, EPI_ISL_443364, EPI_ISL_443365, EPI_ISL_443366, EPI_ISL_443367, EPI_ISL_443368, EPI_ISL_443369, EPI_ISL_443370, EPI_ISL_443371, EPI_ISL_443372, EPI_ISL_443373, EPI_ISL_443374, EPI_ISL_443375, EPI_ISL_443376, EPI_ISL_443377, EPI_ISL_443378, EPI_ISL_443379, EPI_ISL_443380, EPI_ISL_443381, EPI_ISL_443382, EPI_ISL_443383, EPI_ISL_443384, EPI_ISL_443385, EPI_ISL_443386, EPI_ISL_443387, EPI_ISL_443388, EPI_ISL_443389, EPI_ISL_443390, EPI_ISL_443391, EPI_ISL_443392, EPI_ISL_443393, EPI_ISL_443394, EPI_ISL_443395, EPI_ISL_443396, EPI_ISL_443397, EPI_ISL_443398, EPI_ISL_443399, EPI_ISL_443400, EPI_ISL_443401, EPI_ISL_443402, EPI_ISL_443403, EPI_ISL_443404, EPI_ISL_443405, EPI_ISL_443406, EPI_ISL_443407, EPI_ISL_443408, EPI_ISL_443409, EPI_ISL_443410, EPI_ISL_443411, EPI_ISL_443412, EPI_ISL_443413, EPI_ISL_443414, EPI_ISL_443415, EPI_ISL_443416, EPI_ISL_443417, EPI_ISL_443418, EPI_ISL_443419, EPI_ISL_443420, EPI_ISL_443421, EPI_ISL_443422, EPI_ISL_443423, EPI_ISL_443424, EPI_ISL_443425, EPI_ISL_443426, EPI_ISL_443427, EPI_ISL_443428, EPI_ISL_443429, EPI_ISL_443430, EPI_ISL_443431, EPI_ISL_443432, EPI_ISL_443433, EPI_ISL_443434, EPI_ISL_443435, EPI_ISL_443436, EPI_ISL_443437, EPI_ISL_443438, EPI_ISL_443439, EPI_ISL_443440, EPI_ISL_443441, EPI_ISL_443442, EPI_ISL_443443, EPI_ISL_443444, EPI_ISL_443445, EPI_ISL_443446, EPI_ISL_443447, EPI_ISL_443448, EPI_ISL_443449, EPI_ISL_443450, EPI_ISL_443451, EPI_ISL_443452, EPI_ISL_443453, EPI_ISL_443454, EPI_ISL_443455, EPI_ISL_443456, EPI_ISL_443457, EPI_ISL_443458, EPI_ISL_443459, EPI_ISL_443460, EPI_ISL_443461, EPI_ISL_443462, EPI_ISL_443463, EPI_ISL_443464, EPI_ISL_443465, EPI_ISL_443466, EPI_ISL_443467, EPI_ISL_443468, EPI_ISL_443469, EPI_ISL_443470, EPI_ISL_443471, EPI_ISL_443472, EPI_ISL_443473, EPI_ISL_443474, EPI_ISL_443475, EPI_ISL_443476, EPI_ISL_443477, EPI_ISL_443478, EPI_ISL_443479, EPI_ISL_443480, EPI_ISL_443481, EPI_ISL_443482, EPI_ISL_443483, EPI_ISL_443484, EPI_ISL_443485, EPI_ISL_443486, EPI_ISL_443487, EPI_ISL_443488, EPI_ISL_443489, EPI_ISL_443490, EPI_ISL_443491, EPI_ISL_443492, EPI_ISL_443493, EPI_ISL_443494, EPI_ISL_443495, EPI_ISL_443496, EPI_ISL_443497, EPI_ISL_443498, EPI_ISL_443499, EPI_ISL_443500, EPI_ISL_443501, EPI_ISL_443502, EPI_ISL_443503, EPI_ISL_443504, EPI_ISL_443505, EPI_ISL_443506, EPI_ISL_443507, EPI_ISL_443508, EPI_ISL_443509, EPI_ISL_443510, EPI_ISL_443511, EPI_ISL_443512, EPI_ISL_443513, EPI_ISL_443514, EPI_ISL_443515, EPI_ISL_443516, EPI_ISL_443517, EPI_ISL_443518, EPI_ISL_443519, EPI_ISL_443520, EPI_ISL_443521, EPI_ISL_443522, EPI_ISL_443523, EPI_ISL_443524, EPI_ISL_443525, EPI_ISL_443526, EPI_ISL_443527, EPI_ISL_443528, EPI_ISL_443529, EPI_ISL_443530, EPI_ISL_443531, EPI_ISL_443532, EPI_ISL_443533, | | | |

| | | | |
|---|---|---|---|
| EPI_ISL_443534, EPI_ISL_443535, EPI_ISL_443536, EPI_ISL_443537, EPI_ISL_443538, EPI_ISL_443539, EPI_ISL_443540, EPI_ISL_443541, EPI_ISL_443542, EPI_ISL_443543, EPI_ISL_443544, EPI_ISL_443545, EPI_ISL_443546, EPI_ISL_443547, EPI_ISL_443548, EPI_ISL_443549, EPI_ISL_443550, EPI_ISL_443551, EPI_ISL_443552, EPI_ISL_443553, EPI_ISL_443554, EPI_ISL_443555, EPI_ISL_443556, EPI_ISL_443557, EPI_ISL_443558, EPI_ISL_443559, EPI_ISL_443560, EPI_ISL_443561, EPI_ISL_443562, EPI_ISL_443563, EPI_ISL_443564, EPI_ISL_443565, EPI_ISL_443566, EPI_ISL_443567, EPI_ISL_443568, EPI_ISL_443569, EPI_ISL_443570, EPI_ISL_443571, EPI_ISL_443572, EPI_ISL_443573, EPI_ISL_443574, EPI_ISL_443575, EPI_ISL_443576, EPI_ISL_443577, EPI_ISL_443578, EPI_ISL_443579, EPI_ISL_443580, EPI_ISL_443581, EPI_ISL_443582, EPI_ISL_443583, EPI_ISL_443584, EPI_ISL_443585, EPI_ISL_443586, EPI_ISL_443587, EPI_ISL_443588, EPI_ISL_443589, EPI_ISL_443590, EPI_ISL_443591, EPI_ISL_443592, EPI_ISL_443593, EPI_ISL_443594, EPI_ISL_443595, EPI_ISL_443596, EPI_ISL_443597, EPI_ISL_443598, EPI_ISL_443599, EPI_ISL_443600, EPI_ISL_443601, EPI_ISL_443602, EPI_ISL_443603, EPI_ISL_443604, EPI_ISL_443605, EPI_ISL_443606, EPI_ISL_443607, EPI_ISL_443608, EPI_ISL_443609, EPI_ISL_443610, EPI_ISL_443611, EPI_ISL_443612, EPI_ISL_443613, EPI_ISL_443614, EPI_ISL_443615, EPI_ISL_443616, EPI_ISL_443617, EPI_ISL_443618, EPI_ISL_443619, EPI_ISL_443620, EPI_ISL_443621, EPI_ISL_443622, EPI_ISL_443623, EPI_ISL_443624, EPI_ISL_443625, EPI_ISL_443626, EPI_ISL_443627, EPI_ISL_443628, EPI_ISL_443629, EPI_ISL_443630, EPI_ISL_443631, EPI_ISL_443632, EPI_ISL_443633, EPI_ISL_443634, EPI_ISL_443635, EPI_ISL_443636, EPI_ISL_443637, EPI_ISL_443638, EPI_ISL_443639, EPI_ISL_443640, EPI_ISL_443641, EPI_ISL_443642, EPI_ISL_443643, EPI_ISL_443644, EPI_ISL_443645, EPI_ISL_443646, EPI_ISL_443647, EPI_ISL_443648, EPI_ISL_443649, EPI_ISL_443650, EPI_ISL_443651, EPI_ISL_443652, EPI_ISL_443653, EPI_ISL_443654, EPI_ISL_443655, EPI_ISL_443656, EPI_ISL_443657, EPI_ISL_443658, EPI_ISL_443659, EPI_ISL_443660, EPI_ISL_443661, EPI_ISL_443662, EPI_ISL_443663, EPI_ISL_443664, EPI_ISL_443665, EPI_ISL_443666, EPI_ISL_443667, EPI_ISL_443668, EPI_ISL_443669, EPI_ISL_443670, EPI_ISL_443671, EPI_ISL_443672, EPI_ISL_443673, EPI_ISL_443674, EPI_ISL_443675, EPI_ISL_443676, EPI_ISL_443677, EPI_ISL_443678, EPI_ISL_443679, EPI_ISL_443680, EPI_ISL_443681, EPI_ISL_443682, EPI_ISL_443683, EPI_ISL_443684, EPI_ISL_443685 | | | |
| see above | Department of Pathology, University of Cambridge | Wellcome Sanger Institute for the COVID-19 Genomics UK (COG-UK) consortium | Luke W Meredith, M. Estée Török , Myra Hosmillo, William L. Hamilton, Martin D. Curran, Theresa Feltwell, Grant Hall, Anna Yakovleva, Fahad A Khokhar, Charlotte J. Houldcroft, Laura G Caller, Aminu S. Jahun, Sarah L. Caddy, Ian Goodfellow; and Alex Alderton, Roberto Amato, Sonia Goncalves, Ewan Harrison, David K. Jackson, Ian Johnston, Dominic Kwiatkowski, Cordelia Langford, John Sillitoe on behalf of the Wellcome Sanger Institute COVID-19 Surveillance Team (http://www.sanger.ac.uk/covid-team) |
| EPI_ISL_444279, EPI_ISL_444280, EPI_ISL_444281, EPI_ISL_444282, EPI_ISL_444283, EPI_ISL_444284, EPI_ISL_444285, EPI_ISL_444286, EPI_ISL_444287, EPI_ISL_444288, EPI_ISL_444289, EPI_ISL_444290, EPI_ISL_444291, EPI_ISL_444292, EPI_ISL_444293, EPI_ISL_444294, EPI_ISL_444295, EPI_ISL_444296, EPI_ISL_444297, EPI_ISL_444298, EPI_ISL_444299, EPI_ISL_444300, EPI_ISL_444301, EPI_ISL_444302, EPI_ISL_444303, EPI_ISL_444304, EPI_ISL_444305, EPI_ISL_444306, EPI_ISL_444307, EPI_ISL_444308, EPI_ISL_444309, EPI_ISL_444310, EPI_ISL_444311, EPI_ISL_444312 | | | |
| see above | University of Birmingham | COVID-19 Genomics UK (COG-UK) Consortium | Loman Lab: Claire McMurray, Joanne Stockton, Samuel Nicholls, Radoslaw Poplawski, Will Rowe, Josh Quick, Nicholas Loman // UHB Lab: Celina M Whalley, Andrew Bosworth, Charlotte Poxon, Kasun Wanigasooriya, Oliver Pickles, Mike Kidd, Alex Richter, Andrew D Beggs // PHE Heartlands Lab: Husam Osman, Andrew Bosworth |
| EPI_ISL_444313, EPI_ISL_444314, EPI_ISL_444315, EPI_ISL_444316, EPI_ISL_444317, EPI_ISL_444318, EPI_ISL_444319, EPI_ISL_444320, EPI_ISL_444321, EPI_ISL_444322, EPI_ISL_444323, EPI_ISL_444324, EPI_ISL_444325, EPI_ISL_444326, EPI_ISL_444327, EPI_ISL_444328, EPI_ISL_444329, EPI_ISL_444330, EPI_ISL_444331, EPI_ISL_444332, EPI_ISL_444333, EPI_ISL_444334, EPI_ISL_444335, EPI_ISL_444336, EPI_ISL_444337, EPI_ISL_444338, EPI_ISL_444339, EPI_ISL_444340, EPI_ISL_444341, EPI_ISL_444342, EPI_ISL_444344, EPI_ISL_444346, EPI_ISL_444347, EPI_ISL_444348, EPI_ISL_444350, EPI_ISL_444351, EPI_ISL_444352, EPI_ISL_444353, EPI_ISL_444354, EPI_ISL_444356, EPI_ISL_444357, EPI_ISL_444358, EPI_ISL_444359, EPI_ISL_444360, EPI_ISL_444361, EPI_ISL_444362, EPI_ISL_444363, EPI_ISL_444364, EPI_ISL_444365, EPI_ISL_444366, EPI_ISL_444367, EPI_ISL_444368, EPI_ISL_444369, EPI_ISL_444370, EPI_ISL_444371, EPI_ISL_444372, EPI_ISL_444373, EPI_ISL_444374, EPI_ISL_444376, EPI_ISL_444377, EPI_ISL_444378, EPI_ISL_444379, EPI_ISL_444380, EPI_ISL_444381, EPI_ISL_444382, EPI_ISL_444384, EPI_ISL_444385, EPI_ISL_444386, EPI_ISL_444387, EPI_ISL_444388, EPI_ISL_444389, EPI_ISL_444390, EPI_ISL_444391, EPI_ISL_444392, EPI_ISL_444393, EPI_ISL_444394, EPI_ISL_444395, EPI_ISL_444396, EPI_ISL_444397, EPI_ISL_444398, EPI_ISL_444399, EPI_ISL_444400, EPI_ISL_444401, EPI_ISL_444402, EPI_ISL_444403, EPI_ISL_444404, EPI_ISL_444405, EPI_ISL_444406, EPI_ISL_444407, EPI_ISL_444408, EPI_ISL_444410, EPI_ISL_444411, EPI_ISL_444412, EPI_ISL_444413, EPI_ISL_444414, EPI_ISL_444415, EPI_ISL_444416, EPI_ISL_444417, EPI_ISL_444418, EPI_ISL_444419, EPI_ISL_444420, EPI_ISL_444421, EPI_ISL_444422, EPI_ISL_444423, EPI_ISL_444424, EPI_ISL_444425, EPI_ISL_444426, EPI_ISL_444427, EPI_ISL_444428, EPI_ISL_444429, EPI_ISL_444430, EPI_ISL_444431, EPI_ISL_444432, EPI_ISL_444433, EPI_ISL_444435, EPI_ISL_444436, EPI_ISL_444437, EPI_ISL_444438, EPI_ISL_444439, EPI_ISL_444440, EPI_ISL_444441, EPI_ISL_444442, EPI_ISL_444443, EPI_ISL_444446, EPI_ISL_444447, EPI_ISL_444448, EPI_ISL_444449, EPI_ISL_444450, EPI_ISL_444451, EPI_ISL_444452, EPI_ISL_444453 | | | |
| see above | Department of Pathology, University of Cambridge | COVID-19 Genomics UK (COG-UK) Consortium | Luke W Meredith, M. Estée Török , Myra Hosmillo, William L. Hamilton, Martin D. Curran, Theresa Feltwell, Grant Hall, Anna Yakovleva, Fahad A Khokhar, Charlotte J. Houldcroft, Laura G Caller, Aminu S. Jahun, Sarah L. Caddy, Ian Goodfellow |
| EPI_ISL_444467 | B.J. Medical College and Civil hospital | Gujarat Biotechnology Research Centre | Maharshi Pandya, Amit Kanani, Akanksha Verma, Nitin Savaliya, Raghawendra Kumar, Dinesh Kumar, Zuber Saiyed, Dipa Kinariwala, Disha Patel, Binita Aring, Neeta Khandelwal, Geeta Vaghela, Sonia Barve, Bhavesh Modi, Kairavi Joshi, Gaurishankar Shrimali, Nidhi Sood, Pranay Shah, R D Dixit, Snehal Bagatharia, Kamlesh J Upadhyay, Ramesh Pandit, Tejas Shah, Ankit Hinsu, Pritesh Sabara, Apurvasinh Puvar, Janvi Raval, Monika Gandhi, Pinal Trivedi, Pooja P Doshi, Chaitanya Joshi, Madhvi Joshi |
| EPI_ISL_444624 | NYU Langone Health | Departments of Pathology and Medicine, New York University School of Medicine | Maria Aguero-Rosenfeld, Brendan Belovarac, Margaret Black, Ludovic Boytard, John Cadley, Paolo Cotzia, John Chen, Dacia Dimartino, Xiaojun Feng, Tatyana Gindin, Emily Guzman, Adriana Heguy, Megan Hogan, Emily Huang, George Jour, Alireza Khodadadi-Jamayran, Lawrence H. Lin, Raven Luther, Andrew Lytle, Christian Marier, Matthew T. Maurano, Mark J. Mulligan, Peter Meyn, Raquel Ordonez Ciriza, Iman Osman, Jared Pinnell, Vanessa Raabe, Sitharam Ramaswami, Amy Rapkiewicz, Andre M. Ribeiro-dos-Santos, Marie Samanovic-Golden, Antonio Serrano, Guomiao Shen, Matija Snuderl, Theodore Vougiouklakis, Nick Vulpescu, Gael Westby, Paul Zappile, Yutong Zhang |
| EPI_ISL_444994, EPI_ISL_444995, EPI_ISL_444996, EPI_ISL_444997, EPI_ISL_444998, EPI_ISL_444999, EPI_ISL_445000 | Naval Health Research Center | Naval Medical Research Center Biological Defense Research Directorate | Logan Voegtly, Regina Cer, Dessiree Pena-Gomez, Adrian Paskey,Kyle Long, Roger Pan, Melinda Balansay-Ames, Chris Myers, Ewell Hollis, Nathaniel Christy, Kimberly Bishop-Lilly |
| EPI_ISL_445085 | Virology Unit, Agrobiodiversity and Biotechnology Project, CIAT - International Center for Tropical Agriculture | Virology Unit, Agrobiodiversity and Biotechnology Project, CIAT - International Center for Tropical Agriculture | Lopez,D., Parra,B. and Cuellar,W.J. |
| EPI_ISL_445220 | Laborarory for Respiratory Viruses, "Cantacuzino" National Military-Medical Institute for Resararch and Development | Cantacuzino Institute | M.Lazar, L.Ustea, A.Cretu |
| EPI_ISL_445224 | Narhalsan Olskroken VC | The Public Health Agency of Sweden | Mahin Ghoroghi, Oskar Karlsson Lindsjo, Maria Lind Karlberg, Anna-Malin Linde, Olov Svartstrom, Anna Risberg, Theresa Enkirch, Mia Brytting, Karin Tegmark-Wisell |
| EPI_ISL_445241 | Ã-restadsklinikens VC | The Public Health Agency of Sweden | Lisa Kjellberg / Laura Plavitu, Oskar Karlsson Lindsjo, Maria Lind Karlberg, Anna-Malin Linde, Olov Svartstrom, Anna Risberg, Theresa Enkirch, Mia Brytting, Karin Tegmark-Wisell |
| EPI_ISL_445243 | Laboratory for Respiratory Viruses, Cantacuzino National Military-Medical Institute for Research and Development | Cantacuzino Institute | M.Lazar, L.Ustea, A.Cretu |
| EPI_ISL_447039 | B.J. Medical College and Civil hospital | Gujarat Biotechnology Research Centre | Maharshi Pandya, Amit Kanani, Akanksha Verma, Nitin Savaliya, Raghawendra Kumar, Dinesh Kumar, Zuber Saiyed, Dipa Kinariwala, Disha Patel, Binita Aring, Neeta Khandelwal, Geeta Vaghela, Sonia Barve, Bhavesh Modi, Kairavi Joshi, Gaurishankar Shrimali, Nidhi Sood, Pranay Shah, R D Dixit, Snehal Bagatharia, Kamlesh J Upadhyay, Ramesh Pandit, Tejas Shah, Ankit Hinsu, Pritesh Sabara, Apurvasinh Puvar, Janvi Raval, Monika Gandhi, Pinal Trivedi, Bhavya Jindal, Chaitanya Joshi, Madhvi Joshi |
| EPI_ISL_447054 | Cantacuzino National Military-Medical Institute for Research and Development | Cantacuzino Institute | M.Lazar, L.Ustea, A.Cretu |
| EPI_ISL_447474 | Servicio de Microbiologia. Hospital Clinico Universitario de Valencia | Sequencing and Bioinformatics Service and Molecular Epidemiology Research Group. FISABIO-Public Health | Lidia Ruiz Roldan, Neris Garcia-Gonzalez, Inma Galán Vendrell, Sandra Carbo, Loreto Ferrús Abad, Paula Ruiz-Hueso, Mariana Reyes-Prieto, Vicente Soriano Chirona, Ivan Ansari, Llúcia Martínez-Priego, Giuseppe 'Auria, David Navarro, Eliseo Albert, Maria Alma Bracho, Fernando Gonzalez-Candelas |
| EPI_ISL_447478 | Servicio de Microbiologia. Hospital Clinico Universitario de Valencia | Sequencing and Bioinformatics Service and Molecular Epidemiology Research Group. FISABIO-Public Health | Loreto Ferrús Abad, Paula Ruiz-Hueso, Mariana Reyes-Prieto, Vicente Soriano Chirona, Ivan Ansari, Llúcia Martínez-Priego, Giuseppe 'Auria, David Navarro, Eliseo Albert, Maria Alma Bracho, Lidia Ruiz Roldan, Neris Garcia-Gonzalez, Inma Galán Vendrell, Sandra Carbo, Fernando Gonzalez-Candelas |
| EPI_ISL_447483 | Servicio de Microbiologia. Hospital Clinico Universitario de Valencia | Sequencing and Bioinformatics Service and Molecular Epidemiology Research Group. FISABIO-Public Health | Llúcia Martínez-Priego, Giuseppe 'Auria, David Navarro, Eliseo Albert, Maria Alma Bracho, Lidia Ruiz Roldan, Neris Garcia-Gonzalez, Inma Galán Vendrell, Sandra Carbo, Loreto Ferrús Abad, Paula Ruiz-Hueso, Mariana Reyes-Prieto, Vicente Soriano Chirona, Ivan Ansari, Fernando Gonzalez-Candelas |
| EPI_ISL_447488 | Servicio de Microbiologia. Hospital Clinico Universitario de Valencia | Sequencing and Bioinformatics Service and Molecular Epidemiology Research Group. FISABIO-Public Health | Lidia Ruiz Roldan, Neris Garcia-Gonzalez, Inma Galán Vendrell, Sandra Carbo, Loreto Ferrús Abad, Paula Ruiz-Hueso, Mariana Reyes-Prieto, Vicente Soriano Chirona, Ivan Ansari, Llúcia Martínez-Priego, Giuseppe 'Auria, David Navarro, Eliseo Albert, Maria Alma Bracho, Fernando Gonzalez-Candelas |
| EPI_ISL_447492 | Servicio de Microbiologia. Hospital Clinico Universitario de Valencia | Sequencing and Bioinformatics Service and Molecular Epidemiology Research Group. FISABIO-Public Health | Loreto Ferrús Abad, Paula Ruiz-Hueso, Mariana Reyes-Prieto, Vicente Soriano Chirona, Ivan Ansari, Llúcia Martínez-Priego, Giuseppe 'Auria, David Navarro, Eliseo Albert, Maria Alma Bracho, Lidia Ruiz Roldan, Neris Garcia-Gonzalez, Inma Galán Vendrell, Sandra Carbo, Fernando Gonzalez-Candelas |
| EPI_ISL_447497 | Servicio de Microbiologia. Hospital Clinico Universitario de Valencia | Sequencing and Bioinformatics Service and Molecular Epidemiology Research Group. FISABIO-Public Health | Llúcia Martínez-Priego, Giuseppe 'Auria, David Navarro, Eliseo Albert, Maria Alma Bracho, Lidia Ruiz Roldan, Neris Garcia-Gonzalez, Inma Galán Vendrell, Sandra Carbo, Loreto Ferrús Abad, Paula Ruiz-Hueso, Mariana Reyes-Prieto, Vicente Soriano Chirona, Ivan Ansari, Fernando Gonzalez-Candelas |
| EPI_ISL_447502 | Servicio de Microbiologia. Hospital Clinico Universitario de Valencia | Sequencing and Bioinformatics Service and Molecular Epidemiology Research Group. FISABIO-Public Health | Lidia Ruiz Roldan, Neris Garcia-Gonzalez, Inma Galán Vendrell, Sandra Carbo, Loreto Ferrús Abad, Paula Ruiz-Hueso, Mariana Reyes-Prieto, Vicente Soriano Chirona, Ivan Ansari, Llúcia Martínez-Priego, Giuseppe 'Auria, David Navarro, Eliseo Albert, Maria Alma Bracho, Fernando Gonzalez-Candelas |
| EPI_ISL_447506 | Servicio de Microbiologia. Hospital Clinico Universitario de Valencia | Sequencing and Bioinformatics Service and Molecular Epidemiology Research Group. FISABIO-Public Health | Loreto Ferrús Abad, Paula Ruiz-Hueso, Mariana Reyes-Prieto, Vicente Soriano Chirona, Ivan Ansari, Llúcia Martínez-Priego, Giuseppe 'Auria, David Navarro, Eliseo Albert, Maria Alma Bracho, Lidia Ruiz Roldan, Neris Garcia-Gonzalez, Inma Galán Vendrell, Sandra Carbo, Fernando Gonzalez-Candelas |

| | | | |
|---|---|---|---|
| EPI_ISL_447511 | Servicio de Microbiologia. Hospital Clinico Universitario de Valencia | Sequencing and Bioinformatics Service and Molecular Epidemiology Research Group. FISABIO-Public Health | Llúcia Martínez-Priego, Giuseppe 'Auria, David Navarro, Eliseo Albert, Maria Alma Bracho, Lidia Ruiz Roldan, Neris Garcia-Gonzalez, Inma Galán Vendrell, Sandra Carbo, Loreto Ferrús Abad, Paula Ruiz-Hueso, Mariana Reyes-Prieto, Vicente Soriano Chirona, Ivan Ansari, Fernando Gonzalez-Candelas |
| EPI_ISL_447516 | Servicio de Microbiologia. Hospital Clinico Universitario de Valencia | Sequencing and Bioinformatics Service and Molecular Epidemiology Research Group. FISABIO-Public Health | Lidia Ruiz Roldan, Neris Garcia-Gonzalez, Inma Galán Vendrell, Sandra Carbo, Loreto Ferrús Abad, Paula Ruiz-Hueso, Mariana Reyes-Prieto, Vicente Soriano Chirona, Ivan Ansari, Llúcia Martínez-Priego, Giuseppe 'Auria, David Navarro, Eliseo Albert, Maria Alma Bracho, Fernando Gonzalez-Candelas |
| EPI_ISL_447520 | Servicio de Microbiologia. Hospital Clinico Universitario de Valencia | Sequencing and Bioinformatics Service and Molecular Epidemiology Research Group. FISABIO-Public Health | Loreto Ferrús Abad, Paula Ruiz-Hueso, Mariana Reyes-Prieto, Vicente Soriano Chirona, Ivan Ansari, Llúcia Martínez-Priego, Giuseppe 'Auria, David Navarro, Eliseo Albert, Maria Alma Bracho, Lidia Ruiz Roldan, Neris Garcia-Gonzalez, Inma Galán Vendrell, Sandra Carbo, Fernando Gonzalez-Candelas |
| EPI_ISL_447524 | Servicio de Microbiologia. Hospital Clinico Universitario de Valencia | Sequencing and Bioinformatics Service and Molecular Epidemiology Research Group. FISABIO-Public Health | Llúcia Martínez-Priego, Giuseppe 'Auria, David Navarro, Eliseo Albert, Maria Alma Bracho, Lidia Ruiz Roldan, Neris Garcia-Gonzalez, Inma Galán Vendrell, Sandra Carbo, Loreto Ferrús Abad, Paula Ruiz-Hueso, Mariana Reyes-Prieto, Vicente Soriano Chirona, Ivan Ansari, Fernando Gonzalez-Candelas |
| EPI_ISL_447529 | Servicio de Microbiologia. Hospital Clinico Universitario de Valencia | Sequencing and Bioinformatics Service and Molecular Epidemiology Research Group. FISABIO-Public Health | Lidia Ruiz Roldan, Neris Garcia-Gonzalez, Inma Galán Vendrell, Sandra Carbo, Loreto Ferrús Abad, Paula Ruiz-Hueso, Mariana Reyes-Prieto, Vicente Soriano Chirona, Ivan Ansari, Llúcia Martínez-Priego, Giuseppe 'Auria, David Navarro, Eliseo Albert, Maria Alma Bracho, Fernando Gonzalez-Candelas |
| EPI_ISL_447548 | GMERS Medical College and Hospital, Gandhinagar | Gujarat Biotechnology Research Centre | Maharshi Pandya, Amit Kanani, Akanksha Verma, Nitin Savaliya, Raghawendra Kumar, Dinesh Kumar, Zuber Saiyed, Dipa Kinariwala, Disha Patel, Binita Aring, Neeta Khandelwal, Geeta Vaghela, Sonia Barve, Bhavesh Modi, Kairavi Joshi, Gaurishankar Shrimali, Nidhi Sood, Pranay Shah, R D Dixit, Snehal Bagatharia, Kamlesh J Upadhyay, Ramesh Pandit, Tejas Shah, Ankit Hinsu, Pritesh Sabara, Apurvasinh Puvar, Janvi Raval, Monika Gandhi, Pinal Trivedi, Afzal Ansari, Chaitanya Joshi, Madhvi Joshi |
| EPI_ISL_447945, EPI_ISL_447946, EPI_ISL_447947, EPI_ISL_447948, EPI_ISL_447949, EPI_ISL_447950, EPI_ISL_447951, EPI_ISL_447952, EPI_ISL_447953, EPI_ISL_447954, EPI_ISL_447955, EPI_ISL_447956, EPI_ISL_447957, EPI_ISL_447958, EPI_ISL_447959, EPI_ISL_447960, EPI_ISL_447961, EPI_ISL_447962, EPI_ISL_447963, EPI_ISL_447964, EPI_ISL_447965, EPI_ISL_447966, EPI_ISL_447967, EPI_ISL_447968, EPI_ISL_447969, EPI_ISL_447970, EPI_ISL_447971, EPI_ISL_447972, EPI_ISL_447973, EPI_ISL_447974, EPI_ISL_447975, EPI_ISL_447976, EPI_ISL_447977, EPI_ISL_447978, EPI_ISL_447979, EPI_ISL_447980, EPI_ISL_447981, EPI_ISL_447982, EPI_ISL_447983, EPI_ISL_447987, EPI_ISL_447988, EPI_ISL_447989, EPI_ISL_447990, EPI_ISL_447991, EPI_ISL_447992, EPI_ISL_447993, EPI_ISL_447994, EPI_ISL_447995, EPI_ISL_447996, EPI_ISL_447997, EPI_ISL_447998, EPI_ISL_447999, EPI_ISL_448000, EPI_ISL_448001, EPI_ISL_448002, EPI_ISL_448003, EPI_ISL_448004, EPI_ISL_448005, EPI_ISL_448006, EPI_ISL_448007, EPI_ISL_448008, EPI_ISL_448009, EPI_ISL_448010, EPI_ISL_448011, EPI_ISL_448012, EPI_ISL_448013, EPI_ISL_448015, EPI_ISL_448016, EPI_ISL_448017, EPI_ISL_448018, EPI_ISL_448019, EPI_ISL_448020, EPI_ISL_448021, EPI_ISL_448022, EPI_ISL_448023, EPI_ISL_448024, EPI_ISL_448025, EPI_ISL_448026, EPI_ISL_448027, EPI_ISL_448028, EPI_ISL_448029, EPI_ISL_448031, EPI_ISL_448032, EPI_ISL_448033, EPI_ISL_448034, EPI_ISL_448035, EPI_ISL_448036, EPI_ISL_448037, EPI_ISL_448039, EPI_ISL_448040, EPI_ISL_448041, EPI_ISL_448042, EPI_ISL_448043, EPI_ISL_448044, EPI_ISL_448045, EPI_ISL_448046, EPI_ISL_448047, EPI_ISL_448048, EPI_ISL_448049, EPI_ISL_448050, EPI_ISL_448051, EPI_ISL_448052, EPI_ISL_448053, EPI_ISL_448054, EPI_ISL_448055, EPI_ISL_448056, EPI_ISL_448057, EPI_ISL_448058, EPI_ISL_448059, EPI_ISL_448060, EPI_ISL_448061, EPI_ISL_448062, EPI_ISL_448063, EPI_ISL_448064, EPI_ISL_448065, EPI_ISL_448066, EPI_ISL_448067, EPI_ISL_448068, EPI_ISL_448069, EPI_ISL_448070, EPI_ISL_448071, EPI_ISL_448074, EPI_ISL_448075, EPI_ISL_448076, EPI_ISL_448077, EPI_ISL_448078, EPI_ISL_448079, EPI_ISL_448080, EPI_ISL_448081, EPI_ISL_448082, EPI_ISL_448083, EPI_ISL_448084, EPI_ISL_448086, EPI_ISL_448087, EPI_ISL_448088, EPI_ISL_448089, EPI_ISL_448090, EPI_ISL_448092, EPI_ISL_448096, EPI_ISL_448097, EPI_ISL_448098, EPI_ISL_448099, EPI_ISL_448100, EPI_ISL_448101, EPI_ISL_448102, EPI_ISL_448103, EPI_ISL_448104, EPI_ISL_448106, EPI_ISL_448107, EPI_ISL_448108, EPI_ISL_448109, EPI_ISL_448110, EPI_ISL_448111, EPI_ISL_448112, EPI_ISL_448113, EPI_ISL_448114, EPI_ISL_448115 | | | |
| see above | Department of Pathology, University of Cambridge | COVID-19 Genomics UK (COG-UK) Consortium | Luke W Meredith, M. Estée Török , Myra Hosmillo, William L. Hamilton, Martin D. Curran, Theresa Feltwell, Grant Hall, Anna Yakovleva, Fahad A Khokhar, Charlotte J. Houldcroft, Laura G Caller, Aminu S. Jahun, Sarah L. Caddy, Ian Goodfellow |
| EPI_ISL_448222 | Pasteur Insitute Ho Chi Minh City | National Key Laboratory of Gene Technology, Institute of Biotechnology, Vietnam Academy of Science and Technology | Le Tung Lam, Nguyen Trung Hieu, Nguyen Hong Trang, Ho Thi Thuong, Nguyen Thi Ngoc Thao, Huynh Thi Kim Loan, Luu Thuy Tien, Tran Huyen Linh, Pham Duy Quang, Luong Chan Quang, Cao Minh Thang, Nguyen Vu Thuong, Hoang Ha, Chu Hoang Ha, Phan Trong Lan, Truong Nam Hai |
| EPI_ISL_449799 | National Laboratory for Health, Environment and Food | National Laboratory for Health, Environment and Food | Mahnic A., Hedzet S., Janezic S., Duh D., Zavrsnik J., Blazun Vosner H., Rupnik M. |
| EPI_ISL_450193, EPI_ISL_450194, EPI_ISL_450195 | SA Pathology | SA Pathology | Lex Leong |
| EPI_ISL_450442 | The Department of Infectious Disease Prevention and Control, Henan Provincial Center for Disease Control and Prevention | The Department of Infectious Disease Prevention and Control, Henan Provincial Center for Disease Control and Prevention | Li,X., Lu,S., Wu,B., Hu,X., Li,D., Huang,X. and Guo,W. |
| EPI_ISL_450506 | Clinical Laboratory, Hospital Israelita Albert Einstein | Clinical Laboratory, Hospital Israelita Albert Einstein | Malta,F., Amgarten,D., de Oliveira,D.B.L., Araujo,D.B., Machado,R.R.G., Santana,R.A.F., Mangueira,C.L.P., Durigon,E.L. and Pinho,J.R.R. |
| EPI_ISL_450525, EPI_ISL_450526, EPI_ISL_450527, EPI_ISL_450528, EPI_ISL_450529, EPI_ISL_450530 | Hematology Laboratory, Section of Molecular Diagnostics, University Clinical Centre, Medical University of Gdansk | Department of Virology, Faculty of Medicine, University of Helsinki, Helsinki, Finland | Maciej Grzybek, Marlena Robakowska, Aneta Szulc, Olli Vapalahti, Teemu Smura |
| EPI_ISL_450746 | Laboratory of Molecular Biology, Diagnostyka sp. z o.o. | Laboratory of Recombinant Vaccines | Lukasz Rabalski, Anna Piotrowska-Mietelska, Maciej Kosinski, Boguslaw Szewczyk, Krystyna Bienkowska-Szewczyk |
| EPI_ISL_450787 | Pandit Deendayal Upadhyay Government Medical College, Rajkot | Gujarat Biotechnology Research Centre | Manish Pattani, Prakash Modi, Sejul Antala, Zarna Patel, Monika Gandhi, Pinal Trivedi, Maharshi Pandya, Amit Kanani, Nidhi Patel, Nitin Savaliya, Raghawendra Kumar, Dinesh Kumar, Zuber Saiyed, Komal Patel, Labdhi Pandya, Snehal Bagatharia, Ramesh Pandit, Tejas Shah, Ankit Hinsu, Pritesh Sabara, Apurvasinh Puvar, Janvi Raval, Bhavesh Modi, Gaurishankar Shrimali, R D Dixit, A M Kadri, Bhavya Jindal, Chaitanya Joshi, Madhvi Joshi |
| EPI_ISL_450809 | Kungsholmsdoktorn | The Public Health Agency of Sweden | Linus Hammar, Anna-Malin Linde, Maria Lind Karlberg, Oskar Karlsson Lindsjo, Olov Svartstrom, Anna Risberg, Theresa Enkirch, Mia Brytting, Karin Tegmark-Wisell |
| EPI_ISL_450816 | orestadsklinikens VC | The Public Health Agency of Sweden | Lisa Kjellberg / Laura Plavitu, Anna-Malin Linde, Maria Lind Karlberg, Oskar Karlsson Lindsjo, Olov Svartstrom, Anna Risberg, Theresa Enkirch, Mia Brytting, Karin Tegmark-Wisell |
| EPI_ISL_450829 | Narhalsan Sjobo vardcentral | The Public Health Agency of Sweden | Lovisa Hjerten, Anna-Malin Linde, Maria Lind Karlberg, Oskar Karlsson Lindsjo, Olov Svartstrom, Anna Risberg, Theresa Enkirch, Mia Brytting, Karin Tegmark-Wisell |
| EPI_ISL_450830 | Narhalsan Olskroken VC | The Public Health Agency of Sweden | Mahin Ghoroghi, Anna-Malin Linde, Maria Lind Karlberg, Oskar Karlsson Lindsjo, Olov Svartstrom, Anna Risberg, Theresa Enkirch, Mia Brytting, Karin Tegmark-Wisell |
| EPI_ISL_451077, EPI_ISL_451078, EPI_ISL_451079, EPI_ISL_451080, EPI_ISL_451081, EPI_ISL_451082, EPI_ISL_451083, EPI_ISL_451084, EPI_ISL_451085, EPI_ISL_451086, EPI_ISL_451087, EPI_ISL_451088, EPI_ISL_451089, EPI_ISL_451090, EPI_ISL_451091, EPI_ISL_451092, EPI_ISL_451093, EPI_ISL_451094, EPI_ISL_451095, EPI_ISL_451096, EPI_ISL_451097, EPI_ISL_451098, EPI_ISL_451099, EPI_ISL_451100, EPI_ISL_451101, EPI_ISL_451102, EPI_ISL_451103, EPI_ISL_451104, EPI_ISL_451105, EPI_ISL_451106, EPI_ISL_451107, EPI_ISL_451108, EPI_ISL_451109, EPI_ISL_451110, EPI_ISL_451111, EPI_ISL_451112, EPI_ISL_451113, EPI_ISL_451114, EPI_ISL_451115, EPI_ISL_451116, EPI_ISL_451117, EPI_ISL_451118, EPI_ISL_451119, EPI_ISL_451120, EPI_ISL_451121, EPI_ISL_451122, EPI_ISL_451123, EPI_ISL_451124, EPI_ISL_451125, EPI_ISL_451126, EPI_ISL_451127, EPI_ISL_451128, EPI_ISL_451129, EPI_ISL_451130, EPI_ISL_451131, EPI_ISL_451132, EPI_ISL_451133, EPI_ISL_451134, EPI_ISL_451135, EPI_ISL_451136, EPI_ISL_451137, EPI_ISL_451138, EPI_ISL_451139, EPI_ISL_451140, EPI_ISL_451141, EPI_ISL_451142, EPI_ISL_451143, EPI_ISL_451144, EPI_ISL_451145, EPI_ISL_451146, EPI_ISL_451147, EPI_ISL_451148 | | | |
| see above | SA Pathology | SA Pathology | Lex Leong, Chuan Kok Lim, Mark Turra, Ivan Bastian, Geoff Higgins |
| EPI_ISL_451153 | M.P Shah Government Medocal college Jamnagar | Gujarat Biotechnology Research Centre | Maharshi Pandya, Amit Kanani, Nidhi Patel, Nitin Savaliya, Raghawendra Kumar, Dinesh Kumar, Zuber Saiyed, Komal Patel, Labdhi Pandya, Snehal Bagatharia, Ramesh Pandit, Tejas Shah, Ankit Hinsu, Pritesh Sabara, Apurvasinh Puvar, Binita Aring, Janvi Raval, Zarna Patel, Monika Gandhi, Pinal Trivedi, Bhavesh Modi, Gaurishankar Shrimali, R D Dixit, A M Kadri, Fenil Patel, Chaitanya Joshi, Madhvi Joshi, |
| EPI_ISL_451154 | Government Medical College, Vadodara | Gujarat Biotechnology Research Centre | Manish Pattani, Tanuja Javadekar , Maharshi Pandya, Amit Kanani, Nidhi Patel, Nitin Savaliya, Raghawendra Kumar, Dinesh Kumar, Zuber Saiyed, Komal Patel, Labdhi Pandya, Snehal Bagatharia, Ramesh Pandit, Tejas Shah, Ankit Hinsu, Pritesh Sabara, Apurvasinh Puvar, Janvi Raval, Zarna Patel, Monika Gandhi, Pinal Trivedi, Bhavesh Modi, Gaurishankar Shrimali, R D Dixit, A M Kadri, Neelam Nathani, Chaitanya Joshi, Madhvi Joshi |
| EPI_ISL_451164, EPI_ISL_451165, EPI_ISL_451166, EPI_ISL_451167, EPI_ISL_451168, EPI_ISL_451169, EPI_ISL_451170, EPI_ISL_451171, EPI_ISL_451172, EPI_ISL_451173 | Lab voor klinische biologie | Onderzoeksgroep Virologie | Laurens Lambrechts, Nick Vereecke, Marthe Pauwels, Jozefien De Clercq, Bruno Verhasselt, Linos Vandekerckhove, Hans Nauwynck, Sebastiaan Theuns |
| EPI_ISL_451644, EPI_ISL_451645, EPI_ISL_451646, EPI_ISL_451647 | Laboratory of Molecular Biology, Diagnostyka sp. z o.o. | Laboratory of Recombinant Vaccines | Lukasz Rabalski, Anna Piotrowska-Mietelska, Maciej Kosinski, Boguslaw Szewczyk, Krystyna Bienkowska-Szewczyk |
| EPI_ISL_451648, EPI_ISL_451649, EPI_ISL_451650, EPI_ISL_451651, EPI_ISL_451652, EPI_ISL_451653, EPI_ISL_451654 | Hematology Laboratory, Section of Molecular Diagnostics, University Clinical Centre, Medical University of Gdansk | Laboratory of Recombinant Vaccines | Lukasz Rabalski, Adam Sodol, Aneta Szulc, Krzysztof Lewandowski, Ewa Milosz, Marlena Robakowska, Boguslaw Szewczyk, Krystyna Bienkowska-Szewczyk |
| EPI_ISL_451655, EPI_ISL_451656, EPI_ISL_451657, EPI_ISL_451658, | State Sanitary Inspectorate | Laboratory of Recombinant Vaccines | Lukasz Rabalski, Boguslaw Szewczyk, Krystyna Bienkowska-Szewczyk, Jaroslaw Pinkas |

| | | | |
|---|---|---|---|
| EPI_ISL_451659, EPI_ISL_451660, EPI_ISL_451661, EPI_ISL_451662, EPI_ISL_451663, EPI_ISL_451664 | | | |
| EPI_ISL_451971, EPI_ISL_451972, EPI_ISL_451973, EPI_ISL_451974, EPI_ISL_451975, EPI_ISL_451976, EPI_ISL_451977, EPI_ISL_451978, EPI_ISL_451979, EPI_ISL_451980, EPI_ISL_451981, EPI_ISL_451982, EPI_ISL_451983, EPI_ISL_451984, EPI_ISL_451985, EPI_ISL_451986, EPI_ISL_451987 | | | |
| see above | 1. ViroGenetics - BSL3 Laboratory of Virology, Maopolska Centre of Biotechnology, Jagiellonian University; 2. II Department of Internal Medicine, Faculty of Medicine, Jagiellonian University Medical College; 3. DIAGNOSTYKA Ltd. | 1. ViroGenetics - BSL3 Laboratory of Virology, Maopolska Centre of Biotechnology, Jagiellonian University; 2. II Department of Internal Medicine, Faculty of Medicine, Jagiellonian University Medical College. | Marek Sanak, Marcin Surmiak, Monika Gsecka-Czapla, Wojciech Branicki, Pawe P abaj, Marta Rogalska-Kupiec, Jakub Swadba, Krzysztof Pyr |
| EPI_ISL_452236 | VC Sorgenfrimottagningen | The Public Health Agency of Sweden | Lisa Esbjornsson Klemendz, Anna-Malin Linde, Maria Lind Karlberg, Oskar Karlsson Lindsjo, Olov Svartstrom, Anna Risberg, Theresa Enkirch, Mia Brytting, Karin Tegmark-Wisell |
| EPI_ISL_452238 | Narhalsan Sjobo vardcentral | The Public Health Agency of Sweden | Lovisa Hjerten, Anna-Malin Linde, Maria Lind Karlberg, Oskar Karlsson Lindsjo, Olov Svartstrom, Anna Risberg, Theresa Enkirch, Mia Brytting, Karin Tegmark-Wisell |
| EPI_ISL_452857, EPI_ISL_452858, EPI_ISL_452859, EPI_ISL_452860, EPI_ISL_452861, EPI_ISL_452862, EPI_ISL_452863, EPI_ISL_452864, EPI_ISL_452865, EPI_ISL_452866, EPI_ISL_452867, EPI_ISL_452868, EPI_ISL_452869, EPI_ISL_452870, EPI_ISL_452871, EPI_ISL_452872, EPI_ISL_452873, EPI_ISL_452874, EPI_ISL_452875, EPI_ISL_452876, EPI_ISL_452877, EPI_ISL_452878, EPI_ISL_452879, EPI_ISL_452880, EPI_ISL_452881, EPI_ISL_452882, EPI_ISL_452883, EPI_ISL_452884, EPI_ISL_452885, EPI_ISL_452886, EPI_ISL_452887, EPI_ISL_452888, EPI_ISL_452889, EPI_ISL_452890, EPI_ISL_452891, EPI_ISL_452892, EPI_ISL_452893, EPI_ISL_452894, EPI_ISL_452895, EPI_ISL_452896, EPI_ISL_452897, EPI_ISL_452898, EPI_ISL_452899, EPI_ISL_452900, EPI_ISL_452901, EPI_ISL_452903, EPI_ISL_452904, EPI_ISL_452905, EPI_ISL_452906, EPI_ISL_452910, EPI_ISL_452911, EPI_ISL_452912, EPI_ISL_452913, EPI_ISL_452914, EPI_ISL_452915, EPI_ISL_452916, EPI_ISL_452917, EPI_ISL_452918, EPI_ISL_452919, EPI_ISL_452920, EPI_ISL_452921, EPI_ISL_452922, EPI_ISL_452923, EPI_ISL_452924, EPI_ISL_452925, EPI_ISL_452926, EPI_ISL_452927, EPI_ISL_452928, EPI_ISL_452929, EPI_ISL_452930, EPI_ISL_452932, EPI_ISL_452933, EPI_ISL_452934, EPI_ISL_452935, EPI_ISL_452936, EPI_ISL_452937, EPI_ISL_452938, EPI_ISL_452939, EPI_ISL_452940, EPI_ISL_452941, EPI_ISL_452942, EPI_ISL_452943, EPI_ISL_452944, EPI_ISL_452945, EPI_ISL_452946, EPI_ISL_452947, EPI_ISL_452948, EPI_ISL_452949, EPI_ISL_452951, EPI_ISL_452952, EPI_ISL_452953, EPI_ISL_452954, EPI_ISL_452956, EPI_ISL_452957, EPI_ISL_452958, EPI_ISL_452959, EPI_ISL_452960, EPI_ISL_452961, EPI_ISL_452962, EPI_ISL_452963, EPI_ISL_452964, EPI_ISL_452965, EPI_ISL_452966, EPI_ISL_452967, EPI_ISL_452968, EPI_ISL_452969, EPI_ISL_452970, EPI_ISL_452971, EPI_ISL_452972, EPI_ISL_452973, EPI_ISL_452974, EPI_ISL_452975, EPI_ISL_452976, EPI_ISL_452977, EPI_ISL_452978, EPI_ISL_452979, EPI_ISL_452980, EPI_ISL_452981, EPI_ISL_452982, EPI_ISL_452983, EPI_ISL_452984, EPI_ISL_452985, EPI_ISL_452986, EPI_ISL_452987, EPI_ISL_452988, EPI_ISL_452989, EPI_ISL_452990, EPI_ISL_452991, EPI_ISL_452992, EPI_ISL_452993, EPI_ISL_452994, EPI_ISL_452995, EPI_ISL_452996, EPI_ISL_452997, EPI_ISL_452998, EPI_ISL_452999, EPI_ISL_453000, EPI_ISL_453001, EPI_ISL_453002, EPI_ISL_453003, EPI_ISL_453004, EPI_ISL_453005 | | | |
| see above | Department of Pathology, University of Cambridge | COVID-19 Genomics UK (COG-UK) Consortium | Luke W Meredith, M. Estée Török , Myra Hosmillo, William L. Hamilton, Martin D. Curran, Theresa Feltwell, Grant Hall, Anna Yakovleva, Fahad A Khokhar, Charlotte J. Houldcroft, Laura G Caller, Aminu S. Jahun, Sarah L. Caddy, Ian Goodfellow |
| EPI_ISL_454421, EPI_ISL_454422, EPI_ISL_454423, EPI_ISL_454424, EPI_ISL_454425, EPI_ISL_454426, EPI_ISL_454427, EPI_ISL_454428, EPI_ISL_454429, EPI_ISL_454430, EPI_ISL_454431, EPI_ISL_454432, EPI_ISL_454433, EPI_ISL_454434, EPI_ISL_454435 | | | |
| see above | Maryland Department of Health Laboratories Administration | Maryland Department of Health Laboratories Administration | MDH Laboratories Administration |
| EPI_ISL_455468, EPI_ISL_455469, EPI_ISL_455470, EPI_ISL_455471, EPI_ISL_455472, EPI_ISL_455473, EPI_ISL_455474 | Laboratory for Respiratory Viruses, Cantacuzino National Military-Medical Institute for Research and Development | Cantacuzino Institute | M.Lazar, L.Ustea, A.Cretu, Tim Durfee |
| EPI_ISL_455475, EPI_ISL_455476, EPI_ISL_455477, EPI_ISL_455479 | Laboratory for Respiratory Viruses, Cantacuzino National Military-Medical Institute for Research and Development | Cantacuzino Institute | M.Lazar, L.Ustea, A.Cretu, T.Durfee |
| EPI_ISL_455724 | Servicio de Microbiologia. Hospital Clinico Universitario de Valencia | Sequencing and Bioinformatics Service and Molecular Epidemiology Research Group. FISABIO-Public Health, and SeqCOVID-Spain Consortium | Loreto Ferrús Abad, Paula Ruiz-Hueso, Mariana Reyes-Prieto, Vicente Soriano Chirona, Ivan Ansari, Llúcia Martínez-Priego, Giuseppe 'Auria, David Navarro, Eliseo Albert, Maria Alma Bracho, Lidia Ruiz Roldan, Neris Garcia-Gonzalez, Inma Galán Vendrell, Sandra Carbo, Fernando Gonzalez-Candelas |
| EPI_ISL_455732 | Servicio de Microbiologia. Hospital Clinico Universitario de Valencia | Sequencing and Bioinformatics Service and Molecular Epidemiology Research Group. FISABIO-Public Health, and SeqCOVID-Spain Consortium | Lidia Ruiz Roldan, Neris Garcia-Gonzalez, Inma Galán Vendrell, Sandra Carbo, Loreto Ferrús Abad, Paula Ruiz-Hueso, Mariana Reyes-Prieto, Vicente Soriano Chirona, Ivan Ansari, Llúcia Martínez-Priego, Giuseppe 'Auria, David Navarro, Eliseo Albert, Maria Alma Bracho, Fernando Gonzalez-Candelas |
| EPI_ISL_455733 | Servicio de Microbiologia. Hospital Clinico Universitario de Valencia | Sequencing and Bioinformatics Service and Molecular Epidemiology Research Group. FISABIO-Public Health, and SeqCOVID-Spain Consortium | Loreto Ferrús Abad, Paula Ruiz-Hueso, Mariana Reyes-Prieto, Vicente Soriano Chirona, Ivan Ansari, Llúcia Martínez-Priego, Giuseppe 'Auria, David Navarro, Eliseo Albert, Maria Alma Bracho, Lidia Ruiz Roldan, Neris Garcia-Gonzalez, Inma Galán Vendrell, Sandra Carbo, Fernando Gonzalez-Candelas |
| EPI_ISL_455739 | Servicio de Microbiologia. Hospital Clinico Universitario de Valencia | Sequencing and Bioinformatics Service and Molecular Epidemiology Research Group. FISABIO-Public Health, and SeqCOVID-Spain Consortium | Lidia Ruiz Roldan, Neris Garcia-Gonzalez, Inma Galán Vendrell, Sandra Carbo, Loreto Ferrús Abad, Paula Ruiz-Hueso, Mariana Reyes-Prieto, Vicente Soriano Chirona, Ivan Ansari, Llúcia Martínez-Priego, Giuseppe 'Auria, David Navarro, Eliseo Albert, Maria Alma Bracho, Fernando Gonzalez-Candelas |
| EPI_ISL_455744 | Servicio de Microbiologia. Hospital Clinico Universitario de Valencia | Sequencing and Bioinformatics Service and Molecular Epidemiology Research Group. FISABIO-Public Health, and SeqCOVID-Spain Consortium | Llúcia Martínez-Priego, Giuseppe 'Auria, David Navarro, Eliseo Albert, Maria Alma Bracho, Lidia Ruiz Roldan, Neris Garcia-Gonzalez, Inma Galán Vendrell, Sandra Carbo, Loreto Ferrús Abad, Paula Ruiz-Hueso, Mariana Reyes-Prieto, Vicente Soriano Chirona, Ivan Ansari, Fernando Gonzalez-Candelas |
| EPI_ISL_456666, EPI_ISL_456667, EPI_ISL_456668, EPI_ISL_456669, EPI_ISL_456670, EPI_ISL_456671, EPI_ISL_456672, EPI_ISL_456673, EPI_ISL_456674, EPI_ISL_456675, EPI_ISL_456676 | | | |
| see above | University of Birmingham | COVID-19 Genomics UK (COG-UK) Consortium | Loman Lab: Claire McMurray, Joanne Stockton, Samuel Nicholls, Radoslaw Poplawski, Will Rowe, Josh Quick, Nicholas Loman // UHB Lab: Celina M Whalley, Andrew Bosworth, Charlotte Poxon, Kasun Wanigasooriya, Oliver Pickles, Mike Kidd, Alex Richter, Andrew D Beggs // PHE Heartlands Lab: Husam Osman, Andrew Bosworth |
| EPI_ISL_456677, EPI_ISL_456678, EPI_ISL_456679, EPI_ISL_456680, EPI_ISL_456681, EPI_ISL_456682, EPI_ISL_456686, EPI_ISL_456688, EPI_ISL_456690, EPI_ISL_456692, EPI_ISL_456695, EPI_ISL_456696, EPI_ISL_456697, EPI_ISL_456698, EPI_ISL_456699, EPI_ISL_456700, EPI_ISL_456701, EPI_ISL_456702, EPI_ISL_456703, EPI_ISL_456704, EPI_ISL_456705, EPI_ISL_456706, EPI_ISL_456708, EPI_ISL_456709, EPI_ISL_456710, EPI_ISL_456711, EPI_ISL_456712, EPI_ISL_456713, EPI_ISL_456714, EPI_ISL_456715, EPI_ISL_456716, EPI_ISL_456717, EPI_ISL_456718, EPI_ISL_456719, EPI_ISL_456720, EPI_ISL_456721, EPI_ISL_456722, EPI_ISL_456723, EPI_ISL_456724, EPI_ISL_456725, EPI_ISL_456726, EPI_ISL_456727, EPI_ISL_456728, EPI_ISL_456729, EPI_ISL_456730, EPI_ISL_456731, EPI_ISL_456732, EPI_ISL_456733, EPI_ISL_456735, EPI_ISL_456736, EPI_ISL_456737, EPI_ISL_456738, EPI_ISL_456739, EPI_ISL_456740, EPI_ISL_456741, EPI_ISL_456742, EPI_ISL_456743, EPI_ISL_456744, EPI_ISL_456745, EPI_ISL_456746, EPI_ISL_456747, EPI_ISL_456748, EPI_ISL_456749, EPI_ISL_456750, EPI_ISL_456751, EPI_ISL_456752, EPI_ISL_456753, EPI_ISL_456754, EPI_ISL_456755 | | | |
| see above | Department of Pathology, University of Cambridge | COVID-19 Genomics UK (COG-UK) Consortium | Luke W Meredith, M. Estée Török, Myra Hosmillo, William L. Hamilton, Martin D. Curran, Theresa Feltwell, Grant Hall, Anna Yakovleva, Fahad A Khokhar, Charlotte J. Houldcroft, Laura G Caller, Aminu S. Jahun, Sarah L. Caddy, Ian Goodfellow |
| EPI_ISL_458091 | B.J. Medical College and Civil hospital | Gujarat Biotechnology Research Centre | Maharshi Pandya, Amit Kanani, Nidhi Patel, Nitin Savaliya, Raghawendra Kumar, Dinesh Kumar, Zuber Saiyed, Komal Patel, Labdhi Pandya, Snehal Bagatharia, Dhaval Vaghela, Ramesh Patel, Pranay Shah, Kamlesh J Upadhyay, Ramesh Pandit, Tejas Shah, Ankit Hinsu, Pritesh Sabara, Apurvasinh Puvar, Janvi Raval, Zarna Patel, Monika Gandhi, Pinal Trivedi, Bhavya Jindal, Bhavesh Modi, Gaurishankar Shrimali, R D Dixit, A M Kadri, Umang Mishra, Chaitanya Joshi, Madhvi Joshi |
| EPI_ISL_458112 | Gujarat Biotechnology Research Centre | Gujarat Biotechnology Research Centre | Maharshi Pandya, Amit Kanani, Nidhi Patel, Nitin Savaliya, Raghawendra Kumar, Dinesh Kumar, Zuber Saiyed, Komal Patel, Labdhi Pandya, Snehal Bagatharia, Ramesh Pandit, Tejas Shah, Ankit Hinsu, Pritesh Sabara, Apurvasinh Puvar, Janvi Raval, Zarna Patel, Monika Gandhi, Pinal Trivedi, Neha Rajpara, Bhavesh Modi, Gaurishankar Shrimali, R D Dixit, A M Kadri, Umang Mishra, Chaitanya Joshi, Madhvi Joshi, , , , |
| EPI_ISL_458287 | Biosafety Department PCL3 | Biosafety Department PCL3 | Lemriss,S., Souiri,A. and El Kabbaj,S. |
| EPI_ISL_458516, EPI_ISL_458517, EPI_ISL_458518, EPI_ISL_458519, EPI_ISL_458520, EPI_ISL_458521, EPI_ISL_458522, EPI_ISL_458523, EPI_ISL_458524, EPI_ISL_458525, EPI_ISL_458526, EPI_ISL_458527, EPI_ISL_458528, EPI_ISL_458529, EPI_ISL_458530, EPI_ISL_458531, EPI_ISL_458532, EPI_ISL_458533, EPI_ISL_458534, EPI_ISL_458535, EPI_ISL_458536, EPI_ISL_458537, EPI_ISL_458538, EPI_ISL_458539, EPI_ISL_458540, EPI_ISL_458541, EPI_ISL_458542, EPI_ISL_458543, EPI_ISL_458544, EPI_ISL_458545, EPI_ISL_458546, EPI_ISL_458547, EPI_ISL_458548, EPI_ISL_458549, EPI_ISL_458550, EPI_ISL_458551, EPI_ISL_458552, EPI_ISL_458553, EPI_ISL_458554, EPI_ISL_458555, EPI_ISL_458556, EPI_ISL_458557, EPI_ISL_458558, EPI_ISL_458559, EPI_ISL_458560, EPI_ISL_458561, EPI_ISL_458562, EPI_ISL_458563, EPI_ISL_458564, EPI_ISL_458565, EPI_ISL_458566, EPI_ISL_458567, EPI_ISL_458568, EPI_ISL_458569, EPI_ISL_458570 | | | |
| see above | Department of Pathology, University of Cambridge | Wellcome Sanger Institute for the COVID-19 Genomics UK (COG-UK) consortium | Luke W Meredith, M. Estée Török , Myra Hosmillo, William L. Hamilton, Martin D. Curran, Theresa Feltwell, Grant Hall, Anna Yakovleva, Fahad A Khokhar, Charlotte J. Houldcroft, Laura G Caller, Aminu S. Jahun, Sarah L. Caddy, Ian Goodfellow; and Alex Alderton, Roberto Amato, Sonia Goncalves, Ewan Harrison, David K. Jackson, Ian Johnston, Dominic Kwiatkowski, Cordelia Langford, John Sillitoe on behalf of the Wellcome Sanger Institute COVID-19 Surveillance Team (http://www.sanger.ac.uk/covid-team) |
| EPI_ISL_458571 | Department of Pathology, University of Cambridge | Wellcome Sanger Institute for the COVID-19 Genomics UK (COG-UK) Consortium | Luke W Meredith, M. Estée Török , Myra Hosmillo, William L. Hamilton, Martin D. Curran, Theresa Feltwell, Grant Hall, Anna Yakovleva, Fahad A Khokhar, Charlotte J. Houldcroft, Laura G Caller, Aminu S. Jahun, Sarah L. Caddy, Ian Goodfellow; and Alex Alderton, Roberto Amato, Sonia Goncalves, Ewan Harrison, David K. Jackson, Ian Johnston, Dominic Kwiatkowski, Cordelia Langford, John Sillitoe on behalf of the Wellcome Sanger Institute COVID-19 |

| | | | |
|---|---|---|---|
| | | | Surveillance Team |
| EPI_ISL_458572, EPI_ISL_458573, EPI_ISL_458574, EPI_ISL_458575, EPI_ISL_458576, EPI_ISL_458577, EPI_ISL_459166, EPI_ISL_459167, EPI_ISL_459168, EPI_ISL_459169, EPI_ISL_459170, EPI_ISL_459171, EPI_ISL_459172 | | | |
| see above | Department of Pathology, University of Cambridge | Wellcome Sanger Institute for the COVID-19 Genomics UK (COG-UK) consortium | Luke W Meredith, M. Estée Török , Myra Hosmillo, William L. Hamilton, Martin D. Curran, Theresa Feltwell, Grant Hall, Anna Yakovleva, Fahad A Khokhar, Charlotte J. Houldcroft, Laura G Caller, Aminu S. Jahun, Sarah L. Caddy, Ian Goodfellow; and Alex Alderton, Roberto Amato, Sonia Goncalves, Ewan Harrison, David K. Jackson, Ian Johnston, Dominic Kwiatkowski, Cordelia Langford, John Sillitoe on behalf of the Wellcome Sanger Institute COVID-19 Surveillance Team (http://www.sanger.ac.uk/covid-team) |
| EPI_ISL_459173 | Department of Pathology, University of Cambridge | Wellcome Sanger Institute for the COVID-19 Genomics UK (COG-UK) Consortium | Luke W Meredith, M. Estée Török , Myra Hosmillo, William L. Hamilton, Martin D. Curran, Theresa Feltwell, Grant Hall, Anna Yakovleva, Fahad A Khokhar, Charlotte J. Houldcroft, Laura G Caller, Aminu S. Jahun, Sarah L. Caddy, Ian Goodfellow; and Alex Alderton, Roberto Amato, Sonia Goncalves, Ewan Harrison, David K. Jackson, Ian Johnston, Dominic Kwiatkowski, Cordelia Langford, John Sillitoe on behalf of the Wellcome Sanger Institute COVID-19 Surveillance Team |
| EPI_ISL_459174, EPI_ISL_459175, EPI_ISL_459176, EPI_ISL_459177, EPI_ISL_459178, EPI_ISL_459179, EPI_ISL_459180, EPI_ISL_459181, EPI_ISL_459182, EPI_ISL_459183, EPI_ISL_459184, EPI_ISL_459185, EPI_ISL_459186, EPI_ISL_459187, EPI_ISL_459188, EPI_ISL_459189, EPI_ISL_459190, EPI_ISL_459191, EPI_ISL_459192, EPI_ISL_459193, EPI_ISL_459194, EPI_ISL_459195, EPI_ISL_459196, EPI_ISL_459197, EPI_ISL_459198, EPI_ISL_459199, EPI_ISL_459200, EPI_ISL_459201, EPI_ISL_459202, EPI_ISL_459203, EPI_ISL_459204, EPI_ISL_459205, EPI_ISL_459206, EPI_ISL_459207, EPI_ISL_459208, EPI_ISL_459209, EPI_ISL_459210, EPI_ISL_459211, EPI_ISL_459212, EPI_ISL_459213, EPI_ISL_459214, EPI_ISL_459215, EPI_ISL_459216, EPI_ISL_459217, EPI_ISL_459218, EPI_ISL_459219, EPI_ISL_459220, EPI_ISL_459221, EPI_ISL_459222, EPI_ISL_459223, EPI_ISL_459224, EPI_ISL_459225, EPI_ISL_459226, EPI_ISL_459227, EPI_ISL_459228, EPI_ISL_459229, EPI_ISL_459230, EPI_ISL_459231, EPI_ISL_459232, EPI_ISL_459233, EPI_ISL_459234, EPI_ISL_459235, EPI_ISL_459236, EPI_ISL_459237, EPI_ISL_459238, EPI_ISL_459239, EPI_ISL_459240 | | | |
| see above | Department of Pathology, University of Cambridge | Wellcome Sanger Institute for the COVID-19 Genomics UK (COG-UK) consortium | Luke W Meredith, M. Estée Török , Myra Hosmillo, William L. Hamilton, Martin D. Curran, Theresa Feltwell, Grant Hall, Anna Yakovleva, Fahad A Khokhar, Charlotte J. Houldcroft, Laura G Caller, Aminu S. Jahun, Sarah L. Caddy, Ian Goodfellow; and Alex Alderton, Roberto Amato, Sonia Goncalves, Ewan Harrison, David K. Jackson, Ian Johnston, Dominic Kwiatkowski, Cordelia Langford, John Sillitoe on behalf of the Wellcome Sanger Institute COVID-19 Surveillance Team (http://www.sanger.ac.uk/covid-team) |
| EPI_ISL_459241 | Department of Pathology, University of Cambridge | Wellcome Sanger Institute for the COVID-19 Genomics UK (COG-UK) Consortium | Luke W Meredith, M. Estée Török , Myra Hosmillo, William L. Hamilton, Martin D. Curran, Theresa Feltwell, Grant Hall, Anna Yakovleva, Fahad A Khokhar, Charlotte J. Houldcroft, Laura G Caller, Aminu S. Jahun, Sarah L. Caddy, Ian Goodfellow; and Alex Alderton, Roberto Amato, Sonia Goncalves, Ewan Harrison, David K. Jackson, Ian Johnston, Dominic Kwiatkowski, Cordelia Langford, John Sillitoe on behalf of the Wellcome Sanger Institute COVID-19 Surveillance Team |
| EPI_ISL_459242 | Department of Pathology, University of Cambridge | Wellcome Sanger Institute for the COVID-19 Genomics UK (COG-UK) consortium | Luke W Meredith, M. Estée Török , Myra Hosmillo, William L. Hamilton, Martin D. Curran, Theresa Feltwell, Grant Hall, Anna Yakovleva, Fahad A Khokhar, Charlotte J. Houldcroft, Laura G Caller, Aminu S. Jahun, Sarah L. Caddy, Ian Goodfellow; and Alex Alderton, Roberto Amato, Sonia Goncalves, Ewan Harrison, David K. Jackson, Ian Johnston, Dominic Kwiatkowski, Cordelia Langford, John Sillitoe on behalf of the Wellcome Sanger Institute COVID-19 Surveillance Team (http://www.sanger.ac.uk/covid-team) |
| EPI_ISL_459243 | Department of Pathology, University of Cambridge | Wellcome Sanger Institute for the COVID-19 Genomics UK (COG-UK) Consortium | Luke W Meredith, M. Estée Török , Myra Hosmillo, William L. Hamilton, Martin D. Curran, Theresa Feltwell, Grant Hall, Anna Yakovleva, Fahad A Khokhar, Charlotte J. Houldcroft, Laura G Caller, Aminu S. Jahun, Sarah L. Caddy, Ian Goodfellow; and Alex Alderton, Roberto Amato, Sonia Goncalves, Ewan Harrison, David K. Jackson, Ian Johnston, Dominic Kwiatkowski, Cordelia Langford, John Sillitoe on behalf of the Wellcome Sanger Institute COVID-19 Surveillance Team |
| EPI_ISL_459244, EPI_ISL_459245, EPI_ISL_459246, EPI_ISL_459247, EPI_ISL_459248, EPI_ISL_459249, EPI_ISL_459250, EPI_ISL_459251, EPI_ISL_459252, EPI_ISL_459253, EPI_ISL_459254, EPI_ISL_459255, EPI_ISL_459256, EPI_ISL_459257 | | | |
| see above | Department of Pathology, University of Cambridge | Wellcome Sanger Institute for the COVID-19 Genomics UK (COG-UK) consortium | Luke W Meredith, M. Estée Török , Myra Hosmillo, William L. Hamilton, Martin D. Curran, Theresa Feltwell, Grant Hall, Anna Yakovleva, Fahad A Khokhar, Charlotte J. Houldcroft, Laura G Caller, Aminu S. Jahun, Sarah L. Caddy, Ian Goodfellow; and Alex Alderton, Roberto Amato, Sonia Goncalves, Ewan Harrison, David K. Jackson, Ian Johnston, Dominic Kwiatkowski, Cordelia Langford, John Sillitoe on behalf of the Wellcome Sanger Institute COVID-19 Surveillance Team (http://www.sanger.ac.uk/covid-team) |
| EPI_ISL_459258 | Department of Pathology, University of Cambridge | Wellcome Sanger Institute for the COVID-19 Genomics UK (COG-UK) Consortium | Luke W Meredith, M. Estée Török , Myra Hosmillo, William L. Hamilton, Martin D. Curran, Theresa Feltwell, Grant Hall, Anna Yakovleva, Fahad A Khokhar, Charlotte J. Houldcroft, Laura G Caller, Aminu S. Jahun, Sarah L. Caddy, Ian Goodfellow; and Alex Alderton, Roberto Amato, Sonia Goncalves, Ewan Harrison, David K. Jackson, Ian Johnston, Dominic Kwiatkowski, Cordelia Langford, John Sillitoe on behalf of the Wellcome Sanger Institute COVID-19 Surveillance Team |
| EPI_ISL_459259, EPI_ISL_459260, EPI_ISL_459261, EPI_ISL_459262, EPI_ISL_459263, EPI_ISL_459264, EPI_ISL_459265, EPI_ISL_459266, EPI_ISL_459267, EPI_ISL_459268, EPI_ISL_459269, EPI_ISL_459270, EPI_ISL_459271, EPI_ISL_459272, EPI_ISL_459273, EPI_ISL_459274, EPI_ISL_459275, EPI_ISL_459276, EPI_ISL_459277, EPI_ISL_459278, EPI_ISL_459279, EPI_ISL_459280, EPI_ISL_459282, EPI_ISL_459283, EPI_ISL_459284, EPI_ISL_459285, EPI_ISL_459286, EPI_ISL_459287, EPI_ISL_459288, EPI_ISL_459289, EPI_ISL_459290, EPI_ISL_459291, EPI_ISL_459292, EPI_ISL_459293, EPI_ISL_459294, EPI_ISL_459295, EPI_ISL_459296, EPI_ISL_459297, EPI_ISL_459304, EPI_ISL_459305, EPI_ISL_459306, EPI_ISL_459307, EPI_ISL_459308, EPI_ISL_459309, EPI_ISL_459310, EPI_ISL_459311, EPI_ISL_459312, EPI_ISL_459313, EPI_ISL_459314, EPI_ISL_459315, EPI_ISL_459316, EPI_ISL_459317, EPI_ISL_459318, EPI_ISL_459319, EPI_ISL_459320, EPI_ISL_459321, EPI_ISL_459322, EPI_ISL_459323, EPI_ISL_459410, EPI_ISL_459411, EPI_ISL_459412, EPI_ISL_459413, EPI_ISL_459414, EPI_ISL_459415, EPI_ISL_459416, EPI_ISL_459417, EPI_ISL_459418, EPI_ISL_459419, EPI_ISL_459420, EPI_ISL_459421, EPI_ISL_459422, EPI_ISL_459423, EPI_ISL_459424, EPI_ISL_459425, EPI_ISL_459426, EPI_ISL_459427, EPI_ISL_459428, EPI_ISL_459429, EPI_ISL_459430, EPI_ISL_459431, EPI_ISL_459432, EPI_ISL_459433, EPI_ISL_459434, EPI_ISL_459435, EPI_ISL_459436, EPI_ISL_459437, EPI_ISL_459438, EPI_ISL_459439, EPI_ISL_459440, EPI_ISL_459441, EPI_ISL_459442, EPI_ISL_459443, EPI_ISL_459444, EPI_ISL_459445, EPI_ISL_459446, EPI_ISL_459447, EPI_ISL_459448, EPI_ISL_459449, EPI_ISL_459450, EPI_ISL_459451, EPI_ISL_459452, EPI_ISL_459453, EPI_ISL_459454, EPI_ISL_459455, EPI_ISL_459456, EPI_ISL_459457, EPI_ISL_459458, EPI_ISL_459459, EPI_ISL_459460, EPI_ISL_459461, EPI_ISL_459462, EPI_ISL_459463, EPI_ISL_459464, EPI_ISL_459465, EPI_ISL_459466, EPI_ISL_459467, EPI_ISL_459468, EPI_ISL_459469, EPI_ISL_459470, EPI_ISL_459471, EPI_ISL_459472, EPI_ISL_459473, EPI_ISL_459474, EPI_ISL_459475, EPI_ISL_459476, EPI_ISL_459477, EPI_ISL_459478, EPI_ISL_459479, EPI_ISL_459480, EPI_ISL_459481, EPI_ISL_459482, EPI_ISL_459483, EPI_ISL_459484, EPI_ISL_459485, EPI_ISL_459486, EPI_ISL_459487, EPI_ISL_459488, EPI_ISL_459489, EPI_ISL_459490, EPI_ISL_459491, EPI_ISL_459492, EPI_ISL_459493, EPI_ISL_459494, EPI_ISL_459495, EPI_ISL_459496, EPI_ISL_459497, EPI_ISL_459498, EPI_ISL_459499, EPI_ISL_459500, EPI_ISL_459501, EPI_ISL_459502, EPI_ISL_459503, EPI_ISL_459504 | | | |
| see above | Department of Pathology, University of Cambridge | Wellcome Sanger Institute for the COVID-19 Genomics UK (COG-UK) consortium | Luke W Meredith, M. Estée Török , Myra Hosmillo, William L. Hamilton, Martin D. Curran, Theresa Feltwell, Grant Hall, Anna Yakovleva, Fahad A Khokhar, Charlotte J. Houldcroft, Laura G Caller, Aminu S. Jahun, Sarah L. Caddy, Ian Goodfellow; and Alex Alderton, Roberto Amato, Sonia Goncalves, Ewan Harrison, David K. Jackson, Ian Johnston, Dominic Kwiatkowski, Cordelia Langford, John Sillitoe on behalf of the Wellcome Sanger Institute COVID-19 Surveillance Team (http://www.sanger.ac.uk/covid-team) |
| EPI_ISL_460097, EPI_ISL_460098, EPI_ISL_460099, EPI_ISL_460100, EPI_ISL_460101, EPI_ISL_460102, EPI_ISL_460103, EPI_ISL_460104, EPI_ISL_460105, EPI_ISL_460106, EPI_ISL_460107, EPI_ISL_460108, EPI_ISL_460109, EPI_ISL_460110, EPI_ISL_460111, EPI_ISL_460112, EPI_ISL_460113, EPI_ISL_460114, EPI_ISL_460115, EPI_ISL_460116, EPI_ISL_460117, EPI_ISL_460118, EPI_ISL_460119, EPI_ISL_460120, EPI_ISL_460121, EPI_ISL_460122, EPI_ISL_460123, EPI_ISL_460124, EPI_ISL_460125, EPI_ISL_460126, EPI_ISL_460127, EPI_ISL_460128, EPI_ISL_460129, EPI_ISL_460130, EPI_ISL_460131, EPI_ISL_460132, EPI_ISL_460133, EPI_ISL_460134, EPI_ISL_460135, EPI_ISL_460136, EPI_ISL_460137, EPI_ISL_460138, EPI_ISL_460139, EPI_ISL_460140, EPI_ISL_460141, EPI_ISL_460142, EPI_ISL_460143, EPI_ISL_460144, EPI_ISL_460145, EPI_ISL_460146, EPI_ISL_460147, EPI_ISL_460148, EPI_ISL_460149, EPI_ISL_460150, EPI_ISL_460151, EPI_ISL_460152, EPI_ISL_460153, EPI_ISL_460154, EPI_ISL_460155, EPI_ISL_460156, EPI_ISL_460157, EPI_ISL_460158, EPI_ISL_460159, EPI_ISL_460160, EPI_ISL_460161, EPI_ISL_460162, EPI_ISL_460163, EPI_ISL_460164, EPI_ISL_460165, EPI_ISL_460166, EPI_ISL_460167, EPI_ISL_460168, EPI_ISL_460169, EPI_ISL_460170, EPI_ISL_460171, EPI_ISL_460172, EPI_ISL_460173, EPI_ISL_460174, EPI_ISL_460175, EPI_ISL_460176, EPI_ISL_460177, EPI_ISL_460178, EPI_ISL_460179, EPI_ISL_460180, EPI_ISL_460181, EPI_ISL_460182, EPI_ISL_460183, EPI_ISL_460184, EPI_ISL_460185, EPI_ISL_460186, EPI_ISL_460187, EPI_ISL_460188, EPI_ISL_460189, EPI_ISL_460190, EPI_ISL_460191, EPI_ISL_460192, EPI_ISL_460193, EPI_ISL_460194, EPI_ISL_460195, EPI_ISL_460196, EPI_ISL_460197, EPI_ISL_460198, EPI_ISL_460199, EPI_ISL_460200, EPI_ISL_460201, EPI_ISL_460202, EPI_ISL_460203, EPI_ISL_460204, EPI_ISL_460205, EPI_ISL_460206, EPI_ISL_460207, EPI_ISL_460208, EPI_ISL_460209, EPI_ISL_460210, EPI_ISL_460211, EPI_ISL_460212, EPI_ISL_460213, EPI_ISL_460214, EPI_ISL_460215, EPI_ISL_460216, EPI_ISL_460217, EPI_ISL_460218, EPI_ISL_460219, EPI_ISL_460220, EPI_ISL_460221, EPI_ISL_460222, EPI_ISL_460223, EPI_ISL_460224, EPI_ISL_460225, EPI_ISL_460226, EPI_ISL_460227, EPI_ISL_460228, EPI_ISL_460229, EPI_ISL_460230, EPI_ISL_460231, EPI_ISL_460232, EPI_ISL_460233, EPI_ISL_460234, EPI_ISL_460236, EPI_ISL_460237, EPI_ISL_460238, EPI_ISL_460239, EPI_ISL_460240, EPI_ISL_460241, EPI_ISL_460242, EPI_ISL_460243, EPI_ISL_460244, EPI_ISL_460245, EPI_ISL_460246, EPI_ISL_460247, EPI_ISL_460248, EPI_ISL_460249, EPI_ISL_460250, EPI_ISL_460251, EPI_ISL_460252, EPI_ISL_460253, EPI_ISL_460254, EPI_ISL_460255, EPI_ISL_460256, EPI_ISL_460257, EPI_ISL_460258, EPI_ISL_460259, EPI_ISL_460260, EPI_ISL_460261, EPI_ISL_460262, EPI_ISL_460263, EPI_ISL_460264, EPI_ISL_460265, EPI_ISL_460266, EPI_ISL_460267, EPI_ISL_460268, EPI_ISL_460269, EPI_ISL_460270, EPI_ISL_460271, EPI_ISL_460272, EPI_ISL_460273, EPI_ISL_460274, EPI_ISL_460275, EPI_ISL_460276, EPI_ISL_460277, EPI_ISL_460278, EPI_ISL_460279, EPI_ISL_460280, EPI_ISL_460281, EPI_ISL_460282, EPI_ISL_460283, EPI_ISL_460284, EPI_ISL_460285, EPI_ISL_460286, EPI_ISL_460287, EPI_ISL_460288, EPI_ISL_460289, EPI_ISL_460290, EPI_ISL_460291, EPI_ISL_460292, EPI_ISL_460293, EPI_ISL_460294, EPI_ISL_460295, EPI_ISL_460296, EPI_ISL_460297, EPI_ISL_460298, EPI_ISL_460299, EPI_ISL_460300, EPI_ISL_460301, EPI_ISL_460302, EPI_ISL_460303, EPI_ISL_460304, EPI_ISL_460305, EPI_ISL_460306, EPI_ISL_460307, EPI_ISL_460308, EPI_ISL_460309, EPI_ISL_460310, EPI_ISL_460311, EPI_ISL_460312, EPI_ISL_460313, EPI_ISL_460314, EPI_ISL_460315, EPI_ISL_460316, EPI_ISL_460317, EPI_ISL_460318, EPI_ISL_460319, EPI_ISL_460320, EPI_ISL_460321, EPI_ISL_460322, EPI_ISL_460323, EPI_ISL_460324, EPI_ISL_460325, EPI_ISL_460326, EPI_ISL_460327, EPI_ISL_460328, EPI_ISL_460329, EPI_ISL_460330, EPI_ISL_460331, EPI_ISL_460332, EPI_ISL_460333, EPI_ISL_460334, EPI_ISL_460335, EPI_ISL_460336, EPI_ISL_460337, EPI_ISL_460338, EPI_ISL_460339, EPI_ISL_460340, EPI_ISL_460341, EPI_ISL_460342, EPI_ISL_460343, EPI_ISL_460344, EPI_ISL_460345, EPI_ISL_460346, EPI_ISL_460347, EPI_ISL_460348, EPI_ISL_460349, EPI_ISL_460350, EPI_ISL_460351, EPI_ISL_460352, EPI_ISL_460353, EPI_ISL_460354, EPI_ISL_460355, EPI_ISL_460356, EPI_ISL_460357, EPI_ISL_460358, EPI_ISL_460359, EPI_ISL_460360, EPI_ISL_460361, EPI_ISL_460362, EPI_ISL_460363, EPI_ISL_460364, EPI_ISL_460365, EPI_ISL_460366, EPI_ISL_460367, EPI_ISL_460368, EPI_ISL_460369, EPI_ISL_460370, EPI_ISL_460371, EPI_ISL_460372, EPI_ISL_460373, EPI_ISL_460374, EPI_ISL_460375, EPI_ISL_460376, EPI_ISL_460377, EPI_ISL_460378, EPI_ISL_460379, EPI_ISL_460380, EPI_ISL_460381, EPI_ISL_460382, EPI_ISL_460383, EPI_ISL_460384, EPI_ISL_460385, EPI_ISL_460386, EPI_ISL_460388, EPI_ISL_460389, EPI_ISL_460390, EPI_ISL_460391, EPI_ISL_460392, EPI_ISL_460393, EPI_ISL_460394, EPI_ISL_460395, EPI_ISL_460396, EPI_ISL_460397, EPI_ISL_460398, EPI_ISL_460399, EPI_ISL_460400, EPI_ISL_460401, EPI_ISL_460402, EPI_ISL_460403, EPI_ISL_460404, EPI_ISL_460405, EPI_ISL_460406, EPI_ISL_460407, EPI_ISL_460408, EPI_ISL_460409, EPI_ISL_460410, EPI_ISL_460411, EPI_ISL_460412, EPI_ISL_460413, EPI_ISL_460414, EPI_ISL_460415, EPI_ISL_460416, EPI_ISL_460417, EPI_ISL_460418, EPI_ISL_460419, EPI_ISL_460420, EPI_ISL_460421, EPI_ISL_460422, EPI_ISL_460423, EPI_ISL_460424, EPI_ISL_460425, EPI_ISL_460426, EPI_ISL_460427, EPI_ISL_460428, EPI_ISL_460429, EPI_ISL_460430, EPI_ISL_460431, EPI_ISL_460432, EPI_ISL_460433, EPI_ISL_460434, EPI_ISL_460435, EPI_ISL_460436, EPI_ISL_460437, EPI_ISL_460438, EPI_ISL_460439, EPI_ISL_460440, EPI_ISL_460441, EPI_ISL_460442, EPI_ISL_460443, EPI_ISL_460444, EPI_ISL_460445, EPI_ISL_460446, EPI_ISL_460447, EPI_ISL_460448, EPI_ISL_460449, EPI_ISL_460450, EPI_ISL_460451, EPI_ISL_460452, EPI_ISL_460453, EPI_ISL_460454, EPI_ISL_460455, EPI_ISL_460456, EPI_ISL_460457, EPI_ISL_460458, EPI_ISL_460459, EPI_ISL_460460, EPI_ISL_460461, EPI_ISL_460462, EPI_ISL_460463, EPI_ISL_460464, EPI_ISL_460465, EPI_ISL_460466, EPI_ISL_460467, EPI_ISL_460468, EPI_ISL_460469, EPI_ISL_460470, EPI_ISL_460471, EPI_ISL_460472 | | | |
| see above | Massachusetts General Hospital | Infectious Disease Program, Broad Institute of Harvard and MIT | Lemieux,J.E., Siddle,K.J., Shaw,B., Adams,G., Pierce,V., Turbett,S., Anahtar,M., Branda,J., Slater,D., Harris,J., Lin,A.E., Gladden-Young,A., Lagerborg,K., Rudy,M., DeRuff,K., Carter,A., Normandin,E., Bauer,M., Reilly,S., Tomkins-Tinch,C., Loreth,C., Chaluvadi,S., Neumann,A., Cusick,C., Chapman,S.B., Gnirke,A., Flowers,K., Cerrato,F., Birren,B.W., Gallagher,G., Smole,S., Park,D.J., MacInnis,B.L., Ryan,E., LaRocque,R., Rosenberg,E., Sabeti,P.C. |
| EPI_ISL_461546, EPI_ISL_461547, EPI_ISL_461548, EPI_ISL_461549, EPI_ISL_461550, EPI_ISL_461551, EPI_ISL_461552, EPI_ISL_461553, EPI_ISL_461554, EPI_ISL_461555, EPI_ISL_461556, EPI_ISL_461557, EPI_ISL_461558, EPI_ISL_461559, EPI_ISL_461560, EPI_ISL_461561, EPI_ISL_461562, EPI_ISL_461563, EPI_ISL_461565, EPI_ISL_461566, EPI_ISL_461567, EPI_ISL_461568, EPI_ISL_461569, EPI_ISL_461570, EPI_ISL_461571, EPI_ISL_461572, EPI_ISL_461573, EPI_ISL_461574, EPI_ISL_461575, EPI_ISL_461576, EPI_ISL_461577, EPI_ISL_461578, EPI_ISL_461579, EPI_ISL_461580, EPI_ISL_461581, EPI_ISL_461582, EPI_ISL_461583, EPI_ISL_461584, EPI_ISL_461585, EPI_ISL_461586, EPI_ISL_461587, EPI_ISL_461588 | | | |

| | | | |
|---|---|---|---|
| see above | Department of Pathology, University of Cambridge | COVID-19 Genomics UK (COG-UK) Consortium | Luke W Meredith, M. Estée Török, Myra Hosmillo, William L. Hamilton, Martin D. Curran, Theresa Feltwell, Grant Hall, Anna Yakovleva, Fahad A Khokhar, Charlotte J. Houldcroft, Laura G Caller, Aminu S. Jahun, Sarah L. Caddy, Ian Goodfellow |
| EPI_ISL_462276, EPI_ISL_462277, EPI_ISL_462278, EPI_ISL_462279, EPI_ISL_462280, EPI_ISL_462281, EPI_ISL_462282, EPI_ISL_462283, EPI_ISL_462284, EPI_ISL_462285, EPI_ISL_462286, EPI_ISL_462287, EPI_ISL_462288, EPI_ISL_462289, EPI_ISL_462290, EPI_ISL_462291, EPI_ISL_462292, EPI_ISL_462293, EPI_ISL_462294, EPI_ISL_462295, EPI_ISL_462296, EPI_ISL_462297, EPI_ISL_462298, EPI_ISL_462299, EPI_ISL_462300, EPI_ISL_462301, EPI_ISL_462302, EPI_ISL_462303, EPI_ISL_462304, EPI_ISL_462305, EPI_ISL_462306, EPI_ISL_462307, EPI_ISL_462308, EPI_ISL_462309, EPI_ISL_462310, EPI_ISL_462311, EPI_ISL_462312, EPI_ISL_462313, EPI_ISL_462314, EPI_ISL_462315, EPI_ISL_462316, EPI_ISL_462317, EPI_ISL_462318, EPI_ISL_462319, EPI_ISL_462320, EPI_ISL_462321, EPI_ISL_462322, EPI_ISL_462323, EPI_ISL_462324, EPI_ISL_462325, EPI_ISL_462326, EPI_ISL_462327, EPI_ISL_462328, EPI_ISL_462329, EPI_ISL_462330, EPI_ISL_462331, EPI_ISL_462332, EPI_ISL_462333, EPI_ISL_462334, EPI_ISL_462335, EPI_ISL_462336, EPI_ISL_462337, EPI_ISL_462338, EPI_ISL_462339, EPI_ISL_462340, EPI_ISL_462341, EPI_ISL_462342, EPI_ISL_462343, EPI_ISL_462344, EPI_ISL_462345, EPI_ISL_462346, EPI_ISL_462347, EPI_ISL_462348, EPI_ISL_462349, EPI_ISL_462350, EPI_ISL_462351, EPI_ISL_462352, EPI_ISL_462353, EPI_ISL_462354, EPI_ISL_462355, EPI_ISL_462356, EPI_ISL_462357, EPI_ISL_462358, EPI_ISL_462359, EPI_ISL_462360, EPI_ISL_462361, EPI_ISL_462362, EPI_ISL_462363, EPI_ISL_462364, EPI_ISL_462365, EPI_ISL_462366, EPI_ISL_462367, EPI_ISL_462368, EPI_ISL_462369, EPI_ISL_462370, EPI_ISL_462371, EPI_ISL_462372, EPI_ISL_462373, EPI_ISL_462374, EPI_ISL_462375, EPI_ISL_462376, EPI_ISL_462377, EPI_ISL_462378, EPI_ISL_462379, EPI_ISL_462380, EPI_ISL_462381, EPI_ISL_462382, EPI_ISL_462383, EPI_ISL_462384, EPI_ISL_462385, EPI_ISL_462386, EPI_ISL_462387, EPI_ISL_462388, EPI_ISL_462389, EPI_ISL_462390, EPI_ISL_462391, EPI_ISL_462392, EPI_ISL_462393, EPI_ISL_462394, EPI_ISL_462395, EPI_ISL_462396, EPI_ISL_462397, EPI_ISL_462398, EPI_ISL_462399, EPI_ISL_462400, EPI_ISL_462401, EPI_ISL_462402, EPI_ISL_462403, EPI_ISL_462404, EPI_ISL_462405, EPI_ISL_462406, EPI_ISL_462407, EPI_ISL_462408, EPI_ISL_462409, EPI_ISL_462410, EPI_ISL_462411, EPI_ISL_462412, EPI_ISL_462413, EPI_ISL_462414, EPI_ISL_462415, EPI_ISL_462416, EPI_ISL_462417, EPI_ISL_462418, EPI_ISL_462419, EPI_ISL_462420, EPI_ISL_462421, EPI_ISL_462422, EPI_ISL_462423, EPI_ISL_462424, EPI_ISL_462425, EPI_ISL_462426, EPI_ISL_462427, EPI_ISL_462428, EPI_ISL_462429, EPI_ISL_462430, EPI_ISL_462431, EPI_ISL_462432, EPI_ISL_462433 | | | |
| see above | National Public Health Laboratory, National Centre for Infectious Diseases | National Public Health Laboratory, National Centre for Infectious Diseases | Mak TM, Octavia S, Chavatte JM, Cui L, Lin RTP |
| EPI_ISL_462447, EPI_ISL_462448 | Fundació Lluita contra la SIDA (FLSida)/Hospital Universitari Germans Trias i Pujol | IrsiCaixa AIDS Research Lab | Marc Noguera-Julian, Mariona Parera, Maria Pilar Armengol, Marc Corbacho, Maria Ubals, Oriol Mitjà, Lidia Ruiz, Nuria Izquierdo, Jorge Carrillo, Roger Paredes, Julia Blanco, Joaquim Segalés, Bonaventura Clotet |
| EPI_ISL_462449 | Fundació Lluita contra la SIDA (FLSida)/Hospital Universitari Germans Trias i Pujol | IrsiCaixa AIDS Research Lab | Marc Noguera-Julian, Mariona Parera, Maria Pilar Armengol, Marc Corbacho, Maria Ubals, Oriol Mitjà, Lidia Ruiz, Nuria Izquierdo, Jorge Carrillo, Roger Paredes, Julia Blanco, Bonaventura Clotet |
| EPI_ISL_463749 | Pasteur Institute of Iran | Rapid Response Team | Mahboobeh Rafigh, Kayhan Azadmanesh, Tahmineh Jalali, Fatemeh Fotouhi-Chahooki, Mohammad Hassan Pouriayevali, Arash Arashkia, Zahra Ahmadi, Mohammad Sadegh Shams Nosrati, Ali Maleki, Zabihollah Shoja, Sanam Azad-Mazjiri, Mehdi Rohani, Saber Esmaeili, Ahmad Ghasemi, Amir Hesam Nemati, Ahmad Mahmoudi, Zahra Fereydouni, Mahsa Tavakolirad, Tahereh Mohammadi, Sahar Khakifirouz, Mehdi Fazlalipour, Hesam Karimi, Kazem Baesi, Seyed Dawood Mousavi Nasab, Mahmood Barati, Mohammad Reza Asadi Karam, Mehri Habibi, Neda Afzali, Ali Torabi, Azita Eshratkhah mohammadnejad, Seyedeh Sahar Bathaeian, Mohamad Mahdi Mortazavipour, Seyedeh Atefe Hosseini, Farideh niknam oskouei, Zahra Nejatipour, Parastoo Yekta Sanati, Hadiseh Shokouhi Targhi, Mahsa Ghalejoogh, Azam Amirian, Afsaneh Zokaei, Hajarossadat Ghaderi, Elmira Vadaye kheiri, Mina Agharezaei, Akram Abouie Mehrizi, Seyedeh Zahra Moravej, Mostafa Salehi-Vaziri |
| EPI_ISL_467043 | B.J. Medical College and Civil hospital | Gujarat Biotechnology Research Centre | Maharshi Pandya, Nidhi Patel, Nitin Savaliya, Raghawendra Kumar, Dinesh Kumar, Zuber Saiyed, Komal Patel, Labdhi Pandya, Snehal Bagatharia, Pranay Shah, Kamlesh J Upadhyay, Nirav Mungalpara, Tejas Shah, Ankit Hinsu, Pritesh Sabara, Apurvasinh Puvar, Janvi Raval, Zarna Patel, Monika Gandhi, Pinal Trivedi, Akanksha Verma, R D Dixit, A M Kadri, Harsh Bakshi, Chaitanya Joshi, Madhvi Joshi, |
| EPI_ISL_467086, EPI_ISL_467087, EPI_ISL_467088, EPI_ISL_467089, EPI_ISL_467090, EPI_ISL_467091 | Hospital Universitario de Gran Canaria Dr. Negrín | SeqCOVID-SPAIN consortium/IBV(CSIC) | M. Carmen Pérez González, Francisco J. Chamizo López, Ana Bordes Benítez and SeqCOVID-SPAIN consortium |
| EPI_ISL_467184, EPI_ISL_467185, EPI_ISL_467186, EPI_ISL_467187, EPI_ISL_467188, EPI_ISL_467189, EPI_ISL_467190, EPI_ISL_467191, EPI_ISL_467192, EPI_ISL_467193, EPI_ISL_467194, EPI_ISL_467195, EPI_ISL_467196, EPI_ISL_467197, EPI_ISL_467198, EPI_ISL_467199, EPI_ISL_467200, EPI_ISL_467201, EPI_ISL_467202, EPI_ISL_467203, EPI_ISL_467204, EPI_ISL_467205, EPI_ISL_467206, EPI_ISL_467207, EPI_ISL_467208, EPI_ISL_467209, EPI_ISL_467210, EPI_ISL_467211, EPI_ISL_467212, EPI_ISL_467213, EPI_ISL_467214, EPI_ISL_467215, EPI_ISL_467216, EPI_ISL_467217, EPI_ISL_467218, EPI_ISL_467219, EPI_ISL_467220, EPI_ISL_467221, EPI_ISL_467222, EPI_ISL_467223, EPI_ISL_467224, EPI_ISL_467225, EPI_ISL_467226, EPI_ISL_467227, EPI_ISL_467228, EPI_ISL_467229, EPI_ISL_467230, EPI_ISL_467231, EPI_ISL_467232, EPI_ISL_467233, EPI_ISL_467234, EPI_ISL_467235, EPI_ISL_467236, EPI_ISL_467237, EPI_ISL_467238, EPI_ISL_467239, EPI_ISL_467240, EPI_ISL_467241, EPI_ISL_467242, EPI_ISL_467243, EPI_ISL_467244, EPI_ISL_467245, EPI_ISL_467246, EPI_ISL_467247, EPI_ISL_467248, EPI_ISL_467249, EPI_ISL_467250, EPI_ISL_467251, EPI_ISL_467252, EPI_ISL_467253, EPI_ISL_467254, EPI_ISL_467255, EPI_ISL_467256, EPI_ISL_467257, EPI_ISL_467258, EPI_ISL_467259, EPI_ISL_467260, EPI_ISL_467261 | | | |
| see above | Hospital General Universitario Gregorio Marañón | SeqCOVID-SPAIN consortium/IBV(CSIC) | Laura Pérez-Lago, Marta Herranz, Jon Sicilia, Julia Suárez, Pilar Catalán, Patricia Muñoz, Darío García de Viedma and SeqCOVID-SPAIN consortium |
| EPI_ISL_467985, EPI_ISL_467986, EPI_ISL_467987, EPI_ISL_467988, EPI_ISL_467989 | SA Pathology | SA Pathology | Lex Leong, Chuan Kok Lim, Mark Turra, Ivan Bastian, Geoff Higgins |
| EPI_ISL_467990 | SA Pathology | SA Pathology | Leong, LEX, Lim, CK, Turra, M, Bastian, I, Higgins, G |
| EPI_ISL_467991, EPI_ISL_467992, EPI_ISL_467993, EPI_ISL_467994, EPI_ISL_467995, EPI_ISL_467996, EPI_ISL_467997, EPI_ISL_467998, EPI_ISL_467999, EPI_ISL_468000, EPI_ISL_468001, EPI_ISL_468002, EPI_ISL_468003, EPI_ISL_468004, EPI_ISL_468005, EPI_ISL_468006, EPI_ISL_468007, EPI_ISL_468008, EPI_ISL_468009, EPI_ISL_468010, EPI_ISL_468011, EPI_ISL_468012, EPI_ISL_468013, EPI_ISL_468014, EPI_ISL_468015, EPI_ISL_468016, EPI_ISL_468017, EPI_ISL_468018, EPI_ISL_468019, EPI_ISL_468020, EPI_ISL_468021, EPI_ISL_468022, EPI_ISL_468023, EPI_ISL_468024, EPI_ISL_468025, EPI_ISL_468026, EPI_ISL_468027, EPI_ISL_468028, EPI_ISL_468029, EPI_ISL_468030, EPI_ISL_468031, EPI_ISL_468032, EPI_ISL_468033, EPI_ISL_468034, EPI_ISL_468035, EPI_ISL_468036, EPI_ISL_468037, EPI_ISL_468038, EPI_ISL_468039, EPI_ISL_468040, EPI_ISL_468041, EPI_ISL_468042, EPI_ISL_468043 | | | |
| see above | SA Pathology | SA Pathology | Lex Leong, Chuan Kok Lim, Mark Turra, Ivan Bastian, Geoff Higgins |
| EPI_ISL_468134, EPI_ISL_468135, EPI_ISL_468136, EPI_ISL_468137, EPI_ISL_468138, EPI_ISL_468139, EPI_ISL_468140, EPI_ISL_468141, EPI_ISL_468142, EPI_ISL_468143, EPI_ISL_468144, EPI_ISL_468145, EPI_ISL_468146, EPI_ISL_468147, EPI_ISL_468148, EPI_ISL_468149, EPI_ISL_468150, EPI_ISL_468151, EPI_ISL_468152, EPI_ISL_468153, EPI_ISL_468154, EPI_ISL_468155, EPI_ISL_468156, EPI_ISL_468157, EPI_ISL_468158 | | | |
| see above | [Romania, Bucharest] National Institute for Infectious Diseases "Prof. Dr. Matei Bal" | [Romania, Bucharest] National Institute for Infectious Diseases "Prof. Dr. Matei Bal" | Leontina Banica, Marius Cotic, Corina Casangiu, Marius Surleac, Simona Paraschiv |
| EPI_ISL_468727, EPI_ISL_468728, EPI_ISL_468729, EPI_ISL_468730, EPI_ISL_468731, EPI_ISL_468732, EPI_ISL_468733, EPI_ISL_468734, EPI_ISL_468735 | Lab voor klinische biologie | Onderzoeksgroep Virologie | Laurens Lambrechts, Nick Vereecke, Marthe Pauwels, Bruno Verhasselt, Linos Vandekerckhove, Hans Nauwynck, Sebastiaan Theuns |
| EPI_ISL_469032 | Government Medical College, Vadodara | Gujarat Biotechnology Research Centre | Maharshi Pandya, Nidhi Patel, Nitin Savaliya, Raghawendra Kumar, Dinesh Kumar, Zuber Saiyed, Komal Patel, Labdhi Pandya, Snehal Bagatharia, Meenakshi Shah, Neena Doshi, Varsha Godbole, Tejas Shah, Ankit Hinsu, Pritesh Sabara, Apurvasinh Puvar, Janvi Raval, Zarna Patel, Monika Gandhi, Pinal Trivedi, Pragya Sharma, R D Dixit, A M Kadri, Harsh Bakshi, Chaitanya Joshi, Madhvi Joshi |
| EPI_ISL_469080, EPI_ISL_469081, EPI_ISL_469082, EPI_ISL_469083, EPI_ISL_469084, EPI_ISL_469085, EPI_ISL_469086, EPI_ISL_469087, EPI_ISL_469088, EPI_ISL_469089, EPI_ISL_469090, EPI_ISL_469091, EPI_ISL_469092, EPI_ISL_469093, EPI_ISL_469094, EPI_ISL_469095, EPI_ISL_469096, EPI_ISL_469097, EPI_ISL_469098, EPI_ISL_469099, EPI_ISL_469100, EPI_ISL_469101, EPI_ISL_469102, EPI_ISL_469103, EPI_ISL_469104, EPI_ISL_469105, EPI_ISL_469106, EPI_ISL_469107, EPI_ISL_469108, EPI_ISL_469109, EPI_ISL_469110, EPI_ISL_469111, EPI_ISL_469112, EPI_ISL_469113, EPI_ISL_469114, EPI_ISL_469115, EPI_ISL_469117, EPI_ISL_469118, EPI_ISL_469119, EPI_ISL_469120, EPI_ISL_469121, EPI_ISL_469122, EPI_ISL_469123, EPI_ISL_469124, EPI_ISL_469125, EPI_ISL_469126, EPI_ISL_469127, EPI_ISL_469128, EPI_ISL_469129, EPI_ISL_469130, EPI_ISL_469131, EPI_ISL_469132, EPI_ISL_469133, EPI_ISL_469134, EPI_ISL_469135, EPI_ISL_469136, EPI_ISL_469137, EPI_ISL_469138, EPI_ISL_469139, EPI_ISL_469140, EPI_ISL_469141, EPI_ISL_469142, EPI_ISL_469143, EPI_ISL_469144, EPI_ISL_469145, EPI_ISL_469146, EPI_ISL_469147, EPI_ISL_469148, EPI_ISL_469149, EPI_ISL_469150, EPI_ISL_469151, EPI_ISL_469152, EPI_ISL_469153, EPI_ISL_469154, EPI_ISL_469155, EPI_ISL_469274 | | | |
| see above | National Public Health Laboratory, National Centre for Infectious Diseases | National Public Health Laboratory, National Centre for Infectious Diseases | Mak TM, Octavia S, Chavatte JM, Cui L, Lin RTP |
| EPI_ISL_469282 | Service de Virologie Hôpital Saint-Louis | Laboratory of Cell Biology of viral infection, Unit INSERM-U944 | Laurent Meertens, Lucie Bonnet-Madin, Constance Delaugerre, Ali Amara |
| EPI_ISL_469283, EPI_ISL_469284 | Service de Virologie Hôpital Saint-Louis | Laboratory Cell Biology of Viral Infection-INSERM unit 944 | Laurent Meertens, Lucie Bonnet-Madin, Séverine Mercier-Delarue, Maud SALMONA, Constance Delaugerre, Ali Amara |
| EPI_ISL_469891, EPI_ISL_469892, EPI_ISL_469893, EPI_ISL_469894, EPI_ISL_469895, EPI_ISL_469896, EPI_ISL_469897, EPI_ISL_469898, EPI_ISL_469899, EPI_ISL_469900, EPI_ISL_469901, EPI_ISL_469902, EPI_ISL_469903, EPI_ISL_469904, EPI_ISL_469905, EPI_ISL_469906, EPI_ISL_469907, EPI_ISL_469908, EPI_ISL_469909, EPI_ISL_469910, EPI_ISL_469911, EPI_ISL_469912, EPI_ISL_469913, EPI_ISL_470090, EPI_ISL_470091, EPI_ISL_470092, EPI_ISL_470093, EPI_ISL_470094, EPI_ISL_470095, EPI_ISL_470096, EPI_ISL_470097, EPI_ISL_470098, EPI_ISL_470099, EPI_ISL_470100, EPI_ISL_470101, EPI_ISL_470102, EPI_ISL_470103, EPI_ISL_470104, EPI_ISL_470105, EPI_ISL_470106, EPI_ISL_470107, EPI_ISL_470108, EPI_ISL_470109, EPI_ISL_470110, EPI_ISL_470111, EPI_ISL_470112, EPI_ISL_470113, EPI_ISL_470114, EPI_ISL_470115, EPI_ISL_470116, EPI_ISL_470117, EPI_ISL_470118, EPI_ISL_470119, EPI_ISL_470120, EPI_ISL_470121, EPI_ISL_470122, EPI_ISL_470123, EPI_ISL_470124, EPI_ISL_470125, EPI_ISL_470126, EPI_ISL_470127, EPI_ISL_470128, EPI_ISL_470129, EPI_ISL_470130, EPI_ISL_470131, EPI_ISL_470132, EPI_ISL_470133, EPI_ISL_470134, EPI_ISL_470135, EPI_ISL_470136, EPI_ISL_470137, EPI_ISL_470138, EPI_ISL_470139, EPI_ISL_470140, EPI_ISL_470141, EPI_ISL_470142, EPI_ISL_470143, EPI_ISL_470144, EPI_ISL_470145, EPI_ISL_470146, EPI_ISL_470147, EPI_ISL_470148, EPI_ISL_470149, EPI_ISL_470150, EPI_ISL_470151, EPI_ISL_470152, EPI_ISL_470153, EPI_ISL_470154, EPI_ISL_470155, EPI_ISL_470156, EPI_ISL_470157, EPI_ISL_470158, EPI_ISL_470159, EPI_ISL_470160, EPI_ISL_470161, EPI_ISL_470162, EPI_ISL_470163, EPI_ISL_470164, EPI_ISL_470165, EPI_ISL_470166, EPI_ISL_470167, EPI_ISL_470168, EPI_ISL_470169, EPI_ISL_470170, EPI_ISL_470171, EPI_ISL_470172, EPI_ISL_470173, EPI_ISL_470174, EPI_ISL_470175, EPI_ISL_470176, EPI_ISL_470177, EPI_ISL_470178, EPI_ISL_470179, EPI_ISL_470180, EPI_ISL_470181, EPI_ISL_470182, EPI_ISL_470183, EPI_ISL_470184, EPI_ISL_470185, EPI_ISL_470186, EPI_ISL_470187, EPI_ISL_470188, EPI_ISL_470189, EPI_ISL_470190, EPI_ISL_470191, EPI_ISL_470192, EPI_ISL_470193, EPI_ISL_470194, EPI_ISL_470195, EPI_ISL_470196, EPI_ISL_470197, EPI_ISL_470198, EPI_ISL_470199, EPI_ISL_470200, EPI_ISL_470201, EPI_ISL_470202, EPI_ISL_470203, EPI_ISL_470204, EPI_ISL_470205, EPI_ISL_470206, EPI_ISL_470207, EPI_ISL_470208, EPI_ISL_470209, EPI_ISL_470210, EPI_ISL_470211, EPI_ISL_470212, EPI_ISL_470213, EPI_ISL_470214, EPI_ISL_470215, EPI_ISL_470216, EPI_ISL_470217, EPI_ISL_470218, EPI_ISL_470219, EPI_ISL_470220, EPI_ISL_470221, EPI_ISL_470222, EPI_ISL_470223, EPI_ISL_470224, EPI_ISL_470225, EPI_ISL_470226, EPI_ISL_470227, EPI_ISL_470228, EPI_ISL_470229, EPI_ISL_470230, EPI_ISL_470231, EPI_ISL_470232, EPI_ISL_470233, EPI_ISL_470234, EPI_ISL_470235, EPI_ISL_470236, EPI_ISL_470237, EPI_ISL_470238, EPI_ISL_470239, EPI_ISL_470240, EPI_ISL_470241, EPI_ISL_470242, EPI_ISL_470243, EPI_ISL_470244, EPI_ISL_470245, EPI_ISL_470246, | | | |

| | | | |
|---|---|---|---|
| EPI_ISL_470247, EPI_ISL_470248, EPI_ISL_470249, EPI_ISL_470250, EPI_ISL_470251, EPI_ISL_470252, EPI_ISL_470253, EPI_ISL_470254, EPI_ISL_470255, EPI_ISL_470256, EPI_ISL_470257, EPI_ISL_470258, EPI_ISL_470259, EPI_ISL_470260, EPI_ISL_470261, EPI_ISL_470262, EPI_ISL_470263, EPI_ISL_470264, EPI_ISL_470265, EPI_ISL_470266, EPI_ISL_470267, EPI_ISL_470268, EPI_ISL_470269, EPI_ISL_470270, EPI_ISL_470271, EPI_ISL_470272, EPI_ISL_470273, EPI_ISL_470274, EPI_ISL_470275, EPI_ISL_470276, EPI_ISL_470277, EPI_ISL_470278, EPI_ISL_470279, EPI_ISL_470280, EPI_ISL_470281, EPI_ISL_470282, EPI_ISL_470283, EPI_ISL_470284, EPI_ISL_470285, EPI_ISL_470286, EPI_ISL_470287, EPI_ISL_470288, EPI_ISL_470289, EPI_ISL_470290, EPI_ISL_470291, EPI_ISL_470292, EPI_ISL_470293, EPI_ISL_470294, EPI_ISL_470295, EPI_ISL_470296, EPI_ISL_470297, EPI_ISL_470298, EPI_ISL_470299, EPI_ISL_470300, EPI_ISL_470301, EPI_ISL_470302, EPI_ISL_470303, EPI_ISL_470304, EPI_ISL_470305, EPI_ISL_470306, EPI_ISL_470307, EPI_ISL_470308, EPI_ISL_470309, EPI_ISL_470310, EPI_ISL_470311, EPI_ISL_470312, EPI_ISL_470313, EPI_ISL_470314, EPI_ISL_470315, EPI_ISL_470316, EPI_ISL_470317, EPI_ISL_470318, EPI_ISL_470319, EPI_ISL_470320, EPI_ISL_470321, EPI_ISL_470322, EPI_ISL_470323, EPI_ISL_470324, EPI_ISL_470325, EPI_ISL_470326, EPI_ISL_470327, EPI_ISL_470328, EPI_ISL_470329, EPI_ISL_470330, EPI_ISL_470331, EPI_ISL_470332, EPI_ISL_470333, EPI_ISL_470334, EPI_ISL_470335, EPI_ISL_470336, EPI_ISL_470337, EPI_ISL_470338, EPI_ISL_470339, EPI_ISL_470340, EPI_ISL_470341, EPI_ISL_470342, EPI_ISL_470343, EPI_ISL_470344, EPI_ISL_470345, EPI_ISL_470346, EPI_ISL_470347, EPI_ISL_470348, EPI_ISL_470349, EPI_ISL_470350, EPI_ISL_470351, EPI_ISL_470352, EPI_ISL_470353, EPI_ISL_470354, EPI_ISL_470355, EPI_ISL_470358, EPI_ISL_470359, EPI_ISL_470360, EPI_ISL_470361, EPI_ISL_470362, EPI_ISL_470363, EPI_ISL_470364, EPI_ISL_470365, EPI_ISL_470366, EPI_ISL_470367, EPI_ISL_470368, EPI_ISL_470369, EPI_ISL_470370, EPI_ISL_470371, EPI_ISL_470372, EPI_ISL_470373, EPI_ISL_470374, EPI_ISL_470375, EPI_ISL_470376, EPI_ISL_470377, EPI_ISL_470378, EPI_ISL_470379, EPI_ISL_470380, EPI_ISL_470381, EPI_ISL_470382, EPI_ISL_470383, EPI_ISL_470384, EPI_ISL_470385, EPI_ISL_470386, EPI_ISL_470387, EPI_ISL_470388, EPI_ISL_470389, EPI_ISL_470390, EPI_ISL_470391, EPI_ISL_470392, EPI_ISL_470393, EPI_ISL_470394, EPI_ISL_470395, EPI_ISL_470396, EPI_ISL_470397, EPI_ISL_470398, EPI_ISL_470399, EPI_ISL_470400, EPI_ISL_470401, EPI_ISL_470402, EPI_ISL_470403, EPI_ISL_470404, EPI_ISL_470405, EPI_ISL_470406, EPI_ISL_470407, EPI_ISL_470408, EPI_ISL_470409, EPI_ISL_470410, EPI_ISL_470411, EPI_ISL_470412, EPI_ISL_470413, EPI_ISL_470414, EPI_ISL_470415, EPI_ISL_470416, EPI_ISL_470417, EPI_ISL_470418, EPI_ISL_470419, EPI_ISL_470420, EPI_ISL_470421, EPI_ISL_470422, EPI_ISL_470423, EPI_ISL_470424, EPI_ISL_470425, EPI_ISL_470426, EPI_ISL_470427, EPI_ISL_470428, EPI_ISL_470429, EPI_ISL_470430, EPI_ISL_470431, EPI_ISL_470432, EPI_ISL_470433, EPI_ISL_470434, EPI_ISL_470435, EPI_ISL_470436, EPI_ISL_470437, EPI_ISL_470438, EPI_ISL_470439, EPI_ISL_470440, EPI_ISL_470441, EPI_ISL_470442, EPI_ISL_470443, EPI_ISL_470444, EPI_ISL_470445, EPI_ISL_470446, EPI_ISL_470447, EPI_ISL_470448, EPI_ISL_470449, EPI_ISL_470450, EPI_ISL_470451, EPI_ISL_470452, EPI_ISL_470453, EPI_ISL_470454, EPI_ISL_470455, EPI_ISL_470456, EPI_ISL_470457, EPI_ISL_470458, EPI_ISL_470459, EPI_ISL_470460, EPI_ISL_470461, EPI_ISL_470462, EPI_ISL_470463, EPI_ISL_470464, EPI_ISL_470465, EPI_ISL_470466, EPI_ISL_470467, EPI_ISL_470468, EPI_ISL_470469, EPI_ISL_470470, EPI_ISL_470471, EPI_ISL_470472, EPI_ISL_470473, EPI_ISL_470474, EPI_ISL_470475, EPI_ISL_470476, EPI_ISL_470477, EPI_ISL_470478, EPI_ISL_470479, EPI_ISL_470480, EPI_ISL_470481, EPI_ISL_470482, EPI_ISL_470483, EPI_ISL_470484, EPI_ISL_470485, EPI_ISL_470486, EPI_ISL_470487, EPI_ISL_470488, EPI_ISL_470489, EPI_ISL_470490, EPI_ISL_470491, EPI_ISL_470492, EPI_ISL_470493, EPI_ISL_470494, EPI_ISL_470495, EPI_ISL_470496, EPI_ISL_470497, EPI_ISL_470498, EPI_ISL_470499, EPI_ISL_470500, EPI_ISL_470501, EPI_ISL_470502, EPI_ISL_470503, EPI_ISL_470504, EPI_ISL_470505, EPI_ISL_470506, EPI_ISL_470507, EPI_ISL_470508, EPI_ISL_470509, EPI_ISL_470510, EPI_ISL_470511, EPI_ISL_470512, EPI_ISL_470513, EPI_ISL_470514, EPI_ISL_470515, EPI_ISL_470516, EPI_ISL_470517, EPI_ISL_470518, EPI_ISL_470519, EPI_ISL_470520, EPI_ISL_470521, EPI_ISL_470522, EPI_ISL_470523, EPI_ISL_470524, EPI_ISL_470525, EPI_ISL_470526, EPI_ISL_470527, EPI_ISL_470528, EPI_ISL_470531, EPI_ISL_470532, EPI_ISL_470533, EPI_ISL_470535, EPI_ISL_470536, EPI_ISL_470537, EPI_ISL_470538 | | | |
| see above | Department of Pathology, University of Cambridge | Wellcome Sanger Institute for the COVID-19 Genomics UK (COG-UK) consortium | Luke W Meredith, M. Estée Török , Myra Hosmillo, William L. Hamilton, Martin D. Curran, Theresa Feltwell, Grant Hall, Anna Yakovleva, Fahad A Khokhar, Charlotte J. Houldcroft, Laura G Caller, Aminu S. Jahun, Sarah L. Caddy, Ian Goodfellow; and Alex Alderton, Roberto Amato, Sonia Goncalves, Ewan Harrison, David K. Jackson, Ian Johnston, Dominic Kwiatkowski, Cordelia Langford, John Sillitoe on behalf of the Wellcome Sanger Institute COVID-19 Surveillance Team (http://www.sanger.ac.uk/covid-team) |
| EPI_ISL_471416, EPI_ISL_471417, EPI_ISL_471418, EPI_ISL_471419, EPI_ISL_471420, EPI_ISL_471421, EPI_ISL_471422, EPI_ISL_471423, EPI_ISL_471424 | Laboratory for Respiratory Viruses, National Influenza Centre, Cantacuzino National Military-Medical Institute for Research and Development | Cantacuzino Institute | Luiza Ustea, Nicoleta Paraschiv, Tim Durfee, Mihaela Lazar |
| EPI_ISL_471472 | Hospital Universitari Germans Trias i Pujol(HUGTiP)/Fundació Lluita contra la SIDA (FLSida)/IRTA-CReSA | IrsiCaixa AIDS Research Lab | Marc Noguera-Julian, Pilar Armengol, Jordi Rodón, Julia Vergara, Lidia Ruiz, Nuria Izquierdo, Jorge Carrillo, Roger Paredes, Albert Bensaid, Julia Blanco, Joaquim Segalés, Bonaventura Clotet |
| EPI_ISL_473451, EPI_ISL_473452, EPI_ISL_473453, EPI_ISL_473454, EPI_ISL_473455, EPI_ISL_473456, EPI_ISL_473457, EPI_ISL_473458, EPI_ISL_473459, EPI_ISL_473460, EPI_ISL_473461, EPI_ISL_473462, EPI_ISL_473463, EPI_ISL_473464, EPI_ISL_473465, EPI_ISL_473466, EPI_ISL_473467, EPI_ISL_473468, EPI_ISL_473469, EPI_ISL_473470, EPI_ISL_473471, EPI_ISL_473472, EPI_ISL_473473, EPI_ISL_473474, EPI_ISL_473475, EPI_ISL_473476, EPI_ISL_473477, EPI_ISL_473478, EPI_ISL_473479, EPI_ISL_473480, EPI_ISL_473481, EPI_ISL_473482, EPI_ISL_473483, EPI_ISL_473484, EPI_ISL_473485, EPI_ISL_473486, EPI_ISL_473487, EPI_ISL_473488, EPI_ISL_473489, EPI_ISL_473490, EPI_ISL_473491, EPI_ISL_473492, EPI_ISL_473493, EPI_ISL_473495, EPI_ISL_473496, EPI_ISL_473497, EPI_ISL_473498, EPI_ISL_473499, EPI_ISL_473500, EPI_ISL_473501, EPI_ISL_473502, EPI_ISL_473503, EPI_ISL_473504, EPI_ISL_473505, EPI_ISL_473506, EPI_ISL_473507 | | | |
| see above | Department of Pathology, University of Cambridge | COVID-19 Genomics UK (COG-UK) Consortium | Luke W Meredith, M. Estée Török, Myra Hosmillo, William L. Hamilton, Martin D. Curran, Theresa Feltwell, Grant Hall, Anna Yakovleva, Fahad A Khokhar, Charlotte J. Houldcroft, Laura G Caller, Aminu S. Jahun, Sarah L. Caddy, Yasmin Chaudhry, Malte Pinckert, Ian Goodfellow |
| EPI_ISL_474910, EPI_ISL_474911, EPI_ISL_474912, EPI_ISL_474913, EPI_ISL_474914, EPI_ISL_474915, EPI_ISL_474916, EPI_ISL_474917, EPI_ISL_474918 | Hospital Universitario de Gran Canaria Dr. Negrín | SeqCOVID-SPAIN consortium/IBV(CSIC) | M. Carmen Pérez González, Francisco J. Chamizo López, Ana Bordes Benítez and SeqCOVID-SPAIN consortium |
| EPI_ISL_475034 | Department of MicroBiology, Government Medical College, Surat | Gujarat Biotechnology Research Centre | Maharshi Pandya, Nidhi Patel, Nitin Savaliya, Raghawendra Kumar, Dinesh Kumar, Zuber Saiyed, Komal Patel, Labdhi Pandya, Snehal Bagatharia, Naresh Chauhan, Summaiya Mullan, Amit gamit, Pritesh Sabara, Apurvasinh Puvar, Janvi Raval, Zarna Patel, Monika Gandhi, Pinal Trivedi, Akanksha Verma, R D Dixit, A M Kadri, Harsh Bakshi, Chaitanya Joshi, Madhvi Joshi |
| EPI_ISL_475052 | GMERS Medical College & Hospital | Gujarat Biotechnology Research Centre | Maharshi Pandya, Nidhi Patel, Nitin Savaliya, Raghawendra Kumar, Dinesh Kumar, Zuber Saiyed, Komal Patel, Labdhi Pandya, Snehal Bagatharia, Meenakshi Shah, Neena Doshi, Varsha Godbole, Pritesh Sabara, Apurvasinh Puvar, Janvi Raval, Zarna Patel, Monika Gandhi, Pinal Trivedi, Armi Chaudhari, R D Dixit, A M Kadri, Harsh Bakshi, Chaitanya Joshi, Madhvi Joshi |
| EPI_ISL_475060, EPI_ISL_475061, EPI_ISL_475062, EPI_ISL_475063, EPI_ISL_475064, EPI_ISL_475065, EPI_ISL_475066, EPI_ISL_475067, EPI_ISL_475068, EPI_ISL_475069, EPI_ISL_475070 | | | |
| see above | Lab voor klinische biologie | Onderzoeksgroep Virologie | Laurens Lambrechts, Nick Vereecke, Marthe Pauwels, Bruno Verhasselt, Linos Vandekerckhove, Hans Nauwynck, Sebastiaan Theuns |
| EPI_ISL_475937, EPI_ISL_475938, EPI_ISL_475939, EPI_ISL_475940, EPI_ISL_475941, EPI_ISL_475942, EPI_ISL_475943, EPI_ISL_475944, EPI_ISL_475945, EPI_ISL_475946, EPI_ISL_475947, EPI_ISL_475948, EPI_ISL_475949, EPI_ISL_475950, EPI_ISL_475951, EPI_ISL_475952, EPI_ISL_475953, EPI_ISL_475954, EPI_ISL_475955, EPI_ISL_475956, EPI_ISL_475957, EPI_ISL_475958, EPI_ISL_475959, EPI_ISL_475960, EPI_ISL_475961, EPI_ISL_475962, EPI_ISL_475963, EPI_ISL_475964, EPI_ISL_475965, EPI_ISL_475966, EPI_ISL_475967, EPI_ISL_475968, EPI_ISL_475969, EPI_ISL_475970, EPI_ISL_475971, EPI_ISL_475972, EPI_ISL_475973, EPI_ISL_475974, EPI_ISL_475975, EPI_ISL_475976, EPI_ISL_475977, EPI_ISL_475978, EPI_ISL_475979, EPI_ISL_475980, EPI_ISL_475981, EPI_ISL_475982, EPI_ISL_475983, EPI_ISL_475984, EPI_ISL_475985, EPI_ISL_475986, EPI_ISL_475987, EPI_ISL_475988, EPI_ISL_475989, EPI_ISL_475990, EPI_ISL_475991, EPI_ISL_475992, EPI_ISL_475993, EPI_ISL_475994, EPI_ISL_475995, EPI_ISL_475996, EPI_ISL_475997, EPI_ISL_475998, EPI_ISL_475999 | | | |
| see above | National Public Health Laboratory, National Centre for Infectious Diseases | National Public Health Laboratory, National Centre for Infectious Diseases | Mak TM, Octavia S, Chavatte JM, Cui L, Lin RTP |
| EPI_ISL_476860 | GMERS Medical College & Hospital, Gotri, Vadodara | Gujarat Biotechnology Research Centre | Maharshi Pandya, Nidhi Patel, Nitin Savaliya, Raghawendra Kumar, Dinesh Kumar, Zuber Saiyed, Komal Patel, Labdhi Pandya, Afzal Ansari, Nikha Trivedi, Meenakshi Shah, Neena Doshi, Varsha Godbole, Apurvasinh Puvar, Janvi Raval, Zarna Patel, Monika Gandhi, Pinal Trivedi, R D Dixit, A M Kadri, Harsh Bakshi, Chaitanya Joshi, Madhvi Joshi |
| EPI_ISL_476877 | Department of MicroBiology, Government Medical College, Surat | Gujarat Biotechnology Research Centre | Maharshi Pandya, Nidhi Patel, Nitin Savaliya, Raghawendra Kumar, Dinesh Kumar, Zuber Saiyed, Komal Patel, Labdhi Pandya, Afzal Ansari, Nikha Trivedi, Naresh Chauhan, Summaiya Mullan, Amit gamit, Apurvasinh Puvar, Janvi Raval, Zarna Patel, Monika Gandhi, Pinal Trivedi, R D Dixit, A M Kadri, Harsh Bakshi, Chaitanya Joshi, Madhvi Joshi |
| EPI_ISL_477783, EPI_ISL_477784, EPI_ISL_477785, EPI_ISL_477787, EPI_ISL_477788, EPI_ISL_477789, EPI_ISL_477790, EPI_ISL_477791, EPI_ISL_477792, EPI_ISL_477793, EPI_ISL_477794, EPI_ISL_477795, EPI_ISL_477796, EPI_ISL_477797, EPI_ISL_477798, EPI_ISL_477799, EPI_ISL_477800, EPI_ISL_477801, EPI_ISL_477802, EPI_ISL_477803, EPI_ISL_477804, EPI_ISL_477805, EPI_ISL_477806, EPI_ISL_477807, EPI_ISL_477808, EPI_ISL_477809, EPI_ISL_477810, EPI_ISL_477811, EPI_ISL_477812, EPI_ISL_477813, EPI_ISL_477814, EPI_ISL_477815, EPI_ISL_477816, EPI_ISL_477817, EPI_ISL_477818, EPI_ISL_477819 | | | |
| see above | Department of Pathology, University of Cambridge | COVID-19 Genomics UK (COG-UK) Consortium | Luke W Meredith, M. Estée Török, Myra Hosmillo, William L. Hamilton, Martin D. Curran, Theresa Feltwell, Grant Hall, Anna Yakovleva, Fahad A Khokhar, Charlotte J. Houldcroft, Laura G Caller, Aminu S. Jahun, Sarah L. Caddy, Yasmin Chaudhry, Malte Pinckert, Ian Goodfellow |
| EPI_ISL_479574, EPI_ISL_479575, EPI_ISL_479576, EPI_ISL_479577, EPI_ISL_479578, EPI_ISL_479579, EPI_ISL_479580, EPI_ISL_479581, EPI_ISL_479582, EPI_ISL_479583, EPI_ISL_479584, EPI_ISL_479585, EPI_ISL_479586, EPI_ISL_479587, EPI_ISL_479588, EPI_ISL_479589, EPI_ISL_479590, EPI_ISL_479591, EPI_ISL_479592, EPI_ISL_479593, EPI_ISL_479594, EPI_ISL_479595, EPI_ISL_479596, EPI_ISL_479597, EPI_ISL_479598, EPI_ISL_479599, EPI_ISL_479600, EPI_ISL_479601, EPI_ISL_479602, EPI_ISL_479603 | | | |
| see above | National Public Health Laboratory, National Centre for Infectious Diseases | National Public Health Laboratory, National Centre for Infectious Diseases | Mak TM, Octavia S, Zhou Z, Chavatte JM, Cui L, Lin RTP |
| EPI_ISL_479616, EPI_ISL_479617 | Laboratory of Molecular Virology of the International Centre for Genetic Engineering and Biotechnology (ICGEB) | ARGO Open Lab Platform for Genome Sequencing | Licastro, D, Rajasekharan S, Dal Monego S, Segat L, D'Agaro P, Salton F, Confalonieri P, Confalonieri M Marcello A |
| EPI_ISL_479618, EPI_ISL_479619 | Laboratory of Molecular Virology of the International Centre for Genetic Engineering and Biotechnology (ICGEB) | ARGO Open Lab Platform for Genome Sequencing | Licastro, D, Rajasekharan S, Dal Monego S, Segat L, D'Agaro P, Salton F, Confalonieri P, Confalonieri M, Marcello A |
| EPI_ISL_479756, EPI_ISL_479757, EPI_ISL_479758 | National Institute of Hygiene and Epidemiology (NIHE) | National Key Laboratory of Gene Technology, Institute of Biotechnology (IBT) | Le Tung Lam, Nguyen Hong Trang, Ho Thi Thuong, Tran Huyen Linh, Ung Thi Hong Trang, Le Thi Thanh, Nguyen Vu Son, Vuong Duc Cuong, Tran Thu Huong, Pham Thi Hien, Nguyen Phuong Anh, Nguyen Le Khanh Hang, Hoang Vu Mai Phuong, Hoang Ha, Taichiro Takemura, Futoshi Hasebe, Chu Hoang Ha, Le Quynh Mai, Dang Duc Anh, Truong Nam Hai |
| EPI_ISL_479790, EPI_ISL_479791 | Laboratory of Molecular Virology of the International Centre for Genetic Engineering and Biotechnology (ICGEB) | ARGO Open Lab Platform for Genome Sequencing | Licastro, D, Rajasekharan S, Dal Monego S, Segat L, D'Agaro P, Salton F, Confalonieri P, Confalonieri M, Marcello A |
| EPI_ISL_481041, EPI_ISL_481042, EPI_ISL_481043, EPI_ISL_481044, EPI_ISL_481045, EPI_ISL_481046, EPI_ISL_481047, EPI_ISL_481048, EPI_ISL_481049, EPI_ISL_481050, EPI_ISL_481051, EPI_ISL_481052, EPI_ISL_481053, EPI_ISL_481054, EPI_ISL_481055, EPI_ISL_481056, EPI_ISL_481057, EPI_ISL_481058, | | | |

| | | | |
|---|---|---|---|
| EPI_ISL_481059, EPI_ISL_481060, EPI_ISL_481061, EPI_ISL_481062, EPI_ISL_481063, EPI_ISL_481064, EPI_ISL_481065, EPI_ISL_481066, EPI_ISL_481067, EPI_ISL_481068, EPI_ISL_481069, EPI_ISL_481070, EPI_ISL_481071, EPI_ISL_481072, EPI_ISL_481073, EPI_ISL_481074, EPI_ISL_481075, EPI_ISL_481076, EPI_ISL_481077, EPI_ISL_481078, EPI_ISL_481079, EPI_ISL_481080, EPI_ISL_481081, EPI_ISL_481082, EPI_ISL_481083, EPI_ISL_481084, EPI_ISL_481085, EPI_ISL_481086, EPI_ISL_481087, EPI_ISL_481088, EPI_ISL_481089, EPI_ISL_481090, EPI_ISL_481091, EPI_ISL_481092, EPI_ISL_481093, EPI_ISL_481094, EPI_ISL_481095, EPI_ISL_481096, EPI_ISL_481097, EPI_ISL_481098, EPI_ISL_481099, EPI_ISL_481100, EPI_ISL_481101, EPI_ISL_481102, EPI_ISL_481103, EPI_ISL_481104, EPI_ISL_481105, EPI_ISL_481106, EPI_ISL_481107, EPI_ISL_481108, EPI_ISL_481109 | | | |
| see above | Hospital General Universitario Gregorio Marañón | SeqCOVID-SPAIN consortium/IBV(CSIC) | Laura Pérez-Lago, Marta Herranz, Jon Sicilia, Julia Suárez, Pilar Catalán, Patricia Muñoz, Darío García de Viedma and SeqCOVID-SPAIN consortium |
| EPI_ISL_481221, EPI_ISL_481222, EPI_ISL_481223, EPI_ISL_481224, EPI_ISL_481225, EPI_ISL_481226 | Lab voor klinische biologie | Onderzoeksgroep Virologie | Laurens Lambrechts, Nick Vereecke, Marthe Pauwels, Bruno Verhasselt, Linos Vandekerckhove, Hans Nauwynck, Sebastiaan Theuns |
| EPI_ISL_482124, EPI_ISL_482131 | North West London Pathology, Imperial College Healthcare NHS Trust | Wellcome Sanger Institute for the COVID-19 Genomics UK (COG-UK) consortium | Ling Li, Paul Randell, David Muir, Frankie Bolt, Alison Holmes, James Price, Aileen Rowan, Graham Taylor, Anjna Badhan, Carolina Herrera and Alex Alderton, Roberto Amato, Sonia Goncalves, Ewan Harrison, David K. Jackson, Ian Johnston, Dominic Kwiatkowski, Cordelia Langford, John Sillitoe on behalf of the Wellcome Sanger Institute COVID-19 Surveillance Team (http://www.sanger.ac.uk/covid-team) |
| EPI_ISL_483066, EPI_ISL_483067, EPI_ISL_483068, EPI_ISL_483069, EPI_ISL_483070, EPI_ISL_483071, EPI_ISL_483072, EPI_ISL_483073, EPI_ISL_483074, EPI_ISL_483075, EPI_ISL_483076, EPI_ISL_483077, EPI_ISL_483078, EPI_ISL_483079, EPI_ISL_483080, EPI_ISL_483081, EPI_ISL_483082, EPI_ISL_483084, EPI_ISL_483085, EPI_ISL_483086, EPI_ISL_483087, EPI_ISL_483088, EPI_ISL_483089, EPI_ISL_483090, EPI_ISL_483091, EPI_ISL_483092, EPI_ISL_483093, EPI_ISL_483094, EPI_ISL_483095, EPI_ISL_483096, EPI_ISL_483097, EPI_ISL_483098, EPI_ISL_483099, EPI_ISL_483100, EPI_ISL_483101, EPI_ISL_483102, EPI_ISL_483103, EPI_ISL_483104, EPI_ISL_483105, EPI_ISL_483107, EPI_ISL_483108, EPI_ISL_483109, EPI_ISL_483110, EPI_ISL_483111, EPI_ISL_483113, EPI_ISL_483114, EPI_ISL_483115, EPI_ISL_483116, EPI_ISL_483117, EPI_ISL_483118, EPI_ISL_483119, EPI_ISL_483120, EPI_ISL_483121, EPI_ISL_483122, EPI_ISL_483123, EPI_ISL_483124, EPI_ISL_483125, EPI_ISL_483126, EPI_ISL_483127, EPI_ISL_483128, EPI_ISL_483129, EPI_ISL_483130, EPI_ISL_483131, EPI_ISL_483132, EPI_ISL_483133, EPI_ISL_483134, EPI_ISL_483135, EPI_ISL_483136, EPI_ISL_483137, EPI_ISL_483138 | | | |
| see above | SA Pathology | SA Pathology | Lex Leong, Chuan Kok Lim, Mark Turra, Ivan Bastian, Geoff Higgins |
| EPI_ISL_483576, EPI_ISL_483577, EPI_ISL_483578, EPI_ISL_483579, EPI_ISL_483580, EPI_ISL_483581, EPI_ISL_483582, EPI_ISL_483583, EPI_ISL_483584, EPI_ISL_483585, EPI_ISL_483586, EPI_ISL_483587, EPI_ISL_483588, EPI_ISL_483589, EPI_ISL_483590, EPI_ISL_483591, EPI_ISL_483592, EPI_ISL_483593, EPI_ISL_483594, EPI_ISL_483595, EPI_ISL_483596, EPI_ISL_483597, EPI_ISL_483598, EPI_ISL_483599, EPI_ISL_483600, EPI_ISL_483601, EPI_ISL_483602, EPI_ISL_483603, EPI_ISL_483604, EPI_ISL_483605, EPI_ISL_483606, EPI_ISL_483607, EPI_ISL_483608, EPI_ISL_483609, EPI_ISL_483610, EPI_ISL_483611, EPI_ISL_483612, EPI_ISL_483613, EPI_ISL_483614, EPI_ISL_483615, EPI_ISL_483616, EPI_ISL_483617, EPI_ISL_483618, EPI_ISL_483619, EPI_ISL_483620, EPI_ISL_483621 | | | |
| see above | National Public Health Laboratory, National Centre for Infectious Diseases | National Public Health Laboratory, National Centre for Infectious Diseases | Mak TM, Octavia S, Zhou Z, Chavatte JM, Cui L, Lin RTP |
| EPI_ISL_483627, EPI_ISL_483628 | National Institute of Laboratory Medicine and Referral Center | Genomic Research Lab, BCSIR | Mahmuda Yeasmin, Abu Sayeed Mohammad Mahmud, Mohammad Samir Uzzaman, Eshrar Osman, Md. Ahasan Habib, Shahina Akter, Tanjina Akhter Banu, Md. Murshed Hasan Sarkar, Barna Goswami, Iffat Jahan, Md. Saddam Hossain, Tasnim Nafisa, Md. Maruf Ahmed Molla, Asish Kumar Ghosh, A. K. M. Shamsuzzaman, Sheikh Md. Selim Al Din, Utpal Chandra Ray, Salek Ahmed Sajib, Md. Salim Khan |
| EPI_ISL_483831, EPI_ISL_483848 | Pandit Deendayal Upadhyay Government Medical College, Rajkot | Gujarat Biotechnology Research Centre | Maharshi Pandya, Nidhi Patel, Nitin Savaliya, Raghawendra Kumar, Dinesh Kumar, Zuber Saiyed, Komal Patel, Labdhi Pandya, Afzal Ansari, Nikha Trivedi, Gauravi Dhruv, Arti Trivedi, Apurvasinh Puvar, Janvi Raval, Zarna Patel, Monika Gandhi, Pinal Trivedi, R D Dixit, A M Kadri, Harsh Bakshi, Chaitanya Joshi, Madhvi Joshi |
| EPI_ISL_483866 | Department of Microbiology, Government Medical College, Surat | Gujarat Biotechnology Research Centre | Maharshi Pandya, Nidhi Patel, Nitin Savaliya, Raghawendra Kumar, Dinesh Kumar, Zuber Saiyed, Komal Patel, Labdhi Pandya, Afzal Ansari, Nikha Trivedi, Naresh Chauhan, Summaiya Mullan, Amit gamit, Apurvasinh Puvar, Janvi Raval, Zarna Patel, Monika Gandhi, Pinal Trivedi, R D Dixit, A M Kadri, Harsh Bakshi, Chaitanya Joshi, Madhvi Joshi |
| EPI_ISL_483913, EPI_ISL_483914, EPI_ISL_483915 | Department of Pathology, University of Cambridge | COVID-19 Genomics UK (COG-UK) Consortium | Luke W Meredith, M. Estée Török, Myra Hosmillo, William L. Hamilton, Martin D. Curran, Theresa Feltwell, Grant Hall, Anna Yakovleva, Fahad A Khokhar, Charlotte J. Houldcroft, Laura G Caller, Aminu S. Jahun, Sarah L. Caddy, Yasmin Chaudhry, Malte Pinckert, Ian Goodfellow |
| EPI_ISL_486429 | unknown | Clinical Laboratory, Hospital Israelita Albert Einstein | Malta,F., Amgarten,D., Guedes,R.L., Santana,R.A., de Menezes,F.G., Mangueira,C.L. and Pinho,J.R. |
| EPI_ISL_486646, EPI_ISL_486647, EPI_ISL_486651, EPI_ISL_486655, EPI_ISL_486662 | Microbiology, Virology and Biemergency Laboratory-ASST FBF Sacco | Microbiology, Virology and Biemergency Laboratory-ASST FBF Sacco | Mancon A, Comandatore F, Romeri F, Micheli V, Rimoldi SG |
| EPI_ISL_486834 | Suceava County Energency Hospital "Sf. Ioan cel Nou" | SMU Metagenomics lab | Lobiuc Andrei, Antoniadis Panagiotis |
| EPI_ISL_486854 | Emergency County Hospital Suceava | Stefan cel Mare, University Metagenomics lab | Lobiuc Andrei et al. |
| EPI_ISL_486855 | Emergency county Hospital Suceava | "Stefan cel Mare" University Metagenomics Lab | Lobiuc Andrei et al. |
| EPI_ISL_488840, EPI_ISL_488845, EPI_ISL_488847, EPI_ISL_488853, EPI_ISL_488854, EPI_ISL_488855, EPI_ISL_488856, EPI_ISL_488861, EPI_ISL_488863, EPI_ISL_488864, EPI_ISL_488872 | | | |
| see above | Department of Medical Microbiology, Western Sussex Hospitals NHS Foundation Trust, St Richard's Hospital | Wellcome Sanger Institute for the COVID-19 Genomics UK (COG-UK) consortium | Manasa Mutingwende, Sarah Lowdon, Olga Podplomyk, Michelle Erkiert, Jonathan Lewis, Paul Randell and Alex Alderton, Roberto Amato, Sonia Goncalves, Ewan Harrison, David K. Jackson, Ian Johnston, Dominic Kwiatkowski, Cordelia Langford, John Sillitoe on behalf of the Wellcome Sanger Institute COVID-19 Surveillance Team (http://www.sanger.ac.uk/covid-team) |
| EPI_ISL_489381, EPI_ISL_489382, EPI_ISL_489383, EPI_ISL_489384, EPI_ISL_489385, EPI_ISL_489386, EPI_ISL_489387, EPI_ISL_489388, EPI_ISL_489389, EPI_ISL_489390, EPI_ISL_489394, EPI_ISL_489395, EPI_ISL_489397, EPI_ISL_489398, EPI_ISL_489399, EPI_ISL_489400, EPI_ISL_489402, EPI_ISL_489403, EPI_ISL_489404, EPI_ISL_489405, EPI_ISL_489406, EPI_ISL_489409, EPI_ISL_489410, EPI_ISL_489411, EPI_ISL_489412, EPI_ISL_489414, EPI_ISL_489415, EPI_ISL_489416, EPI_ISL_489417, EPI_ISL_489419, EPI_ISL_489420, EPI_ISL_489421, EPI_ISL_489422, EPI_ISL_489423, EPI_ISL_489424, EPI_ISL_489425, EPI_ISL_489426, EPI_ISL_489427, EPI_ISL_489428, EPI_ISL_489430, EPI_ISL_489431, EPI_ISL_489432, EPI_ISL_489433, EPI_ISL_489434, EPI_ISL_489435, EPI_ISL_489436, EPI_ISL_489437, EPI_ISL_489438, EPI_ISL_489439, EPI_ISL_489440, EPI_ISL_489441, EPI_ISL_489443, EPI_ISL_489444, EPI_ISL_489445, EPI_ISL_489446, EPI_ISL_489447, EPI_ISL_489448, EPI_ISL_489450, EPI_ISL_489451, EPI_ISL_489452, EPI_ISL_489453, EPI_ISL_489454, EPI_ISL_489456, EPI_ISL_489457, EPI_ISL_489458, EPI_ISL_489459, EPI_ISL_489460, EPI_ISL_489461, EPI_ISL_489462, EPI_ISL_489463, EPI_ISL_489464, EPI_ISL_489465, EPI_ISL_489466, EPI_ISL_489467, EPI_ISL_489468, EPI_ISL_489469, EPI_ISL_489471, EPI_ISL_489472, EPI_ISL_489473, EPI_ISL_489474, EPI_ISL_489475, EPI_ISL_489476, EPI_ISL_489477, EPI_ISL_489478, EPI_ISL_489479, EPI_ISL_489481, EPI_ISL_489482, EPI_ISL_489483, EPI_ISL_489484, EPI_ISL_489485, EPI_ISL_489486, EPI_ISL_489487, EPI_ISL_489488, EPI_ISL_489490, EPI_ISL_489491, EPI_ISL_489492, EPI_ISL_489493, EPI_ISL_489494, EPI_ISL_489495, EPI_ISL_489496, EPI_ISL_489497, EPI_ISL_489498, EPI_ISL_489499, EPI_ISL_489500, EPI_ISL_489501, EPI_ISL_489502, EPI_ISL_489503, EPI_ISL_489504, EPI_ISL_489505, EPI_ISL_489506, EPI_ISL_489507, EPI_ISL_489508, EPI_ISL_489509, EPI_ISL_489510, EPI_ISL_489511, EPI_ISL_489512, EPI_ISL_489513, EPI_ISL_489514, EPI_ISL_489515, EPI_ISL_489516, EPI_ISL_489517, EPI_ISL_489518, EPI_ISL_489519, EPI_ISL_489520, EPI_ISL_489521, EPI_ISL_489522, EPI_ISL_489524, EPI_ISL_489525, EPI_ISL_489526, EPI_ISL_489527, EPI_ISL_489528, EPI_ISL_489529, EPI_ISL_489531, EPI_ISL_489532, EPI_ISL_489533, EPI_ISL_489534, EPI_ISL_489535, EPI_ISL_489538, EPI_ISL_489539, EPI_ISL_489540, EPI_ISL_489541, EPI_ISL_489542, EPI_ISL_489544, EPI_ISL_489545, EPI_ISL_489546, EPI_ISL_489548, EPI_ISL_489549, EPI_ISL_489550, EPI_ISL_489551, EPI_ISL_489552, EPI_ISL_489553, EPI_ISL_489554, EPI_ISL_489555, EPI_ISL_489556, EPI_ISL_489557, EPI_ISL_489558, EPI_ISL_489559, EPI_ISL_489560, EPI_ISL_489561, EPI_ISL_489562, EPI_ISL_489563, EPI_ISL_489564, EPI_ISL_489565, EPI_ISL_489566, EPI_ISL_489568, EPI_ISL_489569, EPI_ISL_489570, EPI_ISL_489571, EPI_ISL_489572, EPI_ISL_489573, EPI_ISL_489574, EPI_ISL_489575, EPI_ISL_489576, EPI_ISL_489577 | | | |
| see above | Department of Pathology, University of Cambridge | Wellcome Sanger Institute for the COVID-19 Genomics UK (COG-UK) consortium | Luke W Meredith, M. Estée Török , Myra Hosmillo, William L. Hamilton, Martin D. Curran, Theresa Feltwell, Grant Hall, Anna Yakovleva, Fahad A Khokhar, Charlotte J. Houldcroft, Laura G Caller, Aminu S. Jahun, Sarah L. Caddy, Ian Goodfellow; and Alex Alderton, Roberto Amato, Sonia Goncalves, Ewan Harrison, David K. Jackson, Ian Johnston, Dominic Kwiatkowski, Cordelia Langford, John Sillitoe on behalf of the Wellcome Sanger Institute COVID-19 Surveillance Team (http://www.sanger.ac.uk/covid-team) |
| EPI_ISL_490049, EPI_ISL_490050, EPI_ISL_490051, EPI_ISL_490052, EPI_ISL_490053, EPI_ISL_490054, EPI_ISL_490055, EPI_ISL_490056, EPI_ISL_490057, EPI_ISL_490058, EPI_ISL_490059, EPI_ISL_490060, EPI_ISL_490061, EPI_ISL_490062, EPI_ISL_490063, EPI_ISL_490064, EPI_ISL_490065, EPI_ISL_490066, EPI_ISL_490067, EPI_ISL_490068, EPI_ISL_490069, EPI_ISL_490070, EPI_ISL_490071, EPI_ISL_490072, EPI_ISL_490073, EPI_ISL_490074, EPI_ISL_490075, EPI_ISL_490076, EPI_ISL_490077, EPI_ISL_490078, EPI_ISL_490079 | | | |
| see above | National Public Health Laboratory, National Centre for Infectious Diseases | National Public Health Laboratory, National Centre for Infectious Diseases | Mak TM, Octavia S, Zhou Z, Chavatte JM, Cui L, Lin RTP |
| EPI_ISL_490317, EPI_ISL_490318, EPI_ISL_490319, EPI_ISL_490320, EPI_ISL_490321, EPI_ISL_490322, EPI_ISL_490323, EPI_ISL_490324, EPI_ISL_490325, EPI_ISL_490326, EPI_ISL_490327, EPI_ISL_490328, EPI_ISL_490329 | | | |
| see above | Department of Pathology, University of Cambridge | COVID-19 Genomics UK (COG-UK) Consortium | Luke W Meredith, M. Estée Török, Myra Hosmillo, William L. Hamilton, Martin D. Curran, Theresa Feltwell, Grant Hall, Anna Yakovleva, Fahad A Khokhar, Charlotte J. Houldcroft, Laura G Caller, Aminu S. Jahun, Sarah L. Caddy, Yasmin Chaudhry, Malte Pinckert, Ian Goodfellow |
| EPI_ISL_491036, EPI_ISL_491037, EPI_ISL_491038 | Suceava County Emergency Hospital | "Stefan cel Mare" University Metagenomics Lab | Lobiuc Andrei, Antoniadis Panagiotis et al. |
| EPI_ISL_491039, EPI_ISL_491040, EPI_ISL_491041 | Suceava County Emergency Hospital | "Stefan cel Mare" University Metagenomics Lab | Lobiuc Andrei et al. |
| EPI_ISL_491042, EPI_ISL_491043 | Suceava County Emergency Hospital | "Stefan cel Mare" University Metagenomics Lab | Lobiuc Andrei, Antoniadis Panagiotis et al. |
| EPI_ISL_491044 | Suceava County Emergency Hospital | "Stefan cel Mare" University Metagenomics Lab | Lobiuc Andrei et al. |
| EPI_ISL_491045, EPI_ISL_491046, EPI_ISL_491047 | Suceava County Emergency Hospital | "Stefan cel Mare" University Metagenomics Lab | Lobiuc Andrei, Antoniadis Panagiotis et al. |
| EPI_ISL_491048 | Suceava County Emergency Hospital | "Stefan cel Mare" University Metagenomics Lab | Lobiuc Andrei et al. |

| | | | |
|---|---|---|---|
| EPI_ISL_491049 | Suceava County Emergency Hospital | "Stefan cel Mare" University Metagenomics Lab | Lobiuc Andrei, Antoniadis Panagiotis et al. |
| EPI_ISL_491050, EPI_ISL_491051, EPI_ISL_491052, EPI_ISL_491053, EPI_ISL_491054, EPI_ISL_491055, EPI_ISL_491056, EPI_ISL_491057 | Suceava County Emergency Hospital | "Stefan cel Mare" University Metagenomics Lab | Lobiuc Andrei et al. |
| EPI_ISL_491058, EPI_ISL_491059 | Suceava County Emergency Hospital | "Stefan cel Mare" University Metagenomics Lab | Lobiuc Andrei, Antoniadis Panagiotis et al. |
| EPI_ISL_491060, EPI_ISL_491061 | Suceava County Emergency Hospital | "Stefan cel Mare" University Metagenomics Lab | Lobiuc Andrei et al. |
| EPI_ISL_491062, EPI_ISL_491063, EPI_ISL_491064, EPI_ISL_491065 | Suceava County Emergency Hospital | "Stefan cel Mare" University Metagenomics Lab | Lobiuc Andrei, Antoniadis Panagiotis et al. |
| EPI_ISL_491066, EPI_ISL_491067, EPI_ISL_491068, EPI_ISL_491069, EPI_ISL_491070, EPI_ISL_491071, EPI_ISL_491072 | Suceava County Emergency Hospital | "Stefan cel Mare" University Metagenomics Lab | Lobiuc Andrei et al. |
| EPI_ISL_491073 | Suceava County Emergency Hospital | "Stefan cel Mare" University Metagenomics Lab | Lobiuc Andrei, Antoniadis Panagiotis et al. |
| EPI_ISL_491074 | Suceava County Emergency Hospital | "Stefan cel Mare" University Metagenomics Lab | Lobiuc Andrei et al. |
| EPI_ISL_491075 | Suceava County Emergency Hospital | "Stefan cel Mare" University Metagenomics Lab | Lobiuc Andrei, Antoniadis Panagiotis et al. |
| EPI_ISL_491076 | Suceava County Emergency Hospital | "Stefan cel Mare" University Metagenomics Lab | Lobiuc Andrei et al. |
| EPI_ISL_491077 | Suceava County Emergency Hospital | "Stefan cel Mare" University Metagenomics Lab | Lobiuc Andrei, Antoniadis Panagiotis et al. |
| EPI_ISL_491078 | Suceava County Emergency Hospital | "Stefan cel Mare" University Metagenomics Lab | Lobiuc Andrei et al. |
| EPI_ISL_491079, EPI_ISL_491080 | Suceava County Emergency Hospital | "Stefan cel Mare" University Metagenomics Lab | Lobiuc Andrei, Antoniadis Panagiotis et al. |
| EPI_ISL_491081 | Suceava County Emergency Hospital | "Stefan cel Mare" University Metagenomics Lab | Lobiuc Andrei et al. |
| EPI_ISL_491082 | Suceava County Emergency Hospital | "Stefan cel Mare" University Metagenomics Lab | Lobiuc Andrei, Antoniadis Panagiotis et al. |
| EPI_ISL_491083, EPI_ISL_491084 | Suceava County Emergency Hospital | "Stefan cel Mare" University Metagenomics Lab | Lobiuc Andrei et al. |
| EPI_ISL_491085 | Suceava County Emergency Hospital | "Stefan cel Mare" University Metagenomics Lab | Lobiuc Andrei, Antoniadis Panagiotis et al. |
| EPI_ISL_491086 | Suceava County Emergency Hospital | "Stefan cel Mare" University Metagenomics Lab | Lobiuc Andrei et al. |
| EPI_ISL_491087, EPI_ISL_491088 | Suceava County Emergency Hospital | "Stefan cel Mare" University Metagenomics Lab | Lobiuc Andrei, Antoniadis Panagiotis et al. |
| EPI_ISL_491089 | Suceava County Emergency Hospital | "Stefan cel Mare" University Metagenomics Lab | Lobiuc Andrei et al. |
| EPI_ISL_491090 | Suceava County Emergency Hospital | "Stefan cel Mare" University Metagenomics Lab | Lobiuc Andrei, Antoniadis Panagiotis et al. |
| EPI_ISL_491091 | Suceava County Emergency Hospital | "Stefan cel Mare" University Metagenomics Lab | Lobiuc Andrei et al. |
| EPI_ISL_491115 | Cicin-Sain Lab | Cicin-Sain Lab | M. Zeeshan Chaudhry, Kathrin Eschke, Yeonsu Kim, Luka Cicin-Sain |
| EPI_ISL_491465, EPI_ISL_491466 | San Lazaro Hospital | Research Institute for Tropical Medicine | Ma. Angelica Tujan, Othoniel Jan Onza, Francisco Gerardo Polotan, Inez Andrea Medado, Criselda Bautista, Kirstyn Brunker, Edelwisa Mercado, Daria Manalo, Catalino Demetria |
| EPI_ISL_491467, EPI_ISL_491468 | Research Institute for Tropical Medicine | Research Institute for Tropical Medicine | Ma. Angelica Tujan, Othoniel Jan Onza, Francisco Gerardo Polotan, Inez Andrea Medado, Criselda Bautista, Kirstyn Brunker, Edelwisa Mercado, Daria Manalo, Catalino Demetria |
| EPI_ISL_491471 | San Lazaro Hospital | Research Institute for Tropical Medicine | Ma. Angelica Tujan, Othoniel Jan Onza, Francisco Gerardo Polotan, Inez Andrea Medado, Criselda Bautista, Kirstyn Brunker, Edelwisa Mercado, Daria Manalo, Catalino Demetria |
| EPI_ISL_491472 | Cardinal Santos Medical Center | Research Institute for Tropical Medicine | Ma. Angelica Tujan, Othoniel Jan Onza, Francisco Gerardo Polotan, Inez Andrea Medado, Criselda Bautista, Kirstyn Brunker, Edelwisa Mercado, Daria Manalo, Catalino Demetria |
| EPI_ISL_491473 | The Medical City | Research Institute for Tropical Medicine | Ma. Angelica Tujan, Othoniel Jan Onza, Francisco Gerardo Polotan, Inez Andrea Medado, Criselda Bautista, Kirstyn Brunker, Edelwisa Mercado, Daria Manalo, Catalino Demetria |
| EPI_ISL_491474 | Research Institute for Tropical Medicine | Research Institute for Tropical Medicine | Ma. Angelica Tujan, Othoniel Jan Onza, Francisco Gerardo Polotan, Inez Andrea Medado, Criselda Bautista, Kirstyn Brunker, Edelwisa Mercado, Daria Manalo, Catalino Demetria |
| EPI_ISL_491475 | Protacio Hospital | Research Institute for Tropical Medicine | Ma. Angelica Tujan, Othoniel Jan Onza, Francisco Gerardo Polotan, Inez Andrea Medado, Criselda Bautista, Kirstyn Brunker, Edelwisa Mercado, Daria Manalo, Catalino Demetria |
| EPI_ISL_491907, EPI_ISL_491908, EPI_ISL_491909, EPI_ISL_491910, EPI_ISL_491911, EPI_ISL_491912, EPI_ISL_491913, EPI_ISL_491914, EPI_ISL_491915, EPI_ISL_491916, EPI_ISL_491917, EPI_ISL_491918, EPI_ISL_491919, EPI_ISL_491920, EPI_ISL_491921, EPI_ISL_491922, EPI_ISL_491923, EPI_ISL_491924, EPI_ISL_491925, EPI_ISL_491926, EPI_ISL_491927, EPI_ISL_491928, EPI_ISL_491929, EPI_ISL_491930, EPI_ISL_491931, EPI_ISL_491942, EPI_ISL_491943 | | | |
| see above | Naval Infectious Diseases Diagnostic Laboratory | Naval Medical Research Center Biological Defense Research Directorate | Logan Voegtly, Regina Cer, Lindsay Glang, Victor Sugiharto, Francisco Malgon Bautista, Hua Wei Chen, Dessiree Pena-Gomez, Megan Schilling, Adrian Paskey, Kyle Long, Mark Simons, Kimberly Bishop-Lilly |
| EPI_ISL_492109, EPI_ISL_492110, EPI_ISL_492111, EPI_ISL_492112, EPI_ISL_492113, EPI_ISL_492114, EPI_ISL_492115, EPI_ISL_492116, EPI_ISL_492117, EPI_ISL_492118, EPI_ISL_492119, EPI_ISL_492120, EPI_ISL_492121, EPI_ISL_492122, EPI_ISL_492123, EPI_ISL_492124, EPI_ISL_492125, EPI_ISL_492126, EPI_ISL_492127, EPI_ISL_492128, EPI_ISL_492129, EPI_ISL_492130, EPI_ISL_492131, EPI_ISL_492132, EPI_ISL_492133, EPI_ISL_492134, EPI_ISL_492135, EPI_ISL_492136, EPI_ISL_492137, EPI_ISL_492138, EPI_ISL_492139, EPI_ISL_492140, EPI_ISL_492141, EPI_ISL_492142, EPI_ISL_492143, EPI_ISL_492144, EPI_ISL_492145, EPI_ISL_492146, EPI_ISL_492147, EPI_ISL_492148, EPI_ISL_492149, EPI_ISL_492150, EPI_ISL_492151, EPI_ISL_492152, EPI_ISL_492153, EPI_ISL_492154, EPI_ISL_492155, EPI_ISL_492156, EPI_ISL_492157, EPI_ISL_492158, EPI_ISL_492159, EPI_ISL_492160, EPI_ISL_492161, EPI_ISL_492162, EPI_ISL_492163, EPI_ISL_492164, EPI_ISL_492165, EPI_ISL_492166, EPI_ISL_492167, EPI_ISL_492168, EPI_ISL_492169, EPI_ISL_492170, EPI_ISL_492171, EPI_ISL_492172, EPI_ISL_492173, EPI_ISL_492174, EPI_ISL_492175, EPI_ISL_492176, EPI_ISL_492177, EPI_ISL_492178, EPI_ISL_492179, EPI_ISL_492180 | | | |
| see above | SA Pathology | SA Pathology | Lex Leong, Chuan Kok Lim, Mark Turra, Ivan Bastian, Geoff Higgins |
| EPI_ISL_492189, EPI_ISL_492190, EPI_ISL_492191, EPI_ISL_492192, EPI_ISL_492193, EPI_ISL_492194, EPI_ISL_492196, EPI_ISL_492198, EPI_ISL_492199, EPI_ISL_492200, EPI_ISL_492201, EPI_ISL_492204, EPI_ISL_492205, EPI_ISL_492206 | | | |
| see above | Department of Pathology, University of Cambridge | Wellcome Sanger Institute for the COVID-19 Genomics UK (COG-UK) consortium | Luke W Meredith, M. Estée Török , Myra Hosmillo, William L. Hamilton, Martin D. Curran, Theresa Feltwell, Grant Hall, Anna Yakovleva, Fahad A Khokhar, Charlotte J. Houldcroft, Laura G Caller, Aminu S. Jahun, Sarah L. Caddy, Ian Goodfellow; and Alex Alderton, Roberto Amato, Sonia Goncalves, Ewan Harrison, David K. Jackson, Ian Johnston, Dominic Kwiatkowski, Cordelia Langford, John Sillitoe on behalf of the Wellcome Sanger Institute COVID-19 Surveillance Team (http://www.sanger.ac.uk/covid-team) |
| EPI_ISL_492836, EPI_ISL_492838, EPI_ISL_492839, EPI_ISL_492841, EPI_ISL_492842, EPI_ISL_492848, EPI_ISL_492849, EPI_ISL_492853, EPI_ISL_492854, EPI_ISL_492855, EPI_ISL_492857, EPI_ISL_492858, EPI_ISL_492859, EPI_ISL_492860, EPI_ISL_492861, EPI_ISL_492866, EPI_ISL_492867, EPI_ISL_492870, EPI_ISL_492871, EPI_ISL_492873, EPI_ISL_492875, EPI_ISL_492878, EPI_ISL_492881, EPI_ISL_492884, EPI_ISL_492886, EPI_ISL_492889, EPI_ISL_492891, EPI_ISL_492892, EPI_ISL_492893, EPI_ISL_492895, EPI_ISL_492896, EPI_ISL_492897, EPI_ISL_492898, EPI_ISL_492899, EPI_ISL_492900, EPI_ISL_492903, EPI_ISL_492904, EPI_ISL_492907, EPI_ISL_492909, EPI_ISL_492910, EPI_ISL_492912, EPI_ISL_492913, EPI_ISL_492915 | | | |
| see above | Department of Medical Microbiology, Western Sussex Hospitals NHS Foundation Trust, St Richard's Hospital | Wellcome Sanger Institute for the COVID-19 Genomics UK (COG-UK) consortium | Manasa Mutingwende, Sarah Lowdon, Olga Podplomyk, Michelle Erkiert, Jonathan Lewis, Paul Randell and Alex Alderton, Roberto Amato, Sonia Goncalves, Ewan Harrison, David K. Jackson, Ian Johnston, Dominic Kwiatkowski, Cordelia Langford, John Sillitoe on behalf of the Wellcome Sanger Institute COVID-19 Surveillance Team (http://www.sanger.ac.uk/covid-team) |
| EPI_ISL_493390, EPI_ISL_493391, EPI_ISL_493392, EPI_ISL_493393, EPI_ISL_493394, EPI_ISL_493395, EPI_ISL_493396, EPI_ISL_493397, EPI_ISL_493398, EPI_ISL_493399, EPI_ISL_493400, EPI_ISL_493401, EPI_ISL_493402, EPI_ISL_493403, EPI_ISL_493404, EPI_ISL_493405, EPI_ISL_493406, EPI_ISL_493407, EPI_ISL_493408, EPI_ISL_493409, EPI_ISL_493410, EPI_ISL_493411, EPI_ISL_493412, EPI_ISL_493413, EPI_ISL_493414, EPI_ISL_493415, EPI_ISL_493416, EPI_ISL_493417, EPI_ISL_493418, EPI_ISL_493419, EPI_ISL_493420, EPI_ISL_493421, EPI_ISL_493422, EPI_ISL_493423, EPI_ISL_493424, EPI_ISL_493425 | | | |
| see above | National Public Health Laboratory, National Centre for Infectious Diseases | National Public Health Laboratory, National Centre for Infectious Diseases | Mak TM, Octavia S, Zhou Z, Chavatte JM, Cui L, Lin RTP |

| | | | |
|---|---|---|---|
| EPI_ISL_495018 | B.J. Medical College and Civil hospital | Gujarat Biotechnology Research Centre | Maharshi Pandya, Nidhi Patel, Nitin Savaliya, Raghawendra Kumar, Dinesh Kumar, Zuber Saiyed, Komal Patel, Labdhi Pandya, Afzal Ansari, Nikha Trivedi, Pranay Shah, Kamlesh J Upadhyay, Sanjay Kapadia, Apurvasinh Puvar, Janvi Raval, Zarna Patel, Monika Gandhi, Pinal Trivedi, R D Dixit, A M Kadri, Harsh Bakshi, Chaitanya Joshi, Madhvi Joshi |
| EPI_ISL_495038, EPI_ISL_495056 | GMERS Medical College & Hospital, Gotri, Vadodara | Gujarat Biotechnology Research Centre | Maharshi Pandya, Nidhi Patel, Nitin Savaliya, Raghawendra Kumar, Dinesh Kumar, Zuber Saiyed, Komal Patel, Labdhi Pandya, Afzal Ansari, Nikha Trivedi, Meenakshi Shah, Neena Doshi, Varsha Godbole, Apurvasinh Puvar, Janvi Raval, Zarna Patel, Monika Gandhi, Pinal Trivedi, R D Dixit, A M Kadri, Harsh Bakshi, Chaitanya Joshi, Madhvi Joshi |
| EPI_ISL_495081, EPI_ISL_495082, EPI_ISL_495083, EPI_ISL_495084, EPI_ISL_495085, EPI_ISL_495086, EPI_ISL_495087, EPI_ISL_495088, EPI_ISL_495089, EPI_ISL_495090, EPI_ISL_495091, EPI_ISL_495092, EPI_ISL_495093, EPI_ISL_495094, EPI_ISL_495095, EPI_ISL_495096 | | | |
| see above | Department of Medical Microbiology, Western Sussex Hospitals NHS Foundation Trust, St Richard's Hospital | Wellcome Sanger Institute for the COVID-19 Genomics UK (COG-UK) consortium | Manasa Mutingwende, Sarah Lowdon, Olga Podplomyk, Michelle Erkiert, Jonathan Lewis, Paul Randell and Alex Alderton, Roberto Amato, Sonia Goncalves, Ewan Harrison, David K. Jackson, Ian Johnston, Dominic Kwiatkowski, Cordelia Langford, John Sillitoe on behalf of the Wellcome Sanger Institute COVID-19 Surveillance Team (http://www.sanger.ac.uk/covid-team) |
| EPI_ISL_495171 | CSIR-Centre for Cellular and Molecular Biology | CSIR-Centre for Cellular and Molecular Biology | M Soujanya Reddy, Nikhil Hajirnis, Pratheusa Maccha, Namami Gaur, Sakshi Shambhavi, Lamuk Zaveri, Shagufta Khan, Tulasi Nagabandi, Purushotham Vodnala, Payel Mukherjee, Sofia Banu, Priya Singh, Onkar Kulkarni, Dhiviya Vedagiri, Divya Gupta, Vishal Sah, Santosh Kumar Kuncha, Krishnan Harinivas Harshan, Archana Bharadwaj Siva, Karthik Bharadwaj Tallapaka, Zeba Rizvi, Zuberwasim Sayyad, Kakade Aishwarya Arun, Amrutha H C, Ananga Ghosh, Rakesh K Mishra, Divya Tej Sowpati |
| EPI_ISL_495174 | CSIR-Centre for Cellular and Molecular Biology | CSIR-Centre for Cellular and Molecular Biology | M Soujanya Reddy, Nikhil Hajirnis, Pratheusa Maccha, Payel Mukherjee, Sofia Banu, Priya Singh, Onkar Kulkarni,Tulasi Nagabandi, Namami Gaur, Sakshi Shambhavi, Lamuk Zaveri, Shagufta Khan, Purushotham Vodnala, Dhiviya Vedagiri, Divya Gupta, Vishal Sah, Santosh Kumar Kuncha, Krishnan Harinivas Harshan, Archana Bharadwaj Siva, Karthik Bharadwaj Tallapaka,Kezia J Ann, Radhika Khandelwal, Roshan Maku Venkata, Shemin Mansuri, Sonu Uday, Rakesh K Mishra, Divya Tej Sowpati |
| EPI_ISL_495180, EPI_ISL_495212 | CSIR-Centre for Cellular and Molecular Biology | CSIR-Centre for Cellular and Molecular Biology | M Soujanya Reddy, Nikhil Hajirnis, Pratheusa Maccha, Sakshi Shambhavi, Lamuk Zaveri, Shagufta Khan, Namami Gaur, Tulasi Nagabandi, Purushotham Vodnala, Payel Mukherjee, Sofia Banu, Priya Singh,Onkar Kulkarni, Dhiviya Vedagiri, Divya Gupta, Vishal Sah, Santosh Kumar Kuncha, Krishnan Harinivas Harshan, Archana Bharadwaj Siva, Karthik Bharadwaj Tallapaka, G. Aditya Kumar, Koushick Sivakumar, Pooja Ramesh Gupta, Rajan Kumar Jha, Shraddha Vijay Lahoti, Rakesh K Mishra, Divya Tej Sowpati |
| EPI_ISL_495226 | CSIR-Centre for Cellular and Molecular Biology | CSIR-Centre for Cellular and Molecular Biology | M Soujanya Reddy, Nikhil Hajirnis, Pratheusa Maccha, Namami Gaur, Sakshi Shambhavi, Lamuk Zaveri, Shagufta Khan, Tulasi Nagabandi, Purushotham Vodnala, Payel Mukherjee, Sofia Banu, Priya Singh, Onkar Kulkarni, Dhiviya Vedagiri, Divya Gupta, Vishal Sah, Santosh Kumar Kuncha, Krishnan Harinivas Harshan, Archana Bharadwaj Siva, Karthik Bharadwaj Tallapaka, Zeba Rizvi, Zuberwasim Sayyad, Kakade Aishwarya Arun, Amrutha H C, Ananga Ghosh, Rakesh K Mishra, Divya Tej Sowpati |
| EPI_ISL_495240 | CSIR-Centre for Cellular and Molecular Biology | CSIR-Centre for Cellular and Molecular Biology | M Soujanya Reddy, Nikhil Hajirnis, Pratheusa Maccha, Payel Mukherjee, Sofia Banu, Priya Singh, Onkar Kulkarni,Tulasi Nagabandi, Namami Gaur, Sakshi Shambhavi, Lamuk Zaveri, Shagufta Khan, Purushotham Vodnala, Dhiviya Vedagiri, Divya Gupta, Vishal Sah, Santosh Kumar Kuncha, Krishnan Harinivas Harshan, Archana Bharadwaj Siva, Karthik Bharadwaj Tallapaka,Kezia J Ann, Radhika Khandelwal, Roshan Maku Venkata, Shemin Mansuri, Sonu Uday, Rakesh K Mishra, Divya Tej Sowpati |
| EPI_ISL_495253 | CSIR-Centre for Cellular and Molecular Biology | CSIR-Centre for Cellular and Molecular Biology | M Soujanya Reddy, Nikhil Hajirnis, Pratheusa Maccha, Namami Gaur, Sakshi Shambhavi, Lamuk Zaveri, Shagufta Khan, Tulasi Nagabandi, Purushotham Vodnala, Payel Mukherjee, Sofia Banu, Priya Singh, Onkar Kulkarni, Dhiviya Vedagiri, Divya Gupta, Vishal Sah, Santosh Kumar Kuncha, Krishnan Harinivas Harshan, Archana Bharadwaj Siva, Karthik Bharadwaj Tallapaka, Zeba Rizvi, Zuberwasim Sayyad, Kakade Aishwarya Arun, Amrutha H C, Ananga Ghosh, Rakesh K Mishra, Divya Tej Sowpati |
| EPI_ISL_495257 | CSIR-Centre for Cellular and Molecular Biology | CSIR-Centre for Cellular and Molecular Biology | M Soujanya Reddy, Nikhil Hajirnis, Pratheusa Maccha, Payel Mukherjee, Sofia Banu, Priya Singh, Onkar Kulkarni,Tulasi Nagabandi, Namami Gaur, Sakshi Shambhavi, Lamuk Zaveri, Shagufta Khan, Purushotham Vodnala, Dhiviya Vedagiri, Divya Gupta, Vishal Sah, Santosh Kumar Kuncha, Krishnan Harinivas Harshan, Archana Bharadwaj Siva, Karthik Bharadwaj Tallapaka,Kezia J Ann, Radhika Khandelwal, Roshan Maku Venkata, Shemin Mansuri, Sonu Uday, Rakesh K Mishra, Divya Tej Sowpati |
| EPI_ISL_495271 | CSIR-Centre for Cellular and Molecular Biology | CSIR-Centre for Cellular and Molecular Biology | M Soujanya Reddy, Nikhil Hajirnis, Pratheusa Maccha, Sakshi Shambhavi, Lamuk Zaveri, Shagufta Khan, Namami Gaur, Tulasi Nagabandi, Purushotham Vodnala, Payel Mukherjee, Sofia Banu, Priya Singh,Onkar Kulkarni, Dhiviya Vedagiri, Divya Gupta, Vishal Sah, Santosh Kumar Kuncha, Krishnan Harinivas Harshan, Archana Bharadwaj Siva, Karthik Bharadwaj Tallapaka, G. Aditya Kumar, Koushick Sivakumar, Pooja Ramesh Gupta, Rajan Kumar Jha, Shraddha Vijay Lahoti, Rakesh K Mishra, Divya Tej Sowpati |
| EPI_ISL_495272 | CSIR-Centre for Cellular and Molecular Biology | CSIR-Centre for Cellular and Molecular Biology | M Soujanya Reddy, Nikhil Hajirnis, Pratheusa Maccha, Namami Gaur, Sakshi Shambhavi, Lamuk Zaveri, Shagufta Khan, Tulasi Nagabandi, Purushotham Vodnala, Payel Mukherjee, Sofia Banu, Priya Singh, Onkar Kulkarni, Dhiviya Vedagiri, Divya Gupta, Vishal Sah, Santosh Kumar Kuncha, Krishnan Harinivas Harshan, Archana Bharadwaj Siva, Karthik Bharadwaj Tallapaka, Zeba Rizvi, Zuberwasim Sayyad, Kakade Aishwarya Arun, Amrutha H C, Ananga Ghosh, Rakesh K Mishra, Divya Tej Sowpati |
| EPI_ISL_498466, EPI_ISL_498467 | National Institute of Laboratory Medicine and Referral Center | Genomic Research Lab, BCSIR | Mahmuda Yeasmin, Abu Sayeed Mohammad Mahmud, Mohammad Samir Uzzaman, Eshrar Osman, Md. Ahasan Habib, Shahina Akter, Tanjina Akhter Banu, Md. Murshed Hasan Sarkar, Barna Goswami, Iffat Jahan, Md. Saddam Hossain, Tasnim Nafisa, Md. Maruf Ahmed Molla, Asish Kumar Ghosh, A. K. M. Shamsuzzaman, Sheikh Md. Selim Al Din, Utpal Chandra Ray, Salek Ahmed Sajib, Md. Salim Khan |
| EPI_ISL_498559, EPI_ISL_498560, EPI_ISL_498561, EPI_ISL_498562, EPI_ISL_498563 | Laboratory of Molecular Virology International Center for Genetic Engineering and Biotechnology (ICGEB) | ARGO Open Lab Platform for Genome Sequencing | Licastro D, Rajasekharan S, Dal Monego S, Segat L, D'Agaro P, Marcello A |
| EPI_ISL_498564, EPI_ISL_498565, EPI_ISL_498566, EPI_ISL_498567, EPI_ISL_498568, EPI_ISL_498569, EPI_ISL_498570, EPI_ISL_498571, EPI_ISL_498572, EPI_ISL_498573, EPI_ISL_498574, EPI_ISL_498575, EPI_ISL_498576, EPI_ISL_498577, EPI_ISL_498578, EPI_ISL_498579, EPI_ISL_498580, EPI_ISL_498581, EPI_ISL_498582, EPI_ISL_498583, EPI_ISL_498584, EPI_ISL_498585, EPI_ISL_498586, EPI_ISL_498587, EPI_ISL_498588, EPI_ISL_498589, EPI_ISL_498590, EPI_ISL_498591, EPI_ISL_498592, EPI_ISL_498593, EPI_ISL_498594, EPI_ISL_498595, EPI_ISL_498596, EPI_ISL_498597, EPI_ISL_498598, EPI_ISL_498599, EPI_ISL_498600, EPI_ISL_498601, EPI_ISL_498602, EPI_ISL_498603, EPI_ISL_498604, EPI_ISL_498605, EPI_ISL_498606, EPI_ISL_498607, EPI_ISL_498608, EPI_ISL_498609, EPI_ISL_498610, EPI_ISL_498611, EPI_ISL_498612, EPI_ISL_498613, EPI_ISL_498614, EPI_ISL_498615, EPI_ISL_498616, EPI_ISL_498617, EPI_ISL_498618 | | | |
| see above | National Public Health Laboratory, National Centre for Infectious Diseases | National Public Health Laboratory, National Centre for Infectious Diseases | Mak TM, Octavia S, Zhou Z, Chavatte JM, Cui L, Lin RTP |
| EPI_ISL_499853, EPI_ISL_499855, EPI_ISL_499905, EPI_ISL_499908, EPI_ISL_499966, EPI_ISL_499968, EPI_ISL_499969, EPI_ISL_499970, EPI_ISL_499971, EPI_ISL_499972, EPI_ISL_499973 | | | |
| see above | Department of Pathology, University of Cambridge | COVID-19 Genomics UK (COG-UK) Consortium | Luke W Meredith, M. Estée Török, Myra Hosmillo, William L. Hamilton, Martin D. Curran, Theresa Feltwell, Grant Hall, Anna Yakovleva, Fahad A Khokhar, Charlotte J. Houldcroft, Laura G Caller, Aminu S. Jahun, Sarah L. Caddy, Yasmin Chaudhry, Malte Pinckert, Ian Goodfellow |
| EPI_ISL_500460, EPI_ISL_500461, EPI_ISL_500462, EPI_ISL_500463, EPI_ISL_500464, EPI_ISL_500465, EPI_ISL_500466, EPI_ISL_500467, EPI_ISL_500468, EPI_ISL_500469, EPI_ISL_500470, EPI_ISL_500471, EPI_ISL_500472, EPI_ISL_500473, EPI_ISL_500474, EPI_ISL_500475, EPI_ISL_500476, EPI_ISL_500477, EPI_ISL_500478, EPI_ISL_500479, EPI_ISL_500480, EPI_ISL_500481, EPI_ISL_500482, EPI_ISL_500483, EPI_ISL_500484, EPI_ISL_500485, EPI_ISL_500486, EPI_ISL_500865, EPI_ISL_500866, EPI_ISL_500867, EPI_ISL_500868, EPI_ISL_500869, EPI_ISL_500870, EPI_ISL_500871, EPI_ISL_500872, EPI_ISL_500873, EPI_ISL_500874, EPI_ISL_500875 | | | |
| see above | LACEN/PE | WallauLab, Aggeu Magalhaes Institute | Marcelo Henrique Santos Paiva, Duschinka Ribeiro Duarte Guedes, Cássia Docena, Matheus Filgueira Bezerra, Filipe Zimmer Dezordi, Laís Ceschini Machado, Larissa Krokovsky, Elisama Helvecio, Alexandre Freitas da Silva, Luydson Richardson Silva Vasconcelos, Antonio Mauro Rezende, Severino Jefferson Ribeiro da Silva, Kamila Gaudêncio da Silva Sales, Bruna Santos Lima Figueiredo de Sá, Derciliano Lopes da Cruz, Claudio Eduardo Cavalcanti, Armando de Menezes Neto, Caroline Targino Alves da Silva, Renata Pessôa Germano Mendes, Maria Almerice Lopes da Silva, Tiago Gräf, Paola Cristina Resende, Gonzalo Bello, Michelle da Silva Barros, Wheverton Ricardo Correia do Nascimento, Rodrigo Moraes Loyo Arcoverde, Luciane Caroline Albuquerque Bezerra, Sinval Pinto Brandão Filho, Constância Flávia Junqueira Ayres, Gabriel Luz Wallau |
| EPI_ISL_500947 | Department of MicroBiology, Government Medical College, Surat | Gujarat Biotechnology Research Centre | Maharshi Pandya, Nidhi Patel, Nitin Savaliya, Raghawendra Kumar, Dinesh Kumar, Zuber Saiyed, Komal Patel, Labdhi Pandya, Afzal Ansari, Nikha Trivedi, Naresh Chauhan, Summaiya Mullan, Amit gamit, Apurvasinh Puvar, Janvi Raval, Zarna Patel, Monika Gandhi, Pinal Trivedi, R D Dixit, A M Kadri, Harsh Bakshi, Chaitanya Joshi, Madhvi Joshi |
| EPI_ISL_501557, EPI_ISL_501562, EPI_ISL_501602, EPI_ISL_501613 | Department of Medical Microbiology, Western Sussex Hospitals NHS Foundation Trust, St Richard's Hospital | Wellcome Sanger Institute for the COVID-19 Genomics UK (COG-UK) consortium | Manasa Mutingwende, Sarah Lowdon, Olga Podplomyk, Michelle Erkiert, Jonathan Lewis, Paul Randell and Alex Alderton, Roberto Amato, Sonia Goncalves, Ewan Harrison, David K. Jackson, Ian Johnston, Dominic Kwiatkowski, Cordelia Langford, John Sillitoe on behalf of the Wellcome Sanger Institute COVID-19 Surveillance Team (http://www.sanger.ac.uk/covid-team) |

| | | | |
|---|---|---|---|
| EPI_ISL_508122, EPI_ISL_508123, EPI_ISL_508124, EPI_ISL_508125, EPI_ISL_508126, EPI_ISL_508127, EPI_ISL_508128, EPI_ISL_508129, EPI_ISL_508130, EPI_ISL_508131, EPI_ISL_508132, EPI_ISL_508133, EPI_ISL_508134, EPI_ISL_508135, EPI_ISL_508136, EPI_ISL_508137, EPI_ISL_508138, EPI_ISL_508139, EPI_ISL_508140, EPI_ISL_508141, EPI_ISL_508142, EPI_ISL_508143, EPI_ISL_508144, EPI_ISL_508605, EPI_ISL_508606, EPI_ISL_508607 | | | |
| see above | SA Pathology | SA Pathology | Lex Leong, Chuan Kok Lim, Mark Turra, Ivan Bastian, Geoff Higgins |
| EPI_ISL_510103, EPI_ISL_510104, EPI_ISL_510105, EPI_ISL_510106, EPI_ISL_510107, EPI_ISL_510108, EPI_ISL_510109, EPI_ISL_510110, EPI_ISL_510111, EPI_ISL_510112, EPI_ISL_510113, EPI_ISL_510114, EPI_ISL_510115, EPI_ISL_510116, EPI_ISL_510117, EPI_ISL_510118, EPI_ISL_510119, EPI_ISL_510120, EPI_ISL_510121, EPI_ISL_510122, EPI_ISL_510123, EPI_ISL_510124, EPI_ISL_510125, EPI_ISL_510126, EPI_ISL_510127, EPI_ISL_510128, EPI_ISL_510129, EPI_ISL_510130, EPI_ISL_510131, EPI_ISL_510132, EPI_ISL_510133, EPI_ISL_510134, EPI_ISL_510135, EPI_ISL_510136, EPI_ISL_510137, EPI_ISL_510138, EPI_ISL_510139, EPI_ISL_510140, EPI_ISL_510141, EPI_ISL_510142, EPI_ISL_510143, EPI_ISL_510144, EPI_ISL_510145, EPI_ISL_510146, EPI_ISL_510147, EPI_ISL_510148, EPI_ISL_510149, EPI_ISL_510150, EPI_ISL_510151, EPI_ISL_510165, EPI_ISL_510166, EPI_ISL_510167, EPI_ISL_510168, EPI_ISL_510169, EPI_ISL_510170, EPI_ISL_510171, EPI_ISL_510172, EPI_ISL_510173, EPI_ISL_510174, EPI_ISL_510175, EPI_ISL_510176, EPI_ISL_510177, EPI_ISL_510178, EPI_ISL_510179, EPI_ISL_510180, EPI_ISL_510181, EPI_ISL_510182, EPI_ISL_510183, EPI_ISL_510184, EPI_ISL_510185, EPI_ISL_510186, EPI_ISL_510187, EPI_ISL_510188, EPI_ISL_510189, EPI_ISL_510190, EPI_ISL_510191, EPI_ISL_510192, EPI_ISL_510193, EPI_ISL_510194, EPI_ISL_510195, EPI_ISL_510196, EPI_ISL_510197, EPI_ISL_510198, EPI_ISL_510199, EPI_ISL_510200, EPI_ISL_510201, EPI_ISL_510202, EPI_ISL_510203, EPI_ISL_510204, EPI_ISL_510205, EPI_ISL_510206, EPI_ISL_510207, EPI_ISL_510208, EPI_ISL_510209, EPI_ISL_510210, EPI_ISL_510211, EPI_ISL_510212, EPI_ISL_510213, EPI_ISL_510214, EPI_ISL_510215, EPI_ISL_510216, EPI_ISL_510217, EPI_ISL_510218, EPI_ISL_510219, EPI_ISL_510220, EPI_ISL_510221, EPI_ISL_510222, EPI_ISL_510223, EPI_ISL_510224, EPI_ISL_510225, EPI_ISL_510226, EPI_ISL_510227, EPI_ISL_510228, EPI_ISL_510229, EPI_ISL_510230, EPI_ISL_510231, EPI_ISL_510232, EPI_ISL_510233, EPI_ISL_510234, EPI_ISL_510235, EPI_ISL_510236, EPI_ISL_510237, EPI_ISL_510238, EPI_ISL_510239, EPI_ISL_510240, EPI_ISL_510241, EPI_ISL_510242, EPI_ISL_510243, EPI_ISL_510244, EPI_ISL_510245 | | | |
| see above | Hospital General Universitario Gregorio Marañón | SeqCOVID-SPAIN consortium/IBV(CSIC) | Laura Pérez-Lago, Marta Herranz, Jon Sicilia, Julia Suárez, Pilar Catalán, Patricia Muñoz, Darío García de Viedma and SeqCOVID-SPAIN consortium |
| EPI_ISL_510542, EPI_ISL_510543, EPI_ISL_510544 | SA Pathology | SA Pathology | Lex Leong, Chuan Kok Lim, Mark Turra, Ivan Bastian, Geoff Higgins |
| EPI_ISL_511959, EPI_ISL_511970, EPI_ISL_511982 | Department of Pathology, University of Cambridge | Wellcome Sanger Institute for the COVID-19 Genomics UK (COG-UK) consortium | Luke W Meredith, M. Estée Török , Myra Hosmillo, William L. Hamilton, Martin D. Curran, Theresa Feltwell, Grant Hall, Anna Yakovleva, Fahad A Khokhar, Charlotte J. Houldcroft, Laura G Caller, Aminu S. Jahun, Sarah L. Caddy, Ian Goodfellow; and Alex Alderton, Roberto Amato, Sonia Goncalves, Ewan Harrison, David K. Jackson, Ian Johnston, Dominic Kwiatkowski, Cordelia Langford, John Sillitoe on behalf of the Wellcome Sanger Institute COVID-19 Surveillance Team (http://www.sanger.ac.uk/covid-team) |
| EPI_ISL_512060 | B.J. Medical College and Civil hospital, Ahmedabad | Gujarat Biotechnology Research Centre | Maharshi Pandya, Nidhi Patel, Nitin Savaliya, Raghawendra Kumar, Dinesh Kumar, Zuber Saiyed, Komal Patel, Labdhi Pandya, Afzal Ansari, Nikha Trivedi, Pranay Shah, Kamlesh J Upadhyay, Sanjay Kapadia, Apurvasinh Puvar, Janvi Raval, Zarna Patel, Monika Gandhi, Pinal Trivedi, R D Dixit, A M Kadri, Harsh Bakshi, Chaitanya Joshi, Madhvi Joshi |
| EPI_ISL_512077 | Dr. RSS Hospital, Modasa | Gujarat Biotechnology Research Centre | Maharshi Pandya, Nidhi Patel, Nitin Savaliya, Raghawendra Kumar, Dinesh Kumar, Zuber Saiyed, Komal Patel, Labdhi Pandya, Afzal Ansari, Nikha Trivedi, Harsh Chaudhari, Apurvasinh Puvar, Janvi Raval, Zarna Patel, Monika Gandhi, Pinal Trivedi, R D Dixit, A M Kadri, Harsh Bakshi, Chaitanya Joshi, Madhvi Joshi |
| EPI_ISL_512296, EPI_ISL_512297, EPI_ISL_512298, EPI_ISL_512299, EPI_ISL_512300, EPI_ISL_512301, EPI_ISL_512302, EPI_ISL_512303 | Hematology Laboratory, Section of Molecular Diagnostics, University Clinical Centre, Medical University of Gdansk | Department of Virology, Faculty of Medicine, University of Helsinki, Helsinki, Finland | Maciej Grzybek, Marlena Robakowska, Aneta Szulc, Ewa Miosz, Olli Vapalahti, Teemu Smura |
| EPI_ISL_512330, EPI_ISL_512331, EPI_ISL_512332, EPI_ISL_512333, EPI_ISL_512334, EPI_ISL_512335 | Department of Pathology, University of Cambridge | COVID-19 Genomics UK (COG-UK) Consortium | Luke W Meredith, M. Estée Török, Myra Hosmillo, William L. Hamilton, Martin D. Curran, Theresa Feltwell, Grant Hall, Anna Yakovleva, Fahad A Khokhar, Charlotte J. Houldcroft, Laura G Caller, Aminu S. Jahun, Sarah L. Caddy, Yasmin Chaudhry, Malte Pinckert, Ian Goodfellow |
| EPI_ISL_512822, EPI_ISL_512823, EPI_ISL_512824, EPI_ISL_512825, EPI_ISL_512826, EPI_ISL_512827, EPI_ISL_512828, EPI_ISL_512829, EPI_ISL_512830, EPI_ISL_512831, EPI_ISL_512832, EPI_ISL_512833, EPI_ISL_512834, EPI_ISL_512835, EPI_ISL_512836, EPI_ISL_512837, EPI_ISL_512838, EPI_ISL_512839, EPI_ISL_512840, EPI_ISL_512841, EPI_ISL_512842 | | | |
| see above | National Public Health Laboratory, National Centre for Infectious Diseases | National Public Health Laboratory, National Centre for Infectious Diseases | Mak TM, Octavia S, Zhou Z, Chavatte JM, Cui L, Lin RTP |
| EPI_ISL_513911, EPI_ISL_513912, EPI_ISL_513913 | SA Pathology | SA Pathology | Lex Leong, Chuan Kok Lim, Mark Turra, Ivan Bastian, Geoff Higgins |
| EPI_ISL_514131 | Rondônia Central Public Health Laboratory (LACEN/RO), vinculated to State Health Secretariat of Rondônia (SESAU/RO) | Molecular Virology Laboratory of Oswaldo Cruz Foundation of Rondônia | Luan Felipo Botelho-Souza, Felipe Souza Nogueira-Lima, Tárcio Peixoto Roca, Alcione de Oliveira dos Santos, Felipe Gomes Naveca, Adriana Cristina Salvador Maia, Cicileia Correia da Silva, Aline Linhares Ferreira de Melo Mendonça, Celina Aparecida Bertoni Lugtenburg, Camila Flávia Gomes Azzi, Juliana Loca Furtado, Suelen Cavalcante, Rita de Cássia Pontello Rampazzo, Caio Henrique Nemeth Santos, Alice Paula Di Sabatino Guimarães, Jansen Fernandes de Medeiros, Fernando Rodrigues Máximo, Juan Miguel Vilallobos-Salcedo and Deusilene Souza Vieira1 |
| EPI_ISL_514132 | Rondônia Central Public Health Laboratory (LACEN/RO), vinculated to State Health Secretariat of Rondônia (SESAU/RO) | Molecular Virology Laboratory of Oswaldo Cruz Foundation of Rondônia | Luan Felipo Botelho-Souza, Felipe Souza Nogueira-Lima, Tárcio Peixoto Roca, Alcione de Oliveira dos Santos, Felipe Gomes Naveca, Adriana Cristina Salvador Maia, Cicileia Correia da Silva, Aline Linhares Ferreira de Melo Mendonça, Celina Aparecida Bertoni Lugtenburg, Camila Flávia Gomes Azzi, Juliana Loca Furtado, Suelen Cavalcante, Rita de Cássia Pontello Rampazzo, Caio Henrique Nemeth Santos, Alice Paula Di Sabatino Guimarães, Jansen Fernandes de Medeiros, Fernando Rodrigues Máximo, Juan Miguel Vilallobos-Salcedo and Deusilene Souza Vieira. |
| EPI_ISL_514133, EPI_ISL_514134, EPI_ISL_514135, EPI_ISL_514136, EPI_ISL_514137, EPI_ISL_514138 | Rondônia Central Public Health Laboratory (LACEN/RO), vinculated to State Health Secretariat of Rondônia (SESAU/RO) | Molecular Virology Laboratory of Oswaldo Cruz Foundation of Rondônia | Luan Felipo Botelho-Souza, Felipe Souza Nogueira-Lima, Tárcio Peixoto Roca, Alcione de Oliveira dos Santos, Felipe Gomes Naveca, Adriana Cristina Salvador Maia, Cicileia Correia da Silva, Aline Linhares Ferreira de Melo Mendonça, Celina Aparecida Bertoni Lugtenburg, Camila Flávia Gomes Azzi, Juliana Loca Furtado, Suelen Cavalcante, Rita de Cássia Pontello Rampazzo, Caio Henrique Nemeth Santos, Alice Paula Di Sabatino Guimarães, Jansen Fernandes de Medeiros, Fernando Rodrigues Máximo, Juan Miguel Vilallobos-Salcedo and Deusilene Souza Vieira |
| EPI_ISL_514443, EPI_ISL_514444, EPI_ISL_514445, EPI_ISL_514446, EPI_ISL_514447, EPI_ISL_514448, EPI_ISL_514449, EPI_ISL_514450, EPI_ISL_514451 | Department of Pathology, University of Cambridge | COVID-19 Genomics UK (COG-UK) Consortium | Luke W Meredith, M. Estée Török, Myra Hosmillo, William L. Hamilton, Martin D. Curran, Theresa Feltwell, Grant Hall, Anna Yakovleva, Fahad A Khokhar, Charlotte J. Houldcroft, Laura G Caller, Aminu S. Jahun, Sarah L. Caddy, Yasmin Chaudhry, Malte Pinckert, Ian Goodfellow |
| EPI_ISL_514597 | B.J. Medical College and Civil hospital, Ahmedabad | Gujarat Biotechnology Research Centre | Maharshi Pandya, Nidhi Patel, Nitin Savaliya, Raghawendra Kumar, Dinesh Kumar, Zuber Saiyed, Komal Patel, Labdhi Pandya, Afzal Ansari, Nikha Trivedi, Pranay Shah, Kamlesh J Upadhyay, Sanjay Kapadia, Apurvasinh Puvar, Janvi Raval, Zarna Patel, Monika Gandhi, Pinal Trivedi, R D Dixit, A M Kadri, Harsh Bakshi, Chaitanya Joshi, Madhvi Joshi |
| EPI_ISL_514752 | Infectious Disease Control Center, Center for Disease Control and Prevention of PLA | Infectious Disease Control Center, Center for Disease Control and Prevention of PLA | Li, P. |
| EPI_ISL_516193 | Hospital Universitari Germans Trias i Pujol. | IrsiCaixa AIDS Research Lab | Marc Noguera-Julian, Mariona Parera, Maria Pilar Armengol, Marta Massanella, Ester Ballana, Lidia Ruiz, Nuria Izquierdo, Jorge Carrillo, Roger Paredes, Julia Blanco, Joaquim Segalés, Bonaventura Clotet |
| EPI_ISL_516550, EPI_ISL_516551 | SA Pathology | SA Pathology | Lex Leong, Julien Soubrier, Luke Walters, Chuan Kok Lim, Song Gao, Mark Turra, Karin Kassahn, Ivan Bastian, Geoff Higgins |
| EPI_ISL_516807, EPI_ISL_516808, EPI_ISL_516809, EPI_ISL_516810, EPI_ISL_516811, EPI_ISL_516812, EPI_ISL_516813, EPI_ISL_516814, EPI_ISL_516815, EPI_ISL_516816, EPI_ISL_516817, EPI_ISL_516818, EPI_ISL_516819, EPI_ISL_516820, EPI_ISL_516821, EPI_ISL_516822, EPI_ISL_516823, EPI_ISL_516824, EPI_ISL_516825, EPI_ISL_516826, EPI_ISL_516827, EPI_ISL_516828 | | | |
| see above | National Public Health Laboratory, National Centre for Infectious Diseases | National Public Health Laboratory, National Centre for Infectious Diseases | Mak TM, Octavia S, Zhou Z, Cui L, Lin RTP |
| EPI_ISL_516831, EPI_ISL_516832, EPI_ISL_516833, EPI_ISL_516834, EPI_ISL_516835, EPI_ISL_516836, EPI_ISL_516837, EPI_ISL_516838, EPI_ISL_516839, EPI_ISL_516840, EPI_ISL_516841, EPI_ISL_516842, EPI_ISL_516843, EPI_ISL_516844, EPI_ISL_516845, EPI_ISL_516846, EPI_ISL_516847, EPI_ISL_516848, EPI_ISL_516849, EPI_ISL_516850, EPI_ISL_516851, EPI_ISL_516852, EPI_ISL_516853, EPI_ISL_516854, EPI_ISL_516855, EPI_ISL_516856, EPI_ISL_516857, EPI_ISL_516858, EPI_ISL_516859, EPI_ISL_516860, EPI_ISL_516861, EPI_ISL_516862, EPI_ISL_516863, EPI_ISL_516864, EPI_ISL_516865, EPI_ISL_516866, EPI_ISL_516867, EPI_ISL_516868, EPI_ISL_516869, EPI_ISL_516870, EPI_ISL_516871, EPI_ISL_516872, EPI_ISL_516873, EPI_ISL_516874, EPI_ISL_516875, EPI_ISL_516876, EPI_ISL_516877, EPI_ISL_516878, EPI_ISL_516879, EPI_ISL_516880, EPI_ISL_516881, EPI_ISL_516882, EPI_ISL_516883 | | | |
| see above | North West London Pathology, Imperial College Healthcare NHS Trust | Wellcome Sanger Institute for the COVID-19 Genomics UK (COG-UK) consortium | Ling Li, Paul Randell, David Muir, Frankie Bolt, Alison Holmes, James Price, Aileen Rowan, Graham Taylor, Anjna Badhan, Carolina Herrera and Alex Alderton, Roberto Amato, Sonia Goncalves, Ewan Harrison, David K. Jackson, Ian Johnston, Dominic Kwiatkowski, Cordelia Langford, John Sillitoe on behalf of the Wellcome Sanger Institute COVID-19 Surveillance Team (http://www.sanger.ac.uk/covid-team) |
| EPI_ISL_516990, EPI_ISL_516991, EPI_ISL_516992, EPI_ISL_516993, EPI_ISL_516994, EPI_ISL_516995, EPI_ISL_516996, EPI_ISL_516997, EPI_ISL_516998, EPI_ISL_516999, EPI_ISL_517000 | | | |
| see above | Department of Pathology, University of Cambridge | COVID-19 Genomics UK (COG-UK) Consortium | Luke W Meredith, M. Estée Török, Myra Hosmillo, William L. Hamilton, Martin D. Curran, Theresa Feltwell, Grant Hall, Anna Yakovleva, Fahad A Khokhar, |

| | | | |
|---|---|---|---|
| | | | Charlotte J. Houldcroft, Laura G Caller, Aminu S. Jahun, Sarah L. Caddy, Yasmin Chaudhry, Malte Pinckert, Ian Goodfellow |
| EPI_ISL_524493, EPI_ISL_524495, EPI_ISL_524497, EPI_ISL_524498, EPI_ISL_524501, EPI_ISL_524502, EPI_ISL_524503, EPI_ISL_524505, EPI_ISL_524506, EPI_ISL_524508, EPI_ISL_524509, EPI_ISL_524515, EPI_ISL_524521, EPI_ISL_524522, EPI_ISL_524525, EPI_ISL_524528, EPI_ISL_524529, EPI_ISL_524530, EPI_ISL_524531, EPI_ISL_524537, EPI_ISL_524538, EPI_ISL_524541, EPI_ISL_524544, EPI_ISL_524546, EPI_ISL_524548, EPI_ISL_524551, EPI_ISL_524553, EPI_ISL_524558, EPI_ISL_524560, EPI_ISL_524564, EPI_ISL_524568, EPI_ISL_524570, EPI_ISL_524573, EPI_ISL_524574, EPI_ISL_524575, EPI_ISL_524576, EPI_ISL_524577, EPI_ISL_524578, EPI_ISL_524579, EPI_ISL_524580, EPI_ISL_524581, EPI_ISL_524582, EPI_ISL_524583, EPI_ISL_524584, EPI_ISL_524585, EPI_ISL_524586, EPI_ISL_524587, EPI_ISL_524588, EPI_ISL_524589, EPI_ISL_524591, EPI_ISL_524592, EPI_ISL_524593, EPI_ISL_524594, EPI_ISL_524596, EPI_ISL_524597, EPI_ISL_524598, EPI_ISL_524599, EPI_ISL_524600, EPI_ISL_524601, EPI_ISL_524602, EPI_ISL_524603, EPI_ISL_524604, EPI_ISL_524605, EPI_ISL_524606, EPI_ISL_524607, EPI_ISL_524608, EPI_ISL_524609, EPI_ISL_524610, EPI_ISL_524611, EPI_ISL_524612, EPI_ISL_524613, EPI_ISL_524614, EPI_ISL_524615, EPI_ISL_524616, EPI_ISL_524617, EPI_ISL_524618 | | | |
| see above | Department of Pathology, University of Cambridge | Wellcome Sanger Institute for the COVID-19 Genomics UK (COG-UK) consortium | Luke W Meredith, M. Estée Török , Myra Hosmillo, William L. Hamilton, Martin D. Curran, Theresa Feltwell, Grant Hall, Anna Yakovleva, Fahad A Khokhar, Charlotte J. Houldcroft, Laura G Caller, Aminu S. Jahun, Sarah L. Caddy, Ian Goodfellow; and Alex Alderton, Roberto Amato, Sonia Goncalves, Ewan Harrison, David K. Jackson, Ian Johnston, Dominic Kwiatkowski, Cordelia Langford, John Sillitoe on behalf of the Wellcome Sanger Institute COVID-19 Surveillance Team (http://www.sanger.ac.uk/covid-team) |
| EPI_ISL_524619, EPI_ISL_524620, EPI_ISL_524621, EPI_ISL_524622, EPI_ISL_524623, EPI_ISL_524624, EPI_ISL_524625, EPI_ISL_524626, EPI_ISL_524627, EPI_ISL_524628, EPI_ISL_524629, EPI_ISL_524630, EPI_ISL_524631, EPI_ISL_524632, EPI_ISL_524633, EPI_ISL_524634, EPI_ISL_524635, EPI_ISL_524636, EPI_ISL_524637, EPI_ISL_524638, EPI_ISL_524639, EPI_ISL_524640, EPI_ISL_524641, EPI_ISL_524642, EPI_ISL_524643, EPI_ISL_524644, EPI_ISL_524645, EPI_ISL_524646, EPI_ISL_524647, EPI_ISL_524648, EPI_ISL_524649, EPI_ISL_524650, EPI_ISL_524651, EPI_ISL_524652, EPI_ISL_524653, EPI_ISL_524654, EPI_ISL_524655, EPI_ISL_524656, EPI_ISL_524657, EPI_ISL_524658, EPI_ISL_524659, EPI_ISL_524660, EPI_ISL_524661, EPI_ISL_524662, EPI_ISL_524663, EPI_ISL_524664, EPI_ISL_524665, EPI_ISL_524666, EPI_ISL_524667, EPI_ISL_524668, EPI_ISL_524669, EPI_ISL_524670, EPI_ISL_524671, EPI_ISL_524672, EPI_ISL_524673, EPI_ISL_524674, EPI_ISL_524675, EPI_ISL_524676, EPI_ISL_524677, EPI_ISL_524678, EPI_ISL_524679, EPI_ISL_524680, EPI_ISL_524681, EPI_ISL_524682, EPI_ISL_524683, EPI_ISL_524684, EPI_ISL_524685, EPI_ISL_524686, EPI_ISL_524687, EPI_ISL_524688, EPI_ISL_524689, EPI_ISL_524690, EPI_ISL_524691, EPI_ISL_524692, EPI_ISL_524693, EPI_ISL_524694, EPI_ISL_524695, EPI_ISL_524696, EPI_ISL_524697, EPI_ISL_524698, EPI_ISL_524699, EPI_ISL_524700, EPI_ISL_524701, EPI_ISL_524702, EPI_ISL_524703, EPI_ISL_524704, EPI_ISL_524705, EPI_ISL_524706, EPI_ISL_524707, EPI_ISL_524708, EPI_ISL_524709, EPI_ISL_524710, EPI_ISL_524711, EPI_ISL_524712 | | | |
| see above | North West London Pathology, Imperial College Healthcare NHS Trust | Wellcome Sanger Institute for the COVID-19 Genomics UK (COG-UK) consortium | Ling Li, Paul Randell, David Muir, Frankie Bolt, Alison Holmes, James Price, Aileen Rowan, Graham Taylor, Anjna Badhan, Carolina Herrera and Alex Alderton, Roberto Amato, Sonia Goncalves, Ewan Harrison, David K. Jackson, Ian Johnston, Dominic Kwiatkowski, Cordelia Langford, John Sillitoe on behalf of the Wellcome Sanger Institute COVID-19 Surveillance Team (http://www.sanger.ac.uk/covid-team) |
| EPI_ISL_524718 | B.J. Medical College and Civil hospital, Ahmedabad | Gujarat Biotechnology Research Centre | Maharshi Pandya, Nidhi Patel, Nitin Savaliya, Raghawendra Kumar, Dinesh Kumar, Zuber Saiyed, Komal Patel, Labdhi Pandya, Afzal Ansari, Nikha Trivedi, Pranay Shah, Kamlesh J Upadhyay, Sanjay Kapadia, Apurvasinh Puvar, Janvi Raval, Zarna Patel, Monika Gandhi, Pinal Trivedi, R D Dixit, A M Kadri, Harsh Bakshi, Chaitanya Joshi, Madhvi Joshi |
| EPI_ISL_524740 | GMERS Medical College and Hospital, Gandhinagar | Gujarat Biotechnology Research Centre | Maharshi Pandya, Nidhi Patel, Nitin Savaliya, Raghawendra Kumar, Dinesh Kumar, Zuber Saiyed, Komal Patel, Labdhi Pandya, Afzal Ansari, Nikha Trivedi, Seema Bhatt, Gaurishankar Shrimali, Bhavesh Modi, Bharti Rajani, Apurvasinh Puvar, Janvi Raval, Zarna Patel, Monika Gandhi, Pinal Trivedi, R D Dixit, A M Kadri, Harsh Bakshi, Chaitanya Joshi, Madhvi Joshi |
| EPI_ISL_524754 | GMERS Medical College and Hospital, Dharpur, Patan | Gujarat Biotechnology Research Centre | Maharshi Pandya, Nidhi Patel, Nitin Savaliya, Raghawendra Kumar, Dinesh Kumar, Zuber Saiyed, Komal Patel, Labdhi Pandya, Afzal Ansari, Nikha Trivedi, A N Parmar, Apurvasinh Puvar, Janvi Raval, Zarna Patel, Monika Gandhi, Pinal Trivedi, R D Dixit, A M Kadri, Harsh Bakshi, Chaitanya Joshi, Madhvi Joshi |
| EPI_ISL_525495, EPI_ISL_525496 | Laboratory of Molecular Virology of the International Centre for Genetic Engineering and Biotechnology (ICGEB) | ARGO Open Lab Platform for Genome Sequencing | Licastro D, Rajasekharan S, Dal Monego S, Segat L, D'Agaro P, Marcello A |
| EPI_ISL_527359, EPI_ISL_527360, EPI_ISL_527361, EPI_ISL_527362, EPI_ISL_527363, EPI_ISL_527364, EPI_ISL_527365, EPI_ISL_527366, EPI_ISL_527367, EPI_ISL_527368, EPI_ISL_527369, EPI_ISL_527370, EPI_ISL_527371, EPI_ISL_527372, EPI_ISL_527373, EPI_ISL_527374, EPI_ISL_527375, EPI_ISL_527376, EPI_ISL_527377, EPI_ISL_527378, EPI_ISL_527379 | | | |
| see above | National Public Health Laboratory, National Centre for Infectious Diseases | National Public Health Laboratory, National Centre for Infectious Diseases | Mak TM, Octavia S, Zhou Z, Cui L, Lin RTP |
| EPI_ISL_527916, EPI_ISL_527917, EPI_ISL_527918, EPI_ISL_527919, EPI_ISL_527920, EPI_ISL_527921, EPI_ISL_527922, EPI_ISL_527923, EPI_ISL_527924, EPI_ISL_527925, EPI_ISL_527926, EPI_ISL_527927, EPI_ISL_527928, EPI_ISL_527929, EPI_ISL_527930, EPI_ISL_527931, EPI_ISL_527932, EPI_ISL_527933, EPI_ISL_527934, EPI_ISL_527935, EPI_ISL_527936, EPI_ISL_527937, EPI_ISL_527938, EPI_ISL_527939, EPI_ISL_527940, EPI_ISL_527941, EPI_ISL_527942, EPI_ISL_527943, EPI_ISL_527944, EPI_ISL_527945, EPI_ISL_527946, EPI_ISL_527947, EPI_ISL_527948, EPI_ISL_527949, EPI_ISL_527950, EPI_ISL_527951, EPI_ISL_527952, EPI_ISL_527953, EPI_ISL_527954, EPI_ISL_527955, EPI_ISL_527956, EPI_ISL_527957, EPI_ISL_527958, EPI_ISL_527959, EPI_ISL_527960, EPI_ISL_527961, EPI_ISL_527962, EPI_ISL_527963, EPI_ISL_527964, EPI_ISL_527965, EPI_ISL_527966, EPI_ISL_527967, EPI_ISL_527968, EPI_ISL_527969, EPI_ISL_527970, EPI_ISL_527971, EPI_ISL_527972, EPI_ISL_527973, EPI_ISL_527974, EPI_ISL_527975, EPI_ISL_527976, EPI_ISL_527977, EPI_ISL_527978, EPI_ISL_527979, EPI_ISL_527980, EPI_ISL_527981, EPI_ISL_527982, EPI_ISL_527983, EPI_ISL_527984, EPI_ISL_527985, EPI_ISL_527986, EPI_ISL_527987, EPI_ISL_527988, EPI_ISL_527989, EPI_ISL_527990, EPI_ISL_527991, EPI_ISL_527992, EPI_ISL_527993, EPI_ISL_527994, EPI_ISL_527995, EPI_ISL_527996, EPI_ISL_527997, EPI_ISL_527998, EPI_ISL_527999, EPI_ISL_528000, EPI_ISL_528001, EPI_ISL_528002, EPI_ISL_528003, EPI_ISL_528004, EPI_ISL_528005, EPI_ISL_528006, EPI_ISL_528007, EPI_ISL_528008, EPI_ISL_528009, EPI_ISL_528010, EPI_ISL_528011, EPI_ISL_528012, EPI_ISL_528013, EPI_ISL_528014, EPI_ISL_528015, EPI_ISL_528016, EPI_ISL_528017, EPI_ISL_528018, EPI_ISL_528019, EPI_ISL_528020, EPI_ISL_528021, EPI_ISL_528022, EPI_ISL_528023, EPI_ISL_528024, EPI_ISL_528025, EPI_ISL_528026, EPI_ISL_528027, EPI_ISL_528028, EPI_ISL_528029, EPI_ISL_528030, EPI_ISL_528031, EPI_ISL_528032, EPI_ISL_528033, EPI_ISL_528034, EPI_ISL_528035, EPI_ISL_528036, EPI_ISL_528037, EPI_ISL_528038, EPI_ISL_528039, EPI_ISL_528040, EPI_ISL_528041, EPI_ISL_528042, EPI_ISL_528043, EPI_ISL_528044, EPI_ISL_528045, EPI_ISL_528046, EPI_ISL_528047, EPI_ISL_528048, EPI_ISL_528049, EPI_ISL_528050, EPI_ISL_528051, EPI_ISL_528052, EPI_ISL_528053, EPI_ISL_528054, EPI_ISL_528055, EPI_ISL_528056, EPI_ISL_528057, EPI_ISL_528058, EPI_ISL_528059, EPI_ISL_528060, EPI_ISL_528061, EPI_ISL_528062, EPI_ISL_528063, EPI_ISL_528064, EPI_ISL_528065, EPI_ISL_528066, EPI_ISL_528067, EPI_ISL_528068, EPI_ISL_528069, EPI_ISL_528070, EPI_ISL_528071, EPI_ISL_528072, EPI_ISL_528073, EPI_ISL_528074, EPI_ISL_528075, EPI_ISL_528076, EPI_ISL_528077, EPI_ISL_528078, EPI_ISL_528079, EPI_ISL_528080, EPI_ISL_528081, EPI_ISL_528082, EPI_ISL_528083, EPI_ISL_528084, EPI_ISL_528085, EPI_ISL_528086, EPI_ISL_528087, EPI_ISL_528088, EPI_ISL_528089, EPI_ISL_528090, EPI_ISL_528091, EPI_ISL_528092, EPI_ISL_528093, EPI_ISL_528094, EPI_ISL_528095, EPI_ISL_528096, EPI_ISL_528097, EPI_ISL_528098, EPI_ISL_528099, EPI_ISL_528100, EPI_ISL_528101, EPI_ISL_528102, EPI_ISL_528103, EPI_ISL_528104, EPI_ISL_528105, EPI_ISL_528106, EPI_ISL_528107, EPI_ISL_528108, EPI_ISL_528109, EPI_ISL_528110, EPI_ISL_528111, EPI_ISL_528112, EPI_ISL_528113, EPI_ISL_528114, EPI_ISL_528115, EPI_ISL_528116, EPI_ISL_528117, EPI_ISL_528118, EPI_ISL_528119, EPI_ISL_528120, EPI_ISL_528121, EPI_ISL_528122, EPI_ISL_528123, EPI_ISL_528124, EPI_ISL_528125, EPI_ISL_528126, EPI_ISL_528127, EPI_ISL_528128, EPI_ISL_528129, EPI_ISL_528130, EPI_ISL_528131, EPI_ISL_528132, EPI_ISL_528133, EPI_ISL_528134, EPI_ISL_528135, EPI_ISL_528136, EPI_ISL_528137, EPI_ISL_528138, EPI_ISL_528139, EPI_ISL_528140, EPI_ISL_528141, EPI_ISL_528142, EPI_ISL_528143, EPI_ISL_528144, EPI_ISL_528145, EPI_ISL_528146, EPI_ISL_528147, EPI_ISL_528148, EPI_ISL_528149, EPI_ISL_528150, EPI_ISL_528151, EPI_ISL_528152, EPI_ISL_528153, EPI_ISL_528154, EPI_ISL_528155, EPI_ISL_528156, EPI_ISL_528157, EPI_ISL_528158, EPI_ISL_528159, EPI_ISL_528160, EPI_ISL_528161, EPI_ISL_528162, EPI_ISL_528163, EPI_ISL_528164, EPI_ISL_528165, EPI_ISL_528166, EPI_ISL_528167, EPI_ISL_528168, EPI_ISL_528169, EPI_ISL_528170, EPI_ISL_528171, EPI_ISL_528172, EPI_ISL_528173, EPI_ISL_528174, EPI_ISL_528175, EPI_ISL_528176, EPI_ISL_528177, EPI_ISL_528178, EPI_ISL_528179, EPI_ISL_528180, EPI_ISL_528181, EPI_ISL_528182, EPI_ISL_528183, EPI_ISL_528184, EPI_ISL_528185, EPI_ISL_528186, EPI_ISL_528187, EPI_ISL_528188, EPI_ISL_528189, EPI_ISL_528190, EPI_ISL_528191, EPI_ISL_528192, EPI_ISL_528193, EPI_ISL_528194, EPI_ISL_528195, EPI_ISL_528196, EPI_ISL_528197, EPI_ISL_528198, EPI_ISL_528199, EPI_ISL_528200, EPI_ISL_528201, EPI_ISL_528202, EPI_ISL_528203, EPI_ISL_528204, EPI_ISL_528205, EPI_ISL_528206, EPI_ISL_528207, EPI_ISL_528208, EPI_ISL_528209, EPI_ISL_528210, EPI_ISL_528211, EPI_ISL_528212, EPI_ISL_528213, EPI_ISL_528214, EPI_ISL_528215, EPI_ISL_528216, EPI_ISL_528217, EPI_ISL_528218, EPI_ISL_528219, EPI_ISL_528220, EPI_ISL_528221, EPI_ISL_528222, EPI_ISL_528223, EPI_ISL_528224, EPI_ISL_528225, EPI_ISL_528226, EPI_ISL_528227, EPI_ISL_528228, EPI_ISL_528229, EPI_ISL_528230, EPI_ISL_528231, EPI_ISL_528232, EPI_ISL_528233, EPI_ISL_528234, EPI_ISL_528235, EPI_ISL_528236, EPI_ISL_528237, EPI_ISL_528238, EPI_ISL_528239, EPI_ISL_528240, EPI_ISL_528241, EPI_ISL_528242, EPI_ISL_528243, EPI_ISL_528244, EPI_ISL_528245, EPI_ISL_528246, EPI_ISL_528247, EPI_ISL_528248, EPI_ISL_528249, EPI_ISL_528250, EPI_ISL_528251, EPI_ISL_528252, EPI_ISL_528253, EPI_ISL_528254, EPI_ISL_528255, EPI_ISL_528256, EPI_ISL_528257, EPI_ISL_528258, EPI_ISL_528259, EPI_ISL_528260, EPI_ISL_528261, EPI_ISL_528262, EPI_ISL_528263, EPI_ISL_528264, EPI_ISL_528265, EPI_ISL_528266, EPI_ISL_528267, EPI_ISL_528268, EPI_ISL_528269, EPI_ISL_528270, EPI_ISL_528271, EPI_ISL_528272, EPI_ISL_528273, EPI_ISL_528274, EPI_ISL_528275, EPI_ISL_528276, EPI_ISL_528277, EPI_ISL_528278, EPI_ISL_528279, EPI_ISL_528280, EPI_ISL_528281, EPI_ISL_528282, EPI_ISL_528283, EPI_ISL_528284, EPI_ISL_528285, EPI_ISL_528286, EPI_ISL_528287, EPI_ISL_528288, EPI_ISL_528289, EPI_ISL_528290, EPI_ISL_528291, EPI_ISL_528292, EPI_ISL_528293, EPI_ISL_528294, EPI_ISL_528295, EPI_ISL_528296, EPI_ISL_528297, EPI_ISL_528298, EPI_ISL_528299, EPI_ISL_528300, EPI_ISL_528301, EPI_ISL_528302, EPI_ISL_528303, EPI_ISL_528304, EPI_ISL_528305, EPI_ISL_528306, EPI_ISL_528307, EPI_ISL_528308, EPI_ISL_528309, EPI_ISL_528310, EPI_ISL_528311, EPI_ISL_528312, EPI_ISL_528313, EPI_ISL_528314, EPI_ISL_528315, EPI_ISL_528316, EPI_ISL_528317, EPI_ISL_528318, EPI_ISL_528319, EPI_ISL_528320, EPI_ISL_528321, EPI_ISL_528322, EPI_ISL_528323, EPI_ISL_528324, EPI_ISL_528325, EPI_ISL_528326, EPI_ISL_528327, EPI_ISL_528328, EPI_ISL_528329, EPI_ISL_528330, EPI_ISL_528331, EPI_ISL_528332, EPI_ISL_528333, EPI_ISL_528334, EPI_ISL_528335, EPI_ISL_528336, EPI_ISL_528337, EPI_ISL_528338, EPI_ISL_528339, EPI_ISL_528340, EPI_ISL_528341, EPI_ISL_528342, EPI_ISL_528343, EPI_ISL_528344, EPI_ISL_528345, EPI_ISL_528346, EPI_ISL_528347, EPI_ISL_528348, EPI_ISL_528349, EPI_ISL_528350, EPI_ISL_528351, EPI_ISL_528352, EPI_ISL_528353, EPI_ISL_528354, EPI_ISL_528355, EPI_ISL_528356, EPI_ISL_528357, EPI_ISL_528358, EPI_ISL_528359, EPI_ISL_528360, EPI_ISL_528361, EPI_ISL_528362, EPI_ISL_528363, EPI_ISL_528364, EPI_ISL_528365, EPI_ISL_528366, EPI_ISL_528367, EPI_ISL_528368, EPI_ISL_528369, EPI_ISL_528370, EPI_ISL_528371, EPI_ISL_528372, EPI_ISL_528373, EPI_ISL_528374, EPI_ISL_528375, EPI_ISL_528376, EPI_ISL_528377, EPI_ISL_528378, EPI_ISL_528379, EPI_ISL_528380, EPI_ISL_528381 | | | |
| see above | University Hospital Basel, Clinical Virology | University Hospital Basel, Clinical Bacteriology | Madlen Stange, Alfredo Mari, Tim Roloff, Helena MB Seth-Smith, Michael Schweitzer, Myrta Brunner, Karoline Leuzinger, Kirstine K. Soegaard, Alexander Gensch, Sarah Tschudin-Sutter, Simon Fuchs, Julia Bielicki, Hans Pargger, Martin Siegemund, Christian Nickel, Roland Bingisser, Michael Osthoff, Stefano Bassetti, Rita Schneider-Sliwa, Manuel Battegay, Hans Hirsch, Adrian Egli |
| EPI_ISL_528835, EPI_ISL_528867 | CSIR-Centre for Cellular and Molecular Biology | CSIR-Centre for Cellular and Molecular Biology | M Soujanya Reddy, Nikhil Hajirnis, Pratheusa Maccha, Namami Gaur, Sakshi Shambhavi, Lamuk Zaveri, Shagufta Khan, Tulasi Nagabandi, Purushotham Vodnala, Payel Mukherjee, Sofia Banu, Priya Singh, Onkar Kulkarni, Dhiviya Vedagiri, Divya Gupta, Vishal Sah, Santosh Kumar Kuncha, Krishnan Harinivas Harshan, Archana Bharadwaj Siva, Karthik Bharadwaj Tallapaka, Zeba Rizvi, Zuberwasim Sayyad, Kakade Aishwarya Arun, Amrutha H C, Ananga Ghosh, Rakesh K Mishra, Divya Tej Sowpati |
| EPI_ISL_528919 | Ospedale Civile S. Liberatore-Atri | Istituto Zooprofilattico Sperimentale dell'Abruzzo e Molise "G.Caporale" | Lorusso A, Marcacci M, Di Domenico M, Curini V, Ancora M, Cammà C, Rinaldi A, Mangone I, Di Pasquale A, Puglia I, Savini G. |
| EPI_ISL_528920, EPI_ISL_528921 | Presidio Ospedaliero "Santo Spirito"-Pescara | Istituto Zooprofilattico Sperimentale dell'Abruzzo e Molise "G.Caporale" | Lorusso A, Marcacci M, Di Domenico M, Curini V, Ancora M, Cammà C, Rinaldi A, Mangone I, Di Pasquale A, Puglia I, Savini G. |
| EPI_ISL_528922 | Ospedale "Giuseppe Mazzini"-Teramo | Istituto Zooprofilattico Sperimentale dell'Abruzzo e Molise "G.Caporale" | Lorusso A, Marcacci M, Di Domenico M, Curini V, Ancora M, Cammà C, Rinaldi A, Mangone I, Di Pasquale A, Puglia I, Savini G. |
| EPI_ISL_528923 | Ospedale Civile S. Liberatore-Atri | Istituto Zooprofilattico Sperimentale dell'Abruzzo e Molise "G.Caporale" | Lorusso A, Marcacci M, Di Domenico M, Curini V, Ancora M, Cammà C, Rinaldi A, Mangone I, Di Pasquale A, Puglia I, Savini G. |
| EPI_ISL_528924 | Ospedale "Giuseppe Mazzini"-Teramo | Istituto Zooprofilattico Sperimentale dell'Abruzzo e Molise "G.Caporale" | Lorusso A, Marcacci M, Di Domenico M, Curini V, Ancora M, Cammà C, Rinaldi A, Mangone I, Di Pasquale A, Puglia I, Savini G. |
| EPI_ISL_528925 | Ospedale Regionale San Salvatore-L'Aquila | Istituto Zooprofilattico Sperimentale dell'Abruzzo e Molise | Lorusso A, Marcacci M, Di Domenico M, Curini V, Ancora M, Cammà C, Rinaldi A, Mangone I, Di Pasquale A, Puglia I, Savini G. |

| | | | |
|---|---|---|---|
| | | "G.Caporale" | |
| EPI_ISL_528926, EPI_ISL_528927, EPI_ISL_528928 | Ospedale "Giuseppe Mazzini"-Teramo | Istituto Zooprofilattico Sperimentale dell'Abruzzo e Molise "G.Caporale" | Lorusso A, Marcacci M, Di Domenico M, Curini V, Ancora M, Cammà C, Rinaldi A, Mangone I, Di Pasquale A, Puglia I, Savini G. |
| EPI_ISL_528929 | Ospedale Civile S. Liberatore-Atri | Istituto Zooprofilattico Sperimentale dell'Abruzzo e Molise "G.Caporale" | Lorusso A, Marcacci M, Di Domenico M, Curini V, Ancora M, Cammà C, Rinaldi A, Mangone I, Di Pasquale A, Puglia I, Savini G. |
| EPI_ISL_528934, EPI_ISL_528935, EPI_ISL_528936, EPI_ISL_528937, EPI_ISL_528938, EPI_ISL_528939, EPI_ISL_528940, EPI_ISL_528941, EPI_ISL_528942, EPI_ISL_528943, EPI_ISL_528944, EPI_ISL_528945, EPI_ISL_528946, EPI_ISL_528947, EPI_ISL_528948, EPI_ISL_528949 | | | |
| see above | Agenzia di Tutela della Salute di Bergamo | Istituto Zooprofilattico Sperimentale dell'Abruzzo e Molise "G.Caporale" | Lorusso A, Marcacci M, Di Domenico M, Curini V, Ancora M, Cammà C, Rinaldi A, Mangone I, Di Pasquale A, Puglia I, Savini G. |
| EPI_ISL_528990 | Ospedale Civile Maria SS. dello Splendore | Istituto Zooprofilattico Sperimentale dell'Abruzzo e Molise "G.Caporale" | Lorusso A, Marcacci M, Di Domenico M, Curini V, Ancora M, Cammà C, Rinaldi A, Mangone I, Di Pasquale A, Puglia I, Savini G. |
| EPI_ISL_528991, EPI_ISL_528992 | Ospedale SS Annunziata-Sulmona | Istituto Zooprofilattico Sperimentale dell'Abruzzo e Molise "G.Caporale" | Lorusso A, Marcacci M, Di Domenico M, Curini V, Ancora M, Cammà C, Rinaldi A, Mangone I, Di Pasquale A, Puglia I, Savini G. |
| EPI_ISL_528993 | Ospedale Civile S. Liberatore-Atri | Istituto Zooprofilattico Sperimentale dell'Abruzzo e Molise "G.Caporale" | Lorusso A, Marcacci M, Di Domenico M, Curini V, Ancora M, Cammà C, Rinaldi A, Mangone I, Di Pasquale A, Puglia I, Savini G. |
| EPI_ISL_528994, EPI_ISL_528995, EPI_ISL_528996, EPI_ISL_528997, EPI_ISL_528998, EPI_ISL_528999, EPI_ISL_529000, EPI_ISL_529001, EPI_ISL_529002, EPI_ISL_529003, EPI_ISL_529004, EPI_ISL_529005 | | | |
| see above | Servizio di igiene epidemiologia e sanità pubblica (SIESP)-Chieti | Istituto Zooprofilattico Sperimentale dell'Abruzzo e Molise "G.Caporale" | Lorusso A, Marcacci M, Di Domenico M, Curini V, Ancora M, Cammà C, Rinaldi A, Mangone I, Di Pasquale A, Puglia I, Savini G. |
| EPI_ISL_529006 | Servizio di igiene e sanità pubblica (SIESP)-Teramo | Istituto Zooprofilattico Sperimentale dell'Abruzzo e Molise "G.Caporale" | Lorusso A, Marcacci M, Di Domenico M, Curini V, Ancora M, Cammà C, Rinaldi A, Mangone I, Di Pasquale A, Puglia I, Savini G. |
| EPI_ISL_529007 | Ospedale Civile S. Liberatore-Atri | Istituto Zooprofilattico Sperimentale dell'Abruzzo e Molise "G.Caporale" | Lorusso A, Marcacci M, Di Domenico M, Curini V, Ancora M, Cammà C, Rinaldi A, Mangone I, Di Pasquale A, Puglia I, Savini G. |
| EPI_ISL_529008 | Servizio di igiene e sanità pubblica (SIESP)-Teramo | Istituto Zooprofilattico Sperimentale dell'Abruzzo e Molise "G.Caporale" | Lorusso A, Marcacci M, Di Domenico M, Curini V, Ancora M, Cammà C, Rinaldi A, Mangone I, Di Pasquale A, Puglia I, Savini G. |
| EPI_ISL_529009 | Ospedale Civile S. Liberatore-Atri | Istituto Zooprofilattico Sperimentale dell'Abruzzo e Molise "G.Caporale" | Lorusso A, Marcacci M, Di Domenico M, Curini V, Ancora M, Cammà C, Rinaldi A, Mangone I, Di Pasquale A, Puglia I, Savini G. |
| EPI_ISL_529010, EPI_ISL_529011, EPI_ISL_529012 | Servizio Igiene Epidemiologia e Sanità Pubblica (SIESP)-L'Aquila | Istituto Zooprofilattico Sperimentale dell'Abruzzo e Molise "G.Caporale" | Lorusso A, Marcacci M, Di Domenico M, Curini V, Ancora M, Cammà C, Rinaldi A, Mangone I, Di Pasquale A, Puglia I, Savini G. |
| EPI_ISL_529013 | Presidio Ospedaliero "S.Filippo e Nicola"-Avezzano | Istituto Zooprofilattico Sperimentale dell'Abruzzo e Molise "G.Caporale" | Lorusso A, Marcacci M, Di Domenico M, Curini V, Ancora M, Cammà C, Rinaldi A, Mangone I, Di Pasquale A, Puglia I, Savini G. |
| EPI_ISL_529014, EPI_ISL_529015 | Ospedale "Ss. Annunziata" | Istituto Zooprofilattico Sperimentale dell'Abruzzo e Molise "G.Caporale" | Lorusso A, Marcacci M, Di Domenico M, Curini V, Ancora M, Cammà C, Rinaldi A, Mangone I, Di Pasquale A, Puglia I, Savini G. |
| EPI_ISL_529016 | Ospedale SS Annunziata-Sulmona | Istituto Zooprofilattico Sperimentale dell'Abruzzo e Molise "G.Caporale" | Lorusso A, Marcacci M, Di Domenico M, Curini V, Ancora M, Cammà C, Rinaldi A, Mangone I, Di Pasquale A, Puglia I, Savini G. |
| EPI_ISL_529017 | RSA San Raffaele Sulmona | Istituto Zooprofilattico Sperimentale dell'Abruzzo e Molise "G.Caporale" | Lorusso A, Marcacci M, Di Domenico M, Curini V, Ancora M, Cammà C, Rinaldi A, Mangone I, Di Pasquale A, Puglia I, Savini G. |
| EPI_ISL_529018 | Ospedale "Giuseppe Mazzini"-Teramo | Istituto Zooprofilattico Sperimentale dell'Abruzzo e Molise "G.Caporale" | Lorusso A, Marcacci M, Di Domenico M, Curini V, Ancora M, Cammà C, Rinaldi A, Mangone I, Di Pasquale A, Puglia I, Savini G. |
| EPI_ISL_529019 | RSA/RP Villa San Giovanni - Gruppo Edos | Istituto Zooprofilattico Sperimentale dell'Abruzzo e Molise "G.Caporale" | Lorusso A, Marcacci M, Di Domenico M, Curini V, Ancora M, Cammà C, Rinaldi A, Mangone I, Di Pasquale A, Puglia I, Savini G. |
| EPI_ISL_529020, EPI_ISL_529021 | Ospedale Civile S. Liberatore-Atri | Istituto Zooprofilattico Sperimentale dell'Abruzzo e Molise "G.Caporale" | Lorusso A, Marcacci M, Di Domenico M, Curini V, Ancora M, Cammà C, Rinaldi A, Mangone I, Di Pasquale A, Puglia I, Savini G. |
| EPI_ISL_529022 | Ospedale "Ss. Annunziata" | Istituto Zooprofilattico Sperimentale dell'Abruzzo e Molise "G.Caporale" | Lorusso A, Marcacci M, Di Domenico M, Curini V, Ancora M, Cammà C, Rinaldi A, Mangone I, Di Pasquale A, Puglia I, Savini G. |
| EPI_ISL_529023, EPI_ISL_529024, EPI_ISL_529025 | Servizio di igiene epidemiologia e sanità pubblica (SIESP)-Chieti | Istituto Zooprofilattico Sperimentale dell'Abruzzo e Molise "G.Caporale" | Lorusso A, Marcacci M, Di Domenico M, Curini V, Ancora M, Cammà C, Rinaldi A, Mangone I, Di Pasquale A, Puglia I, Savini G. |
| EPI_ISL_529026 | Ospedale "Giuseppe Mazzini"-Teramo | Istituto Zooprofilattico Sperimentale dell'Abruzzo e Molise "G.Caporale" | Lorusso A, Marcacci M, Di Domenico M, Curini V, Ancora M, Cammà C, Rinaldi A, Mangone I, Di Pasquale A, Puglia I, Savini G. |
| EPI_ISL_529224, EPI_ISL_529225, EPI_ISL_529236, EPI_ISL_529257, EPI_ISL_529261, EPI_ISL_529265, EPI_ISL_529269, EPI_ISL_529283, EPI_ISL_529299, EPI_ISL_529445, EPI_ISL_529524, EPI_ISL_529656, EPI_ISL_529657, EPI_ISL_529658, EPI_ISL_529659, EPI_ISL_529660 | | | |
| see above | Department of Pathology, University of Cambridge | COVID-19 Genomics UK (COG-UK) Consortium | Luke W Meredith, M. Estée Török, Myra Hosmillo, William L. Hamilton, Martin D. Curran, Theresa Feltwell, Grant Hall, Anna Yakovleva, Fahad A Khokhar, Charlotte J. Houldcroft, Laura G Caller, Aminu S. Jahun, Sarah L. Caddy, Yasmin Chaudhry, Malte Pinckert, Ian Goodfellow |
| EPI_ISL_530122, EPI_ISL_530123 | University Hospital Basel, Clinical Virology | University Hospital Basel, Clinical Bacteriology | Madlen Stange, Alfredo Mari, Tim Roloff, Helena MB Seth-Smith, Michael Schweitzer, Myrta Brunner, Karoline Leuzinger, Kirstine K. Soegaard, Alexander Gensch, Sarah Tschudin-Sutter, Simon Fuchs, Julia Bielicki, Hans Pargger, Martin Siegemund, Christian Nickel, Roland Bingisser, Michael Osthoff, Stefano Bassetti, Rita Schneider-Sliwa, Manuel Battegay, Hans Hirsch, Adrian Egli |
| EPI_ISL_535575 | Hospital Universitari Germans Trias i Pujol(HUGTiP)/Fundació Lluita contra la SIDA (FLSida) | IrsiCaixa AIDS Research Lab | Marc Noguera-Julian, Mariona Parera, Maria Pilar Armengol, Marta Massanella, Ester Ballana, Lidia Ruiz, Nuria Izquierdo, Jorge Carrillo, Roger Paredes, Julia Blanco, Joaquim Segalés, Bonaventura Clotet |
| EPI_ISL_536398 | Lithuanian University of Health Sciences Hospital, Department of Laboratory Medicine | Lithuanian University of Health Sciences, Molecular cardiology lab. | Lukas Zemaitis, Arnoldas Pautienius, Kamile Tamusauskaite, Dovydas Gecys, Vaiva Lesauskaite, Astra Vitkauskiene |
| EPI_ISL_536412, EPI_ISL_536413, EPI_ISL_536414, EPI_ISL_536415, EPI_ISL_536416, EPI_ISL_536417, EPI_ISL_536418, EPI_ISL_536419, EPI_ISL_536420, EPI_ISL_536421, EPI_ISL_536422, EPI_ISL_536423, EPI_ISL_536424, EPI_ISL_536425, EPI_ISL_536426, EPI_ISL_536427, EPI_ISL_536428, EPI_ISL_536429, EPI_ISL_536430, EPI_ISL_536431, EPI_ISL_536432, EPI_ISL_536433, EPI_ISL_536434, EPI_ISL_536435, EPI_ISL_536436, EPI_ISL_536437, EPI_ISL_536438, EPI_ISL_536439, EPI_ISL_536440, EPI_ISL_536441, EPI_ISL_536442, EPI_ISL_536443, EPI_ISL_536444, EPI_ISL_536445, EPI_ISL_536446, EPI_ISL_536447, EPI_ISL_536448, EPI_ISL_536449, EPI_ISL_536450, EPI_ISL_536451, EPI_ISL_536452, EPI_ISL_536453, EPI_ISL_536454 | | | |
| see above | National Public Health Laboratory, National Centre for Infectious Diseases | National Public Health Laboratory, National Centre for Infectious Diseases | Mak TM, Octavia S, Zhou Z, Cui L, Lin RTP |
| EPI_ISL_537691, EPI_ISL_537692, EPI_ISL_537693, EPI_ISL_537694, EPI_ISL_537695, EPI_ISL_537696, EPI_ISL_537697, EPI_ISL_537698, EPI_ISL_537699, EPI_ISL_537700, EPI_ISL_537701, EPI_ISL_537702, EPI_ISL_537703, EPI_ISL_537704, EPI_ISL_537705, EPI_ISL_537706, EPI_ISL_537707, EPI_ISL_537708, EPI_ISL_537709, EPI_ISL_537710, EPI_ISL_537711, EPI_ISL_537712, EPI_ISL_537713, EPI_ISL_537714, EPI_ISL_537715, EPI_ISL_537716, EPI_ISL_537717, EPI_ISL_537718, EPI_ISL_537719, EPI_ISL_537720, EPI_ISL_537721, EPI_ISL_537722, EPI_ISL_537723, EPI_ISL_537724, EPI_ISL_537725, EPI_ISL_537726, EPI_ISL_537727, EPI_ISL_537728, EPI_ISL_537788, EPI_ISL_537789, EPI_ISL_537790, EPI_ISL_537791, EPI_ISL_537792, EPI_ISL_537793, EPI_ISL_537794, EPI_ISL_537795, EPI_ISL_537796, EPI_ISL_537797, EPI_ISL_537798, EPI_ISL_537799, EPI_ISL_537800, EPI_ISL_537801, EPI_ISL_537802, EPI_ISL_537803, EPI_ISL_537804, EPI_ISL_537805, EPI_ISL_537806, EPI_ISL_537807, EPI_ISL_537808 | | | |
| see above | Hospital Universitario de Gran Canaria Dr. Negrín | SeqCOVID-SPAIN consortium/IBV(CSIC) | M. Carmen Pérez González, Francisco J. Chamizo López, Ana Bordes Benítez and SeqCOVID-SPAIN consortium |
| EPI_ISL_538552 | Hospital Universitari Germans Trias i Pujol(HUGTiP)/Fundació Lluita contra la SIDA (FLSida) | IrsiCaixa AIDS Research Lab | Marc Noguera-Julian, Mariona Parera, Maria Pilar Armengol, Marta Massanella, Ester Ballana, Lidia Ruiz, Nuria Izquierdo, Jorge Carrillo, Roger Paredes, Julia Blanco, Joaquim Segalés, Bonaventura Clotet |
| EPI_ISL_538748, EPI_ISL_538749, EPI_ISL_538750, EPI_ISL_538751, EPI_ISL_538752, EPI_ISL_538753, EPI_ISL_538754, EPI_ISL_538755, EPI_ISL_538756, EPI_ISL_538757, EPI_ISL_538758, EPI_ISL_538759, EPI_ISL_538760, EPI_ISL_538761, EPI_ISL_538762, EPI_ISL_538763, EPI_ISL_538764, EPI_ISL_538765, | | | |

| | | | |
|---|---|---|---|
| EPI_ISL_538766, EPI_ISL_538767, EPI_ISL_538768, EPI_ISL_538769, EPI_ISL_538770, EPI_ISL_538771, EPI_ISL_538772, EPI_ISL_538773, EPI_ISL_538774, EPI_ISL_538775, EPI_ISL_538776, EPI_ISL_538777, EPI_ISL_538778, EPI_ISL_538779, EPI_ISL_538780, EPI_ISL_538781, EPI_ISL_538782, EPI_ISL_538783, EPI_ISL_538784, EPI_ISL_538785, EPI_ISL_538786, EPI_ISL_538787, EPI_ISL_538788, EPI_ISL_538789, EPI_ISL_538790, EPI_ISL_538791, EPI_ISL_538792, EPI_ISL_538793, EPI_ISL_538794, EPI_ISL_538795, EPI_ISL_538796, EPI_ISL_538797, EPI_ISL_538798, EPI_ISL_538799, EPI_ISL_538800, EPI_ISL_538801, EPI_ISL_538802, EPI_ISL_538803, EPI_ISL_538804, EPI_ISL_538805, EPI_ISL_538806, EPI_ISL_538807, EPI_ISL_538808, EPI_ISL_538809, EPI_ISL_538810, EPI_ISL_538812, EPI_ISL_538813, EPI_ISL_538814, EPI_ISL_538815, EPI_ISL_538816, EPI_ISL_538817, EPI_ISL_538819, EPI_ISL_538820, EPI_ISL_538821, EPI_ISL_538822, EPI_ISL_538823, EPI_ISL_538824, EPI_ISL_538825, EPI_ISL_538826, EPI_ISL_538827, EPI_ISL_538828, EPI_ISL_538829, EPI_ISL_538830, EPI_ISL_538831, EPI_ISL_538832, EPI_ISL_538833, EPI_ISL_538834, EPI_ISL_538835, EPI_ISL_538836, EPI_ISL_538837, EPI_ISL_538838, EPI_ISL_538839, EPI_ISL_538840, EPI_ISL_538842, EPI_ISL_538843, EPI_ISL_538844, EPI_ISL_538845, EPI_ISL_538846, EPI_ISL_538847, EPI_ISL_538848, EPI_ISL_538849, EPI_ISL_538850, EPI_ISL_538852, EPI_ISL_538853, EPI_ISL_538854, EPI_ISL_538855, EPI_ISL_538856, EPI_ISL_538857, EPI_ISL_538858, EPI_ISL_538859, EPI_ISL_538860, EPI_ISL_538861, EPI_ISL_538862, EPI_ISL_538864, EPI_ISL_538865, EPI_ISL_538867, EPI_ISL_538868, EPI_ISL_538869, EPI_ISL_538870, EPI_ISL_538871, EPI_ISL_538872, EPI_ISL_538873, EPI_ISL_538874, EPI_ISL_538875, EPI_ISL_538876, EPI_ISL_538877, EPI_ISL_538878, EPI_ISL_538879, EPI_ISL_538880, EPI_ISL_538881, EPI_ISL_538882, EPI_ISL_538883, EPI_ISL_538884, EPI_ISL_538885, EPI_ISL_538886, EPI_ISL_538887, EPI_ISL_538888, EPI_ISL_538889, EPI_ISL_538890, EPI_ISL_538891, EPI_ISL_538892, EPI_ISL_538893, EPI_ISL_538894, EPI_ISL_538895, EPI_ISL_538896, EPI_ISL_538897, EPI_ISL_538898, EPI_ISL_538900, EPI_ISL_538901, EPI_ISL_538902, EPI_ISL_538903, EPI_ISL_538904, EPI_ISL_538905, EPI_ISL_538906, EPI_ISL_538907, EPI_ISL_538908, EPI_ISL_538909, EPI_ISL_538910, EPI_ISL_538911, EPI_ISL_538912, EPI_ISL_538913, EPI_ISL_538914, EPI_ISL_538915, EPI_ISL_538916, EPI_ISL_538918, EPI_ISL_538919, EPI_ISL_538920, EPI_ISL_538921, EPI_ISL_538922, EPI_ISL_538923, EPI_ISL_538924, EPI_ISL_538925, EPI_ISL_538926, EPI_ISL_538927, EPI_ISL_538928, EPI_ISL_538929, EPI_ISL_538930, EPI_ISL_538931, EPI_ISL_538932, EPI_ISL_538933, EPI_ISL_538934, EPI_ISL_538935, EPI_ISL_538936, EPI_ISL_538937, EPI_ISL_538938, EPI_ISL_538939, EPI_ISL_538940, EPI_ISL_538941, EPI_ISL_538942, EPI_ISL_538943, EPI_ISL_538944, EPI_ISL_538946, EPI_ISL_538947, EPI_ISL_538948, EPI_ISL_538949, EPI_ISL_538950, EPI_ISL_538951, EPI_ISL_538952, EPI_ISL_538953, EPI_ISL_538954, EPI_ISL_538955, EPI_ISL_538956, EPI_ISL_538957, EPI_ISL_538958, EPI_ISL_538959, EPI_ISL_538960, EPI_ISL_538961, EPI_ISL_538962, EPI_ISL_538963, EPI_ISL_538964, EPI_ISL_538965, EPI_ISL_538966, EPI_ISL_538967, EPI_ISL_538968, EPI_ISL_538969, EPI_ISL_538970, EPI_ISL_538971, EPI_ISL_538972, EPI_ISL_538973, EPI_ISL_538974, EPI_ISL_538975, EPI_ISL_538976, EPI_ISL_538977, EPI_ISL_538978, EPI_ISL_538979, EPI_ISL_538980, EPI_ISL_538981, EPI_ISL_538982, EPI_ISL_538983, EPI_ISL_538984, EPI_ISL_538985, EPI_ISL_538986, EPI_ISL_538988, EPI_ISL_538989, EPI_ISL_538990, EPI_ISL_538991, EPI_ISL_538992, EPI_ISL_538993, EPI_ISL_538995, EPI_ISL_538996, EPI_ISL_538997, EPI_ISL_538998, EPI_ISL_538999, EPI_ISL_539000, EPI_ISL_539001, EPI_ISL_539002, EPI_ISL_539003, EPI_ISL_539004, EPI_ISL_539005, EPI_ISL_539007, EPI_ISL_539008, EPI_ISL_539009, EPI_ISL_539010, EPI_ISL_539011, EPI_ISL_539012, EPI_ISL_539013, EPI_ISL_539014, EPI_ISL_539015, EPI_ISL_539016, EPI_ISL_539017, EPI_ISL_539018, EPI_ISL_539019, EPI_ISL_539020, EPI_ISL_539021, EPI_ISL_539022, EPI_ISL_539023, EPI_ISL_539024, EPI_ISL_539025, EPI_ISL_539026, EPI_ISL_539027, EPI_ISL_539028, EPI_ISL_539029, EPI_ISL_539031, EPI_ISL_539032, EPI_ISL_539033, EPI_ISL_539034, EPI_ISL_539035, EPI_ISL_539036, EPI_ISL_539037, EPI_ISL_539038, EPI_ISL_539039, EPI_ISL_539040, EPI_ISL_539041, EPI_ISL_539042, EPI_ISL_539043, EPI_ISL_539044, EPI_ISL_539045, EPI_ISL_539046, EPI_ISL_539047, EPI_ISL_539048, EPI_ISL_539049, EPI_ISL_539050, EPI_ISL_539051, EPI_ISL_539052, EPI_ISL_539053, EPI_ISL_539054, EPI_ISL_539055, EPI_ISL_539056, EPI_ISL_539057, EPI_ISL_539058, EPI_ISL_539059, EPI_ISL_539060, EPI_ISL_539061, EPI_ISL_539062, EPI_ISL_539063, EPI_ISL_539064, EPI_ISL_539065, EPI_ISL_539066, EPI_ISL_539067, EPI_ISL_539068, EPI_ISL_539069, EPI_ISL_539071, EPI_ISL_539072, EPI_ISL_539073, EPI_ISL_539074, EPI_ISL_539075, EPI_ISL_539076, EPI_ISL_539077, EPI_ISL_539078, EPI_ISL_539079, EPI_ISL_539080, EPI_ISL_539081, EPI_ISL_539082, EPI_ISL_539083, EPI_ISL_539084, EPI_ISL_539085, EPI_ISL_539086, EPI_ISL_539087, EPI_ISL_539088, EPI_ISL_539089, EPI_ISL_539090, EPI_ISL_539091, EPI_ISL_539092, EPI_ISL_539093, EPI_ISL_539094, EPI_ISL_539095, EPI_ISL_539096, EPI_ISL_539097, EPI_ISL_539098, EPI_ISL_539099, EPI_ISL_539100, EPI_ISL_539101, EPI_ISL_539102, EPI_ISL_539103, EPI_ISL_539104, EPI_ISL_539105, EPI_ISL_539106, EPI_ISL_539107, EPI_ISL_539108, EPI_ISL_539109, EPI_ISL_539110, EPI_ISL_539111, EPI_ISL_539112, EPI_ISL_539113, EPI_ISL_539114, EPI_ISL_539115, EPI_ISL_539118, EPI_ISL_539119, EPI_ISL_539121, EPI_ISL_539122, EPI_ISL_539123, EPI_ISL_539124, EPI_ISL_539125, EPI_ISL_539126, EPI_ISL_539127, EPI_ISL_539128, EPI_ISL_539129, EPI_ISL_539130, EPI_ISL_539131, EPI_ISL_539132, EPI_ISL_539133, EPI_ISL_539134, EPI_ISL_539135, EPI_ISL_539136, EPI_ISL_539137, EPI_ISL_539138, EPI_ISL_539139, EPI_ISL_539140, EPI_ISL_539141, EPI_ISL_539142, EPI_ISL_539144, EPI_ISL_539145, EPI_ISL_539146, EPI_ISL_539147, EPI_ISL_539148, EPI_ISL_539149, EPI_ISL_539151, EPI_ISL_539153, EPI_ISL_539154, EPI_ISL_539155, EPI_ISL_539156, EPI_ISL_539158, EPI_ISL_539159, EPI_ISL_539160, EPI_ISL_539161, EPI_ISL_539162, EPI_ISL_539163, EPI_ISL_539164, EPI_ISL_539165, EPI_ISL_539166, EPI_ISL_539167, EPI_ISL_539168, EPI_ISL_539169, EPI_ISL_539170, EPI_ISL_539171, EPI_ISL_539172, EPI_ISL_539173, EPI_ISL_539174, EPI_ISL_539175, EPI_ISL_539176, EPI_ISL_539177, EPI_ISL_539178, EPI_ISL_539179, EPI_ISL_539180, EPI_ISL_539181, EPI_ISL_539182, EPI_ISL_539183, EPI_ISL_539184, EPI_ISL_539186, EPI_ISL_539187, EPI_ISL_539188, EPI_ISL_539189, EPI_ISL_539190, EPI_ISL_539191, EPI_ISL_539192, EPI_ISL_539193, EPI_ISL_539194, EPI_ISL_539195, EPI_ISL_539196, EPI_ISL_539198, EPI_ISL_539199, EPI_ISL_539200, EPI_ISL_539201, EPI_ISL_539202, EPI_ISL_539203, EPI_ISL_539204, EPI_ISL_539205, EPI_ISL_539206, EPI_ISL_539207, EPI_ISL_539208, EPI_ISL_539209, EPI_ISL_539210, EPI_ISL_539211, EPI_ISL_539212, EPI_ISL_539214, EPI_ISL_539215, EPI_ISL_539216, EPI_ISL_539217, EPI_ISL_539218, EPI_ISL_539219, EPI_ISL_539220 | | | |
| see above | Leeds Teaching Hospitals NHS Trust and Public Health England, National Infection Service (Leeds laboratory) | Wellcome Sanger Institute for the COVID-19 Genomics UK (COG-UK) consortium | Louissa Macfarlane-Smith, Holli Carden, Katherine L. Harper, Antony Hale and Alex Alderton, Roberto Amato, Sonia Goncalves, Ewan Harrison, David K. Jackson, Ian Johnston, Dominic Kwiatkowski, Cordelia Langford, John Sillitoe on behalf of the Wellcome Sanger Institute COVID-19 Surveillance Team |
| EPI_ISL_539619 | CSIR-Centre for Cellular and Molecular Biology | CSIR-Centre for Cellular and Molecular Biology | M Soujanya Reddy, Nikhil Hajirnis, Pratheusa Maccha, Namami Gaur, Sakshi Shambhavi, Lamuk Zaveri, Shagufta Khan, Tulasi Nagabandi, Purushotham Vodnala, Payel Mukherjee, Sofia Banu, Priya Singh, Onkar Kulkarni, Dhiviya Vedagiri, Divya Gupta, Vishal Sah, Santosh Kumar Kuncha, Krishnan Harinivas Harshan, Archana Bharadwaj Siva, Karthik Bharadwaj Tallapaka, Zeba Rizvi, Zuberwasim Sayyad, Kakade Aishwarya Arun, Amrutha H C, Ananga Ghosh, Rakesh K Mishra, Divya Tej Sowpati |
| EPI_ISL_539620 | CSIR-Centre for Cellular and Molecular Biology | CSIR-Centre for Cellular and Molecular Biology | M Soujanya Reddy, Nikhil Hajirnis, Pratheusa Maccha, Payel Mukherjee, Sofia Banu, Priya Singh, Onkar Kulkarni,Tulasi Nagabandi, Namami Gaur, Sakshi Shambhavi, Lamuk Zaveri, Shagufta Khan, Purushotham Vodnala, Dhiviya Vedagiri, Divya Gupta, Vishal Sah, Santosh Kumar Kuncha, Krishnan Harinivas Harshan, Archana Bharadwaj Siva, Karthik Bharadwaj Tallapaka,Kezia J Ann, Radhika Khandelwal, Roshan Maku Venkata, Shemin Mansuri, Sonu Uday, Rakesh K Mishra, Divya Tej Sowpati |
| EPI_ISL_539621 | CSIR-Centre for Cellular and Molecular Biology | CSIR-Centre for Cellular and Molecular Biology | M Soujanya Reddy, Nikhil Hajirnis, Pratheusa Maccha, Sakshi Shambhavi, Lamuk Zaveri, Shagufta Khan, Namami Gaur, Tulasi Nagabandi, Purushotham Vodnala, Payel Mukherjee, Sofia Banu, Priya Singh,Onkar Kulkarni, Dhiviya Vedagiri, Divya Gupta, Vishal Sah, Santosh Kumar Kuncha, Krishnan Harinivas Harshan, Archana Bharadwaj Siva, Karthik Bharadwaj Tallapaka, G. Aditya Kumar, Koushick Sivakumar, Pooja Ramesh Gupta, Rajan Kumar Jha, Shraddha Vijay Lahoti, Rakesh K Mishra, Divya Tej Sowpati |
| EPI_ISL_539649 | CSIR-Centre for Cellular and Molecular Biology | CSIR-Centre for Cellular and Molecular Biology | M Soujanya Reddy, Nikhil Hajirnis, Pratheusa Maccha, Namami Gaur, Sakshi Shambhavi, Lamuk Zaveri, Shagufta Khan, Tulasi Nagabandi, Purushotham Vodnala, Payel Mukherjee, Sofia Banu, Priya Singh, Onkar Kulkarni, Dhiviya Vedagiri, Divya Gupta, Vishal Sah, Santosh Kumar Kuncha, Krishnan Harinivas Harshan, Archana Bharadwaj Siva, Karthik Bharadwaj Tallapaka, Zeba Rizvi, Zuberwasim Sayyad, Kakade Aishwarya Arun, Amrutha H C, Ananga Ghosh, Rakesh K Mishra, Divya Tej Sowpati |
| EPI_ISL_539650 | CSIR-Centre for Cellular and Molecular Biology | CSIR-Centre for Cellular and Molecular Biology | M Soujanya Reddy, Nikhil Hajirnis, Pratheusa Maccha, Payel Mukherjee, Sofia Banu, Priya Singh, Onkar Kulkarni,Tulasi Nagabandi, Namami Gaur, Sakshi Shambhavi, Lamuk Zaveri, Shagufta Khan, Purushotham Vodnala, Dhiviya Vedagiri, Divya Gupta, Vishal Sah, Santosh Kumar Kuncha, Krishnan Harinivas Harshan, Archana Bharadwaj Siva, Karthik Bharadwaj Tallapaka,Kezia J Ann, Radhika Khandelwal, Roshan Maku Venkata, Shemin Mansuri, Sonu Uday, Rakesh K Mishra, Divya Tej Sowpati |
| EPI_ISL_539651 | CSIR-Centre for Cellular and Molecular Biology | CSIR-Centre for Cellular and Molecular Biology | M Soujanya Reddy, Nikhil Hajirnis, Pratheusa Maccha, Sakshi Shambhavi, Lamuk Zaveri, Shagufta Khan, Namami Gaur, Tulasi Nagabandi, Purushotham Vodnala, Payel Mukherjee, Sofia Banu, Priya Singh,Onkar Kulkarni, Dhiviya Vedagiri, Divya Gupta, Vishal Sah, Santosh Kumar Kuncha, Krishnan Harinivas Harshan, Archana Bharadwaj Siva, Karthik Bharadwaj Tallapaka, G. Aditya Kumar, Koushick Sivakumar, Pooja Ramesh Gupta, Rajan Kumar Jha, Shraddha Vijay Lahoti, Rakesh K Mishra, Divya Tej Sowpati |
| EPI_ISL_539679 | CSIR-Centre for Cellular and Molecular Biology | CSIR-Centre for Cellular and Molecular Biology | M Soujanya Reddy, Nikhil Hajirnis, Pratheusa Maccha, Namami Gaur, Sakshi Shambhavi, Lamuk Zaveri, Shagufta Khan, Tulasi Nagabandi, Purushotham Vodnala, Payel Mukherjee, Sofia Banu, Priya Singh, Onkar Kulkarni, Dhiviya Vedagiri, Divya Gupta, Vishal Sah, Santosh Kumar Kuncha, Krishnan Harinivas Harshan, Archana Bharadwaj Siva, Karthik Bharadwaj Tallapaka, Zeba Rizvi, Zuberwasim Sayyad, Kakade Aishwarya Arun, Amrutha H C, Ananga Ghosh, Rakesh K Mishra, Divya Tej Sowpati |
| EPI_ISL_539680 | CSIR-Centre for Cellular and Molecular Biology | CSIR-Centre for Cellular and Molecular Biology | M Soujanya Reddy, Nikhil Hajirnis, Pratheusa Maccha, Payel Mukherjee, Sofia Banu, Priya Singh, Onkar Kulkarni,Tulasi Nagabandi, Namami Gaur, Sakshi Shambhavi, Lamuk Zaveri, Shagufta Khan, Purushotham Vodnala, Dhiviya Vedagiri, Divya Gupta, Vishal Sah, Santosh Kumar Kuncha, Krishnan Harinivas Harshan, Archana Bharadwaj Siva, Karthik Bharadwaj Tallapaka,Kezia J Ann, Radhika Khandelwal, Roshan Maku Venkata, Shemin Mansuri, Sonu Uday, Rakesh K Mishra, Divya Tej Sowpati |
| EPI_ISL_539681 | CSIR-Centre for Cellular and Molecular Biology | CSIR-Centre for Cellular and Molecular Biology | M Soujanya Reddy, Nikhil Hajirnis, Pratheusa Maccha, Sakshi Shambhavi, Lamuk Zaveri, Shagufta Khan, Namami Gaur, Tulasi Nagabandi, Purushotham Vodnala, Payel Mukherjee, Sofia Banu, Priya Singh,Onkar Kulkarni, Dhiviya Vedagiri, Divya Gupta, Vishal Sah, Santosh Kumar Kuncha, Krishnan Harinivas Harshan, Archana Bharadwaj Siva, Karthik Bharadwaj Tallapaka, G. Aditya Kumar, Koushick Sivakumar, Pooja Ramesh Gupta, Rajan Kumar Jha, Shraddha Vijay Lahoti, Rakesh K Mishra, Divya Tej Sowpati |
| EPI_ISL_539709 | CSIR-Centre for Cellular and Molecular Biology | CSIR-Centre for Cellular and Molecular Biology | M Soujanya Reddy, Nikhil Hajirnis, Pratheusa Maccha, Namami Gaur, Sakshi Shambhavi, Lamuk Zaveri, Shagufta Khan, Tulasi Nagabandi, Purushotham Vodnala, Payel Mukherjee, Sofia Banu, Priya Singh, Onkar Kulkarni, Dhiviya Vedagiri, Divya Gupta, Vishal Sah, Santosh Kumar Kuncha, Krishnan Harinivas Harshan, Archana Bharadwaj Siva, Karthik Bharadwaj Tallapaka, Zeba Rizvi, Zuberwasim Sayyad, Kakade Aishwarya Arun, Amrutha H C, Ananga Ghosh, Rakesh K Mishra, Divya Tej Sowpati |
| EPI_ISL_539710 | CSIR-Centre for Cellular and Molecular Biology | CSIR-Centre for Cellular and Molecular Biology | M Soujanya Reddy, Nikhil Hajirnis, Pratheusa Maccha, Payel Mukherjee, Sofia Banu, Priya Singh, Onkar Kulkarni,Tulasi Nagabandi, Namami Gaur, Sakshi Shambhavi, Lamuk Zaveri, Shagufta Khan, Purushotham Vodnala, Dhiviya Vedagiri, Divya Gupta, Vishal Sah, Santosh Kumar Kuncha, Krishnan Harinivas Harshan, Archana Bharadwaj Siva, Karthik Bharadwaj Tallapaka,Kezia J Ann, Radhika Khandelwal, Roshan Maku Venkata, Shemin Mansuri, Sonu Uday, Rakesh K Mishra, Divya Tej Sowpati |
| EPI_ISL_539711 | CSIR-Centre for Cellular and Molecular Biology | CSIR-Centre for Cellular and Molecular Biology | M Soujanya Reddy, Nikhil Hajirnis, Pratheusa Maccha, Sakshi Shambhavi, Lamuk Zaveri, Shagufta Khan, Namami Gaur, Tulasi Nagabandi, Purushotham Vodnala, Payel Mukherjee, Sofia Banu, Priya Singh,Onkar Kulkarni, Dhiviya Vedagiri, Divya Gupta, Vishal Sah, Santosh Kumar Kuncha, Krishnan Harinivas Harshan, Archana Bharadwaj Siva, Karthik Bharadwaj Tallapaka, G. Aditya Kumar, Koushick Sivakumar, Pooja Ramesh Gupta, Rajan Kumar Jha, Shraddha Vijay Lahoti, Rakesh K Mishra, Divya Tej Sowpati |
| EPI_ISL_539739 | CSIR-Centre for Cellular and Molecular Biology | CSIR-Centre for Cellular and Molecular Biology | M Soujanya Reddy, Nikhil Hajirnis, Pratheusa Maccha, Namami Gaur, Sakshi Shambhavi, Lamuk Zaveri, Shagufta Khan, Tulasi Nagabandi, Purushotham Vodnala, Payel Mukherjee, Sofia Banu, Priya Singh, Onkar Kulkarni, Dhiviya Vedagiri, Divya Gupta, Vishal Sah, Santosh Kumar Kuncha, Krishnan Harinivas Harshan, Archana Bharadwaj Siva, Karthik Bharadwaj Tallapaka, Zeba Rizvi, Zuberwasim Sayyad, Kakade Aishwarya Arun, Amrutha H C, Ananga Ghosh, Rakesh K Mishra, Divya Tej Sowpati |
| EPI_ISL_539740 | CSIR-Centre for Cellular and Molecular Biology | CSIR-Centre for Cellular and Molecular Biology | M Soujanya Reddy, Nikhil Hajirnis, Pratheusa Maccha, Payel Mukherjee, Sofia Banu, Priya Singh, Onkar Kulkarni,Tulasi Nagabandi, Namami Gaur, Sakshi Shambhavi, Lamuk Zaveri, Shagufta Khan, Purushotham Vodnala, Dhiviya Vedagiri, Divya Gupta, Vishal Sah, Santosh Kumar Kuncha, Krishnan Harinivas |

| | | | |
|---|---|---|---|
| | | | Harshan, Archana Bharadwaj Siva, Karthik Bharadwaj Tallapaka,Kezia J Ann, Radhika Khandelwal, Roshan Maku Venkata, Shemin Mansuri, Sonu Uday, Rakesh K Mishra, Divya Tej Sowpati |
| EPI_ISL_539741 | CSIR-Centre for Cellular and Molecular Biology | CSIR-Centre for Cellular and Molecular Biology | M Soujanya Reddy, Nikhil Hajirnis, Pratheusa Maccha, Sakshi Shambhavi, Lamuk Zaveri, Shagufta Khan, Namami Gaur, Tulasi Nagabandi, Purushotham Vodnala, Payel Mukherjee, Sofia Banu, Priya Singh,Onkar Kulkarni, Dhiviya Vedagiri, Divya Gupta, Vishal Sah, Santosh Kumar Kuncha, Krishnan Harinivas Harshan, Archana Bharadwaj Siva, Karthik Bharadwaj Tallapaka, G. Aditya Kumar, Koushick Sivakumar, Pooja Ramesh Gupta, Rajan Kumar Jha, Shraddha Vijay Lahoti, Rakesh K Mishra, Divya Tej Sowpati |
| EPI_ISL_541847, EPI_ISL_541848, EPI_ISL_541849, EPI_ISL_541850, EPI_ISL_541851, EPI_ISL_541852, EPI_ISL_541853, EPI_ISL_541854, EPI_ISL_541855, EPI_ISL_541856, EPI_ISL_541857, EPI_ISL_541858, EPI_ISL_541859, EPI_ISL_541860, EPI_ISL_541861, EPI_ISL_541862, EPI_ISL_541863, EPI_ISL_541864, EPI_ISL_541865, EPI_ISL_541866, EPI_ISL_541867, EPI_ISL_541868, EPI_ISL_541869, EPI_ISL_541870, EPI_ISL_541871, EPI_ISL_541872, EPI_ISL_541873, EPI_ISL_541874, EPI_ISL_541875, EPI_ISL_541876, EPI_ISL_541877 | | | |
| see above | Lithuanian University of Health Sciences Hospital, Department of Laboratory Medicine | Lithuanian University of Health Sciences, Laboratory of Molecular Cardiology | Lukas Zemaitis, Arnoldas Pautienius, Kamile Tamusauskaite, Dovydas Gecys, Laura Pareckaite, Vaiva Lesauskaite, Astra Vitkauskiene |
| EPI_ISL_541878, EPI_ISL_541879, EPI_ISL_541880, EPI_ISL_541881, EPI_ISL_541882, EPI_ISL_541883, EPI_ISL_541884, EPI_ISL_541885, EPI_ISL_541886, EPI_ISL_541887, EPI_ISL_541888, EPI_ISL_541889, EPI_ISL_541890, EPI_ISL_541891, EPI_ISL_541892, EPI_ISL_541893, EPI_ISL_541894, EPI_ISL_541895, EPI_ISL_541896, EPI_ISL_541897, EPI_ISL_541898, EPI_ISL_541899, EPI_ISL_541900, EPI_ISL_541901, EPI_ISL_541902, EPI_ISL_541903, EPI_ISL_541904, EPI_ISL_541905, EPI_ISL_541906, EPI_ISL_541907, EPI_ISL_541908, EPI_ISL_541909, EPI_ISL_541910, EPI_ISL_541911, EPI_ISL_541912, EPI_ISL_541913, EPI_ISL_541914, EPI_ISL_541915, EPI_ISL_541916, EPI_ISL_541917, EPI_ISL_541918, EPI_ISL_541919, EPI_ISL_541920, EPI_ISL_541921, EPI_ISL_541922, EPI_ISL_541923, EPI_ISL_541924, EPI_ISL_541925, EPI_ISL_541926, EPI_ISL_541927, EPI_ISL_541928, EPI_ISL_541929, EPI_ISL_541930, EPI_ISL_541931, EPI_ISL_541932, EPI_ISL_541933, EPI_ISL_541934, EPI_ISL_541935, EPI_ISL_541936, EPI_ISL_541937, EPI_ISL_541938, EPI_ISL_541939, EPI_ISL_541940, EPI_ISL_541941 | | | |
| see above | Hospital General Universitario Gregorio Marañón | SeqCOVID-SPAIN consortium/IBV(CSIC) | Laura Pérez-Lago, Marta Herranz, Jon Sicilia, Julia Suárez, Pilar Catalán, Patricia Muñoz, Darío García de Viedma and SeqCOVID-SPAIN consortium |
| EPI_ISL_542486 | Walter Reed National Military Medical Center | Naval Medical Research Center Biological Defense Research Directorate | Logan J. Voegtly, Regina Z. Cer, Lindsay A. Glang, Francisco J. Malagon, Kyle A. Long, Derek Larson, Sterling L. Brodniak, Kimberly A Bishop-Lilly |
| EPI_ISL_548972, EPI_ISL_548973, EPI_ISL_548974, EPI_ISL_548975, EPI_ISL_548976, EPI_ISL_548977, EPI_ISL_548978, EPI_ISL_548979, EPI_ISL_548980, EPI_ISL_548981, EPI_ISL_548982, EPI_ISL_548983, EPI_ISL_548984, EPI_ISL_548985, EPI_ISL_548986, EPI_ISL_548987, EPI_ISL_548988, EPI_ISL_548989, EPI_ISL_548990, EPI_ISL_548991, EPI_ISL_548992, EPI_ISL_548993, EPI_ISL_548994, EPI_ISL_548995, EPI_ISL_548996, EPI_ISL_548997, EPI_ISL_548998, EPI_ISL_548999, EPI_ISL_549000, EPI_ISL_549001, EPI_ISL_549002, EPI_ISL_549003, EPI_ISL_549004, EPI_ISL_549005 | | | |
| see above | National Public Health Laboratory, National Centre for Infectious Diseases | National Public Health Laboratory, National Centre for Infectious Diseases | Mak TM, Octavia S, Zhou Z, Cui L, Lin RTP |
| EPI_ISL_561370, EPI_ISL_561371, EPI_ISL_561372 | Hospital Universitario de Gran Canaria Dr. Negrín | SeqCOVID-SPAIN consortium/IBV(CSIC) | M. Carmen Pérez González, Francisco J. Chamizo López, Ana Bordes Benítez and SeqCOVID-SPAIN consortium |
| EPI_ISL_572334, EPI_ISL_572335, EPI_ISL_572336, EPI_ISL_572337, EPI_ISL_572338, EPI_ISL_572339, EPI_ISL_572340, EPI_ISL_572341, EPI_ISL_572342, EPI_ISL_572343, EPI_ISL_572344, EPI_ISL_572345, EPI_ISL_572346, EPI_ISL_572347, EPI_ISL_572348, EPI_ISL_572349, EPI_ISL_572350, EPI_ISL_572351, EPI_ISL_572352, EPI_ISL_572353, EPI_ISL_572354, EPI_ISL_572355, EPI_ISL_572356, EPI_ISL_572357, EPI_ISL_572358, EPI_ISL_572359, EPI_ISL_572360, EPI_ISL_572361, EPI_ISL_572362, EPI_ISL_572363, EPI_ISL_572364, EPI_ISL_572365, EPI_ISL_572366, EPI_ISL_572367, EPI_ISL_572368, EPI_ISL_572369, EPI_ISL_572370, EPI_ISL_572371, EPI_ISL_572372, EPI_ISL_572373, EPI_ISL_572374, EPI_ISL_572375, EPI_ISL_572376, EPI_ISL_572377, EPI_ISL_572378, EPI_ISL_572379, EPI_ISL_572380, EPI_ISL_572381, EPI_ISL_572382, EPI_ISL_572383, EPI_ISL_572384, EPI_ISL_572385, EPI_ISL_572386, EPI_ISL_572387, EPI_ISL_572388, EPI_ISL_572389, EPI_ISL_572390, EPI_ISL_572391, EPI_ISL_572392, EPI_ISL_572393, EPI_ISL_572394, EPI_ISL_572395, EPI_ISL_572396 | | | |
| see above | LACEN/PE | WallauLab, Aggeu Magalhaes Institute | Marcelo Henrique Santos Paiva, Duschinka Ribeiro Duarte Guedes, Cássia Docena, Matheus Filgueira Bezerra, Filipe Zimmer Dezordi, Laís Ceschini Machado, Larissa Krokovsky, Elisama Helvecio, Alexandre Freitas da Silva, Luydson Richardson Silva Vasconcelos, Antonio Mauro Rezende, Severino Jefferson Ribeiro da Silva, Kamila Gaudêncio da Silva Sales, Bruna Santos Lima Figueiredo de Sá, Derciliano Lopes da Cruz, Claudio Eduardo Cavalcanti, Armando de Menezes Neto, Caroline Targino Alves da Silva, Renata Pessôa Germano Mendes, Maria Almerice Lopes da Silva, Tiago Gräf, Paola Cristina Resende, Gonzalo Bello0, Michelle da Silva Barros, Wheverton Ricardo Correia do Nascimento,, Rodrigo Moraes Loyo Arcoverde, Luciane Caroline Albuquerque Bezerra, Sinval Pinto Brandão Filho, Constância Flávia Junqueira Ayres, Gabriel Luz Wallau |
| EPI_ISL_574292 | Instituto Nacional de Investigación en Salud Pública | Instituto Nacional de Investigación en Salud Pública | Leandro Patino Patino, Domenica de Mora Coloma, Denisse Portugal, Alfredo Bruno Caicedo, Andrés Carrasco, Orson Mestanza, Manuel Gonzalez, Alberto Orlando Narvaez. |
| EPI_ISL_574293 | Instituto Nacional de Investigación en Salud Pública | Instituto Nacional de Investigación en Salud Publica | Leandro Patino Patino, Doménica de Mora Coloma, Denisse Portugal, Alfredo Bruno Caicedo,Andrés Carrasco, Orson Mestanza, Manuel Gonzalez, Alberto Orlando Narvaez. |
| EPI_ISL_574294 | Instituto Nacional de Investigacion en Salud Pública | Instituto Nacional de Investigación en Salud Pública | Leandro Patino Patino, Doménica de Mora Coloma, Denisse Portugal, Alfredo Bruno,Andrés Carrasco , Orson Mestanza , Manuel Gonzalez, Alberto Orlando Narvaez |
| EPI_ISL_578698, EPI_ISL_578704, EPI_ISL_578705, EPI_ISL_578706, EPI_ISL_578707, EPI_ISL_578708, EPI_ISL_578709, EPI_ISL_578710, EPI_ISL_578711, EPI_ISL_578712, EPI_ISL_578713, EPI_ISL_578714, EPI_ISL_578715, EPI_ISL_578716, EPI_ISL_578717, EPI_ISL_578718, EPI_ISL_578719, EPI_ISL_578720, EPI_ISL_578721, EPI_ISL_578722, EPI_ISL_578723, EPI_ISL_578724, EPI_ISL_578725, EPI_ISL_578726, EPI_ISL_578727 | | | |
| see above | LSUHS Emerging Viral Threat Laboratory | Microbial Genome Sequencing Center | Maarten Van Diest, Jeremy P. Kamil, Rona S. Scott, Malgorzata Bienkowska-Haba, Katarzyna Zwolinska, Andrew D. Yurochko, Christopher G. Kevil, Martin J. Sapp, Daniel J. Snyder, Vaughn S. Cooper, John A. Vanchiere |
| EPI_ISL_581668, EPI_ISL_581669, EPI_ISL_581670, EPI_ISL_581671, EPI_ISL_581672, EPI_ISL_581673, EPI_ISL_581674, EPI_ISL_581675, EPI_ISL_581676, EPI_ISL_581677, EPI_ISL_581678, EPI_ISL_581679, EPI_ISL_581680, EPI_ISL_581681, EPI_ISL_581682, EPI_ISL_581683, EPI_ISL_581684, EPI_ISL_581685, EPI_ISL_581686, EPI_ISL_581687, EPI_ISL_581688, EPI_ISL_581689, EPI_ISL_581690, EPI_ISL_581691, EPI_ISL_581692, EPI_ISL_581693, EPI_ISL_581694, EPI_ISL_581695, EPI_ISL_581696, EPI_ISL_581697, EPI_ISL_581698, EPI_ISL_581699, EPI_ISL_581700, EPI_ISL_581701, EPI_ISL_581702, EPI_ISL_581703, EPI_ISL_581704, EPI_ISL_581705, EPI_ISL_581706, EPI_ISL_581707, EPI_ISL_581708, EPI_ISL_581709, EPI_ISL_581710, EPI_ISL_581711, EPI_ISL_581712, EPI_ISL_581713, EPI_ISL_581714, EPI_ISL_581715, EPI_ISL_581716, EPI_ISL_581717, EPI_ISL_581718, EPI_ISL_581719, EPI_ISL_581720, EPI_ISL_581721, EPI_ISL_581722, EPI_ISL_581723, EPI_ISL_581724, EPI_ISL_581725, EPI_ISL_581726, EPI_ISL_581727, EPI_ISL_581728, EPI_ISL_581729, EPI_ISL_581730, EPI_ISL_581731, EPI_ISL_581732, EPI_ISL_581733, EPI_ISL_581734, EPI_ISL_581735, EPI_ISL_581736, EPI_ISL_581737, EPI_ISL_581738, EPI_ISL_581739, EPI_ISL_581740, EPI_ISL_581741, EPI_ISL_581742, EPI_ISL_581743, EPI_ISL_581744, EPI_ISL_581745, EPI_ISL_581746, EPI_ISL_581747, EPI_ISL_581748, EPI_ISL_581749, EPI_ISL_581750, EPI_ISL_581751, EPI_ISL_581752, EPI_ISL_581753, EPI_ISL_581754, EPI_ISL_581755, EPI_ISL_581756, EPI_ISL_581757, EPI_ISL_581758, EPI_ISL_581759, EPI_ISL_581760, EPI_ISL_581761, EPI_ISL_581762, EPI_ISL_581763, EPI_ISL_581764, EPI_ISL_581765, EPI_ISL_581766, EPI_ISL_581767, EPI_ISL_581768, EPI_ISL_581769, EPI_ISL_581770, EPI_ISL_581771, EPI_ISL_581772, EPI_ISL_581773, EPI_ISL_581774, EPI_ISL_581775, EPI_ISL_581776, EPI_ISL_581777, EPI_ISL_581778, EPI_ISL_581779, EPI_ISL_581780, EPI_ISL_581781, EPI_ISL_581782, EPI_ISL_581783, EPI_ISL_581784, EPI_ISL_581785, EPI_ISL_581786, EPI_ISL_581787, EPI_ISL_581788, EPI_ISL_581789, EPI_ISL_581791, EPI_ISL_581792, EPI_ISL_581793, EPI_ISL_581794, EPI_ISL_581795, EPI_ISL_581796, EPI_ISL_581797, EPI_ISL_581798, EPI_ISL_581799, EPI_ISL_581800, EPI_ISL_581801, EPI_ISL_581802, EPI_ISL_581803, EPI_ISL_581804, EPI_ISL_581805, EPI_ISL_581806, EPI_ISL_581807, EPI_ISL_581808, EPI_ISL_581809, EPI_ISL_581810, EPI_ISL_581811, EPI_ISL_581812, EPI_ISL_581813, EPI_ISL_581814, EPI_ISL_581815, EPI_ISL_581816, EPI_ISL_581817, EPI_ISL_581818, EPI_ISL_581819, EPI_ISL_581820, EPI_ISL_581821, EPI_ISL_581822, EPI_ISL_581823, EPI_ISL_581824, EPI_ISL_581825, EPI_ISL_581826, EPI_ISL_581827, EPI_ISL_581828, EPI_ISL_581829, EPI_ISL_581830, EPI_ISL_581831, EPI_ISL_581832, EPI_ISL_581833, EPI_ISL_581834, EPI_ISL_581835, EPI_ISL_581836, EPI_ISL_581837, EPI_ISL_581838, EPI_ISL_581839, EPI_ISL_581840, EPI_ISL_581841, EPI_ISL_581842, EPI_ISL_581843, EPI_ISL_581844, EPI_ISL_581845, EPI_ISL_581846, EPI_ISL_581847, EPI_ISL_581848, EPI_ISL_581849, EPI_ISL_581850, EPI_ISL_581851, EPI_ISL_581852, EPI_ISL_581853, EPI_ISL_581854, EPI_ISL_581855, EPI_ISL_581856, EPI_ISL_581857, EPI_ISL_581858, EPI_ISL_581859, EPI_ISL_581860, EPI_ISL_581861, EPI_ISL_581862, EPI_ISL_581863, EPI_ISL_581864, EPI_ISL_581865, EPI_ISL_581866, EPI_ISL_581867, EPI_ISL_581868, EPI_ISL_581869, EPI_ISL_581870, EPI_ISL_581871, EPI_ISL_581872, EPI_ISL_581873, EPI_ISL_581874, EPI_ISL_581875, EPI_ISL_581876, EPI_ISL_581877, EPI_ISL_581878, EPI_ISL_581879, EPI_ISL_581880, EPI_ISL_581881, EPI_ISL_581882, EPI_ISL_581883, EPI_ISL_581884, EPI_ISL_581885, EPI_ISL_581886, EPI_ISL_581887, EPI_ISL_581888, EPI_ISL_581889, EPI_ISL_581890, EPI_ISL_581891, EPI_ISL_581892, EPI_ISL_581893, EPI_ISL_581894, EPI_ISL_581895, EPI_ISL_581896, EPI_ISL_581897, EPI_ISL_581898, EPI_ISL_581899, EPI_ISL_581900, EPI_ISL_581901, EPI_ISL_581902, EPI_ISL_581903, EPI_ISL_581904, EPI_ISL_581905, EPI_ISL_581906, EPI_ISL_581907, EPI_ISL_581908, EPI_ISL_581909, EPI_ISL_581910, EPI_ISL_581911, EPI_ISL_581912, EPI_ISL_581913, EPI_ISL_581914, EPI_ISL_581915, EPI_ISL_581916, EPI_ISL_581917, EPI_ISL_581918, EPI_ISL_581919, EPI_ISL_581920, EPI_ISL_581921, EPI_ISL_581922, EPI_ISL_581923, EPI_ISL_581924, EPI_ISL_581925, EPI_ISL_581926, EPI_ISL_581927, EPI_ISL_581928, EPI_ISL_581929, EPI_ISL_581930, EPI_ISL_581931, EPI_ISL_581932, EPI_ISL_581933, EPI_ISL_581934, EPI_ISL_581935, EPI_ISL_581936, EPI_ISL_581937, EPI_ISL_581938, EPI_ISL_581939, EPI_ISL_581940, EPI_ISL_581941, EPI_ISL_581942, EPI_ISL_581943, EPI_ISL_581944, EPI_ISL_581945, EPI_ISL_581946, EPI_ISL_581947, EPI_ISL_581948, EPI_ISL_581949, EPI_ISL_581950, EPI_ISL_581951, EPI_ISL_581952, EPI_ISL_581953, EPI_ISL_581954, EPI_ISL_581955, EPI_ISL_581956, EPI_ISL_581957, EPI_ISL_581958, EPI_ISL_581959, EPI_ISL_581960, EPI_ISL_581961, EPI_ISL_581962, EPI_ISL_581963, EPI_ISL_581964, EPI_ISL_581965, EPI_ISL_581966, EPI_ISL_581967, EPI_ISL_581968, EPI_ISL_581969, EPI_ISL_581970, EPI_ISL_581971, EPI_ISL_581972, EPI_ISL_581973, EPI_ISL_581974, EPI_ISL_581975, EPI_ISL_581976, EPI_ISL_581977, EPI_ISL_581978, EPI_ISL_581979, EPI_ISL_581980, EPI_ISL_581981, EPI_ISL_581982, EPI_ISL_581983, EPI_ISL_581984, EPI_ISL_581985, EPI_ISL_581986, EPI_ISL_581987, EPI_ISL_581988, EPI_ISL_581989, EPI_ISL_581990, EPI_ISL_581991, EPI_ISL_581992, EPI_ISL_581993, EPI_ISL_581994, EPI_ISL_581995, EPI_ISL_581996, EPI_ISL_581997, EPI_ISL_581998, EPI_ISL_581999, EPI_ISL_582000, EPI_ISL_582001, EPI_ISL_582002 | | | |
| see above | University Hospital Basel, Clinical Virology | University Hospital Basel, Clinical Bacteriology | Madlen Stange, Alfredo Mari, Tim Roloff, Helena MB Seth-Smith, Michael Schweitzer, Myrta Brunner, Karoline Leuzinger, Kirstine K. Soegaard, Alexander Gensch, Sarah Tschudin-Sutter, Simon Fuchs, Julia Bielicki, Hans Pargger, Martin Siegemund, Christian Nickel, Roland Bingisser, Michael Osthoff, Stefano Bassetti, Rita Schneider-Sliwa, Manuel Battegay, Hans Hirsch, Adrian Egli |
| EPI_ISL_582695, EPI_ISL_582696, EPI_ISL_582697, EPI_ISL_582698, EPI_ISL_582699, EPI_ISL_582700, EPI_ISL_582701, EPI_ISL_582702, EPI_ISL_582703, EPI_ISL_582704, EPI_ISL_582705, EPI_ISL_582706, EPI_ISL_582707, EPI_ISL_582708, EPI_ISL_582709, EPI_ISL_582710, EPI_ISL_582711, EPI_ISL_582712, EPI_ISL_582713, EPI_ISL_582714, EPI_ISL_582715, EPI_ISL_582716, EPI_ISL_582717, EPI_ISL_582718, EPI_ISL_582719, EPI_ISL_582720, EPI_ISL_582721, EPI_ISL_582722, EPI_ISL_582723, EPI_ISL_582724, EPI_ISL_582725, EPI_ISL_582726, EPI_ISL_582727, EPI_ISL_582728, EPI_ISL_582729, EPI_ISL_582730, EPI_ISL_582731, EPI_ISL_582732, EPI_ISL_582733, EPI_ISL_582734, EPI_ISL_582735, EPI_ISL_582736, EPI_ISL_582737, EPI_ISL_582738, EPI_ISL_582739, EPI_ISL_582740, EPI_ISL_582741, EPI_ISL_582742, EPI_ISL_582743, EPI_ISL_582744, EPI_ISL_582745, EPI_ISL_582746, EPI_ISL_582747, EPI_ISL_582748, EPI_ISL_582749, EPI_ISL_582750, EPI_ISL_582751, EPI_ISL_582752, EPI_ISL_582753, EPI_ISL_582754, EPI_ISL_582755, EPI_ISL_582756, EPI_ISL_582757, EPI_ISL_582758, EPI_ISL_582759, EPI_ISL_582760, EPI_ISL_582761, EPI_ISL_582762, EPI_ISL_582763, EPI_ISL_582764, EPI_ISL_582765, EPI_ISL_582766 | | | |
| see above | Servicio de Microbiología, Hospital Universitario Cruces, Bilbao | SeqCOVID-SPAIN consortium/IBV(CSIC) | Maitane Aranzamendi Zaldumbide and SeqCOVID-SPAIN consortium |
| EPI_ISL_586514 | B.J. Medical College and Civil hospital, Ahmedabad | Gujarat Biotechnology Research Centre | Maharshi Pandya, Nidhi Patel, Nitin Savaliya, Raghawendra Kumar, Dinesh Kumar, Zuber Saiyed, Komal Patel, Labdhi Pandya, Afzal Ansari, Nikha Trivedi, Pranay Shah, Kamlesh J Upadhyay, Sanjay Kapadia, Apurvasinh Puvar, Janvi Raval, Zarna Patel, Monika Gandhi, Pinal Trivedi, R D Dixit, A M Kadri, Harsh Bakshi, Chaitanya Joshi, Madhvi Joshi |
| EPI_ISL_586534 | General Hospital,Chhotaudepur | Gujarat Biotechnology Research Centre | Maharshi Pandya, Nidhi Patel, Nitin Savaliya, Raghawendra Kumar, Dinesh Kumar, Zuber Saiyed, Komal Patel, Labdhi Pandya, Afzal Ansari, Nikha Trivedi, Apurvasinh Puvar, Janvi Raval, Zarna Patel, Monika Gandhi, Pinal Trivedi, Yogesh Parmar, Arti Thakur, Dharak Pandya, R D Dixit, A M Kadri, |

| | | | |
|---|---|---|---|
| | | | Harsh Bakshi, Chaitanya Joshi, Madhvi Joshi |
| EPI_ISL_586553 | Government Hospital,Veraval | Gujarat Biotechnology Research Centre | Maharshi Pandya, Nidhi Patel, Nitin Savaliya, Raghawendra Kumar, Dinesh Kumar, Zuber Saiyed, Komal Patel, Labdhi Pandya, Afzal Ansari, Nikha Trivedi, Jignesh Parmar, Jitendra Bamrotia, Apurvasinh Puvar, Janvi Raval, Zarna Patel, Monika Gandhi, Pinal Trivedi, R D Dixit, A M Kadri, Harsh Bakshi, Chaitanya Joshi, Madhvi Joshi |
| EPI_ISL_591076, EPI_ISL_591077, EPI_ISL_591078, EPI_ISL_591079, EPI_ISL_591080, EPI_ISL_591081, EPI_ISL_591082, EPI_ISL_591091, EPI_ISL_591092, EPI_ISL_591093 | Department of Pathology, University of Cambridge | Wellcome Sanger Institute for the COVID-19 Genomics UK (COG-UK) consortium | Luke W Meredith, M. Estée Török , Myra Hosmillo, William L. Hamilton, Martin D. Curran, Theresa Feltwell, Grant Hall, Anna Yakovleva, Fahad A Khokhar, Charlotte J. Houldcroft, Laura G Caller, Aminu S. Jahun, Sarah L. Caddy, Ian Goodfellow; and Alex Alderton, Roberto Amato, Sonia Goncalves, Ewan Harrison, David K. Jackson, Ian Johnston, Dominic Kwiatkowski, Cordelia Langford, John Sillitoe on behalf of the Wellcome Sanger Institute COVID-19 Surveillance Team |
| EPI_ISL_591542, EPI_ISL_591543, EPI_ISL_591544, EPI_ISL_591545, EPI_ISL_591546, EPI_ISL_591547 | CHU Purpan - Laboratoire de Virologie - Institut Fédératif de Biologie | CHU Purpan - Laboratoire de Virologie - Institut Fédératif de Biologie | Latour J., Ranger N., Dubois M., Carcenac R., Harter A., Boyer P., Tremeaux P., Izopet J. |
| EPI_ISL_593478, EPI_ISL_593479, EPI_ISL_593480 | Brigham and Women's Hospital | Jonathan Li Laboratory | Manish C. Choudhary, James Regan, Jonathan Z. Li |
| EPI_ISL_593481, EPI_ISL_593483, EPI_ISL_593486, EPI_ISL_593487, EPI_ISL_593489, EPI_ISL_593490, EPI_ISL_593491, EPI_ISL_593492, EPI_ISL_593493, EPI_ISL_593494, EPI_ISL_593495, EPI_ISL_593496, EPI_ISL_593497, EPI_ISL_593498, EPI_ISL_593499, EPI_ISL_593501, EPI_ISL_593502, EPI_ISL_593503, EPI_ISL_593504, EPI_ISL_593506, EPI_ISL_593507, EPI_ISL_593508, EPI_ISL_593509, EPI_ISL_593511, EPI_ISL_593512, EPI_ISL_593513, EPI_ISL_593514, EPI_ISL_593515, EPI_ISL_593517, EPI_ISL_593518, EPI_ISL_593519, EPI_ISL_593523, EPI_ISL_593524, EPI_ISL_593525, EPI_ISL_593526, EPI_ISL_593527, EPI_ISL_593528, EPI_ISL_593529, EPI_ISL_593531, EPI_ISL_593532, EPI_ISL_593533, EPI_ISL_593534, EPI_ISL_593536, EPI_ISL_593537, EPI_ISL_593538, EPI_ISL_593539, EPI_ISL_593540, EPI_ISL_593541, EPI_ISL_593543, EPI_ISL_593544, EPI_ISL_593545, EPI_ISL_593546, EPI_ISL_593547, EPI_ISL_593548, EPI_ISL_593549, EPI_ISL_593550, EPI_ISL_593551 | | | |
| see above | Eastern Ontario Regional Laboratory Association | McMaster University | Leanne Mortimer, Hooman Derakhshani, Emily Panousis, Ahmed Draia, Jalees Nasir, Robert Slinger, Andrew G. McArthur |
| EPI_ISL_593553, EPI_ISL_593554, EPI_ISL_593555, EPI_ISL_593556, EPI_ISL_593557, EPI_ISL_593558 | Brigham and Women's Hospital | Jonathan Li Laboratory | Manish C. Choudhary, James Regan, Jonathan Z. Li |
| EPI_ISL_593855, EPI_ISL_593857, EPI_ISL_593858, EPI_ISL_593859, EPI_ISL_593860, EPI_ISL_593861, EPI_ISL_593862, EPI_ISL_593863, EPI_ISL_593864, EPI_ISL_593865, EPI_ISL_593866, EPI_ISL_593867, EPI_ISL_593868, EPI_ISL_593869, EPI_ISL_593870, EPI_ISL_593871, EPI_ISL_593872, EPI_ISL_593873, EPI_ISL_593874, EPI_ISL_593884, EPI_ISL_593885, EPI_ISL_593890, EPI_ISL_593891, EPI_ISL_593892, EPI_ISL_593893, EPI_ISL_593894, EPI_ISL_593895, EPI_ISL_593896, EPI_ISL_593897, EPI_ISL_593898, EPI_ISL_593899, EPI_ISL_593900, EPI_ISL_593901 | | | |
| see above | CHU Purpan - Laboratoire de Virologie - Institut Fédératif de Biologie | CHU Purpan - Laboratoire de Virologie - Institut Fédératif de Biologie | Latour J., Ranger N., Dubois M., Carcenac R., Harter A., Boyer P., Tremeaux P., Izopet J. |
| EPI_ISL_600428, EPI_ISL_600430, EPI_ISL_600431, EPI_ISL_600432, EPI_ISL_600433, EPI_ISL_600434, EPI_ISL_600435, EPI_ISL_600436, EPI_ISL_600437, EPI_ISL_600438, EPI_ISL_600439, EPI_ISL_600440, EPI_ISL_600441, EPI_ISL_600442, EPI_ISL_600443, EPI_ISL_600444, EPI_ISL_600445, EPI_ISL_600446, EPI_ISL_600449, EPI_ISL_600451, EPI_ISL_600454, EPI_ISL_600463, EPI_ISL_600468, EPI_ISL_600471, EPI_ISL_600473, EPI_ISL_600476, EPI_ISL_600478, EPI_ISL_600480, EPI_ISL_600484, EPI_ISL_600492, EPI_ISL_600495, EPI_ISL_600498, EPI_ISL_600500, EPI_ISL_600502, EPI_ISL_600505, EPI_ISL_600507, EPI_ISL_600510, EPI_ISL_600512, EPI_ISL_600515, EPI_ISL_600517, EPI_ISL_600519, EPI_ISL_600524, EPI_ISL_600530, EPI_ISL_600533, EPI_ISL_600535, EPI_ISL_600538, EPI_ISL_600541, EPI_ISL_600543, EPI_ISL_600546, EPI_ISL_600549, EPI_ISL_600552, EPI_ISL_600554, EPI_ISL_600559, EPI_ISL_600562, EPI_ISL_600564, EPI_ISL_600567 | | | |
| see above | Institute of Epidemiology Disease Control And Research | Institute for Developing Science and Health Initiatives | Lauren Cowley, Mokibul Hassan Afrad, Sadia Isfat Ara Rahman, Md. Mahfuz-Al-mamun, Firadausi Qadri, Tahmina Shirin |
| EPI_ISL_602577, EPI_ISL_602578, EPI_ISL_602579, EPI_ISL_602580, EPI_ISL_602581 | SA Pathology | SA Pathology | Lex Leong, Julien Soubrier, Chuan Kok Lim, Song Gao, Mark Turra, Karin Kassahn, Ivan Bastian, Geoff Higgins |
| EPI_ISL_603082, EPI_ISL_603083, EPI_ISL_603084, EPI_ISL_603085, EPI_ISL_603086, EPI_ISL_603087, EPI_ISL_603088, EPI_ISL_603089, EPI_ISL_603090, EPI_ISL_603091, EPI_ISL_603092, EPI_ISL_603093, EPI_ISL_603094, EPI_ISL_603095, EPI_ISL_603096, EPI_ISL_603097, EPI_ISL_603098, EPI_ISL_603099, EPI_ISL_603100, EPI_ISL_603101, EPI_ISL_603102, EPI_ISL_603103, EPI_ISL_603104, EPI_ISL_603105, EPI_ISL_603106, EPI_ISL_603107, EPI_ISL_603108, EPI_ISL_603109, EPI_ISL_603110, EPI_ISL_603111, EPI_ISL_603112, EPI_ISL_603113, EPI_ISL_603114, EPI_ISL_603115, EPI_ISL_603116, EPI_ISL_603117, EPI_ISL_603118, EPI_ISL_603119, EPI_ISL_603120, EPI_ISL_603122 | | | |
| see above | Lithuanian University of Health Sciences Hospital, Department of Laboratory Medicine | Lithuanian University of Health Sciences, Molecular cardiology lab. | Lukas Zemaitis, Ingrida Olendrait, Arnoldas Pautienius, Kamile Tamusauskaite, Dovydas Gecys, Laura Pareckaite, Vaiva Lesauskaite, Astra Vitkauskiene |
| EPI_ISL_603123, EPI_ISL_603124, EPI_ISL_603125, EPI_ISL_603126, EPI_ISL_603127, EPI_ISL_603128 | SA Pathology | SA Pathology | Lex Leong, Julien Soubrier, Chuan Kok Lim, Song Gao, Mark Turra, Karin Kassahn, Ivan Bastian, Geoff Higgins |
| EPI_ISL_603217, EPI_ISL_603218, EPI_ISL_603219, EPI_ISL_603220 | CHU Purpan - Laboratoire de Virologie - Institut Fédératif de Biologie | CHU Purpan - Laboratoire de Virologie - Institut Fédératif de Biologie | Latour J., Ranger N., Dubois M., Carcenac R., Harter A., Boyer P., Tremeaux P., Izopet J. |
| EPI_ISL_605929, EPI_ISL_605930 | Department of Infectious Disease Prevention and Control, Henan Provincial Center for Disease Control and Prevention | Department of Infectious Disease Prevention and Control, Henan Provincial Center for Disease Control and Prevention | Li,X., Lu,S., Wu,B., Hu,X., Li,D., Ye,Y., Huang,X., Guo,W. |
| EPI_ISL_609991, EPI_ISL_609997 | INMI Lazzaro Spallanzani IRCCS | INMI Lazzaro Spallanzani IRCCS | M Rueca, B Bartolini, C.E.M Gruber, F Messina, E Giombini, A Di Caro, MR Capobianchi |
| EPI_ISL_622761, EPI_ISL_622762, EPI_ISL_622763, EPI_ISL_622764, EPI_ISL_622765, EPI_ISL_622766 | SA Pathology | SA Pathology | Lex Leong, Julien Soubrier, Chuan Kok Lim, Song Gao, Mark Turra, Karin Kassahn, Ivan Bastian, Geoff Higgins |

We gratefully acknowledge the following Authors from the Originating laboratories responsible for obtaining the specimens, as well as the Submitting laboratories where the genome data were generated and shared via GISAID, on which this research is based.

All Submitters of data may be contacted directly via www.gisaid.org

Authors are sorted alphabetically.

| Accession ID | Originating Laboratory | Submitting Laboratory | Authors |
|---|---|---|---|
| EPI_ISL_403932, EPI_ISL_403933, EPI_ISL_403934, EPI_ISL_403935, EPI_ISL_403936, EPI_ISL_403937 | Guangdong Provincial Center for Diseases Control and Prevention; Guangdong Provincial Public Health | Department of Microbiology, Guangdong Provincial Center for Diseases Control and Prevention | Min Kang, Jie Wu, Jing Lu, Tao Liu, Baisheng Li, Shujiang Mei, Feng Ruan, Lifeng Lin, Changwen Ke, Haojie Zhong, Yingtao Zhang, Lirong Zou, Xuguang Chen, Qi Zhu, Jianpeng Xiao, Jianxiang Geng, Zhe Liu, Jianxiong Hu, Weilin Zeng, Xing Li, Yuhuang Liao, Xiujuan Tang, Songjian Xiao, Ying Wang, Yingchao Song, Xue Zhuang, Lijun Liang, Guanhao He, Huihong Deng, Tie Song, Jianfeng He, Wenjun Ma |
| EPI_ISL_406531 | Guangdong Provincial Center for Diseases Contorl and Prevention; Guangdong Provinical Public Health | Guangdong Provincial Center for Disease Control and Prevention | Min Kang, Jie Wu, Jing Lu, Tao Liu, Baisheng Li, Shujiang Mei, Feng Ruan, Lifeng Lin, Changwen Ke, Haojie Zhong, Yingtao Zhang, Lirong Zou, Xuguang Chen, Qi Zhu, Jianpeng Xiao, Jianxiang Geng, Zhe Liu, Jianxiong Hu, Weilin Zeng, Xing Li, Yuhuang Liao, Xiujuan Tang, Songjian Xiao, Ying Wang, Yingchao Song, Xue Zhuang, Lijun Liang, Guanhao He, Huihong Deng, Tie Song, Jianfeng He, Wenjun Ma |
| EPI_ISL_406533 | Guangdong Provincial Center for Diseases Contorl and Prevention; Guangdong Provinical Public Health | Guangdong Provincial Center for Diseases Control and Prevention | Min Kang, Jie Wu, Jing Lu, Tao Liu, Baisheng Li, Shujiang Mei, Feng Ruan, Lifeng Lin, Changwen Ke, Haojie Zhong, Yingtao Zhang, Lirong Zou, Xuguang Chen, Qi Zhu, Jianpeng Xiao, Jianxiang Geng, Zhe Liu, Jianxiong Hu, Weilin Zeng, Xing Li, Yuhuang Liao, Xiujuan Tang, Songjian Xiao, Ying Wang, Yingchao Song, Xue Zhuang, Lijun Liang, Guanhao He, Huihong Deng, Tie Song, Jianfeng He, Wenjun Ma |
| EPI_ISL_406534, EPI_ISL_406535, EPI_ISL_406536 | Guangdong Provincial Center for Diseases Control and Prevention; Guangdong Provincial Public Health | Guangdong Provincial Center for Diseases Control and Prevention | Min Kang, Jie Wu, Jing Lu, Tao Liu, Baisheng Li, Shujiang Mei, Feng Ruan, Lifeng Lin, Changwen Ke, Haojie Zhong, Yingtao Zhang, Lirong Zou, Xuguang Chen, Qi Zhu, Jianpeng Xiao, Jianxiang Geng, Zhe Liu, Jianxiong Hu, Weilin Zeng, Xing Li, Yuhuang Liao, Xiujuan Tang, Songjian Xiao, Ying Wang, Yingchao Song, Xue Zhuang, Lijun Liang, Guanhao He, Huihong Deng, Tie Song, Jianfeng He, Wenjun Ma |
| EPI_ISL_406538 | Guangdong Provincial Center for Diseases Control and Prevention;Guangdong Provincial Institute of Public Health | Guangdong Provincial Center for Diseases Control and Prevention | Min Kang, Jie Wu, Jing Lu, Tao Liu, Baisheng Li, Shujiang Mei, Feng Ruan, Lifeng Lin, Changwen Ke, Haojie Zhong, Yingtao Zhang, Lirong Zou, Xuguang Chen, Qi Zhu, Jianpeng Xiao, Jianxiang Geng, Zhe Liu, Jianxiong Hu, Weilin Zeng, Xing Li, Yuhuang Liao, Xiujuan Tang, Songjian Xiao, Ying Wang, Yingchao Song, Xue Zhuang, Lijun Liang, Guanhao He, Huihong Deng, Tie Song, Jianfeng He, Wenjun Ma |
| EPI_ISL_406596, EPI_ISL_406597 | Department of Infectious and Tropical Diseases, Bichat Claude Bernard Hospital, Paris | National Reference Center for Viruses of Respiratory Infections, Institut Pasteur, Paris | Mélanie Albert, Marion Barbet, Sylvie Behillil, Méline Bizard, Angela Brisebarre, Flora Donati, Vincent Enouf, Maud Vanpeene, Sylvie van der Werf, Yazdan Yazdanpanah, Xavier Lescure. |
| EPI_ISL_407071 | Respiratory Virus Unit, Microbiology Services Colindale, Public Health England | Respiratory Virus Unit, Microbiology Services Colindale, Public Health England | Monica Galiano, Shahjahan Miah, Richard Myers, Angie Lackenby, Omolola Akinbami, Tiina Talts, Leena Bhaw, Kirstin Edwards, Jonathan Hubb, Joanna Ellis, Maria Zambon |
| EPI_ISL_407073 | Respiratory Virus Unit, Microbiology Services Colindale, Public Health England | Respiratory Virus Unit, Microbiology Services Colindale, Public Health England | Monica Galiano, Shahjahan Miah, Richard Myers, Angie Lackenby, Omolola Akinbami, Tiina Talts, Leena Bhaw, Kirstin Edwards, Jonathan Hubb, Joanna Ellis, Maria Zambon. |
| EPI_ISL_408430 | Department of Infectious and Tropical Diseases, Bichat Claude Bernard Hospital, Paris | National Reference Center for Viruses of Respiratory Infections, Institut Pasteur, Paris | Mélanie Albert, Marion Barbet, Sylvie Behillil, Méline Bizard, Angela Brisebarre, Flora Donati, Vincent Enouf, Maud Vanpeene, Sylvie van der Werf, Yazdan Yazdanpanah, Xavier Lescure |
| EPI_ISL_408431 | Sorbonne Université, Inserm et Assistance Publique-Hôpitaux de Paris (Pitié Salpétrière) | National Reference Center for Viruses of Respiratory Infections, Institut Pasteur, Paris | Mélanie Albert, Marion Barbet, Sylvie Behillil, Méline Bizard, Angela Brisebarre, Flora Donati, Vincent Enouf, Maud Vanpeene, Sylvie van der Werf, Sonia Burrel, Anne-Geneviève Marcelin, Vincent Calvez, David Boutolleau, Elise Klément, Valérie Pourcher, Eric Caumes. |
| EPI_ISL_410545 | INMI Lazzaro Spallanzani IRCCS | Laboratory of Virology, INMI Lazzaro Spallanzani IRCCS | Maria R. Capobianchi, Cesare E. M. Gruber, Martina Rueca, Barbara Bartolini, Francesco Messina, Emanuela Giombini, Francesca Colavita, Concetta Castilletti, Eleonora Lalle, Fabrizio Carletti, Emanuele Nicastri, Giuseppe Ippolito. |
| EPI_ISL_410546 | INMI Lazzaro Spallanzani IRCCS | Laboratory of Virology, INMI Lazzaro Spallanzani IRCCS | Maria R. Capobianchi, Cesare E. M. Gruber, Martina Rueca, Fabrizio Carletti, Barbara Bartolini, Francesco Messina, Emanuela Giombini, Francesca Colavita, Concetta Castilletti, Eleonora Lalle, Emanuele Nicastri, Giuseppe Ippolito. |
| EPI_ISL_410720 | Department of Infectious and Tropical Diseases, Bichat Claude Bernard Hospital, Paris | National Reference Center for Viruses of Respiratory Infections, Institut Pasteur, Paris | Mélanie Albert, Marion Barbet, Sylvie Behillil, Méline Bizard, Angela Brisebarre, Flora Donati, Vincent Enouf, Maud Vanpeene, Sylvie van der Werf, Yazdan Yazdanpanah, Xavier Lescure. |
| EPI_ISL_410984 | Department of Infectious and Tropical Diseases, Bichat Claude Bernard Hospital, Paris | National Reference Center for Viruses of Respiratory Infections, Institut Pasteur, Paris | Mélanie Albert, Marion Barbet, Sylvie Behillil, Méline Bizard, Angela Brisebarre, Flora Donati, Vincent Enouf, Maud Vanpeene, Sylvie van der Werf, Yazdan Yazdanpanah, Xavier Lescure |
| EPI_ISL_412116 | Respiratory Virus Unit, Microbiology Services Colindale, Public Health England | Respiratory Virus Unit, Microbiology Services Colindale, Public Health England | Monica Galiano, Shahjahan Miah, Angie Lackenby, Omolola Akinbami, Tiina Talts, Leena Bhaw, Richard Myers, Steven Platt, Kirstin Edwards, Jonathan Hubb, Joanna Ellis, Maria Zambon |
| EPI_ISL_413490 | Auckland Hospital | Institute of Environmental Science and Research (ESR) | Matt Storey, Xiaoyun Ren, Gary McAuliffe, Sally Roberts, Matthew Blakiston, Erasmus Smit, Lauren Jelly, Joep de Ligt |
| EPI_ISL_414005, EPI_ISL_414006, EPI_ISL_414007, EPI_ISL_414008, EPI_ISL_414009, EPI_ISL_414010, EPI_ISL_414011, EPI_ISL_414012, EPI_ISL_414013, EPI_ISL_414040, EPI_ISL_414041, EPI_ISL_414042, EPI_ISL_414043, EPI_ISL_414044 | | | |
| see above | Respiratory Virus Unit, Microbiology Services Colindale, Public Health England | Respiratory Virus Unit, Microbiology Services Colindale, Public Health England | Monica Galiano, Shahjahan Miah, Angie Lackenby, Omolola Akinbami, Tiina Talts, Leena Bhaw, Richard Myers, Steven Platt, Kirstin Edwards, Jonathan Hubb, Joanna Ellis, Maria Zambon |
| EPI_ISL_414487 | UCD National Virus Reference Laboratory | UCD National Virus Reference Laboratory | Michael Carr, Gabriel Gonzalez, Jonathan Dean, Suzie Coughlan, Alison Murphy, Kevin Byrne, Ken Wolfe, Jeff Connell, Brendan Loftus, Cillian F De Gascun |
| EPI_ISL_414520, EPI_ISL_414521 | Bundeswehr Institute of Microbiology | Bundeswehr Institute of Microbiology | Mathias C Walter, Markus H Antwerpen and Roman Wölfel |
| EPI_ISL_414522, EPI_ISL_414523, EPI_ISL_414524, EPI_ISL_414525, EPI_ISL_414526 | Respiratory Virus Unit, Microbiology Services Colindale, Public Health England | Respiratory Virus Unit, Microbiology Services Colindale, Public Health England | Monica Galiano, Shahjahan Miah, Angie Lackenby, Omolola Akinbami, Tiina Talts, Leena Bhaw, Richard Myers, Steven Platt, Kirstin Edwards, Jonathan Hubb, Joanna Ellis, Maria Zambon |
| EPI_ISL_414584, EPI_ISL_414585, EPI_ISL_414586, EPI_ISL_414587 | UCD National Virus Reference Laboratory | UCD National Virus Reference Laboratory | Michael Carr, Gabriel Gonzalez, Jonathan Dean, Suzie Coughlan, Alison Murphy, Kevin Byrne, Ken Wolfe, Jeff Connell, Brendan Loftus, Cillian F De Gascun |
| EPI_ISL_414588, EPI_ISL_414589, EPI_ISL_414590 | Minnesota Department of Health, Public Health Laboratory | Minnesota Department of Health, Public Health Laboratory | Matt Plumb, Jake Garfin and Xiong Wang |
| EPI_ISL_414600, EPI_ISL_414623 | Laboratoire de Virologie Institut de Virologie - INSERM U 1109 Hôpitaux Universitaires de Strasbourg | National Reference Center for Viruses of Respiratory Infections, Institut Pasteur, Paris | Mélnie Albert, Marion Barbet, Sylvie Behillil, Méline Bizard, Angela Brisebarre, Flora Donati Vincent Enouf, Maud Vanpeene, Sylvie van der Werf, Samira Fafi-Kremer |
| EPI_ISL_414624 | Centre Hositalier Universitaire de Rouen Laboratoire de Virologie | National Reference Center for Viruses of Respiratory Infections, Institut Pasteur, Paris | Mélnie Albert, Marion Barbet, Sylvie Behillil, Méline Bizard, Angela Brisebarre, Flora Donati Vincent Enouf, Maud Vanpeene, Sylvie van der Werf, Jean-Christophe Plantier |
| EPI_ISL_414625 | Centre Hospitalier Régional Universitaire de Nantes Laboratoire de Virologie | National Reference Center for Viruses of Respiratory Infections, Institut Pasteur, Paris | Mélnie Albert, Marion Barbet, Sylvie Behillil, Méline Bizard, Angela Brisebarre, Flora Donati Vincent Enouf, Maud Vanpeene, Sylvie van der Werf, Marianne Coste-Burel |
| EPI_ISL_414626 | unknown | National Reference Center for Viruses of Respiratory Infections, Institut Pasteur, Paris | Mélnie Albert, Marion Barbet, Sylvie Behillil, Méline Bizard, Angela Brisebarre, Flora Donati Vincent Enouf, Maud Vanpeene, Sylvie van der Werf |
| EPI_ISL_414627, EPI_ISL_414628, EPI_ISL_414629, EPI_ISL_414630 | Centre Hospitalier Compiègne Laboratoire de Biologie | National Reference Center for Viruses of Respiratory Infections, Institut Pasteur, Paris | Mélnie Albert, Marion Barbet, Sylvie Behillil, Méline Bizard, Angela Brisebarre, Flora Donati Vincent Enouf, Maud Vanpeene, Sylvie van der Werf, Raulin Olivia |
| EPI_ISL_414631, EPI_ISL_414632 | Hôpital Robert Debré Laboratoire de Virologie | National Reference Center for Viruses of Respiratory Infections, Institut Pasteur, Paris | Mélnie Albert, Marion Barbet, Sylvie Behillil, Méline Bizard, Angela Brisebarre, Flora Donati Vincent Enouf, Maud Vanpeene, Sylvie van der Werf, Laurent Andreoletti |

| | | | |
|---|---|---|---|
| EPI_ISL_414633 | Centre Hospitalier René Dubois Laboratoire de Microbiologie - Bât A | National Reference Center for Viruses of Respiratory Infections, Institut Pasteur, Paris | Mélnie Albert, Marion Barbet, Sylvie Behillil, Méline Bizard, Angela Brisebarre, Flora Donati Vincent Enouf, Maud Vanpeene, Sylvie van der Werf, Pascale Martres |
| EPI_ISL_414634, EPI_ISL_414635, EPI_ISL_414636, EPI_ISL_414637, EPI_ISL_414638 | Centre Hospitalier Compiègne Laboratoire de Biologie | National Reference Center for Viruses of Respiratory Infections, Institut Pasteur, Paris | Mélnie Albert, Marion Barbet, Sylvie Behillil, Méline Bizard, Angela Brisebarre, Flora Donati Vincent Enouf, Maud Vanpeene, Sylvie van der Werf, Raulin Olivia |
| EPI_ISL_414648 | Andersen Lab, The Scripps Research Institute | Andersen Lab, The Scripps Research Institute | Mark Zeller, Catie Anderson, Emily Spender, Sarah Topol, Raphaelle Klitting, Refugio Robles-Sikisaka, Karthik Gangavarapu, Laura Nicholson, Kristian Andersen |
| EPI_ISL_415129, EPI_ISL_415130, EPI_ISL_415131, EPI_ISL_415132, EPI_ISL_415133, EPI_ISL_415134, EPI_ISL_415135, EPI_ISL_415136, EPI_ISL_415137, EPI_ISL_415138, EPI_ISL_415139, EPI_ISL_415140, EPI_ISL_415141, EPI_ISL_415142, EPI_ISL_415143, EPI_ISL_415144, EPI_ISL_415145, EPI_ISL_415146, EPI_ISL_415147, EPI_ISL_415148, EPI_ISL_415149, EPI_ISL_415150 | | | |
| see above | Respiratory Virus Unit, Microbiology Services Colindale, Public Health England | Respiratory Virus Unit, Microbiology Services Colindale, Public Health England | Monica Galiano, Shahjahan Miah, Angie Lackenby, Omolola Akinbami, Tiina Talts, Leena Bhaw, Richard Myers, Steven Platt, Kirstin Edwards, Jonathan Hubb, Joanna Ellis, Maria Zambon |
| EPI_ISL_415629, EPI_ISL_415632, EPI_ISL_415633, EPI_ISL_415634, EPI_ISL_415635, EPI_ISL_415636, EPI_ISL_415637, EPI_ISL_415638, EPI_ISL_415639, EPI_ISL_415640 | Virology Department, Royal Infirmary of Edinburgh, NHS Lothian | Virology Department, Royal Infirmary of Edinburgh, NHS Lothian | McHugh M, Dewar R, O'Toole Á, Rambaut A, Williams TC, Templeton K |
| EPI_ISL_415646 | Department of Virus and Microbiological Special diagnostics, Statens Serum Institut, Copenhagen, Denmark. | ViFU | Morten Rasmussen, Maiken Worsoe Rosenstierne , Anders Fomsgaard |
| EPI_ISL_415647 | Department of Virus and Microbiological Special diagnostics, Statens Serum Institut, Copenhagen, Denmark. | Statens Serum Institute | Morten Rasmussen, Maiken Worsoe Rosenstierne , Anders Fomsgaard |
| EPI_ISL_415648 | Department of Virus and Microbiological Special diagnostics, Statens Serum Institut, Copenhagen, Denmark. | ViFU | Morten Rasmussen, Maiken Worsoe Rosenstierne , Anders Fomsgaard |
| EPI_ISL_415649 | unknown | National Reference Center for Viruses of Respiratory Infections, Institut Pasteur, Paris | Mélnie Albert, Marion Barbet, Sylvie Behillil, Méline Bizard, Angela Brisebarre, Flora Donati Vincent Enouf, Maud Vanpeene, Sylvie van der Werf |
| EPI_ISL_415650 | Hôpital Instruction des Armées - BEGIN | National Reference Center for Viruses of Respiratory Infections, Institut Pasteur, Paris | Mélnie Albert, Marion Barbet, Sylvie Behillil, Méline Bizard, Angela Brisebarre, Flora Donati Vincent Enouf, Maud Vanpeene, Sylvie van der Werf, Christine Bigaillon |
| EPI_ISL_415651, EPI_ISL_415652 | unknown | National Reference Center for Viruses of Respiratory Infections, Institut Pasteur, Paris | Mélnie Albert, Marion Barbet, Sylvie Behillil, Méline Bizard, Angela Brisebarre, Flora Donati Vincent Enouf, Maud Vanpeene, Sylvie van der Werf |
| EPI_ISL_415653, EPI_ISL_415654 | Centre Hospitalier Compiègne Laboratoire de Biologie | National Reference Center for Viruses of Respiratory Infections, Institut Pasteur, Paris | Mélnie Albert, Marion Barbet, Sylvie Behillil, Méline Bizard, Angela Brisebarre, Flora Donati Vincent Enouf, Maud Vanpeene, Sylvie van der Werf, Raulin Olivia |
| EPI_ISL_416140, EPI_ISL_416141, EPI_ISL_416142 | Department of Virus and Microbiological Special diagnostics, Statens Serum Institut, Copenhagen, Denmark. | Statens Serum Institute | Morten Rasmussen, Maiken Worsoe Rosenstierne , Anders Fomsgaard |
| EPI_ISL_416143, EPI_ISL_416144, EPI_ISL_416153 | Department of Virus and Microbiological Special diagnostics, Statens Serum Institut, Copenhagen, Denmark. | ViFU | Morten Rasmussen, Maiken Worsoe Rosenstierne , Anders Fomsgaard |
| EPI_ISL_416457 | Andersen Lab, The Scripps Research Institute | Andersen Lab, The Scripps Research Institute | Mark Zeller, Catie Anderson, Emily Spender, Sarah Topol, Raphaelle Klitting, Refugio Robles-Sikisaka, Karthik Gangavarapu, Laura Nicholson, Kristian Andersen |
| EPI_ISL_416477, EPI_ISL_416478, EPI_ISL_416479 | R. G. Lugar Center for Public Health Research, National Center for Disease Control and Public Health (NCDC) of Georgia. | R. G. Lugar Center for Public Health Research, National Center for Disease Control and Public Health (NCDC) of Georgia. | Marine Murtskhvaladze, Nato Kotaria, Ann Machablishvili, Lela Sabadze, Mari Gavashelidze, Ana Papkiauri, Meri Pantsulaia, Gvantsa Brachveli, Tata Imnadze, Tamar Jashiashvili, Tea Tevdoradze, Ketevan Sidamonidze, Ekaterine Khmaladze, Ekaterine Zhghenti, Roena Sukhiashvili, Mariam Zakalashvili, Lela Urushadze, Magda Dgebuadze, Giorgi Tomashvili, Davit Tsaguria, Ekaterine Zangaladze, Nino Berishvili, Gvantsa Chanturia, Adam Kotorashvili, Maia Alkhazashvili, Irma Burjanadze, Anna Kasradze, Khatuna Zakhashvili, Paata Imnadze, Amiran Gamkrelidze. |
| EPI_ISL_416483 | Servicio de Microbiología. Consorcio Hospital General Universitario de Valencia | Sequencing and Bioinformatics Service and Molecular Epidemiology Research Group. FISABIO-Public Health | Maria Alma Bracho, Maria Dolores Ocete, Concepcion Gimeno, Giuseppe D'Auria, Griselda De Marco, Neris Garcia-Gonzalez, Fernando Gonzalez-Candelas |
| EPI_ISL_416484 | Servicio de Microbiología. Consorcio Hospital General Universitario de Valencia | Sequencing and Bioinformatics Service and Molecular Epidemiology Research Group. FISABIO-Public Health | Maria Dolores Ocete, Concepcion Gimeno, Giuseppe D'Auria, Griselda De Marco, Neris Garcia-Gonzalez, Maria Alma Bracho, Fernando Gonzalez-Candelas |
| EPI_ISL_416493 | CH Jean de Navarre Laboratoire de Biologie | National Reference Center for Viruses of Respiratory Infections, Institut Pasteur, Paris | Mélnie Albert, Marion Barbet, Sylvie Behillil, Méline Bizard, Angela Brisebarre, Flora Donati, Etienne Simon-Lorière, Vincent Enouf, Maud Vanpeene, Sylvie van der Werf |
| EPI_ISL_416494 | Centre Hositalier Universitaire de Rouen Laboratoire de Virologie | National Reference Center for Viruses of Respiratory Infections, Institut Pasteur, Paris | Mélnie Albert, Marion Barbet, Sylvie Behillil, Méline Bizard, Angela Brisebarre, Flora Donati, Etienne Simon-Lorière, Vincent Enouf, Maud Vanpeene, Sylvie van der Werf, Jean-Christophe Plantier |
| EPI_ISL_416495, EPI_ISL_416496, EPI_ISL_416497 | Centre Hospitalier Compiègne Laboratoire de Biologie | National Reference Center for Viruses of Respiratory Infections, Institut Pasteur, Paris | Mélnie Albert, Marion Barbet, Sylvie Behillil, Méline Bizard, Angela Brisebarre, Flora Donati, Etienne Simon-Lorière, Vincent Enouf, Maud Vanpeene, Sylvie van der Werf, Raulin Olivia |
| EPI_ISL_416498 | Institut Médico légal- Hop R. Poincaré | National Reference Center for Viruses of Respiratory Infections, Institut Pasteur, Paris | Mélnie Albert, Marion Barbet, Sylvie Behillil, Méline Bizard, Angela Brisebarre, Flora Donati, Etienne Simon-Lorière, Vincent Enouf, Maud Vanpeene, Sylvie van der Werf |
| EPI_ISL_416499, EPI_ISL_416500 | LABM GH nord Essonne | National Reference Center for Viruses of Respiratory Infections, Institut Pasteur, Paris | Mélnie Albert, Marion Barbet, Sylvie Behillil, Méline Bizard, Angela Brisebarre, Flora Donati, Etienne Simon-Lorière, Vincent Enouf, Maud Vanpeene, Sylvie van der Werf |
| EPI_ISL_416501 | Hopital franco britannique - Service des Urgences | National Reference Center for Viruses of Respiratory Infections, Institut Pasteur, Paris | Mélnie Albert, Marion Barbet, Sylvie Behillil, Méline Bizard, Angela Brisebarre, Flora Donati, Etienne Simon-Lorière, Vincent Enouf, Maud Vanpeene, Sylvie van der Werf |
| EPI_ISL_416502, EPI_ISL_416503, EPI_ISL_416504, EPI_ISL_416505, EPI_ISL_416506, EPI_ISL_416507, EPI_ISL_416508, EPI_ISL_416509, EPI_ISL_416510, EPI_ISL_416511, EPI_ISL_416512, EPI_ISL_416513 | | | |
| see above | CHRU Pontchaillou - Laboratoire de Virologie | National Reference Center for Viruses of Respiratory Infections, Institut Pasteur, Paris | Mélnie Albert, Marion Barbet, Sylvie Behillil, Méline Bizard, Angela Brisebarre, Flora Donati, Etienne Simon-Lorière, Vincent Enouf, Maud Vanpeene, Sylvie van der Werf, Gisèle Lagathu |
| EPI_ISL_416519, EPI_ISL_416526 | Auckland Hospital | Institute of Environmental Science and Research (ESR) | Matt Storey, Xiaoyun Ren, Gary McAuliffe, Sally Roberts, Matthew Blakiston, Erasmus Smit, Lauren Jelly, Joep de Ligt |
| EPI_ISL_416539 | Wellington Hospital | Institute of Environmental Science and Research (ESR) | Matt Storey, Xiaoyun Ren, Craig Thornley, Maxim Bloomfield, Erasmus Smit, Lauren Jelly, Joep de Ligt |
| EPI_ISL_416829 | National Public Health Laboratory | Malaysia Genome Institute | Mohd Noor Mat Isa, Irni Suhayu Sapian, Yusuf Muhammad Noor, Nurhezreen Md Iqbal, Mohd Faizal Abu Bakar, Enizza Kasim, Shamsidar Sopie, Siti Noraini Othman, Azrin Ahmad, Nor Azfa Johari, Norazimah Tajudin, Noorliza Mohamad Noordin, W Afiza W Mohd Arifin, Rehan Shuhada Abu Bakar, Yu Kie Chem, Selvanesan Sengol, Hani Mat Hussin, Shahrul Hisham Zainal Ariffin |
| EPI_ISL_416866 | National Public Health Laboratory | Malaysia Genome Institute | Mohd Noor Mat Isa, Irni Suhayu Sapian, Yusuf Muhammad Noor, Nurhezreen Md Iqbal, Mohd Faizal Abu Bakar, Enizza Kasim, Shamsidar Sopie, Siti Noraini Othman, Azrin Ahmad, Nor Azfa Johari, Norazimah Tajudin, Noorliza Mohamad Noordin, W Afiza W Mohd Arifin, Rehan Shuhada Abu Bakar, Yu Kie Chem, Selvanesan Sengol, Hani Mat Hussin, Shahrul Hisham Zainal Ariffin |
| EPI_ISL_416885, EPI_ISL_416886, EPI_ISL_416907 | National Public Health Laboratory | Malaysia Genome Institute | Mohd Noor Mat Isa, Irni Suhayu Sapian, Yusuf Muhammad Noor, Nurhezreen Md Iqbal, Mohd Faizal Abu Bakar, Enizza Kasim, Shamsidar Sopie, Siti Noraini Othman, Azrin Ahmad, Nor Azfa Johari, Norazimah Tajudin, Noorliza Mohamad Noordin, W Afiza W Mohd Arifin, Rehan Shuhada Abu Bakar, Yu Kie Chem, Selvanesan Sengol, Hani Mat Hussin, Shahrul Hisham Zainal Ariffin |
| EPI_ISL_417189, EPI_ISL_417191, EPI_ISL_417192, EPI_ISL_417194, | Minnesota Department of Health, Public Health Laboratory | Minnesota Department of Health, Public Health Laboratory | Matt Plumb, Jake Garfin and Xiong Wang |

| | | | |
|---|---|---|---|
| EPI_ISL_417196, EPI_ISL_417198 | | | |
| EPI_ISL_417205 | Servicio de Microbiología. Consorcio Hospital General Universitario de Valencia | Sequencing and Bioinformatics Service and Molecular Epidemiology Research Group. FISABIO-Public Health | Maria Alma Bracho, Maria Dolores Ocete, Concepcion Gimeno, Giuseppe D'Auria, Griselda De Marco, Neris Garcia-Gonzalez, Fernando Gonzalez-Candelas |
| EPI_ISL_417206 | Servicio de Microbiologia. Consorcio Hospital General Universitario de Valencia | Sequencing and Bioinformatics Service and Molecular Epidemiology Research Group. FISABIO-Public Health | Maria Alma Bracho, Maria Dolores Ocete, Concepcion Gimeno, Giuseppe D'Auria, Griselda De Marco, Neris Garcia-Gonzalez, Fernando Gonzalez-Candelas |
| EPI_ISL_417213, EPI_ISL_417214, EPI_ISL_417215, EPI_ISL_417216, EPI_ISL_417217, EPI_ISL_417218, EPI_ISL_417219, EPI_ISL_417220, EPI_ISL_417221, EPI_ISL_417222, EPI_ISL_417223, EPI_ISL_417224, EPI_ISL_417225, EPI_ISL_417226, EPI_ISL_417227, EPI_ISL_417228, EPI_ISL_417229, EPI_ISL_417230, EPI_ISL_417231, EPI_ISL_417232, EPI_ISL_417233, EPI_ISL_417234, EPI_ISL_417235, EPI_ISL_417236, EPI_ISL_417237, EPI_ISL_417238, EPI_ISL_417239, EPI_ISL_417240, EPI_ISL_417241, EPI_ISL_417242, EPI_ISL_417243, EPI_ISL_417244, EPI_ISL_417245, EPI_ISL_417246, EPI_ISL_417247, EPI_ISL_417248, EPI_ISL_417249, EPI_ISL_417250, EPI_ISL_417251, EPI_ISL_417252, EPI_ISL_417253, EPI_ISL_417254, EPI_ISL_417255, EPI_ISL_417256, EPI_ISL_417257, EPI_ISL_417258, EPI_ISL_417259, EPI_ISL_417260, EPI_ISL_417261, EPI_ISL_417262, EPI_ISL_417263, EPI_ISL_417264, EPI_ISL_417265, EPI_ISL_417266, EPI_ISL_417267, EPI_ISL_417268, EPI_ISL_417269, EPI_ISL_417270, EPI_ISL_417271, EPI_ISL_417272, EPI_ISL_417273, EPI_ISL_417274, EPI_ISL_417275, EPI_ISL_417276, EPI_ISL_417277, EPI_ISL_417278, EPI_ISL_417279, EPI_ISL_417280, EPI_ISL_417281, EPI_ISL_417282, EPI_ISL_417283, EPI_ISL_417284, EPI_ISL_417285, EPI_ISL_417286, EPI_ISL_417287, EPI_ISL_417288, EPI_ISL_417289, EPI_ISL_417290, EPI_ISL_417291, EPI_ISL_417292, EPI_ISL_417293, EPI_ISL_417294, EPI_ISL_417295, EPI_ISL_417296, EPI_ISL_417297, EPI_ISL_417298, EPI_ISL_417299, EPI_ISL_417300, EPI_ISL_417301, EPI_ISL_417302, EPI_ISL_417303, EPI_ISL_417304, EPI_ISL_417305, EPI_ISL_417306, EPI_ISL_417307, EPI_ISL_417308, EPI_ISL_417309, EPI_ISL_417310, EPI_ISL_417311, EPI_ISL_417312, EPI_ISL_417313, EPI_ISL_417314, EPI_ISL_417315 | | | |
| see above | Respiratory Virus Unit, Microbiology Services Colindale, Public Health England | Respiratory Virus Unit, Microbiology Services Colindale, Public Health England | Monica Galiano, Shahjahan Miah, Angie Lackenby, Omolola Akinbami, Tiina Talts, Leena Bhaw, Richard Myers, Steven Platt, Kirstin Edwards, Jonathan Hubb, Joanna Ellis, Maria Zambon |
| EPI_ISL_417469, EPI_ISL_417470, EPI_ISL_417471, EPI_ISL_417472, EPI_ISL_417473, EPI_ISL_417474, EPI_ISL_417475, EPI_ISL_417476, EPI_ISL_417477, EPI_ISL_417478, EPI_ISL_417479, EPI_ISL_417480, EPI_ISL_417492, EPI_ISL_417493, EPI_ISL_417494, EPI_ISL_417495, EPI_ISL_417496, EPI_ISL_417497, EPI_ISL_417498, EPI_ISL_417499, EPI_ISL_417500, EPI_ISL_417501, EPI_ISL_417502, EPI_ISL_417503 | | | |
| see above | Minnesota Department of Health, Public Health Laboratory | Minnesota Department of Health, Public Health Laboratory | Matt Plumb, Jake Garfin and Xiong Wang |
| EPI_ISL_417877, EPI_ISL_417878, EPI_ISL_417879, EPI_ISL_417880 | Institute of Virology, Biomedical Research Center of the Slovak Academy of Sciences, Bratislava; Public Health Authority of the Slovak Republic, Bratislava | Institute of Virology, Biomedical Research Center of the Slovak Academy of Sciences, Bratislava; Comenius University Science Park, Bratislava | Monika Slávikova, Martina Liková, Sabina Fumaová Havlíková, Juraj Koi, Juraj Kopáek, Elena Tichá, Edita Staroová, Jaroslav Budiš, Werner Krampl, Miroslav Böhmer, Diana Rusáková, Tomáš Szemeš, Boris Klempa |
| EPI_ISL_417921 | INMI Lazzaro Spallanzani IRCCS | Laboratory of Virology, INMI Lazzaro Spallanzani IRCCS | Martina Rueca, Barbara Bartolini, Francesco Messina, Cesare E. M. Gruber, Emanuela Giombini, Maria R. Capobianchi, Fabrizio Carletti, Francesca Colavita, Concetta Castilletti, Eleonora Lalle, Daniele Lapa, Giuseppe Ippolito. |
| EPI_ISL_418218 | Centre Hospitalier Compiègne Laboratoire de Biologie | National Reference Center for Viruses of Respiratory Infections, Institut Pasteur, Paris | Mélanie Albert, Marion Barbet, Sylvie Behillil, Méline Bizard, Angela Brisebarre, Flora Donati, Fabiana Gambaro, Etienne Simon-Lorière, Vincent Enouf, Maud Vanpeene, Sylvie van der Werf, Raulin Olivia |
| EPI_ISL_418219 | CHU - Hôpital Cavale Blanche - Labo. de Virologie | National Reference Center for Viruses of Respiratory Infections, Institut Pasteur, Paris | Mélanie Albert, Marion Barbet, Sylvie Behillil, Méline Bizard, Angela Brisebarre, Flora Donati, Fabiana Gambaro, Etienne Simon-Lorière, Vincent Enouf, Maud Vanpeene, Sylvie van der Werf, Léa Pilorge |
| EPI_ISL_418220, EPI_ISL_418221 | Centre Hospitalier Compiègne Laboratoire de Biologie | National Reference Center for Viruses of Respiratory Infections, Institut Pasteur, Paris | Mélanie Albert, Marion Barbet, Sylvie Behillil, Méline Bizard, Angela Brisebarre, Flora Donati, Fabiana Gambaro, Etienne Simon-Lorière, Vincent Enouf, Maud Vanpeene, Sylvie van der Werf, Raulin Olivia |
| EPI_ISL_418222 | CHRU Bretonneau - Serv. Bacterio-Virol. | National Reference Center for Viruses of Respiratory Infections, Institut Pasteur, Paris | Mélanie Albert, Marion Barbet, Sylvie Behillil, Méline Bizard, Angela Brisebarre, Flora Donati, Fabiana Gambaro, Etienne Simon-Lorière, Vincent Enouf, Maud Vanpeene, Sylvie van der Werf, Julien Marlet |
| EPI_ISL_418223, EPI_ISL_418224, EPI_ISL_418225 | Centre Hospitalier Compiègne Laboratoire de Biologie | National Reference Center for Viruses of Respiratory Infections, Institut Pasteur, Paris | Mélanie Albert, Marion Barbet, Sylvie Behillil, Méline Bizard, Angela Brisebarre, Flora Donati, Fabiana Gambaro, Etienne Simon-Lorière, Vincent Enouf, Maud Vanpeene, Sylvie van der Werf, Raulin Olivia |
| EPI_ISL_418226 | EHPAD - Résidences les Cèdres | National Reference Center for Viruses of Respiratory Infections, Institut Pasteur, Paris | Mélanie Albert, Marion Barbet, Sylvie Behillil, Méline Bizard, Angela Brisebarre, Flora Donati, Etienne Simon-Lorière, Vincent Enouf, Maud Vanpeene, Sylvie van der Werf |
| EPI_ISL_418227, EPI_ISL_418228 | Centre Hospitalier Compiègne Laboratoire de Biologie | National Reference Center for Viruses of Respiratory Infections, Institut Pasteur, Paris | Mélanie Albert, Marion Barbet, Sylvie Behillil, Méline Bizard, Angela Brisebarre, Flora Donati, Etienne Simon-Lorière, Vincent Enouf, Maud Vanpeene, Sylvie van der Werf, Raulin Olivia |
| EPI_ISL_418229 | Hopital franco britannique - Laboratoire | National Reference Center for Viruses of Respiratory Infections, Institut Pasteur, Paris | Mélanie Albert, Marion Barbet, Sylvie Behillil, Méline Bizard, Angela Brisebarre, Flora Donati, Etienne Simon-Lorière, Vincent Enouf, Maud Vanpeene, Sylvie van der Werf, Marianne Asso Bonnet |
| EPI_ISL_418230 | Clinique AVERAY LA BROUSTE, Med. Polyvalente | National Reference Center for Viruses of Respiratory Infections, Institut Pasteur, Paris | Mélanie Albert, Marion Barbet, Sylvie Behillil, Méline Bizard, Angela Brisebarre, Flora Donati, Etienne Simon-Lorière, Vincent Enouf, Maud Vanpeene, Sylvie van der Werf, Elsa Ngwem |
| EPI_ISL_418231 | Centre Hospitalier Compiègne Laboratoire de Biologie | National Reference Center for Viruses of Respiratory Infections, Institut Pasteur, Paris | Mélanie Albert, Marion Barbet, Sylvie Behillil, Méline Bizard, Angela Brisebarre, Flora Donati, Etienne Simon-Lorière, Vincent Enouf, Maud Vanpeene, Sylvie van der Werf, Raulin Olivia |
| EPI_ISL_418232, EPI_ISL_418233 | Service des Urgences | National Reference Center for Viruses of Respiratory Infections, Institut Pasteur, Paris | Mélanie Albert, Marion Barbet, Sylvie Behillil, Méline Bizard, Angela Brisebarre, Flora Donati, Etienne Simon-Lorière, Vincent Enouf, Maud Vanpeene, Sylvie van der Werf, Boubkeur |
| EPI_ISL_418234 | LABM GH nord Essonne | National Reference Center for Viruses of Respiratory Infections, Institut Pasteur, Paris | Mélanie Albert, Marion Barbet, Sylvie Behillil, Méline Bizard, Angela Brisebarre, Flora Donati, Etienne Simon-Lorière, Vincent Enouf, Maud Vanpeene, Sylvie van der Werf, Christine Lambert |
| EPI_ISL_418235 | Cabinet médical | National Reference Center for Viruses of Respiratory Infections, Institut Pasteur, Paris | Mélanie Albert, Marion Barbet, Sylvie Behillil, Méline Bizard, Angela Brisebarre, Flora Donati, Etienne Simon-Lorière, Vincent Enouf, Maud Vanpeene, Sylvie van der Werf |
| EPI_ISL_418236, EPI_ISL_418237, EPI_ISL_418238, EPI_ISL_418239 | Centre Hospitalier Compiègne Laboratoire de Biologie | National Reference Center for Viruses of Respiratory Infections, Institut Pasteur, Paris | Mélanie Albert, Marion Barbet, Sylvie Behillil, Méline Bizard, Angela Brisebarre, Flora Donati, Etienne Simon-Lorière, Vincent Enouf, Maud Vanpeene, Sylvie van der Werf, Raulin Olivia |
| EPI_ISL_418240 | LABM GH nord Essonne | National Reference Center for Viruses of Respiratory Infections, Institut Pasteur, Paris | Mélanie Albert, Marion Barbet, Sylvie Behillil, Méline Bizard, Angela Brisebarre, Flora Donati, Etienne Simon-Lorière, Vincent Enouf, Maud Vanpeene, Sylvie van der Werf, Christine Lambert |
| EPI_ISL_418241, EPI_ISL_418242 | NIC Viral Respiratory Unit - Institut Pasteur of Algeria | National Reference Center for Viruses of Respiratory Infections, Institut Pasteur, Paris | Mélanie Albert, Marion Barbet, Sylvie Behillil, Méline Bizard, Angela Brisebarre, Flora Donati, Etienne Simon-Lorière, Vincent Enouf, Maud Vanpeene, Sylvie van der Werf, Fawzi Derrar |
| EPI_ISL_418263, EPI_ISL_418264, EPI_ISL_418265 | Laboratory of Microbiology, Department of Medicine, National and Kapodistrian University of Athens, Greece | Laboratory of Biology, Department of Medicine, Democritus University of Thrace, Greece | Maria Bampali, Elisavet Gatzidou, Nikolaos Dovrolis, Stavroula Veletza, Nikolaos Spanakis, Ioannis Karakasiliotis |
| EPI_ISL_418516, EPI_ISL_418548, EPI_ISL_418580, EPI_ISL_418581, EPI_ISL_418582, EPI_ISL_418583, EPI_ISL_418584 | UCD National Virus Reference Laboratory | UCD National Virus Reference Laboratory | Michael Carr, Gabriel Gonzalez, Jonathan Dean, Suzie Coughlan, Alison Murphy, Kevin Byrne, Ken Wolfe, Jeff Connell, Brendan Loftus, Cillian F De Gascun |
| EPI_ISL_418667, EPI_ISL_418668, EPI_ISL_418669, EPI_ISL_418670, EPI_ISL_418671, EPI_ISL_418672, EPI_ISL_418673, EPI_ISL_418674, EPI_ISL_418675, EPI_ISL_418676, EPI_ISL_418677, EPI_ISL_418678, EPI_ISL_418679, EPI_ISL_418680, EPI_ISL_418681, EPI_ISL_418682, EPI_ISL_418683, EPI_ISL_418684, EPI_ISL_418685, EPI_ISL_418686, EPI_ISL_418687, EPI_ISL_418688, EPI_ISL_418689, EPI_ISL_418690, EPI_ISL_418691, EPI_ISL_418692, EPI_ISL_418693, EPI_ISL_418694, EPI_ISL_418695, EPI_ISL_418696, EPI_ISL_418697, EPI_ISL_418698, EPI_ISL_418699, EPI_ISL_418700, EPI_ISL_418701, EPI_ISL_418702, EPI_ISL_418703, EPI_ISL_418704, EPI_ISL_418705, EPI_ISL_418706, EPI_ISL_418707, EPI_ISL_418708, EPI_ISL_418709, EPI_ISL_418710, EPI_ISL_418711, EPI_ISL_418712, EPI_ISL_418713, EPI_ISL_418714, EPI_ISL_418715, EPI_ISL_418716, EPI_ISL_418717, EPI_ISL_418718, EPI_ISL_418719, EPI_ISL_418720, EPI_ISL_418721, EPI_ISL_418722, EPI_ISL_418723, EPI_ISL_418724, EPI_ISL_418725, EPI_ISL_418726, EPI_ISL_418727, EPI_ISL_418728, EPI_ISL_418729, EPI_ISL_418730, EPI_ISL_418731, EPI_ISL_418732, EPI_ISL_418733, EPI_ISL_418734, EPI_ISL_418735, EPI_ISL_418736, EPI_ISL_418737, EPI_ISL_418738, EPI_ISL_418739, EPI_ISL_418740, EPI_ISL_418741, EPI_ISL_418742, EPI_ISL_418743, EPI_ISL_418744, EPI_ISL_418745, EPI_ISL_418746, EPI_ISL_418747, EPI_ISL_418748, EPI_ISL_418749, EPI_ISL_418750, EPI_ISL_418751, EPI_ISL_418752, EPI_ISL_418753, EPI_ISL_418754, EPI_ISL_418755, EPI_ISL_418756, EPI_ISL_418757, EPI_ISL_418758, EPI_ISL_418759, EPI_ISL_418760, EPI_ISL_418761, EPI_ISL_418762, EPI_ISL_418763, EPI_ISL_418764, EPI_ISL_418765, EPI_ISL_418766, EPI_ISL_418767, EPI_ISL_418768, EPI_ISL_418769, EPI_ISL_418770 | | | |
| see above | Respiratory Virus Unit, Microbiology Services Colindale, Public Health England | Respiratory Virus Unit, Microbiology Services Colindale, Public Health England | Monica Galiano, Shahjahan Miah, Angie Lackenby, Omolola Akinbami, Tiina Talts, Leena Bhaw, Richard Myers, Steven Platt, Kirstin Edwards, Jonathan Hubb, Joanna Ellis, Maria Zambon |
| EPI_ISL_418968, EPI_ISL_418969, EPI_ISL_418970, EPI_ISL_418971, EPI_ISL_418972, EPI_ISL_418973, EPI_ISL_418974, EPI_ISL_418975, EPI_ISL_418976, EPI_ISL_418977, EPI_ISL_418978, EPI_ISL_418979, EPI_ISL_418980 | | | |
| see above | NYU Langone Health | Department of Pathology and Medicine, New York University School of Medicine | Maria Aguero-Rosenfeld, Margaret Black, John Cadley, Paolo Cotzia, John Chen, Dacia Dimartino, Xiaojun Feng, Adriana Heguy, Megan Hogan, Emily Huang, George Jour, Christian Marier, Matthew T. Maurano, Mark J. Mulligan, Peter Meyn, Jared Pinnell, Sitharam Ramaswami, Amy Rapkiewicz, Marie Samanovic-Golden, Antonio Serrano, Guomiao Shen, Matija Snuderl, Nick Vulpescu, Gael Westby, Paul Zappile, Yutong Zhang |
| EPI_ISL_419388, EPI_ISL_419390, | Minnesota Department of Health, Public Health Laboratory | Minnesota Department of Health, Public Health Laboratory | Matt Plumb, Jake Garfin and Xiong Wang |

| | | | |
|---|---|---|---|
| EPI_ISL_419391, EPI_ISL_419392, EPI_ISL_419393, EPI_ISL_419394, EPI_ISL_419395, EPI_ISL_419396, EPI_ISL_419397 | | | |
| EPI_ISL_419675 | Servicio de Microbiología. Consorcio Hospital General Universitario de Valencia | Sequencing and Bioinformatics Service and Molecular Epidemiology Research Group. FISABIO-Public Health | Maria Alma Bracho, Maria Dolores Ocete, Giuseppe D'Auria, Griselda De Marco, Neris Garcia-Gonzalez, Concepcion Gimeno, Fernando Gonzalez-Candelas |
| EPI_ISL_419676 | Servicio de Microbiología. Consorcio Hospital General Universitario de Valencia | Sequencing and Bioinformatics Service and Molecular Epidemiology Research Group. FISABIO-Public Health | Maria Dolores Ocete, Giuseppe D'Auria, Griselda De Marco, Neris Garcia-Gonzalez, Maria Alma Bracho, Concepcion Gimeno, Fernando Gonzalez-Candelas |
| EPI_ISL_419680 | Servicio de Microbiología. Consorcio Hospital General Universitario de Valencia | Sequencing and Bioinformatics Service and Molecular Epidemiology Research Group. FISABIO-Public Health | Maria Alma Bracho, Maria Dolores Ocete, Giuseppe D'Auria, Griselda De Marco, Neris Garcia-Gonzalez, Concepcion Gimeno, Fernando Gonzalez-Candelas |
| EPI_ISL_419681 | Servicio de Microbiología. Consorcio Hospital General Universitario de Valencia | Sequencing and Bioinformatics Service and Molecular Epidemiology Research Group. FISABIO-Public Health | Maria Dolores Ocete, Giuseppe D'Auria, Griselda De Marco, Neris Garcia-Gonzalez, Maria Alma Bracho, Concepcion Gimeno, Fernando Gonzalez-Candelas |
| EPI_ISL_419685 | Servicio de Microbiología. Consorcio Hospital General Universitario de Valencia | Sequencing and Bioinformatics Service and Molecular Epidemiology Research Group. FISABIO-Public Health | Maria Alma Bracho, Maria Dolores Ocete, Giuseppe D'Auria, Griselda De Marco, Neris Garcia-Gonzalez, Concepcion Gimeno, Fernando Gonzalez-Candelas |
| EPI_ISL_419686 | Servicio de Microbiología. Consorcio Hospital General Universitario de Valencia | Sequencing and Bioinformatics Service and Molecular Epidemiology Research Group. FISABIO-Public Health | Maria Dolores Ocete, Giuseppe D'Auria, Griselda De Marco, Neris Garcia-Gonzalez, Maria Alma Bracho, Concepcion Gimeno, Fernando Gonzalez-Candelas |
| EPI_ISL_419690 | Servicio de Microbiología. Consorcio Hospital General Universitario de Valencia | Sequencing and Bioinformatics Service and Molecular Epidemiology Research Group. FISABIO-Public Health | Maria Alma Bracho, Maria Dolores Ocete, Giuseppe D'Auria, Griselda De Marco, Neris Garcia-Gonzalez, Concepcion Gimeno, Fernando Gonzalez-Candelas |
| EPI_ISL_419696, EPI_ISL_419697, EPI_ISL_419698, EPI_ISL_419699, EPI_ISL_419700, EPI_ISL_419701, EPI_ISL_419702, EPI_ISL_419704, EPI_ISL_419705 | NYU Langone Health | Departments of Pathology and Medicine, New York University School of Medicine | Maria Aguero-Rosenfeld, Margaret Black, John Cadley, Paolo Cotzia, John Chen, Dacia Dimartino, Xiaojun Feng, Adriana Heguy, Megan Hogan, Emily Huang, George Jour, Christian Marier, Matthew T. Maurano, Mark J. Mulligan, Peter Meyn, Jared Pinnell, Sitharam Ramaswami, Amy Rapkiewicz, Marie Samanovic-Golden, Antonio Serrano, Guomiao Shen, Matija Snuderl, Nick Vulpescu, Gael Westby, Paul Zappile, Yutong Zhang |
| EPI_ISL_420037 | NIC Viral Respiratory Unit - Institut Pasteur of Algeria | National Reference Center for Viruses of Respiratory Infections, Institut Pasteur, Paris | Mélanie Albert, Marion Barbet, Sylvie Behillil, Méline Bizard, Angela Brisebarre, Flora Donati, Etienne Simon-Lorière, Vincent Enouf, Maud Vanpeene, Sylvie van der Werf, Fawzi Derrar |
| EPI_ISL_420038 | Sentinelles network | National Reference Center for Viruses of Respiratory Infections, Institut Pasteur, Paris | Mélanie Albert, Marion Barbet, Sylvie Behillil, Méline Bizard, Angela Brisebarre, Flora Donati, Etienne Simon-Lorière, Vincent Enouf, Maud Vanpeene, Sylvie van der Werf |
| EPI_ISL_420039, EPI_ISL_420040 | L'Air du Temps | National Reference Center for Viruses of Respiratory Infections, Institut Pasteur, Paris | Mélanie Albert, Marion Barbet, Sylvie Behillil, Méline Bizard, Angela Brisebarre, Flora Donati, Etienne Simon-Lorière, Vincent Enouf, Maud Vanpeene, Sylvie van der Werf |
| EPI_ISL_420041 | CH Compiègne Laboratoire de Biologie | National Reference Center for Viruses of Respiratory Infections, Institut Pasteur, Paris | Mélanie Albert, Marion Barbet, Sylvie Behillil, Méline Bizard, Angela Brisebarre, Flora Donati, Etienne Simon-Lorière, Vincent Enouf, Maud Vanpeene, Sylvie van der Werf, Raulin Olivia |
| EPI_ISL_420042 | Service de Biologie clinique | National Reference Center for Viruses of Respiratory Infections, Institut Pasteur, Paris | Mélanie Albert, Marion Barbet, Sylvie Behillil, Méline Bizard, Angela Brisebarre, Flora Donati, Etienne Simon-Lorière, Vincent Enouf, Maud Vanpeene, Sylvie van der Werf |
| EPI_ISL_420043 | CMIP | National Reference Center for Viruses of Respiratory Infections, Institut Pasteur, Paris | Mélanie Albert, Marion Barbet, Sylvie Behillil, Méline Bizard, Angela Brisebarre, Flora Donati, Etienne Simon-Lorière, Vincent Enouf, Maud Vanpeene, Sylvie van der Werf |
| EPI_ISL_420044 | CH Jean de Navarre Laboratoire de Biologie | National Reference Center for Viruses of Respiratory Infections, Institut Pasteur, Paris | Mélanie Albert, Marion Barbet, Sylvie Behillil, Méline Bizard, Angela Brisebarre, Flora Donati, Etienne Simon-Lorière, Vincent Enouf, Maud Vanpeene, Sylvie van der Werf |
| EPI_ISL_420045 | Sentinelles network | National Reference Center for Viruses of Respiratory Infections, Institut Pasteur, Paris | Mélanie Albert, Marion Barbet, Sylvie Behillil, Méline Bizard, Angela Brisebarre, Flora Donati, Etienne Simon-Lorière, Vincent Enouf, Maud Vanpeene, Sylvie van der Werf |
| EPI_ISL_420046, EPI_ISL_420047 | Résidence Villa Caroline | National Reference Center for Viruses of Respiratory Infections, Institut Pasteur, Paris | Mélanie Albert, Marion Barbet, Sylvie Behillil, Méline Bizard, Angela Brisebarre, Flora Donati, Etienne Simon-Lorière, Vincent Enouf, Maud Vanpeene, Sylvie van der Werf |
| EPI_ISL_420048 | Service de Biologie Médicale - BP 125 | National Reference Center for Viruses of Respiratory Infections, Institut Pasteur, Paris | Mélanie Albert, Marion Barbet, Sylvie Behillil, Méline Bizard, Angela Brisebarre, Flora Donati, Etienne Simon-Lorière, Vincent Enouf, Maud Vanpeene, Sylvie van der Werf, Christine Lambert |
| EPI_ISL_420049, EPI_ISL_420050 | CH Compiègne Laboratoire de Biologie | National Reference Center for Viruses of Respiratory Infections, Institut Pasteur, Paris | Mélanie Albert, Marion Barbet, Sylvie Behillil, Méline Bizard, Angela Brisebarre, Flora Donati, Etienne Simon-Lorière, Vincent Enouf, Maud Vanpeene, Sylvie van der Werf, Raulin Olivia |
| EPI_ISL_420051 | Résidence Eleusis | National Reference Center for Viruses of Respiratory Infections, Institut Pasteur, Paris | Mélanie Albert, Marion Barbet, Sylvie Behillil, Méline Bizard, Angela Brisebarre, Flora Donati, Etienne Simon-Lorière, Vincent Enouf, Maud Vanpeene, Sylvie van der Werf |
| EPI_ISL_420052 | Résidence les Marines | National Reference Center for Viruses of Respiratory Infections, Institut Pasteur, Paris | Mélanie Albert, Marion Barbet, Sylvie Behillil, Méline Bizard, Angela Brisebarre, Flora Donati, Etienne Simon-Lorière, Vincent Enouf, Maud Vanpeene, Sylvie van der Werf |
| EPI_ISL_420053 | CH Jean de Navarre Laboratoire de Biologie | National Reference Center for Viruses of Respiratory Infections, Institut Pasteur, Paris | Mélanie Albert, Marion Barbet, Sylvie Behillil, Méline Bizard, Angela Brisebarre, Flora Donati, Etienne Simon-Lorière, Vincent Enouf, Maud Vanpeene, Sylvie van der Werf |
| EPI_ISL_420054 | Résidence de maintenon | National Reference Center for Viruses of Respiratory Infections, Institut Pasteur, Paris | Mélanie Albert, Marion Barbet, Sylvie Behillil, Méline Bizard, Angela Brisebarre, Flora Donati, Etienne Simon-Lorière, Vincent Enouf, Maud Vanpeene, Sylvie van der Werf |
| EPI_ISL_420055 | Sentinelles network | National Reference Center for Viruses of Respiratory Infections, Institut Pasteur, Paris | Mélanie Albert, Marion Barbet, Sylvie Behillil, Méline Bizard, Angela Brisebarre, Flora Donati, Etienne Simon-Lorière, Vincent Enouf, Maud Vanpeene, Sylvie van der Werf |
| EPI_ISL_420056, EPI_ISL_420057 | CH Compiègne Laboratoire de Biologie | National Reference Center for Viruses of Respiratory Infections, Institut Pasteur, Paris | Mélanie Albert, Marion Barbet, Sylvie Behillil, Méline Bizard, Angela Brisebarre, Flora Donati, Etienne Simon-Lorière, Vincent Enouf, Maud Vanpeene, Sylvie van der Werf, Raulin Olivia |
| EPI_ISL_420058, EPI_ISL_420059, EPI_ISL_420060 | Service de Biologie Médicale - BP 125 | National Reference Center for Viruses of Respiratory Infections, Institut Pasteur, Paris | Mélanie Albert, Marion Barbet, Sylvie Behillil, Méline Bizard, Angela Brisebarre, Flora Donati, Etienne Simon-Lorière, Vincent Enouf, Maud Vanpeene, Sylvie van der Werf, Christine Lambert |
| EPI_ISL_420061 | CMIP | National Reference Center for Viruses of Respiratory Infections, Institut Pasteur, Paris | Mélanie Albert, Marion Barbet, Sylvie Behillil, Méline Bizard, Angela Brisebarre, Flora Donati, Etienne Simon-Lorière, Vincent Enouf, Maud Vanpeene, Sylvie van der Werf |
| EPI_ISL_420062 | Service de Biologie Médicale - BP 125 | National Reference Center for Viruses of Respiratory Infections, Institut Pasteur, Paris | Mélanie Albert, Marion Barbet, Sylvie Behillil, Méline Bizard, Angela Brisebarre, Flora Donati, Etienne Simon-Lorière, Vincent Enouf, Maud Vanpeene, Sylvie van der Werf, Christine Lambert |
| EPI_ISL_420063 | Labo BM - Site de Juvisy - Hopital Général | National Reference Center for Viruses of Respiratory Infections, Institut Pasteur, Paris | Mélanie Albert, Marion Barbet, Sylvie Behillil, Méline Bizard, Angela Brisebarre, Flora Donati, Etienne Simon-Lorière, Vincent Enouf, Maud Vanpeene, Sylvie van der Werf |
| EPI_ISL_420064 | Service de Biologie Médicale - BP 125 | National Reference Center for Viruses of Respiratory Infections, Institut Pasteur, Paris | Mélanie Albert, Marion Barbet, Sylvie Behillil, Méline Bizard, Angela Brisebarre, Flora Donati, Etienne Simon-Lorière, Vincent Enouf, Maud Vanpeene, Sylvie van der Werf, Christine Lambert |
| EPI_ISL_420115 | Servicio de Microbiología. Consorcio Hospital General Universitario de Valencia | Sequencing and Bioinformatics Service and Molecular Epidemiology Research Group. FISABIO-Public Health | Marta Pla Diaz, Neris Garcia-Gonzalez, Loreto Ferrús Abad, Inma Galán Vendrell, Paula Ruiz-Hueso, Mariana Reyes-Prieto, Vicente Soriano Chirona, Maria Alma Bracho, Griselda De Marco, Beatriz Beamud, Lidia Ruiz Roldan, Maria Dolores Ocete, Llúcia Martínez-Priego, Concepcion Gimeno, Giuseppe D'Auria, Fernando Gonzalez-Candelas |
| EPI_ISL_420120 | Servicio de Microbiología. Consorcio Hospital General Universitario de Valencia | Sequencing and Bioinformatics Service and Molecular Epidemiology Research Group. FISABIO-Public Health | Mariana Reyes-Prieto, Vicente Soriano Chirona, Maria Alma Bracho, Griselda De Marco, Beatriz Beamud, Lidia Ruiz Roldan, Marta Pla Diaz,Neris Garcia-Gonzalez, Loreto Ferrús Abad, Inma Galán Vendrell, Paula Ruiz-Hueso, Maria Dolores Ocete, Llúcia Martínez-Priego, Concepcion Gimeno, |

| | | | |
|---|---|---|---|
| | | | Giuseppe D'Auria, Fernando Gonzalez-Candelas |
| EPI_ISL_420122 | Servicio de Microbiología. Consorcio Hospital General Universitario de Valencia | Sequencing and Bioinformatics Service and Molecular Epidemiology Research Group. FISABIO-Public Health | Maria Alma Bracho, Griselda De Marco, Beatriz Beamud, Lidia Ruiz Roldan, Marta Pla Diaz, Neris Garcia-Gonzalez, Loreto Ferrús Abad, Inma Galán Vendrell, Paula Ruiz-Hueso, Mariana Reyes-Prieto, Vicente Soriano Chirona, Maria Dolores Ocete, Llúcia Martínez-Priego, Concepcion Gimeno, Giuseppe D'Auria, Fernando Gonzalez-Candelas |
| EPI_ISL_420123 | Servicio de Microbiología. Consorcio Hospital General Universitario de Valencia | Sequencing and Bioinformatics Service and Molecular Epidemiology Research Group. FISABIO-Public Health | Maria Dolores Ocete, Maria Alma Bracho, Griselda De Marco, Beatriz Beamud, Lidia Ruiz Roldan, Marta Pla Diaz, Neris Garcia-Gonzalez, Loreto Ferrús Abad, Inma Galán Vendrell, Paula Ruiz-Hueso, Mariana Reyes-Prieto, Vicente Soriano Chirona, Llúcia Martínez-Priego, Concepcion Gimeno, Giuseppe D'Auria, Fernando Gonzalez-Candelas |
| EPI_ISL_420127 | Servicio de Microbiologia. Hospital Clinico Universitario de Valencia | Sequencing and Bioinformatics Service and Molecular Epidemiology Research Group. FISABIO-Public Health | Maria Alma Bracho, Griselda De Marco, Beatriz Beamud, Lidia Ruiz Roldan, Marta Pla Diaz, Neris Garcia-Gonzalez, Loreto Ferrús Abad, Inma Galán Vendrell, Paula Ruiz-Hueso, Mariana Reyes-Prieto, Vicente Soriano Chirona, David Navarro, Llúcia Martínez-Priego, Giuseppe D'Auria, Fernando Gonzalez-Candelas |
| EPI_ISL_420142 | Department for Virology, Molecular Biology and Genome Research, R. G. Lugar Center for Public Health Research, National Center for Disease Control and Public Health (NCDC) of Georgia. | Department for Virology, Molecular Biology and Genome Research, R. G. Lugar Center for Public Health Research, National Center for Disease Control and Public Health (NCDC) of Georgia. | Marine Murtskhvaladze, Ann Machablishvili, Lela Sabadze, Mari Gavashelidze, Ana Papkiauri, Meri Pantsulaia, Gvantsa Brachveli, Tata Imnadze, Tamar Jashiashvili, Tea Tevdoradze, Ketevan Sidamonidze, Ekaterine Khmaladze, Ekaterine Zhghenti, Roena Sukhiashvili, Mariam Zakalashvili, Lela Urushadze, Magda Dgebuadze, Giorgi Tomashvili, Davit Tsaguria, Ekaterine Zangaladze, Nino Berishvili, Gvantsa Chanturia, Adam Kotorashvili, Maia Alkhazashvili, Irma Burjanadze, Anna Kasradze, Khatuna Zakhashvili, Paata Imnadze, Amiran Gamkrelidze. |
| EPI_ISL_420296, EPI_ISL_420297, EPI_ISL_420298, EPI_ISL_420299, EPI_ISL_420300, EPI_ISL_420301, EPI_ISL_420302, EPI_ISL_420307, EPI_ISL_420308, EPI_ISL_420309 | NYU Langone Health | Departments of Pathology and Medicine, New York University School of Medicine | Maria Aguero-Rosenfeld, Margaret Black, John Cadley, Paolo Cotzia, John Chen, Dacia Dimartino, Xiaojun Feng, Adriana Heguy, Megan Hogan, Emily Huang, George Jour, Christian Marier, Matthew T. Maurano, Mark J. Mulligan, Peter Meyn, Jared Pinnell, Sitharam Ramaswami, Amy Rapkiewicz, Marie Samanovic-Golden, Antonio Serrano, Guomiao Shen, Matija Snuderl, Nick Vulpescu, Gael Westby, Paul Zappile, Yutong Zhang |
| EPI_ISL_420457, EPI_ISL_420458, EPI_ISL_420459, EPI_ISL_420460, EPI_ISL_420461, EPI_ISL_420462, EPI_ISL_420463, EPI_ISL_420464, EPI_ISL_420465, EPI_ISL_420466, EPI_ISL_420467, EPI_ISL_420468, EPI_ISL_420469, EPI_ISL_420470, EPI_ISL_420471, EPI_ISL_420472, EPI_ISL_420473, EPI_ISL_420474, EPI_ISL_420475, EPI_ISL_420476, EPI_ISL_420477, EPI_ISL_420478, EPI_ISL_420479, EPI_ISL_420480, EPI_ISL_420481, EPI_ISL_420482, EPI_ISL_420483, EPI_ISL_420484, EPI_ISL_420485, EPI_ISL_420486, EPI_ISL_420487, EPI_ISL_420488, EPI_ISL_420489, EPI_ISL_420490, EPI_ISL_420491, EPI_ISL_420492, EPI_ISL_420493, EPI_ISL_420494, EPI_ISL_420495, EPI_ISL_420496, EPI_ISL_420497, EPI_ISL_420498, EPI_ISL_420499, EPI_ISL_420500, EPI_ISL_420501, EPI_ISL_420502, EPI_ISL_420503, EPI_ISL_420504, EPI_ISL_420505, EPI_ISL_420506, EPI_ISL_420507, EPI_ISL_420508, EPI_ISL_420509, EPI_ISL_420510, EPI_ISL_420511, EPI_ISL_420512, EPI_ISL_420513, EPI_ISL_420514, EPI_ISL_420515, EPI_ISL_420516, EPI_ISL_420517, EPI_ISL_420518, EPI_ISL_420519, EPI_ISL_420520, EPI_ISL_420521, EPI_ISL_420522, EPI_ISL_420523, EPI_ISL_420524, EPI_ISL_420525, EPI_ISL_420526, EPI_ISL_420527, EPI_ISL_420528, EPI_ISL_420529, EPI_ISL_420530, EPI_ISL_420542 | | | |
| see above | Respiratory Virus Unit, Microbiology Services Colindale, Public Health England | Respiratory Virus Unit, Microbiology Services Colindale, Public Health England | Monica Galiano, Shahjahan Miah, Angie Lackenby, Omolola Akinbami, Tiina Talts, Leena Bhaw, Richard Myers, Steven Platt, Kirstin Edwards, Jonathan Hubb, Joanna Ellis, Maria Zambon |
| EPI_ISL_420570, EPI_ISL_420571, EPI_ISL_420572, EPI_ISL_420573, EPI_ISL_420574, EPI_ISL_420575, EPI_ISL_420576, EPI_ISL_420577, EPI_ISL_420578, EPI_ISL_420579, EPI_ISL_420580, EPI_ISL_420581, EPI_ISL_420582, EPI_ISL_420584, EPI_ISL_420585, EPI_ISL_420586, EPI_ISL_420587, EPI_ISL_420588, EPI_ISL_420589, EPI_ISL_420590, EPI_ISL_420591 | | | |
| see above | NYU Langone Health | Departments of Pathology and Medicine, New York University School of Medicine | Maria Aguero-Rosenfeld, Brendan Belovarac, Margaret Black, Ludovic Boytard, John Cadley, Paolo Cotzia, John Chen, Dacia Dimartino, Xiaojun Feng, Tatyana Gindin, Adriana Heguy, Megan Hogan, Emily Huang, George Jour, Andrew Lytle, Christian Marier, Matthew T. Maurano, Mark J. Mulligan, Peter Meyn, Iman Osman, Jared Pinnell, Sitharam Ramaswami, Amy Rapkiewicz, Marie Samanovic-Golden, Antonio Serrano, Guomiao Shen, Matija Snuderl, Theodore Vougiouklakis, Nick Vulpescu, Gael Westby, Paul Zappile, Yutong Zhang |
| EPI_ISL_420631, EPI_ISL_420632, EPI_ISL_420633, EPI_ISL_420634, EPI_ISL_420635, EPI_ISL_420636, EPI_ISL_420637, EPI_ISL_420638, EPI_ISL_420639, EPI_ISL_420640, EPI_ISL_420641, EPI_ISL_420642, EPI_ISL_420643, EPI_ISL_420644, EPI_ISL_420645, EPI_ISL_420646, EPI_ISL_420647, EPI_ISL_420648, EPI_ISL_420649, EPI_ISL_420650, EPI_ISL_420651, EPI_ISL_420652, EPI_ISL_420653, EPI_ISL_420654, EPI_ISL_420655, EPI_ISL_420656, EPI_ISL_420657, EPI_ISL_420658, EPI_ISL_420659, EPI_ISL_420660, EPI_ISL_420661, EPI_ISL_420662, EPI_ISL_420663, EPI_ISL_420664, EPI_ISL_420665, EPI_ISL_420666, EPI_ISL_420667, EPI_ISL_420668, EPI_ISL_420669, EPI_ISL_420670, EPI_ISL_420671, EPI_ISL_420672, EPI_ISL_420673, EPI_ISL_420674, EPI_ISL_420675, EPI_ISL_420676, EPI_ISL_420677, EPI_ISL_420678, EPI_ISL_420679, EPI_ISL_420680, EPI_ISL_420681, EPI_ISL_420682, EPI_ISL_420683, EPI_ISL_420684, EPI_ISL_420685, EPI_ISL_420686, EPI_ISL_420687, EPI_ISL_420688, EPI_ISL_420689, EPI_ISL_420690, EPI_ISL_420691, EPI_ISL_420692, EPI_ISL_420693, EPI_ISL_420694, EPI_ISL_420695, EPI_ISL_420696, EPI_ISL_420697, EPI_ISL_420698, EPI_ISL_420699, EPI_ISL_420700, EPI_ISL_420701, EPI_ISL_420702, EPI_ISL_420703, EPI_ISL_420704, EPI_ISL_420705, EPI_ISL_420706, EPI_ISL_420707, EPI_ISL_420708, EPI_ISL_420709, EPI_ISL_420710, EPI_ISL_420711, EPI_ISL_420712, EPI_ISL_420713, EPI_ISL_420714, EPI_ISL_420715, EPI_ISL_420716, EPI_ISL_420717, EPI_ISL_420718, EPI_ISL_420719, EPI_ISL_420720, EPI_ISL_420721, EPI_ISL_420722, EPI_ISL_420723, EPI_ISL_420724, EPI_ISL_420725, EPI_ISL_420726, EPI_ISL_420727, EPI_ISL_420728, EPI_ISL_420729, EPI_ISL_420730, EPI_ISL_420731, EPI_ISL_420732, EPI_ISL_420733, EPI_ISL_420734, EPI_ISL_420735, EPI_ISL_420736, EPI_ISL_420737, EPI_ISL_420738, EPI_ISL_420739, EPI_ISL_420740, EPI_ISL_420741, EPI_ISL_420742, EPI_ISL_420743, EPI_ISL_420744, EPI_ISL_420745, EPI_ISL_420746, EPI_ISL_420747, EPI_ISL_420748, EPI_ISL_420749, EPI_ISL_420750, EPI_ISL_420751, EPI_ISL_420752, EPI_ISL_420753, EPI_ISL_420754, EPI_ISL_420755, EPI_ISL_420756, EPI_ISL_420757, EPI_ISL_420758, EPI_ISL_420759, EPI_ISL_420760, EPI_ISL_420761, EPI_ISL_420762, EPI_ISL_420763, EPI_ISL_420764, EPI_ISL_420765, EPI_ISL_420766, EPI_ISL_420767, EPI_ISL_420768, EPI_ISL_420769, EPI_ISL_420770, EPI_ISL_420771, EPI_ISL_420772, EPI_ISL_420773, EPI_ISL_420774, EPI_ISL_420775, EPI_ISL_420776, EPI_ISL_420777 | | | |
| see above | Respiratory Virus Unit, Microbiology Services Colindale, Public Health England | Respiratory Virus Unit, Microbiology Services Colindale, Public Health England | Monica Galiano, Shahjahan Miah, Angie Lackenby, Omolola Akinbami, Tiina Talts, Leena Bhaw, Richard Myers, Steven Platt, Kirstin Edwards, Jonathan Hubb, Joanna Ellis, Maria Zambon |
| EPI_ISL_420898, EPI_ISL_420899, EPI_ISL_420900, EPI_ISL_420901, EPI_ISL_420902, EPI_ISL_420903, EPI_ISL_420904, EPI_ISL_420905, EPI_ISL_420906, EPI_ISL_420907, EPI_ISL_420908, EPI_ISL_420909 | | | |
| see above | Max von Pettenkofer Institute, Virology, National Reference Center for Retroviruses, LMU Munich | Laboratory for Functional Genome Analysis, Dept. Genomics, Gene Center of the LMU Munich | Max Muenchhoff, Stefan Krebs, Alexander Graf, Ashok Varadharajan, Oliver Keppler, Helmut Blum |
| EPI_ISL_420910 | Respiratory Virus Unit, Microbiology Services Colindale, Public Health England | Respiratory Virus Unit, Microbiology Services Colindale, Public Health England | Monica Galiano, Shahjahan Miah, Angie Lackenby, Omolola Akinbami, Tiina Talts, Leena Bhaw, Richard Myers, Steven Platt, Kirstin Edwards, Jonathan Hubb, Joanna Ellis, Maria Zambon |
| EPI_ISL_420911, EPI_ISL_420912 | Max von Pettenkofer Institute, Virology, National Reference Center for Retroviruses, LMU Munich | Laboratory for Functional Genome Analysis, Dept. Genomics, Gene Center of the LMU Munich | Max Muenchhoff, Stefan Krebs, Alexander Graf, Ashok Varadharajan, Oliver Keppler, Helmut Blum |
| EPI_ISL_421500 | CH Compiègne Laboratoire de Biologie | National Reference Center for Viruses of Respiratory Infections, Institut Pasteur, Paris | Mélanie Albert, Marion Barbet, Sylvie Behillil, Méline Bizard, Angela Brisebarre, Flora Donati, Etienne Simon-Lorière, Vincent Enouf, Maud Vanpeene, Sylvie van der Werf, Raulin Olivia |
| EPI_ISL_421501 | Service de Biologie Médicale - BP 125 | National Reference Center for Viruses of Respiratory Infections, Institut Pasteur, Paris | Mélanie Albert, Marion Barbet, Sylvie Behillil, Méline Bizard, Angela Brisebarre, Flora Donati, Etienne Simon-Lorière, Vincent Enouf, Maud Vanpeene, Sylvie van der Werf, Christine Lambert |
| EPI_ISL_421502, EPI_ISL_421503 | Parc des Dames | National Reference Center for Viruses of Respiratory Infections, Institut Pasteur, Paris | Mélanie Albert, Marion Barbet, Sylvie Behillil, Méline Bizard, Angela Brisebarre, Flora Donati, Etienne Simon-Lorière, Vincent Enouf, Maud Vanpeene, Sylvie van der Werf |
| EPI_ISL_421504, EPI_ISL_421505, EPI_ISL_421506 | Service de Biologie Médicale - BP 125 | National Reference Center for Viruses of Respiratory Infections, Institut Pasteur, Paris | Mélanie Albert, Marion Barbet, Sylvie Behillil, Méline Bizard, Angela Brisebarre, Flora Donati, Etienne Simon-Lorière, Vincent Enouf, Maud Vanpeene, Sylvie van der Werf, Christine Lambert |
| EPI_ISL_421507, EPI_ISL_421508 | Le Château de Seine-Port | National Reference Center for Viruses of Respiratory Infections, Institut Pasteur, Paris | Mélanie Albert, Marion Barbet, Sylvie Behillil, Méline Bizard, Angela Brisebarre, Flora Donati, Etienne Simon-Lorière, Vincent Enouf, Maud Vanpeene, Sylvie van der Werf |
| EPI_ISL_421509, EPI_ISL_421510, EPI_ISL_421511 | CH Compiègne Laboratoire de Biologie | National Reference Center for Viruses of Respiratory Infections, Institut Pasteur, Paris | Mélanie Albert, Marion Barbet, Sylvie Behillil, Méline Bizard, Angela Brisebarre, Flora Donati, Etienne Simon-Lorière, Vincent Enouf, Maud Vanpeene, Sylvie van der Werf, Raulin Olivia |
| EPI_ISL_421512 | Service de Biologie Médicale - BP 125 | National Reference Center for Viruses of Respiratory Infections, Institut Pasteur, Paris | Mélanie Albert, Marion Barbet, Sylvie Behillil, Méline Bizard, Angela Brisebarre, Flora Donati, Etienne Simon-Lorière, Vincent Enouf, Maud Vanpeene, Sylvie van der Werf, Christine Lambert |
| EPI_ISL_421513 | Service de Biologie clinique | National Reference Center for Viruses of Respiratory Infections, Institut Pasteur, Paris | Mélanie Albert, Marion Barbet, Sylvie Behillil, Méline Bizard, Angela Brisebarre, Flora Donati, Etienne Simon-Lorière, Vincent Enouf, Maud Vanpeene, Sylvie van der Werf, Christine Lambert |
| EPI_ISL_421514 | Sentinelles network | National Reference Center for Viruses of Respiratory Infections, Institut Pasteur, Paris | Mélanie Albert, Marion Barbet, Sylvie Behillil, Méline Bizard, Angela Brisebarre, Flora Donati, Etienne Simon-Lorière, Vincent Enouf, Maud Vanpeene, Sylvie van der Werf |
| EPI_ISL_421577, EPI_ISL_421578, EPI_ISL_421579, EPI_ISL_421580, EPI_ISL_421581, EPI_ISL_421582, EPI_ISL_421583, EPI_ISL_421584, EPI_ISL_421585, EPI_ISL_421586, EPI_ISL_421587, EPI_ISL_421588, EPI_ISL_421589, EPI_ISL_421590, EPI_ISL_421591, EPI_ISL_421592 | | | |
| see above | NYU Langone Health | Departments of Pathology and Medicine, New York University School of Medicine | Maria Aguero-Rosenfeld, Brendan Belovarac, Margaret Black, Ludovic Boytard, John Cadley, Paolo Cotzia, John Chen, Dacia Dimartino, Xiaojun Feng, Tatyana Gindin, Adriana Heguy, Megan Hogan, Emily Huang, George Jour, Andrew Lytle, Christian Marier, Matthew T. Maurano, Mark J. Mulligan, Peter Meyn, Iman Osman, Jared Pinnell, Sitharam Ramaswami, Amy Rapkiewicz, Marie Samanovic-Golden, Antonio Serrano, Guomiao Shen, Matija Snuderl, Theodore Vougiouklakis, Nick Vulpescu, Gael Westby, Paul Zappile, Yutong Zhang |
| EPI_ISL_421683, EPI_ISL_421684, | Minnesota Department of Health, Public Health Laboratory | Minnesota Department of Health, Public Health Laboratory | Matt Plumb, Jacob Garfin, Xiong Wang |

EPI_ISL_421685, EPI_ISL_421686, EPI_ISL_421687, EPI_ISL_421688, EPI_ISL_421689, EPI_ISL_421690, EPI_ISL_421691, EPI_ISL_421692

EPI_ISL_421704, EPI_ISL_421705, EPI_ISL_421706, EPI_ISL_421707, EPI_ISL_421708, EPI_ISL_421709, EPI_ISL_421710, EPI_ISL_421711, EPI_ISL_421712, EPI_ISL_421713, EPI_ISL_421714, EPI_ISL_421715, EPI_ISL_421716, EPI_ISL_421717, EPI_ISL_421718, EPI_ISL_421719, EPI_ISL_421720, EPI_ISL_421721, EPI_ISL_421722, EPI_ISL_421723, EPI_ISL_421724, EPI_ISL_421725, EPI_ISL_421726, EPI_ISL_421727, EPI_ISL_421728, EPI_ISL_421729, EPI_ISL_421730, EPI_ISL_421731, EPI_ISL_421732, EPI_ISL_421733

| | | | |
|---|---|---|---|
| see above | NYU Langone Health | Departments of Pathology and Medicine, New York University School of Medicine | Maria Aguero-Rosenfeld, Brendan Belovarac, Margaret Black, Ludovic Boytard, John Cadley, Paolo Cotzia, John Chen, Dacia Dimartino, Xiaojun Feng, Tatyana Gindin, Adriana Heguy, Megan Hogan, Emily Huang, George Jour, Andrew Lytle, Christian Marier, Matthew T. Maurano, Mark J. Mulligan, Peter Meyn, Iman Osman, Jared Pinnell, Sitharam Ramaswami, Amy Rapkiewicz, Marie Samanovic-Golden, Antonio Serrano, Guomiao Shen, Matija Snuderl, Theodore Vougiouklakis, Nick Vulpescu, Gael Westby, Paul Zappile, Yutong Zhang |

EPI_ISL_421768, EPI_ISL_421769, EPI_ISL_421770, EPI_ISL_421771, EPI_ISL_421772, EPI_ISL_421773, EPI_ISL_421774, EPI_ISL_421775, EPI_ISL_421776, EPI_ISL_421777, EPI_ISL_421778, EPI_ISL_421779, EPI_ISL_421780, EPI_ISL_421781, EPI_ISL_421782, EPI_ISL_421783, EPI_ISL_421784, EPI_ISL_421785, EPI_ISL_421786, EPI_ISL_421787, EPI_ISL_421788, EPI_ISL_421789, EPI_ISL_421790, EPI_ISL_421791, EPI_ISL_421792, EPI_ISL_421793, EPI_ISL_421794, EPI_ISL_421795, EPI_ISL_421796, EPI_ISL_421797, EPI_ISL_421798, EPI_ISL_421799, EPI_ISL_421800, EPI_ISL_421801, EPI_ISL_421802, EPI_ISL_421803, EPI_ISL_421804, EPI_ISL_421805, EPI_ISL_421806, EPI_ISL_421807, EPI_ISL_421808, EPI_ISL_421809, EPI_ISL_421810, EPI_ISL_421811, EPI_ISL_421812, EPI_ISL_421813, EPI_ISL_421814, EPI_ISL_421815, EPI_ISL_421816, EPI_ISL_421817, EPI_ISL_421818, EPI_ISL_421819, EPI_ISL_421820, EPI_ISL_421821, EPI_ISL_421822, EPI_ISL_421823, EPI_ISL_421824, EPI_ISL_421825, EPI_ISL_421826, EPI_ISL_421827, EPI_ISL_421828, EPI_ISL_421829, EPI_ISL_421830, EPI_ISL_421831, EPI_ISL_421832, EPI_ISL_421833, EPI_ISL_421834, EPI_ISL_421835, EPI_ISL_421836, EPI_ISL_421837, EPI_ISL_421838, EPI_ISL_421839, EPI_ISL_421840, EPI_ISL_421841, EPI_ISL_421842, EPI_ISL_421843, EPI_ISL_421844, EPI_ISL_421845, EPI_ISL_421846, EPI_ISL_421847, EPI_ISL_421848, EPI_ISL_421849, EPI_ISL_421850, EPI_ISL_421851, EPI_ISL_421852, EPI_ISL_421853, EPI_ISL_421854, EPI_ISL_421855, EPI_ISL_421856, EPI_ISL_421857, EPI_ISL_421858, EPI_ISL_421859, EPI_ISL_421860, EPI_ISL_421861, EPI_ISL_421862, EPI_ISL_421863, EPI_ISL_421864, EPI_ISL_421865, EPI_ISL_421866, EPI_ISL_421867, EPI_ISL_421868, EPI_ISL_421869, EPI_ISL_421870, EPI_ISL_421871, EPI_ISL_421872, EPI_ISL_421873, EPI_ISL_421874, EPI_ISL_421875, EPI_ISL_421876, EPI_ISL_421877, EPI_ISL_421878, EPI_ISL_421879, EPI_ISL_421880, EPI_ISL_421881, EPI_ISL_421882, EPI_ISL_421883, EPI_ISL_421884, EPI_ISL_421885, EPI_ISL_421886, EPI_ISL_421887, EPI_ISL_421888, EPI_ISL_421889, EPI_ISL_421890, EPI_ISL_421891, EPI_ISL_421892, EPI_ISL_421893, EPI_ISL_421894, EPI_ISL_421895, EPI_ISL_421896, EPI_ISL_421897, EPI_ISL_421898, EPI_ISL_421899, EPI_ISL_421900, EPI_ISL_421901, EPI_ISL_421902, EPI_ISL_421903, EPI_ISL_421904, EPI_ISL_421905, EPI_ISL_421906, EPI_ISL_421907, EPI_ISL_421908, EPI_ISL_421909, EPI_ISL_421910, EPI_ISL_421911, EPI_ISL_421912, EPI_ISL_421913, EPI_ISL_421914, EPI_ISL_421915, EPI_ISL_421916, EPI_ISL_421917, EPI_ISL_421918, EPI_ISL_421919, EPI_ISL_421920, EPI_ISL_421921, EPI_ISL_421922, EPI_ISL_421923, EPI_ISL_421924, EPI_ISL_421925, EPI_ISL_421926, EPI_ISL_421927, EPI_ISL_421928, EPI_ISL_421929, EPI_ISL_421930, EPI_ISL_421931, EPI_ISL_421932, EPI_ISL_421933, EPI_ISL_421934, EPI_ISL_421935, EPI_ISL_421936, EPI_ISL_421937, EPI_ISL_421938, EPI_ISL_421939, EPI_ISL_421940, EPI_ISL_421941, EPI_ISL_421942, EPI_ISL_421943, EPI_ISL_421944, EPI_ISL_421945, EPI_ISL_421946, EPI_ISL_421947, EPI_ISL_421948, EPI_ISL_421949, EPI_ISL_421951, EPI_ISL_421952, EPI_ISL_421953, EPI_ISL_421954, EPI_ISL_421955, EPI_ISL_421956, EPI_ISL_421957, EPI_ISL_421958, EPI_ISL_421959, EPI_ISL_421960, EPI_ISL_421961, EPI_ISL_421962, EPI_ISL_421963, EPI_ISL_421964, EPI_ISL_421965, EPI_ISL_421966, EPI_ISL_421967, EPI_ISL_421968, EPI_ISL_421969, EPI_ISL_421970, EPI_ISL_421971, EPI_ISL_421972, EPI_ISL_421973, EPI_ISL_421974, EPI_ISL_421975, EPI_ISL_421976, EPI_ISL_421977, EPI_ISL_421978, EPI_ISL_421979, EPI_ISL_421980, EPI_ISL_421981, EPI_ISL_421982, EPI_ISL_421983, EPI_ISL_421984, EPI_ISL_421985, EPI_ISL_421986, EPI_ISL_421987, EPI_ISL_421988, EPI_ISL_421989, EPI_ISL_421990, EPI_ISL_421991, EPI_ISL_421992, EPI_ISL_421993, EPI_ISL_421994, EPI_ISL_421995, EPI_ISL_421996, EPI_ISL_421997, EPI_ISL_421998, EPI_ISL_421999, EPI_ISL_422000, EPI_ISL_422001, EPI_ISL_422002, EPI_ISL_422003, EPI_ISL_422004, EPI_ISL_422005, EPI_ISL_422006, EPI_ISL_422007, EPI_ISL_422008, EPI_ISL_422009, EPI_ISL_422010, EPI_ISL_422011, EPI_ISL_422012, EPI_ISL_422436, EPI_ISL_423044, EPI_ISL_423045, EPI_ISL_423046, EPI_ISL_423047, EPI_ISL_423048, EPI_ISL_423049, EPI_ISL_423050, EPI_ISL_423051, EPI_ISL_423052, EPI_ISL_423053, EPI_ISL_423054, EPI_ISL_423055, EPI_ISL_423056, EPI_ISL_423057, EPI_ISL_423058, EPI_ISL_423059, EPI_ISL_423060, EPI_ISL_423061, EPI_ISL_423062, EPI_ISL_423063, EPI_ISL_423064, EPI_ISL_423065, EPI_ISL_423066, EPI_ISL_423067, EPI_ISL_423068, EPI_ISL_423069, EPI_ISL_423070, EPI_ISL_423071, EPI_ISL_423072, EPI_ISL_423073, EPI_ISL_423074, EPI_ISL_423075, EPI_ISL_423076, EPI_ISL_423077, EPI_ISL_423078, EPI_ISL_423079, EPI_ISL_423080, EPI_ISL_423081, EPI_ISL_423082, EPI_ISL_423083, EPI_ISL_423084, EPI_ISL_423085, EPI_ISL_423086, EPI_ISL_423087, EPI_ISL_423088, EPI_ISL_423089, EPI_ISL_423090, EPI_ISL_423091, EPI_ISL_423092, EPI_ISL_423093, EPI_ISL_423094, EPI_ISL_423095, EPI_ISL_423096, EPI_ISL_423097, EPI_ISL_423098, EPI_ISL_423099, EPI_ISL_423100, EPI_ISL_423101, EPI_ISL_423102, EPI_ISL_423103, EPI_ISL_423104, EPI_ISL_423105, EPI_ISL_423106, EPI_ISL_423107, EPI_ISL_423108, EPI_ISL_423109, EPI_ISL_423110, EPI_ISL_423111, EPI_ISL_423112, EPI_ISL_423113, EPI_ISL_423114, EPI_ISL_423115, EPI_ISL_423116, EPI_ISL_423117, EPI_ISL_423118, EPI_ISL_423119, EPI_ISL_423120, EPI_ISL_423121, EPI_ISL_423122, EPI_ISL_423123, EPI_ISL_423124, EPI_ISL_423125, EPI_ISL_423126, EPI_ISL_423127, EPI_ISL_423128, EPI_ISL_423129, EPI_ISL_423130, EPI_ISL_423131, EPI_ISL_423132, EPI_ISL_423133, EPI_ISL_423134, EPI_ISL_423135, EPI_ISL_423136, EPI_ISL_423137, EPI_ISL_423138, EPI_ISL_423139, EPI_ISL_423140, EPI_ISL_423141, EPI_ISL_423142, EPI_ISL_423143, EPI_ISL_423144, EPI_ISL_423145, EPI_ISL_423146, EPI_ISL_423147, EPI_ISL_423148, EPI_ISL_423149, EPI_ISL_423150, EPI_ISL_423151, EPI_ISL_423152, EPI_ISL_423153, EPI_ISL_423154, EPI_ISL_423155, EPI_ISL_423156, EPI_ISL_423157, EPI_ISL_423158, EPI_ISL_423159, EPI_ISL_423160, EPI_ISL_423161, EPI_ISL_423162, EPI_ISL_423163, EPI_ISL_423164, EPI_ISL_423165, EPI_ISL_423166, EPI_ISL_423167, EPI_ISL_423168, EPI_ISL_423169, EPI_ISL_423170, EPI_ISL_423171, EPI_ISL_423172, EPI_ISL_423173, EPI_ISL_423174, EPI_ISL_423175, EPI_ISL_423176, EPI_ISL_423177, EPI_ISL_423178, EPI_ISL_423179, EPI_ISL_423180, EPI_ISL_423181, EPI_ISL_423182, EPI_ISL_423183, EPI_ISL_423184, EPI_ISL_423185, EPI_ISL_423186, EPI_ISL_423187, EPI_ISL_423188, EPI_ISL_423189, EPI_ISL_423190, EPI_ISL_423191, EPI_ISL_423192, EPI_ISL_423193, EPI_ISL_423194, EPI_ISL_423195, EPI_ISL_423196, EPI_ISL_423197, EPI_ISL_423198, EPI_ISL_423199, EPI_ISL_423200, EPI_ISL_423201, EPI_ISL_423202, EPI_ISL_423203, EPI_ISL_423204, EPI_ISL_423205, EPI_ISL_423206, EPI_ISL_423207, EPI_ISL_423208, EPI_ISL_423209, EPI_ISL_423210, EPI_ISL_423211, EPI_ISL_423212, EPI_ISL_423213, EPI_ISL_423214, EPI_ISL_423215, EPI_ISL_423216, EPI_ISL_423217, EPI_ISL_423218, EPI_ISL_423219, EPI_ISL_423220, EPI_ISL_423221, EPI_ISL_423222, EPI_ISL_423223, EPI_ISL_423224, EPI_ISL_423225, EPI_ISL_423226, EPI_ISL_423227, EPI_ISL_423228, EPI_ISL_423229, EPI_ISL_423230, EPI_ISL_423231, EPI_ISL_423232, EPI_ISL_423233, EPI_ISL_423234, EPI_ISL_423235, EPI_ISL_423236, EPI_ISL_423237, EPI_ISL_423238, EPI_ISL_423239, EPI_ISL_423240, EPI_ISL_423241, EPI_ISL_423242, EPI_ISL_423243, EPI_ISL_423244, EPI_ISL_423245, EPI_ISL_423246, EPI_ISL_423247, EPI_ISL_423248, EPI_ISL_423249, EPI_ISL_423250, EPI_ISL_423251, EPI_ISL_423252, EPI_ISL_423253, EPI_ISL_423254, EPI_ISL_423255, EPI_ISL_423256, EPI_ISL_423257, EPI_ISL_423258, EPI_ISL_423259, EPI_ISL_423260, EPI_ISL_423261, EPI_ISL_423262, EPI_ISL_423263, EPI_ISL_423264, EPI_ISL_423265, EPI_ISL_423266, EPI_ISL_423267, EPI_ISL_423268, EPI_ISL_423269, EPI_ISL_423270, EPI_ISL_423271, EPI_ISL_423272, EPI_ISL_423273, EPI_ISL_423274, EPI_ISL_423275, EPI_ISL_423276, EPI_ISL_423277, EPI_ISL_423278, EPI_ISL_423279, EPI_ISL_423280, EPI_ISL_423281, EPI_ISL_423282, EPI_ISL_423283, EPI_ISL_423284, EPI_ISL_423285, EPI_ISL_423286, EPI_ISL_423287, EPI_ISL_423288, EPI_ISL_423289, EPI_ISL_423290, EPI_ISL_423291, EPI_ISL_423292, EPI_ISL_423293, EPI_ISL_423294, EPI_ISL_423295, EPI_ISL_423296, EPI_ISL_423297, EPI_ISL_423298, EPI_ISL_423299, EPI_ISL_423300, EPI_ISL_423301, EPI_ISL_423302, EPI_ISL_423303, EPI_ISL_423304, EPI_ISL_423305, EPI_ISL_423306, EPI_ISL_423307, EPI_ISL_423308, EPI_ISL_423309, EPI_ISL_423310, EPI_ISL_423311, EPI_ISL_423312, EPI_ISL_423313, EPI_ISL_423314, EPI_ISL_423315, EPI_ISL_423316, EPI_ISL_423317, EPI_ISL_423318, EPI_ISL_423319, EPI_ISL_423320, EPI_ISL_423321, EPI_ISL_423322, EPI_ISL_423323, EPI_ISL_423324, EPI_ISL_423325, EPI_ISL_423326, EPI_ISL_423327, EPI_ISL_423328, EPI_ISL_423329, EPI_ISL_423330, EPI_ISL_423331, EPI_ISL_423332, EPI_ISL_423333, EPI_ISL_423334, EPI_ISL_423335, EPI_ISL_423336, EPI_ISL_423337, EPI_ISL_423338, EPI_ISL_423339, EPI_ISL_423340, EPI_ISL_423341, EPI_ISL_423342, EPI_ISL_423343, EPI_ISL_423344, EPI_ISL_423345, EPI_ISL_423346, EPI_ISL_423347, EPI_ISL_423348, EPI_ISL_423349, EPI_ISL_423350, EPI_ISL_423351, EPI_ISL_423352, EPI_ISL_423353, EPI_ISL_423354, EPI_ISL_423355, EPI_ISL_423356, EPI_ISL_423357, EPI_ISL_423358, EPI_ISL_423359, EPI_ISL_423360, EPI_ISL_423361, EPI_ISL_423362, EPI_ISL_423363, EPI_ISL_423364, EPI_ISL_423365, EPI_ISL_423366, EPI_ISL_423367, EPI_ISL_423368, EPI_ISL_423369, EPI_ISL_423370, EPI_ISL_423371, EPI_ISL_423372, EPI_ISL_423373, EPI_ISL_423374, EPI_ISL_423375, EPI_ISL_423376, EPI_ISL_423377, EPI_ISL_423378, EPI_ISL_423379, EPI_ISL_423380, EPI_ISL_423381, EPI_ISL_423382, EPI_ISL_423383, EPI_ISL_423384, EPI_ISL_423385, EPI_ISL_423386, EPI_ISL_423387, EPI_ISL_423388, EPI_ISL_423389, EPI_ISL_423390, EPI_ISL_423391, EPI_ISL_423392, EPI_ISL_423393, EPI_ISL_423394, EPI_ISL_423395, EPI_ISL_423396, EPI_ISL_423397, EPI_ISL_423398, EPI_ISL_423399, EPI_ISL_423400, EPI_ISL_423401, EPI_ISL_423402, EPI_ISL_423403, EPI_ISL_423404, EPI_ISL_423405, EPI_ISL_423406, EPI_ISL_423407, EPI_ISL_423408, EPI_ISL_423409, EPI_ISL_423410, EPI_ISL_423411, EPI_ISL_423412, EPI_ISL_423413, EPI_ISL_423414, EPI_ISL_423415, EPI_ISL_423416, EPI_ISL_423417, EPI_ISL_423418, EPI_ISL_423419, EPI_ISL_423420, EPI_ISL_423421, EPI_ISL_423422, EPI_ISL_423423, EPI_ISL_423424, EPI_ISL_423425, EPI_ISL_423426, EPI_ISL_423427, EPI_ISL_423428, EPI_ISL_423429, EPI_ISL_423430, EPI_ISL_423431, EPI_ISL_423432, EPI_ISL_423433, EPI_ISL_423434, EPI_ISL_423435, EPI_ISL_423436, EPI_ISL_423437, EPI_ISL_423438, EPI_ISL_423439, EPI_ISL_423440, EPI_ISL_423441, EPI_ISL_423442, EPI_ISL_423443, EPI_ISL_423444, EPI_ISL_423445, EPI_ISL_423446, EPI_ISL_423447, EPI_ISL_423448, EPI_ISL_423449, EPI_ISL_423450, EPI_ISL_423451, EPI_ISL_423452, EPI_ISL_423453, EPI_ISL_423454, EPI_ISL_423455, EPI_ISL_423456, EPI_ISL_423457, EPI_ISL_423458, EPI_ISL_423459, EPI_ISL_423460, EPI_ISL_423461, EPI_ISL_423462, EPI_ISL_423463, EPI_ISL_423464, EPI_ISL_423465, EPI_ISL_423466, EPI_ISL_423467, EPI_ISL_423468, EPI_ISL_423469, EPI_ISL_423470, EPI_ISL_423471, EPI_ISL_423472, EPI_ISL_423473, EPI_ISL_423474, EPI_ISL_423475, EPI_ISL_423476, EPI_ISL_423477, EPI_ISL_423478, EPI_ISL_423479, EPI_ISL_423480, EPI_ISL_423481, EPI_ISL_423482, EPI_ISL_423483, EPI_ISL_423484, EPI_ISL_423485, EPI_ISL_423486, EPI_ISL_423487, EPI_ISL_423488, EPI_ISL_423489, EPI_ISL_423490, EPI_ISL_423491, EPI_ISL_423492, EPI_ISL_423493, EPI_ISL_423494, EPI_ISL_423495, EPI_ISL_423496, EPI_ISL_423497, EPI_ISL_423498, EPI_ISL_423499, EPI_ISL_423500, EPI_ISL_423501, EPI_ISL_423502, EPI_ISL_423503, EPI_ISL_423504, EPI_ISL_423505, EPI_ISL_423506, EPI_ISL_423507, EPI_ISL_423508, EPI_ISL_423509, EPI_ISL_423510, EPI_ISL_423511, EPI_ISL_423512, EPI_ISL_423513, EPI_ISL_423514, EPI_ISL_423515, EPI_ISL_423516, EPI_ISL_423517, EPI_ISL_423518, EPI_ISL_423519, EPI_ISL_423520, EPI_ISL_423521, EPI_ISL_423522, EPI_ISL_423523, EPI_ISL_423524, EPI_ISL_423526, EPI_ISL_423527, EPI_ISL_423528, EPI_ISL_423529, EPI_ISL_423530, EPI_ISL_423531, EPI_ISL_423532, EPI_ISL_423533, EPI_ISL_423534, EPI_ISL_423535, EPI_ISL_423536, EPI_ISL_423537, EPI_ISL_423538, EPI_ISL_423539, EPI_ISL_423540, EPI_ISL_423541, EPI_ISL_423542, EPI_ISL_423543, EPI_ISL_423544, EPI_ISL_423545, EPI_ISL_423546, EPI_ISL_423547, EPI_ISL_423548, EPI_ISL_423549, EPI_ISL_423550, EPI_ISL_423551, EPI_ISL_423552, EPI_ISL_423553, EPI_ISL_423554, EPI_ISL_423555, EPI_ISL_423556, EPI_ISL_423557, EPI_ISL_423558, EPI_ISL_423559, EPI_ISL_423560, EPI_ISL_423561, EPI_ISL_423562, EPI_ISL_423563, EPI_ISL_423564, EPI_ISL_423565, EPI_ISL_423566, EPI_ISL_423567, EPI_ISL_423568, EPI_ISL_423569, EPI_ISL_423570, EPI_ISL_423571, EPI_ISL_423572, EPI_ISL_423573, EPI_ISL_423574, EPI_ISL_423575, EPI_ISL_423576, EPI_ISL_423577, EPI_ISL_423578, EPI_ISL_423579, EPI_ISL_423580, EPI_ISL_423581, EPI_ISL_423582, EPI_ISL_423583, EPI_ISL_423584, EPI_ISL_423585, EPI_ISL_423586, EPI_ISL_423587, EPI_ISL_423588, EPI_ISL_423589, EPI_ISL_423590, EPI_ISL_423591, EPI_ISL_423592, EPI_ISL_423593, EPI_ISL_423594, EPI_ISL_423595, EPI_ISL_423596, EPI_ISL_423597, EPI_ISL_423598, EPI_ISL_423599, EPI_ISL_423600, EPI_ISL_423601, EPI_ISL_423602, EPI_ISL_423603, EPI_ISL_423604, EPI_ISL_423605, EPI_ISL_423606, EPI_ISL_423607, EPI_ISL_423608, EPI_ISL_423609, EPI_ISL_423610, EPI_ISL_423611, EPI_ISL_423612, EPI_ISL_423613, EPI_ISL_423614, EPI_ISL_423615, EPI_ISL_423616, EPI_ISL_423617, EPI_ISL_423618, EPI_ISL_423619, EPI_ISL_423620, EPI_ISL_423621, EPI_ISL_423622, EPI_ISL_423623, EPI_ISL_423624, EPI_ISL_423625, EPI_ISL_423626, EPI_ISL_423627, EPI_ISL_423628, EPI_ISL_423629, EPI_ISL_423630, EPI_ISL_423631, EPI_ISL_423632, EPI_ISL_423633, EPI_ISL_423634, EPI_ISL_423635, EPI_ISL_423636, EPI_ISL_423637, EPI_ISL_423638, EPI_ISL_423639, EPI_ISL_423640, EPI_ISL_423641, EPI_ISL_423642, EPI_ISL_423643, EPI_ISL_423644, EPI_ISL_423645, EPI_ISL_423646, EPI_ISL_423647, EPI_ISL_423648, EPI_ISL_423649, EPI_ISL_423650, EPI_ISL_423651, EPI_ISL_423652, EPI_ISL_423653, EPI_ISL_423654, EPI_ISL_423655, EPI_ISL_423656, EPI_ISL_423657, EPI_ISL_423658, EPI_ISL_423659, EPI_ISL_423660, EPI_ISL_423661, EPI_ISL_423662, EPI_ISL_423663, EPI_ISL_423664, EPI_ISL_423665, EPI_ISL_423666, EPI_ISL_423667, EPI_ISL_423668, EPI_ISL_423669, EPI_ISL_423670, EPI_ISL_423671, EPI_ISL_423672, EPI_ISL_423673, EPI_ISL_423674, EPI_ISL_423675, EPI_ISL_423676, EPI_ISL_423677, EPI_ISL_423678, EPI_ISL_423679, EPI_ISL_423680, EPI_ISL_423681, EPI_ISL_423682, EPI_ISL_423683, EPI_ISL_423684, EPI_ISL_423685, EPI_ISL_423686, EPI_ISL_423687, EPI_ISL_423688, EPI_ISL_423689, EPI_ISL_423690, EPI_ISL_423691, EPI_ISL_423692, EPI_ISL_423693, EPI_ISL_423694, EPI_ISL_423695, EPI_ISL_423696, EPI_ISL_423697, EPI_ISL_423698, EPI_ISL_423699, EPI_ISL_423700, EPI_ISL_423701, EPI_ISL_423702, EPI_ISL_423703, EPI_ISL_423704, EPI_ISL_423705, EPI_ISL_423706, EPI_ISL_423707, EPI_ISL_423708, EPI_ISL_423709, EPI_ISL_423710, EPI_ISL_423711, EPI_ISL_423712, EPI_ISL_423713, EPI_ISL_423714, EPI_ISL_423715, EPI_ISL_423716, EPI_ISL_423717, EPI_ISL_423718, EPI_ISL_423719, EPI_ISL_423720, EPI_ISL_423721, EPI_ISL_423722, EPI_ISL_423723, EPI_ISL_423724, EPI_ISL_423725, EPI_ISL_423726, EPI_ISL_423727, EPI_ISL_423728, EPI_ISL_423729, EPI_ISL_423730, EPI_ISL_423731, EPI_ISL_423732, EPI_ISL_423733, EPI_ISL_423734, EPI_ISL_423735, EPI_ISL_423736, EPI_ISL_423737, EPI_ISL_423738, EPI_ISL_423739, EPI_ISL_423740, EPI_ISL_423741, EPI_ISL_423742, EPI_ISL_423743, EPI_ISL_423744, EPI_ISL_423745, EPI_ISL_423746, EPI_ISL_423747, EPI_ISL_423748, EPI_ISL_423749, EPI_ISL_423750, EPI_ISL_423751, EPI_ISL_423752, EPI_ISL_423753, EPI_ISL_423754, EPI_ISL_423755, EPI_ISL_423756, EPI_ISL_423757, EPI_ISL_423758, EPI_ISL_423759, EPI_ISL_423760, EPI_ISL_423761, EPI_ISL_423762, EPI_ISL_423763, EPI_ISL_423764, EPI_ISL_423765, EPI_ISL_423766, EPI_ISL_423767, EPI_ISL_423768, EPI_ISL_423769, EPI_ISL_423770, EPI_ISL_423771, EPI_ISL_423772, EPI_ISL_423773, EPI_ISL_423774, EPI_ISL_423775, EPI_ISL_423776, EPI_ISL_423777, EPI_ISL_423778, EPI_ISL_423779, EPI_ISL_423780, EPI_ISL_423781, EPI_ISL_423782, EPI_ISL_423783, EPI_ISL_423784, EPI_ISL_423785, EPI_ISL_423786, EPI_ISL_423787, EPI_ISL_423788, EPI_ISL_423789, EPI_ISL_423790, EPI_ISL_423791, EPI_ISL_423792, EPI_ISL_423793, EPI_ISL_423794, EPI_ISL_423795, EPI_ISL_423796, EPI_ISL_423797, EPI_ISL_423798, EPI_ISL_423799, EPI_ISL_423800, EPI_ISL_423801, EPI_ISL_423802, EPI_ISL_423803, EPI_ISL_423804, EPI_ISL_423805, EPI_ISL_423806, EPI_ISL_423807, EPI_ISL_423808, EPI_ISL_423809, EPI_ISL_423810, EPI_ISL_423811, EPI_ISL_423812, EPI_ISL_423813, EPI_ISL_423814, EPI_ISL_423815, EPI_ISL_423816, EPI_ISL_423817, EPI_ISL_423818, EPI_ISL_423819, EPI_ISL_423820, EPI_ISL_423821, EPI_ISL_423822, EPI_ISL_423823, EPI_ISL_423824, EPI_ISL_423825, EPI_ISL_423826, EPI_ISL_423827, EPI_ISL_423828, EPI_ISL_423829, EPI_ISL_423830, EPI_ISL_423831, EPI_ISL_423832, EPI_ISL_423833, EPI_ISL_423834, EPI_ISL_423835, EPI_ISL_423836, EPI_ISL_423837, EPI_ISL_423838, EPI_ISL_423839, EPI_ISL_423840, EPI_ISL_423841, EPI_ISL_423842, EPI_ISL_423843, EPI_ISL_423844, EPI_ISL_423845, EPI_ISL_423846, EPI_ISL_423847, EPI_ISL_423848, EPI_ISL_423849, EPI_ISL_423850, EPI_ISL_423851, EPI_ISL_423852, EPI_ISL_423853, EPI_ISL_423854, EPI_ISL_423855, EPI_ISL_423856, EPI_ISL_423857, EPI_ISL_423858, EPI_ISL_423859, EPI_ISL_423860, EPI_ISL_423861, EPI_ISL_423862, EPI_ISL_423863, EPI_ISL_423864, EPI_ISL_423865, EPI_ISL_423866, EPI_ISL_423867, EPI_ISL_423868, EPI_ISL_423869, EPI_ISL_423870, EPI_ISL_423871, EPI_ISL_423872, EPI_ISL_423873, EPI_ISL_423874, EPI_ISL_423875, EPI_ISL_423876, EPI_ISL_423877, EPI_ISL_423878, EPI_ISL_423879, EPI_ISL_423880, EPI_ISL_423881, EPI_ISL_423882, EPI_ISL_423883, EPI_ISL_423884, EPI_ISL_423885, EPI_ISL_423886, EPI_ISL_423887, EPI_ISL_423888, EPI_ISL_423889, EPI_ISL_423890, EPI_ISL_423891, EPI_ISL_423892, EPI_ISL_423893, EPI_ISL_423894, EPI_ISL_423895, EPI_ISL_423896, EPI_ISL_423897, EPI_ISL_423898, EPI_ISL_423899, EPI_ISL_423900, EPI_ISL_423901, EPI_ISL_423902, EPI_ISL_423903, EPI_ISL_423904, EPI_ISL_423905, EPI_ISL_423906, EPI_ISL_423907, EPI_ISL_423908, EPI_ISL_423909, EPI_ISL_423910, EPI_ISL_423911, EPI_ISL_423912, EPI_ISL_423913, EPI_ISL_423914, EPI_ISL_423915, EPI_ISL_423916, EPI_ISL_423917, EPI_ISL_423918, EPI_ISL_423919, EPI_ISL_423920, EPI_ISL_423921, EPI_ISL_423922, EPI_ISL_423923, EPI_ISL_423924, EPI_ISL_423925, EPI_ISL_423926, EPI_ISL_423927, EPI_ISL_423928, EPI_ISL_423929, EPI_ISL_423930, EPI_ISL_423931, EPI_ISL_423932, EPI_ISL_423933, EPI_ISL_423934, EPI_ISL_423935, EPI_ISL_423936, EPI_ISL_423937, EPI_ISL_423938, EPI_ISL_423939, EPI_ISL_423940, EPI_ISL_423941, EPI_ISL_423942, EPI_ISL_423943, EPI_ISL_423944, EPI_ISL_423945, EPI_ISL_423946, EPI_ISL_423947, EPI_ISL_423948, EPI_ISL_423949, EPI_ISL_423950, EPI_ISL_423951, EPI_ISL_423952, EPI_ISL_423953, EPI_ISL_423954, EPI_ISL_423955, EPI_ISL_423956, EPI_ISL_423957, EPI_ISL_423958, EPI_ISL_423959, EPI_ISL_423960, EPI_ISL_423961, EPI_ISL_423962, EPI_ISL_423963, EPI_ISL_423964, EPI_ISL_423965, EPI_ISL_423966, EPI_ISL_423967, EPI_ISL_423968, EPI_ISL_423969, EPI_ISL_423970, EPI_ISL_423971, EPI_ISL_423972, EPI_ISL_423973, EPI_ISL_423974, EPI_ISL_423975, EPI_ISL_423976, EPI_ISL_423977, EPI_ISL_423978, EPI_ISL_423979, EPI_ISL_423980, EPI_ISL_423981, EPI_ISL_423982, EPI_ISL_423983, EPI_ISL_423984, EPI_ISL_423985, EPI_ISL_423986, EPI_ISL_423987, EPI_ISL_423988, EPI_ISL_423989, EPI_ISL_423990, EPI_ISL_423991, EPI_ISL_423992, EPI_ISL_423993, EPI_ISL_423994, EPI_ISL_423995, EPI_ISL_423996, EPI_ISL_423997, EPI_ISL_423998, EPI_ISL_423999, EPI_ISL_424000, EPI_ISL_424001, EPI_ISL_424002, EPI_ISL_424003, EPI_ISL_424004, EPI_ISL_424005, EPI_ISL_424006, EPI_ISL_424007, EPI_ISL_424008, EPI_ISL_424009, EPI_ISL_424010, EPI_ISL_424011, EPI_ISL_424012, EPI_ISL_424013, EPI_ISL_424014, EPI_ISL_424015, EPI_ISL_424016, EPI_ISL_424017, EPI_ISL_424018, EPI_ISL_424019, EPI_ISL_424020, EPI_ISL_424021, EPI_ISL_424022, EPI_ISL_424023, EPI_ISL_424024, EPI_ISL_424025, EPI_ISL_424026, EPI_ISL_424027, EPI_ISL_424028, EPI_ISL_424029, EPI_ISL_424030, EPI_ISL_424031, EPI_ISL_424032, EPI_ISL_424033, EPI_ISL_424034, EPI_ISL_424035, EPI_ISL_424036, EPI_ISL_424037, EPI_ISL_424038, EPI_ISL_424039, EPI_ISL_424040, EPI_ISL_424041, EPI_ISL_424042, EPI_ISL_424043, EPI_ISL_424044, EPI_ISL_424045, EPI_ISL_424046, EPI_ISL_424047, EPI_ISL_424048, EPI_ISL_424049, EPI_ISL_424050, EPI_ISL_424051, EPI_ISL_424052, EPI_ISL_424053, EPI_ISL_424054, EPI_ISL_424055, EPI_ISL_424056, EPI_ISL_424057, EPI_ISL_424058, EPI_ISL_424059, EPI_ISL_424060, EPI_ISL_424061, EPI_ISL_424062, EPI_ISL_424063, EPI_ISL_424064, EPI_ISL_424065, EPI_ISL_424066, EPI_ISL_424067, EPI_ISL_424068, EPI_ISL_424069, EPI_ISL_424070, EPI_ISL_424071, EPI_ISL_424072, EPI_ISL_424073, EPI_ISL_424074, EPI_ISL_424075, EPI_ISL_424076, EPI_ISL_424077, EPI_ISL_424078, EPI_ISL_424079, EPI_ISL_424080, EPI_ISL_424081, EPI_ISL_424082, EPI_ISL_424083, EPI_ISL_424084, EPI_ISL_424085, EPI_ISL_424086, EPI_ISL_424087, EPI_ISL_424088, EPI_ISL_424089, EPI_ISL_424090, EPI_ISL_424091, EPI_ISL_424092, EPI_ISL_424093, EPI_ISL_424094, EPI_ISL_424095, EPI_ISL_424096, EPI_ISL_424097, EPI_ISL_424098, EPI_ISL_424099, EPI_ISL_424100, EPI_ISL_424101, EPI_ISL_424102, EPI_ISL_424103, EPI_ISL_424104, EPI_ISL_424105, EPI_ISL_424106, EPI_ISL_424107, EPI_ISL_424108, EPI_ISL_424109, EPI_ISL_424110, EPI_ISL_424111, EPI_ISL_424112, EPI_ISL_424113, EPI_ISL_424114, EPI_ISL_424115, EPI_ISL_424116, EPI_ISL_424117, EPI_ISL_424118, EPI_ISL_424119, EPI_ISL_424120, EPI_ISL_424121, EPI_ISL_424122, EPI_ISL_424123, EPI_ISL_424124, EPI_ISL_424125, EPI_ISL_424126, EPI_ISL_424127, EPI_ISL_424128, EPI_ISL_424129, EPI_ISL_424130, EPI_ISL_424131, EPI_ISL_424132, EPI_ISL_424133, EPI_ISL_424134, EPI_ISL_424135, EPI_ISL_424136, EPI_ISL_424137, EPI_ISL_424138, EPI_ISL_424139, EPI_ISL_424140, EPI_ISL_424141, EPI_ISL_424142, EPI_ISL_424143, EPI_ISL_424144, EPI_ISL_424145, EPI_ISL_424146, EPI_ISL_424147, EPI_ISL_424148, EPI_ISL_424149, EPI_ISL_424150, EPI_ISL_424151, EPI_ISL_424152, EPI_ISL_424153, EPI_ISL_424154, EPI_ISL_424155, EPI_ISL_424156, EPI_ISL_424157, EPI_ISL_424158, EPI_ISL_424159, EPI_ISL_424160, EPI_ISL_424161, EPI_ISL_424162, EPI_ISL_424163, EPI_ISL_424164, EPI_ISL_424165

| | | | |
|---|---|---|---|
| see above | Respiratory Virus Unit, Microbiology Services Colindale, Public Health England | Respiratory Virus Unit, Microbiology Services Colindale, Public Health England | Monica Galiano, Shahjahan Miah, Angie Lackenby, Omolola Akinbami, Tiina Talts, Leena Bhaw, Richard Myers, Steven Platt, Kirstin Edwards, Jonathan Hubb, Joanna Ellis, Maria Zambon |
| EPI_ISL_424731 | Hospital General Regional No.66, Ciudad Juárez, Chihuahua. | Laboratorio Central de Epidemiología-DLVIE / Laboratorio de Secuenciación-Centro de Instrumentos. Instituto Mexicano del Seguro Social | Muñoz-Medina JE, González-Torres C, Gaytán-Cervantes FJ, López-Martínez I, Hernández-Rivas L, Barrera-Badillo G, Mendieta-Condado E, Garcés-Ayala F, Araiza-Rodríguez A, Ramírez-González JE, Boukadida C, Ávila-Ríos S, Mújica-Sánchez M, Martínez-Orozco JA, Becerril-Vargas E, Vázquez-Pérez JA, Taboada B, Sánchez A, Isa P, Grande R, Vázquez G, Pulido F. López S, Arias C, Ramos-Cervantes P, Ruiz-Palacios GM, Ibarra-González V, Ledesma-Barrientos F, García-Andrade LA, de León-Garduño AP, González-Bonilla CR, Grajales-Muñiz C, Borja-Aburto VH. |
| EPI_ISL_424929, EPI_ISL_424930, EPI_ISL_424931, EPI_ISL_424932, EPI_ISL_424933, EPI_ISL_424934, EPI_ISL_424935, EPI_ISL_424936, EPI_ISL_424937, EPI_ISL_424938, EPI_ISL_424939, EPI_ISL_424940, EPI_ISL_424941, EPI_ISL_424942, EPI_ISL_424943, EPI_ISL_424944, EPI_ISL_424945, EPI_ISL_424946, EPI_ISL_424947, EPI_ISL_424948, EPI_ISL_424949, EPI_ISL_424950, EPI_ISL_424951, EPI_ISL_424952, EPI_ISL_424953, EPI_ISL_424954, EPI_ISL_424955, EPI_ISL_424956, EPI_ISL_424957, EPI_ISL_424958, EPI_ISL_424959, EPI_ISL_424960, EPI_ISL_424961, EPI_ISL_424962, EPI_ISL_424963, EPI_ISL_424964, EPI_ISL_424965, EPI_ISL_424966, EPI_ISL_424967, EPI_ISL_424968 | | | |
| see above | NYU Langone Health | Departments of Pathology and Medicine, New York University School of Medicine | Maria Aguero-Rosenfeld, Brendan Belovarac, Margaret Black, Ludovic Boytard, John Cadley, Paolo Cotzia, John Chen, Dacia Dimartino, Xiaojun Feng, Tatyana Gindin, Adriana Heguy, Megan Hogan, Emily Huang, George Jour, Andrew Lytle, Christian Marier, Matthew T. Maurano, Mark J. Mulligan, Peter Meyn, Iman Osman, Jared Pinnell, Sitharam Ramaswami, Amy Rapkiewicz, Marie Samanovic-Golden, Antonio Serrano, Guomiao Shen, Matija Snuderl, Theodore Vougiouklakis, Nick Vulpescu, Gael Westby, Paul Zappile, Yutong Zhang |
| EPI_ISL_425183 | Servicio de Microbiología. Consorcio Hospital General Universitario de Valencia | Sequencing and Bioinformatics Service and Molecular Epidemiology Research Group. FISABIO-Public Health | Marta Pla Diaz, Neris Garcia-Gonzalez, Inma Galán Vendrell, Sandra Carbo, Loreto Ferrús Abad, Paula Ruiz-Hueso, Mariana Reyes-Prieto, Vicente Soriano Chirona, Ivan Ansari, David Navarro, Maria Alma Bracho, Griselda De Marco, Beatriz Beamud, Lidia Ruiz Roldan, Llúcia Martínez-Priego, Giuseppe D'Auria, Fernando Gonzalez-Candelas |
| EPI_ISL_425189 | Servicio de Microbiología. Consorcio Hospital General Universitario de Valencia | Sequencing and Bioinformatics Service and Molecular Epidemiology Research Group. FISABIO-Public Health | Mariana Reyes-Prieto, Vicente Soriano Chirona, Ivan Ansari, David Navarro, Maria Alma Bracho, Griselda De Marco, Beatriz Beamud, Lidia Ruiz Roldan, Marta Pla Diaz, Neris Garcia-Gonzalez, Inma Galán Vendrell, Sandra Carbo, Loreto Ferrús Abad, Paula Ruiz-Hueso, Llúcia Martínez-Priego, Giuseppe D'Auria, Fernando Gonzalez-Candelas |
| EPI_ISL_425193 | Servicio de Microbiología. Consorcio Hospital General Universitario de Valencia | Sequencing and Bioinformatics Service and Molecular Epidemiology Research Group. FISABIO-Public Health | Maria Alma Bracho, Griselda De Marco, Beatriz Beamud, Lidia Ruiz Roldan, Marta Pla Diaz, Neris Garcia-Gonzalez, Inma Galán Vendrell, Sandra Carbo, Loreto Ferrús Abad, Paula Ruiz-Hueso, Mariana Reyes-Prieto, Vicente Soriano Chirona, Ivan Ansari, David Navarro, Llúcia Martínez-Priego, Giuseppe D'Auria, Fernando Gonzalez-Candelas |
| EPI_ISL_425197 | Servicio de Microbiología. Consorcio Hospital General Universitario de Valencia | Sequencing and Bioinformatics Service and Molecular Epidemiology Research Group. FISABIO-Public Health | Marta Pla Diaz, Neris Garcia-Gonzalez, Inma Galán Vendrell, Sandra Carbo, Loreto Ferrús Abad, Paula Ruiz-Hueso, Mariana Reyes-Prieto, Vicente Soriano Chirona, Ivan Ansari, David Navarro, Maria Alma Bracho, Griselda De Marco, Beatriz Beamud, Lidia Ruiz Roldan, Llúcia Martínez-Priego, Giuseppe D'Auria, Fernando Gonzalez-Candelas |
| EPI_ISL_425201 | Servicio de Microbiologia. Hospital Clinico Universitario de Valencia | Sequencing and Bioinformatics Service and Molecular Epidemiology Research Group. FISABIO-Public Health | Mariana Reyes-Prieto, Vicente Soriano Chirona, Maria Alma Bracho, Griselda De Marco, Beatriz Beamud, Lidia Ruiz Roldan, Marta Pla Diaz,Neris Garcia-Gonzalez, Loreto Ferrús Abad, Maria Dolores Ocete, Llúcia Martínez-Priego, Inma Galán Vendrell, Paula Ruiz-Hueso,Concepcion Gimeno, Giuseppe D'Auria, Fernando Gonzalez-Candelas |
| EPI_ISL_425203 | Servicio de Microbiologia. Hospital Clinico Universitario de Valencia | Sequencing and Bioinformatics Service and Molecular Epidemiology Research Group. FISABIO-Public Health | Maria Alma Bracho, Griselda De Marco, Beatriz Beamud, Lidia Ruiz Roldan, Marta Pla Diaz,Neris Garcia-Gonzalez, Loreto Ferrús Abad, Maria Dolores Ocete, Inma Galán Vendrell, Paula Ruiz-Hueso, Mariana Reyes-Prieto, Vicente Soriano Chirona, Llúcia Martínez-Priego, Concepcion Gimeno, Giuseppe D'Auria, Fernando Gonzalez-Candelas |
| EPI_ISL_425204 | Servicio de Microbiologia. Hospital Clinico Universitario de Valencia | Sequencing and Bioinformatics Service and Molecular Epidemiology Research Group. FISABIO-Public Health | Maria Dolores Ocete, Inma Galán Vendrell, Paula Ruiz-Hueso, Mariana Reyes-Prieto, Vicente Soriano Chirona, Maria Alma Bracho, Griselda De Marco, Beatriz Beamud, Lidia Ruiz Roldan, Marta Pla Diaz,Neris Garcia-Gonzalez, Loreto Ferrús Abad, Llúcia Martínez-Priego, Concepcion Gimeno, Giuseppe D'Auria, Fernando Gonzalez-Candelas |
| EPI_ISL_425208 | Servicio de Microbiologia. Hospital Clinico Universitario de Valencia | Sequencing and Bioinformatics Service and Molecular Epidemiology Research Group. FISABIO-Public Health | Marta Pla Diaz,Neris Garcia-Gonzalez, Loreto Ferrús Abad, Maria Dolores Ocete, Inma Galán Vendrell, Paula Ruiz-Hueso, Mariana Reyes-Prieto, Vicente Soriano Chirona, Maria Alma Bracho, Griselda De Marco, Beatriz Beamud, Lidia Ruiz Roldan, Llúcia Martínez-Priego, Concepcion Gimeno, Giuseppe D'Auria, Fernando Gonzalez-Candelas |
| EPI_ISL_425213 | Servicio de Microbiología. Consorcio Hospital General Universitario de Valencia | Sequencing and Bioinformatics Service and Molecular Epidemiology Research Group. FISABIO-Public Health | Maria Alma Bracho, Griselda De Marco, Beatriz Beamud, Lidia Ruiz Roldan, Marta Pla Diaz, Neris Garcia-Gonzalez, Inma Galán Vendrell, Sandra Carbo, Loreto Ferrús Abad, Paula Ruiz-Hueso, Mariana Reyes-Prieto, Vicente Soriano Chirona, Ivan Ansari, David Navarro, Llúcia Martínez-Priego, Giuseppe D'Auria, Fernando Gonzalez-Candelas |
| EPI_ISL_425217 | Servicio de Microbiología. Consorcio Hospital General Universitario de Valencia | Sequencing and Bioinformatics Service and Molecular Epidemiology Research Group. FISABIO-Public Health | Marta Pla Diaz, Neris Garcia-Gonzalez, Inma Galán Vendrell, Sandra Carbo, Loreto Ferrús Abad, Paula Ruiz-Hueso, Mariana Reyes-Prieto, Vicente Soriano Chirona, Ivan Ansari, David Navarro, Maria Alma Bracho, Griselda De Marco, Beatriz Beamud, Lidia Ruiz Roldan, Llúcia Martínez-Priego, Giuseppe D'Auria, Fernando Gonzalez-Candelas |
| EPI_ISL_425819, EPI_ISL_425820, EPI_ISL_425821, EPI_ISL_425822, EPI_ISL_425823, EPI_ISL_425824, EPI_ISL_425825, EPI_ISL_425826, EPI_ISL_425827, EPI_ISL_425828, EPI_ISL_425829, EPI_ISL_425830, EPI_ISL_425831, EPI_ISL_425832, EPI_ISL_425833, EPI_ISL_425834, EPI_ISL_425835, EPI_ISL_425836, EPI_ISL_425837, EPI_ISL_425838, EPI_ISL_425839, EPI_ISL_425840, EPI_ISL_425841, EPI_ISL_425842, EPI_ISL_425843, EPI_ISL_425844, EPI_ISL_425845, EPI_ISL_425846, EPI_ISL_425847, EPI_ISL_425848, EPI_ISL_425849, EPI_ISL_425850, EPI_ISL_425851, EPI_ISL_425852, EPI_ISL_425853, EPI_ISL_425854, EPI_ISL_425855, EPI_ISL_425856, EPI_ISL_425857, EPI_ISL_425858, EPI_ISL_425859, EPI_ISL_425860, EPI_ISL_425861, EPI_ISL_425862, EPI_ISL_425863, EPI_ISL_425864, EPI_ISL_425865, EPI_ISL_425866, EPI_ISL_425867, EPI_ISL_425868, EPI_ISL_425869, EPI_ISL_425870, EPI_ISL_425871, EPI_ISL_425872, EPI_ISL_425873, EPI_ISL_425874, EPI_ISL_425875, EPI_ISL_425876, EPI_ISL_425877, EPI_ISL_425878, EPI_ISL_425879, EPI_ISL_425880, EPI_ISL_425881, EPI_ISL_425882, EPI_ISL_425883, EPI_ISL_425884, EPI_ISL_425885, EPI_ISL_425886, EPI_ISL_425887, EPI_ISL_425888, EPI_ISL_425889, EPI_ISL_425890, EPI_ISL_425891, EPI_ISL_425892, EPI_ISL_425893, EPI_ISL_425894, EPI_ISL_425895, EPI_ISL_425896, EPI_ISL_425897, EPI_ISL_425898, EPI_ISL_425899, EPI_ISL_425900, EPI_ISL_425901, EPI_ISL_425902, EPI_ISL_425903, EPI_ISL_425904, EPI_ISL_425905, EPI_ISL_425906, EPI_ISL_425907, EPI_ISL_425908, EPI_ISL_425909, EPI_ISL_425910, EPI_ISL_425911, EPI_ISL_425912, EPI_ISL_425913, EPI_ISL_425914, EPI_ISL_425915, EPI_ISL_425916, EPI_ISL_425917, EPI_ISL_425918, EPI_ISL_425919, EPI_ISL_425920, EPI_ISL_425921, EPI_ISL_425922, EPI_ISL_425923, EPI_ISL_425924, EPI_ISL_425925, EPI_ISL_425926, EPI_ISL_425927, EPI_ISL_425928, EPI_ISL_425929, EPI_ISL_425930, EPI_ISL_425931, EPI_ISL_425932, EPI_ISL_425933, EPI_ISL_425934, EPI_ISL_425935, EPI_ISL_425936, EPI_ISL_425937, EPI_ISL_425938, EPI_ISL_425939, EPI_ISL_425940, EPI_ISL_425941, EPI_ISL_425942, EPI_ISL_425943, EPI_ISL_425944, EPI_ISL_425945, EPI_ISL_425946, EPI_ISL_425947, EPI_ISL_425948, EPI_ISL_425949, EPI_ISL_425950, EPI_ISL_425951, EPI_ISL_425952, EPI_ISL_425953, EPI_ISL_425954, EPI_ISL_425955, EPI_ISL_425956, EPI_ISL_425957, EPI_ISL_425958, EPI_ISL_425959, EPI_ISL_425960, EPI_ISL_425961, EPI_ISL_425962, EPI_ISL_425963, EPI_ISL_425964, EPI_ISL_425965, EPI_ISL_425966, EPI_ISL_425967, EPI_ISL_425968, EPI_ISL_425969, EPI_ISL_425970, EPI_ISL_425971, EPI_ISL_425972, EPI_ISL_425973, EPI_ISL_425974, EPI_ISL_425975, EPI_ISL_425976, EPI_ISL_425977, EPI_ISL_425978, EPI_ISL_425979, EPI_ISL_425980, EPI_ISL_425981, EPI_ISL_425982, EPI_ISL_425983, EPI_ISL_425984, EPI_ISL_425985, EPI_ISL_425986, EPI_ISL_425987, EPI_ISL_425988, EPI_ISL_425989, EPI_ISL_425990, EPI_ISL_425991, EPI_ISL_425992, EPI_ISL_425993, EPI_ISL_425994, EPI_ISL_425995, EPI_ISL_425996, EPI_ISL_425997, EPI_ISL_425998, EPI_ISL_425999, EPI_ISL_426000, EPI_ISL_426001, EPI_ISL_426002, EPI_ISL_426003, EPI_ISL_426004, EPI_ISL_426005, EPI_ISL_426006, EPI_ISL_426007, EPI_ISL_426008, EPI_ISL_426009, EPI_ISL_426010, EPI_ISL_426011, EPI_ISL_426012, EPI_ISL_426013, EPI_ISL_426014, EPI_ISL_426015, EPI_ISL_426016, EPI_ISL_426017, EPI_ISL_426018, EPI_ISL_426019, EPI_ISL_426020, EPI_ISL_426021, EPI_ISL_426022, EPI_ISL_426023, EPI_ISL_426024 | | | |
| see above | Virology Department, Royal Infirmary of Edinburgh, NHS Lothian / School of Biological Sciences, University of Edinburgh / Institute of Genetics and Molecular Medicine, University of Edinburgh | COVID-19 Genomics UK (COG-UK) Consortium | McHugh M, Dewar R, Rooke S, Gallagher M, Balcaza C, O'Toole A, Hill V, McCrone JT, Colquhoun R, Yu X, Jackson B, Scher E, Rambaut A, Williams TC, Templeton K |
| EPI_ISL_426617, EPI_ISL_426618, EPI_ISL_426619, EPI_ISL_426620, EPI_ISL_426621, EPI_ISL_426622, EPI_ISL_426623, EPI_ISL_426624, EPI_ISL_426625, EPI_ISL_426626 | NYU Langone Health | Departments of Pathology and Medicine, New York University School of Medicine | Maria Aguero-Rosenfeld, Brendan Belovarac, Margaret Black, Ludovic Boytard, John Cadley, Paolo Cotzia, John Chen, Dacia Dimartino, Xiaojun Feng, Tatyana Gindin, Emily Guzman, Adriana Heguy, Megan Hogan, Emily Huang, George Jour, Andrew Lytle, Christian Marier, Matthew T. Maurano, Mark J. Mulligan, Peter Meyn, Iman Osman, Jared Pinnell, Vanessa Raabe, Sitharam Ramaswami, Amy Rapkiewicz, Marie Samanovic-Golden, Antonio Serrano, Guomiao Shen, Matija Snuderl, Theodore Vougiouklakis, Nick Vulpescu, Gael Westby, Paul Zappile, Yutong Zhang |
| EPI_ISL_427273, EPI_ISL_427274, EPI_ISL_427275, EPI_ISL_427276, EPI_ISL_427277, EPI_ISL_427278, EPI_ISL_427279, EPI_ISL_427280, EPI_ISL_427281, EPI_ISL_427282, EPI_ISL_427283, EPI_ISL_427284, EPI_ISL_427285, EPI_ISL_427286, EPI_ISL_427287 | | | |
| see above | Minnesota Department of Health, Public Health Laboratory | Minnesota Department of Health, Public Health Laboratory | Matt Plumb, Jacob Garfin and Xiong Wang |
| EPI_ISL_427469, EPI_ISL_427470, EPI_ISL_427471, EPI_ISL_427472, EPI_ISL_427473, EPI_ISL_427474, EPI_ISL_427475, EPI_ISL_427476, EPI_ISL_427477, EPI_ISL_427478, EPI_ISL_427479, EPI_ISL_427480, EPI_ISL_427481, EPI_ISL_427482, EPI_ISL_427483, EPI_ISL_427484, EPI_ISL_427485, EPI_ISL_427486, EPI_ISL_427487, EPI_ISL_427488, EPI_ISL_427489, EPI_ISL_427490, EPI_ISL_427491, EPI_ISL_427492, EPI_ISL_427493, EPI_ISL_427494, EPI_ISL_427495, EPI_ISL_427496, EPI_ISL_427497, EPI_ISL_427498, EPI_ISL_427499, EPI_ISL_427500, EPI_ISL_427501, EPI_ISL_427502, EPI_ISL_427503, EPI_ISL_427504, EPI_ISL_427505, EPI_ISL_427506, EPI_ISL_427507, EPI_ISL_427508, EPI_ISL_427509, EPI_ISL_427510, EPI_ISL_427511, EPI_ISL_427512, EPI_ISL_427513, EPI_ISL_427514, EPI_ISL_427515, EPI_ISL_427516, EPI_ISL_427517, EPI_ISL_427518, EPI_ISL_427519, EPI_ISL_427520, EPI_ISL_427521, EPI_ISL_427524, EPI_ISL_427525, EPI_ISL_427627, EPI_ISL_427628, EPI_ISL_427629, EPI_ISL_427630, EPI_ISL_427631, EPI_ISL_427632, EPI_ISL_427633, EPI_ISL_427634, EPI_ISL_427635, EPI_ISL_427636, EPI_ISL_427637, EPI_ISL_427638, EPI_ISL_427639, EPI_ISL_427640, EPI_ISL_427641, EPI_ISL_427642 | | | |
| see above | NYU Langone Health | Departments of Pathology and Medicine, New York University School of Medicine | Maria Aguero-Rosenfeld, Brendan Belovarac, Margaret Black, Ludovic Boytard, John Cadley, Paolo Cotzia, John Chen, Dacia Dimartino, Xiaojun Feng, Tatyana Gindin, Emily Guzman, Adriana Heguy, Megan Hogan, Emily Huang, George Jour, Andrew Lytle, Christian Marier, Matthew T. Maurano, Mark J. Mulligan, Peter Meyn, Iman Osman, Jared Pinnell, Vanessa Raabe, Sitharam Ramaswami, Amy Rapkiewicz, Marie Samanovic-Golden, Antonio Serrano, Guomiao Shen, Matija Snuderl, Theodore Vougiouklakis, Nick Vulpescu, Gael Westby, Paul Zappile, Yutong Zhang |
| EPI_ISL_428232, EPI_ISL_428233, EPI_ISL_428234, EPI_ISL_428235, EPI_ISL_428236 | Hematology Laboratory, Section of Molecular Diagnostics, University Clinical Centre, Medical University of Gdansk | Department of Virology, Faculty of Medicine, University of Helsinki, Helsinki, Finland | Marlena Robakowska, Aneta Szulc, Maciej Grzybek, Olli Vapalahti, Teemu Smura |

| | | | |
|---|---|---|---|
| EPI_ISL_428347 | Service de Biologie Médicale - BP 125 | National Reference Center for Viruses of Respiratory Infections, Institut Pasteur, Paris | Mélanie Albert, Marion Barbet, Sylvie Behillil, Méline Bizard, Angela Brisebarre, Flora Donati, Etienne Simon-Lorière, Vincent Enouf, Maud Vanpeene, Sylvie van der Werf |
| EPI_ISL_428348 | Maison de Santé du Val d'Ormois | National Reference Center for Viruses of Respiratory Infections, Institut Pasteur, Paris | Mélanie Albert, Marion Barbet, Sylvie Behillil, Méline Bizard, Angela Brisebarre, Flora Donati, Etienne Simon-Lorière, Vincent Enouf, Maud Vanpeene, Sylvie van der Werf |
| EPI_ISL_428349 | Service de Biologie Médicale - BP 125 | National Reference Center for Viruses of Respiratory Infections, Institut Pasteur, Paris | Mélanie Albert, Marion Barbet, Sylvie Behillil, Méline Bizard, Angela Brisebarre, Flora Donati, Etienne Simon-Lorière, Vincent Enouf, Maud Vanpeene, Sylvie van der Werf |
| EPI_ISL_428350 | CH Jean de Navarre Laboratoire de Biologie | National Reference Center for Viruses of Respiratory Infections, Institut Pasteur, Paris | Mélanie Albert, Marion Barbet, Sylvie Behillil, Méline Bizard, Angela Brisebarre, Flora Donati, Etienne Simon-Lorière, Vincent Enouf, Maud Vanpeene, Sylvie van der Werf |
| EPI_ISL_428351, EPI_ISL_428352 | GH Nord Essonne Service de Biologie clinique | National Reference Center for Viruses of Respiratory Infections, Institut Pasteur, Paris | Mélanie Albert, Marion Barbet, Sylvie Behillil, Méline Bizard, Angela Brisebarre, Flora Donati, Etienne Simon-Lorière, Vincent Enouf, Maud Vanpeene, Sylvie van der Werf |
| EPI_ISL_428353 | CH Compiègne Laboratoire de Biologie | National Reference Center for Viruses of Respiratory Infections, Institut Pasteur, Paris | Mélanie Albert, Marion Barbet, Sylvie Behillil, Méline Bizard, Angela Brisebarre, Flora Donati, Etienne Simon-Lorière, Vincent Enouf, Maud Vanpeene, Sylvie van der Werf |
| EPI_ISL_428354 | LABM GH nord Essonne de Longjumeau - BP 125 | National Reference Center for Viruses of Respiratory Infections, Institut Pasteur, Paris | Mélanie Albert, Marion Barbet, Sylvie Behillil, Méline Bizard, Angela Brisebarre, Flora Donati, Etienne Simon-Lorière, Vincent Enouf, Maud Vanpeene, Sylvie van der Werf |
| EPI_ISL_428355, EPI_ISL_428356, EPI_ISL_428357 | Institut Médico légal- Hop R. Poincaré | National Reference Center for Viruses of Respiratory Infections, Institut Pasteur, Paris | Mélanie Albert, Marion Barbet, Sylvie Behillil, Méline Bizard, Angela Brisebarre, Flora Donati, Etienne Simon-Lorière, Vincent Enouf, Maud Vanpeene, Sylvie van der Werf |
| EPI_ISL_428358 | CH Jeanne de Navarre Laboratoire de Biologie | National Reference Center for Viruses of Respiratory Infections, Institut Pasteur, Paris | Mélanie Albert, Marion Barbet, Sylvie Behillil, Méline Bizard, Angela Brisebarre, Flora Donati, Etienne Simon-Lorière, Vincent Enouf, Maud Vanpeene, Sylvie van der Werf |
| EPI_ISL_428359, EPI_ISL_428360 | CH Compiègne Laboratoire de Biologie | National Reference Center for Viruses of Respiratory Infections, Institut Pasteur, Paris | Mélanie Albert, Marion Barbet, Sylvie Behillil, Méline Bizard, Angela Brisebarre, Flora Donati, Etienne Simon-Lorière, Vincent Enouf, Maud Vanpeene, Sylvie van der Werf |
| EPI_ISL_428361, EPI_ISL_428362 | LABM GH nord Essonne de Longjumeau - BP 125 | National Reference Center for Viruses of Respiratory Infections, Institut Pasteur, Paris | Mélanie Albert, Marion Barbet, Sylvie Behillil, Méline Bizard, Angela Brisebarre, Flora Donati, Etienne Simon-Lorière, Vincent Enouf, Maud Vanpeene, Sylvie van der Werf |
| EPI_ISL_428363 | GH Nord Essonne Service de Biologie clinique | National Reference Center for Viruses of Respiratory Infections, Institut Pasteur, Paris | Mélanie Albert, Marion Barbet, Sylvie Behillil, Méline Bizard, Angela Brisebarre, Flora Donati, Etienne Simon-Lorière, Vincent Enouf, Maud Vanpeene, Sylvie van der Werf |
| EPI_ISL_428364 | Cabinet Médical | National Reference Center for Viruses of Respiratory Infections, Institut Pasteur, Paris | Mélanie Albert, Marion Barbet, Sylvie Behillil, Méline Bizard, Angela Brisebarre, Flora Donati, Etienne Simon-Lorière, Vincent Enouf, Maud Vanpeene, Sylvie van der Werf |
| EPI_ISL_428365 | LABM GH nord Essonne de Longjumeau - BP 125 | National Reference Center for Viruses of Respiratory Infections, Institut Pasteur, Paris | Mélanie Albert, Marion Barbet, Sylvie Behillil, Méline Bizard, Angela Brisebarre, Flora Donati, Etienne Simon-Lorière, Vincent Enouf, Maud Vanpeene, Sylvie van der Werf |
| EPI_ISL_428366 | CH Jeanne de Navarre Laboratoire de Biologie | National Reference Center for Viruses of Respiratory Infections, Institut Pasteur, Paris | Mélanie Albert, Marion Barbet, Sylvie Behillil, Méline Bizard, Angela Brisebarre, Flora Donati, Etienne Simon-Lorière, Vincent Enouf, Maud Vanpeene, Sylvie van der Werf |
| EPI_ISL_428367 | Cabinet Médical | National Reference Center for Viruses of Respiratory Infections, Institut Pasteur, Paris | Mélanie Albert, Marion Barbet, Sylvie Behillil, Méline Bizard, Angela Brisebarre, Flora Donati, Etienne Simon-Lorière, Vincent Enouf, Maud Vanpeene, Sylvie van der Werf |
| EPI_ISL_428757, EPI_ISL_428758, EPI_ISL_428759, EPI_ISL_428760, EPI_ISL_428761, EPI_ISL_428762, EPI_ISL_428763, EPI_ISL_428764, EPI_ISL_428765, EPI_ISL_428766, EPI_ISL_428767, EPI_ISL_428768, EPI_ISL_428769, EPI_ISL_428770, EPI_ISL_428771, EPI_ISL_428772, EPI_ISL_428773, EPI_ISL_428774, EPI_ISL_428775, EPI_ISL_428776, EPI_ISL_428777, EPI_ISL_428778, EPI_ISL_428779, EPI_ISL_428780, EPI_ISL_428781, EPI_ISL_428782, EPI_ISL_428783, EPI_ISL_428784, EPI_ISL_428785, EPI_ISL_428786, EPI_ISL_428787, EPI_ISL_428788, EPI_ISL_428789, EPI_ISL_428790, EPI_ISL_428791, EPI_ISL_428792, EPI_ISL_428793, EPI_ISL_428794, EPI_ISL_428795, EPI_ISL_428796, EPI_ISL_428797, EPI_ISL_428798, EPI_ISL_428799, EPI_ISL_428800, EPI_ISL_428801, EPI_ISL_428802, EPI_ISL_428803, EPI_ISL_428804, EPI_ISL_428805 | | | |
| see above | NYU Langone Health | Departments of Pathology and Medicine, New York University School of Medicine | Maria Aguero-Rosenfeld, Brendan Belovarac, Margaret Black, Ludovic Boytard, John Cadley, Paolo Cotzia, John Chen, Dacia Dimartino, Xiaojun Feng, Tatyana Gindin, Emily Guzman, Adriana Heguy, Megan Hogan, Emily Huang, George Jour, Andrew Lytle, Christian Marier, Matthew T. Maurano, Mark J. Mulligan, Peter Meyn, Iman Osman, Jared Pinnell, Vanessa Raabe, Sitharam Ramaswami, Amy Rapkiewicz, Marie Samanovic-Golden, Antonio Serrano, Guomiao Shen, Matija Snuderl, Theodore Vougiouklakis, Nick Vulpescu, Gael Westby, Paul Zappile, Yutong Zhang |
| EPI_ISL_429136, EPI_ISL_429137, EPI_ISL_429138, EPI_ISL_429139, EPI_ISL_429140, EPI_ISL_429141, EPI_ISL_429142, EPI_ISL_429143, EPI_ISL_429144, EPI_ISL_429145, EPI_ISL_429146, EPI_ISL_429147, EPI_ISL_429148, EPI_ISL_429150, EPI_ISL_429151, EPI_ISL_429152, EPI_ISL_429153, EPI_ISL_429154, EPI_ISL_429155, EPI_ISL_429156, EPI_ISL_429157, EPI_ISL_429158, EPI_ISL_429160 | | | |
| see above | Klinisk mikrobiologi Orebro | The Public Health Agency of Sweden | Martin Sundqvist, Olov Svartstrom, Maria Lind Karlberg, Anna-Malin Linde, Oskar Karlsson Lindsjo, Anna Risberg, Shaman Muradrasoli, Karin Tegmark-Wisell |
| EPI_ISL_429968 | Centre Hospitalier Compiègne Laboratoire de Biologie | National Reference Center for Viruses of Respiratory Infections, Institut Pasteur, Paris | Mélanie Albert, Marion Barbet, Sylvie Behillil, Méline Bizard, Angela Brisebarre, Flora Donati, Fabiana Gambaro, Etienne Simon-Lorière, Vincent Enouf, Maud Vanpeene, Sylvie van der Werf, Raulin Olivia |
| EPI_ISL_430319, EPI_ISL_430320, EPI_ISL_430321, EPI_ISL_430322, EPI_ISL_430323, EPI_ISL_430324, EPI_ISL_430325, EPI_ISL_430326, EPI_ISL_430327, EPI_ISL_430328, EPI_ISL_430329, EPI_ISL_430330, EPI_ISL_430331, EPI_ISL_430332, EPI_ISL_430333, EPI_ISL_430334, EPI_ISL_430335, EPI_ISL_430336, EPI_ISL_430337, EPI_ISL_430338, EPI_ISL_430339, EPI_ISL_430340, EPI_ISL_430341, EPI_ISL_430342, EPI_ISL_430343, EPI_ISL_430344, EPI_ISL_430345, EPI_ISL_430346, EPI_ISL_430347, EPI_ISL_430348, EPI_ISL_430349, EPI_ISL_430350, EPI_ISL_430351, EPI_ISL_430352, EPI_ISL_430353, EPI_ISL_430354, EPI_ISL_430355, EPI_ISL_430356, EPI_ISL_430357, EPI_ISL_430358, EPI_ISL_430359, EPI_ISL_430360, EPI_ISL_430361, EPI_ISL_430362, EPI_ISL_430363, EPI_ISL_430364, EPI_ISL_430365, EPI_ISL_430366, EPI_ISL_430367, EPI_ISL_430368, EPI_ISL_430369, EPI_ISL_430370, EPI_ISL_430371, EPI_ISL_430372, EPI_ISL_430373, EPI_ISL_430374, EPI_ISL_430375, EPI_ISL_430376, EPI_ISL_430377, EPI_ISL_430378, EPI_ISL_430379, EPI_ISL_430380, EPI_ISL_430381, EPI_ISL_430382, EPI_ISL_430383, EPI_ISL_430384, EPI_ISL_430385, EPI_ISL_430386, EPI_ISL_430387, EPI_ISL_430388, EPI_ISL_430389, EPI_ISL_430390, EPI_ISL_430391, EPI_ISL_430392, EPI_ISL_430393, EPI_ISL_430394, EPI_ISL_430395, EPI_ISL_430396, EPI_ISL_430397, EPI_ISL_430398, EPI_ISL_430399, EPI_ISL_430400, EPI_ISL_430401, EPI_ISL_430402, EPI_ISL_430403, EPI_ISL_430404, EPI_ISL_430405, EPI_ISL_430406, EPI_ISL_430407, EPI_ISL_430408, EPI_ISL_430409, EPI_ISL_430410, EPI_ISL_430411, EPI_ISL_430412, EPI_ISL_430413, EPI_ISL_430414, EPI_ISL_430415, EPI_ISL_430416, EPI_ISL_430417, EPI_ISL_430418, EPI_ISL_430428, EPI_ISL_430430, EPI_ISL_430431, EPI_ISL_430432, EPI_ISL_430433, EPI_ISL_430434 | | | |
| see above | NYU Langone Health | Departments of Pathology and Medicine, New York University School of Medicine | Maria Aguero-Rosenfeld, Brendan Belovarac, Margaret Black, Ludovic Boytard, John Cadley, Paolo Cotzia, John Chen, Dacia Dimartino, Xiaojun Feng, Tatyana Gindin, Emily Guzman, Adriana Heguy, Megan Hogan, Emily Huang, George Jour, Lawrence H. Lin, Raven Luther, Andrew Lytle, Christian Marier, Matthew T. Maurano, Mark J. Mulligan, Peter Meyn, Raquel Ordonez Ciriza, Iman Osman, Jared Pinnell, Vanessa Raabe, Sitharam Ramaswami, Amy Rapkiewicz, Andre M. Ribeiro-dos-Santos, Marie Samanovic-Golden, Antonio Serrano, Guomiao Shen, Matija Snuderl, Theodore Vougiouklakis, Nick Vulpescu, Gael Westby, Paul Zappile, Yutong Zhang |
| EPI_ISL_430456 | Rizal Medical Center | Research Institute for Tropical Medicine | Medado,I.A.P., Bautista,C.T., Onza,O.J.T., Polotan,F.G.M., Brunker, K., Mercado,E.S., Manalo, D.L., Demetria, C.S. |
| EPI_ISL_430793, EPI_ISL_430794 | Laboratorio Análisis Clínicos, Unidad de Servicios Diagnósticos, Swiss Medical Group | Área de Secuenciación del Laboratorio de Virología del Hospital de Niños Dr. Ricardo Gutierrez on behalf of 'Proyecto Argentino Interinstitucional de genomica de SARS-CoV-2' (PAIS Consortium) | Nabaes Jodar, MS; Goya, S; Natale, MI; Lusso, S; Sanchez, O; Guevara, D; Vicario, SM; Mistchenko, AS; Valinotto, LE; Viegas, M. |
| EPI_ISL_430795 | Laboratorio de Virología del Hospital de Niños Dr. Ricardo Gutierrez | Área de Secuenciación del Laboratorio de Virología del Hospital de Niños Dr. Ricardo Gutierrez on behalf of 'Proyecto Argentino Interinstitucional de genomica de SARS-CoV-2' (PAIS Consortium) | Nabaes Jodar, MS; Goya, S; Natale, MI; Lusso, S; Gravis, E; Mistchenko, AS; Valinotto, LE; Viegas, M. |
| EPI_ISL_430796, EPI_ISL_430797, EPI_ISL_430798 | Departamento de Biología y genética molecular, IACA Laboratorios. | Área de Secuenciación del Laboratorio de Virología del Hospital de Niños Dr. Ricardo Gutierrez on behalf of 'Proyecto Argentino Interinstitucional de genomica de SARS-CoV-2' (PAIS Consortium) | Nabaes Jodar, MS; Goya, S; Natale, MI; Lusso, S; Tittarelli, E; Suárez, A; Masciovecchio MV; Streitenberger ER; Mistchenko, AS; Valinotto, LE; Viegas, M. |
| EPI_ISL_430799, EPI_ISL_430800, EPI_ISL_430801 | Laboratorio de Virología del Hospital de Niños Dr. Ricardo Gutierrez | Área de Secuenciación del Laboratorio de Virología del Hospital de Niños Dr. Ricardo Gutierrez on behalf of 'Proyecto Argentino Interinstitucional de genomica de SARS-CoV-2' (PAIS Consortium) | Nabaes Jodar, MS; Goya, S; Natale, MI; Lusso, S; Gravis, E; Mistchenko, AS; Valinotto, LE; Viegas, M. |

| | | | |
|---|---|---|---|
| EPI_ISL_430802 | Departamento de Biología y genética molecular, IACA Laboratorios. | Área de Secuenciación del Laboratorio de Virología del Hospital de Niños Dr. Ricardo Gutierrez on behalf of 'Proyecto Argentino Interinstitucional de genomica de SARS-CoV-2' (PAIS Consortium) | Nabaes Jodar, MS; Goya, S; Natale, MI; Lusso, S; Tittarelli, E; Suárez, A; Masciovecchio MV; Streitenberger ER; Mistchenko, AS; Valinotto, LE; Viegas, M. |
| EPI_ISL_430803, EPI_ISL_430804 | Laboratorio de Virología del Hospital de Niños Dr. Ricardo Gutierrez | Área de Secuenciación del Laboratorio de Virología del Hospital de Niños Dr. Ricardo Gutierrez on behalf of 'Proyecto Argentino Interinstitucional de genomica de SARS-CoV-2' (PAIS Consortium) | Nabaes Jodar, MS; Goya, S; Natale, MI; Lusso, S; Gravis, E; Mistchenko, AS; Valinotto, LE; Viegas, M. |
| EPI_ISL_430805, EPI_ISL_430806 | Departamento de Biología y genética molecular, IACA Laboratorios. | Área de Secuenciación del Laboratorio de Virología del Hospital de Niños Dr. Ricardo Gutierrez on behalf of 'Proyecto Argentino Interinstitucional de genomica de SARS-CoV-2' (PAIS Consortium) | Nabaes Jodar, MS; Goya, S; Natale, MI; Lusso, S; Tittarelli, E; Suárez, A; Masciovecchio MV; Streitenberger ER; Mistchenko, AS; Valinotto, LE; Viegas, M. |
| EPI_ISL_430807 | Laboratorio de Virología del Hospital de Niños Dr. Ricardo Gutierrez | Área de Secuenciación del Laboratorio de Virología del Hospital de Niños Dr. Ricardo Gutierrez on behalf of 'Proyecto Argentino Interinstitucional de genomica de SARS-CoV-2' (PAIS Consortium) | Nabaes Jodar, MS; Goya, S; Natale, MI; Lusso, S; Gravis, E; Mistchenko, AS; Valinotto, LE; Viegas, M. |
| EPI_ISL_430808 | Departamento de Biología y genética molecular, IACA Laboratorios. | Área de Secuenciación del Laboratorio de Virología del Hospital de Niños Dr. Ricardo Gutierrez on behalf of 'Proyecto Argentino Interinstitucional de genomica de SARS-CoV-2' (PAIS Consortium) | Nabaes Jodar, MS; Goya, S; Natale, MI; Lusso, S; Tittarelli, E; Suárez, A; Masciovecchio MV; Streitenberger ER; Mistchenko, AS; Valinotto, LE; Viegas, M. |
| EPI_ISL_430809, EPI_ISL_430810, EPI_ISL_430811, EPI_ISL_430812, EPI_ISL_430813, EPI_ISL_430814, EPI_ISL_430815, EPI_ISL_430816, EPI_ISL_430817, EPI_ISL_430818 | Laboratorio de Virología del Hospital de Niños Dr. Ricardo Gutierrez | Área de Secuenciación del Laboratorio de Virología del Hospital de Niños Dr. Ricardo Gutierrez on behalf of 'Proyecto Argentino Interinstitucional de genomica de SARS-CoV-2' (PAIS Consortium) | Nabaes Jodar, MS; Goya, S; Natale, MI; Lusso, S; Gravis, E; Mistchenko, AS; Valinotto, LE; Viegas, M. |
| EPI_ISL_430819 | Center of Scientific Excellence for Influenza Viruses,National Research Centre (NRC), Egypt. | Center of Scientific Excellence for Influenza Viruses,National Research Centre (NRC), Egypt. | Mohamed Ahmed Ali, Ahmed Kandeil, Ahmed Mostafa, Rabeh El-Shesheny, Mahmoud Shehata, Wael Roshdy, Shymaa Showky Ahmed , Amal Naguib, Nancy M. El Guindy, Mokhtar Gomaa, Ahmed El-Taweel, Ahmed E Kayed, Yassmin Moatasim, Omnia Kutkat, Sara Mahmoud, Mina Kamel, Abo Shama, M Noura, Mohamed El Sayes |
| EPI_ISL_430820 | Center of Scientific Excellence for Influenza Viruses, National Research Centre (NRC), Egypt. | Center of Scientific Excellence for Influenza Viruses, National Research Centre (NRC), Egypt. | Mohamed Ahmed Ali, Ahmed Kandeil, Ahmed Mostafa, Rabeh El-Shesheny, Mahmoud Shehata, Wael Roshdy, Shymaa Showky Ahmed , Amal Naguib, Mokhtar Gomaa, Ahmed El-Taweel, Ahmed E Kayed, Yassmin Moatasim, Omnia Kutkat, Sara Mahmoud, Mina Kamel, Abo Shama, M Noura, Mohamed El Sayes, Nancy M. El Guindy |
| EPI_ISL_430838 | Makati Medical Center | Research Institute for Tropical Medicine | Medado,I.A.P., Bautista,C.T., Onza,O.J.T., Polotan,F.G.M., Brunker, K., Mercado,E.S., Manalo, D.L., Demetria, C.S. |
| EPI_ISL_430839 | Research Institute for Tropical Medicine | Research Institute for Tropical Medicine | Medado,I.A.P., Bautista,C.T., Onza,O.J.T., Polotan,F.G.M., Brunker, K., Mercado,E.S., Manalo, D.L., Demetria, C.S. |
| EPI_ISL_430840 | Veterans Memorial Medical Center | Research Institute for Tropical Medicine | Medado,I.A.P., Bautista,C.T., Onza,O.J.T., Polotan,F.G.M., Brunker, K., Mercado,E.S., Manalo, D.L., Demetria, C.S. |
| EPI_ISL_430843 | Bethany Hospital | Research Institute for Tropical Medicine | Medado,I.A.P., Bautista,C.T., Onza,O.J.T., Polotan,F.G.M., Brunker, K., Mercado,E.S., Manalo, D.L., Demetria, C.S. |
| EPI_ISL_430844 | Lung Center of the Philippines | Research Institute for Tropical Medicine | Medado,I.A.P., Bautista,C.T., Onza,O.J.T., Polotan,F.G.M., Brunker, K., Mercado,E.S., Manalo, D.L., Demetria, C.S. |
| EPI_ISL_430845 | Pasig City General Hospital | Research Institute for Tropical Medicine | Medado,I.A.P., Bautista,C.T., Onza,O.J.T., Polotan,F.G.M., Brunker, K., Mercado,E.S., Manalo, D.L., Demetria, C.S. |
| EPI_ISL_430860, EPI_ISL_430861, EPI_ISL_430863 | Klinisk mikrobiologi Orebro | The Public Health Agency of Sweden | Martin Sundqvist, Oskar Karlsson Lindsjo, Maria Lind Karlberg, Anna-Malin Linde, Olov Svartstrom, Anna Risberg, Shaman Muradrasoli, Karin Tegmark-Wisell |
| EPI_ISL_431101 | Department of Microbiology,Gandhi Medical College and Hospital | Virus Research Laboratory, Department of Zoology, Osmania University,Hyderabad,India | Muttineni Radhakrishna, Nagamani K, Thrilok Chander B, Raja Rao M, Kalyani Putty, Ravikumar P, Sunitha P, Pankaj Singh D, Anand Kumar K, Amit A. Upadhyay, Steven E. Bosinger, Rama Amara |
| EPI_ISL_431102 | Department of MicroBiology,Gandhi Medical College and Hospital,Secendrabad,Hyderabad,India | Department of Microbiology, Gandhi Medical College and Hospital, Secendrabad, Hyderabad | Nagamani K, Muttineni Radhakrishna, Thrilok Chander B, Raja Rao M, Kalyani Putty, Ravikumar P, Sunitha P, Pankaj Singh D, Anand Kumar K, Amit A. Upadhyay, Steven E. Bosinger, Rama Amara |
| EPI_ISL_431103 | Department of Microbiology, Gandhi Medical College and Hospital, Secendrabad, Hyderabad, India | Department of Microbiology, Gandhi Medical College and Hospital, Secendrabad, Hyderabad, India | Nagamani K, Muttineni Radhakrishna, Thrilok Chander B, Raja Rao M, Kalyani Putty, Ravikumar P, Sunitha P, Pankaj Singh D, Anand Kumar K, Amit A. Upadhyay, Steven E. Bosinger, Rama Amara |
| EPI_ISL_432868, EPI_ISL_432869, EPI_ISL_432870, EPI_ISL_432871, EPI_ISL_432872, EPI_ISL_432873, EPI_ISL_432874, EPI_ISL_432875, EPI_ISL_432876, EPI_ISL_432877, EPI_ISL_432878, EPI_ISL_432879, EPI_ISL_432880, EPI_ISL_432881, EPI_ISL_432882, EPI_ISL_432883, EPI_ISL_432884, EPI_ISL_432885, EPI_ISL_432886, EPI_ISL_432887, EPI_ISL_432888, EPI_ISL_432889, EPI_ISL_432890, EPI_ISL_432891, EPI_ISL_432892, EPI_ISL_432893, EPI_ISL_432894, EPI_ISL_432895, EPI_ISL_432896, EPI_ISL_432897, EPI_ISL_432898, EPI_ISL_432899, EPI_ISL_433068, EPI_ISL_433069, EPI_ISL_433070, EPI_ISL_433071, EPI_ISL_433072, EPI_ISL_433073, EPI_ISL_433074, EPI_ISL_433075, EPI_ISL_433076, EPI_ISL_433077, EPI_ISL_433078, EPI_ISL_433079, EPI_ISL_433080, EPI_ISL_433081, EPI_ISL_433082, EPI_ISL_433083, EPI_ISL_433084, EPI_ISL_433085, EPI_ISL_433086, EPI_ISL_433087, EPI_ISL_433088, EPI_ISL_433089, EPI_ISL_433090, EPI_ISL_433091, EPI_ISL_433092, EPI_ISL_433093, EPI_ISL_433094, EPI_ISL_433095, EPI_ISL_433096, EPI_ISL_433097, EPI_ISL_433098, EPI_ISL_433099, EPI_ISL_433100, EPI_ISL_433101, EPI_ISL_433102, EPI_ISL_433103, EPI_ISL_433104, EPI_ISL_433105, EPI_ISL_433106, EPI_ISL_433107, EPI_ISL_433108, EPI_ISL_433109, EPI_ISL_433110, EPI_ISL_433111, EPI_ISL_433112, EPI_ISL_433113, EPI_ISL_433114, EPI_ISL_433115, EPI_ISL_433116, EPI_ISL_433117, EPI_ISL_433118, EPI_ISL_433119, EPI_ISL_433120, EPI_ISL_433121, EPI_ISL_433122, EPI_ISL_433123, EPI_ISL_433124, EPI_ISL_433125, EPI_ISL_433126, EPI_ISL_433127, EPI_ISL_433128, EPI_ISL_433129, EPI_ISL_433130, EPI_ISL_433131, EPI_ISL_433132, EPI_ISL_433133, EPI_ISL_433134, EPI_ISL_433135, EPI_ISL_433136, EPI_ISL_433137, EPI_ISL_433138, EPI_ISL_433139, EPI_ISL_433140, EPI_ISL_433141, EPI_ISL_433142, EPI_ISL_433143, EPI_ISL_433144, EPI_ISL_433145, EPI_ISL_433146, EPI_ISL_433147, EPI_ISL_433148, EPI_ISL_433149, EPI_ISL_433150, EPI_ISL_433151, EPI_ISL_433152, EPI_ISL_433153, EPI_ISL_433154, EPI_ISL_433155, EPI_ISL_433156, EPI_ISL_433157, EPI_ISL_433158, EPI_ISL_433159, EPI_ISL_433160, EPI_ISL_433161, EPI_ISL_433162, EPI_ISL_433163, EPI_ISL_433164, EPI_ISL_433165, EPI_ISL_433166, EPI_ISL_433167, EPI_ISL_433168, EPI_ISL_433169, EPI_ISL_433170, EPI_ISL_433171, EPI_ISL_433172, EPI_ISL_433173, EPI_ISL_433174, EPI_ISL_433175, EPI_ISL_433176, EPI_ISL_433177, EPI_ISL_433178, EPI_ISL_433179, EPI_ISL_433180, EPI_ISL_433181, EPI_ISL_433182, EPI_ISL_433183, EPI_ISL_433184, EPI_ISL_433185, EPI_ISL_433186, EPI_ISL_433187, EPI_ISL_433188, EPI_ISL_433189, EPI_ISL_433190, EPI_ISL_433191, EPI_ISL_433192, EPI_ISL_433193, EPI_ISL_433194, EPI_ISL_433195, EPI_ISL_433196, EPI_ISL_433197, EPI_ISL_433198, EPI_ISL_433199, EPI_ISL_433200, EPI_ISL_433201, EPI_ISL_433202, EPI_ISL_433203, EPI_ISL_433204, EPI_ISL_433205, EPI_ISL_433206, EPI_ISL_433207, EPI_ISL_433208, EPI_ISL_433209, EPI_ISL_433210, EPI_ISL_433211, EPI_ISL_433212, EPI_ISL_433213, EPI_ISL_433214, EPI_ISL_433215, EPI_ISL_433216, EPI_ISL_433217, EPI_ISL_433218, EPI_ISL_433219, EPI_ISL_433220, EPI_ISL_433221, EPI_ISL_433222, EPI_ISL_433223, EPI_ISL_433224, EPI_ISL_433225, EPI_ISL_433226, EPI_ISL_433227, EPI_ISL_433228, EPI_ISL_433229, EPI_ISL_433230, EPI_ISL_433231, EPI_ISL_433232, EPI_ISL_433233, EPI_ISL_433234, EPI_ISL_433235, EPI_ISL_433236, EPI_ISL_433237, EPI_ISL_433238, EPI_ISL_433239, EPI_ISL_433240, EPI_ISL_433241, EPI_ISL_433242, EPI_ISL_433243, EPI_ISL_433244, EPI_ISL_433245, EPI_ISL_433246, EPI_ISL_433247, EPI_ISL_433248, EPI_ISL_433249, EPI_ISL_433250, EPI_ISL_433251, EPI_ISL_433252, EPI_ISL_433253, EPI_ISL_433254, EPI_ISL_433255, EPI_ISL_433256, EPI_ISL_433257, EPI_ISL_433258, EPI_ISL_433259, EPI_ISL_433260, EPI_ISL_433261, EPI_ISL_433262, EPI_ISL_433263, EPI_ISL_433264, EPI_ISL_433265, EPI_ISL_433266, EPI_ISL_433267, EPI_ISL_433374, EPI_ISL_433375, EPI_ISL_433376, EPI_ISL_433377, EPI_ISL_433378, EPI_ISL_433379, EPI_ISL_433380, EPI_ISL_433381, EPI_ISL_433382, EPI_ISL_433383, EPI_ISL_433384, EPI_ISL_433385, EPI_ISL_433386, EPI_ISL_433387, EPI_ISL_433388, EPI_ISL_433389, EPI_ISL_433390, EPI_ISL_433391, EPI_ISL_433392, EPI_ISL_433393, EPI_ISL_433394, EPI_ISL_433395, EPI_ISL_433396, EPI_ISL_433397, EPI_ISL_433398, EPI_ISL_433399, EPI_ISL_433400, EPI_ISL_433401, EPI_ISL_433402, EPI_ISL_433403, EPI_ISL_433404, EPI_ISL_433405, EPI_ISL_433406, EPI_ISL_433407, EPI_ISL_433408, EPI_ISL_433409, EPI_ISL_433410, EPI_ISL_433411, EPI_ISL_433412, EPI_ISL_433413, EPI_ISL_433414, EPI_ISL_433415, EPI_ISL_433416, EPI_ISL_433417, EPI_ISL_433418, EPI_ISL_433419, EPI_ISL_433420, EPI_ISL_433421, EPI_ISL_433422, EPI_ISL_433423, EPI_ISL_433424, EPI_ISL_433425, EPI_ISL_433426, EPI_ISL_433427, EPI_ISL_433428, EPI_ISL_433429, EPI_ISL_433430, EPI_ISL_433431, EPI_ISL_433432, EPI_ISL_433433, EPI_ISL_433434, EPI_ISL_433435, EPI_ISL_433436, EPI_ISL_433437, EPI_ISL_433438, EPI_ISL_433439, EPI_ISL_433440, EPI_ISL_433441, EPI_ISL_433442, EPI_ISL_433443, EPI_ISL_433444, EPI_ISL_433445, EPI_ISL_433446, EPI_ISL_433447, EPI_ISL_433448, EPI_ISL_433449, EPI_ISL_433450, EPI_ISL_433451, EPI_ISL_433452, EPI_ISL_433453, EPI_ISL_433454, EPI_ISL_433455, EPI_ISL_433456, EPI_ISL_433457, EPI_ISL_433458, EPI_ISL_433459, EPI_ISL_433460, EPI_ISL_433461, EPI_ISL_433462, EPI_ISL_433463, EPI_ISL_433464, EPI_ISL_433465 | | | |
| see above | Virology Department, Royal Infirmary of Edinburgh, NHS Lothian / School of Biological Sciences, University of Edinburgh / Institute of Genetics and Molecular Medicine, University of Edinburgh | COVID-19 Genomics UK (COG-UK) Consortium | McHugh M, Dewar R, Rooke S, Gallagher M, Balcaza C, O'Toole A, Hill V, McCrone JT, Colquhoun R, Yu X, Jackson B, Rambaut A, Williams TC, Templeton K |
| EPI_ISL_434651 | Krokoms Halsocentral | The Public Health Agency of Sweden | Martin Ersson, Oskar Karlsson Lindsjo, Maria Lind Karlberg, Anna-Malin Linde, Olov Svartstrom, Anna Risberg, Theresa Enkirch, Mia Brytting, Karin Tegmark-Wisell |
| EPI_ISL_434657 | Kristianstadkliniken | The Public Health Agency of Sweden | Mia Settergren Hammer, Oskar Karlsson Lindsjo, Maria Lind Karlberg, Anna-Malin Linde, Olov Svartstrom, Anna Risberg, Theresa Enkirch, Mia Brytting, Karin Tegmark-Wisell |
| EPI_ISL_434658 | Lundens VC | The Public Health Agency of Sweden | Marita Dagner, Oskar Karlsson Lindsjo, Maria Lind Karlberg, Anna-Malin Linde, Olov Svartstrom, Anna Risberg, Theresa Enkirch, Mia Brytting, Karin Tegmark-Wisell |
| EPI_ISL_434660 | Narhalsan Backa vardcentral | The Public Health Agency of Sweden | Mats Olsson, Oskar Karlsson Lindsjo, Maria Lind Karlberg, Anna-Malin Linde, Olov Svartstrom, Anna Risberg, Theresa Enkirch, Mia Brytting, Karin Tegmark-Wisell |

| | | | |
|---|---|---|---|
| EPI_ISL_434670 | Narhalsan Molnlycke, Barn och ungdomsmedicin | The Public Health Agency of Sweden | Mats Reimer, Oskar Karlsson Lindsjo, Maria Lind Karlberg, Anna-Malin Linde, Olov Svartstrom, Anna Risberg, Theresa Enkirch, Mia Brytting, Karin Tegmark-Wisell |
| EPI_ISL_434674 | Narhalsan Backa vardcentral | The Public Health Agency of Sweden | Mats Olsson, Oskar Karlsson Lindsjo, Maria Lind Karlberg, Anna-Malin Linde, Olov Svartstrom, Anna Risberg, Theresa Enkirch, Mia Brytting, Karin Tegmark-Wisell |
| EPI_ISL_435054 | B.J. Medical College and Civil hospital | Gujarat Biotechnology Research Centre | Monika Gandhi, Pinal Trivedi, Maharshi Pandya, Amit Kanani, Akanksha Verma, Nitin Savaliya, Raghawendra Kumar, Dinesh Kumar, Zuber Saiyed, Dipa Kinariwala, Disha Patel, Binita Aring, Geeta Vaghela, Sonia Barve, Bhavesh Modi, Kairavi Joshi, Gaurishankar Shrimali, Nidhi Sood, Pranay Shah, R D Dixit, Snehal Bagatharia, Kamlesh J Upadhyay, Ramesh Pandit, Tejas Shah, Ankit Hinsu, Pritesh Sabara, Apurvasinh Puvar, Janvi Raval, Priti Pandita, Chaitanya Joshi, Madhvi Joshi |
| EPI_ISL_435475, EPI_ISL_435476, EPI_ISL_435477, EPI_ISL_435478, EPI_ISL_435479, EPI_ISL_435480, EPI_ISL_435483, EPI_ISL_435488, EPI_ISL_435493, EPI_ISL_435494, EPI_ISL_435495, EPI_ISL_435496, EPI_ISL_435497, EPI_ISL_435498, EPI_ISL_435499, EPI_ISL_435500, EPI_ISL_435501, EPI_ISL_435502, EPI_ISL_435503, EPI_ISL_435504, EPI_ISL_435505, EPI_ISL_435506, EPI_ISL_435507, EPI_ISL_435508, EPI_ISL_435509, EPI_ISL_435510, EPI_ISL_435511, EPI_ISL_435512, EPI_ISL_435513, EPI_ISL_435514, EPI_ISL_435515, EPI_ISL_435516, EPI_ISL_435517, EPI_ISL_435518, EPI_ISL_435519, EPI_ISL_435520, EPI_ISL_435521, EPI_ISL_435522, EPI_ISL_435523, EPI_ISL_435524, EPI_ISL_435525, EPI_ISL_435526, EPI_ISL_435527, EPI_ISL_435528, EPI_ISL_435529, EPI_ISL_435530, EPI_ISL_435531, EPI_ISL_435532, EPI_ISL_435533, EPI_ISL_435534, EPI_ISL_435535, EPI_ISL_435536, EPI_ISL_435537, EPI_ISL_435538, EPI_ISL_435539, EPI_ISL_435540, EPI_ISL_435541, EPI_ISL_435542, EPI_ISL_435543, EPI_ISL_435544, EPI_ISL_435545, EPI_ISL_435546, EPI_ISL_435547, EPI_ISL_435548, EPI_ISL_435549, EPI_ISL_435721, EPI_ISL_435722 | | | |
| see above | NYU Langone Health | Departments of Pathology and Medicine, New York University School of Medicine | Maria Aguero-Rosenfeld, Brendan Belovarac, Margaret Black, Ludovic Boytard, John Cadley, Paolo Cotzia, John Chen, Dacia Dimartino, Xiaojun Feng, Tatyana Gindin, Emily Guzman, Adriana Heguy, Megan Hogan, Emily Huang, George Jour, Lawrence H. Lin, Raven Luther, Andrew Lytle, Christian Marier, Matthew T. Maurano, Mark J. Mulligan, Peter Meyn, Raquel Ordonez Ciriza, Iman Osman, Jared Pinnell, Vanessa Raabe, Sitharam Ramaswami, Amy Rapkiewicz, Andre M. Ribeiro-dos-Santos, Marie Samanovic-Golden, Antonio Serrano, Guomiao Shen, Matija Snuderl, Theodore Vougiouklakis, Nick Vulpescu, Gael Westby, Paul Zappile, Yutong Zhang |
| EPI_ISL_436195 | Servicio de Microbiología. Consorcio Hospital General Universitario de Valencia | Sequencing and Bioinformatics Service and Molecular Epidemiology Research Group. FISABIO-Public Health | Maria Dolores Ocete, Inma Galán Vendrell, Paula Ruiz-Hueso, Mariana Reyes-Prieto, Vicente Soriano Chirona, Maria Alma Bracho, Griselda De Marco, Beatriz Beamud, Lidia Ruiz Roldan, Marta Pla Diaz,Neris Garcia-Gonzalez, Loreto Ferrús Abad, Llúcia Martínez-Priego, Concepcion Gimeno, Giuseppe D'Auria, Fernando Gonzalez-Candelas |
| EPI_ISL_436199 | Servicio de Microbiología. Consorcio Hospital General Universitario de Valencia | Sequencing and Bioinformatics Service and Molecular Epidemiology Research Group. FISABIO-Public Health | Marta Pla Diaz,Neris Garcia-Gonzalez, Loreto Ferrús Abad, Maria Dolores Ocete, Inma Galán Vendrell, Paula Ruiz-Hueso, Mariana Reyes-Prieto, Vicente Soriano Chirona, Maria Alma Bracho, Griselda De Marco, Beatriz Beamud, Lidia Ruiz Roldan, Llúcia Martínez-Priego, Concepcion Gimeno, Giuseppe D'Auria, Fernando Gonzalez-Candelas |
| EPI_ISL_436203 | Servicio de Microbiología. Consorcio Hospital General Universitario de Valencia | Sequencing and Bioinformatics Service and Molecular Epidemiology Research Group. FISABIO-Public Health | Maria Dolores Ocete, Inma Galán Vendrell, Paula Ruiz-Hueso, Mariana Reyes-Prieto, Vicente Soriano Chirona, Maria Alma Bracho, Griselda De Marco, Beatriz Beamud, Lidia Ruiz Roldan, Marta Pla Diaz,Neris Garcia-Gonzalez, Loreto Ferrús Abad, Llúcia Martínez-Priego, Concepcion Gimeno, Giuseppe D'Auria, Fernando Gonzalez-Candelas |
| EPI_ISL_436207 | Servicio de Microbiología. Consorcio Hospital General Universitario de Valencia | Sequencing and Bioinformatics Service and Molecular Epidemiology Research Group. FISABIO-Public Health | Marta Pla Diaz,Neris Garcia-Gonzalez, Loreto Ferrús Abad, Maria Dolores Ocete, Inma Galán Vendrell, Paula Ruiz-Hueso, Mariana Reyes-Prieto, Vicente Soriano Chirona, Maria Alma Bracho, Griselda De Marco, Beatriz Beamud, Lidia Ruiz Roldan, Llúcia Martínez-Priego, Concepcion Gimeno, Giuseppe D'Auria, Fernando Gonzalez-Candelas |
| EPI_ISL_436211 | Servicio de Microbiología. Consorcio Hospital General Universitario de Valencia | Sequencing and Bioinformatics Service and Molecular Epidemiology Research Group. FISABIO-Public Health | Maria Dolores Ocete, Inma Galán Vendrell, Paula Ruiz-Hueso, Mariana Reyes-Prieto, Vicente Soriano Chirona, Maria Alma Bracho, Griselda De Marco, Beatriz Beamud, Lidia Ruiz Roldan, Marta Pla Diaz,Neris Garcia-Gonzalez, Loreto Ferrús Abad, Llúcia Martínez-Priego, Concepcion Gimeno, Giuseppe D'Auria, Fernando Gonzalez-Candelas |
| EPI_ISL_436215 | Servicio de Microbiología. Consorcio Hospital General Universitario de Valencia | Sequencing and Bioinformatics Service and Molecular Epidemiology Research Group. FISABIO-Public Health | Marta Pla Diaz,Neris Garcia-Gonzalez, Loreto Ferrús Abad, Maria Dolores Ocete, Inma Galán Vendrell, Paula Ruiz-Hueso, Mariana Reyes-Prieto, Vicente Soriano Chirona, Maria Alma Bracho, Griselda De Marco, Beatriz Beamud, Lidia Ruiz Roldan, Llúcia Martínez-Priego, Concepcion Gimeno, Giuseppe D'Auria, Fernando Gonzalez-Candelas |
| EPI_ISL_436219 | Servicio de Microbiología. Consorcio Hospital General Universitario de Valencia | Sequencing and Bioinformatics Service and Molecular Epidemiology Research Group. FISABIO-Public Health | Maria Dolores Ocete, Inma Galán Vendrell, Paula Ruiz-Hueso, Mariana Reyes-Prieto, Vicente Soriano Chirona, Maria Alma Bracho, Griselda De Marco, Beatriz Beamud, Lidia Ruiz Roldan, Marta Pla Diaz,Neris Garcia-Gonzalez, Loreto Ferrús Abad, Llúcia Martínez-Priego, Concepcion Gimeno, Giuseppe D'Auria, Fernando Gonzalez-Candelas |
| EPI_ISL_436223 | Servicio de Microbiología. Consorcio Hospital General Universitario de Valencia | Sequencing and Bioinformatics Service and Molecular Epidemiology Research Group. FISABIO-Public Health | Marta Pla Diaz,Neris Garcia-Gonzalez, Loreto Ferrús Abad, Maria Dolores Ocete, Inma Galán Vendrell, Paula Ruiz-Hueso, Mariana Reyes-Prieto, Vicente Soriano Chirona, Maria Alma Bracho, Griselda De Marco, Beatriz Beamud, Lidia Ruiz Roldan, Llúcia Martínez-Priego, Concepcion Gimeno, Giuseppe D'Auria, Fernando Gonzalez-Candelas |
| EPI_ISL_436226 | Servicio de Microbiología. Consorcio Hospital General Universitario de Valencia | Sequencing and Bioinformatics Service and Molecular Epidemiology Research Group. FISABIO-Public Health | Maria Dolores Ocete, Inma Galán Vendrell, Paula Ruiz-Hueso, Mariana Reyes-Prieto, Vicente Soriano Chirona, Maria Alma Bracho, Griselda De Marco, Beatriz Beamud, Lidia Ruiz Roldan, Marta Pla Diaz,Neris Garcia-Gonzalez, Loreto Ferrús Abad, Llúcia Martínez-Priego, Concepcion Gimeno, Giuseppe D'Auria, Fernando Gonzalez-Candelas |
| EPI_ISL_436230 | Servicio de Microbiología. Consorcio Hospital General Universitario de Valencia | Sequencing and Bioinformatics Service and Molecular Epidemiology Research Group. FISABIO-Public Health | Marta Pla Diaz,Neris Garcia-Gonzalez, Loreto Ferrús Abad, Maria Dolores Ocete, Inma Galán Vendrell, Paula Ruiz-Hueso, Mariana Reyes-Prieto, Vicente Soriano Chirona, Maria Alma Bracho, Griselda De Marco, Beatriz Beamud, Lidia Ruiz Roldan, Llúcia Martínez-Priego, Concepcion Gimeno, Giuseppe D'Auria, Fernando Gonzalez-Candelas |
| EPI_ISL_436234 | Servicio de Microbiología. Consorcio Hospital General Universitario de Valencia | Sequencing and Bioinformatics Service and Molecular Epidemiology Research Group. FISABIO-Public Health | Maria Dolores Ocete, Inma Galán Vendrell, Paula Ruiz-Hueso, Mariana Reyes-Prieto, Vicente Soriano Chirona, Maria Alma Bracho, Griselda De Marco, Beatriz Beamud, Lidia Ruiz Roldan, Marta Pla Diaz,Neris Garcia-Gonzalez, Loreto Ferrús Abad, Llúcia Martínez-Priego, Concepcion Gimeno, Giuseppe D'Auria, Fernando Gonzalez-Candelas |
| EPI_ISL_436274 | Servicio de Microbiología. Consorcio Hospital General Universitario de Valencia | Sequencing and Bioinformatics Service and Molecular Epidemiology Research Group. FISABIO-Public Health | Marta Pla Diaz,Neris Garcia-Gonzalez, Loreto Ferrús Abad, Maria Dolores Ocete, Inma Galán Vendrell, Paula Ruiz-Hueso, Mariana Reyes-Prieto, Vicente Soriano Chirona, Maria Alma Bracho, Griselda De Marco, Beatriz Beamud, Lidia Ruiz Roldan, Llúcia Martínez-Priego, Concepcion Gimeno, Giuseppe D'Auria, Fernando Gonzalez-Candelas |
| EPI_ISL_436282 | Servicio de Microbiologia. Hospital Clinico Universitario de Valencia | Sequencing and Bioinformatics Service and Molecular Epidemiology Research Group. FISABIO-Public Health | Marta Pla Diaz, Neris Garcia-Gonzalez, Inma Galán Vendrell, Sandra Carbo, Loreto Ferrús Abad, Paula Ruiz-Hueso, Mariana Reyes-Prieto, Vicente Soriano Chirona, Ivan Ansari, David Navarro, Maria Alma Bracho, Griselda De Marco, Beatriz Beamud, Lidia Ruiz Roldan, Llúcia Martínez-Priego, Giuseppe D'Auria, Fernando Gonzalez-Candelas |
| EPI_ISL_436288 | Servicio de Microbiologia. Hospital Clinico Universitario de Valencia | Sequencing and Bioinformatics Service and Molecular Epidemiology Research Group. FISABIO-Public Health | Mariana Reyes-Prieto, Vicente Soriano Chirona, Ivan Ansari, David Navarro, Maria Alma Bracho, Griselda De Marco, Beatriz Beamud, Lidia Ruiz Roldan, Marta Pla Diaz, Neris Garcia-Gonzalez, Inma Galán Vendrell, Sandra Carbo, Loreto Ferrús Abad, Paula Ruiz-Hueso, Llúcia Martínez-Priego, Giuseppe D'Auria, Fernando Gonzalez-Candelas |
| EPI_ISL_436292 | Servicio de Microbiologia. Hospital Clinico Universitario de Valencia | Sequencing and Bioinformatics Service and Molecular Epidemiology Research Group. FISABIO-Public Health | Maria Alma Bracho, Griselda De Marco, Beatriz Beamud, Lidia Ruiz Roldan, Marta Pla Diaz, Neris Garcia-Gonzalez, Inma Galán Vendrell, Sandra Carbo, Loreto Ferrús Abad, Paula Ruiz-Hueso, Mariana Reyes-Prieto, Vicente Soriano Chirona, Ivan Ansari, David Navarro, Llúcia Martínez-Priego, Giuseppe D'Auria, Fernando Gonzalez-Candelas |
| EPI_ISL_436296 | Servicio de Microbiologia. Hospital Clinico Universitario de Valencia | Sequencing and Bioinformatics Service and Molecular Epidemiology Research Group. FISABIO-Public Health | Marta Pla Diaz, Neris Garcia-Gonzalez, Inma Galán Vendrell, Sandra Carbo, Loreto Ferrús Abad, Paula Ruiz-Hueso, Mariana Reyes-Prieto, Vicente Soriano Chirona, Ivan Ansari, David Navarro, Maria Alma Bracho, Griselda De Marco, Beatriz Beamud, Lidia Ruiz Roldan, Llúcia Martínez-Priego, Giuseppe D'Auria, Fernando Gonzalez-Candelas |
| EPI_ISL_436302 | Servicio de Microbiologia. Hospital Clinico Universitario de Valencia | Sequencing and Bioinformatics Service and Molecular Epidemiology Research Group. FISABIO-Public Health | Marta Pla Diaz, Neris Garcia-Gonzalez, Inma Galán Vendrell, Sandra Carbo, Loreto Ferrús Abad, Paula Ruiz-Hueso, Mariana Reyes-Prieto, Vicente Soriano Chirona, Ivan Ansari, David Navarro, Maria Alma Bracho, Griselda De Marco, Beatriz Beamud, Lidia Ruiz Roldan, Llúcia Martínez-Priego, Giuseppe D'Auria, Fernando Gonzalez-Candelas |
| EPI_ISL_436308 | Servicio de Microbiologia. Hospital Clinico Universitario de Valencia | Sequencing and Bioinformatics Service and Molecular Epidemiology Research Group. FISABIO-Public Health | Mariana Reyes-Prieto, Vicente Soriano Chirona, Ivan Ansari, David Navarro, Maria Alma Bracho, Griselda De Marco, Beatriz Beamud, Lidia Ruiz Roldan, Marta Pla Diaz, Neris Garcia-Gonzalez, Inma Galán Vendrell, Sandra Carbo, Loreto Ferrús Abad, Paula Ruiz-Hueso, Llúcia Martínez-Priego, Giuseppe D'Auria, Fernando Gonzalez-Candelas |
| EPI_ISL_436312 | Servicio de Microbiologia. Hospital Clinico Universitario de Valencia | Sequencing and Bioinformatics Service and Molecular Epidemiology Research Group. FISABIO-Public Health | Maria Alma Bracho, Griselda De Marco, Beatriz Beamud, Lidia Ruiz Roldan, Marta Pla Diaz, Neris Garcia-Gonzalez, Inma Galán Vendrell, Sandra Carbo, Loreto Ferrús Abad, Paula Ruiz-Hueso, Mariana Reyes-Prieto, Vicente Soriano Chirona, Ivan Ansari, David Navarro, Llúcia Martínez-Priego, Giuseppe D'Auria, Fernando Gonzalez-Candelas |
| EPI_ISL_436316 | Servicio de Microbiologia. Hospital Clinico Universitario de | Sequencing and Bioinformatics Service and Molecular | Marta Pla Diaz, Neris Garcia-Gonzalez, Inma Galán Vendrell, Sandra Carbo, Loreto Ferrús Abad, Paula Ruiz-Hueso, Mariana Reyes-Prieto, Vicente |

| | | | |
|---|---|---|---|
| | Valencia | Epidemiology Research Group. FISABIO-Public Health | Soriano Chirona, Ivan Ansari, David Navarro, Maria Alma Bracho, Griselda De Marco, Beatriz Beamud, Lidia Ruiz Roldan, Llúcia Martínez-Priego, Giuseppe D'Auria, Fernando Gonzalez-Candelas |
| EPI_ISL_436322 | Servicio de Microbiologia. Hospital Clinico Universitario de Valencia | Sequencing and Bioinformatics Service and Molecular Epidemiology Research Group. FISABIO-Public Health | Marta Pla Diaz, Neris Garcia-Gonzalez, Inma Galán Vendrell, Sandra Carbo, Loreto Ferrús Abad, Paula Ruiz-Hueso, Mariana Reyes-Prieto, Vicente Soriano Chirona, Ivan Ansari, David Navarro, Maria Alma Bracho, Griselda De Marco, Beatriz Beamud, Lidia Ruiz Roldan, Llúcia Martínez-Priego, Giuseppe D'Auria, Fernando Gonzalez-Candelas |
| EPI_ISL_436328 | Servicio de Microbiologia. Hospital Clinico Universitario de Valencia | Sequencing and Bioinformatics Service and Molecular Epidemiology Research Group. FISABIO-Public Health | Mariana Reyes-Prieto, Vicente Soriano Chirona, Ivan Ansari, David Navarro, Maria Alma Bracho, Griselda De Marco, Beatriz Beamud, Lidia Ruiz Roldan, Marta Pla Diaz, Neris Garcia-Gonzalez, Inma Galán Vendrell, Sandra Carbo, Loreto Ferrús Abad, Paula Ruiz-Hueso, Llúcia Martínez-Priego, Giuseppe D'Auria, Fernando Gonzalez-Candelas |
| EPI_ISL_436332 | Servicio de Microbiologia. Hospital Clinico Universitario de Valencia | Sequencing and Bioinformatics Service and Molecular Epidemiology Research Group. FISABIO-Public Health | Maria Alma Bracho, Griselda De Marco, Beatriz Beamud, Lidia Ruiz Roldan, Marta Pla Diaz, Neris Garcia-Gonzalez, Inma Galán Vendrell, Sandra Carbo, Loreto Ferrús Abad, Paula Ruiz-Hueso, Mariana Reyes-Prieto, Vicente Soriano Chirona, Ivan Ansari, David Navarro, Llúcia Martínez-Priego, Giuseppe D'Auria, Fernando Gonzalez-Candelas |
| EPI_ISL_436336 | Servicio de Microbiologia. Hospital Clinico Universitario de Valencia | Sequencing and Bioinformatics Service and Molecular Epidemiology Research Group. FISABIO-Public Health | Marta Pla Diaz, Neris Garcia-Gonzalez, Inma Galán Vendrell, Sandra Carbo, Loreto Ferrús Abad, Paula Ruiz-Hueso, Mariana Reyes-Prieto, Vicente Soriano Chirona, Ivan Ansari, David Navarro, Maria Alma Bracho, Griselda De Marco, Beatriz Beamud, Lidia Ruiz Roldan, Llúcia Martínez-Priego, Giuseppe D'Auria, Fernando Gonzalez-Candelas |
| EPI_ISL_436342 | Servicio de Microbiologia. Hospital Clinico Universitario de Valencia | Sequencing and Bioinformatics Service and Molecular Epidemiology Research Group. FISABIO-Public Health | Marta Pla Diaz, Neris Garcia-Gonzalez, Inma Galán Vendrell, Sandra Carbo, Loreto Ferrús Abad, Paula Ruiz-Hueso, Mariana Reyes-Prieto, Vicente Soriano Chirona, Ivan Ansari, David Navarro, Maria Alma Bracho, Griselda De Marco, Beatriz Beamud, Lidia Ruiz Roldan, Llúcia Martínez-Priego, Giuseppe D'Auria, Fernando Gonzalez-Candelas |
| EPI_ISL_436348 | Servicio de Microbiologia. Hospital Clinico Universitario de Valencia | Sequencing and Bioinformatics Service and Molecular Epidemiology Research Group. FISABIO-Public Health | Mariana Reyes-Prieto, Vicente Soriano Chirona, Ivan Ansari, David Navarro, Maria Alma Bracho, Griselda De Marco, Beatriz Beamud, Lidia Ruiz Roldan, Marta Pla Diaz, Neris Garcia-Gonzalez, Inma Galán Vendrell, Sandra Carbo, Loreto Ferrús Abad, Paula Ruiz-Hueso, Llúcia Martínez-Priego, Giuseppe D'Auria, Fernando Gonzalez-Candelas |
| EPI_ISL_436352 | Servicio de Microbiologia. Hospital Clinico Universitario de Valencia | Sequencing and Bioinformatics Service and Molecular Epidemiology Research Group. FISABIO-Public Health | Maria Alma Bracho, Griselda De Marco, Beatriz Beamud, Lidia Ruiz Roldan, Marta Pla Diaz, Neris Garcia-Gonzalez, Inma Galán Vendrell, Sandra Carbo, Loreto Ferrús Abad, Paula Ruiz-Hueso, Mariana Reyes-Prieto, Vicente Soriano Chirona, Ivan Ansari, David Navarro, Llúcia Martínez-Priego, Giuseppe D'Auria, Fernando Gonzalez-Candelas |
| EPI_ISL_436356, EPI_ISL_436359 | Servicio de Microbiologia. Hospital Clinico Universitario de Valencia | Sequencing and Bioinformatics Service and Molecular Epidemiology Research Group. FISABIO-Public Health | Marta Pla Diaz, Neris Garcia-Gonzalez, Inma Galán Vendrell, Sandra Carbo, Loreto Ferrús Abad, Paula Ruiz-Hueso, Mariana Reyes-Prieto, Vicente Soriano Chirona, Ivan Ansari, David Navarro, Maria Alma Bracho, Griselda De Marco, Beatriz Beamud, Lidia Ruiz Roldan, Llúcia Martínez-Priego, Giuseppe D'Auria, Fernando Gonzalez-Candelas |
| EPI_ISL_436366 | Servicio de Microbiologia. Hospital Clinico Universitario de Valencia | Sequencing and Bioinformatics Service and Molecular Epidemiology Research Group. FISABIO-Public Health | Marta Pla Diaz, Neris Garcia-Gonzalez, Inma Galán Vendrell, Sandra Carbo, Loreto Ferrús Abad, Paula Ruiz-Hueso, Mariana Reyes-Prieto, Vicente Soriano Chirona, Ivan Ansari, David Navarro, Maria Alma Bracho, Griselda De Marco, Beatriz Beamud, Lidia Ruiz Roldan, Llúcia Martínez-Priego, Giuseppe D'Auria, Fernando Gonzalez-Candelas |
| EPI_ISL_436383 | Servicio de Microbiología. Consorcio Hospital General Universitario de Valencia | Sequencing and Bioinformatics Service and Molecular Epidemiology Research Group. FISABIO-Public Health | Maria Dolores Ocete, Inma Galán Vendrell, Paula Ruiz-Hueso, Mariana Reyes-Prieto, Vicente Soriano Chirona, Maria Alma Bracho, Griselda De Marco, Beatriz Beamud, Lidia Ruiz Roldan, Marta Pla Diaz,Neris Garcia-Gonzalez, Loreto Ferrús Abad, Llúcia Martínez-Priego, Concepcion Gimeno, Giuseppe D'Auria, Fernando Gonzalez-Candelas |
| EPI_ISL_436387 | Servicio de Microbiología. Consorcio Hospital General Universitario de Valencia | Sequencing and Bioinformatics Service and Molecular Epidemiology Research Group. FISABIO-Public Health | Marta Pla Diaz,Neris Garcia-Gonzalez, Loreto Ferrús Abad, Maria Dolores Ocete, Inma Galán Vendrell, Paula Ruiz-Hueso, Mariana Reyes-Prieto, Vicente Soriano Chirona, Maria Alma Bracho, Griselda De Marco, Beatriz Beamud, Lidia Ruiz Roldan, Llúcia Martínez-Priego, Concepcion Gimeno, Giuseppe D'Auria, Fernando Gonzalez-Candelas |
| EPI_ISL_436390 | Servicio de Microbiología. Consorcio Hospital General Universitario de Valencia | Sequencing and Bioinformatics Service and Molecular Epidemiology Research Group. FISABIO-Public Health | Maria Dolores Ocete, Inma Galán Vendrell, Paula Ruiz-Hueso, Mariana Reyes-Prieto, Vicente Soriano Chirona, Maria Alma Bracho, Griselda De Marco, Beatriz Beamud, Lidia Ruiz Roldan, Marta Pla Diaz,Neris Garcia-Gonzalez, Loreto Ferrús Abad, Llúcia Martínez-Priego, Concepcion Gimeno, Giuseppe D'Auria, Fernando Gonzalez-Candelas |
| EPI_ISL_436394 | Servicio de Microbiología. Consorcio Hospital General Universitario de Valencia | Sequencing and Bioinformatics Service and Molecular Epidemiology Research Group. FISABIO-Public Health | Marta Pla Diaz,Neris Garcia-Gonzalez, Loreto Ferrús Abad, Maria Dolores Ocete, Inma Galán Vendrell, Paula Ruiz-Hueso, Mariana Reyes-Prieto, Vicente Soriano Chirona, Maria Alma Bracho, Griselda De Marco, Beatriz Beamud, Lidia Ruiz Roldan, Llúcia Martínez-Priego, Concepcion Gimeno, Giuseppe D'Auria, Fernando Gonzalez-Candelas |
| EPI_ISL_436399 | Servicio de Microbiologia. Hospital Clinico Universitario de Valencia | Sequencing and Bioinformatics Service and Molecular Epidemiology Research Group. FISABIO-Public Health | Mariana Reyes-Prieto, Vicente Soriano Chirona, Ivan Ansari, David Navarro, Maria Alma Bracho, Griselda De Marco, Beatriz Beamud, Lidia Ruiz Roldan, Marta Pla Diaz, Neris Garcia-Gonzalez, Inma Galán Vendrell, Sandra Carbo, Loreto Ferrús Abad, Paula Ruiz-Hueso, Llúcia Martínez-Priego, Giuseppe D'Auria, Fernando Gonzalez-Candelas |
| EPI_ISL_436403 | Servicio de Microbiologia. Hospital Clinico Universitario de Valencia | Sequencing and Bioinformatics Service and Molecular Epidemiology Research Group. FISABIO-Public Health | Maria Alma Bracho, Griselda De Marco, Beatriz Beamud, Lidia Ruiz Roldan, Marta Pla Diaz, Neris Garcia-Gonzalez, Inma Galán Vendrell, Sandra Carbo, Loreto Ferrús Abad, Paula Ruiz-Hueso, Mariana Reyes-Prieto, Vicente Soriano Chirona, Ivan Ansari, David Navarro, Llúcia Martínez-Priego, Giuseppe D'Auria, Fernando Gonzalez-Candelas |
| EPI_ISL_436407, EPI_ISL_436410 | Servicio de Microbiologia. Hospital Clinico Universitario de Valencia | Sequencing and Bioinformatics Service and Molecular Epidemiology Research Group. FISABIO-Public Health | Marta Pla Diaz, Neris Garcia-Gonzalez, Inma Galán Vendrell, Sandra Carbo, Loreto Ferrús Abad, Paula Ruiz-Hueso, Mariana Reyes-Prieto, Vicente Soriano Chirona, Ivan Ansari, David Navarro, Maria Alma Bracho, Griselda De Marco, Beatriz Beamud, Lidia Ruiz Roldan, Llúcia Martínez-Priego, Giuseppe D'Auria, Fernando Gonzalez-Candelas |
| EPI_ISL_436733, EPI_ISL_436734, EPI_ISL_436735, EPI_ISL_436736, EPI_ISL_436737, EPI_ISL_436738, EPI_ISL_436739, EPI_ISL_436740, EPI_ISL_436741, EPI_ISL_436742, EPI_ISL_436743 | | | |
| see above | NYU Langone Health | Departments of Pathology and Medicine, New York University School of Medicine | Maria Aguero-Rosenfeld, Brendan Belovarac, Margaret Black, Ludovic Boytard, John Cadley, Paolo Cotzia, John Chen, Dacia Dimartino, Xiaojun Feng, Tatyana Gindin, Emily Guzman, Adriana Heguy, Megan Hogan, Emily Huang, George Jour, Alireza Khodadadi-Jamayran, Lawrence H. Lin, Raven Luther, Andrew Lytle, Christian Marier, Matthew T. Maurano, Mark J. Mulligan, Peter Meyn, Raquel Ordonez Ciriza, Iman Osman, Jared Pinnell, Vanessa Raabe, Sitharam Ramaswami, Amy Rapkiewicz, Andre M. Ribeiro-dos-Santos, Marie Samanovic-Golden, Antonio Serrano, Guomiao Shen, Matija Snuderl, Theodore Vougiouklakis, Nick Vulpescu, Gael Westby, Paul Zappile, Yutong Zhang |
| EPI_ISL_437204, EPI_ISL_437205, EPI_ISL_437206, EPI_ISL_437207, EPI_ISL_437208, EPI_ISL_437209, EPI_ISL_437210, EPI_ISL_437211, EPI_ISL_437212, EPI_ISL_437213, EPI_ISL_437214, EPI_ISL_437215, EPI_ISL_437216, EPI_ISL_437217, EPI_ISL_437218, EPI_ISL_437219, EPI_ISL_437220, EPI_ISL_437221, EPI_ISL_437222, EPI_ISL_437223, EPI_ISL_437224, EPI_ISL_437225, EPI_ISL_437226, EPI_ISL_437227, EPI_ISL_437228, EPI_ISL_437229, EPI_ISL_437230, EPI_ISL_437231, EPI_ISL_437232, EPI_ISL_437233, EPI_ISL_437234, EPI_ISL_437235, EPI_ISL_437236, EPI_ISL_437237, EPI_ISL_437238, EPI_ISL_437239, EPI_ISL_437240, EPI_ISL_437241, EPI_ISL_437242, EPI_ISL_437243, EPI_ISL_437244, EPI_ISL_437245, EPI_ISL_437246, EPI_ISL_437247, EPI_ISL_437248, EPI_ISL_437249, EPI_ISL_437250, EPI_ISL_437251, EPI_ISL_437252, EPI_ISL_437253, EPI_ISL_437254, EPI_ISL_437255, EPI_ISL_437256, EPI_ISL_437257, EPI_ISL_437258, EPI_ISL_437259, EPI_ISL_437260, EPI_ISL_437261, EPI_ISL_437262, EPI_ISL_437263, EPI_ISL_437264, EPI_ISL_437265, EPI_ISL_437266, EPI_ISL_437267, EPI_ISL_437268, EPI_ISL_437269, EPI_ISL_437270, EPI_ISL_437271, EPI_ISL_437272, EPI_ISL_437273, EPI_ISL_437274, EPI_ISL_437275, EPI_ISL_437276, EPI_ISL_437277, EPI_ISL_437278, EPI_ISL_437279, EPI_ISL_437280, EPI_ISL_437281, EPI_ISL_437282, EPI_ISL_437283, EPI_ISL_437284, EPI_ISL_437285, EPI_ISL_437286, EPI_ISL_437287, EPI_ISL_437288, EPI_ISL_437289, EPI_ISL_437290, EPI_ISL_437291, EPI_ISL_437292, EPI_ISL_437293, EPI_ISL_437294, EPI_ISL_437295, EPI_ISL_437296, EPI_ISL_437297, EPI_ISL_437359 | | | |
| see above | Max von Pettenkofer Institute, Virology, National Reference Center for Retroviruses, LMU München | Laboratory for Functional Genome Analysis, Dept. Genomics, Gene Center of the LMU Munich | Max Muenchhoff, Stefan Krebs, Alexander Graf, Oliver Keppler, Helmut Blum |
| EPI_ISL_437360, EPI_ISL_437361, EPI_ISL_437362, EPI_ISL_437363, EPI_ISL_437364, EPI_ISL_437365, EPI_ISL_437366, EPI_ISL_437367, EPI_ISL_437368, EPI_ISL_437369, EPI_ISL_437370, EPI_ISL_437371, EPI_ISL_437372, EPI_ISL_437373, EPI_ISL_437374, EPI_ISL_437375, EPI_ISL_437376, EPI_ISL_437377, EPI_ISL_437378, EPI_ISL_437379, EPI_ISL_437380, EPI_ISL_437381, EPI_ISL_437382, EPI_ISL_437383, EPI_ISL_437384, EPI_ISL_437385, EPI_ISL_437386 | | | |
| see above | Minnesota Department of Health, Public Health Laboratory | Minnesota Department of Health, Public Health Laboratory | Matt Plumb, Jacob Garfin, and Xiong Wang |
| EPI_ISL_438139 | Department of Microbiology,Gandhi Medical College and Hospital,Hyderabad | Virus Research Laboratory, Department of Zoology, Osmania University, Hyderabad, India | Muttineni Radhakrishna, Nagamani K, Thrilok Chander B, Raja Rao M, Kalyani Putty, Ravikumar P, Sunitha P, Pankaj Singh D, Anand Kumar K, Amit A. Upadhyay, Steven Bosinger, Rama Amara |
| EPI_ISL_439144, EPI_ISL_439145, EPI_ISL_439146, EPI_ISL_439147, EPI_ISL_439148, EPI_ISL_439149, EPI_ISL_439150, EPI_ISL_439151, EPI_ISL_439152, EPI_ISL_439153, EPI_ISL_439154, EPI_ISL_439155, EPI_ISL_439156, EPI_ISL_439157, EPI_ISL_439158, EPI_ISL_439159, EPI_ISL_439160, EPI_ISL_439161, EPI_ISL_439162, EPI_ISL_439163, EPI_ISL_439164, EPI_ISL_439165, EPI_ISL_439166, EPI_ISL_439167, EPI_ISL_439168, EPI_ISL_439169, EPI_ISL_439170, EPI_ISL_439171, EPI_ISL_439172, EPI_ISL_439173, EPI_ISL_439174, EPI_ISL_439175, EPI_ISL_439176, EPI_ISL_439177, EPI_ISL_439178, EPI_ISL_439179, EPI_ISL_439180, EPI_ISL_439181, EPI_ISL_439182, EPI_ISL_439183, EPI_ISL_439184, EPI_ISL_439185, EPI_ISL_439186, EPI_ISL_439187, EPI_ISL_439188, EPI_ISL_439189, EPI_ISL_439190, EPI_ISL_439191, EPI_ISL_439192, EPI_ISL_439193, EPI_ISL_439194, EPI_ISL_439195, EPI_ISL_439196, EPI_ISL_439197, EPI_ISL_439198, EPI_ISL_439199, EPI_ISL_439200, EPI_ISL_439201, EPI_ISL_439202, EPI_ISL_439203, EPI_ISL_439204, EPI_ISL_439205, EPI_ISL_439206, EPI_ISL_439207, EPI_ISL_439208, EPI_ISL_439209, EPI_ISL_439210, EPI_ISL_439211, EPI_ISL_439212, EPI_ISL_439213, EPI_ISL_439214, EPI_ISL_439215, EPI_ISL_439216, EPI_ISL_439217, EPI_ISL_439218, EPI_ISL_439219, EPI_ISL_439220, EPI_ISL_439221, EPI_ISL_439222, EPI_ISL_439223, EPI_ISL_439224, EPI_ISL_439225, EPI_ISL_439226, EPI_ISL_439227, EPI_ISL_439228, EPI_ISL_439229, EPI_ISL_439230, EPI_ISL_439231, EPI_ISL_439232, EPI_ISL_439233, EPI_ISL_439234, EPI_ISL_439235, EPI_ISL_439236, EPI_ISL_439237, EPI_ISL_439238, EPI_ISL_439239, EPI_ISL_439240, EPI_ISL_439241, EPI_ISL_439242, EPI_ISL_439243, EPI_ISL_439244, EPI_ISL_439245, EPI_ISL_439246, EPI_ISL_439247, EPI_ISL_439248, EPI_ISL_439249, EPI_ISL_439250, EPI_ISL_439251, | | | |

| | | | |
|---|---|---|---|
| EPI_ISL_439252, EPI_ISL_439253, EPI_ISL_439254, EPI_ISL_439255, EPI_ISL_439256, EPI_ISL_439257, EPI_ISL_439258, EPI_ISL_439259, EPI_ISL_439260, EPI_ISL_439261, EPI_ISL_439262, EPI_ISL_439263, EPI_ISL_439264, EPI_ISL_439265, EPI_ISL_439266, EPI_ISL_439267, EPI_ISL_439268, EPI_ISL_439269, EPI_ISL_439270, EPI_ISL_439271, EPI_ISL_439272, EPI_ISL_439273, EPI_ISL_439274, EPI_ISL_439275, EPI_ISL_439276, EPI_ISL_439277, EPI_ISL_439278, EPI_ISL_439279, EPI_ISL_439280, EPI_ISL_439281, EPI_ISL_439282, EPI_ISL_439283, EPI_ISL_439284, EPI_ISL_439285, EPI_ISL_439286, EPI_ISL_439287, EPI_ISL_439288, EPI_ISL_439289, EPI_ISL_439290, EPI_ISL_439291, EPI_ISL_439292, EPI_ISL_439293, EPI_ISL_439294, EPI_ISL_439295, EPI_ISL_439296, EPI_ISL_439297, EPI_ISL_439298, EPI_ISL_439299, EPI_ISL_439300, EPI_ISL_439301, EPI_ISL_439302, EPI_ISL_439303, EPI_ISL_439304, EPI_ISL_439305, EPI_ISL_439306, EPI_ISL_439307, EPI_ISL_439308, EPI_ISL_439309, EPI_ISL_439310, EPI_ISL_439311, EPI_ISL_439312, EPI_ISL_439313, EPI_ISL_439314, EPI_ISL_439315, EPI_ISL_439316, EPI_ISL_439317, EPI_ISL_439318, EPI_ISL_439319, EPI_ISL_439320, EPI_ISL_439321, EPI_ISL_439322, EPI_ISL_439323, EPI_ISL_439324, EPI_ISL_439325, EPI_ISL_439326, EPI_ISL_439327, EPI_ISL_439328, EPI_ISL_439329, EPI_ISL_439330, EPI_ISL_439331, EPI_ISL_439332, EPI_ISL_439333, EPI_ISL_439334, EPI_ISL_439335, EPI_ISL_439336, EPI_ISL_439337, EPI_ISL_439338, EPI_ISL_439339, EPI_ISL_439340, EPI_ISL_439341, EPI_ISL_439342, EPI_ISL_439343, EPI_ISL_439344, EPI_ISL_439345, EPI_ISL_439346, EPI_ISL_439347, EPI_ISL_439348, EPI_ISL_439349, EPI_ISL_439350, EPI_ISL_439351, EPI_ISL_439352, EPI_ISL_439353, EPI_ISL_439354, EPI_ISL_439355, EPI_ISL_439356, EPI_ISL_439357, EPI_ISL_439358, EPI_ISL_439359, EPI_ISL_439360, EPI_ISL_439361, EPI_ISL_439362, EPI_ISL_439363, EPI_ISL_439364, EPI_ISL_439365, EPI_ISL_439366, EPI_ISL_439367, EPI_ISL_439665, EPI_ISL_439666, EPI_ISL_439667, EPI_ISL_439668, EPI_ISL_439669, EPI_ISL_439670, EPI_ISL_439671 | | | |
| see above | Virology Department, Royal Infirmary of Edinburgh, NHS Lothian / School of Biological Sciences, University of Edinburgh / Institute of Genetics and Molecular Medicine, University of Edinburgh | COVID-19 Genomics UK (COG-UK) Consortium | McHugh M, Dewar R, Rooke S, Gallagher M, Balcaza C, O'ÄôToole Å, Scher E, Hill V, McCrone JT, Colquhoun R, Yu X, Jackson B, Rambaut A, Williams TC, Templeton K |
| EPI_ISL_442044 | Kawsar Human Genetic Research Center | Kawsar Human Genetic Research Center | Mohammad Ali Khosravi, Maryam Abbasalipour Bashash, Sirous Zeinali, Solmaz Sabeghi, Yeganeh Keshvar, Fatemeh Hosseini, Yeganeh Haghdoost |
| EPI_ISL_443258, EPI_ISL_443259 | Résidence Ornano | National Reference Center for Viruses of Respiratory Infections, Institut Pasteur, Paris | Mélanie Albert, Marion Barbet, Sylvie Behillil, Méline Bizard, Angela Brisebarre, Flora Donati, Etienne Simon-Lorière, Vincent Enouf, Maud Vanpeene, Sylvie van der Werf |
| EPI_ISL_443260 | LABM GH nord Essonne de Longjumeau - BP 125 | National Reference Center for Viruses of Respiratory Infections, Institut Pasteur, Paris | Mélanie Albert, Marion Barbet, Sylvie Behillil, Méline Bizard, Angela Brisebarre, Flora Donati, Etienne Simon-Lorière, Vincent Enouf, Maud Vanpeene, Sylvie van der Werf |
| EPI_ISL_443261, EPI_ISL_443262, EPI_ISL_443263, EPI_ISL_443264 | CHU de Dijon - Laboratoire de Virologie | National Reference Center for Viruses of Respiratory Infections, Institut Pasteur, Paris | Mélanie Albert, Marion Barbet, Sylvie Behillil, Méline Bizard, Angela Brisebarre, Flora Donati, Etienne Simon-Lorière, Vincent Enouf, Maud Vanpeene, Sylvie van der Werf, Jean-Baptiste Bour |
| EPI_ISL_443265, EPI_ISL_443266, EPI_ISL_443267, EPI_ISL_443268, EPI_ISL_443269, EPI_ISL_443270, EPI_ISL_443271, EPI_ISL_443272, EPI_ISL_443273, EPI_ISL_443274, EPI_ISL_443275, EPI_ISL_443276, EPI_ISL_443277, EPI_ISL_443278, EPI_ISL_443279, EPI_ISL_443280, EPI_ISL_443281, EPI_ISL_443282, EPI_ISL_443283 | | | |
| see above | CHU - Hôpital Cavale Blanche - Labo. de Virologie | National Reference Center for Viruses of Respiratory Infections, Institut Pasteur, Paris | Mélanie Albert, Marion Barbet, Sylvie Behillil, Méline Bizard, Angela Brisebarre, Flora Donati, Etienne Simon-Lorière, Vincent Enouf, Maud Vanpeene, Sylvie van der Werf, Léa Pilorge |
| EPI_ISL_443284, EPI_ISL_443285, EPI_ISL_443286, EPI_ISL_443287, EPI_ISL_443288 | Laboratoire de Microbiologie - Bât A - CH René Dubois | National Reference Center for Viruses of Respiratory Infections, Institut Pasteur, Paris | Mélanie Albert, Marion Barbet, Sylvie Behillil, Méline Bizard, Angela Brisebarre, Flora Donati, Etienne Simon-Lorière, Vincent Enouf, Maud Vanpeene, Sylvie van der Werf, Pascale Martres |
| EPI_ISL_443289, EPI_ISL_443290, EPI_ISL_443291, EPI_ISL_443292, EPI_ISL_443293, EPI_ISL_443294 | CHRU Pontchaillou - Laboratoire de Virologie | National Reference Center for Viruses of Respiratory Infections, Institut Pasteur, Paris | Mélanie Albert, Marion Barbet, Sylvie Behillil, Méline Bizard, Angela Brisebarre, Flora Donati, Etienne Simon-Lorière, Vincent Enouf, Maud Vanpeene, Sylvie van der Werf, Gisèle Lagathu |
| EPI_ISL_443295, EPI_ISL_443296, EPI_ISL_443297, EPI_ISL_443298, EPI_ISL_443299 | Hôpital Necker - Enfants - Malades Laboratoire de Virologie | National Reference Center for Viruses of Respiratory Infections, Institut Pasteur, Paris | Mélanie Albert, Marion Barbet, Sylvie Behillil, Méline Bizard, Angela Brisebarre, Flora Donati, Etienne Simon-Lorière, Vincent Enouf, Maud Vanpeene, Sylvie van der Werf, Marianne Leruez-Ville |
| EPI_ISL_443300, EPI_ISL_443301, EPI_ISL_443302 | Cabinet Médical | National Reference Center for Viruses of Respiratory Infections, Institut Pasteur, Paris | Mélanie Albert, Marion Barbet, Sylvie Behillil, Méline Bizard, Angela Brisebarre, Flora Donati, Etienne Simon-Lorière, Vincent Enouf, Maud Vanpeene, Sylvie van der Werf |
| EPI_ISL_443303 | Résidence Les Marines | National Reference Center for Viruses of Respiratory Infections, Institut Pasteur, Paris | Mélanie Albert, Marion Barbet, Sylvie Behillil, Méline Bizard, Angela Brisebarre, Flora Donati, Etienne Simon-Lorière, Vincent Enouf, Maud Vanpeene, Sylvie van der Werf |
| EPI_ISL_443304 | Résidence Esterel | National Reference Center for Viruses of Respiratory Infections, Institut Pasteur, Paris | Mélanie Albert, Marion Barbet, Sylvie Behillil, Méline Bizard, Angela Brisebarre, Flora Donati, Etienne Simon-Lorière, Vincent Enouf, Maud Vanpeene, Sylvie van der Werf |
| EPI_ISL_443305 | LABM GH nord Essonne de Longjumeau - BP 125 | National Reference Center for Viruses of Respiratory Infections, Institut Pasteur, Paris | Mélanie Albert, Marion Barbet, Sylvie Behillil, Méline Bizard, Angela Brisebarre, Flora Donati, Etienne Simon-Lorière, Vincent Enouf, Maud Vanpeene, Sylvie van der Werf |
| EPI_ISL_443306 | Cabinet Médical | National Reference Center for Viruses of Respiratory Infections, Institut Pasteur, Paris | Mélanie Albert, Marion Barbet, Sylvie Behillil, Méline Bizard, Angela Brisebarre, Flora Donati, Etienne Simon-Lorière, Vincent Enouf, Maud Vanpeene, Sylvie van der Werf |
| EPI_ISL_443307 | La Villa Papyri | National Reference Center for Viruses of Respiratory Infections, Institut Pasteur, Paris | Mélanie Albert, Marion Barbet, Sylvie Behillil, Méline Bizard, Angela Brisebarre, Flora Donati, Etienne Simon-Lorière, Vincent Enouf, Maud Vanpeene, Sylvie van der Werf |
| EPI_ISL_443308 | Plaisance | National Reference Center for Viruses of Respiratory Infections, Institut Pasteur, Paris | Mélanie Albert, Marion Barbet, Sylvie Behillil, Méline Bizard, Angela Brisebarre, Flora Donati, Etienne Simon-Lorière, Vincent Enouf, Maud Vanpeene, Sylvie van der Werf |
| EPI_ISL_443309 | CH Compiègne Laboratoire de Biologie | National Reference Center for Viruses of Respiratory Infections, Institut Pasteur, Paris | Mélanie Albert, Marion Barbet, Sylvie Behillil, Méline Bizard, Angela Brisebarre, Flora Donati, Etienne Simon-Lorière, Vincent Enouf, Maud Vanpeene, Sylvie van der Werf |
| EPI_ISL_443310 | Centre de santé Filieris | National Reference Center for Viruses of Respiratory Infections, Institut Pasteur, Paris | Mélanie Albert, Marion Barbet, Sylvie Behillil, Méline Bizard, Angela Brisebarre, Flora Donati, Etienne Simon-Lorière, Vincent Enouf, Maud Vanpeene, Sylvie van der Werf |
| EPI_ISL_443311, EPI_ISL_443312, EPI_ISL_443313 | Cabinet Médical | National Reference Center for Viruses of Respiratory Infections, Institut Pasteur, Paris | Mélanie Albert, Marion Barbet, Sylvie Behillil, Méline Bizard, Angela Brisebarre, Flora Donati, Etienne Simon-Lorière, Vincent Enouf, Maud Vanpeene, Sylvie van der Werf |
| EPI_ISL_443314 | LABM GH nord Essonne de Longjumeau - BP 125 | National Reference Center for Viruses of Respiratory Infections, Institut Pasteur, Paris | Mélanie Albert, Marion Barbet, Sylvie Behillil, Méline Bizard, Angela Brisebarre, Flora Donati, Etienne Simon-Lorière, Vincent Enouf, Maud Vanpeene, Sylvie van der Werf |
| EPI_ISL_443315 | Château de la Source | National Reference Center for Viruses of Respiratory Infections, Institut Pasteur, Paris | Mélanie Albert, Marion Barbet, Sylvie Behillil, Méline Bizard, Angela Brisebarre, Flora Donati, Etienne Simon-Lorière, Vincent Enouf, Maud Vanpeene, Sylvie van der Werf |
| EPI_ISL_443316 | CH Compiègne Laboratoire de Biologie | National Reference Center for Viruses of Respiratory Infections, Institut Pasteur, Paris | Mélanie Albert, Marion Barbet, Sylvie Behillil, Méline Bizard, Angela Brisebarre, Flora Donati, Etienne Simon-Lorière, Vincent Enouf, Maud Vanpeene, Sylvie van der Werf, Olivia Raulin |
| EPI_ISL_443317 | Cabinet Médical | National Reference Center for Viruses of Respiratory Infections, Institut Pasteur, Paris | Mélanie Albert, Marion Barbet, Sylvie Behillil, Méline Bizard, Angela Brisebarre, Flora Donati, Etienne Simon-Lorière, Vincent Enouf, Maud Vanpeene, Sylvie van der Werf |
| EPI_ISL_444465 | B.J. Medical College and Civil hospital | Gujarat Biotechnology Research Centre | Monika Gandhi, Pinal Trivedi, Maharshi Pandya, Amit Kanani, Akanksha Verma, Nitin Savaliya, Raghawendra Kumar, Dinesh Kumar, Zuber Saiyed, Dipa Kinariwala, Disha Patel, Binita Aring, Neeta Khandelwal, Geeta Vaghela, Sonia Barve, Bhavesh Modi, Kairavi Joshi, Gaurishankar Shrimali, Nidhi Sood, Pranay Shah, R D Dixit, Snehal Bagatharia, Kamlesh J Upadhyay, Ramesh Pandit, Tejas Shah, Ankit Hinsu, Pritesh Sabara, Apurvasinh Puvar, Janvi Raval, Anjali Rajwar, Chaitanya Joshi, Madhvi Joshi |
| EPI_ISL_444494, EPI_ISL_444495, EPI_ISL_444496, EPI_ISL_444497, EPI_ISL_444498, EPI_ISL_444499, EPI_ISL_444500, EPI_ISL_444501, EPI_ISL_444502, EPI_ISL_444503, EPI_ISL_444504, EPI_ISL_444505, EPI_ISL_444506, EPI_ISL_444507, EPI_ISL_444508, EPI_ISL_444509, EPI_ISL_444510, EPI_ISL_444511, EPI_ISL_444512, EPI_ISL_444513, EPI_ISL_444514, EPI_ISL_444515, EPI_ISL_444516 | | | |
| see above | Laboratoire de microbiologie, Hopital de Verdun | Smith Laboratory, Centre de Recherche CHU Sainte-Justine | Martin Smith, Marieke Rozendaal, Ivan Pavlov |
| EPI_ISL_444613, EPI_ISL_444614, EPI_ISL_444615, EPI_ISL_444616, EPI_ISL_444617, EPI_ISL_444618, EPI_ISL_444619, EPI_ISL_444620, EPI_ISL_444621, EPI_ISL_444622, EPI_ISL_444623, EPI_ISL_444625, EPI_ISL_444626, EPI_ISL_444627, EPI_ISL_444628, EPI_ISL_444629, EPI_ISL_444630, EPI_ISL_444631, EPI_ISL_444632, EPI_ISL_444633, EPI_ISL_444634, EPI_ISL_444635, EPI_ISL_444636, EPI_ISL_444637, EPI_ISL_444638, EPI_ISL_444639, EPI_ISL_444640, EPI_ISL_444641, EPI_ISL_444642, EPI_ISL_444643, EPI_ISL_444644, EPI_ISL_444645, EPI_ISL_444646, EPI_ISL_444647, EPI_ISL_444648, EPI_ISL_444649, EPI_ISL_444650, EPI_ISL_444651, EPI_ISL_444652, EPI_ISL_444653, EPI_ISL_444654, EPI_ISL_444655, EPI_ISL_444656, EPI_ISL_444657, EPI_ISL_444658, EPI_ISL_444659, EPI_ISL_444660, EPI_ISL_444661, EPI_ISL_444662, EPI_ISL_444663, EPI_ISL_444664, EPI_ISL_444665, EPI_ISL_444666, EPI_ISL_444667, EPI_ISL_444668, EPI_ISL_444669, EPI_ISL_444670, EPI_ISL_444671, EPI_ISL_444673, EPI_ISL_444674, EPI_ISL_444675, EPI_ISL_444676, EPI_ISL_444677, EPI_ISL_444678, EPI_ISL_444679, EPI_ISL_444680, EPI_ISL_444681, EPI_ISL_444682, EPI_ISL_444683, EPI_ISL_444684, EPI_ISL_444685, EPI_ISL_444686, EPI_ISL_444687, EPI_ISL_444688, EPI_ISL_444689, EPI_ISL_444690, EPI_ISL_444691, EPI_ISL_444692, EPI_ISL_444693, EPI_ISL_444694, EPI_ISL_444695, EPI_ISL_444696, EPI_ISL_444697, EPI_ISL_444698, EPI_ISL_444699, EPI_ISL_444700, EPI_ISL_444701, EPI_ISL_444702, EPI_ISL_444703, EPI_ISL_444704, EPI_ISL_444705, EPI_ISL_444706, EPI_ISL_444707, EPI_ISL_444708, EPI_ISL_444709, EPI_ISL_444710, EPI_ISL_444711, EPI_ISL_444712, EPI_ISL_444713, EPI_ISL_444714, EPI_ISL_444715, EPI_ISL_444716, EPI_ISL_444717, EPI_ISL_444718, EPI_ISL_444719, EPI_ISL_444720, EPI_ISL_444721, EPI_ISL_444722, EPI_ISL_444723, EPI_ISL_444724, EPI_ISL_444725, EPI_ISL_444726, EPI_ISL_444727, EPI_ISL_444728, EPI_ISL_444729, EPI_ISL_444730, EPI_ISL_444731, EPI_ISL_444732, EPI_ISL_444733, EPI_ISL_444734, EPI_ISL_444735, EPI_ISL_444736, EPI_ISL_444737, EPI_ISL_444738, EPI_ISL_444739, EPI_ISL_444740, | | | |

| | | | |
|---|---|---|---|
| EPI_ISL_444741, EPI_ISL_444742, EPI_ISL_444743, EPI_ISL_444744, EPI_ISL_444745, EPI_ISL_444746, EPI_ISL_444747, EPI_ISL_444748, EPI_ISL_444749, EPI_ISL_444750, EPI_ISL_444751, EPI_ISL_444752, EPI_ISL_444753, EPI_ISL_444754, EPI_ISL_444755, EPI_ISL_444756, EPI_ISL_444757, EPI_ISL_444758, EPI_ISL_444759, EPI_ISL_444760, EPI_ISL_444761, EPI_ISL_444762, EPI_ISL_444763, EPI_ISL_444764, EPI_ISL_444765, EPI_ISL_444766, EPI_ISL_444767, EPI_ISL_444768, EPI_ISL_444769, EPI_ISL_444770, EPI_ISL_444771, EPI_ISL_444772, EPI_ISL_444773, EPI_ISL_444774, EPI_ISL_444775, EPI_ISL_444776, EPI_ISL_444777, EPI_ISL_444778, EPI_ISL_444779, EPI_ISL_444780, EPI_ISL_444781, EPI_ISL_444782, EPI_ISL_444783, EPI_ISL_444784, EPI_ISL_444785, EPI_ISL_444786, EPI_ISL_444787, EPI_ISL_444788, EPI_ISL_444789, EPI_ISL_444790, EPI_ISL_444791, EPI_ISL_444792 | | | |
| see above | NYU Langone Health | Departments of Pathology and Medicine, New York University School of Medicine | Maria Aguero-Rosenfeld, Brendan Belovarac, Margaret Black, Ludovic Boytard, John Cadley, Paolo Cotzia, John Chen, Dacia Dimartino, Xiaojun Feng, Tatyana Gindin, Emily Guzman, Adriana Heguy, Megan Hogan, Emily Huang, George Jour, Alireza Khodadadi-Jamayran, Lawrence H. Lin, Raven Luther, Andrew Lytle, Christian Marier, Matthew T. Maurano, Mark J. Mulligan, Peter Meyn, Raquel Ordonez Ciriza, Iman Osman, Jared Pinnell, Vanessa Raabe, Sitharam Ramaswami, Amy Rapkiewicz, Andre M. Ribeiro-dos-Santos, Marie Samanovic-Golden, Antonio Serrano, Guomiao Shen, Matija Snuderl, Theodore Vougiouklakis, Nick Vulpescu, Gael Westby, Paul Zappile, Yutong Zhang |
| EPI_ISL_445213 | DNA Solution Ltd | DNA Solution Ltd | Md. Imran Khan, Kazi Nadim Hasan, Abu Sufian, Mohammed Nafiz Imtiaz Polol, Abdul Khaleque, Mizanur Rahman, MSM Chowdhury, Hasan Ul Haider, Mamudul Hasan Razu, Mala Khan, Mohammad Fazle Alam Rabbi |
| EPI_ISL_445214, EPI_ISL_445215, EPI_ISL_445216, EPI_ISL_445217 | DNA Solution Ltd. | DNA Solution Ltd. | Md. Imran Khan, Kazi Nadim Hasan, Abu Sufian, Mohammed Nafiz Imtiaz Polol, Abdul Khaleque, Mizanur Rahman, MSM Chowdhury, Hasan Ul Haider, Mamudul Hasan Razu, Mala Khan, Mohammad Fazle Alam Rabbi |
| EPI_ISL_445229, EPI_ISL_445236 | Narhalsan Backa vardcentral | The Public Health Agency of Sweden | Mats Olsson, Oskar Karlsson Lindsjo, Maria Lind Karlberg, Anna-Malin Linde, Olov Svartstrom, Anna Risberg, Theresa Enkirch, Mia Brytting, Karin Tegmark-Wisell |
| EPI_ISL_445237 | Narhalsan Molnlycke, Barn och ungdomsmedicin | The Public Health Agency of Sweden | Mats Reimer, Oskar Karlsson Lindsjo, Maria Lind Karlberg, Anna-Malin Linde, Olov Svartstrom, Anna Risberg, Theresa Enkirch, Mia Brytting, Karin Tegmark-Wisell |
| EPI_ISL_445242 | Jokkmokks Halsocentral | The Public Health Agency of Sweden | Markus Beland, Oskar Karlsson Lindsjo, Maria Lind Karlberg, Anna-Malin Linde, Olov Svartstrom, Anna Risberg, Theresa Enkirch, Mia Brytting, Karin Tegmark-Wisell |
| EPI_ISL_445244 | Akbiomed lab | Tejgaon College bmb lab | Md.Abdul kaium,Md.Easin Arafat |
| EPI_ISL_447037 | B.J. Medical College and Civil hospital | Gujarat Biotechnology Research Centre | Monika Gandhi, Pinal Trivedi, Maharshi Pandya, Amit Kanani, Akanksha Verma, Nitin Savaliya, Raghawendra Kumar, Dinesh Kumar, Zuber Saiyed, Dipa Kinariwala, Disha Patel, Binita Aring, Neeta Khandelwal, Geeta Vaghela, Sonia Barve, Bhavesh Modi, Kairavi Joshi, Gaurishankar Shrimali, Nidhi Sood, Pranay Shah, R D Dixit, Snehal Bagatharia, Kamlesh J Upadhyay, Ramesh Pandit, Tejas Shah, Ankit Hinsu, Pritesh Sabara, Apurvasinh Puvar, Janvi Raval, Neelam Nathani, Chaitanya Joshi, Madhvi Joshi |
| EPI_ISL_447055 | Department for Virology, Molecular Biology and Genome Research, R. G. Lugar Center for Public Health Research, National Center for Disease Control and Public Health (NCDC) of Georgia. | Department for Virology, Molecular Biology and Genome Research, R. G. Lugar Center for Public Health Research, National Center for Disease Control and Public Health (NCDC) of Georgia. | Meri Pantsulaia, Gvantsa Brachveli, Giorgi Tomashvili, Gvantsa Chanturia, Ann Machablishvili, Nato Kotaria, Marine Murtskhvaladze, Lela Sabadze, Mari Gavashelidze, Ana Papkiauri, Tata Imnadze, Tamar Jashiashvili, Tea Tevdoradze, Ketevan Sidamonidze, Ekaterine Khmaladze, Ekaterine Zhghenti, Roena Sukhiashvili, Mariam Zakalashvili, Lela Urushadze, Magda Dgebuadze, Davit Tsaguria, Ekaterine Zangaladze, Nino Berishvili, Adam Kotorashvili, Maia Alkhazashvili, Irma Burjanadze, Anna Kasradze, Khatuna Zakhashvili, Paata Imnadze, Amiran Gamkrelidze. |
| EPI_ISL_447472 | Servicio de Microbiologia. Hospital Clinico Universitario de Valencia | Sequencing and Bioinformatics Service and Molecular Epidemiology Research Group. FISABIO-Public Health | Maria Alma Bracho, Griselda De Marco, Lidia Ruiz Roldan, Neris Garcia-Gonzalez, Inma Galán Vendrell, Sandra Carbo, Loreto Ferrús Abad, Paula Ruiz-Hueso, Mariana Reyes-Prieto, Vicente Soriano Chirona, Ivan Ansari, Llúcia Martínez-Priego, Giuseppe 'Auria, David Navarro, Eliseo Albert, Fernando Gonzalez-Candelas |
| EPI_ISL_447480 | Servicio de Microbiologia. Hospital Clinico Universitario de Valencia | Sequencing and Bioinformatics Service and Molecular Epidemiology Research Group. FISABIO-Public Health | Mariana Reyes-Prieto, Vicente Soriano Chirona, Ivan Ansari, Llúcia Martínez-Priego, Giuseppe 'Auria, David Navarro, Eliseo Albert, Maria Alma Bracho, Lidia Ruiz Roldan, Neris Garcia-Gonzalez, Inma Galán Vendrell, Sandra Carbo, Loreto Ferrús Abad, Paula Ruiz-Hueso, Fernando Gonzalez-Candelas |
| EPI_ISL_447486 | Servicio de Microbiologia. Hospital Clinico Universitario de Valencia | Sequencing and Bioinformatics Service and Molecular Epidemiology Research Group. FISABIO-Public Health | Maria Alma Bracho, Griselda De Marco, Lidia Ruiz Roldan, Neris Garcia-Gonzalez, Inma Galán Vendrell, Sandra Carbo, Loreto Ferrús Abad, Paula Ruiz-Hueso, Mariana Reyes-Prieto, Vicente Soriano Chirona, Ivan Ansari, Llúcia Martínez-Priego, Giuseppe 'Auria, David Navarro, Eliseo Albert, Fernando Gonzalez-Candelas |
| EPI_ISL_447494 | Servicio de Microbiologia. Hospital Clinico Universitario de Valencia | Sequencing and Bioinformatics Service and Molecular Epidemiology Research Group. FISABIO-Public Health | Mariana Reyes-Prieto, Vicente Soriano Chirona, Ivan Ansari, Llúcia Martínez-Priego, Giuseppe 'Auria, David Navarro, Eliseo Albert, Maria Alma Bracho, Lidia Ruiz Roldan, Neris Garcia-Gonzalez, Inma Galán Vendrell, Sandra Carbo, Loreto Ferrús Abad, Paula Ruiz-Hueso, Fernando Gonzalez-Candelas |
| EPI_ISL_447500 | Servicio de Microbiologia. Hospital Clinico Universitario de Valencia | Sequencing and Bioinformatics Service and Molecular Epidemiology Research Group. FISABIO-Public Health | Maria Alma Bracho, Griselda De Marco, Lidia Ruiz Roldan, Neris Garcia-Gonzalez, Inma Galán Vendrell, Sandra Carbo, Loreto Ferrús Abad, Paula Ruiz-Hueso, Mariana Reyes-Prieto, Vicente Soriano Chirona, Ivan Ansari, Llúcia Martínez-Priego, Giuseppe 'Auria, David Navarro, Eliseo Albert, Fernando Gonzalez-Candelas |
| EPI_ISL_447508 | Servicio de Microbiologia. Hospital Clinico Universitario de Valencia | Sequencing and Bioinformatics Service and Molecular Epidemiology Research Group. FISABIO-Public Health | Mariana Reyes-Prieto, Vicente Soriano Chirona, Ivan Ansari, Llúcia Martínez-Priego, Giuseppe 'Auria, David Navarro, Eliseo Albert, Maria Alma Bracho, Lidia Ruiz Roldan, Neris Garcia-Gonzalez, Inma Galán Vendrell, Sandra Carbo, Loreto Ferrús Abad, Paula Ruiz-Hueso, Fernando Gonzalez-Candelas |
| EPI_ISL_447514, EPI_ISL_447527 | Servicio de Microbiologia. Hospital Clinico Universitario de Valencia | Sequencing and Bioinformatics Service and Molecular Epidemiology Research Group. FISABIO-Public Health | Maria Alma Bracho, Griselda De Marco, Lidia Ruiz Roldan, Neris Garcia-Gonzalez, Inma Galán Vendrell, Sandra Carbo, Loreto Ferrús Abad, Paula Ruiz-Hueso, Mariana Reyes-Prieto, Vicente Soriano Chirona, Ivan Ansari, Llúcia Martínez-Priego, Giuseppe 'Auria, David Navarro, Eliseo Albert, Fernando Gonzalez-Candelas |
| EPI_ISL_447546 | Gujarat Biotechnology Research Centre | Gujarat Biotechnology Research Centre | Monika Gandhi, Pinal Trivedi, Maharshi Pandya, Amit Kanani, Akanksha Verma, Nitin Savaliya, Raghawendra Kumar, Dinesh Kumar, Zuber Saiyed, Dipa Kinariwala, Disha Patel, Binita Aring, Neeta Khandelwal, Geeta Vaghela, Sonia Barve, Bhavesh Modi, Kairavi Joshi, Gaurishankar Shrimali, Nidhi Sood, Pranay Shah, R D Dixit, Snehal Bagatharia, Kamlesh J Upadhyay, Ramesh Pandit, Tejas Shah, Ankit Hinsu, Pritesh Sabara, Apurvasinh Puvar, Janvi Raval, Priti Pandita, Chaitanya Joshi, Madhvi Joshi |
| EPI_ISL_447574, EPI_ISL_447576, EPI_ISL_447577, EPI_ISL_447578, EPI_ISL_447853 | CSIR-Centre for Cellular and Molecular Biology | CSIR-Centre for Cellular and Molecular Biology | Namami Gaur, Sakshi Shambhavi, Lamuk Zaveri, Shagufta Khan, Tulasi Nagabandi, Purushotham Vodnala, Payel Mukherjee, Sofia Banu, Priya Singh, Dhiviya Vedagiri, Divya Gupta, Vishal Sah, Santosh Kumar Kuncha, Krishnan Harinivas Harshan, Archana Bharadwaj Siva, Karthik Bharadwaj Tallapaka, Rakesh K Mishra, Divya Tej Sowpati |
| EPI_ISL_447904 | National Institute of Biotechnology | National Institute of Biotechnology | Md. Moniruzzaman, Mohammad Uzzal Hossain, Md. Nazrul Islam, Md. Hadisur Rahman, Irfan Ahmed, Tahia Anan Rahman, Arittra Bhattacharjee, Md. Ruhul Amin, Asif Rashid, Chaman Ara Keya, Keshob Chandra Das, Md. Salimullah |
| EPI_ISL_449259, EPI_ISL_449260, EPI_ISL_449261, EPI_ISL_449262, EPI_ISL_449263, EPI_ISL_449264, EPI_ISL_449265, EPI_ISL_449266, EPI_ISL_449267, EPI_ISL_449268, EPI_ISL_449269, EPI_ISL_449270, EPI_ISL_449271, EPI_ISL_449272, EPI_ISL_449273, EPI_ISL_449274, EPI_ISL_449275, EPI_ISL_449276, EPI_ISL_449277, EPI_ISL_449278, EPI_ISL_449279, EPI_ISL_449280, EPI_ISL_449281, EPI_ISL_449282, EPI_ISL_449283, EPI_ISL_449284, EPI_ISL_449285, EPI_ISL_449286, EPI_ISL_449287, EPI_ISL_449288, EPI_ISL_449289, EPI_ISL_449290, EPI_ISL_449291, EPI_ISL_449292, EPI_ISL_449293, EPI_ISL_449294, EPI_ISL_449295, EPI_ISL_449296, EPI_ISL_449297, EPI_ISL_449298, EPI_ISL_449299, EPI_ISL_449300, EPI_ISL_449301, EPI_ISL_449302, EPI_ISL_449303, EPI_ISL_449304, EPI_ISL_449305, EPI_ISL_449306, EPI_ISL_449307, EPI_ISL_449308, EPI_ISL_449309, EPI_ISL_449310, EPI_ISL_449311, EPI_ISL_449312, EPI_ISL_449313, EPI_ISL_449314, EPI_ISL_449315, EPI_ISL_449316, EPI_ISL_449317, EPI_ISL_449318, EPI_ISL_449319, EPI_ISL_449320, EPI_ISL_449321, EPI_ISL_449322, EPI_ISL_449323, EPI_ISL_449324, EPI_ISL_449325, EPI_ISL_449326, EPI_ISL_449327, EPI_ISL_449328, EPI_ISL_449329 | | | |
| see above | Virology Department, Royal Infirmary of Edinburgh, NHS Lothian / School of Biological Sciences, University of Edinburgh / Institute of Genetics and Molecular Medicine, University of Edinburgh | COVID-19 Genomics UK (COG-UK) Consortium | McHugh M, Dewar R, Rooke S, Gallagher M, Balcaza C, O'Toole Á, Scher E, Hill V, McCrone JT, Colquhoun R, Yu X, Jackson B, Rambaut A, Williams TC, Templeton K |
| EPI_ISL_450329 | CSIR-Centre for Cellular and Molecular Biology | CSIR-Centre for Cellular and Molecular Biology | Namami Gaur, Sakshi Shambhavi, Lamuk Zaveri, Shagufta Khan, Tulasi Nagabandi, Purushotham Vodnala, Payel Mukherjee, Sofia Banu, Priya Singh, Dhiviya Vedagiri, Divya Gupta, Vishal Sah, Santosh Kumar Kuncha, Krishnan Harinivas Harshan, Archana Bharadwaj Siva, Karthik Bharadwaj Tallapaka, Sonu Uday, Sudipta Mondal, Annapoorna P Karthyayani, Debabrata Jana, Debrya Saha, Gokulan C G, Gunjan Purohit, Hanuman Tulashiram Kale, Pankaj Kumar, Prachand Issarapu, Preethi Jampala Rakesh K Mishra, Divya Tej Sowpati |
| EPI_ISL_450341 | Bangladesh Institute of Tropical & Infectious Diseases, COVID-19 Testing Laboratory | Basic and Applied Research on Jute Project | Md. Sabbir Hossain, Rasel Ahmed, Shah Md Tamim Kabir, Emdadul Mannan Emdad, Md. Nazmul Haq Rony, Eaftekhar Ahmed Rana, Paritous Kumar Biswas, M A Hassan Chowdhury, Md. Shakeel Ahmed, Md. Samiul Haque, Md. Monjurul Alam, Md. Sharifur Rahman, A S M Anwarul Huq, Md. Shahidul Islam, Goutam Buddha Das, AMAM Zonaed Siddiki |
| EPI_ISL_450343, EPI_ISL_450345 | Bangladesh Institute of Tropical & Infectious Diseases, COVID-19 Testing Laboratory | Basic and Applied Research on Jute Project | Md. Sabbir Hossain, Rasel Ahmed, Shah Md Tamim Kabir, Emdadul Mannan Emdad, Md. Nazmul Haq Rony, Eaftekhar Ahmed Rana, Paritous Kumar Biswas, M A Hassan Chowdhury, Md. Shakeel Ahmed, Md. Samiul Haque, Md. Monjurul Alam, Md. Sharifur Rahman, A S M Anwarul Huq, Md. Shahidul Islam, Goutam Buddha Das, AMAM Zonaed Siddiki |
| EPI_ISL_450393, EPI_ISL_450394, EPI_ISL_450395, EPI_ISL_450396, EPI_ISL_450397, EPI_ISL_450398, | NYU Langone Health | Departments of Pathology and Medicine, New York University School of Medicine | Maria Aguero-Rosenfeld, Brendan Belovarac, Margaret Black, Ludovic Boytard, John Cadley, Paolo Cotzia, John Chen, Dacia Dimartino, Xiaojun Feng, Tatyana Gindin, Emily Guzman, Adriana Heguy, Megan Hogan, Emily Huang, George Jour, Alireza Khodadadi-Jamayran, Lawrence H. Lin, Raven Luther, Andrew Lytle, Christian Marier, Matthew T. Maurano, Mark J. Mulligan, Peter Meyn, Raquel Ordonez Ciriza, Iman Osman, Jared Pinnell, Vanessa Raabe, |

| | | | |
|---|---|---|---|
| EPI_ISL_450399, EPI_ISL_450400, EPI_ISL_450401, EPI_ISL_450402 | | | Sitharam Ramaswami, Amy Rapkiewicz, Andre M. Ribeiro-dos-Santos, Marie Samanovic-Golden, Antonio Serrano, Guomiao Shen, Matija Snuderl, Theodore Vougiouklakis, Nick Vulpescu, Gael Westby, Paul Zappile, Yutong Zhang |
| EPI_ISL_450638, EPI_ISL_450639, EPI_ISL_450640, EPI_ISL_450641, EPI_ISL_450642, EPI_ISL_450643, EPI_ISL_450644, EPI_ISL_450645, EPI_ISL_450646, EPI_ISL_450647, EPI_ISL_450648, EPI_ISL_450649, EPI_ISL_450650, EPI_ISL_450651, EPI_ISL_450652, EPI_ISL_450653 | | | |
| see above | Laboratoire de microbiologie, Hopital de Verdun | Smith Laboratory, Centre de Recherche CHU Sainte-Justine | Martin Smith, Marieke Rozendaal, Ivan Pavlov |
| EPI_ISL_450748, EPI_ISL_450749, EPI_ISL_450750, EPI_ISL_450751, EPI_ISL_450752, EPI_ISL_450753, EPI_ISL_450754, EPI_ISL_450755, EPI_ISL_450756, EPI_ISL_450757, EPI_ISL_450758, EPI_ISL_450759, EPI_ISL_450760, EPI_ISL_450761, EPI_ISL_450762, EPI_ISL_450763, EPI_ISL_450764, EPI_ISL_450765, EPI_ISL_450766, EPI_ISL_450767, EPI_ISL_450768, EPI_ISL_450769, EPI_ISL_450770, EPI_ISL_450771, EPI_ISL_450772, EPI_ISL_450773, EPI_ISL_450774, EPI_ISL_450775, EPI_ISL_450776, EPI_ISL_450777, EPI_ISL_450778, EPI_ISL_450779, EPI_ISL_450780 | | | |
| see above | Minnesota Department of Health, Public Health Laboratory | Minnesota Department of Health, Public Health Laboratory | Matt Plumb, Jacob Garfin, and Xiong Wang |
| EPI_ISL_450815 | Narhalsan Molnlycke, Barn och ungdomsmedicin | The Public Health Agency of Sweden | Mats Reimer, Anna-Malin Linde, Maria Lind Karlberg, Oskar Karlsson Lindsjo, Olov Svartstrom, Anna Risberg, Theresa Enkirch, Mia Brytting, Karin Tegmark-Wisell |
| EPI_ISL_450817 | Narhalsan Backa vardcentral | The Public Health Agency of Sweden | Mats Olsson, Anna-Malin Linde, Maria Lind Karlberg, Oskar Karlsson Lindsjo, Olov Svartstrom, Anna Risberg, Theresa Enkirch, Mia Brytting, Karin Tegmark-Wisell |
| EPI_ISL_450820 | Narhalsan Molnlycke, Barn och ungdomsmedicin | The Public Health Agency of Sweden | Mats Reimer, Anna-Malin Linde, Maria Lind Karlberg, Oskar Karlsson Lindsjo, Olov Svartstrom, Anna Risberg, Theresa Enkirch, Mia Brytting, Karin Tegmark-Wisell |
| EPI_ISL_450823, EPI_ISL_450825 | Narhalsan Backa vardcentral | The Public Health Agency of Sweden | Mats Olsson, Anna-Malin Linde, Maria Lind Karlberg, Oskar Karlsson Lindsjo, Olov Svartstrom, Anna Risberg, Theresa Enkirch, Mia Brytting, Karin Tegmark-Wisell |
| EPI_ISL_451151 | M.P Shah Government Medocal college Jamnagar | Gujarat Biotechnology Research Centre | Monika Gandhi, Pinal Trivedi, Maharshi Pandya, Amit Kanani, Nidhi Patel, Nitin Savaliya, Raghawendra Kumar, Dinesh Kumar, Zuber Saiyed, Komal Patel, Labdhi Pandya, Snehal Bagatharia, Ramesh Pandit, Tejas Shah, Ankit Hinsu, Pritesh Sabara, Apurvasinh Puvar, Binita Aring, Janvi Raval, Zarna Patel, Bhavesh Modi, Gaurishankar Shrimali, R D Dixit, A M Kadri, Neha Rajpara, Chaitanya Joshi, Madhvi Joshi, |
| EPI_ISL_451299, EPI_ISL_451301, EPI_ISL_451303 | Laboratory of Virology, INMI Lazzaro Spallanzani IRCCS | Laboratory of Virology, INMI Lazzaro Spallanzani IRCCS | Martina Rueca, Cesare E.M. Gruber, Barbara Bartolini, Francesco Messina, Antonino Di Caro, Maria R. Capobianchi, Giuseppe Ippolito |
| EPI_ISL_451401, EPI_ISL_451402, EPI_ISL_451403, EPI_ISL_451404, EPI_ISL_451405, EPI_ISL_451406, EPI_ISL_451407, EPI_ISL_451408, EPI_ISL_451409, EPI_ISL_451410, EPI_ISL_451411, EPI_ISL_451412, EPI_ISL_451413, EPI_ISL_451414, EPI_ISL_451415, EPI_ISL_451416, EPI_ISL_451417, EPI_ISL_451418, EPI_ISL_451419, EPI_ISL_451420, EPI_ISL_451421, EPI_ISL_451422, EPI_ISL_451423, EPI_ISL_451424, EPI_ISL_451425, EPI_ISL_451426, EPI_ISL_451427, EPI_ISL_451428, EPI_ISL_451429, EPI_ISL_451430, EPI_ISL_451431, EPI_ISL_451432, EPI_ISL_451433, EPI_ISL_451434, EPI_ISL_451435, EPI_ISL_451436, EPI_ISL_451437, EPI_ISL_451438, EPI_ISL_451439, EPI_ISL_451440, EPI_ISL_451441, EPI_ISL_451442, EPI_ISL_451443, EPI_ISL_451444, EPI_ISL_451445, EPI_ISL_451446, EPI_ISL_451447, EPI_ISL_451448, EPI_ISL_451449, EPI_ISL_451450, EPI_ISL_451451, EPI_ISL_451452, EPI_ISL_451453, EPI_ISL_451454, EPI_ISL_451455, EPI_ISL_451456, EPI_ISL_451457, EPI_ISL_451458, EPI_ISL_451459, EPI_ISL_451460, EPI_ISL_451461, EPI_ISL_451462, EPI_ISL_451463, EPI_ISL_451464, EPI_ISL_451465, EPI_ISL_451466, EPI_ISL_451467, EPI_ISL_451468, EPI_ISL_451469, EPI_ISL_451470, EPI_ISL_451471, EPI_ISL_451472, EPI_ISL_451473, EPI_ISL_451474, EPI_ISL_451475, EPI_ISL_451476, EPI_ISL_451477, EPI_ISL_451478, EPI_ISL_451479, EPI_ISL_451665 | | | |
| see above | NYU Langone Health | Departments of Pathology and Medicine, New York University School of Medicine | Maria Aguero-Rosenfeld, Brendan Belovarac, Margaret Black, Ludovic Boytard, John Cadley, Paolo Cotzia, John Chen, Dacia Dimartino, Xiaojun Feng, Tatyana Gindin, Emily Guzman, Adriana Heguy, Megan Hogan, Emily Huang, George Jour, Alireza Khodadadi-Jamayran, Lawrence H. Lin, Raven Luther, Andrew Lytle, Christian Marier, Matthew T. Maurano, Mark J. Mulligan, Peter Meyn, Raquel Ordonez Ciriza, Iman Osman, Jared Pinnell, Vanessa Raabe, Sitharam Ramaswami, Amy Rapkiewicz, Andre M. Ribeiro-dos-Santos, Marie Samanovic-Golden, Antonio Serrano, Guomiao Shen, Matija Snuderl, Theodore Vougiouklakis, Nick Vulpescu, Gael Westby, Paul Zappile, Yutong Zhang |
| EPI_ISL_451936, EPI_ISL_451937, EPI_ISL_451938, EPI_ISL_451939, EPI_ISL_451940, EPI_ISL_451941, EPI_ISL_451942, EPI_ISL_451943, EPI_ISL_451944, EPI_ISL_451945, EPI_ISL_451946, EPI_ISL_451947, EPI_ISL_452103, EPI_ISL_452104 | | | |
| see above | Max von Pettenkofer Institute, Virology, National Reference Center for Retroviruses, LMU München | Laboratory for Functional Genome Analysis, Dept. Genomics, Gene Center of the LMU Munich | Max Muenchhoff, Stefan Krebs, Alexander Graf, Oliver Keppler, Helmut Blum |
| EPI_ISL_452229, EPI_ISL_452230, EPI_ISL_452235, EPI_ISL_452239 | Narhalsan Backa vardcentral | The Public Health Agency of Sweden | Mats Olsson, Anna-Malin Linde, Maria Lind Karlberg, Oskar Karlsson Lindsjo, Olov Svartstrom, Anna Risberg, Theresa Enkirch, Mia Brytting, Karin Tegmark-Wisell |
| EPI_ISL_452472, EPI_ISL_452473, EPI_ISL_452474, EPI_ISL_452475, EPI_ISL_452476, EPI_ISL_452477, EPI_ISL_452478, EPI_ISL_452479, EPI_ISL_452480, EPI_ISL_452481, EPI_ISL_452482, EPI_ISL_452483, EPI_ISL_452484, EPI_ISL_452485, EPI_ISL_452486, EPI_ISL_452487, EPI_ISL_452488, EPI_ISL_452489, EPI_ISL_452490, EPI_ISL_452491, EPI_ISL_452492, EPI_ISL_452493, EPI_ISL_452494, EPI_ISL_452495, EPI_ISL_452496, EPI_ISL_452497, EPI_ISL_452498, EPI_ISL_452499, EPI_ISL_452500, EPI_ISL_452501, EPI_ISL_452502, EPI_ISL_452503, EPI_ISL_452504, EPI_ISL_452505, EPI_ISL_452506, EPI_ISL_452507, EPI_ISL_452508, EPI_ISL_452509, EPI_ISL_452510, EPI_ISL_452511, EPI_ISL_452512, EPI_ISL_452513, EPI_ISL_452514, EPI_ISL_452515, EPI_ISL_452516, EPI_ISL_452517, EPI_ISL_452518, EPI_ISL_452519, EPI_ISL_452520, EPI_ISL_452521, EPI_ISL_452522, EPI_ISL_452523, EPI_ISL_452524, EPI_ISL_452525, EPI_ISL_452526, EPI_ISL_452527, EPI_ISL_452528, EPI_ISL_452529, EPI_ISL_452530, EPI_ISL_452531, EPI_ISL_452532, EPI_ISL_452533, EPI_ISL_452534, EPI_ISL_452535, EPI_ISL_452536, EPI_ISL_452537, EPI_ISL_452538, EPI_ISL_452539, EPI_ISL_452540, EPI_ISL_452541, EPI_ISL_452542, EPI_ISL_452543 | | | |
| see above | Clínica Universidad de Navarra. Servicio de Enfermedades Infecciosas y Microbiología clínica | SeqCOVID-SPAIN consortium/IBV(CSIC) | Mirian Fernández-Alonso, Jose Luis del Pozo and SeqCOVID-SPAIN consortium |
| EPI_ISL_453099, EPI_ISL_453100, EPI_ISL_453101, EPI_ISL_453102, EPI_ISL_453103, EPI_ISL_453104, EPI_ISL_453105, EPI_ISL_453106, EPI_ISL_453107, EPI_ISL_453108, EPI_ISL_453109, EPI_ISL_453110, EPI_ISL_453111, EPI_ISL_453112, EPI_ISL_453113, EPI_ISL_453114, EPI_ISL_453115, EPI_ISL_453116, EPI_ISL_453117, EPI_ISL_453118, EPI_ISL_453119, EPI_ISL_453120, EPI_ISL_453121, EPI_ISL_453122, EPI_ISL_453123, EPI_ISL_453124, EPI_ISL_453125, EPI_ISL_453126, EPI_ISL_453127, EPI_ISL_453128, EPI_ISL_453129, EPI_ISL_453130, EPI_ISL_453131, EPI_ISL_453132, EPI_ISL_453133, EPI_ISL_453134, EPI_ISL_453135, EPI_ISL_453136, EPI_ISL_453137, EPI_ISL_453138, EPI_ISL_453139, EPI_ISL_453140, EPI_ISL_453141, EPI_ISL_453142, EPI_ISL_453143, EPI_ISL_453144, EPI_ISL_453145, EPI_ISL_453146, EPI_ISL_453147, EPI_ISL_453148, EPI_ISL_453149, EPI_ISL_453150, EPI_ISL_453151, EPI_ISL_453152, EPI_ISL_453153, EPI_ISL_453154, EPI_ISL_453155, EPI_ISL_453156, EPI_ISL_453157, EPI_ISL_453158, EPI_ISL_453159, EPI_ISL_453160, EPI_ISL_453161, EPI_ISL_453162, EPI_ISL_453163, EPI_ISL_453164, EPI_ISL_453165, EPI_ISL_453166, EPI_ISL_453167, EPI_ISL_453168, EPI_ISL_453169, EPI_ISL_453170, EPI_ISL_453171, EPI_ISL_453172, EPI_ISL_453173, EPI_ISL_453174, EPI_ISL_453175, EPI_ISL_453176, EPI_ISL_453177, EPI_ISL_453178, EPI_ISL_453179, EPI_ISL_453180, EPI_ISL_453181, EPI_ISL_453182, EPI_ISL_453183, EPI_ISL_453184, EPI_ISL_453185, EPI_ISL_453186, EPI_ISL_453187, EPI_ISL_453188, EPI_ISL_453189, EPI_ISL_453190, EPI_ISL_453191, EPI_ISL_453192, EPI_ISL_453193, EPI_ISL_453194 | | | |
| see above | Virology Department, Royal Infirmary of Edinburgh, NHS Lothian / School of Biological Sciences, University of Edinburgh / Institute of Genetics and Molecular Medicine, University of Edinburgh | COVID-19 Genomics UK (COG-UK) Consortium | McHugh M, Dewar R, Rooke S, Gallagher M, Balcaza C, O'Toole Á, Scher E, Hill V, McCrone JT, Colquhoun R, Yu X, Jackson B, Rambaut A, Williams TC, Templeton K |
| EPI_ISL_454353 | Microbial Genome Sequencing Center; Microbial Genomic Epidemiology Laboratory | Microbial Genomic Epidemiology Laboratory, University of Pittsburgh | Mustapha M. Mustapha, Jane W. Marsh, Dan Snyder, Marissa P. Griffith, Stephanie L. Mitchell, Vatsala R. Srinivasa, Kady D. Waggle, Chinelo Ezeonwuku, Vaughn S. Cooper, Lee H. Harrison |
| EPI_ISL_454354 | UPMC Clinical Microbiology Laboratory | Microbial Genome Sequencing Center; Microbial Genomic Epidemiology Laboratory, University of Pittsburgh | Mustapha M. Mustapha, Jane W. Marsh, Dan Snyder, Marissa P. Griffith, Stephanie L. Mitchell, Vatsala R. Srinivasa, Kady D. Waggle, Chinelo Ezeonwuku, Vaughn S. Cooper, Lee H. Harrison |
| EPI_ISL_454355, EPI_ISL_454356, EPI_ISL_454357, EPI_ISL_454358, EPI_ISL_454359, EPI_ISL_454360, EPI_ISL_454361, EPI_ISL_454362, EPI_ISL_454363, EPI_ISL_454364, EPI_ISL_454365, EPI_ISL_454366, EPI_ISL_454367, EPI_ISL_454368, EPI_ISL_454370, EPI_ISL_454371, EPI_ISL_454372, EPI_ISL_454373, EPI_ISL_454374, EPI_ISL_454375, EPI_ISL_454376, EPI_ISL_454378, EPI_ISL_454379, EPI_ISL_454380, EPI_ISL_454381, EPI_ISL_454382, EPI_ISL_454383, EPI_ISL_454384, EPI_ISL_454387, EPI_ISL_454399, EPI_ISL_454401, EPI_ISL_454402, EPI_ISL_454403, EPI_ISL_454404, EPI_ISL_454406, EPI_ISL_454407, EPI_ISL_454408, EPI_ISL_454409, EPI_ISL_454410, EPI_ISL_454411 | | | |
| see above | UPMC Clinical Microbiology Laboratory | Microbial Genome Sequencing Center, Microbial Genomic Epidemiological Laboratory | Mustapha M. Mustapha, Jane W. Marsh, Dan Snyder, Marissa P. Griffith, Stephanie L. Mitchell, Vatsala R. Srinivasa, Kady D. Waggle, Chinelo Ezeonwuku, Vaughn S. Cooper, Lee H. Harrison |
| EPI_ISL_455026 | Government Medical College, Vadodara | Gujarat Biotechnology Research Centre | Monika Gandhi, Pinal Trivedi, Maharshi Pandya, Amit Kanani, Nidhi Patel, Nitin Savaliya, Raghawendra Kumar, Dinesh Kumar, Zuber Saiyed, Komal Patel, Labdhi Pandya, Snehal Bagatharia, Tanuja Javadekar , R N Daveshwar, Ramesh Pandit, Tejas Shah, Ankit Hinsu, Pritesh Sabara, Apurvasinh Puvar, Janvi Raval, Zarna Patel, Priti Pandita, Bhavesh Modi, Gaurishankar Shrimali, R D Dixit, A M Kadri, Umang Mishra, Chaitanya Joshi, Madhvi Joshi, |
| EPI_ISL_455107 | Narhalsan Mellerud | The Public Health Agency of Sweden | Maria Nykvist, Anna-Malin Linde, Maria Lind Karlberg, Oskar Karlsson Lindsjo, Olov Svartstrom, Anna Risberg, Theresa Enkirch, Mia Brytting, Karin Tegmark-Wisell |
| EPI_ISL_455439 | Instituto de Diagnostico y Referencia Epidemiologicos (INDRE) | Instituto de Diagnostico y Referencia Epidemiologicos (INDRE) | Mendieta-Condado Edgar, Araiza-Rodriguez Adnan, Garces-Ayala Fabiola, Rodriguez-Maldonado Abril, Wong-Arambula Claudia, Barrera-Badillo Gisela, Hernandez-Rivas Lucia, Lopez-Martinez Irma, Taboada Ramirez Blanca, Ramirez-Gonzalez Ernesto. |
| EPI_ISL_455454 | Instituto de Diagnostico y Referencia Epidemiologicos (INDRE) | Instituto de Diagnostico y Referencia Epidemiologicos (INDRE) | Mendieta-Condado Edgar, Araiza-Rodriguez Adnan, Garces-Ayala Fabiola, , Rodriguez-Maldonado Abril, Wong-Arambula Claudia, Barrera-Badillo Gisela, Taboada Ramírez Blanca, Ramirez-Gonzalez Ernesto, Hernandez-Rivas Lucia, Lopez-Martinez Irma. |
| EPI_ISL_455719 | T.C. Salk Bakanl Adyaman l Salk Müdürlüü Adyaman Eitim Ve Aratrma Hastanesi | VETAL Animal Health Products Company, BSL3+ Production Laboratuary /Turkey | Mehmet Turgut, Muhittin Önderci, Fatma Nilay Tutak, Abidin Ercan Yonucu, Murat Dönen, Haluk Uluca, Fethiye Sevimli, O. Ugur Sezerman |
| EPI_ISL_455728 | Servicio de Microbiologia. Hospital Clinico Universitario de Valencia | Sequencing and Bioinformatics Service and Molecular Epidemiology Research Group. FISABIO-Public Health, and | Maria Alma Bracho, Griselda De Marco, Lidia Ruiz Roldan, Neris Garcia-Gonzalez, Inma Galán Vendrell, Sandra Carbo, Loreto Ferrús Abad, Paula Ruiz-Hueso, Mariana Reyes-Prieto, Vicente Soriano Chirona, Ivan Ansari, Llúcia Martínez-Priego, Giuseppe 'Auria, David Navarro, Eliseo Albert, Fernando |

| | | | |
|---|---|---|---|
| | | SeqCOVID-Spain Consortium | Gonzalez-Candelas |
| EPI_ISL_455730, EPI_ISL_455731 | Servicio de Microbiologia. Hospital Clinico Universitario de Valencia | Sequencing and Bioinformatics Service and Molecular Epidemiology Research Group. FISABIO-Public Health, and SeqCOVID-Spain Consortium | Mariana Reyes-Prieto, Vicente Soriano Chirona, Ivan Ansari, Llúcia Martínez-Priego, Giuseppe 'Auria, David Navarro, Eliseo Albert, Maria Alma Bracho, Lidia Ruiz Roldan, Neris Garcia-Gonzalez, Inma Galán Vendrell, Sandra Carbo, Loreto Ferrús Abad, Paula Ruiz-Hueso, Fernando Gonzalez-Candelas |
| EPI_ISL_455738 | Servicio de Microbiologia. Hospital Clinico Universitario de Valencia | Sequencing and Bioinformatics Service and Molecular Epidemiology Research Group. FISABIO-Public Health, and SeqCOVID-Spain Consortium | Maria Alma Bracho, Griselda De Marco, Lidia Ruiz Roldan, Neris Garcia-Gonzalez, Inma Galán Vendrell, Sandra Carbo, Loreto Ferrús Abad, Paula Ruiz-Hueso, Mariana Reyes-Prieto, Vicente Soriano Chirona, Ivan Ansari, Llúcia Martínez-Priego, Giuseppe 'Auria, David Navarro, Eliseo Albert, Fernando Gonzalez-Candelas |
| EPI_ISL_455790, EPI_ISL_455791 | Institute for Medical Research, Infectious Disease Research Centre, National Institutes of Health, Ministry of Health Malaysia | Malaysia Genome Institute | Mohd Noor Mat Isa, Irni Suhayu Sapian, Yusuf Muhammad Noor, Jeyanthi Suppiah, Nurhezreen Md Iqbal, Enizza Kasim, Zarina Mohd Zawawi, Siti Noraini Othman, Mohd Faizal Abu Bakar, Shamsidar Sopie, Azrin Ahmad, Ravindran Thayan, Norazah Ahmad, Tahir Aris, Shahrul Hisham Zainal Ariffin |
| EPI_ISL_455792, EPI_ISL_455793 | Institute for Medical Research, Infectious Disease Research Centre, National Institutes of Health, Ministry of Health Malaysia | Malaysia Genome Institute | Mohd Noor Mat Isa, Irni Suhayu Sapian, Yusuf Muhammad Noor, Jeyanthi Suppiah, Nurhezreen Md Iqbal, Enizza Kasim, Zarina Mohd Zawawi, Siti Noraini Othman, Mohd Faizal Abu Bakar, Shamsidar Sopie, Azrin Ahmad, Ravindran Thayan, Norazah Ahmad, Tahir Aris, Shahrul Hisham Zainal Ariffin |
| EPI_ISL_456004, EPI_ISL_456005, EPI_ISL_456006, EPI_ISL_456007, EPI_ISL_456008, EPI_ISL_456009, EPI_ISL_456010, EPI_ISL_456011, EPI_ISL_456012, EPI_ISL_456013, EPI_ISL_456014, EPI_ISL_456015, EPI_ISL_456016, EPI_ISL_456017, EPI_ISL_456018, EPI_ISL_456019, EPI_ISL_456020, EPI_ISL_456021, EPI_ISL_456022, EPI_ISL_456023, EPI_ISL_456024, EPI_ISL_456025, EPI_ISL_456026, EPI_ISL_456027, EPI_ISL_456028, EPI_ISL_456029, EPI_ISL_456030, EPI_ISL_456031, EPI_ISL_456032, EPI_ISL_456033, EPI_ISL_456034, EPI_ISL_456035, EPI_ISL_456036, EPI_ISL_456037, EPI_ISL_456038, EPI_ISL_456039, EPI_ISL_456040, EPI_ISL_456041, EPI_ISL_456042, EPI_ISL_456043, EPI_ISL_456044, EPI_ISL_456045, EPI_ISL_456046, EPI_ISL_456047, EPI_ISL_456048, EPI_ISL_456049, EPI_ISL_456050, EPI_ISL_456051, EPI_ISL_456052, EPI_ISL_456053, EPI_ISL_456054, EPI_ISL_456055, EPI_ISL_456056, EPI_ISL_456057, EPI_ISL_456058, EPI_ISL_456059, EPI_ISL_456060, EPI_ISL_456061, EPI_ISL_456062, EPI_ISL_456063, EPI_ISL_456064, EPI_ISL_456065, EPI_ISL_456066, EPI_ISL_456067, EPI_ISL_456068, EPI_ISL_456069, EPI_ISL_456070, EPI_ISL_456107, EPI_ISL_456108, EPI_ISL_456109, EPI_ISL_456110, EPI_ISL_456111, EPI_ISL_456112, EPI_ISL_456113, EPI_ISL_456114, EPI_ISL_456115 | | | |
| see above | NYU Langone Health | Departments of Pathology and Medicine, New York University School of Medicine | Maria Aguero-Rosenfeld, Brendan Belovarac, Margaret Black, Ludovic Boytard, John Cadley, Paolo Cotzia, John Chen, Dacia Dimartino, Xiaojun Feng, Tatyana Gindin, Emily Guzman, Adriana Heguy, Megan Hogan, Emily Huang, George Jour, Alireza Khodadadi-Jamayran, Lawrence H. Lin, Raven Luther, Andrew Lytle, Christian Marier, Matthew T. Maurano, Mark J. Mulligan, Peter Meyn, Raquel Ordonez Ciriza, Iman Osman, Jared Pinnell, Vanessa Raabe, Sitharam Ramaswami, Amy Rapkiewicz, Andre M. Ribeiro-dos-Santos, Marie Samanovic-Golden, Antonio Serrano, Guomiao Shen, Matija Snuderl, Theodore Vougiouklakis, Nick Vulpescu, Gael Westby, Paul Zappile, Yutong Zhang |
| EPI_ISL_456157 | LabPLUS | Institute of Environmental Science and Research (ESR) | Matt Storey, Xiaoyun Ren, Anja Werno, Antje van der Linden, Arlo Upton, Chris Mansell, David Hammer, Dragana Drinkovic, Erasmus Smit, Gary McAuliffe, Hana Sofia Andersson, James Ussher, Jill Sherwood, Josh Freeman, Julia Howard, Juliet Elvy, Mary DeAlmeida, Matt Blakiston, Matthew Rogers, Max Bloomfield, Michael Addidle, Michelle Balm, Sally Roberts, Sarah Jefferies, Sharmini Muttaiyah, Susan Morpeth, Susan Taylor, Timothy Blackmore, Vani Sathyendran, Veronica Playle, Virginia Hope, Erasmus Smit, Lauren Jelly, Joep de Ligt |
| EPI_ISL_456158, EPI_ISL_456159 | Southern Community Labs Dunedin | Institute of Environmental Science and Research (ESR) | Matt Storey, Xiaoyun Ren, Anja Werno, Antje van der Linden, Arlo Upton, Chris Mansell, David Hammer, Dragana Drinkovic, Erasmus Smit, Gary McAuliffe, Hana Sofia Andersson, James Ussher, Jill Sherwood, Josh Freeman, Julia Howard, Juliet Elvy, Mary DeAlmeida, Matt Blakiston, Matthew Rogers, Max Bloomfield, Michael Addidle, Michelle Balm, Sally Roberts, Sarah Jefferies, Sharmini Muttaiyah, Susan Morpeth, Susan Taylor, Timothy Blackmore, Vani Sathyendran, Veronica Playle, Virginia Hope, Erasmus Smit, Lauren Jelly, Joep de Ligt |
| EPI_ISL_456160 | Waikato Hospital | Institute of Environmental Science and Research (ESR) | Matt Storey, Xiaoyun Ren, Anja Werno, Antje van der Linden, Arlo Upton, Chris Mansell, David Hammer, Dragana Drinkovic, Erasmus Smit, Gary McAuliffe, Hana Sofia Andersson, James Ussher, Jill Sherwood, Josh Freeman, Julia Howard, Juliet Elvy, Mary DeAlmeida, Matt Blakiston, Matthew Rogers, Max Bloomfield, Michael Addidle, Michelle Balm, Sally Roberts, Sarah Jefferies, Sharmini Muttaiyah, Susan Morpeth, Susan Taylor, Timothy Blackmore, Vani Sathyendran, Veronica Playle, Virginia Hope, Erasmus Smit, Lauren Jelly, Joep de Ligt |
| EPI_ISL_456161 | Wellington SCL | Institute of Environmental Science and Research (ESR) | Matt Storey, Xiaoyun Ren, Anja Werno, Antje van der Linden, Arlo Upton, Chris Mansell, David Hammer, Dragana Drinkovic, Erasmus Smit, Gary McAuliffe, Hana Sofia Andersson, James Ussher, Jill Sherwood, Josh Freeman, Julia Howard, Juliet Elvy, Mary DeAlmeida, Matt Blakiston, Matthew Rogers, Max Bloomfield, Michael Addidle, Michelle Balm, Sally Roberts, Sarah Jefferies, Sharmini Muttaiyah, Susan Morpeth, Susan Taylor, Timothy Blackmore, Vani Sathyendran, Veronica Playle, Virginia Hope, Erasmus Smit, Lauren Jelly, Joep de Ligt |
| EPI_ISL_456162 | Southern Community Labs Dunedin | Institute of Environmental Science and Research (ESR) | Matt Storey, Xiaoyun Ren, Anja Werno, Antje van der Linden, Arlo Upton, Chris Mansell, David Hammer, Dragana Drinkovic, Erasmus Smit, Gary McAuliffe, Hana Sofia Andersson, James Ussher, Jill Sherwood, Josh Freeman, Julia Howard, Juliet Elvy, Mary DeAlmeida, Matt Blakiston, Matthew Rogers, Max Bloomfield, Michael Addidle, Michelle Balm, Sally Roberts, Sarah Jefferies, Sharmini Muttaiyah, Susan Morpeth, Susan Taylor, Timothy Blackmore, Vani Sathyendran, Veronica Playle, Virginia Hope, Erasmus Smit, Lauren Jelly, Joep de Ligt |
| EPI_ISL_456163 | PathLab Bay of Plenty | Institute of Environmental Science and Research (ESR) | Matt Storey, Xiaoyun Ren, Anja Werno, Antje van der Linden, Arlo Upton, Chris Mansell, David Hammer, Dragana Drinkovic, Erasmus Smit, Gary McAuliffe, Hana Sofia Andersson, James Ussher, Jill Sherwood, Josh Freeman, Julia Howard, Juliet Elvy, Mary DeAlmeida, Matt Blakiston, Matthew Rogers, Max Bloomfield, Michael Addidle, Michelle Balm, Sally Roberts, Sarah Jefferies, Sharmini Muttaiyah, Susan Morpeth, Susan Taylor, Timothy Blackmore, Vani Sathyendran, Veronica Playle, Virginia Hope, Erasmus Smit, Lauren Jelly, Joep de Ligt |
| EPI_ISL_456164, EPI_ISL_456165, EPI_ISL_456166 | Wellington SCL | Institute of Environmental Science and Research (ESR) | Matt Storey, Xiaoyun Ren, Anja Werno, Antje van der Linden, Arlo Upton, Chris Mansell, David Hammer, Dragana Drinkovic, Erasmus Smit, Gary McAuliffe, Hana Sofia Andersson, James Ussher, Jill Sherwood, Josh Freeman, Julia Howard, Juliet Elvy, Mary DeAlmeida, Matt Blakiston, Matthew Rogers, Max Bloomfield, Michael Addidle, Michelle Balm, Sally Roberts, Sarah Jefferies, Sharmini Muttaiyah, Susan Morpeth, Susan Taylor, Timothy Blackmore, Vani Sathyendran, Veronica Playle, Virginia Hope, Erasmus Smit, Lauren Jelly, Joep de Ligt |
| EPI_ISL_456167 | Southern Community Labs Dunedin | Institute of Environmental Science and Research (ESR) | Matt Storey, Xiaoyun Ren, Anja Werno, Antje van der Linden, Arlo Upton, Chris Mansell, David Hammer, Dragana Drinkovic, Erasmus Smit, Gary McAuliffe, Hana Sofia Andersson, James Ussher, Jill Sherwood, Josh Freeman, Julia Howard, Juliet Elvy, Mary DeAlmeida, Matt Blakiston, Matthew Rogers, Max Bloomfield, Michael Addidle, Michelle Balm, Sally Roberts, Sarah Jefferies, Sharmini Muttaiyah, Susan Morpeth, Susan Taylor, Timothy Blackmore, Vani Sathyendran, Veronica Playle, Virginia Hope, Erasmus Smit, Lauren Jelly, Joep de Ligt |
| EPI_ISL_456168 | Waikato Hospital | Institute of Environmental Science and Research (ESR) | Matt Storey, Xiaoyun Ren, Anja Werno, Antje van der Linden, Arlo Upton, Chris Mansell, David Hammer, Dragana Drinkovic, Erasmus Smit, Gary McAuliffe, Hana Sofia Andersson, James Ussher, Jill Sherwood, Josh Freeman, Julia Howard, Juliet Elvy, Mary DeAlmeida, Matt Blakiston, Matthew Rogers, Max Bloomfield, Michael Addidle, Michelle Balm, Sally Roberts, Sarah Jefferies, Sharmini Muttaiyah, Susan Morpeth, Susan Taylor, Timothy Blackmore, Vani Sathyendran, Veronica Playle, Virginia Hope, Erasmus Smit, Lauren Jelly, Joep de Ligt |
| EPI_ISL_456169, EPI_ISL_456170 | Wellington SCL | Institute of Environmental Science and Research (ESR) | Matt Storey, Xiaoyun Ren, Anja Werno, Antje van der Linden, Arlo Upton, Chris Mansell, David Hammer, Dragana Drinkovic, Erasmus Smit, Gary McAuliffe, Hana Sofia Andersson, James Ussher, Jill Sherwood, Josh Freeman, Julia Howard, Juliet Elvy, Mary DeAlmeida, Matt Blakiston, Matthew Rogers, Max Bloomfield, Michael Addidle, Michelle Balm, Sally Roberts, Sarah Jefferies, Sharmini Muttaiyah, Susan Morpeth, Susan Taylor, Timothy Blackmore, Vani Sathyendran, Veronica Playle, Virginia Hope, Erasmus Smit, Lauren Jelly, Joep de Ligt |
| EPI_ISL_456171 | PathLab Bay of Plenty | Institute of Environmental Science and Research (ESR) | Matt Storey, Xiaoyun Ren, Anja Werno, Antje van der Linden, Arlo Upton, Chris Mansell, David Hammer, Dragana Drinkovic, Erasmus Smit, Gary McAuliffe, Hana Sofia Andersson, James Ussher, Jill Sherwood, Josh Freeman, Julia Howard, Juliet Elvy, Mary DeAlmeida, Matt Blakiston, Matthew Rogers, Max Bloomfield, Michael Addidle, Michelle Balm, Sally Roberts, Sarah Jefferies, Sharmini Muttaiyah, Susan Morpeth, Susan Taylor, Timothy Blackmore, Vani Sathyendran, Veronica Playle, Virginia Hope, Erasmus Smit, Lauren Jelly, Joep de Ligt |
| EPI_ISL_456172, EPI_ISL_456173, EPI_ISL_456174, EPI_ISL_456175, EPI_ISL_456176, EPI_ISL_456177, EPI_ISL_456178, EPI_ISL_456179, EPI_ISL_456180 | LabPLUS | Institute of Environmental Science and Research (ESR) | Matt Storey, Xiaoyun Ren, Anja Werno, Antje van der Linden, Arlo Upton, Chris Mansell, David Hammer, Dragana Drinkovic, Erasmus Smit, Gary McAuliffe, Hana Sofia Andersson, James Ussher, Jill Sherwood, Josh Freeman, Julia Howard, Juliet Elvy, Mary DeAlmeida, Matt Blakiston, Matthew Rogers, Max Bloomfield, Michael Addidle, Michelle Balm, Sally Roberts, Sarah Jefferies, Sharmini Muttaiyah, Susan Morpeth, Susan Taylor, Timothy Blackmore, Vani Sathyendran, Veronica Playle, Virginia Hope, Erasmus Smit, Lauren Jelly, Joep de Ligt |
| EPI_ISL_456181 | Southern Community Labs Dunedin | Institute of Environmental Science and Research (ESR) | Matt Storey, Xiaoyun Ren, Anja Werno, Antje van der Linden, Arlo Upton, Chris Mansell, David Hammer, Dragana Drinkovic, Erasmus Smit, Gary McAuliffe, Hana Sofia Andersson, James Ussher, Jill Sherwood, Josh Freeman, Julia Howard, Juliet Elvy, Mary DeAlmeida, Matt Blakiston, Matthew Rogers, Max Bloomfield, Michael Addidle, Michelle Balm, Sally Roberts, Sarah Jefferies, Sharmini Muttaiyah, Susan Morpeth, Susan Taylor, Timothy Blackmore, Vani Sathyendran, Veronica Playle, Virginia Hope, Erasmus Smit, Lauren Jelly, Joep de Ligt |
| EPI_ISL_456182, EPI_ISL_456183, | Wellington SCL | Institute of Environmental Science and Research (ESR) | Matt Storey, Xiaoyun Ren, Anja Werno, Antje van der Linden, Arlo Upton, Chris Mansell, David Hammer, Dragana Drinkovic, Erasmus Smit, Gary |

| | | | |
|---|---|---|---|
| EPI_ISL_456184, EPI_ISL_456185, EPI_ISL_456186, EPI_ISL_456187, EPI_ISL_456188 | | | McAuliffe, Hana Sofia Andersson, James Ussher, Jill Sherwood, Josh Freeman, Julia Howard, Juliet Elvy, Mary DeAlmeida, Matt Blakiston, Matthew Rogers, Max Bloomfield, Michael Addidle, Michelle Balm, Sally Roberts, Sarah Jefferies, Sharmini Muttaiyah, Susan Morpeth, Susan Taylor, Timothy Blackmore, Vani Sathyendran, Veronica Playle, Virginia Hope, Erasmus Smit, Lauren Jelly, Joep de Ligt |
| EPI_ISL_456189, EPI_ISL_456190, EPI_ISL_456191, EPI_ISL_456192, EPI_ISL_456193 | Waikato Hospital | Institute of Environmental Science and Research (ESR) | Matt Storey, Xiaoyun Ren, Anja Werno, Antje van der Linden, Arlo Upton, Chris Mansell, David Hammer, Dragana Drinkovic, Erasmus Smit, Gary McAuliffe, Hana Sofia Andersson, James Ussher, Jill Sherwood, Josh Freeman, Julia Howard, Juliet Elvy, Mary DeAlmeida, Matt Blakiston, Matthew Rogers, Max Bloomfield, Michael Addidle, Michelle Balm, Sally Roberts, Sarah Jefferies, Sharmini Muttaiyah, Susan Morpeth, Susan Taylor, Timothy Blackmore, Vani Sathyendran, Veronica Playle, Virginia Hope, Erasmus Smit, Lauren Jelly, Joep de Ligt |
| EPI_ISL_456194, EPI_ISL_456195 | PathLab Bay of Plenty | Institute of Environmental Science and Research (ESR) | Matt Storey, Xiaoyun Ren, Anja Werno, Antje van der Linden, Arlo Upton, Chris Mansell, David Hammer, Dragana Drinkovic, Erasmus Smit, Gary McAuliffe, Hana Sofia Andersson, James Ussher, Jill Sherwood, Josh Freeman, Julia Howard, Juliet Elvy, Mary DeAlmeida, Matt Blakiston, Matthew Rogers, Max Bloomfield, Michael Addidle, Michelle Balm, Sally Roberts, Sarah Jefferies, Sharmini Muttaiyah, Susan Morpeth, Susan Taylor, Timothy Blackmore, Vani Sathyendran, Veronica Playle, Virginia Hope, Erasmus Smit, Lauren Jelly, Joep de Ligt |
| EPI_ISL_456196, EPI_ISL_456197, EPI_ISL_456198, EPI_ISL_456199, EPI_ISL_456200, EPI_ISL_456201, EPI_ISL_456202, EPI_ISL_456203, EPI_ISL_456204, EPI_ISL_456205, EPI_ISL_456206, EPI_ISL_456207, EPI_ISL_456208, EPI_ISL_456209, EPI_ISL_456210, EPI_ISL_456211, EPI_ISL_456212 | | | |
| see above | LabPLUS | Institute of Environmental Science and Research (ESR) | Matt Storey, Xiaoyun Ren, Anja Werno, Antje van der Linden, Arlo Upton, Chris Mansell, David Hammer, Dragana Drinkovic, Erasmus Smit, Gary McAuliffe, Hana Sofia Andersson, James Ussher, Jill Sherwood, Josh Freeman, Julia Howard, Juliet Elvy, Mary DeAlmeida, Matt Blakiston, Matthew Rogers, Max Bloomfield, Michael Addidle, Michelle Balm, Sally Roberts, Sarah Jefferies, Sharmini Muttaiyah, Susan Morpeth, Susan Taylor, Timothy Blackmore, Vani Sathyendran, Veronica Playle, Virginia Hope, Erasmus Smit, Lauren Jelly, Joep de Ligt |
| EPI_ISL_456213, EPI_ISL_456214 | Middlemore Hospital | Institute of Environmental Science and Research (ESR) | Matt Storey, Xiaoyun Ren, Anja Werno, Antje van der Linden, Arlo Upton, Chris Mansell, David Hammer, Dragana Drinkovic, Erasmus Smit, Gary McAuliffe, Hana Sofia Andersson, James Ussher, Jill Sherwood, Josh Freeman, Julia Howard, Juliet Elvy, Mary DeAlmeida, Matt Blakiston, Matthew Rogers, Max Bloomfield, Michael Addidle, Michelle Balm, Sally Roberts, Sarah Jefferies, Sharmini Muttaiyah, Susan Morpeth, Susan Taylor, Timothy Blackmore, Vani Sathyendran, Veronica Playle, Virginia Hope, Erasmus Smit, Lauren Jelly, Joep de Ligt |
| EPI_ISL_456215 | PathLab Bay of Plenty | Institute of Environmental Science and Research (ESR) | Matt Storey, Xiaoyun Ren, Anja Werno, Antje van der Linden, Arlo Upton, Chris Mansell, David Hammer, Dragana Drinkovic, Erasmus Smit, Gary McAuliffe, Hana Sofia Andersson, James Ussher, Jill Sherwood, Josh Freeman, Julia Howard, Juliet Elvy, Mary DeAlmeida, Matt Blakiston, Matthew Rogers, Max Bloomfield, Michael Addidle, Michelle Balm, Sally Roberts, Sarah Jefferies, Sharmini Muttaiyah, Susan Morpeth, Susan Taylor, Timothy Blackmore, Vani Sathyendran, Veronica Playle, Virginia Hope, Erasmus Smit, Lauren Jelly, Joep de Ligt |
| EPI_ISL_456216, EPI_ISL_456217, EPI_ISL_456218, EPI_ISL_456219, EPI_ISL_456220 | Southern Community Labs Dunedin | Institute of Environmental Science and Research (ESR) | Matt Storey, Xiaoyun Ren, Anja Werno, Antje van der Linden, Arlo Upton, Chris Mansell, David Hammer, Dragana Drinkovic, Erasmus Smit, Gary McAuliffe, Hana Sofia Andersson, James Ussher, Jill Sherwood, Josh Freeman, Julia Howard, Juliet Elvy, Mary DeAlmeida, Matt Blakiston, Matthew Rogers, Max Bloomfield, Michael Addidle, Michelle Balm, Sally Roberts, Sarah Jefferies, Sharmini Muttaiyah, Susan Morpeth, Susan Taylor, Timothy Blackmore, Vani Sathyendran, Veronica Playle, Virginia Hope, Erasmus Smit, Lauren Jelly, Joep de Ligt |
| EPI_ISL_456221, EPI_ISL_456222, EPI_ISL_456223, EPI_ISL_456224 | PathLab Bay of Plenty | Institute of Environmental Science and Research (ESR) | Matt Storey, Xiaoyun Ren, Anja Werno, Antje van der Linden, Arlo Upton, Chris Mansell, David Hammer, Dragana Drinkovic, Erasmus Smit, Gary McAuliffe, Hana Sofia Andersson, James Ussher, Jill Sherwood, Josh Freeman, Julia Howard, Juliet Elvy, Mary DeAlmeida, Matt Blakiston, Matthew Rogers, Max Bloomfield, Michael Addidle, Michelle Balm, Sally Roberts, Sarah Jefferies, Sharmini Muttaiyah, Susan Morpeth, Susan Taylor, Timothy Blackmore, Vani Sathyendran, Veronica Playle, Virginia Hope, Erasmus Smit, Lauren Jelly, Joep de Ligt |
| EPI_ISL_456225, EPI_ISL_456226, EPI_ISL_456227, EPI_ISL_456228, EPI_ISL_456229, EPI_ISL_456230, EPI_ISL_456231, EPI_ISL_456232, EPI_ISL_456233, EPI_ISL_456234, EPI_ISL_456235, EPI_ISL_456236, EPI_ISL_456237, EPI_ISL_456238, EPI_ISL_456239, EPI_ISL_456240 | | | |
| see above | Southern Community Labs Dunedin | Institute of Environmental Science and Research (ESR) | Matt Storey, Xiaoyun Ren, Anja Werno, Antje van der Linden, Arlo Upton, Chris Mansell, David Hammer, Dragana Drinkovic, Erasmus Smit, Gary McAuliffe, Hana Sofia Andersson, James Ussher, Jill Sherwood, Josh Freeman, Julia Howard, Juliet Elvy, Mary DeAlmeida, Matt Blakiston, Matthew Rogers, Max Bloomfield, Michael Addidle, Michelle Balm, Sally Roberts, Sarah Jefferies, Sharmini Muttaiyah, Susan Morpeth, Susan Taylor, Timothy Blackmore, Vani Sathyendran, Veronica Playle, Virginia Hope, Erasmus Smit, Lauren Jelly, Joep de Ligt |
| EPI_ISL_456241 | Waikato Hospital | Institute of Environmental Science and Research (ESR) | Matt Storey, Xiaoyun Ren, Anja Werno, Antje van der Linden, Arlo Upton, Chris Mansell, David Hammer, Dragana Drinkovic, Erasmus Smit, Gary McAuliffe, Hana Sofia Andersson, James Ussher, Jill Sherwood, Josh Freeman, Julia Howard, Juliet Elvy, Mary DeAlmeida, Matt Blakiston, Matthew Rogers, Max Bloomfield, Michael Addidle, Michelle Balm, Sally Roberts, Sarah Jefferies, Sharmini Muttaiyah, Susan Morpeth, Susan Taylor, Timothy Blackmore, Vani Sathyendran, Veronica Playle, Virginia Hope, Erasmus Smit, Lauren Jelly, Joep de Ligt |
| EPI_ISL_456242, EPI_ISL_456243, EPI_ISL_456244, EPI_ISL_456245, EPI_ISL_456246, EPI_ISL_456247, EPI_ISL_456248, EPI_ISL_456249, EPI_ISL_456250, EPI_ISL_456251, EPI_ISL_456252, EPI_ISL_456253, EPI_ISL_456254 | | | |
| see above | Southern Community Labs Dunedin | Institute of Environmental Science and Research (ESR) | Matt Storey, Xiaoyun Ren, Anja Werno, Antje van der Linden, Arlo Upton, Chris Mansell, David Hammer, Dragana Drinkovic, Erasmus Smit, Gary McAuliffe, Hana Sofia Andersson, James Ussher, Jill Sherwood, Josh Freeman, Julia Howard, Juliet Elvy, Mary DeAlmeida, Matt Blakiston, Matthew Rogers, Max Bloomfield, Michael Addidle, Michelle Balm, Sally Roberts, Sarah Jefferies, Sharmini Muttaiyah, Susan Morpeth, Susan Taylor, Timothy Blackmore, Vani Sathyendran, Veronica Playle, Virginia Hope, Erasmus Smit, Lauren Jelly, Joep de Ligt |
| EPI_ISL_456255, EPI_ISL_456256, EPI_ISL_456257, EPI_ISL_456258 | Middlemore Hospital | Institute of Environmental Science and Research (ESR) | Matt Storey, Xiaoyun Ren, Anja Werno, Antje van der Linden, Arlo Upton, Chris Mansell, David Hammer, Dragana Drinkovic, Erasmus Smit, Gary McAuliffe, Hana Sofia Andersson, James Ussher, Jill Sherwood, Josh Freeman, Julia Howard, Juliet Elvy, Mary DeAlmeida, Matt Blakiston, Matthew Rogers, Max Bloomfield, Michael Addidle, Michelle Balm, Sally Roberts, Sarah Jefferies, Sharmini Muttaiyah, Susan Morpeth, Susan Taylor, Timothy Blackmore, Vani Sathyendran, Veronica Playle, Virginia Hope, Erasmus Smit, Lauren Jelly, Joep de Ligt |
| EPI_ISL_456259, EPI_ISL_456260 | Waikato Hospital | Institute of Environmental Science and Research (ESR) | Matt Storey, Xiaoyun Ren, Anja Werno, Antje van der Linden, Arlo Upton, Chris Mansell, David Hammer, Dragana Drinkovic, Erasmus Smit, Gary McAuliffe, Hana Sofia Andersson, James Ussher, Jill Sherwood, Josh Freeman, Julia Howard, Juliet Elvy, Mary DeAlmeida, Matt Blakiston, Matthew Rogers, Max Bloomfield, Michael Addidle, Michelle Balm, Sally Roberts, Sarah Jefferies, Sharmini Muttaiyah, Susan Morpeth, Susan Taylor, Timothy Blackmore, Vani Sathyendran, Veronica Playle, Virginia Hope, Erasmus Smit, Lauren Jelly, Joep de Ligt |
| EPI_ISL_456261, EPI_ISL_456262, EPI_ISL_456263, EPI_ISL_456264, EPI_ISL_456265, EPI_ISL_456266, EPI_ISL_456267, EPI_ISL_456268, EPI_ISL_456269, EPI_ISL_456270, EPI_ISL_456271, EPI_ISL_456272, EPI_ISL_456273, EPI_ISL_456274, EPI_ISL_456275, EPI_ISL_456276, EPI_ISL_456277, EPI_ISL_456278, EPI_ISL_456279, EPI_ISL_456280, EPI_ISL_456281, EPI_ISL_456282, EPI_ISL_456283, EPI_ISL_456284, EPI_ISL_456285, EPI_ISL_456286, EPI_ISL_456287, EPI_ISL_456288, EPI_ISL_456289, EPI_ISL_456290, EPI_ISL_456291, EPI_ISL_456292, EPI_ISL_456293 | | | |
| see above | Southern Community Labs Dunedin | Institute of Environmental Science and Research (ESR) | Matt Storey, Xiaoyun Ren, Anja Werno, Antje van der Linden, Arlo Upton, Chris Mansell, David Hammer, Dragana Drinkovic, Erasmus Smit, Gary McAuliffe, Hana Sofia Andersson, James Ussher, Jill Sherwood, Josh Freeman, Julia Howard, Juliet Elvy, Mary DeAlmeida, Matt Blakiston, Matthew Rogers, Max Bloomfield, Michael Addidle, Michelle Balm, Sally Roberts, Sarah Jefferies, Sharmini Muttaiyah, Susan Morpeth, Susan Taylor, Timothy Blackmore, Vani Sathyendran, Veronica Playle, Virginia Hope, Erasmus Smit, Lauren Jelly, Joep de Ligt |
| EPI_ISL_456294 | North Shore Hospital | Institute of Environmental Science and Research (ESR) | Matt Storey, Xiaoyun Ren, Anja Werno, Antje van der Linden, Arlo Upton, Chris Mansell, David Hammer, Dragana Drinkovic, Erasmus Smit, Gary McAuliffe, Hana Sofia Andersson, James Ussher, Jill Sherwood, Josh Freeman, Julia Howard, Juliet Elvy, Mary DeAlmeida, Matt Blakiston, Matthew Rogers, Max Bloomfield, Michael Addidle, Michelle Balm, Sally Roberts, Sarah Jefferies, Sharmini Muttaiyah, Susan Morpeth, Susan Taylor, Timothy Blackmore, Vani Sathyendran, Veronica Playle, Virginia Hope, Erasmus Smit, Lauren Jelly, Joep de Ligt |
| EPI_ISL_456295, EPI_ISL_456296, EPI_ISL_456297, EPI_ISL_456298, EPI_ISL_456299 | Southern Community Labs Dunedin | Institute of Environmental Science and Research (ESR) | Matt Storey, Xiaoyun Ren, Anja Werno, Antje van der Linden, Arlo Upton, Chris Mansell, David Hammer, Dragana Drinkovic, Erasmus Smit, Gary McAuliffe, Hana Sofia Andersson, James Ussher, Jill Sherwood, Josh Freeman, Julia Howard, Juliet Elvy, Mary DeAlmeida, Matt Blakiston, Matthew Rogers, Max Bloomfield, Michael Addidle, Michelle Balm, Sally Roberts, Sarah Jefferies, Sharmini Muttaiyah, Susan Morpeth, Susan Taylor, Timothy Blackmore, Vani Sathyendran, Veronica Playle, Virginia Hope, Erasmus Smit, Lauren Jelly, Joep de Ligt |
| EPI_ISL_456300 | LabPLUS | Institute of Environmental Science and Research (ESR) | Matt Storey, Xiaoyun Ren, Anja Werno, Antje van der Linden, Arlo Upton, Chris Mansell, David Hammer, Dragana Drinkovic, Erasmus Smit, Gary McAuliffe, Hana Sofia Andersson, James Ussher, Jill Sherwood, Josh Freeman, Julia Howard, Juliet Elvy, Mary DeAlmeida, Matt Blakiston, Matthew Rogers, Max Bloomfield, Michael Addidle, Michelle Balm, Sally Roberts, Sarah Jefferies, Sharmini Muttaiyah, Susan Morpeth, Susan Taylor, Timothy Blackmore, Vani Sathyendran, Veronica Playle, Virginia Hope, Erasmus Smit, Lauren Jelly, Joep de Ligt |
| EPI_ISL_456301 | PathLab Bay of Plenty | Institute of Environmental Science and Research (ESR) | Matt Storey, Xiaoyun Ren, Anja Werno, Antje van der Linden, Arlo Upton, Chris Mansell, David Hammer, Dragana Drinkovic, Erasmus Smit, Gary McAuliffe, Hana Sofia Andersson, James Ussher, Jill Sherwood, Josh Freeman, Julia Howard, Juliet Elvy, Mary DeAlmeida, Matt Blakiston, Matthew Rogers, Max Bloomfield, Michael Addidle, Michelle Balm, Sally Roberts, Sarah Jefferies, Sharmini Muttaiyah, Susan Morpeth, Susan Taylor, Timothy Blackmore, Vani Sathyendran, Veronica Playle, Virginia Hope, Erasmus Smit, Lauren Jelly, Joep de Ligt |

| | | | |
|---|---|---|---|
| EPI_ISL_456302, EPI_ISL_456303, EPI_ISL_456304, EPI_ISL_456305, EPI_ISL_456306, EPI_ISL_456307, EPI_ISL_456308, EPI_ISL_456309, EPI_ISL_456310, EPI_ISL_456311, EPI_ISL_456312, EPI_ISL_456313, EPI_ISL_456314, EPI_ISL_456315, EPI_ISL_456316, EPI_ISL_456317, EPI_ISL_456318 | | | |
| see above | Southern Community Labs Dunedin | Institute of Environmental Science and Research (ESR) | Matt Storey, Xiaoyun Ren, Anja Werno, Antje van der Linden, Arlo Upton, Chris Mansell, David Hammer, Dragana Drinkovic, Erasmus Smit, Gary McAuliffe, Hana Sofia Andersson, James Ussher, Jill Sherwood, Josh Freeman, Julia Howard, Juliet Elvy, Mary DeAlmeida, Matt Blakiston, Matthew Rogers, Max Bloomfield, Michael Addidle, Michelle Balm, Sally Roberts, Sarah Jefferies, Sharmini Muttaiyah, Susan Morpeth, Susan Taylor, Timothy Blackmore, Vani Sathyendran, Veronica Playle, Virginia Hope, Erasmus Smit, Lauren Jelly, Joep de Ligt |
| EPI_ISL_456319, EPI_ISL_456320 | Wellington SCL | Institute of Environmental Science and Research (ESR) | Matt Storey, Xiaoyun Ren, Anja Werno, Antje van der Linden, Arlo Upton, Chris Mansell, David Hammer, Dragana Drinkovic, Erasmus Smit, Gary McAuliffe, Hana Sofia Andersson, James Ussher, Jill Sherwood, Josh Freeman, Julia Howard, Juliet Elvy, Mary DeAlmeida, Matt Blakiston, Matthew Rogers, Max Bloomfield, Michael Addidle, Michelle Balm, Sally Roberts, Sarah Jefferies, Sharmini Muttaiyah, Susan Morpeth, Susan Taylor, Timothy Blackmore, Vani Sathyendran, Veronica Playle, Virginia Hope, Erasmus Smit, Lauren Jelly, Joep de Ligt |
| EPI_ISL_456321, EPI_ISL_456322 | Canterbury Health Laboratories | Institute of Environmental Science and Research (ESR) | Matt Storey, Xiaoyun Ren, Anja Werno, Antje van der Linden, Arlo Upton, Chris Mansell, David Hammer, Dragana Drinkovic, Erasmus Smit, Gary McAuliffe, Hana Sofia Andersson, James Ussher, Jill Sherwood, Josh Freeman, Julia Howard, Juliet Elvy, Mary DeAlmeida, Matt Blakiston, Matthew Rogers, Max Bloomfield, Michael Addidle, Michelle Balm, Sally Roberts, Sarah Jefferies, Sharmini Muttaiyah, Susan Morpeth, Susan Taylor, Timothy Blackmore, Vani Sathyendran, Veronica Playle, Virginia Hope, Erasmus Smit, Lauren Jelly, Joep de Ligt |
| EPI_ISL_456323 | North Shore Hospital | Institute of Environmental Science and Research (ESR) | Matt Storey, Xiaoyun Ren, Anja Werno, Antje van der Linden, Arlo Upton, Chris Mansell, David Hammer, Dragana Drinkovic, Erasmus Smit, Gary McAuliffe, Hana Sofia Andersson, James Ussher, Jill Sherwood, Josh Freeman, Julia Howard, Juliet Elvy, Mary DeAlmeida, Matt Blakiston, Matthew Rogers, Max Bloomfield, Michael Addidle, Michelle Balm, Sally Roberts, Sarah Jefferies, Sharmini Muttaiyah, Susan Morpeth, Susan Taylor, Timothy Blackmore, Vani Sathyendran, Veronica Playle, Virginia Hope, Erasmus Smit, Lauren Jelly, Joep de Ligt |
| EPI_ISL_456324, EPI_ISL_456325, EPI_ISL_456326, EPI_ISL_456327, EPI_ISL_456328, EPI_ISL_456329, EPI_ISL_456330, EPI_ISL_456331, EPI_ISL_456332, EPI_ISL_456333, EPI_ISL_456334, EPI_ISL_456335, EPI_ISL_456336, EPI_ISL_456337, EPI_ISL_456338, EPI_ISL_456339, EPI_ISL_456340, EPI_ISL_456341, EPI_ISL_456342, EPI_ISL_456343 | | | |
| see above | Wellington SCL | Institute of Environmental Science and Research (ESR) | Matt Storey, Xiaoyun Ren, Anja Werno, Antje van der Linden, Arlo Upton, Chris Mansell, David Hammer, Dragana Drinkovic, Erasmus Smit, Gary McAuliffe, Hana Sofia Andersson, James Ussher, Jill Sherwood, Josh Freeman, Julia Howard, Juliet Elvy, Mary DeAlmeida, Matt Blakiston, Matthew Rogers, Max Bloomfield, Michael Addidle, Michelle Balm, Sally Roberts, Sarah Jefferies, Sharmini Muttaiyah, Susan Morpeth, Susan Taylor, Timothy Blackmore, Vani Sathyendran, Veronica Playle, Virginia Hope, Erasmus Smit, Lauren Jelly, Joep de Ligt |
| EPI_ISL_456344, EPI_ISL_456345, EPI_ISL_456346, EPI_ISL_456347 | Southern Community Labs Dunedin | Institute of Environmental Science and Research (ESR) | Matt Storey, Xiaoyun Ren, Anja Werno, Antje van der Linden, Arlo Upton, Chris Mansell, David Hammer, Dragana Drinkovic, Erasmus Smit, Gary McAuliffe, Hana Sofia Andersson, James Ussher, Jill Sherwood, Josh Freeman, Julia Howard, Juliet Elvy, Mary DeAlmeida, Matt Blakiston, Matthew Rogers, Max Bloomfield, Michael Addidle, Michelle Balm, Sally Roberts, Sarah Jefferies, Sharmini Muttaiyah, Susan Morpeth, Susan Taylor, Timothy Blackmore, Vani Sathyendran, Veronica Playle, Virginia Hope, Erasmus Smit, Lauren Jelly, Joep de Ligt |
| EPI_ISL_456348, EPI_ISL_456349, EPI_ISL_456350, EPI_ISL_456351, EPI_ISL_456352, EPI_ISL_456353, EPI_ISL_456354, EPI_ISL_456355, EPI_ISL_456356, EPI_ISL_456357, EPI_ISL_456358, EPI_ISL_456359, EPI_ISL_456360, EPI_ISL_456361, EPI_ISL_456362, EPI_ISL_456363, EPI_ISL_456364, EPI_ISL_456365, EPI_ISL_456366, EPI_ISL_456367, EPI_ISL_456368, EPI_ISL_456369, EPI_ISL_456370, EPI_ISL_456371, EPI_ISL_456372, EPI_ISL_456373, EPI_ISL_456374, EPI_ISL_456375 | | | |
| see above | Canterbury Health Laboratories | Institute of Environmental Science and Research (ESR) | Matt Storey, Xiaoyun Ren, Anja Werno, Antje van der Linden, Arlo Upton, Chris Mansell, David Hammer, Dragana Drinkovic, Erasmus Smit, Gary McAuliffe, Hana Sofia Andersson, James Ussher, Jill Sherwood, Josh Freeman, Julia Howard, Juliet Elvy, Mary DeAlmeida, Matt Blakiston, Matthew Rogers, Max Bloomfield, Michael Addidle, Michelle Balm, Sally Roberts, Sarah Jefferies, Sharmini Muttaiyah, Susan Morpeth, Susan Taylor, Timothy Blackmore, Vani Sathyendran, Veronica Playle, Virginia Hope, Erasmus Smit, Lauren Jelly, Joep de Ligt |
| EPI_ISL_456376 | Middlemore Hospital | Institute of Environmental Science and Research (ESR) | Matt Storey, Xiaoyun Ren, Anja Werno, Antje van der Linden, Arlo Upton, Chris Mansell, David Hammer, Dragana Drinkovic, Erasmus Smit, Gary McAuliffe, Hana Sofia Andersson, James Ussher, Jill Sherwood, Josh Freeman, Julia Howard, Juliet Elvy, Mary DeAlmeida, Matt Blakiston, Matthew Rogers, Max Bloomfield, Michael Addidle, Michelle Balm, Sally Roberts, Sarah Jefferies, Sharmini Muttaiyah, Susan Morpeth, Susan Taylor, Timothy Blackmore, Vani Sathyendran, Veronica Playle, Virginia Hope, Erasmus Smit, Lauren Jelly, Joep de Ligt |
| EPI_ISL_456377, EPI_ISL_456378 | MedLab Central Ltd | Institute of Environmental Science and Research (ESR) | Matt Storey, Xiaoyun Ren, Anja Werno, Antje van der Linden, Arlo Upton, Chris Mansell, David Hammer, Dragana Drinkovic, Erasmus Smit, Gary McAuliffe, Hana Sofia Andersson, James Ussher, Jill Sherwood, Josh Freeman, Julia Howard, Juliet Elvy, Mary DeAlmeida, Matt Blakiston, Matthew Rogers, Max Bloomfield, Michael Addidle, Michelle Balm, Sally Roberts, Sarah Jefferies, Sharmini Muttaiyah, Susan Morpeth, Susan Taylor, Timothy Blackmore, Vani Sathyendran, Veronica Playle, Virginia Hope, Erasmus Smit, Lauren Jelly, Joep de Ligt |
| EPI_ISL_456379, EPI_ISL_456380, EPI_ISL_456381, EPI_ISL_456382, EPI_ISL_456383, EPI_ISL_456384 | LabPLUS | Institute of Environmental Science and Research (ESR) | Matt Storey, Xiaoyun Ren, Anja Werno, Antje van der Linden, Arlo Upton, Chris Mansell, David Hammer, Dragana Drinkovic, Erasmus Smit, Gary McAuliffe, Hana Sofia Andersson, James Ussher, Jill Sherwood, Josh Freeman, Julia Howard, Juliet Elvy, Mary DeAlmeida, Matt Blakiston, Matthew Rogers, Max Bloomfield, Michael Addidle, Michelle Balm, Sally Roberts, Sarah Jefferies, Sharmini Muttaiyah, Susan Morpeth, Susan Taylor, Timothy Blackmore, Vani Sathyendran, Veronica Playle, Virginia Hope, Erasmus Smit, Lauren Jelly, Joep de Ligt |
| EPI_ISL_456385, EPI_ISL_456386, EPI_ISL_456387 | Middlemore Hospital | Institute of Environmental Science and Research (ESR) | Matt Storey, Xiaoyun Ren, Anja Werno, Antje van der Linden, Arlo Upton, Chris Mansell, David Hammer, Dragana Drinkovic, Erasmus Smit, Gary McAuliffe, Hana Sofia Andersson, James Ussher, Jill Sherwood, Josh Freeman, Julia Howard, Juliet Elvy, Mary DeAlmeida, Matt Blakiston, Matthew Rogers, Max Bloomfield, Michael Addidle, Michelle Balm, Sally Roberts, Sarah Jefferies, Sharmini Muttaiyah, Susan Morpeth, Susan Taylor, Timothy Blackmore, Vani Sathyendran, Veronica Playle, Virginia Hope, Erasmus Smit, Lauren Jelly, Joep de Ligt |
| EPI_ISL_456388, EPI_ISL_456389, EPI_ISL_456390, EPI_ISL_456391, EPI_ISL_456392, EPI_ISL_456393, EPI_ISL_456394 | LabPLUS | Institute of Environmental Science and Research (ESR) | Matt Storey, Xiaoyun Ren, Anja Werno, Antje van der Linden, Arlo Upton, Chris Mansell, David Hammer, Dragana Drinkovic, Erasmus Smit, Gary McAuliffe, Hana Sofia Andersson, James Ussher, Jill Sherwood, Josh Freeman, Julia Howard, Juliet Elvy, Mary DeAlmeida, Matt Blakiston, Matthew Rogers, Max Bloomfield, Michael Addidle, Michelle Balm, Sally Roberts, Sarah Jefferies, Sharmini Muttaiyah, Susan Morpeth, Susan Taylor, Timothy Blackmore, Vani Sathyendran, Veronica Playle, Virginia Hope, Erasmus Smit, Lauren Jelly, Joep de Ligt |
| EPI_ISL_456395, EPI_ISL_456396, EPI_ISL_456397 | North Shore Hospital | Institute of Environmental Science and Research (ESR) | Matt Storey, Xiaoyun Ren, Anja Werno, Antje van der Linden, Arlo Upton, Chris Mansell, David Hammer, Dragana Drinkovic, Erasmus Smit, Gary McAuliffe, Hana Sofia Andersson, James Ussher, Jill Sherwood, Josh Freeman, Julia Howard, Juliet Elvy, Mary DeAlmeida, Matt Blakiston, Matthew Rogers, Max Bloomfield, Michael Addidle, Michelle Balm, Sally Roberts, Sarah Jefferies, Sharmini Muttaiyah, Susan Morpeth, Susan Taylor, Timothy Blackmore, Vani Sathyendran, Veronica Playle, Virginia Hope, Erasmus Smit, Lauren Jelly, Joep de Ligt |
| EPI_ISL_456398, EPI_ISL_456399, EPI_ISL_456400, EPI_ISL_456401, EPI_ISL_456402 | Wellington SCL | Institute of Environmental Science and Research (ESR) | Matt Storey, Xiaoyun Ren, Anja Werno, Antje van der Linden, Arlo Upton, Chris Mansell, David Hammer, Dragana Drinkovic, Erasmus Smit, Gary McAuliffe, Hana Sofia Andersson, James Ussher, Jill Sherwood, Josh Freeman, Julia Howard, Juliet Elvy, Mary DeAlmeida, Matt Blakiston, Matthew Rogers, Max Bloomfield, Michael Addidle, Michelle Balm, Sally Roberts, Sarah Jefferies, Sharmini Muttaiyah, Susan Morpeth, Susan Taylor, Timothy Blackmore, Vani Sathyendran, Veronica Playle, Virginia Hope, Erasmus Smit, Lauren Jelly, Joep de Ligt |
| EPI_ISL_456403 | Middlemore Hospital | Institute of Environmental Science and Research (ESR) | Matt Storey, Xiaoyun Ren, Anja Werno, Antje van der Linden, Arlo Upton, Chris Mansell, David Hammer, Dragana Drinkovic, Erasmus Smit, Gary McAuliffe, Hana Sofia Andersson, James Ussher, Jill Sherwood, Josh Freeman, Julia Howard, Juliet Elvy, Mary DeAlmeida, Matt Blakiston, Matthew Rogers, Max Bloomfield, Michael Addidle, Michelle Balm, Sally Roberts, Sarah Jefferies, Sharmini Muttaiyah, Susan Morpeth, Susan Taylor, Timothy Blackmore, Vani Sathyendran, Veronica Playle, Virginia Hope, Erasmus Smit, Lauren Jelly, Joep de Ligt |
| EPI_ISL_456890, EPI_ISL_456891, EPI_ISL_456892, EPI_ISL_456893, EPI_ISL_456894, EPI_ISL_456895, EPI_ISL_456896, EPI_ISL_456897, EPI_ISL_456898, EPI_ISL_456899, EPI_ISL_456900, EPI_ISL_456901, EPI_ISL_456902, EPI_ISL_456903, EPI_ISL_456904, EPI_ISL_456905, EPI_ISL_456906, EPI_ISL_456907, EPI_ISL_456908, EPI_ISL_456909, EPI_ISL_456910, EPI_ISL_456911, EPI_ISL_456912, EPI_ISL_456913, EPI_ISL_456914, EPI_ISL_456915, EPI_ISL_456916, EPI_ISL_456917, EPI_ISL_456918, EPI_ISL_456919, EPI_ISL_456920, EPI_ISL_456921, EPI_ISL_456922, EPI_ISL_456923, EPI_ISL_456924, EPI_ISL_456925, EPI_ISL_456926, EPI_ISL_456927, EPI_ISL_456928, EPI_ISL_456929, EPI_ISL_456930, EPI_ISL_456931, EPI_ISL_456932, EPI_ISL_456933, EPI_ISL_456934, EPI_ISL_456935, EPI_ISL_456936, EPI_ISL_456937, EPI_ISL_456938, EPI_ISL_456939, EPI_ISL_456940, EPI_ISL_456941, EPI_ISL_456942, EPI_ISL_456943, EPI_ISL_456944, EPI_ISL_456945, EPI_ISL_456946, EPI_ISL_456947, EPI_ISL_456948, EPI_ISL_456949, EPI_ISL_456950, EPI_ISL_456951, EPI_ISL_456952, EPI_ISL_456953, EPI_ISL_456954, EPI_ISL_456955, EPI_ISL_456956, EPI_ISL_456957, EPI_ISL_456958, EPI_ISL_456959, EPI_ISL_456960, EPI_ISL_456961, EPI_ISL_456962, EPI_ISL_456963, EPI_ISL_456964, EPI_ISL_456965, EPI_ISL_456966, EPI_ISL_456967, EPI_ISL_456968, EPI_ISL_456969, EPI_ISL_456970, EPI_ISL_456971, EPI_ISL_456972, EPI_ISL_456973, EPI_ISL_456974, EPI_ISL_456975, EPI_ISL_456976, EPI_ISL_456977, EPI_ISL_456978, EPI_ISL_456979, EPI_ISL_456980, EPI_ISL_456981, EPI_ISL_456982, EPI_ISL_456983, EPI_ISL_456984, EPI_ISL_456985, EPI_ISL_456986, EPI_ISL_456987, EPI_ISL_456988, EPI_ISL_456989, EPI_ISL_456990, EPI_ISL_456991, EPI_ISL_456992, EPI_ISL_456993, EPI_ISL_456994, EPI_ISL_456995, EPI_ISL_456996, EPI_ISL_456997, EPI_ISL_456998, EPI_ISL_456999, EPI_ISL_457000, EPI_ISL_457001, EPI_ISL_457002, EPI_ISL_457003, EPI_ISL_457004, EPI_ISL_457005, EPI_ISL_457006, EPI_ISL_457007, EPI_ISL_457008, EPI_ISL_457009, EPI_ISL_457010, EPI_ISL_457011, EPI_ISL_457012, EPI_ISL_457013, EPI_ISL_457014, EPI_ISL_457015, EPI_ISL_457016, EPI_ISL_457017, EPI_ISL_457018, EPI_ISL_457019, EPI_ISL_457020, EPI_ISL_457021, EPI_ISL_457022, EPI_ISL_457023 | | | |
| see above | Virology Department, Royal Infirmary of Edinburgh, NHS Lothian / School of Biological Sciences, University of Edinburgh / Institute of Genetics and Molecular Medicine, University of Edinburgh | COVID-19 Genomics UK (COG-UK) Consortium | McHugh M, Dewar R, Rooke S, Gallagher M, Balcaza C, O'Toole Á, Scher E, Hill V, McCrone JT, Colquhoun R, Yu X, Jackson B, Rambaut A, Williams TC, Templeton K |

| | | | |
|---|---|---|---|
| EPI_ISL_457824 | Bezmialem Vakif University, Dept Microbiology, Medical School, Fatih, Istanbul, Turkey | Bezmialem Vakif University, Medical School & Beykoz Institute of Life Sciences & Biotechnology | Mehmet Z. Doymaz, Merve Kalkan, Nesibe Cetin, Elif Karaaslan, Bilge Sumbul, Filiz Guney |
| EPI_ISL_457940, EPI_ISL_457941, EPI_ISL_457942, EPI_ISL_457943, EPI_ISL_457944, EPI_ISL_457945, EPI_ISL_457946, EPI_ISL_457947, EPI_ISL_457948, EPI_ISL_457949, EPI_ISL_457950, EPI_ISL_457951, EPI_ISL_457952, EPI_ISL_457953, EPI_ISL_457954, EPI_ISL_457955, EPI_ISL_457956, EPI_ISL_457957, EPI_ISL_457958, EPI_ISL_457959, EPI_ISL_457960, EPI_ISL_457961, EPI_ISL_457962, EPI_ISL_457963, EPI_ISL_457964, EPI_ISL_457965, EPI_ISL_457966, EPI_ISL_457967, EPI_ISL_457968, EPI_ISL_457969, EPI_ISL_457970, EPI_ISL_457971, EPI_ISL_457972, EPI_ISL_457973 | | | |
| see above | Laboratorio de Biología Molecular Asociación Española Primera en Salud | Departments of Pathology and Medicine, New York University School of Medicine | Maria Victoria Elizondo, Maria Noel Zubillaga, Gonzalo Manrique, Paul Zappile, Gael Westby, Matthew T Maurano, Christian Marier, Adriana Heguy |
| EPI_ISL_457999 | Centre For Biotechnology Research and Development | Centre For Biotechnology Research and Development | Matoke-Muhia,D., Symeker,S.L., Muuo,S.N., Ochwoto,M., Zablon,J.O., Kimotho,J., Waruhiu,C.N. and Michuki,G.N. |
| EPI_ISL_458017, EPI_ISL_458018, EPI_ISL_458019, EPI_ISL_458020, EPI_ISL_458021 | NYU Langone Health | Departments of Pathology and Medicine, New York University School of Medicine | Maria Aguero-Rosenfeld, Brendan Belovarac, Margaret Black, Ludovic Boytard, John Cadley, Paolo Cotzia, John Chen, Dacia Dimartino, Xiaojun Feng, Tatyana Gindin, Emily Guzman, Adriana Heguy, Megan Hogan, Emily Huang, George Jour, Alireza Khodadadi-Jamayran, Lawrence H. Lin, Raven Luther, Andrew Lytle, Christian Marier, Matthew T. Maurano, Mark J. Mulligan, Peter Meyn, Raquel Ordonez Ciriza, Iman Osman, Jared Pinnell, Vanessa Raabe, Sitharam Ramaswami, Amy Rapkiewicz, Andre M. Ribeiro-dos-Santos, Marie Samanovic-Golden, Antonio Serrano, Guomiao Shen, Matija Snuderl, Theodore Vougiouklakis, Nick Vulpescu, Gael Westby, Paul Zappile, Yutong Zhang |
| EPI_ISL_458049 | CSIR-Centre for Cellular and Molecular Biology | CSIR-Centre for Cellular and Molecular Biology | Namami Gaur, Sakshi Shambhavi, Lamuk Zaveri, Shagufta Khan, Tulasi Nagabandi, Purushotham Vodnala, Payel Mukherjee, Sofia Banu, Priya Singh, Dhiviya Vedagiri, Divya Gupta, Vishal Sah, Santosh Kumar Kuncha, Krishnan Harinivas Harshan, Archana Bharadwaj Siva, Karthik Bharadwaj Tallapaka, Zeba Rizvi, Zuberwasim Sayyad, Kakade Aishwarya Arun, Amrutha H C, Ananga Ghosh, Rakesh K Mishra, Divya Tej Sowpati |
| EPI_ISL_458055 | CSIR-Centre for Cellular and Molecular Biology | CSIR-Centre for Cellular and Molecular Biology | Namami Gaur, Sakshi Shambhavi, Lamuk Zaveri, Shagufta Khan, Tulasi Nagabandi, Purushotham Vodnala, Payel Mukherjee, Sofia Banu, Priya Singh, Dhiviya Vedagiri, Divya Gupta, Vishal Sah, Santosh Kumar Kuncha, Krishnan Harinivas Harshan, Archana Bharadwaj Siva, Karthik Bharadwaj Tallapaka, Nikhil Hajirnis, Pratheusa Maccha, M Soujanya Reddy,G. Aditya Kumar, Koushick Sivakumar, Rakesh K Mishra, Divya Tej Sowpati |
| EPI_ISL_458061 | CSIR-Centre for Cellular and Molecular Biology | CSIR-Centre for Cellular and Molecular Biology | Namami Gaur, Sakshi Shambhavi, Lamuk Zaveri, Shagufta Khan, Tulasi Nagabandi, Purushotham Vodnala, Payel Mukherjee, Sofia Banu, Priya Singh, Dhiviya Vedagiri, Divya Gupta, Vishal Sah, Santosh Kumar Kuncha, Krishnan Harinivas Harshan, Archana Bharadwaj Siva, Karthik Bharadwaj Tallapaka,Kezia J Ann, Radhika Khandelwal, Roshan Maku Venkata, Shemin Mansuri, Sonu Uday, Rakesh K Mishra, Divya Tej Sowpati |
| EPI_ISL_458110 | Gujarat Biotechnology Research Centre | Gujarat Biotechnology Research Centre | Monika Gandhi, Pinal Trivedi, Maharshi Pandya, Amit Kanani, Nidhi Patel, Nitin Savaliya, Raghawendra Kumar, Dinesh Kumar, Zuber Saiyed, Komal Patel, Labdhi Pandya, Snehal Bagatharia, Ramesh Pandit, Tejas Shah, Ankit Hinsu, Pritesh Sabara, Apurvasinh Puvar, Janvi Raval, Zarna Patel, Priti Pandita, Bhavesh Modi, Gaurishankar Shrimali, R D Dixit, A M Kadri, Umang Mishra, Chaitanya Joshi, Madhvi Joshi, , , , |
| EPI_ISL_458133 | National Institute of Biotechnology | Bioinformatics Division, National Institute of Biotechnology | Mohammad Uzzal Hossain, Md. Moniruzzaman, Md. Salim Khan, Md. Nazrul Islam, Md. Hadisur Rahman, Arittra Bhattacharjee, Md. Ruhul Amin, Asif Rashid, Chaman Ara Keya, Keshob Chandra Das, Md. Salimullah |
| EPI_ISL_459965, EPI_ISL_459966, EPI_ISL_459967, EPI_ISL_459968, EPI_ISL_459969, EPI_ISL_459970, EPI_ISL_459971, EPI_ISL_459972, EPI_ISL_459973, EPI_ISL_459974, EPI_ISL_459975, EPI_ISL_459976, EPI_ISL_459977, EPI_ISL_459978, EPI_ISL_459979, EPI_ISL_459980, EPI_ISL_459981, EPI_ISL_459982, EPI_ISL_459983, EPI_ISL_459984 | | | |
| see above | Institut Pasteur du Maroc | Institut Pasteur du Maroc | Marion Barbet, Sylvie Behillil, Méline Bizard, Angela Brisebarre, Camille Capel, Etienne Simon-Lorière, Vincent Enouf, Maud Vanpeene, Sylvie van der Werf, Latifa Anga, Abdellah Faouzi, Anass Abbad, Mjid Eloualid, Jalal Nourlil, Anderrahmane Maaroufi |
| EPI_ISL_460045, EPI_ISL_460046, EPI_ISL_460047, EPI_ISL_460048, EPI_ISL_460049, EPI_ISL_460050, EPI_ISL_460051, EPI_ISL_460052, EPI_ISL_460053, EPI_ISL_460054, EPI_ISL_460055, EPI_ISL_460056, EPI_ISL_460057, EPI_ISL_460058, EPI_ISL_460059, EPI_ISL_460060, EPI_ISL_460061, EPI_ISL_460062, EPI_ISL_460063, EPI_ISL_460064, EPI_ISL_460065, EPI_ISL_460066, EPI_ISL_460067, EPI_ISL_460068, EPI_ISL_460069, EPI_ISL_460070, EPI_ISL_460071, EPI_ISL_460072, EPI_ISL_460073, EPI_ISL_460074, EPI_ISL_460075, EPI_ISL_460076, EPI_ISL_460077, EPI_ISL_460078 | | | |
| see above | Minnesota Department of Health, Public Health Laboratory | Minnesota Department of Health, Public Health Laboratory | Matt Plumb, Jacob Garfin, and Xiong Wang |
| EPI_ISL_460082 | Molecular Virology Unit, Fondazione IRCCS Policlinico San Matteo , Pavia | Laboratory of Virology, INMI Lazzaro Spallanzani IRCCS | Martina Rueca, Cesare E.M. Gruber, Antonio Piralla, Antonino Di Caro, Barbara Bartolini, Maria R. Capobianchi, Fausto Baldanti |
| EPI_ISL_460083 | Molecular Virology Unit, Fondazione IRCCS Policlinico San Matteo , Pavia | Laboratory of Virology, INMI Lazzaro Spallanzani IRCCS | Martina Rueca, Antonino Di Caro, Cesare E.M. Gruber, Barbara Bartolini, Fausto Baldanti, Antonio Piralla, Maria R. Capobianchi |
| EPI_ISL_460086 | Molecular Virology Unit, Fondazione IRCCS Policlinico San Matteo , Pavia | Laboratory of Virology, INMI Lazzaro Spallanzani IRCCS | Maria R. Capobianchi, Fausto Baldanti, Antonio Piralla, Antonino Di Caro, Barbara Bartolini, Cesare E.M. Gruber, Martina Rueca |
| EPI_ISL_460088 | Molecular Virology Unit, Fondazione IRCCS Policlinico San Matteo , Pavia | Laboratory of Virology, INMI Lazzaro Spallanzani IRCCS | Martina Rueca, Barbara Bartolini, Fausto Baldanti, Maria R. Capobianchi, Cesare E.M. Gruber, Antonino Di Caro, Antonio Piralla |
| EPI_ISL_460093 | Molecular Virology Unit, Fondazione IRCCS Policlinico San Matteo , Pavia | Laboratory of Virology, INMI Lazzaro Spallanzani IRCCS | Maria R. Capobianchi, Antonio Piralla, Antonino Di Caro, Fausto Baldanti, Martina Rueca, Cesare E.M. Gruber, Barbara Bartolini |
| EPI_ISL_460603 | NYU Langone Health | Departments of Pathology and Medicine, New York University School of Medicine | Maria Aguero-Rosenfeld, Brendan Belovarac, Margaret Black, Ludovic Boytard, John Cadley, Paolo Cotzia, John Chen, Dacia Dimartino, Xiaojun Feng, Tatyana Gindin, Emily Guzman, Adriana Heguy, Megan Hogan, Emily Huang, George Jour, Alireza Khodadadi-Jamayran, Lawrence H. Lin, Raven Luther, Andrew Lytle, Christian Marier, Matthew T. Maurano, Mark J. Mulligan, Peter Meyn, Raquel Ordonez Ciriza, Iman Osman, Jared Pinnell, Vanessa Raabe, Sitharam Ramaswami, Amy Rapkiewicz, Andre M. Ribeiro-dos-Santos, Marie Samanovic-Golden, Antonio Serrano, Guomiao Shen, Matija Snuderl, Theodore Vougiouklakis, Nick Vulpescu, Gael Westby, Paul Zappile, Yutong Zhang |
| EPI_ISL_461706, EPI_ISL_461707, EPI_ISL_461708, EPI_ISL_461709, EPI_ISL_461710, EPI_ISL_461711, EPI_ISL_461712, EPI_ISL_461713, EPI_ISL_461714, EPI_ISL_461715, EPI_ISL_461716, EPI_ISL_461717, EPI_ISL_461718, EPI_ISL_461719, EPI_ISL_461720, EPI_ISL_461721, EPI_ISL_461722, EPI_ISL_461723, EPI_ISL_461724, EPI_ISL_461725, EPI_ISL_461726, EPI_ISL_461727, EPI_ISL_461728, EPI_ISL_461729, EPI_ISL_461730, EPI_ISL_461731, EPI_ISL_461732, EPI_ISL_461733, EPI_ISL_461734, EPI_ISL_461735, EPI_ISL_461736, EPI_ISL_461737, EPI_ISL_461738, EPI_ISL_461739, EPI_ISL_461740, EPI_ISL_461741, EPI_ISL_461742, EPI_ISL_461743, EPI_ISL_461744, EPI_ISL_461745, EPI_ISL_461746, EPI_ISL_461747, EPI_ISL_461748, EPI_ISL_461749, EPI_ISL_461750, EPI_ISL_461751, EPI_ISL_461752, EPI_ISL_461753, EPI_ISL_461754, EPI_ISL_461755, EPI_ISL_461756, EPI_ISL_461757, EPI_ISL_461758, EPI_ISL_461759, EPI_ISL_461760, EPI_ISL_461761, EPI_ISL_461762 | | | |
| see above | Virology Department, Royal Infirmary of Edinburgh, NHS Lothian / School of Biological Sciences, University of Edinburgh / Institute of Genetics and Molecular Medicine, University of Edinburgh | COVID-19 Genomics UK (COG-UK) Consortium | McHugh M, Dewar R, Rooke S, Gallagher M, Balcaza C, O'Toole Á, Scher E, Hill V, McCrone JT, Colquhoun R, Yu X, Jackson B, Rambaut A, Williams TC, Templeton K |
| EPI_ISL_462845, EPI_ISL_462846, EPI_ISL_462847, EPI_ISL_462848, EPI_ISL_462849, EPI_ISL_462850, EPI_ISL_462851, EPI_ISL_462852, EPI_ISL_462853, EPI_ISL_462854, EPI_ISL_462855, EPI_ISL_462856, EPI_ISL_462857, EPI_ISL_462858, EPI_ISL_462859, EPI_ISL_462860, EPI_ISL_462861, EPI_ISL_462862, EPI_ISL_462863, EPI_ISL_462864, EPI_ISL_462865, EPI_ISL_462866, EPI_ISL_462867, EPI_ISL_462868, EPI_ISL_462869, EPI_ISL_462870, EPI_ISL_462871, EPI_ISL_462872, EPI_ISL_462873, EPI_ISL_462874, EPI_ISL_462875, EPI_ISL_462876, EPI_ISL_462877, EPI_ISL_462878, EPI_ISL_462879, EPI_ISL_462880, EPI_ISL_462881, EPI_ISL_462882, EPI_ISL_462883, EPI_ISL_462884, EPI_ISL_462885, EPI_ISL_462886, EPI_ISL_462887, EPI_ISL_462888, EPI_ISL_462889, EPI_ISL_462890, EPI_ISL_462891, EPI_ISL_462892, EPI_ISL_462893, EPI_ISL_462894, EPI_ISL_462895, EPI_ISL_462896, EPI_ISL_462897, EPI_ISL_462898, EPI_ISL_462899, EPI_ISL_462900, EPI_ISL_462901, EPI_ISL_462902, EPI_ISL_462903, EPI_ISL_462904, EPI_ISL_462905, EPI_ISL_462906, EPI_ISL_462907, EPI_ISL_462908, EPI_ISL_462909, EPI_ISL_462910, EPI_ISL_462911 | | | |
| see above | Minnesota Department of Health, Public Health Laboratory | Minnesota Department of Health, Public Health Laboratory | Matt Plumb, Jacob Garfin, and Xiong Wang |
| EPI_ISL_463903, EPI_ISL_463904, EPI_ISL_463905, EPI_ISL_463906, EPI_ISL_463907, EPI_ISL_463908, EPI_ISL_463909, EPI_ISL_463910, EPI_ISL_463911, EPI_ISL_463912, EPI_ISL_463913, EPI_ISL_463914, EPI_ISL_463915, EPI_ISL_463916, EPI_ISL_463917, EPI_ISL_463918, EPI_ISL_463919, EPI_ISL_463920, EPI_ISL_463921, EPI_ISL_463922, EPI_ISL_463923, EPI_ISL_463924, EPI_ISL_463925, EPI_ISL_463926, EPI_ISL_463927, EPI_ISL_463928, EPI_ISL_463929, EPI_ISL_463930, EPI_ISL_463931, EPI_ISL_463932, EPI_ISL_463933, EPI_ISL_463934, EPI_ISL_463935, EPI_ISL_463936, EPI_ISL_463937, EPI_ISL_463938, EPI_ISL_463939, EPI_ISL_463940, EPI_ISL_463941, EPI_ISL_463942, EPI_ISL_463943, EPI_ISL_463944, EPI_ISL_463945, EPI_ISL_463946, EPI_ISL_463947, EPI_ISL_463948, EPI_ISL_463949, EPI_ISL_463950, EPI_ISL_463951, EPI_ISL_463952, EPI_ISL_463953, EPI_ISL_463954, EPI_ISL_463955, EPI_ISL_463956, EPI_ISL_463957, EPI_ISL_463958, EPI_ISL_463959, EPI_ISL_463960, EPI_ISL_463961, EPI_ISL_463962, EPI_ISL_463963, EPI_ISL_463964, EPI_ISL_463965, EPI_ISL_463966, EPI_ISL_463967, EPI_ISL_463968, EPI_ISL_463969 | | | |
| see above | Laboratoire de microbiologie, Hopital de Verdun | Smith Laboratory, Centre de Recherche CHU Sainte-Justine | Martin Smith, Marieke Rozendaal, Ivan Pavlov |
| EPI_ISL_464161, EPI_ISL_464162 | National Institute of Laboratory Medicine and Referral Center | Genomic Research Lab, BCSIR | Md. Ahasan Habib, Abu Sayeed Mohammad Mahmud, Mohammad Samir Uzzaman, Eshrar Osman, Shahina Akter, Tanjina Akhter Banu, Md. Murshed Hasan Sarker, Barna Goswami, Iffat Jahan, Md. Saddam Hossain, Tasnim Nafisa, Md. Maruf Ahmed Molla, Mahmuda Yeasmin, Asish Kumar Ghosh, Arifa Akram, A. K. M. Shamsuzzaman, Sheikh Md. Selim Al Din, Utpal Chandra Ray, Salek Ahmed Sajib, Md. Salim Khan |
| EPI_ISL_466874, EPI_ISL_466875, EPI_ISL_466876, EPI_ISL_466877, EPI_ISL_466878, EPI_ISL_466879, EPI_ISL_466880, EPI_ISL_466881, EPI_ISL_466882, EPI_ISL_466883, EPI_ISL_466884, EPI_ISL_466885, EPI_ISL_466886, EPI_ISL_466887, EPI_ISL_466888, EPI_ISL_466889, EPI_ISL_466890, EPI_ISL_466891, EPI_ISL_466892, EPI_ISL_466893, EPI_ISL_466894, EPI_ISL_466895, EPI_ISL_466896, EPI_ISL_466897, EPI_ISL_466898, EPI_ISL_466899, EPI_ISL_466900, EPI_ISL_466901, EPI_ISL_466902, EPI_ISL_466903, EPI_ISL_466904, EPI_ISL_466905, EPI_ISL_466906, EPI_ISL_466907, EPI_ISL_466908, EPI_ISL_466909, EPI_ISL_466910, EPI_ISL_466911, EPI_ISL_466912, EPI_ISL_466913, EPI_ISL_466914, EPI_ISL_466915, EPI_ISL_466916, EPI_ISL_466917, EPI_ISL_466918, EPI_ISL_466919, EPI_ISL_466920, EPI_ISL_466921, EPI_ISL_466922, EPI_ISL_466923, EPI_ISL_466924, EPI_ISL_466925 | | | |
| see above | Max von Pettenkofer Institute, Virology, National Reference | Laboratory for Functional Genome Analysis, Dept. Genomics, | Max Muenchhoff, Stefan Krebs, Alexander Graf, Oliver Keppler, Helmut Blum |

| | | | |
|---|---|---|---|
| | Center for Retroviruses, LMU München | Gene Center of the LMU Munich | |
| EPI_ISL_467039 | Government Medical College, Vadodara | Gujarat Biotechnology Research Centre | Meenakshi Shah, Neena Doshi, Varsha Godbole, Tejas Shah, Ankit Hinsu, Pritesh Sabara, Apurvasinh Puvar, Janvi Raval, Zarna Patel, Monika Gandhi, Pinal Trivedi, Maharshi Pandya, Nidhi Patel, Nitin Savaliya, Raghawendra Kumar, Dinesh Kumar, Zuber Saiyed, Komal Patel, Labdhi Pandya, Snehal Bagatharia, Armi Chaudhari, R D Dixit, A M Kadri, Harsh Bakshi, Chaitanya Joshi, Madhvi Joshi, |
| EPI_ISL_467041 | B.J. Medical College and Civil hospital | Gujarat Biotechnology Research Centre | Monika Gandhi, Pinal Trivedi, Maharshi Pandya, Nidhi Patel, Nitin Savaliya, Raghawendra Kumar, Dinesh Kumar, Zuber Saiyed, Komal Patel, Labdhi Pandya, Snehal Bagatharia, Pranay Shah, Kamlesh J Upadhyay, Nirav Mungalpara, Tejas Shah, Ankit Hinsu, Pritesh Sabara, Apurvasinh Puvar, Janvi Raval, Zarna Patel, Priyanka P Vatsa, R D Dixit, A M Kadri, Harsh Bakshi, Chaitanya Joshi, Madhvi Joshi, |
| EPI_ISL_467059, EPI_ISL_467060, EPI_ISL_467061, EPI_ISL_467062, EPI_ISL_467063 | Hospital Universitario Virgen de las Nieves de Granada-SAS | SeqCOVID-SPAIN consortium/IBV(CSIC) | Mercedes Pérez Ruiz, Sara Sanbonmatsu Gámez, Irene Pedrosa Corral, José M. Navarro-Marí and SeqCOVID-SPAIN consortium |
| EPI_ISL_467377, EPI_ISL_467378, EPI_ISL_467379, EPI_ISL_467380, EPI_ISL_467381, EPI_ISL_467382, EPI_ISL_467383, EPI_ISL_467384, EPI_ISL_467385, EPI_ISL_467386, EPI_ISL_467387, EPI_ISL_467388, EPI_ISL_467389, EPI_ISL_467390, EPI_ISL_467391, EPI_ISL_467392, EPI_ISL_467393, EPI_ISL_467394, EPI_ISL_467395, EPI_ISL_467396, EPI_ISL_467397, EPI_ISL_467398, EPI_ISL_467399, EPI_ISL_467400, EPI_ISL_467401, EPI_ISL_467402, EPI_ISL_467403, EPI_ISL_467405, EPI_ISL_467406, EPI_ISL_467407, EPI_ISL_467408, EPI_ISL_467409, EPI_ISL_467410, EPI_ISL_467411, EPI_ISL_467412, EPI_ISL_467413, EPI_ISL_467414, EPI_ISL_467415, EPI_ISL_467416, EPI_ISL_467417, EPI_ISL_467418, EPI_ISL_467419, EPI_ISL_467420, EPI_ISL_467421, EPI_ISL_467422 | | | |
| see above | NYU Langone Health | Departments of Pathology and Medicine, New York University School of Medicine | Maria Aguero-Rosenfeld, Brendan Belovarac, Margaret Black, Ludovic Boytard, John Cadley, Paolo Cotzia, John Chen, Dacia Dimartino, Xiaojun Feng, Tatyana Gindin, Emily Guzman, Adriana Heguy, Megan Hogan, Emily Huang, George Jour, Alireza Khodadadi-Jamayran, Lawrence H. Lin, Raven Luther, Andrew Lytle, Christian Marier, Matthew T. Maurano, Mark J. Mulligan, Peter Meyn, Raquel Ordonez Ciriza, Iman Osman, Jared Pinnell, Vanessa Raabe, Sitharam Ramaswami, Amy Rapkiewicz, Andre M. Ribeiro-dos-Santos, Marie Samanovic-Golden, Antonio Serrano, Guomiao Shen, Matija Snuderl, Theodore Vougiouklakis, Nick Vulpescu, Gael Westby, Paul Zappile, Yutong Zhang |
| EPI_ISL_468159, EPI_ISL_468160 | unknown | Department of Virology, Public Health Laboratories Division, National Institute of Health | Massab Umair, Aamer Ikram, Muhammad Salman, Adnan Khurshid, Nazish Badar, Shannon Whitmer, John Klena |
| EPI_ISL_468161 | Department of Virology, Public Health Laboratories Division, National Institute of Health | Department of Virology, Public Health Laboratories Division, National Institute of Health | Massab Umair, Aamer Ikram, Muhammad Salman, Adnan Khurshid, Nazish Badar, Shannon Whitmer, John Klena |
| EPI_ISL_468162 | unknown | Department of Virology, Public Health Laboratories Division, National Institute of Health | Massab Umair, Aamer Ikram, Muhammad Salman, Adnan Khurshid, Nazish Badar, Shannon Whitmer, John Klena |
| EPI_ISL_468163 | Department of Virology, Public Health Laboratories Division, National Institute of Health | Department of Virology, Public Health Laboratories Division, National Institute of Health | Massab Umair, Aamer Ikram, Muhammad Salman, Adnan Khurshid, Nazish Badar, Shannon Whitmer, John Klena |
| EPI_ISL_468761, EPI_ISL_468762, EPI_ISL_468763, EPI_ISL_468764 | Centro de Investigación Biomédica de La Rioja - Hospital San Pedro Logroño | SeqCOVID-SPAIN consortium/IBV(CSIC) | María de Toro, José Manuel Azcona Gutiérrez, María Pilar Bea Escudero, Miriam Blasco Alberdi and SeqCOVID-SPAIN consortium |
| EPI_ISL_469017 | LNR National Reference Laboratory, Mohammed VI University of Health Sciences | Medical Biotechnology Laboratory, Rabat Medical and Pharmacy School, Mohammed The Vth University in Rabat | Meriem LAAMARTI, Souad KARTTI, Rokaia LAAMRTI , M.W. CHEMAO-ELFIHRI, Loubna ALLAM, Mouna OUADGHIRI, Imane SMYEJ, Jalila RAHOUI, Houda BENRAHMA, Jalil El Atar, Idrissa Diawara, Rachid EL JAOUDI, Laila SBABOU, Chakib NEJJARI, Saaid AMZAZI, Rachid MENTAG, Lahcen BELYAMANI and Azeddine IBRAHIMI |
| EPI_ISL_469030 | Government Medical College, Vadodara | Gujarat Biotechnology Research Centre | Monika Gandhi, Pinal Trivedi, Maharshi Pandya, Nidhi Patel, Nitin Savaliya, Raghawendra Kumar, Dinesh Kumar, Zuber Saiyed, Komal Patel, Labdhi Pandya, Snehal Bagatharia, Meenakshi Shah, Neena Doshi, Varsha Godbole, Tejas Shah, Ankit Hinsu, Pritesh Sabara, Apurvasinh Puvar, Janvi Raval, Zarna Patel, Armi Chaudhari, R D Dixit, A M Kadri, Harsh Bakshi, Chaitanya Joshi, Madhvi Joshi |
| EPI_ISL_469041 | GMERS Medical College & Hospital | Gujarat Biotechnology Research Centre | Meenakshi Shah, Neena Doshi, Varsha Godbole, Tejas Shah, Ankit Hinsu, Pritesh Sabara, Apurvasinh Puvar, Janvi Raval, Zarna Patel, Monika Gandhi, Pinal Trivedi, Maharshi Pandya, Nidhi Patel, Nitin Savaliya, Raghawendra Kumar, Dinesh Kumar, Zuber Saiyed, Komal Patel, Labdhi Pandya, Snehal Bagatharia, Neelam Nathani, R D Dixit, A M Kadri, Harsh Bakshi, Chaitanya Joshi, Madhvi Joshi |
| EPI_ISL_469049, EPI_ISL_469051, EPI_ISL_469052, EPI_ISL_469053, EPI_ISL_469054 | LNR National Reference Laboratory, Mohammed VI University of Health Sciences | Medical Biotechnology Laboratory, Rabat Medical and Pharmacy School, Mohammed The Vth University in Rabat | Meriem LAAMARTI, Souad KARTTI, Rokaia LAAMRTI , M.W. CHEMAO-ELFIHRI, Loubna ALLAM, Mouna OUADGHIRI, Imane SMYEJ, Jalila RAHOUI, Houda BENRAHMA, Jalil El Atar, Idrissa Diawara, Rachid EL JAOUDI, Laila SBABOU, Chakib NEJJARI, Saaid AMZAZI, Rachid MENTAG, Lahcen BELYAMANI and Azeddine IBRAHIMI |
| EPI_ISL_469298 | National Institute of Laboratory Medicine and Referral Center | Genomic Research Lab, BCSIR | Md. Murshed Hasan Sarkar, Abu Sayeed Mohammad Mahmud, Mohammad Samir Uzzaman, Eshrar Osman, Md. Ahasan Habib, Shahina Akter, Tanjina Akhter Banu, Barna Goswami, Iffat Jahan, Md. Saddam Hossain, Tasnim Nafisa, Md. Maruf Ahmed Molla, Mahmuda Yeasmin, Asish Kumar Ghosh, Bayzid Bin Monir, A. K. M. Shamsuzzaman, Sheikh Md. Selim Al Din, Utpal Chandra Ray, Salek Ahmed Sajib, Md. Salim Khan |
| EPI_ISL_470747, EPI_ISL_470748, EPI_ISL_470749, EPI_ISL_470750, EPI_ISL_470751, EPI_ISL_470752, EPI_ISL_470753, EPI_ISL_470754, EPI_ISL_470755, EPI_ISL_470756, EPI_ISL_470757, EPI_ISL_470758, EPI_ISL_470759, EPI_ISL_470760, EPI_ISL_470761, EPI_ISL_470762, EPI_ISL_470763, EPI_ISL_470764, EPI_ISL_470765, EPI_ISL_470766, EPI_ISL_470767, EPI_ISL_470768, EPI_ISL_470769, EPI_ISL_470770, EPI_ISL_470771, EPI_ISL_470772, EPI_ISL_470773, EPI_ISL_470774, EPI_ISL_470775, EPI_ISL_470776, EPI_ISL_470777, EPI_ISL_470778, EPI_ISL_470779, EPI_ISL_470780, EPI_ISL_470781, EPI_ISL_470782, EPI_ISL_470783, EPI_ISL_470784, EPI_ISL_470785, EPI_ISL_470786, EPI_ISL_470787, EPI_ISL_470788, EPI_ISL_470789 | | | |
| see above | Minnesota Department of Health, Public Health Laboratory | Minnesota Department of Health, Public Health Laboratory | Matt Plumb, Jacob Garfin, and Xiong Wang |
| EPI_ISL_470790, EPI_ISL_470791, EPI_ISL_470792, EPI_ISL_470793, EPI_ISL_470794, EPI_ISL_470795, EPI_ISL_470796, EPI_ISL_470797, EPI_ISL_470798, EPI_ISL_470799, EPI_ISL_470800 | | | |
| see above | M Health Fairview | Minnesota Department of Health, Public Health Laboratory | Matt Plumb, Jacob Garfin, and Xiong Wang |
| EPI_ISL_471528 | The National Institute of Public Health | State Veterinary Institute Prague and The National Institute of Public Health | Nagy,A;Jirincova,H;Novakova,L;Trnka,D;Vecerova,J |
| EPI_ISL_471529 | Department for Virology, Molecular Biology and Genome Research, R. G. Lugar Center for Public Health Research, National Center for Disease Control and Public Health (NCDC) of Georgia. | Department for Virology, Molecular Biology and Genome Research, R. G. Lugar Center for Public Health Research, National Center for Disease Control and Public Health (NCDC) of Georgia. | Meri Pantsulaia, Gvantsa Brachveli, Giorgi Tomashvili, Gvantsa Chanturia, Ann Machablishvili, Nato Kotaria, Marine Murtskhvaladze, Lela Sabadze, Mari Gavashelidze, Ana Papkiauri, Gvantsa Brachveli, Tata Imnadze, Tamar Jashiashvili, Tea Tevdoradze, Ketevan Sidamonidze, Ekaterine Khmaladze, Ekaterine Zhghenti, Roena Sukhiashvili, Mariam Zakalashvili, Lela Urushadze, Magda Dgebuadze, Davit Tsaguria, Ekaterine Zangaladze, Nino Berishvili, Adam Kotorashvili, Maia Alkhazashvili, Irma Burjanadze, Anna Kasradze, Khatuna Zakhashvili, Paata Imnadze, Amiran Gamkrelidze. |
| EPI_ISL_471530, EPI_ISL_471540, EPI_ISL_471544, EPI_ISL_471547, EPI_ISL_471550, EPI_ISL_471553, EPI_ISL_471555 | The National Institute of Public Health | State Veterinary Institute Prague and The National Institute of Public Health | Nagy,A;Jirincova,H;Novakova,L;Trnka,D;Vecerova,J |
| EPI_ISL_471591 | CSIR-Centre for Cellular and Molecular Biology | CSIR-Centre for Cellular and Molecular Biology | Namami Gaur, Sakshi Shambhavi, Lamuk Zaveri, Shagufta Khan, Tulasi Nagabandi, Purushotham Vodnala, Payel Mukherjee, Sofia Banu, Priya Singh, Dhiviya Vedagiri, Divya Gupta, Vishal Sah, Santosh Kumar Kuncha, Krishnan Harinivas Harshan, Archana Bharadwaj Siva, Karthik Bharadwaj Tallapaka, Zeba Rizvi, Zuberwasim Sayyad, Kakade Aishwarya Arun, Amrutha H C, Ananga Ghosh, Rakesh K Mishra, Divya Tej Sowpati |
| EPI_ISL_471592 | CSIR-Centre for Cellular and Molecular Biology | CSIR-Centre for Cellular and Molecular Biology | Namami Gaur, Sakshi Shambhavi, Lamuk Zaveri, Shagufta Khan, Tulasi Nagabandi, Purushotham Vodnala, Payel Mukherjee, Sofia Banu, Priya Singh, Dhiviya Vedagiri, Divya Gupta, Vishal Sah, Santosh Kumar Kuncha, Krishnan Harinivas Harshan, Archana Bharadwaj Siva, Karthik Bharadwaj Tallapaka, Nikhil Hajirnis, Pratheusa Maccha, M Soujanya Reddy,G. Aditya Kumar, Koushick Sivakumar, Rakesh K Mishra, Divya Tej Sowpati |
| EPI_ISL_471593 | CSIR-Centre for Cellular and Molecular Biology | CSIR-Centre for Cellular and Molecular Biology | Namami Gaur, Sakshi Shambhavi, Lamuk Zaveri, Shagufta Khan, Tulasi Nagabandi, Purushotham Vodnala, Payel Mukherjee, Sofia Banu, Priya Singh, Dhiviya Vedagiri, Divya Gupta, Vishal Sah, Santosh Kumar Kuncha, Krishnan Harinivas Harshan, Archana Bharadwaj Siva, Karthik Bharadwaj Tallapaka, Zeba Rizvi, Zuberwasim Sayyad, Kakade Aishwarya Arun, Amrutha H C, Ananga Ghosh, Rakesh K Mishra, Divya Tej Sowpati |
| EPI_ISL_471594 | CSIR-Centre for Cellular and Molecular Biology | CSIR-Centre for Cellular and Molecular Biology | Namami Gaur, Sakshi Shambhavi, Lamuk Zaveri, Shagufta Khan, Tulasi Nagabandi, Purushotham Vodnala, Payel Mukherjee, Sofia Banu, Priya Singh, Dhiviya Vedagiri, Divya Gupta, Vishal Sah, Santosh Kumar Kuncha, Krishnan Harinivas Harshan, Archana Bharadwaj Siva, Karthik Bharadwaj Tallapaka,Kezia J Ann, Radhika Khandelwal, Roshan Maku Venkata, Shemin Mansuri, Sonu Uday, Rakesh K Mishra, Divya Tej Sowpati |
| EPI_ISL_471619 | CSIR-Centre for Cellular and Molecular Biology | CSIR-Centre for Cellular and Molecular Biology | Namami Gaur, Sakshi Shambhavi, Lamuk Zaveri, Shagufta Khan, Tulasi Nagabandi, Purushotham Vodnala, Payel Mukherjee, Sofia Banu, Priya Singh, Dhiviya Vedagiri, Divya Gupta, Vishal Sah, Santosh Kumar Kuncha, Krishnan Harinivas Harshan, Archana Bharadwaj Siva, Karthik Bharadwaj Tallapaka, Zeba Rizvi, Zuberwasim Sayyad, Kakade Aishwarya Arun, Amrutha H C, Ananga Ghosh, Rakesh K Mishra, Divya Tej Sowpati |

| | | | |
|---|---|---|---|
| EPI_ISL_471620 | CSIR-Centre for Cellular and Molecular Biology | CSIR-Centre for Cellular and Molecular Biology | Namami Gaur, Sakshi Shambhavi, Lamuk Zaveri, Shagufta Khan, Tulasi Nagabandi, Purushotham Vodnala, Payel Mukherjee, Sofia Banu, Priya Singh, Dhiviya Vedagiri, Divya Gupta, Vishal Sah, Santosh Kumar Kuncha, Krishnan Harinivas Harshan, Archana Bharadwaj Siva, Karthik Bharadwaj Tallapaka, Nikhil Hajirnis, Pratheusa Maccha, M Soujanya Reddy,G. Aditya Kumar, Koushick Sivakumar, Rakesh K Mishra, Divya Tej Sowpati |
| EPI_ISL_471621 | CSIR-Centre for Cellular and Molecular Biology | CSIR-Centre for Cellular and Molecular Biology | Namami Gaur, Sakshi Shambhavi, Lamuk Zaveri, Shagufta Khan, Tulasi Nagabandi, Purushotham Vodnala, Payel Mukherjee, Sofia Banu, Priya Singh, Dhiviya Vedagiri, Divya Gupta, Vishal Sah, Santosh Kumar Kuncha, Krishnan Harinivas Harshan, Archana Bharadwaj Siva, Karthik Bharadwaj Tallapaka, Zeba Rizvi, Zuberwasim Sayyad, Kakade Aishwarya Arun, Amrutha H C, Ananga Ghosh, Rakesh K Mishra, Divya Tej Sowpati |
| EPI_ISL_473783, EPI_ISL_473784, EPI_ISL_473785, EPI_ISL_473786, EPI_ISL_473787, EPI_ISL_473788, EPI_ISL_473789, EPI_ISL_473790, EPI_ISL_473791, EPI_ISL_473792, EPI_ISL_473793, EPI_ISL_473794, EPI_ISL_473795, EPI_ISL_473796, EPI_ISL_473797, EPI_ISL_473798, EPI_ISL_473799, EPI_ISL_473800, EPI_ISL_473801, EPI_ISL_473802, EPI_ISL_473803, EPI_ISL_473804, EPI_ISL_473805, EPI_ISL_473806, EPI_ISL_473807, EPI_ISL_473808, EPI_ISL_473809, EPI_ISL_473810, EPI_ISL_473811, EPI_ISL_473812, EPI_ISL_473813, EPI_ISL_473814, EPI_ISL_473815, EPI_ISL_473816, EPI_ISL_473817, EPI_ISL_473818, EPI_ISL_473819, EPI_ISL_473820, EPI_ISL_473821, EPI_ISL_473822, EPI_ISL_473823, EPI_ISL_473824, EPI_ISL_473825, EPI_ISL_473826, EPI_ISL_473827, EPI_ISL_473828, EPI_ISL_473829, EPI_ISL_473830, EPI_ISL_473831, EPI_ISL_473832, EPI_ISL_473833, EPI_ISL_473834, EPI_ISL_473835, EPI_ISL_473836, EPI_ISL_473837, EPI_ISL_473838, EPI_ISL_473839, EPI_ISL_473840, EPI_ISL_473841, EPI_ISL_473842, EPI_ISL_473843, EPI_ISL_473844, EPI_ISL_473845, EPI_ISL_473846, EPI_ISL_473847, EPI_ISL_473848, EPI_ISL_473849, EPI_ISL_473850, EPI_ISL_473851, EPI_ISL_473852, EPI_ISL_473853, EPI_ISL_473854, EPI_ISL_473855, EPI_ISL_473856, EPI_ISL_473857, EPI_ISL_473858, EPI_ISL_473859, EPI_ISL_473860, EPI_ISL_473861, EPI_ISL_473862, EPI_ISL_473863, EPI_ISL_473864, EPI_ISL_473865, EPI_ISL_473866, EPI_ISL_473867, EPI_ISL_473868, EPI_ISL_473869, EPI_ISL_473870, EPI_ISL_473871, EPI_ISL_473872, EPI_ISL_473873, EPI_ISL_473874, EPI_ISL_473875, EPI_ISL_473876, EPI_ISL_473877, EPI_ISL_473878, EPI_ISL_473879, EPI_ISL_473880, EPI_ISL_473881, EPI_ISL_473882, EPI_ISL_473883, EPI_ISL_473884, EPI_ISL_473885, EPI_ISL_473886, EPI_ISL_473887, EPI_ISL_473888, EPI_ISL_473889, EPI_ISL_473890, EPI_ISL_473891, EPI_ISL_473892, EPI_ISL_473893, EPI_ISL_473894, EPI_ISL_473895, EPI_ISL_473896, EPI_ISL_473897, EPI_ISL_473898, EPI_ISL_473899, EPI_ISL_473900, EPI_ISL_473901, EPI_ISL_473902, EPI_ISL_473903, EPI_ISL_473904, EPI_ISL_473905, EPI_ISL_473906, EPI_ISL_473907, EPI_ISL_473908, EPI_ISL_473909, EPI_ISL_473910, EPI_ISL_473911, EPI_ISL_473912, EPI_ISL_473913, EPI_ISL_473914, EPI_ISL_473915, EPI_ISL_473916, EPI_ISL_473917, EPI_ISL_473918, EPI_ISL_473919, EPI_ISL_473920, EPI_ISL_473921, EPI_ISL_473922, EPI_ISL_473923, EPI_ISL_473924, EPI_ISL_473925, EPI_ISL_473926, EPI_ISL_473927, EPI_ISL_473928, EPI_ISL_473929, EPI_ISL_473930, EPI_ISL_473931, EPI_ISL_473932, EPI_ISL_473933, EPI_ISL_473934, EPI_ISL_473935, EPI_ISL_473936, EPI_ISL_473937, EPI_ISL_473938, EPI_ISL_473939, EPI_ISL_473940, EPI_ISL_473941, EPI_ISL_473942, EPI_ISL_473943, EPI_ISL_473944, EPI_ISL_473945, EPI_ISL_473946, EPI_ISL_473947, EPI_ISL_473948, EPI_ISL_473949, EPI_ISL_473950, EPI_ISL_473951, EPI_ISL_473952, EPI_ISL_473953, EPI_ISL_473954, EPI_ISL_473955 | | | |
| see above | Virology Department, Royal Infirmary of Edinburgh, NHS Lothian / School of Biological Sciences, University of Edinburgh / Institute of Genetics and Molecular Medicine, University of Edinburgh | COVID-19 Genomics UK (COG-UK) Consortium | McHugh M, Dewar R, Rooke S, Gallagher M, Balcaza C, O'Toole Á, Scher E, Hill V, McCrone JT, Colquhoun R, Yu X, Jackson B, Rambaut A, Williams TC, Templeton K |
| EPI_ISL_474797, EPI_ISL_474832, EPI_ISL_474833, EPI_ISL_474834, EPI_ISL_474835, EPI_ISL_474836, EPI_ISL_474841, EPI_ISL_474842, EPI_ISL_474843, EPI_ISL_474844, EPI_ISL_474845, EPI_ISL_474846, EPI_ISL_474850, EPI_ISL_474851, EPI_ISL_474852, EPI_ISL_474854, EPI_ISL_474855, EPI_ISL_474856, EPI_ISL_474857, EPI_ISL_474858, EPI_ISL_474859, EPI_ISL_474860, EPI_ISL_474861, EPI_ISL_474862, EPI_ISL_474863, EPI_ISL_474864, EPI_ISL_474865, EPI_ISL_474866, EPI_ISL_474867, EPI_ISL_474868, EPI_ISL_474869, EPI_ISL_474870, EPI_ISL_474871, EPI_ISL_474872, EPI_ISL_474873, EPI_ISL_474874, EPI_ISL_474875, EPI_ISL_474876, EPI_ISL_474877, EPI_ISL_474878, EPI_ISL_474879, EPI_ISL_474880, EPI_ISL_474881, EPI_ISL_474882, EPI_ISL_474883, EPI_ISL_474884, EPI_ISL_474885, EPI_ISL_474886, EPI_ISL_474887, EPI_ISL_474888, EPI_ISL_474889, EPI_ISL_474890, EPI_ISL_474891, EPI_ISL_474892, EPI_ISL_474893, EPI_ISL_474894, EPI_ISL_474895, EPI_ISL_474896, EPI_ISL_474897, EPI_ISL_474898, EPI_ISL_474899, EPI_ISL_474900, EPI_ISL_474907, EPI_ISL_474908, EPI_ISL_474909, EPI_ISL_474920, EPI_ISL_474922, EPI_ISL_474923, EPI_ISL_474924, EPI_ISL_474925, EPI_ISL_474926, EPI_ISL_474927, EPI_ISL_474928, EPI_ISL_474929, EPI_ISL_474930, EPI_ISL_474931, EPI_ISL_474932, EPI_ISL_474934, EPI_ISL_474935, EPI_ISL_474936, EPI_ISL_474937, EPI_ISL_474938, EPI_ISL_474939, EPI_ISL_474942, EPI_ISL_474943, EPI_ISL_474944, EPI_ISL_474948, EPI_ISL_474949, EPI_ISL_474950, EPI_ISL_474957 | | | |
| see above | Hospital Universitario Virgen de las Nieves de Granada-SAS | SeqCOVID-SPAIN consortium/IBV(CSIC) | Mercedes Pérez Ruiz, Sara Sanbonmatsu Gámez, Irene Pedrosa Corral, José M. Navarro-Marí and SeqCOVID-SPAIN consortium |
| EPI_ISL_475032 | Department of MicroBiology, Government Medical College, Surat | Gujarat Biotechnology Research Centre | Monika Gandhi, Pinal Trivedi, Maharshi Pandya, Nidhi Patel, Nitin Savaliya, Raghawendra Kumar, Dinesh Kumar, Zuber Saiyed, Komal Patel, Labdhi Pandya, Snehal Bagatharia, Naresh Chauhan, Summaiya Mullan, Amit gamit, Pritesh Sabara, Apurvasinh Puvar, Janvi Raval, Zarna Patel, Priyanka P Vatsa, R D Dixit, A M Kadri, Harsh Bakshi, Chaitanya Joshi, Madhvi Joshi |
| EPI_ISL_475050 | GMERS Medical College & Hospital | Gujarat Biotechnology Research Centre | Monika Gandhi, Pinal Trivedi, Maharshi Pandya, Nidhi Patel, Nitin Savaliya, Raghawendra Kumar, Dinesh Kumar, Zuber Saiyed, Komal Patel, Labdhi Pandya, Snehal Bagatharia, Meenakshi Shah, Neena Doshi, Varsha Godbole, Pritesh Sabara, Apurvasinh Puvar, Janvi Raval, Zarna Patel, Fenil Patel, R D Dixit, A M Kadri, Harsh Bakshi, Chaitanya Joshi, Madhvi Joshi |
| EPI_ISL_475083, EPI_ISL_475084 | National Institute of Laboratory Medicine and Referral Center | Genomic Research Lab, BCSIR | Md. Murshed Hasan Sarkar, Abu Sayeed Mohammad Mahmud, Mohammad Samir Uzzaman, Eshrar Osman, Md. Ahasan Habib, Shahina Akter, Tanjina Akhter Banu, Barna Goswami, Iffat Jahan, Md. Saddam Hossain, Tasnim Nafisa, Md. Maruf Ahmed Molla, Mahmuda Yeasmin, Asish Kumar Ghosh, Bayzid Bin Monir, A. K. M. Shamsuzzaman, Sheikh Md. Selim Al Din, Utpal Chandra Ray, Salek Ahmed Sajib, Md. Salim Khan |
| EPI_ISL_475169 | National Institute of Laboratory Medicine and Referral Center | Genomic Research Lab, BCSIR | Md. Saddam Hossain, Abu Sayeed Mohammad Mahmud, Mohammad Samir Uzzaman, Eshrar Osman, Md. Ahasan Habib, Shahina Akter, Tanjina Akhter Banu, Md. Murshed Hasan Sarkar, Barna Goswami, Iffat Jahan, Tasnim Nafisa, Md. Maruf Ahmed Molla, Mahmuda Yeasmin, Asish Kumar Ghosh, Bayzid Bin Monir, A. K. M. Shamsuzzaman, Sheikh Md. Selim Al Din, Utpal Chandra Ray, Salek Ahmed Sajib, Md. Salim Khan |
| EPI_ISL_475573 | Genome Center | Genome Center | Md. Shazid Hasan, Hassan M. Al-Emran, Ovinu Kibria Islam, A. S. M. Rubayet- Ul- Alam, Selina Akter, Shireen Nigar, Md. Tanvir Islam, Pravas Chandra Roy, Shovon Lal Sarkar, Md. Nazmul Hasan, Tanay Chakrovarty, Md. Ali Ahasan Setu, Sourav Dutta, Ruhul Amin, Md. Iqbal Kabir Jahid, Md. Anwar Hossain |
| EPI_ISL_475755 | National Institute of Laboratory Medicine and Referral Center | Genomic Research Lab, BCSIR | Md. Murshed Hasan Sarkar, Abu Sayeed Mohammad Mahmud, Mohammad Samir Uzzaman, Eshrar Osman, Md. Ahasan Habib, Shahina Akter, Tanjina Akhter Banu, Barna Goswami, Iffat Jahan, Md. Saddam Hossain, Tasnim Nafisa, Md. Maruf Ahmed Molla, Mahmuda Yeasmin, Asish Kumar Ghosh, Arifa Akram, A. K. M. Shamsuzzaman, Sheikh Md. Selim Al Din, Utpal Chandra Ray, Salek Ahmed Sajib, Md. Salim Khan |
| EPI_ISL_475759 | National Institute of Laboratory Medicine and Referral Center | Genomic Research Lab, BCSIR | Md. Saddam Hossain, Abu Sayeed Mohammad Mahmud, Mohammad Samir Uzzaman, Eshrar Osman, Md. Ahasan Habib, Shahina Akter, Tanjina Akhter Banu, Md. Murshed Hasan Sarkar, Barna Goswami, Iffat Jahan, Tasnim Nafisa, Md. Maruf Ahmed Molla, Mahmuda Yeasmin, Asish Kumar Ghosh, Arifa Akram, A. K. M. Shamsuzzaman, Sheikh Md. Selim Al Din, Utpal Chandra Ray, Salek Ahmed Sajib, Md. Salim Khan |
| EPI_ISL_476067 | The National Institute of Public Health | State Veterinary Institute Prague and The National Institute of Public Health | Nagy,A;Jirincova,H;Novakova,L;Trnka,D;Vecerova,J |
| EPI_ISL_476498, EPI_ISL_476499, EPI_ISL_476500, EPI_ISL_476501, EPI_ISL_476502, EPI_ISL_476503, EPI_ISL_476504, EPI_ISL_476505, EPI_ISL_476506, EPI_ISL_476507, EPI_ISL_476508, EPI_ISL_476509, EPI_ISL_476510, EPI_ISL_476511, EPI_ISL_476512, EPI_ISL_476513 | | | |
| see above | Laboratoire de microbiologie, Hopital de Verdun | Smith Laboratory, Centre de Recherche CHU Sainte-Justine | Martin Smith, Marieke Rozendaal, Ivan Pavlov |
| EPI_ISL_476706, EPI_ISL_476707, EPI_ISL_476708, EPI_ISL_476709, EPI_ISL_476710, EPI_ISL_476711, EPI_ISL_476712, EPI_ISL_476713, EPI_ISL_476714, EPI_ISL_476715, EPI_ISL_476716, EPI_ISL_476717, EPI_ISL_476718, EPI_ISL_476719, EPI_ISL_476720, EPI_ISL_476721, EPI_ISL_476722, EPI_ISL_476723, EPI_ISL_476724, EPI_ISL_476725, EPI_ISL_476726, EPI_ISL_476727, EPI_ISL_476728, EPI_ISL_476729, EPI_ISL_476730, EPI_ISL_476731, EPI_ISL_476732, EPI_ISL_476733, EPI_ISL_476734, EPI_ISL_476735, EPI_ISL_476736, EPI_ISL_476737, EPI_ISL_476738, EPI_ISL_476739, EPI_ISL_476740, EPI_ISL_476741, EPI_ISL_476742, EPI_ISL_476743, EPI_ISL_476744, EPI_ISL_476745, EPI_ISL_476746, EPI_ISL_476747, EPI_ISL_476748, EPI_ISL_476749, EPI_ISL_476750, EPI_ISL_476751, EPI_ISL_476752, EPI_ISL_476753, EPI_ISL_476754, EPI_ISL_476755, EPI_ISL_476756, EPI_ISL_476757, EPI_ISL_476758, EPI_ISL_476759, EPI_ISL_476760, EPI_ISL_476761, EPI_ISL_476762, EPI_ISL_476763, EPI_ISL_476764, EPI_ISL_476765, EPI_ISL_476766 | | | |
| see above | Minnesota Department of Health, Public Health Laboratory | Minnesota Department of Health, Public Health Laboratory | Matt Plumb, Jacob Garfin, and Xiong Wang |
| EPI_ISL_476835 | National Influenza Centre for Northern Greece | National Influenza Centre for Northern Greece | Maria Christoforidi |
| EPI_ISL_476836 | National Influenza Centre for Nothern Greece | National Influenza Centre for Nothern Greece | Maria Christoforidi |
| EPI_ISL_476837, EPI_ISL_476838, EPI_ISL_476839, EPI_ISL_476841, EPI_ISL_476843, EPI_ISL_476845, EPI_ISL_476847, EPI_ISL_476851 | National Influenza Centre for Northern Greece | National Influenza Centre for Nothern Greece | Maria Christoforidi |
| EPI_ISL_476858 | GMERS Medical College & Hospital, Gotri, Vadodara | Gujarat Biotechnology Research Centre | Monika Gandhi, Pinal Trivedi, Maharshi Pandya, Nidhi Patel, Nitin Savaliya, Raghawendra Kumar, Dinesh Kumar, Zuber Saiyed, Komal Patel, Labdhi Pandya, Afzal Ansari, Nikha Trivedi, Meenakshi Shah, Neena Doshi, Varsha Godbole, Apurvasinh Puvar, Janvi Raval, Zarna Patel, R D Dixit, A M Kadri, Harsh Bakshi, Chaitanya Joshi, Madhvi Joshi |
| EPI_ISL_476897 | University of South Carolina Functional Genomics Core | University of South Carolina Functional Genomics Core | Michael Shtutman |
| EPI_ISL_477183 | Department of MicroBiology, Government Medical College, Surat | Gujarat Biotechnology Research Centre | Monika Gandhi, Pinal Trivedi, Maharshi Pandya, Nidhi Patel, Nitin Savaliya, Raghawendra Kumar, Dinesh Kumar, Zuber Saiyed, Komal Patel, Labdhi Pandya, Afzal Ansari, Nikha Trivedi, Naresh Chauhan, Summaiya Mullan, Amit gamit, Apurvasinh Puvar, Janvi Raval, Zarna Patel, R D Dixit, A M Kadri, Harsh Bakshi, Chaitanya Joshi, Madhvi Joshi |
| EPI_ISL_477272, EPI_ISL_477273, EPI_ISL_477274, EPI_ISL_477275, EPI_ISL_477276 | Mayo Clinic & Mayo Clinic Laboratories | Minnesota Department of Health, Public Health Laboratory | Matt Plumb, Jacob Garfin, Kelly Pung, and Xiong Wang |
| EPI_ISL_477277, EPI_ISL_477278, EPI_ISL_477279, EPI_ISL_477280, EPI_ISL_477281, EPI_ISL_477282, EPI_ISL_477283, EPI_ISL_477284, EPI_ISL_477285, EPI_ISL_477286, EPI_ISL_477287, EPI_ISL_477288, EPI_ISL_477289, EPI_ISL_477290 | | | |
| see above | M Health Fairview | Minnesota Department of Health, Public Health Laboratory | Matt Plumb, Jacob Garfin, Kelly Pung, and Xiong Wang |

| | | | |
|---|---|---|---|
| EPI_ISL_477291, EPI_ISL_477292, EPI_ISL_477293, EPI_ISL_477294, EPI_ISL_477295, EPI_ISL_477296, EPI_ISL_477297, EPI_ISL_477298, EPI_ISL_477299, EPI_ISL_477300, EPI_ISL_477301, EPI_ISL_477302, EPI_ISL_477303, EPI_ISL_477304, EPI_ISL_477305, EPI_ISL_477306, EPI_ISL_477307, EPI_ISL_477308, EPI_ISL_477309, EPI_ISL_477310, EPI_ISL_477311, EPI_ISL_477312 | | | |
| see above | Mayo Clinic & Mayo Clinic Laboratories | Minnesota Department of Health, Public Health Laboratory | Matt Plumb, Jacob Garfin, Kelly Pung, and Xiong Wang |
| EPI_ISL_478163, EPI_ISL_478164, EPI_ISL_478165, EPI_ISL_478166, EPI_ISL_478167, EPI_ISL_478168, EPI_ISL_478169, EPI_ISL_478170, EPI_ISL_478171, EPI_ISL_478172, EPI_ISL_478173, EPI_ISL_478174, EPI_ISL_478175, EPI_ISL_478176, EPI_ISL_478177, EPI_ISL_478178, EPI_ISL_478179, EPI_ISL_478180, EPI_ISL_478181, EPI_ISL_478182, EPI_ISL_478183, EPI_ISL_478184, EPI_ISL_478185, EPI_ISL_478186, EPI_ISL_478187, EPI_ISL_478188, EPI_ISL_478189, EPI_ISL_478190, EPI_ISL_478191, EPI_ISL_478192, EPI_ISL_478193, EPI_ISL_478194, EPI_ISL_478195, EPI_ISL_478196, EPI_ISL_478197, EPI_ISL_478198, EPI_ISL_478199, EPI_ISL_478200, EPI_ISL_478201, EPI_ISL_478202, EPI_ISL_478203, EPI_ISL_478204, EPI_ISL_478205, EPI_ISL_478206, EPI_ISL_478207, EPI_ISL_478208, EPI_ISL_478209, EPI_ISL_478210, EPI_ISL_478211, EPI_ISL_478212, EPI_ISL_478213, EPI_ISL_478214, EPI_ISL_478215, EPI_ISL_478216, EPI_ISL_478217, EPI_ISL_478218, EPI_ISL_478219, EPI_ISL_478220, EPI_ISL_478221, EPI_ISL_478222, EPI_ISL_478223, EPI_ISL_478224, EPI_ISL_478225, EPI_ISL_478226, EPI_ISL_478227, EPI_ISL_478228, EPI_ISL_478229, EPI_ISL_478230, EPI_ISL_478231, EPI_ISL_478232, EPI_ISL_478233, EPI_ISL_478234, EPI_ISL_478235, EPI_ISL_478236, EPI_ISL_478237, EPI_ISL_478238, EPI_ISL_478239, EPI_ISL_478240, EPI_ISL_478241, EPI_ISL_478242, EPI_ISL_478243, EPI_ISL_478244, EPI_ISL_478245, EPI_ISL_478246, EPI_ISL_478247, EPI_ISL_478248, EPI_ISL_478249, EPI_ISL_478250, EPI_ISL_478251, EPI_ISL_478252, EPI_ISL_478253, EPI_ISL_478254, EPI_ISL_478255, EPI_ISL_478256, EPI_ISL_478257, EPI_ISL_478258, EPI_ISL_478259, EPI_ISL_478260, EPI_ISL_478261, EPI_ISL_478262, EPI_ISL_478263, EPI_ISL_478264, EPI_ISL_478265, EPI_ISL_478266, EPI_ISL_478267, EPI_ISL_478268, EPI_ISL_478269, EPI_ISL_478270, EPI_ISL_478271, EPI_ISL_478272, EPI_ISL_478273 | | | |
| see above | Virology Department, Royal Infirmary of Edinburgh, NHS Lothian / School of Biological Sciences, University of Edinburgh / Institute of Genetics and Molecular Medicine, University of Edinburgh | COVID-19 Genomics UK (COG-UK) Consortium | McHugh M, Dewar R, Rooke S, Gallagher M, Balcaza C, O'Toole Á, Scher E, Hill V, McCrone JT, Colquhoun R, Yu X, Jackson B, Rambaut A, Williams TC, Templeton K |
| EPI_ISL_480414, EPI_ISL_480415, EPI_ISL_480416, EPI_ISL_480417, EPI_ISL_480418 | National Institute of Laboratory Medicine and Referral Center | Bangladesh Council of Scientific and Industrial Research | Md. Saddam Hossain, Abu Sayeed Mohammad Mahmud, Mohammad Samir Uzzaman, Eshrar Osman, Md. Ahasan Habib, Shahina Akter, Tanjina Akhter Banu, Md. Murshed Hasan Sarkar, Barna Goswami, Iffat Jahan, Tasnim Nafisa, Md. Maruf Ahmed Molla, Mahmuda Yeasmin, Asish Kumar Ghosh, Shahjahan Siddike, A. K. M. Shamsuzzaman, Sheikh Md. Selim Al Din, Utpal Chandra Ray, Salek Ahmed Sajib, Md. Salim Khan |
| EPI_ISL_480419, EPI_ISL_480420, EPI_ISL_480421, EPI_ISL_480424, EPI_ISL_480425 | National Institute of Laboratory Medicine and Referral Center | Bangladesh Council of Scientific and Industrial Research | Md. Murshed Hasan Sarkar, Abu Sayeed Mohammad Mahmud, Mohammad Samir Uzzaman, Eshrar Osman, Md. Ahasan Habib, Shahina Akter, Tanjina Akhter Banu, Barna Goswami, Iffat Jahan, Md. Saddam Hossain, Tasnim Nafisa, Md. Maruf Ahmed Molla, Mahmuda Yeasmin, Asish Kumar Ghosh, Shahjahan Siddike, A. K. M. Shamsuzzaman, Sheikh Md. Selim Al Din, Utpal Chandra Ray, Salek Ahmed Sajib, Md. Salim Khan |
| EPI_ISL_480428, EPI_ISL_480429, EPI_ISL_480431, EPI_ISL_480432, EPI_ISL_480433, EPI_ISL_480434, EPI_ISL_480435, EPI_ISL_480436, EPI_ISL_480437, EPI_ISL_480438 | Laboratorio de Biología Molecular Asociación Española Primera en Salud | Departments of Pathology and Medicine, New York University School of Medicine | Maria Victoria Elizondo, Maria Noel Zubillaga, Gonzalo Manrique, Paul Zappile, Gael Westby, Matthew T Maurano, Christian Marier, Adriana Heguy |
| EPI_ISL_480445 | National Institute of Laboratory Medicine and Referral Center | Genomic Research Lab, BCSIR | Md. Ahasan Habib, Abu Sayeed Mohammad Mahmud, Mohammad Samir Uzzaman, Eshrar Osman, , Shahina Akter, Tanjina Akhter Banu, Md. Murshed Hasan Sarkar, Barna Goswami, Iffat Jahan, Md. Saddam Hossain, Tasnim Nafisa, Md. Maruf Ahmed Molla, Mahmuda Yeasmin, Asish Kumar Ghosh, Shahjahan Siddike, A. K. M. Shamsuzzaman, Sheikh Md. Selim Al Din, Utpal Chandra Ray, Salek Ahmed Sajib, Md. Salim Khan |
| EPI_ISL_481241 | M Health Fairview | Minnesota Department of Health, Public Health Laboratory | Matt Plumb, Jacob Garfin, Kelly Pung, and Xiong Wang |
| EPI_ISL_481242 | Mayo Clinic & Mayo Clinic Laboratories | Minnesota Department of Health, Public Health Laboratory | Matt Plumb, Jacob Garfin, Kelly Pung, and Xiong Wang |
| EPI_ISL_481253, EPI_ISL_481256, EPI_ISL_481262, EPI_ISL_481264 | Robert Koch Institute, National Reference center for Influenza, Berlin, Germany | Robert Koch Institute, Bioinformatics MF1, Berlin, Germany | Marianne Wedde, Oliver Drechsel, Andrea Thuermer, Rene Kmiecinski, Ralf Duerrwald, Thorsten Wolff, Stephan Fuchs, Max v. Kleist |
| EPI_ISL_482488 | National Institute of Laboratory Medicine and Referral Center | Genomic Research Lab, BCSIR | Md. Murshed Hasan Sarkar, Abu Sayeed Mohammad Mahmud, Mohammad Samir Uzzaman, Eshrar Osman, Md. Ahasan Habib, Shahina Akter, Tanjina Akhter Banu, Barna Goswami, Iffat Jahan, Md. Saddam Hossain, Tasnim Nafisa, Md. Maruf Ahmed Molla, Mahmuda Yeasmin, Asish Kumar Ghosh, Shahjahan Siddike, A. K. M. Shamsuzzaman, Sheikh Md. Selim Al Din, Utpal Chandra Ray, Salek Ahmed Sajib, Md. Salim Khan |
| EPI_ISL_482489 | National Institute of Laboratory Medicine and Referral Center | Genomic Research Lab, BCSIR | Md. Ahasan Habib, Abu Sayeed Mohammad Mahmud, Mohammad Samir Uzzaman, Eshrar Osman, Shahina Akter, Tanjina Akhter Banu, Md. Murshed Hasan Sarkar, Barna Goswami, Iffat Jahan, Md. Saddam Hossain, Tasnim Nafisa, Md. Maruf Ahmed Molla, Mahmuda Yeasmin, Asish Kumar Ghosh, Shahjahan Siddike, A. K. M. Shamsuzzaman, Sheikh Md. Selim Al Din, Utpal Chandra Ray, Salek Ahmed Sajib, Md. Salim Khan |
| EPI_ISL_482732, EPI_ISL_482733, EPI_ISL_482734, EPI_ISL_482735, EPI_ISL_482736, EPI_ISL_482737, EPI_ISL_482738, EPI_ISL_482739, EPI_ISL_482740 | LNR National Reference Laboratory, Mohammed VI University of Health Sciences | Medical Biotechnology Laboratory, Rabat Medical and Pharmacy School, Mohammed The Vth University in Rabat | Meriem LAAMARTI, Souad KARTTI, Rokia LAAMARTI , M.W. CHEMAO-ELFIHRI, Loubna ALLAM, Mouna OUADGHIRI, Imane SMYEJ, Jalila RAHOUI, Houda BENRAHMA, Jalil El ATAR, Idrissa DIAWARA, Rachid EL JAOUDI, Laila SBABOU, Chakib NEJJARI, Saaid AMZAZI, Rachid MENTAG, Lahcen BELYAMANI and Azeddine IBRAHIMI |
| EPI_ISL_482946, EPI_ISL_482947, EPI_ISL_482948, EPI_ISL_482949, EPI_ISL_482950, EPI_ISL_482951, EPI_ISL_482952, EPI_ISL_482953, EPI_ISL_482954, EPI_ISL_482955, EPI_ISL_482956, EPI_ISL_482957, EPI_ISL_482958, EPI_ISL_482959, EPI_ISL_482960, EPI_ISL_482961, EPI_ISL_482962, EPI_ISL_482963, EPI_ISL_482964, EPI_ISL_482965, EPI_ISL_482966 | | | |
| see above | Minnesota Department of Health, Public Health Laboratory | Minnesota Department of Health, Public Health Laboratory | Matt Plumb, Jacob Garfin, and Xiong Wang |
| EPI_ISL_482967, EPI_ISL_482968, EPI_ISL_482969, EPI_ISL_482970, EPI_ISL_482971, EPI_ISL_482972, EPI_ISL_482973, EPI_ISL_482974, EPI_ISL_482975, EPI_ISL_482976, EPI_ISL_482977, EPI_ISL_482978, EPI_ISL_482979, EPI_ISL_482980, EPI_ISL_482981, EPI_ISL_482982, EPI_ISL_482983, EPI_ISL_482984, EPI_ISL_482985, EPI_ISL_482986, EPI_ISL_482987 | | | |
| see above | Mayo Clinic & Mayo Clinic Laboratories | Minnesota Department of Health, Public Health Laboratory | Matt Plumb, Jacob Garfin, and Xiong Wang |
| EPI_ISL_482988, EPI_ISL_482989, EPI_ISL_482990, EPI_ISL_482991, EPI_ISL_482992, EPI_ISL_482993, EPI_ISL_482994, EPI_ISL_482995, EPI_ISL_482996, EPI_ISL_482997, EPI_ISL_482998, EPI_ISL_482999, EPI_ISL_483000, EPI_ISL_483001, EPI_ISL_483002, EPI_ISL_483003, EPI_ISL_483004, EPI_ISL_483005, EPI_ISL_483006, EPI_ISL_483007, EPI_ISL_483008, EPI_ISL_483009, EPI_ISL_483010, EPI_ISL_483011, EPI_ISL_483012, EPI_ISL_483013, EPI_ISL_483014, EPI_ISL_483015, EPI_ISL_483016, EPI_ISL_483017 | | | |
| see above | Minnesota Department of Health, Public Health Laboratory | Minnesota Department of Health, Public Health Laboratory | Matt Plumb, Jacob Garfin, and Xiong Wang |
| EPI_ISL_483566 | Clinical Microbiology Laboratory- Basurto University Hospital | Biocruces-Bizkaia | Mikel J. Urrutikoetxea-Gutierrez, Ana Belén Belén de la Hoz, Matxalen Vidal-García, Mº Carmen Nieto Toboso, Estibaliz Ugalde-Zarraga, José Luis Díaz de Tuesta del Arco |
| EPI_ISL_483570 | Clinical Microbiology Laboratory- Basurto University Hospita | Biocruces-Bizkaia | Mikel J. Urrutikoetxea-Gutierrez, Ana Belén Belén de la Hoz, Matxalen Vidal-García, Mº Carmen Nieto Toboso, Estibaliz Ugalde-Zarraga, José Luis Díaz de Tuesta del Arco |
| EPI_ISL_483571, EPI_ISL_483572, EPI_ISL_483573 | Clinical Microbiology Laboratory- Basurto University Hospital | Biocruces-Bizkaia | Mikel J. Urrutikoetxea-Gutierrez, Ana Belén Belén de la Hoz, Matxalen Vidal-García, Mº Carmen Nieto Toboso, Estibaliz Ugalde-Zarraga, José Luis Díaz de Tuesta del Arco |
| EPI_ISL_483624 | National Institute of Laboratory Medicine and Referral Center | Genomic Research Lab, BCSIR | Md. Maruf Ahmed Molla, Abu Sayeed Mohammad Mahmud, Mohammad Samir Uzzaman, Eshrar Osman, Md. Ahasan Habib, Shahina Akter, Tanjina Akhter Banu, Md. Murshed Hasan Sarkar, Barna Goswami, Iffat Jahan, Md. Saddam Hossain, Tasnim Nafisa, Mahmuda Yeasmin, Asish Kumar Ghosh, A. K. M. Shamsuzzaman, Sheikh Md. Selim Al Din, Utpal Chandra Ray, Salek Ahmed Sajib, Md. Salim Khan |
| EPI_ISL_483626 | National Institute of Laboratory Medicine and Referral Center | Genomic Research Lab, BCSIR | Md. Maruf Ahmed Molla, Abu Sayeed Mohammad Mahmud, Mohammad Samir Uzzaman, Eshrar Osman, Md. Ahasan Habib, Shahina Akter, Tanjina Akhter Banu, Md. Murshed Hasan Sarkar, Barna Goswami, Iffat Jahan, Md. Saddam Hossain, Tasnim Nafisa, Mahmuda Yeasmin, Asish Kumar Ghosh, A. K. M. Shamsuzzaman, Sheikh Md. Selim Al Din, Utpal Chandra Ray, Salek Ahmed Sajib, Md. Salim Khan |
| EPI_ISL_483631, EPI_ISL_483632 | National Institute of Laboratory Medicine and Referral Center | Genomic Research Lab, BCSIR | Md. Ahasan Habib, Abu Sayeed Mohammad Mahmud, Mohammad Samir Uzzaman, Eshrar Osman, Shahina Akter, Tanjina Akhter Banu, Md. Murshed Hasan Sarkar, Barna Goswami, Iffat Jahan, Md. Saddam Hossain, Tasnim Nafisa, Md. Maruf Ahmed Molla, Mahmuda Yeasmin, Asish Kumar Ghosh, A. K. M. Shamsuzzaman, Sheikh Md. Selim Al Din, Utpal Chandra Ray, Salek Ahmed Sajib, Md. Salim Khan |
| EPI_ISL_483645, EPI_ISL_483646, EPI_ISL_483647 | National Institute of Laboratory Medicine and Referral Center | Genomic Research Lab, BCSIR | Md. Saddam Hossain, Abu Sayeed Mohammad Mahmud, Mohammad Samir Uzzaman, Eshrar Osman, Md. Ahasan Habib, Shahina Akter, Tanjina Akhter Banu, Md. Murshed Hasan Sarkar, Barna Goswami, Iffat Jahan, Tasnim Nafisa, Md. Maruf Ahmed Molla, Mahmuda Yeasmin, Asish Kumar Ghosh, A. K. M. Shamsuzzaman, Sheikh Md. Selim Al Din, Utpal Chandra Ray, Salek Ahmed Sajib, Md. Salim Khan |
| EPI_ISL_483686, EPI_ISL_483687 | National Institute of Laboratory Medicine and Referral Center | Genomic Research Lab, BCSIR | Md. Murshed Hasan Sarkar, Abu Sayeed Mohammad Mahmud, Mohammad Samir Uzzaman, Eshrar Osman, Md. Ahasan Habib, Shahina Akter, Tanjina Akhter Banu, Barna Goswami, Iffat Jahan, Md. Saddam Hossain, Tasnim Nafisa, Md. Maruf Ahmed Molla, Mahmuda Yeasmin, Asish Kumar Ghosh, A. K. M. Shamsuzzaman, Sheikh Md. Selim Al Din, Utpal Chandra Ray, Salek Ahmed Sajib, Md. Salim Khan |

| | | | |
|---|---|---|---|
| EPI_ISL_483688 | Genomic Research Lab, BCSIR | Genomic Research Lab, BCSIR | Md. Murshed Hasan Sarkar, Abu Sayeed Mohammad Mahmud, Mohammad Samir Uzzaman, Eshrar Osman, Md. Ahasan Habib, Shahina Akter, Tanjina Akhter Banu, Barna Goswami, Iffat Jahan, Md. Saddam Hossain, Tasnim Nafisa, Md. Maruf Ahmed Molla, Mahmuda Yeasmin, Asish Kumar Ghosh, A. K. M. Shamsuzzaman, Sheikh Md. Selim Al Din, Utpal Chandra Ray, Salek Ahmed Sajib, Md. Salim Khan |
| EPI_ISL_483689, EPI_ISL_483690, EPI_ISL_483691, EPI_ISL_483692 | National Institute of Laboratory Medicine and Referral Center | Genomic Research Lab, BCSIR | Md. Murshed Hasan Sarkar, Abu Sayeed Mohammad Mahmud, Mohammad Samir Uzzaman, Eshrar Osman, Md. Ahasan Habib, Shahina Akter, Tanjina Akhter Banu, Barna Goswami, Iffat Jahan, Md. Saddam Hossain, Tasnim Nafisa, Md. Maruf Ahmed Molla, Mahmuda Yeasmin, Asish Kumar Ghosh, A. K. M. Shamsuzzaman, Sheikh Md. Selim Al Din, Utpal Chandra Ray, Salek Ahmed Sajib, Md. Salim Khan |
| EPI_ISL_483825 | GMERS Medical College Himmatnagar | Gujarat Biotechnology Research Centre | Mayur Gandhi, Apurvasinh Puvar, Janvi Raval, Zarna Patel, Monika Gandhi, Pinal Trivedi, Maharshi Pandya, Nidhi Patel, Nitin Savaliya, Raghawendra Kumar, Dinesh Kumar, Zuber Saiyed, Komal Patel, Labdhi Pandya, Afzal Ansari, Nikha Trivedi, Himanshu Khatri, R D Dixit, A M Kadri, Harsh Bakshi, Chaitanya Joshi, Madhvi Joshi |
| EPI_ISL_483829, EPI_ISL_483846 | Pandit Deendayal Upadhyay Government Medical College, Rajkot | Gujarat Biotechnology Research Centre | Monika Gandhi, Pinal Trivedi, Maharshi Pandya, Nidhi Patel, Nitin Savaliya, Raghawendra Kumar, Dinesh Kumar, Zuber Saiyed, Komal Patel, Labdhi Pandya, Afzal Ansari, Nikha Trivedi, Gauravi Dhruv, Arti Trivedi, Apurvasinh Puvar, Janvi Raval, Zarna Patel, R D Dixit, A M Kadri, Harsh Bakshi, Chaitanya Joshi, Madhvi Joshi |
| EPI_ISL_483864 | Department of Microbiology, Government Medical College, Surat | Gujarat Biotechnology Research Centre | Monika Gandhi, Pinal Trivedi, Maharshi Pandya, Nidhi Patel, Nitin Savaliya, Raghawendra Kumar, Dinesh Kumar, Zuber Saiyed, Komal Patel, Labdhi Pandya, Afzal Ansari, Nikha Trivedi, Naresh Chauhan, Summaiya Mullan, Amit gamit, Apurvasinh Puvar, Janvi Raval, Zarna Patel, R D Dixit, A M Kadri, Harsh Bakshi, Chaitanya Joshi, Madhvi Joshi |
| EPI_ISL_484680, EPI_ISL_484681, EPI_ISL_484682, EPI_ISL_484683, EPI_ISL_484684 | Virology Department, Royal Infirmary of Edinburgh, NHS Lothian / School of Biological Sciences, University of Edinburgh / Institute of Genetics and Molecular Medicine, University of Edinburgh | COVID-19 Genomics UK (COG-UK) Consortium | McHugh M, Dewar R, Rooke S, Gallagher M, Balcaza C, O'Toole Á, Scher E, Hill V, McCrone JT, Colquhoun R, Yu X, Jackson B, Rambaut A, Williams TC, Templeton K |
| EPI_ISL_484692 | Minnesota Department of Health, Public Health Laboratory | Minnesota Department of Health, Public Health Laboratory | Matt Plumb, Jacob Garfin, and Xiong Wang |
| EPI_ISL_486648, EPI_ISL_486652, EPI_ISL_486656, EPI_ISL_486659, EPI_ISL_486663 | Microbiology, Virology and Biemergency Laboratory-ASST FBF Sacco | Microbiology, Virology and Biemergency Laboratory-ASST FBF Sacco | Micheli V, Comandatore F, Romeri F, Mancon A, Rimoldi SG |
| EPI_ISL_486665 | Microbiology, Virology and Biemergency Laboratory-ASST FBF Sacco | Microbiology, Virology and Biemergency Laboratory-ASST FBF Sacco | Micheli V, Rimoldi SG, Comandatore F, Mancon A, Romeri F |
| EPI_ISL_486876 | Clinical Microbiology Laboratory- Basurto University Hospital | Biocruces-Bizkaia | Mikel J. Urrutikoetxea-Gutierrez, Ana Belén Belén de la Hoz, Matxalen Vidal-García, Mº Carmen Nieto Toboso, Estibaliz Ugalde-Zarraga, José Luis Díaz de Tuesta del Arco |
| EPI_ISL_487362 | National Institute of Laboratory Medicine and Referral Center | Genomic Research Lab, BCSIR | Md. Ahasan Habib, Abu Sayeed Mohammad Mahmud, Mohammad Samir Uzzaman, Eshrar Osman, Shahina Akter, Tanjina Akhter Banu, Md. Murshed Hasan Sarkar, Barna Goswami, Iffat Jahan, Md. Saddam Hossain, Tasnim Nafisa, Md. Maruf Ahmed Molla, Mahmuda Yeasmin, Asish Kumar Ghosh, A. K. M. Shamsuzzaman, Sheikh Md. Selim Al Din, Utpal Chandra Ray, Salek Ahmed Sajib, Md. Salim Khan |
| EPI_ISL_487375, EPI_ISL_487376 | National Institute of Laboratory Medicine and Referral Center | Genomic Research Lab, BCSIR | Md. Saddam Hossain, Abu Sayeed Mohammad Mahmud, Mohammad Samir Uzzaman, Eshrar Osman, Md. Ahasan Habib, Shahina Akter, Tanjina Akhter Banu, Md. Murshed Hasan Sarkar, Barna Goswami, Iffat Jahan, Tasnim Nafisa, Md. Maruf Ahmed Molla, Mahmuda Yeasmin, Asish Kumar Ghosh, A. K. M. Shamsuzzaman, Sheikh Md. Selim Al Din, Utpal Chandra Ray, Salek Ahmed Sajib, Md. Salim Khan |
| EPI_ISL_487378, EPI_ISL_487380, EPI_ISL_487382, EPI_ISL_487383, EPI_ISL_487384, EPI_ISL_487385 | National Institute of Laboratory Medicine and Referral Center | Genomic Research Lab, BCSIR | Md. Murshed Hasan Sarkar, Abu Sayeed Mohammad Mahmud, Mohammad Samir Uzzaman, Eshrar Osman, Md. Ahasan Habib, Shahina Akter, Tanjina Akhter Banu, Barna Goswami, Iffat Jahan, Md. Saddam Hossain, Tasnim Nafisa, Md. Maruf Ahmed Molla, Mahmuda Yeasmin, Asish Kumar Ghosh, A. K. M. Shamsuzzaman, Sheikh Md. Selim Al Din, Utpal Chandra Ray, Salek Ahmed Sajib, Md. Salim Khan |
| EPI_ISL_487647, EPI_ISL_487648, EPI_ISL_487649, EPI_ISL_487650, EPI_ISL_487651, EPI_ISL_487652, EPI_ISL_487653, EPI_ISL_487654, EPI_ISL_487655, EPI_ISL_487656, EPI_ISL_487657, EPI_ISL_487658, EPI_ISL_487659, EPI_ISL_487660, EPI_ISL_487661, EPI_ISL_487662, EPI_ISL_487663, EPI_ISL_487664, EPI_ISL_487665, EPI_ISL_487666, EPI_ISL_487667, EPI_ISL_487668, EPI_ISL_487669, EPI_ISL_487670, EPI_ISL_487671, EPI_ISL_487672, EPI_ISL_487673, EPI_ISL_487674, EPI_ISL_487675, EPI_ISL_487676, EPI_ISL_487677, EPI_ISL_487678, EPI_ISL_487679, EPI_ISL_487680, EPI_ISL_487681, EPI_ISL_487682, EPI_ISL_487683, EPI_ISL_487684, EPI_ISL_487685, EPI_ISL_487686, EPI_ISL_487687, EPI_ISL_487688, EPI_ISL_487689, EPI_ISL_487690, EPI_ISL_487691, EPI_ISL_487692, EPI_ISL_487693, EPI_ISL_487694, EPI_ISL_487695, EPI_ISL_487696, EPI_ISL_487697, EPI_ISL_487698, EPI_ISL_487699, EPI_ISL_487700, EPI_ISL_487701, EPI_ISL_487702, EPI_ISL_487703, EPI_ISL_487704, EPI_ISL_487705, EPI_ISL_487706, EPI_ISL_487707, EPI_ISL_487708, EPI_ISL_487709, EPI_ISL_487710, EPI_ISL_487711, EPI_ISL_487712, EPI_ISL_487713, EPI_ISL_487714, EPI_ISL_487715, EPI_ISL_487716, EPI_ISL_487717, EPI_ISL_487718, EPI_ISL_487719, EPI_ISL_487720, EPI_ISL_487721, EPI_ISL_487722, EPI_ISL_487723, EPI_ISL_487724, EPI_ISL_487725, EPI_ISL_487726, EPI_ISL_487727, EPI_ISL_487728, EPI_ISL_487729, EPI_ISL_487730, EPI_ISL_487731, EPI_ISL_487732, EPI_ISL_487733, EPI_ISL_487734, EPI_ISL_487735, EPI_ISL_487736, EPI_ISL_487737, EPI_ISL_487738, EPI_ISL_487739, EPI_ISL_487740, EPI_ISL_487741, EPI_ISL_487742, EPI_ISL_487743, EPI_ISL_487744, EPI_ISL_487745, EPI_ISL_487746, EPI_ISL_487747, EPI_ISL_487748, EPI_ISL_487749, EPI_ISL_487750, EPI_ISL_487751, EPI_ISL_487752, EPI_ISL_487753, EPI_ISL_487754, EPI_ISL_487755, EPI_ISL_487756, EPI_ISL_487757, EPI_ISL_487758, EPI_ISL_487759, EPI_ISL_487760, EPI_ISL_487761, EPI_ISL_487762, EPI_ISL_487763, EPI_ISL_487764, EPI_ISL_487765, EPI_ISL_487766, EPI_ISL_487768, EPI_ISL_487769, EPI_ISL_487770, EPI_ISL_487771, EPI_ISL_487772, EPI_ISL_487773, EPI_ISL_487774, EPI_ISL_487775, EPI_ISL_487776, EPI_ISL_487777, EPI_ISL_487778, EPI_ISL_487779, EPI_ISL_487780, EPI_ISL_487781, EPI_ISL_487782, EPI_ISL_487783, EPI_ISL_487784, EPI_ISL_487785, EPI_ISL_487786, EPI_ISL_487787, EPI_ISL_487788, EPI_ISL_487789, EPI_ISL_487790, EPI_ISL_487791, EPI_ISL_487792, EPI_ISL_487793, EPI_ISL_487794, EPI_ISL_487795, EPI_ISL_487796, EPI_ISL_487797, EPI_ISL_487798, EPI_ISL_487799, EPI_ISL_487800, EPI_ISL_487801, EPI_ISL_487802, EPI_ISL_487803, EPI_ISL_487804, EPI_ISL_487805, EPI_ISL_487806, EPI_ISL_487807, EPI_ISL_487808, EPI_ISL_487809, EPI_ISL_487810, EPI_ISL_487811, EPI_ISL_487812, EPI_ISL_487813, EPI_ISL_487814, EPI_ISL_487815, EPI_ISL_487816, EPI_ISL_487817, EPI_ISL_487818, EPI_ISL_487819, EPI_ISL_487820, EPI_ISL_487821, EPI_ISL_487822, EPI_ISL_487823, EPI_ISL_487824, EPI_ISL_487825, EPI_ISL_487826, EPI_ISL_487827, EPI_ISL_487828, EPI_ISL_487829, EPI_ISL_487830, EPI_ISL_487831, EPI_ISL_487832, EPI_ISL_487833, EPI_ISL_487834, EPI_ISL_487835, EPI_ISL_487836, EPI_ISL_487837, EPI_ISL_487838, EPI_ISL_487839, EPI_ISL_487840, EPI_ISL_487841, EPI_ISL_487842, EPI_ISL_487843, EPI_ISL_487844, EPI_ISL_487845, EPI_ISL_487846, EPI_ISL_487847, EPI_ISL_487848, EPI_ISL_487849, EPI_ISL_487850, EPI_ISL_487851, EPI_ISL_487852, EPI_ISL_487853, EPI_ISL_487854, EPI_ISL_487855, EPI_ISL_487856, EPI_ISL_487857, EPI_ISL_487858, EPI_ISL_487859, EPI_ISL_487860, EPI_ISL_487861, EPI_ISL_487862, EPI_ISL_487863, EPI_ISL_487864, EPI_ISL_487865, EPI_ISL_487866, EPI_ISL_487867, EPI_ISL_487868, EPI_ISL_487869, EPI_ISL_487870, EPI_ISL_487871, EPI_ISL_487872, EPI_ISL_487873, EPI_ISL_487874, EPI_ISL_487875, EPI_ISL_487876, EPI_ISL_487877, EPI_ISL_487878, EPI_ISL_487879, EPI_ISL_487880, EPI_ISL_487881, EPI_ISL_487882, EPI_ISL_487883, EPI_ISL_487884, EPI_ISL_487885, EPI_ISL_487886, EPI_ISL_487887, EPI_ISL_487888, EPI_ISL_487889, EPI_ISL_487890, EPI_ISL_487891, EPI_ISL_487892, EPI_ISL_487893, EPI_ISL_487894, EPI_ISL_487895, EPI_ISL_487896, EPI_ISL_487897, EPI_ISL_487898, EPI_ISL_487899, EPI_ISL_487900, EPI_ISL_487901, EPI_ISL_487902, EPI_ISL_487903, EPI_ISL_487904, EPI_ISL_487905, EPI_ISL_487906, EPI_ISL_487907, EPI_ISL_487908, EPI_ISL_487909, EPI_ISL_487910, EPI_ISL_487911, EPI_ISL_487912, EPI_ISL_487913, EPI_ISL_487914, EPI_ISL_487915, EPI_ISL_487916, EPI_ISL_487917, EPI_ISL_487918, EPI_ISL_487919, EPI_ISL_487920, EPI_ISL_487921, EPI_ISL_487922, EPI_ISL_487923, EPI_ISL_487924, EPI_ISL_487925, EPI_ISL_487926, EPI_ISL_487927, EPI_ISL_487928, EPI_ISL_487929, EPI_ISL_487930, EPI_ISL_487931, EPI_ISL_487932, EPI_ISL_487933, EPI_ISL_487934, EPI_ISL_487935, EPI_ISL_487936, EPI_ISL_487937, EPI_ISL_487938, EPI_ISL_487939, EPI_ISL_487940, EPI_ISL_487941, EPI_ISL_487942, EPI_ISL_487943, EPI_ISL_487944, EPI_ISL_487945, EPI_ISL_487946, EPI_ISL_487947, EPI_ISL_487948, EPI_ISL_487949, EPI_ISL_487950, EPI_ISL_487951, EPI_ISL_487952, EPI_ISL_487953, EPI_ISL_487954, EPI_ISL_487955, EPI_ISL_487956, EPI_ISL_487957, EPI_ISL_487958, EPI_ISL_487959, EPI_ISL_487960, EPI_ISL_487961, EPI_ISL_487962, EPI_ISL_487963, EPI_ISL_487964, EPI_ISL_487965, EPI_ISL_487966, EPI_ISL_487967, EPI_ISL_487968, EPI_ISL_487969, EPI_ISL_487970, EPI_ISL_487971, EPI_ISL_487972, EPI_ISL_487973, EPI_ISL_487974, EPI_ISL_487975, EPI_ISL_487976, EPI_ISL_487977, EPI_ISL_487978, EPI_ISL_487979, EPI_ISL_487980, EPI_ISL_487981, EPI_ISL_487982, EPI_ISL_487983, EPI_ISL_487984, EPI_ISL_487985, EPI_ISL_487986, EPI_ISL_487987, EPI_ISL_487988, EPI_ISL_487989, EPI_ISL_487990, EPI_ISL_487991, EPI_ISL_487992, EPI_ISL_487993, EPI_ISL_487994, EPI_ISL_487996, EPI_ISL_487997, EPI_ISL_487998, EPI_ISL_487999, EPI_ISL_488000, EPI_ISL_488001, EPI_ISL_488002, EPI_ISL_488879, EPI_ISL_488880, EPI_ISL_488881, EPI_ISL_488882, EPI_ISL_488883, EPI_ISL_488884, EPI_ISL_488885, EPI_ISL_488886, EPI_ISL_488887, EPI_ISL_488888, EPI_ISL_488889, EPI_ISL_488890, EPI_ISL_488891, EPI_ISL_488892, EPI_ISL_488893, EPI_ISL_488894, EPI_ISL_488895, EPI_ISL_488896, EPI_ISL_488897, EPI_ISL_488898, EPI_ISL_488899, EPI_ISL_488900, EPI_ISL_488901, EPI_ISL_488902, EPI_ISL_488903, EPI_ISL_488904, EPI_ISL_488905, EPI_ISL_488906, EPI_ISL_488907, EPI_ISL_488908, EPI_ISL_488909, EPI_ISL_488910, EPI_ISL_488911, EPI_ISL_488912, EPI_ISL_488913, EPI_ISL_488914, EPI_ISL_488915, EPI_ISL_488916, EPI_ISL_488917, EPI_ISL_488918, EPI_ISL_488919, EPI_ISL_488920, EPI_ISL_488921, EPI_ISL_488922, EPI_ISL_488923, EPI_ISL_488924, EPI_ISL_488925, EPI_ISL_488926, EPI_ISL_488927, EPI_ISL_488928, EPI_ISL_488929, EPI_ISL_488930, EPI_ISL_488931, EPI_ISL_488932, EPI_ISL_488933, EPI_ISL_488934, EPI_ISL_488935, EPI_ISL_488936, EPI_ISL_488937, EPI_ISL_488938, EPI_ISL_488939, EPI_ISL_488940, EPI_ISL_488941, EPI_ISL_488942, EPI_ISL_488943, EPI_ISL_488944, EPI_ISL_488945, EPI_ISL_488946, EPI_ISL_488947, EPI_ISL_488948, EPI_ISL_488949, EPI_ISL_488950, EPI_ISL_488951, EPI_ISL_488952, EPI_ISL_488953, EPI_ISL_488954, EPI_ISL_488955, EPI_ISL_488956, EPI_ISL_488957, EPI_ISL_488958, EPI_ISL_488959, EPI_ISL_488960, EPI_ISL_488961, EPI_ISL_488962, EPI_ISL_488963, EPI_ISL_488964, EPI_ISL_488965, EPI_ISL_488966, EPI_ISL_488967, EPI_ISL_488968, EPI_ISL_488969, EPI_ISL_488970, EPI_ISL_488971, EPI_ISL_488972, EPI_ISL_488973, EPI_ISL_488974, EPI_ISL_488975, EPI_ISL_488976, EPI_ISL_488977, EPI_ISL_488978, EPI_ISL_488979, EPI_ISL_488980, EPI_ISL_488981, EPI_ISL_488982, EPI_ISL_488983, EPI_ISL_488984, EPI_ISL_488985, EPI_ISL_488986, EPI_ISL_488987, EPI_ISL_488988, EPI_ISL_488989, EPI_ISL_488990, EPI_ISL_488991, EPI_ISL_488992, EPI_ISL_488993, EPI_ISL_488994, EPI_ISL_488995, EPI_ISL_488996, EPI_ISL_488997, EPI_ISL_488998, EPI_ISL_488999, EPI_ISL_489000, EPI_ISL_489001, EPI_ISL_489002, EPI_ISL_489003, EPI_ISL_489004, EPI_ISL_489005, EPI_ISL_489006, EPI_ISL_489007, EPI_ISL_489008, EPI_ISL_489009, EPI_ISL_489010, EPI_ISL_489011, EPI_ISL_489012, EPI_ISL_489013, EPI_ISL_489014, EPI_ISL_489015, EPI_ISL_489016, EPI_ISL_489017, EPI_ISL_489018, EPI_ISL_489019, EPI_ISL_489020, EPI_ISL_489021, EPI_ISL_489022, EPI_ISL_489023, EPI_ISL_489024, EPI_ISL_489025, EPI_ISL_489026, EPI_ISL_489027, EPI_ISL_489028, EPI_ISL_489029, EPI_ISL_489030, EPI_ISL_489031, EPI_ISL_489032, EPI_ISL_489033, EPI_ISL_489034, EPI_ISL_489035, EPI_ISL_489036, EPI_ISL_489037, EPI_ISL_489038, EPI_ISL_489039, EPI_ISL_489040, EPI_ISL_489041, EPI_ISL_489042, EPI_ISL_489044, EPI_ISL_489045, EPI_ISL_489046, EPI_ISL_489047, EPI_ISL_489048, EPI_ISL_489049, EPI_ISL_489050, EPI_ISL_489051, EPI_ISL_489052, EPI_ISL_489053, EPI_ISL_489054, EPI_ISL_489055, EPI_ISL_489056, EPI_ISL_489057, EPI_ISL_489058, EPI_ISL_489059, EPI_ISL_489060, EPI_ISL_489061, EPI_ISL_489062, EPI_ISL_489063 | | | |
| see above | Virology Department, Royal Infirmary of Edinburgh, NHS Lothian / School of Biological Sciences, University of Edinburgh | Wellcome Sanger Institute for the COVID-19 Genomics UK (COG-UK) consortium | McHugh M, Dewar R, Rooke S, O'Toole Á, Scher E, Hill V, McCrone JT, Colquhoun R, Yu X, Jackson B, Rambaut A, Templeton K and Alex Alderton, Roberto Amato, Sonia Goncalves, Ewan Harrison, David K. Jackson, Ian Johnston, Dominic Kwiatkowski, Cordelia Langford, John Sillitoe on behalf of the Wellcome Sanger Institute COVID-19 Surveillance Team (http://www.sanger.ac.uk/covid-team) |
| EPI_ISL_489708, EPI_ISL_489709 | The National Institute of Public Health | The National Institute of Public Health and State Veterinary Institute Prague | Nagy,A;Jirincova,H;Novakova,L;Trnka,D;Vecerova,J |
| EPI_ISL_489833, EPI_ISL_489834, EPI_ISL_489835 | Clinical Microbiology Laboratory- Basurto University Hospital | Biocruces-Bizkaia | Mikel J. Urrutikoetxea-Gutierrez, Ana Belén Belén de la Hoz, Matxalen Vidal-García, Mº Carmen Nieto Toboso, Estibaliz Ugalde-Zarraga, José Luis Díaz de Tuesta del Arco |
| EPI_ISL_490109, EPI_ISL_490110, EPI_ISL_490111 | National Institute of Laboratory Medicine and Referral Center | Genomic Research Lab, BCSIR | Md. Murshed Hasan Sarkar, Abu Sayeed Mohammad Mahmud, Mohammad Samir Uzzaman, Eshrar Osman, Md. Ahasan Habib, Shahina Akter, Tanjina Akhter Banu, Barna Goswami, Iffat Jahan, Md. Saddam Hossain, Tasnim Nafisa, Md. Maruf Ahmed Molla, Mahmuda Yeasmin, Asish Kumar Ghosh, A. K. M. Shamsuzzaman, Sheikh Md. Selim Al Din, Utpal Chandra Ray, Salek Ahmed Sajib, Md. Salim Khan |
| EPI_ISL_490112 | The National Institute of Public Health | The National Institute of Public Health and State Veterinary Institute Prague | Nagy,A;Jirincova,H;Novakova,L;Trnka,D;Vecerova,J |

| | | | |
|---|---|---|---|
| EPI_ISL_490165 | National Institute of Laboratory Medicine and Referral Center | Genomic Research Lab, BCSIR | Md. Saddam Hossain, Abu Sayeed Mohammad Mahmud, Mohammad Samir Uzzaman, Eshrar Osman, Md. Ahasan Habib, Shahina Akter, Tanjina Akhter Banu, Md. Murshed Hasan Sarkar, Barna Goswami, Iffat Jahan, Tasnim Nafisa, Md. Maruf Ahmed Molla, Mahmuda Yeasmin, Asish Kumar Ghosh, A. K. M. Shamsuzzaman, Sheikh Md. Selim Al Din, Utpal Chandra Ray, Salek Ahmed Sajib, Md. Salim Khan |
| EPI_ISL_490202, EPI_ISL_490203, EPI_ISL_490204, EPI_ISL_490977 | Clinical Microbiology Laboratory- Basurto University Hospital | Biocruces-Bizkaia | Mikel J. Urrutikoetxea-Gutierrez, Ana Belén Belén de la Hoz, Matxalen Vidal-García, Mº Carmen Nieto Toboso, Estibaliz Ugalde-Zarraga, José Luis Díaz de Tuesta del Arco |
| EPI_ISL_491092 | The National Institute of Public Health | State Veterinary Institute Prague | Nagy,A;Jirincova,H;Novakova,L;Trnka,D;Vecerova,J |
| EPI_ISL_491093, EPI_ISL_491094, EPI_ISL_491095 | The National Institute of Public Health | The National Institute of Public Health and State Veterinary Institute Prague | Nagy,A; Jirincova,H; Novakova,L; Trnka,D; Vecerova,J |
| EPI_ISL_491117, EPI_ISL_491118, EPI_ISL_491119 | The National Institute of Public Health | The National Institute of Public Health and State Veterinary Institute Prague | Nagy,A;Jirincova,H;Novakova,L;Trnka,D;Vecerova,J |
| EPI_ISL_491476 | BSL3 Lab, Pendik Veterinary Control Enstitue | Genomic Laboratory (GLAB), Istanbul Technical University | Mustafa HASOKSUZ, Fahriye SARAC, Osman ERGANIS, Serdar UZAR, Hakan ENUL, Cumhur ADIAY, Ahmet SAIT, Orbay SAYI, Kadir YESILBAG, Oguz KARABEY |
| EPI_ISL_491494, EPI_ISL_491495, EPI_ISL_491496, EPI_ISL_491497, EPI_ISL_491498, EPI_ISL_491499, EPI_ISL_491500, EPI_ISL_491501, EPI_ISL_491502, EPI_ISL_491503, EPI_ISL_491504, EPI_ISL_491505, EPI_ISL_491506, EPI_ISL_491507, EPI_ISL_491508, EPI_ISL_491509, EPI_ISL_491510, EPI_ISL_491511, EPI_ISL_491512, EPI_ISL_491513, EPI_ISL_491514, EPI_ISL_491515, EPI_ISL_491516, EPI_ISL_491517, EPI_ISL_491518, EPI_ISL_491519, EPI_ISL_491520, EPI_ISL_491521, EPI_ISL_491522, EPI_ISL_491523, EPI_ISL_491524, EPI_ISL_491525, EPI_ISL_491526, EPI_ISL_491527, EPI_ISL_491528, EPI_ISL_491529, EPI_ISL_491530, EPI_ISL_491531, EPI_ISL_491532, EPI_ISL_491533, EPI_ISL_491534, EPI_ISL_491535, EPI_ISL_491536, EPI_ISL_491537, EPI_ISL_491538, EPI_ISL_491539, EPI_ISL_491540, EPI_ISL_491541, EPI_ISL_491542, EPI_ISL_491543, EPI_ISL_491544, EPI_ISL_491545, EPI_ISL_491546, EPI_ISL_491547, EPI_ISL_491548, EPI_ISL_491549, EPI_ISL_491550, EPI_ISL_491551, EPI_ISL_491552, EPI_ISL_491553, EPI_ISL_491554, EPI_ISL_491555, EPI_ISL_491556, EPI_ISL_491557, EPI_ISL_491558, EPI_ISL_491559, EPI_ISL_491560, EPI_ISL_491561, EPI_ISL_491562, EPI_ISL_491563, EPI_ISL_491564, EPI_ISL_491565, EPI_ISL_491566, EPI_ISL_491567, EPI_ISL_491568, EPI_ISL_491569, EPI_ISL_491570, EPI_ISL_491571, EPI_ISL_491573, EPI_ISL_491574, EPI_ISL_491575, EPI_ISL_491576, EPI_ISL_491577, EPI_ISL_491578, EPI_ISL_491579, EPI_ISL_491580, EPI_ISL_491581, EPI_ISL_491582, EPI_ISL_491583, EPI_ISL_491584, EPI_ISL_491585, EPI_ISL_491586, EPI_ISL_491587, EPI_ISL_491588, EPI_ISL_491589, EPI_ISL_491590, EPI_ISL_491591, EPI_ISL_491592, EPI_ISL_491593, EPI_ISL_491594, EPI_ISL_491595, EPI_ISL_491596, EPI_ISL_491597, EPI_ISL_491598, EPI_ISL_491599, EPI_ISL_491600, EPI_ISL_491601, EPI_ISL_491602, EPI_ISL_491603, EPI_ISL_491604, EPI_ISL_491605, EPI_ISL_491606, EPI_ISL_491607, EPI_ISL_491609, EPI_ISL_491610, EPI_ISL_491612, EPI_ISL_491613, EPI_ISL_491614, EPI_ISL_491615, EPI_ISL_491617, EPI_ISL_491618, EPI_ISL_491619, EPI_ISL_491620, EPI_ISL_491621, EPI_ISL_491622, EPI_ISL_491623, EPI_ISL_491624, EPI_ISL_491625, EPI_ISL_491626, EPI_ISL_491627, EPI_ISL_491628, EPI_ISL_491629, EPI_ISL_491630, EPI_ISL_491631, EPI_ISL_491632, EPI_ISL_491633, EPI_ISL_491634, EPI_ISL_491635, EPI_ISL_491636, EPI_ISL_491637, EPI_ISL_491638, EPI_ISL_491639, EPI_ISL_491640, EPI_ISL_491641, EPI_ISL_491642, EPI_ISL_491643, EPI_ISL_491644, EPI_ISL_491645, EPI_ISL_491646, EPI_ISL_491647, EPI_ISL_491648, EPI_ISL_491649, EPI_ISL_491650, EPI_ISL_491651, EPI_ISL_491652, EPI_ISL_491653, EPI_ISL_491654, EPI_ISL_491656, EPI_ISL_491657, EPI_ISL_491658, EPI_ISL_491659, EPI_ISL_491660, EPI_ISL_491661, EPI_ISL_491662, EPI_ISL_491663, EPI_ISL_491664, EPI_ISL_491665, EPI_ISL_491666, EPI_ISL_491667, EPI_ISL_491668, EPI_ISL_491669, EPI_ISL_491670, EPI_ISL_491671, EPI_ISL_491672, EPI_ISL_491673, EPI_ISL_491674, EPI_ISL_491675, EPI_ISL_491676, EPI_ISL_491677, EPI_ISL_491678, EPI_ISL_491679, EPI_ISL_491680, EPI_ISL_491681, EPI_ISL_491682, EPI_ISL_491683, EPI_ISL_491684, EPI_ISL_491685, EPI_ISL_491686, EPI_ISL_491687, EPI_ISL_491688, EPI_ISL_491689, EPI_ISL_491690, EPI_ISL_491691, EPI_ISL_491692, EPI_ISL_491693, EPI_ISL_491694, EPI_ISL_491695, EPI_ISL_491696, EPI_ISL_491697, EPI_ISL_491698, EPI_ISL_491699, EPI_ISL_491700 | | | |
| see above | Virology Department, Royal Infirmary of Edinburgh, NHS Lothian / School of Biological Sciences, University of Edinburgh | Wellcome Sanger Institute for the COVID-19 Genomics UK (COG-UK) consortium | McHugh M, Dewar R, Rooke S, O'Toole Á, Scher E, Hill V, McCrone JT, Colquhoun R, Yu X, Jackson B, Rambaut A, Templeton K and Alex Alderton, Roberto Amato, Sonia Goncalves, Ewan Harrison, David K. Jackson, Ian Johnston, Dominic Kwiatkowski, Cordelia Langford, John Sillitoe on behalf of the Wellcome Sanger Institute COVID-19 Surveillance Team (http://www.sanger.ac.uk/covid-team) |
| EPI_ISL_493328, EPI_ISL_493331 | INMI Lazzaro Spallanzani IRCCS | INMI Lazzaro Spallanzani IRCCS | Martina Rueca, Cesare E.M. Gruber, Barbara Bartolini, Francesco Messina, Maria R. Capobianchi, Antonino Di Caro |
| EPI_ISL_493898, EPI_ISL_493899, EPI_ISL_493900, EPI_ISL_493901, EPI_ISL_493902, EPI_ISL_493903, EPI_ISL_493904, EPI_ISL_493905, EPI_ISL_493906, EPI_ISL_493907, EPI_ISL_493908, EPI_ISL_493909, EPI_ISL_493910, EPI_ISL_493911, EPI_ISL_493912, EPI_ISL_493913, EPI_ISL_493914, EPI_ISL_493915, EPI_ISL_493916, EPI_ISL_493917, EPI_ISL_493918, EPI_ISL_493919, EPI_ISL_493920, EPI_ISL_493921, EPI_ISL_493922, EPI_ISL_493923, EPI_ISL_493924, EPI_ISL_493925, EPI_ISL_493926, EPI_ISL_493927, EPI_ISL_493928, EPI_ISL_493929, EPI_ISL_493930, EPI_ISL_493931, EPI_ISL_493932, EPI_ISL_493933, EPI_ISL_493934, EPI_ISL_493935, EPI_ISL_493936, EPI_ISL_493937, EPI_ISL_493938, EPI_ISL_493939, EPI_ISL_493940, EPI_ISL_493941, EPI_ISL_493942, EPI_ISL_493943, EPI_ISL_493944, EPI_ISL_493945, EPI_ISL_493946, EPI_ISL_493947, EPI_ISL_493948, EPI_ISL_493949, EPI_ISL_493950, EPI_ISL_493951, EPI_ISL_493952, EPI_ISL_493953, EPI_ISL_493954, EPI_ISL_493955, EPI_ISL_493956, EPI_ISL_493957, EPI_ISL_493958, EPI_ISL_493959, EPI_ISL_493960, EPI_ISL_493961, EPI_ISL_493962, EPI_ISL_493963, EPI_ISL_493964, EPI_ISL_493965, EPI_ISL_493966, EPI_ISL_493967, EPI_ISL_493968, EPI_ISL_493969, EPI_ISL_493970, EPI_ISL_493971, EPI_ISL_493972, EPI_ISL_493973, EPI_ISL_493974 | | | |
| see above | Virology Department, Royal Infirmary of Edinburgh, NHS Lothian / School of Biological Sciences, University of Edinburgh / Institute of Genetics and Molecular Medicine, University of Edinburgh | COVID-19 Genomics UK (COG-UK) Consortium | McHugh M, Dewar R, Rooke S, Gallagher M, Balcaza C, O'Toole Á, Scher E, Hill V, McCrone JT, Colquhoun R, Yu X, Jackson B, Rambaut A, Williams TC, Templeton K |
| EPI_ISL_495016 | B.J. Medical College and Civil hospital | Gujarat Biotechnology Research Centre | Monika Gandhi, Pinal Trivedi, Maharshi Pandya, Nidhi Patel, Nitin Savaliya, Raghawendra Kumar, Dinesh Kumar, Zuber Saiyed, Komal Patel, Labdhi Pandya, Afzal Ansari, Nikha Trivedi, Pranay Shah, Kamlesh J Upadhyay, Sanjay Kapadia, Apurvasinh Puvar, Janvi Raval, Zarna Patel, R D Dixit, A M Kadri, Harsh Bakshi, Chaitanya Joshi, Madhvi Joshi |
| EPI_ISL_495030 | GMERS Medical College & Hospital, Gotri, Vadodara | Gujarat Biotechnology Research Centre | Meenakshi Shah, Neena Doshi, Varsha Godbole, Apurvasinh Puvar, Janvi Raval, Zarna Patel, Monika Gandhi, Pinal Trivedi, Maharshi Pandya, Nidhi Patel, Nitin Savaliya, Raghawendra Kumar, Dinesh Kumar, Zuber Saiyed, Komal Patel, Labdhi Pandya, Afzal Ansari, Nikha Trivedi, R D Dixit, A M Kadri, Harsh Bakshi, Chaitanya Joshi, Madhvi Joshi |
| EPI_ISL_495036 | GMERS Medical College & Hospital, Gotri, Vadodara | Gujarat Biotechnology Research Centre | Monika Gandhi, Pinal Trivedi, Maharshi Pandya, Nidhi Patel, Nitin Savaliya, Raghawendra Kumar, Dinesh Kumar, Zuber Saiyed, Komal Patel, Labdhi Pandya, Afzal Ansari, Nikha Trivedi, Meenakshi Shah, Neena Doshi, Varsha Godbole, Apurvasinh Puvar, Janvi Raval, Zarna Patel, R D Dixit, A M Kadri, Harsh Bakshi, Chaitanya Joshi, Madhvi Joshi |
| EPI_ISL_495048 | GMERS Medical College & Hospital, Gotri, Vadodara | Gujarat Biotechnology Research Centre | Meenakshi Shah, Neena Doshi, Varsha Godbole, Apurvasinh Puvar, Janvi Raval, Zarna Patel, Monika Gandhi, Pinal Trivedi, Maharshi Pandya, Nidhi Patel, Nitin Savaliya, Raghawendra Kumar, Dinesh Kumar, Zuber Saiyed, Komal Patel, Labdhi Pandya, Afzal Ansari, Nikha Trivedi, R D Dixit, A M Kadri, Harsh Bakshi, Chaitanya Joshi, Madhvi Joshi |
| EPI_ISL_495054 | GMERS Medical College & Hospital, Gotri, Vadodara | Gujarat Biotechnology Research Centre | Monika Gandhi, Pinal Trivedi, Maharshi Pandya, Nidhi Patel, Nitin Savaliya, Raghawendra Kumar, Dinesh Kumar, Zuber Saiyed, Komal Patel, Labdhi Pandya, Afzal Ansari, Nikha Trivedi, Meenakshi Shah, Neena Doshi, Varsha Godbole, Apurvasinh Puvar, Janvi Raval, Zarna Patel, R D Dixit, A M Kadri, Harsh Bakshi, Chaitanya Joshi, Madhvi Joshi |
| EPI_ISL_495073 | Department of MicroBiology, Government Medical College, Surat | Gujarat Biotechnology Research Centre | Monika Gandhi, Pinal Trivedi, Maharshi Pandya, Nidhi Patel, Nitin Savaliya, Raghawendra Kumar, Dinesh Kumar, Zuber Saiyed, Komal Patel, Labdhi Pandya, Afzal Ansari, Nikha Trivedi, Naresh Chauhan, Summaiya Mullan, Amit gamit, Apurvasinh Puvar, Janvi Raval, Zarna Patel, R D Dixit, A M Kadri, Harsh Bakshi, Chaitanya Joshi, Madhvi Joshi |
| EPI_ISL_495183 | CSIR-Centre for Cellular and Molecular Biology | CSIR-Centre for Cellular and Molecular Biology | Namami Gaur, Sakshi Shambhavi, Lamuk Zaveri, Shagufta Khan, Nikhil Hajirnis, M Soujanya Reddy, Pratheusa Maccha, Tulasi Nagabandi, Purushotham Vodnala, Payel Mukherjee, Sofia Banu, Priya Singh, Onkar Kulkarni, Dhiviya Vedagiri, Divya Gupta, Vishal Sah, Santosh Kumar Kuncha, Krishnan Harinivas Harshan, Archana Bharadwaj Siva, Karthik Bharadwaj Tallapaka, Zeba Rizvi, Zuberwasim Sayyad, Kakade Aishwarya Arun, Amrutha H C, Ananga Ghosh, Rakesh K Mishra, Divya Tej Sowpati |
| EPI_ISL_495184 | CSIR-Centre for Cellular and Molecular Biology | CSIR-Centre for Cellular and Molecular Biology | Namami Gaur, Sakshi Shambhavi, Lamuk Zaveri, Shagufta Khan, Nikhil Hajirnis, M Soujanya Reddy, Pratheusa Maccha, Tulasi Nagabandi, Purushotham Vodnala, Payel Mukherjee, Sofia Banu, Priya Singh,Onkar Kulkarni, Dhiviya Vedagiri, Divya Gupta, Vishal Sah, Santosh Kumar Kuncha, Krishnan Harinivas Harshan, Archana Bharadwaj Siva, Karthik Bharadwaj Tallapaka, Zeba Rizvi, Zuberwasim Sayyad, Kakade Aishwarya Arun, Amrutha H C, Ananga Ghosh, Rakesh K Mishra, Divya Tej Sowpati |
| EPI_ISL_495193 | CSIR-Centre for Cellular and Molecular Biology | CSIR-Centre for Cellular and Molecular Biology | Namami Gaur, Sakshi Shambhavi, Lamuk Zaveri, Shagufta Khan, Nikhil Hajirnis, M Soujanya Reddy, Pratheusa Maccha, Tulasi Nagabandi, Purushotham Vodnala, Payel Mukherjee, Sofia Banu, Priya Singh, Onkar Kulkarni, Dhiviya Vedagiri, Divya Gupta, Vishal Sah, Santosh Kumar Kuncha, Krishnan Harinivas Harshan, Archana Bharadwaj Siva, Karthik Bharadwaj Tallapaka, Zeba Rizvi, Zuberwasim Sayyad, Kakade Aishwarya Arun, Amrutha H C, Ananga Ghosh, Rakesh K Mishra, Divya Tej Sowpati |
| EPI_ISL_495207 | CSIR-Centre for Cellular and Molecular Biology | CSIR-Centre for Cellular and Molecular Biology | Namami Gaur, Sakshi Shambhavi, Lamuk Zaveri, Shagufta Khan, Nikhil Hajirnis, M Soujanya Reddy, Pratheusa Maccha, Tulasi Nagabandi, Purushotham Vodnala, Payel Mukherjee, Sofia Banu, Priya Singh, Onkar Kulkarni, Dhiviya Vedagiri, Divya Gupta, Vishal Sah, Santosh Kumar Kuncha, Krishnan Harinivas Harshan, Archana Bharadwaj Siva, Karthik Bharadwaj Tallapaka,G. Aditya Kumar, Koushick Sivakumar, Rakesh K Mishra, Divya Tej Sowpati |
| EPI_ISL_495270 | CSIR-Centre for Cellular and Molecular Biology | CSIR-Centre for Cellular and Molecular Biology | Namami Gaur, Sakshi Shambhavi, Lamuk Zaveri, Shagufta Khan, Nikhil Hajirnis, M Soujanya Reddy, Pratheusa Maccha, Tulasi Nagabandi, Purushotham Vodnala, Payel Mukherjee, Sofia Banu, Priya Singh, Onkar Kulkarni, Dhiviya Vedagiri, Divya Gupta, Vishal Sah, Santosh Kumar Kuncha, Krishnan Harinivas Harshan, Archana Bharadwaj Siva, Karthik Bharadwaj Tallapaka, Zeba Rizvi, Zuberwasim Sayyad, Kakade Aishwarya Arun, Amrutha H C, Ananga Ghosh, Rakesh K Mishra, Divya Tej Sowpati |
| EPI_ISL_495411, EPI_ISL_495412, EPI_ISL_495413, EPI_ISL_495414, EPI_ISL_495415, EPI_ISL_495416, EPI_ISL_495417, EPI_ISL_495418, EPI_ISL_495419, EPI_ISL_495420, EPI_ISL_495421, EPI_ISL_495422, EPI_ISL_495423, EPI_ISL_495424, EPI_ISL_495425, EPI_ISL_495426, EPI_ISL_495427, EPI_ISL_495428, EPI_ISL_495429, EPI_ISL_495430, EPI_ISL_495431, EPI_ISL_495432, EPI_ISL_495433, EPI_ISL_495434, EPI_ISL_495435, EPI_ISL_495436, EPI_ISL_495437, EPI_ISL_495438, EPI_ISL_495439, EPI_ISL_495440, EPI_ISL_495441, EPI_ISL_495442, EPI_ISL_495443, EPI_ISL_495444, EPI_ISL_495445, EPI_ISL_495446, | | | |

| | | | |
|---|---|---|---|
| EPI_ISL_495447, EPI_ISL_495448, EPI_ISL_495449, EPI_ISL_495450, EPI_ISL_495451, EPI_ISL_495452, EPI_ISL_495453, EPI_ISL_495454, EPI_ISL_495455, EPI_ISL_495456, EPI_ISL_495457 | | | |
| see above | Kafkas University, Faculty of Medicine, Department of Medical Microbiology | Kafkas University, Faculty of Medicine, Department of Medical Microbiology | Murat Karamese, Didem Ozgur, E. Ediz Tutuncu |
| EPI_ISL_495597, EPI_ISL_495598, EPI_ISL_495599, EPI_ISL_495600, EPI_ISL_495601, EPI_ISL_495602, EPI_ISL_495603, EPI_ISL_495604, EPI_ISL_495605, EPI_ISL_495606, EPI_ISL_495607, EPI_ISL_495608, EPI_ISL_495609 | | | |
| see above | Mayo Clinic & Mayo Clinic Laboratories | Minnesota Department of Health, Public Health Laboratory | Matt Plumb, Jacob Garfin, and Xiong Wang |
| EPI_ISL_495610, EPI_ISL_495611, EPI_ISL_495612, EPI_ISL_495613, EPI_ISL_495614, EPI_ISL_495615, EPI_ISL_495616, EPI_ISL_495617, EPI_ISL_495618, EPI_ISL_495619, EPI_ISL_495620, EPI_ISL_495621, EPI_ISL_495622, EPI_ISL_495623, EPI_ISL_495624, EPI_ISL_495625, EPI_ISL_495626, EPI_ISL_495627, EPI_ISL_495628, EPI_ISL_496916, EPI_ISL_496917 | | | |
| see above | Minnesota Department of Health, Public Health Laboratory | Minnesota Department of Health, Public Health Laboratory | Matt Plumb, Jacob Garfin, and Xiong Wang |
| EPI_ISL_496918 | Mayo Clinic & Mayo Clinic Laboratories | Minnesota Department of Health, Public Health Laboratory | Matt Plumb, Jacob Garfin, and Xiong Wang |
| EPI_ISL_496919, EPI_ISL_496920, EPI_ISL_496921, EPI_ISL_496922, EPI_ISL_496923, EPI_ISL_496924, EPI_ISL_496925, EPI_ISL_496926, EPI_ISL_496927, EPI_ISL_496928, EPI_ISL_496929, EPI_ISL_496930 | | | |
| see above | Minnesota Department of Health, Public Health Laboratory | Minnesota Department of Health, Public Health Laboratory | Matt Plumb, Jacob Garfin, and Xiong Wang |
| EPI_ISL_498418, EPI_ISL_498419 | National Institute of Laboratory Medicine and Referral Center | Genomic Research Lab, BCSIR | Md. Maruf Ahmed Molla, Abu Sayeed Mohammad Mahmud, Mohammad Samir Uzzaman, Eshrar Osman, Md. Ahasan Habib, Shahina Akter, Tanjina Akhter Banu, Md. Murshed Hasan Sarkar, Barna Goswami, Iffat Jahan, Md. Saddam Hossain, Tasnim Nafisa, Mahmuda Yeasmin, Asish Kumar Ghosh, A. K. M. Shamsuzzaman, Sheikh Md. Selim Al Din, Utpal Chandra Ray, Salek Ahmed Sajib, Md. Salim Khan |
| EPI_ISL_498783, EPI_ISL_498784, EPI_ISL_498785, EPI_ISL_498786, EPI_ISL_498787, EPI_ISL_498788, EPI_ISL_498789, EPI_ISL_498790, EPI_ISL_498791, EPI_ISL_498792 | National Institute of Laboratory Medicine and Referral Center | Genomic Research Lab, BCSIR | Md. Saddam Hossain, Abu Sayeed Mohammad Mahmud, Mohammad Samir Uzzaman, Eshrar Osman, Md. Ahasan Habib, Shahina Akter, Tanjina Akhter Banu, Md. Murshed Hasan Sarkar, Barna Goswami, Iffat Jahan, Tasnim Nafisa, Md. Maruf Ahmed Molla, Mahmuda Yeasmin, Asish Kumar Ghosh, A. K. M. Shamsuzzaman, Sheikh Md. Selim Al Din, Utpal Chandra Ray, Salek Ahmed Sajib, Md. Salim Khan |
| EPI_ISL_498793, EPI_ISL_498794, EPI_ISL_498795, EPI_ISL_498796, EPI_ISL_498797, EPI_ISL_498800, EPI_ISL_498801, EPI_ISL_498802, EPI_ISL_498803, EPI_ISL_498804, EPI_ISL_498805, EPI_ISL_498806 | | | |
| see above | National Institute of Laboratory Medicine and Referral Center | Genomic Research Lab, BCSIR | Md. Murshed Hasan Sarkar, Abu Sayeed Mohammad Mahmud, Mohammad Samir Uzzaman, Eshrar Osman, Md. Ahasan Habib, Shahina Akter, Tanjina Akhter Banu, Barna Goswami, Iffat Jahan, Md. Saddam Hossain, Tasnim Nafisa, Md. Maruf Ahmed Molla, Mahmuda Yeasmin, Asish Kumar Ghosh, A. K. M. Shamsuzzaman, Sheikh Md. Selim Al Din, Utpal Chandra Ray, Salek Ahmed Sajib, Md. Salim Khan |
| EPI_ISL_500160, EPI_ISL_500161, EPI_ISL_500163, EPI_ISL_500164, EPI_ISL_500165, EPI_ISL_500168, EPI_ISL_500169, EPI_ISL_500170, EPI_ISL_500171, EPI_ISL_500172, EPI_ISL_500173, EPI_ISL_500174, EPI_ISL_500175, EPI_ISL_500176, EPI_ISL_500177, EPI_ISL_500178, EPI_ISL_500179, EPI_ISL_500180, EPI_ISL_500181, EPI_ISL_500182, EPI_ISL_500183, EPI_ISL_500184, EPI_ISL_500185, EPI_ISL_500186, EPI_ISL_500187, EPI_ISL_500188, EPI_ISL_500189, EPI_ISL_500190, EPI_ISL_500191, EPI_ISL_500192, EPI_ISL_500193, EPI_ISL_500194, EPI_ISL_500195, EPI_ISL_500196, EPI_ISL_500197, EPI_ISL_500198, EPI_ISL_500199, EPI_ISL_500200, EPI_ISL_500201, EPI_ISL_500202, EPI_ISL_500203, EPI_ISL_500204, EPI_ISL_500205, EPI_ISL_500206, EPI_ISL_500208, EPI_ISL_500210, EPI_ISL_500211, EPI_ISL_500212, EPI_ISL_500213, EPI_ISL_500214, EPI_ISL_500215, EPI_ISL_500216, EPI_ISL_500217, EPI_ISL_500220, EPI_ISL_500222, EPI_ISL_500223, EPI_ISL_500224, EPI_ISL_500225, EPI_ISL_500226, EPI_ISL_500227 | | | |
| see above | Hospital Universitario Virgen de las Nieves de Granada-SAS | SeqCOVID-SPAIN consortium/IBV(CSIC) | Mercedes Pérez Ruiz, Sara Sanbonmatsu Gámez, Irene Pedrosa Corral, José M. Navarro-Marí and SeqCOVID-SPAIN consortium |
| EPI_ISL_500369, EPI_ISL_500370, EPI_ISL_500371, EPI_ISL_500372, EPI_ISL_500374, EPI_ISL_500375, EPI_ISL_500376, EPI_ISL_500377, EPI_ISL_500378, EPI_ISL_500379, EPI_ISL_500380, EPI_ISL_500381, EPI_ISL_500383, EPI_ISL_500385, EPI_ISL_500386, EPI_ISL_500387, EPI_ISL_500388, EPI_ISL_500389, EPI_ISL_500390, EPI_ISL_500391, EPI_ISL_500392, EPI_ISL_500393, EPI_ISL_500394, EPI_ISL_500395, EPI_ISL_500396, EPI_ISL_500397, EPI_ISL_500398, EPI_ISL_500399, EPI_ISL_500400, EPI_ISL_500401, EPI_ISL_500402, EPI_ISL_500403, EPI_ISL_500404, EPI_ISL_500405, EPI_ISL_500406, EPI_ISL_500407, EPI_ISL_500408, EPI_ISL_500409, EPI_ISL_500410, EPI_ISL_500411, EPI_ISL_500412, EPI_ISL_500413, EPI_ISL_500414, EPI_ISL_500415, EPI_ISL_500416, EPI_ISL_500417, EPI_ISL_500418, EPI_ISL_500419, EPI_ISL_500420, EPI_ISL_500421, EPI_ISL_500422, EPI_ISL_500423, EPI_ISL_500424, EPI_ISL_500425, EPI_ISL_500426, EPI_ISL_500427, EPI_ISL_500428, EPI_ISL_500429, EPI_ISL_500430, EPI_ISL_500431, EPI_ISL_500432, EPI_ISL_500433, EPI_ISL_500434, EPI_ISL_500435, EPI_ISL_500436, EPI_ISL_500437, EPI_ISL_500438, EPI_ISL_500439, EPI_ISL_500440, EPI_ISL_500441, EPI_ISL_500442, EPI_ISL_500443, EPI_ISL_500444, EPI_ISL_500445, EPI_ISL_500446, EPI_ISL_500447, EPI_ISL_500448, EPI_ISL_500449, EPI_ISL_500450, EPI_ISL_500451, EPI_ISL_500452, EPI_ISL_500453, EPI_ISL_500454, EPI_ISL_500455, EPI_ISL_500456, EPI_ISL_500457, EPI_ISL_500458 | | | |
| see above | Centro de Investigación Biomédica de La Rioja - Hospital San Pedro Logroño | SeqCOVID-SPAIN consortium/IBV(CSIC) | María de Toro, José Manuel Azcona Gutiérrez, María Pilar Bea Escudero, Miriam Blasco Alberdi and SeqCOVID-SPAIN consortium |
| EPI_ISL_500573, EPI_ISL_500574, EPI_ISL_500575, EPI_ISL_500576, EPI_ISL_500577, EPI_ISL_500578, EPI_ISL_500579, EPI_ISL_500580, EPI_ISL_500581, EPI_ISL_500582, EPI_ISL_500583, EPI_ISL_500584, EPI_ISL_500585, EPI_ISL_500586, EPI_ISL_500587, EPI_ISL_500588, EPI_ISL_500589, EPI_ISL_500590, EPI_ISL_500591, EPI_ISL_500592, EPI_ISL_500593, EPI_ISL_500594, EPI_ISL_500595 | | | |
| see above | National Virus Reference Laboratory | National Virus Reference Laboratory | Michael Carr, Gabriel Gonzalez, Jonathan Dean, Suzie Coughlan, Cillian F De Gascun |
| EPI_ISL_500716 | BSL3 Lab Pendik Veterinary Control Institute | Department of Medicinal Genetics, Bursa Uluda University, Faculty of medicine By Sehime Gülsün Temel, Adem Alemdar, Kadir Yeilba | Mustafa HASOKSUZ, Fahriye SARAC, Osman ERGANIS, Serdar UZAR, Hakan ENUL, Cumhur ADIAY, Ahmet SAIT, Orbay SAYI, Kadir YESILBAG, Oguz KARABEY |
| EPI_ISL_501252, EPI_ISL_501253, EPI_ISL_501254, EPI_ISL_501255, EPI_ISL_501256, EPI_ISL_501257, EPI_ISL_501258, EPI_ISL_501259, EPI_ISL_501260, EPI_ISL_501261, EPI_ISL_501262, EPI_ISL_501263, EPI_ISL_501264, EPI_ISL_501265, EPI_ISL_501266, EPI_ISL_501267, EPI_ISL_501268, EPI_ISL_501269, EPI_ISL_501270, EPI_ISL_501271, EPI_ISL_501272, EPI_ISL_501273, EPI_ISL_501274 | | | |
| see above | National Virus Reference Laboratory | National Virus Reference Laboratory | Michael Carr, Gabriel Gonzalez, Jonathan Dean, Suzie Coughlan, Cillian F De Gascun |
| EPI_ISL_501632 | Virology Department, Royal Infirmary of Edinburgh, NHS Lothian / School of Biological Sciences, University of Edinburgh | Wellcome Sanger Institute for the COVID-19 Genomics UK (COG-UK) consortium | McHugh M, Dewar R, Rooke S, O'Toole Á, Scher E, Hill V, McCrone JT, Colquhoun R, Yu X, Jackson B, Rambaut A, Templeton K and Alex Alderton, Roberto Amato, Sonia Goncalves, Ewan Harrison, David K. Jackson, Ian Johnston, Dominic Kwiatkowski, Cordelia Langford, John Sillitoe on behalf of the Wellcome Sanger Institute COVID-19 Surveillance Team (http://www.sanger.ac.uk/covid-team) |
| EPI_ISL_503958, EPI_ISL_504040 | National Institute of Laboratory Medicine and Referral Center | Genomic Research Lab, BCSIR | Md. Ahasan Habib, Abu Sayeed Mohammad Mahmud, Mohammad Samir Uzzaman, Eshrar Osman, Shahina Akter, Tanjina Akhter Banu, Md. Murshed Hasan Sarkar, Barna Goswami, Iffat Jahan, Md. Saddam Hossain, Tarannum Taznin, Tasnim Nafisa, Md. Maruf Ahmed Molla, Mahmuda Yeasmin, Asish Kumar Ghosh, A. K. M. Shamsuzzaman, Sheikh Md. Selim Al Din, Utpal Chandra Ray, Salek Ahmed Sajib, Md. Salim Khan |
| EPI_ISL_507180, EPI_ISL_507181, EPI_ISL_507182, EPI_ISL_507183, EPI_ISL_507184, EPI_ISL_507185, EPI_ISL_507186, EPI_ISL_507187, EPI_ISL_507188, EPI_ISL_507189, EPI_ISL_507190, EPI_ISL_507191, EPI_ISL_507192, EPI_ISL_507193, EPI_ISL_507194, EPI_ISL_507195, EPI_ISL_507196, EPI_ISL_507197, EPI_ISL_507198, EPI_ISL_507199, EPI_ISL_507200, EPI_ISL_507201 | | | |
| see above | Virology Department, Royal Infirmary of Edinburgh, NHS Lothian / School of Biological Sciences, University of Edinburgh / Institute of Genetics and Molecular Medicine, University of Edinburgh | COVID-19 Genomics UK (COG-UK) Consortium | McHugh M, Dewar R, Rooke S, Gallagher M, Balcaza C, O'Toole Á, Scher E, Hill V, McCrone JT, Colquhoun R, Yu X, Jackson B, Rambaut A, Williams TC, Templeton K |
| EPI_ISL_507934, EPI_ISL_507935, EPI_ISL_507936, EPI_ISL_507937, EPI_ISL_507938, EPI_ISL_507939, EPI_ISL_507940, EPI_ISL_507941, EPI_ISL_507942, EPI_ISL_507943, EPI_ISL_507944, EPI_ISL_507945, EPI_ISL_507946, EPI_ISL_507947, EPI_ISL_507948, EPI_ISL_507949, EPI_ISL_507950, EPI_ISL_507951, EPI_ISL_507952, EPI_ISL_507953, EPI_ISL_507954, EPI_ISL_507955, EPI_ISL_507956 | | | |
| see above | Minnesota Department of Health, Public Health Laboratory | Minnesota Department of Health, Public Health Laboratory | Matt Plumb, Jacob Garfin, and Xiong Wang |
| EPI_ISL_507957, EPI_ISL_507958, EPI_ISL_507959, EPI_ISL_507960, EPI_ISL_507961 | Mayo Clinic & Mayo Clinic Laboratories | Minnesota Department of Health, Public Health Laboratory | Matt Plumb, Jacob Garfin, and Xiong Wang |
| EPI_ISL_507962 | Children's Hospitals and Clinics of Minnesota | Minnesota Department of Health, Public Health Laboratory | Matt Plumb, Jacob Garfin, and Xiong Wang |
| EPI_ISL_507963, EPI_ISL_507964, EPI_ISL_507965, EPI_ISL_507966, EPI_ISL_507967, EPI_ISL_507968, EPI_ISL_507969, EPI_ISL_507970, EPI_ISL_507971, EPI_ISL_507972 | Avera Mckennan Laboratory | Minnesota Department of Health, Public Health Laboratory | Matt Plumb, Jacob Garfin, and Xiong Wang |
| EPI_ISL_507973, EPI_ISL_507974, EPI_ISL_507975, EPI_ISL_507976, EPI_ISL_507977, EPI_ISL_507978, EPI_ISL_507979, EPI_ISL_507980, EPI_ISL_507981, EPI_ISL_507982, EPI_ISL_507983 | | | |
| see above | Minnesota Department of Health, Public Health Laboratory | Minnesota Department of Health, Public Health Laboratory | Matt Plumb, Jacob Garfin, and Xiong Wang |
| EPI_ISL_508865, EPI_ISL_508867, | Division of Infectious Diseases and Hospital Epidemiology, | Institute of Medical Virology, University of Zurich | Maryam Zaheri, Verena Kufner, Dana Weissberg, Jürg Böni, Silvana K. Rampini, Peter W. Schreiber, Irene A. Abela, Hugo Sax, Aline Wolfensberger, |

| | | | |
|---|---|---|---|
| EPI_ISL_508869, EPI_ISL_509222 | University Hospital Zürich | | Michael Huber |
| EPI_ISL_509686 | M Health Fairview | Minnesota Department of Health, Public Health Laboratory | Matt Plumb, Jacob Garfin, and Xiong Wang |
| EPI_ISL_510424, EPI_ISL_510425, EPI_ISL_510426, EPI_ISL_510427, EPI_ISL_510428, EPI_ISL_510429, EPI_ISL_510430, EPI_ISL_510431, EPI_ISL_510432, EPI_ISL_510433, EPI_ISL_510434, EPI_ISL_510435, EPI_ISL_510436, EPI_ISL_510437, EPI_ISL_510438, EPI_ISL_510439, EPI_ISL_510440, EPI_ISL_510441, EPI_ISL_510442, EPI_ISL_510443, EPI_ISL_510444, EPI_ISL_510445, EPI_ISL_510446, EPI_ISL_510447, EPI_ISL_510448, EPI_ISL_510449, EPI_ISL_510450, EPI_ISL_510451, EPI_ISL_510452, EPI_ISL_510453, EPI_ISL_510454, EPI_ISL_510455, EPI_ISL_510456, EPI_ISL_510457, EPI_ISL_510458, EPI_ISL_510459, EPI_ISL_510460 | | | |
| see above | Hospital Universitario Virgen de las Nieves de Granada-SAS | SeqCOVID-SPAIN consortium/IBV(CSIC) | Mercedes Pérez Ruiz, Sara Sanbonmatsu Gámez, Irene Pedrosa Corral, José M. Navarro-Marí and SeqCOVID-SPAIN consortium |
| EPI_ISL_511879 | Laboratorium Kesehatan Provinsi Jawa Barat | Molecular Genetics Laboratory-Faculty of Medicine-Universitas Padjadjaran; School of Life Sciences and Technology & School of Pharmacy-Institut Teknologi Bandung; Laboratorium Kesehatan Provinsi Jawa Barat | Marselina Irasonia Tan, Yunia Sribudiani, Catur Riani, Azzania Fibriani, Husna Nugrahapraja, Tarwadi, Ema Rahmawati, Hesti Lina Wiraswati, Lia Faridah, Savira Ekawardhani, Ryan Bayusantika Ristandi, Rifky Waluyajati Rachman, Cut Nur Cinthia Alamanda, Hammam Riza, Soni Solistia Wirawan, Agung Eru Wibowo |
| EPI_ISL_512058 | B.J. Medical College and Civil hospital, Ahmedabad | Gujarat Biotechnology Research Centre | Monika Gandhi, Pinal Trivedi, Maharshi Pandya, Nidhi Patel, Nitin Savaliya, Raghawendra Kumar, Dinesh Kumar, Zuber Saiyed, Komal Patel, Labdhi Pandya, Afzal Ansari, Nikha Trivedi, Pranay Shah, Kamlesh J Upadhyay, Sanjay Kapadia, Apurvasinh Puvar, Janvi Raval, Zarna Patel, R D Dixit, A M Kadri, Harsh Bakshi, Chaitanya Joshi, Madhvi Joshi |
| EPI_ISL_512075 | Dr. RSS Hospital, Modasa | Gujarat Biotechnology Research Centre | Monika Gandhi, Pinal Trivedi, Maharshi Pandya, Nidhi Patel, Nitin Savaliya, Raghawendra Kumar, Dinesh Kumar, Zuber Saiyed, Komal Patel, Labdhi Pandya, Afzal Ansari, Nikha Trivedi, Harsh Chaudhari, Apurvasinh Puvar, Janvi Raval, Zarna Patel, R D Dixit, A M Kadri, Harsh Bakshi, Chaitanya Joshi, Madhvi Joshi |
| EPI_ISL_512087, EPI_ISL_512088, EPI_ISL_512089, EPI_ISL_512090, EPI_ISL_512091, EPI_ISL_512092, EPI_ISL_512093, EPI_ISL_512094, EPI_ISL_512095, EPI_ISL_512096, EPI_ISL_512097, EPI_ISL_512098, EPI_ISL_512099, EPI_ISL_512100, EPI_ISL_512101, EPI_ISL_512102, EPI_ISL_512103, EPI_ISL_512104, EPI_ISL_512105, EPI_ISL_512106 | | | |
| see above | National Virus Reference Laboratory | National Virus Reference Laboratory | Michael Carr, Gabriel Gonzalez, Jonathan Dean, Aditi Chaturvedi, Suzie Coughlan, Cillian F De Gascun |
| EPI_ISL_512844 | Department of Medical Research | DMR_Myanmar | Myat Htut Nyunt, Hnin Ohnmar Soe, Kay Thi Aye, Wah Wah Aung,Yi Yi Kyaw, Aung Kyaw Kyaw, Theingi Win Myat, Phyu Win Ei, Aung Zaw Latt, Nan Aye Thida Oo, Lai Lai San, Su Mon Win, Ni Ni Zaw, Htin Lin, Hlaing Myat Thu, Zaw Than Htun |
| EPI_ISL_513314 | The National Institute of Public Health | The National Institute of Public Health and State Veterinary Institute Prague | Nagy,A;Jirincova,H;Novakova,L;Trnka,D;Vecerova,J |
| EPI_ISL_513414, EPI_ISL_513415, EPI_ISL_513416, EPI_ISL_513417, EPI_ISL_513418, EPI_ISL_513419, EPI_ISL_513420, EPI_ISL_513421, EPI_ISL_513422, EPI_ISL_513423, EPI_ISL_513424, EPI_ISL_513425, EPI_ISL_513426, EPI_ISL_513427, EPI_ISL_513428, EPI_ISL_513429, EPI_ISL_513430, EPI_ISL_513431, EPI_ISL_513432, EPI_ISL_513433, EPI_ISL_513434, EPI_ISL_513435, EPI_ISL_513436, EPI_ISL_513437, EPI_ISL_513438, EPI_ISL_513439, EPI_ISL_513440, EPI_ISL_513441, EPI_ISL_513442, EPI_ISL_513443, EPI_ISL_513444, EPI_ISL_513445, EPI_ISL_513446, EPI_ISL_513447, EPI_ISL_513448, EPI_ISL_513449, EPI_ISL_513450, EPI_ISL_513451, EPI_ISL_513452, EPI_ISL_513453, EPI_ISL_513454, EPI_ISL_513455, EPI_ISL_513456, EPI_ISL_513457, EPI_ISL_513458, EPI_ISL_513459, EPI_ISL_513460, EPI_ISL_513461, EPI_ISL_513462, EPI_ISL_513463, EPI_ISL_513464, EPI_ISL_513465, EPI_ISL_513466, EPI_ISL_513467, EPI_ISL_513468, EPI_ISL_513469, EPI_ISL_513470, EPI_ISL_513471, EPI_ISL_513472, EPI_ISL_513473, EPI_ISL_513474, EPI_ISL_513475, EPI_ISL_513476, EPI_ISL_513477, EPI_ISL_513478, EPI_ISL_513479, EPI_ISL_513480, EPI_ISL_513481, EPI_ISL_513482, EPI_ISL_513483, EPI_ISL_513484, EPI_ISL_513485, EPI_ISL_513486, EPI_ISL_513487, EPI_ISL_513488, EPI_ISL_513489, EPI_ISL_513490, EPI_ISL_513491, EPI_ISL_513492, EPI_ISL_513493, EPI_ISL_513494, EPI_ISL_513495, EPI_ISL_513496, EPI_ISL_513497, EPI_ISL_513498, EPI_ISL_513499, EPI_ISL_513500, EPI_ISL_513501, EPI_ISL_513502, EPI_ISL_513503, EPI_ISL_513504, EPI_ISL_513505, EPI_ISL_513506, EPI_ISL_513507, EPI_ISL_513508, EPI_ISL_513509, EPI_ISL_513510 | | | |
| see above | Maine HETL | Tewhey Lab, The Jackson Laboratory | Matluk,N., Dewey,H., Barter,M., Lynch,R., Munger,H. and Tewhey,R. |
| EPI_ISL_513511, EPI_ISL_513512 | The National Institute of Public Health | The National Institute of Public Health and State Veterinary Institute Prague | Nagy,A;Jirincova,H;Novakova,L;Trnka,D;Vecerova,J |
| EPI_ISL_514129, EPI_ISL_514130 | National Institute of Laboratory Medicine and Referral Center | Genomic Research Lab, BCSIR | Md. Murshed Hasan Sarkar, Abu Sayeed Mohammad Mahmud, Mohammad Samir Uzzaman, Eshrar Osman, Md. Ahashan Habib, Shahina Akter, Tanjina Akhter Banu, Barna Goswami, Iffat Jahan, Md. Saddam Hossain, Tasnim Nafisa, Md. Maruf Ahmed Molla, Mahmuda Yeasmin, Asish Kumar Ghosh, A. K. M. Shamsuzzaman, Sheikh Md. Selim Al Din, Utpal Chandra Ray, Salek Ahmed Sajib, Md. Salim Khan |
| EPI_ISL_514233, EPI_ISL_514234, EPI_ISL_514235, EPI_ISL_514236 | National Institute of Laboratory Medicine and Referral Center | Genomic Research Lab, BCSIR | Md. Murshed Hasan Sarkar, Abu Sayeed Mohammad Mahmud, Mohammad Samir Uzzaman, Eshrar Osman, Md. Ahashan Habib, Shahina Akter, Tanjina Akhter Banu, Barna Goswami, Iffat Jahan, Md. Saddam Hossain, Tasnim Nafisa, Md. Maruf Ahmed Molla, Mahmuda Yeasmin, Asish Kumar Ghosh, A. K. M. Shamsuzzaman, Sheikh Md. Selim Al Din, Utpal Chandra Ray, Salek Ahmed Sajib, Md. Salim Khan |
| EPI_ISL_514237, EPI_ISL_514238, EPI_ISL_514239, EPI_ISL_514240, EPI_ISL_514241 | National Institute of Laboratory Medicine and Referral Center | Genomic Research Lab, BCSIR | Md. Saddam Hossain, Abu Sayeed Mohammad Mahmud, Mohammad Samir Uzzaman, Eshrar Osman, Md. Ahashan Habib, Shahina Akter, Tanjina Akhter Banu, Md. Murshed Hasan Sarkar, Barna Goswami, Iffat Jahan, Tasnim Nafisa, Md. Maruf Ahmed Molla, Mahmuda Yeasmin, Asish Kumar Ghosh, A. K. M. Shamsuzzaman, Sheikh Md. Selim Al Din, Utpal Chandra Ray, Salek Ahmed Sajib, Md. Salim Khan |
| EPI_ISL_514438, EPI_ISL_514439 | Tewhey Lab, The Jackson Laboratory | Tewhey Lab, The Jackson Laboratory | Matluk,N., Dewey,H., Barter,M., Lynch,R., Munger,H., Tewhey,R. |
| EPI_ISL_514516, EPI_ISL_514517, EPI_ISL_514518, EPI_ISL_514519, EPI_ISL_514520, EPI_ISL_514521, EPI_ISL_514522, EPI_ISL_514523, EPI_ISL_514524, EPI_ISL_514525, EPI_ISL_514526, EPI_ISL_514527, EPI_ISL_514528, EPI_ISL_514529, EPI_ISL_514530, EPI_ISL_514531, EPI_ISL_514532, EPI_ISL_514533, EPI_ISL_514534, EPI_ISL_514535, EPI_ISL_514536, EPI_ISL_514537, EPI_ISL_514538, EPI_ISL_514539, EPI_ISL_514540, EPI_ISL_514541, EPI_ISL_514542, EPI_ISL_514543, EPI_ISL_514544, EPI_ISL_514545, EPI_ISL_514546, EPI_ISL_514547, EPI_ISL_514548, EPI_ISL_514549, EPI_ISL_514550, EPI_ISL_514551, EPI_ISL_514552, EPI_ISL_514553, EPI_ISL_514554, EPI_ISL_514555, EPI_ISL_514556, EPI_ISL_514557, EPI_ISL_514558, EPI_ISL_514559, EPI_ISL_514560, EPI_ISL_514561, EPI_ISL_514562 | | | |
| see above | Virology Department, Royal Infirmary of Edinburgh, NHS Lothian / School of Biological Sciences, University of Edinburgh / Institute of Genetics and Molecular Medicine, University of Edinburgh | COVID-19 Genomics UK (COG-UK) Consortium | McHugh M, Dewar R, Rooke S, Gallagher M, Balcaza C, O'Toole Á, Scher E, Hill V, McCrone JT, Colquhoun R, Yu X, Jackson B, Rambaut A, Williams TC, Templeton K |
| EPI_ISL_514595 | B.J. Medical College and Civil hospital, Ahmedabad | Gujarat Biotechnology Research Centre | Monika Gandhi, Pinal Trivedi, Maharshi Pandya, Nidhi Patel, Nitin Savaliya, Raghawendra Kumar, Dinesh Kumar, Zuber Saiyed, Komal Patel, Labdhi Pandya, Afzal Ansari, Nikha Trivedi, Pranay Shah, Kamlesh J Upadhyay, Sanjay Kapadia, Apurvasinh Puvar, Janvi Raval, Zarna Patel, R D Dixit, A M Kadri, Harsh Bakshi, Chaitanya Joshi, Madhvi Joshi |
| EPI_ISL_514610 | GMERS Medical College & Hospital, Gotri, Vadodara | Gujarat Biotechnology Research Centre | Meenakshi Shah, Neena Doshi, Varsha Godbole, Apurvasinh Puvar, Janvi Raval, Zarna Patel, Monika Gandhi, Pinal Trivedi, Maharshi Pandya, Nidhi Patel, Nitin Savaliya, Raghawendra Kumar, Dinesh Kumar, Zuber Saiyed, Komal Patel, Labdhi Pandya, Afzal Ansari, Nikha Trivedi, Pranay Shah, Kamlesh J Upadhyay, Sanjay Kapadia, R D Dixit, A M Kadri, Harsh Bakshi, Chaitanya Joshi, Madhvi Joshi, |
| EPI_ISL_514616, EPI_ISL_514617, EPI_ISL_514618 | Minnesota Department of Health, Public Health Laboratory | Minnesota Department of Health, Public Health Laboratory | Matt Plumb, Jacob Garfin, and Xiong Wang |
| EPI_ISL_514619, EPI_ISL_514620, EPI_ISL_514621, EPI_ISL_514622, EPI_ISL_514623, EPI_ISL_514624, EPI_ISL_514625, EPI_ISL_514626, EPI_ISL_514627, EPI_ISL_514628 | Mayo Clinic & Mayo Clinic Laboratories | Minnesota Department of Health, Public Health Laboratory | Matt Plumb, Jacob Garfin, and Xiong Wang |
| EPI_ISL_514629, EPI_ISL_514630, EPI_ISL_514631, EPI_ISL_514632, EPI_ISL_514633, EPI_ISL_514634, EPI_ISL_514635 | M Health Fairview | Minnesota Department of Health, Public Health Laboratory | Matt Plumb, Jacob Garfin, and Xiong Wang |
| EPI_ISL_514636 | Mayo Clinic & Mayo Clinic Laboratories | Minnesota Department of Health, Public Health Laboratory | Matt Plumb, Jacob Garfin, and Xiong Wang |
| EPI_ISL_514637 | M Health Fairview St. Joseph's Hospital | Minnesota Department of Health, Public Health Laboratory | Matt Plumb, Jacob Garfin, and Xiong Wang |
| EPI_ISL_514638, EPI_ISL_514639, EPI_ISL_514640 | Mayo Clinic & Mayo Clinic Laboratories | Minnesota Department of Health, Public Health Laboratory | Matt Plumb, Jacob Garfin, and Xiong Wang |
| EPI_ISL_514641 | St. Luke's Hospital | Minnesota Department of Health, Public Health Laboratory | Matt Plumb, Jacob Garfin, and Xiong Wang |
| EPI_ISL_514642 | Essentia Health-St. Mary's Medical Center | Minnesota Department of Health, Public Health Laboratory | Matt Plumb, Jacob Garfin, and Xiong Wang |
| EPI_ISL_514643 | M Health Fairview St. Joseph's Hospital | Minnesota Department of Health, Public Health Laboratory | Matt Plumb, Jacob Garfin, and Xiong Wang |

| | | | |
|---|---|---|---|
| EPI_ISL_514644 | Mayo Clinic & Mayo Clinic Laboratories | Minnesota Department of Health, Public Health Laboratory | Matt Plumb, Jacob Garfin, and Xiong Wang |
| EPI_ISL_514645 | Avera Mckennan Laboratory | Minnesota Department of Health, Public Health Laboratory | Matt Plumb, Jacob Garfin, and Xiong Wang |
| EPI_ISL_514646 | St. Luke's Hospital | Minnesota Department of Health, Public Health Laboratory | Matt Plumb, Jacob Garfin, and Xiong Wang |
| EPI_ISL_514647 | Allina Health Laboratory | Minnesota Department of Health, Public Health Laboratory | Matt Plumb, Jacob Garfin, and Xiong Wang |
| EPI_ISL_514648 | Essentia Health-St. Joseph's Medical Center | Minnesota Department of Health, Public Health Laboratory | Matt Plumb, Jacob Garfin, and Xiong Wang |
| EPI_ISL_514649 | Allina Health Laboratory | Minnesota Department of Health, Public Health Laboratory | Matt Plumb, Jacob Garfin, and Xiong Wang |
| EPI_ISL_514650 | M Health Fairview St. Joseph's Hospital | Minnesota Department of Health, Public Health Laboratory | Matt Plumb, Jacob Garfin, and Xiong Wang |
| EPI_ISL_514651 | Allina Health Laboratory | Minnesota Department of Health, Public Health Laboratory | Matt Plumb, Jacob Garfin, and Xiong Wang |
| EPI_ISL_514652 | M Health Fairview St. Joseph's Hospital | Minnesota Department of Health, Public Health Laboratory | Matt Plumb, Jacob Garfin, and Xiong Wang |
| EPI_ISL_514653, EPI_ISL_514654, EPI_ISL_514655, EPI_ISL_514656 | Mayo Clinic & Mayo Clinic Laboratories | Minnesota Department of Health, Public Health Laboratory | Matt Plumb, Jacob Garfin, and Xiong Wang |
| EPI_ISL_514657 | M Health Fairview St. Joseph's Hospital | Minnesota Department of Health, Public Health Laboratory | Matt Plumb, Jacob Garfin, and Xiong Wang |
| EPI_ISL_514658, EPI_ISL_514659, EPI_ISL_514660, EPI_ISL_514661, EPI_ISL_514662, EPI_ISL_514663, EPI_ISL_514664, EPI_ISL_514665, EPI_ISL_514666, EPI_ISL_514667, EPI_ISL_514668, EPI_ISL_514669, EPI_ISL_514670, EPI_ISL_514671, EPI_ISL_514672 | | | |
| see above | Minnesota Department of Health, Public Health Laboratory | Minnesota Department of Health, Public Health Laboratory | Matt Plumb, Jacob Garfin, and Xiong Wang |
| EPI_ISL_515197, EPI_ISL_515198, EPI_ISL_515199, EPI_ISL_515200, EPI_ISL_515201, EPI_ISL_515202, EPI_ISL_515203, EPI_ISL_515204, EPI_ISL_515205, EPI_ISL_515206, EPI_ISL_515207, EPI_ISL_515208, EPI_ISL_515209, EPI_ISL_515210, EPI_ISL_515211, EPI_ISL_515212, EPI_ISL_515213, EPI_ISL_515214, EPI_ISL_515215, EPI_ISL_515216, EPI_ISL_515217, EPI_ISL_515218, EPI_ISL_515219, EPI_ISL_515220, EPI_ISL_515221, EPI_ISL_515222, EPI_ISL_515223, EPI_ISL_515224, EPI_ISL_515225, EPI_ISL_515226, EPI_ISL_515227, EPI_ISL_515228, EPI_ISL_515229, EPI_ISL_515230, EPI_ISL_515231, EPI_ISL_515232, EPI_ISL_515233, EPI_ISL_515234, EPI_ISL_515235, EPI_ISL_515236, EPI_ISL_515237, EPI_ISL_515238, EPI_ISL_515239, EPI_ISL_515240, EPI_ISL_515241, EPI_ISL_515242, EPI_ISL_515243, EPI_ISL_515244, EPI_ISL_515245, EPI_ISL_515246 | | | |
| see above | Laboratoire de microbiologie, Hopital de Verdun | Smith Laboratory, Centre de Recherche CHU Sainte-Justine | Martin Smith, Marieke Rozendaal, Ivan Pavlov |
| EPI_ISL_515262 | Minnesota Department of Health, Public Health Laboratory | Minnesota Department of Health, Public Health Laboratory | Matt Plumb, Jacob Garfin, and Xiong Wang |
| EPI_ISL_515263, EPI_ISL_515264, EPI_ISL_515265, EPI_ISL_515266 | Mayo Clinic & Mayo Clinic Laboratories | Minnesota Department of Health, Public Health Laboratory | Matt Plumb, Jacob Garfin, and Xiong Wang |
| EPI_ISL_515267, EPI_ISL_515268 | M Health Fairview | Minnesota Department of Health, Public Health Laboratory | Matt Plumb, Jacob Garfin, and Xiong Wang |
| EPI_ISL_515269 | Minnesota Department of Health, Public Health Laboratory | Minnesota Department of Health, Public Health Laboratory | Matt Plumb, Jacob Garfin, and Xiong Wang |
| EPI_ISL_517583, EPI_ISL_517584, EPI_ISL_517585, EPI_ISL_517586, EPI_ISL_517587, EPI_ISL_517588, EPI_ISL_517589 | Virology Department, Royal Infirmary of Edinburgh, NHS Lothian / School of Biological Sciences, University of Edinburgh / Institute of Genetics and Molecular Medicine, University of Edinburgh | COVID-19 Genomics UK (COG-UK) Consortium | McHugh M, Dewar R, Rooke S, Gallagher M, Balcaza C, O'Toole Á, Scher E, Hill V, McCrone JT, Colquhoun R, Yu X, Jackson B, Rambaut A, Williams TC, Templeton K |
| EPI_ISL_518855 | Minnesota Department of Health, Public Health Laboratory | Minnesota Department of Health, Public Health Laboratory | Matt Plumb, Jacob Garfin, and Xiong Wang |
| EPI_ISL_518856, EPI_ISL_518857, EPI_ISL_518858, EPI_ISL_518859, EPI_ISL_518860, EPI_ISL_518861, EPI_ISL_518862, EPI_ISL_518863, EPI_ISL_518864, EPI_ISL_518865, EPI_ISL_518866, EPI_ISL_518867, EPI_ISL_518868, EPI_ISL_518869, EPI_ISL_518870, EPI_ISL_518871, EPI_ISL_518872, EPI_ISL_518873, EPI_ISL_518874, EPI_ISL_518875, EPI_ISL_518876, EPI_ISL_518877, EPI_ISL_518878, EPI_ISL_518879, EPI_ISL_518880, EPI_ISL_518881, EPI_ISL_518882, EPI_ISL_518883, EPI_ISL_518884, EPI_ISL_518885, EPI_ISL_518886, EPI_ISL_518887, EPI_ISL_518888, EPI_ISL_518889, EPI_ISL_518890, EPI_ISL_518891, EPI_ISL_518892, EPI_ISL_518893, EPI_ISL_518894, EPI_ISL_518895, EPI_ISL_518896, EPI_ISL_518897, EPI_ISL_518898, EPI_ISL_518899, EPI_ISL_518900, EPI_ISL_521153 | | | |
| see above | Mayo Clinic & Mayo Clinic Laboratories | Minnesota Department of Health, Public Health Laboratory | Matt Plumb, Jacob Garfin, and Xiong Wang |
| EPI_ISL_521860, EPI_ISL_521861, EPI_ISL_521862 | Royal Darwin Hospital Pathology | MDU-PHL | Meumann, E., Caly L., Seemann T., Sait, M., Schultz M., Druce J., Sherry, N. |
| EPI_ISL_522824, EPI_ISL_522825, EPI_ISL_522826, EPI_ISL_522827, EPI_ISL_522828, EPI_ISL_522829, EPI_ISL_522830, EPI_ISL_522831, EPI_ISL_522832, EPI_ISL_522833, EPI_ISL_522834, EPI_ISL_522835, EPI_ISL_522836, EPI_ISL_522837, EPI_ISL_522838, EPI_ISL_522839, EPI_ISL_522840, EPI_ISL_522841, EPI_ISL_522842, EPI_ISL_522843, EPI_ISL_522844, EPI_ISL_522845, EPI_ISL_522846, EPI_ISL_522847, EPI_ISL_522848, EPI_ISL_522849, EPI_ISL_522850, EPI_ISL_522851, EPI_ISL_522852, EPI_ISL_522853, EPI_ISL_522854, EPI_ISL_522869, EPI_ISL_522870, EPI_ISL_522871 | | | |
| see above | Maryland Department of Health | Maryland Department of Health | Maryland Department of Health Laboratories Administration |
| EPI_ISL_523927, EPI_ISL_523928, EPI_ISL_523929 | Center of Medical Microbiology, Virology, and Hospital Hygiene, University of Duesseldorf | Center of Medical Microbiology, Virology, and Hospital Hygiene, University of Duesseldorf | Maximilian Damagnez, Alexander Dilthey, Torsten Houwaart, Malte Kohns Vasconcelos, Marek Korencak, Jessica Nicolai, Klaus Pfeffer, Hendrik Streeck, Daniel Strelow, Jörg Timm, Andreas Walker, Tobias Wienemann |
| EPI_ISL_523930 | Center of Medical Microbiology, Virology, and Hospital Hygiene, University of Duesseldorf | Universitätstr.1 40225 Düsseldorf Germany | Maximilian Damagnez, Alexander Dilthey, Torsten Houwaart, Malte Kohns Vasconcelos, Marek Korencak, Jessica Nicolai, Klaus Pfeffer, Hendrik Streeck, Daniel Strelow, Jörg Timm, Andreas Walker, Tobias Wienemann |
| EPI_ISL_523931, EPI_ISL_523932, EPI_ISL_523933, EPI_ISL_523934, EPI_ISL_523935, EPI_ISL_523936, EPI_ISL_523937, EPI_ISL_523938, EPI_ISL_523939, EPI_ISL_523940, EPI_ISL_523941, EPI_ISL_523942, EPI_ISL_523943, EPI_ISL_523944, EPI_ISL_523945, EPI_ISL_523946, EPI_ISL_523947, EPI_ISL_523948, EPI_ISL_523949 | | | |
| see above | Center of Medical Microbiology, Virology, and Hospital Hygiene, University of Duesseldorf | Center of Medical Microbiology, Virology, and Hospital Hygiene, University of Duesseldorf | Maximilian Damagnez, Alexander Dilthey, Torsten Houwaart, Malte Kohns Vasconcelos, Marek Korencak, Jessica Nicolai, Klaus Pfeffer, Hendrik Streeck, Daniel Strelow, Jörg Timm, Andreas Walker, Tobias Wienemann |
| EPI_ISL_523950 | Center of Medical Microbiology, Virology, and Hospital Hygiene, University of Duesseldorf | Center of Medical Microbiology, Virology, and Hospital Hygiene, Heinrich Heine University Düsseldorf | Maximilian Damagnez, Alexander Dilthey, Torsten Houwaart, Malte Kohns Vasconcelos, Marek Korencak, Jessica Nicolai, Klaus Pfeffer, Hendrik Streeck, Daniel Strelow, Jörg Timm, Andreas Walker, Tobias Wienemann |
| EPI_ISL_524716 | B.J. Medical College and Civil hospital, Ahmedabad | Gujarat Biotechnology Research Centre | Monika Gandhi, Pinal Trivedi, Maharshi Pandya, Nidhi Patel, Nitin Savaliya, Raghawendra Kumar, Dinesh Kumar, Zuber Saiyed, Komal Patel, Labdhi Pandya, Afzal Ansari, Nikha Trivedi, Pranay Shah, Kamlesh J Upadhyay, Sanjay Kapadia, Apurvasinh Puvar, Janvi Raval, Zarna Patel, R D Dixit, A M Kadri, Harsh Bakshi, Chaitanya Joshi, Madhvi Joshi |
| EPI_ISL_524730 | GMERS Medical College & Hospital, Gotri, Vadodara | Gujarat Biotechnology Research Centre | Meenakshi Shah, Neena Doshi, Varsha Godbole, Apurvasinh Puvar, Janvi Raval, Zarna Patel, Monika Gandhi, Pinal Trivedi, Maharshi Pandya, Nidhi Patel, Nitin Savaliya, Raghawendra Kumar, Dinesh Kumar, Zuber Saiyed, Komal Patel, Labdhi Pandya, Afzal Ansari, Nikha Trivedi, R D Dixit, A M Kadri, Harsh Bakshi, Chaitanya Joshi, Madhvi Joshi |
| EPI_ISL_524738 | GMERS Medical College and Hospital, Gandhinagar | Gujarat Biotechnology Research Centre | Monika Gandhi, Pinal Trivedi, Maharshi Pandya, Nidhi Patel, Nitin Savaliya, Raghawendra Kumar, Dinesh Kumar, Zuber Saiyed, Komal Patel, Labdhi Pandya, Afzal Ansari, Nikha Trivedi, Seema Bhatt, Gaurishankar Shrimali, Bhavesh Modi, Bharti Rajani, Apurvasinh Puvar, Janvi Raval, Zarna Patel, R D Dixit, A M Kadri, Harsh Bakshi, Chaitanya Joshi, Madhvi Joshi |
| EPI_ISL_524752 | GMERS Medical College and Hospital, Dharpur, Patan | Gujarat Biotechnology Research Centre | Monika Gandhi, Pinal Trivedi, Maharshi Pandya, Nidhi Patel, Nitin Savaliya, Raghawendra Kumar, Dinesh Kumar, Zuber Saiyed, Komal Patel, Labdhi Pandya, Afzal Ansari, Nikha Trivedi, A N Parmar, Apurvasinh Puvar, Janvi Raval, Zarna Patel, R D Dixit, A M Kadri, Harsh Bakshi, Chaitanya Joshi, Madhvi Joshi |
| EPI_ISL_524873, EPI_ISL_524874, EPI_ISL_524875, EPI_ISL_524876, EPI_ISL_524877, EPI_ISL_524878, EPI_ISL_524879, EPI_ISL_524880, EPI_ISL_524881, EPI_ISL_524882, EPI_ISL_524883, EPI_ISL_524884, EPI_ISL_524885, EPI_ISL_524886, EPI_ISL_524887, EPI_ISL_524888, EPI_ISL_524889, EPI_ISL_524890, EPI_ISL_524891, EPI_ISL_524892, EPI_ISL_524893, EPI_ISL_524894, EPI_ISL_524895, EPI_ISL_524896, EPI_ISL_524897, EPI_ISL_524898, EPI_ISL_524899, EPI_ISL_524900, EPI_ISL_524901, EPI_ISL_524902, EPI_ISL_524903 | | | |
| see above | MD PHL | MD PHL | Maryland Department of Health Laboratories Administration |
| EPI_ISL_525118 | Maryland Department of Health Laboratory | Maryland Department of Health Laboratory | Maryland Department of Health Laboratories Administration |
| EPI_ISL_525352, EPI_ISL_525353, EPI_ISL_525354, EPI_ISL_525355, EPI_ISL_525356, EPI_ISL_525357, EPI_ISL_525358, EPI_ISL_525359, EPI_ISL_525360, EPI_ISL_525361, EPI_ISL_525362, EPI_ISL_525363, EPI_ISL_525364, EPI_ISL_525365, EPI_ISL_525366, EPI_ISL_525367, EPI_ISL_525368, EPI_ISL_525369, EPI_ISL_525370, EPI_ISL_525371, EPI_ISL_525372, EPI_ISL_525373, EPI_ISL_525374, EPI_ISL_525375, EPI_ISL_525376, EPI_ISL_525377, EPI_ISL_525378, EPI_ISL_525379, EPI_ISL_525380, EPI_ISL_525381, EPI_ISL_525382, EPI_ISL_525383, EPI_ISL_525384, EPI_ISL_525385, EPI_ISL_525386, EPI_ISL_525387, EPI_ISL_525388, EPI_ISL_525389, EPI_ISL_525390, EPI_ISL_525391, EPI_ISL_525392, EPI_ISL_525393, EPI_ISL_525394, EPI_ISL_525395, EPI_ISL_525396, EPI_ISL_525397, EPI_ISL_525398, EPI_ISL_525399, EPI_ISL_525400, EPI_ISL_525401, EPI_ISL_525402, EPI_ISL_525403, EPI_ISL_525404, EPI_ISL_525405, EPI_ISL_525406, EPI_ISL_525407, EPI_ISL_525408, EPI_ISL_525409, EPI_ISL_525410, EPI_ISL_525411, EPI_ISL_525412, EPI_ISL_525413, EPI_ISL_525414, EPI_ISL_525415, EPI_ISL_525416, EPI_ISL_525417, EPI_ISL_525418 | | | |
| see above | National Virus Reference Laboratory | National Virus Reference Laboratory | Michael Carr, Gabriel Gonzalez, Jonathan Dean, Aditi Chaturvedi, Suzie Coughlan, Cillian F De Gascun |

| | | | |
|---|---|---|---|
| EPI_ISL_525422 | B.J. Medical College and Civil hospital, Ahmedabad | Gujarat Biotechnology Research Centre | Monika Gandhi, Pinal Trivedi, Maharshi Pandya, Nidhi Patel, Nitin Savaliya, Raghawendra Kumar, Dinesh Kumar, Zuber Saiyed, Komal Patel, Labdhi Pandya, Afzal Ansari, Nikha Trivedi, Pranay Shah, Kamlesh J Upadhyay, Sanjay Kapadia, Apurvasinh Puvar, Janvi Raval, Zarna Patel, R D Dixit, A M Kadri, Harsh Bakshi, Chaitanya Joshi, Madhvi Joshi |
| EPI_ISL_526459, EPI_ISL_526460, EPI_ISL_526461, EPI_ISL_526462, EPI_ISL_526463, EPI_ISL_526464, EPI_ISL_526465, EPI_ISL_526466, EPI_ISL_526467, EPI_ISL_526468, EPI_ISL_526469, EPI_ISL_526470, EPI_ISL_526471, EPI_ISL_526472, EPI_ISL_526473, EPI_ISL_526474, EPI_ISL_526475, EPI_ISL_526476, EPI_ISL_526477, EPI_ISL_526478, EPI_ISL_526479, EPI_ISL_526480, EPI_ISL_526481, EPI_ISL_526482, EPI_ISL_526483, EPI_ISL_526484, EPI_ISL_526485, EPI_ISL_526486, EPI_ISL_526487, EPI_ISL_526488, EPI_ISL_526489, EPI_ISL_526490, EPI_ISL_526491, EPI_ISL_526492, EPI_ISL_526493, EPI_ISL_526494, EPI_ISL_526495, EPI_ISL_526496, EPI_ISL_526497, EPI_ISL_526498, EPI_ISL_526499, EPI_ISL_526500, EPI_ISL_526501, EPI_ISL_526502, EPI_ISL_526503, EPI_ISL_526504, EPI_ISL_526505, EPI_ISL_526506, EPI_ISL_526507, EPI_ISL_526508, EPI_ISL_526509, EPI_ISL_526510, EPI_ISL_526511, EPI_ISL_526512, EPI_ISL_526513, EPI_ISL_526514, EPI_ISL_526515, EPI_ISL_526516, EPI_ISL_526517, EPI_ISL_526518, EPI_ISL_526519, EPI_ISL_526520, EPI_ISL_526521, EPI_ISL_526522, EPI_ISL_526523, EPI_ISL_526524, EPI_ISL_526525, EPI_ISL_526526, EPI_ISL_526527, EPI_ISL_526528, EPI_ISL_526529, EPI_ISL_526530, EPI_ISL_526531, EPI_ISL_526532, EPI_ISL_526533, EPI_ISL_526534, EPI_ISL_526535, EPI_ISL_526536 | | | |
| see above | Virology Department, Royal Infirmary of Edinburgh, NHS Lothian / School of Biological Sciences, University of Edinburgh / Institute of Genetics and Molecular Medicine, University of Edinburgh | COVID-19 Genomics UK (COG-UK) Consortium | McHugh M, Dewar R, Rooke S, Gallagher M, Balcaza C, O'Toole Á, Scher E, Hill V, McCrone JT, Colquhoun R, Yu X, Jackson B, Rambaut A, Williams TC, Templeton K |
| EPI_ISL_526935, EPI_ISL_526936, EPI_ISL_526937, EPI_ISL_526938, EPI_ISL_526939, EPI_ISL_526940, EPI_ISL_526941, EPI_ISL_526942, EPI_ISL_526943, EPI_ISL_526944, EPI_ISL_526945, EPI_ISL_526946, EPI_ISL_526947, EPI_ISL_526948 | | | |
| see above | Faroese National Reference Laboratory for Fish and Animal Diseases | Faroese National Reference Laboratory for Fish and Animal Diseases | Maria Marjunardóttir Dahl, Petra Elisabeth Petersen, Debes Hammershaimb Christiansen |
| EPI_ISL_527573, EPI_ISL_527574, EPI_ISL_527575, EPI_ISL_527576, EPI_ISL_527577, EPI_ISL_527578, EPI_ISL_527579, EPI_ISL_527580, EPI_ISL_527581, EPI_ISL_527582, EPI_ISL_527583, EPI_ISL_527584, EPI_ISL_527585, EPI_ISL_527586, EPI_ISL_527587, EPI_ISL_527588, EPI_ISL_527589, EPI_ISL_527590, EPI_ISL_527591, EPI_ISL_527592, EPI_ISL_527593, EPI_ISL_527594, EPI_ISL_527595, EPI_ISL_527596, EPI_ISL_527597, EPI_ISL_527598, EPI_ISL_527599, EPI_ISL_527600, EPI_ISL_527601, EPI_ISL_527602, EPI_ISL_527603, EPI_ISL_527604, EPI_ISL_527605, EPI_ISL_527606, EPI_ISL_527607, EPI_ISL_527608, EPI_ISL_527609, EPI_ISL_527610, EPI_ISL_527611, EPI_ISL_527612, EPI_ISL_527613, EPI_ISL_527614, EPI_ISL_527615, EPI_ISL_527616, EPI_ISL_527617, EPI_ISL_527618, EPI_ISL_527619, EPI_ISL_527620, EPI_ISL_527621, EPI_ISL_527622, EPI_ISL_527623, EPI_ISL_527624, EPI_ISL_527625, EPI_ISL_527626, EPI_ISL_527627, EPI_ISL_527628, EPI_ISL_527629 | | | |
| see above | Minnesota Department of Health, Public Health Laboratory | Minnesota Department of Health, Public Health Laboratory | Matt Plumb, Jacob Garfin, and Xiong Wang |
| EPI_ISL_527630, EPI_ISL_527631 | Mayo Clinic & Mayo Clinic Laboratories | Minnesota Department of Health, Public Health Laboratory | Matt Plumb, Jacob Garfin, and Xiong Wang |
| EPI_ISL_528440, EPI_ISL_528441, EPI_ISL_528442, EPI_ISL_528443, EPI_ISL_528444, EPI_ISL_528445, EPI_ISL_528446, EPI_ISL_528447, EPI_ISL_528448, EPI_ISL_528449, EPI_ISL_528450, EPI_ISL_528451, EPI_ISL_528452, EPI_ISL_528453, EPI_ISL_528454, EPI_ISL_528455, EPI_ISL_528456, EPI_ISL_528457, EPI_ISL_528458, EPI_ISL_528459, EPI_ISL_528460, EPI_ISL_528461, EPI_ISL_528462, EPI_ISL_528463, EPI_ISL_528464, EPI_ISL_528465, EPI_ISL_528466, EPI_ISL_528467, EPI_ISL_528468, EPI_ISL_528469, EPI_ISL_528470, EPI_ISL_528471, EPI_ISL_528472, EPI_ISL_528473, EPI_ISL_528474, EPI_ISL_528475, EPI_ISL_528476, EPI_ISL_528477, EPI_ISL_528478, EPI_ISL_528479, EPI_ISL_528480, EPI_ISL_528481, EPI_ISL_528482, EPI_ISL_528483, EPI_ISL_528484 | | | |
| see above | National Virus Reference Laboratory | National Virus Reference Laboratory | Michael Carr, Gabriel Gonzalez, Jonathan Dean, Suzie Coughlan, Cillian F De Gascun |
| EPI_ISL_528738, EPI_ISL_528739, EPI_ISL_528740, EPI_ISL_528741 | Malaysia Genome Institute | Malaysia Genome Institute | Mohd Noor Mat Isa, Irni Suhayu Sapian, Yusuf Muhammad Noor, Nurhezreen Md Iqbal, Mohd Faizal Abu Bakar, Enizza Kasim, Shamsidar Sopie, Siti Noraini Othman, Azrin Ahmad, Nor Azfa Johari, Shahrul Hisham Zainal Ariffin |
| EPI_ISL_528742 | Malaysia Genome Institute | Malaysia Genome Institute | Mohd Noor Mat Isa, Irni Suhayu Sapian, Gan Han Ming, Yusuf Muhammad Noor, Tan Ju Lin, Nurhezreen Md Iqbal, Mohd Faizal Abu Bakar, Enizza Kasim, Shamsidar Sopie, Siti Noraini Othman, Azrin Ahmad, Nor Azfa Johari, Shahrul Hisham Zainal Ariffin |
| EPI_ISL_528743, EPI_ISL_528744 | Malaysia Genome Institute | Malaysia Genome Institute | Mohd Noor Mat Isa, Irni Suhayu Sapian, Yusuf Muhammad Noor, Nurhezreen Md Iqbal, Mohd Faizal Abu Bakar, Enizza Kasim, Shamsidar Sopie, Siti Noraini Othman, Azrin Ahmad, Nor Azfa Johari, Shahrul Hisham Zainal Ariffin |
| EPI_ISL_528745 | Laboratorium Kesehatan Provinsi Jawa Barat | School of Life Sciences and Technology & School of Pharmacy-Institut Teknologi Bandung; Molecular Genetics Laboratory-Faculty of Medicine-Universitas Padjadjaran; Laboratorium Kesehatan Provinsi Jawa Barat | Marselina Irasonia Tan, Yunia Sribudiani, Catur Riani, Azzania Fibriani, Husna Nugrahapraja, Tarwadi, Ema Rahmawati, Savira Ekawardhani, Hesti Lina Wiraswati, Ryan Bayusantika Ristandi, Rifky Waluyajati Rachman, Cut Nur Cinthia Alamanda, Lia Faridah, Tri Hanggono Achmad, Mas Rizky A.A. Syamsunarno, Fensi Amalina, Hammam Riza, Sony Solistia Wirawan, Agung Eru Wibowo, Irvan Faizal |
| EPI_ISL_528750 | Santo Borromeus Hospital | School of Life Sciences and Technology & School of Pharmacy-Institut Teknologi Bandung; Molecular Genetics Laboratory-Faculty of Medicine-Universitas Padjadjaran; Laboratorium Kesehatan Provinsi Jawa Barat | Marselina Irasonia Tan, Yunia Sribudiani, Catur Riani, Azzania Fibriani, Husna Nugrahapraja, Tarwadi, Ema Rahmawati, Savira Ekawardhani, Hesti Lina Wiraswati, Ryan Bayusantika Ristandi, Rifky Waluyajati Rachman, Cut Nur Cinthia Alamanda, Lia Faridah, Miftahul Faridl, Karimatu Khoirunnisa, Hammam Riza, Sony Solistia Wirawan, Agung Eru Wibowo, Irvan Faizal |
| EPI_ISL_528843, EPI_ISL_528844, EPI_ISL_528846, EPI_ISL_528852 | CSIR-Centre for Cellular and Molecular Biology | CSIR-Centre for Cellular and Molecular Biology | Namami Gaur, Sakshi Shambhavi, Lamuk Zaveri, Shagufta Khan, Nikhil Hajirnis, M Soujanya Reddy, Pratheusa Maccha, Tulasi Nagabandi, Purushotham Vodnala, Payel Mukherjee, Sofia Banu, Priya Singh, Onkar Kulkarni, Dhiviya Vedagiri, Divya Gupta, Vishal Sah, Santosh Kumar Kuncha, Krishnan Harinivas Harshan, Archana Bharadwaj Siva, Karthik Bharadwaj Tallapaka, Zeba Rizvi, Zuberwasim Sayyad, Kakade Aishwarya Arun, Amrutha H C, Ananga Ghosh, Rakesh K Mishra, Divya Tej Sowpati |
| EPI_ISL_529031 | Central Molecular Microbiology Laboratory, Clinical and Chemical Pathology Department, Faculty of Medicine, CAIRO UNIVERSITY | Next Generation Sequencing Reference Laboratory, Faculty of Medicine, Cairo University and The Center for Genome and Microbiome Research, Faculty of Pharmacy, CAIRO UNIVERSITY | May Sherif Soliman, May Abdelfattah, Ramy Karam Aziz |
| EPI_ISL_529032 | Central Molecular Microbiology Laboratory and Next Generation Sequencing Reference Laboratory, Clinical and Chemical Pathology Department, Faculty of Medicine, CAIRO UNIVERSITY | Next Generation Sequencing Reference Laboratory, Faculty of Medicine, CAIRO UNIVERSITY and The Center for Genome and Microbiome Research, Faculty of Pharmacy, CAIRO UNIVERSITY | May Sherif Soliman, May Abdelfattah, Ramy Karam Aziz |
| EPI_ISL_529285, EPI_ISL_529317, EPI_ISL_529318, EPI_ISL_529478, EPI_ISL_529689, EPI_ISL_529690, EPI_ISL_529691, EPI_ISL_529692, EPI_ISL_529693, EPI_ISL_529694, EPI_ISL_529695, EPI_ISL_529696 | | | |
| see above | Virology Department, Royal Infirmary of Edinburgh, NHS Lothian / School of Biological Sciences, University of Edinburgh / Institute of Genetics and Molecular Medicine, University of Edinburgh | COVID-19 Genomics UK (COG-UK) Consortium | McHugh M, Dewar R, Rooke S, Gallagher M, Balcaza C, O'Toole Á, Scher E, Hill V, McCrone JT, Colquhoun R, Yu X, Jackson B, Rambaut A, Williams TC, Templeton K |
| EPI_ISL_530025, EPI_ISL_530026, EPI_ISL_530027, EPI_ISL_530028, EPI_ISL_530029, EPI_ISL_530030, EPI_ISL_530031, EPI_ISL_530032, EPI_ISL_530033, EPI_ISL_530034, EPI_ISL_530035, EPI_ISL_530036, EPI_ISL_530037, EPI_ISL_530038, EPI_ISL_530039, EPI_ISL_530040, EPI_ISL_530041, EPI_ISL_530042, EPI_ISL_530043, EPI_ISL_530070, EPI_ISL_530071, EPI_ISL_530072, EPI_ISL_530073, EPI_ISL_530074, EPI_ISL_530075, EPI_ISL_530076, EPI_ISL_530077, EPI_ISL_530078, EPI_ISL_530079, EPI_ISL_530080, EPI_ISL_530081, EPI_ISL_530082, EPI_ISL_530083, EPI_ISL_530084, EPI_ISL_530085, EPI_ISL_530086, EPI_ISL_530087, EPI_ISL_530088, EPI_ISL_530089, EPI_ISL_530090, EPI_ISL_530091, EPI_ISL_530092, EPI_ISL_530093 | | | |
| see above | Hospital Universitario La Paz | Hospital Universitario La Paz | María Rodríguez, Elias Dahdouh, Sara González, Raúl Recio, Fernando Lázaro, Esther Viedma, Natalia Stella, Julio García, Juan Carlos Galán, Rafael Cantón, Mª Dolores Folgueira, Rafael Delgado, Jesús Mingorance |
| EPI_ISL_530114, EPI_ISL_530115, EPI_ISL_530116, EPI_ISL_530117, EPI_ISL_530118, EPI_ISL_530119 | Hospital Universitario Ramón y Cajal | Hospital Universitario La Paz | María Rodríguez, Elias Dahdouh, Sara González, Raúl Recio, Fernando Lázaro, Esther Viedma, Natalia Stella, Julio García, Juan Carlos Galán, Rafael Cantón, Mª Dolores Folgueira, Rafael Delgado, Jesús Mingorance |
| EPI_ISL_530163, EPI_ISL_530164, EPI_ISL_530165, EPI_ISL_530166, EPI_ISL_530167, EPI_ISL_530168 | M Health Fairview | Minnesota Department of Health, Public Health Laboratory | Matt Plumb, Jacob Garfin, and Xiong Wang |
| EPI_ISL_530169, EPI_ISL_530170, EPI_ISL_530171 | Hennepin County Medical Center | Minnesota Department of Health, Public Health Laboratory | Matt Plumb, Jacob Garfin, and Xiong Wang |
| EPI_ISL_530172, EPI_ISL_530173, EPI_ISL_530174, EPI_ISL_530175, EPI_ISL_530176, EPI_ISL_530177, EPI_ISL_530178, EPI_ISL_530179, EPI_ISL_530180, EPI_ISL_530181, EPI_ISL_530182, EPI_ISL_530183, EPI_ISL_530184, EPI_ISL_530185, EPI_ISL_530186, EPI_ISL_530187, EPI_ISL_530188, EPI_ISL_530189, EPI_ISL_530190, EPI_ISL_530191, EPI_ISL_530192, EPI_ISL_530193, EPI_ISL_530194, EPI_ISL_530195, EPI_ISL_530196, EPI_ISL_530197, EPI_ISL_530198, EPI_ISL_530199, EPI_ISL_530200, EPI_ISL_530201, EPI_ISL_530202, EPI_ISL_530203, EPI_ISL_530204, EPI_ISL_530205, EPI_ISL_530206, EPI_ISL_530207, EPI_ISL_530208, EPI_ISL_530209, EPI_ISL_530210, EPI_ISL_530211, EPI_ISL_530212, EPI_ISL_530213, EPI_ISL_530214, EPI_ISL_530215, EPI_ISL_530216, EPI_ISL_530217, EPI_ISL_530218, EPI_ISL_530219, EPI_ISL_530220, EPI_ISL_530221, EPI_ISL_530222, EPI_ISL_530223, EPI_ISL_530224 | | | |
| see above | Minnesota Department of Health, Public Health Laboratory | Minnesota Department of Health, Public Health Laboratory | Matt Plumb, Jacob Garfin, and Xiong Wang |
| EPI_ISL_530347, EPI_ISL_530348, | The National Institute of Public Health | State Veterinary Institute Prague | Nagy,A;Jirincova,H;Novakova,L;Trnka,D;Vecerova,J |

| | | | |
|---|---|---|---|
| EPI_ISL_530349, EPI_ISL_530350 | | | |
| EPI_ISL_532659, EPI_ISL_532683, EPI_ISL_532689, EPI_ISL_532705, EPI_ISL_532926, EPI_ISL_532978, EPI_ISL_532983, EPI_ISL_532990, EPI_ISL_532992, EPI_ISL_532994 | Virology Department, Royal Infirmary of Edinburgh, NHS Lothian / School of Biological Sciences, University of Edinburgh | Wellcome Sanger Institute for the COVID-19 Genomics UK (COG-UK) consortium | McHugh M, Dewar R, Rooke S, O'Toole Á, Scher E, Hill V, McCrone JT, Colquhoun R, Yu X, Jackson B, Rambaut A, Templeton K and Alex Alderton, Roberto Amato, Sonia Goncalves, Ewan Harrison, David K. Jackson, Ian Johnston, Dominic Kwiatkowski, Cordelia Langford, John Sillitoe |
| EPI_ISL_534217 | M Health Fairview | Minnesota Department of Health, Public Health Laboratory | Matt Plumb, Jacob Garfin, and Xiong Wang |
| EPI_ISL_534330, EPI_ISL_534331, EPI_ISL_534332, EPI_ISL_534333 | Hospital Universitario La Paz | Hospital Universitario La Paz | María Rodríguez, Elias Dahdouh, Sara González, Raúl Recio, Fernando Lázaro, Esther Viedma, Natalia Stella, Julio García, Juan Carlos Galán, Rafael Cantón, Ma Dolores Folgueira, Rafael Delgado, Jesús Mingorance |
| EPI_ISL_534334 | Hospital Universitario Ramón y Cajal | Hospital Universitario La Paz | María Rodríguez, Elias Dahdouh, Sara González, Raúl Recio, Fernando Lázaro, Esther Viedma, Natalia Stella, Julio García, Juan Carlos Galán, Rafael Cantón, Ma Dolores Folgueira, Rafael Delgado, Jesús Mingorance |
| EPI_ISL_534335 | LabPLUS | Institute of Environmental Science and Research (ESR) | Matt Storey, Xiaoyun Ren, Anja Werno, Antje van der Linden, Arlo Upton, Chris Mansell, David Hammer, Dragana Drinkovic, Gary McAuliffe, Hana Sofia Andersson, James Ussher, Jill Sherwood, Josh Freeman, Julia Howard, Juliet Elvy, Mary DeAlmeida, Matt Blakiston, Matthew Rogers, Max Bloomfield, Michael Addidle, Michelle Balm, Sally Roberts, Sarah Jefferies, Sharmini Muttaiyah, Susan Morpeth, Susan Taylor, Timothy Blackmore, Vani Sathyendran, Veronica Playle, Virginia Hope, Erasmus Smit, Lauren Jelly, Joep de Ligt |
| EPI_ISL_534699, EPI_ISL_534700, EPI_ISL_534701, EPI_ISL_534702, EPI_ISL_534703, EPI_ISL_534704, EPI_ISL_534705, EPI_ISL_534706, EPI_ISL_534707, EPI_ISL_534708, EPI_ISL_534709, EPI_ISL_534710, EPI_ISL_534711, EPI_ISL_534712, EPI_ISL_534713, EPI_ISL_534714, EPI_ISL_534715, EPI_ISL_534716 | | | |
| see above | MD PHL | MD PHL | Maryland Department of Health Laboratories Administration |
| EPI_ISL_537382, EPI_ISL_537383, EPI_ISL_537384, EPI_ISL_537385, EPI_ISL_537386, EPI_ISL_537387, EPI_ISL_537388, EPI_ISL_537389, EPI_ISL_537390, EPI_ISL_537391, EPI_ISL_537392, EPI_ISL_537393, EPI_ISL_537394, EPI_ISL_537395, EPI_ISL_537396, EPI_ISL_537397, EPI_ISL_537398, EPI_ISL_537399, EPI_ISL_537400, EPI_ISL_537401, EPI_ISL_537402, EPI_ISL_537403, EPI_ISL_537404, EPI_ISL_537405, EPI_ISL_537406, EPI_ISL_537407, EPI_ISL_537408, EPI_ISL_537409, EPI_ISL_537410, EPI_ISL_537411, EPI_ISL_537412, EPI_ISL_537413, EPI_ISL_537414, EPI_ISL_537415, EPI_ISL_537416, EPI_ISL_537417, EPI_ISL_537418, EPI_ISL_537419, EPI_ISL_537420, EPI_ISL_537421, EPI_ISL_537422, EPI_ISL_537423, EPI_ISL_537424, EPI_ISL_537425, EPI_ISL_537426, EPI_ISL_537427, EPI_ISL_537428, EPI_ISL_537429, EPI_ISL_537430, EPI_ISL_537431, EPI_ISL_537432, EPI_ISL_537433, EPI_ISL_537434, EPI_ISL_537435, EPI_ISL_537436, EPI_ISL_537437, EPI_ISL_537438, EPI_ISL_537439, EPI_ISL_537440, EPI_ISL_537441, EPI_ISL_537442, EPI_ISL_537443, EPI_ISL_537444, EPI_ISL_537445, EPI_ISL_537446, EPI_ISL_537447, EPI_ISL_537448, EPI_ISL_537449, EPI_ISL_537450, EPI_ISL_537451, EPI_ISL_537452, EPI_ISL_537453, EPI_ISL_537454, EPI_ISL_537455, EPI_ISL_537456, EPI_ISL_537457, EPI_ISL_537458, EPI_ISL_537459, EPI_ISL_537460, EPI_ISL_537461, EPI_ISL_537462, EPI_ISL_537463, EPI_ISL_537464, EPI_ISL_537465, EPI_ISL_537466 | | | |
| see above | Centro de Investigación Biomédica de La Rioja - Hospital San Pedro Logroño | SeqCOVID-SPAIN consortium/IBV(CSIC) | María de Toro, José Manuel Azcona Gutiérrez, María Pilar Bea Escudero, Miriam Blasco Alberdi and SeqCOVID-SPAIN consortium |
| EPI_ISL_537809, EPI_ISL_537810 | Hospital Universitario Marqués de Valdecilla (Santander), Servicio de Microbiología | SeqCOVID-SPAIN consortium/IBV(CSIC) | Mónica Gozalo Margüello, María Eliecer Cano García, Jose Manuel Méndez Legaza, Daniel Pablo Marcos, Jesús Rodríguez Lozano, María Siller Ruiz and SeqCOVID-SPAIN consortium |
| EPI_ISL_537811, EPI_ISL_537812, EPI_ISL_537813, EPI_ISL_537814, EPI_ISL_537815, EPI_ISL_537816, EPI_ISL_537817, EPI_ISL_537818, EPI_ISL_537819, EPI_ISL_537820, EPI_ISL_537821, EPI_ISL_537822, EPI_ISL_537823, EPI_ISL_537824, EPI_ISL_537825, EPI_ISL_537826, EPI_ISL_537827, EPI_ISL_537828, EPI_ISL_537829, EPI_ISL_537830, EPI_ISL_537831, EPI_ISL_537832, EPI_ISL_537833, EPI_ISL_537834, EPI_ISL_537835, EPI_ISL_537836, EPI_ISL_537837, EPI_ISL_537838, EPI_ISL_537839, EPI_ISL_537840, EPI_ISL_537841, EPI_ISL_537842, EPI_ISL_537843, EPI_ISL_537844, EPI_ISL_537845, EPI_ISL_537846, EPI_ISL_537847, EPI_ISL_537848, EPI_ISL_537849, EPI_ISL_537850, EPI_ISL_537851, EPI_ISL_537852, EPI_ISL_537853, EPI_ISL_537854, EPI_ISL_537855, EPI_ISL_537856, EPI_ISL_537857, EPI_ISL_537858, EPI_ISL_537859, EPI_ISL_537860, EPI_ISL_537861, EPI_ISL_537862, EPI_ISL_537863, EPI_ISL_537864, EPI_ISL_537865, EPI_ISL_537866, EPI_ISL_537867, EPI_ISL_537868, EPI_ISL_537869, EPI_ISL_537870, EPI_ISL_537871, EPI_ISL_537872, EPI_ISL_537873 | | | |
| see above | Centro de Investigación Biomédica de La Rioja - Hospital San Pedro Logroño | SeqCOVID-SPAIN consortium/IBV(CSIC) | María de Toro, José Manuel Azcona Gutiérrez, María Pilar Bea Escudero, Miriam Blasco Alberdi and SeqCOVID-SPAIN consortium |
| EPI_ISL_538003, EPI_ISL_538004, EPI_ISL_538005, EPI_ISL_538006, EPI_ISL_538007, EPI_ISL_538008, EPI_ISL_538009, EPI_ISL_538010, EPI_ISL_538011, EPI_ISL_538012, EPI_ISL_538013, EPI_ISL_538014, EPI_ISL_538015, EPI_ISL_538016, EPI_ISL_538017, EPI_ISL_538018, EPI_ISL_538019, EPI_ISL_538020, EPI_ISL_538021, EPI_ISL_538022, EPI_ISL_538023, EPI_ISL_538024, EPI_ISL_538025, EPI_ISL_538026, EPI_ISL_538027, EPI_ISL_538028, EPI_ISL_538029, EPI_ISL_538030, EPI_ISL_538031, EPI_ISL_538032, EPI_ISL_538033, EPI_ISL_538034, EPI_ISL_538035, EPI_ISL_538036, EPI_ISL_538037, EPI_ISL_538038, EPI_ISL_538039, EPI_ISL_538040, EPI_ISL_538041, EPI_ISL_538042, EPI_ISL_538043, EPI_ISL_538044, EPI_ISL_538045, EPI_ISL_538046, EPI_ISL_538047, EPI_ISL_538048, EPI_ISL_538049, EPI_ISL_538050, EPI_ISL_538051, EPI_ISL_538052, EPI_ISL_538053, EPI_ISL_538054 | | | |
| see above | Hospital Universitari i Politècnic La Fe de València | SeqCOVID-SPAIN consortium/IBV(CSIC) | María Dolores Gómez Ruiz, Eva González Barbera, Ana Gil Brusola, Salvador Giner Almaraz, José Luis López Hontangas and SeqCOVID-SPAIN consortium |
| EPI_ISL_538082, EPI_ISL_538083, EPI_ISL_538084, EPI_ISL_538085, EPI_ISL_538086, EPI_ISL_538087, EPI_ISL_538088, EPI_ISL_538089, EPI_ISL_538090, EPI_ISL_538091, EPI_ISL_538092, EPI_ISL_538093, EPI_ISL_538094, EPI_ISL_538095, EPI_ISL_538096, EPI_ISL_538097, EPI_ISL_538098, EPI_ISL_538099, EPI_ISL_538100, EPI_ISL_538101, EPI_ISL_538102, EPI_ISL_538103, EPI_ISL_538104, EPI_ISL_538105, EPI_ISL_538106, EPI_ISL_538107, EPI_ISL_538108, EPI_ISL_538109, EPI_ISL_538110, EPI_ISL_538111, EPI_ISL_538112, EPI_ISL_538113, EPI_ISL_538114, EPI_ISL_538115, EPI_ISL_538116, EPI_ISL_538117 | | | |
| see above | Clínica Universidad de Navarra. Servicio de Enfermedades Infecciosas y Microbiología clínica | SeqCOVID-SPAIN consortium/IBV(CSIC) | Mirian Fernández-Alonso, Jose Luis del Pozo and SeqCOVID-SPAIN consortium |
| EPI_ISL_538265, EPI_ISL_538266, EPI_ISL_538267, EPI_ISL_538268, EPI_ISL_538269, EPI_ISL_538270, EPI_ISL_538271, EPI_ISL_538272, EPI_ISL_538273, EPI_ISL_538274 | Maryland Public Health Laboratory | Maryland Public Health Laboratory | Maryland Department of Health Laboratories Administration |
| EPI_ISL_538608, EPI_ISL_538609, EPI_ISL_538610, EPI_ISL_538611, EPI_ISL_538612 | Clínica Universidad de Navarra. Servicio de Enfermedades Infecciosas y Microbiología clínica | SeqCOVID-SPAIN consortium/IBV(CSIC) | Mirian Fernández-Alonso, Jose Luis del Pozo and SeqCOVID-SPAIN consortium |
| EPI_ISL_538672, EPI_ISL_538673, EPI_ISL_538674, EPI_ISL_538675, EPI_ISL_538676, EPI_ISL_538677, EPI_ISL_538678, EPI_ISL_538679, EPI_ISL_538680, EPI_ISL_538681, EPI_ISL_538682, EPI_ISL_538683, EPI_ISL_538684, EPI_ISL_538685, EPI_ISL_538686, EPI_ISL_538687, EPI_ISL_538688, EPI_ISL_538689 | | | |
| see above | Hospital Universitario Virgen de las Nieves de Granada-SAS | SeqCOVID-SPAIN consortium/IBV(CSIC) | Mercedes Pérez Ruiz, Sara Sanbonmatsu Gámez, Irene Pedrosa Corral, José M. Navarro-Marí and SeqCOVID-SPAIN consortium |
| EPI_ISL_538690, EPI_ISL_538691, EPI_ISL_538692, EPI_ISL_538693, EPI_ISL_538694, EPI_ISL_538695, EPI_ISL_538696, EPI_ISL_538697, EPI_ISL_538698, EPI_ISL_538699, EPI_ISL_538700, EPI_ISL_538701, EPI_ISL_538702, EPI_ISL_538703, EPI_ISL_538704, EPI_ISL_538705, EPI_ISL_538706, EPI_ISL_538707, EPI_ISL_538708, EPI_ISL_538709, EPI_ISL_538710, EPI_ISL_538711, EPI_ISL_538712, EPI_ISL_538713, EPI_ISL_538714, EPI_ISL_538715, EPI_ISL_538716, EPI_ISL_538717, EPI_ISL_538718, EPI_ISL_538719, EPI_ISL_538720, EPI_ISL_538721, EPI_ISL_538722, EPI_ISL_538723, EPI_ISL_538724, EPI_ISL_538725, EPI_ISL_538726, EPI_ISL_538727, EPI_ISL_538728, EPI_ISL_538729, EPI_ISL_538730, EPI_ISL_538731, EPI_ISL_538732, EPI_ISL_538733, EPI_ISL_538734, EPI_ISL_538735, EPI_ISL_538736, EPI_ISL_538737, EPI_ISL_538738, EPI_ISL_538739, EPI_ISL_538740, EPI_ISL_538741, EPI_ISL_538742, EPI_ISL_538743, EPI_ISL_538744, EPI_ISL_538745, EPI_ISL_538746, EPI_ISL_538747 | | | |
| see above | Hospital Universitario Marqués de Valdecilla - IDIVAL (Santander, Cantabria) | SeqCOVID-SPAIN consortium/IBV(CSIC) | María Eliecer Cano García, Mónica Gozalo Margüello, Jose Manuel Méndez Legaza, Daniel Pablo Marcos, Jesús Rodríguez Rodríguez, María Siller Ruiz and SeqCOVID-SPAIN consortium |
| EPI_ISL_539221, EPI_ISL_539222, EPI_ISL_539223, EPI_ISL_539224, EPI_ISL_539225, EPI_ISL_539226, EPI_ISL_539227, EPI_ISL_539228, EPI_ISL_539229, EPI_ISL_539230, EPI_ISL_539231, EPI_ISL_539232, EPI_ISL_539233, EPI_ISL_539235, EPI_ISL_539236, EPI_ISL_539237, EPI_ISL_539238, EPI_ISL_539239, EPI_ISL_539243, EPI_ISL_539244 | | | |
| see above | Servicio de Microbiologia. Hospital Clinico Universitario de Valencia | SeqCOVID-SPAIN consortium/IBV(CSIC) | Maria Dolores Ocete, Inma Galán Vendrell, Paula Ruiz-Hueso, Mariana Reyes-Prieto, Vicente Soriano Chirona, Maria Alma Bracho, Griselda De Marco, Beatriz Beamud, Lidia Ruiz Roldan, Marta Pla Diaz,Neris Garcia-Gonzalez, Loreto Ferrús Abad, Llúcia Martínez-Priego, Concepcion Gimeno, Giuseppe D'Auria, Fernando Gonzalez-Candelas and SeqCOVID-SPAIN consortium |
| EPI_ISL_539245 | Hospital Universitari i Politècnic La Fe de València | SeqCOVID-SPAIN consortium/IBV(CSIC) | María Dolores Gómez Ruiz, Eva González Barbera, Ana Gil Brusola, Salvador Giner Almaraz, José Luis López Hontangas and SeqCOVID-SPAIN consortium |
| EPI_ISL_539288 | Servicio de Microbiologia. Hospital Clinico Universitario de Valencia | SeqCOVID-SPAIN consortium/IBV(CSIC) | Maria Dolores Ocete, Inma Galán Vendrell, Paula Ruiz-Hueso, Mariana Reyes-Prieto, Vicente Soriano Chirona, Maria Alma Bracho, Griselda De Marco, Beatriz Beamud, Lidia Ruiz Roldan, Marta Pla Diaz,Neris Garcia-Gonzalez, Loreto Ferrús Abad, Llúcia Martínez-Priego, Concepcion Gimeno, Giuseppe D'Auria, Fernando Gonzalez-Candelas and SeqCOVID-SPAIN consortium |
| EPI_ISL_539493, EPI_ISL_539494 | The National Institute of Public Health | State Veterinary Institute Prague | Nagy,A;Jirincova,H;Novakova,L;Trnka,D;Vecerova,J |
| EPI_ISL_539577, EPI_ISL_539578, EPI_ISL_539579, EPI_ISL_539580, EPI_ISL_539581, EPI_ISL_539582, EPI_ISL_539583, EPI_ISL_539584, EPI_ISL_539585, EPI_ISL_539586, EPI_ISL_539587, EPI_ISL_539588, EPI_ISL_539589, EPI_ISL_539590, EPI_ISL_539591, EPI_ISL_539592, EPI_ISL_539593, EPI_ISL_539594, EPI_ISL_539595, EPI_ISL_539596, EPI_ISL_539597, EPI_ISL_539598, EPI_ISL_539599, EPI_ISL_539600, EPI_ISL_539601, EPI_ISL_539602, EPI_ISL_539603, EPI_ISL_539604, EPI_ISL_539605, EPI_ISL_539606, EPI_ISL_539607, EPI_ISL_539608, EPI_ISL_539609, EPI_ISL_539610, EPI_ISL_539611, EPI_ISL_539612, EPI_ISL_539613, EPI_ISL_539614, EPI_ISL_539615 | | | |
| see above | ZOTZ KLIMAS MVZ Düsseldorf-Centrum GbR ÜBAG für Labormedizin, Genetik, Zytologie, Pathologie | Center of Medical Microbiology, Virology, and Hospital Hygiene, University of Duesseldorf | Maximilian Damagnez, Alexander Dilthey, Ashley-Jane Duplessis, Patrick Finzer, Katrin Hoffmann, Torsten Houwaart, Malte Kohns Vasconcelos, Marek Korencak, Nadine Lübke, Jessica Nicolai, Klaus Pfeffer, Daniel Strelow, Jörg Timm, Andreas Walker, Tobias Wienemann, Rainer Zotz |

| | | | |
|---|---|---|---|
| EPI_ISL_539652 | CSIR-Centre for Cellular and Molecular Biology | CSIR-Centre for Cellular and Molecular Biology | Namami Gaur, Sakshi Shambhavi, Lamuk Zaveri, Shagufta Khan, Nikhil Hajirnis, M Soujanya Reddy, Pratheusa Maccha, Tulasi Nagabandi, Purushotham Vodnala, Payel Mukherjee, Sofia Banu, Priya Singh, Onkar Kulkarni, Dhiviya Vedagiri, Divya Gupta, Vishal Sah, Santosh Kumar Kuncha, Krishnan Harinivas Harshan, Archana Bharadwaj Siva, Karthik Bharadwaj Tallapaka, Zeba Rizvi, Zuberwasim Sayyad, Kakade Aishwarya Arun, Amrutha H C, Ananga Ghosh, Rakesh K Mishra, Divya Tej Sowpati |
| EPI_ISL_539653 | CSIR-Centre for Cellular and Molecular Biology | CSIR-Centre for Cellular and Molecular Biology | Namami Gaur, Sakshi Shambhavi, Lamuk Zaveri, Shagufta Khan, Nikhil Hajirnis, M Soujanya Reddy, Pratheusa Maccha, Tulasi Nagabandi, Purushotham Vodnala, Payel Mukherjee, Sofia Banu, Priya Singh, Onkar Kulkarni, Dhiviya Vedagiri, Divya Gupta, Vishal Sah, Santosh Kumar Kuncha, Krishnan Harinivas Harshan, Archana Bharadwaj Siva, Karthik Bharadwaj Tallapaka,G. Aditya Kumar, Koushick Sivakumar, Rakesh K Mishra, Divya Tej Sowpati |
| EPI_ISL_539654 | CSIR-Centre for Cellular and Molecular Biology | CSIR-Centre for Cellular and Molecular Biology | Namami Gaur, Sakshi Shambhavi, Lamuk Zaveri, Shagufta Khan, Nikhil Hajirnis, M Soujanya Reddy, Pratheusa Maccha, Tulasi Nagabandi, Purushotham Vodnala, Payel Mukherjee, Sofia Banu, Priya Singh,Onkar Kulkarni, Dhiviya Vedagiri, Divya Gupta, Vishal Sah, Santosh Kumar Kuncha, Krishnan Harinivas Harshan, Archana Bharadwaj Siva, Karthik Bharadwaj Tallapaka, Zeba Rizvi, Zuberwasim Sayyad, Kakade Aishwarya Arun, Amrutha H C, Ananga Ghosh, Rakesh K Mishra, Divya Tej Sowpati |
| EPI_ISL_539682 | CSIR-Centre for Cellular and Molecular Biology | CSIR-Centre for Cellular and Molecular Biology | Namami Gaur, Sakshi Shambhavi, Lamuk Zaveri, Shagufta Khan, Nikhil Hajirnis, M Soujanya Reddy, Pratheusa Maccha, Tulasi Nagabandi, Purushotham Vodnala, Payel Mukherjee, Sofia Banu, Priya Singh, Onkar Kulkarni, Dhiviya Vedagiri, Divya Gupta, Vishal Sah, Santosh Kumar Kuncha, Krishnan Harinivas Harshan, Archana Bharadwaj Siva, Karthik Bharadwaj Tallapaka, Zeba Rizvi, Zuberwasim Sayyad, Kakade Aishwarya Arun, Amrutha H C, Ananga Ghosh, Rakesh K Mishra, Divya Tej Sowpati |
| EPI_ISL_539683 | CSIR-Centre for Cellular and Molecular Biology | CSIR-Centre for Cellular and Molecular Biology | Namami Gaur, Sakshi Shambhavi, Lamuk Zaveri, Shagufta Khan, Nikhil Hajirnis, M Soujanya Reddy, Pratheusa Maccha, Tulasi Nagabandi, Purushotham Vodnala, Payel Mukherjee, Sofia Banu, Priya Singh, Onkar Kulkarni, Dhiviya Vedagiri, Divya Gupta, Vishal Sah, Santosh Kumar Kuncha, Krishnan Harinivas Harshan, Archana Bharadwaj Siva, Karthik Bharadwaj Tallapaka,G. Aditya Kumar, Koushick Sivakumar, Rakesh K Mishra, Divya Tej Sowpati |
| EPI_ISL_539684 | CSIR-Centre for Cellular and Molecular Biology | CSIR-Centre for Cellular and Molecular Biology | Namami Gaur, Sakshi Shambhavi, Lamuk Zaveri, Shagufta Khan, Nikhil Hajirnis, M Soujanya Reddy, Pratheusa Maccha, Tulasi Nagabandi, Purushotham Vodnala, Payel Mukherjee, Sofia Banu, Priya Singh,Onkar Kulkarni, Dhiviya Vedagiri, Divya Gupta, Vishal Sah, Santosh Kumar Kuncha, Krishnan Harinivas Harshan, Archana Bharadwaj Siva, Karthik Bharadwaj Tallapaka, Zeba Rizvi, Zuberwasim Sayyad, Kakade Aishwarya Arun, Amrutha H C, Ananga Ghosh, Rakesh K Mishra, Divya Tej Sowpati |
| EPI_ISL_539712 | CSIR-Centre for Cellular and Molecular Biology | CSIR-Centre for Cellular and Molecular Biology | Namami Gaur, Sakshi Shambhavi, Lamuk Zaveri, Shagufta Khan, Nikhil Hajirnis, M Soujanya Reddy, Pratheusa Maccha, Tulasi Nagabandi, Purushotham Vodnala, Payel Mukherjee, Sofia Banu, Priya Singh, Onkar Kulkarni, Dhiviya Vedagiri, Divya Gupta, Vishal Sah, Santosh Kumar Kuncha, Krishnan Harinivas Harshan, Archana Bharadwaj Siva, Karthik Bharadwaj Tallapaka, Zeba Rizvi, Zuberwasim Sayyad, Kakade Aishwarya Arun, Amrutha H C, Ananga Ghosh, Rakesh K Mishra, Divya Tej Sowpati |
| EPI_ISL_539713 | CSIR-Centre for Cellular and Molecular Biology | CSIR-Centre for Cellular and Molecular Biology | Namami Gaur, Sakshi Shambhavi, Lamuk Zaveri, Shagufta Khan, Nikhil Hajirnis, M Soujanya Reddy, Pratheusa Maccha, Tulasi Nagabandi, Purushotham Vodnala, Payel Mukherjee, Sofia Banu, Priya Singh, Onkar Kulkarni, Dhiviya Vedagiri, Divya Gupta, Vishal Sah, Santosh Kumar Kuncha, Krishnan Harinivas Harshan, Archana Bharadwaj Siva, Karthik Bharadwaj Tallapaka,G. Aditya Kumar, Koushick Sivakumar, Rakesh K Mishra, Divya Tej Sowpati |
| EPI_ISL_539714 | CSIR-Centre for Cellular and Molecular Biology | CSIR-Centre for Cellular and Molecular Biology | Namami Gaur, Sakshi Shambhavi, Lamuk Zaveri, Shagufta Khan, Nikhil Hajirnis, M Soujanya Reddy, Pratheusa Maccha, Tulasi Nagabandi, Purushotham Vodnala, Payel Mukherjee, Sofia Banu, Priya Singh,Onkar Kulkarni, Dhiviya Vedagiri, Divya Gupta, Vishal Sah, Santosh Kumar Kuncha, Krishnan Harinivas Harshan, Archana Bharadwaj Siva, Karthik Bharadwaj Tallapaka, Zeba Rizvi, Zuberwasim Sayyad, Kakade Aishwarya Arun, Amrutha H C, Ananga Ghosh, Rakesh K Mishra, Divya Tej Sowpati |
| EPI_ISL_539742 | CSIR-Centre for Cellular and Molecular Biology | CSIR-Centre for Cellular and Molecular Biology | Namami Gaur, Sakshi Shambhavi, Lamuk Zaveri, Shagufta Khan, Nikhil Hajirnis, M Soujanya Reddy, Pratheusa Maccha, Tulasi Nagabandi, Purushotham Vodnala, Payel Mukherjee, Sofia Banu, Priya Singh, Onkar Kulkarni, Dhiviya Vedagiri, Divya Gupta, Vishal Sah, Santosh Kumar Kuncha, Krishnan Harinivas Harshan, Archana Bharadwaj Siva, Karthik Bharadwaj Tallapaka, Zeba Rizvi, Zuberwasim Sayyad, Kakade Aishwarya Arun, Amrutha H C, Ananga Ghosh, Rakesh K Mishra, Divya Tej Sowpati |
| EPI_ISL_539743 | CSIR-Centre for Cellular and Molecular Biology | CSIR-Centre for Cellular and Molecular Biology | Namami Gaur, Sakshi Shambhavi, Lamuk Zaveri, Shagufta Khan, Nikhil Hajirnis, M Soujanya Reddy, Pratheusa Maccha, Tulasi Nagabandi, Purushotham Vodnala, Payel Mukherjee, Sofia Banu, Priya Singh, Onkar Kulkarni, Dhiviya Vedagiri, Divya Gupta, Vishal Sah, Santosh Kumar Kuncha, Krishnan Harinivas Harshan, Archana Bharadwaj Siva, Karthik Bharadwaj Tallapaka,G. Aditya Kumar, Koushick Sivakumar, Rakesh K Mishra, Divya Tej Sowpati |
| EPI_ISL_539744 | CSIR-Centre for Cellular and Molecular Biology | CSIR-Centre for Cellular and Molecular Biology | Namami Gaur, Sakshi Shambhavi, Lamuk Zaveri, Shagufta Khan, Nikhil Hajirnis, M Soujanya Reddy, Pratheusa Maccha, Tulasi Nagabandi, Purushotham Vodnala, Payel Mukherjee, Sofia Banu, Priya Singh,Onkar Kulkarni, Dhiviya Vedagiri, Divya Gupta, Vishal Sah, Santosh Kumar Kuncha, Krishnan Harinivas Harshan, Archana Bharadwaj Siva, Karthik Bharadwaj Tallapaka, Zeba Rizvi, Zuberwasim Sayyad, Kakade Aishwarya Arun, Amrutha H C, Ananga Ghosh, Rakesh K Mishra, Divya Tej Sowpati |
| EPI_ISL_539776 | National Institute of Public Health (Czech Republic) | State Veterinary Institute Prague | Nagy, A; Jirincova, H; Novakova, L; Trnka, D; Vecerova, J. |
| EPI_ISL_539777 | The National Institute of Public Health | State Veterinary Institute Prague | Nagy,A;Jirincova,H;Novakova,L;Trnka,D;Vecerova,J |
| EPI_ISL_539778, EPI_ISL_539779, EPI_ISL_539780, EPI_ISL_539781, EPI_ISL_539782 | National Institute of Public Health (Czech Republic) | State Veterinary Institute Prague | Nagy, A; Jirincova, H; Novakova, L; Trnka, D; Vecerova, J. |
| EPI_ISL_539824, EPI_ISL_539825, EPI_ISL_539826, EPI_ISL_539827, EPI_ISL_539828, EPI_ISL_539829, EPI_ISL_539830, EPI_ISL_539831, EPI_ISL_539832, EPI_ISL_539833, EPI_ISL_539834, EPI_ISL_539835, EPI_ISL_539836, EPI_ISL_539837, EPI_ISL_539838, EPI_ISL_539839, EPI_ISL_539840, EPI_ISL_539841 | | | |
| see above | Minnesota Department of Health, Public Health Laboratory | Minnesota Department of Health, Public Health Laboratory | Matt Plumb, Jacob Garfin, and Xiong Wang |
| EPI_ISL_539842, EPI_ISL_539843, EPI_ISL_539844, EPI_ISL_539845, EPI_ISL_539846, EPI_ISL_539847, EPI_ISL_539848, EPI_ISL_539849 | Mayo Clinic & Mayo Clinic Laboratories | Minnesota Department of Health, Public Health Laboratory | Matt Plumb, Jacob Garfin, and Xiong Wang |
| EPI_ISL_540871, EPI_ISL_540872, EPI_ISL_540873, EPI_ISL_540874, EPI_ISL_540875, EPI_ISL_540876, EPI_ISL_540877, EPI_ISL_540878, EPI_ISL_540879, EPI_ISL_540880, EPI_ISL_540881, EPI_ISL_540882, EPI_ISL_540883, EPI_ISL_540884, EPI_ISL_540885, EPI_ISL_540886, EPI_ISL_540887, EPI_ISL_540888, EPI_ISL_540889, EPI_ISL_540890, EPI_ISL_540891, EPI_ISL_540892, EPI_ISL_540893 | | | |
| see above | Virology Department, Royal Infirmary of Edinburgh, NHS Lothian / School of Biological Sciences, University of Edinburgh / Institute of Genetics and Molecular Medicine, University of Edinburgh | COVID-19 Genomics UK (COG-UK) Consortium | McHugh M, Dewar R, Rooke S, Gallagher M, Balcaza C, O'Toole Á, Scher E, Hill V, McCrone JT, Colquhoun R, Yu X, Jackson B, Rambaut A, Williams TC, Templeton K |
| EPI_ISL_540993, EPI_ISL_540994, EPI_ISL_540995, EPI_ISL_540996, EPI_ISL_540997, EPI_ISL_540998, EPI_ISL_540999, EPI_ISL_541000, EPI_ISL_541001, EPI_ISL_541002, EPI_ISL_541003, EPI_ISL_541004, EPI_ISL_541005, EPI_ISL_541006, EPI_ISL_541007 | | | |
| see above | Health and Environmental Research Institute of Gwangju Metropolitan city | Health and Environmental Research Institute of Gwangju Metropolitan city | Min Ji Kim, Ji-eun Lee |
| EPI_ISL_541082 | The National Institute of Public Health | State Veterinary Institute Prague | Nagy,A;Jirincova,H;Novakova,L;Trnka,D;Vecerova,J |
| EPI_ISL_541138, EPI_ISL_541332, EPI_ISL_541333, EPI_ISL_541334 | The National Institute of Public Health | State Veterinary Institute Prague | Nagy,A; Jirincova,H; Novakova,L; Trnka,D; Vecerova,J |
| EPI_ISL_541335 | The National Institute of Public Health | Sídlištní 136/24 165 03, Prague Czech Republic | Nagy,A; Jirincova,H; Novakova,L; Trnka,D; Vecerova,J |
| EPI_ISL_541336, EPI_ISL_541337 | The National Institute of Public Health | State Veterinary Institute Prague | Nagy,A; Jirincova,H; Novakova,L; Trnka,D; Vecerova,J |
| EPI_ISL_541942 | Health and Environmental Research Institute of Gwangju | Health & Environment Institute of Gwangju | Min Ji Kim, Ji-eun Lee |
| EPI_ISL_545580, EPI_ISL_545954, EPI_ISL_545956 | The National Institute of Public Health | State Veterinary Institute Prague | Nagy,A;Jirincova,H;Novakova,L;Trnka,D;Vecerova,J |
| EPI_ISL_545957, EPI_ISL_546436, | The National Institute of Public Health | State Veterinary Institute Prague | Nagy,A; Jirincova,H; Novakova,L; Trnka,D; Vecerova,J |

| | | | |
|---|---|---|---|
| EPI_ISL_546935, EPI_ISL_546936 | | | |
| EPI_ISL_547966, EPI_ISL_547967, EPI_ISL_547968 | The National Institute of Public Health | State Veterinary Institute Prague | Nagy,A;Jirincova,H;Novakova,L;Trnka,D;Vecerova,J |
| EPI_ISL_548949, EPI_ISL_548950, EPI_ISL_548951, EPI_ISL_548952, EPI_ISL_548953, EPI_ISL_548954, EPI_ISL_548955, EPI_ISL_548956, EPI_ISL_548957, EPI_ISL_548958, EPI_ISL_548959, EPI_ISL_548960, EPI_ISL_548961, EPI_ISL_549025 | | | |
| see above | Max von Pettenkofer Institute, Virology, National Reference Center for Retroviruses, LMU München | Laboratory for Functional Genome Analysis, Dept. Genomics, Gene Center of the LMU Munich | Max Muenchhoff, Stefan Krebs, Alexander Graf, Oliver Keppler, Helmut Blum |
| EPI_ISL_559971, EPI_ISL_559972, EPI_ISL_559973, EPI_ISL_559974, EPI_ISL_559975, EPI_ISL_559976, EPI_ISL_559977, EPI_ISL_559978, EPI_ISL_559979, EPI_ISL_559980, EPI_ISL_559981, EPI_ISL_559982 | | | |
| see above | Virology Department, Royal Infirmary of Edinburgh, NHS Lothian / School of Biological Sciences, University of Edinburgh / Institute of Genetics and Molecular Medicine, University of Edinburgh | COVID-19 Genomics UK (COG-UK) Consortium | McHugh M, Dewar R, Rooke S, Gallagher M, Balcaza C, O'Toole Á, Scher E, Hill V, McCrone JT, Colquhoun R, Yu X, Jackson B, Rambaut A, Williams TC, Templeton K |
| EPI_ISL_560408, EPI_ISL_560409, EPI_ISL_560410, EPI_ISL_560411, EPI_ISL_560412 | The National Institute of Public Health | State Veterinary Institute Prague | Nagy, A; Jirincova, H; Novakova, L; Trnka, D; Vecerova, J |
| EPI_ISL_560743, EPI_ISL_560744, EPI_ISL_560745, EPI_ISL_560746, EPI_ISL_560747, EPI_ISL_560748, EPI_ISL_560749, EPI_ISL_560750, EPI_ISL_560751, EPI_ISL_560752, EPI_ISL_560753, EPI_ISL_560754, EPI_ISL_560755, EPI_ISL_560756, EPI_ISL_560757, EPI_ISL_560758, EPI_ISL_560759, EPI_ISL_560760, EPI_ISL_560761, EPI_ISL_560762, EPI_ISL_560763, EPI_ISL_560764, EPI_ISL_560765, EPI_ISL_560766, EPI_ISL_560767, EPI_ISL_560768, EPI_ISL_560769, EPI_ISL_560770, EPI_ISL_560771, EPI_ISL_560772, EPI_ISL_560773, EPI_ISL_560774, EPI_ISL_560775, EPI_ISL_560776, EPI_ISL_560777, EPI_ISL_560778, EPI_ISL_560779, EPI_ISL_560780, EPI_ISL_560781, EPI_ISL_560782, EPI_ISL_560783, EPI_ISL_560784, EPI_ISL_560785, EPI_ISL_560786, EPI_ISL_560787, EPI_ISL_560788, EPI_ISL_560789, EPI_ISL_560790, EPI_ISL_560791 | | | |
| see above | Minnesota Department of Health, Public Health Laboratory | Minnesota Department of Health, Public Health Laboratory | Matt Plumb, Jacob Garfin, and Xiong Wang |
| EPI_ISL_560792, EPI_ISL_560793, EPI_ISL_560794, EPI_ISL_560795, EPI_ISL_560796, EPI_ISL_560797 | Mayo Clinic & Mayo Clinic Laboratories | Minnesota Department of Health, Public Health Laboratory | Matt Plumb, Jacob Garfin, and Xiong Wang |
| EPI_ISL_560798, EPI_ISL_560799, EPI_ISL_560800, EPI_ISL_560801, EPI_ISL_560802, EPI_ISL_560803, EPI_ISL_560804, EPI_ISL_560805 | M Health Fairview | Minnesota Department of Health, Public Health Laboratory | Matt Plumb, Jacob Garfin, and Xiong Wang |
| EPI_ISL_560806, EPI_ISL_560807, EPI_ISL_560808, EPI_ISL_560809, EPI_ISL_560810, EPI_ISL_560811, EPI_ISL_560812, EPI_ISL_560813, EPI_ISL_560814, EPI_ISL_560815, EPI_ISL_560816, EPI_ISL_560817, EPI_ISL_560818, EPI_ISL_560819, EPI_ISL_560820, EPI_ISL_560821, EPI_ISL_560822, EPI_ISL_560823, EPI_ISL_560824, EPI_ISL_560825, EPI_ISL_560826, EPI_ISL_560827, EPI_ISL_560828, EPI_ISL_560829, EPI_ISL_560830, EPI_ISL_560831, EPI_ISL_560832 | | | |
| see above | Maryland Public Health Laboratory | Maryland Public Health Laboratory | Maryland Department of Health Laboratories Administration |
| EPI_ISL_568577 | The National Institute of Public Health | State Veterinary Institute Prague | Nagy, A; Jirincova, H; Novakova, L; Trnka,D; Vecerova, J |
| EPI_ISL_568873, EPI_ISL_568874, EPI_ISL_568875 | Malaysia Genome Institute | Malaysia Genome Institute | Mohd Noor Mat Isa, Irni Suhayu Sapian, Yusuf Muhammad Noor, Nurhezreen Md Iqbal, Mohd Faizal Abu Bakar, Enizza Kasim, Shamsidar Sopie, Siti Noraini Othman, Azrin Ahmad, Nor Azfa Johari, Shahrul Hisham Zainal Ariffin |
| EPI_ISL_569609 | Bennet County Hospital | South Dakota Public Health Laboratory | Matt Plumb, Jacob Garfin, Xiong Wang, and Chris Carlson |
| EPI_ISL_569610 | Avera McKennan Hospital | South Dakota Public Health Laboratory | Matt Plumb, Jacob Garfin, Xiong Wang, and Chris Carlson |
| EPI_ISL_569611 | Sioux Falls Urgent Care | South Dakota Public Health Laboratory | Matt Plumb, Jacob Garfin, Xiong Wang, and Chris Carlson |
| EPI_ISL_569612, EPI_ISL_569613 | Cheyenne River Health Center | South Dakota Public Health Laboratory | Matt Plumb, Jacob Garfin, Xiong Wang, and Chris Carlson |
| EPI_ISL_569614 | IHS Fort Yates Hospital | South Dakota Public Health Laboratory | Matt Plumb, Jacob Garfin, Xiong Wang, and Chris Carlson |
| EPI_ISL_569615 | Quick Care Huron | South Dakota Public Health Laboratory | Matt Plumb, Jacob Garfin, Xiong Wang, and Chris Carlson |
| EPI_ISL_569616 | Aurora County Clinic | South Dakota Public Health Laboratory | Matt Plumb, Jacob Garfin, Xiong Wang, and Chris Carlson |
| EPI_ISL_569617 | SD Urban Indian Health Pierre | South Dakota Public Health Laboratory | Matt Plumb, Jacob Garfin, Xiong Wang, and Chris Carlson |
| EPI_ISL_569618, EPI_ISL_569619 | Bethel Lutheran Home | South Dakota Public Health Laboratory | Matt Plumb, Jacob Garfin, Xiong Wang, and Chris Carlson |
| EPI_ISL_569620 | McCrossan Boys Ranch | South Dakota Public Health Laboratory | Matt Plumb, Jacob Garfin, Xiong Wang, and Chris Carlson |
| EPI_ISL_569621 | Quick Care Watertown | South Dakota Public Health Laboratory | Matt Plumb, Jacob Garfin, Xiong Wang, and Chris Carlson |
| EPI_ISL_569622 | Woodrow Wilson Keeble Memorial Healthcare | South Dakota Public Health Laboratory | Matt Plumb, Jacob Garfin, Xiong Wang, and Chris Carlson |
| EPI_ISL_569623 | Quick Care Watertown | South Dakota Public Health Laboratory | Matt Plumb, Jacob Garfin, Xiong Wang, and Chris Carlson |
| EPI_ISL_569624 | Avera McKennan Hospital | South Dakota Public Health Laboratory | Matt Plumb, Jacob Garfin, Xiong Wang, and Chris Carlson |
| EPI_ISL_569865, EPI_ISL_569866, EPI_ISL_569867, EPI_ISL_569868, EPI_ISL_569869, EPI_ISL_569870, EPI_ISL_569871, EPI_ISL_569872, EPI_ISL_569873, EPI_ISL_569874, EPI_ISL_569875, EPI_ISL_569876, EPI_ISL_569877, EPI_ISL_569878, EPI_ISL_569879, EPI_ISL_569880, EPI_ISL_569881, EPI_ISL_569882, EPI_ISL_569883, EPI_ISL_569884, EPI_ISL_569885, EPI_ISL_569886 | | | |
| see above | Amedeo di savoia | Crosetto lab, Karolinska Institutet, SciLifeLab | Michele Simonetti, Maria Grazia Milia, Luuk Harbers, Ning Zhang, Anna Sapino, Valeria Ghisetti, Nicola Crosetto |
| EPI_ISL_572320, EPI_ISL_572321, EPI_ISL_572322, EPI_ISL_572323, EPI_ISL_572324 | IZSM | IZSM | Maurizio Viscardi, Lorena Cardillo, Giovanna Fusco |
| EPI_ISL_572330, EPI_ISL_572331, EPI_ISL_572333, EPI_ISL_572397 | Institute for Virology, University Hospital Duesseldorf, Medical Faculty, Heinrich-Heine-University Duesseldorf | Institute for Virology, University Hospital Duesseldorf, Medical Faculty, Heinrich-Heine-University Duesseldorf | Maximilian Damagnez, Verena Keitel, Björn Jensen, Nadine Lübke, Lisa Müller, Philipp Ostermann, Tina Senff, Ortwin Adams, Philipp Albrecht, Gerald Antoch, Johannes Bode, Edwin Bölke, Saskia Elben, Torsten Feldt, Johannes C. Fischer, , Anselm Kunstein, Caroline Klindt, Alexander Killer, Tom Lüdde, Annemarie Mohring, Jennifer Neubert, Heiner Schaal, Ansgar Schulz, Jörg Timm, Andreas Walker |
| EPI_ISL_572411, EPI_ISL_572413, EPI_ISL_572414, EPI_ISL_572416, EPI_ISL_572421, EPI_ISL_572425, EPI_ISL_572427, EPI_ISL_572435, EPI_ISL_572442, EPI_ISL_572483, EPI_ISL_572490, EPI_ISL_572514, EPI_ISL_572535, EPI_ISL_572536, EPI_ISL_572537, EPI_ISL_572538, EPI_ISL_572539, EPI_ISL_572542, EPI_ISL_572543, EPI_ISL_572544, EPI_ISL_572573, EPI_ISL_572590, EPI_ISL_572593, EPI_ISL_572599, EPI_ISL_572608, EPI_ISL_572610, EPI_ISL_572617, EPI_ISL_572689, EPI_ISL_572692, EPI_ISL_572707, EPI_ISL_572756, EPI_ISL_572760, EPI_ISL_572762, EPI_ISL_572774, EPI_ISL_572776, EPI_ISL_572777, EPI_ISL_572801, EPI_ISL_572803, EPI_ISL_572821, EPI_ISL_572834, EPI_ISL_572839, EPI_ISL_572871, EPI_ISL_572874, EPI_ISL_572955 | | | |
| see above | Virology Department, Royal Infirmary of Edinburgh, NHS Lothian / School of Biological Sciences, University of Edinburgh / Institute of Genetics and Molecular Medicine, University of Edinburgh | COVID-19 Genomics UK (COG-UK) Consortium | McHugh M, Dewar R, Rooke S, Gallagher M, Balcaza C, O'Toole Á, Scher E, Hill V, McCrone JT, Colquhoun R, Yu X, Jackson B, Rambaut A, Williams TC, Templeton K |
| EPI_ISL_573152 | Institute for Virology, University Hospital Duesseldorf, Medical Faculty, Heinrich-Heine-University Duesseldorf | Institute for Virology, University Hospital Duesseldorf, Medical Faculty, Heinrich-Heine-University Duesseldorf | Maximilian Damagnez, Verena Keitel, Björn Jensen, Nadine Lübke, Lisa Müller, Philipp Ostermann, Tina Senff, Ortwin Adams, Philipp Albrecht, Gerald Antoch, Johannes Bode, Edwin Bölke, Saskia Elben, Torsten Feldt, Johannes C. Fischer, , Anselm Kunstein, Caroline Klindt, Alexander Killer, Tom Lüdde, Annemarie Mohring, Jennifer Neubert, Heiner Schaal, Ansgar Schulz, Jörg Timm, Andreas Walker |
| EPI_ISL_573247, EPI_ISL_573248, EPI_ISL_573249, EPI_ISL_573250, EPI_ISL_573251, EPI_ISL_573252, EPI_ISL_573794, EPI_ISL_573795, EPI_ISL_573796, EPI_ISL_573797, EPI_ISL_573798 | | | |
| see above | Virology Department, Royal Infirmary of Edinburgh, NHS Lothian / School of Biological Sciences, University of Edinburgh / Institute of Genetics and Molecular Medicine, University of Edinburgh | COVID-19 Genomics UK (COG-UK) Consortium | McHugh M, Dewar R, Rooke S, Gallagher M, Balcaza C, O'Toole Á, Scher E, Hill V, McCrone JT, Colquhoun R, Yu X, Jackson B, Rambaut A, Williams TC, Templeton K |

| | | | |
|---|---|---|---|
| EPI_ISL_574259 | Institute for Virology, University Hospital Duesseldorf, Medical Faculty, Heinrich-Heine-University Duesseldorf | Institute for Virology, University Hospital Duesseldorf, Medical Faculty, Heinrich-Heine-University Duesseldorf | Maximilian Damagnez, Verena Keitel, Björn Jensen, Nadine Lübke, Lisa Müller, Philipp Ostermann, Tina Senff, Ortwin Adams, Philipp Albrecht, Gerald Antoch, Johannes Bode, Edwin Bölke, Saskia Elben, Torsten Feldt, Johannes C. Fischer, , Anselm Kunstein, Caroline Klindt, Alexander Killer, Tom Lüdde, Annemarie Mohring, Jennifer Neubert, Heiner Schaal, Ansgar Schulz, Jörg Timm, Andreas Walker |
| EPI_ISL_574790, EPI_ISL_574791, EPI_ISL_574792, EPI_ISL_574793, EPI_ISL_574794, EPI_ISL_574795, EPI_ISL_574796, EPI_ISL_574797, EPI_ISL_574798, EPI_ISL_574799, EPI_ISL_574800, EPI_ISL_574801, EPI_ISL_574802, EPI_ISL_574803, EPI_ISL_574804, EPI_ISL_574805, EPI_ISL_574806, EPI_ISL_574807, EPI_ISL_574808, EPI_ISL_574809, EPI_ISL_574810, EPI_ISL_574811, EPI_ISL_574812, EPI_ISL_574813, EPI_ISL_574814, EPI_ISL_574815, EPI_ISL_574816, EPI_ISL_574817, EPI_ISL_574818, EPI_ISL_574819, EPI_ISL_574820, EPI_ISL_574821, EPI_ISL_574822, EPI_ISL_574823, EPI_ISL_574824, EPI_ISL_574825, EPI_ISL_574826, EPI_ISL_574827, EPI_ISL_574828, EPI_ISL_574829, EPI_ISL_574830, EPI_ISL_574831, EPI_ISL_574832, EPI_ISL_574833, EPI_ISL_574834, EPI_ISL_574835, EPI_ISL_574836, EPI_ISL_574837, EPI_ISL_574838, EPI_ISL_574839, EPI_ISL_574840, EPI_ISL_574841, EPI_ISL_574842, EPI_ISL_574843, EPI_ISL_574844, EPI_ISL_574845, EPI_ISL_574846, EPI_ISL_574847, EPI_ISL_574848, EPI_ISL_574849, EPI_ISL_574850, EPI_ISL_574851, EPI_ISL_574852, EPI_ISL_574853, EPI_ISL_574854, EPI_ISL_574855, EPI_ISL_574856, EPI_ISL_574857, EPI_ISL_574858, EPI_ISL_574859, EPI_ISL_574860, EPI_ISL_574861, EPI_ISL_574862 | | | |
| see above | Institute for Infectious Diseases, University of Bern | Institute for Infectious Diseases, University of Bern | Michel C Koch, Christian Baumann, Miguel A Terrazos Miani, Cora Sägesser, Stephen L Leib, Peter Keller, Franziska Suter-Riniker, Alban Ramette |
| EPI_ISL_576228, EPI_ISL_576229, EPI_ISL_576230, EPI_ISL_576231, EPI_ISL_576232, EPI_ISL_576233, EPI_ISL_576234, EPI_ISL_576235, EPI_ISL_576236, EPI_ISL_576237, EPI_ISL_576238, EPI_ISL_576239, EPI_ISL_576240, EPI_ISL_576241, EPI_ISL_576242, EPI_ISL_576243, EPI_ISL_576244, EPI_ISL_576245, EPI_ISL_576246, EPI_ISL_576247, EPI_ISL_576248, EPI_ISL_576249, EPI_ISL_576250, EPI_ISL_576251, EPI_ISL_576252, EPI_ISL_576253, EPI_ISL_576254, EPI_ISL_576255, EPI_ISL_576256 | | | |
| see above | Minnesota Department of Health, Public Health Laboratory | Minnesota Department of Health, Public Health Laboratory | Matt Plumb, Jacob Garfin, Alexandra Lorentz, and Xiong Wang |
| EPI_ISL_577347, EPI_ISL_577348, EPI_ISL_577349, EPI_ISL_577350, EPI_ISL_577351, EPI_ISL_577352, EPI_ISL_577353, EPI_ISL_577354, EPI_ISL_577355, EPI_ISL_577356, EPI_ISL_577357, EPI_ISL_577358 | | | |
| see above | Virology Department, Royal Infirmary of Edinburgh, NHS Lothian / School of Biological Sciences, University of Edinburgh / Institute of Genetics and Molecular Medicine, University of Edinburgh | COVID-19 Genomics UK (COG-UK) Consortium | McHugh M, Dewar R, Rooke S, Gallagher M, Balcaza C, O'Toole Á, Scher E, Hill V, McCrone JT, Colquhoun R, Yu X, Jackson B, Rambaut A, Williams TC, Templeton K |
| EPI_ISL_577623, EPI_ISL_577624, EPI_ISL_577625, EPI_ISL_577626, EPI_ISL_577627, EPI_ISL_577628, EPI_ISL_577629, EPI_ISL_577630, EPI_ISL_577631, EPI_ISL_577632, EPI_ISL_577633, EPI_ISL_577634, EPI_ISL_577635, EPI_ISL_577636, EPI_ISL_577637, EPI_ISL_577638 | | | |
| see above | The National Institute of Public Health | State Veterinary Institute Prague | Nagy,A;Jirincova,H;Novakova,L;Trnka,D;Vecerova,J |
| EPI_ISL_578202, EPI_ISL_578203, EPI_ISL_578204, EPI_ISL_578205, EPI_ISL_578206, EPI_ISL_578207, EPI_ISL_578208, EPI_ISL_578209, EPI_ISL_578210, EPI_ISL_578211, EPI_ISL_578212, EPI_ISL_578213, EPI_ISL_578214, EPI_ISL_578215, EPI_ISL_578216, EPI_ISL_578217, EPI_ISL_578218, EPI_ISL_578219, EPI_ISL_578220, EPI_ISL_578221, EPI_ISL_578222, EPI_ISL_578223, EPI_ISL_578224, EPI_ISL_578225, EPI_ISL_578226, EPI_ISL_578227, EPI_ISL_578228, EPI_ISL_578229, EPI_ISL_578230, EPI_ISL_578231, EPI_ISL_578232, EPI_ISL_578233, EPI_ISL_578234, EPI_ISL_578235, EPI_ISL_578236, EPI_ISL_578237, EPI_ISL_578238, EPI_ISL_578239, EPI_ISL_578240, EPI_ISL_578241, EPI_ISL_578242, EPI_ISL_578243, EPI_ISL_578244, EPI_ISL_578245, EPI_ISL_578246, EPI_ISL_578247, EPI_ISL_578248, EPI_ISL_578249, EPI_ISL_578250, EPI_ISL_578251, EPI_ISL_578252, EPI_ISL_578253, EPI_ISL_578256, EPI_ISL_578257, EPI_ISL_578258, EPI_ISL_578259, EPI_ISL_578260, EPI_ISL_578261, EPI_ISL_578262, EPI_ISL_578263, EPI_ISL_578264, EPI_ISL_578265, EPI_ISL_578266, EPI_ISL_578267, EPI_ISL_578268, EPI_ISL_578269, EPI_ISL_578270, EPI_ISL_578271, EPI_ISL_578272, EPI_ISL_578273, EPI_ISL_578274, EPI_ISL_578275, EPI_ISL_578276, EPI_ISL_578277, EPI_ISL_578278, EPI_ISL_578279, EPI_ISL_578280, EPI_ISL_578281, EPI_ISL_578282, EPI_ISL_578283, EPI_ISL_578284, EPI_ISL_578285, EPI_ISL_578286, EPI_ISL_578287, EPI_ISL_578288, EPI_ISL_578289, EPI_ISL_578290, EPI_ISL_578291, EPI_ISL_578292, EPI_ISL_578293, EPI_ISL_578294, EPI_ISL_578295, EPI_ISL_578296, EPI_ISL_578297, EPI_ISL_578298, EPI_ISL_578299, EPI_ISL_578300, EPI_ISL_578301, EPI_ISL_578302, EPI_ISL_578303, EPI_ISL_578304, EPI_ISL_578305, EPI_ISL_578306, EPI_ISL_578307, EPI_ISL_578308, EPI_ISL_578309, EPI_ISL_578310, EPI_ISL_578311, EPI_ISL_578312, EPI_ISL_578313, EPI_ISL_578314, EPI_ISL_578315, EPI_ISL_578316, EPI_ISL_578317, EPI_ISL_578318, EPI_ISL_578319, EPI_ISL_578320, EPI_ISL_578321, EPI_ISL_578322, EPI_ISL_578323, EPI_ISL_578324, EPI_ISL_578325, EPI_ISL_578326, EPI_ISL_578327, EPI_ISL_578328, EPI_ISL_578329, EPI_ISL_578330, EPI_ISL_578331, EPI_ISL_578332, EPI_ISL_578333, EPI_ISL_578334 | | | |
| see above | National Virus Reference Laboratory | National Virus Reference Laboratory | Michael Carr, Gabriel Gonzalez, Jonathan Dean, Suzie Coughlan, Cillian F De Gascun |
| EPI_ISL_582091, EPI_ISL_582092, EPI_ISL_582093, EPI_ISL_582094, EPI_ISL_582095, EPI_ISL_582096, EPI_ISL_582097, EPI_ISL_582098, EPI_ISL_582099, EPI_ISL_582100, EPI_ISL_582101, EPI_ISL_582102, EPI_ISL_582103, EPI_ISL_582104, EPI_ISL_582105, EPI_ISL_582106, EPI_ISL_582107, EPI_ISL_582108, EPI_ISL_582109 | | | |
| see above | Hospital Universitario Marqués de Valdecilla - IDIVAL (Santander, Cantabria) | SeqCOVID-SPAIN consortium/IBV(CSIC) | María Eliecer Cano García, Mónica Gozalo Margüello, Jose Manuel Méndez Legaza, Daniel Pablo Marcos, Jesús Rodríguez Rodríguez, María Siller Ruiz and SeqCOVID-SPAIN consortium |
| EPI_ISL_582123 | UOC Microbiologia e Virologia, Azienda Ospedaliera Universitaria Senese, Siena, Italy | Dipartimento di Biotecnologie Mediche | Maria Grazia Cusi, David Pinzauti, Claudia Gandolfo, Gabriele Anichini, Gianni Pozzi, Francesco Santoro |
| EPI_ISL_582124 | Malaysia Genome Institute | Malaysia Genome Institute | Mohd Noor Mat Isa, Irni Suhayu Sapian, Yusuf Muhammad Noor, Nurhezreen Md Iqbal, Mohd Faizal Abu Bakar, Enizza Kasim, Shamsidar Sopie, Siti Noraini Othman, Azrin Ahmad, Nor Azfa Johari, Shahrul Hisham Zainal Ariffin |
| EPI_ISL_583954, EPI_ISL_583955, EPI_ISL_583956, EPI_ISL_583957, EPI_ISL_583958, EPI_ISL_583959, EPI_ISL_583960, EPI_ISL_583961, EPI_ISL_583962, EPI_ISL_583963, EPI_ISL_583964, EPI_ISL_583965, EPI_ISL_583966, EPI_ISL_583967 | | | |
| see above | UOC Microbiologia e Virologia, Azienda Ospedaliera Universitaria Senese, Siena, Italy | Dipartimento di Biotecnologie Mediche | Maria Grazia Cusi, David Pinzauti, Claudia Gandolfo, Gabriele Anichini, Gianni Pozzi, Francesco Santoro |
| EPI_ISL_584069, EPI_ISL_584071, EPI_ISL_584072 | IZSM | IZSM | Maurizio Viscardi, Lorena Cardillo, Giovanna Fusco |
| EPI_ISL_584073, EPI_ISL_584074, EPI_ISL_584075, EPI_ISL_584076, EPI_ISL_584077, EPI_ISL_584078, EPI_ISL_584079, EPI_ISL_584080, EPI_ISL_584081, EPI_ISL_584082 | The National Institute of Public Health | State Veterinary Institute Prague | Nagy,A;Jirincova,H;Novakova,L;Trnka,D;Vecerova,J |
| EPI_ISL_585426, EPI_ISL_585427, EPI_ISL_585428, EPI_ISL_585429, EPI_ISL_585430, EPI_ISL_585431, EPI_ISL_585432, EPI_ISL_585433, EPI_ISL_585434, EPI_ISL_585435, EPI_ISL_585436, EPI_ISL_585437, EPI_ISL_585438, EPI_ISL_585439, EPI_ISL_585440, EPI_ISL_585441, EPI_ISL_585442, EPI_ISL_585443, EPI_ISL_585444, EPI_ISL_585445, EPI_ISL_585446, EPI_ISL_585447, EPI_ISL_585448, EPI_ISL_585449, EPI_ISL_585450, EPI_ISL_585451, EPI_ISL_585452, EPI_ISL_585453, EPI_ISL_585454, EPI_ISL_585455, EPI_ISL_585456, EPI_ISL_585457, EPI_ISL_585458, EPI_ISL_585459, EPI_ISL_585460, EPI_ISL_585461, EPI_ISL_585462, EPI_ISL_585463, EPI_ISL_585464, EPI_ISL_585465, EPI_ISL_585466, EPI_ISL_585467, EPI_ISL_585468, EPI_ISL_585469, EPI_ISL_585470, EPI_ISL_585471, EPI_ISL_585472, EPI_ISL_585473, EPI_ISL_585474, EPI_ISL_585475, EPI_ISL_585476, EPI_ISL_585477, EPI_ISL_585478, EPI_ISL_585479, EPI_ISL_585480, EPI_ISL_585481, EPI_ISL_585482, EPI_ISL_585483, EPI_ISL_585484, EPI_ISL_585485, EPI_ISL_585486, EPI_ISL_585487, EPI_ISL_585488, EPI_ISL_585489, EPI_ISL_585490, EPI_ISL_585491, EPI_ISL_585492, EPI_ISL_585493, EPI_ISL_585494, EPI_ISL_585495, EPI_ISL_585496, EPI_ISL_585497, EPI_ISL_585498, EPI_ISL_585499, EPI_ISL_585500, EPI_ISL_585501 | | | |
| see above | Virology Department, Royal Infirmary of Edinburgh, NHS Lothian / School of Biological Sciences, University of Edinburgh / Institute of Genetics and Molecular Medicine, University of Edinburgh | COVID-19 Genomics UK (COG-UK) Consortium | McHugh M, Dewar R, Rooke S, Gallagher M, Balcaza C, O'Toole Á, Scher E, Hill V, McCrone JT, Colquhoun R, Yu X, Jackson B, Rambaut A, Williams TC, Templeton K |
| EPI_ISL_586529 | B.J. Medical College and Civil hospital, Ahmedabad | Gujarat Biotechnology Research Centre | Monika Gandhi, Pinal Trivedi, Maharshi Pandya, Nidhi Patel, Nitin Savaliya, Raghawendra Kumar, Dinesh Kumar, Zuber Saiyed, Komal Patel, Labdhi Pandya, Afzal Ansari, Nikha Trivedi, Pranay Shah, Kamlesh J Upadhyay, Sanjay Kapadia, Apurvasinh Puvar, Janvi Raval, Zarna Patel, R D Dixit, A M Kadri, Harsh Bakshi, Chaitanya Joshi, Madhvi Joshi |
| EPI_ISL_586537 | Zydus Medical Hospital,Dahod | Gujarat Biotechnology Research Centre | Mohit Desai, Nitin Savaliya, Raghawendra Kumar, Dinesh Kumar, Zuber Saiyed, Komal Patel, Labdhi Pandya, Afzal Ansari, Nikha Trivedi, Apurvasinh Puvar, Janvi Raval, Zarna Patel, Monika Gandhi, Pinal Trivedi, Maharshi Pandya, Nidhi Patel, Kalpesh Labana, R D Dixit, A M Kadri, Harsh Bakshi, Chaitanya Joshi, Madhvi Joshi |
| EPI_ISL_586551 | Government Hospital,Veraval | Gujarat Biotechnology Research Centre | Monika Gandhi, Pinal Trivedi, Maharshi Pandya, Nidhi Patel, Nitin Savaliya, Raghawendra Kumar, Dinesh Kumar, Zuber Saiyed, Komal Patel, Labdhi Pandya, Afzal Ansari, Nikha Trivedi, Jignesh Parmar, Jitendra Bamrotia, Apurvasinh Puvar, Janvi Raval, Zarna Patel, R D Dixit, A M Kadri, Harsh Bakshi, Chaitanya Joshi, Madhvi Joshi |
| EPI_ISL_590693 | INMI Lazzaro Spallanzani IRCCS | INMI Lazzaro Spallanzani IRCCS | Martina Rueca, Barbara Bartolini, Cesare E.M. Gruber, Francesco Messina, Emanuela Giombini, Beatrice Valli, Eleonora Lalle, Simone Lanini, Francesco Vairo, Maria R. Capobianchi, Antonino Di Caro |
| EPI_ISL_590697 | INMI Lazzaro Spallanzani IRCCS | INMI Lazzaro Spallanzani IRCCS | Martina Rueca, Cesare E.M. Gruber, Barbara Bartolini, Francesco Messina, Emanuela Giombini, Beatrice Valli, Eleonora Lalle, Simone Lanini, Francesco Vairo, Antonino Di Caro, Maria R. Capobianchi |
| EPI_ISL_590753, EPI_ISL_590754, EPI_ISL_590755, EPI_ISL_590756, EPI_ISL_590757, EPI_ISL_590758, EPI_ISL_590759, EPI_ISL_590760, EPI_ISL_590761, EPI_ISL_590762, EPI_ISL_590763, EPI_ISL_590764, EPI_ISL_590765, EPI_ISL_590766, EPI_ISL_590767, EPI_ISL_590768, EPI_ISL_590769, EPI_ISL_590770, EPI_ISL_590771, EPI_ISL_590772, EPI_ISL_590773, EPI_ISL_590774, EPI_ISL_590775, EPI_ISL_590776, EPI_ISL_590777, EPI_ISL_590778, EPI_ISL_590779, EPI_ISL_590780, EPI_ISL_590781, EPI_ISL_590782, EPI_ISL_590783, EPI_ISL_590784, EPI_ISL_590785, EPI_ISL_590786, EPI_ISL_590787, EPI_ISL_590788, EPI_ISL_590789, EPI_ISL_590790, EPI_ISL_590791, EPI_ISL_590792, EPI_ISL_590793, EPI_ISL_590794, EPI_ISL_590795, EPI_ISL_590796, EPI_ISL_590797, EPI_ISL_590798, EPI_ISL_590799, EPI_ISL_590800, EPI_ISL_590801, EPI_ISL_590802, EPI_ISL_590803, EPI_ISL_590804, EPI_ISL_590805, EPI_ISL_590806, EPI_ISL_590807, EPI_ISL_590808, EPI_ISL_590809, EPI_ISL_590810, EPI_ISL_590811, EPI_ISL_590812, EPI_ISL_590813, EPI_ISL_590814, EPI_ISL_590815, EPI_ISL_590816, EPI_ISL_590817, EPI_ISL_590818, EPI_ISL_590819, EPI_ISL_590820, EPI_ISL_590821, EPI_ISL_590822 | | | |
| see above | Minnesota Department of Health, Public Health Laboratory | Minnesota Department of Health, Public Health Laboratory | Matt Plumb, Jacob Garfin, Alexandra Lorentz, and Xiong Wang |

| | | | |
|---|---|---|---|
| EPI_ISL_590823, EPI_ISL_590824, EPI_ISL_590825, EPI_ISL_590826, EPI_ISL_590827, EPI_ISL_590828, EPI_ISL_590829, EPI_ISL_590830, EPI_ISL_590831, EPI_ISL_590832, EPI_ISL_590833, EPI_ISL_590834, EPI_ISL_590835, EPI_ISL_590836 | | | |
| see above | Institute of Medical Virology, University of Zurich | Institute of Medical Virology, University of Zurich | Marie O. Pohl, Idoia Busnadiego, Verena Kufner, Stefan Schmutz, Maryam Zaheri, Irene Abela, Alexandra Trkola, Michael Huber, Silke Stertz, Benjamin G. Hale |
| EPI_ISL_590837 | Minnesota Department of Health, Public Health Laboratory | Minnesota Department of Health, Public Health Laboratory | Matt Plumb, Jacob Garfin, Alexandra Lorentz, and Xiong Wang |
| EPI_ISL_591022, EPI_ISL_591023, EPI_ISL_591024, EPI_ISL_591025, EPI_ISL_591026, EPI_ISL_591027, EPI_ISL_591028, EPI_ISL_591029, EPI_ISL_591030, EPI_ISL_591031, EPI_ISL_591032, EPI_ISL_591033, EPI_ISL_591034, EPI_ISL_591035, EPI_ISL_591036 | | | |
| see above | MD PHL | MD PHL | Maryland Department of Health Laboratories Administration |
| EPI_ISL_594816, EPI_ISL_594817, EPI_ISL_594818, EPI_ISL_594819, EPI_ISL_594820, EPI_ISL_594821, EPI_ISL_594822, EPI_ISL_594823, EPI_ISL_594824, EPI_ISL_594825, EPI_ISL_594826, EPI_ISL_594827, EPI_ISL_594828, EPI_ISL_594829, EPI_ISL_594830, EPI_ISL_594831, EPI_ISL_594832, EPI_ISL_594833, EPI_ISL_594834, EPI_ISL_594835, EPI_ISL_594836, EPI_ISL_594837, EPI_ISL_594838, EPI_ISL_594839, EPI_ISL_594840, EPI_ISL_594841, EPI_ISL_594842, EPI_ISL_594843, EPI_ISL_594844, EPI_ISL_594845, EPI_ISL_594846, EPI_ISL_594847, EPI_ISL_594848 | | | |
| see above | Virology Department, Royal Infirmary of Edinburgh, NHS Lothian / School of Biological Sciences, University of Edinburgh / Institute of Genetics and Molecular Medicine, University of Edinburgh | COVID-19 Genomics UK (COG-UK) Consortium | McHugh M, Dewar R, Rooke S, Gallagher M, Balcaza C, O'Toole Á, Scher E, Hill V, McCrone JT, Colquhoun R, Yu X, Jackson B, Rambaut A, Williams TC, Templeton K |
| EPI_ISL_596627, EPI_ISL_596628, EPI_ISL_596629, EPI_ISL_596630, EPI_ISL_596631, EPI_ISL_596632, EPI_ISL_596633, EPI_ISL_596634, EPI_ISL_596635, EPI_ISL_596636, EPI_ISL_596637, EPI_ISL_596638, EPI_ISL_596639, EPI_ISL_596640, EPI_ISL_596641, EPI_ISL_596642, EPI_ISL_596643, EPI_ISL_596644, EPI_ISL_596645, EPI_ISL_596646, EPI_ISL_596647, EPI_ISL_596648, EPI_ISL_596649, EPI_ISL_596650, EPI_ISL_596651, EPI_ISL_596652, EPI_ISL_596653, EPI_ISL_596654, EPI_ISL_596655, EPI_ISL_596656, EPI_ISL_596657, EPI_ISL_596658, EPI_ISL_596659, EPI_ISL_596660, EPI_ISL_596661, EPI_ISL_596662, EPI_ISL_596663, EPI_ISL_596664, EPI_ISL_596665, EPI_ISL_596666, EPI_ISL_596667, EPI_ISL_596668, EPI_ISL_596669, EPI_ISL_596670, EPI_ISL_596671, EPI_ISL_596672, EPI_ISL_596673, EPI_ISL_596674, EPI_ISL_596675, EPI_ISL_596676 | | | |
| see above | St.Vincent's University Hospital | St.Vincent's University Hospital | Mary Lucey, Guerrino Macori, Niamh Mullane, Una Sutton-Fitzpatrick, Gabriel Gonzalez, Suzie Coughlan, Aisling Purcell, Lynda Fenelon, Séamus Fanning, Kirsten Schaffer |
| EPI_ISL_596889, EPI_ISL_596890, EPI_ISL_596891, EPI_ISL_596892, EPI_ISL_596893, EPI_ISL_596894, EPI_ISL_596895, EPI_ISL_596896, EPI_ISL_596897, EPI_ISL_596898, EPI_ISL_596899, EPI_ISL_596900, EPI_ISL_596901, EPI_ISL_596902, EPI_ISL_596903, EPI_ISL_596904, EPI_ISL_596905, EPI_ISL_596906, EPI_ISL_596907, EPI_ISL_596908, EPI_ISL_596909, EPI_ISL_596910, EPI_ISL_596911, EPI_ISL_596912, EPI_ISL_596913, EPI_ISL_596914, EPI_ISL_596915, EPI_ISL_596916, EPI_ISL_596917, EPI_ISL_596918, EPI_ISL_596919, EPI_ISL_596920, EPI_ISL_596921, EPI_ISL_596922, EPI_ISL_596923, EPI_ISL_596924, EPI_ISL_596925, EPI_ISL_596926, EPI_ISL_596927 | | | |
| see above | National Virus Reference Laboratory | National Virus Reference Laboratory | Michael Carr, Gabriel Gonzalez, Jonathan Dean, Daniel Hare, Cillian F De Gascun |
| EPI_ISL_602258, EPI_ISL_602259, EPI_ISL_602260, EPI_ISL_602261, EPI_ISL_602262, EPI_ISL_602263, EPI_ISL_602264, EPI_ISL_602265, EPI_ISL_602266, EPI_ISL_602267, EPI_ISL_602268, EPI_ISL_602269, EPI_ISL_602270, EPI_ISL_602271, EPI_ISL_602272, EPI_ISL_602273, EPI_ISL_602274, EPI_ISL_602275, EPI_ISL_602276, EPI_ISL_602277 | | | |
| see above | MD PHL | MD PHL | Maryland Department of Health Laboratories Administration |
| EPI_ISL_602304 | Istituto Zooprofilattico Sperimentale del Mezzogiorno | U.O. Diagnostica Virologica Dip. Sanità Animale IZSM | Maurizio Viscardi, Lorena Cardillo , e Giovanna Fusco |
| EPI_ISL_602510, EPI_ISL_602511, EPI_ISL_602512, EPI_ISL_602513, EPI_ISL_602514, EPI_ISL_602515, EPI_ISL_602516, EPI_ISL_602517 | Center of Medical Microbiology, Virology, and Hospital Hygiene, University of Duesseldorf | Center of Medical Microbiology, Virology, and Hospital Hygiene, University of Duesseldorf | Maximilian Damagnez, Alexander Dilthey, Torsten Houwaart, Lisanna Hülse, Malte Kohns Vasconcelos, Marek Korencak, Nadine Lübke, Jessica Nicolai, Klaus Pfeffer, Hendrik Streeck, Daniel Strelow, Jörg Timm, Andreas Walker, Tobias Wienemann |
| EPI_ISL_602533, EPI_ISL_602534, EPI_ISL_602535, EPI_ISL_602536, EPI_ISL_602537, EPI_ISL_602538, EPI_ISL_602539, EPI_ISL_602540, EPI_ISL_602541, EPI_ISL_602542, EPI_ISL_602543, EPI_ISL_602544, EPI_ISL_602545, EPI_ISL_602546, EPI_ISL_602547, EPI_ISL_602548, EPI_ISL_602549, EPI_ISL_602550, EPI_ISL_602551, EPI_ISL_602552 | | | |
| see above | Center of Medical Microbiology, Virology, and Hospital Hygiene, University of Duesseldorf | Center of Medical Microbiology, Virology, and Hospital Hygiene, University of Duesseldorf | Maximilian Damagnez, Alexander Dilthey, Torsten Houwaart, Malte Kohns Vasconcelos, Marek Korencak, Nadine Lübke, Jessica Nicolai, Klaus Pfeffer, Hendrik Streeck, Daniel Strelow, Jörg Timm, Andreas Walker, Tobias Wienemann |
| EPI_ISL_602956, EPI_ISL_602957, EPI_ISL_602958, EPI_ISL_602959, EPI_ISL_602960, EPI_ISL_602961, EPI_ISL_602962, EPI_ISL_602963, EPI_ISL_602964, EPI_ISL_602965, EPI_ISL_602966, EPI_ISL_602967, EPI_ISL_602968, EPI_ISL_602969, EPI_ISL_602970, EPI_ISL_602971, EPI_ISL_602972, EPI_ISL_602973, EPI_ISL_602974, EPI_ISL_602975, EPI_ISL_602976, EPI_ISL_602977, EPI_ISL_602978, EPI_ISL_602979, EPI_ISL_602980, EPI_ISL_602981, EPI_ISL_602982, EPI_ISL_602983, EPI_ISL_602984, EPI_ISL_602985, EPI_ISL_602986, EPI_ISL_602987, EPI_ISL_602988, EPI_ISL_602989, EPI_ISL_602990, EPI_ISL_602991, EPI_ISL_602992, EPI_ISL_602993, EPI_ISL_602994, EPI_ISL_602995, EPI_ISL_602996, EPI_ISL_602997, EPI_ISL_602998, EPI_ISL_602999 | | | |
| see above | Minnesota Department of Health, Public Health Laboratory | Minnesota Department of Health, Public Health Laboratory | Matt Plumb, Jacob Garfin, Alexandra Lorentz, and Xiong Wang |
| EPI_ISL_603000, EPI_ISL_603001, EPI_ISL_603002, EPI_ISL_603003 | Sanford South University Medical Center | Minnesota Department of Health, Public Health Laboratory | Matt Plumb, Jacob Garfin, Alexandra Lorentz, and Xiong Wang |
| EPI_ISL_603004, EPI_ISL_603005 | Essentia Health-St. Mary's Medical Center | Minnesota Department of Health, Public Health Laboratory | Matt Plumb, Jacob Garfin, Alexandra Lorentz, and Xiong Wang |
| EPI_ISL_603006, EPI_ISL_603007, EPI_ISL_603008, EPI_ISL_603009 | Minnesota Department of Health, Public Health Laboratory | Minnesota Department of Health, Public Health Laboratory | Matt Plumb, Jacob Garfin, Alexandra Lorentz, and Xiong Wang |
| EPI_ISL_603010, EPI_ISL_603011 | Mayo Clinic & Mayo Clinic Laboratories | Minnesota Department of Health, Public Health Laboratory | Matt Plumb, Jacob Garfin, Alexandra Lorentz, and Xiong Wang |
| EPI_ISL_603012, EPI_ISL_603013, EPI_ISL_603014 | Essentia Health-St. Mary's Medical Center | Minnesota Department of Health, Public Health Laboratory | Matt Plumb, Jacob Garfin, Alexandra Lorentz, and Xiong Wang |
| EPI_ISL_603015 | Minnesota Department of Health, Public Health Laboratory | Minnesota Department of Health, Public Health Laboratory | Matt Plumb, Jacob Garfin, Alexandra Lorentz, and Xiong Wang |
| EPI_ISL_603016, EPI_ISL_603017 | Essentia Health-St. Mary's Medical Center | Minnesota Department of Health, Public Health Laboratory | Matt Plumb, Jacob Garfin, Alexandra Lorentz, and Xiong Wang |
| EPI_ISL_603018, EPI_ISL_603019, EPI_ISL_603020 | Minnesota Department of Health, Public Health Laboratory | Minnesota Department of Health, Public Health Laboratory | Matt Plumb, Jacob Garfin, Alexandra Lorentz, and Xiong Wang |
| EPI_ISL_603138 | INMI Lazzaro Spallanzani IRCCS | INMI Lazzaro Spallanzani IRCCS | Martina Rueca, Francesco Messina, Barbara Bartolini, Cesare E.M. Gruber, Emanuela Giombini, Simone Lanini, Antonino Di Caro, Maria R. Capobianchi |
| EPI_ISL_603139 | INMI Lazzaro Spallanzani IRCCS | INMI Lazzaro Spallanzani IRCCS | Martina Rueca, Cesare E.M. Gruber, Barbara Bartolini, Francesco Messina, Emanuela Giombini, Simone Lanini, Antonino Di Caro, Maria R. Capobianchi |
| EPI_ISL_603140 | INMI Lazzaro Spallanzani IRCCS | INMI Lazzaro Spallanzani IRCCS | Martina Rueca, Cesare E.M. Gruber, Francesco Messina, Barbara Bartolini, Emanuela Giombini, Simone Lanini, Antonino Di Caro, Maria R. Capobianchi |
| EPI_ISL_603143, EPI_ISL_603154 | INMI Lazzaro Spallanzani IRCCS | INMI Lazzaro Spallanzani IRCCS | Martina Rueca, Francesco Messina, Cesare E.M. Gruber, Barbara Bartolini, Emanuela Giombini, Simone Lanini, Antonino Di Caro, Maria R. Capobianchi |
| EPI_ISL_603155 | INMI Lazzaro Spallanzani IRCCS | INMI Lazzaro Spallanzani IRCCS | Martina Rueca, Francesco Messina, Barbara Bartolini, Cesare E.M. Gruber, Simone Lanini, Emanuela Giombini, Maria R. Capobianchi, Antonino Di Caro |
| EPI_ISL_603165 | INMI Lazzaro Spallanzani IRCCS | INMI Lazzaro Spallanzani IRCCS | Martina Rueca, Francesco Messina, Barbara Bartolini, Cesare E.M. Gruber, Emanuela Giombini, Simone Lanini, Fulvia Pimpinelli, Maria R. Capobianchi, Antonino Di Caro |
| EPI_ISL_603166 | INMI Lazzaro Spallanzani IRCCS | INMI Lazzaro Spallanzani IRCCS | Martina Rueca, Cesare E.M. Gruber, Barbara Bartolini, Francesco Messina, Simone Lanini, Emanuela Giombini, Fulvia Pimpinelli, Maria R. Capobianchi, Antonino Di Caro |
| EPI_ISL_603167 | INMI Lazzaro Spallanzani IRCCS | INMI Lazzaro Spallanzani IRCCS | Martina Rueca, Cesare E.M. Gruber, Francesco Messina, Barbara Bartolini, Simone Lanini, Emanuela Giombini, Fulvia Pimpinelli, Maria R. Capobianchi, Antonino Di Caro |
| EPI_ISL_603168 | INMI Lazzaro Spallanzani IRCCS | INMI Lazzaro Spallanzani IRCCS | Martina Rueca, Barbara Bartolini, Cesare E.M. Gruber, Simone Lanini, Emanuela Giombini, Francesco Messina, Fulvia Pimpinelli, Antonino Di Caro, Maria R. Capobianchi |
| EPI_ISL_603170 | INMI Lazzaro Spallanzani IRCCS | INMI Lazzaro Spallanzani IRCCS | Martina Rueca, Barbara Bartolini, Cesare E.M. Gruber, Francesco Messina, Simone Lanini, Emanuela Giombini, Maria R. Capobianchi, Antonino Di Caro |
| EPI_ISL_603178 | INMI Lazzaro Spallanzani IRCCS | INMI Lazzaro Spallanzani IRCCS | Martina Rueca, Francesco Messina, Cesare E.M. Gruber, Barbara Bartolini, Simone Lanini, Emanuela Giombini, Antonino Di Caro, Maria R. Capobianchi |
| EPI_ISL_603182 | INMI Lazzaro Spallanzani IRCCS | INMI Lazzaro Spallanzani IRCCS | Martina Rueca, Barbara Bartolini, Cesare E.M. Gruber, Francesco Messina, Emanuela Giombini, Simone Lanini, Maria R. Capobianchi, Antonino Di Caro |
| EPI_ISL_603184 | INMI Lazzaro Spallanzani IRCCS | INMI Lazzaro Spallanzani IRCCS | Martina Rueca, Barbara Bartolini, Francesco Messina, Cesare E.M. Gruber, Emanuela Giombini, Simone Lanini, Maria R. Capobianchi, Antonino Di Caro |
| EPI_ISL_603223, EPI_ISL_603224, EPI_ISL_603225 | National Institute of Laboratory Medicine and Referral Center | Genomic Research Lab, BCSIR | Md. Murshed Hasan Sarkar, Abu Sayeed Mohammad Mahmud, Mohammad Samir Uzzaman, Eshrar Osman, Md. Ahashan Habib, Shahina Akter, Tanjina Akhter Banu, Barna Goswami, Iffat Jahan, Md. Saddam Hossain, Tasnim Nafisa, Md. Maruf Ahmed Molla, Mahmuda Yeasmin, Asish Kumar Ghosh, A. K. |

| | | | |
|---|---|---|---|
| | | | M. Shamsuzzaman, Monira Parveen, Md. Masum Hossain Arif, Md. Salim Khan |
| EPI_ISL_603246, EPI_ISL_603247 | National Institute of Laboratory Medicine and Referral Center | Genomic Research Lab, BCSIR | Md. Saddam Hossain, Abu Sayeed Mohammad Mahmud, Mohammad Samir Uzzaman, Eshrar Osman, Md. Ahashan Habib, Shahina Akter, Tanjina Akhter Banu, Md. Murshed Hasan Sarkar, Barna Goswami, Iffat Jahan, Tasnim Nafisa, Md. Maruf Ahmed Molla, Mahmuda Yeasmin, Asish Kumar Ghosh, A. K. M. Shamsuzzaman, Monira Parveen, Md. Masum Hossain Arif, Md. Salim Khan |
| EPI_ISL_603249, EPI_ISL_603250 | National Institute of Laboratory Medicine and Referral Center | Genomic Research Lab, BCSIR | Md. Ahashan Habib, Abu Sayeed Mohammad Mahmud, Mohammad Samir Uzzaman, Eshrar Osman, Shahina Akter, Tanjina Akhter Banu, Md. Murshed Hasan Sarkar, Barna Goswami, Iffat Jahan, Md. Saddam Hossain, Tasnim Nafisa, Md. Maruf Ahmed Molla, Mahmuda Yeasmin, Asish Kumar Ghosh, A. K. M. Shamsuzzaman, Monira Parveen, Md. Masum Hossain Arif, Md. Salim Khan |
| EPI_ISL_605783 | Genome Center | Genome Center | Md. Shazid Hasan, Hassan M. Al-Emran, Ovinu Kibria Islam, A. S. M. Rubayet- Ul- Alam, Selina Akter, Shireen Nigar, Md. Tanvir Islam, Pravas Chandra Roy, Shovon Lal Sarkar, Najmuj Sakib, S. M. Tanjil Shah, Md. Iqbal Kabir Jahid, Md. Anwar Hossain |
| EPI_ISL_610245 | Virology, National Institute for Biological Standards and Control | Virology, National Institute for Biological Standards and Control | Mee,E.T., Bentley,E., Routley,S., Mate,R., Fritzsche,M. and Mattiuzzo,G. |
| EPI_ISL_611544, EPI_ISL_611549, EPI_ISL_611685, EPI_ISL_611707, EPI_ISL_611842, EPI_ISL_611850, EPI_ISL_611894, EPI_ISL_611924, EPI_ISL_611989, EPI_ISL_612037, EPI_ISL_612074, EPI_ISL_612126, EPI_ISL_612127, EPI_ISL_612128, EPI_ISL_612129, EPI_ISL_612187, EPI_ISL_612188, EPI_ISL_612189, EPI_ISL_612314, EPI_ISL_612315, EPI_ISL_612316, EPI_ISL_612317, EPI_ISL_612318, EPI_ISL_612319, EPI_ISL_612320, EPI_ISL_612321, EPI_ISL_612322, EPI_ISL_612323, EPI_ISL_612324, EPI_ISL_612325, EPI_ISL_612326, EPI_ISL_612327, EPI_ISL_612328, EPI_ISL_612329, EPI_ISL_612330, EPI_ISL_612331, EPI_ISL_612332, EPI_ISL_612333, EPI_ISL_612334, EPI_ISL_612335, EPI_ISL_612336, EPI_ISL_612337, EPI_ISL_612338, EPI_ISL_612339, EPI_ISL_612340, EPI_ISL_612341, EPI_ISL_612342, EPI_ISL_612343, EPI_ISL_612344, EPI_ISL_612345, EPI_ISL_612346, EPI_ISL_612347, EPI_ISL_612348, EPI_ISL_612349, EPI_ISL_612350, EPI_ISL_612351, EPI_ISL_612352, EPI_ISL_612353, EPI_ISL_612354, EPI_ISL_612355, EPI_ISL_612356, EPI_ISL_612357, EPI_ISL_612358, EPI_ISL_612359, EPI_ISL_612360, EPI_ISL_612361, EPI_ISL_612362, EPI_ISL_612363, EPI_ISL_612364, EPI_ISL_612365, EPI_ISL_612366, EPI_ISL_612367, EPI_ISL_612368, EPI_ISL_612369, EPI_ISL_612370, EPI_ISL_612371, EPI_ISL_612372, EPI_ISL_612373, EPI_ISL_612374 | | | |
| see above | Virology Department, Royal Infirmary of Edinburgh, NHS Lothian / School of Biological Sciences, University of Edinburgh / Institute of Genetics and Molecular Medicine, University of Edinburgh | COVID-19 Genomics UK (COG-UK) Consortium | McHugh M, Dewar R, Rooke S, Gallagher M, Balcaza C, O'Toole Á, Scher E, Hill V, McCrone JT, Colquhoun R, Yu X, Jackson B, Rambaut A, Williams TC, Templeton K |
| EPI_ISL_613419, EPI_ISL_613420, EPI_ISL_613421, EPI_ISL_613422, EPI_ISL_613423, EPI_ISL_613424, EPI_ISL_613425, EPI_ISL_613426, EPI_ISL_613427, EPI_ISL_613428, EPI_ISL_613429, EPI_ISL_613430, EPI_ISL_613431, EPI_ISL_613432, EPI_ISL_613433, EPI_ISL_613434, EPI_ISL_613435, EPI_ISL_613436, EPI_ISL_613437, EPI_ISL_613438, EPI_ISL_613439, EPI_ISL_613440, EPI_ISL_613441, EPI_ISL_613442, EPI_ISL_613443, EPI_ISL_613444, EPI_ISL_613445, EPI_ISL_613446 | | | |
| see above | Institut Pasteur de la Guadeloupe | Institut Pasteur de la Guadeloupe | Marion Barbet, Sylvie Behillil, Méline Bizard, Angela Brisebarre, Camille Capel, Etienne Simon-Lorière, Vincent Enouf, Maud Vanpeene, Sylvie van der Werf, Stéphanie Guyomard, Sébastien Breurec, Antoine Talarmin |
| EPI_ISL_613447, EPI_ISL_613448, EPI_ISL_613449, EPI_ISL_613450, EPI_ISL_613451, EPI_ISL_613452, EPI_ISL_613453, EPI_ISL_613454, EPI_ISL_613455, EPI_ISL_613456 | Institut Pasteur de la Guadeloupe | Institut Pasteur de la Guadeloupe | Marion Barbet, Sylvie Behillil, Méline Bizard, Angela Brisebarre, Camille Capel, Etienne Simon-Lorière, Vincent Enouf, Maud Vanpeene, Sylvie van der Werf, Stéphanie Guyomard, Sébastien Breurec, Radjin Steingrover, Cherina Fleming, Antoine Talarmin |
| EPI_ISL_613545, EPI_ISL_613546, EPI_ISL_613547, EPI_ISL_613548, EPI_ISL_613549, EPI_ISL_613550, EPI_ISL_613551, EPI_ISL_613552, EPI_ISL_613553, EPI_ISL_613554, EPI_ISL_613555, EPI_ISL_613556, EPI_ISL_613557, EPI_ISL_613558, EPI_ISL_613559 | | | |
| see above | CHRU Pontchaillou - Laboratoire de Virologie 2, rue Henri Le Guilloux | National Reference Center for Viruses of Respiratory Infections, Institut Pasteur, Paris | Marion Barbet, Sylvie Behillil, Méline Bizard, Angela Brisebarre, Camille Capel, Etienne Simon-Lorière, Vincent Enouf, Maud Vanpeene, Sylvie van der Werf, Gisèle Lagathu |
| EPI_ISL_614265, EPI_ISL_614266, EPI_ISL_614267, EPI_ISL_614268, EPI_ISL_614269, EPI_ISL_614270, EPI_ISL_614271, EPI_ISL_614272, EPI_ISL_614273, EPI_ISL_614274, EPI_ISL_614275, EPI_ISL_614276, EPI_ISL_614277, EPI_ISL_614278, EPI_ISL_614279, EPI_ISL_614280 | | | |
| see above | Eurofins | National Reference Center for Viruses of Respiratory Infections, Institut Pasteur, Paris | Marion Barbet, Sylvie Behillil, Méline Bizard, Angela Brisebarre, Camille Capel, Etienne Simon-Lorière, Vincent Enouf, Maud Vanpeene, Sylvie van der Werf |
| EPI_ISL_614281 | CHRU Pontchaillou - Laboratoire de Virologie 2, rue Henri Le Guilloux | National Reference Center for Viruses of Respiratory Infections, Institut Pasteur, Paris | Marion Barbet, Sylvie Behillil, Méline Bizard, Angela Brisebarre, Camille Capel, Etienne Simon-Lorière, Vincent Enouf, Maud Vanpeene, Sylvie van der Werf, Gisèle Lagathu |
| EPI_ISL_614282, EPI_ISL_614283, EPI_ISL_614284, EPI_ISL_614285, EPI_ISL_614286, EPI_ISL_614287, EPI_ISL_614288, EPI_ISL_614289, EPI_ISL_614290, EPI_ISL_614291, EPI_ISL_614292, EPI_ISL_614293 | | | |
| see above | General practitioner | National Reference Center for Viruses of Respiratory Infections, Institut Pasteur, Paris | Marion Barbet, Sylvie Behillil, Méline Bizard, Angela Brisebarre, Camille Capel, Etienne Simon-Lorière, Vincent Enouf, Maud Vanpeene, Sylvie van der Werf |
| EPI_ISL_614295, EPI_ISL_614296, EPI_ISL_614297, EPI_ISL_614298, EPI_ISL_614299, EPI_ISL_614300, EPI_ISL_614301, EPI_ISL_614302, EPI_ISL_614303, EPI_ISL_614304, EPI_ISL_614305, EPI_ISL_614306, EPI_ISL_614307, EPI_ISL_614308, EPI_ISL_614309, EPI_ISL_614310, EPI_ISL_614311, EPI_ISL_614312, EPI_ISL_614313, EPI_ISL_614314, EPI_ISL_614315, EPI_ISL_614316, EPI_ISL_614317 | | | |
| see above | Faroese National Reference Laboratory for Fish and Animal Diseases | Faroese National Reference Laboratory for Fish and Animal Diseases | Maria Marjunardóttir Dahl, Petra Elisabeth Petersen, Debes Hammershaimb Christiansen |
| EPI_ISL_623096, EPI_ISL_623097, EPI_ISL_623103 | General practitioner | National Reference Center for Viruses of Respiratory Infections, Institut Pasteur, Paris | Marion Barbet, Sylvie Behillil, Méline Bizard, Angela Brisebarre, Camille Capel, Etienne Simon-Lorière, Vincent Enouf, Maud Vanpeene, Sylvie van der Werf |
| EPI_ISL_626570, EPI_ISL_626571, EPI_ISL_626572, EPI_ISL_626573, EPI_ISL_626574, EPI_ISL_626575, EPI_ISL_626576, EPI_ISL_626577, EPI_ISL_626578, EPI_ISL_626579, EPI_ISL_626580, EPI_ISL_626581, EPI_ISL_626582, EPI_ISL_626583, EPI_ISL_626584, EPI_ISL_626585, EPI_ISL_626586, EPI_ISL_626587, EPI_ISL_626588, EPI_ISL_626589, EPI_ISL_626590, EPI_ISL_626591, EPI_ISL_626592, EPI_ISL_626593, EPI_ISL_626594, EPI_ISL_626595, EPI_ISL_626596, EPI_ISL_626597, EPI_ISL_626598, EPI_ISL_626599, EPI_ISL_626600, EPI_ISL_626601, EPI_ISL_626602, EPI_ISL_626603, EPI_ISL_626604, EPI_ISL_626605, EPI_ISL_626606, EPI_ISL_626607, EPI_ISL_626608, EPI_ISL_626609, EPI_ISL_626610, EPI_ISL_626611, EPI_ISL_626612, EPI_ISL_626613, EPI_ISL_626614, EPI_ISL_626615, EPI_ISL_626616, EPI_ISL_626617, EPI_ISL_626618, EPI_ISL_626619, EPI_ISL_626620, EPI_ISL_626621, EPI_ISL_626622, EPI_ISL_626623, EPI_ISL_626624, EPI_ISL_626625 | | | |
| see above | The National Institute of Public Health | State Veterinary Institute Prague | Nagy,A;Jirincova,H;Novakova,L;Trnka,D;Vecerova,J |
| EPI_ISL_629036, EPI_ISL_629037, EPI_ISL_629038, EPI_ISL_629039, EPI_ISL_629040, EPI_ISL_629041, EPI_ISL_629042, EPI_ISL_629043, EPI_ISL_629044, EPI_ISL_629045, EPI_ISL_629046, EPI_ISL_629047, EPI_ISL_629048, EPI_ISL_629049, EPI_ISL_629050, EPI_ISL_629051, EPI_ISL_629052, EPI_ISL_629053, EPI_ISL_629054, EPI_ISL_629055, EPI_ISL_629056, EPI_ISL_629057, EPI_ISL_629058, EPI_ISL_629059, EPI_ISL_629060, EPI_ISL_629061, EPI_ISL_629062, EPI_ISL_629063, EPI_ISL_629064, EPI_ISL_629065, EPI_ISL_629066, EPI_ISL_629067, EPI_ISL_629068, EPI_ISL_629069, EPI_ISL_629070, EPI_ISL_629071, EPI_ISL_629072, EPI_ISL_629073, EPI_ISL_629074, EPI_ISL_629075, EPI_ISL_629076, EPI_ISL_629077, EPI_ISL_629078, EPI_ISL_629079, EPI_ISL_629080 | | | |
| see above | Maryland Public Health Laboratory | Maryland Public Health Laboratory | Maryland Department of Health Laboratories Administration |
| EPI_ISL_631304, EPI_ISL_631305, EPI_ISL_631306, EPI_ISL_631307, EPI_ISL_631308, EPI_ISL_631309, EPI_ISL_631310, EPI_ISL_631311, EPI_ISL_631312, EPI_ISL_631313, EPI_ISL_631314, EPI_ISL_631315, EPI_ISL_631316, EPI_ISL_631317, EPI_ISL_631318, EPI_ISL_631319, EPI_ISL_631320, EPI_ISL_631344, EPI_ISL_631345, EPI_ISL_631346, EPI_ISL_631347, EPI_ISL_631348, EPI_ISL_631349, EPI_ISL_631350, EPI_ISL_631351, EPI_ISL_631352, EPI_ISL_631353, EPI_ISL_631354, EPI_ISL_631355, EPI_ISL_631356, EPI_ISL_631357, EPI_ISL_631358, EPI_ISL_631359, EPI_ISL_631360, EPI_ISL_631361, EPI_ISL_631362, EPI_ISL_631363, EPI_ISL_631364, EPI_ISL_631365, EPI_ISL_631366, EPI_ISL_631367, EPI_ISL_631368, EPI_ISL_631369, EPI_ISL_631370, EPI_ISL_631371, EPI_ISL_631372, EPI_ISL_631373, EPI_ISL_631374, EPI_ISL_631375, EPI_ISL_631376 | | | |
| see above | ZOTZ KLIMAS MVZ Düsseldorf-Centrum GbR ÜBAG für Labormedizin, Genetik, Zytologie, Pathologie | Center of Medical Microbiology, Virology, and Hospital Hygiene, University of Duesseldorf | Maximilian Damagnez, Alexander Dilthey, Ashley-Jane Duplessis, Patrick Finzer, Katrin Hoffmann, Torsten Houwaart, Lisanna Hülse, Malte Kohns Vasconcelos, Marek Korencak, Nadine Lübke, Jessica Nicolai, Klaus Pfeffer, Daniel Strelow, Jörg Timm, Andreas Walker, Tobias Wienemann, Rainer Zotz |
| EPI_ISL_631377, EPI_ISL_631378, EPI_ISL_631379, EPI_ISL_631380, EPI_ISL_631381, EPI_ISL_631382, EPI_ISL_631383, EPI_ISL_631384, EPI_ISL_631385, EPI_ISL_631386 | University Hospital Cologne | Center of Medical Microbiology, Virology, and Hospital Hygiene, University of Duesseldorf | Maximilian Damagnez, Alexander Dilthey, Ashley-Jane Duplessis, Eva Heger, Torsten Houwaart, Rolf Kaiser, Florian Klein, Elena Knops, Malte Kohns Vasconcelos, Jessica Nicolai, Klaus Pfeffer, Gibran Rubio Quintanares, Saleta Sierra-Aragón, Daniel Strelow, Jörg Timm, Andreas Walker, Tobias Wienemann |
| EPI_ISL_631387 | ZOTZ KLIMAS MVZ Düsseldorf-Centrum GbR ÜBAG für Labormedizin, Genetik, Zytologie, Pathologie | Center of Medical Microbiology, Virology, and Hospital Hygiene, University of Duesseldorf | Maximilian Damagnez, Alexander Dilthey, Ashley-Jane Duplessis, Patrick Finzer, Katrin Hoffmann, Torsten Houwaart, Lisanna Hülse, Malte Kohns Vasconcelos, Marek Korencak, Nadine Lübke, Jessica Nicolai, Klaus Pfeffer, Daniel Strelow, Jörg Timm, Andreas Walker, Tobias Wienemann, Rainer Zotz |
| EPI_ISL_631388 | University Hospital Cologne | Center of Medical Microbiology, Virology, and Hospital Hygiene, University of Duesseldorf | Maximilian Damagnez, Alexander Dilthey, Ashley-Jane Duplessis, Eva Heger, Torsten Houwaart, Rolf Kaiser, Florian Klein, Elena Knops, Malte Kohns Vasconcelos, Jessica Nicolai, Klaus Pfeffer, Gibran Rubio Quintanares, Saleta Sierra-Aragón, Daniel Strelow, Jörg Timm, Andreas Walker, Tobias Wienemann |

We gratefully acknowledge the following Authors from the Originating laboratories responsible for obtaining the specimens, as well as the Submitting laboratories where the genome data were generated and shared via GISAID, on which this research is based.

All Submitters of data may be contacted directly via www.gisaid.org

Authors are sorted alphabetically.

| Accession ID | Originating Laboratory | Submitting Laboratory | Authors |
|---|---|---|---|
| EPI_ISL_402124, EPI_ISL_402127, EPI_ISL_402128, EPI_ISL_402129, EPI_ISL_402130 | Wuhan Jinyintan Hospital | Wuhan Institute of Virology, Chinese Academy of Sciences | Peng Zhou, Xing-Lou Yang, Ding-Yu Zhang, Lei Zhang, Yan Zhu, Hao-Rui Si, Zhengli Shi |
| EPI_ISL_403962, EPI_ISL_403963 | Bamrasnaradura Hospital | 1. Department of Medical Sciences, Ministry of Public Health, Thailand 2. Thai Red Cross Emerging Infectious Diseases - Health Science Centre 3. Department of Disease Control, Ministry of Public Health, Thailand | Pilailuk,Okada; Siripaporn,Phuygun; Thanutsapa,Thanadachakul; Supaporn,Wacharapluesadee; Sittiporn,Parnmen; Warawan,Wongboot; Sunthareeya,Waicharoen; Rome,Buathong; Malinee,Chittaganpitch; Nanthawan,Mekha |
| EPI_ISL_404895 | Providence Regional Medical Center | Division of Viral Diseases, Centers for Disease Control and Prevention | Queen,K., Tao,Y., Li,Y., Paden,C.R., Lu,X., Zhang,J., Gerber,S.I., Lindstrom,S., Tong,S. |
| EPI_ISL_410713, EPI_ISL_410714, EPI_ISL_410715 | National Public Health Laboratory, National Centre for Infectious Diseases | National Public Health Laboratory, National Centre for Infectious Diseases | Octavia S, Mak TM, Cui L, Lin RTP |
| EPI_ISL_410716 | National Public Health Laboratory, National Centre for Infectious Diseases | National Centre for Infectious Diseases, National Centre for Infectious Diseases | Octavia S, Mak TM, Cui L, Lin RTP |
| EPI_ISL_410719 | National Public Health Laboratory | National Public Health Laboratory | Octavia S, Mak TM, Cui L, Lin RTP |
| EPI_ISL_411218 | Department of Infectious and Tropical Diseases, Bichat Claude Bernard Hospital, Paris | Laboratoire Virpath, CIRI U111, UCBL1, INSERM, CNRS, ENS Lyon | Olivier Terrier, Aurélien Traversier, Julien Fouret, Yazdan Yazdanpanah, Xavier Lescure, Catherine Legras-Lachuer, Alexandre Gaymard, Bruno Lina, Manuel Rosa-Calatrava |
| EPI_ISL_411219, EPI_ISL_411220 | Department of Infectious and Tropical Diseases, Bichat Claude Bernard Hospital, Paris | Laboratoire Virpath, CIRI U111, UCBL1, INSERM, CNRS, ENS Lyon | Olivier Terrier, Aurélien Traversier, Julien Fouret, Yazdan Yazdanpanah, Xavier Lescure, Alexandre Gaymard, Bruno Lina, Manuel Rosa-Calatrava |
| EPI_ISL_412973 | Department of Infectious Diseases, Istituto Superiore di Sanità, Roma , Italy | Virology Laboratory, Scientific Department, Army Medical Center | Paola Stefanelli, Stefano Fiore, Antonella Marchi, Eleonora Benedetti, Concetta Fabiani, Giovanni Faggioni, Antonella Fortunato, Riccardo De Santis, Silvia Fillo, Anna Anselmo, Andrea Ciammaruconi, Stefano Palomba, Florigio Lista |
| EPI_ISL_412974 | Department of Infectious Diseases, Istituto Superiore di Sanità, Rome, Italy | Virology Laboratory, Scientific Department, Army Medical Center | Paola Stefanelli, Stefano Fiore, Antonella Marchi, Eleonora Benedetti, Concetta Fabiani, Giovanni Faggioni, Antonella Fortunato, Silvia Fillo, Riccardo De Santis, Andrea Ciammaruconi, Giancarlo Petralito, Filippo Molinari, Florigio Lista |
| EPI_ISL_413025 | Harborview Medical Center | UW Virology Lab | Pavitra Roychoudhury, Arun Nalla, Hong Xie, Keith Jerome, Alexander Greninger |
| EPI_ISL_413455 | Washington State Public Health Lab | University of Washington Virology Lab | Pavitra Roychoudhury, Arun Nalla, Hong Xie, Keith Jerome, Alexander Greninger |
| EPI_ISL_413457, EPI_ISL_413458 | Washington State Public Health Lab | UW Virology Lab | Pavitra Roychoudhury, Arun Nalla, Hong Xie, Keith Jerome, Alexander Greninger |
| EPI_ISL_413486 | Valley Medical Center | University of Washington Virology Lab | Pavitra Roychoudhury, Arun Nalla, Hong Xie, Keith Jerome, Alexander Greninger |
| EPI_ISL_413487 | Harborview Medical Center | University of Washington Virology Lab | Pavitra Roychoudhury, Arun Nalla, Hong Xie, Keith Jerome, Alexander Greninger |
| EPI_ISL_413488 | Center of Medical Microbiology, Virology, and Hospital Hygiene, University of Duesseldorf | Center of Medical Microbiology, Virology, and Hospital Hygiene, University of Duesseldorf | Ortwin Adams, Marcel Andree, Alexander Dilthey, Torsten Feldt, Sandra Hauka, Torsten Houwaart, Björn-Erik Jensen, Detlef Kindgen-Milles, Malte Kohns Vasconcelos, Klaus Pfeffer, Tina Senff, Daniel Strelow, Jörg Timm, Andreas Walker, Tobias Wienemann |
| EPI_ISL_413489 | Laboratorio di Microbiologia e Virologia, Università Vita-Salute San Raffaele, Milano | Laboratorio di Microbiologia e Virologia, Università Vita-Salute San Raffaele, Milano | R.A Diotti, E. Criscuolo, M. Castelli, V. Caputo, R. Ferrarese, M. Sampaolo, E. Boeri, I. Negri, V. Amato, G. Lo Raso, C. Di Resta, R. Burioni, M. Clementi, N. Mancini & N. Clementi |
| EPI_ISL_413522 | Indian Council of Medical Research - National Institute of Virology | National Influenza Center, Indian Council of Medical Research - National Institute of Virology | Potdar V, Yadav PD, Choudhary ML, Shete-Aich A |
| EPI_ISL_413523 | Indian Council of Medical Research-National Institute of Virology | National Influenza Center, Indian Council of Medical Research-National Institute of Virology | Potdar V, Yadav PD, Choudhary ML, Shete-Aich A |
| EPI_ISL_413550 | Centre for Human and Zoonotic Virology (CHAZVY), College of Medicine University of Lagos/Lagos University Teaching Hospital (LUTH), part of the Laboratory Network of the Nigeria Centre for Disease Control (NCDC) | African Centre of Excellence for Genomics of Infectious Diseases (ACEGID), Redeemer's University, Ede, Osun State, Nigeria | Oluniyi P.E., Ajogbasile F.V., Kayode A., Oguzie J., Folarin O.A., Ihekweazu C. Happi C.T. |
| EPI_ISL_413562, EPI_ISL_413563, EPI_ISL_413601, EPI_ISL_413649, EPI_ISL_413650, EPI_ISL_413651, EPI_ISL_413652, EPI_ISL_413653 | UW Virology Lab | UW Virology Lab | Pavitra Roychoudhury, Hong Xie, Keith Jerome, Alexander Greninger |
| EPI_ISL_414045 | LACEN RJ - Laboratório Central de Saúde Pública Noel Nutels | Instituto Oswaldo Cruz FIOCRUZ - Laboratory of Respiratory Viruses and Measles (LVRS) | Paola Resende, Alisson Fabri, Joilson Xavier, Sunando Roy, Fernando Motta, Aline Mattos, Milene Miranda, Cristiana Garcia, Braulia Caetano, Maria Ogrzewalska, Jonathan Lopes, Luciana Appolinario, Maria Nóbrega, Marilda Siqueira |
| EPI_ISL_414363, EPI_ISL_414364, EPI_ISL_414365, EPI_ISL_414366, EPI_ISL_414367, EPI_ISL_414368, EPI_ISL_414369 | UW Virology Lab | UW Virology Lab | Pavitra Roychoudhury, Hong Xie, Keith Jerome, Alexander Greninger |
| EPI_ISL_414497, EPI_ISL_414498, EPI_ISL_414499, EPI_ISL_414504, EPI_ISL_414505, EPI_ISL_414506, EPI_ISL_414507, EPI_ISL_414508, EPI_ISL_414509, EPI_ISL_414574 | Center of Medical Microbiology, Virology, and Hospital Hygiene, University of Duesseldorf | Center of Medical Microbiology, Virology, and Hospital Hygiene, University of Duesseldorf | Ortwin Adams, Marcel Andree, Alexander Dilthey, Torsten Feldt, Sandra Hauka, Torsten Houwaart, Björn-Erik Jensen, Detlef Kindgen-Milles, Malte Kohns Vasconcelos, Klaus Pfeffer, Tina Senff, Daniel Strelow, Jörg Timm, Andreas Walker, Tobias Wienemann |
| EPI_ISL_414591, EPI_ISL_414592, EPI_ISL_414593, EPI_ISL_414594, EPI_ISL_414595, EPI_ISL_414596, EPI_ISL_414597, EPI_ISL_414616, EPI_ISL_414617, EPI_ISL_414618, EPI_ISL_414619, EPI_ISL_414620, EPI_ISL_414621, EPI_ISL_414622 | | | |
| see above | UW Virology Lab | UW Virology Lab | Pavitra Roychoudhury, Hong Xie, Keith Jerome, Alexander Greninger |
| EPI_ISL_414647 | Viral Respiratory Lab, National Institute for Biomedical Research (INRB) | Pathogen Sequencing Lab, National Institute for Biomedical Research (INRB) | Placide Mbala-Kingebeni, Edith Nkwembe, Eddy Kinganda-Lusamaki, Amuri Aziza, Catherine Pratt, Matthias Pauthner, Josh Quick, Allison Black, James Hadfield, Trevor Bedford, Ian Goodfellow, Nick Loman, Kristian Andersen, Michael Wiley, Steve Ahuka-Mundeke, Jean-Jacques Muyembe Tamfum |
| EPI_ISL_415105 | Laboratório Central de Saúde Pública Professor Gonçalo Moniz - LACEN/BA | Instituto Oswaldo Cruz FIOCRUZ - Laboratory of Respiratory Viruses and Measles (LVRS) | Paola Resende, Allison Fabri, Joilson Xavier, Sunando Roy, Fernando Motta, Aline Mattos, Milene Miranda, Cristiana Garcia, Braulia Caetano, Maria Ogrzewalska, Jonathan Lopes, Luciana Appolinario, Maria Nóbrega, Marilda Siqueira |
| EPI_ISL_415128 | LACEN/ES - Laboratório Central de Saúde Pública do | Instituto Oswaldo Cruz FIOCRUZ - Laboratory of Respiratory | Paola Resende, Allison Fabri, Joilson Xavier, Sunando Roy, Fernando Motta, Aline Mattos, Milene Miranda, Cristiana Garcia, Braulia Caetano, Maria |

| | | | |
|---|---|---|---|
| | Espírito Santo | Viruses and Measles (LVRS) | Ogrzewalska, Jonathan Lopes, Luciana Appolinario, Maria Nóbrega, Marilda Siqueira |
| EPI_ISL_415591, EPI_ISL_415592, EPI_ISL_415593, EPI_ISL_415594, EPI_ISL_415595, EPI_ISL_415596, EPI_ISL_415597, EPI_ISL_415598, EPI_ISL_415599, EPI_ISL_415600, EPI_ISL_415601, EPI_ISL_415602, EPI_ISL_415603, EPI_ISL_415604, EPI_ISL_415605, EPI_ISL_415606, EPI_ISL_415607, EPI_ISL_415608, EPI_ISL_415609, EPI_ISL_415610, EPI_ISL_415611, EPI_ISL_415612, EPI_ISL_415613, EPI_ISL_415614, EPI_ISL_415615, EPI_ISL_415616, EPI_ISL_415617, EPI_ISL_415618, EPI_ISL_415619, EPI_ISL_415620, EPI_ISL_415621, EPI_ISL_415622, EPI_ISL_415623, EPI_ISL_415624, EPI_ISL_415625, EPI_ISL_415626, EPI_ISL_415627 | | | |
| see above | UW Virology Lab | UW Virology Lab | Pavitra Roychoudhury, Hong Xie, Keith Jerome, Alexander Greninger |
| EPI_ISL_415641, EPI_ISL_415642, EPI_ISL_415643, EPI_ISL_415644 | R. G. Lugar Center for Public Health Research, National Center for Disease Control and Public Health (NCDC) of Georgia. | R. G. Lugar Center for Public Health Research, National Center for Disease Control and Public Health (NCDC) of Georgia. | Nato Kotaria, Marine Murtskhvaladze, Ann Machablishvili, Lela Sabadze, Mari Gavashelidze, Ana Papkiauri, Meri Pantsulaia, Gvantsa Brachveli, Tata Imnadze, Tamar Jashiashvili, Tea Tevdoradze, Ketevan Sidamonidze, Ekaterine Khmaladze, Ekaterine Zhghenti, Roena Sukhiashvili, Mariam Zakalashvili, Lela Urushadze, Magda Dgebuadze, Giorgi Tomashvili, Davit Tsaguria, Ekaterine Zangaladze, Nino Berishvili, Gvantsa Chanturia, Adam Kotorashvili, Maia Alkhazashvili, Irma Burjanadze, Anna Kasradze, Khatuna Zakhashvili, Paata Imnadze, Amiran Gamkrelidze. |
| EPI_ISL_415658, EPI_ISL_415660, EPI_ISL_415661 | Laboratory of Molecular Virology, Pontificia Universidad Católica de Chile | MSHS Pathogen Surveillance Program | Rafael A. Medina, Pablo Vial, Tamara Garcia, Eileen Serrano, Ana Silvia Gonzalez-Reiche, Zenab Khan, Mitchell Sullivan, Ajay Obla, Matthew Hernandez, Hala Alshammary, Juan Soto, Shwetha Sridhar Hara, Ying-Chih Wang, Melissa Smith, Robert Sebra, Viviana Simon, Harm van Bakel |
| EPI_ISL_416426 | Virological Research Group, Szentágothai Research Centre, University of Pécs | Bioinformatics Research Group, Szentágothai Research Centre, University of Pécs | Péter Urbán, Endre Gábor Tóth, Gábor Kemenesi, Róbert Herczeg, Attila Gyenesei, Ferenc Jakab |
| EPI_ISL_416433, EPI_ISL_416434, EPI_ISL_416435, EPI_ISL_416436, EPI_ISL_416437, EPI_ISL_416438, EPI_ISL_416439, EPI_ISL_416440, EPI_ISL_416441, EPI_ISL_416442, EPI_ISL_416443, EPI_ISL_416444, EPI_ISL_416445, EPI_ISL_416446, EPI_ISL_416447, EPI_ISL_416448, EPI_ISL_416449, EPI_ISL_416450, EPI_ISL_416451, EPI_ISL_416452, EPI_ISL_416453, EPI_ISL_416454, EPI_ISL_416455, EPI_ISL_416456 | | | |
| see above | UW Virology Lab | UW Virology Lab | Pavitra Roychoudhury, Hong Xie, Keith Jerome, Alexander Greninger |
| EPI_ISL_416486 | Servicio de Microbiología. Consorcio Hospital General Universitario de Valencia | Sequencing and Bioinformatics Service and Molecular Epidemiology Research Group. FISABIO-Public Health | Neris Garcia-Gonzalez, Maria Alma Bracho, Maria Dolores Ocete, Concepcion Gimeno, Giuseppe D'Auria, Griselda De Marco, Fernando Gonzalez-Candelas |
| EPI_ISL_416635, EPI_ISL_416636, EPI_ISL_416637, EPI_ISL_416638, EPI_ISL_416639, EPI_ISL_416640, EPI_ISL_416641, EPI_ISL_416642, EPI_ISL_416643, EPI_ISL_416644, EPI_ISL_416645, EPI_ISL_416646, EPI_ISL_416647, EPI_ISL_416648, EPI_ISL_416649, EPI_ISL_416650, EPI_ISL_416651, EPI_ISL_416652, EPI_ISL_416653, EPI_ISL_416654, EPI_ISL_416655, EPI_ISL_416656, EPI_ISL_416657, EPI_ISL_416658, EPI_ISL_416659, EPI_ISL_416660, EPI_ISL_416661, EPI_ISL_416662, EPI_ISL_416663, EPI_ISL_416664, EPI_ISL_416665, EPI_ISL_416666, EPI_ISL_416667, EPI_ISL_416668, EPI_ISL_416669, EPI_ISL_416670, EPI_ISL_416671, EPI_ISL_416673, EPI_ISL_416674, EPI_ISL_416675, EPI_ISL_416676, EPI_ISL_416677, EPI_ISL_416678, EPI_ISL_416679, EPI_ISL_416680, EPI_ISL_416681, EPI_ISL_416682, EPI_ISL_416683, EPI_ISL_416684, EPI_ISL_416685, EPI_ISL_416686, EPI_ISL_416687, EPI_ISL_416688, EPI_ISL_416689, EPI_ISL_416690, EPI_ISL_416691, EPI_ISL_416692, EPI_ISL_416693, EPI_ISL_416694, EPI_ISL_416695, EPI_ISL_416696, EPI_ISL_416697, EPI_ISL_416698, EPI_ISL_416699, EPI_ISL_416700, EPI_ISL_416701, EPI_ISL_416702, EPI_ISL_416703, EPI_ISL_416704, EPI_ISL_416705, EPI_ISL_416706, EPI_ISL_416707, EPI_ISL_416708, EPI_ISL_416709, EPI_ISL_416710, EPI_ISL_416711, EPI_ISL_416712, EPI_ISL_416713, EPI_ISL_416714, EPI_ISL_416715, EPI_ISL_416716, EPI_ISL_416717, EPI_ISL_416718, EPI_ISL_416719, EPI_ISL_416721, EPI_ISL_416722, EPI_ISL_416723, EPI_ISL_416724, EPI_ISL_416725, EPI_ISL_416726, EPI_ISL_416727, EPI_ISL_416728, EPI_ISL_416729 | | | |
| see above | UW Virology Lab | UW Virology Lab | Pavitra Roychoudhury, Hong Xie, Keith Jerome, Alexander Greninger |
| EPI_ISL_416744 | Virological Research Group, Szentágothai Research Centre | Bioinformatics Research Group, Szentágothai Research Centre | Péter Urbán, Endre Gábor Tóth, Gábor Kemenesi, Róbert Herczeg, Attila Gyenesei, Ferenc Jakab |
| EPI_ISL_417341, EPI_ISL_417342, EPI_ISL_417343, EPI_ISL_417344, EPI_ISL_417345, EPI_ISL_417346, EPI_ISL_417347, EPI_ISL_417348, EPI_ISL_417349, EPI_ISL_417350, EPI_ISL_417351, EPI_ISL_417352, EPI_ISL_417353, EPI_ISL_417354, EPI_ISL_417355, EPI_ISL_417356, EPI_ISL_417357, EPI_ISL_417358, EPI_ISL_417359, EPI_ISL_417360, EPI_ISL_417361, EPI_ISL_417362, EPI_ISL_417363, EPI_ISL_417364, EPI_ISL_417365, EPI_ISL_417366, EPI_ISL_417367, EPI_ISL_417368, EPI_ISL_417369, EPI_ISL_417370, EPI_ISL_417371, EPI_ISL_417372, EPI_ISL_417373, EPI_ISL_417374, EPI_ISL_417375, EPI_ISL_417376, EPI_ISL_417377, EPI_ISL_417378, EPI_ISL_417379, EPI_ISL_417380, EPI_ISL_417381, EPI_ISL_417382 | | | |
| see above | UW Virology Lab | UW Virology Lab | Pavitra Roychoudhury, Hong Xie, Keith Jerome, Alexander Greninger |
| EPI_ISL_417394, EPI_ISL_417411 | Centre for Infectious Diseases and Microbiology Public Health | NSW Health Pathology - Institute of Clinical Pathology and Medical Research; Westmead Hospital; University of Sydney | O'Sullivan MV, Sintchenko V, Chen SC, Maddocks S, Kok J, Dwyer DE, Rockett R, Eden J-S, Lam C, Gray K, Timms V, Gall M, Arnott A, Sadsad R, Carter I, Rahman H and Holmes EC for the 2019-nCoV Study Group |
| EPI_ISL_417433, EPI_ISL_417434, EPI_ISL_417435, EPI_ISL_417436, EPI_ISL_417437, EPI_ISL_417438, EPI_ISL_417439, EPI_ISL_417440, EPI_ISL_417441, EPI_ISL_417442 | Viral Respiratory Lab, National Institute for Biomedical Research (INRB) | Pathogen Sequencing Lab, National Institute for Biomedical Research (INRB) | Placide Mbala-Kingebeni, Edith Nkwembe, Eddy Kinganda-Lusamaki, Amuri Aziza, Catherine Pratt, Matthias Pauthner, Josh Quick, Allison Black, James Hadfield, Trevor Bedford, Ian Goodfellow, Nick Loman, Kristian Andersen, Michael Wiley, Steve Ahuka-Mundeke, Jean-Jacques Muyembe Tamfum |
| EPI_ISL_417443 | State Key Laboratory for Emerging Infectious Diseases Department of Microbiology Li Ka Shing Faculty of Medicine The University of Hong Kong | State Key Laboratory for Emerging Infectious Diseases Department of Microbiology Li Ka Shing Faculty of Medicine The University of Hong Kong | Pui Wang, Siu-Ying Lau, Shaofeng Deng, Bobo Wing-Yee Mok, Wenjun Song, Kwok-Yung Yuen, Honglin Chen |
| EPI_ISL_417448, EPI_ISL_417449, EPI_ISL_417450, EPI_ISL_417451, EPI_ISL_417452, EPI_ISL_417453, EPI_ISL_417454, EPI_ISL_417455, EPI_ISL_417456 | UW Virology Lab | UW Virology Lab | Pavitra Roychoudhury, Hong Xie, Keith Jerome, Alexander Greninger |
| EPI_ISL_417457, EPI_ISL_417458, EPI_ISL_417459, EPI_ISL_417460, EPI_ISL_417461, EPI_ISL_417462, EPI_ISL_417463, EPI_ISL_417464, EPI_ISL_417465, EPI_ISL_417466, EPI_ISL_417467, EPI_ISL_417468 | | | |
| see above | Center of Medical Microbiology, Virology, and Hospital Hygiene, University of Duesseldorf | Center of Medical Microbiology, Virology, and Hospital Hygiene, University of Duesseldorf | Ortwin Adams, Marcel Andree, Alexander Dilthey, Torsten Feldt, Sandra Hauka, Torsten Houwaart, Björn-Erik Jensen, Detlef Kindgen-Milles, Malte Kohns Vasconcelos, Klaus Pfeffer, Tina Senff, Daniel Strelow, Jörg Timm, Andreas Walker, Tobias Wienemann |
| EPI_ISL_417941, EPI_ISL_417942, EPI_ISL_417944, EPI_ISL_417946, EPI_ISL_417947, EPI_ISL_417948, EPI_ISL_417950, EPI_ISL_417955 | Viral Respiratory Lab, National Institute for Biomedical Research (INRB) | Pathogen Sequencing Lab, National Institute for Biomedical Research (INRB) | Placide Mbala-Kingebeni, Edith Nkwembe, Eddy Kinganda-Lusamaki, Amuri Aziza, Catherine Pratt, Matthias Pauthner, Josh Quick, Allison Black, James Hadfield, Trevor Bedford, Ian Goodfellow, Nick Loman, Kristian Andersen, Michael Wiley, Steve Ahuka-Mundeke, Jean-Jacques Muyembe Tamfum |
| EPI_ISL_418028, EPI_ISL_418029, EPI_ISL_418030, EPI_ISL_418031, EPI_ISL_418032, EPI_ISL_418033, EPI_ISL_418034, EPI_ISL_418035, EPI_ISL_418036, EPI_ISL_418037, EPI_ISL_418038, EPI_ISL_418039, EPI_ISL_418040, EPI_ISL_418041, EPI_ISL_418042, EPI_ISL_418043, EPI_ISL_418044, EPI_ISL_418045, EPI_ISL_418046, EPI_ISL_418047, EPI_ISL_418048, EPI_ISL_418049, EPI_ISL_418050, EPI_ISL_418051, EPI_ISL_418052, EPI_ISL_418053, EPI_ISL_418054, EPI_ISL_418055, EPI_ISL_418056, EPI_ISL_418057, EPI_ISL_418058, EPI_ISL_418059, EPI_ISL_418060, EPI_ISL_418061, EPI_ISL_418062, EPI_ISL_418063, EPI_ISL_418064, EPI_ISL_418065, EPI_ISL_418067, EPI_ISL_418068, EPI_ISL_418069, EPI_ISL_418070, EPI_ISL_418071, EPI_ISL_418072, EPI_ISL_418073, EPI_ISL_418074, EPI_ISL_418075, EPI_ISL_418076, EPI_ISL_418077, EPI_ISL_418078, EPI_ISL_418079, EPI_ISL_418080, EPI_ISL_418081, EPI_ISL_418082 | | | |
| see above | UW Virology Lab | UW Virology Lab | Pavitra Roychoudhury, Hong Xie, Keith Jerome, Alexander Greninger |
| EPI_ISL_418183 | Virological Research Group, Szentágothai Research Centre | Bioinformatics Research Group, Szentágothai Research Centre | Péter Urbán, Endre Gábor Tóth, Gábor Kemenesi, Róbert Herczeg, Attila Gyenesei, Ferenc Jakab |
| EPI_ISL_418206, EPI_ISL_418207, EPI_ISL_418208, EPI_ISL_418209, EPI_ISL_418210, EPI_ISL_418211 | Institut Pasteur Dakar | Institut Pasteur de Dakar | Ndongo Dia, Ousmane Faye, Amadou Alpha Sall |
| EPI_ISL_418212 | Institut Pasteur Dakar | Institut Pasteur de Dakar | Ndongo Dia, Ousmane Faye, Amadou Alpha sall |
| EPI_ISL_418213, EPI_ISL_418214 | Institut Pasteur Dakar | Institut Pasteur de Dakar | Ndongo Dia, Ousmane Faye, Amadou Alpha Sall |
| EPI_ISL_418215 | Instirut Pasteur Dakar | Institut Pasteur de Dakar | Ndongo Dia, Ousmane Faye, Amadou Alpha Sall |
| EPI_ISL_418216, EPI_ISL_418217 | Institut Pasteur Dakar | Institut Pasteur de Dakar | Ndongo Dia, Ousmane Faye, Amadou Alpha Sall |
| EPI_ISL_418267 | Microbiology and Immunology department, Pasteur institute in Ho Chi Minh city | Microbiology and Immunology department, Pasteur institute in Ho Chi Minh city | Nguyen,H.T., Cao,T.M., Pham,H.T.T., Vu,N.P.H., Dao,M.H., Huynh,L.T.K., Nguyen,L.T., Nguyen,N.T., Nguyen,T.T.N., Nguyen,A.H., Luong,Q.C., Nguyen,T.V., Tran,K.C., Pham,Q.D., Tran,T., Hoang,C.Q., Nguyen,T.T., Le,H.Q., Phung,T.M., Vo,T.N.A., Nguyen,S.N., Pham,D.T., Nguyen,T.V. and Phan,L.T. |
| EPI_ISL_418866, EPI_ISL_418867, EPI_ISL_418868, EPI_ISL_418869, EPI_ISL_418870, EPI_ISL_418871, EPI_ISL_418872, EPI_ISL_418873, EPI_ISL_418874, EPI_ISL_418875, EPI_ISL_418876, EPI_ISL_418877, EPI_ISL_418878, EPI_ISL_418879, EPI_ISL_418880, EPI_ISL_418881, EPI_ISL_418882, EPI_ISL_418883, EPI_ISL_418884, EPI_ISL_418885, EPI_ISL_418886, EPI_ISL_418887, EPI_ISL_418888, EPI_ISL_418889, EPI_ISL_418890, EPI_ISL_418891, EPI_ISL_418892, EPI_ISL_418893, EPI_ISL_418894, EPI_ISL_418895, EPI_ISL_418896, EPI_ISL_418897, EPI_ISL_418898, EPI_ISL_418899, EPI_ISL_418900, EPI_ISL_418901, EPI_ISL_418902, EPI_ISL_418903, EPI_ISL_418904, EPI_ISL_418905, EPI_ISL_418906, EPI_ISL_418907, EPI_ISL_418908, EPI_ISL_418909, EPI_ISL_418910, EPI_ISL_418911, EPI_ISL_418912, EPI_ISL_418913, EPI_ISL_418914, EPI_ISL_418915, EPI_ISL_418916, EPI_ISL_418917, EPI_ISL_418918, EPI_ISL_418919, EPI_ISL_418920, EPI_ISL_418921, EPI_ISL_418922, EPI_ISL_418923, EPI_ISL_418924, EPI_ISL_418926, EPI_ISL_418927, EPI_ISL_418928, EPI_ISL_418929, EPI_ISL_418930, EPI_ISL_418931, EPI_ISL_418932, EPI_ISL_418933, EPI_ISL_418934, EPI_ISL_418935, EPI_ISL_418936, EPI_ISL_418937, EPI_ISL_418938, EPI_ISL_418939, EPI_ISL_418940, EPI_ISL_418942, EPI_ISL_418943, EPI_ISL_418944, EPI_ISL_418945, EPI_ISL_418946, EPI_ISL_418947, EPI_ISL_418948, EPI_ISL_418949, EPI_ISL_418950, EPI_ISL_418951, EPI_ISL_418952, EPI_ISL_418953, EPI_ISL_418954, EPI_ISL_418955 | | | |

| | | | |
|---|---|---|---|
| see above | UW Virology Lab | UW Virology Lab | Pavitra Roychoudhury, Hong Xie, Keith Jerome, Alexander Greninger |
| EPI_ISL_419264, EPI_ISL_419266 | Lab voor klinische biologie | Onderzoeksgroep Virologie | Nick Vereecke, Laurens Lambrechts, Marthe Pauwels, Basiel Cole, Bruno Verhasselt, Linos Vandekerckhove, Hans Nauwynck, Sebastiaan Theuns |
| EPI_ISL_419541, EPI_ISL_419542, EPI_ISL_419543, EPI_ISL_419544, EPI_ISL_419545, EPI_ISL_419546, EPI_ISL_419547, EPI_ISL_419548, EPI_ISL_419549, EPI_ISL_419550, EPI_ISL_419551, EPI_ISL_419552 | | | |
| see above | Center of Medical Microbiology, Virology, and Hospital Hygiene, University of Duesseldorf | Center of Medical Microbiology, Virology, and Hospital Hygiene, University of Duesseldorf | Ortwin Adams, Marcel Andree, Alexander Dilthey, Torsten Feldt, Sandra Hauka, Torsten Houwaart, Björn-Erik Jensen, Detlef Kindgen-Milles, Malte Kohns Vasconcelos, Klaus Pfeffer, Tina Senff, Daniel Strelow, Jörg Timm, Andreas Walker, Tobias Wienemann |
| EPI_ISL_419679, EPI_ISL_419684, EPI_ISL_419689 | Servicio de Microbiología. Consorcio Hospital General Universitario de Valencia | Sequencing and Bioinformatics Service and Molecular Epidemiology Research Group. FISABIO-Public Health | Neris Garcia-Gonzalez, Maria Alma Bracho, Maria Dolores Ocete, Giuseppe D'Auria, Griselda De Marco, Concepcion Gimeno, Fernando Gonzalez-Candelas |
| EPI_ISL_420030, EPI_ISL_420031, EPI_ISL_420032, EPI_ISL_420033, EPI_ISL_420034, EPI_ISL_420035 | Viral Respiratory Lab, National Institute for Biomedical Research (INRB) | Pathogen Sequencing Lab, National Institute for Biomedical Research (INRB) | Placide Mbala-Kingebeni, Edith Nkwembe, Eddy Kinganda-Lusamaki, Amuri Aziza, Catherine Pratt, Matthias Pauthner, Josh Quick, Allison Black, James Hadfield, Trevor Bedford, Ian Goodfellow, Nick Loman, Kristian Andersen, Michael Wiley, Steve Ahuka-Mundeke, Jean-Jacques Muyembe Tamfum |
| EPI_ISL_420069, EPI_ISL_420070, EPI_ISL_420071 | Institut Pasteur Dakar | Institut Pasteur de Dakar | Ndongo Dia, Moussa Moise Diagne, Mamadou Diop, Ousmane Faye, Amadou Alpha Sall |
| EPI_ISL_420072, EPI_ISL_420073, EPI_ISL_420074 | Institut Pasteur Dakar | Institut Pasteur de Dakar | Ndongo Dia, Moussa Moise Diagne, Mamadou Diop, Ousmane Faye , Amadou Alpha Sall |
| EPI_ISL_420075 | Institut pasteur Dakar | Institut Pasteur de Dakar | Ndongo Dia, Moussa Moise Diagne, Mamadou Diop, Ousmane Faye , Amadou Alpha Sall |
| EPI_ISL_420076 | Institut Pasteur Dakar | Institut Pasteur de Dakar | Ndongo Dia, Moussa Moise Diagne, Mamadou Diop, Ousmane Faye , Ndongo Dia |
| EPI_ISL_420077, EPI_ISL_420078, EPI_ISL_420079 | Institut Pasteur Dakar | Institut Pasteur de Dakar | Ndongo Dia, Moussa Moise Diagne, Mamadou Diop, Ousmane Faye , Amadou Alpha Sall |
| EPI_ISL_420116 | Servicio de Microbiología. Consorcio Hospital General Universitario de Valencia | Sequencing and Bioinformatics Service and Molecular Epidemiology Research Group. FISABIO-Public Health | Neris Garcia-Gonzalez, Loreto Ferrús Abad, Inma Galán Vendrell, Paula Ruiz-Hueso, Mariana Reyes-Prieto, Vicente Soriano Chirona, Maria Alma Bracho, Griselda De Marco, Beatriz Beamud, Lidia Ruiz Roldan, Marta Pla Diaz, Maria Dolores Ocete, Llúcia Martínez-Priego, Concepcion Gimeno, Giuseppe D'Auria, Fernando Gonzalez-Candelas |
| EPI_ISL_420119 | Servicio de Microbiología. Consorcio Hospital General Universitario de Valencia | Sequencing and Bioinformatics Service and Molecular Epidemiology Research Group. FISABIO-Public Health | Paula Ruiz-Hueso, Mariana Reyes-Prieto, Vicente Soriano Chirona, Maria Alma Bracho, Griselda De Marco, Beatriz Beamud, Lidia Ruiz Roldan, Marta Pla Diaz,Neris Garcia-Gonzalez, Loreto Ferrús Abad, Inma Galán Vendrell, Maria Dolores Ocete, Llúcia Martínez-Priego, Concepcion Gimeno, Giuseppe D'Auria, Fernando Gonzalez-Candelas |
| EPI_ISL_420131 | Servicio de Microbiologia. Hospital Clinico Universitario de Valencia | Sequencing and Bioinformatics Service and Molecular Epidemiology Research Group. FISABIO-Public Health | Paula Ruiz-Hueso,Loreto Ferrús Abad, Maria Alma Bracho, Griselda De Marco, Beatriz Beamud, Sandra Carbo, Lidia Ruiz Roldan, Marta Pla Diaz, Neris Garcia-Gonzalez, Inma Galán Vendrell, Mariana Reyes-Prieto, Vicente Soriano Chirona, Ivan Ansari, David Navarro, Llúcia Martínez-Priego, Giuseppe D'Auria, Fernando Gonzalez-Candelas |
| EPI_ISL_420140 | Department for Virology, Molecular Biology and Genome Research, R. G. Lugar Center for Public Health Research, National Center for Disease Control and Public Health (NCDC) of Georgia. | Department for Virology, Molecular Biology and Genome Research, R. G. Lugar Center for Public Health Research, National Center for Disease Control and Public Health (NCDC) of Georgia. | Nato Kotaria, Marine Murtskhvaladze, Ann Machablishvili, Lela Sabadze, Mari Gavashelidze, Ana Papkiauri, Meri Pantsulaia, Gvantsa Brachveli, Tata Imnadze, Tamar Jashiashvili, Tea Tevdoradze, Ketevan Sidamonidze, Ekaterine Khmaladze, Ekaterine Zhghenti, Roena Sukhiashvili, Mariam Zakalashvili, Lela Urushadze, Magda Dgebuadze, Giorgi Tomashvili, Davit Tsaguria, Ekaterine Zangaladze, Nino Berishvili, Gvantsa Chanturia, Adam Kotorashvili, Maia Alkhazashvili, Irma Burjanadze, Anna Kasradze, Khatuna Zakhashvili, Paata Imnadze, Amiran Gamkrelidze |
| EPI_ISL_420543, EPI_ISL_420545, EPI_ISL_420547, EPI_ISL_420549, EPI_ISL_420551, EPI_ISL_420553, EPI_ISL_420555 | National Influenza Center, Indian Council of Medical Research - National Institute of Virology | Indian Council of Medical Research-National Institute of Virology, Microbial Containment Complex | Pragya D. Yadav. Savita Patil, Varsha Potdar, Prasad Sarkale, Dimpal A. Nyayanit, Gajanan Sapkal, Anita M. Shete, Atanu Basu, Lalit Dar, M Choudhary, Amita Jain, Bharati Malhotra, Pranita Gawande, Sarah Cherian, Priya Abraham |
| EPI_ISL_420838, EPI_ISL_420839, EPI_ISL_420840, EPI_ISL_420841, EPI_ISL_420845, EPI_ISL_420847, EPI_ISL_420849, EPI_ISL_420851, EPI_ISL_420852, EPI_ISL_420853, EPI_ISL_420854 | | | |
| see above | Viral Respiratory Lab, National Institute for Biomedical Research (INRB) | Pathogen Sequencing Lab, National Institute for Biomedical Research (INRB) | Placide Mbala-Kingebeni, Edith Nkwembe, Eddy Kinganda-Lusamaki, Amuri Aziza, Catherine Pratt, Matthias Pauthner, Josh Quick, Allison Black, James Hadfield, Trevor Bedford, Ian Goodfellow, Nick Loman, Kristian Andersen, Michael Wiley, Steve Ahuka-Mundeke, Jean-Jacques Muyembe Tamfum |
| EPI_ISL_421662, EPI_ISL_421663, EPI_ISL_421664, EPI_ISL_421665, EPI_ISL_421666, EPI_ISL_421667, EPI_ISL_421668, EPI_ISL_421669, EPI_ISL_421670, EPI_ISL_421671, EPI_ISL_421672 | | | |
| see above | National Influenza Center, Indian Council of Medical Research - National Institute of Virology | Indian Council of Medical Research-National Institute of Virology, Microbial Containment Complex | Pragya D. Yadav, Varsha Potdar, Savita Patil, Dimpal A. Nyayanit, Triparna Majumdar, Manohar. L. Chaudhary, Gururaj Deshpande, Padinjaremattathil Thankappan Ullas, Anita Shete-Aich, Hitesh Dighe, Sreelekshmy Mohandas, Gajanan Sapkal, Atanu Basu, Amita Jain, Bharti Malhotra, Deepika Chaudhary, Sarah Cherian, Priya Abraham |
| EPI_ISL_422961, EPI_ISL_422962, EPI_ISL_422963, EPI_ISL_422964, EPI_ISL_422965, EPI_ISL_422966, EPI_ISL_422967, EPI_ISL_422969, EPI_ISL_422970, EPI_ISL_422971, EPI_ISL_422972, EPI_ISL_422973, EPI_ISL_422974, EPI_ISL_422975, EPI_ISL_422976, EPI_ISL_422977, EPI_ISL_422978, EPI_ISL_422979, EPI_ISL_422980, EPI_ISL_422981, EPI_ISL_422982, EPI_ISL_422983, EPI_ISL_422984, EPI_ISL_422985, EPI_ISL_422986, EPI_ISL_422987, EPI_ISL_422988, EPI_ISL_422989, EPI_ISL_422990, EPI_ISL_422991, EPI_ISL_422992, EPI_ISL_422993, EPI_ISL_422994, EPI_ISL_422995, EPI_ISL_422996, EPI_ISL_422997, EPI_ISL_422998, EPI_ISL_422999, EPI_ISL_423000, EPI_ISL_423001, EPI_ISL_423002, EPI_ISL_423003, EPI_ISL_423004, EPI_ISL_423005, EPI_ISL_423006, EPI_ISL_423007, EPI_ISL_423008, EPI_ISL_423009, EPI_ISL_423010, EPI_ISL_423011, EPI_ISL_423012, EPI_ISL_423013, EPI_ISL_423014, EPI_ISL_423015, EPI_ISL_423016, EPI_ISL_423017, EPI_ISL_423018, EPI_ISL_423019, EPI_ISL_423020, EPI_ISL_423021, EPI_ISL_423022, EPI_ISL_423024, EPI_ISL_423025, EPI_ISL_423026, EPI_ISL_423028, EPI_ISL_423029, EPI_ISL_423030, EPI_ISL_423031, EPI_ISL_423032, EPI_ISL_423033, EPI_ISL_424166, EPI_ISL_424167, EPI_ISL_424168, EPI_ISL_424169, EPI_ISL_424170, EPI_ISL_424171, EPI_ISL_424172, EPI_ISL_424173, EPI_ISL_424174, EPI_ISL_424175, EPI_ISL_424176, EPI_ISL_424177, EPI_ISL_424178, EPI_ISL_424179, EPI_ISL_424180, EPI_ISL_424181, EPI_ISL_424182, EPI_ISL_424183, EPI_ISL_424184, EPI_ISL_424185, EPI_ISL_424186, EPI_ISL_424187, EPI_ISL_424188, EPI_ISL_424190, EPI_ISL_424191, EPI_ISL_424192, EPI_ISL_424193, EPI_ISL_424194, EPI_ISL_424195, EPI_ISL_424196, EPI_ISL_424197, EPI_ISL_424198, EPI_ISL_424199, EPI_ISL_424200, EPI_ISL_424201, EPI_ISL_424202, EPI_ISL_424203, EPI_ISL_424204, EPI_ISL_424205, EPI_ISL_424206, EPI_ISL_424207, EPI_ISL_424208, EPI_ISL_424209, EPI_ISL_424210, EPI_ISL_424211, EPI_ISL_424212, EPI_ISL_424213, EPI_ISL_424214, EPI_ISL_424215, EPI_ISL_424216, EPI_ISL_424217, EPI_ISL_424218, EPI_ISL_424219, EPI_ISL_424220, EPI_ISL_424221, EPI_ISL_424222, EPI_ISL_424223, EPI_ISL_424224, EPI_ISL_424225, EPI_ISL_424226, EPI_ISL_424227, EPI_ISL_424228, EPI_ISL_424229, EPI_ISL_424230, EPI_ISL_424231, EPI_ISL_424232, EPI_ISL_424233, EPI_ISL_424234, EPI_ISL_424235, EPI_ISL_424236, EPI_ISL_424237, EPI_ISL_424238, EPI_ISL_424239, EPI_ISL_424240, EPI_ISL_424241, EPI_ISL_424242, EPI_ISL_424243, EPI_ISL_424244, EPI_ISL_424245, EPI_ISL_424246, EPI_ISL_424247, EPI_ISL_424249, EPI_ISL_424250, EPI_ISL_424251, EPI_ISL_424252, EPI_ISL_424253, EPI_ISL_424254, EPI_ISL_424255, EPI_ISL_424256, EPI_ISL_424257, EPI_ISL_424258, EPI_ISL_424259, EPI_ISL_424260, EPI_ISL_424261, EPI_ISL_424262, EPI_ISL_424263, EPI_ISL_424264, EPI_ISL_424265, EPI_ISL_424266, EPI_ISL_424267, EPI_ISL_424268, EPI_ISL_424269, EPI_ISL_424270, EPI_ISL_424271, EPI_ISL_424272, EPI_ISL_424273, EPI_ISL_424275, EPI_ISL_424276, EPI_ISL_424277, EPI_ISL_424278, EPI_ISL_424279, EPI_ISL_424280, EPI_ISL_424281, EPI_ISL_424282, EPI_ISL_424283, EPI_ISL_424284, EPI_ISL_424285, EPI_ISL_424286, EPI_ISL_424287, EPI_ISL_424288, EPI_ISL_424289, EPI_ISL_424290, EPI_ISL_424291, EPI_ISL_424292, EPI_ISL_424293, EPI_ISL_424294, EPI_ISL_424295, EPI_ISL_424296, EPI_ISL_424297, EPI_ISL_424298, EPI_ISL_424299, EPI_ISL_424300, EPI_ISL_424301, EPI_ISL_424302, EPI_ISL_424303, EPI_ISL_424304, EPI_ISL_424305, EPI_ISL_424306, EPI_ISL_424307, EPI_ISL_424308, EPI_ISL_424309, EPI_ISL_424310, EPI_ISL_424311, EPI_ISL_424312, EPI_ISL_424313, EPI_ISL_424314, EPI_ISL_424315, EPI_ISL_424316, EPI_ISL_424317, EPI_ISL_424318, EPI_ISL_424319, EPI_ISL_424320, EPI_ISL_424321, EPI_ISL_424322, EPI_ISL_424323, EPI_ISL_424324, EPI_ISL_424325, EPI_ISL_424326, EPI_ISL_424328, EPI_ISL_424329, EPI_ISL_424330, EPI_ISL_424331, EPI_ISL_424332, EPI_ISL_424333, EPI_ISL_424334, EPI_ISL_424335, EPI_ISL_424336, EPI_ISL_424337, EPI_ISL_424338, EPI_ISL_424339, EPI_ISL_424340, EPI_ISL_424341 | | | |
| see above | UW Virology Lab | UW Virology Lab | Pavitra Roychoudhury, Hong Xie, Keith Jerome, Alexander Greninger |
| EPI_ISL_424361, EPI_ISL_424362, EPI_ISL_424363, EPI_ISL_424364, EPI_ISL_424365 | National Influenza Center, Indian Council of Medical Research - National Institute of Virology | Indian Council of Medical Research-National Institute of Virology, Microbial Containment Complex | Pragya D. Yadav, Varsha Potdar, Savita Patil, Dimpal A. Nyayanit, Triparna Majumdar, Manohar. L. Chaudhary, Gururaj Deshpande, Padinjaremattathil Thankappan Ullas, Anita Shete-Aich, Hitesh Dighe, Sreelekshmy Mohandas, Gajanan Sapkal, Atanu Basu, Amita Jain, Bharti Malhotra, Deepika Chaudhary, Sarah Cherian, Priya Abraham |
| EPI_ISL_424668, EPI_ISL_424669, EPI_ISL_424671 | Arizona State University Health Services | Arizona State University | Rabia Maqsood, LaRinda A. Holland, Emily A. Kaelin, Bereket Estifanos, Nicholas J. Mellor, Jason Steel, Lily I. Wu, Arvind Varsani, Rolf U. Halden, Brenda G. Hogue, Matthew Scotch, Efrem S. Lim |
| EPI_ISL_425051, EPI_ISL_425052, EPI_ISL_425053, EPI_ISL_425054, EPI_ISL_425055, EPI_ISL_425059, EPI_ISL_425060, EPI_ISL_425061 | Lab voor klinische biologie | Onderzoeksgroep Virologie | Nick Vereecke, Laurens Lambrechts, Marthe Pauwels, Basiel Cole, Bruno Verhasselt, Linos Vandekerckhove, Hans Nauwynck, Sebastiaan Theuns |
| EPI_ISL_425120, EPI_ISL_425121, EPI_ISL_425122, EPI_ISL_425123, EPI_ISL_425124, EPI_ISL_425125, EPI_ISL_425126, EPI_ISL_425127, EPI_ISL_425128 | Center of Medical Microbiology, Virology, and Hospital Hygiene, University of Duesseldorf | Center of Medical Microbiology, Virology, and Hospital Hygiene, University of Duesseldorf | Ortwin Adams, Marcel Andree, Alexander Dilthey, Torsten Feldt, Sandra Hauka, Torsten Houwaart, Björn-Erik Jensen, Detlef Kindgen-Milles, Malte Kohns Vasconcelos, Klaus Pfeffer, Tina Senff, Daniel Strelow, Jörg Timm, Andreas Walker, Tobias Wienemann |
| EPI_ISL_425129, EPI_ISL_425130, EPI_ISL_425131, EPI_ISL_425132, EPI_ISL_425133, EPI_ISL_425134, | Center of Medical Microbiology, Virology, and Hospital Hygiene, University of Duesseldorf | Center of Medical Microbiology, Virology, and Hospital Hygiene, University of Duesseldorf | Ortwin Adams, Marcel Andree, Alexander Dilthey, Torsten Feldt, Sandra Hauka, Torsten Houwaart, Bjorn-Erik Jensen, Detlef Kindgen-Milles, Malte Kohns Vasconcelos, Klaus Pfeffer, Tina Senff, Daniel Strelow, Jorg Timm, Andreas Walker, Tobias Wienemann |

| | | | |
|---|---|---|---|
| EPI_ISL_425135, EPI_ISL_425136, EPI_ISL_425137 | | | |
| EPI_ISL_425138, EPI_ISL_425139, EPI_ISL_425140 | Center of Medical Microbiology, Virology, and Hospital Hygiene, University of Duesseldorf | Center of Medical Microbiology, Virology, and Hospital Hygiene, University of Duesseldorf | Ortwin Adams, Marcel Andree, Alexander Dilthey, Torsten Feldt, Sandra Hauka, Torsten Houwaart, Björn-Erik Jensen, Detlef Kindgen-Milles, Malte Kohns Vasconcelos, Klaus Pfeffer, Tina Senff, Daniel Strelow, Jörg Timm,Andreas Walker, Tobias Wienemann |
| EPI_ISL_425184 | Servicio de Microbiología. Consorcio Hospital General Universitario de Valencia | Sequencing and Bioinformatics Service and Molecular Epidemiology Research Group. FISABIO-Public Health | Neris Garcia-Gonzalez, Inma Galán Vendrell, Sandra Carbo, Loreto Ferrús Abad, Paula Ruiz-Hueso, Mariana Reyes-Prieto, Vicente Soriano Chirona, Ivan Ansari, David Navarro, Maria Alma Bracho, Griselda De Marco, Beatriz Beamud, Lidia Ruiz Roldan, Marta Pla Diaz, Llúcia Martínez-Priego, Giuseppe D'Auria, Fernando Gonzalez-Candelas |
| EPI_ISL_425188 | Servicio de Microbiología. Consorcio Hospital General Universitario de Valencia | Sequencing and Bioinformatics Service and Molecular Epidemiology Research Group. FISABIO-Public Health | Paula Ruiz-Hueso, Mariana Reyes-Prieto, Vicente Soriano Chirona, Ivan Ansari, David Navarro, Maria Alma Bracho, Griselda De Marco, Beatriz Beamud, Lidia Ruiz Roldan, Marta Pla Diaz, Neris Garcia-Gonzalez, Inma Galán Vendrell, Sandra Carbo, Loreto Ferrús Abad, Llúcia Martínez-Priego, Giuseppe D'Auria, Fernando Gonzalez-Candelas |
| EPI_ISL_425198 | Servicio de Microbiología. Consorcio Hospital General Universitario de Valencia | Sequencing and Bioinformatics Service and Molecular Epidemiology Research Group. FISABIO-Public Health | Neris Garcia-Gonzalez, Inma Galán Vendrell, Sandra Carbo, Loreto Ferrús Abad, Paula Ruiz-Hueso, Mariana Reyes-Prieto, Vicente Soriano Chirona, Ivan Ansari, David Navarro, Maria Alma Bracho, Griselda De Marco, Beatriz Beamud, Lidia Ruiz Roldan, Marta Pla Diaz, Llúcia Martínez-Priego, Giuseppe D'Auria, Fernando Gonzalez-Candelas |
| EPI_ISL_425200 | Servicio de Microbiologia. Hospital Clinico Universitario de Valencia | Sequencing and Bioinformatics Service and Molecular Epidemiology Research Group. FISABIO-Public Health | Paula Ruiz-Hueso, Mariana Reyes-Prieto, Vicente Soriano Chirona, Maria Alma Bracho, Griselda De Marco, Beatriz Beamud, Lidia Ruiz Roldan, Marta Pla Diaz,Neris Garcia-Gonzalez, Loreto Ferrús Abad, Maria Dolores Ocete, Llúcia Martínez-Priego, Inma Galán Vendrell, Concepcion Gimeno, Giuseppe D'Auria, Fernando Gonzalez-Candelas |
| EPI_ISL_425209 | Servicio de Microbiologia. Hospital Clinico Universitario de Valencia | Sequencing and Bioinformatics Service and Molecular Epidemiology Research Group. FISABIO-Public Health | Neris Garcia-Gonzalez, Loreto Ferrús Abad, Maria Dolores Ocete, Inma Galán Vendrell, Paula Ruiz-Hueso, Mariana Reyes-Prieto, Vicente Soriano Chirona, Maria Alma Bracho, Griselda De Marco, Beatriz Beamud, Lidia Ruiz Roldan, Marta Pla Diaz, Llúcia Martínez-Priego, Concepcion Gimeno, Giuseppe D'Auria, Fernando Gonzalez-Candelas |
| EPI_ISL_425218, EPI_ISL_425219 | Servicio de Microbiología. Consorcio Hospital General Universitario de Valencia | Sequencing and Bioinformatics Service and Molecular Epidemiology Research Group. FISABIO-Public Health | Neris Garcia-Gonzalez, Inma Galán Vendrell, Sandra Carbo, Loreto Ferrús Abad, Paula Ruiz-Hueso, Mariana Reyes-Prieto, Vicente Soriano Chirona, Ivan Ansari, David Navarro, Maria Alma Bracho, Griselda De Marco, Beatriz Beamud, Lidia Ruiz Roldan, Marta Pla Diaz, Llúcia Martínez-Priego, Giuseppe D'Auria, Fernando Gonzalez-Candelas |
| EPI_ISL_425223 | Servicio de Microbiología. Consorcio Hospital General Universitario de Valencia | Sequencing and Bioinformatics Service and Molecular Epidemiology Research Group. FISABIO-Public Health | Paula Ruiz-Hueso, Mariana Reyes-Prieto, Vicente Soriano Chirona, Ivan Ansari, David Navarro, Maria Alma Bracho, Griselda De Marco, Beatriz Beamud, Lidia Ruiz Roldan, Marta Pla Diaz, Neris Garcia-Gonzalez, Inma Galán Vendrell, Sandra Carbo, Loreto Ferrús Abad, Llúcia Martínez-Priego, Giuseppe D'Auria, Fernando Gonzalez-Candelas |
| EPI_ISL_426052, EPI_ISL_426053, EPI_ISL_426054, EPI_ISL_426055, EPI_ISL_426056, EPI_ISL_426057, EPI_ISL_426058, EPI_ISL_426059, EPI_ISL_426060, EPI_ISL_426061, EPI_ISL_426062, EPI_ISL_426063, EPI_ISL_426064, EPI_ISL_426065, EPI_ISL_426066, EPI_ISL_426067, EPI_ISL_426068, EPI_ISL_426069, EPI_ISL_426070, EPI_ISL_426071, EPI_ISL_426072, EPI_ISL_426073, EPI_ISL_426074, EPI_ISL_426075, EPI_ISL_426076, EPI_ISL_426077, EPI_ISL_426078, EPI_ISL_426079, EPI_ISL_426080, EPI_ISL_426081, EPI_ISL_426082, EPI_ISL_426083, EPI_ISL_426084, EPI_ISL_426085, EPI_ISL_426086, EPI_ISL_426087, EPI_ISL_426088, EPI_ISL_426089, EPI_ISL_426090, EPI_ISL_426091, EPI_ISL_426092, EPI_ISL_426093, EPI_ISL_426094, EPI_ISL_426095, EPI_ISL_426096, EPI_ISL_426097, EPI_ISL_426098, EPI_ISL_426099, EPI_ISL_426100, EPI_ISL_426101, EPI_ISL_426102, EPI_ISL_426103, EPI_ISL_426104, EPI_ISL_426105, EPI_ISL_426106, EPI_ISL_426107, EPI_ISL_426108, EPI_ISL_426109, EPI_ISL_426110, EPI_ISL_426111, EPI_ISL_426112, EPI_ISL_426113, EPI_ISL_426114, EPI_ISL_426115, EPI_ISL_426116, EPI_ISL_426117, EPI_ISL_426118, EPI_ISL_426119, EPI_ISL_426120, EPI_ISL_426121, EPI_ISL_426122, EPI_ISL_426123, EPI_ISL_426124, EPI_ISL_426125, EPI_ISL_426126, EPI_ISL_426127, EPI_ISL_426128, EPI_ISL_426129, EPI_ISL_426130, EPI_ISL_426131, EPI_ISL_426132, EPI_ISL_426133, EPI_ISL_426135, EPI_ISL_426136, EPI_ISL_426137 | | | |
| see above | UW Virology Lab | UW Virology Lab | Pavitra Roychoudhury, Hong Xie, Keith Jerome, Alexander Greninger |
| EPI_ISL_426179 | National Influenza Center, Indian Council of Medical Research - National Institute of Virology | Indian Council of Medical Research-National Institute of Virology, Microbial Containment Complex | Pragya D. Yadav, Varsha Potdar, Savita Patil, Dimpal A. Nyayanit, Triparna Majumdar, Manohar. L. Chaudhary, Gururaj Deshpande, Padinjaremattathil Thankappan Ullas, Anita Shete-Aich, Hitesh Dighe, Sreelekshmy Mohandas, Gajanan Sapkal, Atanu Basu, Amita Jain, Bharti Malhotra, Deepika Chaudhary, Sarah Cherian, Priya Abraham |
| EPI_ISL_426883, EPI_ISL_426884, EPI_ISL_426885, EPI_ISL_426886, EPI_ISL_426887, EPI_ISL_426888, EPI_ISL_426889, EPI_ISL_426890, EPI_ISL_426891, EPI_ISL_426892, EPI_ISL_426893, EPI_ISL_426894, EPI_ISL_426895, EPI_ISL_426896, EPI_ISL_426897 | | | |
| see above | Motol University Hospital | Institute of Applied Biotechnologies a.s. | Petr Brož, Jan Geryk, Petr Klempt, Martin Kašný, Adam Novotný, Kateina Kvapilová, Pavel Devínek, Petr Kvapil, Milan Macek |
| EPI_ISL_427161, EPI_ISL_427162, EPI_ISL_427163, EPI_ISL_427164, EPI_ISL_427165, EPI_ISL_427166, EPI_ISL_427167, EPI_ISL_427168, EPI_ISL_427169, EPI_ISL_427170, EPI_ISL_427171, EPI_ISL_427172, EPI_ISL_427173, EPI_ISL_427174, EPI_ISL_427175, EPI_ISL_427176, EPI_ISL_427177, EPI_ISL_427178, EPI_ISL_427179, EPI_ISL_427180, EPI_ISL_427181, EPI_ISL_427182, EPI_ISL_427183, EPI_ISL_427184, EPI_ISL_427185, EPI_ISL_427186, EPI_ISL_427187, EPI_ISL_427188, EPI_ISL_427189, EPI_ISL_427190, EPI_ISL_427191, EPI_ISL_427192, EPI_ISL_427193, EPI_ISL_427194, EPI_ISL_427195, EPI_ISL_427196, EPI_ISL_427197, EPI_ISL_427198, EPI_ISL_427199, EPI_ISL_427200, EPI_ISL_427201, EPI_ISL_427202, EPI_ISL_427203, EPI_ISL_427204, EPI_ISL_427205, EPI_ISL_427206, EPI_ISL_427207, EPI_ISL_427208, EPI_ISL_427209, EPI_ISL_427210, EPI_ISL_427211, EPI_ISL_427212, EPI_ISL_427213, EPI_ISL_427214, EPI_ISL_427215, EPI_ISL_427216, EPI_ISL_427217, EPI_ISL_427218, EPI_ISL_427219, EPI_ISL_427220, EPI_ISL_427221, EPI_ISL_427222, EPI_ISL_427223, EPI_ISL_427224, EPI_ISL_427225, EPI_ISL_427226, EPI_ISL_427227, EPI_ISL_427228, EPI_ISL_427229, EPI_ISL_427230, EPI_ISL_427231, EPI_ISL_427232, EPI_ISL_427233, EPI_ISL_427234, EPI_ISL_427235, EPI_ISL_427236, EPI_ISL_427237, EPI_ISL_427238, EPI_ISL_427239, EPI_ISL_427240, EPI_ISL_427241, EPI_ISL_427242, EPI_ISL_427243, EPI_ISL_427244, EPI_ISL_427245, EPI_ISL_427246, EPI_ISL_427247, EPI_ISL_427248, EPI_ISL_427249, EPI_ISL_427250, EPI_ISL_427251, EPI_ISL_427252, EPI_ISL_427253, EPI_ISL_427254, EPI_ISL_427255, EPI_ISL_427256, EPI_ISL_427257, EPI_ISL_427258, EPI_ISL_427259, EPI_ISL_427260, EPI_ISL_427261, EPI_ISL_427262, EPI_ISL_427263, EPI_ISL_427264, EPI_ISL_427265, EPI_ISL_427266, EPI_ISL_427268, EPI_ISL_427269, EPI_ISL_427270 | | | |
| see above | UW Virology Lab | UW Virology Lab | Pavitra Roychoudhury, Hong Xie, Keith Jerome, Alexander Greninger |
| EPI_ISL_427292 | LACEN-AL - Laboratorio Central de Alagoas | Instituto Oswaldo Cruz FIOCRUZ - Laboratory of Respiratory Viruses and Measles (LVRS) | Paola Resende, Fernando Motta, Luciana Appolinario, Sunando Roy, Aline Mattos, Milene Miranda, Cristiana Garcia, Braulia Caetano, Maria Ogrzewalska, Priscila Born, Jonathan Lopes, Marilda Siqueira |
| EPI_ISL_427293 | LACEN-BA - Laboratório Central de Saúde Pública Professor Gonçalo Moniz | Instituto Oswaldo Cruz FIOCRUZ - Laboratory of Respiratory Viruses and Measles (LVRS) | Paola Resende, Fernando Motta, Luciana Appolinario, Sunando Roy, Aline Mattos, Milene Miranda, Cristiana Garcia, Braulia Caetano, Maria Ogrzewalska, Priscila Born, Jonathan Lopes, Marilda Siqueira |
| EPI_ISL_427294, EPI_ISL_427295, EPI_ISL_427296, EPI_ISL_427297, EPI_ISL_427298, EPI_ISL_427299, EPI_ISL_427300, EPI_ISL_427301, EPI_ISL_427302, EPI_ISL_427303, EPI_ISL_427304 | | | |
| see above | Instituto Oswaldo Cruz FIOCRUZ - Laboratory of Respiratory Viruses and Measles (LVRS) | Instituto Oswaldo Cruz FIOCRUZ - Laboratory of Respiratory Viruses and Measles (LVRS) | Paola Resende, Fernando Motta, Luciana Appolinario, Sunando Roy, Aline Mattos, Milene Miranda, Cristiana Garcia, Braulia Caetano, Maria Ogrzewalska, Priscila Born, Jonathan Lopes, Marilda Siqueira |
| EPI_ISL_427305, EPI_ISL_427306 | LACEN-SC - Laboratorio Central de Santa Catarina | Instituto Oswaldo Cruz FIOCRUZ - Laboratory of Respiratory Viruses and Measles (LVRS) | Paola Resende, Fernando Motta, Luciana Appolinario, Sunando Roy, Aline Mattos, Milene Miranda, Cristiana Garcia, Braulia Caetano, Maria Ogrzewalska, Priscila Born, Jonathan Lopes, Marilda Siqueira |
| EPI_ISL_428860, EPI_ISL_428861 | State Research Center of Virology and Biotechnology VECTOR, Department of Collection of Microorganisms | State Research Center of Virology and Biotechnology VECTOR, Department of Collection of Microorganisms | Oleg V. Pyankov, Sergey A. Bodnev, Tatyana V. Tregubchak, Alexander N. Shvalov, Elena V. Gavrilova, Rinat A. Maksyutov |
| EPI_ISL_428862, EPI_ISL_428863 | State Research Center of Virology and Biotechnology VECTOR, Department of Collection of Microorganisms | State Research Center of Virology and Biotechnology VECTOR, Department of Collection of Microorganisms | Oleg V. Pyankov, Sergey A. Bodnev, Anastasiya M. Smirnova, Anastasiya A. Nazarenko, Tatyana V. Tregubchak, Alexander N. Shvalov, Elena V. Gavrilova, Rinat A. Maksyutov |
| EPI_ISL_428864, EPI_ISL_428865, EPI_ISL_428866, EPI_ISL_428867, EPI_ISL_428868, EPI_ISL_428869, EPI_ISL_428870, EPI_ISL_428871, EPI_ISL_428882, EPI_ISL_428883, EPI_ISL_428884, EPI_ISL_428885, EPI_ISL_428886, EPI_ISL_428887, EPI_ISL_428888, EPI_ISL_428889, EPI_ISL_428890, EPI_ISL_428891, EPI_ISL_428892, EPI_ISL_428893, EPI_ISL_428894, EPI_ISL_428906, EPI_ISL_428907, EPI_ISL_428908, EPI_ISL_428909, EPI_ISL_428910, EPI_ISL_428911, EPI_ISL_428912, EPI_ISL_428913, EPI_ISL_428914, EPI_ISL_428915, EPI_ISL_428916 | | | |
| see above | State Research Center of Virology and Biotechnology VECTOR, Department of Collection of Microorganisms | State Research Center of Virology and Biotechnology VECTOR, Department of Collection of Microorganisms | Oleg V. Pyankov, Sergey A. Bodnev, Tatyana V. Tregubchak, Alexander N. Shvalov, Elena V. Gavrilova, Rinat A. Maksyutov |
| EPI_ISL_429128, EPI_ISL_429129, EPI_ISL_429130, EPI_ISL_429131, EPI_ISL_429132, EPI_ISL_429133, EPI_ISL_429134, EPI_ISL_429135 | Laboratoriemedicin | The Public Health Agency of Sweden | Olov Svartstrom, Maria Lind Karlberg, Anna-Malin Linde, Oskar Karlsson Lindsjo, Anna Risberg, Shaman Muradrasoli, Karin Tegmark-Wisell |
| EPI_ISL_429161, EPI_ISL_429162, EPI_ISL_429163 | The Public Health Agency of Sweden | The Public Health Agency of Sweden | Olov Svartstrom, Maria Lind Karlberg, Anna-Malin Linde, Oskar Karlsson Lindsjo, Anna Risberg, Shaman Muradrasoli, Karin Tegmark-Wisell |
| EPI_ISL_429254, EPI_ISL_429255, EPI_ISL_429258, EPI_ISL_429259 | Viral Respiratory Lab, National Institute for Biomedical Research (INRB) | Pathogen Sequencing Lab, National Institute for Biomedical Research (INRB) | Placide Mbala-Kingebeni, Edith Nkwembe, Eddy Kinganda-Lusamaki, Amuri Aziza, Catherine Pratt, Matthias Pauthner, Josh Quick, Allison Black, James Hadfield, Trevor Bedford, Ian Goodfellow, Nick Loman, Kristian Andersen, Michael Wiley, Steve Ahuka-Mundeke, Jean-Jacques Muyembe Tamfum |
| EPI_ISL_429597, EPI_ISL_429598, EPI_ISL_429599, EPI_ISL_429600, EPI_ISL_429601, EPI_ISL_429602, EPI_ISL_429603, EPI_ISL_429604, EPI_ISL_429605, EPI_ISL_429606, EPI_ISL_429607, EPI_ISL_429608, EPI_ISL_429609, EPI_ISL_429610, EPI_ISL_429611, EPI_ISL_429612, EPI_ISL_429613, EPI_ISL_429614, EPI_ISL_429615, EPI_ISL_429616, EPI_ISL_429617, EPI_ISL_429618, EPI_ISL_429619, EPI_ISL_429620, EPI_ISL_429621, EPI_ISL_429622, EPI_ISL_429623, EPI_ISL_429624, EPI_ISL_429625, EPI_ISL_429626, EPI_ISL_429627, EPI_ISL_429628, EPI_ISL_429629, EPI_ISL_429630, EPI_ISL_429631, EPI_ISL_429632, EPI_ISL_429633, EPI_ISL_429634, EPI_ISL_429635, EPI_ISL_429636, EPI_ISL_429637, EPI_ISL_429638, EPI_ISL_429639, EPI_ISL_429640, EPI_ISL_429641, EPI_ISL_429642, EPI_ISL_429643, EPI_ISL_429644, EPI_ISL_429645, EPI_ISL_429647, EPI_ISL_429648, EPI_ISL_429649, EPI_ISL_429650, EPI_ISL_429651, EPI_ISL_429652, EPI_ISL_429653, EPI_ISL_429654, EPI_ISL_429655, EPI_ISL_429656 | | | |

| | | | |
|---|---|---|---|
| see above | UW Virology Lab | UW Virology Lab | Pavitra Roychoudhury, Hong Xie, Keith Jerome, Alexander Greninger |
| EPI_ISL_430722, EPI_ISL_430723, EPI_ISL_430724, EPI_ISL_430725, EPI_ISL_430726, EPI_ISL_430727, EPI_ISL_430728, EPI_ISL_430729, EPI_ISL_430730, EPI_ISL_430731, EPI_ISL_430732, EPI_ISL_430733, EPI_ISL_430734, EPI_ISL_430735, EPI_ISL_430736, EPI_ISL_430737, EPI_ISL_430738, EPI_ISL_430739, EPI_ISL_430740, EPI_ISL_430741, EPI_ISL_430742 | | | |
| see above | Chinese PLA Institute for Disease Control and Prevention | Chinese PLA Institute for Disease Control and Prevention | Peng LiJinhui Li, Lizhong Li |
| EPI_ISL_430837 | n/a | Thai National Influenza Center, Department of medical Science, Ministry of Public Health, Thailand | Pilailuk,Okada; Siripaporn,Phuygun; Thanutsapa,Thanadachakul;Sittiporn,Parnmen;Warawan,Wongboot;Sunthareeya,Waicharoen; Malinee,Chittaganpitch |
| EPI_ISL_430841 | Praram 9 Hospital | National Institute of Health. Department of medical Sciences, Ministry of Public Health, Thailand | Pilailuk,Okada; Siripaporn,Phuygun; Thanutsapa,Thanadachakul; Sittiporn,Parnmen;Warawan,Wongboot; Sunthareeya,Waicharoen; Malinee,Chittaganpitch |
| EPI_ISL_430842 | Central chest Institute of Thailand | National Institute of Health. Department of medical Sciences, Ministry of Public Health, Thailand | Pilailuk,Okada; Siripaporn,Phuygun; Thanutsapa,Thanadachakul; Sittiporn,Parnmen;Warawan,Wongboot; Sunthareeya,Waicharoen; Malinee,Chittaganpitch |
| EPI_ISL_430856, EPI_ISL_430857, EPI_ISL_430858, EPI_ISL_430859 | Laboratoriemedicin | The Public Health Agency of Sweden | Oskar Karlsson Lindsjo, Maria Lind Karlberg, Anna-Malin Linde, Olov Svartstrom, Anna Risberg, Shaman Muradrasoli, Karin Tegmark-Wisell |
| EPI_ISL_430862, EPI_ISL_430864 | The Public Health Agency of Sweden | The Public Health Agency of Sweden | Oskar Karlsson Lindsjo, Maria Lind Karlberg, Anna-Malin Linde, Olov Svartstrom, Anna Risberg, Shaman Muradrasoli, Karin Tegmark-Wisell |
| EPI_ISL_430867, EPI_ISL_430868, EPI_ISL_430869, EPI_ISL_430870, EPI_ISL_430871, EPI_ISL_430872, EPI_ISL_430873, EPI_ISL_430874, EPI_ISL_430875, EPI_ISL_430876, EPI_ISL_430877, EPI_ISL_430878, EPI_ISL_430879, EPI_ISL_430880, EPI_ISL_430881, EPI_ISL_430882, EPI_ISL_430883, EPI_ISL_430884, EPI_ISL_430885, EPI_ISL_430886, EPI_ISL_430887, EPI_ISL_430888, EPI_ISL_430889, EPI_ISL_430890, EPI_ISL_430891, EPI_ISL_430892, EPI_ISL_430893, EPI_ISL_430894, EPI_ISL_430895, EPI_ISL_430897, EPI_ISL_430898, EPI_ISL_430899, EPI_ISL_430900, EPI_ISL_430901, EPI_ISL_430902, EPI_ISL_430903, EPI_ISL_430904, EPI_ISL_430905, EPI_ISL_430906, EPI_ISL_430907, EPI_ISL_430908, EPI_ISL_430909, EPI_ISL_430910, EPI_ISL_430911, EPI_ISL_430912, EPI_ISL_430913, EPI_ISL_430914, EPI_ISL_430915, EPI_ISL_430916, EPI_ISL_430917, EPI_ISL_430918, EPI_ISL_430919, EPI_ISL_430920, EPI_ISL_430921, EPI_ISL_430922, EPI_ISL_430923, EPI_ISL_430924, EPI_ISL_430925, EPI_ISL_430926, EPI_ISL_430927, EPI_ISL_430928, EPI_ISL_430929, EPI_ISL_430930, EPI_ISL_430931, EPI_ISL_430932, EPI_ISL_430933, EPI_ISL_430934, EPI_ISL_430935, EPI_ISL_430936, EPI_ISL_430937, EPI_ISL_430938, EPI_ISL_430939, EPI_ISL_430940, EPI_ISL_430941, EPI_ISL_430942, EPI_ISL_430943, EPI_ISL_430944, EPI_ISL_430945, EPI_ISL_430946, EPI_ISL_430947, EPI_ISL_430948, EPI_ISL_430949, EPI_ISL_430950, EPI_ISL_430951, EPI_ISL_430952, EPI_ISL_430953, EPI_ISL_430954, EPI_ISL_430955, EPI_ISL_430956, EPI_ISL_430957, EPI_ISL_430958, EPI_ISL_430959, EPI_ISL_430960, EPI_ISL_430961, EPI_ISL_430962, EPI_ISL_430963, EPI_ISL_430964, EPI_ISL_430965, EPI_ISL_430966, EPI_ISL_430967, EPI_ISL_430968, EPI_ISL_430969, EPI_ISL_430970, EPI_ISL_430971, EPI_ISL_430972, EPI_ISL_430973, EPI_ISL_430974, EPI_ISL_430975, EPI_ISL_430976, EPI_ISL_430977, EPI_ISL_430978, EPI_ISL_430979, EPI_ISL_430980 | | | |
| see above | UW Virology Lab | UW Virology Lab | Pavitra Roychoudhury, Hong Xie, Keith Jerome, Alexander Greninger |
| EPI_ISL_431011, EPI_ISL_431012 | Viral Respiratory Lab, National Institute for Biomedical Research (INRB) | Pathogen Sequencing Lab, National Institute for Biomedical Research (INRB) | Placide Mbala-Kingebeni, Edith Nkwembe, Eddy Kinganda-Lusamaki, Amuri Aziza, Francisca Muyembe Mawete, Catherine Pratt, Matthias Pauthner, Josh Quick, Allison Black, James Hadfield, Trevor Bedford, Ian Goodfellow, Andrew Rambaut, Nick Loman, Kristian Andersen, Michael Wiley, Steve Ahuka-Mundeke, Jean-Jacques Muyembe Tamfum |
| EPI_ISL_434356, EPI_ISL_434357, EPI_ISL_434358, EPI_ISL_434359, EPI_ISL_434360, EPI_ISL_434361, EPI_ISL_434362, EPI_ISL_434363, EPI_ISL_434364, EPI_ISL_434365 | Lab voor klinische biologie | Onderzoeksgroep Virologie | Nick Vereecke, Laurens Lambrechts, Marthe Pauwels, Jozefien De Clercq, Bruno Verhasselt, Linos Vandekerckhove, Hans Nauwynck, Sebastiaan Theuns |
| EPI_ISL_434366, EPI_ISL_434367, EPI_ISL_434368, EPI_ISL_434369, EPI_ISL_434370, EPI_ISL_434371 | Hospital AZ Rivierenland | Institute of Tropical Medicine | Philippe Selhorst, Colin Anthony |
| EPI_ISL_434372, EPI_ISL_434373, EPI_ISL_434374, EPI_ISL_434375, EPI_ISL_434376, EPI_ISL_434377, EPI_ISL_434378, EPI_ISL_434379, EPI_ISL_434380, EPI_ISL_434381, EPI_ISL_434382, EPI_ISL_434383 | | | |
| see above | Hospital AZ Rivierenland | Institute of Tropical Medicine | Philippe Selhorst, Colin Anthony, |
| EPI_ISL_434384, EPI_ISL_434385, EPI_ISL_434386 | Hospital AZ Rivierenland | Institute of Tropical Medicine | Philippe Selhorst, Colin Anthony |
| EPI_ISL_434586, EPI_ISL_434587, EPI_ISL_434588, EPI_ISL_434638, EPI_ISL_434639, EPI_ISL_434640 | Johns Hopkins Hospital Department of Pathology | Johns Hopkins Hospital Department of Pathology | Peter M. Thielen, Thomas Mehoke, Shirlee Wohl, Srividya Ramakrishnan, Oluwaseun Nwulia-Falade, Amanda Ernlund, Melanie Kirsche, Paul Morris, Norah Sadowski, Nídiá Trovao, Victoria Gniazdowski, Michael Schatz, Stuart C. Ray, Winston Timp, Heba Mostafa |
| EPI_ISL_434641, EPI_ISL_434642 | Laboratoriemedicin | The Public Health Agency of Sweden | Oskar Karlsson Lindsjo, Maria Lind Karlberg, Anna-Malin Linde, Olov Svartstrom, Anna Risberg, Shaman Muradrasoli, Karin Tegmark-Wisell |
| EPI_ISL_434678, EPI_ISL_434679, EPI_ISL_434680, EPI_ISL_434681 | Viral Respiratory Lab, National Institute for Biomedical Research (INRB) | Pathogen Sequencing Lab, National Institute for Biomedical Research (INRB) | Placide Mbala-Kingebeni; Edith Nkwembe; Eddy Kinganda-Lusamaki; Amuri Aziza; Francisca Muyembe Mawete; Catherine Pratt; Matthias Pauthner; Josh Quick; Allison Black; James Hadfield; Trevor Bedford; Ian Goodfellow; Andrew Rambaut; Nick Loman; Kristian Andersen; Michael Wiley; Steve Ahuka-Mundeke; Jean-Jacques Muyembe Tamfum |
| EPI_ISL_434682, EPI_ISL_434683, EPI_ISL_434684, EPI_ISL_434685, EPI_ISL_434686, EPI_ISL_434687, EPI_ISL_434688, EPI_ISL_434689, EPI_ISL_434690, EPI_ISL_434691 | Johns Hopkins Hospital Department of Pathology | Johns Hopkins Hospital Department of Pathology | Peter M. Thielen, Thomas Mehoke, Shirlee Wohl, Srividya Ramakrishnan, Melanie Kirsche, Amanda Ernlund, Oluwaseun Falade-Nwulia, Timothy Gilpatrick, Paul Morris, Norah Sadowski, Nídiá Trovao, Victoria Gniazdowski, Michael Schatz, Stuart C. Ray, Winston Timp, Heba Mostafa |
| EPI_ISL_434692, EPI_ISL_434693, EPI_ISL_434694 | Bamrasnaradura hospital | National Institute of Health. Department of medical Sciences, Ministry of Public Health, Thailand | Pilailuk,Okada; Siripaporn,Phuygun; Thanutsapa,Thanadachakul; Sittiporn,Parnmen;Warawan,Wongboot; Sunthareeya,Waicharoen; Malinee,Chittaganpitch |
| EPI_ISL_434695 | unknown | National Institute of Health. Department of medical Sciences, Ministry of Public Health, Thailand | Pilailuk,Okada; Siripaporn,Phuygun; Thanutsapa,Thanadachakul; Sittiporn,Parnmen;Warawan,Wongboot; Sunthareeya,Waicharoen; Malinee,Chittaganpitch |
| EPI_ISL_434696 | Bamrasnaradura hospital | National Institute of Health. Department of medical Sciences, Ministry of Public Health, Thailand | Pilailuk,Okada; Siripaporn,Phuygun; Thanutsapa,Thanadachakul; Sittiporn,Parnmen;Warawan,Wongboot; Sunthareeya,Waicharoen; Malinee,Chittaganpitch |
| EPI_ISL_434697 | unknown | National Institute of Health. Department of medical Sciences, Ministry of Public Health, Thailand | Pilailuk,Okada; Siripaporn,Phuygun; Thanutsapa,Thanadachakul; Sittiporn,Parnmen;Warawan,Wongboot; Sunthareeya,Waicharoen; Malinee,Chittaganpitch |
| EPI_ISL_434698, EPI_ISL_434699 | Praram 9 Hospital | National Institute of Health. Department of medical Sciences, Ministry of Public Health, Thailand | Pilailuk,Okada; Siripaporn,Phuygun; Thanutsapa,Thanadachakul; Sittiporn,Parnmen;Warawan,Wongboot; Sunthareeya,Waicharoen; Malinee,Chittaganpitch |
| EPI_ISL_434700 | Ramkhamhaeng Hospital | National Institute of Health. Department of medical Sciences, Ministry of Public Health, Thailand | Pilailuk,Okada; Siripaporn,Phuygun; Thanutsapa,Thanadachakul; Sittiporn,Parnmen;Warawan,Wongboot; Sunthareeya,Waicharoen; Malinee,Chittaganpitch |
| EPI_ISL_434701 | Panyanunthaphikkhu Chonprathan Medical Center (PCMC) | National Institute of Health. Department of medical Sciences, Ministry of Public Health, Thailand | Pilailuk,Okada; Siripaporn,Phuygun; Thanutsapa,Thanadachakul; Sittiporn,Parnmen;Warawan,Wongboot; Sunthareeya,Waicharoen; Malinee,Chittaganpitch |
| EPI_ISL_434702, EPI_ISL_434703, EPI_ISL_434704, EPI_ISL_434705, EPI_ISL_434706 | Praram 9 Hospital | National Institute of Health. Department of medical Sciences, Ministry of Public Health, Thailand | Pilailuk,Okada; Siripaporn,Phuygun; Thanutsapa,Thanadachakul; Sittiporn,Parnmen;Warawan,Wongboot; Sunthareeya,Waicharoen; Malinee,Chittaganpitch |
| EPI_ISL_434707 | Thammasat University Hospital | National Institute of Health. Department of medical Sciences, Ministry of Public Health, Thailand | Pilailuk,Okada; Siripaporn,Phuygun; Thanutsapa,Thanadachakul; Sittiporn,Parnmen;Warawan,Wongboot; Sunthareeya,Waicharoen; Malinee,Chittaganpitch |
| EPI_ISL_434708 | unknown | National Institute of Health. Department of medical Sciences, Ministry of Public Health, Thailand | Pilailuk,Okada; Siripaporn,Phuygun; Thanutsapa,Thanadachakul; Sittiporn,Parnmen;Warawan,Wongboot; Sunthareeya,Waicharoen; Malinee,Chittaganpitch |
| EPI_ISL_434710, EPI_ISL_434711, EPI_ISL_435032, EPI_ISL_435033 | Viral Respiratory Lab, National Institute for Biomedical Research (INRB) | Pathogen Sequencing Lab, National Institute for Biomedical Research (INRB) | Placide Mbala-Kingebeni, Edith Nkwembe, Eddy Kinganda-Lusamaki, Adrienne Amuri Aziza, Francisca Muyembe Mawete, Catherine Pratt, Matthias Pauthner, Josh Quick, Allison Black, James Hadfield, Trevor Bedford, Ian Goodfellow, Andrew Rambaut, Nick Loman, Kristian Andersen, Michael Wiley, Steve Ahuka-Mundeke, Jean-Jacques Muyembe Tamfum |

| | | | |
|---|---|---|---|
| EPI_ISL_435049 | B.J. Medical College and Civil hospital | Gujarat Biotechnology Research Centre | Pinal Trivedi, Maharshi Pandya, Amit Kanani, Akanksha Verma, Nitin Savaliya, Raghawendra Kumar, Dinesh Kumar, Zuber Saiyed, Dipa Kinariwala, Disha Patel, Binita Aring, Geeta Vaghela, Sonia Barve, Bhavesh Modi, Kairavi Joshi, Gaurishankar Shrimali, Nidhi Sood, Pranay Shah, R D Dixit, Snehal Bagatharia, Kamlesh J Upadhyay, Ramesh Pandit, Tejas Shah, Ankit Hinsu, Pritesh Sabara, Apurvasinh Puvar, Janvi Raval, Monika Gandhi, Neha Rajpara, Chaitanya Joshi, Madhvi Joshi |
| EPI_ISL_435051 | B.J. Medical College and Civil hospital | Gujarat Biotechnology Research Centre | Pritesh Sabara, Apurvasinh Puvar, Janvi Raval, Monika Gandhi, Pinal Trivedi, Maharshi Pandya, Amit Kanani, Akanksha Verma, Nitin Savaliya, Raghawendra Kumar, Dinesh Kumar, Zuber Saiyed, Dipa Kinariwala, Disha Patel, Binita Aring, Geeta Vaghela, Sonia Barve, Bhavesh Modi, Kairavi Joshi, Gaurishankar Shrimali, Nidhi Sood, Pranay Shah, R D Dixit, Snehal Bagatharia, Kamlesh J Upadhyay, Ramesh Pandit, Tejas Shah, Ankit Hinsu, Vasudha Sharma, Chaitanya Joshi, Madhvi Joshi |
| EPI_ISL_435060, EPI_ISL_435061, EPI_ISL_435062, EPI_ISL_435063, EPI_ISL_435064, EPI_ISL_435065, EPI_ISL_435066, EPI_ISL_435067, EPI_ISL_435068, EPI_ISL_435069, EPI_ISL_435070, EPI_ISL_435071, EPI_ISL_435072, EPI_ISL_435073, EPI_ISL_435074, EPI_ISL_435075, EPI_ISL_435076, EPI_ISL_435077, EPI_ISL_435078, EPI_ISL_435079, EPI_ISL_435080, EPI_ISL_435081, EPI_ISL_435082, EPI_ISL_435083, EPI_ISL_435084, EPI_ISL_435085, EPI_ISL_435086, EPI_ISL_435087, EPI_ISL_435088, EPI_ISL_435089, EPI_ISL_435090, EPI_ISL_435091, EPI_ISL_435092, EPI_ISL_435093, EPI_ISL_435094, EPI_ISL_435095, EPI_ISL_435096, EPI_ISL_435097, EPI_ISL_435098, EPI_ISL_435099, EPI_ISL_435100, EPI_ISL_435101, EPI_ISL_435102, EPI_ISL_435103, EPI_ISL_435104, EPI_ISL_435105, EPI_ISL_435106, EPI_ISL_435107, EPI_ISL_435108, EPI_ISL_435109, EPI_ISL_435110, EPI_ISL_435111, EPI_ISL_435112 | | | |
| see above | National Centre for Disease control (NCDC), CSIR-Institute of Genomics and Integrative Biology (CSIR-IGIB) | NCDC/CSIR-IGIB | Pramod Kumar, Rajesh Pandey, Pooja Sharma, Mahesh Dhar, Vivekanand A, Bharathram Uppili, Himanshu Vashisht, Saruchi Wadhwa, Nishu Tyagi, Uma Sharma, Priyanka Singh, Hemlata Lall, Meena Datta, Poonam Gupta, Nidhi Saini, Aarti Tewari, Bibhash Nandi, Dhirendra Kumar, Satyabrata Bag, Varun Jaiswal, Hema Gogia, Preeti Madan, Simrita Singh, Prateek Singh, Debasis Dash, Mitali Mukerji, Manju Bala, Sandhya Kabra, Sujeet Singh, Mohammed Faruq, Anurag Agrawal, Partha Rakshit |
| EPI_ISL_435113, EPI_ISL_435114, EPI_ISL_435116, EPI_ISL_435117, EPI_ISL_435118 | Viral Respiratory Lab, National Institute for Biomedical Research (INRB) | Pathogen Sequencing Lab, National Institute for Biomedical Research (INRB) | Placide Mbala-Kingebeni, Edith Nkwembe, Eddy Kinganda-Lusamaki, Adrienne Amuri Aziza, Francisca Muyembe Mawete, Catherine Pratt, Matthias Pauthner, Josh Quick, Allison Black, James Hadfield, Trevor Bedford, Ian Goodfellow, Andrew Rambaut, Nick Loman, Kristian Andersen, Michael Wiley, Steve Ahuka-Mundeke, Jean-Jacques Muyembe Tamfum |
| EPI_ISL_435156, EPI_ISL_435157, EPI_ISL_435158, EPI_ISL_435159, EPI_ISL_435160, EPI_ISL_435161, EPI_ISL_435162, EPI_ISL_435163, EPI_ISL_435164, EPI_ISL_435165, EPI_ISL_435166, EPI_ISL_435167, EPI_ISL_435168 | | | |
| see above | Viral Respiratory Lab, National Institute for Biomedical Research (INRB) | Pathogen Sequencing Lab, National Institute for Biomedical Research (INRB) | Placide Mbala-Kingebeni, Edith Nkwembe, Eddy Kinganda-Lusamaki, Amuri Aziza, Francisca Muyembe Mawete, Catherine Pratt, Matthias Pauthner, Josh Quick, Allison Black, James Hadfield, Trevor Bedford, Ian Goodfellow, Andrew Rambaut, Nick Loman, Kristian Andersen, Michael Wiley, Steve Ahuka-Mundeke, Jean-Jacques Muyembe Tamfum |
| EPI_ISL_435303 | National Hospital of Tropical Diseases | Oxford University Clinical Research Unit, Hanoi, Vietnam | Nguyen Thi Tam, Van Dinh Trang, Nguyen Thu Trang, Nguyen Thi Ngoc Diep, Le Nguyen Minh Hoa, Pham Ngoc Thach, H.Rogier van Doorn, on behalf of the OUCRU COVID-19 research group |
| EPI_ISL_435305, EPI_ISL_435308, EPI_ISL_435310, EPI_ISL_435311, EPI_ISL_435312, EPI_ISL_435313, EPI_ISL_435314, EPI_ISL_435315, EPI_ISL_435316, EPI_ISL_435317 | National Hospital of Tropical Diseases | Oxford University Clinical Research Unit, Hanoi, Vietnam | Nguyen Thi Tam, Van Dinh Trang, Nguyen Thu Trang, Nguyen Thi Ngoc Diep, Le Nguyen Minh Hoa, Pham Ngoc Thach, H. Rogier van Doorn, on behalf of the OUCRU COVID-19 research group |
| EPI_ISL_435403, EPI_ISL_435404, EPI_ISL_435405, EPI_ISL_435406, EPI_ISL_435407, EPI_ISL_435408, EPI_ISL_435409, EPI_ISL_435410, EPI_ISL_435411, EPI_ISL_435412, EPI_ISL_435413, EPI_ISL_435414, EPI_ISL_435415, EPI_ISL_435416, EPI_ISL_435417, EPI_ISL_435418, EPI_ISL_435419, EPI_ISL_435420, EPI_ISL_435421, EPI_ISL_435422, EPI_ISL_435423, EPI_ISL_435424, EPI_ISL_435425, EPI_ISL_435426, EPI_ISL_435427, EPI_ISL_435428, EPI_ISL_435429, EPI_ISL_435430, EPI_ISL_435431 | | | |
| see above | Virological Research Group, Szentágothai Research Centre | Bioinformatics Research Group, Szentágothai Research Centre | Péter Urbán, Endre Gábor Tóth, Gábor Kemenesi, Róbert Herczeg, Attila Gyenesei, Ferenc Jakab |
| EPI_ISL_436194 | Viral Respiratory Lab, National Institute for Biomedical Research (INRB) | Pathogen Sequencing Lab, National Institute for Biomedical Research (INRB) | Placide Mbala-Kingebeni, Edith Nkwembe, Eddy Kinganda-Lusamaki, Amuri Aziza, Francisca Muyembe Mawete, Catherine Pratt, Matthias Pauthner, Josh Quick, Allison Black, James Hadfield, Trevor Bedford, Ian Goodfellow, Andrew Rambaut, Nick Loman, Kristian Andersen, Michael Wiley, Steve Ahuka-Mundeke, Jean-Jacques Muyembe Tamfum |
| EPI_ISL_436200, EPI_ISL_436208, EPI_ISL_436216, EPI_ISL_436224, EPI_ISL_436231, EPI_ISL_436275, EPI_ISL_436277 | Servicio de Microbiología. Consorcio Hospital General Universitario de Valencia | Sequencing and Bioinformatics Service and Molecular Epidemiology Research Group. FISABIO-Public Health | Neris Garcia-Gonzalez, Loreto Ferrús Abad, Maria Dolores Ocete, Inma Galán Vendrell, Paula Ruiz-Hueso, Mariana Reyes-Prieto, Vicente Soriano Chirona, Maria Alma Bracho, Griselda De Marco, Beatriz Beamud, Lidia Ruiz Roldan, Marta Pla Diaz, Llúcia Martínez-Priego, Concepcion Gimeno, Giuseppe D'Auria, Fernando Gonzalez-Candelas |
| EPI_ISL_436283 | Servicio de Microbiologia. Hospital Clinico Universitario de Valencia | Sequencing and Bioinformatics Service and Molecular Epidemiology Research Group. FISABIO-Public Health | Neris Garcia-Gonzalez, Inma Galán Vendrell, Sandra Carbo, Loreto Ferrús Abad, Paula Ruiz-Hueso, Mariana Reyes-Prieto, Vicente Soriano Chirona, Ivan Ansari, David Navarro, Maria Alma Bracho, Griselda De Marco, Beatriz Beamud, Lidia Ruiz Roldan, Marta Pla Diaz, Llúcia Martínez-Priego, Giuseppe D'Auria, Fernando Gonzalez-Candelas |
| EPI_ISL_436287 | Servicio de Microbiologia. Hospital Clinico Universitario de Valencia | Sequencing and Bioinformatics Service and Molecular Epidemiology Research Group. FISABIO-Public Health | Paula Ruiz-Hueso, Mariana Reyes-Prieto, Vicente Soriano Chirona, Ivan Ansari, David Navarro, Maria Alma Bracho, Griselda De Marco, Beatriz Beamud, Lidia Ruiz Roldan, Marta Pla Diaz, Neris Garcia-Gonzalez, Inma Galán Vendrell, Sandra Carbo, Loreto Ferrús Abad, Llúcia Martínez-Priego, Giuseppe D'Auria, Fernando Gonzalez-Candelas |
| EPI_ISL_436297, EPI_ISL_436303 | Servicio de Microbiologia. Hospital Clinico Universitario de Valencia | Sequencing and Bioinformatics Service and Molecular Epidemiology Research Group. FISABIO-Public Health | Neris Garcia-Gonzalez, Inma Galán Vendrell, Sandra Carbo, Loreto Ferrús Abad, Paula Ruiz-Hueso, Mariana Reyes-Prieto, Vicente Soriano Chirona, Ivan Ansari, David Navarro, Maria Alma Bracho, Griselda De Marco, Beatriz Beamud, Lidia Ruiz Roldan, Marta Pla Diaz, Llúcia Martínez-Priego, Giuseppe D'Auria, Fernando Gonzalez-Candelas |
| EPI_ISL_436307 | Servicio de Microbiologia. Hospital Clinico Universitario de Valencia | Sequencing and Bioinformatics Service and Molecular Epidemiology Research Group. FISABIO-Public Health | Paula Ruiz-Hueso, Mariana Reyes-Prieto, Vicente Soriano Chirona, Ivan Ansari, David Navarro, Maria Alma Bracho, Griselda De Marco, Beatriz Beamud, Lidia Ruiz Roldan, Marta Pla Diaz, Neris Garcia-Gonzalez, Inma Galán Vendrell, Sandra Carbo, Loreto Ferrús Abad, Llúcia Martínez-Priego, Giuseppe D'Auria, Fernando Gonzalez-Candelas |
| EPI_ISL_436317, EPI_ISL_436323 | Servicio de Microbiologia. Hospital Clinico Universitario de Valencia | Sequencing and Bioinformatics Service and Molecular Epidemiology Research Group. FISABIO-Public Health | Neris Garcia-Gonzalez, Inma Galán Vendrell, Sandra Carbo, Loreto Ferrús Abad, Paula Ruiz-Hueso, Mariana Reyes-Prieto, Vicente Soriano Chirona, Ivan Ansari, David Navarro, Maria Alma Bracho, Griselda De Marco, Beatriz Beamud, Lidia Ruiz Roldan, Marta Pla Diaz, Llúcia Martínez-Priego, Giuseppe D'Auria, Fernando Gonzalez-Candelas |
| EPI_ISL_436327 | Servicio de Microbiologia. Hospital Clinico Universitario de Valencia | Sequencing and Bioinformatics Service and Molecular Epidemiology Research Group. FISABIO-Public Health | Paula Ruiz-Hueso, Mariana Reyes-Prieto, Vicente Soriano Chirona, Ivan Ansari, David Navarro, Maria Alma Bracho, Griselda De Marco, Beatriz Beamud, Lidia Ruiz Roldan, Marta Pla Diaz, Neris Garcia-Gonzalez, Inma Galán Vendrell, Sandra Carbo, Loreto Ferrús Abad, Llúcia Martínez-Priego, Giuseppe D'Auria, Fernando Gonzalez-Candelas |
| EPI_ISL_436337, EPI_ISL_436343 | Servicio de Microbiologia. Hospital Clinico Universitario de Valencia | Sequencing and Bioinformatics Service and Molecular Epidemiology Research Group. FISABIO-Public Health | Neris Garcia-Gonzalez, Inma Galán Vendrell, Sandra Carbo, Loreto Ferrús Abad, Paula Ruiz-Hueso, Mariana Reyes-Prieto, Vicente Soriano Chirona, Ivan Ansari, David Navarro, Maria Alma Bracho, Griselda De Marco, Beatriz Beamud, Lidia Ruiz Roldan, Marta Pla Diaz, Llúcia Martínez-Priego, Giuseppe D'Auria, Fernando Gonzalez-Candelas |
| EPI_ISL_436347 | Servicio de Microbiologia. Hospital Clinico Universitario de Valencia | Sequencing and Bioinformatics Service and Molecular Epidemiology Research Group. FISABIO-Public Health | Paula Ruiz-Hueso, Mariana Reyes-Prieto, Vicente Soriano Chirona, Ivan Ansari, David Navarro, Maria Alma Bracho, Griselda De Marco, Beatriz Beamud, Lidia Ruiz Roldan, Marta Pla Diaz, Neris Garcia-Gonzalez, Inma Galán Vendrell, Sandra Carbo, Loreto Ferrús Abad, Llúcia Martínez-Priego, Giuseppe D'Auria, Fernando Gonzalez-Candelas |
| EPI_ISL_436357, EPI_ISL_436360 | Servicio de Microbiologia. Hospital Clinico Universitario de Valencia | Sequencing and Bioinformatics Service and Molecular Epidemiology Research Group. FISABIO-Public Health | Neris Garcia-Gonzalez, Inma Galán Vendrell, Sandra Carbo, Loreto Ferrús Abad, Paula Ruiz-Hueso, Mariana Reyes-Prieto, Vicente Soriano Chirona, Ivan Ansari, David Navarro, Maria Alma Bracho, Griselda De Marco, Beatriz Beamud, Lidia Ruiz Roldan, Marta Pla Diaz, Llúcia Martínez-Priego, Giuseppe D'Auria, Fernando Gonzalez-Candelas |
| EPI_ISL_436361 | Servicio de Microbiología. Consorcio Hospital General Universitario de Valencia | Sequencing and Bioinformatics Service and Molecular Epidemiology Research Group. FISABIO-Public Health | Neris Garcia-Gonzalez, Loreto Ferrús Abad, Maria Dolores Ocete, Inma Galán Vendrell, Paula Ruiz-Hueso, Mariana Reyes-Prieto, Vicente Soriano Chirona, Maria Alma Bracho, Griselda De Marco, Beatriz Beamud, Lidia Ruiz Roldan, Marta Pla Diaz, Llúcia Martínez-Priego, Concepcion Gimeno, Giuseppe D'Auria, Fernando Gonzalez-Candelas |
| EPI_ISL_436367 | Servicio de Microbiologia. Hospital Clinico Universitario de Valencia | Sequencing and Bioinformatics Service and Molecular Epidemiology Research Group. FISABIO-Public Health | Neris Garcia-Gonzalez, Inma Galán Vendrell, Sandra Carbo, Loreto Ferrús Abad, Paula Ruiz-Hueso, Mariana Reyes-Prieto, Vicente Soriano Chirona, Ivan Ansari, David Navarro, Maria Alma Bracho, Griselda De Marco, Beatriz Beamud, Lidia Ruiz Roldan, Marta Pla Diaz, Llúcia Martínez-Priego, Giuseppe D'Auria, Fernando Gonzalez-Candelas |
| EPI_ISL_436371 | Servicio de Microbiologia. Hospital Clinico Universitario de Valencia | Sequencing and Bioinformatics Service and Molecular Epidemiology Research Group. FISABIO-Public Health | Paula Ruiz-Hueso, Mariana Reyes-Prieto, Vicente Soriano Chirona, Ivan Ansari, David Navarro, Maria Alma Bracho, Griselda De Marco, Beatriz Beamud, Lidia Ruiz Roldan, Marta Pla Diaz, Neris Garcia-Gonzalez, Inma Galán Vendrell, Sandra Carbo, Loreto Ferrús Abad, Llúcia Martínez-Priego, Giuseppe |

| | | | |
|---|---|---|---|
| | | | D'Auria, Fernando Gonzalez-Candelas |
| EPI_ISL_436388, EPI_ISL_436395 | Servicio de Microbiología. Consorcio Hospital General Universitario de Valencia | Sequencing and Bioinformatics Service and Molecular Epidemiology Research Group. FISABIO-Public Health | Neris Garcia-Gonzalez, Loreto Ferrús Abad, Maria Dolores Ocete, Inma Galán Vendrell, Paula Ruiz-Hueso, Mariana Reyes-Prieto, Vicente Soriano Chirona, Maria Alma Bracho, Griselda De Marco, Beatriz Beamud, Lidia Ruiz Roldan, Marta Pla Diaz, Llúcia Martínez-Priego, Concepcion Gimeno, Giuseppe D'Auria, Fernando Gonzalez-Candelas |
| EPI_ISL_436398 | Servicio de Microbiologia. Hospital Clinico Universitario de Valencia | Sequencing and Bioinformatics Service and Molecular Epidemiology Research Group. FISABIO-Public Health | Paula Ruiz-Hueso, Mariana Reyes-Prieto, Vicente Soriano Chirona, Ivan Ansari, David Navarro, Maria Alma Bracho, Griselda De Marco, Beatriz Beamud, Lidia Ruiz Roldan, Marta Pla Diaz, Neris Garcia-Gonzalez, Inma Galán Vendrell, Sandra Carbo, Loreto Ferrús Abad, Llúcia Martínez-Priego, Giuseppe D'Auria, Fernando Gonzalez-Candelas |
| EPI_ISL_436408, EPI_ISL_436411 | Servicio de Microbiologia. Hospital Clinico Universitario de Valencia | Sequencing and Bioinformatics Service and Molecular Epidemiology Research Group. FISABIO-Public Health | Neris Garcia-Gonzalez, Inma Galán Vendrell, Sandra Carbo, Loreto Ferrús Abad, Paula Ruiz-Hueso, Mariana Reyes-Prieto, Vicente Soriano Chirona, Ivan Ansari, David Navarro, Maria Alma Bracho, Griselda De Marco, Beatriz Beamud, Lidia Ruiz Roldan, Marta Pla Diaz, Llúcia Martínez-Priego, Giuseppe D'Auria, Fernando Gonzalez-Candelas |
| EPI_ISL_436412 | Viral Respiratory Lab, National Institute for Biomedical Research (INRB) | Pathogen Sequencing Lab, National Institute for Biomedical Research (INRB) | Placide Mbala-Kingebeni, Edith Nkwembe, Eddy Kinganda-Lusamaki, Amuri Aziza, Francisca Muyembe Mawete, Catherine Pratt, Matthias Pauthner, Josh Quick, Allison Black, James Hadfield, Trevor Bedford, Ian Goodfellow, Andrew Rambaut, Nick Loman, Kristian Andersen, Michael Wiley, Steve Ahuka-Mundeke, Jean-Jacques Muyembe Tamfum |
| EPI_ISL_436413, EPI_ISL_436414, EPI_ISL_436415, EPI_ISL_436417, EPI_ISL_436418, EPI_ISL_436419, EPI_ISL_436420, EPI_ISL_436421, EPI_ISL_436422, EPI_ISL_436423, EPI_ISL_436424, EPI_ISL_436425, EPI_ISL_436426, EPI_ISL_436427, EPI_ISL_436428, EPI_ISL_436429, EPI_ISL_436430, EPI_ISL_436431, EPI_ISL_436432, EPI_ISL_436433, EPI_ISL_436434, EPI_ISL_436435, EPI_ISL_436436, EPI_ISL_436437, EPI_ISL_436438, EPI_ISL_436439, EPI_ISL_436440, EPI_ISL_436441, EPI_ISL_436443, EPI_ISL_436444, EPI_ISL_436445, EPI_ISL_436446, EPI_ISL_436447, EPI_ISL_436448, EPI_ISL_436449, EPI_ISL_436450, EPI_ISL_436451, EPI_ISL_436452, EPI_ISL_436453, EPI_ISL_436454, EPI_ISL_436455, EPI_ISL_436456, EPI_ISL_436457, EPI_ISL_436458, EPI_ISL_436459, EPI_ISL_436460, EPI_ISL_436461, EPI_ISL_436462, EPI_ISL_436463 | | | |
| see above | National Centre for Disease control (NCDC) | NCDC/CSIR-IGIB | Pramod Kumar#, Rajesh Pandey#, Pooja Sharma, Mahesh S Dhar, Vivekanand A, Bharathram Uppili, Himanshu Vashisht, Saruchi Wadhwa, Nishu Tyagi, Uma Sharma, Priyanka Singh, Hemlata Lall, Meena Datta, Poonam Gupta, Nidhi Saini, Aarti Tewari, Bibhash Nandi, Dhirendra Kumar, Satyabrata Bag, Varun Jaiswal, Hema Gogia, Preeti Madan, Simrita Singh, Prateek Singh, Debasis Dash, Mitali Mukerji, Manju Bala, Sandhya Kabra, Sujeet Singh, Mohammed Faruq, Anurag Agrawal*, Partha Rakshit* |
| EPI_ISL_437193, EPI_ISL_437194, EPI_ISL_437195, EPI_ISL_437196, EPI_ISL_437337, EPI_ISL_437338, EPI_ISL_437339, EPI_ISL_437340, EPI_ISL_437341, EPI_ISL_437342, EPI_ISL_437343, EPI_ISL_437344, EPI_ISL_437345, EPI_ISL_437346, EPI_ISL_437347, EPI_ISL_437348, EPI_ISL_437350, EPI_ISL_437351, EPI_ISL_437352, EPI_ISL_437353, EPI_ISL_437354, EPI_ISL_437355, EPI_ISL_437356, EPI_ISL_437357, EPI_ISL_437358 | | | |
| see above | Viral Respiratory Lab, National Institute for Biomedical Research (INRB) | Pathogen Sequencing Lab, National Institute for Biomedical Research (INRB) | Placide Mbala-Kingebeni, Edith Nkwembe, Eddy Kinganda-Lusamaki, Amuri Aziza, Francisca Muyembe Mawete, Catherine Pratt, Matthias Pauthner, Josh Quick, Allison Black, James Hadfield, Trevor Bedford, Ian Goodfellow, Andrew Rambaut, Nick Loman, Kristian Andersen, Michael Wiley, Steve Ahuka-Mundeke, Jean-Jacques Muyembe Tamfum |
| EPI_ISL_437440 | Department of MicroBiology, Government Medical College, Surat | Gujarat Biotechnology Research Centre | Nitin Savaliya, Raghawendra Kumar, Dinesh Kumar, Zuber Saiyed, Dipa Kinariwala, Disha Patel, Binita Aring, Neeta Khandelwal, Geeta Vaghela, Sonia Barve, Bhavesh Modi, Kairavi Joshi, Gaurishankar Shrimali, Nidhi Sood, Pranay Shah, R D Dixit, Snehal Bagatharia, Kamlesh J Upadhyay, Ramesh Pandit, Tejas Shah, Ankit Hinsu, Pritesh Sabara, Apurvasinh Puvar, Janvi Raval, Monika Gandhi, Pinal Trivedi, Maharshi Pandya, Amit Kanani, Akanksha Verma, Bhavya Jindal, Chaitanya Joshi, Madhvi Joshi |
| EPI_ISL_437447 | B.J. Medical College and Civil hospital | Gujarat Biotechnology Research Centre | Neeta Khandelwal, Geeta Vaghela, Sonia Barve, Bhavesh Modi, Kairavi Joshi, Gaurishankar Shrimali, Nidhi Sood, Pranay Shah, R D Dixit, Snehal Bagatharia, Kamlesh J Upadhyay, Ramesh Pandit, Tejas Shah, Ankit Hinsu, Pritesh Sabara, Apurvasinh Puvar, Janvi Raval, Monika Gandhi, Pinal Trivedi, Maharshi Pandya, Amit Kanani, Akanksha Verma, Nitin Savaliya, Raghawendra Kumar, Dinesh Kumar, Zuber Saiyed, Dipa Kinariwala, Disha Patel, Binita Aring, Neha Rajpara, Chaitanya Joshi, Madhvi Joshi |
| EPI_ISL_437453 | B.J. Medical College and Civil hospital | Gujarat Biotechnology Research Centre | Nidhi Sood, Pranay Shah, R D Dixit, Snehal Bagatharia, Kamlesh J Upadhyay, Ramesh Pandit, Tejas Shah, Ankit Hinsu, Pritesh Sabara, Apurvasinh Puvar, Janvi Raval, Monika Gandhi, Pinal Trivedi, Maharshi Pandya, Amit Kanani, Akanksha Verma, Nitin Savaliya, Raghawendra Kumar, Dinesh Kumar, Zuber Saiyed, Dipa Kinariwala, Disha Patel, Binita Aring, Neeta Khandelwal, Geeta Vaghela, Sonia Barve, Bhavesh Modi, Kairavi Joshi, Gaurishankar Shrimali, Chaitanya Joshi, Dipeshwari Shewale, Madhvi Joshi |
| EPI_ISL_437454 | B.J. Medical College and Civil hospital | Gujarat Biotechnology Research Centre | Pranay Shah, R D Dixit, Snehal Bagatharia, Kamlesh J Upadhyay, Ramesh Pandit, Tejas Shah, Ankit Hinsu, Pritesh Sabara, Apurvasinh Puvar, Janvi Raval, Monika Gandhi, Pinal Trivedi, Maharshi Pandya, Amit Kanani, Akanksha Verma, Nitin Savaliya, Raghawendra Kumar, Dinesh Kumar, Zuber Saiyed, Dipa Kinariwala, Disha Patel, Binita Aring, Neeta Khandelwal, Geeta Vaghela, Sonia Barve, Bhavesh Modi, Kairavi Joshi, Gaurishankar Shrimali, Nidhi Sood, Chaitanya Joshi, Sharmistha Majumdar, Madhvi Joshi |
| EPI_ISL_437803, EPI_ISL_437804, EPI_ISL_437805, EPI_ISL_437806, EPI_ISL_437807, EPI_ISL_437808, EPI_ISL_437809, EPI_ISL_437810, EPI_ISL_437811, EPI_ISL_437812, EPI_ISL_437813, EPI_ISL_437814, EPI_ISL_437815, EPI_ISL_437816, EPI_ISL_437817, EPI_ISL_437818, EPI_ISL_437819, EPI_ISL_437820, EPI_ISL_437821, EPI_ISL_437822, EPI_ISL_437823, EPI_ISL_437824, EPI_ISL_437825, EPI_ISL_437826, EPI_ISL_437827, EPI_ISL_437828, EPI_ISL_437829, EPI_ISL_437830, EPI_ISL_437831, EPI_ISL_437832, EPI_ISL_437833, EPI_ISL_437834, EPI_ISL_437835, EPI_ISL_437836, EPI_ISL_437837, EPI_ISL_437838, EPI_ISL_437839, EPI_ISL_437840, EPI_ISL_437841, EPI_ISL_437842, EPI_ISL_437843, EPI_ISL_437844, EPI_ISL_437845, EPI_ISL_437846, EPI_ISL_437847, EPI_ISL_437848, EPI_ISL_437849, EPI_ISL_437850, EPI_ISL_437851, EPI_ISL_437852, EPI_ISL_437853, EPI_ISL_437854, EPI_ISL_437855, EPI_ISL_437856, EPI_ISL_437857, EPI_ISL_437858, EPI_ISL_437859, EPI_ISL_437860, EPI_ISL_437861, EPI_ISL_437862, EPI_ISL_437863, EPI_ISL_437864, EPI_ISL_437865, EPI_ISL_437866, EPI_ISL_437868, EPI_ISL_437869, EPI_ISL_437870, EPI_ISL_437871, EPI_ISL_437872 | | | |
| see above | UW Virology Lab | UW Virology Lab | Pavitra Roychoudhury, Hong Xie, Keith Jerome, Alexander Greninger |
| EPI_ISL_438222, EPI_ISL_438223, EPI_ISL_438224, EPI_ISL_438225, EPI_ISL_438226, EPI_ISL_438227, EPI_ISL_438228, EPI_ISL_438229, EPI_ISL_438230, EPI_ISL_438231, EPI_ISL_438232, EPI_ISL_438233, EPI_ISL_438234 | | | |
| see above | Johns Hopkins Hospital Department of Pathology | Johns Hopkins Hospital Department of Pathology | Peter M. Thielen, Thomas Mehoke, Shirlee Wohl, Srividya Ramakrishnan, Melanie Kirsche,_Amanda Ernlund,_Oluwaseun Falade-Nwulia, Timothy Gilpatrick, Paul Morris, Norah Sadowski, N_di_ Trovao, Victoria Gniazdowski, Michael Schatz, Stuart C. Ray, Winston Timp, Heba Mostafa |
| EPI_ISL_438235, EPI_ISL_438236, EPI_ISL_438237, EPI_ISL_438238, EPI_ISL_438239, EPI_ISL_438240, EPI_ISL_438241, EPI_ISL_438242, EPI_ISL_438243, EPI_ISL_438244, EPI_ISL_438245, EPI_ISL_438246, EPI_ISL_438247 | | | |
| see above | Johns Hopkins Hospital Department of Pathology | Johns Hopkins Hospital Department of Pathology | Peter M. Thielen, Thomas Mehoke, Shirlee Wohl, Srividya Ramakrishnan, Melanie Kirsche, Amanda Ernlund, Oluwaseun Falade-Nwulia, Timothy Gilpatrick, Paul Morris, Norah Sadowski, Nidia Trovao, Victoria Gniazdowski, Michael Schatz, Stuart C. Ray, Winston Timp, Heba Mostafa |
| EPI_ISL_443184, EPI_ISL_443185, EPI_ISL_443186 | UW Virology Lab | UW Virology Lab | Pavitra Roychoudhury, Hong Xie, Keith Jerome, Alexander Greninger |
| EPI_ISL_444456 | B.J. Medical College and Civil hospital | Gujarat Biotechnology Research Centre | R D Dixit, Snehal Bagatharia, Kamlesh J Upadhyay, Ramesh Pandit, Tejas Shah, Ankit Hinsu, Pritesh Sabara, Apurvasinh Puvar, Janvi Raval, Monika Gandhi, Pinal Trivedi, Maharshi Pandya, Amit Kanani, Akanksha Verma, Nitin Savaliya, Raghawendra Kumar, Dinesh Kumar, Zuber Saiyed, Dipa Kinariwala, Disha Patel, Binita Aring, Neeta Khandelwal, Geeta Vaghela, Sonia Barve, Bhavesh Modi, Kairavi Joshi, Gaurishankar Shrimali, Nidhi Sood, Pranay Shah, Pooja P Doshi, Chaitanya Joshi, Madhvi Joshi |
| EPI_ISL_444462 | B.J. Medical College and Civil hospital | Gujarat Biotechnology Research Centre | Pritesh Sabara, Apurvasinh Puvar, Janvi Raval, Monika Gandhi, Pinal Trivedi, Maharshi Pandya, Amit Kanani, Akanksha Verma, Nitin Savaliya, Raghawendra Kumar, Dinesh Kumar, Zuber Saiyed, Dipa Kinariwala, Disha Patel, Binita Aring, Neeta Khandelwal, Geeta Vaghela, Sonia Barve, Bhavesh Modi, Kairavi Joshi, Gaurishankar Shrimali, Nidhi Sood, Pranay Shah, R D Dixit, Snehal Bagatharia, Kamlesh J Upadhyay, Ramesh Pandit, Tejas Shah, Ankit Hinsu, Armi Chaudhari, Chaitanya Joshi, Madhvi Joshi |
| EPI_ISL_444466 | B.J. Medical College and Civil hospital | Gujarat Biotechnology Research Centre | Pinal Trivedi, Maharshi Pandya, Amit Kanani, Akanksha Verma, Nitin Savaliya, Raghawendra Kumar, Dinesh Kumar, Zuber Saiyed, Dipa Kinariwala, Disha Patel, Binita Aring, Neeta Khandelwal, Geeta Vaghela, Sonia Barve, Bhavesh Modi, Kairavi Joshi, Gaurishankar Shrimali, Nidhi Sood, Pranay Shah, R D Dixit, Snehal Bagatharia, Kamlesh J Upadhyay, Ramesh Pandit, Tejas Shah, Ankit Hinsu, Pritesh Sabara, Apurvasinh Puvar, Janvi Raval, Monika Gandhi, Sharmistha Majumdar, Chaitanya Joshi, Madhvi Joshi |
| EPI_ISL_444470 | B.J. Medical College and Civil hospital | Gujarat Biotechnology Research Centre | Nitin Savaliya, Raghawendra Kumar, Dinesh Kumar, Zuber Saiyed, Dipa Kinariwala, Disha Patel, Binita Aring, Neeta Khandelwal, Geeta Vaghela, Sonia Barve, Bhavesh Modi, Kairavi Joshi, Gaurishankar Shrimali, Nidhi Sood, Pranay Shah, R D Dixit, Snehal Bagatharia, Kamlesh J Upadhyay, Ramesh Pandit, Tejas Shah, Ankit Hinsu, Pritesh Sabara, Apurvasinh Puvar, Janvi Raval, Monika Gandhi, Pinal Trivedi, Maharshi Pandya, Amit Kanani, Akanksha Verma, Neha Rajpara, Chaitanya Joshi, Madhvi Joshi |
| EPI_ISL_444477 | B.J. Medical College and Civil hospital | Gujarat Biotechnology Research Centre | Neeta Khandelwal, Geeta Vaghela, Sonia Barve, Bhavesh Modi, Kairavi Joshi, Gaurishankar Shrimali, Nidhi Sood, Pranay Shah, R D Dixit, Snehal Bagatharia, Kamlesh J Upadhyay, Ramesh Pandit, Tejas Shah, Ankit Hinsu, Pritesh Sabara, Apurvasinh Puvar, Janvi Raval, Monika Gandhi, Pinal Trivedi, Maharshi Pandya, Amit Kanani, Akanksha Verma, Nitin Savaliya, Raghawendra Kumar, Dinesh Kumar, Zuber Saiyed, Dipa Kinariwala, Disha Patel, Binita Aring, Sharmistha Majumdar, Chaitanya Joshi, Madhvi Joshi |
| EPI_ISL_444483 | Gujarat Biotechnology Research Centre | Gujarat Biotechnology Research Centre | Nidhi Sood, Pranay Shah, R D Dixit, Snehal Bagatharia, Kamlesh J Upadhyay, Ramesh Pandit, Tejas Shah, Ankit Hinsu, Pritesh Sabara, Apurvasinh Puvar, |

| | | | |
|---|---|---|---|
| | | | Janvi Raval, Monika Gandhi, Pinal Trivedi, Maharshi Pandya, Amit Kanani, Akanksha Verma, Nitin Savaliya, Raghawendra Kumar, Dinesh Kumar, Zuber Saiyed, Dipa Kinariwala, Disha Patel, Binita Aring, Neeta Khandelwal, Geeta Vaghela, Sonia Barve, Bhavesh Modi, Kairavi Joshi, Gaurishankar Shrimali, Neelam Nathani, Chaitanya Joshi, Madhvi Joshi |
| EPI_ISL_444484 | Gujarat Biotechnology Research Centre | Gujarat Biotechnology Research Centre | Pranay Shah, R D Dixit, Snehal Bagatharia, Kamlesh J Upadhyay, Ramesh Pandit, Tejas Shah, Ankit Hinsu, Pritesh Sabara, Apurvasinh Puvar, Janvi Raval, Monika Gandhi, Pinal Trivedi, Maharshi Pandya, Amit Kanani, Akanksha Verma, Nitin Savaliya, Raghawendra Kumar, Dinesh Kumar, Zuber Saiyed, Dipa Kinariwala, Disha Patel, Binita Aring, Neeta Khandelwal, Geeta Vaghela, Sonia Barve, Bhavesh Modi, Kairavi Joshi, Gaurishankar Shrimali, Nidhi Sood, Armi Chaudhari, Chaitanya Joshi, Madhvi Joshi |
| EPI_ISL_444485 | Gujarat Biotechnology Research Centre | Gujarat Biotechnology Research Centre | R D Dixit, Snehal Bagatharia, Kamlesh J Upadhyay, Ramesh Pandit, Tejas Shah, Ankit Hinsu, Pritesh Sabara, Apurvasinh Puvar, Janvi Raval, Monika Gandhi, Pinal Trivedi, Maharshi Pandya, Amit Kanani, Akanksha Verma, Nitin Savaliya, Raghawendra Kumar, Dinesh Kumar, Zuber Saiyed, Dipa Kinariwala, Disha Patel, Binita Aring, Neeta Khandelwal, Geeta Vaghela, Sonia Barve, Bhavesh Modi, Kairavi Joshi, Gaurishankar Shrimali, Nidhi Sood, Pranay Shah, Bhavya Jindal, Chaitanya Joshi, Madhvi Joshi |
| EPI_ISL_444493 | Departamento de Laboratorios de Salud Publica (DLSP, Division Epidemiologia, Ministerio de Salud Publica) | Facultad de Ciencias (Sección Genética Evolutiva, Sección Virología). | Panzera,Y., Delfraro,A., Ramos,N., Frabasile,S., Calleros,L., Techera,C., Grecco,S., Fuques,E., Goni,N., Coppola,L., Ramos,V., Chiparelli,H., Arbiza,J. and Perez,R. |
| EPI_ISL_447034 | B.J. Medical College and Civil hospital | Gujarat Biotechnology Research Centre | Pritesh Sabara, Apurvasinh Puvar, Janvi Raval, Monika Gandhi, Pinal Trivedi, Maharshi Pandya, Amit Kanani, Akanksha Verma, Nitin Savaliya, Raghawendra Kumar, Dinesh Kumar, Zuber Saiyed, Dipa Kinariwala, Disha Patel, Binita Aring, Neeta Khandelwal, Geeta Vaghela, Sonia Barve, Bhavesh Modi, Kairavi Joshi, Gaurishankar Shrimali, Nidhi Sood, Pranay Shah, R D Dixit, Snehal Bagatharia, Kamlesh J Upadhyay, Ramesh Pandit, Tejas Shah, Ankit Hinsu, Priti Pandita, Chaitanya Joshi, Madhvi Joshi |
| EPI_ISL_447038 | B.J. Medical College and Civil hospital | Gujarat Biotechnology Research Centre | Pinal Trivedi, Maharshi Pandya, Amit Kanani, Akanksha Verma, Nitin Savaliya, Raghawendra Kumar, Dinesh Kumar, Zuber Saiyed, Dipa Kinariwala, Disha Patel, Binita Aring, Neeta Khandelwal, Geeta Vaghela, Sonia Barve, Bhavesh Modi, Kairavi Joshi, Gaurishankar Shrimali, Nidhi Sood, Pranay Shah, R D Dixit, Snehal Bagatharia, Kamlesh J Upadhyay, Ramesh Pandit, Tejas Shah, Ankit Hinsu, Pritesh Sabara, Apurvasinh Puvar, Janvi Raval, Monika Gandhi, Armi Chaudhari, Chaitanya Joshi, Madhvi Joshi |
| EPI_ISL_447042 | B.J. Medical College and Civil hospital | Gujarat Biotechnology Research Centre | Nitin Savaliya, Raghawendra Kumar, Dinesh Kumar, Zuber Saiyed, Dipa Kinariwala, Disha Patel, Binita Aring, Neeta Khandelwal, Geeta Vaghela, Sonia Barve, Bhavesh Modi, Kairavi Joshi, Gaurishankar Shrimali, Nidhi Sood, Pranay Shah, R D Dixit, Snehal Bagatharia, Kamlesh J Upadhyay, Ramesh Pandit, Tejas Shah, Ankit Hinsu, Pritesh Sabara, Apurvasinh Puvar, Janvi Raval, Monika Gandhi, Pinal Trivedi, Maharshi Pandya, Amit Kanani, Akanksha Verma, Pooja P Doshi, Chaitanya Joshi, Madhvi Joshi |
| EPI_ISL_447049 | GMERS Medical College and Hospital, Gandhinagar | Gujarat Biotechnology Research Centre | Neeta Khandelwal, Geeta Vaghela, Sonia Barve, Bhavesh Modi, Kairavi Joshi, Gaurishankar Shrimali, Nidhi Sood, Pranay Shah, R D Dixit, Snehal Bagatharia, Kamlesh J Upadhyay, Ramesh Pandit, Tejas Shah, Ankit Hinsu, Pritesh Sabara, Apurvasinh Puvar, Janvi Raval, Monika Gandhi, Pinal Trivedi, Maharshi Pandya, Amit Kanani, Akanksha Verma, Nitin Savaliya, Raghawendra Kumar, Dinesh Kumar, Zuber Saiyed, Dipa Kinariwala, Disha Patel, Binita Aring, Bhavya Jindal, Chaitanya Joshi, Madhvi Joshi |
| EPI_ISL_447230, EPI_ISL_447231, EPI_ISL_447232, EPI_ISL_447233, EPI_ISL_447234, EPI_ISL_447235, EPI_ISL_447236, EPI_ISL_447237, EPI_ISL_447238, EPI_ISL_447239, EPI_ISL_447240, EPI_ISL_447241, EPI_ISL_447242, EPI_ISL_447243, EPI_ISL_447244, EPI_ISL_447245, EPI_ISL_447246, EPI_ISL_447247, EPI_ISL_447248, EPI_ISL_447249 | | | |
| see above | Viral Respiratory Lab, National Institute for Biomedical Research (INRB) | Pathogen Sequencing Lab, National Institute for Biomedical Research (INRB) | Placide Mbala-Kingebeni, Edith Nkwembe, Eddy Kinganda-Lusamaki, Amuri Aziza, Francisca Muyembe Mawete, Catherine Pratt, Matthias Pauthner, Josh Quick, Allison Black, James Hadfield, Trevor Bedford, Ian Goodfellow, Andrew Rambaut, Nick Loman, Kristian Andersen, Michael Wiley, Steve Ahuka-Mundeke, Jean-Jacques Muyembe Tamfum |
| EPI_ISL_447250, EPI_ISL_447251 | Central Virology Laboratory | Central Virology Laboratory | Neta Zuckerman, Efrat Bucris, Oran Erster, Danit Sofer, Orna Mor, Ella Mendelson, Michal Mandelboim |
| EPI_ISL_447475 | Servicio de Microbiologia. Hospital Clinico Universitario de Valencia | Sequencing and Bioinformatics Service and Molecular Epidemiology Research Group. FISABIO-Public Health | Neris Garcia-Gonzalez, Inma Galán Vendrell, Sandra Carbo, Loreto Ferrús Abad, Paula Ruiz-Hueso, Mariana Reyes-Prieto, Vicente Soriano Chirona, Ivan Ansari, Llúcia Martínez-Priego, Giuseppe 'Auria, David Navarro, Eliseo Albert, Maria Alma Bracho, Lidia Ruiz Roldan, Fernando Gonzalez-Candelas |
| EPI_ISL_447479 | Servicio de Microbiologia. Hospital Clinico Universitario de Valencia | Sequencing and Bioinformatics Service and Molecular Epidemiology Research Group. FISABIO-Public Health | Paula Ruiz-Hueso, Mariana Reyes-Prieto, Vicente Soriano Chirona, Ivan Ansari, Llúcia Martínez-Priego, Giuseppe 'Auria, David Navarro, Eliseo Albert, Maria Alma Bracho, Lidia Ruiz Roldan, Neris Garcia-Gonzalez, Inma Galán Vendrell, Sandra Carbo, Loreto Ferrús Abad, Fernando Gonzalez-Candelas |
| EPI_ISL_447489 | Servicio de Microbiologia. Hospital Clinico Universitario de Valencia | Sequencing and Bioinformatics Service and Molecular Epidemiology Research Group. FISABIO-Public Health | Neris Garcia-Gonzalez, Inma Galán Vendrell, Sandra Carbo, Loreto Ferrús Abad, Paula Ruiz-Hueso, Mariana Reyes-Prieto, Vicente Soriano Chirona, Ivan Ansari, Llúcia Martínez-Priego, Giuseppe 'Auria, David Navarro, Eliseo Albert, Maria Alma Bracho, Lidia Ruiz Roldan, Fernando Gonzalez-Candelas |
| EPI_ISL_447493 | Servicio de Microbiologia. Hospital Clinico Universitario de Valencia | Sequencing and Bioinformatics Service and Molecular Epidemiology Research Group. FISABIO-Public Health | Paula Ruiz-Hueso, Mariana Reyes-Prieto, Vicente Soriano Chirona, Ivan Ansari, Llúcia Martínez-Priego, Giuseppe 'Auria, David Navarro, Eliseo Albert, Maria Alma Bracho, Lidia Ruiz Roldan, Neris Garcia-Gonzalez, Inma Galán Vendrell, Sandra Carbo, Loreto Ferrús Abad, Fernando Gonzalez-Candelas |
| EPI_ISL_447503 | Servicio de Microbiologia. Hospital Clinico Universitario de Valencia | Sequencing and Bioinformatics Service and Molecular Epidemiology Research Group. FISABIO-Public Health | Neris Garcia-Gonzalez, Inma Galán Vendrell, Sandra Carbo, Loreto Ferrús Abad, Paula Ruiz-Hueso, Mariana Reyes-Prieto, Vicente Soriano Chirona, Ivan Ansari, Llúcia Martínez-Priego, Giuseppe 'Auria, David Navarro, Eliseo Albert, Maria Alma Bracho, Lidia Ruiz Roldan, Fernando Gonzalez-Candelas |
| EPI_ISL_447507 | Servicio de Microbiologia. Hospital Clinico Universitario de Valencia | Sequencing and Bioinformatics Service and Molecular Epidemiology Research Group. FISABIO-Public Health | Paula Ruiz-Hueso, Mariana Reyes-Prieto, Vicente Soriano Chirona, Ivan Ansari, Llúcia Martínez-Priego, Giuseppe 'Auria, David Navarro, Eliseo Albert, Maria Alma Bracho, Lidia Ruiz Roldan, Neris Garcia-Gonzalez, Inma Galán Vendrell, Sandra Carbo, Loreto Ferrús Abad, Fernando Gonzalez-Candelas |
| EPI_ISL_447517 | Servicio de Microbiologia. Hospital Clinico Universitario de Valencia | Sequencing and Bioinformatics Service and Molecular Epidemiology Research Group. FISABIO-Public Health | Neris Garcia-Gonzalez, Inma Galán Vendrell, Sandra Carbo, Loreto Ferrús Abad, Paula Ruiz-Hueso, Mariana Reyes-Prieto, Vicente Soriano Chirona, Ivan Ansari, Llúcia Martínez-Priego, Giuseppe 'Auria, David Navarro, Eliseo Albert, Maria Alma Bracho, Lidia Ruiz Roldan, Fernando Gonzalez-Candelas |
| EPI_ISL_447521 | Servicio de Microbiologia. Hospital Clinico Universitario de Valencia | Sequencing and Bioinformatics Service and Molecular Epidemiology Research Group. FISABIO-Public Health | Paula Ruiz-Hueso, Mariana Reyes-Prieto, Vicente Soriano Chirona, Ivan Ansari, Llúcia Martínez-Priego, Giuseppe 'Auria, David Navarro, Eliseo Albert, Maria Alma Bracho, Lidia Ruiz Roldan, Neris Garcia-Gonzalez, Inma Galán Vendrell, Sandra Carbo, Loreto Ferrús Abad, Fernando Gonzalez-Candelas |
| EPI_ISL_447530 | Servicio de Microbiologia. Hospital Clinico Universitario de Valencia | Sequencing and Bioinformatics Service and Molecular Epidemiology Research Group. FISABIO-Public Health | Neris Garcia-Gonzalez, Inma Galán Vendrell, Sandra Carbo, Loreto Ferrús Abad, Paula Ruiz-Hueso, Mariana Reyes-Prieto, Vicente Soriano Chirona, Ivan Ansari, Llúcia Martínez-Priego, Giuseppe 'Auria, David Navarro, Eliseo Albert, Maria Alma Bracho, Lidia Ruiz Roldan, Fernando Gonzalez-Candelas |
| EPI_ISL_447535 | Gujarat Biotechnology Research Centre | Gujarat Biotechnology Research Centre | Nidhi Sood, Pranay Shah, R D Dixit, Snehal Bagatharia, Kamlesh J Upadhyay, Ramesh Pandit, Tejas Shah, Ankit Hinsu, Pritesh Sabara, Apurvasinh Puvar, Janvi Raval, Monika Gandhi, Pinal Trivedi, Maharshi Pandya, Amit Kanani, Akanksha Verma, Nitin Savaliya, Raghawendra Kumar, Dinesh Kumar, Zuber Saiyed, Dipa Kinariwala, Disha Patel, Binita Aring, Neeta Khandelwal, Geeta Vaghela, Sonia Barve, Bhavesh Modi, Kairavi Joshi, Gaurishankar Shrimali, Priti Pandita, Chaitanya Joshi, Madhvi Joshi |
| EPI_ISL_447536 | Gujarat Biotechnology Research Centre | Gujarat Biotechnology Research Centre | Pranay Shah, R D Dixit, Snehal Bagatharia, Kamlesh J Upadhyay, Ramesh Pandit, Tejas Shah, Ankit Hinsu, Pritesh Sabara, Apurvasinh Puvar, Janvi Raval, Monika Gandhi, Pinal Trivedi, Maharshi Pandya, Amit Kanani, Akanksha Verma, Nitin Savaliya, Raghawendra Kumar, Dinesh Kumar, Zuber Saiyed, Dipa Kinariwala, Disha Patel, Binita Aring, Neeta Khandelwal, Geeta Vaghela, Sonia Barve, Bhavesh Modi, Kairavi Joshi, Gaurishankar Shrimali, Nidhi Sood, Neha Rajpara, Chaitanya Joshi, Madhvi Joshi |
| EPI_ISL_447537 | Gujarat Biotechnology Research Centre | Gujarat Biotechnology Research Centre | R D Dixit, Snehal Bagatharia, Kamlesh J Upadhyay, Ramesh Pandit, Tejas Shah, Ankit Hinsu, Pritesh Sabara, Apurvasinh Puvar, Janvi Raval, Monika Gandhi, Pinal Trivedi, Maharshi Pandya, Amit Kanani, Akanksha Verma, Nitin Savaliya, Raghawendra Kumar, Dinesh Kumar, Zuber Saiyed, Dipa Kinariwala, Disha Patel, Binita Aring, Neeta Khandelwal, Geeta Vaghela, Sonia Barve, Bhavesh Modi, Kairavi Joshi, Gaurishankar Shrimali, Nidhi Sood, Pranay Shah, Afzal Ansari, Chaitanya Joshi, Madhvi Joshi |
| EPI_ISL_447543 | Gujarat Biotechnology Research Centre | Gujarat Biotechnology Research Centre | Pritesh Sabara, Apurvasinh Puvar, Janvi Raval, Monika Gandhi, Pinal Trivedi, Maharshi Pandya, Amit Kanani, Akanksha Verma, Nitin Savaliya, Raghawendra Kumar, Dinesh Kumar, Zuber Saiyed, Dipa Kinariwala, Disha Patel, Binita Aring, Neeta Khandelwal, Geeta Vaghela, Sonia Barve, Bhavesh Modi, Kairavi Joshi, Gaurishankar Shrimali, Nidhi Sood, Pranay Shah, R D Dixit, Snehal Bagatharia, Kamlesh J Upadhyay, Ramesh Pandit, Tejas Shah, Ankit Hinsu, Sharmistha Majumdar, Chaitanya Joshi, Madhvi Joshi |
| EPI_ISL_447547 | GMERS Medical College and Hospital, Gandhinagar | Gujarat Biotechnology Research Centre | Pinal Trivedi, Maharshi Pandya, Amit Kanani, Akanksha Verma, Nitin Savaliya, Raghawendra Kumar, Dinesh Kumar, Zuber Saiyed, Dipa Kinariwala, Disha Patel, Binita Aring, Neeta Khandelwal, Geeta Vaghela, Sonia Barve, Bhavesh Modi, Kairavi Joshi, Gaurishankar Shrimali, Nidhi Sood, Pranay Shah, R D Dixit, Snehal Bagatharia, Kamlesh J Upadhyay, Ramesh Pandit, Tejas Shah, Ankit Hinsu, Pritesh Sabara, Apurvasinh Puvar, Janvi Raval, Monika Gandhi, Neha Rajpara, Chaitanya Joshi, Madhvi Joshi |
| EPI_ISL_447551 | GMERS Medical College and Hospital, Gandhinagar | Gujarat Biotechnology Research Centre | Nitin Savaliya, Raghawendra Kumar, Dinesh Kumar, Zuber Saiyed, Dipa Kinariwala, Disha Patel, Binita Aring, Neeta Khandelwal, Geeta Vaghela, Sonia Barve, Bhavesh Modi, Kairavi Joshi, Gaurishankar Shrimali, Nidhi Sood, Pranay Shah, R D Dixit, Snehal Bagatharia, Kamlesh J Upadhyay, Ramesh Pandit, |

| | | | |
|---|---|---|---|
| | | | Tejas Shah, Ankit Hinsu, Pritesh Sabara, Apurvasinh Puvar, Janvi Raval, Monika Gandhi, Pinal Trivedi, Maharshi Pandya, Amit Kanani, Akanksha Verma, Bhavya Jindal, Chaitanya Joshi, Madhvi Joshi |
| EPI_ISL_447557, EPI_ISL_447558, EPI_ISL_447560, EPI_ISL_447567 | CSIR-Centre for Cellular and Molecular Biology | CSIR-Centre for Cellular and Molecular Biology | Payel Mukherjee, Sofia Banu, Priya Singh, Dhiviya Vedagiri, Divya Gupta, Vishal Sah, Santosh Kumar Kuncha, Krishnan Harinivas Harshan, Archana Bharadwaj Siva, Karthik Bharadwaj Tallapaka, Shagufta Khan, Lamuk Zaveri, Namami Gaur, Sakshi Shambhavi, Tulasi Nagabandi, Purushotham Vodnala, Rakesh K Mishra, Divya Tej Sowpati |
| EPI_ISL_447596, EPI_ISL_447597, EPI_ISL_447598, EPI_ISL_447599, EPI_ISL_447600, EPI_ISL_447601, EPI_ISL_447602, EPI_ISL_447603, EPI_ISL_447604, EPI_ISL_447605, EPI_ISL_447606, EPI_ISL_447607 | | | |
| see above | Viral Respiratory Lab, National Institute for Biomedical Research (INRB) | Pathogen Sequencing Lab, National Institute for Biomedical Research (INRB) | Placide Mbala-Kingebeni, Edith Nkwembe, Eddy Kinganda-Lusamaki, Amuri Aziza, Francisca Muyembe Mawete, Catherine Pratt, Matthias Pauthner, Josh Quick, Allison Black, James Hadfield, Trevor Bedford, Ian Goodfellow, Andrew Rambaut, Nick Loman, Kristian Andersen, Michael Wiley, Steve Ahuka-Mundeke, Jean-Jacques Muyembe Tamfum |
| EPI_ISL_447847, EPI_ISL_447854, EPI_ISL_447859, EPI_ISL_447862, EPI_ISL_447863, EPI_ISL_447864 | CSIR-Centre for Cellular and Molecular Biology | CSIR-Centre for Cellular and Molecular Biology | Payel Mukherjee, Sofia Banu, Priya Singh, Dhiviya Vedagiri, Divya Gupta, Vishal Sah, Santosh Kumar Kuncha, Krishnan Harinivas Harshan, Archana Bharadwaj Siva, Karthik Bharadwaj Tallapaka, Shagufta Khan, Lamuk Zaveri, Namami Gaur, Sakshi Shambhavi, Tulasi Nagabandi, Purushotham Vodnala, Rakesh K Mishra, Divya Tej Sowpati |
| EPI_ISL_447909, EPI_ISL_447910, EPI_ISL_447911, EPI_ISL_447912, EPI_ISL_447913, EPI_ISL_447914, EPI_ISL_447915, EPI_ISL_447916, EPI_ISL_447917, EPI_ISL_447918, EPI_ISL_447919, EPI_ISL_447920, EPI_ISL_447921 | | | |
| see above | n/a | National Institute of Health. Department of medical Sciences, Ministry of Public Health, Thailand | Pilailuk,Okada; Siripaporn,Phuygun; Thanutsapa,Thanadachakul; Sittiporn,Parnmen;Warawan,Wongboot; Sunthareeya,Waicharoen; Malinee,Chittaganpitch |
| EPI_ISL_450196 | bumrungrad international hospital | National Institute of Health. Department of medical Sciences, Ministry of Public Health, Thailand | Pilailuk,Okada; Siripaporn,Phuygun; Thanutsapa,Thanadachakul; Sittiporn,Parnmen;Warawan,Wongboot; Sunthareeya,Waicharoen; Malinee,Chittaganpitch |
| EPI_ISL_450197 | National Institute of Health. Department of medical Sciences, Ministry of Public Health, Thailand | National Institute of Health. Department of medical Sciences, Ministry of Public Health, Thailand | Pilailuk,Okada; Siripaporn,Phuygun; Thanutsapa,Thanadachakul; Sittiporn,Parnmen;Warawan,Wongboot; Sunthareeya,Waicharoen; Malinee,Chittaganpitch |
| EPI_ISL_450326, EPI_ISL_450332 | CSIR-Centre for Cellular and Molecular Biology | CSIR-Centre for Cellular and Molecular Biology | Payel Mukherjee, Sofia Banu, Priya Singh, Dhiviya Vedagiri, Divya Gupta, Vishal Sah, Santosh Kumar Kuncha, Krishnan Harinivas Harshan, Archana Bharadwaj Siva, Karthik Bharadwaj Tallapaka, Shagufta Khan, Lamuk Zaveri, Namami Gaur, Sakshi Shambhavi, Tulasi Nagabandi, Purushotham Vodnala,G. Aditya Kumar, Koushick Sivakumar, Pooja Ramesh Gupta, Rajan Kumar Jha, Shraddha Vijay Lahoti, Deepak Kumar, Devi Prasad Vijayashankara, Disha Nanda, Divya Das, Jotin Gogoi, Manish Bhattacharjee, Rakesh K Mishra, Divya Tej Sowpati |
| EPI_ISL_450724, EPI_ISL_450725, EPI_ISL_450726, EPI_ISL_450727, EPI_ISL_450728, EPI_ISL_450729, EPI_ISL_450730, EPI_ISL_450731, EPI_ISL_450732, EPI_ISL_450734, EPI_ISL_450735, EPI_ISL_450736, EPI_ISL_450737 | | | |
| see above | Hospital AZ Rivierenland | Institute of Tropical Medicine | Philippe Selhorst, Colin Anthony |
| EPI_ISL_450738, EPI_ISL_450739, EPI_ISL_450740, EPI_ISL_450741, EPI_ISL_450742, EPI_ISL_450743, EPI_ISL_450744, EPI_ISL_450745 | OUCRU/HTD | OUCRU/HTD | Nguyen Van Vinh Chau, Nguyen Thi Thu Hong, Nguyen Thi Han Ny, Le Nguyen Truc Nhu, Nghiem My Ngoc, Vo Thanh Lam, Nguyen Thanh Dung, Lam Minh Yen, Ngo Ngoc Quang Minh, Le Manh Hung, Nguyen Tri Dung, Dinh Nguyen Huy Man, Lam Anh Nguyet, Tran Chanh Xuan, Tran Tinh Hien, Nguyen Thanh Phong, Tran Nguyen Hoang Tu, Tran Tan Thanh, Nguyen Thanh Truong, Nguyen Tan Binh, Tang Chi Thuong, Guy Thwaites, and Le Van Tan, for OUCRU COVID-19 research group* |
| EPI_ISL_450785 | Pandit Deendayal Upadhyay Government Medical College, Rajkot | Gujarat Biotechnology Research Centre | Prakash Modi, Sejul Antala, Manish Pattani, Apurvasinh Puvar, Janvi Raval, Zarna Patel, Monika Gandhi, Pinal Trivedi, Maharshi Pandya, Amit Kanani, Nidhi Patel, Nitin Savaliya, Raghawendra Kumar, Dinesh Kumar, Zuber Saiyed, Komal Patel, Labdhi Pandya, Snehal Bagatharia, Ramesh Pandit, Tejas Shah, Ankit Hinsu, Pritesh Sabara, Bhavesh Modi, Gaurishankar Shrimali, R D Dixit, A M Kadri, Neelam Nathani, Chaitanya Joshi, Madhvi Joshi |
| EPI_ISL_450813 | Bla Kustens halsocentral | The Public Health Agency of Sweden | Olof Norrby, Anna-Malin Linde, Maria Lind Karlberg, Oskar Karlsson Lindsjo, Olov Svartstrom, Anna Risberg, Theresa Enkirch, Mia Brytting, Karin Tegmark-Wisell |
| EPI_ISL_450832 | Byjorden vardcentral | The Public Health Agency of Sweden | Pernilla Brunman, Anna-Malin Linde, Maria Lind Karlberg, Oskar Karlsson Lindsjo, Olov Svartstrom, Anna Risberg, Theresa Enkirch, Mia Brytting, Karin Tegmark-Wisell |
| EPI_ISL_451152 | M.P Shah Government Medocal college Jamnagar | Gujarat Biotechnology Research Centre | Pinal Trivedi, Maharshi Pandya, Amit Kanani, Nidhi Patel, Nitin Savaliya, Raghawendra Kumar, Dinesh Kumar, Zuber Saiyed, Komal Patel, Labdhi Pandya, Snehal Bagatharia, Ramesh Pandit, Tejas Shah, Ankit Hinsu, Pritesh Sabara, Apurvasinh Puvar, Binita Aring, Janvi Raval, Zarna Patel, Monika Gandhi, Bhavesh Modi, Gaurishankar Shrimali, R D Dixit, A M Kadri, Afzal Ansari, Chaitanya Joshi, Madhvi Joshi, |
| EPI_ISL_451157 | Government Medical College, Vadodara | Gujarat Biotechnology Research Centre | Nidhi Patel, Nitin Savaliya, Raghawendra Kumar, Dinesh Kumar, Zuber Saiyed, Komal Patel, Labdhi Pandya, Snehal Bagatharia, Ramesh Pandit, Tejas Shah, Ankit Hinsu, Pritesh Sabara, Apurvasinh Puvar, Janvi Raval, Zarna Patel, Monika Gandhi, Pinal Trivedi, Maharshi Pandya, Manish Pattani, Tanuja Javadekar , Amit Kanani, Bhavesh Modi, Gaurishankar Shrimali, R D Dixit, A M Kadri, Camellia Chakraborty, Chaitanya Joshi, Madhvi Joshi |
| EPI_ISL_451158 | Government Medical College, Vadodara | Gujarat Biotechnology Research Centre | Nitin Savaliya, Raghawendra Kumar, Dinesh Kumar, Zuber Saiyed, Komal Patel, Labdhi Pandya, Snehal Bagatharia, Ramesh Pandit, Tejas Shah, Ankit Hinsu, Pritesh Sabara, Apurvasinh Puvar, Janvi Raval, Zarna Patel, Monika Gandhi, Pinal Trivedi, Maharshi Pandya, Manish Pattani, Tanuja Javadekar , Amit Kanani, Nidhi Patel, Bhavesh Modi, Gaurishankar Shrimali, R D Dixit, A M Kadri, Siddhant Kumar, Chaitanya Joshi, Madhvi Joshi |
| EPI_ISL_451174, EPI_ISL_451175, EPI_ISL_451176, EPI_ISL_451177, EPI_ISL_451178, EPI_ISL_451179, EPI_ISL_451180, EPI_ISL_451181, EPI_ISL_451182 | Lab voor klinische biologie | Onderzoeksgroep Virologie | Nick Vereecke, Laurens Lambrechts, Marthe Pauwels, Jozefien De Clercq, Bruno Verhasselt, Linos Vandekerckhove, Hans Nauwynck, Sebastiaan Theuns |
| EPI_ISL_451935 | CUB Hopital Erasme Laboratoire d'Anatomie Pathologique | CUB Hopital Erasme Laboratoire d'Anatomie Pathologique | Prof. Isabelle Salmon, Dr. Nicky D'Haene |
| EPI_ISL_451960 | VI-US Virgin Islands Department of Health | Centers for Disease Control and Prevention | Queen,K., Li,Y., Montmayeur,A., Tao,Y., Zhang,J., Uehara,A., Paden,C.R., Marine,R., Wang,H., Padilla,J., Lee,J., Weiner,Z., Bankamp,B. and Tong,S. |
| EPI_ISL_451961 | Istituto Zooprofilattico Sperimentale Puglia e Basilicata; Dipartimento di Bioscienze, Biotecnologie e Biofarmaceutica dell'Università degli Studi di Bari "A.Moro"; Istituto di Biomembrane. Bioenergetica e Biotecnologie Molecolari del Consiglio Nazionale delle Ricerche di Bari | Beaconlab (Bioinformatics Evolution and Comparative Genomics lab), Dept of Biosciences, University of Milan | Parisi A.,Pesole G., Manzari C., Chiara M. |
| EPI_ISL_451962 | Istituto Zooprofilattico Sperimentale Puglia e Basilicata; Dipartimento di Bioscienze, Biotecnologie e Biofarmaceutica dell'Università degli Studi di Bari "A.Moro"; Istituto di Biomembrane. Bioenergetica e Biotecnologie Molecolari del Consiglio Nazionale delle Ricerche di Bari | Beaconlab (Bioinformatics, Evolution and Comparative Genomics lab), Dept of Biosciences, University on Milan | Parisi A.,Pesole G., Manzari C., Chiara M. |
| EPI_ISL_452150 | CUB Hopital Erasme Laboratoire d'Anatomie Pathologique | CUB Hopital Erasme Laboratoire d'Anatomie Pathologique | Prof. Isabelle Salmon, Dr Nicky D'Haene |
| EPI_ISL_452151 | CUB Hopital Erasme Laboratoire d'Anatomie Pathologique | CUB Hopital Erasme Laboratoire d'Anatomie Pathologique | Prof. Isabelle Salmon, Dr.Nikcy D'Haene |
| EPI_ISL_452152 | CUB Hopital Erasme Laboratoire d'Anatomie Pathologique | CUB Hopital Erasme Laboratoire d'Anatomie Pathologique | Prof. Isabelle Salmon, Dr Nicky D'Haene |
| EPI_ISL_452180 | Center of Medical Microbiology, Virology, and Hospital Hygiene, University of Duesseldorf | Center of Medical Microbiology, Virology, and Hospital Hygiene, University of Duesseldorf | Ortwin Adams, Marcel Andree, Alexander Dilthey, Torsten Feldt, Sandra Hauka, Torsten Houwaart, Björn-ErikJensen, Detlef Kindgen-Milles, Malte Kohns Vasconcelos, Klaus Pfeffer, Tina Senff, Daniel Strelow, Jörg Timm, Andreas Walker, Tobias Wienemann |
| EPI_ISL_452192, EPI_ISL_452193, EPI_ISL_452194, EPI_ISL_452195, EPI_ISL_452196, EPI_ISL_452197, EPI_ISL_452198, EPI_ISL_452199, EPI_ISL_452200, EPI_ISL_452201, EPI_ISL_452202, EPI_ISL_452203, EPI_ISL_452204, EPI_ISL_452205, EPI_ISL_452206, EPI_ISL_452207, EPI_ISL_452208, EPI_ISL_452209, EPI_ISL_452210, EPI_ISL_452211, EPI_ISL_452212, EPI_ISL_452213, EPI_ISL_452214, EPI_ISL_452215, EPI_ISL_452216, EPI_ISL_452217, EPI_ISL_454521, EPI_ISL_454522, EPI_ISL_454523, EPI_ISL_454524, EPI_ISL_454525, EPI_ISL_454526, EPI_ISL_454527, EPI_ISL_454528, EPI_ISL_454529, EPI_ISL_454530, EPI_ISL_454531, EPI_ISL_454532, EPI_ISL_454533, EPI_ISL_454534, EPI_ISL_454536, EPI_ISL_454537, EPI_ISL_454538, EPI_ISL_454539, EPI_ISL_454540, EPI_ISL_454541, EPI_ISL_454542, EPI_ISL_454543, EPI_ISL_454544, EPI_ISL_454546, EPI_ISL_454547, EPI_ISL_454548, EPI_ISL_454549, EPI_ISL_454550, EPI_ISL_454551, EPI_ISL_454552, EPI_ISL_454553, EPI_ISL_454554, EPI_ISL_454555, EPI_ISL_454556, EPI_ISL_454557, EPI_ISL_454558, EPI_ISL_454559, EPI_ISL_454560, EPI_ISL_454561, EPI_ISL_454562, EPI_ISL_454563, EPI_ISL_454564, EPI_ISL_454565, EPI_ISL_454566, EPI_ISL_454567, EPI_ISL_454568, EPI_ISL_454569, EPI_ISL_454570 | | | |
| see above | NIV Influenza | NIV Influenza | Potdar V |

| | | | |
|---|---|---|---|
| EPI_ISL_455016 | Pandit Deendayal Upadhyay Government Medical College, Rajkot | Gujarat Biotechnology Research Centre | Prakash Modi, Sejul Antala, Manish Pattani, Ramesh Pandit, Tejas Shah, Ankit Hinsu, Pritesh Sabara, Apurvasinh Puvar, Janvi Raval, Zarna Patel, Monika Gandhi, Pinal Trivedi, Maharshi Pandya, Amit Kanani, Nidhi Patel, Nitin Savaliya, Raghawendra Kumar, Dinesh Kumar, Zuber Saiyed, Komal Patel, Labdhi Pandya, Snehal Bagatharia, Afzal Ansari, Bhavesh Modi, Gaurishankar Shrimali, R D Dixit, A M Kadri, Umang Mishra, Chaitanya Joshi, Madhvi Joshi |
| EPI_ISL_455018 | Government Medical College, Vadodara | Gujarat Biotechnology Research Centre | R N Daveshwar, Ramesh Pandit, Tejas Shah, Ankit Hinsu, Pritesh Sabara, Apurvasinh Puvar, Janvi Raval, Zarna Patel, Monika Gandhi, Pinal Trivedi, Maharshi Pandya, Amit Kanani, Nidhi Patel, Nitin Savaliya, Raghawendra Kumar, Dinesh Kumar, Zuber Saiyed, Komal Patel, Labdhi Pandya, Snehal Bagatharia, Tanuja Javadekar , Neelam Nathani, Bhavesh Modi, Gaurishankar Shrimali, R D Dixit, A M Kadri, Umang Mishra, Chaitanya Joshi, Madhvi Joshi, |
| EPI_ISL_455022 | Government Medical College, Vadodara | Gujarat Biotechnology Research Centre | Pritesh Sabara, Apurvasinh Puvar, Janvi Raval, Zarna Patel, Monika Gandhi, Pinal Trivedi, Maharshi Pandya, Amit Kanani, Nidhi Patel, Nitin Savaliya, Raghawendra Kumar, Dinesh Kumar, Zuber Saiyed, Komal Patel, Labdhi Pandya, Snehal Bagatharia, Tanuja Javadekar , R N Daveshwar, Ramesh Pandit, Tejas Shah, Ankit Hinsu, Siddhant Kumar, Bhavesh Modi, Gaurishankar Shrimali, R D Dixit, A M Kadri, Umang Mishra, Chaitanya Joshi, Madhvi Joshi, |
| EPI_ISL_455027 | Government Medical College, Vadodara | Gujarat Biotechnology Research Centre | Pinal Trivedi, Maharshi Pandya, Amit Kanani, Nidhi Patel, Nitin Savaliya, Raghawendra Kumar, Dinesh Kumar, Zuber Saiyed, Komal Patel, Labdhi Pandya, Snehal Bagatharia, Tanuja Javadekar , R N Daveshwar, Ramesh Pandit, Tejas Shah, Ankit Hinsu, Pritesh Sabara, Apurvasinh Puvar, Janvi Raval, Zarna Patel, Monika Gandhi, Pragya Sharma, Bhavesh Modi, Gaurishankar Shrimali, R D Dixit, A M Kadri, Umang Mishra, Chaitanya Joshi, Madhvi Joshi, |
| EPI_ISL_455312 | Microbiology Unit, Department of Pathology & Laboratory Medicine, IIUM Medical Centre | SEA Microbiome Unit, Faculty of Industrial Sciences & Technology, Universiti Malaysia Pahang | Norhidayah Binti Kamarudin, Ahmad Hafiz Bin Zulkifly, Hajar Fauzan Ahmad, Muhammad Adam Lee Abdullah, Mohd Fazli Farida Asras, Ahmad Mahfuz Gazali, Mohd Nazli Bin Kamarulzaman, IIUM Medical Centre Covid19 Taskforce, UMP Covid19 Team |
| EPI_ISL_455362 | Nigeria Centre for Disease Control (NCDC) | African Centre of Excellence for Genomics of Infectious Diseases (ACEGID), Redeemer's University, Ede, Osun State, Nigeria | Oluniyi P.E., Ajogbasile F.V., Kayode A., Olawoye I., Uwanibe J., Oguzie J., Olumade T., Folarin O.A., Ihekweazu C., Happi C.T. |
| EPI_ISL_455412, EPI_ISL_455413 | Nigeria Centre for Disease Control (NCDC) | African Centre of Excellence for Genomics of Infectious Diseases (ACEGID), Redeemer's University, Ede, Osun State, Nigeria | Oluniyi P.E., Ajogbasile F.V., Kayode A., Oguzie J., Olawoye I., Uwanibe J., Olumade T., Folarin O.A., Ihekweazu C., Happi C.T. |
| EPI_ISL_455415 | Nigeria Centre for Disease Control (NCDC) | African Centre of Excellence for Genomics of Infectious Diseases (ACEGID), Redeemer's University, Ede, Osun State, Nigeria | Oluniyi P.E., Ajogbasile F.V., Kayode A., Oguzie J., Olawoye I., Uwanibe J., Olumade T., Folarin O.A., Ihekweazu C., Happi C.T. |
| EPI_ISL_455418, EPI_ISL_455419 | Nigeria Centre for Disease Control (NCDC) | African Centre of Excellence for Genomics of Infectious Diseases (ACEGID), Redeemer's University, Ede, Osun State, Nigeria | Oluniyi P.E., Ajogbasile F.V., Kayode A., Oguzie J., Olawoye I., Uwanibe J., Olumade T., Folarin O.A., Ihekweazu C., Happi C.T. |
| EPI_ISL_455422 | Nigeria Centre for Disease Control | African Centre of Excellence for Genomics of Infectious Diseases (ACEGID), Redeemer's University, Ede, Osun State, Nigeria | Oluniyi P.E., Ajogbasile F.V., Kayode A., Oguzie J., Olawoye I., Uwanibe J., Olumade T., Folarin O.A., Ihekweazu C., Happi C.T. |
| EPI_ISL_455423, EPI_ISL_455424 | Nigeria Centre for Disease Control (NCDC) | African Centre of Excellence for Genomics of Infectious Diseases (ACEGID), Redeemer's University, Ede, Osun State, Nigeria | Oluniyi P.E., Ajogbasile F.V., Kayode A., Oguzie J., Olawoye I., Uwanibe J., Olumade T., Folarin O.A., Ihekweazu C., Happi C.T. |
| EPI_ISL_455426 | Nigeria Centre for Disease Control | African Centre of Excellence for Genomics of Infectious Diseases (ACEGID), Redeemer's University, Ede, Osun State, Nigeria | Oluniyi P.E., Ajogbasile F.V., Kayode A., Oguzie J., Olawoye I., Uwanibe J., Olumade T., Folarin O.A., Ihekweazu C., Happi C.T. |
| EPI_ISL_455427, EPI_ISL_455429, EPI_ISL_455431 | Nigeria Centre for Disease Control (NCDC) | African Centre of Excellence for Genomics of Infectious Diseases (ACEGID), Redeemer's University, Ede, Osun State, Nigeria | Oluniyi P.E., Ajogbasile F.V., Kayode A., Oguzie J., Olawoye I., Uwanibe J., Olumade T., Folarin O.A., Ihekweazu C., Happi C.T. |
| EPI_ISL_455583 | Central Chest Institute of Thailand | National Institute of Health. Department of medical Sciences, Ministry of Public Health, Thailand | Pilailuk,Okada; Siripaporn,Phuygun; Thanutsapa,Thanadachakul; Sittiporn,Parnmen;Warawan,Wongboot; Sunthareeya,Waicharoen; Malinee,Chittaganpitch |
| EPI_ISL_455584 | National Institute of Health. Department of medical Sciences, Ministry of Public Health, Thailand | National Institute of Health. Department of medical Sciences, Ministry of Public Health, Thailand | Pilailuk,Okada; Siripaporn,Phuygun; Thanutsapa,Thanadachakul; Sittiporn,Parnmen;Warawan,Wongboot; Sunthareeya,Waicharoen; Malinee,Chittaganpitch |
| EPI_ISL_455585 | Phramongkutklao Hospital | National Institute of Health. Department of medical Sciences, Ministry of Public Health, Thailand | Pilailuk,Okada; Siripaporn,Phuygun; Thanutsapa,Thanadachakul; Sittiporn,Parnmen;Warawan,Wongboot; Sunthareeya,Waicharoen; Malinee,Chittaganpitch |
| EPI_ISL_455586 | Siriraj hospital | National Institute of Health. Department of medical Sciences, Ministry of Public Health, Thailand | Pilailuk,Okada; Siripaporn,Phuygun; Thanutsapa,Thanadachakul; Sittiporn,Parnmen;Warawan,Wongboot; Sunthareeya,Waicharoen; Malinee,Chittaganpitch |
| EPI_ISL_455587 | H.R.H. Maha Chakri Sirindhorn Medical Center | National Institute of Health. Department of medical Sciences, Ministry of Public Health, Thailand | Pilailuk,Okada; Siripaporn,Phuygun; Thanutsapa,Thanadachakul; Sittiporn,Parnmen;Warawan,Wongboot; Sunthareeya,Waicharoen; Malinee,Chittaganpitch |
| EPI_ISL_455588 | Trang Hospital | National Institute of Health. Department of medical Sciences, Ministry of Public Health, Thailand | Pilailuk,Okada; Siripaporn,Phuygun; Thanutsapa,Thanadachakul; Sittiporn,Parnmen;Warawan,Wongboot; Sunthareeya,Waicharoen; Malinee,Chittaganpitch |
| EPI_ISL_455589, EPI_ISL_455590, EPI_ISL_455591, EPI_ISL_455592, EPI_ISL_455593 | National Institute of Health. Department of medical Sciences, Ministry of Public Health, Thailand | National Institute of Health. Department of medical Sciences, Ministry of Public Health, Thailand | Pilailuk,Okada; Siripaporn,Phuygun; Thanutsapa,Thanadachakul; Sittiporn,Parnmen;Warawan,Wongboot; Sunthareeya,Waicharoen; Malinee,Chittaganpitch |
| EPI_ISL_455594 | Central Chest Institute of Thailand | National Institute of Health. Department of medical Sciences, Ministry of Public Health, Thailand | Pilailuk,Okada; Siripaporn,Phuygun; Thanutsapa,Thanadachakul; Sittiporn,Parnmen;Warawan,Wongboot; Sunthareeya,Waicharoen; Malinee,Chittaganpitch |
| EPI_ISL_455604 | Ramkhamhaeng Hospital | National Institute of Health. Department of medical Sciences, Ministry of Public Health, Thailand | Pilailuk,Okada; Siripaporn,Phuygun; Thanutsapa,Thanadachakul; Sittiporn,Parnmen;Warawan,Wongboot; Sunthareeya,Waicharoen; Malinee,Chittaganpitch |
| EPI_ISL_455605 | Panyananthaphikkhu Chonprathan Medical Center | National Institute of Health. Department of medical Sciences, Ministry of Public Health, Thailand | Pilailuk,Okada; Siripaporn,Phuygun; Thanutsapa,Thanadachakul; Sittiporn,Parnmen;Warawan,Wongboot; Sunthareeya,Waicharoen; Malinee,Chittaganpitch |
| EPI_ISL_455606, EPI_ISL_455607 | Praram 9 Hospital | National Institute of Health. Department of medical Sciences, Ministry of Public Health, Thailand | Pilailuk,Okada; Siripaporn,Phuygun; Thanutsapa,Thanadachakul; Sittiporn,Parnmen;Warawan,Wongboot; Sunthareeya,Waicharoen; Malinee,Chittaganpitch |
| EPI_ISL_455608 | Phramongkutklao Hospital | National Institute of Health. Department of medical Sciences, Ministry of Public Health, Thailand | Pilailuk,Okada; Siripaporn,Phuygun; Thanutsapa,Thanadachakul; Sittiporn,Parnmen;Warawan,Wongboot; Sunthareeya,Waicharoen; Malinee,Chittaganpitch |
| EPI_ISL_455624 | National Institute of Health. Department of medical Sciences, Ministry of Public Health, Thailand | National Institute of Health. Department of medical Sciences, Ministry of Public Health, Thailand | Pilailuk,Okada; Siripaporn,Phuygun; Thanutsapa,Thanadachakul; Sittiporn,Parnmen;Warawan,Wongboot; Sunthareeya,Waicharoen; Malinee,Chittaganpitch |
| EPI_ISL_455694, EPI_ISL_455695, EPI_ISL_455696, EPI_ISL_455697, EPI_ISL_455698, EPI_ISL_455699, EPI_ISL_455700, EPI_ISL_455701, EPI_ISL_455702, EPI_ISL_455703, EPI_ISL_455704, EPI_ISL_455705, EPI_ISL_455706, EPI_ISL_455707, EPI_ISL_455708, EPI_ISL_455709, EPI_ISL_455710, EPI_ISL_455711, EPI_ISL_455712 | | | |
| see above | National Hospital of Tropical Diseases | Oxford University Clinical Research Unit, Hanoi, Vietnam | Nguyen Thi Tam, Van Dinh Trang, Nguyen Thu Trang, Nguyen Thi Ngoc Diep, Le Nguyen Minh Hoa, Pham Ngoc Thach, H. Rogier van Doorn, on behalf of the OUCRU COVID-19 research group |
| EPI_ISL_455713, EPI_ISL_455714, EPI_ISL_455715, EPI_ISL_455716, EPI_ISL_455717, EPI_ISL_455718 | National Hospital of Tropical Diseases | Oxford University Clinical Research Unit, Hanoi, Vietnam | Nguyen Thi Tam, Van Dinh Trang, Nguyen Thi Hong Thuong, Vu Thi Ngoc Bich, Nguyen Thu Trang, Nguyen Thi Ngoc Diep, Le Nguyen Minh Hoa, Pham Ngoc Thach, H. Rogier van Doorn, on behalf of the OUCRU COVID-19 research group |
| EPI_ISL_455740 | Servicio de Microbiologia. Hospital Clinico Universitario de | Sequencing and Bioinformatics Service and Molecular | Paula Ruiz-Hueso, Mariana Reyes-Prieto, Vicente Soriano Chirona, Ivan Ansari, Llúcia Martínez-Priego, Giuseppe 'Auria, David Navarro, Eliseo Albert, Maria |

| | | | |
|---|---|---|---|
| | Valencia | Epidemiology Research Group. FISABIO-Public Health, and SeqCOVID-Spain Consortium | Alma Bracho, Lidia Ruiz Roldan, Neris Garcia-Gonzalez, Inma Galán Vendrell, Sandra Carbo, Loreto Ferrús Abad, Fernando Gonzalez-Candelas |
| EPI_ISL_455742 | Servicio de Microbiologia. Hospital Clinico Universitario de Valencia | Sequencing and Bioinformatics Service and Molecular Epidemiology Research Group. FISABIO-Public Health, and SeqCOVID-Spain Consortium | Neris Garcia-Gonzalez, Inma Galán Vendrell, Sandra Carbo, Loreto Ferrús Abad, Paula Ruiz-Hueso, Mariana Reyes-Prieto, Vicente Soriano Chirona, Ivan Ansari, Llúcia Martínez-Priego, Giuseppe 'Auria, David Navarro, Eliseo Albert, Maria Alma Bracho, Lidia Ruiz Roldan, Fernando Gonzalez-Candelas |
| EPI_ISL_456071, EPI_ISL_456072, EPI_ISL_456073, EPI_ISL_456074, EPI_ISL_456075 | Laboratory of Respiratory Viruses and Measles, Oswaldo Cruz Institute, FIOCRUZ | Laboratory of Respiratory Viruses and Measles, Oswaldo Cruz Institute, FIOCRUZ | Paola Resende, Luciana Appolinario, Fernando Motta, Aline Mattos, Milene Miranda, Cristiana Garcia, Braulia Caetano, Maria Ogrzewalska, Jonathan Lopes, Marilda Siqueira |
| EPI_ISL_456076, EPI_ISL_456077 | LACEN RJ - Laboratório Central de Saúde Pública Noel Nutels | Laboratory of Respiratory Viruses and Measles, Oswaldo Cruz Institute, FIOCRUZ | Paola Resende, Luciana Appolinario, Fernando Motta, Aline Mattos, Milene Miranda, Cristiana Garcia, Braulia Caetano, Maria Ogrzewalska, Jonathan Lopes, Marilda Siqueira |
| EPI_ISL_456079, EPI_ISL_456080, EPI_ISL_456081 | Laboratory of Respiratory Viruses and Measles, Oswaldo Cruz Institute, FIOCRUZ | Laboratory of Respiratory Viruses and Measles, Oswaldo Cruz Institute, FIOCRUZ | Paola Resende, Luciana Appolinario, Fernando Motta, Aline Mattos, Milene Miranda, Cristiana Garcia, Braulia Caetano, Maria Ogrzewalska, Jonathan Lopes, Marilda Siqueira |
| EPI_ISL_456082, EPI_ISL_456083 | LACEN RJ - Laboratório Central de Saúde Pública Noel Nutels | Laboratory of Respiratory Viruses and Measles, Oswaldo Cruz Institute, FIOCRUZ | Paola Resende, Luciana Appolinario, Fernando Motta, Aline Mattos, Milene Miranda, Cristiana Garcia, Braulia Caetano, Maria Ogrzewalska, Jonathan Lopes, Marilda Siqueira |
| EPI_ISL_456084, EPI_ISL_456085, EPI_ISL_456086, EPI_ISL_456087 | Laboratory of Respiratory Viruses and Measles, Oswaldo Cruz Institute, FIOCRUZ | Laboratory of Respiratory Viruses and Measles, Oswaldo Cruz Institute, FIOCRUZ | Paola Resende, Luciana Appolinario, Fernando Motta, Aline Mattos, Milene Miranda, Cristiana Garcia, Braulia Caetano, Maria Ogrzewalska, Jonathan Lopes, Marilda Siqueira |
| EPI_ISL_456088 | LACEN RJ - Laboratório Central de Saúde Pública Noel Nutels | Laboratory of Respiratory Viruses and Measles, Oswaldo Cruz Institute, FIOCRUZ | Paola Resende, Luciana Appolinario, Fernando Motta, Aline Mattos, Milene Miranda, Cristiana Garcia, Braulia Caetano, Maria Ogrzewalska, Jonathan Lopes, Marilda Siqueira |
| EPI_ISL_456089, EPI_ISL_456090, EPI_ISL_456091, EPI_ISL_456092, EPI_ISL_456093, EPI_ISL_456094, EPI_ISL_456095, EPI_ISL_456096, EPI_ISL_456097, EPI_ISL_456098, EPI_ISL_456099, EPI_ISL_456100, EPI_ISL_456101, EPI_ISL_456102, EPI_ISL_456103, EPI_ISL_456104, EPI_ISL_456105, EPI_ISL_456106 | | | |
| see above | Laboratory of Respiratory Viruses and Measles, Oswaldo Cruz Institute, FIOCRUZ | Laboratory of Respiratory Viruses and Measles, Oswaldo Cruz Institute, FIOCRUZ | Paola Resende, Luciana Appolinario, Fernando Motta, Aline Mattos, Milene Miranda, Cristiana Garcia, Braulia Caetano, Maria Ogrzewalska, Jonathan Lopes, Marilda Siqueira |
| EPI_ISL_457699, EPI_ISL_457700 | Department of Infectious Diseases, Istituto Superiore di Sanità, Roma , Italy | Army Medical and Veterinary Research Center | Paola Stefanelli, Alessandra Lo Presti, Stefano Fiore, Antonella Marchi, Eleonora Benedetti, Concetta Fabiani Silvia Fillo, Giovanni Faggioni, Riccardo De Sanctis, Antonella Fortunato, Anna Anselmo, Francesco Giordani, Vanessa Vera Fain, Nino D'Amore, Florigio Lista |
| EPI_ISL_457721, EPI_ISL_457724, EPI_ISL_457728, EPI_ISL_457732, EPI_ISL_457736, EPI_ISL_457749 | Department of Infectious Diseases, Istituto Superiore di Sanità, Roma , Italy | Army Medical and Veterinary Research Center | Paola Stefanelli, Alessandra Lo Presti, Stefano Fiore, Antonella Marchi, Eleonora Benedetti, Concetta Fabiani Silvia Fillo, Giovanni Faggioni, Riccardo De Sanctis, Antonella Fortunato, Anna Anselmo, Francesco Giordani, Vanessa Vera Fain, Nino D'Amore, Florigio Lista |
| EPI_ISL_457750 | Centogene AG | Centogene AG | Prof. Dr. Peter Bauer, Dr. Krishna Kumar Kandaswamy |
| EPI_ISL_457751, EPI_ISL_457752, EPI_ISL_457753, EPI_ISL_457754, EPI_ISL_457755, EPI_ISL_457756, EPI_ISL_457757, EPI_ISL_457758, EPI_ISL_457759, EPI_ISL_457760, EPI_ISL_457761, EPI_ISL_457762, EPI_ISL_457763, EPI_ISL_457764, EPI_ISL_457765, EPI_ISL_457766, EPI_ISL_457767, EPI_ISL_457768, EPI_ISL_457769, EPI_ISL_457770, EPI_ISL_457771, EPI_ISL_457772, EPI_ISL_457773, EPI_ISL_457774, EPI_ISL_457775, EPI_ISL_457776, EPI_ISL_457777, EPI_ISL_457778, EPI_ISL_457779, EPI_ISL_457780, EPI_ISL_457781, EPI_ISL_457782, EPI_ISL_457783, EPI_ISL_457784, EPI_ISL_457785, EPI_ISL_457786, EPI_ISL_457787, EPI_ISL_457788, EPI_ISL_457789, EPI_ISL_457790, EPI_ISL_457791, EPI_ISL_457792, EPI_ISL_457793, EPI_ISL_457794, EPI_ISL_457795, EPI_ISL_457796, EPI_ISL_457797, EPI_ISL_457798, EPI_ISL_457799, EPI_ISL_457800, EPI_ISL_457801, EPI_ISL_457802, EPI_ISL_457803, EPI_ISL_457804, EPI_ISL_457805, EPI_ISL_457806, EPI_ISL_457807, EPI_ISL_457808, EPI_ISL_457809, EPI_ISL_457810, EPI_ISL_457811, EPI_ISL_457812, EPI_ISL_457813, EPI_ISL_457814, EPI_ISL_457815, EPI_ISL_457816, EPI_ISL_457817, EPI_ISL_457818, EPI_ISL_457819, EPI_ISL_457820, EPI_ISL_457821, EPI_ISL_457822, EPI_ISL_457823 | | | |
| see above | Johns Hopkins Hospital Department of Pathology | Johns Hopkins Hospital Department of Pathology | Peter M. Thielen, Thomas Mehoke, Shirlee Wohl, Srividya Ramakrishnan, Melanie Kirsche, Amanda Ernlund, Craig Howser, Kristina Zudock, Oluwaseun Falade-Nwulia, Norah Sadowski, Paul Morris, Mark Hopkins, Yunfan Fan, Nidia Trovao, Victoria Gniazdowski, Michael C. Schatz, Stuart C. Ray, Winston Timp, Heba H. Mostafa |
| EPI_ISL_458045 | CSIR-Centre for Cellular and Molecular Biology | CSIR-Centre for Cellular and Molecular Biology | Payel Mukherjee, Sofia Banu, Priya Singh, Dhiviya Vedagiri, Divya Gupta, Vishal Sah, Santosh Kumar Kuncha, Krishnan Harinivas Harshan, Archana Bharadwaj Siva, Karthik Bharadwaj Tallapaka, Shagufta Khan, Lamuk Zaveri, Namami Gaur, Sakshi Shambhavi, Tulasi Nagabandi, Purushotham Vodnala, G. Aditya Kumar, Koushick Sivakumar, Pooja Ramesh Gupta, Rajan Kumar Jha, Shraddha Vijay Lahoti, Rakesh K Mishra, Divya Tej Sowpati |
| EPI_ISL_458051 | CSIR-Centre for Cellular and Molecular Biology | CSIR-Centre for Cellular and Molecular Biology | Payel Mukherjee, Sofia Banu, Priya Singh, Dhiviya Vedagiri, Divya Gupta, Vishal Sah, Santosh Kumar Kuncha, Krishnan Harinivas Harshan, Archana Bharadwaj Siva, Karthik Bharadwaj Tallapaka, Shagufta Khan, Lamuk Zaveri, Namami Gaur, Sakshi Shambhavi, Tulasi Nagabandi, Purushotham Vodnala, Gokulan C G, Gunjan Purohit, Hanuman Tulashiram Kale, Pankaj Kumar, Prachand Issarapu, Rakesh K Mishra, Divya Tej Sowpati |
| EPI_ISL_458057 | CSIR-Centre for Cellular and Molecular Biology | CSIR-Centre for Cellular and Molecular Biology | Payel Mukherjee, Sofia Banu, Priya Singh, Dhiviya Vedagiri, Divya Gupta, Vishal Sah, Santosh Kumar Kuncha, Krishnan Harinivas Harshan, Archana Bharadwaj Siva, Karthik Bharadwaj Tallapaka, Shagufta Khan, Lamuk Zaveri, Namami Gaur, Sakshi Shambhavi, Tulasi Nagabandi, Purushotham Vodnala,Deepak Kumar, Devi Prasad Vijayashankar, Disha Nanda, Divya Das, Jotin Gogoi, Manish Bhattacharjee, Rakesh K Mishra, Divya Tej Sowpati |
| EPI_ISL_458062 | CSIR-Centre for Cellular and Molecular Biology | CSIR-Centre for Cellular and Molecular Biology | Payel Mukherjee, Sofia Banu, Priya Singh, Dhiviya Vedagiri, Divya Gupta, Vishal Sah, Santosh Kumar Kuncha, Krishnan Harinivas Harshan, Archana Bharadwaj Siva, Karthik Bharadwaj Tallapaka, Shagufta Khan, Lamuk Zaveri, Namami Gaur, Sakshi Shambhavi, Tulasi Nagabandi, Purushotham Vodnala, Rakesh K Mishra, Sonu Uday, Sudipta Mondal, Annapoorna P Karthyayani, Debabrata Jana, Debrya Saha, Divya Tej Sowpati |
| EPI_ISL_458084 | Laboratorio Biologia Molecolare Sars Cov2 - UOC Laboratorio Analisi - Servizio Medicina di Laboratorio, Ospedale "San Francesco" - ATS-ASSL Nuoro | Laboratorio specialistico UOC Ematologia - Ospedale "San Francesco" - ATS-ASSL Nuoro | Piras Giovanna, Fancello Tatiana, Asproni Rosanna, Fiamma Maura, Monne Maria Itria, Toja Alessandro, Sanna Filomena, Floris Anna Rita, Sulis Vincenzo, Palmas Angelo Domenico, Casu Gavino, Lo Maglio Iana, Mameli Giuseppe. |
| EPI_ISL_458085 | Laboratorio Biologia Molecolare Sars Cov2 - UOC Laboratorio Analisi - Servizio Medicina di Laboratorio , Ospedale "San Francesco" - ATS- ASSL Nuoro | Laboratorio specialistico UOC Ematologia - Ospedale "San Francesco" - ATS-ASSL Nuoro | Piras Giovanna, Fancello Tatiana, Asproni Rosanna, Fiamma Maura, Monne Maria Itria, Toja Alessandro, Sanna Filomena, Floris Anna Rita, Sulis Vincenzo, Palmas Angelo Domenico, Casu Gavino, Lo Maglio Iana, Mameli Giuseppe. |
| EPI_ISL_458088 | B.J. Medical College and Civil hospital | Gujarat Biotechnology Research Centre | Pranay Shah, Kamlesh J Upadhyay, Ramesh Pandit, Tejas Shah, Ankit Hinsu, Pritesh Sabara, Apurvasinh Puvar, Janvi Raval, Zarna Patel, Monika Gandhi, Pinal Trivedi, Maharshi Pandya, Amit Kanani, Nidhi Patel, Nitin Savaliya, Raghawendra Kumar, Dinesh Kumar, Zuber Saiyed, Komal Patel, Labdhi Pandya, Snehal Bagatharia, Dhaval Vaghela, Ramesh Patel, Fenil Patel, Bhavesh Modi, Gaurishankar Shrimali, R D Dixit, A M Kadri, Umang Mishra, Chaitanya Joshi, Madhvi Joshi |
| EPI_ISL_458089 | B.J. Medical College and Civil hospital | Gujarat Biotechnology Research Centre | Pranay Shah, Kamlesh J Upadhyay, Ramesh Pandit, Tejas Shah, Ankit Hinsu, Pritesh Sabara, Apurvasinh Puvar, Janvi Raval, Zarna Patel, Monika Gandhi, Pinal Trivedi, Maharshi Pandya, Amit Kanani, Nidhi Patel, Nitin Savaliya, Raghawendra Kumar, Dinesh Kumar, Zuber Saiyed, Komal Patel, Labdhi Pandya, Snehal Bagatharia, Dhaval Vaghela, Ramesh Patel, Neelam Nathani, Bhavesh Modi, Gaurishankar Shrimali, R D Dixit, A M Kadri, Umang Mishra, Chaitanya Joshi, Madhvi Joshi |
| EPI_ISL_458093 | B.J. Medical College and Civil hospital | Gujarat Biotechnology Research Centre | Nidhi Patel, Nitin Savaliya, Raghawendra Kumar, Dinesh Kumar, Zuber Saiyed, Komal Patel, Labdhi Pandya, Snehal Bagatharia, Dhaval Vaghela, Ramesh Patel, Pranay Shah, Kamlesh J Upadhyay, Ramesh Pandit, Tejas Shah, Ankit Hinsu, Pritesh Sabara, Apurvasinh Puvar, Janvi Raval, Zarna Patel, Monika Gandhi, Pinal Trivedi, Maharshi Pandya, Amit Kanani, Siddhant Kumar, Bhavesh Modi, Gaurishankar Shrimali, R D Dixit, A M Kadri, Umang Mishra, Chaitanya Joshi, Madhvi Joshi |
| EPI_ISL_458094 | B.J. Medical College and Civil hospital | Gujarat Biotechnology Research Centre | Nitin Savaliya, Raghawendra Kumar, Dinesh Kumar, Zuber Saiyed, Komal Patel, Labdhi Pandya, Snehal Bagatharia, Dhaval Vaghela, Ramesh Patel, Pranay Shah, Kamlesh J Upadhyay, Ramesh Pandit, Tejas Shah, Ankit Hinsu, Pritesh Sabara, Apurvasinh Puvar, Janvi Raval, Zarna Patel, Monika Gandhi, Pinal Trivedi, Maharshi Pandya, Amit Kanani, Nidhi Patel, Priyanka P Vatsa, Bhavesh Modi, Gaurishankar Shrimali, R D Dixit, A M Kadri, Umang Mishra, Chaitanya Joshi, Madhvi Joshi |
| EPI_ISL_458106 | Gujarat Biotechnology Research Centre | Gujarat Biotechnology Research Centre | Pritesh Sabara, Apurvasinh Puvar, Janvi Raval, Zarna Patel, Monika Gandhi, Pinal Trivedi, Maharshi Pandya, Amit Kanani, Nidhi Patel, Nitin Savaliya, Raghawendra Kumar, Dinesh Kumar, Zuber Saiyed, Komal Patel, Labdhi Pandya, Snehal Bagatharia, Ramesh Pandit, Tejas Shah, Ankit Hinsu, Siddhant Kumar, Bhavesh Modi, Gaurishankar Shrimali, R D Dixit, A M Kadri, Umang Mishra, Chaitanya Joshi, Madhvi Joshi, , , , , |
| EPI_ISL_458111 | Gujarat Biotechnology Research Centre | Gujarat Biotechnology Research Centre | Pinal Trivedi, Maharshi Pandya, Amit Kanani, Nidhi Patel, Nitin Savaliya, Raghawendra Kumar, Dinesh Kumar, Zuber Saiyed, Komal Patel, Labdhi Pandya, Snehal Bagatharia, Ramesh Pandit, Tejas Shah, Ankit Hinsu, Pritesh Sabara, Apurvasinh Puvar, Janvi Raval, Zarna Patel, Monika Gandhi, Pragya Sharma, |

| | | | |
|---|---|---|---|
| | | | Bhavesh Modi, Gaurishankar Shrimali, R D Dixit, A M Kadri, Umang Mishra, Chaitanya Joshi, Madhvi Joshi, , , , , |
| EPI_ISL_459962, EPI_ISL_459963, EPI_ISL_459964 | Centogene AG | Centogene AG | Prof. Dr. Peter Bauer, Dr. Krishna Kumar Kandaswamy |
| EPI_ISL_460617, EPI_ISL_460618, EPI_ISL_460619 | unknown | Physiology | Pence,S., Caykara,B., Pence,H.H., Tekin,S., Yiyit,N., Cevher Keskin,B., Kara,A. |
| EPI_ISL_460621, EPI_ISL_460622, EPI_ISL_460623, EPI_ISL_460624, EPI_ISL_460625, EPI_ISL_460626, EPI_ISL_460627, EPI_ISL_460628, EPI_ISL_460629, EPI_ISL_460630, EPI_ISL_460631, EPI_ISL_460632, EPI_ISL_460633, EPI_ISL_460634, EPI_ISL_461399, EPI_ISL_461400, EPI_ISL_461401, EPI_ISL_461402, EPI_ISL_461403, EPI_ISL_461404, EPI_ISL_461405, EPI_ISL_461406, EPI_ISL_461407, EPI_ISL_461408, EPI_ISL_461409, EPI_ISL_461410, EPI_ISL_461411, EPI_ISL_461412, EPI_ISL_461413, EPI_ISL_461414, EPI_ISL_461415, EPI_ISL_461416, EPI_ISL_461417, EPI_ISL_461418, EPI_ISL_461419, EPI_ISL_461420, EPI_ISL_461421, EPI_ISL_461422, EPI_ISL_461423, EPI_ISL_461424, EPI_ISL_461425, EPI_ISL_461426, EPI_ISL_461427, EPI_ISL_461428, EPI_ISL_461429, EPI_ISL_461430, EPI_ISL_461431, EPI_ISL_461432, EPI_ISL_461433, EPI_ISL_461434, EPI_ISL_461435, EPI_ISL_461436, EPI_ISL_461437, EPI_ISL_461438, EPI_ISL_461439, EPI_ISL_461440, EPI_ISL_461441, EPI_ISL_461442, EPI_ISL_461443, EPI_ISL_461444, EPI_ISL_461445, EPI_ISL_461446, EPI_ISL_461447, EPI_ISL_461448, EPI_ISL_461449, EPI_ISL_461450, EPI_ISL_461451, EPI_ISL_461452, EPI_ISL_461453, EPI_ISL_461454, EPI_ISL_461455, EPI_ISL_461456, EPI_ISL_461457, EPI_ISL_461458, EPI_ISL_461459, EPI_ISL_461460, EPI_ISL_461461, EPI_ISL_461462, EPI_ISL_461463, EPI_ISL_461464, EPI_ISL_461465, EPI_ISL_461466, EPI_ISL_461467, EPI_ISL_461468, EPI_ISL_461469, EPI_ISL_461470, EPI_ISL_461471, EPI_ISL_461472, EPI_ISL_461473, EPI_ISL_461474, EPI_ISL_461475, EPI_ISL_461476, EPI_ISL_461477 | | | |
| see above | UW Virology Lab | UW Virology Lab | Pavitra Roychoudhury, Amin Addetia, Hong Xie, Lasata Shrestha, Truong Nguyen, Meei-Li Huang, Keith Jerome, Alexander Greninger |
| EPI_ISL_461479 | Government Medical College, Vadodara | Gujarat Biotechnology Research Centre | Neelam Nathani, Nitin Savaliya, Raghawendra Kumar, Dinesh Kumar, Zuber Saiyed, Komal Patel, Labdhi Pandya, Snehal Bagatharia, Tanuja Javadekar , R N Daveshwar, Tejas Shah, Ankit Hinsu, Pritesh Sabara, Apurvasinh Puvar, Janvi Raval, Zarna Patel, Monika Gandhi, Pinal Trivedi, Maharshi Pandya, Nidhi Patel, R D Dixit, A M Kadri, Harsh Bakshi, Chaitanya Joshi, Madhvi Joshi, |
| EPI_ISL_461484 | B.J. Medical College and Civil hospital | Gujarat Biotechnology Research Centre | Priyanka P Vatsa, Labdhi Pandya, Snehal Bagatharia, Pranay Shah, Kamlesh J Upadhyay, Tejas Shah, Ankit Hinsu, Pritesh Sabara, Apurvasinh Puvar, Janvi Raval, Zarna Patel, Monika Gandhi, Pinal Trivedi, Maharshi Pandya, Nidhi Patel, Nitin Savaliya, Raghawendra Kumar, Dinesh Kumar, Zuber Saiyed, Komal Patel, R D Dixit, A M Kadri, Harsh Bakshi, Chaitanya Joshi, Madhvi Joshi, |
| EPI_ISL_461485 | B.J. Medical College and Civil hospital | Gujarat Biotechnology Research Centre | Pooja P Doshi, Snehal Bagatharia, Pranay Shah, Kamlesh J Upadhyay, Tejas Shah, Ankit Hinsu, Pritesh Sabara, Apurvasinh Puvar, Janvi Raval, Zarna Patel, Monika Gandhi, Pinal Trivedi, Maharshi Pandya, Nidhi Patel, Nitin Savaliya, Raghawendra Kumar, Dinesh Kumar, Zuber Saiyed, Komal Patel, Labdhi Pandya, R D Dixit, A M Kadri, Harsh Bakshi, Chaitanya Joshi, Madhvi Joshi, |
| EPI_ISL_461487 | B.J. Medical College and Civil hospital | Gujarat Biotechnology Research Centre | Priti Pandita, Kamlesh J Upadhyay, Tejas Shah, Ankit Hinsu, Pritesh Sabara, Apurvasinh Puvar, Janvi Raval, Zarna Patel, Monika Gandhi, Pinal Trivedi, Maharshi Pandya, Nidhi Patel, Nitin Savaliya, Raghawendra Kumar, Dinesh Kumar, Zuber Saiyed, Komal Patel, Labdhi Pandya, Snehal Bagatharia, Pranay Shah, R D Dixit, A M Kadri, Harsh Bakshi, Chaitanya Joshi, Madhvi Joshi, |
| EPI_ISL_461488 | B.J. Medical College and Civil hospital | Gujarat Biotechnology Research Centre | Pragya Sharma, Tejas Shah, Ankit Hinsu, Pritesh Sabara, Apurvasinh Puvar, Janvi Raval, Zarna Patel, Monika Gandhi, Pinal Trivedi, Maharshi Pandya, Nidhi Patel, Nitin Savaliya, Raghawendra Kumar, Dinesh Kumar, Zuber Saiyed, Komal Patel, Labdhi Pandya, Snehal Bagatharia, Pranay Shah, Kamlesh J Upadhyay, R D Dixit, A M Kadri, Harsh Bakshi, Chaitanya Joshi, Madhvi Joshi, |
| EPI_ISL_461489 | B.J. Medical College and Civil hospital | Gujarat Biotechnology Research Centre | Neha Rajpara, Ankit Hinsu, Pritesh Sabara, Apurvasinh Puvar, Janvi Raval, Zarna Patel, Monika Gandhi, Pinal Trivedi, Maharshi Pandya, Nidhi Patel, Nitin Savaliya, Raghawendra Kumar, Dinesh Kumar, Zuber Saiyed, Komal Patel, Labdhi Pandya, Snehal Bagatharia, Pranay Shah, Kamlesh J Upadhyay, Tejas Shah, R D Dixit, A M Kadri, Harsh Bakshi, Chaitanya Joshi, Madhvi Joshi, |
| EPI_ISL_461492 | B.J. Medical College and Civil hospital | Gujarat Biotechnology Research Centre | Neelam Nathani, Janvi Raval, Zarna Patel, Monika Gandhi, Pinal Trivedi, Maharshi Pandya, Nidhi Patel, Nitin Savaliya, Raghawendra Kumar, Dinesh Kumar, Zuber Saiyed, Komal Patel, Labdhi Pandya, Snehal Bagatharia, Pranay Shah, Kamlesh J Upadhyay, Tejas Shah, Ankit Hinsu, Pritesh Sabara, Apurvasinh Puvar, R D Dixit, A M Kadri, Harsh Bakshi, Chaitanya Joshi, Madhvi Joshi, |
| EPI_ISL_461497 | B.J. Medical College and Civil hospital | Gujarat Biotechnology Research Centre | Priyanka P Vatsa, Nidhi Patel, Nitin Savaliya, Raghawendra Kumar, Dinesh Kumar, Zuber Saiyed, Komal Patel, Labdhi Pandya, Snehal Bagatharia, Pranay Shah, Kamlesh J Upadhyay, Tejas Shah, Ankit Hinsu, Pritesh Sabara, Apurvasinh Puvar, Janvi Raval, Zarna Patel, Monika Gandhi, Pinal Trivedi, Maharshi Pandya, R D Dixit, A M Kadri, Harsh Bakshi, Chaitanya Joshi, Madhvi Joshi, |
| EPI_ISL_461498 | B.J. Medical College and Civil hospital | Gujarat Biotechnology Research Centre | Pooja P Doshi, Nitin Savaliya, Raghawendra Kumar, Dinesh Kumar, Zuber Saiyed, Komal Patel, Labdhi Pandya, Snehal Bagatharia, Pranay Shah, Kamlesh J Upadhyay, Tejas Shah, Ankit Hinsu, Pritesh Sabara, Apurvasinh Puvar, Janvi Raval, Zarna Patel, Monika Gandhi, Pinal Trivedi, Maharshi Pandya, Nidhi Patel, R D Dixit, A M Kadri, Harsh Bakshi, Chaitanya Joshi, Madhvi Joshi, |
| EPI_ISL_461500 | B.J. Medical College and Civil hospital | Gujarat Biotechnology Research Centre | Priti Pandita, Dinesh Kumar, Zuber Saiyed, Komal Patel, Labdhi Pandya, Snehal Bagatharia, Pranay Shah, Kamlesh J Upadhyay, Tejas Shah, Ankit Hinsu, Pritesh Sabara, Apurvasinh Puvar, Janvi Raval, Zarna Patel, Monika Gandhi, Pinal Trivedi, Maharshi Pandya, Nidhi Patel, Nitin Savaliya, Raghawendra Kumar, R D Dixit, A M Kadri, Harsh Bakshi, Chaitanya Joshi, Madhvi Joshi, |
| EPI_ISL_461501 | B.J. Medical College and Civil hospital | Gujarat Biotechnology Research Centre | Pragya Sharma, Zuber Saiyed, Komal Patel, Labdhi Pandya, Snehal Bagatharia, Pranay Shah, Kamlesh J Upadhyay, Tejas Shah, Ankit Hinsu, Pritesh Sabara, Apurvasinh Puvar, Janvi Raval, Zarna Patel, Monika Gandhi, Pinal Trivedi, Maharshi Pandya, Nidhi Patel, Nitin Savaliya, Raghawendra Kumar, Dinesh Kumar, R D Dixit, A M Kadri, Harsh Bakshi, Chaitanya Joshi, Madhvi Joshi, |
| EPI_ISL_461502 | B.J. Medical College and Civil hospital | Gujarat Biotechnology Research Centre | Neha Rajpara, Komal Patel, Labdhi Pandya, Snehal Bagatharia, Pranay Shah, Kamlesh J Upadhyay, Tejas Shah, Ankit Hinsu, Pritesh Sabara, Apurvasinh Puvar, Janvi Raval, Zarna Patel, Monika Gandhi, Pinal Trivedi, Maharshi Pandya, Nidhi Patel, Nitin Savaliya, Raghawendra Kumar, Dinesh Kumar, Zuber Saiyed, R D Dixit, A M Kadri, Harsh Bakshi, Chaitanya Joshi, Madhvi Joshi, |
| EPI_ISL_461505 | B.J. Medical College and Civil hospital | Gujarat Biotechnology Research Centre | Pranay Shah, Kamlesh J Upadhyay, Tejas Shah, Ankit Hinsu, Pritesh Sabara, Apurvasinh Puvar, Janvi Raval, Zarna Patel, Monika Gandhi, Pinal Trivedi, Maharshi Pandya, Nidhi Patel, Nitin Savaliya, Raghawendra Kumar, Dinesh Kumar, Zuber Saiyed, Komal Patel, Labdhi Pandya, Snehal Bagatharia, Neelam Nathani, R D Dixit, A M Kadri, Harsh Bakshi, Chaitanya Joshi, Madhvi Joshi, |
| EPI_ISL_462085, EPI_ISL_462086, EPI_ISL_462087, EPI_ISL_462088, EPI_ISL_462089 | Singapore General Hospital | Department of Microbiology | Nurdyana Abdul Rahman, Kun Lee Lim, Chenhao Li, Kian Sing Chan, Lynette Oon, Kern Rei Chng, Niranjan Nagarajan, Karrie Ko |
| EPI_ISL_464168, EPI_ISL_464169, EPI_ISL_464170, EPI_ISL_464171, EPI_ISL_464172, EPI_ISL_464173, EPI_ISL_464174, EPI_ISL_464175, EPI_ISL_464176, EPI_ISL_464177, EPI_ISL_464178, EPI_ISL_464179, EPI_ISL_464180, EPI_ISL_464181, EPI_ISL_464182, EPI_ISL_464183, EPI_ISL_464184, EPI_ISL_464185, EPI_ISL_464186, EPI_ISL_464187, EPI_ISL_464188, EPI_ISL_464189, EPI_ISL_464190, EPI_ISL_464191, EPI_ISL_464192, EPI_ISL_464193, EPI_ISL_464194, EPI_ISL_464195, EPI_ISL_464196, EPI_ISL_464197, EPI_ISL_464198, EPI_ISL_464199, EPI_ISL_464200, EPI_ISL_464201, EPI_ISL_464202, EPI_ISL_464203, EPI_ISL_464204, EPI_ISL_464205, EPI_ISL_464206, EPI_ISL_464207, EPI_ISL_464208, EPI_ISL_464209, EPI_ISL_464210, EPI_ISL_464211, EPI_ISL_464212, EPI_ISL_464213, EPI_ISL_464214, EPI_ISL_464215, EPI_ISL_464216, EPI_ISL_464217, EPI_ISL_464218, EPI_ISL_464219, EPI_ISL_464220, EPI_ISL_464221, EPI_ISL_464222, EPI_ISL_464223, EPI_ISL_464224, EPI_ISL_464225, EPI_ISL_464226, EPI_ISL_464227, EPI_ISL_464228, EPI_ISL_464229, EPI_ISL_464230, EPI_ISL_464231, EPI_ISL_464232, EPI_ISL_464233, EPI_ISL_464234, EPI_ISL_464235, EPI_ISL_464236, EPI_ISL_464237, EPI_ISL_464238, EPI_ISL_464239, EPI_ISL_464240, EPI_ISL_464241, EPI_ISL_464242, EPI_ISL_464243, EPI_ISL_464244, EPI_ISL_464245, EPI_ISL_464246, EPI_ISL_464247, EPI_ISL_464248, EPI_ISL_464249, EPI_ISL_464250, EPI_ISL_464251, EPI_ISL_464252, EPI_ISL_464253, EPI_ISL_464254, EPI_ISL_464255, EPI_ISL_464256, EPI_ISL_464258, EPI_ISL_464259, EPI_ISL_464260, EPI_ISL_464261, EPI_ISL_464262, EPI_ISL_464263, EPI_ISL_464264, EPI_ISL_464265, EPI_ISL_464266, EPI_ISL_464267, EPI_ISL_464268, EPI_ISL_464269, EPI_ISL_464270, EPI_ISL_464271, EPI_ISL_464272, EPI_ISL_464273, EPI_ISL_464274, EPI_ISL_464275, EPI_ISL_464276, EPI_ISL_464277, EPI_ISL_464278, EPI_ISL_464279, EPI_ISL_464280, EPI_ISL_464281, EPI_ISL_464282, EPI_ISL_464283, EPI_ISL_464284, EPI_ISL_464285, EPI_ISL_464286, EPI_ISL_464287, EPI_ISL_464288, EPI_ISL_464289, EPI_ISL_464290, EPI_ISL_464291, EPI_ISL_464292, EPI_ISL_464293, EPI_ISL_464294, EPI_ISL_464295, EPI_ISL_464296, EPI_ISL_464297, EPI_ISL_464298, EPI_ISL_464299, EPI_ISL_464300, EPI_ISL_464301, EPI_ISL_464302, EPI_ISL_464303, EPI_ISL_464304, EPI_ISL_464305, EPI_ISL_464306, EPI_ISL_464307, EPI_ISL_464308, EPI_ISL_464309, EPI_ISL_464310, EPI_ISL_464311, EPI_ISL_464312, EPI_ISL_464313, EPI_ISL_464314, EPI_ISL_464315, EPI_ISL_464316, EPI_ISL_464317, EPI_ISL_464318, EPI_ISL_464319, EPI_ISL_464320, EPI_ISL_464321, EPI_ISL_464322, EPI_ISL_464323, EPI_ISL_464324, EPI_ISL_464325, EPI_ISL_464326, EPI_ISL_464327, EPI_ISL_464328, EPI_ISL_464329, EPI_ISL_464330, EPI_ISL_464331, EPI_ISL_464332, EPI_ISL_464333, EPI_ISL_464334, EPI_ISL_464335, EPI_ISL_464336, EPI_ISL_464337, EPI_ISL_464338, EPI_ISL_464339, EPI_ISL_464340, EPI_ISL_464341, EPI_ISL_464342, EPI_ISL_464343, EPI_ISL_464344, EPI_ISL_464345, EPI_ISL_464346, EPI_ISL_464347, EPI_ISL_464348, EPI_ISL_464349, EPI_ISL_464350, EPI_ISL_464351, EPI_ISL_464352, EPI_ISL_464353, EPI_ISL_464354, EPI_ISL_464355, EPI_ISL_464356, EPI_ISL_464357, EPI_ISL_464358, EPI_ISL_464359, EPI_ISL_464360, EPI_ISL_464361, EPI_ISL_464362, EPI_ISL_464363, EPI_ISL_464364, EPI_ISL_464365, EPI_ISL_464366, EPI_ISL_464367, EPI_ISL_464368, EPI_ISL_464369, EPI_ISL_464370, EPI_ISL_464371, EPI_ISL_464372, EPI_ISL_464373, EPI_ISL_464374, EPI_ISL_464375, EPI_ISL_464376, EPI_ISL_464377, EPI_ISL_464378, EPI_ISL_464379, EPI_ISL_464380, EPI_ISL_464381, EPI_ISL_464382, EPI_ISL_464383, EPI_ISL_464384, EPI_ISL_464385, EPI_ISL_464386, EPI_ISL_464387, EPI_ISL_464388, EPI_ISL_464389, EPI_ISL_464390, EPI_ISL_464391, EPI_ISL_464392, EPI_ISL_464393, EPI_ISL_464394, EPI_ISL_464395, EPI_ISL_464396, EPI_ISL_464397, EPI_ISL_464398, EPI_ISL_464399, EPI_ISL_464400, EPI_ISL_464401, EPI_ISL_464402, EPI_ISL_464403, EPI_ISL_464404, EPI_ISL_464405, EPI_ISL_464406, EPI_ISL_464407, EPI_ISL_464408, EPI_ISL_464409, EPI_ISL_464410, EPI_ISL_464411, EPI_ISL_464412, EPI_ISL_464413, EPI_ISL_464414, EPI_ISL_464415, EPI_ISL_464416, EPI_ISL_464417, EPI_ISL_464418, EPI_ISL_464419, EPI_ISL_464420, EPI_ISL_464421, EPI_ISL_464422, EPI_ISL_464423, EPI_ISL_464424, EPI_ISL_464425, EPI_ISL_464426, EPI_ISL_464427, EPI_ISL_464428, EPI_ISL_464429, EPI_ISL_464430, EPI_ISL_464431, EPI_ISL_464432, EPI_ISL_464433, EPI_ISL_464434, EPI_ISL_464435, EPI_ISL_464436, EPI_ISL_464437, EPI_ISL_464438, EPI_ISL_464439, EPI_ISL_464440, EPI_ISL_464441, EPI_ISL_464442, EPI_ISL_464443, EPI_ISL_464444, EPI_ISL_464445, EPI_ISL_464446, EPI_ISL_464447, EPI_ISL_464448, EPI_ISL_464449, EPI_ISL_464450, EPI_ISL_464451, EPI_ISL_464452, EPI_ISL_464453, EPI_ISL_464454, EPI_ISL_464455, EPI_ISL_464456, EPI_ISL_464457, EPI_ISL_464458, EPI_ISL_464459, EPI_ISL_464460, EPI_ISL_464461, EPI_ISL_464462, EPI_ISL_464463, EPI_ISL_464464, EPI_ISL_464465, EPI_ISL_464466, EPI_ISL_464467, EPI_ISL_464468, EPI_ISL_464469, EPI_ISL_464470, EPI_ISL_464471, EPI_ISL_464472, EPI_ISL_464473, EPI_ISL_464474, EPI_ISL_464475, EPI_ISL_464476, EPI_ISL_464477, EPI_ISL_464478, EPI_ISL_464479, EPI_ISL_464480, EPI_ISL_464481, EPI_ISL_464482, EPI_ISL_464483, EPI_ISL_464484, EPI_ISL_464485, EPI_ISL_464486, EPI_ISL_464487, EPI_ISL_464488, EPI_ISL_464489, EPI_ISL_464490, EPI_ISL_464491, EPI_ISL_464492, EPI_ISL_464493, EPI_ISL_464494, EPI_ISL_464495, EPI_ISL_464496, EPI_ISL_464497, EPI_ISL_464498, EPI_ISL_464499, EPI_ISL_464500, EPI_ISL_464501, EPI_ISL_464502, EPI_ISL_464503, EPI_ISL_464504, EPI_ISL_464505, EPI_ISL_464506, EPI_ISL_464507, EPI_ISL_464508, EPI_ISL_464509, EPI_ISL_464510, EPI_ISL_464511, EPI_ISL_464512, EPI_ISL_464513, EPI_ISL_464514, EPI_ISL_464515, EPI_ISL_464516, EPI_ISL_464517, EPI_ISL_464518, EPI_ISL_464519, EPI_ISL_464520, EPI_ISL_464521, EPI_ISL_464522, EPI_ISL_464523, EPI_ISL_464524, EPI_ISL_464525, EPI_ISL_464526, EPI_ISL_464527, EPI_ISL_464528, EPI_ISL_464529, EPI_ISL_464530, EPI_ISL_464531, EPI_ISL_464532, EPI_ISL_464533, EPI_ISL_464534, EPI_ISL_464535, EPI_ISL_464536, EPI_ISL_464537, EPI_ISL_464538, EPI_ISL_464539, EPI_ISL_464540, EPI_ISL_464541, EPI_ISL_464542, EPI_ISL_464543, EPI_ISL_464544, EPI_ISL_464545, EPI_ISL_464546, EPI_ISL_464547, EPI_ISL_464548, EPI_ISL_464549, EPI_ISL_464550, EPI_ISL_464551, EPI_ISL_464552, EPI_ISL_464553, EPI_ISL_464554, EPI_ISL_464555, EPI_ISL_464556, EPI_ISL_464557, EPI_ISL_464558, EPI_ISL_464559, EPI_ISL_464560, EPI_ISL_464561, EPI_ISL_464562, EPI_ISL_464563, EPI_ISL_464564, EPI_ISL_464565, EPI_ISL_464566, EPI_ISL_464567, EPI_ISL_464568, EPI_ISL_464569, EPI_ISL_464570, EPI_ISL_464571, EPI_ISL_464572, EPI_ISL_464573, EPI_ISL_464574, EPI_ISL_464575, EPI_ISL_464576, EPI_ISL_464577, EPI_ISL_464578, EPI_ISL_464579, EPI_ISL_464580, EPI_ISL_464581, EPI_ISL_464582, EPI_ISL_464583, EPI_ISL_464584, EPI_ISL_464585, EPI_ISL_464586, EPI_ISL_464587, EPI_ISL_464588, EPI_ISL_464589, EPI_ISL_464590, EPI_ISL_464591, EPI_ISL_464592, EPI_ISL_464593, EPI_ISL_464594, EPI_ISL_464595, EPI_ISL_464596, EPI_ISL_464597, EPI_ISL_464598, EPI_ISL_464599, EPI_ISL_464600, | | | |

EPI_ISL_464601, EPI_ISL_464602, EPI_ISL_464603, EPI_ISL_464604, EPI_ISL_464605, EPI_ISL_464606, EPI_ISL_464607, EPI_ISL_464608, EPI_ISL_464609, EPI_ISL_464610, EPI_ISL_464611, EPI_ISL_464612, EPI_ISL_464613, EPI_ISL_464614, EPI_ISL_464615, EPI_ISL_464616, EPI_ISL_464617, EPI_ISL_464618,
EPI_ISL_464619, EPI_ISL_464620, EPI_ISL_464621, EPI_ISL_464622, EPI_ISL_464623, EPI_ISL_464624, EPI_ISL_464625, EPI_ISL_464626, EPI_ISL_464627, EPI_ISL_464628, EPI_ISL_464629, EPI_ISL_464630, EPI_ISL_464631, EPI_ISL_464632, EPI_ISL_464633, EPI_ISL_464634, EPI_ISL_464635, EPI_ISL_464636,
EPI_ISL_464637, EPI_ISL_464638, EPI_ISL_464639, EPI_ISL_464640, EPI_ISL_464641, EPI_ISL_464642, EPI_ISL_464643, EPI_ISL_464644, EPI_ISL_464645, EPI_ISL_464646, EPI_ISL_464647, EPI_ISL_464648, EPI_ISL_464649, EPI_ISL_464650, EPI_ISL_464651, EPI_ISL_464652, EPI_ISL_464653, EPI_ISL_464654,
EPI_ISL_464655, EPI_ISL_464656, EPI_ISL_464657, EPI_ISL_464658, EPI_ISL_464659, EPI_ISL_464660, EPI_ISL_464661, EPI_ISL_464662, EPI_ISL_464663, EPI_ISL_464664, EPI_ISL_464665, EPI_ISL_464666, EPI_ISL_464667, EPI_ISL_464668, EPI_ISL_464669, EPI_ISL_464670, EPI_ISL_464671, EPI_ISL_464672,
EPI_ISL_464673, EPI_ISL_464674, EPI_ISL_464675, EPI_ISL_464676, EPI_ISL_464677, EPI_ISL_464678, EPI_ISL_464679, EPI_ISL_464680, EPI_ISL_464681, EPI_ISL_464682, EPI_ISL_464683, EPI_ISL_464684, EPI_ISL_464685, EPI_ISL_464686, EPI_ISL_464687, EPI_ISL_464688, EPI_ISL_464689, EPI_ISL_464690,
EPI_ISL_464691, EPI_ISL_464692, EPI_ISL_464693, EPI_ISL_464694, EPI_ISL_464695, EPI_ISL_464696, EPI_ISL_464697, EPI_ISL_464698, EPI_ISL_464699, EPI_ISL_464700, EPI_ISL_464701, EPI_ISL_464702, EPI_ISL_464703, EPI_ISL_464704, EPI_ISL_464705, EPI_ISL_464706, EPI_ISL_464707, EPI_ISL_464708,
EPI_ISL_464709, EPI_ISL_464710, EPI_ISL_464711, EPI_ISL_464712, EPI_ISL_464713, EPI_ISL_464714, EPI_ISL_464715, EPI_ISL_464716, EPI_ISL_464717, EPI_ISL_464718, EPI_ISL_464719, EPI_ISL_464720, EPI_ISL_464721, EPI_ISL_464722, EPI_ISL_464723, EPI_ISL_464724, EPI_ISL_464725, EPI_ISL_464726,
EPI_ISL_464727, EPI_ISL_464728, EPI_ISL_464729, EPI_ISL_464730, EPI_ISL_464731, EPI_ISL_464732, EPI_ISL_464733, EPI_ISL_464734, EPI_ISL_464735, EPI_ISL_464736, EPI_ISL_464737, EPI_ISL_464738, EPI_ISL_464739, EPI_ISL_464740, EPI_ISL_464741, EPI_ISL_464742, EPI_ISL_464743, EPI_ISL_464744,
EPI_ISL_464745, EPI_ISL_464746, EPI_ISL_464747, EPI_ISL_464748, EPI_ISL_464749, EPI_ISL_464750, EPI_ISL_464751, EPI_ISL_464752, EPI_ISL_464753, EPI_ISL_464754, EPI_ISL_464755, EPI_ISL_464756, EPI_ISL_464757, EPI_ISL_464758, EPI_ISL_464759, EPI_ISL_464760, EPI_ISL_464761, EPI_ISL_464762,
EPI_ISL_464763, EPI_ISL_464764, EPI_ISL_464765, EPI_ISL_464766, EPI_ISL_464767, EPI_ISL_464768, EPI_ISL_464769, EPI_ISL_464770, EPI_ISL_464771, EPI_ISL_464772, EPI_ISL_464773, EPI_ISL_464774, EPI_ISL_464775, EPI_ISL_464776, EPI_ISL_464777, EPI_ISL_464778, EPI_ISL_464779, EPI_ISL_464780,
EPI_ISL_464781, EPI_ISL_464782, EPI_ISL_464783, EPI_ISL_464784, EPI_ISL_464785, EPI_ISL_464786, EPI_ISL_464787, EPI_ISL_464788, EPI_ISL_464789, EPI_ISL_464790, EPI_ISL_464791, EPI_ISL_464792, EPI_ISL_464793, EPI_ISL_464794, EPI_ISL_464795, EPI_ISL_464796, EPI_ISL_464797, EPI_ISL_464798,
EPI_ISL_464799, EPI_ISL_464800, EPI_ISL_464801, EPI_ISL_464802, EPI_ISL_464803, EPI_ISL_464804, EPI_ISL_464805, EPI_ISL_464806, EPI_ISL_464807, EPI_ISL_464808, EPI_ISL_464809, EPI_ISL_464810, EPI_ISL_464811, EPI_ISL_464812, EPI_ISL_464813, EPI_ISL_464814, EPI_ISL_464815, EPI_ISL_464816,
EPI_ISL_464817, EPI_ISL_464818, EPI_ISL_464819, EPI_ISL_464820, EPI_ISL_464821, EPI_ISL_464822, EPI_ISL_464823, EPI_ISL_464824, EPI_ISL_464825, EPI_ISL_464826, EPI_ISL_464827, EPI_ISL_464828, EPI_ISL_464829, EPI_ISL_464830, EPI_ISL_464831, EPI_ISL_464832, EPI_ISL_464833, EPI_ISL_464834,
EPI_ISL_464835, EPI_ISL_464836, EPI_ISL_464837, EPI_ISL_464838, EPI_ISL_464839, EPI_ISL_464840, EPI_ISL_464841, EPI_ISL_464842, EPI_ISL_464843, EPI_ISL_464844, EPI_ISL_464845, EPI_ISL_464846, EPI_ISL_464847, EPI_ISL_464848, EPI_ISL_464849, EPI_ISL_464850, EPI_ISL_464851, EPI_ISL_464852,
EPI_ISL_464853, EPI_ISL_464854, EPI_ISL_464855, EPI_ISL_464856, EPI_ISL_464857, EPI_ISL_464858, EPI_ISL_464859, EPI_ISL_464860, EPI_ISL_464861, EPI_ISL_464862, EPI_ISL_464863, EPI_ISL_464864, EPI_ISL_464865, EPI_ISL_464866, EPI_ISL_464867, EPI_ISL_464868, EPI_ISL_464869, EPI_ISL_464870,
EPI_ISL_464871, EPI_ISL_464872, EPI_ISL_464873, EPI_ISL_464874, EPI_ISL_464875, EPI_ISL_464876, EPI_ISL_464877, EPI_ISL_464878, EPI_ISL_464879, EPI_ISL_464880, EPI_ISL_464881, EPI_ISL_464882, EPI_ISL_464883, EPI_ISL_464884, EPI_ISL_464885, EPI_ISL_464886, EPI_ISL_464887, EPI_ISL_464888,
EPI_ISL_464889, EPI_ISL_464890, EPI_ISL_464891, EPI_ISL_464892, EPI_ISL_464893, EPI_ISL_464894, EPI_ISL_464895, EPI_ISL_464896, EPI_ISL_464897, EPI_ISL_464898, EPI_ISL_464899, EPI_ISL_464900, EPI_ISL_464901, EPI_ISL_464902, EPI_ISL_464903, EPI_ISL_464904, EPI_ISL_464905, EPI_ISL_464906,
EPI_ISL_464907, EPI_ISL_464908, EPI_ISL_464909, EPI_ISL_464910, EPI_ISL_464911, EPI_ISL_464912, EPI_ISL_464913, EPI_ISL_464914, EPI_ISL_464915, EPI_ISL_464916, EPI_ISL_464917, EPI_ISL_464918, EPI_ISL_464919, EPI_ISL_464920, EPI_ISL_464921, EPI_ISL_464922, EPI_ISL_464923, EPI_ISL_464924,
EPI_ISL_464925, EPI_ISL_464926, EPI_ISL_464927, EPI_ISL_464928, EPI_ISL_464929, EPI_ISL_464930, EPI_ISL_464931, EPI_ISL_464932, EPI_ISL_464933, EPI_ISL_464934, EPI_ISL_464935, EPI_ISL_464936, EPI_ISL_464937, EPI_ISL_464938, EPI_ISL_464939, EPI_ISL_464940, EPI_ISL_464941, EPI_ISL_464942,
EPI_ISL_464943, EPI_ISL_464944, EPI_ISL_464945, EPI_ISL_464946, EPI_ISL_464947, EPI_ISL_464948, EPI_ISL_464949, EPI_ISL_464950, EPI_ISL_464951, EPI_ISL_464952, EPI_ISL_464953, EPI_ISL_464954, EPI_ISL_464955, EPI_ISL_464956, EPI_ISL_464957, EPI_ISL_464958, EPI_ISL_464959, EPI_ISL_464960,
EPI_ISL_464961, EPI_ISL_464962, EPI_ISL_464963, EPI_ISL_464964, EPI_ISL_464965, EPI_ISL_464966, EPI_ISL_464967, EPI_ISL_464968, EPI_ISL_464969, EPI_ISL_464970, EPI_ISL_464971, EPI_ISL_464972, EPI_ISL_464973, EPI_ISL_464974, EPI_ISL_464975, EPI_ISL_464976, EPI_ISL_464977, EPI_ISL_464978,
EPI_ISL_464979, EPI_ISL_464980, EPI_ISL_464981, EPI_ISL_464982, EPI_ISL_464983, EPI_ISL_464984, EPI_ISL_464985, EPI_ISL_464986, EPI_ISL_464987, EPI_ISL_464988, EPI_ISL_464989, EPI_ISL_464990, EPI_ISL_464991, EPI_ISL_464992, EPI_ISL_464993, EPI_ISL_464994, EPI_ISL_464995, EPI_ISL_464996,
EPI_ISL_464997, EPI_ISL_464998, EPI_ISL_464999, EPI_ISL_465000, EPI_ISL_465001, EPI_ISL_465002, EPI_ISL_465003, EPI_ISL_465004, EPI_ISL_465005, EPI_ISL_465006, EPI_ISL_465007, EPI_ISL_465008, EPI_ISL_465009, EPI_ISL_465010, EPI_ISL_465011, EPI_ISL_465012, EPI_ISL_465013, EPI_ISL_465014,
EPI_ISL_465015, EPI_ISL_465016, EPI_ISL_465017, EPI_ISL_465018, EPI_ISL_465019, EPI_ISL_465020, EPI_ISL_465021, EPI_ISL_465022, EPI_ISL_465023, EPI_ISL_465024, EPI_ISL_465025, EPI_ISL_465026, EPI_ISL_465027, EPI_ISL_465028, EPI_ISL_465029, EPI_ISL_465030, EPI_ISL_465031, EPI_ISL_465032,
EPI_ISL_465033, EPI_ISL_465034, EPI_ISL_465035, EPI_ISL_465036, EPI_ISL_465037, EPI_ISL_465038, EPI_ISL_465039, EPI_ISL_465040, EPI_ISL_465041, EPI_ISL_465042, EPI_ISL_465043, EPI_ISL_465044, EPI_ISL_465045, EPI_ISL_465046, EPI_ISL_465047, EPI_ISL_465048, EPI_ISL_465049, EPI_ISL_465050,
EPI_ISL_465051, EPI_ISL_465052, EPI_ISL_465053, EPI_ISL_465054, EPI_ISL_465055, EPI_ISL_465056, EPI_ISL_465057, EPI_ISL_465058, EPI_ISL_465059, EPI_ISL_465060, EPI_ISL_465061, EPI_ISL_465062, EPI_ISL_465063, EPI_ISL_465064, EPI_ISL_465065, EPI_ISL_465066, EPI_ISL_465067, EPI_ISL_465068,
EPI_ISL_465069, EPI_ISL_465070, EPI_ISL_465071, EPI_ISL_465072, EPI_ISL_465073, EPI_ISL_465074, EPI_ISL_465075, EPI_ISL_465076, EPI_ISL_465077, EPI_ISL_465078, EPI_ISL_465079, EPI_ISL_465080, EPI_ISL_465081, EPI_ISL_465082, EPI_ISL_465083, EPI_ISL_465084, EPI_ISL_465085, EPI_ISL_465086,
EPI_ISL_465087, EPI_ISL_465088, EPI_ISL_465089, EPI_ISL_465090, EPI_ISL_465091, EPI_ISL_465092, EPI_ISL_465093, EPI_ISL_465094, EPI_ISL_465095, EPI_ISL_465096, EPI_ISL_465097, EPI_ISL_465098, EPI_ISL_465099, EPI_ISL_465100, EPI_ISL_465101, EPI_ISL_465102, EPI_ISL_465103, EPI_ISL_465104,
EPI_ISL_465105, EPI_ISL_465106, EPI_ISL_465107, EPI_ISL_465108, EPI_ISL_465109, EPI_ISL_465110, EPI_ISL_465111, EPI_ISL_465112, EPI_ISL_465113, EPI_ISL_465114, EPI_ISL_465115, EPI_ISL_465116, EPI_ISL_465117, EPI_ISL_465118, EPI_ISL_465119, EPI_ISL_465120, EPI_ISL_465121, EPI_ISL_465122,
EPI_ISL_465123, EPI_ISL_465124, EPI_ISL_465125, EPI_ISL_465126, EPI_ISL_465127, EPI_ISL_465128, EPI_ISL_465129, EPI_ISL_465130, EPI_ISL_465131, EPI_ISL_465132, EPI_ISL_465133, EPI_ISL_465134, EPI_ISL_465135, EPI_ISL_465136, EPI_ISL_465137, EPI_ISL_465138, EPI_ISL_465139, EPI_ISL_465140,
EPI_ISL_465141, EPI_ISL_465142, EPI_ISL_465143, EPI_ISL_465144, EPI_ISL_465145, EPI_ISL_465146, EPI_ISL_465147, EPI_ISL_465148, EPI_ISL_465149, EPI_ISL_465150, EPI_ISL_465151, EPI_ISL_465152, EPI_ISL_465153, EPI_ISL_465154, EPI_ISL_465155, EPI_ISL_465156, EPI_ISL_465157, EPI_ISL_465158,
EPI_ISL_465159, EPI_ISL_465160, EPI_ISL_465161, EPI_ISL_465162, EPI_ISL_465165, EPI_ISL_465166, EPI_ISL_465167, EPI_ISL_465168, EPI_ISL_465169, EPI_ISL_465170, EPI_ISL_465171, EPI_ISL_465172, EPI_ISL_465173, EPI_ISL_465174, EPI_ISL_465175, EPI_ISL_465176, EPI_ISL_465177, EPI_ISL_465178,
EPI_ISL_465179, EPI_ISL_465180, EPI_ISL_465181, EPI_ISL_465182, EPI_ISL_465183, EPI_ISL_465184, EPI_ISL_465185, EPI_ISL_465186, EPI_ISL_465187, EPI_ISL_465188, EPI_ISL_465189, EPI_ISL_465190, EPI_ISL_465191, EPI_ISL_465192, EPI_ISL_465193, EPI_ISL_465194, EPI_ISL_465195, EPI_ISL_465196,
EPI_ISL_465197, EPI_ISL_465198, EPI_ISL_465199, EPI_ISL_465200, EPI_ISL_465201, EPI_ISL_465202, EPI_ISL_465203, EPI_ISL_465204, EPI_ISL_465205, EPI_ISL_465206, EPI_ISL_465207, EPI_ISL_465208, EPI_ISL_465210, EPI_ISL_465211, EPI_ISL_465212, EPI_ISL_465213, EPI_ISL_465214, EPI_ISL_465215,
EPI_ISL_465216, EPI_ISL_465217, EPI_ISL_465218, EPI_ISL_465219, EPI_ISL_465220, EPI_ISL_465221, EPI_ISL_465222, EPI_ISL_465223, EPI_ISL_465224, EPI_ISL_465225, EPI_ISL_465226, EPI_ISL_465227, EPI_ISL_465228, EPI_ISL_465229, EPI_ISL_465230, EPI_ISL_465231, EPI_ISL_465232, EPI_ISL_465233,
EPI_ISL_465234, EPI_ISL_465235, EPI_ISL_465236, EPI_ISL_465237, EPI_ISL_465238, EPI_ISL_465239, EPI_ISL_465240, EPI_ISL_465241, EPI_ISL_465242, EPI_ISL_465243, EPI_ISL_465244, EPI_ISL_465245, EPI_ISL_465246, EPI_ISL_465247, EPI_ISL_465248, EPI_ISL_465249, EPI_ISL_465250, EPI_ISL_465251,
EPI_ISL_465252, EPI_ISL_465253, EPI_ISL_465254, EPI_ISL_465255, EPI_ISL_465256, EPI_ISL_465257, EPI_ISL_465258, EPI_ISL_465259, EPI_ISL_465260, EPI_ISL_465261, EPI_ISL_465262, EPI_ISL_465263, EPI_ISL_465264, EPI_ISL_465265, EPI_ISL_465266, EPI_ISL_465267, EPI_ISL_465268, EPI_ISL_465269,
EPI_ISL_465270, EPI_ISL_465271, EPI_ISL_465272, EPI_ISL_465273, EPI_ISL_465274, EPI_ISL_465275, EPI_ISL_465276, EPI_ISL_465277, EPI_ISL_465278, EPI_ISL_465279, EPI_ISL_465280, EPI_ISL_465281, EPI_ISL_465282, EPI_ISL_465283, EPI_ISL_465284, EPI_ISL_465285, EPI_ISL_465286, EPI_ISL_465287,
EPI_ISL_465288, EPI_ISL_465289, EPI_ISL_465290, EPI_ISL_465291, EPI_ISL_465292, EPI_ISL_465293, EPI_ISL_465294, EPI_ISL_465295, EPI_ISL_465296, EPI_ISL_465297, EPI_ISL_465298, EPI_ISL_465299, EPI_ISL_465300, EPI_ISL_465301, EPI_ISL_465302, EPI_ISL_465303, EPI_ISL_465304, EPI_ISL_465305,
EPI_ISL_465306, EPI_ISL_465307, EPI_ISL_465308, EPI_ISL_465309, EPI_ISL_465310, EPI_ISL_465311, EPI_ISL_465312, EPI_ISL_465313, EPI_ISL_465314, EPI_ISL_465315, EPI_ISL_465316, EPI_ISL_465317, EPI_ISL_465318, EPI_ISL_465319, EPI_ISL_465320, EPI_ISL_465321, EPI_ISL_465322, EPI_ISL_465323,
EPI_ISL_465324, EPI_ISL_465325, EPI_ISL_465326, EPI_ISL_465327, EPI_ISL_465328, EPI_ISL_465331, EPI_ISL_465332, EPI_ISL_465333, EPI_ISL_465334, EPI_ISL_465335, EPI_ISL_465336, EPI_ISL_465337, EPI_ISL_465338, EPI_ISL_465339, EPI_ISL_465340, EPI_ISL_465341, EPI_ISL_465342, EPI_ISL_465343,
EPI_ISL_465344, EPI_ISL_465345, EPI_ISL_465346, EPI_ISL_465347, EPI_ISL_465348, EPI_ISL_465349, EPI_ISL_465350, EPI_ISL_465351, EPI_ISL_465352, EPI_ISL_465353, EPI_ISL_465354, EPI_ISL_465355, EPI_ISL_465356, EPI_ISL_465357, EPI_ISL_465358, EPI_ISL_465359, EPI_ISL_465360, EPI_ISL_465361,
EPI_ISL_465362, EPI_ISL_465363, EPI_ISL_465364, EPI_ISL_465365, EPI_ISL_465366, EPI_ISL_465367, EPI_ISL_465368, EPI_ISL_465369, EPI_ISL_465370, EPI_ISL_465371, EPI_ISL_465372, EPI_ISL_465373, EPI_ISL_465374, EPI_ISL_465375, EPI_ISL_465376, EPI_ISL_465377, EPI_ISL_465378, EPI_ISL_465379,
EPI_ISL_465381, EPI_ISL_465382, EPI_ISL_465383, EPI_ISL_465384, EPI_ISL_465385, EPI_ISL_465386, EPI_ISL_465387, EPI_ISL_465388, EPI_ISL_465389, EPI_ISL_465390, EPI_ISL_465391, EPI_ISL_465392, EPI_ISL_465393, EPI_ISL_465394, EPI_ISL_465395, EPI_ISL_465396, EPI_ISL_465397, EPI_ISL_465398,
EPI_ISL_465399, EPI_ISL_465400, EPI_ISL_465402, EPI_ISL_465403, EPI_ISL_465404, EPI_ISL_465405, EPI_ISL_465406, EPI_ISL_465407, EPI_ISL_465408, EPI_ISL_465409, EPI_ISL_465410, EPI_ISL_465411, EPI_ISL_465412, EPI_ISL_465413, EPI_ISL_465414, EPI_ISL_465415, EPI_ISL_465416, EPI_ISL_465417,
EPI_ISL_465418, EPI_ISL_465419, EPI_ISL_465420, EPI_ISL_465421, EPI_ISL_465422, EPI_ISL_465423, EPI_ISL_465424, EPI_ISL_465425, EPI_ISL_465426, EPI_ISL_465427, EPI_ISL_465428, EPI_ISL_465429, EPI_ISL_465430, EPI_ISL_465431, EPI_ISL_465432, EPI_ISL_465433, EPI_ISL_465434, EPI_ISL_465435,
EPI_ISL_465436, EPI_ISL_465437, EPI_ISL_465438, EPI_ISL_465439, EPI_ISL_465440, EPI_ISL_465441, EPI_ISL_465442, EPI_ISL_465443, EPI_ISL_465444, EPI_ISL_465445, EPI_ISL_465446, EPI_ISL_465447, EPI_ISL_465448, EPI_ISL_465449, EPI_ISL_465450, EPI_ISL_465451, EPI_ISL_465452, EPI_ISL_465453,
EPI_ISL_465454, EPI_ISL_465455, EPI_ISL_465456, EPI_ISL_465457, EPI_ISL_465458, EPI_ISL_465459, EPI_ISL_465460, EPI_ISL_465461, EPI_ISL_465462, EPI_ISL_465464, EPI_ISL_465466, EPI_ISL_465467, EPI_ISL_465468, EPI_ISL_465469, EPI_ISL_465470, EPI_ISL_465471, EPI_ISL_465472, EPI_ISL_465473,
EPI_ISL_465474, EPI_ISL_465475, EPI_ISL_465476, EPI_ISL_465477, EPI_ISL_465478, EPI_ISL_465479, EPI_ISL_465480, EPI_ISL_465481, EPI_ISL_465482, EPI_ISL_465484, EPI_ISL_465485, EPI_ISL_465486, EPI_ISL_465487, EPI_ISL_465488, EPI_ISL_465489, EPI_ISL_465490, EPI_ISL_465491, EPI_ISL_465492,
EPI_ISL_465493, EPI_ISL_465494, EPI_ISL_465495, EPI_ISL_465496, EPI_ISL_465497, EPI_ISL_465498, EPI_ISL_465499, EPI_ISL_465500, EPI_ISL_465501, EPI_ISL_465502, EPI_ISL_465503, EPI_ISL_465504, EPI_ISL_465505, EPI_ISL_465506, EPI_ISL_465507, EPI_ISL_465508, EPI_ISL_465509, EPI_ISL_465510,
EPI_ISL_465511, EPI_ISL_465512, EPI_ISL_465513, EPI_ISL_465514, EPI_ISL_465515, EPI_ISL_465516, EPI_ISL_465517, EPI_ISL_465519, EPI_ISL_465520, EPI_ISL_465521, EPI_ISL_465522, EPI_ISL_465523, EPI_ISL_465524, EPI_ISL_465525, EPI_ISL_465526, EPI_ISL_465527, EPI_ISL_465528, EPI_ISL_465529,
EPI_ISL_465530, EPI_ISL_465531, EPI_ISL_465532, EPI_ISL_465533, EPI_ISL_465534, EPI_ISL_465535, EPI_ISL_465536, EPI_ISL_465537, EPI_ISL_465538, EPI_ISL_465539, EPI_ISL_465540, EPI_ISL_465541, EPI_ISL_465542, EPI_ISL_465543, EPI_ISL_465544, EPI_ISL_465545, EPI_ISL_465546, EPI_ISL_465547,
EPI_ISL_465548, EPI_ISL_465549, EPI_ISL_465550, EPI_ISL_465551, EPI_ISL_465552, EPI_ISL_465553, EPI_ISL_465554, EPI_ISL_465555, EPI_ISL_465556, EPI_ISL_465557, EPI_ISL_465558, EPI_ISL_465559, EPI_ISL_465560, EPI_ISL_465561, EPI_ISL_465562, EPI_ISL_465563, EPI_ISL_465564, EPI_ISL_465565,
EPI_ISL_465566, EPI_ISL_465567, EPI_ISL_465568, EPI_ISL_465569, EPI_ISL_465570, EPI_ISL_465571, EPI_ISL_465572, EPI_ISL_465573, EPI_ISL_465574, EPI_ISL_465575, EPI_ISL_465576, EPI_ISL_465577, EPI_ISL_465578, EPI_ISL_465579, EPI_ISL_465580, EPI_ISL_465581, EPI_ISL_465582, EPI_ISL_465583,
EPI_ISL_465584, EPI_ISL_465585, EPI_ISL_465586, EPI_ISL_465587, EPI_ISL_465588, EPI_ISL_465589, EPI_ISL_465590, EPI_ISL_465591, EPI_ISL_465592, EPI_ISL_465593, EPI_ISL_465594, EPI_ISL_465595, EPI_ISL_465596, EPI_ISL_465597, EPI_ISL_465598, EPI_ISL_465599, EPI_ISL_465600, EPI_ISL_465601,
EPI_ISL_465602, EPI_ISL_465603, EPI_ISL_465604, EPI_ISL_465605, EPI_ISL_465606, EPI_ISL_465607, EPI_ISL_465608, EPI_ISL_465609, EPI_ISL_465610, EPI_ISL_465611, EPI_ISL_465612, EPI_ISL_465613, EPI_ISL_465614, EPI_ISL_465615, EPI_ISL_465616, EPI_ISL_465617, EPI_ISL_465618, EPI_ISL_465619,
EPI_ISL_465620, EPI_ISL_465621, EPI_ISL_465622, EPI_ISL_465623, EPI_ISL_465624, EPI_ISL_465625, EPI_ISL_465626, EPI_ISL_465627, EPI_ISL_465628, EPI_ISL_465629, EPI_ISL_465630, EPI_ISL_465631, EPI_ISL_465632, EPI_ISL_465633, EPI_ISL_465634, EPI_ISL_465636, EPI_ISL_465637, EPI_ISL_465638,
EPI_ISL_465639, EPI_ISL_465640, EPI_ISL_465641, EPI_ISL_465642, EPI_ISL_465643, EPI_ISL_465645, EPI_ISL_465648, EPI_ISL_465649, EPI_ISL_465650, EPI_ISL_465651, EPI_ISL_465652, EPI_ISL_465653, EPI_ISL_465654, EPI_ISL_465655, EPI_ISL_465656, EPI_ISL_465657, EPI_ISL_465658, EPI_ISL_465659,
EPI_ISL_465660, EPI_ISL_465661, EPI_ISL_465662, EPI_ISL_465663, EPI_ISL_465664, EPI_ISL_465665, EPI_ISL_465666, EPI_ISL_465667, EPI_ISL_465668, EPI_ISL_465669, EPI_ISL_465670, EPI_ISL_465671, EPI_ISL_465672, EPI_ISL_465673, EPI_ISL_465674, EPI_ISL_465675, EPI_ISL_465676, EPI_ISL_465677,
EPI_ISL_465678, EPI_ISL_465704, EPI_ISL_465705, EPI_ISL_465706, EPI_ISL_465707, EPI_ISL_465708, EPI_ISL_465709, EPI_ISL_465710, EPI_ISL_465711, EPI_ISL_465712, EPI_ISL_465713, EPI_ISL_465714, EPI_ISL_465715, EPI_ISL_465716, EPI_ISL_465717, EPI_ISL_465718, EPI_ISL_465719, EPI_ISL_465720,
EPI_ISL_465721, EPI_ISL_465722, EPI_ISL_465723, EPI_ISL_465724, EPI_ISL_465725, EPI_ISL_465726, EPI_ISL_465727, EPI_ISL_465728, EPI_ISL_465729, EPI_ISL_465730, EPI_ISL_465731, EPI_ISL_465732, EPI_ISL_465733, EPI_ISL_465734, EPI_ISL_465735, EPI_ISL_465736, EPI_ISL_465737, EPI_ISL_465738,
EPI_ISL_465739, EPI_ISL_465740, EPI_ISL_465741, EPI_ISL_465742, EPI_ISL_465743, EPI_ISL_465744, EPI_ISL_465745, EPI_ISL_465746, EPI_ISL_465747, EPI_ISL_465748, EPI_ISL_465749, EPI_ISL_465750, EPI_ISL_465751, EPI_ISL_465752, EPI_ISL_465753, EPI_ISL_465754, EPI_ISL_465755, EPI_ISL_465756,
EPI_ISL_465757, EPI_ISL_465758, EPI_ISL_465759, EPI_ISL_465760, EPI_ISL_465761, EPI_ISL_465762, EPI_ISL_465763, EPI_ISL_465764, EPI_ISL_465765, EPI_ISL_465766, EPI_ISL_465767, EPI_ISL_465768, EPI_ISL_465769, EPI_ISL_465770, EPI_ISL_465771, EPI_ISL_465772, EPI_ISL_465773, EPI_ISL_465774,
EPI_ISL_465775, EPI_ISL_465776, EPI_ISL_465777, EPI_ISL_465778, EPI_ISL_465779, EPI_ISL_465780, EPI_ISL_465781, EPI_ISL_465782, EPI_ISL_465783, EPI_ISL_465784, EPI_ISL_465785, EPI_ISL_465786, EPI_ISL_465787, EPI_ISL_465788, EPI_ISL_465789, EPI_ISL_465790, EPI_ISL_465791, EPI_ISL_465792,
EPI_ISL_465793, EPI_ISL_465794, EPI_ISL_465795, EPI_ISL_465796, EPI_ISL_465797, EPI_ISL_465798, EPI_ISL_465799, EPI_ISL_465800, EPI_ISL_465801, EPI_ISL_465802, EPI_ISL_465803, EPI_ISL_465804, EPI_ISL_465805, EPI_ISL_465806, EPI_ISL_465807, EPI_ISL_465808, EPI_ISL_465809, EPI_ISL_465810,
EPI_ISL_465811, EPI_ISL_465812, EPI_ISL_465813, EPI_ISL_465814, EPI_ISL_465815, EPI_ISL_465816, EPI_ISL_465817, EPI_ISL_465818, EPI_ISL_465819, EPI_ISL_465820, EPI_ISL_465821, EPI_ISL_465822, EPI_ISL_465823, EPI_ISL_465824, EPI_ISL_465825, EPI_ISL_465826, EPI_ISL_465827, EPI_ISL_465828,
EPI_ISL_465829, EPI_ISL_465830, EPI_ISL_465831, EPI_ISL_465832, EPI_ISL_465833, EPI_ISL_465834, EPI_ISL_465835, EPI_ISL_465836, EPI_ISL_465837, EPI_ISL_465838, EPI_ISL_465839, EPI_ISL_465840, EPI_ISL_465841, EPI_ISL_465842, EPI_ISL_465843, EPI_ISL_465844, EPI_ISL_465845, EPI_ISL_465846,
EPI_ISL_465847, EPI_ISL_465848, EPI_ISL_465849, EPI_ISL_465850, EPI_ISL_465851, EPI_ISL_465852, EPI_ISL_465853, EPI_ISL_465854, EPI_ISL_465855, EPI_ISL_465856, EPI_ISL_465857, EPI_ISL_465858, EPI_ISL_465859, EPI_ISL_465860, EPI_ISL_465861, EPI_ISL_465862, EPI_ISL_465863, EPI_ISL_465864,
EPI_ISL_465865, EPI_ISL_465866, EPI_ISL_465867, EPI_ISL_465868, EPI_ISL_465869, EPI_ISL_465870, EPI_ISL_465871, EPI_ISL_465872, EPI_ISL_465873, EPI_ISL_465874, EPI_ISL_465875, EPI_ISL_465876, EPI_ISL_465877, EPI_ISL_465878, EPI_ISL_465879, EPI_ISL_465880, EPI_ISL_465881, EPI_ISL_465882,
EPI_ISL_465883, EPI_ISL_465884, EPI_ISL_465885, EPI_ISL_465886, EPI_ISL_465887, EPI_ISL_465888, EPI_ISL_465889, EPI_ISL_465890, EPI_ISL_465891, EPI_ISL_465892, EPI_ISL_465893, EPI_ISL_465894, EPI_ISL_465895, EPI_ISL_465896, EPI_ISL_465897, EPI_ISL_465898, EPI_ISL_465899, EPI_ISL_465900,
EPI_ISL_465901, EPI_ISL_465902, EPI_ISL_465903, EPI_ISL_465904, EPI_ISL_465905, EPI_ISL_465906, EPI_ISL_465907, EPI_ISL_465908, EPI_ISL_465909, EPI_ISL_465910, EPI_ISL_465911, EPI_ISL_465912, EPI_ISL_465913, EPI_ISL_465914, EPI_ISL_465915, EPI_ISL_465916, EPI_ISL_465917, EPI_ISL_465918,
EPI_ISL_465919, EPI_ISL_465920, EPI_ISL_465921, EPI_ISL_465922, EPI_ISL_465923, EPI_ISL_465924, EPI_ISL_465925, EPI_ISL_465926, EPI_ISL_465927, EPI_ISL_465928, EPI_ISL_465929, EPI_ISL_465930, EPI_ISL_465931, EPI_ISL_465932, EPI_ISL_465933, EPI_ISL_465934, EPI_ISL_465935, EPI_ISL_465936,
EPI_ISL_465937, EPI_ISL_465938, EPI_ISL_465939, EPI_ISL_465940, EPI_ISL_465941, EPI_ISL_465942, EPI_ISL_465943, EPI_ISL_465944, EPI_ISL_465945, EPI_ISL_465946, EPI_ISL_465947, EPI_ISL_465948, EPI_ISL_465949, EPI_ISL_465950, EPI_ISL_465951, EPI_ISL_465952, EPI_ISL_465953, EPI_ISL_465954,
EPI_ISL_465955, EPI_ISL_465956, EPI_ISL_465957, EPI_ISL_465958, EPI_ISL_465959, EPI_ISL_465960, EPI_ISL_465961, EPI_ISL_465962, EPI_ISL_465963, EPI_ISL_465964, EPI_ISL_465965, EPI_ISL_465966, EPI_ISL_465967, EPI_ISL_465968, EPI_ISL_465969, EPI_ISL_465970, EPI_ISL_465971, EPI_ISL_465972,
EPI_ISL_465973, EPI_ISL_465974, EPI_ISL_465975, EPI_ISL_465976, EPI_ISL_465977, EPI_ISL_465978, EPI_ISL_465979, EPI_ISL_465980, EPI_ISL_465981, EPI_ISL_465982, EPI_ISL_465983, EPI_ISL_465984, EPI_ISL_465985, EPI_ISL_465986, EPI_ISL_465987, EPI_ISL_465988, EPI_ISL_465989, EPI_ISL_465990,
EPI_ISL_465991, EPI_ISL_465992, EPI_ISL_465993, EPI_ISL_465994, EPI_ISL_465995, EPI_ISL_465996, EPI_ISL_465997, EPI_ISL_465998, EPI_ISL_465999, EPI_ISL_466000, EPI_ISL_466001, EPI_ISL_466002, EPI_ISL_466003, EPI_ISL_466004, EPI_ISL_466005, EPI_ISL_466006, EPI_ISL_466007, EPI_ISL_466008,
EPI_ISL_466009, EPI_ISL_466010, EPI_ISL_466011, EPI_ISL_466012, EPI_ISL_466013, EPI_ISL_466014, EPI_ISL_466015, EPI_ISL_466016, EPI_ISL_466017, EPI_ISL_466018, EPI_ISL_466019, EPI_ISL_466020, EPI_ISL_466021, EPI_ISL_466022, EPI_ISL_466023, EPI_ISL_466024, EPI_ISL_466025, EPI_ISL_466026,
EPI_ISL_466027, EPI_ISL_466028, EPI_ISL_466029, EPI_ISL_466030, EPI_ISL_466031, EPI_ISL_466032, EPI_ISL_466033, EPI_ISL_466034, EPI_ISL_466035, EPI_ISL_466036, EPI_ISL_466037, EPI_ISL_466038, EPI_ISL_466039, EPI_ISL_466040, EPI_ISL_466041, EPI_ISL_466042, EPI_ISL_466043, EPI_ISL_466044,
EPI_ISL_466045, EPI_ISL_466046, EPI_ISL_466047, EPI_ISL_466048, EPI_ISL_466049, EPI_ISL_466050, EPI_ISL_466051, EPI_ISL_466052, EPI_ISL_466053, EPI_ISL_466054, EPI_ISL_466055, EPI_ISL_466056, EPI_ISL_466057, EPI_ISL_466058, EPI_ISL_466059, EPI_ISL_466060, EPI_ISL_466061, EPI_ISL_466062,
EPI_ISL_466063, EPI_ISL_466064, EPI_ISL_466065, EPI_ISL_466066, EPI_ISL_466067, EPI_ISL_466068, EPI_ISL_466069, EPI_ISL_466070, EPI_ISL_466071, EPI_ISL_466072, EPI_ISL_466073, EPI_ISL_466074, EPI_ISL_466075, EPI_ISL_466076, EPI_ISL_466077, EPI_ISL_466078, EPI_ISL_466079, EPI_ISL_466080,
EPI_ISL_466081, EPI_ISL_466082, EPI_ISL_466083, EPI_ISL_466084, EPI_ISL_466085, EPI_ISL_466086, EPI_ISL_466087, EPI_ISL_466088, EPI_ISL_466089, EPI_ISL_466090, EPI_ISL_466091, EPI_ISL_466092, EPI_ISL_466093, EPI_ISL_466094, EPI_ISL_466095, EPI_ISL_466096, EPI_ISL_466097, EPI_ISL_466098,
EPI_ISL_466099, EPI_ISL_466100, EPI_ISL_466101, EPI_ISL_466102, EPI_ISL_466103, EPI_ISL_466104, EPI_ISL_466105, EPI_ISL_466106, EPI_ISL_466107, EPI_ISL_466108, EPI_ISL_466109, EPI_ISL_466110, EPI_ISL_466111, EPI_ISL_466112, EPI_ISL_466113, EPI_ISL_466114, EPI_ISL_466115, EPI_ISL_466116,
EPI_ISL_466117, EPI_ISL_466118, EPI_ISL_466119, EPI_ISL_466120, EPI_ISL_466121, EPI_ISL_466122, EPI_ISL_466123, EPI_ISL_466124, EPI_ISL_466125, EPI_ISL_466126, EPI_ISL_466127, EPI_ISL_466128, EPI_ISL_466129, EPI_ISL_466130, EPI_ISL_466131, EPI_ISL_466132, EPI_ISL_466133, EPI_ISL_466134,
EPI_ISL_466135, EPI_ISL_466136, EPI_ISL_466137, EPI_ISL_466138, EPI_ISL_466139, EPI_ISL_466140, EPI_ISL_466141, EPI_ISL_466142, EPI_ISL_466143, EPI_ISL_466144, EPI_ISL_466145, EPI_ISL_466146, EPI_ISL_466147, EPI_ISL_466148, EPI_ISL_466149, EPI_ISL_466150, EPI_ISL_466151, EPI_ISL_466152,
EPI_ISL_466153, EPI_ISL_466154, EPI_ISL_466155, EPI_ISL_466156, EPI_ISL_466157, EPI_ISL_466158, EPI_ISL_466159, EPI_ISL_466160, EPI_ISL_466161, EPI_ISL_466162, EPI_ISL_466163, EPI_ISL_466164, EPI_ISL_466165, EPI_ISL_466166, EPI_ISL_466167, EPI_ISL_466168, EPI_ISL_466169, EPI_ISL_466170,
EPI_ISL_466171, EPI_ISL_466172, EPI_ISL_466173, EPI_ISL_466174, EPI_ISL_466175, EPI_ISL_466176, EPI_ISL_466177, EPI_ISL_466178, EPI_ISL_466179, EPI_ISL_466180, EPI_ISL_466181, EPI_ISL_466182, EPI_ISL_466184, EPI_ISL_466185, EPI_ISL_466186, EPI_ISL_466187, EPI_ISL_466188, EPI_ISL_466189,
EPI_ISL_466190, EPI_ISL_466191, EPI_ISL_466192, EPI_ISL_466193, EPI_ISL_466194, EPI_ISL_466195, EPI_ISL_466196, EPI_ISL_466197, EPI_ISL_466198, EPI_ISL_466199, EPI_ISL_466200, EPI_ISL_466201, EPI_ISL_466202, EPI_ISL_466203, EPI_ISL_466204, EPI_ISL_466205, EPI_ISL_466206, EPI_ISL_466207,

| | | | |
|---|---|---|---|
| EPI_ISL_466208, EPI_ISL_466209, EPI_ISL_466210, EPI_ISL_466211, EPI_ISL_466212, EPI_ISL_466213, EPI_ISL_466214, EPI_ISL_466215, EPI_ISL_466216, EPI_ISL_466217, EPI_ISL_466218, EPI_ISL_466219, EPI_ISL_466220, EPI_ISL_466221, EPI_ISL_466222, EPI_ISL_466223, EPI_ISL_466224, EPI_ISL_466225, EPI_ISL_466226, EPI_ISL_466227, EPI_ISL_466228, EPI_ISL_466229, EPI_ISL_466230, EPI_ISL_466231, EPI_ISL_466232, EPI_ISL_466233, EPI_ISL_466234, EPI_ISL_466235, EPI_ISL_466237, EPI_ISL_466238, EPI_ISL_466239, EPI_ISL_466240, EPI_ISL_466241, EPI_ISL_466242, EPI_ISL_466243, EPI_ISL_466244, EPI_ISL_466245, EPI_ISL_466246, EPI_ISL_466247, EPI_ISL_466248, EPI_ISL_466249, EPI_ISL_466250, EPI_ISL_466251, EPI_ISL_466252, EPI_ISL_466253, EPI_ISL_466254, EPI_ISL_466255, EPI_ISL_466256, EPI_ISL_466257, EPI_ISL_466258, EPI_ISL_466259, EPI_ISL_466260, EPI_ISL_466261, EPI_ISL_466262, EPI_ISL_466263, EPI_ISL_466264, EPI_ISL_466265, EPI_ISL_466266, EPI_ISL_466267, EPI_ISL_466269, EPI_ISL_466270, EPI_ISL_466271, EPI_ISL_466272, EPI_ISL_466273, EPI_ISL_466274, EPI_ISL_466275, EPI_ISL_466276, EPI_ISL_466277, EPI_ISL_466278, EPI_ISL_466279, EPI_ISL_466280, EPI_ISL_466281, EPI_ISL_466282, EPI_ISL_466283, EPI_ISL_466284, EPI_ISL_466285, EPI_ISL_466286, EPI_ISL_466287, EPI_ISL_466288, EPI_ISL_466289, EPI_ISL_466290, EPI_ISL_466291, EPI_ISL_466292, EPI_ISL_466293, EPI_ISL_466294, EPI_ISL_466295, EPI_ISL_466296, EPI_ISL_466297, EPI_ISL_466298, EPI_ISL_466299, EPI_ISL_466300, EPI_ISL_466301, EPI_ISL_466302, EPI_ISL_466303, EPI_ISL_466304, EPI_ISL_466305, EPI_ISL_466307, EPI_ISL_466308, EPI_ISL_466309, EPI_ISL_466310, EPI_ISL_466311, EPI_ISL_466312, EPI_ISL_466313, EPI_ISL_466314, EPI_ISL_466315, EPI_ISL_466316, EPI_ISL_466317, EPI_ISL_466318, EPI_ISL_466319, EPI_ISL_466320, EPI_ISL_466321, EPI_ISL_466322, EPI_ISL_466323, EPI_ISL_466324, EPI_ISL_466325, EPI_ISL_466326, EPI_ISL_466327, EPI_ISL_466328, EPI_ISL_466329, EPI_ISL_466330, EPI_ISL_466331, EPI_ISL_466332, EPI_ISL_466333, EPI_ISL_466334, EPI_ISL_466335, EPI_ISL_466336, EPI_ISL_466337, EPI_ISL_466338, EPI_ISL_466339, EPI_ISL_466340, EPI_ISL_466341, EPI_ISL_466342, EPI_ISL_466343, EPI_ISL_466344, EPI_ISL_466345, EPI_ISL_466346, EPI_ISL_466347, EPI_ISL_466348, EPI_ISL_466349, EPI_ISL_466350, EPI_ISL_466351, EPI_ISL_466352, EPI_ISL_466354, EPI_ISL_466355, EPI_ISL_466356, EPI_ISL_466357, EPI_ISL_466358, EPI_ISL_466359, EPI_ISL_466360, EPI_ISL_466361, EPI_ISL_466362, EPI_ISL_466363, EPI_ISL_466364, EPI_ISL_466365, EPI_ISL_466366, EPI_ISL_466367, EPI_ISL_466368, EPI_ISL_466369, EPI_ISL_466370, EPI_ISL_466371, EPI_ISL_466372, EPI_ISL_466373, EPI_ISL_466374, EPI_ISL_466375, EPI_ISL_466376, EPI_ISL_466377, EPI_ISL_466378, EPI_ISL_466379, EPI_ISL_466380, EPI_ISL_466381, EPI_ISL_466382, EPI_ISL_466383, EPI_ISL_466384, EPI_ISL_466385, EPI_ISL_466386, EPI_ISL_466387, EPI_ISL_466389, EPI_ISL_466390, EPI_ISL_466391, EPI_ISL_466392, EPI_ISL_466393, EPI_ISL_466394, EPI_ISL_466395, EPI_ISL_466396, EPI_ISL_466397, EPI_ISL_466398, EPI_ISL_466399, EPI_ISL_466400, EPI_ISL_466401, EPI_ISL_466402, EPI_ISL_466403, EPI_ISL_466404, EPI_ISL_466405, EPI_ISL_466406, EPI_ISL_466407, EPI_ISL_466408, EPI_ISL_466409, EPI_ISL_466410, EPI_ISL_466411, EPI_ISL_466412, EPI_ISL_466413, EPI_ISL_466414, EPI_ISL_466415, EPI_ISL_466416, EPI_ISL_466417, EPI_ISL_466418, EPI_ISL_466419, EPI_ISL_466420, EPI_ISL_466421, EPI_ISL_466422, EPI_ISL_466423, EPI_ISL_466424, EPI_ISL_466425, EPI_ISL_466426, EPI_ISL_466427, EPI_ISL_466428, EPI_ISL_466429, EPI_ISL_466430, EPI_ISL_466431, EPI_ISL_466432, EPI_ISL_466433, EPI_ISL_466434, EPI_ISL_466435, EPI_ISL_466436, EPI_ISL_466437, EPI_ISL_466438, EPI_ISL_466439, EPI_ISL_466440, EPI_ISL_466441, EPI_ISL_466442, EPI_ISL_466444, EPI_ISL_466445, EPI_ISL_466446, EPI_ISL_466447, EPI_ISL_466448, EPI_ISL_466449, EPI_ISL_466450, EPI_ISL_466451, EPI_ISL_466452, EPI_ISL_466453, EPI_ISL_466454, EPI_ISL_466455, EPI_ISL_466456, EPI_ISL_466457, EPI_ISL_466458, EPI_ISL_466459, EPI_ISL_466460, EPI_ISL_466461, EPI_ISL_466462, EPI_ISL_466463, EPI_ISL_466464, EPI_ISL_466465, EPI_ISL_466466, EPI_ISL_466467, EPI_ISL_466468, EPI_ISL_466469, EPI_ISL_466470, EPI_ISL_466471, EPI_ISL_466472, EPI_ISL_466473, EPI_ISL_466474, EPI_ISL_466475, EPI_ISL_466476, EPI_ISL_466477, EPI_ISL_466478, EPI_ISL_466479, EPI_ISL_466480, EPI_ISL_466481, EPI_ISL_466482, EPI_ISL_466483, EPI_ISL_466484, EPI_ISL_466485, EPI_ISL_466486, EPI_ISL_466487, EPI_ISL_466488, EPI_ISL_466489, EPI_ISL_466490, EPI_ISL_466491, EPI_ISL_466492, EPI_ISL_466493, EPI_ISL_466494, EPI_ISL_466495, EPI_ISL_466496, EPI_ISL_466497, EPI_ISL_466498, EPI_ISL_466499, EPI_ISL_466500, EPI_ISL_466501, EPI_ISL_466502, EPI_ISL_466503, EPI_ISL_466504, EPI_ISL_466505, EPI_ISL_466506, EPI_ISL_466507, EPI_ISL_466508, EPI_ISL_466509, EPI_ISL_466510, EPI_ISL_466511, EPI_ISL_466512, EPI_ISL_466513, EPI_ISL_466514, EPI_ISL_466515, EPI_ISL_466516, EPI_ISL_466517, EPI_ISL_466518, EPI_ISL_466519, EPI_ISL_466520, EPI_ISL_466522, EPI_ISL_466523, EPI_ISL_466524, EPI_ISL_466525, EPI_ISL_466526, EPI_ISL_466527, EPI_ISL_466528, EPI_ISL_466529, EPI_ISL_466530, EPI_ISL_466531, EPI_ISL_466532, EPI_ISL_466533, EPI_ISL_466534, EPI_ISL_466535, EPI_ISL_466536, EPI_ISL_466537, EPI_ISL_466538, EPI_ISL_466539, EPI_ISL_466540, EPI_ISL_466541, EPI_ISL_466542, EPI_ISL_466543, EPI_ISL_466544, EPI_ISL_466545, EPI_ISL_466546, EPI_ISL_466547, EPI_ISL_466548, EPI_ISL_466549, EPI_ISL_466550, EPI_ISL_466551, EPI_ISL_466552, EPI_ISL_466554, EPI_ISL_466555, EPI_ISL_466556, EPI_ISL_466557, EPI_ISL_466558, EPI_ISL_466559, EPI_ISL_466560, EPI_ISL_466561, EPI_ISL_466562, EPI_ISL_466563, EPI_ISL_466564, EPI_ISL_466565, EPI_ISL_466566, EPI_ISL_466567, EPI_ISL_466568, EPI_ISL_466569, EPI_ISL_466570, EPI_ISL_466571, EPI_ISL_466572, EPI_ISL_466573, EPI_ISL_466574, EPI_ISL_466575, EPI_ISL_466576, EPI_ISL_466577, EPI_ISL_466578, EPI_ISL_466579, EPI_ISL_466580, EPI_ISL_466581, EPI_ISL_466582, EPI_ISL_466583, EPI_ISL_466584, EPI_ISL_466585, EPI_ISL_466586, EPI_ISL_466587, EPI_ISL_466588, EPI_ISL_466589, EPI_ISL_466590, EPI_ISL_466591, EPI_ISL_466592, EPI_ISL_466593, EPI_ISL_466594, EPI_ISL_466595, EPI_ISL_466596, EPI_ISL_466597, EPI_ISL_466598, EPI_ISL_466599, EPI_ISL_466600, EPI_ISL_466601, EPI_ISL_466602, EPI_ISL_466603, EPI_ISL_466604, EPI_ISL_466605, EPI_ISL_466606, EPI_ISL_466607, EPI_ISL_466608, EPI_ISL_466609, EPI_ISL_466610, EPI_ISL_466611, EPI_ISL_466612, EPI_ISL_466613, EPI_ISL_466614, EPI_ISL_466615, EPI_ISL_466616, EPI_ISL_466617, EPI_ISL_466618, EPI_ISL_466619, EPI_ISL_466620, EPI_ISL_466621, EPI_ISL_466622, EPI_ISL_466623, EPI_ISL_466624, EPI_ISL_466625 | | | |
| see above | Respiratory Virus Unit, Microbiology Services Colindale, Public Health England | Respiratory Virus Unit, Microbiology Services Colindale, Public Health England | PHE Covid Sequencing Team |
| EPI_ISL_467035 | GMERS Medical College and Hospital, Gandhinagar | Gujarat Biotechnology Research Centre | Pritesh Sabara, Apurvasinh Puvar, Janvi Raval, Zarna Patel, Monika Gandhi, Pinal Trivedi, Maharshi Pandya, Nidhi Patel, Nitin Savaliya, Raghawendra Kumar, Dinesh Kumar, Zuber Saiyed, Komal Patel, Labdhi Pandya, Snehal Bagatharia, Seema Bhatt, Gaurishankar Shrimali, Bhavesh Modi, Bharti Rajani, Tejas Shah, Ankit Hinsu, Neha Rajpara, R D Dixit, A M Kadri, Harsh Bakshi, Chaitanya Joshi, Madhvi Joshi |
| EPI_ISL_467040 | Government Medical College, Vadodara | Gujarat Biotechnology Research Centre | Neena Doshi, Varsha Godbole, Tejas Shah, Ankit Hinsu, Pritesh Sabara, Apurvasinh Puvar, Janvi Raval, Zarna Patel, Monika Gandhi, Pinal Trivedi, Maharshi Pandya, Nidhi Patel, Nitin Savaliya, Raghawendra Kumar, Dinesh Kumar, Zuber Saiyed, Komal Patel, Labdhi Pandya, Snehal Bagatharia, Meenakshi Shah, Bhavya Jindal, R D Dixit, A M Kadri, Harsh Bakshi, Chaitanya Joshi, Madhvi Joshi, |
| EPI_ISL_467042 | B.J. Medical College and Civil hospital | Gujarat Biotechnology Research Centre | Pinal Trivedi, Maharshi Pandya, Nidhi Patel, Nitin Savaliya, Raghawendra Kumar, Dinesh Kumar, Zuber Saiyed, Komal Patel, Labdhi Pandya, Snehal Bagatharia, Pranay Shah, Kamlesh J Upadhyay, Nirav Mungalpara, Tejas Shah, Ankit Hinsu, Pritesh Sabara, Apurvasinh Puvar, Janvi Raval, Zarna Patel, Monika Gandhi, Pooja P Doshi, R D Dixit, A M Kadri, Harsh Bakshi, Chaitanya Joshi, Madhvi Joshi, |
| EPI_ISL_467044 | B.J. Medical College and Civil hospital | Gujarat Biotechnology Research Centre | Nidhi Patel, Nitin Savaliya, Raghawendra Kumar, Dinesh Kumar, Zuber Saiyed, Komal Patel, Labdhi Pandya, Snehal Bagatharia, Pranay Shah, Kamlesh J Upadhyay, Nirav Mungalpara, Tejas Shah, Ankit Hinsu, Pritesh Sabara, Apurvasinh Puvar, Janvi Raval, Zarna Patel, Monika Gandhi, Pinal Trivedi, Maharshi Pandya, Priti Pandita, R D Dixit, A M Kadri, Harsh Bakshi, Chaitanya Joshi, Madhvi Joshi, |
| EPI_ISL_467045 | B.J. Medical College and Civil hospital | Gujarat Biotechnology Research Centre | Nitin Savaliya, Raghawendra Kumar, Dinesh Kumar, Zuber Saiyed, Komal Patel, Labdhi Pandya, Snehal Bagatharia, Pranay Shah, Kamlesh J Upadhyay, Nirav Mungalpara, Tejas Shah, Ankit Hinsu, Pritesh Sabara, Apurvasinh Puvar, Janvi Raval, Zarna Patel, Monika Gandhi, Pinal Trivedi, Maharshi Pandya, Nidhi Patel, Pragya Sharma, R D Dixit, A M Kadri, Harsh Bakshi, Chaitanya Joshi, Madhvi Joshi, |
| EPI_ISL_467052 | B.J. Medical College and Civil hospital | Gujarat Biotechnology Research Centre | Pranay Shah, Kamlesh J Upadhyay, Nirav Mungalpara, Tejas Shah, Ankit Hinsu, Pritesh Sabara, Apurvasinh Puvar, Janvi Raval, Zarna Patel, Monika Gandhi, Pinal Trivedi, Maharshi Pandya, Nidhi Patel, Nitin Savaliya, Raghawendra Kumar, Dinesh Kumar, Zuber Saiyed, Komal Patel, Labdhi Pandya, Snehal Bagatharia, Priyanka P Vatsa, R D Dixit, A M Kadri, Harsh Bakshi, Chaitanya Joshi, Madhvi Joshi, |
| EPI_ISL_467054 | B.J. Medical College and Civil hospital | Gujarat Biotechnology Research Centre | Nirav Mungalpara, Tejas Shah, Ankit Hinsu, Pritesh Sabara, Apurvasinh Puvar, Janvi Raval, Zarna Patel, Monika Gandhi, Pinal Trivedi, Maharshi Pandya, Nidhi Patel, Nitin Savaliya, Raghawendra Kumar, Dinesh Kumar, Zuber Saiyed, Komal Patel, Labdhi Pandya, Snehal Bagatharia, Pranay Shah, Kamlesh J Upadhyay, Akanksha Verma, R D Dixit, A M Kadri, Harsh Bakshi, Chaitanya Joshi, Madhvi Joshi, |
| EPI_ISL_467344, EPI_ISL_467345, EPI_ISL_467346, EPI_ISL_467347, EPI_ISL_467348, EPI_ISL_467349, EPI_ISL_467350, EPI_ISL_467351, EPI_ISL_467352, EPI_ISL_467353, EPI_ISL_467354, EPI_ISL_467355, EPI_ISL_467356, EPI_ISL_467357, EPI_ISL_467358, EPI_ISL_467359, EPI_ISL_467360, EPI_ISL_467361, EPI_ISL_467362, EPI_ISL_467363, EPI_ISL_467364, EPI_ISL_467365, EPI_ISL_467366, EPI_ISL_467367, EPI_ISL_467368, EPI_ISL_467369, EPI_ISL_467370, EPI_ISL_467371 | | | |
| see above | Laboratory of Respiratory Viruses and Measles, Oswaldo Cruz Institute, FIOCRUZ | Laboratory of Respiratory Viruses and Measles, Oswaldo Cruz Institute, FIOCRUZ | Paola Resende, Luciana Appolinario, Fernando Motta, Anna Carolina Paixão, Ana Carolina Mendonça, Aline Mattos, Milene Miranda, Cristiana Garcia, Braulia Caetano, Maria Ogrzewalska, Jonathan Lopes, Marilda Siqueira |
| EPI_ISL_467372, EPI_ISL_467373 | Arizona State University Health Services | Arizona State University | Peter T. Skidmore, Rabia Maqsood, LaRinda A. Holland, Emily A. Kaelin, Lily I. Wu, Arvind Varsani, Rolf U. Halden, Brenda G. Hogue, Matthew Scotch, Efrem S. Lim |
| EPI_ISL_467666 | Virology lab, NIC, NCCD, Ulaanbaatar, Mongolia | National Centre for Communicable Deseases (NCCD) | Naranzul Ts,Darmaa B,Bayasgalan N,Ankhbayar S,Tsogtbaatar B, Erdene-Ochir Ts,Nymadawa P |
| EPI_ISL_468066 | Physiology, Istanbul Medeniyet University | Physiology, Istanbul Medeniyet University | Pence,S., Caykara,B., Pence,H.H., Tekin,S., Yiyit,N., Cevher Keskin,B. and Kara,A. |
| EPI_ISL_468726 | unknown | Department of Microbiology | Peng,H., Tang,H., Jiang,L., Qi,Z., Zhao,P. |
| EPI_ISL_468736, EPI_ISL_468737, EPI_ISL_468738, EPI_ISL_468739, EPI_ISL_468740, EPI_ISL_468741, EPI_ISL_468742, EPI_ISL_468743, EPI_ISL_468744, EPI_ISL_468745, EPI_ISL_468746 | | | |
| see above | Lab voor klinische biologie | Onderzoeksgroep Virologie | Nick Vereecke, Laurens Lambrechts, Marthe Pauwels, Bruno Verhasselt, Linos Vandekerckhove, Hans Nauwynck, Sebastiaan Theuns |
| EPI_ISL_468914, EPI_ISL_469016, EPI_ISL_469018, EPI_ISL_469019, EPI_ISL_469020, EPI_ISL_469021, EPI_ISL_469022 | Istituto Zooprofilattico Sperimentale Puglia e Basilicata; Dipartimento di Bioscienze, Biotecnologie e Biofarmaceutica dell'Università degli Studi di Bari "A.Moro"; Istituto di Biomembrane. Bioenergetica e Biotecnologie Molecolari del Consiglio Nazionale delle Ricerche di Bari | Beaconlab (Bioinformatics, Evolution and Comparative Genomics lab), Dept of Biosciences, University on Milan | Parisi A.,Pesole G., Manzari C., Chiara M. |
| EPI_ISL_469023 | Istituto Zooprofilattico Sperimentale Puglia e Basilicata; Dipartimento di Bioscienze, Biotecnologie e Biofarmaceutica dell'Università degli Studi di Bari "A.Moro"; Istituto di Biomembrane. Bioenergetica e Biotecnologie Molecolari del Consiglio Nazionale delle Ricerche di Bari | Beaconlab (Bioinformatics, Evolution and Comparative Genomics lab), Dept of Biosciences, University on Milan | Parisi A.,Pesole G., Manzari C., Chiara M |
| EPI_ISL_469026 | B.J. Medical College and Civil hospital | Gujarat Biotechnology Research Centre | Pritesh Sabara, Apurvasinh Puvar, Janvi Raval, Zarna Patel, Monika Gandhi, Pinal Trivedi, Maharshi Pandya, Nidhi Patel, Nitin Savaliya, Raghawendra Kumar, Dinesh Kumar, Zuber Saiyed, Komal Patel, Labdhi Pandya, Snehal Bagatharia, Pranay Shah, Kamlesh J Upadhyay, Nirav Mungalpara, Tejas Shah, Ankit Hinsu, Neha Rajpara, R D Dixit, A M Kadri, Harsh Bakshi, Chaitanya Joshi, Madhvi Joshi |
| EPI_ISL_469031 | Government Medical College, Vadodara | Gujarat Biotechnology Research Centre | Pinal Trivedi, Maharshi Pandya, Nidhi Patel, Nitin Savaliya, Raghawendra Kumar, Dinesh Kumar, Zuber Saiyed, Komal Patel, Labdhi Pandya, Snehal Bagatharia, Meenakshi Shah, Neena Doshi, Varsha Godbole, Tejas Shah, Ankit Hinsu, Pritesh Sabara, Apurvasinh Puvar, Janvi Raval, Zarna Patel, Monika Gandhi, Bhavya Jindal, R D Dixit, A M Kadri, Harsh Bakshi, Chaitanya Joshi, Madhvi Joshi |
| EPI_ISL_469033 | Government Medical College, Vadodara | Gujarat Biotechnology Research Centre | Nidhi Patel, Nitin Savaliya, Raghawendra Kumar, Dinesh Kumar, Zuber Saiyed, Komal Patel, Labdhi Pandya, Snehal Bagatharia, Meenakshi Shah, Neena Doshi, Varsha Godbole, Tejas Shah, Ankit Hinsu, Pritesh Sabara, Apurvasinh Puvar, Janvi Raval, Zarna Patel, Monika Gandhi, Pinal Trivedi, Maharshi Pandya, Priyanka P Vatsa, R D Dixit, A M Kadri, Harsh Bakshi, Chaitanya Joshi, Madhvi Joshi |

| | | | |
|---|---|---|---|
| EPI_ISL_469034 | Government Medical College, Vadodara | Gujarat Biotechnology Research Centre | Nitin Savaliya, Raghawendra Kumar, Dinesh Kumar, Zuber Saiyed, Komal Patel, Labdhi Pandya, Snehal Bagatharia, Meenakshi Shah, Neena Doshi, Varsha Godbole, Tejas Shah, Ankit Hinsu, Pritesh Sabara, Apurvasinh Puvar, Janvi Raval, Zarna Patel, Monika Gandhi, Pinal Trivedi, Maharshi Pandya, Nidhi Patel, Pooja P Doshi, R D Dixit, A M Kadri, Harsh Bakshi, Chaitanya Joshi, Madhvi Joshi |
| EPI_ISL_469042 | GMERS Medical College & Hospital | Gujarat Biotechnology Research Centre | Neena Doshi, Varsha Godbole, Tejas Shah, Ankit Hinsu, Pritesh Sabara, Apurvasinh Puvar, Janvi Raval, Zarna Patel, Monika Gandhi, Pinal Trivedi, Maharshi Pandya, Nidhi Patel, Nitin Savaliya, Raghawendra Kumar, Dinesh Kumar, Zuber Saiyed, Komal Patel, Labdhi Pandya, Snehal Bagatharia, Meenakshi Shah, Armi Chaudhari, R D Dixit, A M Kadri, Harsh Bakshi, Chaitanya Joshi, Madhvi Joshi |
| EPI_ISL_469048 | Banas Medical College and Research Institute | Gujarat Biotechnology Research Centre | Radhika Khara, Sunil R Joshi, Viren s Doshi, Zarna Patel, Monika Gandhi, Pinal Trivedi, Maharshi Pandya, Nidhi Patel, Nitin Savaliya, Raghawendra Kumar, Dinesh Kumar, Zuber Saiyed, Komal Patel, Labdhi Pandya, Snehal Bagatharia, Tejas Shah, Ankit Hinsu, Pritesh Sabara, Apurvasinh Puvar, Janvi Raval, Priti Pandita, R D Dixit, A M Kadri, Harsh Bakshi, Chaitanya Joshi, Madhvi Joshi |
| EPI_ISL_469055, EPI_ISL_469056 | Jourcentralen | The Public Health Agency of Sweden | Oskar Karlsson Lindsjo, Maria Lind Karlberg, Mattias Haukland, Reza Advani, Olov Svartstrom, Anna-Malin Linde, Sandra Broddesson, Petra Edquist, Shamam Muradrasoli, Anna Risberg, Karin Tegmark-Wisell |
| EPI_ISL_469057 | Inger Landgren | The Public Health Agency of Sweden | Oskar Karlsson Lindsjo, Maria Lind Karlberg, Mattias Haukland, Reza Advani, Olov Svartstrom, Anna-Malin Linde, Sandra Broddesson, Petra Edquist, Shamam Muradrasoli, Anna Risberg, Karin Tegmark-Wisell |
| EPI_ISL_469058 | Narhalsan Sjobo vardcentral | The Public Health Agency of Sweden | Oskar Karlsson Lindsjo, Maria Lind Karlberg, Mattias Haukland, Reza Advani, Olov Svartstrom, Anna-Malin Linde, Sandra Broddesson, Petra Edquist, Shamam Muradrasoli, Anna Risberg, Karin Tegmark-Wisell |
| EPI_ISL_469059 | Hovas Askim Familjelakare och BVC | The Public Health Agency of Sweden | Oskar Karlsson Lindsjo, Maria Lind Karlberg, Mattias Haukland, Reza Advani, Olov Svartstrom, Anna-Malin Linde, Sandra Broddesson, Petra Edquist, Shamam Muradrasoli, Anna Risberg, Karin Tegmark-Wisell |
| EPI_ISL_469060, EPI_ISL_469061 | Narhalsan Sjobo vardcentral | The Public Health Agency of Sweden | Oskar Karlsson Lindsjo, Maria Lind Karlberg, Mattias Haukland, Reza Advani, Olov Svartstrom, Anna-Malin Linde, Sandra Broddesson, Petra Edquist, Shamam Muradrasoli, Anna Risberg, Karin Tegmark-Wisell |
| EPI_ISL_469062 | Huddinge VC | The Public Health Agency of Sweden | Oskar Karlsson Lindsjo, Maria Lind Karlberg, Mattias Haukland, Reza Advani, Olov Svartstrom, Anna-Malin Linde, Sandra Broddesson, Petra Edquist, Shamam Muradrasoli, Anna Risberg, Karin Tegmark-Wisell |
| EPI_ISL_469063 | Ulltuna Vardcentral | The Public Health Agency of Sweden | Oskar Karlsson Lindsjo, Maria Lind Karlberg, Mattias Haukland, Reza Advani, Olov Svartstrom, Anna-Malin Linde, Sandra Broddesson, Petra Edquist, Shamam Muradrasoli, Anna Risberg, Karin Tegmark-Wisell |
| EPI_ISL_469064, EPI_ISL_469065 | Huddinge VC | The Public Health Agency of Sweden | Oskar Karlsson Lindsjo, Maria Lind Karlberg, Mattias Haukland, Reza Advani, Olov Svartstrom, Anna-Malin Linde, Sandra Broddesson, Petra Edquist, Shamam Muradrasoli, Anna Risberg, Karin Tegmark-Wisell |
| EPI_ISL_469066 | Surbrunns VC | The Public Health Agency of Sweden | Oskar Karlsson Lindsjo, Maria Lind Karlberg, Mattias Haukland, Reza Advani, Olov Svartstrom, Anna-Malin Linde, Sandra Broddesson, Petra Edquist, Shamam Muradrasoli, Anna Risberg, Karin Tegmark-Wisell |
| EPI_ISL_469067 | Kungsholmsdoktorn | The Public Health Agency of Sweden | Oskar Karlsson Lindsjo, Maria Lind Karlberg, Mattias Haukland, Reza Advani, Olov Svartstrom, Anna-Malin Linde, Sandra Broddesson, Petra Edquist, Shamam Muradrasoli, Anna Risberg, Karin Tegmark-Wisell |
| EPI_ISL_469068 | Hovas Askim Familjelakare och BVC | The Public Health Agency of Sweden | Oskar Karlsson Lindsjo, Maria Lind Karlberg, Mattias Haukland, Reza Advani, Olov Svartstrom, Anna-Malin Linde, Sandra Broddesson, Petra Edquist, Shamam Muradrasoli, Anna Risberg, Karin Tegmark-Wisell |
| EPI_ISL_469069 | Narhalsan Olskroken VC | The Public Health Agency of Sweden | Oskar Karlsson Lindsjo, Maria Lind Karlberg, Mattias Haukland, Reza Advani, Olov Svartstrom, Anna-Malin Linde, Sandra Broddesson, Petra Edquist, Shamam Muradrasoli, Anna Risberg, Karin Tegmark-Wisell |
| EPI_ISL_469070 | Surbrunns VC | The Public Health Agency of Sweden | Oskar Karlsson Lindsjo, Maria Lind Karlberg, Mattias Haukland, Reza Advani, Olov Svartstrom, Anna-Malin Linde, Sandra Broddesson, Petra Edquist, Shamam Muradrasoli, Anna Risberg, Karin Tegmark-Wisell |
| EPI_ISL_469071 | Wasterlakarna | The Public Health Agency of Sweden | Oskar Karlsson Lindsjo, Maria Lind Karlberg, Mattias Haukland, Reza Advani, Olov Svartstrom, Anna-Malin Linde, Sandra Broddesson, Petra Edquist, Shamam Muradrasoli, Anna Risberg, Karin Tegmark-Wisell |
| EPI_ISL_469072 | Ulltuna Vardcentral | The Public Health Agency of Sweden | Oskar Karlsson Lindsjo, Maria Lind Karlberg, Mattias Haukland, Reza Advani, Olov Svartstrom, Anna-Malin Linde, Sandra Broddesson, Petra Edquist, Shamam Muradrasoli, Anna Risberg, Karin Tegmark-Wisell |
| EPI_ISL_469073, EPI_ISL_469074 | Halmstad klinisk mikrobiologi | The Public Health Agency of Sweden | Oskar Karlsson Lindsjo, Maria Lind Karlberg, Mattias Haukland, Reza Advani, Olov Svartstrom, Anna-Malin Linde, Sandra Broddesson, Petra Edquist, Shamam Muradrasoli, Anna Risberg, Karin Tegmark-Wisell |
| EPI_ISL_469075 | Karolinska Universitetslaboratoriet | The Public Health Agency of Sweden | Oskar Karlsson Lindsjo, Maria Lind Karlberg, Mattias Haukland, Reza Advani, Olov Svartstrom, Anna-Malin Linde, Sandra Broddesson, Petra Edquist, Shamam Muradrasoli, Anna Risberg, Karin Tegmark-Wisell |
| EPI_ISL_469076 | Uppsala klinisk mikrobiologi | The Public Health Agency of Sweden | Oskar Karlsson Lindsjo, Maria Lind Karlberg, Mattias Haukland, Reza Advani, Olov Svartstrom, Anna-Malin Linde, Sandra Broddesson, Petra Edquist, Shamam Muradrasoli, Anna Risberg, Karin Tegmark-Wisell |
| EPI_ISL_469077, EPI_ISL_469078, EPI_ISL_469079 | Karolinska Universitetslaboratoriet | The Public Health Agency of Sweden | Oskar Karlsson Lindsjo, Maria Lind Karlberg, Mattias Haukland, Reza Advani, Olov Svartstrom, Anna-Malin Linde, Sandra Broddesson, Petra Edquist, Shamam Muradrasoli, Anna Risberg, Karin Tegmark-Wisell |
| EPI_ISL_469253 | Second Military Medical University, Department of Microbiology | Second Military Medical University, Department of Microbiology | Peng,H., Tang,H., Jiang,L., Qi,Z. and Zhao,P. |
| EPI_ISL_471396, EPI_ISL_471397, EPI_ISL_471398, EPI_ISL_471399, EPI_ISL_471400, EPI_ISL_471401, EPI_ISL_471402, EPI_ISL_471403, EPI_ISL_471404, EPI_ISL_471405, EPI_ISL_471406, EPI_ISL_471407, EPI_ISL_471408, EPI_ISL_471409, EPI_ISL_471410, EPI_ISL_471411, EPI_ISL_471412, EPI_ISL_471413, EPI_ISL_471414, EPI_ISL_471415 | | | |
| see above | Viral Respiratory Lab, National Institute for Biomedical Research (INRB) | Pathogen Sequencing Lab, National Institute for Biomedical Research (INRB) | Placide Mbala-Kingebeni, Edith Nkwembe, Eddy Kinganda-Lusamaki, Amuri Aziza, Francisca Muyembe Mawete, Catherine Pratt, Matthias Pauthner, Josh Quick, Allison Black, James Hadfield, Trevor Bedford, Ian Goodfellow, Andrew Rambaut, Nick Loman, Kristian Andersen, Michael Wiley, Steve Ahuka-Mundeke, Jean-Jacques Muyembe Tamfum |
| EPI_ISL_471510, EPI_ISL_471511, EPI_ISL_471512, EPI_ISL_471513, EPI_ISL_471514, EPI_ISL_471515, EPI_ISL_471516, EPI_ISL_471517, EPI_ISL_471518, EPI_ISL_471519, EPI_ISL_471520, EPI_ISL_471521, EPI_ISL_471522, EPI_ISL_471523, EPI_ISL_471524, EPI_ISL_471525, EPI_ISL_471526, EPI_ISL_471527 | | | |
| see above | Respiratory Virus Unit, Microbiology Services Colindale, Public Health England | Respiratory Virus Unit, Microbiology Services Colindale, Public Health England | PHE Covid Sequencing Team |
| EPI_ISL_471595 | CSIR-Centre for Cellular and Molecular Biology | CSIR-Centre for Cellular and Molecular Biology | Payel Mukherjee, Sofia Banu, Priya Singh, Dhiviya Vedagiri, Divya Gupta, Vishal Sah, Santosh Kumar Kuncha, Krishnan Harinivas Harshan, Archana Bharadwaj Siva, Karthik Bharadwaj Tallapaka, Shagufta Khan, Lamuk Zaveri, Namami Gaur, Sakshi Shambhavi, Tulasi Nagabandi, Purushotham Vodnala, G. Aditya Kumar, Koushick Sivakumar, Pooja Ramesh Gupta, Rajan Kumar Jha, Shraddha Vijay Lahoti, Rakesh K Mishra, Divya Tej Sowpati |
| EPI_ISL_471596 | CSIR-Centre for Cellular and Molecular Biology | CSIR-Centre for Cellular and Molecular Biology | Payel Mukherjee, Sofia Banu, Priya Singh, Dhiviya Vedagiri, Divya Gupta, Vishal Sah, Santosh Kumar Kuncha, Krishnan Harinivas Harshan, Archana Bharadwaj Siva, Karthik Bharadwaj Tallapaka, Shagufta Khan, Lamuk Zaveri, Namami Gaur, Sakshi Shambhavi, Tulasi Nagabandi, Purushotham Vodnala, Gokulan C G, Gunjan Purohit, Hanuman Tulashiram Kale, Pankaj Kumar, Prachand Issarapu, Rakesh K Mishra, Divya Tej Sowpati |
| EPI_ISL_471597 | CSIR-Centre for Cellular and Molecular Biology | CSIR-Centre for Cellular and Molecular Biology | Payel Mukherjee, Sofia Banu, Priya Singh, Dhiviya Vedagiri, Divya Gupta, Vishal Sah, Santosh Kumar Kuncha, Krishnan Harinivas Harshan, Archana Bharadwaj Siva, Karthik Bharadwaj Tallapaka, Shagufta Khan, Lamuk Zaveri, Namami Gaur, Sakshi Shambhavi, Tulasi Nagabandi, Purushotham Vodnala, Rakesh K Mishra, Sonu Uday, Sudipta Mondal, Annapoorna P Karthyayani, Debabrata Jana, Debrya Saha, Divya Tej Sowpati |
| EPI_ISL_471598 | CSIR-Centre for Cellular and Molecular Biology | CSIR-Centre for Cellular and Molecular Biology | Payel Mukherjee, Sofia Banu, Priya Singh, Dhiviya Vedagiri, Divya Gupta, Vishal Sah, Santosh Kumar Kuncha, Krishnan Harinivas Harshan, Archana Bharadwaj Siva, Karthik Bharadwaj Tallapaka, Shagufta Khan, Lamuk Zaveri, Namami Gaur, Sakshi Shambhavi, Tulasi Nagabandi, Purushotham Vodnala,Deepak Kumar, Devi Prasad Vijayashankar, Disha Nanda, Divya Das, Jotin Gogoi, Manish Bhattacharjee, Rakesh K Mishra, Divya Tej Sowpati |
| EPI_ISL_471622 | CSIR-Centre for Cellular and Molecular Biology | CSIR-Centre for Cellular and Molecular Biology | Namami Gaur, Sakshi Shambhavi, Lamuk Zaveri, Shagufta Khan, Tulasi Nagabandi, Purushotham Vodnala, Payel Mukherjee, Sofia Banu, Priya Singh, Dhiviya Vedagiri, Divya Gupta, Vishal Sah, Santosh Kumar Kuncha, Krishnan Harinivas Harshan, Archana Bharadwaj Siva, Karthik Bharadwaj Tallapaka,Kezia J Ann, Radhika Khandelwal, Roshan Maku Venkata, Shemin Mansuri, Sonu Uday, Rakesh K Mishra, Divya Tej Sowpati |

| | | | |
|---|---|---|---|
| EPI_ISL_471623 | CSIR-Centre for Cellular and Molecular Biology | CSIR-Centre for Cellular and Molecular Biology | Payel Mukherjee, Sofia Banu, Priya Singh, Dhiviya Vedagiri, Divya Gupta, Vishal Sah, Santosh Kumar Kuncha, Krishnan Harinivas Harshan, Archana Bharadwaj Siva, Karthik Bharadwaj Tallapaka, Shagufta Khan, Lamuk Zaveri, Namami Gaur, Sakshi Shambhavi, Tulasi Nagabandi, Purushotham Vodnala, G. Aditya Kumar, Koushick Sivakumar, Pooja Ramesh Gupta, Rajan Kumar Jha, Shraddha Vijay Lahoti, Rakesh K Mishra, Divya Tej Sowpati |
| EPI_ISL_471624 | CSIR-Centre for Cellular and Molecular Biology | CSIR-Centre for Cellular and Molecular Biology | Payel Mukherjee, Sofia Banu, Priya Singh, Dhiviya Vedagiri, Divya Gupta, Vishal Sah, Santosh Kumar Kuncha, Krishnan Harinivas Harshan, Archana Bharadwaj Siva, Karthik Bharadwaj Tallapaka, Shagufta Khan, Lamuk Zaveri, Namami Gaur, Sakshi Shambhavi, Tulasi Nagabandi, Purushotham Vodnala, Gokulan C G, Gunjan Purohit, Hanuman Tulashiram Kale, Pankaj Kumar, Prachand Issarapu, Rakesh K Mishra, Divya Tej Sowpati |
| EPI_ISL_471625 | CSIR-Centre for Cellular and Molecular Biology | CSIR-Centre for Cellular and Molecular Biology | Payel Mukherjee, Sofia Banu, Priya Singh, Dhiviya Vedagiri, Divya Gupta, Vishal Sah, Santosh Kumar Kuncha, Krishnan Harinivas Harshan, Archana Bharadwaj Siva, Karthik Bharadwaj Tallapaka, Shagufta Khan, Lamuk Zaveri, Namami Gaur, Sakshi Shambhavi, Tulasi Nagabandi, Purushotham Vodnala, Rakesh K Mishra, Sonu Uday, Sudipta Mondal, Annapoorna P Karthyayani, Debabrata Jana, Debrya Saha, Divya Tej Sowpati |
| EPI_ISL_471626 | CSIR-Centre for Cellular and Molecular Biology | CSIR-Centre for Cellular and Molecular Biology | Payel Mukherjee, Sofia Banu, Priya Singh, Dhiviya Vedagiri, Divya Gupta, Vishal Sah, Santosh Kumar Kuncha, Krishnan Harinivas Harshan, Archana Bharadwaj Siva, Karthik Bharadwaj Tallapaka, Shagufta Khan, Lamuk Zaveri, Namami Gaur, Sakshi Shambhavi, Tulasi Nagabandi, Purushotham Vodnala,Deepak Kumar, Devi Prasad Vijayashankar, Disha Nanda, Divya Das, Jotin Gogoi, Manish Bhattacharjee, Rakesh K Mishra, Divya Tej Sowpati |
| EPI_ISL_472430, EPI_ISL_472431 | Queens Medical Centre, Clinical Microbiology Department / DeepSeq Nottingham | COVID-19 Genomics UK (COG-UK) Consortium | Nichola Duckworth, Tim Sloan, Sarah Walsh, Jonathan Ball, Patrick McClure, Joeseph Chappell, Nadine Holmes, Matthew Carlisle, Christopher Moore, Fei Sang, Johnny Debebe, Victoria Wright, Matthew Loose |
| EPI_ISL_474958 | Israeli Central Virology laboratory | Israel Central Virology laboratory | Neta Zuckerman, Efrat Dahan Bucris, Oran Erster, Ella Mendelson, Michal Mandelboim |
| EPI_ISL_474959, EPI_ISL_474960, EPI_ISL_474961, EPI_ISL_474962, EPI_ISL_474963, EPI_ISL_474964, EPI_ISL_474965, EPI_ISL_474966, EPI_ISL_474967, EPI_ISL_474968, EPI_ISL_474969, EPI_ISL_474970, EPI_ISL_474971, EPI_ISL_474972, EPI_ISL_474973, EPI_ISL_474974, EPI_ISL_474975, EPI_ISL_474976, EPI_ISL_474977, EPI_ISL_474978, EPI_ISL_474979, EPI_ISL_474980, EPI_ISL_474981, EPI_ISL_474982, EPI_ISL_474983, EPI_ISL_474984, EPI_ISL_474985, EPI_ISL_474986, EPI_ISL_474987, EPI_ISL_474988, EPI_ISL_474989, EPI_ISL_474990, EPI_ISL_474991, EPI_ISL_474992, EPI_ISL_474993, EPI_ISL_474994, EPI_ISL_474996, EPI_ISL_474997, EPI_ISL_474998, EPI_ISL_474999, EPI_ISL_475000, EPI_ISL_475001, EPI_ISL_475002, EPI_ISL_475003, EPI_ISL_475004, EPI_ISL_475005, EPI_ISL_475006, EPI_ISL_475007, EPI_ISL_475008, EPI_ISL_475009, EPI_ISL_475010, EPI_ISL_475011, EPI_ISL_475012, EPI_ISL_475013, EPI_ISL_475014, EPI_ISL_475015, EPI_ISL_475016, EPI_ISL_475017, EPI_ISL_475018, EPI_ISL_475019, EPI_ISL_475020, EPI_ISL_475021, EPI_ISL_475022, EPI_ISL_475023, EPI_ISL_475024, EPI_ISL_475025 | | | |
| see above | Israel Central Virology laboratory | Israel Central Virology laboratory | Neta Zuckerman, Efrat Dahan Bucris, Oran Erster, Ella Mendelson, Michal Mandelboim |
| EPI_ISL_475028 | Banas Medical College and Research Institute | Gujarat Biotechnology Research Centre | Pritesh Sabara, Apurvasinh Puvar, Janvi Raval, Zarna Patel, Monika Gandhi, Pinal Trivedi, Maharshi Pandya, Nidhi Patel, Nitin Savaliya, Raghawendra Kumar, Dinesh Kumar, Zuber Saiyed, Komal Patel, Labdhi Pandya, Snehal Bagatharia, Radhika Khara, Sunil R Joshi, Viren s Doshi, Fenil Patel, R D Dixit, A M Kadri, Harsh Bakshi, Chaitanya Joshi, Madhvi Joshi |
| EPI_ISL_475033 | Department of MicroBiology, Government Medical College, Surat | Gujarat Biotechnology Research Centre | Pinal Trivedi, Maharshi Pandya, Nidhi Patel, Nitin Savaliya, Raghawendra Kumar, Dinesh Kumar, Zuber Saiyed, Komal Patel, Labdhi Pandya, Snehal Bagatharia, Naresh Chauhan, Summaiya Mullan, Amit gamit, Pritesh Sabara, Apurvasinh Puvar, Janvi Raval, Zarna Patel, Monika Gandhi, Pooja P Doshi, R D Dixit, A M Kadri, Harsh Bakshi, Chaitanya Joshi, Madhvi Joshi |
| EPI_ISL_475035 | Department of MicroBiology, Government Medical College, Surat | Gujarat Biotechnology Research Centre | Nidhi Patel, Nitin Savaliya, Raghawendra Kumar, Dinesh Kumar, Zuber Saiyed, Komal Patel, Labdhi Pandya, Snehal Bagatharia, Naresh Chauhan, Summaiya Mullan, Amit gamit, Pritesh Sabara, Apurvasinh Puvar, Janvi Raval, Zarna Patel, Monika Gandhi, Pinal Trivedi, Maharshi Pandya, Priti Pandita, R D Dixit, A M Kadri, Harsh Bakshi, Chaitanya Joshi, Madhvi Joshi |
| EPI_ISL_475036 | Department of MicroBiology, Government Medical College, Surat | Gujarat Biotechnology Research Centre | Nitin Savaliya, Raghawendra Kumar, Dinesh Kumar, Zuber Saiyed, Komal Patel, Labdhi Pandya, Snehal Bagatharia, Naresh Chauhan, Summaiya Mullan, Amit gamit, Pritesh Sabara, Apurvasinh Puvar, Janvi Raval, Zarna Patel, Monika Gandhi, Pinal Trivedi, Maharshi Pandya, Nidhi Patel, Pragya Sharma, R D Dixit, A M Kadri, Harsh Bakshi, Chaitanya Joshi, Madhvi Joshi |
| EPI_ISL_475043 | Department of MicroBiology, Government Medical College, Surat | Gujarat Biotechnology Research Centre | Naresh Chauhan, Summaiya Mullan, Amit gamit, Pritesh Sabara, Apurvasinh Puvar, Janvi Raval, Zarna Patel, Monika Gandhi, Pinal Trivedi, Maharshi Pandya, Nidhi Patel, Nitin Savaliya, Raghawendra Kumar, Dinesh Kumar, Zuber Saiyed, Komal Patel, Labdhi Pandya, Snehal Bagatharia, Priyanka P Vatsa, R D Dixit, A M Kadri, Harsh Bakshi, Chaitanya Joshi, Madhvi Joshi |
| EPI_ISL_475046 | Department of MicroBiology, Government Medical College, Surat | Gujarat Biotechnology Research Centre | Pritesh Sabara, Apurvasinh Puvar, Janvi Raval, Zarna Patel, Monika Gandhi, Pinal Trivedi, Maharshi Pandya, Nidhi Patel, Nitin Savaliya, Raghawendra Kumar, Dinesh Kumar, Zuber Saiyed, Komal Patel, Labdhi Pandya, Snehal Bagatharia, Naresh Chauhan, Summaiya Mullan, Amit gamit, Priti Pandita, R D Dixit, A M Kadri, Harsh Bakshi, Chaitanya Joshi, Madhvi Joshi |
| EPI_ISL_475051 | GMERS Medical College & Hospital | Gujarat Biotechnology Research Centre | Pinal Trivedi, Maharshi Pandya, Nidhi Patel, Nitin Savaliya, Raghawendra Kumar, Dinesh Kumar, Zuber Saiyed, Komal Patel, Labdhi Pandya, Snehal Bagatharia, Meenakshi Shah, Neena Doshi, Varsha Godbole, Pritesh Sabara, Apurvasinh Puvar, Janvi Raval, Zarna Patel, Monika Gandhi, Neelam Nathani, R D Dixit, A M Kadri, Harsh Bakshi, Chaitanya Joshi, Madhvi Joshi |
| EPI_ISL_475053 | GMERS Medical College & Hospital | Gujarat Biotechnology Research Centre | Nidhi Patel, Nitin Savaliya, Raghawendra Kumar, Dinesh Kumar, Zuber Saiyed, Komal Patel, Labdhi Pandya, Snehal Bagatharia, Meenakshi Shah, Neena Doshi, Varsha Godbole, Pritesh Sabara, Apurvasinh Puvar, Janvi Raval, Zarna Patel, Monika Gandhi, Pinal Trivedi, Maharshi Pandya, Bhavya Jindal, R D Dixit, A M Kadri, Harsh Bakshi, Chaitanya Joshi, Madhvi Joshi |
| EPI_ISL_475054 | GMERS Medical College & Hospital | Gujarat Biotechnology Research Centre | Nitin Savaliya, Raghawendra Kumar, Dinesh Kumar, Zuber Saiyed, Komal Patel, Labdhi Pandya, Snehal Bagatharia, Meenakshi Shah, Neena Doshi, Varsha Godbole, Pritesh Sabara, Apurvasinh Puvar, Janvi Raval, Zarna Patel, Monika Gandhi, Pinal Trivedi, Maharshi Pandya, Nidhi Patel, Priyanka P Vatsa, R D Dixit, A M Kadri, Harsh Bakshi, Chaitanya Joshi, Madhvi Joshi |
| EPI_ISL_475071, EPI_ISL_475072, EPI_ISL_475073, EPI_ISL_475074, EPI_ISL_475075, EPI_ISL_475076, EPI_ISL_475077, EPI_ISL_475078, EPI_ISL_475079, EPI_ISL_475080, EPI_ISL_475081, EPI_ISL_475082 | | | |
| see above | Lab voor klinische biologie | Onderzoeksgroep Virologie | Nick Vereecke, Laurens Lambrechts, Marthe Pauwels, Bruno Verhasselt, Linos Vandekerckhove, Hans Nauwynck, Sebastiaan Theuns |
| EPI_ISL_475085 | Skovde/Unilabs | The Public Health Agency of Sweden | Oskar Karlsson Lindsjo, Maria Lind Karlberg, Mattias Haukland, Reza Advani, Olov Svartstrom, Anna-Malin Linde, Sandra Broddesson, Petra Edquist, Shamam Muradrasoli, Anna Risberg, Karin Tegmark-Wisell |
| EPI_ISL_475086, EPI_ISL_475087, EPI_ISL_475088, EPI_ISL_475089, EPI_ISL_475090, EPI_ISL_475091 | Karolinska Universitetslaboratoriet | The Public Health Agency of Sweden | Oskar Karlsson Lindsjo, Maria Lind Karlberg, Mattias Haukland, Reza Advani, Olov Svartstrom, Anna-Malin Linde, Sandra Broddesson, Petra Edquist, Shamam Muradrasoli, Anna Risberg, Karin Tegmark-Wisell |
| EPI_ISL_475092 | Skovde/Unilabs | The Public Health Agency of Sweden | Oskar Karlsson Lindsjo, Maria Lind Karlberg, Mattias Haukland, Reza Advani, Olov Svartstrom, Anna-Malin Linde, Sandra Broddesson, Petra Edquist, Shamam Muradrasoli, Anna Risberg, Karin Tegmark-Wisell |
| EPI_ISL_475093, EPI_ISL_475094 | Halmstad klinisk mikrobiologi | The Public Health Agency of Sweden | Oskar Karlsson Lindsjo, Maria Lind Karlberg, Mattias Haukland, Reza Advani, Olov Svartstrom, Anna-Malin Linde, Sandra Broddesson, Petra Edquist, Shamam Muradrasoli, Anna Risberg, Karin Tegmark-Wisell |
| EPI_ISL_475095 | Karolinska Universitetslaboratoriet | The Public Health Agency of Sweden | Oskar Karlsson Lindsjo, Maria Lind Karlberg, Mattias Haukland, Reza Advani, Olov Svartstrom, Anna-Malin Linde, Sandra Broddesson, Petra Edquist, Shamam Muradrasoli, Anna Risberg, Karin Tegmark-Wisell |
| EPI_ISL_475096, EPI_ISL_475097, EPI_ISL_475098 | Halmstad klinisk mikrobiologi | The Public Health Agency of Sweden | Oskar Karlsson Lindsjo, Maria Lind Karlberg, Mattias Haukland, Reza Advani, Olov Svartstrom, Anna-Malin Linde, Sandra Broddesson, Petra Edquist, Shamam Muradrasoli, Anna Risberg, Karin Tegmark-Wisell |
| EPI_ISL_475099 | Skovde/Unilabs | The Public Health Agency of Sweden | Oskar Karlsson Lindsjo, Maria Lind Karlberg, Mattias Haukland, Reza Advani, Olov Svartstrom, Anna-Malin Linde, Sandra Broddesson, Petra Edquist, Shamam Muradrasoli, Anna Risberg, Karin Tegmark-Wisell |
| EPI_ISL_475100 | Halmstad klinisk mikrobiologi | The Public Health Agency of Sweden | Oskar Karlsson Lindsjo, Maria Lind Karlberg, Mattias Haukland, Reza Advani, Olov Svartstrom, Anna-Malin Linde, Sandra Broddesson, Petra Edquist, Shamam Muradrasoli, Anna Risberg, Karin Tegmark-Wisell |
| EPI_ISL_475101, EPI_ISL_475102, EPI_ISL_475103, EPI_ISL_475104 | Klinisk Mikrobiologi | The Public Health Agency of Sweden | Oskar Karlsson Lindsjo, Maria Lind Karlberg, Mattias Haukland, Reza Advani, Olov Svartstrom, Anna-Malin Linde, Sandra Broddesson, Petra Edquist, Shamam Muradrasoli, Anna Risberg, Karin Tegmark-Wisell |
| EPI_ISL_475105, EPI_ISL_475106, EPI_ISL_475107, EPI_ISL_475108, EPI_ISL_475109 | Skovde/Unilabs | The Public Health Agency of Sweden | Oskar Karlsson Lindsjo, Maria Lind Karlberg, Mattias Haukland, Reza Advani, Olov Svartstrom, Anna-Malin Linde, Sandra Broddesson, Petra Edquist, Shamam Muradrasoli, Anna Risberg, Karin Tegmark-Wisell |
| EPI_ISL_475110 | Gavle klinisk mikrobiologi | The Public Health Agency of Sweden | Oskar Karlsson Lindsjo, Maria Lind Karlberg, Mattias Haukland, Reza Advani, Olov Svartstrom, Anna-Malin Linde, Sandra Broddesson, Petra Edquist, Shamam Muradrasoli, Anna Risberg, Karin Tegmark-Wisell |

| | | | |
|---|---|---|---|
| EPI_ISL_475111, EPI_ISL_475112, EPI_ISL_475113 | Skovde/Unilabs | The Public Health Agency of Sweden | Oskar Karlsson Lindsjo, Maria Lind Karlberg, Mattias Haukland, Reza Advani, Olov Svartstrom, Anna-Malin Linde, Sandra Broddesson, Petra Edquist, Shamam Muradrasoli, Anna Risberg, Karin Tegmark-Wisell |
| EPI_ISL_475114 | Halmstad klinisk mikrobiologi | The Public Health Agency of Sweden | Oskar Karlsson Lindsjo, Maria Lind Karlberg, Mattias Haukland, Reza Advani, Olov Svartstrom, Anna-Malin Linde, Sandra Broddesson, Petra Edquist, Shamam Muradrasoli, Anna Risberg, Karin Tegmark-Wisell |
| EPI_ISL_475115 | Gavle klinisk mikrobiologi | The Public Health Agency of Sweden | Oskar Karlsson Lindsjo, Maria Lind Karlberg, Mattias Haukland, Reza Advani, Olov Svartstrom, Anna-Malin Linde, Sandra Broddesson, Petra Edquist, Shamam Muradrasoli, Anna Risberg, Karin Tegmark-Wisell |
| EPI_ISL_475116, EPI_ISL_475117 | Halmstad klinisk mikrobiologi | The Public Health Agency of Sweden | Oskar Karlsson Lindsjo, Maria Lind Karlberg, Mattias Haukland, Reza Advani, Olov Svartstrom, Anna-Malin Linde, Sandra Broddesson, Petra Edquist, Shamam Muradrasoli, Anna Risberg, Karin Tegmark-Wisell |
| EPI_ISL_475118, EPI_ISL_475119 | Uppsala klinisk mikrobiologi | The Public Health Agency of Sweden | Oskar Karlsson Lindsjo, Maria Lind Karlberg, Mattias Haukland, Reza Advani, Olov Svartstrom, Anna-Malin Linde, Sandra Broddesson, Petra Edquist, Shamam Muradrasoli, Anna Risberg, Karin Tegmark-Wisell |
| EPI_ISL_475120, EPI_ISL_475121 | Halmstad klinisk mikrobiologi | The Public Health Agency of Sweden | Oskar Karlsson Lindsjo, Maria Lind Karlberg, Mattias Haukland, Reza Advani, Olov Svartstrom, Anna-Malin Linde, Sandra Broddesson, Petra Edquist, Shamam Muradrasoli, Anna Risberg, Karin Tegmark-Wisell |
| EPI_ISL_475122 | Umea klinisk mikrobiologi | The Public Health Agency of Sweden | Oskar Karlsson Lindsjo, Maria Lind Karlberg, Mattias Haukland, Reza Advani, Olov Svartstrom, Anna-Malin Linde, Sandra Broddesson, Petra Edquist, Shamam Muradrasoli, Anna Risberg, Karin Tegmark-Wisell |
| EPI_ISL_475123 | Halmstad klinisk mikrobiologi | The Public Health Agency of Sweden | Oskar Karlsson Lindsjo, Maria Lind Karlberg, Mattias Haukland, Reza Advani, Olov Svartstrom, Anna-Malin Linde, Sandra Broddesson, Petra Edquist, Shamam Muradrasoli, Anna Risberg, Karin Tegmark-Wisell |
| EPI_ISL_475124 | Ostersund klinisk mikrobiologi | The Public Health Agency of Sweden | Oskar Karlsson Lindsjo, Maria Lind Karlberg, Mattias Haukland, Reza Advani, Olov Svartstrom, Anna-Malin Linde, Sandra Broddesson, Petra Edquist, Shamam Muradrasoli, Anna Risberg, Karin Tegmark-Wisell |
| EPI_ISL_475125 | Halmstad klinisk mikrobiologi | The Public Health Agency of Sweden | Oskar Karlsson Lindsjo, Maria Lind Karlberg, Mattias Haukland, Reza Advani, Olov Svartstrom, Anna-Malin Linde, Sandra Broddesson, Petra Edquist, Shamam Muradrasoli, Anna Risberg, Karin Tegmark-Wisell |
| EPI_ISL_475126, EPI_ISL_475127 | Umea klinisk mikrobiologi | The Public Health Agency of Sweden | Oskar Karlsson Lindsjo, Maria Lind Karlberg, Mattias Haukland, Reza Advani, Olov Svartstrom, Anna-Malin Linde, Sandra Broddesson, Petra Edquist, Shamam Muradrasoli, Anna Risberg, Karin Tegmark-Wisell |
| EPI_ISL_475128, EPI_ISL_475129, EPI_ISL_475130, EPI_ISL_475131, EPI_ISL_475132, EPI_ISL_475133, EPI_ISL_475134, EPI_ISL_475135 | Orebro klinisk mikrobiologi | The Public Health Agency of Sweden | Oskar Karlsson Lindsjo, Maria Lind Karlberg, Mattias Haukland, Reza Advani, Olov Svartstrom, Anna-Malin Linde, Sandra Broddesson, Petra Edquist, Shamam Muradrasoli, Anna Risberg, Karin Tegmark-Wisell |
| EPI_ISL_475136 | Karolinska Universitetslaboratoriet | The Public Health Agency of Sweden | Oskar Karlsson Lindsjo, Maria Lind Karlberg, Mattias Haukland, Reza Advani, Olov Svartstrom, Anna-Malin Linde, Sandra Broddesson, Petra Edquist, Shamam Muradrasoli, Anna Risberg, Karin Tegmark-Wisell |
| EPI_ISL_475137, EPI_ISL_475138 | Kalmar klinisk mikrobiologi | The Public Health Agency of Sweden | Oskar Karlsson Lindsjo, Maria Lind Karlberg, Mattias Haukland, Reza Advani, Olov Svartstrom, Anna-Malin Linde, Sandra Broddesson, Petra Edquist, Shamam Muradrasoli, Anna Risberg, Karin Tegmark-Wisell |
| EPI_ISL_475139 | Orebro klinisk mikrobiologi | The Public Health Agency of Sweden | Oskar Karlsson Lindsjo, Maria Lind Karlberg, Mattias Haukland, Reza Advani, Olov Svartstrom, Anna-Malin Linde, Sandra Broddesson, Petra Edquist, Shamam Muradrasoli, Anna Risberg, Karin Tegmark-Wisell |
| EPI_ISL_475140 | Skovde/Unilabs | The Public Health Agency of Sweden | Oskar Karlsson Lindsjo, Maria Lind Karlberg, Mattias Haukland, Reza Advani, Olov Svartstrom, Anna-Malin Linde, Sandra Broddesson, Petra Edquist, Shamam Muradrasoli, Anna Risberg, Karin Tegmark-Wisell |
| EPI_ISL_475141, EPI_ISL_475142 | Karolinska Universitetslaboratoriet | The Public Health Agency of Sweden | Oskar Karlsson Lindsjo, Maria Lind Karlberg, Mattias Haukland, Reza Advani, Olov Svartstrom, Anna-Malin Linde, Sandra Broddesson, Petra Edquist, Shamam Muradrasoli, Anna Risberg, Karin Tegmark-Wisell |
| EPI_ISL_475143 | Umea klinisk mikrobiologi | The Public Health Agency of Sweden | Oskar Karlsson Lindsjo, Maria Lind Karlberg, Mattias Haukland, Reza Advani, Olov Svartstrom, Anna-Malin Linde, Sandra Broddesson, Petra Edquist, Shamam Muradrasoli, Anna Risberg, Karin Tegmark-Wisell |
| EPI_ISL_475144, EPI_ISL_475145, EPI_ISL_475146, EPI_ISL_475147, EPI_ISL_475148 | Klinisk mikrobiologi Vasternorrland | The Public Health Agency of Sweden | Oskar Karlsson Lindsjo, Maria Lind Karlberg, Mattias Haukland, Reza Advani, Olov Svartstrom, Anna-Malin Linde, Sandra Broddesson, Petra Edquist, Shamam Muradrasoli, Anna Risberg, Karin Tegmark-Wisell |
| EPI_ISL_475149, EPI_ISL_475150 | Karolinska Universitetslaboratoriet | The Public Health Agency of Sweden | Oskar Karlsson Lindsjo, Maria Lind Karlberg, Mattias Haukland, Reza Advani, Olov Svartstrom, Anna-Malin Linde, Sandra Broddesson, Petra Edquist, Shamam Muradrasoli, Anna Risberg, Karin Tegmark-Wisell |
| EPI_ISL_475151 | Uppsala klinisk mikrobiologi | The Public Health Agency of Sweden | Oskar Karlsson Lindsjo, Maria Lind Karlberg, Mattias Haukland, Reza Advani, Olov Svartstrom, Anna-Malin Linde, Sandra Broddesson, Petra Edquist, Shamam Muradrasoli, Anna Risberg, Karin Tegmark-Wisell |
| EPI_ISL_475152 | Folkhalsomyndigheten | The Public Health Agency of Sweden | Oskar Karlsson Lindsjo, Maria Lind Karlberg, Mattias Haukland, Reza Advani, Olov Svartstrom, Anna-Malin Linde, Sandra Broddesson, Petra Edquist, Shamam Muradrasoli, Anna Risberg, Karin Tegmark-Wisell |
| EPI_ISL_475153, EPI_ISL_475154, EPI_ISL_475155 | Klinisk mikrobiologi Vasternorrland | The Public Health Agency of Sweden | Oskar Karlsson Lindsjo, Maria Lind Karlberg, Mattias Haukland, Reza Advani, Olov Svartstrom, Anna-Malin Linde, Sandra Broddesson, Petra Edquist, Shamam Muradrasoli, Anna Risberg, Karin Tegmark-Wisell |
| EPI_ISL_475156, EPI_ISL_475157, EPI_ISL_475158, EPI_ISL_475159 | Halmstad klinisk mikrobiologi | The Public Health Agency of Sweden | Oskar Karlsson Lindsjo, Maria Lind Karlberg, Mattias Haukland, Reza Advani, Olov Svartstrom, Anna-Malin Linde, Sandra Broddesson, Petra Edquist, Shamam Muradrasoli, Anna Risberg, Karin Tegmark-Wisell |
| EPI_ISL_475160, EPI_ISL_475161, EPI_ISL_475162, EPI_ISL_475163 | Klinisk mikrobiologi Vasternorrland | The Public Health Agency of Sweden | Oskar Karlsson Lindsjo, Maria Lind Karlberg, Mattias Haukland, Reza Advani, Olov Svartstrom, Anna-Malin Linde, Sandra Broddesson, Petra Edquist, Shamam Muradrasoli, Anna Risberg, Karin Tegmark-Wisell |
| EPI_ISL_475164 | Halmstad klinisk mikrobiologi | The Public Health Agency of Sweden | Oskar Karlsson Lindsjo, Maria Lind Karlberg, Mattias Haukland, Reza Advani, Olov Svartstrom, Anna-Malin Linde, Sandra Broddesson, Petra Edquist, Shamam Muradrasoli, Anna Risberg, Karin Tegmark-Wisell |
| EPI_ISL_475511 | Orestadsklinikens VC | The Public Health Agency of Sweden | Oskar Karlsson Lindsjo, Maria Lind Karlberg, Mattias Haukland, Reza Advani, Olov Svartstrom, Anna-Malin Linde, Sandra Broddesson, Mia Brytting, Anna Risberg, Karin Tegmark-Wisell |
| EPI_ISL_475512 | Din Klinik | The Public Health Agency of Sweden | Oskar Karlsson Lindsjo, Maria Lind Karlberg, Mattias Haukland, Reza Advani, Olov Svartstrom, Anna-Malin Linde, Sandra Broddesson, Mia Brytting, Anna Risberg, Karin Tegmark-Wisell |
| EPI_ISL_475513 | Huddinge VC | The Public Health Agency of Sweden | Oskar Karlsson Lindsjo, Maria Lind Karlberg, Mattias Haukland, Reza Advani, Olov Svartstrom, Anna-Malin Linde, Sandra Broddesson, Mia Brytting, Anna Risberg, Karin Tegmark-Wisell |
| EPI_ISL_475514 | Uppsala Narakut Aleris | The Public Health Agency of Sweden | Oskar Karlsson Lindsjo, Maria Lind Karlberg, Mattias Haukland, Reza Advani, Olov Svartstrom, Anna-Malin Linde, Sandra Broddesson, Mia Brytting, Anna Risberg, Karin Tegmark-Wisell |
| EPI_ISL_475515 | Lakargruppen | The Public Health Agency of Sweden | Oskar Karlsson Lindsjo, Maria Lind Karlberg, Mattias Haukland, Reza Advani, Olov Svartstrom, Anna-Malin Linde, Sandra Broddesson, Mia Brytting, Anna Risberg, Karin Tegmark-Wisell |
| EPI_ISL_475516, EPI_ISL_475517 | Uppsala Narakut Aleris | The Public Health Agency of Sweden | Oskar Karlsson Lindsjo, Maria Lind Karlberg, Mattias Haukland, Reza Advani, Olov Svartstrom, Anna-Malin Linde, Sandra Broddesson, Mia Brytting, Anna Risberg, Karin Tegmark-Wisell |
| EPI_ISL_475518 | Trollbackens VC | The Public Health Agency of Sweden | Oskar Karlsson Lindsjo, Maria Lind Karlberg, Mattias Haukland, Reza Advani, Olov Svartstrom, Anna-Malin Linde, Sandra Broddesson, Mia Brytting, Anna Risberg, Karin Tegmark-Wisell |
| EPI_ISL_475519 | Orsa VC | The Public Health Agency of Sweden | Oskar Karlsson Lindsjo, Maria Lind Karlberg, Mattias Haukland, Reza Advani, Olov Svartstrom, Anna-Malin Linde, Sandra Broddesson, Mia Brytting, Anna Risberg, Karin Tegmark-Wisell |

| | | | |
|---|---|---|---|
| EPI_ISL_475520 | Vardcentralen Brinken | The Public Health Agency of Sweden | Oskar Karlsson Lindsjo, Maria Lind Karlberg, Mattias Haukland, Reza Advani, Olov Svartstrom, Anna-Malin Linde, Sandra Broddesson, Mia Brytting, Anna Risberg, Karin Tegmark-Wisell |
| EPI_ISL_475521 | Ulltuna Vardcentral | The Public Health Agency of Sweden | Oskar Karlsson Lindsjo, Maria Lind Karlberg, Mattias Haukland, Reza Advani, Olov Svartstrom, Anna-Malin Linde, Sandra Broddesson, Mia Brytting, Anna Risberg, Karin Tegmark-Wisell |
| EPI_ISL_475522, EPI_ISL_475523 | Huddinge VC | The Public Health Agency of Sweden | Oskar Karlsson Lindsjo, Maria Lind Karlberg, Mattias Haukland, Reza Advani, Olov Svartstrom, Anna-Malin Linde, Sandra Broddesson, Mia Brytting, Anna Risberg, Karin Tegmark-Wisell |
| EPI_ISL_475524 | Narhalsan Sjobo vardcentral | The Public Health Agency of Sweden | Oskar Karlsson Lindsjo, Maria Lind Karlberg, Mattias Haukland, Reza Advani, Olov Svartstrom, Anna-Malin Linde, Sandra Broddesson, Mia Brytting, Anna Risberg, Karin Tegmark-Wisell |
| EPI_ISL_475525 | Huddinge VC | The Public Health Agency of Sweden | Oskar Karlsson Lindsjo, Maria Lind Karlberg, Mattias Haukland, Reza Advani, Olov Svartstrom, Anna-Malin Linde, Sandra Broddesson, Mia Brytting, Anna Risberg, Karin Tegmark-Wisell |
| EPI_ISL_475526, EPI_ISL_475527 | Uppsala Narakut Aleris | The Public Health Agency of Sweden | Oskar Karlsson Lindsjo, Maria Lind Karlberg, Mattias Haukland, Reza Advani, Olov Svartstrom, Anna-Malin Linde, Sandra Broddesson, Mia Brytting, Anna Risberg, Karin Tegmark-Wisell |
| EPI_ISL_475528 | Omtanken Grimmered | The Public Health Agency of Sweden | Oskar Karlsson Lindsjo, Maria Lind Karlberg, Mattias Haukland, Reza Advani, Olov Svartstrom, Anna-Malin Linde, Sandra Broddesson, Mia Brytting, Anna Risberg, Karin Tegmark-Wisell |
| EPI_ISL_475529, EPI_ISL_475530, EPI_ISL_475531, EPI_ISL_475532 | Kungsors VC | The Public Health Agency of Sweden | Oskar Karlsson Lindsjo, Maria Lind Karlberg, Mattias Haukland, Reza Advani, Olov Svartstrom, Anna-Malin Linde, Sandra Broddesson, Mia Brytting, Anna Risberg, Karin Tegmark-Wisell |
| EPI_ISL_475533, EPI_ISL_475534, EPI_ISL_475535 | Omtanken Grimmered | The Public Health Agency of Sweden | Oskar Karlsson Lindsjo, Maria Lind Karlberg, Mattias Haukland, Reza Advani, Olov Svartstrom, Anna-Malin Linde, Sandra Broddesson, Mia Brytting, Anna Risberg, Karin Tegmark-Wisell |
| EPI_ISL_475536 | Follinge Halsocentral | The Public Health Agency of Sweden | Oskar Karlsson Lindsjo, Maria Lind Karlberg, Mattias Haukland, Reza Advani, Olov Svartstrom, Anna-Malin Linde, Sandra Broddesson, Mia Brytting, Anna Risberg, Karin Tegmark-Wisell |
| EPI_ISL_475537 | Narhalsan Oden VC | The Public Health Agency of Sweden | Oskar Karlsson Lindsjo, Maria Lind Karlberg, Mattias Haukland, Reza Advani, Olov Svartstrom, Anna-Malin Linde, Sandra Broddesson, Mia Brytting, Anna Risberg, Karin Tegmark-Wisell |
| EPI_ISL_475538 | Omtanken Grimmered | The Public Health Agency of Sweden | Oskar Karlsson Lindsjo, Maria Lind Karlberg, Mattias Haukland, Reza Advani, Olov Svartstrom, Anna-Malin Linde, Sandra Broddesson, Mia Brytting, Anna Risberg, Karin Tegmark-Wisell |
| EPI_ISL_475539 | Narhalsan Sjobo vardcentral | The Public Health Agency of Sweden | Oskar Karlsson Lindsjo, Maria Lind Karlberg, Mattias Haukland, Reza Advani, Olov Svartstrom, Anna-Malin Linde, Sandra Broddesson, Mia Brytting, Anna Risberg, Karin Tegmark-Wisell |
| EPI_ISL_475540 | Bla Kustens halsocentral | The Public Health Agency of Sweden | Oskar Karlsson Lindsjo, Maria Lind Karlberg, Mattias Haukland, Reza Advani, Olov Svartstrom, Anna-Malin Linde, Sandra Broddesson, Mia Brytting, Anna Risberg, Karin Tegmark-Wisell |
| EPI_ISL_475541 | Follinge Halsocentral | The Public Health Agency of Sweden | Oskar Karlsson Lindsjo, Maria Lind Karlberg, Mattias Haukland, Reza Advani, Olov Svartstrom, Anna-Malin Linde, Sandra Broddesson, Mia Brytting, Anna Risberg, Karin Tegmark-Wisell |
| EPI_ISL_475542 | Kungsholmsdoktorn | The Public Health Agency of Sweden | Oskar Karlsson Lindsjo, Maria Lind Karlberg, Mattias Haukland, Reza Advani, Olov Svartstrom, Anna-Malin Linde, Sandra Broddesson, Mia Brytting, Anna Risberg, Karin Tegmark-Wisell |
| EPI_ISL_475543 | Surbrunns VC | The Public Health Agency of Sweden | Oskar Karlsson Lindsjo, Maria Lind Karlberg, Mattias Haukland, Reza Advani, Olov Svartstrom, Anna-Malin Linde, Sandra Broddesson, Mia Brytting, Anna Risberg, Karin Tegmark-Wisell |
| EPI_ISL_475544, EPI_ISL_475545, EPI_ISL_475546, EPI_ISL_475547 | Karolinska Universitetslaboratoriet | The Public Health Agency of Sweden | Oskar Karlsson Lindsjo, Maria Lind Karlberg, Mattias Haukland, Reza Advani, Olov Svartstrom, Anna-Malin Linde, Sandra Broddesson, Shaman Muradrasoli, Anna Risberg, Karin Tegmark-Wisell |
| EPI_ISL_475548 | Halmstad klinisk mikrobiologi | The Public Health Agency of Sweden | Oskar Karlsson Lindsjo, Maria Lind Karlberg, Mattias Haukland, Reza Advani, Olov Svartstrom, Anna-Malin Linde, Sandra Broddesson, Shaman Muradrasoli, Anna Risberg, Karin Tegmark-Wisell |
| EPI_ISL_475549, EPI_ISL_475550 | Skovde/Unilabs | The Public Health Agency of Sweden | Oskar Karlsson Lindsjo, Maria Lind Karlberg, Mattias Haukland, Reza Advani, Olov Svartstrom, Anna-Malin Linde, Sandra Broddesson, Shaman Muradrasoli, Anna Risberg, Karin Tegmark-Wisell |
| EPI_ISL_475551, EPI_ISL_475552 | Karolinska Universitetslaboratoriet | The Public Health Agency of Sweden | Oskar Karlsson Lindsjo, Maria Lind Karlberg, Mattias Haukland, Reza Advani, Olov Svartstrom, Anna-Malin Linde, Sandra Broddesson, Shaman Muradrasoli, Anna Risberg, Karin Tegmark-Wisell |
| EPI_ISL_475553 | Halmstad klinisk mikrobiologi | The Public Health Agency of Sweden | Oskar Karlsson Lindsjo, Maria Lind Karlberg, Mattias Haukland, Reza Advani, Olov Svartstrom, Anna-Malin Linde, Sandra Broddesson, Shaman Muradrasoli, Anna Risberg, Karin Tegmark-Wisell |
| EPI_ISL_475554, EPI_ISL_475555 | Skovde/Unilabs | The Public Health Agency of Sweden | Oskar Karlsson Lindsjo, Maria Lind Karlberg, Mattias Haukland, Reza Advani, Olov Svartstrom, Anna-Malin Linde, Sandra Broddesson, Shaman Muradrasoli, Anna Risberg, Karin Tegmark-Wisell |
| EPI_ISL_475556, EPI_ISL_475557 | Halmstad klinisk mikrobiologi | The Public Health Agency of Sweden | Oskar Karlsson Lindsjo, Maria Lind Karlberg, Mattias Haukland, Reza Advani, Olov Svartstrom, Anna-Malin Linde, Sandra Broddesson, Shaman Muradrasoli, Anna Risberg, Karin Tegmark-Wisell |
| EPI_ISL_475558, EPI_ISL_475559, EPI_ISL_475560, EPI_ISL_475561 | Karolinska Universitetslaboratoriet | The Public Health Agency of Sweden | Oskar Karlsson Lindsjo, Maria Lind Karlberg, Mattias Haukland, Reza Advani, Olov Svartstrom, Anna-Malin Linde, Sandra Broddesson, Shaman Muradrasoli, Anna Risberg, Karin Tegmark-Wisell |
| EPI_ISL_475562, EPI_ISL_475563 | Din Klinik | The Public Health Agency of Sweden | Oskar Karlsson Lindsjo, Maria Lind Karlberg, Mattias Haukland, Reza Advani, Olov Svartstrom, Anna-Malin Linde, Sandra Broddesson, Mia Brytting, Anna Risberg, Karin Tegmark-Wisell |
| EPI_ISL_475564 | Surbrunns VC | The Public Health Agency of Sweden | Oskar Karlsson Lindsjo, Maria Lind Karlberg, Mattias Haukland, Reza Advani, Olov Svartstrom, Anna-Malin Linde, Sandra Broddesson, Mia Brytting, Anna Risberg, Karin Tegmark-Wisell |
| EPI_ISL_475565 | Narhalsan Backa vardcentral | The Public Health Agency of Sweden | Oskar Karlsson Lindsjo, Maria Lind Karlberg, Mattias Haukland, Reza Advani, Olov Svartstrom, Anna-Malin Linde, Sandra Broddesson, Mia Brytting, Anna Risberg, Karin Tegmark-Wisell |
| EPI_ISL_475566 | Vardcentralen Brinken | The Public Health Agency of Sweden | Oskar Karlsson Lindsjo, Maria Lind Karlberg, Mattias Haukland, Reza Advani, Olov Svartstrom, Anna-Malin Linde, Sandra Broddesson, Mia Brytting, Anna Risberg, Karin Tegmark-Wisell |
| EPI_ISL_475567 | Huddinge VC | The Public Health Agency of Sweden | Oskar Karlsson Lindsjo, Maria Lind Karlberg, Mattias Haukland, Reza Advani, Olov Svartstrom, Anna-Malin Linde, Sandra Broddesson, Mia Brytting, Anna Risberg, Karin Tegmark-Wisell |
| EPI_ISL_475568 | Kungsors VC | The Public Health Agency of Sweden | Oskar Karlsson Lindsjo, Maria Lind Karlberg, Mattias Haukland, Reza Advani, Olov Svartstrom, Anna-Malin Linde, Sandra Broddesson, Mia Brytting, Anna Risberg, Karin Tegmark-Wisell |
| EPI_ISL_475569 | Kungsholmsdoktorn | The Public Health Agency of Sweden | Oskar Karlsson Lindsjo, Maria Lind Karlberg, Mattias Haukland, Reza Advani, Olov Svartstrom, Anna-Malin Linde, Sandra Broddesson, Mia Brytting, Anna Risberg, Karin Tegmark-Wisell |
| EPI_ISL_476135 | Achima Care Fristadens VC | The Public Health Agency of Sweden | Oskar Karlsson Lindsjo, Maria Lind Karlberg, Mattias Haukland, Reza Advani, Olov Svartstrom, Anna-Malin Linde, Sandra Broddesson, Petra Edquist, Mia Brytting, Anna Risberg, Karin Tegmark-Wisell |
| EPI_ISL_476136 | Surbrunns VC | The Public Health Agency of Sweden | Oskar Karlsson Lindsjo, Maria Lind Karlberg, Mattias Haukland, Reza Advani, Olov Svartstrom, Anna-Malin Linde, Sandra Broddesson, Petra Edquist, Mia Brytting, Anna Risberg, Karin Tegmark-Wisell |
| EPI_ISL_476137 | Wasterlakarna | The Public Health Agency of Sweden | Oskar Karlsson Lindsjo, Maria Lind Karlberg, Mattias Haukland, Reza Advani, Olov Svartstrom, Anna-Malin Linde, Sandra Broddesson, Petra Edquist, Mia Brytting, Anna Risberg, Karin Tegmark-Wisell |

| | | | |
|---|---|---|---|
| EPI_ISL_476138 | Ulltuna Vardcentral | The Public Health Agency of Sweden | Oskar Karlsson Lindsjo, Maria Lind Karlberg, Mattias Haukland, Reza Advani, Olov Svartstrom, Anna-Malin Linde, Sandra Broddesson, Petra Edquist, Mia Brytting, Anna Risberg, Karin Tegmark-Wisell |
| EPI_ISL_476139 | Folkhalsomyndigheten | The Public Health Agency of Sweden | Oskar Karlsson Lindsjo, Maria Lind Karlberg, Mattias Haukland, Reza Advani, Olov Svartstrom, Anna-Malin Linde, Sandra Broddesson, Petra Edquist, Shamam Muradrasoli, Anna Risberg, Karin Tegmark-Wisell |
| EPI_ISL_476140, EPI_ISL_476141, EPI_ISL_476142 | Klinisk Mikrobiologi | The Public Health Agency of Sweden | Oskar Karlsson Lindsjo, Maria Lind Karlberg, Mattias Haukland, Reza Advani, Olov Svartstrom, Anna-Malin Linde, Sandra Broddesson, Petra Edquist, Shamam Muradrasoli, Anna Risberg, Karin Tegmark-Wisell |
| EPI_ISL_476143, EPI_ISL_476144 | Skovde/Unilabs | The Public Health Agency of Sweden | Oskar Karlsson Lindsjo, Maria Lind Karlberg, Mattias Haukland, Reza Advani, Olov Svartstrom, Anna-Malin Linde, Sandra Broddesson, Petra Edquist, Shamam Muradrasoli, Anna Risberg, Karin Tegmark-Wisell |
| EPI_ISL_476145, EPI_ISL_476146, EPI_ISL_476147 | Ostersund klinisk mikrobiologi | The Public Health Agency of Sweden | Oskar Karlsson Lindsjo, Maria Lind Karlberg, Mattias Haukland, Reza Advani, Olov Svartstrom, Anna-Malin Linde, Sandra Broddesson, Petra Edquist, Shamam Muradrasoli, Anna Risberg, Karin Tegmark-Wisell |
| EPI_ISL_476148, EPI_ISL_476149 | Institut Pasteur Dakar | Institut Pasteur de Dakar | Ndongo Dia, Moussa Moise Diagne, Mamadou Diop, Ousmane Faye, Amadou Alpha Sall |
| EPI_ISL_476150 | Institut Pasteur Dakar | Institut Pasteur de Dakar | Ndongo Dia, Moussa Moise Diagne, Mamadou diop, Ousmane Faye, Amadou Alpha Sall |
| EPI_ISL_476151 | Institut Pasteur Dakar | Institut Pasteur de Dakar | Ndongo Dia, Moussa Moise Diagne, Mamadou Diop, Ousmane faye, Amadou Alpha Sall |
| EPI_ISL_476491, EPI_ISL_476492 | Institut Pasteur Dakar | Institut Pasteur de Dakar | Ndongo Dia, Moussa Moise Diagne, Mamadou Diop, Ousmane Faye, Amadou Alpha Sall |
| EPI_ISL_476493 | Institut Pasteur Dakar | Institut Pasteur de Dakar | Ndongo Dia, Moussa Moise Diagne, Mamadou Diop, Ousmane Faye, Amadou alpha Sall |
| EPI_ISL_476494 | Institut Pasteur Dakar | Institut Pasteur de Dakar | Ndongo Dia, Moussa Moise Diagne, Mamadou Diop, Ousmane Faye, Amadou Alpha Sall |
| EPI_ISL_476495, EPI_ISL_476497 | Institut Pasteur Dakar | Institut Pasteur de Dakar | Ndongo Dia, Moussa Moise Diagne, Mamadou Diop, Ousmane Faye, Amadou alpha Sall |
| EPI_ISL_476514 | Institut Pasteur Dakar | Institut Pasteur de Dakar | Ndongo Dia, Moussa Moise Diagne, Mamadou Diop, Ousmane Faye, Amadou Alpha Sall |
| EPI_ISL_476515 | Institut Pasteur Dakar | Institut Pasteur de Dakar | Ndongo Dia, Moussa Moise Diagne, Mamadou diop, Ousmane Faye, Amadou alpha Sall |
| EPI_ISL_476516 | Institut Pasteur Dakar | Institut Pasteur de Dakar | Ndongo Dia, Moussa Moise Diagne, mamadou Diop, Ousmane Faye, Amadou Alpha Sall |
| EPI_ISL_476558, EPI_ISL_476560, EPI_ISL_476562 | Institut Pasteur Dakar | Institut Pasteur de Dakar | Ndongo Dia, Moussa Moise Diagne, Mamadou Diop, Ousmane Faye, Amadou Alpha Sall |
| EPI_ISL_476564 | Institut Pasteur Dakar | Institut Pasteur de Dakar | Ndongo Dia, Moussa Moise Diagne, Mamadou diop, Ousmane Faye, Amadou alpha Sall |
| EPI_ISL_476566 | Institut pasteur Dakar | Institut Pasteur de Dakar | Ndongo Dia, Moussa Moise Diagne, Mamadou Diop, Ousmane Faye, Amadou Alpha Sall |
| EPI_ISL_476569 | Institut Pasteur Dakar | Institut Pasteur de Dakar | Ndongo Dia, Moussa Moise, Mamadou Diop, Ousmane Faye, Amadou Alpha Sall |
| EPI_ISL_476570, EPI_ISL_476572, EPI_ISL_476574 | Institut Pasteur Dakar | Institut Pasteur de Dakar | Ndongo Dia, Moussa Moise Diagne, Mamadou Diop, Ousmane Faye, Amadou Alpha Sall |
| EPI_ISL_476859 | GMERS Medical College & Hospital, Gotri, Vadodara | Gujarat Biotechnology Research Centre | Pinal Trivedi, Maharshi Pandya, Nidhi Patel, Nitin Savaliya, Raghawendra Kumar, Dinesh Kumar, Zuber Saiyed, Komal Patel, Labdhi Pandya, Afzal Ansari, Nikha Trivedi, Meenakshi Shah, Neena Doshi, Varsha Godbole, Apurvasinh Puvar, Janvi Raval, Zarna Patel, Monika Gandhi, R D Dixit, A M Kadri, Harsh Bakshi, Chaitanya Joshi, Madhvi Joshi |
| EPI_ISL_476861 | GMERS Medical College & Hospital, Gotri, Vadodara | Gujarat Biotechnology Research Centre | Nidhi Patel, Nitin Savaliya, Raghawendra Kumar, Dinesh Kumar, Zuber Saiyed, Komal Patel, Labdhi Pandya, Afzal Ansari, Nikha Trivedi, Meenakshi Shah, Neena Doshi, Varsha Godbole, Apurvasinh Puvar, Janvi Raval, Zarna Patel, Monika Gandhi, Pinal Trivedi, Maharshi Pandya, R D Dixit, A M Kadri, Harsh Bakshi, Chaitanya Joshi, Madhvi Joshi |
| EPI_ISL_476862 | GMERS Medical College & Hospital, Gotri, Vadodara | Gujarat Biotechnology Research Centre | Nitin Savaliya, Raghawendra Kumar, Dinesh Kumar, Zuber Saiyed, Komal Patel, Labdhi Pandya, Afzal Ansari, Nikha Trivedi, Meenakshi Shah, Neena Doshi, Varsha Godbole, Apurvasinh Puvar, Janvi Raval, Zarna Patel, Monika Gandhi, Pinal Trivedi, Maharshi Pandya, Nidhi Patel, R D Dixit, A M Kadri, Harsh Bakshi, Chaitanya Joshi, Madhvi Joshi |
| EPI_ISL_476869 | Department of MicroBiology, Government Medical College, Surat | Gujarat Biotechnology Research Centre | Nikha Trivedi, Naresh Chauhan, Summaiya Mullan, Amit gamit, Apurvasinh Puvar, Janvi Raval, Zarna Patel, Monika Gandhi, Pinal Trivedi, Maharshi Pandya, Nidhi Patel, Nitin Savaliya, Raghawendra Kumar, Dinesh Kumar, Zuber Saiyed, Komal Patel, Labdhi Pandya, Afzal Ansari, R D Dixit, A M Kadri, Harsh Bakshi, Chaitanya Joshi, Madhvi Joshi |
| EPI_ISL_476870 | Department of MicroBiology, Government Medical College, Surat | Gujarat Biotechnology Research Centre | Naresh Chauhan, Summaiya Mullan, Amit gamit, Apurvasinh Puvar, Janvi Raval, Zarna Patel, Monika Gandhi, Pinal Trivedi, Maharshi Pandya, Nidhi Patel, Nitin Savaliya, Raghawendra Kumar, Dinesh Kumar, Zuber Saiyed, Komal Patel, Labdhi Pandya, Afzal Ansari, Nikha Trivedi, R D Dixit, A M Kadri, Harsh Bakshi, Chaitanya Joshi, Madhvi Joshi |
| EPI_ISL_476876 | Department of MicroBiology, Government Medical College, Surat | Gujarat Biotechnology Research Centre | Pinal Trivedi, Maharshi Pandya, Nidhi Patel, Nitin Savaliya, Raghawendra Kumar, Dinesh Kumar, Zuber Saiyed, Komal Patel, Labdhi Pandya, Afzal Ansari, Nikha Trivedi, Naresh Chauhan, Summaiya Mullan, Amit gamit, Apurvasinh Puvar, Janvi Raval, Zarna Patel, Monika Gandhi, R D Dixit, A M Kadri, Harsh Bakshi, Chaitanya Joshi, Madhvi Joshi |
| EPI_ISL_476878 | Department of MicroBiology, Government Medical College, Surat | Gujarat Biotechnology Research Centre | Nidhi Patel, Nitin Savaliya, Raghawendra Kumar, Dinesh Kumar, Zuber Saiyed, Komal Patel, Labdhi Pandya, Afzal Ansari, Nikha Trivedi, Naresh Chauhan, Summaiya Mullan, Amit gamit, Apurvasinh Puvar, Janvi Raval, Zarna Patel, Monika Gandhi, Pinal Trivedi, Maharshi Pandya, R D Dixit, A M Kadri, Harsh Bakshi, Chaitanya Joshi, Madhvi Joshi |
| EPI_ISL_476879 | Department of MicroBiology, Government Medical College, Surat | Gujarat Biotechnology Research Centre | Nitin Savaliya, Raghawendra Kumar, Dinesh Kumar, Zuber Saiyed, Komal Patel, Labdhi Pandya, Afzal Ansari, Nikha Trivedi, Naresh Chauhan, Summaiya Mullan, Amit gamit, Apurvasinh Puvar, Janvi Raval, Zarna Patel, Monika Gandhi, Pinal Trivedi, Maharshi Pandya, Nidhi Patel, R D Dixit, A M Kadri, Harsh Bakshi, Chaitanya Joshi, Madhvi Joshi |
| EPI_ISL_476900, EPI_ISL_476902, EPI_ISL_476903, EPI_ISL_476904, EPI_ISL_476905, EPI_ISL_476906, EPI_ISL_476907, EPI_ISL_476908, EPI_ISL_476909, EPI_ISL_476910, EPI_ISL_476911, EPI_ISL_476912, EPI_ISL_476914, EPI_ISL_476915, EPI_ISL_476916, EPI_ISL_476917, EPI_ISL_476918, EPI_ISL_476919, EPI_ISL_476920, EPI_ISL_476921, EPI_ISL_476922, EPI_ISL_476923, EPI_ISL_476924, EPI_ISL_476925, EPI_ISL_476926, EPI_ISL_476927, EPI_ISL_476928, EPI_ISL_476929, EPI_ISL_476930, EPI_ISL_476931, EPI_ISL_476932, EPI_ISL_476933, EPI_ISL_476935, EPI_ISL_476936, EPI_ISL_476937, EPI_ISL_476938, EPI_ISL_476939, EPI_ISL_476940 | | | |
| see above | UW Virology Lab | UW Virology Lab | Pavitra Roychoudhury, Hong Xie, Lasata Shrestha, Amin Addetia, Truong Nguyen, Victoria M Rachleff, Meei-Li Huang, Keith R Jerome, Alexander Greninger |
| EPI_ISL_477141, EPI_ISL_477142, EPI_ISL_477143, EPI_ISL_477144, EPI_ISL_477145, EPI_ISL_477146, EPI_ISL_477147, EPI_ISL_477148, EPI_ISL_477149, EPI_ISL_477150, EPI_ISL_477151, EPI_ISL_477152, EPI_ISL_477153, EPI_ISL_477154, EPI_ISL_477155, EPI_ISL_477156, EPI_ISL_477157, EPI_ISL_477158, EPI_ISL_477159 | | | |
| see above | Institut Pasteur Dakar | Institut Pasteur de Dakar | Ndongo Dia, Moussa Moise Diagne, Mamadou Diop, Mamadou Malado Jallow, Marie Henriette Dior Ndione, Safietou Sankhe, Ousmane Faye, Amadou Alpha Sall. |
| EPI_ISL_477193, EPI_ISL_477194 | Istituto Zooprofilattico Sperimentale Puglia e Basilicata; | Beaconlab (Bioinformatics, Evolution and Comparative Genomics lab), Dept of Biosciences, University on Mila | Parisi A.,Pesole G., Manzari C., Chiara M. |
| EPI_ISL_477195, EPI_ISL_477196, EPI_ISL_477197, EPI_ISL_477198, EPI_ISL_477199, EPI_ISL_477200, EPI_ISL_477201 | Istituto Zooprofilattico Sperimentale Puglia e Basilicata; | Beaconlab (Bioinformatics, Evolution and Comparative Genomics lab), Dept of Biosciences, University on Milan | Parisi A.,Pesole G., Manzari C., Chiara M. |
| EPI_ISL_477202, EPI_ISL_477203 | Istituto Zooprofilattico Sperimentale Puglia e Basilicata; | Beaconlab (Bioinformatics, Evolution and Comparative Genomics lab), Dept of Biosciences, University on Mila | Parisi A.,Pesole G., Manzari C., Chiara M. |
| EPI_ISL_477673, EPI_ISL_477674, EPI_ISL_477675, EPI_ISL_477676, EPI_ISL_477677, EPI_ISL_477678, EPI_ISL_477679, EPI_ISL_477680, EPI_ISL_477681, EPI_ISL_477682, EPI_ISL_477683, EPI_ISL_477684, EPI_ISL_477685, EPI_ISL_477686, EPI_ISL_477687, EPI_ISL_477689, EPI_ISL_477690, EPI_ISL_477691, EPI_ISL_477692, EPI_ISL_477693, EPI_ISL_477694, EPI_ISL_477695, EPI_ISL_477696, EPI_ISL_477697, EPI_ISL_477698, EPI_ISL_477699, EPI_ISL_477700, EPI_ISL_477701, EPI_ISL_477702, EPI_ISL_477703, EPI_ISL_477704, EPI_ISL_477705, EPI_ISL_477706, EPI_ISL_477707, EPI_ISL_477708, EPI_ISL_477709, EPI_ISL_477710, EPI_ISL_477711, EPI_ISL_477712, EPI_ISL_477713, EPI_ISL_477714, EPI_ISL_477715, EPI_ISL_477716, EPI_ISL_477717, EPI_ISL_477718, EPI_ISL_477719, EPI_ISL_477720, EPI_ISL_477721, EPI_ISL_477723, EPI_ISL_477724 | | | |

| | | | |
|---|---|---|---|
| see above | UW Virology Lab | UW Virology Lab | Pavitra Roychoudhury, Hong Xie, Lasata Shrestha, Amin Addetia, Truong Nguyen, Victoria M Rachleff, Meei-Li Huang, Keith R Jerome, Alexander Greninger |
| EPI_ISL_479493, EPI_ISL_479494, EPI_ISL_479495, EPI_ISL_479496, EPI_ISL_479497, EPI_ISL_479498, EPI_ISL_479499, EPI_ISL_479500, EPI_ISL_479501, EPI_ISL_479502, EPI_ISL_479503, EPI_ISL_479504, EPI_ISL_479505, EPI_ISL_479506, EPI_ISL_479507, EPI_ISL_479508, EPI_ISL_479509, EPI_ISL_479510, EPI_ISL_479511, EPI_ISL_479512, EPI_ISL_479513, EPI_ISL_479514, EPI_ISL_479515, EPI_ISL_479516, EPI_ISL_479517, EPI_ISL_479518, EPI_ISL_479519, EPI_ISL_479520, EPI_ISL_479521, EPI_ISL_479522, EPI_ISL_479523, EPI_ISL_479524, EPI_ISL_479525, EPI_ISL_479526, EPI_ISL_479527, EPI_ISL_479528, EPI_ISL_479529, EPI_ISL_479530, EPI_ISL_479531, EPI_ISL_479532, EPI_ISL_479533, EPI_ISL_479534, EPI_ISL_479535, EPI_ISL_479536, EPI_ISL_479537, EPI_ISL_479538, EPI_ISL_479539, EPI_ISL_479540, EPI_ISL_479541, EPI_ISL_479542, EPI_ISL_479543, EPI_ISL_479544, EPI_ISL_479545, EPI_ISL_479546, EPI_ISL_479547, EPI_ISL_479548, EPI_ISL_479549, EPI_ISL_479550, EPI_ISL_479551, EPI_ISL_479552, EPI_ISL_479553, EPI_ISL_479554, EPI_ISL_479555, EPI_ISL_479556, EPI_ISL_479557, EPI_ISL_479558, EPI_ISL_479559, EPI_ISL_479560, EPI_ISL_479561, EPI_ISL_479562, EPI_ISL_479563, EPI_ISL_479564, EPI_ISL_479565, EPI_ISL_479566, EPI_ISL_479567, EPI_ISL_479568, EPI_ISL_479569, EPI_ISL_479570, EPI_ISL_479571, EPI_ISL_479572, EPI_ISL_479573, EPI_ISL_479657, EPI_ISL_479658, EPI_ISL_479659, EPI_ISL_479661 | | | |
| see above | NIV Influenza | NIV Influenza | Potdar V |
| EPI_ISL_480554, EPI_ISL_480556, EPI_ISL_480782, EPI_ISL_480783, EPI_ISL_480786, EPI_ISL_480787, EPI_ISL_480788, EPI_ISL_480789, EPI_ISL_481220 | Institut Pasteur Dakar | Institut Pasteur de Dakar | Ndongo Dia, Moussa Moise Diagne, Mamadou Diop, Marie Henriette Dior Ndione, Mamadou Malado Jallow, Safietou Sanke, Ousmane Faye, Amadou Alpha Sall. |
| EPI_ISL_481227, EPI_ISL_481228, EPI_ISL_481229, EPI_ISL_481230, EPI_ISL_481231, EPI_ISL_481232, EPI_ISL_481233 | Lab voor klinische biologie | Onderzoeksgroep Virologie | Nick Vereecke, Laurens Lambrechts, Marthe Pauwels, Bruno Verhasselt, Linos Vandekerckhove, Hans Nauwynck, Sebastiaan Theuns |
| EPI_ISL_481234, EPI_ISL_481235, EPI_ISL_481236, EPI_ISL_481237, EPI_ISL_481238, EPI_ISL_481239, EPI_ISL_481240, EPI_ISL_481243 | Institut Pasteur Dakar | Institut Pasteur de Dakar | Ndongo Dia, Moussa Moise Diagne, Mamadou Diop, Marie Henriette Dior Ndione, Mamadou Malado Jallow, Safietou Sanke, Ousmane Faye, Amadou Alpha Sall. |
| EPI_ISL_481483 | Department for Virology, Molecular Biology and Genome Research, R. G. Lugar Center for Public Health Research, National Center for Disease Control and Public Health (NCDC) of Georgia. | Department for Virology, Molecular Biology and Genome Research, R. G. Lugar Center for Public Health Research, National Center for Disease Control and Public Health (NCDC) of Georgia. | Nino Berishvili, Tata Imnadze, Giorgi Tomashvili, Ana Papkiauri, Meri Pantsulaia, Gvantsa Brachveli, Gvantsa Chanturia, Ann Machablishvili, Nato Kotaria, Marine Murtskhvaladze, Lela Sabadze, Mari Gavashelidze, Tamar Jashiashvili, Tea Tevdoradze, Ketevan Sidamonidze, Ekaterine Khmaladze, Ekaterine Zhghenti, Roena Sukhiashvili, Mariam Zakalashvili, Lela Urushadze, Magda Dgebuadze, Davit Tsaguria, Ekaterine Zangaladze, Adam Kotorashvili, Maia Alkhazashvili, Irma Burjanadze, Anna Kasradze, Khatuna Zakhashvili, Paata Imnadze, Amiran Gamkrelidze. |
| EPI_ISL_482491, EPI_ISL_482492, EPI_ISL_482493, EPI_ISL_482494, EPI_ISL_482495, EPI_ISL_482496, EPI_ISL_482497, EPI_ISL_482498, EPI_ISL_482499, EPI_ISL_482500, EPI_ISL_482501, EPI_ISL_482502, EPI_ISL_482503, EPI_ISL_482505, EPI_ISL_482506, EPI_ISL_482507, EPI_ISL_482508, EPI_ISL_482509, EPI_ISL_482510, EPI_ISL_482511, EPI_ISL_482512, EPI_ISL_482514, EPI_ISL_482515, EPI_ISL_482516, EPI_ISL_482517, EPI_ISL_482518, EPI_ISL_482519, EPI_ISL_482520, EPI_ISL_482521, EPI_ISL_482522, EPI_ISL_482523, EPI_ISL_482524, EPI_ISL_482525, EPI_ISL_482526, EPI_ISL_482527, EPI_ISL_482528, EPI_ISL_482529, EPI_ISL_482530, EPI_ISL_482531, EPI_ISL_482532, EPI_ISL_482533, EPI_ISL_482534, EPI_ISL_482535, EPI_ISL_482536, EPI_ISL_482537, EPI_ISL_482538, EPI_ISL_482539, EPI_ISL_482540, EPI_ISL_482541, EPI_ISL_482542, EPI_ISL_482543, EPI_ISL_482544, EPI_ISL_482545, EPI_ISL_482546, EPI_ISL_482547, EPI_ISL_482548, EPI_ISL_482549, EPI_ISL_482550, EPI_ISL_482551, EPI_ISL_482552, EPI_ISL_482553, EPI_ISL_482554, EPI_ISL_482555, EPI_ISL_482556, EPI_ISL_482557, EPI_ISL_482558, EPI_ISL_482559, EPI_ISL_482560, EPI_ISL_482561, EPI_ISL_482562, EPI_ISL_482563, EPI_ISL_482564, EPI_ISL_482565, EPI_ISL_482566, EPI_ISL_482567, EPI_ISL_482568, EPI_ISL_482569, EPI_ISL_482570, EPI_ISL_482571, EPI_ISL_482572, EPI_ISL_482573, EPI_ISL_482574, EPI_ISL_482587, EPI_ISL_482588, EPI_ISL_482589, EPI_ISL_482590, EPI_ISL_482591, EPI_ISL_482592, EPI_ISL_482593, EPI_ISL_482594, EPI_ISL_482595, EPI_ISL_482596, EPI_ISL_482597, EPI_ISL_482598, EPI_ISL_482599, EPI_ISL_482600, EPI_ISL_482601, EPI_ISL_482602, EPI_ISL_482603, EPI_ISL_482604, EPI_ISL_482605, EPI_ISL_482606, EPI_ISL_482607, EPI_ISL_482608, EPI_ISL_482609, EPI_ISL_482610, EPI_ISL_482611, EPI_ISL_482612, EPI_ISL_482613, EPI_ISL_482614, EPI_ISL_482615, EPI_ISL_482616, EPI_ISL_482617, EPI_ISL_482618, EPI_ISL_482619, EPI_ISL_482620, EPI_ISL_482621, EPI_ISL_482622, EPI_ISL_482623, EPI_ISL_482624, EPI_ISL_482625, EPI_ISL_482626, EPI_ISL_482627, EPI_ISL_482628, EPI_ISL_482629, EPI_ISL_482630, EPI_ISL_482631, EPI_ISL_482632, EPI_ISL_482634, EPI_ISL_482635, EPI_ISL_482636, EPI_ISL_482637, EPI_ISL_482639, EPI_ISL_482640, EPI_ISL_482641, EPI_ISL_482642, EPI_ISL_482643, EPI_ISL_482644, EPI_ISL_482645, EPI_ISL_482646, EPI_ISL_482647, EPI_ISL_482648, EPI_ISL_482649, EPI_ISL_482650, EPI_ISL_482651, EPI_ISL_482652, EPI_ISL_482653, EPI_ISL_482654, EPI_ISL_482655, EPI_ISL_482656, EPI_ISL_482657, EPI_ISL_482658, EPI_ISL_482659, EPI_ISL_482660, EPI_ISL_482661, EPI_ISL_482662, EPI_ISL_482663, EPI_ISL_482664, EPI_ISL_482665, EPI_ISL_482666, EPI_ISL_482667, EPI_ISL_482668, EPI_ISL_482669, EPI_ISL_482670, EPI_ISL_482671 | | | |
| see above | National Centre for Disease control (NCDC) | NCDC/CSIR-IGIB | Pramod Kumar#, Rajesh Pandey#, Pooja Sharma, Mahesh S Dhar, Vivekanand A, Bharathram Uppili, Robin Marwal, Radhakrishanan VS, Saruchi Wadhwa, Nishu Tyagi, Uma Sharma, Priyanka Singh, Hemlata Lall, Meena Datta, Varun Jaiswal, Hema Gogia, Preeti Madan, Prateek Singh, Debasis Dash, Mitali Mukerji, Sandhya Kabra, Sujeet Singh, Mohammed Faruq, Anurag Agrawal*, Partha Rakshit* |
| EPI_ISL_482672, EPI_ISL_482673, EPI_ISL_482674, EPI_ISL_482675, EPI_ISL_482676, EPI_ISL_482677, EPI_ISL_482678, EPI_ISL_482679, EPI_ISL_482680, EPI_ISL_482681, EPI_ISL_482682, EPI_ISL_482683, EPI_ISL_482684, EPI_ISL_482685, EPI_ISL_482686, EPI_ISL_482687, EPI_ISL_482688, EPI_ISL_482689, EPI_ISL_482690, EPI_ISL_482691, EPI_ISL_482692, EPI_ISL_482693, EPI_ISL_482694, EPI_ISL_482695, EPI_ISL_482696, EPI_ISL_482697, EPI_ISL_482698, EPI_ISL_482699 | | | |
| see above | Singapore General Hospital | Department of Microbiology | Nurdyana Abdul Rahman, Kun Lee Lim, Chenhao Li, Kian Sing Chan, Lynette Oon, Kern Rei Chng, Niranjan Nagarajan, Karrie Ko |
| EPI_ISL_482874, EPI_ISL_482875, EPI_ISL_482876, EPI_ISL_482877, EPI_ISL_482878 | Institut Pasteur Dakar | Institut Pasteur de Dakar | Ndongo Dia, Moussa Moise Diagne, Mamadou Diop, Marie Henriette Dior Ndione, Mamadou malado Jallow, Safietou Sankhe, Ousmane Faye, Amadou Alpha Sall. |
| EPI_ISL_483701, EPI_ISL_483704, EPI_ISL_483706, EPI_ISL_483708, EPI_ISL_483709, EPI_ISL_483711, EPI_ISL_483712, EPI_ISL_483713, EPI_ISL_483714, EPI_ISL_483715, EPI_ISL_483716, EPI_ISL_483717, EPI_ISL_483718, EPI_ISL_483719, EPI_ISL_483720, EPI_ISL_483721, EPI_ISL_483722, EPI_ISL_483723, EPI_ISL_483724, EPI_ISL_483725 | | | |
| see above | Israel Central Virology laboratory | Israel Central Virology laboratory | Neta Zuckerman, Efrat Dahan Bucris, Oran Erster, Ella Mendelson, Michal Mandelboim |
| EPI_ISL_483823 | GMERS Medical College Himmatnagar | Gujarat Biotechnology Research Centre | Nikha Trivedi, Himanshu Khatri, Mayur Gandhi, Apurvasinh Puvar, Janvi Raval, Zarna Patel, Monika Gandhi, Pinal Trivedi, Maharshi Pandya, Nidhi Patel, Nitin Savaliya, Raghawendra Kumar, Dinesh Kumar, Zuber Saiyed, Komal Patel, Labdhi Pandya, Afzal Ansari, R D Dixit, A M Kadri, Harsh Bakshi, Chaitanya Joshi, Madhvi Joshi |
| EPI_ISL_483830 | Pandit Deendayal Upadhyay Government Medical College, Rajkot | Gujarat Biotechnology Research Centre | Pinal Trivedi, Maharshi Pandya, Nidhi Patel, Nitin Savaliya, Raghawendra Kumar, Dinesh Kumar, Zuber Saiyed, Komal Patel, Labdhi Pandya, Afzal Ansari, Nikha Trivedi, Gauravi Dhruv, Arti Trivedi, Apurvasinh Puvar, Janvi Raval, Zarna Patel, Monika Gandhi, R D Dixit, A M Kadri, Harsh Bakshi, Chaitanya Joshi, Madhvi Joshi |
| EPI_ISL_483832 | Pandit Deendayal Upadhyay Government Medical College, Rajkot | Gujarat Biotechnology Research Centre | Nidhi Patel, Nitin Savaliya, Raghawendra Kumar, Dinesh Kumar, Zuber Saiyed, Komal Patel, Labdhi Pandya, Afzal Ansari, Nikha Trivedi, Gauravi Dhruv, Arti Trivedi, Apurvasinh Puvar, Janvi Raval, Zarna Patel, Monika Gandhi, Pinal Trivedi, Maharshi Pandya, R D Dixit, A M Kadri, Harsh Bakshi, Chaitanya Joshi, Madhvi Joshi |
| EPI_ISL_483833 | Pandit Deendayal Upadhyay Government Medical College, Rajkot | Gujarat Biotechnology Research Centre | Nitin Savaliya, Raghawendra Kumar, Dinesh Kumar, Zuber Saiyed, Komal Patel, Labdhi Pandya, Afzal Ansari, Nikha Trivedi, Gauravi Dhruv, Arti Trivedi, Apurvasinh Puvar, Janvi Raval, Zarna Patel, Monika Gandhi, Pinal Trivedi, Maharshi Pandya, Nidhi Patel, R D Dixit, A M Kadri, Harsh Bakshi, Chaitanya Joshi, Madhvi Joshi |
| EPI_ISL_483840 | Pandit Deendayal Upadhyay Government Medical College, Rajkot | Gujarat Biotechnology Research Centre | Nikha Trivedi, Gauravi Dhruv, Arti Trivedi, Apurvasinh Puvar, Janvi Raval, Zarna Patel, Monika Gandhi, Pinal Trivedi, Maharshi Pandya, Nidhi Patel, Nitin Savaliya, Raghawendra Kumar, Dinesh Kumar, Zuber Saiyed, Komal Patel, Labdhi Pandya, Afzal Ansari, R D Dixit, A M Kadri, Harsh Bakshi, Chaitanya Joshi, Madhvi Joshi |
| EPI_ISL_483847 | Pandit Deendayal Upadhyay Government Medical College, Rajkot | Gujarat Biotechnology Research Centre | Pinal Trivedi, Maharshi Pandya, Nidhi Patel, Nitin Savaliya, Raghawendra Kumar, Dinesh Kumar, Zuber Saiyed, Komal Patel, Labdhi Pandya, Afzal Ansari, Nikha Trivedi, Gauravi Dhruv, Arti Trivedi, Apurvasinh Puvar, Janvi Raval, Zarna Patel, Monika Gandhi, R D Dixit, A M Kadri, Harsh Bakshi, Chaitanya Joshi, Madhvi Joshi |
| EPI_ISL_483849 | Pandit Deendayal Upadhyay Government Medical College, Rajkot | Gujarat Biotechnology Research Centre | Nidhi Patel, Nitin Savaliya, Raghawendra Kumar, Dinesh Kumar, Zuber Saiyed, Komal Patel, Labdhi Pandya, Afzal Ansari, Nikha Trivedi, Gauravi Dhruv, Arti Trivedi, Apurvasinh Puvar, Janvi Raval, Zarna Patel, Monika Gandhi, Pinal Trivedi, Maharshi Pandya, R D Dixit, A M Kadri, Harsh Bakshi, Chaitanya Joshi, Madhvi Joshi |
| EPI_ISL_483850 | Department of Microbiology, Government Medical College, Surat | Gujarat Biotechnology Research Centre | Nitin Savaliya, Raghawendra Kumar, Dinesh Kumar, Zuber Saiyed, Komal Patel, Labdhi Pandya, Afzal Ansari, Nikha Trivedi, Naresh Chauhan, Summaiya Mullan, Amit gamit, Apurvasinh Puvar, Janvi Raval, Zarna Patel, Monika Gandhi, Pinal Trivedi, Maharshi Pandya, Nidhi Patel, R D Dixit, A M Kadri, Harsh Bakshi, Chaitanya Joshi, Madhvi Joshi |
| EPI_ISL_483857 | Department of Microbiology, Government Medical College, Surat | Gujarat Biotechnology Research Centre | Nikha Trivedi, Naresh Chauhan, Summaiya Mullan, Amit gamit, Apurvasinh Puvar, Janvi Raval, Zarna Patel, Monika Gandhi, Pinal Trivedi, Maharshi Pandya, Nidhi Patel, Nitin Savaliya, Raghawendra Kumar, Dinesh Kumar, Zuber Saiyed, Komal Patel, Labdhi Pandya, Afzal Ansari, R D Dixit, A M Kadri, Harsh Bakshi, Chaitanya Joshi, Madhvi Joshi |
| EPI_ISL_483858 | Department of Microbiology, Government Medical College, Surat | Gujarat Biotechnology Research Centre | Naresh Chauhan, Summaiya Mullan, Amit gamit, Apurvasinh Puvar, Janvi Raval, Zarna Patel, Monika Gandhi, Pinal Trivedi, Maharshi Pandya, Nidhi Patel, Nitin Savaliya, Raghawendra Kumar, Dinesh Kumar, Zuber Saiyed, Komal Patel, Labdhi Pandya, Afzal Ansari, Nikha Trivedi, R D Dixit, A M Kadri, Harsh Bakshi, Chaitanya Joshi, Madhvi Joshi |

| | | | |
|---|---|---|---|
| EPI_ISL_483865 | Department of Microbiology, Government Medical College, Surat | Gujarat Biotechnology Research Centre | Pinal Trivedi, Maharshi Pandya, Nidhi Patel, Nitin Savaliya, Raghawendra Kumar, Dinesh Kumar, Zuber Saiyed, Komal Patel, Labdhi Pandya, Afzal Ansari, Nikha Trivedi, Naresh Chauhan, Summaiya Mullan, Amit gamit, Apurvasinh Puvar, Janvi Raval, Zarna Patel, Monika Gandhi, R D Dixit, A M Kadri, Harsh Bakshi, Chaitanya Joshi, Madhvi Joshi |
| EPI_ISL_483867 | Department of Microbiology, Government Medical College, Surat | Gujarat Biotechnology Research Centre | Nidhi Patel, Nitin Savaliya, Raghawendra Kumar, Dinesh Kumar, Zuber Saiyed, Komal Patel, Labdhi Pandya, Afzal Ansari, Nikha Trivedi, Naresh Chauhan, Summaiya Mullan, Amit gamit, Apurvasinh Puvar, Janvi Raval, Zarna Patel, Monika Gandhi, Pinal Trivedi, Maharshi Pandya, R D Dixit, A M Kadri, Harsh Bakshi, Chaitanya Joshi, Madhvi Joshi |
| EPI_ISL_483868 | Department of Microbiology, Government Medical College, Surat | Gujarat Biotechnology Research Centre | Nitin Savaliya, Raghawendra Kumar, Dinesh Kumar, Zuber Saiyed, Komal Patel, Labdhi Pandya, Afzal Ansari, Nikha Trivedi, Naresh Chauhan, Summaiya Mullan, Amit gamit, Apurvasinh Puvar, Janvi Raval, Zarna Patel, Monika Gandhi, Pinal Trivedi, Maharshi Pandya, Nidhi Patel, R D Dixit, A M Kadri, Harsh Bakshi, Chaitanya Joshi, Madhvi Joshi |
| EPI_ISL_483875 | Department of Microbiology, Government Medical College, Surat | Gujarat Biotechnology Research Centre | Nikha Trivedi, Naresh Chauhan, Summaiya Mullan, Amit gamit, Apurvasinh Puvar, Janvi Raval, Zarna Patel, Monika Gandhi, Pinal Trivedi, Maharshi Pandya, Nidhi Patel, Nitin Savaliya, Raghawendra Kumar, Dinesh Kumar, Zuber Saiyed, Komal Patel, Labdhi Pandya, Afzal Ansari, R D Dixit, A M Kadri, Harsh Bakshi, Chaitanya Joshi, Madhvi Joshi |
| EPI_ISL_483876 | Department of Microbiology, Government Medical College, Surat | Gujarat Biotechnology Research Centre | Naresh Chauhan, Summaiya Mullan, Amit gamit, Apurvasinh Puvar, Janvi Raval, Zarna Patel, Monika Gandhi, Pinal Trivedi, Maharshi Pandya, Nidhi Patel, Nitin Savaliya, Raghawendra Kumar, Dinesh Kumar, Zuber Saiyed, Komal Patel, Labdhi Pandya, Afzal Ansari, Nikha Trivedi, R D Dixit, A M Kadri, Harsh Bakshi, Chaitanya Joshi, Madhvi Joshi |
| EPI_ISL_484354, EPI_ISL_484355, EPI_ISL_484356, EPI_ISL_484357, EPI_ISL_484358, EPI_ISL_484359, EPI_ISL_484360, EPI_ISL_484361, EPI_ISL_484362, EPI_ISL_484363, EPI_ISL_484364, EPI_ISL_484365, EPI_ISL_484366, EPI_ISL_484367, EPI_ISL_484368, EPI_ISL_484369, EPI_ISL_484370, EPI_ISL_484371, EPI_ISL_484372, EPI_ISL_484373, EPI_ISL_484392, EPI_ISL_484393, EPI_ISL_484394, EPI_ISL_484395, EPI_ISL_484396, EPI_ISL_484397, EPI_ISL_484398, EPI_ISL_484399, EPI_ISL_484400, EPI_ISL_484401, EPI_ISL_484402, EPI_ISL_484403, EPI_ISL_484404, EPI_ISL_484405, EPI_ISL_484406 | | | |
| see above | Lincolnshire Hospitals and DeepSeq Nottingham | COVID-19 Genomics UK (COG-UK) Consortium | Nichola Duckworth, Tim Sloan, Sarah Walsh, Jonathan Ball, Patrick McClure, Joeseph Chappell, Nadine Holmes, Matthew Carlisle, Christopher Moore, Fei Sang, Johnny Debebe, Victoria Wright, Matthew Loose |
| EPI_ISL_485604, EPI_ISL_485605, EPI_ISL_485606, EPI_ISL_485607, EPI_ISL_485608, EPI_ISL_485609, EPI_ISL_485610, EPI_ISL_485611 | Respiratory Virus Unit, Microbiology Services Colindale, Public Health England | Respiratory Virus Unit, Microbiology Services Colindale, Public Health England | PHE Covid Sequencing Team |
| EPI_ISL_485635, EPI_ISL_485708, EPI_ISL_485710, EPI_ISL_485711 | Institut Pasteur Dakar | Institut Pasteur de Dakar | Ndongo Dia, Moussa Moise Diagne, Mamadou diop, Marie Henriette Dior Ndione, Mamadou Malado Jallow, Safietou Sanke, Ousmane Faye, Amadou Alpha Sall. |
| EPI_ISL_485712 | Institut Pasteur | Institut Pasteur de Dakar | Ndongo Dia, Moussa Moise Diagne, Mamadou diop, Marie Henriette Dior Ndione, Mamadou Malado Jallow, Safietou Sanke, Ousmane Faye, Amadou Alpha Sall. |
| EPI_ISL_485713, EPI_ISL_485715, EPI_ISL_485716, EPI_ISL_485717 | Institut Pasteur Dakar | Institut Pasteur de Dakar | Ndongo Dia, Moussa Moise Diagne, Mamadou diop, Marie Henriette Dior Ndione, Mamadou Malado Jallow, Safietou Sanke, Ousmane Faye, Amadou Alpha Sall. |
| EPI_ISL_485917, EPI_ISL_485918, EPI_ISL_485919, EPI_ISL_485920, EPI_ISL_485921, EPI_ISL_485922, EPI_ISL_485923, EPI_ISL_485924, EPI_ISL_485925, EPI_ISL_485926, EPI_ISL_485927, EPI_ISL_485928, EPI_ISL_485929, EPI_ISL_485930, EPI_ISL_485931, EPI_ISL_485932, EPI_ISL_485933, EPI_ISL_485934, EPI_ISL_485936, EPI_ISL_485937, EPI_ISL_485938, EPI_ISL_485939, EPI_ISL_485940, EPI_ISL_485941, EPI_ISL_485942, EPI_ISL_485944, EPI_ISL_485945, EPI_ISL_485946, EPI_ISL_485947, EPI_ISL_485948, EPI_ISL_485949, EPI_ISL_485950, EPI_ISL_485951, EPI_ISL_485952, EPI_ISL_485953, EPI_ISL_485954, EPI_ISL_485955, EPI_ISL_485956, EPI_ISL_485957, EPI_ISL_485958, EPI_ISL_485959, EPI_ISL_485960, EPI_ISL_485961, EPI_ISL_485963, EPI_ISL_485964, EPI_ISL_485967, EPI_ISL_485968, EPI_ISL_485969, EPI_ISL_485972, EPI_ISL_485973, EPI_ISL_485974, EPI_ISL_485975, EPI_ISL_485976, EPI_ISL_485977, EPI_ISL_485978, EPI_ISL_485979, EPI_ISL_485981, EPI_ISL_485982, EPI_ISL_485983, EPI_ISL_485984, EPI_ISL_485985, EPI_ISL_485986, EPI_ISL_485987, EPI_ISL_485988, EPI_ISL_485989, EPI_ISL_485990, EPI_ISL_485991, EPI_ISL_485992, EPI_ISL_485993, EPI_ISL_485994, EPI_ISL_485995, EPI_ISL_485996, EPI_ISL_485997, EPI_ISL_485998, EPI_ISL_485999, EPI_ISL_486000, EPI_ISL_486001, EPI_ISL_486002, EPI_ISL_486003, EPI_ISL_486004, EPI_ISL_486005, EPI_ISL_486006, EPI_ISL_486007, EPI_ISL_486008, EPI_ISL_486009, EPI_ISL_486010, EPI_ISL_486011, EPI_ISL_486012, EPI_ISL_486013, EPI_ISL_486014, EPI_ISL_486015, EPI_ISL_486016, EPI_ISL_486017, EPI_ISL_486018, EPI_ISL_486019, EPI_ISL_486020, EPI_ISL_486021, EPI_ISL_486022, EPI_ISL_486023, EPI_ISL_486024, EPI_ISL_486025, EPI_ISL_486026, EPI_ISL_486027, EPI_ISL_486028, EPI_ISL_486029, EPI_ISL_486030, EPI_ISL_486032, EPI_ISL_486033, EPI_ISL_486034, EPI_ISL_486035, EPI_ISL_486036, EPI_ISL_486037, EPI_ISL_486038, EPI_ISL_486039, EPI_ISL_486040, EPI_ISL_486041, EPI_ISL_486042, EPI_ISL_486043, EPI_ISL_486044, EPI_ISL_486046, EPI_ISL_486047, EPI_ISL_486048, EPI_ISL_486049, EPI_ISL_486050, EPI_ISL_486051, EPI_ISL_486052, EPI_ISL_486053, EPI_ISL_486054, EPI_ISL_486055, EPI_ISL_486056, EPI_ISL_486057, EPI_ISL_486058, EPI_ISL_486059, EPI_ISL_486060, EPI_ISL_486061, EPI_ISL_486062, EPI_ISL_486063, EPI_ISL_486064, EPI_ISL_486065, EPI_ISL_486066, EPI_ISL_486067, EPI_ISL_486068, EPI_ISL_486069, EPI_ISL_486070, EPI_ISL_486071, EPI_ISL_486072, EPI_ISL_486074, EPI_ISL_486075, EPI_ISL_486076, EPI_ISL_486077, EPI_ISL_486078, EPI_ISL_486079, EPI_ISL_486080, EPI_ISL_486081, EPI_ISL_486082, EPI_ISL_486083, EPI_ISL_486084, EPI_ISL_486085, EPI_ISL_486086, EPI_ISL_486087, EPI_ISL_486088, EPI_ISL_486089, EPI_ISL_486093, EPI_ISL_486094, EPI_ISL_486095, EPI_ISL_486097, EPI_ISL_486098, EPI_ISL_486099, EPI_ISL_486102, EPI_ISL_486103, EPI_ISL_486105, EPI_ISL_486106, EPI_ISL_486108, EPI_ISL_486109, EPI_ISL_486110, EPI_ISL_486112, EPI_ISL_486113, EPI_ISL_486114 | | | |
| see above | UW Virology Lab | UW Virology Lab | Pavitra Roychoudhury, Hong Xie, Lasata Shrestha, Amin Addetia, Truong Nguyen, Victoria M Rachleff, Meei-Li Huang, Keith R Jerome, Alexander Greninger |
| EPI_ISL_486857, EPI_ISL_486858, EPI_ISL_486859, EPI_ISL_486860, EPI_ISL_486861, EPI_ISL_486862, EPI_ISL_486863, EPI_ISL_486864, EPI_ISL_486865, EPI_ISL_486866, EPI_ISL_486867, EPI_ISL_486868, EPI_ISL_486869, EPI_ISL_486870, EPI_ISL_486871, EPI_ISL_486872, EPI_ISL_486873 | | | |
| see above | Institut Pasteur Dakar | Institut Pasteur de Dakar | Ndongo Dia, Moussa Moise Diagne, Mamadou Diop, Marie Henriette Dior Ndione, Mamadou Malado Jallow, Safietou Sanke, Ousmane Faye, Amadou Alpha Sall. |
| EPI_ISL_487089, EPI_ISL_487091, EPI_ISL_487092, EPI_ISL_487096, EPI_ISL_487098, EPI_ISL_487099, EPI_ISL_487101, EPI_ISL_487102, EPI_ISL_487103, EPI_ISL_487104, EPI_ISL_487105, EPI_ISL_487106, EPI_ISL_487107, EPI_ISL_487108, EPI_ISL_487109, EPI_ISL_487110, EPI_ISL_487111, EPI_ISL_487112 | | | |
| see above | Nigeria Centre for Disease Control (NCDC) | African Centre of Excellence for Genomics of Infectious Diseases (ACEGID), Redeemer's University, Ede, Osun State, Nigeria | Oluniyi P.E., Ajogbasile F.V., Kayode A., Oguzie J., Olawoye I., Uwanibe J., Olumade T., Folarin O.A., Ihekweazu C., Happi C.T. |
| EPI_ISL_487113 | Nigeria Centre for Disease Control (NCDC) | Redeemer's University, ACEGID | Oluniyi P.E., Ajogbasile F.V., Kayode A., Oguzie J., Olawoye I., Uwanibe J., Olumade T., Folarin O.A., Ihekweazu C., Happi C.T. |
| EPI_ISL_487192 | Viral Respiratory Lab, National institute for Biomedical Research (INRB) | Pathogen Sequencing Lab, National Institute for Biomedical Research (INRB) | Placide Mbala-Kingebeni, Edith Nkwembe, Eddy Kinganda-Lusamaki, Amuri Aziza, Francisca Muyembe-Mawete, Emmanuel Lokilo-Lofiko, Catherine Pratt, Matthias Pauthner, Josh Quick, Allison Black, James Hadfield, Trevor Bedford, Ian Goodfellow, Andrew Rambault, Nick Loman, Kristian Andersen, Michael Wiley, Steve Ahuka-Mundeke, Jean-Jacques Muyembe Tamfum. |
| EPI_ISL_487365, EPI_ISL_487369 | Viral Respiratory Lab, National Institute for Biomedical Research (INRB) | Pathogen Sequencing Lab, National Institute for Biomedical Research (INRB) | Placide Mbala-Kingebeni, Edith Nkwembe, Eddy Kinganda-Lusamaki, Amuri Aziza, Francisca Muyembe-Mawete, Emmanuel Lokilo-Lofiko, Catherine Pratt, Matthias Pauthner, Josh Quick, Allison Black, James Hadfield, Trevor Bedford, Ian Goodfellow, Andrew Rambault, Nick Loman, Kristian Andersen, Michael Wiley, Steve Ahuka-Mundeke, Jean-Jacques Muyembe Tamfum |
| EPI_ISL_487432, EPI_ISL_487433, EPI_ISL_487434, EPI_ISL_487435, EPI_ISL_487436 | Queen Astrid Military Hospital | Institute of Tropical Medicine | Philippe Selhorst, Colin Anthony |
| EPI_ISL_490224, EPI_ISL_490225, EPI_ISL_490226, EPI_ISL_490227, EPI_ISL_490228, EPI_ISL_490229, EPI_ISL_490230, EPI_ISL_490231, EPI_ISL_490232, EPI_ISL_490233, EPI_ISL_490234, EPI_ISL_490235, EPI_ISL_490236, EPI_ISL_490237, EPI_ISL_490238, EPI_ISL_490239, EPI_ISL_490240, EPI_ISL_490241, EPI_ISL_490242, EPI_ISL_490243, EPI_ISL_490244, EPI_ISL_490245, EPI_ISL_490246, EPI_ISL_490247, EPI_ISL_490248, EPI_ISL_490249, EPI_ISL_490250, EPI_ISL_490251, EPI_ISL_490252, EPI_ISL_490253, EPI_ISL_490254 | | | |
| see above | Respiratory Virus Unit, Microbiology Services Colindale, Public Health England | Respiratory Virus Unit, Microbiology Services Colindale, Public Health England | PHE Covid Sequencing Team |
| EPI_ISL_490563, EPI_ISL_490564, EPI_ISL_490565, EPI_ISL_490566, EPI_ISL_490567, EPI_ISL_490568, EPI_ISL_490569, EPI_ISL_490570, EPI_ISL_490571, EPI_ISL_490572, EPI_ISL_490573, EPI_ISL_490574, EPI_ISL_490575, EPI_ISL_490576, EPI_ISL_490577, EPI_ISL_490578, EPI_ISL_490579, EPI_ISL_490580 | | | |
| see above | Lincolnshire Hospitals and DeepSeq Nottingham | COVID-19 Genomics UK (COG-UK) Consortium | Nichola Duckworth, Tim Sloan, Sarah Walsh, Jonathan Ball, Patrick McClure, Joeseph Chappell, Nadine Holmes, Matthew Carlisle, Christopher Moore, Fei Sang, Johnny Debebe, Victoria Wright, Matthew Loose |
| EPI_ISL_490978 | Mayo Clinic Laboratories | UW Virology Lab | Pavitra Roychoudhury, Hong Xie, Lasata Shrestha, Amin Addetia, Truong Nguyen, Victoria M Rachleff, Meei-Li Huang, Keith R Jerome, Alexander Greninger |
| EPI_ISL_490979, EPI_ISL_490980, EPI_ISL_490982, EPI_ISL_490983, EPI_ISL_490984, EPI_ISL_490985, EPI_ISL_490986, EPI_ISL_490987 | UW Virology Lab | UW Virology Lab | Pavitra Roychoudhury, Hong Xie, Lasata Shrestha, Amin Addetia, Truong Nguyen, Victoria M Rachleff, Meei-Li Huang, Keith R Jerome, Alexander Greninger |
| EPI_ISL_490988 | Mayo Clinic Laboratories | UW Virology Lab | Pavitra Roychoudhury, Hong Xie, Lasata Shrestha, Amin Addetia, Truong Nguyen, Victoria M Rachleff, Meei-Li Huang, Keith R Jerome, Alexander Greninger |
| EPI_ISL_490989, EPI_ISL_490990, EPI_ISL_490991, EPI_ISL_490992, EPI_ISL_490993, EPI_ISL_490994, | UW Virology Lab | UW Virology Lab | Pavitra Roychoudhury, Hong Xie, Lasata Shrestha, Amin Addetia, Truong Nguyen, Victoria M Rachleff, Meei-Li Huang, Keith R Jerome, Alexander Greninger |

| | | | |
|---|---|---|---|
| EPI_ISL_490995, EPI_ISL_490996 | | | |
| EPI_ISL_490997, EPI_ISL_490998, EPI_ISL_490999, EPI_ISL_491000 | Mayo Clinic Laboratories | UW Virology Lab | Pavitra Roychoudhury, Hong Xie, Lasata Shrestha, Amin Addetia, Truong Nguyen, Victoria M Rachleff, Meei-Li Huang, Keith R Jerome, Alexander Greninger |
| EPI_ISL_491001, EPI_ISL_491002, EPI_ISL_491003 | UW Virology Lab | UW Virology Lab | Pavitra Roychoudhury, Hong Xie, Lasata Shrestha, Amin Addetia, Truong Nguyen, Victoria M Rachleff, Meei-Li Huang, Keith R Jerome, Alexander Greninger |
| EPI_ISL_491004 | Mayo Clinic Laboratories | UW Virology Lab | Pavitra Roychoudhury, Hong Xie, Lasata Shrestha, Amin Addetia, Truong Nguyen, Victoria M Rachleff, Meei-Li Huang, Keith R Jerome, Alexander Greninger |
| EPI_ISL_491005, EPI_ISL_491006, EPI_ISL_491007, EPI_ISL_491009, EPI_ISL_491010, EPI_ISL_491011, EPI_ISL_491012, EPI_ISL_491013, EPI_ISL_491014, EPI_ISL_491016, EPI_ISL_491017, EPI_ISL_491018, EPI_ISL_491019, EPI_ISL_491020, EPI_ISL_491021, EPI_ISL_491022, EPI_ISL_491023, EPI_ISL_491024, EPI_ISL_491025, EPI_ISL_491026, EPI_ISL_491027, EPI_ISL_491028, EPI_ISL_491029, EPI_ISL_491030, EPI_ISL_491031, EPI_ISL_491032, EPI_ISL_491033, EPI_ISL_491034 | | | |
| see above | UW Virology Lab | UW Virology Lab | Pavitra Roychoudhury, Hong Xie, Lasata Shrestha, Amin Addetia, Truong Nguyen, Victoria M Rachleff, Meei-Li Huang, Keith R Jerome, Alexander Greninger |
| EPI_ISL_491709, EPI_ISL_491710, EPI_ISL_491711, EPI_ISL_491712, EPI_ISL_491713, EPI_ISL_491714, EPI_ISL_491715, EPI_ISL_491716, EPI_ISL_491717, EPI_ISL_491718, EPI_ISL_491719, EPI_ISL_491720, EPI_ISL_491721, EPI_ISL_491722, EPI_ISL_491723, EPI_ISL_491724, EPI_ISL_491725, EPI_ISL_491726, EPI_ISL_491727, EPI_ISL_491728, EPI_ISL_491729, EPI_ISL_491730, EPI_ISL_491731, EPI_ISL_491732, EPI_ISL_491733, EPI_ISL_491734, EPI_ISL_491735 | | | |
| see above | Respiratory Virus Unit, Microbiology Services Colindale, Public Health England | Respiratory Virus Unit, Microbiology Services Colindale, Public Health England | PHE Covid Sequencing Team |
| EPI_ISL_492181, EPI_ISL_492182 | University of Arkansas for Medical Sciences (UAMS) | Department of Biomedical Informatics, University of Arkansas for Medical Sciences (UAMS) | Piroon Jenjaroenpun, David W Ussery, Thidathip Wongsurawat |
| EPI_ISL_493001 | Wyoming Public Health Laboratory | Wyoming Public Health Laboratory | Noah Hull, Rob Christensen, Jim Mildenberger, Joel Sevinsky, Cari Sloma, and Wanda Manley |
| EPI_ISL_493002, EPI_ISL_493003 | Respiratory Virus Unit, Microbiology Services Colindale, Public Health England | Respiratory Virus Unit, Microbiology Services Colindale, Public Health England | PHE Covid Sequencing Team |
| EPI_ISL_493061, EPI_ISL_493062, EPI_ISL_493063, EPI_ISL_493064, EPI_ISL_493125, EPI_ISL_493126, EPI_ISL_493127, EPI_ISL_493128 | Wyoming Public Health Laboratory | Wyoming Public Health Laboratory | Noah Hull, Rob Christensen, Jim Mildenberger, Joel Sevinsky, Cari Sloma, and Wanda Manley |
| EPI_ISL_493549, EPI_ISL_493550, EPI_ISL_493551, EPI_ISL_493552, EPI_ISL_493553, EPI_ISL_493554, EPI_ISL_493555, EPI_ISL_493556, EPI_ISL_493557, EPI_ISL_493558, EPI_ISL_493559, EPI_ISL_493560, EPI_ISL_493561, EPI_ISL_493562, EPI_ISL_493563, EPI_ISL_493564, EPI_ISL_493565, EPI_ISL_493566, EPI_ISL_493567, EPI_ISL_493568, EPI_ISL_493569, EPI_ISL_493570, EPI_ISL_493571, EPI_ISL_493572, EPI_ISL_493573, EPI_ISL_493574, EPI_ISL_493575, EPI_ISL_493576, EPI_ISL_493577, EPI_ISL_493578, EPI_ISL_493579, EPI_ISL_493580, EPI_ISL_493581, EPI_ISL_493582, EPI_ISL_493583, EPI_ISL_493584, EPI_ISL_493585, EPI_ISL_493586, EPI_ISL_493587, EPI_ISL_493588, EPI_ISL_493589, EPI_ISL_493590, EPI_ISL_493591, EPI_ISL_493592, EPI_ISL_493593, EPI_ISL_493594, EPI_ISL_493595, EPI_ISL_493596, EPI_ISL_493597, EPI_ISL_493598, EPI_ISL_493599, EPI_ISL_493600, EPI_ISL_493603, EPI_ISL_493604, EPI_ISL_493605, EPI_ISL_493610, EPI_ISL_493611, EPI_ISL_493612, EPI_ISL_493613 | | | |
| see above | Lincolnshire Hospitals and DeepSeq Nottingham | COVID-19 Genomics UK (COG-UK) Consortium | Nichola Duckworth, Tim Sloan, Sarah Walsh, Jonathan Ball, Patrick McClure, Joeseph Chappell, Nadine Holmes, Matthew Carlisle, Christopher Moore, Fei Sang, Johnny Debebe, Victoria Wright, Matthew Loose |
| EPI_ISL_495017 | B.J. Medical College and Civil hospital | Gujarat Biotechnology Research Centre | Pinal Trivedi, Maharshi Pandya, Nidhi Patel, Nitin Savaliya, Raghawendra Kumar, Dinesh Kumar, Zuber Saiyed, Komal Patel, Labdhi Pandya, Afzal Ansari, Nikha Trivedi, Pranay Shah, Kamlesh J Upadhyay, Sanjay Kapadia, Apurvasinh Puvar, Janvi Raval, Zarna Patel, Monika Gandhi, R D Dixit, A M Kadri, Harsh Bakshi, Chaitanya Joshi, Madhvi Joshi |
| EPI_ISL_495019 | B.J. Medical College and Civil hospital | Gujarat Biotechnology Research Centre | Nidhi Patel, Nitin Savaliya, Raghawendra Kumar, Dinesh Kumar, Zuber Saiyed, Komal Patel, Labdhi Pandya, Afzal Ansari, Nikha Trivedi, Pranay Shah, Kamlesh J Upadhyay, Sanjay Kapadia, Apurvasinh Puvar, Janvi Raval, Zarna Patel, Monika Gandhi, Pinal Trivedi, Maharshi Pandya, R D Dixit, A M Kadri, Harsh Bakshi, Chaitanya Joshi, Madhvi Joshi |
| EPI_ISL_495023 | Government Medical College, Bhavnagar | Gujarat Biotechnology Research Centre | Nitin Savaliya, Raghawendra Kumar, Dinesh Kumar, Zuber Saiyed, Komal Patel, Labdhi Pandya, Afzal Ansari, Nikha Trivedi, Kairavi Desai, Saklin Malek, Shirish Patel, Apurvasinh Puvar, Janvi Raval, Zarna Patel, Monika Gandhi, Pinal Trivedi, Maharshi Pandya, Nidhi Patel, R D Dixit, A M Kadri, Harsh Bakshi, Chaitanya Joshi, Madhvi Joshi |
| EPI_ISL_495029 | GMERS Medical College & Hospital, Gotri, Vadodara | Gujarat Biotechnology Research Centre | Nikha Trivedi, Meenakshi Shah, Neena Doshi, Varsha Godbole, Apurvasinh Puvar, Janvi Raval, Zarna Patel, Monika Gandhi, Pinal Trivedi, Maharshi Pandya, Nidhi Patel, Nitin Savaliya, Raghawendra Kumar, Dinesh Kumar, Zuber Saiyed, Komal Patel, Labdhi Pandya, Afzal Ansari, R D Dixit, A M Kadri, Harsh Bakshi, Chaitanya Joshi, Madhvi Joshi |
| EPI_ISL_495031 | GMERS Medical College & Hospital, Gotri, Vadodara | Gujarat Biotechnology Research Centre | Neena Doshi, Varsha Godbole, Apurvasinh Puvar, Janvi Raval, Zarna Patel, Monika Gandhi, Pinal Trivedi, Maharshi Pandya, Nidhi Patel, Nitin Savaliya, Raghawendra Kumar, Dinesh Kumar, Zuber Saiyed, Komal Patel, Labdhi Pandya, Afzal Ansari, Nikha Trivedi, Meenakshi Shah, R D Dixit, A M Kadri, Harsh Bakshi, Chaitanya Joshi, Madhvi Joshi |
| EPI_ISL_495037 | GMERS Medical College & Hospital, Gotri, Vadodara | Gujarat Biotechnology Research Centre | Pinal Trivedi, Maharshi Pandya, Nidhi Patel, Nitin Savaliya, Raghawendra Kumar, Dinesh Kumar, Zuber Saiyed, Komal Patel, Labdhi Pandya, Afzal Ansari, Nikha Trivedi, Meenakshi Shah, Neena Doshi, Varsha Godbole, Apurvasinh Puvar, Janvi Raval, Zarna Patel, Monika Gandhi, R D Dixit, A M Kadri, Harsh Bakshi, Chaitanya Joshi, Madhvi Joshi |
| EPI_ISL_495039 | GMERS Medical College & Hospital, Gotri, Vadodara | Gujarat Biotechnology Research Centre | Nidhi Patel, Nitin Savaliya, Raghawendra Kumar, Dinesh Kumar, Zuber Saiyed, Komal Patel, Labdhi Pandya, Afzal Ansari, Nikha Trivedi, Meenakshi Shah, Neena Doshi, Varsha Godbole, Apurvasinh Puvar, Janvi Raval, Zarna Patel, Monika Gandhi, Pinal Trivedi, Maharshi Pandya, R D Dixit, A M Kadri, Harsh Bakshi, Chaitanya Joshi, Madhvi Joshi |
| EPI_ISL_495040 | GMERS Medical College & Hospital, Gotri, Vadodara | Gujarat Biotechnology Research Centre | Nitin Savaliya, Raghawendra Kumar, Dinesh Kumar, Zuber Saiyed, Komal Patel, Labdhi Pandya, Afzal Ansari, Nikha Trivedi, Meenakshi Shah, Neena Doshi, Varsha Godbole, Apurvasinh Puvar, Janvi Raval, Zarna Patel, Monika Gandhi, Pinal Trivedi, Maharshi Pandya, Nidhi Patel, R D Dixit, A M Kadri, Harsh Bakshi, Chaitanya Joshi, Madhvi Joshi |
| EPI_ISL_495047 | GMERS Medical College & Hospital, Gotri, Vadodara | Gujarat Biotechnology Research Centre | Nikha Trivedi, Meenakshi Shah, Neena Doshi, Varsha Godbole, Apurvasinh Puvar, Janvi Raval, Zarna Patel, Monika Gandhi, Pinal Trivedi, Maharshi Pandya, Nidhi Patel, Nitin Savaliya, Raghawendra Kumar, Dinesh Kumar, Zuber Saiyed, Komal Patel, Labdhi Pandya, Afzal Ansari, R D Dixit, A M Kadri, Harsh Bakshi, Chaitanya Joshi, Madhvi Joshi |
| EPI_ISL_495049 | GMERS Medical College & Hospital, Gotri, Vadodara | Gujarat Biotechnology Research Centre | Neena Doshi, Varsha Godbole, Apurvasinh Puvar, Janvi Raval, Zarna Patel, Monika Gandhi, Pinal Trivedi, Maharshi Pandya, Nidhi Patel, Nitin Savaliya, Raghawendra Kumar, Dinesh Kumar, Zuber Saiyed, Komal Patel, Labdhi Pandya, Afzal Ansari, Nikha Trivedi, Meenakshi Shah, R D Dixit, A M Kadri, Harsh Bakshi, Chaitanya Joshi, Madhvi Joshi |
| EPI_ISL_495055 | GMERS Medical College & Hospital, Gotri, Vadodara | Gujarat Biotechnology Research Centre | Pinal Trivedi, Maharshi Pandya, Nidhi Patel, Nitin Savaliya, Raghawendra Kumar, Dinesh Kumar, Zuber Saiyed, Komal Patel, Labdhi Pandya, Afzal Ansari, Nikha Trivedi, Meenakshi Shah, Neena Doshi, Varsha Godbole, Apurvasinh Puvar, Janvi Raval, Zarna Patel, Monika Gandhi, R D Dixit, A M Kadri, Harsh Bakshi, Chaitanya Joshi, Madhvi Joshi |
| EPI_ISL_495057 | GMERS Medical College & Hospital, Gotri, Vadodara | Gujarat Biotechnology Research Centre | Nidhi Patel, Nitin Savaliya, Raghawendra Kumar, Dinesh Kumar, Zuber Saiyed, Komal Patel, Labdhi Pandya, Afzal Ansari, Nikha Trivedi, Meenakshi Shah, Neena Doshi, Varsha Godbole, Apurvasinh Puvar, Janvi Raval, Zarna Patel, Monika Gandhi, Pinal Trivedi, Maharshi Pandya, R D Dixit, A M Kadri, Harsh Bakshi, Chaitanya Joshi, Madhvi Joshi |
| EPI_ISL_495058 | GMERS Medical College & Hospital, Gotri, Vadodara | Gujarat Biotechnology Research Centre | Nitin Savaliya, Raghawendra Kumar, Dinesh Kumar, Zuber Saiyed, Komal Patel, Labdhi Pandya, Afzal Ansari, Nikha Trivedi, Meenakshi Shah, Neena Doshi, Varsha Godbole, Apurvasinh Puvar, Janvi Raval, Zarna Patel, Monika Gandhi, Pinal Trivedi, Maharshi Pandya, Nidhi Patel, R D Dixit, A M Kadri, Harsh Bakshi, Chaitanya Joshi, Madhvi Joshi |
| EPI_ISL_495065 | GAIMS & G K General Hospital | Gujarat Biotechnology Research Centre | Nikha Trivedi, Babulal Babhoria, Hitesh Assudani, Apurvasinh Puvar, Janvi Raval, Zarna Patel, Monika Gandhi, Pinal Trivedi, Maharshi Pandya, Nidhi Patel, Nitin Savaliya, Raghawendra Kumar, Dinesh Kumar, Zuber Saiyed, Komal Patel, Labdhi Pandya, Afzal Ansari, R D Dixit, A M Kadri, Harsh Bakshi, Chaitanya Joshi, Madhvi Joshi, |
| EPI_ISL_495074 | Department of MicroBiology, Government Medical College, Surat | Gujarat Biotechnology Research Centre | Pinal Trivedi, Maharshi Pandya, Nidhi Patel, Nitin Savaliya, Raghawendra Kumar, Dinesh Kumar, Zuber Saiyed, Komal Patel, Labdhi Pandya, Afzal Ansari, Nikha Trivedi, Naresh Chauhan, Summaiya Mullan, Amit gamit, Apurvasinh Puvar, Janvi Raval, Zarna Patel, Monika Gandhi, R D Dixit, A M Kadri, Harsh Bakshi, Chaitanya Joshi, Madhvi Joshi |

| | | | |
|---|---|---|---|
| EPI_ISL_495075 | Department of MicroBiology, Government Medical College, Surat | Gujarat Biotechnology Research Centre | Nidhi Patel, Nitin Savaliya, Raghawendra Kumar, Dinesh Kumar, Zuber Saiyed, Komal Patel, Labdhi Pandya, Afzal Ansari, Nikha Trivedi, Naresh Chauhan, Summaiya Mullan, Amit gamit, Apurvasinh Puvar, Janvi Raval, Zarna Patel, Monika Gandhi, Pinal Trivedi, Maharshi Pandya, R D Dixit, A M Kadri, Harsh Bakshi, Chaitanya Joshi, Madhvi Joshi |
| EPI_ISL_495076 | Department of MicroBiology, Government Medical College, Surat | Gujarat Biotechnology Research Centre | Nitin Savaliya, Raghawendra Kumar, Dinesh Kumar, Zuber Saiyed, Komal Patel, Labdhi Pandya, Afzal Ansari, Nikha Trivedi, Naresh Chauhan, Summaiya Mullan, Amit gamit, Apurvasinh Puvar, Janvi Raval, Zarna Patel, Monika Gandhi, Pinal Trivedi, Maharshi Pandya, Nidhi Patel, R D Dixit, A M Kadri, Harsh Bakshi, Chaitanya Joshi, Madhvi Joshi |
| EPI_ISL_495080 | Department of MicroBiology, Government Medical College, Surat | Gujarat Biotechnology Research Centre | Nikha Trivedi, Naresh Chauhan, Summaiya Mullan, Amit gamit, Apurvasinh Puvar, Janvi Raval, Zarna Patel, Monika Gandhi, Pinal Trivedi, Maharshi Pandya, Nidhi Patel, Nitin Savaliya, Raghawendra Kumar, Dinesh Kumar, Zuber Saiyed, Komal Patel, Labdhi Pandya, Afzal Ansari, R D Dixit, A M Kadri, Harsh Bakshi, Chaitanya Joshi, Madhvi Joshi |
| EPI_ISL_495161 | CSIR-Centre for Cellular and Molecular Biology | CSIR-Centre for Cellular and Molecular Biology | Onkar Kulkarni,Sofia Banu, Payel Mukherjee, Priya Singh, Dhiviya Vedagiri, Divya Gupta, Vishal Sah, Santosh Kumar Kuncha, Krishnan Harinivas Harshan, Archana Bharadwaj Siva, Karthik Bharadwaj Tallapaka, Shagufta Khan, Lamuk Zaveri,Nikhil Hajirnis, M Soujanya Reddy, Pratheusa Maccha, Namami Gaur, Sakshi Shambhavi, Tulasi Nagabandi, Purushotham Vodnala, Deepak Kumar, Devi Prasad Vijayashankar, Disha Nanda, Divya Das, Jotin Gogoi, Manish Bhattacharjee, Rakesh K Mishra, Divya Tej Sowpati |
| EPI_ISL_495162 | CSIR-Centre for Cellular and Molecular Biology | CSIR-Centre for Cellular and Molecular Biology | Onkar Kulkarni, Payel Mukherjee, Sofia Banu, Priya Singh, Dhiviya Vedagiri, Divya Gupta, Vishal Sah, Santosh Kumar Kuncha, Krishnan Harinivas Harshan, Archana Bharadwaj Siva, Karthik Bharadwaj Tallapaka, Shagufta Khan, Lamuk Zaveri,Nikhil Hajirnis, M Soujanya Reddy, Pratheusa Maccha, Namami Gaur, Sakshi Shambhavi, Tulasi Nagabandi, Purushotham Vodnala, Deepak Kumar, Devi Prasad Vijayashankar, Disha Nanda, Divya Das, Jotin Gogoi, Manish Bhattacharjee, Rakesh K Mishra, Divya Tej Sowpati |
| EPI_ISL_495163 | CSIR-Centre for Cellular and Molecular Biology | CSIR-Centre for Cellular and Molecular Biology | Onkar Kulkarni, Payel Mukherjee, Sofia Banu, Priya Singh, Dhiviya Vedagiri, Divya Gupta, Vishal Sah, Santosh Kumar Kuncha, Krishnan Harinivas Harshan, Archana Bharadwaj Siva, Karthik Bharadwaj Tallapaka, Shagufta Khan, Lamuk Zaveri, Nikhil Hajirnis, M Soujanya Reddy, Pratheusa Maccha, Namami Gaur, Sakshi Shambhavi, Tulasi Nagabandi, Purushotham Vodnala, Deepak Kumar, Devi Prasad Vijayashankar, Disha Nanda, Divya Das, Jotin Gogoi, Manish Bhattacharjee, Rakesh K Mishra, Divya Tej Sowpati |
| EPI_ISL_495164 | CSIR-Centre for Cellular and Molecular Biology | CSIR-Centre for Cellular and Molecular Biology | Onkar Kulkarni, Payel Mukherjee, Sofia Banu, Priya Singh, Dhiviya Vedagiri, Divya Gupta, Vishal Sah, Santosh Kumar Kuncha, Krishnan Harinivas Harshan, Archana Bharadwaj Siva, Karthik Bharadwaj Tallapaka, Shagufta Khan, Lamuk Zaveri, Namami Gaur, Sakshi Shambhavi, Nikhil Hajirnis, M Soujanya Reddy, Pratheusa Maccha, Tulasi Nagabandi, Purushotham Vodnala,Preethi Jampala, Sharada Ravi Iyer, Sulagana Mukherjee, Swetha Sundar, Peddapuvala Sai Uday Kiran, Rakesh K Mishra, Divya Tej Sowpati |
| EPI_ISL_495165 | CSIR-Centre for Cellular and Molecular Biology | CSIR-Centre for Cellular and Molecular Biology | Onkar Kulkarni, Payel Mukherjee, Sofia Banu, Priya Singh, Dhiviya Vedagiri, Divya Gupta, Vishal Sah, Santosh Kumar Kuncha, Krishnan Harinivas Harshan, Archana Bharadwaj Siva, Karthik Bharadwaj Tallapaka, Shagufta Khan, Lamuk Zaveri, Namami Gaur, Sakshi Shambhavi,Nikhil Hajirnis, M Soujanya Reddy, Pratheusa Maccha, Tulasi Nagabandi, Purushotham Vodnala,Preethi Jampala, Sharada Ravi Iyer, Sulagana Mukherjee, Swetha Sundar, Peddapuvala Sai Uday Kiran, Rakesh K Mishra, Divya Tej Sowpati |
| EPI_ISL_495166 | CSIR-Centre for Cellular and Molecular Biology | CSIR-Centre for Cellular and Molecular Biology | Onkar Kulkarni,Sofia Banu, Payel Mukherjee, Priya Singh, Dhiviya Vedagiri, Divya Gupta, Vishal Sah, Santosh Kumar Kuncha, Krishnan Harinivas Harshan, Archana Bharadwaj Siva, Karthik Bharadwaj Tallapaka, Shagufta Khan, Lamuk Zaveri, Namami Gaur, Nikhil Hajirnis, M Soujanya Reddy, Pratheusa Maccha,Sakshi Shambhavi, Tulasi Nagabandi, Purushotham Vodnala, Deepak Kumar, Devi Prasad Vijayashankar, Disha Nanda, Divya Das, Jotin Gogoi, Manish Bhattacharjee, Rakesh K Mishra, Divya Tej Sowpati |
| EPI_ISL_495167 | CSIR-Centre for Cellular and Molecular Biology | CSIR-Centre for Cellular and Molecular Biology | Onkar Kulkarni, Payel Mukherjee, Sofia Banu, Priya Singh, Dhiviya Vedagiri, Divya Gupta, Vishal Sah, Santosh Kumar Kuncha, Krishnan Harinivas Harshan, Archana Bharadwaj Siva, Karthik Bharadwaj Tallapaka, Shagufta Khan, Lamuk Zaveri, Namami Gaur, Sakshi Shambhavi, Nikhil Hajirnis, M Soujanya Reddy, Pratheusa Maccha,Tulasi Nagabandi, Purushotham Vodnala, Deepak Kumar, Devi Prasad Vijayashankar, Disha Nanda, Divya Das, Jotin Gogoi, Manish Bhattacharjee, Rakesh K Mishra, Divya Tej Sowpati |
| EPI_ISL_495169 | CSIR-Centre for Cellular and Molecular Biology | CSIR-Centre for Cellular and Molecular Biology | Onkar Kulkarni, Payel Mukherjee, Sofia Banu, Priya Singh, Dhiviya Vedagiri, Divya Gupta, Vishal Sah, Santosh Kumar Kuncha, Krishnan Harinivas Harshan, Archana Bharadwaj Siva, Karthik Bharadwaj Tallapaka, Shagufta Khan, Lamuk Zaveri, Namami Gaur,Nikhil Hajirnis, M Soujanya Reddy, Pratheusa Maccha, Sakshi Shambhavi, Tulasi Nagabandi, Purushotham Vodnala, Deepak Kumar, Devi Prasad Vijayashankar, Disha Nanda, Divya Das, Jotin Gogoi, Manish Bhattacharjee, Rakesh K Mishra, Divya Tej Sowpati |
| EPI_ISL_495170 | CSIR-Centre for Cellular and Molecular Biology | CSIR-Centre for Cellular and Molecular Biology | Onkar Kulkarni, Tulasi Nagabandi, Namami Gaur, Sakshi Shambhavi, Lamuk Zaveri,Nikhil Hajirnis, M Soujanya Reddy, Pratheusa Maccha, Shagufta Khan, Purushotham Vodnala, Payel Mukherjee, Sofia Banu, Priya Singh, Onkar Kulkarni, Dhiviya Vedagiri, Divya Gupta, Vishal Sah, Santosh Kumar Kuncha, Krishnan Harinivas Harshan, Archana Bharadwaj Siva, Karthik Bharadwaj Tallapaka,G. Aditya Kumar, Koushick Sivakumar, Pooja Ramesh Gupta, Rajan Kumar Jha, Shraddha Vijay Lahoti, Rakesh K Mishra, Divya Tej Sowpati |
| EPI_ISL_495172 | CSIR-Centre for Cellular and Molecular Biology | CSIR-Centre for Cellular and Molecular Biology | Onkar Kulkarni, Tulasi Nagabandi, Namami Gaur, Sakshi Shambhavi, Lamuk Zaveri, Nikhil Hajirnis, M Soujanya Reddy, Pratheusa Maccha, Shagufta Khan, Purushotham Vodnala, Payel Mukherjee, Sofia Banu, Priya Singh,Onkar Kulkarni, Dhiviya Vedagiri, Divya Gupta, Vishal Sah, Santosh Kumar Kuncha, Krishnan Harinivas Harshan, Archana Bharadwaj Siva, Karthik Bharadwaj Tallapaka,Kezia J Ann, Radhika Khandelwal, Roshan Maku Venkata, Shemin Mansuri, Sonu Uday, Rakesh K Mishra, Divya Tej Sowpati |
| EPI_ISL_495173 | CSIR-Centre for Cellular and Molecular Biology | CSIR-Centre for Cellular and Molecular Biology | Onkar Kulkarni, Lamuk Zaveri, Nikhil Hajirnis, M Soujanya Reddy, Pratheusa Maccha, Tulasi Nagabandi, Namami Gaur, Sakshi Shambhavi,Shagufta Khan, Purushotham Vodnala, Payel Mukherjee, Sofia Banu, Priya Singh, Dhiviya Vedagiri, Divya Gupta, Vishal Sah, Santosh Kumar Kuncha, Krishnan Harinivas Harshan, Archana Bharadwaj Siva, Karthik Bharadwaj Tallapaka,G. Aditya Kumar, Koushick Sivakumar, Pooja Ramesh Gupta, Rajan Kumar Jha, Shraddha Vijay Lahoti, Rakesh K Mishra, Divya Tej Sowpati |
| EPI_ISL_495175 | CSIR-Centre for Cellular and Molecular Biology | CSIR-Centre for Cellular and Molecular Biology | Onkar Kulkarni, Lamuk Zaveri, Tulasi Nagabandi, Namami Gaur, Sakshi Shambhavi,Shagufta Khan, Nikhil Hajirnis, M Soujanya Reddy, Pratheusa Maccha, Purushotham Vodnala, Payel Mukherjee, Sofia Banu, Priya Singh, Dhiviya Vedagiri, Divya Gupta, Vishal Sah, Santosh Kumar Kuncha, Krishnan Harinivas Harshan, Archana Bharadwaj Siva, Karthik Bharadwaj Tallapaka,G. Aditya Kumar, Koushick Sivakumar, Pooja Ramesh Gupta, Rajan Kumar Jha, Shraddha Vijay Lahoti, Rakesh K Mishra, Divya Tej Sowpati |
| EPI_ISL_495178 | CSIR-Centre for Cellular and Molecular Biology | CSIR-Centre for Cellular and Molecular Biology | Payel Mukherjee, Sofia Banu, Priya Singh, Onkar Kulkarni, Dhiviya Vedagiri, Divya Gupta, Vishal Sah, Santosh Kumar Kuncha, Krishnan Harinivas Harshan, Archana Bharadwaj Siva, Karthik Bharadwaj Tallapaka, Shagufta Khan, Lamuk Zaveri, Nikhil Hajirnis, M Soujanya Reddy, Pratheusa Maccha, Namami Gaur, Sakshi Shambhavi, Tulasi Nagabandi, Purushotham Vodnala, Rakesh K Mishra, Sonu Uday, Sudipta Mondal, Annapoorna P Karthyayani, Debabrata Jana, Debrya Saha, Divya Tej Sowpati |
| EPI_ISL_495179 | CSIR-Centre for Cellular and Molecular Biology | CSIR-Centre for Cellular and Molecular Biology | Pratheusa Maccha, Sakshi Shambhavi, Lamuk Zaveri, Shagufta Khan, Namami Gaur, Nikhil Hajirnis, M Soujanya Reddy, Tulasi Nagabandi, Purushotham Vodnala, Payel Mukherjee, Sofia Banu, Priya Singh,Onkar Kulkarni, Dhiviya Vedagiri, Divya Gupta, Vishal Sah, Santosh Kumar Kuncha, Krishnan Harinivas Harshan, Archana Bharadwaj Siva, Karthik Bharadwaj Tallapaka, G. Aditya Kumar, Koushick Sivakumar,Disha Nanda, Divya Das, Jotin Gogoi, Manish Bhattacharjee, Ravi Prasad Mukku, Rakesh K Mishra, Divya Tej Sowpati |
| EPI_ISL_495181 | CSIR-Centre for Cellular and Molecular Biology | CSIR-Centre for Cellular and Molecular Biology | Pratheusa Maccha, Sofia Banu, Payel Mukherjee, Priya Singh, Onkar Kulkarni, Dhiviya Vedagiri, Divya Gupta, Vishal Sah, Santosh Kumar Kuncha, Krishnan Harinivas Harshan, Archana Bharadwaj Siva, Karthik Bharadwaj Tallapaka, Shagufta Khan, Lamuk Zaveri, Namami Gaur, Sakshi Shambhavi, Nikhil Hajirnis, M Soujanya Reddy, Tulasi Nagabandi, Purushotham Vodnala,Preethi Jampala, Sharada Ravi Iyer, Sulagana Mukherjee, Swetha Sundar, Peddapuvala Sai Uday Kiran, Rakesh K Mishra, Divya Tej Sowpati |
| EPI_ISL_495188 | CSIR-Centre for Cellular and Molecular Biology | CSIR-Centre for Cellular and Molecular Biology | Nikhil Hajirnis, M Soujanya Reddy, Pratheusa Maccha, Payel Mukherjee, Sofia Banu, Priya Singh,Onkar Kulkarni, Dhiviya Vedagiri, Divya Gupta, Vishal Sah, Santosh Kumar Kuncha, Krishnan Harinivas Harshan, Archana Bharadwaj Siva, Karthik Bharadwaj Tallapaka, Shagufta Khan, Lamuk Zaveri, Namami Gaur, Sakshi Shambhavi, Tulasi Nagabandi, Purushotham Vodnala,Deepak Kumar, Devi Prasad Vijayashankar, Disha Nanda, Divya Das, Jotin Gogoi, Manish Bhattacharjee, Rakesh K Mishra, Divya Tej Sowpati |
| EPI_ISL_495191 | CSIR-Centre for Cellular and Molecular Biology | CSIR-Centre for Cellular and Molecular Biology | Nikhil Hajirnis, M Soujanya Reddy, Pratheusa Maccha, Lamuk Zaveri, Shagufta Khan, Namami Gaur, Sakshi Shambhavi, Tulasi Nagabandi, Purushotham Vodnala, Payel Mukherjee, Sofia Banu, Priya Singh, Onkar Kulkarni, Dhiviya Vedagiri, Divya Gupta, Vishal Sah, Santosh Kumar Kuncha, Krishnan Harinivas Harshan, Archana Bharadwaj Siva, Karthik Bharadwaj Tallapaka,Zeba Rizvi, Zuberwasim Sayyad, Kakade Aishwarya Arun, Amrutha H C, Ananga Ghosh, Rakesh K Mishra, Divya Tej Sowpati |

| | | | |
|---|---|---|---|
| EPI_ISL_495195 | CSIR-Centre for Cellular and Molecular Biology | CSIR-Centre for Cellular and Molecular Biology | Nikhil Hajirnis, M Soujanya Reddy, Pratheusa Maccha, Namami Gaur, Sakshi Shambhavi, Lamuk Zaveri, Shagufta Khan, Tulasi Nagabandi, Purushotham Vodnala, Payel Mukherjee, Sofia Banu, Priya Singh,Onkar Kulkarni, Dhiviya Vedagiri, Divya Gupta, Vishal Sah, Santosh Kumar Kuncha, Krishnan Harinivas Harshan, Archana Bharadwaj Siva, Karthik Bharadwaj Tallapaka,Kezia J Ann, Radhika Khandelwal, Roshan Maku Venkata, Shemin Mansuri, Sonu Uday, Rakesh K Mishra, Divya Tej Sowpati |
| EPI_ISL_495196 | CSIR-Centre for Cellular and Molecular Biology | CSIR-Centre for Cellular and Molecular Biology | Payel Mukherjee, Sofia Banu, Priya Singh, Onkar Kulkarni, Dhiviya Vedagiri, Divya Gupta, Vishal Sah, Santosh Kumar Kuncha, Krishnan Harinivas Harshan, Archana Bharadwaj Siva, Karthik Bharadwaj Tallapaka, Shagufta Khan, Lamuk Zaveri, Nikhil Hajirnis, M Soujanya Reddy, Pratheusa Maccha, Namami Gaur, Sakshi Shambhavi, Tulasi Nagabandi, Purushotham Vodnala, G. Aditya Kumar, Koushick Sivakumar, Pooja Ramesh Gupta, Rajan Kumar Jha, Shraddha Vijay Lahoti, Rakesh K Mishra, Divya Tej Sowpati |
| EPI_ISL_495199 | CSIR-Centre for Cellular and Molecular Biology | CSIR-Centre for Cellular and Molecular Biology | Pratheusa Maccha, Sakshi Shambhavi, Lamuk Zaveri, Shagufta Khan, Namami Gaur, Nikhil Hajirnis, M Soujanya Reddy, Tulasi Nagabandi, Purushotham Vodnala, Payel Mukherjee, Sofia Banu, Priya Singh,Onkar Kulkarni, Dhiviya Vedagiri, Divya Gupta, Vishal Sah, Santosh Kumar Kuncha, Krishnan Harinivas Harshan, Archana Bharadwaj Siva, Karthik Bharadwaj Tallapaka, G. Aditya Kumar, Koushick Sivakumar,Disha Nanda, Divya Das, Jotin Gogoi, Manish Bhattacharjee, Ravi Prasad Mukku, Rakesh K Mishra, Divya Tej Sowpati |
| EPI_ISL_495204 | CSIR-Centre for Cellular and Molecular Biology | CSIR-Centre for Cellular and Molecular Biology | Pratheusa Maccha, Shagufta Khan, Lamuk Zaveri, Namami Gaur, Sakshi Shambhavi, Tulasi Nagabandi, Nikhil Hajirnis, M Soujanya Reddy, Purushotham Vodnala, Payel Mukherjee, Sofia Banu, Priya Singh, Onkar Kulkarni, Dhiviya Vedagiri, Divya Gupta, Vishal Sah, Santosh Kumar Kuncha, Krishnan Harinivas Harshan, Archana Bharadwaj Siva, Karthik Bharadwaj Tallapaka, Disha Nanda, Divya Das, Jotin Gogoi, Manish Bhattacharjee, Ravi Prasad Mukku, Rakesh K Mishra, Divya Tej Sowpati |
| EPI_ISL_495209 | CSIR-Centre for Cellular and Molecular Biology | CSIR-Centre for Cellular and Molecular Biology | Payel Mukherjee, Sofia Banu, Priya Singh, Onkar Kulkarni, Dhiviya Vedagiri, Divya Gupta, Vishal Sah, Santosh Kumar Kuncha, Krishnan Harinivas Harshan, Archana Bharadwaj Siva, Karthik Bharadwaj Tallapaka, Shagufta Khan, Lamuk Zaveri, Nikhil Hajirnis, M Soujanya Reddy, Pratheusa Maccha, Namami Gaur, Sakshi Shambhavi, Tulasi Nagabandi, Purushotham Vodnala, Gokulan C G, Gunjan Purohit, Hanuman Tulashiram Kale, Pankaj Kumar, Prachand Issarapu, Rakesh K Mishra, Divya Tej Sowpati |
| EPI_ISL_495216 | CSIR-Centre for Cellular and Molecular Biology | CSIR-Centre for Cellular and Molecular Biology | Nikhil Hajirnis, M Soujanya Reddy, Pratheusa Maccha, Lamuk Zaveri, Shagufta Khan, Namami Gaur, Sakshi Shambhavi, Tulasi Nagabandi, Purushotham Vodnala, Payel Mukherjee, Sofia Banu, Priya Singh, Onkar Kulkarni, Dhiviya Vedagiri, Divya Gupta, Vishal Sah, Santosh Kumar Kuncha, Krishnan Harinivas Harshan, Archana Bharadwaj Siva, Karthik Bharadwaj Tallapaka,Zeba Rizvi, Zuberwasim Sayyad, Kakade Aishwarya Arun, Amrutha H C, Ananga Ghosh, Rakesh K Mishra, Divya Tej Sowpati |
| EPI_ISL_495217 | CSIR-Centre for Cellular and Molecular Biology | CSIR-Centre for Cellular and Molecular Biology | Nikhil Hajirnis, M Soujanya Reddy, Pratheusa Maccha, Namami Gaur, Sakshi Shambhavi, Lamuk Zaveri, Shagufta Khan, Tulasi Nagabandi, Purushotham Vodnala, Payel Mukherjee, Sofia Banu, Priya Singh,Onkar Kulkarni, Dhiviya Vedagiri, Divya Gupta, Vishal Sah, Santosh Kumar Kuncha, Krishnan Harinivas Harshan, Archana Bharadwaj Siva, Karthik Bharadwaj Tallapaka,Kezia J Ann, Radhika Khandelwal, Roshan Maku Venkata, Shemin Mansuri, Sonu Uday, Rakesh K Mishra, Divya Tej Sowpati |
| EPI_ISL_495222 | CSIR-Centre for Cellular and Molecular Biology | CSIR-Centre for Cellular and Molecular Biology | Pratheusa Maccha, Shagufta Khan, Lamuk Zaveri, Namami Gaur, Sakshi Shambhavi, Tulasi Nagabandi, Nikhil Hajirnis, M Soujanya Reddy, Purushotham Vodnala, Payel Mukherjee, Sofia Banu, Priya Singh, Onkar Kulkarni, Dhiviya Vedagiri, Divya Gupta, Vishal Sah, Santosh Kumar Kuncha, Krishnan Harinivas Harshan, Archana Bharadwaj Siva, Karthik Bharadwaj Tallapaka, Disha Nanda, Divya Das, Jotin Gogoi, Manish Bhattacharjee, Ravi Prasad Mukku, Rakesh K Mishra, Divya Tej Sowpati |
| EPI_ISL_495223 | CSIR-Centre for Cellular and Molecular Biology | CSIR-Centre for Cellular and Molecular Biology | Nikhil Hajirnis, M Soujanya Reddy, Pratheusa Maccha, Payel Mukherjee, Sofia Banu, Priya Singh,Onkar Kulkarni, Dhiviya Vedagiri, Divya Gupta, Vishal Sah, Santosh Kumar Kuncha, Krishnan Harinivas Harshan, Archana Bharadwaj Siva, Karthik Bharadwaj Tallapaka, Shagufta Khan, Lamuk Zaveri, Namami Gaur, Sakshi Shambhavi, Tulasi Nagabandi, Purushotham Vodnala,Deepak Kumar, Devi Prasad Vijayashankar, Disha Nanda, Divya Das, Jotin Gogoi, Manish Bhattacharjee, Rakesh K Mishra, Divya Tej Sowpati |
| EPI_ISL_495224 | CSIR-Centre for Cellular and Molecular Biology | CSIR-Centre for Cellular and Molecular Biology | Namami Gaur, Sakshi Shambhavi, Lamuk Zaveri, Shagufta Khan, Nikhil Hajirnis, M Soujanya Reddy, Pratheusa Maccha, Tulasi Nagabandi, Purushotham Vodnala, Payel Mukherjee, Sofia Banu, Priya Singh, Onkar Kulkarni, Dhiviya Vedagiri, Divya Gupta, Vishal Sah, Santosh Kumar Kuncha, Krishnan Harinivas Harshan, Archana Bharadwaj Siva, Karthik Bharadwaj Tallapaka,G. Aditya Kumar, Koushick Sivakumar, Rakesh K Mishra, Divya Tej Sowpati |
| EPI_ISL_495225 | CSIR-Centre for Cellular and Molecular Biology | CSIR-Centre for Cellular and Molecular Biology | Pratheusa Maccha, Sakshi Shambhavi, Lamuk Zaveri, Shagufta Khan, Namami Gaur, Nikhil Hajirnis, M Soujanya Reddy, Tulasi Nagabandi, Purushotham Vodnala, Payel Mukherjee, Sofia Banu, Priya Singh,Onkar Kulkarni, Dhiviya Vedagiri, Divya Gupta, Vishal Sah, Santosh Kumar Kuncha, Krishnan Harinivas Harshan, Archana Bharadwaj Siva, Karthik Bharadwaj Tallapaka, G. Aditya Kumar, Koushick Sivakumar,Disha Nanda, Divya Das, Jotin Gogoi, Manish Bhattacharjee, Ravi Prasad Mukku, Rakesh K Mishra, Divya Tej Sowpati |
| EPI_ISL_495227 | CSIR-Centre for Cellular and Molecular Biology | CSIR-Centre for Cellular and Molecular Biology | Payel Mukherjee, Sofia Banu, Priya Singh, Onkar Kulkarni, Dhiviya Vedagiri, Divya Gupta, Vishal Sah, Santosh Kumar Kuncha, Krishnan Harinivas Harshan, Archana Bharadwaj Siva, Karthik Bharadwaj Tallapaka, Shagufta Khan, Lamuk Zaveri, Nikhil Hajirnis, M Soujanya Reddy, Pratheusa Maccha, Namami Gaur, Sakshi Shambhavi, Tulasi Nagabandi, Purushotham Vodnala, Gokulan C G, Gunjan Purohit, Hanuman Tulashiram Kale, Pankaj Kumar, Prachand Issarapu, Rakesh K Mishra, Divya Tej Sowpati |
| EPI_ISL_495228 | CSIR-Centre for Cellular and Molecular Biology | CSIR-Centre for Cellular and Molecular Biology | Payel Mukherjee, Sofia Banu, Priya Singh, Onkar Kulkarni, Dhiviya Vedagiri, Divya Gupta, Vishal Sah, Santosh Kumar Kuncha, Krishnan Harinivas Harshan, Archana Bharadwaj Siva, Karthik Bharadwaj Tallapaka, Shagufta Khan, Lamuk Zaveri, Nikhil Hajirnis, M Soujanya Reddy, Pratheusa Maccha, Namami Gaur, Sakshi Shambhavi, Tulasi Nagabandi, Purushotham Vodnala, Rakesh K Mishra, Sonu Uday, Sudipta Mondal, Annapoorna P Karthyayani, Debabrata Jana, Debrya Saha, Divya Tej Sowpati |
| EPI_ISL_495229 | CSIR-Centre for Cellular and Molecular Biology | CSIR-Centre for Cellular and Molecular Biology | Namami Gaur, Sakshi Shambhavi, Lamuk Zaveri, Shagufta Khan, Nikhil Hajirnis, M Soujanya Reddy, Pratheusa Maccha, Tulasi Nagabandi, Purushotham Vodnala, Payel Mukherjee, Sofia Banu, Priya Singh,Onkar Kulkarni, Dhiviya Vedagiri, Divya Gupta, Vishal Sah, Santosh Kumar Kuncha, Krishnan Harinivas Harshan, Archana Bharadwaj Siva, Karthik Bharadwaj Tallapaka, Zeba Rizvi, Zuberwasim Sayyad, Kakade Aishwarya Arun, Amrutha H C, Ananga Ghosh, Rakesh K Mishra, Divya Tej Sowpati |
| EPI_ISL_495231 | CSIR-Centre for Cellular and Molecular Biology | CSIR-Centre for Cellular and Molecular Biology | Namami Gaur, Sakshi Shambhavi, Lamuk Zaveri, Shagufta Khan, Nikhil Hajirnis, M Soujanya Reddy, Pratheusa Maccha, Tulasi Nagabandi, Purushotham Vodnala, Payel Mukherjee, Sofia Banu, Priya Singh, Onkar Kulkarni, Dhiviya Vedagiri, Divya Gupta, Vishal Sah, Santosh Kumar Kuncha, Krishnan Harinivas Harshan, Archana Bharadwaj Siva, Karthik Bharadwaj Tallapaka, Zeba Rizvi, Zuberwasim Sayyad, Kakade Aishwarya Arun, Amrutha H C, Ananga Ghosh, Rakesh K Mishra, Divya Tej Sowpati |
| EPI_ISL_495236 | CSIR-Centre for Cellular and Molecular Biology | CSIR-Centre for Cellular and Molecular Biology | Nikhil Hajirnis, M Soujanya Reddy, Pratheusa Maccha, Lamuk Zaveri, Shagufta Khan, Namami Gaur, Sakshi Shambhavi, Tulasi Nagabandi, Purushotham Vodnala, Payel Mukherjee, Sofia Banu, Priya Singh, Onkar Kulkarni, Dhiviya Vedagiri, Divya Gupta, Vishal Sah, Santosh Kumar Kuncha, Krishnan Harinivas Harshan, Archana Bharadwaj Siva, Karthik Bharadwaj Tallapaka,Zeba Rizvi, Zuberwasim Sayyad, Kakade Aishwarya Arun, Amrutha H C, Ananga Ghosh, Rakesh K Mishra, Divya Tej Sowpati |
| EPI_ISL_495237 | CSIR-Centre for Cellular and Molecular Biology | CSIR-Centre for Cellular and Molecular Biology | Payel Mukherjee, Sofia Banu, Priya Singh, Onkar Kulkarni, Dhiviya Vedagiri, Divya Gupta, Vishal Sah, Santosh Kumar Kuncha, Krishnan Harinivas Harshan, Archana Bharadwaj Siva, Karthik Bharadwaj Tallapaka, Shagufta Khan, Lamuk Zaveri, Nikhil Hajirnis, M Soujanya Reddy, Pratheusa Maccha, Namami Gaur, Sakshi Shambhavi, Tulasi Nagabandi, Purushotham Vodnala, Rakesh K Mishra, Sonu Uday, Sudipta Mondal, Annapoorna P Karthyayani, Debabrata Jana, Debrya Saha, Divya Tej Sowpati |
| EPI_ISL_495238 | CSIR-Centre for Cellular and Molecular Biology | CSIR-Centre for Cellular and Molecular Biology | Pratheusa Maccha, Sakshi Shambhavi, Lamuk Zaveri, Shagufta Khan, Namami Gaur, Nikhil Hajirnis, M Soujanya Reddy, Tulasi Nagabandi, Purushotham Vodnala, Payel Mukherjee, Sofia Banu, Priya Singh,Onkar Kulkarni, Dhiviya Vedagiri, Divya Gupta, Vishal Sah, Santosh Kumar Kuncha, Krishnan Harinivas Harshan, Archana Bharadwaj Siva, Karthik Bharadwaj Tallapaka, G. Aditya Kumar, Koushick Sivakumar,Disha Nanda, Divya Das, Jotin Gogoi, Manish Bhattacharjee, Ravi Prasad Mukku, Rakesh K Mishra, Divya Tej Sowpati |
| EPI_ISL_495239 | CSIR-Centre for Cellular and Molecular Biology | CSIR-Centre for Cellular and Molecular Biology | Nikhil Hajirnis, M Soujanya Reddy, Pratheusa Maccha, Payel Mukherjee, Sofia Banu, Priya Singh,Onkar Kulkarni, Dhiviya Vedagiri, Divya Gupta, Vishal Sah, Santosh Kumar Kuncha, Krishnan Harinivas Harshan, Archana Bharadwaj Siva, Karthik Bharadwaj Tallapaka, Shagufta Khan, Lamuk Zaveri, Namami Gaur, Sakshi Shambhavi, Tulasi Nagabandi, Purushotham Vodnala,Deepak Kumar, Devi Prasad Vijayashankar, Disha Nanda, Divya Das, Jotin Gogoi, Manish Bhattacharjee, Rakesh K Mishra, Divya Tej Sowpati |
| EPI_ISL_495244 | CSIR-Centre for Cellular and Molecular Biology | CSIR-Centre for Cellular and Molecular Biology | Payel Mukherjee, Sofia Banu, Priya Singh, Onkar Kulkarni, Dhiviya Vedagiri, Divya Gupta, Vishal Sah, Santosh Kumar Kuncha, Krishnan Harinivas Harshan, Archana Bharadwaj Siva, Karthik Bharadwaj Tallapaka, Shagufta Khan, Lamuk Zaveri, Nikhil Hajirnis, M Soujanya Reddy, Pratheusa Maccha, Namami Gaur, Sakshi Shambhavi, Tulasi Nagabandi, Purushotham Vodnala, G. Aditya Kumar, Koushick Sivakumar, Pooja Ramesh Gupta, Rajan Kumar Jha, Shraddha Vijay |

| | | | |
|---|---|---|---|
| | | | Lahoti, Rakesh K Mishra, Divya Tej Sowpati |
| EPI_ISL_495245, EPI_ISL_495249 | CSIR-Centre for Cellular and Molecular Biology | CSIR-Centre for Cellular and Molecular Biology | Payel Mukherjee, Sofia Banu, Priya Singh, Onkar Kulkarni, Dhiviya Vedagiri, Divya Gupta, Vishal Sah, Santosh Kumar Kuncha, Krishnan Harinivas Harshan, Archana Bharadwaj Siva, Karthik Bharadwaj Tallapaka, Shagufta Khan, Lamuk Zaveri, Nikhil Hajirnis, M Soujanya Reddy, Pratheusa Maccha, Namami Gaur, Sakshi Shambhavi, Tulasi Nagabandi, Purushotham Vodnala, Gokulan C G, Gunjan Purohit, Hanuman Tulashiram Kale, Pankaj Kumar, Prachand Issarapu, Rakesh K Mishra, Divya Tej Sowpati |
| EPI_ISL_495251 | CSIR-Centre for Cellular and Molecular Biology | CSIR-Centre for Cellular and Molecular Biology | Pratheusa Maccha, Shagufta Khan, Lamuk Zaveri, Namami Gaur, Sakshi Shambhavi, Tulasi Nagabandi, Nikhil Hajirnis, M Soujanya Reddy, Purushotham Vodnala, Payel Mukherjee, Sofia Banu, Priya Singh,Onkar Kulkarni, Dhiviya Vedagiri, Divya Gupta, Vishal Sah, Santosh Kumar Kuncha, Krishnan Harinivas Harshan, Archana Bharadwaj Siva, Karthik Bharadwaj Tallapaka, Disha Nanda, Divya Das, Jotin Gogoi, Manish Bhattacharjee, Ravi Prasad Mukku, Rakesh K Mishra, Divya Tej Sowpati |
| EPI_ISL_495252 | CSIR-Centre for Cellular and Molecular Biology | CSIR-Centre for Cellular and Molecular Biology | Payel Mukherjee, Sofia Banu, Priya Singh, Onkar Kulkarni, Dhiviya Vedagiri, Divya Gupta, Vishal Sah, Santosh Kumar Kuncha, Krishnan Harinivas Harshan, Archana Bharadwaj Siva, Karthik Bharadwaj Tallapaka, Shagufta Khan, Lamuk Zaveri, Nikhil Hajirnis, M Soujanya Reddy, Pratheusa Maccha, Namami Gaur, Sakshi Shambhavi, Tulasi Nagabandi, Purushotham Vodnala, G. Aditya Kumar, Koushick Sivakumar, Pooja Ramesh Gupta, Rajan Kumar Jha, Shraddha Vijay Lahoti, Rakesh K Mishra, Divya Tej Sowpati |
| EPI_ISL_495254 | CSIR-Centre for Cellular and Molecular Biology | CSIR-Centre for Cellular and Molecular Biology | Pratheusa Maccha, Sofia Banu, Payel Mukherjee, Priya Singh, Onkar Kulkarni, Dhiviya Vedagiri, Divya Gupta, Vishal Sah, Santosh Kumar Kuncha, Krishnan Harinivas Harshan, Archana Bharadwaj Siva, Karthik Bharadwaj Tallapaka, Shagufta Khan, Lamuk Zaveri, Namami Gaur, Sakshi Shambhavi, Nikhil Hajirnis, M Soujanya Reddy, Tulasi Nagabandi, Purushotham Vodnala,Preethi Jampala, Sharada Ravi Iyer, Sulagana Mukherjee, Swetha Sundar, Peddapuvala Sai Uday Kiran, Rakesh K Mishra, Divya Tej Sowpati |
| EPI_ISL_495259 | CSIR-Centre for Cellular and Molecular Biology | CSIR-Centre for Cellular and Molecular Biology | Nikhil Hajirnis, M Soujanya Reddy, Pratheusa Maccha, Namami Gaur, Sakshi Shambhavi, Lamuk Zaveri, Shagufta Khan, Tulasi Nagabandi, Purushotham Vodnala, Payel Mukherjee, Sofia Banu, Priya Singh,Onkar Kulkarni, Dhiviya Vedagiri, Divya Gupta, Vishal Sah, Santosh Kumar Kuncha, Krishnan Harinivas Harshan, Archana Bharadwaj Siva, Karthik Bharadwaj Tallapaka,Kezia J Ann, Radhika Khandelwal, Roshan Maku Venkata, Shemin Mansuri, Sonu Uday, Rakesh K Mishra, Divya Tej Sowpati |
| EPI_ISL_495260 | CSIR-Centre for Cellular and Molecular Biology | CSIR-Centre for Cellular and Molecular Biology | Namami Gaur, Sakshi Shambhavi, Lamuk Zaveri, Shagufta Khan, Nikhil Hajirnis, M Soujanya Reddy, Pratheusa Maccha, Tulasi Nagabandi, Purushotham Vodnala, Payel Mukherjee, Sofia Banu, Priya Singh, Onkar Kulkarni, Dhiviya Vedagiri, Divya Gupta, Vishal Sah, Santosh Kumar Kuncha, Krishnan Harinivas Harshan, Archana Bharadwaj Siva, Karthik Bharadwaj Tallapaka,G. Aditya Kumar, Koushick Sivakumar, Rakesh K Mishra, Divya Tej Sowpati |
| EPI_ISL_495261 | CSIR-Centre for Cellular and Molecular Biology | CSIR-Centre for Cellular and Molecular Biology | Nikhil Hajirnis, M Soujanya Reddy, Pratheusa Maccha, Lamuk Zaveri, Shagufta Khan, Namami Gaur, Sakshi Shambhavi, Tulasi Nagabandi, Purushotham Vodnala, Payel Mukherjee, Sofia Banu, Priya Singh, Onkar Kulkarni, Dhiviya Vedagiri, Divya Gupta, Vishal Sah, Santosh Kumar Kuncha, Krishnan Harinivas Harshan, Archana Bharadwaj Siva, Karthik Bharadwaj Tallapaka,Zeba Rizvi, Zuberwasim Sayyad, Kakade Aishwarya Arun, Amrutha H C, Ananga Ghosh, Rakesh K Mishra, Divya Tej Sowpati |
| EPI_ISL_495266 | CSIR-Centre for Cellular and Molecular Biology | CSIR-Centre for Cellular and Molecular Biology | Namami Gaur, Sakshi Shambhavi, Lamuk Zaveri, Shagufta Khan, Nikhil Hajirnis, M Soujanya Reddy, Pratheusa Maccha, Tulasi Nagabandi, Purushotham Vodnala, Payel Mukherjee, Sofia Banu, Priya Singh,Onkar Kulkarni, Dhiviya Vedagiri, Divya Gupta, Vishal Sah, Santosh Kumar Kuncha, Krishnan Harinivas Harshan, Archana Bharadwaj Siva, Karthik Bharadwaj Tallapaka, Zeba Rizvi, Zuberwasim Sayyad, Kakade Aishwarya Arun, Amrutha H C, Ananga Ghosh, Rakesh K Mishra, Divya Tej Sowpati |
| EPI_ISL_495273 | CSIR-Centre for Cellular and Molecular Biology | CSIR-Centre for Cellular and Molecular Biology | Pratheusa Maccha, Sofia Banu, Payel Mukherjee, Priya Singh, Onkar Kulkarni, Dhiviya Vedagiri, Divya Gupta, Vishal Sah, Santosh Kumar Kuncha, Krishnan Harinivas Harshan, Archana Bharadwaj Siva, Karthik Bharadwaj Tallapaka, Shagufta Khan, Lamuk Zaveri, Namami Gaur, Sakshi Shambhavi, Nikhil Hajirnis, M Soujanya Reddy, Tulasi Nagabandi, Purushotham Vodnala,Preethi Jampala, Sharada Ravi Iyer, Sulagana Mukherjee, Swetha Sundar, Peddapuvala Sai Uday Kiran, Rakesh K Mishra, Divya Tej Sowpati |
| EPI_ISL_495629, EPI_ISL_495630, EPI_ISL_495631, EPI_ISL_495632, EPI_ISL_495633, EPI_ISL_495634, EPI_ISL_495635, EPI_ISL_495636, EPI_ISL_495637, EPI_ISL_495638, EPI_ISL_495639, EPI_ISL_495640, EPI_ISL_495641, EPI_ISL_496491, EPI_ISL_496492, EPI_ISL_496493, EPI_ISL_496494, EPI_ISL_496495, EPI_ISL_496496, EPI_ISL_496497, EPI_ISL_496498, EPI_ISL_496499, EPI_ISL_496500, EPI_ISL_496501, EPI_ISL_496502, EPI_ISL_496503, EPI_ISL_496504, EPI_ISL_496505, EPI_ISL_496506, EPI_ISL_496507, EPI_ISL_496508, EPI_ISL_496509, EPI_ISL_496510, EPI_ISL_496511, EPI_ISL_496512, EPI_ISL_496513, EPI_ISL_496514, EPI_ISL_496515, EPI_ISL_496516, EPI_ISL_496517 | | | |
| see above | Viral Respiratory Lab, National Institute for Biomedical Research (INRB) | Pathogen Sequencing Lab, National Institute for Biomedical Research (INRB) | Placide Mbala-Kingebeni, Edith Nkwembe, Eddy Kinganda-Lusamaki, Amuri Aziza, Francisca Muyembe Mawete, Emmanuel Lokilo Lofiko, Catherine Pratt, Matthias Pauthner, Josh Quick, Allison Black, James Hadfield, Trevor Bedford, Ian Goodfellow, Andrew Rambaut, Nick Loman, Kristian Andersen, Michael Wiley, Steve Ahuka-Mundeke, Jean-Jacques Muyembe Tamfum |
| EPI_ISL_498171, EPI_ISL_498172, EPI_ISL_498173, EPI_ISL_498174, EPI_ISL_498175, EPI_ISL_498176, EPI_ISL_498177, EPI_ISL_498178, EPI_ISL_498179, EPI_ISL_498180, EPI_ISL_498181, EPI_ISL_498182, EPI_ISL_498183, EPI_ISL_498184, EPI_ISL_498185, EPI_ISL_498186, EPI_ISL_498187, EPI_ISL_498188, EPI_ISL_498189, EPI_ISL_498190, EPI_ISL_498191, EPI_ISL_498192 | | | |
| see above | OUCRU | OUCRU | Nguyen Van Vinh Chau, Nguyen Thi Thu Hong, Nguyen Thi Han Ny, Le Nguyen Truc Nhu, Nghiem My Ngoc, Vo Thanh Lam, Nguyen Thanh Dung, Lam Minh Yen, Ngo Ngoc Quang Minh, Le Manh Hung, Nguyen Tri Dung, Dinh Nguyen Huy Man, Lam Anh Nguyet, Tran Chanh Xuan, Tran Tinh Hien, Nguyen Thanh Phong, Tran Nguyen Hoang Tu, Tran Tan Thanh, Nguyen Thanh Truong, Nguyen Tan Binh, Tang Chi Thuong, Guy Thwaites, and Le Van Tan, for OUCRU COVID-19 research group* |
| EPI_ISL_498229, EPI_ISL_498230, EPI_ISL_498231, EPI_ISL_498232, EPI_ISL_498233, EPI_ISL_498234, EPI_ISL_498235, EPI_ISL_498236, EPI_ISL_498237, EPI_ISL_498238, EPI_ISL_498239, EPI_ISL_498240, EPI_ISL_498241, EPI_ISL_498242, EPI_ISL_498243, EPI_ISL_498244, EPI_ISL_498245, EPI_ISL_498246, EPI_ISL_498247, EPI_ISL_498248, EPI_ISL_498249, EPI_ISL_498250, EPI_ISL_498251, EPI_ISL_498252 | | | |
| see above | Institut Pasteur de Dakar | Institut Pasteur de Dakar | Ndongo Dia, Moussa Moise Diagne, Mamadou Diop, Marie Henriette Dior Ndione, Mamadou Malado Jallow, Safietou Sankhe Mbengue, Ousmane Faye, Amadou Alpha Sall. |
| EPI_ISL_500500, EPI_ISL_500501, EPI_ISL_500502 | University of Washington Virology Lab | University of Washington Virology Lab | Pavitra Roychoudhury, Hong Xie, Lasata Shrestha, Amin Addetia, Truong Nguyen, Victoria M Rachleff, Meei-Li Huang, Keith R Jerome, Alexander Greninger |
| EPI_ISL_500503, EPI_ISL_500504, EPI_ISL_500505, EPI_ISL_500506, EPI_ISL_500507, EPI_ISL_500508, EPI_ISL_500509, EPI_ISL_500510, EPI_ISL_500511, EPI_ISL_500512, EPI_ISL_500513, EPI_ISL_500514, EPI_ISL_500515, EPI_ISL_500516, EPI_ISL_500517, EPI_ISL_500518, EPI_ISL_500519, EPI_ISL_500520, EPI_ISL_500521, EPI_ISL_500522, EPI_ISL_500523, EPI_ISL_500524, EPI_ISL_500525, EPI_ISL_500526, EPI_ISL_500527, EPI_ISL_500528, EPI_ISL_500529, EPI_ISL_500530, EPI_ISL_500531, EPI_ISL_500532, EPI_ISL_500533, EPI_ISL_500534, EPI_ISL_500535, EPI_ISL_500536, EPI_ISL_500537, EPI_ISL_500538 | | | |
| see above | Mayo Clinic Laboratories | University of Washington Virology Lab | Pavitra Roychoudhury, Hong Xie, Lasata Shrestha, Amin Addetia, Truong Nguyen, Victoria M Rachleff, Meei-Li Huang, Keith R Jerome, Alexander Greninger |
| EPI_ISL_500539, EPI_ISL_500540, EPI_ISL_500541, EPI_ISL_500542, EPI_ISL_500543, EPI_ISL_500544, EPI_ISL_500545, EPI_ISL_500546, EPI_ISL_500547, EPI_ISL_500548, EPI_ISL_500549, EPI_ISL_500550, EPI_ISL_500551, EPI_ISL_500552, EPI_ISL_500553, EPI_ISL_500554, EPI_ISL_500555, EPI_ISL_500556, EPI_ISL_500557, EPI_ISL_500558, EPI_ISL_500559, EPI_ISL_500560, EPI_ISL_500561, EPI_ISL_500562, EPI_ISL_500563, EPI_ISL_500564, EPI_ISL_500565, EPI_ISL_500566, EPI_ISL_500567, EPI_ISL_500568, EPI_ISL_500569, EPI_ISL_500570, EPI_ISL_500571, EPI_ISL_500572 | | | |
| see above | Singapore General Hospital | Department of Microbiology | Nurdyana Abdul Rahman, Kun Lee Lim, Chenhao Li, Kian Sing Chan, Lynette Oon, Kern Rei Chng, Niranjan Nagarajan, Karrie Ko |
| EPI_ISL_500707, EPI_ISL_500708, EPI_ISL_500709, EPI_ISL_500710, EPI_ISL_500711, EPI_ISL_500712, EPI_ISL_500713, EPI_ISL_500714, EPI_ISL_500715, EPI_ISL_500719, EPI_ISL_500951, EPI_ISL_500952, EPI_ISL_500953 | | | |
| see above | Respiratory Virus Unit, Microbiology Services Colindale, Public Health England | Respiratory Virus Unit, Microbiology Services Colindale, Public Health England | PHE Covid Sequencing Team |
| EPI_ISL_501072, EPI_ISL_501073, EPI_ISL_501074, EPI_ISL_501075, EPI_ISL_501076, EPI_ISL_501077, EPI_ISL_501078, EPI_ISL_501079, EPI_ISL_501080, EPI_ISL_501081, EPI_ISL_501082 | | | |
| see above | Mayo Clinic Laboratories | University of Washington Virology Lab | Pavitra Roychoudhury, Hong Xie, Lasata Shrestha, Amin Addetia, Truong Nguyen, Victoria M Rachleff, Meei-Li Huang, Keith R Jerome, Alexander Greninger |
| EPI_ISL_501083, EPI_ISL_501084, EPI_ISL_501085, EPI_ISL_501086, EPI_ISL_501087, EPI_ISL_501088, EPI_ISL_501089, EPI_ISL_501090, EPI_ISL_501091, EPI_ISL_501092, EPI_ISL_501093, EPI_ISL_501094, EPI_ISL_501095, EPI_ISL_501096, EPI_ISL_501097, EPI_ISL_501098, EPI_ISL_501099, EPI_ISL_501100, EPI_ISL_501101, EPI_ISL_501102, EPI_ISL_501103, EPI_ISL_501104, EPI_ISL_501105, EPI_ISL_501106, EPI_ISL_501107, EPI_ISL_501108, EPI_ISL_501109, EPI_ISL_501110, EPI_ISL_501111, EPI_ISL_501112, EPI_ISL_501113, EPI_ISL_501114, EPI_ISL_501115, EPI_ISL_501116, EPI_ISL_501117, EPI_ISL_501118, EPI_ISL_501120, EPI_ISL_501121, EPI_ISL_501122, EPI_ISL_501123, EPI_ISL_501124, EPI_ISL_501125, EPI_ISL_501126, EPI_ISL_501127, EPI_ISL_501128, EPI_ISL_501129, EPI_ISL_501130, EPI_ISL_501131, EPI_ISL_501132, EPI_ISL_501133, EPI_ISL_501134, EPI_ISL_501135, EPI_ISL_501136, EPI_ISL_501137, EPI_ISL_501138, EPI_ISL_501139, EPI_ISL_501140, EPI_ISL_501141, EPI_ISL_501142, EPI_ISL_501143, EPI_ISL_501144, EPI_ISL_501145, EPI_ISL_501146, EPI_ISL_501147, EPI_ISL_501148, EPI_ISL_501149, EPI_ISL_501150, EPI_ISL_501151, EPI_ISL_501152, EPI_ISL_501153, EPI_ISL_501154, EPI_ISL_501155, EPI_ISL_501156, EPI_ISL_501157, EPI_ISL_501158, EPI_ISL_501159, EPI_ISL_501160, EPI_ISL_501161, EPI_ISL_501162, EPI_ISL_501163, EPI_ISL_501164 | | | |
| see above | University of Washington Virology Lab | University of Washington Virology Lab | Pavitra Roychoudhury, Hong Xie, Lasata Shrestha, Amin Addetia, Truong Nguyen, Victoria M Rachleff, Meei-Li Huang, Keith R Jerome, Alexander Greninger |
| EPI_ISL_501165 | Mayo Clinic Laboratories | University of Washington Virology Lab | Pavitra Roychoudhury, Hong Xie, Lasata Shrestha, Amin Addetia, Truong Nguyen, Victoria M Rachleff, Meei-Li Huang, Keith R Jerome, Alexander Greninger |
| EPI_ISL_509372, EPI_ISL_509373, EPI_ISL_509374, EPI_ISL_509375, EPI_ISL_509376, EPI_ISL_509377, EPI_ISL_509378, EPI_ISL_509379, EPI_ISL_509380, EPI_ISL_509381, EPI_ISL_509382, EPI_ISL_509383, EPI_ISL_509384, EPI_ISL_509385, EPI_ISL_509386, EPI_ISL_509387, EPI_ISL_509388, EPI_ISL_509389, | | | |

EPI_ISL_509390, EPI_ISL_509391, EPI_ISL_509392, EPI_ISL_509393, EPI_ISL_509394, EPI_ISL_509395

| | | | |
|---|---|---|---|
| see above | Singapore General Hospital | Department of Microbiology | Nurdyana Abdul Rahman, Kun Lee Lim, Chenhao Li, Kian Sing Chan, Lynette Oon, Kern Rei Chng, Niranjan Nagarajan, Karrie Ko |
| EPI_ISL_509413, EPI_ISL_509414, EPI_ISL_509415, EPI_ISL_509416, EPI_ISL_509417 | Acibadem Labcell Cellular Therapy Laboratory | Acibadem Mehmet Ali Aydinlar University School of Medicine, Medical Genetics Department | Ozden Hatirnaz Ng, Sezer Akyoney, Ilayda Sahin, Gunseli Bayram Akcapinar, Ozkan Ozdemir, Derya Dilek Kancagi, Gozde Sir Karakus, Bulut Yurtsever, Cihan Tastan, Ercument Ovali, Ugur Ozbek |
| EPI_ISL_509436, EPI_ISL_509437, EPI_ISL_509438, EPI_ISL_509439, EPI_ISL_509440, EPI_ISL_509441, EPI_ISL_509442, EPI_ISL_509443 | The Princess Alexandra Hospital | Wellcome Sanger Institute for the COVID-19 Genomics UK (COG-UK) consortium | Nick Levene, Louise Lopez, Lynn Monaghan, Jessica Scott, Claudia McCrea and Alex Alderton, Roberto Amato, Sonia Goncalves, Ewan Harrison, David K. Jackson, Ian Johnston, Dominic Kwiatkowski, Cordelia Langford, John Sillitoe on behalf of the Wellcome Sanger Institute COVID-19 Surveillance Team (http://www.sanger.ac.uk/covid-team) |
| EPI_ISL_510689 | Hospital Universitari Germans Trias i Pujol(HUGTiP)/Fundació Lluita contra la SIDA (FLSida)/IRTA-CReSA | IrsiCaixa AIDS Research Lab | Pilar Armengol, Marc Noguera-Julian, Jordi Rodón, Julia Vergara, Lidia Ruiz, Nuria Izquierdo, Jorge Carrillo, Roger Paredes, Albert Bensaid, Julia Blanco, Joaquim Segalés, Bonaventura Clotet |
| EPI_ISL_510810, EPI_ISL_510811, EPI_ISL_510812, EPI_ISL_510813, EPI_ISL_510814, EPI_ISL_510815, EPI_ISL_510816 | NA | The Public Health Agency of Sweden | Oskar Karlsson Lindsjo, Maria Lind Karlberg, Mattias Haukland, Reza Advani, Olov Svartstrom, Anna-Malin Linde, Sandra Broddesson, Petra Edquist, Mia Brytting, Anna Risberg, Karin Tegmark-Wisell |
| EPI_ISL_510817 | Orebro klinisk mikrobiologi | The Public Health Agency of Sweden | Oskar Karlsson Lindsjo, Maria Lind Karlberg, Mattias Haukland, Reza Advani, Olov Svartstrom, Anna-Malin Linde, Sandra Broddesson, Petra Edquist, Mia Brytting, Anna Risberg, Karin Tegmark-Wisell |
| EPI_ISL_510818, EPI_ISL_510819 | Klinisk mikrobiologi Vasternorrland | The Public Health Agency of Sweden | Oskar Karlsson Lindsjo, Maria Lind Karlberg, Mattias Haukland, Reza Advani, Olov Svartstrom, Anna-Malin Linde, Sandra Broddesson, Petra Edquist, Mia Brytting, Anna Risberg, Karin Tegmark-Wisell |
| EPI_ISL_510820, EPI_ISL_510821, EPI_ISL_510822, EPI_ISL_510823, EPI_ISL_510824 | Klinisk mikrobiologi centralsjukhuset Karlstad | The Public Health Agency of Sweden | Oskar Karlsson Lindsjo, Maria Lind Karlberg, Mattias Haukland, Reza Advani, Olov Svartstrom, Anna-Malin Linde, Sandra Broddesson, Petra Edquist, Mia Brytting, Anna Risberg, Karin Tegmark-Wisell |
| EPI_ISL_510825, EPI_ISL_510826 | Kalmar klinisk mikrobiologi | The Public Health Agency of Sweden | Oskar Karlsson Lindsjo, Maria Lind Karlberg, Mattias Haukland, Reza Advani, Olov Svartstrom, Anna-Malin Linde, Sandra Broddesson, Petra Edquist, Mia Brytting, Anna Risberg, Karin Tegmark-Wisell |
| EPI_ISL_510827, EPI_ISL_510828, EPI_ISL_510829, EPI_ISL_510830 | Unilabs Eskilstuna | The Public Health Agency of Sweden | Oskar Karlsson Lindsjo, Maria Lind Karlberg, Mattias Haukland, Reza Advani, Olov Svartstrom, Anna-Malin Linde, Sandra Broddesson, Petra Edquist, Mia Brytting, Anna Risberg, Karin Tegmark-Wisell |
| EPI_ISL_510831 | Halmstad klinisk mikrobiologi | The Public Health Agency of Sweden | Oskar Karlsson Lindsjo, Maria Lind Karlberg, Mattias Haukland, Reza Advani, Olov Svartstrom, Anna-Malin Linde, Sandra Broddesson, Petra Edquist, Mia Brytting, Anna Risberg, Karin Tegmark-Wisell |
| EPI_ISL_510832, EPI_ISL_510833 | Kalmar klinisk mikrobiologi | The Public Health Agency of Sweden | Oskar Karlsson Lindsjo, Maria Lind Karlberg, Mattias Haukland, Reza Advani, Olov Svartstrom, Anna-Malin Linde, Sandra Broddesson, Petra Edquist, Mia Brytting, Anna Risberg, Karin Tegmark-Wisell |
| EPI_ISL_510834, EPI_ISL_510835 | Karolinska Universitetslaboratoriet | The Public Health Agency of Sweden | Oskar Karlsson Lindsjo, Maria Lind Karlberg, Mattias Haukland, Reza Advani, Olov Svartstrom, Anna-Malin Linde, Sandra Broddesson, Petra Edquist, Mia Brytting, Anna Risberg, Karin Tegmark-Wisell |
| EPI_ISL_510836, EPI_ISL_510837 | Kalmar klinisk mikrobiologi | The Public Health Agency of Sweden | Oskar Karlsson Lindsjo, Maria Lind Karlberg, Mattias Haukland, Reza Advani, Olov Svartstrom, Anna-Malin Linde, Sandra Broddesson, Petra Edquist, Mia Brytting, Anna Risberg, Karin Tegmark-Wisell |
| EPI_ISL_510838, EPI_ISL_510839, EPI_ISL_510840 | Karolinska Universitetslaboratoriet | The Public Health Agency of Sweden | Oskar Karlsson Lindsjo, Maria Lind Karlberg, Mattias Haukland, Reza Advani, Olov Svartstrom, Anna-Malin Linde, Sandra Broddesson, Petra Edquist, Mia Brytting, Anna Risberg, Karin Tegmark-Wisell |
| EPI_ISL_510841, EPI_ISL_510842, EPI_ISL_510843, EPI_ISL_510844 | Klinisk mikrobiologi NAL Trollhattan | The Public Health Agency of Sweden | Oskar Karlsson Lindsjo, Maria Lind Karlberg, Mattias Haukland, Reza Advani, Olov Svartstrom, Anna-Malin Linde, Sandra Broddesson, Petra Edquist, Mia Brytting, Anna Risberg, Karin Tegmark-Wisell |
| EPI_ISL_510845, EPI_ISL_510846, EPI_ISL_510847, EPI_ISL_510848, EPI_ISL_510849, EPI_ISL_510850, EPI_ISL_510851 | Skovde/Unilabs | The Public Health Agency of Sweden | Oskar Karlsson Lindsjo, Maria Lind Karlberg, Mattias Haukland, Reza Advani, Olov Svartstrom, Anna-Malin Linde, Sandra Broddesson, Petra Edquist, Mia Brytting, Anna Risberg, Karin Tegmark-Wisell |
| EPI_ISL_510852 | Karolinska Universitetslaboratoriet | The Public Health Agency of Sweden | Oskar Karlsson Lindsjo, Maria Lind Karlberg, Mattias Haukland, Reza Advani, Olov Svartstrom, Anna-Malin Linde, Sandra Broddesson, Petra Edquist, Mia Brytting, Anna Risberg, Karin Tegmark-Wisell |
| EPI_ISL_510853, EPI_ISL_510854, EPI_ISL_510855, EPI_ISL_510856, EPI_ISL_510857, EPI_ISL_510858 | Klinisk mikrobiologi Vasternorrland | The Public Health Agency of Sweden | Oskar Karlsson Lindsjo, Maria Lind Karlberg, Mattias Haukland, Reza Advani, Olov Svartstrom, Anna-Malin Linde, Sandra Broddesson, Petra Edquist, Mia Brytting, Anna Risberg, Karin Tegmark-Wisell |
| EPI_ISL_510859 | Kalmar klinisk mikrobiologi | The Public Health Agency of Sweden | Oskar Karlsson Lindsjo, Maria Lind Karlberg, Mattias Haukland, Reza Advani, Olov Svartstrom, Anna-Malin Linde, Sandra Broddesson, Petra Edquist, Mia Brytting, Anna Risberg, Karin Tegmark-Wisell |
| EPI_ISL_510860, EPI_ISL_510861, EPI_ISL_510862, EPI_ISL_510863 | Karolinska Universitetslaboratoriet | The Public Health Agency of Sweden | Oskar Karlsson Lindsjo, Maria Lind Karlberg, Mattias Haukland, Reza Advani, Olov Svartstrom, Anna-Malin Linde, Sandra Broddesson, Petra Edquist, Mia Brytting, Anna Risberg, Karin Tegmark-Wisell |
| EPI_ISL_510864, EPI_ISL_510865 | Kalmar klinisk mikrobiologi | The Public Health Agency of Sweden | Oskar Karlsson Lindsjo, Maria Lind Karlberg, Mattias Haukland, Reza Advani, Olov Svartstrom, Anna-Malin Linde, Sandra Broddesson, Petra Edquist, Mia Brytting, Anna Risberg, Karin Tegmark-Wisell |
| EPI_ISL_510866 | Halmstad klinisk mikrobiologi | The Public Health Agency of Sweden | Oskar Karlsson Lindsjo, Maria Lind Karlberg, Mattias Haukland, Reza Advani, Olov Svartstrom, Anna-Malin Linde, Sandra Broddesson, Petra Edquist, Mia Brytting, Anna Risberg, Karin Tegmark-Wisell |
| EPI_ISL_510867 | Klinisk mikrobiologi centralsjukhuset Karlstad | The Public Health Agency of Sweden | Oskar Karlsson Lindsjo, Maria Lind Karlberg, Mattias Haukland, Reza Advani, Olov Svartstrom, Anna-Malin Linde, Sandra Broddesson, Petra Edquist, Mia Brytting, Anna Risberg, Karin Tegmark-Wisell |
| EPI_ISL_510868, EPI_ISL_510869, EPI_ISL_510870 | Klinisk mikrobiologi NAL Trollhattan | The Public Health Agency of Sweden | Oskar Karlsson Lindsjo, Maria Lind Karlberg, Mattias Haukland, Reza Advani, Olov Svartstrom, Anna-Malin Linde, Sandra Broddesson, Petra Edquist, Mia Brytting, Anna Risberg, Karin Tegmark-Wisell |
| EPI_ISL_510871, EPI_ISL_510872, EPI_ISL_510873, EPI_ISL_510874, EPI_ISL_510875, EPI_ISL_510876, EPI_ISL_510877 | Klinsisk mikrobiologi Linkoping | The Public Health Agency of Sweden | Oskar Karlsson Lindsjo, Maria Lind Karlberg, Mattias Haukland, Reza Advani, Olov Svartstrom, Anna-Malin Linde, Sandra Broddesson, Petra Edquist, Mia Brytting, Anna Risberg, Karin Tegmark-Wisell |
| EPI_ISL_511852, EPI_ISL_511853, EPI_ISL_511854, EPI_ISL_511855, EPI_ISL_511856, EPI_ISL_511857, EPI_ISL_511858, EPI_ISL_511859, EPI_ISL_511860, EPI_ISL_511861 | UW Virology Lab | UW Virology Lab | Pavitra Roychoudhury, Amin Addetia, Hong Xie, Lasata Shrestha, Truong Nguyen, Meei-Li Huang, Keith Jerome, Alexander Greninger |

EPI_ISL_511863, EPI_ISL_511865, EPI_ISL_511866, EPI_ISL_511867, EPI_ISL_511868, EPI_ISL_511869, EPI_ISL_511870, EPI_ISL_511871, EPI_ISL_511872, EPI_ISL_511873, EPI_ISL_511874, EPI_ISL_511875, EPI_ISL_511876, EPI_ISL_511877

| | | | |
|---|---|---|---|
| see above | Johns Hopkins Hospital Department of Pathology | Johns Hopkins Hospital Department of Pathology | Peter M. Thielen, Thomas Mehoke, Shirlee Wohl, Srividya Ramakrishnan, Melanie Kirsche, Amanda Ernlund, Craig Howser, Kristina Zudock, Oluwaseun Falade-Nwulia, Norah Sadowski, Paul Morris, Mark Hopkins, Yunfan Fan, Nidia Trovao, Victoria Gniazdowski, Michael C. Schatz, Stuart C. Ray, Winston Timp, Heba H. Mostafa |

| | | | |
|---|---|---|---|
| EPI_ISL_511891, EPI_ISL_511892, EPI_ISL_511893, EPI_ISL_511894, EPI_ISL_511895, EPI_ISL_511896, EPI_ISL_511897, EPI_ISL_511898 | National Hospital of Tropical Diseases | Oxford University Clinical Research Unit, Hanoi, Vietnam | Nguyen Thi Tam, Van Dinh Trang, Nguyen Thi Hong Thuong, Vu Thi Ngoc Bich, Nguyen Thu Trang, Nguyen Thi Ngoc Diep, Le Nguyen Minh Hoa, Pham Ngoc Thach, H. Rogier van Doorn, on behalf of the OUCRU COVID-19 research group |
| EPI_ISL_512059 | B.J. Medical College and Civil hospital, Ahmedabad | Gujarat Biotechnology Research Centre | Pinal Trivedi, Maharshi Pandya, Nidhi Patel, Nitin Savaliya, Raghawendra Kumar, Dinesh Kumar, Zuber Saiyed, Komal Patel, Labdhi Pandya, Afzal Ansari, Nikha Trivedi, Pranay Shah, Kamlesh J Upadhyay, Sanjay Kapadia, Apurvasinh Puvar, Janvi Raval, Zarna Patel, Monika Gandhi, R D Dixit, A M Kadri, Harsh Bakshi, Chaitanya Joshi, Madhvi Joshi |
| EPI_ISL_512061 | B.J. Medical College and Civil hospital, Ahmedabad | Gujarat Biotechnology Research Centre | Nidhi Patel, Nitin Savaliya, Raghawendra Kumar, Dinesh Kumar, Zuber Saiyed, Komal Patel, Labdhi Pandya, Afzal Ansari, Nikha Trivedi, Pranay Shah, Kamlesh J Upadhyay, Sanjay Kapadia, Apurvasinh Puvar, Janvi Raval, Zarna Patel, Monika Gandhi, Pinal Trivedi, Maharshi Pandya, R D Dixit, A M Kadri, Harsh Bakshi, Chaitanya Joshi, Madhvi Joshi |
| EPI_ISL_512062 | B.J. Medical College and Civil hospital, Ahmedabad | Gujarat Biotechnology Research Centre | Nitin Savaliya, Raghawendra Kumar, Dinesh Kumar, Zuber Saiyed, Komal Patel, Labdhi Pandya, Afzal Ansari, Nikha Trivedi, Pranay Shah, Kamlesh J Upadhyay, Sanjay Kapadia, Apurvasinh Puvar, Janvi Raval, Zarna Patel, Monika Gandhi, Pinal Trivedi, Maharshi Pandya, Nidhi Patel, R D Dixit, A M Kadri, Harsh Bakshi, Chaitanya Joshi, Madhvi Joshi |
| EPI_ISL_512069 | Sardar Vallabhbhai Patel Institute of Medical Sciences & Research | Gujarat Biotechnology Research Centre | Nikha Trivedi, Pranay Shah, Kamlesh J Upadhyay, Sanjay Kapadia, Apurvasinh Puvar, Janvi Raval, Zarna Patel, Monika Gandhi, Pinal Trivedi, Maharshi Pandya, Nidhi Patel, Nitin Savaliya, Raghawendra Kumar, Dinesh Kumar, Zuber Saiyed, Komal Patel, Labdhi Pandya, Afzal Ansari, R D Dixit, A M Kadri, Harsh Bakshi, Chaitanya Joshi, Madhvi Joshi |
| EPI_ISL_512070 | Department of MicroBiology, Government Medical College, Surat | Gujarat Biotechnology Research Centre | Naresh Chauhan, Summaiya Mullan, Amit gamit, Apurvasinh Puvar, Janvi Raval, Zarna Patel, Monika Gandhi, Pinal Trivedi, Maharshi Pandya, Nidhi Patel, Nitin Savaliya, Raghawendra Kumar, Dinesh Kumar, Zuber Saiyed, Komal Patel, Labdhi Pandya, Afzal Ansari, Nikha Trivedi, R D Dixit, A M Kadri, Harsh Bakshi, Chaitanya Joshi, Madhvi Joshi |
| EPI_ISL_512076 | Dr. RSS Hospital, Modasa | Gujarat Biotechnology Research Centre | Pinal Trivedi, Maharshi Pandya, Nidhi Patel, Nitin Savaliya, Raghawendra Kumar, Dinesh Kumar, Zuber Saiyed, Komal Patel, Labdhi Pandya, Afzal Ansari, Nikha Trivedi, Harsh Chaudhari, Apurvasinh Puvar, Janvi Raval, Zarna Patel, Monika Gandhi, R D Dixit, A M Kadri, Harsh Bakshi, Chaitanya Joshi, Madhvi Joshi |
| EPI_ISL_512086 | UW Virology lab | UW Virology lab | Pavitra Roychoudhury, Amin Addetia, Hong Xie, Lasata Shrestha, Truong Nguyen, Meei-Li Huang, Keith Jerome, Alexander Greninger |
| EPI_ISL_512597, EPI_ISL_512598, EPI_ISL_512599, EPI_ISL_512600, EPI_ISL_512601, EPI_ISL_512602, EPI_ISL_512603, EPI_ISL_512604, EPI_ISL_512605, EPI_ISL_512606, EPI_ISL_512607, EPI_ISL_512608, EPI_ISL_512609, EPI_ISL_512610, EPI_ISL_512611, EPI_ISL_512612, EPI_ISL_512613, EPI_ISL_512614, EPI_ISL_512615, EPI_ISL_512616, EPI_ISL_512617, EPI_ISL_512618, EPI_ISL_512619, EPI_ISL_512620, EPI_ISL_512621, EPI_ISL_512622, EPI_ISL_512623, EPI_ISL_512624, EPI_ISL_512625, EPI_ISL_512626, EPI_ISL_512627, EPI_ISL_512628, EPI_ISL_512629, EPI_ISL_512630, EPI_ISL_512631, EPI_ISL_512632, EPI_ISL_512633, EPI_ISL_512634, EPI_ISL_512635, EPI_ISL_512636, EPI_ISL_512637, EPI_ISL_512639, EPI_ISL_512640, EPI_ISL_512641, EPI_ISL_512642, EPI_ISL_512643, EPI_ISL_512644 | | | |
| see above | National Laboratory for Influenza/Virology reference laboratory, Public Health Center of the Ministry of Health of Ukraine | Respiratory Virus Unit, Microbiology Services Colindale, Public Health England | PHE Covid Sequencing Team, Dr. Iryna Demchyshyna |
| EPI_ISL_512712, EPI_ISL_512713, EPI_ISL_512714, EPI_ISL_512715, EPI_ISL_512716, EPI_ISL_512717, EPI_ISL_512718, EPI_ISL_512719, EPI_ISL_512720, EPI_ISL_512721, EPI_ISL_512722, EPI_ISL_512723, EPI_ISL_512724, EPI_ISL_512725, EPI_ISL_512726, EPI_ISL_512727, EPI_ISL_512728, EPI_ISL_512729, EPI_ISL_512730, EPI_ISL_512731, EPI_ISL_512732, EPI_ISL_512733, EPI_ISL_512734, EPI_ISL_512735, EPI_ISL_512736, EPI_ISL_512737, EPI_ISL_512738, EPI_ISL_512739, EPI_ISL_512741, EPI_ISL_512742, EPI_ISL_512743, EPI_ISL_512744, EPI_ISL_512745, EPI_ISL_512746, EPI_ISL_512747, EPI_ISL_512748, EPI_ISL_512749, EPI_ISL_512750, EPI_ISL_512751, EPI_ISL_512752, EPI_ISL_512753, EPI_ISL_512754, EPI_ISL_512755, EPI_ISL_512756, EPI_ISL_512757, EPI_ISL_512758, EPI_ISL_512759, EPI_ISL_512760, EPI_ISL_512761, EPI_ISL_512762, EPI_ISL_512763, EPI_ISL_512764, EPI_ISL_512765 | | | |
| see above | PathWest Laboratory Medicine WA | PathWest Laboratory Medicine WA Microbial Surveillance Unit | PathWest Laboratory Medicine WA Microbial Surveillance Unit |
| EPI_ISL_512810 | National Laboratory for Influenza/Virology reference laboratory, Public Health Center of the Ministry of Health of Ukraine | Respiratory Virus Unit, Microbiology Services Colindale, Public Health England | PHE Covid Sequencing Team, Dr. Iryna Demchyshyna |
| EPI_ISL_512874, EPI_ISL_512875, EPI_ISL_512876, EPI_ISL_512877, EPI_ISL_512878, EPI_ISL_512879, EPI_ISL_512880, EPI_ISL_512881, EPI_ISL_512882, EPI_ISL_512883, EPI_ISL_512884, EPI_ISL_512885, EPI_ISL_512886, EPI_ISL_512887, EPI_ISL_512888, EPI_ISL_512889, EPI_ISL_512890, EPI_ISL_512891, EPI_ISL_512892, EPI_ISL_512893, EPI_ISL_512894, EPI_ISL_512895, EPI_ISL_512896, EPI_ISL_512897, EPI_ISL_512898, EPI_ISL_512899, EPI_ISL_512900, EPI_ISL_512901, EPI_ISL_512902, EPI_ISL_512903, EPI_ISL_513064, EPI_ISL_513065, EPI_ISL_513066, EPI_ISL_513067, EPI_ISL_513068, EPI_ISL_513069, EPI_ISL_513070, EPI_ISL_513071, EPI_ISL_513072, EPI_ISL_513073, EPI_ISL_513074, EPI_ISL_513075, EPI_ISL_513076, EPI_ISL_513208, EPI_ISL_513209, EPI_ISL_513210, EPI_ISL_513211, EPI_ISL_513212, EPI_ISL_513213 | | | |
| see above | Pathogen Genomics Lab King Abdullah University of Science and Technology(KAUST) | Pathogen Genomics Lab King Abdullah University of Science and Technology(KAUST) | Raeece Naeem, Rahul P Salunke, Sharif Hala, Sara Mfarrej, Amit Kumar Subudhi, Fadwa Alofi, Fathia Ben Rached, Afrah Alsomali, Asim Khogeer, Ahmad Bakur Mahmoud, Anwar Hashem, Naif Almontashiri, Arnab Pain |
| EPI_ISL_513589, EPI_ISL_513590, EPI_ISL_513591, EPI_ISL_513592, EPI_ISL_513593, EPI_ISL_513594, EPI_ISL_513595, EPI_ISL_513596, EPI_ISL_513597, EPI_ISL_513598, EPI_ISL_513599, EPI_ISL_513600, EPI_ISL_513601, EPI_ISL_513602, EPI_ISL_513603, EPI_ISL_513604, EPI_ISL_513605, EPI_ISL_513606, EPI_ISL_513607, EPI_ISL_513608, EPI_ISL_513609, EPI_ISL_513610, EPI_ISL_513611, EPI_ISL_513612, EPI_ISL_513613, EPI_ISL_513614, EPI_ISL_513615, EPI_ISL_513616, EPI_ISL_513617, EPI_ISL_513618, EPI_ISL_513619, EPI_ISL_513620, EPI_ISL_513621, EPI_ISL_513622, EPI_ISL_513623, EPI_ISL_513624, EPI_ISL_513625, EPI_ISL_513626, EPI_ISL_513627, EPI_ISL_513628, EPI_ISL_513629, EPI_ISL_513630, EPI_ISL_513631 | | | |
| see above | Viral Respiratory Lab, National Institute for Biomedical Research (INRB) | Pathogen Sequencing Lab, National Institute for Biomedical Research (INRB) | Placide Mbala-Kingebeni, Edith Nkwembe, Eddy Kinganda-Lusamaki, Amuri Aziza, Francisca Muyembe Mawete, Emmanuel Lokilo Lofiko, Catherine Pratt, Matthias Pauthner, Josh Quick, Allison Black, James Hadfield, Trevor Bedford, Ian Goodfellow, Andrew Rambaut, Nick Loman, Kristian Andersen, Michael Wiley, Steve Ahuka-Mundeke, Jean-Jacques Muyembe Tamfum |
| EPI_ISL_513632, EPI_ISL_513633, EPI_ISL_513634, EPI_ISL_513635, EPI_ISL_513636, EPI_ISL_513637, EPI_ISL_513638 | University of Washington Virology Lab | University of Washington Virology Lab | Pavitra Roychoudhury, Hong Xie, Lasata Shrestha, Amin Addetia, Truong Nguyen, Victoria M Rachleff, Meei-Li Huang, Keith R Jerome, Alexander Greninger |
| EPI_ISL_514103, EPI_ISL_514104, EPI_ISL_514105, EPI_ISL_514106, EPI_ISL_514107, EPI_ISL_514108, EPI_ISL_514109, EPI_ISL_514110, EPI_ISL_514111, EPI_ISL_514112, EPI_ISL_514113, EPI_ISL_514114, EPI_ISL_514115, EPI_ISL_514116, EPI_ISL_514117, EPI_ISL_514118, EPI_ISL_514119, EPI_ISL_514120, EPI_ISL_514121, EPI_ISL_514122, EPI_ISL_514123, EPI_ISL_514124, EPI_ISL_514125 | | | |
| see above | Viral Respiratory Lab, National Institute for Biomedical Research (INRB) | Pathogen Sequencing Lab, National Institute for Biomedical Research (INRB) | Placide Mbala-Kingebeni, Edith Nkwembe, Eddy Kinganda-Lusamaki, Amuri Aziza, Francisca Muyembe Mawete, Emmanuel Lokilo Lofiko, Catherine Pratt, Matthias Pauthner, Josh Quick, Allison Black, James Hadfield, Trevor Bedford, Ian Goodfellow, Andrew Rambaut, Nick Loman, Kristian Andersen, Michael Wiley, Steve Ahuka-Mundeke, Jean-Jacques Muyembe Tamfum |
| EPI_ISL_514267, EPI_ISL_514268, EPI_ISL_514269, EPI_ISL_514270, EPI_ISL_514271, EPI_ISL_514272, EPI_ISL_514273, EPI_ISL_514274 | Israel Central Virology laboratory | Israel Central Virology laboratory | Neta Zuckerman, Efrat Dahan Bucris, Oran Erster, Ella Mendelson, Michal Mandelboim |
| EPI_ISL_514275 | Israeli Central Virology laboratory | Israel Central Virology laboratory | Neta Zuckerman, Efrat Dahan Bucris, Oran Erster, Ella Mendelson, Michal Mandelboim |
| EPI_ISL_514276, EPI_ISL_514277, EPI_ISL_514278, EPI_ISL_514279, EPI_ISL_514280, EPI_ISL_514281, EPI_ISL_514282, EPI_ISL_514283, EPI_ISL_514284, EPI_ISL_514285, EPI_ISL_514286, EPI_ISL_514287, EPI_ISL_514288, EPI_ISL_514289, EPI_ISL_514290, EPI_ISL_514291, EPI_ISL_514292, EPI_ISL_514293, EPI_ISL_514294, EPI_ISL_514295, EPI_ISL_514296, EPI_ISL_514297, EPI_ISL_514298, EPI_ISL_514299, EPI_ISL_514300, EPI_ISL_514301, EPI_ISL_514302, EPI_ISL_514303, EPI_ISL_514304, EPI_ISL_514305, EPI_ISL_514306, EPI_ISL_514307, EPI_ISL_514308, EPI_ISL_514309, EPI_ISL_514310, EPI_ISL_514311, EPI_ISL_514312, EPI_ISL_514313, EPI_ISL_514314, EPI_ISL_514315, EPI_ISL_514316 | | | |
| see above | Israel Central Virology laboratory | Israel Central Virology laboratory | Neta Zuckerman, Efrat Dahan Bucris, Oran Erster, Ella Mendelson, Michal Mandelboim |
| EPI_ISL_514343, EPI_ISL_514344, EPI_ISL_514345, EPI_ISL_514346, EPI_ISL_514347, EPI_ISL_514348, EPI_ISL_514349, EPI_ISL_514350, EPI_ISL_514351, EPI_ISL_514352, EPI_ISL_514353 | | | |
| see above | Respiratory Virus Unit, Microbiology Services Colindale, Public Health England | Respiratory Virus Unit, Microbiology Services Colindale, Public Health England | PHE Covid Sequencing Team |
| EPI_ISL_514354 | General Hospital - Prilep | Research Center for Genetic Engineering and Biotechnology "Georgi D. Efremov" , Macedonian Academy of Sciences and Arts | RCGEB - MASA |
| EPI_ISL_514437 | Dr. RSS Hospital, Modasa | Gujarat Biotechnology Research Centre | Nidhi Patel, Nitin Savaliya, Raghawendra Kumar, Dinesh Kumar, Zuber Saiyed, Komal Patel, Labdhi Pandya, Afzal Ansari, Nikha Trivedi, Harsh Chaudhari, Apurvasinh Puvar, Janvi Raval, Zarna Patel, Monika Gandhi, Pinal Trivedi, Maharshi Pandya, R D Dixit, A M Kadri, Harsh Bakshi, Chaitanya Joshi, Madhvi |

| | | | |
|---|---|---|---|
| | | | Joshi |
| EPI_ISL_514581 | B.J. Medical College and Civil hospital, Ahmedabad | Gujarat Biotechnology Research Centre | Nitin Savaliya, Raghawendra Kumar, Dinesh Kumar, Zuber Saiyed, Komal Patel, Labdhi Pandya, Afzal Ansari, Nikha Trivedi, Pranay Shah, Kamlesh J Upadhyay, Sanjay Kapadia, Apurvasinh Puvar, Janvi Raval, Zarna Patel, Monika Gandhi, Pinal Trivedi, Maharshi Pandya, Nidhi Patel, R D Dixit, A M Kadri, Harsh Bakshi, Chaitanya Joshi, Madhvi Joshi |
| EPI_ISL_514588 | B.J. Medical College and Civil hospital, Ahmedabad | Gujarat Biotechnology Research Centre | Nikha Trivedi, Pranay Shah, Kamlesh J Upadhyay, Sanjay Kapadia, Apurvasinh Puvar, Janvi Raval, Zarna Patel, Monika Gandhi, Pinal Trivedi, Maharshi Pandya, Nidhi Patel, Nitin Savaliya, Raghawendra Kumar, Dinesh Kumar, Zuber Saiyed, Komal Patel, Labdhi Pandya, Afzal Ansari, R D Dixit, A M Kadri, Harsh Bakshi, Chaitanya Joshi, Madhvi Joshi |
| EPI_ISL_514589 | B.J. Medical College and Civil hospital, Ahmedabad | Gujarat Biotechnology Research Centre | Pranay Shah, Kamlesh J Upadhyay, Sanjay Kapadia, Apurvasinh Puvar, Janvi Raval, Zarna Patel, Monika Gandhi, Pinal Trivedi, Maharshi Pandya, Nidhi Patel, Nitin Savaliya, Raghawendra Kumar, Dinesh Kumar, Zuber Saiyed, Komal Patel, Labdhi Pandya, Afzal Ansari, Nikha Trivedi, R D Dixit, A M Kadri, Harsh Bakshi, Chaitanya Joshi, Madhvi Joshi |
| EPI_ISL_514596 | B.J. Medical College and Civil hospital, Ahmedabad | Gujarat Biotechnology Research Centre | Pinal Trivedi, Maharshi Pandya, Nidhi Patel, Nitin Savaliya, Raghawendra Kumar, Dinesh Kumar, Zuber Saiyed, Komal Patel, Labdhi Pandya, Afzal Ansari, Nikha Trivedi, Pranay Shah, Kamlesh J Upadhyay, Sanjay Kapadia, Apurvasinh Puvar, Janvi Raval, Zarna Patel, Monika Gandhi, R D Dixit, A M Kadri, Harsh Bakshi, Chaitanya Joshi, Madhvi Joshi |
| EPI_ISL_514598 | B.J. Medical College and Civil hospital, Ahmedabad | Gujarat Biotechnology Research Centre | Nidhi Patel, Nitin Savaliya, Raghawendra Kumar, Dinesh Kumar, Zuber Saiyed, Komal Patel, Labdhi Pandya, Afzal Ansari, Nikha Trivedi, Pranay Shah, Kamlesh J Upadhyay, Sanjay Kapadia, Apurvasinh Puvar, Janvi Raval, Zarna Patel, Monika Gandhi, Pinal Trivedi, Maharshi Pandya, R D Dixit, A M Kadri, Harsh Bakshi, Chaitanya Joshi, Madhvi Joshi |
| EPI_ISL_514599 | B.J. Medical College and Civil hospital, Ahmedabad | Gujarat Biotechnology Research Centre | Nitin Savaliya, Raghawendra Kumar, Dinesh Kumar, Zuber Saiyed, Komal Patel, Labdhi Pandya, Afzal Ansari, Nikha Trivedi, Pranay Shah, Kamlesh J Upadhyay, Sanjay Kapadia, Apurvasinh Puvar, Janvi Raval, Zarna Patel, Monika Gandhi, Pinal Trivedi, Maharshi Pandya, Nidhi Patel, R D Dixit, A M Kadri, Harsh Bakshi, Chaitanya Joshi, Madhvi Joshi |
| EPI_ISL_514606 | B.J. Medical College and Civil hospital, Ahmedabad | Gujarat Biotechnology Research Centre | Nikha Trivedi, Pranay Shah, Kamlesh J Upadhyay, Sanjay Kapadia, Apurvasinh Puvar, Janvi Raval, Zarna Patel, Monika Gandhi, Pinal Trivedi, Maharshi Pandya, Nidhi Patel, Nitin Savaliya, Raghawendra Kumar, Dinesh Kumar, Zuber Saiyed, Komal Patel, Labdhi Pandya, Afzal Ansari, R D Dixit, A M Kadri, Harsh Bakshi, Chaitanya Joshi, Madhvi Joshi |
| EPI_ISL_514607 | B.J. Medical College and Civil hospital, Ahmedabad | Gujarat Biotechnology Research Centre | Pranay Shah, Kamlesh J Upadhyay, Sanjay Kapadia, Apurvasinh Puvar, Janvi Raval, Zarna Patel, Monika Gandhi, Pinal Trivedi, Maharshi Pandya, Nidhi Patel, Nitin Savaliya, Raghawendra Kumar, Dinesh Kumar, Zuber Saiyed, Komal Patel, Labdhi Pandya, Afzal Ansari, Nikha Trivedi, R D Dixit, A M Kadri, Harsh Bakshi, Chaitanya Joshi, Madhvi Joshi |
| EPI_ISL_515082, EPI_ISL_515083, EPI_ISL_515084, EPI_ISL_515085, EPI_ISL_515086, EPI_ISL_515087, EPI_ISL_515088, EPI_ISL_515089, EPI_ISL_515090, EPI_ISL_515091, EPI_ISL_515092, EPI_ISL_515093, EPI_ISL_515094, EPI_ISL_515095, EPI_ISL_515096, EPI_ISL_515097, EPI_ISL_515098, EPI_ISL_515099, EPI_ISL_515100, EPI_ISL_515101, EPI_ISL_515102, EPI_ISL_515103, EPI_ISL_515104, EPI_ISL_515105, EPI_ISL_515106, EPI_ISL_515107, EPI_ISL_515108, EPI_ISL_515109, EPI_ISL_515110, EPI_ISL_515111, EPI_ISL_515112 | | | |
| see above | Department of Biochemistry, Cell and Molecular Biology | WACCBIP, University of Ghana | Ngoi,J.M., Quashie,P., Morang'a,C.M., Amuzu,D.S., Adu,B., Kumordjie,S., Eshun,M., Boatemaa,L., Magnussen,V., Kotey,E., Tei-Maya,F., Arjarquah,A., Mutungi,J.K., Bediako,Y., Asante,I., Bonney,E., Kyei,G.B., Bonney,K., Amenga-Etego,L.N., Anang,A.K., Awandare,G.A., Ampofo,W. |
| EPI_ISL_515270, EPI_ISL_515271, EPI_ISL_515272, EPI_ISL_515273, EPI_ISL_515274, EPI_ISL_515275, EPI_ISL_515276, EPI_ISL_515277, EPI_ISL_515278, EPI_ISL_515279, EPI_ISL_515280, EPI_ISL_515281, EPI_ISL_515282, EPI_ISL_515283, EPI_ISL_515284, EPI_ISL_515285, EPI_ISL_515286 | | | |
| see above | University of Washington Virology Lab | University of Washington Virology Lab | Pavitra Roychoudhury, Hong Xie, Lasata Shrestha, Amin Addetia, Truong Nguyen, Victoria M Rachleff, Meei-Li Huang, Keith R Jerome, Alexander Greninger |
| EPI_ISL_516413 | General Hospital - Kumanovo | Research Center for Genetic Engineering and Biotechnology "Georgi D. Efremov" , Macedonian Academy of Sciences and Arts | RCGEB - MASA |
| EPI_ISL_516414, EPI_ISL_516415, EPI_ISL_516416, EPI_ISL_516417, EPI_ISL_516418, EPI_ISL_516419, EPI_ISL_516420, EPI_ISL_516421, EPI_ISL_516422, EPI_ISL_516423, EPI_ISL_516424, EPI_ISL_516425 | | | |
| see above | Center for public health - Skopje | Research Center for Genetic Engineering and Biotechnology "Georgi D. Efremov" , Macedonian Academy of Sciences and Arts | RCGEB - MASA |
| EPI_ISL_516426, EPI_ISL_516427, EPI_ISL_516428, EPI_ISL_516429, EPI_ISL_516430, EPI_ISL_516431 | Clinical Hospital - Shtip | Research Center for Genetic Engineering and Biotechnology "Georgi D. Efremov" , Macedonian Academy of Sciences and Arts | RCGEB - MASA |
| EPI_ISL_516650 | Institute of Microbiology, Universidad San Francisco de Quito | Institute of Microbiology, Universidad San Francisco de Quito | Prado-Vivar, Sully Márquez, Juan José Guadalupe, Monica Becerra-Wong, Bernardo Gutiérrez, Verónica Barragán, Patricio Rojas-Silva, Gabriel Trueba, Michelle Grunauer, Paúl Cárdenas |
| EPI_ISL_516651 | Institute of Microbiology, Universidad San Francisco de Quito | Institute of Microbiology, Universidad San Francisco de Quito | Prado-Vivar, Sully Márquez, Juan José Guadalupe, Monica Becerra-Wong, Bernardo Gutiérrez, Nabih Dahik, Carlos Mena, Eulalia Pazmiño, Carolina Pacheco, DamarisSandoya, Verónica Barragán, Patricio Rojas-Silva, Gabriel Trueba, Michelle Grunauer, Paúl Cárdenas |
| EPI_ISL_516652 | Institute of Microbiology, Universidad San Francisco de Quito | Institute of Microbiology, Universidad San Francisco de Quito | Prado-Vivar, Sully Márquez, Juan José Guadalupe, Monica Becerra-Wong, Bernardo Gutiérrez, Nabih Dahik, Carlos Mena, Edy Quizhpe, Yomara Napa, Verónica Barragán, Patricio Rojas-Silva, Gabriel Trueba, Michelle Grunauer, Paúl Cárdenas |
| EPI_ISL_516884, EPI_ISL_516885, EPI_ISL_516886, EPI_ISL_516887, EPI_ISL_516888, EPI_ISL_516889, EPI_ISL_516890, EPI_ISL_516891 | Israel Central Virology laboratory | Israel Central Virology laboratory | Neta Zuckerman, Efrat Dahan Bucris, Oran Erster, Ella Mendelson, Michal Mandelboim |
| EPI_ISL_516892 | Israeli Central Virology laboratory | Israel Central Virology laboratory | Neta Zuckerman, Efrat Dahan Bucris, Oran Erster, Ella Mendelson, Michal Mandelboim |
| EPI_ISL_516893, EPI_ISL_516894, EPI_ISL_516895, EPI_ISL_516896, EPI_ISL_516897, EPI_ISL_516898, EPI_ISL_516899, EPI_ISL_516900, EPI_ISL_516901, EPI_ISL_516902, EPI_ISL_516903, EPI_ISL_516904, EPI_ISL_516905, EPI_ISL_516906, EPI_ISL_516907, EPI_ISL_516908, EPI_ISL_516909, EPI_ISL_516910, EPI_ISL_516911, EPI_ISL_516912, EPI_ISL_516913, EPI_ISL_516914, EPI_ISL_516915, EPI_ISL_516916, EPI_ISL_516917, EPI_ISL_516918, EPI_ISL_516919, EPI_ISL_516920, EPI_ISL_516921 | | | |
| see above | Israel Central Virology laboratory | Israel Central Virology laboratory | Neta Zuckerman, Efrat Dahan Bucris, Oran Erster, Ella Mendelson, Michal Mandelboim |
| EPI_ISL_516940, EPI_ISL_516941, EPI_ISL_516942, EPI_ISL_516943, EPI_ISL_516944, EPI_ISL_516945, EPI_ISL_516946, EPI_ISL_516947, EPI_ISL_516948, EPI_ISL_516949, EPI_ISL_516950, EPI_ISL_516951, EPI_ISL_516952, EPI_ISL_516953, EPI_ISL_516954, EPI_ISL_516955, EPI_ISL_516956, EPI_ISL_516957, EPI_ISL_516958, EPI_ISL_516959, EPI_ISL_516960, EPI_ISL_516961, EPI_ISL_516962, EPI_ISL_516963, EPI_ISL_516964, EPI_ISL_516965, EPI_ISL_516966, EPI_ISL_516967, EPI_ISL_516968, EPI_ISL_516969, EPI_ISL_516970, EPI_ISL_516971, EPI_ISL_516972, EPI_ISL_516973, EPI_ISL_516974, EPI_ISL_516975, EPI_ISL_516976, EPI_ISL_516977, EPI_ISL_516978, EPI_ISL_516979, EPI_ISL_516980, EPI_ISL_516981, EPI_ISL_516982, EPI_ISL_516983, EPI_ISL_516984, EPI_ISL_516985, EPI_ISL_516986 | | | |
| see above | King Georges Medical University | CSIR-National Botanical Research Institute | Priti Prasad, Shantanu Prakash, Kishan Sahu, Babita Singh, Suruchi Shukla, Hricha Mishra, Danish Nasar Khan , Om Prakash, MLB Bhatt, SK Barik, Mehar H.Asif,Samir V. Sawant,Amita Jain, Sumit Kr. Bag |
| EPI_ISL_517996, EPI_ISL_517997, EPI_ISL_517998, EPI_ISL_517999, EPI_ISL_518000, EPI_ISL_518001, EPI_ISL_518002, EPI_ISL_518003, EPI_ISL_518004, EPI_ISL_518005, EPI_ISL_518006, EPI_ISL_518007, EPI_ISL_518008, EPI_ISL_518009, EPI_ISL_518010, EPI_ISL_518011, EPI_ISL_518012, EPI_ISL_518013, EPI_ISL_518014, EPI_ISL_518015, EPI_ISL_518016, EPI_ISL_518017 | | | |
| see above | Singapore General Hospital | Department of Microbiology | Nurdyana Abdul Rahman, Kun Lee Lim, Chenhao Li, Kian Sing Chan, Lynette Oon, Kern Rei Chng, Niranjan Nagarajan, Karrie Ko |
| EPI_ISL_523951 | Respiratory Virus Unit, Microbiology Services Colindale, Public Health England | Respiratory Virus Unit, Microbiology Services Colindale, Public Health England | PHE Covid Sequencing Team |
| EPI_ISL_524445, EPI_ISL_524446, EPI_ISL_524447, EPI_ISL_524448, EPI_ISL_524449 | Singapore General Hospital | Department of Microbiology | Nurdyana Abdul Rahman, Kun Lee Lim, Chenhao Li, Kian Sing Chan, Lynette Oon, Kern Rei Chng, Niranjan Nagarajan, Karrie Ko |
| EPI_ISL_524717 | B.J. Medical College and Civil hospital, Ahmedabad | Gujarat Biotechnology Research Centre | Pinal Trivedi, Maharshi Pandya, Nidhi Patel, Nitin Savaliya, Raghawendra Kumar, Dinesh Kumar, Zuber Saiyed, Komal Patel, Labdhi Pandya, Afzal Ansari, Nikha Trivedi, Pranay Shah, Kamlesh J Upadhyay, Sanjay Kapadia, Apurvasinh Puvar, Janvi Raval, Zarna Patel, Monika Gandhi, R D Dixit, A M Kadri, Harsh Bakshi, Chaitanya Joshi, Madhvi Joshi |
| EPI_ISL_524719 | B.J. Medical College and Civil hospital, Ahmedabad | Gujarat Biotechnology Research Centre | Nidhi Patel, Nitin Savaliya, Raghawendra Kumar, Dinesh Kumar, Zuber Saiyed, Komal Patel, Labdhi Pandya, Afzal Ansari, Nikha Trivedi, Pranay Shah, |

| | | | |
|---|---|---|---|
| | | | Kamlesh J Upadhyay, Sanjay Kapadia, Apurvasinh Puvar, Janvi Raval, Zarna Patel, Monika Gandhi, Pinal Trivedi, Maharshi Pandya, R D Dixit, A M Kadri, Harsh Bakshi, Chaitanya Joshi, Madhvi Joshi |
| EPI_ISL_524720 | B.J. Medical College and Civil hospital, Ahmedabad | Gujarat Biotechnology Research Centre | Nitin Savaliya, Raghawendra Kumar, Dinesh Kumar, Zuber Saiyed, Komal Patel, Labdhi Pandya, Afzal Ansari, Nikha Trivedi, Pranay Shah, Kamlesh J Upadhyay, Sanjay Kapadia, Apurvasinh Puvar, Janvi Raval, Zarna Patel, Monika Gandhi, Pinal Trivedi, Maharshi Pandya, Nidhi Patel, R D Dixit, A M Kadri, Harsh Bakshi, Chaitanya Joshi, Madhvi Joshi |
| EPI_ISL_524727 | B.J. Medical College and Civil hospital, Ahmedabad | Gujarat Biotechnology Research Centre | Nikha Trivedi, Pranay Shah, Kamlesh J Upadhyay, Sanjay Kapadia, Apurvasinh Puvar, Janvi Raval, Zarna Patel, Monika Gandhi, Pinal Trivedi, Maharshi Pandya, Nidhi Patel, Nitin Savaliya, Raghawendra Kumar, Dinesh Kumar, Zuber Saiyed, Komal Patel, Labdhi Pandya, Afzal Ansari, R D Dixit, A M Kadri, Harsh Bakshi, Chaitanya Joshi, Madhvi Joshi |
| EPI_ISL_524728 | B.J. Medical College and Civil hospital, Ahmedabad | Gujarat Biotechnology Research Centre | Pranay Shah, Kamlesh J Upadhyay, Sanjay Kapadia, Apurvasinh Puvar, Janvi Raval, Zarna Patel, Monika Gandhi, Pinal Trivedi, Maharshi Pandya, Nidhi Patel, Nitin Savaliya, Raghawendra Kumar, Dinesh Kumar, Zuber Saiyed, Komal Patel, Labdhi Pandya, Afzal Ansari, Nikha Trivedi, R D Dixit, A M Kadri, Harsh Bakshi, Chaitanya Joshi, Madhvi Joshi |
| EPI_ISL_524731 | GMERS Medical College & Hospital, Gotri, Vadodara | Gujarat Biotechnology Research Centre | Neena Doshi, Varsha Godbole, Apurvasinh Puvar, Janvi Raval, Zarna Patel, Monika Gandhi, Pinal Trivedi, Maharshi Pandya, Nidhi Patel, Nitin Savaliya, Raghawendra Kumar, Dinesh Kumar, Zuber Saiyed, Komal Patel, Labdhi Pandya, Afzal Ansari, Nikha Trivedi, Meenakshi Shah, R D Dixit, A M Kadri, Harsh Bakshi, Chaitanya Joshi, Madhvi Joshi |
| EPI_ISL_524739 | GMERS Medical College and Hospital, Gandhinagar | Gujarat Biotechnology Research Centre | Pinal Trivedi, Maharshi Pandya, Nidhi Patel, Nitin Savaliya, Raghawendra Kumar, Dinesh Kumar, Zuber Saiyed, Komal Patel, Labdhi Pandya, Afzal Ansari, Nikha Trivedi, Seema Bhatt, Gaurishankar Shrimali, Bhavesh Modi, Bharti Rajani, Apurvasinh Puvar, Janvi Raval, Zarna Patel, Monika Gandhi, R D Dixit, A M Kadri, Harsh Bakshi, Chaitanya Joshi, Madhvi Joshi |
| EPI_ISL_524741 | GMERS Medical College and Hospital, Gandhinagar | Gujarat Biotechnology Research Centre | Nidhi Patel, Nitin Savaliya, Raghawendra Kumar, Dinesh Kumar, Zuber Saiyed, Komal Patel, Labdhi Pandya, Afzal Ansari, Nikha Trivedi, Seema Bhatt, Gaurishankar Shrimali, Bhavesh Modi, Bharti Rajani, Apurvasinh Puvar, Janvi Raval, Zarna Patel, Monika Gandhi, Pinal Trivedi, Maharshi Pandya, R D Dixit, A M Kadri, Harsh Bakshi, Chaitanya Joshi, Madhvi Joshi |
| EPI_ISL_524748 | Sir M P Shah Government Medical College, Jamnagar | Gujarat Biotechnology Research Centre | Nikha Trivedi, Apurvasinh Puvar, Janvi Raval, Binita Aring, Zarna Patel, Monika Gandhi, Pinal Trivedi, Maharshi Pandya, Nidhi Patel, Nitin Savaliya, Raghawendra Kumar, Dinesh Kumar, Zuber Saiyed, Komal Patel, Labdhi Pandya, Afzal Ansari, R D Dixit, A M Kadri, Harsh Bakshi, Chaitanya Joshi, Madhvi Joshi |
| EPI_ISL_524753 | GMERS Medical College and Hospital, Dharpur, Patan | Gujarat Biotechnology Research Centre | Pinal Trivedi, Maharshi Pandya, Nidhi Patel, Nitin Savaliya, Raghawendra Kumar, Dinesh Kumar, Zuber Saiyed, Komal Patel, Labdhi Pandya, Afzal Ansari, Nikha Trivedi, A N Parmar, Apurvasinh Puvar, Janvi Raval, Zarna Patel, Monika Gandhi, R D Dixit, A M Kadri, Harsh Bakshi, Chaitanya Joshi, Madhvi Joshi |
| EPI_ISL_524755 | GMERS Medical College and Hospital, Dharpur, Patan | Gujarat Biotechnology Research Centre | Nidhi Patel, Nitin Savaliya, Raghawendra Kumar, Dinesh Kumar, Zuber Saiyed, Komal Patel, Labdhi Pandya, Afzal Ansari, Nikha Trivedi, A N Parmar, Apurvasinh Puvar, Janvi Raval, Zarna Patel, Monika Gandhi, Pinal Trivedi, Maharshi Pandya, R D Dixit, A M Kadri, Harsh Bakshi, Chaitanya Joshi, Madhvi Joshi |
| EPI_ISL_524756 | GMERS Medical College and Hospital, Dharpur, Patan | Gujarat Biotechnology Research Centre | Nitin Savaliya, Raghawendra Kumar, Dinesh Kumar, Zuber Saiyed, Komal Patel, Labdhi Pandya, Afzal Ansari, Nikha Trivedi, A N Parmar, Apurvasinh Puvar, Janvi Raval, Zarna Patel, Monika Gandhi, Pinal Trivedi, Maharshi Pandya, Nidhi Patel, R D Dixit, A M Kadri, Harsh Bakshi, Chaitanya Joshi, Madhvi Joshi |
| EPI_ISL_524763 | GMERS Medical College and Hospital, Dharpur, Patan | Gujarat Biotechnology Research Centre | Nikha Trivedi, A N Parmar, Apurvasinh Puvar, Janvi Raval, Zarna Patel, Monika Gandhi, Pinal Trivedi, Maharshi Pandya, Nidhi Patel, Nitin Savaliya, Raghawendra Kumar, Dinesh Kumar, Zuber Saiyed, Komal Patel, Labdhi Pandya, Afzal Ansari, R D Dixit, A M Kadri, Harsh Bakshi, Chaitanya Joshi, Madhvi Joshi |
| EPI_ISL_525419 | GMERS Medical College and Hospital, Gandhinagar | Gujarat Biotechnology Research Centre | Nitin Savaliya, Raghawendra Kumar, Dinesh Kumar, Zuber Saiyed, Komal Patel, Labdhi Pandya, Afzal Ansari, Nikha Trivedi, Seema Bhatt, Gaurishankar Shrimali, Bhavesh Modi, Bharti Rajani, Apurvasinh Puvar, Janvi Raval, Zarna Patel, Monika Gandhi, Pinal Trivedi, Maharshi Pandya, Nidhi Patel, R D Dixit, A M Kadri, Harsh Bakshi, Chaitanya Joshi, Madhvi Joshi |
| EPI_ISL_525553, EPI_ISL_525554, EPI_ISL_525555, EPI_ISL_525556, EPI_ISL_525557, EPI_ISL_525558, EPI_ISL_525559, EPI_ISL_525560, EPI_ISL_525561, EPI_ISL_525562, EPI_ISL_525563, EPI_ISL_525564, EPI_ISL_525565, EPI_ISL_525566, EPI_ISL_525567, EPI_ISL_525568, EPI_ISL_525569, EPI_ISL_525570, EPI_ISL_525571, EPI_ISL_525572, EPI_ISL_525573, EPI_ISL_525574 | | | |
| see above | Istituto Zooprofilattico Sperimentale Puglia e Basilicata; Dipartimento di Bioscienze, Biotecnologie e Biofarmaceutica dell'Università degli Studi di Bari "A.Moro"; Istituto di Biomembrane. Bioenergetica e Biotecnologie Molecolari del Consiglio Nazionale delle Ricerche di Bari | Beaconlab (Bioinformatics, Evolution and Comparative Genomics lab), Dept of Biosciences, University on Milan | Parisi A.,Pesole G., Manzari C., Chiara M |
| EPI_ISL_526431, EPI_ISL_526432 | Lincolnshire Hospitals and DeepSeq Nottingham | COVID-19 Genomics UK (COG-UK) Consortium | Nichola Duckworth, Tim Sloan, Sarah Walsh, Jonathan Ball, Patrick McClure, Joeseph Chappell, Nadine Holmes, Matthew Carlisle, Christopher Moore, Fei Sang, Johnny Debebe, Victoria Wright, Matthew Loose |
| EPI_ISL_526537, EPI_ISL_526538, EPI_ISL_526539, EPI_ISL_526540, EPI_ISL_526541, EPI_ISL_526543, EPI_ISL_526544, EPI_ISL_526545, EPI_ISL_526546, EPI_ISL_526547, EPI_ISL_526548, EPI_ISL_526550, EPI_ISL_526754, EPI_ISL_526755, EPI_ISL_526756, EPI_ISL_526757, EPI_ISL_526758, EPI_ISL_526759, EPI_ISL_526760, EPI_ISL_526761, EPI_ISL_526762, EPI_ISL_526763, EPI_ISL_526764, EPI_ISL_526765, EPI_ISL_526766, EPI_ISL_526767, EPI_ISL_526768, EPI_ISL_526769, EPI_ISL_526770, EPI_ISL_526771, EPI_ISL_526772, EPI_ISL_526773, EPI_ISL_527257, EPI_ISL_527329, EPI_ISL_527330, EPI_ISL_527331, EPI_ISL_527332, EPI_ISL_527333, EPI_ISL_527334, EPI_ISL_527335, EPI_ISL_527336, EPI_ISL_527337, EPI_ISL_527338, EPI_ISL_527339, EPI_ISL_527340, EPI_ISL_527341, EPI_ISL_527342, EPI_ISL_527343, EPI_ISL_527344, EPI_ISL_527345, EPI_ISL_527346, EPI_ISL_527347, EPI_ISL_527348, EPI_ISL_527349, EPI_ISL_527350, EPI_ISL_527351, EPI_ISL_527352, EPI_ISL_527353, EPI_ISL_527354, EPI_ISL_527355, EPI_ISL_527356, EPI_ISL_527357, EPI_ISL_527358 | | | |
| see above | Respiratory Virus Unit, Microbiology Services Colindale, Public Health England | Respiratory Virus Unit, Microbiology Services Colindale, Public Health England | PHE Covid Sequencing Team |
| EPI_ISL_527380 | Istituto Zooprofilattico Sperimentale Puglia e Basilicata; Dipartimento di Bioscienze, Biotecnologie e Biofarmaceutica dell'Università degli Studi di Bari "A.Moro"; Istituto di Biomembrane. Bioenergetica e Biotecnologie Molecolari del Consiglio Nazionale delle Ricerche di Bari | Beaconlab (Bioinformatics, Evolution and Comparative Genomics lab), Dept of Biosciences, University on Milan | Parisi A.,Pesole G., Manzari C., Chiara M |
| EPI_ISL_527489, EPI_ISL_527490, EPI_ISL_527491, EPI_ISL_527492, EPI_ISL_527493, EPI_ISL_527494, EPI_ISL_527495, EPI_ISL_527496, EPI_ISL_527497, EPI_ISL_527498, EPI_ISL_527499, EPI_ISL_527500, EPI_ISL_527501, EPI_ISL_527502, EPI_ISL_527503, EPI_ISL_527504, EPI_ISL_527505, EPI_ISL_527506, EPI_ISL_527507, EPI_ISL_527508, EPI_ISL_527509, EPI_ISL_527510, EPI_ISL_527511, EPI_ISL_527512, EPI_ISL_527513, EPI_ISL_527514, EPI_ISL_527515, EPI_ISL_527516, EPI_ISL_527517, EPI_ISL_527518, EPI_ISL_527519, EPI_ISL_527520, EPI_ISL_527521, EPI_ISL_527522, EPI_ISL_527523, EPI_ISL_527524, EPI_ISL_527525, EPI_ISL_527526, EPI_ISL_527527, EPI_ISL_527528, EPI_ISL_527529, EPI_ISL_527530, EPI_ISL_527531, EPI_ISL_527532, EPI_ISL_527533, EPI_ISL_527534, EPI_ISL_527535, EPI_ISL_527536, EPI_ISL_527537, EPI_ISL_527538, EPI_ISL_527539, EPI_ISL_527540, EPI_ISL_527541, EPI_ISL_527542, EPI_ISL_527543, EPI_ISL_527544, EPI_ISL_527545, EPI_ISL_527546, EPI_ISL_527547, EPI_ISL_527548, EPI_ISL_527549, EPI_ISL_527550, EPI_ISL_527551, EPI_ISL_527552, EPI_ISL_527553, EPI_ISL_527554, EPI_ISL_527555, EPI_ISL_527556, EPI_ISL_527557, EPI_ISL_527558, EPI_ISL_527559, EPI_ISL_527560, EPI_ISL_527561, EPI_ISL_527562, EPI_ISL_527563, EPI_ISL_527564, EPI_ISL_527565, EPI_ISL_527566, EPI_ISL_527567, EPI_ISL_527568, EPI_ISL_527569, EPI_ISL_527570, EPI_ISL_527571, EPI_ISL_527572 | | | |
| see above | Viral Respiratory Lab, National Institute for Biomedical Research (INRB) | Pathogen Sequencing Lab, National Institute for Biomedical Research (INRB) | Placide Mbala-Kingebeni, Edith Nkwembe, Eddy Kinganda-Lusamaki, Amuri Aziza, Francisca Muyembe Mawete, Emmanuel Lokilo Lofiko, Catherine Pratt, Matthias Pauthner, Josh Quick, Allison Black, James Hadfield, Trevor Bedford, Ian Goodfellow, Andrew Rambaut, Nick Loman, Kristian Andersen, Michael Wiley, Steve Ahuka-Mundeke, Jean-Jacques Muyembe Tamfum |
| EPI_ISL_527872, EPI_ISL_527873, EPI_ISL_527874, EPI_ISL_527875, EPI_ISL_527876, EPI_ISL_527877, EPI_ISL_527878, EPI_ISL_527879, EPI_ISL_527880, EPI_ISL_527881, EPI_ISL_527882, EPI_ISL_527883, EPI_ISL_527884, EPI_ISL_527885, EPI_ISL_527886, EPI_ISL_527887, EPI_ISL_527888, EPI_ISL_527889, EPI_ISL_527890, EPI_ISL_527891, EPI_ISL_527892, EPI_ISL_527893, EPI_ISL_527895, EPI_ISL_527896, EPI_ISL_527898, EPI_ISL_527899, EPI_ISL_527900, EPI_ISL_527901, EPI_ISL_527902, EPI_ISL_527903, EPI_ISL_527904, EPI_ISL_527905, EPI_ISL_527906, EPI_ISL_527909, EPI_ISL_527910, EPI_ISL_527911, EPI_ISL_527912, EPI_ISL_527914, EPI_ISL_527915 | | | |
| see above | Nigeria Centre for Disease Control (NCDC) | African Centre of Excellence for Genomics of Infectious Diseases (ACEGID), Redeemer's University, Ede, Osun State, Nigeria | Oluniyi P.E. et al |
| EPI_ISL_528388, EPI_ISL_528389, EPI_ISL_528390, EPI_ISL_528391, EPI_ISL_528392, EPI_ISL_528393, EPI_ISL_528394, EPI_ISL_528395, EPI_ISL_528398, EPI_ISL_528399, EPI_ISL_528400, EPI_ISL_528401, EPI_ISL_528402, EPI_ISL_528403, EPI_ISL_528404, EPI_ISL_528405, EPI_ISL_528406, EPI_ISL_528407, EPI_ISL_528408, EPI_ISL_528409, EPI_ISL_528410, EPI_ISL_528411, EPI_ISL_528412, EPI_ISL_528413, EPI_ISL_528414, EPI_ISL_528415, EPI_ISL_528416, EPI_ISL_528417, EPI_ISL_528418, EPI_ISL_528430, EPI_ISL_528431, EPI_ISL_528432, EPI_ISL_528433, EPI_ISL_528434, EPI_ISL_528435, EPI_ISL_528437, EPI_ISL_528438, EPI_ISL_528439 | | | |
| see above | Respiratory Virus Unit, Microbiology Services Colindale, Public Health England | Respiratory Virus Unit, Microbiology Services Colindale, Public Health England | PHE Covid Sequencing Team |

| | | | |
|---|---|---|---|
| EPI_ISL_528827 | CSIR-Centre for Cellular and Molecular Biology | CSIR-Centre for Cellular and Molecular Biology | Namami Gaur, Sakshi Shambhavi, Lamuk Zaveri, Shagufta Khan, Nikhil Hajirnis, M Soujanya Reddy, Pratheusa Maccha, Tulasi Nagabandi, Purushotham Vodnala, Payel Mukherjee, Sofia Banu, Priya Singh, Onkar Kulkarni, Dhiviya Vedagiri, Divya Gupta, Vishal Sah, Santosh Kumar Kuncha, Krishnan Harinivas Harshan, Archana Bharadwaj Siva, Karthik Bharadwaj Tallapaka, Zeba Rizvi, Zuberwasim Sayyad, Kakade Aishwarya Arun, Amrutha H C, Ananga Ghosh, Rakesh K Mishra, Divya Tej Sowpati |
| EPI_ISL_528834 | CSIR-Centre for Cellular and Molecular Biology | CSIR-Centre for Cellular and Molecular Biology | Pratheusa Maccha, Sakshi Shambhavi, Lamuk Zaveri, Shagufta Khan, Namami Gaur, Nikhil Hajirnis, M Soujanya Reddy, Tulasi Nagabandi, Purushotham Vodnala, Payel Mukherjee, Sofia Banu, Priya Singh,Onkar Kulkarni, Dhiviya Vedagiri, Divya Gupta, Vishal Sah, Santosh Kumar Kuncha, Krishnan Harinivas Harshan, Archana Bharadwaj Siva, Karthik Bharadwaj Tallapaka, G. Aditya Kumar, Koushick Sivakumar,Disha Nanda, Divya Das, Jotin Gogoi, Manish Bhattacharjee, Ravi Prasad Mukku, Rakesh K Mishra, Divya Tej Sowpati |
| EPI_ISL_528838 | CSIR-Centre for Cellular and Molecular Biology | CSIR-Centre for Cellular and Molecular Biology | Payel Mukherjee, Sofia Banu, Priya Singh, Onkar Kulkarni, Dhiviya Vedagiri, Divya Gupta, Vishal Sah, Santosh Kumar Kuncha, Krishnan Harinivas Harshan, Archana Bharadwaj Siva, Karthik Bharadwaj Tallapaka, Shagufta Khan, Lamuk Zaveri, Nikhil Hajirnis, M Soujanya Reddy, Pratheusa Maccha, Namami Gaur, Sakshi Shambhavi, Tulasi Nagabandi, Purushotham Vodnala, Rakesh K Mishra, Sonu Uday, Sudipta Mondal, Annapoorna P Karthyayani, Debabrata Jana, Debrya Saha, Divya Tej Sowpati |
| EPI_ISL_528839 | CSIR-Centre for Cellular and Molecular Biology | CSIR-Centre for Cellular and Molecular Biology | Pratheusa Maccha, Sakshi Shambhavi, Lamuk Zaveri, Shagufta Khan, Namami Gaur, Nikhil Hajirnis, M Soujanya Reddy, Tulasi Nagabandi, Purushotham Vodnala, Payel Mukherjee, Sofia Banu, Priya Singh,Onkar Kulkarni, Dhiviya Vedagiri, Divya Gupta, Vishal Sah, Santosh Kumar Kuncha, Krishnan Harinivas Harshan, Archana Bharadwaj Siva, Karthik Bharadwaj Tallapaka, G. Aditya Kumar, Koushick Sivakumar,Disha Nanda, Divya Das, Jotin Gogoi, Manish Bhattacharjee, Ravi Prasad Mukku, Rakesh K Mishra, Divya Tej Sowpati |
| EPI_ISL_528841, EPI_ISL_528842, EPI_ISL_528845 | CSIR-Centre for Cellular and Molecular Biology | CSIR-Centre for Cellular and Molecular Biology | Payel Mukherjee, Sofia Banu, Priya Singh, Onkar Kulkarni, Dhiviya Vedagiri, Divya Gupta, Vishal Sah, Santosh Kumar Kuncha, Krishnan Harinivas Harshan, Archana Bharadwaj Siva, Karthik Bharadwaj Tallapaka, Shagufta Khan, Lamuk Zaveri, Nikhil Hajirnis, M Soujanya Reddy, Pratheusa Maccha, Namami Gaur, Sakshi Shambhavi, Tulasi Nagabandi, Purushotham Vodnala, Rakesh K Mishra, Sonu Uday, Sudipta Mondal, Annapoorna P Karthyayani, Debabrata Jana, Debrya Saha, Divya Tej Sowpati |
| EPI_ISL_528849 | CSIR-Centre for Cellular and Molecular Biology | CSIR-Centre for Cellular and Molecular Biology | Nikhil Hajirnis, M Soujanya Reddy, Pratheusa Maccha, Lamuk Zaveri, Shagufta Khan, Namami Gaur, Sakshi Shambhavi, Tulasi Nagabandi, Purushotham Vodnala, Payel Mukherjee, Sofia Banu, Priya Singh, Onkar Kulkarni, Dhiviya Vedagiri, Divya Gupta, Vishal Sah, Santosh Kumar Kuncha, Krishnan Harinivas Harshan, Archana Bharadwaj Siva, Karthik Bharadwaj Tallapaka,Zeba Rizvi, Zuberwasim Sayyad, Kakade Aishwarya Arun, Amrutha H C, Ananga Ghosh, Rakesh K Mishra, Divya Tej Sowpati |
| EPI_ISL_528851, EPI_ISL_528854 | CSIR-Centre for Cellular and Molecular Biology | CSIR-Centre for Cellular and Molecular Biology | Payel Mukherjee, Sofia Banu, Priya Singh, Onkar Kulkarni, Dhiviya Vedagiri, Divya Gupta, Vishal Sah, Santosh Kumar Kuncha, Krishnan Harinivas Harshan, Archana Bharadwaj Siva, Karthik Bharadwaj Tallapaka, Shagufta Khan, Lamuk Zaveri, Nikhil Hajirnis, M Soujanya Reddy, Pratheusa Maccha, Namami Gaur, Sakshi Shambhavi, Tulasi Nagabandi, Purushotham Vodnala, Rakesh K Mishra, Sonu Uday, Sudipta Mondal, Annapoorna P Karthyayani, Debabrata Jana, Debrya Saha, Divya Tej Sowpati |
| EPI_ISL_528855 | CSIR-Centre for Cellular and Molecular Biology | CSIR-Centre for Cellular and Molecular Biology | Nikhil Hajirnis, M Soujanya Reddy, Pratheusa Maccha, Lamuk Zaveri, Shagufta Khan, Namami Gaur, Sakshi Shambhavi, Tulasi Nagabandi, Purushotham Vodnala, Payel Mukherjee, Sofia Banu, Priya Singh, Onkar Kulkarni, Dhiviya Vedagiri, Divya Gupta, Vishal Sah, Santosh Kumar Kuncha, Krishnan Harinivas Harshan, Archana Bharadwaj Siva, Karthik Bharadwaj Tallapaka,Zeba Rizvi, Zuberwasim Sayyad, Kakade Aishwarya Arun, Amrutha H C, Ananga Ghosh, Rakesh K Mishra, Divya Tej Sowpati |
| EPI_ISL_528856 | CSIR-Centre for Cellular and Molecular Biology | CSIR-Centre for Cellular and Molecular Biology | Pratheusa Maccha, Sakshi Shambhavi, Lamuk Zaveri, Shagufta Khan, Namami Gaur, Nikhil Hajirnis, M Soujanya Reddy, Tulasi Nagabandi, Purushotham Vodnala, Payel Mukherjee, Sofia Banu, Priya Singh,Onkar Kulkarni, Dhiviya Vedagiri, Divya Gupta, Vishal Sah, Santosh Kumar Kuncha, Krishnan Harinivas Harshan, Archana Bharadwaj Siva, Karthik Bharadwaj Tallapaka, G. Aditya Kumar, Koushick Sivakumar,Disha Nanda, Divya Das, Jotin Gogoi, Manish Bhattacharjee, Ravi Prasad Mukku, Rakesh K Mishra, Divya Tej Sowpati |
| EPI_ISL_528857 | CSIR-Centre for Cellular and Molecular Biology | CSIR-Centre for Cellular and Molecular Biology | Nikhil Hajirnis, M Soujanya Reddy, Pratheusa Maccha, Lamuk Zaveri, Shagufta Khan, Namami Gaur, Sakshi Shambhavi, Tulasi Nagabandi, Purushotham Vodnala, Payel Mukherjee, Sofia Banu, Priya Singh, Onkar Kulkarni, Dhiviya Vedagiri, Divya Gupta, Vishal Sah, Santosh Kumar Kuncha, Krishnan Harinivas Harshan, Archana Bharadwaj Siva, Karthik Bharadwaj Tallapaka,Zeba Rizvi, Zuberwasim Sayyad, Kakade Aishwarya Arun, Amrutha H C, Ananga Ghosh, Rakesh K Mishra, Divya Tej Sowpati |
| EPI_ISL_528858, EPI_ISL_528859 | CSIR-Centre for Cellular and Molecular Biology | CSIR-Centre for Cellular and Molecular Biology | Pratheusa Maccha, Sakshi Shambhavi, Lamuk Zaveri, Shagufta Khan, Namami Gaur, Nikhil Hajirnis, M Soujanya Reddy, Tulasi Nagabandi, Purushotham Vodnala, Payel Mukherjee, Sofia Banu, Priya Singh,Onkar Kulkarni, Dhiviya Vedagiri, Divya Gupta, Vishal Sah, Santosh Kumar Kuncha, Krishnan Harinivas Harshan, Archana Bharadwaj Siva, Karthik Bharadwaj Tallapaka, G. Aditya Kumar, Koushick Sivakumar,Disha Nanda, Divya Das, Jotin Gogoi, Manish Bhattacharjee, Ravi Prasad Mukku, Rakesh K Mishra, Divya Tej Sowpati |
| EPI_ISL_528863 | CSIR-Centre for Cellular and Molecular Biology | CSIR-Centre for Cellular and Molecular Biology | Payel Mukherjee, Sofia Banu, Priya Singh, Onkar Kulkarni, Dhiviya Vedagiri, Divya Gupta, Vishal Sah, Santosh Kumar Kuncha, Krishnan Harinivas Harshan, Archana Bharadwaj Siva, Karthik Bharadwaj Tallapaka, Shagufta Khan, Lamuk Zaveri, Nikhil Hajirnis, M Soujanya Reddy, Pratheusa Maccha, Namami Gaur, Sakshi Shambhavi, Tulasi Nagabandi, Purushotham Vodnala, Rakesh K Mishra, Sonu Uday, Sudipta Mondal, Annapoorna P Karthyayani, Debabrata Jana, Debrya Saha, Divya Tej Sowpati |
| EPI_ISL_528930, EPI_ISL_528932, EPI_ISL_528933, EPI_ISL_528950, EPI_ISL_529027, EPI_ISL_529028 | Respiratory Virus Unit, Microbiology Services Colindale, Public Health England | Respiratory Virus Unit, Microbiology Services Colindale, Public Health England | PHE Covid Sequencing Team |
| EPI_ISL_529065, EPI_ISL_529066, EPI_ISL_529067, EPI_ISL_529068, EPI_ISL_529069, EPI_ISL_529070, EPI_ISL_529071, EPI_ISL_529072, EPI_ISL_529073, EPI_ISL_529074, EPI_ISL_529075, EPI_ISL_529076, EPI_ISL_529077, EPI_ISL_529078, EPI_ISL_529079 | | | |
| see above | Laboratorio de Referencia Nacional de Virus Respiratorios, Instituto Nacional de Salud Peru | Laboratorio de Genómica Microbiana, Universidad Peruana Cayetano Heredia | Pablo Tsukayama, Alejandra Dávila-Barclay, Luis González, Pedro E. Romero, Brenda Ayzanoa, Janet Huancachoque, Pool Marcos, Maribel Huaringa |
| EPI_ISL_529151, EPI_ISL_529152, EPI_ISL_529153 | Department of Immunology, The Scripps Research Institute | Andersen lab at Scripps Research | Quigley, M., Stefanski, E., Mchardy, I. with SEARCH Alliance San Diego |
| EPI_ISL_534717, EPI_ISL_534718, EPI_ISL_534719, EPI_ISL_534720, EPI_ISL_534721, EPI_ISL_534722, EPI_ISL_534723, EPI_ISL_534724, EPI_ISL_534725, EPI_ISL_534726, EPI_ISL_534727, EPI_ISL_534728, EPI_ISL_534729, EPI_ISL_534730 | | | |
| see above | Respiratory Virus Unit, Microbiology Services Colindale, Public Health England | Respiratory Virus Unit, Microbiology Services Colindale, Public Health England | PHE Covid Sequencing Team |
| EPI_ISL_537608 | Servicio de Microbiología. Consorcio Hospital General Universitario de Valencia | SeqCOVID-SPAIN consortium/IBV(CSIC) | Paula Ruiz-Hueso, Mariana Reyes-Prieto, Vicente Soriano Chirona, Ivan Ansari, David Navarro, Maria Alma Bracho, Griselda De Marco, Beatriz Beamud, Lidia Ruiz Roldan, Marta Pla Diaz, Neris Garcia-Gonzalez, Inma Galán Vendrell, Sandra Carbo, Loreto Ferrús Abad, Llúcia Martínez-Priego, Giuseppe D'Auria, Fernando Gonzalez-Candelas and SeqCOVID-SPAIN consortium |
| EPI_ISL_538321, EPI_ISL_538322, EPI_ISL_538323, EPI_ISL_538324, EPI_ISL_538325, EPI_ISL_538326, EPI_ISL_538327, EPI_ISL_538328, EPI_ISL_538329, EPI_ISL_538330, EPI_ISL_538331, EPI_ISL_538332, EPI_ISL_538333, EPI_ISL_538334, EPI_ISL_538335, EPI_ISL_538336, EPI_ISL_538337, EPI_ISL_538338, EPI_ISL_538339, EPI_ISL_538340, EPI_ISL_538341, EPI_ISL_538342, EPI_ISL_538343, EPI_ISL_538344, EPI_ISL_538345, EPI_ISL_538346, EPI_ISL_538347, EPI_ISL_538348, EPI_ISL_538349, EPI_ISL_538350, EPI_ISL_538351, EPI_ISL_538352, EPI_ISL_538353, EPI_ISL_538354, EPI_ISL_538355, EPI_ISL_538356, EPI_ISL_538357, EPI_ISL_538358, EPI_ISL_538359, EPI_ISL_538360, EPI_ISL_538361, EPI_ISL_538362, EPI_ISL_538363, EPI_ISL_538364, EPI_ISL_538365, EPI_ISL_538366, EPI_ISL_538367, EPI_ISL_538368, EPI_ISL_538369, EPI_ISL_538370, EPI_ISL_538371, EPI_ISL_538372, EPI_ISL_538373, EPI_ISL_538374, EPI_ISL_538375, EPI_ISL_538376, EPI_ISL_538377, EPI_ISL_538378, EPI_ISL_538379, EPI_ISL_538380, EPI_ISL_538381, EPI_ISL_538382, EPI_ISL_538383, EPI_ISL_538384, EPI_ISL_538385, EPI_ISL_538386 | | | |
| see above | Kingston Health Sciences Centre / Queen's University | Ontario Institute for Cancer Research | Prameet M. Sheth, Calvin Sjaarda, Robert Colautti, Katya Douchant, Ilinca Lungu, Bernard Lam, Paul Krzyzanowski, Michael Laszloffy, Lawrence E Heisler, Richard de Borja, Jared T. Simpson |
| EPI_ISL_538504, EPI_ISL_538505 | National Institute of Health Research and Development | National Institute of Health Research and Development | Pawestri, HA; Subangkit; Puspa, KD; Nugraha, AA; Ikawati, HD; Pangesti, KNA; Soekarso, T; Susilarini, NK; Hariastuti, NI; Nikmah, UA; Mursinah; Febriyani, A; Herman, R; Susanti, N; Herna; Febriyanti, T; Nurhadi, M; Paisal; Ramadhany, R; Agustiningsih; Kurniawati, J; Kipuw, NL; Muna, F; Indalau, IL; Adam, K; Wibowo, HA; Rizki, A; Puspandary, N; Setiawaty,V. |
| EPI_ISL_538506, EPI_ISL_538507 | Balai Penelitian dan Pengembangan Biomedis Papua | National Institute of Health Research and Development | Pawestri, HA; Subangkit; Puspa, KD; Nugraha, AA; Ikawati, HD; Pangesti, KNA; Soekarso, T; Paisal; Oktavian, A; Hutapea, HML; Setiawaty,V. |
| EPI_ISL_538508, EPI_ISL_538509 | National Institute of Health Research and Development | National Institute of Health Research and Development | Pawestri, HA; Subangkit; Puspa, KD; Nugraha, AA; Ikawati, HD; Pangesti, KNA; Soekarso, T; Susilarini, NK; Hariastuti, NI; Nikmah, UA; Mursinah; Febriyani, A; Herman, R; Susanti, N; Herna; Febriyanti, T; Nurhadi, M; Paisal; Ramadhany, R; Agustiningsih; Kurniawati, J; Kipuw, NL; Muna, F; Indalau, IL; Adam, K; Wibowo, HA; Rizki, A; Puspandary, N; Setiawaty,V. |

| | | | |
|---|---|---|---|
| EPI_ISL_538510 | RSUD Ulin Banjarmasin South Kalimantan | National Institute of Health Research and Development | Pawestri, HA; Subangkit; Puspa, KD; Nugraha, AA; Ikawati, HD; Pangesti, KNA; Soekarso, T; Paisal; Pasaribu, M; Setiawaty,V. |
| EPI_ISL_538512 | Balai Penelitian dan Pengembangan Biomedis Papua | National Institute of Health Research and Development | Pawestri, HA; Subangkit; Puspa, KD; Nugraha, AA; Ikawati, HD; Pangesti, KNA; Soekarso, T; Paisal; Pasaribu, M; Setiawaty,V. |
| EPI_ISL_539234, EPI_ISL_539240, EPI_ISL_539241, EPI_ISL_539242 | Servicio de Microbiología. Consorcio Hospital General Universitario de Valencia | SeqCOVID-SPAIN consortium/IBV(CSIC) | Paula Ruiz-Hueso, Mariana Reyes-Prieto, Vicente Soriano Chirona, Ivan Ansari, David Navarro, Maria Alma Bracho, Griselda De Marco, Beatriz Beamud, Lidia Ruiz Roldan, Marta Pla Diaz, Neris Garcia-Gonzalez, Inma Galán Vendrell, Sandra Carbo, Loreto Ferrús Abad, Llúcia Martínez-Priego, Giuseppe D'Auria, Fernando Gonzalez-Candelas and SeqCOVID-SPAIN consortium |
| EPI_ISL_539246, EPI_ISL_539247, EPI_ISL_539248, EPI_ISL_539249, EPI_ISL_539250, EPI_ISL_539251, EPI_ISL_539252, EPI_ISL_539253, EPI_ISL_539254, EPI_ISL_539255, EPI_ISL_539256, EPI_ISL_539257, EPI_ISL_539258, EPI_ISL_539260, EPI_ISL_539261, EPI_ISL_539262, EPI_ISL_539263, EPI_ISL_539264, EPI_ISL_539265, EPI_ISL_539266, EPI_ISL_539267, EPI_ISL_539268, EPI_ISL_539269, EPI_ISL_539270, EPI_ISL_539271, EPI_ISL_539272, EPI_ISL_539273, EPI_ISL_539274, EPI_ISL_539275, EPI_ISL_539276, EPI_ISL_539277, EPI_ISL_539278, EPI_ISL_539279, EPI_ISL_539280, EPI_ISL_539281, EPI_ISL_539282, EPI_ISL_539283 | | | |
| see above | Hospital Universitario de La Ribera (Alzira, València) | SeqCOVID-SPAIN consortium/IBV(CSIC) | Olalla Martínez Macias, Julia González and SeqCOVID-SPAIN consortium |
| EPI_ISL_539289, EPI_ISL_539290, EPI_ISL_539291, EPI_ISL_539292, EPI_ISL_539293, EPI_ISL_539294, EPI_ISL_539295, EPI_ISL_539296, EPI_ISL_539297, EPI_ISL_539298 | Servicio de Microbiología. Consorcio Hospital General Universitario de Valencia | SeqCOVID-SPAIN consortium/IBV(CSIC) | Paula Ruiz-Hueso, Mariana Reyes-Prieto, Vicente Soriano Chirona, Ivan Ansari, David Navarro, Maria Alma Bracho, Griselda De Marco, Beatriz Beamud, Lidia Ruiz Roldan, Marta Pla Diaz, Neris Garcia-Gonzalez, Inma Galán Vendrell, Sandra Carbo, Loreto Ferrús Abad, Llúcia Martínez-Priego, Giuseppe D'Auria, Fernando Gonzalez-Candelas and SeqCOVID-SPAIN consortium |
| EPI_ISL_539333, EPI_ISL_539334, EPI_ISL_539335, EPI_ISL_539336, EPI_ISL_539337, EPI_ISL_539338, EPI_ISL_539339 | Institute of Disease Control and Prevention, People's Liberation Army | Institute of Disease Control and Prevention, People's Liberation Army | Qiu,S., Li,P. |
| EPI_ISL_539622 | CSIR-Centre for Cellular and Molecular Biology | CSIR-Centre for Cellular and Molecular Biology | Namami Gaur, Sakshi Shambhavi, Lamuk Zaveri, Shagufta Khan, Nikhil Hajirnis, M Soujanya Reddy, Pratheusa Maccha, Tulasi Nagabandi, Purushotham Vodnala, Payel Mukherjee, Sofia Banu, Priya Singh, Onkar Kulkarni, Dhiviya Vedagiri, Divya Gupta, Vishal Sah, Santosh Kumar Kuncha, Krishnan Harinivas Harshan, Archana Bharadwaj Siva, Karthik Bharadwaj Tallapaka, Zeba Rizvi, Zuberwasim Sayyad, Kakade Aishwarya Arun, Amrutha H C, Ananga Ghosh, Rakesh K Mishra, Divya Tej Sowpati |
| EPI_ISL_539623 | CSIR-Centre for Cellular and Molecular Biology | CSIR-Centre for Cellular and Molecular Biology | Namami Gaur, Sakshi Shambhavi, Lamuk Zaveri, Shagufta Khan, Nikhil Hajirnis, M Soujanya Reddy, Pratheusa Maccha, Tulasi Nagabandi, Purushotham Vodnala, Payel Mukherjee, Sofia Banu, Priya Singh, Onkar Kulkarni, Dhiviya Vedagiri, Divya Gupta, Vishal Sah, Santosh Kumar Kuncha, Krishnan Harinivas Harshan, Archana Bharadwaj Siva, Karthik Bharadwaj Tallapaka,G. Aditya Kumar, Koushick Sivakumar, Rakesh K Mishra, Divya Tej Sowpati |
| EPI_ISL_539624 | CSIR-Centre for Cellular and Molecular Biology | CSIR-Centre for Cellular and Molecular Biology | Namami Gaur, Sakshi Shambhavi, Lamuk Zaveri, Shagufta Khan, Nikhil Hajirnis, M Soujanya Reddy, Pratheusa Maccha, Tulasi Nagabandi, Purushotham Vodnala, Payel Mukherjee, Sofia Banu, Priya Singh,Onkar Kulkarni, Dhiviya Vedagiri, Divya Gupta, Vishal Sah, Santosh Kumar Kuncha, Krishnan Harinivas Harshan, Archana Bharadwaj Siva, Karthik Bharadwaj Tallapaka, Zeba Rizvi, Zuberwasim Sayyad, Kakade Aishwarya Arun, Amrutha H C, Ananga Ghosh, Rakesh K Mishra, Divya Tej Sowpati |
| EPI_ISL_539625 | CSIR-Centre for Cellular and Molecular Biology | CSIR-Centre for Cellular and Molecular Biology | Nikhil Hajirnis, M Soujanya Reddy, Pratheusa Maccha, Lamuk Zaveri, Shagufta Khan, Namami Gaur, Sakshi Shambhavi, Tulasi Nagabandi, Purushotham Vodnala, Payel Mukherjee, Sofia Banu, Priya Singh, Onkar Kulkarni, Dhiviya Vedagiri, Divya Gupta, Vishal Sah, Santosh Kumar Kuncha, Krishnan Harinivas Harshan, Archana Bharadwaj Siva, Karthik Bharadwaj Tallapaka,Zeba Rizvi, Zuberwasim Sayyad, Kakade Aishwarya Arun, Amrutha H C, Ananga Ghosh, Rakesh K Mishra, Divya Tej Sowpati |
| EPI_ISL_539626 | CSIR-Centre for Cellular and Molecular Biology | CSIR-Centre for Cellular and Molecular Biology | Nikhil Hajirnis, M Soujanya Reddy, Pratheusa Maccha, Namami Gaur, Sakshi Shambhavi, Lamuk Zaveri, Shagufta Khan, Tulasi Nagabandi, Purushotham Vodnala, Payel Mukherjee, Sofia Banu, Priya Singh,Onkar Kulkarni, Dhiviya Vedagiri, Divya Gupta, Vishal Sah, Santosh Kumar Kuncha, Krishnan Harinivas Harshan, Archana Bharadwaj Siva, Karthik Bharadwaj Tallapaka,Kezia J Ann, Radhika Khandelwal, Roshan Maku Venkata, Shemin Mansuri, Sonu Uday, Rakesh K Mishra, Divya Tej Sowpati |
| EPI_ISL_539627 | CSIR-Centre for Cellular and Molecular Biology | CSIR-Centre for Cellular and Molecular Biology | Nikhil Hajirnis, M Soujanya Reddy, Pratheusa Maccha, Payel Mukherjee, Sofia Banu, Priya Singh,Onkar Kulkarni, Dhiviya Vedagiri, Divya Gupta, Vishal Sah, Santosh Kumar Kuncha, Krishnan Harinivas Harshan, Archana Bharadwaj Siva, Karthik Bharadwaj Tallapaka, Shagufta Khan, Lamuk Zaveri, Namami Gaur, Sakshi Shambhavi, Tulasi Nagabandi, Purushotham Vodnala,Deepak Kumar, Devi Prasad Vijayashankar, Disha Nanda, Divya Das, Jotin Gogoi, Manish Bhattacharjee, Rakesh K Mishra, Divya Tej Sowpati |
| EPI_ISL_539628 | CSIR-Centre for Cellular and Molecular Biology | CSIR-Centre for Cellular and Molecular Biology | Payel Mukherjee, Sofia Banu, Priya Singh, Onkar Kulkarni, Dhiviya Vedagiri, Divya Gupta, Vishal Sah, Santosh Kumar Kuncha, Krishnan Harinivas Harshan, Archana Bharadwaj Siva, Karthik Bharadwaj Tallapaka, Shagufta Khan, Lamuk Zaveri, Nikhil Hajirnis, M Soujanya Reddy, Pratheusa Maccha, Namami Gaur, Sakshi Shambhavi, Tulasi Nagabandi, Purushotham Vodnala, G. Aditya Kumar, Koushick Sivakumar, Pooja Ramesh Gupta, Rajan Kumar Jha, Shraddha Vijay Lahoti, Rakesh K Mishra, Divya Tej Sowpati |
| EPI_ISL_539629 | CSIR-Centre for Cellular and Molecular Biology | CSIR-Centre for Cellular and Molecular Biology | Payel Mukherjee, Sofia Banu, Priya Singh, Onkar Kulkarni, Dhiviya Vedagiri, Divya Gupta, Vishal Sah, Santosh Kumar Kuncha, Krishnan Harinivas Harshan, Archana Bharadwaj Siva, Karthik Bharadwaj Tallapaka, Shagufta Khan, Lamuk Zaveri, Nikhil Hajirnis, M Soujanya Reddy, Pratheusa Maccha, Namami Gaur, Sakshi Shambhavi, Tulasi Nagabandi, Purushotham Vodnala, Gokulan C G, Gunjan Purohit, Hanuman Tulashiram Kale, Pankaj Kumar, Prachand Issarapu, Rakesh K Mishra, Divya Tej Sowpati |
| EPI_ISL_539630 | CSIR-Centre for Cellular and Molecular Biology | CSIR-Centre for Cellular and Molecular Biology | Payel Mukherjee, Sofia Banu, Priya Singh, Onkar Kulkarni, Dhiviya Vedagiri, Divya Gupta, Vishal Sah, Santosh Kumar Kuncha, Krishnan Harinivas Harshan, Archana Bharadwaj Siva, Karthik Bharadwaj Tallapaka, Shagufta Khan, Lamuk Zaveri, Nikhil Hajirnis, M Soujanya Reddy, Pratheusa Maccha, Namami Gaur, Sakshi Shambhavi, Tulasi Nagabandi, Purushotham Vodnala, Rakesh K Mishra, Sonu Uday, Sudipta Mondal, Annapoorna P Karthyayani, Debabrata Jana, Debrya Saha, Divya Tej Sowpati |
| EPI_ISL_539631 | CSIR-Centre for Cellular and Molecular Biology | CSIR-Centre for Cellular and Molecular Biology | Pratheusa Maccha, Sakshi Shambhavi, Lamuk Zaveri, Shagufta Khan, Namami Gaur, Nikhil Hajirnis, M Soujanya Reddy, Tulasi Nagabandi, Purushotham Vodnala, Payel Mukherjee, Sofia Banu, Priya Singh,Onkar Kulkarni, Dhiviya Vedagiri, Divya Gupta, Vishal Sah, Santosh Kumar Kuncha, Krishnan Harinivas Harshan, Archana Bharadwaj Siva, Karthik Bharadwaj Tallapaka, G. Aditya Kumar, Koushick Sivakumar,Disha Nanda, Divya Das, Jotin Gogoi, Manish Bhattacharjee, Ravi Prasad Mukku, Rakesh K Mishra, Divya Tej Sowpati |
| EPI_ISL_539632 | CSIR-Centre for Cellular and Molecular Biology | CSIR-Centre for Cellular and Molecular Biology | Pratheusa Maccha, Shagufta Khan, Lamuk Zaveri, Namami Gaur, Sakshi Shambhavi, Tulasi Nagabandi, Nikhil Hajirnis, M Soujanya Reddy, Purushotham Vodnala, Payel Mukherjee, Sofia Banu, Priya Singh, Onkar Kulkarni, Dhiviya Vedagiri, Divya Gupta, Vishal Sah, Santosh Kumar Kuncha, Krishnan Harinivas Harshan, Archana Bharadwaj Siva, Karthik Bharadwaj Tallapaka, Disha Nanda, Divya Das, Jotin Gogoi, Manish Bhattacharjee, Ravi Prasad Mukku, Rakesh K Mishra, Divya Tej Sowpati |
| EPI_ISL_539633 | CSIR-Centre for Cellular and Molecular Biology | CSIR-Centre for Cellular and Molecular Biology | Pratheusa Maccha, Sofia Banu, Payel Mukherjee, Priya Singh, Onkar Kulkarni, Dhiviya Vedagiri, Divya Gupta, Vishal Sah, Santosh Kumar Kuncha, Krishnan Harinivas Harshan, Archana Bharadwaj Siva, Karthik Bharadwaj Tallapaka, Shagufta Khan, Lamuk Zaveri, Namami Gaur, Sakshi Shambhavi, Nikhil Hajirnis, M Soujanya Reddy, Tulasi Nagabandi, Purushotham Vodnala,Preethi Jampala, Sharada Ravi Iyer, Sulagana Mukherjee, Swetha Sundar, Peddapuvala Sai Uday Kiran, Rakesh K Mishra, Divya Tej Sowpati |
| EPI_ISL_539655 | CSIR-Centre for Cellular and Molecular Biology | CSIR-Centre for Cellular and Molecular Biology | Nikhil Hajirnis, M Soujanya Reddy, Pratheusa Maccha, Lamuk Zaveri, Shagufta Khan, Namami Gaur, Sakshi Shambhavi, Tulasi Nagabandi, Purushotham Vodnala, Payel Mukherjee, Sofia Banu, Priya Singh, Onkar Kulkarni, Dhiviya Vedagiri, Divya Gupta, Vishal Sah, Santosh Kumar Kuncha, Krishnan Harinivas Harshan, Archana Bharadwaj Siva, Karthik Bharadwaj Tallapaka,Zeba Rizvi, Zuberwasim Sayyad, Kakade Aishwarya Arun, Amrutha H C, Ananga Ghosh, Rakesh K Mishra, Divya Tej Sowpati |
| EPI_ISL_539656 | CSIR-Centre for Cellular and Molecular Biology | CSIR-Centre for Cellular and Molecular Biology | Nikhil Hajirnis, M Soujanya Reddy, Pratheusa Maccha, Namami Gaur, Sakshi Shambhavi, Lamuk Zaveri, Shagufta Khan, Tulasi Nagabandi, Purushotham Vodnala, Payel Mukherjee, Sofia Banu, Priya Singh,Onkar Kulkarni, Dhiviya Vedagiri, Divya Gupta, Vishal Sah, Santosh Kumar Kuncha, Krishnan Harinivas Harshan, Archana Bharadwaj Siva, Karthik Bharadwaj Tallapaka,Kezia J Ann, Radhika Khandelwal, Roshan Maku Venkata, Shemin Mansuri, Sonu Uday, Rakesh K Mishra, Divya Tej Sowpati |
| EPI_ISL_539657 | CSIR-Centre for Cellular and Molecular Biology | CSIR-Centre for Cellular and Molecular Biology | Nikhil Hajirnis, M Soujanya Reddy, Pratheusa Maccha, Payel Mukherjee, Sofia Banu, Priya Singh,Onkar Kulkarni, Dhiviya Vedagiri, Divya Gupta, Vishal Sah, Santosh Kumar Kuncha, Krishnan Harinivas Harshan, Archana Bharadwaj Siva, Karthik Bharadwaj Tallapaka, Shagufta Khan, Lamuk Zaveri, Namami Gaur, Sakshi Shambhavi, Tulasi Nagabandi, Purushotham Vodnala,Deepak Kumar, Devi Prasad Vijayashankar, Disha Nanda, Divya Das, Jotin Gogoi, Manish Bhattacharjee, Rakesh K Mishra, Divya Tej Sowpati |

| | | | |
|---|---|---|---|
| EPI_ISL_539658 | CSIR-Centre for Cellular and Molecular Biology | CSIR-Centre for Cellular and Molecular Biology | Payel Mukherjee, Sofia Banu, Priya Singh, Onkar Kulkarni, Dhiviya Vedagiri, Divya Gupta, Vishal Sah, Santosh Kumar Kuncha, Krishnan Harinivas Harshan, Archana Bharadwaj Siva, Karthik Bharadwaj Tallapaka, Shagufta Khan, Lamuk Zaveri, Nikhil Hajirnis, M Soujanya Reddy, Pratheusa Maccha, Namami Gaur, Sakshi Shambhavi, Tulasi Nagabandi, Purushotham Vodnala, G. Aditya Kumar, Koushick Sivakumar, Pooja Ramesh Gupta, Rajan Kumar Jha, Shraddha Vijay Lahoti, Rakesh K Mishra, Divya Tej Sowpati |
| EPI_ISL_539659 | CSIR-Centre for Cellular and Molecular Biology | CSIR-Centre for Cellular and Molecular Biology | Payel Mukherjee, Sofia Banu, Priya Singh, Onkar Kulkarni, Dhiviya Vedagiri, Divya Gupta, Vishal Sah, Santosh Kumar Kuncha, Krishnan Harinivas Harshan, Archana Bharadwaj Siva, Karthik Bharadwaj Tallapaka, Shagufta Khan, Lamuk Zaveri, Nikhil Hajirnis, M Soujanya Reddy, Pratheusa Maccha, Namami Gaur, Sakshi Shambhavi, Tulasi Nagabandi, Purushotham Vodnala, Gokulan C G, Gunjan Purohit, Hanuman Tulashiram Kale, Pankaj Kumar, Prachand Issarapu, Rakesh K Mishra, Divya Tej Sowpati |
| EPI_ISL_539660 | CSIR-Centre for Cellular and Molecular Biology | CSIR-Centre for Cellular and Molecular Biology | Payel Mukherjee, Sofia Banu, Priya Singh, Onkar Kulkarni, Dhiviya Vedagiri, Divya Gupta, Vishal Sah, Santosh Kumar Kuncha, Krishnan Harinivas Harshan, Archana Bharadwaj Siva, Karthik Bharadwaj Tallapaka, Shagufta Khan, Lamuk Zaveri, Nikhil Hajirnis, M Soujanya Reddy, Pratheusa Maccha, Namami Gaur, Sakshi Shambhavi, Tulasi Nagabandi, Purushotham Vodnala, Rakesh K Mishra, Sonu Uday, Sudipta Mondal, Annapoorna P Karthyayani, Debabrata Jana, Debrya Saha, Divya Tej Sowpati |
| EPI_ISL_539661 | CSIR-Centre for Cellular and Molecular Biology | CSIR-Centre for Cellular and Molecular Biology | Pratheusa Maccha, Sakshi Shambhavi, Lamuk Zaveri, Shagufta Khan, Namami Gaur, Nikhil Hajirnis, M Soujanya Reddy, Tulasi Nagabandi, Purushotham Vodnala, Payel Mukherjee, Sofia Banu, Priya Singh,Onkar Kulkarni, Dhiviya Vedagiri, Divya Gupta, Vishal Sah, Santosh Kumar Kuncha, Krishnan Harinivas Harshan, Archana Bharadwaj Siva, Karthik Bharadwaj Tallapaka, G. Aditya Kumar, Koushick Sivakumar,Disha Nanda, Divya Das, Jotin Gogoi, Manish Bhattacharjee, Ravi Prasad Mukku, Rakesh K Mishra, Divya Tej Sowpati |
| EPI_ISL_539662 | CSIR-Centre for Cellular and Molecular Biology | CSIR-Centre for Cellular and Molecular Biology | Pratheusa Maccha, Shagufta Khan, Lamuk Zaveri, Namami Gaur, Sakshi Shambhavi, Tulasi Nagabandi, Nikhil Hajirnis, M Soujanya Reddy, Purushotham Vodnala, Payel Mukherjee, Sofia Banu, Priya Singh, Onkar Kulkarni, Dhiviya Vedagiri, Divya Gupta, Vishal Sah, Santosh Kumar Kuncha, Krishnan Harinivas Harshan, Archana Bharadwaj Siva, Karthik Bharadwaj Tallapaka, Disha Nanda, Divya Das, Jotin Gogoi, Manish Bhattacharjee, Ravi Prasad Mukku, Rakesh K Mishra, Divya Tej Sowpati |
| EPI_ISL_539663 | CSIR-Centre for Cellular and Molecular Biology | CSIR-Centre for Cellular and Molecular Biology | Pratheusa Maccha, Sofia Banu, Payel Mukherjee, Priya Singh, Onkar Kulkarni, Dhiviya Vedagiri, Divya Gupta, Vishal Sah, Santosh Kumar Kuncha, Krishnan Harinivas Harshan, Archana Bharadwaj Siva, Karthik Bharadwaj Tallapaka, Shagufta Khan, Lamuk Zaveri, Namami Gaur, Sakshi Shambhavi, Nikhil Hajirnis, M Soujanya Reddy, Tulasi Nagabandi, Purushotham Vodnala,Preethi Jampala, Sharada Ravi Iyer, Sulagana Mukherjee, Swetha Sundar, Peddapuvala Sai Uday Kiran, Rakesh K Mishra, Divya Tej Sowpati |
| EPI_ISL_539685 | CSIR-Centre for Cellular and Molecular Biology | CSIR-Centre for Cellular and Molecular Biology | Nikhil Hajirnis, M Soujanya Reddy, Pratheusa Maccha, Lamuk Zaveri, Shagufta Khan, Namami Gaur, Sakshi Shambhavi, Tulasi Nagabandi, Purushotham Vodnala, Payel Mukherjee, Sofia Banu, Priya Singh, Onkar Kulkarni, Dhiviya Vedagiri, Divya Gupta, Vishal Sah, Santosh Kumar Kuncha, Krishnan Harinivas Harshan, Archana Bharadwaj Siva, Karthik Bharadwaj Tallapaka,Zeba Rizvi, Zuberwasim Sayyad, Kakade Aishwarya Arun, Amrutha H C, Ananga Ghosh, Rakesh K Mishra, Divya Tej Sowpati |
| EPI_ISL_539686 | CSIR-Centre for Cellular and Molecular Biology | CSIR-Centre for Cellular and Molecular Biology | Nikhil Hajirnis, M Soujanya Reddy, Pratheusa Maccha, Namami Gaur, Sakshi Shambhavi, Lamuk Zaveri, Shagufta Khan, Tulasi Nagabandi, Purushotham Vodnala, Payel Mukherjee, Sofia Banu, Priya Singh,Onkar Kulkarni, Dhiviya Vedagiri, Divya Gupta, Vishal Sah, Santosh Kumar Kuncha, Krishnan Harinivas Harshan, Archana Bharadwaj Siva, Karthik Bharadwaj Tallapaka,Kezia J Ann, Radhika Khandelwal, Roshan Maku Venkata, Shemin Mansuri, Sonu Uday, Rakesh K Mishra, Divya Tej Sowpati |
| EPI_ISL_539687 | CSIR-Centre for Cellular and Molecular Biology | CSIR-Centre for Cellular and Molecular Biology | Nikhil Hajirnis, M Soujanya Reddy, Pratheusa Maccha, Payel Mukherjee, Sofia Banu, Priya Singh,Onkar Kulkarni, Dhiviya Vedagiri, Divya Gupta, Vishal Sah, Santosh Kumar Kuncha, Krishnan Harinivas Harshan, Archana Bharadwaj Siva, Karthik Bharadwaj Tallapaka, Shagufta Khan, Lamuk Zaveri, Namami Gaur, Sakshi Shambhavi, Tulasi Nagabandi, Purushotham Vodnala,Deepak Kumar, Devi Prasad Vijayashankar, Disha Nanda, Divya Das, Jotin Gogoi, Manish Bhattacharjee, Rakesh K Mishra, Divya Tej Sowpati |
| EPI_ISL_539688 | CSIR-Centre for Cellular and Molecular Biology | CSIR-Centre for Cellular and Molecular Biology | Payel Mukherjee, Sofia Banu, Priya Singh, Onkar Kulkarni, Dhiviya Vedagiri, Divya Gupta, Vishal Sah, Santosh Kumar Kuncha, Krishnan Harinivas Harshan, Archana Bharadwaj Siva, Karthik Bharadwaj Tallapaka, Shagufta Khan, Lamuk Zaveri, Nikhil Hajirnis, M Soujanya Reddy, Pratheusa Maccha, Namami Gaur, Sakshi Shambhavi, Tulasi Nagabandi, Purushotham Vodnala, G. Aditya Kumar, Koushick Sivakumar, Pooja Ramesh Gupta, Rajan Kumar Jha, Shraddha Vijay Lahoti, Rakesh K Mishra, Divya Tej Sowpati |
| EPI_ISL_539689 | CSIR-Centre for Cellular and Molecular Biology | CSIR-Centre for Cellular and Molecular Biology | Payel Mukherjee, Sofia Banu, Priya Singh, Onkar Kulkarni, Dhiviya Vedagiri, Divya Gupta, Vishal Sah, Santosh Kumar Kuncha, Krishnan Harinivas Harshan, Archana Bharadwaj Siva, Karthik Bharadwaj Tallapaka, Shagufta Khan, Lamuk Zaveri, Nikhil Hajirnis, M Soujanya Reddy, Pratheusa Maccha, Namami Gaur, Sakshi Shambhavi, Tulasi Nagabandi, Purushotham Vodnala, Gokulan C G, Gunjan Purohit, Hanuman Tulashiram Kale, Pankaj Kumar, Prachand Issarapu, Rakesh K Mishra, Divya Tej Sowpati |
| EPI_ISL_539690 | CSIR-Centre for Cellular and Molecular Biology | CSIR-Centre for Cellular and Molecular Biology | Payel Mukherjee, Sofia Banu, Priya Singh, Onkar Kulkarni, Dhiviya Vedagiri, Divya Gupta, Vishal Sah, Santosh Kumar Kuncha, Krishnan Harinivas Harshan, Archana Bharadwaj Siva, Karthik Bharadwaj Tallapaka, Shagufta Khan, Lamuk Zaveri, Nikhil Hajirnis, M Soujanya Reddy, Pratheusa Maccha, Namami Gaur, Sakshi Shambhavi, Tulasi Nagabandi, Purushotham Vodnala, Rakesh K Mishra, Sonu Uday, Sudipta Mondal, Annapoorna P Karthyayani, Debabrata Jana, Debrya Saha, Divya Tej Sowpati |
| EPI_ISL_539691 | CSIR-Centre for Cellular and Molecular Biology | CSIR-Centre for Cellular and Molecular Biology | Pratheusa Maccha, Sakshi Shambhavi, Lamuk Zaveri, Shagufta Khan, Namami Gaur, Nikhil Hajirnis, M Soujanya Reddy, Tulasi Nagabandi, Purushotham Vodnala, Payel Mukherjee, Sofia Banu, Priya Singh,Onkar Kulkarni, Dhiviya Vedagiri, Divya Gupta, Vishal Sah, Santosh Kumar Kuncha, Krishnan Harinivas Harshan, Archana Bharadwaj Siva, Karthik Bharadwaj Tallapaka, G. Aditya Kumar, Koushick Sivakumar,Disha Nanda, Divya Das, Jotin Gogoi, Manish Bhattacharjee, Ravi Prasad Mukku, Rakesh K Mishra, Divya Tej Sowpati |
| EPI_ISL_539692 | CSIR-Centre for Cellular and Molecular Biology | CSIR-Centre for Cellular and Molecular Biology | Pratheusa Maccha, Shagufta Khan, Lamuk Zaveri, Namami Gaur, Sakshi Shambhavi, Tulasi Nagabandi, Nikhil Hajirnis, M Soujanya Reddy, Purushotham Vodnala, Payel Mukherjee, Sofia Banu, Priya Singh, Onkar Kulkarni, Dhiviya Vedagiri, Divya Gupta, Vishal Sah, Santosh Kumar Kuncha, Krishnan Harinivas Harshan, Archana Bharadwaj Siva, Karthik Bharadwaj Tallapaka, Disha Nanda, Divya Das, Jotin Gogoi, Manish Bhattacharjee, Ravi Prasad Mukku, Rakesh K Mishra, Divya Tej Sowpati |
| EPI_ISL_539693 | CSIR-Centre for Cellular and Molecular Biology | CSIR-Centre for Cellular and Molecular Biology | Pratheusa Maccha, Sofia Banu, Payel Mukherjee, Priya Singh, Onkar Kulkarni, Dhiviya Vedagiri, Divya Gupta, Vishal Sah, Santosh Kumar Kuncha, Krishnan Harinivas Harshan, Archana Bharadwaj Siva, Karthik Bharadwaj Tallapaka, Shagufta Khan, Lamuk Zaveri, Namami Gaur, Sakshi Shambhavi, Nikhil Hajirnis, M Soujanya Reddy, Tulasi Nagabandi, Purushotham Vodnala,Preethi Jampala, Sharada Ravi Iyer, Sulagana Mukherjee, Swetha Sundar, Peddapuvala Sai Uday Kiran, Rakesh K Mishra, Divya Tej Sowpati |
| EPI_ISL_539715 | CSIR-Centre for Cellular and Molecular Biology | CSIR-Centre for Cellular and Molecular Biology | Nikhil Hajirnis, M Soujanya Reddy, Pratheusa Maccha, Lamuk Zaveri, Shagufta Khan, Namami Gaur, Sakshi Shambhavi, Tulasi Nagabandi, Purushotham Vodnala, Payel Mukherjee, Sofia Banu, Priya Singh, Onkar Kulkarni, Dhiviya Vedagiri, Divya Gupta, Vishal Sah, Santosh Kumar Kuncha, Krishnan Harinivas Harshan, Archana Bharadwaj Siva, Karthik Bharadwaj Tallapaka,Zeba Rizvi, Zuberwasim Sayyad, Kakade Aishwarya Arun, Amrutha H C, Ananga Ghosh, Rakesh K Mishra, Divya Tej Sowpati |
| EPI_ISL_539716 | CSIR-Centre for Cellular and Molecular Biology | CSIR-Centre for Cellular and Molecular Biology | Nikhil Hajirnis, M Soujanya Reddy, Pratheusa Maccha, Namami Gaur, Sakshi Shambhavi, Lamuk Zaveri, Shagufta Khan, Tulasi Nagabandi, Purushotham Vodnala, Payel Mukherjee, Sofia Banu, Priya Singh,Onkar Kulkarni, Dhiviya Vedagiri, Divya Gupta, Vishal Sah, Santosh Kumar Kuncha, Krishnan Harinivas Harshan, Archana Bharadwaj Siva, Karthik Bharadwaj Tallapaka,Kezia J Ann, Radhika Khandelwal, Roshan Maku Venkata, Shemin Mansuri, Sonu Uday, Rakesh K Mishra, Divya Tej Sowpati |
| EPI_ISL_539717 | CSIR-Centre for Cellular and Molecular Biology | CSIR-Centre for Cellular and Molecular Biology | Nikhil Hajirnis, M Soujanya Reddy, Pratheusa Maccha, Payel Mukherjee, Sofia Banu, Priya Singh,Onkar Kulkarni, Dhiviya Vedagiri, Divya Gupta, Vishal Sah, Santosh Kumar Kuncha, Krishnan Harinivas Harshan, Archana Bharadwaj Siva, Karthik Bharadwaj Tallapaka, Shagufta Khan, Lamuk Zaveri, Namami Gaur, Sakshi Shambhavi, Tulasi Nagabandi, Purushotham Vodnala,Deepak Kumar, Devi Prasad Vijayashankar, Disha Nanda, Divya Das, Jotin Gogoi, Manish Bhattacharjee, Rakesh K Mishra, Divya Tej Sowpati |
| EPI_ISL_539718 | CSIR-Centre for Cellular and Molecular Biology | CSIR-Centre for Cellular and Molecular Biology | Payel Mukherjee, Sofia Banu, Priya Singh, Onkar Kulkarni, Dhiviya Vedagiri, Divya Gupta, Vishal Sah, Santosh Kumar Kuncha, Krishnan Harinivas Harshan, Archana Bharadwaj Siva, Karthik Bharadwaj Tallapaka, Shagufta Khan, Lamuk Zaveri, Nikhil Hajirnis, M Soujanya Reddy, Pratheusa Maccha, Namami Gaur, Sakshi Shambhavi, Tulasi Nagabandi, Purushotham Vodnala, G. Aditya Kumar, Koushick Sivakumar, Pooja Ramesh Gupta, Rajan Kumar Jha, Shraddha Vijay Lahoti, Rakesh K Mishra, Divya Tej Sowpati |
| EPI_ISL_539719 | CSIR-Centre for Cellular and Molecular Biology | CSIR-Centre for Cellular and Molecular Biology | Payel Mukherjee, Sofia Banu, Priya Singh, Onkar Kulkarni, Dhiviya Vedagiri, Divya Gupta, Vishal Sah, Santosh Kumar Kuncha, Krishnan Harinivas Harshan, Archana Bharadwaj Siva, Karthik Bharadwaj Tallapaka, Shagufta Khan, Lamuk Zaveri, Nikhil Hajirnis, M Soujanya Reddy, Pratheusa Maccha, Namami Gaur, |

| | | | |
|---|---|---|---|
| | | | Sakshi Shambhavi, Tulasi Nagabandi, Purushotham Vodnala, Gokulan C G, Gunjan Purohit, Hanuman Tulashiram Kale, Pankaj Kumar, Prachand Issarapu, Rakesh K Mishra, Divya Tej Sowpati |
| EPI_ISL_539720 | CSIR-Centre for Cellular and Molecular Biology | CSIR-Centre for Cellular and Molecular Biology | Payel Mukherjee, Sofia Banu, Priya Singh, Onkar Kulkarni, Dhiviya Vedagiri, Divya Gupta, Vishal Sah, Santosh Kumar Kuncha, Krishnan Harinivas Harshan, Archana Bharadwaj Siva, Karthik Bharadwaj Tallapaka, Shagufta Khan, Lamuk Zaveri, Nikhil Hajirnis, M Soujanya Reddy, Pratheusa Maccha, Namami Gaur, Sakshi Shambhavi, Tulasi Nagabandi, Purushotham Vodnala, Rakesh K Mishra, Sonu Uday, Sudipta Mondal, Annapoorna P Karthyayani, Debabrata Jana, Debrya Saha, Divya Tej Sowpati |
| EPI_ISL_539721 | CSIR-Centre for Cellular and Molecular Biology | CSIR-Centre for Cellular and Molecular Biology | Pratheusa Maccha, Sakshi Shambhavi, Lamuk Zaveri, Shagufta Khan, Namami Gaur, Nikhil Hajirnis, M Soujanya Reddy, Tulasi Nagabandi, Purushotham Vodnala, Payel Mukherjee, Sofia Banu, Priya Singh,Onkar Kulkarni, Dhiviya Vedagiri, Divya Gupta, Vishal Sah, Santosh Kumar Kuncha, Krishnan Harinivas Harshan, Archana Bharadwaj Siva, Karthik Bharadwaj Tallapaka, G. Aditya Kumar, Koushick Sivakumar,Disha Nanda, Divya Das, Jotin Gogoi, Manish Bhattacharjee, Ravi Prasad Mukku, Rakesh K Mishra, Divya Tej Sowpati |
| EPI_ISL_539722 | CSIR-Centre for Cellular and Molecular Biology | CSIR-Centre for Cellular and Molecular Biology | Pratheusa Maccha, Shagufta Khan, Lamuk Zaveri, Namami Gaur, Sakshi Shambhavi, Tulasi Nagabandi, Nikhil Hajirnis, M Soujanya Reddy, Purushotham Vodnala, Payel Mukherjee, Sofia Banu, Priya Singh, Onkar Kulkarni, Dhiviya Vedagiri, Divya Gupta, Vishal Sah, Santosh Kumar Kuncha, Krishnan Harinivas Harshan, Archana Bharadwaj Siva, Karthik Bharadwaj Tallapaka, Disha Nanda, Divya Das, Jotin Gogoi, Manish Bhattacharjee, Ravi Prasad Mukku, Rakesh K Mishra, Divya Tej Sowpati |
| EPI_ISL_539723 | CSIR-Centre for Cellular and Molecular Biology | CSIR-Centre for Cellular and Molecular Biology | Pratheusa Maccha, Sofia Banu, Payel Mukherjee, Priya Singh, Onkar Kulkarni, Dhiviya Vedagiri, Divya Gupta, Vishal Sah, Santosh Kumar Kuncha, Krishnan Harinivas Harshan, Archana Bharadwaj Siva, Karthik Bharadwaj Tallapaka, Shagufta Khan, Lamuk Zaveri, Namami Gaur, Sakshi Shambhavi, Nikhil Hajirnis, M Soujanya Reddy, Tulasi Nagabandi, Purushotham Vodnala,Preethi Jampala, Sharada Ravi Iyer, Sulagana Mukherjee, Swetha Sundar, Peddapuvala Sai Uday Kiran, Rakesh K Mishra, Divya Tej Sowpati |
| EPI_ISL_539745 | CSIR-Centre for Cellular and Molecular Biology | CSIR-Centre for Cellular and Molecular Biology | Nikhil Hajirnis, M Soujanya Reddy, Pratheusa Maccha, Lamuk Zaveri, Shagufta Khan, Namami Gaur, Sakshi Shambhavi, Tulasi Nagabandi, Purushotham Vodnala, Payel Mukherjee, Sofia Banu, Priya Singh, Onkar Kulkarni, Dhiviya Vedagiri, Divya Gupta, Vishal Sah, Santosh Kumar Kuncha, Krishnan Harinivas Harshan, Archana Bharadwaj Siva, Karthik Bharadwaj Tallapaka,Zeba Rizvi, Zuberwasim Sayyad, Kakade Aishwarya Arun, Amrutha H C, Ananga Ghosh, Rakesh K Mishra, Divya Tej Sowpati |
| EPI_ISL_539746 | CSIR-Centre for Cellular and Molecular Biology | CSIR-Centre for Cellular and Molecular Biology | Nikhil Hajirnis, M Soujanya Reddy, Pratheusa Maccha, Namami Gaur, Sakshi Shambhavi, Lamuk Zaveri, Shagufta Khan, Tulasi Nagabandi, Purushotham Vodnala, Payel Mukherjee, Sofia Banu, Priya Singh,Onkar Kulkarni, Dhiviya Vedagiri, Divya Gupta, Vishal Sah, Santosh Kumar Kuncha, Krishnan Harinivas Harshan, Archana Bharadwaj Siva, Karthik Bharadwaj Tallapaka,Kezia J Ann, Radhika Khandelwal, Roshan Maku Venkata, Shemin Mansuri, Sonu Uday, Rakesh K Mishra, Divya Tej Sowpati |
| EPI_ISL_539747 | CSIR-Centre for Cellular and Molecular Biology | CSIR-Centre for Cellular and Molecular Biology | Nikhil Hajirnis, M Soujanya Reddy, Pratheusa Maccha, Payel Mukherjee, Sofia Banu, Priya Singh,Onkar Kulkarni, Dhiviya Vedagiri, Divya Gupta, Vishal Sah, Santosh Kumar Kuncha, Krishnan Harinivas Harshan, Archana Bharadwaj Siva, Karthik Bharadwaj Tallapaka, Shagufta Khan, Lamuk Zaveri, Namami Gaur, Sakshi Shambhavi, Tulasi Nagabandi, Purushotham Vodnala,Deepak Kumar, Devi Prasad Vijayashankar, Disha Nanda, Divya Das, Jotin Gogoi, Manish Bhattacharjee, Rakesh K Mishra, Divya Tej Sowpati |
| EPI_ISL_539748 | CSIR-Centre for Cellular and Molecular Biology | CSIR-Centre for Cellular and Molecular Biology | Payel Mukherjee, Sofia Banu, Priya Singh, Onkar Kulkarni, Dhiviya Vedagiri, Divya Gupta, Vishal Sah, Santosh Kumar Kuncha, Krishnan Harinivas Harshan, Archana Bharadwaj Siva, Karthik Bharadwaj Tallapaka, Shagufta Khan, Lamuk Zaveri, Nikhil Hajirnis, M Soujanya Reddy, Pratheusa Maccha, Namami Gaur, Sakshi Shambhavi, Tulasi Nagabandi, Purushotham Vodnala, G. Aditya Kumar, Koushick Sivakumar, Pooja Ramesh Gupta, Rajan Kumar Jha, Shraddha Vijay Lahoti, Rakesh K Mishra, Divya Tej Sowpati |
| EPI_ISL_539749 | CSIR-Centre for Cellular and Molecular Biology | CSIR-Centre for Cellular and Molecular Biology | Payel Mukherjee, Sofia Banu, Priya Singh, Onkar Kulkarni, Dhiviya Vedagiri, Divya Gupta, Vishal Sah, Santosh Kumar Kuncha, Krishnan Harinivas Harshan, Archana Bharadwaj Siva, Karthik Bharadwaj Tallapaka, Shagufta Khan, Lamuk Zaveri, Nikhil Hajirnis, M Soujanya Reddy, Pratheusa Maccha, Namami Gaur, Sakshi Shambhavi, Tulasi Nagabandi, Purushotham Vodnala, Gokulan C G, Gunjan Purohit, Hanuman Tulashiram Kale, Pankaj Kumar, Prachand Issarapu, Rakesh K Mishra, Divya Tej Sowpati |
| EPI_ISL_539750 | CSIR-Centre for Cellular and Molecular Biology | CSIR-Centre for Cellular and Molecular Biology | Payel Mukherjee, Sofia Banu, Priya Singh, Onkar Kulkarni, Dhiviya Vedagiri, Divya Gupta, Vishal Sah, Santosh Kumar Kuncha, Krishnan Harinivas Harshan, Archana Bharadwaj Siva, Karthik Bharadwaj Tallapaka, Shagufta Khan, Lamuk Zaveri, Nikhil Hajirnis, M Soujanya Reddy, Pratheusa Maccha, Namami Gaur, Sakshi Shambhavi, Tulasi Nagabandi, Purushotham Vodnala, Rakesh K Mishra, Sonu Uday, Sudipta Mondal, Annapoorna P Karthyayani, Debabrata Jana, Debrya Saha, Divya Tej Sowpati |
| EPI_ISL_539751 | CSIR-Centre for Cellular and Molecular Biology | CSIR-Centre for Cellular and Molecular Biology | Pratheusa Maccha, Sakshi Shambhavi, Lamuk Zaveri, Shagufta Khan, Namami Gaur, Nikhil Hajirnis, M Soujanya Reddy, Tulasi Nagabandi, Purushotham Vodnala, Payel Mukherjee, Sofia Banu, Priya Singh,Onkar Kulkarni, Dhiviya Vedagiri, Divya Gupta, Vishal Sah, Santosh Kumar Kuncha, Krishnan Harinivas Harshan, Archana Bharadwaj Siva, Karthik Bharadwaj Tallapaka, G. Aditya Kumar, Koushick Sivakumar,Disha Nanda, Divya Das, Jotin Gogoi, Manish Bhattacharjee, Ravi Prasad Mukku, Rakesh K Mishra, Divya Tej Sowpati |
| EPI_ISL_539752 | CSIR-Centre for Cellular and Molecular Biology | CSIR-Centre for Cellular and Molecular Biology | Pratheusa Maccha, Shagufta Khan, Lamuk Zaveri, Namami Gaur, Sakshi Shambhavi, Tulasi Nagabandi, Nikhil Hajirnis, M Soujanya Reddy, Purushotham Vodnala, Payel Mukherjee, Sofia Banu, Priya Singh, Onkar Kulkarni, Dhiviya Vedagiri, Divya Gupta, Vishal Sah, Santosh Kumar Kuncha, Krishnan Harinivas Harshan, Archana Bharadwaj Siva, Karthik Bharadwaj Tallapaka, Disha Nanda, Divya Das, Jotin Gogoi, Manish Bhattacharjee, Ravi Prasad Mukku, Rakesh K Mishra, Divya Tej Sowpati |
| EPI_ISL_539753 | CSIR-Centre for Cellular and Molecular Biology | CSIR-Centre for Cellular and Molecular Biology | Pratheusa Maccha, Sofia Banu, Payel Mukherjee, Priya Singh, Onkar Kulkarni, Dhiviya Vedagiri, Divya Gupta, Vishal Sah, Santosh Kumar Kuncha, Krishnan Harinivas Harshan, Archana Bharadwaj Siva, Karthik Bharadwaj Tallapaka, Shagufta Khan, Lamuk Zaveri, Namami Gaur, Sakshi Shambhavi, Nikhil Hajirnis, M Soujanya Reddy, Tulasi Nagabandi, Purushotham Vodnala,Preethi Jampala, Sharada Ravi Iyer, Sulagana Mukherjee, Swetha Sundar, Peddapuvala Sai Uday Kiran, Rakesh K Mishra, Divya Tej Sowpati |
| EPI_ISL_539771 | CSIR-Centre for Cellular and Molecular Biology | CSIR-Centre for Cellular and Molecular Biology | Pratheusa Maccha, Sakshi Shambhavi, Lamuk Zaveri, Shagufta Khan, Namami Gaur, Nikhil Hajirnis, M Soujanya Reddy, Tulasi Nagabandi, Purushotham Vodnala, Payel Mukherjee, Sofia Banu, Priya Singh,Onkar Kulkarni, Dhiviya Vedagiri, Divya Gupta, Vishal Sah, Santosh Kumar Kuncha, Krishnan Harinivas Harshan, Archana Bharadwaj Siva, Karthik Bharadwaj Tallapaka, G. Aditya Kumar, Koushick Sivakumar,Disha Nanda, Divya Das, Jotin Gogoi, Manish Bhattacharjee, Ravi Prasad Mukku, Rakesh K Mishra, Divya Tej Sowpati |
| EPI_ISL_539772 | CSIR-Centre for Cellular and Molecular Biology | CSIR-Centre for Cellular and Molecular Biology | Pratheusa Maccha, Shagufta Khan, Lamuk Zaveri, Namami Gaur, Sakshi Shambhavi, Tulasi Nagabandi, Nikhil Hajirnis, M Soujanya Reddy, Purushotham Vodnala, Payel Mukherjee, Sofia Banu, Priya Singh, Onkar Kulkarni, Dhiviya Vedagiri, Divya Gupta, Vishal Sah, Santosh Kumar Kuncha, Krishnan Harinivas Harshan, Archana Bharadwaj Siva, Karthik Bharadwaj Tallapaka, Disha Nanda, Divya Das, Jotin Gogoi, Manish Bhattacharjee, Ravi Prasad Mukku, Rakesh K Mishra, Divya Tej Sowpati |
| EPI_ISL_539773 | CSIR-Centre for Cellular and Molecular Biology | CSIR-Centre for Cellular and Molecular Biology | Payel Mukherjee, Sofia Banu, Priya Singh, Onkar Kulkarni, Dhiviya Vedagiri, Divya Gupta, Vishal Sah, Santosh Kumar Kuncha, Krishnan Harinivas Harshan, Archana Bharadwaj Siva, Karthik Bharadwaj Tallapaka, Shagufta Khan, Lamuk Zaveri, Nikhil Hajirnis, M Soujanya Reddy, Pratheusa Maccha, Namami Gaur, Sakshi Shambhavi, Tulasi Nagabandi, Purushotham Vodnala, Gokulan C G, Gunjan Purohit, Hanuman Tulashiram Kale, Pankaj Kumar, Prachand Issarapu, Rakesh K Mishra, Divya Tej Sowpati |
| EPI_ISL_539774 | CSIR-Centre for Cellular and Molecular Biology | CSIR-Centre for Cellular and Molecular Biology | Payel Mukherjee, Sofia Banu, Priya Singh, Onkar Kulkarni, Dhiviya Vedagiri, Divya Gupta, Vishal Sah, Santosh Kumar Kuncha, Krishnan Harinivas Harshan, Archana Bharadwaj Siva, Karthik Bharadwaj Tallapaka, Shagufta Khan, Lamuk Zaveri, Nikhil Hajirnis, M Soujanya Reddy, Pratheusa Maccha, Namami Gaur, Sakshi Shambhavi, Tulasi Nagabandi, Purushotham Vodnala, Rakesh K Mishra, Sonu Uday, Sudipta Mondal, Annapoorna P Karthyayani, Debabrata Jana, Debrya Saha, Divya Tej Sowpati |
| EPI_ISL_539775 | CSIR-Centre for Cellular and Molecular Biology | CSIR-Centre for Cellular and Molecular Biology | Payel Mukherjee, Sofia Banu, Priya Singh, Onkar Kulkarni, Dhiviya Vedagiri, Divya Gupta, Vishal Sah, Santosh Kumar Kuncha, Krishnan Harinivas Harshan, Archana Bharadwaj Siva, Karthik Bharadwaj Tallapaka, Shagufta Khan, Lamuk Zaveri, Nikhil Hajirnis, M Soujanya Reddy, Pratheusa Maccha, Namami Gaur, Sakshi Shambhavi, Tulasi Nagabandi, Purushotham Vodnala, Gokulan C G, Gunjan Purohit, Hanuman Tulashiram Kale, Pankaj Kumar, Prachand Issarapu, Rakesh K Mishra, Divya Tej Sowpati |
| EPI_ISL_539794, EPI_ISL_539795, EPI_ISL_539796, EPI_ISL_539797, | Wyoming Public Health Laboratory | Wyoming Public Health Laboratory | Noah Hull, Rob Christensen, Jim Mildenberger, Joel Sevinsky, Cari Sloma, and Wanda Manley |

| | | | |
|---|---|---|---|
| EPI_ISL_539798, EPI_ISL_539799, EPI_ISL_539800, EPI_ISL_539801, EPI_ISL_539802 | | | |
| EPI_ISL_539880 | Respiratory Virus Unit, Microbiology Services Colindale, Public Health England | Respiratory Virus Unit, Microbiology Services Colindale, Public Health England | PHE Covid Sequencing Team |
| EPI_ISL_540421, EPI_ISL_540422, EPI_ISL_540921 | Wyoming Public Health Laboratory | Wyoming Public Health Laboratory | Noah Hull, Rob Christensen, Jim Mildenberger, Joel Sevinsky, Cari Sloma, and Wanda Manley |
| EPI_ISL_540923, EPI_ISL_540924, EPI_ISL_540925, EPI_ISL_540926, EPI_ISL_540927, EPI_ISL_540928, EPI_ISL_540929, EPI_ISL_540930, EPI_ISL_540931, EPI_ISL_540932, EPI_ISL_540933, EPI_ISL_540934, EPI_ISL_540935, EPI_ISL_540936, EPI_ISL_540937, EPI_ISL_540938, EPI_ISL_540939, EPI_ISL_540940, EPI_ISL_540941, EPI_ISL_540942, EPI_ISL_540943, EPI_ISL_540944, EPI_ISL_540945, EPI_ISL_540946, EPI_ISL_540947, EPI_ISL_540948, EPI_ISL_540949, EPI_ISL_540950, EPI_ISL_540951, EPI_ISL_540952, EPI_ISL_540953, EPI_ISL_540954, EPI_ISL_540955, EPI_ISL_540956, EPI_ISL_540957, EPI_ISL_540958, EPI_ISL_540959, EPI_ISL_540960, EPI_ISL_540961, EPI_ISL_540962, EPI_ISL_540963, EPI_ISL_540964, EPI_ISL_540965, EPI_ISL_540966, EPI_ISL_540967, EPI_ISL_540968, EPI_ISL_540969, EPI_ISL_540970, EPI_ISL_540971, EPI_ISL_540972, EPI_ISL_540973, EPI_ISL_540974, EPI_ISL_540975, EPI_ISL_540976, EPI_ISL_540977, EPI_ISL_540978, EPI_ISL_540979, EPI_ISL_540980, EPI_ISL_540981, EPI_ISL_540982, EPI_ISL_540983, EPI_ISL_540984, EPI_ISL_540985, EPI_ISL_540986, EPI_ISL_540987, EPI_ISL_540988, EPI_ISL_540989, EPI_ISL_540990, EPI_ISL_540991, EPI_ISL_540992 | | | |
| see above | Laboratorio de Referencia Nacional de Virus Respiratorios, Instituto Nacional de Salud Peru | Laboratorio de Genómica Microbiana, Universidad Peruana Cayetano Heredia | Pablo Tsukayama, Alejandra Dávila-Barclay, Luis González, Pedro E. Romero, Brenda Ayzanoa, Janet Huancachoque, Pool Marcos, Maribel Huaringa, Camila Castillo-Vilcahuaman, Guillermo Salvatierra |
| EPI_ISL_541340, EPI_ISL_541341, EPI_ISL_541342, EPI_ISL_541343, EPI_ISL_541344, EPI_ISL_541345, EPI_ISL_541346 | LACEN/PR | Laboratory of Respiratory Viruses and Measles, Oswaldo Cruz Institute, FIOCRUZ | Paola Resende, Luciana Appolinario, Fernando Motta, Anna Carolina Paixão, Ana Carolina Mendonça, Jonathan Lopes, Irina Riediger, Maria do Carmo Debur, Marilda Siqueira |
| EPI_ISL_541347, EPI_ISL_541348, EPI_ISL_541349, EPI_ISL_541350, EPI_ISL_541351, EPI_ISL_541352, EPI_ISL_541353, EPI_ISL_541354, EPI_ISL_541355, EPI_ISL_541356, EPI_ISL_541357, EPI_ISL_541358, EPI_ISL_541359, EPI_ISL_541360, EPI_ISL_541361, EPI_ISL_541362, EPI_ISL_541363, EPI_ISL_541364, EPI_ISL_541365, EPI_ISL_541366, EPI_ISL_541367, EPI_ISL_541368, EPI_ISL_541369 | | | |
| see above | Laboratory of Respiratory Viruses and Measles, Oswaldo Cruz Institute, FIOCRUZ | Laboratory of Respiratory Viruses and Measles, Oswaldo Cruz Institute, FIOCRUZ | Paola Resende, Luciana Appolinario, Fernando Motta, Anna Carolina Paixão, Ana Carolina Mendonça, Jonathan Lopes, Marilda Siqueira |
| EPI_ISL_541370, EPI_ISL_541371 | LACEN/SC | Laboratory of Respiratory Viruses and Measles, Oswaldo Cruz Institute, FIOCRUZ | Paola Resende, Luciana Appolinario, Fernando Motta, Anna Carolina Paixão, Ana Carolina Mendonça, Jonathan Lopes, Sandra Bianchini, Marilda Siqueira |
| EPI_ISL_541372, EPI_ISL_541373, EPI_ISL_541374, EPI_ISL_541375, EPI_ISL_541376, EPI_ISL_541377, EPI_ISL_541378, EPI_ISL_541379, EPI_ISL_541380, EPI_ISL_541381, EPI_ISL_541382, EPI_ISL_541383, EPI_ISL_541384, EPI_ISL_541385, EPI_ISL_541386, EPI_ISL_541387, EPI_ISL_541388, EPI_ISL_541389, EPI_ISL_541390, EPI_ISL_541391, EPI_ISL_541392, EPI_ISL_541393, EPI_ISL_541394, EPI_ISL_541395, EPI_ISL_541396 | | | |
| see above | LACEN/SE | Laboratory of Respiratory Viruses and Measles, Oswaldo Cruz Institute, FIOCRUZ | Paola Resende, Luciana Appolinario, Fernando Motta, Anna Carolina Paixão, Ana Carolina Mendonça, Jonathan Lopes, Clioma Santos, Marilda Siqueira |
| EPI_ISL_541681, EPI_ISL_541682, EPI_ISL_541683, EPI_ISL_541684, EPI_ISL_541685, EPI_ISL_541687, EPI_ISL_541688, EPI_ISL_541689, EPI_ISL_541690, EPI_ISL_541691, EPI_ISL_541692, EPI_ISL_541693, EPI_ISL_541694, EPI_ISL_541695, EPI_ISL_541696, EPI_ISL_541697, EPI_ISL_541698, EPI_ISL_541699, EPI_ISL_541700, EPI_ISL_541701, EPI_ISL_541702, EPI_ISL_541703, EPI_ISL_541704, EPI_ISL_541705, EPI_ISL_541706, EPI_ISL_541707, EPI_ISL_541708, EPI_ISL_541710, EPI_ISL_541711, EPI_ISL_541712, EPI_ISL_541713, EPI_ISL_541714, EPI_ISL_541715, EPI_ISL_541716, EPI_ISL_541717, EPI_ISL_541718, EPI_ISL_541719, EPI_ISL_541720, EPI_ISL_541721, EPI_ISL_541722, EPI_ISL_541723, EPI_ISL_541724, EPI_ISL_541725, EPI_ISL_541726, EPI_ISL_541727, EPI_ISL_541728, EPI_ISL_541729, EPI_ISL_541730, EPI_ISL_541731, EPI_ISL_541732, EPI_ISL_541733, EPI_ISL_541734, EPI_ISL_541735, EPI_ISL_541736, EPI_ISL_541737, EPI_ISL_541738, EPI_ISL_541739, EPI_ISL_541740, EPI_ISL_541741, EPI_ISL_541742, EPI_ISL_541743, EPI_ISL_541744, EPI_ISL_541745, EPI_ISL_541746, EPI_ISL_541747, EPI_ISL_541748, EPI_ISL_541749, EPI_ISL_541750, EPI_ISL_541751 | | | |
| see above | National Institute of Virology, NIV Influenza | National Institute of Virology, NIV Influenza | Potdar V |
| EPI_ISL_547433, EPI_ISL_547434, EPI_ISL_547435, EPI_ISL_547436, EPI_ISL_547437, EPI_ISL_547438, EPI_ISL_547439, EPI_ISL_547440, EPI_ISL_547441, EPI_ISL_547442, EPI_ISL_547443, EPI_ISL_547444 | | | |
| see above | Microbiology, Department of Pathology, St. Bernard's Hospital, Gibraltar Health Authority | Respiratory Virus Unit, Microbiology Services Colindale, Public Health England | PHE Covid Sequencing Team, Dr Nicholas Cortes (Gibraltar), Charlotte Gillborn-Jones (Gibraltar) |
| EPI_ISL_547965 | Laboratorio Biologia Molecolare SarsCov2 UOC LAboratorio Analisi SErvizio Medicina di Laboratorio Ospedale San Francesco ATS-ASSL Nuoro | Laboratorio Specialistico UOC Ematologia Ospedale San Francesco - ATS ASSL NUORO | Piras Giovanna, Asproni Rosanna, Monne Maria Itria, Fancello Tatiana,Fiamma Maura,Toja Alessandro, Sanna Filomena, Floris Anna Rita, Sulis Vincenzo, Palmas Angelo Domenico, Casu Gavino, Lo Maglio Iana, Mameli Giuseppe. |
| EPI_ISL_549352, EPI_ISL_549356, EPI_ISL_549362, EPI_ISL_549365, EPI_ISL_549371, EPI_ISL_549372, EPI_ISL_549375, EPI_ISL_549381, EPI_ISL_549387, EPI_ISL_549517, EPI_ISL_549518, EPI_ISL_549519, EPI_ISL_549520, EPI_ISL_549521, EPI_ISL_549522 | | | |
| see above | Lincolnshire Hospitals and DeepSeq Nottingham | COVID-19 Genomics UK (COG-UK) Consortium | Nichola Duckworth, Tim Sloan, Sarah Walsh, Jonathan Ball, Patrick McClure, Joeseph Chappell, Nadine Holmes, Matthew Carlisle, Christopher Moore, Fei Sang, Johnny Debebe, Victoria Wright, Matthew Loose |
| EPI_ISL_566045, EPI_ISL_566046, EPI_ISL_566047, EPI_ISL_566048, EPI_ISL_566049, EPI_ISL_566050, EPI_ISL_566051, EPI_ISL_566052, EPI_ISL_566053, EPI_ISL_566054, EPI_ISL_566055, EPI_ISL_566056, EPI_ISL_566057, EPI_ISL_566058, EPI_ISL_566059, EPI_ISL_566060, EPI_ISL_566061, EPI_ISL_566062, EPI_ISL_566063, EPI_ISL_566064, EPI_ISL_566065, EPI_ISL_566066, EPI_ISL_566067, EPI_ISL_566068, EPI_ISL_566069, EPI_ISL_566070, EPI_ISL_566071, EPI_ISL_566072, EPI_ISL_566073, EPI_ISL_566074, EPI_ISL_566075, EPI_ISL_566076 | | | |
| see above | Respiratory Virus Unit, Microbiology Services Colindale, Public Health England | Respiratory Virus Unit, Microbiology Services Colindale, Public Health England | PHE Covid Sequencing Team |
| EPI_ISL_568510, EPI_ISL_568511, EPI_ISL_568512, EPI_ISL_568513, EPI_ISL_568514, EPI_ISL_568515, EPI_ISL_568516, EPI_ISL_568517, EPI_ISL_568518, EPI_ISL_568519, EPI_ISL_568520, EPI_ISL_568521, EPI_ISL_568522, EPI_ISL_568523, EPI_ISL_568524, EPI_ISL_568525, EPI_ISL_568526, EPI_ISL_568527, EPI_ISL_568528, EPI_ISL_568529, EPI_ISL_568530, EPI_ISL_568531, EPI_ISL_568532, EPI_ISL_568533, EPI_ISL_568534, EPI_ISL_568535, EPI_ISL_568536, EPI_ISL_568537, EPI_ISL_568538, EPI_ISL_568539, EPI_ISL_568540, EPI_ISL_568541, EPI_ISL_568542, EPI_ISL_568543, EPI_ISL_568544, EPI_ISL_568545, EPI_ISL_568546, EPI_ISL_568547, EPI_ISL_568548, EPI_ISL_568549, EPI_ISL_568550, EPI_ISL_568551, EPI_ISL_568552, EPI_ISL_568553, EPI_ISL_568554, EPI_ISL_568555 | | | |
| see above | Laboratorio de Referencia Nacional de Virus Respiratorios, Instituto Nacional de Salud Peru | Laboratorio de Genómica Microbiana, Universidad Peruana Cayetano Heredia | Pablo Tsukayama, Alejandra Dávila-Barclay, Luis González, Pedro E. Romero, Brenda Ayzanoa, Janet Huancachoque, Pool Marcos, Maribel Huaringa, Camila Castillo-Vilcahuaman, Guillermo Salvatierra |
| EPI_ISL_570027, EPI_ISL_570028, EPI_ISL_570029, EPI_ISL_570030, EPI_ISL_570031, EPI_ISL_570032, EPI_ISL_570033, EPI_ISL_570034, EPI_ISL_570035, EPI_ISL_570036, EPI_ISL_570037, EPI_ISL_570038, EPI_ISL_570039, EPI_ISL_570040, EPI_ISL_570041, EPI_ISL_570042, EPI_ISL_570043, EPI_ISL_570044, EPI_ISL_570045, EPI_ISL_570046, EPI_ISL_570047, EPI_ISL_570048, EPI_ISL_570049, EPI_ISL_570050, EPI_ISL_570051, EPI_ISL_570052, EPI_ISL_570053, EPI_ISL_570054, EPI_ISL_570055, EPI_ISL_570056, EPI_ISL_570057, EPI_ISL_570058, EPI_ISL_570059, EPI_ISL_570060, EPI_ISL_570061, EPI_ISL_570062, EPI_ISL_570063, EPI_ISL_570064, EPI_ISL_570065, EPI_ISL_570066, EPI_ISL_570067, EPI_ISL_570068, EPI_ISL_570069, EPI_ISL_570070, EPI_ISL_570071, EPI_ISL_570072, EPI_ISL_570073, EPI_ISL_570074, EPI_ISL_570075, EPI_ISL_570076, EPI_ISL_570077, EPI_ISL_570078, EPI_ISL_570079, EPI_ISL_570080, EPI_ISL_570081, EPI_ISL_570082, EPI_ISL_570083, EPI_ISL_570084, EPI_ISL_570085, EPI_ISL_570086, EPI_ISL_570087, EPI_ISL_570088, EPI_ISL_570089, EPI_ISL_570090, EPI_ISL_570091, EPI_ISL_570092, EPI_ISL_570093, EPI_ISL_570094, EPI_ISL_570095, EPI_ISL_570096, EPI_ISL_570097, EPI_ISL_570098, EPI_ISL_570099, EPI_ISL_570100, EPI_ISL_570101, EPI_ISL_570102, EPI_ISL_570103, EPI_ISL_570104, EPI_ISL_570105, EPI_ISL_570106, EPI_ISL_570107, EPI_ISL_570108, EPI_ISL_570109, EPI_ISL_570110, EPI_ISL_570111, EPI_ISL_570112, EPI_ISL_570113, EPI_ISL_570114, EPI_ISL_570115, EPI_ISL_570116, EPI_ISL_570117, EPI_ISL_570118, EPI_ISL_570119, EPI_ISL_570120, EPI_ISL_570121, EPI_ISL_570122, EPI_ISL_570123, EPI_ISL_570124, EPI_ISL_570125, EPI_ISL_570126, EPI_ISL_570127, EPI_ISL_570128, EPI_ISL_570129, EPI_ISL_570130, EPI_ISL_570131, EPI_ISL_570132, EPI_ISL_570133, EPI_ISL_570134, EPI_ISL_570135, EPI_ISL_570136, EPI_ISL_570137, EPI_ISL_570138, EPI_ISL_570139, EPI_ISL_570140, EPI_ISL_570141, EPI_ISL_570142, EPI_ISL_570143, EPI_ISL_570144, EPI_ISL_570145, EPI_ISL_570146, EPI_ISL_570147, EPI_ISL_570148, EPI_ISL_570149, EPI_ISL_570150, EPI_ISL_570151, EPI_ISL_570152, EPI_ISL_570153, EPI_ISL_570154, EPI_ISL_570155, EPI_ISL_570156, EPI_ISL_570157, EPI_ISL_570158, EPI_ISL_570159, EPI_ISL_570160, EPI_ISL_570161, EPI_ISL_570162, EPI_ISL_570163, EPI_ISL_570164, EPI_ISL_570165, EPI_ISL_570166, EPI_ISL_570167, EPI_ISL_570168, EPI_ISL_570169, EPI_ISL_570170, EPI_ISL_570171, EPI_ISL_570172, EPI_ISL_570173, EPI_ISL_570174, EPI_ISL_570175, EPI_ISL_570176, EPI_ISL_570177, EPI_ISL_570178, EPI_ISL_570179, EPI_ISL_570180, EPI_ISL_570181, EPI_ISL_570182, EPI_ISL_570184, EPI_ISL_570185, EPI_ISL_570186, EPI_ISL_570187, EPI_ISL_570188, EPI_ISL_570189, EPI_ISL_570190, EPI_ISL_570191, EPI_ISL_570192, EPI_ISL_570193, EPI_ISL_570194, EPI_ISL_570195, EPI_ISL_570196, EPI_ISL_570197, EPI_ISL_570198, EPI_ISL_570199, EPI_ISL_570200, EPI_ISL_570201, EPI_ISL_570202, EPI_ISL_570203, EPI_ISL_570204, EPI_ISL_570205, EPI_ISL_570206, EPI_ISL_570207, EPI_ISL_570208, EPI_ISL_570209, EPI_ISL_570210, EPI_ISL_570211, EPI_ISL_570212, EPI_ISL_570213, EPI_ISL_570214, EPI_ISL_570215, EPI_ISL_570216, EPI_ISL_570217, EPI_ISL_570218, EPI_ISL_570219, EPI_ISL_570220, EPI_ISL_570221, EPI_ISL_570222, EPI_ISL_570223, EPI_ISL_570224, EPI_ISL_570225, EPI_ISL_570226, EPI_ISL_570227, EPI_ISL_570228, EPI_ISL_570229, EPI_ISL_570230, EPI_ISL_570231, EPI_ISL_570232, EPI_ISL_570233, EPI_ISL_570234, EPI_ISL_570235, EPI_ISL_570236, EPI_ISL_570237, EPI_ISL_570238, EPI_ISL_570239, EPI_ISL_570240, EPI_ISL_570241, EPI_ISL_570242, EPI_ISL_570243, EPI_ISL_570244, EPI_ISL_570245, EPI_ISL_570246, EPI_ISL_570247, EPI_ISL_570248, EPI_ISL_570249, EPI_ISL_570250, EPI_ISL_570251, EPI_ISL_570252, EPI_ISL_570253, EPI_ISL_570254, EPI_ISL_570255, EPI_ISL_570256, EPI_ISL_570257, EPI_ISL_570258, EPI_ISL_570259, EPI_ISL_570260, EPI_ISL_570261, EPI_ISL_570262, EPI_ISL_570263, EPI_ISL_570264, EPI_ISL_570265, EPI_ISL_570266, EPI_ISL_570267, EPI_ISL_570268, EPI_ISL_570269, EPI_ISL_570270, EPI_ISL_570271, EPI_ISL_570272, EPI_ISL_570273, EPI_ISL_570274, EPI_ISL_570275, EPI_ISL_570276, EPI_ISL_570277, EPI_ISL_570278, EPI_ISL_570279, EPI_ISL_570280, EPI_ISL_570281, EPI_ISL_570282, EPI_ISL_570283, EPI_ISL_570284, EPI_ISL_570285, EPI_ISL_570286, EPI_ISL_570287, EPI_ISL_570288, EPI_ISL_570289, EPI_ISL_570290, EPI_ISL_570291, EPI_ISL_570292, EPI_ISL_570293, EPI_ISL_570294, EPI_ISL_570295, EPI_ISL_570296, EPI_ISL_570297, EPI_ISL_570298, EPI_ISL_570299, EPI_ISL_570300, EPI_ISL_570301, EPI_ISL_570302, EPI_ISL_570303, EPI_ISL_570304, EPI_ISL_570305, EPI_ISL_570306, EPI_ISL_570307, EPI_ISL_570308, EPI_ISL_570309, EPI_ISL_570310, EPI_ISL_570311, EPI_ISL_570312, EPI_ISL_570313, EPI_ISL_570314, EPI_ISL_570315, EPI_ISL_570316, EPI_ISL_570317, EPI_ISL_570318, EPI_ISL_570319, EPI_ISL_570320, EPI_ISL_570321, EPI_ISL_570322, EPI_ISL_570323, EPI_ISL_570324, EPI_ISL_570325, EPI_ISL_570326, EPI_ISL_570327, EPI_ISL_570328, EPI_ISL_570329, EPI_ISL_570330, EPI_ISL_570331, EPI_ISL_570332, EPI_ISL_570333, EPI_ISL_570334, EPI_ISL_570335, EPI_ISL_570336, EPI_ISL_570337, EPI_ISL_570338, EPI_ISL_570339, EPI_ISL_570340, EPI_ISL_570341, EPI_ISL_570342, EPI_ISL_570343, EPI_ISL_570344, EPI_ISL_570345, EPI_ISL_570346, EPI_ISL_570347, EPI_ISL_570348, EPI_ISL_570349, EPI_ISL_570350, EPI_ISL_570351, EPI_ISL_570352, EPI_ISL_570353, EPI_ISL_570354, EPI_ISL_570355, EPI_ISL_570356, EPI_ISL_570357, EPI_ISL_570358, EPI_ISL_570359, EPI_ISL_570360, EPI_ISL_570361, EPI_ISL_570362, EPI_ISL_570363, EPI_ISL_570364, EPI_ISL_570365, EPI_ISL_570366, EPI_ISL_570367, EPI_ISL_570368, EPI_ISL_570369, EPI_ISL_570370, EPI_ISL_570371, EPI_ISL_570372, EPI_ISL_570373, EPI_ISL_570374, EPI_ISL_570375, EPI_ISL_570376, EPI_ISL_570377, EPI_ISL_570378, EPI_ISL_570379, EPI_ISL_570380, EPI_ISL_570381, EPI_ISL_570382, EPI_ISL_570383, EPI_ISL_570384, EPI_ISL_570385, EPI_ISL_570386, EPI_ISL_570387, EPI_ISL_570388, EPI_ISL_570389, EPI_ISL_570390, EPI_ISL_570391, EPI_ISL_570392, EPI_ISL_570393, EPI_ISL_570394, EPI_ISL_570395, EPI_ISL_570396, EPI_ISL_570397, EPI_ISL_570398, EPI_ISL_570399, EPI_ISL_570400, EPI_ISL_570401, EPI_ISL_570402, EPI_ISL_570403, EPI_ISL_570404, EPI_ISL_570405, EPI_ISL_570406, EPI_ISL_570407, EPI_ISL_570408, EPI_ISL_570409, EPI_ISL_570410, EPI_ISL_570411, EPI_ISL_570412, EPI_ISL_570413, EPI_ISL_570414, EPI_ISL_570415, EPI_ISL_570416, EPI_ISL_570417, EPI_ISL_570418, EPI_ISL_570419, EPI_ISL_570420, EPI_ISL_570421, EPI_ISL_570422, EPI_ISL_570423, EPI_ISL_570424, EPI_ISL_570425, EPI_ISL_570426, EPI_ISL_570427, EPI_ISL_570428, EPI_ISL_570429, EPI_ISL_570430, EPI_ISL_570431, EPI_ISL_570432, EPI_ISL_570433, EPI_ISL_570434, EPI_ISL_570435, EPI_ISL_570436, EPI_ISL_570437, EPI_ISL_570438, EPI_ISL_570439, EPI_ISL_570440, EPI_ISL_570441, EPI_ISL_570442, EPI_ISL_570443, EPI_ISL_570444, EPI_ISL_570445, EPI_ISL_570446, EPI_ISL_570447, EPI_ISL_570448, EPI_ISL_570449, EPI_ISL_570450, EPI_ISL_570451, EPI_ISL_570452, EPI_ISL_570453, EPI_ISL_570454, EPI_ISL_570455, EPI_ISL_570456, EPI_ISL_570457, EPI_ISL_570458, EPI_ISL_570459, EPI_ISL_570460, EPI_ISL_570461, EPI_ISL_570462, EPI_ISL_570463, EPI_ISL_570464, EPI_ISL_570465, EPI_ISL_570466, EPI_ISL_570467, EPI_ISL_570468, EPI_ISL_570469, EPI_ISL_570470, EPI_ISL_570471, EPI_ISL_570472, EPI_ISL_570473, EPI_ISL_570474, EPI_ISL_570475, EPI_ISL_570476, EPI_ISL_570477, EPI_ISL_570478, EPI_ISL_570479, EPI_ISL_570480, EPI_ISL_570481, EPI_ISL_570482, EPI_ISL_570483, EPI_ISL_570484, EPI_ISL_570485, EPI_ISL_570486, EPI_ISL_570487, EPI_ISL_570488, EPI_ISL_570489, EPI_ISL_570490, EPI_ISL_570491, EPI_ISL_570492, EPI_ISL_570493, EPI_ISL_570494, EPI_ISL_570495, | | | |

| | | | |
|---|---|---|---|
| EPI_ISL_570496, EPI_ISL_570497, EPI_ISL_570498, EPI_ISL_570499, EPI_ISL_570500, EPI_ISL_570501, EPI_ISL_570502, EPI_ISL_570503, EPI_ISL_570504, EPI_ISL_570505, EPI_ISL_570506, EPI_ISL_570507, EPI_ISL_570508, EPI_ISL_570509, EPI_ISL_570510, EPI_ISL_570511, EPI_ISL_570512, EPI_ISL_570513, EPI_ISL_570514, EPI_ISL_570515, EPI_ISL_570516, EPI_ISL_570517, EPI_ISL_570518, EPI_ISL_570519, EPI_ISL_570520, EPI_ISL_570521, EPI_ISL_570522, EPI_ISL_570523, EPI_ISL_570524, EPI_ISL_570525, EPI_ISL_570526, EPI_ISL_570527, EPI_ISL_570528, EPI_ISL_570529, EPI_ISL_570530, EPI_ISL_570531, EPI_ISL_570532, EPI_ISL_570533, EPI_ISL_570534, EPI_ISL_570535, EPI_ISL_570536, EPI_ISL_570537, EPI_ISL_570538, EPI_ISL_570539, EPI_ISL_570540, EPI_ISL_570541, EPI_ISL_570542, EPI_ISL_570543, EPI_ISL_570544, EPI_ISL_570545, EPI_ISL_570546, EPI_ISL_570547, EPI_ISL_570548, EPI_ISL_570549, EPI_ISL_570550, EPI_ISL_570551, EPI_ISL_570552, EPI_ISL_570553, EPI_ISL_570554, EPI_ISL_570555, EPI_ISL_570556, EPI_ISL_570557, EPI_ISL_570558, EPI_ISL_570559, EPI_ISL_570560, EPI_ISL_570561, EPI_ISL_570562, EPI_ISL_570563, EPI_ISL_570564, EPI_ISL_570565, EPI_ISL_570566, EPI_ISL_570567, EPI_ISL_570568, EPI_ISL_570569, EPI_ISL_570570, EPI_ISL_570571, EPI_ISL_570572, EPI_ISL_570573, EPI_ISL_570574, EPI_ISL_570575, EPI_ISL_570576, EPI_ISL_570577, EPI_ISL_570578, EPI_ISL_570579, EPI_ISL_570580, EPI_ISL_570581, EPI_ISL_570582, EPI_ISL_570583, EPI_ISL_570584, EPI_ISL_570585, EPI_ISL_570586, EPI_ISL_570587, EPI_ISL_570588, EPI_ISL_570589, EPI_ISL_570590, EPI_ISL_570591, EPI_ISL_570592, EPI_ISL_570593, EPI_ISL_570594, EPI_ISL_570595, EPI_ISL_570596, EPI_ISL_570597, EPI_ISL_570598, EPI_ISL_570599, EPI_ISL_570600, EPI_ISL_570601, EPI_ISL_570602, EPI_ISL_570603, EPI_ISL_570604, EPI_ISL_570605, EPI_ISL_570606, EPI_ISL_570607, EPI_ISL_570608, EPI_ISL_570609, EPI_ISL_570610, EPI_ISL_570611, EPI_ISL_570612, EPI_ISL_570613, EPI_ISL_570614, EPI_ISL_570615, EPI_ISL_570616, EPI_ISL_570617, EPI_ISL_570618, EPI_ISL_570619, EPI_ISL_570620, EPI_ISL_570621, EPI_ISL_570622, EPI_ISL_570623, EPI_ISL_570624, EPI_ISL_570625, EPI_ISL_570626, EPI_ISL_570627, EPI_ISL_570628, EPI_ISL_570629, EPI_ISL_570630, EPI_ISL_570631, EPI_ISL_570632, EPI_ISL_570633, EPI_ISL_570634, EPI_ISL_570635, EPI_ISL_570636, EPI_ISL_570637, EPI_ISL_570638, EPI_ISL_570639, EPI_ISL_570640, EPI_ISL_570641, EPI_ISL_570642, EPI_ISL_570643, EPI_ISL_570644, EPI_ISL_570645, EPI_ISL_570646, EPI_ISL_570647, EPI_ISL_570648, EPI_ISL_570649, EPI_ISL_570650, EPI_ISL_570651, EPI_ISL_570652, EPI_ISL_570653, EPI_ISL_570654, EPI_ISL_570655, EPI_ISL_570656, EPI_ISL_570657, EPI_ISL_570658, EPI_ISL_570659, EPI_ISL_570660, EPI_ISL_570661, EPI_ISL_570662, EPI_ISL_570663, EPI_ISL_570664, EPI_ISL_570665, EPI_ISL_570666, EPI_ISL_570667, EPI_ISL_570668, EPI_ISL_570669, EPI_ISL_570670, EPI_ISL_570671, EPI_ISL_570672, EPI_ISL_570673, EPI_ISL_570674, EPI_ISL_570675, EPI_ISL_570676, EPI_ISL_570677, EPI_ISL_570678, EPI_ISL_570679, EPI_ISL_570680, EPI_ISL_570681, EPI_ISL_570682, EPI_ISL_570683, EPI_ISL_570684, EPI_ISL_570685, EPI_ISL_570686, EPI_ISL_570687, EPI_ISL_570688, EPI_ISL_570689, EPI_ISL_570690, EPI_ISL_570691, EPI_ISL_570692, EPI_ISL_570693, EPI_ISL_570694, EPI_ISL_570695, EPI_ISL_570696, EPI_ISL_570697, EPI_ISL_570698, EPI_ISL_570699, EPI_ISL_570700, EPI_ISL_570701, EPI_ISL_570702, EPI_ISL_570703, EPI_ISL_570704, EPI_ISL_570705, EPI_ISL_570706, EPI_ISL_570707, EPI_ISL_570708, EPI_ISL_570709, EPI_ISL_570710, EPI_ISL_570711, EPI_ISL_570712, EPI_ISL_570713, EPI_ISL_570714, EPI_ISL_570715, EPI_ISL_570716, EPI_ISL_570717, EPI_ISL_570718, EPI_ISL_570719, EPI_ISL_570720, EPI_ISL_570721, EPI_ISL_570722, EPI_ISL_570723, EPI_ISL_570724, EPI_ISL_570725, EPI_ISL_570726, EPI_ISL_570727, EPI_ISL_570728, EPI_ISL_570729, EPI_ISL_570730, EPI_ISL_570731, EPI_ISL_570732, EPI_ISL_570733, EPI_ISL_570734, EPI_ISL_570735, EPI_ISL_570736, EPI_ISL_570737, EPI_ISL_570738, EPI_ISL_570739, EPI_ISL_570740, EPI_ISL_570741, EPI_ISL_570742, EPI_ISL_570743, EPI_ISL_570744, EPI_ISL_570745, EPI_ISL_570746, EPI_ISL_570747, EPI_ISL_570748, EPI_ISL_570749, EPI_ISL_570750, EPI_ISL_570751, EPI_ISL_570752, EPI_ISL_570753, EPI_ISL_570754, EPI_ISL_570755, EPI_ISL_570756, EPI_ISL_570757, EPI_ISL_570758, EPI_ISL_570759, EPI_ISL_570760, EPI_ISL_570761, EPI_ISL_570762, EPI_ISL_570763, EPI_ISL_570764, EPI_ISL_570765, EPI_ISL_570766, EPI_ISL_570767, EPI_ISL_570768, EPI_ISL_570769, EPI_ISL_570770, EPI_ISL_570771, EPI_ISL_570772, EPI_ISL_570773, EPI_ISL_570774, EPI_ISL_570775, EPI_ISL_570776, EPI_ISL_570777, EPI_ISL_570778, EPI_ISL_570779, EPI_ISL_570780, EPI_ISL_570781, EPI_ISL_570782, EPI_ISL_570783, EPI_ISL_570784, EPI_ISL_570785, EPI_ISL_570786, EPI_ISL_570787, EPI_ISL_570788, EPI_ISL_570789, EPI_ISL_570790, EPI_ISL_570791, EPI_ISL_570792, EPI_ISL_570793, EPI_ISL_570794, EPI_ISL_570795, EPI_ISL_570796, EPI_ISL_570797, EPI_ISL_570798, EPI_ISL_570799, EPI_ISL_570800, EPI_ISL_570801, EPI_ISL_570802, EPI_ISL_570803, EPI_ISL_570804, EPI_ISL_570805, EPI_ISL_570806, EPI_ISL_570807, EPI_ISL_570808, EPI_ISL_570809, EPI_ISL_570810, EPI_ISL_570811, EPI_ISL_570812, EPI_ISL_570813, EPI_ISL_570814, EPI_ISL_570815, EPI_ISL_570816, EPI_ISL_570817, EPI_ISL_570818, EPI_ISL_570819, EPI_ISL_570820, EPI_ISL_570821, EPI_ISL_570822, EPI_ISL_570823, EPI_ISL_570824, EPI_ISL_570825, EPI_ISL_570826, EPI_ISL_570827, EPI_ISL_570828, EPI_ISL_570829, EPI_ISL_570830, EPI_ISL_570831, EPI_ISL_570832, EPI_ISL_570833, EPI_ISL_570834, EPI_ISL_570835, EPI_ISL_570836, EPI_ISL_570837, EPI_ISL_570838, EPI_ISL_570839, EPI_ISL_570840, EPI_ISL_570841, EPI_ISL_570842, EPI_ISL_570843, EPI_ISL_570844, EPI_ISL_570846, EPI_ISL_570847, EPI_ISL_570848, EPI_ISL_570849, EPI_ISL_570850, EPI_ISL_570851, EPI_ISL_570852, EPI_ISL_570853, EPI_ISL_570854, EPI_ISL_570855, EPI_ISL_570856, EPI_ISL_570857, EPI_ISL_570858, EPI_ISL_570859, EPI_ISL_570860, EPI_ISL_570861, EPI_ISL_570862, EPI_ISL_570863, EPI_ISL_570864, EPI_ISL_570865, EPI_ISL_570866, EPI_ISL_570867, EPI_ISL_570868, EPI_ISL_570869, EPI_ISL_570870, EPI_ISL_570871, EPI_ISL_570872, EPI_ISL_570873, EPI_ISL_570874, EPI_ISL_570875, EPI_ISL_570876, EPI_ISL_570877, EPI_ISL_570878, EPI_ISL_570879, EPI_ISL_570880, EPI_ISL_570881, EPI_ISL_570882, EPI_ISL_570883, EPI_ISL_570884, EPI_ISL_570885, EPI_ISL_570886, EPI_ISL_570887, EPI_ISL_570888, EPI_ISL_570889, EPI_ISL_570890, EPI_ISL_570891, EPI_ISL_570892, EPI_ISL_570893, EPI_ISL_570894, EPI_ISL_570895, EPI_ISL_570896, EPI_ISL_570897, EPI_ISL_570898, EPI_ISL_570899, EPI_ISL_570900, EPI_ISL_570901, EPI_ISL_570902, EPI_ISL_570903, EPI_ISL_570904, EPI_ISL_570905, EPI_ISL_570906, EPI_ISL_570907, EPI_ISL_570908, EPI_ISL_570909, EPI_ISL_570910, EPI_ISL_570911, EPI_ISL_570912, EPI_ISL_570913, EPI_ISL_570914, EPI_ISL_570915, EPI_ISL_570916, EPI_ISL_570917, EPI_ISL_570918, EPI_ISL_570919, EPI_ISL_570920, EPI_ISL_570921, EPI_ISL_570922, EPI_ISL_570923, EPI_ISL_570924, EPI_ISL_570925, EPI_ISL_570926, EPI_ISL_570927, EPI_ISL_570928, EPI_ISL_570929, EPI_ISL_570930, EPI_ISL_570931, EPI_ISL_570932, EPI_ISL_570933, EPI_ISL_570934, EPI_ISL_570935, EPI_ISL_570936, EPI_ISL_570937, EPI_ISL_570938, EPI_ISL_570939, EPI_ISL_570940, EPI_ISL_570941, EPI_ISL_570942, EPI_ISL_570943, EPI_ISL_570944, EPI_ISL_570945, EPI_ISL_570946, EPI_ISL_570947, EPI_ISL_570948, EPI_ISL_570949, EPI_ISL_570950, EPI_ISL_570951, EPI_ISL_570952, EPI_ISL_570953, EPI_ISL_570954, EPI_ISL_570955, EPI_ISL_570956, EPI_ISL_570957, EPI_ISL_570958, EPI_ISL_570959, EPI_ISL_570960, EPI_ISL_570961, EPI_ISL_570962, EPI_ISL_570963, EPI_ISL_570964, EPI_ISL_570965, EPI_ISL_570966, EPI_ISL_570967, EPI_ISL_570968, EPI_ISL_570969, EPI_ISL_570970, EPI_ISL_570971, EPI_ISL_570972, EPI_ISL_570973, EPI_ISL_570974, EPI_ISL_570975, EPI_ISL_570976, EPI_ISL_570977, EPI_ISL_570978, EPI_ISL_570979, EPI_ISL_570980, EPI_ISL_570981, EPI_ISL_570982, EPI_ISL_570983, EPI_ISL_570984, EPI_ISL_570985, EPI_ISL_570986, EPI_ISL_570987, EPI_ISL_570988, EPI_ISL_570989, EPI_ISL_570990, EPI_ISL_570991, EPI_ISL_570992, EPI_ISL_570993, EPI_ISL_570994, EPI_ISL_570995, EPI_ISL_570996, EPI_ISL_570997, EPI_ISL_570998, EPI_ISL_570999, EPI_ISL_571000, EPI_ISL_571001, EPI_ISL_571002, EPI_ISL_571003, EPI_ISL_571004, EPI_ISL_571005, EPI_ISL_571006, EPI_ISL_571007, EPI_ISL_571008, EPI_ISL_571009, EPI_ISL_571010, EPI_ISL_571011, EPI_ISL_571012, EPI_ISL_571013, EPI_ISL_571014, EPI_ISL_571015 | | | |
| see above | UW Virology Lab | UW Virology Lab | Pavitra Roychoudhury, Hong Xie, Lasata Shrestha, Amin Addetia, Victoria M Rachleff, Meei-Li Huang, Keith R Jerome, Alexander Greninger |
| EPI_ISL_572507, EPI_ISL_572508, EPI_ISL_572510, EPI_ISL_572527, EPI_ISL_572705, EPI_ISL_572938, EPI_ISL_572943, EPI_ISL_572944, EPI_ISL_573437, EPI_ISL_573438, EPI_ISL_573439, EPI_ISL_573440, EPI_ISL_573441, EPI_ISL_573442, EPI_ISL_573443, EPI_ISL_573444, EPI_ISL_573445, EPI_ISL_573446, EPI_ISL_573447, EPI_ISL_573448, EPI_ISL_573449 | | | |
| see above | Lincolnshire Hospitals and DeepSeq Nottingham | COVID-19 Genomics UK (COG-UK) Consortium | Nichola Duckworth, Tim Sloan, Sarah Walsh, Jonathan Ball, Patrick McClure, Joeseph Chappell, Nadine Holmes, Matthew Carlisle, Christopher Moore, Fei Sang, Johnny Debebe, Victoria Wright, Matthew Loose |
| EPI_ISL_574433 | National Institute of Health Research and Development | National Institute of Health Research and Development | Pawestri,HA;Subangkit;Puspa,KD;Nugraha,AA;Ikawati,HD;Pangesti, KNA;Soekarso,T;Susilarini,NK;Hariastuti,NI;Nikmah,UA;Mursinah;Febriyani,A;Herman,R;Susanti,N;Herna;Febriyanti,T; Nurhadi,M; Paisal;Ramadhany,R;Agustiningsih;Kurniawati,J;Kipuw,NL;Muna,F;Indalau,IL;Adam,K;Wibowo,HA;Rizki,A;Puspandari,N;Setiawaty,V |
| EPI_ISL_574697, EPI_ISL_574698, EPI_ISL_574699, EPI_ISL_574700, EPI_ISL_574701, EPI_ISL_574702, EPI_ISL_574703, EPI_ISL_574704 | Respiratory Virus Unit, Microbiology Services Colindale, Public Health England | Respiratory Virus Unit, Microbiology Services Colindale, Public Health England | PHE Covid Sequencing Team |
| EPI_ISL_575332, EPI_ISL_575333, EPI_ISL_575334 | Israel Central Virology laboratory | Israel Central Virology laboratory | Neta Zuckerman, Efrat Dahan Bucris, Oran Erster, Ella Mendelson, Michal Mandelboim |
| EPI_ISL_576127 | Oklahoma Animal Disease Diagnostic Laboratory, Oklahoma State University | Oklahoma Animal Disease Diagnostic Laboratory, Oklahoma State University | Narayanan,S., Ritchey,J.C., Patil,G., Narasaraju,T., More,S., Malayer,J.R., Saliki,J.T., Kaul,A., Ramachandran,A. |
| EPI_ISL_576386 | National Institute of Health Research and Development | National Institute of Health Research and Development | Pawestri,HA;Subangkit;Puspa,KD;Nugraha,AA;Ikawati,HD;Pangesti, KNA;Soekarso,T;Susilarini,NK;Hariastuti,NI;Nikmah,UA;Mursinah;Febriyani,A;Herman,R;Susanti,N;Herna;Febriyanti,T; Nurhadi,M; Paisal;Ramadhany,R;Agustiningsih;Kurniawati,J;Kipuw,NL;Muna,F;Indalau,IL;Adam,K;Wibowo,HA;Rizki,A;Puspandari,N;Setiawaty,V |
| EPI_ISL_576399, EPI_ISL_576400, EPI_ISL_576401, EPI_ISL_576402, EPI_ISL_576403, EPI_ISL_576404, EPI_ISL_576405, EPI_ISL_576407, EPI_ISL_576408, EPI_ISL_576409, EPI_ISL_576410, EPI_ISL_576411, EPI_ISL_576412, EPI_ISL_576414, EPI_ISL_576415, EPI_ISL_576416, EPI_ISL_576417, EPI_ISL_576418, EPI_ISL_576419, EPI_ISL_576420, EPI_ISL_576422, EPI_ISL_576423, EPI_ISL_576424, EPI_ISL_576425, EPI_ISL_576426, EPI_ISL_576427, EPI_ISL_576428, EPI_ISL_576429, EPI_ISL_576430, EPI_ISL_576431, EPI_ISL_576432, EPI_ISL_576433, EPI_ISL_576434, EPI_ISL_576435, EPI_ISL_576436, EPI_ISL_576437, EPI_ISL_576438, EPI_ISL_576439, EPI_ISL_576440, EPI_ISL_576441, EPI_ISL_576442, EPI_ISL_576443, EPI_ISL_576444, EPI_ISL_576445, EPI_ISL_576446, EPI_ISL_576447, EPI_ISL_576448, EPI_ISL_576449, EPI_ISL_576450, EPI_ISL_576451, EPI_ISL_576452, EPI_ISL_576453, EPI_ISL_576454, EPI_ISL_576455, EPI_ISL_576456, EPI_ISL_576457, EPI_ISL_576458, EPI_ISL_576459, EPI_ISL_576460, EPI_ISL_576461, EPI_ISL_576462, EPI_ISL_576463, EPI_ISL_576464, EPI_ISL_576465, EPI_ISL_576466, EPI_ISL_576467, EPI_ISL_576468, EPI_ISL_576469, EPI_ISL_576470, EPI_ISL_576471, EPI_ISL_576472, EPI_ISL_576473, EPI_ISL_576474, EPI_ISL_576475, EPI_ISL_576476, EPI_ISL_576477, EPI_ISL_576478, EPI_ISL_576479, EPI_ISL_576480, EPI_ISL_576481, EPI_ISL_576482, EPI_ISL_576483, EPI_ISL_576484, EPI_ISL_576485, EPI_ISL_576486, EPI_ISL_576487, EPI_ISL_576488, EPI_ISL_576489, EPI_ISL_576490, EPI_ISL_576491, EPI_ISL_576492, EPI_ISL_576493, EPI_ISL_576494, EPI_ISL_576495, EPI_ISL_576496, EPI_ISL_576497, EPI_ISL_576498, EPI_ISL_576499, EPI_ISL_576500, EPI_ISL_576501, EPI_ISL_576502, EPI_ISL_576503, EPI_ISL_576504, EPI_ISL_576505, EPI_ISL_576506, EPI_ISL_576507, EPI_ISL_576508, EPI_ISL_576509, EPI_ISL_576510, EPI_ISL_576511, EPI_ISL_576512, EPI_ISL_576513, EPI_ISL_576514, EPI_ISL_576515, EPI_ISL_576516, EPI_ISL_576517, EPI_ISL_576557, EPI_ISL_576558, EPI_ISL_576559, EPI_ISL_576560, EPI_ISL_576561, EPI_ISL_576562, EPI_ISL_576563, EPI_ISL_576564 | | | |
| see above | UW Virology Lab | UW Virology Lab | Pavitra Roychoudhury, Hong Xie, Lasata Shrestha, Amin Addetia, Victoria M Rachleff, Meei-Li Huang, Keith R Jerome, Alexander Greninger |
| EPI_ISL_577639, EPI_ISL_577640, EPI_ISL_577641, EPI_ISL_577642, EPI_ISL_577643, EPI_ISL_577644, EPI_ISL_577645, EPI_ISL_577646, EPI_ISL_577647, EPI_ISL_577648, EPI_ISL_577649, EPI_ISL_577650, EPI_ISL_577651, EPI_ISL_577652, EPI_ISL_577653, EPI_ISL_577654, EPI_ISL_577655, EPI_ISL_577656, EPI_ISL_577657, EPI_ISL_577658, EPI_ISL_577659, EPI_ISL_577660, EPI_ISL_577661, EPI_ISL_577662, EPI_ISL_577663, EPI_ISL_577664, EPI_ISL_577665, EPI_ISL_577666, EPI_ISL_577667, EPI_ISL_577668, EPI_ISL_577669, EPI_ISL_577670, EPI_ISL_577671, EPI_ISL_577672, EPI_ISL_577673, EPI_ISL_577675, EPI_ISL_577676, EPI_ISL_577677, EPI_ISL_577678, EPI_ISL_577679, EPI_ISL_577680, EPI_ISL_577681, EPI_ISL_577682, EPI_ISL_577683, EPI_ISL_577684, EPI_ISL_577685, EPI_ISL_577686, EPI_ISL_577687, EPI_ISL_577688, EPI_ISL_577689, EPI_ISL_577690, EPI_ISL_577691, EPI_ISL_577692, EPI_ISL_577693, EPI_ISL_577694, EPI_ISL_577695, EPI_ISL_577696, EPI_ISL_577697, EPI_ISL_577698, EPI_ISL_577699, EPI_ISL_577700, EPI_ISL_577701, EPI_ISL_577702, EPI_ISL_577704, EPI_ISL_577705, EPI_ISL_577706, EPI_ISL_577707, EPI_ISL_577708, EPI_ISL_577709, EPI_ISL_577710, EPI_ISL_577711, EPI_ISL_577712, EPI_ISL_577713, EPI_ISL_577716, EPI_ISL_577717, EPI_ISL_577720, EPI_ISL_577721, EPI_ISL_577722, EPI_ISL_577723, EPI_ISL_577724, EPI_ISL_577725, EPI_ISL_577726, EPI_ISL_577727, EPI_ISL_577728, EPI_ISL_577729, EPI_ISL_577730, EPI_ISL_577731 | | | |
| see above | NIV Influenza | NIV Influenza | Potdar V |
| EPI_ISL_582029, EPI_ISL_582030 | Biology Department, College of Science, Al-Muthanna University | International Centre for Genetic Engineering and Biotechnology (ICGEB) and ARGO Open Lab Platform | Nihad Al-Rashedi, Danilo Licastro, Sreejith Rajasekharan, Simeone Dal Monego, Alessandro Marcello |
| EPI_ISL_582127, EPI_ISL_582128, EPI_ISL_582129, EPI_ISL_582130, EPI_ISL_582131, EPI_ISL_582132 | Antwerp University Hospital | Institute of Tropical Medicine | Philippe Selhorst, Colin Anthony |
| EPI_ISL_582241 | NIV Influenza | NIV Influenza | Potdar V |
| EPI_ISL_583893, EPI_ISL_583894, EPI_ISL_583895, EPI_ISL_583896, EPI_ISL_583897, EPI_ISL_583898, EPI_ISL_583899, EPI_ISL_583900 | Singapore General Hospital | Department of Microbiology | Nurdyana Abdul Rahman, Kun Lee Lim, Chenhao Li, Sui Sin Goh, Kenneth Xin Long Chan, Kian Sing Chan, Lynette Oon, Kern Rei Chng, Niranjan Nagarajan, Karrie Ko |
| EPI_ISL_583968, EPI_ISL_583969, EPI_ISL_583970, EPI_ISL_583971, EPI_ISL_583972, EPI_ISL_583973, EPI_ISL_583974, EPI_ISL_583975, EPI_ISL_583976, EPI_ISL_583977, EPI_ISL_583978, EPI_ISL_583979, EPI_ISL_583980, EPI_ISL_583981, EPI_ISL_583982, EPI_ISL_583983, EPI_ISL_583984, EPI_ISL_583985, EPI_ISL_583986, EPI_ISL_583987, EPI_ISL_583988, EPI_ISL_583989, EPI_ISL_583990, EPI_ISL_583991, EPI_ISL_583992, EPI_ISL_583993, EPI_ISL_584061, EPI_ISL_584066 | | | |
| see above | Respiratory Virus Unit, Microbiology Services Colindale, Public Health England | Respiratory Virus Unit, Microbiology Services Colindale, Public Health England | PHE Covid Sequencing Team |
| EPI_ISL_586513 | B.J. Medical College and Civil hospital, Ahmedabad | Gujarat Biotechnology Research Centre | Pinal Trivedi, Maharshi Pandya, Nidhi Patel, Nitin Savaliya, Raghawendra Kumar, Dinesh Kumar, Zuber Saiyed, Komal Patel, Labdhi Pandya, Afzal Ansari, Nikha Trivedi, Pranay Shah, Kamlesh J Upadhyay, Sanjay Kapadia, Apurvasinh Puvar, Janvi Raval, Zarna Patel, Monika Gandhi, R D Dixit, A M Kadri, Harsh |

| | | | |
|---|---|---|---|
| | | | Bakshi, Chaitanya Joshi, Madhvi Joshi |
| EPI_ISL_586515 | B.J. Medical College and Civil hospital, Ahmedabad | Gujarat Biotechnology Research Centre | Nidhi Patel, Nitin Savaliya, Raghawendra Kumar, Dinesh Kumar, Zuber Saiyed, Komal Patel, Labdhi Pandya, Afzal Ansari, Nikha Trivedi, Pranay Shah, Kamlesh J Upadhyay, Sanjay Kapadia, Apurvasinh Puvar, Janvi Raval, Zarna Patel, Monika Gandhi, Pinal Trivedi, Maharshi Pandya, R D Dixit, A M Kadri, Harsh Bakshi, Chaitanya Joshi, Madhvi Joshi |
| EPI_ISL_586516 | B.J. Medical College and Civil hospital, Ahmedabad | Gujarat Biotechnology Research Centre | Nitin Savaliya, Raghawendra Kumar, Dinesh Kumar, Zuber Saiyed, Komal Patel, Labdhi Pandya, Afzal Ansari, Nikha Trivedi, Pranay Shah, Kamlesh J Upadhyay, Sanjay Kapadia, Apurvasinh Puvar, Janvi Raval, Zarna Patel, Monika Gandhi, Pinal Trivedi, Maharshi Pandya, Nidhi Patel, R D Dixit, A M Kadri, Harsh Bakshi, Chaitanya Joshi, Madhvi Joshi |
| EPI_ISL_586523 | B.J. Medical College and Civil hospital, Ahmedabad | Gujarat Biotechnology Research Centre | Nikha Trivedi, Pranay Shah, Kamlesh J Upadhyay, Sanjay Kapadia, Apurvasinh Puvar, Janvi Raval, Zarna Patel, Monika Gandhi, Pinal Trivedi, Maharshi Pandya, Nidhi Patel, Nitin Savaliya, Raghawendra Kumar, Dinesh Kumar, Zuber Saiyed, Komal Patel, Labdhi Pandya, Afzal Ansari, R D Dixit, A M Kadri, Harsh Bakshi, Chaitanya Joshi, Madhvi Joshi |
| EPI_ISL_586524 | B.J. Medical College and Civil hospital, Ahmedabad | Gujarat Biotechnology Research Centre | Pranay Shah, Kamlesh J Upadhyay, Sanjay Kapadia, Apurvasinh Puvar, Janvi Raval, Zarna Patel, Monika Gandhi, Pinal Trivedi, Maharshi Pandya, Nidhi Patel, Nitin Savaliya, Raghawendra Kumar, Dinesh Kumar, Zuber Saiyed, Komal Patel, Labdhi Pandya, Afzal Ansari, Nikha Trivedi, R D Dixit, A M Kadri, Harsh Bakshi, Chaitanya Joshi, Madhvi Joshi |
| EPI_ISL_586530 | B.J. Medical College and Civil hospital, Ahmedabad | Gujarat Biotechnology Research Centre | Pinal Trivedi, Maharshi Pandya, Nidhi Patel, Nitin Savaliya, Raghawendra Kumar, Dinesh Kumar, Zuber Saiyed, Komal Patel, Labdhi Pandya, Afzal Ansari, Nikha Trivedi, Pranay Shah, Kamlesh J Upadhyay, Sanjay Kapadia, Apurvasinh Puvar, Janvi Raval, Zarna Patel, Monika Gandhi, R D Dixit, A M Kadri, Harsh Bakshi, Chaitanya Joshi, Madhvi Joshi |
| EPI_ISL_586535 | General Hospital,Chhotaudepur | Gujarat Biotechnology Research Centre | Nidhi Patel, Nitin Savaliya, Raghawendra Kumar, Dinesh Kumar, Zuber Saiyed, Komal Patel, Labdhi Pandya, Afzal Ansari, Nikha Trivedi, Apurvasinh Puvar, Janvi Raval, Zarna Patel, Monika Gandhi, Pinal Trivedi, Yogesh Parmar, Arti Thakur, Dharak Pandya, Maharshi Pandya, R D Dixit, A M Kadri, Harsh Bakshi, Chaitanya Joshi, Madhvi Joshi |
| EPI_ISL_586538 | Zydus Medical Hospital,Dahod | Gujarat Biotechnology Research Centre | Nitin Savaliya, Raghawendra Kumar, Dinesh Kumar, Zuber Saiyed, Komal Patel, Labdhi Pandya, Afzal Ansari, Nikha Trivedi, Apurvasinh Puvar, Janvi Raval, Zarna Patel, Monika Gandhi, Pinal Trivedi, Maharshi Pandya, Nidhi Patel, Kalpesh Labana, Mohit Desai, R D Dixit, A M Kadri, Harsh Bakshi, Chaitanya Joshi, Madhvi Joshi |
| EPI_ISL_586545 | Government General Hospital, Jam Khambhaliya, Devbhoomi Dwarka | Gujarat Biotechnology Research Centre | Nikha Trivedi, Harish Matani, Apurvasinh Puvar, Janvi Raval, Zarna Patel, Monika Gandhi, Pinal Trivedi, Maharshi Pandya, Nidhi Patel, Nitin Savaliya, Raghawendra Kumar, Dinesh Kumar, Zuber Saiyed, Komal Patel, Labdhi Pandya, Afzal Ansari, R D Dixit, A M Kadri, Harsh Bakshi, Chaitanya Joshi, Madhvi Joshi |
| EPI_ISL_586546 | General Hospital,Godhra | Gujarat Biotechnology Research Centre | Neela Katara, Pinal Gandhi, Sangeeta Kumar, Apurvasinh Puvar, Janvi Raval, Zarna Patel, Monika Gandhi, Pinal Trivedi, Maharshi Pandya, Nidhi Patel, Nitin Savaliya, Raghawendra Kumar, Dinesh Kumar, Zuber Saiyed, Komal Patel, Labdhi Pandya, Afzal Ansari, Nikha Trivedi, R D Dixit, A M Kadri, Harsh Bakshi, Chaitanya Joshi, Madhvi Joshi |
| EPI_ISL_586547 | General Hospital,Godhra | Gujarat Biotechnology Research Centre | Pinal Gandhi, Sangeeta Kumar, Apurvasinh Puvar, Janvi Raval, Zarna Patel, Monika Gandhi, Pinal Trivedi, Maharshi Pandya, Nidhi Patel, Nitin Savaliya, Raghawendra Kumar, Dinesh Kumar, Zuber Saiyed, Komal Patel, Labdhi Pandya, Afzal Ansari, Nikha Trivedi, Neela Katara, R D Dixit, A M Kadri, Harsh Bakshi, Chaitanya Joshi, Madhvi Joshi |
| EPI_ISL_586552 | Government Hospital,Veraval | Gujarat Biotechnology Research Centre | Pinal Trivedi, Maharshi Pandya, Nidhi Patel, Nitin Savaliya, Raghawendra Kumar, Dinesh Kumar, Zuber Saiyed, Komal Patel, Labdhi Pandya, Afzal Ansari, Nikha Trivedi, Jignesh Parmar, Jitendra Bamrotia, Apurvasinh Puvar, Janvi Raval, Zarna Patel, Monika Gandhi, R D Dixit, A M Kadri, Harsh Bakshi, Chaitanya Joshi, Madhvi Joshi |
| EPI_ISL_586554 | Government Hospital,Veraval | Gujarat Biotechnology Research Centre | Nidhi Patel, Nitin Savaliya, Raghawendra Kumar, Dinesh Kumar, Zuber Saiyed, Komal Patel, Labdhi Pandya, Afzal Ansari, Nikha Trivedi, Jignesh Parmar, Jitendra Bamrotia, Apurvasinh Puvar, Janvi Raval, Zarna Patel, Monika Gandhi, Pinal Trivedi, Maharshi Pandya, R D Dixit, A M Kadri, Harsh Bakshi, Chaitanya Joshi, Madhvi Joshi |
| EPI_ISL_586555 | CCC,Veraval | Gujarat Biotechnology Research Centre | Nitin Savaliya, Raghawendra Kumar, Dinesh Kumar, Zuber Saiyed, Komal Patel, Labdhi Pandya, Afzal Ansari, Nikha Trivedi, Jignesh Parmar, Jitendra Bamrotia, Apurvasinh Puvar, Janvi Raval, Zarna Patel, Monika Gandhi, Pinal Trivedi, Maharshi Pandya, Nidhi Patel, R D Dixit, A M Kadri, Harsh Bakshi, Chaitanya Joshi, Madhvi Joshi |
| EPI_ISL_586562 | GMERS GMERS Medical College and Hospital, Junagadh | Gujarat Biotechnology Research Centre | Nikha Trivedi, Bhavesh Bagda , Ravi Dedaniya, Apurvasinh Puvar, Janvi Raval, Zarna Patel, Monika Gandhi, Pinal Trivedi, Maharshi Pandya, Nidhi Patel, Nitin Savaliya, Raghawendra Kumar, Dinesh Kumar, Zuber Saiyed, Komal Patel, Labdhi Pandya, Afzal Ansari, R D Dixit, A M Kadri, Harsh Bakshi, Chaitanya Joshi, Madhvi Joshi |
| EPI_ISL_591083, EPI_ISL_591084, EPI_ISL_591085, EPI_ISL_591086, EPI_ISL_591087, EPI_ISL_591088, EPI_ISL_591089 | Viral Respiratory Lab, National Institute for Biomedical Research (INRB) | Pathogen Sequencing Lab, National Institute for Biomedical Research (INRB) | Placide Mbala-Kingebeni, Edith Nkwembe, Eddy Kinganda-Lusamaki, Amuri Aziza, Francisca Muyembe Mawete, Emmanuel Lokilo Lofiko, Jean Claude Makangara, Catherine Pratt, Matthias Pauthner, Josh Quick, Allison Black, James Hadfield, Trevor Bedford, Ian Goodfellow, Andrew Rambaut, Nick Loman, Kristian Andersen, Michael Wiley, Steve Ahuka-Mundeke, Jean-Jacques Muyembe Tamfum |
| EPI_ISL_593804, EPI_ISL_593805, EPI_ISL_593810, EPI_ISL_593824, EPI_ISL_593825, EPI_ISL_593829, EPI_ISL_593831, EPI_ISL_593834, EPI_ISL_593837, EPI_ISL_593843, EPI_ISL_593846, EPI_ISL_593848, EPI_ISL_593850, EPI_ISL_593998 | | | |
| see above | Respiratory Virus Unit, Microbiology Services Colindale, Public Health England | Respiratory Virus Unit, Microbiology Services Colindale, Public Health England | PHE Covid Sequencing Team |
| EPI_ISL_594166, EPI_ISL_594167, EPI_ISL_594168, EPI_ISL_594169, EPI_ISL_594170, EPI_ISL_594171, EPI_ISL_594172, EPI_ISL_594173, EPI_ISL_594174, EPI_ISL_594175, EPI_ISL_594176, EPI_ISL_594179 | | | |
| see above | PathWest Laboratory Medicine WA | PathWest Laboratory Medicine WA Microbial Surveillance Unit | PathWest Laboratory Medicine WA Microbial Surveillance Unit |
| EPI_ISL_596500, EPI_ISL_596501, EPI_ISL_596502, EPI_ISL_596503, EPI_ISL_596504, EPI_ISL_596505, EPI_ISL_596506, EPI_ISL_596507, EPI_ISL_596508, EPI_ISL_596509, EPI_ISL_596510, EPI_ISL_596511, EPI_ISL_596512, EPI_ISL_596513, EPI_ISL_596514, EPI_ISL_596515, EPI_ISL_596516, EPI_ISL_596517, EPI_ISL_596518, EPI_ISL_596519, EPI_ISL_596520, EPI_ISL_596521, EPI_ISL_596522, EPI_ISL_596523, EPI_ISL_596525, EPI_ISL_596526, EPI_ISL_596527, EPI_ISL_596528, EPI_ISL_596529, EPI_ISL_596530, EPI_ISL_596531, EPI_ISL_596532, EPI_ISL_596533, EPI_ISL_596534, EPI_ISL_596535, EPI_ISL_596536, EPI_ISL_596537, EPI_ISL_596538, EPI_ISL_596542, EPI_ISL_596543, EPI_ISL_596544, EPI_ISL_596545, EPI_ISL_596546, EPI_ISL_596547, EPI_ISL_596548, EPI_ISL_596549, EPI_ISL_596550, EPI_ISL_596551, EPI_ISL_596552, EPI_ISL_596553, EPI_ISL_596554, EPI_ISL_596555, EPI_ISL_596556, EPI_ISL_596557, EPI_ISL_596558, EPI_ISL_596559, EPI_ISL_596560, EPI_ISL_596561, EPI_ISL_596562, EPI_ISL_596563, EPI_ISL_596564, EPI_ISL_596565, EPI_ISL_596567, EPI_ISL_596568 | | | |
| see above | Palestinian Ministry of Health | Molecular Genetics Lab | Nouar Qutob, Zaidoun Salah, Damien Richard, Hisham Darwish, Husam Sallam, Issa Shtayeh, Osama Najjar, Mahmoud Ruzayqat, Dana Najjar, Francois Balloux, Lucy van Dorp |
| EPI_ISL_596677, EPI_ISL_596678, EPI_ISL_596679, EPI_ISL_596680, EPI_ISL_596681, EPI_ISL_596682, EPI_ISL_596683, EPI_ISL_596684, EPI_ISL_596685, EPI_ISL_596686, EPI_ISL_596687, EPI_ISL_596688, EPI_ISL_596689, EPI_ISL_596690, EPI_ISL_596691, EPI_ISL_596692, EPI_ISL_596693, EPI_ISL_596694, EPI_ISL_596695, EPI_ISL_596696, EPI_ISL_596697, EPI_ISL_596698, EPI_ISL_596699, EPI_ISL_596700, EPI_ISL_596701, EPI_ISL_596702, EPI_ISL_596703, EPI_ISL_596704, EPI_ISL_596705, EPI_ISL_596706, EPI_ISL_596707, EPI_ISL_596708, EPI_ISL_596709, EPI_ISL_596710, EPI_ISL_596711, EPI_ISL_596712, EPI_ISL_596713, EPI_ISL_596714, EPI_ISL_596715, EPI_ISL_596716, EPI_ISL_596717, EPI_ISL_596718, EPI_ISL_596719, EPI_ISL_596720, EPI_ISL_596721, EPI_ISL_596722, EPI_ISL_596723, EPI_ISL_596724, EPI_ISL_596725, EPI_ISL_596726, EPI_ISL_596727, EPI_ISL_596728, EPI_ISL_596729, EPI_ISL_596730, EPI_ISL_596731, EPI_ISL_596732, EPI_ISL_596733, EPI_ISL_596734, EPI_ISL_596735, EPI_ISL_596736, EPI_ISL_596737, EPI_ISL_596738, EPI_ISL_596739, EPI_ISL_596740, EPI_ISL_596741, EPI_ISL_596742, EPI_ISL_596743, EPI_ISL_596744, EPI_ISL_596745, EPI_ISL_596746, EPI_ISL_596747, EPI_ISL_596748, EPI_ISL_596749, EPI_ISL_596750, EPI_ISL_596751, EPI_ISL_596752, EPI_ISL_596753, EPI_ISL_596754, EPI_ISL_596755, EPI_ISL_596756, EPI_ISL_596757, EPI_ISL_596758, EPI_ISL_596759, EPI_ISL_596760, EPI_ISL_596761, EPI_ISL_596762, EPI_ISL_596763, EPI_ISL_596764, EPI_ISL_596765, EPI_ISL_596766, EPI_ISL_596767, EPI_ISL_596768, EPI_ISL_596769, EPI_ISL_596770, EPI_ISL_596771, EPI_ISL_596772, EPI_ISL_596773, EPI_ISL_596774, EPI_ISL_596775, EPI_ISL_596776, EPI_ISL_596777, EPI_ISL_596778, EPI_ISL_596779, EPI_ISL_596780, EPI_ISL_596781, EPI_ISL_596782, EPI_ISL_596783, EPI_ISL_596784, EPI_ISL_596785, EPI_ISL_596786, EPI_ISL_596787, EPI_ISL_596788, EPI_ISL_596789, EPI_ISL_596790, EPI_ISL_596792, EPI_ISL_596793, EPI_ISL_596794, EPI_ISL_596795, EPI_ISL_596796, EPI_ISL_596798, EPI_ISL_596799, EPI_ISL_596800, EPI_ISL_596802, EPI_ISL_596803, EPI_ISL_596805, EPI_ISL_596806, EPI_ISL_596807, EPI_ISL_596809, EPI_ISL_596810, EPI_ISL_596811, EPI_ISL_596812, EPI_ISL_596815, EPI_ISL_596816, EPI_ISL_596819, EPI_ISL_596820, EPI_ISL_596821, EPI_ISL_596822, EPI_ISL_596823, EPI_ISL_596824, EPI_ISL_596825, EPI_ISL_596826, EPI_ISL_596827, EPI_ISL_596828, EPI_ISL_596829, EPI_ISL_596830, EPI_ISL_596831, EPI_ISL_596832, EPI_ISL_596833, EPI_ISL_596834, EPI_ISL_596835, EPI_ISL_596836, EPI_ISL_596837, EPI_ISL_596838, EPI_ISL_596839, EPI_ISL_596840, EPI_ISL_596841, EPI_ISL_596842, EPI_ISL_596843, EPI_ISL_596844, EPI_ISL_596845, EPI_ISL_596846, EPI_ISL_596847, EPI_ISL_596848, EPI_ISL_596849, EPI_ISL_596850, EPI_ISL_596851, EPI_ISL_596852, EPI_ISL_596853, EPI_ISL_596854, EPI_ISL_596855, EPI_ISL_596856, EPI_ISL_596857, EPI_ISL_596858, EPI_ISL_596859, EPI_ISL_596860, EPI_ISL_596861, EPI_ISL_596862, EPI_ISL_596863, EPI_ISL_596864, EPI_ISL_596865, EPI_ISL_596866, EPI_ISL_596867, EPI_ISL_596868, EPI_ISL_596869, EPI_ISL_596870, EPI_ISL_596871, EPI_ISL_596872, EPI_ISL_596873, EPI_ISL_596874, EPI_ISL_596875, EPI_ISL_596876, EPI_ISL_596877, EPI_ISL_596879, EPI_ISL_596880, EPI_ISL_596881, EPI_ISL_596882, EPI_ISL_596884, EPI_ISL_596885, EPI_ISL_596886 | | | |
| see above | PathWest Laboratory Medicine WA | PathWest Laboratory Medicine WA Microbial Surveillance Unit | PathWest Laboratory Medicine WA Microbial Surveillance Unit |
| EPI_ISL_602462, EPI_ISL_602463, EPI_ISL_602464, EPI_ISL_602465, EPI_ISL_602466, EPI_ISL_602467, EPI_ISL_602468, EPI_ISL_602469, EPI_ISL_602470, EPI_ISL_602471, EPI_ISL_602472, EPI_ISL_602473, EPI_ISL_602474, EPI_ISL_602475, EPI_ISL_602476, EPI_ISL_602477, EPI_ISL_602478, EPI_ISL_602479, EPI_ISL_602480, EPI_ISL_602481, EPI_ISL_602482, EPI_ISL_602483, EPI_ISL_602484, EPI_ISL_602485, EPI_ISL_602486, EPI_ISL_602487, EPI_ISL_602488, EPI_ISL_602489, EPI_ISL_602490, EPI_ISL_602491, EPI_ISL_602492, EPI_ISL_602493, EPI_ISL_602494, EPI_ISL_602495, EPI_ISL_602496, EPI_ISL_602497, | | | |

| | | | |
|---|---|---|---|
| EPI_ISL_602498, EPI_ISL_602499, EPI_ISL_602500, EPI_ISL_602501, EPI_ISL_602502, EPI_ISL_602503, EPI_ISL_602504, EPI_ISL_602505, EPI_ISL_602506, EPI_ISL_602507, EPI_ISL_602508, EPI_ISL_602509, EPI_ISL_602518, EPI_ISL_602519, EPI_ISL_602520, EPI_ISL_602521, EPI_ISL_602522, EPI_ISL_602523, EPI_ISL_602524, EPI_ISL_602525, EPI_ISL_602526, EPI_ISL_602527, EPI_ISL_602528, EPI_ISL_602529, EPI_ISL_602530, EPI_ISL_602531, EPI_ISL_602532 | | | |
| see above | Institute for Virology, University Hospital Essen | Center of Medical Microbiology, Virology, and Hospital Hygiene, University of Duesseldorf | Olympia E. Anastasiou, Ulf Dittmer, Maximilian Damagnez, Alexander Dilthey, Torsten Houwaart, Lisanna Hülse, Malte Kohns Vasconcelos, Nadine Lübke, Jessica Nicolai, Klaus Pfeffer, Daniel Strelow, Jörg Timm, Andreas Walker, Tobias Wienemann |
| EPI_ISL_603188, EPI_ISL_603189, EPI_ISL_603190, EPI_ISL_603191, EPI_ISL_603192, EPI_ISL_603193, EPI_ISL_603194, EPI_ISL_603195, EPI_ISL_603196, EPI_ISL_603197, EPI_ISL_603198, EPI_ISL_603199, EPI_ISL_603200, EPI_ISL_603201, EPI_ISL_603202, EPI_ISL_603203, EPI_ISL_603204, EPI_ISL_603205, EPI_ISL_603206, EPI_ISL_603207, EPI_ISL_603208, EPI_ISL_603209, EPI_ISL_603210, EPI_ISL_603211, EPI_ISL_603212, EPI_ISL_603213, EPI_ISL_603214, EPI_ISL_603215 | | | |
| see above | Respiratory Virus Unit, Microbiology Services Colindale, Public Health England | Respiratory Virus Unit, Microbiology Services Colindale, Public Health England | PHE Covid Sequencing Team |
| EPI_ISL_603251, EPI_ISL_603252, EPI_ISL_603253, EPI_ISL_603254, EPI_ISL_603255, EPI_ISL_603256, EPI_ISL_603257, EPI_ISL_603258, EPI_ISL_603259, EPI_ISL_603260, EPI_ISL_603261, EPI_ISL_603262, EPI_ISL_603264, EPI_ISL_603265 | | | |
| see above | UW Virology Lab | UW Virology Lab | Pavitra Roychoudhury, Hong Xie, Lasata Shrestha, Meei-Li Huang, Keith R Jerome, Alexander Greninger |
| EPI_ISL_605825, EPI_ISL_605826, EPI_ISL_605827, EPI_ISL_605828, EPI_ISL_605829, EPI_ISL_605830, EPI_ISL_605831, EPI_ISL_605832, EPI_ISL_605833, EPI_ISL_605834, EPI_ISL_605835, EPI_ISL_605836, EPI_ISL_605837, EPI_ISL_605838, EPI_ISL_605839, EPI_ISL_605840, EPI_ISL_605841, EPI_ISL_605842, EPI_ISL_605843, EPI_ISL_605844, EPI_ISL_605845, EPI_ISL_605846, EPI_ISL_605847, EPI_ISL_605848, EPI_ISL_605849, EPI_ISL_605850, EPI_ISL_605851, EPI_ISL_605852, EPI_ISL_605853, EPI_ISL_605854, EPI_ISL_605855, EPI_ISL_605856, EPI_ISL_605857, EPI_ISL_605859, EPI_ISL_605860, EPI_ISL_605861, EPI_ISL_605862, EPI_ISL_605863, EPI_ISL_605864, EPI_ISL_605865, EPI_ISL_605866, EPI_ISL_605867, EPI_ISL_605868, EPI_ISL_605869, EPI_ISL_605870, EPI_ISL_605873, EPI_ISL_605877, EPI_ISL_605879, EPI_ISL_605880, EPI_ISL_605881 | | | |
| see above | PathWest Laboratory Medicine WA | PathWest Laboratory Medicine WA Microbial Surveillance Unit | PathWest Laboratory Medicine WA Microbial Surveillance Unit |
| EPI_ISL_609827, EPI_ISL_609828, EPI_ISL_609829, EPI_ISL_609830, EPI_ISL_609831, EPI_ISL_609832, EPI_ISL_609833, EPI_ISL_609834, EPI_ISL_609835, EPI_ISL_609836, EPI_ISL_609837, EPI_ISL_609838, EPI_ISL_609839, EPI_ISL_609840, EPI_ISL_609841, EPI_ISL_609842, EPI_ISL_609843, EPI_ISL_609844, EPI_ISL_609845, EPI_ISL_609846, EPI_ISL_609847, EPI_ISL_609848, EPI_ISL_609849, EPI_ISL_609850, EPI_ISL_609851, EPI_ISL_609852, EPI_ISL_609853, EPI_ISL_609854, EPI_ISL_609855, EPI_ISL_609856, EPI_ISL_609857, EPI_ISL_609858, EPI_ISL_609859, EPI_ISL_609860, EPI_ISL_609861, EPI_ISL_609862, EPI_ISL_609863, EPI_ISL_609864, EPI_ISL_609865, EPI_ISL_609866, EPI_ISL_609867, EPI_ISL_609868, EPI_ISL_609869, EPI_ISL_609870, EPI_ISL_609871, EPI_ISL_609872, EPI_ISL_609873, EPI_ISL_609874, EPI_ISL_609875, EPI_ISL_609876, EPI_ISL_609877, EPI_ISL_609878, EPI_ISL_609879, EPI_ISL_609880, EPI_ISL_609881, EPI_ISL_609882, EPI_ISL_609883, EPI_ISL_609884, EPI_ISL_609885, EPI_ISL_609886, EPI_ISL_609887, EPI_ISL_609888, EPI_ISL_609889, EPI_ISL_609890, EPI_ISL_609891, EPI_ISL_609892, EPI_ISL_609893, EPI_ISL_609894, EPI_ISL_609895, EPI_ISL_609896, EPI_ISL_609897, EPI_ISL_609898, EPI_ISL_609899, EPI_ISL_609900, EPI_ISL_609901, EPI_ISL_609902, EPI_ISL_609903, EPI_ISL_609904, EPI_ISL_609905, EPI_ISL_609906, EPI_ISL_609907, EPI_ISL_609908, EPI_ISL_609909 | | | |
| see above | Respiratory Virus Unit, Microbiology Services Colindale, Public Health England | Respiratory Virus Unit, Microbiology Services Colindale, Public Health England | PHE Covid Sequencing Team |
| EPI_ISL_610152, EPI_ISL_610153, EPI_ISL_610154 | Singapore General Hospital | Department of Microbiology | Nurdyana Abdul Rahman, Kun Lee Lim, Chenhao Li, Sui Sin Goh, Kenneth Xin Long Chan, Kian Sing Chan, Lynette Oon, Kern Rei Chng, Niranjan Nagarajan, Karrie Ko |
| EPI_ISL_610283, EPI_ISL_610284, EPI_ISL_610285 | UW Virology Lab | UW Virology Lab | Pavitra Roychoudhury, Hong Xie, Lasata Shrestha, Meei-Li Huang, Keith R Jerome, Alexander Greninger |
| EPI_ISL_613459, EPI_ISL_613460 | Microbiology, Koc University | Microbiology, Koc University | Ozer,B., Nurtop,E., Kuskucu,M.A., Dogan,O., Can,F. |
| EPI_ISL_613461, EPI_ISL_613462 | Microbiology, Koc University | Microbiology, Koc University | Nurtop,E., Ozer,B., Kuskucu,M.A., Dogan,O., Can,F. |
| EPI_ISL_613463, EPI_ISL_613464, EPI_ISL_613465, EPI_ISL_613466, EPI_ISL_613467, EPI_ISL_613468, EPI_ISL_613469, EPI_ISL_613470, EPI_ISL_613471, EPI_ISL_613472, EPI_ISL_613473, EPI_ISL_613474, EPI_ISL_613475, EPI_ISL_613476, EPI_ISL_613477, EPI_ISL_613478, EPI_ISL_613479, EPI_ISL_613480, EPI_ISL_613481, EPI_ISL_613482, EPI_ISL_613483, EPI_ISL_613484, EPI_ISL_613485, EPI_ISL_613486, EPI_ISL_613487, EPI_ISL_613488, EPI_ISL_613489, EPI_ISL_613490, EPI_ISL_613491, EPI_ISL_613492, EPI_ISL_613493, EPI_ISL_613494, EPI_ISL_613495, EPI_ISL_613496, EPI_ISL_613497, EPI_ISL_613498, EPI_ISL_613499, EPI_ISL_613500, EPI_ISL_613501, EPI_ISL_613502, EPI_ISL_613503, EPI_ISL_613504, EPI_ISL_613505, EPI_ISL_613506, EPI_ISL_613507, EPI_ISL_613508, EPI_ISL_613509, EPI_ISL_613510, EPI_ISL_613511, EPI_ISL_613512, EPI_ISL_613513, EPI_ISL_613514, EPI_ISL_613515, EPI_ISL_613516, EPI_ISL_613517, EPI_ISL_613518, EPI_ISL_613519, EPI_ISL_613520, EPI_ISL_613521, EPI_ISL_613522, EPI_ISL_613523, EPI_ISL_613524, EPI_ISL_613525, EPI_ISL_613526, EPI_ISL_613527, EPI_ISL_613528, EPI_ISL_613529, EPI_ISL_613530, EPI_ISL_613531, EPI_ISL_613532, EPI_ISL_613533, EPI_ISL_613534, EPI_ISL_613535, EPI_ISL_613536, EPI_ISL_613537, EPI_ISL_613538, EPI_ISL_613539, EPI_ISL_613540 | | | |
| see above | Public Health Laboratory - Infectious Disease Lab, Minnesota Department of Health Infectious Disease Laboratory Submission Group | Minnesota Department of Health, Public Health Laboratory | Plumb,M., Garfin,J., Lorentz,A., Wang,X. |
| EPI_ISL_613560 | Laboratorio Biologia Molecolare Sars Cov2 - UOC Laboratorio Analisi - Servizio Medicina di Laboratorio, Ospedale "San Francesco" - ATS-ASSL Nuoro Via Manniron 1, 08100 Nuoro | Laboratorio specialistico UOC Ematologia - Ospedale "San Francesco" - ATS-ASSL Nuoro Nuoro | Piras Giovanna, Fancello Tatiana, Asproni Rosanna, Fiamma Maura, Monne Maria Itria, Toja Alessandro, Sanna Filomena, Floris Anna Rita, Sulis Vincenzo, Palmas Angelo Domenico, Casu Gavino, Lo Maglio Iana, Mameli Giuseppe |
| EPI_ISL_613565 | Respiratory Virus Unit, Microbiology Services Colindale, Public Health England | Respiratory Virus Unit, Microbiology Services Colindale, Public Health England | PHE Covid Sequencing Team |
| EPI_ISL_613566, EPI_ISL_613567 | Microbiology, Department of Pathology, St. Bernard's Hospital, Gibraltar Health Authority | Respiratory Virus Unit, Microbiology Services Colindale, Public Health England | PHE Covid Sequencing Team, Dr Nicholas Cortes (Gibraltar), Charlotte Gillborn-Jones (Gibraltar) |
| EPI_ISL_613568, EPI_ISL_613569 | Respiratory Virus Unit, Microbiology Services Colindale, Public Health England | Respiratory Virus Unit, Microbiology Services Colindale, Public Health England | PHE Covid Sequencing Team |
| EPI_ISL_613570 | Microbiology, Department of Pathology, St. Bernard's Hospital, Gibraltar Health Authority | Respiratory Virus Unit, Microbiology Services Colindale, Public Health England | PHE Covid Sequencing Team, Dr Nicholas Cortes (Gibraltar), Charlotte Gillborn-Jones (Gibraltar) |
| EPI_ISL_613571, EPI_ISL_613572, EPI_ISL_613573, EPI_ISL_613574, EPI_ISL_613575, EPI_ISL_613576, EPI_ISL_613577, EPI_ISL_613578, EPI_ISL_613579, EPI_ISL_613580, EPI_ISL_613581, EPI_ISL_613582, EPI_ISL_613583, EPI_ISL_613584, EPI_ISL_613585, EPI_ISL_613586, EPI_ISL_613588, EPI_ISL_613589, EPI_ISL_613590, EPI_ISL_613591, EPI_ISL_613592, EPI_ISL_613593, EPI_ISL_613594, EPI_ISL_613595, EPI_ISL_613596, EPI_ISL_613597, EPI_ISL_613598, EPI_ISL_613599, EPI_ISL_613600, EPI_ISL_613601, EPI_ISL_613602, EPI_ISL_613603, EPI_ISL_613604, EPI_ISL_613605, EPI_ISL_613606, EPI_ISL_613607, EPI_ISL_613608, EPI_ISL_613609, EPI_ISL_613610, EPI_ISL_613611, EPI_ISL_613612, EPI_ISL_613613, EPI_ISL_613614, EPI_ISL_613615, EPI_ISL_613616, EPI_ISL_613617, EPI_ISL_613618, EPI_ISL_613619, EPI_ISL_613620, EPI_ISL_613621, EPI_ISL_613622, EPI_ISL_613623, EPI_ISL_613624, EPI_ISL_613625, EPI_ISL_613626, EPI_ISL_613627, EPI_ISL_613628, EPI_ISL_613629, EPI_ISL_613630, EPI_ISL_613631, EPI_ISL_613632, EPI_ISL_613633, EPI_ISL_613634, EPI_ISL_613635 | | | |
| see above | Respiratory Virus Unit, Microbiology Services Colindale, Public Health England | Respiratory Virus Unit, Microbiology Services Colindale, Public Health England | PHE Covid Sequencing Team |
| EPI_ISL_613636, EPI_ISL_613637, EPI_ISL_613638, EPI_ISL_613639, EPI_ISL_613640, EPI_ISL_613641, EPI_ISL_613642, EPI_ISL_613643, EPI_ISL_613644, EPI_ISL_613645, EPI_ISL_613646, EPI_ISL_613647, EPI_ISL_613648, EPI_ISL_613649, EPI_ISL_613650, EPI_ISL_613651, EPI_ISL_613652, EPI_ISL_613653, EPI_ISL_613654, EPI_ISL_613655, EPI_ISL_613656, EPI_ISL_613657, EPI_ISL_613658, EPI_ISL_613659, EPI_ISL_613660, EPI_ISL_613661, EPI_ISL_613662, EPI_ISL_613663, EPI_ISL_613664, EPI_ISL_613665, EPI_ISL_613666, EPI_ISL_613667, EPI_ISL_613668, EPI_ISL_613669, EPI_ISL_613670, EPI_ISL_613671, EPI_ISL_613672, EPI_ISL_613673, EPI_ISL_613674, EPI_ISL_613675, EPI_ISL_613676, EPI_ISL_613677, EPI_ISL_613678, EPI_ISL_613679, EPI_ISL_613680, EPI_ISL_613681, EPI_ISL_613682, EPI_ISL_613683, EPI_ISL_613684, EPI_ISL_613685, EPI_ISL_613686, EPI_ISL_613687, EPI_ISL_613688, EPI_ISL_613689, EPI_ISL_613690, EPI_ISL_613691, EPI_ISL_613692, EPI_ISL_613693, EPI_ISL_613694, EPI_ISL_613695, EPI_ISL_613696, EPI_ISL_613697, EPI_ISL_613698, EPI_ISL_613699, EPI_ISL_613700, EPI_ISL_613701, EPI_ISL_613702, EPI_ISL_613703, EPI_ISL_613704, EPI_ISL_613705 | | | |
| see above | Microbiology, Department of Pathology, St. Bernard's Hospital, Gibraltar Health Authority | Respiratory Virus Unit, Microbiology Services Colindale, Public Health England | PHE Covid Sequencing Team, Dr Nicholas Cortes (Gibraltar), Charlotte Gillborn-Jones (Gibraltar) |
| EPI_ISL_613953, EPI_ISL_613955 | Laboratorio Biologia Molecolare Sars Cov2 - UOC Laboratorio Analisi - Servizio Medicina di Laboratorio, Ospedale "San Francesco" - ATS-ASSL Nuoro | Laboratorio specialistico UOC Ematologia - Ospedale "San Francesco" - ATS-ASSL Nuoro | Piras Giovanna, Fancello Tatiana, Asproni Rosanna, Fiamma Maura, Monne Maria Itria, Toja Alessandro, Sanna Filomena, Floris Anna Rita, Sulis Vincenzo, Palmas Angelo Domenico, Casu Gavino, Lo Maglio Iana, Mameli Giuseppe |
| EPI_ISL_613957 | Respiratory Virus Unit, Microbiology Services Colindale, Public Health England | Respiratory Virus Unit, Microbiology Services Colindale, Public Health England | PHE Covid Sequencing Team |
| EPI_ISL_613958, EPI_ISL_613960 | Microbiology, Department of Pathology, St. Bernard's Hospital, Gibraltar Health Authority | Respiratory Virus Unit, Microbiology Services Colindale, Public Health England | PHE Covid Sequencing Team, Dr Nicholas Cortes (Gibraltar), Charlotte Gillborn-Jones (Gibraltar) |
| EPI_ISL_614396, EPI_ISL_614397, EPI_ISL_614398, EPI_ISL_614889 | Laboratorio Biologia Molecolare Sars Cov2 - UOC Laboratorio Analisi - Servizio Medicina di Laboratorio, Ospedale "San Francesco" - ATS-ASSL Nuoro | Laboratorio specialistico UOC Ematologia - Ospedale "San Francesco" - ATS-ASSL Nuoro | Piras Giovanna, Fancello Tatiana, Asproni Rosanna, Fiamma Maura, Monne Maria Itria, Toja Alessandro, Sanna Filomena, Floris Anna Rita, Sulis Vincenzo, Palmas Angelo Domenico, Casu Gavino, Lo Maglio Iana, Mameli Giuseppe |
| EPI_ISL_614981, EPI_ISL_614982, EPI_ISL_614983, EPI_ISL_614984, EPI_ISL_614985, EPI_ISL_614986, EPI_ISL_614987, EPI_ISL_614988, EPI_ISL_614989, EPI_ISL_614990, EPI_ISL_614991, EPI_ISL_614992, EPI_ISL_614993, EPI_ISL_614994, EPI_ISL_614995, EPI_ISL_614996, EPI_ISL_614997, EPI_ISL_614998, EPI_ISL_614999, EPI_ISL_615000, EPI_ISL_615001, EPI_ISL_615002, EPI_ISL_615003, EPI_ISL_615005, EPI_ISL_615006, EPI_ISL_615007, EPI_ISL_615008, EPI_ISL_615009, EPI_ISL_615010, EPI_ISL_615011, EPI_ISL_615012, EPI_ISL_615013, EPI_ISL_615014, EPI_ISL_615015, EPI_ISL_615016, EPI_ISL_615017, EPI_ISL_615018, EPI_ISL_615019, EPI_ISL_615020, EPI_ISL_615021, EPI_ISL_615022, EPI_ISL_615023, EPI_ISL_615024, EPI_ISL_615025, EPI_ISL_615026, EPI_ISL_615027, EPI_ISL_615028, EPI_ISL_615029, EPI_ISL_615030, EPI_ISL_615031, EPI_ISL_615032, EPI_ISL_615033, EPI_ISL_615034, EPI_ISL_615035, | | | |

| | | | |
|---|---|---|---|
| EPI_ISL_615036, EPI_ISL_615037, EPI_ISL_615038, EPI_ISL_615039, EPI_ISL_615040, EPI_ISL_615041, EPI_ISL_615042, EPI_ISL_615043, EPI_ISL_615044, EPI_ISL_615045, EPI_ISL_615046, EPI_ISL_615047, EPI_ISL_615048, EPI_ISL_615049, EPI_ISL_615050, EPI_ISL_615051, EPI_ISL_615052, EPI_ISL_615053, EPI_ISL_615054, EPI_ISL_615055, EPI_ISL_615056, EPI_ISL_615057, EPI_ISL_615058, EPI_ISL_615059, EPI_ISL_615060, EPI_ISL_615061, EPI_ISL_615062 | | | |
| see above | Microbiology, Department of Pathology, St. Bernard's Hospital, Gibraltar Health Authority | Respiratory Virus Unit, Microbiology Services Colindale, Public Health England | PHE Covid Sequencing Team, Dr Nicholas Cortes (Gibraltar), Charlotte Gillborn-Jones (Gibraltar) |
| EPI_ISL_622833, EPI_ISL_622834, EPI_ISL_622835, EPI_ISL_622836, EPI_ISL_622837, EPI_ISL_622838, EPI_ISL_622839, EPI_ISL_622840, EPI_ISL_622841, EPI_ISL_622842, EPI_ISL_622843, EPI_ISL_622844, EPI_ISL_622845, EPI_ISL_622846, EPI_ISL_622847, EPI_ISL_622848, EPI_ISL_622849, EPI_ISL_622850, EPI_ISL_622851, EPI_ISL_622852, EPI_ISL_622853, EPI_ISL_622854, EPI_ISL_622855, EPI_ISL_622856, EPI_ISL_622857, EPI_ISL_622858, EPI_ISL_622859, EPI_ISL_622860, EPI_ISL_622861, EPI_ISL_622862, EPI_ISL_622863, EPI_ISL_622864, EPI_ISL_622865, EPI_ISL_622866, EPI_ISL_622867, EPI_ISL_622868, EPI_ISL_622869, EPI_ISL_622870, EPI_ISL_622871, EPI_ISL_622872, EPI_ISL_622873, EPI_ISL_622874, EPI_ISL_622875, EPI_ISL_622876, EPI_ISL_622877, EPI_ISL_622878, EPI_ISL_622880, EPI_ISL_622881, EPI_ISL_622882, EPI_ISL_622883, EPI_ISL_622884, EPI_ISL_622885, EPI_ISL_622886, EPI_ISL_622887, EPI_ISL_622888, EPI_ISL_622889, EPI_ISL_622890, EPI_ISL_622891 | | | |
| see above | Respiratory Virus Unit, Microbiology Services Colindale, Public Health England | Respiratory Virus Unit, Microbiology Services Colindale, Public Health England | PHE Covid Sequencing Team |
| EPI_ISL_626211, EPI_ISL_626212, EPI_ISL_626213, EPI_ISL_626214, EPI_ISL_626215, EPI_ISL_626216, EPI_ISL_626217, EPI_ISL_626218, EPI_ISL_626219, EPI_ISL_626220, EPI_ISL_626221, EPI_ISL_626222, EPI_ISL_626223, EPI_ISL_626224, EPI_ISL_626225, EPI_ISL_626226, EPI_ISL_626227, EPI_ISL_626228, EPI_ISL_626229, EPI_ISL_626230 | | | |
| see above | Institute for Virology, University Hospital Essen | Center of Medical Microbiology, Virology, and Hospital Hygiene, University of Duesseldorf | Olympia E. Anastasiou, Ulf Dittmer, Maximilian Damagnez, Alexander Dilthey, Torsten Houwaart, Lisanna Hülse, Malte Kohns Vasconcelos, Nadine Lübke, Jessica Nicolai, Klaus Pfeffer, Daniel Strelow, Jörg Timm, Andreas Walker, Tobias Wienemann |
| EPI_ISL_626566 | National Centre for Communication Disease (NCCD) National Influenza Center | National Centre for Communication Disease (NCCD) National Influenza Center | Naranzul Ts,Darmaa B,Bayasgalan N,Ankhbayar S,Tsogtbaatar B,Erden-Ochir Ts,Nymadawa P,Chang-Seon Song |
| EPI_ISL_626567, EPI_ISL_626568, EPI_ISL_626569, EPI_ISL_626626, EPI_ISL_626627 | National Centre for Communicable Disease (NCCD) National Influenza Center | National Centre for Communicable Disease (NCCD) National Influenza Center | Naranzul Ts,Darmaa B,Bayasgalan N,Ankhbayar S, Nyamkhuu D, Erden-Ochir Ts,Nymadawa P Chang-Seon Song |
| EPI_ISL_632796 | Respiratory Virus Unit, Microbiology Services Colindale, Public Health England | Respiratory Virus Unit, Microbiology Services Colindale, Public Health England | PHE Covid Sequencing Team |
| EPI_ISL_632797, EPI_ISL_632798, EPI_ISL_632799, EPI_ISL_632800, EPI_ISL_632801, EPI_ISL_632802, EPI_ISL_632803, EPI_ISL_632804, EPI_ISL_632805, EPI_ISL_632806, EPI_ISL_632807, EPI_ISL_632808, EPI_ISL_632809, EPI_ISL_632810, EPI_ISL_632811, EPI_ISL_632812, EPI_ISL_632813, EPI_ISL_632815, EPI_ISL_632816, EPI_ISL_632817, EPI_ISL_632818, EPI_ISL_632819, EPI_ISL_632820, EPI_ISL_632821, EPI_ISL_632822 | | | |
| see above | Microbiology, Department of Pathology, St. Bernard's Hospital, Gibraltar Health Authority | Respiratory Virus Unit, Microbiology Services Colindale, Public Health England | PHE Covid Sequencing Team, Dr Nicholas Cortes (Gibraltar), Charlotte Gillborn-Jones (Gibraltar) |

We gratefully acknowledge the following Authors from the Originating laboratories responsible for obtaining the specimens, as well as the Submitting laboratories where the genome data were generated and shared via GISAID, on which this research is based.

All Submitters of data may be contacted directly via www.gisaid.org

Authors are sorted alphabetically.

| Accession ID | Originating Laboratory | Submitting Laboratory | Authors |
|---|---|---|---|
| EPI_ISL_408976 | Centre for Infectious Diseases and Microbiology Laboratory Services | NSW Health Pathology - Institute of Clinical Pathology and Medical Research; Westmead Hospital; University of Sydney | Rockett R, Sadsad R, Eden J-S, Carter I, Rahman H, Holmes EC, O'Sullivan MV, Sintchenko V, Chen SC, Maddocks S, Kok J and Dwyer DE for the 2019-nCoV Study Group* |
| EPI_ISL_410301 | National Influenza Centre, National Public Health Laboratory, Kathmandu, Nepal | The University of Hong Kong | Ranjit Sah , Runa Jha, Daniel Chu, Haogao Gu, Malik Peiris, Anup Bastola, Alfonso J. Rodriguez-Morales, Bibek Kumar Lal, Basu Dev Pandey, Leo Poon |
| EPI_ISL_412972 | Instituto Nacional de Enfermedades Respiratorias | Instituto de Diagnostico y Referencia Epidemiologicos (INDRE) | Ramirez-Gonzalez Ernesto, Garces-Ayala Fabiola, Araiza-Rodriguez Adnan, Mendieta-Condado Edgar, Rodriguez-Maldonado Abril, Wong-Arambula Claudia, Vazquez-Perez Joel, Martinez Arturo, Boukadida Celia, Munoz-Medina Esteban, Sanchez Alejandro, Isa Pavel, Taboada Blanca, Lopez Susana, Arias Carlos, Barrera-Badillo Gisela, Hernandez-Rivas Lucia, Lopez-Martinez Irma |
| EPI_ISL_413594 | Centre for Infectious Diseases and Microbiology Laboratory Services | NSW Health Pathology - Institute of Clinical Pathology and Medical Research; Westmead Hospital; University of Sydney | Rockett R, Eden J-S, Lam C, Gray K, Timms, V, Gall, M, Alicia, A, Carter I, Rahman H, Holmes EC, , O'Sullivan MV, Sintchenko V, Chen SC, Maddocks S, Kok J and Dwyer DE for the 2019-nCoV Study Group* |
| EPI_ISL_413595 | Centre for Infectious Diseases and Microbiology Laboratory Services | NSW Health Pathology - Institute of Clinical Pathology and Medical Research; Westmead Hospital; University of Sydney | Rockett R, Eden J-S, Lam C, Gray K, Timms, V, Gall, M, Carter I, Rahman H, Holmes EC, O'Sullivan MV, Sintchenko V, Chen SC, Maddocks S, Kok J and Dwyer DE for the 2019-nCoV Study Group* |
| EPI_ISL_413596 | Centre for Infectious Diseases and Microbiology - Public Health | NSW Health Pathology - Institute of Clinical Pathology and Medical Research; Westmead Hospital; University of Sydney | Rockett R, Eden J-S, Lam C, Gray K, Timms, V, Gall, M, Carter I, Rahman H, Holmes EC, O'Sullivan MV, Sintchenko V, Chen SC, Maddocks S, Kok J and Dwyer DE for the 2019-nCoV Study Group* |
| EPI_ISL_413647 | Centro Hospital do Porto, E.P.E. - H. Geral de Santo Antonio | Instituto Nacional de Saude (INSA) | Raquel Guiomar, Inês Costa, Pedro Pechirra, Joana Mendonça, Luís Vieira, Helena Ramos, Joana Isidro, Vítor Borges, João Paulo Gomes |
| EPI_ISL_413648 | Centro Hospitalar e Universitário de Sao Joao, Porto | Instituto Nacional de Saude (INSA) | Raquel Guiomar, Inês Costa, Pedro Pechirra, Joana Mendonça, Luís Vieira, João Tiago Guimarães, Joana Isidro, Vítor Borges, João Paulo Gomes |
| EPI_ISL_417383 | Centre for Infectious Diseases and Microbiology Public Health | NSW Health Pathology - Institute of Clinical Pathology and Medical Research; Westmead Hospital; University of Sydney | Rockett R, Eden J-S, Lam C, Gray K, Timms V, Gall M, Arnott A, Sadsad R, Carter I, Rahman H, Holmes EC, O'Sullivan MV, Sintchenko V, Chen SC, Maddocks S, Kok J and Dwyer DE for the 2019-nCoV Study Group |
| EPI_ISL_417392 | Centre for Infectious Diseases and Microbiology Public Health | NSW Health Pathology - Institute of Clinical Pathology and Medical Research; Westmead Hospital; University of Sydney | Rahman H, Holmes EC, O'Sullivan MV, Sintchenko V, Chen SC, Maddocks S, Kok J, Dwyer DE, Rockett R, Eden J-S, Lam C, Gray K, Timms V, Gall M, Arnott A, Sadsad R and Carter I for the 2019-nCoV Study Group |
| EPI_ISL_417400 | Centre for Infectious Diseases and Microbiology Public Health | NSW Health Pathology - Institute of Clinical Pathology and Medical Research; Westmead Hospital; University of Sydney | Rockett R, Eden J-S, Lam C, Gray K, Timms V, Gall M, Arnott A, Sadsad R, Carter I, Rahman H, Holmes EC, O'Sullivan MV, Sintchenko V, Chen SC, Maddocks S, Kok J and Dwyer DE for the 2019-nCoV Study Group |
| EPI_ISL_417409 | Centre for Infectious Diseases and Microbiology Public Health | NSW Health Pathology - Institute of Clinical Pathology and Medical Research; Westmead Hospital; University of Sydney | Rahman H, Holmes EC, O'Sullivan MV, Sintchenko V, Chen SC, Maddocks S, Kok J, Dwyer DE, Rockett R, Eden J-S, Lam C, Gray K, Timms V, Gall M, Arnott A, Sadsad R and Carter I for the 2019-nCoV Study Group |
| EPI_ISL_426414 | Sir M P Shah Government Medical College | Gujarat Biotechnology Research Centre | Ramesh Pandit, Tejas Shah, Ankit Hinsu, Pritesh Sabara, Apurvasinh Puvar, Janvi Raval, Monika Gandhi, Pinal Trivedi, Maharshi Pandya, Amit Kanani, Akanksha Verma, Nitin Savaliya, Raghawendra Kumar, Dinesh Kumar, Zubair Saiyed, Dipa Kinariwala, Disha Patel, Binita Aring, Geeta Vaghela, Sonia Barve, Bhavesh Modi, Kairavi Joshi, Nidhi Sood, Pranay Shah, R D Dixit, Snehal Bagatharia, Madhvi Joshi, Chaitanya Joshi |
| EPI_ISL_426415 | Sir M P Shah Government Medical College, Jamnagar | Gujarat Biotechnology Research Centre, Gandhinagar | Ramesh Pandit, Tejas Shah, Ankit Hinsu, Pritesh Sabara, Apurvasinh Puvar, Janvi Raval, Monika Gandhi, Pinal Trivedi, Maharshi Pandya, Amit Kanani, Akanksha Verma, Nitin Savaliya, Raghvendra Kumar, Dinesh Kumar, Zuber Saiyed, Dipa Kinariwala, Disha Patel, Binita Aring, Geeta Vaghela, Sonia Barve, Bhavesh Modi, Kairavi Joshi, Nidhi Sood, Pranay Shah, R D Dixit, Snehal Bagatharia, Madhvi Joshi, Chaitanya Joshi |
| EPI_ISL_427644, EPI_ISL_427653, EPI_ISL_427657, EPI_ISL_427660, EPI_ISL_427681 | Centre for Infectious Diseases and Microbiology Public Health | NSW Health Pathology - Institute of Clinical Pathology and Medical Research; Westmead Hospital; University of Sydney | Rockett R, Lam C, Gray K, Timms V, Gall M, Arnott A, Sadsad R, Draper J, Sim E, Bachmann N, Carter I, Holmes EC, O'Sullivan MV, Byun R, Sintchenko V, Chen SC, Eden JS, Maddocks S, Kok J, Propenko M, Sorrell T, Chang S, Basile K, Dwyer DE for the 2019-nCoV Study Group |
| EPI_ISL_427711 | ACT Pathology, The Canberra Hospital | NSW Health Pathology - Institute of Clinical Pathology and Medical Research; Westmead Hospital; University of Sydney | Rockett R, Lam C, Gray K, Timms V, Gall M, Arnott A, Sadsad R, Draper J, Sim E, Bachmann N, Carter I, Holmes EC, O'Sullivan MV, Byun R, Sintchenko V, Chen SC, Eden JS, Maddocks S, Kok J, Propenko M, Sorrell T, Chang S, Basile K, Dwyer DE for the 2019-nCoV Study Group |
| EPI_ISL_427714, EPI_ISL_427726, EPI_ISL_427728, EPI_ISL_427737, EPI_ISL_427750, EPI_ISL_427765, EPI_ISL_427780, EPI_ISL_427789, EPI_ISL_427791, EPI_ISL_427798, EPI_ISL_427801 | | | |
| see above | Centre for Infectious Diseases and Microbiology Public Health | NSW Health Pathology - Institute of Clinical Pathology and Medical Research; Westmead Hospital; University of Sydney | Rockett R, Lam C, Gray K, Timms V, Gall M, Arnott A, Sadsad R, Draper J, Sim E, Bachmann N, Carter I, Holmes EC, O'Sullivan MV, Byun R, Sintchenko V, Chen SC, Eden JS, Maddocks S, Kok J, Propenko M, Sorrell T, Chang S, Basile K, Dwyer DE for the 2019-nCoV Study Group |
| EPI_ISL_428695, EPI_ISL_428696, EPI_ISL_428697, EPI_ISL_428698, EPI_ISL_428699 | Hospital Universitario 12 de Octubre | Hospital Universitario 12 de Octubre | Raúl Recio, Sara González, Elias Dahdouh, Fernando Lázaro, Esther Viedma, Natalia Stella, Julio García, Juan Carlos Galán, Rafael Cantón, Mª Dolores Folgueira, Rafael Delgado, Jesús Mingorance |
| EPI_ISL_429262, EPI_ISL_429263, EPI_ISL_429264, EPI_ISL_429265, EPI_ISL_429266, EPI_ISL_429267, EPI_ISL_429268, EPI_ISL_429269, EPI_ISL_429270, EPI_ISL_429271, EPI_ISL_429272, EPI_ISL_429273, EPI_ISL_429274, EPI_ISL_429275, EPI_ISL_429276, EPI_ISL_429277, EPI_ISL_429278, EPI_ISL_429279, EPI_ISL_429280, EPI_ISL_429281, EPI_ISL_429282, EPI_ISL_429283, EPI_ISL_429284, EPI_ISL_429285, EPI_ISL_429286, EPI_ISL_429287, EPI_ISL_429288, EPI_ISL_429289, EPI_ISL_429290, EPI_ISL_429291, EPI_ISL_429292, EPI_ISL_429293, EPI_ISL_429294, EPI_ISL_429295, EPI_ISL_429296, EPI_ISL_429297, EPI_ISL_429298, EPI_ISL_429299, EPI_ISL_429300, EPI_ISL_429301, EPI_ISL_429302, EPI_ISL_429303, EPI_ISL_429304, EPI_ISL_429305, EPI_ISL_429306, EPI_ISL_429307, EPI_ISL_429308, EPI_ISL_429309, EPI_ISL_429310, EPI_ISL_429311, EPI_ISL_429312, EPI_ISL_429313, EPI_ISL_429314, EPI_ISL_429315, EPI_ISL_429316, EPI_ISL_429317, EPI_ISL_429318, EPI_ISL_429320, EPI_ISL_429321, EPI_ISL_429322, EPI_ISL_429323, EPI_ISL_429324, EPI_ISL_429325, EPI_ISL_429326, EPI_ISL_429327, EPI_ISL_429328, EPI_ISL_429329, EPI_ISL_429330, EPI_ISL_429331, EPI_ISL_429332 | | | |
| see above | Department of Clinical Microbiology, Copenhagen University Hospital, Hvidovre, Kettegaard Alle 30, 2650 Hvidovre. | Albertsen lab, Department of Chemistry and Bioscience, Aalborg University, Denmark | Rasmus Kirkegaard |
| EPI_ISL_429333, EPI_ISL_429334, EPI_ISL_429335, EPI_ISL_429336, EPI_ISL_429337, EPI_ISL_429338, EPI_ISL_429339, EPI_ISL_429340, EPI_ISL_429341, EPI_ISL_429342, EPI_ISL_429343, EPI_ISL_429344, EPI_ISL_429345, EPI_ISL_429346, EPI_ISL_429347, EPI_ISL_429348, EPI_ISL_429349, EPI_ISL_429350, EPI_ISL_429351, EPI_ISL_429352, EPI_ISL_429353, EPI_ISL_429354, EPI_ISL_429355, EPI_ISL_429356, EPI_ISL_429357, EPI_ISL_429358, EPI_ISL_429359, EPI_ISL_429360, EPI_ISL_429361, EPI_ISL_429362, EPI_ISL_429363, EPI_ISL_429364, EPI_ISL_429365, EPI_ISL_429366, EPI_ISL_429367, EPI_ISL_429368, EPI_ISL_429369, EPI_ISL_429370, EPI_ISL_429371, EPI_ISL_429372, EPI_ISL_429373, EPI_ISL_429374, EPI_ISL_429375, EPI_ISL_429376, EPI_ISL_429377, EPI_ISL_429378, EPI_ISL_429379, EPI_ISL_429380, EPI_ISL_429381, EPI_ISL_429382, EPI_ISL_429383, EPI_ISL_429384, EPI_ISL_429385, EPI_ISL_429386, EPI_ISL_429387, EPI_ISL_429388, EPI_ISL_429389, EPI_ISL_429390, EPI_ISL_429391, EPI_ISL_429392, EPI_ISL_429393, EPI_ISL_429394, EPI_ISL_429395, EPI_ISL_429396, EPI_ISL_429397, EPI_ISL_429398, EPI_ISL_429399, EPI_ISL_429400, EPI_ISL_429401, EPI_ISL_429402, EPI_ISL_429403, EPI_ISL_429404, EPI_ISL_429405, EPI_ISL_429406, EPI_ISL_429407, EPI_ISL_429408, EPI_ISL_429409, EPI_ISL_429410, EPI_ISL_429411, EPI_ISL_429412, EPI_ISL_429413, EPI_ISL_429414, EPI_ISL_429415, EPI_ISL_429416, EPI_ISL_429417, EPI_ISL_429418, EPI_ISL_429419, EPI_ISL_429420, EPI_ISL_429421, EPI_ISL_429422, EPI_ISL_429423, EPI_ISL_429424, EPI_ISL_429425, EPI_ISL_429426, EPI_ISL_429427, EPI_ISL_429428, EPI_ISL_429429, EPI_ISL_429430, EPI_ISL_429431, EPI_ISL_429432, EPI_ISL_429433, EPI_ISL_429434, EPI_ISL_429435, EPI_ISL_429436, EPI_ISL_429437, EPI_ISL_429438, EPI_ISL_429439, EPI_ISL_429440, EPI_ISL_429441, EPI_ISL_429442, EPI_ISL_429443, EPI_ISL_429444, EPI_ISL_429445, EPI_ISL_429446, EPI_ISL_429447, EPI_ISL_429448, EPI_ISL_429449, EPI_ISL_429450, EPI_ISL_429451, EPI_ISL_429452, EPI_ISL_429453, EPI_ISL_429454, EPI_ISL_429455, EPI_ISL_429456, EPI_ISL_429457, EPI_ISL_429458, EPI_ISL_429459, EPI_ISL_429460, EPI_ISL_429461, EPI_ISL_429462, EPI_ISL_429463, EPI_ISL_429464, EPI_ISL_429465, EPI_ISL_429466, EPI_ISL_429467, EPI_ISL_429468, EPI_ISL_429469, EPI_ISL_429470, EPI_ISL_429471, EPI_ISL_429472, EPI_ISL_429473, EPI_ISL_429474, EPI_ISL_429475, EPI_ISL_429476, EPI_ISL_429477, EPI_ISL_429478, EPI_ISL_429479, EPI_ISL_429480, EPI_ISL_429481, EPI_ISL_429482, EPI_ISL_429483, EPI_ISL_429484, EPI_ISL_429485, EPI_ISL_429486, EPI_ISL_429487, EPI_ISL_429488, EPI_ISL_429489, EPI_ISL_429490, EPI_ISL_429491, EPI_ISL_429492, EPI_ISL_429493, EPI_ISL_429494, EPI_ISL_429495, EPI_ISL_429496, EPI_ISL_429497, EPI_ISL_429498, EPI_ISL_429499, EPI_ISL_429500, EPI_ISL_429501, EPI_ISL_429502, EPI_ISL_429503, EPI_ISL_429504, EPI_ISL_429505, EPI_ISL_429506, EPI_ISL_429507, EPI_ISL_429508, EPI_ISL_429509, EPI_ISL_429510, EPI_ISL_429511, EPI_ISL_429512, EPI_ISL_429513, EPI_ISL_429514, EPI_ISL_429515, EPI_ISL_429516, EPI_ISL_429517, EPI_ISL_429518, EPI_ISL_429519, EPI_ISL_429520, EPI_ISL_429521, EPI_ISL_429522, EPI_ISL_429523, EPI_ISL_429524, EPI_ISL_429525, EPI_ISL_429526, EPI_ISL_429527, EPI_ISL_429528, EPI_ISL_429529, EPI_ISL_429530, EPI_ISL_429531, EPI_ISL_429532, EPI_ISL_429533, EPI_ISL_429534, EPI_ISL_429535, EPI_ISL_429536, EPI_ISL_429537, EPI_ISL_429538, EPI_ISL_429539, EPI_ISL_429540, EPI_ISL_429541, EPI_ISL_429542, EPI_ISL_429543, EPI_ISL_429544, EPI_ISL_429545, EPI_ISL_429546, EPI_ISL_429547, EPI_ISL_429548, EPI_ISL_429549, EPI_ISL_429550, EPI_ISL_429551, EPI_ISL_429552, EPI_ISL_429553, EPI_ISL_429554, EPI_ISL_429555, EPI_ISL_429556, EPI_ISL_429557, EPI_ISL_429558, EPI_ISL_429559, EPI_ISL_429560, EPI_ISL_429561, EPI_ISL_429562, EPI_ISL_429563, EPI_ISL_429564, EPI_ISL_429565, EPI_ISL_429566, EPI_ISL_429567, EPI_ISL_429568, EPI_ISL_429569, EPI_ISL_429570, EPI_ISL_429571, EPI_ISL_429572, EPI_ISL_429573, EPI_ISL_429574, EPI_ISL_429575, EPI_ISL_429576, EPI_ISL_429577, EPI_ISL_429578, EPI_ISL_429579, EPI_ISL_429580, EPI_ISL_429581, EPI_ISL_429582, EPI_ISL_429583, EPI_ISL_429584, EPI_ISL_429585, EPI_ISL_429586, EPI_ISL_429587, EPI_ISL_429588, EPI_ISL_429589, EPI_ISL_429590 | | | |
| see above | Department of Virus and Microbiological Special Diagnostics, Statens Serum Institut, Copenhagen, Denmark, Artillerivej 5, 2300 Copenahgen S | Albertsen lab, Department of Chemistry and Bioscience, Aalborg University, Denmark | Rasmus Kirkegaard |

| | | | |
|---|---|---|---|
| EPI_ISL_435550, EPI_ISL_435551, EPI_ISL_435552, EPI_ISL_435553, EPI_ISL_435554 | LSUHS Emerging Viral Threat Laboratory | Microbial Genome Sequencing Center | Rona S. Scott, Jeremy P. Kamil, John A. Vanchiere, Camille F. Abshire, Abida Siddiqa, Byeong-Jae Lee, Chan-ki Min, Md Maksudul Alam, Monica Gestal-Cartele, Edna Ondari, Adam Greer, Malgorzata Bienkowska-Haba, Katarzyna Zwolinska, Jason M. Bodily, Andrew D. Yurochko, Paul M. Weinberger, Christopher G. Kevil, Martin J. Sapp, Daniel J. Snyder, Vaughn S. Cooper |
| EPI_ISL_436962, EPI_ISL_436963, EPI_ISL_436964, EPI_ISL_436965, EPI_ISL_436966, EPI_ISL_436967, EPI_ISL_436968, EPI_ISL_436969, EPI_ISL_436970, EPI_ISL_436971, EPI_ISL_436972, EPI_ISL_436973, EPI_ISL_436974, EPI_ISL_436975, EPI_ISL_436976, EPI_ISL_436977, EPI_ISL_436978, EPI_ISL_436979, EPI_ISL_436980, EPI_ISL_436981, EPI_ISL_436982, EPI_ISL_436983, EPI_ISL_436984, EPI_ISL_436985, EPI_ISL_436986, EPI_ISL_436987, EPI_ISL_436988, EPI_ISL_436989, EPI_ISL_436990, EPI_ISL_436991, EPI_ISL_436992, EPI_ISL_436993, EPI_ISL_436994, EPI_ISL_436995, EPI_ISL_436996, EPI_ISL_436997, EPI_ISL_436998, EPI_ISL_436999, EPI_ISL_437000, EPI_ISL_437001, EPI_ISL_437002, EPI_ISL_437003, EPI_ISL_437004, EPI_ISL_437005, EPI_ISL_437006, EPI_ISL_437007, EPI_ISL_437008, EPI_ISL_437009, EPI_ISL_437010, EPI_ISL_437011, EPI_ISL_437012, EPI_ISL_437013, EPI_ISL_437014, EPI_ISL_437015, EPI_ISL_437016, EPI_ISL_437017, EPI_ISL_437018, EPI_ISL_437019, EPI_ISL_437020, EPI_ISL_437021, EPI_ISL_437022, EPI_ISL_437023, EPI_ISL_437024, EPI_ISL_437025, EPI_ISL_437026, EPI_ISL_437027, EPI_ISL_437028, EPI_ISL_437029, EPI_ISL_437030, EPI_ISL_437031, EPI_ISL_437032, EPI_ISL_437033, EPI_ISL_437034, EPI_ISL_437035, EPI_ISL_437036, EPI_ISL_437037, EPI_ISL_437038, EPI_ISL_437039, EPI_ISL_437040, EPI_ISL_437041, EPI_ISL_437042 | | | |
| see above | Department of Virus and Microbiological Special Diagnostics, Statens Serum Institut, Copenhagen, Denmark, Artillerivej 5, 2300 Copenahgen S | Albertsen lab, Department of Chemistry and Bioscience, Aalborg University, Denmark | Rasmus Kirkegaard |
| EPI_ISL_437441 | Department of MicroBiology, Government Medical College, Surat | Gujarat Biotechnology Research Centre | Raghawendra Kumar, Dinesh Kumar, Zuber Saiyed, Dipa Kinariwala, Disha Patel, Binita Aring, Neeta Khandelwal, Geeta Vaghela, Sonia Barve, Bhavesh Modi, Kairavi Joshi, Gaurishankar Shrimali, Nidhi Sood, Pranay Shah, R D Dixit, Snehal Bagatharia, Kamlesh J Upadhyay, Ramesh Pandit, Tejas Shah, Ankit Hinsu, Pritesh Sabara, Apurvasinh Puvar, Janvi Raval, Monika Gandhi, Pinal Trivedi, Maharshi Pandya, Amit Kanani, Akanksha Verma, Nitin Savaliya, Anjali Rajwar, Chaitanya Joshi, Madhvi Joshi |
| EPI_ISL_437602, EPI_ISL_437603, EPI_ISL_437604, EPI_ISL_437605, EPI_ISL_437606, EPI_ISL_437607, EPI_ISL_437608, EPI_ISL_437609, EPI_ISL_437610, EPI_ISL_437611, EPI_ISL_437612, EPI_ISL_437613, EPI_ISL_437614, EPI_ISL_437615, EPI_ISL_437616, EPI_ISL_437617, EPI_ISL_437618, EPI_ISL_437619, EPI_ISL_437620, EPI_ISL_437621, EPI_ISL_437622, EPI_ISL_437623, EPI_ISL_437624 | | | |
| see above | unknown | Faculty of Medicine | Rodpan,A., Joyjinda,Y., Wacharapluesadee,S., Buathong,R., Ghai,S., Petcharat,S., Bunprakob,S., Sirichan,N., Prasithsirikul,W., Mungaomklang,A., Plipat,T. and Hemachudha,T. |
| EPI_ISL_437627, EPI_ISL_437628, EPI_ISL_437629, EPI_ISL_437630, EPI_ISL_437631, EPI_ISL_437632, EPI_ISL_437633, EPI_ISL_437634, EPI_ISL_437635, EPI_ISL_437636, EPI_ISL_437637, EPI_ISL_437638, EPI_ISL_437639, EPI_ISL_437640, EPI_ISL_437641, EPI_ISL_437642, EPI_ISL_437643, EPI_ISL_437644, EPI_ISL_437645, EPI_ISL_437646, EPI_ISL_437647, EPI_ISL_437648, EPI_ISL_437649, EPI_ISL_437650, EPI_ISL_437651, EPI_ISL_437652, EPI_ISL_437653, EPI_ISL_437654, EPI_ISL_437655, EPI_ISL_437656, EPI_ISL_437657, EPI_ISL_437658, EPI_ISL_437659, EPI_ISL_437660, EPI_ISL_437661, EPI_ISL_437662, EPI_ISL_437663, EPI_ISL_437664, EPI_ISL_437665, EPI_ISL_437666, EPI_ISL_437667, EPI_ISL_437668, EPI_ISL_437669, EPI_ISL_437670, EPI_ISL_437671, EPI_ISL_437672, EPI_ISL_437673, EPI_ISL_437674, EPI_ISL_437675, EPI_ISL_437676, EPI_ISL_437677, EPI_ISL_437678, EPI_ISL_437679, EPI_ISL_437680, EPI_ISL_437681, EPI_ISL_437682, EPI_ISL_437683 | | | |
| see above | Department of Virus and Microbiological Special Diagnostics, Statens Serum Institut, Copenhagen, Denmark, Artillerivej 5, 2300 Copenahgen S | Albertsen lab, Department of Chemistry and Bioscience, Aalborg University, Denmark | Rasmus Kirkegaard |
| EPI_ISL_438138 | Department of Microbiology,Gandhi Medical College and Hospital | Department of Microbiology, Gandhi Medical College and Hospital Secendrabad, Hyderbad, India | Raja Rao Mesipogu, Muttineni Radhakrishna, Nagamani K, Thrilok Chander B, Kalyani Putty, Ravikumar P, Sunitha P, Pankaj Singh D, Anand Kumar K, Amit A. Upadhyay, Steven Bosinger, Rama Amara |
| EPI_ISL_444459 | B.J. Medical College and Civil hospital | Gujarat Biotechnology Research Centre | Ramesh Pandit, Tejas Shah, Ankit Hinsu, Pritesh Sabara, Apurvasinh Puvar, Janvi Raval, Monika Gandhi, Pinal Trivedi, Maharshi Pandya, Amit Kanani, Akanksha Verma, Nitin Savaliya, Raghawendra Kumar, Dinesh Kumar, Zuber Saiyed, Dipa Kinariwala, Disha Patel, Binita Aring, Neeta Khandelwal, Geeta Vaghela, Sonia Barve, Bhavesh Modi, Kairavi Joshi, Gaurishankar Shrimali, Nidhi Sood, Pranay Shah, R D Dixit, Snehal Bagatharia, Kamlesh J Upadhyay, Neha Rajpara, Chaitanya Joshi, Madhvi Joshi |
| EPI_ISL_444471 | B.J. Medical College and Civil hospital | Gujarat Biotechnology Research Centre | Raghawendra Kumar, Dinesh Kumar, Zuber Saiyed, Dipa Kinariwala, Disha Patel, Binita Aring, Neeta Khandelwal, Geeta Vaghela, Sonia Barve, Bhavesh Modi, Kairavi Joshi, Gaurishankar Shrimali, Nidhi Sood, Pranay Shah, R D Dixit, Snehal Bagatharia, Kamlesh J Upadhyay, Ramesh Pandit, Tejas Shah, Ankit Hinsu, Pritesh Sabara, Apurvasinh Puvar, Janvi Raval, Monika Gandhi, Pinal Trivedi, Maharshi Pandya, Amit Kanani, Akanksha Verma, Nitin Savaliya, Afzal Ansari, Chaitanya Joshi, Madhvi Joshi |
| EPI_ISL_444520, EPI_ISL_444521, EPI_ISL_444522, EPI_ISL_444523, EPI_ISL_444524, EPI_ISL_444525, EPI_ISL_444526, EPI_ISL_444527, EPI_ISL_444528, EPI_ISL_444529, EPI_ISL_444530, EPI_ISL_444531, EPI_ISL_444532, EPI_ISL_444533, EPI_ISL_444534, EPI_ISL_444535, EPI_ISL_444536, EPI_ISL_444537, EPI_ISL_444538, EPI_ISL_444539, EPI_ISL_444540, EPI_ISL_444541, EPI_ISL_444542, EPI_ISL_444543, EPI_ISL_444544, EPI_ISL_444545, EPI_ISL_444546, EPI_ISL_444547, EPI_ISL_444548, EPI_ISL_444549, EPI_ISL_444550, EPI_ISL_444551, EPI_ISL_444552, EPI_ISL_444553, EPI_ISL_444554, EPI_ISL_444555, EPI_ISL_444556, EPI_ISL_444557, EPI_ISL_444558, EPI_ISL_444559, EPI_ISL_444560, EPI_ISL_444561, EPI_ISL_444562, EPI_ISL_444563, EPI_ISL_444564, EPI_ISL_444565, EPI_ISL_444566, EPI_ISL_444567, EPI_ISL_444568, EPI_ISL_444569, EPI_ISL_444570, EPI_ISL_444571, EPI_ISL_444572, EPI_ISL_444573, EPI_ISL_444574, EPI_ISL_444575, EPI_ISL_444576, EPI_ISL_444577, EPI_ISL_444578, EPI_ISL_444579, EPI_ISL_444580, EPI_ISL_444581, EPI_ISL_444582, EPI_ISL_444583, EPI_ISL_444584, EPI_ISL_444585, EPI_ISL_444586, EPI_ISL_444587, EPI_ISL_444588, EPI_ISL_444589, EPI_ISL_444590, EPI_ISL_444591, EPI_ISL_444592, EPI_ISL_444593, EPI_ISL_444594, EPI_ISL_444595, EPI_ISL_444596, EPI_ISL_444597, EPI_ISL_444598, EPI_ISL_444599, EPI_ISL_444600, EPI_ISL_444601, EPI_ISL_444602, EPI_ISL_444603, EPI_ISL_444604, EPI_ISL_444605, EPI_ISL_444606, EPI_ISL_444607, EPI_ISL_444608, EPI_ISL_444609 | | | |
| see above | Northwestern Memorial Hospital | Ozer Lab | Ramon Lorenzo-Redondo, Hannah H. Nam, Scott C. Roberts, Lacy M. Simons, Chad J. Achenbach, Lawrence J. Jennings, Chao Qi, Alan R. Hauser, Michael G. Ison, Judd F. Hultquist, Egon A. Ozer |
| EPI_ISL_444817, EPI_ISL_444818, EPI_ISL_444819, EPI_ISL_444820, EPI_ISL_444821, EPI_ISL_444822, EPI_ISL_444823, EPI_ISL_444824, EPI_ISL_444825, EPI_ISL_444826, EPI_ISL_444827, EPI_ISL_444828, EPI_ISL_444829, EPI_ISL_444830, EPI_ISL_444831, EPI_ISL_444832, EPI_ISL_444833, EPI_ISL_444834, EPI_ISL_444835, EPI_ISL_444836, EPI_ISL_444837, EPI_ISL_444838, EPI_ISL_444839, EPI_ISL_444840, EPI_ISL_444841, EPI_ISL_444842, EPI_ISL_444843, EPI_ISL_444844, EPI_ISL_444845, EPI_ISL_444846, EPI_ISL_444847, EPI_ISL_444848, EPI_ISL_444849, EPI_ISL_444850, EPI_ISL_444851, EPI_ISL_444852, EPI_ISL_444853, EPI_ISL_444854, EPI_ISL_444855, EPI_ISL_444856, EPI_ISL_444857, EPI_ISL_444858, EPI_ISL_444859, EPI_ISL_444860, EPI_ISL_444861, EPI_ISL_444862, EPI_ISL_444863, EPI_ISL_444864, EPI_ISL_444865, EPI_ISL_444866, EPI_ISL_444867, EPI_ISL_444868, EPI_ISL_444869, EPI_ISL_444870, EPI_ISL_444871, EPI_ISL_444872, EPI_ISL_444873, EPI_ISL_444874, EPI_ISL_444875, EPI_ISL_444876, EPI_ISL_444877, EPI_ISL_444878, EPI_ISL_444879, EPI_ISL_444880, EPI_ISL_444881, EPI_ISL_444882, EPI_ISL_444883, EPI_ISL_444884, EPI_ISL_444885, EPI_ISL_444886, EPI_ISL_444887, EPI_ISL_444888, EPI_ISL_444889, EPI_ISL_444890, EPI_ISL_444891, EPI_ISL_444892, EPI_ISL_444893, EPI_ISL_444894, EPI_ISL_444895, EPI_ISL_444896, EPI_ISL_444897, EPI_ISL_444898, EPI_ISL_444899, EPI_ISL_444900, EPI_ISL_444901, EPI_ISL_444902, EPI_ISL_444903, EPI_ISL_444904, EPI_ISL_444905, EPI_ISL_444906, EPI_ISL_444907, EPI_ISL_444908, EPI_ISL_444909, EPI_ISL_444910, EPI_ISL_444911, EPI_ISL_444912, EPI_ISL_444913, EPI_ISL_444914, EPI_ISL_444915, EPI_ISL_444916, EPI_ISL_444917, EPI_ISL_444918, EPI_ISL_444919, EPI_ISL_444920, EPI_ISL_444921, EPI_ISL_444922, EPI_ISL_444923, EPI_ISL_444924, EPI_ISL_444925, EPI_ISL_444926, EPI_ISL_444927, EPI_ISL_444928, EPI_ISL_444929, EPI_ISL_444930, EPI_ISL_444931, EPI_ISL_444932, EPI_ISL_444933, EPI_ISL_444934, EPI_ISL_444935, EPI_ISL_444936, EPI_ISL_444937, EPI_ISL_444938, EPI_ISL_444939, EPI_ISL_444940, EPI_ISL_444941, EPI_ISL_444942, EPI_ISL_444943, EPI_ISL_444944, EPI_ISL_444945, EPI_ISL_444946, EPI_ISL_444947, EPI_ISL_444948, EPI_ISL_444949, EPI_ISL_444950, EPI_ISL_444951, EPI_ISL_444952, EPI_ISL_444953, EPI_ISL_444954, EPI_ISL_444955, EPI_ISL_444956, EPI_ISL_444957, EPI_ISL_444958, EPI_ISL_444959, EPI_ISL_444960, EPI_ISL_444961, EPI_ISL_444962, EPI_ISL_444963, EPI_ISL_444964, EPI_ISL_444965, EPI_ISL_444966, EPI_ISL_444967, EPI_ISL_444968 | | | |
| see above | Department of Virus and Microbiological Special Diagnostics, Statens Serum Institut, Copenhagen, Denmark, Artillerivej 5, 2300 Copenahgen S | Albertsen lab, Department of Chemistry and Bioscience, Aalborg University, Denmark | Rasmus Kirkegaard |
| EPI_ISL_447031 | B.J. Medical College and Civil hospital | Gujarat Biotechnology Research Centre | Ramesh Pandit, Tejas Shah, Ankit Hinsu, Pritesh Sabara, Apurvasinh Puvar, Janvi Raval, Monika Gandhi, Pinal Trivedi, Maharshi Pandya, Amit Kanani, Akanksha Verma, Nitin Savaliya, Raghawendra Kumar, Dinesh Kumar, Zuber Saiyed, Dipa Kinariwala, Disha Patel, Binita Aring, Neeta Khandelwal, Geeta Vaghela, Sonia Barve, Bhavesh Modi, Kairavi Joshi, Gaurishankar Shrimali, Nidhi Sood, Pranay Shah, R D Dixit, Snehal Bagatharia, Kamlesh J Upadhyay, Sharmistha Majumdar, Chaitanya Joshi, Madhvi Joshi |
| EPI_ISL_447043 | B.J. Medical College and Civil hospital | Gujarat Biotechnology Research Centre | Raghawendra Kumar, Dinesh Kumar, Zuber Saiyed, Dipa Kinariwala, Disha Patel, Binita Aring, Neeta Khandelwal, Geeta Vaghela, Sonia Barve, Bhavesh Modi, Kairavi Joshi, Gaurishankar Shrimali, Nidhi Sood, Pranay Shah, R D Dixit, Snehal Bagatharia, Kamlesh J Upadhyay, Ramesh Pandit, Tejas Shah, Ankit Hinsu, Pritesh Sabara, Apurvasinh Puvar, Janvi Raval, Monika Gandhi, Pinal Trivedi, Maharshi Pandya, Amit Kanani, Akanksha Verma, Nitin Savaliya, Nidhi Patel, Chaitanya Joshi, Madhvi Joshi |
| EPI_ISL_447540 | Gujarat Biotechnology Research Centre | Gujarat Biotechnology Research Centre | Ramesh Pandit, Tejas Shah, Ankit Hinsu, Pritesh Sabara, Apurvasinh Puvar, Janvi Raval, Monika Gandhi, Pinal Trivedi, Maharshi Pandya, Amit Kanani, Akanksha Verma, Nitin Savaliya, Raghawendra Kumar, Dinesh Kumar, Zuber Saiyed, Dipa Kinariwala, Disha Patel, Binita Aring, Neeta Khandelwal, Geeta Vaghela, Sonia Barve, Bhavesh Modi, Kairavi Joshi, Gaurishankar Shrimali, Nidhi Sood, Pranay Shah, R D Dixit, Snehal Bagatharia, Kamlesh J Upadhyay, Bhavya Jindal, Chaitanya Joshi, Madhvi Joshi |
| EPI_ISL_447552 | GMERS Medical College and Hospital, Gandhinagar | Gujarat Biotechnology Research Centre | Raghawendra Kumar, Dinesh Kumar, Zuber Saiyed, Dipa Kinariwala, Disha Patel, Binita Aring, Neeta Khandelwal, Geeta Vaghela, Sonia Barve, Bhavesh Modi, Kairavi Joshi, Gaurishankar Shrimali, Nidhi Sood, Pranay Shah, R D Dixit, Snehal Bagatharia, Kamlesh J Upadhyay, Ramesh Pandit, Tejas Shah, Ankit Hinsu, Pritesh Sabara, Apurvasinh Puvar, Janvi Raval, Monika Gandhi, Pinal Trivedi, Maharshi Pandya, Amit Kanani, Akanksha Verma, Nitin Savaliya, Anjali Rajwar, Chaitanya Joshi, Madhvi Joshi |
| EPI_ISL_450339, EPI_ISL_450340, EPI_ISL_450342, EPI_ISL_450344 | Bangladesh Institute of Tropical & Infectious Diseases, COVID-19 Testing Laboratory | Basic and Applied Research on Jute Project | Rasel Ahmed, Md. Sabbir Hossain, Shah Md Tamim Kabir, Emdadul Mannan Emdad, Md. Nazmul Haq Rony, Eaftekhar Ahmed Rana, Paritous Kumar Biswas, M A Hassan Chowdhury, Md. Shakeel Ahmed, Md. Samiul Haque, Md. Monjurul Alam, Md. Sharifur Rahman, A S M Anwarul Huq, Md. Shahidul Islam, Goutam Buddha Das, AMAM Zonaed Siddiki |
| EPI_ISL_450508, EPI_ISL_450509, EPI_ISL_450510, EPI_ISL_450511, EPI_ISL_450512, EPI_ISL_450513, | Rafik Hariri University Hospital | Rafik Hariri University Hospital | Rita Feghali |

| | | | |
|---|---|---|---|
| EPI_ISL_450514, EPI_ISL_450515, EPI_ISL_450516, EPI_ISL_450517 | | | |
| EPI_ISL_451159 | Government Medical College, Vadodara | Gujarat Biotechnology Research Centre | Raghawendra Kumar, Dinesh Kumar, Zuber Saiyed, Komal Patel, Labdhi Pandya, Snehal Bagatharia, Ramesh Pandit, Tejas Shah, Ankit Hinsu, Pritesh Sabara, Apurvasinh Puvar, Janvi Raval, Zarna Patel, Monika Gandhi, Pinal Trivedi, Maharshi Pandya, Manish Pattani, Tanuja Javadekar , Amit Kanani, Nidhi Patel, Nitin Savaliya, Bhavesh Modi, Gaurishankar Shrimali, R D Dixit, A M Kadri, Sharmistha Majumdar, Chaitanya Joshi, Madhvi Joshi |
| EPI_ISL_451219, EPI_ISL_451220, EPI_ISL_451221, EPI_ISL_451222, EPI_ISL_451223, EPI_ISL_451224, EPI_ISL_451225, EPI_ISL_451226, EPI_ISL_451227, EPI_ISL_451228, EPI_ISL_451229 | | | |
| see above | LSUHS Emerging Viral Threat Laboratory | Microbial Genome Sequencing Center | Rona S. Scott, Jeremy P. Kamil, John A. Vanchiere, Camille F. Abshire, Abida Siddiqa, Byeong-Jae Lee, Chan-ki Min, Md Maksudul Alam, Monica Gestal-Cartele, Edna Ondari, Adam Greer, Malgorzata Bienkowska-Haba, Katarzyna Zwolinska, Jason M. Bodily, Andrew D. Yurochko, Paul M. Weinberger, Christopher G. Kevil, Martin J. Sapp, Daniel J. Snyder, Vaughn S. Cooper |
| EPI_ISL_451988, EPI_ISL_451989, EPI_ISL_451990, EPI_ISL_451991, EPI_ISL_451992, EPI_ISL_451993, EPI_ISL_451994, EPI_ISL_451995, EPI_ISL_451996, EPI_ISL_451997, EPI_ISL_451998, EPI_ISL_451999, EPI_ISL_452000, EPI_ISL_452001, EPI_ISL_452002, EPI_ISL_452003, EPI_ISL_452004, EPI_ISL_452005, EPI_ISL_452006, EPI_ISL_452007, EPI_ISL_452008, EPI_ISL_452009, EPI_ISL_452010, EPI_ISL_452011, EPI_ISL_452012, EPI_ISL_452013, EPI_ISL_452014, EPI_ISL_452015, EPI_ISL_452016, EPI_ISL_452017, EPI_ISL_452018, EPI_ISL_452019, EPI_ISL_452020, EPI_ISL_452021, EPI_ISL_452022, EPI_ISL_452023, EPI_ISL_452024, EPI_ISL_452025, EPI_ISL_452026, EPI_ISL_452027, EPI_ISL_452028, EPI_ISL_452029, EPI_ISL_452030, EPI_ISL_452031, EPI_ISL_452032, EPI_ISL_452033, EPI_ISL_452034, EPI_ISL_452035, EPI_ISL_452036, EPI_ISL_452037, EPI_ISL_452038, EPI_ISL_452039, EPI_ISL_452040, EPI_ISL_452041, EPI_ISL_452042, EPI_ISL_452043, EPI_ISL_452044, EPI_ISL_452045, EPI_ISL_452046, EPI_ISL_452047, EPI_ISL_452048, EPI_ISL_452049, EPI_ISL_452050, EPI_ISL_452051, EPI_ISL_452052, EPI_ISL_452053, EPI_ISL_452054, EPI_ISL_452055, EPI_ISL_452056, EPI_ISL_452057, EPI_ISL_452058, EPI_ISL_452059, EPI_ISL_452060, EPI_ISL_452061, EPI_ISL_452062, EPI_ISL_452063, EPI_ISL_452064, EPI_ISL_452065, EPI_ISL_452066, EPI_ISL_452067, EPI_ISL_452068, EPI_ISL_452069, EPI_ISL_452070, EPI_ISL_452071, EPI_ISL_452072, EPI_ISL_452073, EPI_ISL_452074, EPI_ISL_452075, EPI_ISL_452076, EPI_ISL_452077, EPI_ISL_452078, EPI_ISL_452079, EPI_ISL_452080, EPI_ISL_452081, EPI_ISL_452082, EPI_ISL_452083, EPI_ISL_452084, EPI_ISL_452085, EPI_ISL_452086, EPI_ISL_452087, EPI_ISL_452088, EPI_ISL_452089, EPI_ISL_452090, EPI_ISL_452091, EPI_ISL_452092, EPI_ISL_452093, EPI_ISL_452094, EPI_ISL_452095, EPI_ISL_452096, EPI_ISL_452097, EPI_ISL_452098, EPI_ISL_452099, EPI_ISL_452100 | | | |
| see above | Department of Clinical Microbiology, Copenhagen University Hospital, Hvidovre, Kettegaard Alle 30, 2650 Hvidovre. | Albertsen lab, Department of Chemistry and Bioscience, Aalborg University, Denmark | Rasmus Kirkegaard |
| EPI_ISL_452101, EPI_ISL_452102 | Department of Virus and Microbiological Special Diagnostics, Statens Serum Institut, Copenhagen, Denmark, Artillerivej 5, 2300 Copenahgen S | Albertsen lab, Department of Chemistry and Bioscience, Aalborg University, Denmark | Rasmus Kirkegaard |
| EPI_ISL_452139 | Instituto de Diagnostico y Referencia Epidemiologicos (INDRE) | Instituto de diagnóstico y Referencia Epidemiologicos (INDRE) | Ramirez-Gonzalez Ernesto, Garces-Ayala Fabiola, Araiza-Rodriguez Adnan, Mendieta-Condado Edgar, Rodriguez-Maldonado Abril, Wong-Arambula Claudia, Barrera-Badillo Gisela, Hernandez-Rivas Lucia, Lopez-Martinez Irma |
| EPI_ISL_452141 | Instituto de Diagnostico y Referencia Epidemiologicos (INDRE) | Instituto de diagnóstico y Referencia Epidemiologicos (INDRE) | Ramirez-Gonzalez Ernesto, Garces-Ayala Fabiola, Araiza-Rodriguez Adnan, Mendieta-Condado Edgar, Rodriguez-Maldonado Abril, Wong-Arambula Claudia, Barrera-Badillo Gisela, Hernandez-Rivas Lucia, Lopez-Martinez Irma. |
| EPI_ISL_454420 | Rafik Hariri University Hospital | Rafik Hariri University Hospital | Rita Feghali |
| EPI_ISL_455019 | Government Medical College, Vadodara | Gujarat Biotechnology Research Centre | Ramesh Pandit, Tejas Shah, Ankit Hinsu, Pritesh Sabara, Apurvasinh Puvar, Janvi Raval, Zarna Patel, Monika Gandhi, Pinal Trivedi, Maharshi Pandya, Amit Kanani, Nidhi Patel, Nitin Savaliya, Raghawendra Kumar, Dinesh Kumar, Zuber Saiyed, Komal Patel, Labdhi Pandya, Snehal Bagatharia, Tanuja Javadekar , R N Daveshwar, Armi Chaudhari, Bhavesh Modi, Gaurishankar Shrimali, R D Dixit, A M Kadri, Umang Mishra, Chaitanya Joshi, Madhvi Joshi, |
| EPI_ISL_455455 | Instituto de Diagnostico y Referencia Epidemiologicos (INDRE) | Instituto de Diagnostico y Referencia Epidemiologicos (INDRE) | Rodriguez-Maldonado Abril, Mendieta-Condado Edgar, Araiza-Rodriguez Adnan, Garces-Ayala Fabiola. Taboada Ramirez Blanca. Ramirez-Gonzalez Ernesto, Barrera-Badillo Gisela, Hernandez-Rivas Lucia, Lopez-Martinez Irma, Wong-Arambula Claudia. |
| EPI_ISL_455456 | Instituto de Diagnostico y Referencia Epidemiologicos (INDRE) | Instituto de Diagnostico y Referencia Epidemiologicos (INDRE) | Rodriguez-Maldonado Abril, Mendieta-Condado Edgar, Araiza-Rodriguez Adnan, Garces-Ayala Fabiola. Taboada Ramírez Blanca. Ramirez-Gonzalez Ernesto, , Barrera-Badillo Gisela, Hernandez-Rivas Lucia, Lopez-Martinez Irma, Wong-Arambula Claudia. |
| EPI_ISL_456000, EPI_ISL_456001, EPI_ISL_456002 | LSUHS Emerging Viral Threat Laboratory | Microbial Genome Sequencing Center | Rona S. Scott, Jeremy P. Kamil, John A. Vanchiere, Camille F. Abshire, Abida Siddiqa, Byeong-Jae Lee, Chan-ki Min, Md Maksudul Alam, Monica Gestal-Cartele, Edna Ondari, Adam Greer, Malgorzata Bienkowska-Haba, Katarzyna Zwolinska, Michelle M. Arnold, Jason M. Bodily, Andrew D. Yurochko, Paul M. Weinberger, Christopher G. Kevil, Martin J. Sapp, Daniel J. Snyder, Vaughn S. Cooper |
| EPI_ISL_456656 | American Type Culture Collection Inc. (ATCC) | American Type Culture Collection Inc. (ATCC) | Riojas, M.A., Frank,A.M., Puthuveetil, N.P., Benton,B., Peiris, J.S.M., Chu, D.K.W., King,S.P., Flores, B., Parker, M. and Rashid, S. |
| EPI_ISL_458083 | Adi Husada Undaan Hospital | Institute of Tropical Disease, Universitas Airlangga | Rima R Prasetya, Krisnoadi Rahardjo, Aldise M Nastri, Jezzy R Dewantari, Irawati Marga, Gatot Soegiarto, Laksmi Wulandari, Retno A Setyoningrum, Resti Yudhawati, Yohko K Shimizu, Mitsuhiro Nishimura, Yasuko Mori, Soetjipto, Kazufumi Shimizu, Maria I Lusida |
| EPI_ISL_458087 | B.J. Medical College and Civil hospital | Gujarat Biotechnology Research Centre | Ramesh Patel, Pranay Shah, Kamlesh J Upadhyay, Ramesh Pandit, Tejas Shah, Ankit Hinsu, Pritesh Sabara, Apurvasinh Puvar, Janvi Raval, Zarna Patel, Monika Gandhi, Pinal Trivedi, Maharshi Pandya, Amit Kanani, Nidhi Patel, Nitin Savaliya, Raghawendra Kumar, Dinesh Kumar, Zuber Saiyed, Komal Patel, Labdhi Pandya, Snehal Bagatharia, Dhaval Vaghela, Afzal Ansari, Bhavesh Modi, Gaurishankar Shrimali, R D Dixit, A M Kadri, Umang Mishra, Chaitanya Joshi, Madhvi Joshi |
| EPI_ISL_458095 | B.J. Medical College and Civil hospital | Gujarat Biotechnology Research Centre | Raghawendra Kumar, Dinesh Kumar, Zuber Saiyed, Komal Patel, Labdhi Pandya, Snehal Bagatharia, Dhaval Vaghela, Ramesh Patel, Pranay Shah, Kamlesh J Upadhyay, Ramesh Pandit, Tejas Shah, Ankit Hinsu, Pritesh Sabara, Apurvasinh Puvar, Janvi Raval, Zarna Patel, Monika Gandhi, Pinal Trivedi, Maharshi Pandya, Amit Kanani, Nidhi Patel, Nitin Savaliya, Pooja P Doshi, Bhavesh Modi, Gaurishankar Shrimali, R D Dixit, A M Kadri, Umang Mishra, Chaitanya Joshi, Madhvi Joshi |
| EPI_ISL_458102 | B.J. Medical College and Civil hospital | Gujarat Biotechnology Research Centre | Ramesh Patel, Pranay Shah, Kamlesh J Upadhyay, Ramesh Pandit, Tejas Shah, Ankit Hinsu, Pritesh Sabara, Apurvasinh Puvar, Janvi Raval, Zarna Patel, Monika Gandhi, Pinal Trivedi, Maharshi Pandya, Amit Kanani, Nidhi Patel, Nitin Savaliya, Raghawendra Kumar, Dinesh Kumar, Zuber Saiyed, Komal Patel, Labdhi Pandya, Snehal Bagatharia, Dhaval Vaghela, Neelam Nathani, Bhavesh Modi, Gaurishankar Shrimali, R D Dixit, A M Kadri, Umang Mishra, Chaitanya Joshi, Madhvi Joshi |
| EPI_ISL_458103 | Gujarat Biotechnology Research Centre | Gujarat Biotechnology Research Centre | Ramesh Pandit, Tejas Shah, Ankit Hinsu, Pritesh Sabara, Apurvasinh Puvar, Janvi Raval, Zarna Patel, Monika Gandhi, Pinal Trivedi, Maharshi Pandya, Amit Kanani, Nidhi Patel, Nitin Savaliya, Raghawendra Kumar, Dinesh Kumar, Zuber Saiyed, Komal Patel, Labdhi Pandya, Snehal Bagatharia, Armi Chaudhari, Bhavesh Modi, Gaurishankar Shrimali, R D Dixit, A M Kadri, Umang Mishra, Chaitanya Joshi, Madhvi Joshi, , , , , |
| EPI_ISL_459911 | Devki Devi Foundation, a unit of Max Healthcare | CSIR-IGIB/Max | Rajesh Pandey#, Samreen Siddiqui, Pooja Sharma, Bansidhar Tarai, Vivekanand A, Bharathram Uppili, Saruchi Wadhwa, Nishu Tyagi, Mitali Mukerji, Poonam Das, Sujeet Jha, Mohammed Faruq, Vinita Jha, Anurag Agrawal |
| EPI_ISL_459912, EPI_ISL_459913, EPI_ISL_459914, EPI_ISL_459915, EPI_ISL_459916, EPI_ISL_459917, EPI_ISL_459918, EPI_ISL_459919, EPI_ISL_459920, EPI_ISL_459921, EPI_ISL_459922, EPI_ISL_459923, EPI_ISL_459924, EPI_ISL_459925, EPI_ISL_459931, EPI_ISL_459932, EPI_ISL_459933, EPI_ISL_459934, EPI_ISL_459935, EPI_ISL_459936, EPI_ISL_459937, EPI_ISL_459938, EPI_ISL_459939, EPI_ISL_459940, EPI_ISL_459941, EPI_ISL_459942, EPI_ISL_459943, EPI_ISL_459944, EPI_ISL_459945, EPI_ISL_459946, EPI_ISL_459947, EPI_ISL_459948, EPI_ISL_459949, EPI_ISL_459950, EPI_ISL_459951, EPI_ISL_459952 | | | |
| see above | Devki Devi Foundation, a unit of Max Healthcare | CSIR-IGIB/Max | Rajesh Pandey#, Samreen Siddiqui, Pooja Sharma, Bansidhar Tarai, Vivekanand A, Bharathram Uppili, Saruchi Wadhwa, Nishu Tyagi, Mitali Mukerji, Bansidhar Tarai, Poonam Das, Sujeet Jha, Mohammed Faruq, Vinita Jha, Anurag Agrawal |
| EPI_ISL_460635 | UHCW Pathology / University of Warwick | University of Warwick, for the COVID-19 Genomics (COG) UK Consortium | Richard Stark, Chrystala Constantinidou, Meera Unnikrishnan, Laura Baxter, Jeff Cheng, Grace Taylor-Joyce, Hannah Elizabeth Bridgewater, Lucy Frost, Sarojini Pandey, Paul Brown, Tauqeer Alam, Sascha Ott, Dimitris Grammatopoulos |
| EPI_ISL_463995, EPI_ISL_463996, EPI_ISL_463997, EPI_ISL_463998, EPI_ISL_463999, EPI_ISL_464000, EPI_ISL_464001, EPI_ISL_464002, EPI_ISL_464003, EPI_ISL_464004, EPI_ISL_464005, EPI_ISL_464006, EPI_ISL_464007, EPI_ISL_464008, EPI_ISL_464009, EPI_ISL_464010, EPI_ISL_464011, EPI_ISL_464012, EPI_ISL_464013, EPI_ISL_464014, EPI_ISL_464015, EPI_ISL_464016, EPI_ISL_464017, EPI_ISL_464018, EPI_ISL_464019, EPI_ISL_464020, EPI_ISL_464021, EPI_ISL_464022, EPI_ISL_464023, EPI_ISL_464024, EPI_ISL_464025, EPI_ISL_464026, EPI_ISL_464027, EPI_ISL_464028, EPI_ISL_464029, EPI_ISL_464030, EPI_ISL_464031, EPI_ISL_464032, EPI_ISL_464033, EPI_ISL_464034, EPI_ISL_464035, EPI_ISL_464036, EPI_ISL_464037, EPI_ISL_464038, EPI_ISL_464039, EPI_ISL_464040, EPI_ISL_464041, EPI_ISL_464042, EPI_ISL_464043, EPI_ISL_464044, EPI_ISL_464045, EPI_ISL_464046, EPI_ISL_464047, EPI_ISL_464048, EPI_ISL_464049, EPI_ISL_464050, EPI_ISL_464051, EPI_ISL_464052, EPI_ISL_464053, EPI_ISL_464054, EPI_ISL_464055, EPI_ISL_464056, EPI_ISL_464057, EPI_ISL_464058, EPI_ISL_464059, EPI_ISL_464060, EPI_ISL_464061, EPI_ISL_464062, EPI_ISL_464063, EPI_ISL_464064 | | | |
| see above | Unity Health Toronto | Ontario Institute for Cancer Research | Ramzi Fattouh,Larissa M. Matukas,Mark Downing,Annette Gower,Karel Boissinot,Samira Mubareka,TIBDN,Ilinca Lungu,Bernard Lam,Jeremy Johns,Paul Krzyzanowski,Richard de Borja,Philip Zuzarte,Jared Simpson |
| EPI_ISL_467046 | B.J. Medical College and Civil hospital | Gujarat Biotechnology Research Centre | Raghawendra Kumar, Dinesh Kumar, Zuber Saiyed, Komal Patel, Labdhi Pandya, Snehal Bagatharia, Pranay Shah, Kamlesh J Upadhyay, Nirav Mungalpara, Tejas Shah, Ankit Hinsu, Pritesh Sabara, Apurvasinh Puvar, Janvi Raval, Zarna Patel, Monika Gandhi, Pinal Trivedi, Maharshi Pandya, Nidhi Patel, Nitin Savaliya, Neha Rajpara, R D Dixit, A M Kadri, Harsh Bakshi, Chaitanya Joshi, Madhvi Joshi, |
| EPI_ISL_467300 | General Hospital "Abdulah Nakas" | Alea Genetic Center | Rijad Konjhodzic; Lana Salihefendic; Teufik Goletic; Sead Jazic; Dino Pecar; Nihad Fejzic; Damir Marjanovic; Enis Kandic |
| EPI_ISL_468063, EPI_ISL_468064, | unknown | Computer Science and Engineering | Rouchka,E.C., Chariker,J.H., Chung,D., Ramirez,J., Palmer,K.E., Lasnik,A.B., Carrico,R., Arnold,F.W., Adcock,R.S., Zhang,M., Alejandro,B., Wolf,L.A., |

| | | | |
|---|---|---|---|
| EPI_ISL_468065 | | | Hwang,J.Y., Park,J.W., Waigel,S., Zacharias,W. |
| EPI_ISL_468701, EPI_ISL_468702, EPI_ISL_468703, EPI_ISL_468704, EPI_ISL_468705, EPI_ISL_468706, EPI_ISL_468707, EPI_ISL_468708, EPI_ISL_468709, EPI_ISL_468710, EPI_ISL_468711, EPI_ISL_468712, EPI_ISL_468713, EPI_ISL_468714, EPI_ISL_468715, EPI_ISL_468716, EPI_ISL_468717 | | | |
| see above | Ochsner Health | Bioinfoexperts, LLC | Rebecca Rose, Amy Feehan, David J. Nolan, Sissy Cross, David Moraga Amador, Tong Yang, Luke Caruso, Wayra Navia, Lydia Von Borstel, Xiao Hui Zhou, Julia-Garcia-Diaz, Susanna L. Lamers |
| EPI_ISL_469035 | Government Medical College, Vadodara | Gujarat Biotechnology Research Centre | Raghawendra Kumar, Dinesh Kumar, Zuber Saiyed, Komal Patel, Labdhi Pandya, Snehal Bagatharia, Meenakshi Shah, Neena Doshi, Varsha Godbole, Tejas Shah, Ankit Hinsu, Pritesh Sabara, Apurvasinh Puvar, Janvi Raval, Zarna Patel, Monika Gandhi, Pinal Trivedi, Maharshi Pandya, Nidhi Patel, Nitin Savaliya, Akanksha Verma, R D Dixit, A M Kadri, Harsh Bakshi, Chaitanya Joshi, Madhvi Joshi |
| EPI_ISL_475037 | Department of MicroBiology, Government Medical College, Surat | Gujarat Biotechnology Research Centre | Raghawendra Kumar, Dinesh Kumar, Zuber Saiyed, Komal Patel, Labdhi Pandya, Snehal Bagatharia, Naresh Chauhan, Summaiya Mullan, Amit gamit, Pritesh Sabara, Apurvasinh Puvar, Janvi Raval, Zarna Patel, Monika Gandhi, Pinal Trivedi, Maharshi Pandya, Nidhi Patel, Nitin Savaliya, Neha Rajpara, R D Dixit, A M Kadri, Harsh Bakshi, Chaitanya Joshi, Madhvi Joshi |
| EPI_ISL_475055 | GMERS Medical College & Hospital | Gujarat Biotechnology Research Centre | Raghawendra Kumar, Dinesh Kumar, Zuber Saiyed, Komal Patel, Labdhi Pandya, Snehal Bagatharia, Meenakshi Shah, Neena Doshi, Varsha Godbole, Pritesh Sabara, Apurvasinh Puvar, Janvi Raval, Zarna Patel, Monika Gandhi, Pinal Trivedi, Maharshi Pandya, Nidhi Patel, Nitin Savaliya, Pooja P Doshi, R D Dixit, A M Kadri, Harsh Bakshi, Chaitanya Joshi, Madhvi Joshi |
| EPI_ISL_476561 | Hospital Garrahan | Héritas | Roberta Crespo, Dalmacio Pereyra, Mauricio Grisolia, Cristian Rohr, Andrea Mangano, Maria Florencia Fernandez, Fabian Fay, Martin Vazquez |
| EPI_ISL_476863 | GMERS Medical College and Hospital, Gandhinagar | Gujarat Biotechnology Research Centre | Raghawendra Kumar, Dinesh Kumar, Zuber Saiyed, Komal Patel, Labdhi Pandya, Afzal Ansari, Nikha Trivedi, Seema Bhatt, Gaurishankar Shrimali, Bhavesh Modi, Bharti Rajani, Apurvasinh Puvar, Janvi Raval, Zarna Patel, Monika Gandhi, Pinal Trivedi, Maharshi Pandya, Nidhi Patel, Nitin Savaliya, R D Dixit, A M Kadri, Harsh Bakshi, Chaitanya Joshi, Madhvi Joshi |
| EPI_ISL_476880 | Department of MicroBiology, Government Medical College, Surat | Gujarat Biotechnology Research Centre | Raghawendra Kumar, Dinesh Kumar, Zuber Saiyed, Komal Patel, Labdhi Pandya, Afzal Ansari, Nikha Trivedi, Naresh Chauhan, Summaiya Mullan, Amit gamit, Apurvasinh Puvar, Janvi Raval, Zarna Patel, Monika Gandhi, Pinal Trivedi, Maharshi Pandya, Nidhi Patel, Nitin Savaliya, R D Dixit, A M Kadri, Harsh Bakshi, Chaitanya Joshi, Madhvi Joshi |
| EPI_ISL_479663, EPI_ISL_479664, EPI_ISL_479665, EPI_ISL_479666, EPI_ISL_479667, EPI_ISL_479668, EPI_ISL_479669, EPI_ISL_479670, EPI_ISL_479671, EPI_ISL_479672, EPI_ISL_479673, EPI_ISL_479674, EPI_ISL_479675 | | | |
| see above | Center for Genomics and System Biology, New York University | Center for Genomics and System Biology, New York University | Roder,A., Banakis,S., Johnson,K., Khalfan,M., Borenstein,E.S., Samanovic,M., Cornelius,A., Herati,R., Ulrich,R., Fleming,A., Kottkamp,A., Raabe,V., Mulligan,M.J., Gresham,D., Ghedin,E. |
| EPI_ISL_481283 | Center for Genomics and System Biology, New York University | Center for Genomics and System Biology, New York University | Roder,A., Banakis,S., Johnson,K., Khalfan,M., Borenstein,E.S., Samanovic,M., Cornelius,A., Herati,R., Ulrich,R., Fleming,A., Kottkamp,A., Raabe,V., Mulligan,M.J., Gresham,D. and Ghedin,E. |
| EPI_ISL_483834 | Pandit Deendayal Upadhyay Government Medical College, Rajkot | Gujarat Biotechnology Research Centre | Raghawendra Kumar, Dinesh Kumar, Zuber Saiyed, Komal Patel, Labdhi Pandya, Afzal Ansari, Nikha Trivedi, Gauravi Dhruv, Arti Trivedi, Apurvasinh Puvar, Janvi Raval, Zarna Patel, Monika Gandhi, Pinal Trivedi, Maharshi Pandya, Nidhi Patel, Nitin Savaliya, R D Dixit, A M Kadri, Harsh Bakshi, Chaitanya Joshi, Madhvi Joshi |
| EPI_ISL_483851, EPI_ISL_483869 | Department of Microbiology, Government Medical College, Surat | Gujarat Biotechnology Research Centre | Raghawendra Kumar, Dinesh Kumar, Zuber Saiyed, Komal Patel, Labdhi Pandya, Afzal Ansari, Nikha Trivedi, Naresh Chauhan, Summaiya Mullan, Amit gamit, Apurvasinh Puvar, Janvi Raval, Zarna Patel, Monika Gandhi, Pinal Trivedi, Maharshi Pandya, Nidhi Patel, Nitin Savaliya, R D Dixit, A M Kadri, Harsh Bakshi, Chaitanya Joshi, Madhvi Joshi |
| EPI_ISL_485003, EPI_ISL_485004, EPI_ISL_485005, EPI_ISL_485006, EPI_ISL_485007, EPI_ISL_485008, EPI_ISL_485009, EPI_ISL_485010, EPI_ISL_485011, EPI_ISL_485012, EPI_ISL_485013, EPI_ISL_485014, EPI_ISL_485015, EPI_ISL_485016, EPI_ISL_485017, EPI_ISL_485018, EPI_ISL_485019, EPI_ISL_485020, EPI_ISL_485021, EPI_ISL_485022, EPI_ISL_485023, EPI_ISL_485024, EPI_ISL_485025, EPI_ISL_485026, EPI_ISL_485027, EPI_ISL_485028, EPI_ISL_485029, EPI_ISL_485030, EPI_ISL_485031, EPI_ISL_485032, EPI_ISL_485033, EPI_ISL_485034, EPI_ISL_485035, EPI_ISL_485036, EPI_ISL_485037, EPI_ISL_485038, EPI_ISL_485039, EPI_ISL_485040, EPI_ISL_485041, EPI_ISL_485042, EPI_ISL_485043, EPI_ISL_485044, EPI_ISL_485045, EPI_ISL_485046, EPI_ISL_485047, EPI_ISL_485048, EPI_ISL_485049, EPI_ISL_485050, EPI_ISL_485051, EPI_ISL_485052, EPI_ISL_485053, EPI_ISL_485054, EPI_ISL_485055, EPI_ISL_485056, EPI_ISL_485057, EPI_ISL_485058, EPI_ISL_485059, EPI_ISL_485060, EPI_ISL_485061, EPI_ISL_485062, EPI_ISL_485063, EPI_ISL_485064, EPI_ISL_485065, EPI_ISL_485066, EPI_ISL_485067, EPI_ISL_485068, EPI_ISL_485069, EPI_ISL_485070, EPI_ISL_485071, EPI_ISL_485072, EPI_ISL_485073, EPI_ISL_485074, EPI_ISL_485075, EPI_ISL_485076, EPI_ISL_485077, EPI_ISL_485078, EPI_ISL_485079, EPI_ISL_485080, EPI_ISL_485081, EPI_ISL_485082, EPI_ISL_485083, EPI_ISL_485084, EPI_ISL_485085, EPI_ISL_485086, EPI_ISL_485087, EPI_ISL_485088, EPI_ISL_485089, EPI_ISL_485090, EPI_ISL_485091, EPI_ISL_485092, EPI_ISL_485093, EPI_ISL_485094, EPI_ISL_485095, EPI_ISL_485096, EPI_ISL_485097, EPI_ISL_485098, EPI_ISL_485099, EPI_ISL_485100, EPI_ISL_485101, EPI_ISL_485102, EPI_ISL_485103, EPI_ISL_485104, EPI_ISL_485105, EPI_ISL_485106, EPI_ISL_485107, EPI_ISL_485108, EPI_ISL_485109, EPI_ISL_485110, EPI_ISL_485111, EPI_ISL_485112, EPI_ISL_485113, EPI_ISL_485114, EPI_ISL_485115, EPI_ISL_485116, EPI_ISL_485117, EPI_ISL_485118, EPI_ISL_485119, EPI_ISL_485120, EPI_ISL_485121, EPI_ISL_485122, EPI_ISL_485123, EPI_ISL_485124, EPI_ISL_485125, EPI_ISL_485126, EPI_ISL_485127, EPI_ISL_485128, EPI_ISL_485129, EPI_ISL_485130, EPI_ISL_485131, EPI_ISL_485132, EPI_ISL_485133, EPI_ISL_485134, EPI_ISL_485135, EPI_ISL_485136, EPI_ISL_485137, EPI_ISL_485138, EPI_ISL_485139, EPI_ISL_485140, EPI_ISL_485141, EPI_ISL_485142, EPI_ISL_485143, EPI_ISL_485144, EPI_ISL_485145, EPI_ISL_485146, EPI_ISL_485147, EPI_ISL_485148, EPI_ISL_485149, EPI_ISL_485150, EPI_ISL_485151, EPI_ISL_485152, EPI_ISL_485153, EPI_ISL_485154, EPI_ISL_485155, EPI_ISL_485156, EPI_ISL_485157, EPI_ISL_485158, EPI_ISL_485159, EPI_ISL_485160, EPI_ISL_485161, EPI_ISL_485162, EPI_ISL_485163, EPI_ISL_485164, EPI_ISL_485165, EPI_ISL_485166, EPI_ISL_485167, EPI_ISL_485168, EPI_ISL_485169, EPI_ISL_485170, EPI_ISL_485171, EPI_ISL_485172, EPI_ISL_485173, EPI_ISL_485174, EPI_ISL_485175, EPI_ISL_485176, EPI_ISL_485177, EPI_ISL_485178, EPI_ISL_485179, EPI_ISL_485180, EPI_ISL_485181, EPI_ISL_485182, EPI_ISL_485183, EPI_ISL_485184, EPI_ISL_485185, EPI_ISL_485186, EPI_ISL_485187, EPI_ISL_485188, EPI_ISL_485189, EPI_ISL_485190, EPI_ISL_485191, EPI_ISL_485192, EPI_ISL_485193, EPI_ISL_485194, EPI_ISL_485195, EPI_ISL_485196, EPI_ISL_485197, EPI_ISL_485198, EPI_ISL_485199, EPI_ISL_485200, EPI_ISL_485201, EPI_ISL_485202, EPI_ISL_485203, EPI_ISL_485204, EPI_ISL_485205, EPI_ISL_485206, EPI_ISL_485207, EPI_ISL_485208, EPI_ISL_485209, EPI_ISL_485210, EPI_ISL_485211, EPI_ISL_485212, EPI_ISL_485213, EPI_ISL_485214, EPI_ISL_485215, EPI_ISL_485216, EPI_ISL_485217, EPI_ISL_485218, EPI_ISL_485219, EPI_ISL_485220, EPI_ISL_485221, EPI_ISL_485222, EPI_ISL_485223, EPI_ISL_485224, EPI_ISL_485225, EPI_ISL_485226, EPI_ISL_485227, EPI_ISL_485228, EPI_ISL_485229, EPI_ISL_485230, EPI_ISL_485231, EPI_ISL_485232, EPI_ISL_485233, EPI_ISL_485234, EPI_ISL_485235, EPI_ISL_485236, EPI_ISL_485237, EPI_ISL_485238, EPI_ISL_485239, EPI_ISL_485240, EPI_ISL_485241, EPI_ISL_485242, EPI_ISL_485243, EPI_ISL_485244, EPI_ISL_485245, EPI_ISL_485246, EPI_ISL_485247, EPI_ISL_485248, EPI_ISL_485249, EPI_ISL_485250, EPI_ISL_485251, EPI_ISL_485252, EPI_ISL_485253, EPI_ISL_485254, EPI_ISL_485255, EPI_ISL_485256, EPI_ISL_485257, EPI_ISL_485258, EPI_ISL_485259, EPI_ISL_485260, EPI_ISL_485261, EPI_ISL_485262, EPI_ISL_485263, EPI_ISL_485264, EPI_ISL_485265, EPI_ISL_485266, EPI_ISL_485267, EPI_ISL_485268, EPI_ISL_485269, EPI_ISL_485270, EPI_ISL_485271, EPI_ISL_485272, EPI_ISL_485273, EPI_ISL_485274, EPI_ISL_485275, EPI_ISL_485276, EPI_ISL_485277, EPI_ISL_485278, EPI_ISL_485279, EPI_ISL_485280, EPI_ISL_485281, EPI_ISL_485282, EPI_ISL_485283, EPI_ISL_485284, EPI_ISL_485285, EPI_ISL_485286, EPI_ISL_485287, EPI_ISL_485288, EPI_ISL_485289, EPI_ISL_485290, EPI_ISL_485291, EPI_ISL_485292, EPI_ISL_485293, EPI_ISL_485294, EPI_ISL_485295, EPI_ISL_485296, EPI_ISL_485297, EPI_ISL_485298, EPI_ISL_485299, EPI_ISL_485300, EPI_ISL_485301, EPI_ISL_485302, EPI_ISL_485303, EPI_ISL_485304, EPI_ISL_485305, EPI_ISL_485306, EPI_ISL_485307, EPI_ISL_485308, EPI_ISL_485309, EPI_ISL_485310, EPI_ISL_485311, EPI_ISL_485312, EPI_ISL_485313, EPI_ISL_485314, EPI_ISL_485315, EPI_ISL_485316, EPI_ISL_485317, EPI_ISL_485318, EPI_ISL_485319, EPI_ISL_485320, EPI_ISL_485321, EPI_ISL_485322, EPI_ISL_485323, EPI_ISL_485324, EPI_ISL_485325, EPI_ISL_485326, EPI_ISL_485327, EPI_ISL_485328, EPI_ISL_485329, EPI_ISL_485330, EPI_ISL_485331, EPI_ISL_485332, EPI_ISL_485333, EPI_ISL_485334, EPI_ISL_485335, EPI_ISL_485336, EPI_ISL_485337, EPI_ISL_485338, EPI_ISL_485339, EPI_ISL_485340, EPI_ISL_485341, EPI_ISL_485342, EPI_ISL_485343, EPI_ISL_485344, EPI_ISL_485345, EPI_ISL_485346, EPI_ISL_485347, EPI_ISL_485348, EPI_ISL_485349, EPI_ISL_485350, EPI_ISL_485351, EPI_ISL_485352, EPI_ISL_485353, EPI_ISL_485354, EPI_ISL_485355, EPI_ISL_485356, EPI_ISL_485357, EPI_ISL_485358, EPI_ISL_485359, EPI_ISL_485360, EPI_ISL_485361, EPI_ISL_485362, EPI_ISL_485363, EPI_ISL_485364, EPI_ISL_485365, EPI_ISL_485366, EPI_ISL_485367, EPI_ISL_485368, EPI_ISL_485369, EPI_ISL_485370, EPI_ISL_485371, EPI_ISL_485372, EPI_ISL_485373, EPI_ISL_485374, EPI_ISL_485375, EPI_ISL_485376, EPI_ISL_485377, EPI_ISL_485378, EPI_ISL_485379, EPI_ISL_485380, EPI_ISL_485381, EPI_ISL_485382, EPI_ISL_485383, EPI_ISL_485384, EPI_ISL_485385, EPI_ISL_485386, EPI_ISL_485387, EPI_ISL_485873, EPI_ISL_485874, EPI_ISL_485875, EPI_ISL_485876, EPI_ISL_485877, EPI_ISL_485878, EPI_ISL_485879, EPI_ISL_485880, EPI_ISL_485881, EPI_ISL_485882, EPI_ISL_485883, EPI_ISL_485884, EPI_ISL_485885, EPI_ISL_485886, EPI_ISL_485887, EPI_ISL_485888, EPI_ISL_485889, EPI_ISL_485890, EPI_ISL_485891, EPI_ISL_485892, EPI_ISL_485893, EPI_ISL_485894, EPI_ISL_485895, EPI_ISL_485896, EPI_ISL_485897, EPI_ISL_485898, EPI_ISL_485899, EPI_ISL_485900, EPI_ISL_485901, EPI_ISL_485902, EPI_ISL_485903, EPI_ISL_485904, EPI_ISL_485905, EPI_ISL_485906 | | | |
| see above | River Road Testing Lab | Ginkgo Bioworks Clinical Laboratory | Rebecca C. Christofferson, Stephania A. Cormier, Luan V. Dinh, E. Handly Mayton, Hollis R. O'Neil, Thaya Stoufflet, Malaika Mckenzie-Bennett, James McGann, Jim Griffin, Keith Robison, Alex Plocik, Becky Schilling, Rebecca Littlefield, Michelle Spencer, Birgitte Simen |
| EPI_ISL_486649 | Microbiology, Virology and Biemergency Laboratory-ASST FBF Sacco | Microbiology, Virology and Biemergency Laboratory-ASST FBF Sacco | Rimoldi SG, Comandatore F, Romeri F, Mancon A, Micheli V |
| EPI_ISL_486650 | Microbiology, Virology and Biemergency Laboratory-ASST FBF Sacco | Microbiology, Virology and Biemergency Laboratory-ASST FBF Sacco | Romeri F, Comandatore F, Mancon A, Micheli V, Rimoldi SG |
| EPI_ISL_486653, EPI_ISL_486657 | Microbiology, Virology and Biemergency Laboratory-ASST FBF Sacco | Microbiology, Virology and Biemergency Laboratory-ASST FBF Sacco | Rimoldi SG, Comandatore F, Romeri F, Mancon A, Micheli V |
| EPI_ISL_486658 | Microbiology, Virology and Biemergency Laboratory-ASST FBF Sacco | Microbiology, Virology and Biemergency Laboratory-ASST FBF Sacco | Romeri F, Comandatore F, Mancon A, Micheli V, Rimoldi SG |
| EPI_ISL_486660 | Microbiology, Virology and Biemergency Laboratory-ASST FBF Sacco | Microbiology, Virology and Biemergency Laboratory-ASST FBF Sacco | Rimoldi SG, Comandatore F, Romeri F, Mancon A, Micheli V |
| EPI_ISL_486661 | Microbiology, Virology and Biemergency Laboratory-ASST FBF Sacco | Microbiology, Virology and Biemergency Laboratory-ASST FBF Sacco | Romeri F, Comandatore F, Mancon A, Micheli V, Rimoldi SG |
| EPI_ISL_486664 | Microbiology, Virology and Biemergency Laboratory-ASST FBF Sacco | Microbiology, Virology and Biemergency Laboratory-ASST FBF Sacco | Rimoldi SG, Comandatore F, Romeri F, Mancon A, Micheli V |
| EPI_ISL_495024 | Government Medical College, Bhavnagar | Gujarat Biotechnology Research Centre | Raghawendra Kumar, Dinesh Kumar, Zuber Saiyed, Komal Patel, Labdhi Pandya, Afzal Ansari, Nikha Trivedi, Kairavi Desai, Saklin Malek, Shirish Patel, Apurvasinh Puvar, Janvi Raval, Zarna Patel, Monika Gandhi, Pinal Trivedi, Maharshi Pandya, Nidhi Patel, Nitin Savaliya, R D Dixit, A M Kadri, Harsh Bakshi, Chaitanya Joshi, Madhvi Joshi |
| EPI_ISL_495041, EPI_ISL_495059 | GMERS Medical College & Hospital, Gotri, Vadodara | Gujarat Biotechnology Research Centre | Raghawendra Kumar, Dinesh Kumar, Zuber Saiyed, Komal Patel, Labdhi Pandya, Afzal Ansari, Nikha Trivedi, Meenakshi Shah, Neena Doshi, Varsha |

| | | | |
|---|---|---|---|
| | | | Godbole, Apurvasinh Puvar, Janvi Raval, Zarna Patel, Monika Gandhi, Pinal Trivedi, Maharshi Pandya, Nidhi Patel, Nitin Savaliya, R D Dixit, A M Kadri, Harsh Bakshi, Chaitanya Joshi, Madhvi Joshi |
| EPI_ISL_495077 | Department of MicroBiology, Government Medical College, Surat | Gujarat Biotechnology Research Centre | Raghawendra Kumar, Dinesh Kumar, Zuber Saiyed, Komal Patel, Labdhi Pandya, Afzal Ansari, Nikha Trivedi, Naresh Chauhan, Summaiya Mullan, Amit gamit, Apurvasinh Puvar, Janvi Raval, Zarna Patel, Monika Gandhi, Pinal Trivedi, Maharshi Pandya, Nidhi Patel, Nitin Savaliya, R D Dixit, A M Kadri, Harsh Bakshi, Chaitanya Joshi, Madhvi Joshi |
| EPI_ISL_498695, EPI_ISL_498696, EPI_ISL_498697, EPI_ISL_498698, EPI_ISL_498699, EPI_ISL_498700, EPI_ISL_498701, EPI_ISL_498702, EPI_ISL_498703, EPI_ISL_498704, EPI_ISL_498705, EPI_ISL_498706, EPI_ISL_498707, EPI_ISL_498708, EPI_ISL_498709, EPI_ISL_498710, EPI_ISL_498711, EPI_ISL_498712, EPI_ISL_498713, EPI_ISL_498714, EPI_ISL_498715, EPI_ISL_498716, EPI_ISL_498717, EPI_ISL_498718, EPI_ISL_498719, EPI_ISL_498720, EPI_ISL_498721, EPI_ISL_498722, EPI_ISL_498723, EPI_ISL_498724, EPI_ISL_498725, EPI_ISL_498726, EPI_ISL_498727, EPI_ISL_498728, EPI_ISL_498729, EPI_ISL_498730, EPI_ISL_498731, EPI_ISL_498732, EPI_ISL_498733, EPI_ISL_498734, EPI_ISL_498735, EPI_ISL_498736, EPI_ISL_498737, EPI_ISL_498738, EPI_ISL_498739, EPI_ISL_498740, EPI_ISL_498741, EPI_ISL_498742, EPI_ISL_498743, EPI_ISL_498744, EPI_ISL_498745, EPI_ISL_498746, EPI_ISL_498747 | | | |
| see above | Quest Diagnostics | Quest Diagnostics | Rosenthal,S.H., Gerasimova,A., Kagan,R.M. and Owen, R. |
| EPI_ISL_500459 | Quest Diagnostics | Quest Diagnostics | Rosenthal,S.H., Gerasimova,A., Kagan,R.M., Owen, R. AND Lacbawan, F. |
| EPI_ISL_500596, EPI_ISL_500597, EPI_ISL_500598, EPI_ISL_500599, EPI_ISL_500600, EPI_ISL_500601, EPI_ISL_500602, EPI_ISL_500603, EPI_ISL_500604, EPI_ISL_500605, EPI_ISL_500606, EPI_ISL_500607, EPI_ISL_500608, EPI_ISL_500609, EPI_ISL_500610, EPI_ISL_500611, EPI_ISL_500612, EPI_ISL_500613, EPI_ISL_500614, EPI_ISL_500615, EPI_ISL_500616, EPI_ISL_500617, EPI_ISL_500618, EPI_ISL_500619, EPI_ISL_500620, EPI_ISL_500621, EPI_ISL_500622, EPI_ISL_500623, EPI_ISL_500624, EPI_ISL_500625, EPI_ISL_500626, EPI_ISL_500627, EPI_ISL_500628, EPI_ISL_500629, EPI_ISL_500630, EPI_ISL_500631, EPI_ISL_500632, EPI_ISL_500633, EPI_ISL_500634, EPI_ISL_500635, EPI_ISL_500636, EPI_ISL_500637, EPI_ISL_500638, EPI_ISL_500639, EPI_ISL_500640, EPI_ISL_500641, EPI_ISL_500642, EPI_ISL_500643, EPI_ISL_500644, EPI_ISL_500645, EPI_ISL_500646, EPI_ISL_500647, EPI_ISL_500648, EPI_ISL_500649, EPI_ISL_500650, EPI_ISL_500651, EPI_ISL_500652, EPI_ISL_500653, EPI_ISL_500654, EPI_ISL_500655, EPI_ISL_500656, EPI_ISL_500657, EPI_ISL_500658, EPI_ISL_500659, EPI_ISL_500660, EPI_ISL_500661, EPI_ISL_500662, EPI_ISL_500663, EPI_ISL_500664, EPI_ISL_500665, EPI_ISL_500666, EPI_ISL_500667, EPI_ISL_500668, EPI_ISL_500669, EPI_ISL_500670, EPI_ISL_500671, EPI_ISL_500672, EPI_ISL_500673, EPI_ISL_500674, EPI_ISL_500675, EPI_ISL_500676, EPI_ISL_500677, EPI_ISL_500678, EPI_ISL_500679, EPI_ISL_500680, EPI_ISL_500681, EPI_ISL_500682, EPI_ISL_500683, EPI_ISL_500684, EPI_ISL_500685, EPI_ISL_500686, EPI_ISL_500687, EPI_ISL_500688, EPI_ISL_500689, EPI_ISL_500690, EPI_ISL_500691, EPI_ISL_500692, EPI_ISL_500693, EPI_ISL_500694, EPI_ISL_500695, EPI_ISL_500696, EPI_ISL_500697, EPI_ISL_500698, EPI_ISL_500699, EPI_ISL_500700, EPI_ISL_500701, EPI_ISL_500702, EPI_ISL_500703, EPI_ISL_500704, EPI_ISL_500705, EPI_ISL_500706 | | | |
| see above | Area of Virology, Serology and Virology Division (SAViD), New South Wales Health Pathology Randwick | Area of Virology, Serology and Virology Division (SAViD), New South Wales Health Pathology Randwick | Rawlinson, W. |
| EPI_ISL_500717 | Area of Virology, Serology and Virology Division (SAViD), New South Wales Health Pathology Randwick | Area of Virology, Serology and Virology Division (SAViD), New South Wales Health Pathology Randwick | Rawlinson, W |
| EPI_ISL_509492, EPI_ISL_509493, EPI_ISL_509494, EPI_ISL_509495, EPI_ISL_509496, EPI_ISL_509497, EPI_ISL_509498, EPI_ISL_509499, EPI_ISL_509500, EPI_ISL_509501, EPI_ISL_509502, EPI_ISL_509503, EPI_ISL_509504, EPI_ISL_509505, EPI_ISL_509506, EPI_ISL_509507, EPI_ISL_509508, EPI_ISL_509509, EPI_ISL_509510, EPI_ISL_509511, EPI_ISL_509512, EPI_ISL_509513, EPI_ISL_509514, EPI_ISL_509515, EPI_ISL_509516, EPI_ISL_509517, EPI_ISL_509518, EPI_ISL_509519, EPI_ISL_509520, EPI_ISL_509521, EPI_ISL_509522, EPI_ISL_509523 | | | |
| see above | Area of Virology, Serology and Virology Division (SAViD), New South Wales Health Pathology Randwick | Area of Virology, Serology and Virology Division (SAViD), New South Wales Health Pathology Randwick | Rawlinson, W. |
| EPI_ISL_512063 | B.J. Medical College and Civil hospital, Ahmedabad | Gujarat Biotechnology Research Centre | Raghawendra Kumar, Dinesh Kumar, Zuber Saiyed, Komal Patel, Labdhi Pandya, Afzal Ansari, Nikha Trivedi, Pranay Shah, Kamlesh J Upadhyay, Sanjay Kapadia, Apurvasinh Puvar, Janvi Raval, Zarna Patel, Monika Gandhi, Pinal Trivedi, Maharshi Pandya, Nidhi Patel, Nitin Savaliya, R D Dixit, A M Kadri, Harsh Bakshi, Chaitanya Joshi, Madhvi Joshi |
| EPI_ISL_512873 | Centre Pasteur of Cameroun | Virology Service, Centre Pasteur of Cameroun | Richard Njouom and Serge Alain SADEUH-Mba |
| EPI_ISL_513027, EPI_ISL_513028, EPI_ISL_513029, EPI_ISL_513030, EPI_ISL_513031, EPI_ISL_513032, EPI_ISL_513033, EPI_ISL_513034, EPI_ISL_513035, EPI_ISL_513036, EPI_ISL_513037, EPI_ISL_513038, EPI_ISL_513039, EPI_ISL_513040, EPI_ISL_513041, EPI_ISL_513042, EPI_ISL_513043, EPI_ISL_513044, EPI_ISL_513045, EPI_ISL_513046, EPI_ISL_513047, EPI_ISL_513048, EPI_ISL_513049, EPI_ISL_513050, EPI_ISL_513051, EPI_ISL_513052, EPI_ISL_513053, EPI_ISL_513054, EPI_ISL_513055, EPI_ISL_513056, EPI_ISL_513057, EPI_ISL_513058, EPI_ISL_513059, EPI_ISL_513060, EPI_ISL_513061, EPI_ISL_513062, EPI_ISL_513063, EPI_ISL_513198, EPI_ISL_513199, EPI_ISL_513200, EPI_ISL_513201, EPI_ISL_513202, EPI_ISL_513203, EPI_ISL_513204, EPI_ISL_513205, EPI_ISL_513206, EPI_ISL_513207 | | | |
| see above | Pathogen Genomics Lab King Abdullah University of Science and Technology(KAUST) | Pathogen Genomics Lab King Abdullah University of Science and Technology(KAUST) | Rahul P Salunke, Sharif Hala, Raeece Naeem, Sara Mfarrej, Amit Kumar Subudhi, Amanda Ooi, Luke Esau, Fadwa Alofi, Fathia Ben Rached, Afrah Alsomali, Asim Khogeer, Ahmad Bakur Mahmoud, Anwar Hashem, Naif Almontashiri, Arnab Pain |
| EPI_ISL_514582, EPI_ISL_514600 | B.J. Medical College and Civil hospital, Ahmedabad | Gujarat Biotechnology Research Centre | Raghawendra Kumar, Dinesh Kumar, Zuber Saiyed, Komal Patel, Labdhi Pandya, Afzal Ansari, Nikha Trivedi, Pranay Shah, Kamlesh J Upadhyay, Sanjay Kapadia, Apurvasinh Puvar, Janvi Raval, Zarna Patel, Monika Gandhi, Pinal Trivedi, Maharshi Pandya, Nidhi Patel, Nitin Savaliya, R D Dixit, A M Kadri, Harsh Bakshi, Chaitanya Joshi, Madhvi Joshi |
| EPI_ISL_514673 | Nevada State Public Health Laboratory | Nevada State Public Health Laboratory | Richard Tillet, Joel R. Sevinsky, Paul Hartley, Heather Kerwin, David Jackson, Subhash C. Verma, Cyprian Rosetto, Andrew Gorzalski, Chris Laverdure, Natalie Crawford, Stephanie Van Hooser, and Mark Pandori |
| EPI_ISL_514674 | Nevada State Public Health Laboratory | Nevada State Public Health Laboratory | Richard Tillet, Joel R. Sevinsky, Paul Hartley, Heather Kerwin, David Jackson, Subhash C. Verma, Cyprian Rossetto, Andrew Gorzalski, Chris Laverdure, Natalie Crawford, Stephanie Van Hooser, and Mark Pandori |
| EPI_ISL_515250, EPI_ISL_515251, EPI_ISL_515252, EPI_ISL_515253, EPI_ISL_515254, EPI_ISL_515255, EPI_ISL_515256, EPI_ISL_515257, EPI_ISL_515258, EPI_ISL_515259, EPI_ISL_515260, EPI_ISL_515261 | | | |
| see above | Texas Department of State Health Services | Texas Department of State Health Services | Rashmi Tuladhar, Bonnie Oh, Cara Akrout, Jenny Zhang, Maliha Rahman, Anita Pokharel, Myong Koag, Chun Wang, Rachel Lee, Grace Kubin |
| EPI_ISL_515293, EPI_ISL_515294, EPI_ISL_515295, EPI_ISL_515296, EPI_ISL_515297, EPI_ISL_515298, EPI_ISL_515299, EPI_ISL_515300, EPI_ISL_515301, EPI_ISL_515302, EPI_ISL_515303, EPI_ISL_515304, EPI_ISL_515305, EPI_ISL_515306, EPI_ISL_515307, EPI_ISL_515308, EPI_ISL_515309, EPI_ISL_515310, EPI_ISL_515311, EPI_ISL_515312, EPI_ISL_515313, EPI_ISL_515314, EPI_ISL_515315, EPI_ISL_515316, EPI_ISL_515317, EPI_ISL_515318, EPI_ISL_515319, EPI_ISL_515320, EPI_ISL_515321, EPI_ISL_515322, EPI_ISL_515323, EPI_ISL_515324, EPI_ISL_515325, EPI_ISL_515326, EPI_ISL_515327, EPI_ISL_515328, EPI_ISL_515329, EPI_ISL_515330, EPI_ISL_515331, EPI_ISL_515332, EPI_ISL_515333, EPI_ISL_515334, EPI_ISL_515335, EPI_ISL_515336, EPI_ISL_515337, EPI_ISL_515338, EPI_ISL_515339, EPI_ISL_515340, EPI_ISL_515341, EPI_ISL_515342, EPI_ISL_515343, EPI_ISL_515344, EPI_ISL_515345, EPI_ISL_515346, EPI_ISL_515347, EPI_ISL_515348, EPI_ISL_515349, EPI_ISL_515350, EPI_ISL_515351, EPI_ISL_515352, EPI_ISL_515353, EPI_ISL_515354, EPI_ISL_515355, EPI_ISL_515356, EPI_ISL_515357, EPI_ISL_515358, EPI_ISL_515359, EPI_ISL_515360, EPI_ISL_515361, EPI_ISL_515362, EPI_ISL_515363, EPI_ISL_515364, EPI_ISL_515365, EPI_ISL_515366, EPI_ISL_515367, EPI_ISL_515368, EPI_ISL_515369, EPI_ISL_515370, EPI_ISL_515371, EPI_ISL_515372, EPI_ISL_515373, EPI_ISL_515374, EPI_ISL_515375, EPI_ISL_515376, EPI_ISL_515377, EPI_ISL_515378, EPI_ISL_515379, EPI_ISL_515380, EPI_ISL_515381, EPI_ISL_515382, EPI_ISL_515383, EPI_ISL_515384, EPI_ISL_515385, EPI_ISL_515386, EPI_ISL_515387, EPI_ISL_515388, EPI_ISL_515389, EPI_ISL_515390, EPI_ISL_515391, EPI_ISL_515392, EPI_ISL_515393, EPI_ISL_515394, EPI_ISL_515395, EPI_ISL_515397, EPI_ISL_515399, EPI_ISL_515400, EPI_ISL_515401, EPI_ISL_515402, EPI_ISL_515403, EPI_ISL_515404, EPI_ISL_515405, EPI_ISL_515406, EPI_ISL_515407, EPI_ISL_515408, EPI_ISL_515409, EPI_ISL_515410, EPI_ISL_515411, EPI_ISL_515412, EPI_ISL_515413, EPI_ISL_515414, EPI_ISL_515415, EPI_ISL_515416, EPI_ISL_515417, EPI_ISL_515418, EPI_ISL_515419, EPI_ISL_515420, EPI_ISL_515421, EPI_ISL_515422, EPI_ISL_515423, EPI_ISL_515424, EPI_ISL_515425, EPI_ISL_515426, EPI_ISL_515427, EPI_ISL_515428, EPI_ISL_515429, EPI_ISL_515430, EPI_ISL_515431, EPI_ISL_515432, EPI_ISL_515433, EPI_ISL_515434, EPI_ISL_515435, EPI_ISL_515436, EPI_ISL_515437, EPI_ISL_515438, EPI_ISL_515439, EPI_ISL_515440, EPI_ISL_515441, EPI_ISL_515442, EPI_ISL_515443, EPI_ISL_515444, EPI_ISL_515445, EPI_ISL_515446, EPI_ISL_515447, EPI_ISL_515448, EPI_ISL_515449, EPI_ISL_515450, EPI_ISL_515451, EPI_ISL_515452, EPI_ISL_515453, EPI_ISL_515454, EPI_ISL_515455, EPI_ISL_515456, EPI_ISL_515457, EPI_ISL_515458, EPI_ISL_515459, EPI_ISL_515460, EPI_ISL_515461, EPI_ISL_515462 | | | |
| see above | Nevada State Public Health Laboratory | Nevada State Public Health Laboratory | Richard Tillett, Joel R. Sevinsky, Paul Hartley, Heather Kerwin, David Jackson, Subhash C. Verma, Cyprian Rossetto, Andrew Gorzalski, Chris Laverdure, Natalie Crawford, Stephanie Van Hooser, and Mark Pandori |
| EPI_ISL_517960, EPI_ISL_517961, EPI_ISL_517962, EPI_ISL_517963, EPI_ISL_517964, EPI_ISL_517965, EPI_ISL_517966, EPI_ISL_517967, EPI_ISL_517968, EPI_ISL_517969, EPI_ISL_517970, EPI_ISL_517971, EPI_ISL_517972, EPI_ISL_517974, EPI_ISL_517975, EPI_ISL_517976, EPI_ISL_517977, EPI_ISL_517978, EPI_ISL_517979, EPI_ISL_517981, EPI_ISL_517982, EPI_ISL_517983, EPI_ISL_517984, EPI_ISL_517985, EPI_ISL_517986, EPI_ISL_517987, EPI_ISL_517989, EPI_ISL_517990, EPI_ISL_517991, EPI_ISL_517992, EPI_ISL_517993, EPI_ISL_517994, EPI_ISL_517995, EPI_ISL_522351, EPI_ISL_522352, EPI_ISL_522353, EPI_ISL_522354, EPI_ISL_522355, EPI_ISL_522356, EPI_ISL_522357, EPI_ISL_522358, EPI_ISL_522359, EPI_ISL_522360, EPI_ISL_522361, EPI_ISL_522362, EPI_ISL_522363, EPI_ISL_522364, EPI_ISL_522365, EPI_ISL_522366, EPI_ISL_522367, EPI_ISL_522368, EPI_ISL_522369, EPI_ISL_522370, EPI_ISL_522371, EPI_ISL_522372, EPI_ISL_522373, EPI_ISL_522374, EPI_ISL_522375, EPI_ISL_522376, EPI_ISL_522377, EPI_ISL_522378, EPI_ISL_522379, EPI_ISL_522380, EPI_ISL_522381, EPI_ISL_522382, EPI_ISL_522383, EPI_ISL_522384, EPI_ISL_522385, EPI_ISL_522386, EPI_ISL_522387, EPI_ISL_522388, EPI_ISL_522389, EPI_ISL_522390, EPI_ISL_522391, EPI_ISL_522392, EPI_ISL_522393, EPI_ISL_522394, EPI_ISL_522395, EPI_ISL_522943, EPI_ISL_522944, EPI_ISL_522945, EPI_ISL_522946, EPI_ISL_522947, EPI_ISL_522948, EPI_ISL_522949, EPI_ISL_522950, EPI_ISL_522951, EPI_ISL_522952, EPI_ISL_522953, EPI_ISL_522954, EPI_ISL_522955, EPI_ISL_522956, EPI_ISL_522957, EPI_ISL_522958, EPI_ISL_522959, EPI_ISL_522960, EPI_ISL_522961, EPI_ISL_522962, EPI_ISL_522963, EPI_ISL_522964, EPI_ISL_522965, EPI_ISL_522966, EPI_ISL_522967, EPI_ISL_522968, EPI_ISL_522969, EPI_ISL_522970, EPI_ISL_522971, EPI_ISL_522972, EPI_ISL_522973, EPI_ISL_522974, EPI_ISL_522975, EPI_ISL_522976, EPI_ISL_522977 | | | |
| see above | Texas Department of State Health Services | Texas Department of State Health Services | Rashmi Tuladhar, Bonnie Oh, Cara Akrout, Jenny Zhang, Maliha Rahman, Anita Pokharel, Myong Koag, Chun Wang, Rachel Lee, Grace Kubin |
| EPI_ISL_523811, EPI_ISL_523812 | Universidad Iberoamericana, Instituto de Medicina Tropical & Salud Global | International Centre for Genetic Engineering and Biotechnology (ICGEB) and ARGO Open Lab Platform | Robert Paulino-Ramirez, Eileen Riego, Alejandro Vallejo Degaudenzi, Victor Virgilio Calderon, Leandro Tapia, Danilo Licastro, Simeone Dal Monego, Sreejith Rajasekharan and Alessandro Marcello. |
| EPI_ISL_524053, EPI_ISL_524054, EPI_ISL_524055, EPI_ISL_524056, EPI_ISL_524057, EPI_ISL_524058, EPI_ISL_524059, EPI_ISL_524060, EPI_ISL_524061, EPI_ISL_524062, EPI_ISL_524063, EPI_ISL_524064, EPI_ISL_524065, EPI_ISL_524066, EPI_ISL_524067, EPI_ISL_524068, EPI_ISL_524069, EPI_ISL_524070, EPI_ISL_524071, EPI_ISL_524072, EPI_ISL_524073 | | | |
| see above | Texas Department of State Health Services | Texas Department of State Health Services | Rashmi Tuladhar, Bonnie Oh, Cara Akrout, Jenny Zhang, Maliha Rahman, Anita Pokharel, Myong Koag, Chun Wang, Rachel Lee, Grace Kubin |
| EPI_ISL_524437, EPI_ISL_524438, EPI_ISL_524439, EPI_ISL_524440, EPI_ISL_524441, EPI_ISL_524442, EPI_ISL_524443 | University of Washington Virology Lab | Laboratory Medicine, University of Washington | Roychoudhury,P., Greninger,A., Jerome,K. |
| EPI_ISL_524444 | University of Washington Virology Lab | Fred Hutchinson Cancer Research Center | Roychoudhury,P., Greninger,A., Jerome,K. |

| | | | |
|---|---|---|---|
| EPI_ISL_524721 | B.J. Medical College and Civil hospital, Ahmedabad | Gujarat Biotechnology Research Centre | Raghawendra Kumar, Dinesh Kumar, Zuber Saiyed, Komal Patel, Labdhi Pandya, Afzal Ansari, Nikha Trivedi, Pranay Shah, Kamlesh J Upadhyay, Sanjay Kapadia, Apurvasinh Puvar, Janvi Raval, Zarna Patel, Monika Gandhi, Pinal Trivedi, Maharshi Pandya, Nidhi Patel, Nitin Savaliya, R D Dixit, A M Kadri, Harsh Bakshi, Chaitanya Joshi, Madhvi Joshi |
| EPI_ISL_524757 | GMERS Medical College and Hospital, Dharpur, Patan | Gujarat Biotechnology Research Centre | Raghawendra Kumar, Dinesh Kumar, Zuber Saiyed, Komal Patel, Labdhi Pandya, Afzal Ansari, Nikha Trivedi, A N Parmar, Apurvasinh Puvar, Janvi Raval, Zarna Patel, Monika Gandhi, Pinal Trivedi, Maharshi Pandya, Nidhi Patel, Nitin Savaliya, R D Dixit, A M Kadri, Harsh Bakshi, Chaitanya Joshi, Madhvi Joshi |
| EPI_ISL_525420 | GMERS Medical College and Hospital, Gandhinagar | Gujarat Biotechnology Research Centre | Raghawendra Kumar, Dinesh Kumar, Zuber Saiyed, Komal Patel, Labdhi Pandya, Afzal Ansari, Nikha Trivedi, Seema Bhatt, Gaurishankar Shrimali, Bhavesh Modi, Bharti Rajani, Apurvasinh Puvar, Janvi Raval, Zarna Patel, Monika Gandhi, Pinal Trivedi, Maharshi Pandya, Nidhi Patel, Nitin Savaliya, R D Dixit, A M Kadri, Harsh Bakshi, Chaitanya Joshi, Madhvi Joshi |
| EPI_ISL_525467 | Universidad Iberoamericana | International Centre for Genetic Engineering and Biotechnology (ICGEB) and ARGO Open Lab Platform | Robert Paulino-Ramirez, Eileen Riego, Alejandro Vallejo Degaudenzi, Victor Virgilio Calderon, Leandro Tapia, Patricia Leon, Danilo Licastro, Simeone Dal Monego, Sreejith Rajasekharan and Alessandro Marcello. |
| EPI_ISL_525468 | Universidad Iberoamericana | International Centre for Genetic Engineering and Biotechnology (ICGEB) and ARGO Open Lab Platform | Robert Paulino-Ramirez, Eileen Riego, Alejandro Vallejo Degaudenzi, Victor Virgilio Calderon, Leandro Tapia,Patricia Leon,Danilo Licastro, Simeone Dal Monego, Sreejith Rajasekharan and Alessandro Marcello. |
| EPI_ISL_525469 | Universidad Iberoamericana | International Centre for Genetic Engineering and Biotechnology (ICGEB) and ARGO Open Lab Platform | Robert Paulino-Ramirez, Eileen Riego, Alejandro Vallejo Degaudenzi, Victor Virgilio Calderon, Leandro Tapia, Patricia Leon, Danilo Licastro, Simeone Dal Monego, Sreejith Rajasekharan and Alessandro Marcello. |
| EPI_ISL_526253, EPI_ISL_526254, EPI_ISL_526255, EPI_ISL_526256, EPI_ISL_526257, EPI_ISL_526258, EPI_ISL_526259, EPI_ISL_526260, EPI_ISL_526261, EPI_ISL_526262, EPI_ISL_526263, EPI_ISL_526264, EPI_ISL_526265, EPI_ISL_526266, EPI_ISL_526267, EPI_ISL_526268, EPI_ISL_526269, EPI_ISL_526270, EPI_ISL_526271, EPI_ISL_526272, EPI_ISL_526273, EPI_ISL_526274, EPI_ISL_526275, EPI_ISL_526276, EPI_ISL_526277, EPI_ISL_526278, EPI_ISL_526279, EPI_ISL_526280, EPI_ISL_526281, EPI_ISL_526282, EPI_ISL_526283, EPI_ISL_526284, EPI_ISL_526285, EPI_ISL_526286 | | | |
| see above | Unity Health Toronto | Ontario Institute for Cancer Research | Ramzi Fattouh, Larissa M. Matukas, Mark Downing, Annette Gower, Karel Boissinot, Samira Mubareka, TIBDN, Ilinca Lungu, Bernard Lam, Jeremy Johns, Paul Krzyzanowski, Richard de Borja, Felicia Vincelli, Philip Zuzarte, Jared Simpson |
| EPI_ISL_527008, EPI_ISL_527009, EPI_ISL_527010, EPI_ISL_527011, EPI_ISL_527012, EPI_ISL_527013, EPI_ISL_527014, EPI_ISL_527015, EPI_ISL_527016, EPI_ISL_527017, EPI_ISL_527018, EPI_ISL_527019, EPI_ISL_527020, EPI_ISL_527021, EPI_ISL_527022, EPI_ISL_527023, EPI_ISL_527024, EPI_ISL_527025, EPI_ISL_527026, EPI_ISL_527027, EPI_ISL_527028, EPI_ISL_527029, EPI_ISL_527030, EPI_ISL_527031, EPI_ISL_527032, EPI_ISL_527033, EPI_ISL_527034, EPI_ISL_527035, EPI_ISL_527036, EPI_ISL_527037, EPI_ISL_527038, EPI_ISL_527039, EPI_ISL_527040, EPI_ISL_527041, EPI_ISL_527042, EPI_ISL_527043, EPI_ISL_527044, EPI_ISL_527045, EPI_ISL_527046, EPI_ISL_527047, EPI_ISL_527048, EPI_ISL_527049, EPI_ISL_527050, EPI_ISL_527051, EPI_ISL_527052, EPI_ISL_527053, EPI_ISL_527054, EPI_ISL_527055, EPI_ISL_527056, EPI_ISL_527057, EPI_ISL_527058, EPI_ISL_527059, EPI_ISL_527060, EPI_ISL_527061, EPI_ISL_527062, EPI_ISL_527063, EPI_ISL_527064, EPI_ISL_527179, EPI_ISL_527180 | | | |
| see above | Area of Virology, Serology and Virology Division (SAViD), New South Wales Health Pathology Randwick | Area of Virology, Serology and Virology Division (SAViD), New South Wales Health Pathology Randwick | Rawlinson, W. |
| EPI_ISL_527400 | Area of Virology, Serology and Virology Division (SAViD), New South Wales Health Pathology Randwick | Area of Virology, Serology and Virology Division (SAViD), New South Wales Health Pathology Randwick | Rawlinson, W., Deveson, I., Bull, R. |
| EPI_ISL_528421, EPI_ISL_528422, EPI_ISL_528423 | TNMC & BYL NAIR CH. HOSPITAL | Institute of Genomics and Integrative Biology - Council of Scientific and Industrial Research | Rajesh Pandey, Jayanthi Shastri, Akshay Kanakan, Vivekanand A, Janani Srinivasa Vasudevan, Ranjeet Maurya, Sachee Agrawal, Nirjhar Chatterjee, Swapneil Parikh, Manish Pathak, Subrat Thanapati, Jasmina Savak, Suresh Poojari, Mahesh Sangar, Amol Borse, Shweta Kawankar, Vasil Nachan, Mayuresh Vishwanathan, Shruthi Sachidanandan, Shrutika Pophale, Utkarsha Yelve |
| EPI_ISL_528425, EPI_ISL_528426 | P. D. Hinduja Hospital and Medical Research Centre | Institute of Genomics and Integrative Biology - Council of Scientific and Industrial Research | Rajesh Pandey, Jayanthi Shastri, Akshay Kanakan, Vivekanand A, Janani Srinivasa Vasudevan, Ranjeet Maurya, Sachee Agrawal, Nirjhar Chatterjee, Swapneil Parikh, Manish Pathak, Subrat Thanapati, Jasmina Savak, Suresh Poojari, Mahesh Sangar, Amol Borse, Shweta Kawankar, Vasil Nachan, Mayuresh Vishwanathan, Shruthi Sachidanandan, Shrutika Pophale, Utkarsha Yelve |
| EPI_ISL_528809, EPI_ISL_528814, EPI_ISL_528816, EPI_ISL_528819, EPI_ISL_528820 | Department of Medicine, Gandhi hospital, Hyderabad | CSIR-Centre for Cellular and Molecular Biology | Rajarao Mesipogu , Thrilok Chander Bingi ,Vinayasekhar Aedula,Tulasi Nagabandi, Namami Gaur, Sakshi Shambhavi, Lamuk Zaveri, Shagufta Khan, Nikhil Hajirnis, M Soujanya Reddy, Pratheusa Maccha, Purushotham Vodnala, Payel Mukherjee, Sofia Banu, Priya Singh, Onkar Kulkarni, Dhiviya Vedagiri, Divya Gupta, Vishal Sah, Santosh Kumar Kuncha, Krishnan Harinivas Harshan, Archana Bharadwaj Siva, Karthik Bharadwaj Tallapaka,G. Aditya Kumar, Koushick Sivakumar, Pooja Ramesh Gupta, Rajan Kumar Jha, Shraddha Vijay Lahoti, Rakesh K Mishra, Divya Tej Sowpati |
| EPI_ISL_529155, EPI_ISL_529156, EPI_ISL_529157, EPI_ISL_529158, EPI_ISL_529159 | University of Washington, Laboratory Medicine | University of Washington, Laboratory Medicine | Roychoudhury,P., Greninger,A., Jerome,K. |
| EPI_ISL_529160, EPI_ISL_529161 | University of Washington, Laboratory Medicine | University of Washington, Laboratory Medicine | Roychoudhury, P., Greninger, A., Jerome, K. |
| EPI_ISL_529170, EPI_ISL_529200 | University of Washington, Laboratory Medicine | University of Washington, Laboratory Medicine | Roychoudhury,P., Greninger,A., Jerome,K. |
| EPI_ISL_529208, EPI_ISL_529209, EPI_ISL_529210, EPI_ISL_529211, EPI_ISL_529212 | Laboratory Medicine, University of Washington | University of Washington, Laboratory Medicine | Roychoudhury,P., Greninger,A., Jerome,K. |
| EPI_ISL_529963 | Universitas Airlangga Hospital | Institute of Tropical Disease, Universitas Airlangga | Rima R Prasetya, Krisnoadi Rahardjo, Aldise M Nastri, Jezzy R Dewantari, Nasronudin, Gatot Soegiarto, Laksmi Wulandari, Retno A Setyoningrum, Resti Yudhawati, Yohko K Shimizu, Mitsuhiro Nishimura, Yasuko Mori, Soetjipto, Kazufumi Shimizu, Maria I Lusida |
| EPI_ISL_529972, EPI_ISL_529973, EPI_ISL_529974, EPI_ISL_529975, EPI_ISL_529976, EPI_ISL_529977, EPI_ISL_529978, EPI_ISL_529979, EPI_ISL_529980, EPI_ISL_529981, EPI_ISL_529982, EPI_ISL_529983, EPI_ISL_529984, EPI_ISL_529985, EPI_ISL_529986, EPI_ISL_529987, EPI_ISL_529988 | | | |
| see above | Hospital Universitario 12 de Octubre | Hospital Universitario 12 de Octubre | Raúl Recio, Sara González, Esther Viedma, Elias Dahdouh, Fernando Lázaro, Natalia Stella, Julio García, Juan Carlos Galán, Rafael Cantón, Mª Dolores Folgueira, Rafael Delgado, Jesús Mingorance |
| EPI_ISL_530094, EPI_ISL_530095, EPI_ISL_530096, EPI_ISL_530097, EPI_ISL_530098, EPI_ISL_530099 | Hospital Universitario Ramón y Cajal | Hospital Universitario 12 de Octubre | Raúl Recio, Sara González, Elias Dahdouh, Fernando Lázaro, Esther Viedma, Natalia Stella, Julio García, Juan Carlos Galán, Rafael Cantón, Mª Dolores Folgueira, Rafael Delgado, Jesús Mingorance |
| EPI_ISL_530100, EPI_ISL_530101, EPI_ISL_530102, EPI_ISL_530103, EPI_ISL_530104, EPI_ISL_530105, EPI_ISL_530106, EPI_ISL_530107, EPI_ISL_530108, EPI_ISL_530109, EPI_ISL_530110, EPI_ISL_530111, EPI_ISL_530112, EPI_ISL_530113 | | | |
| see above | Hospital Universitario Ramón y Cajal | Hospital Universitario La Paz | Raúl Recio, Sara González, Elias Dahdouh, Fernando Lázaro, Esther Viedma, Natalia Stella, Julio García, Juan Carlos Galán, Rafael Cantón, Mª Dolores Folgueira, Rafael Delgado, Jesús Mingorance |
| EPI_ISL_530334, EPI_ISL_530335, EPI_ISL_530336, EPI_ISL_530337, EPI_ISL_530338, EPI_ISL_530339, EPI_ISL_530340 | Area of Virology, Serology and Virology Division (SAViD), New South Wales Health Pathology Randwick | Area of Virology, Serology and Virology Division (SAViD), New South Wales Health Pathology Randwick | Rawlinson, W., Deveson, I., Bull, R., Van Hal, S. |
| EPI_ISL_534260, EPI_ISL_534261, EPI_ISL_534262, EPI_ISL_534263, EPI_ISL_534264, EPI_ISL_534265, EPI_ISL_534266, EPI_ISL_534267, EPI_ISL_534268, EPI_ISL_534269, EPI_ISL_534270, EPI_ISL_534271, EPI_ISL_534272, EPI_ISL_534273, EPI_ISL_534274, EPI_ISL_534275, EPI_ISL_534276, EPI_ISL_534277, EPI_ISL_534278, EPI_ISL_534279, EPI_ISL_534280, EPI_ISL_534281, EPI_ISL_534282, EPI_ISL_534283, EPI_ISL_534284, EPI_ISL_534285, EPI_ISL_534286, EPI_ISL_534287, EPI_ISL_534288, EPI_ISL_534289, EPI_ISL_534290, EPI_ISL_534291, EPI_ISL_534292, EPI_ISL_534293, EPI_ISL_534294, EPI_ISL_534295, EPI_ISL_534296, EPI_ISL_534297, EPI_ISL_534298, EPI_ISL_534299, EPI_ISL_534300, EPI_ISL_534301, EPI_ISL_534302 | | | |
| see above | Texas Department of State Health Services | Texas Department of State Health Services | Rashmi Tuladhar, Bonnie Oh, Jenny Zhang, Maliha Rahman, Anita Pokharel, Myong Koag, Chun Wang, Rachel Lee, Grace Kubin |
| EPI_ISL_534327, EPI_ISL_534328 | Hospital Universitario 12 de Octubre | Hospital Universitario 12 de Octubre | Raúl Recio, Sara González, Esther Viedma, Elias Dahdouh, Fernando Lázaro, Natalia Stella, Julio García, Juan Carlos Galán, Rafael Cantón, Ma Dolores Folgueira, Rafael Delgado, Jesús Mingorance |
| EPI_ISL_537286, EPI_ISL_537287 | Area of Virology, Serology and Virology Division (SAViD), New South Wales Health Pathology Randwick | Area of Virology, Serology and Virology Division (SAViD), New South Wales Health Pathology Randwick | Rawlinson, W., Deveson, I., Bull, R., Van Hal, S. |
| EPI_ISL_537946, EPI_ISL_537947, EPI_ISL_537948, EPI_ISL_537949, EPI_ISL_537950, EPI_ISL_537951, EPI_ISL_537952, EPI_ISL_537953, EPI_ISL_537954, EPI_ISL_537955, EPI_ISL_537956, EPI_ISL_537957, EPI_ISL_537958, EPI_ISL_537959, EPI_ISL_537960, EPI_ISL_537961, EPI_ISL_537962, EPI_ISL_537963, EPI_ISL_537964, EPI_ISL_537965, EPI_ISL_537966, EPI_ISL_537967, EPI_ISL_537968, EPI_ISL_537969, EPI_ISL_537970, EPI_ISL_537971, EPI_ISL_537972, EPI_ISL_537973, EPI_ISL_537974, EPI_ISL_537975, EPI_ISL_537976, EPI_ISL_537977, EPI_ISL_537978, EPI_ISL_537979, EPI_ISL_537980, EPI_ISL_537981, EPI_ISL_537982, EPI_ISL_537983, EPI_ISL_537984, EPI_ISL_537985, EPI_ISL_537986, EPI_ISL_537987, EPI_ISL_537988, EPI_ISL_537989, EPI_ISL_537990, EPI_ISL_537991, EPI_ISL_537992, EPI_ISL_537993, EPI_ISL_537994, EPI_ISL_537995, EPI_ISL_538613, EPI_ISL_538614, EPI_ISL_538615, EPI_ISL_538616, EPI_ISL_538617, EPI_ISL_538618, EPI_ISL_538619, EPI_ISL_538620, EPI_ISL_538621, EPI_ISL_538622, EPI_ISL_538623, EPI_ISL_538624, EPI_ISL_538625, EPI_ISL_538626, EPI_ISL_538627, EPI_ISL_538628, EPI_ISL_538629, EPI_ISL_538630, EPI_ISL_538631, EPI_ISL_538632, EPI_ISL_538633, EPI_ISL_538634, | | | |

EPI_ISL_538635, EPI_ISL_538636, EPI_ISL_538637, EPI_ISL_538638, EPI_ISL_538639, EPI_ISL_538640, EPI_ISL_538641, EPI_ISL_538642, EPI_ISL_538643, EPI_ISL_538644, EPI_ISL_538645, EPI_ISL_538646, EPI_ISL_538647, EPI_ISL_538648, EPI_ISL_538649, EPI_ISL_538650, EPI_ISL_538651, EPI_ISL_538652, EPI_ISL_538653, EPI_ISL_538654, EPI_ISL_538655, EPI_ISL_538656, EPI_ISL_538657, EPI_ISL_538658, EPI_ISL_538659, EPI_ISL_538660, EPI_ISL_538661, EPI_ISL_538662, EPI_ISL_538663, EPI_ISL_538664, EPI_ISL_538665, EPI_ISL_538666, EPI_ISL_538667, EPI_ISL_538668, EPI_ISL_538669, EPI_ISL_538670, EPI_ISL_538671, EPI_ISL_539284, EPI_ISL_539285, EPI_ISL_539286, EPI_ISL_539287

| | | | |
|---|---|---|---|
| see above | Servicio de Microbiología. Hospital General Universitario de Castellón | SeqCOVID-SPAIN consortium/IBV(CSIC) | Rosario Moreno, María Dolores Tirado and SeqCOVID-SPAIN consortium |
| EPI_ISL_539327 | Area of Virology, Serology and Virology Division (SAViD), New South Wales Health Pathology Randwick | Area of Virology, Serology and Virology Division (SAViD), New South Wales Health Pathology Randwick | Rawlinson, W., Bull, R., Deveson, I., Van Hal, S. |

EPI_ISL_541971, EPI_ISL_541972, EPI_ISL_541973, EPI_ISL_541974, EPI_ISL_541975, EPI_ISL_541976, EPI_ISL_541977, EPI_ISL_541978, EPI_ISL_541979, EPI_ISL_541980, EPI_ISL_541981, EPI_ISL_541982, EPI_ISL_541983, EPI_ISL_541984, EPI_ISL_541985, EPI_ISL_541986, EPI_ISL_541987, EPI_ISL_541988, EPI_ISL_541989, EPI_ISL_541990, EPI_ISL_541991, EPI_ISL_541992, EPI_ISL_541993, EPI_ISL_541994, EPI_ISL_541995, EPI_ISL_541996, EPI_ISL_541997, EPI_ISL_541998, EPI_ISL_541999, EPI_ISL_542000, EPI_ISL_542001, EPI_ISL_542002, EPI_ISL_542003, EPI_ISL_542004, EPI_ISL_542005, EPI_ISL_542006, EPI_ISL_542007, EPI_ISL_542008, EPI_ISL_542009, EPI_ISL_542010, EPI_ISL_542011

| | | | |
|---|---|---|---|
| see above | Texas Department of State Health Services | Texas Department of State Health Services | Rashmi Tuladhar, Bonnie Oh, Jenny Zhang, Maliha Rahman, Anita Pokharel, Myong Koag, Chun Wang, Rachel Lee, Grace Kubin |

EPI_ISL_542444, EPI_ISL_542445, EPI_ISL_542446, EPI_ISL_542447, EPI_ISL_542448, EPI_ISL_542449, EPI_ISL_542450, EPI_ISL_542451, EPI_ISL_542452, EPI_ISL_542453, EPI_ISL_542454, EPI_ISL_542455, EPI_ISL_542456, EPI_ISL_542457, EPI_ISL_542458, EPI_ISL_542459, EPI_ISL_542460, EPI_ISL_542461, EPI_ISL_542462, EPI_ISL_542463, EPI_ISL_542464, EPI_ISL_542465, EPI_ISL_542466, EPI_ISL_542467, EPI_ISL_542468, EPI_ISL_542469, EPI_ISL_542470, EPI_ISL_542471, EPI_ISL_542472, EPI_ISL_542473, EPI_ISL_542474, EPI_ISL_542475, EPI_ISL_542476, EPI_ISL_542477, EPI_ISL_542478, EPI_ISL_542479, EPI_ISL_542480, EPI_ISL_542481, EPI_ISL_542482, EPI_ISL_542483, EPI_ISL_542484, EPI_ISL_542485

| | | | |
|---|---|---|---|
| see above | Texas Department of State Health Services | Texas Department of State Health Services | Rashmi Tuladhar, Bonnie Oh,Jenny Zhang, Maliha Rahman, Anita Pokharel, Myong Koag, Chun Wang, Rachel Lee, Grace Kubin |

EPI_ISL_542487, EPI_ISL_542488, EPI_ISL_542489, EPI_ISL_542490, EPI_ISL_542491, EPI_ISL_542492, EPI_ISL_542493, EPI_ISL_542494, EPI_ISL_542495, EPI_ISL_542496, EPI_ISL_542497, EPI_ISL_542498, EPI_ISL_542500, EPI_ISL_542501, EPI_ISL_542502, EPI_ISL_542503, EPI_ISL_542504, EPI_ISL_542505,
EPI_ISL_542506, EPI_ISL_542507, EPI_ISL_542509, EPI_ISL_542510, EPI_ISL_542511, EPI_ISL_542512, EPI_ISL_542513, EPI_ISL_542514, EPI_ISL_542515, EPI_ISL_542516, EPI_ISL_542517, EPI_ISL_542518, EPI_ISL_542519, EPI_ISL_542520, EPI_ISL_542521, EPI_ISL_542522, EPI_ISL_542524, EPI_ISL_542526,
EPI_ISL_542528, EPI_ISL_542530, EPI_ISL_542532, EPI_ISL_542533, EPI_ISL_542534, EPI_ISL_542535, EPI_ISL_542536, EPI_ISL_542537, EPI_ISL_542539, EPI_ISL_542542, EPI_ISL_542543, EPI_ISL_542544, EPI_ISL_542546, EPI_ISL_542547, EPI_ISL_542549, EPI_ISL_542551, EPI_ISL_542553, EPI_ISL_542556,
EPI_ISL_542557, EPI_ISL_542561, EPI_ISL_542562, EPI_ISL_542564, EPI_ISL_542565, EPI_ISL_542568, EPI_ISL_542570, EPI_ISL_542571, EPI_ISL_542572, EPI_ISL_542573, EPI_ISL_542574, EPI_ISL_542575, EPI_ISL_542576, EPI_ISL_542577, EPI_ISL_542578, EPI_ISL_542579, EPI_ISL_542580, EPI_ISL_542581,
EPI_ISL_542582, EPI_ISL_542583, EPI_ISL_542584, EPI_ISL_542586, EPI_ISL_542587, EPI_ISL_542588, EPI_ISL_542589, EPI_ISL_542590, EPI_ISL_542592, EPI_ISL_542593, EPI_ISL_542594, EPI_ISL_542595, EPI_ISL_542596, EPI_ISL_542597, EPI_ISL_542598, EPI_ISL_542599, EPI_ISL_542600, EPI_ISL_542601,
EPI_ISL_542602, EPI_ISL_542603, EPI_ISL_542604, EPI_ISL_542605, EPI_ISL_542606, EPI_ISL_542607, EPI_ISL_542609, EPI_ISL_542610, EPI_ISL_542611, EPI_ISL_542612, EPI_ISL_542613, EPI_ISL_542614, EPI_ISL_542615, EPI_ISL_542617, EPI_ISL_542618, EPI_ISL_542620, EPI_ISL_542621, EPI_ISL_542622,
EPI_ISL_542623, EPI_ISL_542624, EPI_ISL_542626, EPI_ISL_542628, EPI_ISL_542629, EPI_ISL_543284, EPI_ISL_543285, EPI_ISL_543292, EPI_ISL_543294, EPI_ISL_543298, EPI_ISL_543301, EPI_ISL_543303, EPI_ISL_543304, EPI_ISL_543305, EPI_ISL_543306, EPI_ISL_543308, EPI_ISL_543311, EPI_ISL_543312,
EPI_ISL_543313, EPI_ISL_543314, EPI_ISL_543318, EPI_ISL_543319, EPI_ISL_543320, EPI_ISL_543321, EPI_ISL_543322, EPI_ISL_543324, EPI_ISL_543325, EPI_ISL_543326, EPI_ISL_543328, EPI_ISL_543329, EPI_ISL_543332, EPI_ISL_543335, EPI_ISL_543336, EPI_ISL_543338, EPI_ISL_543341, EPI_ISL_543343,
EPI_ISL_543346, EPI_ISL_543348, EPI_ISL_543354, EPI_ISL_543358, EPI_ISL_543359, EPI_ISL_543360, EPI_ISL_543364, EPI_ISL_543367, EPI_ISL_543368, EPI_ISL_543369, EPI_ISL_543371, EPI_ISL_543372, EPI_ISL_543374, EPI_ISL_543375, EPI_ISL_543382, EPI_ISL_543383, EPI_ISL_543386, EPI_ISL_543387,
EPI_ISL_543433, EPI_ISL_543434, EPI_ISL_543435, EPI_ISL_543438, EPI_ISL_543439, EPI_ISL_543440, EPI_ISL_543441, EPI_ISL_543442, EPI_ISL_543456, EPI_ISL_543460, EPI_ISL_543461, EPI_ISL_543469, EPI_ISL_543470, EPI_ISL_543471, EPI_ISL_543474, EPI_ISL_543476, EPI_ISL_543478, EPI_ISL_543479,
EPI_ISL_543487, EPI_ISL_543490, EPI_ISL_543491, EPI_ISL_543493, EPI_ISL_543495, EPI_ISL_543497, EPI_ISL_543498, EPI_ISL_543502, EPI_ISL_543504, EPI_ISL_543506, EPI_ISL_543508, EPI_ISL_543509, EPI_ISL_543511, EPI_ISL_543512, EPI_ISL_543513, EPI_ISL_543515, EPI_ISL_543519, EPI_ISL_543521,
EPI_ISL_543524, EPI_ISL_543525, EPI_ISL_543527, EPI_ISL_543529, EPI_ISL_543530, EPI_ISL_543533, EPI_ISL_543534, EPI_ISL_543537, EPI_ISL_543538, EPI_ISL_543540, EPI_ISL_543541, EPI_ISL_543544, EPI_ISL_543545, EPI_ISL_543546, EPI_ISL_543547, EPI_ISL_543549, EPI_ISL_543551, EPI_ISL_543552,
EPI_ISL_543553, EPI_ISL_543555, EPI_ISL_543558, EPI_ISL_543565, EPI_ISL_543567, EPI_ISL_543568, EPI_ISL_543569, EPI_ISL_543571, EPI_ISL_543572, EPI_ISL_543574, EPI_ISL_543575, EPI_ISL_543582, EPI_ISL_543583, EPI_ISL_543585, EPI_ISL_543590, EPI_ISL_543591, EPI_ISL_543597, EPI_ISL_543601,
EPI_ISL_543602, EPI_ISL_543605, EPI_ISL_543606, EPI_ISL_543607, EPI_ISL_543608, EPI_ISL_543611, EPI_ISL_543612, EPI_ISL_543615, EPI_ISL_543619, EPI_ISL_543621, EPI_ISL_543631, EPI_ISL_543635, EPI_ISL_543655, EPI_ISL_543658, EPI_ISL_543662, EPI_ISL_543668, EPI_ISL_543671, EPI_ISL_543672,
EPI_ISL_543673, EPI_ISL_543674, EPI_ISL_543677, EPI_ISL_543687, EPI_ISL_543691, EPI_ISL_543692, EPI_ISL_543695, EPI_ISL_543696, EPI_ISL_543697, EPI_ISL_543698, EPI_ISL_543699, EPI_ISL_543700, EPI_ISL_543703, EPI_ISL_543710, EPI_ISL_543711, EPI_ISL_543712, EPI_ISL_543715, EPI_ISL_543717,
EPI_ISL_543723, EPI_ISL_543724, EPI_ISL_543725, EPI_ISL_543729, EPI_ISL_543734, EPI_ISL_543735, EPI_ISL_543737, EPI_ISL_543744, EPI_ISL_543753, EPI_ISL_543755, EPI_ISL_543756, EPI_ISL_543759, EPI_ISL_543762, EPI_ISL_543763, EPI_ISL_543767, EPI_ISL_543768, EPI_ISL_543770, EPI_ISL_543773,
EPI_ISL_543776, EPI_ISL_543786, EPI_ISL_544240, EPI_ISL_544242, EPI_ISL_544243, EPI_ISL_544245, EPI_ISL_544246, EPI_ISL_544247, EPI_ISL_544249, EPI_ISL_544251, EPI_ISL_544255, EPI_ISL_544259, EPI_ISL_544264, EPI_ISL_544265, EPI_ISL_544268, EPI_ISL_544271, EPI_ISL_544275, EPI_ISL_544278,
EPI_ISL_544279, EPI_ISL_544281, EPI_ISL_544282, EPI_ISL_544283, EPI_ISL_544285, EPI_ISL_544296, EPI_ISL_544309, EPI_ISL_544311, EPI_ISL_544314, EPI_ISL_544318, EPI_ISL_544325, EPI_ISL_544332, EPI_ISL_544339, EPI_ISL_544345, EPI_ISL_544350, EPI_ISL_544351, EPI_ISL_544352, EPI_ISL_544353,
EPI_ISL_544354, EPI_ISL_544355, EPI_ISL_544356, EPI_ISL_544357, EPI_ISL_544359, EPI_ISL_544360, EPI_ISL_544362, EPI_ISL_544365, EPI_ISL_544366, EPI_ISL_544367, EPI_ISL_544368, EPI_ISL_544369, EPI_ISL_544370, EPI_ISL_544371, EPI_ISL_544372, EPI_ISL_544373, EPI_ISL_544374, EPI_ISL_544376,
EPI_ISL_544377, EPI_ISL_544381, EPI_ISL_544382, EPI_ISL_544384, EPI_ISL_544386, EPI_ISL_544387, EPI_ISL_544389, EPI_ISL_544390, EPI_ISL_544391, EPI_ISL_544392, EPI_ISL_544393, EPI_ISL_544394, EPI_ISL_544395, EPI_ISL_544396, EPI_ISL_544397, EPI_ISL_544398, EPI_ISL_544402, EPI_ISL_544404,
EPI_ISL_544406, EPI_ISL_544407, EPI_ISL_544408, EPI_ISL_544409, EPI_ISL_544410, EPI_ISL_544412, EPI_ISL_544415, EPI_ISL_544416, EPI_ISL_544421, EPI_ISL_544422, EPI_ISL_544423, EPI_ISL_544424, EPI_ISL_544425, EPI_ISL_544429, EPI_ISL_544430, EPI_ISL_544433, EPI_ISL_544434, EPI_ISL_544439,
EPI_ISL_544440, EPI_ISL_544454, EPI_ISL_544456, EPI_ISL_544457, EPI_ISL_544460, EPI_ISL_544461, EPI_ISL_544462, EPI_ISL_544465, EPI_ISL_544466, EPI_ISL_544469, EPI_ISL_544470, EPI_ISL_544471, EPI_ISL_544473, EPI_ISL_544474, EPI_ISL_544475, EPI_ISL_544476, EPI_ISL_544478, EPI_ISL_544479,
EPI_ISL_544480, EPI_ISL_544481, EPI_ISL_544482, EPI_ISL_544483, EPI_ISL_544486, EPI_ISL_544487, EPI_ISL_544489, EPI_ISL_544491, EPI_ISL_544492, EPI_ISL_544495, EPI_ISL_544496, EPI_ISL_544497, EPI_ISL_544498, EPI_ISL_544501, EPI_ISL_544502, EPI_ISL_544503, EPI_ISL_544504, EPI_ISL_544505,
EPI_ISL_544506, EPI_ISL_544507, EPI_ISL_544508, EPI_ISL_544511, EPI_ISL_544512, EPI_ISL_544514, EPI_ISL_544517, EPI_ISL_544518, EPI_ISL_544519, EPI_ISL_544522, EPI_ISL_544523, EPI_ISL_544524, EPI_ISL_544527, EPI_ISL_544528, EPI_ISL_544532, EPI_ISL_544533, EPI_ISL_544534, EPI_ISL_544538,
EPI_ISL_544539, EPI_ISL_544543, EPI_ISL_544545, EPI_ISL_544548, EPI_ISL_544551, EPI_ISL_544552, EPI_ISL_544558, EPI_ISL_544561, EPI_ISL_544562, EPI_ISL_544565, EPI_ISL_544568, EPI_ISL_544569, EPI_ISL_544572, EPI_ISL_544575, EPI_ISL_544576, EPI_ISL_544578, EPI_ISL_544579, EPI_ISL_544580,
EPI_ISL_544585, EPI_ISL_544588, EPI_ISL_544595, EPI_ISL_544601, EPI_ISL_544604, EPI_ISL_544607, EPI_ISL_544611, EPI_ISL_544613, EPI_ISL_544620, EPI_ISL_544623, EPI_ISL_544630, EPI_ISL_544635, EPI_ISL_544639, EPI_ISL_544641, EPI_ISL_544642, EPI_ISL_544645, EPI_ISL_544647, EPI_ISL_544649,
EPI_ISL_544651, EPI_ISL_544656, EPI_ISL_544658, EPI_ISL_544659, EPI_ISL_544661, EPI_ISL_544664, EPI_ISL_544673, EPI_ISL_544676, EPI_ISL_544677, EPI_ISL_544680, EPI_ISL_544681, EPI_ISL_544682, EPI_ISL_544683, EPI_ISL_544684, EPI_ISL_544687, EPI_ISL_544695, EPI_ISL_544697, EPI_ISL_544698,
EPI_ISL_544700, EPI_ISL_544702, EPI_ISL_544703, EPI_ISL_544705, EPI_ISL_544706, EPI_ISL_544707, EPI_ISL_544708, EPI_ISL_544711, EPI_ISL_544713, EPI_ISL_544714, EPI_ISL_544715, EPI_ISL_544716, EPI_ISL_544721, EPI_ISL_544722, EPI_ISL_544723, EPI_ISL_544724, EPI_ISL_544725, EPI_ISL_544726,
EPI_ISL_544727, EPI_ISL_544730, EPI_ISL_544733, EPI_ISL_544734, EPI_ISL_544737, EPI_ISL_544738, EPI_ISL_544741, EPI_ISL_544742, EPI_ISL_544748, EPI_ISL_544755, EPI_ISL_544756, EPI_ISL_544759, EPI_ISL_544760, EPI_ISL_544761, EPI_ISL_544762, EPI_ISL_544764, EPI_ISL_544766, EPI_ISL_544767,
EPI_ISL_544772, EPI_ISL_544773, EPI_ISL_544775, EPI_ISL_544781, EPI_ISL_544784, EPI_ISL_544795, EPI_ISL_544796, EPI_ISL_544797, EPI_ISL_544798, EPI_ISL_544799, EPI_ISL_544800, EPI_ISL_544805, EPI_ISL_544806, EPI_ISL_544807, EPI_ISL_544812, EPI_ISL_544815, EPI_ISL_544820, EPI_ISL_544822,
EPI_ISL_544830, EPI_ISL_544835, EPI_ISL_544838, EPI_ISL_544839, EPI_ISL_544840, EPI_ISL_544841, EPI_ISL_544848, EPI_ISL_544850, EPI_ISL_544852, EPI_ISL_544853, EPI_ISL_544867, EPI_ISL_544868, EPI_ISL_544873, EPI_ISL_544875, EPI_ISL_544876, EPI_ISL_544879, EPI_ISL_544882, EPI_ISL_544883,
EPI_ISL_544884, EPI_ISL_544886, EPI_ISL_544889, EPI_ISL_544893, EPI_ISL_544896, EPI_ISL_544904, EPI_ISL_544905, EPI_ISL_544906, EPI_ISL_544907, EPI_ISL_544910, EPI_ISL_544912, EPI_ISL_544914, EPI_ISL_544915, EPI_ISL_544916, EPI_ISL_544917, EPI_ISL_544921, EPI_ISL_544922, EPI_ISL_544923,
EPI_ISL_544925, EPI_ISL_544928, EPI_ISL_544929, EPI_ISL_544932, EPI_ISL_544935, EPI_ISL_544936, EPI_ISL_544939, EPI_ISL_544950, EPI_ISL_545262, EPI_ISL_545582, EPI_ISL_545587, EPI_ISL_545589, EPI_ISL_545591, EPI_ISL_545595, EPI_ISL_545596, EPI_ISL_545600, EPI_ISL_545601, EPI_ISL_545606,
EPI_ISL_545608, EPI_ISL_545610, EPI_ISL_545611, EPI_ISL_545614, EPI_ISL_545615, EPI_ISL_545619, EPI_ISL_545621, EPI_ISL_545625, EPI_ISL_545626, EPI_ISL_545627, EPI_ISL_545633, EPI_ISL_545636, EPI_ISL_545637, EPI_ISL_545638, EPI_ISL_545640, EPI_ISL_545644, EPI_ISL_545645, EPI_ISL_545647,
EPI_ISL_545650, EPI_ISL_545651, EPI_ISL_545653, EPI_ISL_545656, EPI_ISL_545658, EPI_ISL_545659, EPI_ISL_545660, EPI_ISL_545661, EPI_ISL_545662, EPI_ISL_545664, EPI_ISL_545666, EPI_ISL_545667, EPI_ISL_545668, EPI_ISL_545669, EPI_ISL_545670, EPI_ISL_545671, EPI_ISL_545672, EPI_ISL_545673,
EPI_ISL_545675, EPI_ISL_545676, EPI_ISL_545677, EPI_ISL_545678, EPI_ISL_545679, EPI_ISL_545680, EPI_ISL_545681, EPI_ISL_545682, EPI_ISL_545687, EPI_ISL_545689, EPI_ISL_545691, EPI_ISL_545692, EPI_ISL_545695, EPI_ISL_545697, EPI_ISL_545698, EPI_ISL_545700, EPI_ISL_545707, EPI_ISL_545708,
EPI_ISL_545714, EPI_ISL_545719, EPI_ISL_545720, EPI_ISL_545722, EPI_ISL_545724, EPI_ISL_545728, EPI_ISL_545732, EPI_ISL_545734, EPI_ISL_545739, EPI_ISL_545744, EPI_ISL_545745, EPI_ISL_545748, EPI_ISL_545751, EPI_ISL_545752, EPI_ISL_545757, EPI_ISL_545758, EPI_ISL_545759, EPI_ISL_545760,
EPI_ISL_545766, EPI_ISL_545767, EPI_ISL_545769, EPI_ISL_545770, EPI_ISL_545771, EPI_ISL_545772, EPI_ISL_545776, EPI_ISL_545777, EPI_ISL_545778, EPI_ISL_545780, EPI_ISL_545781, EPI_ISL_545782, EPI_ISL_545783, EPI_ISL_545784, EPI_ISL_545785, EPI_ISL_545786, EPI_ISL_545788, EPI_ISL_545789,
EPI_ISL_545790, EPI_ISL_545791, EPI_ISL_545793, EPI_ISL_545794, EPI_ISL_545796, EPI_ISL_545797, EPI_ISL_545798, EPI_ISL_545800, EPI_ISL_545801, EPI_ISL_545802, EPI_ISL_545803, EPI_ISL_545804, EPI_ISL_545805, EPI_ISL_545807, EPI_ISL_545809, EPI_ISL_545811, EPI_ISL_545812, EPI_ISL_545816,
EPI_ISL_545817, EPI_ISL_545819, EPI_ISL_545822, EPI_ISL_545825, EPI_ISL_545960, EPI_ISL_545961, EPI_ISL_545963, EPI_ISL_545964, EPI_ISL_545965, EPI_ISL_545966, EPI_ISL_545967, EPI_ISL_545968, EPI_ISL_545977, EPI_ISL_545981, EPI_ISL_545982, EPI_ISL_545983, EPI_ISL_545987, EPI_ISL_545988,
EPI_ISL_545989, EPI_ISL_545994, EPI_ISL_545995, EPI_ISL_545999, EPI_ISL_546000, EPI_ISL_546005, EPI_ISL_546006, EPI_ISL_546007, EPI_ISL_546008, EPI_ISL_546009, EPI_ISL_546010, EPI_ISL_546011, EPI_ISL_546014, EPI_ISL_546017, EPI_ISL_546019, EPI_ISL_546020, EPI_ISL_546022, EPI_ISL_546025,
EPI_ISL_546026, EPI_ISL_546027, EPI_ISL_546028, EPI_ISL_546033, EPI_ISL_546036, EPI_ISL_546037, EPI_ISL_546038, EPI_ISL_546039, EPI_ISL_546040, EPI_ISL_546041, EPI_ISL_546042, EPI_ISL_546043, EPI_ISL_546046, EPI_ISL_546047, EPI_ISL_546049, EPI_ISL_546050, EPI_ISL_546052, EPI_ISL_546053,
EPI_ISL_546054, EPI_ISL_546055, EPI_ISL_546056, EPI_ISL_546058, EPI_ISL_546059, EPI_ISL_546060, EPI_ISL_546061, EPI_ISL_546062, EPI_ISL_546065, EPI_ISL_546066, EPI_ISL_546067, EPI_ISL_546069, EPI_ISL_546070, EPI_ISL_546071, EPI_ISL_546072, EPI_ISL_546080, EPI_ISL_546082, EPI_ISL_546083,
EPI_ISL_546084, EPI_ISL_546088, EPI_ISL_546091, EPI_ISL_546093, EPI_ISL_546096, EPI_ISL_546097, EPI_ISL_546098, EPI_ISL_546099, EPI_ISL_546100, EPI_ISL_546102, EPI_ISL_546105, EPI_ISL_546106, EPI_ISL_546108, EPI_ISL_546109, EPI_ISL_546111, EPI_ISL_546112, EPI_ISL_546113, EPI_ISL_546114,
EPI_ISL_546115, EPI_ISL_546116, EPI_ISL_546117, EPI_ISL_546119, EPI_ISL_546120, EPI_ISL_546121, EPI_ISL_546122, EPI_ISL_546123, EPI_ISL_546128, EPI_ISL_546129, EPI_ISL_546132, EPI_ISL_546136, EPI_ISL_546137, EPI_ISL_546138, EPI_ISL_546139, EPI_ISL_546146, EPI_ISL_546147, EPI_ISL_546148,
EPI_ISL_546178, EPI_ISL_546179, EPI_ISL_546181, EPI_ISL_546182, EPI_ISL_546183, EPI_ISL_546184, EPI_ISL_546188, EPI_ISL_546190, EPI_ISL_546193, EPI_ISL_546194, EPI_ISL_546196, EPI_ISL_546197, EPI_ISL_546200, EPI_ISL_546201, EPI_ISL_546207, EPI_ISL_546209, EPI_ISL_546213, EPI_ISL_546216,
EPI_ISL_546217, EPI_ISL_546218, EPI_ISL_546219, EPI_ISL_546220, EPI_ISL_546221, EPI_ISL_546222, EPI_ISL_546225, EPI_ISL_546227, EPI_ISL_546231, EPI_ISL_546232, EPI_ISL_546233, EPI_ISL_546234, EPI_ISL_546235, EPI_ISL_546236, EPI_ISL_546237, EPI_ISL_546240, EPI_ISL_546241, EPI_ISL_546242,
EPI_ISL_546243, EPI_ISL_546244, EPI_ISL_546245, EPI_ISL_546247, EPI_ISL_546248, EPI_ISL_546251, EPI_ISL_546252, EPI_ISL_546253, EPI_ISL_546254, EPI_ISL_546255, EPI_ISL_546256, EPI_ISL_546257, EPI_ISL_546258, EPI_ISL_546262, EPI_ISL_546264, EPI_ISL_546271, EPI_ISL_546288, EPI_ISL_546290,
EPI_ISL_546294, EPI_ISL_546295, EPI_ISL_546297, EPI_ISL_546299, EPI_ISL_546302, EPI_ISL_546305, EPI_ISL_546308, EPI_ISL_546310, EPI_ISL_546311, EPI_ISL_546312, EPI_ISL_546313, EPI_ISL_546315, EPI_ISL_546316, EPI_ISL_546319, EPI_ISL_546322, EPI_ISL_546324, EPI_ISL_546326, EPI_ISL_546328,
EPI_ISL_546332, EPI_ISL_546333, EPI_ISL_546334, EPI_ISL_546335, EPI_ISL_546336, EPI_ISL_546337, EPI_ISL_546342, EPI_ISL_546352, EPI_ISL_546361, EPI_ISL_546365, EPI_ISL_546368, EPI_ISL_546372, EPI_ISL_546373, EPI_ISL_546379, EPI_ISL_546385, EPI_ISL_546386, EPI_ISL_546390, EPI_ISL_546392,
EPI_ISL_546397, EPI_ISL_546400, EPI_ISL_546414, EPI_ISL_546423, EPI_ISL_546426, EPI_ISL_546437, EPI_ISL_546438, EPI_ISL_546440, EPI_ISL_546443, EPI_ISL_546447, EPI_ISL_546448, EPI_ISL_546450, EPI_ISL_546451, EPI_ISL_546454, EPI_ISL_546464, EPI_ISL_546465, EPI_ISL_546466, EPI_ISL_546468,
EPI_ISL_546469, EPI_ISL_546475, EPI_ISL_546476, EPI_ISL_546477, EPI_ISL_546479, EPI_ISL_546480, EPI_ISL_546482, EPI_ISL_546484, EPI_ISL_546486, EPI_ISL_546487, EPI_ISL_546488, EPI_ISL_546489, EPI_ISL_546494, EPI_ISL_546495, EPI_ISL_546498, EPI_ISL_546500, EPI_ISL_546502, EPI_ISL_546503,
EPI_ISL_546504, EPI_ISL_546505, EPI_ISL_546506, EPI_ISL_546507, EPI_ISL_546509, EPI_ISL_546510, EPI_ISL_546511, EPI_ISL_546512, EPI_ISL_546515, EPI_ISL_546517, EPI_ISL_546518, EPI_ISL_546519, EPI_ISL_546522, EPI_ISL_546524, EPI_ISL_546525, EPI_ISL_546526, EPI_ISL_546527, EPI_ISL_546529,
EPI_ISL_546531, EPI_ISL_546541, EPI_ISL_546547, EPI_ISL_546555, EPI_ISL_546556, EPI_ISL_546557, EPI_ISL_546563, EPI_ISL_546564, EPI_ISL_546565, EPI_ISL_546566, EPI_ISL_546567, EPI_ISL_546569, EPI_ISL_546570, EPI_ISL_546571, EPI_ISL_546574, EPI_ISL_546575, EPI_ISL_546585, EPI_ISL_546586,
EPI_ISL_546593, EPI_ISL_546594, EPI_ISL_546598, EPI_ISL_546599, EPI_ISL_546604, EPI_ISL_546607, EPI_ISL_546610, EPI_ISL_546613, EPI_ISL_546615, EPI_ISL_546617, EPI_ISL_546619, EPI_ISL_546621, EPI_ISL_546623, EPI_ISL_546624, EPI_ISL_546631, EPI_ISL_546632, EPI_ISL_546636, EPI_ISL_546639,
EPI_ISL_546650, EPI_ISL_546653, EPI_ISL_546656, EPI_ISL_546657, EPI_ISL_546659, EPI_ISL_546660, EPI_ISL_546667, EPI_ISL_546669, EPI_ISL_546670, EPI_ISL_546671, EPI_ISL_546672, EPI_ISL_546675, EPI_ISL_546677, EPI_ISL_546678, EPI_ISL_546680, EPI_ISL_546681, EPI_ISL_546683, EPI_ISL_546687,
EPI_ISL_546688, EPI_ISL_546689, EPI_ISL_546692, EPI_ISL_546700, EPI_ISL_546704, EPI_ISL_546706, EPI_ISL_546707, EPI_ISL_546709, EPI_ISL_546710, EPI_ISL_546711, EPI_ISL_546712, EPI_ISL_546714, EPI_ISL_546719, EPI_ISL_546722, EPI_ISL_546723, EPI_ISL_546728, EPI_ISL_546729, EPI_ISL_546730,
EPI_ISL_546731, EPI_ISL_546733, EPI_ISL_546734, EPI_ISL_546736, EPI_ISL_546738, EPI_ISL_546740, EPI_ISL_546741, EPI_ISL_546742, EPI_ISL_546743, EPI_ISL_546744, EPI_ISL_546747, EPI_ISL_546749, EPI_ISL_546750, EPI_ISL_546753, EPI_ISL_546754, EPI_ISL_546771, EPI_ISL_546775, EPI_ISL_546786,
EPI_ISL_546790, EPI_ISL_546792, EPI_ISL_546793, EPI_ISL_546794, EPI_ISL_546795, EPI_ISL_546797, EPI_ISL_546808, EPI_ISL_546810, EPI_ISL_546811, EPI_ISL_546815, EPI_ISL_546816, EPI_ISL_546819, EPI_ISL_546824, EPI_ISL_546825, EPI_ISL_546831, EPI_ISL_546833, EPI_ISL_546834, EPI_ISL_546837,
EPI_ISL_546841, EPI_ISL_546842, EPI_ISL_546862, EPI_ISL_546864, EPI_ISL_546866, EPI_ISL_546868, EPI_ISL_546869, EPI_ISL_546871, EPI_ISL_546879, EPI_ISL_546880, EPI_ISL_546885, EPI_ISL_546889, EPI_ISL_546893, EPI_ISL_546895, EPI_ISL_546900, EPI_ISL_546902, EPI_ISL_546904, EPI_ISL_546905,
EPI_ISL_546907, EPI_ISL_546908, EPI_ISL_546909, EPI_ISL_546911, EPI_ISL_546913, EPI_ISL_546914, EPI_ISL_546940, EPI_ISL_546944, EPI_ISL_546949, EPI_ISL_546950, EPI_ISL_546952, EPI_ISL_546955, EPI_ISL_546956, EPI_ISL_546957, EPI_ISL_546967, EPI_ISL_546968, EPI_ISL_546971, EPI_ISL_546973,
EPI_ISL_546974, EPI_ISL_546979, EPI_ISL_546980, EPI_ISL_546983, EPI_ISL_546984, EPI_ISL_546988, EPI_ISL_546989, EPI_ISL_546990, EPI_ISL_546991, EPI_ISL_546992, EPI_ISL_546995, EPI_ISL_546997, EPI_ISL_546999, EPI_ISL_547001, EPI_ISL_547002, EPI_ISL_547003, EPI_ISL_547004, EPI_ISL_547007,
EPI_ISL_547008, EPI_ISL_547009, EPI_ISL_547010, EPI_ISL_547011, EPI_ISL_547013, EPI_ISL_547014, EPI_ISL_547015, EPI_ISL_547016, EPI_ISL_547020, EPI_ISL_547021, EPI_ISL_547022, EPI_ISL_547027, EPI_ISL_547028, EPI_ISL_547029, EPI_ISL_547034, EPI_ISL_547037, EPI_ISL_547040, EPI_ISL_547046,
EPI_ISL_547048, EPI_ISL_547056, EPI_ISL_547069, EPI_ISL_547071, EPI_ISL_547074, EPI_ISL_547077, EPI_ISL_547079, EPI_ISL_547087, EPI_ISL_547092, EPI_ISL_547095, EPI_ISL_547097, EPI_ISL_547100, EPI_ISL_547104, EPI_ISL_547106, EPI_ISL_547107, EPI_ISL_547109, EPI_ISL_547111, EPI_ISL_547118,
EPI_ISL_547119, EPI_ISL_547122, EPI_ISL_547126, EPI_ISL_547132, EPI_ISL_547139, EPI_ISL_547140, EPI_ISL_547156, EPI_ISL_547159, EPI_ISL_547160, EPI_ISL_547162, EPI_ISL_547163, EPI_ISL_547165, EPI_ISL_547166, EPI_ISL_547168, EPI_ISL_547169, EPI_ISL_547170, EPI_ISL_547172, EPI_ISL_547173,
EPI_ISL_547175, EPI_ISL_547176, EPI_ISL_547177, EPI_ISL_547178, EPI_ISL_547179, EPI_ISL_547180, EPI_ISL_547181, EPI_ISL_547182, EPI_ISL_547183, EPI_ISL_547184, EPI_ISL_547188, EPI_ISL_547191, EPI_ISL_547195, EPI_ISL_547200, EPI_ISL_547201, EPI_ISL_547202, EPI_ISL_547204, EPI_ISL_547205,
EPI_ISL_547209, EPI_ISL_547213, EPI_ISL_547217, EPI_ISL_547222, EPI_ISL_547228, EPI_ISL_547237, EPI_ISL_547245, EPI_ISL_547255, EPI_ISL_547269, EPI_ISL_547280, EPI_ISL_547281, EPI_ISL_547283, EPI_ISL_547286, EPI_ISL_547287, EPI_ISL_547288, EPI_ISL_547294, EPI_ISL_547295, EPI_ISL_547296,
EPI_ISL_547297, EPI_ISL_547298, EPI_ISL_547299, EPI_ISL_547300, EPI_ISL_547301, EPI_ISL_547302, EPI_ISL_547306, EPI_ISL_547310, EPI_ISL_547311, EPI_ISL_547312, EPI_ISL_547313, EPI_ISL_547314, EPI_ISL_547315, EPI_ISL_547318, EPI_ISL_547323, EPI_ISL_547324, EPI_ISL_547328, EPI_ISL_547333,
EPI_ISL_547336, EPI_ISL_547340, EPI_ISL_547344, EPI_ISL_547347, EPI_ISL_547348, EPI_ISL_547350, EPI_ISL_547352, EPI_ISL_547353, EPI_ISL_547354, EPI_ISL_547357, EPI_ISL_547359, EPI_ISL_547363, EPI_ISL_547372, EPI_ISL_547373, EPI_ISL_547375, EPI_ISL_547376, EPI_ISL_547378, EPI_ISL_547380,

| | | | |
|---|---|---|---|
| EPI_ISL_547381, EPI_ISL_547383, EPI_ISL_547384, EPI_ISL_547388, EPI_ISL_547390, EPI_ISL_547394, EPI_ISL_547395, EPI_ISL_547397, EPI_ISL_547399, EPI_ISL_547402, EPI_ISL_547405, EPI_ISL_547408, EPI_ISL_547409, EPI_ISL_547410, EPI_ISL_547411, EPI_ISL_547413, EPI_ISL_547416, EPI_ISL_547420, EPI_ISL_547421, EPI_ISL_547424, EPI_ISL_547426, EPI_ISL_547427, EPI_ISL_547431, EPI_ISL_547432 | | | |
| see above | Houston Methodist Hospital | Houston Methodist Hospital | S. Wesley Long, Randall J. Olsen, Paul A. Christensen, David W. Bernard, James J. Davis, Maulik Shukla, Marcus Nguyen, Matthew Ojeda Saavedra, Concepcion C. Cantu, Prasanti Yerramilli, Layne Pruitt, Sishir Subedi, Hung-Che Kuo, Heather Hendrickson, Ghazaleh Eskandari, Hoang A. T. Nguyen, J. Hunter Long, Muthiah Kumaraswami, Jule Goike, Daniel Boutz, Jimmy Gollihar, Jason S. McLellan, Chia-Wei Chou, Kamyab Javanmardi, Ilya J. Finkelstein, and James M. Musser |
| EPI_ISL_551835, EPI_ISL_551836, EPI_ISL_551837, EPI_ISL_551838, EPI_ISL_551839, EPI_ISL_551840, EPI_ISL_551841, EPI_ISL_551842, EPI_ISL_551843, EPI_ISL_551844, EPI_ISL_551845, EPI_ISL_551846, EPI_ISL_551847, EPI_ISL_551848, EPI_ISL_551849, EPI_ISL_551851, EPI_ISL_551852, EPI_ISL_551853, EPI_ISL_551854, EPI_ISL_551855, EPI_ISL_551856, EPI_ISL_551857, EPI_ISL_551858, EPI_ISL_551859, EPI_ISL_551860, EPI_ISL_551861, EPI_ISL_551862, EPI_ISL_551863, EPI_ISL_551864, EPI_ISL_551865, EPI_ISL_551866, EPI_ISL_551867, EPI_ISL_551868, EPI_ISL_551869, EPI_ISL_551870, EPI_ISL_551871, EPI_ISL_551872, EPI_ISL_551873, EPI_ISL_551874, EPI_ISL_551875, EPI_ISL_551876, EPI_ISL_551877, EPI_ISL_551878, EPI_ISL_551879, EPI_ISL_551880, EPI_ISL_551881, EPI_ISL_551882, EPI_ISL_551883, EPI_ISL_551884, EPI_ISL_551885, EPI_ISL_551886, EPI_ISL_551887, EPI_ISL_551888, EPI_ISL_551889, EPI_ISL_551890, EPI_ISL_551891, EPI_ISL_551892, EPI_ISL_551893, EPI_ISL_551894, EPI_ISL_551895, EPI_ISL_551896, EPI_ISL_551897, EPI_ISL_551898, EPI_ISL_551899, EPI_ISL_551900, EPI_ISL_551901, EPI_ISL_551902, EPI_ISL_551903, EPI_ISL_551904, EPI_ISL_551905, EPI_ISL_551906, EPI_ISL_551907, EPI_ISL_551908, EPI_ISL_551909, EPI_ISL_551910, EPI_ISL_551911, EPI_ISL_551912, EPI_ISL_551913, EPI_ISL_551914, EPI_ISL_551915, EPI_ISL_551916, EPI_ISL_551917, EPI_ISL_551918, EPI_ISL_551919, EPI_ISL_551920, EPI_ISL_551921, EPI_ISL_551925, EPI_ISL_551932, EPI_ISL_551951, EPI_ISL_551953, EPI_ISL_551967, EPI_ISL_551980, EPI_ISL_551983, EPI_ISL_551990, EPI_ISL_551994, EPI_ISL_552001, EPI_ISL_552005, EPI_ISL_552008, EPI_ISL_552009, EPI_ISL_552010, EPI_ISL_552017, EPI_ISL_552051, EPI_ISL_552065, EPI_ISL_552073, EPI_ISL_552091, EPI_ISL_552094, EPI_ISL_552098, EPI_ISL_552105, EPI_ISL_552124, EPI_ISL_552151, EPI_ISL_552166, EPI_ISL_552170, EPI_ISL_552173, EPI_ISL_552180, EPI_ISL_552185, EPI_ISL_555843, EPI_ISL_555844, EPI_ISL_555864, EPI_ISL_555868, EPI_ISL_555875, EPI_ISL_555910, EPI_ISL_555939, EPI_ISL_555945, EPI_ISL_555948, EPI_ISL_555955, EPI_ISL_555973, EPI_ISL_555976, EPI_ISL_555993, EPI_ISL_556012, EPI_ISL_556021, EPI_ISL_556022, EPI_ISL_556026, EPI_ISL_556039, EPI_ISL_556041, EPI_ISL_556047, EPI_ISL_556058, EPI_ISL_556063, EPI_ISL_556066, EPI_ISL_556069, EPI_ISL_556305, EPI_ISL_556309, EPI_ISL_556317, EPI_ISL_556318, EPI_ISL_556321, EPI_ISL_556324, EPI_ISL_556338, EPI_ISL_556344, EPI_ISL_556349, EPI_ISL_556351, EPI_ISL_556357, EPI_ISL_556425, EPI_ISL_556434, EPI_ISL_556446, EPI_ISL_556459, EPI_ISL_556471, EPI_ISL_556482, EPI_ISL_556488, EPI_ISL_556494, EPI_ISL_556500, EPI_ISL_556508, EPI_ISL_556509, EPI_ISL_556516, EPI_ISL_556517, EPI_ISL_556518, EPI_ISL_556519, EPI_ISL_556520, EPI_ISL_556521, EPI_ISL_556522, EPI_ISL_556523, EPI_ISL_556524, EPI_ISL_556525, EPI_ISL_556526, EPI_ISL_556527, EPI_ISL_556528, EPI_ISL_556529, EPI_ISL_556530, EPI_ISL_556531, EPI_ISL_556532, EPI_ISL_556533, EPI_ISL_556534, EPI_ISL_556535, EPI_ISL_556536, EPI_ISL_556537, EPI_ISL_556538, EPI_ISL_556539, EPI_ISL_556540, EPI_ISL_556541, EPI_ISL_556542, EPI_ISL_556543, EPI_ISL_556544, EPI_ISL_556545, EPI_ISL_556546, EPI_ISL_556547, EPI_ISL_556548, EPI_ISL_556549, EPI_ISL_556550, EPI_ISL_556551, EPI_ISL_556552, EPI_ISL_556553, EPI_ISL_556554, EPI_ISL_556555, EPI_ISL_556556, EPI_ISL_556557, EPI_ISL_556558, EPI_ISL_556559, EPI_ISL_556560, EPI_ISL_556561, EPI_ISL_556562, EPI_ISL_556563, EPI_ISL_556564, EPI_ISL_556565, EPI_ISL_556566, EPI_ISL_556567, EPI_ISL_556568, EPI_ISL_556569, EPI_ISL_556570, EPI_ISL_556571, EPI_ISL_556572, EPI_ISL_556573, EPI_ISL_556574, EPI_ISL_556575, EPI_ISL_556639, EPI_ISL_556640, EPI_ISL_556641, EPI_ISL_556642, EPI_ISL_556643 | | | |
| see above | Lighthouse Lab in Cambridge | Wellcome Sanger Institute for the COVID-19 Genomics UK (COG-UK) consortium | Rob Howes, The Lighthouse Lab in Cambridge and Alex Alderton, Roberto Amato, Sonia Goncalves, Ewan Harrison, David K. Jackson, Ian Johnston, Dominic Kwiatkowski, Cordelia Langford, John Sillitoe on behalf of the Wellcome Sanger Institute COVID-19 Surveillance Team |
| EPI_ISL_556644 | Lighthouse Lab in Cambridge | Wellcome Sanger Institute for the COVID-19 Genomics UK (COG-UK) consortium | Rob Howes, The Lighthouse Lab in Cambridge and Alex Alderton, Roberto Amato, Sonia Goncalves, Ewan Harrison, David K. Jackson, Ian Johnston, Dominic Kwiatkowski, Cordelia Langford, John Sillitoe on behalf of the Wellcome Sanger Institute COVID-19 Surveillance Team (http://www.sanger.ac.uk/covid-team) |
| EPI_ISL_556645, EPI_ISL_556646, EPI_ISL_556647, EPI_ISL_556648, EPI_ISL_556649, EPI_ISL_556650, EPI_ISL_556651, EPI_ISL_556652, EPI_ISL_556653, EPI_ISL_556654, EPI_ISL_556655, EPI_ISL_556656, EPI_ISL_556657, EPI_ISL_556658, EPI_ISL_556659, EPI_ISL_556660, EPI_ISL_556661, EPI_ISL_556662, EPI_ISL_556663, EPI_ISL_556664, EPI_ISL_556665, EPI_ISL_556666, EPI_ISL_556667, EPI_ISL_556668, EPI_ISL_556669, EPI_ISL_556670, EPI_ISL_556671, EPI_ISL_556672, EPI_ISL_556673, EPI_ISL_556674, EPI_ISL_556675, EPI_ISL_556676, EPI_ISL_556677, EPI_ISL_556678, EPI_ISL_556679, EPI_ISL_556680, EPI_ISL_556681 | | | |
| see above | Lighthouse Lab in Cambridge | Wellcome Sanger Institute for the COVID-19 Genomics UK (COG-UK) consortium | Rob Howes, The Lighthouse Lab in Cambridge and Alex Alderton, Roberto Amato, Sonia Goncalves, Ewan Harrison, David K. Jackson, Ian Johnston, Dominic Kwiatkowski, Cordelia Langford, John Sillitoe on behalf of the Wellcome Sanger Institute COVID-19 Surveillance Team |
| EPI_ISL_556682 | Lighthouse Lab in Cambridge | Wellcome Sanger Institute for the COVID-19 Genomics UK (COG-UK) consortium | Rob Howes, The Lighthouse Lab in Cambridge and Alex Alderton, Roberto Amato, Sonia Goncalves, Ewan Harrison, David K. Jackson, Ian Johnston, Dominic Kwiatkowski, Cordelia Langford, John Sillitoe on behalf of the Wellcome Sanger Institute COVID-19 Surveillance Team (http://www.sanger.ac.uk/covid-team) |
| EPI_ISL_556683, EPI_ISL_556684, EPI_ISL_556685, EPI_ISL_556686, EPI_ISL_556687, EPI_ISL_556688, EPI_ISL_556689, EPI_ISL_556690, EPI_ISL_556691, EPI_ISL_556692, EPI_ISL_556693, EPI_ISL_556694, EPI_ISL_556695, EPI_ISL_556696, EPI_ISL_556697, EPI_ISL_556699, EPI_ISL_556700, EPI_ISL_556701, EPI_ISL_556702, EPI_ISL_556703, EPI_ISL_556704, EPI_ISL_556705, EPI_ISL_556706, EPI_ISL_556707, EPI_ISL_556708, EPI_ISL_556709, EPI_ISL_556710, EPI_ISL_556711, EPI_ISL_556712, EPI_ISL_556713, EPI_ISL_556714, EPI_ISL_556715, EPI_ISL_556716, EPI_ISL_556717, EPI_ISL_556718, EPI_ISL_556719, EPI_ISL_556720, EPI_ISL_556721, EPI_ISL_556722, EPI_ISL_556723, EPI_ISL_556724, EPI_ISL_556725, EPI_ISL_556726, EPI_ISL_556727, EPI_ISL_556728, EPI_ISL_556729, EPI_ISL_556730, EPI_ISL_556731, EPI_ISL_556732, EPI_ISL_556733, EPI_ISL_556734, EPI_ISL_556735, EPI_ISL_556736, EPI_ISL_556737, EPI_ISL_556738, EPI_ISL_556739, EPI_ISL_556740, EPI_ISL_556741, EPI_ISL_556742, EPI_ISL_556743, EPI_ISL_556744, EPI_ISL_556745, EPI_ISL_556746, EPI_ISL_556747, EPI_ISL_556748, EPI_ISL_556749, EPI_ISL_556750 | | | |
| see above | Lighthouse Lab in Cambridge | Wellcome Sanger Institute for the COVID-19 Genomics UK (COG-UK) consortium | Rob Howes, The Lighthouse Lab in Cambridge and Alex Alderton, Roberto Amato, Sonia Goncalves, Ewan Harrison, David K. Jackson, Ian Johnston, Dominic Kwiatkowski, Cordelia Langford, John Sillitoe on behalf of the Wellcome Sanger Institute COVID-19 Surveillance Team |
| EPI_ISL_556751 | Lighthouse Lab in Cambridge | Wellcome Sanger Institute for the COVID-19 Genomics UK (COG-UK) consortium | Rob Howes, The Lighthouse Lab in Cambridge and Alex Alderton, Roberto Amato, Sonia Goncalves, Ewan Harrison, David K. Jackson, Ian Johnston, Dominic Kwiatkowski, Cordelia Langford, John Sillitoe on behalf of the Wellcome Sanger Institute COVID-19 Surveillance Team (http://www.sanger.ac.uk/covid-team) |
| EPI_ISL_556752, EPI_ISL_556753, EPI_ISL_556754, EPI_ISL_556755, EPI_ISL_556756, EPI_ISL_556757, EPI_ISL_556758, EPI_ISL_556759, EPI_ISL_556760 | Lighthouse Lab in Cambridge | Wellcome Sanger Institute for the COVID-19 Genomics UK (COG-UK) consortium | Rob Howes, The Lighthouse Lab in Cambridge and Alex Alderton, Roberto Amato, Sonia Goncalves, Ewan Harrison, David K. Jackson, Ian Johnston, Dominic Kwiatkowski, Cordelia Langford, John Sillitoe on behalf of the Wellcome Sanger Institute COVID-19 Surveillance Team |
| EPI_ISL_556761 | Lighthouse Lab in Cambridge | Wellcome Sanger Institute for the COVID-19 Genomics UK (COG-UK) consortium | Rob Howes, The Lighthouse Lab in Cambridge and Alex Alderton, Roberto Amato, Sonia Goncalves, Ewan Harrison, David K. Jackson, Ian Johnston, Dominic Kwiatkowski, Cordelia Langford, John Sillitoe on behalf of the Wellcome Sanger Institute COVID-19 Surveillance Team (http://www.sanger.ac.uk/covid-team) |
| EPI_ISL_556762, EPI_ISL_556763, EPI_ISL_556764, EPI_ISL_556765, EPI_ISL_556766, EPI_ISL_556767, EPI_ISL_556768, EPI_ISL_556769, EPI_ISL_556771, EPI_ISL_556772, EPI_ISL_556773, EPI_ISL_556774, EPI_ISL_556775, EPI_ISL_556776, EPI_ISL_556777, EPI_ISL_556778, EPI_ISL_556779, EPI_ISL_556780, EPI_ISL_556781, EPI_ISL_556782, EPI_ISL_556783, EPI_ISL_556784, EPI_ISL_556785, EPI_ISL_556786 | | | |
| see above | Lighthouse Lab in Cambridge | Wellcome Sanger Institute for the COVID-19 Genomics UK (COG-UK) consortium | Rob Howes, The Lighthouse Lab in Cambridge and Alex Alderton, Roberto Amato, Sonia Goncalves, Ewan Harrison, David K. Jackson, Ian Johnston, Dominic Kwiatkowski, Cordelia Langford, John Sillitoe on behalf of the Wellcome Sanger Institute COVID-19 Surveillance Team |
| EPI_ISL_556787 | Lighthouse Lab in Cambridge | Wellcome Sanger Institute for the COVID-19 Genomics UK (COG-UK) consortium | Rob Howes, The Lighthouse Lab in Cambridge and Alex Alderton, Roberto Amato, Sonia Goncalves, Ewan Harrison, David K. Jackson, Ian Johnston, Dominic Kwiatkowski, Cordelia Langford, John Sillitoe on behalf of the Wellcome Sanger Institute COVID-19 Surveillance Team (http://www.sanger.ac.uk/covid-team) |
| EPI_ISL_556788, EPI_ISL_556789, EPI_ISL_556790, EPI_ISL_556791, EPI_ISL_556792, EPI_ISL_556793, EPI_ISL_556794, EPI_ISL_556795, EPI_ISL_556796, EPI_ISL_556797, EPI_ISL_556798, EPI_ISL_556799, EPI_ISL_556800, EPI_ISL_556801, EPI_ISL_556802, EPI_ISL_556803, EPI_ISL_556804, EPI_ISL_556805, EPI_ISL_556806, EPI_ISL_556807, EPI_ISL_556808, EPI_ISL_556809, EPI_ISL_556810, EPI_ISL_556811, EPI_ISL_556812, EPI_ISL_556813, EPI_ISL_556814, EPI_ISL_556815, EPI_ISL_556816, EPI_ISL_556817, EPI_ISL_556818, EPI_ISL_556819, EPI_ISL_556820, EPI_ISL_556821, EPI_ISL_556822, EPI_ISL_556823, EPI_ISL_556824, EPI_ISL_556825, EPI_ISL_556826, EPI_ISL_556827, EPI_ISL_556828, EPI_ISL_556829, EPI_ISL_556830, EPI_ISL_556831, EPI_ISL_556832, EPI_ISL_556833, EPI_ISL_556834, EPI_ISL_556835, EPI_ISL_556836, EPI_ISL_556837, EPI_ISL_556838, EPI_ISL_556839, EPI_ISL_556840, EPI_ISL_556841, EPI_ISL_556842, EPI_ISL_556843, EPI_ISL_556844, EPI_ISL_556845, EPI_ISL_556846, EPI_ISL_556847, EPI_ISL_556848, EPI_ISL_556849, EPI_ISL_556850, EPI_ISL_556851, EPI_ISL_556852, EPI_ISL_556853, EPI_ISL_556854, EPI_ISL_556855, EPI_ISL_556856, EPI_ISL_556857, EPI_ISL_556858, EPI_ISL_556859, EPI_ISL_556860, EPI_ISL_556861, EPI_ISL_556862, EPI_ISL_556863, EPI_ISL_556864, EPI_ISL_556865, EPI_ISL_556866, EPI_ISL_556867, EPI_ISL_556868, EPI_ISL_556869, EPI_ISL_556870, EPI_ISL_556871, EPI_ISL_556872, EPI_ISL_556873, EPI_ISL_556874, EPI_ISL_556875, EPI_ISL_556876, EPI_ISL_556877, EPI_ISL_556878, EPI_ISL_556879, EPI_ISL_556880, EPI_ISL_556881, EPI_ISL_556882 | | | |
| see above | Lighthouse Lab in Cambridge | Wellcome Sanger Institute for the COVID-19 Genomics UK (COG-UK) consortium | Rob Howes, The Lighthouse Lab in Cambridge and Alex Alderton, Roberto Amato, Sonia Goncalves, Ewan Harrison, David K. Jackson, Ian Johnston, Dominic Kwiatkowski, Cordelia Langford, John Sillitoe on behalf of the Wellcome Sanger Institute COVID-19 Surveillance Team |
| EPI_ISL_556883 | Lighthouse Lab in Cambridge | Wellcome Sanger Institute for the COVID-19 Genomics UK (COG-UK) consortium | Rob Howes, The Lighthouse Lab in Cambridge and Alex Alderton, Roberto Amato, Sonia Goncalves, Ewan Harrison, David K. Jackson, Ian Johnston, Dominic Kwiatkowski, Cordelia Langford, John Sillitoe on behalf of the Wellcome Sanger Institute COVID-19 Surveillance Team (http://www.sanger.ac.uk/covid-team) |
| EPI_ISL_556884, EPI_ISL_556885, EPI_ISL_556886, EPI_ISL_556888, EPI_ISL_556889, EPI_ISL_556890, EPI_ISL_556891, EPI_ISL_556892, EPI_ISL_556893 | Lighthouse Lab in Cambridge | Wellcome Sanger Institute for the COVID-19 Genomics UK (COG-UK) consortium | Rob Howes, The Lighthouse Lab in Cambridge and Alex Alderton, Roberto Amato, Sonia Goncalves, Ewan Harrison, David K. Jackson, Ian Johnston, Dominic Kwiatkowski, Cordelia Langford, John Sillitoe on behalf of the Wellcome Sanger Institute COVID-19 Surveillance Team |
| EPI_ISL_556894 | Lighthouse Lab in Cambridge | Wellcome Sanger Institute for the COVID-19 Genomics UK (COG-UK) consortium | Rob Howes, The Lighthouse Lab in Cambridge and Alex Alderton, Roberto Amato, Sonia Goncalves, Ewan Harrison, David K. Jackson, Ian Johnston, Dominic Kwiatkowski, Cordelia Langford, John Sillitoe on behalf of the Wellcome Sanger Institute COVID-19 Surveillance Team (http://www.sanger.ac.uk/covid-team) |
| EPI_ISL_556895, EPI_ISL_556896, EPI_ISL_556897, EPI_ISL_556898, | Lighthouse Lab in Cambridge | Wellcome Sanger Institute for the COVID-19 Genomics UK (COG-UK) consortium | Rob Howes, The Lighthouse Lab in Cambridge and Alex Alderton, Roberto Amato, Sonia Goncalves, Ewan Harrison, David K. Jackson, Ian Johnston, Dominic Kwiatkowski, Cordelia Langford, John Sillitoe on behalf of the Wellcome Sanger Institute COVID-19 Surveillance Team |

| | | | |
|---|---|---|---|
| EPI_ISL_556899, EPI_ISL_556900, EPI_ISL_556901, EPI_ISL_556902, EPI_ISL_556903 | | | |
| EPI_ISL_556904 | Lighthouse Lab in Cambridge | Wellcome Sanger Institute for the COVID-19 Genomics UK (COG-UK) consortium | Rob Howes, The Lighthouse Lab in Cambridge and Alex Alderton, Roberto Amato, Sonia Goncalves, Ewan Harrison, David K. Jackson, Ian Johnston, Dominic Kwiatkowski, Cordelia Langford, John Sillitoe on behalf of the Wellcome Sanger Institute COVID-19 Surveillance Team (http://www.sanger.ac.uk/covid-team) |
| EPI_ISL_556905, EPI_ISL_556906, EPI_ISL_556907, EPI_ISL_556908, EPI_ISL_556909, EPI_ISL_556910, EPI_ISL_556911, EPI_ISL_556912, EPI_ISL_556913, EPI_ISL_556914, EPI_ISL_556915, EPI_ISL_556916, EPI_ISL_556917 | | | |
| see above | Lighthouse Lab in Cambridge | Wellcome Sanger Institute for the COVID-19 Genomics UK (COG-UK) consortium | Rob Howes, The Lighthouse Lab in Cambridge and Alex Alderton, Roberto Amato, Sonia Goncalves, Ewan Harrison, David K. Jackson, Ian Johnston, Dominic Kwiatkowski, Cordelia Langford, John Sillitoe on behalf of the Wellcome Sanger Institute COVID-19 Surveillance Team |
| EPI_ISL_556918 | Lighthouse Lab in Cambridge | Wellcome Sanger Institute for the COVID-19 Genomics UK (COG-UK) consortium | Rob Howes, The Lighthouse Lab in Cambridge and Alex Alderton, Roberto Amato, Sonia Goncalves, Ewan Harrison, David K. Jackson, Ian Johnston, Dominic Kwiatkowski, Cordelia Langford, John Sillitoe on behalf of the Wellcome Sanger Institute COVID-19 Surveillance Team (http://www.sanger.ac.uk/covid-team) |
| EPI_ISL_556919, EPI_ISL_556920, EPI_ISL_556921, EPI_ISL_556922, EPI_ISL_556923, EPI_ISL_556924, EPI_ISL_556925, EPI_ISL_556926, EPI_ISL_556927, EPI_ISL_556928 | Lighthouse Lab in Cambridge | Wellcome Sanger Institute for the COVID-19 Genomics UK (COG-UK) consortium | Rob Howes, The Lighthouse Lab in Cambridge and Alex Alderton, Roberto Amato, Sonia Goncalves, Ewan Harrison, David K. Jackson, Ian Johnston, Dominic Kwiatkowski, Cordelia Langford, John Sillitoe on behalf of the Wellcome Sanger Institute COVID-19 Surveillance Team |
| EPI_ISL_556929 | Lighthouse Lab in Cambridge | Wellcome Sanger Institute for the COVID-19 Genomics UK (COG-UK) consortium | Rob Howes, The Lighthouse Lab in Cambridge and Alex Alderton, Roberto Amato, Sonia Goncalves, Ewan Harrison, David K. Jackson, Ian Johnston, Dominic Kwiatkowski, Cordelia Langford, John Sillitoe on behalf of the Wellcome Sanger Institute COVID-19 Surveillance Team (http://www.sanger.ac.uk/covid-team) |
| EPI_ISL_556930, EPI_ISL_556931, EPI_ISL_556932, EPI_ISL_556933, EPI_ISL_556934, EPI_ISL_556935, EPI_ISL_556936, EPI_ISL_556937, EPI_ISL_556938, EPI_ISL_556939, EPI_ISL_556941, EPI_ISL_556942, EPI_ISL_556943, EPI_ISL_556944, EPI_ISL_556945, EPI_ISL_556946, EPI_ISL_556947, EPI_ISL_556948, EPI_ISL_556949, EPI_ISL_556950, EPI_ISL_556951, EPI_ISL_556952, EPI_ISL_556953, EPI_ISL_556954, EPI_ISL_556955, EPI_ISL_556956, EPI_ISL_556957, EPI_ISL_556958, EPI_ISL_556959, EPI_ISL_556960, EPI_ISL_556961, EPI_ISL_556962 | | | |
| see above | Lighthouse Lab in Cambridge | Wellcome Sanger Institute for the COVID-19 Genomics UK (COG-UK) consortium | Rob Howes, The Lighthouse Lab in Cambridge and Alex Alderton, Roberto Amato, Sonia Goncalves, Ewan Harrison, David K. Jackson, Ian Johnston, Dominic Kwiatkowski, Cordelia Langford, John Sillitoe on behalf of the Wellcome Sanger Institute COVID-19 Surveillance Team |
| EPI_ISL_556963 | Lighthouse Lab in Cambridge | Wellcome Sanger Institute for the COVID-19 Genomics UK (COG-UK) consortium | Rob Howes, The Lighthouse Lab in Cambridge and Alex Alderton, Roberto Amato, Sonia Goncalves, Ewan Harrison, David K. Jackson, Ian Johnston, Dominic Kwiatkowski, Cordelia Langford, John Sillitoe on behalf of the Wellcome Sanger Institute COVID-19 Surveillance Team (http://www.sanger.ac.uk/covid-team) |
| EPI_ISL_556964, EPI_ISL_556965, EPI_ISL_556966, EPI_ISL_556967, EPI_ISL_556968, EPI_ISL_556969, EPI_ISL_556970, EPI_ISL_556971, EPI_ISL_556972, EPI_ISL_556973, EPI_ISL_556974, EPI_ISL_556975, EPI_ISL_556976, EPI_ISL_556977, EPI_ISL_556978, EPI_ISL_556979, EPI_ISL_556980, EPI_ISL_556981, EPI_ISL_556982, EPI_ISL_556983 | | | |
| see above | Lighthouse Lab in Cambridge | Wellcome Sanger Institute for the COVID-19 Genomics UK (COG-UK) consortium | Rob Howes, The Lighthouse Lab in Cambridge and Alex Alderton, Roberto Amato, Sonia Goncalves, Ewan Harrison, David K. Jackson, Ian Johnston, Dominic Kwiatkowski, Cordelia Langford, John Sillitoe on behalf of the Wellcome Sanger Institute COVID-19 Surveillance Team |
| EPI_ISL_556984 | Lighthouse Lab in Cambridge | Wellcome Sanger Institute for the COVID-19 Genomics UK (COG-UK) consortium | Rob Howes, The Lighthouse Lab in Cambridge and Alex Alderton, Roberto Amato, Sonia Goncalves, Ewan Harrison, David K. Jackson, Ian Johnston, Dominic Kwiatkowski, Cordelia Langford, John Sillitoe on behalf of the Wellcome Sanger Institute COVID-19 Surveillance Team (http://www.sanger.ac.uk/covid-team) |
| EPI_ISL_560926, EPI_ISL_560927, EPI_ISL_560928, EPI_ISL_560929, EPI_ISL_560930, EPI_ISL_560931, EPI_ISL_560932, EPI_ISL_560933, EPI_ISL_560934, EPI_ISL_560935, EPI_ISL_560936, EPI_ISL_560937, EPI_ISL_560938, EPI_ISL_560939, EPI_ISL_560940, EPI_ISL_560941, EPI_ISL_560942, EPI_ISL_560943, EPI_ISL_560944, EPI_ISL_560945, EPI_ISL_560946, EPI_ISL_560947, EPI_ISL_560948, EPI_ISL_560949, EPI_ISL_560950, EPI_ISL_560951, EPI_ISL_560952, EPI_ISL_560953, EPI_ISL_560954, EPI_ISL_560955, EPI_ISL_560956, EPI_ISL_560957, EPI_ISL_560958, EPI_ISL_560959, EPI_ISL_560960, EPI_ISL_560961, EPI_ISL_560962, EPI_ISL_560963, EPI_ISL_560964, EPI_ISL_560965, EPI_ISL_560966, EPI_ISL_560967, EPI_ISL_560968, EPI_ISL_560969, EPI_ISL_560970 | | | |
| see above | Texas Department of State Health Services | Texas Department of State Health Services | Rashmi Tuladhar, Bonnie Oh, Jenny Zhang, Maliha Rahman, Anita Pokharel, Myong Koag, Chun Wang, Rachel Lee, Grace Kubin |
| EPI_ISL_560991 | RSUD Dr. Soetomo | Institute of Tropical Disease, Universitas Airlangga | Rima R Prasetya, Krisnoadi Rahardjo, Aldise M Nastri, Jezzy R Dewantari, Joni Wahyuhadi, Gatot Soegiarto, Laksmi Wulandari, Retno A Setyoningrum, Resti Yudhawati, Yohko K Shimizu, Mitsuhiro Nishimura, Yasuko Mori, Soetjipto, Kazufumi Shimizu, Maria I Lusida |
| EPI_ISL_566423, EPI_ISL_566428, EPI_ISL_566429, EPI_ISL_566444, EPI_ISL_566445, EPI_ISL_566453, EPI_ISL_566456, EPI_ISL_566457, EPI_ISL_566467, EPI_ISL_566473, EPI_ISL_566492, EPI_ISL_566493, EPI_ISL_566495, EPI_ISL_566500, EPI_ISL_566508, EPI_ISL_566518, EPI_ISL_566519, EPI_ISL_566524, EPI_ISL_566528, EPI_ISL_566534, EPI_ISL_566537, EPI_ISL_566543, EPI_ISL_566544, EPI_ISL_566567, EPI_ISL_566576, EPI_ISL_566586, EPI_ISL_566589, EPI_ISL_566610, EPI_ISL_566611, EPI_ISL_566625, EPI_ISL_566631, EPI_ISL_566638, EPI_ISL_566659, EPI_ISL_566661, EPI_ISL_566662, EPI_ISL_566666, EPI_ISL_566672, EPI_ISL_566674, EPI_ISL_568385, EPI_ISL_568436, EPI_ISL_568453, EPI_ISL_568454, EPI_ISL_568458, EPI_ISL_568461, EPI_ISL_568472, EPI_ISL_568473 | | | |
| see above | Lighthouse Lab in Cambridge | Wellcome Sanger Institute for the COVID-19 Genomics UK (COG-UK) consortium | Rob Howes, The Lighthouse Lab in Cambridge and Alex Alderton, Roberto Amato, Sonia Goncalves, Ewan Harrison, David K. Jackson, Ian Johnston, Dominic Kwiatkowski, Cordelia Langford, John Sillitoe on behalf of the Wellcome Sanger Institute COVID-19 Surveillance Team |
| EPI_ISL_569689, EPI_ISL_569690, EPI_ISL_569691, EPI_ISL_569692, EPI_ISL_569693, EPI_ISL_569694, EPI_ISL_569695, EPI_ISL_569696, EPI_ISL_569697, EPI_ISL_569698, EPI_ISL_569699, EPI_ISL_569700, EPI_ISL_569701, EPI_ISL_569702, EPI_ISL_569703, EPI_ISL_569704, EPI_ISL_569705, EPI_ISL_569706, EPI_ISL_569707, EPI_ISL_569708, EPI_ISL_569709, EPI_ISL_569710, EPI_ISL_569711, EPI_ISL_569712, EPI_ISL_569713, EPI_ISL_569714, EPI_ISL_569715, EPI_ISL_569716, EPI_ISL_569717, EPI_ISL_569718, EPI_ISL_569719, EPI_ISL_569720, EPI_ISL_569721, EPI_ISL_569722, EPI_ISL_569723, EPI_ISL_569724, EPI_ISL_569725, EPI_ISL_569726, EPI_ISL_569727, EPI_ISL_569728, EPI_ISL_569729, EPI_ISL_569730, EPI_ISL_569731, EPI_ISL_569732, EPI_ISL_569733, EPI_ISL_569734 | | | |
| see above | Texas Department of State Health Services | Texas Department of State Health Services | Rashmi Tuladhar, Bonnie Oh,Jenny Zhang, Maliha Rahman, Anita Pokharel, Myong Koag, Chun Wang, Rachel Lee, Grace Kubin |
| EPI_ISL_569858 | Area of Virology, Serology and Virology Division (SAViD), New South Wales Health Pathology Randwick | Area of Virology, Serology and Virology Division (SAViD), New South Wales Health Pathology Randwick | Rawlinson, W., Deveson, I., Bull, R., Van Hal, S. |
| EPI_ISL_569949, EPI_ISL_569950, EPI_ISL_569951, EPI_ISL_569952, EPI_ISL_569953, EPI_ISL_569954, EPI_ISL_569955, EPI_ISL_569956, EPI_ISL_569957, EPI_ISL_569958, EPI_ISL_569959, EPI_ISL_569960, EPI_ISL_569961, EPI_ISL_569962, EPI_ISL_569963, EPI_ISL_569964, EPI_ISL_569965, EPI_ISL_569966, EPI_ISL_569967, EPI_ISL_569968, EPI_ISL_569969, EPI_ISL_569970, EPI_ISL_569971, EPI_ISL_569972, EPI_ISL_569973, EPI_ISL_569974, EPI_ISL_569975, EPI_ISL_569976, EPI_ISL_569977, EPI_ISL_569978, EPI_ISL_569979, EPI_ISL_569980, EPI_ISL_569981, EPI_ISL_569982, EPI_ISL_569983, EPI_ISL_569984, EPI_ISL_569985, EPI_ISL_569986, EPI_ISL_569987, EPI_ISL_569988, EPI_ISL_569989, EPI_ISL_569990, EPI_ISL_569991, EPI_ISL_569992, EPI_ISL_569993, EPI_ISL_569994, EPI_ISL_569995, EPI_ISL_569996, EPI_ISL_569997, EPI_ISL_569998, EPI_ISL_569999, EPI_ISL_570000, EPI_ISL_570001, EPI_ISL_570002, EPI_ISL_570003, EPI_ISL_570004, EPI_ISL_570005, EPI_ISL_570006, EPI_ISL_570007, EPI_ISL_570008, EPI_ISL_570009, EPI_ISL_570010, EPI_ISL_570011, EPI_ISL_570012, EPI_ISL_570013, EPI_ISL_570014, EPI_ISL_570015, EPI_ISL_570016, EPI_ISL_570017, EPI_ISL_570018, EPI_ISL_570019, EPI_ISL_570020, EPI_ISL_570021, EPI_ISL_570022, EPI_ISL_570023, EPI_ISL_570024, EPI_ISL_570025, EPI_ISL_570026 | | | |
| see above | Unity Health Toronto | Ontario Institute for Cancer Research | Ramzi Fattouh, Larissa M. Matukas, Yan Chen,Mark Downing, Trina Otterman, Karel Boissinot, Wai Sum Siu, Zhi Cui, Le Luu, Samira Mubareka, TIBDN, Ilinca Lungu, Bernard Lam, Jeremy Johns, Paul Krzyzanowski, Richard de Borja, Felicia Vincelli, Philip Zuzarte, Jared T. Simpson |
| EPI_ISL_571016, EPI_ISL_571017, EPI_ISL_571018, EPI_ISL_571019, EPI_ISL_571020, EPI_ISL_571021, EPI_ISL_571022, EPI_ISL_571023, EPI_ISL_571024, EPI_ISL_571025, EPI_ISL_571026, EPI_ISL_571027, EPI_ISL_571028, EPI_ISL_571029, EPI_ISL_571030, EPI_ISL_571031, EPI_ISL_571032, EPI_ISL_571033, EPI_ISL_571034, EPI_ISL_571035, EPI_ISL_571036, EPI_ISL_571037, EPI_ISL_571038, EPI_ISL_571039, EPI_ISL_571040, EPI_ISL_571041, EPI_ISL_571042, EPI_ISL_571043, EPI_ISL_571044, EPI_ISL_571045, EPI_ISL_571046, EPI_ISL_571047, EPI_ISL_571048, EPI_ISL_571049, EPI_ISL_571050, EPI_ISL_571051, EPI_ISL_571052, EPI_ISL_571053, EPI_ISL_571054, EPI_ISL_571055, EPI_ISL_571056, EPI_ISL_571057, EPI_ISL_571058, EPI_ISL_571059, EPI_ISL_571060, EPI_ISL_571061, EPI_ISL_571062, EPI_ISL_571063, EPI_ISL_571064, EPI_ISL_571065, EPI_ISL_571066, EPI_ISL_571067, EPI_ISL_571068, EPI_ISL_571069, EPI_ISL_571070, EPI_ISL_571071, EPI_ISL_571072, EPI_ISL_571073, EPI_ISL_571074, EPI_ISL_571075, EPI_ISL_571076, EPI_ISL_571077, EPI_ISL_571078, EPI_ISL_571079, EPI_ISL_571080, EPI_ISL_571081, EPI_ISL_571082, EPI_ISL_571083, EPI_ISL_571084, EPI_ISL_571085, EPI_ISL_571086, EPI_ISL_571087, EPI_ISL_571088, EPI_ISL_571089, EPI_ISL_571090, EPI_ISL_571091, EPI_ISL_571092, EPI_ISL_571093, EPI_ISL_571094, EPI_ISL_571095, EPI_ISL_571096, EPI_ISL_571097, EPI_ISL_571098, EPI_ISL_571099, EPI_ISL_571100, EPI_ISL_571101, EPI_ISL_571102, EPI_ISL_571103, EPI_ISL_571104, EPI_ISL_571105, EPI_ISL_571106, EPI_ISL_571107, EPI_ISL_571108, EPI_ISL_571109, EPI_ISL_571110, EPI_ISL_571111, EPI_ISL_571112, EPI_ISL_571113, EPI_ISL_571114, EPI_ISL_571115, EPI_ISL_571116, EPI_ISL_571117, EPI_ISL_571118, EPI_ISL_571119, EPI_ISL_571120, EPI_ISL_571121, EPI_ISL_571122, EPI_ISL_571123, EPI_ISL_571124, EPI_ISL_571125, EPI_ISL_571126, EPI_ISL_571127, EPI_ISL_571128, EPI_ISL_571129, EPI_ISL_571130, EPI_ISL_571131, EPI_ISL_571132, EPI_ISL_571133, EPI_ISL_571134, EPI_ISL_571135, EPI_ISL_571136, EPI_ISL_571137, EPI_ISL_571138, EPI_ISL_571139, EPI_ISL_571140, EPI_ISL_571141, EPI_ISL_571142, EPI_ISL_571143, EPI_ISL_571144, EPI_ISL_571145, EPI_ISL_571146, EPI_ISL_571147, EPI_ISL_571148, EPI_ISL_571149, EPI_ISL_571150, EPI_ISL_571151, EPI_ISL_571152, EPI_ISL_571153, EPI_ISL_571154, EPI_ISL_571155, EPI_ISL_571156, EPI_ISL_571157, EPI_ISL_571158, EPI_ISL_571159, EPI_ISL_571160, EPI_ISL_571161, EPI_ISL_571162, EPI_ISL_571163, EPI_ISL_571164, EPI_ISL_571165, EPI_ISL_571166, EPI_ISL_571167, EPI_ISL_571168, EPI_ISL_571169, EPI_ISL_571170, EPI_ISL_571171, EPI_ISL_571172, EPI_ISL_571173, EPI_ISL_571174, EPI_ISL_571175, EPI_ISL_571176, EPI_ISL_571177, EPI_ISL_571178, EPI_ISL_571179, EPI_ISL_571180, EPI_ISL_571181, EPI_ISL_571182, EPI_ISL_571183, EPI_ISL_571184, EPI_ISL_571185, EPI_ISL_571186, EPI_ISL_571187, EPI_ISL_571188, EPI_ISL_571189, EPI_ISL_571190, EPI_ISL_571191, EPI_ISL_571192, EPI_ISL_571193, EPI_ISL_571194, EPI_ISL_571195, EPI_ISL_571196, EPI_ISL_571197, EPI_ISL_571198, EPI_ISL_571199, EPI_ISL_571200, EPI_ISL_571201, EPI_ISL_571202, EPI_ISL_571203, EPI_ISL_571204, EPI_ISL_571205, EPI_ISL_571206, EPI_ISL_571207, EPI_ISL_571208, EPI_ISL_571209, EPI_ISL_571210, EPI_ISL_571211, EPI_ISL_571212, EPI_ISL_571213, EPI_ISL_571214, EPI_ISL_571215, EPI_ISL_571216, EPI_ISL_571217, EPI_ISL_571218, EPI_ISL_571219, EPI_ISL_571220, EPI_ISL_571221, EPI_ISL_571222, EPI_ISL_571223, EPI_ISL_571224, EPI_ISL_571225, EPI_ISL_571226, EPI_ISL_571227, EPI_ISL_571228, EPI_ISL_571229, EPI_ISL_571230, EPI_ISL_571231, EPI_ISL_571232, EPI_ISL_571233, EPI_ISL_571234, EPI_ISL_571235, EPI_ISL_571236, EPI_ISL_571237, EPI_ISL_571238, EPI_ISL_571239, EPI_ISL_571240, EPI_ISL_571241, EPI_ISL_571242, EPI_ISL_571243, EPI_ISL_571244, EPI_ISL_571245, EPI_ISL_571246, EPI_ISL_571247, EPI_ISL_571248, EPI_ISL_571249, EPI_ISL_571250, EPI_ISL_571251, EPI_ISL_571252, EPI_ISL_571253, EPI_ISL_571254, EPI_ISL_571255, EPI_ISL_571256, EPI_ISL_571257, EPI_ISL_571258, EPI_ISL_571259, EPI_ISL_571260, EPI_ISL_571261, EPI_ISL_571262, EPI_ISL_571263, EPI_ISL_571264, EPI_ISL_571265, EPI_ISL_571266, EPI_ISL_571267, EPI_ISL_571268, EPI_ISL_571269, EPI_ISL_571270, EPI_ISL_571271, EPI_ISL_571272, EPI_ISL_571273, EPI_ISL_571274, EPI_ISL_571275, EPI_ISL_571276, EPI_ISL_571277, EPI_ISL_571278, EPI_ISL_571279, EPI_ISL_571280, EPI_ISL_571281, EPI_ISL_571282, EPI_ISL_571283, EPI_ISL_571284, EPI_ISL_571285, EPI_ISL_571286, EPI_ISL_571287, EPI_ISL_571288, EPI_ISL_571289, EPI_ISL_571290, EPI_ISL_571291, EPI_ISL_571292, EPI_ISL_571293, EPI_ISL_571294, EPI_ISL_571295, EPI_ISL_571296, EPI_ISL_571297, EPI_ISL_571298, EPI_ISL_571299, EPI_ISL_571300, EPI_ISL_571301, EPI_ISL_571302, EPI_ISL_571303, EPI_ISL_571304, EPI_ISL_571305, EPI_ISL_571306, EPI_ISL_571307, EPI_ISL_571308, EPI_ISL_571309, EPI_ISL_571310, EPI_ISL_571311, EPI_ISL_571312, EPI_ISL_571313, EPI_ISL_571314, EPI_ISL_571315, EPI_ISL_571316, EPI_ISL_571317, EPI_ISL_571318, EPI_ISL_571319, EPI_ISL_571320, EPI_ISL_571321, EPI_ISL_571322, EPI_ISL_571323, EPI_ISL_571324, EPI_ISL_571325, EPI_ISL_571326, EPI_ISL_571327, EPI_ISL_571328, EPI_ISL_571329, EPI_ISL_571330, EPI_ISL_571331, EPI_ISL_571332, EPI_ISL_571333, EPI_ISL_571334, EPI_ISL_571335, EPI_ISL_571336, EPI_ISL_571337, EPI_ISL_571338, EPI_ISL_571339, | | | |

| | | | |
|---|---|---|---|
| EPI_ISL_571340, EPI_ISL_571341, EPI_ISL_571342, EPI_ISL_571343, EPI_ISL_571344, EPI_ISL_571345, EPI_ISL_571346, EPI_ISL_571347, EPI_ISL_571348, EPI_ISL_571349, EPI_ISL_571350, EPI_ISL_571351, EPI_ISL_571352, EPI_ISL_571353, EPI_ISL_571354, EPI_ISL_571355, EPI_ISL_571356, EPI_ISL_571357, EPI_ISL_571358, EPI_ISL_571359, EPI_ISL_571360, EPI_ISL_571361, EPI_ISL_571362, EPI_ISL_571363, EPI_ISL_571364, EPI_ISL_571365, EPI_ISL_571366, EPI_ISL_571367, EPI_ISL_571368, EPI_ISL_571369, EPI_ISL_571370, EPI_ISL_571371, EPI_ISL_571372, EPI_ISL_571373, EPI_ISL_571374, EPI_ISL_571375, EPI_ISL_571376, EPI_ISL_571377, EPI_ISL_571378, EPI_ISL_571379, EPI_ISL_571380, EPI_ISL_571381, EPI_ISL_571382, EPI_ISL_571383, EPI_ISL_571384, EPI_ISL_571385, EPI_ISL_571386, EPI_ISL_571387, EPI_ISL_571388, EPI_ISL_571389, EPI_ISL_571390, EPI_ISL_571391, EPI_ISL_571392, EPI_ISL_571393, EPI_ISL_571394, EPI_ISL_571395, EPI_ISL_571396, EPI_ISL_571397, EPI_ISL_571398, EPI_ISL_571399, EPI_ISL_571400, EPI_ISL_571401, EPI_ISL_571402, EPI_ISL_571403, EPI_ISL_571404, EPI_ISL_571405, EPI_ISL_571406, EPI_ISL_571407, EPI_ISL_571408, EPI_ISL_571409, EPI_ISL_571410, EPI_ISL_571411, EPI_ISL_571412, EPI_ISL_571413, EPI_ISL_571414, EPI_ISL_571415, EPI_ISL_571416, EPI_ISL_571417, EPI_ISL_571418, EPI_ISL_571419, EPI_ISL_571420, EPI_ISL_571421, EPI_ISL_571422, EPI_ISL_571423, EPI_ISL_571424, EPI_ISL_571425, EPI_ISL_571426, EPI_ISL_571427, EPI_ISL_571428, EPI_ISL_571429, EPI_ISL_571430, EPI_ISL_571431, EPI_ISL_571432, EPI_ISL_571433, EPI_ISL_571434, EPI_ISL_571435, EPI_ISL_571436, EPI_ISL_571437, EPI_ISL_571438, EPI_ISL_571439, EPI_ISL_571440, EPI_ISL_571441, EPI_ISL_571442, EPI_ISL_571443, EPI_ISL_571444, EPI_ISL_571445, EPI_ISL_571446, EPI_ISL_571447, EPI_ISL_571448, EPI_ISL_571449, EPI_ISL_571450, EPI_ISL_571451, EPI_ISL_571452, EPI_ISL_571453, EPI_ISL_571454, EPI_ISL_571455, EPI_ISL_571456, EPI_ISL_571457, EPI_ISL_571458, EPI_ISL_571459, EPI_ISL_571460, EPI_ISL_571461, EPI_ISL_571462, EPI_ISL_571463, EPI_ISL_571464, EPI_ISL_571465, EPI_ISL_571466, EPI_ISL_571467, EPI_ISL_571468, EPI_ISL_571469, EPI_ISL_571470, EPI_ISL_571471, EPI_ISL_571472, EPI_ISL_571473, EPI_ISL_571474, EPI_ISL_571475, EPI_ISL_571476, EPI_ISL_571477, EPI_ISL_571478, EPI_ISL_571479, EPI_ISL_571480, EPI_ISL_571481, EPI_ISL_571482, EPI_ISL_571483, EPI_ISL_571484, EPI_ISL_571485, EPI_ISL_571486, EPI_ISL_571487, EPI_ISL_571488, EPI_ISL_571489, EPI_ISL_571490, EPI_ISL_571491, EPI_ISL_571492, EPI_ISL_571493, EPI_ISL_571494, EPI_ISL_571495, EPI_ISL_571496, EPI_ISL_571497, EPI_ISL_571498, EPI_ISL_571499, EPI_ISL_571500, EPI_ISL_571501, EPI_ISL_571502, EPI_ISL_571503, EPI_ISL_571504, EPI_ISL_571505, EPI_ISL_571506, EPI_ISL_571507, EPI_ISL_571508, EPI_ISL_571509, EPI_ISL_571510, EPI_ISL_571511, EPI_ISL_571512, EPI_ISL_571513, EPI_ISL_571514, EPI_ISL_571515, EPI_ISL_571516, EPI_ISL_571517, EPI_ISL_571518, EPI_ISL_571519, EPI_ISL_571520, EPI_ISL_571521, EPI_ISL_571522, EPI_ISL_571523, EPI_ISL_571524, EPI_ISL_571525, EPI_ISL_571526, EPI_ISL_571527, EPI_ISL_571528, EPI_ISL_571529, EPI_ISL_571530, EPI_ISL_571531, EPI_ISL_571532, EPI_ISL_571533, EPI_ISL_571534, EPI_ISL_571535, EPI_ISL_571536, EPI_ISL_571537, EPI_ISL_571538, EPI_ISL_571539, EPI_ISL_571540, EPI_ISL_571541, EPI_ISL_571542, EPI_ISL_571543, EPI_ISL_571544, EPI_ISL_571545, EPI_ISL_571546, EPI_ISL_571547, EPI_ISL_571548, EPI_ISL_571549, EPI_ISL_571550, EPI_ISL_571551, EPI_ISL_571552, EPI_ISL_571553, EPI_ISL_571554, EPI_ISL_571555, EPI_ISL_571556, EPI_ISL_571557, EPI_ISL_571558, EPI_ISL_571559, EPI_ISL_571560, EPI_ISL_571561, EPI_ISL_571562, EPI_ISL_571563, EPI_ISL_571564, EPI_ISL_571565, EPI_ISL_571566, EPI_ISL_571567, EPI_ISL_571568, EPI_ISL_571569, EPI_ISL_571570, EPI_ISL_571571, EPI_ISL_571572, EPI_ISL_571573, EPI_ISL_571574, EPI_ISL_571575, EPI_ISL_571576, EPI_ISL_571577, EPI_ISL_571578, EPI_ISL_571579, EPI_ISL_571580, EPI_ISL_571581, EPI_ISL_571582, EPI_ISL_571583, EPI_ISL_571584, EPI_ISL_571585, EPI_ISL_571586, EPI_ISL_571587, EPI_ISL_571588, EPI_ISL_571589, EPI_ISL_571590, EPI_ISL_571591, EPI_ISL_571592, EPI_ISL_571593, EPI_ISL_571594, EPI_ISL_571595, EPI_ISL_571596, EPI_ISL_571597, EPI_ISL_571598, EPI_ISL_571599, EPI_ISL_571600, EPI_ISL_571601, EPI_ISL_571602, EPI_ISL_571603, EPI_ISL_571604, EPI_ISL_571605, EPI_ISL_571606, EPI_ISL_571607, EPI_ISL_571608, EPI_ISL_571609, EPI_ISL_571610, EPI_ISL_571611, EPI_ISL_571612, EPI_ISL_571613, EPI_ISL_571614, EPI_ISL_571615, EPI_ISL_571616, EPI_ISL_571617, EPI_ISL_571618, EPI_ISL_571619, EPI_ISL_571620, EPI_ISL_571621, EPI_ISL_571622, EPI_ISL_571623, EPI_ISL_571624, EPI_ISL_571625, EPI_ISL_571626, EPI_ISL_571627, EPI_ISL_571628, EPI_ISL_571629, EPI_ISL_571630, EPI_ISL_571631, EPI_ISL_571632, EPI_ISL_571633, EPI_ISL_571634, EPI_ISL_571635, EPI_ISL_571636, EPI_ISL_571637, EPI_ISL_571638, EPI_ISL_571639, EPI_ISL_571640, EPI_ISL_571641, EPI_ISL_571642, EPI_ISL_571643, EPI_ISL_571644, EPI_ISL_571645, EPI_ISL_571646, EPI_ISL_571647, EPI_ISL_571648, EPI_ISL_571649, EPI_ISL_571650, EPI_ISL_571651, EPI_ISL_571652, EPI_ISL_571653, EPI_ISL_571654, EPI_ISL_571655, EPI_ISL_571656, EPI_ISL_571657, EPI_ISL_571658, EPI_ISL_571659, EPI_ISL_571660, EPI_ISL_571661, EPI_ISL_571662, EPI_ISL_571663, EPI_ISL_571664, EPI_ISL_571665, EPI_ISL_571666, EPI_ISL_571667, EPI_ISL_571668, EPI_ISL_571669, EPI_ISL_571670, EPI_ISL_571671, EPI_ISL_571672, EPI_ISL_571673, EPI_ISL_571674, EPI_ISL_571675, EPI_ISL_571676, EPI_ISL_571677, EPI_ISL_571678, EPI_ISL_571679, EPI_ISL_571680, EPI_ISL_571681, EPI_ISL_571682, EPI_ISL_571683, EPI_ISL_571684, EPI_ISL_571685, EPI_ISL_571686, EPI_ISL_571687, EPI_ISL_571688, EPI_ISL_571689, EPI_ISL_571690, EPI_ISL_571691, EPI_ISL_571692, EPI_ISL_571693, EPI_ISL_571694, EPI_ISL_571695, EPI_ISL_571696, EPI_ISL_571697, EPI_ISL_571698, EPI_ISL_571699, EPI_ISL_571700, EPI_ISL_571701, EPI_ISL_571702, EPI_ISL_571703, EPI_ISL_571704, EPI_ISL_571705, EPI_ISL_571706, EPI_ISL_571707, EPI_ISL_571708, EPI_ISL_571709, EPI_ISL_571710, EPI_ISL_571711, EPI_ISL_571712, EPI_ISL_571713, EPI_ISL_571714, EPI_ISL_571715, EPI_ISL_571716, EPI_ISL_571717, EPI_ISL_571718, EPI_ISL_571719, EPI_ISL_571720, EPI_ISL_571721, EPI_ISL_571722, EPI_ISL_571723, EPI_ISL_571724, EPI_ISL_571725, EPI_ISL_571726, EPI_ISL_571727, EPI_ISL_571728, EPI_ISL_571729, EPI_ISL_571730, EPI_ISL_571731, EPI_ISL_571732, EPI_ISL_571733, EPI_ISL_571734, EPI_ISL_571735, EPI_ISL_571736, EPI_ISL_571737, EPI_ISL_571738, EPI_ISL_571739, EPI_ISL_571740, EPI_ISL_571741, EPI_ISL_571742, EPI_ISL_571743, EPI_ISL_571744, EPI_ISL_571745, EPI_ISL_571746, EPI_ISL_571747, EPI_ISL_571748, EPI_ISL_571749, EPI_ISL_571750, EPI_ISL_571751, EPI_ISL_571752, EPI_ISL_571753, EPI_ISL_571754, EPI_ISL_571755, EPI_ISL_571756, EPI_ISL_571757, EPI_ISL_571758, EPI_ISL_571759, EPI_ISL_571760, EPI_ISL_571761, EPI_ISL_571762, EPI_ISL_571763, EPI_ISL_571764, EPI_ISL_571765, EPI_ISL_571766, EPI_ISL_571767, EPI_ISL_571768, EPI_ISL_571769, EPI_ISL_571770, EPI_ISL_571771, EPI_ISL_571772, EPI_ISL_571773, EPI_ISL_571774, EPI_ISL_571775, EPI_ISL_571776, EPI_ISL_571777, EPI_ISL_571778, EPI_ISL_571779, EPI_ISL_571780, EPI_ISL_571781, EPI_ISL_571782, EPI_ISL_571783, EPI_ISL_571784, EPI_ISL_571785, EPI_ISL_571786, EPI_ISL_571787, EPI_ISL_571788, EPI_ISL_571789, EPI_ISL_571790, EPI_ISL_571791, EPI_ISL_571792, EPI_ISL_571793, EPI_ISL_571794, EPI_ISL_571795, EPI_ISL_571796, EPI_ISL_571797, EPI_ISL_571798, EPI_ISL_571799, EPI_ISL_571800, EPI_ISL_571801, EPI_ISL_571802, EPI_ISL_571803, EPI_ISL_571804, EPI_ISL_571805, EPI_ISL_571806, EPI_ISL_571807, EPI_ISL_571808, EPI_ISL_571809, EPI_ISL_571810, EPI_ISL_571811, EPI_ISL_571812, EPI_ISL_571813, EPI_ISL_571814, EPI_ISL_571815, EPI_ISL_571816, EPI_ISL_571817, EPI_ISL_571818, EPI_ISL_571819, EPI_ISL_571820, EPI_ISL_571821, EPI_ISL_571822, EPI_ISL_571823, EPI_ISL_571824, EPI_ISL_571825, EPI_ISL_571826, EPI_ISL_571827, EPI_ISL_571828, EPI_ISL_571829, EPI_ISL_571830, EPI_ISL_571831, EPI_ISL_571832, EPI_ISL_571833, EPI_ISL_571834, EPI_ISL_571835, EPI_ISL_571836, EPI_ISL_571837, EPI_ISL_571838, EPI_ISL_571839, EPI_ISL_571840, EPI_ISL_571841, EPI_ISL_571842, EPI_ISL_571843, EPI_ISL_571844, EPI_ISL_571845, EPI_ISL_571846, EPI_ISL_571847, EPI_ISL_571848, EPI_ISL_571849, EPI_ISL_571850, EPI_ISL_571851, EPI_ISL_571852, EPI_ISL_571853, EPI_ISL_571854, EPI_ISL_571855, EPI_ISL_571856, EPI_ISL_571857, EPI_ISL_571858, EPI_ISL_571859, EPI_ISL_571860, EPI_ISL_571861, EPI_ISL_571862, EPI_ISL_571863, EPI_ISL_571864, EPI_ISL_571865, EPI_ISL_571866, EPI_ISL_571867, EPI_ISL_571868, EPI_ISL_571869, EPI_ISL_571870, EPI_ISL_571871, EPI_ISL_571872, EPI_ISL_571873, EPI_ISL_571874, EPI_ISL_571875, EPI_ISL_571876, EPI_ISL_571877, EPI_ISL_571878, EPI_ISL_571879, EPI_ISL_571880, EPI_ISL_571881, EPI_ISL_571882, EPI_ISL_571883, EPI_ISL_571884, EPI_ISL_571885, EPI_ISL_571886, EPI_ISL_571887, EPI_ISL_571888, EPI_ISL_571889, EPI_ISL_571890, EPI_ISL_571891, EPI_ISL_571892, EPI_ISL_571893, EPI_ISL_571894, EPI_ISL_571895, EPI_ISL_571896, EPI_ISL_571897, EPI_ISL_571898, EPI_ISL_571899, EPI_ISL_571900, EPI_ISL_571901, EPI_ISL_571902, EPI_ISL_571903, EPI_ISL_571904, EPI_ISL_571905, EPI_ISL_571906, EPI_ISL_571907, EPI_ISL_571908, EPI_ISL_571909, EPI_ISL_571910, EPI_ISL_571911, EPI_ISL_571912, EPI_ISL_571913, EPI_ISL_571914, EPI_ISL_571915, EPI_ISL_571916, EPI_ISL_571917, EPI_ISL_571918, EPI_ISL_571919, EPI_ISL_571920, EPI_ISL_571921, EPI_ISL_571922, EPI_ISL_571923, EPI_ISL_571924, EPI_ISL_571925, EPI_ISL_571926, EPI_ISL_571927, EPI_ISL_571928, EPI_ISL_571929, EPI_ISL_571930, EPI_ISL_571931, EPI_ISL_571932, EPI_ISL_571933, EPI_ISL_571934, EPI_ISL_571935, EPI_ISL_571936, EPI_ISL_571937, EPI_ISL_571938, EPI_ISL_571939, EPI_ISL_571940, EPI_ISL_571941, EPI_ISL_571942, EPI_ISL_571943, EPI_ISL_571944, EPI_ISL_571945, EPI_ISL_571946, EPI_ISL_571947, EPI_ISL_571948, EPI_ISL_571949, EPI_ISL_571950, EPI_ISL_571951, EPI_ISL_571952, EPI_ISL_571953, EPI_ISL_571954, EPI_ISL_571955, EPI_ISL_571956, EPI_ISL_571957, EPI_ISL_571958, EPI_ISL_571959, EPI_ISL_571960, EPI_ISL_571961, EPI_ISL_571962, EPI_ISL_571963, EPI_ISL_571964, EPI_ISL_571965, EPI_ISL_571966, EPI_ISL_571967, EPI_ISL_571968, EPI_ISL_571969, EPI_ISL_571970, EPI_ISL_571971, EPI_ISL_571972, EPI_ISL_571973, EPI_ISL_571974, EPI_ISL_571975, EPI_ISL_571976, EPI_ISL_571977, EPI_ISL_571978, EPI_ISL_571979, EPI_ISL_571980, EPI_ISL_571981, EPI_ISL_571982, EPI_ISL_571983, EPI_ISL_571984, EPI_ISL_571985, EPI_ISL_571986, EPI_ISL_571987, EPI_ISL_571988, EPI_ISL_571989, EPI_ISL_571990, EPI_ISL_571991, EPI_ISL_571992, EPI_ISL_571993, EPI_ISL_571994, EPI_ISL_571995, EPI_ISL_571996, EPI_ISL_571997, EPI_ISL_571998, EPI_ISL_571999, EPI_ISL_572000, EPI_ISL_572001, EPI_ISL_572002, EPI_ISL_572003, EPI_ISL_572004, EPI_ISL_572005, EPI_ISL_572006, EPI_ISL_572007, EPI_ISL_572008, EPI_ISL_572009, EPI_ISL_572010, EPI_ISL_572011, EPI_ISL_572012, EPI_ISL_572013, EPI_ISL_572014, EPI_ISL_572015, EPI_ISL_572016, EPI_ISL_572017, EPI_ISL_572018, EPI_ISL_572019, EPI_ISL_572020, EPI_ISL_572021, EPI_ISL_572022, EPI_ISL_572023, EPI_ISL_572024, EPI_ISL_572025, EPI_ISL_572026, EPI_ISL_572027, EPI_ISL_572028, EPI_ISL_572029, EPI_ISL_572030, EPI_ISL_572031, EPI_ISL_572032, EPI_ISL_572033, EPI_ISL_572034, EPI_ISL_572035, EPI_ISL_572036, EPI_ISL_572037, EPI_ISL_572038, EPI_ISL_572039, EPI_ISL_572040, EPI_ISL_572041, EPI_ISL_572042, EPI_ISL_572043, EPI_ISL_572044, EPI_ISL_572045, EPI_ISL_572046, EPI_ISL_572047, EPI_ISL_572048, EPI_ISL_572049, EPI_ISL_572050, EPI_ISL_572051, EPI_ISL_572052, EPI_ISL_572053, EPI_ISL_572054, EPI_ISL_572055, EPI_ISL_572056, EPI_ISL_572057, EPI_ISL_572058, EPI_ISL_572059, EPI_ISL_572060, EPI_ISL_572061, EPI_ISL_572062, EPI_ISL_572063, EPI_ISL_572064, EPI_ISL_572065, EPI_ISL_572066, EPI_ISL_572067, EPI_ISL_572068, EPI_ISL_572069, EPI_ISL_572070, EPI_ISL_572071, EPI_ISL_572072, EPI_ISL_572073, EPI_ISL_572074, EPI_ISL_572075, EPI_ISL_572076, EPI_ISL_572077, EPI_ISL_572078, EPI_ISL_572079, EPI_ISL_572080, EPI_ISL_572081, EPI_ISL_572082, EPI_ISL_572083, EPI_ISL_572084, EPI_ISL_572085, EPI_ISL_572086, EPI_ISL_572087, EPI_ISL_572088, EPI_ISL_572089, EPI_ISL_572090, EPI_ISL_572091, EPI_ISL_572092, EPI_ISL_572093, EPI_ISL_572094, EPI_ISL_572095, EPI_ISL_572096, EPI_ISL_572097, EPI_ISL_572098, EPI_ISL_572099, EPI_ISL_572100, EPI_ISL_572101, EPI_ISL_572102, EPI_ISL_572103, EPI_ISL_572104, EPI_ISL_572105, EPI_ISL_572106, EPI_ISL_572107, EPI_ISL_572108, EPI_ISL_572109, EPI_ISL_572110, EPI_ISL_572111, EPI_ISL_572112, EPI_ISL_572113, EPI_ISL_572114, EPI_ISL_572115, EPI_ISL_572116, EPI_ISL_572117, EPI_ISL_572118, EPI_ISL_572119, EPI_ISL_572120, EPI_ISL_572121, EPI_ISL_572122, EPI_ISL_572123, EPI_ISL_572124, EPI_ISL_572125, EPI_ISL_572126, EPI_ISL_572127, EPI_ISL_572128, EPI_ISL_572129, EPI_ISL_572130, EPI_ISL_572131, EPI_ISL_572132, EPI_ISL_572133, EPI_ISL_572134, EPI_ISL_572135, EPI_ISL_572136, EPI_ISL_572137, EPI_ISL_572138, EPI_ISL_572139, EPI_ISL_572140, EPI_ISL_572141, EPI_ISL_572142, EPI_ISL_572143, EPI_ISL_572144, EPI_ISL_572145, EPI_ISL_572146, EPI_ISL_572147, EPI_ISL_572148, EPI_ISL_572149, EPI_ISL_572150, EPI_ISL_572151, EPI_ISL_572152, EPI_ISL_572153, EPI_ISL_572154, EPI_ISL_572155, EPI_ISL_572156, EPI_ISL_572157, EPI_ISL_572158, EPI_ISL_572159, EPI_ISL_572160, EPI_ISL_572161, EPI_ISL_572162, EPI_ISL_572163, EPI_ISL_572164, EPI_ISL_572165, EPI_ISL_572166, EPI_ISL_572167, EPI_ISL_572168, EPI_ISL_572169, EPI_ISL_572170, EPI_ISL_572171, EPI_ISL_572172, EPI_ISL_572173, EPI_ISL_572174, EPI_ISL_572175, EPI_ISL_572176, EPI_ISL_572177, EPI_ISL_572178, EPI_ISL_572179, EPI_ISL_572180, EPI_ISL_572181, EPI_ISL_572182, EPI_ISL_572183, EPI_ISL_572184, EPI_ISL_572185 | | | |
| see above | Quest Diagnostics | Quest Diagnostics | Rosenthal,S.H., Gerasimova,A., Kagan,R.M., Anderson, B., Grover, D., Livingston, K.E., Hua, M., Liu Y., Shalhout, D.F., Owen, R., Lacbawan, F. |
| EPI_ISL_574301, EPI_ISL_574302, EPI_ISL_574303, EPI_ISL_574304, EPI_ISL_574305, EPI_ISL_574306, EPI_ISL_574307, EPI_ISL_574308, EPI_ISL_574309, EPI_ISL_574310, EPI_ISL_574311, EPI_ISL_574312, EPI_ISL_574313, EPI_ISL_574314, EPI_ISL_574315, EPI_ISL_574316, EPI_ISL_574317, EPI_ISL_574318, EPI_ISL_574319 | | | |
| see above | LSUHS Emerging Viral Threat Laboratory | Microbial Genome Sequencing Center | Rona S. Scott, Jeremy P. Kamil, Maarten Van Diest, Malgorzata Bienkowska-Haba, Katarzyna Zwolinska, Andrew D. Yurochko, Christopher G. Kevil, Martin J. Sapp, Daniel J. Snyder, Vaughn S. Cooper, John A. Vanchiere |
| EPI_ISL_574434, EPI_ISL_574435, EPI_ISL_574436, EPI_ISL_574437, EPI_ISL_574438, EPI_ISL_574439, EPI_ISL_574440, EPI_ISL_574441, EPI_ISL_574442, EPI_ISL_574443, EPI_ISL_574444, EPI_ISL_574445, EPI_ISL_574446, EPI_ISL_574447, EPI_ISL_574448, EPI_ISL_574449, EPI_ISL_574450, EPI_ISL_574451, EPI_ISL_574452, EPI_ISL_574453, EPI_ISL_574454, EPI_ISL_574455, EPI_ISL_574456, EPI_ISL_574457, EPI_ISL_574458, EPI_ISL_574459, EPI_ISL_574460, EPI_ISL_574461, EPI_ISL_574462, EPI_ISL_574463, EPI_ISL_574464, EPI_ISL_574465, EPI_ISL_574466, EPI_ISL_574467, EPI_ISL_574468, EPI_ISL_574469, EPI_ISL_574470, EPI_ISL_574471, EPI_ISL_574472, EPI_ISL_574473, EPI_ISL_574474, EPI_ISL_574475, EPI_ISL_574476, EPI_ISL_574477, EPI_ISL_574478 | | | |
| see above | Texas Department of State Health Services | Texas Department of State Health Services | Rashmi Tuladhar, Bonnie Oh, Jenny Zhang, Maliha Rahman, Anita Pokharel, Myong Koag, Chun Wang, Rachel Lee, Grace Kubin |
| EPI_ISL_575329, EPI_ISL_575330 | National Institute for Viral Disease Control and Prevention, China CDC | National Institute for Viral Disease Control and Prevention, China CDC | Rongbao Gao, Kang Xiao, Qinqin Song, ZhiQiang Xia, Dong Xia, Juan Song, Haijun Du, Yuan He, Shuai Pang, Xuancheng Lu, Guizhen Wu, Geogia Fu Gao, Jun Han |
| EPI_ISL_576281, EPI_ISL_576282, EPI_ISL_576283, EPI_ISL_576284, EPI_ISL_576285, EPI_ISL_576286, EPI_ISL_576287, EPI_ISL_576288, EPI_ISL_576289, EPI_ISL_576290, EPI_ISL_576291, EPI_ISL_576292, EPI_ISL_576293, EPI_ISL_576294, EPI_ISL_576295, EPI_ISL_576296, EPI_ISL_576297, EPI_ISL_576298, EPI_ISL_576299, EPI_ISL_576300, EPI_ISL_576301, EPI_ISL_576302, EPI_ISL_576303, EPI_ISL_576304, EPI_ISL_576305, EPI_ISL_576306, EPI_ISL_576307, EPI_ISL_576308, EPI_ISL_576309, EPI_ISL_576310, EPI_ISL_576311, EPI_ISL_576312, EPI_ISL_576313, EPI_ISL_576314, EPI_ISL_576315, EPI_ISL_576316, EPI_ISL_576317, EPI_ISL_576318, EPI_ISL_576319, EPI_ISL_576320, EPI_ISL_576321, EPI_ISL_576322, EPI_ISL_576323, EPI_ISL_576324, EPI_ISL_576325 | | | |
| see above | Texas Department of State Health Services | Texas Department of State Health Services | Rashmi Tuladhar, Bonnie Oh, Jenny Zhang, Maliha Rahman, Anita Pokharel, Myong Koag, Chun Wang, Rachel Lee, Grace Kubin |
| EPI_ISL_576326, EPI_ISL_576327, EPI_ISL_576328, EPI_ISL_576329, EPI_ISL_576330, EPI_ISL_576331, EPI_ISL_576332, EPI_ISL_576333, EPI_ISL_576334, EPI_ISL_576335, EPI_ISL_576336, EPI_ISL_576337, EPI_ISL_576338, EPI_ISL_576339, EPI_ISL_576340, EPI_ISL_576341, EPI_ISL_576342, EPI_ISL_576343, EPI_ISL_576344, EPI_ISL_576345, EPI_ISL_576346, EPI_ISL_576347, EPI_ISL_576348, EPI_ISL_576349, EPI_ISL_576350, EPI_ISL_576351, EPI_ISL_576352, EPI_ISL_576353, EPI_ISL_576354, EPI_ISL_576355, EPI_ISL_576356, EPI_ISL_576357, EPI_ISL_576358, EPI_ISL_576359, EPI_ISL_576360, EPI_ISL_576361, EPI_ISL_576362, EPI_ISL_576568 | | | |
| see above | Texas Department of State Health Services | Texas Department of State Health Services | Rashmi Tuladhar, Bonnie Oh, Mayela Pedrueza, Jenny Zhang, Maliha Rahman, Anita Pokharel, Myong Koag, Chun Wang, Rachel Lee, Grace Kubin |
| EPI_ISL_577597 | Area of Virology, Serology and Virology Division (SAViD), New South Wales Health Pathology Randwick | Area of Virology, Serology and Virology Division (SAViD), New South Wales Health Pathology Randwick | Rawlinson, W., Bull, R., Deveson, I. |
| EPI_ISL_577604 | Area of Virology, Serology and Virology Division (SAViD), New South Wales Health Pathology Randwick | Area of Virology, Serology and Virology Division (SAViD), New South Wales Health Pathology Randwick | Rawlinson, W., Deveson, I., Bull, R., Van Hal, S. |
| EPI_ISL_578728, EPI_ISL_578729, EPI_ISL_578730, EPI_ISL_578731, EPI_ISL_578732, EPI_ISL_578733, EPI_ISL_578734, EPI_ISL_578735, EPI_ISL_578736, EPI_ISL_578737, EPI_ISL_578738, EPI_ISL_578739, EPI_ISL_578740, EPI_ISL_578741, EPI_ISL_578742, EPI_ISL_578743, EPI_ISL_578744, EPI_ISL_578745, EPI_ISL_578746, EPI_ISL_578747, EPI_ISL_578748, EPI_ISL_578749, EPI_ISL_578750, EPI_ISL_578751, EPI_ISL_578752 | | | |
| see above | LSUHS Emerging Viral Threat Laboratory | Microbial Genome Sequencing Center | Rona S. Scott, Jeremy P. Kamil, Maarten Van Diest, Malgorzata Bienkowska-Haba, Katarzyna Zwolinska, Andrew D. Yurochko, Christopher G. Kevil, Martin J. Sapp, Daniel J. Snyder, Vaughn S. Cooper, John A. Vanchiere |
| EPI_ISL_580282, EPI_ISL_580283, EPI_ISL_580286, EPI_ISL_580287, EPI_ISL_580299, EPI_ISL_580300, EPI_ISL_580301, EPI_ISL_580305, EPI_ISL_580313, EPI_ISL_580322, EPI_ISL_580325, EPI_ISL_580327, EPI_ISL_580333, EPI_ISL_580338, EPI_ISL_580340, EPI_ISL_580341, EPI_ISL_580353, EPI_ISL_580354, EPI_ISL_580362, EPI_ISL_580364, EPI_ISL_580367, EPI_ISL_580368, EPI_ISL_580369, EPI_ISL_580371, EPI_ISL_580373, EPI_ISL_580385, EPI_ISL_580391, EPI_ISL_580393, EPI_ISL_580394, EPI_ISL_580396, EPI_ISL_580400, EPI_ISL_580414, EPI_ISL_580418, EPI_ISL_580421, EPI_ISL_580435, EPI_ISL_580444, EPI_ISL_580449, EPI_ISL_580453, EPI_ISL_580459, EPI_ISL_580465, EPI_ISL_580467, EPI_ISL_580471, EPI_ISL_580478, EPI_ISL_580479, EPI_ISL_580480, EPI_ISL_580481, EPI_ISL_580485, EPI_ISL_580486, EPI_ISL_580495, EPI_ISL_580499, EPI_ISL_580502, EPI_ISL_580508, EPI_ISL_580509, EPI_ISL_580511, EPI_ISL_580515, EPI_ISL_580532, EPI_ISL_580533, EPI_ISL_580536, EPI_ISL_580550, EPI_ISL_580557, EPI_ISL_580559, EPI_ISL_580564, EPI_ISL_580565, EPI_ISL_580567, EPI_ISL_580569, EPI_ISL_580573, EPI_ISL_580576, EPI_ISL_580577, EPI_ISL_580579, EPI_ISL_580582, EPI_ISL_580586, EPI_ISL_580587, EPI_ISL_580593, EPI_ISL_580603, EPI_ISL_580607, EPI_ISL_580610, EPI_ISL_580618, EPI_ISL_580621, EPI_ISL_580625, EPI_ISL_580626, EPI_ISL_581394, EPI_ISL_581396 | | | |

| | | | |
|---|---|---|---|
| see above | Lighthouse Lab in Cambridge | Wellcome Sanger Institute for the COVID-19 Genomics UK (COG-UK) consortium | Rob Howes, The Lighthouse Lab in Cambridge and Alex Alderton, Roberto Amato, Sonia Goncalves, Ewan Harrison, David K. Jackson, Ian Johnston, Dominic Kwiatkowski, Cordelia Langford, John Sillitoe on behalf of the Wellcome Sanger Institute COVID-19 Surveillance Team |
| EPI_ISL_583917 | Servicio de Microbiología. Hospital General Universitario de Castellón | SeqCOVID-SPAIN consortium/IBV (CSIC) | Rosario Moreno, María Dolores Tirado and SeqCOVID-SPAIN consortium |
| EPI_ISL_583918, EPI_ISL_583919, EPI_ISL_583920, EPI_ISL_583921, EPI_ISL_583922, EPI_ISL_583923, EPI_ISL_583924, EPI_ISL_583925, EPI_ISL_583926, EPI_ISL_583927, EPI_ISL_583928, EPI_ISL_583929, EPI_ISL_583930, EPI_ISL_583931, EPI_ISL_583932, EPI_ISL_583933, EPI_ISL_583934, EPI_ISL_583935, EPI_ISL_583936, EPI_ISL_583937, EPI_ISL_583938, EPI_ISL_583939, EPI_ISL_583940, EPI_ISL_583941, EPI_ISL_583942, EPI_ISL_583943, EPI_ISL_583944, EPI_ISL_583945, EPI_ISL_583946, EPI_ISL_583947, EPI_ISL_583948, EPI_ISL_583949, EPI_ISL_583950, EPI_ISL_583951, EPI_ISL_583952 | | | |
| see above | Servicio de Microbiología. Hospital General Universitario de Castellón | SeqCOVID-SPAIN consortium/IBV(CSIC) | Rosario Moreno, María Dolores Tirado and SeqCOVID-SPAIN consortium |
| EPI_ISL_584123, EPI_ISL_584124, EPI_ISL_584125, EPI_ISL_584126, EPI_ISL_584127, EPI_ISL_584128, EPI_ISL_584129, EPI_ISL_584130, EPI_ISL_584131, EPI_ISL_584132, EPI_ISL_584133, EPI_ISL_584134, EPI_ISL_584135, EPI_ISL_584136, EPI_ISL_584137, EPI_ISL_584138, EPI_ISL_584139, EPI_ISL_584140, EPI_ISL_584141, EPI_ISL_584142, EPI_ISL_584143, EPI_ISL_584144, EPI_ISL_584145, EPI_ISL_584146, EPI_ISL_584147, EPI_ISL_584148, EPI_ISL_584149, EPI_ISL_584150, EPI_ISL_584151, EPI_ISL_584152, EPI_ISL_584153, EPI_ISL_584154, EPI_ISL_584155, EPI_ISL_584156, EPI_ISL_584157, EPI_ISL_584158, EPI_ISL_584159, EPI_ISL_584160, EPI_ISL_584161, EPI_ISL_584162, EPI_ISL_584163, EPI_ISL_584164 | | | |
| see above | Texas Department of State Health Services | Texas Department of State Health Services | Rashmi Tuladhar, Bonnie Oh, Jenny Zhang, Maliha Rahman, Anita Pokharel, Mayela Pedrueza, Myong Koag, Chun Wang, Rachel Lee, Grace Kubin |
| EPI_ISL_584440, EPI_ISL_584441, EPI_ISL_584442, EPI_ISL_584443, EPI_ISL_584444, EPI_ISL_584445, EPI_ISL_584446, EPI_ISL_584447, EPI_ISL_584448, EPI_ISL_584449, EPI_ISL_584450, EPI_ISL_584451, EPI_ISL_584452, EPI_ISL_584453, EPI_ISL_584454, EPI_ISL_584455, EPI_ISL_584456, EPI_ISL_584457, EPI_ISL_584458, EPI_ISL_584459, EPI_ISL_584460, EPI_ISL_584461, EPI_ISL_584462, EPI_ISL_584463, EPI_ISL_584464, EPI_ISL_584465, EPI_ISL_584466, EPI_ISL_584467, EPI_ISL_584468, EPI_ISL_584469, EPI_ISL_584470, EPI_ISL_584471, EPI_ISL_584472, EPI_ISL_584473, EPI_ISL_584474, EPI_ISL_584475, EPI_ISL_584476, EPI_ISL_584477, EPI_ISL_584478, EPI_ISL_584479, EPI_ISL_584480, EPI_ISL_584481, EPI_ISL_584482, EPI_ISL_584483, EPI_ISL_584484, EPI_ISL_584485, EPI_ISL_584486, EPI_ISL_584487, EPI_ISL_584488, EPI_ISL_584489, EPI_ISL_584490, EPI_ISL_584491, EPI_ISL_584492, EPI_ISL_584493, EPI_ISL_584494, EPI_ISL_584495, EPI_ISL_584496, EPI_ISL_584497, EPI_ISL_584498, EPI_ISL_584499, EPI_ISL_584500, EPI_ISL_584501, EPI_ISL_584502, EPI_ISL_584503, EPI_ISL_584504, EPI_ISL_584505, EPI_ISL_584506, EPI_ISL_584507, EPI_ISL_584508, EPI_ISL_584509, EPI_ISL_584510, EPI_ISL_584511, EPI_ISL_584512, EPI_ISL_584513, EPI_ISL_584514, EPI_ISL_584515, EPI_ISL_584516, EPI_ISL_584517, EPI_ISL_584518, EPI_ISL_584519, EPI_ISL_584520, EPI_ISL_584521, EPI_ISL_584522, EPI_ISL_584523, EPI_ISL_584524, EPI_ISL_584525, EPI_ISL_584526, EPI_ISL_584527, EPI_ISL_584528, EPI_ISL_584529, EPI_ISL_584530, EPI_ISL_584531, EPI_ISL_584532, EPI_ISL_584533, EPI_ISL_584534, EPI_ISL_584535, EPI_ISL_584536, EPI_ISL_584537, EPI_ISL_584538, EPI_ISL_584539, EPI_ISL_584540, EPI_ISL_584541, EPI_ISL_584542, EPI_ISL_584543, EPI_ISL_584544, EPI_ISL_584545, EPI_ISL_584546, EPI_ISL_584547, EPI_ISL_584548, EPI_ISL_584549, EPI_ISL_584550, EPI_ISL_584551, EPI_ISL_584552, EPI_ISL_584553, EPI_ISL_584554, EPI_ISL_584555, EPI_ISL_584556, EPI_ISL_584557, EPI_ISL_584558, EPI_ISL_584559, EPI_ISL_584560, EPI_ISL_584561, EPI_ISL_584562, EPI_ISL_584563, EPI_ISL_584564, EPI_ISL_584565, EPI_ISL_584566, EPI_ISL_584567, EPI_ISL_584568, EPI_ISL_584569, EPI_ISL_584570, EPI_ISL_584571, EPI_ISL_584572, EPI_ISL_584573, EPI_ISL_584574, EPI_ISL_584575, EPI_ISL_584576, EPI_ISL_584577, EPI_ISL_584578, EPI_ISL_584579, EPI_ISL_584580, EPI_ISL_584581, EPI_ISL_584582, EPI_ISL_584583, EPI_ISL_584584, EPI_ISL_584585, EPI_ISL_584586, EPI_ISL_584587, EPI_ISL_584588, EPI_ISL_584589, EPI_ISL_584590 | | | |
| see above | UHCW / University of Warwick | COVID-19 Genomics UK (COG-UK) Consortium | Richard Stark, Chrystala Constantinidou, Meera Unnikrishnan, Laura Baxter, Jeff Cheng, Grace Taylor-Joyce, Hannah Elizabeth Bridgewater, Lucy Frost, Sarojini Pandey, Paul Brown, Tauqeer Alam, Sascha Ott, Dimitris Grammatopoulos |
| EPI_ISL_586517 | B.J. Medical College and Civil hospital, Ahmedabad | Gujarat Biotechnology Research Centre | Raghawendra Kumar, Dinesh Kumar, Zuber Saiyed, Komal Patel, Labdhi Pandya, Afzal Ansari, Nikha Trivedi, Pranay Shah, Kamlesh J Upadhyay, Sanjay Kapadia, Apurvasinh Puvar, Janvi Raval, Zarna Patel, Monika Gandhi, Pinal Trivedi, Maharshi Pandya, Nidhi Patel, Nitin Savaliya, R D Dixit, A M Kadri, Harsh Bakshi, Chaitanya Joshi, Madhvi Joshi |
| EPI_ISL_586539 | Zydus Medical Hospital,Dahod | Gujarat Biotechnology Research Centre | Raghawendra Kumar, Dinesh Kumar, Zuber Saiyed, Komal Patel, Labdhi Pandya, Afzal Ansari, Nikha Trivedi, Apurvasinh Puvar, Janvi Raval, Zarna Patel, Monika Gandhi, Pinal Trivedi, Maharshi Pandya, Nidhi Patel, Kalpesh Labana, Mohit Desai, Nitin Savaliya, R D Dixit, A M Kadri, Harsh Bakshi, Chaitanya Joshi, Madhvi Joshi |
| EPI_ISL_586556 | CCC,Veraval | Gujarat Biotechnology Research Centre | Raghawendra Kumar, Dinesh Kumar, Zuber Saiyed, Komal Patel, Labdhi Pandya, Afzal Ansari, Nikha Trivedi, Jignesh Parmar, Jitendra Bamrotia, Apurvasinh Puvar, Janvi Raval, Zarna Patel, Monika Gandhi, Pinal Trivedi, Maharshi Pandya, Nidhi Patel, Nitin Savaliya, R D Dixit, A M Kadri, Harsh Bakshi, Chaitanya Joshi, Madhvi Joshi |
| EPI_ISL_586564 | GMERS GMERS Medical College and Hospital, Junagadh | Gujarat Biotechnology Research Centre | Ravi Dedaniya, Apurvasinh Puvar, Janvi Raval, Zarna Patel, Monika Gandhi, Pinal Trivedi, Maharshi Pandya, Nidhi Patel, Nitin Savaliya, Raghawendra Kumar, Dinesh Kumar, Zuber Saiyed, Komal Patel, Labdhi Pandya, Afzal Ansari, Nikha Trivedi, Bhavesh Bagda , R D Dixit, A M Kadri, Harsh Bakshi, Chaitanya Joshi, Madhvi Joshi |
| EPI_ISL_586571 | Area of Virology, Serology and Virology Division (SAViD), New South Wales Health Pathology Randwick | Area of Virology, Serology and Virology Division (SAViD), New South Wales Health Pathology Randwick | Rawlinson, W., Deveson, I., Bull, R. |
| EPI_ISL_590173 | Lighthouse Lab in Cambridge | Wellcome Sanger Institute for the COVID-19 Genomics UK (COG-UK) consortium | Rob Howes, The Lighthouse Lab in Cambridge and Alex Alderton, Roberto Amato, Sonia Goncalves, Ewan Harrison, David K. Jackson, Ian Johnston, Dominic Kwiatkowski, Cordelia Langford, John Sillitoe on behalf of the Wellcome Sanger Institute COVID-19 Surveillance Team (http://www.sanger.ac.uk/covid-team) |
| EPI_ISL_590838, EPI_ISL_590839, EPI_ISL_590840, EPI_ISL_590841, EPI_ISL_590842, EPI_ISL_590843, EPI_ISL_590844, EPI_ISL_590845, EPI_ISL_590846, EPI_ISL_590847, EPI_ISL_590848, EPI_ISL_590849, EPI_ISL_590850, EPI_ISL_590851, EPI_ISL_590852, EPI_ISL_590853, EPI_ISL_590854, EPI_ISL_590855, EPI_ISL_590856, EPI_ISL_590857, EPI_ISL_590858, EPI_ISL_590859, EPI_ISL_590860, EPI_ISL_590861, EPI_ISL_590862, EPI_ISL_590863, EPI_ISL_590864, EPI_ISL_590865, EPI_ISL_590866, EPI_ISL_590867, EPI_ISL_590868, EPI_ISL_590869, EPI_ISL_590870, EPI_ISL_590871, EPI_ISL_590872, EPI_ISL_590873, EPI_ISL_590874, EPI_ISL_590875, EPI_ISL_593559, EPI_ISL_593560, EPI_ISL_593561, EPI_ISL_593562, EPI_ISL_593563, EPI_ISL_593564, EPI_ISL_593565, EPI_ISL_593566, EPI_ISL_593567, EPI_ISL_593568, EPI_ISL_593569, EPI_ISL_593570, EPI_ISL_593571, EPI_ISL_593572, EPI_ISL_593573, EPI_ISL_593574, EPI_ISL_593575, EPI_ISL_593576, EPI_ISL_593577, EPI_ISL_593578, EPI_ISL_593579, EPI_ISL_593580, EPI_ISL_593581, EPI_ISL_593582, EPI_ISL_593583, EPI_ISL_593584, EPI_ISL_593585, EPI_ISL_593586, EPI_ISL_593587, EPI_ISL_593588, EPI_ISL_593589, EPI_ISL_593590, EPI_ISL_593591, EPI_ISL_593592, EPI_ISL_593593, EPI_ISL_593594, EPI_ISL_593595 | | | |
| see above | Texas Department of State Health Services | Texas Department of State Health Services | Rashmi Tuladhar, Bonnie Oh, Jenny Zhang, Maliha Rahman, Anita Pokharel, Myong Koag, Chung Wang, Rachel Lee, Grace Kubin, Mayela Pedrueza |
| EPI_ISL_596928, EPI_ISL_596929, EPI_ISL_596930, EPI_ISL_596931, EPI_ISL_596932, EPI_ISL_596933, EPI_ISL_596934, EPI_ISL_596935, EPI_ISL_596936, EPI_ISL_596937, EPI_ISL_596938, EPI_ISL_596939, EPI_ISL_596940, EPI_ISL_596941, EPI_ISL_596942, EPI_ISL_596943, EPI_ISL_596944, EPI_ISL_596945, EPI_ISL_596946, EPI_ISL_596947, EPI_ISL_596948, EPI_ISL_596949, EPI_ISL_596950, EPI_ISL_596951, EPI_ISL_596952, EPI_ISL_596953, EPI_ISL_596954, EPI_ISL_596955, EPI_ISL_596956, EPI_ISL_596957, EPI_ISL_596958, EPI_ISL_596959, EPI_ISL_596960, EPI_ISL_596961, EPI_ISL_596962, EPI_ISL_596963, EPI_ISL_596964, EPI_ISL_596965, EPI_ISL_596966, EPI_ISL_596967, EPI_ISL_596968, EPI_ISL_596970, EPI_ISL_596971, EPI_ISL_596972, EPI_ISL_596973, EPI_ISL_596974, EPI_ISL_596975, EPI_ISL_596976, EPI_ISL_596977, EPI_ISL_596978, EPI_ISL_596979, EPI_ISL_596980, EPI_ISL_596981, EPI_ISL_596982, EPI_ISL_596983, EPI_ISL_596984, EPI_ISL_596985, EPI_ISL_596986, EPI_ISL_596987, EPI_ISL_596988, EPI_ISL_596989, EPI_ISL_596990, EPI_ISL_596991, EPI_ISL_596992, EPI_ISL_596993, EPI_ISL_596994, EPI_ISL_596995, EPI_ISL_596996, EPI_ISL_596997, EPI_ISL_596998, EPI_ISL_596999, EPI_ISL_597000, EPI_ISL_597001, EPI_ISL_597002, EPI_ISL_597003, EPI_ISL_597004, EPI_ISL_597005, EPI_ISL_597006, EPI_ISL_597007, EPI_ISL_597008, EPI_ISL_597009, EPI_ISL_597010, EPI_ISL_597011, EPI_ISL_597012, EPI_ISL_597013, EPI_ISL_597014, EPI_ISL_597015, EPI_ISL_597016, EPI_ISL_597017, EPI_ISL_597018, EPI_ISL_597019, EPI_ISL_597020, EPI_ISL_597021, EPI_ISL_597022, EPI_ISL_597023, EPI_ISL_597024, EPI_ISL_597025, EPI_ISL_597026, EPI_ISL_597027, EPI_ISL_597028, EPI_ISL_597029, EPI_ISL_597030, EPI_ISL_597031, EPI_ISL_597032, EPI_ISL_597033, EPI_ISL_597034, EPI_ISL_597035, EPI_ISL_597036, EPI_ISL_597037, EPI_ISL_597038, EPI_ISL_597039, EPI_ISL_597040, EPI_ISL_597041, EPI_ISL_597042, EPI_ISL_597043, EPI_ISL_597044, EPI_ISL_597045, EPI_ISL_597046, EPI_ISL_597047, EPI_ISL_597048, EPI_ISL_597049, EPI_ISL_597050, EPI_ISL_597051, EPI_ISL_597052, EPI_ISL_597053, EPI_ISL_597054, EPI_ISL_597055, EPI_ISL_597056, EPI_ISL_597057, EPI_ISL_597058, EPI_ISL_597059, EPI_ISL_597060, EPI_ISL_597061, EPI_ISL_597062, EPI_ISL_597063, EPI_ISL_597064, EPI_ISL_597065, EPI_ISL_597066, EPI_ISL_597067, EPI_ISL_597068, EPI_ISL_597069, EPI_ISL_597070, EPI_ISL_597071, EPI_ISL_597072, EPI_ISL_597073, EPI_ISL_597075, EPI_ISL_597076, EPI_ISL_597077, EPI_ISL_597078, EPI_ISL_597079, EPI_ISL_597080, EPI_ISL_597081, EPI_ISL_597082, EPI_ISL_597083, EPI_ISL_597084, EPI_ISL_597085, EPI_ISL_597086, EPI_ISL_597087, EPI_ISL_597088, EPI_ISL_597089, EPI_ISL_597090, EPI_ISL_597091, EPI_ISL_597092, EPI_ISL_597093, EPI_ISL_597094, EPI_ISL_597095, EPI_ISL_597096, EPI_ISL_597097, EPI_ISL_597098, EPI_ISL_597100, EPI_ISL_597101, EPI_ISL_597102, EPI_ISL_597103, EPI_ISL_597104, EPI_ISL_597105, EPI_ISL_597106, EPI_ISL_597107, EPI_ISL_597108, EPI_ISL_597109, EPI_ISL_597110, EPI_ISL_597111, EPI_ISL_597112, EPI_ISL_597113, EPI_ISL_597114, EPI_ISL_597115, EPI_ISL_597116, EPI_ISL_597117, EPI_ISL_597118, EPI_ISL_597119, EPI_ISL_597120, EPI_ISL_597121, EPI_ISL_597122, EPI_ISL_597123, EPI_ISL_597124, EPI_ISL_597125, EPI_ISL_597126, EPI_ISL_597127, EPI_ISL_597128, EPI_ISL_597129, EPI_ISL_597130, EPI_ISL_597131, EPI_ISL_597132, EPI_ISL_597133, EPI_ISL_597134, EPI_ISL_597135, EPI_ISL_597136, EPI_ISL_597137, EPI_ISL_597138, EPI_ISL_597139, EPI_ISL_597140, EPI_ISL_597141, EPI_ISL_597142, EPI_ISL_597143, EPI_ISL_597144, EPI_ISL_597145, EPI_ISL_597146, EPI_ISL_597147, EPI_ISL_597148, EPI_ISL_597149, EPI_ISL_597150, EPI_ISL_597151, EPI_ISL_597152, EPI_ISL_597153, EPI_ISL_597154, EPI_ISL_597155, EPI_ISL_597156, EPI_ISL_597157, EPI_ISL_597158, EPI_ISL_597159, EPI_ISL_597160, EPI_ISL_597161, EPI_ISL_597162, EPI_ISL_597163, EPI_ISL_597164, EPI_ISL_597165, EPI_ISL_597166, EPI_ISL_597167, EPI_ISL_597168, EPI_ISL_597169, EPI_ISL_597170, EPI_ISL_597171, EPI_ISL_597172, EPI_ISL_597173, EPI_ISL_597174, EPI_ISL_597175, EPI_ISL_597176, EPI_ISL_597177, EPI_ISL_597178, EPI_ISL_597179, EPI_ISL_597180, EPI_ISL_597181, EPI_ISL_597182, EPI_ISL_597183, EPI_ISL_597184, EPI_ISL_597185, EPI_ISL_597186, EPI_ISL_597187, EPI_ISL_597188, EPI_ISL_597190, EPI_ISL_597191, EPI_ISL_597192, EPI_ISL_597193, EPI_ISL_597194, EPI_ISL_597195, EPI_ISL_597196, EPI_ISL_597197, EPI_ISL_597198, EPI_ISL_597199, EPI_ISL_597200, EPI_ISL_597201, EPI_ISL_597202, EPI_ISL_597203, EPI_ISL_597204, EPI_ISL_597205, EPI_ISL_597206, EPI_ISL_597207, EPI_ISL_597208, EPI_ISL_597209, EPI_ISL_597210, EPI_ISL_597211, EPI_ISL_597212, EPI_ISL_597213, EPI_ISL_597214, EPI_ISL_597215, EPI_ISL_597216, EPI_ISL_597217, EPI_ISL_597218, EPI_ISL_597219, EPI_ISL_597220, EPI_ISL_597221, EPI_ISL_597222, EPI_ISL_597223, EPI_ISL_597224, EPI_ISL_597225, EPI_ISL_597226, EPI_ISL_597227, EPI_ISL_597228, EPI_ISL_597229, EPI_ISL_597230, EPI_ISL_597231, EPI_ISL_597232, EPI_ISL_597233, EPI_ISL_597237, EPI_ISL_597238, EPI_ISL_597239, EPI_ISL_597241, EPI_ISL_597243, EPI_ISL_597246, EPI_ISL_597247, EPI_ISL_597248, EPI_ISL_597250, EPI_ISL_597251, EPI_ISL_597253, EPI_ISL_597254, EPI_ISL_597255, EPI_ISL_597256, EPI_ISL_597257, EPI_ISL_597258, EPI_ISL_597259, EPI_ISL_597262, EPI_ISL_597264, EPI_ISL_597265, EPI_ISL_597266, EPI_ISL_597269, EPI_ISL_597270, EPI_ISL_597271, EPI_ISL_597273, EPI_ISL_597274, EPI_ISL_597279, EPI_ISL_597282, EPI_ISL_597283, EPI_ISL_597284, EPI_ISL_597285, EPI_ISL_597286, EPI_ISL_597287, EPI_ISL_597289, EPI_ISL_597293, EPI_ISL_597294, EPI_ISL_597295, EPI_ISL_597297, EPI_ISL_597300, EPI_ISL_597301, EPI_ISL_597304, EPI_ISL_597305, EPI_ISL_597306, EPI_ISL_597307, EPI_ISL_597308, EPI_ISL_597309, EPI_ISL_597310, EPI_ISL_597316, EPI_ISL_597317, EPI_ISL_597319, EPI_ISL_597320, EPI_ISL_597321, EPI_ISL_597323, EPI_ISL_597324, EPI_ISL_597325, EPI_ISL_597327, EPI_ISL_597331, EPI_ISL_597335, EPI_ISL_597337, EPI_ISL_597340, EPI_ISL_597341, EPI_ISL_597342, EPI_ISL_597343, EPI_ISL_597344, EPI_ISL_597345, EPI_ISL_597346, EPI_ISL_597349, EPI_ISL_597350, EPI_ISL_597354, EPI_ISL_597358, EPI_ISL_597361, EPI_ISL_597365, EPI_ISL_597366, EPI_ISL_597367, EPI_ISL_597368, EPI_ISL_597371, EPI_ISL_597372, EPI_ISL_597373, EPI_ISL_597374, EPI_ISL_597375, EPI_ISL_597377, EPI_ISL_597378, EPI_ISL_597380, EPI_ISL_597381, EPI_ISL_597388, EPI_ISL_597390, EPI_ISL_597392, EPI_ISL_597396, EPI_ISL_597397, EPI_ISL_597398, EPI_ISL_597399, EPI_ISL_597401, EPI_ISL_597402, EPI_ISL_597403, EPI_ISL_597405, EPI_ISL_597412, EPI_ISL_597414, EPI_ISL_597418, EPI_ISL_597424, EPI_ISL_597425, EPI_ISL_597426, EPI_ISL_597427, EPI_ISL_597428, EPI_ISL_597432, EPI_ISL_597435, EPI_ISL_597437, EPI_ISL_597438, EPI_ISL_597441, EPI_ISL_597445, EPI_ISL_597446, EPI_ISL_597449, EPI_ISL_597450, EPI_ISL_597451, EPI_ISL_597453, EPI_ISL_597458, EPI_ISL_597459, EPI_ISL_597460, EPI_ISL_597461, EPI_ISL_597462, EPI_ISL_597464, EPI_ISL_597466, EPI_ISL_597467, EPI_ISL_597468, EPI_ISL_597469, EPI_ISL_597470, EPI_ISL_597472, EPI_ISL_597473, EPI_ISL_597476, EPI_ISL_597477, EPI_ISL_597478, EPI_ISL_597479, EPI_ISL_597480, EPI_ISL_597483, EPI_ISL_597487, EPI_ISL_597488, EPI_ISL_597489, EPI_ISL_597490, EPI_ISL_597494, EPI_ISL_597495, EPI_ISL_597497, EPI_ISL_597502, EPI_ISL_597504, EPI_ISL_597505, EPI_ISL_597507, EPI_ISL_597508, EPI_ISL_597510, EPI_ISL_597512, EPI_ISL_597517, EPI_ISL_597519, EPI_ISL_597520, EPI_ISL_601493, EPI_ISL_601495, EPI_ISL_601499, EPI_ISL_601501, EPI_ISL_601506, EPI_ISL_601515, EPI_ISL_601522, EPI_ISL_601530, EPI_ISL_601541, EPI_ISL_601547, EPI_ISL_601549, EPI_ISL_601550, EPI_ISL_601555, EPI_ISL_601567, EPI_ISL_601569, EPI_ISL_601571, EPI_ISL_601578, EPI_ISL_601579, EPI_ISL_601580, EPI_ISL_601585, EPI_ISL_601588, EPI_ISL_601589, EPI_ISL_601590, EPI_ISL_601591, EPI_ISL_601594, EPI_ISL_601604, EPI_ISL_601606, EPI_ISL_601611, EPI_ISL_601615, EPI_ISL_601617, EPI_ISL_601618, EPI_ISL_601619, EPI_ISL_601622, EPI_ISL_601624, EPI_ISL_601627, EPI_ISL_601628, EPI_ISL_601634, EPI_ISL_601639, EPI_ISL_601641, EPI_ISL_601648, EPI_ISL_601650, EPI_ISL_601657, EPI_ISL_601663, EPI_ISL_601664, EPI_ISL_601666, EPI_ISL_601668, EPI_ISL_601673, EPI_ISL_601682, EPI_ISL_601685, EPI_ISL_601690, EPI_ISL_601691, EPI_ISL_601700, EPI_ISL_601704, EPI_ISL_601709, EPI_ISL_601715, EPI_ISL_601728, EPI_ISL_601730, EPI_ISL_601733, EPI_ISL_601740, EPI_ISL_601747, EPI_ISL_601748, EPI_ISL_601764, EPI_ISL_601765, EPI_ISL_601766, EPI_ISL_601768, EPI_ISL_601771, EPI_ISL_601778, EPI_ISL_601783, EPI_ISL_601785, EPI_ISL_602154 | | | |
| see above | Lighthouse Lab in Cambridge | Wellcome Sanger Institute for the COVID-19 Genomics UK (COG-UK) consortium | Rob Howes, The Lighthouse Lab in Cambridge and Alex Alderton, Roberto Amato, Sonia Goncalves, Ewan Harrison, David K. Jackson, Ian Johnston, Dominic Kwiatkowski, Cordelia Langford, John Sillitoe on behalf of the Wellcome Sanger Institute COVID-19 Surveillance Team (http://www.sanger.ac.uk/covid-team) |

EPI_ISL_602210, EPI_ISL_602211, EPI_ISL_602212, EPI_ISL_602213, EPI_ISL_602214, EPI_ISL_602215, EPI_ISL_602216, EPI_ISL_602217, EPI_ISL_602218, EPI_ISL_602219, EPI_ISL_602220, EPI_ISL_602221, EPI_ISL_602222, EPI_ISL_602223, EPI_ISL_602224, EPI_ISL_602225, EPI_ISL_602226, EPI_ISL_602227,
EPI_ISL_602228, EPI_ISL_602229, EPI_ISL_602230, EPI_ISL_602231, EPI_ISL_602232, EPI_ISL_602233, EPI_ISL_602234, EPI_ISL_602235, EPI_ISL_602236, EPI_ISL_602237, EPI_ISL_602238, EPI_ISL_602239, EPI_ISL_602240, EPI_ISL_602241, EPI_ISL_602242, EPI_ISL_602243, EPI_ISL_602244, EPI_ISL_602245,
EPI_ISL_602246, EPI_ISL_602247, EPI_ISL_602248, EPI_ISL_602249, EPI_ISL_602250, EPI_ISL_602251, EPI_ISL_602252, EPI_ISL_602253, EPI_ISL_602254, EPI_ISL_602255

| see above | Texas Department of State Health Services | Texas Department of State Health Services | Rashmi Tuladhar, Bonnie Oh, Jenny Zhang, Maliha Rahman, Anita Pokharel, Myong Koag, Chung Wang, Rachel Lee, Grace Kubin, Mayela Pedrueza |
|---|---|---|---|

EPI_ISL_603759, EPI_ISL_603760, EPI_ISL_603761, EPI_ISL_603762, EPI_ISL_603763, EPI_ISL_603764, EPI_ISL_603765, EPI_ISL_603766, EPI_ISL_603767, EPI_ISL_603768, EPI_ISL_603769, EPI_ISL_603770, EPI_ISL_603771, EPI_ISL_603772, EPI_ISL_603773, EPI_ISL_603774, EPI_ISL_603775, EPI_ISL_603776,
EPI_ISL_603777, EPI_ISL_603778, EPI_ISL_603779, EPI_ISL_603780, EPI_ISL_603781, EPI_ISL_603782, EPI_ISL_603783, EPI_ISL_603784, EPI_ISL_603785, EPI_ISL_603786, EPI_ISL_603787, EPI_ISL_603788, EPI_ISL_603789, EPI_ISL_603790, EPI_ISL_603791, EPI_ISL_603792, EPI_ISL_603793, EPI_ISL_603794,
EPI_ISL_603795, EPI_ISL_603796, EPI_ISL_603797, EPI_ISL_603798, EPI_ISL_603799, EPI_ISL_603800, EPI_ISL_603801, EPI_ISL_603802, EPI_ISL_603803, EPI_ISL_603804, EPI_ISL_603805, EPI_ISL_603806, EPI_ISL_603807, EPI_ISL_603808, EPI_ISL_603809, EPI_ISL_603810, EPI_ISL_603811, EPI_ISL_603812,
EPI_ISL_603813, EPI_ISL_603814, EPI_ISL_603815, EPI_ISL_603816, EPI_ISL_603817, EPI_ISL_603818, EPI_ISL_603819, EPI_ISL_603820, EPI_ISL_603821, EPI_ISL_603822, EPI_ISL_603823, EPI_ISL_603824, EPI_ISL_603825, EPI_ISL_603826, EPI_ISL_603827, EPI_ISL_603828, EPI_ISL_603829, EPI_ISL_603830,
EPI_ISL_603831, EPI_ISL_603832, EPI_ISL_603833, EPI_ISL_603834, EPI_ISL_603835, EPI_ISL_603836, EPI_ISL_603837, EPI_ISL_603838, EPI_ISL_603839, EPI_ISL_603840, EPI_ISL_603841, EPI_ISL_603842, EPI_ISL_603843, EPI_ISL_603844, EPI_ISL_603845, EPI_ISL_603846, EPI_ISL_603847, EPI_ISL_603848,
EPI_ISL_603849, EPI_ISL_603850, EPI_ISL_603851, EPI_ISL_603852, EPI_ISL_603853, EPI_ISL_603854, EPI_ISL_603855, EPI_ISL_603856, EPI_ISL_603857, EPI_ISL_603858, EPI_ISL_603859, EPI_ISL_603860, EPI_ISL_603861, EPI_ISL_603862, EPI_ISL_603863, EPI_ISL_603864, EPI_ISL_603865, EPI_ISL_603866,
EPI_ISL_603867, EPI_ISL_603868, EPI_ISL_603869, EPI_ISL_603870, EPI_ISL_603871, EPI_ISL_603872, EPI_ISL_603873, EPI_ISL_603874, EPI_ISL_603875, EPI_ISL_603876, EPI_ISL_603877, EPI_ISL_603878, EPI_ISL_603879, EPI_ISL_603880, EPI_ISL_603881, EPI_ISL_603882, EPI_ISL_603883, EPI_ISL_603884,
EPI_ISL_603885, EPI_ISL_603886, EPI_ISL_603887, EPI_ISL_603888, EPI_ISL_603889, EPI_ISL_603890, EPI_ISL_603891, EPI_ISL_603892, EPI_ISL_603893, EPI_ISL_603894, EPI_ISL_603895, EPI_ISL_603896, EPI_ISL_603897, EPI_ISL_603898, EPI_ISL_603899, EPI_ISL_603900, EPI_ISL_603901, EPI_ISL_603902,
EPI_ISL_603903, EPI_ISL_603904, EPI_ISL_603905, EPI_ISL_603906, EPI_ISL_603907, EPI_ISL_603908, EPI_ISL_603909, EPI_ISL_603910, EPI_ISL_603911, EPI_ISL_603912, EPI_ISL_603913, EPI_ISL_603914, EPI_ISL_603915, EPI_ISL_603916, EPI_ISL_603917, EPI_ISL_603918, EPI_ISL_603919, EPI_ISL_603920,
EPI_ISL_603921, EPI_ISL_603922, EPI_ISL_603923, EPI_ISL_603924, EPI_ISL_603925, EPI_ISL_603926, EPI_ISL_603927, EPI_ISL_603928, EPI_ISL_603929, EPI_ISL_603930, EPI_ISL_603931, EPI_ISL_603932, EPI_ISL_603933, EPI_ISL_603934, EPI_ISL_603935, EPI_ISL_603936, EPI_ISL_603937, EPI_ISL_603938,
EPI_ISL_603939, EPI_ISL_603940, EPI_ISL_603941, EPI_ISL_603942, EPI_ISL_603943, EPI_ISL_603944, EPI_ISL_603945, EPI_ISL_603946, EPI_ISL_603947, EPI_ISL_603948, EPI_ISL_603949, EPI_ISL_603950, EPI_ISL_603951, EPI_ISL_603952, EPI_ISL_603953, EPI_ISL_603954, EPI_ISL_603955, EPI_ISL_603956,
EPI_ISL_603957, EPI_ISL_603958, EPI_ISL_603959, EPI_ISL_603960, EPI_ISL_603961, EPI_ISL_603962, EPI_ISL_603963, EPI_ISL_603964, EPI_ISL_603965, EPI_ISL_603966, EPI_ISL_603967, EPI_ISL_603968, EPI_ISL_603969, EPI_ISL_603970, EPI_ISL_603971, EPI_ISL_603972, EPI_ISL_603973, EPI_ISL_603974,
EPI_ISL_603975, EPI_ISL_603976, EPI_ISL_603977, EPI_ISL_603978, EPI_ISL_603979, EPI_ISL_603980, EPI_ISL_603981, EPI_ISL_603982, EPI_ISL_603983, EPI_ISL_603984, EPI_ISL_603985, EPI_ISL_603986, EPI_ISL_603987, EPI_ISL_603988, EPI_ISL_603989, EPI_ISL_603990, EPI_ISL_603991, EPI_ISL_603992,
EPI_ISL_603993, EPI_ISL_603994, EPI_ISL_603995, EPI_ISL_603996, EPI_ISL_603997, EPI_ISL_603998, EPI_ISL_603999, EPI_ISL_604000, EPI_ISL_604001, EPI_ISL_604002, EPI_ISL_604003, EPI_ISL_604004, EPI_ISL_604005, EPI_ISL_604006, EPI_ISL_604007, EPI_ISL_604008, EPI_ISL_604009, EPI_ISL_604010,
EPI_ISL_604011, EPI_ISL_604012, EPI_ISL_604013, EPI_ISL_604014, EPI_ISL_604015, EPI_ISL_604016, EPI_ISL_604017, EPI_ISL_604018, EPI_ISL_604019, EPI_ISL_604020, EPI_ISL_604021, EPI_ISL_604022, EPI_ISL_604023, EPI_ISL_604024, EPI_ISL_604025, EPI_ISL_604026, EPI_ISL_604027, EPI_ISL_604028,
EPI_ISL_604029, EPI_ISL_604030, EPI_ISL_604031, EPI_ISL_604032, EPI_ISL_604033, EPI_ISL_604034, EPI_ISL_604035, EPI_ISL_604036, EPI_ISL_604037, EPI_ISL_604038, EPI_ISL_604039, EPI_ISL_604040, EPI_ISL_604041, EPI_ISL_604042, EPI_ISL_604043, EPI_ISL_604044, EPI_ISL_604045, EPI_ISL_604046,
EPI_ISL_604047, EPI_ISL_604048, EPI_ISL_604049, EPI_ISL_604050, EPI_ISL_604051, EPI_ISL_604052, EPI_ISL_604053, EPI_ISL_604054, EPI_ISL_604055, EPI_ISL_604056, EPI_ISL_604057, EPI_ISL_604058, EPI_ISL_604059, EPI_ISL_604060, EPI_ISL_604061, EPI_ISL_604062, EPI_ISL_604063, EPI_ISL_604064,
EPI_ISL_604065, EPI_ISL_604066, EPI_ISL_604067, EPI_ISL_604068, EPI_ISL_604069, EPI_ISL_604070, EPI_ISL_604071, EPI_ISL_604072, EPI_ISL_604073, EPI_ISL_604074, EPI_ISL_604075, EPI_ISL_604076, EPI_ISL_604077, EPI_ISL_604078, EPI_ISL_604079, EPI_ISL_604080, EPI_ISL_604081, EPI_ISL_604082,
EPI_ISL_604083, EPI_ISL_604084, EPI_ISL_604085, EPI_ISL_604086, EPI_ISL_604087, EPI_ISL_604088, EPI_ISL_604089, EPI_ISL_604090, EPI_ISL_604091, EPI_ISL_604092, EPI_ISL_604093, EPI_ISL_604094, EPI_ISL_604095, EPI_ISL_604096, EPI_ISL_604097, EPI_ISL_604098, EPI_ISL_604099, EPI_ISL_604100,
EPI_ISL_604101, EPI_ISL_604102, EPI_ISL_604103, EPI_ISL_604104, EPI_ISL_604105, EPI_ISL_604106, EPI_ISL_604107, EPI_ISL_604108, EPI_ISL_604109, EPI_ISL_604110, EPI_ISL_604111, EPI_ISL_604112, EPI_ISL_604113, EPI_ISL_604114, EPI_ISL_604115, EPI_ISL_604116, EPI_ISL_604117, EPI_ISL_604118,
EPI_ISL_604119, EPI_ISL_604120, EPI_ISL_604121, EPI_ISL_604122, EPI_ISL_604123, EPI_ISL_604124, EPI_ISL_604125, EPI_ISL_604126, EPI_ISL_604127, EPI_ISL_604128, EPI_ISL_604129, EPI_ISL_604130, EPI_ISL_604131, EPI_ISL_604132, EPI_ISL_604133, EPI_ISL_604134, EPI_ISL_604135, EPI_ISL_604136,
EPI_ISL_604137, EPI_ISL_604138, EPI_ISL_604139, EPI_ISL_604140, EPI_ISL_604141, EPI_ISL_604142, EPI_ISL_604143, EPI_ISL_604144, EPI_ISL_604145, EPI_ISL_604146, EPI_ISL_604147, EPI_ISL_604148, EPI_ISL_604149, EPI_ISL_604150, EPI_ISL_604151, EPI_ISL_604152, EPI_ISL_604153, EPI_ISL_604154,
EPI_ISL_604155, EPI_ISL_604156, EPI_ISL_604157, EPI_ISL_604158, EPI_ISL_604159, EPI_ISL_604160, EPI_ISL_604161, EPI_ISL_604162, EPI_ISL_604163, EPI_ISL_604164, EPI_ISL_604165, EPI_ISL_604166, EPI_ISL_604167, EPI_ISL_604168, EPI_ISL_604169, EPI_ISL_604170, EPI_ISL_604171, EPI_ISL_604172,
EPI_ISL_604173, EPI_ISL_604174, EPI_ISL_604175, EPI_ISL_604176, EPI_ISL_604177, EPI_ISL_604178, EPI_ISL_604179, EPI_ISL_604180, EPI_ISL_604181, EPI_ISL_604182, EPI_ISL_604183, EPI_ISL_604184, EPI_ISL_604185, EPI_ISL_604186, EPI_ISL_604187, EPI_ISL_604188, EPI_ISL_604189, EPI_ISL_604190,
EPI_ISL_604191, EPI_ISL_604192, EPI_ISL_604193, EPI_ISL_604194, EPI_ISL_604195, EPI_ISL_604196, EPI_ISL_604197, EPI_ISL_604198, EPI_ISL_604199, EPI_ISL_604200, EPI_ISL_604201, EPI_ISL_604202, EPI_ISL_604203, EPI_ISL_604204, EPI_ISL_604205, EPI_ISL_604206, EPI_ISL_604207, EPI_ISL_604208,
EPI_ISL_604209, EPI_ISL_604210, EPI_ISL_604211, EPI_ISL_604212, EPI_ISL_604213, EPI_ISL_604214, EPI_ISL_604215, EPI_ISL_604216, EPI_ISL_604217, EPI_ISL_604218, EPI_ISL_604219, EPI_ISL_604220, EPI_ISL_604221, EPI_ISL_604222, EPI_ISL_604223, EPI_ISL_604224, EPI_ISL_604225, EPI_ISL_604226,
EPI_ISL_604227, EPI_ISL_604228, EPI_ISL_604229, EPI_ISL_604230, EPI_ISL_604231, EPI_ISL_604232, EPI_ISL_604233, EPI_ISL_604234, EPI_ISL_604235, EPI_ISL_604236, EPI_ISL_604237, EPI_ISL_604238, EPI_ISL_604239, EPI_ISL_604240, EPI_ISL_604241, EPI_ISL_604242, EPI_ISL_604243, EPI_ISL_604244,
EPI_ISL_604245, EPI_ISL_604246, EPI_ISL_604247, EPI_ISL_604248, EPI_ISL_604249, EPI_ISL_604250, EPI_ISL_604251, EPI_ISL_604252, EPI_ISL_604253, EPI_ISL_604254, EPI_ISL_604255, EPI_ISL_604256, EPI_ISL_604257, EPI_ISL_604258, EPI_ISL_604259, EPI_ISL_604260, EPI_ISL_604261, EPI_ISL_604262,
EPI_ISL_604263, EPI_ISL_604264, EPI_ISL_604265, EPI_ISL_604266, EPI_ISL_604267, EPI_ISL_604268, EPI_ISL_604269, EPI_ISL_604270, EPI_ISL_604271, EPI_ISL_604272, EPI_ISL_604273, EPI_ISL_604274, EPI_ISL_604275, EPI_ISL_604276, EPI_ISL_604277, EPI_ISL_604278, EPI_ISL_604279, EPI_ISL_604280,
EPI_ISL_604281, EPI_ISL_604282, EPI_ISL_604283, EPI_ISL_604284, EPI_ISL_604285, EPI_ISL_604286, EPI_ISL_604287, EPI_ISL_604288, EPI_ISL_604289, EPI_ISL_604290, EPI_ISL_604291, EPI_ISL_604292, EPI_ISL_604293, EPI_ISL_604294, EPI_ISL_604295, EPI_ISL_604296, EPI_ISL_604297, EPI_ISL_604298,
EPI_ISL_604299, EPI_ISL_604300, EPI_ISL_604301, EPI_ISL_604302, EPI_ISL_604303, EPI_ISL_604304, EPI_ISL_604305, EPI_ISL_604306, EPI_ISL_604307, EPI_ISL_604308, EPI_ISL_604309, EPI_ISL_604310, EPI_ISL_604311, EPI_ISL_604312, EPI_ISL_604313, EPI_ISL_604314, EPI_ISL_604315, EPI_ISL_604316,
EPI_ISL_604317, EPI_ISL_604318, EPI_ISL_604319, EPI_ISL_604320, EPI_ISL_604321, EPI_ISL_604322, EPI_ISL_604323, EPI_ISL_604324, EPI_ISL_604325, EPI_ISL_604326, EPI_ISL_604327, EPI_ISL_604328, EPI_ISL_604329, EPI_ISL_604330, EPI_ISL_604331, EPI_ISL_604332, EPI_ISL_604333, EPI_ISL_604334,
EPI_ISL_604335, EPI_ISL_604336, EPI_ISL_604337, EPI_ISL_604338, EPI_ISL_604339, EPI_ISL_604340, EPI_ISL_604341, EPI_ISL_604342, EPI_ISL_604343, EPI_ISL_604344, EPI_ISL_604345, EPI_ISL_604346, EPI_ISL_604347, EPI_ISL_604348, EPI_ISL_604349, EPI_ISL_604350, EPI_ISL_604351, EPI_ISL_604352,
EPI_ISL_604353, EPI_ISL_604354, EPI_ISL_604355, EPI_ISL_604356, EPI_ISL_604357, EPI_ISL_604358, EPI_ISL_604359, EPI_ISL_604360, EPI_ISL_604361, EPI_ISL_604362, EPI_ISL_604363, EPI_ISL_604364, EPI_ISL_604365, EPI_ISL_604366, EPI_ISL_604367, EPI_ISL_604368, EPI_ISL_604369, EPI_ISL_604370,
EPI_ISL_604371, EPI_ISL_604372, EPI_ISL_604373, EPI_ISL_604374, EPI_ISL_604375, EPI_ISL_604376, EPI_ISL_604377, EPI_ISL_604378, EPI_ISL_604379, EPI_ISL_604380, EPI_ISL_604381, EPI_ISL_604382, EPI_ISL_604383, EPI_ISL_604384, EPI_ISL_604385, EPI_ISL_604386, EPI_ISL_604387, EPI_ISL_604388,
EPI_ISL_604389, EPI_ISL_604390, EPI_ISL_604391, EPI_ISL_604392, EPI_ISL_604393, EPI_ISL_604394, EPI_ISL_604395, EPI_ISL_604396, EPI_ISL_604397, EPI_ISL_604398, EPI_ISL_604399, EPI_ISL_604400, EPI_ISL_604401, EPI_ISL_604402, EPI_ISL_604403, EPI_ISL_604404, EPI_ISL_604405, EPI_ISL_604406,
EPI_ISL_604407, EPI_ISL_604408, EPI_ISL_604409, EPI_ISL_604410, EPI_ISL_604411, EPI_ISL_604412, EPI_ISL_604413, EPI_ISL_604414, EPI_ISL_604415, EPI_ISL_604416, EPI_ISL_604417, EPI_ISL_604418, EPI_ISL_604419, EPI_ISL_604420, EPI_ISL_604421, EPI_ISL_604422, EPI_ISL_604423, EPI_ISL_604424,
EPI_ISL_604425, EPI_ISL_604426, EPI_ISL_604427, EPI_ISL_604428, EPI_ISL_604429, EPI_ISL_604430, EPI_ISL_604431, EPI_ISL_604432, EPI_ISL_604433, EPI_ISL_604434, EPI_ISL_604435, EPI_ISL_604436, EPI_ISL_604437, EPI_ISL_604438, EPI_ISL_604439, EPI_ISL_604440, EPI_ISL_604441, EPI_ISL_604442,
EPI_ISL_604443, EPI_ISL_604444, EPI_ISL_604445, EPI_ISL_604446, EPI_ISL_604447, EPI_ISL_604448, EPI_ISL_604449, EPI_ISL_604450, EPI_ISL_604451, EPI_ISL_604452, EPI_ISL_604453, EPI_ISL_604454, EPI_ISL_604455, EPI_ISL_604456, EPI_ISL_604457, EPI_ISL_604458, EPI_ISL_604459, EPI_ISL_604460,
EPI_ISL_604461, EPI_ISL_604462, EPI_ISL_604463, EPI_ISL_604464, EPI_ISL_604465, EPI_ISL_604466, EPI_ISL_604467, EPI_ISL_604468, EPI_ISL_604469, EPI_ISL_604470, EPI_ISL_604471, EPI_ISL_604472, EPI_ISL_604473, EPI_ISL_604474, EPI_ISL_604475, EPI_ISL_604476, EPI_ISL_604477, EPI_ISL_604478,
EPI_ISL_604479, EPI_ISL_604480, EPI_ISL_604481, EPI_ISL_604482, EPI_ISL_604483, EPI_ISL_604484, EPI_ISL_604485, EPI_ISL_604486, EPI_ISL_604487, EPI_ISL_604488, EPI_ISL_604489, EPI_ISL_604490, EPI_ISL_604491, EPI_ISL_604492, EPI_ISL_604493, EPI_ISL_604494, EPI_ISL_604495, EPI_ISL_604496,
EPI_ISL_604497, EPI_ISL_604498, EPI_ISL_604499, EPI_ISL_604500, EPI_ISL_604501, EPI_ISL_604502, EPI_ISL_604503, EPI_ISL_604504, EPI_ISL_604505, EPI_ISL_604506, EPI_ISL_604507, EPI_ISL_604508, EPI_ISL_604509, EPI_ISL_604510, EPI_ISL_604511, EPI_ISL_604512, EPI_ISL_604513, EPI_ISL_604514,
EPI_ISL_604515, EPI_ISL_604516, EPI_ISL_604517, EPI_ISL_604518, EPI_ISL_604519, EPI_ISL_604520, EPI_ISL_604521, EPI_ISL_604522, EPI_ISL_604523, EPI_ISL_604524, EPI_ISL_604525, EPI_ISL_604526, EPI_ISL_604527, EPI_ISL_604528, EPI_ISL_604529, EPI_ISL_604530, EPI_ISL_604531, EPI_ISL_604532,
EPI_ISL_604533, EPI_ISL_604534, EPI_ISL_604535, EPI_ISL_604536, EPI_ISL_604537, EPI_ISL_604538, EPI_ISL_604539, EPI_ISL_604540, EPI_ISL_604541, EPI_ISL_604542, EPI_ISL_604543, EPI_ISL_604544, EPI_ISL_604545, EPI_ISL_604546, EPI_ISL_604547, EPI_ISL_604548, EPI_ISL_604549, EPI_ISL_604550,
EPI_ISL_604551, EPI_ISL_604552, EPI_ISL_604553, EPI_ISL_604554, EPI_ISL_604555, EPI_ISL_604556, EPI_ISL_604557, EPI_ISL_604558, EPI_ISL_604559, EPI_ISL_604560, EPI_ISL_604561, EPI_ISL_604562, EPI_ISL_604563, EPI_ISL_604564, EPI_ISL_604565, EPI_ISL_604566, EPI_ISL_604567, EPI_ISL_604568,
EPI_ISL_604569, EPI_ISL_604570, EPI_ISL_604571, EPI_ISL_604572, EPI_ISL_604573, EPI_ISL_604574, EPI_ISL_604575, EPI_ISL_604576, EPI_ISL_604577, EPI_ISL_604578, EPI_ISL_604579, EPI_ISL_604580, EPI_ISL_604581, EPI_ISL_604582, EPI_ISL_604583, EPI_ISL_604584, EPI_ISL_604585, EPI_ISL_604586,
EPI_ISL_604587, EPI_ISL_604588, EPI_ISL_604589, EPI_ISL_604590, EPI_ISL_604591, EPI_ISL_604592, EPI_ISL_604593, EPI_ISL_604594, EPI_ISL_604595, EPI_ISL_604596, EPI_ISL_604597, EPI_ISL_604598, EPI_ISL_604599, EPI_ISL_604600, EPI_ISL_604601, EPI_ISL_604602, EPI_ISL_604603, EPI_ISL_604604,
EPI_ISL_604605, EPI_ISL_604606, EPI_ISL_604607, EPI_ISL_604608, EPI_ISL_604609, EPI_ISL_604610, EPI_ISL_604611, EPI_ISL_604612, EPI_ISL_604613, EPI_ISL_604614, EPI_ISL_604615, EPI_ISL_604616, EPI_ISL_604617, EPI_ISL_604618, EPI_ISL_604619, EPI_ISL_604620, EPI_ISL_604621, EPI_ISL_604622,
EPI_ISL_604623, EPI_ISL_604624, EPI_ISL_604625, EPI_ISL_604626, EPI_ISL_604627, EPI_ISL_604628, EPI_ISL_604629, EPI_ISL_604630, EPI_ISL_604631, EPI_ISL_604632, EPI_ISL_604633, EPI_ISL_604634, EPI_ISL_604635, EPI_ISL_604636, EPI_ISL_604637, EPI_ISL_604638, EPI_ISL_604639, EPI_ISL_604640,
EPI_ISL_604641, EPI_ISL_604642, EPI_ISL_604643, EPI_ISL_604644, EPI_ISL_604645, EPI_ISL_604646, EPI_ISL_604647, EPI_ISL_604648, EPI_ISL_604649, EPI_ISL_604650, EPI_ISL_604651, EPI_ISL_604652, EPI_ISL_604653, EPI_ISL_604654, EPI_ISL_604655, EPI_ISL_604656, EPI_ISL_604657, EPI_ISL_604658,
EPI_ISL_604659, EPI_ISL_604660, EPI_ISL_604661, EPI_ISL_604662, EPI_ISL_604663, EPI_ISL_604664, EPI_ISL_604665, EPI_ISL_604666, EPI_ISL_604667, EPI_ISL_604668, EPI_ISL_604669, EPI_ISL_604670, EPI_ISL_604671, EPI_ISL_604672, EPI_ISL_604673, EPI_ISL_604674, EPI_ISL_604675, EPI_ISL_604676,
EPI_ISL_604677, EPI_ISL_604678, EPI_ISL_604679, EPI_ISL_604680, EPI_ISL_604681, EPI_ISL_604682, EPI_ISL_604683, EPI_ISL_604684, EPI_ISL_604685, EPI_ISL_604686, EPI_ISL_604687, EPI_ISL_604688, EPI_ISL_604689, EPI_ISL_604690, EPI_ISL_604691, EPI_ISL_604692, EPI_ISL_604693, EPI_ISL_604694,
EPI_ISL_604695, EPI_ISL_604696, EPI_ISL_604697, EPI_ISL_604698, EPI_ISL_604699, EPI_ISL_604700, EPI_ISL_604701, EPI_ISL_604702, EPI_ISL_604703, EPI_ISL_604704, EPI_ISL_604705, EPI_ISL_604706, EPI_ISL_604707, EPI_ISL_604708, EPI_ISL_604709, EPI_ISL_604710, EPI_ISL_604711, EPI_ISL_604712,
EPI_ISL_604713, EPI_ISL_604714, EPI_ISL_604715, EPI_ISL_604716, EPI_ISL_604717, EPI_ISL_604718, EPI_ISL_604719, EPI_ISL_604720, EPI_ISL_604721, EPI_ISL_604722, EPI_ISL_604723, EPI_ISL_604724, EPI_ISL_604725, EPI_ISL_604726, EPI_ISL_604727, EPI_ISL_604728, EPI_ISL_604729, EPI_ISL_604730,
EPI_ISL_604731, EPI_ISL_604732, EPI_ISL_604733, EPI_ISL_604734, EPI_ISL_604735, EPI_ISL_604736, EPI_ISL_604737, EPI_ISL_604738, EPI_ISL_604739, EPI_ISL_604740, EPI_ISL_604741, EPI_ISL_604742, EPI_ISL_604743, EPI_ISL_604744, EPI_ISL_604745, EPI_ISL_604746, EPI_ISL_604747, EPI_ISL_604748,
EPI_ISL_604749, EPI_ISL_604750, EPI_ISL_604751, EPI_ISL_604752, EPI_ISL_604753, EPI_ISL_604754, EPI_ISL_604755, EPI_ISL_604756, EPI_ISL_604757, EPI_ISL_604758, EPI_ISL_604759, EPI_ISL_604760, EPI_ISL_604761, EPI_ISL_604762, EPI_ISL_604763, EPI_ISL_604764, EPI_ISL_604765, EPI_ISL_604766,
EPI_ISL_604767, EPI_ISL_604768, EPI_ISL_604769, EPI_ISL_604770, EPI_ISL_604771, EPI_ISL_604772, EPI_ISL_604773, EPI_ISL_604774, EPI_ISL_604775, EPI_ISL_604776, EPI_ISL_604777, EPI_ISL_604778, EPI_ISL_604779, EPI_ISL_604780, EPI_ISL_604781, EPI_ISL_604782, EPI_ISL_604783, EPI_ISL_604784,
EPI_ISL_604785, EPI_ISL_604786, EPI_ISL_604787, EPI_ISL_604788, EPI_ISL_604789, EPI_ISL_604790, EPI_ISL_604791, EPI_ISL_604792, EPI_ISL_604793, EPI_ISL_604794, EPI_ISL_604795, EPI_ISL_604796, EPI_ISL_604797, EPI_ISL_604798, EPI_ISL_604799, EPI_ISL_604800, EPI_ISL_604801, EPI_ISL_604802,
EPI_ISL_604803, EPI_ISL_604804, EPI_ISL_604805, EPI_ISL_604806, EPI_ISL_604807, EPI_ISL_604808, EPI_ISL_604809, EPI_ISL_604810, EPI_ISL_604811, EPI_ISL_604812, EPI_ISL_604813, EPI_ISL_604814, EPI_ISL_604815, EPI_ISL_604816, EPI_ISL_604817, EPI_ISL_604818, EPI_ISL_604819, EPI_ISL_604820,
EPI_ISL_604821, EPI_ISL_604822, EPI_ISL_604823, EPI_ISL_604824, EPI_ISL_604825, EPI_ISL_604826, EPI_ISL_604827, EPI_ISL_604828, EPI_ISL_604829, EPI_ISL_604830, EPI_ISL_604831, EPI_ISL_604832, EPI_ISL_604833, EPI_ISL_604834, EPI_ISL_604835, EPI_ISL_604836, EPI_ISL_604837, EPI_ISL_604838,
EPI_ISL_604839, EPI_ISL_604840, EPI_ISL_604841, EPI_ISL_604842, EPI_ISL_604843, EPI_ISL_604844, EPI_ISL_604845, EPI_ISL_604846, EPI_ISL_604847, EPI_ISL_604848, EPI_ISL_604849, EPI_ISL_604850, EPI_ISL_604851, EPI_ISL_604852, EPI_ISL_604853, EPI_ISL_604854, EPI_ISL_604855, EPI_ISL_604856,
EPI_ISL_604857, EPI_ISL_604858, EPI_ISL_604859, EPI_ISL_604860, EPI_ISL_604861, EPI_ISL_604862, EPI_ISL_604863, EPI_ISL_604864, EPI_ISL_604865, EPI_ISL_604866, EPI_ISL_604867, EPI_ISL_604868, EPI_ISL_604869, EPI_ISL_604870, EPI_ISL_604871, EPI_ISL_604872, EPI_ISL_604873, EPI_ISL_604874,
EPI_ISL_604875, EPI_ISL_604876, EPI_ISL_604877, EPI_ISL_604878, EPI_ISL_604879, EPI_ISL_604880, EPI_ISL_604881, EPI_ISL_604882, EPI_ISL_604883, EPI_ISL_604884, EPI_ISL_604885, EPI_ISL_604886, EPI_ISL_604887, EPI_ISL_604888, EPI_ISL_604889, EPI_ISL_604890, EPI_ISL_604891, EPI_ISL_604892,
EPI_ISL_604893, EPI_ISL_604894, EPI_ISL_604895, EPI_ISL_604896, EPI_ISL_604897, EPI_ISL_604898, EPI_ISL_604899, EPI_ISL_604900, EPI_ISL_604901, EPI_ISL_604902, EPI_ISL_604903, EPI_ISL_604904, EPI_ISL_604905, EPI_ISL_604906, EPI_ISL_604907, EPI_ISL_604908, EPI_ISL_604909, EPI_ISL_604910,
EPI_ISL_604911, EPI_ISL_604912, EPI_ISL_604913, EPI_ISL_604914, EPI_ISL_604915, EPI_ISL_604916, EPI_ISL_604917, EPI_ISL_604918, EPI_ISL_604919, EPI_ISL_604920, EPI_ISL_604921, EPI_ISL_604922, EPI_ISL_604923, EPI_ISL_604924, EPI_ISL_604925, EPI_ISL_604926, EPI_ISL_604927, EPI_ISL_604928,
EPI_ISL_604929, EPI_ISL_604930, EPI_ISL_604931, EPI_ISL_604932, EPI_ISL_604933, EPI_ISL_604934, EPI_ISL_604935, EPI_ISL_604936, EPI_ISL_604937, EPI_ISL_604938, EPI_ISL_604939, EPI_ISL_604940, EPI_ISL_604941, EPI_ISL_604942, EPI_ISL_604943, EPI_ISL_604944, EPI_ISL_604945, EPI_ISL_604946,
EPI_ISL_604947, EPI_ISL_604948, EPI_ISL_604949, EPI_ISL_604950, EPI_ISL_604951, EPI_ISL_604952, EPI_ISL_604953, EPI_ISL_604954, EPI_ISL_604955, EPI_ISL_604956, EPI_ISL_604957, EPI_ISL_604958, EPI_ISL_604959, EPI_ISL_604960, EPI_ISL_604961, EPI_ISL_604962, EPI_ISL_604963, EPI_ISL_604964,
EPI_ISL_604965, EPI_ISL_604966, EPI_ISL_604967, EPI_ISL_604968, EPI_ISL_604969, EPI_ISL_604970, EPI_ISL_604971, EPI_ISL_604972, EPI_ISL_604973, EPI_ISL_604974, EPI_ISL_604975, EPI_ISL_604976, EPI_ISL_604977, EPI_ISL_604978, EPI_ISL_604979, EPI_ISL_604980, EPI_ISL_604981, EPI_ISL_604982,
EPI_ISL_604983, EPI_ISL_604984, EPI_ISL_604985, EPI_ISL_604986, EPI_ISL_604987, EPI_ISL_604988, EPI_ISL_604989, EPI_ISL_604990, EPI_ISL_604991, EPI_ISL_604992, EPI_ISL_604993, EPI_ISL_604994, EPI_ISL_604995, EPI_ISL_604996, EPI_ISL_604997, EPI_ISL_604998, EPI_ISL_604999, EPI_ISL_605000,
EPI_ISL_605001, EPI_ISL_605002, EPI_ISL_605003, EPI_ISL_605004, EPI_ISL_605005, EPI_ISL_605006, EPI_ISL_605007, EPI_ISL_605008, EPI_ISL_605009, EPI_ISL_605010, EPI_ISL_605011, EPI_ISL_605012, EPI_ISL_605013, EPI_ISL_605014, EPI_ISL_605015, EPI_ISL_605016, EPI_ISL_605017, EPI_ISL_605018,
EPI_ISL_605019, EPI_ISL_605020, EPI_ISL_605021, EPI_ISL_605022, EPI_ISL_605023, EPI_ISL_605024, EPI_ISL_605025, EPI_ISL_605026, EPI_ISL_605027, EPI_ISL_605028, EPI_ISL_605029, EPI_ISL_605030, EPI_ISL_605031, EPI_ISL_605032, EPI_ISL_605033, EPI_ISL_605034, EPI_ISL_605035, EPI_ISL_605036,
EPI_ISL_605037, EPI_ISL_605038, EPI_ISL_605039, EPI_ISL_605040, EPI_ISL_605041, EPI_ISL_605042, EPI_ISL_605043, EPI_ISL_605044, EPI_ISL_605045, EPI_ISL_605046, EPI_ISL_605047, EPI_ISL_605048, EPI_ISL_605049, EPI_ISL_605050, EPI_ISL_605051, EPI_ISL_605052, EPI_ISL_605053, EPI_ISL_605054,
EPI_ISL_605055, EPI_ISL_605056, EPI_ISL_605057

| see above | Quest Diagnostics | Quest Diagnostics | Rosenthal,S.H., Gerasimova,A., Kagan,R.M., Anderson, B., Grover, D., Livingston, K.E., Hua, M., Liu Y., Shalhout, D.F., Owen, R., Lacbawan, F. |
|---|---|---|---|
| EPI_ISL_605780, EPI_ISL_605781, EPI_ISL_605782 | CEIRS Data Processing and Coordinating Center, St. Jude Center of Excellence for Influenza Research and Surveillance (CEIRS) | CEIRS Data Processing and Coordinating Center, St. Jude Center of Excellence for Influenza Research and Surveillance (CEIRS) | Roshdy,W.H., Kayed,A.E., Naguib,A., Kamel,M.N., El-Taweel,A., El-Shesheny,R., Kandeil,A., Mostafa,A., Shehata,M., Gomaa,M., Mahmoud,S.H., Moatasim,Y., Kutkat,O., Mahrous,N., El-Sayes,M., Showky,S., El-Guindy,N.M., Webby,R., Kayali,G., Ali,M.A. |

EPI_ISL_608348, EPI_ISL_608349, EPI_ISL_608353, EPI_ISL_608354, EPI_ISL_608357, EPI_ISL_608359, EPI_ISL_608360, EPI_ISL_608363, EPI_ISL_608368, EPI_ISL_608370, EPI_ISL_608373, EPI_ISL_608381, EPI_ISL_608382, EPI_ISL_608388, EPI_ISL_608389, EPI_ISL_608391, EPI_ISL_608392, EPI_ISL_608393,
EPI_ISL_608394, EPI_ISL_608395, EPI_ISL_608396, EPI_ISL_608397, EPI_ISL_608398, EPI_ISL_608400, EPI_ISL_608402, EPI_ISL_608404, EPI_ISL_608407, EPI_ISL_608408, EPI_ISL_608410, EPI_ISL_608411, EPI_ISL_608412, EPI_ISL_608416, EPI_ISL_608417, EPI_ISL_608423, EPI_ISL_608424, EPI_ISL_608426,
EPI_ISL_608427, EPI_ISL_608428, EPI_ISL_608429, EPI_ISL_608430, EPI_ISL_608432, EPI_ISL_608433, EPI_ISL_608436, EPI_ISL_608437, EPI_ISL_608439, EPI_ISL_608442, EPI_ISL_608446, EPI_ISL_608447, EPI_ISL_608449, EPI_ISL_608450, EPI_ISL_608451, EPI_ISL_608456, EPI_ISL_608458, EPI_ISL_608459,
EPI_ISL_608463, EPI_ISL_608464, EPI_ISL_608466, EPI_ISL_608467, EPI_ISL_608471, EPI_ISL_608472, EPI_ISL_608475, EPI_ISL_608476, EPI_ISL_608477, EPI_ISL_608478, EPI_ISL_608480, EPI_ISL_608482, EPI_ISL_608484, EPI_ISL_608485, EPI_ISL_608486, EPI_ISL_608490, EPI_ISL_608492, EPI_ISL_608493,

EPI_ISL_608494, EPI_ISL_608495, EPI_ISL_608497, EPI_ISL_608498, EPI_ISL_608501, EPI_ISL_608504, EPI_ISL_608506, EPI_ISL_608507, EPI_ISL_608508, EPI_ISL_608509, EPI_ISL_608510, EPI_ISL_608514, EPI_ISL_608515, EPI_ISL_608521, EPI_ISL_608522, EPI_ISL_608523, EPI_ISL_608524, EPI_ISL_608525, EPI_ISL_608527, EPI_ISL_608529, EPI_ISL_608530, EPI_ISL_608531, EPI_ISL_608532, EPI_ISL_608535, EPI_ISL_608536, EPI_ISL_608537, EPI_ISL_608538, EPI_ISL_608540, EPI_ISL_608544, EPI_ISL_608545, EPI_ISL_608546, EPI_ISL_608547, EPI_ISL_608550, EPI_ISL_608552, EPI_ISL_608557, EPI_ISL_608558, EPI_ISL_608559, EPI_ISL_608560, EPI_ISL_608561, EPI_ISL_608565, EPI_ISL_608566, EPI_ISL_608568, EPI_ISL_608570, EPI_ISL_608571, EPI_ISL_608572, EPI_ISL_608573, EPI_ISL_608576, EPI_ISL_608577, EPI_ISL_608578, EPI_ISL_608585, EPI_ISL_608588, EPI_ISL_608596, EPI_ISL_608597, EPI_ISL_608598, EPI_ISL_608602, EPI_ISL_608604, EPI_ISL_608607, EPI_ISL_608608, EPI_ISL_608611, EPI_ISL_608612, EPI_ISL_608613, EPI_ISL_608615, EPI_ISL_608616, EPI_ISL_608618, EPI_ISL_608619, EPI_ISL_608620, EPI_ISL_608624, EPI_ISL_608626, EPI_ISL_608627, EPI_ISL_608628, EPI_ISL_608629, EPI_ISL_608631, EPI_ISL_608632, EPI_ISL_608633, EPI_ISL_608636, EPI_ISL_608637, EPI_ISL_608638, EPI_ISL_608641, EPI_ISL_608642, EPI_ISL_608643, EPI_ISL_608644, EPI_ISL_608645, EPI_ISL_608650, EPI_ISL_608651, EPI_ISL_608653, EPI_ISL_608654, EPI_ISL_608656, EPI_ISL_608660, EPI_ISL_608664, EPI_ISL_608666, EPI_ISL_608671, EPI_ISL_608673, EPI_ISL_608678, EPI_ISL_608679, EPI_ISL_608682, EPI_ISL_608683, EPI_ISL_608688, EPI_ISL_608689, EPI_ISL_608693, EPI_ISL_608695, EPI_ISL_608696, EPI_ISL_608845, EPI_ISL_608852, EPI_ISL_608867, EPI_ISL_608868, EPI_ISL_608873, EPI_ISL_608875, EPI_ISL_608876, EPI_ISL_608879, EPI_ISL_608881, EPI_ISL_608892, EPI_ISL_608902, EPI_ISL_608906, EPI_ISL_608908, EPI_ISL_608910, EPI_ISL_608912, EPI_ISL_608918, EPI_ISL_608923, EPI_ISL_608924, EPI_ISL_608927, EPI_ISL_608935, EPI_ISL_608937, EPI_ISL_608939, EPI_ISL_608943, EPI_ISL_608947, EPI_ISL_608949, EPI_ISL_608953, EPI_ISL_608958, EPI_ISL_608959, EPI_ISL_608960, EPI_ISL_608965, EPI_ISL_608967, EPI_ISL_608968, EPI_ISL_608969, EPI_ISL_608974, EPI_ISL_608982, EPI_ISL_608987, EPI_ISL_608990, EPI_ISL_608996, EPI_ISL_608997, EPI_ISL_609003, EPI_ISL_609005, EPI_ISL_609014, EPI_ISL_609018, EPI_ISL_609023, EPI_ISL_609025, EPI_ISL_609033, EPI_ISL_609037, EPI_ISL_609045, EPI_ISL_609048, EPI_ISL_609050, EPI_ISL_609054, EPI_ISL_609067, EPI_ISL_609072, EPI_ISL_609077, EPI_ISL_609081, EPI_ISL_609082, EPI_ISL_609086, EPI_ISL_609088, EPI_ISL_609091, EPI_ISL_609097, EPI_ISL_609099, EPI_ISL_609100, EPI_ISL_609101, EPI_ISL_609111, EPI_ISL_609113, EPI_ISL_609114, EPI_ISL_609118, EPI_ISL_609119, EPI_ISL_609122, EPI_ISL_609123, EPI_ISL_609129, EPI_ISL_609131, EPI_ISL_609142, EPI_ISL_609143, EPI_ISL_609145, EPI_ISL_609156, EPI_ISL_609158, EPI_ISL_609159, EPI_ISL_609160, EPI_ISL_609161, EPI_ISL_609162, EPI_ISL_609163, EPI_ISL_609164, EPI_ISL_609165, EPI_ISL_609166, EPI_ISL_609167, EPI_ISL_609168, EPI_ISL_609169, EPI_ISL_609170, EPI_ISL_609171, EPI_ISL_609172, EPI_ISL_609173, EPI_ISL_609174, EPI_ISL_609175, EPI_ISL_609176, EPI_ISL_609177, EPI_ISL_609180, EPI_ISL_609181, EPI_ISL_609183, EPI_ISL_609188, EPI_ISL_609193, EPI_ISL_609195, EPI_ISL_609196, EPI_ISL_609200, EPI_ISL_609201, EPI_ISL_609203, EPI_ISL_609204, EPI_ISL_609205, EPI_ISL_609308, EPI_ISL_609309, EPI_ISL_609310, EPI_ISL_609311, EPI_ISL_609312, EPI_ISL_609313, EPI_ISL_609314, EPI_ISL_609315, EPI_ISL_609316, EPI_ISL_609317, EPI_ISL_609318, EPI_ISL_609319, EPI_ISL_609320, EPI_ISL_609321, EPI_ISL_609322, EPI_ISL_609323, EPI_ISL_609324, EPI_ISL_609325, EPI_ISL_609326, EPI_ISL_609327, EPI_ISL_609328, EPI_ISL_609329, EPI_ISL_609330, EPI_ISL_609331, EPI_ISL_609332, EPI_ISL_609333, EPI_ISL_609334, EPI_ISL_609335, EPI_ISL_609336, EPI_ISL_609337, EPI_ISL_609338, EPI_ISL_609339, EPI_ISL_609340, EPI_ISL_609341, EPI_ISL_609342, EPI_ISL_609343, EPI_ISL_609344, EPI_ISL_609345, EPI_ISL_609346, EPI_ISL_609347, EPI_ISL_609348, EPI_ISL_609349, EPI_ISL_609350, EPI_ISL_609351, EPI_ISL_609352, EPI_ISL_609353, EPI_ISL_609354, EPI_ISL_609355, EPI_ISL_609356, EPI_ISL_609357, EPI_ISL_609358, EPI_ISL_609359, EPI_ISL_609360, EPI_ISL_609361, EPI_ISL_609362, EPI_ISL_609363, EPI_ISL_609364, EPI_ISL_609365, EPI_ISL_609366, EPI_ISL_609367, EPI_ISL_609368, EPI_ISL_609369, EPI_ISL_609370, EPI_ISL_609371, EPI_ISL_609372, EPI_ISL_609373, EPI_ISL_609374, EPI_ISL_609375, EPI_ISL_609376, EPI_ISL_609377, EPI_ISL_609379, EPI_ISL_609380, EPI_ISL_609381, EPI_ISL_609382, EPI_ISL_609383, EPI_ISL_609384, EPI_ISL_609385, EPI_ISL_609386, EPI_ISL_609387, EPI_ISL_609388, EPI_ISL_609389, EPI_ISL_609390, EPI_ISL_609391, EPI_ISL_609392, EPI_ISL_609394, EPI_ISL_609395, EPI_ISL_609396, EPI_ISL_609397, EPI_ISL_609398, EPI_ISL_609399, EPI_ISL_609400, EPI_ISL_609401, EPI_ISL_609402, EPI_ISL_609403, EPI_ISL_609404, EPI_ISL_609405, EPI_ISL_609406, EPI_ISL_609407, EPI_ISL_609408, EPI_ISL_609409, EPI_ISL_609410, EPI_ISL_609411, EPI_ISL_609412, EPI_ISL_609413, EPI_ISL_609414, EPI_ISL_609415, EPI_ISL_609416, EPI_ISL_609417, EPI_ISL_609418, EPI_ISL_609419, EPI_ISL_609420, EPI_ISL_609421, EPI_ISL_609422, EPI_ISL_609423, EPI_ISL_609424, EPI_ISL_609425, EPI_ISL_609426, EPI_ISL_609427, EPI_ISL_609428, EPI_ISL_609429, EPI_ISL_609430, EPI_ISL_609431, EPI_ISL_609432, EPI_ISL_609433, EPI_ISL_609434, EPI_ISL_609435, EPI_ISL_609436, EPI_ISL_609437, EPI_ISL_609438, EPI_ISL_609439, EPI_ISL_609440, EPI_ISL_609441, EPI_ISL_609442, EPI_ISL_609443, EPI_ISL_609444, EPI_ISL_609445, EPI_ISL_609446, EPI_ISL_609447, EPI_ISL_609448, EPI_ISL_609449, EPI_ISL_609451, EPI_ISL_609452, EPI_ISL_609453, EPI_ISL_609454, EPI_ISL_609455, EPI_ISL_609456, EPI_ISL_609457, EPI_ISL_609458, EPI_ISL_609459, EPI_ISL_609460, EPI_ISL_609461, EPI_ISL_609462, EPI_ISL_609463, EPI_ISL_609464, EPI_ISL_609465, EPI_ISL_609466, EPI_ISL_609467, EPI_ISL_609468, EPI_ISL_609469, EPI_ISL_609470, EPI_ISL_609471, EPI_ISL_609473, EPI_ISL_609474, EPI_ISL_609475, EPI_ISL_609476, EPI_ISL_609477, EPI_ISL_609478, EPI_ISL_609479, EPI_ISL_609480, EPI_ISL_609481, EPI_ISL_609482, EPI_ISL_609483, EPI_ISL_609484, EPI_ISL_609485, EPI_ISL_609486, EPI_ISL_609487, EPI_ISL_609488, EPI_ISL_609489, EPI_ISL_609490, EPI_ISL_609491, EPI_ISL_609492, EPI_ISL_609493, EPI_ISL_609494, EPI_ISL_609495, EPI_ISL_609496, EPI_ISL_609497, EPI_ISL_609498, EPI_ISL_609499, EPI_ISL_609500, EPI_ISL_609501, EPI_ISL_609502, EPI_ISL_609503, EPI_ISL_609504, EPI_ISL_609505, EPI_ISL_609506, EPI_ISL_609507, EPI_ISL_609508, EPI_ISL_609509, EPI_ISL_609510, EPI_ISL_609511, EPI_ISL_609512, EPI_ISL_609513, EPI_ISL_609514, EPI_ISL_609515, EPI_ISL_609516, EPI_ISL_609517, EPI_ISL_609518, EPI_ISL_609519, EPI_ISL_609520, EPI_ISL_609521, EPI_ISL_609522, EPI_ISL_609523, EPI_ISL_609524, EPI_ISL_609525, EPI_ISL_609526, EPI_ISL_609528, EPI_ISL_609529, EPI_ISL_609530, EPI_ISL_609531, EPI_ISL_609532, EPI_ISL_609533, EPI_ISL_609535, EPI_ISL_609536, EPI_ISL_609537, EPI_ISL_609538, EPI_ISL_609539, EPI_ISL_609540, EPI_ISL_609541, EPI_ISL_609542, EPI_ISL_609543, EPI_ISL_609544, EPI_ISL_609545, EPI_ISL_609546, EPI_ISL_609547, EPI_ISL_609548, EPI_ISL_609549, EPI_ISL_609550, EPI_ISL_609551, EPI_ISL_609552, EPI_ISL_609553, EPI_ISL_609554, EPI_ISL_609555, EPI_ISL_609556, EPI_ISL_609557, EPI_ISL_609558, EPI_ISL_609559, EPI_ISL_609560, EPI_ISL_609561, EPI_ISL_609562, EPI_ISL_609563, EPI_ISL_609564, EPI_ISL_609565, EPI_ISL_609566, EPI_ISL_609567, EPI_ISL_609568, EPI_ISL_609569, EPI_ISL_609570, EPI_ISL_609571, EPI_ISL_609572, EPI_ISL_609573, EPI_ISL_609574, EPI_ISL_609575, EPI_ISL_609576, EPI_ISL_609577, EPI_ISL_609578, EPI_ISL_609579, EPI_ISL_609580, EPI_ISL_609581, EPI_ISL_609582, EPI_ISL_609583, EPI_ISL_609584, EPI_ISL_609585, EPI_ISL_609586, EPI_ISL_609587, EPI_ISL_609588, EPI_ISL_609589, EPI_ISL_609590, EPI_ISL_609591, EPI_ISL_609592, EPI_ISL_609593, EPI_ISL_609594, EPI_ISL_609595, EPI_ISL_609596, EPI_ISL_609597, EPI_ISL_609598, EPI_ISL_609599, EPI_ISL_609601, EPI_ISL_609602, EPI_ISL_609603, EPI_ISL_609604, EPI_ISL_609605, EPI_ISL_609606, EPI_ISL_609607, EPI_ISL_609608, EPI_ISL_609609, EPI_ISL_609610, EPI_ISL_609611, EPI_ISL_609612, EPI_ISL_609613, EPI_ISL_609614, EPI_ISL_609615, EPI_ISL_609616, EPI_ISL_609617, EPI_ISL_609618, EPI_ISL_609619, EPI_ISL_609620, EPI_ISL_609621, EPI_ISL_609622, EPI_ISL_609623, EPI_ISL_609624, EPI_ISL_609625, EPI_ISL_609626, EPI_ISL_609627, EPI_ISL_609628, EPI_ISL_609629, EPI_ISL_609630, EPI_ISL_609631, EPI_ISL_609632, EPI_ISL_609633, EPI_ISL_609634, EPI_ISL_609635, EPI_ISL_609636, EPI_ISL_609637, EPI_ISL_609638, EPI_ISL_609639, EPI_ISL_609640, EPI_ISL_609641, EPI_ISL_609642, EPI_ISL_609643, EPI_ISL_609644, EPI_ISL_609645, EPI_ISL_609646, EPI_ISL_609647, EPI_ISL_609648, EPI_ISL_609649, EPI_ISL_609650, EPI_ISL_609651, EPI_ISL_609652, EPI_ISL_609653, EPI_ISL_609654, EPI_ISL_609655, EPI_ISL_609656, EPI_ISL_609657, EPI_ISL_609658, EPI_ISL_609659, EPI_ISL_609660, EPI_ISL_609661, EPI_ISL_609662, EPI_ISL_609663, EPI_ISL_609664, EPI_ISL_609665, EPI_ISL_609666, EPI_ISL_609667, EPI_ISL_609668, EPI_ISL_609669, EPI_ISL_609670, EPI_ISL_609671, EPI_ISL_609672, EPI_ISL_609673, EPI_ISL_609674, EPI_ISL_609675, EPI_ISL_609676, EPI_ISL_609677, EPI_ISL_609678, EPI_ISL_609679, EPI_ISL_609680, EPI_ISL_609681, EPI_ISL_609682, EPI_ISL_609683, EPI_ISL_609684, EPI_ISL_609685, EPI_ISL_609686, EPI_ISL_609687, EPI_ISL_609688, EPI_ISL_609689, EPI_ISL_609690, EPI_ISL_609691, EPI_ISL_609692, EPI_ISL_609693, EPI_ISL_609694, EPI_ISL_609695, EPI_ISL_609696, EPI_ISL_609697, EPI_ISL_609698, EPI_ISL_609699, EPI_ISL_609700, EPI_ISL_609701, EPI_ISL_609702, EPI_ISL_609703, EPI_ISL_609704, EPI_ISL_609705, EPI_ISL_609706, EPI_ISL_609707, EPI_ISL_609708, EPI_ISL_609709, EPI_ISL_609710, EPI_ISL_609711, EPI_ISL_609712, EPI_ISL_609713, EPI_ISL_609714, EPI_ISL_609715, EPI_ISL_609716, EPI_ISL_609717, EPI_ISL_609718, EPI_ISL_609719, EPI_ISL_609720, EPI_ISL_609721, EPI_ISL_609722, EPI_ISL_609723, EPI_ISL_609724, EPI_ISL_609725, EPI_ISL_609726, EPI_ISL_609727, EPI_ISL_609728, EPI_ISL_609729, EPI_ISL_609730, EPI_ISL_609731, EPI_ISL_609732, EPI_ISL_609733, EPI_ISL_609734, EPI_ISL_609735, EPI_ISL_609736, EPI_ISL_609737, EPI_ISL_609738, EPI_ISL_609739, EPI_ISL_609740, EPI_ISL_609741, EPI_ISL_609742, EPI_ISL_609743, EPI_ISL_609744, EPI_ISL_609745, EPI_ISL_609747, EPI_ISL_609748, EPI_ISL_609749, EPI_ISL_609750, EPI_ISL_609751, EPI_ISL_609752, EPI_ISL_609753, EPI_ISL_609754, EPI_ISL_609755, EPI_ISL_609756, EPI_ISL_609757, EPI_ISL_609758, EPI_ISL_609759, EPI_ISL_609760, EPI_ISL_609761, EPI_ISL_609762, EPI_ISL_609763, EPI_ISL_609764, EPI_ISL_609765, EPI_ISL_609766, EPI_ISL_609767, EPI_ISL_609768, EPI_ISL_609769, EPI_ISL_609770, EPI_ISL_609771, EPI_ISL_609772, EPI_ISL_609773, EPI_ISL_609774, EPI_ISL_609775, EPI_ISL_609776, EPI_ISL_609777, EPI_ISL_609778, EPI_ISL_609779, EPI_ISL_609780, EPI_ISL_609781, EPI_ISL_609782, EPI_ISL_609783, EPI_ISL_609784, EPI_ISL_609785, EPI_ISL_609786, EPI_ISL_609787, EPI_ISL_609788, EPI_ISL_609789, EPI_ISL_609790, EPI_ISL_609791, EPI_ISL_609792, EPI_ISL_609793, EPI_ISL_609794, EPI_ISL_609795, EPI_ISL_609796, EPI_ISL_609797, EPI_ISL_609798, EPI_ISL_609799, EPI_ISL_609800

| | | | |
|---|---|---|---|
| see above | Lighthouse Lab in Cambridge | Wellcome Sanger Institute for the COVID-19 Genomics UK (COG-UK) consortium | Rob Howes, The Lighthouse Lab in Cambridge and Alex Alderton, Roberto Amato, Sonia Goncalves, Ewan Harrison, David K. Jackson, Ian Johnston, Dominic Kwiatkowski, Cordelia Langford, John Sillitoe on behalf of the Wellcome Sanger Institute COVID-19 Surveillance Team |

EPI_ISL_609801, EPI_ISL_609802, EPI_ISL_609803, EPI_ISL_609804, EPI_ISL_609805, EPI_ISL_609806, EPI_ISL_609807, EPI_ISL_609808, EPI_ISL_609809, EPI_ISL_609810, EPI_ISL_609811, EPI_ISL_609812, EPI_ISL_609813, EPI_ISL_609814, EPI_ISL_609815, EPI_ISL_609816, EPI_ISL_609817, EPI_ISL_609818, EPI_ISL_609819, EPI_ISL_609820, EPI_ISL_609821, EPI_ISL_609822, EPI_ISL_609823, EPI_ISL_609824, EPI_ISL_609825, EPI_ISL_609826

| | | | |
|---|---|---|---|
| see above | Unity Health Toronto | Ontario Institute for Cancer Research | Ramzi Fattouh, Larissa M. Matukas, Yan Chen,Mark Downing, Trina Otterman, Karel Boissinot, Wai Sum Siu, Zhi Cui, Le Luu, Samira Mubareka, TIBDN, Ilinca Lungu, Bernard Lam, Jeremy Johns, Paul Krzyzanowski, Richard de Borja, Felicia Vincelli, Philip Zuzarte, Jared T. Simpson |

EPI_ISL_610000, EPI_ISL_610001, EPI_ISL_610002, EPI_ISL_610003, EPI_ISL_610004, EPI_ISL_610005, EPI_ISL_610006, EPI_ISL_610007, EPI_ISL_610008, EPI_ISL_610009, EPI_ISL_610010, EPI_ISL_610011, EPI_ISL_610012, EPI_ISL_610013, EPI_ISL_610014, EPI_ISL_610015, EPI_ISL_610016, EPI_ISL_610017, EPI_ISL_610018, EPI_ISL_610019, EPI_ISL_610020, EPI_ISL_610021, EPI_ISL_610022, EPI_ISL_610023, EPI_ISL_610024, EPI_ISL_610025, EPI_ISL_610026, EPI_ISL_610027, EPI_ISL_610028, EPI_ISL_610029, EPI_ISL_610030, EPI_ISL_610031, EPI_ISL_610032, EPI_ISL_610033, EPI_ISL_610034, EPI_ISL_610035, EPI_ISL_610036, EPI_ISL_610037, EPI_ISL_610038, EPI_ISL_610039, EPI_ISL_610040, EPI_ISL_610041, EPI_ISL_610042, EPI_ISL_610043, EPI_ISL_610044, EPI_ISL_610045, EPI_ISL_610046, EPI_ISL_610047

| | | | |
|---|---|---|---|
| see above | Texas Department of State Health Services | Texas Department of State Health Services | Rashmi Tuladhar, Bonnie Oh, Jenny Zhang, Maliha Rahman, Anita Pokharel, Myong Koag, Chung Wang, Rachel Lee, Grace Kubin, Mayela Pedrueza |

EPI_ISL_610629, EPI_ISL_610630, EPI_ISL_610631, EPI_ISL_610632, EPI_ISL_610633, EPI_ISL_610634, EPI_ISL_610635, EPI_ISL_610636, EPI_ISL_610637, EPI_ISL_610638, EPI_ISL_610639, EPI_ISL_610640, EPI_ISL_610641, EPI_ISL_610642, EPI_ISL_610643, EPI_ISL_610644, EPI_ISL_610645, EPI_ISL_610646, EPI_ISL_610647, EPI_ISL_610648, EPI_ISL_610649, EPI_ISL_610650, EPI_ISL_610651, EPI_ISL_610652, EPI_ISL_610653, EPI_ISL_610654, EPI_ISL_610655, EPI_ISL_610656, EPI_ISL_610657, EPI_ISL_610658, EPI_ISL_610659, EPI_ISL_610660, EPI_ISL_610661, EPI_ISL_610662, EPI_ISL_610663, EPI_ISL_610664, EPI_ISL_610665, EPI_ISL_610666, EPI_ISL_610667, EPI_ISL_610668, EPI_ISL_610669, EPI_ISL_610670, EPI_ISL_610671, EPI_ISL_610672, EPI_ISL_610673, EPI_ISL_610674, EPI_ISL_610675, EPI_ISL_610676, EPI_ISL_610677, EPI_ISL_610678, EPI_ISL_610679, EPI_ISL_610680, EPI_ISL_610681, EPI_ISL_610682, EPI_ISL_610683, EPI_ISL_610684, EPI_ISL_610685, EPI_ISL_610686, EPI_ISL_610687, EPI_ISL_610688, EPI_ISL_610689, EPI_ISL_610690, EPI_ISL_610691, EPI_ISL_610692, EPI_ISL_610693, EPI_ISL_610694, EPI_ISL_610695, EPI_ISL_610696, EPI_ISL_610697, EPI_ISL_610698, EPI_ISL_610699, EPI_ISL_610700, EPI_ISL_610701, EPI_ISL_610703, EPI_ISL_610704, EPI_ISL_610705, EPI_ISL_610706, EPI_ISL_610707, EPI_ISL_610708, EPI_ISL_610709, EPI_ISL_610710, EPI_ISL_610711, EPI_ISL_610712, EPI_ISL_610713, EPI_ISL_610714, EPI_ISL_610715, EPI_ISL_610716, EPI_ISL_610717, EPI_ISL_610718, EPI_ISL_610719, EPI_ISL_610720, EPI_ISL_610721, EPI_ISL_610722, EPI_ISL_610723, EPI_ISL_610724, EPI_ISL_610725, EPI_ISL_610726, EPI_ISL_610727, EPI_ISL_610728, EPI_ISL_610729, EPI_ISL_610730, EPI_ISL_610731, EPI_ISL_610732, EPI_ISL_610733, EPI_ISL_610734, EPI_ISL_610735, EPI_ISL_610736, EPI_ISL_610737, EPI_ISL_610738, EPI_ISL_610739, EPI_ISL_610740, EPI_ISL_610741, EPI_ISL_610742, EPI_ISL_610743, EPI_ISL_610744, EPI_ISL_610745, EPI_ISL_610746, EPI_ISL_610747, EPI_ISL_610748, EPI_ISL_610749, EPI_ISL_610750, EPI_ISL_610751, EPI_ISL_610752, EPI_ISL_610753, EPI_ISL_610754, EPI_ISL_610755, EPI_ISL_610756, EPI_ISL_610757, EPI_ISL_610758, EPI_ISL_610759, EPI_ISL_610760, EPI_ISL_610761, EPI_ISL_610762, EPI_ISL_610763, EPI_ISL_610764, EPI_ISL_610765, EPI_ISL_610766, EPI_ISL_610767, EPI_ISL_610768, EPI_ISL_610769, EPI_ISL_610770, EPI_ISL_610771, EPI_ISL_610772, EPI_ISL_610773, EPI_ISL_610774, EPI_ISL_610775, EPI_ISL_610776, EPI_ISL_610777, EPI_ISL_610778, EPI_ISL_610780, EPI_ISL_610781, EPI_ISL_610782, EPI_ISL_610783, EPI_ISL_610784, EPI_ISL_610785

| | | | |
|---|---|---|---|
| see above | Lighthouse Lab in Cambridge | Wellcome Sanger Institute for the COVID-19 Genomics UK (COG-UK) consortium | Rob Howes, The Lighthouse Lab in Cambridge and Alex Alderton, Roberto Amato, Sonia Goncalves, Ewan Harrison, David K. Jackson, Ian Johnston, Dominic Kwiatkowski, Cordelia Langford, John Sillitoe on behalf of the Wellcome Sanger Institute COVID-19 Surveillance Team |

EPI_ISL_610786, EPI_ISL_610787, EPI_ISL_610788, EPI_ISL_610789, EPI_ISL_610790, EPI_ISL_610791, EPI_ISL_610792, EPI_ISL_610793, EPI_ISL_610794, EPI_ISL_610795, EPI_ISL_610796, EPI_ISL_610797, EPI_ISL_610798, EPI_ISL_610799, EPI_ISL_610800, EPI_ISL_610801, EPI_ISL_610802, EPI_ISL_610803, EPI_ISL_610804, EPI_ISL_610805, EPI_ISL_610806, EPI_ISL_610807, EPI_ISL_610808, EPI_ISL_610809, EPI_ISL_610810, EPI_ISL_610811, EPI_ISL_610812, EPI_ISL_610813, EPI_ISL_610814, EPI_ISL_610815, EPI_ISL_610816, EPI_ISL_610817, EPI_ISL_610818, EPI_ISL_610819, EPI_ISL_610820, EPI_ISL_610821, EPI_ISL_610822, EPI_ISL_610823, EPI_ISL_610824, EPI_ISL_610825, EPI_ISL_610826, EPI_ISL_610827, EPI_ISL_610828, EPI_ISL_610829, EPI_ISL_610830, EPI_ISL_610831, EPI_ISL_610832, EPI_ISL_610833, EPI_ISL_610834, EPI_ISL_610835, EPI_ISL_610836, EPI_ISL_610837, EPI_ISL_610838, EPI_ISL_610839, EPI_ISL_610840, EPI_ISL_610841, EPI_ISL_610842, EPI_ISL_610844, EPI_ISL_610845, EPI_ISL_610846, EPI_ISL_610847, EPI_ISL_610848, EPI_ISL_610849, EPI_ISL_610850, EPI_ISL_610851, EPI_ISL_610852, EPI_ISL_610853, EPI_ISL_610854, EPI_ISL_610855, EPI_ISL_610856, EPI_ISL_610857, EPI_ISL_610858, EPI_ISL_610859, EPI_ISL_610860, EPI_ISL_610861, EPI_ISL_610862, EPI_ISL_610863, EPI_ISL_610864, EPI_ISL_610865, EPI_ISL_610866, EPI_ISL_610867, EPI_ISL_610868, EPI_ISL_610869, EPI_ISL_610870, EPI_ISL_610871, EPI_ISL_610872, EPI_ISL_610873, EPI_ISL_610874, EPI_ISL_610875, EPI_ISL_610876, EPI_ISL_610877, EPI_ISL_610878, EPI_ISL_610879, EPI_ISL_610880, EPI_ISL_610881, EPI_ISL_610882, EPI_ISL_610883, EPI_ISL_610884, EPI_ISL_610885, EPI_ISL_610886, EPI_ISL_610887, EPI_ISL_610888, EPI_ISL_610889, EPI_ISL_610890, EPI_ISL_610891, EPI_ISL_610892, EPI_ISL_610893, EPI_ISL_610894, EPI_ISL_610895, EPI_ISL_610896, EPI_ISL_610897, EPI_ISL_610898, EPI_ISL_610899, EPI_ISL_610900, EPI_ISL_610901, EPI_ISL_610902, EPI_ISL_610903, EPI_ISL_610904, EPI_ISL_610905, EPI_ISL_610906, EPI_ISL_610907, EPI_ISL_610908, EPI_ISL_610909, EPI_ISL_610910, EPI_ISL_610911, EPI_ISL_610912, EPI_ISL_610913, EPI_ISL_610914, EPI_ISL_610915, EPI_ISL_610916, EPI_ISL_610917, EPI_ISL_610918, EPI_ISL_610919, EPI_ISL_610920, EPI_ISL_610921

| | | | |
|---|---|---|---|
| see above | Lighthouse Lab in Cambridge | Wellcome Sanger Institute for the COVID-19 Genomics UK (COG-UK) consortium | Rob Howes, The Lighthouse Lab in Cambridge and Alex Alderton, Roberto Amato, Sonia Goncalves, Ewan Harrison, David K. Jackson, Ian Johnston, Dominic Kwiatkowski, Cordelia Langford, John Sillitoe on behalf of the Wellcome Sanger Institute COVID-19 Surveillance Team (http://www.sanger.ac.uk/covid-team) |

EPI_ISL_613966, EPI_ISL_613967, EPI_ISL_613968, EPI_ISL_613969, EPI_ISL_613970, EPI_ISL_613971, EPI_ISL_613972, EPI_ISL_613973, EPI_ISL_613974, EPI_ISL_613975, EPI_ISL_613976, EPI_ISL_613977, EPI_ISL_613978, EPI_ISL_613979, EPI_ISL_613980, EPI_ISL_613981, EPI_ISL_613982, EPI_ISL_613983, EPI_ISL_613984, EPI_ISL_613985, EPI_ISL_613986, EPI_ISL_613987, EPI_ISL_613988, EPI_ISL_613989, EPI_ISL_613990, EPI_ISL_613991, EPI_ISL_613992, EPI_ISL_613993, EPI_ISL_613994, EPI_ISL_613995, EPI_ISL_613996, EPI_ISL_613997, EPI_ISL_613998, EPI_ISL_613999, EPI_ISL_614000, EPI_ISL_614001, EPI_ISL_614002, EPI_ISL_614003, EPI_ISL_614004, EPI_ISL_614005, EPI_ISL_614006, EPI_ISL_614007, EPI_ISL_614008, EPI_ISL_614009, EPI_ISL_614010, EPI_ISL_615122, EPI_ISL_615123, EPI_ISL_615124, EPI_ISL_615125, EPI_ISL_615126, EPI_ISL_615127, EPI_ISL_615128, EPI_ISL_615129, EPI_ISL_615130, EPI_ISL_615131, EPI_ISL_615132, EPI_ISL_615133, EPI_ISL_615134, EPI_ISL_615135, EPI_ISL_615136, EPI_ISL_615137, EPI_ISL_615138, EPI_ISL_615139, EPI_ISL_615140, EPI_ISL_615141, EPI_ISL_615142, EPI_ISL_615143, EPI_ISL_615144, EPI_ISL_615145, EPI_ISL_615146, EPI_ISL_615147, EPI_ISL_615148, EPI_ISL_615149, EPI_ISL_615150, EPI_ISL_615151, EPI_ISL_615152, EPI_ISL_615153, EPI_ISL_615154, EPI_ISL_615155, EPI_ISL_615156, EPI_ISL_615157, EPI_ISL_615158, EPI_ISL_615159, EPI_ISL_615160, EPI_ISL_615161, EPI_ISL_615162, EPI_ISL_615163, EPI_ISL_615164, EPI_ISL_615165

| | | | |
|---|---|---|---|
| see above | Texas Department of State Health Services | Texas Department of State Health Services | Rashmi Tuladhar, Bonnie Oh, Jenny Zhang, Maliha Rahman, Anita Pokharel, Myong Koag, Chung Wang, Rachel Lee, Grace Kubin, Mayela Pedrueza |

EPI_ISL_623220, EPI_ISL_623222, EPI_ISL_623234, EPI_ISL_623245, EPI_ISL_623247, EPI_ISL_623248, EPI_ISL_623256, EPI_ISL_623268, EPI_ISL_623273, EPI_ISL_623276, EPI_ISL_623281, EPI_ISL_623286, EPI_ISL_623288, EPI_ISL_623301, EPI_ISL_623308, EPI_ISL_623312, EPI_ISL_623321, EPI_ISL_623323, EPI_ISL_623330, EPI_ISL_623333, EPI_ISL_623334, EPI_ISL_623337, EPI_ISL_623343, EPI_ISL_623344, EPI_ISL_623345, EPI_ISL_623346, EPI_ISL_623347, EPI_ISL_623359, EPI_ISL_623360, EPI_ISL_623363, EPI_ISL_623364, EPI_ISL_623366, EPI_ISL_623370, EPI_ISL_623385, EPI_ISL_623386, EPI_ISL_623390, EPI_ISL_623392, EPI_ISL_623393, EPI_ISL_623397, EPI_ISL_623398, EPI_ISL_623404, EPI_ISL_623409, EPI_ISL_623411, EPI_ISL_623424, EPI_ISL_623425, EPI_ISL_623426, EPI_ISL_623427, EPI_ISL_623433, EPI_ISL_623437, EPI_ISL_623440, EPI_ISL_623444, EPI_ISL_623445, EPI_ISL_623448, EPI_ISL_623449, EPI_ISL_623451, EPI_ISL_623474, EPI_ISL_623475, EPI_ISL_623482, EPI_ISL_623485, EPI_ISL_623488, EPI_ISL_623489, EPI_ISL_623490, EPI_ISL_623494, EPI_ISL_623495, EPI_ISL_623520, EPI_ISL_623527, EPI_ISL_623528, EPI_ISL_623531, EPI_ISL_623532, EPI_ISL_623537, EPI_ISL_623539, EPI_ISL_623544,

EPI_ISL_623545, EPI_ISL_623547, EPI_ISL_623548, EPI_ISL_623550, EPI_ISL_623551, EPI_ISL_623555, EPI_ISL_623556, EPI_ISL_623557, EPI_ISL_623558, EPI_ISL_623559, EPI_ISL_623560, EPI_ISL_623562, EPI_ISL_623567, EPI_ISL_623568, EPI_ISL_623569, EPI_ISL_623570, EPI_ISL_623572, EPI_ISL_623574, EPI_ISL_623575, EPI_ISL_623576, EPI_ISL_623578, EPI_ISL_623581, EPI_ISL_623584, EPI_ISL_623585, EPI_ISL_623587, EPI_ISL_623592, EPI_ISL_623593, EPI_ISL_623595, EPI_ISL_623600, EPI_ISL_623605, EPI_ISL_623608, EPI_ISL_623609, EPI_ISL_623611, EPI_ISL_623612, EPI_ISL_623616, EPI_ISL_623617, EPI_ISL_623619, EPI_ISL_623626, EPI_ISL_623630, EPI_ISL_623633, EPI_ISL_623639, EPI_ISL_623640, EPI_ISL_623641, EPI_ISL_623643, EPI_ISL_623644, EPI_ISL_623645, EPI_ISL_623647, EPI_ISL_623650, EPI_ISL_623651, EPI_ISL_623652, EPI_ISL_623653, EPI_ISL_623654, EPI_ISL_623655, EPI_ISL_623657, EPI_ISL_623658, EPI_ISL_623661, EPI_ISL_623662, EPI_ISL_623663, EPI_ISL_623665, EPI_ISL_623666, EPI_ISL_623667, EPI_ISL_623674, EPI_ISL_623675, EPI_ISL_623679, EPI_ISL_623680, EPI_ISL_623683, EPI_ISL_623688, EPI_ISL_623690, EPI_ISL_623694, EPI_ISL_623700, EPI_ISL_623702, EPI_ISL_623703, EPI_ISL_623710, EPI_ISL_623712, EPI_ISL_623716, EPI_ISL_623717, EPI_ISL_623721, EPI_ISL_623724, EPI_ISL_623725, EPI_ISL_623726, EPI_ISL_623731, EPI_ISL_623733, EPI_ISL_623735, EPI_ISL_623736, EPI_ISL_623740, EPI_ISL_623741, EPI_ISL_623742, EPI_ISL_623746, EPI_ISL_623749, EPI_ISL_623750, EPI_ISL_623751, EPI_ISL_623754, EPI_ISL_623758, EPI_ISL_623759, EPI_ISL_623762, EPI_ISL_623763, EPI_ISL_623764, EPI_ISL_623772, EPI_ISL_623774, EPI_ISL_623775, EPI_ISL_623776, EPI_ISL_623785, EPI_ISL_623788, EPI_ISL_623790, EPI_ISL_623791, EPI_ISL_623793, EPI_ISL_623794, EPI_ISL_623796, EPI_ISL_623797, EPI_ISL_623798, EPI_ISL_623799, EPI_ISL_623800, EPI_ISL_623802, EPI_ISL_623806, EPI_ISL_623808, EPI_ISL_623809, EPI_ISL_623810, EPI_ISL_623811, EPI_ISL_623812, EPI_ISL_623815, EPI_ISL_623816, EPI_ISL_623822, EPI_ISL_623823, EPI_ISL_623828, EPI_ISL_623829, EPI_ISL_623831, EPI_ISL_623832, EPI_ISL_623833, EPI_ISL_623834, EPI_ISL_623838, EPI_ISL_623840, EPI_ISL_623841, EPI_ISL_623843, EPI_ISL_623844, EPI_ISL_623846, EPI_ISL_623847, EPI_ISL_623848, EPI_ISL_623849, EPI_ISL_623854, EPI_ISL_623855, EPI_ISL_623856, EPI_ISL_623857, EPI_ISL_625281, EPI_ISL_625282, EPI_ISL_625283, EPI_ISL_625284, EPI_ISL_625285, EPI_ISL_625286, EPI_ISL_625287, EPI_ISL_625288, EPI_ISL_625290, EPI_ISL_625291, EPI_ISL_625292, EPI_ISL_625293, EPI_ISL_625294, EPI_ISL_625295, EPI_ISL_625296, EPI_ISL_625297, EPI_ISL_625298, EPI_ISL_625299, EPI_ISL_625300, EPI_ISL_625301, EPI_ISL_625302, EPI_ISL_625303, EPI_ISL_625304, EPI_ISL_625305, EPI_ISL_625306, EPI_ISL_625307, EPI_ISL_625308, EPI_ISL_625309, EPI_ISL_625310, EPI_ISL_625311, EPI_ISL_625313, EPI_ISL_625314, EPI_ISL_625432, EPI_ISL_625437, EPI_ISL_625438, EPI_ISL_625439, EPI_ISL_625440, EPI_ISL_625441, EPI_ISL_625442, EPI_ISL_625443, EPI_ISL_625444, EPI_ISL_625445, EPI_ISL_625446, EPI_ISL_625447, EPI_ISL_625448, EPI_ISL_625449, EPI_ISL_625450, EPI_ISL_625451, EPI_ISL_625452, EPI_ISL_625453

| | | | |
|---|---|---|---|
| see above | Lighthouse Lab in Cambridge | Wellcome Sanger Institute for the COVID-19 Genomics UK (COG-UK) consortium | Rob Howes, The Lighthouse Lab in Cambridge and Alex Alderton, Roberto Amato, Sonia Goncalves, Ewan Harrison, David K. Jackson, Ian Johnston, Dominic Kwiatkowski, Cordelia Langford, John Sillitoe on behalf of the Wellcome Sanger Institute COVID-19 Surveillance Team (http://www.sanger.ac.uk/covid-team) |

EPI_ISL_626357, EPI_ISL_626358, EPI_ISL_626359, EPI_ISL_626360, EPI_ISL_626361, EPI_ISL_626362, EPI_ISL_626363, EPI_ISL_626364, EPI_ISL_626365, EPI_ISL_626366, EPI_ISL_626367, EPI_ISL_626368, EPI_ISL_626369, EPI_ISL_626370, EPI_ISL_626371, EPI_ISL_626372, EPI_ISL_626373, EPI_ISL_626374, EPI_ISL_626375, EPI_ISL_626376, EPI_ISL_626377, EPI_ISL_626378, EPI_ISL_626379, EPI_ISL_626380, EPI_ISL_626381, EPI_ISL_626382, EPI_ISL_626383, EPI_ISL_626384, EPI_ISL_626385, EPI_ISL_626386, EPI_ISL_626387, EPI_ISL_626388, EPI_ISL_626389, EPI_ISL_626390, EPI_ISL_626391, EPI_ISL_626392, EPI_ISL_626393, EPI_ISL_626394, EPI_ISL_626395, EPI_ISL_626396, EPI_ISL_626397, EPI_ISL_626398, EPI_ISL_626399, EPI_ISL_626400, EPI_ISL_626401, EPI_ISL_626402, EPI_ISL_626403, EPI_ISL_626404, EPI_ISL_626405, EPI_ISL_626406, EPI_ISL_626407, EPI_ISL_626408, EPI_ISL_626409, EPI_ISL_626410, EPI_ISL_626411, EPI_ISL_626412, EPI_ISL_626413, EPI_ISL_626414, EPI_ISL_626415, EPI_ISL_626416, EPI_ISL_626417, EPI_ISL_626418, EPI_ISL_626419, EPI_ISL_626420, EPI_ISL_626421, EPI_ISL_626422, EPI_ISL_626423, EPI_ISL_626424, EPI_ISL_626425, EPI_ISL_626426, EPI_ISL_626427, EPI_ISL_626428, EPI_ISL_626429, EPI_ISL_626430, EPI_ISL_626431, EPI_ISL_626432, EPI_ISL_626433, EPI_ISL_626434, EPI_ISL_626435, EPI_ISL_626436, EPI_ISL_626437, EPI_ISL_626438, EPI_ISL_626439, EPI_ISL_626440, EPI_ISL_626441, EPI_ISL_626442, EPI_ISL_626443, EPI_ISL_626444, EPI_ISL_626445, EPI_ISL_626446, EPI_ISL_626447, EPI_ISL_626448, EPI_ISL_626449, EPI_ISL_626450, EPI_ISL_626451, EPI_ISL_626452, EPI_ISL_626453, EPI_ISL_626454, EPI_ISL_626455, EPI_ISL_626456, EPI_ISL_626457, EPI_ISL_626458, EPI_ISL_626459, EPI_ISL_626460, EPI_ISL_626461, EPI_ISL_626462, EPI_ISL_626463, EPI_ISL_626464, EPI_ISL_626465, EPI_ISL_626466, EPI_ISL_626467, EPI_ISL_626468, EPI_ISL_626469, EPI_ISL_626470, EPI_ISL_626471, EPI_ISL_626472, EPI_ISL_626473, EPI_ISL_626474, EPI_ISL_626475, EPI_ISL_626476, EPI_ISL_626477, EPI_ISL_626478, EPI_ISL_626479, EPI_ISL_626480, EPI_ISL_626481, EPI_ISL_626482, EPI_ISL_626483, EPI_ISL_626484, EPI_ISL_626485, EPI_ISL_626486, EPI_ISL_626487, EPI_ISL_626488, EPI_ISL_626489, EPI_ISL_626490, EPI_ISL_626491, EPI_ISL_626492, EPI_ISL_626493, EPI_ISL_626494, EPI_ISL_626495, EPI_ISL_626496, EPI_ISL_626497, EPI_ISL_626498, EPI_ISL_626499, EPI_ISL_626500, EPI_ISL_626501, EPI_ISL_626502, EPI_ISL_626503, EPI_ISL_626504, EPI_ISL_626505, EPI_ISL_626506, EPI_ISL_626507, EPI_ISL_626508, EPI_ISL_626509, EPI_ISL_626510, EPI_ISL_626511, EPI_ISL_626512, EPI_ISL_626513, EPI_ISL_626514, EPI_ISL_626515, EPI_ISL_626516, EPI_ISL_626517, EPI_ISL_626518, EPI_ISL_626519, EPI_ISL_626520, EPI_ISL_626521, EPI_ISL_626522, EPI_ISL_626523, EPI_ISL_626524, EPI_ISL_626525, EPI_ISL_626526, EPI_ISL_626527, EPI_ISL_626528, EPI_ISL_626529, EPI_ISL_626530, EPI_ISL_626531, EPI_ISL_626532, EPI_ISL_626533, EPI_ISL_626534, EPI_ISL_626535, EPI_ISL_626536, EPI_ISL_626537, EPI_ISL_626538, EPI_ISL_626539, EPI_ISL_626540, EPI_ISL_626541, EPI_ISL_626542, EPI_ISL_626543, EPI_ISL_626544, EPI_ISL_626545, EPI_ISL_626546, EPI_ISL_626547, EPI_ISL_626548

| | | | |
|---|---|---|---|
| see above | Northwestern Memorial Hospital | Ozer Lab | Ramon Lorenzo-Redondo, Hannah H. Nam, Scott C. Roberts, Lacy M. Simons, Chad J. Achenbach, Lawrence J. Jennings, Chao Qi, Alan R. Hauser, Michael G. Ison, Judd F. Hultquist, Egon A. Ozer |

EPI_ISL_628701, EPI_ISL_628702, EPI_ISL_628703, EPI_ISL_628704, EPI_ISL_628705, EPI_ISL_628706, EPI_ISL_628707, EPI_ISL_628708, EPI_ISL_628709, EPI_ISL_628710, EPI_ISL_628711, EPI_ISL_628712, EPI_ISL_628713, EPI_ISL_628714, EPI_ISL_628715, EPI_ISL_628716, EPI_ISL_628717, EPI_ISL_628718, EPI_ISL_628719, EPI_ISL_628720, EPI_ISL_628721, EPI_ISL_628722, EPI_ISL_628723, EPI_ISL_628724, EPI_ISL_628725, EPI_ISL_628726, EPI_ISL_628727, EPI_ISL_628728, EPI_ISL_628729, EPI_ISL_628730, EPI_ISL_628731, EPI_ISL_628732, EPI_ISL_628733, EPI_ISL_628734, EPI_ISL_628735, EPI_ISL_628736, EPI_ISL_628737, EPI_ISL_628738, EPI_ISL_628739, EPI_ISL_628740, EPI_ISL_628741, EPI_ISL_628742, EPI_ISL_628743, EPI_ISL_628744

| | | | |
|---|---|---|---|
| see above | Texas Department of State Health Services | Texas Department of State Health Services | Rashmi Tuladhar, Bonnie Oh, Jenny Zhang, Maliha Rahman, Anita Pokharel, Myong Koag, Chung Wang, Rachel Lee, Grace Kubin, Mayela Pedrueza |

EPI_ISL_630972, EPI_ISL_630977, EPI_ISL_630979, EPI_ISL_630984, EPI_ISL_630988, EPI_ISL_630996, EPI_ISL_630997, EPI_ISL_631001, EPI_ISL_631003, EPI_ISL_631005, EPI_ISL_631008, EPI_ISL_631016, EPI_ISL_631017, EPI_ISL_631025, EPI_ISL_631028, EPI_ISL_631038, EPI_ISL_631039, EPI_ISL_631045, EPI_ISL_631053, EPI_ISL_631055, EPI_ISL_631062, EPI_ISL_631063, EPI_ISL_631069, EPI_ISL_631078, EPI_ISL_631080, EPI_ISL_631094, EPI_ISL_631101, EPI_ISL_631105, EPI_ISL_631109, EPI_ISL_631117, EPI_ISL_631118, EPI_ISL_631123, EPI_ISL_631129, EPI_ISL_631132, EPI_ISL_631135, EPI_ISL_631137, EPI_ISL_631142, EPI_ISL_631144, EPI_ISL_631148, EPI_ISL_631149, EPI_ISL_631154, EPI_ISL_631155, EPI_ISL_631160, EPI_ISL_631164, EPI_ISL_631171, EPI_ISL_631174, EPI_ISL_631177, EPI_ISL_631180, EPI_ISL_631181, EPI_ISL_631185, EPI_ISL_631191, EPI_ISL_631192, EPI_ISL_631194, EPI_ISL_631199, EPI_ISL_631204, EPI_ISL_631208, EPI_ISL_631219, EPI_ISL_631220, EPI_ISL_631221, EPI_ISL_631224, EPI_ISL_631225, EPI_ISL_631235, EPI_ISL_631237, EPI_ISL_631245, EPI_ISL_631254, EPI_ISL_631257, EPI_ISL_631262, EPI_ISL_631266, EPI_ISL_631268, EPI_ISL_631270, EPI_ISL_631271, EPI_ISL_631272, EPI_ISL_631273, EPI_ISL_631274

| | | | |
|---|---|---|---|
| see above | Lighthouse Lab in Cambridge | Wellcome Sanger Institute for the COVID-19 Genomics UK (COG-UK) consortium | Rob Howes, The Lighthouse Lab in Cambridge and Alex Alderton, Roberto Amato, Sonia Goncalves, Ewan Harrison, David K. Jackson, Ian Johnston, Dominic Kwiatkowski, Cordelia Langford, John Sillitoe on behalf of the Wellcome Sanger Institute COVID-19 Surveillance Team |

EPI_ISL_631640, EPI_ISL_631641, EPI_ISL_631642, EPI_ISL_631643, EPI_ISL_631644, EPI_ISL_631645, EPI_ISL_631646, EPI_ISL_631647, EPI_ISL_631648, EPI_ISL_631649, EPI_ISL_631650, EPI_ISL_631651, EPI_ISL_631652, EPI_ISL_631653, EPI_ISL_631654, EPI_ISL_631655, EPI_ISL_631656, EPI_ISL_631657, EPI_ISL_631658, EPI_ISL_631659, EPI_ISL_631660, EPI_ISL_631661, EPI_ISL_631662, EPI_ISL_631663, EPI_ISL_631664, EPI_ISL_631665, EPI_ISL_631666, EPI_ISL_631667, EPI_ISL_631668, EPI_ISL_631669, EPI_ISL_631670, EPI_ISL_631671, EPI_ISL_631672, EPI_ISL_631673, EPI_ISL_631674, EPI_ISL_631675, EPI_ISL_631676, EPI_ISL_631677, EPI_ISL_631678, EPI_ISL_631679, EPI_ISL_631680, EPI_ISL_631681, EPI_ISL_631682, EPI_ISL_631683, EPI_ISL_631684

| | | | |
|---|---|---|---|
| see above | Texas Department of State Health Services | Texas Department of State Health Services | Rashmi Tuladhar, Bonnie Oh, Jenny Zhang, Maliha Rahman, Anita Pokharel, Myong Koag, Chung Wang, Rachel Lee, Grace Kubin, Mayela Pedrueza |

We gratefully acknowledge the following Authors from the Originating laboratories responsible for obtaining the specimens, as well as the Submitting laboratories where the genome data were generated and shared via GISAID, on which this research is based.

All Submitters of data may be contacted directly via www.gisaid.org

Authors are sorted alphabetically.

| Accession ID | Originating Laboratory | Submitting Laboratory | Authors |
|---|---|---|---|
| EPI_ISL_417390, EPI_ISL_417407 | Centre for Infectious Diseases and Microbiology Public Health | NSW Health Pathology - Institute of Clinical Pathology and Medical Research; Westmead Hospital; University of Sydney | Sadsad R, Carter I, Rahman H, Holmes EC, O'Sullivan MV, Sintchenko V, Chen SC, Maddocks S, Kok J, Dwyer DE, Rockett R, Eden J-S, Lam C, Gray K, Timms V, Gall M and Arnott A for the 2019-nCoV Study Group |
| EPI_ISL_420294, EPI_ISL_420295 | Institute of Microbiology and Immunology, Faculty of Medicine, University of Ljubljana | Institute of Microbiology and Immunology, Faculty of Medicine, University of Ljubljana | Samo Zakotnik, Tomaž Mark Zorec, Lucijan Skubic, Miša Korva, Mario Poljak, Tatjana Avši - Županc |
| EPI_ISL_421517 | Servicio de Microbiologia. Hospital Clinico Universitario de Valencia | Sequencing and Bioinformatics Service and Molecular Epidemiology Research Group. FISABIO-Public Health | Sandra Carbo, Loreto Ferrús Abad, Maria Alma Bracho, Griselda De Marco, Beatriz Beamud, Lidia Ruiz Roldan, Marta Pla Diaz, Neris Garcia-Gonzalez, Inma Galán Vendrell, Paula Ruiz-Hueso, Mariana Reyes-Prieto, Vicente Soriano Chirona, Ivan Ansari, David Navarro, Llúcia Martínez-Priego, Giuseppe D'Auria, Fernando Gonzalez-Candelas |
| EPI_ISL_425186, EPI_ISL_425221 | Servicio de Microbiología. Consorcio Hospital General Universitario de Valencia | Sequencing and Bioinformatics Service and Molecular Epidemiology Research Group. FISABIO-Public Health | Sandra Carbo, Loreto Ferrús Abad, Paula Ruiz-Hueso, Mariana Reyes-Prieto, Vicente Soriano Chirona, Ivan Ansari, David Navarro, Maria Alma Bracho, Griselda De Marco, Beatriz Beamud, Lidia Ruiz Roldan, Marta Pla Diaz, Neris Garcia-Gonzalez, Inma Galán Vendrell, Llúcia Martínez-Priego, Giuseppe D'Auria, Fernando Gonzalez-Candelas |
| EPI_ISL_427654, EPI_ISL_427669, EPI_ISL_427685, EPI_ISL_427705, EPI_ISL_427740, EPI_ISL_427747, EPI_ISL_427754, EPI_ISL_427755, EPI_ISL_427760, EPI_ISL_427775, EPI_ISL_427776, EPI_ISL_427777, EPI_ISL_427779, EPI_ISL_427781, EPI_ISL_427790, EPI_ISL_427795 | | | |
| see above | Centre for Infectious Diseases and Microbiology Public Health | NSW Health Pathology - Institute of Clinical Pathology and Medical Research; Westmead Hospital; University of Sydney | Sadsad R, Draper J, Sim E, Bachmann N, Rockett R, Lam C, Gray K, Timms V, Gall M, Arnott A, Carter I, Holmes EC, O'Sullivan MV, Byun R, Sintchenko V, Chen SC, Eden JS, Maddocks S, Kok J, Propenko M, Sorrell T, Chang S, Basile K, Dwyer DE for the 2019-nCoV Study Group |
| EPI_ISL_428683, EPI_ISL_428684, EPI_ISL_428685, EPI_ISL_428686, EPI_ISL_428687, EPI_ISL_428688, EPI_ISL_428689, EPI_ISL_428690, EPI_ISL_428691, EPI_ISL_428692, EPI_ISL_428693, EPI_ISL_428694 | | | |
| see above | Hospital Universitario 12 de Octubre | Hospital Universitario 12 de Octubre | Sara González, Raúl Recio,Elias Dahdouh, Fernando Lázaro, Esther Viedma, Natalia Stella, Julio García, Juan Carlos Galán, Rafael Cantón, Mª Dolores Folgueira, Rafael Delgado, Jesús Mingorance |
| EPI_ISL_429990 | Rady's Childrens Hospital | Andersen lab at Scripps Research | SEARCH Alliance San Diego with Christina Clarke, Michelle Vanderpool, Teresa Mueller, Denise Malicki |
| EPI_ISL_429991, EPI_ISL_430016 | Andersen lab at Scripps Research | Andersen lab at Scripps Research | SEARCH Alliance San Diego |
| EPI_ISL_430718 | Hospital Universitario 12 de Octubre | Hospital Universitario 12 de Octubre | Sara González, Raúl Recio,Elias Dahdouh, Fernando Lázaro, Esther Viedma, Natalia Stella, Julio García, Juan Carlos Galán, Rafael Cantón, Mª Dolores Folgueira, Rafael Delgado, Jesús Mingorance |
| EPI_ISL_434647, EPI_ISL_434648 | Victoria Vard och Halsa | The Public Health Agency of Sweden | Sarah Henriksson, Oskar Karlsson Lindsjo, Maria Lind Karlberg, Anna-Malin Linde, Olov Svartstrom, Anna Risberg, Theresa Enkirch, Mia Brytting, Karin Tegmark-Wisell |
| EPI_ISL_434712, EPI_ISL_434713, EPI_ISL_434714, EPI_ISL_434715, EPI_ISL_434716, EPI_ISL_434717, EPI_ISL_434718, EPI_ISL_434719, EPI_ISL_434720, EPI_ISL_434721, EPI_ISL_434722, EPI_ISL_434723, EPI_ISL_434724, EPI_ISL_434725, EPI_ISL_434726, EPI_ISL_434727, EPI_ISL_434728, EPI_ISL_434729, EPI_ISL_434730, EPI_ISL_434731, EPI_ISL_434732, EPI_ISL_434733, EPI_ISL_434734, EPI_ISL_434735, EPI_ISL_434736, EPI_ISL_434737, EPI_ISL_434738, EPI_ISL_434739, EPI_ISL_434740, EPI_ISL_434741, EPI_ISL_434742, EPI_ISL_434743, EPI_ISL_434744, EPI_ISL_434745, EPI_ISL_434746, EPI_ISL_434747, EPI_ISL_434748, EPI_ISL_434749, EPI_ISL_434750, EPI_ISL_434751, EPI_ISL_434752, EPI_ISL_434753, EPI_ISL_434754, EPI_ISL_434755, EPI_ISL_434756, EPI_ISL_434757, EPI_ISL_434758, EPI_ISL_434759, EPI_ISL_434760, EPI_ISL_434761, EPI_ISL_434762, EPI_ISL_434763, EPI_ISL_434764, EPI_ISL_434765, EPI_ISL_434766, EPI_ISL_434767, EPI_ISL_434768, EPI_ISL_434769, EPI_ISL_434770, EPI_ISL_434771, EPI_ISL_434772, EPI_ISL_434773, EPI_ISL_434774, EPI_ISL_434775, EPI_ISL_434776, EPI_ISL_434777, EPI_ISL_434778, EPI_ISL_434779, EPI_ISL_434780, EPI_ISL_434781, EPI_ISL_434782, EPI_ISL_434783, EPI_ISL_434784, EPI_ISL_434785, EPI_ISL_434786, EPI_ISL_434787, EPI_ISL_434788, EPI_ISL_434789, EPI_ISL_434790, EPI_ISL_434791, EPI_ISL_434792, EPI_ISL_434793, EPI_ISL_434794, EPI_ISL_434795, EPI_ISL_434796, EPI_ISL_434797, EPI_ISL_434798, EPI_ISL_434799, EPI_ISL_434800, EPI_ISL_434801, EPI_ISL_434802, EPI_ISL_434803, EPI_ISL_434804, EPI_ISL_434805, EPI_ISL_434806, EPI_ISL_434807, EPI_ISL_434808, EPI_ISL_434809, EPI_ISL_434810, EPI_ISL_434811, EPI_ISL_434812, EPI_ISL_434813, EPI_ISL_434814, EPI_ISL_434815, EPI_ISL_434816, EPI_ISL_434817, EPI_ISL_434818, EPI_ISL_434819, EPI_ISL_434820, EPI_ISL_434821, EPI_ISL_434822, EPI_ISL_434823, EPI_ISL_434824, EPI_ISL_434825, EPI_ISL_434826, EPI_ISL_434827, EPI_ISL_434828, EPI_ISL_434829, EPI_ISL_434830, EPI_ISL_434831, EPI_ISL_434832, EPI_ISL_434833, EPI_ISL_434834, EPI_ISL_434835, EPI_ISL_434836, EPI_ISL_434837, EPI_ISL_434838, EPI_ISL_434839, EPI_ISL_434840, EPI_ISL_434841, EPI_ISL_434842, EPI_ISL_434843, EPI_ISL_434844, EPI_ISL_434845, EPI_ISL_434846, EPI_ISL_434847, EPI_ISL_434848, EPI_ISL_434849, EPI_ISL_434850, EPI_ISL_434851, EPI_ISL_434852, EPI_ISL_434853, EPI_ISL_434854, EPI_ISL_434855, EPI_ISL_434856, EPI_ISL_434857, EPI_ISL_434858, EPI_ISL_434859, EPI_ISL_434860, EPI_ISL_434861, EPI_ISL_434862, EPI_ISL_434863, EPI_ISL_434864, EPI_ISL_434865, EPI_ISL_434866, EPI_ISL_434867, EPI_ISL_434868, EPI_ISL_434869, EPI_ISL_434870, EPI_ISL_434871, EPI_ISL_434872, EPI_ISL_434873, EPI_ISL_434874, EPI_ISL_434875, EPI_ISL_434876, EPI_ISL_434877, EPI_ISL_434878, EPI_ISL_434879, EPI_ISL_434880, EPI_ISL_434881, EPI_ISL_434882, EPI_ISL_434883, EPI_ISL_434884, EPI_ISL_434885, EPI_ISL_434886, EPI_ISL_434887, EPI_ISL_434888, EPI_ISL_434889, EPI_ISL_434890, EPI_ISL_434891, EPI_ISL_434892, EPI_ISL_434893, EPI_ISL_434894, EPI_ISL_434895, EPI_ISL_434896, EPI_ISL_434897, EPI_ISL_434898, EPI_ISL_434899, EPI_ISL_434900, EPI_ISL_434901, EPI_ISL_434902, EPI_ISL_434903, EPI_ISL_434904, EPI_ISL_434905, EPI_ISL_434906, EPI_ISL_434907, EPI_ISL_434908, EPI_ISL_434909, EPI_ISL_434910, EPI_ISL_434911, EPI_ISL_434912, EPI_ISL_434913, EPI_ISL_434914, EPI_ISL_434915, EPI_ISL_434916, EPI_ISL_434917, EPI_ISL_434918, EPI_ISL_434919, EPI_ISL_434920, EPI_ISL_434921, EPI_ISL_434922, EPI_ISL_434923, EPI_ISL_434924, EPI_ISL_434925, EPI_ISL_434926, EPI_ISL_434927, EPI_ISL_434928, EPI_ISL_434929, EPI_ISL_434930, EPI_ISL_434931, EPI_ISL_434932, EPI_ISL_434933, EPI_ISL_434934, EPI_ISL_434935, EPI_ISL_434936, EPI_ISL_434937, EPI_ISL_434938, EPI_ISL_434939, EPI_ISL_434940, EPI_ISL_434941, EPI_ISL_434942, EPI_ISL_434943, EPI_ISL_434944, EPI_ISL_434945, EPI_ISL_434946, EPI_ISL_434947, EPI_ISL_434948, EPI_ISL_434949, EPI_ISL_434950, EPI_ISL_434951, EPI_ISL_434952, EPI_ISL_434953, EPI_ISL_434954, EPI_ISL_434955, EPI_ISL_434956, EPI_ISL_434957, EPI_ISL_434958, EPI_ISL_434959, EPI_ISL_434960, EPI_ISL_434961, EPI_ISL_434962, EPI_ISL_434963, EPI_ISL_434964, EPI_ISL_434965, EPI_ISL_434966, EPI_ISL_434967, EPI_ISL_434968, EPI_ISL_434969, EPI_ISL_434970, EPI_ISL_434971, EPI_ISL_434972, EPI_ISL_434973, EPI_ISL_434974, EPI_ISL_434975, EPI_ISL_434976, EPI_ISL_434977, EPI_ISL_434978, EPI_ISL_434979, EPI_ISL_434980, EPI_ISL_434981, EPI_ISL_434982, EPI_ISL_434983, EPI_ISL_434984, EPI_ISL_434985, EPI_ISL_434986, EPI_ISL_434987, EPI_ISL_434988, EPI_ISL_434989, EPI_ISL_434990, EPI_ISL_434991, EPI_ISL_434992, EPI_ISL_434993, EPI_ISL_434994, EPI_ISL_434995, EPI_ISL_434996, EPI_ISL_434997, EPI_ISL_434998, EPI_ISL_434999, EPI_ISL_435000, EPI_ISL_435001, EPI_ISL_435002, EPI_ISL_435003, EPI_ISL_435004, EPI_ISL_435005, EPI_ISL_435006, EPI_ISL_435007, EPI_ISL_435008, EPI_ISL_435009, EPI_ISL_435010, EPI_ISL_435011, EPI_ISL_435012, EPI_ISL_435013, EPI_ISL_435014, EPI_ISL_435015, EPI_ISL_435016, EPI_ISL_435017, EPI_ISL_435018, EPI_ISL_435019, EPI_ISL_435020, EPI_ISL_435021, EPI_ISL_435022, EPI_ISL_435023, EPI_ISL_435024, EPI_ISL_435025, EPI_ISL_435026, EPI_ISL_435027, EPI_ISL_435028, EPI_ISL_435029, EPI_ISL_435030, EPI_ISL_435031 | | | |
| see above | Houston Methodist Hospital | Houston Methodist Hospital | S. Wesley Long, Randall J. Olsen, Paul A. Christensen, David W. Bernard, James J. Davis, Maulik Shukla, Marcus Nguyen, Matthew Ojeda Saavedra, Concepcion C. Cantu, Prasanti Yerramilli, Layne Pruitt, Sishir Subedi, Heather Hendrickson, Ghazaleh Eskandari, Muthiah Kumaraswami, Jason S. McLellan, Hakon Jonsson, Kari Stefansson, and James M. Musser |
| EPI_ISL_435473, EPI_ISL_435474 | Rady's Childrens Hospital | Andersen lab at Scripps Research | SEARCH Alliance San Diego |
| EPI_ISL_436285, EPI_ISL_436305, EPI_ISL_436325, EPI_ISL_436345, EPI_ISL_436369, EPI_ISL_436396 | Servicio de Microbiologia. Hospital Clinico Universitario de Valencia | Sequencing and Bioinformatics Service and Molecular Epidemiology Research Group. FISABIO-Public Health | Sandra Carbo, Loreto Ferrús Abad, Paula Ruiz-Hueso, Mariana Reyes-Prieto, Vicente Soriano Chirona, Ivan Ansari, David Navarro, Maria Alma Bracho, Griselda De Marco, Beatriz Beamud, Lidia Ruiz Roldan, Marta Pla Diaz, Neris Garcia-Gonzalez, Inma Galán Vendrell, Llúcia Martínez-Priego, Giuseppe D'Auria, Fernando Gonzalez-Candelas |
| EPI_ISL_436508, EPI_ISL_436509, EPI_ISL_436510, EPI_ISL_436511, EPI_ISL_436513, EPI_ISL_436541, EPI_ISL_436542, EPI_ISL_436543, EPI_ISL_436544, EPI_ISL_436545, EPI_ISL_436546, EPI_ISL_436547, EPI_ISL_436548, EPI_ISL_436549, EPI_ISL_436550, EPI_ISL_436551, EPI_ISL_436552, EPI_ISL_436553, EPI_ISL_436554, EPI_ISL_436555, EPI_ISL_436558, EPI_ISL_436559, EPI_ISL_436560, EPI_ISL_436562 | | | |
| see above | Florida Bureau of Public Health Laboratories | Florida Bureau of Public Health Laboratories | Sarah Schmedes, Jason Blanton |
| EPI_ISL_437481, EPI_ISL_437482, EPI_ISL_437483, EPI_ISL_437484, EPI_ISL_437485, EPI_ISL_437486, EPI_ISL_437487, EPI_ISL_437488, EPI_ISL_437489, EPI_ISL_437490, EPI_ISL_437491, EPI_ISL_437492, EPI_ISL_437493, EPI_ISL_437494, EPI_ISL_437495, EPI_ISL_437496, EPI_ISL_437497 | | | |
| see above | Pathogen Genomics Lab King Abdullah University of Science and Technology(KAUST) | Pathogen Genomics Lab King Abdullah University of Science and Technology(KAUST) | Sara Mfarrej,Raeece Naeem,Sharif Hala,Amit Subudhi,Fathia Rached,Arnab Pain |
| EPI_ISL_437549, EPI_ISL_437550, EPI_ISL_437551, EPI_ISL_437552, EPI_ISL_437553, EPI_ISL_437554, EPI_ISL_437555, EPI_ISL_437556, EPI_ISL_437557, EPI_ISL_437559, EPI_ISL_437560, EPI_ISL_437561, EPI_ISL_437562, EPI_ISL_437563, EPI_ISL_437564, EPI_ISL_437565, EPI_ISL_437566, EPI_ISL_437567, EPI_ISL_437568, EPI_ISL_437569, EPI_ISL_437570, EPI_ISL_437571, EPI_ISL_437572, EPI_ISL_437573, EPI_ISL_437574, EPI_ISL_437575, EPI_ISL_437576, EPI_ISL_437577, EPI_ISL_437578, EPI_ISL_437579, EPI_ISL_437580, EPI_ISL_437581, EPI_ISL_437582, EPI_ISL_437583, EPI_ISL_437584, EPI_ISL_437585, EPI_ISL_437586, EPI_ISL_437587, EPI_ISL_437588, EPI_ISL_437589, EPI_ISL_437590, EPI_ISL_437591, EPI_ISL_437592, EPI_ISL_437593, EPI_ISL_437594, EPI_ISL_437595, EPI_ISL_437596, EPI_ISL_437598, EPI_ISL_437599, EPI_ISL_437600 | | | |
| see above | Scripps Medical Laboratory | Andersen lab at Scripps Research | SEARCH Alliance San Diego with Michael Quigley, Ellen Stefanski, Ian Mchardy |
| EPI_ISL_439672, EPI_ISL_439673, EPI_ISL_439674, EPI_ISL_439675, EPI_ISL_439676, EPI_ISL_439677, EPI_ISL_439678, EPI_ISL_439679, EPI_ISL_439680, EPI_ISL_439681, EPI_ISL_439682, EPI_ISL_439683, EPI_ISL_439684, EPI_ISL_439685, EPI_ISL_439686, EPI_ISL_439687, EPI_ISL_439688, EPI_ISL_439689, EPI_ISL_439690, EPI_ISL_439691, EPI_ISL_439692, EPI_ISL_439693, EPI_ISL_439694, EPI_ISL_439695, EPI_ISL_439696, EPI_ISL_439697, EPI_ISL_439698, EPI_ISL_439699, EPI_ISL_439700, EPI_ISL_439701, EPI_ISL_439702, EPI_ISL_439703, EPI_ISL_439704, EPI_ISL_439705, EPI_ISL_439706, EPI_ISL_439707, EPI_ISL_439708, EPI_ISL_439709, EPI_ISL_439710, EPI_ISL_439711, EPI_ISL_439712, EPI_ISL_439713, EPI_ISL_439714, EPI_ISL_439715, EPI_ISL_439716, EPI_ISL_439717, EPI_ISL_439718, EPI_ISL_439719, EPI_ISL_439720, EPI_ISL_439721, EPI_ISL_439722, EPI_ISL_439723, EPI_ISL_439724, EPI_ISL_439725, EPI_ISL_439726, EPI_ISL_439727, EPI_ISL_439728, EPI_ISL_439729, EPI_ISL_439730, EPI_ISL_439731, EPI_ISL_439732, EPI_ISL_439733, EPI_ISL_439734, EPI_ISL_439735, EPI_ISL_439736, EPI_ISL_439737, EPI_ISL_439738, EPI_ISL_439739, EPI_ISL_439740, EPI_ISL_439741, EPI_ISL_439742, EPI_ISL_439743, EPI_ISL_439744, EPI_ISL_439745, EPI_ISL_439746, EPI_ISL_439747, EPI_ISL_439748, EPI_ISL_439749, EPI_ISL_439750, EPI_ISL_439751, EPI_ISL_439752, EPI_ISL_439753, EPI_ISL_439754, EPI_ISL_439755, EPI_ISL_439756, EPI_ISL_439757, EPI_ISL_439758, EPI_ISL_439759, EPI_ISL_439760, EPI_ISL_439761, EPI_ISL_439762, EPI_ISL_439763, EPI_ISL_439764, EPI_ISL_439765, EPI_ISL_439766, EPI_ISL_439767, EPI_ISL_439768, EPI_ISL_439769, EPI_ISL_439770, EPI_ISL_439771, EPI_ISL_439772, EPI_ISL_439773, EPI_ISL_439774, EPI_ISL_439775, EPI_ISL_439776, EPI_ISL_439777, EPI_ISL_439778, EPI_ISL_439779, EPI_ISL_439780, EPI_ISL_439781, EPI_ISL_439782, EPI_ISL_439783, EPI_ISL_439784, EPI_ISL_439785, EPI_ISL_439786, EPI_ISL_439787, EPI_ISL_439788, EPI_ISL_439789, EPI_ISL_439790, EPI_ISL_439791, EPI_ISL_439792, EPI_ISL_439793, EPI_ISL_439794, EPI_ISL_439795, EPI_ISL_439796, EPI_ISL_439797, | | | |

| | | | |
|---|---|---|---|
| EPI_ISL_439798, EPI_ISL_439799, EPI_ISL_439800, EPI_ISL_439801, EPI_ISL_439802, EPI_ISL_439803, EPI_ISL_439804, EPI_ISL_439805, EPI_ISL_439806, EPI_ISL_439807, EPI_ISL_439808, EPI_ISL_439809, EPI_ISL_439810, EPI_ISL_439811, EPI_ISL_439812, EPI_ISL_439813, EPI_ISL_439814, EPI_ISL_439815, EPI_ISL_439816, EPI_ISL_439817, EPI_ISL_439818, EPI_ISL_439819, EPI_ISL_439820, EPI_ISL_439821, EPI_ISL_439822, EPI_ISL_439823, EPI_ISL_439824, EPI_ISL_439825, EPI_ISL_439826, EPI_ISL_439827, EPI_ISL_439828, EPI_ISL_439829, EPI_ISL_439830, EPI_ISL_439831, EPI_ISL_439832, EPI_ISL_439833, EPI_ISL_439834, EPI_ISL_439835, EPI_ISL_439836, EPI_ISL_439837, EPI_ISL_439838, EPI_ISL_439839, EPI_ISL_439840, EPI_ISL_439841, EPI_ISL_439842, EPI_ISL_439843, EPI_ISL_439844, EPI_ISL_439845, EPI_ISL_439846, EPI_ISL_439847, EPI_ISL_439848, EPI_ISL_439849, EPI_ISL_439850, EPI_ISL_439851, EPI_ISL_439852, EPI_ISL_439853, EPI_ISL_439854, EPI_ISL_439855, EPI_ISL_439856, EPI_ISL_439857, EPI_ISL_439858, EPI_ISL_439859, EPI_ISL_439860, EPI_ISL_439861, EPI_ISL_439862, EPI_ISL_440852, EPI_ISL_440853, EPI_ISL_440854, EPI_ISL_440855, EPI_ISL_440856, EPI_ISL_440857, EPI_ISL_440858, EPI_ISL_440859, EPI_ISL_440860, EPI_ISL_440861, EPI_ISL_440862, EPI_ISL_440863, EPI_ISL_440864, EPI_ISL_440865, EPI_ISL_440866, EPI_ISL_440867, EPI_ISL_440868, EPI_ISL_440869, EPI_ISL_440870, EPI_ISL_440871, EPI_ISL_440872, EPI_ISL_440873, EPI_ISL_440874, EPI_ISL_440875, EPI_ISL_440876, EPI_ISL_440877, EPI_ISL_440878, EPI_ISL_440879, EPI_ISL_440880, EPI_ISL_440881, EPI_ISL_440882, EPI_ISL_440883, EPI_ISL_440884, EPI_ISL_440885, EPI_ISL_440886, EPI_ISL_440887, EPI_ISL_440888, EPI_ISL_440889, EPI_ISL_440890, EPI_ISL_440891, EPI_ISL_440892, EPI_ISL_440893, EPI_ISL_440894, EPI_ISL_440895, EPI_ISL_440896, EPI_ISL_440897, EPI_ISL_440898, EPI_ISL_440899, EPI_ISL_440900, EPI_ISL_440901, EPI_ISL_440902, EPI_ISL_440903, EPI_ISL_440904, EPI_ISL_440905, EPI_ISL_440906, EPI_ISL_440907, EPI_ISL_440908, EPI_ISL_440909, EPI_ISL_440910, EPI_ISL_440911, EPI_ISL_440912, EPI_ISL_440913, EPI_ISL_440914, EPI_ISL_440915, EPI_ISL_440916, EPI_ISL_440917, EPI_ISL_440918, EPI_ISL_440919, EPI_ISL_440920, EPI_ISL_440921, EPI_ISL_440922, EPI_ISL_440923, EPI_ISL_440924, EPI_ISL_440925, EPI_ISL_440926, EPI_ISL_440927, EPI_ISL_440928, EPI_ISL_440929, EPI_ISL_440930, EPI_ISL_440931, EPI_ISL_440932, EPI_ISL_440933, EPI_ISL_440934, EPI_ISL_440935, EPI_ISL_440936, EPI_ISL_440937, EPI_ISL_440938, EPI_ISL_440939, EPI_ISL_440940, EPI_ISL_440941, EPI_ISL_440942, EPI_ISL_440943, EPI_ISL_440944, EPI_ISL_440945, EPI_ISL_440946, EPI_ISL_440947, EPI_ISL_440948, EPI_ISL_440949, EPI_ISL_440950 | | | |
| see above | Liverpool Clinical Laboratories | COVID-19 Genomics UK (COG-UK) Consortium | Sam Haldenby, Anita Lucaci, Steve Paterson, Julian Hiscox, Alistair Darby, M Almsaud, A Alrezaihi, Muhannad Alruwaili, Stuart D Armstrong, Jones Benjamin , Eleanor G Bentley, Anu Chawla, Jordan J Clark, Angela Cowell, Richard Eccles, Isabel Garca-Dorival, Matthew Gemmell, Alessandro Gerada, PKF Gilmore, Richard Gregory, Ximeng Han, Catherine Hartley, Margaret Hughes, Miren Iturriza-Gomara, James Johnson, L Luu, Jenifer Manson , Charlotte Nelson, Elaine O'ÂôToole, Cassie Olateju, Rebekah Penrice-Randal¬†, Lucille Rainbow, N.P Randle, Trevor Ian Robinson, Parul Sharma, Ghada T Shawli, James P Stewart , Neil Swainston, Ecaterina Vamos, Joanne Watts, Mark Whitehead |
| EPI_ISL_445019, EPI_ISL_445024, EPI_ISL_445030, EPI_ISL_445035, EPI_ISL_445036, EPI_ISL_445041, EPI_ISL_445043, EPI_ISL_445044, EPI_ISL_445045, EPI_ISL_445046, EPI_ISL_445047, EPI_ISL_445048, EPI_ISL_445049 | | | |
| see above | Florida Bureau of Public Health Laboratories | Florida Bureau of Public Health Laboratories | Sarah Schmedes, Jason Blanton |
| EPI_ISL_445094, EPI_ISL_445095, EPI_ISL_445096, EPI_ISL_445097, EPI_ISL_445098, EPI_ISL_445100, EPI_ISL_445101, EPI_ISL_445102, EPI_ISL_445103, EPI_ISL_445104, EPI_ISL_445105, EPI_ISL_445106, EPI_ISL_445107, EPI_ISL_445108, EPI_ISL_445109, EPI_ISL_445110, EPI_ISL_445111, EPI_ISL_445113, EPI_ISL_445114, EPI_ISL_445115, EPI_ISL_445116, EPI_ISL_445117 | | | |
| see above | UC San Diego Center for Advanced Laboratory Medicine | Andersen lab at Scripps Research | SEARCH Alliance San Diego with David Pride, Ji H Shin |
| EPI_ISL_445118 | Rady's Childrens Hospital | Andersen lab at Scripps Research | SEARCH Alliance San Diego |
| EPI_ISL_445164, EPI_ISL_445166, EPI_ISL_445167, EPI_ISL_445168 | Scripps Medical Laboratory | Andersen lab at Scripps Research | SEARCH Alliance San Diego with Michael Quigley, Ellen Stefanski, Ian Mchardy |
| EPI_ISL_445223 | Victoria Vard och Halsa | The Public Health Agency of Sweden | Sarah Henriksson, Oskar Karlsson Lindsjo, Maria Lind Karlberg, Anna-Malin Linde, Olov Svartstrom, Anna Risberg, Theresa Enkirch, Mia Brytting, Karin Tegmark-Wisell |
| EPI_ISL_447477, EPI_ISL_447491, EPI_ISL_447505, EPI_ISL_447519 | Servicio de Microbiologia. Hospital Clinico Universitario de Valencia | Sequencing and Bioinformatics Service and Molecular Epidemiology Research Group. FISABIO-Public Health | Sandra Carbo, Loreto Ferrús Abad, Paula Ruiz-Hueso, Mariana Reyes-Prieto, Vicente Soriano Chirona, Ivan Ansari, Llúcia Martínez-Priego, Giuseppe 'Auria, David Navarro, Eliseo Albert, Maria Alma Bracho, Lidia Ruiz Roldan, Neris Garcia-Gonzalez, Inma Galán Vendrell, Fernando Gonzalez-Candelas |
| EPI_ISL_447563, EPI_ISL_447573, EPI_ISL_447581, EPI_ISL_447856, EPI_ISL_447857, EPI_ISL_447858 | CSIR-Centre for Cellular and Molecular Biology | CSIR-Centre for Cellular and Molecular Biology | Sakshi Shambhavi, Lamuk Zaveri, Shagufta Khan, Namami Gaur, Tulasi Nagabandi, Purushotham Vodnala, Payel Mukherjee, Sofia Banu, Priya Singh, Dhiviya Vedagiri, Divya Gupta, Vishal Sah, Santosh Kumar Kuncha, Krishnan Harinivas Harshan, Archana Bharadwaj Siva, Karthik Bharadwaj Tallapaka, Rakesh K Mishra, Divya Tej Sowpati |
| EPI_ISL_449330, EPI_ISL_449331, EPI_ISL_449332, EPI_ISL_449333, EPI_ISL_449334, EPI_ISL_449335, EPI_ISL_449336, EPI_ISL_449337, EPI_ISL_449338, EPI_ISL_449339, EPI_ISL_449340, EPI_ISL_449341, EPI_ISL_449342, EPI_ISL_449343, EPI_ISL_449344, EPI_ISL_449345, EPI_ISL_449346, EPI_ISL_449347, EPI_ISL_449348, EPI_ISL_449349, EPI_ISL_449350, EPI_ISL_449351, EPI_ISL_449352, EPI_ISL_449353, EPI_ISL_449354, EPI_ISL_449355, EPI_ISL_449356, EPI_ISL_449357, EPI_ISL_449358, EPI_ISL_449359, EPI_ISL_449360, EPI_ISL_449361, EPI_ISL_449362, EPI_ISL_449363, EPI_ISL_449364, EPI_ISL_449365, EPI_ISL_449366, EPI_ISL_449367, EPI_ISL_449368, EPI_ISL_449369, EPI_ISL_449370, EPI_ISL_449371, EPI_ISL_449372, EPI_ISL_449373, EPI_ISL_449374, EPI_ISL_449375, EPI_ISL_449376, EPI_ISL_449377, EPI_ISL_449378, EPI_ISL_449379, EPI_ISL_449380, EPI_ISL_449381, EPI_ISL_449382, EPI_ISL_449383, EPI_ISL_449384, EPI_ISL_449385, EPI_ISL_449386, EPI_ISL_449387, EPI_ISL_449388, EPI_ISL_449389, EPI_ISL_449390, EPI_ISL_449391, EPI_ISL_449392, EPI_ISL_449393, EPI_ISL_449394, EPI_ISL_449395, EPI_ISL_449396, EPI_ISL_449397, EPI_ISL_449398, EPI_ISL_449399, EPI_ISL_449400, EPI_ISL_449401, EPI_ISL_449402, EPI_ISL_449403, EPI_ISL_449404, EPI_ISL_449405, EPI_ISL_449406, EPI_ISL_449407, EPI_ISL_449408, EPI_ISL_449409, EPI_ISL_449410, EPI_ISL_449411, EPI_ISL_449412, EPI_ISL_449413, EPI_ISL_449414, EPI_ISL_449415, EPI_ISL_449416, EPI_ISL_449417, EPI_ISL_449418, EPI_ISL_449419, EPI_ISL_449420, EPI_ISL_449421, EPI_ISL_449422, EPI_ISL_449423, EPI_ISL_449424, EPI_ISL_449425, EPI_ISL_449426, EPI_ISL_449427, EPI_ISL_449428, EPI_ISL_449429, EPI_ISL_449430, EPI_ISL_449431, EPI_ISL_449432, EPI_ISL_449433, EPI_ISL_449434, EPI_ISL_449435, EPI_ISL_449436, EPI_ISL_449437, EPI_ISL_449438, EPI_ISL_449439, EPI_ISL_449440, EPI_ISL_449441, EPI_ISL_449442, EPI_ISL_449443, EPI_ISL_449444, EPI_ISL_449445, EPI_ISL_449446, EPI_ISL_449447, EPI_ISL_449448, EPI_ISL_449449, EPI_ISL_449450, EPI_ISL_449451, EPI_ISL_449452, EPI_ISL_449453, EPI_ISL_449454, EPI_ISL_449455, EPI_ISL_449456, EPI_ISL_449457, EPI_ISL_449458, EPI_ISL_449459, EPI_ISL_449460, EPI_ISL_449461, EPI_ISL_449462, EPI_ISL_449463, EPI_ISL_449464, EPI_ISL_449465, EPI_ISL_449466, EPI_ISL_449467, EPI_ISL_449468, EPI_ISL_449469, EPI_ISL_449470, EPI_ISL_449471, EPI_ISL_449472, EPI_ISL_449473, EPI_ISL_449474, EPI_ISL_449475, EPI_ISL_449488, EPI_ISL_449489, EPI_ISL_449490, EPI_ISL_449491, EPI_ISL_449492, EPI_ISL_449493, EPI_ISL_449494, EPI_ISL_449495, EPI_ISL_449496, EPI_ISL_449497, EPI_ISL_449498, EPI_ISL_449499, EPI_ISL_449500, EPI_ISL_449501, EPI_ISL_449502, EPI_ISL_449503, EPI_ISL_449504, EPI_ISL_449505, EPI_ISL_449506, EPI_ISL_449507, EPI_ISL_449508, EPI_ISL_449509, EPI_ISL_449510, EPI_ISL_449511, EPI_ISL_449512, EPI_ISL_449513, EPI_ISL_449514, EPI_ISL_449515, EPI_ISL_449516, EPI_ISL_449517, EPI_ISL_449518, EPI_ISL_449519, EPI_ISL_449520, EPI_ISL_449521, EPI_ISL_449522, EPI_ISL_449523, EPI_ISL_449524, EPI_ISL_449525, EPI_ISL_449526, EPI_ISL_449527, EPI_ISL_449528, EPI_ISL_449529, EPI_ISL_449530, EPI_ISL_449531, EPI_ISL_449532, EPI_ISL_449533, EPI_ISL_449534, EPI_ISL_449535, EPI_ISL_449536, EPI_ISL_449537, EPI_ISL_449538, EPI_ISL_449539, EPI_ISL_449540, EPI_ISL_449541, EPI_ISL_449542, EPI_ISL_449543, EPI_ISL_449544, EPI_ISL_449545, EPI_ISL_449546, EPI_ISL_449547, EPI_ISL_449548, EPI_ISL_449549, EPI_ISL_449550, EPI_ISL_449551, EPI_ISL_449552, EPI_ISL_449553, EPI_ISL_449554, EPI_ISL_449555, EPI_ISL_449556, EPI_ISL_449557, EPI_ISL_449558, EPI_ISL_449559, EPI_ISL_449560, EPI_ISL_449561, EPI_ISL_449562, EPI_ISL_449563, EPI_ISL_449564, EPI_ISL_449565, EPI_ISL_449566, EPI_ISL_449567, EPI_ISL_449568, EPI_ISL_449569, EPI_ISL_449570, EPI_ISL_449571, EPI_ISL_449572, EPI_ISL_449573, EPI_ISL_449574, EPI_ISL_449575, EPI_ISL_449576, EPI_ISL_449577, EPI_ISL_449578, EPI_ISL_449579, EPI_ISL_449580, EPI_ISL_449581, EPI_ISL_449582, EPI_ISL_449583, EPI_ISL_449584, EPI_ISL_449585, EPI_ISL_449586, EPI_ISL_449587, EPI_ISL_449588, EPI_ISL_449589, EPI_ISL_449590, EPI_ISL_449591, EPI_ISL_449592, EPI_ISL_449593, EPI_ISL_449594, EPI_ISL_449595, EPI_ISL_449596, EPI_ISL_449597, EPI_ISL_449598, EPI_ISL_449599, EPI_ISL_449600, EPI_ISL_449601, EPI_ISL_449602, EPI_ISL_449603, EPI_ISL_449604, EPI_ISL_449605, EPI_ISL_449606, EPI_ISL_449607, EPI_ISL_449608, EPI_ISL_449609, EPI_ISL_449610, EPI_ISL_449611, EPI_ISL_449612, EPI_ISL_449613, EPI_ISL_449614, EPI_ISL_449615, EPI_ISL_449616, EPI_ISL_449617, EPI_ISL_449618, EPI_ISL_449619, EPI_ISL_449620, EPI_ISL_449621, EPI_ISL_449622, EPI_ISL_449623, EPI_ISL_449624, EPI_ISL_449625 | | | |
| see above | Liverpool Clinical Laboratories | COVID-19 Genomics UK (COG-UK) Consortium | Sam Haldenby, Anita Lucaci, Steve Paterson, Julian Hiscox, Alistair Darby, M Almsaud, A Alrezaihi, Muhannad Alruwaili, Stuart D Armstrong, Jones Benjamin , Eleanor G Bentley, Anu Chawla, Jordan J Clark, Angela Cowell, Richard Eccles, Isabel García-Dorival, Matthew Gemmell, Alessandro Gerada, PKF Gilmore, Richard Gregory, Ximeng Han, Catherine Hartley, Margaret Hughes, Miren Iturriza-Gomara, James Johnson, L Luu, Jenifer Manson , Charlotte Nelson, Elaine O'Toole, Cassie Olateju, Rebekah Penrice-Randal , Lucille Rainbow, N.P Randle, Trevor Ian Robinson, Parul Sharma, Ghada T Shawli, James P Stewart , Neil Swainston, Ecaterina Vamos, Joanne Watts, Mark Whitehead |
| EPI_ISL_450190 | Rady's Childrens Hospital | Andersen lab at Scripps Research | SEARCH Alliance San Diego |
| EPI_ISL_450191, EPI_ISL_450192 | Scripps Medical Laboratory | Andersen lab at Scripps Research | SEARCH Alliance San Diego with Michael Quigley, Ellen Stefanski, Ian Mchardy |
| EPI_ISL_450302, EPI_ISL_450303 | Centre hospitalier Anna-Laberge | Laboratoire de santé publique du Québec | Sandrine Moreira, Ioannis Ragoussis, Guillaume Bourque, Jesse Shapiro, Mark Lathrop and Michel Roger on behalf of the CoVSeQ research group (http://covseq.ca/researchgroup) |
| EPI_ISL_450304, EPI_ISL_450305 | Hôpital Charles-LeMoyne | Laboratoire de santé publique du Québec | Sandrine Moreira, Ioannis Ragoussis, Guillaume Bourque, Jesse Shapiro, Mark Lathrop and Michel Roger on behalf of the CoVSeQ research group (http://covseq.ca/researchgroup) |
| EPI_ISL_450306 | CSSS Haut-Richelieu/Rouville (Hôpital) | Laboratoire de santé publique du Québec | Sandrine Moreira, Ioannis Ragoussis, Guillaume Bourque, Jesse Shapiro, Mark Lathrop and Michel Roger on behalf of the CoVSeQ research group (http://covseq.ca/researchgroup) |
| EPI_ISL_450307 | Hôpital du Suroît | Laboratoire de santé publique du Québec | Sandrine Moreira, Ioannis Ragoussis, Guillaume Bourque, Jesse Shapiro, Mark Lathrop and Michel Roger on behalf of the CoVSeQ research group (http://covseq.ca/researchgroup) |
| EPI_ISL_450308 | Hôpital Charles-LeMoyne | Laboratoire de santé publique du Québec | Sandrine Moreira, Ioannis Ragoussis, Guillaume Bourque, Jesse Shapiro, Mark Lathrop and Michel Roger on behalf of the CoVSeQ research group (http://covseq.ca/researchgroup) |
| EPI_ISL_450309 | Hôpital Pierre-Boucher | Laboratoire de santé publique du Québec | Sandrine Moreira, Ioannis Ragoussis, Guillaume Bourque, Jesse Shapiro, Mark Lathrop and Michel Roger on behalf of the CoVSeQ research group (http://covseq.ca/researchgroup) |
| EPI_ISL_450310 | Hôpital Charles-LeMoyne | Laboratoire de santé publique du Québec | Sandrine Moreira, Ioannis Ragoussis, Guillaume Bourque, Jesse Shapiro, Mark Lathrop and Michel Roger on behalf of the CoVSeQ research group (http://covseq.ca/researchgroup) |
| EPI_ISL_450311, EPI_ISL_450312, EPI_ISL_450313 | Hôpital du Suroît | Laboratoire de santé publique du Québec | Sandrine Moreira, Ioannis Ragoussis, Guillaume Bourque, Jesse Shapiro, Mark Lathrop and Michel Roger on behalf of the CoVSeQ research group (http://covseq.ca/researchgroup) |
| EPI_ISL_450314, EPI_ISL_450315 | Hôpital Pierre-Boucher | Laboratoire de santé publique du Québec | Sandrine Moreira, Ioannis Ragoussis, Guillaume Bourque, Jesse Shapiro, Mark Lathrop and Michel Roger on behalf of the CoVSeQ research group (http://covseq.ca/researchgroup) |
| EPI_ISL_450316 | Hôpital Charles-LeMoyne | Laboratoire de santé publique du Québec | Sandrine Moreira, Ioannis Ragoussis, Guillaume Bourque, Jesse Shapiro, Mark Lathrop and Michel Roger on behalf of the CoVSeQ research group (http://covseq.ca/researchgroup) |
| EPI_ISL_450317, EPI_ISL_450318 | Hôpital Pierre-Boucher | Laboratoire de santé publique du Québec | Sandrine Moreira, Ioannis Ragoussis, Guillaume Bourque, Jesse Shapiro, Mark Lathrop and Michel Roger on behalf of the CoVSeQ research group (http://covseq.ca/researchgroup) |

| | | | |
|---|---|---|---|
| EPI_ISL_450330 | CSIR-Centre for Cellular and Molecular Biology | CSIR-Centre for Cellular and Molecular Biology | Sakshi Shambhavi, Lamuk Zaveri, Shagufta Khan, Namami Gaur, Tulasi Nagabandi, Purushotham Vodnala, Payel Mukherjee, Sofia Banu, Priya Singh, Dhiviya Vedagiri, Divya Gupta, Vishal Sah, Santosh Kumar Kuncha, Krishnan Harinivas Harshan, Archana Bharadwaj Siva, Karthik Bharadwaj Tallapaka,Preethi Jampala, Sharada Ravi Iyer, Sulagana Mukherjee, Swetha Sundar, Peddapuvala Sai Uday Kiran, Umesh Kumar, Unis Ahmad Bhat, Ajay Sarawagi, Priyanka Pant, Rajkanwar Nathawat, Nikhil Hajirnis, Pratheusa Maccha, M Soujanya Reddy Rakesh K Mishra, Divya Tej Sowpati |
| EPI_ISL_450782 | Government Medical College-Bhavnagar | Gujarat Biotechnology Research Centre | Saklain Malek, Shirish Patel, Kairavi Desai, Tejas Shah, Ankit Hinsu, Pritesh Sabara, Apurvasinh Puvar, Janvi Raval, Zarna Patel, Monika Gandhi, Pinal Trivedi, Maharshi Pandya, Amit Kanani, Nidhi Patel, Nitin Savaliya, Raghawendra Kumar, Dinesh Kumar, Zuber Saiyed, Komal Patel, Labdhi Pandya, Snehal Bagatharia, Ramesh Pandit, Bhavesh Modi, Gaurishankar Shrimali, R D Dixit, A M Kadri, Priti Pandita, Chaitanya Joshi, Madhvi Joshi |
| EPI_ISL_450807 | Victoria Vard och Halsa | The Public Health Agency of Sweden | Sarah Henriksson, Anna-Malin Linde, Maria Lind Karlberg, Oskar Karlsson Lindsjo, Olov Svartstrom, Anna Risberg, Theresa Enkirch, Mia Brytting, Karin Tegmark-Wisell |
| EPI_ISL_450844, EPI_ISL_450847, EPI_ISL_450849, EPI_ISL_450850, EPI_ISL_450851, EPI_ISL_450857, EPI_ISL_450864, EPI_ISL_450865, EPI_ISL_450869, EPI_ISL_450871 | Florida Bureau of Public Health Laboratories | Florida Bureau of Public Health Laboratories | Sarah Schmedes, Jason Blanton |
| EPI_ISL_450873 | Evandro Chagas Institute | Evandro Chagas Institute | Santos, M.C.; Silva, A.M.; Junior, W.D.C.; Barbagelata, L.S.; Ferreira, J.A.; Sousa, E.M.A.; da Silva, P.S.; Martins, L.C.; Sousa Junior, E.C.; Viana, G.M.R |
| EPI_ISL_450874 | Evandro Chagas Institute | Evandro Chagas Institute | Santos, M.C.; Silva, A.M.; Junior, W.D.C.; Barbagelata, L.S.; Ferreira, J.A.; Sousa, E.M.A.; da Silva, P.S.; Martins, L.C.;Sousa Junior, E.C.;Viana, G.M.R |
| EPI_ISL_451400 | Laboratoire de Recherche et d'Analyse Médicale de la Gendarmerie Royale | Laboratoire de Recherche et d'Analyse Médicale de la Gendarmerie Royale | Sanaâ LEMRISS, Amal SOUIRI, Saâd EL KABBAJ |
| EPI_ISL_452233 | Victoria Vard och Halsa | The Public Health Agency of Sweden | Sarah Henriksson, Anna-Malin Linde, Maria Lind Karlberg, Oskar Karlsson Lindsjo, Olov Svartstrom, Anna Risberg, Theresa Enkirch, Mia Brytting, Karin Tegmark-Wisell |
| EPI_ISL_452453, EPI_ISL_452454, EPI_ISL_452455, EPI_ISL_452456, EPI_ISL_452457, EPI_ISL_452458, EPI_ISL_452459, EPI_ISL_452460, EPI_ISL_452461, EPI_ISL_452462, EPI_ISL_452463, EPI_ISL_452464, EPI_ISL_452465, EPI_ISL_452466, EPI_ISL_452467, EPI_ISL_452468, EPI_ISL_452469, EPI_ISL_452470, EPI_ISL_452471 | | | |
| see above | Hospital Universitario Puerta del Mar de Cádiz - INIBICA | SeqCOVID-SPAIN consortium/IBV(CSIC) | Salud Rodríguez-Pallares, Fátima-Galán-Sánchez, Manuel Rodrí-guez-Iglesias and SeqCOVID-SPAIN consortium |
| EPI_ISL_453195, EPI_ISL_453196, EPI_ISL_453197, EPI_ISL_453198, EPI_ISL_453199, EPI_ISL_453200, EPI_ISL_453201, EPI_ISL_453202, EPI_ISL_453203, EPI_ISL_453204, EPI_ISL_453205, EPI_ISL_453206, EPI_ISL_453207, EPI_ISL_453208, EPI_ISL_453209, EPI_ISL_453210, EPI_ISL_453211, EPI_ISL_453212, EPI_ISL_453213, EPI_ISL_453214, EPI_ISL_453215, EPI_ISL_453216, EPI_ISL_453217, EPI_ISL_453218, EPI_ISL_453219, EPI_ISL_453220, EPI_ISL_453221, EPI_ISL_453222, EPI_ISL_453223, EPI_ISL_453224, EPI_ISL_453225, EPI_ISL_453226, EPI_ISL_453227, EPI_ISL_453228, EPI_ISL_453229, EPI_ISL_453230, EPI_ISL_453231, EPI_ISL_453232, EPI_ISL_453233, EPI_ISL_453234, EPI_ISL_453235, EPI_ISL_453236, EPI_ISL_453237, EPI_ISL_453238, EPI_ISL_453239, EPI_ISL_453240, EPI_ISL_453241, EPI_ISL_453242, EPI_ISL_453243, EPI_ISL_453244, EPI_ISL_453245, EPI_ISL_453246, EPI_ISL_453247, EPI_ISL_453248, EPI_ISL_453249, EPI_ISL_453250, EPI_ISL_453251, EPI_ISL_453252, EPI_ISL_453253, EPI_ISL_453254, EPI_ISL_453255, EPI_ISL_453256, EPI_ISL_453257, EPI_ISL_453258, EPI_ISL_453259, EPI_ISL_453260, EPI_ISL_453261, EPI_ISL_453262, EPI_ISL_453263, EPI_ISL_453264, EPI_ISL_453265, EPI_ISL_453266, EPI_ISL_453267, EPI_ISL_453268, EPI_ISL_453269, EPI_ISL_453270, EPI_ISL_453271, EPI_ISL_453272, EPI_ISL_453273, EPI_ISL_453274, EPI_ISL_453275, EPI_ISL_453276, EPI_ISL_453277, EPI_ISL_453278, EPI_ISL_453279, EPI_ISL_453280, EPI_ISL_453281, EPI_ISL_453282, EPI_ISL_453283, EPI_ISL_453284, EPI_ISL_453285, EPI_ISL_453286, EPI_ISL_453287, EPI_ISL_453288, EPI_ISL_453289, EPI_ISL_453290, EPI_ISL_453291, EPI_ISL_453292, EPI_ISL_453293, EPI_ISL_453294, EPI_ISL_453295, EPI_ISL_453296, EPI_ISL_453297, EPI_ISL_453298, EPI_ISL_453299, EPI_ISL_453300, EPI_ISL_453301, EPI_ISL_453302, EPI_ISL_453303, EPI_ISL_453304, EPI_ISL_453305, EPI_ISL_453306, EPI_ISL_453307, EPI_ISL_453308, EPI_ISL_453309, EPI_ISL_453310, EPI_ISL_453311, EPI_ISL_453312, EPI_ISL_453313, EPI_ISL_453314, EPI_ISL_453315, EPI_ISL_453316, EPI_ISL_453317, EPI_ISL_453318, EPI_ISL_453319, EPI_ISL_453320, EPI_ISL_453321, EPI_ISL_453322, EPI_ISL_453323, EPI_ISL_453324, EPI_ISL_453325, EPI_ISL_453326, EPI_ISL_453327, EPI_ISL_453328, EPI_ISL_453329, EPI_ISL_453330, EPI_ISL_453331, EPI_ISL_453332, EPI_ISL_453333, EPI_ISL_453334, EPI_ISL_453335, EPI_ISL_453336, EPI_ISL_453337, EPI_ISL_453338, EPI_ISL_453339, EPI_ISL_453340, EPI_ISL_453341, EPI_ISL_453342, EPI_ISL_453343, EPI_ISL_453344, EPI_ISL_453345, EPI_ISL_453346, EPI_ISL_453347, EPI_ISL_453348, EPI_ISL_453349, EPI_ISL_453350, EPI_ISL_453351, EPI_ISL_453352, EPI_ISL_453353, EPI_ISL_453354, EPI_ISL_453355, EPI_ISL_453356, EPI_ISL_453357, EPI_ISL_453358, EPI_ISL_453359, EPI_ISL_453360, EPI_ISL_453361, EPI_ISL_453362, EPI_ISL_453363, EPI_ISL_453364, EPI_ISL_453365, EPI_ISL_453366, EPI_ISL_453367, EPI_ISL_453368, EPI_ISL_453369, EPI_ISL_453370, EPI_ISL_453371, EPI_ISL_453372, EPI_ISL_453373, EPI_ISL_453374, EPI_ISL_453375, EPI_ISL_453376, EPI_ISL_453377, EPI_ISL_453378, EPI_ISL_453379, EPI_ISL_453380, EPI_ISL_453381, EPI_ISL_453382, EPI_ISL_453383, EPI_ISL_453384, EPI_ISL_453385, EPI_ISL_453386, EPI_ISL_453387, EPI_ISL_453388, EPI_ISL_453389, EPI_ISL_453390, EPI_ISL_453391, EPI_ISL_453392, EPI_ISL_453393, EPI_ISL_453394, EPI_ISL_453395, EPI_ISL_453396, EPI_ISL_453397, EPI_ISL_453398, EPI_ISL_453399, EPI_ISL_453400, EPI_ISL_453401, EPI_ISL_453402, EPI_ISL_453403, EPI_ISL_453404, EPI_ISL_453405, EPI_ISL_453406, EPI_ISL_453407, EPI_ISL_453408, EPI_ISL_453409, EPI_ISL_453410, EPI_ISL_453411, EPI_ISL_453412, EPI_ISL_453413, EPI_ISL_453414, EPI_ISL_453415, EPI_ISL_453416, EPI_ISL_453417, EPI_ISL_453418, EPI_ISL_453419, EPI_ISL_453420, EPI_ISL_453421, EPI_ISL_453422, EPI_ISL_453423, EPI_ISL_453424, EPI_ISL_453425, EPI_ISL_453426, EPI_ISL_453427, EPI_ISL_453428, EPI_ISL_453429, EPI_ISL_453430, EPI_ISL_453431, EPI_ISL_453432, EPI_ISL_453433, EPI_ISL_453434, EPI_ISL_453435, EPI_ISL_453436, EPI_ISL_453437, EPI_ISL_453438, EPI_ISL_453439, EPI_ISL_453440, EPI_ISL_453441, EPI_ISL_453442, EPI_ISL_453443, EPI_ISL_453444, EPI_ISL_453445, EPI_ISL_453446, EPI_ISL_453447, EPI_ISL_453448, EPI_ISL_453449, EPI_ISL_453450, EPI_ISL_453451, EPI_ISL_453452, EPI_ISL_453453, EPI_ISL_453454, EPI_ISL_453455, EPI_ISL_453456, EPI_ISL_453457, EPI_ISL_453458, EPI_ISL_453459 | | | |
| see above | Liverpool Clinical Laboratories | COVID-19 Genomics UK (COG-UK) Consortium | Sam Haldenby, Anita Lucaci, Steve Paterson, Julian Hiscox, Alistair Darby, M Almsaud, A Alrezaihi, Muhannad Alruwaili, Stuart D Armstrong, Jones Benjamin , Eleanor G Bentley, Anu Chawla, Jordan J Clark, Angela Cowell, Richard Eccles, Isabel García-Dorival, Matthew Gemmell, Alessandro Gerada, PKF Gilmore, Richard Gregory, Ximeng Han, Catherine Hartley, Margaret Hughes, Miren Iturriza-Gomara, James Johnson, L Luu, Jenifer Manson , Charlotte Nelson, Elaine O'Toole, Cassie Olateju, Rebekah Penrice-Randal , Lucille Rainbow, N.P Randle, Trevor Ian Robinson, Parul Sharma, Ghada T Shawli, James P Stewart , Neil Swainston, Ecaterina Vamos, Joanne Watts, Mark Whitehead |
| EPI_ISL_455101 | Jourcentralen | The Public Health Agency of Sweden | Salvatore Ascione, Anna-Malin Linde, Maria Lind Karlberg, Oskar Karlsson Lindsjo, Olov Svartstrom, Anna Risberg, Theresa Enkirch, Mia Brytting, Karin Tegmark-Wisell |
| EPI_ISL_455721, EPI_ISL_455729 | Servicio de Microbiologia. Hospital Clinico Universitario de Valencia | Sequencing and Bioinformatics Service and Molecular Epidemiology Research Group. FISABIO-Public Health, and SeqCOVID-Spain Consortium | Sandra Carbo, Loreto Ferrús Abad, Paula Ruiz-Hueso, Mariana Reyes-Prieto, Vicente Soriano Chirona, Ivan Ansari, Llúcia Martínez-Priego, Giuseppe 'Auria, David Navarro, Eliseo Albert, Maria Alma Bracho, Lidia Ruiz Roldan, Neris Garcia-Gonzalez, Inma Galán Vendrell, Fernando Gonzalez-Candelas |
| EPI_ISL_457259, EPI_ISL_457260, EPI_ISL_457261, EPI_ISL_457262, EPI_ISL_457263, EPI_ISL_457264, EPI_ISL_457265, EPI_ISL_457266, EPI_ISL_457267, EPI_ISL_457268, EPI_ISL_457269 | | | |
| see above | Liverpool Clinical Laboratories | COVID-19 Genomics UK (COG-UK) Consortium | Sam Haldenby, Anita Lucaci, Steve Paterson, Julian Hiscox, Alistair Darby, M Almsaud, A Alrezaihi, Muhannad Alruwaili, Stuart D Armstrong, Jones Benjamin, Eleanor G Bentley, Anu Chawla, Jordan J Clark, Angela Cowell, Richard Eccles, Isabel García-Dorival, Matthew Gemmell, Alessandro Gerada, PKF Gilmore, Richard Gregory, Ximeng Han, Catherine Hartley, Margaret Hughes, Miren Iturriza-Gomara, James Johnson, L Luu, Jenifer Manson, Charlotte Nelson, Elaine O'Toole, Cassie Olateju, Rebekah Penrice-Randal , Lucille Rainbow, N.P Randle, Trevor Ian Robinson, Parul Sharma, Ghada T Shawli, James P Stewart, Neil Swainston, Ecaterina Vamos, Joanne Watts, Mark Whitehead |
| EPI_ISL_457701, EPI_ISL_457704, EPI_ISL_457706, EPI_ISL_457937, EPI_ISL_457938, EPI_ISL_457939, EPI_ISL_457974, EPI_ISL_457975, EPI_ISL_457976, EPI_ISL_457977, EPI_ISL_457978, EPI_ISL_457979, EPI_ISL_457980, EPI_ISL_457985, EPI_ISL_457986, EPI_ISL_457987, EPI_ISL_457988, EPI_ISL_457989, EPI_ISL_457990, EPI_ISL_457991, EPI_ISL_457992, EPI_ISL_457993, EPI_ISL_457994, EPI_ISL_457995, EPI_ISL_457996, EPI_ISL_457997, EPI_ISL_457998 | | | |
| see above | Oman-NIC | Oman-NIC | Samira Al-Maruqi, Fahad Zadjali, Amina Al Jardani, Khulood Al-Mammary, Hanan Al-kindi, Fatma BaAlawi, Hamida AL Barwani, Zeyana AL-Dahmani, Intisar Al-Shukri, Aisha Al-Busaidi, Aisha Al-Amri, Ahlam Al-Amri, Mohammed Al-Tobi, Samiha Al Kharusi, Abdulla Balkhair |
| EPI_ISL_458070 | CSIR-Centre for Cellular and Molecular Biology | CSIR-Centre for Cellular and Molecular Biology | Sakshi Shambhavi, Lamuk Zaveri, Shagufta Khan, Namami Gaur, Tulasi Nagabandi, Purushotham Vodnala, Payel Mukherjee, Sofia Banu, Priya Singh, Dhiviya Vedagiri, Divya Gupta, Vishal Sah, Santosh Kumar Kuncha, Krishnan Harinivas Harshan, Archana Bharadwaj Siva, Karthik Bharadwaj Tallapaka,Nikhil Hajirnis, Pratheusa Maccha, M Soujanya Reddy,G. Aditya Kumar, Koushick Sivakumar,Disha Nanda, Divya Das, Jotin Gogoi, Manish Bhattacharjee, Ravi Prasad Mukku, Rakesh K Mishra, Divya Tej Sowpati |
| EPI_ISL_458071 | CSIR-Centre for Cellular and Molecular Biology | CSIR-Centre for Cellular and Molecular Biology | Sakshi Shambhavi, Lamuk Zaveri, Shagufta Khan, Namami Gaur, Tulasi Nagabandi, Purushotham Vodnala, Payel Mukherjee, Sofia Banu, Priya Singh, Dhiviya Vedagiri, Divya Gupta, Vishal Sah, Santosh Kumar Kuncha, Krishnan Harinivas Harshan, Archana Bharadwaj Siva, Karthik Bharadwaj Tallapaka,Nikhil Hajirnis, Pratheusa Maccha, M Soujanya Reddy,G. Aditya Kumar, Koushick Sivakumar, Rakesh K Mishra, Divya Tej Sowpati |
| EPI_ISL_458080 | CSIR-Centre for Cellular and Molecular Biology | CSIR-Centre for Cellular and Molecular Biology | Sakshi Shambhavi, Lamuk Zaveri, Shagufta Khan, Namami Gaur, Tulasi Nagabandi, Purushotham Vodnala, Payel Mukherjee, Sofia Banu, Priya Singh, Dhiviya Vedagiri, Divya Gupta, Vishal Sah, Santosh Kumar Kuncha, Krishnan Harinivas Harshan, Archana Bharadwaj Siva, Karthik Bharadwaj Tallapaka, G. Aditya Kumar, Koushick Sivakumar, Pooja Ramesh Gupta, Rajan Kumar Jha, Shraddha Vijay Lahoti, Rakesh K Mishra, Divya Tej Sowpati |
| EPI_ISL_458138, EPI_ISL_458139, EPI_ISL_458140, EPI_ISL_458141, EPI_ISL_458142, EPI_ISL_458143, EPI_ISL_458144, EPI_ISL_458145, EPI_ISL_458146, EPI_ISL_458147, EPI_ISL_458148, EPI_ISL_458149 | | | |
| see above | Evandro Chagas Institute | Evandro Chagas Institute | Santos, M.C.; Silva, A.M.; Junior, W.D.C.; Barbagelata, L.S.; Ferreira, J.A.; Sousa, E.M.A.; da Silva, P.S.; Resque, H.R; Martins, L.C.; Sousa Junior, E.C.;Viana, G.M.R |
| EPI_ISL_458237, EPI_ISL_458238, EPI_ISL_458239, EPI_ISL_458240, EPI_ISL_458241, EPI_ISL_458242, EPI_ISL_458243, EPI_ISL_458244, EPI_ISL_458245, EPI_ISL_458246, EPI_ISL_458247, EPI_ISL_458248, EPI_ISL_458249, EPI_ISL_458250, EPI_ISL_458251, EPI_ISL_458252, EPI_ISL_458253, EPI_ISL_458254, EPI_ISL_458255, EPI_ISL_458256, EPI_ISL_458257, EPI_ISL_458258, EPI_ISL_458259, EPI_ISL_458260, EPI_ISL_458261, EPI_ISL_458262, EPI_ISL_458263, EPI_ISL_458264, EPI_ISL_458265, EPI_ISL_458266, EPI_ISL_458267, EPI_ISL_458268, EPI_ISL_458269, EPI_ISL_458270, EPI_ISL_458271, EPI_ISL_458272, EPI_ISL_458273, EPI_ISL_458274, EPI_ISL_458275, EPI_ISL_458276, EPI_ISL_458277, EPI_ISL_458278, EPI_ISL_458279, EPI_ISL_458280, EPI_ISL_458281, EPI_ISL_458282, EPI_ISL_458283, EPI_ISL_458284 | | | |
| see above | Scripps Medical Laboratory | Andersen lab at Scripps Research | SEARCH Alliance San Diego with Michael Quigley, Ellen Stefanski, Ian Mchardy |

| | | | |
|---|---|---|---|
| EPI_ISL_458298 | CSIR-Centre for Cellular and Molecular Biology | CSIR-Centre for Cellular and Molecular Biology | Sakshi Shambhavi, Lamuk Zaveri, Shagufta Khan, Namami Gaur, Tulasi Nagabandi, Purushotham Vodnala, Payel Mukherjee, Sofia Banu, Priya Singh, Dhiviya Vedagiri, Divya Gupta, Vishal Sah, Santosh Kumar Kuncha, Krishnan Harinivas Harshan, Archana Bharadwaj Siva, Karthik Bharadwaj Tallapaka, Deepak Kumar, Devi Prasad Vijayashankar, Disha Nanda, Divya Das, Jotin Gogoi, Manish Bhattacharjee, Rakesh K Mishra, Divya Tej Sowpati |
| EPI_ISL_465679, EPI_ISL_465680 | Hôpital du Suroît | Laboratoire de santé publique du Québec | Sandrine Moreira, Ioannis Ragoussis, Guillaume Bourque, Jesse Shapiro, Mark Lathrop and Michel Roger on behalf of the CoVSeQ research group (http://covseq.ca/researchgroup) |
| EPI_ISL_465681, EPI_ISL_465682 | Hôpital Charles-LeMoyne | Laboratoire de santé publique du Québec | Sandrine Moreira, Ioannis Ragoussis, Guillaume Bourque, Jesse Shapiro, Mark Lathrop and Michel Roger on behalf of the CoVSeQ research group (http://covseq.ca/researchgroup) |
| EPI_ISL_465683, EPI_ISL_465684 | Hôpital de Maria | Laboratoire de santé publique du Québec | Sandrine Moreira, Ioannis Ragoussis, Guillaume Bourque, Jesse Shapiro, Mark Lathrop and Michel Roger on behalf of the CoVSeQ research group (http://covseq.ca/researchgroup) |
| EPI_ISL_465685 | CSSS Haut-Richelieu/Rouville (Hôpital) | Laboratoire de santé publique du Québec | Sandrine Moreira, Ioannis Ragoussis, Guillaume Bourque, Jesse Shapiro, Mark Lathrop and Michel Roger on behalf of the CoVSeQ research group (http://covseq.ca/researchgroup) |
| EPI_ISL_465686 | Hôpital de Hull | Laboratoire de santé publique du Québec | Sandrine Moreira, Ioannis Ragoussis, Guillaume Bourque, Jesse Shapiro, Mark Lathrop and Michel Roger on behalf of the CoVSeQ research group (http://covseq.ca/researchgroup) |
| EPI_ISL_465687 | CSSS Haut-Richelieu/Rouville (Hôpital) | Laboratoire de santé publique du Québec | Sandrine Moreira, Ioannis Ragoussis, Guillaume Bourque, Jesse Shapiro, Mark Lathrop and Michel Roger on behalf of the CoVSeQ research group (http://covseq.ca/researchgroup) |
| EPI_ISL_465688, EPI_ISL_465689, EPI_ISL_465690 | Centre hospitalier Anna-Laberge | Laboratoire de santé publique du Québec | Sandrine Moreira, Ioannis Ragoussis, Guillaume Bourque, Jesse Shapiro, Mark Lathrop and Michel Roger on behalf of the CoVSeQ research group (http://covseq.ca/researchgroup) |
| EPI_ISL_465691 | Hôpital de Gatineau | Laboratoire de santé publique du Québec | Sandrine Moreira, Ioannis Ragoussis, Guillaume Bourque, Jesse Shapiro, Mark Lathrop and Michel Roger on behalf of the CoVSeQ research group (http://covseq.ca/researchgroup) |
| EPI_ISL_465692 | CSSS Haut-Richelieu/Rouville (Hôpital) | Laboratoire de santé publique du Québec | Sandrine Moreira, Ioannis Ragoussis, Guillaume Bourque, Jesse Shapiro, Mark Lathrop and Michel Roger on behalf of the CoVSeQ research group (http://covseq.ca/researchgroup) |
| EPI_ISL_465693 | Hôpital du Suroît | Laboratoire de santé publique du Québec | Sandrine Moreira, Ioannis Ragoussis, Guillaume Bourque, Jesse Shapiro, Mark Lathrop and Michel Roger on behalf of the CoVSeQ research group (http://covseq.ca/researchgroup) |
| EPI_ISL_465694 | Hôpital Charles-LeMoyne | Laboratoire de santé publique du Québec | Sandrine Moreira, Ioannis Ragoussis, Guillaume Bourque, Jesse Shapiro, Mark Lathrop and Michel Roger on behalf of the CoVSeQ research group (http://covseq.ca/researchgroup) |
| EPI_ISL_465695, EPI_ISL_465696 | Hôpital Pierre-Boucher | Laboratoire de santé publique du Québec | Sandrine Moreira, Ioannis Ragoussis, Guillaume Bourque, Jesse Shapiro, Mark Lathrop and Michel Roger on behalf of the CoVSeQ research group (http://covseq.ca/researchgroup) |
| EPI_ISL_465697 | Centre de santé Inuulitsivik | Laboratoire de santé publique du Québec | Sandrine Moreira, Ioannis Ragoussis, Guillaume Bourque, Jesse Shapiro, Mark Lathrop and Michel Roger on behalf of the CoVSeQ research group (http://covseq.ca/researchgroup) |
| EPI_ISL_465698, EPI_ISL_465699 | Hôpital Charles-LeMoyne | Laboratoire de santé publique du Québec | Sandrine Moreira, Ioannis Ragoussis, Guillaume Bourque, Jesse Shapiro, Mark Lathrop and Michel Roger on behalf of the CoVSeQ research group (http://covseq.ca/researchgroup) |
| EPI_ISL_465700 | Hôpital Pierre-Boucher | Laboratoire de santé publique du Québec | Sandrine Moreira, Ioannis Ragoussis, Guillaume Bourque, Jesse Shapiro, Mark Lathrop and Michel Roger on behalf of the CoVSeQ research group (http://covseq.ca/researchgroup) |
| EPI_ISL_465701 | Hôpital du Suroît | Laboratoire de santé publique du Québec | Sandrine Moreira, Ioannis Ragoussis, Guillaume Bourque, Jesse Shapiro, Mark Lathrop and Michel Roger on behalf of the CoVSeQ research group (http://covseq.ca/researchgroup) |
| EPI_ISL_465702, EPI_ISL_465703 | Hôpital Pierre-Boucher | Laboratoire de santé publique du Québec | Sandrine Moreira, Ioannis Ragoussis, Guillaume Bourque, Jesse Shapiro, Mark Lathrop and Michel Roger on behalf of the CoVSeQ research group (http://covseq.ca/researchgroup) |
| EPI_ISL_467064, EPI_ISL_467065, EPI_ISL_467066, EPI_ISL_467068, EPI_ISL_467069, EPI_ISL_467070, EPI_ISL_467071, EPI_ISL_467072, EPI_ISL_467073, EPI_ISL_467074, EPI_ISL_467075, EPI_ISL_467076, EPI_ISL_467077, EPI_ISL_467078, EPI_ISL_467079, EPI_ISL_467080, EPI_ISL_467081, EPI_ISL_467082, EPI_ISL_467083, EPI_ISL_467084, EPI_ISL_467085 | | | |
| see above | Hospital Universitario Puerta del Mar de Cádiz - INIBICA | SeqCOVID-SPAIN consortium/IBV(CSIC) | Salud Rodríguez-Pallares, Fátima-Galán-Sánchez, Manuel Rodríguez-Iglesias and SeqCOVID-SPAIN consortium |
| EPI_ISL_467299 | Research and Medical Analysis Laboratory of Gendarmerie Royale | Research and Medical Analysis Laboratory of Gendarmerie Royale | Sanaâ LEMRISS Amal SOUIRI Hicham EL OSSMANI Saâd EL Kabbaj |
| EPI_ISL_467950, EPI_ISL_467951, EPI_ISL_467952, EPI_ISL_467953, EPI_ISL_467954, EPI_ISL_467955, EPI_ISL_467956, EPI_ISL_467957, EPI_ISL_467958, EPI_ISL_467959, EPI_ISL_467960, EPI_ISL_467961, EPI_ISL_467962 | | | |
| see above | San Diego County Public Health Laboratory | Andersen lab at Scripps Research | SEARCH Alliance San Diego with Tracy Basler, Jovan Shephard, Brett Austin |
| EPI_ISL_467963 | San Diego County Public Health Laboratory | Andersen lab at Scripps Research | SEARCH Alliance San Diego with Michael Quigley, Ellen Stefanski, Ian Mchardy |
| EPI_ISL_467964, EPI_ISL_467965, EPI_ISL_467966 | San Diego County Public Health Laboratory | Andersen lab at Scripps Research | SEARCH Alliance San Diego with Tracy Basler, Jovan Shephard, Brett Austin |
| EPI_ISL_467967 | Scripps Medical Laboratory | Andersen lab at Scripps Research | SEARCH Alliance San Diego with Tracy Basler, Jovan Shephard, Brett Austin |
| EPI_ISL_467968, EPI_ISL_467969, EPI_ISL_467970, EPI_ISL_467971 | San Diego County Public Health Laboratory | Andersen lab at Scripps Research | SEARCH Alliance San Diego with Tracy Basler, Jovan Shephard, Brett Austin |
| EPI_ISL_467972 | Scripps Medical Laboratory | Andersen lab at Scripps Research | SEARCH Alliance San Diego with Michael Quigley, Ellen Stefanski, Ian Mchardy |
| EPI_ISL_467973, EPI_ISL_467974, EPI_ISL_467975 | San Diego County Public Health Laboratory | Andersen lab at Scripps Research | SEARCH Alliance San Diego with Tracy Basler, Jovan Shephard, Brett Austin |
| EPI_ISL_467976 | Rady's Childrens Hospital | Andersen lab at Scripps Research | SEARCH Alliance San Diego |
| EPI_ISL_467977, EPI_ISL_467978, EPI_ISL_467979, EPI_ISL_467980 | San Diego County Public Health Laboratory | Andersen lab at Scripps Research | SEARCH Alliance San Diego with Tracy Basler, Jovan Shephard, Brett Austin |
| EPI_ISL_467981 | Rady's Childrens Hospital | Andersen lab at Scripps Research | SEARCH Alliance San Diego |
| EPI_ISL_467982, EPI_ISL_467983 | San Diego County Public Health Laboratory | Andersen lab at Scripps Research | SEARCH Alliance San Diego with Tracy Basler, Jovan Shephard, Brett Austin |
| EPI_ISL_467984 | Rady's Childrens Hospital | Andersen lab at Scripps Research | SEARCH Alliance San Diego |
| EPI_ISL_471599 | CSIR-Centre for Cellular and Molecular Biology | CSIR-Centre for Cellular and Molecular Biology | Sakshi Shambhavi, Lamuk Zaveri, Shagufta Khan, Namami Gaur, Tulasi Nagabandi, Purushotham Vodnala, Payel Mukherjee, Sofia Banu, Priya Singh, Dhiviya Vedagiri, Divya Gupta, Vishal Sah, Santosh Kumar Kuncha, Krishnan Harinivas Harshan, Archana Bharadwaj Siva, Karthik Bharadwaj Tallapaka, Deepak Kumar, Devi Prasad Vijayashankar, Disha Nanda, Divya Das, Jotin Gogoi, Manish Bhattacharjee, Rakesh K Mishra, Divya Tej Sowpati |
| EPI_ISL_471600 | CSIR-Centre for Cellular and Molecular Biology | CSIR-Centre for Cellular and Molecular Biology | Sakshi Shambhavi, Lamuk Zaveri, Shagufta Khan, Namami Gaur, Tulasi Nagabandi, Purushotham Vodnala, Payel Mukherjee, Sofia Banu, Priya Singh, Dhiviya Vedagiri, Divya Gupta, Vishal Sah, Santosh Kumar Kuncha, Krishnan Harinivas Harshan, Archana Bharadwaj Siva, Karthik Bharadwaj Tallapaka, G. Aditya Kumar, Koushick Sivakumar, Pooja Ramesh Gupta, Rajan Kumar Jha, Shraddha Vijay Lahoti, Rakesh K Mishra, Divya Tej Sowpati |
| EPI_ISL_471601 | CSIR-Centre for Cellular and Molecular Biology | CSIR-Centre for Cellular and Molecular Biology | Sakshi Shambhavi, Lamuk Zaveri, Shagufta Khan, Namami Gaur, Tulasi Nagabandi, Purushotham Vodnala, Payel Mukherjee, Sofia Banu, Priya Singh, Dhiviya Vedagiri, Divya Gupta, Vishal Sah, Santosh Kumar Kuncha, Krishnan Harinivas Harshan, Archana Bharadwaj Siva, Karthik Bharadwaj Tallapaka,Nikhil Hajirnis, Pratheusa Maccha, M Soujanya Reddy,G. Aditya Kumar, Koushick Sivakumar, Rakesh K Mishra, Divya Tej Sowpati |
| EPI_ISL_471602 | CSIR-Centre for Cellular and Molecular Biology | CSIR-Centre for Cellular and Molecular Biology | Sakshi Shambhavi, Lamuk Zaveri, Shagufta Khan, Namami Gaur, Tulasi Nagabandi, Purushotham Vodnala, Payel Mukherjee, Sofia Banu, Priya Singh, |

| | | | |
|---|---|---|---|
| | | | Dhiviya Vedagiri, Divya Gupta, Vishal Sah, Santosh Kumar Kuncha, Krishnan Harinivas Harshan, Archana Bharadwaj Siva, Karthik Bharadwaj Tallapaka,Nikhil Hajirnis, Pratheusa Maccha, M Soujanya Reddy,G. Aditya Kumar, Koushick Sivakumar,Disha Nanda, Divya Das, Jotin Gogoi, Manish Bhattacharjee, Ravi Prasad Mukku, Rakesh K Mishra, Divya Tej Sowpati |
| EPI_ISL_471627 | CSIR-Centre for Cellular and Molecular Biology | CSIR-Centre for Cellular and Molecular Biology | Sakshi Shambhavi, Lamuk Zaveri, Shagufta Khan, Namami Gaur, Tulasi Nagabandi, Purushotham Vodnala, Payel Mukherjee, Sofia Banu, Priya Singh, Dhiviya Vedagiri, Divya Gupta, Vishal Sah, Santosh Kumar Kuncha, Krishnan Harinivas Harshan, Archana Bharadwaj Siva, Karthik Bharadwaj Tallapaka, Deepak Kumar, Devi Prasad Vijayashankar, Disha Nanda, Divya Das, Jotin Gogoi, Manish Bhattacharjee, Rakesh K Mishra, Divya Tej Sowpati |
| EPI_ISL_471628 | CSIR-Centre for Cellular and Molecular Biology | CSIR-Centre for Cellular and Molecular Biology | Sakshi Shambhavi, Lamuk Zaveri, Shagufta Khan, Namami Gaur, Tulasi Nagabandi, Purushotham Vodnala, Payel Mukherjee, Sofia Banu, Priya Singh, Dhiviya Vedagiri, Divya Gupta, Vishal Sah, Santosh Kumar Kuncha, Krishnan Harinivas Harshan, Archana Bharadwaj Siva, Karthik Bharadwaj Tallapaka, G. Aditya Kumar, Koushick Sivakumar, Pooja Ramesh Gupta, Rajan Kumar Jha, Shraddha Vijay Lahoti, Rakesh K Mishra, Divya Tej Sowpati |
| EPI_ISL_471629 | CSIR-Centre for Cellular and Molecular Biology | CSIR-Centre for Cellular and Molecular Biology | Sakshi Shambhavi, Lamuk Zaveri, Shagufta Khan, Namami Gaur, Tulasi Nagabandi, Purushotham Vodnala, Payel Mukherjee, Sofia Banu, Priya Singh, Dhiviya Vedagiri, Divya Gupta, Vishal Sah, Santosh Kumar Kuncha, Krishnan Harinivas Harshan, Archana Bharadwaj Siva, Karthik Bharadwaj Tallapaka,Nikhil Hajirnis, Pratheusa Maccha, M Soujanya Reddy,G. Aditya Kumar, Koushick Sivakumar, Rakesh K Mishra, Divya Tej Sowpati |
| EPI_ISL_471630 | CSIR-Centre for Cellular and Molecular Biology | CSIR-Centre for Cellular and Molecular Biology | Sakshi Shambhavi, Lamuk Zaveri, Shagufta Khan, Namami Gaur, Tulasi Nagabandi, Purushotham Vodnala, Payel Mukherjee, Sofia Banu, Priya Singh, Dhiviya Vedagiri, Divya Gupta, Vishal Sah, Santosh Kumar Kuncha, Krishnan Harinivas Harshan, Archana Bharadwaj Siva, Karthik Bharadwaj Tallapaka,Nikhil Hajirnis, Pratheusa Maccha, M Soujanya Reddy,G. Aditya Kumar, Koushick Sivakumar,Disha Nanda, Divya Das, Jotin Gogoi, Manish Bhattacharjee, Ravi Prasad Mukku, Rakesh K Mishra, Divya Tej Sowpati |
| EPI_ISL_471993, EPI_ISL_471994, EPI_ISL_471995, EPI_ISL_471996, EPI_ISL_471997, EPI_ISL_471998, EPI_ISL_471999, EPI_ISL_472000, EPI_ISL_472001, EPI_ISL_472002, EPI_ISL_472003, EPI_ISL_472004, EPI_ISL_472005, EPI_ISL_472006, EPI_ISL_472007, EPI_ISL_472008, EPI_ISL_472009, EPI_ISL_472010, EPI_ISL_472011, EPI_ISL_472012, EPI_ISL_472013, EPI_ISL_472014, EPI_ISL_472015, EPI_ISL_472016, EPI_ISL_472017, EPI_ISL_472018, EPI_ISL_472019, EPI_ISL_472020, EPI_ISL_472021, EPI_ISL_472022, EPI_ISL_472023, EPI_ISL_472024, EPI_ISL_472025, EPI_ISL_472026, EPI_ISL_472027, EPI_ISL_472028, EPI_ISL_472029, EPI_ISL_472030, EPI_ISL_472031, EPI_ISL_472032, EPI_ISL_472033, EPI_ISL_472034, EPI_ISL_472035, EPI_ISL_472036, EPI_ISL_472037, EPI_ISL_472038, EPI_ISL_472039, EPI_ISL_472040, EPI_ISL_472041, EPI_ISL_472042, EPI_ISL_472043, EPI_ISL_472044, EPI_ISL_472045, EPI_ISL_472046, EPI_ISL_472047, EPI_ISL_472048, EPI_ISL_472049, EPI_ISL_472050, EPI_ISL_472051, EPI_ISL_472052, EPI_ISL_472053, EPI_ISL_472054, EPI_ISL_472055, EPI_ISL_472056, EPI_ISL_472057, EPI_ISL_472058, EPI_ISL_472059, EPI_ISL_472060, EPI_ISL_472061, EPI_ISL_472062, EPI_ISL_472063, EPI_ISL_472064, EPI_ISL_472065, EPI_ISL_472066, EPI_ISL_472067, EPI_ISL_472068, EPI_ISL_472069, EPI_ISL_472070, EPI_ISL_472071, EPI_ISL_472072, EPI_ISL_472073, EPI_ISL_472074, EPI_ISL_472075, EPI_ISL_472076, EPI_ISL_472077, EPI_ISL_472078, EPI_ISL_472079, EPI_ISL_472080, EPI_ISL_472081, EPI_ISL_472082, EPI_ISL_472083, EPI_ISL_472084, EPI_ISL_472085, EPI_ISL_472086, EPI_ISL_472087, EPI_ISL_472088, EPI_ISL_472089, EPI_ISL_472090, EPI_ISL_472091, EPI_ISL_472092, EPI_ISL_472093, EPI_ISL_472094, EPI_ISL_472095, EPI_ISL_472096, EPI_ISL_472097, EPI_ISL_472098, EPI_ISL_472099, EPI_ISL_472100, EPI_ISL_472101, EPI_ISL_472102, EPI_ISL_472103, EPI_ISL_472104, EPI_ISL_472105, EPI_ISL_472106, EPI_ISL_472107, EPI_ISL_472108, EPI_ISL_472109, EPI_ISL_472110, EPI_ISL_472111, EPI_ISL_472112, EPI_ISL_472113, EPI_ISL_472114, EPI_ISL_472115, EPI_ISL_472116, EPI_ISL_472117, EPI_ISL_472118, EPI_ISL_472119, EPI_ISL_472120, EPI_ISL_472121, EPI_ISL_472122, EPI_ISL_472123, EPI_ISL_472124, EPI_ISL_472125, EPI_ISL_472126, EPI_ISL_472127, EPI_ISL_472128, EPI_ISL_472129, EPI_ISL_472130, EPI_ISL_472131, EPI_ISL_472132, EPI_ISL_472133, EPI_ISL_472134, EPI_ISL_472135 | | | |
| see above | Liverpool Clinical Laboratories | COVID-19 Genomics UK (COG-UK) Consortium | Sam Haldenby, Anita Lucaci, Steve Paterson, Julian Hiscox, Alistair Darby, M Almsaud, A Alrezaihi, Muhannad Alruwaili, Stuart D Armstrong, Jones Benjamin, Eleanor G Bentley, Anu Chawla, Jordan J Clark, Angela Cowell, Richard Eccles, Isabel García-Dorival, Matthew Gemmell, Alessandro Gerada, PKF Gilmore, Richard Gregory, Ximeng Han, Catherine Hartley, Margaret Hughes, Miren Iturriza-Gomara, James Johnson, L Luu, Jenifer Manson, Charlotte Nelson, Elaine O'Toole, Cassie Olateju, Rebekah Penrice-Randal , Lucille Rainbow, N.P Randle, Trevor Ian Robinson, Parul Sharma, Ghada T Shawli, James P Stewart, Neil Swainston, Ecaterina Vamos, Joanne Watts, Mark Whitehead |
| EPI_ISL_476024 | Laboratoire de Recherche et d'Analyses Médicales de la Gendarmerie Royale | Laboratoire de Recherche et d'Analyses Médicales de la Gendarmerie Royale | Sanaâ Lemriss, Amal SOUIRI, Nabil Lemzaoui, Omar Mestoui, Mohamed Labioui, Nabil Ouaariba, Ayoub Jibjibe, Mahmoud Yartaoui, Mohamed Chahmi, Marouane El Rhouila, Samiha Sellak, Nadia Kandoussi, Saâd El Kabbaj. |
| EPI_ISL_476025 | Laboratoire de Recherche et d'Analyses Médicales de la Gendarmerie Royale | Laboratoire de Recherche et d'Analyses Médicales de la Gendarmerie Royale | Sanaâ LEMRISS, Amal Souiri, Saâd EL KABBAJ |
| EPI_ISL_476026 | Laboratoire de Recherche et d'Analyses Médicales de la Gendarmerie Royale | Laboratoire de Recherche et d'Analyses Médicales de la Gendarmerie Royale | Sanaâ Lemriss, Amal SOUIRI, Saâd EL KABBAJ |
| EPI_ISL_476171, EPI_ISL_476172, EPI_ISL_476173, EPI_ISL_476174, EPI_ISL_476175, EPI_ISL_476176, EPI_ISL_476177, EPI_ISL_476178, EPI_ISL_476179, EPI_ISL_476180, EPI_ISL_476181, EPI_ISL_476182, EPI_ISL_476183, EPI_ISL_476184, EPI_ISL_476185, EPI_ISL_476186, EPI_ISL_476187, EPI_ISL_476188, EPI_ISL_476189, EPI_ISL_476190, EPI_ISL_476191, EPI_ISL_476192, EPI_ISL_476193, EPI_ISL_476194, EPI_ISL_476195, EPI_ISL_476196, EPI_ISL_476197, EPI_ISL_476198, EPI_ISL_476199, EPI_ISL_476200, EPI_ISL_476201, EPI_ISL_476202 | | | |
| see above | DB Diagnósticos do Brasil | Instituto de Medicina Tropical da Univesidade de São Paulo | Samples: Nelson Gaburo Jr; Sequencing: Ingra Morales Claro, Jaqueline Goes de Jesus, Erika Regina Manuli, Flavia Cristina da Silva Sales, Thais de Moura Coletti, Camila Alves Maia da Silva, Mariana Severo Ramundo, Giula Magalhaes Ferreira, Darlan da Silva Candido, Julien Theze, Nuno Faria, Ester Sabino |
| EPI_ISL_476203, EPI_ISL_476204, EPI_ISL_476205, EPI_ISL_476206, EPI_ISL_476207, EPI_ISL_476208 | Hospital da Clínicas da Faculdade de Medicina da Universidade de São Paulo | Instituto de Medicina Tropical da Univesidade de São Paulo | Samples: Ingra Morales Claro, Erika Regina Manuli, Cecilia Salete Alencar, Carolina S. Lazar, Silvia F. Costa; Sequencing: Ingra Morales Claro, Jaqueline Goes de Jesus, Erika Regina Manuli, Flavia Cristina da Silva Sales, Thais de Moura Coletti, Camila Alves Maia da Silva, Mariana Severo Ramundo, Giula Magalhaes Ferreira, Darlan da Silva Candido, Julien Theze, Nuno Faria, Ester Sabino |
| EPI_ISL_476209, EPI_ISL_476210, EPI_ISL_476211, EPI_ISL_476212, EPI_ISL_476213, EPI_ISL_476214, EPI_ISL_476215, EPI_ISL_476216 | DB Diagnósticos do Brasil | Instituto de Medicina Tropical da Univesidade de São Paulo | Samples: Nelson Gaburo Jr; Sequencing: Ingra Morales Claro, Jaqueline Goes de Jesus, Erika Regina Manuli, Flavia Cristina da Silva Sales, Thais de Moura Coletti, Camila Alves Maia da Silva, Mariana Severo Ramundo, Giula Magalhaes Ferreira, Darlan da Silva Candido, Julien Theze, Nuno Faria, Ester Sabino |
| EPI_ISL_476217 | Hospital da Clínicas da Faculdade de Medicina da Universidade de São Paulo | Instituto de Medicina Tropical da Univesidade de São Paulo | Samples: Ingra Morales Claro, Erika Regina Manuli, Cecilia Salete Alencar, Carolina S. Lazar, Silvia F. Costa; Sequencing: Ingra Morales Claro, Jaqueline Goes de Jesus, Erika Regina Manuli, Flavia Cristina da Silva Sales, Thais de Moura Coletti, Camila Alves Maia da Silva, Mariana Severo Ramundo, Giula Magalhaes Ferreira, Darlan da Silva Candido, Julien Theze, Nuno Faria, Ester Sabino |
| EPI_ISL_476218, EPI_ISL_476219 | DB Diagnósticos do Brasil | Instituto de Medicina Tropical da Univesidade de São Paulo | Samples: Nelson Gaburo Jr; Sequencing: Ingra Morales Claro, Jaqueline Goes de Jesus, Erika Regina Manuli, Flavia Cristina da Silva Sales, Thais de Moura Coletti, Camila Alves Maia da Silva, Mariana Severo Ramundo, Giula Magalhaes Ferreira, Darlan da Silva Candido, Julien Theze, Nuno Faria, Ester Sabino |
| EPI_ISL_476220, EPI_ISL_476221 | Laboratory Fleury | Instituto de Medicina Tropical da Univesidade de São Paulo | Samples: Celso Granato; Sequencing: Ingra Morales Claro, Jaqueline Goes de Jesus, Erika Regina Manuli, Flavia Cristina da Silva Sales, Thais de Moura Coletti, Camila Alves Maia da Silva, Mariana Severo Ramundo, Giula Magalhaes Ferreira, Darlan da Silva Candido, Julien Theze, Nuno Faria, Ester Sabino |
| EPI_ISL_476222, EPI_ISL_476223 | Hospital da Clínicas da Faculdade de Medicina da Universidade de São Paulo | Instituto de Medicina Tropical da Univesidade de São Paulo | Samples: Ingra Morales Claro, Erika Regina Manuli, Cecilia Salete Alencar, Carolina S. Lazar, Silvia F. Costa; Sequencing: Ingra Morales Claro, Jaqueline Goes de Jesus, Erika Regina Manuli, Flavia Cristina da Silva Sales, Thais de Moura Coletti, Camila Alves Maia da Silva, Mariana Severo Ramundo, Giula Magalhaes Ferreira, Darlan da Silva Candido, Julien Theze, Nuno Faria, Ester Sabino |
| EPI_ISL_476224, EPI_ISL_476225, EPI_ISL_476226, EPI_ISL_476227, EPI_ISL_476228, EPI_ISL_476229 | DB Diagnósticos do Brasil | Instituto de Medicina Tropical da Univesidade de São Paulo | Samples: Nelson Gaburo Jr; Sequencing: Ingra Morales Claro, Jaqueline Goes de Jesus, Erika Regina Manuli, Flavia Cristina da Silva Sales, Thais de Moura Coletti, Camila Alves Maia da Silva, Mariana Severo Ramundo, Giula Magalhaes Ferreira, Darlan da Silva Candido, Julien Theze, Nuno Faria, Ester Sabino |
| EPI_ISL_476230, EPI_ISL_476231 | Laboratory Fleury | Instituto de Medicina Tropical da Univesidade de São Paulo | Samples: Celso Granato; Sequencing: Ingra Morales Claro, Jaqueline Goes de Jesus, Erika Regina Manuli, Flavia Cristina da Silva Sales, Thais de Moura Coletti, Camila Alves Maia da Silva, Mariana Severo Ramundo, Giula Magalhaes Ferreira, Darlan da Silva Candido, Julien Theze, Nuno Faria, Ester Sabino |
| EPI_ISL_476232 | DB Diagnósticos do Brasil | Instituto de Medicina Tropical da Univesidade de São Paulo | Samples: Nelson Gaburo Jr; Sequencing: Ingra Morales Claro, Jaqueline Goes de Jesus, Erika Regina Manuli, Flavia Cristina da Silva Sales, Thais de Moura Coletti, Camila Alves Maia da Silva, Mariana Severo Ramundo, Giula Magalhaes Ferreira, Darlan da Silva Candido, Julien Theze, Nuno Faria, Ester Sabino |
| EPI_ISL_476233, EPI_ISL_476234, EPI_ISL_476235, EPI_ISL_476236 | Laboratory Fleury | Instituto de Medicina Tropical da Univesidade de São Paulo | Samples: Celso Granato; Sequencing: Ingra Morales Claro, Jaqueline Goes de Jesus, Erika Regina Manuli, Flavia Cristina da Silva Sales, Thais de Moura Coletti, Camila Alves Maia da Silva, Mariana Severo Ramundo, Giula Magalhaes Ferreira, Darlan da Silva Candido, Julien Theze, Nuno Faria, Ester Sabino |
| EPI_ISL_476237, EPI_ISL_476238, EPI_ISL_476239, EPI_ISL_476240, EPI_ISL_476241, EPI_ISL_476242, EPI_ISL_476243, EPI_ISL_476244, EPI_ISL_476245, EPI_ISL_476246, EPI_ISL_476247, EPI_ISL_476248, EPI_ISL_476249, EPI_ISL_476250, EPI_ISL_476251, EPI_ISL_476252, EPI_ISL_476253, EPI_ISL_476254, EPI_ISL_476255, EPI_ISL_476256, EPI_ISL_476257, EPI_ISL_476258, EPI_ISL_476259, EPI_ISL_476260, EPI_ISL_476261, EPI_ISL_476262, EPI_ISL_476263, EPI_ISL_476264, EPI_ISL_476265, EPI_ISL_476266, EPI_ISL_476267, EPI_ISL_476268, EPI_ISL_476269, EPI_ISL_476270, EPI_ISL_476271, EPI_ISL_476272, EPI_ISL_476273, EPI_ISL_476274, EPI_ISL_476275, EPI_ISL_476276, EPI_ISL_476277 | | | |
| see above | Hospital da Clínicas da Faculdade de Medicina da Universidade de São Paulo | Instituto de Medicina Tropical da Univesidade de São Paulo | Samples: Ingra Morales Claro, Erika Regina Manuli, Cecilia Salete Alencar, Carolina S. Lazar, Silvia F. Costa; Sequencing: Ingra Morales Claro, Jaqueline Goes de Jesus, Erika Regina Manuli, Flavia Cristina da Silva Sales, Thais de Moura Coletti, Camila Alves Maia da Silva, Mariana Severo Ramundo, Giula |

| | | | |
|---|---|---|---|
| | | | Magalhaes Ferreira, Darlan da Silva Candido, Julien Theze, Nuno Faria, Ester Sabino |
| EPI_ISL_476278, EPI_ISL_476279, EPI_ISL_476280, EPI_ISL_476281, EPI_ISL_476282, EPI_ISL_476283, EPI_ISL_476284, EPI_ISL_476285, EPI_ISL_476286, EPI_ISL_476287, EPI_ISL_476288, EPI_ISL_476289, EPI_ISL_476290, EPI_ISL_476291, EPI_ISL_476292, EPI_ISL_476293, EPI_ISL_476294, EPI_ISL_476295, EPI_ISL_476296, EPI_ISL_476297, EPI_ISL_476298, EPI_ISL_476299, EPI_ISL_476300, EPI_ISL_476301, EPI_ISL_476302, EPI_ISL_476303, EPI_ISL_476304, EPI_ISL_476305, EPI_ISL_476306, EPI_ISL_476307, EPI_ISL_476308, EPI_ISL_476309, EPI_ISL_476310, EPI_ISL_476311, EPI_ISL_476312, EPI_ISL_476313, EPI_ISL_476314, EPI_ISL_476315, EPI_ISL_476316, EPI_ISL_476317, EPI_ISL_476318, EPI_ISL_476319, EPI_ISL_476320, EPI_ISL_476321, EPI_ISL_476322, EPI_ISL_476323, EPI_ISL_476324, EPI_ISL_476325, EPI_ISL_476326, EPI_ISL_476327, EPI_ISL_476328, EPI_ISL_476329, EPI_ISL_476330, EPI_ISL_476331, EPI_ISL_476332, EPI_ISL_476333, EPI_ISL_476334, EPI_ISL_476335, EPI_ISL_476336, EPI_ISL_476350, EPI_ISL_476351, EPI_ISL_476352, EPI_ISL_476353, EPI_ISL_476354, EPI_ISL_476355, EPI_ISL_476356, EPI_ISL_476357, EPI_ISL_476358, EPI_ISL_476359, EPI_ISL_476360, EPI_ISL_476361, EPI_ISL_476362, EPI_ISL_476363, EPI_ISL_476364, EPI_ISL_476365, EPI_ISL_476366, EPI_ISL_476367, EPI_ISL_476368, EPI_ISL_476369, EPI_ISL_476370, EPI_ISL_476371 | | | |
| see above | DB Diagnósticos do Brasil | Instituto de Medicina Tropical da Univesidade de São Paulo | Samples: Nelson Gaburo Jr; Sequencing: Ingra Morales Claro, Jaqueline Goes de Jesus, Erika Regina Manuli, Flavia Cristina da Silva Sales, Thais de Moura Coletti, Camila Alves Maia da Silva, Mariana Severo Ramundo, Giula Magalhaes Ferreira, Darlan da Silva Candido, Julien Theze, Nuno Faria, Ester Sabino |
| EPI_ISL_476372, EPI_ISL_476373, EPI_ISL_476374, EPI_ISL_476375, EPI_ISL_476376, EPI_ISL_476377, EPI_ISL_476378, EPI_ISL_476379, EPI_ISL_476380, EPI_ISL_476381, EPI_ISL_476382, EPI_ISL_476383, EPI_ISL_476384, EPI_ISL_476385, EPI_ISL_476386 | | | |
| see above | Hospital da Clínicas da Faculdade de Medicina da Universidade de São Paulo | Instituto de Medicina Tropical da Univesidade de São Paulo | Samples: Ingra Morales Claro, Erika Regina Manuli, Cecilia Salete Alencar, Carolina S. Lazar, Silvia F. Costa; Sequencing: Ingra Morales Claro, Jaqueline Goes de Jesus, Erika Regina Manuli, Flavia Cristina da Silva Sales, Thais de Moura Coletti, Camila Alves Maia da Silva, Mariana Severo Ramundo, Giula Magalhaes Ferreira, Darlan da Silva Candido, Julien Theze, Nuno Faria, Ester Sabino |
| EPI_ISL_476403, EPI_ISL_476404, EPI_ISL_476405 | Laboratory DASA | Instituto de Medicina Tropical da Univesidade de São Paulo | Samples: José Eduardo Levi; Sequencing: Ingra Morales Claro, Jaqueline Goes de Jesus, Erika Regina Manuli, Flavia Cristina da Silva Sales, Thais de Moura Coletti, Camila Alves Maia da Silva, Mariana Severo Ramundo, Giula Magalhaes Ferreira, Darlan da Silva Candido, Julien Theze, Nuno Faria, Ester Sabino |
| EPI_ISL_476406, EPI_ISL_476407, EPI_ISL_476426 | Laboratory Fleury | Instituto de Medicina Tropical da Univesidade de São Paulo | Samples: Celso Granato; Sequencing: Ingra Morales Claro, Jaqueline Goes de Jesus, Erika Regina Manuli, Flavia Cristina da Silva Sales, Thais de Moura Coletti, Camila Alves Maia da Silva, Mariana Severo Ramundo, Giula Magalhaes Ferreira, Darlan da Silva Candido, Julien Theze, Nuno Faria, Ester Sabino |
| EPI_ISL_476428, EPI_ISL_476429, EPI_ISL_476430, EPI_ISL_476431, EPI_ISL_476432, EPI_ISL_476433, EPI_ISL_476434, EPI_ISL_476435, EPI_ISL_476436, EPI_ISL_476437, EPI_ISL_476438, EPI_ISL_476439, EPI_ISL_476440, EPI_ISL_476441, EPI_ISL_476442, EPI_ISL_476443, EPI_ISL_476444, EPI_ISL_476445, EPI_ISL_476446, EPI_ISL_476447, EPI_ISL_476448, EPI_ISL_476449, EPI_ISL_476450, EPI_ISL_476451, EPI_ISL_476452, EPI_ISL_476453, EPI_ISL_476454, EPI_ISL_476455, EPI_ISL_476456, EPI_ISL_476457, EPI_ISL_476458, EPI_ISL_476459, EPI_ISL_476460, EPI_ISL_476461, EPI_ISL_476462, EPI_ISL_476463, EPI_ISL_476464, EPI_ISL_476465, EPI_ISL_476466, EPI_ISL_476467, EPI_ISL_476468, EPI_ISL_476469, EPI_ISL_476470, EPI_ISL_476471, EPI_ISL_476472, EPI_ISL_476473, EPI_ISL_476474, EPI_ISL_476475, EPI_ISL_476476, EPI_ISL_476477, EPI_ISL_476478, EPI_ISL_476479, EPI_ISL_476480, EPI_ISL_476481, EPI_ISL_476482, EPI_ISL_476483, EPI_ISL_476484, EPI_ISL_476485, EPI_ISL_476486, EPI_ISL_476487, EPI_ISL_476488, EPI_ISL_476489, EPI_ISL_476490 | | | |
| see above | Hospital da Clínicas da Faculdade de Medicina da Universidade de São Paulo | Instituto de Medicina Tropical da Univesidade de São Paulo | Samples: Ingra Morales Claro, Erika Regina Manuli, Cecilia Salete Alencar, Carolina S. Lazar, Silvia F. Costa; Sequencing: Ingra Morales Claro, Jaqueline Goes de Jesus, Erika Regina Manuli, Flavia Cristina da Silva Sales, Thais de Moura Coletti, Camila Alves Maia da Silva, Mariana Severo Ramundo, Giula Magalhaes Ferreira, Darlan da Silva Candido, Julien Theze, Nuno Faria, Ester Sabino |
| EPI_ISL_478372, EPI_ISL_478373, EPI_ISL_478374, EPI_ISL_478375, EPI_ISL_478376, EPI_ISL_478377, EPI_ISL_478378, EPI_ISL_478379, EPI_ISL_478380, EPI_ISL_478381, EPI_ISL_478382, EPI_ISL_478383, EPI_ISL_478384, EPI_ISL_478385, EPI_ISL_478386, EPI_ISL_478387, EPI_ISL_478388, EPI_ISL_478389, EPI_ISL_478390, EPI_ISL_478391, EPI_ISL_478392, EPI_ISL_478393, EPI_ISL_478394, EPI_ISL_478395, EPI_ISL_478396, EPI_ISL_478397, EPI_ISL_478398, EPI_ISL_478399, EPI_ISL_478400, EPI_ISL_478401, EPI_ISL_478402, EPI_ISL_478403 | | | |
| see above | Liverpool Clinical Laboratories | COVID-19 Genomics UK (COG-UK) Consortium | Sam Haldenby, Anita Lucaci, Steve Paterson, Julian Hiscox, Alistair Darby, M Almsaud, A Alrezaihi, Muhannad Alruwaili, Stuart D Armstrong, Jones Benjamin, Eleanor G Bentley, Anu Chawla, Jordan J Clark, Angela Cowell, Richard Eccles, Isabel García-Dorival, Matthew Gemmell, Alessandro Gerada, PKF Gilmore, Richard Gregory, Ximeng Han, Catherine Hartley, Margaret Hughes, Miren Iturriza-Gomara, James Johnson, L Luu, Jenifer Manson, Charlotte Nelson, Elaine O'Toole, Cassie Olateju, Rebekah Penrice-Randal , Lucille Rainbow, N.P Randle, Trevor Ian Robinson, Parul Sharma, Ghada T Shawli, James P Stewart, Neil Swainston, Ecaterina Vamos, Joanne Watts, Mark Whitehead |
| EPI_ISL_480791, EPI_ISL_480796, EPI_ISL_480801, EPI_ISL_480806, EPI_ISL_480808, EPI_ISL_480811, EPI_ISL_480812, EPI_ISL_480814, EPI_ISL_480815, EPI_ISL_480816, EPI_ISL_480817, EPI_ISL_480818, EPI_ISL_480821, EPI_ISL_480822, EPI_ISL_480823, EPI_ISL_480824, EPI_ISL_480825, EPI_ISL_480826, EPI_ISL_480827, EPI_ISL_480828, EPI_ISL_480829, EPI_ISL_480830, EPI_ISL_480831, EPI_ISL_480832, EPI_ISL_480833, EPI_ISL_480834, EPI_ISL_480836, EPI_ISL_480837, EPI_ISL_480838, EPI_ISL_480839, EPI_ISL_480840, EPI_ISL_480841, EPI_ISL_480842, EPI_ISL_480844, EPI_ISL_480845, EPI_ISL_480847, EPI_ISL_480850, EPI_ISL_480851, EPI_ISL_480852, EPI_ISL_480853, EPI_ISL_480855, EPI_ISL_480856, EPI_ISL_480857, EPI_ISL_480858, EPI_ISL_480859, EPI_ISL_480861, EPI_ISL_480862, EPI_ISL_480863, EPI_ISL_480866, EPI_ISL_480870, EPI_ISL_480871, EPI_ISL_480872, EPI_ISL_480873, EPI_ISL_480874, EPI_ISL_480875, EPI_ISL_480876, EPI_ISL_480877, EPI_ISL_480879, EPI_ISL_480880, EPI_ISL_480883, EPI_ISL_480921, EPI_ISL_480922, EPI_ISL_480925, EPI_ISL_480926, EPI_ISL_480927, EPI_ISL_480928, EPI_ISL_480933, EPI_ISL_480934, EPI_ISL_480935, EPI_ISL_480936, EPI_ISL_480938, EPI_ISL_480940, EPI_ISL_480942, EPI_ISL_480944, EPI_ISL_480945, EPI_ISL_480950, EPI_ISL_480951 | | | |
| see above | Florida Bureau of Public Health Laboratories | Florida Bureau of Public Health Laboratories | Sarah Schmedes, Jason Blanton |
| EPI_ISL_483158, EPI_ISL_483159, EPI_ISL_483160, EPI_ISL_483161, EPI_ISL_483162, EPI_ISL_483163, EPI_ISL_483164 | San Diego County Public Health Laboratory | Andersen lab at Scripps Research | SEARCH Alliance San Diego with Tracy Basler, Jovan Shephard, Brett Austin |
| EPI_ISL_483165, EPI_ISL_483166, EPI_ISL_483167, EPI_ISL_483168, EPI_ISL_483169, EPI_ISL_483170, EPI_ISL_483171, EPI_ISL_483172, EPI_ISL_483173, EPI_ISL_483174, EPI_ISL_483175, EPI_ISL_483176, EPI_ISL_483177, EPI_ISL_483178, EPI_ISL_483179, EPI_ISL_483180, EPI_ISL_483181, EPI_ISL_483182, EPI_ISL_483183, EPI_ISL_483184, EPI_ISL_483185, EPI_ISL_483186, EPI_ISL_483187, EPI_ISL_483188, EPI_ISL_483189, EPI_ISL_483190, EPI_ISL_483191, EPI_ISL_483192, EPI_ISL_483193, EPI_ISL_483194, EPI_ISL_483195, EPI_ISL_483196, EPI_ISL_483197, EPI_ISL_483198, EPI_ISL_483199, EPI_ISL_483200, EPI_ISL_483201, EPI_ISL_483202, EPI_ISL_483203, EPI_ISL_483204, EPI_ISL_483205, EPI_ISL_483206, EPI_ISL_483207, EPI_ISL_483208, EPI_ISL_483209, EPI_ISL_483210, EPI_ISL_483211, EPI_ISL_483212, EPI_ISL_483213, EPI_ISL_483214, EPI_ISL_483215, EPI_ISL_483216, EPI_ISL_483217, EPI_ISL_483218, EPI_ISL_483219, EPI_ISL_483220, EPI_ISL_483221, EPI_ISL_483222, EPI_ISL_483223, EPI_ISL_483224, EPI_ISL_483225, EPI_ISL_483226, EPI_ISL_483227, EPI_ISL_483228, EPI_ISL_483229, EPI_ISL_483230, EPI_ISL_483231, EPI_ISL_483232, EPI_ISL_483233, EPI_ISL_483234, EPI_ISL_483235, EPI_ISL_483236, EPI_ISL_483237, EPI_ISL_483238, EPI_ISL_483239, EPI_ISL_483240, EPI_ISL_483241, EPI_ISL_483242, EPI_ISL_483243, EPI_ISL_483244, EPI_ISL_483245, EPI_ISL_483246, EPI_ISL_483247, EPI_ISL_483248, EPI_ISL_483249, EPI_ISL_483250, EPI_ISL_483251, EPI_ISL_483252, EPI_ISL_483253, EPI_ISL_483254, EPI_ISL_483255, EPI_ISL_483256, EPI_ISL_483257, EPI_ISL_483258, EPI_ISL_483259, EPI_ISL_483260, EPI_ISL_483261, EPI_ISL_483262, EPI_ISL_483263, EPI_ISL_483264, EPI_ISL_483265, EPI_ISL_483266, EPI_ISL_483267, EPI_ISL_483268, EPI_ISL_483269, EPI_ISL_483270, EPI_ISL_483271, EPI_ISL_483272, EPI_ISL_483273, EPI_ISL_483274, EPI_ISL_483275, EPI_ISL_483276, EPI_ISL_483277, EPI_ISL_483278, EPI_ISL_483279, EPI_ISL_483280, EPI_ISL_483281, EPI_ISL_483282, EPI_ISL_483283, EPI_ISL_483284, EPI_ISL_483285, EPI_ISL_483286, EPI_ISL_483287, EPI_ISL_483288, EPI_ISL_483289, EPI_ISL_483290, EPI_ISL_483291, EPI_ISL_483292, EPI_ISL_483293, EPI_ISL_483294, EPI_ISL_483295, EPI_ISL_483296, EPI_ISL_483297, EPI_ISL_483298, EPI_ISL_483299, EPI_ISL_483300, EPI_ISL_483301, EPI_ISL_483302, EPI_ISL_483303, EPI_ISL_483304, EPI_ISL_483305, EPI_ISL_483306, EPI_ISL_483307, EPI_ISL_483308, EPI_ISL_483309, EPI_ISL_483310, EPI_ISL_483311, EPI_ISL_483312, EPI_ISL_483313, EPI_ISL_483314, EPI_ISL_483315, EPI_ISL_483316, EPI_ISL_483317, EPI_ISL_483318, EPI_ISL_483319, EPI_ISL_483320, EPI_ISL_483321, EPI_ISL_483322, EPI_ISL_483323, EPI_ISL_483324, EPI_ISL_483325, EPI_ISL_483326, EPI_ISL_483327, EPI_ISL_483328, EPI_ISL_483329, EPI_ISL_483330, EPI_ISL_483331, EPI_ISL_483332, EPI_ISL_483333, EPI_ISL_483334, EPI_ISL_483335, EPI_ISL_483336, EPI_ISL_483337, EPI_ISL_483338, EPI_ISL_483339, EPI_ISL_483340, EPI_ISL_483341, EPI_ISL_483342, EPI_ISL_483343, EPI_ISL_483344, EPI_ISL_483345, EPI_ISL_483346, EPI_ISL_483347, EPI_ISL_483348, EPI_ISL_483349, EPI_ISL_483350, EPI_ISL_483351, EPI_ISL_483352, EPI_ISL_483353, EPI_ISL_483354, EPI_ISL_483355, EPI_ISL_483356, EPI_ISL_483357, EPI_ISL_483358, EPI_ISL_483359, EPI_ISL_483360, EPI_ISL_483361, EPI_ISL_483362, EPI_ISL_483363, EPI_ISL_483364, EPI_ISL_483365, EPI_ISL_483366, EPI_ISL_483367, EPI_ISL_483368, EPI_ISL_483369, EPI_ISL_483370, EPI_ISL_483371, EPI_ISL_483372, EPI_ISL_483373, EPI_ISL_483374, EPI_ISL_483375, EPI_ISL_483376, EPI_ISL_483377, EPI_ISL_483378, EPI_ISL_483379, EPI_ISL_483380, EPI_ISL_483381, EPI_ISL_483382, EPI_ISL_483383, EPI_ISL_483384, EPI_ISL_483385, EPI_ISL_483386, EPI_ISL_483387, EPI_ISL_483388, EPI_ISL_483389, EPI_ISL_483390, EPI_ISL_483391, EPI_ISL_483392, EPI_ISL_483393, EPI_ISL_483394, EPI_ISL_483395, EPI_ISL_483396, EPI_ISL_483397, EPI_ISL_483403, EPI_ISL_483404, EPI_ISL_483405, EPI_ISL_483406, EPI_ISL_483407, EPI_ISL_483408, EPI_ISL_483409, EPI_ISL_483410, EPI_ISL_483411, EPI_ISL_483412, EPI_ISL_483413, EPI_ISL_483414, EPI_ISL_483415, EPI_ISL_483416, EPI_ISL_483417, EPI_ISL_483418, EPI_ISL_483419, EPI_ISL_483420, EPI_ISL_483421, EPI_ISL_483422, EPI_ISL_483423, EPI_ISL_483424, EPI_ISL_483425, EPI_ISL_483426, EPI_ISL_483427, EPI_ISL_483428, EPI_ISL_483429, EPI_ISL_483430, EPI_ISL_483431, EPI_ISL_483432, EPI_ISL_483433, EPI_ISL_483434, EPI_ISL_483435, EPI_ISL_483436, EPI_ISL_483437, EPI_ISL_483438, EPI_ISL_483439, EPI_ISL_483440, EPI_ISL_483441, EPI_ISL_483442, EPI_ISL_483443, EPI_ISL_483444, EPI_ISL_483445, EPI_ISL_483446, EPI_ISL_483447, EPI_ISL_483448, EPI_ISL_483449, EPI_ISL_483450, EPI_ISL_483451, EPI_ISL_483452, EPI_ISL_483453, EPI_ISL_483454, EPI_ISL_483455, EPI_ISL_483456, EPI_ISL_483457, EPI_ISL_483458, EPI_ISL_483459, EPI_ISL_483460, EPI_ISL_483461, EPI_ISL_483462, EPI_ISL_483463, EPI_ISL_483464, EPI_ISL_483465, EPI_ISL_483466, EPI_ISL_483467, EPI_ISL_483468, EPI_ISL_483469, EPI_ISL_483470, EPI_ISL_483471, EPI_ISL_483472, EPI_ISL_483473, EPI_ISL_483474, EPI_ISL_483475, EPI_ISL_483476, EPI_ISL_483477, EPI_ISL_483478, EPI_ISL_483479, EPI_ISL_483480, EPI_ISL_483481, EPI_ISL_483482, EPI_ISL_483483, EPI_ISL_483484, EPI_ISL_483485, EPI_ISL_483486, EPI_ISL_483487, EPI_ISL_483488, EPI_ISL_483489, EPI_ISL_483490, EPI_ISL_483491, EPI_ISL_483492, EPI_ISL_483493, EPI_ISL_483494, EPI_ISL_483495, EPI_ISL_483496, EPI_ISL_483497, EPI_ISL_483498, EPI_ISL_483499, EPI_ISL_483500, EPI_ISL_483501, EPI_ISL_483502, EPI_ISL_483503, EPI_ISL_483504, EPI_ISL_483505, EPI_ISL_483506, EPI_ISL_483507, EPI_ISL_483508, EPI_ISL_483509, EPI_ISL_483510, EPI_ISL_483511, EPI_ISL_483512, EPI_ISL_483513, EPI_ISL_483514, EPI_ISL_483515, EPI_ISL_483516, EPI_ISL_483517, EPI_ISL_483518, EPI_ISL_483519, EPI_ISL_483520, EPI_ISL_483521, EPI_ISL_483522, EPI_ISL_483523, EPI_ISL_483524, EPI_ISL_483525, EPI_ISL_483526, EPI_ISL_483527, EPI_ISL_483528, EPI_ISL_483529, EPI_ISL_483530 | | | |
| see above | UC San Diego Center for Advanced Laboratory Medicine | Andersen lab at Scripps Research | SEARCH Alliance San Diego with David Pride, Ji H Shin |
| EPI_ISL_483531, EPI_ISL_483532, EPI_ISL_483533, EPI_ISL_483534, EPI_ISL_483535, EPI_ISL_483536, EPI_ISL_483537, EPI_ISL_483538, EPI_ISL_483539, EPI_ISL_483540, EPI_ISL_483541 | | | |
| see above | San Diego County Public Health Laboratory | Andersen lab at Scripps Research | SEARCH Alliance San Diego with Tracy Basler, Jovan Shephard, Brett Austin |
| EPI_ISL_484254, EPI_ISL_484255, EPI_ISL_484256, EPI_ISL_484257, EPI_ISL_484258, EPI_ISL_484259, EPI_ISL_484260, EPI_ISL_484261, EPI_ISL_484262, EPI_ISL_484263 | Liverpool Clinical Laboratories | COVID-19 Genomics UK (COG-UK) Consortium | Sam Haldenby, Anita Lucaci, Steve Paterson, Julian Hiscox, Alistair Darby, M Almsaud, A Alrezaihi, Muhannad Alruwaili, Stuart D Armstrong, Jones Benjamin, Eleanor G Bentley, Anu Chawla, Jordan J Clark, Angela Cowell, Richard Eccles, Isabel García-Dorival, Matthew Gemmell, Alessandro Gerada, PKF Gilmore, Richard Gregory, Ximeng Han, Catherine Hartley, Margaret Hughes, Miren Iturriza-Gomara, James Johnson, L Luu, Jenifer Manson, Charlotte Nelson, Elaine O'Toole, Cassie Olateju, Rebekah Penrice-Randal , Lucille Rainbow, N.P Randle, Trevor Ian Robinson, Parul Sharma, Ghada T Shawli, James P Stewart, Neil Swainston, Ecaterina Vamos, Joanne Watts, Mark Whitehead |
| EPI_ISL_487231 | Oklahoma State University Diagnostic Laboratory-Oklahoma Animal Disease Diagnostic Laboratory | Oklahoma Animal Disease Diagnostic Laboratory | Sai Narayanan, Teluguakula Narasaraju, John Ritchey, Girish Patil, Sunil More, Jerry Saliki, Anil Kaul, Akhilesh Ramachandran |
| EPI_ISL_489710, EPI_ISL_489711, EPI_ISL_489712, EPI_ISL_489716, EPI_ISL_489717, EPI_ISL_489718, EPI_ISL_489720, EPI_ISL_489722, EPI_ISL_489723, EPI_ISL_489724, EPI_ISL_489725, EPI_ISL_489727, EPI_ISL_489728, EPI_ISL_489729, EPI_ISL_489730, EPI_ISL_489731, EPI_ISL_489733, EPI_ISL_489734, EPI_ISL_489737, EPI_ISL_489739, EPI_ISL_489745, EPI_ISL_489747, EPI_ISL_489752, EPI_ISL_489756, EPI_ISL_489758, EPI_ISL_489759, EPI_ISL_489760, EPI_ISL_489761, EPI_ISL_489762, EPI_ISL_489763, EPI_ISL_489764, EPI_ISL_489765, EPI_ISL_489767, EPI_ISL_489769, EPI_ISL_489770, EPI_ISL_489772, EPI_ISL_489773, EPI_ISL_489775, EPI_ISL_489776, EPI_ISL_489777, EPI_ISL_489778, EPI_ISL_489779, EPI_ISL_489780, EPI_ISL_489781, EPI_ISL_489782, EPI_ISL_489783, EPI_ISL_489784, EPI_ISL_489785, EPI_ISL_489786, EPI_ISL_489787, EPI_ISL_489788, EPI_ISL_489789, EPI_ISL_489790, EPI_ISL_489791, | | | |

| | | | |
|---|---|---|---|
| EPI_ISL_489792, EPI_ISL_489793, EPI_ISL_489794, EPI_ISL_489795, EPI_ISL_489796, EPI_ISL_489797, EPI_ISL_489798, EPI_ISL_489799, EPI_ISL_489800, EPI_ISL_489801, EPI_ISL_489802, EPI_ISL_489803, EPI_ISL_489804, EPI_ISL_489805, EPI_ISL_489806, EPI_ISL_489807, EPI_ISL_489808, EPI_ISL_489809, EPI_ISL_489810, EPI_ISL_489811, EPI_ISL_489812, EPI_ISL_489813, EPI_ISL_489814, EPI_ISL_489815, EPI_ISL_489816, EPI_ISL_489817, EPI_ISL_489818, EPI_ISL_489819, EPI_ISL_489820, EPI_ISL_489821, EPI_ISL_489822, EPI_ISL_489823, EPI_ISL_489824, EPI_ISL_489825, EPI_ISL_489826, EPI_ISL_489827, EPI_ISL_489828, EPI_ISL_489829, EPI_ISL_489830, EPI_ISL_489831, EPI_ISL_489832 | | | |
| see above | Florida Bureau of Public Health Laboratories | Florida Bureau of Public Health Laboratories | Sarah Schmedes, Jason Blanton |
| EPI_ISL_490331, EPI_ISL_490332, EPI_ISL_490333, EPI_ISL_490334, EPI_ISL_490335, EPI_ISL_490336, EPI_ISL_490337, EPI_ISL_490338, EPI_ISL_490339, EPI_ISL_490340, EPI_ISL_490341, EPI_ISL_490342, EPI_ISL_490343, EPI_ISL_490344, EPI_ISL_490345, EPI_ISL_490346, EPI_ISL_490347, EPI_ISL_490348, EPI_ISL_490349, EPI_ISL_490350, EPI_ISL_490351, EPI_ISL_490352, EPI_ISL_490353, EPI_ISL_490354, EPI_ISL_490355, EPI_ISL_490356, EPI_ISL_490357, EPI_ISL_490359, EPI_ISL_490360, EPI_ISL_490361, EPI_ISL_490362, EPI_ISL_490363, EPI_ISL_490364, EPI_ISL_490365, EPI_ISL_490366, EPI_ISL_490367, EPI_ISL_490368, EPI_ISL_490369, EPI_ISL_490370, EPI_ISL_490371, EPI_ISL_490372, EPI_ISL_490373, EPI_ISL_490374, EPI_ISL_490375, EPI_ISL_490376, EPI_ISL_490377, EPI_ISL_490378, EPI_ISL_490379, EPI_ISL_490380, EPI_ISL_490381, EPI_ISL_490382, EPI_ISL_490383, EPI_ISL_490384, EPI_ISL_490385, EPI_ISL_490386, EPI_ISL_490387, EPI_ISL_490388, EPI_ISL_490389, EPI_ISL_490390, EPI_ISL_490391, EPI_ISL_490392, EPI_ISL_490393, EPI_ISL_490394, EPI_ISL_490395, EPI_ISL_490396, EPI_ISL_490397, EPI_ISL_490398, EPI_ISL_490399, EPI_ISL_490400, EPI_ISL_490401, EPI_ISL_490402, EPI_ISL_490403, EPI_ISL_490404, EPI_ISL_490405, EPI_ISL_490406, EPI_ISL_490407, EPI_ISL_490408, EPI_ISL_490409, EPI_ISL_490410, EPI_ISL_490411, EPI_ISL_490412, EPI_ISL_490413, EPI_ISL_490414, EPI_ISL_490415, EPI_ISL_490416, EPI_ISL_490417, EPI_ISL_490418, EPI_ISL_490419, EPI_ISL_490420 | | | |
| see above | Liverpool Clinical Laboratories | COVID-19 Genomics UK (COG-UK) Consortium | Sam Haldenby, Anita Lucaci, Steve Paterson, Julian Hiscox, Alistair Darby, M Almsaud, A Alrezaihi, Muhannad Alruwaili, Stuart D Armstrong, Jones Benjamin, Eleanor G Bentley, Anu Chawla, Jordan J Clark, Angela Cowell, Richard Eccles, Isabel García-Dorival, Matthew Gemmell, Alessandro Gerada, PKF Gilmore, Richard Gregory, Ximeng Han, Catherine Hartley, Margaret Hughes, Miren Iturriza-Gomara, James Johnson, L Luu, Jenifer Manson, Charlotte Nelson, Elaine O'Toole, Cassie Olateju, Rebekah Penrice-Randal , Lucille Rainbow, N.P Randle, Trevor Ian Robinson, Parul Sharma, Ghada T Shawli, James P Stewart, Neil Swainston, Ecaterina Vamos, Joanne Watts, Mark Whitehead |
| EPI_ISL_491116, EPI_ISL_491121, EPI_ISL_491122, EPI_ISL_491123, EPI_ISL_491124, EPI_ISL_491125, EPI_ISL_491126, EPI_ISL_491127, EPI_ISL_491128, EPI_ISL_491129, EPI_ISL_491130, EPI_ISL_491131, EPI_ISL_491132 | | | |
| see above | Oman-National Influenza Center | Biotechnology & OMICs Laboratory | Samira Al-Mahruqi, Abdul Latif Khan, Samiha Al-Kharusi, Adil Khan , Ahmed Al-Rawahi, Sajjad Asaf, Amina Al-Jardani, Hanan Al-Kindi, Intisar Al-Shukri, Ahlam Al-Amri, Aisha Al-Amri, Aisha Al-Busaidi, Adil Al-Wahaibi, Seif Al-Abri, Ahmed Al-Harrasi |
| EPI_ISL_491133, EPI_ISL_491134, EPI_ISL_491135, EPI_ISL_491136, EPI_ISL_491137, EPI_ISL_491138, EPI_ISL_491139, EPI_ISL_491140, EPI_ISL_491141, EPI_ISL_491142, EPI_ISL_491143, EPI_ISL_491144, EPI_ISL_491145 | | | |
| see above | Oman-National Influenza Center | Biotechnology & OMICs Laboratory | Samiha Al-Kharusi, Sajjad Asaf, Abdul Latif Khan, Samira Al-Mahruqi, Adil Khan, Ahmed Al-Rawahi, Amina Al-Jardani, Hanan Al-Kindi, Intisar Al-Shukri, Ahlam Al-Amri, Aisha Al-Amri, Aisha Al-Busaidi, Adil Al-Wahaibi, Seif Al-Abri, Ahmed Al-Harrasi |
| EPI_ISL_491159, EPI_ISL_491160, EPI_ISL_491161, EPI_ISL_491162, EPI_ISL_491163, EPI_ISL_491164, EPI_ISL_491165, EPI_ISL_491166, EPI_ISL_491167, EPI_ISL_491168, EPI_ISL_491169, EPI_ISL_491170, EPI_ISL_491171 | | | |
| see above | Oman-National Influenza Center | Biotechnology & OMICs Laboratory | Sajjad Asaf, Samiha Al-Kharusi, Ahmed Al-Harrasi, Samira Al-Mahruqi, Adil Khan, Ahmed Al-Rawahi, Abdul Latif Khan, Amina Al-Jardani, Hanan Al-Kindi, Intisar Al-Shukri, Ahlam Al-Amri, Aisha Al-Amri, Aisha Al-Busaidi, Adil Al-Wahaibi, Seif Al-Abri. |
| EPI_ISL_492065 | Oman-National Influenza Center | Department of Microbiology and Immunology-SQUH Department of Microbiology and Immunology, Sultan Qaboos University Hopsital, P.O 35, Postal code 123 | Samira Al-Maruqi, Fahad Zadjali, Amina Al Jardani, Khulood Al-Mammary, Hanan Al-kindi, Fatma BaAlawi, Hamida AL Barwani, Zeyana AL-Dahmani, Intisar Al-Shukri, Azza Al-Rashdi, Samiha Al Kharusi, Abdulla Balkhair |
| EPI_ISL_494372, EPI_ISL_494373, EPI_ISL_494374, EPI_ISL_494375, EPI_ISL_494376, EPI_ISL_494377, EPI_ISL_494378, EPI_ISL_494379, EPI_ISL_494380, EPI_ISL_494381, EPI_ISL_494382, EPI_ISL_494383, EPI_ISL_494384, EPI_ISL_494385, EPI_ISL_494386, EPI_ISL_494387, EPI_ISL_494388, EPI_ISL_494389, EPI_ISL_494390, EPI_ISL_494391, EPI_ISL_494392, EPI_ISL_494393, EPI_ISL_494394, EPI_ISL_494395, EPI_ISL_494396, EPI_ISL_494397, EPI_ISL_494398, EPI_ISL_494399, EPI_ISL_494400, EPI_ISL_494401, EPI_ISL_494402, EPI_ISL_494403, EPI_ISL_494404, EPI_ISL_494405, EPI_ISL_494406, EPI_ISL_494407, EPI_ISL_494408, EPI_ISL_494409, EPI_ISL_494410, EPI_ISL_494411, EPI_ISL_494412, EPI_ISL_494413, EPI_ISL_494414, EPI_ISL_494415, EPI_ISL_494416, EPI_ISL_494417, EPI_ISL_494418, EPI_ISL_494419, EPI_ISL_494420, EPI_ISL_494421, EPI_ISL_494422, EPI_ISL_494423, EPI_ISL_494424, EPI_ISL_494425, EPI_ISL_494426, EPI_ISL_494427, EPI_ISL_494428, EPI_ISL_494429, EPI_ISL_494430, EPI_ISL_494431, EPI_ISL_494432, EPI_ISL_494433, EPI_ISL_494434, EPI_ISL_494435, EPI_ISL_494436, EPI_ISL_494437, EPI_ISL_494438, EPI_ISL_494439, EPI_ISL_494440, EPI_ISL_494441, EPI_ISL_494442, EPI_ISL_494443, EPI_ISL_494444, EPI_ISL_494445, EPI_ISL_494446, EPI_ISL_494447, EPI_ISL_494448, EPI_ISL_494449, EPI_ISL_494450, EPI_ISL_494451, EPI_ISL_494452, EPI_ISL_494453, EPI_ISL_494454, EPI_ISL_494455, EPI_ISL_494456, EPI_ISL_494457, EPI_ISL_494458, EPI_ISL_494459, EPI_ISL_494460, EPI_ISL_494461, EPI_ISL_494462, EPI_ISL_494463, EPI_ISL_494464, EPI_ISL_494465, EPI_ISL_494466, EPI_ISL_494467, EPI_ISL_494468, EPI_ISL_494469, EPI_ISL_494470, EPI_ISL_494471, EPI_ISL_494472, EPI_ISL_494473, EPI_ISL_494474, EPI_ISL_494475, EPI_ISL_494476, EPI_ISL_494477, EPI_ISL_494478, EPI_ISL_494479, EPI_ISL_494480, EPI_ISL_494481, EPI_ISL_494482, EPI_ISL_494483, EPI_ISL_494484, EPI_ISL_494485, EPI_ISL_494486, EPI_ISL_494487, EPI_ISL_494488, EPI_ISL_494489, EPI_ISL_494490, EPI_ISL_494491, EPI_ISL_494492, EPI_ISL_494493, EPI_ISL_494494, EPI_ISL_494495, EPI_ISL_494496, EPI_ISL_494497, EPI_ISL_494498, EPI_ISL_494499, EPI_ISL_494500, EPI_ISL_494501, EPI_ISL_494502, EPI_ISL_494503, EPI_ISL_494568, EPI_ISL_494569 | | | |
| see above | San Diego County Public Health Laboratory | Andersen lab at Scripps Research | SEARCH Alliance San Diego with Tracy Basler, Jovan Shephard, Brett Austin |
| EPI_ISL_494570 | Rady's Childrens Hospital | Andersen lab at Scripps Research | SEARCH Alliance San Diego |
| EPI_ISL_494571, EPI_ISL_494572, EPI_ISL_494573, EPI_ISL_494574, EPI_ISL_494575, EPI_ISL_494576, EPI_ISL_494577, EPI_ISL_494578, EPI_ISL_494579, EPI_ISL_494580, EPI_ISL_494581, EPI_ISL_494582, EPI_ISL_494583, EPI_ISL_494584, EPI_ISL_494585, EPI_ISL_494586, EPI_ISL_494587, EPI_ISL_494588, EPI_ISL_494589, EPI_ISL_494590, EPI_ISL_494591, EPI_ISL_494592, EPI_ISL_494593, EPI_ISL_494594, EPI_ISL_494595 | | | |
| see above | San Diego County Public Health Laboratory | Andersen lab at Scripps Research | SEARCH Alliance San Diego with Tracy Basler, Jovan Shephard, Brett Austin |
| EPI_ISL_494596, EPI_ISL_494597, EPI_ISL_494598, EPI_ISL_494599, EPI_ISL_494600, EPI_ISL_494601, EPI_ISL_494602, EPI_ISL_494603, EPI_ISL_494604, EPI_ISL_494605, EPI_ISL_494606, EPI_ISL_494607, EPI_ISL_494608, EPI_ISL_494609, EPI_ISL_494610, EPI_ISL_494611, EPI_ISL_494612, EPI_ISL_494613, EPI_ISL_494614, EPI_ISL_494615, EPI_ISL_494616, EPI_ISL_494617, EPI_ISL_494618, EPI_ISL_494619 | | | |
| see above | Scripps Medical Laboratory | Andersen lab at Scripps Research | SEARCH Alliance San Diego with Michael Quigley, Ellen Stefanski, Ian Mchardy |
| EPI_ISL_494620, EPI_ISL_494621, EPI_ISL_494622, EPI_ISL_494623, EPI_ISL_494624, EPI_ISL_494625, EPI_ISL_494626, EPI_ISL_494627, EPI_ISL_494628, EPI_ISL_494629, EPI_ISL_494630, EPI_ISL_494631, EPI_ISL_494632, EPI_ISL_494633, EPI_ISL_494634 | | | |
| see above | San Diego County Public Health Laboratory | Andersen lab at Scripps Research | SEARCH Alliance San Diego with Tracy Basler, Jovan Shephard, Brett Austin |
| EPI_ISL_494635, EPI_ISL_494636, EPI_ISL_494637, EPI_ISL_494638, EPI_ISL_494639, EPI_ISL_494640, EPI_ISL_494641, EPI_ISL_494642, EPI_ISL_494643, EPI_ISL_494644, EPI_ISL_494645, EPI_ISL_494646, EPI_ISL_494647, EPI_ISL_494648, EPI_ISL_494649, EPI_ISL_494650, EPI_ISL_494651, EPI_ISL_494652, EPI_ISL_494653, EPI_ISL_494654, EPI_ISL_494655, EPI_ISL_494656, EPI_ISL_494657, EPI_ISL_494658, EPI_ISL_494659, EPI_ISL_494660, EPI_ISL_494661, EPI_ISL_494662, EPI_ISL_494663, EPI_ISL_494664, EPI_ISL_494665, EPI_ISL_494666, EPI_ISL_494667, EPI_ISL_494668, EPI_ISL_494669, EPI_ISL_494670, EPI_ISL_494671, EPI_ISL_494672, EPI_ISL_494673, EPI_ISL_494674, EPI_ISL_494675, EPI_ISL_494676, EPI_ISL_494677, EPI_ISL_494678, EPI_ISL_494679, EPI_ISL_494680, EPI_ISL_494681, EPI_ISL_494682, EPI_ISL_494683, EPI_ISL_494684, EPI_ISL_494685, EPI_ISL_494686, EPI_ISL_494687, EPI_ISL_494688, EPI_ISL_494689, EPI_ISL_494690, EPI_ISL_494691, EPI_ISL_494692, EPI_ISL_494693, EPI_ISL_494694, EPI_ISL_494695, EPI_ISL_494696, EPI_ISL_494697, EPI_ISL_494698, EPI_ISL_494699, EPI_ISL_494700, EPI_ISL_494701, EPI_ISL_494702, EPI_ISL_494703, EPI_ISL_494704, EPI_ISL_494705, EPI_ISL_494706, EPI_ISL_494707, EPI_ISL_494708, EPI_ISL_494709, EPI_ISL_494710, EPI_ISL_494711, EPI_ISL_494712, EPI_ISL_494713 | | | |
| see above | Scripps Medical Laboratory | Andersen lab at Scripps Research | SEARCH Alliance San Diego with Michael Quigley, Ellen Stefanski, Ian Mchardy |
| EPI_ISL_494714, EPI_ISL_494715, EPI_ISL_494716, EPI_ISL_494717, EPI_ISL_494718, EPI_ISL_494719, EPI_ISL_494720, EPI_ISL_494721, EPI_ISL_494722, EPI_ISL_494723, EPI_ISL_494724, EPI_ISL_494725, EPI_ISL_494726, EPI_ISL_494727, EPI_ISL_494728, EPI_ISL_494729, EPI_ISL_494730, EPI_ISL_494731, EPI_ISL_494732, EPI_ISL_494733, EPI_ISL_494734, EPI_ISL_494735, EPI_ISL_494736, EPI_ISL_494737, EPI_ISL_494738, EPI_ISL_494739, EPI_ISL_494740, EPI_ISL_494741, EPI_ISL_494742, EPI_ISL_494743, EPI_ISL_494744, EPI_ISL_494745, EPI_ISL_494746 | | | |
| see above | San Diego County Public Health Laboratory | Andersen lab at Scripps Research | SEARCH Alliance San Diego with Tracy Basler, Jovan Shephard, Brett Austin |
| EPI_ISL_495021 | Government Medical College, Bhavnagar | Gujarat Biotechnology Research Centre | Saklin Malek, Shirish Patel, Kairavi Desai, Raghawendra Kumar, Dinesh Kumar, Zuber Saiyed, Komal Patel, Labdhi Pandya, Afzal Ansari, Nikha Trivedi, Apurvasinh Puvar, Janvi Raval, Zarna Patel, Monika Gandhi, Pinal Trivedi, Maharshi Pandya, Nidhi Patel, Nitin Savaliya, R D Dixit, A M Kadri, Harsh Bakshi, Chaitanya Joshi, Madhvi Joshi |
| EPI_ISL_495177 | CSIR-Centre for Cellular and Molecular Biology | CSIR-Centre for Cellular and Molecular Biology | Sakshi Shambhavi, Lamuk Zaveri, Shagufta Khan, Namami Gaur, Nikhil Hajirnis, M Soujanya Reddy, Pratheusa Maccha,Tulasi Nagabandi, Purushotham Vodnala, Payel Mukherjee, Sofia Banu, Priya Singh,Onkar Kulkarni, Dhiviya Vedagiri, Divya Gupta, Vishal Sah, Santosh Kumar Kuncha, Krishnan Harinivas Harshan, Archana Bharadwaj Siva, Karthik Bharadwaj Tallapaka, G. Aditya Kumar, Koushick Sivakumar, Pooja Ramesh Gupta, Rajan Kumar Jha, Shraddha Vijay Lahoti, Rakesh K Mishra, Divya Tej Sowpati |
| EPI_ISL_495182 | CSIR-Centre for Cellular and Molecular Biology | CSIR-Centre for Cellular and Molecular Biology | Sakshi Shambhavi, Lamuk Zaveri, Shagufta Khan, Namami Gaur, Nikhil Hajirnis, M Soujanya Reddy, Pratheusa Maccha, Tulasi Nagabandi, Purushotham Vodnala, Payel Mukherjee, Sofia Banu, Priya Singh, Onkar Kulkarni, Dhiviya Vedagiri, Divya Gupta, Vishal Sah, Santosh Kumar Kuncha, Krishnan Harinivas Harshan, Archana Bharadwaj Siva, Karthik Bharadwaj Tallapaka, Deepak Kumar, Devi Prasad Vijayashankar, Disha Nanda, Divya Das, Jotin Gogoi, Manish Bhattacharjee, Rakesh K Mishra, Divya Tej Sowpati |
| EPI_ISL_495190 | CSIR-Centre for Cellular and Molecular Biology | CSIR-Centre for Cellular and Molecular Biology | Sakshi Shambhavi, Lamuk Zaveri, Shagufta Khan, Nikhil Hajirnis, M Soujanya Reddy, Pratheusa Maccha, Namami Gaur, Tulasi Nagabandi, Purushotham Vodnala, Payel Mukherjee, Sofia Banu, Priya Singh, Onkar Kulkarni, Dhiviya Vedagiri, Divya Gupta, Vishal Sah, Santosh Kumar Kuncha, Krishnan Harinivas Harshan, Archana Bharadwaj Siva, Karthik Bharadwaj Tallapaka,G. Aditya Kumar, Koushick Sivakumar, Rakesh K Mishra, Divya Tej Sowpati |
| EPI_ISL_495201 | CSIR-Centre for Cellular and Molecular Biology | CSIR-Centre for Cellular and Molecular Biology | Sakshi Shambhavi, Lamuk Zaveri, Shagufta Khan, Namami Gaur, Nikhil Hajirnis, M Soujanya Reddy, Pratheusa Maccha,Tulasi Nagabandi, Purushotham Vodnala, Payel Mukherjee, Sofia Banu, Priya Singh,Onkar Kulkarni, Dhiviya Vedagiri, Divya Gupta, Vishal Sah, Santosh Kumar Kuncha, Krishnan Harinivas Harshan, Archana Bharadwaj Siva, Karthik Bharadwaj Tallapaka, G. Aditya Kumar, Koushick Sivakumar, Pooja Ramesh Gupta, Rajan Kumar Jha, Shraddha Vijay Lahoti, Rakesh K Mishra, Divya Tej Sowpati |
| EPI_ISL_495213 | CSIR-Centre for Cellular and Molecular Biology | CSIR-Centre for Cellular and Molecular Biology | Sakshi Shambhavi, Lamuk Zaveri, Shagufta Khan, Nikhil Hajirnis, M Soujanya Reddy, Pratheusa Maccha, Namami Gaur, Tulasi Nagabandi, Purushotham Vodnala, Payel Mukherjee, Sofia Banu, Priya Singh, Onkar Kulkarni, Dhiviya Vedagiri, Divya Gupta, Vishal Sah, Santosh Kumar Kuncha, Krishnan |

| | | | |
|---|---|---|---|
| | | | Harinivas Harshan, Archana Bharadwaj Siva, Karthik Bharadwaj Tallapaka,G. Aditya Kumar, Koushick Sivakumar, Rakesh K Mishra, Divya Tej Sowpati |
| EPI_ISL_495218 | CSIR-Centre for Cellular and Molecular Biology | CSIR-Centre for Cellular and Molecular Biology | Sakshi Shambhavi, Lamuk Zaveri, Shagufta Khan, Namami Gaur, Nikhil Hajirnis, M Soujanya Reddy, Pratheusa Maccha, Tulasi Nagabandi, Purushotham Vodnala, Payel Mukherjee, Sofia Banu, Priya Singh, Onkar Kulkarni, Dhiviya Vedagiri, Divya Gupta, Vishal Sah, Santosh Kumar Kuncha, Krishnan Harinivas Harshan, Archana Bharadwaj Siva, Karthik Bharadwaj Tallapaka, Deepak Kumar, Devi Prasad Vijayashankar, Disha Nanda, Divya Das, Jotin Gogoi, Manish Bhattacharjee, Rakesh K Mishra, Divya Tej Sowpati |
| EPI_ISL_495248 | CSIR-Centre for Cellular and Molecular Biology | CSIR-Centre for Cellular and Molecular Biology | Sakshi Shambhavi, Lamuk Zaveri, Shagufta Khan, Nikhil Hajirnis, M Soujanya Reddy, Pratheusa Maccha, Namami Gaur, Tulasi Nagabandi, Purushotham Vodnala, Payel Mukherjee, Sofia Banu, Priya Singh, Onkar Kulkarni, Dhiviya Vedagiri, Divya Gupta, Vishal Sah, Santosh Kumar Kuncha, Krishnan Harinivas Harshan, Archana Bharadwaj Siva, Karthik Bharadwaj Tallapaka,G. Aditya Kumar, Koushick Sivakumar, Rakesh K Mishra, Divya Tej Sowpati |
| EPI_ISL_495256, EPI_ISL_495264 | CSIR-Centre for Cellular and Molecular Biology | CSIR-Centre for Cellular and Molecular Biology | Sakshi Shambhavi, Lamuk Zaveri, Shagufta Khan, Namami Gaur, Nikhil Hajirnis, M Soujanya Reddy, Pratheusa Maccha, Tulasi Nagabandi, Purushotham Vodnala, Payel Mukherjee, Sofia Banu, Priya Singh, Onkar Kulkarni, Dhiviya Vedagiri, Divya Gupta, Vishal Sah, Santosh Kumar Kuncha, Krishnan Harinivas Harshan, Archana Bharadwaj Siva, Karthik Bharadwaj Tallapaka, Deepak Kumar, Devi Prasad Vijayashankar, Disha Nanda, Divya Das, Jotin Gogoi, Manish Bhattacharjee, Rakesh K Mishra, Divya Tej Sowpati |
| EPI_ISL_495269 | CSIR-Centre for Cellular and Molecular Biology | CSIR-Centre for Cellular and Molecular Biology | Sakshi Shambhavi, Lamuk Zaveri, Shagufta Khan, Namami Gaur, Nikhil Hajirnis, M Soujanya Reddy, Pratheusa Maccha,Tulasi Nagabandi, Purushotham Vodnala, Payel Mukherjee, Sofia Banu, Priya Singh,Onkar Kulkarni, Dhiviya Vedagiri, Divya Gupta, Vishal Sah, Santosh Kumar Kuncha, Krishnan Harinivas Harshan, Archana Bharadwaj Siva, Karthik Bharadwaj Tallapaka, G. Aditya Kumar, Koushick Sivakumar, Pooja Ramesh Gupta, Rajan Kumar Jha, Shraddha Vijay Lahoti, Rakesh K Mishra, Divya Tej Sowpati |
| EPI_ISL_495330, EPI_ISL_495331, EPI_ISL_495332, EPI_ISL_495333, EPI_ISL_495334, EPI_ISL_495335, EPI_ISL_495336, EPI_ISL_495337, EPI_ISL_495338, EPI_ISL_495339, EPI_ISL_495340, EPI_ISL_495341, EPI_ISL_495342, EPI_ISL_495343, EPI_ISL_495344, EPI_ISL_495345, EPI_ISL_495346, EPI_ISL_495347, EPI_ISL_495348, EPI_ISL_495349, EPI_ISL_495350, EPI_ISL_495351, EPI_ISL_495352, EPI_ISL_495353, EPI_ISL_495354, EPI_ISL_495355, EPI_ISL_495356, EPI_ISL_495357, EPI_ISL_495358, EPI_ISL_495359, EPI_ISL_495360, EPI_ISL_495361, EPI_ISL_495362, EPI_ISL_495363, EPI_ISL_495364, EPI_ISL_495365, EPI_ISL_495366, EPI_ISL_495367, EPI_ISL_495368, EPI_ISL_495369, EPI_ISL_495370, EPI_ISL_495371, EPI_ISL_495372, EPI_ISL_495373, EPI_ISL_495374, EPI_ISL_495375, EPI_ISL_495376, EPI_ISL_495377, EPI_ISL_495378, EPI_ISL_495379, EPI_ISL_495380, EPI_ISL_495381, EPI_ISL_495382, EPI_ISL_495383, EPI_ISL_495384, EPI_ISL_495385, EPI_ISL_495386, EPI_ISL_495387, EPI_ISL_495388, EPI_ISL_495389, EPI_ISL_495390, EPI_ISL_495391, EPI_ISL_495392, EPI_ISL_495393, EPI_ISL_495394, EPI_ISL_495395, EPI_ISL_495396, EPI_ISL_495397, EPI_ISL_495398, EPI_ISL_495399, EPI_ISL_495400, EPI_ISL_495401, EPI_ISL_495402, EPI_ISL_495403, EPI_ISL_495404, EPI_ISL_495405, EPI_ISL_495406, EPI_ISL_495407, EPI_ISL_495408, EPI_ISL_495409, EPI_ISL_495410 | | | |
| see above | Florida Bureau of Public Health Laboratories | Florida Bureau of Public Health Laboratories | Sarah Schmedes, Jason Blanton |
| EPI_ISL_496339, EPI_ISL_496340, EPI_ISL_496341, EPI_ISL_496342, EPI_ISL_496343, EPI_ISL_496344, EPI_ISL_496345, EPI_ISL_496346, EPI_ISL_496347, EPI_ISL_496348, EPI_ISL_496349, EPI_ISL_496350, EPI_ISL_496351, EPI_ISL_496352, EPI_ISL_496353, EPI_ISL_496354, EPI_ISL_496355, EPI_ISL_496356, EPI_ISL_496357, EPI_ISL_496358, EPI_ISL_496359, EPI_ISL_496360, EPI_ISL_496361, EPI_ISL_496362, EPI_ISL_496363, EPI_ISL_496364, EPI_ISL_496365, EPI_ISL_496366, EPI_ISL_496367, EPI_ISL_496368, EPI_ISL_496369, EPI_ISL_496370, EPI_ISL_496371, EPI_ISL_496372, EPI_ISL_496373, EPI_ISL_496374, EPI_ISL_496375, EPI_ISL_496376 | | | |
| see above | Infectolab | Andersen lab at Scripps Research | SEARCH Alliance San Diego with Samuel Navarro Alvarez, Carlos A. Cota Haros, Octavio Renteria Pacheco |
| EPI_ISL_499083 | Instituto de Virologia "Dr. J. M. Vanella", Facultad de Ciencias Medicas, Universidad Nacional de Cordoba. Laboratorio Central de la Provincia de Cordoba, Argentina. Ministerio de Salud de la provincia de Cordoba, Argentina. | Laboratorio de Virología, Hospital de Niños Ricardo Gutiérrez, CABA, Argentina. | Sandra Gallego, Brenda Konigheim, Sebastian Blanco, Lorena Spinsanti, Javier Aguilar, Adrian Diaz, Gonzalo Castro, Gabriela Barbas, Mercedes Nabaes, Stephanie Goya, Monica Natale, Silvina Lusso, Mariana Viegas. |
| EPI_ISL_499460, EPI_ISL_499461, EPI_ISL_499462, EPI_ISL_499463, EPI_ISL_499464, EPI_ISL_499465, EPI_ISL_499466, EPI_ISL_499467, EPI_ISL_499468, EPI_ISL_499469, EPI_ISL_499470, EPI_ISL_499471, EPI_ISL_499473, EPI_ISL_499475, EPI_ISL_499476, EPI_ISL_499477, EPI_ISL_499480, EPI_ISL_499481, EPI_ISL_499483, EPI_ISL_499484, EPI_ISL_499486, EPI_ISL_499487, EPI_ISL_499488, EPI_ISL_499489, EPI_ISL_499491, EPI_ISL_499492, EPI_ISL_499496, EPI_ISL_499497, EPI_ISL_499498, EPI_ISL_499499, EPI_ISL_499500, EPI_ISL_499501, EPI_ISL_499509, EPI_ISL_499511, EPI_ISL_499512, EPI_ISL_499513, EPI_ISL_499514, EPI_ISL_499516, EPI_ISL_499517, EPI_ISL_499518, EPI_ISL_499519, EPI_ISL_499520, EPI_ISL_499521, EPI_ISL_499522, EPI_ISL_499523, EPI_ISL_499524, EPI_ISL_499525, EPI_ISL_499526, EPI_ISL_499527, EPI_ISL_499528, EPI_ISL_499529, EPI_ISL_499530, EPI_ISL_499531, EPI_ISL_499532, EPI_ISL_499533, EPI_ISL_499534, EPI_ISL_499535, EPI_ISL_499536, EPI_ISL_499537, EPI_ISL_499538, EPI_ISL_499539, EPI_ISL_499540, EPI_ISL_499541, EPI_ISL_499542, EPI_ISL_499543, EPI_ISL_499544, EPI_ISL_499545, EPI_ISL_499557, EPI_ISL_499564, EPI_ISL_499565, EPI_ISL_499566, EPI_ISL_499567, EPI_ISL_499569, EPI_ISL_499570, EPI_ISL_499571, EPI_ISL_499573, EPI_ISL_499574, EPI_ISL_499575, EPI_ISL_499576, EPI_ISL_499577, EPI_ISL_499578, EPI_ISL_499579, EPI_ISL_499580, EPI_ISL_499581, EPI_ISL_499582, EPI_ISL_499583, EPI_ISL_499584, EPI_ISL_499585, EPI_ISL_499586, EPI_ISL_499589, EPI_ISL_499590, EPI_ISL_499593, EPI_ISL_499594, EPI_ISL_499600, EPI_ISL_499604, EPI_ISL_499605, EPI_ISL_499607, EPI_ISL_499608, EPI_ISL_499611, EPI_ISL_499612, EPI_ISL_499613, EPI_ISL_499614, EPI_ISL_499615, EPI_ISL_499616, EPI_ISL_499617, EPI_ISL_499618, EPI_ISL_499619, EPI_ISL_499620, EPI_ISL_499621, EPI_ISL_499622, EPI_ISL_499624, EPI_ISL_499625, EPI_ISL_499627, EPI_ISL_499628, EPI_ISL_499629, EPI_ISL_499630, EPI_ISL_499632, EPI_ISL_499633, EPI_ISL_499634, EPI_ISL_499635, EPI_ISL_499636, EPI_ISL_499637, EPI_ISL_499638, EPI_ISL_499639, EPI_ISL_499640, EPI_ISL_499641, EPI_ISL_499642, EPI_ISL_499643, EPI_ISL_499644, EPI_ISL_499645, EPI_ISL_499646, EPI_ISL_499647, EPI_ISL_499648, EPI_ISL_499649, EPI_ISL_499650, EPI_ISL_499651, EPI_ISL_499652, EPI_ISL_499653, EPI_ISL_499654, EPI_ISL_499655, EPI_ISL_499656, EPI_ISL_499657, EPI_ISL_499658, EPI_ISL_499659, EPI_ISL_499660, EPI_ISL_499661, EPI_ISL_499662, EPI_ISL_499663, EPI_ISL_499664, EPI_ISL_499665, EPI_ISL_499666, EPI_ISL_499667, EPI_ISL_499668, EPI_ISL_499669, EPI_ISL_499670, EPI_ISL_499671, EPI_ISL_499672, EPI_ISL_499673, EPI_ISL_499674, EPI_ISL_499675, EPI_ISL_499676, EPI_ISL_499677, EPI_ISL_499678, EPI_ISL_499679, EPI_ISL_499680, EPI_ISL_499681, EPI_ISL_499682, EPI_ISL_499683, EPI_ISL_499684, EPI_ISL_499685, EPI_ISL_499686, EPI_ISL_499687, EPI_ISL_499688, EPI_ISL_499689, EPI_ISL_499690, EPI_ISL_499691, EPI_ISL_499692, EPI_ISL_499693, EPI_ISL_499694, EPI_ISL_499695, EPI_ISL_499696, EPI_ISL_499697, EPI_ISL_499698, EPI_ISL_499699, EPI_ISL_499700, EPI_ISL_499701, EPI_ISL_499702, EPI_ISL_499703, EPI_ISL_499704, EPI_ISL_499705, EPI_ISL_499706, EPI_ISL_499707, EPI_ISL_499708, EPI_ISL_499709, EPI_ISL_499710, EPI_ISL_499711, EPI_ISL_499712, EPI_ISL_499713, EPI_ISL_499714, EPI_ISL_499715, EPI_ISL_499716, EPI_ISL_499717, EPI_ISL_499718, EPI_ISL_499719, EPI_ISL_499720, EPI_ISL_499721, EPI_ISL_499722, EPI_ISL_499723, EPI_ISL_499724, EPI_ISL_499725, EPI_ISL_499726, EPI_ISL_499727, EPI_ISL_499728, EPI_ISL_499729, EPI_ISL_499730, EPI_ISL_499731, EPI_ISL_499732, EPI_ISL_499733, EPI_ISL_499734, EPI_ISL_499735, EPI_ISL_499736, EPI_ISL_499737, EPI_ISL_499738, EPI_ISL_499739, EPI_ISL_499740, EPI_ISL_499741, EPI_ISL_499742, EPI_ISL_499743, EPI_ISL_499744, EPI_ISL_499745, EPI_ISL_499746, EPI_ISL_499747, EPI_ISL_499748, EPI_ISL_499749, EPI_ISL_499750, EPI_ISL_499751, EPI_ISL_499752, EPI_ISL_499753, EPI_ISL_499754, EPI_ISL_499755, EPI_ISL_499756, EPI_ISL_499757, EPI_ISL_499758, EPI_ISL_499759, EPI_ISL_499760, EPI_ISL_499761, EPI_ISL_499762, EPI_ISL_499763, EPI_ISL_499764, EPI_ISL_499765, EPI_ISL_499766, EPI_ISL_499767, EPI_ISL_499768, EPI_ISL_499769, EPI_ISL_499770, EPI_ISL_499771, EPI_ISL_499772, EPI_ISL_499773, EPI_ISL_499774, EPI_ISL_499775, EPI_ISL_499776, EPI_ISL_499811, EPI_ISL_499812, EPI_ISL_499813, EPI_ISL_499814, EPI_ISL_499815, EPI_ISL_499816, EPI_ISL_499817, EPI_ISL_499818, EPI_ISL_499819, EPI_ISL_499820, EPI_ISL_499821, EPI_ISL_499822, EPI_ISL_499823, EPI_ISL_499824, EPI_ISL_499825, EPI_ISL_499826, EPI_ISL_499827, EPI_ISL_499831, EPI_ISL_499832, EPI_ISL_499833, EPI_ISL_499834, EPI_ISL_499835, EPI_ISL_499836, EPI_ISL_499837, EPI_ISL_499838, EPI_ISL_499840, EPI_ISL_499841, EPI_ISL_499842, EPI_ISL_499843, EPI_ISL_499844, EPI_ISL_499845, EPI_ISL_499846, EPI_ISL_499847, EPI_ISL_499848, EPI_ISL_499849, EPI_ISL_499850, EPI_ISL_499854, EPI_ISL_499856, EPI_ISL_499857, EPI_ISL_499859, EPI_ISL_499860, EPI_ISL_499862, EPI_ISL_499863, EPI_ISL_499864, EPI_ISL_499865, EPI_ISL_499866, EPI_ISL_499867, EPI_ISL_499868, EPI_ISL_499870, EPI_ISL_499871, EPI_ISL_499872, EPI_ISL_499873, EPI_ISL_499874, EPI_ISL_499875, EPI_ISL_499876, EPI_ISL_499877, EPI_ISL_499878, EPI_ISL_499879, EPI_ISL_499880, EPI_ISL_499881, EPI_ISL_499882, EPI_ISL_499883, EPI_ISL_499884, EPI_ISL_499885, EPI_ISL_499886, EPI_ISL_499887, EPI_ISL_499888, EPI_ISL_499889, EPI_ISL_499890, EPI_ISL_499891, EPI_ISL_499892, EPI_ISL_499893, EPI_ISL_499896, EPI_ISL_499898, EPI_ISL_499899, EPI_ISL_499900, EPI_ISL_499901, EPI_ISL_499902, EPI_ISL_499903, EPI_ISL_499904, EPI_ISL_499906, EPI_ISL_499907, EPI_ISL_499909, EPI_ISL_499911, EPI_ISL_499914, EPI_ISL_499915, EPI_ISL_499916, EPI_ISL_499917, EPI_ISL_499918, EPI_ISL_499919, EPI_ISL_499920, EPI_ISL_499921, EPI_ISL_499922, EPI_ISL_499923, EPI_ISL_499924, EPI_ISL_499925, EPI_ISL_499926, EPI_ISL_499927, EPI_ISL_499928, EPI_ISL_499929, EPI_ISL_499930, EPI_ISL_499931, EPI_ISL_499932, EPI_ISL_499934, EPI_ISL_499935, EPI_ISL_499936, EPI_ISL_499937, EPI_ISL_499938, EPI_ISL_499939, EPI_ISL_499940, EPI_ISL_499941, EPI_ISL_499942, EPI_ISL_499943, EPI_ISL_499944, EPI_ISL_499945, EPI_ISL_499946, EPI_ISL_499948, EPI_ISL_499949, EPI_ISL_499951, EPI_ISL_499952, EPI_ISL_499953, EPI_ISL_499954, EPI_ISL_499955, EPI_ISL_499956, EPI_ISL_499957, EPI_ISL_499958, EPI_ISL_499959, EPI_ISL_499960, EPI_ISL_499961, EPI_ISL_499962, EPI_ISL_499963, EPI_ISL_499964, EPI_ISL_499965, EPI_ISL_499967, EPI_ISL_499984, EPI_ISL_499985, EPI_ISL_499986, EPI_ISL_499987, EPI_ISL_499988, EPI_ISL_499989, EPI_ISL_499990, EPI_ISL_499991, EPI_ISL_499992, EPI_ISL_499993, EPI_ISL_499994, EPI_ISL_499995, EPI_ISL_499996, EPI_ISL_499997, EPI_ISL_499998, EPI_ISL_499999, EPI_ISL_500000, EPI_ISL_500001, EPI_ISL_500002, EPI_ISL_500003, EPI_ISL_500004, EPI_ISL_500005, EPI_ISL_500006, EPI_ISL_500007, EPI_ISL_500008, EPI_ISL_500009, EPI_ISL_500010, EPI_ISL_500011, EPI_ISL_500012, EPI_ISL_500013, EPI_ISL_500014, EPI_ISL_500015, EPI_ISL_500016, EPI_ISL_500017, EPI_ISL_500018, EPI_ISL_500019, EPI_ISL_500020, EPI_ISL_500021, EPI_ISL_500022, EPI_ISL_500023, EPI_ISL_500024, EPI_ISL_500025, EPI_ISL_500026, EPI_ISL_500027, EPI_ISL_500028, EPI_ISL_500029, EPI_ISL_500030, EPI_ISL_500031, EPI_ISL_500032, EPI_ISL_500033, EPI_ISL_500034, EPI_ISL_500035, EPI_ISL_500036, EPI_ISL_500037, EPI_ISL_500038, EPI_ISL_500039, EPI_ISL_500040, EPI_ISL_500041, EPI_ISL_500042, EPI_ISL_500043, EPI_ISL_500044, EPI_ISL_500045, EPI_ISL_500046, EPI_ISL_500047, EPI_ISL_500048, EPI_ISL_500049, EPI_ISL_500050, EPI_ISL_500051, EPI_ISL_500052, EPI_ISL_500053, EPI_ISL_500054, EPI_ISL_500055, EPI_ISL_500056, EPI_ISL_500057, EPI_ISL_500058, EPI_ISL_500059, EPI_ISL_500060, EPI_ISL_500061, EPI_ISL_500062, EPI_ISL_500063, EPI_ISL_500064, EPI_ISL_500065, EPI_ISL_500066, EPI_ISL_500067, EPI_ISL_500068, EPI_ISL_500069, EPI_ISL_500070, EPI_ISL_500071, EPI_ISL_500072, EPI_ISL_500073, EPI_ISL_500074, EPI_ISL_500075, EPI_ISL_500076, EPI_ISL_500077, EPI_ISL_500078, EPI_ISL_500079, EPI_ISL_500080, EPI_ISL_500081, EPI_ISL_500082, EPI_ISL_500083, EPI_ISL_500084, EPI_ISL_500085, EPI_ISL_500086, EPI_ISL_500087, EPI_ISL_500088, EPI_ISL_500089, EPI_ISL_500090, EPI_ISL_500091, EPI_ISL_500092, EPI_ISL_500093, EPI_ISL_500094, EPI_ISL_500095, EPI_ISL_500096, EPI_ISL_500097, EPI_ISL_500098, EPI_ISL_500099, EPI_ISL_500100, EPI_ISL_500101, EPI_ISL_500102, EPI_ISL_500103, EPI_ISL_500104, EPI_ISL_500105, EPI_ISL_500106, EPI_ISL_500107, EPI_ISL_500108, EPI_ISL_500109, EPI_ISL_500110, EPI_ISL_500111, EPI_ISL_500112, EPI_ISL_500113, EPI_ISL_500114, EPI_ISL_500115, EPI_ISL_500116, EPI_ISL_500117, EPI_ISL_500118, EPI_ISL_500119, EPI_ISL_500120, EPI_ISL_500121, EPI_ISL_500122, EPI_ISL_500123, EPI_ISL_500124, EPI_ISL_500125, EPI_ISL_500126, EPI_ISL_500127, EPI_ISL_500128, EPI_ISL_500129, EPI_ISL_500130, EPI_ISL_500131, EPI_ISL_500132, EPI_ISL_500133, EPI_ISL_500134, EPI_ISL_500135, EPI_ISL_500136, EPI_ISL_500137, EPI_ISL_500138, EPI_ISL_500139, EPI_ISL_500140, EPI_ISL_500141, EPI_ISL_500142, EPI_ISL_500143, EPI_ISL_500144, EPI_ISL_500145, EPI_ISL_500146, EPI_ISL_500147, EPI_ISL_500148, EPI_ISL_500149, EPI_ISL_500150, EPI_ISL_500151, EPI_ISL_500152, EPI_ISL_500153, EPI_ISL_500154, EPI_ISL_500155, EPI_ISL_500156 | | | |
| see above | Liverpool Clinical Laboratories | COVID-19 Genomics UK (COG-UK) Consortium | Sam Haldenby, Anita Lucaci, Steve Paterson, Julian Hiscox, Alistair Darby, M Almsaud, A Alrezaihi, Muhannad Alruwaili, Stuart D Armstrong, Jones Benjamin, Eleanor G Bentley, Anu Chawla, Jordan J Clark, Angela Cowell, Richard Eccles, Isabel García-Dorival, Matthew Gemmell, Alessandro Gerada, PKF Gilmore, Richard Gregory, Ximeng Han, Catherine Hartley, Margaret Hughes, Miren Iturriza-Gomara, James Johnson, L Luu, Jenifer Manson, Charlotte Nelson, Elaine O'Toole, Cassie Olateju, Rebekah Penrice-Randal , Lucille Rainbow, N.P Randle, Trevor Ian Robinson, Parul Sharma, Ghada T Shawli, James P Stewart, Neil Swainston, Ecaterina Vamos, Joanne Watts, Mark Whitehead |
| EPI_ISL_501614, EPI_ISL_501615, EPI_ISL_501616, EPI_ISL_501617, EPI_ISL_501618, EPI_ISL_501619, EPI_ISL_501620, EPI_ISL_501621, EPI_ISL_501622, EPI_ISL_501623, EPI_ISL_501624, EPI_ISL_501625, EPI_ISL_501626, EPI_ISL_501627, EPI_ISL_501628, EPI_ISL_501629, EPI_ISL_501630, EPI_ISL_501631 | | | |
| see above | Lab Microbiology, Pathology Department, William Harvey Hospital | Wellcome Sanger Institute for the COVID-19 Genomics UK (COG-UK) consortium | Samuel Moses, Hannah Lowe, Felicity Ryan and Alex Alderton, Roberto Amato, Sonia Goncalves, Ewan Harrison, David K. Jackson, Ian Johnston, Dominic Kwiatkowski, Cordelia Langford, John Sillitoe on behalf of the Wellcome Sanger Institute COVID-19 Surveillance Team (http://www.sanger.ac.uk/covid-team) |
| EPI_ISL_508708, EPI_ISL_508709, EPI_ISL_508710, EPI_ISL_508711, EPI_ISL_508712, EPI_ISL_508713, EPI_ISL_508714, EPI_ISL_508715, EPI_ISL_508716, EPI_ISL_508717, EPI_ISL_508718, EPI_ISL_508719, EPI_ISL_508720, EPI_ISL_508721, EPI_ISL_508723, EPI_ISL_508724, EPI_ISL_508725, EPI_ISL_508726, EPI_ISL_508727, EPI_ISL_508728, EPI_ISL_508729, EPI_ISL_508730, EPI_ISL_508731, EPI_ISL_508732, EPI_ISL_508733, EPI_ISL_508734, EPI_ISL_508735, EPI_ISL_508736, EPI_ISL_508737, EPI_ISL_508738, EPI_ISL_508739, EPI_ISL_508740, EPI_ISL_508741, EPI_ISL_508742, EPI_ISL_508743, EPI_ISL_508744, EPI_ISL_508745, EPI_ISL_508746, EPI_ISL_508747, EPI_ISL_508748, EPI_ISL_508749, EPI_ISL_508750, EPI_ISL_508751, EPI_ISL_508752, EPI_ISL_508753, EPI_ISL_508754, EPI_ISL_508755, EPI_ISL_508756, EPI_ISL_508757, EPI_ISL_508758, EPI_ISL_508759, EPI_ISL_508760, EPI_ISL_508761, EPI_ISL_508762, EPI_ISL_508763, EPI_ISL_508765, EPI_ISL_508766, EPI_ISL_508767, EPI_ISL_508768, EPI_ISL_508769, EPI_ISL_508770, EPI_ISL_508771, EPI_ISL_508772, EPI_ISL_508773, EPI_ISL_508774, EPI_ISL_508775, EPI_ISL_508776, EPI_ISL_508777, EPI_ISL_508778, EPI_ISL_508779, EPI_ISL_508780, EPI_ISL_508781, EPI_ISL_508782, EPI_ISL_508783, EPI_ISL_508784, EPI_ISL_508785, EPI_ISL_508786, EPI_ISL_508787, EPI_ISL_508789, EPI_ISL_508806, EPI_ISL_508808, EPI_ISL_508809, EPI_ISL_508811, EPI_ISL_508812, EPI_ISL_508813, EPI_ISL_508814, EPI_ISL_508815, EPI_ISL_508816, EPI_ISL_508818, EPI_ISL_508819, EPI_ISL_508820, EPI_ISL_508823, EPI_ISL_508824, EPI_ISL_508825, EPI_ISL_508826, EPI_ISL_508827, EPI_ISL_508829, EPI_ISL_508830, EPI_ISL_508831, EPI_ISL_508832, EPI_ISL_508833, EPI_ISL_508834, EPI_ISL_508835, EPI_ISL_508836, EPI_ISL_508842, EPI_ISL_508843, EPI_ISL_508844, EPI_ISL_508845, EPI_ISL_508846, EPI_ISL_508847, EPI_ISL_508848, EPI_ISL_508852, EPI_ISL_508853, EPI_ISL_508854, EPI_ISL_508855, EPI_ISL_508856, EPI_ISL_508857, EPI_ISL_508859, EPI_ISL_508861, EPI_ISL_509720, EPI_ISL_509723, EPI_ISL_509729, EPI_ISL_509733, EPI_ISL_509737, EPI_ISL_509743, EPI_ISL_509744, EPI_ISL_509746, EPI_ISL_509749, EPI_ISL_509750, EPI_ISL_509755, EPI_ISL_509777, EPI_ISL_509779, EPI_ISL_509782, EPI_ISL_509785, EPI_ISL_509790, EPI_ISL_509796, EPI_ISL_509797, EPI_ISL_509798 | | | |
| see above | Florida Bureau of Public Health Laboratories | Florida Bureau of Public Health Laboratories | Sarah Schmedes, Jason Blanton |

| | | | |
|---|---|---|---|
| EPI_ISL_512158, EPI_ISL_512159, EPI_ISL_512160, EPI_ISL_512161, EPI_ISL_512162, EPI_ISL_512163, EPI_ISL_512164, EPI_ISL_512165, EPI_ISL_512166, EPI_ISL_512167, EPI_ISL_512168, EPI_ISL_512169, EPI_ISL_512170, EPI_ISL_512171, EPI_ISL_512172, EPI_ISL_512173, EPI_ISL_512174, EPI_ISL_512175, EPI_ISL_512176, EPI_ISL_512177, EPI_ISL_512178, EPI_ISL_512179, EPI_ISL_512180, EPI_ISL_512181, EPI_ISL_512182, EPI_ISL_512183, EPI_ISL_512184, EPI_ISL_512185, EPI_ISL_512186, EPI_ISL_512187, EPI_ISL_512188, EPI_ISL_512189, EPI_ISL_512190, EPI_ISL_512191, EPI_ISL_512192, EPI_ISL_512193, EPI_ISL_512194, EPI_ISL_512195, EPI_ISL_512196, EPI_ISL_512197, EPI_ISL_512198, EPI_ISL_512199, EPI_ISL_512200, EPI_ISL_512201, EPI_ISL_512202, EPI_ISL_512203, EPI_ISL_512204, EPI_ISL_512205, EPI_ISL_512206, EPI_ISL_512207, EPI_ISL_512208, EPI_ISL_512209, EPI_ISL_512210, EPI_ISL_512211, EPI_ISL_512212, EPI_ISL_512213, EPI_ISL_512214, EPI_ISL_512215, EPI_ISL_512216, EPI_ISL_512217, EPI_ISL_512218, EPI_ISL_512219, EPI_ISL_512220, EPI_ISL_512221, EPI_ISL_512222, EPI_ISL_512223, EPI_ISL_512224, EPI_ISL_512225, EPI_ISL_512226, EPI_ISL_512227, EPI_ISL_512228, EPI_ISL_512229, EPI_ISL_512230, EPI_ISL_512231, EPI_ISL_512232, EPI_ISL_512233, EPI_ISL_512234, EPI_ISL_512235, EPI_ISL_512236, EPI_ISL_512237, EPI_ISL_512238, EPI_ISL_512239, EPI_ISL_512240, EPI_ISL_512241, EPI_ISL_512242, EPI_ISL_512243, EPI_ISL_512244, EPI_ISL_512245, EPI_ISL_512246, EPI_ISL_512247, EPI_ISL_512248, EPI_ISL_512249, EPI_ISL_512250, EPI_ISL_512251, EPI_ISL_512252, EPI_ISL_512253, EPI_ISL_512254, EPI_ISL_512255, EPI_ISL_512256, EPI_ISL_512257, EPI_ISL_512258, EPI_ISL_512259, EPI_ISL_512260, EPI_ISL_512261, EPI_ISL_512262, EPI_ISL_512263, EPI_ISL_512264, EPI_ISL_512265, EPI_ISL_512266, EPI_ISL_512267, EPI_ISL_512268, EPI_ISL_512269, EPI_ISL_512270, EPI_ISL_512271, EPI_ISL_512272, EPI_ISL_512273 | | | |
| see above | San Diego County Public Health Laboratory | Andersen lab at Scripps Research | SEARCH Alliance San Diego with Tracy Basler, Jovan Shephard, Brett Austin |
| EPI_ISL_512342, EPI_ISL_512343, EPI_ISL_512344, EPI_ISL_512345, EPI_ISL_512346, EPI_ISL_512347 | Liverpool Clinical Laboratories | COVID-19 Genomics UK (COG-UK) Consortium | Sam Haldenby, Anita Lucaci, Steve Paterson, Julian Hiscox, Alistair Darby, M Almsaud, A Alrezaihi, Muhannad Alruwaili, Stuart D Armstrong, Jones Benjamin, Eleanor G Bentley, Anu Chawla, Jordan J Clark, Angela Cowell, Richard Eccles, Isabel García-Dorival, Matthew Gemmell, Alessandro Gerada, PKF Gilmore, Richard Gregory, Ximeng Han, Catherine Hartley, Margaret Hughes, Miren Iturriza-Gomara, James Johnson, L Luu, Jenifer Manson, Charlotte Nelson, Elaine O'Toole, Cassie Olateju, Rebekah Penrice-Randal , Lucille Rainbow, N.P Randle, Trevor Ian Robinson, Parul Sharma, Ghada T Shawli, James P Stewart, Neil Swainston, Ecaterina Vamos, Joanne Watts, Mark Whitehead |
| EPI_ISL_512548, EPI_ISL_512550, EPI_ISL_512551, EPI_ISL_512553, EPI_ISL_512554, EPI_ISL_512559, EPI_ISL_512569, EPI_ISL_512570, EPI_ISL_512574, EPI_ISL_512576, EPI_ISL_512577, EPI_ISL_512579, EPI_ISL_512580, EPI_ISL_512581, EPI_ISL_512583, EPI_ISL_512584, EPI_ISL_512585, EPI_ISL_512586, EPI_ISL_512587, EPI_ISL_512588, EPI_ISL_512589, EPI_ISL_512591, EPI_ISL_512592, EPI_ISL_512593, EPI_ISL_512596, EPI_ISL_512676, EPI_ISL_512677 | | | |
| see above | Florida Bureau of Public Health Laboratories | Florida Bureau of Public Health Laboratories | Sarah Schmedes, Jason Blanton |
| EPI_ISL_512946, EPI_ISL_512947, EPI_ISL_512948, EPI_ISL_512949, EPI_ISL_512950, EPI_ISL_512951, EPI_ISL_512952, EPI_ISL_512953, EPI_ISL_512954, EPI_ISL_512955, EPI_ISL_512956, EPI_ISL_512957, EPI_ISL_512958, EPI_ISL_512959, EPI_ISL_512960, EPI_ISL_512961, EPI_ISL_512962, EPI_ISL_512963, EPI_ISL_512964, EPI_ISL_512965, EPI_ISL_512966, EPI_ISL_512967, EPI_ISL_512968, EPI_ISL_512969, EPI_ISL_512970, EPI_ISL_512971, EPI_ISL_512973, EPI_ISL_512974, EPI_ISL_512975, EPI_ISL_512976, EPI_ISL_512977, EPI_ISL_512978, EPI_ISL_512979, EPI_ISL_512980, EPI_ISL_512981, EPI_ISL_512982, EPI_ISL_512983, EPI_ISL_512984, EPI_ISL_512985, EPI_ISL_512986, EPI_ISL_512987, EPI_ISL_512988, EPI_ISL_513168, EPI_ISL_513169, EPI_ISL_513170, EPI_ISL_513171, EPI_ISL_513172, EPI_ISL_513173, EPI_ISL_513174 | | | |
| see above | Pathogen Genomics Lab King Abdullah University of Science and Technology(KAUST) | Pathogen Genomics Lab King Abdullah University of Science and Technology(KAUST) | Sara Mfarrej, Raeece Naeem, Rahul P Salunke, Sharif Hala, Fadwa Alofi, Amit Kumar Subudhi, Fathia Ben Rached, Afrah Alsomali, Jumana Taha, Abdulaziz Alahmadi, Asim Khogeer, Nashwa Al-khotani, Anwar Hashem, Naif Almontashiri, Arnab Pain |
| EPI_ISL_513585, EPI_ISL_513586, EPI_ISL_513587, EPI_ISL_513588 | Microbiology, Pathology, Kettering General Hospital | Wellcome Sanger Institute for the COVID-19 Genomics UK (COG-UK) consortium | Sahar El-dirdiri, Anita Kenyon, Thomas Davis and Alex Alderton, Roberto Amato, Sonia Goncalves, Ewan Harrison, David K. Jackson, Ian Johnston, Dominic Kwiatkowski, Cordelia Langford, John Sillitoe on behalf of the Wellcome Sanger Institute COVID-19 Surveillance Team (http://www.sanger.ac.uk/covid-team) |
| EPI_ISL_514139, EPI_ISL_514141, EPI_ISL_514144, EPI_ISL_514147, EPI_ISL_514148, EPI_ISL_514149, EPI_ISL_514150, EPI_ISL_514151, EPI_ISL_514154, EPI_ISL_514158, EPI_ISL_514164, EPI_ISL_514165, EPI_ISL_514166, EPI_ISL_514167, EPI_ISL_514170, EPI_ISL_514172, EPI_ISL_514173, EPI_ISL_514175, EPI_ISL_514182, EPI_ISL_514190, EPI_ISL_514191, EPI_ISL_514192, EPI_ISL_514193, EPI_ISL_514195, EPI_ISL_514196, EPI_ISL_514198, EPI_ISL_514199, EPI_ISL_514206, EPI_ISL_514208, EPI_ISL_514209, EPI_ISL_514210, EPI_ISL_514215, EPI_ISL_514217, EPI_ISL_514218 | | | |
| see above | Florida Bureau of Public Health Laboratories | Florida Bureau of Public Health Laboratories | Sarah Schmedes, Jason Blanton |
| EPI_ISL_514591 | B.J. Medical College and Civil hospital, Ahmedabad | Gujarat Biotechnology Research Centre | Sanjay Kapadia,Apurvasinh Puvar,Janvi Raval,Zarna Patel, Monika Gandhi, Pinal Trivedi, Maharshi Pandya, Nidhi Patel, Nitin Savaliya, Raghawendra Kumar, Dinesh Kumar, Zuber Saiyed, Komal Patel, Labdhi Pandya, Afzal Ansari, Nikha Trivedi, Pranay Shah, Kamlesh J Upadhyay, R D Dixit, A M Kadri, Harsh Bakshi, Chaitanya Joshi, Madhvi Joshi |
| EPI_ISL_514609 | B.J. Medical College and Civil hospital, Ahmedabad | Gujarat Biotechnology Research Centre | Sanjay Kapadia, Apurvasinh Puvar, Janvi Raval, Zarna Patel, Monika Gandhi, Pinal Trivedi, Maharshi Pandya, Nidhi Patel, Nitin Savaliya, Raghawendra Kumar, Dinesh Kumar, Zuber Saiyed, Komal Patel, Labdhi Pandya, Afzal Ansari, Nikha Trivedi, Pranay Shah, Kamlesh J Upadhyay, R D Dixit, A M Kadri, Harsh Bakshi, Chaitanya Joshi, Madhvi Joshi |
| EPI_ISL_517001, EPI_ISL_517002, EPI_ISL_517003, EPI_ISL_517004, EPI_ISL_517005, EPI_ISL_517006, EPI_ISL_517007, EPI_ISL_517008, EPI_ISL_517009, EPI_ISL_517010, EPI_ISL_517011, EPI_ISL_517012, EPI_ISL_517013, EPI_ISL_517014, EPI_ISL_517015, EPI_ISL_517016, EPI_ISL_517017, EPI_ISL_517018, EPI_ISL_517019, EPI_ISL_517020, EPI_ISL_517021, EPI_ISL_517022, EPI_ISL_517023, EPI_ISL_517024, EPI_ISL_517025, EPI_ISL_517026, EPI_ISL_517027, EPI_ISL_517028, EPI_ISL_517029, EPI_ISL_517030, EPI_ISL_517031, EPI_ISL_517032, EPI_ISL_517033, EPI_ISL_517034, EPI_ISL_517035, EPI_ISL_517036, EPI_ISL_517037, EPI_ISL_517038, EPI_ISL_517039, EPI_ISL_517040, EPI_ISL_517041, EPI_ISL_517042, EPI_ISL_517043, EPI_ISL_517044, EPI_ISL_517045, EPI_ISL_517046, EPI_ISL_517047, EPI_ISL_517048, EPI_ISL_517049, EPI_ISL_517050, EPI_ISL_517051, EPI_ISL_517052, EPI_ISL_517053, EPI_ISL_517054, EPI_ISL_517055, EPI_ISL_517056, EPI_ISL_517057, EPI_ISL_517058, EPI_ISL_517059, EPI_ISL_517060, EPI_ISL_517061, EPI_ISL_517062, EPI_ISL_517063, EPI_ISL_517064, EPI_ISL_517065, EPI_ISL_517066, EPI_ISL_517067, EPI_ISL_517068, EPI_ISL_517069, EPI_ISL_517070, EPI_ISL_517071, EPI_ISL_517072, EPI_ISL_517073, EPI_ISL_517074, EPI_ISL_517075, EPI_ISL_517076, EPI_ISL_517077, EPI_ISL_517078, EPI_ISL_517079, EPI_ISL_517080, EPI_ISL_517081, EPI_ISL_517082, EPI_ISL_517083, EPI_ISL_517084, EPI_ISL_517085, EPI_ISL_517086, EPI_ISL_517087, EPI_ISL_517088, EPI_ISL_517089, EPI_ISL_517090, EPI_ISL_517091, EPI_ISL_517092, EPI_ISL_517093, EPI_ISL_517094, EPI_ISL_517095, EPI_ISL_517096, EPI_ISL_517097, EPI_ISL_517098, EPI_ISL_517099, EPI_ISL_517100, EPI_ISL_517101, EPI_ISL_517102, EPI_ISL_517103, EPI_ISL_517104, EPI_ISL_517105, EPI_ISL_517106, EPI_ISL_517107, EPI_ISL_517108, EPI_ISL_517109, EPI_ISL_517110, EPI_ISL_517111, EPI_ISL_517112, EPI_ISL_517113, EPI_ISL_517114, EPI_ISL_517115, EPI_ISL_517116, EPI_ISL_517117, EPI_ISL_517118, EPI_ISL_517119, EPI_ISL_517120, EPI_ISL_517121, EPI_ISL_517122, EPI_ISL_517123, EPI_ISL_517124, EPI_ISL_517125, EPI_ISL_517126, EPI_ISL_517127, EPI_ISL_517128, EPI_ISL_517129, EPI_ISL_517130, EPI_ISL_517131, EPI_ISL_517132, EPI_ISL_517133, EPI_ISL_517134, EPI_ISL_517135, EPI_ISL_517136, EPI_ISL_517137, EPI_ISL_517138, EPI_ISL_517139, EPI_ISL_517140, EPI_ISL_517141, EPI_ISL_517142, EPI_ISL_517143, EPI_ISL_517144, EPI_ISL_517145, EPI_ISL_517146, EPI_ISL_517147, EPI_ISL_517148, EPI_ISL_517149, EPI_ISL_517150, EPI_ISL_517151, EPI_ISL_517152, EPI_ISL_517153, EPI_ISL_517154, EPI_ISL_517155, EPI_ISL_517156, EPI_ISL_517157, EPI_ISL_517158, EPI_ISL_517159, EPI_ISL_517160, EPI_ISL_517161, EPI_ISL_517162, EPI_ISL_517163, EPI_ISL_517164, EPI_ISL_517165, EPI_ISL_517166, EPI_ISL_517167, EPI_ISL_517168, EPI_ISL_517169, EPI_ISL_517170, EPI_ISL_517171, EPI_ISL_517172, EPI_ISL_517173, EPI_ISL_517174, EPI_ISL_517175, EPI_ISL_517176, EPI_ISL_517177, EPI_ISL_517178, EPI_ISL_517179, EPI_ISL_517180, EPI_ISL_517181, EPI_ISL_517182, EPI_ISL_517183, EPI_ISL_517184, EPI_ISL_517185, EPI_ISL_517186, EPI_ISL_517187, EPI_ISL_517188, EPI_ISL_517189, EPI_ISL_517190, EPI_ISL_517191, EPI_ISL_517192, EPI_ISL_517193, EPI_ISL_517194, EPI_ISL_517195, EPI_ISL_517196, EPI_ISL_517197, EPI_ISL_517198, EPI_ISL_517199, EPI_ISL_517200, EPI_ISL_517201, EPI_ISL_517202, EPI_ISL_517203, EPI_ISL_517204, EPI_ISL_517205, EPI_ISL_517206, EPI_ISL_517207, EPI_ISL_517208, EPI_ISL_517209, EPI_ISL_517210, EPI_ISL_517211, EPI_ISL_517212, EPI_ISL_517213, EPI_ISL_517214, EPI_ISL_517215, EPI_ISL_517216, EPI_ISL_517217, EPI_ISL_517218, EPI_ISL_517219, EPI_ISL_517220, EPI_ISL_517221, EPI_ISL_517222, EPI_ISL_517223, EPI_ISL_517224, EPI_ISL_517225, EPI_ISL_517226, EPI_ISL_517227, EPI_ISL_517228, EPI_ISL_517229, EPI_ISL_517230, EPI_ISL_517231, EPI_ISL_517232, EPI_ISL_517233, EPI_ISL_517234, EPI_ISL_517235, EPI_ISL_517236, EPI_ISL_517237, EPI_ISL_517238, EPI_ISL_517239, EPI_ISL_517240, EPI_ISL_517241, EPI_ISL_517242, EPI_ISL_517243, EPI_ISL_517244, EPI_ISL_517245, EPI_ISL_517246, EPI_ISL_517247, EPI_ISL_517248, EPI_ISL_517249, EPI_ISL_517250, EPI_ISL_517251, EPI_ISL_517252, EPI_ISL_517253, EPI_ISL_517254, EPI_ISL_517255, EPI_ISL_517256, EPI_ISL_517257, EPI_ISL_517258, EPI_ISL_517259, EPI_ISL_517260, EPI_ISL_517261, EPI_ISL_517262, EPI_ISL_517263, EPI_ISL_517264, EPI_ISL_517265, EPI_ISL_517266, EPI_ISL_517267, EPI_ISL_517268, EPI_ISL_517269, EPI_ISL_517270, EPI_ISL_517271, EPI_ISL_517272, EPI_ISL_517273, EPI_ISL_517274, EPI_ISL_517275, EPI_ISL_517276, EPI_ISL_517277, EPI_ISL_517278, EPI_ISL_517279, EPI_ISL_517280, EPI_ISL_517281, EPI_ISL_517282, EPI_ISL_517283, EPI_ISL_517284, EPI_ISL_517285, EPI_ISL_517286, EPI_ISL_517287, EPI_ISL_517288, EPI_ISL_517289, EPI_ISL_517290, EPI_ISL_517291, EPI_ISL_517292, EPI_ISL_517293, EPI_ISL_517294, EPI_ISL_517295, EPI_ISL_517296, EPI_ISL_517297, EPI_ISL_517298, EPI_ISL_517299, EPI_ISL_517300, EPI_ISL_517301, EPI_ISL_517302, EPI_ISL_517303, EPI_ISL_517304, EPI_ISL_517305, EPI_ISL_517306, EPI_ISL_517307, EPI_ISL_517308, EPI_ISL_517309, EPI_ISL_517310, EPI_ISL_517311, EPI_ISL_517312, EPI_ISL_517313, EPI_ISL_517314, EPI_ISL_517315, EPI_ISL_517316, EPI_ISL_517317, EPI_ISL_517318, EPI_ISL_517319, EPI_ISL_517320, EPI_ISL_517321, EPI_ISL_517322, EPI_ISL_517323, EPI_ISL_517324, EPI_ISL_517325, EPI_ISL_517326, EPI_ISL_517327, EPI_ISL_517328, EPI_ISL_517329, EPI_ISL_517330, EPI_ISL_517331, EPI_ISL_517332, EPI_ISL_517333, EPI_ISL_517334, EPI_ISL_517335, EPI_ISL_517336, EPI_ISL_517337, EPI_ISL_517338, EPI_ISL_517339, EPI_ISL_517340, EPI_ISL_517341, EPI_ISL_517342, EPI_ISL_517343, EPI_ISL_517344, EPI_ISL_517345, EPI_ISL_517346, EPI_ISL_517347, EPI_ISL_517348, EPI_ISL_517349, EPI_ISL_517350, EPI_ISL_517351, EPI_ISL_517352, EPI_ISL_517353, EPI_ISL_517354, EPI_ISL_517355, EPI_ISL_517356, EPI_ISL_517357, EPI_ISL_517358, EPI_ISL_517359, EPI_ISL_517360, EPI_ISL_517361, EPI_ISL_517362, EPI_ISL_517363, EPI_ISL_517364, EPI_ISL_517365, EPI_ISL_517366, EPI_ISL_517367, EPI_ISL_517368, EPI_ISL_517369, EPI_ISL_517370, EPI_ISL_517371, EPI_ISL_517372, EPI_ISL_517373, EPI_ISL_517374, EPI_ISL_517375, EPI_ISL_517376, EPI_ISL_517377, EPI_ISL_517378, EPI_ISL_517379, EPI_ISL_517380, EPI_ISL_517381, EPI_ISL_517382, EPI_ISL_517383, EPI_ISL_517384, EPI_ISL_517385, EPI_ISL_517386, EPI_ISL_517387, EPI_ISL_517388, EPI_ISL_517389, EPI_ISL_517390, EPI_ISL_517391, EPI_ISL_517392, EPI_ISL_517393, EPI_ISL_517394, EPI_ISL_517395, EPI_ISL_517396, EPI_ISL_517397, EPI_ISL_517398, EPI_ISL_517399, EPI_ISL_517400, EPI_ISL_517401, EPI_ISL_517402, EPI_ISL_517403, EPI_ISL_517404, EPI_ISL_517405, EPI_ISL_517406, EPI_ISL_517407, EPI_ISL_517408, EPI_ISL_517409, EPI_ISL_517410, EPI_ISL_517411, EPI_ISL_517412, EPI_ISL_517413, EPI_ISL_517414, EPI_ISL_517415, EPI_ISL_517416, EPI_ISL_517417, EPI_ISL_517418, EPI_ISL_517419, EPI_ISL_517420, EPI_ISL_517421, EPI_ISL_517422, EPI_ISL_517423, EPI_ISL_517424, EPI_ISL_517425, EPI_ISL_517426, EPI_ISL_517427, EPI_ISL_517428, EPI_ISL_517429, EPI_ISL_517430, EPI_ISL_517431, EPI_ISL_517432, EPI_ISL_517433, EPI_ISL_517434, EPI_ISL_517435, EPI_ISL_517436, EPI_ISL_517437, EPI_ISL_517438, EPI_ISL_517439, EPI_ISL_517440, EPI_ISL_517441, EPI_ISL_517442, EPI_ISL_517443, EPI_ISL_517444, EPI_ISL_517445, EPI_ISL_517446, EPI_ISL_517447, EPI_ISL_517448, EPI_ISL_517449, EPI_ISL_517450, EPI_ISL_517451, EPI_ISL_517452, EPI_ISL_517453, EPI_ISL_517454, EPI_ISL_517455, EPI_ISL_517456, EPI_ISL_517457, EPI_ISL_517458, EPI_ISL_517459, EPI_ISL_517460, EPI_ISL_517461, EPI_ISL_517462, EPI_ISL_517463, EPI_ISL_517464, EPI_ISL_517465, EPI_ISL_517466, EPI_ISL_517467, EPI_ISL_517468, EPI_ISL_517469, EPI_ISL_517470, EPI_ISL_517471, EPI_ISL_517472, EPI_ISL_517473, EPI_ISL_517474, EPI_ISL_517475, EPI_ISL_517476, EPI_ISL_517477, EPI_ISL_517478, EPI_ISL_517479, EPI_ISL_517480, EPI_ISL_517481, EPI_ISL_517482, EPI_ISL_517483, EPI_ISL_517484, EPI_ISL_517485, EPI_ISL_517486, EPI_ISL_517487, EPI_ISL_517488, EPI_ISL_517489, EPI_ISL_517490, EPI_ISL_517491, EPI_ISL_517492, EPI_ISL_517493, EPI_ISL_517494, EPI_ISL_517495, EPI_ISL_517496, EPI_ISL_517497, EPI_ISL_517498, EPI_ISL_517499, EPI_ISL_517500, EPI_ISL_517501, EPI_ISL_517502, EPI_ISL_517503, EPI_ISL_517504, EPI_ISL_517505, EPI_ISL_517506, EPI_ISL_517507, EPI_ISL_517508, EPI_ISL_517509 | | | |
| see above | Liverpool Clinical Laboratories | COVID-19 Genomics UK (COG-UK) Consortium | Sam Haldenby, Anita Lucaci, Steve Paterson, Julian Hiscox, Alistair Darby, M Almsaud, A Alrezaihi, Muhannad Alruwaili, Stuart D Armstrong, Jones Benjamin, Eleanor G Bentley, Anu Chawla, Jordan J Clark, Angela Cowell, Richard Eccles, Isabel García-Dorival, Matthew Gemmell, Alessandro Gerada, PKF Gilmore, Richard Gregory, Ximeng Han, Catherine Hartley, Margaret Hughes, Miren Iturriza-Gomara, James Johnson, L Luu, Jenifer Manson, Charlotte Nelson, Elaine O'Toole, Cassie Olateju, Rebekah Penrice-Randal , Lucille Rainbow, N.P Randle, Trevor Ian Robinson, Parul Sharma, Ghada T Shawli, James P Stewart, Neil Swainston, Ecaterina Vamos, Joanne Watts, Mark Whitehead |
| EPI_ISL_517771, EPI_ISL_517773, EPI_ISL_517774, EPI_ISL_517775, EPI_ISL_517776, EPI_ISL_517779, EPI_ISL_517781, EPI_ISL_517782, EPI_ISL_517783, EPI_ISL_517784, EPI_ISL_517786, EPI_ISL_517787, EPI_ISL_517789, EPI_ISL_517793, EPI_ISL_517796, EPI_ISL_517797, EPI_ISL_517798, EPI_ISL_517799, EPI_ISL_517800, EPI_ISL_517804, EPI_ISL_517806, EPI_ISL_517809, EPI_ISL_517810, EPI_ISL_517811, EPI_ISL_517818, EPI_ISL_517819, EPI_ISL_517820, EPI_ISL_517821, EPI_ISL_517822, EPI_ISL_517826 | | | |
| see above | Florida Bureau of Public Health Laboratories | Florida Bureau of Public Health Laboratories | Sarah Schmedes, Jason Blanton |
| EPI_ISL_518821, EPI_ISL_518822, EPI_ISL_518823, EPI_ISL_518825 | Oman-National Influenza Center | Biotechnology & OMICs Laboratory, Natural & Medical Sciences Research Center, University of Nizwa | Samira Al-Mahruqi, Abdul Latif Khan, Samiha Al-Kharusi, Adil Khan , Ahmed Al-Rawahi, Sajjad Asaf, Amina Al-Jardani, Hanan Al-Kindi, Intisar Al-Shukri, Adil Al-Wahaibi, Seif Al-Abri, Ahmed Al-Harrasi |
| EPI_ISL_518826, EPI_ISL_518827, EPI_ISL_518828, EPI_ISL_518829, EPI_ISL_518830, EPI_ISL_518831, EPI_ISL_518832 | Oman-National Influenza Center | Biotechnology & OMICs Laboratory, Natural & Medical Sciences Research Center, University of Nizwa | Sajjad Asaf, Samiha Al-Kharusi, Ahmed Al-Harrasi, Samira Al-Mahruqi, Adil Khan, Ahmed Al-Rawahi, Abdul Latif Khan, Amina Al-Jardani, Hanan Al-Kindi, Intisar Al-Shukri, Ahlam Al-Amri, Aisha Al-Amri, Aisha Al-Busaidi, Adil Al-Wahaibi, Seif Al-Abri. |
| EPI_ISL_518833, EPI_ISL_518834, EPI_ISL_518835, EPI_ISL_518836 | Oman-National Influenza Center | Biotechnology & OMICs Laboratory, Natural & Medical Sciences Research Center, University of Nizwa | Samira Al-Mahruqi, Abdul Latif Khan, Samiha Al-Kharusi, Adil Khan , Ahmed Al-Rawahi, Sajjad Asaf, Amina Al-Jardani, Hanan Al-Kindi, Intisar Al-Shukri, Adil Al-Wahaibi, Seif Al-Abri, Ahmed Al-Harrasi |

| | | | |
|---|---|---|---|
| EPI_ISL_518837, EPI_ISL_518838, EPI_ISL_518839, EPI_ISL_518840, EPI_ISL_518841, EPI_ISL_518842, EPI_ISL_518843, EPI_ISL_518844, EPI_ISL_518845 | Oman-National Influenza Center | Biotechnology & OMICs Laboratory, Natural & Medical Sciences Research Center, University of Nizwa | Samiha Al-Kharusi, Sajjad Asaf, Abdul Latif Khan, Samira Al-Mahruqi, Adil Khan, Ahmed Al-Rawahi, Amina Al-Jardani, Hanan Al-Kindi, Intisar Al-Shukri, Ahlam Al-Amri, Aisha Al-Amri, Aisha Al-Busaidi, Adil Al-Wahaibi, Seif Al-Abri, Ahmed Al-Harrasi |
| EPI_ISL_524435 | Department of Immunology, The Scripps Research Institute | Department of Immunology, The Scripps Research Institute | SEARCH Alliance,S.D., Pride,D., Shin,J.H. |
| EPI_ISL_524783, EPI_ISL_524784, EPI_ISL_524785, EPI_ISL_524786, EPI_ISL_524787, EPI_ISL_524788, EPI_ISL_524789, EPI_ISL_524790, EPI_ISL_524791, EPI_ISL_524792, EPI_ISL_524793, EPI_ISL_524794, EPI_ISL_524795, EPI_ISL_524796, EPI_ISL_524797, EPI_ISL_524798, EPI_ISL_524799, EPI_ISL_524800, EPI_ISL_524801 | | | |
| see above | Evandro Chagas Institute | Evandro Chagas Institute | Santos, M.C.; Silva, A.M.; Junior, W.D.C.; Barbagelata, L.S.; Ferreira, J.A.; Sousa, E.M.A.; da Silva, P.S.; Resque, H.R; Martins, L.C.; Sousa Junior, E.C.;Viana, G.M.R |
| EPI_ISL_525423 | Oman-National Influenza Center | Biotechnology & OMICs Laboratory | Sajjad Asaf, Samiha Al-Kharusi, Ahmed Al-Harrasi, Samira Al-Mahruqi, Adil Khan, Ahmed Al-Rawahi, Abdul Latif Khan, Amina Al-Jardani, Hanan Al-Kindi, Intisar Al-Shukri, Ahlam Al-Amri, Aisha Al-Amri, Aisha Al-Busaidi, Adil Al-Wahaibi, Seif Al-Abri. |
| EPI_ISL_525424 | Oman-National Influenza Center | Biotechnology & OMICs Laboratory | Samira Al-Mahruqi, Abdul Latif Khan, Samiha Al-Kharusi, Adil Khan , Ahmed Al-Rawahi, Sajjad Asaf, Amina Al-Jardani, Hanan Al-Kindi, Intisar Al-Shukri, Ahlam Al-Amri, Aisha Al-Amri, Aisha Al-Busaidi, Adil Al-Wahaibi, Seif Al-Abri, Ahmed Al-Harrasi |
| EPI_ISL_525425 | Oman-National Influenza Center | Biotechnology & OMICs Laboratory | Samira Al-Mahruqi, Abdul Latif Khan, Samiha Al-Kharusi, Adil Khan , Ahmed Al-Rawahi, Sajjad Asaf, Amina Al-Jardani, Hanan Al-Kindi, Intisar Al-Shukri, Adil Al-Wahaibi, Seif Al-Abri, Ahmed Al-Harrasi |
| EPI_ISL_525426, EPI_ISL_525427, EPI_ISL_525428 | Oman-National Influenza Center | Biotechnology & OMICs Laboratory | Sajjad Asaf, Samiha Al-Kharusi, Ahmed Al-Harrasi, Samira Al-Mahruqi, Adil Khan, Ahmed Al-Rawahi, Abdul Latif Khan, Amina Al-Jardani, Hanan Al-Kindi, Intisar Al-Shukri, Ahlam Al-Amri, Aisha Al-Amri, Aisha Al-Busaidi, Adil Al-Wahaibi, Seif Al-Abri. |
| EPI_ISL_525429 | Oman-National Influenza Center | Biotechnology & OMICs Laboratory | Samira Al-Mahruqi, Abdul Latif Khan, Samiha Al-Kharusi, Adil Khan , Ahmed Al-Rawahi, Sajjad Asaf, Amina Al-Jardani, Hanan Al-Kindi, Intisar Al-Shukri, Adil Al-Wahaibi, Seif Al-Abri, Ahmed Al-Harrasi |
| EPI_ISL_525466 | Oman-National Influenza Center | Biotechnology & OMICs Laboratory | Samiha Al-Kharusi, Sajjad Asaf, Abdul Latif Khan, Samira Al-Mahruqi, Adil Khan, Ahmed Al-Rawahi, Amina Al-Jardani, Hanan Al-Kindi, Intisar Al-Shukri, Aisha Al-Busaidi, Adil Al-Wahaibi, Seif Al-Abri, Ahmed Al-Harrasi |
| EPI_ISL_526367, EPI_ISL_526368, EPI_ISL_526369, EPI_ISL_526370, EPI_ISL_526371, EPI_ISL_526372, EPI_ISL_526373, EPI_ISL_526374, EPI_ISL_526375, EPI_ISL_526376, EPI_ISL_526377, EPI_ISL_526378, EPI_ISL_526379, EPI_ISL_526380, EPI_ISL_526381, EPI_ISL_526382, EPI_ISL_526383, EPI_ISL_526384, EPI_ISL_526385, EPI_ISL_526386, EPI_ISL_526387, EPI_ISL_526388, EPI_ISL_526389, EPI_ISL_526390, EPI_ISL_526391, EPI_ISL_526392, EPI_ISL_526393, EPI_ISL_526394 | | | |
| see above | Liverpool Clinical Laboratories | COVID-19 Genomics UK (COG-UK) Consortium | Sam Haldenby, Anita Lucaci, Steve Paterson, Julian Hiscox, Alistair Darby, M Almsaud, A Alrezaihi, Muhannad Alruwaili, Stuart D Armstrong, Jones Benjamin, Eleanor G Bentley, Anu Chawla, Jordan J Clark, Angela Cowell, Richard Eccles, Isabel García-Dorival, Matthew Gemmell, Alessandro Gerada, PKF Gilmore, Richard Gregory, Ximeng Han, Catherine Hartley, Margaret Hughes, Miren Iturriza-Gomara, James Johnson, L Luu, Jenifer Manson, Charlotte Nelson, Elaine O'Toole, Cassie Olateju, Rebekah Penrice-Randal , Lucille Rainbow, N.P Randle, Trevor Ian Robinson, Parul Sharma, Ghada T Shawli, James P Stewart, Neil Swainston, Ecaterina Vamos, Joanne Watts, Mark Whitehead |
| EPI_ISL_526558, EPI_ISL_526560, EPI_ISL_526561, EPI_ISL_526562, EPI_ISL_526563, EPI_ISL_526564, EPI_ISL_526565, EPI_ISL_526566, EPI_ISL_526567, EPI_ISL_526568, EPI_ISL_526570, EPI_ISL_526574, EPI_ISL_526576, EPI_ISL_526577, EPI_ISL_526578, EPI_ISL_526579, EPI_ISL_526580, EPI_ISL_526581, EPI_ISL_526582, EPI_ISL_526587, EPI_ISL_526588, EPI_ISL_526601, EPI_ISL_526604, EPI_ISL_526609, EPI_ISL_526612, EPI_ISL_526615, EPI_ISL_526634, EPI_ISL_526636, EPI_ISL_526640, EPI_ISL_526641, EPI_ISL_526642, EPI_ISL_526647, EPI_ISL_526648, EPI_ISL_526651, EPI_ISL_526654, EPI_ISL_526655, EPI_ISL_526656 | | | |
| see above | Florida Bureau of Public Health Laboratories | Florida Bureau of Public Health Laboratories | Sarah Schmedes, Jason Blanton |
| EPI_ISL_528386 | Viral vaccines, VSVRI- Veterinary serum and vaccine research institute | Viral vaccines, VSVRI- Veterinary serum and vaccine research institute | Saleh,A.A., Saad,M.A. |
| EPI_ISL_528829, EPI_ISL_528837, EPI_ISL_528861, EPI_ISL_528866 | CSIR-Centre for Cellular and Molecular Biology | CSIR-Centre for Cellular and Molecular Biology | Sakshi Shambhavi, Lamuk Zaveri, Shagufta Khan, Namami Gaur, Nikhil Hajirnis, M Soujanya Reddy, Pratheusa Maccha, Tulasi Nagabandi, Purushotham Vodnala, Payel Mukherjee, Sofia Banu, Priya Singh, Onkar Kulkarni, Dhiviya Vedagiri, Divya Gupta, Vishal Sah, Santosh Kumar Kuncha, Krishnan Harinivas Harshan, Archana Bharadwaj Siva, Karthik Bharadwaj Tallapaka, Deepak Kumar, Devi Prasad Vijayashankar, Disha Nanda, Divya Das, Jotin Gogoi, Manish Bhattacharjee, Rakesh K Mishra, Divya Tej Sowpati |
| EPI_ISL_529254, EPI_ISL_529258, EPI_ISL_529259, EPI_ISL_529260, EPI_ISL_529263, EPI_ISL_529270, EPI_ISL_529271, EPI_ISL_529274, EPI_ISL_529280, EPI_ISL_529284, EPI_ISL_529287, EPI_ISL_529293, EPI_ISL_529295, EPI_ISL_529296, EPI_ISL_529302, EPI_ISL_529304, EPI_ISL_529305, EPI_ISL_529466, EPI_ISL_529484, EPI_ISL_529490, EPI_ISL_529498, EPI_ISL_529509, EPI_ISL_529510, EPI_ISL_529513, EPI_ISL_529516 | | | |
| see above | Liverpool Clinical Laboratories | COVID-19 Genomics UK (COG-UK) Consortium | Sam Haldenby, Anita Lucaci, Steve Paterson, Julian Hiscox, Alistair Darby, M Almsaud, A Alrezaihi, Muhannad Alruwaili, Stuart D Armstrong, Jones Benjamin, Eleanor G Bentley, Anu Chawla, Jordan J Clark, Angela Cowell, Richard Eccles, Isabel García-Dorival, Matthew Gemmell, Alessandro Gerada, PKF Gilmore, Richard Gregory, Ximeng Han, Catherine Hartley, Margaret Hughes, Miren Iturriza-Gomara, James Johnson, L Luu, Jenifer Manson, Charlotte Nelson, Elaine O'Toole, Cassie Olateju, Rebekah Penrice-Randal , Lucille Rainbow, N.P Randle, Trevor Ian Robinson, Parul Sharma, Ghada T Shawli, James P Stewart, Neil Swainston, Ecaterina Vamos, Joanne Watts, Mark Whitehead |
| EPI_ISL_529989, EPI_ISL_529990, EPI_ISL_529991, EPI_ISL_529992, EPI_ISL_529993, EPI_ISL_529994, EPI_ISL_529995, EPI_ISL_529996, EPI_ISL_529997, EPI_ISL_529998, EPI_ISL_529999, EPI_ISL_530000, EPI_ISL_530001, EPI_ISL_530002, EPI_ISL_530003, EPI_ISL_530004, EPI_ISL_530005, EPI_ISL_530006 | | | |
| see above | Hospital Universitario 12 de Octubre | Hospital Universitario 12 de Octubre | Sara González, Esther Viedma, Raúl Recio, Elias Dahdouh, Fernando Lázaro, Natalia Stella, Julio García, Juan Carlos Galán, Rafael Cantón, Mª Dolores Folgueira, Rafael Delgado, Jesús Mingorance |
| EPI_ISL_534732, EPI_ISL_534733, EPI_ISL_534734, EPI_ISL_534735, EPI_ISL_534736, EPI_ISL_534737, EPI_ISL_534738, EPI_ISL_534739, EPI_ISL_534740, EPI_ISL_534741, EPI_ISL_534742, EPI_ISL_534743, EPI_ISL_534744, EPI_ISL_534745, EPI_ISL_534746, EPI_ISL_534747, EPI_ISL_534748, EPI_ISL_534749, EPI_ISL_534750, EPI_ISL_534751, EPI_ISL_534752, EPI_ISL_534753, EPI_ISL_534754, EPI_ISL_534755 | | | |
| see above | Liverpool Clinical Laboratories | COVID-19 Genomics UK (COG-UK) Consortium | Sam Haldenby, Anita Lucaci, Steve Paterson, Julian Hiscox, Alistair Darby, M Almsaud, A Alrezaihi, Muhannad Alruwaili, Stuart D Armstrong, Jones Benjamin, Eleanor G Bentley, Anu Chawla, Jordan J Clark, Angela Cowell, Richard Eccles, Isabel García-Dorival, Matthew Gemmell, Alessandro Gerada, PKF Gilmore, Richard Gregory, Ximeng Han, Catherine Hartley, Margaret Hughes, Miren Iturriza-Gomara, James Johnson, L Luu, Jenifer Manson, Charlotte Nelson, Elaine O'Toole, Cassie Olateju, Rebekah Penrice-Randal , Lucille Rainbow, N.P Randle, Trevor Ian Robinson, Parul Sharma, Ghada T Shawli, James P Stewart, Neil Swainston, Ecaterina Vamos, Joanne Watts, Mark Whitehead |
| EPI_ISL_535361, EPI_ISL_535362, EPI_ISL_535363, EPI_ISL_535364 | Oklahoma Animal Disease Diagnostic Laboratory | Oklahoma Animal Disease Diagnostic Laboratory | Sai Narayanan, John C Ritchey, Girish Patil, Teluguakula Narasaraju, Sunil More, Jerry Malayer, Jeremiah Saliki, Anil Kaul, Akhilesh Ramachandran |
| EPI_ISL_535716 | Hôpital de Verdun | Laboratoire de santé publique du Québec | Sandrine Moreira, Ioannis Ragoussis, Guillaume Bourque, Jesse Shapiro, Mark Lathrop and Michel Roger |
| EPI_ISL_535717 | Hôpital Honoré-Mercier | Laboratoire de santé publique du Québec | Sandrine Moreira, Ioannis Ragoussis, Guillaume Bourque, Jesse Shapiro, Mark Lathrop and Michel Roger |
| EPI_ISL_535718 | Hôpital Pierre-Boucher | Laboratoire de santé publique du Québec | Sandrine Moreira, Ioannis Ragoussis, Guillaume Bourque, Jesse Shapiro, Mark Lathrop and Michel Roger |
| EPI_ISL_535719, EPI_ISL_535720 | Hôpital général Juif | Laboratoire de santé publique du Québec | Sandrine Moreira, Ioannis Ragoussis, Guillaume Bourque, Jesse Shapiro, Mark Lathrop and Michel Roger |
| EPI_ISL_535721 | Hôpital Notre-Dame | Laboratoire de santé publique du Québec | Sandrine Moreira, Ioannis Ragoussis, Guillaume Bourque, Jesse Shapiro, Mark Lathrop and Michel Roger |
| EPI_ISL_535722 | Centre de SSS de Trois-Rivières | Laboratoire de santé publique du Québec | Sandrine Moreira, Ioannis Ragoussis, Guillaume Bourque, Jesse Shapiro, Mark Lathrop and Michel Roger |
| EPI_ISL_535723 | Hôpital de Verdun | Laboratoire de santé publique du Québec | Sandrine Moreira, Ioannis Ragoussis, Guillaume Bourque, Jesse Shapiro, Mark Lathrop and Michel Roger |
| EPI_ISL_535724 | Centre de SSS La Pommeraie | Laboratoire de santé publique du Québec | Sandrine Moreira, Ioannis Ragoussis, Guillaume Bourque, Jesse Shapiro, Mark Lathrop and Michel Roger |
| EPI_ISL_535725 | Hôtel-Dieu de Lévis | Laboratoire de santé publique du Québec | Sandrine Moreira, Ioannis Ragoussis, Guillaume Bourque, Jesse Shapiro, Mark Lathrop and Michel Roger |
| EPI_ISL_535726 | CHUM - Microbiologie - Hôpital Saint-Luc | Laboratoire de santé publique du Québec | Sandrine Moreira, Ioannis Ragoussis, Guillaume Bourque, Jesse Shapiro, Mark Lathrop and Michel Roger |
| EPI_ISL_535727 | Centre de SSS de la Haute-Yamaska | Laboratoire de santé publique du Québec | Sandrine Moreira, Ioannis Ragoussis, Guillaume Bourque, Jesse Shapiro, Mark Lathrop and Michel Roger |
| EPI_ISL_535728 | Hôpital de Verdun | Laboratoire de santé publique du Québec | Sandrine Moreira, Ioannis Ragoussis, Guillaume Bourque, Jesse Shapiro, Mark Lathrop and Michel Roger |

| | | | |
|---|---|---|---|
| EPI_ISL_535729, EPI_ISL_535730 | CHUM - Microbiologie - Hôpital Saint-Luc | Laboratoire de santé publique du Québec | Sandrine Moreira, Ioannis Ragoussis, Guillaume Bourque, Jesse Shapiro, Mark Lathrop and Michel Roger |
| EPI_ISL_535731 | Hôpital Pierre-Le Gardeur | Laboratoire de santé publique du Québec | Sandrine Moreira, Ioannis Ragoussis, Guillaume Bourque, Jesse Shapiro, Mark Lathrop and Michel Roger |
| EPI_ISL_535732 | CHU Sainte-Justine | Laboratoire de santé publique du Québec | Sandrine Moreira, Ioannis Ragoussis, Guillaume Bourque, Jesse Shapiro, Mark Lathrop and Michel Roger |
| EPI_ISL_535733, EPI_ISL_535734 | Hôpital Pierre-Boucher | Laboratoire de santé publique du Québec | Sandrine Moreira, Ioannis Ragoussis, Guillaume Bourque, Jesse Shapiro, Mark Lathrop and Michel Roger |
| EPI_ISL_535735 | CHUM - Microbiologie - Hôpital Saint-Luc | Laboratoire de santé publique du Québec | Sandrine Moreira, Ioannis Ragoussis, Guillaume Bourque, Jesse Shapiro, Mark Lathrop and Michel Roger |
| EPI_ISL_535736, EPI_ISL_535737 | CHUL-LABO MULTI / MICRO | Laboratoire de santé publique du Québec | Sandrine Moreira, Ioannis Ragoussis, Guillaume Bourque, Jesse Shapiro, Mark Lathrop and Michel Roger |
| EPI_ISL_535738 | Hôpital Charles-LeMoyne | Laboratoire de santé publique du Québec | Sandrine Moreira, Ioannis Ragoussis, Guillaume Bourque, Jesse Shapiro, Mark Lathrop and Michel Roger |
| EPI_ISL_535739 | Hôpital de l'Enfant-Jésus | Laboratoire de santé publique du Québec | Sandrine Moreira, Ioannis Ragoussis, Guillaume Bourque, Jesse Shapiro, Mark Lathrop and Michel Roger |
| EPI_ISL_535740 | CHUM - Microbiologie - Hôpital Saint-Luc | Laboratoire de santé publique du Québec | Sandrine Moreira, Ioannis Ragoussis, Guillaume Bourque, Jesse Shapiro, Mark Lathrop and Michel Roger |
| EPI_ISL_535741 | CHUL-LABO MULTI / MICRO | Laboratoire de santé publique du Québec | Sandrine Moreira, Ioannis Ragoussis, Guillaume Bourque, Jesse Shapiro, Mark Lathrop and Michel Roger |
| EPI_ISL_535742, EPI_ISL_535743 | Centre de SSS La Pommeraie | Laboratoire de santé publique du Québec | Sandrine Moreira, Ioannis Ragoussis, Guillaume Bourque, Jesse Shapiro, Mark Lathrop and Michel Roger |
| EPI_ISL_535744 | Hôpital de Lasalle | Laboratoire de santé publique du Québec | Sandrine Moreira, Ioannis Ragoussis, Guillaume Bourque, Jesse Shapiro, Mark Lathrop and Michel Roger |
| EPI_ISL_535745 | Hôpital Laurentien | Laboratoire de santé publique du Québec | Sandrine Moreira, Ioannis Ragoussis, Guillaume Bourque, Jesse Shapiro, Mark Lathrop and Michel Roger |
| EPI_ISL_535746 | Centre hospitalier de St-Mary | Laboratoire de santé publique du Québec | Sandrine Moreira, Ioannis Ragoussis, Guillaume Bourque, Jesse Shapiro, Mark Lathrop and Michel Roger |
| EPI_ISL_535747, EPI_ISL_535748 | Centre de SSS de Trois-Rivières | Laboratoire de santé publique du Québec | Sandrine Moreira, Ioannis Ragoussis, Guillaume Bourque, Jesse Shapiro, Mark Lathrop and Michel Roger |
| EPI_ISL_535749 | CUSM-Site Glen-LAB Microbiologie | Laboratoire de santé publique du Québec | Sandrine Moreira, Ioannis Ragoussis, Guillaume Bourque, Jesse Shapiro, Mark Lathrop and Michel Roger |
| EPI_ISL_535750, EPI_ISL_535751 | Hôtel-Dieu de Lévis | Laboratoire de santé publique du Québec | Sandrine Moreira, Ioannis Ragoussis, Guillaume Bourque, Jesse Shapiro, Mark Lathrop and Michel Roger |
| EPI_ISL_535752 | Hôpital Pierre-Le Gardeur | Laboratoire de santé publique du Québec | Sandrine Moreira, Ioannis Ragoussis, Guillaume Bourque, Jesse Shapiro, Mark Lathrop and Michel Roger |
| EPI_ISL_535753, EPI_ISL_535754 | CHUM - Microbiologie - Hôpital Saint-Luc | Laboratoire de santé publique du Québec | Sandrine Moreira, Ioannis Ragoussis, Guillaume Bourque, Jesse Shapiro, Mark Lathrop and Michel Roger |
| EPI_ISL_535755 | Hôpital Maisonneuve-Rosemont | Laboratoire de santé publique du Québec | Sandrine Moreira, Ioannis Ragoussis, Guillaume Bourque, Jesse Shapiro, Mark Lathrop and Michel Roger |
| EPI_ISL_535756, EPI_ISL_535757 | Hôpital général Juif | Laboratoire de santé publique du Québec | Sandrine Moreira, Ioannis Ragoussis, Guillaume Bourque, Jesse Shapiro, Mark Lathrop and Michel Roger |
| EPI_ISL_535758, EPI_ISL_535759 | Hôtel-Dieu de Lévis | Laboratoire de santé publique du Québec | Sandrine Moreira, Ioannis Ragoussis, Guillaume Bourque, Jesse Shapiro, Mark Lathrop and Michel Roger |
| EPI_ISL_535760 | CHUM - Microbiologie - Hôpital Saint-Luc | Laboratoire de santé publique du Québec | Sandrine Moreira, Ioannis Ragoussis, Guillaume Bourque, Jesse Shapiro, Mark Lathrop and Michel Roger |
| EPI_ISL_535761 | CUSM-Site Glen-LAB Microbiologie | Laboratoire de santé publique du Québec | Sandrine Moreira, Ioannis Ragoussis, Guillaume Bourque, Jesse Shapiro, Mark Lathrop and Michel Roger |
| EPI_ISL_535762 | Hôpital Pierre-Le Gardeur | Laboratoire de santé publique du Québec | Sandrine Moreira, Ioannis Ragoussis, Guillaume Bourque, Jesse Shapiro, Mark Lathrop and Michel Roger |
| EPI_ISL_535763 | Hôpital Pierre-Boucher | Laboratoire de santé publique du Québec | Sandrine Moreira, Ioannis Ragoussis, Guillaume Bourque, Jesse Shapiro, Mark Lathrop and Michel Roger |
| EPI_ISL_535764, EPI_ISL_535765 | CUSM-Site Glen-LAB Microbiologie | Laboratoire de santé publique du Québec | Sandrine Moreira, Ioannis Ragoussis, Guillaume Bourque, Jesse Shapiro, Mark Lathrop and Michel Roger |
| EPI_ISL_535766 | Hôpital Pierre-Le Gardeur | Laboratoire de santé publique du Québec | Sandrine Moreira, Ioannis Ragoussis, Guillaume Bourque, Jesse Shapiro, Mark Lathrop and Michel Roger |
| EPI_ISL_535767, EPI_ISL_535768 | Centre hospitalier de St-Mary | Laboratoire de santé publique du Québec | Sandrine Moreira, Ioannis Ragoussis, Guillaume Bourque, Jesse Shapiro, Mark Lathrop and Michel Roger |
| EPI_ISL_535769 | Hôpital général Juif | Laboratoire de santé publique du Québec | Sandrine Moreira, Ioannis Ragoussis, Guillaume Bourque, Jesse Shapiro, Mark Lathrop and Michel Roger |
| EPI_ISL_535770, EPI_ISL_535771 | Hôpital Pierre-Le Gardeur | Laboratoire de santé publique du Québec | Sandrine Moreira, Ioannis Ragoussis, Guillaume Bourque, Jesse Shapiro, Mark Lathrop and Michel Roger |
| EPI_ISL_535772, EPI_ISL_535773, EPI_ISL_535774, EPI_ISL_535775 | CHUL-LABO MULTI / MICRO | Laboratoire de santé publique du Québec | Sandrine Moreira, Ioannis Ragoussis, Guillaume Bourque, Jesse Shapiro, Mark Lathrop and Michel Roger |
| EPI_ISL_535776, EPI_ISL_535777, EPI_ISL_535778, EPI_ISL_535779, EPI_ISL_535780 | Centre de SSS de la Haute-Yamaska | Laboratoire de santé publique du Québec | Sandrine Moreira, Ioannis Ragoussis, Guillaume Bourque, Jesse Shapiro, Mark Lathrop and Michel Roger |
| EPI_ISL_535781, EPI_ISL_535782 | CHUL-LABO MULTI / MICRO | Laboratoire de santé publique du Québec | Sandrine Moreira, Ioannis Ragoussis, Guillaume Bourque, Jesse Shapiro, Mark Lathrop and Michel Roger |
| EPI_ISL_535783 | CHUM - Microbiologie - Hôpital Saint-Luc | Laboratoire de santé publique du Québec | Sandrine Moreira, Ioannis Ragoussis, Guillaume Bourque, Jesse Shapiro, Mark Lathrop and Michel Roger |
| EPI_ISL_535784, EPI_ISL_535785 | CHUL-LABO MULTI / MICRO | Laboratoire de santé publique du Québec | Sandrine Moreira, Ioannis Ragoussis, Guillaume Bourque, Jesse Shapiro, Mark Lathrop and Michel Roger |
| EPI_ISL_535786 | Hôpital Pierre-Boucher | Laboratoire de santé publique du Québec | Sandrine Moreira, Ioannis Ragoussis, Guillaume Bourque, Jesse Shapiro, Mark Lathrop and Michel Roger |
| EPI_ISL_535787, EPI_ISL_535788 | Centre Hospitalier Régional de Lanaudière | Laboratoire de santé publique du Québec | Sandrine Moreira, Ioannis Ragoussis, Guillaume Bourque, Jesse Shapiro, Mark Lathrop and Michel Roger |
| EPI_ISL_535789 | CHUM - Microbiologie - Hôpital Saint-Luc | Laboratoire de santé publique du Québec | Sandrine Moreira, Ioannis Ragoussis, Guillaume Bourque, Jesse Shapiro, Mark Lathrop and Michel Roger |
| EPI_ISL_535790 | Centre Hospitalier Régional de Lanaudière | Laboratoire de santé publique du Québec | Sandrine Moreira, Ioannis Ragoussis, Guillaume Bourque, Jesse Shapiro, Mark Lathrop and Michel Roger |
| EPI_ISL_535791 | Hôpital régional de Saint-Jérôme | Laboratoire de santé publique du Québec | Sandrine Moreira, Ioannis Ragoussis, Guillaume Bourque, Jesse Shapiro, Mark Lathrop and Michel Roger |
| EPI_ISL_535792 | Hôpital Fleurimont | Laboratoire de santé publique du Québec | Sandrine Moreira, Ioannis Ragoussis, Guillaume Bourque, Jesse Shapiro, Mark Lathrop and Michel Roger |
| EPI_ISL_535793 | Hôpital de Saint-Eustache | Laboratoire de santé publique du Québec | Sandrine Moreira, Ioannis Ragoussis, Guillaume Bourque, Jesse Shapiro, Mark Lathrop and Michel Roger |
| EPI_ISL_535794 | Hôpital de Hull | Laboratoire de santé publique du Québec | Sandrine Moreira, Ioannis Ragoussis, Guillaume Bourque, Jesse Shapiro, Mark Lathrop and Michel Roger |
| EPI_ISL_535795 | Hôpital Notre-Dame | Laboratoire de santé publique du Québec | Sandrine Moreira, Ioannis Ragoussis, Guillaume Bourque, Jesse Shapiro, Mark Lathrop and Michel Roger |
| EPI_ISL_535796 | Hôpital Charles-LeMoyne | Laboratoire de santé publique du Québec | Sandrine Moreira, Ioannis Ragoussis, Guillaume Bourque, Jesse Shapiro, Mark Lathrop and Michel Roger |
| EPI_ISL_535797 | Hôpital Honoré-Mercier | Laboratoire de santé publique du Québec | Sandrine Moreira, Ioannis Ragoussis, Guillaume Bourque, Jesse Shapiro, Mark Lathrop and Michel Roger |
| EPI_ISL_535798 | Hôpital régional de Saint-Jérôme | Laboratoire de santé publique du Québec | Sandrine Moreira, Ioannis Ragoussis, Guillaume Bourque, Jesse Shapiro, Mark Lathrop and Michel Roger |
| EPI_ISL_535799, EPI_ISL_535800 | CSSS Haut-Richelieu/Rouville (Hôpital) | Laboratoire de santé publique du Québec | Sandrine Moreira, Ioannis Ragoussis, Guillaume Bourque, Jesse Shapiro, Mark Lathrop and Michel Roger |
| EPI_ISL_535801 | CHU Sainte-Justine | Laboratoire de santé publique du Québec | Sandrine Moreira, Ioannis Ragoussis, Guillaume Bourque, Jesse Shapiro, Mark Lathrop and Michel Roger |
| EPI_ISL_535802 | CHUM - Microbiologie - Hôpital Saint-Luc | Laboratoire de santé publique du Québec | Sandrine Moreira, Ioannis Ragoussis, Guillaume Bourque, Jesse Shapiro, Mark Lathrop and Michel Roger |
| EPI_ISL_535803, EPI_ISL_535804, EPI_ISL_535805 | Centre de SSS de la Haute-Yamaska | Laboratoire de santé publique du Québec | Sandrine Moreira, Ioannis Ragoussis, Guillaume Bourque, Jesse Shapiro, Mark Lathrop and Michel Roger |
| EPI_ISL_535806 | CHUM - Microbiologie - Hôpital Saint-Luc | Laboratoire de santé publique du Québec | Sandrine Moreira, Ioannis Ragoussis, Guillaume Bourque, Jesse Shapiro, Mark Lathrop and Michel Roger |
| EPI_ISL_535807 | Centre hospitalier régional du Grand Portage | Laboratoire de santé publique du Québec | Sandrine Moreira, Ioannis Ragoussis, Guillaume Bourque, Jesse Shapiro, Mark Lathrop and Michel Roger |
| EPI_ISL_535808 | Centre de santé Chibougamau | Laboratoire de santé publique du Québec | Sandrine Moreira, Ioannis Ragoussis, Guillaume Bourque, Jesse Shapiro, Mark Lathrop and Michel Roger |
| EPI_ISL_535809 | Hôpital de Hull | Laboratoire de santé publique du Québec | Sandrine Moreira, Ioannis Ragoussis, Guillaume Bourque, Jesse Shapiro, Mark Lathrop and Michel Roger |

| | | | |
|---|---|---|---|
| EPI_ISL_535810 | Hôpital Sainte-Croix | Laboratoire de santé publique du Québec | Sandrine Moreira, Ioannis Ragoussis, Guillaume Bourque, Jesse Shapiro, Mark Lathrop and Michel Roger |
| EPI_ISL_535811 | CUSM-Site Glen-LAB Microbiologie | Laboratoire de santé publique du Québec | Sandrine Moreira, Ioannis Ragoussis, Guillaume Bourque, Jesse Shapiro, Mark Lathrop and Michel Roger |
| EPI_ISL_535812 | Hôpital du Sacré-Coeur de Montréal | Laboratoire de santé publique du Québec | Sandrine Moreira, Ioannis Ragoussis, Guillaume Bourque, Jesse Shapiro, Mark Lathrop and Michel Roger |
| EPI_ISL_535813 | CSSS Haut-Richelieu/Rouville (Hôpital) | Laboratoire de santé publique du Québec | Sandrine Moreira, Ioannis Ragoussis, Guillaume Bourque, Jesse Shapiro, Mark Lathrop and Michel Roger |
| EPI_ISL_535814 | CHUM - Microbiologie - Hôpital Saint-Luc | Laboratoire de santé publique du Québec | Sandrine Moreira, Ioannis Ragoussis, Guillaume Bourque, Jesse Shapiro, Mark Lathrop and Michel Roger |
| EPI_ISL_535815 | CSSS Haut-Richelieu/Rouville (Hôpital) | Laboratoire de santé publique du Québec | Sandrine Moreira, Ioannis Ragoussis, Guillaume Bourque, Jesse Shapiro, Mark Lathrop and Michel Roger |
| EPI_ISL_535816 | Centre de SSS de la Haute-Yamaska | Laboratoire de santé publique du Québec | Sandrine Moreira, Ioannis Ragoussis, Guillaume Bourque, Jesse Shapiro, Mark Lathrop and Michel Roger |
| EPI_ISL_535817 | Hôpital Maisonneuve-Rosemont | Laboratoire de santé publique du Québec | Sandrine Moreira, Ioannis Ragoussis, Guillaume Bourque, Jesse Shapiro, Mark Lathrop and Michel Roger |
| EPI_ISL_535818 | Hôpital Pierre-Boucher | Laboratoire de santé publique du Québec | Sandrine Moreira, Ioannis Ragoussis, Guillaume Bourque, Jesse Shapiro, Mark Lathrop and Michel Roger |
| EPI_ISL_535819 | Hôpital Charles-LeMoyne | Laboratoire de santé publique du Québec | Sandrine Moreira, Ioannis Ragoussis, Guillaume Bourque, Jesse Shapiro, Mark Lathrop and Michel Roger |
| EPI_ISL_535820, EPI_ISL_535821 | Hôtel-Dieu de Lévis | Laboratoire de santé publique du Québec | Sandrine Moreira, Ioannis Ragoussis, Guillaume Bourque, Jesse Shapiro, Mark Lathrop and Michel Roger |
| EPI_ISL_535822, EPI_ISL_535823, EPI_ISL_535824, EPI_ISL_535825 | CHUL-LABO MULTI / MICRO | Laboratoire de santé publique du Québec | Sandrine Moreira, Ioannis Ragoussis, Guillaume Bourque, Jesse Shapiro, Mark Lathrop and Michel Roger |
| EPI_ISL_535826, EPI_ISL_535827 | CHUM - Microbiologie - Hôpital Saint-Luc | Laboratoire de santé publique du Québec | Sandrine Moreira, Ioannis Ragoussis, Guillaume Bourque, Jesse Shapiro, Mark Lathrop and Michel Roger |
| EPI_ISL_535828 | CHUL-LABO MULTI / MICRO | Laboratoire de santé publique du Québec | Sandrine Moreira, Ioannis Ragoussis, Guillaume Bourque, Jesse Shapiro, Mark Lathrop and Michel Roger |
| EPI_ISL_535829 | Hôpital de Chicoutimi | Laboratoire de santé publique du Québec | Sandrine Moreira, Ioannis Ragoussis, Guillaume Bourque, Jesse Shapiro, Mark Lathrop and Michel Roger |
| EPI_ISL_535830 | CHUM - Microbiologie - Hôpital Saint-Luc | Laboratoire de santé publique du Québec | Sandrine Moreira, Ioannis Ragoussis, Guillaume Bourque, Jesse Shapiro, Mark Lathrop and Michel Roger |
| EPI_ISL_535831 | Hôpital Maisonneuve-Rosemont | Laboratoire de santé publique du Québec | Sandrine Moreira, Ioannis Ragoussis, Guillaume Bourque, Jesse Shapiro, Mark Lathrop and Michel Roger |
| EPI_ISL_535832 | CHUM - Microbiologie - Hôpital Saint-Luc | Laboratoire de santé publique du Québec | Sandrine Moreira, Ioannis Ragoussis, Guillaume Bourque, Jesse Shapiro, Mark Lathrop and Michel Roger |
| EPI_ISL_535833, EPI_ISL_535834, EPI_ISL_535835, EPI_ISL_535836 | Hôtel-Dieu de Lévis | Laboratoire de santé publique du Québec | Sandrine Moreira, Ioannis Ragoussis, Guillaume Bourque, Jesse Shapiro, Mark Lathrop and Michel Roger |
| EPI_ISL_535837, EPI_ISL_535838 | Hôpital régional de Saint-Jérôme | Laboratoire de santé publique du Québec | Sandrine Moreira, Ioannis Ragoussis, Guillaume Bourque, Jesse Shapiro, Mark Lathrop and Michel Roger |
| EPI_ISL_535839 | Hôpital Cité de la Santé | Laboratoire de santé publique du Québec | Sandrine Moreira, Ioannis Ragoussis, Guillaume Bourque, Jesse Shapiro, Mark Lathrop and Michel Roger |
| EPI_ISL_535840 | Hôpital Pierre-Boucher | Laboratoire de santé publique du Québec | Sandrine Moreira, Ioannis Ragoussis, Guillaume Bourque, Jesse Shapiro, Mark Lathrop and Michel Roger |
| EPI_ISL_535841 | CSSS Haut-Richelieu/Rouville (Hôpital) | Laboratoire de santé publique du Québec | Sandrine Moreira, Ioannis Ragoussis, Guillaume Bourque, Jesse Shapiro, Mark Lathrop and Michel Roger |
| EPI_ISL_535842, EPI_ISL_535843, EPI_ISL_535844 | Centre Hospitalier Régional de Lanaudière | Laboratoire de santé publique du Québec | Sandrine Moreira, Ioannis Ragoussis, Guillaume Bourque, Jesse Shapiro, Mark Lathrop and Michel Roger |
| EPI_ISL_535845 | Centre de SSS D'Arthabaska-et-de-l'érable - Hôtel-Dieu | Laboratoire de santé publique du Québec | Sandrine Moreira, Ioannis Ragoussis, Guillaume Bourque, Jesse Shapiro, Mark Lathrop and Michel Roger |
| EPI_ISL_535846 | CSSS Haut-Richelieu/Rouville (Hôpital) | Laboratoire de santé publique du Québec | Sandrine Moreira, Ioannis Ragoussis, Guillaume Bourque, Jesse Shapiro, Mark Lathrop and Michel Roger |
| EPI_ISL_535847 | Hôpital du Suroît | Laboratoire de santé publique du Québec | Sandrine Moreira, Ioannis Ragoussis, Guillaume Bourque, Jesse Shapiro, Mark Lathrop and Michel Roger |
| EPI_ISL_535848 | Hôpital Charles-LeMoyne | Laboratoire de santé publique du Québec | Sandrine Moreira, Ioannis Ragoussis, Guillaume Bourque, Jesse Shapiro, Mark Lathrop and Michel Roger |
| EPI_ISL_535849 | Centre de SSS de Trois-Rivières | Laboratoire de santé publique du Québec | Sandrine Moreira, Ioannis Ragoussis, Guillaume Bourque, Jesse Shapiro, Mark Lathrop and Michel Roger |
| EPI_ISL_535850 | Hôpital Pierre-Le Gardeur | Laboratoire de santé publique du Québec | Sandrine Moreira, Ioannis Ragoussis, Guillaume Bourque, Jesse Shapiro, Mark Lathrop and Michel Roger |
| EPI_ISL_535851 | Hôpital Charles-LeMoyne | Laboratoire de santé publique du Québec | Sandrine Moreira, Ioannis Ragoussis, Guillaume Bourque, Jesse Shapiro, Mark Lathrop and Michel Roger |
| EPI_ISL_535852 | Hôpital général du Lakeshore | Laboratoire de santé publique du Québec | Sandrine Moreira, Ioannis Ragoussis, Guillaume Bourque, Jesse Shapiro, Mark Lathrop and Michel Roger |
| EPI_ISL_535853 | Hôpital Charles-LeMoyne | Laboratoire de santé publique du Québec | Sandrine Moreira, Ioannis Ragoussis, Guillaume Bourque, Jesse Shapiro, Mark Lathrop and Michel Roger |
| EPI_ISL_535854, EPI_ISL_535855 | Centre de SSS de la Haute-Yamaska | Laboratoire de santé publique du Québec | Sandrine Moreira, Ioannis Ragoussis, Guillaume Bourque, Jesse Shapiro, Mark Lathrop and Michel Roger |
| EPI_ISL_535856 | Hôpital de Verdun | Laboratoire de santé publique du Québec | Sandrine Moreira, Ioannis Ragoussis, Guillaume Bourque, Jesse Shapiro, Mark Lathrop and Michel Roger |
| EPI_ISL_535857 | Hôpital Pierre-Boucher | Laboratoire de santé publique du Québec | Sandrine Moreira, Ioannis Ragoussis, Guillaume Bourque, Jesse Shapiro, Mark Lathrop and Michel Roger |
| EPI_ISL_535858 | Hôpital de Gatineau | Laboratoire de santé publique du Québec | Sandrine Moreira, Ioannis Ragoussis, Guillaume Bourque, Jesse Shapiro, Mark Lathrop and Michel Roger |
| EPI_ISL_535859, EPI_ISL_535860 | Hôpital Sainte-Croix | Laboratoire de santé publique du Québec | Sandrine Moreira, Ioannis Ragoussis, Guillaume Bourque, Jesse Shapiro, Mark Lathrop and Michel Roger |
| EPI_ISL_535861, EPI_ISL_535862, EPI_ISL_535863 | Hôpital Pierre-Le Gardeur | Laboratoire de santé publique du Québec | Sandrine Moreira, Ioannis Ragoussis, Guillaume Bourque, Jesse Shapiro, Mark Lathrop and Michel Roger |
| EPI_ISL_535864 | CSSS Haut-Richelieu/Rouville (Hôpital) | Laboratoire de santé publique du Québec | Sandrine Moreira, Ioannis Ragoussis, Guillaume Bourque, Jesse Shapiro, Mark Lathrop and Michel Roger |
| EPI_ISL_535865 | Hôpital de Saint-Eustache | Laboratoire de santé publique du Québec | Sandrine Moreira, Ioannis Ragoussis, Guillaume Bourque, Jesse Shapiro, Mark Lathrop and Michel Roger |
| EPI_ISL_535866 | Hôpital Charles-LeMoyne | Laboratoire de santé publique du Québec | Sandrine Moreira, Ioannis Ragoussis, Guillaume Bourque, Jesse Shapiro, Mark Lathrop and Michel Roger |
| EPI_ISL_535867 | Hôpital de Saint-Eustache | Laboratoire de santé publique du Québec | Sandrine Moreira, Ioannis Ragoussis, Guillaume Bourque, Jesse Shapiro, Mark Lathrop and Michel Roger |
| EPI_ISL_535868 | Hôpital du Suroît | Laboratoire de santé publique du Québec | Sandrine Moreira, Ioannis Ragoussis, Guillaume Bourque, Jesse Shapiro, Mark Lathrop and Michel Roger |
| EPI_ISL_535869 | Hôpital Charles-LeMoyne | Laboratoire de santé publique du Québec | Sandrine Moreira, Ioannis Ragoussis, Guillaume Bourque, Jesse Shapiro, Mark Lathrop and Michel Roger |
| EPI_ISL_535870, EPI_ISL_535871 | Centre Hospitalier Régional de Lanaudière | Laboratoire de santé publique du Québec | Sandrine Moreira, Ioannis Ragoussis, Guillaume Bourque, Jesse Shapiro, Mark Lathrop and Michel Roger |
| EPI_ISL_535872, EPI_ISL_535873 | Centre hospitalier régional du Grand Portage | Laboratoire de santé publique du Québec | Sandrine Moreira, Ioannis Ragoussis, Guillaume Bourque, Jesse Shapiro, Mark Lathrop and Michel Roger |
| EPI_ISL_535874 | Hôpital Charles-LeMoyne | Laboratoire de santé publique du Québec | Sandrine Moreira, Ioannis Ragoussis, Guillaume Bourque, Jesse Shapiro, Mark Lathrop and Michel Roger |
| EPI_ISL_535875 | Hôpital général du Lakeshore | Laboratoire de santé publique du Québec | Sandrine Moreira, Ioannis Ragoussis, Guillaume Bourque, Jesse Shapiro, Mark Lathrop and Michel Roger |
| EPI_ISL_535876, EPI_ISL_535877 | CSSS Haut-Richelieu/Rouville (Hôpital) | Laboratoire de santé publique du Québec | Sandrine Moreira, Ioannis Ragoussis, Guillaume Bourque, Jesse Shapiro, Mark Lathrop and Michel Roger |
| EPI_ISL_535878 | Hôpital de Saint-Eustache | Laboratoire de santé publique du Québec | Sandrine Moreira, Ioannis Ragoussis, Guillaume Bourque, Jesse Shapiro, Mark Lathrop and Michel Roger |
| EPI_ISL_535879 | Hôpital Pierre-Boucher | Laboratoire de santé publique du Québec | Sandrine Moreira, Ioannis Ragoussis, Guillaume Bourque, Jesse Shapiro, Mark Lathrop and Michel Roger |
| EPI_ISL_535880 | Hôpital du Suroît | Laboratoire de santé publique du Québec | Sandrine Moreira, Ioannis Ragoussis, Guillaume Bourque, Jesse Shapiro, Mark Lathrop and Michel Roger |
| EPI_ISL_535881, EPI_ISL_535882, EPI_ISL_535883 | Hôpital Pierre-Le Gardeur | Laboratoire de santé publique du Québec | Sandrine Moreira, Ioannis Ragoussis, Guillaume Bourque, Jesse Shapiro, Mark Lathrop and Michel Roger |
| EPI_ISL_535884 | CSSS Haut-Richelieu/Rouville (Hôpital) | Laboratoire de santé publique du Québec | Sandrine Moreira, Ioannis Ragoussis, Guillaume Bourque, Jesse Shapiro, Mark Lathrop and Michel Roger |

| | | | |
|---|---|---|---|
| EPI_ISL_535885 | Hôpital de Maria | Laboratoire de santé publique du Québec | Sandrine Moreira, Ioannis Ragoussis, Guillaume Bourque, Jesse Shapiro, Mark Lathrop and Michel Roger |
| EPI_ISL_535886, EPI_ISL_535887 | Hôpital de Hull | Laboratoire de santé publique du Québec | Sandrine Moreira, Ioannis Ragoussis, Guillaume Bourque, Jesse Shapiro, Mark Lathrop and Michel Roger |
| EPI_ISL_535888 | Hôpital Charles-LeMoyne | Laboratoire de santé publique du Québec | Sandrine Moreira, Ioannis Ragoussis, Guillaume Bourque, Jesse Shapiro, Mark Lathrop and Michel Roger |
| EPI_ISL_535889 | Hôpital de Saint-Eustache | Laboratoire de santé publique du Québec | Sandrine Moreira, Ioannis Ragoussis, Guillaume Bourque, Jesse Shapiro, Mark Lathrop and Michel Roger |
| EPI_ISL_535890, EPI_ISL_535891 | Hôpital Charles-LeMoyne | Laboratoire de santé publique du Québec | Sandrine Moreira, Ioannis Ragoussis, Guillaume Bourque, Jesse Shapiro, Mark Lathrop and Michel Roger |
| EPI_ISL_535892, EPI_ISL_535893 | Hôpital Pierre-Le Gardeur | Laboratoire de santé publique du Québec | Sandrine Moreira, Ioannis Ragoussis, Guillaume Bourque, Jesse Shapiro, Mark Lathrop and Michel Roger |
| EPI_ISL_535894, EPI_ISL_535895, EPI_ISL_535896 | Centre Hospitalier Régional de Lanaudière | Laboratoire de santé publique du Québec | Sandrine Moreira, Ioannis Ragoussis, Guillaume Bourque, Jesse Shapiro, Mark Lathrop and Michel Roger |
| EPI_ISL_535897, EPI_ISL_535898, EPI_ISL_535899, EPI_ISL_535900, EPI_ISL_535901 | Hôpital Sainte-Croix | Laboratoire de santé publique du Québec | Sandrine Moreira, Ioannis Ragoussis, Guillaume Bourque, Jesse Shapiro, Mark Lathrop and Michel Roger |
| EPI_ISL_535902 | CSSS Haut-Richelieu/Rouville (Hôpital) | Laboratoire de santé publique du Québec | Sandrine Moreira, Ioannis Ragoussis, Guillaume Bourque, Jesse Shapiro, Mark Lathrop and Michel Roger |
| EPI_ISL_535903, EPI_ISL_535904 | Hôtel-Dieu de Sorel | Laboratoire de santé publique du Québec | Sandrine Moreira, Ioannis Ragoussis, Guillaume Bourque, Jesse Shapiro, Mark Lathrop and Michel Roger |
| EPI_ISL_535905, EPI_ISL_535906 | Hôpital de Chicoutimi | Laboratoire de santé publique du Québec | Sandrine Moreira, Ioannis Ragoussis, Guillaume Bourque, Jesse Shapiro, Mark Lathrop and Michel Roger |
| EPI_ISL_535907 | Centre Hospitalier Régional de Lanaudière | Laboratoire de santé publique du Québec | Sandrine Moreira, Ioannis Ragoussis, Guillaume Bourque, Jesse Shapiro, Mark Lathrop and Michel Roger |
| EPI_ISL_535908, EPI_ISL_535909 | Hôpital Pierre-Boucher | Laboratoire de santé publique du Québec | Sandrine Moreira, Ioannis Ragoussis, Guillaume Bourque, Jesse Shapiro, Mark Lathrop and Michel Roger |
| EPI_ISL_535910 | CSSS Haut-Richelieu/Rouville (Hôpital) | Laboratoire de santé publique du Québec | Sandrine Moreira, Ioannis Ragoussis, Guillaume Bourque, Jesse Shapiro, Mark Lathrop and Michel Roger |
| EPI_ISL_535911 | Hôpital de Hull | Laboratoire de santé publique du Québec | Sandrine Moreira, Ioannis Ragoussis, Guillaume Bourque, Jesse Shapiro, Mark Lathrop and Michel Roger |
| EPI_ISL_535912 | Hôpital Honoré-Mercier | Laboratoire de santé publique du Québec | Sandrine Moreira, Ioannis Ragoussis, Guillaume Bourque, Jesse Shapiro, Mark Lathrop and Michel Roger |
| EPI_ISL_535913 | Hôpital du Suroît | Laboratoire de santé publique du Québec | Sandrine Moreira, Ioannis Ragoussis, Guillaume Bourque, Jesse Shapiro, Mark Lathrop and Michel Roger |
| EPI_ISL_535914 | Hôpital du Centre-de-la-Mauricie | Laboratoire de santé publique du Québec | Sandrine Moreira, Ioannis Ragoussis, Guillaume Bourque, Jesse Shapiro, Mark Lathrop and Michel Roger |
| EPI_ISL_535915 | Hôpital Honoré-Mercier | Laboratoire de santé publique du Québec | Sandrine Moreira, Ioannis Ragoussis, Guillaume Bourque, Jesse Shapiro, Mark Lathrop and Michel Roger |
| EPI_ISL_535916 | Centre de SSS de Trois-Rivières | Laboratoire de santé publique du Québec | Sandrine Moreira, Ioannis Ragoussis, Guillaume Bourque, Jesse Shapiro, Mark Lathrop and Michel Roger |
| EPI_ISL_535917, EPI_ISL_535918, EPI_ISL_535919 | Hôpital du Suroît | Laboratoire de santé publique du Québec | Sandrine Moreira, Ioannis Ragoussis, Guillaume Bourque, Jesse Shapiro, Mark Lathrop and Michel Roger |
| EPI_ISL_535920, EPI_ISL_535921, EPI_ISL_535922, EPI_ISL_535923 | Hôpital Charles-LeMoyne | Laboratoire de santé publique du Québec | Sandrine Moreira, Ioannis Ragoussis, Guillaume Bourque, Jesse Shapiro, Mark Lathrop and Michel Roger |
| EPI_ISL_535924 | Hôpital de Saint-Eustache | Laboratoire de santé publique du Québec | Sandrine Moreira, Ioannis Ragoussis, Guillaume Bourque, Jesse Shapiro, Mark Lathrop and Michel Roger |
| EPI_ISL_535925 | CSSS Haut-Richelieu/Rouville (Hôpital) | Laboratoire de santé publique du Québec | Sandrine Moreira, Ioannis Ragoussis, Guillaume Bourque, Jesse Shapiro, Mark Lathrop and Michel Roger |
| EPI_ISL_535926 | Hôpital de Saint-Eustache | Laboratoire de santé publique du Québec | Sandrine Moreira, Ioannis Ragoussis, Guillaume Bourque, Jesse Shapiro, Mark Lathrop and Michel Roger |
| EPI_ISL_535927 | Hôpital Charles-LeMoyne | Laboratoire de santé publique du Québec | Sandrine Moreira, Ioannis Ragoussis, Guillaume Bourque, Jesse Shapiro, Mark Lathrop and Michel Roger |
| EPI_ISL_535928 | CSSS Haut-Richelieu/Rouville (Hôpital) | Laboratoire de santé publique du Québec | Sandrine Moreira, Ioannis Ragoussis, Guillaume Bourque, Jesse Shapiro, Mark Lathrop and Michel Roger |
| EPI_ISL_535929, EPI_ISL_535930 | Hôpital de Saint-Eustache | Laboratoire de santé publique du Québec | Sandrine Moreira, Ioannis Ragoussis, Guillaume Bourque, Jesse Shapiro, Mark Lathrop and Michel Roger |
| EPI_ISL_535931, EPI_ISL_535932, EPI_ISL_535933, EPI_ISL_535934, EPI_ISL_535935 | Centre Hospitalier Régional de Lanaudière | Laboratoire de santé publique du Québec | Sandrine Moreira, Ioannis Ragoussis, Guillaume Bourque, Jesse Shapiro, Mark Lathrop and Michel Roger |
| EPI_ISL_535936 | Hôpital Pierre-Le Gardeur | Laboratoire de santé publique du Québec | Sandrine Moreira, Ioannis Ragoussis, Guillaume Bourque, Jesse Shapiro, Mark Lathrop and Michel Roger |
| EPI_ISL_535937 | Hôpital du Centre-de-la-Mauricie | Laboratoire de santé publique du Québec | Sandrine Moreira, Ioannis Ragoussis, Guillaume Bourque, Jesse Shapiro, Mark Lathrop and Michel Roger |
| EPI_ISL_535938 | Hôpital Pierre-Le Gardeur | Laboratoire de santé publique du Québec | Sandrine Moreira, Ioannis Ragoussis, Guillaume Bourque, Jesse Shapiro, Mark Lathrop and Michel Roger |
| EPI_ISL_535939, EPI_ISL_535940 | Centre de SSS D'Arthabaska-et-de-l'érable - Hôtel-Dieu | Laboratoire de santé publique du Québec | Sandrine Moreira, Ioannis Ragoussis, Guillaume Bourque, Jesse Shapiro, Mark Lathrop and Michel Roger |
| EPI_ISL_535941, EPI_ISL_535942, EPI_ISL_535943 | Hôpital de Saint-Eustache | Laboratoire de santé publique du Québec | Sandrine Moreira, Ioannis Ragoussis, Guillaume Bourque, Jesse Shapiro, Mark Lathrop and Michel Roger |
| EPI_ISL_535944 | Hôpital Charles-LeMoyne | Laboratoire de santé publique du Québec | Sandrine Moreira, Ioannis Ragoussis, Guillaume Bourque, Jesse Shapiro, Mark Lathrop and Michel Roger |
| EPI_ISL_535945 | Hôpital de Mont-Laurier | Laboratoire de santé publique du Québec | Sandrine Moreira, Ioannis Ragoussis, Guillaume Bourque, Jesse Shapiro, Mark Lathrop and Michel Roger |
| EPI_ISL_535946 | CSSS Haut-Richelieu/Rouville (Hôpital) | Laboratoire de santé publique du Québec | Sandrine Moreira, Ioannis Ragoussis, Guillaume Bourque, Jesse Shapiro, Mark Lathrop and Michel Roger |
| EPI_ISL_535947 | Hôpital de Saint-Eustache | Laboratoire de santé publique du Québec | Sandrine Moreira, Ioannis Ragoussis, Guillaume Bourque, Jesse Shapiro, Mark Lathrop and Michel Roger |
| EPI_ISL_535948 | Hôpital Pierre-Boucher | Laboratoire de santé publique du Québec | Sandrine Moreira, Ioannis Ragoussis, Guillaume Bourque, Jesse Shapiro, Mark Lathrop and Michel Roger |
| EPI_ISL_535949, EPI_ISL_535950 | Hôpital Charles-LeMoyne | Laboratoire de santé publique du Québec | Sandrine Moreira, Ioannis Ragoussis, Guillaume Bourque, Jesse Shapiro, Mark Lathrop and Michel Roger |
| EPI_ISL_535951 | Hôpital de Saint-Eustache | Laboratoire de santé publique du Québec | Sandrine Moreira, Ioannis Ragoussis, Guillaume Bourque, Jesse Shapiro, Mark Lathrop and Michel Roger |
| EPI_ISL_535952, EPI_ISL_535953 | Hôpital Charles-LeMoyne | Laboratoire de santé publique du Québec | Sandrine Moreira, Ioannis Ragoussis, Guillaume Bourque, Jesse Shapiro, Mark Lathrop and Michel Roger |
| EPI_ISL_535954 | Hôpital Pierre-Le Gardeur | Laboratoire de santé publique du Québec | Sandrine Moreira, Ioannis Ragoussis, Guillaume Bourque, Jesse Shapiro, Mark Lathrop and Michel Roger |
| EPI_ISL_535955 | Centre Hospitalier Régional de Lanaudière | Laboratoire de santé publique du Québec | Sandrine Moreira, Ioannis Ragoussis, Guillaume Bourque, Jesse Shapiro, Mark Lathrop and Michel Roger |
| EPI_ISL_535956 | Hôpital Pierre-Le Gardeur | Laboratoire de santé publique du Québec | Sandrine Moreira, Ioannis Ragoussis, Guillaume Bourque, Jesse Shapiro, Mark Lathrop and Michel Roger |
| EPI_ISL_535957 | Hôpital Charles-LeMoyne | Laboratoire de santé publique du Québec | Sandrine Moreira, Ioannis Ragoussis, Guillaume Bourque, Jesse Shapiro, Mark Lathrop and Michel Roger |
| EPI_ISL_535958, EPI_ISL_535959 | Centre hospitalier régional du Grand Portage | Laboratoire de santé publique du Québec | Sandrine Moreira, Ioannis Ragoussis, Guillaume Bourque, Jesse Shapiro, Mark Lathrop and Michel Roger |
| EPI_ISL_535960 | Centre hospitalier Anna-Laberge | Laboratoire de santé publique du Québec | Sandrine Moreira, Ioannis Ragoussis, Guillaume Bourque, Jesse Shapiro, Mark Lathrop and Michel Roger |
| EPI_ISL_535961, EPI_ISL_535962 | Hôpital Charles-LeMoyne | Laboratoire de santé publique du Québec | Sandrine Moreira, Ioannis Ragoussis, Guillaume Bourque, Jesse Shapiro, Mark Lathrop and Michel Roger |
| EPI_ISL_535963 | Hôpital d'Alma | Laboratoire de santé publique du Québec | Sandrine Moreira, Ioannis Ragoussis, Guillaume Bourque, Jesse Shapiro, Mark Lathrop and Michel Roger |
| EPI_ISL_535964, EPI_ISL_535965 | Hôpital de Chicoutimi | Laboratoire de santé publique du Québec | Sandrine Moreira, Ioannis Ragoussis, Guillaume Bourque, Jesse Shapiro, Mark Lathrop and Michel Roger |
| EPI_ISL_535966 | Hôpital Charles-LeMoyne | Laboratoire de santé publique du Québec | Sandrine Moreira, Ioannis Ragoussis, Guillaume Bourque, Jesse Shapiro, Mark Lathrop and Michel Roger |
| EPI_ISL_535967 | CSSS Haut-Richelieu/Rouville (Hôpital) | Laboratoire de santé publique du Québec | Sandrine Moreira, Ioannis Ragoussis, Guillaume Bourque, Jesse Shapiro, Mark Lathrop and Michel Roger |

| | | | |
|---|---|---|---|
| EPI_ISL_535968 | Hôpital du Centre-de-la-Mauricie | Laboratoire de santé publique du Québec | Sandrine Moreira, Ioannis Ragoussis, Guillaume Bourque, Jesse Shapiro, Mark Lathrop and Michel Roger |
| EPI_ISL_535969 | Hôpital Sainte-Croix | Laboratoire de santé publique du Québec | Sandrine Moreira, Ioannis Ragoussis, Guillaume Bourque, Jesse Shapiro, Mark Lathrop and Michel Roger |
| EPI_ISL_535970 | CSSS de la Minganie | Laboratoire de santé publique du Québec | Sandrine Moreira, Ioannis Ragoussis, Guillaume Bourque, Jesse Shapiro, Mark Lathrop and Michel Roger |
| EPI_ISL_535971 | Hôpital du Centre-de-la-Mauricie | Laboratoire de santé publique du Québec | Sandrine Moreira, Ioannis Ragoussis, Guillaume Bourque, Jesse Shapiro, Mark Lathrop and Michel Roger |
| EPI_ISL_535972, EPI_ISL_535973 | Hôpital Charles-LeMoyne | Laboratoire de santé publique du Québec | Sandrine Moreira, Ioannis Ragoussis, Guillaume Bourque, Jesse Shapiro, Mark Lathrop and Michel Roger |
| EPI_ISL_535974, EPI_ISL_535975 | Hôpital du Centre-de-la-Mauricie | Laboratoire de santé publique du Québec | Sandrine Moreira, Ioannis Ragoussis, Guillaume Bourque, Jesse Shapiro, Mark Lathrop and Michel Roger |
| EPI_ISL_535976, EPI_ISL_535977 | Centre de SSS de Trois-Rivières | Laboratoire de santé publique du Québec | Sandrine Moreira, Ioannis Ragoussis, Guillaume Bourque, Jesse Shapiro, Mark Lathrop and Michel Roger |
| EPI_ISL_535978 | Hôpital du Centre-de-la-Mauricie | Laboratoire de santé publique du Québec | Sandrine Moreira, Ioannis Ragoussis, Guillaume Bourque, Jesse Shapiro, Mark Lathrop and Michel Roger |
| EPI_ISL_535979 | Centre de SSS de Trois-Rivières | Laboratoire de santé publique du Québec | Sandrine Moreira, Ioannis Ragoussis, Guillaume Bourque, Jesse Shapiro, Mark Lathrop and Michel Roger |
| EPI_ISL_535980, EPI_ISL_535981 | Centre hospitalier Anna-Laberge | Laboratoire de santé publique du Québec | Sandrine Moreira, Ioannis Ragoussis, Guillaume Bourque, Jesse Shapiro, Mark Lathrop and Michel Roger |
| EPI_ISL_535982 | Hôpital de Saint-Eustache | Laboratoire de santé publique du Québec | Sandrine Moreira, Ioannis Ragoussis, Guillaume Bourque, Jesse Shapiro, Mark Lathrop and Michel Roger |
| EPI_ISL_535983 | Hôpital Charles-LeMoyne | Laboratoire de santé publique du Québec | Sandrine Moreira, Ioannis Ragoussis, Guillaume Bourque, Jesse Shapiro, Mark Lathrop and Michel Roger |
| EPI_ISL_535984 | CSSS Haut-Richelieu/Rouville (Hôpital) | Laboratoire de santé publique du Québec | Sandrine Moreira, Ioannis Ragoussis, Guillaume Bourque, Jesse Shapiro, Mark Lathrop and Michel Roger |
| EPI_ISL_535985, EPI_ISL_535986, EPI_ISL_535987 | Hôpital de Rouyn-Noranda | Laboratoire de santé publique du Québec | Sandrine Moreira, Ioannis Ragoussis, Guillaume Bourque, Jesse Shapiro, Mark Lathrop and Michel Roger |
| EPI_ISL_535988 | Hôpital du Suroît | Laboratoire de santé publique du Québec | Sandrine Moreira, Ioannis Ragoussis, Guillaume Bourque, Jesse Shapiro, Mark Lathrop and Michel Roger |
| EPI_ISL_535989 | Hôpital de Hull | Laboratoire de santé publique du Québec | Sandrine Moreira, Ioannis Ragoussis, Guillaume Bourque, Jesse Shapiro, Mark Lathrop and Michel Roger |
| EPI_ISL_535990 | Hôtel-Dieu de Sorel | Laboratoire de santé publique du Québec | Sandrine Moreira, Ioannis Ragoussis, Guillaume Bourque, Jesse Shapiro, Mark Lathrop and Michel Roger |
| EPI_ISL_535991 | Hôpital et Centre d'Hébergement de Sept-Iles | Laboratoire de santé publique du Québec | Sandrine Moreira, Ioannis Ragoussis, Guillaume Bourque, Jesse Shapiro, Mark Lathrop and Michel Roger |
| EPI_ISL_535992 | Hôtel-Dieu de Sorel | Laboratoire de santé publique du Québec | Sandrine Moreira, Ioannis Ragoussis, Guillaume Bourque, Jesse Shapiro, Mark Lathrop and Michel Roger |
| EPI_ISL_535993 | Hôpital de Hull | Laboratoire de santé publique du Québec | Sandrine Moreira, Ioannis Ragoussis, Guillaume Bourque, Jesse Shapiro, Mark Lathrop and Michel Roger |
| EPI_ISL_535994 | Hôpital Laurentien | Laboratoire de santé publique du Québec | Sandrine Moreira, Ioannis Ragoussis, Guillaume Bourque, Jesse Shapiro, Mark Lathrop and Michel Roger |
| EPI_ISL_535995 | Hôpital de Hull | Laboratoire de santé publique du Québec | Sandrine Moreira, Ioannis Ragoussis, Guillaume Bourque, Jesse Shapiro, Mark Lathrop and Michel Roger |
| EPI_ISL_535996, EPI_ISL_535997, EPI_ISL_535998, EPI_ISL_535999 | Hôpital du Centre-de-la-Mauricie | Laboratoire de santé publique du Québec | Sandrine Moreira, Ioannis Ragoussis, Guillaume Bourque, Jesse Shapiro, Mark Lathrop and Michel Roger |
| EPI_ISL_536000 | Centre hospitalier Anna-Laberge | Laboratoire de santé publique du Québec | Sandrine Moreira, Ioannis Ragoussis, Guillaume Bourque, Jesse Shapiro, Mark Lathrop and Michel Roger |
| EPI_ISL_536001 | Hôpital du Centre-de-la-Mauricie | Laboratoire de santé publique du Québec | Sandrine Moreira, Ioannis Ragoussis, Guillaume Bourque, Jesse Shapiro, Mark Lathrop and Michel Roger |
| EPI_ISL_536002 | Centre Hospitalier Régional de Lanaudière | Laboratoire de santé publique du Québec | Sandrine Moreira, Ioannis Ragoussis, Guillaume Bourque, Jesse Shapiro, Mark Lathrop and Michel Roger |
| EPI_ISL_536003 | Hôpital Sainte-Croix | Laboratoire de santé publique du Québec | Sandrine Moreira, Ioannis Ragoussis, Guillaume Bourque, Jesse Shapiro, Mark Lathrop and Michel Roger |
| EPI_ISL_536004 | Centre hospitalier Anna-Laberge | Laboratoire de santé publique du Québec | Sandrine Moreira, Ioannis Ragoussis, Guillaume Bourque, Jesse Shapiro, Mark Lathrop and Michel Roger |
| EPI_ISL_536005, EPI_ISL_536006 | Centre Hospitalier Régional de Lanaudière | Laboratoire de santé publique du Québec | Sandrine Moreira, Ioannis Ragoussis, Guillaume Bourque, Jesse Shapiro, Mark Lathrop and Michel Roger |
| EPI_ISL_536007 | Centre de SSS de Trois-Rivières | Laboratoire de santé publique du Québec | Sandrine Moreira, Ioannis Ragoussis, Guillaume Bourque, Jesse Shapiro, Mark Lathrop and Michel Roger |
| EPI_ISL_536008 | Hôpital de Saint-Eustache | Laboratoire de santé publique du Québec | Sandrine Moreira, Ioannis Ragoussis, Guillaume Bourque, Jesse Shapiro, Mark Lathrop and Michel Roger |
| EPI_ISL_536009 | Centre Hospitalier Régional de Lanaudière | Laboratoire de santé publique du Québec | Sandrine Moreira, Ioannis Ragoussis, Guillaume Bourque, Jesse Shapiro, Mark Lathrop and Michel Roger |
| EPI_ISL_536010, EPI_ISL_536011 | Hôpital Sainte-Croix | Laboratoire de santé publique du Québec | Sandrine Moreira, Ioannis Ragoussis, Guillaume Bourque, Jesse Shapiro, Mark Lathrop and Michel Roger |
| EPI_ISL_536012 | Centre Hospitalier Régional de Lanaudière | Laboratoire de santé publique du Québec | Sandrine Moreira, Ioannis Ragoussis, Guillaume Bourque, Jesse Shapiro, Mark Lathrop and Michel Roger |
| EPI_ISL_536013 | Hôpital Honoré-Mercier | Laboratoire de santé publique du Québec | Sandrine Moreira, Ioannis Ragoussis, Guillaume Bourque, Jesse Shapiro, Mark Lathrop and Michel Roger |
| EPI_ISL_536014, EPI_ISL_536015, EPI_ISL_536016 | Centre Hospitalier Régional de Lanaudière | Laboratoire de santé publique du Québec | Sandrine Moreira, Ioannis Ragoussis, Guillaume Bourque, Jesse Shapiro, Mark Lathrop and Michel Roger |
| EPI_ISL_536017, EPI_ISL_536018 | Hôpital Charles-LeMoyne | Laboratoire de santé publique du Québec | Sandrine Moreira, Ioannis Ragoussis, Guillaume Bourque, Jesse Shapiro, Mark Lathrop and Michel Roger |
| EPI_ISL_536019 | Hôpital de Chicoutimi | Laboratoire de santé publique du Québec | Sandrine Moreira, Ioannis Ragoussis, Guillaume Bourque, Jesse Shapiro, Mark Lathrop and Michel Roger |
| EPI_ISL_536020 | Hôpital Charles-LeMoyne | Laboratoire de santé publique du Québec | Sandrine Moreira, Ioannis Ragoussis, Guillaume Bourque, Jesse Shapiro, Mark Lathrop and Michel Roger |
| EPI_ISL_536021 | Hôpital Pierre-Boucher | Laboratoire de santé publique du Québec | Sandrine Moreira, Ioannis Ragoussis, Guillaume Bourque, Jesse Shapiro, Mark Lathrop and Michel Roger |
| EPI_ISL_536022 | Hôpital régional de Saint-Jérôme | Laboratoire de santé publique du Québec | Sandrine Moreira, Ioannis Ragoussis, Guillaume Bourque, Jesse Shapiro, Mark Lathrop and Michel Roger |
| EPI_ISL_536023 | Centre hospitalier Anna-Laberge | Laboratoire de santé publique du Québec | Sandrine Moreira, Ioannis Ragoussis, Guillaume Bourque, Jesse Shapiro, Mark Lathrop and Michel Roger |
| EPI_ISL_536024, EPI_ISL_536025 | Hôpital Charles-LeMoyne | Laboratoire de santé publique du Québec | Sandrine Moreira, Ioannis Ragoussis, Guillaume Bourque, Jesse Shapiro, Mark Lathrop and Michel Roger |
| EPI_ISL_536026 | Hôpital du Suroît | Laboratoire de santé publique du Québec | Sandrine Moreira, Ioannis Ragoussis, Guillaume Bourque, Jesse Shapiro, Mark Lathrop and Michel Roger |
| EPI_ISL_536027, EPI_ISL_536028 | CSSS Haut-Richelieu/Rouville (Hôpital) | Laboratoire de santé publique du Québec | Sandrine Moreira, Ioannis Ragoussis, Guillaume Bourque, Jesse Shapiro, Mark Lathrop and Michel Roger |
| EPI_ISL_536029 | Hôpital Honoré-Mercier | Laboratoire de santé publique du Québec | Sandrine Moreira, Ioannis Ragoussis, Guillaume Bourque, Jesse Shapiro, Mark Lathrop and Michel Roger |
| EPI_ISL_536030 | Hôpital du Centre-de-la-Mauricie | Laboratoire de santé publique du Québec | Sandrine Moreira, Ioannis Ragoussis, Guillaume Bourque, Jesse Shapiro, Mark Lathrop and Michel Roger |
| EPI_ISL_536031 | Centre de SSS de Trois-Rivières | Laboratoire de santé publique du Québec | Sandrine Moreira, Ioannis Ragoussis, Guillaume Bourque, Jesse Shapiro, Mark Lathrop and Michel Roger |
| EPI_ISL_536032, EPI_ISL_536033 | Hôpital du Centre-de-la-Mauricie | Laboratoire de santé publique du Québec | Sandrine Moreira, Ioannis Ragoussis, Guillaume Bourque, Jesse Shapiro, Mark Lathrop and Michel Roger |
| EPI_ISL_536034, EPI_ISL_536035, EPI_ISL_536036 | Centre de SSS de Trois-Rivières | Laboratoire de santé publique du Québec | Sandrine Moreira, Ioannis Ragoussis, Guillaume Bourque, Jesse Shapiro, Mark Lathrop and Michel Roger |
| EPI_ISL_536037 | Hôpital du Centre-de-la-Mauricie | Laboratoire de santé publique du Québec | Sandrine Moreira, Ioannis Ragoussis, Guillaume Bourque, Jesse Shapiro, Mark Lathrop and Michel Roger |
| EPI_ISL_536038 | Centre de SSS de Trois-Rivières | Laboratoire de santé publique du Québec | Sandrine Moreira, Ioannis Ragoussis, Guillaume Bourque, Jesse Shapiro, Mark Lathrop and Michel Roger |
| EPI_ISL_536039 | Hôpital Pierre-Boucher | Laboratoire de santé publique du Québec | Sandrine Moreira, Ioannis Ragoussis, Guillaume Bourque, Jesse Shapiro, Mark Lathrop and Michel Roger |
| EPI_ISL_536040 | CSSS Haut-Richelieu/Rouville (Hôpital) | Laboratoire de santé publique du Québec | Sandrine Moreira, Ioannis Ragoussis, Guillaume Bourque, Jesse Shapiro, Mark Lathrop and Michel Roger |
| EPI_ISL_536041 | Hôpital Pierre-Boucher | Laboratoire de santé publique du Québec | Sandrine Moreira, Ioannis Ragoussis, Guillaume Bourque, Jesse Shapiro, Mark Lathrop and Michel Roger |
| EPI_ISL_536042 | CSSS Haut-Richelieu/Rouville (Hôpital) | Laboratoire de santé publique du Québec | Sandrine Moreira, Ioannis Ragoussis, Guillaume Bourque, Jesse Shapiro, Mark Lathrop and Michel Roger |

| | | | |
|---|---|---|---|
| EPI_ISL_536043 | Centre hospitalier Anna-Laberge | Laboratoire de santé publique du Québec | Sandrine Moreira, Ioannis Ragoussis, Guillaume Bourque, Jesse Shapiro, Mark Lathrop and Michel Roger |
| EPI_ISL_536044 | CSSS Haut-Richelieu/Rouville (Hôpital) | Laboratoire de santé publique du Québec | Sandrine Moreira, Ioannis Ragoussis, Guillaume Bourque, Jesse Shapiro, Mark Lathrop and Michel Roger |
| EPI_ISL_536045 | Centre hospitalier Anna-Laberge | Laboratoire de santé publique du Québec | Sandrine Moreira, Ioannis Ragoussis, Guillaume Bourque, Jesse Shapiro, Mark Lathrop and Michel Roger |
| EPI_ISL_536046 | Hôpital d'Alma | Laboratoire de santé publique du Québec | Sandrine Moreira, Ioannis Ragoussis, Guillaume Bourque, Jesse Shapiro, Mark Lathrop and Michel Roger |
| EPI_ISL_536047, EPI_ISL_536048 | Centre Hospitalier Régional de Lanaudière | Laboratoire de santé publique du Québec | Sandrine Moreira, Ioannis Ragoussis, Guillaume Bourque, Jesse Shapiro, Mark Lathrop and Michel Roger |
| EPI_ISL_536049 | Hôpital de Saint-Eustache | Laboratoire de santé publique du Québec | Sandrine Moreira, Ioannis Ragoussis, Guillaume Bourque, Jesse Shapiro, Mark Lathrop and Michel Roger |
| EPI_ISL_536050 | Centre Hospitalier Régional de Lanaudière | Laboratoire de santé publique du Québec | Sandrine Moreira, Ioannis Ragoussis, Guillaume Bourque, Jesse Shapiro, Mark Lathrop and Michel Roger |
| EPI_ISL_536051 | Hôpital de Hull | Laboratoire de santé publique du Québec | Sandrine Moreira, Ioannis Ragoussis, Guillaume Bourque, Jesse Shapiro, Mark Lathrop and Michel Roger |
| EPI_ISL_536052, EPI_ISL_536053, EPI_ISL_536054 | Centre Hospitalier Régional de Lanaudière | Laboratoire de santé publique du Québec | Sandrine Moreira, Ioannis Ragoussis, Guillaume Bourque, Jesse Shapiro, Mark Lathrop and Michel Roger |
| EPI_ISL_536055 | Hôpital Honoré-Mercier | Laboratoire de santé publique du Québec | Sandrine Moreira, Ioannis Ragoussis, Guillaume Bourque, Jesse Shapiro, Mark Lathrop and Michel Roger |
| EPI_ISL_536056, EPI_ISL_536057 | Hôpital Sainte-Croix | Laboratoire de santé publique du Québec | Sandrine Moreira, Ioannis Ragoussis, Guillaume Bourque, Jesse Shapiro, Mark Lathrop and Michel Roger |
| EPI_ISL_536058 | Hôpital de Saint-Eustache | Laboratoire de santé publique du Québec | Sandrine Moreira, Ioannis Ragoussis, Guillaume Bourque, Jesse Shapiro, Mark Lathrop and Michel Roger |
| EPI_ISL_536059, EPI_ISL_536060 | Centre Hospitalier Régional de Lanaudière | Laboratoire de santé publique du Québec | Sandrine Moreira, Ioannis Ragoussis, Guillaume Bourque, Jesse Shapiro, Mark Lathrop and Michel Roger |
| EPI_ISL_536061, EPI_ISL_536062 | Hôpital Pierre-Le Gardeur | Laboratoire de santé publique du Québec | Sandrine Moreira, Ioannis Ragoussis, Guillaume Bourque, Jesse Shapiro, Mark Lathrop and Michel Roger |
| EPI_ISL_536063 | Centre Hospitalier Régional de Lanaudière | Laboratoire de santé publique du Québec | Sandrine Moreira, Ioannis Ragoussis, Guillaume Bourque, Jesse Shapiro, Mark Lathrop and Michel Roger |
| EPI_ISL_536064, EPI_ISL_536065, EPI_ISL_536066, EPI_ISL_536067, EPI_ISL_536068 | Hôpital Pierre-Le Gardeur | Laboratoire de santé publique du Québec | Sandrine Moreira, Ioannis Ragoussis, Guillaume Bourque, Jesse Shapiro, Mark Lathrop and Michel Roger |
| EPI_ISL_536069 | Hôpital de Saint-Eustache | Laboratoire de santé publique du Québec | Sandrine Moreira, Ioannis Ragoussis, Guillaume Bourque, Jesse Shapiro, Mark Lathrop and Michel Roger |
| EPI_ISL_536070 | Hôpital Pierre-Le Gardeur | Laboratoire de santé publique du Québec | Sandrine Moreira, Ioannis Ragoussis, Guillaume Bourque, Jesse Shapiro, Mark Lathrop and Michel Roger |
| EPI_ISL_536071, EPI_ISL_536072 | Hôpital de Saint-Eustache | Laboratoire de santé publique du Québec | Sandrine Moreira, Ioannis Ragoussis, Guillaume Bourque, Jesse Shapiro, Mark Lathrop and Michel Roger |
| EPI_ISL_536073, EPI_ISL_536074 | Centre Hospitalier Régional de Lanaudière | Laboratoire de santé publique du Québec | Sandrine Moreira, Ioannis Ragoussis, Guillaume Bourque, Jesse Shapiro, Mark Lathrop and Michel Roger |
| EPI_ISL_536075 | Hôpital Charles-LeMoyne | Laboratoire de santé publique du Québec | Sandrine Moreira, Ioannis Ragoussis, Guillaume Bourque, Jesse Shapiro, Mark Lathrop and Michel Roger |
| EPI_ISL_536076 | Centre Hospitalier Régional de Lanaudière | Laboratoire de santé publique du Québec | Sandrine Moreira, Ioannis Ragoussis, Guillaume Bourque, Jesse Shapiro, Mark Lathrop and Michel Roger |
| EPI_ISL_536077, EPI_ISL_536078 | Hôpital Charles-LeMoyne | Laboratoire de santé publique du Québec | Sandrine Moreira, Ioannis Ragoussis, Guillaume Bourque, Jesse Shapiro, Mark Lathrop and Michel Roger |
| EPI_ISL_536079 | Centre Hospitalier Régional de Lanaudière | Laboratoire de santé publique du Québec | Sandrine Moreira, Ioannis Ragoussis, Guillaume Bourque, Jesse Shapiro, Mark Lathrop and Michel Roger |
| EPI_ISL_536080 | Centre hospitalier Anna-Laberge | Laboratoire de santé publique du Québec | Sandrine Moreira, Ioannis Ragoussis, Guillaume Bourque, Jesse Shapiro, Mark Lathrop and Michel Roger |
| EPI_ISL_536081, EPI_ISL_536082 | Hôpital Charles-LeMoyne | Laboratoire de santé publique du Québec | Sandrine Moreira, Ioannis Ragoussis, Guillaume Bourque, Jesse Shapiro, Mark Lathrop and Michel Roger |
| EPI_ISL_536083, EPI_ISL_536084, EPI_ISL_536085, EPI_ISL_536086 | Hôpital Pierre-Le Gardeur | Laboratoire de santé publique du Québec | Sandrine Moreira, Ioannis Ragoussis, Guillaume Bourque, Jesse Shapiro, Mark Lathrop and Michel Roger |
| EPI_ISL_536087, EPI_ISL_536088, EPI_ISL_536089, EPI_ISL_536090, EPI_ISL_536091 | Hôpital Charles-LeMoyne | Laboratoire de santé publique du Québec | Sandrine Moreira, Ioannis Ragoussis, Guillaume Bourque, Jesse Shapiro, Mark Lathrop and Michel Roger |
| EPI_ISL_536092 | Hôpital Pierre-Boucher | Laboratoire de santé publique du Québec | Sandrine Moreira, Ioannis Ragoussis, Guillaume Bourque, Jesse Shapiro, Mark Lathrop and Michel Roger |
| EPI_ISL_536093, EPI_ISL_536094 | Hôpital Charles-LeMoyne | Laboratoire de santé publique du Québec | Sandrine Moreira, Ioannis Ragoussis, Guillaume Bourque, Jesse Shapiro, Mark Lathrop and Michel Roger |
| EPI_ISL_536095 | Hôpital du Suroît | Laboratoire de santé publique du Québec | Sandrine Moreira, Ioannis Ragoussis, Guillaume Bourque, Jesse Shapiro, Mark Lathrop and Michel Roger |
| EPI_ISL_536096 | Hôpital Pierre-Boucher | Laboratoire de santé publique du Québec | Sandrine Moreira, Ioannis Ragoussis, Guillaume Bourque, Jesse Shapiro, Mark Lathrop and Michel Roger |
| EPI_ISL_536097, EPI_ISL_536098 | Hôpital du Suroît | Laboratoire de santé publique du Québec | Sandrine Moreira, Ioannis Ragoussis, Guillaume Bourque, Jesse Shapiro, Mark Lathrop and Michel Roger |
| EPI_ISL_536099 | Hôpital Pierre-Boucher | Laboratoire de santé publique du Québec | Sandrine Moreira, Ioannis Ragoussis, Guillaume Bourque, Jesse Shapiro, Mark Lathrop and Michel Roger |
| EPI_ISL_536100 | Hôpital Charles-LeMoyne | Laboratoire de santé publique du Québec | Sandrine Moreira, Ioannis Ragoussis, Guillaume Bourque, Jesse Shapiro, Mark Lathrop and Michel Roger |
| EPI_ISL_536101 | Hôpital, CLSC et Centre d'Hébergement de Roberval | Laboratoire de santé publique du Québec | Sandrine Moreira, Ioannis Ragoussis, Guillaume Bourque, Jesse Shapiro, Mark Lathrop and Michel Roger |
| EPI_ISL_536102 | Hôpital du Suroît | Laboratoire de santé publique du Québec | Sandrine Moreira, Ioannis Ragoussis, Guillaume Bourque, Jesse Shapiro, Mark Lathrop and Michel Roger |
| EPI_ISL_536103 | Hôpital Pierre-Boucher | Laboratoire de santé publique du Québec | Sandrine Moreira, Ioannis Ragoussis, Guillaume Bourque, Jesse Shapiro, Mark Lathrop and Michel Roger |
| EPI_ISL_536104 | Centre hospitalier Anna-Laberge | Laboratoire de santé publique du Québec | Sandrine Moreira, Ioannis Ragoussis, Guillaume Bourque, Jesse Shapiro, Mark Lathrop and Michel Roger |
| EPI_ISL_536105 | Hôpital du Suroît | Laboratoire de santé publique du Québec | Sandrine Moreira, Ioannis Ragoussis, Guillaume Bourque, Jesse Shapiro, Mark Lathrop and Michel Roger |
| EPI_ISL_536106 | Hôpital régional de Rimouski | Laboratoire de santé publique du Québec | Sandrine Moreira, Ioannis Ragoussis, Guillaume Bourque, Jesse Shapiro, Mark Lathrop and Michel Roger |
| EPI_ISL_536107 | Hôpital de Chicoutimi | Laboratoire de santé publique du Québec | Sandrine Moreira, Ioannis Ragoussis, Guillaume Bourque, Jesse Shapiro, Mark Lathrop and Michel Roger |
| EPI_ISL_536108 | Hôpital Le Royer | Laboratoire de santé publique du Québec | Sandrine Moreira, Ioannis Ragoussis, Guillaume Bourque, Jesse Shapiro, Mark Lathrop and Michel Roger |
| EPI_ISL_536109 | CSSS de Port-Cartier | Laboratoire de santé publique du Québec | Sandrine Moreira, Ioannis Ragoussis, Guillaume Bourque, Jesse Shapiro, Mark Lathrop and Michel Roger |
| EPI_ISL_536110 | CSSS Haut-Richelieu/Rouville (Hôpital) | Laboratoire de santé publique du Québec | Sandrine Moreira, Ioannis Ragoussis, Guillaume Bourque, Jesse Shapiro, Mark Lathrop and Michel Roger |
| EPI_ISL_536111, EPI_ISL_536112 | Hôpital de Maria | Laboratoire de santé publique du Québec | Sandrine Moreira, Ioannis Ragoussis, Guillaume Bourque, Jesse Shapiro, Mark Lathrop and Michel Roger |
| EPI_ISL_536113, EPI_ISL_536114 | Hôpital et Centre d'Hébergement de Sept-Iles | Laboratoire de santé publique du Québec | Sandrine Moreira, Ioannis Ragoussis, Guillaume Bourque, Jesse Shapiro, Mark Lathrop and Michel Roger |
| EPI_ISL_536115 | Centre hospitalier régional du Grand Portage | Laboratoire de santé publique du Québec | Sandrine Moreira, Ioannis Ragoussis, Guillaume Bourque, Jesse Shapiro, Mark Lathrop and Michel Roger |
| EPI_ISL_536116 | Hôtel-Dieu de Sorel | Laboratoire de santé publique du Québec | Sandrine Moreira, Ioannis Ragoussis, Guillaume Bourque, Jesse Shapiro, Mark Lathrop and Michel Roger |
| EPI_ISL_536117, EPI_ISL_536118 | Hôpital de Hull | Laboratoire de santé publique du Québec | Sandrine Moreira, Ioannis Ragoussis, Guillaume Bourque, Jesse Shapiro, Mark Lathrop and Michel Roger |
| EPI_ISL_536119, EPI_ISL_536120 | Centre Hospitalier Régional de Lanaudière | Laboratoire de santé publique du Québec | Sandrine Moreira, Ioannis Ragoussis, Guillaume Bourque, Jesse Shapiro, Mark Lathrop and Michel Roger |
| EPI_ISL_536121 | Hôtel-Dieu de Sorel | Laboratoire de santé publique du Québec | Sandrine Moreira, Ioannis Ragoussis, Guillaume Bourque, Jesse Shapiro, Mark Lathrop and Michel Roger |
| EPI_ISL_536122 | Hôpital de Hull | Laboratoire de santé publique du Québec | Sandrine Moreira, Ioannis Ragoussis, Guillaume Bourque, Jesse Shapiro, Mark Lathrop and Michel Roger |
| EPI_ISL_536123 | Centre Hospitalier Régional de Lanaudière | Laboratoire de santé publique du Québec | Sandrine Moreira, Ioannis Ragoussis, Guillaume Bourque, Jesse Shapiro, Mark Lathrop and Michel Roger |
| EPI_ISL_536124 | Hôpital de Hull | Laboratoire de santé publique du Québec | Sandrine Moreira, Ioannis Ragoussis, Guillaume Bourque, Jesse Shapiro, Mark Lathrop and Michel Roger |

| | | | |
|---|---|---|---|
| EPI_ISL_536125 | Hôpital Pierre-Le Gardeur | Laboratoire de santé publique du Québec | Sandrine Moreira, Ioannis Ragoussis, Guillaume Bourque, Jesse Shapiro, Mark Lathrop and Michel Roger |
| EPI_ISL_536126 | Centre Hospitalier Régional de Lanaudière | Laboratoire de santé publique du Québec | Sandrine Moreira, Ioannis Ragoussis, Guillaume Bourque, Jesse Shapiro, Mark Lathrop and Michel Roger |
| EPI_ISL_536127 | Hôpital de Hull | Laboratoire de santé publique du Québec | Sandrine Moreira, Ioannis Ragoussis, Guillaume Bourque, Jesse Shapiro, Mark Lathrop and Michel Roger |
| EPI_ISL_536128 | Hôpital Pierre-Le Gardeur | Laboratoire de santé publique du Québec | Sandrine Moreira, Ioannis Ragoussis, Guillaume Bourque, Jesse Shapiro, Mark Lathrop and Michel Roger |
| EPI_ISL_536129, EPI_ISL_536130 | Centre Hospitalier Régional de Lanaudière | Laboratoire de santé publique du Québec | Sandrine Moreira, Ioannis Ragoussis, Guillaume Bourque, Jesse Shapiro, Mark Lathrop and Michel Roger |
| EPI_ISL_536131 | Hôpital Pierre-Le Gardeur | Laboratoire de santé publique du Québec | Sandrine Moreira, Ioannis Ragoussis, Guillaume Bourque, Jesse Shapiro, Mark Lathrop and Michel Roger |
| EPI_ISL_536132 | Hôpital de Hull | Laboratoire de santé publique du Québec | Sandrine Moreira, Ioannis Ragoussis, Guillaume Bourque, Jesse Shapiro, Mark Lathrop and Michel Roger |
| EPI_ISL_536133, EPI_ISL_536134 | Centre Hospitalier Régional de Lanaudière | Laboratoire de santé publique du Québec | Sandrine Moreira, Ioannis Ragoussis, Guillaume Bourque, Jesse Shapiro, Mark Lathrop and Michel Roger |
| EPI_ISL_536135, EPI_ISL_536136 | Hôpital de Hull | Laboratoire de santé publique du Québec | Sandrine Moreira, Ioannis Ragoussis, Guillaume Bourque, Jesse Shapiro, Mark Lathrop and Michel Roger |
| EPI_ISL_536137 | Hôpital Pierre-Le Gardeur | Laboratoire de santé publique du Québec | Sandrine Moreira, Ioannis Ragoussis, Guillaume Bourque, Jesse Shapiro, Mark Lathrop and Michel Roger |
| EPI_ISL_536138 | Centre Hospitalier Régional de Lanaudière | Laboratoire de santé publique du Québec | Sandrine Moreira, Ioannis Ragoussis, Guillaume Bourque, Jesse Shapiro, Mark Lathrop and Michel Roger |
| EPI_ISL_536139, EPI_ISL_536140 | Hôpital Pierre-Le Gardeur | Laboratoire de santé publique du Québec | Sandrine Moreira, Ioannis Ragoussis, Guillaume Bourque, Jesse Shapiro, Mark Lathrop and Michel Roger |
| EPI_ISL_536141, EPI_ISL_536142 | Centre hospitalier Anna-Laberge | Laboratoire de santé publique du Québec | Sandrine Moreira, Ioannis Ragoussis, Guillaume Bourque, Jesse Shapiro, Mark Lathrop and Michel Roger |
| EPI_ISL_536143, EPI_ISL_536144 | Hôpital Pierre-Le Gardeur | Laboratoire de santé publique du Québec | Sandrine Moreira, Ioannis Ragoussis, Guillaume Bourque, Jesse Shapiro, Mark Lathrop and Michel Roger |
| EPI_ISL_536145 | Centre hospitalier Anna-Laberge | Laboratoire de santé publique du Québec | Sandrine Moreira, Ioannis Ragoussis, Guillaume Bourque, Jesse Shapiro, Mark Lathrop and Michel Roger |
| EPI_ISL_536146, EPI_ISL_536147, EPI_ISL_536148, EPI_ISL_536149 | Hôpital Charles-LeMoyne | Laboratoire de santé publique du Québec | Sandrine Moreira, Ioannis Ragoussis, Guillaume Bourque, Jesse Shapiro, Mark Lathrop and Michel Roger |
| EPI_ISL_536150 | CSSS Haut-Richelieu/Rouville (Hôpital) | Laboratoire de santé publique du Québec | Sandrine Moreira, Ioannis Ragoussis, Guillaume Bourque, Jesse Shapiro, Mark Lathrop and Michel Roger |
| EPI_ISL_536151 | Centre Hospitalier Régional de Lanaudière | Laboratoire de santé publique du Québec | Sandrine Moreira, Ioannis Ragoussis, Guillaume Bourque, Jesse Shapiro, Mark Lathrop and Michel Roger |
| EPI_ISL_536152 | Hôpital Pierre-Le Gardeur | Laboratoire de santé publique du Québec | Sandrine Moreira, Ioannis Ragoussis, Guillaume Bourque, Jesse Shapiro, Mark Lathrop and Michel Roger |
| EPI_ISL_536153, EPI_ISL_536154 | Centre Hospitalier Régional de Lanaudière | Laboratoire de santé publique du Québec | Sandrine Moreira, Ioannis Ragoussis, Guillaume Bourque, Jesse Shapiro, Mark Lathrop and Michel Roger |
| EPI_ISL_536155, EPI_ISL_536156 | Centre hospitalier Anna-Laberge | Laboratoire de santé publique du Québec | Sandrine Moreira, Ioannis Ragoussis, Guillaume Bourque, Jesse Shapiro, Mark Lathrop and Michel Roger |
| EPI_ISL_536157 | Hôpital Pierre-Le Gardeur | Laboratoire de santé publique du Québec | Sandrine Moreira, Ioannis Ragoussis, Guillaume Bourque, Jesse Shapiro, Mark Lathrop and Michel Roger |
| EPI_ISL_536158 | CSSS Haut-Richelieu/Rouville (Hôpital) | Laboratoire de santé publique du Québec | Sandrine Moreira, Ioannis Ragoussis, Guillaume Bourque, Jesse Shapiro, Mark Lathrop and Michel Roger |
| EPI_ISL_536159 | Centre hospitalier Anna-Laberge | Laboratoire de santé publique du Québec | Sandrine Moreira, Ioannis Ragoussis, Guillaume Bourque, Jesse Shapiro, Mark Lathrop and Michel Roger |
| EPI_ISL_536160, EPI_ISL_536161 | CSSS Haut-Richelieu/Rouville (Hôpital) | Laboratoire de santé publique du Québec | Sandrine Moreira, Ioannis Ragoussis, Guillaume Bourque, Jesse Shapiro, Mark Lathrop and Michel Roger |
| EPI_ISL_536162 | Centre hospitalier Anna-Laberge | Laboratoire de santé publique du Québec | Sandrine Moreira, Ioannis Ragoussis, Guillaume Bourque, Jesse Shapiro, Mark Lathrop and Michel Roger |
| EPI_ISL_536163 | Hôpital du Suroît | Laboratoire de santé publique du Québec | Sandrine Moreira, Ioannis Ragoussis, Guillaume Bourque, Jesse Shapiro, Mark Lathrop and Michel Roger |
| EPI_ISL_536164 | CSSS Haut-Richelieu/Rouville (Hôpital) | Laboratoire de santé publique du Québec | Sandrine Moreira, Ioannis Ragoussis, Guillaume Bourque, Jesse Shapiro, Mark Lathrop and Michel Roger |
| EPI_ISL_536165 | Hôpital du Suroît | Laboratoire de santé publique du Québec | Sandrine Moreira, Ioannis Ragoussis, Guillaume Bourque, Jesse Shapiro, Mark Lathrop and Michel Roger |
| EPI_ISL_536166 | CSSS Haut-Richelieu/Rouville (Hôpital) | Laboratoire de santé publique du Québec | Sandrine Moreira, Ioannis Ragoussis, Guillaume Bourque, Jesse Shapiro, Mark Lathrop and Michel Roger |
| EPI_ISL_536167 | Hôpital du Suroît | Laboratoire de santé publique du Québec | Sandrine Moreira, Ioannis Ragoussis, Guillaume Bourque, Jesse Shapiro, Mark Lathrop and Michel Roger |
| EPI_ISL_536168, EPI_ISL_536169 | Hôpital de Hull | Laboratoire de santé publique du Québec | Sandrine Moreira, Ioannis Ragoussis, Guillaume Bourque, Jesse Shapiro, Mark Lathrop and Michel Roger |
| EPI_ISL_536170 | Hôpital du Suroît | Laboratoire de santé publique du Québec | Sandrine Moreira, Ioannis Ragoussis, Guillaume Bourque, Jesse Shapiro, Mark Lathrop and Michel Roger |
| EPI_ISL_536171 | Hôpital de Papineau | Laboratoire de santé publique du Québec | Sandrine Moreira, Ioannis Ragoussis, Guillaume Bourque, Jesse Shapiro, Mark Lathrop and Michel Roger |
| EPI_ISL_536172 | CSSS Haut-Richelieu/Rouville (Hôpital) | Laboratoire de santé publique du Québec | Sandrine Moreira, Ioannis Ragoussis, Guillaume Bourque, Jesse Shapiro, Mark Lathrop and Michel Roger |
| EPI_ISL_536173, EPI_ISL_536174, EPI_ISL_536175, EPI_ISL_536176 | Hôpital Pierre-Boucher | Laboratoire de santé publique du Québec | Sandrine Moreira, Ioannis Ragoussis, Guillaume Bourque, Jesse Shapiro, Mark Lathrop and Michel Roger |
| EPI_ISL_536177 | Hôpital Honoré-Mercier | Laboratoire de santé publique du Québec | Sandrine Moreira, Ioannis Ragoussis, Guillaume Bourque, Jesse Shapiro, Mark Lathrop and Michel Roger |
| EPI_ISL_536178, EPI_ISL_536179 | Hôpital Pierre-Boucher | Laboratoire de santé publique du Québec | Sandrine Moreira, Ioannis Ragoussis, Guillaume Bourque, Jesse Shapiro, Mark Lathrop and Michel Roger |
| EPI_ISL_536180 | Hôpital Honoré-Mercier | Laboratoire de santé publique du Québec | Sandrine Moreira, Ioannis Ragoussis, Guillaume Bourque, Jesse Shapiro, Mark Lathrop and Michel Roger |
| EPI_ISL_536181 | CSSS Haut-Richelieu/Rouville (Hôpital) | Laboratoire de santé publique du Québec | Sandrine Moreira, Ioannis Ragoussis, Guillaume Bourque, Jesse Shapiro, Mark Lathrop and Michel Roger |
| EPI_ISL_536182, EPI_ISL_536183, EPI_ISL_536184 | Centre hospitalier Anna-Laberge | Laboratoire de santé publique du Québec | Sandrine Moreira, Ioannis Ragoussis, Guillaume Bourque, Jesse Shapiro, Mark Lathrop and Michel Roger |
| EPI_ISL_536185, EPI_ISL_536186, EPI_ISL_536187 | Hôpital Charles-LeMoyne | Laboratoire de santé publique du Québec | Sandrine Moreira, Ioannis Ragoussis, Guillaume Bourque, Jesse Shapiro, Mark Lathrop and Michel Roger |
| EPI_ISL_536188, EPI_ISL_536189 | Hôpital Pierre-Boucher | Laboratoire de santé publique du Québec | Sandrine Moreira, Ioannis Ragoussis, Guillaume Bourque, Jesse Shapiro, Mark Lathrop and Michel Roger |
| EPI_ISL_536190 | Hôpital du Suroît | Laboratoire de santé publique du Québec | Sandrine Moreira, Ioannis Ragoussis, Guillaume Bourque, Jesse Shapiro, Mark Lathrop and Michel Roger |
| EPI_ISL_536191 | Hôpital Charles-LeMoyne | Laboratoire de santé publique du Québec | Sandrine Moreira, Ioannis Ragoussis, Guillaume Bourque, Jesse Shapiro, Mark Lathrop and Michel Roger |
| EPI_ISL_536192, EPI_ISL_536193 | Hôpital du Suroît | Laboratoire de santé publique du Québec | Sandrine Moreira, Ioannis Ragoussis, Guillaume Bourque, Jesse Shapiro, Mark Lathrop and Michel Roger |
| EPI_ISL_536194 | Hôpital Charles-LeMoyne | Laboratoire de santé publique du Québec | Sandrine Moreira, Ioannis Ragoussis, Guillaume Bourque, Jesse Shapiro, Mark Lathrop and Michel Roger |
| EPI_ISL_536195, EPI_ISL_536196 | Hôpital Pierre-Boucher | Laboratoire de santé publique du Québec | Sandrine Moreira, Ioannis Ragoussis, Guillaume Bourque, Jesse Shapiro, Mark Lathrop and Michel Roger |
| EPI_ISL_536197 | CSSS Haut-Richelieu/Rouville (Hôpital) | Laboratoire de santé publique du Québec | Sandrine Moreira, Ioannis Ragoussis, Guillaume Bourque, Jesse Shapiro, Mark Lathrop and Michel Roger |
| EPI_ISL_536198 | Hôpital du Suroît | Laboratoire de santé publique du Québec | Sandrine Moreira, Ioannis Ragoussis, Guillaume Bourque, Jesse Shapiro, Mark Lathrop and Michel Roger |
| EPI_ISL_536199 | Centre hospitalier régional du Grand Portage | Laboratoire de santé publique du Québec | Sandrine Moreira, Ioannis Ragoussis, Guillaume Bourque, Jesse Shapiro, Mark Lathrop and Michel Roger |
| EPI_ISL_536200, EPI_ISL_536201, EPI_ISL_536202 | Hôpital du Suroît | Laboratoire de santé publique du Québec | Sandrine Moreira, Ioannis Ragoussis, Guillaume Bourque, Jesse Shapiro, Mark Lathrop and Michel Roger |
| EPI_ISL_536203 | Hôpital Charles-LeMoyne | Laboratoire de santé publique du Québec | Sandrine Moreira, Ioannis Ragoussis, Guillaume Bourque, Jesse Shapiro, Mark Lathrop and Michel Roger |
| EPI_ISL_536204 | Centre hospitalier régional du Grand Portage | Laboratoire de santé publique du Québec | Sandrine Moreira, Ioannis Ragoussis, Guillaume Bourque, Jesse Shapiro, Mark Lathrop and Michel Roger |
| EPI_ISL_536205 | Hôpital régional de Rimouski | Laboratoire de santé publique du Québec | Sandrine Moreira, Ioannis Ragoussis, Guillaume Bourque, Jesse Shapiro, Mark Lathrop and Michel Roger |

| | | | |
|---|---|---|---|
| EPI_ISL_536206 | CSSS de la Minganie | Laboratoire de santé publique du Québec | Sandrine Moreira, Ioannis Ragoussis, Guillaume Bourque, Jesse Shapiro, Mark Lathrop and Michel Roger |
| EPI_ISL_536207, EPI_ISL_536208, EPI_ISL_536209, EPI_ISL_536210 | Hôpital de Hull | Laboratoire de santé publique du Québec | Sandrine Moreira, Ioannis Ragoussis, Guillaume Bourque, Jesse Shapiro, Mark Lathrop and Michel Roger |
| EPI_ISL_536211, EPI_ISL_536212 | Centre hospitalier Anna-Laberge | Laboratoire de santé publique du Québec | Sandrine Moreira, Ioannis Ragoussis, Guillaume Bourque, Jesse Shapiro, Mark Lathrop and Michel Roger |
| EPI_ISL_536213 | CSSS Haut-Richelieu/Rouville (Hôpital) | Laboratoire de santé publique du Québec | Sandrine Moreira, Ioannis Ragoussis, Guillaume Bourque, Jesse Shapiro, Mark Lathrop and Michel Roger |
| EPI_ISL_536214 | Hôpital Charles-LeMoyne | Laboratoire de santé publique du Québec | Sandrine Moreira, Ioannis Ragoussis, Guillaume Bourque, Jesse Shapiro, Mark Lathrop and Michel Roger |
| EPI_ISL_536215, EPI_ISL_536216, EPI_ISL_536217 | CSSS Haut-Richelieu/Rouville (Hôpital) | Laboratoire de santé publique du Québec | Sandrine Moreira, Ioannis Ragoussis, Guillaume Bourque, Jesse Shapiro, Mark Lathrop and Michel Roger |
| EPI_ISL_536218, EPI_ISL_536219, EPI_ISL_536220, EPI_ISL_536221 | Centre hospitalier Anna-Laberge | Laboratoire de santé publique du Québec | Sandrine Moreira, Ioannis Ragoussis, Guillaume Bourque, Jesse Shapiro, Mark Lathrop and Michel Roger |
| EPI_ISL_536222 | Conseil Cri de la SSS de la Baie-James | Laboratoire de santé publique du Québec | Sandrine Moreira, Ioannis Ragoussis, Guillaume Bourque, Jesse Shapiro, Mark Lathrop and Michel Roger |
| EPI_ISL_536223 | Hôpital Charles-LeMoyne | Laboratoire de santé publique du Québec | Sandrine Moreira, Ioannis Ragoussis, Guillaume Bourque, Jesse Shapiro, Mark Lathrop and Michel Roger |
| EPI_ISL_536224 | Centre hospitalier Anna-Laberge | Laboratoire de santé publique du Québec | Sandrine Moreira, Ioannis Ragoussis, Guillaume Bourque, Jesse Shapiro, Mark Lathrop and Michel Roger |
| EPI_ISL_536225 | Hôpital Charles-LeMoyne | Laboratoire de santé publique du Québec | Sandrine Moreira, Ioannis Ragoussis, Guillaume Bourque, Jesse Shapiro, Mark Lathrop and Michel Roger |
| EPI_ISL_536226, EPI_ISL_536227, EPI_ISL_536228 | Hôpital du Suroît | Laboratoire de santé publique du Québec | Sandrine Moreira, Ioannis Ragoussis, Guillaume Bourque, Jesse Shapiro, Mark Lathrop and Michel Roger |
| EPI_ISL_536229 | Hôpital Pierre-Boucher | Laboratoire de santé publique du Québec | Sandrine Moreira, Ioannis Ragoussis, Guillaume Bourque, Jesse Shapiro, Mark Lathrop and Michel Roger |
| EPI_ISL_536230 | Hôpital de Hull | Laboratoire de santé publique du Québec | Sandrine Moreira, Ioannis Ragoussis, Guillaume Bourque, Jesse Shapiro, Mark Lathrop and Michel Roger |
| EPI_ISL_536231 | Hôpital Pierre-Boucher | Laboratoire de santé publique du Québec | Sandrine Moreira, Ioannis Ragoussis, Guillaume Bourque, Jesse Shapiro, Mark Lathrop and Michel Roger |
| EPI_ISL_536232 | Centre hospitalier Anna-Laberge | Laboratoire de santé publique du Québec | Sandrine Moreira, Ioannis Ragoussis, Guillaume Bourque, Jesse Shapiro, Mark Lathrop and Michel Roger |
| EPI_ISL_536233 | Hôpital Charles-LeMoyne | Laboratoire de santé publique du Québec | Sandrine Moreira, Ioannis Ragoussis, Guillaume Bourque, Jesse Shapiro, Mark Lathrop and Michel Roger |
| EPI_ISL_536234 | Hôpital Pierre-Boucher | Laboratoire de santé publique du Québec | Sandrine Moreira, Ioannis Ragoussis, Guillaume Bourque, Jesse Shapiro, Mark Lathrop and Michel Roger |
| EPI_ISL_536235 | CSSS Haut-Richelieu/Rouville (Hôpital) | Laboratoire de santé publique du Québec | Sandrine Moreira, Ioannis Ragoussis, Guillaume Bourque, Jesse Shapiro, Mark Lathrop and Michel Roger |
| EPI_ISL_536236 | Hôpital Pierre-Boucher | Laboratoire de santé publique du Québec | Sandrine Moreira, Ioannis Ragoussis, Guillaume Bourque, Jesse Shapiro, Mark Lathrop and Michel Roger |
| EPI_ISL_536237 | CSSS Haut-Richelieu/Rouville (Hôpital) | Laboratoire de santé publique du Québec | Sandrine Moreira, Ioannis Ragoussis, Guillaume Bourque, Jesse Shapiro, Mark Lathrop and Michel Roger |
| EPI_ISL_536238, EPI_ISL_536239 | Hôpital de Hull | Laboratoire de santé publique du Québec | Sandrine Moreira, Ioannis Ragoussis, Guillaume Bourque, Jesse Shapiro, Mark Lathrop and Michel Roger |
| EPI_ISL_536240, EPI_ISL_536241 | Centre hospitalier Anna-Laberge | Laboratoire de santé publique du Québec | Sandrine Moreira, Ioannis Ragoussis, Guillaume Bourque, Jesse Shapiro, Mark Lathrop and Michel Roger |
| EPI_ISL_536242 | Hôpital de Hull | Laboratoire de santé publique du Québec | Sandrine Moreira, Ioannis Ragoussis, Guillaume Bourque, Jesse Shapiro, Mark Lathrop and Michel Roger |
| EPI_ISL_536243 | CSSS Haut-Richelieu/Rouville (Hôpital) | Laboratoire de santé publique du Québec | Sandrine Moreira, Ioannis Ragoussis, Guillaume Bourque, Jesse Shapiro, Mark Lathrop and Michel Roger |
| EPI_ISL_536244 | Hôpital de Hull | Laboratoire de santé publique du Québec | Sandrine Moreira, Ioannis Ragoussis, Guillaume Bourque, Jesse Shapiro, Mark Lathrop and Michel Roger |
| EPI_ISL_536245 | Hôpital Pierre-Boucher | Laboratoire de santé publique du Québec | Sandrine Moreira, Ioannis Ragoussis, Guillaume Bourque, Jesse Shapiro, Mark Lathrop and Michel Roger |
| EPI_ISL_536246 | Hôpital du Centre-de-la-Mauricie | Laboratoire de santé publique du Québec | Sandrine Moreira, Ioannis Ragoussis, Guillaume Bourque, Jesse Shapiro, Mark Lathrop and Michel Roger |
| EPI_ISL_536247 | Hôpital Charles-LeMoyne | Laboratoire de santé publique du Québec | Sandrine Moreira, Ioannis Ragoussis, Guillaume Bourque, Jesse Shapiro, Mark Lathrop and Michel Roger |
| EPI_ISL_536248, EPI_ISL_536249, EPI_ISL_536250, EPI_ISL_536251 | Hôpital du Centre-de-la-Mauricie | Laboratoire de santé publique du Québec | Sandrine Moreira, Ioannis Ragoussis, Guillaume Bourque, Jesse Shapiro, Mark Lathrop and Michel Roger |
| EPI_ISL_536252 | Hôpital Sainte-Croix | Laboratoire de santé publique du Québec | Sandrine Moreira, Ioannis Ragoussis, Guillaume Bourque, Jesse Shapiro, Mark Lathrop and Michel Roger |
| EPI_ISL_536253 | Hôpital Pierre-Boucher | Laboratoire de santé publique du Québec | Sandrine Moreira, Ioannis Ragoussis, Guillaume Bourque, Jesse Shapiro, Mark Lathrop and Michel Roger |
| EPI_ISL_536254 | Hôpital du Suroît | Laboratoire de santé publique du Québec | Sandrine Moreira, Ioannis Ragoussis, Guillaume Bourque, Jesse Shapiro, Mark Lathrop and Michel Roger |
| EPI_ISL_536255 | Centre hospitalier Anna-Laberge | Laboratoire de santé publique du Québec | Sandrine Moreira, Ioannis Ragoussis, Guillaume Bourque, Jesse Shapiro, Mark Lathrop and Michel Roger |
| EPI_ISL_536256 | CSSS Haut-Richelieu/Rouville (Hôpital) | Laboratoire de santé publique du Québec | Sandrine Moreira, Ioannis Ragoussis, Guillaume Bourque, Jesse Shapiro, Mark Lathrop and Michel Roger |
| EPI_ISL_536257 | Hôpital du Suroît | Laboratoire de santé publique du Québec | Sandrine Moreira, Ioannis Ragoussis, Guillaume Bourque, Jesse Shapiro, Mark Lathrop and Michel Roger |
| EPI_ISL_536258 | CSSS Haut-Richelieu/Rouville (Hôpital) | Laboratoire de santé publique du Québec | Sandrine Moreira, Ioannis Ragoussis, Guillaume Bourque, Jesse Shapiro, Mark Lathrop and Michel Roger |
| EPI_ISL_536259 | Hôpital du Suroît | Laboratoire de santé publique du Québec | Sandrine Moreira, Ioannis Ragoussis, Guillaume Bourque, Jesse Shapiro, Mark Lathrop and Michel Roger |
| EPI_ISL_536260 | CSSS de la Minganie | Laboratoire de santé publique du Québec | Sandrine Moreira, Ioannis Ragoussis, Guillaume Bourque, Jesse Shapiro, Mark Lathrop and Michel Roger |
| EPI_ISL_536261 | Hôpital du Suroît | Laboratoire de santé publique du Québec | Sandrine Moreira, Ioannis Ragoussis, Guillaume Bourque, Jesse Shapiro, Mark Lathrop and Michel Roger |
| EPI_ISL_536262 | Hôpital de Maria | Laboratoire de santé publique du Québec | Sandrine Moreira, Ioannis Ragoussis, Guillaume Bourque, Jesse Shapiro, Mark Lathrop and Michel Roger |
| EPI_ISL_536263, EPI_ISL_536264 | Point de Service de Témiscaming-et-Kipawa | Laboratoire de santé publique du Québec | Sandrine Moreira, Ioannis Ragoussis, Guillaume Bourque, Jesse Shapiro, Mark Lathrop and Michel Roger |
| EPI_ISL_536265 | Hôpital de Chandler | Laboratoire de santé publique du Québec | Sandrine Moreira, Ioannis Ragoussis, Guillaume Bourque, Jesse Shapiro, Mark Lathrop and Michel Roger |
| EPI_ISL_536266 | Hôpital de Gatineau | Laboratoire de santé publique du Québec | Sandrine Moreira, Ioannis Ragoussis, Guillaume Bourque, Jesse Shapiro, Mark Lathrop and Michel Roger |
| EPI_ISL_536267 | Hôpital de Hull | Laboratoire de santé publique du Québec | Sandrine Moreira, Ioannis Ragoussis, Guillaume Bourque, Jesse Shapiro, Mark Lathrop and Michel Roger |
| EPI_ISL_536268, EPI_ISL_536269, EPI_ISL_536270 | Hôpital Charles-LeMoyne | Laboratoire de santé publique du Québec | Sandrine Moreira, Ioannis Ragoussis, Guillaume Bourque, Jesse Shapiro, Mark Lathrop and Michel Roger |
| EPI_ISL_536271 | Hôpital Honoré-Mercier | Laboratoire de santé publique du Québec | Sandrine Moreira, Ioannis Ragoussis, Guillaume Bourque, Jesse Shapiro, Mark Lathrop and Michel Roger |
| EPI_ISL_536272 | CSSS Haut-Richelieu/Rouville (Hôpital) | Laboratoire de santé publique du Québec | Sandrine Moreira, Ioannis Ragoussis, Guillaume Bourque, Jesse Shapiro, Mark Lathrop and Michel Roger |
| EPI_ISL_536273, EPI_ISL_536274 | Hôpital du Suroît | Laboratoire de santé publique du Québec | Sandrine Moreira, Ioannis Ragoussis, Guillaume Bourque, Jesse Shapiro, Mark Lathrop and Michel Roger |
| EPI_ISL_536275 | Hôpital de Hull | Laboratoire de santé publique du Québec | Sandrine Moreira, Ioannis Ragoussis, Guillaume Bourque, Jesse Shapiro, Mark Lathrop and Michel Roger |
| EPI_ISL_536276, EPI_ISL_536277 | Hôpital Pierre-Boucher | Laboratoire de santé publique du Québec | Sandrine Moreira, Ioannis Ragoussis, Guillaume Bourque, Jesse Shapiro, Mark Lathrop and Michel Roger |
| EPI_ISL_536278 | CSSS Haut-Richelieu/Rouville (Hôpital) | Laboratoire de santé publique du Québec | Sandrine Moreira, Ioannis Ragoussis, Guillaume Bourque, Jesse Shapiro, Mark Lathrop and Michel Roger |
| EPI_ISL_536279, EPI_ISL_536280, EPI_ISL_536281, EPI_ISL_536282 | Hôpital Charles-LeMoyne | Laboratoire de santé publique du Québec | Sandrine Moreira, Ioannis Ragoussis, Guillaume Bourque, Jesse Shapiro, Mark Lathrop and Michel Roger |
| EPI_ISL_536283 | Centre hospitalier Anna-Laberge | Laboratoire de santé publique du Québec | Sandrine Moreira, Ioannis Ragoussis, Guillaume Bourque, Jesse Shapiro, Mark Lathrop and Michel Roger |

| | | | |
|---|---|---|---|
| EPI_ISL_536284, EPI_ISL_536285 | Hôpital Charles-LeMoyne | Laboratoire de santé publique du Québec | Sandrine Moreira, Ioannis Ragoussis, Guillaume Bourque, Jesse Shapiro, Mark Lathrop and Michel Roger |
| EPI_ISL_536286 | CSSS Haut-Richelieu/Rouville (Hôpital) | Laboratoire de santé publique du Québec | Sandrine Moreira, Ioannis Ragoussis, Guillaume Bourque, Jesse Shapiro, Mark Lathrop and Michel Roger |
| EPI_ISL_536287 | Hôpital du Suroît | Laboratoire de santé publique du Québec | Sandrine Moreira, Ioannis Ragoussis, Guillaume Bourque, Jesse Shapiro, Mark Lathrop and Michel Roger |
| EPI_ISL_536288 | Conseil Cri de la SSS de la Baie-James | Laboratoire de santé publique du Québec | Sandrine Moreira, Ioannis Ragoussis, Guillaume Bourque, Jesse Shapiro, Mark Lathrop and Michel Roger |
| EPI_ISL_536289 | Hôpital de Maria | Laboratoire de santé publique du Québec | Sandrine Moreira, Ioannis Ragoussis, Guillaume Bourque, Jesse Shapiro, Mark Lathrop and Michel Roger |
| EPI_ISL_536290 | Hôpital Charles-LeMoyne | Laboratoire de santé publique du Québec | Sandrine Moreira, Ioannis Ragoussis, Guillaume Bourque, Jesse Shapiro, Mark Lathrop and Michel Roger |
| EPI_ISL_536291 | Hôpital de Hull | Laboratoire de santé publique du Québec | Sandrine Moreira, Ioannis Ragoussis, Guillaume Bourque, Jesse Shapiro, Mark Lathrop and Michel Roger |
| EPI_ISL_536292, EPI_ISL_536293, EPI_ISL_536294 | Hôpital Charles-LeMoyne | Laboratoire de santé publique du Québec | Sandrine Moreira, Ioannis Ragoussis, Guillaume Bourque, Jesse Shapiro, Mark Lathrop and Michel Roger |
| EPI_ISL_536295 | Hôpital Pierre-Boucher | Laboratoire de santé publique du Québec | Sandrine Moreira, Ioannis Ragoussis, Guillaume Bourque, Jesse Shapiro, Mark Lathrop and Michel Roger |
| EPI_ISL_536296, EPI_ISL_536297, EPI_ISL_536298 | Hôpital Charles-LeMoyne | Laboratoire de santé publique du Québec | Sandrine Moreira, Ioannis Ragoussis, Guillaume Bourque, Jesse Shapiro, Mark Lathrop and Michel Roger |
| EPI_ISL_536299 | CSSS Haut-Richelieu/Rouville (Hôpital) | Laboratoire de santé publique du Québec | Sandrine Moreira, Ioannis Ragoussis, Guillaume Bourque, Jesse Shapiro, Mark Lathrop and Michel Roger |
| EPI_ISL_536300 | Hôpital Pierre-Boucher | Laboratoire de santé publique du Québec | Sandrine Moreira, Ioannis Ragoussis, Guillaume Bourque, Jesse Shapiro, Mark Lathrop and Michel Roger |
| EPI_ISL_536301, EPI_ISL_536302 | CSSS Haut-Richelieu/Rouville (Hôpital) | Laboratoire de santé publique du Québec | Sandrine Moreira, Ioannis Ragoussis, Guillaume Bourque, Jesse Shapiro, Mark Lathrop and Michel Roger |
| EPI_ISL_536303 | Hôpital Pierre-Boucher | Laboratoire de santé publique du Québec | Sandrine Moreira, Ioannis Ragoussis, Guillaume Bourque, Jesse Shapiro, Mark Lathrop and Michel Roger |
| EPI_ISL_536304 | CSSS Haut-Richelieu/Rouville (Hôpital) | Laboratoire de santé publique du Québec | Sandrine Moreira, Ioannis Ragoussis, Guillaume Bourque, Jesse Shapiro, Mark Lathrop and Michel Roger |
| EPI_ISL_536305, EPI_ISL_536306, EPI_ISL_536307, EPI_ISL_536308, EPI_ISL_536309 | Hôpital Pierre-Boucher | Laboratoire de santé publique du Québec | Sandrine Moreira, Ioannis Ragoussis, Guillaume Bourque, Jesse Shapiro, Mark Lathrop and Michel Roger |
| EPI_ISL_536310 | CLSC et Centre d'Hébergement la Petite-Nation | Laboratoire de santé publique du Québec | Sandrine Moreira, Ioannis Ragoussis, Guillaume Bourque, Jesse Shapiro, Mark Lathrop and Michel Roger |
| EPI_ISL_536311 | Hôpital de Gatineau | Laboratoire de santé publique du Québec | Sandrine Moreira, Ioannis Ragoussis, Guillaume Bourque, Jesse Shapiro, Mark Lathrop and Michel Roger |
| EPI_ISL_536312 | Hôpital de Hull | Laboratoire de santé publique du Québec | Sandrine Moreira, Ioannis Ragoussis, Guillaume Bourque, Jesse Shapiro, Mark Lathrop and Michel Roger |
| EPI_ISL_536313, EPI_ISL_536314 | Hôpital Charles-LeMoyne | Laboratoire de santé publique du Québec | Sandrine Moreira, Ioannis Ragoussis, Guillaume Bourque, Jesse Shapiro, Mark Lathrop and Michel Roger |
| EPI_ISL_536315 | Centre hospitalier Anna-Laberge | Laboratoire de santé publique du Québec | Sandrine Moreira, Ioannis Ragoussis, Guillaume Bourque, Jesse Shapiro, Mark Lathrop and Michel Roger |
| EPI_ISL_536316, EPI_ISL_536317 | Hôpital Charles-LeMoyne | Laboratoire de santé publique du Québec | Sandrine Moreira, Ioannis Ragoussis, Guillaume Bourque, Jesse Shapiro, Mark Lathrop and Michel Roger |
| EPI_ISL_536318, EPI_ISL_536319, EPI_ISL_536320 | Hôpital du Suroît | Laboratoire de santé publique du Québec | Sandrine Moreira, Ioannis Ragoussis, Guillaume Bourque, Jesse Shapiro, Mark Lathrop and Michel Roger |
| EPI_ISL_536321 | CSSS Haut-Richelieu/Rouville (Hôpital) | Laboratoire de santé publique du Québec | Sandrine Moreira, Ioannis Ragoussis, Guillaume Bourque, Jesse Shapiro, Mark Lathrop and Michel Roger |
| EPI_ISL_536322, EPI_ISL_536323, EPI_ISL_536324 | Hôpital Pierre-Boucher | Laboratoire de santé publique du Québec | Sandrine Moreira, Ioannis Ragoussis, Guillaume Bourque, Jesse Shapiro, Mark Lathrop and Michel Roger |
| EPI_ISL_536325 | Centre hospitalier Anna-Laberge | Laboratoire de santé publique du Québec | Sandrine Moreira, Ioannis Ragoussis, Guillaume Bourque, Jesse Shapiro, Mark Lathrop and Michel Roger |
| EPI_ISL_536326, EPI_ISL_536327, EPI_ISL_536328 | Hôpital Charles-LeMoyne | Laboratoire de santé publique du Québec | Sandrine Moreira, Ioannis Ragoussis, Guillaume Bourque, Jesse Shapiro, Mark Lathrop and Michel Roger |
| EPI_ISL_536329 | Centre hospitalier Anna-Laberge | Laboratoire de santé publique du Québec | Sandrine Moreira, Ioannis Ragoussis, Guillaume Bourque, Jesse Shapiro, Mark Lathrop and Michel Roger |
| EPI_ISL_536330 | Hôpital Honoré-Mercier | Laboratoire de santé publique du Québec | Sandrine Moreira, Ioannis Ragoussis, Guillaume Bourque, Jesse Shapiro, Mark Lathrop and Michel Roger |
| EPI_ISL_536331 | Hôpital Charles-LeMoyne | Laboratoire de santé publique du Québec | Sandrine Moreira, Ioannis Ragoussis, Guillaume Bourque, Jesse Shapiro, Mark Lathrop and Michel Roger |
| EPI_ISL_536332, EPI_ISL_536333, EPI_ISL_536334, EPI_ISL_536335 | Hôpital Pierre-Boucher | Laboratoire de santé publique du Québec | Sandrine Moreira, Ioannis Ragoussis, Guillaume Bourque, Jesse Shapiro, Mark Lathrop and Michel Roger |
| EPI_ISL_536336 | Hôpital Charles-LeMoyne | Laboratoire de santé publique du Québec | Sandrine Moreira, Ioannis Ragoussis, Guillaume Bourque, Jesse Shapiro, Mark Lathrop and Michel Roger |
| EPI_ISL_536337 | CUSM-Site Glen-LAB Microbiologie | Laboratoire de santé publique du Québec | Sandrine Moreira, Ioannis Ragoussis, Guillaume Bourque, Jesse Shapiro, Mark Lathrop and Michel Roger |
| EPI_ISL_536338, EPI_ISL_536339, EPI_ISL_536340, EPI_ISL_536341, EPI_ISL_536342, EPI_ISL_536343 | Hôpital Charles-LeMoyne | Laboratoire de santé publique du Québec | Sandrine Moreira, Ioannis Ragoussis, Guillaume Bourque, Jesse Shapiro, Mark Lathrop and Michel Roger |
| EPI_ISL_536344, EPI_ISL_536345 | CSSS Haut-Richelieu/Rouville (Hôpital) | Laboratoire de santé publique du Québec | Sandrine Moreira, Ioannis Ragoussis, Guillaume Bourque, Jesse Shapiro, Mark Lathrop and Michel Roger |
| EPI_ISL_536346, EPI_ISL_536347 | Hôpital Pierre-Boucher | Laboratoire de santé publique du Québec | Sandrine Moreira, Ioannis Ragoussis, Guillaume Bourque, Jesse Shapiro, Mark Lathrop and Michel Roger |
| EPI_ISL_536348 | Centre hospitalier Anna-Laberge | Laboratoire de santé publique du Québec | Sandrine Moreira, Ioannis Ragoussis, Guillaume Bourque, Jesse Shapiro, Mark Lathrop and Michel Roger |
| EPI_ISL_536349 | Hôpital du Suroît | Laboratoire de santé publique du Québec | Sandrine Moreira, Ioannis Ragoussis, Guillaume Bourque, Jesse Shapiro, Mark Lathrop and Michel Roger |
| EPI_ISL_536350 | Hôpital Charles-LeMoyne | Laboratoire de santé publique du Québec | Sandrine Moreira, Ioannis Ragoussis, Guillaume Bourque, Jesse Shapiro, Mark Lathrop and Michel Roger |
| EPI_ISL_536351, EPI_ISL_536352 | Hôpital Pierre-Boucher | Laboratoire de santé publique du Québec | Sandrine Moreira, Ioannis Ragoussis, Guillaume Bourque, Jesse Shapiro, Mark Lathrop and Michel Roger |
| EPI_ISL_536353 | Centre de santé Inuulitsivik | Laboratoire de santé publique du Québec | Sandrine Moreira, Ioannis Ragoussis, Guillaume Bourque, Jesse Shapiro, Mark Lathrop and Michel Roger |
| EPI_ISL_536354, EPI_ISL_536355, EPI_ISL_536356, EPI_ISL_536357, EPI_ISL_536358, EPI_ISL_536359 | Hôpital du Suroît | Laboratoire de santé publique du Québec | Sandrine Moreira, Ioannis Ragoussis, Guillaume Bourque, Jesse Shapiro, Mark Lathrop and Michel Roger |
| EPI_ISL_536360 | Hôpital Pierre-Boucher | Laboratoire de santé publique du Québec | Sandrine Moreira, Ioannis Ragoussis, Guillaume Bourque, Jesse Shapiro, Mark Lathrop and Michel Roger |
| EPI_ISL_536361 | CSSS Haut-Richelieu/Rouville (Hôpital) | Laboratoire de santé publique du Québec | Sandrine Moreira, Ioannis Ragoussis, Guillaume Bourque, Jesse Shapiro, Mark Lathrop and Michel Roger |
| EPI_ISL_536362 | Hôpital du Suroît | Laboratoire de santé publique du Québec | Sandrine Moreira, Ioannis Ragoussis, Guillaume Bourque, Jesse Shapiro, Mark Lathrop and Michel Roger |
| EPI_ISL_536363, EPI_ISL_536364 | Hôpital de Hull | Laboratoire de santé publique du Québec | Sandrine Moreira, Ioannis Ragoussis, Guillaume Bourque, Jesse Shapiro, Mark Lathrop and Michel Roger |
| EPI_ISL_536365, EPI_ISL_536366 | CUSM-Site Glen-LAB Microbiologie | Laboratoire de santé publique du Québec | Sandrine Moreira, Ioannis Ragoussis, Guillaume Bourque, Jesse Shapiro, Mark Lathrop and Michel Roger |
| EPI_ISL_536367 | Hôpital de Hull | Laboratoire de santé publique du Québec | Sandrine Moreira, Ioannis Ragoussis, Guillaume Bourque, Jesse Shapiro, Mark Lathrop and Michel Roger |
| EPI_ISL_536368, EPI_ISL_536369, EPI_ISL_536370 | Centre hospitalier Anna-Laberge | Laboratoire de santé publique du Québec | Sandrine Moreira, Ioannis Ragoussis, Guillaume Bourque, Jesse Shapiro, Mark Lathrop and Michel Roger |
| EPI_ISL_536371, EPI_ISL_536372, | Hôpital Charles-LeMoyne | Laboratoire de santé publique du Québec | Sandrine Moreira, Ioannis Ragoussis, Guillaume Bourque, Jesse Shapiro, Mark Lathrop and Michel Roger |

| | | | |
|---|---|---|---|
| EPI_ISL_536373 | | | |
| EPI_ISL_536374, EPI_ISL_536375 | Hôpital Pierre-Boucher | Laboratoire de santé publique du Québec | Sandrine Moreira, Ioannis Ragoussis, Guillaume Bourque, Jesse Shapiro, Mark Lathrop and Michel Roger |
| EPI_ISL_536376, EPI_ISL_536377, EPI_ISL_536378 | Centre hospitalier Anna-Laberge | Laboratoire de santé publique du Québec | Sandrine Moreira, Ioannis Ragoussis, Guillaume Bourque, Jesse Shapiro, Mark Lathrop and Michel Roger |
| EPI_ISL_536379 | Hôpital Honoré-Mercier | Laboratoire de santé publique du Québec | Sandrine Moreira, Ioannis Ragoussis, Guillaume Bourque, Jesse Shapiro, Mark Lathrop and Michel Roger |
| EPI_ISL_536380 | Hôpital Pierre-Boucher | Laboratoire de santé publique du Québec | Sandrine Moreira, Ioannis Ragoussis, Guillaume Bourque, Jesse Shapiro, Mark Lathrop and Michel Roger |
| EPI_ISL_536381 | Centre Hospitalier Régional de Lanaudière | Laboratoire de santé publique du Québec | Sandrine Moreira, Ioannis Ragoussis, Guillaume Bourque, Jesse Shapiro, Mark Lathrop and Michel Roger |
| EPI_ISL_536382, EPI_ISL_536383 | CSSS Haut-Richelieu/Rouville (Hôpital) | Laboratoire de santé publique du Québec | Sandrine Moreira, Ioannis Ragoussis, Guillaume Bourque, Jesse Shapiro, Mark Lathrop and Michel Roger |
| EPI_ISL_536384 | Hôpital de Gatineau | Laboratoire de santé publique du Québec | Sandrine Moreira, Ioannis Ragoussis, Guillaume Bourque, Jesse Shapiro, Mark Lathrop and Michel Roger |
| EPI_ISL_536385, EPI_ISL_536386, EPI_ISL_536387 | Hôpital de Hull | Laboratoire de santé publique du Québec | Sandrine Moreira, Ioannis Ragoussis, Guillaume Bourque, Jesse Shapiro, Mark Lathrop and Michel Roger |
| EPI_ISL_536388, EPI_ISL_536389 | Hôpital Pierre-Boucher | Laboratoire de santé publique du Québec | Sandrine Moreira, Ioannis Ragoussis, Guillaume Bourque, Jesse Shapiro, Mark Lathrop and Michel Roger |
| EPI_ISL_536390 | Hôpital du Suroît | Laboratoire de santé publique du Québec | Sandrine Moreira, Ioannis Ragoussis, Guillaume Bourque, Jesse Shapiro, Mark Lathrop and Michel Roger |
| EPI_ISL_536391 | Hôpital Pierre-Boucher | Laboratoire de santé publique du Québec | Sandrine Moreira, Ioannis Ragoussis, Guillaume Bourque, Jesse Shapiro, Mark Lathrop and Michel Roger |
| EPI_ISL_536392, EPI_ISL_536393, EPI_ISL_536394 | Hôpital de Hull | Laboratoire de santé publique du Québec | Sandrine Moreira, Ioannis Ragoussis, Guillaume Bourque, Jesse Shapiro, Mark Lathrop and Michel Roger |
| EPI_ISL_536395 | Hôpital Pierre-Boucher | Laboratoire de santé publique du Québec | Sandrine Moreira, Ioannis Ragoussis, Guillaume Bourque, Jesse Shapiro, Mark Lathrop and Michel Roger |
| EPI_ISL_536396 | Hôpital de Hull | Laboratoire de santé publique du Québec | Sandrine Moreira, Ioannis Ragoussis, Guillaume Bourque, Jesse Shapiro, Mark Lathrop and Michel Roger |
| EPI_ISL_536397 | Hôpital Pierre-Boucher | Laboratoire de santé publique du Québec | Sandrine Moreira, Ioannis Ragoussis, Guillaume Bourque, Jesse Shapiro, Mark Lathrop and Michel Roger |
| EPI_ISL_539634 | CSIR-Centre for Cellular and Molecular Biology | CSIR-Centre for Cellular and Molecular Biology | Sakshi Shambhavi, Lamuk Zaveri, Shagufta Khan, Namami Gaur, Nikhil Hajirnis, M Soujanya Reddy, Pratheusa Maccha, Tulasi Nagabandi, Purushotham Vodnala, Payel Mukherjee, Sofia Banu, Priya Singh, Onkar Kulkarni, Dhiviya Vedagiri, Divya Gupta, Vishal Sah, Santosh Kumar Kuncha, Krishnan Harinivas Harshan, Archana Bharadwaj Siva, Karthik Bharadwaj Tallapaka, Deepak Kumar, Devi Prasad Vijayashankar, Disha Nanda, Divya Das, Jotin Gogoi, Manish Bhattacharjee, Rakesh K Mishra, Divya Tej Sowpati |
| EPI_ISL_539635 | CSIR-Centre for Cellular and Molecular Biology | CSIR-Centre for Cellular and Molecular Biology | Sakshi Shambhavi, Lamuk Zaveri, Shagufta Khan, Namami Gaur, Nikhil Hajirnis, M Soujanya Reddy, Pratheusa Maccha,Tulasi Nagabandi, Purushotham Vodnala, Payel Mukherjee, Sofia Banu, Priya Singh,Onkar Kulkarni, Dhiviya Vedagiri, Divya Gupta, Vishal Sah, Santosh Kumar Kuncha, Krishnan Harinivas Harshan, Archana Bharadwaj Siva, Karthik Bharadwaj Tallapaka, G. Aditya Kumar, Koushick Sivakumar, Pooja Ramesh Gupta, Rajan Kumar Jha, Shraddha Vijay Lahoti, Rakesh K Mishra, Divya Tej Sowpati |
| EPI_ISL_539636 | CSIR-Centre for Cellular and Molecular Biology | CSIR-Centre for Cellular and Molecular Biology | Sakshi Shambhavi, Lamuk Zaveri, Shagufta Khan, Nikhil Hajirnis, M Soujanya Reddy, Pratheusa Maccha, Namami Gaur, Tulasi Nagabandi, Purushotham Vodnala, Payel Mukherjee, Sofia Banu, Priya Singh, Onkar Kulkarni, Dhiviya Vedagiri, Divya Gupta, Vishal Sah, Santosh Kumar Kuncha, Krishnan Harinivas Harshan, Archana Bharadwaj Siva, Karthik Bharadwaj Tallapaka,G. Aditya Kumar, Koushick Sivakumar, Rakesh K Mishra, Divya Tej Sowpati |
| EPI_ISL_539664 | CSIR-Centre for Cellular and Molecular Biology | CSIR-Centre for Cellular and Molecular Biology | Sakshi Shambhavi, Lamuk Zaveri, Shagufta Khan, Namami Gaur, Nikhil Hajirnis, M Soujanya Reddy, Pratheusa Maccha, Tulasi Nagabandi, Purushotham Vodnala, Payel Mukherjee, Sofia Banu, Priya Singh, Onkar Kulkarni, Dhiviya Vedagiri, Divya Gupta, Vishal Sah, Santosh Kumar Kuncha, Krishnan Harinivas Harshan, Archana Bharadwaj Siva, Karthik Bharadwaj Tallapaka, Deepak Kumar, Devi Prasad Vijayashankar, Disha Nanda, Divya Das, Jotin Gogoi, Manish Bhattacharjee, Rakesh K Mishra, Divya Tej Sowpati |
| EPI_ISL_539665 | CSIR-Centre for Cellular and Molecular Biology | CSIR-Centre for Cellular and Molecular Biology | Sakshi Shambhavi, Lamuk Zaveri, Shagufta Khan, Namami Gaur, Nikhil Hajirnis, M Soujanya Reddy, Pratheusa Maccha,Tulasi Nagabandi, Purushotham Vodnala, Payel Mukherjee, Sofia Banu, Priya Singh,Onkar Kulkarni, Dhiviya Vedagiri, Divya Gupta, Vishal Sah, Santosh Kumar Kuncha, Krishnan Harinivas Harshan, Archana Bharadwaj Siva, Karthik Bharadwaj Tallapaka, G. Aditya Kumar, Koushick Sivakumar, Pooja Ramesh Gupta, Rajan Kumar Jha, Shraddha Vijay Lahoti, Rakesh K Mishra, Divya Tej Sowpati |
| EPI_ISL_539666 | CSIR-Centre for Cellular and Molecular Biology | CSIR-Centre for Cellular and Molecular Biology | Sakshi Shambhavi, Lamuk Zaveri, Shagufta Khan, Nikhil Hajirnis, M Soujanya Reddy, Pratheusa Maccha, Namami Gaur, Tulasi Nagabandi, Purushotham Vodnala, Payel Mukherjee, Sofia Banu, Priya Singh, Onkar Kulkarni, Dhiviya Vedagiri, Divya Gupta, Vishal Sah, Santosh Kumar Kuncha, Krishnan Harinivas Harshan, Archana Bharadwaj Siva, Karthik Bharadwaj Tallapaka,G. Aditya Kumar, Koushick Sivakumar, Rakesh K Mishra, Divya Tej Sowpati |
| EPI_ISL_539694 | CSIR-Centre for Cellular and Molecular Biology | CSIR-Centre for Cellular and Molecular Biology | Sakshi Shambhavi, Lamuk Zaveri, Shagufta Khan, Namami Gaur, Nikhil Hajirnis, M Soujanya Reddy, Pratheusa Maccha, Tulasi Nagabandi, Purushotham Vodnala, Payel Mukherjee, Sofia Banu, Priya Singh, Onkar Kulkarni, Dhiviya Vedagiri, Divya Gupta, Vishal Sah, Santosh Kumar Kuncha, Krishnan Harinivas Harshan, Archana Bharadwaj Siva, Karthik Bharadwaj Tallapaka, Deepak Kumar, Devi Prasad Vijayashankar, Disha Nanda, Divya Das, Jotin Gogoi, Manish Bhattacharjee, Rakesh K Mishra, Divya Tej Sowpati |
| EPI_ISL_539695 | CSIR-Centre for Cellular and Molecular Biology | CSIR-Centre for Cellular and Molecular Biology | Sakshi Shambhavi, Lamuk Zaveri, Shagufta Khan, Namami Gaur, Nikhil Hajirnis, M Soujanya Reddy, Pratheusa Maccha,Tulasi Nagabandi, Purushotham Vodnala, Payel Mukherjee, Sofia Banu, Priya Singh,Onkar Kulkarni, Dhiviya Vedagiri, Divya Gupta, Vishal Sah, Santosh Kumar Kuncha, Krishnan Harinivas Harshan, Archana Bharadwaj Siva, Karthik Bharadwaj Tallapaka, G. Aditya Kumar, Koushick Sivakumar, Pooja Ramesh Gupta, Rajan Kumar Jha, Shraddha Vijay Lahoti, Rakesh K Mishra, Divya Tej Sowpati |
| EPI_ISL_539696 | CSIR-Centre for Cellular and Molecular Biology | CSIR-Centre for Cellular and Molecular Biology | Sakshi Shambhavi, Lamuk Zaveri, Shagufta Khan, Nikhil Hajirnis, M Soujanya Reddy, Pratheusa Maccha, Namami Gaur, Tulasi Nagabandi, Purushotham Vodnala, Payel Mukherjee, Sofia Banu, Priya Singh, Onkar Kulkarni, Dhiviya Vedagiri, Divya Gupta, Vishal Sah, Santosh Kumar Kuncha, Krishnan Harinivas Harshan, Archana Bharadwaj Siva, Karthik Bharadwaj Tallapaka,G. Aditya Kumar, Koushick Sivakumar, Rakesh K Mishra, Divya Tej Sowpati |
| EPI_ISL_539724 | CSIR-Centre for Cellular and Molecular Biology | CSIR-Centre for Cellular and Molecular Biology | Sakshi Shambhavi, Lamuk Zaveri, Shagufta Khan, Namami Gaur, Nikhil Hajirnis, M Soujanya Reddy, Pratheusa Maccha, Tulasi Nagabandi, Purushotham Vodnala, Payel Mukherjee, Sofia Banu, Priya Singh, Onkar Kulkarni, Dhiviya Vedagiri, Divya Gupta, Vishal Sah, Santosh Kumar Kuncha, Krishnan Harinivas Harshan, Archana Bharadwaj Siva, Karthik Bharadwaj Tallapaka, Deepak Kumar, Devi Prasad Vijayashankar, Disha Nanda, Divya Das, Jotin Gogoi, Manish Bhattacharjee, Rakesh K Mishra, Divya Tej Sowpati |
| EPI_ISL_539725 | CSIR-Centre for Cellular and Molecular Biology | CSIR-Centre for Cellular and Molecular Biology | Sakshi Shambhavi, Lamuk Zaveri, Shagufta Khan, Namami Gaur, Nikhil Hajirnis, M Soujanya Reddy, Pratheusa Maccha,Tulasi Nagabandi, Purushotham Vodnala, Payel Mukherjee, Sofia Banu, Priya Singh,Onkar Kulkarni, Dhiviya Vedagiri, Divya Gupta, Vishal Sah, Santosh Kumar Kuncha, Krishnan Harinivas Harshan, Archana Bharadwaj Siva, Karthik Bharadwaj Tallapaka, G. Aditya Kumar, Koushick Sivakumar, Pooja Ramesh Gupta, Rajan Kumar Jha, Shraddha Vijay Lahoti, Rakesh K Mishra, Divya Tej Sowpati |
| EPI_ISL_539726 | CSIR-Centre for Cellular and Molecular Biology | CSIR-Centre for Cellular and Molecular Biology | Sakshi Shambhavi, Lamuk Zaveri, Shagufta Khan, Nikhil Hajirnis, M Soujanya Reddy, Pratheusa Maccha, Namami Gaur, Tulasi Nagabandi, Purushotham Vodnala, Payel Mukherjee, Sofia Banu, Priya Singh, Onkar Kulkarni, Dhiviya Vedagiri, Divya Gupta, Vishal Sah, Santosh Kumar Kuncha, Krishnan Harinivas Harshan, Archana Bharadwaj Siva, Karthik Bharadwaj Tallapaka,G. Aditya Kumar, Koushick Sivakumar, Rakesh K Mishra, Divya Tej Sowpati |
| EPI_ISL_539754 | CSIR-Centre for Cellular and Molecular Biology | CSIR-Centre for Cellular and Molecular Biology | Sakshi Shambhavi, Lamuk Zaveri, Shagufta Khan, Namami Gaur, Nikhil Hajirnis, M Soujanya Reddy, Pratheusa Maccha, Tulasi Nagabandi, Purushotham Vodnala, Payel Mukherjee, Sofia Banu, Priya Singh, Onkar Kulkarni, Dhiviya Vedagiri, Divya Gupta, Vishal Sah, Santosh Kumar Kuncha, Krishnan Harinivas Harshan, Archana Bharadwaj Siva, Karthik Bharadwaj Tallapaka, Deepak Kumar, Devi Prasad Vijayashankar, Disha Nanda, Divya Das, Jotin Gogoi, Manish Bhattacharjee, Rakesh K Mishra, Divya Tej Sowpati |
| EPI_ISL_539755 | CSIR-Centre for Cellular and Molecular Biology | CSIR-Centre for Cellular and Molecular Biology | Sakshi Shambhavi, Lamuk Zaveri, Shagufta Khan, Namami Gaur, Nikhil Hajirnis, M Soujanya Reddy, Pratheusa Maccha,Tulasi Nagabandi, Purushotham Vodnala, Payel Mukherjee, Sofia Banu, Priya Singh,Onkar Kulkarni, Dhiviya Vedagiri, Divya Gupta, Vishal Sah, Santosh Kumar Kuncha, Krishnan Harinivas Harshan, Archana Bharadwaj Siva, Karthik Bharadwaj Tallapaka, G. Aditya Kumar, Koushick Sivakumar, Pooja Ramesh Gupta, Rajan Kumar Jha, Shraddha Vijay Lahoti, Rakesh K Mishra, Divya Tej Sowpati |
| EPI_ISL_539756 | CSIR-Centre for Cellular and Molecular Biology | CSIR-Centre for Cellular and Molecular Biology | Sakshi Shambhavi, Lamuk Zaveri, Shagufta Khan, Nikhil Hajirnis, M Soujanya Reddy, Pratheusa Maccha, Namami Gaur, Tulasi Nagabandi, Purushotham Vodnala, Payel Mukherjee, Sofia Banu, Priya Singh, Onkar Kulkarni, Dhiviya Vedagiri, Divya Gupta, Vishal Sah, Santosh Kumar Kuncha, Krishnan |

| | | | |
|---|---|---|---|
| | | | Harinivas Harshan, Archana Bharadwaj Siva, Karthik Bharadwaj Tallapaka,G. Aditya Kumar, Koushick Sivakumar, Rakesh K Mishra, Divya Tej Sowpati |
| EPI_ISL_540584, EPI_ISL_540585, EPI_ISL_540586, EPI_ISL_540587, EPI_ISL_540588, EPI_ISL_540589, EPI_ISL_540590, EPI_ISL_540591, EPI_ISL_540592, EPI_ISL_540593, EPI_ISL_540594, EPI_ISL_540595, EPI_ISL_540596, EPI_ISL_540597, EPI_ISL_540598, EPI_ISL_540599, EPI_ISL_540600, EPI_ISL_540601, EPI_ISL_540602, EPI_ISL_540603, EPI_ISL_540604, EPI_ISL_540605, EPI_ISL_540606, EPI_ISL_540607, EPI_ISL_540608, EPI_ISL_540609, EPI_ISL_540610, EPI_ISL_540611, EPI_ISL_540612, EPI_ISL_540613, EPI_ISL_540614, EPI_ISL_540615, EPI_ISL_540616, EPI_ISL_540617, EPI_ISL_540618, EPI_ISL_540619, EPI_ISL_540620, EPI_ISL_540621, EPI_ISL_540622, EPI_ISL_540623, EPI_ISL_540624, EPI_ISL_540625, EPI_ISL_540626, EPI_ISL_540627, EPI_ISL_540628, EPI_ISL_540629, EPI_ISL_540630, EPI_ISL_540631, EPI_ISL_540632, EPI_ISL_540633, EPI_ISL_540634, EPI_ISL_540635, EPI_ISL_540636, EPI_ISL_540637, EPI_ISL_540638, EPI_ISL_540639 | | | |
| see above | Liverpool Clinical Laboratories | COVID-19 Genomics UK (COG-UK) Consortium | Sam Haldenby, Anita Lucaci, Steve Paterson, Julian Hiscox, Alistair Darby, M Almsaud, A Alrezaihi, Muhannad Alruwaili, Stuart D Armstrong, Jones Benjamin, Eleanor G Bentley, Anu Chawla, Jordan J Clark, Angela Cowell, Richard Eccles, Isabel García-Dorival, Matthew Gemmell, Alessandro Gerada, PKF Gilmore, Richard Gregory, Ximeng Han, Catherine Hartley, Margaret Hughes, Miren Iturriza-Gomara, James Johnson, L Luu, Jenifer Manson, Charlotte Nelson, Elaine O'Toole, Cassie Olateju, Rebekah Penrice-Randal , Lucille Rainbow, N.P Randle, Trevor Ian Robinson, Parul Sharma, Ghada T Shawli, James P Stewart, Neil Swainston, Ecaterina Vamos, Joanne Watts, Mark Whitehead |
| EPI_ISL_541338 | Florida Bureau of Public Health Laboratories | Florida Bureau of Public Health Laboratories | Sarah Schmedes, Jason Blanton |
| EPI_ISL_542499, EPI_ISL_542508, EPI_ISL_542523, EPI_ISL_542525, EPI_ISL_542527, EPI_ISL_542529, EPI_ISL_542531, EPI_ISL_542538, EPI_ISL_542540, EPI_ISL_542541, EPI_ISL_542545, EPI_ISL_542548, EPI_ISL_542550, EPI_ISL_542552, EPI_ISL_542554, EPI_ISL_542555, EPI_ISL_542558, EPI_ISL_542559, EPI_ISL_542560, EPI_ISL_542563, EPI_ISL_542566, EPI_ISL_542567, EPI_ISL_542569, EPI_ISL_542585, EPI_ISL_542591, EPI_ISL_542608, EPI_ISL_542616, EPI_ISL_542619, EPI_ISL_542625, EPI_ISL_542627, EPI_ISL_542630, EPI_ISL_542631, EPI_ISL_542632, EPI_ISL_542633, EPI_ISL_542634, EPI_ISL_542635, EPI_ISL_542636, EPI_ISL_542637, EPI_ISL_542638, EPI_ISL_542639, EPI_ISL_542640, EPI_ISL_542641, EPI_ISL_542642, EPI_ISL_542643, EPI_ISL_542644, EPI_ISL_542645, EPI_ISL_542646, EPI_ISL_542647, EPI_ISL_542648, EPI_ISL_542649, EPI_ISL_542650, EPI_ISL_542651, EPI_ISL_542652, EPI_ISL_542653, EPI_ISL_542654, EPI_ISL_542655, EPI_ISL_542656, EPI_ISL_542657, EPI_ISL_542658, EPI_ISL_542659, EPI_ISL_542660, EPI_ISL_542661, EPI_ISL_542662, EPI_ISL_542663, EPI_ISL_542664, EPI_ISL_542665, EPI_ISL_542666, EPI_ISL_542667, EPI_ISL_542668, EPI_ISL_542669, EPI_ISL_542670, EPI_ISL_542671, EPI_ISL_542672, EPI_ISL_542673, EPI_ISL_542674, EPI_ISL_542675, EPI_ISL_542676, EPI_ISL_542677, EPI_ISL_542678, EPI_ISL_542679, EPI_ISL_542680, EPI_ISL_542681, EPI_ISL_542682, EPI_ISL_542683, EPI_ISL_542684, EPI_ISL_542685, EPI_ISL_542686, EPI_ISL_542687, EPI_ISL_542688, EPI_ISL_542689, EPI_ISL_542690, EPI_ISL_542691, EPI_ISL_542692, EPI_ISL_542693, EPI_ISL_542694, EPI_ISL_542695, EPI_ISL_542696, EPI_ISL_542697, EPI_ISL_542698, EPI_ISL_542699, EPI_ISL_542700, EPI_ISL_542701, EPI_ISL_542702, EPI_ISL_542703, EPI_ISL_542704, EPI_ISL_542705, EPI_ISL_542706, EPI_ISL_542707, EPI_ISL_542708, EPI_ISL_542709, EPI_ISL_542710, EPI_ISL_542711, EPI_ISL_542712, EPI_ISL_542713, EPI_ISL_542714, EPI_ISL_542715, EPI_ISL_542716, EPI_ISL_542717, EPI_ISL_542718, EPI_ISL_542719, EPI_ISL_542720, EPI_ISL_542721, EPI_ISL_542722, EPI_ISL_542723, EPI_ISL_542724, EPI_ISL_542725, EPI_ISL_542726, EPI_ISL_542727, EPI_ISL_542728, EPI_ISL_542729, EPI_ISL_542730, EPI_ISL_542731, EPI_ISL_542732, EPI_ISL_542733, EPI_ISL_542734, EPI_ISL_542735, EPI_ISL_542736, EPI_ISL_542737, EPI_ISL_542738, EPI_ISL_542739, EPI_ISL_542740, EPI_ISL_542741, EPI_ISL_542742, EPI_ISL_542743, EPI_ISL_542744, EPI_ISL_542745, EPI_ISL_542746, EPI_ISL_542747, EPI_ISL_542748, EPI_ISL_542749, EPI_ISL_542750, EPI_ISL_542751, EPI_ISL_542752, EPI_ISL_542753, EPI_ISL_542754, EPI_ISL_542755, EPI_ISL_542756, EPI_ISL_542757, EPI_ISL_542758, EPI_ISL_542759, EPI_ISL_542760, EPI_ISL_542761, EPI_ISL_542762, EPI_ISL_542763, EPI_ISL_542764, EPI_ISL_542765, EPI_ISL_542766, EPI_ISL_542767, EPI_ISL_542768, EPI_ISL_542769, EPI_ISL_542770, EPI_ISL_542771, EPI_ISL_542772, EPI_ISL_542773, EPI_ISL_542774, EPI_ISL_542775, EPI_ISL_542776, EPI_ISL_542777, EPI_ISL_542778, EPI_ISL_542779, EPI_ISL_542780, EPI_ISL_542781, EPI_ISL_542782, EPI_ISL_542783, EPI_ISL_542784, EPI_ISL_542785, EPI_ISL_542786, EPI_ISL_542787, EPI_ISL_542788, EPI_ISL_542789, EPI_ISL_542790, EPI_ISL_542791, EPI_ISL_542792, EPI_ISL_542793, EPI_ISL_542794, EPI_ISL_542795, EPI_ISL_542796, EPI_ISL_542797, EPI_ISL_542798, EPI_ISL_542799, EPI_ISL_542800, EPI_ISL_542801, EPI_ISL_542802, EPI_ISL_542803, EPI_ISL_542804, EPI_ISL_542805, EPI_ISL_542806, EPI_ISL_542807, EPI_ISL_542808, EPI_ISL_542809, EPI_ISL_542810, EPI_ISL_542811, EPI_ISL_542812, EPI_ISL_542813, EPI_ISL_542814, EPI_ISL_542815, EPI_ISL_542816, EPI_ISL_542817, EPI_ISL_542818, EPI_ISL_542819, EPI_ISL_542820, EPI_ISL_542821, EPI_ISL_542822, EPI_ISL_542823, EPI_ISL_542824, EPI_ISL_542825, EPI_ISL_542826, EPI_ISL_542827, EPI_ISL_542828, EPI_ISL_542829, EPI_ISL_542830, EPI_ISL_542831, EPI_ISL_542832, EPI_ISL_542833, EPI_ISL_542834, EPI_ISL_542835, EPI_ISL_542836, EPI_ISL_542837, EPI_ISL_542838, EPI_ISL_542839, EPI_ISL_542840, EPI_ISL_542841, EPI_ISL_542842, EPI_ISL_542843, EPI_ISL_542844, EPI_ISL_542845, EPI_ISL_542846, EPI_ISL_542847, EPI_ISL_542848, EPI_ISL_542849, EPI_ISL_542850, EPI_ISL_542851, EPI_ISL_542852, EPI_ISL_542853, EPI_ISL_542854, EPI_ISL_542855, EPI_ISL_542856, EPI_ISL_542857, EPI_ISL_542858, EPI_ISL_542859, EPI_ISL_542860, EPI_ISL_542861, EPI_ISL_542862, EPI_ISL_542863, EPI_ISL_542864, EPI_ISL_542865, EPI_ISL_542866, EPI_ISL_542867, EPI_ISL_542868, EPI_ISL_542869, EPI_ISL_542870, EPI_ISL_542871, EPI_ISL_542872, EPI_ISL_542873, EPI_ISL_542874, EPI_ISL_542875, EPI_ISL_542876, EPI_ISL_542877, EPI_ISL_542878, EPI_ISL_542879, EPI_ISL_542880, EPI_ISL_542881, EPI_ISL_542882, EPI_ISL_542883, EPI_ISL_542884, EPI_ISL_542885, EPI_ISL_542886, EPI_ISL_542887, EPI_ISL_542888, EPI_ISL_542889, EPI_ISL_542890, EPI_ISL_542891, EPI_ISL_542892, EPI_ISL_542893, EPI_ISL_542894, EPI_ISL_542895, EPI_ISL_542896, EPI_ISL_542897, EPI_ISL_542898, EPI_ISL_542899, EPI_ISL_542900, EPI_ISL_542901, EPI_ISL_542902, EPI_ISL_542903, EPI_ISL_542904, EPI_ISL_542905, EPI_ISL_542906, EPI_ISL_542907, EPI_ISL_542908, EPI_ISL_542909, EPI_ISL_542910, EPI_ISL_542911, EPI_ISL_542912, EPI_ISL_542913, EPI_ISL_542914, EPI_ISL_542915, EPI_ISL_542916, EPI_ISL_542917, EPI_ISL_542918, EPI_ISL_542919, EPI_ISL_542920, EPI_ISL_542921, EPI_ISL_542922, EPI_ISL_542923, EPI_ISL_542924, EPI_ISL_542925, EPI_ISL_542926, EPI_ISL_542927, EPI_ISL_542928, EPI_ISL_542929, EPI_ISL_542930, EPI_ISL_542931, EPI_ISL_542932, EPI_ISL_542933, EPI_ISL_542934, EPI_ISL_543007, EPI_ISL_543008, EPI_ISL_543009, EPI_ISL_543010, EPI_ISL_543011, EPI_ISL_543012, EPI_ISL_543013, EPI_ISL_543014, EPI_ISL_543015, EPI_ISL_543016, EPI_ISL_543017, EPI_ISL_543018, EPI_ISL_543019, EPI_ISL_543020, EPI_ISL_543021, EPI_ISL_543022, EPI_ISL_543023, EPI_ISL_543024, EPI_ISL_543025, EPI_ISL_543026, EPI_ISL_543027, EPI_ISL_543028, EPI_ISL_543029, EPI_ISL_543030, EPI_ISL_543031, EPI_ISL_543032, EPI_ISL_543033, EPI_ISL_543034, EPI_ISL_543035, EPI_ISL_543036, EPI_ISL_543037, EPI_ISL_543038, EPI_ISL_543039, EPI_ISL_543040, EPI_ISL_543041, EPI_ISL_543042, EPI_ISL_543043, EPI_ISL_543044, EPI_ISL_543045, EPI_ISL_543046, EPI_ISL_543047, EPI_ISL_543048, EPI_ISL_543049, EPI_ISL_543050, EPI_ISL_543051, EPI_ISL_543052, EPI_ISL_543053, EPI_ISL_543054, EPI_ISL_543055, EPI_ISL_543056, EPI_ISL_543057, EPI_ISL_543058, EPI_ISL_543059, EPI_ISL_543060, EPI_ISL_543061, EPI_ISL_543062, EPI_ISL_543063, EPI_ISL_543064, EPI_ISL_543065, EPI_ISL_543066, EPI_ISL_543067, EPI_ISL_543068, EPI_ISL_543069, EPI_ISL_543070, EPI_ISL_543071, EPI_ISL_543072, EPI_ISL_543073, EPI_ISL_543074, EPI_ISL_543075, EPI_ISL_543076, EPI_ISL_543077, EPI_ISL_543078, EPI_ISL_543079, EPI_ISL_543080, EPI_ISL_543081, EPI_ISL_543082, EPI_ISL_543083, EPI_ISL_543084, EPI_ISL_543085, EPI_ISL_543086, EPI_ISL_543087, EPI_ISL_543088, EPI_ISL_543089, EPI_ISL_543090, EPI_ISL_543091, EPI_ISL_543092, EPI_ISL_543093, EPI_ISL_543094, EPI_ISL_543095, EPI_ISL_543096, EPI_ISL_543097, EPI_ISL_543098, EPI_ISL_543099, EPI_ISL_543100, EPI_ISL_543101, EPI_ISL_543102, EPI_ISL_543103, EPI_ISL_543104, EPI_ISL_543105, EPI_ISL_543106, EPI_ISL_543107, EPI_ISL_543108, EPI_ISL_543109, EPI_ISL_543110, EPI_ISL_543111, EPI_ISL_543112, EPI_ISL_543113, EPI_ISL_543114, EPI_ISL_543115, EPI_ISL_543116, EPI_ISL_543117, EPI_ISL_543118, EPI_ISL_543119, EPI_ISL_543120, EPI_ISL_543121, EPI_ISL_543122, EPI_ISL_543123, EPI_ISL_543124, EPI_ISL_543125, EPI_ISL_543126, EPI_ISL_543127, EPI_ISL_543128, EPI_ISL_543129, EPI_ISL_543130, EPI_ISL_543131, EPI_ISL_543132, EPI_ISL_543133, EPI_ISL_543134, EPI_ISL_543135, EPI_ISL_543136, EPI_ISL_543137, EPI_ISL_543138, EPI_ISL_543139, EPI_ISL_543140, EPI_ISL_543141, EPI_ISL_543142, EPI_ISL_543143, EPI_ISL_543144, EPI_ISL_543145, EPI_ISL_543146, EPI_ISL_543147, EPI_ISL_543148, EPI_ISL_543149, EPI_ISL_543150, EPI_ISL_543151, EPI_ISL_543152, EPI_ISL_543153, EPI_ISL_543154, EPI_ISL_543155, EPI_ISL_543156, EPI_ISL_543157, EPI_ISL_543158, EPI_ISL_543159, EPI_ISL_543160, EPI_ISL_543161, EPI_ISL_543162, EPI_ISL_543163, EPI_ISL_543164, EPI_ISL_543165, EPI_ISL_543166, EPI_ISL_543167, EPI_ISL_543168, EPI_ISL_543169, EPI_ISL_543170, EPI_ISL_543171, EPI_ISL_543172, EPI_ISL_543173, EPI_ISL_543174, EPI_ISL_543175, EPI_ISL_543176, EPI_ISL_543177, EPI_ISL_543178, EPI_ISL_543179, EPI_ISL_543180, EPI_ISL_543181, EPI_ISL_543182, EPI_ISL_543183, EPI_ISL_543184, EPI_ISL_543185, EPI_ISL_543186, EPI_ISL_543187, EPI_ISL_543188, EPI_ISL_543189, EPI_ISL_543190, EPI_ISL_543191, EPI_ISL_543192, EPI_ISL_543193, EPI_ISL_543194, EPI_ISL_543195, EPI_ISL_543196, EPI_ISL_543197, EPI_ISL_543198, EPI_ISL_543199, EPI_ISL_543200, EPI_ISL_543201, EPI_ISL_543202, EPI_ISL_543203, EPI_ISL_543204, EPI_ISL_543205, EPI_ISL_543206, EPI_ISL_543207, EPI_ISL_543208, EPI_ISL_543209, EPI_ISL_543210, EPI_ISL_543211, EPI_ISL_543212, EPI_ISL_543213, EPI_ISL_543214, EPI_ISL_543215, EPI_ISL_543216, EPI_ISL_543217, EPI_ISL_543218, EPI_ISL_543219, EPI_ISL_543220, EPI_ISL_543221, EPI_ISL_543222, EPI_ISL_543223, EPI_ISL_543224, EPI_ISL_543225, EPI_ISL_543226, EPI_ISL_543227, EPI_ISL_543228, EPI_ISL_543229, EPI_ISL_543230, EPI_ISL_543231, EPI_ISL_543232, EPI_ISL_543233, EPI_ISL_543234, EPI_ISL_543235, EPI_ISL_543236, EPI_ISL_543237, EPI_ISL_543238, EPI_ISL_543239, EPI_ISL_543240, EPI_ISL_543241, EPI_ISL_543242, EPI_ISL_543243, EPI_ISL_543244, EPI_ISL_543245, EPI_ISL_543246, EPI_ISL_543247, EPI_ISL_543248, EPI_ISL_543249, EPI_ISL_543250, EPI_ISL_543251, EPI_ISL_543252, EPI_ISL_543253, EPI_ISL_543254, EPI_ISL_543255, EPI_ISL_543256, EPI_ISL_543257, EPI_ISL_543258, EPI_ISL_543259, EPI_ISL_543260, EPI_ISL_543261, EPI_ISL_543262, EPI_ISL_543263, EPI_ISL_543264, EPI_ISL_543265, EPI_ISL_543266, EPI_ISL_543267, EPI_ISL_543268, EPI_ISL_543269, EPI_ISL_543270, EPI_ISL_543271, EPI_ISL_543272, EPI_ISL_543273, EPI_ISL_543274, EPI_ISL_543275, EPI_ISL_543276, EPI_ISL_543277, EPI_ISL_543278, EPI_ISL_543279, EPI_ISL_543280, EPI_ISL_543281, EPI_ISL_543282, EPI_ISL_543283, EPI_ISL_543286, EPI_ISL_543287, EPI_ISL_543288, EPI_ISL_543289, EPI_ISL_543290, EPI_ISL_543291, EPI_ISL_543293, EPI_ISL_543295, EPI_ISL_543296, EPI_ISL_543297, EPI_ISL_543299, EPI_ISL_543300, EPI_ISL_543302, EPI_ISL_543307, EPI_ISL_543309, EPI_ISL_543310, EPI_ISL_543315, EPI_ISL_543316, EPI_ISL_543317, EPI_ISL_543323, EPI_ISL_543327, EPI_ISL_543330, EPI_ISL_543331, EPI_ISL_543333, EPI_ISL_543334, EPI_ISL_543337, EPI_ISL_543339, EPI_ISL_543340, EPI_ISL_543342, EPI_ISL_543344, EPI_ISL_543345, EPI_ISL_543347, EPI_ISL_543349, EPI_ISL_543350, EPI_ISL_543351, EPI_ISL_543352, EPI_ISL_543353, EPI_ISL_543355, EPI_ISL_543356, EPI_ISL_543357, EPI_ISL_543361, EPI_ISL_543362, EPI_ISL_543363, EPI_ISL_543365, EPI_ISL_543366, EPI_ISL_543370, EPI_ISL_543373, EPI_ISL_543376, EPI_ISL_543377, EPI_ISL_543378, EPI_ISL_543379, EPI_ISL_543380, EPI_ISL_543381, EPI_ISL_543384, EPI_ISL_543385, EPI_ISL_543388, EPI_ISL_543389, EPI_ISL_543390, EPI_ISL_543391, EPI_ISL_543392, EPI_ISL_543393, EPI_ISL_543394, EPI_ISL_543395, EPI_ISL_543396, EPI_ISL_543397, EPI_ISL_543398, EPI_ISL_543399, EPI_ISL_543400, EPI_ISL_543401, EPI_ISL_543402, EPI_ISL_543403, EPI_ISL_543404, EPI_ISL_543405, EPI_ISL_543406, EPI_ISL_543407, EPI_ISL_543408, EPI_ISL_543409, EPI_ISL_543410, EPI_ISL_543411, EPI_ISL_543412, EPI_ISL_543413, EPI_ISL_543414, EPI_ISL_543415, EPI_ISL_543416, EPI_ISL_543417, EPI_ISL_543418, EPI_ISL_543419, EPI_ISL_543420, EPI_ISL_543421, EPI_ISL_543422, EPI_ISL_543423, EPI_ISL_543424, EPI_ISL_543425, EPI_ISL_543426, EPI_ISL_543427, EPI_ISL_543428, EPI_ISL_543429, EPI_ISL_543430, EPI_ISL_543431, EPI_ISL_543432, EPI_ISL_543436, EPI_ISL_543437, EPI_ISL_543443, EPI_ISL_543444, EPI_ISL_543445, EPI_ISL_543446, EPI_ISL_543447, EPI_ISL_543448, EPI_ISL_543449, EPI_ISL_543450, EPI_ISL_543451, EPI_ISL_543452, EPI_ISL_543453, EPI_ISL_543454, EPI_ISL_543455, EPI_ISL_543457, EPI_ISL_543458, EPI_ISL_543459, EPI_ISL_543462, EPI_ISL_543463, EPI_ISL_543464, EPI_ISL_543465, EPI_ISL_543466, EPI_ISL_543467, EPI_ISL_543468, EPI_ISL_543472, EPI_ISL_543473, EPI_ISL_543475, EPI_ISL_543477, EPI_ISL_543480, EPI_ISL_543481, EPI_ISL_543482, EPI_ISL_543483, EPI_ISL_543484, EPI_ISL_543485, EPI_ISL_543486, EPI_ISL_543488, EPI_ISL_543489, EPI_ISL_543492, EPI_ISL_543494, EPI_ISL_543496, EPI_ISL_543499, EPI_ISL_543500, EPI_ISL_543501, EPI_ISL_543503, EPI_ISL_543505, EPI_ISL_543507, EPI_ISL_543510, EPI_ISL_543514, EPI_ISL_543516, EPI_ISL_543517, EPI_ISL_543518, EPI_ISL_543520, EPI_ISL_543522, EPI_ISL_543523, EPI_ISL_543526, EPI_ISL_543528, EPI_ISL_543531, EPI_ISL_543532, EPI_ISL_543535, EPI_ISL_543536, EPI_ISL_543539, EPI_ISL_543542, EPI_ISL_543543, EPI_ISL_543548, EPI_ISL_543550, EPI_ISL_543554, EPI_ISL_543556, EPI_ISL_543557, EPI_ISL_543559, EPI_ISL_543560, EPI_ISL_543561, EPI_ISL_543562, EPI_ISL_543563, EPI_ISL_543564, EPI_ISL_543566, EPI_ISL_543570, EPI_ISL_543573, EPI_ISL_543576, EPI_ISL_543577, EPI_ISL_543578, EPI_ISL_543579, EPI_ISL_543580, EPI_ISL_543581, EPI_ISL_543584, EPI_ISL_543586, EPI_ISL_543587, EPI_ISL_543588, EPI_ISL_543589, EPI_ISL_543592, EPI_ISL_543593, EPI_ISL_543594, EPI_ISL_543595, EPI_ISL_543596, EPI_ISL_543598, EPI_ISL_543599, EPI_ISL_543600, EPI_ISL_543603, EPI_ISL_543604, EPI_ISL_543609, EPI_ISL_543610, EPI_ISL_543613, EPI_ISL_543614, EPI_ISL_543616, EPI_ISL_543617, EPI_ISL_543618, EPI_ISL_543620, EPI_ISL_543622, EPI_ISL_543623, EPI_ISL_543624, EPI_ISL_543625, EPI_ISL_543626, EPI_ISL_543627, EPI_ISL_543628, EPI_ISL_543629, EPI_ISL_543630, EPI_ISL_543632, EPI_ISL_543633, EPI_ISL_543634, EPI_ISL_543636, EPI_ISL_543637, EPI_ISL_543638, EPI_ISL_543639, EPI_ISL_543640, EPI_ISL_543641, EPI_ISL_543642, EPI_ISL_543643, EPI_ISL_543644, EPI_ISL_543645, EPI_ISL_543646, EPI_ISL_543647, EPI_ISL_543648, EPI_ISL_543649, EPI_ISL_543650, EPI_ISL_543651, EPI_ISL_543652, EPI_ISL_543653, EPI_ISL_543654, EPI_ISL_543656, EPI_ISL_543657, EPI_ISL_543659, EPI_ISL_543660, EPI_ISL_543661, EPI_ISL_543663, EPI_ISL_543664, EPI_ISL_543665, EPI_ISL_543666, EPI_ISL_543667, EPI_ISL_543669, EPI_ISL_543670, EPI_ISL_543675, EPI_ISL_543676, EPI_ISL_543678, EPI_ISL_543679, EPI_ISL_543680, EPI_ISL_543681, EPI_ISL_543682, EPI_ISL_543683, EPI_ISL_543684, EPI_ISL_543685, EPI_ISL_543686, EPI_ISL_543688, EPI_ISL_543689, EPI_ISL_543690, EPI_ISL_543693, EPI_ISL_543694, EPI_ISL_543701, EPI_ISL_543702, EPI_ISL_543704, EPI_ISL_543705, EPI_ISL_543706, EPI_ISL_543707, EPI_ISL_543708, EPI_ISL_543709, EPI_ISL_543713, EPI_ISL_543714, EPI_ISL_543716, EPI_ISL_543718, EPI_ISL_543719, EPI_ISL_543720, EPI_ISL_543721, EPI_ISL_543722, EPI_ISL_543726, EPI_ISL_543727, EPI_ISL_543728, EPI_ISL_543730, EPI_ISL_543731, EPI_ISL_543732, EPI_ISL_543733, EPI_ISL_543736, EPI_ISL_543738, EPI_ISL_543739, EPI_ISL_543740, EPI_ISL_543741, EPI_ISL_543742, EPI_ISL_543743, EPI_ISL_543745, EPI_ISL_543746, EPI_ISL_543747, EPI_ISL_543748, EPI_ISL_543749, EPI_ISL_543750, EPI_ISL_543751, EPI_ISL_543752, EPI_ISL_543754, EPI_ISL_543757, EPI_ISL_543758, EPI_ISL_543760, EPI_ISL_543761, EPI_ISL_543764, EPI_ISL_543765, EPI_ISL_543766, EPI_ISL_543769, EPI_ISL_543771, EPI_ISL_543772, EPI_ISL_543774, EPI_ISL_543775, EPI_ISL_543777, EPI_ISL_543778, EPI_ISL_543779, EPI_ISL_543780, EPI_ISL_543781, EPI_ISL_543782, EPI_ISL_543783, EPI_ISL_543784, EPI_ISL_543785, EPI_ISL_543787, EPI_ISL_543788, EPI_ISL_543789, EPI_ISL_543790, EPI_ISL_543791, EPI_ISL_543792, EPI_ISL_543793, EPI_ISL_543794, EPI_ISL_543795, EPI_ISL_543796, EPI_ISL_543797, EPI_ISL_543798, EPI_ISL_543799, EPI_ISL_543800, EPI_ISL_543801, EPI_ISL_543802, EPI_ISL_543803, EPI_ISL_543804, EPI_ISL_543805, EPI_ISL_543806, EPI_ISL_543807, EPI_ISL_543808, EPI_ISL_543809, EPI_ISL_543810, EPI_ISL_543811, EPI_ISL_543812, EPI_ISL_543813, EPI_ISL_543814, EPI_ISL_543815, EPI_ISL_543816, EPI_ISL_543817, EPI_ISL_543818, EPI_ISL_543819, EPI_ISL_543820, EPI_ISL_543821, EPI_ISL_543822, EPI_ISL_543823, EPI_ISL_543824, EPI_ISL_543825, EPI_ISL_543826, EPI_ISL_543827, EPI_ISL_543828, EPI_ISL_543829, EPI_ISL_543830, EPI_ISL_543831, EPI_ISL_543832, EPI_ISL_543833, EPI_ISL_543834, EPI_ISL_543835, EPI_ISL_543836, EPI_ISL_543837, EPI_ISL_543838, EPI_ISL_543839, EPI_ISL_543840, EPI_ISL_543841, EPI_ISL_543842, EPI_ISL_543843, EPI_ISL_543844, EPI_ISL_543845, EPI_ISL_543846, EPI_ISL_543847, EPI_ISL_543848, EPI_ISL_543849, EPI_ISL_543850, EPI_ISL_543851, EPI_ISL_543852, EPI_ISL_543853, EPI_ISL_543854, EPI_ISL_543855, EPI_ISL_543856, EPI_ISL_543857, EPI_ISL_543858, EPI_ISL_543859, EPI_ISL_543860, EPI_ISL_543861, EPI_ISL_543862, EPI_ISL_543863, EPI_ISL_543864, EPI_ISL_543865, EPI_ISL_543866, EPI_ISL_543867, EPI_ISL_543868, EPI_ISL_543869, EPI_ISL_543870, EPI_ISL_543871, EPI_ISL_543872, EPI_ISL_543873, EPI_ISL_543874, EPI_ISL_543875, EPI_ISL_543876, EPI_ISL_543877, EPI_ISL_543878, EPI_ISL_543879, EPI_ISL_543880, EPI_ISL_543881, EPI_ISL_543882, EPI_ISL_543883, EPI_ISL_543884, EPI_ISL_543885, EPI_ISL_543886, EPI_ISL_543887, EPI_ISL_543888, EPI_ISL_543889, EPI_ISL_543890, EPI_ISL_543891, EPI_ISL_543892, EPI_ISL_543893, EPI_ISL_543894, EPI_ISL_543895, EPI_ISL_543896, EPI_ISL_543897, EPI_ISL_543898, EPI_ISL_543899, EPI_ISL_543900, EPI_ISL_543901, EPI_ISL_543902, EPI_ISL_543903, EPI_ISL_543904, EPI_ISL_543905, EPI_ISL_543906, EPI_ISL_543907, EPI_ISL_543908, EPI_ISL_543909, EPI_ISL_543910, EPI_ISL_543911, EPI_ISL_543912, EPI_ISL_543913, EPI_ISL_543914, EPI_ISL_543915, EPI_ISL_543916, EPI_ISL_543917, EPI_ISL_543918, EPI_ISL_543919, EPI_ISL_543920, EPI_ISL_543921, EPI_ISL_543922, EPI_ISL_543923, EPI_ISL_543924, EPI_ISL_543925, EPI_ISL_543926, EPI_ISL_543927, EPI_ISL_543928, EPI_ISL_543929, EPI_ISL_543930, EPI_ISL_543931, EPI_ISL_543932, EPI_ISL_543933, EPI_ISL_543934, EPI_ISL_543935, EPI_ISL_543936, EPI_ISL_543937, EPI_ISL_543938, EPI_ISL_543939, EPI_ISL_543940, EPI_ISL_543941, EPI_ISL_543942, EPI_ISL_543943, EPI_ISL_543944, EPI_ISL_543945, EPI_ISL_543946, EPI_ISL_543947, EPI_ISL_543948, EPI_ISL_543949, EPI_ISL_543950, EPI_ISL_543951, EPI_ISL_543952, EPI_ISL_543953, EPI_ISL_543954, EPI_ISL_543955, EPI_ISL_543956, EPI_ISL_543957, EPI_ISL_543958, EPI_ISL_543959, EPI_ISL_543960, EPI_ISL_543961, EPI_ISL_543962, EPI_ISL_543963, EPI_ISL_543964, EPI_ISL_543965, EPI_ISL_543966, EPI_ISL_543967, EPI_ISL_543968, EPI_ISL_543969, EPI_ISL_543970, EPI_ISL_543971, EPI_ISL_543972, EPI_ISL_543973, EPI_ISL_543974, EPI_ISL_543975, EPI_ISL_543976, EPI_ISL_543977, EPI_ISL_543978, EPI_ISL_543979, EPI_ISL_543980, EPI_ISL_543981, EPI_ISL_543982, EPI_ISL_543983, EPI_ISL_543984, EPI_ISL_543985, EPI_ISL_543986, EPI_ISL_543987, EPI_ISL_543988, EPI_ISL_543989, EPI_ISL_543990, EPI_ISL_543991, EPI_ISL_543992, EPI_ISL_543993, EPI_ISL_543994, EPI_ISL_543995, EPI_ISL_543996, EPI_ISL_543997, EPI_ISL_543998, EPI_ISL_543999, EPI_ISL_544000, EPI_ISL_544001, EPI_ISL_544002, EPI_ISL_544003, EPI_ISL_544004, EPI_ISL_544005, EPI_ISL_544006, EPI_ISL_544007, EPI_ISL_544008, EPI_ISL_544009, EPI_ISL_544010, EPI_ISL_544011, EPI_ISL_544012, EPI_ISL_544013, EPI_ISL_544014, EPI_ISL_544015, EPI_ISL_544016, EPI_ISL_544017, EPI_ISL_544018, EPI_ISL_544019, EPI_ISL_544020, EPI_ISL_544021, EPI_ISL_544022, EPI_ISL_544023, EPI_ISL_544024, EPI_ISL_544025, EPI_ISL_544026, EPI_ISL_544027, EPI_ISL_544028, EPI_ISL_544029, EPI_ISL_544030, EPI_ISL_544031, EPI_ISL_544032, EPI_ISL_544033, EPI_ISL_544034, EPI_ISL_544035, EPI_ISL_544036, EPI_ISL_544037, EPI_ISL_544038, EPI_ISL_544039, EPI_ISL_544040, EPI_ISL_544041, EPI_ISL_544042, EPI_ISL_544043, EPI_ISL_544044, EPI_ISL_544045, EPI_ISL_544046, EPI_ISL_544047, EPI_ISL_544048, EPI_ISL_544049, EPI_ISL_544050, EPI_ISL_544051, EPI_ISL_544052, EPI_ISL_544053, EPI_ISL_544054, EPI_ISL_544055, EPI_ISL_544056, EPI_ISL_544057, EPI_ISL_544058, EPI_ISL_544059, EPI_ISL_544060, EPI_ISL_544061, EPI_ISL_544062, EPI_ISL_544063, EPI_ISL_544064, EPI_ISL_544065, EPI_ISL_544066, EPI_ISL_544067, EPI_ISL_544068, EPI_ISL_544069, EPI_ISL_544070, EPI_ISL_544071, EPI_ISL_544072, EPI_ISL_544073, EPI_ISL_544074, EPI_ISL_544075, EPI_ISL_544076, EPI_ISL_544077, EPI_ISL_544078, EPI_ISL_544079, EPI_ISL_544080, EPI_ISL_544081, EPI_ISL_544082, EPI_ISL_544083, EPI_ISL_544084, EPI_ISL_544085, EPI_ISL_544086, EPI_ISL_544087, EPI_ISL_544088, EPI_ISL_544089, EPI_ISL_544090, EPI_ISL_544091, EPI_ISL_544092, EPI_ISL_544093, EPI_ISL_544094, EPI_ISL_544095, EPI_ISL_544096, EPI_ISL_544097, EPI_ISL_544098, EPI_ISL_544099, EPI_ISL_544100, EPI_ISL_544101, EPI_ISL_544102, EPI_ISL_544103, EPI_ISL_544104, EPI_ISL_544105, EPI_ISL_544106, EPI_ISL_544107, EPI_ISL_544108, EPI_ISL_544109, EPI_ISL_544110, EPI_ISL_544111, EPI_ISL_544112, EPI_ISL_544113, EPI_ISL_544114, EPI_ISL_544115, EPI_ISL_544116, EPI_ISL_544117, EPI_ISL_544118, EPI_ISL_544119, EPI_ISL_544120, EPI_ISL_544121, EPI_ISL_544122, EPI_ISL_544123, EPI_ISL_544124, EPI_ISL_544125, EPI_ISL_544126, EPI_ISL_544127, EPI_ISL_544128, EPI_ISL_544129, EPI_ISL_544130, EPI_ISL_544131, EPI_ISL_544132, EPI_ISL_544133, EPI_ISL_544134, EPI_ISL_544135, EPI_ISL_544136, EPI_ISL_544137, EPI_ISL_544138, EPI_ISL_544139, EPI_ISL_544140, EPI_ISL_544141, EPI_ISL_544142, EPI_ISL_544143, EPI_ISL_544144, EPI_ISL_544145, EPI_ISL_544146, EPI_ISL_544147, EPI_ISL_544148, EPI_ISL_544149, EPI_ISL_544150, EPI_ISL_544151, EPI_ISL_544152, EPI_ISL_544153, EPI_ISL_544154, EPI_ISL_544155, EPI_ISL_544156, EPI_ISL_544157, EPI_ISL_544158, EPI_ISL_544159, EPI_ISL_544160, EPI_ISL_544161, EPI_ISL_544162, EPI_ISL_544163, EPI_ISL_544164, EPI_ISL_544165, EPI_ISL_544166, EPI_ISL_544167, EPI_ISL_544168, EPI_ISL_544169, EPI_ISL_544170, EPI_ISL_544171, EPI_ISL_544172, EPI_ISL_544173, EPI_ISL_544174, EPI_ISL_544175, EPI_ISL_544176, EPI_ISL_544177, EPI_ISL_544178, EPI_ISL_544179, EPI_ISL_544180, | | | |

EPI_ISL_544181, EPI_ISL_544182, EPI_ISL_544183, EPI_ISL_544184, EPI_ISL_544185, EPI_ISL_544186, EPI_ISL_544187, EPI_ISL_544188, EPI_ISL_544189, EPI_ISL_544190, EPI_ISL_544191, EPI_ISL_544192, EPI_ISL_544193, EPI_ISL_544194, EPI_ISL_544195, EPI_ISL_544196, EPI_ISL_544197, EPI_ISL_544198, EPI_ISL_544199, EPI_ISL_544200, EPI_ISL_544201, EPI_ISL_544202, EPI_ISL_544203, EPI_ISL_544204, EPI_ISL_544205, EPI_ISL_544206, EPI_ISL_544207, EPI_ISL_544208, EPI_ISL_544209, EPI_ISL_544210, EPI_ISL_544211, EPI_ISL_544212, EPI_ISL_544213, EPI_ISL_544214, EPI_ISL_544215, EPI_ISL_544216, EPI_ISL_544217, EPI_ISL_544218, EPI_ISL_544219, EPI_ISL_544220, EPI_ISL_544221, EPI_ISL_544222, EPI_ISL_544223, EPI_ISL_544224, EPI_ISL_544225, EPI_ISL_544226, EPI_ISL_544227, EPI_ISL_544228, EPI_ISL_544229, EPI_ISL_544230, EPI_ISL_544231, EPI_ISL_544232, EPI_ISL_544233, EPI_ISL_544234, EPI_ISL_544235, EPI_ISL_544236, EPI_ISL_544237, EPI_ISL_544238, EPI_ISL_544239, EPI_ISL_544241, EPI_ISL_544244, EPI_ISL_544248, EPI_ISL_544250, EPI_ISL_544252, EPI_ISL_544253, EPI_ISL_544254, EPI_ISL_544256, EPI_ISL_544257, EPI_ISL_544258, EPI_ISL_544260, EPI_ISL_544261, EPI_ISL_544262, EPI_ISL_544263, EPI_ISL_544266, EPI_ISL_544267, EPI_ISL_544269, EPI_ISL_544270, EPI_ISL_544272, EPI_ISL_544273, EPI_ISL_544274, EPI_ISL_544276, EPI_ISL_544277, EPI_ISL_544280, EPI_ISL_544284, EPI_ISL_544286, EPI_ISL_544287, EPI_ISL_544288, EPI_ISL_544289, EPI_ISL_544290, EPI_ISL_544291, EPI_ISL_544292, EPI_ISL_544293, EPI_ISL_544294, EPI_ISL_544295, EPI_ISL_544297, EPI_ISL_544298, EPI_ISL_544299, EPI_ISL_544300, EPI_ISL_544301, EPI_ISL_544302, EPI_ISL_544303, EPI_ISL_544304, EPI_ISL_544305, EPI_ISL_544306, EPI_ISL_544307, EPI_ISL_544308, EPI_ISL_544310, EPI_ISL_544312, EPI_ISL_544313, EPI_ISL_544315, EPI_ISL_544316, EPI_ISL_544317, EPI_ISL_544319, EPI_ISL_544320, EPI_ISL_544321, EPI_ISL_544322, EPI_ISL_544323, EPI_ISL_544324, EPI_ISL_544326, EPI_ISL_544327, EPI_ISL_544328, EPI_ISL_544329, EPI_ISL_544330, EPI_ISL_544331, EPI_ISL_544333, EPI_ISL_544334, EPI_ISL_544335, EPI_ISL_544336, EPI_ISL_544337, EPI_ISL_544338, EPI_ISL_544340, EPI_ISL_544341, EPI_ISL_544342, EPI_ISL_544343, EPI_ISL_544344, EPI_ISL_544346, EPI_ISL_544347, EPI_ISL_544348, EPI_ISL_544349, EPI_ISL_544358, EPI_ISL_544361, EPI_ISL_544363, EPI_ISL_544364, EPI_ISL_544375, EPI_ISL_544378, EPI_ISL_544379, EPI_ISL_544380, EPI_ISL_544383, EPI_ISL_544385, EPI_ISL_544388, EPI_ISL_544399, EPI_ISL_544400, EPI_ISL_544401, EPI_ISL_544403, EPI_ISL_544405, EPI_ISL_544411, EPI_ISL_544413, EPI_ISL_544414, EPI_ISL_544417, EPI_ISL_544418, EPI_ISL_544419, EPI_ISL_544420, EPI_ISL_544426, EPI_ISL_544427, EPI_ISL_544428, EPI_ISL_544431, EPI_ISL_544432, EPI_ISL_544435, EPI_ISL_544436, EPI_ISL_544437, EPI_ISL_544438, EPI_ISL_544441, EPI_ISL_544442, EPI_ISL_544443, EPI_ISL_544444, EPI_ISL_544445, EPI_ISL_544446, EPI_ISL_544447, EPI_ISL_544448, EPI_ISL_544449, EPI_ISL_544450, EPI_ISL_544451, EPI_ISL_544452, EPI_ISL_544453, EPI_ISL_544455, EPI_ISL_544458, EPI_ISL_544459, EPI_ISL_544463, EPI_ISL_544464, EPI_ISL_544467, EPI_ISL_544468, EPI_ISL_544472, EPI_ISL_544477, EPI_ISL_544484, EPI_ISL_544485, EPI_ISL_544488, EPI_ISL_544490, EPI_ISL_544493, EPI_ISL_544494, EPI_ISL_544499, EPI_ISL_544500, EPI_ISL_544509, EPI_ISL_544510, EPI_ISL_544513, EPI_ISL_544515, EPI_ISL_544516, EPI_ISL_544520, EPI_ISL_544521, EPI_ISL_544525, EPI_ISL_544526, EPI_ISL_544529, EPI_ISL_544530, EPI_ISL_544531, EPI_ISL_544535, EPI_ISL_544536, EPI_ISL_544537, EPI_ISL_544540, EPI_ISL_544541, EPI_ISL_544542, EPI_ISL_544544, EPI_ISL_544546, EPI_ISL_544547, EPI_ISL_544549, EPI_ISL_544550, EPI_ISL_544553, EPI_ISL_544554, EPI_ISL_544555, EPI_ISL_544556, EPI_ISL_544557, EPI_ISL_544559, EPI_ISL_544560, EPI_ISL_544563, EPI_ISL_544564, EPI_ISL_544566, EPI_ISL_544567, EPI_ISL_544570, EPI_ISL_544571, EPI_ISL_544573, EPI_ISL_544574, EPI_ISL_544577, EPI_ISL_544581, EPI_ISL_544582, EPI_ISL_544583, EPI_ISL_544584, EPI_ISL_544586, EPI_ISL_544587, EPI_ISL_544589, EPI_ISL_544590, EPI_ISL_544591, EPI_ISL_544592, EPI_ISL_544593, EPI_ISL_544594, EPI_ISL_544596, EPI_ISL_544597, EPI_ISL_544598, EPI_ISL_544599, EPI_ISL_544600, EPI_ISL_544602, EPI_ISL_544603, EPI_ISL_544605, EPI_ISL_544606, EPI_ISL_544608, EPI_ISL_544609, EPI_ISL_544610, EPI_ISL_544612, EPI_ISL_544614, EPI_ISL_544615, EPI_ISL_544616, EPI_ISL_544617, EPI_ISL_544618, EPI_ISL_544619, EPI_ISL_544621, EPI_ISL_544622, EPI_ISL_544624, EPI_ISL_544625, EPI_ISL_544626, EPI_ISL_544627, EPI_ISL_544628, EPI_ISL_544629, EPI_ISL_544631, EPI_ISL_544632, EPI_ISL_544633, EPI_ISL_544634, EPI_ISL_544636, EPI_ISL_544637, EPI_ISL_544638, EPI_ISL_544640, EPI_ISL_544643, EPI_ISL_544644, EPI_ISL_544646, EPI_ISL_544648, EPI_ISL_544650, EPI_ISL_544652, EPI_ISL_544653, EPI_ISL_544654, EPI_ISL_544655, EPI_ISL_544657, EPI_ISL_544660, EPI_ISL_544662, EPI_ISL_544663, EPI_ISL_544665, EPI_ISL_544666, EPI_ISL_544667, EPI_ISL_544668, EPI_ISL_544669, EPI_ISL_544670, EPI_ISL_544671, EPI_ISL_544672, EPI_ISL_544674, EPI_ISL_544675, EPI_ISL_544678, EPI_ISL_544679, EPI_ISL_544685, EPI_ISL_544686, EPI_ISL_544688, EPI_ISL_544689, EPI_ISL_544690, EPI_ISL_544691, EPI_ISL_544692, EPI_ISL_544693, EPI_ISL_544694, EPI_ISL_544696, EPI_ISL_544699, EPI_ISL_544701, EPI_ISL_544704, EPI_ISL_544709, EPI_ISL_544710, EPI_ISL_544712, EPI_ISL_544717, EPI_ISL_544718, EPI_ISL_544719, EPI_ISL_544720, EPI_ISL_544728, EPI_ISL_544729, EPI_ISL_544731, EPI_ISL_544732, EPI_ISL_544735, EPI_ISL_544736, EPI_ISL_544739, EPI_ISL_544740, EPI_ISL_544743, EPI_ISL_544744, EPI_ISL_544745, EPI_ISL_544746, EPI_ISL_544747, EPI_ISL_544749, EPI_ISL_544750, EPI_ISL_544751, EPI_ISL_544752, EPI_ISL_544753, EPI_ISL_544754, EPI_ISL_544757, EPI_ISL_544758, EPI_ISL_544763, EPI_ISL_544765, EPI_ISL_544768, EPI_ISL_544769, EPI_ISL_544770, EPI_ISL_544771, EPI_ISL_544774, EPI_ISL_544776, EPI_ISL_544777, EPI_ISL_544778, EPI_ISL_544779, EPI_ISL_544780, EPI_ISL_544782, EPI_ISL_544783, EPI_ISL_544785, EPI_ISL_544786, EPI_ISL_544787, EPI_ISL_544788, EPI_ISL_544789, EPI_ISL_544790, EPI_ISL_544791, EPI_ISL_544792, EPI_ISL_544793, EPI_ISL_544794, EPI_ISL_544801, EPI_ISL_544802, EPI_ISL_544803, EPI_ISL_544804, EPI_ISL_544808, EPI_ISL_544809, EPI_ISL_544810, EPI_ISL_544811, EPI_ISL_544813, EPI_ISL_544814, EPI_ISL_544816, EPI_ISL_544817, EPI_ISL_544818, EPI_ISL_544819, EPI_ISL_544821, EPI_ISL_544823, EPI_ISL_544824, EPI_ISL_544825, EPI_ISL_544826, EPI_ISL_544827, EPI_ISL_544828, EPI_ISL_544829, EPI_ISL_544831, EPI_ISL_544832, EPI_ISL_544833, EPI_ISL_544834, EPI_ISL_544836, EPI_ISL_544837, EPI_ISL_544842, EPI_ISL_544843, EPI_ISL_544844, EPI_ISL_544845, EPI_ISL_544846, EPI_ISL_544847, EPI_ISL_544849, EPI_ISL_544851, EPI_ISL_544854, EPI_ISL_544855, EPI_ISL_544856, EPI_ISL_544857, EPI_ISL_544858, EPI_ISL_544859, EPI_ISL_544860, EPI_ISL_544861, EPI_ISL_544862, EPI_ISL_544863, EPI_ISL_544864, EPI_ISL_544865, EPI_ISL_544866, EPI_ISL_544869, EPI_ISL_544870, EPI_ISL_544871, EPI_ISL_544872, EPI_ISL_544874, EPI_ISL_544877, EPI_ISL_544878, EPI_ISL_544880, EPI_ISL_544881, EPI_ISL_544885, EPI_ISL_544887, EPI_ISL_544888, EPI_ISL_544890, EPI_ISL_544891, EPI_ISL_544892, EPI_ISL_544894, EPI_ISL_544895, EPI_ISL_544897, EPI_ISL_544898, EPI_ISL_544899, EPI_ISL_544900, EPI_ISL_544901, EPI_ISL_544902, EPI_ISL_544903, EPI_ISL_544908, EPI_ISL_544909, EPI_ISL_544911, EPI_ISL_544913, EPI_ISL_544918, EPI_ISL_544919, EPI_ISL_544920, EPI_ISL_544924, EPI_ISL_544926, EPI_ISL_544927, EPI_ISL_544930, EPI_ISL_544931, EPI_ISL_544933, EPI_ISL_544934, EPI_ISL_544937, EPI_ISL_544938, EPI_ISL_544940, EPI_ISL_544941, EPI_ISL_544942, EPI_ISL_544943, EPI_ISL_544944, EPI_ISL_544945, EPI_ISL_544946, EPI_ISL_544947, EPI_ISL_544948, EPI_ISL_544949, EPI_ISL_544951, EPI_ISL_544952, EPI_ISL_544953, EPI_ISL_544954, EPI_ISL_545051, EPI_ISL_545052, EPI_ISL_545053, EPI_ISL_545054, EPI_ISL_545055, EPI_ISL_545056, EPI_ISL_545057, EPI_ISL_545058, EPI_ISL_545059, EPI_ISL_545060, EPI_ISL_545061, EPI_ISL_545062, EPI_ISL_545063, EPI_ISL_545064, EPI_ISL_545065, EPI_ISL_545066, EPI_ISL_545067, EPI_ISL_545068, EPI_ISL_545069, EPI_ISL_545070, EPI_ISL_545071, EPI_ISL_545072, EPI_ISL_545073, EPI_ISL_545074, EPI_ISL_545075, EPI_ISL_545076, EPI_ISL_545077, EPI_ISL_545078, EPI_ISL_545079, EPI_ISL_545080, EPI_ISL_545081, EPI_ISL_545082, EPI_ISL_545083, EPI_ISL_545084, EPI_ISL_545085, EPI_ISL_545086, EPI_ISL_545087, EPI_ISL_545088, EPI_ISL_545089, EPI_ISL_545090, EPI_ISL_545091, EPI_ISL_545092, EPI_ISL_545093, EPI_ISL_545094, EPI_ISL_545095, EPI_ISL_545096, EPI_ISL_545097, EPI_ISL_545098, EPI_ISL_545099, EPI_ISL_545100, EPI_ISL_545101, EPI_ISL_545102, EPI_ISL_545103, EPI_ISL_545104, EPI_ISL_545105, EPI_ISL_545106, EPI_ISL_545107, EPI_ISL_545108, EPI_ISL_545109, EPI_ISL_545110, EPI_ISL_545111, EPI_ISL_545112, EPI_ISL_545113, EPI_ISL_545114, EPI_ISL_545115, EPI_ISL_545116, EPI_ISL_545117, EPI_ISL_545118, EPI_ISL_545119, EPI_ISL_545120, EPI_ISL_545121, EPI_ISL_545122, EPI_ISL_545123, EPI_ISL_545124, EPI_ISL_545125, EPI_ISL_545126, EPI_ISL_545127, EPI_ISL_545128, EPI_ISL_545129, EPI_ISL_545130, EPI_ISL_545131, EPI_ISL_545132, EPI_ISL_545133, EPI_ISL_545134, EPI_ISL_545135, EPI_ISL_545136, EPI_ISL_545137, EPI_ISL_545138, EPI_ISL_545139, EPI_ISL_545140, EPI_ISL_545141, EPI_ISL_545142, EPI_ISL_545143, EPI_ISL_545144, EPI_ISL_545145, EPI_ISL_545146, EPI_ISL_545147, EPI_ISL_545148, EPI_ISL_545149, EPI_ISL_545150, EPI_ISL_545151, EPI_ISL_545152, EPI_ISL_545153, EPI_ISL_545154, EPI_ISL_545155, EPI_ISL_545156, EPI_ISL_545157, EPI_ISL_545158, EPI_ISL_545159, EPI_ISL_545160, EPI_ISL_545161, EPI_ISL_545162, EPI_ISL_545163, EPI_ISL_545164, EPI_ISL_545165, EPI_ISL_545166, EPI_ISL_545167, EPI_ISL_545168, EPI_ISL_545169, EPI_ISL_545170, EPI_ISL_545171, EPI_ISL_545172, EPI_ISL_545173, EPI_ISL_545174, EPI_ISL_545175, EPI_ISL_545176, EPI_ISL_545177, EPI_ISL_545178, EPI_ISL_545179, EPI_ISL_545180, EPI_ISL_545181, EPI_ISL_545182, EPI_ISL_545183, EPI_ISL_545184, EPI_ISL_545185, EPI_ISL_545186, EPI_ISL_545187, EPI_ISL_545188, EPI_ISL_545189, EPI_ISL_545190, EPI_ISL_545191, EPI_ISL_545192, EPI_ISL_545193, EPI_ISL_545194, EPI_ISL_545195, EPI_ISL_545196, EPI_ISL_545197, EPI_ISL_545198, EPI_ISL_545199, EPI_ISL_545200, EPI_ISL_545201, EPI_ISL_545202, EPI_ISL_545203, EPI_ISL_545204, EPI_ISL_545205, EPI_ISL_545206, EPI_ISL_545207, EPI_ISL_545208, EPI_ISL_545209, EPI_ISL_545210, EPI_ISL_545211, EPI_ISL_545212, EPI_ISL_545213, EPI_ISL_545214, EPI_ISL_545215, EPI_ISL_545216, EPI_ISL_545217, EPI_ISL_545218, EPI_ISL_545219, EPI_ISL_545220, EPI_ISL_545221, EPI_ISL_545222, EPI_ISL_545223, EPI_ISL_545224, EPI_ISL_545225, EPI_ISL_545226, EPI_ISL_545227, EPI_ISL_545228, EPI_ISL_545229, EPI_ISL_545230, EPI_ISL_545231, EPI_ISL_545232, EPI_ISL_545233, EPI_ISL_545234, EPI_ISL_545235, EPI_ISL_545236, EPI_ISL_545237, EPI_ISL_545238, EPI_ISL_545239, EPI_ISL_545240, EPI_ISL_545241, EPI_ISL_545242, EPI_ISL_545243, EPI_ISL_545244, EPI_ISL_545245, EPI_ISL_545246, EPI_ISL_545247, EPI_ISL_545248, EPI_ISL_545249, EPI_ISL_545250, EPI_ISL_545251, EPI_ISL_545252, EPI_ISL_545253, EPI_ISL_545254, EPI_ISL_545255, EPI_ISL_545256, EPI_ISL_545257, EPI_ISL_545258, EPI_ISL_545259, EPI_ISL_545260, EPI_ISL_545261, EPI_ISL_545263, EPI_ISL_545264, EPI_ISL_545265, EPI_ISL_545266, EPI_ISL_545267, EPI_ISL_545268, EPI_ISL_545269, EPI_ISL_545270, EPI_ISL_545271, EPI_ISL_545272, EPI_ISL_545273, EPI_ISL_545274, EPI_ISL_545275, EPI_ISL_545276, EPI_ISL_545277, EPI_ISL_545278, EPI_ISL_545279, EPI_ISL_545280, EPI_ISL_545281, EPI_ISL_545282, EPI_ISL_545283, EPI_ISL_545284, EPI_ISL_545285, EPI_ISL_545286, EPI_ISL_545287, EPI_ISL_545288, EPI_ISL_545289, EPI_ISL_545290, EPI_ISL_545291, EPI_ISL_545292, EPI_ISL_545293, EPI_ISL_545294, EPI_ISL_545295, EPI_ISL_545296, EPI_ISL_545297, EPI_ISL_545298, EPI_ISL_545299, EPI_ISL_545300, EPI_ISL_545301, EPI_ISL_545302, EPI_ISL_545303, EPI_ISL_545304, EPI_ISL_545305, EPI_ISL_545306, EPI_ISL_545307, EPI_ISL_545308, EPI_ISL_545309, EPI_ISL_545310, EPI_ISL_545311, EPI_ISL_545312, EPI_ISL_545313, EPI_ISL_545314, EPI_ISL_545315, EPI_ISL_545316, EPI_ISL_545317, EPI_ISL_545318, EPI_ISL_545319, EPI_ISL_545320, EPI_ISL_545321, EPI_ISL_545322, EPI_ISL_545323, EPI_ISL_545324, EPI_ISL_545325, EPI_ISL_545326, EPI_ISL_545327, EPI_ISL_545328, EPI_ISL_545329, EPI_ISL_545330, EPI_ISL_545331, EPI_ISL_545332, EPI_ISL_545333, EPI_ISL_545334, EPI_ISL_545335, EPI_ISL_545336, EPI_ISL_545337, EPI_ISL_545338, EPI_ISL_545339, EPI_ISL_545340, EPI_ISL_545341, EPI_ISL_545342, EPI_ISL_545343, EPI_ISL_545344, EPI_ISL_545345, EPI_ISL_545346, EPI_ISL_545347, EPI_ISL_545348, EPI_ISL_545349, EPI_ISL_545350, EPI_ISL_545351, EPI_ISL_545352, EPI_ISL_545353, EPI_ISL_545354, EPI_ISL_545355, EPI_ISL_545356, EPI_ISL_545357, EPI_ISL_545358, EPI_ISL_545359, EPI_ISL_545360, EPI_ISL_545361, EPI_ISL_545362, EPI_ISL_545363, EPI_ISL_545364, EPI_ISL_545365, EPI_ISL_545366, EPI_ISL_545367, EPI_ISL_545368, EPI_ISL_545369, EPI_ISL_545370, EPI_ISL_545371, EPI_ISL_545372, EPI_ISL_545373, EPI_ISL_545374, EPI_ISL_545375, EPI_ISL_545376, EPI_ISL_545377, EPI_ISL_545378, EPI_ISL_545379, EPI_ISL_545380, EPI_ISL_545381, EPI_ISL_545382, EPI_ISL_545383, EPI_ISL_545384, EPI_ISL_545385, EPI_ISL_545386, EPI_ISL_545387, EPI_ISL_545388, EPI_ISL_545389, EPI_ISL_545390, EPI_ISL_545391, EPI_ISL_545392, EPI_ISL_545393, EPI_ISL_545394, EPI_ISL_545395, EPI_ISL_545396, EPI_ISL_545397, EPI_ISL_545398, EPI_ISL_545399, EPI_ISL_545400, EPI_ISL_545401, EPI_ISL_545402, EPI_ISL_545403, EPI_ISL_545404, EPI_ISL_545405, EPI_ISL_545406, EPI_ISL_545407, EPI_ISL_545408, EPI_ISL_545409, EPI_ISL_545410, EPI_ISL_545411, EPI_ISL_545412, EPI_ISL_545413, EPI_ISL_545414, EPI_ISL_545415, EPI_ISL_545416, EPI_ISL_545417, EPI_ISL_545418, EPI_ISL_545419, EPI_ISL_545420, EPI_ISL_545421, EPI_ISL_545422, EPI_ISL_545423, EPI_ISL_545424, EPI_ISL_545425, EPI_ISL_545426, EPI_ISL_545427, EPI_ISL_545428, EPI_ISL_545429, EPI_ISL_545430, EPI_ISL_545431, EPI_ISL_545432, EPI_ISL_545433, EPI_ISL_545434, EPI_ISL_545435, EPI_ISL_545436, EPI_ISL_545437, EPI_ISL_545438, EPI_ISL_545439, EPI_ISL_545440, EPI_ISL_545441, EPI_ISL_545442, EPI_ISL_545443, EPI_ISL_545444, EPI_ISL_545445, EPI_ISL_545446, EPI_ISL_545447, EPI_ISL_545448, EPI_ISL_545449, EPI_ISL_545450, EPI_ISL_545451, EPI_ISL_545452, EPI_ISL_545453, EPI_ISL_545454, EPI_ISL_545455, EPI_ISL_545456, EPI_ISL_545457, EPI_ISL_545458, EPI_ISL_545459, EPI_ISL_545460, EPI_ISL_545461, EPI_ISL_545462, EPI_ISL_545463, EPI_ISL_545464, EPI_ISL_545465, EPI_ISL_545466, EPI_ISL_545467, EPI_ISL_545468, EPI_ISL_545469, EPI_ISL_545470, EPI_ISL_545471, EPI_ISL_545472, EPI_ISL_545473, EPI_ISL_545474, EPI_ISL_545475, EPI_ISL_545476, EPI_ISL_545477, EPI_ISL_545478, EPI_ISL_545479, EPI_ISL_545480, EPI_ISL_545481, EPI_ISL_545482, EPI_ISL_545483, EPI_ISL_545484, EPI_ISL_545485, EPI_ISL_545486, EPI_ISL_545487, EPI_ISL_545488, EPI_ISL_545489, EPI_ISL_545490, EPI_ISL_545491, EPI_ISL_545492, EPI_ISL_545493, EPI_ISL_545494, EPI_ISL_545495, EPI_ISL_545496, EPI_ISL_545497, EPI_ISL_545498, EPI_ISL_545499, EPI_ISL_545500, EPI_ISL_545501, EPI_ISL_545502, EPI_ISL_545503, EPI_ISL_545504, EPI_ISL_545505, EPI_ISL_545506, EPI_ISL_545507, EPI_ISL_545508, EPI_ISL_545509, EPI_ISL_545510, EPI_ISL_545511, EPI_ISL_545512, EPI_ISL_545513, EPI_ISL_545514, EPI_ISL_545515, EPI_ISL_545516, EPI_ISL_545517, EPI_ISL_545518, EPI_ISL_545519, EPI_ISL_545520, EPI_ISL_545521, EPI_ISL_545522, EPI_ISL_545523, EPI_ISL_545524, EPI_ISL_545525, EPI_ISL_545526, EPI_ISL_545527, EPI_ISL_545528, EPI_ISL_545529, EPI_ISL_545530, EPI_ISL_545531, EPI_ISL_545532, EPI_ISL_545533, EPI_ISL_545534, EPI_ISL_545535, EPI_ISL_545536, EPI_ISL_545537, EPI_ISL_545538, EPI_ISL_545539, EPI_ISL_545540, EPI_ISL_545541, EPI_ISL_545542, EPI_ISL_545543, EPI_ISL_545544, EPI_ISL_545545, EPI_ISL_545546, EPI_ISL_545547, EPI_ISL_545548, EPI_ISL_545549, EPI_ISL_545550, EPI_ISL_545551, EPI_ISL_545552, EPI_ISL_545553, EPI_ISL_545554, EPI_ISL_545555, EPI_ISL_545556, EPI_ISL_545557, EPI_ISL_545558, EPI_ISL_545559, EPI_ISL_545560, EPI_ISL_545561, EPI_ISL_545562, EPI_ISL_545563, EPI_ISL_545564, EPI_ISL_545565, EPI_ISL_545566, EPI_ISL_545567, EPI_ISL_545568, EPI_ISL_545569, EPI_ISL_545570, EPI_ISL_545571, EPI_ISL_545572, EPI_ISL_545573, EPI_ISL_545574, EPI_ISL_545581, EPI_ISL_545583, EPI_ISL_545584, EPI_ISL_545585, EPI_ISL_545586, EPI_ISL_545588, EPI_ISL_545590, EPI_ISL_545592, EPI_ISL_545593, EPI_ISL_545594, EPI_ISL_545597, EPI_ISL_545598, EPI_ISL_545599, EPI_ISL_545602, EPI_ISL_545603, EPI_ISL_545604, EPI_ISL_545605, EPI_ISL_545607, EPI_ISL_545609, EPI_ISL_545612, EPI_ISL_545613, EPI_ISL_545616, EPI_ISL_545617, EPI_ISL_545618, EPI_ISL_545620, EPI_ISL_545622, EPI_ISL_545623, EPI_ISL_545624, EPI_ISL_545628, EPI_ISL_545629, EPI_ISL_545630, EPI_ISL_545631, EPI_ISL_545632, EPI_ISL_545634, EPI_ISL_545635, EPI_ISL_545639, EPI_ISL_545641, EPI_ISL_545642, EPI_ISL_545643, EPI_ISL_545646, EPI_ISL_545648, EPI_ISL_545649, EPI_ISL_545652, EPI_ISL_545654, EPI_ISL_545655, EPI_ISL_545657, EPI_ISL_545663, EPI_ISL_545665, EPI_ISL_545674, EPI_ISL_545683, EPI_ISL_545684, EPI_ISL_545685, EPI_ISL_545686, EPI_ISL_545688, EPI_ISL_545690, EPI_ISL_545693, EPI_ISL_545694, EPI_ISL_545696, EPI_ISL_545699, EPI_ISL_545701, EPI_ISL_545702, EPI_ISL_545703, EPI_ISL_545704, EPI_ISL_545705, EPI_ISL_545706, EPI_ISL_545709, EPI_ISL_545710, EPI_ISL_545711, EPI_ISL_545712, EPI_ISL_545713, EPI_ISL_545715, EPI_ISL_545716, EPI_ISL_545717, EPI_ISL_545718, EPI_ISL_545721, EPI_ISL_545723, EPI_ISL_545725, EPI_ISL_545726, EPI_ISL_545727, EPI_ISL_545729, EPI_ISL_545730, EPI_ISL_545731, EPI_ISL_545733, EPI_ISL_545735, EPI_ISL_545736, EPI_ISL_545737, EPI_ISL_545738, EPI_ISL_545740, EPI_ISL_545741, EPI_ISL_545742, EPI_ISL_545743, EPI_ISL_545746, EPI_ISL_545747, EPI_ISL_545749, EPI_ISL_545750, EPI_ISL_545753, EPI_ISL_545754, EPI_ISL_545755, EPI_ISL_545756, EPI_ISL_545761, EPI_ISL_545762, EPI_ISL_545763, EPI_ISL_545764, EPI_ISL_545765, EPI_ISL_545768, EPI_ISL_545773, EPI_ISL_545774, EPI_ISL_545775, EPI_ISL_545779, EPI_ISL_545787, EPI_ISL_545792, EPI_ISL_545795, EPI_ISL_545799, EPI_ISL_545806, EPI_ISL_545808, EPI_ISL_545810, EPI_ISL_545813, EPI_ISL_545814, EPI_ISL_545815, EPI_ISL_545818, EPI_ISL_545820, EPI_ISL_545821, EPI_ISL_545823, EPI_ISL_545824, EPI_ISL_545826, EPI_ISL_545827, EPI_ISL_545828, EPI_ISL_545829, EPI_ISL_545830, EPI_ISL_545831, EPI_ISL_545832, EPI_ISL_545833, EPI_ISL_545834, EPI_ISL_545835, EPI_ISL_545836, EPI_ISL_545837, EPI_ISL_545838, EPI_ISL_545839, EPI_ISL_545840, EPI_ISL_545841, EPI_ISL_545842, EPI_ISL_545843, EPI_ISL_545844, EPI_ISL_545845, EPI_ISL_545846, EPI_ISL_545847, EPI_ISL_545848, EPI_ISL_545849, EPI_ISL_545850, EPI_ISL_545851, EPI_ISL_545852, EPI_ISL_545853, EPI_ISL_545854, EPI_ISL_545855, EPI_ISL_545856, EPI_ISL_545857, EPI_ISL_545858, EPI_ISL_545859, EPI_ISL_545860, EPI_ISL_545861, EPI_ISL_545862, EPI_ISL_545863, EPI_ISL_545864, EPI_ISL_545865, EPI_ISL_545866, EPI_ISL_545867, EPI_ISL_545868, EPI_ISL_545869, EPI_ISL_545870, EPI_ISL_545871, EPI_ISL_545872, EPI_ISL_545873, EPI_ISL_545874, EPI_ISL_545875, EPI_ISL_545876, EPI_ISL_545877, EPI_ISL_545878, EPI_ISL_545879, EPI_ISL_545880, EPI_ISL_545881, EPI_ISL_545882, EPI_ISL_545883, EPI_ISL_545884, EPI_ISL_545885, EPI_ISL_545886, EPI_ISL_545887, EPI_ISL_545888, EPI_ISL_545889, EPI_ISL_545890, EPI_ISL_545891, EPI_ISL_545892, EPI_ISL_545893, EPI_ISL_545894, EPI_ISL_545895, EPI_ISL_545896, EPI_ISL_545897, EPI_ISL_545898, EPI_ISL_545899, EPI_ISL_545900, EPI_ISL_545901, EPI_ISL_545902, EPI_ISL_545903, EPI_ISL_545904, EPI_ISL_545905, EPI_ISL_545906, EPI_ISL_545907, EPI_ISL_545908, EPI_ISL_545909, EPI_ISL_545910, EPI_ISL_545911, EPI_ISL_545912, EPI_ISL_545913, EPI_ISL_545914, EPI_ISL_545915, EPI_ISL_545916, EPI_ISL_545917, EPI_ISL_545918, EPI_ISL_545919, EPI_ISL_545920, EPI_ISL_545921, EPI_ISL_545922, EPI_ISL_545923, EPI_ISL_545924, EPI_ISL_545925, EPI_ISL_545926, EPI_ISL_545927, EPI_ISL_545928, EPI_ISL_545929, EPI_ISL_545930, EPI_ISL_545931, EPI_ISL_545932, EPI_ISL_545933, EPI_ISL_545934, EPI_ISL_545935, EPI_ISL_545936, EPI_ISL_545937, EPI_ISL_545938, EPI_ISL_545939, EPI_ISL_545940, EPI_ISL_545941, EPI_ISL_545942, EPI_ISL_545943, EPI_ISL_545944, EPI_ISL_545945, EPI_ISL_545946, EPI_ISL_545947, EPI_ISL_545948, EPI_ISL_545949, EPI_ISL_545950, EPI_ISL_545951, EPI_ISL_545952, EPI_ISL_545953, EPI_ISL_545958, EPI_ISL_545959, EPI_ISL_545962, EPI_ISL_545969, EPI_ISL_545970, EPI_ISL_545971, EPI_ISL_545972, EPI_ISL_545973, EPI_ISL_545974, EPI_ISL_545975, EPI_ISL_545976, EPI_ISL_545978, EPI_ISL_545979, EPI_ISL_545980, EPI_ISL_545984, EPI_ISL_545985, EPI_ISL_545986, EPI_ISL_545990, EPI_ISL_545991, EPI_ISL_545992, EPI_ISL_545993, EPI_ISL_545996, EPI_ISL_545997, EPI_ISL_545998, EPI_ISL_546001, EPI_ISL_546002, EPI_ISL_546003, EPI_ISL_546004, EPI_ISL_546012, EPI_ISL_546013, EPI_ISL_546015, EPI_ISL_546016, EPI_ISL_546018, EPI_ISL_546021, EPI_ISL_546023, EPI_ISL_546024, EPI_ISL_546029, EPI_ISL_546030, EPI_ISL_546031, EPI_ISL_546032, EPI_ISL_546034, EPI_ISL_546035, EPI_ISL_546044, EPI_ISL_546045, EPI_ISL_546048, EPI_ISL_546051, EPI_ISL_546057, EPI_ISL_546063, EPI_ISL_546064, EPI_ISL_546068, EPI_ISL_546073, EPI_ISL_546074, EPI_ISL_546075, EPI_ISL_546076, EPI_ISL_546077, EPI_ISL_546078, EPI_ISL_546079, EPI_ISL_546081, EPI_ISL_546085, EPI_ISL_546086, EPI_ISL_546087, EPI_ISL_546089, EPI_ISL_546090, EPI_ISL_546092, EPI_ISL_546094, EPI_ISL_546095, EPI_ISL_546101, EPI_ISL_546103, EPI_ISL_546104, EPI_ISL_546107, EPI_ISL_546110, EPI_ISL_546118, EPI_ISL_546124, EPI_ISL_546125, EPI_ISL_546126, EPI_ISL_546127, EPI_ISL_546130, EPI_ISL_546131, EPI_ISL_546133, EPI_ISL_546134, EPI_ISL_546135, EPI_ISL_546140, EPI_ISL_546141, EPI_ISL_546142, EPI_ISL_546143, EPI_ISL_546144, EPI_ISL_546145, EPI_ISL_546149, EPI_ISL_546150, EPI_ISL_546151, EPI_ISL_546152, EPI_ISL_546153, EPI_ISL_546154, EPI_ISL_546155, EPI_ISL_546156, EPI_ISL_546157, EPI_ISL_546158, EPI_ISL_546159, EPI_ISL_546160, EPI_ISL_546161, EPI_ISL_546162, EPI_ISL_546163, EPI_ISL_546164, EPI_ISL_546165, EPI_ISL_546166, EPI_ISL_546167, EPI_ISL_546168, EPI_ISL_546169, EPI_ISL_546170, EPI_ISL_546171, EPI_ISL_546172, EPI_ISL_546173, EPI_ISL_546174, EPI_ISL_546175, EPI_ISL_546176, EPI_ISL_546177, EPI_ISL_546180, EPI_ISL_546185, EPI_ISL_546186, EPI_ISL_546187, EPI_ISL_546189, EPI_ISL_546191, EPI_ISL_546192, EPI_ISL_546195, EPI_ISL_546198, EPI_ISL_546199, EPI_ISL_546202, EPI_ISL_546203, EPI_ISL_546204, EPI_ISL_546205, EPI_ISL_546206, EPI_ISL_546208, EPI_ISL_546210, EPI_ISL_546211, EPI_ISL_546212, EPI_ISL_546214, EPI_ISL_546215, EPI_ISL_546223, EPI_ISL_546224, EPI_ISL_546226, EPI_ISL_546228, EPI_ISL_546229, EPI_ISL_546230, EPI_ISL_546238, EPI_ISL_546239, EPI_ISL_546246, EPI_ISL_546249, EPI_ISL_546250, EPI_ISL_546259, EPI_ISL_546260, EPI_ISL_546261, EPI_ISL_546263, EPI_ISL_546265, EPI_ISL_546266, EPI_ISL_546267, EPI_ISL_546268, EPI_ISL_546269, EPI_ISL_546270, EPI_ISL_546272, EPI_ISL_546273, EPI_ISL_546274, EPI_ISL_546275, EPI_ISL_546276, EPI_ISL_546277, EPI_ISL_546278, EPI_ISL_546279, EPI_ISL_546280, EPI_ISL_546281, EPI_ISL_546282, EPI_ISL_546283, EPI_ISL_546284, EPI_ISL_546285, EPI_ISL_546286, EPI_ISL_546287, EPI_ISL_546289, EPI_ISL_546291, EPI_ISL_546292, EPI_ISL_546293, EPI_ISL_546296, EPI_ISL_546298, EPI_ISL_546300, EPI_ISL_546301, EPI_ISL_546303, EPI_ISL_546304, EPI_ISL_546306, EPI_ISL_546307, EPI_ISL_546309, EPI_ISL_546314, EPI_ISL_546317, EPI_ISL_546318, EPI_ISL_546320, EPI_ISL_546321, EPI_ISL_546323, EPI_ISL_546325, EPI_ISL_546327, EPI_ISL_546329, EPI_ISL_546330, EPI_ISL_546331, EPI_ISL_546338, EPI_ISL_546339, EPI_ISL_546340, EPI_ISL_546341, EPI_ISL_546343, EPI_ISL_546344, EPI_ISL_546345, EPI_ISL_546346, EPI_ISL_546347, EPI_ISL_546348, EPI_ISL_546349, EPI_ISL_546350, EPI_ISL_546351, EPI_ISL_546353, EPI_ISL_546354, EPI_ISL_546355, EPI_ISL_546356, EPI_ISL_546357, EPI_ISL_546358, EPI_ISL_546359, EPI_ISL_546360, EPI_ISL_546362, EPI_ISL_546363, EPI_ISL_546364, EPI_ISL_546366, EPI_ISL_546367, EPI_ISL_546369, EPI_ISL_546370, EPI_ISL_546371, EPI_ISL_546374, EPI_ISL_546375, EPI_ISL_546376, EPI_ISL_546377, EPI_ISL_546378, EPI_ISL_546380, EPI_ISL_546381, EPI_ISL_546382, EPI_ISL_546383, EPI_ISL_546384, EPI_ISL_546387, EPI_ISL_546388, EPI_ISL_546389, EPI_ISL_546391, EPI_ISL_546393, EPI_ISL_546394, EPI_ISL_546395, EPI_ISL_546396, EPI_ISL_546398, EPI_ISL_546399, EPI_ISL_546401, EPI_ISL_546402, EPI_ISL_546403, EPI_ISL_546404, EPI_ISL_546405, EPI_ISL_546406, EPI_ISL_546407, EPI_ISL_546408, EPI_ISL_546409, EPI_ISL_546410, EPI_ISL_546411, EPI_ISL_546412, EPI_ISL_546413, EPI_ISL_546415, EPI_ISL_546416, EPI_ISL_546417, EPI_ISL_546418, EPI_ISL_546419, EPI_ISL_546420, EPI_ISL_546421, EPI_ISL_546422, EPI_ISL_546424, EPI_ISL_546425, EPI_ISL_546427, EPI_ISL_546428, EPI_ISL_546429, EPI_ISL_546430, EPI_ISL_546431, EPI_ISL_546432, EPI_ISL_546433, EPI_ISL_546434, EPI_ISL_546435, EPI_ISL_546439, EPI_ISL_546441, EPI_ISL_546442, EPI_ISL_546444, EPI_ISL_546445, EPI_ISL_546446, EPI_ISL_546449, EPI_ISL_546452, EPI_ISL_546453, EPI_ISL_546455, EPI_ISL_546456, EPI_ISL_546457, EPI_ISL_546458, EPI_ISL_546459, EPI_ISL_546460, EPI_ISL_546461, EPI_ISL_546462, EPI_ISL_546463, EPI_ISL_546467, EPI_ISL_546470, EPI_ISL_546471, EPI_ISL_546472, EPI_ISL_546473, EPI_ISL_546474, EPI_ISL_546478, EPI_ISL_546481, EPI_ISL_546483, EPI_ISL_546485, EPI_ISL_546490, EPI_ISL_546491, EPI_ISL_546492, EPI_ISL_546493,

| | | | |
|---|---|---|---|
| EPI_ISL_546496, EPI_ISL_546497, EPI_ISL_546499, EPI_ISL_546501, EPI_ISL_546508, EPI_ISL_546513, EPI_ISL_546514, EPI_ISL_546516, EPI_ISL_546520, EPI_ISL_546521, EPI_ISL_546523, EPI_ISL_546528, EPI_ISL_546530, EPI_ISL_546532, EPI_ISL_546533, EPI_ISL_546534, EPI_ISL_546535, EPI_ISL_546536, EPI_ISL_546537, EPI_ISL_546538, EPI_ISL_546539, EPI_ISL_546540, EPI_ISL_546542, EPI_ISL_546543, EPI_ISL_546544, EPI_ISL_546545, EPI_ISL_546546, EPI_ISL_546548, EPI_ISL_546549, EPI_ISL_546550, EPI_ISL_546551, EPI_ISL_546552, EPI_ISL_546553, EPI_ISL_546554, EPI_ISL_546558, EPI_ISL_546559, EPI_ISL_546560, EPI_ISL_546561, EPI_ISL_546562, EPI_ISL_546568, EPI_ISL_546572, EPI_ISL_546573, EPI_ISL_546576, EPI_ISL_546577, EPI_ISL_546578, EPI_ISL_546579, EPI_ISL_546580, EPI_ISL_546581, EPI_ISL_546582, EPI_ISL_546583, EPI_ISL_546584, EPI_ISL_546587, EPI_ISL_546588, EPI_ISL_546589, EPI_ISL_546590, EPI_ISL_546591, EPI_ISL_546592, EPI_ISL_546595, EPI_ISL_546596, EPI_ISL_546597, EPI_ISL_546600, EPI_ISL_546601, EPI_ISL_546602, EPI_ISL_546603, EPI_ISL_546605, EPI_ISL_546606, EPI_ISL_546608, EPI_ISL_546609, EPI_ISL_546611, EPI_ISL_546612, EPI_ISL_546614, EPI_ISL_546616, EPI_ISL_546618, EPI_ISL_546620, EPI_ISL_546622, EPI_ISL_546625, EPI_ISL_546626, EPI_ISL_546627, EPI_ISL_546628, EPI_ISL_546629, EPI_ISL_546630, EPI_ISL_546633, EPI_ISL_546634, EPI_ISL_546635, EPI_ISL_546637, EPI_ISL_546638, EPI_ISL_546640, EPI_ISL_546641, EPI_ISL_546642, EPI_ISL_546643, EPI_ISL_546644, EPI_ISL_546645, EPI_ISL_546646, EPI_ISL_546647, EPI_ISL_546648, EPI_ISL_546649, EPI_ISL_546651, EPI_ISL_546652, EPI_ISL_546654, EPI_ISL_546655, EPI_ISL_546658, EPI_ISL_546661, EPI_ISL_546662, EPI_ISL_546663, EPI_ISL_546664, EPI_ISL_546665, EPI_ISL_546666, EPI_ISL_546668, EPI_ISL_546673, EPI_ISL_546674, EPI_ISL_546676, EPI_ISL_546679, EPI_ISL_546682, EPI_ISL_546684, EPI_ISL_546685, EPI_ISL_546686, EPI_ISL_546690, EPI_ISL_546691, EPI_ISL_546693, EPI_ISL_546694, EPI_ISL_546695, EPI_ISL_546696, EPI_ISL_546697, EPI_ISL_546698, EPI_ISL_546699, EPI_ISL_546701, EPI_ISL_546702, EPI_ISL_546703, EPI_ISL_546705, EPI_ISL_546708, EPI_ISL_546713, EPI_ISL_546715, EPI_ISL_546716, EPI_ISL_546717, EPI_ISL_546718, EPI_ISL_546720, EPI_ISL_546721, EPI_ISL_546724, EPI_ISL_546725, EPI_ISL_546726, EPI_ISL_546727, EPI_ISL_546732, EPI_ISL_546735, EPI_ISL_546737, EPI_ISL_546739, EPI_ISL_546745, EPI_ISL_546746, EPI_ISL_546748, EPI_ISL_546751, EPI_ISL_546752, EPI_ISL_546755, EPI_ISL_546756, EPI_ISL_546757, EPI_ISL_546758, EPI_ISL_546759, EPI_ISL_546760, EPI_ISL_546761, EPI_ISL_546762, EPI_ISL_546763, EPI_ISL_546764, EPI_ISL_546765, EPI_ISL_546766, EPI_ISL_546767, EPI_ISL_546768, EPI_ISL_546769, EPI_ISL_546770, EPI_ISL_546772, EPI_ISL_546773, EPI_ISL_546774, EPI_ISL_546776, EPI_ISL_546777, EPI_ISL_546778, EPI_ISL_546779, EPI_ISL_546780, EPI_ISL_546781, EPI_ISL_546782, EPI_ISL_546783, EPI_ISL_546784, EPI_ISL_546785, EPI_ISL_546787, EPI_ISL_546788, EPI_ISL_546789, EPI_ISL_546791, EPI_ISL_546796, EPI_ISL_546798, EPI_ISL_546799, EPI_ISL_546800, EPI_ISL_546801, EPI_ISL_546802, EPI_ISL_546803, EPI_ISL_546804, EPI_ISL_546805, EPI_ISL_546806, EPI_ISL_546807, EPI_ISL_546809, EPI_ISL_546812, EPI_ISL_546813, EPI_ISL_546814, EPI_ISL_546817, EPI_ISL_546818, EPI_ISL_546820, EPI_ISL_546821, EPI_ISL_546822, EPI_ISL_546823, EPI_ISL_546826, EPI_ISL_546827, EPI_ISL_546828, EPI_ISL_546829, EPI_ISL_546830, EPI_ISL_546832, EPI_ISL_546835, EPI_ISL_546836, EPI_ISL_546838, EPI_ISL_546839, EPI_ISL_546840, EPI_ISL_546843, EPI_ISL_546844, EPI_ISL_546845, EPI_ISL_546846, EPI_ISL_546847, EPI_ISL_546848, EPI_ISL_546849, EPI_ISL_546850, EPI_ISL_546851, EPI_ISL_546852, EPI_ISL_546853, EPI_ISL_546854, EPI_ISL_546855, EPI_ISL_546856, EPI_ISL_546857, EPI_ISL_546858, EPI_ISL_546859, EPI_ISL_546860, EPI_ISL_546861, EPI_ISL_546863, EPI_ISL_546865, EPI_ISL_546867, EPI_ISL_546870, EPI_ISL_546872, EPI_ISL_546873, EPI_ISL_546874, EPI_ISL_546875, EPI_ISL_546876, EPI_ISL_546877, EPI_ISL_546878, EPI_ISL_546881, EPI_ISL_546882, EPI_ISL_546883, EPI_ISL_546884, EPI_ISL_546886, EPI_ISL_546887, EPI_ISL_546888, EPI_ISL_546890, EPI_ISL_546891, EPI_ISL_546892, EPI_ISL_546894, EPI_ISL_546896, EPI_ISL_546897, EPI_ISL_546898, EPI_ISL_546899, EPI_ISL_546901, EPI_ISL_546903, EPI_ISL_546906, EPI_ISL_546910, EPI_ISL_546912, EPI_ISL_546915, EPI_ISL_546916, EPI_ISL_546917, EPI_ISL_546918, EPI_ISL_546919, EPI_ISL_546920, EPI_ISL_546921, EPI_ISL_546922, EPI_ISL_546923, EPI_ISL_546924, EPI_ISL_546925, EPI_ISL_546926, EPI_ISL_546927, EPI_ISL_546928, EPI_ISL_546929, EPI_ISL_546930, EPI_ISL_546931, EPI_ISL_546932, EPI_ISL_546933, EPI_ISL_546934, EPI_ISL_546937, EPI_ISL_546938, EPI_ISL_546939, EPI_ISL_546941, EPI_ISL_546942, EPI_ISL_546943, EPI_ISL_546945, EPI_ISL_546946, EPI_ISL_546947, EPI_ISL_546948, EPI_ISL_546951, EPI_ISL_546953, EPI_ISL_546954, EPI_ISL_546958, EPI_ISL_546959, EPI_ISL_546960, EPI_ISL_546961, EPI_ISL_546962, EPI_ISL_546963, EPI_ISL_546964, EPI_ISL_546965, EPI_ISL_546966, EPI_ISL_546969, EPI_ISL_546970, EPI_ISL_546972, EPI_ISL_546975, EPI_ISL_546976, EPI_ISL_546977, EPI_ISL_546978, EPI_ISL_546981, EPI_ISL_546982, EPI_ISL_546985, EPI_ISL_546986, EPI_ISL_546987, EPI_ISL_546993, EPI_ISL_546994, EPI_ISL_546996, EPI_ISL_546998, EPI_ISL_547000, EPI_ISL_547005, EPI_ISL_547006, EPI_ISL_547012, EPI_ISL_547017, EPI_ISL_547018, EPI_ISL_547019, EPI_ISL_547023, EPI_ISL_547024, EPI_ISL_547025, EPI_ISL_547026, EPI_ISL_547030, EPI_ISL_547031, EPI_ISL_547032, EPI_ISL_547033, EPI_ISL_547035, EPI_ISL_547036, EPI_ISL_547038, EPI_ISL_547039, EPI_ISL_547041, EPI_ISL_547042, EPI_ISL_547043, EPI_ISL_547044, EPI_ISL_547045, EPI_ISL_547047, EPI_ISL_547049, EPI_ISL_547050, EPI_ISL_547051, EPI_ISL_547052, EPI_ISL_547053, EPI_ISL_547054, EPI_ISL_547055, EPI_ISL_547057, EPI_ISL_547058, EPI_ISL_547059, EPI_ISL_547060, EPI_ISL_547061, EPI_ISL_547062, EPI_ISL_547063, EPI_ISL_547064, EPI_ISL_547065, EPI_ISL_547066, EPI_ISL_547067, EPI_ISL_547068, EPI_ISL_547070, EPI_ISL_547072, EPI_ISL_547073, EPI_ISL_547075, EPI_ISL_547076, EPI_ISL_547078, EPI_ISL_547080, EPI_ISL_547081, EPI_ISL_547082, EPI_ISL_547083, EPI_ISL_547084, EPI_ISL_547085, EPI_ISL_547086, EPI_ISL_547088, EPI_ISL_547089, EPI_ISL_547090, EPI_ISL_547091, EPI_ISL_547093, EPI_ISL_547094, EPI_ISL_547096, EPI_ISL_547098, EPI_ISL_547099, EPI_ISL_547101, EPI_ISL_547102, EPI_ISL_547103, EPI_ISL_547105, EPI_ISL_547108, EPI_ISL_547110, EPI_ISL_547112, EPI_ISL_547113, EPI_ISL_547114, EPI_ISL_547115, EPI_ISL_547116, EPI_ISL_547117, EPI_ISL_547120, EPI_ISL_547121, EPI_ISL_547123, EPI_ISL_547124, EPI_ISL_547125, EPI_ISL_547127, EPI_ISL_547128, EPI_ISL_547129, EPI_ISL_547130, EPI_ISL_547131, EPI_ISL_547133, EPI_ISL_547134, EPI_ISL_547135, EPI_ISL_547136, EPI_ISL_547137, EPI_ISL_547138, EPI_ISL_547141, EPI_ISL_547142, EPI_ISL_547143, EPI_ISL_547144, EPI_ISL_547145, EPI_ISL_547146, EPI_ISL_547147, EPI_ISL_547148, EPI_ISL_547149, EPI_ISL_547150, EPI_ISL_547151, EPI_ISL_547152, EPI_ISL_547153, EPI_ISL_547154, EPI_ISL_547155, EPI_ISL_547157, EPI_ISL_547158, EPI_ISL_547161, EPI_ISL_547164, EPI_ISL_547167, EPI_ISL_547171, EPI_ISL_547174, EPI_ISL_547185, EPI_ISL_547186, EPI_ISL_547187, EPI_ISL_547189, EPI_ISL_547190, EPI_ISL_547192, EPI_ISL_547193, EPI_ISL_547194, EPI_ISL_547196, EPI_ISL_547197, EPI_ISL_547198, EPI_ISL_547199, EPI_ISL_547203, EPI_ISL_547206, EPI_ISL_547207, EPI_ISL_547208, EPI_ISL_547210, EPI_ISL_547211, EPI_ISL_547212, EPI_ISL_547214, EPI_ISL_547215, EPI_ISL_547216, EPI_ISL_547218, EPI_ISL_547219, EPI_ISL_547220, EPI_ISL_547221, EPI_ISL_547223, EPI_ISL_547224, EPI_ISL_547225, EPI_ISL_547226, EPI_ISL_547227, EPI_ISL_547229, EPI_ISL_547230, EPI_ISL_547231, EPI_ISL_547232, EPI_ISL_547233, EPI_ISL_547234, EPI_ISL_547235, EPI_ISL_547236, EPI_ISL_547238, EPI_ISL_547239, EPI_ISL_547240, EPI_ISL_547241, EPI_ISL_547242, EPI_ISL_547243, EPI_ISL_547244, EPI_ISL_547246, EPI_ISL_547247, EPI_ISL_547248, EPI_ISL_547249, EPI_ISL_547250, EPI_ISL_547251, EPI_ISL_547252, EPI_ISL_547253, EPI_ISL_547254, EPI_ISL_547256, EPI_ISL_547257, EPI_ISL_547258, EPI_ISL_547259, EPI_ISL_547260, EPI_ISL_547261, EPI_ISL_547262, EPI_ISL_547263, EPI_ISL_547264, EPI_ISL_547265, EPI_ISL_547266, EPI_ISL_547267, EPI_ISL_547268, EPI_ISL_547270, EPI_ISL_547271, EPI_ISL_547272, EPI_ISL_547273, EPI_ISL_547274, EPI_ISL_547275, EPI_ISL_547276, EPI_ISL_547277, EPI_ISL_547278, EPI_ISL_547279, EPI_ISL_547282, EPI_ISL_547284, EPI_ISL_547285, EPI_ISL_547289, EPI_ISL_547290, EPI_ISL_547291, EPI_ISL_547292, EPI_ISL_547293, EPI_ISL_547303, EPI_ISL_547304, EPI_ISL_547305, EPI_ISL_547307, EPI_ISL_547308, EPI_ISL_547309, EPI_ISL_547316, EPI_ISL_547317, EPI_ISL_547319, EPI_ISL_547320, EPI_ISL_547321, EPI_ISL_547322, EPI_ISL_547325, EPI_ISL_547326, EPI_ISL_547327, EPI_ISL_547329, EPI_ISL_547330, EPI_ISL_547331, EPI_ISL_547332, EPI_ISL_547334, EPI_ISL_547335, EPI_ISL_547337, EPI_ISL_547338, EPI_ISL_547339, EPI_ISL_547341, EPI_ISL_547342, EPI_ISL_547343, EPI_ISL_547345, EPI_ISL_547346, EPI_ISL_547349, EPI_ISL_547351, EPI_ISL_547355, EPI_ISL_547356, EPI_ISL_547358, EPI_ISL_547360, EPI_ISL_547361, EPI_ISL_547362, EPI_ISL_547364, EPI_ISL_547365, EPI_ISL_547366, EPI_ISL_547367, EPI_ISL_547368, EPI_ISL_547369, EPI_ISL_547370, EPI_ISL_547371, EPI_ISL_547374, EPI_ISL_547377, EPI_ISL_547379, EPI_ISL_547382, EPI_ISL_547385, EPI_ISL_547386, EPI_ISL_547387, EPI_ISL_547389, EPI_ISL_547391, EPI_ISL_547392, EPI_ISL_547393, EPI_ISL_547396, EPI_ISL_547398, EPI_ISL_547400, EPI_ISL_547401, EPI_ISL_547403, EPI_ISL_547404, EPI_ISL_547406, EPI_ISL_547407, EPI_ISL_547412, EPI_ISL_547414, EPI_ISL_547415, EPI_ISL_547417, EPI_ISL_547418, EPI_ISL_547419, EPI_ISL_547422, EPI_ISL_547423, EPI_ISL_547425, EPI_ISL_547428, EPI_ISL_547429, EPI_ISL_547430 | | | |
| see above | Houston Methodist Hospital | Houston Methodist Hospital | S. Wesley Long, Randall J. Olsen, Paul A. Christensen, David W. Bernard, James J. Davis, Maulik Shukla, Marcus Nguyen, Matthew Ojeda Saavedra, Concepcion C. Cantu, Prasanti Yerramilli, Layne Pruitt, Sishir Subedi, Hung-Che Kuo, Heather Hendrickson, Ghazaleh Eskandari, Hoang A. T. Nguyen, J. Hunter Long, Muthiah Kumaraswami, Jule Goike, Daniel Boutz, Jimmy Gollihar, Jason S. McLellan, Chia-Wei Chou, Kamyab Javanmardi, Ilya J. Finkelstein, and James M. Musser |
| EPI_ISL_549178, EPI_ISL_549179, EPI_ISL_549180, EPI_ISL_549181, EPI_ISL_549182, EPI_ISL_549183, EPI_ISL_549185, EPI_ISL_549186, EPI_ISL_549187, EPI_ISL_549188, EPI_ISL_549189, EPI_ISL_549190, EPI_ISL_549191, EPI_ISL_549196, EPI_ISL_549198, EPI_ISL_549209, EPI_ISL_549211, EPI_ISL_549212, EPI_ISL_549221, EPI_ISL_549228, EPI_ISL_549229, EPI_ISL_549230, EPI_ISL_549232, EPI_ISL_549233, EPI_ISL_549234, EPI_ISL_549235, EPI_ISL_549241, EPI_ISL_549242, EPI_ISL_549243, EPI_ISL_549244, EPI_ISL_549246, EPI_ISL_549252, EPI_ISL_549254, EPI_ISL_549256, EPI_ISL_549257, EPI_ISL_549260, EPI_ISL_549263, EPI_ISL_549265, EPI_ISL_549270 | | | |
| see above | Florida Bureau of Public Health Laboratories | Florida Bureau of Public Health Laboratories | Sarah Schmedes, Jason Blanton |
| EPI_ISL_549354, EPI_ISL_549358 | Liverpool Clinical Laboratories | COVID-19 Genomics UK (COG-UK) Consortium | Sam Haldenby, Anita Lucaci, Steve Paterson, Julian Hiscox, Alistair Darby, M Almsaud, A Alrezaihi, Muhannad Alruwaili, Stuart D Armstrong, Jones Benjamin, Eleanor G Bentley, Anu Chawla, Jordan J Clark, Angela Cowell, Richard Eccles, Isabel García-Dorival, Matthew Gemmell, Alessandro Gerada, PKF Gilmore, Richard Gregory, Ximeng Han, Catherine Hartley, Margaret Hughes, Miren Iturriza-Gomara, James Johnson, L Luu, Jenifer Manson, Charlotte Nelson, Elaine O'Toole, Cassie Olateju, Rebekah Penrice-Randal , Lucille Rainbow, N.P Randle, Trevor Ian Robinson, Parul Sharma, Ghada T Shawli, James P Stewart, Neil Swainston, Ecaterina Vamos, Joanne Watts, Mark Whitehead |
| EPI_ISL_560413, EPI_ISL_560414 | Centre de SSS D'Arthabaska-et-de-l'érable - Hôtel-Dieu | Laboratoire de santé publique du Québec | Sandrine Moreira, Ioannis Ragoussis, Guillaume Bourque, Jesse Shapiro, Mark Lathrop and Michel Roger on behalf of the CoVSeQ research group |
| EPI_ISL_560415 | Centre Hospitalier Régional de Lanaudière | Laboratoire de santé publique du Québec | Sandrine Moreira, Ioannis Ragoussis, Guillaume Bourque, Jesse Shapiro, Mark Lathrop and Michel Roger on behalf of the CoVSeQ research group |
| EPI_ISL_568587, EPI_ISL_568588, EPI_ISL_568589, EPI_ISL_568592, EPI_ISL_568593, EPI_ISL_568594, EPI_ISL_568596, EPI_ISL_568599, EPI_ISL_568600, EPI_ISL_568601, EPI_ISL_568602, EPI_ISL_568603, EPI_ISL_568604, EPI_ISL_568605, EPI_ISL_568606, EPI_ISL_568607, EPI_ISL_568608, EPI_ISL_568609, EPI_ISL_568610, EPI_ISL_568611, EPI_ISL_568612, EPI_ISL_568613, EPI_ISL_568614, EPI_ISL_568615, EPI_ISL_568616, EPI_ISL_568617, EPI_ISL_568618, EPI_ISL_568619, EPI_ISL_568620, EPI_ISL_568621, EPI_ISL_568622, EPI_ISL_568623, EPI_ISL_568624, EPI_ISL_568625, EPI_ISL_568626, EPI_ISL_568627, EPI_ISL_568628, EPI_ISL_568629, EPI_ISL_568630, EPI_ISL_568631, EPI_ISL_568632, EPI_ISL_568633, EPI_ISL_568634, EPI_ISL_568635, EPI_ISL_568636, EPI_ISL_568637, EPI_ISL_568638, EPI_ISL_568639, EPI_ISL_568640, EPI_ISL_568641, EPI_ISL_568642, EPI_ISL_568643, EPI_ISL_568644, EPI_ISL_568645, EPI_ISL_568646, EPI_ISL_568647, EPI_ISL_568648, EPI_ISL_568649, EPI_ISL_568650, EPI_ISL_568651, EPI_ISL_568652, EPI_ISL_568653, EPI_ISL_568654, EPI_ISL_568655, EPI_ISL_568656, EPI_ISL_568657, EPI_ISL_568658, EPI_ISL_568659, EPI_ISL_568660, EPI_ISL_568661, EPI_ISL_568662, EPI_ISL_568663, EPI_ISL_568664, EPI_ISL_568665, EPI_ISL_568666, EPI_ISL_568667, EPI_ISL_568668, EPI_ISL_568669, EPI_ISL_568670, EPI_ISL_568671, EPI_ISL_568672, EPI_ISL_568673, EPI_ISL_568674, EPI_ISL_568675, EPI_ISL_568676, EPI_ISL_568677, EPI_ISL_568678, EPI_ISL_568679, EPI_ISL_568680, EPI_ISL_568681, EPI_ISL_568682, EPI_ISL_568683, EPI_ISL_568684, EPI_ISL_568685 | | | |
| see above | Florida Bureau of Public Health Laboratories | Florida Bureau of Public Health Laboratories | Sarah Schmedes, Jason Blanton |
| EPI_ISL_572402, EPI_ISL_572404, EPI_ISL_572426, EPI_ISL_572429, EPI_ISL_572431, EPI_ISL_572436, EPI_ISL_572443, EPI_ISL_572445, EPI_ISL_572451, EPI_ISL_572466, EPI_ISL_572473, EPI_ISL_572478, EPI_ISL_572479, EPI_ISL_572500, EPI_ISL_572502, EPI_ISL_572506, EPI_ISL_572511, EPI_ISL_572552, EPI_ISL_572554, EPI_ISL_572557, EPI_ISL_572562, EPI_ISL_572578, EPI_ISL_572584, EPI_ISL_572588, EPI_ISL_572601, EPI_ISL_572604, EPI_ISL_572650, EPI_ISL_572669, EPI_ISL_572677, EPI_ISL_572679, EPI_ISL_572682, EPI_ISL_572687, EPI_ISL_572688, EPI_ISL_572691, EPI_ISL_572695, EPI_ISL_572696, EPI_ISL_572710, EPI_ISL_572733, EPI_ISL_572734, EPI_ISL_572736, EPI_ISL_572737, EPI_ISL_572738, EPI_ISL_572739, EPI_ISL_572740, EPI_ISL_572748, EPI_ISL_572750, EPI_ISL_572761, EPI_ISL_572769, EPI_ISL_572771, EPI_ISL_572773, EPI_ISL_572813, EPI_ISL_572815, EPI_ISL_572817, EPI_ISL_572822, EPI_ISL_572824, EPI_ISL_572827, EPI_ISL_572843, EPI_ISL_572904, EPI_ISL_572905, EPI_ISL_572906, EPI_ISL_572907, EPI_ISL_572908, EPI_ISL_573236, EPI_ISL_573237, EPI_ISL_573238, EPI_ISL_573239, EPI_ISL_573240, EPI_ISL_573241, EPI_ISL_573242, EPI_ISL_573243, EPI_ISL_573244, EPI_ISL_573245, EPI_ISL_573246, EPI_ISL_576961, EPI_ISL_576962, EPI_ISL_576963, EPI_ISL_576964, EPI_ISL_576965, EPI_ISL_576966, EPI_ISL_576967, EPI_ISL_576968, EPI_ISL_576969, EPI_ISL_576970, EPI_ISL_576971, EPI_ISL_576972, EPI_ISL_576973, EPI_ISL_576974, EPI_ISL_576975, EPI_ISL_576976, EPI_ISL_576977, EPI_ISL_576978, EPI_ISL_576979, EPI_ISL_576980, EPI_ISL_576981, EPI_ISL_576982, EPI_ISL_576983, EPI_ISL_584591, EPI_ISL_584592, EPI_ISL_584593, EPI_ISL_584594, EPI_ISL_584595, EPI_ISL_584596, EPI_ISL_584597, EPI_ISL_584598, EPI_ISL_584599, EPI_ISL_584600, EPI_ISL_584601, EPI_ISL_584602, EPI_ISL_584603, EPI_ISL_584604, EPI_ISL_584605, EPI_ISL_584606, EPI_ISL_584607, EPI_ISL_584608, EPI_ISL_584609, EPI_ISL_584610, EPI_ISL_584611, EPI_ISL_584612, EPI_ISL_584613, EPI_ISL_584614, EPI_ISL_584615, EPI_ISL_584616, EPI_ISL_584617, EPI_ISL_584618, EPI_ISL_584619, EPI_ISL_584620, EPI_ISL_584621 | | | |
| see above | Liverpool Clinical Laboratories | COVID-19 Genomics UK (COG-UK) Consortium | Sam Haldenby, Anita Lucaci, Steve Paterson, Julian Hiscox, Alistair Darby, M Almsaud, A Alrezaihi, Muhannad Alruwaili, Stuart D Armstrong, Jones Benjamin, Eleanor G Bentley, Anu Chawla, Jordan J Clark, Angela Cowell, Richard Eccles, Isabel García-Dorival, Matthew Gemmell, Alessandro Gerada, PKF Gilmore, Richard Gregory, Ximeng Han, Catherine Hartley, Margaret Hughes, Miren Iturriza-Gomara, James Johnson, L Luu, Jenifer Manson, Charlotte Nelson, Elaine O'Toole, Cassie Olateju, Rebekah Penrice-Randal , Lucille Rainbow, N.P Randle, Trevor Ian Robinson, Parul Sharma, Ghada T Shawli, James P Stewart, Neil Swainston, Ecaterina Vamos, Joanne Watts, Mark Whitehead |
| EPI_ISL_586526 | B.J. Medical College and Civil hospital, Ahmedabad | Gujarat Biotechnology Research Centre | Sanjay Kapadia, Apurvasinh Puvar, Janvi Raval, Zarna Patel, Monika Gandhi, Pinal Trivedi, Maharshi Pandya, Nidhi Patel, Nitin Savaliya, Raghawendra Kumar, Dinesh Kumar, Zuber Saiyed, Komal Patel, Labdhi Pandya, Afzal Ansari, Nikha Trivedi, Pranay Shah, Kamlesh J Upadhyay, R D Dixit, A M Kadri, Harsh Bakshi, Chaitanya Joshi, Madhvi Joshi |
| EPI_ISL_586548 | General Hospital,Godhra | Gujarat Biotechnology Research Centre | Sangeeta Kumar, Apurvasinh Puvar, Janvi Raval, Zarna Patel, Monika Gandhi, Pinal Trivedi, Maharshi Pandya, Nidhi Patel, Nitin Savaliya, Raghawendra Kumar, Dinesh Kumar, Zuber Saiyed, Komal Patel, Labdhi Pandya, Afzal Ansari, Nikha Trivedi, Neela Katara, Pinal Gandhi, R D Dixit, A M Kadri, Harsh Bakshi, Chaitanya Joshi, Madhvi Joshi |
| EPI_ISL_594277, EPI_ISL_594278, EPI_ISL_594279, EPI_ISL_594280, EPI_ISL_594281, EPI_ISL_594282, EPI_ISL_594283, EPI_ISL_594284, EPI_ISL_594285, EPI_ISL_594286, EPI_ISL_594287, EPI_ISL_594288, EPI_ISL_594289, EPI_ISL_594290, EPI_ISL_594291, EPI_ISL_594292, EPI_ISL_594293, EPI_ISL_594294, EPI_ISL_594295, EPI_ISL_594296, EPI_ISL_594297, EPI_ISL_594298, EPI_ISL_594299, EPI_ISL_594300, EPI_ISL_594301, EPI_ISL_594302, EPI_ISL_594303, EPI_ISL_594304, EPI_ISL_594305, EPI_ISL_594306, EPI_ISL_594307, EPI_ISL_594308, EPI_ISL_594309, EPI_ISL_594310, EPI_ISL_594311, EPI_ISL_594312, EPI_ISL_594313, EPI_ISL_594314, EPI_ISL_594315, EPI_ISL_594316, EPI_ISL_594317, EPI_ISL_594318, EPI_ISL_594319, EPI_ISL_594320, EPI_ISL_594321, EPI_ISL_594322, EPI_ISL_594323, EPI_ISL_594324, EPI_ISL_594325, EPI_ISL_594326, EPI_ISL_594327, EPI_ISL_594328, EPI_ISL_594329, EPI_ISL_594330, EPI_ISL_594331, EPI_ISL_594332, EPI_ISL_594333, EPI_ISL_594334, EPI_ISL_594335, EPI_ISL_594336, EPI_ISL_594337, EPI_ISL_594338, EPI_ISL_594339, EPI_ISL_594340, EPI_ISL_594341, EPI_ISL_594342, EPI_ISL_594343, EPI_ISL_594344, EPI_ISL_594345, EPI_ISL_594346, EPI_ISL_594347, EPI_ISL_594348, | | | |

| | | | |
|---|---|---|---|
| EPI_ISL_594349, EPI_ISL_594350, EPI_ISL_594351, EPI_ISL_594352, EPI_ISL_594353, EPI_ISL_594354, EPI_ISL_594355, EPI_ISL_594356, EPI_ISL_594357, EPI_ISL_594358, EPI_ISL_594359, EPI_ISL_594360, EPI_ISL_594361, EPI_ISL_594362, EPI_ISL_594363, EPI_ISL_594364, EPI_ISL_594365, EPI_ISL_594366, EPI_ISL_594367, EPI_ISL_594368, EPI_ISL_594369, EPI_ISL_594370, EPI_ISL_594371, EPI_ISL_594372, EPI_ISL_594373, EPI_ISL_594374, EPI_ISL_594375, EPI_ISL_594376, EPI_ISL_594377, EPI_ISL_594378, EPI_ISL_594379, EPI_ISL_594380, EPI_ISL_594381, EPI_ISL_594382, EPI_ISL_594383, EPI_ISL_594384, EPI_ISL_594385, EPI_ISL_594386, EPI_ISL_594387, EPI_ISL_594389, EPI_ISL_594390, EPI_ISL_594391, EPI_ISL_594392, EPI_ISL_594393, EPI_ISL_594466 | | | |
| see above | Florida Bureau of Public Health Laboratories | Florida Bureau of Public Health Laboratories | Sarah Schmedes, Jason Blanton |
| EPI_ISL_594849, EPI_ISL_594850, EPI_ISL_594851, EPI_ISL_594852, EPI_ISL_594853, EPI_ISL_594854, EPI_ISL_594855, EPI_ISL_594856, EPI_ISL_594857, EPI_ISL_594858, EPI_ISL_594859, EPI_ISL_594860, EPI_ISL_595799, EPI_ISL_595800, EPI_ISL_595801, EPI_ISL_595822 | | | |
| see above | Liverpool Clinical Laboratories | COVID-19 Genomics UK (COG-UK) Consortium | Sam Haldenby, Anita Lucaci, Steve Paterson, Julian Hiscox, Alistair Darby, M Almsaud, A Alrezaihi, Muhannad Alruwaili, Stuart D Armstrong, Jones Benjamin, Eleanor G Bentley, Anu Chawla, Jordan J Clark, Angela Cowell, Richard Eccles, Isabel García-Dorival, Matthew Gemmell, Alessandro Gerada, PKF Gilmore, Richard Gregory, Ximeng Han, Catherine Hartley, Margaret Hughes, Miren Iturriza-Gomara, James Johnson, L Luu, Jenifer Manson, Charlotte Nelson, Elaine O'Toole, Cassie Olateju, Rebekah Penrice-Randal , Lucille Rainbow, N.P Randle, Trevor Ian Robinson, Parul Sharma, Ghada T Shawli, James P Stewart, Neil Swainston, Ecaterina Vamos, Joanne Watts, Mark Whitehead |
| EPI_ISL_605058, EPI_ISL_605059, EPI_ISL_605060, EPI_ISL_605061, EPI_ISL_605062, EPI_ISL_605063, EPI_ISL_605064, EPI_ISL_605065, EPI_ISL_605066, EPI_ISL_605067, EPI_ISL_605068, EPI_ISL_605069, EPI_ISL_605070, EPI_ISL_605071, EPI_ISL_605072, EPI_ISL_605073, EPI_ISL_605074, EPI_ISL_605075, EPI_ISL_605076, EPI_ISL_605077, EPI_ISL_605078, EPI_ISL_605079, EPI_ISL_605080 | | | |
| see above | National Virus Reference Laboratory | Irish Coronavirus Sequencing Consortium - Helixworks | Sachin Chalapati, Conor Crosbie, Nimesh Pinnamaneni |
| EPI_ISL_610227, EPI_ISL_610228, EPI_ISL_610229, EPI_ISL_610230, EPI_ISL_610231, EPI_ISL_610232, EPI_ISL_610233, EPI_ISL_610234, EPI_ISL_610235, EPI_ISL_610236 | Molecular diagnostic laboratory of Federal Budget Institution of Science "Central Research Institute of Epidemiology" of The Federal Service on Customers' Rights Protection and Human Well-being Surveillance | Group of Genomics and Postgenomic Technologies of Central Research Institute of Epidemilology | Samoilov AE, Kaptelova VV, Valdokhina AV, Bulanenko VP, Speranskaya AS, Tivanova EV, Shipulina OY, Akimkin VG |
| EPI_ISL_610237, EPI_ISL_610238 | Molecular diagnostic laboratory of Federal Budget Institution of Science "Central Research Institute of Epidemiology" of The Federal Service on Customers' Rights Protection and Human Well-being Surveillance | Group of Genomics and Postgenomic Technologies of Central Research Institute of Epidemilology | Samoilov AE, Kaptelova VV, Bukharina AY, Speranskaya AS, Tivanova EV, Shipulina OY, Akimkin VG |
| EPI_ISL_610239, EPI_ISL_610240, EPI_ISL_610241, EPI_ISL_610242, EPI_ISL_610243, EPI_ISL_610244 | Molecular diagnostic laboratory of Federal Budget Institution of Science "Central Research Institute of Epidemiology" of The Federal Service on Customers' Rights Protection and Human Well-being Surveillance | Group of Genomics and Postgenomic Technologies of Central Research Institute of Epidemilology | Samoilov AE, Kaptelova VV, Valdokhina AV, Bulanenko VP, Speranskaya AS, Tivanova EV, Shipulina OY, Akimkin VG |
| EPI_ISL_611514, EPI_ISL_611515, EPI_ISL_611516, EPI_ISL_611517, EPI_ISL_611521, EPI_ISL_611522, EPI_ISL_611542, EPI_ISL_611543, EPI_ISL_611565, EPI_ISL_611567, EPI_ISL_611568, EPI_ISL_611569, EPI_ISL_611570, EPI_ISL_611571, EPI_ISL_611575, EPI_ISL_611579, EPI_ISL_611580, EPI_ISL_611600, EPI_ISL_611603, EPI_ISL_611613, EPI_ISL_611621, EPI_ISL_611627, EPI_ISL_611629, EPI_ISL_611630, EPI_ISL_611632, EPI_ISL_611634, EPI_ISL_611635, EPI_ISL_611642, EPI_ISL_611653, EPI_ISL_611654, EPI_ISL_611658, EPI_ISL_611664, EPI_ISL_611667, EPI_ISL_611672, EPI_ISL_611675, EPI_ISL_611694, EPI_ISL_611695, EPI_ISL_611721, EPI_ISL_611723, EPI_ISL_611724, EPI_ISL_611727, EPI_ISL_611728, EPI_ISL_611729, EPI_ISL_611730, EPI_ISL_611731, EPI_ISL_611734, EPI_ISL_611736, EPI_ISL_611737, EPI_ISL_611739, EPI_ISL_611741, EPI_ISL_611743, EPI_ISL_611750, EPI_ISL_611764, EPI_ISL_611765, EPI_ISL_611784, EPI_ISL_611785, EPI_ISL_611786, EPI_ISL_611790, EPI_ISL_611796, EPI_ISL_611811, EPI_ISL_611815, EPI_ISL_611821, EPI_ISL_611827, EPI_ISL_611845, EPI_ISL_611851, EPI_ISL_611853, EPI_ISL_611857, EPI_ISL_611885, EPI_ISL_611889, EPI_ISL_611891, EPI_ISL_611892, EPI_ISL_611893, EPI_ISL_611895, EPI_ISL_611897, EPI_ISL_611898, EPI_ISL_611899, EPI_ISL_611908, EPI_ISL_611910, EPI_ISL_611911, EPI_ISL_611914, EPI_ISL_611915, EPI_ISL_611918, EPI_ISL_611921, EPI_ISL_611929, EPI_ISL_611974, EPI_ISL_611977, EPI_ISL_611978, EPI_ISL_611981, EPI_ISL_611984, EPI_ISL_611985, EPI_ISL_611991, EPI_ISL_611995, EPI_ISL_611999, EPI_ISL_612001, EPI_ISL_612003, EPI_ISL_612009, EPI_ISL_612010, EPI_ISL_612016, EPI_ISL_612017, EPI_ISL_612019, EPI_ISL_612024, EPI_ISL_612035, EPI_ISL_612036, EPI_ISL_612039, EPI_ISL_612041, EPI_ISL_612042, EPI_ISL_612043, EPI_ISL_612044, EPI_ISL_612046, EPI_ISL_612051, EPI_ISL_612059, EPI_ISL_612060, EPI_ISL_612061, EPI_ISL_612064, EPI_ISL_612065, EPI_ISL_612066, EPI_ISL_612070, EPI_ISL_612071, EPI_ISL_612072, EPI_ISL_612075, EPI_ISL_612080, EPI_ISL_612088, EPI_ISL_612101, EPI_ISL_612108, EPI_ISL_612111, EPI_ISL_612112, EPI_ISL_612114, EPI_ISL_612115, EPI_ISL_612190, EPI_ISL_612191, EPI_ISL_612401, EPI_ISL_612402, EPI_ISL_612403, EPI_ISL_612404, EPI_ISL_612405, EPI_ISL_612406, EPI_ISL_612407, EPI_ISL_612408, EPI_ISL_612409, EPI_ISL_612410, EPI_ISL_612411, EPI_ISL_612412, EPI_ISL_612413, EPI_ISL_612414, EPI_ISL_612415, EPI_ISL_612416, EPI_ISL_612417, EPI_ISL_612418, EPI_ISL_612419, EPI_ISL_612420, EPI_ISL_612421, EPI_ISL_612422, EPI_ISL_612423, EPI_ISL_612424, EPI_ISL_612425, EPI_ISL_612426, EPI_ISL_612427, EPI_ISL_612428, EPI_ISL_612429, EPI_ISL_612430, EPI_ISL_612431, EPI_ISL_612432, EPI_ISL_612433, EPI_ISL_612434, EPI_ISL_612435, EPI_ISL_612436, EPI_ISL_612437, EPI_ISL_612438, EPI_ISL_612439, EPI_ISL_612440, EPI_ISL_612441, EPI_ISL_612442, EPI_ISL_612443, EPI_ISL_612444, EPI_ISL_612445, EPI_ISL_612446, EPI_ISL_612447, EPI_ISL_612448, EPI_ISL_612449 | | | |
| see above | Liverpool Clinical Laboratories | COVID-19 Genomics UK (COG-UK) Consortium | Sam Haldenby, Anita Lucaci, Steve Paterson, Julian Hiscox, Alistair Darby, M Almsaud, A Alrezaihi, Muhannad Alruwaili, Stuart D Armstrong, Jones Benjamin, Eleanor G Bentley, Anu Chawla, Jordan J Clark, Angela Cowell, Richard Eccles, Isabel García-Dorival, Matthew Gemmell, Alessandro Gerada, PKF Gilmore, Richard Gregory, Ximeng Han, Catherine Hartley, Margaret Hughes, Miren Iturriza-Gomara, James Johnson, L Luu, Jenifer Manson, Charlotte Nelson, Elaine O'Toole, Cassie Olateju, Rebekah Penrice-Randal , Lucille Rainbow, N.P Randle, Trevor Ian Robinson, Parul Sharma, Ghada T Shawli, James P Stewart, Neil Swainston, Ecaterina Vamos, Joanne Watts, Mark Whitehead |
| EPI_ISL_613712, EPI_ISL_613713, EPI_ISL_613714, EPI_ISL_613715, EPI_ISL_613716, EPI_ISL_613717, EPI_ISL_613718, EPI_ISL_613719, EPI_ISL_613720, EPI_ISL_613721, EPI_ISL_613722, EPI_ISL_613723, EPI_ISL_613725, EPI_ISL_613726, EPI_ISL_613727, EPI_ISL_613728, EPI_ISL_613729, EPI_ISL_613730, EPI_ISL_613731, EPI_ISL_613732, EPI_ISL_613733, EPI_ISL_613734, EPI_ISL_613735, EPI_ISL_613736, EPI_ISL_613737, EPI_ISL_613738, EPI_ISL_613740, EPI_ISL_613741, EPI_ISL_613742, EPI_ISL_613743, EPI_ISL_613744, EPI_ISL_613745, EPI_ISL_613746, EPI_ISL_613748, EPI_ISL_613749, EPI_ISL_613750, EPI_ISL_613752, EPI_ISL_613753, EPI_ISL_613754, EPI_ISL_613755, EPI_ISL_613757, EPI_ISL_613759, EPI_ISL_613761, EPI_ISL_613762, EPI_ISL_613763, EPI_ISL_613764, EPI_ISL_613765, EPI_ISL_613767, EPI_ISL_613768, EPI_ISL_613769, EPI_ISL_613773, EPI_ISL_613774, EPI_ISL_613775, EPI_ISL_613776, EPI_ISL_613777, EPI_ISL_613778, EPI_ISL_613779, EPI_ISL_613780, EPI_ISL_613781, EPI_ISL_613782, EPI_ISL_613783, EPI_ISL_613784, EPI_ISL_613785, EPI_ISL_613786, EPI_ISL_613787, EPI_ISL_613788, EPI_ISL_613789, EPI_ISL_613790, EPI_ISL_613791, EPI_ISL_613792, EPI_ISL_613793, EPI_ISL_613794, EPI_ISL_613795, EPI_ISL_613796, EPI_ISL_613797, EPI_ISL_613798, EPI_ISL_613799, EPI_ISL_613801, EPI_ISL_613802, EPI_ISL_613803, EPI_ISL_613804, EPI_ISL_613805, EPI_ISL_613806, EPI_ISL_613807, EPI_ISL_613808, EPI_ISL_613809, EPI_ISL_613810, EPI_ISL_613811, EPI_ISL_613812, EPI_ISL_613813, EPI_ISL_613814, EPI_ISL_613815, EPI_ISL_613816, EPI_ISL_613817, EPI_ISL_613818, EPI_ISL_613819, EPI_ISL_613820, EPI_ISL_613821, EPI_ISL_613822, EPI_ISL_613823, EPI_ISL_613824, EPI_ISL_613825, EPI_ISL_613826, EPI_ISL_613827, EPI_ISL_613828, EPI_ISL_613832, EPI_ISL_613833, EPI_ISL_613835, EPI_ISL_613836, EPI_ISL_613839, EPI_ISL_613844, EPI_ISL_613845, EPI_ISL_613846, EPI_ISL_613848, EPI_ISL_613855, EPI_ISL_613860, EPI_ISL_613869, EPI_ISL_613870, EPI_ISL_613872, EPI_ISL_613873, EPI_ISL_613879, EPI_ISL_613891, EPI_ISL_613892, EPI_ISL_613899, EPI_ISL_613911, EPI_ISL_613926, EPI_ISL_613941 | | | |
| see above | Florida Bureau of Public Health Laboratories | Florida Bureau of Public Health Laboratories | Sarah Schmedes, Jason Blanton |
| EPI_ISL_626661, EPI_ISL_626665, EPI_ISL_626694, EPI_ISL_626697, EPI_ISL_626709, EPI_ISL_626712, EPI_ISL_626714, EPI_ISL_626736, EPI_ISL_626742, EPI_ISL_626760, EPI_ISL_626766, EPI_ISL_626789, EPI_ISL_626807, EPI_ISL_626815, EPI_ISL_626832, EPI_ISL_626866, EPI_ISL_626905, EPI_ISL_626994, EPI_ISL_627004, EPI_ISL_627017, EPI_ISL_627106, EPI_ISL_627107, EPI_ISL_627109, EPI_ISL_627123, EPI_ISL_627125, EPI_ISL_627127, EPI_ISL_627131, EPI_ISL_627132, EPI_ISL_627133, EPI_ISL_627135, EPI_ISL_627136, EPI_ISL_627137, EPI_ISL_627138, EPI_ISL_627139, EPI_ISL_627141, EPI_ISL_627152, EPI_ISL_627153, EPI_ISL_627156, EPI_ISL_627162, EPI_ISL_627171, EPI_ISL_627174, EPI_ISL_627176, EPI_ISL_627177, EPI_ISL_627178, EPI_ISL_627179, EPI_ISL_627181, EPI_ISL_627183, EPI_ISL_627185, EPI_ISL_627188, EPI_ISL_627445, EPI_ISL_627446, EPI_ISL_627447, EPI_ISL_627448, EPI_ISL_627449 | | | |
| see above | Liverpool Clinical Laboratories | COVID-19 Genomics UK (COG-UK) Consortium | Sam Haldenby, Anita Lucaci, Steve Paterson, Julian Hiscox, Alistair Darby, M Almsaud, A Alrezaihi, Muhannad Alruwaili, Stuart D Armstrong, Jones Benjamin, Eleanor G Bentley, Anu Chawla, Jordan J Clark, Angela Cowell, Richard Eccles, Isabel García-Dorival, Matthew Gemmell, Alessandro Gerada, PKF Gilmore, Richard Gregory, Ximeng Han, Catherine Hartley, Margaret Hughes, Miren Iturriza-Gomara, James Johnson, L Luu, Jenifer Manson, Charlotte Nelson, Elaine O'Toole, Cassie Olateju, Rebekah Penrice-Randal , Lucille Rainbow, N.P Randle, Trevor Ian Robinson, Parul Sharma, Ghada T Shawli, James P Stewart, Neil Swainston, Ecaterina Vamos, Joanne Watts, Mark Whitehead |

We gratefully acknowledge the following Authors from the Originating laboratories responsible for obtaining the specimens, as well as the Submitting laboratories where the genome data were generated and shared via GISAID, on which this research is based.

All Submitters of data may be contacted directly via www.gisaid.org

Authors are sorted alphabetically.

| Accession ID | Originating Laboratory | Submitting Laboratory | Authors |
|---|---|---|---|
| EPI_ISL_419714, EPI_ISL_419715, EPI_ISL_419716, EPI_ISL_419717, EPI_ISL_419718, EPI_ISL_419719, EPI_ISL_419720, EPI_ISL_419721, EPI_ISL_419722, EPI_ISL_419723, EPI_ISL_419724, EPI_ISL_419725, EPI_ISL_419726, EPI_ISL_419727, EPI_ISL_419728, EPI_ISL_419729, EPI_ISL_419730, EPI_ISL_419731, EPI_ISL_419732, EPI_ISL_419823, EPI_ISL_419824, EPI_ISL_419825, EPI_ISL_419827, EPI_ISL_419828, EPI_ISL_419829, EPI_ISL_419830, EPI_ISL_419980, EPI_ISL_419981, EPI_ISL_419999, EPI_ISL_420000, EPI_ISL_420001, EPI_ISL_420002, EPI_ISL_420003, EPI_ISL_420006, EPI_ISL_420007, EPI_ISL_420008, EPI_ISL_420009, EPI_ISL_420010, EPI_ISL_420011, EPI_ISL_420012, EPI_ISL_420013, EPI_ISL_420014, EPI_ISL_420015, EPI_ISL_420016, EPI_ISL_420017, EPI_ISL_426905, EPI_ISL_426906, EPI_ISL_426907, EPI_ISL_426908, EPI_ISL_426909, EPI_ISL_426910, EPI_ISL_426911, EPI_ISL_426912, EPI_ISL_426913, EPI_ISL_426914, EPI_ISL_426915, EPI_ISL_426916, EPI_ISL_426917, EPI_ISL_426918, EPI_ISL_426919, EPI_ISL_426920, EPI_ISL_426921, EPI_ISL_426922, EPI_ISL_427054, EPI_ISL_427055, EPI_ISL_427056, EPI_ISL_427057, EPI_ISL_427058, EPI_ISL_427059, EPI_ISL_427060, EPI_ISL_427061, EPI_ISL_427062, EPI_ISL_427063, EPI_ISL_427064, EPI_ISL_427065, EPI_ISL_427066, EPI_ISL_427067, EPI_ISL_427068, EPI_ISL_427069, EPI_ISL_427070, EPI_ISL_427071, EPI_ISL_427072, EPI_ISL_427073, EPI_ISL_427074, EPI_ISL_427075, EPI_ISL_427076, EPI_ISL_427077, EPI_ISL_427078, EPI_ISL_427133, EPI_ISL_427148, EPI_ISL_427149 | | | |
| see above | Microbiological Diagnostic Unit Public Health Laboratory | Microbiological Diagnostic Unit Public Health Laboratory | Seemann T., Schultz M., Sait, M., Sherry, N. |
| EPI_ISL_428855 | MRCG at LSHTM Geomics lab | MRCG at LSHTM Genomics lab | Sesay et al |
| EPI_ISL_428856 | MRCG at LSHTM Genomics Lab | MRCG at LSHTM Genomics lab | Sesay et al |
| EPI_ISL_428857 | MRCG at LSHTM Genomics lab | MRCG at LSHTM Genomics lab | Sesay et al |
| EPI_ISL_428872, EPI_ISL_428873, EPI_ISL_428874, EPI_ISL_428875 | State Research Center of Virology and Biotechnology VECTOR, Department of Collection of Microorganisms | State Research Center of Virology and Biotechnology VECTOR, Department of Collection of Microorganisms | Sergey A. Bodnev, Oleg V. Pyankov, Tatyana V. Tregubchak, Alexander N. Shvalov, Elena V. Gavrilova, Rinat A. Maksyutov |
| EPI_ISL_428876, EPI_ISL_428877 | State Research Center of Virology and Biotechnology VECTOR, Department of Collection of Microorganisms | State Research Center of Virology and Biotechnology VECTOR, Department of Collection of Microorganisms | Sergey A. Bodnev, Oleg V. Pyankov, Anastasiya M. Smirnova, Anastasiya A. Nazarenko, Tatyana V. Tregubchak, Alexander N. Shvalov, Elena V. Gavrilova, Rinat A. Maksyutov |
| EPI_ISL_428878, EPI_ISL_428879, EPI_ISL_428880, EPI_ISL_428881, EPI_ISL_428895, EPI_ISL_428896, EPI_ISL_428897, EPI_ISL_428898, EPI_ISL_428899, EPI_ISL_428900, EPI_ISL_428901, EPI_ISL_428902, EPI_ISL_428903, EPI_ISL_428904, EPI_ISL_428905, EPI_ISL_428917, EPI_ISL_428918, EPI_ISL_428919, EPI_ISL_428920, EPI_ISL_428921, EPI_ISL_428922, EPI_ISL_428923, EPI_ISL_428924 | | | |
| see above | State Research Center of Virology and Biotechnology VECTOR, Department of Collection of Microorganisms | State Research Center of Virology and Biotechnology VECTOR, Department of Collection of Microorganisms | Sergey A. Bodnev, Oleg V. Pyankov, Tatyana V. Tregubchak, Alexander N. Shvalov, Elena V. Gavrilova, Rinat A. Maksyutov |
| EPI_ISL_430470, EPI_ISL_430471, EPI_ISL_430472, EPI_ISL_430639, EPI_ISL_430640, EPI_ISL_430641, EPI_ISL_430642, EPI_ISL_430643, EPI_ISL_430644, EPI_ISL_430645, EPI_ISL_430646, EPI_ISL_430647, EPI_ISL_430648, EPI_ISL_430649, EPI_ISL_430650, EPI_ISL_430651, EPI_ISL_430652, EPI_ISL_430653, EPI_ISL_430654, EPI_ISL_430655, EPI_ISL_430656, EPI_ISL_430657, EPI_ISL_430658, EPI_ISL_430659, EPI_ISL_430660, EPI_ISL_430661, EPI_ISL_430662, EPI_ISL_430663, EPI_ISL_430664, EPI_ISL_430665, EPI_ISL_430666, EPI_ISL_430667, EPI_ISL_430668, EPI_ISL_430669, EPI_ISL_430670, EPI_ISL_430671, EPI_ISL_430672, EPI_ISL_430673, EPI_ISL_430674, EPI_ISL_430675, EPI_ISL_430676, EPI_ISL_430677, EPI_ISL_430678, EPI_ISL_430679, EPI_ISL_430680, EPI_ISL_430681, EPI_ISL_430682, EPI_ISL_430683, EPI_ISL_430684, EPI_ISL_430685, EPI_ISL_430686, EPI_ISL_430687, EPI_ISL_430715, EPI_ISL_430716, EPI_ISL_430717 | | | |
| see above | Microbiological Diagnostic Unit Public Health Laboratory | Microbiological Diagnostic Unit Public Health Laboratory | Seemann T., Schultz M., Sait, M., Sherry, N. |
| EPI_ISL_436505, EPI_ISL_436506, EPI_ISL_436507, EPI_ISL_436512, EPI_ISL_436514, EPI_ISL_436515, EPI_ISL_436516, EPI_ISL_436517, EPI_ISL_436518, EPI_ISL_436519, EPI_ISL_436520, EPI_ISL_436521, EPI_ISL_436522, EPI_ISL_436523, EPI_ISL_436524, EPI_ISL_436525, EPI_ISL_436526, EPI_ISL_436527, EPI_ISL_436528, EPI_ISL_436529, EPI_ISL_436530, EPI_ISL_436531, EPI_ISL_436532, EPI_ISL_436533, EPI_ISL_436534, EPI_ISL_436535, EPI_ISL_436536, EPI_ISL_436537, EPI_ISL_436538, EPI_ISL_436539, EPI_ISL_436540, EPI_ISL_436556, EPI_ISL_436563 | | | |
| see above | Florida Bureau of Public Health Laboratories | Florida Bureau of Public Health Laboratories | Sarah Schmedes, Jason Blanton |
| EPI_ISL_437912 | Child Health Research Foundation | Child Health Research Lab | Senjuti Saha, Roly Malaker, Md Saiful Islam Sajib, Md Hasanuzzaman, Md Hafizur Rahman, Md Shahidul Islam, Zabed B Ahmed, Maksuda Islam, Samir K Saha |
| EPI_ISL_440951, EPI_ISL_440952, EPI_ISL_440953, EPI_ISL_440954, EPI_ISL_440955, EPI_ISL_440956, EPI_ISL_440957, EPI_ISL_440958, EPI_ISL_440959, EPI_ISL_440960, EPI_ISL_440961, EPI_ISL_440962, EPI_ISL_440963, EPI_ISL_440964, EPI_ISL_440965, EPI_ISL_440966, EPI_ISL_440967, EPI_ISL_440968, EPI_ISL_440969, EPI_ISL_440970, EPI_ISL_440971, EPI_ISL_440972, EPI_ISL_440973, EPI_ISL_440974, EPI_ISL_440975, EPI_ISL_440976, EPI_ISL_440977, EPI_ISL_440978, EPI_ISL_440979, EPI_ISL_440980, EPI_ISL_440981, EPI_ISL_440982, EPI_ISL_440983, EPI_ISL_440984, EPI_ISL_440985, EPI_ISL_440986, EPI_ISL_440987, EPI_ISL_440988, EPI_ISL_440989, EPI_ISL_440990, EPI_ISL_440991, EPI_ISL_440992, EPI_ISL_440993, EPI_ISL_440994, EPI_ISL_440995, EPI_ISL_440996, EPI_ISL_440997, EPI_ISL_440998, EPI_ISL_440999, EPI_ISL_441000, EPI_ISL_441001, EPI_ISL_441002, EPI_ISL_441003, EPI_ISL_441004, EPI_ISL_441005, EPI_ISL_441006, EPI_ISL_441007, EPI_ISL_441008, EPI_ISL_441009, EPI_ISL_441010, EPI_ISL_441011, EPI_ISL_441012, EPI_ISL_441013, EPI_ISL_441014, EPI_ISL_441015, EPI_ISL_441016, EPI_ISL_441017, EPI_ISL_441018, EPI_ISL_441019, EPI_ISL_441020, EPI_ISL_441021, EPI_ISL_441022, EPI_ISL_441023, EPI_ISL_441024, EPI_ISL_441025, EPI_ISL_441026, EPI_ISL_441027, EPI_ISL_441028, EPI_ISL_441029, EPI_ISL_441030, EPI_ISL_441031, EPI_ISL_441032, EPI_ISL_441033, EPI_ISL_441034, EPI_ISL_441035, EPI_ISL_441036, EPI_ISL_441037, EPI_ISL_441038, EPI_ISL_441039, EPI_ISL_441040, EPI_ISL_441041, EPI_ISL_441042, EPI_ISL_441043, EPI_ISL_441044, EPI_ISL_441045, EPI_ISL_441046, EPI_ISL_441047, EPI_ISL_441048, EPI_ISL_441049, EPI_ISL_441050, EPI_ISL_441051, EPI_ISL_441350, EPI_ISL_441351, EPI_ISL_441352, EPI_ISL_441353, EPI_ISL_441354, EPI_ISL_444079, EPI_ISL_444080, EPI_ISL_444081, EPI_ISL_444082, EPI_ISL_444083, EPI_ISL_444084, EPI_ISL_444085, EPI_ISL_444086, EPI_ISL_444087, EPI_ISL_444088, EPI_ISL_444089, EPI_ISL_444090, EPI_ISL_444091, EPI_ISL_444092, EPI_ISL_444093, EPI_ISL_444094, EPI_ISL_444095, EPI_ISL_444096, EPI_ISL_444097, EPI_ISL_444098, EPI_ISL_444099, EPI_ISL_444100, EPI_ISL_444101, EPI_ISL_444102, EPI_ISL_444103, EPI_ISL_444104, EPI_ISL_444105, EPI_ISL_444106, EPI_ISL_444107, EPI_ISL_444108, EPI_ISL_444109, EPI_ISL_444110, EPI_ISL_444111, EPI_ISL_444112, EPI_ISL_444113, EPI_ISL_444114, EPI_ISL_444115, EPI_ISL_444116, EPI_ISL_444117, EPI_ISL_444118, EPI_ISL_444119, EPI_ISL_444120, EPI_ISL_444121, EPI_ISL_444122, EPI_ISL_444123, EPI_ISL_444124, EPI_ISL_444125, EPI_ISL_444126, EPI_ISL_444127, EPI_ISL_444128, EPI_ISL_444129, EPI_ISL_444130, EPI_ISL_444131, EPI_ISL_444132, EPI_ISL_444133, EPI_ISL_444134, EPI_ISL_444136, EPI_ISL_444137, EPI_ISL_444138, EPI_ISL_444139, EPI_ISL_444140, EPI_ISL_444141, EPI_ISL_444142, EPI_ISL_444143, EPI_ISL_444144, EPI_ISL_444145, EPI_ISL_444146, EPI_ISL_444147, EPI_ISL_444148, EPI_ISL_444149, EPI_ISL_444150, EPI_ISL_444151, EPI_ISL_444152, EPI_ISL_444153, EPI_ISL_444154, EPI_ISL_444155, EPI_ISL_444156, EPI_ISL_444157, EPI_ISL_444158, EPI_ISL_444159, EPI_ISL_444160, EPI_ISL_444161, EPI_ISL_444162, EPI_ISL_444163, EPI_ISL_444164, EPI_ISL_444165, EPI_ISL_444166, EPI_ISL_444167, EPI_ISL_444168, EPI_ISL_444169, EPI_ISL_444170, EPI_ISL_444171, EPI_ISL_444172, EPI_ISL_444173, EPI_ISL_444174, EPI_ISL_444175, EPI_ISL_444176, EPI_ISL_444177, EPI_ISL_444178, EPI_ISL_444179, EPI_ISL_444180, EPI_ISL_444181, EPI_ISL_444182, EPI_ISL_444183, EPI_ISL_444184, EPI_ISL_444185, EPI_ISL_444186, EPI_ISL_444187, EPI_ISL_444188, EPI_ISL_444189, EPI_ISL_444190, EPI_ISL_444191, EPI_ISL_444192, EPI_ISL_444193, EPI_ISL_444194, EPI_ISL_444195, EPI_ISL_444196, EPI_ISL_444197, EPI_ISL_444198, EPI_ISL_444199, EPI_ISL_444200, EPI_ISL_444201, EPI_ISL_444202, EPI_ISL_444203, EPI_ISL_444204, EPI_ISL_444205, EPI_ISL_444206, EPI_ISL_444207, EPI_ISL_444208, EPI_ISL_444209, EPI_ISL_444210, EPI_ISL_444211, EPI_ISL_444212, EPI_ISL_444213, EPI_ISL_444214, EPI_ISL_444215, EPI_ISL_444216, EPI_ISL_444217, EPI_ISL_444218, EPI_ISL_444219, EPI_ISL_444220, EPI_ISL_444221, EPI_ISL_444222, EPI_ISL_444223, EPI_ISL_444224, EPI_ISL_444225, EPI_ISL_444226, EPI_ISL_444227, EPI_ISL_444228, EPI_ISL_444229, EPI_ISL_444230, EPI_ISL_444231, EPI_ISL_444232, EPI_ISL_444233, EPI_ISL_444234, EPI_ISL_444235, EPI_ISL_444236, EPI_ISL_444237, EPI_ISL_444238, EPI_ISL_444239, EPI_ISL_444240, EPI_ISL_444241, EPI_ISL_444242, EPI_ISL_444243, EPI_ISL_444244, EPI_ISL_444245, EPI_ISL_444246, EPI_ISL_444247, EPI_ISL_444248, EPI_ISL_444249, EPI_ISL_444250, EPI_ISL_444251, EPI_ISL_444252, EPI_ISL_444253, EPI_ISL_444254, EPI_ISL_444255, EPI_ISL_444256, EPI_ISL_444257, EPI_ISL_444258, EPI_ISL_444259, EPI_ISL_444260, EPI_ISL_444261, EPI_ISL_444262, EPI_ISL_444263, EPI_ISL_444264, EPI_ISL_444265, EPI_ISL_444266, EPI_ISL_444267, EPI_ISL_444268, EPI_ISL_444269, EPI_ISL_444270, EPI_ISL_444271, EPI_ISL_444272 | | | |
| see above | University College London, Great Ormond Street Hospital for Children NHS Foundation Trust, Imperial College Healthcare NHS Trust | COVID-19 Genomics UK (COG-UK) Consortium | Sergi Castellano, Rachel Williams, Mark Kristiansen, Paola Resende Silva, Sunando Roy, Tony Brooks, Helena Tutill, Paola Niola, Patricia Dyal, Charlotte Williams, Leysa Forrest, Yasmin Panchbhaya, Jacqueline Findlay, Sam Weeks, Julianne Brown, Kathryn Harris, Paul Randell, James Price, Alison Holmes, Judith Breuer |
| EPI_ISL_445003, EPI_ISL_445004, EPI_ISL_445005, EPI_ISL_445006, EPI_ISL_445007, EPI_ISL_445008, EPI_ISL_445009, EPI_ISL_445010, EPI_ISL_445011, EPI_ISL_445012, EPI_ISL_445013, EPI_ISL_445014, EPI_ISL_445015, EPI_ISL_445016, EPI_ISL_445017, EPI_ISL_445018, EPI_ISL_445020, EPI_ISL_445021, EPI_ISL_445022, EPI_ISL_445023, EPI_ISL_445025, EPI_ISL_445026, EPI_ISL_445027, EPI_ISL_445028, EPI_ISL_445029, EPI_ISL_445031, EPI_ISL_445032, EPI_ISL_445033, EPI_ISL_445034, EPI_ISL_445037, EPI_ISL_445038, EPI_ISL_445039, EPI_ISL_445040, EPI_ISL_445042, EPI_ISL_445050 | | | |
| see above | Florida Bureau of Public Health Laboratories | Florida Bureau of Public Health Laboratories | Sarah Schmedes, Jason Blanton |
| EPI_ISL_447561, EPI_ISL_447562, EPI_ISL_447568, EPI_ISL_447569, EPI_ISL_447849, EPI_ISL_447850 | CSIR-Centre for Cellular and Molecular Biology | CSIR-Centre for Cellular and Molecular Biology | Shagufta Khan, Lamuk Zaveri, Namami Gaur, Sakshi Shambhavi, Tulasi Nagabandi, Purushotham Vodnala, Payel Mukherjee, Sofia Banu, Priya Singh, Dhiviya Vedagiri, Divya Gupta, Vishal Sah, Santosh Kumar Kuncha, Krishnan Harinivas Harshan, Archana Bharadwaj Siva, Karthik Bharadwaj Tallapaka, Rakesh K Mishra, Divya Tej Sowpati |
| EPI_ISL_449626, EPI_ISL_449627, EPI_ISL_449628, EPI_ISL_449629, EPI_ISL_449630, EPI_ISL_449631, EPI_ISL_449632, EPI_ISL_449633, EPI_ISL_449634, EPI_ISL_449635, EPI_ISL_449636, EPI_ISL_449637, EPI_ISL_449638, EPI_ISL_449639, EPI_ISL_449640, EPI_ISL_449641, EPI_ISL_449642, EPI_ISL_449643, EPI_ISL_449644, EPI_ISL_449645, EPI_ISL_449646, EPI_ISL_449647, EPI_ISL_449648, EPI_ISL_449649, EPI_ISL_449650, EPI_ISL_449651, EPI_ISL_449652, EPI_ISL_449653, EPI_ISL_449654, EPI_ISL_449655, EPI_ISL_449656, EPI_ISL_449657, EPI_ISL_449658, EPI_ISL_449659, EPI_ISL_449660, EPI_ISL_449661, EPI_ISL_449662, EPI_ISL_449663, EPI_ISL_449664, EPI_ISL_449665, EPI_ISL_449666, EPI_ISL_449667, EPI_ISL_449668, EPI_ISL_449669, EPI_ISL_449670, EPI_ISL_449671, EPI_ISL_449672, EPI_ISL_449673, EPI_ISL_449674, EPI_ISL_449675, EPI_ISL_449676, EPI_ISL_449677, EPI_ISL_449678, EPI_ISL_449679, EPI_ISL_449680, EPI_ISL_449681, EPI_ISL_449682, EPI_ISL_449683, EPI_ISL_449684, EPI_ISL_449685, EPI_ISL_449686, EPI_ISL_449687, EPI_ISL_449688, EPI_ISL_449689, EPI_ISL_449690, EPI_ISL_449691, EPI_ISL_449692, EPI_ISL_449693, EPI_ISL_449694, EPI_ISL_449695, EPI_ISL_449696, EPI_ISL_449697, EPI_ISL_449698, EPI_ISL_449699, EPI_ISL_449700, EPI_ISL_449701, EPI_ISL_449702, EPI_ISL_449703, EPI_ISL_449704, EPI_ISL_449705, EPI_ISL_449706, EPI_ISL_449707, EPI_ISL_449708, EPI_ISL_449709, EPI_ISL_449710, EPI_ISL_449711, EPI_ISL_449712, EPI_ISL_449713, EPI_ISL_449714, EPI_ISL_449715, EPI_ISL_449716, EPI_ISL_449717, EPI_ISL_449718, EPI_ISL_449719, EPI_ISL_449720, EPI_ISL_449721, EPI_ISL_449722, EPI_ISL_449723, EPI_ISL_449724, EPI_ISL_449725, EPI_ISL_449726, EPI_ISL_449727, EPI_ISL_449728, EPI_ISL_449729 | | | |
| see above | University College London, Great Ormond Street Hospital for Children NHS Foundation Trust, Imperial College Healthcare NHS Trust | COVID-19 Genomics UK (COG-UK) Consortium | Sergi Castellano, Rachel Williams, Mark Kristiansen, Paola Resende Silva, Sunando Roy, Tony Brooks, Helena Tutill, Paola Niola, Patricia Dyal, Charlotte Williams, Leysa Forrest, Yasmin Panchbhaya, Jacqueline Findlay, Sam Weeks, Julianne Brown, Kathryn Harris, Paul Randell, James Price, Alison Holmes, Judith Breuer |
| EPI_ISL_450212, EPI_ISL_450213, EPI_ISL_450214, EPI_ISL_450215, | unknown | Microbiological Diagnostic Unit Public Health Laboratory (MDU-PHL) and Victorian Infectious Disease Reference | Seemann,T., Lane,C.R., Sherry,N.L., Duchene,S., Goncalves da Silva,A., Caly,L., Sait,M., Ballard,S.A., Horan,K., Schultz,M.B., Hoang,T., Easton,M., Dougal,S., Stinear,T.P., Druce,J., Catton,M., Sutton,B., van Diemen,A., Alpren,C., Williamson,D.A., Howden,B.P. |

| | | | |
|---|---|---|---|
| EPI_ISL_450216 | | Laboratory (VIDRL) | |
| EPI_ISL_450328 | CSIR-Centre for Cellular and Molecular Biology | CSIR-Centre for Cellular and Molecular Biology | Shagufta Khan, Lamuk Zaveri, Namami Gaur, Sakshi Shambhavi, Tulasi Nagabandi, Purushotham Vodnala, Payel Mukherjee, Sofia Banu, Priya Singh, Dhiviya Vedagiri, Divya Gupta, Vishal Sah, Santosh Kumar Kuncha, Krishnan Harinivas Harshan, Archana Bharadwaj Siva, Karthik Bharadwaj Tallapaka, Zeba Rizvi, Zuberwasim Sayyad, Kakade Aishwarya Arun, Amrutha H C, Ananga Ghosh, Kezia J Ann, Radhika Khandelwal, Roshan Maku Venkata, Shemin Mansuri, Sonu Uday, Sudipta Mondal, Rakesh K Mishra, Divya Tej Sowpati |
| EPI_ISL_450786 | Pandit Deendayal Upadhyay Government Medical College, Rajkot | Gujarat Biotechnology Research Centre | Sejul Antala, Manish Pattani, Prakash Modi, Janvi Raval, Zarna Patel, Monika Gandhi, Pinal Trivedi, Maharshi Pandya, Amit Kanani, Nidhi Patel, Nitin Savaliya, Raghawendra Kumar, Dinesh Kumar, Zuber Saiyed, Komal Patel, Labdhi Pandya, Snehal Bagatharia, Ramesh Pandit, Tejas Shah, Ankit Hinsu, Pritesh Sabara, Apurvasinh Puvar, Bhavesh Modi, Gaurishankar Shrimali, R D Dixit, A M Kadri, Armi Chaudhari, Chaitanya Joshi, Madhvi Joshi |
| EPI_ISL_450845, EPI_ISL_450846, EPI_ISL_450848, EPI_ISL_450852, EPI_ISL_450853, EPI_ISL_450854, EPI_ISL_450855, EPI_ISL_450856, EPI_ISL_450858, EPI_ISL_450859, EPI_ISL_450860, EPI_ISL_450861, EPI_ISL_450862, EPI_ISL_450863, EPI_ISL_450866, EPI_ISL_450867, EPI_ISL_450868, EPI_ISL_450870, EPI_ISL_450872 | | | |
| see above | Florida Bureau of Public Health Laboratories | Florida Bureau of Public Health Laboratories | Sarah Schmedes, Jason Blanton |
| EPI_ISL_453460, EPI_ISL_453461, EPI_ISL_453462, EPI_ISL_453463, EPI_ISL_453464 | University College London, Great Ormond Street Hospital for Children NHS Foundation Trust, Imperial College Healthcare NHS Trust | COVID-19 Genomics UK (COG-UK) Consortium | Sergi Castellano, Rachel Williams, Mark Kristiansen, Paola Resende Silva, Sunando Roy, Tony Brooks, Helena Tutill, Paola Niola, Patricia Dyal, Charlotte Williams, Leysa Forrest, Yasmin Panchbhaya, Jacqueline Findlay, Sam Weeks, Julianne Brown, Kathryn Harris, Paul Randell, James Price, Alison Holmes, Judith Breuer |
| EPI_ISL_454858, EPI_ISL_454859, EPI_ISL_454860, EPI_ISL_454861, EPI_ISL_454862, EPI_ISL_454863, EPI_ISL_454864, EPI_ISL_454865, EPI_ISL_454866, EPI_ISL_454867 | Translational Health Science and Technology Institute -ESIC medical college and hospital, Faridabad | THSTI Bioassay laboratory | Saurabh Kumar, Jigme Wangchuk, Anil Kumar Pandey, Asim Das, Guruprasad R. Medigeshi |
| EPI_ISL_456478, EPI_ISL_456479, EPI_ISL_456594, EPI_ISL_456595 | Microbiological Diagnostic Unit Public Health Laboratory | Microbiological Diagnostic Unit Public Health Laboratory, The Peter Doherty Institute for Infection and Immunity | Seemann T., Schultz M., Sait, M., Sherry, N. |
| EPI_ISL_457270, EPI_ISL_457271, EPI_ISL_457272, EPI_ISL_457273, EPI_ISL_457274, EPI_ISL_457275, EPI_ISL_457276, EPI_ISL_457277, EPI_ISL_457278, EPI_ISL_457279, EPI_ISL_457280, EPI_ISL_457281, EPI_ISL_457282, EPI_ISL_457283, EPI_ISL_457284, EPI_ISL_457285, EPI_ISL_457286, EPI_ISL_457287, EPI_ISL_457288, EPI_ISL_457289, EPI_ISL_457290, EPI_ISL_457291, EPI_ISL_457292, EPI_ISL_457293, EPI_ISL_457294, EPI_ISL_457295, EPI_ISL_457296, EPI_ISL_457297, EPI_ISL_457298, EPI_ISL_457299, EPI_ISL_457300, EPI_ISL_457301, EPI_ISL_457302 | | | |
| see above | University College London, Great Ormond Street Hospital for Children NHS Foundation Trust, Imperial College Healthcare NHS Trust | COVID-19 Genomics UK (COG-UK) Consortium | Sergi Castellano, Rachel Williams, Mark Kristiansen, Paola Resende Silva, Sunando Roy, Tony Brooks, Helena Tutill, Paola Niola, Patricia Dyal, Charlotte Williams, Leysa Forrest, Yasmin Panchbhaya, Jacqueline Findlay, Sam Weeks, Julianne Brown, Kathryn Harris, Paul Randell, James Price, Alison Holmes, Judith Breuer |
| EPI_ISL_458047 | CSIR-Centre for Cellular and Molecular Biology | CSIR-Centre for Cellular and Molecular Biology | Shagufta Khan, Lamuk Zaveri, Namami Gaur, Sakshi Shambhavi, Tulasi Nagabandi, Purushotham Vodnala, Payel Mukherjee, Sofia Banu, Priya Singh, Dhiviya Vedagiri, Divya Gupta, Vishal Sah, Santosh Kumar Kuncha, Krishnan Harinivas Harshan, Archana Bharadwaj Siva, Karthik Bharadwaj Tallapaka, Disha Nanda, Divya Das, Jotin Gogoi, Manish Bhattacharjee, Ravi Prasad Mukku, Rakesh K Mishra, Divya Tej Sowpati |
| EPI_ISL_458053 | CSIR-Centre for Cellular and Molecular Biology | CSIR-Centre for Cellular and Molecular Biology | Shagufta Khan, Lamuk Zaveri, Namami Gaur, Sakshi Shambhavi, Tulasi Nagabandi, Purushotham Vodnala, Payel Mukherjee, Sofia Banu, Priya Singh, Dhiviya Vedagiri, Divya Gupta, Vishal Sah, Santosh Kumar Kuncha, Krishnan Harinivas Harshan, Archana Bharadwaj Siva, Karthik Bharadwaj Tallapaka,Umesh Kumar, Unis Ahmad Bhat, Ajay Sarawagi, Priyanka Pant, Rajkanwar Nathawat, Rakesh K Mishra, Divya Tej Sowpati |
| EPI_ISL_458059 | CSIR-Centre for Cellular and Molecular Biology | CSIR-Centre for Cellular and Molecular Biology | Shagufta Khan, Lamuk Zaveri, Namami Gaur, Sakshi Shambhavi, Tulasi Nagabandi, Purushotham Vodnala, Payel Mukherjee, Sofia Banu, Priya Singh, Dhiviya Vedagiri, Divya Gupta, Vishal Sah, Santosh Kumar Kuncha, Krishnan Harinivas Harshan, Archana Bharadwaj Siva, Karthik Bharadwaj Tallapaka, Renu Sudhakar, Somesh Gorde, Gangumala Srinivas Reddy, Sujoy Deb, Swati Bayyana, Rakesh K Mishra, Divya Tej Sowpati |
| EPI_ISL_458064 | CSIR-Centre for Cellular and Molecular Biology | CSIR-Centre for Cellular and Molecular Biology | Shagufta Khan, Lamuk Zaveri, Namami Gaur, Sakshi Shambhavi, Tulasi Nagabandi, Purushotham Vodnala, Payel Mukherjee, Sofia Banu, Priya Singh, Dhiviya Vedagiri, Divya Gupta, Vishal Sah, Santosh Kumar Kuncha, Krishnan Harinivas Harshan, Archana Bharadwaj Siva, Karthik Bharadwaj Tallapaka,Preethi Jampala, Sharada Ravi Iyer, Sulagana Mukherjee, Swetha Sundar, Peddapuvala Sai Uday Kiran Rakesh K Mishra, Divya Tej Sowpati |
| EPI_ISL_461763, EPI_ISL_461764, EPI_ISL_461765, EPI_ISL_461766, EPI_ISL_461767, EPI_ISL_461768, EPI_ISL_461769, EPI_ISL_461770, EPI_ISL_461771, EPI_ISL_461772 | University College London, Great Ormond Street Hospital for Children NHS Foundation Trust, Imperial College Healthcare NHS Trust | COVID-19 Genomics UK (COG-UK) Consortium | Sergi Castellano, Rachel Williams, Mark Kristiansen, Paola Resende Silva, Sunando Roy, Tony Brooks, Helena Tutill, Paola Niola, Patricia Dyal, Charlotte Williams, Leysa Forrest, Yasmin Panchbhaya, Jacqueline Findlay, Sam Weeks, Julianne Brown, Kathryn Harris, Paul Randell, James Price, Alison Holmes, Judith Breuer |
| EPI_ISL_462092 | National Institute of Laboratory Medicine and Referral Center | Genomic Research Lab, BCSIR | Shahina Akter, Abu Sayeed Mohammad Mahmud, Mohammad Samir Uzzaman, Eshrar Osman, Md. Ahasan Habib, Tanjina Akhter Banu, Barna Goswami, Iffat Jahan, Md. Saddam Hossain, Tasnim Nafisa, Md. Maruf Ahmed Molla, Mahmuda Yeasmin, Asish Kumar Ghos, Bayzid Bin Monir, Arifa Akram, Sheikh Md. Selim Al Din, Salek Ahmed Sajib, Utpal Chandra Ray, Md. Salim Khan |
| EPI_ISL_464159, EPI_ISL_464160 | National Institute of Laboratory Medicine and Referral Center | Genomic Research Lab, BCSIR | Shahina Akter, Abu Sayeed Mohammad Mahmud, Mohammad Samir Uzzaman, Eshrar Osman, Md. Ahasan Habib, Tanjina Akhter Banu, Md. Murshed Hasan Sarker, Barna Goswami, Iffat Jahan, Md. Saddam Hossain, Tasnim Nafisa, Md. Maruf Ahmed Molla, Mahmuda Yeasmin, Asish Kumar Ghosh, Arifa Akram, A. K. M. Shamsuzzaman, Sheikh Md. Selim Al Din, Utpal Chandra Ray, Salek Ahmed Sajib, Md. Salim Khan |
| EPI_ISL_467029 | GMERS Medical College and Hospital, Gandhinagar | Gujarat Biotechnology Research Centre | Seema Bhatt, Gaurishankar Shrimali, Bhavesh Modi, Bharti Rajani, Tejas Shah, Ankit Hinsu, Pritesh Sabara, Apurvasinh Puvar, Janvi Raval, Zarna Patel, Monika Gandhi, Pinal Trivedi, Maharshi Pandya, Nidhi Patel, Nitin Savaliya, Raghawendra Kumar, Dinesh Kumar, Zuber Saiyed, Komal Patel, Labdhi Pandya, Snehal Bagatharia, Bhavya Jindal, R D Dixit, A M Kadri, Harsh Bakshi, Chaitanya Joshi, Madhvi Joshi |
| EPI_ISL_467667 | Florida Bureau of Public Health Laboratories | Florida Bureau of Public Health Laboratories | Schmedes,S. and Blanton,J. |
| EPI_ISL_468070, EPI_ISL_468071, EPI_ISL_468072, EPI_ISL_468073 | Child Health Research Foundation | Child Health Research Foundation | Senjuti Saha, Roly Malaker, Md Saiful Islam Sajib, Hafizur Rahman, Maksuda Islam, Samir K Saha |
| EPI_ISL_468074, EPI_ISL_468075, EPI_ISL_468076, EPI_ISL_468077, EPI_ISL_468078 | Child Health Research Foundation | Child Health Research Foundation | Senjuti Saha, Roly Malaker, Md Saiful Islam Sajib, Hafizur Rahman, Afroza Akter Tanni, Syed Muktadir Al Sium, Maksuda Islam, Samir K Saha |
| EPI_ISL_471158, EPI_ISL_471159, EPI_ISL_471160, EPI_ISL_471161, EPI_ISL_471162, EPI_ISL_471163, EPI_ISL_471164, EPI_ISL_471165, EPI_ISL_471166, EPI_ISL_471167, EPI_ISL_471168, EPI_ISL_471169, EPI_ISL_471170, EPI_ISL_471171 | | | |
| see above | MRCG at LSHTM Genomics lab | MRCG at LSHTM Genomics lab | Sesay et al |
| EPI_ISL_471603 | CSIR-Centre for Cellular and Molecular Biology | CSIR-Centre for Cellular and Molecular Biology | Shagufta Khan, Lamuk Zaveri, Namami Gaur, Sakshi Shambhavi, Tulasi Nagabandi, Purushotham Vodnala, Payel Mukherjee, Sofia Banu, Priya Singh, Dhiviya Vedagiri, Divya Gupta, Vishal Sah, Santosh Kumar Kuncha, Krishnan Harinivas Harshan, Archana Bharadwaj Siva, Karthik Bharadwaj Tallapaka, Disha Nanda, Divya Das, Jotin Gogoi, Manish Bhattacharjee, Ravi Prasad Mukku, Rakesh K Mishra, Divya Tej Sowpati |
| EPI_ISL_471604 | CSIR-Centre for Cellular and Molecular Biology | CSIR-Centre for Cellular and Molecular Biology | Shagufta Khan, Lamuk Zaveri, Namami Gaur, Sakshi Shambhavi, Tulasi Nagabandi, Purushotham Vodnala, Payel Mukherjee, Sofia Banu, Priya Singh, Dhiviya Vedagiri, Divya Gupta, Vishal Sah, Santosh Kumar Kuncha, Krishnan Harinivas Harshan, Archana Bharadwaj Siva, Karthik Bharadwaj Tallapaka, Renu Sudhakar, Somesh Gorde, Gangumala Srinivas Reddy, Sujoy Deb, Swati Bayyana, Rakesh K Mishra, Divya Tej Sowpati |
| EPI_ISL_471605 | CSIR-Centre for Cellular and Molecular Biology | CSIR-Centre for Cellular and Molecular Biology | Shagufta Khan, Lamuk Zaveri, Namami Gaur, Sakshi Shambhavi, Tulasi Nagabandi, Purushotham Vodnala, Payel Mukherjee, Sofia Banu, Priya Singh, Dhiviya Vedagiri, Divya Gupta, Vishal Sah, Santosh Kumar Kuncha, Krishnan Harinivas Harshan, Archana Bharadwaj Siva, Karthik Bharadwaj Tallapaka,Preethi Jampala, Sharada Ravi Iyer, Sulagana Mukherjee, Swetha Sundar, Peddapuvala Sai Uday Kiran Rakesh K Mishra, Divya Tej Sowpati |
| EPI_ISL_471606 | CSIR-Centre for Cellular and Molecular Biology | CSIR-Centre for Cellular and Molecular Biology | Shagufta Khan, Lamuk Zaveri, Namami Gaur, Sakshi Shambhavi, Tulasi Nagabandi, Purushotham Vodnala, Payel Mukherjee, Sofia Banu, Priya Singh, Dhiviya Vedagiri, Divya Gupta, Vishal Sah, Santosh Kumar Kuncha, Krishnan Harinivas Harshan, Archana Bharadwaj Siva, Karthik Bharadwaj Tallapaka,Umesh Kumar, Unis Ahmad Bhat, Ajay Sarawagi, Priyanka Pant, Rajkanwar Nathawat, Rakesh K Mishra, Divya Tej Sowpati |
| EPI_ISL_471631 | CSIR-Centre for Cellular and Molecular Biology | CSIR-Centre for Cellular and Molecular Biology | Shagufta Khan, Lamuk Zaveri, Namami Gaur, Sakshi Shambhavi, Tulasi Nagabandi, Purushotham Vodnala, Payel Mukherjee, Sofia Banu, Priya Singh, Dhiviya Vedagiri, Divya Gupta, Vishal Sah, Santosh Kumar Kuncha, Krishnan Harinivas Harshan, Archana Bharadwaj Siva, Karthik Bharadwaj Tallapaka, |

| | | | |
|---|---|---|---|
| | | | Disha Nanda, Divya Das, Jotin Gogoi, Manish Bhattacharjee, Ravi Prasad Mukku, Rakesh K Mishra, Divya Tej Sowpati |
| EPI_ISL_471632 | CSIR-Centre for Cellular and Molecular Biology | CSIR-Centre for Cellular and Molecular Biology | Shagufta Khan, Lamuk Zaveri, Namami Gaur, Sakshi Shambhavi, Tulasi Nagabandi, Purushotham Vodnala, Payel Mukherjee, Sofia Banu, Priya Singh, Dhiviya Vedagiri, Divya Gupta, Vishal Sah, Santosh Kumar Kuncha, Krishnan Harinivas Harshan, Archana Bharadwaj Siva, Karthik Bharadwaj Tallapaka, Renu Sudhakar, Somesh Gorde, Gangumala Srinivas Reddy, Sujoy Deb, Swati Bayyana, Rakesh K Mishra, Divya Tej Sowpati |
| EPI_ISL_471633 | CSIR-Centre for Cellular and Molecular Biology | CSIR-Centre for Cellular and Molecular Biology | Shagufta Khan, Lamuk Zaveri, Namami Gaur, Sakshi Shambhavi, Tulasi Nagabandi, Purushotham Vodnala, Payel Mukherjee, Sofia Banu, Priya Singh, Dhiviya Vedagiri, Divya Gupta, Vishal Sah, Santosh Kumar Kuncha, Krishnan Harinivas Harshan, Archana Bharadwaj Siva, Karthik Bharadwaj Tallapaka,Preethi Jampala, Sharada Ravi Iyer, Sulagana Mukherjee, Swetha Sundar, Peddapuvala Sai Uday Kiran Rakesh K Mishra, Divya Tej Sowpati |
| EPI_ISL_471634 | CSIR-Centre for Cellular and Molecular Biology | CSIR-Centre for Cellular and Molecular Biology | Shagufta Khan, Lamuk Zaveri, Namami Gaur, Sakshi Shambhavi, Tulasi Nagabandi, Purushotham Vodnala, Payel Mukherjee, Sofia Banu, Priya Singh, Dhiviya Vedagiri, Divya Gupta, Vishal Sah, Santosh Kumar Kuncha, Krishnan Harinivas Harshan, Archana Bharadwaj Siva, Karthik Bharadwaj Tallapaka,Umesh Kumar, Unis Ahmad Bhat, Ajay Sarawagi, Priyanka Pant, Rajkanwar Nathawat, Rakesh K Mishra, Divya Tej Sowpati |
| EPI_ISL_475754 | National Institute of Laboratory Medicine and Referral Center | Genomic Research Lab, BCSIR | Shahina Akter, Abu Sayeed Mohammad Mahmud, Mohammad Samir Uzzaman, Eshrar Osman, Md. Ahasan Habib, Tanjina Akhter Banu, Md. Murshed Hasan Sarkar, Barna Goswami, Iffat Jahan, Md. Saddam Hossain, Tasnim Nafisa, Md. Maruf Ahmed Molla, Mahmuda Yeasmin, Asish Kumar Ghosh, Arifa Akram, A. K. M. Shamsuzzaman, Sheikh Md. Selim Al Din, Utpal Chandra Ray, Salek Ahmed Sajib, Md. Salim Khan |
| EPI_ISL_476832 | Medical Biology Department, Kocaeli University | Medical Genetics Department, Kocaeli University | Savli H, Cine N, Sunnetci-Akkoyunlu D, Eren-Keskin S, Ilgazli A, Akhan S, Karadenizli A, Kasap M, Sayan M, Akpinar G, Canturk NZ. |
| EPI_ISL_477125, EPI_ISL_477126, EPI_ISL_477127, EPI_ISL_477128, EPI_ISL_477129, EPI_ISL_477130, EPI_ISL_477131, EPI_ISL_477132, EPI_ISL_477133, EPI_ISL_477134, EPI_ISL_477135, EPI_ISL_477136, EPI_ISL_477137, EPI_ISL_477138, EPI_ISL_477139, EPI_ISL_477140 | | | |
| see above | Child Health Research Foundation | Child Health Research Foundation | Senjuti Saha, Md Saiful Islam Sajib, Roly Malaker, Md Hafizur Rahman, Afroza Akter Tanni, Syed Muktadir Al Sium, Maksuda Islam, Samir K Saha |
| EPI_ISL_478404, EPI_ISL_478405, EPI_ISL_478406, EPI_ISL_478407, EPI_ISL_478408, EPI_ISL_478409, EPI_ISL_478410, EPI_ISL_478411, EPI_ISL_478412, EPI_ISL_478413, EPI_ISL_478414, EPI_ISL_478415, EPI_ISL_478416, EPI_ISL_478417, EPI_ISL_478418, EPI_ISL_478419, EPI_ISL_478420, EPI_ISL_478421, EPI_ISL_478422, EPI_ISL_478423, EPI_ISL_478424, EPI_ISL_478425, EPI_ISL_478426, EPI_ISL_478427, EPI_ISL_478428, EPI_ISL_478429, EPI_ISL_478430, EPI_ISL_478431, EPI_ISL_478432, EPI_ISL_478433, EPI_ISL_478434, EPI_ISL_478435, EPI_ISL_478436, EPI_ISL_478437, EPI_ISL_478438, EPI_ISL_478439, EPI_ISL_478440, EPI_ISL_478441, EPI_ISL_478442, EPI_ISL_478443, EPI_ISL_478444, EPI_ISL_478445, EPI_ISL_478446, EPI_ISL_478447, EPI_ISL_478448, EPI_ISL_478449, EPI_ISL_478450, EPI_ISL_478451, EPI_ISL_478452, EPI_ISL_478453, EPI_ISL_478454, EPI_ISL_478455, EPI_ISL_478456, EPI_ISL_478457, EPI_ISL_478458, EPI_ISL_478459, EPI_ISL_478460, EPI_ISL_478461, EPI_ISL_478462, EPI_ISL_478463, EPI_ISL_478464, EPI_ISL_478465, EPI_ISL_478466, EPI_ISL_478467, EPI_ISL_478468, EPI_ISL_478469, EPI_ISL_478470, EPI_ISL_478471, EPI_ISL_478472 | | | |
| see above | University College London, Great Ormond Street Hospital for Children NHS Foundation Trust, Imperial College Healthcare NHS Trust | COVID-19 Genomics UK (COG-UK) Consortium | Sergi Castellano, Rachel Williams, Mark Kristiansen, Paola Resende Silva, Sunando Roy, Tony Brooks, Helena Tutill, Paola Niola, Patricia Dyal, Charlotte Williams, Leysa Forrest, Yasmin Panchbhaya, Jacqueline Findlay, Samuel Weeks, Julianne Brown, Kathryn Harris, Paul Randell, James Price, Alison Holmes, Judith Breuer |
| EPI_ISL_479662 | College of Veterinary Medicine, Chungnam National University | College of Veterinary Medicine, Chungnam National University | Seo,S. |
| EPI_ISL_480558, EPI_ISL_480559, EPI_ISL_480560, EPI_ISL_480561, EPI_ISL_480562, EPI_ISL_480587, EPI_ISL_480612, EPI_ISL_480613, EPI_ISL_480614, EPI_ISL_480615, EPI_ISL_480616, EPI_ISL_480617, EPI_ISL_480618, EPI_ISL_480620, EPI_ISL_480687, EPI_ISL_480745, EPI_ISL_480746, EPI_ISL_480747, EPI_ISL_480748, EPI_ISL_480749, EPI_ISL_480750, EPI_ISL_480751, EPI_ISL_480752, EPI_ISL_480753, EPI_ISL_480754, EPI_ISL_480755, EPI_ISL_480756, EPI_ISL_480757, EPI_ISL_480758, EPI_ISL_480759, EPI_ISL_480760, EPI_ISL_480761, EPI_ISL_480762, EPI_ISL_480763, EPI_ISL_480767, EPI_ISL_480768, EPI_ISL_480769, EPI_ISL_480770, EPI_ISL_480771, EPI_ISL_480772, EPI_ISL_480773, EPI_ISL_480774, EPI_ISL_480776, EPI_ISL_480777 | | | |
| see above | Microbiological Diagnostic Unit - Public Health Laboratory (MDU-PHL) | MDU-PHL | Seemann T., Schultz M., Sait, M., Sherry, N. |
| EPI_ISL_480790, EPI_ISL_480792, EPI_ISL_480793, EPI_ISL_480794, EPI_ISL_480795, EPI_ISL_480798, EPI_ISL_480799, EPI_ISL_480800, EPI_ISL_480802, EPI_ISL_480803, EPI_ISL_480804, EPI_ISL_480805, EPI_ISL_480807, EPI_ISL_480809, EPI_ISL_480810, EPI_ISL_480813, EPI_ISL_480819, EPI_ISL_480820, EPI_ISL_480835, EPI_ISL_480843, EPI_ISL_480846, EPI_ISL_480848, EPI_ISL_480849, EPI_ISL_480854, EPI_ISL_480860, EPI_ISL_480864, EPI_ISL_480865, EPI_ISL_480867, EPI_ISL_480868, EPI_ISL_480869, EPI_ISL_480878, EPI_ISL_480881, EPI_ISL_480882, EPI_ISL_480884, EPI_ISL_480885, EPI_ISL_480886, EPI_ISL_480887, EPI_ISL_480888, EPI_ISL_480889, EPI_ISL_480890, EPI_ISL_480891, EPI_ISL_480892, EPI_ISL_480893, EPI_ISL_480894, EPI_ISL_480895, EPI_ISL_480896, EPI_ISL_480897, EPI_ISL_480898, EPI_ISL_480899, EPI_ISL_480900, EPI_ISL_480901, EPI_ISL_480902, EPI_ISL_480903, EPI_ISL_480904, EPI_ISL_480905, EPI_ISL_480906, EPI_ISL_480907, EPI_ISL_480908, EPI_ISL_480909, EPI_ISL_480910, EPI_ISL_480911, EPI_ISL_480912, EPI_ISL_480913, EPI_ISL_480914, EPI_ISL_480915, EPI_ISL_480916, EPI_ISL_480917, EPI_ISL_480918, EPI_ISL_480919, EPI_ISL_480920, EPI_ISL_480923, EPI_ISL_480924, EPI_ISL_480929, EPI_ISL_480930, EPI_ISL_480931, EPI_ISL_480932, EPI_ISL_480937, EPI_ISL_480939, EPI_ISL_480941, EPI_ISL_480943, EPI_ISL_480946, EPI_ISL_480947, EPI_ISL_480948, EPI_ISL_480949 | | | |
| see above | Florida Bureau of Public Health Laboratories | Florida Bureau of Public Health Laboratories | Sarah Schmedes, Jason Blanton |
| EPI_ISL_482160, EPI_ISL_482161, EPI_ISL_482162, EPI_ISL_482163, EPI_ISL_482164, EPI_ISL_482165, EPI_ISL_482166, EPI_ISL_482167, EPI_ISL_482168, EPI_ISL_482169, EPI_ISL_482170, EPI_ISL_482171, EPI_ISL_482172, EPI_ISL_482173, EPI_ISL_482174, EPI_ISL_482175, EPI_ISL_482176, EPI_ISL_482177, EPI_ISL_482178, EPI_ISL_482179, EPI_ISL_482180, EPI_ISL_482181, EPI_ISL_482182, EPI_ISL_482183, EPI_ISL_482184, EPI_ISL_482185, EPI_ISL_482186, EPI_ISL_482187, EPI_ISL_482188, EPI_ISL_482189, EPI_ISL_482190, EPI_ISL_482191, EPI_ISL_482192, EPI_ISL_482193, EPI_ISL_482194, EPI_ISL_482195, EPI_ISL_482196, EPI_ISL_482197, EPI_ISL_482198, EPI_ISL_482199, EPI_ISL_482200, EPI_ISL_482201, EPI_ISL_482202, EPI_ISL_482203, EPI_ISL_482204, EPI_ISL_482205, EPI_ISL_482206, EPI_ISL_482207, EPI_ISL_482208, EPI_ISL_482209, EPI_ISL_482210, EPI_ISL_482211, EPI_ISL_482212, EPI_ISL_482213, EPI_ISL_482214, EPI_ISL_482215, EPI_ISL_482216, EPI_ISL_482217, EPI_ISL_482218 | | | |
| see above | University College London, Great Ormond Street Hospital for Children NHS Foundation Trust, Imperial College Healthcare NHS Trust | Wellcome Sanger Institute for the COVID-19 Genomics UK (COG-UK) consortium | Sergi Castellano, Rachel Williams, Mark Kristiansen, Paola Resende Silva, Sunando Roy, Tony Brooks, Helena Tutill, Paola Niola, Patricia Dyal, Charlotte Williams, Leysa Forrest, Yasmin Panchbhaya, Jacqueline Findlay, Sam Weeks, Julianne Brown, Kathryn Harris, Paul Randell, James Price, Alison Holmes, Judith Breuer and Alex Alderton, Roberto Amato, Sonia Goncalves, Ewan Harrison, David K. Jackson, Ian Johnston, Dominic Kwiatkowski, Cordelia Langford, John Sillitoe on behalf of the Wellcome Sanger Institute COVID-19 Surveillance Team (http://www.sanger.ac.uk/covid-team) |
| EPI_ISL_483633 | National Institute of Laboratory Medicine and Referral Center | Genomic Research Lab, BCSIR | Shahina Akter, Abu Sayeed Mohammad Mahmud, Mohammad Samir Uzzaman, Eshrar Osman, Md. Ahasan Habib, Tanjina Akhter Banu, Md. Murshed Hasan Sarkar, Barna Goswami, Iffat Jahan, Md. Saddam Hossain, Tasnim Nafisa, Md. Maruf Ahmed Molla, Mahmuda Yeasmin, Asish Kumar Ghosh, A. K. M. Shamsuzzaman, Sheikh Md. Selim Al Din, Utpal Chandra Ray, Salek Ahmed Sajib, Md. Salim Khan |
| EPI_ISL_486852 | CDRI/SGPGI | CSIR-CDRI/SGPGI | Saumya Sarkar, Dharam Veer Singh, Rahul Vishvkarma, Ujjala Ghoshal, Uday Ghoshal, Ravishankar Ramachandran, Tapas Kumar Kundu, Rajender Singh |
| EPI_ISL_486853 | CSIR-CDRI/SGPGI | CSIR-CDRI/SGPGI | Saumya Sarkar, Dharam Veer Singh, Rahul Vishvkarma, Ujjala Ghoshal, Uday Ghoshal, Ravishankar Ramachandran, Tapas Kumar Kundu, Rajender Singh |
| EPI_ISL_487366 | National Institute of Laboratory Medicine and Referral Center | Genomic Research Lab, BCSIR | Shahina Akter, Abu Sayeed Mohammad Mahmud, Mohammad Samir Uzzaman, Eshrar Osman, Md. Ahasan Habib, Tanjina Akhter Banu, Md. Murshed Hasan Sarkar, Barna Goswami, Iffat Jahan, Md. Saddam Hossain, Tasnim Nafisa, Md. Maruf Ahmed Molla, Mahmuda Yeasmin, Asish Kumar Ghosh, A. K. M. Shamsuzzaman, Sheikh Md. Selim Al Din, Utpal Chandra Ray, Salek Ahmed Sajib, Md. Salim Khan |
| EPI_ISL_487525, EPI_ISL_487529, EPI_ISL_487534, EPI_ISL_487537, EPI_ISL_487540, EPI_ISL_487546, EPI_ISL_487550, EPI_ISL_487556, EPI_ISL_487557, EPI_ISL_487558, EPI_ISL_487559, EPI_ISL_487562, EPI_ISL_487564, EPI_ISL_487568, EPI_ISL_487572, EPI_ISL_487578, EPI_ISL_487581, EPI_ISL_487582, EPI_ISL_487584, EPI_ISL_487585, EPI_ISL_487589, EPI_ISL_487595, EPI_ISL_487597, EPI_ISL_487600, EPI_ISL_487606, EPI_ISL_487607, EPI_ISL_487614, EPI_ISL_487615, EPI_ISL_487616, EPI_ISL_487621, EPI_ISL_487625, EPI_ISL_487626, EPI_ISL_487627, EPI_ISL_487628, EPI_ISL_487630, EPI_ISL_487631, EPI_ISL_487632, EPI_ISL_487633, EPI_ISL_487635, EPI_ISL_487636, EPI_ISL_487637, EPI_ISL_487638, EPI_ISL_487639, EPI_ISL_487640, EPI_ISL_487641, EPI_ISL_487642, EPI_ISL_487643, EPI_ISL_487644, EPI_ISL_487645, EPI_ISL_487646 | | | |
| see above | University College London, Great Ormond Street Hospital for Children NHS Foundation Trust, Imperial College Healthcare NHS Trust | Wellcome Sanger Institute for the COVID-19 Genomics UK (COG-UK) consortium | Sergi Castellano, Rachel Williams, Mark Kristiansen, Paola Resende Silva, Sunando Roy, Tony Brooks, Helena Tutill, Paola Niola, Patricia Dyal, Charlotte Williams, Leysa Forrest, Yasmin Panchbhaya, Jacqueline Findlay, Sam Weeks, Julianne Brown, Kathryn Harris, Paul Randell, James Price, Alison Holmes, Judith Breuer and Alex Alderton, Roberto Amato, Sonia Goncalves, Ewan Harrison, David K. Jackson, Ian Johnston, Dominic Kwiatkowski, Cordelia Langford, John Sillitoe on behalf of the Wellcome Sanger Institute COVID-19 Surveillance Team (http://www.sanger.ac.uk/covid-team) |
| EPI_ISL_489713, EPI_ISL_489714, EPI_ISL_489715, EPI_ISL_489719, EPI_ISL_489721, EPI_ISL_489726, EPI_ISL_489732, EPI_ISL_489735, EPI_ISL_489736, EPI_ISL_489738, EPI_ISL_489740, EPI_ISL_489741, EPI_ISL_489742, EPI_ISL_489743, EPI_ISL_489744, EPI_ISL_489746, EPI_ISL_489748, EPI_ISL_489749, EPI_ISL_489750, EPI_ISL_489751, EPI_ISL_489753, EPI_ISL_489754, EPI_ISL_489755, EPI_ISL_489757, EPI_ISL_489766, EPI_ISL_489768, EPI_ISL_489771, EPI_ISL_489774 | | | |
| see above | Florida Bureau of Public Health Laboratories | Florida Bureau of Public Health Laboratories | Sarah Schmedes, Jason Blanton |
| EPI_ISL_490113 | National Institute of Laboratory Medicine and Referral Center | Genomic Research Lab, BCSIR | Shahina Akter, Abu Sayeed Mohammad Mahmud, Mohammad Samir Uzzaman, Eshrar Osman, Md. Ahasan Habib, Tanjina Akhter Banu, Md. Murshed Hasan Sarkar, Barna Goswami, Iffat Jahan, Md. Saddam Hossain, Tasnim Nafisa, Md. Maruf Ahmed Molla, Mahmuda Yeasmin, Asish Kumar Ghosh, A. K. M. Shamsuzzaman, Sheikh Md. Selim Al Din, Utpal Chandra Ray, Salek Ahmed Sajib, Md. Salim Khan |
| EPI_ISL_492028, EPI_ISL_492029, EPI_ISL_492030, EPI_ISL_492031 | Child Health Research Foundation | Child Health Research Foundation | Senjuti Saha, Md Saiful Islam Sajib, Roly Malaker, Md Hafizur Rahman, Afroza Akter Tanni, Syed Muktadir Al Sium, Maksuda Islam, Samir K Saha |
| EPI_ISL_492496, EPI_ISL_492498, EPI_ISL_492499, EPI_ISL_492500, EPI_ISL_492501, EPI_ISL_492502, EPI_ISL_492503, EPI_ISL_492504, EPI_ISL_492747, EPI_ISL_492749, EPI_ISL_492751, EPI_ISL_492752, EPI_ISL_492753, EPI_ISL_492754, EPI_ISL_492755, EPI_ISL_492756, EPI_ISL_492757, EPI_ISL_492759, EPI_ISL_492760, EPI_ISL_492761, EPI_ISL_492762, EPI_ISL_492763 | | | |
| see above | University College London, Great Ormond Street Hospital for Children NHS Foundation Trust, Imperial College Healthcare | Wellcome Sanger Institute for the COVID-19 Genomics UK (COG-UK) consortium | Sergi Castellano, Rachel Williams, Mark Kristiansen, Paola Resende Silva, Sunando Roy, Tony Brooks, Helena Tutill, Paola Niola, Patricia Dyal, Charlotte Williams, Leysa Forrest, Yasmin Panchbhaya, Jacqueline Findlay, Sam Weeks, Julianne Brown, Kathryn Harris, Paul Randell, James Price, Alison Holmes, |

| | | | |
|---|---|---|---|
| | NHS Trust | | Judith Breuer and Alex Alderton, Roberto Amato, Sonia Goncalves, Ewan Harrison, David K. Jackson, Ian Johnston, Dominic Kwiatkowski, Cordelia Langford, John Sillitoe on behalf of the Wellcome Sanger Institute COVID-19 Surveillance Team (http://www.sanger.ac.uk/covid-team) |
| EPI_ISL_495185, EPI_ISL_495186 | CSIR-Centre for Cellular and Molecular Biology | CSIR-Centre for Cellular and Molecular Biology | Shagufta Khan, Lamuk Zaveri, Namami Gaur, Sakshi Shambhavi, Nikhil Hajirnis, M Soujanya Reddy, Pratheusa Maccha, Tulasi Nagabandi, Purushotham Vodnala, Payel Mukherjee, Sofia Banu, Priya Singh, Onkar Kulkarni, Dhiviya Vedagiri, Divya Gupta, Vishal Sah, Santosh Kumar Kuncha, Krishnan Harinivas Harshan, Archana Bharadwaj Siva, Karthik Bharadwaj Tallapaka, Renu Sudhakar, Somesh Gorde, Gangumala Srinivas Reddy, Sujoy Deb, Swati Bayyana, Rakesh K Mishra, Divya Tej Sowpati |
| EPI_ISL_495187 | CSIR-Centre for Cellular and Molecular Biology | CSIR-Centre for Cellular and Molecular Biology | Shagufta Khan, Lamuk Zaveri, Namami Gaur, Sakshi Shambhavi, Nikhil Hajirnis, M Soujanya Reddy, Pratheusa Maccha,Tulasi Nagabandi, Purushotham Vodnala, Payel Mukherjee, Sofia Banu, Priya Singh, Onkar Kulkarni, Dhiviya Vedagiri, Divya Gupta, Vishal Sah, Santosh Kumar Kuncha, Krishnan Harinivas Harshan, Archana Bharadwaj Siva, Karthik Bharadwaj Tallapaka,Umesh Kumar, Unis Ahmad Bhat, Ajay Sarawagi, Priyanka Pant, Rajkanwar Nathawat, Rakesh K Mishra, Divya Tej Sowpati |
| EPI_ISL_495202 | CSIR-Centre for Cellular and Molecular Biology | CSIR-Centre for Cellular and Molecular Biology | Shagufta Khan, Lamuk Zaveri, Namami Gaur, Sakshi Shambhavi, Nikhil Hajirnis, M Soujanya Reddy, Pratheusa Maccha, Tulasi Nagabandi, Purushotham Vodnala, Payel Mukherjee, Sofia Banu, Priya Singh, Onkar Kulkarni, Dhiviya Vedagiri, Divya Gupta, Vishal Sah, Santosh Kumar Kuncha, Krishnan Harinivas Harshan, Archana Bharadwaj Siva, Karthik Bharadwaj Tallapaka, Renu Sudhakar, Somesh Gorde, Gangumala Srinivas Reddy, Sujoy Deb, Swati Bayyana, Rakesh K Mishra, Divya Tej Sowpati |
| EPI_ISL_495205 | CSIR-Centre for Cellular and Molecular Biology | CSIR-Centre for Cellular and Molecular Biology | Shagufta Khan, Lamuk Zaveri, Namami Gaur, Sakshi Shambhavi, Nikhil Hajirnis, M Soujanya Reddy, Pratheusa Maccha, Tulasi Nagabandi, Purushotham Vodnala, Payel Mukherjee, Sofia Banu, Priya Singh, Onkar Kulkarni, Dhiviya Vedagiri, Divya Gupta, Vishal Sah, Santosh Kumar Kuncha, Krishnan Harinivas Harshan, Archana Bharadwaj Siva, Karthik Bharadwaj Tallapaka,Preethi Jampala, Sharada Ravi Iyer, Sulagana Mukherjee, Swetha Sundar, Peddapuvala Sai Uday Kiran Rakesh K Mishra, Divya Tej Sowpati |
| EPI_ISL_495206 | CSIR-Centre for Cellular and Molecular Biology | CSIR-Centre for Cellular and Molecular Biology | Shagufta Khan, Lamuk Zaveri, Namami Gaur, Sakshi Shambhavi, Nikhil Hajirnis, M Soujanya Reddy, Pratheusa Maccha,Tulasi Nagabandi, Purushotham Vodnala, Payel Mukherjee, Sofia Banu, Priya Singh, Onkar Kulkarni, Dhiviya Vedagiri, Divya Gupta, Vishal Sah, Santosh Kumar Kuncha, Krishnan Harinivas Harshan, Archana Bharadwaj Siva, Karthik Bharadwaj Tallapaka,Umesh Kumar, Unis Ahmad Bhat, Ajay Sarawagi, Priyanka Pant, Rajkanwar Nathawat, Rakesh K Mishra, Divya Tej Sowpati |
| EPI_ISL_495208 | CSIR-Centre for Cellular and Molecular Biology | CSIR-Centre for Cellular and Molecular Biology | Shagufta Khan, Lamuk Zaveri, Namami Gaur, Sakshi Shambhavi, Nikhil Hajirnis, M Soujanya Reddy, Pratheusa Maccha, Tulasi Nagabandi, Purushotham Vodnala, Payel Mukherjee, Sofia Banu, Priya Singh, Onkar Kulkarni, Dhiviya Vedagiri, Divya Gupta, Vishal Sah, Santosh Kumar Kuncha, Krishnan Harinivas Harshan, Archana Bharadwaj Siva, Karthik Bharadwaj Tallapaka, Renu Sudhakar, Somesh Gorde, Gangumala Srinivas Reddy, Sujoy Deb, Swati Bayyana, Rakesh K Mishra, Divya Tej Sowpati |
| EPI_ISL_495211, EPI_ISL_495220 | CSIR-Centre for Cellular and Molecular Biology | CSIR-Centre for Cellular and Molecular Biology | Shagufta Khan, Lamuk Zaveri, Namami Gaur, Sakshi Shambhavi, Nikhil Hajirnis, M Soujanya Reddy, Pratheusa Maccha, Tulasi Nagabandi, Purushotham Vodnala, Payel Mukherjee, Sofia Banu, Priya Singh, Onkar Kulkarni, Dhiviya Vedagiri, Divya Gupta, Vishal Sah, Santosh Kumar Kuncha, Krishnan Harinivas Harshan, Archana Bharadwaj Siva, Karthik Bharadwaj Tallapaka,Preethi Jampala, Sharada Ravi Iyer, Sulagana Mukherjee, Swetha Sundar, Peddapuvala Sai Uday Kiran Rakesh K Mishra, Divya Tej Sowpati |
| EPI_ISL_495241 | CSIR-Centre for Cellular and Molecular Biology | CSIR-Centre for Cellular and Molecular Biology | Shagufta Khan, Lamuk Zaveri, Namami Gaur, Sakshi Shambhavi, Nikhil Hajirnis, M Soujanya Reddy, Pratheusa Maccha,Tulasi Nagabandi, Purushotham Vodnala, Payel Mukherjee, Sofia Banu, Priya Singh, Onkar Kulkarni, Dhiviya Vedagiri, Divya Gupta, Vishal Sah, Santosh Kumar Kuncha, Krishnan Harinivas Harshan, Archana Bharadwaj Siva, Karthik Bharadwaj Tallapaka,Umesh Kumar, Unis Ahmad Bhat, Ajay Sarawagi, Priyanka Pant, Rajkanwar Nathawat, Rakesh K Mishra, Divya Tej Sowpati |
| EPI_ISL_498227 | National Institute of Laboratory Medicine and Referral Center | Genomic Research Lab, BCSIR | Shahina Akter, Abu Sayeed Mohammad Mahmud, Mohammad Samir Uzzaman, Eshrar Osman, Md. Ahasan Habib, Tanjina Akhter Banu, Md. Murshed Hasan Sarkar, Barna Goswami, Iffat Jahan, Md. Saddam Hossain, Tasnim Nafisa, Md. Maruf Ahmed Molla, Mahmuda Yeasmin, Asish Kumar Ghosh, A. K. M. Shamsuzzaman, Sheikh Md. Selim Al Din, Utpal Chandra Ray, Salek Ahmed Sajib, Md. Salim Khan |
| EPI_ISL_508722, EPI_ISL_508788, EPI_ISL_508790, EPI_ISL_508791, EPI_ISL_508792, EPI_ISL_508793, EPI_ISL_508794, EPI_ISL_508795, EPI_ISL_508796, EPI_ISL_508797, EPI_ISL_508798, EPI_ISL_508799, EPI_ISL_508800, EPI_ISL_508801, EPI_ISL_508802, EPI_ISL_508803, EPI_ISL_508804, EPI_ISL_508805, EPI_ISL_508807, EPI_ISL_508817, EPI_ISL_508821, EPI_ISL_508822, EPI_ISL_508828, EPI_ISL_508837, EPI_ISL_508838, EPI_ISL_508839, EPI_ISL_508840, EPI_ISL_508849, EPI_ISL_508850, EPI_ISL_508851, EPI_ISL_508858, EPI_ISL_508860, EPI_ISL_509716, EPI_ISL_509717, EPI_ISL_509718, EPI_ISL_509719, EPI_ISL_509721, EPI_ISL_509722, EPI_ISL_509724, EPI_ISL_509725, EPI_ISL_509726, EPI_ISL_509727, EPI_ISL_509728, EPI_ISL_509730, EPI_ISL_509731, EPI_ISL_509732, EPI_ISL_509734, EPI_ISL_509735, EPI_ISL_509736, EPI_ISL_509738, EPI_ISL_509739, EPI_ISL_509740, EPI_ISL_509741, EPI_ISL_509742, EPI_ISL_509745, EPI_ISL_509747, EPI_ISL_509748, EPI_ISL_509751, EPI_ISL_509752, EPI_ISL_509753, EPI_ISL_509754, EPI_ISL_509757, EPI_ISL_509758, EPI_ISL_509759, EPI_ISL_509760, EPI_ISL_509761, EPI_ISL_509762, EPI_ISL_509763, EPI_ISL_509764, EPI_ISL_509765, EPI_ISL_509766, EPI_ISL_509767, EPI_ISL_509768, EPI_ISL_509769, EPI_ISL_509770, EPI_ISL_509771, EPI_ISL_509772, EPI_ISL_509773, EPI_ISL_509774, EPI_ISL_509775, EPI_ISL_509776, EPI_ISL_509778, EPI_ISL_509780, EPI_ISL_509781, EPI_ISL_509783, EPI_ISL_509784, EPI_ISL_509786, EPI_ISL_509787, EPI_ISL_509788, EPI_ISL_509789, EPI_ISL_509791, EPI_ISL_509792, EPI_ISL_509793, EPI_ISL_509794, EPI_ISL_509795 | | | |
| see above | Florida Bureau of Public Health Laboratories | Florida Bureau of Public Health Laboratories | Sarah Schmedes, Jason Blanton |
| EPI_ISL_510526 | Biological prevention, army | Biological prevention, army | Seadawy, M.G., Shamel,M.D., Harty,B.S., Elhoseny,M.M. and Gad,A.F. |
| EPI_ISL_510532 | Biological prevention, army | Biological prevention, army | Seadawy,M.G., ELnabrawy,H.A., Shamel,M.D., Elhoseiny,M.F., Gad,A.F., Hassan,W.A., Raouf,A.A., Harty,B.E., ElGohary,A.A., Karam,M.A., Amer,k.E., Elnakeeb,M.A., Elnagdy,T.A., Ali,M.A., Kandeil,A.M. and Soliman,Y.A. |
| EPI_ISL_512107, EPI_ISL_512547, EPI_ISL_512549, EPI_ISL_512552, EPI_ISL_512555, EPI_ISL_512556, EPI_ISL_512557, EPI_ISL_512558, EPI_ISL_512560, EPI_ISL_512561, EPI_ISL_512562, EPI_ISL_512563, EPI_ISL_512564, EPI_ISL_512565, EPI_ISL_512566, EPI_ISL_512567, EPI_ISL_512568, EPI_ISL_512571, EPI_ISL_512572, EPI_ISL_512573, EPI_ISL_512575, EPI_ISL_512578, EPI_ISL_512582, EPI_ISL_512590, EPI_ISL_512594, EPI_ISL_512595, EPI_ISL_514140, EPI_ISL_514143, EPI_ISL_514145, EPI_ISL_514146, EPI_ISL_514152, EPI_ISL_514153, EPI_ISL_514155, EPI_ISL_514156, EPI_ISL_514157, EPI_ISL_514159, EPI_ISL_514160, EPI_ISL_514161, EPI_ISL_514162, EPI_ISL_514163, EPI_ISL_514168, EPI_ISL_514169, EPI_ISL_514171, EPI_ISL_514174, EPI_ISL_514176, EPI_ISL_514177, EPI_ISL_514178, EPI_ISL_514179, EPI_ISL_514180, EPI_ISL_514181, EPI_ISL_514183, EPI_ISL_514184, EPI_ISL_514185, EPI_ISL_514186, EPI_ISL_514187, EPI_ISL_514188, EPI_ISL_514189, EPI_ISL_514194, EPI_ISL_514197, EPI_ISL_514200, EPI_ISL_514201, EPI_ISL_514202, EPI_ISL_514203, EPI_ISL_514204, EPI_ISL_514205, EPI_ISL_514207, EPI_ISL_514211, EPI_ISL_514212, EPI_ISL_514213, EPI_ISL_514214, EPI_ISL_514216, EPI_ISL_514219, EPI_ISL_514220, EPI_ISL_514221, EPI_ISL_514222, EPI_ISL_514223, EPI_ISL_514224, EPI_ISL_514225 | | | |
| see above | Florida Bureau of Public Health Laboratories | Florida Bureau of Public Health Laboratories | Sarah Schmedes, Jason Blanton |
| EPI_ISL_514611 | GMERS Medical College and Hospital, Gandhinagar | Gujarat Biotechnology Research Centre | Seema Bhatt, Gaurishankar Shrimali, Bhavesh Modi, Bharti Rajani, Apurvasinh Puvar, Janvi Raval, Zarna Patel, Monika Gandhi, Pinal Trivedi, Maharshi Pandya, Nidhi Patel, Nitin Savaliya, Raghawendra Kumar, Dinesh Kumar, Zuber Saiyed, Komal Patel, Labdhi Pandya, Afzal Ansari, Nikha Trivedi, Pranay Shah, Kamlesh J Upadhyay, Sanjay Kapadia, R D Dixit, A M Kadri, Harsh Bakshi, Chaitanya Joshi, Madhvi Joshi |
| EPI_ISL_514753 | Yaftabad Hospital, COVID Lab Center | University of Tabriz | Shahabzadeh,Z., Hosseinzadeh Gharajeh,N., Hashemian,S.M. and Barati,O. |
| EPI_ISL_517772, EPI_ISL_517777, EPI_ISL_517778, EPI_ISL_517780, EPI_ISL_517785, EPI_ISL_517788, EPI_ISL_517790, EPI_ISL_517791, EPI_ISL_517792, EPI_ISL_517794, EPI_ISL_517795, EPI_ISL_517801, EPI_ISL_517802, EPI_ISL_517803, EPI_ISL_517805, EPI_ISL_517807, EPI_ISL_517808, EPI_ISL_517812, EPI_ISL_517813, EPI_ISL_517814, EPI_ISL_517815, EPI_ISL_517816, EPI_ISL_517817, EPI_ISL_517823, EPI_ISL_517824, EPI_ISL_517825, EPI_ISL_517827, EPI_ISL_517828, EPI_ISL_517829, EPI_ISL_517830, EPI_ISL_517831, EPI_ISL_517832, EPI_ISL_517833, EPI_ISL_517834, EPI_ISL_517835, EPI_ISL_517836, EPI_ISL_517837, EPI_ISL_517838, EPI_ISL_517839, EPI_ISL_517840, EPI_ISL_517841, EPI_ISL_517842, EPI_ISL_517843, EPI_ISL_517844, EPI_ISL_517845, EPI_ISL_517846, EPI_ISL_517847, EPI_ISL_517848, EPI_ISL_517849, EPI_ISL_517850, EPI_ISL_517851, EPI_ISL_517852, EPI_ISL_517853, EPI_ISL_517854, EPI_ISL_517855, EPI_ISL_517856, EPI_ISL_517857, EPI_ISL_517858, EPI_ISL_517859, EPI_ISL_517860, EPI_ISL_517861, EPI_ISL_517862, EPI_ISL_517863, EPI_ISL_517864, EPI_ISL_517865, EPI_ISL_517866, EPI_ISL_517867, EPI_ISL_517868, EPI_ISL_517869, EPI_ISL_517870, EPI_ISL_517871, EPI_ISL_517872, EPI_ISL_517873, EPI_ISL_517874, EPI_ISL_517875, EPI_ISL_517876, EPI_ISL_517877, EPI_ISL_517878, EPI_ISL_517879, EPI_ISL_517880, EPI_ISL_517881, EPI_ISL_517882, EPI_ISL_517883, EPI_ISL_517884, EPI_ISL_517885, EPI_ISL_517886, EPI_ISL_517887, EPI_ISL_517888, EPI_ISL_517889, EPI_ISL_517890, EPI_ISL_517891, EPI_ISL_517892, EPI_ISL_517893, EPI_ISL_517894, EPI_ISL_517895, EPI_ISL_517896, EPI_ISL_517897, EPI_ISL_517898, EPI_ISL_517899, EPI_ISL_517900, EPI_ISL_517901, EPI_ISL_517902, EPI_ISL_517903, EPI_ISL_517904, EPI_ISL_517905, EPI_ISL_517906, EPI_ISL_517907, EPI_ISL_517908, EPI_ISL_517909, EPI_ISL_517910, EPI_ISL_517911, EPI_ISL_517912, EPI_ISL_517913, EPI_ISL_517914, EPI_ISL_517915, EPI_ISL_517916, EPI_ISL_517917, EPI_ISL_517918, EPI_ISL_517919, EPI_ISL_517920, EPI_ISL_517921, EPI_ISL_517922, EPI_ISL_517923, EPI_ISL_517924, EPI_ISL_517925, EPI_ISL_517926, EPI_ISL_517927, EPI_ISL_517928, EPI_ISL_517929, EPI_ISL_517930, EPI_ISL_517931, EPI_ISL_517932, EPI_ISL_517933, EPI_ISL_517934, EPI_ISL_517936, EPI_ISL_517937, EPI_ISL_517938, EPI_ISL_517939, EPI_ISL_517940, EPI_ISL_517941, EPI_ISL_517942, EPI_ISL_517943, EPI_ISL_517944, EPI_ISL_517945, EPI_ISL_517946, EPI_ISL_517947, EPI_ISL_517948, EPI_ISL_517949, EPI_ISL_517950, EPI_ISL_517951, EPI_ISL_517952, EPI_ISL_517953, EPI_ISL_517954, EPI_ISL_517955, EPI_ISL_517956, EPI_ISL_517957 | | | |
| see above | Florida Bureau of Public Health Laboratories | Florida Bureau of Public Health Laboratories | Sarah Schmedes, Jason Blanton |
| EPI_ISL_518063, EPI_ISL_518064, EPI_ISL_518065, EPI_ISL_518066, EPI_ISL_518067, EPI_ISL_518068, EPI_ISL_518069, EPI_ISL_518070, EPI_ISL_518071, EPI_ISL_518072, EPI_ISL_518073, EPI_ISL_518074, EPI_ISL_518075, EPI_ISL_518076, EPI_ISL_518077, EPI_ISL_518078, EPI_ISL_518079, EPI_ISL_518080, EPI_ISL_518093, EPI_ISL_518094, EPI_ISL_518095, EPI_ISL_518096, EPI_ISL_518097, EPI_ISL_518231, EPI_ISL_518232, EPI_ISL_518233, EPI_ISL_518234, EPI_ISL_518235, EPI_ISL_518236, EPI_ISL_518237, EPI_ISL_518241, EPI_ISL_518242, EPI_ISL_518243, EPI_ISL_518250, EPI_ISL_518251, EPI_ISL_518252, EPI_ISL_518253, EPI_ISL_518254, EPI_ISL_518255, EPI_ISL_518256, EPI_ISL_518257, EPI_ISL_518258, EPI_ISL_518259, EPI_ISL_518260, EPI_ISL_518261, EPI_ISL_518262, EPI_ISL_518263, EPI_ISL_518264, EPI_ISL_518265, EPI_ISL_518266, EPI_ISL_518267, EPI_ISL_518268, EPI_ISL_518269, EPI_ISL_518270, EPI_ISL_518271, EPI_ISL_518272, EPI_ISL_518273, EPI_ISL_518274, EPI_ISL_518275, EPI_ISL_518276, EPI_ISL_518277, EPI_ISL_518278, EPI_ISL_518279, EPI_ISL_518280, EPI_ISL_518281, EPI_ISL_518282, EPI_ISL_518283, EPI_ISL_518285, EPI_ISL_518286, EPI_ISL_518287, EPI_ISL_518288, EPI_ISL_518289, EPI_ISL_518290, EPI_ISL_518292, EPI_ISL_518293, EPI_ISL_518294, EPI_ISL_518295, EPI_ISL_518296, EPI_ISL_518297, EPI_ISL_518298, EPI_ISL_518299, EPI_ISL_518300, EPI_ISL_518301, EPI_ISL_518302, EPI_ISL_518303, EPI_ISL_518304, EPI_ISL_518305, EPI_ISL_518306, EPI_ISL_518307, EPI_ISL_518308, EPI_ISL_518309, EPI_ISL_518310, EPI_ISL_518311, EPI_ISL_518312, EPI_ISL_518313, EPI_ISL_518314, EPI_ISL_518315, EPI_ISL_518316, EPI_ISL_518317, EPI_ISL_518318, EPI_ISL_518319, EPI_ISL_518320, EPI_ISL_518321, EPI_ISL_518322, EPI_ISL_518323, EPI_ISL_518324, EPI_ISL_518325, EPI_ISL_518326, EPI_ISL_518327, EPI_ISL_518328, EPI_ISL_518329, EPI_ISL_518330, EPI_ISL_518331, EPI_ISL_518332, EPI_ISL_518333, EPI_ISL_518334, EPI_ISL_518335, EPI_ISL_518336, EPI_ISL_518337, EPI_ISL_518339, EPI_ISL_518340, EPI_ISL_518341, EPI_ISL_518342, EPI_ISL_518343, EPI_ISL_518344, EPI_ISL_518345, EPI_ISL_518346, EPI_ISL_518347, EPI_ISL_518348, EPI_ISL_518349, EPI_ISL_518350, EPI_ISL_518351, EPI_ISL_518352, EPI_ISL_518353, EPI_ISL_518354, EPI_ISL_518355, EPI_ISL_518356, EPI_ISL_518357, EPI_ISL_518358, EPI_ISL_518359, EPI_ISL_518360, EPI_ISL_518361, EPI_ISL_518362, EPI_ISL_518363, EPI_ISL_518364, EPI_ISL_518365, EPI_ISL_518366, EPI_ISL_518367, EPI_ISL_518368, EPI_ISL_518369, EPI_ISL_518370, EPI_ISL_518371, EPI_ISL_518372, EPI_ISL_518373, EPI_ISL_518374, EPI_ISL_518375, EPI_ISL_518376, EPI_ISL_518377, EPI_ISL_518378, EPI_ISL_518379, EPI_ISL_518380, EPI_ISL_518383, EPI_ISL_518384, EPI_ISL_518385, EPI_ISL_518386, EPI_ISL_518387, EPI_ISL_518402, EPI_ISL_518403, EPI_ISL_518404, EPI_ISL_518405, EPI_ISL_518406, EPI_ISL_518407, EPI_ISL_518408, EPI_ISL_518409, EPI_ISL_518410, EPI_ISL_518411, EPI_ISL_518412, EPI_ISL_518413, EPI_ISL_518414, EPI_ISL_518415, | | | |

EPI_ISL_518416, EPI_ISL_518417, EPI_ISL_518418, EPI_ISL_518419, EPI_ISL_518420, EPI_ISL_518421, EPI_ISL_518422, EPI_ISL_518423, EPI_ISL_518472, EPI_ISL_518473, EPI_ISL_518474, EPI_ISL_518475, EPI_ISL_518476, EPI_ISL_518477, EPI_ISL_518478, EPI_ISL_518479, EPI_ISL_518480, EPI_ISL_518481, EPI_ISL_518482, EPI_ISL_518483, EPI_ISL_518484, EPI_ISL_518485, EPI_ISL_518486, EPI_ISL_518487, EPI_ISL_518488, EPI_ISL_518489, EPI_ISL_518490, EPI_ISL_518491, EPI_ISL_518492, EPI_ISL_518493, EPI_ISL_518494, EPI_ISL_518495, EPI_ISL_518496, EPI_ISL_518497, EPI_ISL_518498, EPI_ISL_518499, EPI_ISL_518500, EPI_ISL_518501, EPI_ISL_518502, EPI_ISL_518503, EPI_ISL_518504, EPI_ISL_518505, EPI_ISL_518507, EPI_ISL_518508, EPI_ISL_518509, EPI_ISL_518510, EPI_ISL_518511, EPI_ISL_518512, EPI_ISL_518513, EPI_ISL_518514, EPI_ISL_518515, EPI_ISL_518516, EPI_ISL_518517, EPI_ISL_518518, EPI_ISL_518519, EPI_ISL_518520, EPI_ISL_518521, EPI_ISL_518522, EPI_ISL_518523, EPI_ISL_518524, EPI_ISL_518525, EPI_ISL_518526, EPI_ISL_518527, EPI_ISL_518528, EPI_ISL_518529, EPI_ISL_518530, EPI_ISL_518531, EPI_ISL_518532, EPI_ISL_518533, EPI_ISL_518534, EPI_ISL_518535, EPI_ISL_518536, EPI_ISL_518537, EPI_ISL_518538, EPI_ISL_518539, EPI_ISL_518540, EPI_ISL_518541, EPI_ISL_518542, EPI_ISL_518543, EPI_ISL_518544, EPI_ISL_518545, EPI_ISL_518546, EPI_ISL_518547, EPI_ISL_518548, EPI_ISL_518549, EPI_ISL_518550, EPI_ISL_518551, EPI_ISL_518552, EPI_ISL_518553, EPI_ISL_518554, EPI_ISL_518555, EPI_ISL_518556, EPI_ISL_518557, EPI_ISL_518559, EPI_ISL_518560, EPI_ISL_518561, EPI_ISL_518562, EPI_ISL_518563, EPI_ISL_518564, EPI_ISL_518565, EPI_ISL_518566, EPI_ISL_518567, EPI_ISL_518568, EPI_ISL_518569, EPI_ISL_518570, EPI_ISL_518571, EPI_ISL_518572, EPI_ISL_518573, EPI_ISL_518574, EPI_ISL_518575, EPI_ISL_518576, EPI_ISL_518577, EPI_ISL_518578, EPI_ISL_518579, EPI_ISL_518580, EPI_ISL_518581, EPI_ISL_518582, EPI_ISL_518583, EPI_ISL_518584, EPI_ISL_518585, EPI_ISL_518586, EPI_ISL_518588, EPI_ISL_518589, EPI_ISL_518590, EPI_ISL_518592, EPI_ISL_518593, EPI_ISL_518594, EPI_ISL_518595, EPI_ISL_518596, EPI_ISL_518597, EPI_ISL_518598, EPI_ISL_518600, EPI_ISL_518601, EPI_ISL_518602, EPI_ISL_518603, EPI_ISL_518604, EPI_ISL_518605, EPI_ISL_518606, EPI_ISL_518607, EPI_ISL_518608, EPI_ISL_518609, EPI_ISL_518610, EPI_ISL_518611, EPI_ISL_518613, EPI_ISL_518614, EPI_ISL_518619, EPI_ISL_518621, EPI_ISL_518622, EPI_ISL_518623, EPI_ISL_518624, EPI_ISL_518626, EPI_ISL_518627, EPI_ISL_518628, EPI_ISL_518629, EPI_ISL_518630, EPI_ISL_518631, EPI_ISL_518632, EPI_ISL_518633, EPI_ISL_518634, EPI_ISL_518635, EPI_ISL_518636, EPI_ISL_518637, EPI_ISL_518638, EPI_ISL_518639, EPI_ISL_518640, EPI_ISL_518641, EPI_ISL_518642, EPI_ISL_518643, EPI_ISL_518644, EPI_ISL_518645, EPI_ISL_518646, EPI_ISL_518647, EPI_ISL_518648, EPI_ISL_518649, EPI_ISL_518650

| | | | |
|---|---|---|---|
| see above | Microbiological Diagnostic Unit - Public Health Laboratory (MDU-PHL) | MDU-PHL | Seemann T., Schultz M., Sait, M., Sherry, N. |
| EPI_ISL_518651 | Microbiological Diagnostic Unit - Public Health Laboratory (MDU-PHL) | Microbiological Diagnostic Unit - Public Health Laboratory (MDU-PHL) | Seemann T., Schultz M., Sait, M., Sherry, N. |

EPI_ISL_518652, EPI_ISL_518653, EPI_ISL_518654, EPI_ISL_518655, EPI_ISL_518656, EPI_ISL_518657, EPI_ISL_518658, EPI_ISL_518659, EPI_ISL_518660, EPI_ISL_518661, EPI_ISL_518662, EPI_ISL_518663, EPI_ISL_518664, EPI_ISL_518665, EPI_ISL_518666, EPI_ISL_518667, EPI_ISL_518668, EPI_ISL_518669, EPI_ISL_518670, EPI_ISL_518671, EPI_ISL_518672, EPI_ISL_518673, EPI_ISL_518674, EPI_ISL_518675, EPI_ISL_518676, EPI_ISL_518677, EPI_ISL_518678, EPI_ISL_518679, EPI_ISL_518680, EPI_ISL_518681, EPI_ISL_518682, EPI_ISL_518683, EPI_ISL_518684, EPI_ISL_518685, EPI_ISL_518686, EPI_ISL_518687, EPI_ISL_518688, EPI_ISL_518689, EPI_ISL_518690, EPI_ISL_518691, EPI_ISL_518692, EPI_ISL_518693, EPI_ISL_518694, EPI_ISL_518695, EPI_ISL_518696, EPI_ISL_518697, EPI_ISL_518698, EPI_ISL_518699, EPI_ISL_518700, EPI_ISL_518701, EPI_ISL_518703, EPI_ISL_518704, EPI_ISL_518705, EPI_ISL_518706, EPI_ISL_518707, EPI_ISL_518708, EPI_ISL_518711, EPI_ISL_518712, EPI_ISL_518713, EPI_ISL_518714, EPI_ISL_518715, EPI_ISL_518716, EPI_ISL_518717, EPI_ISL_518718, EPI_ISL_518719, EPI_ISL_518720, EPI_ISL_518721, EPI_ISL_518722, EPI_ISL_518725, EPI_ISL_518726, EPI_ISL_518727, EPI_ISL_518728, EPI_ISL_518729, EPI_ISL_518730, EPI_ISL_518731, EPI_ISL_518732, EPI_ISL_518733, EPI_ISL_518735, EPI_ISL_518736, EPI_ISL_518745, EPI_ISL_518746, EPI_ISL_518747, EPI_ISL_518748, EPI_ISL_518749, EPI_ISL_518750, EPI_ISL_518751, EPI_ISL_518752, EPI_ISL_518753, EPI_ISL_518754, EPI_ISL_518755, EPI_ISL_518756, EPI_ISL_518757, EPI_ISL_518758, EPI_ISL_518759, EPI_ISL_518760, EPI_ISL_518761, EPI_ISL_518762, EPI_ISL_518763, EPI_ISL_518764, EPI_ISL_518765, EPI_ISL_518766, EPI_ISL_518767, EPI_ISL_518768, EPI_ISL_518769, EPI_ISL_518770, EPI_ISL_518771, EPI_ISL_518772, EPI_ISL_518773, EPI_ISL_518774, EPI_ISL_518775, EPI_ISL_518776, EPI_ISL_518777, EPI_ISL_518778, EPI_ISL_518779, EPI_ISL_518780, EPI_ISL_518781, EPI_ISL_518782, EPI_ISL_518783, EPI_ISL_518784, EPI_ISL_518785, EPI_ISL_518786, EPI_ISL_518787, EPI_ISL_518788, EPI_ISL_518789, EPI_ISL_518790, EPI_ISL_518791, EPI_ISL_518792, EPI_ISL_518793, EPI_ISL_518794, EPI_ISL_518795, EPI_ISL_518796, EPI_ISL_518901, EPI_ISL_518902, EPI_ISL_518904, EPI_ISL_518906, EPI_ISL_518907, EPI_ISL_518908, EPI_ISL_518909, EPI_ISL_518910, EPI_ISL_518911, EPI_ISL_518912, EPI_ISL_518913, EPI_ISL_518914, EPI_ISL_518915, EPI_ISL_518917, EPI_ISL_518918, EPI_ISL_518919, EPI_ISL_518920, EPI_ISL_518921, EPI_ISL_518922, EPI_ISL_518923, EPI_ISL_518924, EPI_ISL_518925, EPI_ISL_518926, EPI_ISL_518927, EPI_ISL_518928, EPI_ISL_518929, EPI_ISL_518930, EPI_ISL_518931, EPI_ISL_518933, EPI_ISL_518939, EPI_ISL_518940, EPI_ISL_518941, EPI_ISL_518942, EPI_ISL_518943, EPI_ISL_518944, EPI_ISL_518945, EPI_ISL_518946, EPI_ISL_518947, EPI_ISL_518948, EPI_ISL_518949, EPI_ISL_518950, EPI_ISL_518951, EPI_ISL_518952, EPI_ISL_518953, EPI_ISL_518954, EPI_ISL_518955, EPI_ISL_518956, EPI_ISL_518957, EPI_ISL_518958, EPI_ISL_518959, EPI_ISL_518960, EPI_ISL_518961, EPI_ISL_518962, EPI_ISL_518963, EPI_ISL_518964, EPI_ISL_518965, EPI_ISL_518966, EPI_ISL_518967, EPI_ISL_518968, EPI_ISL_518969, EPI_ISL_518970, EPI_ISL_518971, EPI_ISL_518972, EPI_ISL_518973, EPI_ISL_518975, EPI_ISL_518977, EPI_ISL_518978, EPI_ISL_518979, EPI_ISL_518980, EPI_ISL_519052, EPI_ISL_519054, EPI_ISL_519055, EPI_ISL_519056, EPI_ISL_519057, EPI_ISL_519059, EPI_ISL_519060, EPI_ISL_519061, EPI_ISL_519062, EPI_ISL_519063, EPI_ISL_519064, EPI_ISL_519065, EPI_ISL_519066, EPI_ISL_519067, EPI_ISL_519068, EPI_ISL_519069, EPI_ISL_519070, EPI_ISL_519071, EPI_ISL_519072, EPI_ISL_519073, EPI_ISL_519074, EPI_ISL_519075, EPI_ISL_519076, EPI_ISL_519077, EPI_ISL_519078, EPI_ISL_519079, EPI_ISL_519080, EPI_ISL_519081, EPI_ISL_519082, EPI_ISL_519083, EPI_ISL_519084, EPI_ISL_519085, EPI_ISL_519086, EPI_ISL_519087, EPI_ISL_519088, EPI_ISL_519089, EPI_ISL_519090, EPI_ISL_519091, EPI_ISL_519092, EPI_ISL_519093, EPI_ISL_519094, EPI_ISL_519095, EPI_ISL_519096, EPI_ISL_519097, EPI_ISL_519098, EPI_ISL_519099, EPI_ISL_519100, EPI_ISL_519101, EPI_ISL_519102, EPI_ISL_519104, EPI_ISL_519105, EPI_ISL_519106, EPI_ISL_519107, EPI_ISL_519108, EPI_ISL_519109, EPI_ISL_519110, EPI_ISL_519111, EPI_ISL_519112, EPI_ISL_519113, EPI_ISL_519114, EPI_ISL_519115, EPI_ISL_519116, EPI_ISL_519117, EPI_ISL_519118, EPI_ISL_519119, EPI_ISL_519120, EPI_ISL_519121, EPI_ISL_519122, EPI_ISL_519123, EPI_ISL_519124, EPI_ISL_519125, EPI_ISL_519126, EPI_ISL_519127, EPI_ISL_519128, EPI_ISL_519129, EPI_ISL_519130, EPI_ISL_519131, EPI_ISL_519132, EPI_ISL_519133, EPI_ISL_519134, EPI_ISL_519135, EPI_ISL_519136, EPI_ISL_519137, EPI_ISL_519138, EPI_ISL_519139, EPI_ISL_519140, EPI_ISL_519141, EPI_ISL_519142, EPI_ISL_519143, EPI_ISL_519144, EPI_ISL_519145, EPI_ISL_519146, EPI_ISL_519147, EPI_ISL_519148, EPI_ISL_519149, EPI_ISL_519150, EPI_ISL_519151, EPI_ISL_519152, EPI_ISL_519153, EPI_ISL_519154, EPI_ISL_519155, EPI_ISL_519156, EPI_ISL_519157, EPI_ISL_519158, EPI_ISL_519159, EPI_ISL_519160, EPI_ISL_519161, EPI_ISL_519162, EPI_ISL_519163, EPI_ISL_519164, EPI_ISL_519165, EPI_ISL_519166, EPI_ISL_519168, EPI_ISL_519169, EPI_ISL_519170, EPI_ISL_519171, EPI_ISL_519173, EPI_ISL_519174, EPI_ISL_519175, EPI_ISL_519176, EPI_ISL_519177, EPI_ISL_519178, EPI_ISL_519179, EPI_ISL_519180, EPI_ISL_519181, EPI_ISL_519182, EPI_ISL_519183, EPI_ISL_519184, EPI_ISL_519185, EPI_ISL_519186, EPI_ISL_519187, EPI_ISL_519188, EPI_ISL_519189, EPI_ISL_519190, EPI_ISL_519191, EPI_ISL_519192, EPI_ISL_519193, EPI_ISL_519194, EPI_ISL_519195, EPI_ISL_519196, EPI_ISL_519197, EPI_ISL_519198, EPI_ISL_519199, EPI_ISL_519200, EPI_ISL_519201, EPI_ISL_519202, EPI_ISL_519203, EPI_ISL_519204, EPI_ISL_519205, EPI_ISL_519206, EPI_ISL_519207, EPI_ISL_519208, EPI_ISL_519209, EPI_ISL_519210, EPI_ISL_519211, EPI_ISL_519212, EPI_ISL_519213, EPI_ISL_519214, EPI_ISL_519215, EPI_ISL_519216, EPI_ISL_519217, EPI_ISL_519218, EPI_ISL_519219, EPI_ISL_519220, EPI_ISL_519221, EPI_ISL_519222, EPI_ISL_519223, EPI_ISL_519226, EPI_ISL_519228, EPI_ISL_519230, EPI_ISL_519231, EPI_ISL_519232, EPI_ISL_519233, EPI_ISL_519234, EPI_ISL_519235, EPI_ISL_519236, EPI_ISL_519238, EPI_ISL_519239, EPI_ISL_519240, EPI_ISL_519241, EPI_ISL_519242, EPI_ISL_519243, EPI_ISL_519244, EPI_ISL_519245, EPI_ISL_519246, EPI_ISL_519247, EPI_ISL_519248, EPI_ISL_519249, EPI_ISL_519250, EPI_ISL_519251, EPI_ISL_519252, EPI_ISL_519253, EPI_ISL_519254, EPI_ISL_519255, EPI_ISL_519327, EPI_ISL_519328, EPI_ISL_519329, EPI_ISL_519330, EPI_ISL_519333, EPI_ISL_519334, EPI_ISL_519335, EPI_ISL_519336, EPI_ISL_519337, EPI_ISL_519338, EPI_ISL_519339, EPI_ISL_519340, EPI_ISL_519341, EPI_ISL_519342, EPI_ISL_519343, EPI_ISL_519344, EPI_ISL_519345, EPI_ISL_519346, EPI_ISL_519347, EPI_ISL_519348, EPI_ISL_519349, EPI_ISL_519350, EPI_ISL_519351, EPI_ISL_519352, EPI_ISL_519353, EPI_ISL_519354, EPI_ISL_519355, EPI_ISL_519356, EPI_ISL_519357, EPI_ISL_519358, EPI_ISL_519359, EPI_ISL_519360, EPI_ISL_519361, EPI_ISL_519362, EPI_ISL_519363, EPI_ISL_519364, EPI_ISL_519365, EPI_ISL_519366, EPI_ISL_519367, EPI_ISL_519368, EPI_ISL_519369, EPI_ISL_519371, EPI_ISL_519372, EPI_ISL_519373, EPI_ISL_519374, EPI_ISL_519375, EPI_ISL_519376, EPI_ISL_519377, EPI_ISL_519378, EPI_ISL_519379, EPI_ISL_519380, EPI_ISL_519381, EPI_ISL_519382, EPI_ISL_519383, EPI_ISL_519384, EPI_ISL_519386, EPI_ISL_519387, EPI_ISL_519388, EPI_ISL_519389, EPI_ISL_519390, EPI_ISL_519391, EPI_ISL_519392, EPI_ISL_519393, EPI_ISL_519394, EPI_ISL_519395, EPI_ISL_519396, EPI_ISL_519397, EPI_ISL_519398, EPI_ISL_519399, EPI_ISL_519400, EPI_ISL_519401, EPI_ISL_519402, EPI_ISL_519403, EPI_ISL_519404, EPI_ISL_519405, EPI_ISL_519406, EPI_ISL_519407, EPI_ISL_519408, EPI_ISL_519409, EPI_ISL_519410, EPI_ISL_519411, EPI_ISL_519412, EPI_ISL_519413, EPI_ISL_519414, EPI_ISL_519415, EPI_ISL_519416, EPI_ISL_519417, EPI_ISL_519418, EPI_ISL_519419, EPI_ISL_519420, EPI_ISL_519421, EPI_ISL_519422, EPI_ISL_519423, EPI_ISL_519424, EPI_ISL_519425, EPI_ISL_519426, EPI_ISL_519428, EPI_ISL_519429, EPI_ISL_519430, EPI_ISL_519432, EPI_ISL_519433, EPI_ISL_519434, EPI_ISL_519435, EPI_ISL_519436, EPI_ISL_519437, EPI_ISL_519438, EPI_ISL_519439, EPI_ISL_519440, EPI_ISL_519441, EPI_ISL_519442, EPI_ISL_519443, EPI_ISL_519445, EPI_ISL_519446, EPI_ISL_519449, EPI_ISL_519450, EPI_ISL_519451, EPI_ISL_519453, EPI_ISL_519454, EPI_ISL_519455, EPI_ISL_519456, EPI_ISL_519457, EPI_ISL_519458, EPI_ISL_519459, EPI_ISL_519460, EPI_ISL_519461, EPI_ISL_519462, EPI_ISL_519463, EPI_ISL_519464, EPI_ISL_519465, EPI_ISL_519466, EPI_ISL_519467, EPI_ISL_519468, EPI_ISL_519469, EPI_ISL_519470, EPI_ISL_519471, EPI_ISL_519472, EPI_ISL_519473, EPI_ISL_519474, EPI_ISL_519475, EPI_ISL_519476, EPI_ISL_519477, EPI_ISL_519478, EPI_ISL_519479, EPI_ISL_519480, EPI_ISL_519481, EPI_ISL_519482, EPI_ISL_519483, EPI_ISL_519484, EPI_ISL_519485, EPI_ISL_519486, EPI_ISL_519487, EPI_ISL_519488, EPI_ISL_519489, EPI_ISL_519490, EPI_ISL_519491, EPI_ISL_519492, EPI_ISL_519493, EPI_ISL_519494, EPI_ISL_519495, EPI_ISL_519496, EPI_ISL_519498, EPI_ISL_519501, EPI_ISL_519503, EPI_ISL_519504, EPI_ISL_519505, EPI_ISL_519506, EPI_ISL_519507, EPI_ISL_519508, EPI_ISL_519509, EPI_ISL_519511, EPI_ISL_519512, EPI_ISL_519513, EPI_ISL_519515, EPI_ISL_519517, EPI_ISL_519518, EPI_ISL_519522, EPI_ISL_519523, EPI_ISL_519524, EPI_ISL_519525, EPI_ISL_519526, EPI_ISL_519527, EPI_ISL_519528, EPI_ISL_519529, EPI_ISL_519530, EPI_ISL_519531, EPI_ISL_519532, EPI_ISL_519533, EPI_ISL_519534, EPI_ISL_519535, EPI_ISL_519536, EPI_ISL_519537, EPI_ISL_519538, EPI_ISL_519539, EPI_ISL_519540, EPI_ISL_519541, EPI_ISL_519542, EPI_ISL_519543, EPI_ISL_519547, EPI_ISL_519548, EPI_ISL_519549, EPI_ISL_519550, EPI_ISL_519551, EPI_ISL_519552, EPI_ISL_519553, EPI_ISL_519554, EPI_ISL_519555, EPI_ISL_519556, EPI_ISL_519557, EPI_ISL_519558, EPI_ISL_519559, EPI_ISL_519561, EPI_ISL_519562, EPI_ISL_519563, EPI_ISL_519564, EPI_ISL_519565, EPI_ISL_519566, EPI_ISL_519567, EPI_ISL_519568, EPI_ISL_519569, EPI_ISL_519571, EPI_ISL_519572, EPI_ISL_519573, EPI_ISL_519574, EPI_ISL_519575, EPI_ISL_519576, EPI_ISL_519577, EPI_ISL_519578, EPI_ISL_519579, EPI_ISL_519580, EPI_ISL_519581, EPI_ISL_519582, EPI_ISL_519583, EPI_ISL_519584, EPI_ISL_519585, EPI_ISL_519586, EPI_ISL_519587, EPI_ISL_519588, EPI_ISL_519589, EPI_ISL_519590, EPI_ISL_519591, EPI_ISL_519592, EPI_ISL_519593, EPI_ISL_519594, EPI_ISL_519595, EPI_ISL_519596, EPI_ISL_519597, EPI_ISL_519598, EPI_ISL_519599, EPI_ISL_519600, EPI_ISL_519601, EPI_ISL_519602, EPI_ISL_519603, EPI_ISL_519604, EPI_ISL_519605, EPI_ISL_519607, EPI_ISL_519608, EPI_ISL_519609, EPI_ISL_519610, EPI_ISL_519611, EPI_ISL_519612, EPI_ISL_519613, EPI_ISL_519614, EPI_ISL_519615, EPI_ISL_519616, EPI_ISL_519617, EPI_ISL_519618, EPI_ISL_519619, EPI_ISL_519620, EPI_ISL_519621, EPI_ISL_519622, EPI_ISL_519623, EPI_ISL_519624, EPI_ISL_519625, EPI_ISL_519626, EPI_ISL_519627, EPI_ISL_519628, EPI_ISL_519629, EPI_ISL_519630, EPI_ISL_519631, EPI_ISL_519632, EPI_ISL_519633, EPI_ISL_519634, EPI_ISL_519635, EPI_ISL_519636, EPI_ISL_519637, EPI_ISL_519638, EPI_ISL_519639, EPI_ISL_519640, EPI_ISL_519641, EPI_ISL_519642, EPI_ISL_519643, EPI_ISL_519645, EPI_ISL_519646, EPI_ISL_519647, EPI_ISL_519648, EPI_ISL_519649, EPI_ISL_519650, EPI_ISL_519651, EPI_ISL_519652, EPI_ISL_519653, EPI_ISL_519654, EPI_ISL_519655, EPI_ISL_519656, EPI_ISL_519657, EPI_ISL_519658, EPI_ISL_519659, EPI_ISL_519660, EPI_ISL_519661, EPI_ISL_519662, EPI_ISL_519663, EPI_ISL_519664, EPI_ISL_519665, EPI_ISL_519666, EPI_ISL_519667, EPI_ISL_519668, EPI_ISL_519669, EPI_ISL_519670, EPI_ISL_519671, EPI_ISL_519672, EPI_ISL_519673, EPI_ISL_519674, EPI_ISL_519675, EPI_ISL_519676, EPI_ISL_519677, EPI_ISL_519678, EPI_ISL_519679, EPI_ISL_519680, EPI_ISL_519681, EPI_ISL_519682, EPI_ISL_519683, EPI_ISL_519684, EPI_ISL_519685, EPI_ISL_519686, EPI_ISL_519687, EPI_ISL_519688, EPI_ISL_519689, EPI_ISL_519690, EPI_ISL_519691, EPI_ISL_519692, EPI_ISL_519693, EPI_ISL_519694, EPI_ISL_519695, EPI_ISL_519696, EPI_ISL_519697, EPI_ISL_519698, EPI_ISL_519699, EPI_ISL_519700, EPI_ISL_519701, EPI_ISL_519702, EPI_ISL_519703, EPI_ISL_519704, EPI_ISL_519705, EPI_ISL_519706, EPI_ISL_519707, EPI_ISL_519708, EPI_ISL_519709, EPI_ISL_519710, EPI_ISL_519711, EPI_ISL_519712, EPI_ISL_519713, EPI_ISL_519714, EPI_ISL_519715, EPI_ISL_519716, EPI_ISL_519717, EPI_ISL_519718, EPI_ISL_519719, EPI_ISL_519720, EPI_ISL_519721, EPI_ISL_519722, EPI_ISL_519723, EPI_ISL_519724, EPI_ISL_519725, EPI_ISL_519726, EPI_ISL_519727, EPI_ISL_519728, EPI_ISL_519729, EPI_ISL_519730, EPI_ISL_519731, EPI_ISL_519732, EPI_ISL_519733, EPI_ISL_519734, EPI_ISL_519735, EPI_ISL_519736, EPI_ISL_519737, EPI_ISL_519738, EPI_ISL_519739, EPI_ISL_519740, EPI_ISL_519741, EPI_ISL_519742, EPI_ISL_519743, EPI_ISL_519744, EPI_ISL_519745, EPI_ISL_519746, EPI_ISL_519747, EPI_ISL_519748, EPI_ISL_519749, EPI_ISL_519750, EPI_ISL_519751, EPI_ISL_519752, EPI_ISL_519753, EPI_ISL_519754, EPI_ISL_519755, EPI_ISL_519756, EPI_ISL_519757, EPI_ISL_519758, EPI_ISL_519759, EPI_ISL_519760, EPI_ISL_519761, EPI_ISL_519762, EPI_ISL_519763, EPI_ISL_519764, EPI_ISL_519765, EPI_ISL_519766, EPI_ISL_519767, EPI_ISL_519770, EPI_ISL_519771, EPI_ISL_519773, EPI_ISL_519780, EPI_ISL_519781, EPI_ISL_519782, EPI_ISL_519783, EPI_ISL_519784, EPI_ISL_519785, EPI_ISL_519786, EPI_ISL_519787, EPI_ISL_519788, EPI_ISL_519789, EPI_ISL_519790, EPI_ISL_519791, EPI_ISL_519792, EPI_ISL_519793, EPI_ISL_519794, EPI_ISL_519795, EPI_ISL_519797, EPI_ISL_519798, EPI_ISL_519799, EPI_ISL_519800, EPI_ISL_519801, EPI_ISL_519802, EPI_ISL_519803, EPI_ISL_519804, EPI_ISL_519805, EPI_ISL_519806, EPI_ISL_519807, EPI_ISL_519808, EPI_ISL_519810, EPI_ISL_519811, EPI_ISL_519812, EPI_ISL_519813, EPI_ISL_519815, EPI_ISL_519816, EPI_ISL_519817, EPI_ISL_519818, EPI_ISL_519819, EPI_ISL_519820, EPI_ISL_519821, EPI_ISL_519822, EPI_ISL_519823, EPI_ISL_519824, EPI_ISL_519825, EPI_ISL_519826, EPI_ISL_519827, EPI_ISL_519828, EPI_ISL_519829, EPI_ISL_519830, EPI_ISL_519831, EPI_ISL_519832, EPI_ISL_519833, EPI_ISL_519834, EPI_ISL_519835, EPI_ISL_519836, EPI_ISL_519837, EPI_ISL_519838, EPI_ISL_519839, EPI_ISL_519840, EPI_ISL_519841, EPI_ISL_519842, EPI_ISL_519843, EPI_ISL_519844, EPI_ISL_519845, EPI_ISL_519846, EPI_ISL_519847, EPI_ISL_519848, EPI_ISL_519849, EPI_ISL_519850, EPI_ISL_519851, EPI_ISL_519852, EPI_ISL_519853, EPI_ISL_519854, EPI_ISL_519855, EPI_ISL_519856, EPI_ISL_519857, EPI_ISL_519858, EPI_ISL_519859, EPI_ISL_519860, EPI_ISL_519861, EPI_ISL_519862, EPI_ISL_519863, EPI_ISL_519864, EPI_ISL_519865, EPI_ISL_519866, EPI_ISL_519867, EPI_ISL_519868, EPI_ISL_519869, EPI_ISL_519870, EPI_ISL_519871, EPI_ISL_519872, EPI_ISL_519873, EPI_ISL_519874, EPI_ISL_519875, EPI_ISL_519876, EPI_ISL_519877, EPI_ISL_519878, EPI_ISL_519879, EPI_ISL_519880, EPI_ISL_519881, EPI_ISL_519882, EPI_ISL_519883, EPI_ISL_519884, EPI_ISL_519885, EPI_ISL_519886, EPI_ISL_519887, EPI_ISL_519888, EPI_ISL_519889, EPI_ISL_519890, EPI_ISL_519891, EPI_ISL_519892, EPI_ISL_519893, EPI_ISL_519894

| | | | |
|---|---|---|---|
| see above | Microbiological Diagnostic Unit - Public Health Laboratory (MDU-PHL) | MDU-PHL | Seemann T., Schultz M., Sait, M., Sherry, N. |
| EPI_ISL_519895 | Microbiological Diagnostic Unit - Public Health Laboratory (MDU-PHL) | Microbiological Diagnostic Unit - Public Health Laboratory (MDU-PHL) | Seemann T., Schultz M., Sait, M., Sherry, N. |

EPI_ISL_519896, EPI_ISL_519897, EPI_ISL_519898, EPI_ISL_519899, EPI_ISL_519900, EPI_ISL_519901, EPI_ISL_519902, EPI_ISL_519903, EPI_ISL_519904, EPI_ISL_519905, EPI_ISL_519906, EPI_ISL_519907, EPI_ISL_519908, EPI_ISL_519909, EPI_ISL_519910, EPI_ISL_519911, EPI_ISL_519913, EPI_ISL_519914, EPI_ISL_519915, EPI_ISL_519916, EPI_ISL_519917, EPI_ISL_519918, EPI_ISL_519919, EPI_ISL_519920, EPI_ISL_519921, EPI_ISL_519922, EPI_ISL_519923, EPI_ISL_519924, EPI_ISL_519925, EPI_ISL_519926, EPI_ISL_519927, EPI_ISL_519928, EPI_ISL_519929, EPI_ISL_519930, EPI_ISL_519931, EPI_ISL_519932, EPI_ISL_519933, EPI_ISL_519934, EPI_ISL_519935, EPI_ISL_519936, EPI_ISL_519937, EPI_ISL_519938, EPI_ISL_519939, EPI_ISL_519940, EPI_ISL_519941, EPI_ISL_519942, EPI_ISL_519943, EPI_ISL_519944, EPI_ISL_519945, EPI_ISL_519946, EPI_ISL_519947, EPI_ISL_519951, EPI_ISL_519952, EPI_ISL_519953, EPI_ISL_519954, EPI_ISL_519955, EPI_ISL_519956, EPI_ISL_519957, EPI_ISL_519958, EPI_ISL_519963, EPI_ISL_519964, EPI_ISL_519965, EPI_ISL_519966, EPI_ISL_519967, EPI_ISL_519968, EPI_ISL_519969, EPI_ISL_519970, EPI_ISL_519971, EPI_ISL_519972, EPI_ISL_519973, EPI_ISL_519974, EPI_ISL_519975, EPI_ISL_519976, EPI_ISL_519977, EPI_ISL_519978, EPI_ISL_519979, EPI_ISL_519980, EPI_ISL_519983, EPI_ISL_519984, EPI_ISL_519985, EPI_ISL_519986, EPI_ISL_519987, EPI_ISL_519988, EPI_ISL_519989, EPI_ISL_519990, EPI_ISL_519992, EPI_ISL_519993, EPI_ISL_519994, EPI_ISL_519995, EPI_ISL_519996, EPI_ISL_519997, EPI_ISL_519998, EPI_ISL_519999, EPI_ISL_520000, EPI_ISL_520002, EPI_ISL_520003, EPI_ISL_520004, EPI_ISL_520005, EPI_ISL_520007, EPI_ISL_520008, EPI_ISL_520009, EPI_ISL_520010, EPI_ISL_520011, EPI_ISL_520012, EPI_ISL_520013, EPI_ISL_520014, EPI_ISL_520015, EPI_ISL_520016, EPI_ISL_520017, EPI_ISL_520018, EPI_ISL_520019, EPI_ISL_520020, EPI_ISL_520021, EPI_ISL_520022, EPI_ISL_520023, EPI_ISL_520024, EPI_ISL_520025, EPI_ISL_520026, EPI_ISL_520027, EPI_ISL_520028, EPI_ISL_520029, EPI_ISL_520030, EPI_ISL_520031, EPI_ISL_520041, EPI_ISL_520042, EPI_ISL_520043, EPI_ISL_520044, EPI_ISL_520045, EPI_ISL_520046, EPI_ISL_520047, EPI_ISL_520048, EPI_ISL_520049, EPI_ISL_520050, EPI_ISL_520051, EPI_ISL_520052, EPI_ISL_520053, EPI_ISL_520054, EPI_ISL_520055, EPI_ISL_520057, EPI_ISL_520058, EPI_ISL_520059, EPI_ISL_520061, EPI_ISL_520062, EPI_ISL_520063, EPI_ISL_520066, EPI_ISL_520067, EPI_ISL_520068, EPI_ISL_520069, EPI_ISL_520070, EPI_ISL_520072, EPI_ISL_520073, EPI_ISL_520074, EPI_ISL_520075, EPI_ISL_520077, EPI_ISL_520079, EPI_ISL_520080, EPI_ISL_520081, EPI_ISL_520082, EPI_ISL_520083, EPI_ISL_520085, EPI_ISL_520086, EPI_ISL_520087, EPI_ISL_520089, EPI_ISL_520090, EPI_ISL_520091, EPI_ISL_520092, EPI_ISL_520093, EPI_ISL_520094, EPI_ISL_520095, EPI_ISL_520096, EPI_ISL_520097, EPI_ISL_520098, EPI_ISL_520099, EPI_ISL_520100, EPI_ISL_520101, EPI_ISL_520102, EPI_ISL_520103, EPI_ISL_520104, EPI_ISL_520105, EPI_ISL_520106, EPI_ISL_520107, EPI_ISL_520108, EPI_ISL_520109, EPI_ISL_520110, EPI_ISL_520111, EPI_ISL_520112, EPI_ISL_520113, EPI_ISL_520114, EPI_ISL_520115, EPI_ISL_520116, EPI_ISL_520117, EPI_ISL_520118, EPI_ISL_520119, EPI_ISL_520121, EPI_ISL_520122, EPI_ISL_520123, EPI_ISL_520124, EPI_ISL_520125, EPI_ISL_520126, EPI_ISL_520127, EPI_ISL_520128, EPI_ISL_520129, EPI_ISL_520130, EPI_ISL_520131, EPI_ISL_520132, EPI_ISL_520133, EPI_ISL_520134, EPI_ISL_520135, EPI_ISL_520136, EPI_ISL_520137, EPI_ISL_520138, EPI_ISL_520139, EPI_ISL_520140, EPI_ISL_520141, EPI_ISL_520142, EPI_ISL_520143, EPI_ISL_520144, EPI_ISL_520145, EPI_ISL_520146, EPI_ISL_520147, EPI_ISL_520148, EPI_ISL_520149, EPI_ISL_520150, EPI_ISL_520151, EPI_ISL_520152, EPI_ISL_520153, EPI_ISL_520154, EPI_ISL_520155, EPI_ISL_520156, EPI_ISL_520157, EPI_ISL_520158, EPI_ISL_520159, EPI_ISL_520160, EPI_ISL_520161, EPI_ISL_520162, EPI_ISL_520163, EPI_ISL_520164, EPI_ISL_520165, EPI_ISL_520166, EPI_ISL_520167, EPI_ISL_520168, EPI_ISL_520169, EPI_ISL_520170, EPI_ISL_520171, EPI_ISL_520172, EPI_ISL_520173, EPI_ISL_520174, EPI_ISL_520175, EPI_ISL_520176, EPI_ISL_520177, EPI_ISL_520178,

| | | | |
|---|---|---|---|
| EPI_ISL_520179, EPI_ISL_520180, EPI_ISL_520215, EPI_ISL_520216, EPI_ISL_520217, EPI_ISL_520218, EPI_ISL_520219, EPI_ISL_520220, EPI_ISL_520221, EPI_ISL_520222, EPI_ISL_520223, EPI_ISL_520224, EPI_ISL_520225, EPI_ISL_520226, EPI_ISL_520227, EPI_ISL_520228, EPI_ISL_520229, EPI_ISL_520230, EPI_ISL_520231, EPI_ISL_520233, EPI_ISL_520234, EPI_ISL_520235, EPI_ISL_520236, EPI_ISL_520237, EPI_ISL_520238, EPI_ISL_520239, EPI_ISL_520240, EPI_ISL_520241, EPI_ISL_520242, EPI_ISL_520243, EPI_ISL_520244, EPI_ISL_520245, EPI_ISL_520246, EPI_ISL_520247, EPI_ISL_520248, EPI_ISL_520392, EPI_ISL_520393, EPI_ISL_520394, EPI_ISL_520395, EPI_ISL_520396, EPI_ISL_520397, EPI_ISL_520398, EPI_ISL_520399, EPI_ISL_520400, EPI_ISL_520401, EPI_ISL_520402, EPI_ISL_520403, EPI_ISL_520404, EPI_ISL_520405, EPI_ISL_520406, EPI_ISL_520407, EPI_ISL_520408, EPI_ISL_520409, EPI_ISL_520410, EPI_ISL_520411, EPI_ISL_520412, EPI_ISL_520413, EPI_ISL_520414, EPI_ISL_520415, EPI_ISL_520416, EPI_ISL_520417, EPI_ISL_520418, EPI_ISL_520419, EPI_ISL_520420, EPI_ISL_520421, EPI_ISL_520422, EPI_ISL_520423, EPI_ISL_520424, EPI_ISL_520425, EPI_ISL_520426, EPI_ISL_520427, EPI_ISL_520428, EPI_ISL_520429, EPI_ISL_520430, EPI_ISL_520431, EPI_ISL_520432, EPI_ISL_520433, EPI_ISL_520434, EPI_ISL_520435, EPI_ISL_520436, EPI_ISL_520437, EPI_ISL_520438, EPI_ISL_520439, EPI_ISL_520440, EPI_ISL_520441, EPI_ISL_520442, EPI_ISL_520443, EPI_ISL_520444, EPI_ISL_520445, EPI_ISL_520446, EPI_ISL_520447, EPI_ISL_520448, EPI_ISL_520449, EPI_ISL_520450, EPI_ISL_520451, EPI_ISL_520452, EPI_ISL_520453, EPI_ISL_520454, EPI_ISL_520455, EPI_ISL_520456, EPI_ISL_520457, EPI_ISL_520458, EPI_ISL_520459, EPI_ISL_520460, EPI_ISL_520461, EPI_ISL_520462, EPI_ISL_520463, EPI_ISL_520464, EPI_ISL_520465, EPI_ISL_520466, EPI_ISL_520467, EPI_ISL_520468, EPI_ISL_520469, EPI_ISL_520470, EPI_ISL_520471, EPI_ISL_520472, EPI_ISL_520473, EPI_ISL_520474, EPI_ISL_520475, EPI_ISL_520476, EPI_ISL_520477, EPI_ISL_520478, EPI_ISL_520479, EPI_ISL_520480, EPI_ISL_520481, EPI_ISL_520482, EPI_ISL_520483, EPI_ISL_520484, EPI_ISL_520485, EPI_ISL_520486, EPI_ISL_520487, EPI_ISL_520488, EPI_ISL_520489, EPI_ISL_520490, EPI_ISL_520491, EPI_ISL_520492, EPI_ISL_520493, EPI_ISL_520494, EPI_ISL_520495, EPI_ISL_520496, EPI_ISL_520497, EPI_ISL_520498, EPI_ISL_520499, EPI_ISL_520500, EPI_ISL_520501, EPI_ISL_520502, EPI_ISL_520503, EPI_ISL_520504, EPI_ISL_520505, EPI_ISL_520506, EPI_ISL_520507, EPI_ISL_520508, EPI_ISL_520509, EPI_ISL_520510, EPI_ISL_520511, EPI_ISL_520512, EPI_ISL_520513, EPI_ISL_520514, EPI_ISL_520515, EPI_ISL_520516, EPI_ISL_520517, EPI_ISL_520518, EPI_ISL_520519, EPI_ISL_520520, EPI_ISL_520521, EPI_ISL_520522, EPI_ISL_520523, EPI_ISL_520524, EPI_ISL_520525, EPI_ISL_520526, EPI_ISL_520527, EPI_ISL_520528, EPI_ISL_520529, EPI_ISL_520530, EPI_ISL_520531, EPI_ISL_520532, EPI_ISL_520533, EPI_ISL_520534, EPI_ISL_520535, EPI_ISL_520536, EPI_ISL_520537, EPI_ISL_520538, EPI_ISL_520539, EPI_ISL_520540, EPI_ISL_520541, EPI_ISL_520542, EPI_ISL_520543, EPI_ISL_520544, EPI_ISL_520545, EPI_ISL_520546, EPI_ISL_520547, EPI_ISL_520548, EPI_ISL_520549, EPI_ISL_520550, EPI_ISL_520551, EPI_ISL_520552, EPI_ISL_520553, EPI_ISL_520554, EPI_ISL_520555, EPI_ISL_520556, EPI_ISL_520557, EPI_ISL_520558, EPI_ISL_520559, EPI_ISL_520560, EPI_ISL_520561, EPI_ISL_520563, EPI_ISL_520564, EPI_ISL_520565, EPI_ISL_520566, EPI_ISL_520567, EPI_ISL_520568, EPI_ISL_520569, EPI_ISL_520570, EPI_ISL_520571, EPI_ISL_520572, EPI_ISL_520573, EPI_ISL_520574, EPI_ISL_520575, EPI_ISL_520576, EPI_ISL_520577, EPI_ISL_520578, EPI_ISL_520579, EPI_ISL_520580, EPI_ISL_520581, EPI_ISL_520582, EPI_ISL_520583, EPI_ISL_520584, EPI_ISL_520585, EPI_ISL_520586, EPI_ISL_520587, EPI_ISL_520588, EPI_ISL_520589, EPI_ISL_520590, EPI_ISL_520591, EPI_ISL_520592, EPI_ISL_520593, EPI_ISL_520594, EPI_ISL_520595, EPI_ISL_520596, EPI_ISL_520597, EPI_ISL_520598, EPI_ISL_520599, EPI_ISL_520600, EPI_ISL_520601, EPI_ISL_520602, EPI_ISL_520603 | | | |
| see above | Microbiological Diagnostic Unit - Public Health Laboratory (MDU-PHL) | MDU-PHL | Seemann T., Schultz M., Sait, M., Sherry, N. |
| EPI_ISL_520604, EPI_ISL_520605 | Microbiological Diagnostic Unit - Public Health Laboratory (MDU-PHL) | Microbiological Diagnostic Unit - Public Health Laboratory (MDU-PHL) | Seemann T., Schultz M., Sait, M., Sherry, N. |
| EPI_ISL_520606, EPI_ISL_520607, EPI_ISL_520608, EPI_ISL_520609, EPI_ISL_520610, EPI_ISL_520611, EPI_ISL_520612, EPI_ISL_520613, EPI_ISL_520614, EPI_ISL_520615, EPI_ISL_520616, EPI_ISL_520617, EPI_ISL_520618, EPI_ISL_520619, EPI_ISL_520620, EPI_ISL_520621, EPI_ISL_520623, EPI_ISL_520624, EPI_ISL_520625, EPI_ISL_520626, EPI_ISL_520627, EPI_ISL_520628, EPI_ISL_520629, EPI_ISL_520630, EPI_ISL_520631, EPI_ISL_520632, EPI_ISL_520633, EPI_ISL_520634, EPI_ISL_520635, EPI_ISL_520636, EPI_ISL_520637, EPI_ISL_520638, EPI_ISL_520639, EPI_ISL_520640, EPI_ISL_520641, EPI_ISL_520642, EPI_ISL_520643 | | | |
| see above | Microbiological Diagnostic Unit - Public Health Laboratory (MDU-PHL) | MDU-PHL | Seemann T., Schultz M., Sait, M., Sherry, N. |
| EPI_ISL_520644 | Microbiological Diagnostic Unit - Public Health Laboratory (MDU-PHL) | Microbiological Diagnostic Unit - Public Health Laboratory (MDU-PHL) | Seemann T., Schultz M., Sait, M., Sherry, N. |
| EPI_ISL_520645, EPI_ISL_520646, EPI_ISL_520647, EPI_ISL_520648, EPI_ISL_520649, EPI_ISL_520650, EPI_ISL_520651, EPI_ISL_520652, EPI_ISL_520653, EPI_ISL_520654, EPI_ISL_520655, EPI_ISL_520656, EPI_ISL_520657, EPI_ISL_520658, EPI_ISL_520659, EPI_ISL_520660, EPI_ISL_520661, EPI_ISL_520662, EPI_ISL_520765, EPI_ISL_520767, EPI_ISL_520768, EPI_ISL_520769, EPI_ISL_520770, EPI_ISL_520771, EPI_ISL_520773, EPI_ISL_520774, EPI_ISL_520776, EPI_ISL_520777, EPI_ISL_520778, EPI_ISL_520779, EPI_ISL_520780, EPI_ISL_520781, EPI_ISL_520782, EPI_ISL_520783, EPI_ISL_520784, EPI_ISL_520785, EPI_ISL_520786, EPI_ISL_520787, EPI_ISL_520788, EPI_ISL_520844, EPI_ISL_520845, EPI_ISL_520846, EPI_ISL_520847, EPI_ISL_520848, EPI_ISL_520849, EPI_ISL_520850, EPI_ISL_520851, EPI_ISL_520852, EPI_ISL_520853, EPI_ISL_520854, EPI_ISL_520855, EPI_ISL_520856, EPI_ISL_520857, EPI_ISL_520858, EPI_ISL_520859, EPI_ISL_520860, EPI_ISL_520861, EPI_ISL_520862, EPI_ISL_520863, EPI_ISL_520929, EPI_ISL_520930, EPI_ISL_520931, EPI_ISL_520932, EPI_ISL_520933, EPI_ISL_520934, EPI_ISL_520935, EPI_ISL_520936, EPI_ISL_520937, EPI_ISL_520938, EPI_ISL_520939, EPI_ISL_520940, EPI_ISL_520941, EPI_ISL_520942, EPI_ISL_520943, EPI_ISL_520944, EPI_ISL_520945, EPI_ISL_520946, EPI_ISL_520947, EPI_ISL_520948, EPI_ISL_520949, EPI_ISL_520950, EPI_ISL_520951, EPI_ISL_520952, EPI_ISL_520953, EPI_ISL_520954, EPI_ISL_520955, EPI_ISL_520956, EPI_ISL_520957, EPI_ISL_520958, EPI_ISL_520959, EPI_ISL_520960, EPI_ISL_520961, EPI_ISL_520962, EPI_ISL_520963, EPI_ISL_520964, EPI_ISL_520965, EPI_ISL_520966, EPI_ISL_520967, EPI_ISL_520968, EPI_ISL_520969, EPI_ISL_520970, EPI_ISL_520971, EPI_ISL_520972, EPI_ISL_520973, EPI_ISL_520974, EPI_ISL_520975, EPI_ISL_520976, EPI_ISL_520977, EPI_ISL_520978, EPI_ISL_520979, EPI_ISL_520980, EPI_ISL_520981, EPI_ISL_520982, EPI_ISL_520983, EPI_ISL_520984, EPI_ISL_520985, EPI_ISL_520986, EPI_ISL_520987, EPI_ISL_520988, EPI_ISL_520989, EPI_ISL_520990, EPI_ISL_520991, EPI_ISL_520992, EPI_ISL_520993, EPI_ISL_520994, EPI_ISL_520995, EPI_ISL_520996, EPI_ISL_520997, EPI_ISL_520998, EPI_ISL_520999, EPI_ISL_521000, EPI_ISL_521001, EPI_ISL_521002, EPI_ISL_521003, EPI_ISL_521004, EPI_ISL_521005, EPI_ISL_521006, EPI_ISL_521007, EPI_ISL_521008, EPI_ISL_521009, EPI_ISL_521010, EPI_ISL_521011, EPI_ISL_521012, EPI_ISL_521013, EPI_ISL_521014, EPI_ISL_521015, EPI_ISL_521016, EPI_ISL_521017, EPI_ISL_521018, EPI_ISL_521019, EPI_ISL_521020, EPI_ISL_521021, EPI_ISL_521022, EPI_ISL_521023, EPI_ISL_521024, EPI_ISL_521025, EPI_ISL_521026, EPI_ISL_521027, EPI_ISL_521028, EPI_ISL_521029, EPI_ISL_521030, EPI_ISL_521031, EPI_ISL_521032, EPI_ISL_521033, EPI_ISL_521034, EPI_ISL_521035, EPI_ISL_521036, EPI_ISL_521037, EPI_ISL_521038, EPI_ISL_521039, EPI_ISL_521040, EPI_ISL_521041, EPI_ISL_521042, EPI_ISL_521043, EPI_ISL_521044, EPI_ISL_521045, EPI_ISL_521046, EPI_ISL_521047, EPI_ISL_521048, EPI_ISL_521049, EPI_ISL_521050, EPI_ISL_521051, EPI_ISL_521052, EPI_ISL_521053, EPI_ISL_521054, EPI_ISL_521055, EPI_ISL_521056, EPI_ISL_521057, EPI_ISL_521058, EPI_ISL_521059, EPI_ISL_521060, EPI_ISL_521061, EPI_ISL_521062, EPI_ISL_521063, EPI_ISL_521064, EPI_ISL_521065, EPI_ISL_521066, EPI_ISL_521067, EPI_ISL_521068, EPI_ISL_521069, EPI_ISL_521070, EPI_ISL_521071, EPI_ISL_521072, EPI_ISL_521073, EPI_ISL_521074, EPI_ISL_521075, EPI_ISL_521076, EPI_ISL_521077, EPI_ISL_521078, EPI_ISL_521079, EPI_ISL_521080, EPI_ISL_521081, EPI_ISL_521082, EPI_ISL_521083, EPI_ISL_521084, EPI_ISL_521085, EPI_ISL_521086, EPI_ISL_521089, EPI_ISL_521090, EPI_ISL_521091, EPI_ISL_521092, EPI_ISL_521093, EPI_ISL_521094, EPI_ISL_521095, EPI_ISL_521096, EPI_ISL_521097, EPI_ISL_521098, EPI_ISL_521099, EPI_ISL_521100, EPI_ISL_521101, EPI_ISL_521154, EPI_ISL_521156, EPI_ISL_521157, EPI_ISL_521158, EPI_ISL_521159, EPI_ISL_521160, EPI_ISL_521161, EPI_ISL_521162, EPI_ISL_521163, EPI_ISL_521164, EPI_ISL_521165, EPI_ISL_521166, EPI_ISL_521167, EPI_ISL_521168, EPI_ISL_521170, EPI_ISL_521171, EPI_ISL_521172, EPI_ISL_521173, EPI_ISL_521174, EPI_ISL_521175, EPI_ISL_521176, EPI_ISL_521177, EPI_ISL_521178, EPI_ISL_521179, EPI_ISL_521180, EPI_ISL_521181, EPI_ISL_521182, EPI_ISL_521183, EPI_ISL_521184, EPI_ISL_521185, EPI_ISL_521186, EPI_ISL_521187, EPI_ISL_521188, EPI_ISL_521189, EPI_ISL_521190, EPI_ISL_521191, EPI_ISL_521192, EPI_ISL_521193, EPI_ISL_521194, EPI_ISL_521195, EPI_ISL_521196, EPI_ISL_521197, EPI_ISL_521198, EPI_ISL_521199, EPI_ISL_521200, EPI_ISL_521201, EPI_ISL_521202, EPI_ISL_521203, EPI_ISL_521204, EPI_ISL_521205, EPI_ISL_521206, EPI_ISL_521207, EPI_ISL_521208, EPI_ISL_521209, EPI_ISL_521210, EPI_ISL_521211, EPI_ISL_521212, EPI_ISL_521213, EPI_ISL_521214, EPI_ISL_521215, EPI_ISL_521216, EPI_ISL_521217, EPI_ISL_521218, EPI_ISL_521219, EPI_ISL_521220, EPI_ISL_521221, EPI_ISL_521222, EPI_ISL_521223, EPI_ISL_521226, EPI_ISL_521227, EPI_ISL_521228, EPI_ISL_521229, EPI_ISL_521230, EPI_ISL_521231, EPI_ISL_521232, EPI_ISL_521233, EPI_ISL_521234, EPI_ISL_521235, EPI_ISL_521236, EPI_ISL_521237, EPI_ISL_521238, EPI_ISL_521239, EPI_ISL_521240, EPI_ISL_521241, EPI_ISL_521242, EPI_ISL_521243, EPI_ISL_521244, EPI_ISL_521247, EPI_ISL_521248, EPI_ISL_521251, EPI_ISL_521252, EPI_ISL_521253, EPI_ISL_521254, EPI_ISL_521255, EPI_ISL_521256, EPI_ISL_521257, EPI_ISL_521258, EPI_ISL_521259, EPI_ISL_521260, EPI_ISL_521261, EPI_ISL_521262, EPI_ISL_521263, EPI_ISL_521264, EPI_ISL_521266, EPI_ISL_521267, EPI_ISL_521268, EPI_ISL_521269, EPI_ISL_521278, EPI_ISL_521279, EPI_ISL_521280, EPI_ISL_521281, EPI_ISL_521282, EPI_ISL_521283, EPI_ISL_521284, EPI_ISL_521285, EPI_ISL_521286, EPI_ISL_521287, EPI_ISL_521288, EPI_ISL_521289, EPI_ISL_521290, EPI_ISL_521291, EPI_ISL_521292, EPI_ISL_521293, EPI_ISL_521294, EPI_ISL_521295, EPI_ISL_521296, EPI_ISL_521297, EPI_ISL_521298, EPI_ISL_521299, EPI_ISL_521300, EPI_ISL_521301, EPI_ISL_521302, EPI_ISL_521303, EPI_ISL_521304, EPI_ISL_521305, EPI_ISL_521306, EPI_ISL_521307, EPI_ISL_521308, EPI_ISL_521309, EPI_ISL_521310, EPI_ISL_521311, EPI_ISL_521312, EPI_ISL_521313, EPI_ISL_521314, EPI_ISL_521450, EPI_ISL_521452, EPI_ISL_521453, EPI_ISL_521454, EPI_ISL_521455, EPI_ISL_521456, EPI_ISL_521457, EPI_ISL_521458, EPI_ISL_521459, EPI_ISL_521460, EPI_ISL_521461, EPI_ISL_521562, EPI_ISL_521563, EPI_ISL_521564, EPI_ISL_521565, EPI_ISL_521566, EPI_ISL_521567, EPI_ISL_521568, EPI_ISL_521569, EPI_ISL_521570, EPI_ISL_521571, EPI_ISL_521572, EPI_ISL_521573, EPI_ISL_521574, EPI_ISL_521575, EPI_ISL_521576, EPI_ISL_521577, EPI_ISL_521578, EPI_ISL_521579, EPI_ISL_521580, EPI_ISL_521581, EPI_ISL_521582, EPI_ISL_521583, EPI_ISL_521585, EPI_ISL_521586, EPI_ISL_521587, EPI_ISL_521588, EPI_ISL_521589, EPI_ISL_521590, EPI_ISL_521591, EPI_ISL_521592, EPI_ISL_521593, EPI_ISL_521594, EPI_ISL_521595, EPI_ISL_521596, EPI_ISL_521597, EPI_ISL_521598, EPI_ISL_521599, EPI_ISL_521600, EPI_ISL_521601, EPI_ISL_521602, EPI_ISL_521603, EPI_ISL_521604, EPI_ISL_521605, EPI_ISL_521606, EPI_ISL_521607, EPI_ISL_521608, EPI_ISL_521609, EPI_ISL_521610, EPI_ISL_521611, EPI_ISL_521612, EPI_ISL_521613, EPI_ISL_521614, EPI_ISL_521615, EPI_ISL_521616, EPI_ISL_521617, EPI_ISL_521618, EPI_ISL_521619, EPI_ISL_521620, EPI_ISL_521621, EPI_ISL_521622, EPI_ISL_521623, EPI_ISL_521624, EPI_ISL_521625, EPI_ISL_521626, EPI_ISL_521627, EPI_ISL_521628, EPI_ISL_521629, EPI_ISL_521630, EPI_ISL_521631, EPI_ISL_521632, EPI_ISL_521633, EPI_ISL_521634, EPI_ISL_521635, EPI_ISL_521636, EPI_ISL_521637, EPI_ISL_521638, EPI_ISL_521639, EPI_ISL_521640, EPI_ISL_521641, EPI_ISL_521642, EPI_ISL_521643, EPI_ISL_521644, EPI_ISL_521645, EPI_ISL_521646, EPI_ISL_521647, EPI_ISL_521648, EPI_ISL_521649, EPI_ISL_521650, EPI_ISL_521651, EPI_ISL_521652, EPI_ISL_521653, EPI_ISL_521654, EPI_ISL_521655, EPI_ISL_521656, EPI_ISL_521657, EPI_ISL_521658, EPI_ISL_521659, EPI_ISL_521660, EPI_ISL_521661, EPI_ISL_521662, EPI_ISL_521663, EPI_ISL_521664, EPI_ISL_521665, EPI_ISL_521666, EPI_ISL_521667, EPI_ISL_521668, EPI_ISL_521669, EPI_ISL_521670, EPI_ISL_521671, EPI_ISL_521672, EPI_ISL_521673, EPI_ISL_521674, EPI_ISL_521675, EPI_ISL_521676, EPI_ISL_521677, EPI_ISL_521678, EPI_ISL_521679, EPI_ISL_521680, EPI_ISL_521681, EPI_ISL_521682, EPI_ISL_521683, EPI_ISL_521684, EPI_ISL_521685, EPI_ISL_521686, EPI_ISL_521687, EPI_ISL_521688, EPI_ISL_521689, EPI_ISL_521690, EPI_ISL_521691, EPI_ISL_521692, EPI_ISL_521693, EPI_ISL_521694, EPI_ISL_521695, EPI_ISL_521696, EPI_ISL_521697, EPI_ISL_521698, EPI_ISL_521699, EPI_ISL_521700, EPI_ISL_521701, EPI_ISL_521702, EPI_ISL_521703, EPI_ISL_521704, EPI_ISL_521705, EPI_ISL_521706, EPI_ISL_521707, EPI_ISL_521708, EPI_ISL_521709, EPI_ISL_521710, EPI_ISL_521711, EPI_ISL_521712, EPI_ISL_521713, EPI_ISL_521714, EPI_ISL_521715, EPI_ISL_521716, EPI_ISL_521717, EPI_ISL_521718, EPI_ISL_521719, EPI_ISL_521720, EPI_ISL_521721, EPI_ISL_521722, EPI_ISL_521723, EPI_ISL_521724, EPI_ISL_521725, EPI_ISL_521726, EPI_ISL_521727, EPI_ISL_521728, EPI_ISL_521729, EPI_ISL_521730, EPI_ISL_521731, EPI_ISL_521732, EPI_ISL_521733, EPI_ISL_521734, EPI_ISL_521735, EPI_ISL_521736, EPI_ISL_521737, EPI_ISL_521738, EPI_ISL_521739, EPI_ISL_521740, EPI_ISL_521741, EPI_ISL_521742, EPI_ISL_521743, EPI_ISL_521744, EPI_ISL_521745, EPI_ISL_521746, EPI_ISL_521747, EPI_ISL_521748, EPI_ISL_521749, EPI_ISL_521750, EPI_ISL_521751, EPI_ISL_521752, EPI_ISL_521753, EPI_ISL_521754, EPI_ISL_521755, EPI_ISL_521756, EPI_ISL_521757, EPI_ISL_521758, EPI_ISL_521759, EPI_ISL_521760, EPI_ISL_521761, EPI_ISL_521762, EPI_ISL_521763, EPI_ISL_521764, EPI_ISL_521765, EPI_ISL_521766, EPI_ISL_521767, EPI_ISL_521768, EPI_ISL_521769, EPI_ISL_521770, EPI_ISL_521771, EPI_ISL_521772, EPI_ISL_521773, EPI_ISL_521774, EPI_ISL_521775, EPI_ISL_521776, EPI_ISL_521777, EPI_ISL_521778, EPI_ISL_521779, EPI_ISL_521780, EPI_ISL_521781, EPI_ISL_521782, EPI_ISL_521783, EPI_ISL_521784, EPI_ISL_521785, EPI_ISL_521786, EPI_ISL_521787, EPI_ISL_521788, EPI_ISL_521789, EPI_ISL_521790, EPI_ISL_521791, EPI_ISL_521792, EPI_ISL_521793, EPI_ISL_521794, EPI_ISL_521795, EPI_ISL_521796, EPI_ISL_521797, EPI_ISL_521798, EPI_ISL_521799, EPI_ISL_521800, EPI_ISL_521801, EPI_ISL_521802, EPI_ISL_521803, EPI_ISL_521804, EPI_ISL_521805, EPI_ISL_521806, EPI_ISL_521807, EPI_ISL_521808, EPI_ISL_521809, EPI_ISL_521810, EPI_ISL_521811, EPI_ISL_521812, EPI_ISL_521813, EPI_ISL_521814, EPI_ISL_521815, EPI_ISL_521816, EPI_ISL_521818, EPI_ISL_521819, EPI_ISL_521820, EPI_ISL_521822, EPI_ISL_521823, EPI_ISL_521824, EPI_ISL_521825, EPI_ISL_521826, EPI_ISL_521827, EPI_ISL_521828, EPI_ISL_521829, EPI_ISL_521830, EPI_ISL_521831, EPI_ISL_521832, EPI_ISL_521833, EPI_ISL_521834, EPI_ISL_521835, EPI_ISL_521836, EPI_ISL_521837, EPI_ISL_521838, EPI_ISL_521839, EPI_ISL_521887, EPI_ISL_521888, EPI_ISL_521906, EPI_ISL_521907, EPI_ISL_521908, EPI_ISL_521909, EPI_ISL_521910, EPI_ISL_521923, EPI_ISL_521926, EPI_ISL_521927, EPI_ISL_521928, EPI_ISL_521929, EPI_ISL_521930, EPI_ISL_521931, EPI_ISL_521932, EPI_ISL_521933, EPI_ISL_521934, EPI_ISL_521935, EPI_ISL_521939, EPI_ISL_521940, EPI_ISL_521941, EPI_ISL_521942, EPI_ISL_521943, EPI_ISL_521944, EPI_ISL_521972, EPI_ISL_521973, EPI_ISL_521974, EPI_ISL_521989, EPI_ISL_521990, EPI_ISL_521991, EPI_ISL_521992, EPI_ISL_521993, EPI_ISL_521994, EPI_ISL_521995, EPI_ISL_521996, EPI_ISL_521997, EPI_ISL_521998, EPI_ISL_521999, EPI_ISL_522000, EPI_ISL_522001, EPI_ISL_522002, EPI_ISL_522003, EPI_ISL_522004, EPI_ISL_522005, EPI_ISL_522006, EPI_ISL_522007, EPI_ISL_522008, EPI_ISL_522009, EPI_ISL_522010, EPI_ISL_522011, EPI_ISL_522012, EPI_ISL_522013, EPI_ISL_522014, EPI_ISL_522015, EPI_ISL_522016, EPI_ISL_522017, EPI_ISL_522018, EPI_ISL_522019, EPI_ISL_522020, EPI_ISL_522021, EPI_ISL_522022, EPI_ISL_522023, EPI_ISL_522024, EPI_ISL_522025, EPI_ISL_522026, EPI_ISL_522027, EPI_ISL_522028, EPI_ISL_522029, EPI_ISL_522030, EPI_ISL_522031, EPI_ISL_522032, EPI_ISL_522033, EPI_ISL_522034, EPI_ISL_522035, EPI_ISL_522036, EPI_ISL_522037, EPI_ISL_522038, EPI_ISL_522039, EPI_ISL_522040, EPI_ISL_522041, EPI_ISL_522042, EPI_ISL_522043, EPI_ISL_522044, EPI_ISL_522045, EPI_ISL_522046, EPI_ISL_522047, EPI_ISL_522048, EPI_ISL_522049, EPI_ISL_522050, EPI_ISL_522051, EPI_ISL_522052, EPI_ISL_522053, EPI_ISL_522054, EPI_ISL_522111, EPI_ISL_522112, EPI_ISL_522113, EPI_ISL_522114, EPI_ISL_522115, EPI_ISL_522116, EPI_ISL_522117, EPI_ISL_522118, EPI_ISL_522188, EPI_ISL_522189, EPI_ISL_522190, EPI_ISL_522191, EPI_ISL_522192, EPI_ISL_522193, EPI_ISL_522195, EPI_ISL_522196, EPI_ISL_522197, EPI_ISL_522198, EPI_ISL_522199, EPI_ISL_522200, EPI_ISL_522201, EPI_ISL_522202, EPI_ISL_522203, EPI_ISL_522204, EPI_ISL_522205, EPI_ISL_522206, EPI_ISL_522207, EPI_ISL_522208, EPI_ISL_522209, EPI_ISL_522210, EPI_ISL_522211, EPI_ISL_522212, EPI_ISL_522213, EPI_ISL_522214, EPI_ISL_522215, EPI_ISL_522216, EPI_ISL_522217, EPI_ISL_522218, EPI_ISL_522219, EPI_ISL_522220, EPI_ISL_522221, EPI_ISL_522222, EPI_ISL_522223, EPI_ISL_522224, EPI_ISL_522225, EPI_ISL_522226, EPI_ISL_522227, EPI_ISL_522228, EPI_ISL_522229, EPI_ISL_522230, EPI_ISL_522231, EPI_ISL_522232, EPI_ISL_522243, EPI_ISL_522244, EPI_ISL_522245, EPI_ISL_522246, EPI_ISL_522247, EPI_ISL_522248, EPI_ISL_522249, EPI_ISL_522250, EPI_ISL_522251, EPI_ISL_522252, EPI_ISL_522253 | | | |
| see above | Microbiological Diagnostic Unit - Public Health Laboratory (MDU-PHL) | MDU-PHL | Seemann T., Schultz M., Sait, M., Sherry, N. |
| EPI_ISL_526553, EPI_ISL_526554, EPI_ISL_526555, EPI_ISL_526556, EPI_ISL_526557, EPI_ISL_526559, EPI_ISL_526569, EPI_ISL_526571, EPI_ISL_526572, EPI_ISL_526573, EPI_ISL_526575, EPI_ISL_526583, EPI_ISL_526585, EPI_ISL_526586, EPI_ISL_526589, EPI_ISL_526590, EPI_ISL_526591, EPI_ISL_526592, EPI_ISL_526593, EPI_ISL_526594, EPI_ISL_526595, EPI_ISL_526596, EPI_ISL_526597, EPI_ISL_526598, EPI_ISL_526599, EPI_ISL_526600, EPI_ISL_526602, EPI_ISL_526603, EPI_ISL_526605, EPI_ISL_526606, EPI_ISL_526607, EPI_ISL_526608, EPI_ISL_526610, EPI_ISL_526611, EPI_ISL_526613, EPI_ISL_526614, EPI_ISL_526616, EPI_ISL_526617, EPI_ISL_526618, EPI_ISL_526619, EPI_ISL_526620, EPI_ISL_526621, EPI_ISL_526622, EPI_ISL_526623, EPI_ISL_526624, EPI_ISL_526625, EPI_ISL_526626, EPI_ISL_526627, EPI_ISL_526628, EPI_ISL_526629, EPI_ISL_526630, EPI_ISL_526631, EPI_ISL_526632, EPI_ISL_526633, EPI_ISL_526635, EPI_ISL_526637, EPI_ISL_526638, EPI_ISL_526639, EPI_ISL_526643, EPI_ISL_526644, EPI_ISL_526645, EPI_ISL_526646, EPI_ISL_526649, EPI_ISL_526650, EPI_ISL_526652, EPI_ISL_526653, EPI_ISL_526657, EPI_ISL_526658, EPI_ISL_526659, EPI_ISL_526660, EPI_ISL_526661, EPI_ISL_526662, EPI_ISL_526663, EPI_ISL_526664, EPI_ISL_526665, EPI_ISL_526666, EPI_ISL_526667, EPI_ISL_526668, EPI_ISL_526669, EPI_ISL_526670, EPI_ISL_526671, EPI_ISL_526672, EPI_ISL_526673, EPI_ISL_526674, EPI_ISL_526675, EPI_ISL_526676, EPI_ISL_526677, EPI_ISL_526678, EPI_ISL_526679, EPI_ISL_526680, EPI_ISL_526681, EPI_ISL_526682, EPI_ISL_526683, EPI_ISL_526684, EPI_ISL_526685 | | | |
| see above | Florida Bureau of Public Health Laboratories | Florida Bureau of Public Health Laboratories | Sarah Schmedes, Jason Blanton |
| EPI_ISL_526975, EPI_ISL_526976, EPI_ISL_526977, EPI_ISL_526978, EPI_ISL_526979, EPI_ISL_526980, EPI_ISL_526981, EPI_ISL_526982, EPI_ISL_526983, EPI_ISL_526984, EPI_ISL_526985, EPI_ISL_526986, EPI_ISL_526987, EPI_ISL_526988, EPI_ISL_526989, EPI_ISL_526990, EPI_ISL_526991, EPI_ISL_526992, EPI_ISL_526993, EPI_ISL_526994, EPI_ISL_526995, EPI_ISL_526996 | | | |

| | | | |
|---|---|---|---|
| see above | Biological prevention, army | Biological prevention, army | Seadawy, M.G., Gad, A.F., Harty, B.E., Elhosieny, M.F., Shamel, M.D. |
| EPI_ISL_526997, EPI_ISL_526998 | Biological prevention, army | Biological prevention, army | Seadawy, M.G., Harty,B.E., Gad,A.F., Elhoseiny,M.F., Shamel,M.D., Shabaan,A.E., Ageez,A.M. |
| EPI_ISL_526999, EPI_ISL_527000, EPI_ISL_527001 | Biological prevention, army | Biological prevention, army | Seadawy,M.G., Harty,B.E., Gad,A.F., Elhoseiny,M.F., Shamel,M.D., Shabaan,A.E., Ageez,A.M. |
| EPI_ISL_527002 | Biological prevention, army | Biological prevention, army | Seadawy, M.G., Harty,B.E., Gad,A.F., Elhoseiny,M.F., Shamel,M.D., Shabaan,A.E., Ageez,A.M. |
| EPI_ISL_527003, EPI_ISL_527004, EPI_ISL_527005, EPI_ISL_527006 | Biological prevention, army | Biological prevention, army | Seadawy,M.G., Harty,B.E., Gad,A.F., Elhoseiny,M.F., Shamel,M.D., Shabaan,A.E., Ageez,A.M. |
| EPI_ISL_527007 | Biological Prevention, Army | Biological Prevention, Army | Seadawy, M.G., Harty,B.E., Gad,A.F., Elhoseiny,M.F., Shamel,M.D., Shabaan,A.E., Ageez,A.M. |
| EPI_ISL_528382, EPI_ISL_528383, EPI_ISL_528384, EPI_ISL_528385 | Translational Health Science and Technology Institute -ESIC medical college and hospital, Faridabad | THSTI Bioassay laboratory | Saurabh Kumar, Jigme Wangchuk, Anil Kumar Pandey, Asim Das, Guruprasad R. Medigeshi |
| EPI_ISL_528823, EPI_ISL_528825, EPI_ISL_528826, EPI_ISL_528847, EPI_ISL_528865, EPI_ISL_528868, EPI_ISL_528869, EPI_ISL_539637 | CSIR-Centre for Cellular and Molecular Biology | CSIR-Centre for Cellular and Molecular Biology | Shagufta Khan, Lamuk Zaveri, Namami Gaur, Sakshi Shambhavi, Nikhil Hajirnis, M Soujanya Reddy, Pratheusa Maccha, Tulasi Nagabandi, Purushotham Vodnala, Payel Mukherjee, Sofia Banu, Priya Singh, Onkar Kulkarni, Dhiviya Vedagiri, Divya Gupta, Vishal Sah, Santosh Kumar Kuncha, Krishnan Harinivas Harshan, Archana Bharadwaj Siva, Karthik Bharadwaj Tallapaka, Renu Sudhakar, Somesh Gorde, Gangumala Srinivas Reddy, Sujoy Deb, Swati Bayyana, Rakesh K Mishra, Divya Tej Sowpati |
| EPI_ISL_539638 | CSIR-Centre for Cellular and Molecular Biology | CSIR-Centre for Cellular and Molecular Biology | Shagufta Khan, Lamuk Zaveri, Namami Gaur, Sakshi Shambhavi, Nikhil Hajirnis, M Soujanya Reddy, Pratheusa Maccha, Tulasi Nagabandi, Purushotham Vodnala, Payel Mukherjee, Sofia Banu, Priya Singh, Onkar Kulkarni, Dhiviya Vedagiri, Divya Gupta, Vishal Sah, Santosh Kumar Kuncha, Krishnan Harinivas Harshan, Archana Bharadwaj Siva, Karthik Bharadwaj Tallapaka,Preethi Jampala, Sharada Ravi Iyer, Sulagana Mukherjee, Swetha Sundar, Peddapuvala Sai Uday Kiran Rakesh K Mishra, Divya Tej Sowpati |
| EPI_ISL_539639 | CSIR-Centre for Cellular and Molecular Biology | CSIR-Centre for Cellular and Molecular Biology | Shagufta Khan, Lamuk Zaveri, Namami Gaur, Sakshi Shambhavi, Nikhil Hajirnis, M Soujanya Reddy, Pratheusa Maccha,Tulasi Nagabandi, Purushotham Vodnala, Payel Mukherjee, Sofia Banu, Priya Singh, Onkar Kulkarni, Dhiviya Vedagiri, Divya Gupta, Vishal Sah, Santosh Kumar Kuncha, Krishnan Harinivas Harshan, Archana Bharadwaj Siva, Karthik Bharadwaj Tallapaka,Umesh Kumar, Unis Ahmad Bhat, Ajay Sarawagi, Priyanka Pant, Rajkanwar Nathawat, Rakesh K Mishra, Divya Tej Sowpati |
| EPI_ISL_539667 | CSIR-Centre for Cellular and Molecular Biology | CSIR-Centre for Cellular and Molecular Biology | Shagufta Khan, Lamuk Zaveri, Namami Gaur, Sakshi Shambhavi, Nikhil Hajirnis, M Soujanya Reddy, Pratheusa Maccha, Tulasi Nagabandi, Purushotham Vodnala, Payel Mukherjee, Sofia Banu, Priya Singh, Onkar Kulkarni, Dhiviya Vedagiri, Divya Gupta, Vishal Sah, Santosh Kumar Kuncha, Krishnan Harinivas Harshan, Archana Bharadwaj Siva, Karthik Bharadwaj Tallapaka, Renu Sudhakar, Somesh Gorde, Gangumala Srinivas Reddy, Sujoy Deb, Swati Bayyana, Rakesh K Mishra, Divya Tej Sowpati |
| EPI_ISL_539668 | CSIR-Centre for Cellular and Molecular Biology | CSIR-Centre for Cellular and Molecular Biology | Shagufta Khan, Lamuk Zaveri, Namami Gaur, Sakshi Shambhavi, Nikhil Hajirnis, M Soujanya Reddy, Pratheusa Maccha, Tulasi Nagabandi, Purushotham Vodnala, Payel Mukherjee, Sofia Banu, Priya Singh, Onkar Kulkarni, Dhiviya Vedagiri, Divya Gupta, Vishal Sah, Santosh Kumar Kuncha, Krishnan Harinivas Harshan, Archana Bharadwaj Siva, Karthik Bharadwaj Tallapaka,Preethi Jampala, Sharada Ravi Iyer, Sulagana Mukherjee, Swetha Sundar, Peddapuvala Sai Uday Kiran Rakesh K Mishra, Divya Tej Sowpati |
| EPI_ISL_539669 | CSIR-Centre for Cellular and Molecular Biology | CSIR-Centre for Cellular and Molecular Biology | Shagufta Khan, Lamuk Zaveri, Namami Gaur, Sakshi Shambhavi, Nikhil Hajirnis, M Soujanya Reddy, Pratheusa Maccha,Tulasi Nagabandi, Purushotham Vodnala, Payel Mukherjee, Sofia Banu, Priya Singh, Onkar Kulkarni, Dhiviya Vedagiri, Divya Gupta, Vishal Sah, Santosh Kumar Kuncha, Krishnan Harinivas Harshan, Archana Bharadwaj Siva, Karthik Bharadwaj Tallapaka,Umesh Kumar, Unis Ahmad Bhat, Ajay Sarawagi, Priyanka Pant, Rajkanwar Nathawat, Rakesh K Mishra, Divya Tej Sowpati |
| EPI_ISL_539697 | CSIR-Centre for Cellular and Molecular Biology | CSIR-Centre for Cellular and Molecular Biology | Shagufta Khan, Lamuk Zaveri, Namami Gaur, Sakshi Shambhavi, Nikhil Hajirnis, M Soujanya Reddy, Pratheusa Maccha, Tulasi Nagabandi, Purushotham Vodnala, Payel Mukherjee, Sofia Banu, Priya Singh, Onkar Kulkarni, Dhiviya Vedagiri, Divya Gupta, Vishal Sah, Santosh Kumar Kuncha, Krishnan Harinivas Harshan, Archana Bharadwaj Siva, Karthik Bharadwaj Tallapaka, Renu Sudhakar, Somesh Gorde, Gangumala Srinivas Reddy, Sujoy Deb, Swati Bayyana, Rakesh K Mishra, Divya Tej Sowpati |
| EPI_ISL_539698 | CSIR-Centre for Cellular and Molecular Biology | CSIR-Centre for Cellular and Molecular Biology | Shagufta Khan, Lamuk Zaveri, Namami Gaur, Sakshi Shambhavi, Nikhil Hajirnis, M Soujanya Reddy, Pratheusa Maccha, Tulasi Nagabandi, Purushotham Vodnala, Payel Mukherjee, Sofia Banu, Priya Singh, Onkar Kulkarni, Dhiviya Vedagiri, Divya Gupta, Vishal Sah, Santosh Kumar Kuncha, Krishnan Harinivas Harshan, Archana Bharadwaj Siva, Karthik Bharadwaj Tallapaka,Preethi Jampala, Sharada Ravi Iyer, Sulagana Mukherjee, Swetha Sundar, Peddapuvala Sai Uday Kiran Rakesh K Mishra, Divya Tej Sowpati |
| EPI_ISL_539699 | CSIR-Centre for Cellular and Molecular Biology | CSIR-Centre for Cellular and Molecular Biology | Shagufta Khan, Lamuk Zaveri, Namami Gaur, Sakshi Shambhavi, Nikhil Hajirnis, M Soujanya Reddy, Pratheusa Maccha,Tulasi Nagabandi, Purushotham Vodnala, Payel Mukherjee, Sofia Banu, Priya Singh, Onkar Kulkarni, Dhiviya Vedagiri, Divya Gupta, Vishal Sah, Santosh Kumar Kuncha, Krishnan Harinivas Harshan, Archana Bharadwaj Siva, Karthik Bharadwaj Tallapaka,Umesh Kumar, Unis Ahmad Bhat, Ajay Sarawagi, Priyanka Pant, Rajkanwar Nathawat, Rakesh K Mishra, Divya Tej Sowpati |
| EPI_ISL_539727 | CSIR-Centre for Cellular and Molecular Biology | CSIR-Centre for Cellular and Molecular Biology | Shagufta Khan, Lamuk Zaveri, Namami Gaur, Sakshi Shambhavi, Nikhil Hajirnis, M Soujanya Reddy, Pratheusa Maccha, Tulasi Nagabandi, Purushotham Vodnala, Payel Mukherjee, Sofia Banu, Priya Singh, Onkar Kulkarni, Dhiviya Vedagiri, Divya Gupta, Vishal Sah, Santosh Kumar Kuncha, Krishnan Harinivas Harshan, Archana Bharadwaj Siva, Karthik Bharadwaj Tallapaka, Renu Sudhakar, Somesh Gorde, Gangumala Srinivas Reddy, Sujoy Deb, Swati Bayyana, Rakesh K Mishra, Divya Tej Sowpati |
| EPI_ISL_539728 | CSIR-Centre for Cellular and Molecular Biology | CSIR-Centre for Cellular and Molecular Biology | Shagufta Khan, Lamuk Zaveri, Namami Gaur, Sakshi Shambhavi, Nikhil Hajirnis, M Soujanya Reddy, Pratheusa Maccha, Tulasi Nagabandi, Purushotham Vodnala, Payel Mukherjee, Sofia Banu, Priya Singh, Onkar Kulkarni, Dhiviya Vedagiri, Divya Gupta, Vishal Sah, Santosh Kumar Kuncha, Krishnan Harinivas Harshan, Archana Bharadwaj Siva, Karthik Bharadwaj Tallapaka,Preethi Jampala, Sharada Ravi Iyer, Sulagana Mukherjee, Swetha Sundar, Peddapuvala Sai Uday Kiran Rakesh K Mishra, Divya Tej Sowpati |
| EPI_ISL_539729 | CSIR-Centre for Cellular and Molecular Biology | CSIR-Centre for Cellular and Molecular Biology | Shagufta Khan, Lamuk Zaveri, Namami Gaur, Sakshi Shambhavi, Nikhil Hajirnis, M Soujanya Reddy, Pratheusa Maccha,Tulasi Nagabandi, Purushotham Vodnala, Payel Mukherjee, Sofia Banu, Priya Singh, Onkar Kulkarni, Dhiviya Vedagiri, Divya Gupta, Vishal Sah, Santosh Kumar Kuncha, Krishnan Harinivas Harshan, Archana Bharadwaj Siva, Karthik Bharadwaj Tallapaka,Umesh Kumar, Unis Ahmad Bhat, Ajay Sarawagi, Priyanka Pant, Rajkanwar Nathawat, Rakesh K Mishra, Divya Tej Sowpati |
| EPI_ISL_539757 | CSIR-Centre for Cellular and Molecular Biology | CSIR-Centre for Cellular and Molecular Biology | Shagufta Khan, Lamuk Zaveri, Namami Gaur, Sakshi Shambhavi, Nikhil Hajirnis, M Soujanya Reddy, Pratheusa Maccha, Tulasi Nagabandi, Purushotham Vodnala, Payel Mukherjee, Sofia Banu, Priya Singh, Onkar Kulkarni, Dhiviya Vedagiri, Divya Gupta, Vishal Sah, Santosh Kumar Kuncha, Krishnan Harinivas Harshan, Archana Bharadwaj Siva, Karthik Bharadwaj Tallapaka, Renu Sudhakar, Somesh Gorde, Gangumala Srinivas Reddy, Sujoy Deb, Swati Bayyana, Rakesh K Mishra, Divya Tej Sowpati |
| EPI_ISL_539758 | CSIR-Centre for Cellular and Molecular Biology | CSIR-Centre for Cellular and Molecular Biology | Shagufta Khan, Lamuk Zaveri, Namami Gaur, Sakshi Shambhavi, Nikhil Hajirnis, M Soujanya Reddy, Pratheusa Maccha, Tulasi Nagabandi, Purushotham Vodnala, Payel Mukherjee, Sofia Banu, Priya Singh, Onkar Kulkarni, Dhiviya Vedagiri, Divya Gupta, Vishal Sah, Santosh Kumar Kuncha, Krishnan Harinivas Harshan, Archana Bharadwaj Siva, Karthik Bharadwaj Tallapaka,Preethi Jampala, Sharada Ravi Iyer, Sulagana Mukherjee, Swetha Sundar, Peddapuvala Sai Uday Kiran Rakesh K Mishra, Divya Tej Sowpati |
| EPI_ISL_539759 | CSIR-Centre for Cellular and Molecular Biology | CSIR-Centre for Cellular and Molecular Biology | Shagufta Khan, Lamuk Zaveri, Namami Gaur, Sakshi Shambhavi, Nikhil Hajirnis, M Soujanya Reddy, Pratheusa Maccha,Tulasi Nagabandi, Purushotham Vodnala, Payel Mukherjee, Sofia Banu, Priya Singh, Onkar Kulkarni, Dhiviya Vedagiri, Divya Gupta, Vishal Sah, Santosh Kumar Kuncha, Krishnan Harinivas Harshan, Archana Bharadwaj Siva, Karthik Bharadwaj Tallapaka,Umesh Kumar, Unis Ahmad Bhat, Ajay Sarawagi, Priyanka Pant, Rajkanwar Nathawat, Rakesh K Mishra, Divya Tej Sowpati |

EPI_ISL_541139, EPI_ISL_541140, EPI_ISL_541141, EPI_ISL_541142, EPI_ISL_541143, EPI_ISL_541144, EPI_ISL_541145, EPI_ISL_541146, EPI_ISL_541147, EPI_ISL_541148, EPI_ISL_541149, EPI_ISL_541150, EPI_ISL_541151, EPI_ISL_541152, EPI_ISL_541153, EPI_ISL_541154, EPI_ISL_541155, EPI_ISL_541156, EPI_ISL_541157, EPI_ISL_541158, EPI_ISL_541159, EPI_ISL_541160, EPI_ISL_541161, EPI_ISL_541162, EPI_ISL_541163, EPI_ISL_541164, EPI_ISL_541165, EPI_ISL_541166, EPI_ISL_541167, EPI_ISL_541168, EPI_ISL_541169, EPI_ISL_541170, EPI_ISL_541171, EPI_ISL_541172, EPI_ISL_541173, EPI_ISL_541174, EPI_ISL_541175, EPI_ISL_541176, EPI_ISL_541177, EPI_ISL_541178, EPI_ISL_541179, EPI_ISL_541180, EPI_ISL_541181, EPI_ISL_541182, EPI_ISL_541183, EPI_ISL_541184, EPI_ISL_541185, EPI_ISL_541186, EPI_ISL_541187, EPI_ISL_541188, EPI_ISL_541189, EPI_ISL_541190, EPI_ISL_541191, EPI_ISL_541192, EPI_ISL_541193, EPI_ISL_541194, EPI_ISL_541195, EPI_ISL_541196, EPI_ISL_541197, EPI_ISL_541198, EPI_ISL_541199, EPI_ISL_541200, EPI_ISL_541201, EPI_ISL_541202, EPI_ISL_541203, EPI_ISL_541204, EPI_ISL_541205, EPI_ISL_541206, EPI_ISL_541207, EPI_ISL_541208, EPI_ISL_541209, EPI_ISL_541210, EPI_ISL_541211, EPI_ISL_541212, EPI_ISL_541213, EPI_ISL_541214, EPI_ISL_541215, EPI_ISL_541216, EPI_ISL_541217, EPI_ISL_541218, EPI_ISL_541219, EPI_ISL_541220, EPI_ISL_541221, EPI_ISL_541222, EPI_ISL_541223, EPI_ISL_541224, EPI_ISL_541225, EPI_ISL_541226, EPI_ISL_541227, EPI_ISL_541228, EPI_ISL_541229, EPI_ISL_541230, EPI_ISL_541231, EPI_ISL_541232, EPI_ISL_541233, EPI_ISL_541234, EPI_ISL_541235, EPI_ISL_541236, EPI_ISL_541237, EPI_ISL_541238, EPI_ISL_541239, EPI_ISL_541240, EPI_ISL_541241, EPI_ISL_541242, EPI_ISL_541243, EPI_ISL_541244, EPI_ISL_541245, EPI_ISL_541246,

EPI_ISL_541247, EPI_ISL_541248, EPI_ISL_541249, EPI_ISL_541250, EPI_ISL_541251, EPI_ISL_541252, EPI_ISL_541253, EPI_ISL_541254, EPI_ISL_541255, EPI_ISL_541256, EPI_ISL_541257, EPI_ISL_541258, EPI_ISL_541259, EPI_ISL_541260, EPI_ISL_541261, EPI_ISL_541262, EPI_ISL_541263, EPI_ISL_541264,
EPI_ISL_541265, EPI_ISL_541266, EPI_ISL_541267, EPI_ISL_541268, EPI_ISL_541269, EPI_ISL_541270, EPI_ISL_541271, EPI_ISL_541272, EPI_ISL_541273, EPI_ISL_541274, EPI_ISL_541275, EPI_ISL_541276, EPI_ISL_541277, EPI_ISL_541278, EPI_ISL_541279, EPI_ISL_541280, EPI_ISL_541281, EPI_ISL_541282,
EPI_ISL_541283, EPI_ISL_541284, EPI_ISL_541285, EPI_ISL_541286, EPI_ISL_541287, EPI_ISL_541288, EPI_ISL_541289, EPI_ISL_541290, EPI_ISL_541291, EPI_ISL_541292, EPI_ISL_541293, EPI_ISL_541294, EPI_ISL_541295, EPI_ISL_541296, EPI_ISL_541297, EPI_ISL_541298, EPI_ISL_541299, EPI_ISL_541300,
EPI_ISL_541301, EPI_ISL_541302, EPI_ISL_541303, EPI_ISL_541304, EPI_ISL_541305, EPI_ISL_541306, EPI_ISL_541307, EPI_ISL_541308, EPI_ISL_541309, EPI_ISL_541310, EPI_ISL_541311, EPI_ISL_541312, EPI_ISL_541313, EPI_ISL_541314, EPI_ISL_541315, EPI_ISL_541316, EPI_ISL_541317, EPI_ISL_541318,
EPI_ISL_541319, EPI_ISL_541320, EPI_ISL_541321, EPI_ISL_541322, EPI_ISL_541323, EPI_ISL_541324, EPI_ISL_541325, EPI_ISL_541326, EPI_ISL_541327, EPI_ISL_541328, EPI_ISL_541329, EPI_ISL_541330, EPI_ISL_541331

| | | | |
|---|---|---|---|
| see above | Florida Bureau of Public Health Laboratories, Florida Department of Health | Florida Bureau of Public Health Laboratories, Florida Department of Health | Schmedes,S., Blanton,J. |

EPI_ISL_549184, EPI_ISL_549192, EPI_ISL_549193, EPI_ISL_549194, EPI_ISL_549195, EPI_ISL_549197, EPI_ISL_549199, EPI_ISL_549200, EPI_ISL_549201, EPI_ISL_549202, EPI_ISL_549203, EPI_ISL_549204, EPI_ISL_549205, EPI_ISL_549206, EPI_ISL_549207, EPI_ISL_549208, EPI_ISL_549210, EPI_ISL_549213,
EPI_ISL_549214, EPI_ISL_549215, EPI_ISL_549216, EPI_ISL_549217, EPI_ISL_549218, EPI_ISL_549219, EPI_ISL_549220, EPI_ISL_549222, EPI_ISL_549223, EPI_ISL_549224, EPI_ISL_549225, EPI_ISL_549226, EPI_ISL_549227, EPI_ISL_549231, EPI_ISL_549236, EPI_ISL_549237, EPI_ISL_549238, EPI_ISL_549239,
EPI_ISL_549240, EPI_ISL_549245, EPI_ISL_549247, EPI_ISL_549248, EPI_ISL_549249, EPI_ISL_549250, EPI_ISL_549251, EPI_ISL_549253, EPI_ISL_549255, EPI_ISL_549258, EPI_ISL_549259, EPI_ISL_549261, EPI_ISL_549262, EPI_ISL_549264, EPI_ISL_549266, EPI_ISL_549267, EPI_ISL_549268, EPI_ISL_549269

| | | | |
|---|---|---|---|
| see above | Florida Bureau of Public Health Laboratories | Florida Bureau of Public Health Laboratories | Sarah Schmedes, Jason Blanton |

EPI_ISL_561378, EPI_ISL_561379, EPI_ISL_561381, EPI_ISL_561382, EPI_ISL_561383, EPI_ISL_561384, EPI_ISL_561385, EPI_ISL_561386, EPI_ISL_561387, EPI_ISL_561389, EPI_ISL_561390, EPI_ISL_561391, EPI_ISL_561393, EPI_ISL_561394, EPI_ISL_561395, EPI_ISL_561396, EPI_ISL_561398, EPI_ISL_561399,
EPI_ISL_561400, EPI_ISL_561401, EPI_ISL_561402, EPI_ISL_561405, EPI_ISL_561406, EPI_ISL_561408, EPI_ISL_561410, EPI_ISL_561411, EPI_ISL_561412, EPI_ISL_561413, EPI_ISL_561415, EPI_ISL_561417, EPI_ISL_561418, EPI_ISL_561419, EPI_ISL_561422, EPI_ISL_561423, EPI_ISL_561424, EPI_ISL_561427,
EPI_ISL_561430, EPI_ISL_561431, EPI_ISL_561432, EPI_ISL_561435, EPI_ISL_561436, EPI_ISL_561437, EPI_ISL_561438, EPI_ISL_561439, EPI_ISL_561441, EPI_ISL_561443, EPI_ISL_561444, EPI_ISL_561445, EPI_ISL_561446, EPI_ISL_561447, EPI_ISL_561448, EPI_ISL_561449, EPI_ISL_561451, EPI_ISL_561452,
EPI_ISL_561454, EPI_ISL_561455, EPI_ISL_561456, EPI_ISL_561459, EPI_ISL_561460, EPI_ISL_561461, EPI_ISL_561463, EPI_ISL_561464, EPI_ISL_561466, EPI_ISL_561467, EPI_ISL_561469, EPI_ISL_561471, EPI_ISL_561472, EPI_ISL_561473, EPI_ISL_561474, EPI_ISL_561475, EPI_ISL_561476, EPI_ISL_561477,
EPI_ISL_561478, EPI_ISL_561479, EPI_ISL_561480, EPI_ISL_561481, EPI_ISL_561482, EPI_ISL_561483, EPI_ISL_561484, EPI_ISL_561485, EPI_ISL_561486, EPI_ISL_561487, EPI_ISL_561488, EPI_ISL_561489, EPI_ISL_561490, EPI_ISL_561492, EPI_ISL_561493, EPI_ISL_561494, EPI_ISL_561495, EPI_ISL_561496,
EPI_ISL_561497, EPI_ISL_561499, EPI_ISL_561500, EPI_ISL_561501, EPI_ISL_561502, EPI_ISL_561503, EPI_ISL_561504, EPI_ISL_561505, EPI_ISL_561506, EPI_ISL_561507, EPI_ISL_561509, EPI_ISL_561510, EPI_ISL_561511, EPI_ISL_561512, EPI_ISL_561513, EPI_ISL_561516, EPI_ISL_561518, EPI_ISL_561520,
EPI_ISL_561521, EPI_ISL_561522, EPI_ISL_561525, EPI_ISL_561526, EPI_ISL_561527, EPI_ISL_561528, EPI_ISL_561529, EPI_ISL_561530, EPI_ISL_561534, EPI_ISL_561536, EPI_ISL_561537, EPI_ISL_561538, EPI_ISL_561539, EPI_ISL_561540, EPI_ISL_561542, EPI_ISL_561543, EPI_ISL_561545, EPI_ISL_561546,
EPI_ISL_561547, EPI_ISL_561548, EPI_ISL_561549, EPI_ISL_561550, EPI_ISL_561551, EPI_ISL_561552, EPI_ISL_561553, EPI_ISL_561554, EPI_ISL_561555, EPI_ISL_561556, EPI_ISL_561558, EPI_ISL_561560, EPI_ISL_561561, EPI_ISL_561562, EPI_ISL_561563, EPI_ISL_561564, EPI_ISL_561566, EPI_ISL_561567,
EPI_ISL_561568, EPI_ISL_561569, EPI_ISL_561570, EPI_ISL_561571, EPI_ISL_561572, EPI_ISL_561573, EPI_ISL_561574, EPI_ISL_561575, EPI_ISL_561576, EPI_ISL_561577, EPI_ISL_561579, EPI_ISL_561580, EPI_ISL_561581, EPI_ISL_561583, EPI_ISL_561585, EPI_ISL_561587, EPI_ISL_561588, EPI_ISL_561589,
EPI_ISL_561592, EPI_ISL_561593, EPI_ISL_561594, EPI_ISL_561595, EPI_ISL_561596, EPI_ISL_561597, EPI_ISL_561598, EPI_ISL_561599, EPI_ISL_561600, EPI_ISL_561601, EPI_ISL_561602, EPI_ISL_561603, EPI_ISL_561604, EPI_ISL_561605, EPI_ISL_561606, EPI_ISL_561607, EPI_ISL_561608, EPI_ISL_561609,
EPI_ISL_561610, EPI_ISL_561611, EPI_ISL_561615, EPI_ISL_561616, EPI_ISL_561617, EPI_ISL_561618, EPI_ISL_561619, EPI_ISL_561620, EPI_ISL_561623, EPI_ISL_561624, EPI_ISL_561626, EPI_ISL_561627, EPI_ISL_561628, EPI_ISL_561631, EPI_ISL_561632, EPI_ISL_561634, EPI_ISL_561635, EPI_ISL_561636,
EPI_ISL_561637, EPI_ISL_561638, EPI_ISL_561639, EPI_ISL_561640, EPI_ISL_561641, EPI_ISL_561642, EPI_ISL_561643, EPI_ISL_561644, EPI_ISL_561645, EPI_ISL_561646, EPI_ISL_561647, EPI_ISL_561648, EPI_ISL_561649, EPI_ISL_561650, EPI_ISL_561654, EPI_ISL_561658, EPI_ISL_561659, EPI_ISL_561660,
EPI_ISL_561662, EPI_ISL_561663, EPI_ISL_561664, EPI_ISL_561665, EPI_ISL_561667, EPI_ISL_561668, EPI_ISL_561669, EPI_ISL_561670, EPI_ISL_561672, EPI_ISL_561675, EPI_ISL_561676, EPI_ISL_561677, EPI_ISL_561678, EPI_ISL_561680, EPI_ISL_561681, EPI_ISL_561683, EPI_ISL_561685, EPI_ISL_561687,
EPI_ISL_561688, EPI_ISL_561689, EPI_ISL_561691, EPI_ISL_561692, EPI_ISL_561693, EPI_ISL_561694, EPI_ISL_561695, EPI_ISL_561696, EPI_ISL_561698, EPI_ISL_561699, EPI_ISL_561700, EPI_ISL_561701, EPI_ISL_561702, EPI_ISL_561703, EPI_ISL_561704, EPI_ISL_561705, EPI_ISL_561706, EPI_ISL_561708,
EPI_ISL_561709, EPI_ISL_561710, EPI_ISL_561712, EPI_ISL_561713, EPI_ISL_561714, EPI_ISL_561716, EPI_ISL_561717, EPI_ISL_561718, EPI_ISL_561719, EPI_ISL_561720, EPI_ISL_561722, EPI_ISL_561723, EPI_ISL_561724, EPI_ISL_561725, EPI_ISL_561727, EPI_ISL_561729, EPI_ISL_561730, EPI_ISL_561731,
EPI_ISL_561732, EPI_ISL_561733, EPI_ISL_561734, EPI_ISL_561735, EPI_ISL_561736, EPI_ISL_561737, EPI_ISL_561738, EPI_ISL_561739, EPI_ISL_561740, EPI_ISL_561743, EPI_ISL_561744, EPI_ISL_561745, EPI_ISL_561746, EPI_ISL_561747, EPI_ISL_561748, EPI_ISL_561749, EPI_ISL_561751, EPI_ISL_561753,
EPI_ISL_561754, EPI_ISL_561756, EPI_ISL_561757, EPI_ISL_561758, EPI_ISL_561759, EPI_ISL_561760, EPI_ISL_561761, EPI_ISL_561762, EPI_ISL_561763, EPI_ISL_561767, EPI_ISL_561768, EPI_ISL_561769, EPI_ISL_561770, EPI_ISL_561771, EPI_ISL_561772, EPI_ISL_561773, EPI_ISL_561774, EPI_ISL_561775,
EPI_ISL_561776, EPI_ISL_561779, EPI_ISL_561780, EPI_ISL_561781, EPI_ISL_561782, EPI_ISL_561783, EPI_ISL_561784, EPI_ISL_561785, EPI_ISL_561786, EPI_ISL_561789, EPI_ISL_561790, EPI_ISL_561792, EPI_ISL_561794, EPI_ISL_561795, EPI_ISL_561796, EPI_ISL_561797, EPI_ISL_561798, EPI_ISL_561801,
EPI_ISL_561802, EPI_ISL_561804, EPI_ISL_561805, EPI_ISL_561806, EPI_ISL_561807, EPI_ISL_561809, EPI_ISL_561810, EPI_ISL_561811, EPI_ISL_561812, EPI_ISL_561813, EPI_ISL_561814, EPI_ISL_561815, EPI_ISL_561817, EPI_ISL_561818, EPI_ISL_561819, EPI_ISL_561820, EPI_ISL_561821, EPI_ISL_561822,
EPI_ISL_561823, EPI_ISL_561824, EPI_ISL_561826, EPI_ISL_561827, EPI_ISL_561828, EPI_ISL_561829, EPI_ISL_561830, EPI_ISL_561831, EPI_ISL_561832, EPI_ISL_561833, EPI_ISL_561834, EPI_ISL_561835, EPI_ISL_561836, EPI_ISL_561837, EPI_ISL_561838, EPI_ISL_561839, EPI_ISL_561841, EPI_ISL_561842,
EPI_ISL_561845, EPI_ISL_561846, EPI_ISL_561847, EPI_ISL_561848, EPI_ISL_561849, EPI_ISL_561850, EPI_ISL_561851, EPI_ISL_561853, EPI_ISL_561854, EPI_ISL_561855, EPI_ISL_561857, EPI_ISL_561859, EPI_ISL_561860, EPI_ISL_561861, EPI_ISL_561862, EPI_ISL_561864, EPI_ISL_561865, EPI_ISL_561867,
EPI_ISL_561868, EPI_ISL_561869, EPI_ISL_561870, EPI_ISL_561871, EPI_ISL_561872, EPI_ISL_561873, EPI_ISL_561874, EPI_ISL_561875, EPI_ISL_561876, EPI_ISL_561877, EPI_ISL_561878, EPI_ISL_561880, EPI_ISL_561881, EPI_ISL_561883, EPI_ISL_561884, EPI_ISL_561885, EPI_ISL_561886, EPI_ISL_561887,
EPI_ISL_561888, EPI_ISL_561889, EPI_ISL_561890, EPI_ISL_561891, EPI_ISL_561892, EPI_ISL_561895, EPI_ISL_561896, EPI_ISL_561897, EPI_ISL_561898, EPI_ISL_561899, EPI_ISL_561900, EPI_ISL_561901, EPI_ISL_561902, EPI_ISL_561903, EPI_ISL_561904, EPI_ISL_561905, EPI_ISL_561906, EPI_ISL_561908,
EPI_ISL_561909, EPI_ISL_561912, EPI_ISL_561913, EPI_ISL_561914, EPI_ISL_561915, EPI_ISL_561916, EPI_ISL_561919, EPI_ISL_561920, EPI_ISL_561921, EPI_ISL_561922, EPI_ISL_561923, EPI_ISL_561924, EPI_ISL_561925, EPI_ISL_561926, EPI_ISL_561927, EPI_ISL_561929, EPI_ISL_561930, EPI_ISL_561931,
EPI_ISL_561932, EPI_ISL_561933, EPI_ISL_561934, EPI_ISL_561937, EPI_ISL_561938, EPI_ISL_561939, EPI_ISL_561940, EPI_ISL_561941, EPI_ISL_561942, EPI_ISL_561943, EPI_ISL_561944, EPI_ISL_561945, EPI_ISL_561947, EPI_ISL_561948, EPI_ISL_561949, EPI_ISL_561950, EPI_ISL_561951, EPI_ISL_561952,
EPI_ISL_561953, EPI_ISL_561954, EPI_ISL_561955, EPI_ISL_561956, EPI_ISL_561958, EPI_ISL_561959, EPI_ISL_561960, EPI_ISL_561961, EPI_ISL_561962, EPI_ISL_561963, EPI_ISL_561964, EPI_ISL_561965, EPI_ISL_561966, EPI_ISL_561967, EPI_ISL_561968, EPI_ISL_561969, EPI_ISL_561970, EPI_ISL_561971,
EPI_ISL_561972, EPI_ISL_561973, EPI_ISL_561974, EPI_ISL_561976, EPI_ISL_561977, EPI_ISL_561978, EPI_ISL_561979, EPI_ISL_561980, EPI_ISL_561981, EPI_ISL_561982, EPI_ISL_561983, EPI_ISL_561984, EPI_ISL_561985, EPI_ISL_561989, EPI_ISL_561990, EPI_ISL_561992, EPI_ISL_561993, EPI_ISL_561995,
EPI_ISL_561996, EPI_ISL_561997, EPI_ISL_561998, EPI_ISL_561999, EPI_ISL_562000, EPI_ISL_562001, EPI_ISL_562002, EPI_ISL_562003, EPI_ISL_562004, EPI_ISL_562005, EPI_ISL_562006, EPI_ISL_562007, EPI_ISL_562008, EPI_ISL_562009, EPI_ISL_562010, EPI_ISL_562013, EPI_ISL_562014, EPI_ISL_562015,
EPI_ISL_562017, EPI_ISL_562018, EPI_ISL_562019, EPI_ISL_562020, EPI_ISL_562022, EPI_ISL_562024, EPI_ISL_562025, EPI_ISL_562026, EPI_ISL_562027, EPI_ISL_562028, EPI_ISL_562029, EPI_ISL_562030, EPI_ISL_562031, EPI_ISL_562032, EPI_ISL_562034, EPI_ISL_562035, EPI_ISL_562036, EPI_ISL_562037,
EPI_ISL_562038, EPI_ISL_562039, EPI_ISL_562040, EPI_ISL_562041, EPI_ISL_562042, EPI_ISL_562043, EPI_ISL_562044, EPI_ISL_562045, EPI_ISL_562046, EPI_ISL_562047, EPI_ISL_562049, EPI_ISL_562050, EPI_ISL_562051, EPI_ISL_562052, EPI_ISL_562053, EPI_ISL_562054, EPI_ISL_562058, EPI_ISL_562059,
EPI_ISL_562060, EPI_ISL_562061, EPI_ISL_562062, EPI_ISL_562063, EPI_ISL_562064, EPI_ISL_562065, EPI_ISL_562071, EPI_ISL_562073, EPI_ISL_562074, EPI_ISL_562075, EPI_ISL_562077, EPI_ISL_562078, EPI_ISL_562079, EPI_ISL_562082, EPI_ISL_562083, EPI_ISL_562084, EPI_ISL_562086, EPI_ISL_562087,
EPI_ISL_562088, EPI_ISL_562089, EPI_ISL_562090, EPI_ISL_562091, EPI_ISL_562092, EPI_ISL_562093, EPI_ISL_562094, EPI_ISL_562095, EPI_ISL_562096, EPI_ISL_562097, EPI_ISL_562098, EPI_ISL_562099, EPI_ISL_562100, EPI_ISL_562101, EPI_ISL_562102, EPI_ISL_562103, EPI_ISL_562107, EPI_ISL_562108,
EPI_ISL_562111, EPI_ISL_562112, EPI_ISL_562113, EPI_ISL_562114, EPI_ISL_562116, EPI_ISL_562117, EPI_ISL_562118, EPI_ISL_562119, EPI_ISL_562120, EPI_ISL_562121, EPI_ISL_562122, EPI_ISL_562124, EPI_ISL_562125, EPI_ISL_562126, EPI_ISL_562127, EPI_ISL_562128, EPI_ISL_562129, EPI_ISL_562130,
EPI_ISL_562131, EPI_ISL_562132, EPI_ISL_562133, EPI_ISL_562134, EPI_ISL_562135, EPI_ISL_562137, EPI_ISL_562139, EPI_ISL_562140, EPI_ISL_562141, EPI_ISL_562142, EPI_ISL_562143, EPI_ISL_562144, EPI_ISL_562146, EPI_ISL_562147, EPI_ISL_562148, EPI_ISL_562149, EPI_ISL_562150, EPI_ISL_562151,
EPI_ISL_562153, EPI_ISL_562154, EPI_ISL_562156, EPI_ISL_562159, EPI_ISL_562160, EPI_ISL_562161, EPI_ISL_562162, EPI_ISL_562165, EPI_ISL_562167, EPI_ISL_562168, EPI_ISL_562169, EPI_ISL_562170, EPI_ISL_562171, EPI_ISL_562172, EPI_ISL_562173, EPI_ISL_562174, EPI_ISL_562176, EPI_ISL_562177,
EPI_ISL_562178, EPI_ISL_562180, EPI_ISL_562181, EPI_ISL_562183, EPI_ISL_562185, EPI_ISL_562186, EPI_ISL_562187, EPI_ISL_562188, EPI_ISL_562190, EPI_ISL_562191, EPI_ISL_562192, EPI_ISL_562193, EPI_ISL_562195, EPI_ISL_562196, EPI_ISL_562198, EPI_ISL_562199, EPI_ISL_562201, EPI_ISL_562203,
EPI_ISL_562204, EPI_ISL_562205, EPI_ISL_562206, EPI_ISL_562210, EPI_ISL_562212, EPI_ISL_562214, EPI_ISL_562215, EPI_ISL_562218, EPI_ISL_562219, EPI_ISL_562220, EPI_ISL_562221, EPI_ISL_562222, EPI_ISL_562223, EPI_ISL_562226, EPI_ISL_562227, EPI_ISL_562230, EPI_ISL_562232, EPI_ISL_562233,
EPI_ISL_562234, EPI_ISL_562235, EPI_ISL_562236, EPI_ISL_562238, EPI_ISL_562240, EPI_ISL_562241, EPI_ISL_562243, EPI_ISL_562246, EPI_ISL_562247, EPI_ISL_562248, EPI_ISL_562249, EPI_ISL_562251, EPI_ISL_562252, EPI_ISL_562253, EPI_ISL_562254, EPI_ISL_562255, EPI_ISL_562256, EPI_ISL_562257,
EPI_ISL_562260, EPI_ISL_562262, EPI_ISL_562263, EPI_ISL_562265, EPI_ISL_562268, EPI_ISL_562270, EPI_ISL_562271, EPI_ISL_562276, EPI_ISL_562277, EPI_ISL_562279, EPI_ISL_562281, EPI_ISL_562283, EPI_ISL_562284, EPI_ISL_562285, EPI_ISL_562287, EPI_ISL_562291, EPI_ISL_562292, EPI_ISL_562293,
EPI_ISL_562295, EPI_ISL_562300, EPI_ISL_562301, EPI_ISL_562302, EPI_ISL_562303, EPI_ISL_562305, EPI_ISL_562306, EPI_ISL_562307, EPI_ISL_562309, EPI_ISL_562310, EPI_ISL_562311, EPI_ISL_562313, EPI_ISL_562315, EPI_ISL_562316, EPI_ISL_562318, EPI_ISL_562319, EPI_ISL_562320, EPI_ISL_562323,
EPI_ISL_562324, EPI_ISL_562325, EPI_ISL_562327, EPI_ISL_562329, EPI_ISL_562330, EPI_ISL_562332, EPI_ISL_562334, EPI_ISL_562336, EPI_ISL_562337, EPI_ISL_562338, EPI_ISL_562339, EPI_ISL_562340, EPI_ISL_562341, EPI_ISL_562343, EPI_ISL_562344, EPI_ISL_562345, EPI_ISL_562346, EPI_ISL_562349,
EPI_ISL_562352, EPI_ISL_562353, EPI_ISL_562355, EPI_ISL_562358, EPI_ISL_562359, EPI_ISL_562360, EPI_ISL_562361, EPI_ISL_562362, EPI_ISL_562364, EPI_ISL_562365, EPI_ISL_562367, EPI_ISL_562368, EPI_ISL_562369, EPI_ISL_562370, EPI_ISL_562371, EPI_ISL_562372, EPI_ISL_562373, EPI_ISL_562374,
EPI_ISL_562378, EPI_ISL_562380, EPI_ISL_562382, EPI_ISL_562383, EPI_ISL_562384, EPI_ISL_562386, EPI_ISL_562387, EPI_ISL_562388, EPI_ISL_562392, EPI_ISL_562394, EPI_ISL_562395, EPI_ISL_562397, EPI_ISL_562398, EPI_ISL_562399, EPI_ISL_562402, EPI_ISL_562403, EPI_ISL_562404, EPI_ISL_562405,
EPI_ISL_562407, EPI_ISL_562408, EPI_ISL_562409, EPI_ISL_562411, EPI_ISL_562416, EPI_ISL_562417, EPI_ISL_562418, EPI_ISL_562419, EPI_ISL_562420, EPI_ISL_562421, EPI_ISL_562422, EPI_ISL_562423, EPI_ISL_562424, EPI_ISL_562425, EPI_ISL_562426, EPI_ISL_562427, EPI_ISL_562428, EPI_ISL_562429,
EPI_ISL_562432, EPI_ISL_562433, EPI_ISL_562434, EPI_ISL_562436, EPI_ISL_562438, EPI_ISL_562444, EPI_ISL_562449, EPI_ISL_562450, EPI_ISL_562453, EPI_ISL_562454, EPI_ISL_562456, EPI_ISL_562457, EPI_ISL_562461, EPI_ISL_562465, EPI_ISL_562468, EPI_ISL_562469, EPI_ISL_562470, EPI_ISL_562471,
EPI_ISL_562473, EPI_ISL_562475, EPI_ISL_562477, EPI_ISL_562478, EPI_ISL_562480, EPI_ISL_562481, EPI_ISL_562485, EPI_ISL_562489, EPI_ISL_562490, EPI_ISL_562492, EPI_ISL_562493, EPI_ISL_562494, EPI_ISL_562495, EPI_ISL_562496, EPI_ISL_562497, EPI_ISL_562498, EPI_ISL_562499, EPI_ISL_562500,
EPI_ISL_562501, EPI_ISL_562504, EPI_ISL_562506, EPI_ISL_562507, EPI_ISL_562510, EPI_ISL_562511, EPI_ISL_562512, EPI_ISL_562515, EPI_ISL_562516, EPI_ISL_562519, EPI_ISL_562520, EPI_ISL_562521, EPI_ISL_562522, EPI_ISL_562523, EPI_ISL_562525, EPI_ISL_562528, EPI_ISL_562529, EPI_ISL_562530,
EPI_ISL_562532, EPI_ISL_562533, EPI_ISL_562534, EPI_ISL_562537, EPI_ISL_562541, EPI_ISL_562543, EPI_ISL_562544, EPI_ISL_562547, EPI_ISL_562549, EPI_ISL_562550, EPI_ISL_562551, EPI_ISL_562554, EPI_ISL_562555, EPI_ISL_562556, EPI_ISL_562560, EPI_ISL_562561, EPI_ISL_562562, EPI_ISL_562564,
EPI_ISL_562565, EPI_ISL_562566, EPI_ISL_562567, EPI_ISL_562568, EPI_ISL_562569, EPI_ISL_562570, EPI_ISL_562571, EPI_ISL_562572, EPI_ISL_562573, EPI_ISL_562574, EPI_ISL_562575, EPI_ISL_562576, EPI_ISL_562577, EPI_ISL_562578, EPI_ISL_562580, EPI_ISL_562581, EPI_ISL_562582, EPI_ISL_562584,
EPI_ISL_562591, EPI_ISL_562592, EPI_ISL_562594, EPI_ISL_562596, EPI_ISL_562597, EPI_ISL_562598, EPI_ISL_562599, EPI_ISL_562600, EPI_ISL_562601, EPI_ISL_562602, EPI_ISL_562603, EPI_ISL_562604, EPI_ISL_562605, EPI_ISL_562606, EPI_ISL_562607, EPI_ISL_562608, EPI_ISL_562609, EPI_ISL_562610,
EPI_ISL_562611, EPI_ISL_562612, EPI_ISL_562613, EPI_ISL_562614, EPI_ISL_562615, EPI_ISL_562616, EPI_ISL_562617, EPI_ISL_562618, EPI_ISL_562619, EPI_ISL_562620, EPI_ISL_562621, EPI_ISL_562622, EPI_ISL_562623, EPI_ISL_562624, EPI_ISL_562625, EPI_ISL_562626, EPI_ISL_562627, EPI_ISL_562629,
EPI_ISL_562631, EPI_ISL_562632, EPI_ISL_562633, EPI_ISL_562634, EPI_ISL_562635, EPI_ISL_562636, EPI_ISL_562637, EPI_ISL_562638, EPI_ISL_562639, EPI_ISL_562641, EPI_ISL_562642, EPI_ISL_562643, EPI_ISL_562644, EPI_ISL_562647, EPI_ISL_562648, EPI_ISL_562649, EPI_ISL_562650, EPI_ISL_562651,
EPI_ISL_562652, EPI_ISL_562653, EPI_ISL_562654, EPI_ISL_562655, EPI_ISL_562656, EPI_ISL_562657, EPI_ISL_562660, EPI_ISL_562661, EPI_ISL_562662, EPI_ISL_562663, EPI_ISL_562664, EPI_ISL_562665, EPI_ISL_562666, EPI_ISL_562667, EPI_ISL_562668, EPI_ISL_562670, EPI_ISL_562672, EPI_ISL_562673,
EPI_ISL_562674, EPI_ISL_562675, EPI_ISL_562676, EPI_ISL_562677, EPI_ISL_562678, EPI_ISL_562679, EPI_ISL_562680, EPI_ISL_562681, EPI_ISL_562682, EPI_ISL_562683, EPI_ISL_562684, EPI_ISL_562685, EPI_ISL_562686, EPI_ISL_562687, EPI_ISL_562688, EPI_ISL_562689, EPI_ISL_562690, EPI_ISL_562691,
EPI_ISL_562692, EPI_ISL_562693, EPI_ISL_562694, EPI_ISL_562695, EPI_ISL_562698, EPI_ISL_562699, EPI_ISL_562701, EPI_ISL_562702, EPI_ISL_562703, EPI_ISL_562704, EPI_ISL_562706, EPI_ISL_562707, EPI_ISL_562708, EPI_ISL_562709, EPI_ISL_562710, EPI_ISL_562711, EPI_ISL_562712, EPI_ISL_562713,
EPI_ISL_562714, EPI_ISL_562715, EPI_ISL_562716, EPI_ISL_562717, EPI_ISL_562718, EPI_ISL_562719, EPI_ISL_562720, EPI_ISL_562721, EPI_ISL_562722, EPI_ISL_562723, EPI_ISL_562724, EPI_ISL_562725, EPI_ISL_562726, EPI_ISL_562727, EPI_ISL_562728, EPI_ISL_562729, EPI_ISL_562730, EPI_ISL_562731,
EPI_ISL_562732, EPI_ISL_562733, EPI_ISL_562734, EPI_ISL_562735, EPI_ISL_562736, EPI_ISL_562737, EPI_ISL_562738, EPI_ISL_562739, EPI_ISL_562740, EPI_ISL_562741, EPI_ISL_562742, EPI_ISL_562743, EPI_ISL_562744, EPI_ISL_562745, EPI_ISL_562746, EPI_ISL_562747, EPI_ISL_562748, EPI_ISL_562749,
EPI_ISL_562750, EPI_ISL_562751, EPI_ISL_562752, EPI_ISL_562753, EPI_ISL_562755, EPI_ISL_562756, EPI_ISL_562757, EPI_ISL_562758, EPI_ISL_562759, EPI_ISL_562760, EPI_ISL_562761, EPI_ISL_562762, EPI_ISL_562763, EPI_ISL_562764, EPI_ISL_562765, EPI_ISL_562766, EPI_ISL_562767, EPI_ISL_562768,
EPI_ISL_562769, EPI_ISL_562770, EPI_ISL_562771, EPI_ISL_562772, EPI_ISL_562773, EPI_ISL_562774, EPI_ISL_562775, EPI_ISL_562776, EPI_ISL_562777, EPI_ISL_562778, EPI_ISL_562779, EPI_ISL_562780, EPI_ISL_562781, EPI_ISL_562782, EPI_ISL_562783, EPI_ISL_562784, EPI_ISL_562785, EPI_ISL_562786,
EPI_ISL_562787, EPI_ISL_562788, EPI_ISL_562789, EPI_ISL_562790, EPI_ISL_562793, EPI_ISL_562794, EPI_ISL_562795, EPI_ISL_562796, EPI_ISL_562797, EPI_ISL_562798, EPI_ISL_562799, EPI_ISL_562800, EPI_ISL_562801, EPI_ISL_562802, EPI_ISL_562810, EPI_ISL_562811, EPI_ISL_562812, EPI_ISL_562813,
EPI_ISL_562814, EPI_ISL_562815, EPI_ISL_562816, EPI_ISL_562817, EPI_ISL_562818, EPI_ISL_562819, EPI_ISL_562820, EPI_ISL_562821, EPI_ISL_562822, EPI_ISL_562823, EPI_ISL_562824, EPI_ISL_562825, EPI_ISL_562826, EPI_ISL_562827, EPI_ISL_562828, EPI_ISL_562829, EPI_ISL_562830, EPI_ISL_562831,
EPI_ISL_562832, EPI_ISL_562833, EPI_ISL_562834, EPI_ISL_562835, EPI_ISL_562836, EPI_ISL_562837, EPI_ISL_562846, EPI_ISL_562847, EPI_ISL_562848, EPI_ISL_562849, EPI_ISL_562850, EPI_ISL_562851, EPI_ISL_562852, EPI_ISL_562853, EPI_ISL_562854, EPI_ISL_562855, EPI_ISL_562856, EPI_ISL_562857,
EPI_ISL_562858, EPI_ISL_562859, EPI_ISL_562860, EPI_ISL_562861, EPI_ISL_562862, EPI_ISL_562863, EPI_ISL_562864, EPI_ISL_562865, EPI_ISL_562866, EPI_ISL_562867, EPI_ISL_562868, EPI_ISL_562869, EPI_ISL_562870, EPI_ISL_562871, EPI_ISL_562872, EPI_ISL_562873, EPI_ISL_562874, EPI_ISL_562875,
EPI_ISL_562876, EPI_ISL_562877, EPI_ISL_562878, EPI_ISL_562879, EPI_ISL_562880, EPI_ISL_562881, EPI_ISL_562882, EPI_ISL_562883, EPI_ISL_562884, EPI_ISL_562885, EPI_ISL_562886, EPI_ISL_562887, EPI_ISL_562888, EPI_ISL_562889, EPI_ISL_562890, EPI_ISL_562891, EPI_ISL_562892, EPI_ISL_562893,
EPI_ISL_562894, EPI_ISL_562895, EPI_ISL_562896, EPI_ISL_562897, EPI_ISL_562898, EPI_ISL_562899, EPI_ISL_562900, EPI_ISL_562901, EPI_ISL_562902, EPI_ISL_562903, EPI_ISL_562904, EPI_ISL_562905, EPI_ISL_562907, EPI_ISL_562908, EPI_ISL_562909, EPI_ISL_562914, EPI_ISL_562915, EPI_ISL_562916,
EPI_ISL_562917, EPI_ISL_562918, EPI_ISL_562919, EPI_ISL_562920, EPI_ISL_562921, EPI_ISL_562922, EPI_ISL_562923, EPI_ISL_562925, EPI_ISL_562926, EPI_ISL_562927, EPI_ISL_562928, EPI_ISL_562930, EPI_ISL_562931, EPI_ISL_562932, EPI_ISL_562933, EPI_ISL_562934, EPI_ISL_562935, EPI_ISL_562937,
EPI_ISL_562938, EPI_ISL_562939, EPI_ISL_562940, EPI_ISL_562941, EPI_ISL_562942, EPI_ISL_562943, EPI_ISL_562944, EPI_ISL_562945, EPI_ISL_562947, EPI_ISL_562948, EPI_ISL_562949, EPI_ISL_562950, EPI_ISL_562952, EPI_ISL_562953, EPI_ISL_562954, EPI_ISL_562955, EPI_ISL_562956, EPI_ISL_562957,
EPI_ISL_562958, EPI_ISL_562960, EPI_ISL_562962, EPI_ISL_562963, EPI_ISL_562964, EPI_ISL_562965, EPI_ISL_562966, EPI_ISL_562967, EPI_ISL_562968, EPI_ISL_562969, EPI_ISL_562970, EPI_ISL_562971, EPI_ISL_562972, EPI_ISL_562973, EPI_ISL_562974, EPI_ISL_562975, EPI_ISL_562976, EPI_ISL_562977,
EPI_ISL_562978, EPI_ISL_562979, EPI_ISL_562980, EPI_ISL_562981, EPI_ISL_562982, EPI_ISL_562983, EPI_ISL_562984, EPI_ISL_562985, EPI_ISL_562986, EPI_ISL_562987, EPI_ISL_562988, EPI_ISL_562989, EPI_ISL_562990, EPI_ISL_562991, EPI_ISL_562992, EPI_ISL_562993, EPI_ISL_562994, EPI_ISL_562995,
EPI_ISL_562996, EPI_ISL_562997, EPI_ISL_562998, EPI_ISL_562999, EPI_ISL_563001, EPI_ISL_563002, EPI_ISL_563003, EPI_ISL_563004, EPI_ISL_563005, EPI_ISL_563007, EPI_ISL_563008, EPI_ISL_563009, EPI_ISL_563011, EPI_ISL_563013, EPI_ISL_563014, EPI_ISL_563015, EPI_ISL_563016, EPI_ISL_563018,
EPI_ISL_563019, EPI_ISL_563020, EPI_ISL_563021, EPI_ISL_563022, EPI_ISL_563024, EPI_ISL_563025, EPI_ISL_563026, EPI_ISL_563027, EPI_ISL_563028, EPI_ISL_563029, EPI_ISL_563030, EPI_ISL_563031, EPI_ISL_563032, EPI_ISL_563033, EPI_ISL_563034, EPI_ISL_563035, EPI_ISL_563036, EPI_ISL_563037,
EPI_ISL_563038, EPI_ISL_563039, EPI_ISL_563040, EPI_ISL_563042, EPI_ISL_563043, EPI_ISL_563044, EPI_ISL_563045, EPI_ISL_563046, EPI_ISL_563047, EPI_ISL_563048, EPI_ISL_563049, EPI_ISL_563050, EPI_ISL_563051, EPI_ISL_563053, EPI_ISL_563054, EPI_ISL_563055, EPI_ISL_563056, EPI_ISL_563057,
EPI_ISL_563058, EPI_ISL_563059, EPI_ISL_563060, EPI_ISL_563061, EPI_ISL_563062, EPI_ISL_563063, EPI_ISL_563064, EPI_ISL_563065, EPI_ISL_563067, EPI_ISL_563068, EPI_ISL_563069, EPI_ISL_563070, EPI_ISL_563071, EPI_ISL_563072, EPI_ISL_563074, EPI_ISL_563075, EPI_ISL_563076, EPI_ISL_563077,
EPI_ISL_563078, EPI_ISL_563079, EPI_ISL_563080, EPI_ISL_563081, EPI_ISL_563082, EPI_ISL_563083, EPI_ISL_563084, EPI_ISL_563085, EPI_ISL_563086, EPI_ISL_563087, EPI_ISL_563088, EPI_ISL_563089, EPI_ISL_563090, EPI_ISL_563091, EPI_ISL_563092, EPI_ISL_563093, EPI_ISL_563094, EPI_ISL_563095,

EPI_ISL_563096, EPI_ISL_563097, EPI_ISL_563098, EPI_ISL_563099, EPI_ISL_563100, EPI_ISL_563101, EPI_ISL_563102, EPI_ISL_563103, EPI_ISL_563104, EPI_ISL_563105, EPI_ISL_563106, EPI_ISL_563107, EPI_ISL_563108, EPI_ISL_563109, EPI_ISL_563110, EPI_ISL_563111, EPI_ISL_563113, EPI_ISL_563117,
EPI_ISL_563118, EPI_ISL_563119, EPI_ISL_563120, EPI_ISL_563121, EPI_ISL_563122, EPI_ISL_563123, EPI_ISL_563124, EPI_ISL_563125, EPI_ISL_563126, EPI_ISL_563127, EPI_ISL_563128, EPI_ISL_563129, EPI_ISL_563130, EPI_ISL_563134, EPI_ISL_563135, EPI_ISL_563145, EPI_ISL_563146, EPI_ISL_563147,
EPI_ISL_563148, EPI_ISL_563149, EPI_ISL_563150, EPI_ISL_563151, EPI_ISL_563152, EPI_ISL_563153, EPI_ISL_563154, EPI_ISL_563155, EPI_ISL_563156, EPI_ISL_563157, EPI_ISL_563158, EPI_ISL_563159, EPI_ISL_563160, EPI_ISL_563161, EPI_ISL_563162, EPI_ISL_563163, EPI_ISL_563164, EPI_ISL_563165,
EPI_ISL_563166, EPI_ISL_563167, EPI_ISL_563168, EPI_ISL_563170, EPI_ISL_563171, EPI_ISL_563172, EPI_ISL_563174, EPI_ISL_563175, EPI_ISL_563176, EPI_ISL_563177, EPI_ISL_563178, EPI_ISL_563179, EPI_ISL_563180, EPI_ISL_563181, EPI_ISL_563182, EPI_ISL_563183, EPI_ISL_563184, EPI_ISL_563185,
EPI_ISL_563187, EPI_ISL_563188, EPI_ISL_563190, EPI_ISL_563191, EPI_ISL_563192, EPI_ISL_563193, EPI_ISL_563194, EPI_ISL_563195, EPI_ISL_563198, EPI_ISL_563199, EPI_ISL_563200, EPI_ISL_563202, EPI_ISL_563204, EPI_ISL_563205, EPI_ISL_563206, EPI_ISL_563208, EPI_ISL_563209, EPI_ISL_563210,
EPI_ISL_563211, EPI_ISL_563212, EPI_ISL_563213, EPI_ISL_563214, EPI_ISL_563215, EPI_ISL_563216, EPI_ISL_563218, EPI_ISL_563221, EPI_ISL_563222, EPI_ISL_563224, EPI_ISL_563226, EPI_ISL_563227, EPI_ISL_563228, EPI_ISL_563229, EPI_ISL_563230, EPI_ISL_563231, EPI_ISL_563232, EPI_ISL_563233,
EPI_ISL_563234, EPI_ISL_563235, EPI_ISL_563236, EPI_ISL_563237, EPI_ISL_563238, EPI_ISL_563239, EPI_ISL_563240, EPI_ISL_563241, EPI_ISL_563243, EPI_ISL_563245, EPI_ISL_563246, EPI_ISL_563247, EPI_ISL_563248, EPI_ISL_563249, EPI_ISL_563250, EPI_ISL_563251, EPI_ISL_563252, EPI_ISL_563253,
EPI_ISL_563254, EPI_ISL_563255, EPI_ISL_563256, EPI_ISL_563257, EPI_ISL_563258, EPI_ISL_563259, EPI_ISL_563260, EPI_ISL_563261, EPI_ISL_563262, EPI_ISL_563263, EPI_ISL_563264, EPI_ISL_563265, EPI_ISL_563266, EPI_ISL_563267, EPI_ISL_563268, EPI_ISL_563269, EPI_ISL_563270, EPI_ISL_563271,
EPI_ISL_563272, EPI_ISL_563273, EPI_ISL_563274, EPI_ISL_563275, EPI_ISL_563276, EPI_ISL_563277, EPI_ISL_563278, EPI_ISL_563279, EPI_ISL_563280, EPI_ISL_563281, EPI_ISL_563282, EPI_ISL_563283, EPI_ISL_563284, EPI_ISL_563285, EPI_ISL_563286, EPI_ISL_563287, EPI_ISL_563288, EPI_ISL_563289,
EPI_ISL_563290, EPI_ISL_563291, EPI_ISL_563292, EPI_ISL_563293, EPI_ISL_563294, EPI_ISL_563295, EPI_ISL_563296, EPI_ISL_563297, EPI_ISL_563298, EPI_ISL_563299, EPI_ISL_563300, EPI_ISL_563301, EPI_ISL_563302, EPI_ISL_563303, EPI_ISL_563304, EPI_ISL_563305, EPI_ISL_563306, EPI_ISL_563307,
EPI_ISL_563308, EPI_ISL_563309, EPI_ISL_563310, EPI_ISL_563311, EPI_ISL_563312, EPI_ISL_563313, EPI_ISL_563314, EPI_ISL_563315, EPI_ISL_563316, EPI_ISL_563317, EPI_ISL_563318, EPI_ISL_563319, EPI_ISL_563320, EPI_ISL_563321, EPI_ISL_563322, EPI_ISL_563323, EPI_ISL_563324, EPI_ISL_563325,
EPI_ISL_563326, EPI_ISL_563327, EPI_ISL_563328, EPI_ISL_563329, EPI_ISL_563330, EPI_ISL_563331, EPI_ISL_563332, EPI_ISL_563333, EPI_ISL_563334, EPI_ISL_563335, EPI_ISL_563336, EPI_ISL_563337, EPI_ISL_563338, EPI_ISL_563339, EPI_ISL_563340, EPI_ISL_563341, EPI_ISL_563342, EPI_ISL_563343,
EPI_ISL_563344, EPI_ISL_563345, EPI_ISL_563346, EPI_ISL_563347, EPI_ISL_563348, EPI_ISL_563349, EPI_ISL_563350, EPI_ISL_563351, EPI_ISL_563352, EPI_ISL_563353, EPI_ISL_563354, EPI_ISL_563355, EPI_ISL_563356, EPI_ISL_563357, EPI_ISL_563358, EPI_ISL_563359, EPI_ISL_563360, EPI_ISL_563361,
EPI_ISL_563362, EPI_ISL_563363, EPI_ISL_563364, EPI_ISL_563365, EPI_ISL_563367, EPI_ISL_563368, EPI_ISL_563369, EPI_ISL_563370, EPI_ISL_563371, EPI_ISL_563372, EPI_ISL_563373, EPI_ISL_563374, EPI_ISL_563375, EPI_ISL_563376, EPI_ISL_563377, EPI_ISL_563378, EPI_ISL_563379, EPI_ISL_563380,
EPI_ISL_563381, EPI_ISL_563382, EPI_ISL_563383, EPI_ISL_563384, EPI_ISL_563385, EPI_ISL_563386, EPI_ISL_563387, EPI_ISL_563388, EPI_ISL_563389, EPI_ISL_563390, EPI_ISL_563391, EPI_ISL_563392, EPI_ISL_563393, EPI_ISL_563394, EPI_ISL_563395, EPI_ISL_563396, EPI_ISL_563397, EPI_ISL_563398,
EPI_ISL_563399, EPI_ISL_563400, EPI_ISL_563401, EPI_ISL_563402, EPI_ISL_563403, EPI_ISL_563404, EPI_ISL_563405, EPI_ISL_563406, EPI_ISL_563424, EPI_ISL_563425, EPI_ISL_563426, EPI_ISL_563427, EPI_ISL_563428, EPI_ISL_563429, EPI_ISL_563430, EPI_ISL_563431, EPI_ISL_563432, EPI_ISL_563433,
EPI_ISL_563434, EPI_ISL_563435, EPI_ISL_563436, EPI_ISL_563437, EPI_ISL_563438, EPI_ISL_563439, EPI_ISL_563440, EPI_ISL_563441, EPI_ISL_563442, EPI_ISL_563443, EPI_ISL_563444, EPI_ISL_563445, EPI_ISL_563446, EPI_ISL_563447, EPI_ISL_563448, EPI_ISL_563449, EPI_ISL_563450, EPI_ISL_563451,
EPI_ISL_563452, EPI_ISL_563453, EPI_ISL_563454, EPI_ISL_563455, EPI_ISL_563456, EPI_ISL_563457, EPI_ISL_563458, EPI_ISL_563459, EPI_ISL_563460, EPI_ISL_563461, EPI_ISL_563462, EPI_ISL_563463, EPI_ISL_563464, EPI_ISL_563465, EPI_ISL_563466, EPI_ISL_563467, EPI_ISL_563468, EPI_ISL_563469,
EPI_ISL_563470, EPI_ISL_563471, EPI_ISL_563472, EPI_ISL_563473, EPI_ISL_563474, EPI_ISL_563475, EPI_ISL_563476, EPI_ISL_563477, EPI_ISL_563478, EPI_ISL_563479, EPI_ISL_563480, EPI_ISL_563481, EPI_ISL_563482, EPI_ISL_563483, EPI_ISL_563484, EPI_ISL_563485, EPI_ISL_563486, EPI_ISL_563487,
EPI_ISL_563488, EPI_ISL_563489, EPI_ISL_563490, EPI_ISL_563491, EPI_ISL_563492, EPI_ISL_563493, EPI_ISL_563494, EPI_ISL_563495, EPI_ISL_563496, EPI_ISL_563500, EPI_ISL_563501, EPI_ISL_563502, EPI_ISL_563503, EPI_ISL_563504, EPI_ISL_563505, EPI_ISL_563506, EPI_ISL_563507, EPI_ISL_563508,
EPI_ISL_563509, EPI_ISL_563510, EPI_ISL_563511, EPI_ISL_563512, EPI_ISL_563513, EPI_ISL_563514, EPI_ISL_563515, EPI_ISL_563516, EPI_ISL_563517, EPI_ISL_563518, EPI_ISL_563519, EPI_ISL_563520, EPI_ISL_563521, EPI_ISL_563522, EPI_ISL_563523, EPI_ISL_563524, EPI_ISL_563525, EPI_ISL_563526,
EPI_ISL_563527, EPI_ISL_563528, EPI_ISL_563529, EPI_ISL_563530, EPI_ISL_563531, EPI_ISL_563532, EPI_ISL_563533, EPI_ISL_563534, EPI_ISL_563535, EPI_ISL_563536, EPI_ISL_563537, EPI_ISL_563538, EPI_ISL_563539, EPI_ISL_563540, EPI_ISL_563541, EPI_ISL_563542, EPI_ISL_563543, EPI_ISL_563544,
EPI_ISL_563545, EPI_ISL_563546, EPI_ISL_563547, EPI_ISL_563548, EPI_ISL_563549, EPI_ISL_563550, EPI_ISL_563552, EPI_ISL_563553, EPI_ISL_563554, EPI_ISL_563555, EPI_ISL_563556, EPI_ISL_563557, EPI_ISL_563558, EPI_ISL_563559, EPI_ISL_563560, EPI_ISL_563561, EPI_ISL_563562, EPI_ISL_563563,
EPI_ISL_563564, EPI_ISL_563565, EPI_ISL_563566, EPI_ISL_563567, EPI_ISL_563568, EPI_ISL_563569, EPI_ISL_563570, EPI_ISL_563571, EPI_ISL_563572, EPI_ISL_563573, EPI_ISL_563574, EPI_ISL_563575, EPI_ISL_563576, EPI_ISL_563577, EPI_ISL_563578, EPI_ISL_563579, EPI_ISL_563580, EPI_ISL_563581,
EPI_ISL_563582, EPI_ISL_563583, EPI_ISL_563584, EPI_ISL_563585, EPI_ISL_563586, EPI_ISL_563587, EPI_ISL_563588, EPI_ISL_563589, EPI_ISL_563590, EPI_ISL_563591, EPI_ISL_563592, EPI_ISL_563593, EPI_ISL_563594, EPI_ISL_563595, EPI_ISL_563596, EPI_ISL_563597, EPI_ISL_563598, EPI_ISL_563599,
EPI_ISL_563600, EPI_ISL_563601, EPI_ISL_563602, EPI_ISL_563603, EPI_ISL_563604, EPI_ISL_563605, EPI_ISL_563606, EPI_ISL_563607, EPI_ISL_563608, EPI_ISL_563610, EPI_ISL_563611, EPI_ISL_563612, EPI_ISL_563613, EPI_ISL_563614, EPI_ISL_563615, EPI_ISL_563616, EPI_ISL_563617, EPI_ISL_563618,
EPI_ISL_563619, EPI_ISL_563620, EPI_ISL_563621, EPI_ISL_563622, EPI_ISL_563623, EPI_ISL_563624, EPI_ISL_563625, EPI_ISL_563626, EPI_ISL_563627, EPI_ISL_563628, EPI_ISL_563629, EPI_ISL_563630, EPI_ISL_563631, EPI_ISL_563632, EPI_ISL_563633, EPI_ISL_563634, EPI_ISL_563635, EPI_ISL_563636,
EPI_ISL_563637, EPI_ISL_563638, EPI_ISL_563639, EPI_ISL_563640, EPI_ISL_563641, EPI_ISL_563642, EPI_ISL_563643, EPI_ISL_563644, EPI_ISL_563645, EPI_ISL_563646, EPI_ISL_563647, EPI_ISL_563648, EPI_ISL_563649, EPI_ISL_563650, EPI_ISL_563651, EPI_ISL_563652, EPI_ISL_563653, EPI_ISL_563654,
EPI_ISL_563655, EPI_ISL_563656, EPI_ISL_563657, EPI_ISL_563658, EPI_ISL_563659, EPI_ISL_563660, EPI_ISL_563661, EPI_ISL_563662, EPI_ISL_563663, EPI_ISL_563664, EPI_ISL_563665, EPI_ISL_563666, EPI_ISL_563667, EPI_ISL_563668, EPI_ISL_563669, EPI_ISL_563670, EPI_ISL_563671, EPI_ISL_563672,
EPI_ISL_563673, EPI_ISL_563674, EPI_ISL_563675, EPI_ISL_563676, EPI_ISL_563677, EPI_ISL_563678, EPI_ISL_563679, EPI_ISL_563680, EPI_ISL_563681, EPI_ISL_563682, EPI_ISL_563683, EPI_ISL_563684, EPI_ISL_563685, EPI_ISL_563686, EPI_ISL_563687, EPI_ISL_563688, EPI_ISL_563689, EPI_ISL_563690,
EPI_ISL_563691, EPI_ISL_563692, EPI_ISL_563693, EPI_ISL_563694, EPI_ISL_563695, EPI_ISL_563696, EPI_ISL_563697, EPI_ISL_563698, EPI_ISL_563699, EPI_ISL_563700, EPI_ISL_563701, EPI_ISL_563702, EPI_ISL_563703, EPI_ISL_563704, EPI_ISL_563705, EPI_ISL_563706, EPI_ISL_563707, EPI_ISL_563708,
EPI_ISL_563709, EPI_ISL_563710, EPI_ISL_563711, EPI_ISL_563712, EPI_ISL_563713, EPI_ISL_563714, EPI_ISL_563715, EPI_ISL_563716, EPI_ISL_563717, EPI_ISL_563718, EPI_ISL_563719, EPI_ISL_563720, EPI_ISL_563724, EPI_ISL_563725, EPI_ISL_563726, EPI_ISL_563727, EPI_ISL_563728, EPI_ISL_563729,
EPI_ISL_563730, EPI_ISL_563731, EPI_ISL_563732, EPI_ISL_563733, EPI_ISL_563734, EPI_ISL_563735, EPI_ISL_563736, EPI_ISL_563737, EPI_ISL_563738, EPI_ISL_563739, EPI_ISL_563740, EPI_ISL_563741, EPI_ISL_563742, EPI_ISL_563743, EPI_ISL_563744, EPI_ISL_563745, EPI_ISL_563746, EPI_ISL_563747,
EPI_ISL_563748, EPI_ISL_563749, EPI_ISL_563750, EPI_ISL_563751, EPI_ISL_563752, EPI_ISL_563753, EPI_ISL_563754, EPI_ISL_563755, EPI_ISL_563756, EPI_ISL_563757, EPI_ISL_563758, EPI_ISL_563759, EPI_ISL_563760, EPI_ISL_563761, EPI_ISL_563762, EPI_ISL_563763, EPI_ISL_563764, EPI_ISL_563765,
EPI_ISL_563766, EPI_ISL_563767, EPI_ISL_563768, EPI_ISL_563769, EPI_ISL_563770, EPI_ISL_563771, EPI_ISL_563772, EPI_ISL_563773, EPI_ISL_563774, EPI_ISL_563775, EPI_ISL_563776, EPI_ISL_563777, EPI_ISL_563778, EPI_ISL_563779, EPI_ISL_563780, EPI_ISL_563781, EPI_ISL_563782, EPI_ISL_563783,
EPI_ISL_563784, EPI_ISL_563785, EPI_ISL_563786, EPI_ISL_563787, EPI_ISL_563788, EPI_ISL_563789, EPI_ISL_563791, EPI_ISL_563793, EPI_ISL_563794, EPI_ISL_563795, EPI_ISL_563796, EPI_ISL_563797, EPI_ISL_563798, EPI_ISL_563799, EPI_ISL_563800, EPI_ISL_563801, EPI_ISL_563802, EPI_ISL_563803,
EPI_ISL_563804, EPI_ISL_563805, EPI_ISL_563806, EPI_ISL_563807, EPI_ISL_563809, EPI_ISL_563810, EPI_ISL_563812, EPI_ISL_563813, EPI_ISL_563814, EPI_ISL_563815, EPI_ISL_563816, EPI_ISL_563818, EPI_ISL_563819, EPI_ISL_563821, EPI_ISL_563822, EPI_ISL_563823, EPI_ISL_563824, EPI_ISL_563825,
EPI_ISL_563826, EPI_ISL_563827, EPI_ISL_563828, EPI_ISL_563829, EPI_ISL_563830, EPI_ISL_563831, EPI_ISL_563833, EPI_ISL_563834, EPI_ISL_563835, EPI_ISL_563836, EPI_ISL_563837, EPI_ISL_563838, EPI_ISL_563839, EPI_ISL_563841, EPI_ISL_563842, EPI_ISL_563843, EPI_ISL_563844, EPI_ISL_563845,
EPI_ISL_563846, EPI_ISL_563847, EPI_ISL_563848, EPI_ISL_563849, EPI_ISL_563850, EPI_ISL_563851, EPI_ISL_563852, EPI_ISL_563853, EPI_ISL_563854, EPI_ISL_563855, EPI_ISL_563856, EPI_ISL_563857, EPI_ISL_563858, EPI_ISL_563859, EPI_ISL_563860, EPI_ISL_563861, EPI_ISL_563862, EPI_ISL_563863,
EPI_ISL_563864, EPI_ISL_563865, EPI_ISL_563867, EPI_ISL_563868, EPI_ISL_563869, EPI_ISL_563870, EPI_ISL_563871, EPI_ISL_563872, EPI_ISL_563873, EPI_ISL_563874, EPI_ISL_563875, EPI_ISL_563876, EPI_ISL_563877, EPI_ISL_563878, EPI_ISL_563879, EPI_ISL_563881, EPI_ISL_563882, EPI_ISL_563883,
EPI_ISL_563884, EPI_ISL_563885, EPI_ISL_563886, EPI_ISL_563887, EPI_ISL_563888, EPI_ISL_563889, EPI_ISL_563890, EPI_ISL_563891, EPI_ISL_563892, EPI_ISL_563893, EPI_ISL_563894, EPI_ISL_563895, EPI_ISL_563896, EPI_ISL_563897, EPI_ISL_563898, EPI_ISL_563899, EPI_ISL_563900, EPI_ISL_563901,
EPI_ISL_563902, EPI_ISL_563903, EPI_ISL_563904, EPI_ISL_563905, EPI_ISL_563906, EPI_ISL_563907, EPI_ISL_563908, EPI_ISL_563909, EPI_ISL_563910, EPI_ISL_563911, EPI_ISL_563912, EPI_ISL_563913, EPI_ISL_563914, EPI_ISL_563915, EPI_ISL_563916, EPI_ISL_563917, EPI_ISL_563918, EPI_ISL_563919,
EPI_ISL_563920, EPI_ISL_563921, EPI_ISL_563922, EPI_ISL_563923, EPI_ISL_563924, EPI_ISL_563926, EPI_ISL_563927, EPI_ISL_563928, EPI_ISL_563929, EPI_ISL_563930, EPI_ISL_563931, EPI_ISL_563932, EPI_ISL_563933, EPI_ISL_563934, EPI_ISL_563935, EPI_ISL_563936, EPI_ISL_563937, EPI_ISL_563939,
EPI_ISL_563940, EPI_ISL_563941, EPI_ISL_563942, EPI_ISL_563943, EPI_ISL_563944, EPI_ISL_563945, EPI_ISL_563946, EPI_ISL_563947, EPI_ISL_563948, EPI_ISL_563949, EPI_ISL_563950, EPI_ISL_563951, EPI_ISL_563952, EPI_ISL_563953, EPI_ISL_563954, EPI_ISL_563955, EPI_ISL_563956, EPI_ISL_563957,
EPI_ISL_563958, EPI_ISL_563959, EPI_ISL_563960, EPI_ISL_563961, EPI_ISL_563962, EPI_ISL_563963, EPI_ISL_563964, EPI_ISL_563965, EPI_ISL_563966, EPI_ISL_563967, EPI_ISL_563968, EPI_ISL_563969, EPI_ISL_563970, EPI_ISL_563971, EPI_ISL_563972, EPI_ISL_563976, EPI_ISL_563979, EPI_ISL_563980,
EPI_ISL_563981, EPI_ISL_563982, EPI_ISL_563983, EPI_ISL_563984, EPI_ISL_563985, EPI_ISL_563986, EPI_ISL_563989, EPI_ISL_563990, EPI_ISL_563991, EPI_ISL_563992, EPI_ISL_563993, EPI_ISL_563995, EPI_ISL_563996, EPI_ISL_564050, EPI_ISL_564051, EPI_ISL_564052, EPI_ISL_564053, EPI_ISL_564054,
EPI_ISL_564111, EPI_ISL_564112, EPI_ISL_564113, EPI_ISL_564114, EPI_ISL_564115, EPI_ISL_564116, EPI_ISL_564117, EPI_ISL_564118, EPI_ISL_564119, EPI_ISL_564120, EPI_ISL_564121, EPI_ISL_564122, EPI_ISL_564123, EPI_ISL_564124, EPI_ISL_564125, EPI_ISL_564126, EPI_ISL_564127, EPI_ISL_564128,
EPI_ISL_564129, EPI_ISL_564130, EPI_ISL_564131, EPI_ISL_564132, EPI_ISL_564133, EPI_ISL_564134, EPI_ISL_564135, EPI_ISL_564136, EPI_ISL_564137, EPI_ISL_564138, EPI_ISL_564139, EPI_ISL_564140, EPI_ISL_564141, EPI_ISL_564142, EPI_ISL_564143, EPI_ISL_564144, EPI_ISL_564146, EPI_ISL_564147,
EPI_ISL_564148, EPI_ISL_564149, EPI_ISL_564150, EPI_ISL_564151, EPI_ISL_564152, EPI_ISL_564153, EPI_ISL_564154, EPI_ISL_564155, EPI_ISL_564156, EPI_ISL_564157, EPI_ISL_564158, EPI_ISL_564159, EPI_ISL_564160, EPI_ISL_564161, EPI_ISL_564162, EPI_ISL_564163, EPI_ISL_564164, EPI_ISL_564165,
EPI_ISL_564166, EPI_ISL_564167, EPI_ISL_564168, EPI_ISL_564169, EPI_ISL_564170, EPI_ISL_564171, EPI_ISL_564172, EPI_ISL_564173, EPI_ISL_564174, EPI_ISL_564175, EPI_ISL_564176, EPI_ISL_564177, EPI_ISL_564178, EPI_ISL_564179, EPI_ISL_564255, EPI_ISL_564256, EPI_ISL_564257, EPI_ISL_564258,
EPI_ISL_564259, EPI_ISL_564260, EPI_ISL_564261, EPI_ISL_564262, EPI_ISL_564263, EPI_ISL_564264, EPI_ISL_564265, EPI_ISL_564266, EPI_ISL_564267, EPI_ISL_564268, EPI_ISL_564269, EPI_ISL_564271, EPI_ISL_564272, EPI_ISL_564273, EPI_ISL_564274, EPI_ISL_564275, EPI_ISL_564276, EPI_ISL_564277,
EPI_ISL_564278, EPI_ISL_564279, EPI_ISL_564280, EPI_ISL_564281, EPI_ISL_564282, EPI_ISL_564283, EPI_ISL_564284, EPI_ISL_564285, EPI_ISL_564286, EPI_ISL_564287, EPI_ISL_564288, EPI_ISL_564289, EPI_ISL_564290, EPI_ISL_564291, EPI_ISL_564292, EPI_ISL_564293, EPI_ISL_564294, EPI_ISL_564295,
EPI_ISL_564296, EPI_ISL_564297, EPI_ISL_564298, EPI_ISL_564299, EPI_ISL_564300, EPI_ISL_564301, EPI_ISL_564302, EPI_ISL_564303, EPI_ISL_564304, EPI_ISL_564305, EPI_ISL_564306, EPI_ISL_564307, EPI_ISL_564308, EPI_ISL_564309, EPI_ISL_564310, EPI_ISL_564311, EPI_ISL_564312, EPI_ISL_564313,
EPI_ISL_564314, EPI_ISL_564315, EPI_ISL_564316, EPI_ISL_564317, EPI_ISL_564351, EPI_ISL_564398, EPI_ISL_564399, EPI_ISL_564400, EPI_ISL_564401, EPI_ISL_564402, EPI_ISL_564403, EPI_ISL_564405, EPI_ISL_564406, EPI_ISL_564407, EPI_ISL_564408, EPI_ISL_564409, EPI_ISL_564410, EPI_ISL_564411,
EPI_ISL_564412, EPI_ISL_564413, EPI_ISL_564414, EPI_ISL_564415, EPI_ISL_564416, EPI_ISL_564417, EPI_ISL_564418, EPI_ISL_564419, EPI_ISL_564420, EPI_ISL_564421, EPI_ISL_564422, EPI_ISL_564423, EPI_ISL_564424, EPI_ISL_564425, EPI_ISL_564426, EPI_ISL_564427, EPI_ISL_564428, EPI_ISL_564429,
EPI_ISL_564430, EPI_ISL_564431, EPI_ISL_564432, EPI_ISL_564433, EPI_ISL_564434, EPI_ISL_564435, EPI_ISL_564436, EPI_ISL_564437, EPI_ISL_564438, EPI_ISL_564439, EPI_ISL_564440, EPI_ISL_564441, EPI_ISL_564442, EPI_ISL_564443, EPI_ISL_564444, EPI_ISL_564445, EPI_ISL_564446, EPI_ISL_564447,
EPI_ISL_564448, EPI_ISL_564449, EPI_ISL_564450, EPI_ISL_564451, EPI_ISL_564452, EPI_ISL_564453, EPI_ISL_564454, EPI_ISL_564471, EPI_ISL_564472, EPI_ISL_564473, EPI_ISL_564474, EPI_ISL_564475, EPI_ISL_564476, EPI_ISL_564477, EPI_ISL_564478, EPI_ISL_564479, EPI_ISL_564480, EPI_ISL_564481,
EPI_ISL_564482, EPI_ISL_564483, EPI_ISL_564484, EPI_ISL_564485, EPI_ISL_564486, EPI_ISL_564487, EPI_ISL_564488, EPI_ISL_564489, EPI_ISL_564490, EPI_ISL_564494, EPI_ISL_564495, EPI_ISL_564497, EPI_ISL_564503, EPI_ISL_564504, EPI_ISL_564505, EPI_ISL_564506, EPI_ISL_564507, EPI_ISL_564508,
EPI_ISL_564509, EPI_ISL_564510, EPI_ISL_564511, EPI_ISL_564512, EPI_ISL_564513, EPI_ISL_564514, EPI_ISL_564515, EPI_ISL_564516, EPI_ISL_564517, EPI_ISL_564518, EPI_ISL_564519, EPI_ISL_564520, EPI_ISL_564522, EPI_ISL_564523, EPI_ISL_564524, EPI_ISL_564525, EPI_ISL_564526, EPI_ISL_564527,
EPI_ISL_564528, EPI_ISL_564529, EPI_ISL_564530, EPI_ISL_564531, EPI_ISL_564532, EPI_ISL_564533, EPI_ISL_564534, EPI_ISL_564535, EPI_ISL_564536, EPI_ISL_564537, EPI_ISL_564538, EPI_ISL_564539, EPI_ISL_564540, EPI_ISL_564541, EPI_ISL_564542, EPI_ISL_564543, EPI_ISL_564545, EPI_ISL_564565,
EPI_ISL_564567, EPI_ISL_564568, EPI_ISL_564569, EPI_ISL_564570, EPI_ISL_564571, EPI_ISL_564572, EPI_ISL_564573, EPI_ISL_564574, EPI_ISL_564575, EPI_ISL_564576, EPI_ISL_564577, EPI_ISL_564578, EPI_ISL_564579, EPI_ISL_564580, EPI_ISL_564581, EPI_ISL_564582, EPI_ISL_564583, EPI_ISL_564584,
EPI_ISL_564585, EPI_ISL_564586, EPI_ISL_564587, EPI_ISL_564588, EPI_ISL_564589, EPI_ISL_564590, EPI_ISL_564591, EPI_ISL_564592, EPI_ISL_564593, EPI_ISL_564594, EPI_ISL_564595, EPI_ISL_564596, EPI_ISL_564597, EPI_ISL_564598, EPI_ISL_564599, EPI_ISL_564600, EPI_ISL_564601, EPI_ISL_564602,
EPI_ISL_564603, EPI_ISL_564604, EPI_ISL_564605, EPI_ISL_564606, EPI_ISL_564607, EPI_ISL_564608, EPI_ISL_564609, EPI_ISL_564610, EPI_ISL_564611, EPI_ISL_564612, EPI_ISL_564613, EPI_ISL_564614, EPI_ISL_564615, EPI_ISL_564616, EPI_ISL_564617, EPI_ISL_564618, EPI_ISL_564619, EPI_ISL_564620,
EPI_ISL_564621, EPI_ISL_564622, EPI_ISL_564623, EPI_ISL_564624, EPI_ISL_564625, EPI_ISL_564626, EPI_ISL_564627, EPI_ISL_564628, EPI_ISL_564629, EPI_ISL_564630, EPI_ISL_564631, EPI_ISL_564632, EPI_ISL_564633, EPI_ISL_564634, EPI_ISL_564635, EPI_ISL_564636, EPI_ISL_564637, EPI_ISL_564638,
EPI_ISL_564639, EPI_ISL_564640, EPI_ISL_564641, EPI_ISL_564642, EPI_ISL_564643, EPI_ISL_564644, EPI_ISL_564645, EPI_ISL_564646, EPI_ISL_564647, EPI_ISL_564648, EPI_ISL_564649, EPI_ISL_564650, EPI_ISL_564651, EPI_ISL_564652, EPI_ISL_564653, EPI_ISL_564654, EPI_ISL_564655, EPI_ISL_564656,
EPI_ISL_564657, EPI_ISL_564658, EPI_ISL_564659, EPI_ISL_564660, EPI_ISL_564661, EPI_ISL_564662, EPI_ISL_564663, EPI_ISL_564664, EPI_ISL_564665, EPI_ISL_564666, EPI_ISL_564667, EPI_ISL_564668, EPI_ISL_564669, EPI_ISL_564670, EPI_ISL_564671, EPI_ISL_564672, EPI_ISL_564673, EPI_ISL_564674,
EPI_ISL_564675, EPI_ISL_564676, EPI_ISL_564677, EPI_ISL_564678, EPI_ISL_564679, EPI_ISL_564680, EPI_ISL_564681, EPI_ISL_564682, EPI_ISL_564683, EPI_ISL_564684, EPI_ISL_564685, EPI_ISL_564686, EPI_ISL_564687, EPI_ISL_564688, EPI_ISL_564689, EPI_ISL_564690, EPI_ISL_564691, EPI_ISL_564692,
EPI_ISL_564693, EPI_ISL_564694, EPI_ISL_564695, EPI_ISL_564696, EPI_ISL_564697, EPI_ISL_564698, EPI_ISL_564699, EPI_ISL_564700, EPI_ISL_564701, EPI_ISL_564702, EPI_ISL_564703, EPI_ISL_564704, EPI_ISL_564705, EPI_ISL_564706, EPI_ISL_564707, EPI_ISL_564708, EPI_ISL_564709, EPI_ISL_564710,
EPI_ISL_564711, EPI_ISL_564712, EPI_ISL_564713, EPI_ISL_564714, EPI_ISL_564715, EPI_ISL_564716, EPI_ISL_564717, EPI_ISL_564718, EPI_ISL_564719, EPI_ISL_564720, EPI_ISL_564721, EPI_ISL_564722, EPI_ISL_564723, EPI_ISL_564724, EPI_ISL_564725, EPI_ISL_564726, EPI_ISL_564727, EPI_ISL_564728,
EPI_ISL_564729, EPI_ISL_564730, EPI_ISL_564731, EPI_ISL_564732, EPI_ISL_564733, EPI_ISL_564734, EPI_ISL_564735, EPI_ISL_564736, EPI_ISL_564737, EPI_ISL_564738, EPI_ISL_564739, EPI_ISL_564740, EPI_ISL_564741, EPI_ISL_564742, EPI_ISL_564743, EPI_ISL_564744, EPI_ISL_564745, EPI_ISL_564746,
EPI_ISL_564747, EPI_ISL_564748, EPI_ISL_564749, EPI_ISL_564750, EPI_ISL_564751, EPI_ISL_564752, EPI_ISL_564753, EPI_ISL_564754, EPI_ISL_564755, EPI_ISL_564756, EPI_ISL_564757, EPI_ISL_564758, EPI_ISL_564759, EPI_ISL_564760, EPI_ISL_564761, EPI_ISL_564762, EPI_ISL_564763, EPI_ISL_564764,
EPI_ISL_564765, EPI_ISL_564766, EPI_ISL_564767, EPI_ISL_564768, EPI_ISL_564769, EPI_ISL_564770, EPI_ISL_564771, EPI_ISL_564772, EPI_ISL_564773, EPI_ISL_564774, EPI_ISL_564775, EPI_ISL_564776, EPI_ISL_564777, EPI_ISL_564778, EPI_ISL_564779, EPI_ISL_564780, EPI_ISL_564781, EPI_ISL_564782,
EPI_ISL_564783, EPI_ISL_564784, EPI_ISL_564785, EPI_ISL_564786, EPI_ISL_564787, EPI_ISL_564788, EPI_ISL_564789, EPI_ISL_564790, EPI_ISL_564791, EPI_ISL_564792, EPI_ISL_564793, EPI_ISL_564794, EPI_ISL_564795, EPI_ISL_564796, EPI_ISL_564797, EPI_ISL_564798, EPI_ISL_564799, EPI_ISL_564800,
EPI_ISL_564801, EPI_ISL_564802, EPI_ISL_564803, EPI_ISL_564804, EPI_ISL_564805, EPI_ISL_564806, EPI_ISL_564807, EPI_ISL_564808, EPI_ISL_564809, EPI_ISL_564810, EPI_ISL_564811, EPI_ISL_564812, EPI_ISL_564813, EPI_ISL_564814, EPI_ISL_564815, EPI_ISL_564816, EPI_ISL_564817, EPI_ISL_564818,
EPI_ISL_564819, EPI_ISL_564820, EPI_ISL_564821, EPI_ISL_564822, EPI_ISL_564823, EPI_ISL_564824, EPI_ISL_564825, EPI_ISL_564826, EPI_ISL_564827, EPI_ISL_564828, EPI_ISL_564829, EPI_ISL_564830, EPI_ISL_564831, EPI_ISL_564832, EPI_ISL_564833, EPI_ISL_564834, EPI_ISL_564835, EPI_ISL_564836,
EPI_ISL_564837, EPI_ISL_564838, EPI_ISL_564839, EPI_ISL_564840, EPI_ISL_564841, EPI_ISL_564842, EPI_ISL_564843, EPI_ISL_564844, EPI_ISL_564845, EPI_ISL_564846, EPI_ISL_564847, EPI_ISL_564848, EPI_ISL_564849, EPI_ISL_564850, EPI_ISL_564851, EPI_ISL_564852, EPI_ISL_564853, EPI_ISL_564854,
EPI_ISL_564855, EPI_ISL_564856, EPI_ISL_564857, EPI_ISL_564858, EPI_ISL_564859, EPI_ISL_564860, EPI_ISL_564861, EPI_ISL_564862, EPI_ISL_564863, EPI_ISL_564864, EPI_ISL_564865, EPI_ISL_564866, EPI_ISL_564867, EPI_ISL_564868, EPI_ISL_564869, EPI_ISL_564870, EPI_ISL_564871, EPI_ISL_564872,
EPI_ISL_564873, EPI_ISL_564874, EPI_ISL_564875, EPI_ISL_564876, EPI_ISL_564877, EPI_ISL_564878, EPI_ISL_564879, EPI_ISL_564880, EPI_ISL_564881, EPI_ISL_564882, EPI_ISL_564883, EPI_ISL_564884, EPI_ISL_564885, EPI_ISL_564886, EPI_ISL_564887, EPI_ISL_564888, EPI_ISL_564889, EPI_ISL_564890,
EPI_ISL_564891, EPI_ISL_564892, EPI_ISL_564893, EPI_ISL_564894, EPI_ISL_564895, EPI_ISL_564896, EPI_ISL_564897, EPI_ISL_564898, EPI_ISL_564899, EPI_ISL_564900, EPI_ISL_564901, EPI_ISL_564902, EPI_ISL_564903, EPI_ISL_564904, EPI_ISL_564906, EPI_ISL_564907, EPI_ISL_564908, EPI_ISL_564909,
EPI_ISL_564910, EPI_ISL_564911, EPI_ISL_564912, EPI_ISL_564913, EPI_ISL_564914, EPI_ISL_564915, EPI_ISL_564916, EPI_ISL_564917, EPI_ISL_564918, EPI_ISL_564919, EPI_ISL_564920, EPI_ISL_564921, EPI_ISL_564922, EPI_ISL_564923, EPI_ISL_564924, EPI_ISL_564925, EPI_ISL_564926, EPI_ISL_564927,
EPI_ISL_564928, EPI_ISL_564929, EPI_ISL_564930, EPI_ISL_564931, EPI_ISL_564932, EPI_ISL_564933, EPI_ISL_564934, EPI_ISL_564935, EPI_ISL_564936, EPI_ISL_564937, EPI_ISL_564938, EPI_ISL_564939, EPI_ISL_564940, EPI_ISL_564941, EPI_ISL_564942, EPI_ISL_564943, EPI_ISL_564944, EPI_ISL_564945,
EPI_ISL_564946, EPI_ISL_564947, EPI_ISL_564948, EPI_ISL_564949, EPI_ISL_564950, EPI_ISL_564951, EPI_ISL_564952, EPI_ISL_564953, EPI_ISL_564954, EPI_ISL_564955, EPI_ISL_564956, EPI_ISL_564957, EPI_ISL_564958, EPI_ISL_564959, EPI_ISL_564960, EPI_ISL_564961, EPI_ISL_564962, EPI_ISL_564963,
EPI_ISL_564964, EPI_ISL_564965, EPI_ISL_564966, EPI_ISL_564967, EPI_ISL_564968, EPI_ISL_564969, EPI_ISL_564970, EPI_ISL_564971, EPI_ISL_564972, EPI_ISL_564973, EPI_ISL_564974, EPI_ISL_564975, EPI_ISL_564976, EPI_ISL_564977, EPI_ISL_564978, EPI_ISL_564979, EPI_ISL_564980, EPI_ISL_564981,
EPI_ISL_564982, EPI_ISL_564983, EPI_ISL_564984, EPI_ISL_564985, EPI_ISL_564986, EPI_ISL_564987, EPI_ISL_564988, EPI_ISL_564989, EPI_ISL_564990, EPI_ISL_564991, EPI_ISL_564992, EPI_ISL_564993, EPI_ISL_564994, EPI_ISL_564995, EPI_ISL_564996, EPI_ISL_564997, EPI_ISL_564998, EPI_ISL_564999,
EPI_ISL_565000, EPI_ISL_565001, EPI_ISL_565002, EPI_ISL_565003, EPI_ISL_565008, EPI_ISL_565009, EPI_ISL_565010, EPI_ISL_565011, EPI_ISL_565012, EPI_ISL_565013, EPI_ISL_565014, EPI_ISL_565015, EPI_ISL_565016, EPI_ISL_565017, EPI_ISL_565018, EPI_ISL_565019, EPI_ISL_565020, EPI_ISL_565021,
EPI_ISL_565022, EPI_ISL_565023, EPI_ISL_565024, EPI_ISL_565025, EPI_ISL_565026, EPI_ISL_565027, EPI_ISL_565028, EPI_ISL_565029, EPI_ISL_565030, EPI_ISL_565031, EPI_ISL_565032, EPI_ISL_565033, EPI_ISL_565035, EPI_ISL_565036, EPI_ISL_565037, EPI_ISL_565038, EPI_ISL_565039, EPI_ISL_565042,
EPI_ISL_565043, EPI_ISL_565044, EPI_ISL_565045, EPI_ISL_565046, EPI_ISL_565047, EPI_ISL_565048, EPI_ISL_565049, EPI_ISL_565050, EPI_ISL_565051, EPI_ISL_565052, EPI_ISL_565053, EPI_ISL_565054, EPI_ISL_565055, EPI_ISL_565056, EPI_ISL_565057, EPI_ISL_565058, EPI_ISL_565059, EPI_ISL_565060,

| | | | |
|---|---|---|---|
| EPI_ISL_565061, EPI_ISL_565062, EPI_ISL_565063, EPI_ISL_565064, EPI_ISL_565065, EPI_ISL_565066, EPI_ISL_565067, EPI_ISL_565068, EPI_ISL_565069, EPI_ISL_565070, EPI_ISL_565071, EPI_ISL_565072, EPI_ISL_565073, EPI_ISL_565074, EPI_ISL_565075, EPI_ISL_565076, EPI_ISL_565077, EPI_ISL_565078, EPI_ISL_565079, EPI_ISL_565080, EPI_ISL_565081, EPI_ISL_565082, EPI_ISL_565083, EPI_ISL_565084, EPI_ISL_565085, EPI_ISL_565086, EPI_ISL_565087, EPI_ISL_565088, EPI_ISL_565089, EPI_ISL_565090, EPI_ISL_565091, EPI_ISL_565092, EPI_ISL_565093, EPI_ISL_565094, EPI_ISL_565095, EPI_ISL_565096, EPI_ISL_565097, EPI_ISL_565098, EPI_ISL_565099, EPI_ISL_565100, EPI_ISL_565101, EPI_ISL_565102, EPI_ISL_565103, EPI_ISL_565104, EPI_ISL_565105, EPI_ISL_565106, EPI_ISL_565107, EPI_ISL_565108, EPI_ISL_565109, EPI_ISL_565111, EPI_ISL_565112, EPI_ISL_565113, EPI_ISL_565114, EPI_ISL_565115, EPI_ISL_565116, EPI_ISL_565117, EPI_ISL_565118, EPI_ISL_565119, EPI_ISL_565120, EPI_ISL_565121, EPI_ISL_565122, EPI_ISL_565123, EPI_ISL_565124, EPI_ISL_565125, EPI_ISL_565126, EPI_ISL_565127, EPI_ISL_565128, EPI_ISL_565129, EPI_ISL_565130, EPI_ISL_565131, EPI_ISL_565132, EPI_ISL_565133, EPI_ISL_565134, EPI_ISL_565135, EPI_ISL_565223, EPI_ISL_565224, EPI_ISL_565225, EPI_ISL_565226, EPI_ISL_565227, EPI_ISL_565228, EPI_ISL_565229, EPI_ISL_565230, EPI_ISL_565231, EPI_ISL_565232, EPI_ISL_565233, EPI_ISL_565234, EPI_ISL_565235, EPI_ISL_565236, EPI_ISL_565237, EPI_ISL_565238, EPI_ISL_565239, EPI_ISL_565240, EPI_ISL_565241, EPI_ISL_565242, EPI_ISL_565244, EPI_ISL_565245, EPI_ISL_565246, EPI_ISL_565247, EPI_ISL_565248, EPI_ISL_565249, EPI_ISL_565250, EPI_ISL_565251, EPI_ISL_565252, EPI_ISL_565253, EPI_ISL_565254, EPI_ISL_565255, EPI_ISL_565256, EPI_ISL_565257, EPI_ISL_565258, EPI_ISL_565259, EPI_ISL_565260, EPI_ISL_565261, EPI_ISL_565262, EPI_ISL_565263, EPI_ISL_565265, EPI_ISL_565266, EPI_ISL_565267, EPI_ISL_565268, EPI_ISL_565269, EPI_ISL_565270, EPI_ISL_565271, EPI_ISL_565272, EPI_ISL_565273, EPI_ISL_565274, EPI_ISL_565275, EPI_ISL_565276, EPI_ISL_565280, EPI_ISL_565281, EPI_ISL_565282, EPI_ISL_565283, EPI_ISL_565284, EPI_ISL_565285, EPI_ISL_565286, EPI_ISL_565287, EPI_ISL_565288, EPI_ISL_565290, EPI_ISL_565291, EPI_ISL_565292, EPI_ISL_565293, EPI_ISL_565294, EPI_ISL_565295, EPI_ISL_565296, EPI_ISL_565297, EPI_ISL_565298, EPI_ISL_565299, EPI_ISL_565300, EPI_ISL_565301, EPI_ISL_565302, EPI_ISL_565303, EPI_ISL_565304, EPI_ISL_565305, EPI_ISL_565306, EPI_ISL_565309, EPI_ISL_565315, EPI_ISL_565316, EPI_ISL_565317, EPI_ISL_565318, EPI_ISL_565319, EPI_ISL_565320, EPI_ISL_565321, EPI_ISL_565322, EPI_ISL_565323, EPI_ISL_565324, EPI_ISL_565325, EPI_ISL_565326, EPI_ISL_565327, EPI_ISL_565328, EPI_ISL_565329, EPI_ISL_565330, EPI_ISL_565331, EPI_ISL_565332, EPI_ISL_565333, EPI_ISL_565334, EPI_ISL_565335, EPI_ISL_565336, EPI_ISL_565337, EPI_ISL_565338, EPI_ISL_565339, EPI_ISL_565340, EPI_ISL_565341, EPI_ISL_565342, EPI_ISL_565343, EPI_ISL_565344, EPI_ISL_565345, EPI_ISL_565346, EPI_ISL_565347, EPI_ISL_565348, EPI_ISL_565349, EPI_ISL_565350, EPI_ISL_565351, EPI_ISL_565352, EPI_ISL_565353, EPI_ISL_565354, EPI_ISL_565355, EPI_ISL_565356, EPI_ISL_565357, EPI_ISL_565358, EPI_ISL_565359, EPI_ISL_565360, EPI_ISL_565361, EPI_ISL_565362, EPI_ISL_565363, EPI_ISL_565364, EPI_ISL_565365, EPI_ISL_565366, EPI_ISL_565367, EPI_ISL_565368, EPI_ISL_565369, EPI_ISL_565370, EPI_ISL_565371, EPI_ISL_565372, EPI_ISL_565373, EPI_ISL_565374, EPI_ISL_565375, EPI_ISL_565376, EPI_ISL_565377, EPI_ISL_565378, EPI_ISL_565379, EPI_ISL_565380, EPI_ISL_565381, EPI_ISL_565383, EPI_ISL_565384, EPI_ISL_565386, EPI_ISL_565387, EPI_ISL_565388, EPI_ISL_565389, EPI_ISL_565390, EPI_ISL_565391, EPI_ISL_565392, EPI_ISL_565393, EPI_ISL_565397, EPI_ISL_565398, EPI_ISL_565400, EPI_ISL_565405, EPI_ISL_565406, EPI_ISL_565407, EPI_ISL_565408, EPI_ISL_565410, EPI_ISL_565411, EPI_ISL_565412, EPI_ISL_565413, EPI_ISL_565414, EPI_ISL_565416, EPI_ISL_565417, EPI_ISL_565418, EPI_ISL_565419, EPI_ISL_565420, EPI_ISL_565421, EPI_ISL_565422, EPI_ISL_565423, EPI_ISL_565424, EPI_ISL_565425, EPI_ISL_565426, EPI_ISL_565427, EPI_ISL_565428, EPI_ISL_565429, EPI_ISL_565430, EPI_ISL_565431, EPI_ISL_565432, EPI_ISL_565433, EPI_ISL_565434, EPI_ISL_565435, EPI_ISL_565436, EPI_ISL_565437, EPI_ISL_565438, EPI_ISL_565439, EPI_ISL_565440, EPI_ISL_565441, EPI_ISL_565442, EPI_ISL_565443, EPI_ISL_565444, EPI_ISL_565445, EPI_ISL_565446, EPI_ISL_565447, EPI_ISL_565448, EPI_ISL_565449, EPI_ISL_565450, EPI_ISL_565451, EPI_ISL_565452, EPI_ISL_565453, EPI_ISL_565454, EPI_ISL_565455, EPI_ISL_565456, EPI_ISL_565457, EPI_ISL_565458, EPI_ISL_565460, EPI_ISL_565462, EPI_ISL_565468, EPI_ISL_565470, EPI_ISL_565471, EPI_ISL_565472, EPI_ISL_565473, EPI_ISL_565474, EPI_ISL_565475, EPI_ISL_565476, EPI_ISL_565477, EPI_ISL_565478, EPI_ISL_565484, EPI_ISL_565492, EPI_ISL_565493, EPI_ISL_565494, EPI_ISL_565495, EPI_ISL_565496, EPI_ISL_565497, EPI_ISL_565498, EPI_ISL_565499, EPI_ISL_565500, EPI_ISL_565501, EPI_ISL_565502, EPI_ISL_565503, EPI_ISL_565504, EPI_ISL_565505, EPI_ISL_565506, EPI_ISL_565507, EPI_ISL_565508, EPI_ISL_565510, EPI_ISL_565511, EPI_ISL_565512, EPI_ISL_565513, EPI_ISL_565514, EPI_ISL_565515, EPI_ISL_565516, EPI_ISL_565517, EPI_ISL_565518, EPI_ISL_565519, EPI_ISL_565520, EPI_ISL_565521, EPI_ISL_565522, EPI_ISL_565523, EPI_ISL_565524, EPI_ISL_565525, EPI_ISL_565526, EPI_ISL_565527, EPI_ISL_565528, EPI_ISL_565529, EPI_ISL_565530, EPI_ISL_565531, EPI_ISL_565532, EPI_ISL_565533, EPI_ISL_565534, EPI_ISL_565535, EPI_ISL_565536, EPI_ISL_565537, EPI_ISL_565538, EPI_ISL_565539, EPI_ISL_565540, EPI_ISL_565541, EPI_ISL_565542, EPI_ISL_565543, EPI_ISL_565544, EPI_ISL_565545, EPI_ISL_565546, EPI_ISL_565547, EPI_ISL_565548, EPI_ISL_565549, EPI_ISL_565550, EPI_ISL_565551, EPI_ISL_565552, EPI_ISL_565553, EPI_ISL_565554, EPI_ISL_565555, EPI_ISL_565556, EPI_ISL_565557, EPI_ISL_565558, EPI_ISL_565559, EPI_ISL_565560, EPI_ISL_565561, EPI_ISL_565562, EPI_ISL_565563, EPI_ISL_565564, EPI_ISL_565565, EPI_ISL_565567, EPI_ISL_565568, EPI_ISL_565569, EPI_ISL_565570, EPI_ISL_565571, EPI_ISL_565572, EPI_ISL_565573, EPI_ISL_565574, EPI_ISL_565575, EPI_ISL_565576, EPI_ISL_565577, EPI_ISL_565578, EPI_ISL_565579, EPI_ISL_565580, EPI_ISL_565581, EPI_ISL_565582, EPI_ISL_565583, EPI_ISL_565584, EPI_ISL_565585, EPI_ISL_565586, EPI_ISL_565587, EPI_ISL_565588, EPI_ISL_565589, EPI_ISL_565590, EPI_ISL_565591, EPI_ISL_565592, EPI_ISL_565593, EPI_ISL_565594, EPI_ISL_565595, EPI_ISL_565596, EPI_ISL_565597, EPI_ISL_565598, EPI_ISL_565599, EPI_ISL_565600, EPI_ISL_565601, EPI_ISL_565602, EPI_ISL_565603, EPI_ISL_565604, EPI_ISL_565605, EPI_ISL_565606, EPI_ISL_565607, EPI_ISL_565608, EPI_ISL_565609, EPI_ISL_565610, EPI_ISL_565611, EPI_ISL_565612, EPI_ISL_565613, EPI_ISL_565614, EPI_ISL_565615, EPI_ISL_565616, EPI_ISL_565617, EPI_ISL_565618, EPI_ISL_565619, EPI_ISL_565620, EPI_ISL_565621, EPI_ISL_565622, EPI_ISL_565625, EPI_ISL_565626, EPI_ISL_565627, EPI_ISL_565628, EPI_ISL_565629, EPI_ISL_565630, EPI_ISL_565631, EPI_ISL_565632, EPI_ISL_565633, EPI_ISL_565634, EPI_ISL_565635, EPI_ISL_565636, EPI_ISL_565637, EPI_ISL_565638, EPI_ISL_565639, EPI_ISL_565640, EPI_ISL_565641, EPI_ISL_565642, EPI_ISL_565643, EPI_ISL_565644, EPI_ISL_565645, EPI_ISL_565646, EPI_ISL_565647, EPI_ISL_565648, EPI_ISL_565649, EPI_ISL_565650, EPI_ISL_565651, EPI_ISL_565652, EPI_ISL_565653, EPI_ISL_565654, EPI_ISL_565655, EPI_ISL_565656, EPI_ISL_565657, EPI_ISL_565658, EPI_ISL_565659, EPI_ISL_565660, EPI_ISL_565661, EPI_ISL_565662, EPI_ISL_565663, EPI_ISL_565664, EPI_ISL_565665, EPI_ISL_565666, EPI_ISL_565667, EPI_ISL_565668, EPI_ISL_565669, EPI_ISL_565670, EPI_ISL_565671, EPI_ISL_565672, EPI_ISL_565673, EPI_ISL_565674, EPI_ISL_565675, EPI_ISL_565676, EPI_ISL_565677, EPI_ISL_565678, EPI_ISL_565679, EPI_ISL_565680, EPI_ISL_565681, EPI_ISL_565682, EPI_ISL_565683, EPI_ISL_565684, EPI_ISL_565685, EPI_ISL_565686, EPI_ISL_565687, EPI_ISL_565688, EPI_ISL_565689, EPI_ISL_565690, EPI_ISL_565691, EPI_ISL_565692, EPI_ISL_565693, EPI_ISL_565694, EPI_ISL_565695, EPI_ISL_565696, EPI_ISL_565697, EPI_ISL_565698, EPI_ISL_565699, EPI_ISL_565700, EPI_ISL_565701, EPI_ISL_565702, EPI_ISL_565703, EPI_ISL_565704, EPI_ISL_565705, EPI_ISL_565706, EPI_ISL_565707, EPI_ISL_565708, EPI_ISL_565709, EPI_ISL_565710, EPI_ISL_565711, EPI_ISL_565712, EPI_ISL_565713, EPI_ISL_565714, EPI_ISL_565715, EPI_ISL_565716, EPI_ISL_565717, EPI_ISL_565718, EPI_ISL_565719, EPI_ISL_565720, EPI_ISL_565721, EPI_ISL_565722, EPI_ISL_565723, EPI_ISL_565724, EPI_ISL_565725, EPI_ISL_565726, EPI_ISL_565727, EPI_ISL_565728, EPI_ISL_565729, EPI_ISL_565743, EPI_ISL_565745, EPI_ISL_565746, EPI_ISL_565747, EPI_ISL_565748, EPI_ISL_565749, EPI_ISL_565750, EPI_ISL_565751, EPI_ISL_565752, EPI_ISL_565753, EPI_ISL_565754, EPI_ISL_565755, EPI_ISL_565756, EPI_ISL_565757, EPI_ISL_565758, EPI_ISL_565759, EPI_ISL_565760, EPI_ISL_565761, EPI_ISL_565762, EPI_ISL_565763, EPI_ISL_565764, EPI_ISL_565765, EPI_ISL_565766, EPI_ISL_565767, EPI_ISL_565768, EPI_ISL_565769, EPI_ISL_565770, EPI_ISL_565771, EPI_ISL_565772, EPI_ISL_565773, EPI_ISL_565774, EPI_ISL_565775, EPI_ISL_565776, EPI_ISL_565777, EPI_ISL_565778, EPI_ISL_565779, EPI_ISL_565780, EPI_ISL_565781, EPI_ISL_565782, EPI_ISL_565783, EPI_ISL_565784, EPI_ISL_565785, EPI_ISL_565786, EPI_ISL_565787, EPI_ISL_565788, EPI_ISL_565789, EPI_ISL_565790, EPI_ISL_565791, EPI_ISL_565792, EPI_ISL_565793, EPI_ISL_565794, EPI_ISL_565795, EPI_ISL_565796, EPI_ISL_565797, EPI_ISL_565798, EPI_ISL_565799, EPI_ISL_565800, EPI_ISL_565801, EPI_ISL_565802, EPI_ISL_565803, EPI_ISL_565804, EPI_ISL_565805, EPI_ISL_565806, EPI_ISL_565807, EPI_ISL_565808, EPI_ISL_565809, EPI_ISL_565810, EPI_ISL_565811, EPI_ISL_565812, EPI_ISL_565813, EPI_ISL_565814, EPI_ISL_565815, EPI_ISL_565816, EPI_ISL_565817, EPI_ISL_565818, EPI_ISL_565819, EPI_ISL_565820, EPI_ISL_565821, EPI_ISL_565822, EPI_ISL_565823, EPI_ISL_565824, EPI_ISL_565825, EPI_ISL_565826, EPI_ISL_565827, EPI_ISL_565828, EPI_ISL_565829 | | | |
| see above | Microbiological Diagnostic Unit - Public Health Laboratory (MDU-PHL) | MDU-PHL | Seemann, T., Schultz M. B., Sait, M., Sherry, N. |
| EPI_ISL_568590, EPI_ISL_568591, EPI_ISL_568595, EPI_ISL_568597, EPI_ISL_568598, EPI_ISL_568876 | Florida Bureau of Public Health Laboratories | Florida Bureau of Public Health Laboratories | Sarah Schmedes, Jason Blanton |
| EPI_ISL_572684, EPI_ISL_572685, EPI_ISL_572709, EPI_ISL_572725, EPI_ISL_572770, EPI_ISL_572800, EPI_ISL_572807, EPI_ISL_572808, EPI_ISL_572809, EPI_ISL_572828, EPI_ISL_572829, EPI_ISL_572837, EPI_ISL_572838, EPI_ISL_572957, EPI_ISL_573478, EPI_ISL_573479, EPI_ISL_573480, EPI_ISL_573481, EPI_ISL_573482, EPI_ISL_573483, EPI_ISL_573484, EPI_ISL_573485, EPI_ISL_573486, EPI_ISL_573487, EPI_ISL_573488, EPI_ISL_573489, EPI_ISL_573490, EPI_ISL_573491, EPI_ISL_573492, EPI_ISL_573493, EPI_ISL_573494, EPI_ISL_573495, EPI_ISL_573496, EPI_ISL_573497, EPI_ISL_573498, EPI_ISL_573499, EPI_ISL_573500, EPI_ISL_573501, EPI_ISL_573502, EPI_ISL_573503, EPI_ISL_573504, EPI_ISL_573505, EPI_ISL_573506, EPI_ISL_573507, EPI_ISL_573508, EPI_ISL_573509, EPI_ISL_573510, EPI_ISL_573511, EPI_ISL_573512, EPI_ISL_573513, EPI_ISL_573514, EPI_ISL_573515, EPI_ISL_573516, EPI_ISL_573517, EPI_ISL_573518, EPI_ISL_573519, EPI_ISL_573520, EPI_ISL_573521, EPI_ISL_573522, EPI_ISL_573523, EPI_ISL_573524, EPI_ISL_573525, EPI_ISL_573652, EPI_ISL_573653, EPI_ISL_573654, EPI_ISL_573655, EPI_ISL_573656, EPI_ISL_573657, EPI_ISL_573658, EPI_ISL_573659, EPI_ISL_573660, EPI_ISL_573661, EPI_ISL_573662, EPI_ISL_573663, EPI_ISL_573664, EPI_ISL_573665, EPI_ISL_573666, EPI_ISL_573667, EPI_ISL_573668, EPI_ISL_573669, EPI_ISL_573670, EPI_ISL_573671, EPI_ISL_573672, EPI_ISL_573673, EPI_ISL_573674, EPI_ISL_573675, EPI_ISL_573676, EPI_ISL_573677, EPI_ISL_573678, EPI_ISL_573679, EPI_ISL_573680, EPI_ISL_573681, EPI_ISL_573682, EPI_ISL_573683, EPI_ISL_573684, EPI_ISL_573685, EPI_ISL_573764, EPI_ISL_573800, EPI_ISL_573801, EPI_ISL_573802, EPI_ISL_573803, EPI_ISL_573804, EPI_ISL_573805, EPI_ISL_573856, EPI_ISL_573857, EPI_ISL_573858, EPI_ISL_573859, EPI_ISL_573860, EPI_ISL_573861, EPI_ISL_573862, EPI_ISL_573863, EPI_ISL_573864, EPI_ISL_573865, EPI_ISL_574258, EPI_ISL_576984, EPI_ISL_576985, EPI_ISL_584670, EPI_ISL_584671, EPI_ISL_584672, EPI_ISL_584673, EPI_ISL_584674, EPI_ISL_584675, EPI_ISL_584676, EPI_ISL_584677, EPI_ISL_584678, EPI_ISL_584679, EPI_ISL_584811, EPI_ISL_584812, EPI_ISL_584813, EPI_ISL_584814, EPI_ISL_584815, EPI_ISL_584816, EPI_ISL_584817, EPI_ISL_584818, EPI_ISL_584819, EPI_ISL_584820, EPI_ISL_584821, EPI_ISL_584822, EPI_ISL_584823, EPI_ISL_584824, EPI_ISL_584825, EPI_ISL_584826, EPI_ISL_584827, EPI_ISL_584828, EPI_ISL_584829, EPI_ISL_584830, EPI_ISL_584831, EPI_ISL_584832, EPI_ISL_584833, EPI_ISL_584834, EPI_ISL_584835, EPI_ISL_584836, EPI_ISL_584837, EPI_ISL_584838, EPI_ISL_584839, EPI_ISL_584840, EPI_ISL_584841, EPI_ISL_584842, EPI_ISL_584843, EPI_ISL_584844, EPI_ISL_584845, EPI_ISL_584846, EPI_ISL_585502, EPI_ISL_585503 | | | |
| see above | University College London, Great Ormond Street Hospital for Children NHS Foundation Trust, Imperial College Healthcare NHS Trust | COVID-19 Genomics UK (COG-UK) Consortium | Sergi Castellano, Rachel Williams, Mark Kristiansen, Paola Resende Silva, Sunando Roy, Tony Brooks, Helena Tutill, Paola Niola, Patricia Dyal, Charlotte Williams, Leysa Forrest, Yasmin Panchbhaya, Jacqueline Findlay, Samuel Weeks, Julianne Brown, Kathryn Harris, Paul Randell, James Price, Alison Holmes, Judith Breuer |
| EPI_ISL_591070 | College of Veterinary Medicine, Chungnam National University | College of Veterinary Medicine, Chungnam National University | Seo,S.H. and Jang,Y. |
| EPI_ISL_591550, EPI_ISL_591551, EPI_ISL_591552, EPI_ISL_591553, EPI_ISL_591554, EPI_ISL_591555, EPI_ISL_591556, EPI_ISL_591557, EPI_ISL_591559, EPI_ISL_591560, EPI_ISL_591563, EPI_ISL_591564, EPI_ISL_591566, EPI_ISL_591567, EPI_ISL_591568, EPI_ISL_591569, EPI_ISL_591570, EPI_ISL_591572, EPI_ISL_591575, EPI_ISL_591576, EPI_ISL_591577, EPI_ISL_591579, EPI_ISL_591582, EPI_ISL_591583, EPI_ISL_591586, EPI_ISL_591588, EPI_ISL_591589, EPI_ISL_591593, EPI_ISL_591594, EPI_ISL_591596, EPI_ISL_591597, EPI_ISL_591598, EPI_ISL_591600, EPI_ISL_591601, EPI_ISL_591602, EPI_ISL_591604, EPI_ISL_591605, EPI_ISL_591606, EPI_ISL_591607, EPI_ISL_591608, EPI_ISL_591610, EPI_ISL_591611, EPI_ISL_591612, EPI_ISL_591614, EPI_ISL_591616, EPI_ISL_591617, EPI_ISL_591618, EPI_ISL_591620, EPI_ISL_591622, EPI_ISL_591624, EPI_ISL_591625, EPI_ISL_591627, EPI_ISL_591628, EPI_ISL_591629, EPI_ISL_591630, EPI_ISL_591631, EPI_ISL_591632, EPI_ISL_591633, EPI_ISL_591634, EPI_ISL_591635, EPI_ISL_591636, EPI_ISL_591637, EPI_ISL_591639, EPI_ISL_591641, EPI_ISL_591642, EPI_ISL_591644, EPI_ISL_591646, EPI_ISL_591647, EPI_ISL_591648, EPI_ISL_591649, EPI_ISL_591650, EPI_ISL_591651, EPI_ISL_591655, EPI_ISL_591660, EPI_ISL_591661, EPI_ISL_591662, EPI_ISL_591663, EPI_ISL_591664, EPI_ISL_591665, EPI_ISL_591668, EPI_ISL_591671, EPI_ISL_591673, EPI_ISL_591674, EPI_ISL_591675, EPI_ISL_591676, EPI_ISL_591677, EPI_ISL_591678, EPI_ISL_591680, EPI_ISL_591682, EPI_ISL_591683, EPI_ISL_591685, EPI_ISL_591686, EPI_ISL_591687, EPI_ISL_591689, EPI_ISL_591690, EPI_ISL_591691, EPI_ISL_591692, EPI_ISL_591693, EPI_ISL_591694, EPI_ISL_591695, EPI_ISL_591696, EPI_ISL_591697, EPI_ISL_591699, EPI_ISL_591701, EPI_ISL_591702, EPI_ISL_591705, EPI_ISL_591709, EPI_ISL_591710, EPI_ISL_591711, EPI_ISL_591712, EPI_ISL_591713, EPI_ISL_591714, EPI_ISL_591716, EPI_ISL_591717, EPI_ISL_591718, EPI_ISL_591720, EPI_ISL_591722, EPI_ISL_591724, EPI_ISL_591731, EPI_ISL_591733, EPI_ISL_591735, EPI_ISL_591736, EPI_ISL_591738, EPI_ISL_591741, EPI_ISL_591742, EPI_ISL_591745, EPI_ISL_591748, EPI_ISL_591750, EPI_ISL_591751, EPI_ISL_591752, EPI_ISL_591753, EPI_ISL_591757, EPI_ISL_591759, EPI_ISL_591762, EPI_ISL_591764, EPI_ISL_591765, EPI_ISL_591766, EPI_ISL_591768, EPI_ISL_591776, EPI_ISL_591777, EPI_ISL_591780, EPI_ISL_591781, EPI_ISL_591784, EPI_ISL_591788, EPI_ISL_591793, EPI_ISL_591794, EPI_ISL_591795, EPI_ISL_591796, EPI_ISL_591797, EPI_ISL_591798, EPI_ISL_591801, EPI_ISL_591802, EPI_ISL_591803, EPI_ISL_591804, EPI_ISL_591805, EPI_ISL_591806, EPI_ISL_591808, EPI_ISL_591809, EPI_ISL_591810, EPI_ISL_591811, EPI_ISL_591812, EPI_ISL_591813, EPI_ISL_591814, EPI_ISL_591815, EPI_ISL_591816, EPI_ISL_591817, EPI_ISL_591818, EPI_ISL_591819, EPI_ISL_591820, EPI_ISL_591821, EPI_ISL_591823, EPI_ISL_591826, EPI_ISL_591827, EPI_ISL_591828, EPI_ISL_591829, EPI_ISL_591830, EPI_ISL_591831, EPI_ISL_591832, EPI_ISL_591833, EPI_ISL_591834, EPI_ISL_591835, EPI_ISL_591836, EPI_ISL_591837, EPI_ISL_591838, EPI_ISL_591839, EPI_ISL_591840, EPI_ISL_591841, EPI_ISL_591842, EPI_ISL_591843, EPI_ISL_591844, EPI_ISL_591845, EPI_ISL_591846, EPI_ISL_591847, EPI_ISL_591848, EPI_ISL_591853, EPI_ISL_591905, EPI_ISL_591906, EPI_ISL_591910, EPI_ISL_591911, EPI_ISL_591913, EPI_ISL_591914, EPI_ISL_591915, EPI_ISL_591916, EPI_ISL_591918, EPI_ISL_591920, EPI_ISL_591921, EPI_ISL_591922, EPI_ISL_591923, EPI_ISL_591924, EPI_ISL_591926, EPI_ISL_591927, EPI_ISL_591929, EPI_ISL_591930, EPI_ISL_591931, EPI_ISL_591933, EPI_ISL_591934, EPI_ISL_591935, EPI_ISL_591936, EPI_ISL_591937, EPI_ISL_591938, EPI_ISL_591939, EPI_ISL_591961, EPI_ISL_591962, EPI_ISL_591963, EPI_ISL_591964, EPI_ISL_591965, EPI_ISL_591966, EPI_ISL_591967, EPI_ISL_591969, EPI_ISL_591970, EPI_ISL_591971, EPI_ISL_591972, EPI_ISL_591973, EPI_ISL_591974, EPI_ISL_591975, EPI_ISL_591976, EPI_ISL_591977, EPI_ISL_591978, EPI_ISL_591979, EPI_ISL_591980, EPI_ISL_591981, EPI_ISL_591983, EPI_ISL_591984, EPI_ISL_591985, EPI_ISL_591986, EPI_ISL_591987, EPI_ISL_591988, EPI_ISL_591989, EPI_ISL_591990, EPI_ISL_591991, EPI_ISL_591992, EPI_ISL_591993, EPI_ISL_591994, EPI_ISL_591995, EPI_ISL_591996, EPI_ISL_591997, EPI_ISL_591998, EPI_ISL_591999, EPI_ISL_592000, EPI_ISL_592001, EPI_ISL_592002, EPI_ISL_592003, EPI_ISL_592004, EPI_ISL_592005, EPI_ISL_592006, EPI_ISL_592007, EPI_ISL_592008, EPI_ISL_592009, EPI_ISL_592010, EPI_ISL_592011, EPI_ISL_592012, EPI_ISL_592014, EPI_ISL_592015, EPI_ISL_592016, EPI_ISL_592017, EPI_ISL_592018, EPI_ISL_592019, EPI_ISL_592020, EPI_ISL_592021, EPI_ISL_592022, EPI_ISL_592023, EPI_ISL_592024, EPI_ISL_592025, EPI_ISL_592026, EPI_ISL_592027, EPI_ISL_592029, EPI_ISL_592030, EPI_ISL_592031, EPI_ISL_592032, EPI_ISL_592033, EPI_ISL_592034, EPI_ISL_592035, EPI_ISL_592037, EPI_ISL_592038, EPI_ISL_592039, EPI_ISL_592040, EPI_ISL_592041, EPI_ISL_592042, EPI_ISL_592043, EPI_ISL_592044, EPI_ISL_592045, EPI_ISL_592046, EPI_ISL_592047, EPI_ISL_592048, EPI_ISL_592049, EPI_ISL_592051, EPI_ISL_592052, EPI_ISL_592053, EPI_ISL_592054, EPI_ISL_592055, EPI_ISL_592056, EPI_ISL_592057, EPI_ISL_592058, EPI_ISL_592059, EPI_ISL_592060, EPI_ISL_592061, EPI_ISL_592062, EPI_ISL_592063, EPI_ISL_592064, EPI_ISL_592065, EPI_ISL_592066, EPI_ISL_592067, EPI_ISL_592068, EPI_ISL_592070, EPI_ISL_592071, EPI_ISL_592072, EPI_ISL_592073, EPI_ISL_592074, EPI_ISL_592075, EPI_ISL_592076, EPI_ISL_592077, EPI_ISL_592078, EPI_ISL_592079, EPI_ISL_592080, EPI_ISL_592081, EPI_ISL_592082, EPI_ISL_592083, EPI_ISL_592084, EPI_ISL_592085, EPI_ISL_592086, EPI_ISL_592087, EPI_ISL_592088, EPI_ISL_592089, EPI_ISL_592090, EPI_ISL_592091, EPI_ISL_592092, EPI_ISL_592093, EPI_ISL_592094, EPI_ISL_592095, EPI_ISL_592096, EPI_ISL_592097, EPI_ISL_592098, EPI_ISL_592099, EPI_ISL_592100, EPI_ISL_592101, EPI_ISL_592102, EPI_ISL_592103, EPI_ISL_592106, EPI_ISL_592107, EPI_ISL_592108, EPI_ISL_592109, EPI_ISL_592110, EPI_ISL_592111, EPI_ISL_592112, EPI_ISL_592113, EPI_ISL_592114, EPI_ISL_592115, EPI_ISL_592116, EPI_ISL_592117, EPI_ISL_592118, EPI_ISL_592119, EPI_ISL_592120, EPI_ISL_592121, EPI_ISL_592122, EPI_ISL_592123, EPI_ISL_592124, EPI_ISL_592125, EPI_ISL_592126, EPI_ISL_592127, EPI_ISL_592128, EPI_ISL_592129, EPI_ISL_592130, EPI_ISL_592131, EPI_ISL_592132, EPI_ISL_592133, EPI_ISL_592134, EPI_ISL_592135, EPI_ISL_592196, EPI_ISL_592197, EPI_ISL_592198, EPI_ISL_592199, EPI_ISL_592200, EPI_ISL_592201, EPI_ISL_592202, EPI_ISL_592203, EPI_ISL_592204, EPI_ISL_592205, EPI_ISL_592206, EPI_ISL_592207, EPI_ISL_592208, EPI_ISL_592209, EPI_ISL_592210, EPI_ISL_592211, EPI_ISL_592212, EPI_ISL_592213, EPI_ISL_592214, EPI_ISL_592215, EPI_ISL_592216, EPI_ISL_592217, EPI_ISL_592218, EPI_ISL_592219, EPI_ISL_592220, EPI_ISL_592221, EPI_ISL_592222, EPI_ISL_592223, EPI_ISL_592224, EPI_ISL_592225, EPI_ISL_592226, EPI_ISL_592227, EPI_ISL_592228, EPI_ISL_592229, EPI_ISL_592230, EPI_ISL_592231, EPI_ISL_592232, EPI_ISL_592233, EPI_ISL_592234, EPI_ISL_592235, EPI_ISL_592236, EPI_ISL_592237, EPI_ISL_592238, EPI_ISL_592239, EPI_ISL_592240, EPI_ISL_592241, EPI_ISL_592242, EPI_ISL_592243, EPI_ISL_592244, EPI_ISL_592245, EPI_ISL_592246, EPI_ISL_592247, EPI_ISL_592248, EPI_ISL_592249, EPI_ISL_592250, EPI_ISL_592251, EPI_ISL_592252, EPI_ISL_592253, EPI_ISL_592254, EPI_ISL_592255, EPI_ISL_592256, EPI_ISL_592257, EPI_ISL_592258, EPI_ISL_592259, EPI_ISL_592260, EPI_ISL_592261, EPI_ISL_592262, EPI_ISL_592263, EPI_ISL_592264, EPI_ISL_592265, EPI_ISL_592266, EPI_ISL_592267, EPI_ISL_592268, EPI_ISL_592269, EPI_ISL_592270, EPI_ISL_592271, EPI_ISL_592272, EPI_ISL_592273, EPI_ISL_592274, EPI_ISL_592275, EPI_ISL_592276, EPI_ISL_592277, EPI_ISL_592278, EPI_ISL_592279, EPI_ISL_592280, EPI_ISL_592281, EPI_ISL_592282, EPI_ISL_592284, EPI_ISL_592285, EPI_ISL_592286, EPI_ISL_592287, EPI_ISL_592288, EPI_ISL_592289, EPI_ISL_592290, EPI_ISL_592291, EPI_ISL_592292, EPI_ISL_592293, EPI_ISL_592294, EPI_ISL_592295, EPI_ISL_592296, EPI_ISL_592297, EPI_ISL_592298, EPI_ISL_592299, EPI_ISL_592300, EPI_ISL_592301, EPI_ISL_592302, EPI_ISL_592303, EPI_ISL_592304, EPI_ISL_592305, EPI_ISL_592306, EPI_ISL_592307, EPI_ISL_592308, EPI_ISL_592309, EPI_ISL_592310, EPI_ISL_592311, EPI_ISL_592312, EPI_ISL_592313, EPI_ISL_592314, EPI_ISL_592315, EPI_ISL_592316, EPI_ISL_592317, EPI_ISL_592318, EPI_ISL_592319, EPI_ISL_592320, EPI_ISL_592321, EPI_ISL_592322, EPI_ISL_592323, EPI_ISL_592324, EPI_ISL_592325, EPI_ISL_592326, EPI_ISL_592327, EPI_ISL_592328, EPI_ISL_592329, EPI_ISL_592330, EPI_ISL_592331, | | | |

| | | | |
|---|---|---|---|
| EPI_ISL_592332, EPI_ISL_592333, EPI_ISL_592334, EPI_ISL_592335, EPI_ISL_592336, EPI_ISL_592337, EPI_ISL_592338, EPI_ISL_592339, EPI_ISL_592340, EPI_ISL_592341, EPI_ISL_592342, EPI_ISL_592343, EPI_ISL_592344, EPI_ISL_592345, EPI_ISL_592346, EPI_ISL_592347, EPI_ISL_592348, EPI_ISL_592349, EPI_ISL_592350, EPI_ISL_592351, EPI_ISL_592352, EPI_ISL_592353, EPI_ISL_592354, EPI_ISL_592355, EPI_ISL_592356, EPI_ISL_592357, EPI_ISL_592358, EPI_ISL_592359, EPI_ISL_592360, EPI_ISL_592361, EPI_ISL_592362, EPI_ISL_592363, EPI_ISL_592364, EPI_ISL_592365, EPI_ISL_592366, EPI_ISL_592367, EPI_ISL_592368, EPI_ISL_592369, EPI_ISL_592370, EPI_ISL_592371, EPI_ISL_592372, EPI_ISL_592373, EPI_ISL_592374, EPI_ISL_592375, EPI_ISL_592376, EPI_ISL_592377, EPI_ISL_592378, EPI_ISL_592379, EPI_ISL_592381, EPI_ISL_592382, EPI_ISL_592384, EPI_ISL_592385, EPI_ISL_592386, EPI_ISL_592387, EPI_ISL_592388, EPI_ISL_592389, EPI_ISL_592390, EPI_ISL_592391, EPI_ISL_592393, EPI_ISL_592394, EPI_ISL_592395, EPI_ISL_592396, EPI_ISL_592397, EPI_ISL_592398, EPI_ISL_592399, EPI_ISL_592400, EPI_ISL_592401, EPI_ISL_592402, EPI_ISL_592403, EPI_ISL_592404, EPI_ISL_592405, EPI_ISL_592406, EPI_ISL_592407, EPI_ISL_592408, EPI_ISL_592409, EPI_ISL_592410, EPI_ISL_592411, EPI_ISL_592412, EPI_ISL_592413, EPI_ISL_592414, EPI_ISL_592416, EPI_ISL_592417, EPI_ISL_592418, EPI_ISL_592419, EPI_ISL_592420, EPI_ISL_592421, EPI_ISL_592422, EPI_ISL_592423, EPI_ISL_592424, EPI_ISL_592425, EPI_ISL_592426, EPI_ISL_592427, EPI_ISL_592428, EPI_ISL_592429, EPI_ISL_592430, EPI_ISL_592431, EPI_ISL_592432, EPI_ISL_592433, EPI_ISL_592434, EPI_ISL_592435, EPI_ISL_592436, EPI_ISL_592437, EPI_ISL_592438, EPI_ISL_592439, EPI_ISL_592440, EPI_ISL_592441, EPI_ISL_592442, EPI_ISL_592443, EPI_ISL_592444, EPI_ISL_592445, EPI_ISL_592446, EPI_ISL_592447, EPI_ISL_592448, EPI_ISL_592449, EPI_ISL_592450, EPI_ISL_592451, EPI_ISL_592452, EPI_ISL_592453, EPI_ISL_592454, EPI_ISL_592455, EPI_ISL_592456, EPI_ISL_592457, EPI_ISL_592458, EPI_ISL_592459, EPI_ISL_592460, EPI_ISL_592461, EPI_ISL_592462, EPI_ISL_592463, EPI_ISL_592464, EPI_ISL_592465, EPI_ISL_592466, EPI_ISL_592467, EPI_ISL_592468, EPI_ISL_592469, EPI_ISL_592470, EPI_ISL_592471, EPI_ISL_592472, EPI_ISL_592473, EPI_ISL_592474, EPI_ISL_592475, EPI_ISL_592476, EPI_ISL_592477, EPI_ISL_592478, EPI_ISL_592479, EPI_ISL_592480, EPI_ISL_592481, EPI_ISL_592482, EPI_ISL_592483, EPI_ISL_592484, EPI_ISL_592487, EPI_ISL_592510, EPI_ISL_592517, EPI_ISL_592518, EPI_ISL_592519, EPI_ISL_592520, EPI_ISL_592521, EPI_ISL_592522, EPI_ISL_592523, EPI_ISL_592524, EPI_ISL_592525, EPI_ISL_592526, EPI_ISL_592527, EPI_ISL_592528, EPI_ISL_592529, EPI_ISL_592530, EPI_ISL_592536, EPI_ISL_592538, EPI_ISL_592539, EPI_ISL_592540, EPI_ISL_592541, EPI_ISL_592542, EPI_ISL_592543, EPI_ISL_592544, EPI_ISL_592545, EPI_ISL_592546, EPI_ISL_592547, EPI_ISL_592549, EPI_ISL_592550, EPI_ISL_592551, EPI_ISL_592552, EPI_ISL_592553, EPI_ISL_592554, EPI_ISL_592555, EPI_ISL_592556, EPI_ISL_592557, EPI_ISL_592558, EPI_ISL_592559, EPI_ISL_592560, EPI_ISL_592561, EPI_ISL_592562, EPI_ISL_592563, EPI_ISL_592565, EPI_ISL_592566, EPI_ISL_592567, EPI_ISL_592569, EPI_ISL_592570, EPI_ISL_592571, EPI_ISL_592572, EPI_ISL_592573, EPI_ISL_592574, EPI_ISL_592575, EPI_ISL_592576, EPI_ISL_592579, EPI_ISL_592580, EPI_ISL_592581, EPI_ISL_592582, EPI_ISL_592583, EPI_ISL_592584, EPI_ISL_592585, EPI_ISL_592586, EPI_ISL_592587, EPI_ISL_592588, EPI_ISL_592589, EPI_ISL_592591, EPI_ISL_592592, EPI_ISL_592593, EPI_ISL_592594, EPI_ISL_592595, EPI_ISL_592596, EPI_ISL_592598, EPI_ISL_592599, EPI_ISL_592601, EPI_ISL_592602, EPI_ISL_592603, EPI_ISL_592604, EPI_ISL_592605, EPI_ISL_592606, EPI_ISL_592607, EPI_ISL_592608, EPI_ISL_592609, EPI_ISL_592611, EPI_ISL_592612, EPI_ISL_592613, EPI_ISL_592614, EPI_ISL_592615, EPI_ISL_592617, EPI_ISL_592618, EPI_ISL_592619, EPI_ISL_592620, EPI_ISL_592621, EPI_ISL_592622, EPI_ISL_592623, EPI_ISL_592624, EPI_ISL_592625, EPI_ISL_592626, EPI_ISL_592627, EPI_ISL_592628, EPI_ISL_592630, EPI_ISL_592631, EPI_ISL_592632, EPI_ISL_592633, EPI_ISL_592634, EPI_ISL_592635, EPI_ISL_592636, EPI_ISL_592637, EPI_ISL_592638, EPI_ISL_592639, EPI_ISL_592640, EPI_ISL_592641, EPI_ISL_592642, EPI_ISL_592643, EPI_ISL_592644, EPI_ISL_592645, EPI_ISL_592646, EPI_ISL_592647, EPI_ISL_592648, EPI_ISL_592650, EPI_ISL_592651, EPI_ISL_592652, EPI_ISL_592653, EPI_ISL_592654, EPI_ISL_592655, EPI_ISL_592656, EPI_ISL_592657, EPI_ISL_592658, EPI_ISL_592659, EPI_ISL_592662, EPI_ISL_592663, EPI_ISL_592664, EPI_ISL_592665, EPI_ISL_592666, EPI_ISL_592667, EPI_ISL_592669, EPI_ISL_592670, EPI_ISL_592671, EPI_ISL_592672, EPI_ISL_592673, EPI_ISL_592674, EPI_ISL_592675, EPI_ISL_592676, EPI_ISL_592677, EPI_ISL_592678, EPI_ISL_592679, EPI_ISL_592680, EPI_ISL_592681, EPI_ISL_592682, EPI_ISL_592683, EPI_ISL_592684, EPI_ISL_592685, EPI_ISL_592686, EPI_ISL_592688, EPI_ISL_592689, EPI_ISL_592690, EPI_ISL_592691, EPI_ISL_592692, EPI_ISL_592693, EPI_ISL_592694, EPI_ISL_592695, EPI_ISL_592696, EPI_ISL_592697, EPI_ISL_592698, EPI_ISL_592699, EPI_ISL_592700, EPI_ISL_592701, EPI_ISL_592702, EPI_ISL_592703, EPI_ISL_592704, EPI_ISL_592705, EPI_ISL_592706, EPI_ISL_592708, EPI_ISL_592709, EPI_ISL_592710, EPI_ISL_592711, EPI_ISL_592712, EPI_ISL_592713, EPI_ISL_592714, EPI_ISL_592715, EPI_ISL_592717, EPI_ISL_592718, EPI_ISL_592719, EPI_ISL_592721, EPI_ISL_592722, EPI_ISL_592723, EPI_ISL_592724, EPI_ISL_592725, EPI_ISL_592726, EPI_ISL_592727, EPI_ISL_592728, EPI_ISL_592729, EPI_ISL_592731, EPI_ISL_592732, EPI_ISL_592733, EPI_ISL_592734, EPI_ISL_592735, EPI_ISL_592736, EPI_ISL_592737, EPI_ISL_592738, EPI_ISL_592739, EPI_ISL_592740, EPI_ISL_592741, EPI_ISL_592742, EPI_ISL_592743, EPI_ISL_592744, EPI_ISL_592745, EPI_ISL_592746, EPI_ISL_592747, EPI_ISL_592748, EPI_ISL_592749, EPI_ISL_592750, EPI_ISL_592751, EPI_ISL_592752, EPI_ISL_592754, EPI_ISL_592755, EPI_ISL_592756, EPI_ISL_592757, EPI_ISL_592758, EPI_ISL_592759, EPI_ISL_592760, EPI_ISL_592761, EPI_ISL_592762, EPI_ISL_592763, EPI_ISL_592764, EPI_ISL_592765, EPI_ISL_592766, EPI_ISL_592767, EPI_ISL_592768, EPI_ISL_592769, EPI_ISL_592773, EPI_ISL_592774, EPI_ISL_592775, EPI_ISL_592776, EPI_ISL_592777, EPI_ISL_592778, EPI_ISL_592779, EPI_ISL_592780, EPI_ISL_592784, EPI_ISL_592786, EPI_ISL_592787, EPI_ISL_592789, EPI_ISL_592791, EPI_ISL_592792, EPI_ISL_592793, EPI_ISL_592794, EPI_ISL_592796, EPI_ISL_592797, EPI_ISL_592799, EPI_ISL_592800, EPI_ISL_592801, EPI_ISL_592802, EPI_ISL_592803, EPI_ISL_592805, EPI_ISL_592806, EPI_ISL_592807, EPI_ISL_592808, EPI_ISL_592809, EPI_ISL_592810, EPI_ISL_592811, EPI_ISL_592812, EPI_ISL_592813, EPI_ISL_592815, EPI_ISL_592816, EPI_ISL_592817, EPI_ISL_592818, EPI_ISL_592819, EPI_ISL_592822, EPI_ISL_592823, EPI_ISL_592825, EPI_ISL_592826, EPI_ISL_592827, EPI_ISL_592829, EPI_ISL_592830, EPI_ISL_592831, EPI_ISL_592832, EPI_ISL_592833, EPI_ISL_592834, EPI_ISL_592835, EPI_ISL_592838, EPI_ISL_592839, EPI_ISL_592840, EPI_ISL_592841, EPI_ISL_592842, EPI_ISL_592843, EPI_ISL_592844, EPI_ISL_592845, EPI_ISL_592846, EPI_ISL_592847, EPI_ISL_592848, EPI_ISL_592849, EPI_ISL_592850, EPI_ISL_592851, EPI_ISL_592852, EPI_ISL_592853, EPI_ISL_592854, EPI_ISL_592855, EPI_ISL_592856, EPI_ISL_592857, EPI_ISL_592861, EPI_ISL_592866, EPI_ISL_592867, EPI_ISL_592868, EPI_ISL_592869, EPI_ISL_592870, EPI_ISL_592871, EPI_ISL_592872, EPI_ISL_592873, EPI_ISL_592874, EPI_ISL_592875, EPI_ISL_592876, EPI_ISL_592877, EPI_ISL_592882, EPI_ISL_592883, EPI_ISL_592884, EPI_ISL_592885, EPI_ISL_592886, EPI_ISL_592887, EPI_ISL_592889, EPI_ISL_592890, EPI_ISL_592891, EPI_ISL_592892, EPI_ISL_592893, EPI_ISL_592894, EPI_ISL_592895, EPI_ISL_592896, EPI_ISL_592897, EPI_ISL_592898, EPI_ISL_592899, EPI_ISL_592900, EPI_ISL_592901, EPI_ISL_592902, EPI_ISL_592904, EPI_ISL_592905, EPI_ISL_592906, EPI_ISL_592908, EPI_ISL_592909, EPI_ISL_592910, EPI_ISL_592911, EPI_ISL_592912, EPI_ISL_592913, EPI_ISL_592914, EPI_ISL_592915, EPI_ISL_592916, EPI_ISL_592917, EPI_ISL_592918, EPI_ISL_592919, EPI_ISL_592920, EPI_ISL_592921, EPI_ISL_592922, EPI_ISL_592923, EPI_ISL_592924, EPI_ISL_592925, EPI_ISL_592926, EPI_ISL_592927, EPI_ISL_592928, EPI_ISL_592929, EPI_ISL_592930, EPI_ISL_592931, EPI_ISL_592932, EPI_ISL_592933, EPI_ISL_592934, EPI_ISL_592935, EPI_ISL_592936, EPI_ISL_592937, EPI_ISL_592938, EPI_ISL_592939, EPI_ISL_592940, EPI_ISL_592941, EPI_ISL_592942, EPI_ISL_592943, EPI_ISL_592944, EPI_ISL_592945, EPI_ISL_592946, EPI_ISL_592947, EPI_ISL_592948, EPI_ISL_592949, EPI_ISL_592950, EPI_ISL_592951, EPI_ISL_592952, EPI_ISL_592953, EPI_ISL_592954, EPI_ISL_592955, EPI_ISL_592956, EPI_ISL_592957, EPI_ISL_592959, EPI_ISL_592960, EPI_ISL_592961, EPI_ISL_592962, EPI_ISL_592963, EPI_ISL_592964, EPI_ISL_592965, EPI_ISL_592966, EPI_ISL_592967, EPI_ISL_592968, EPI_ISL_592986, EPI_ISL_592987, EPI_ISL_592988, EPI_ISL_592989, EPI_ISL_592990, EPI_ISL_592991, EPI_ISL_592992, EPI_ISL_592993, EPI_ISL_592994, EPI_ISL_592995, EPI_ISL_592996, EPI_ISL_592997, EPI_ISL_592998, EPI_ISL_592999, EPI_ISL_593000, EPI_ISL_593001, EPI_ISL_593002, EPI_ISL_593008, EPI_ISL_593009, EPI_ISL_593010, EPI_ISL_593011, EPI_ISL_593012, EPI_ISL_593013, EPI_ISL_593014, EPI_ISL_593015, EPI_ISL_593016, EPI_ISL_593017, EPI_ISL_593018, EPI_ISL_593019, EPI_ISL_593020, EPI_ISL_593021, EPI_ISL_593022, EPI_ISL_593023, EPI_ISL_593024, EPI_ISL_593025, EPI_ISL_593026, EPI_ISL_593027, EPI_ISL_593028, EPI_ISL_593029, EPI_ISL_593030, EPI_ISL_593031, EPI_ISL_593032, EPI_ISL_593034, EPI_ISL_593036, EPI_ISL_593037, EPI_ISL_593038, EPI_ISL_593039, EPI_ISL_593040, EPI_ISL_593041, EPI_ISL_593042, EPI_ISL_593043, EPI_ISL_593044, EPI_ISL_593045, EPI_ISL_593046, EPI_ISL_593047, EPI_ISL_593049, EPI_ISL_593050, EPI_ISL_593051, EPI_ISL_593052, EPI_ISL_593053, EPI_ISL_593054, EPI_ISL_593055, EPI_ISL_593056, EPI_ISL_593057, EPI_ISL_593058, EPI_ISL_593059, EPI_ISL_593060, EPI_ISL_593061, EPI_ISL_593062, EPI_ISL_593063, EPI_ISL_593064, EPI_ISL_593065, EPI_ISL_593066, EPI_ISL_593067, EPI_ISL_593068, EPI_ISL_593069, EPI_ISL_593070, EPI_ISL_593071, EPI_ISL_593072, EPI_ISL_593073, EPI_ISL_593074, EPI_ISL_593075, EPI_ISL_593076, EPI_ISL_593077, EPI_ISL_593078, EPI_ISL_593079, EPI_ISL_593080, EPI_ISL_593081, EPI_ISL_593082, EPI_ISL_593083, EPI_ISL_593084, EPI_ISL_593085, EPI_ISL_593086, EPI_ISL_593087, EPI_ISL_593088, EPI_ISL_593089, EPI_ISL_593090, EPI_ISL_593091, EPI_ISL_593092, EPI_ISL_593093, EPI_ISL_593094, EPI_ISL_593095, EPI_ISL_593096, EPI_ISL_593097, EPI_ISL_593098, EPI_ISL_593099, EPI_ISL_593100, EPI_ISL_593101, EPI_ISL_593102, EPI_ISL_593103, EPI_ISL_593104, EPI_ISL_593105, EPI_ISL_593106, EPI_ISL_593107, EPI_ISL_593108, EPI_ISL_593109, EPI_ISL_593110, EPI_ISL_593111, EPI_ISL_593112, EPI_ISL_593113, EPI_ISL_593114, EPI_ISL_593115, EPI_ISL_593116, EPI_ISL_593117, EPI_ISL_593118, EPI_ISL_593119, EPI_ISL_593120, EPI_ISL_593121, EPI_ISL_593122, EPI_ISL_593123, EPI_ISL_593124, EPI_ISL_593129, EPI_ISL_593130, EPI_ISL_593131, EPI_ISL_593132, EPI_ISL_593133, EPI_ISL_593134, EPI_ISL_593135, EPI_ISL_593136, EPI_ISL_593137, EPI_ISL_593138, EPI_ISL_593139, EPI_ISL_593140, EPI_ISL_593141, EPI_ISL_593142, EPI_ISL_593143, EPI_ISL_593144, EPI_ISL_593145, EPI_ISL_593146, EPI_ISL_593147, EPI_ISL_593148, EPI_ISL_593149, EPI_ISL_593150, EPI_ISL_593151, EPI_ISL_593152, EPI_ISL_593153, EPI_ISL_593154, EPI_ISL_593155, EPI_ISL_593156, EPI_ISL_593157, EPI_ISL_593158, EPI_ISL_593159, EPI_ISL_593160, EPI_ISL_593161, EPI_ISL_593162, EPI_ISL_593163, EPI_ISL_593164, EPI_ISL_593165, EPI_ISL_593166, EPI_ISL_593167, EPI_ISL_593168, EPI_ISL_593169, EPI_ISL_593170, EPI_ISL_593171, EPI_ISL_593172, EPI_ISL_593173, EPI_ISL_593174, EPI_ISL_593175, EPI_ISL_593176, EPI_ISL_593177, EPI_ISL_593178, EPI_ISL_593179, EPI_ISL_593180, EPI_ISL_593181, EPI_ISL_593182, EPI_ISL_593183, EPI_ISL_593184, EPI_ISL_593185, EPI_ISL_593186, EPI_ISL_593187, EPI_ISL_593188, EPI_ISL_593189, EPI_ISL_593190, EPI_ISL_593191, EPI_ISL_593192, EPI_ISL_593193, EPI_ISL_593194, EPI_ISL_593195, EPI_ISL_593196, EPI_ISL_593197, EPI_ISL_593198, EPI_ISL_593199, EPI_ISL_593200, EPI_ISL_593201, EPI_ISL_593202, EPI_ISL_593203, EPI_ISL_593204, EPI_ISL_593205, EPI_ISL_593206, EPI_ISL_593207, EPI_ISL_593208, EPI_ISL_593209, EPI_ISL_593210, EPI_ISL_593211, EPI_ISL_593212, EPI_ISL_593213, EPI_ISL_593214, EPI_ISL_593217, EPI_ISL_593218, EPI_ISL_593219, EPI_ISL_593220, EPI_ISL_593221, EPI_ISL_593222, EPI_ISL_593223, EPI_ISL_593224, EPI_ISL_593225, EPI_ISL_593226, EPI_ISL_593227, EPI_ISL_593228, EPI_ISL_593229, EPI_ISL_593230, EPI_ISL_593231, EPI_ISL_593232, EPI_ISL_593233, EPI_ISL_593234, EPI_ISL_593235, EPI_ISL_593236, EPI_ISL_593237, EPI_ISL_593238, EPI_ISL_593239, EPI_ISL_593240, EPI_ISL_593241, EPI_ISL_593242, EPI_ISL_593243, EPI_ISL_593244, EPI_ISL_593245, EPI_ISL_593246, EPI_ISL_593247, EPI_ISL_593248, EPI_ISL_593249, EPI_ISL_593250, EPI_ISL_593251, EPI_ISL_593252, EPI_ISL_593253, EPI_ISL_593254, EPI_ISL_593255, EPI_ISL_593257, EPI_ISL_593258, EPI_ISL_593259, EPI_ISL_593260, EPI_ISL_593261, EPI_ISL_593262, EPI_ISL_593263, EPI_ISL_593264, EPI_ISL_593265, EPI_ISL_593266, EPI_ISL_593267, EPI_ISL_593268, EPI_ISL_593269, EPI_ISL_593270, EPI_ISL_593271, EPI_ISL_593272, EPI_ISL_593273, EPI_ISL_593274, EPI_ISL_593275, EPI_ISL_593276, EPI_ISL_593277, EPI_ISL_593278, EPI_ISL_593279, EPI_ISL_593280, EPI_ISL_593281, EPI_ISL_593282, EPI_ISL_593283, EPI_ISL_593284, EPI_ISL_593285, EPI_ISL_593286, EPI_ISL_593287, EPI_ISL_593288, EPI_ISL_593289, EPI_ISL_593290, EPI_ISL_593291, EPI_ISL_593292, EPI_ISL_593293, EPI_ISL_593295, EPI_ISL_593296, EPI_ISL_593297, EPI_ISL_593298, EPI_ISL_593299, EPI_ISL_593300, EPI_ISL_593301, EPI_ISL_593302, EPI_ISL_593303, EPI_ISL_593304, EPI_ISL_593305, EPI_ISL_593306, EPI_ISL_593307, EPI_ISL_593308, EPI_ISL_593309, EPI_ISL_593310, EPI_ISL_593311, EPI_ISL_593312, EPI_ISL_593313, EPI_ISL_593314, EPI_ISL_593315, EPI_ISL_593316, EPI_ISL_593317, EPI_ISL_593318, EPI_ISL_593319, EPI_ISL_593320, EPI_ISL_593321, EPI_ISL_593322, EPI_ISL_593323, EPI_ISL_593324, EPI_ISL_593325, EPI_ISL_593326, EPI_ISL_593327, EPI_ISL_593328, EPI_ISL_593330, EPI_ISL_593331, EPI_ISL_593332, EPI_ISL_593333, EPI_ISL_593334, EPI_ISL_593335, EPI_ISL_593336, EPI_ISL_593337, EPI_ISL_593338, EPI_ISL_593339, EPI_ISL_593340, EPI_ISL_593341, EPI_ISL_593342, EPI_ISL_593343, EPI_ISL_593344, EPI_ISL_593345, EPI_ISL_593346, EPI_ISL_593347, EPI_ISL_593348, EPI_ISL_593349, EPI_ISL_593350, EPI_ISL_593351, EPI_ISL_593352, EPI_ISL_593353, EPI_ISL_593354, EPI_ISL_593355, EPI_ISL_593356, EPI_ISL_593357, EPI_ISL_593358, EPI_ISL_593359, EPI_ISL_593360, EPI_ISL_593361, EPI_ISL_593362, EPI_ISL_593363, EPI_ISL_593364, EPI_ISL_593365, EPI_ISL_593366, EPI_ISL_593367, EPI_ISL_593368, EPI_ISL_593369, EPI_ISL_593370, EPI_ISL_593371, EPI_ISL_593372, EPI_ISL_593373, EPI_ISL_593374, EPI_ISL_593375, EPI_ISL_593376, EPI_ISL_593377, EPI_ISL_593378, EPI_ISL_593379, EPI_ISL_593380, EPI_ISL_593381, EPI_ISL_593382, EPI_ISL_593383, EPI_ISL_593384, EPI_ISL_593385, EPI_ISL_593386, EPI_ISL_593387, EPI_ISL_593388, EPI_ISL_593389, EPI_ISL_593390, EPI_ISL_593391, EPI_ISL_593392, EPI_ISL_593393, EPI_ISL_593394, EPI_ISL_593395, EPI_ISL_593396, EPI_ISL_593397, EPI_ISL_593398, EPI_ISL_593399, EPI_ISL_593400, EPI_ISL_593401, EPI_ISL_593402, EPI_ISL_593403, EPI_ISL_593404, EPI_ISL_593405, EPI_ISL_593406, EPI_ISL_593407, EPI_ISL_593408, EPI_ISL_593409, EPI_ISL_593410, EPI_ISL_593413, EPI_ISL_593414, EPI_ISL_593415, EPI_ISL_593416, EPI_ISL_593418, EPI_ISL_593419, EPI_ISL_593420, EPI_ISL_593421, EPI_ISL_593427, EPI_ISL_593428, EPI_ISL_593429, EPI_ISL_593431, EPI_ISL_593432, EPI_ISL_593434, EPI_ISL_593435, EPI_ISL_593437, EPI_ISL_593438, EPI_ISL_593439, EPI_ISL_593440, EPI_ISL_593441, EPI_ISL_593443, EPI_ISL_593444, EPI_ISL_593445, EPI_ISL_593446, EPI_ISL_593447, EPI_ISL_593448, EPI_ISL_593450, EPI_ISL_593451, EPI_ISL_593452, EPI_ISL_593453, EPI_ISL_593454, EPI_ISL_593455, EPI_ISL_593456, EPI_ISL_593457, EPI_ISL_593458, EPI_ISL_593459, EPI_ISL_593460, EPI_ISL_593461, EPI_ISL_593462, EPI_ISL_593463, EPI_ISL_593464, EPI_ISL_593465, EPI_ISL_593466, EPI_ISL_593467, EPI_ISL_593468, EPI_ISL_593469, EPI_ISL_593470, EPI_ISL_593471, EPI_ISL_593472, EPI_ISL_593473, EPI_ISL_593474, EPI_ISL_593475 | | | |
| see above | Microbiological Diagnostic Unit - Public Health Laboratory (MDU-PHL) | MDU-PHL | Seemann T., Schultz, M. B., Sait, M., Sherry, N. |
| EPI_ISL_594388 | Florida Bureau of Public Health Laboratories | Florida Bureau of Public Health Laboratories | Sarah Schmedes, Jason Blanton |
| EPI_ISL_594861, EPI_ISL_594862, EPI_ISL_594863, EPI_ISL_594864, EPI_ISL_594865, EPI_ISL_594866, EPI_ISL_594867, EPI_ISL_594868, EPI_ISL_594869, EPI_ISL_594870, EPI_ISL_594871, EPI_ISL_594872, EPI_ISL_594873, EPI_ISL_594874, EPI_ISL_594875, EPI_ISL_594876, EPI_ISL_594877, EPI_ISL_594878, EPI_ISL_594879, EPI_ISL_594880, EPI_ISL_594881, EPI_ISL_594882, EPI_ISL_594883, EPI_ISL_594884, EPI_ISL_594885, EPI_ISL_594886, EPI_ISL_594887, EPI_ISL_594888 | | | |
| see above | University College London, Great Ormond Street Hospital for Children NHS Foundation Trust, Imperial College Healthcare NHS Trust | COVID-19 Genomics UK (COG-UK) Consortium | Sergi Castellano, Rachel Williams, Mark Kristiansen, Paola Resende Silva, Sunando Roy, Tony Brooks, Helena Tutill, Paola Niola, Patricia Dyal, Charlotte Williams, Leysa Forrest, Yasmin Panchbhaya, Jacqueline Findlay, Samuel Weeks, Julianne Brown, Kathryn Harris, Paul Randell, James Price, Alison Holmes, Judith Breuer |
| EPI_ISL_603040, EPI_ISL_603041, EPI_ISL_603042, EPI_ISL_603043, EPI_ISL_603044, EPI_ISL_603045, EPI_ISL_603046, EPI_ISL_603047, EPI_ISL_603048 | MDU-PHL, The Peter Doherty Institute for Infection and Immunity | MDU-PHL, The Peter Doherty Institute for Infection and Immunity | Seemann,T., Caly,L., Sait,M., Schultz,M.B., Druce,J., Sherry,N. |
| EPI_ISL_603238, EPI_ISL_603239 | National Institute of Laboratory Medicine and Referral Center | Genomic Research Lab, BCSIR | Shahina Akter, Abu Sayeed Mohammad Mahmud, Mohammad Samir Uzzaman, Eshrar Osman, Md. Ahashan Habib, Tanjina Akhter Banu, Md. Murshed Hasan Sarkar, Barna Goswami, Iffat Jahan, Md. Saddam Hossain, Tasnim Nafisa, Md. Maruf Ahmed Molla, Mahmuda Yeasmin, Asish Kumar Ghosh, A. K. M. Shamsuzzaman, Monira Parveen, Md. Masum Hossain Arif, Md. Salim Khan |
| EPI_ISL_611523, EPI_ISL_611527, EPI_ISL_611558, EPI_ISL_611559, EPI_ISL_611595, EPI_ISL_611596, EPI_ISL_611597, EPI_ISL_611622, EPI_ISL_611623, EPI_ISL_611665, EPI_ISL_611822, EPI_ISL_611823, EPI_ISL_611996, EPI_ISL_612000, EPI_ISL_612014, EPI_ISL_612078, EPI_ISL_612079, EPI_ISL_612103, EPI_ISL_612105, EPI_ISL_612131, EPI_ISL_612198, EPI_ISL_612450, EPI_ISL_612451, EPI_ISL_612452, EPI_ISL_612453, EPI_ISL_612454, EPI_ISL_612455, EPI_ISL_612456, EPI_ISL_612457 | | | |
| see above | University College London, Great Ormond Street Hospital for Children NHS Foundation Trust, Imperial College Healthcare NHS Trust | COVID-19 Genomics UK (COG-UK) Consortium | Sergi Castellano, Rachel Williams, Mark Kristiansen, Paola Resende Silva, Sunando Roy, Tony Brooks, Helena Tutill, Paola Niola, Patricia Dyal, Charlotte Williams, Leysa Forrest, Yasmin Panchbhaya, Jacqueline Findlay, Samuel Weeks, Julianne Brown, Kathryn Harris, Paul Randell, James Price, Alison Holmes, Judith Breuer |
| EPI_ISL_613724, EPI_ISL_613739, EPI_ISL_613747, EPI_ISL_613751, EPI_ISL_613756, EPI_ISL_613758, EPI_ISL_613760, EPI_ISL_613766, EPI_ISL_613770, EPI_ISL_613771, EPI_ISL_613772, EPI_ISL_613800, EPI_ISL_613829, EPI_ISL_613830, EPI_ISL_613831, EPI_ISL_613834, EPI_ISL_613837, EPI_ISL_613838, EPI_ISL_613840, EPI_ISL_613841, EPI_ISL_613842, EPI_ISL_613843, EPI_ISL_613847, EPI_ISL_613849, EPI_ISL_613850, EPI_ISL_613851, EPI_ISL_613852, EPI_ISL_613853, EPI_ISL_613854, EPI_ISL_613856, EPI_ISL_613857, EPI_ISL_613858, EPI_ISL_613859, EPI_ISL_613861, EPI_ISL_613862, EPI_ISL_613863, EPI_ISL_613864, EPI_ISL_613865, EPI_ISL_613866, EPI_ISL_613867, EPI_ISL_613868, EPI_ISL_613871, EPI_ISL_613874, EPI_ISL_613875, EPI_ISL_613876, EPI_ISL_613877, EPI_ISL_613878, EPI_ISL_613880, EPI_ISL_613881, EPI_ISL_613882, EPI_ISL_613883, EPI_ISL_613884, EPI_ISL_613885, EPI_ISL_613886, EPI_ISL_613887, EPI_ISL_613888, EPI_ISL_613889, EPI_ISL_613890, EPI_ISL_613893, EPI_ISL_613894, EPI_ISL_613895, EPI_ISL_613896, EPI_ISL_613897, EPI_ISL_613898, EPI_ISL_613900, EPI_ISL_613901, EPI_ISL_613902, EPI_ISL_613903, EPI_ISL_613904, EPI_ISL_613905, EPI_ISL_613906, EPI_ISL_613907, | | | |

| | | | |
|---|---|---|---|
| EPI_ISL_613908, EPI_ISL_613909, EPI_ISL_613910, EPI_ISL_613912, EPI_ISL_613913, EPI_ISL_613914, EPI_ISL_613915, EPI_ISL_613916, EPI_ISL_613917, EPI_ISL_613918, EPI_ISL_613919, EPI_ISL_613920, EPI_ISL_613921, EPI_ISL_613922, EPI_ISL_613923, EPI_ISL_613924, EPI_ISL_613925, EPI_ISL_613927, EPI_ISL_613928, EPI_ISL_613929, EPI_ISL_613930, EPI_ISL_613931, EPI_ISL_613932, EPI_ISL_613933, EPI_ISL_613934, EPI_ISL_613935, EPI_ISL_613936, EPI_ISL_613937, EPI_ISL_613938, EPI_ISL_613939, EPI_ISL_613940, EPI_ISL_613942, EPI_ISL_613943, EPI_ISL_613944, EPI_ISL_613945, EPI_ISL_613946, EPI_ISL_613947, EPI_ISL_613948, EPI_ISL_613949, EPI_ISL_613950, EPI_ISL_614247 | | | |
| see above | Florida Bureau of Public Health Laboratories | Florida Bureau of Public Health Laboratories | Sarah Schmedes, Jason Blanton |
| EPI_ISL_625457, EPI_ISL_625458, EPI_ISL_625459, EPI_ISL_625460, EPI_ISL_625461, EPI_ISL_625462, EPI_ISL_625463, EPI_ISL_625464, EPI_ISL_625465, EPI_ISL_625466, EPI_ISL_625467, EPI_ISL_625468, EPI_ISL_625469, EPI_ISL_625470, EPI_ISL_625471, EPI_ISL_625472, EPI_ISL_625473, EPI_ISL_625474, EPI_ISL_625475, EPI_ISL_625476, EPI_ISL_625477 | | | |
| see above | Child Health Research Foundation | Child Health Research Foundation | Senjuti Saha, Md Saiful Islam Sajib, Nikkon Sarkar, Syed Muktadir Al Sium, Afroza Akter Tanni, Roly Malaker, Arif Mohammad Tanmoy, Md Hafizur Rahman, Samir K Saha |
| EPI_ISL_627082, EPI_ISL_627104, EPI_ISL_627108, EPI_ISL_627110, EPI_ISL_627175, EPI_ISL_627211, EPI_ISL_627450, EPI_ISL_627451, EPI_ISL_627452, EPI_ISL_627453, EPI_ISL_627454, EPI_ISL_627455, EPI_ISL_627456, EPI_ISL_627457, EPI_ISL_627458, EPI_ISL_627459, EPI_ISL_627460, EPI_ISL_627461, EPI_ISL_627462, EPI_ISL_627463, EPI_ISL_627464, EPI_ISL_627465, EPI_ISL_627466, EPI_ISL_627467, EPI_ISL_627468, EPI_ISL_627469, EPI_ISL_627470, EPI_ISL_627471, EPI_ISL_627472, EPI_ISL_627473, EPI_ISL_627474, EPI_ISL_627475, EPI_ISL_627476, EPI_ISL_627477, EPI_ISL_627478, EPI_ISL_627479, EPI_ISL_627480, EPI_ISL_627481, EPI_ISL_627491 | | | |
| see above | University College London, Great Ormond Street Hospital for Children NHS Foundation Trust, Imperial College Healthcare NHS Trust | COVID-19 Genomics UK (COG-UK) Consortium | Sergi Castellano, Rachel Williams, Mark Kristiansen, Paola Resende Silva, Sunando Roy, Tony Brooks, Helena Tutill, Paola Niola, Patricia Dyal, Charlotte Williams, Leysa Forrest, Yasmin Panchbhaya, Jacqueline Findlay, Samuel Weeks, Julianne Brown, Kathryn Harris, Paul Randell, James Price, Alison Holmes, Judith Breuer |

We gratefully acknowledge the following Authors from the Originating laboratories responsible for obtaining the specimens, as well as the Submitting laboratories where the genome data were generated and shared via GISAID, on which this research is based.

All Submitters of data may be contacted directly via www.gisaid.org

Authors are sorted alphabetically.

| Accession ID | Originating Laboratory | Submitting Laboratory | Authors |
|---|---|---|---|
| EPI_ISL_407079 | Lapland Central Hospital | Department of Virology, University of Helsinki and Helsinki University Hospital, Helsinki, Finland | Teemu Smura, Suvi Kuivanen, Hannimari Kallio-Kokko, Olli Vapalahti |
| EPI_ISL_408489, EPI_ISL_410218 | Department of Laboratory Medicine, National Taiwan University Hospital | Microbial Genomics Core Lab, National Taiwan University Centers of Genomic and Precision Medicine | Shiou-Hwei Yeh, You-Yu Lin, Ya-Yun Lai, Chiao-Ling Li, Shan-Chwen Chang, Pei-Jer Chen, Sui-Yuan Chang |
| EPI_ISL_412966 | Technology Centre, Guangzhou Customs | Technology Centre, Guangzhou Customs | Shi,Y., Sun,J., Zheng,K., Huang,J. and Zhao,J. |
| EPI_ISL_412967 | Technology Centre, Guangzhou Customs | Technology Centre, Guangzhou Customs | Shi,Y., Zheng,K., Sun,J., Huang,J., Zhu,A., Zhuang,Z., Dai,J., Chen,Z., Sun,F., Zhang,Z., Li,X. and Wang,Y. |
| EPI_ISL_412971 | HUS Diagnostiikkakeskus, Hallinto | Department of Virology Faculty of Medicine, Medicum University of Helsinki | Teemu Smura, Suvi Kuivanen, Hannimari Kallio-Kokko, Olli Vapalahti |
| EPI_ISL_413015 | Public Health Ontario Laboratory | National Microbiology Laboratory | Shari Tyson, Anna Majer, Erika Landry, Morag Graham, Grace Seo, Philip Mabon, Natalie Knox, Adrian Zetner, Samira Mubareka, Rob Kozak, Jocelyne Lew, Darryl Falzarano, Gerdts Volker, Jonathan Gubbay, Stephanie Booth, Guillaume Poliquin, Tom Graefenhan, Matthew Gilmour, Nathalie Bastien, Yan Li, Timothy Booth |
| EPI_ISL_413019, EPI_ISL_413020 | Department of Internal Medicine, Triemli Hospital | Institute of Medical Virology, University of Zurich | Stefan Schmutz, Maryam Zaheri, Verena Kufner, Patrick Redli, Fiona Steiner, Jon Huder, Riccarda Capaul, Andrea Zbinden, Jürg Böni, Michael Huber, Gerhard Eich, Alexandra Trkola |
| EPI_ISL_413021 | Klinik Hirslanden Zurich | Institute of Medical Virology, University of Zurich | Stefan Schmutz, Maryam Zaheri, Verena Kufner, Gabriela Ziltener, Patrick Redli, Fiona Steiner, Jon Huder, Riccarda Capaul, Andrea Zbinden, Jürg Böni, Michael Huber, Christian Ruef, Alexandra Trkola |
| EPI_ISL_413022, EPI_ISL_413023, EPI_ISL_413024 | Division of Infectious Diseases, University Hospital Zurich | Institute of Medical Virology, University of Zurich | Stefan Schmutz, Maryam Zaheri, Verena Kufner, Gabriela Ziltener, Patrick Redli, Fiona Steiner, Jon Huder, Riccarda Capaul, Andrea Zbinden, Jürg Böni, Michael Huber, Roberto Speck, Alexandra Trkola |
| EPI_ISL_413592 | Department of Laboratory Medicine, National Taiwan University Hospital | Microbial Genomics Core Lab, National Taiwan University Centers of Genomic and Precision Medicine | Shiou-Hwei Yeh, You-Yu Lin, Ya-Yun Lai, Chiao-Ling Li, Shan-Chwen Chang, Pei-Jer Chen, Sui-Yuan Chang |
| EPI_ISL_413602, EPI_ISL_413603, EPI_ISL_413604, EPI_ISL_413605, EPI_ISL_414640, EPI_ISL_414641, EPI_ISL_414642, EPI_ISL_414643, EPI_ISL_414644, EPI_ISL_414645, EPI_ISL_414646 | | | |
| see above | Department of Virology and Immunology, University of Helsinki and Helsinki University Hospital, Huslab Finland | Department of Virology, Faculty of Medicine, University of Helsinki, Helsinki, Finland | Teemu Smura, Hannimari Kallio-Kokko, Olli Vapalahti |
| EPI_ISL_414949 | Regional Virus Laboratory, Belfast | Public Health Wales Microbiology Cardiff | Tanya Curran, Conall McCaughey, Catherine Moore, Joanne Watkins, Sally Corden, Tom Connor |
| EPI_ISL_416316, EPI_ISL_416317, EPI_ISL_416318, EPI_ISL_416319, EPI_ISL_416320, EPI_ISL_416321, EPI_ISL_416322, EPI_ISL_416323, EPI_ISL_416324, EPI_ISL_416325, EPI_ISL_416326, EPI_ISL_416327, EPI_ISL_416328, EPI_ISL_416329, EPI_ISL_416330, EPI_ISL_416331, EPI_ISL_416332, EPI_ISL_416333, EPI_ISL_416334, EPI_ISL_416335, EPI_ISL_416336, EPI_ISL_416337, EPI_ISL_416338, EPI_ISL_416339, EPI_ISL_416340, EPI_ISL_416341, EPI_ISL_416342, EPI_ISL_416343, EPI_ISL_416344, EPI_ISL_416345, EPI_ISL_416346, EPI_ISL_416347, EPI_ISL_416348, EPI_ISL_416349, EPI_ISL_416350, EPI_ISL_416351, EPI_ISL_416352, EPI_ISL_416353, EPI_ISL_416354, EPI_ISL_416355, EPI_ISL_416356, EPI_ISL_416357, EPI_ISL_416358, EPI_ISL_416359, EPI_ISL_416360, EPI_ISL_416361, EPI_ISL_416362, EPI_ISL_416363, EPI_ISL_416364, EPI_ISL_416365, EPI_ISL_416366, EPI_ISL_416367, EPI_ISL_416368, EPI_ISL_416369, EPI_ISL_416370, EPI_ISL_416371, EPI_ISL_416372, EPI_ISL_416373, EPI_ISL_416374, EPI_ISL_416375, EPI_ISL_416376, EPI_ISL_416377, EPI_ISL_416378, EPI_ISL_416379, EPI_ISL_416380, EPI_ISL_416381, EPI_ISL_416382, EPI_ISL_416383, EPI_ISL_416384, EPI_ISL_416385, EPI_ISL_416386, EPI_ISL_416387, EPI_ISL_416388, EPI_ISL_416389, EPI_ISL_416390, EPI_ISL_416391, EPI_ISL_416392, EPI_ISL_416393, EPI_ISL_416394, EPI_ISL_416395, EPI_ISL_416396, EPI_ISL_416397, EPI_ISL_416398, EPI_ISL_416399, EPI_ISL_416400, EPI_ISL_416401, EPI_ISL_416402, EPI_ISL_416403, EPI_ISL_416404, EPI_ISL_416405, EPI_ISL_416406, EPI_ISL_416407, EPI_ISL_416408, EPI_ISL_416409 | | | |
| see above | Shanghai Public Health Clinical Center, Shanghai Medical College, Fudan University | National Research Center for Translational Medicine (Shanghai), Ruijin Hospital affiliated to Shanghai Jiao Tong University School of Medicine & Shanghai Public Health Clinical Center | Shengyue Wang, Xiaonan Zhang, Gang Lu, Yun Tan, Yun Ling, Hongzhou Lu, Saijuan Chen |
| EPI_ISL_417395, EPI_ISL_417412 | Centre for Infectious Diseases and Microbiology Public Health | NSW Health Pathology - Institute of Clinical Pathology and Medical Research; Westmead Hospital; University of Sydney | Sintchenko V, Chen SC, Maddocks S, Kok J, Dwyer DE, Rockett R, Eden J-S, Lam C, Gray K, Timms V, Gall M, Arnott A, Sadsad R, Carter I, Rahman H, Holmes EC and O'Sullivan MV for the 2019-nCoV Study Group |
| EPI_ISL_417482 | Institute of Microbiology, Universidad San Francisco de Quito | Institute of Microbiology, Universidad San Francisco de Quito | Sully Márquez, Belén Prado-Vivar, Juan José Guadalupe, Bernardo Gutiérrez, Manuel Jibaja, Milton Tobar, Verónica Barragán, Patricio Rojas-Silva, Gabriel Trueba, Michelle Grunauer, Paúl Cárdenas |
| EPI_ISL_417931, EPI_ISL_417932, EPI_ISL_417933, EPI_ISL_417935, EPI_ISL_417937, EPI_ISL_417938, EPI_ISL_417939 | UCSF Clinical Microbiology Laboratory | Chan-Zuckerberg Biohub | Shaun Arevalo, Josh Batson, Olga Botvinnik, Gloria Castaneda, Angela Detweiler, David Dynerman, Samantha Hao, Jack Kamm, Amy Kistler, G. Renuka Kumar, Chaz Langelier, Lucy Li, Steve Miller, Lusajo Mwakibete, Norma Neff, Angela Pisco, Maira Phelps, Michelle Tan, Chunyu Zhao |
| EPI_ISL_418385, EPI_ISL_418386, EPI_ISL_418387, EPI_ISL_418388, EPI_ISL_418389, EPI_ISL_418390, EPI_ISL_418391, EPI_ISL_418392, EPI_ISL_418393, EPI_ISL_418394, EPI_ISL_418395, EPI_ISL_418396, EPI_ISL_418397, EPI_ISL_418398, EPI_ISL_418399, EPI_ISL_418400, EPI_ISL_418401, EPI_ISL_418402, EPI_ISL_418403, EPI_ISL_418404, EPI_ISL_418405, EPI_ISL_418406, EPI_ISL_418407, EPI_ISL_418408, EPI_ISL_418409, EPI_ISL_418410, EPI_ISL_418411 | | | |
| see above | Department of Virology and Immunology, University of Helsinki and Helsinki University Hospital, Huslab Finland | Department of Virology, Faculty of Medicine, University of Helsinki, Helsinki, Finland | Teemu Smura, Hannimari Kallio-Kokko, Olli Vapalahti |
| EPI_ISL_419313 | Molecular Biology and Biotechnology Lab II | Molecular Biology and Biotechnology Lab II | Tayyaba Zainab, Sana Shamshad, Azka Noureen, Aimen Malik, Muhammad Javaid Asad, Kumail Ali Rizvi |
| EPI_ISL_422407, EPI_ISL_422408, EPI_ISL_422409, EPI_ISL_422410, EPI_ISL_422411, EPI_ISL_422412, EPI_ISL_422413, EPI_ISL_422414, EPI_ISL_422415, EPI_ISL_422416, EPI_ISL_422417, EPI_ISL_422418, EPI_ISL_422419, EPI_ISL_422420, EPI_ISL_422421, EPI_ISL_422422 | | | |
| see above | Department of Laboratory Medicine, National Taiwan University Hospital | Microbial Genomics Core Lab, National Taiwan University Centers of Genomic and Precision Medicine | Shiou-Hwei Yeh, You-Yu Lin, Ya-Yun Lai, Chiao-Ling Li, Shan-Chwen Chang, Pei-Jer Chen, Sui-Yuan Chang |
| EPI_ISL_422565 | Institute of Microbiology Universidad San Francisco de Quito | Institute of Microbiology Universidad San Francisco de Quito | Sully Marquez, Belen Prado-Vivar, Juan Jose Guadalupe, Bernardo Gutierrez, Francisco Mora, Juan Gaviria, Alejandra Ramones, Franklin Espinoza, Edison Ligña, Jorge Reyes, Patricio Rojas-Silva, Veronica Barragan, Gabriel Trueba, Michelle Grunauer, Paul Cardenas |
| EPI_ISL_424366 | Vaccine Research, Development and Application Center, Erciyes University | Gen Era Diagnostics Inc. | Shaikh Terkis Islam Pavel, Hazel Yetiskin, Gunsu Aydin, Can Holyavkin, Muhammet Ali Uygut, Zehra B Dursun, lhami Celik, Alper Iseri, Aykut Ozdarendeli |
| EPI_ISL_425023, EPI_ISL_425024 | Respiratory Virus Unit, Microbiology Services Colindale, Public Health England | Respiratory Virus Unit, Microbiology Services Colindale, Public Health England | Steven Platt, Shahjahan Miah, Angie Lackenby, Omolola Akinbami, Tiina Talts, Leena Bhaw, Richard Myers, Monica Galiano, Kirstin Edwards, Jonathan Hubb, Joanna Ellis, Maria Zambon |
| EPI_ISL_427658, EPI_ISL_427661, EPI_ISL_427688, EPI_ISL_427699, EPI_ISL_427709, EPI_ISL_427710, EPI_ISL_427712 | Centre for Infectious Diseases and Microbiology Public Health | NSW Health Pathology - Institute of Clinical Pathology and Medical Research; Westmead Hospital; University of Sydney | Sim E, Bachmann N, Rockett R, Lam C, Gray K, Timms V, Gall M, Arnott A, Sadsad R, Draper J, Carter I, Holmes EC, O'Sullivan MV, Byun R, Sintchenko V, Chen SC, Eden JS, Maddocks S, Kok J, Propenko M, Sorrell T, Chang S, Basile K, Dwyer DE for the 2019-nCoV Study Group |
| EPI_ISL_427717 | ACT Pathology, The Canberra Hospital | NSW Health Pathology - Institute of Clinical Pathology and Medical Research; Westmead Hospital; University of Sydney | Sim E, Bachmann N, Rockett R, Lam C, Gray K, Timms V, Gall M, Arnott A, Sadsad R, Draper J, Carter I, Holmes EC, O'Sullivan MV, Byun R, Sintchenko V, Chen SC, Eden JS, Maddocks S, Kok J, Propenko M, Sorrell T, Chang S, Basile K, Dwyer DE for the 2019-nCoV Study Group |

| | | | |
|---|---|---|---|
| EPI_ISL_427720 | Centre for Infectious Diseases and Microbiology Public Health | NSW Health Pathology - Institute of Clinical Pathology and Medical Research; Westmead Hospital; University of Sydney | Sim E, Bachmann N, Rockett R, Lam C, Gray K, Timms V, Gall M, Arnott A, Sadsad R, Draper J, Carter I, Holmes EC, O'Sullivan MV, Byun R, Sintchenko V, Chen SC, Eden JS, Maddocks S, Kok J, Propenko M, Sorrell T, Chang S, Basile K, Dwyer DE for the 2019-nCoV Study Group |
| EPI_ISL_427721 | ACT Pathology, The Canberra Hospital | NSW Health Pathology - Institute of Clinical Pathology and Medical Research; Westmead Hospital; University of Sydney | Sim E, Bachmann N, Rockett R, Lam C, Gray K, Timms V, Gall M, Arnott A, Sadsad R, Draper J, Carter I, Holmes EC, O'Sullivan MV, Byun R, Sintchenko V, Chen SC, Eden JS, Maddocks S, Kok J, Propenko M, Sorrell T, Chang S, Basile K, Dwyer DE for the 2019-nCoV Study Group |
| EPI_ISL_427743, EPI_ISL_427759, EPI_ISL_427783, EPI_ISL_427786, EPI_ISL_427788, EPI_ISL_427802 | Centre for Infectious Diseases and Microbiology Public Health | NSW Health Pathology - Institute of Clinical Pathology and Medical Research; Westmead Hospital; University of Sydney | Sim E, Bachmann N, Rockett R, Lam C, Gray K, Timms V, Gall M, Arnott A, Sadsad R, Draper J, Carter I, Holmes EC, O'Sullivan MV, Byun R, Sintchenko V, Chen SC, Eden JS, Maddocks S, Kok J, Propenko M, Sorrell T, Chang S, Basile K, Dwyer DE for the 2019-nCoV Study Group |
| EPI_ISL_429664, EPI_ISL_429665, EPI_ISL_429666, EPI_ISL_429667, EPI_ISL_429668, EPI_ISL_429669, EPI_ISL_429670, EPI_ISL_429671, EPI_ISL_429672, EPI_ISL_429673, EPI_ISL_429674, EPI_ISL_429675, EPI_ISL_429676, EPI_ISL_429677, EPI_ISL_429678, EPI_ISL_429679, EPI_ISL_429680, EPI_ISL_429681, EPI_ISL_429682, EPI_ISL_429683, EPI_ISL_429684, EPI_ISL_429685, EPI_ISL_429686, EPI_ISL_429687, EPI_ISL_429688, EPI_ISL_429689, EPI_ISL_429690, EPI_ISL_429691, EPI_ISL_429692, EPI_ISL_429693, EPI_ISL_429694, EPI_ISL_429695, EPI_ISL_429696, EPI_ISL_429697, EPI_ISL_429698, EPI_ISL_429699, EPI_ISL_429700, EPI_ISL_429701, EPI_ISL_429702, EPI_ISL_429703 | | | |
| see above | Central Public Health Laboratory/Octávio Magalhães Institute (IOM) from the Ezequiel Dias Foundation (FUNED) | Instituto Octávio Magalhães / Fundação Ezequiel Dias (IOM/Funed) | Talita Adelino, Joilson Xavier, Marta Giovanetti, Vagner Fonseca, Marcos Vinícius Silva, Luiz Carlos Junior Alcantara, Marluce Aparecida Assunção Oliveira |
| EPI_ISL_430439 | Institute for Medical Research, Infectious Disease Research Centre, National Institutes of Health, Ministry of Health Malaysia | Institute for Medical Research Infectious Disease Research Centre, National Institutes of Health, Ministry of Health Malaysia | Suppiah.J, Mohd-Zawawi.Z, Kalyanasundram.J, Azizan.M-A, Mat-Sharani.S, Hisham.H-A, Tan.L-P, Abdul-Wahid.M-Z, Mohd-Zain.R, Ahmad.N, Thayan.R |
| EPI_ISL_430440 | Institute for Medical Research, Infectious Disease Research Centre, National Institutes of Health, Ministry of Health Malaysia | Institute for Medical Research, Infectious Disease Research Centre, National Institutes of Health, Ministry of Health Malaysia | Suppiah.J, Mohd-Zawawi.Z, Kalyanasundram.J, Azizan.M-A, Mat-Sharani.S, Hisham.H-A, Tan.L-P, Abdul-Wahid.M-Z, Tengku-Abd-Rashid.T-R, Mohd-Zain.R, Ahmad.N, Thayan.R |
| EPI_ISL_430441, EPI_ISL_430442, EPI_ISL_430443, EPI_ISL_430444 | Institute for Medical Research, Infectious Disease Research Centre, National Institutes of Health, Ministry of Health Malaysia | Institute for Medical Research, Infectious Disease Research Centre, National Institutes of Health, Ministry of Health Malaysia | Suppiah.J, Mohd-Zawawi.Z, Kalyanasundram.J, Azizan.M-A, Mat-Sharani.S, Hisham.H-A, Tan.L-P, Abdul-Wahid.M-Z, Tengku-Rogayah.TAR, Mohd-Zain.R, Ahmad.N, Thayan.R |
| EPI_ISL_435045, EPI_ISL_435046, EPI_ISL_435047, EPI_ISL_435048 | Laboratory of Applied Genetics | RSE "National Center for Biotechnology" | Alexandr Shevtsov, Ilyas Akhmetollayev, Viktoriya Lutsay, Asylulan Amirgazin, Ruslan Kalendar, Yerlan Ramanculov |
| EPI_ISL_435055 | Gujarat Biotechnology Research Centre | Gujarat Biotechnology Research Centre | Tejas Shah, Ankit Hinsu, Pritesh Sabara, Apurvasinh Puvar, Janvi Raval, Monika Gandhi, Pinal Trivedi, Maharshi Pandya, Amit Kanani, Akanksha Verma, Nitin Savaliya, Raghawendra Kumar, Dinesh Kumar, Zuber Saiyed, Dipa Kinariwala, Disha Patel, Binita Aring, Geeta Vaghela, Sonia Barve, Bhavesh Modi, Kairavi Joshi, Gaurishankar Shrimali, Nidhi Sood, Pranay Shah, R D Dixit, Snehal Bagatharia, Kamlesh J Upadhyay, Ramesh Pandit, Anjali Rajwal, Chaitanya Joshi, Madhvi Joshi |
| EPI_ISL_437449 | B.J. Medical College and Civil hospital | Gujarat Biotechnology Research Centre | Sonia Barve, Bhavesh Modi, Kairavi Joshi, Gaurishankar Shrimali, Nidhi Sood, Pranay Shah, R D Dixit, Snehal Bagatharia, Kamlesh J Upadhyay, Ramesh Pandit, Tejas Shah, Ankit Hinsu, Pritesh Sabara, Apurvasinh Puvar, Janvi Raval, Monika Gandhi, Pinal Trivedi, Maharshi Pandya, Amit Kanani, Akanksha Verma, Nitin Savaliya, Raghawendra Kumar, Dinesh Kumar, Zuber Saiyed, Dipa Kinariwala, Disha Patel, Binita Aring, Neeta Khandelwal, Geeta Vaghela, Neelam Nathani, Chaitanya Joshi, Madhvi Joshi |
| EPI_ISL_437459, EPI_ISL_437460, EPI_ISL_437461, EPI_ISL_437462, EPI_ISL_437463, EPI_ISL_437464, EPI_ISL_437465, EPI_ISL_437466, EPI_ISL_437467, EPI_ISL_437468, EPI_ISL_437469, EPI_ISL_437470, EPI_ISL_437471, EPI_ISL_437472, EPI_ISL_437473, EPI_ISL_437474, EPI_ISL_437475, EPI_ISL_437477, EPI_ISL_437479 | | | |
| see above | Pathogen Genomics Lab King Abdullah University of Science and Technology(KAUST) | Pathogen Genomics Lab King Abdullah University of Science and Technology(KAUST) | Sharif Hala,Raeece Naeem,Sara Mfarrej,Arnab Pain |
| EPI_ISL_437691, EPI_ISL_437692, EPI_ISL_437693, EPI_ISL_437694, EPI_ISL_437695, EPI_ISL_437696, EPI_ISL_437697, EPI_ISL_437698, EPI_ISL_437699, EPI_ISL_437700, EPI_ISL_437701, EPI_ISL_437702, EPI_ISL_437703, EPI_ISL_437704, EPI_ISL_437705, EPI_ISL_437706, EPI_ISL_437707, EPI_ISL_437708, EPI_ISL_437709, EPI_ISL_437710, EPI_ISL_437711, EPI_ISL_437712, EPI_ISL_437713, EPI_ISL_437714, EPI_ISL_437715, EPI_ISL_437716, EPI_ISL_437717, EPI_ISL_437718, EPI_ISL_437719, EPI_ISL_437720, EPI_ISL_437721, EPI_ISL_437722, EPI_ISL_437723, EPI_ISL_437724, EPI_ISL_437725, EPI_ISL_437726, EPI_ISL_437727, EPI_ISL_437728, EPI_ISL_437729, EPI_ISL_437730, EPI_ISL_437731, EPI_ISL_437732, EPI_ISL_437733, EPI_ISL_437734, EPI_ISL_437735, EPI_ISL_437736, EPI_ISL_437737, EPI_ISL_437738, EPI_ISL_437739, EPI_ISL_437740, EPI_ISL_437741, EPI_ISL_437742, EPI_ISL_437743, EPI_ISL_437744, EPI_ISL_437745, EPI_ISL_437746, EPI_ISL_437747, EPI_ISL_437748, EPI_ISL_437749, EPI_ISL_437750, EPI_ISL_437751, EPI_ISL_437752, EPI_ISL_437753, EPI_ISL_437754, EPI_ISL_437755, EPI_ISL_437756, EPI_ISL_437757, EPI_ISL_437758, EPI_ISL_437759, EPI_ISL_437760, EPI_ISL_437761, EPI_ISL_437762 | | | |
| see above | Pathogen Genomics Lab King Abdullah University of Science and Technology(KAUST) | Pathogen Genomics Lab King Abdullah University of Science and Technology(KAUST) | Sharif Hala,Fadwa Alofi,Afrah Alsomali, Asim Khogeer, Sara Mfarrej, Khaled Algithami,Raeece Naeem, Amit Kumar Subudhi,Fathia Ben-Rached, Rahul Salunke, Anwar Hashem, Naif Almontashiri, Arnab Pain |
| EPI_ISL_439953, EPI_ISL_439957, EPI_ISL_439959, EPI_ISL_439960, EPI_ISL_439962, EPI_ISL_439964, EPI_ISL_439965, EPI_ISL_439966, EPI_ISL_439967, EPI_ISL_439969, EPI_ISL_439971, EPI_ISL_439972, EPI_ISL_439973, EPI_ISL_439974, EPI_ISL_439976, EPI_ISL_439978, EPI_ISL_439979, EPI_ISL_439981, EPI_ISL_439982, EPI_ISL_439983, EPI_ISL_439984, EPI_ISL_439985, EPI_ISL_439986, EPI_ISL_439988, EPI_ISL_439992, EPI_ISL_439998, EPI_ISL_440000, EPI_ISL_440001, EPI_ISL_440002, EPI_ISL_440005, EPI_ISL_440006, EPI_ISL_440008, EPI_ISL_440013, EPI_ISL_440016, EPI_ISL_440018, EPI_ISL_440020, EPI_ISL_440021, EPI_ISL_440022, EPI_ISL_440023, EPI_ISL_440026, EPI_ISL_440027, EPI_ISL_440028, EPI_ISL_440029, EPI_ISL_440031, EPI_ISL_440032, EPI_ISL_440033, EPI_ISL_440035, EPI_ISL_440036, EPI_ISL_440037, EPI_ISL_440038, EPI_ISL_440039, EPI_ISL_440042, EPI_ISL_440043, EPI_ISL_440050, EPI_ISL_440051, EPI_ISL_440052, EPI_ISL_440054, EPI_ISL_440056, EPI_ISL_440058, EPI_ISL_440063, EPI_ISL_440065, EPI_ISL_440068, EPI_ISL_440070, EPI_ISL_440071, EPI_ISL_440081, EPI_ISL_440084, EPI_ISL_440087, EPI_ISL_440091, EPI_ISL_440092, EPI_ISL_440093, EPI_ISL_440094, EPI_ISL_440095, EPI_ISL_440098, EPI_ISL_440100, EPI_ISL_440101, EPI_ISL_440102, EPI_ISL_440103, EPI_ISL_440105, EPI_ISL_440106, EPI_ISL_440113, EPI_ISL_440114, EPI_ISL_440115, EPI_ISL_440116, EPI_ISL_440118, EPI_ISL_440119, EPI_ISL_440123, EPI_ISL_440124, EPI_ISL_440126, EPI_ISL_440127, EPI_ISL_440128, EPI_ISL_440130, EPI_ISL_440131, EPI_ISL_440134, EPI_ISL_440135, EPI_ISL_440136, EPI_ISL_440137, EPI_ISL_440139, EPI_ISL_440140, EPI_ISL_440141, EPI_ISL_440142, EPI_ISL_440145, EPI_ISL_440148, EPI_ISL_440149, EPI_ISL_440151, EPI_ISL_440153, EPI_ISL_440154, EPI_ISL_440156, EPI_ISL_440157, EPI_ISL_440158, EPI_ISL_440160, EPI_ISL_440162, EPI_ISL_440165, EPI_ISL_440167, EPI_ISL_440168, EPI_ISL_440172, EPI_ISL_440174, EPI_ISL_440175, EPI_ISL_440178, EPI_ISL_440187, EPI_ISL_440189, EPI_ISL_440190, EPI_ISL_440191, EPI_ISL_440192, EPI_ISL_440194, EPI_ISL_440196, EPI_ISL_440198, EPI_ISL_440199, EPI_ISL_440200, EPI_ISL_440201, EPI_ISL_440204, EPI_ISL_440205, EPI_ISL_440206, EPI_ISL_440210, EPI_ISL_440212, EPI_ISL_440214, EPI_ISL_440215, EPI_ISL_440216, EPI_ISL_440218, EPI_ISL_440219, EPI_ISL_440220, EPI_ISL_440221, EPI_ISL_440222, EPI_ISL_440224, EPI_ISL_440225, EPI_ISL_440227, EPI_ISL_440228, EPI_ISL_440229, EPI_ISL_440232, EPI_ISL_440233, EPI_ISL_440235, EPI_ISL_440236, EPI_ISL_440238, EPI_ISL_440241, EPI_ISL_440242, EPI_ISL_440243, EPI_ISL_440245, EPI_ISL_440248, EPI_ISL_440249, EPI_ISL_440250, EPI_ISL_440251, EPI_ISL_440252, EPI_ISL_440253, EPI_ISL_440255, EPI_ISL_440256, EPI_ISL_440262, EPI_ISL_440266, EPI_ISL_440269, EPI_ISL_440272, EPI_ISL_440273, EPI_ISL_440274, EPI_ISL_440275, EPI_ISL_440276, EPI_ISL_440277, EPI_ISL_440286, EPI_ISL_440291, EPI_ISL_440292, EPI_ISL_440294, EPI_ISL_440298, EPI_ISL_440299, EPI_ISL_440623, EPI_ISL_440624, EPI_ISL_440625, EPI_ISL_440626, EPI_ISL_440627, EPI_ISL_440628, EPI_ISL_440629, EPI_ISL_440630, EPI_ISL_440631, EPI_ISL_440632, EPI_ISL_440633, EPI_ISL_440634, EPI_ISL_440635, EPI_ISL_440636, EPI_ISL_440637, EPI_ISL_440638, EPI_ISL_440639, EPI_ISL_440640, EPI_ISL_440641, EPI_ISL_440642, EPI_ISL_440643, EPI_ISL_440644, EPI_ISL_440645, EPI_ISL_440646, EPI_ISL_440647, EPI_ISL_440648, EPI_ISL_440649, EPI_ISL_440650, EPI_ISL_440651, EPI_ISL_440652, EPI_ISL_440653, EPI_ISL_440654, EPI_ISL_440655, EPI_ISL_440656, EPI_ISL_440657, EPI_ISL_440658, EPI_ISL_440659, EPI_ISL_440660, EPI_ISL_440661, EPI_ISL_440662, EPI_ISL_440663, EPI_ISL_440664, EPI_ISL_440665, EPI_ISL_440666, EPI_ISL_440667, EPI_ISL_440668, EPI_ISL_440669, EPI_ISL_440670, EPI_ISL_440671, EPI_ISL_440672, EPI_ISL_440673, EPI_ISL_440674, EPI_ISL_440675, EPI_ISL_440676, EPI_ISL_440677, EPI_ISL_440678, EPI_ISL_440679, EPI_ISL_440680, EPI_ISL_440681, EPI_ISL_440682, EPI_ISL_440683, EPI_ISL_440684, EPI_ISL_440685, EPI_ISL_440686, EPI_ISL_440687, EPI_ISL_440688, EPI_ISL_440689, EPI_ISL_440690, EPI_ISL_440691, EPI_ISL_440692, EPI_ISL_440693, EPI_ISL_440694, EPI_ISL_440695, EPI_ISL_440696, EPI_ISL_440697, EPI_ISL_440698, EPI_ISL_440699, EPI_ISL_440700, EPI_ISL_440701, EPI_ISL_440702, EPI_ISL_440703, EPI_ISL_440704, EPI_ISL_440705, EPI_ISL_440706, EPI_ISL_440707, EPI_ISL_440708, EPI_ISL_440709, EPI_ISL_440710, EPI_ISL_440711, EPI_ISL_440712, EPI_ISL_440713, EPI_ISL_440714, EPI_ISL_440715, EPI_ISL_440716, EPI_ISL_440717, EPI_ISL_440718, EPI_ISL_440719, EPI_ISL_440720, EPI_ISL_440721, EPI_ISL_440722, EPI_ISL_440723, EPI_ISL_440724, EPI_ISL_440725, EPI_ISL_440726, EPI_ISL_440727, EPI_ISL_440728, EPI_ISL_440729, EPI_ISL_440730, EPI_ISL_440731, EPI_ISL_440732, EPI_ISL_440733, EPI_ISL_440734, EPI_ISL_440735, EPI_ISL_440736, EPI_ISL_440737, EPI_ISL_440738, EPI_ISL_440739, EPI_ISL_440740, EPI_ISL_440741, EPI_ISL_440742, EPI_ISL_440743, EPI_ISL_440744, EPI_ISL_440745, EPI_ISL_440746, EPI_ISL_440747, EPI_ISL_440748, EPI_ISL_440749, EPI_ISL_440750, EPI_ISL_440751, EPI_ISL_440752, EPI_ISL_440753, EPI_ISL_440754, EPI_ISL_440755, EPI_ISL_440756, EPI_ISL_440757, EPI_ISL_440758, EPI_ISL_440759, EPI_ISL_440760, EPI_ISL_440761, EPI_ISL_440762, EPI_ISL_440763, EPI_ISL_440764, EPI_ISL_440765, EPI_ISL_440766, EPI_ISL_440767, EPI_ISL_440768, EPI_ISL_440769, EPI_ISL_440770, EPI_ISL_440771, EPI_ISL_440772, EPI_ISL_440773, EPI_ISL_440774, EPI_ISL_440775, EPI_ISL_440776, EPI_ISL_440777, EPI_ISL_440778, EPI_ISL_440779, EPI_ISL_440780, EPI_ISL_440781, EPI_ISL_440782, EPI_ISL_440783, EPI_ISL_440784, EPI_ISL_440785, EPI_ISL_440786, EPI_ISL_440787, EPI_ISL_440788, EPI_ISL_440789, EPI_ISL_440790, EPI_ISL_440791, EPI_ISL_440792, EPI_ISL_440793, EPI_ISL_440794, EPI_ISL_440795, EPI_ISL_440796, EPI_ISL_440797, EPI_ISL_440798, EPI_ISL_440799, EPI_ISL_440800, EPI_ISL_440801, EPI_ISL_440802, EPI_ISL_440803, EPI_ISL_440804, EPI_ISL_440805, EPI_ISL_440806, EPI_ISL_440807, EPI_ISL_440808, EPI_ISL_440809 | | | |
| see above | PHE South West Regional Laboratory, National Infection Service | Wellcome Sanger Institute for the COVID-19 Genomics UK (COG-UK) consortium | Stephanie Hutchings, Hannah Pymont, Dr Peter Muir, Barry Vipond, Rich Hopes, Alex Alderton, Roberto Amato, Sonia Goncalves, Ewan Harrison, David K. Jackson, Ian Johnston, Dominic Kwiatkowski, Cordelia Langford, John Sillitoe on behalf of the Wellcome Sanger Institute COVID-19 Surveillance Team (http://www.sanger.ac.uk/covid-team) |
| EPI_ISL_442523 | Pasteur Institute of Iran | Kawsar Human Genetic Research Company | Sirous Zeinali, Mohammad Ali Khosravi,Maryam Abbasalipour Bashash, Sanaz Mostafavi Jabbari, Maraym Firoozi, Sormeh Pourtavakoli, Elmira Khateri, Razieh Zeinali and Fahimeh Hoseini |
| EPI_ISL_442624, EPI_ISL_442625, EPI_ISL_442626, EPI_ISL_442627, EPI_ISL_442628, EPI_ISL_442629, EPI_ISL_442630, EPI_ISL_442631, EPI_ISL_442632, EPI_ISL_442633, EPI_ISL_442634, EPI_ISL_442635, EPI_ISL_442636, EPI_ISL_442637, EPI_ISL_442638, EPI_ISL_442639, EPI_ISL_442640, EPI_ISL_442641, EPI_ISL_442642, EPI_ISL_442643, EPI_ISL_442644, EPI_ISL_442645, EPI_ISL_442646, EPI_ISL_442647, EPI_ISL_442648, EPI_ISL_442649, EPI_ISL_442650, EPI_ISL_442651, EPI_ISL_442652, EPI_ISL_442653, EPI_ISL_442654, EPI_ISL_442655, EPI_ISL_442656, EPI_ISL_442657, EPI_ISL_442658, EPI_ISL_442659, EPI_ISL_442660, EPI_ISL_442661, EPI_ISL_442662, EPI_ISL_442663, EPI_ISL_442664, EPI_ISL_442665, EPI_ISL_442666, EPI_ISL_442667, EPI_ISL_442668, EPI_ISL_442669, EPI_ISL_442670, EPI_ISL_442671, EPI_ISL_442672, EPI_ISL_442673, EPI_ISL_442674, EPI_ISL_442675, EPI_ISL_442676, EPI_ISL_442677, EPI_ISL_442678, EPI_ISL_442679, EPI_ISL_442680, EPI_ISL_442681, EPI_ISL_442682, EPI_ISL_442683, EPI_ISL_442684, EPI_ISL_442685, EPI_ISL_442686, EPI_ISL_442687, EPI_ISL_442688, EPI_ISL_442689, EPI_ISL_442690, EPI_ISL_442691, EPI_ISL_442692, EPI_ISL_442693, EPI_ISL_442694, EPI_ISL_442695, EPI_ISL_442696, EPI_ISL_442697, EPI_ISL_442698, EPI_ISL_442699, EPI_ISL_442700, EPI_ISL_442701, EPI_ISL_442702, EPI_ISL_442703, EPI_ISL_442704, EPI_ISL_442705, EPI_ISL_442706, EPI_ISL_442707, EPI_ISL_442708, EPI_ISL_442709, EPI_ISL_442710, EPI_ISL_442711, EPI_ISL_442712, EPI_ISL_442713, EPI_ISL_442714, EPI_ISL_442715, EPI_ISL_442716, EPI_ISL_442717, EPI_ISL_442718, EPI_ISL_442719, EPI_ISL_442720, EPI_ISL_442721, EPI_ISL_442722, EPI_ISL_442723, EPI_ISL_442724, EPI_ISL_442725, EPI_ISL_442726, EPI_ISL_442727, EPI_ISL_442728, EPI_ISL_442729, EPI_ISL_442730, EPI_ISL_442731, EPI_ISL_442732, EPI_ISL_442733, EPI_ISL_442734, EPI_ISL_442735, EPI_ISL_442736, EPI_ISL_442737, EPI_ISL_442738, EPI_ISL_442739, EPI_ISL_442740, EPI_ISL_442741, EPI_ISL_442742, EPI_ISL_442743, EPI_ISL_442744, EPI_ISL_442745, EPI_ISL_442746, EPI_ISL_442747, EPI_ISL_442748, EPI_ISL_442749, EPI_ISL_442750, EPI_ISL_442751, EPI_ISL_442752, EPI_ISL_442753, EPI_ISL_442754, EPI_ISL_442755, EPI_ISL_442756, EPI_ISL_442757, EPI_ISL_442758, EPI_ISL_442759, EPI_ISL_442760, EPI_ISL_442761, EPI_ISL_442762, EPI_ISL_442763, EPI_ISL_442764, EPI_ISL_442765, EPI_ISL_442766, EPI_ISL_442767, EPI_ISL_442768, EPI_ISL_442769, EPI_ISL_442770, EPI_ISL_442771, EPI_ISL_442772, EPI_ISL_442773, EPI_ISL_442774, EPI_ISL_442775, EPI_ISL_442776, EPI_ISL_442777, EPI_ISL_442778, EPI_ISL_442779, EPI_ISL_442780, EPI_ISL_442781, EPI_ISL_442782, EPI_ISL_442783, EPI_ISL_442784, EPI_ISL_442785, EPI_ISL_442786, EPI_ISL_442787, EPI_ISL_442788, EPI_ISL_442789, EPI_ISL_442790, EPI_ISL_442791, EPI_ISL_442792, EPI_ISL_442793, EPI_ISL_442794, EPI_ISL_442795, EPI_ISL_442796, EPI_ISL_442797, EPI_ISL_442798, EPI_ISL_442799, EPI_ISL_442800, EPI_ISL_442801, EPI_ISL_442802, EPI_ISL_442803, EPI_ISL_442804, EPI_ISL_442805, EPI_ISL_442806, EPI_ISL_442807, EPI_ISL_442808, EPI_ISL_442809, EPI_ISL_442810 | | | |

| | | | |
|---|---|---|---|
| see above | PHE South West Regional Laboratory, National Infection Service | Wellcome Sanger Institute for the COVID-19 Genomics UK (COG-UK) consortium | Stephanie Hutchings, Hannah Pymont, Dr Peter Muir, Barry Vipond, Rich Hopes, Alex Alderton, Roberto Amato, Sonia Goncalves, Ewan Harrison, David K. Jackson, Ian Johnston, Dominic Kwiatkowski, Cordelia Langford, John Sillitoe on behalf of the Wellcome Sanger Institute COVID-19 Surveillance Team (http://www.sanger.ac.uk/covid-team) |
| EPI_ISL_443686, EPI_ISL_443687, EPI_ISL_443688, EPI_ISL_443689, EPI_ISL_443690, EPI_ISL_443691, EPI_ISL_443692, EPI_ISL_443693, EPI_ISL_443694, EPI_ISL_443695, EPI_ISL_443696, EPI_ISL_443697, EPI_ISL_443698, EPI_ISL_443699, EPI_ISL_443700, EPI_ISL_443701, EPI_ISL_443702, EPI_ISL_443703, EPI_ISL_443704, EPI_ISL_443705, EPI_ISL_443706, EPI_ISL_443707, EPI_ISL_443708, EPI_ISL_443709, EPI_ISL_443710, EPI_ISL_443711, EPI_ISL_443712, EPI_ISL_443713, EPI_ISL_443714, EPI_ISL_443715, EPI_ISL_443716, EPI_ISL_443717, EPI_ISL_443718, EPI_ISL_443719, EPI_ISL_443720, EPI_ISL_443721, EPI_ISL_443722, EPI_ISL_443723, EPI_ISL_443724, EPI_ISL_443725, EPI_ISL_443726, EPI_ISL_443727, EPI_ISL_443728, EPI_ISL_443729, EPI_ISL_443730, EPI_ISL_443731, EPI_ISL_443732, EPI_ISL_443733, EPI_ISL_443734, EPI_ISL_443735, EPI_ISL_443736, EPI_ISL_443737, EPI_ISL_443738, EPI_ISL_443739, EPI_ISL_443740, EPI_ISL_443741, EPI_ISL_443742, EPI_ISL_443743, EPI_ISL_443744, EPI_ISL_443745, EPI_ISL_443746, EPI_ISL_443747, EPI_ISL_443748, EPI_ISL_443749, EPI_ISL_443750, EPI_ISL_443751, EPI_ISL_443752, EPI_ISL_443753, EPI_ISL_443754, EPI_ISL_443755, EPI_ISL_443756, EPI_ISL_443757, EPI_ISL_443758, EPI_ISL_443759, EPI_ISL_443760, EPI_ISL_443761, EPI_ISL_443762, EPI_ISL_443763, EPI_ISL_443764, EPI_ISL_443765, EPI_ISL_443766, EPI_ISL_443767, EPI_ISL_443768, EPI_ISL_443769, EPI_ISL_443770, EPI_ISL_443771, EPI_ISL_443772, EPI_ISL_443773, EPI_ISL_443774, EPI_ISL_443775, EPI_ISL_443776, EPI_ISL_443777, EPI_ISL_443778, EPI_ISL_443779, EPI_ISL_443780, EPI_ISL_443781, EPI_ISL_443782, EPI_ISL_443783, EPI_ISL_443784, EPI_ISL_443785, EPI_ISL_443786, EPI_ISL_443787, EPI_ISL_443788, EPI_ISL_443789, EPI_ISL_443790, EPI_ISL_443791, EPI_ISL_443792, EPI_ISL_443793, EPI_ISL_443794, EPI_ISL_443795, EPI_ISL_443796, EPI_ISL_443797, EPI_ISL_443798, EPI_ISL_443799, EPI_ISL_443800, EPI_ISL_443801, EPI_ISL_443802, EPI_ISL_443803, EPI_ISL_443804, EPI_ISL_443805, EPI_ISL_443806, EPI_ISL_443807, EPI_ISL_443808, EPI_ISL_443809, EPI_ISL_443810, EPI_ISL_443811, EPI_ISL_443812, EPI_ISL_443813, EPI_ISL_443814, EPI_ISL_443815, EPI_ISL_443816, EPI_ISL_443817, EPI_ISL_443818, EPI_ISL_443819, EPI_ISL_443820, EPI_ISL_443821, EPI_ISL_443822, EPI_ISL_443823, EPI_ISL_443824, EPI_ISL_443825, EPI_ISL_443826, EPI_ISL_443827, EPI_ISL_443828, EPI_ISL_443829, EPI_ISL_443830, EPI_ISL_443831, EPI_ISL_443832, EPI_ISL_443833, EPI_ISL_443834, EPI_ISL_443835, EPI_ISL_443836, EPI_ISL_443837, EPI_ISL_443838, EPI_ISL_443839, EPI_ISL_443840, EPI_ISL_443841, EPI_ISL_443842, EPI_ISL_443843, EPI_ISL_443844, EPI_ISL_443845, EPI_ISL_443846, EPI_ISL_443847, EPI_ISL_443848, EPI_ISL_443849, EPI_ISL_443850, EPI_ISL_443851, EPI_ISL_443852, EPI_ISL_443853, EPI_ISL_443854, EPI_ISL_443855, EPI_ISL_443856, EPI_ISL_443857, EPI_ISL_443858, EPI_ISL_443859, EPI_ISL_443860, EPI_ISL_443861, EPI_ISL_443862, EPI_ISL_443863, EPI_ISL_443864, EPI_ISL_443865, EPI_ISL_443866, EPI_ISL_443867, EPI_ISL_443868, EPI_ISL_443869, EPI_ISL_443870, EPI_ISL_443871, EPI_ISL_443872, EPI_ISL_443873, EPI_ISL_443874, EPI_ISL_443875, EPI_ISL_443876, EPI_ISL_443877, EPI_ISL_443878, EPI_ISL_443879, EPI_ISL_443880, EPI_ISL_443881, EPI_ISL_443882, EPI_ISL_443883, EPI_ISL_443884, EPI_ISL_443885, EPI_ISL_443886, EPI_ISL_443887, EPI_ISL_443888, EPI_ISL_443889, EPI_ISL_443890, EPI_ISL_443891, EPI_ISL_443892, EPI_ISL_443893, EPI_ISL_443894, EPI_ISL_443895, EPI_ISL_443896, EPI_ISL_443897, EPI_ISL_443898, EPI_ISL_443899, EPI_ISL_443900, EPI_ISL_443901, EPI_ISL_443902, EPI_ISL_443903, EPI_ISL_443904, EPI_ISL_443905, EPI_ISL_443906, EPI_ISL_443907, EPI_ISL_443908, EPI_ISL_443909, EPI_ISL_443910, EPI_ISL_443911, EPI_ISL_443912, EPI_ISL_443913, EPI_ISL_443914, EPI_ISL_443915, EPI_ISL_443916, EPI_ISL_443917, EPI_ISL_443918, EPI_ISL_443919, EPI_ISL_443920, EPI_ISL_443921, EPI_ISL_443922, EPI_ISL_443923, EPI_ISL_443924, EPI_ISL_443925, EPI_ISL_443926, EPI_ISL_443927, EPI_ISL_443928, EPI_ISL_443929, EPI_ISL_443930, EPI_ISL_443931, EPI_ISL_443932, EPI_ISL_443933, EPI_ISL_443934, EPI_ISL_443935, EPI_ISL_443936, EPI_ISL_443937, EPI_ISL_443938, EPI_ISL_443939, EPI_ISL_443940, EPI_ISL_443941, EPI_ISL_443942, EPI_ISL_443943, EPI_ISL_443944, EPI_ISL_443945, EPI_ISL_443946, EPI_ISL_443947, EPI_ISL_443948, EPI_ISL_443949, EPI_ISL_443950, EPI_ISL_443951, EPI_ISL_443952, EPI_ISL_443953, EPI_ISL_443954, EPI_ISL_443955, EPI_ISL_443956, EPI_ISL_443957, EPI_ISL_443958, EPI_ISL_443959, EPI_ISL_443960, EPI_ISL_443961, EPI_ISL_443962, EPI_ISL_443963, EPI_ISL_443964, EPI_ISL_443965, EPI_ISL_443966, EPI_ISL_443967, EPI_ISL_443968, EPI_ISL_443969, EPI_ISL_443970, EPI_ISL_443971, EPI_ISL_443972, EPI_ISL_443973, EPI_ISL_443974, EPI_ISL_443975, EPI_ISL_443976, EPI_ISL_443977, EPI_ISL_443978, EPI_ISL_443979, EPI_ISL_443980, EPI_ISL_443981, EPI_ISL_443982, EPI_ISL_443983, EPI_ISL_443984, EPI_ISL_443985, EPI_ISL_443986, EPI_ISL_443987, EPI_ISL_443988, EPI_ISL_443989, EPI_ISL_443990, EPI_ISL_443991, EPI_ISL_443992, EPI_ISL_443993, EPI_ISL_443994, EPI_ISL_443995, EPI_ISL_443996, EPI_ISL_443997, EPI_ISL_443998, EPI_ISL_443999, EPI_ISL_444000, EPI_ISL_444001, EPI_ISL_444002, EPI_ISL_444003, EPI_ISL_444004, EPI_ISL_444005, EPI_ISL_444006, EPI_ISL_444007, EPI_ISL_444008, EPI_ISL_444009, EPI_ISL_444010, EPI_ISL_444011, EPI_ISL_444012, EPI_ISL_444013, EPI_ISL_444014, EPI_ISL_444015, EPI_ISL_444016, EPI_ISL_444017, EPI_ISL_444018, EPI_ISL_444019, EPI_ISL_444020, EPI_ISL_444021 | | | |
| see above | PHE South West Regional Laboratory, National Infection Service | Wellcome Sanger Institute for the COVID-19 Genomics UK (COG-UK) consortium | Stephanie Hutchings, Hannah Pymont, Dr Peter Muir, Barry Vipond, Rich Hopes; and Alex Alderton, Roberto Amato, Sonia Goncalves, Ewan Harrison, David K. Jackson, Ian Johnston, Dominic Kwiatkowski, Cordelia Langford, John Sillitoe on behalf of the Wellcome Sanger Institute COVID-19 Surveillance Team (http://www.sanger.ac.uk/covid-team) |
| EPI_ISL_444273 | State Key Laboratory of Respiratory Disease, National Clinical Research Center for Respiratory Disease, Guangzhou Institute of Respiratory Health, the First Affiliated Hospital of Guangzhou Medical University | State Key Laboratory of Respiratory Disease, National Clinical Research Center for Respiratory Disease, Guangzhou Institute of Respiratory Health, the First Affiliated Hospital of Guangzhou Medical University | Sun,J., Shi,Y., Zheng,K., Huang,J. and Zhao,J. |
| EPI_ISL_444457 | B.J. Medical College and Civil hospital | Gujarat Biotechnology Research Centre | Snehal Bagatharia, Kamlesh J Upadhyay, Ramesh Pandit, Tejas Shah, Ankit Hinsu, Pritesh Sabara, Apurvasinh Puvar, Janvi Raval, Monika Gandhi, Pinal Trivedi, Maharshi Pandya, Amit Kanani, Akanksha Verma, Nitin Savaliya, Raghawendra Kumar, Dinesh Kumar, Zuber Saiyed, Dipa Kinariwala, Disha Patel, Binita Aring, Neeta Khandelwal, Geeta Vaghela, Sonia Barve, Bhavesh Modi, Kairavi Joshi, Gaurishankar Shrimali, Nidhi Sood, Pranay Shah, R D Dixit, Nidhi Patel, Chaitanya Joshi, Madhvi Joshi |
| EPI_ISL_444460 | B.J. Medical College and Civil hospital | Gujarat Biotechnology Research Centre | Tejas Shah, Ankit Hinsu, Pritesh Sabara, Apurvasinh Puvar, Janvi Raval, Monika Gandhi, Pinal Trivedi, Maharshi Pandya, Amit Kanani, Akanksha Verma, Nitin Savaliya, Raghawendra Kumar, Dinesh Kumar, Zuber Saiyed, Dipa Kinariwala, Disha Patel, Binita Aring, Neeta Khandelwal, Geeta Vaghela, Sonia Barve, Bhavesh Modi, Kairavi Joshi, Gaurishankar Shrimali, Nidhi Sood, Pranay Shah, R D Dixit, Snehal Bagatharia, Kamlesh J Upadhyay, Ramesh Pandit, Afzal Ansari, Chaitanya Joshi, Madhvi Joshi |
| EPI_ISL_444479 | B.J. Medical College and Civil hospital | Gujarat Biotechnology Research Centre | Sonia Barve, Bhavesh Modi, Kairavi Joshi, Gaurishankar Shrimali, Nidhi Sood, Pranay Shah, R D Dixit, Snehal Bagatharia, Kamlesh J Upadhyay, Ramesh Pandit, Tejas Shah, Ankit Hinsu, Pritesh Sabara, Apurvasinh Puvar, Janvi Raval, Monika Gandhi, Pinal Trivedi, Maharshi Pandya, Amit Kanani, Akanksha Verma, Nitin Savaliya, Raghawendra Kumar, Dinesh Kumar, Zuber Saiyed, Dipa Kinariwala, Disha Patel, Binita Aring, Neeta Khandelwal, Geeta Vaghela, Nidhi Patel, Chaitanya Joshi, Madhvi Joshi |
| EPI_ISL_444486 | Gujarat Biotechnology Research Centre | Gujarat Biotechnology Research Centre | Snehal Bagatharia, Kamlesh J Upadhyay, Ramesh Pandit, Tejas Shah, Ankit Hinsu, Pritesh Sabara, Apurvasinh Puvar, Janvi Raval, Monika Gandhi, Pinal Trivedi, Maharshi Pandya, Amit Kanani, Akanksha Verma, Nitin Savaliya, Raghawendra Kumar, Dinesh Kumar, Zuber Saiyed, Dipa Kinariwala, Disha Patel, Binita Aring, Neeta Khandelwal, Geeta Vaghela, Sonia Barve, Bhavesh Modi, Kairavi Joshi, Gaurishankar Shrimali, Nidhi Sood, Pranay Shah, R D Dixit, Dipeshwari Shewale, Chaitanya Joshi, Madhvi Joshi |
| EPI_ISL_447032 | B.J. Medical College and Civil hospital | Gujarat Biotechnology Research Centre | Tejas Shah, Ankit Hinsu, Pritesh Sabara, Apurvasinh Puvar, Janvi Raval, Monika Gandhi, Pinal Trivedi, Maharshi Pandya, Amit Kanani, Akanksha Verma, Nitin Savaliya, Raghawendra Kumar, Dinesh Kumar, Zuber Saiyed, Dipa Kinariwala, Disha Patel, Binita Aring, Neeta Khandelwal, Geeta Vaghela, Sonia Barve, Bhavesh Modi, Kairavi Joshi, Gaurishankar Shrimali, Nidhi Sood, Pranay Shah, R D Dixit, Snehal Bagatharia, Ramesh Pandit, Kamlesh J Upadhyay, Pooja P Doshi, Chaitanya Joshi, Madhvi Joshi |
| EPI_ISL_447051 | GMERS Medical College and Hospital, Gandhinagar | Gujarat Biotechnology Research Centre | Sonia Barve, Bhavesh Modi, Kairavi Joshi, Gaurishankar Shrimali, Nidhi Sood, Pranay Shah, R D Dixit, Snehal Bagatharia, Kamlesh J Upadhyay, Ramesh Pandit, Tejas Shah, Ankit Hinsu, Pritesh Sabara, Apurvasinh Puvar, Janvi Raval, Monika Gandhi, Pinal Trivedi, Maharshi Pandya, Amit Kanani, Akanksha Verma, Nitin Savaliya, Raghawendra Kumar, Dinesh Kumar, Zuber Saiyed, Dipa Kinariwala, Disha Patel, Binita Aring, Neeta Khandelwal, Geeta Vaghela, Anjali Rajwar, Chaitanya Joshi, Madhvi Joshi |
| EPI_ISL_447258, EPI_ISL_447259, EPI_ISL_447260, EPI_ISL_447261, EPI_ISL_447262, EPI_ISL_447263, EPI_ISL_447264, EPI_ISL_447265, EPI_ISL_447266, EPI_ISL_447267, EPI_ISL_447268, EPI_ISL_447269, EPI_ISL_447270, EPI_ISL_447271, EPI_ISL_447272, EPI_ISL_447273, EPI_ISL_447274, EPI_ISL_447275, EPI_ISL_447276, EPI_ISL_447277, EPI_ISL_447278, EPI_ISL_447279, EPI_ISL_447280 | | | |
| see above | Microbiology laboratory, Assuta Ashdod University-Affiliated Hospital | Stern Lab | Stern Lab |
| EPI_ISL_447281, EPI_ISL_447282, EPI_ISL_447283, EPI_ISL_447284, EPI_ISL_447285, EPI_ISL_447286, EPI_ISL_447287, EPI_ISL_447288, EPI_ISL_447289, EPI_ISL_447290, EPI_ISL_447291, EPI_ISL_447292, EPI_ISL_447293, EPI_ISL_447294, EPI_ISL_447295, EPI_ISL_447296, EPI_ISL_447297, EPI_ISL_447298, EPI_ISL_447299, EPI_ISL_447300, EPI_ISL_447301, EPI_ISL_447302, EPI_ISL_447303, EPI_ISL_447304, EPI_ISL_447305, EPI_ISL_447306, EPI_ISL_447307, EPI_ISL_447308, EPI_ISL_447309, EPI_ISL_447310 | | | |
| see above | Microbiology Division, Barzilai University Medical Center | Stern Lab | Stern Lab |
| EPI_ISL_447311, EPI_ISL_447312, EPI_ISL_447313, EPI_ISL_447314, EPI_ISL_447315, EPI_ISL_447316, EPI_ISL_447317, EPI_ISL_447318, EPI_ISL_447319, EPI_ISL_447320, EPI_ISL_447321, EPI_ISL_447322, EPI_ISL_447323, EPI_ISL_447324, EPI_ISL_447325, EPI_ISL_447326, EPI_ISL_447327, EPI_ISL_447328, EPI_ISL_447329, EPI_ISL_447330 | | | |
| see above | Clinical Virology Laboratory, Soroka Medical Center and the Faculty of Health Sciences, Ben-Gurion University of the Negev | Stern Lab | Stern Lab |
| EPI_ISL_447331, EPI_ISL_447332, EPI_ISL_447333, EPI_ISL_447334, EPI_ISL_447335, EPI_ISL_447336, EPI_ISL_447337, EPI_ISL_447338, EPI_ISL_447339, EPI_ISL_447340, EPI_ISL_447341, EPI_ISL_447342, EPI_ISL_447343, EPI_ISL_447344, EPI_ISL_447345, EPI_ISL_447346, EPI_ISL_447347, EPI_ISL_447348, EPI_ISL_447349, EPI_ISL_447350, EPI_ISL_447351, EPI_ISL_447352, EPI_ISL_447353, EPI_ISL_447354, EPI_ISL_447355, EPI_ISL_447356, EPI_ISL_447357, EPI_ISL_447358, EPI_ISL_447359, EPI_ISL_447360, EPI_ISL_447361, EPI_ISL_447362, EPI_ISL_447363, EPI_ISL_447364, EPI_ISL_447365, EPI_ISL_447366, EPI_ISL_447367, EPI_ISL_447368, EPI_ISL_447369, EPI_ISL_447370, EPI_ISL_447371, EPI_ISL_447372, EPI_ISL_447373, EPI_ISL_447374, EPI_ISL_447375, EPI_ISL_447376, EPI_ISL_447377, EPI_ISL_447378, EPI_ISL_447379, EPI_ISL_447380, EPI_ISL_447381, EPI_ISL_447382 | | | |
| see above | Clinical Virology Unit, Hadassah Hebrew University Medical Center | Stern Lab | Stern Lab |
| EPI_ISL_447383, EPI_ISL_447384, EPI_ISL_447385, EPI_ISL_447386, EPI_ISL_447387, EPI_ISL_447388, EPI_ISL_447389, EPI_ISL_447390, EPI_ISL_447391, EPI_ISL_447392, EPI_ISL_447393, EPI_ISL_447394, EPI_ISL_447395, EPI_ISL_447396, EPI_ISL_447397, EPI_ISL_447398, EPI_ISL_447399, EPI_ISL_447400, EPI_ISL_447401, EPI_ISL_447402, EPI_ISL_447403, EPI_ISL_447404, EPI_ISL_447405, EPI_ISL_447406 | | | |
| see above | Clinical Microbiology Laboratory, The Baruch Padeh Medical Center, Poriya | Stern Lab | Stern Lab |
| EPI_ISL_447407, EPI_ISL_447408, EPI_ISL_447409, EPI_ISL_447410, EPI_ISL_447411, EPI_ISL_447412, | Clinical Virology Unit, Hadassah Hebrew University Medical Center | Stern Lab | Stern Lab |

| | | | |
|---|---|---|---|
| EPI_ISL_447413, EPI_ISL_447414, EPI_ISL_447415, EPI_ISL_447416 | | | |
| EPI_ISL_447417, EPI_ISL_447418 | Clinical Microbiology Laboratory, The Baruch Padeh Medical Center, Poriya | Stern Lab | Stern Lab |
| EPI_ISL_447419, EPI_ISL_447420, EPI_ISL_447421, EPI_ISL_447422, EPI_ISL_447423, EPI_ISL_447424, EPI_ISL_447425, EPI_ISL_447426, EPI_ISL_447427, EPI_ISL_447428, EPI_ISL_447429, EPI_ISL_447430, EPI_ISL_447431, EPI_ISL_447432, EPI_ISL_447433, EPI_ISL_447434, EPI_ISL_447435, EPI_ISL_447436, EPI_ISL_447437, EPI_ISL_447438, EPI_ISL_447439, EPI_ISL_447440, EPI_ISL_447441, EPI_ISL_447442, EPI_ISL_447443, EPI_ISL_447444, EPI_ISL_447445, EPI_ISL_447446, EPI_ISL_447447, EPI_ISL_447448, EPI_ISL_447449, EPI_ISL_447450, EPI_ISL_447451, EPI_ISL_447452, EPI_ISL_447453, EPI_ISL_447454, EPI_ISL_447455, EPI_ISL_447456, EPI_ISL_447457, EPI_ISL_447458, EPI_ISL_447459, EPI_ISL_447460, EPI_ISL_447461, EPI_ISL_447462, EPI_ISL_447463, EPI_ISL_447464, EPI_ISL_447465, EPI_ISL_447466, EPI_ISL_447467, EPI_ISL_447468, EPI_ISL_447469 | | | |
| see above | Clinical Microbiology Laboratory, Sheba Medical Center | Stern Lab | Stern Lab |
| EPI_ISL_447538 | Gujarat Biotechnology Research Centre | Gujarat Biotechnology Research Centre | Snehal Bagatharia, Kamlesh J Upadhyay, Ramesh Pandit, Tejas Shah, Ankit Hinsu, Pritesh Sabara, Apurvasinh Puvar, Janvi Raval, Monika Gandhi, Pinal Trivedi, Maharshi Pandya, Amit Kanani, Akanksha Verma, Nitin Savaliya, Raghawendra Kumar, Dinesh Kumar, Zuber Saiyed, Dipa Kinariwala, Disha Patel, Binita Aring, Neeta Khandelwal, Geeta Vaghela, Sonia Barve, Bhavesh Modi, Kairavi Joshi, Gaurishankar Shrimali, Nidhi Sood, Pranay Shah, R D Dixit, Neelam Nathani, Chaitanya Joshi, Madhvi Joshi |
| EPI_ISL_447541 | Gujarat Biotechnology Research Centre | Gujarat Biotechnology Research Centre | Tejas Shah, Ankit Hinsu, Pritesh Sabara, Apurvasinh Puvar, Janvi Raval, Monika Gandhi, Pinal Trivedi, Maharshi Pandya, Amit Kanani, Akanksha Verma, Nitin Savaliya, Raghawendra Kumar, Dinesh Kumar, Zuber Saiyed, Dipa Kinariwala, Disha Patel, Binita Aring, Neeta Khandelwal, Geeta Vaghela, Sonia Barve, Bhavesh Modi, Kairavi Joshi, Gaurishankar Shrimali, Nidhi Sood, Pranay Shah, R D Dixit, Snehal Bagatharia, Kamlesh J Upadhyay, Ramesh Pandit, Anjali Rajwar, Chaitanya Joshi, Madhvi Joshi |
| EPI_ISL_447556, EPI_ISL_447559, EPI_ISL_447564, EPI_ISL_447565, EPI_ISL_447566, EPI_ISL_447575 | CSIR-Centre for Cellular and Molecular Biology | CSIR-Centre for Cellular and Molecular Biology | Sofia Banu, Payel Mukherjee, Priya Singh, Dhiviya Vedagiri, Divya Gupta, Vishal Sah, Santosh Kumar Kuncha, Krishnan Harinivas Harshan, Archana Bharadwaj Siva, Karthik Bharadwaj Tallapaka, Shagufta Khan, Lamuk Zaveri, Namami Gaur, Sakshi Shambhavi, Tulasi Nagabandi, Purushotham Vodnala, Rakesh K Mishra, Divya Tej Sowpati |
| EPI_ISL_447614, EPI_ISL_447615, EPI_ISL_447616, EPI_ISL_447617, EPI_ISL_447618, EPI_ISL_447619, EPI_ISL_447620, EPI_ISL_447621, EPI_ISL_447622 | Department of Laboratory Medicine, National Taiwan University Hospital | Microbial Genomics Core Lab, National Taiwan University Centers of Genomic and Precision Medicine | Shiou-Hwei Yeh, You-Yu Lin, Ya-Yun Lai, Chiao-Ling Li, Shan-Chwen Chang, Pei-Jer Chen, Sui-Yuan Chang |
| EPI_ISL_447848, EPI_ISL_447865, EPI_ISL_447866 | CSIR-Centre for Cellular and Molecular Biology | CSIR-Centre for Cellular and Molecular Biology | Sofia Banu, Payel Mukherjee, Priya Singh, Dhiviya Vedagiri, Divya Gupta, Vishal Sah, Santosh Kumar Kuncha, Krishnan Harinivas Harshan, Archana Bharadwaj Siva, Karthik Bharadwaj Tallapaka, Shagufta Khan, Lamuk Zaveri, Namami Gaur, Sakshi Shambhavi, Tulasi Nagabandi, Purushotham Vodnala, Rakesh K Mishra, Divya Tej Sowpati |
| EPI_ISL_448449, EPI_ISL_448450, EPI_ISL_448451, EPI_ISL_448452, EPI_ISL_448453, EPI_ISL_448454, EPI_ISL_448455, EPI_ISL_448456, EPI_ISL_448457, EPI_ISL_448458, EPI_ISL_448459, EPI_ISL_448460, EPI_ISL_448461, EPI_ISL_448462, EPI_ISL_448463, EPI_ISL_448464, EPI_ISL_448465, EPI_ISL_448466, EPI_ISL_448467, EPI_ISL_448468, EPI_ISL_448469, EPI_ISL_448470, EPI_ISL_448471, EPI_ISL_448472, EPI_ISL_448473, EPI_ISL_448474, EPI_ISL_448475, EPI_ISL_448476, EPI_ISL_448477, EPI_ISL_448478, EPI_ISL_448479, EPI_ISL_448480, EPI_ISL_448481, EPI_ISL_448482, EPI_ISL_448483, EPI_ISL_448484, EPI_ISL_448485, EPI_ISL_448486, EPI_ISL_448487, EPI_ISL_448488, EPI_ISL_448489, EPI_ISL_448490, EPI_ISL_448491, EPI_ISL_448492, EPI_ISL_448493, EPI_ISL_448494, EPI_ISL_448495, EPI_ISL_448496, EPI_ISL_448497, EPI_ISL_448498, EPI_ISL_448499, EPI_ISL_448500, EPI_ISL_448501, EPI_ISL_448502, EPI_ISL_448503, EPI_ISL_448504, EPI_ISL_448505, EPI_ISL_448506, EPI_ISL_448507, EPI_ISL_448508, EPI_ISL_448509, EPI_ISL_448510, EPI_ISL_448511, EPI_ISL_448512, EPI_ISL_448513, EPI_ISL_448514, EPI_ISL_448515, EPI_ISL_448516, EPI_ISL_448517, EPI_ISL_448518, EPI_ISL_448519, EPI_ISL_448520, EPI_ISL_448521, EPI_ISL_448522, EPI_ISL_448523, EPI_ISL_448524, EPI_ISL_448525, EPI_ISL_448526, EPI_ISL_448527, EPI_ISL_448528, EPI_ISL_448529, EPI_ISL_448530, EPI_ISL_448531, EPI_ISL_448532, EPI_ISL_448533, EPI_ISL_448534, EPI_ISL_448535, EPI_ISL_448536, EPI_ISL_448537, EPI_ISL_448538, EPI_ISL_448539, EPI_ISL_448540, EPI_ISL_448541, EPI_ISL_448542, EPI_ISL_448543, EPI_ISL_448544, EPI_ISL_448545, EPI_ISL_448546, EPI_ISL_448547, EPI_ISL_448549, EPI_ISL_448551, EPI_ISL_448553, EPI_ISL_448555, EPI_ISL_448557, EPI_ISL_448559, EPI_ISL_448561, EPI_ISL_448564, EPI_ISL_448565, EPI_ISL_448774, EPI_ISL_448775, EPI_ISL_448776, EPI_ISL_448777, EPI_ISL_448778, EPI_ISL_448779, EPI_ISL_448780, EPI_ISL_448781, EPI_ISL_448782, EPI_ISL_448783, EPI_ISL_448784, EPI_ISL_448785, EPI_ISL_448786, EPI_ISL_448787, EPI_ISL_448788, EPI_ISL_448789, EPI_ISL_448790, EPI_ISL_448791, EPI_ISL_448792, EPI_ISL_448793, EPI_ISL_448794, EPI_ISL_448795, EPI_ISL_448796, EPI_ISL_448797, EPI_ISL_448798, EPI_ISL_448799, EPI_ISL_448800, EPI_ISL_448801, EPI_ISL_448802, EPI_ISL_448803, EPI_ISL_448804, EPI_ISL_448805, EPI_ISL_448806, EPI_ISL_448807, EPI_ISL_448808, EPI_ISL_448809, EPI_ISL_448810, EPI_ISL_448811, EPI_ISL_448812, EPI_ISL_448813, EPI_ISL_448814, EPI_ISL_448815, EPI_ISL_448816, EPI_ISL_448817, EPI_ISL_448818, EPI_ISL_448819, EPI_ISL_448820, EPI_ISL_448821, EPI_ISL_448822, EPI_ISL_448823, EPI_ISL_448824, EPI_ISL_448825 | | | |
| see above | Oxford Viromics, NDM, University of Oxford; Oxford University Hospitals; Basingstoke and North Hampshire Hospital | COVID-19 Genomics UK (COG-UK) Consortium | Tanya Golubchik, David Bonsall, George Macintyre, Amy Trebes, Mariateresa de Cesare, Catrin Moore, Alex Mobbs, Anita Justice, Robert Shaw, Monique Andersson, Emma Wise, Nathan Moore, Jessica Lynch, Nick Cortes, Stephen Kidd, David Buck, John Todd, Christophe Fraser |
| EPI_ISL_450294 | Institute of Human Genetics, Polish Academy of Sciences Sanitary and Epidemiological Station in Pozna | Institute of Human Genetics, Polish Academy of Sciences | Szymon Hryhorowicz, Adam Ustaszewski, Emilia Lis, Marta Kaczmarek-Ry, Micha Witt, Andrzej Pawski |
| EPI_ISL_450295 | Institute of Human Genetics, Polish Academy of Sciences; Sanitary and Epidemiological Station in Pozna | Institute of Human Genetics, Polish Academy of Sciences | Szymon Hryhorowicz, Adam Ustaszewski, Emilia Lis, Marta Kaczmarek-Ry, Micha Witt, Andrzej Pawski |
| EPI_ISL_450327 | CSIR-Centre for Cellular and Molecular Biology | CSIR-Centre for Cellular and Molecular Biology | Sofia Banu, Payel Mukherjee, Priya Singh, Dhiviya Vedagiri, Divya Gupta, Vishal Sah, Santosh Kumar Kuncha, Krishnan Harinivas Harshan, Archana Bharadwaj Siva, Karthik Bharadwaj Tallapaka, Shagufta Khan, Lamuk Zaveri, Namami Gaur, Sakshi Shambhavi, Tulasi Nagabandi, Purushotham Vodnala, Disha Nanda, Divya Das, Jotin Gogoi, Manish Bhattacharjee, Ravi Prasad Mukku, Renu Sudhakar, Somesh Gorde, Gangumala Srinivas Reddy, Sujoy Deb, Swati Bayyana, Zeba Rizvi, Rakesh K Mishra, Divya Tej Sowpati |
| EPI_ISL_450338 | Institute of Human Genetics, Polish Academy of Sciences; Sanitary and Epidemiological Station in Pozna | Institute of Human Genetics, Polish Academy of Sciences | Szymon Hryhorowicz, Adam Ustaszewski, Emilia Lis, Marta Kaczmarek-Ry, Micha Witt, Andrzej Pawski |
| EPI_ISL_450404, EPI_ISL_450405 | unknown | School of Public Health, The University of Hong Kong | Sit,T.H.S., Brackman,C.J., Sims,L.D., Tsang,D.N.C., Chu,D.K.W., Perera,R.A.P.M., Poon,L.L.M. and Peiris,M. |
| EPI_ISL_450489 | Wuhan Institute of Virology, Chinese Academy of Sciences | Wuhan Institute of Virology, Chinese Academy of Sciences | Si,H., Zhu,Y., Lin,H., Xie,S., Shi,Z. and Zhou,P. |
| EPI_ISL_450500, EPI_ISL_450501, EPI_ISL_450502, EPI_ISL_450503, EPI_ISL_450504 | unknown | CAS Key Laboratory of Special Pathogens and Biosafety and Center for Emerging Infectious Diseases | Si,H., Zhu,Y., Lin,H., Xie,S., Shi,Z., Zhou,P. |
| EPI_ISL_450783 | Government Medical College-Bhavnagar | Gujarat Biotechnology Research Centre | Shirish Patel, Kairavi Desai, Saklain Malek, Ankit Hinsu, Pritesh Sabara, Apurvasinh Puvar, Janvi Raval, Zarna Patel, Monika Gandhi, Pinal Trivedi, Maharshi Pandya, Amit Kanani, Nidhi Patel, Nitin Savaliya, Raghawendra Kumar, Dinesh Kumar, Zuber Saiyed, Komal Patel, Labdhi Pandya, Snehal Bagatharia, Ramesh Pandit, Tejas Shah, Bhavesh Modi, Gaurishankar Shrimali, R D Dixit, A M Kadri, Neha Rajpara, Chaitanya Joshi, Madhvi Joshi |
| EPI_ISL_450839 | COVID-19 Laboratory Centre for Advanced Research in Sciences (CARS), University of Dhaka, Dhaka-1000, Bangladesh | DNA Solution Ltd | Sharif Akhteruzzaman, Zeba Islam Seraj, Nazmul Ahsan, Md Imdadul Hoque, MA Malek, Shahryar Nabi, Sabrina Moriom Elius, ABM Khademul Islam, Richard Malo, Imran Khan, Abu Sufian, Sabita Rezwana Rahman, Habibul Bari Shozib, Mamun Ahmed, AHM Nurun Nabi, Mohammad Riazul Islam, Md Mizanur Rahman, Md Ismail Hosen, Latiful Bari, Gazi Nurun Nahar, Haseena Khan, M Anwar Hossain. |
| EPI_ISL_450840 | COVID-19 Laboratory | DNA Solution Ltd. L-5 | Sharif Akhteruzzaman, Zeba Islam Seraj, Nazmul Ahsan, Md Imdadul Hoque, MA Malek, Shahryar Nabi, Sabrina Moriom Elius, ABM Khademul Islam, Richard Malo, Imran Khan, Abu Sufian, Sabita Rezwana Rahman, Habibul Bari Shozib, Mamun Ahmed, AHM Nurun Nabi, Mohammad Riazul Islam, Md Mizanur Rahman, Md Ismail Hosen, Latiful Bari, Gazi Nurun Nahar, Haseena Khan, M Anwar Hossain. |
| EPI_ISL_450841 | COVID-19 Laboratory | DNA Solution Ltd | Sharif Akhteruzzaman, Zeba Islam Seraj, Nazmul Ahsan, Md Imdadul Hoque, MA Malek, Shahryar Nabi, Sabrina Moriom Elius, ABM Khademul Islam, Richard Malo, Imran Khan, Abu Sufian, Sabita Rezwana Rahman, Habibul Bari Shozib, Mamun Ahmed, AHM Nurun Nabi, Mohammad Riazul Islam, Md Mizanur Rahman, Md Ismail Hosen, Latiful Bari, Gazi Nurun Nahar, Haseena Khan, M Anwar Hossain. |
| EPI_ISL_450842, EPI_ISL_450843 | COVID-19 Laboratory | DNA Solution Ltd. | Sharif Akhteruzzaman, Zeba Islam Seraj, Nazmul Ahsan, Md Imdadul Hoque, MA Malek, Shahryar Nabi, Sabrina Moriom Elius, ABM Khademul Islam, Richard Malo, Imran Khan, Abu Sufian, Sabita Rezwana Rahman, Habibul Bari Shozib, Mamun Ahmed, AHM Nurun Nabi, Mohammad Riazul Islam, Md Mizanur Rahman, Md Ismail Hosen, Latiful Bari, Gazi Nurun Nahar, Haseena Khan, M Anwar Hossain. |
| EPI_ISL_451155 | Government Medical College, Vadodara | Gujarat Biotechnology Research Centre | Tanuja Javadekar , Manish Pattani, Amit Kanani, Nidhi Patel, Nitin Savaliya, Raghawendra Kumar, Dinesh Kumar, Zuber Saiyed, Komal Patel, Labdhi Pandya, Snehal Bagatharia, Ramesh Pandit, Tejas Shah, Ankit Hinsu, Pritesh Sabara, Apurvasinh Puvar, Janvi Raval, Zarna Patel, Monika Gandhi, Pinal Trivedi, Maharshi Pandya, Bhavesh Modi, Gaurishankar Shrimali, R D Dixit, A M Kadri, Armi Chaudhari, Chaitanya Joshi, Madhvi Joshi |
| EPI_ISL_451958 | Jamil-ur-Rahman Center for Genome Research, Dr. Panjwani Center for Molecular Medicine and Drug Research, International Center for Chemical and Biological Sciences, University of Karachi | Jamil-ur-Rahman Center for Genome Research, Dr. Panjwani Center for Molecular Medicine and Drug Research, International Center for Chemical and Biological Sciences, University of Karachi | Shakeel,M., Raza,S.A., Khan,S., Khan,B.A., Zahid,M., Qureshi,M.A.and Khan,I.A |

| | | | |
|---|---|---|---|
| EPI_ISL_452249, EPI_ISL_452250, EPI_ISL_452251, EPI_ISL_452252, EPI_ISL_452253, EPI_ISL_452254, EPI_ISL_452255, EPI_ISL_452256, EPI_ISL_452257, EPI_ISL_452258 | Respiratory Virus Unit, Microbiology Services Colindale, Public Health England | Respiratory Virus Unit, Microbiology Services Colindale, Public Health England | Steven Platt, Shahjahan Miah, Angie Lackenby, Omolola Akinbami, Tina Talts, Leena Bhaw, Richard Myers, Monica Galiano, Kirstin Edwards, Jonathan Hubb, Joanna Ellis, Maria Zambon |
| EPI_ISL_452327, EPI_ISL_452328, EPI_ISL_452329, EPI_ISL_452330, EPI_ISL_452331, EPI_ISL_452332, EPI_ISL_452333, EPI_ISL_452334, EPI_ISL_452335, EPI_ISL_452336, EPI_ISL_452337, EPI_ISL_452338, EPI_ISL_452339, EPI_ISL_452340, EPI_ISL_452341, EPI_ISL_452342, EPI_ISL_452343, EPI_ISL_452344, EPI_ISL_452345, EPI_ISL_452346, EPI_ISL_452347, EPI_ISL_452348, EPI_ISL_452349, EPI_ISL_452350, EPI_ISL_452351, EPI_ISL_452352, EPI_ISL_452353, EPI_ISL_452354, EPI_ISL_452355, EPI_ISL_452356, EPI_ISL_452357, EPI_ISL_452358, EPI_ISL_452359, EPI_ISL_452360, EPI_ISL_452361, EPI_ISL_452362, EPI_ISL_452363, EPI_ISL_452364 | | | |
| see above | Laboratory of Infectious Diseases Center of Beijing Ditan Hospital | Laboratory of Infectious Diseases Center of Beijing Ditan Hospital | Siyuan Yang, Chengjie Jie, Fengting Yu, Yunxia Tang, Liting Yan, Linghang Wang |
| EPI_ISL_452692, EPI_ISL_452693, EPI_ISL_452694, EPI_ISL_452695, EPI_ISL_452696, EPI_ISL_452697, EPI_ISL_452698, EPI_ISL_452699, EPI_ISL_452700, EPI_ISL_452701, EPI_ISL_452702, EPI_ISL_452703, EPI_ISL_452704, EPI_ISL_452705, EPI_ISL_452706, EPI_ISL_452707, EPI_ISL_452708, EPI_ISL_452709, EPI_ISL_452710, EPI_ISL_452711, EPI_ISL_452712, EPI_ISL_452713, EPI_ISL_452714, EPI_ISL_452715, EPI_ISL_452716, EPI_ISL_452717, EPI_ISL_452718, EPI_ISL_452719, EPI_ISL_452720, EPI_ISL_452721, EPI_ISL_452722, EPI_ISL_452723, EPI_ISL_452724, EPI_ISL_452725, EPI_ISL_452726, EPI_ISL_452727, EPI_ISL_452728, EPI_ISL_452729, EPI_ISL_452730, EPI_ISL_452731, EPI_ISL_452732, EPI_ISL_452733, EPI_ISL_452734, EPI_ISL_452735, EPI_ISL_452736, EPI_ISL_452737, EPI_ISL_452738, EPI_ISL_452739, EPI_ISL_452740, EPI_ISL_452741, EPI_ISL_452742, EPI_ISL_452743, EPI_ISL_452744, EPI_ISL_452745, EPI_ISL_452746, EPI_ISL_452747, EPI_ISL_452748, EPI_ISL_452749, EPI_ISL_452750, EPI_ISL_452751, EPI_ISL_452752, EPI_ISL_452753, EPI_ISL_452754, EPI_ISL_452755, EPI_ISL_452756, EPI_ISL_452757, EPI_ISL_452758, EPI_ISL_452759, EPI_ISL_452760, EPI_ISL_452761, EPI_ISL_452762, EPI_ISL_452763, EPI_ISL_452764, EPI_ISL_452765, EPI_ISL_452766, EPI_ISL_452767, EPI_ISL_452768, EPI_ISL_452769, EPI_ISL_452770, EPI_ISL_452771, EPI_ISL_452772, EPI_ISL_452773, EPI_ISL_452774, EPI_ISL_452775, EPI_ISL_452776, EPI_ISL_452777, EPI_ISL_452778, EPI_ISL_452779, EPI_ISL_452780, EPI_ISL_452781, EPI_ISL_452782, EPI_ISL_452783, EPI_ISL_452784, EPI_ISL_452785, EPI_ISL_452786 | | | |
| see above | Hospital Universitario Araba. Vitoria-Gasteiz, | SeqCOVID-SPAIN consortium/IBV(CSIC) | Silvia Hernáez Crespo, Carmen Gómez González, Amaia Aguirre Quiñonero, Marina Fernández Torres, Maria Rosario Almela Ferrer, Maria Concepción Lecaroz Agara, Andrés Canut Blasco and SeqCOVID-SPAIN consortium |
| EPI_ISL_454416 | Department of Medical Microbiology, Leiden University Medical Center | Department of Medical Microbiology, Leiden University Medical Center | Snijder,E.J., Ogando,N.S., Zevenhoven,J.C., Dalebout,T.J., de Vries,J.J. and Sidorov,I. |
| EPI_ISL_454830, EPI_ISL_454831, EPI_ISL_454832, EPI_ISL_454833 | SMS Medical College, Jaipur | CSIR Institute of Genomics and Integrative Biology | Sudhir Bhandari, Rahul Bhoyar, Mohammed Imran, Mohit Divakar, Disha Sharma, Anshul Kumar, Bani Jolly, Rahul Sahlot, Abhinav Jain, Paras Sehgal, Gyan Ranjan, Vinod Scaria, Sridhar Sivasubbu, Sandeep K Mathur |
| EPI_ISL_455015 | Pandit Deendayal Upadhyay Government Medical College, Rajkot | Gujarat Biotechnology Research Centre | Snehal Bagatharia, Prakash Modi, Sejul Antala, Manish Pattani, Ramesh Pandit, Tejas Shah, Ankit Hinsu, Pritesh Sabara, Apurvasinh Puvar, Janvi Raval, Zarna Patel, Monika Gandhi, Pinal Trivedi, Maharshi Pandya, Amit Kanani, Nidhi Patel, Nitin Savaliya, Raghawendra Kumar, Dinesh Kumar, Zuber Saiyed, Komal Patel, Labdhi Pandya, Neha Rajpara, Bhavesh Modi, Gaurishankar Shrimali, R D Dixit, A M Kadri, Umang Mishra, Chaitanya Joshi, Madhvi Joshi |
| EPI_ISL_455017 | Government Medical College, Vadodara | Gujarat Biotechnology Research Centre | Tanuja Javadekar , R N Daveshwar, Ramesh Pandit, Tejas Shah, Ankit Hinsu, Pritesh Sabara, Apurvasinh Puvar, Janvi Raval, Zarna Patel, Monika Gandhi, Pinal Trivedi, Maharshi Pandya, Amit Kanani, Nidhi Patel, Nitin Savaliya, Raghawendra Kumar, Dinesh Kumar, Zuber Saiyed, Komal Patel, Labdhi Pandya, Snehal Bagatharia, Fenil Patel, Bhavesh Modi, Gaurishankar Shrimali, R D Dixit, A M Kadri, Umang Mishra, Chaitanya Joshi, Madhvi Joshi, |
| EPI_ISL_455020 | Government Medical College, Vadodara | Gujarat Biotechnology Research Centre | Tejas Shah, Ankit Hinsu, Pritesh Sabara, Apurvasinh Puvar, Janvi Raval, Zarna Patel, Monika Gandhi, Pinal Trivedi, Maharshi Pandya, Amit Kanani, Nidhi Patel, Nitin Savaliya, Raghawendra Kumar, Dinesh Kumar, Zuber Saiyed, Komal Patel, Labdhi Pandya, Snehal Bagatharia, Tanuja Javadekar , R N Daveshwar, Ramesh Pandit, Bhavya Jindal, Bhavesh Modi, Gaurishankar Shrimali, R D Dixit, A M Kadri, Umang Mishra, Chaitanya Joshi, Madhvi Joshi, |
| EPI_ISL_455108 | Smedby HC | The Public Health Agency of Sweden | Susanne Brunby, Anna-Malin Linde, Maria Lind Karlberg, Oskar Karlsson Lindsjo, Olov Svartstrom, Anna Risberg, Theresa Enkirch, Mia Brytting, Karin Tegmark-Wisell |
| EPI_ISL_455308, EPI_ISL_455309, EPI_ISL_455310, EPI_ISL_455311 | REGIONAL VRDL,ICMR-RMRC BBSR | Immunogenomics group, Institute of Life Sciences, Bhubaneswar | Sunil Raghav, Jyotirmayee Turuk, Arup Ghosh, Atimukta Jha, Viplov K. Biswas, Swati Madhulika, Manasi Priyadarshini, Shuchi Smita, Jaya Singh Khastri, Rupesh Dash, Soma Chattopadhyay, Ghulam Hussain Syed, Shanti Senapati, Tushar K. Beuria, Debdutta Bhattacharya, Rajeeb Swain, Punit Prasad, COVID-19 team of ILS & RMRC, Orissa COVID-19 study group, DBT's PAN-INDIA 1000 SARS-CoV2 RNA genome sequencing consortium, Sanghamitra Pati, Ajay Parida |
| EPI_ISL_455432, EPI_ISL_455434 | Instituto de Diagnostico y Referencia Epidemiologicos (INDRE) | Instituto de Diagnostico y Referencia Epidemiologicos (INDRE) | Taboada Ramírez Blanca. Ramirez-Gonzalez Ernesto, Garces-Ayala Fabiola, Araiza-Rodriguez Adnan, Mendieta-Condado Edgar, Rodriguez-Maldonado Abril, Wong-Arambula Claudia, Barrera-Badillo Gisela, Hernandez-Rivas Lucia, Lopez-Martinez Irma. |
| EPI_ISL_455478 | REGIONAL VRDL,ICMR-RMRC BBSR | Immunogenomics group, Institute of Life Sciences, Bhubaneswar | Sunil Raghav, Jyotirmayee Turuk, Arup Ghosh, Atimukta Jha, Viplov K. Biswas, Swati Madhulika, Manasi Priyadarshini, Shuchi Smita, Jaya Singh Khastri, Rupesh Dash, Soma Chattopadhyay, Ghulam Hussain Syed, Shanti Senapati, Tushar K. Beuria, Debdutta Bhattacharya, Rajeeb Swain, Punit Prasad, COVID-19 team of ILS & RMRC, Orissa COVID-19 study group, DBT's PAN-INDIA 1000 SARS-CoV2 RNA genome sequencing consortium, Sanghamitra Pati, Ajay Parida |
| EPI_ISL_455609, EPI_ISL_455610, EPI_ISL_455611, EPI_ISL_455612, EPI_ISL_455613, EPI_ISL_455614, EPI_ISL_455615, EPI_ISL_455616, EPI_ISL_455617, EPI_ISL_455618, EPI_ISL_455619, EPI_ISL_455620, EPI_ISL_455621, EPI_ISL_455622, EPI_ISL_455623 | | | |
| see above | Ochsner Health | Bioinfoexperts, LLC | Susanna L. Lamers, David J. Nolan, Rebecca Rose, Sissy Cross, David Moraga Amador, Tong Yang, Luke Caruso, Wayra Navia, Lydia Von Borstel, Xiao Hui Zhou, Amy Feehan, Julia-Garcia-Diaz |
| EPI_ISL_455749, EPI_ISL_455750, EPI_ISL_455751, EPI_ISL_455752, EPI_ISL_455753, EPI_ISL_455754, EPI_ISL_455755, EPI_ISL_455756, EPI_ISL_455757, EPI_ISL_455758, EPI_ISL_455759, EPI_ISL_455760, EPI_ISL_455761, EPI_ISL_455762, EPI_ISL_455763, EPI_ISL_455764, EPI_ISL_455765, EPI_ISL_455766, EPI_ISL_455767, EPI_ISL_455768, EPI_ISL_455769, EPI_ISL_455770, EPI_ISL_455771, EPI_ISL_455772, EPI_ISL_455773, EPI_ISL_455774, EPI_ISL_455775, EPI_ISL_455776, EPI_ISL_455777, EPI_ISL_455778, EPI_ISL_455779, EPI_ISL_455780, EPI_ISL_455781, EPI_ISL_455782, EPI_ISL_455783, EPI_ISL_455784, EPI_ISL_455785, EPI_ISL_455786, EPI_ISL_455787, EPI_ISL_455788 | | | |
| see above | REGIONAL VRDL,ICMR-RMRC BBSR | Immunogenomics lab, Institute of Life Sciences, Bhubaneswar | Sunil Raghav, Jyotirmayee Turuk, Arup Ghosh, Atimukta Jha, Viplov K. Biswas, Swati Madhulika, Manasi Priyadarshini, Shuchi Smita, Jaya Singh Khastri, Rupesh Dash, Soma Chattopadhyay, Ghulam Hussain Syed, Shanti Senapati, Tushar K. Beuria, Debdutta Bhattacharya, Rajeeb Swain, Punit Prasad, COVID-19 team of ILS & RMRC, Orissa COVID-19 study group, DBT's PAN-INDIA 1000 SARS-CoV2 RNA genome sequencing consortium, Sanghamitra Pati, Ajay Parida |
| EPI_ISL_456597, EPI_ISL_456598, EPI_ISL_456600, EPI_ISL_456601, EPI_ISL_456602, EPI_ISL_456603, EPI_ISL_456605, EPI_ISL_456606, EPI_ISL_456607, EPI_ISL_456608, EPI_ISL_456609, EPI_ISL_456610, EPI_ISL_456611, EPI_ISL_456612 | | | |
| see above | National Health Laboratory, Timor-Leste | Microbiological Diagnostic Unit Public Health Laboratory, The Peter Doherty Institute for Infection and Immunity | Soares da Silva, E., Dolores de Jesus da Costa, M., Salles de Sousa, A., Jayanti Pereira Tilman, A., Antonia da Costa, E., Barreto, I., Marr, I., Wapling, J., Francis, J., Ximenes, J., Canisia, D., Freeman, K., Dakh, F., Douglas, N., Baird, R., Caly, L., Seemann, T., Sait, M., Schultz, M., Sherry, N. |
| EPI_ISL_457825 | Army Medical Research Center - Scientific Department | Army Medical and Veterinary Research Center | Silvia Fillo, Giovanni Faggioni, Riccardo De Sanctis, Antonella Fortunato, Anna Anselmo, Francesco Giordani, Vanessa Vera Fain, Nino D'Amore, Florigio Lista |
| EPI_ISL_457826 | Army Medical Center - Scientific Department | Army Medical and Veterinary Research Center | Silvia Fillo, Giovanni Faggioni, Riccardo De Sanctis, Antonella Fortunato, Anna Anselmo, Francesco Giordani, Vanessa Vera Fain, Nino D'Amore, Florigio Lista |
| EPI_ISL_458046 | CSIR-Centre for Cellular and Molecular Biology | CSIR-Centre for Cellular and Molecular Biology | Sofia Banu, Payel Mukherjee, Priya Singh, Dhiviya Vedagiri, Divya Gupta, Vishal Sah, Santosh Kumar Kuncha, Krishnan Harinivas Harshan, Archana Bharadwaj Siva, Karthik Bharadwaj Tallapaka, Shagufta Khan, Lamuk Zaveri, Namami Gaur, Sakshi Shambhavi, Tulasi Nagabandi, Purushotham Vodnala, Deepak Kumar, Devi Prasad Vijayashankar, Disha Nanda, Divya Das, Jotin Gogoi, Manish Bhattacharjee, Rakesh K Mishra, Divya Tej Sowpati |
| EPI_ISL_458052 | CSIR-Centre for Cellular and Molecular Biology | CSIR-Centre for Cellular and Molecular Biology | Sofia Banu, Payel Mukherjee, Priya Singh, Dhiviya Vedagiri, Divya Gupta, Vishal Sah, Santosh Kumar Kuncha, Krishnan Harinivas Harshan, Archana Bharadwaj Siva, Karthik Bharadwaj Tallapaka, Shagufta Khan, Lamuk Zaveri, Namami Gaur, Sakshi Shambhavi, Tulasi Nagabandi, Purushotham Vodnala,Preethi Jampala, Sharada Ravi Iyer, Sulagana Mukherjee, Swetha Sundar, Peddapuvala Sai Uday Kiran, Rakesh K Mishra, Divya Tej Sowpati |
| EPI_ISL_458058 | CSIR-Centre for Cellular and Molecular Biology | CSIR-Centre for Cellular and Molecular Biology | Sofia Banu, Payel Mukherjee, Priya Singh, Dhiviya Vedagiri, Divya Gupta, Vishal Sah, Santosh Kumar Kuncha, Krishnan Harinivas Harshan, Archana Bharadwaj Siva, Karthik Bharadwaj Tallapaka, Shagufta Khan, Lamuk Zaveri, Namami Gaur, Sakshi Shambhavi, Tulasi Nagabandi, Purushotham Vodnala, Disha Nanda, Divya Das, Jotin Gogoi, Manish Bhattacharjee, Ravi Prasad Mukku, Rakesh K Mishra, Divya Tej Sowpati |
| EPI_ISL_458063 | CSIR-Centre for Cellular and Molecular Biology | CSIR-Centre for Cellular and Molecular Biology | Sofia Banu, Payel Mukherjee, Priya Singh, Dhiviya Vedagiri, Divya Gupta, Vishal Sah, Santosh Kumar Kuncha, Krishnan Harinivas Harshan, Archana Bharadwaj Siva, Karthik Bharadwaj Tallapaka, Shagufta Khan, Lamuk Zaveri, Namami Gaur, Sakshi Shambhavi, Tulasi Nagabandi, Purushotham Vodnala, Gokulan C G, Gunjan Purohit, Hanuman Tulashiram Kale, Pankaj Kumar, Prachand Issarapu, Rakesh K Mishra, Divya Tej Sowpati |
| EPI_ISL_458066, EPI_ISL_458067, EPI_ISL_458068, EPI_ISL_458069 | Osmania Medical College | CSIR-Centre for Cellular and Molecular Biology | Shashikala Reddy, Mahboob Khan,Payel Mukherjee, Sofia Banu, Priya Singh, Dhiviya Vedagiri, Divya Gupta, Vishal Sah, Santosh Kumar Kuncha, Krishnan Harinivas Harshan, Archana Bharadwaj Siva, Karthik Bharadwaj Tallapaka, Shagufta Khan, Lamuk Zaveri, Namami Gaur, Sakshi Shambhavi, |

| | | | |
|---|---|---|---|
| | | | Tulasi Nagabandi, Purushotham Vodnala, Rakesh K Mishra, Divya Tej Sowpati |
| EPI_ISL_458100 | B.J. Medical College and Civil hospital | Gujarat Biotechnology Research Centre | Snehal Bagatharia, Dhaval Vaghela, Ramesh Patel, Pranay Shah, Kamlesh J Upadhyay, Ramesh Pandit, Tejas Shah, Ankit Hinsu, Pritesh Sabara, Apurvasinh Puvar, Janvi Raval, Zarna Patel, Monika Gandhi, Pinal Trivedi, Maharshi Pandya, Amit Kanani, Nidhi Patel, Nitin Savaliya, Raghawendra Kumar, Dinesh Kumar, Zuber Saiyed, Komal Patel, Labdhi Pandya, Afzal Ansari, Bhavesh Modi, Gaurishankar Shrimali, R D Dixit, A M Kadri, Umang Mishra, Chaitanya Joshi, Madhvi Joshi |
| EPI_ISL_458104 | Gujarat Biotechnology Research Centre | Gujarat Biotechnology Research Centre | Tejas Shah, Ankit Hinsu, Pritesh Sabara, Apurvasinh Puvar, Janvi Raval, Zarna Patel, Monika Gandhi, Pinal Trivedi, Maharshi Pandya, Amit Kanani, Nidhi Patel, Nitin Savaliya, Raghawendra Kumar, Dinesh Kumar, Zuber Saiyed, Komal Patel, Labdhi Pandya, Snehal Bagatharia, Ramesh Pandit, Bhavya Jindal, Bhavesh Modi, Gaurishankar Shrimali, R D Dixit, A M Kadri, Umang Mishra, Chaitanya Joshi, Madhvi Joshi, , , , |
| EPI_ISL_458299, EPI_ISL_458300, EPI_ISL_458301, EPI_ISL_458302, EPI_ISL_458303, EPI_ISL_458304, EPI_ISL_458305, EPI_ISL_458306, EPI_ISL_458307, EPI_ISL_458308, EPI_ISL_458309, EPI_ISL_458310, EPI_ISL_458311, EPI_ISL_458312, EPI_ISL_458313, EPI_ISL_458314, EPI_ISL_458315, EPI_ISL_458316, EPI_ISL_458317, EPI_ISL_458318, EPI_ISL_458319, EPI_ISL_458320, EPI_ISL_458321, EPI_ISL_458322, EPI_ISL_458323, EPI_ISL_458324, EPI_ISL_458325, EPI_ISL_458326, EPI_ISL_458327, EPI_ISL_458328, EPI_ISL_458329, EPI_ISL_458330, EPI_ISL_458331, EPI_ISL_458332, EPI_ISL_458333, EPI_ISL_458334, EPI_ISL_458335, EPI_ISL_458336, EPI_ISL_458337, EPI_ISL_458338, EPI_ISL_458339, EPI_ISL_458340, EPI_ISL_458341, EPI_ISL_458342, EPI_ISL_458343, EPI_ISL_458344, EPI_ISL_458345, EPI_ISL_458346, EPI_ISL_458347, EPI_ISL_458348, EPI_ISL_458349, EPI_ISL_458350, EPI_ISL_458351, EPI_ISL_458352, EPI_ISL_458353, EPI_ISL_458354, EPI_ISL_458355, EPI_ISL_458356, EPI_ISL_458357, EPI_ISL_458358, EPI_ISL_458359, EPI_ISL_458360, EPI_ISL_458361, EPI_ISL_458362, EPI_ISL_458363, EPI_ISL_458364, EPI_ISL_458365, EPI_ISL_458366, EPI_ISL_458367, EPI_ISL_458368, EPI_ISL_458369, EPI_ISL_458370, EPI_ISL_458371, EPI_ISL_458372, EPI_ISL_458373, EPI_ISL_458374, EPI_ISL_458375, EPI_ISL_458376, EPI_ISL_458377, EPI_ISL_458378, EPI_ISL_458379, EPI_ISL_458380, EPI_ISL_458381, EPI_ISL_458382, EPI_ISL_458383, EPI_ISL_458384, EPI_ISL_458385, EPI_ISL_458386, EPI_ISL_458387, EPI_ISL_458388, EPI_ISL_458389, EPI_ISL_458390, EPI_ISL_458391, EPI_ISL_458392, EPI_ISL_458393, EPI_ISL_458394, EPI_ISL_458395, EPI_ISL_458396, EPI_ISL_458397, EPI_ISL_458398, EPI_ISL_458399, EPI_ISL_458400, EPI_ISL_458401, EPI_ISL_458402, EPI_ISL_458403, EPI_ISL_458404, EPI_ISL_458405, EPI_ISL_458406, EPI_ISL_458407, EPI_ISL_458408, EPI_ISL_458409, EPI_ISL_458410, EPI_ISL_458411, EPI_ISL_458412, EPI_ISL_458413, EPI_ISL_458414, EPI_ISL_458415, EPI_ISL_458416, EPI_ISL_458417, EPI_ISL_458418, EPI_ISL_458419, EPI_ISL_458420, EPI_ISL_458421, EPI_ISL_458422, EPI_ISL_458423, EPI_ISL_458424, EPI_ISL_458425, EPI_ISL_458426, EPI_ISL_458427, EPI_ISL_458428, EPI_ISL_458429, EPI_ISL_458430, EPI_ISL_458431, EPI_ISL_458432, EPI_ISL_458433, EPI_ISL_458434, EPI_ISL_458435, EPI_ISL_458436, EPI_ISL_458437, EPI_ISL_458438, EPI_ISL_458439, EPI_ISL_458440, EPI_ISL_458441, EPI_ISL_458442, EPI_ISL_458443, EPI_ISL_458444, EPI_ISL_458445, EPI_ISL_458446, EPI_ISL_458447, EPI_ISL_458448, EPI_ISL_458449, EPI_ISL_458450, EPI_ISL_458451, EPI_ISL_458452, EPI_ISL_458453, EPI_ISL_458454, EPI_ISL_458455, EPI_ISL_458456, EPI_ISL_458457, EPI_ISL_458458, EPI_ISL_458459, EPI_ISL_458460, EPI_ISL_458461, EPI_ISL_458462, EPI_ISL_458463, EPI_ISL_458464, EPI_ISL_458465, EPI_ISL_458466, EPI_ISL_458467, EPI_ISL_458468, EPI_ISL_458469, EPI_ISL_458470, EPI_ISL_458471, EPI_ISL_458472, EPI_ISL_458473, EPI_ISL_458474, EPI_ISL_458475, EPI_ISL_458476, EPI_ISL_458477, EPI_ISL_458478, EPI_ISL_458479, EPI_ISL_458480, EPI_ISL_458481, EPI_ISL_458482, EPI_ISL_458483, EPI_ISL_458484, EPI_ISL_458485, EPI_ISL_458486, EPI_ISL_458487, EPI_ISL_458488, EPI_ISL_458489, EPI_ISL_458490 | | | |
| see above | PHE South West Regional Laboratory, National Infection Service | Wellcome Sanger Institute for the COVID-19 Genomics UK (COG-UK) consortium | Stephanie Hutchings, Hannah Pymont, Dr Peter Muir, Barry Vipond, Rich Hopes; and Alex Alderton, Roberto Amato, Sonia Goncalves, Ewan Harrison, David K. Jackson, Ian Johnston, Dominic Kwiatkowski, Cordelia Langford, John Sillitoe on behalf of the Wellcome Sanger Institute COVID-19 Surveillance Team (http://www.sanger.ac.uk/covid-team) |
| EPI_ISL_458491 | PHE South West Regional Laboratory, National Infection Service | Wellcome Sanger Institute for the COVID-19 Genomics UK (COG-UK) Consortium | Stephanie Hutchings, Hannah Pymont, Dr Peter Muir, Barry Vipond, Rich Hopes; and Alex Alderton, Roberto Amato, Sonia Goncalves, Ewan Harrison, David K. Jackson, Ian Johnston, Dominic Kwiatkowski, Cordelia Langford, John Sillitoe on behalf of the Wellcome Sanger Institute COVID-19 Surveillance Team |
| EPI_ISL_458492, EPI_ISL_458493, EPI_ISL_458494, EPI_ISL_458495, EPI_ISL_458496, EPI_ISL_458497, EPI_ISL_458498, EPI_ISL_458499, EPI_ISL_458500, EPI_ISL_458501, EPI_ISL_458502, EPI_ISL_458503, EPI_ISL_458504, EPI_ISL_458505, EPI_ISL_458506, EPI_ISL_458507, EPI_ISL_458508, EPI_ISL_458509, EPI_ISL_458510, EPI_ISL_458511, EPI_ISL_458512, EPI_ISL_458513, EPI_ISL_458514, EPI_ISL_458515, EPI_ISL_458719, EPI_ISL_458720, EPI_ISL_458721, EPI_ISL_458722, EPI_ISL_458723, EPI_ISL_458724, EPI_ISL_458725, EPI_ISL_458726, EPI_ISL_458727, EPI_ISL_458728, EPI_ISL_458729, EPI_ISL_458730, EPI_ISL_458731, EPI_ISL_458732, EPI_ISL_458733, EPI_ISL_458734, EPI_ISL_458735, EPI_ISL_458736, EPI_ISL_458737, EPI_ISL_458738, EPI_ISL_458739, EPI_ISL_458740, EPI_ISL_458741, EPI_ISL_458742, EPI_ISL_458743, EPI_ISL_458744, EPI_ISL_458745, EPI_ISL_458746, EPI_ISL_458747, EPI_ISL_458748, EPI_ISL_458749, EPI_ISL_458750, EPI_ISL_458751, EPI_ISL_458752, EPI_ISL_458753, EPI_ISL_458754, EPI_ISL_458755, EPI_ISL_458756, EPI_ISL_458757, EPI_ISL_458758, EPI_ISL_458759, EPI_ISL_458760, EPI_ISL_458761, EPI_ISL_458762, EPI_ISL_458763, EPI_ISL_458764, EPI_ISL_458765, EPI_ISL_458766, EPI_ISL_458767, EPI_ISL_458768, EPI_ISL_458769, EPI_ISL_458770, EPI_ISL_458771, EPI_ISL_458772, EPI_ISL_458773, EPI_ISL_458774, EPI_ISL_458775, EPI_ISL_458776, EPI_ISL_458777, EPI_ISL_458778, EPI_ISL_458779, EPI_ISL_458780, EPI_ISL_458781, EPI_ISL_458782, EPI_ISL_458783, EPI_ISL_458784, EPI_ISL_458785 | | | |
| see above | PHE South West Regional Laboratory, National Infection Service | Wellcome Sanger Institute for the COVID-19 Genomics UK (COG-UK) consortium | Stephanie Hutchings, Hannah Pymont, Dr Peter Muir, Barry Vipond, Rich Hopes; and Alex Alderton, Roberto Amato, Sonia Goncalves, Ewan Harrison, David K. Jackson, Ian Johnston, Dominic Kwiatkowski, Cordelia Langford, John Sillitoe on behalf of the Wellcome Sanger Institute COVID-19 Surveillance Team (http://www.sanger.ac.uk/covid-team) |
| EPI_ISL_458786 | PHE South West Regional Laboratory, National Infection Service | Wellcome Sanger Institute for the COVID-19 Genomics UK (COG-UK) Consortium | Stephanie Hutchings, Hannah Pymont, Dr Peter Muir, Barry Vipond, Rich Hopes; and Alex Alderton, Roberto Amato, Sonia Goncalves, Ewan Harrison, David K. Jackson, Ian Johnston, Dominic Kwiatkowski, Cordelia Langford, John Sillitoe on behalf of the Wellcome Sanger Institute COVID-19 Surveillance Team |
| EPI_ISL_458787, EPI_ISL_458788, EPI_ISL_458789, EPI_ISL_458790, EPI_ISL_458791, EPI_ISL_458792, EPI_ISL_458793, EPI_ISL_458794, EPI_ISL_458795, EPI_ISL_458796, EPI_ISL_458797, EPI_ISL_458798, EPI_ISL_458799 | | | |
| see above | PHE South West Regional Laboratory, National Infection Service | Wellcome Sanger Institute for the COVID-19 Genomics UK (COG-UK) consortium | Stephanie Hutchings, Hannah Pymont, Dr Peter Muir, Barry Vipond, Rich Hopes; and Alex Alderton, Roberto Amato, Sonia Goncalves, Ewan Harrison, David K. Jackson, Ian Johnston, Dominic Kwiatkowski, Cordelia Langford, John Sillitoe on behalf of the Wellcome Sanger Institute COVID-19 Surveillance Team (http://www.sanger.ac.uk/covid-team) |
| EPI_ISL_458800 | PHE South West Regional Laboratory, National Infection Service | Wellcome Sanger Institute for the COVID-19 Genomics UK (COG-UK) Consortium | Stephanie Hutchings, Hannah Pymont, Dr Peter Muir, Barry Vipond, Rich Hopes; and Alex Alderton, Roberto Amato, Sonia Goncalves, Ewan Harrison, David K. Jackson, Ian Johnston, Dominic Kwiatkowski, Cordelia Langford, John Sillitoe on behalf of the Wellcome Sanger Institute COVID-19 Surveillance Team |
| EPI_ISL_458801, EPI_ISL_458802, EPI_ISL_458803, EPI_ISL_458804, EPI_ISL_458805, EPI_ISL_458806, EPI_ISL_458807, EPI_ISL_458808, EPI_ISL_458809, EPI_ISL_458810, EPI_ISL_458811, EPI_ISL_458812, EPI_ISL_458813, EPI_ISL_458814, EPI_ISL_458815, EPI_ISL_458816, EPI_ISL_458817, EPI_ISL_458818, EPI_ISL_458819, EPI_ISL_458820, EPI_ISL_458821, EPI_ISL_458822, EPI_ISL_458823, EPI_ISL_458824, EPI_ISL_458825, EPI_ISL_458826, EPI_ISL_458827, EPI_ISL_458828, EPI_ISL_458829, EPI_ISL_458830, EPI_ISL_458831, EPI_ISL_458832, EPI_ISL_458833, EPI_ISL_458834, EPI_ISL_458835, EPI_ISL_458836, EPI_ISL_458837, EPI_ISL_458838, EPI_ISL_458839, EPI_ISL_458840, EPI_ISL_458841, EPI_ISL_458842, EPI_ISL_458843, EPI_ISL_458844, EPI_ISL_458845, EPI_ISL_458846, EPI_ISL_458847, EPI_ISL_458848, EPI_ISL_458849, EPI_ISL_458850, EPI_ISL_458851, EPI_ISL_458852, EPI_ISL_458853, EPI_ISL_458854, EPI_ISL_458855, EPI_ISL_458856, EPI_ISL_458857, EPI_ISL_458858, EPI_ISL_458859, EPI_ISL_458860, EPI_ISL_458861, EPI_ISL_458862, EPI_ISL_458863, EPI_ISL_458864, EPI_ISL_458865, EPI_ISL_458866, EPI_ISL_458867, EPI_ISL_458868, EPI_ISL_458869, EPI_ISL_458870, EPI_ISL_458871 | | | |
| see above | PHE South West Regional Laboratory, National Infection Service | Wellcome Sanger Institute for the COVID-19 Genomics UK (COG-UK) consortium | Stephanie Hutchings, Hannah Pymont, Dr Peter Muir, Barry Vipond, Rich Hopes; and Alex Alderton, Roberto Amato, Sonia Goncalves, Ewan Harrison, David K. Jackson, Ian Johnston, Dominic Kwiatkowski, Cordelia Langford, John Sillitoe on behalf of the Wellcome Sanger Institute COVID-19 Surveillance Team (http://www.sanger.ac.uk/covid-team) |
| EPI_ISL_458872 | PHE South West Regional Laboratory, National Infection Service | Wellcome Sanger Institute for the COVID-19 Genomics UK (COG-UK) Consortium | Stephanie Hutchings, Hannah Pymont, Dr Peter Muir, Barry Vipond, Rich Hopes; and Alex Alderton, Roberto Amato, Sonia Goncalves, Ewan Harrison, David K. Jackson, Ian Johnston, Dominic Kwiatkowski, Cordelia Langford, John Sillitoe on behalf of the Wellcome Sanger Institute COVID-19 Surveillance Team |
| EPI_ISL_458873, EPI_ISL_458874, EPI_ISL_458875, EPI_ISL_458876, EPI_ISL_458877, EPI_ISL_458878, EPI_ISL_458879, EPI_ISL_458880, EPI_ISL_458881, EPI_ISL_458882, EPI_ISL_458883, EPI_ISL_458884, EPI_ISL_458885, EPI_ISL_458886, EPI_ISL_458887, EPI_ISL_458888, EPI_ISL_458889, EPI_ISL_458890, EPI_ISL_458891, EPI_ISL_458892, EPI_ISL_458893, EPI_ISL_458894, EPI_ISL_458895, EPI_ISL_458896, EPI_ISL_458897, EPI_ISL_458898, EPI_ISL_458899, EPI_ISL_458900, EPI_ISL_458901, EPI_ISL_458902, EPI_ISL_458903, EPI_ISL_458904, EPI_ISL_458905, EPI_ISL_458906, EPI_ISL_458907, EPI_ISL_458908, EPI_ISL_458909, EPI_ISL_458910, EPI_ISL_458911, EPI_ISL_458912, EPI_ISL_458913, EPI_ISL_458914, EPI_ISL_458915, EPI_ISL_458916, EPI_ISL_458917, EPI_ISL_458918, EPI_ISL_458919, EPI_ISL_458920, EPI_ISL_458921, EPI_ISL_458922, EPI_ISL_458923, EPI_ISL_458924, EPI_ISL_458925, EPI_ISL_458926, EPI_ISL_458927, EPI_ISL_458928, EPI_ISL_458929, EPI_ISL_458930, EPI_ISL_458931, EPI_ISL_458932, EPI_ISL_458933, EPI_ISL_458934, EPI_ISL_458935, EPI_ISL_458936, EPI_ISL_458937, EPI_ISL_458938, EPI_ISL_458939, EPI_ISL_458940, EPI_ISL_458941, EPI_ISL_458942, EPI_ISL_458943, EPI_ISL_458944, EPI_ISL_458945, EPI_ISL_458946, EPI_ISL_458947, EPI_ISL_458948, EPI_ISL_458949, EPI_ISL_458950, EPI_ISL_458951, EPI_ISL_458952, EPI_ISL_458953, EPI_ISL_458954, EPI_ISL_458955, EPI_ISL_458956, EPI_ISL_458957, EPI_ISL_458958, EPI_ISL_458959, EPI_ISL_458960, EPI_ISL_458961, EPI_ISL_458962, EPI_ISL_458963, EPI_ISL_458964, EPI_ISL_458965, EPI_ISL_458966, EPI_ISL_458967, EPI_ISL_458968, EPI_ISL_458969, EPI_ISL_458970, EPI_ISL_458971, EPI_ISL_458972, EPI_ISL_458973, EPI_ISL_458974, EPI_ISL_458975, EPI_ISL_458976, EPI_ISL_458977, EPI_ISL_458978, EPI_ISL_458979, EPI_ISL_458980, EPI_ISL_458981, EPI_ISL_458982, EPI_ISL_458983, EPI_ISL_458984, EPI_ISL_458985, EPI_ISL_458986, EPI_ISL_458987, EPI_ISL_458988, EPI_ISL_458989, EPI_ISL_458990, EPI_ISL_458991, EPI_ISL_458992, EPI_ISL_458993, EPI_ISL_458994, EPI_ISL_458995, EPI_ISL_458996, EPI_ISL_458997, EPI_ISL_458998, EPI_ISL_458999, EPI_ISL_459000, EPI_ISL_459001, EPI_ISL_459002, EPI_ISL_459003, EPI_ISL_459004, EPI_ISL_459005, EPI_ISL_459006, EPI_ISL_459007, EPI_ISL_459008, EPI_ISL_459009, EPI_ISL_459010, EPI_ISL_459011, EPI_ISL_459012, EPI_ISL_459013, EPI_ISL_459014, EPI_ISL_459015, EPI_ISL_459016, EPI_ISL_459017, EPI_ISL_459018, EPI_ISL_459019, EPI_ISL_459020, EPI_ISL_459021, EPI_ISL_459022, EPI_ISL_459023, EPI_ISL_459024, EPI_ISL_459025, EPI_ISL_459026, EPI_ISL_459027, EPI_ISL_459028, EPI_ISL_459029, EPI_ISL_459030, EPI_ISL_459031, EPI_ISL_459032, EPI_ISL_459033, EPI_ISL_459034, EPI_ISL_459035, EPI_ISL_459036, EPI_ISL_459037, EPI_ISL_459038 | | | |
| see above | PHE South West Regional Laboratory, National Infection Service | Wellcome Sanger Institute for the COVID-19 Genomics UK (COG-UK) consortium | Stephanie Hutchings, Hannah Pymont, Dr Peter Muir, Barry Vipond, Rich Hopes; and Alex Alderton, Roberto Amato, Sonia Goncalves, Ewan Harrison, David K. Jackson, Ian Johnston, Dominic Kwiatkowski, Cordelia Langford, John Sillitoe on behalf of the Wellcome Sanger Institute COVID-19 Surveillance Team (http://www.sanger.ac.uk/covid-team) |
| EPI_ISL_459039 | PHE South West Regional Laboratory, National Infection Service | Wellcome Sanger Institute for the COVID-19 Genomics UK (COG-UK) Consortium | Stephanie Hutchings, Hannah Pymont, Dr Peter Muir, Barry Vipond, Rich Hopes; and Alex Alderton, Roberto Amato, Sonia Goncalves, Ewan Harrison, David K. Jackson, Ian Johnston, Dominic Kwiatkowski, Cordelia Langford, John Sillitoe on behalf of the Wellcome Sanger Institute COVID-19 Surveillance Team |
| EPI_ISL_459040, EPI_ISL_459041, EPI_ISL_459042, EPI_ISL_459043, EPI_ISL_459044, EPI_ISL_459045, EPI_ISL_459046, EPI_ISL_459047, EPI_ISL_459048, EPI_ISL_459049, EPI_ISL_459050, EPI_ISL_459051, EPI_ISL_459052, EPI_ISL_459053, EPI_ISL_459054, EPI_ISL_459055, EPI_ISL_459056, EPI_ISL_459057, EPI_ISL_459058, EPI_ISL_459059, EPI_ISL_459060, EPI_ISL_459061, EPI_ISL_459062, EPI_ISL_459063, EPI_ISL_459064, EPI_ISL_459065, EPI_ISL_459066, EPI_ISL_459067, EPI_ISL_459068, EPI_ISL_459069, EPI_ISL_459070, EPI_ISL_459071, EPI_ISL_459072, EPI_ISL_459073, EPI_ISL_459074, EPI_ISL_459075, EPI_ISL_459076, EPI_ISL_459077, EPI_ISL_459078, EPI_ISL_459079, EPI_ISL_459080, EPI_ISL_459081, EPI_ISL_459082, EPI_ISL_459083, EPI_ISL_459084, EPI_ISL_459085, EPI_ISL_459086, EPI_ISL_459087, EPI_ISL_459088, EPI_ISL_459089, EPI_ISL_459090, EPI_ISL_459091, EPI_ISL_459092, EPI_ISL_459093, EPI_ISL_459094, EPI_ISL_459725, EPI_ISL_459726, EPI_ISL_459727 | | | |
| see above | PHE South West Regional Laboratory, National Infection Service | Wellcome Sanger Institute for the COVID-19 Genomics UK (COG-UK) consortium | Stephanie Hutchings, Hannah Pymont, Dr Peter Muir, Barry Vipond, Rich Hopes; and Alex Alderton, Roberto Amato, Sonia Goncalves, Ewan Harrison, David K. Jackson, Ian Johnston, Dominic Kwiatkowski, Cordelia Langford, John Sillitoe on behalf of the Wellcome Sanger Institute COVID-19 Surveillance Team (http://www.sanger.ac.uk/covid-team) |

| | | | |
|---|---|---|---|
| EPI_ISL_459866, EPI_ISL_459867, EPI_ISL_459868, EPI_ISL_459869, EPI_ISL_459871, EPI_ISL_459872, EPI_ISL_459873, EPI_ISL_459874, EPI_ISL_459875, EPI_ISL_459877, EPI_ISL_459878, EPI_ISL_459879, EPI_ISL_459880, EPI_ISL_459881, EPI_ISL_459882, EPI_ISL_459883, EPI_ISL_459884, EPI_ISL_459885, EPI_ISL_459886, EPI_ISL_459887, EPI_ISL_459888, EPI_ISL_459889, EPI_ISL_459890, EPI_ISL_459891, EPI_ISL_459892 | | | |
| see above | Kingston Health Sciences Center | Queen's Genomics Lab at Ongwanada (Q-GLO) | Sjaarda CP, Rustom N, Huang D, Perez-Patrigeon S, Hudson ML, Wong H,Guan H, Ayub M, Soares CN, Colautti R, Evans GA, Sheth P |
| EPI_ISL_459953 | Institute for Medical Research, Infectious Disease Research Centre, National Institutes of Health, Ministry of Health Malaysia | Institute for Medical Research Infectious Disease Research Centre, National Institutes of Health, Ministry of Health Malaysia | Suppiah J, Mohd-Zawawi Z, Kamel KA, Ellan K, Kalyanasundram J, Mohd-Zain R, Thayan R |
| EPI_ISL_459954, EPI_ISL_459955, EPI_ISL_459956 | Institute for Medical Research, Infectious Disease Research Centre, National Institutes of Health, Ministry of Health Malaysia | Institute for Medical Research, Infectious Disease Research Centre, National Institutes of Health, Ministry of Health Malaysia | Suppiah J, Mohd-Zawawi Z, Kamel KA, Ellan K, Kalyanasundram J, Mohd-Zain R, Thayan R |
| EPI_ISL_459957 | Institute for Medical Research, Infectious Disease Research Centre, National Institutes of Health, Minis | Institute for Medical Research, Infectious Disease Research Centre, National Institutes of Health, Minis | Suppiah J, Mohd-Zawawi Z, Kamel KA, Ellan K, Kalyanasundram J, Mohd-Zain R, Thayan R |
| EPI_ISL_459958, EPI_ISL_459959, EPI_ISL_459960, EPI_ISL_459961 | Respiratory Virus Unit, Microbiology Services Colindale, Public Health England | Respiratory Virus Unit, Microbiology Services Colindale, Public Health England | Steven Platt, Shahjahan Miah, Angie Lackenby, Omolola Akinbami, Tina Talts, Leena Bhaw, Richard Myers, Monica Galiano, Kirstin Edwards, Jonathan Hubb, Joanna Ellis, Maria Zambon |
| EPI_ISL_460096, EPI_ISL_460604, EPI_ISL_460605 | Molecular diagnostic laboratory of Federal Budget Institution of Science "Central Research Institute of Epidemiology" of The Federal Service on Customers' Rights Protection and Human Well-being Surveillance | Group of Genomics and Postgenomic Technologies of Central Research Institute of Epidemilology | Speranskaya AS, Kaptelova VV, Samoilov AE, Korneenko EV, Tivanova EV, Shipulina OY, Akimkin VG |
| EPI_ISL_461504 | B.J. Medical College and Civil hospital | Gujarat Biotechnology Research Centre | Snehal Bagatharia, Pranay Shah, Kamlesh J Upadhyay, Tejas Shah, Ankit Hinsu, Pritesh Sabara, Apurvasinh Puvar, Janvi Raval, Zarna Patel, Monika Gandhi, Pinal Trivedi, Maharshi Pandya, Nidhi Patel, Nitin Savaliya, Raghawendra Kumar, Dinesh Kumar, Zuber Saiyed, Komal Patel, Labdhi Pandya, Fenil Patel, R D Dixit, A M Kadri, Harsh Bakshi, Chaitanya Joshi, Madhvi Joshi, |
| EPI_ISL_462149, EPI_ISL_462150 | Molecular diagnostic laboratory of Federal Budget Institution of Science "Central Research Institute of Epidemiology" of The Federal Service on Customers' Rights Protection and Human Well-being Surveillance | Group of Genomics and Postgenomic Technologies of Central Research Institute of Epidemilology | Speranskaya AS, Kaptelova VV, Samoilov AE, Korneenko EV, Sizova TV, Tivanova EV, Shipulina OY, Akimkin VG |
| EPI_ISL_462480 | Institute of Human Genetics, Polish Academy of Sciences | Institute of Human Genetics, Polish Academy of Sciences | Szymon Hryhorowicz, Adam Ustaszewski, Emilia Lis, Marta Kaczmarek-Ry, Micha Witt, Andrzej Pawski |
| EPI_ISL_462990 | University Clinical Centre of the Republic of Srpska | University of Sarajevo, Veterinary Faculty | Teufik, G., Šejla, G., Toni, E., Maja, T., Mirsada, H., Aida, K., Alma, Š.A. |
| EPI_ISL_463007 | Department of Laboratory Medicine, National Taiwan University Hospital | Microbial Genomics Core Lab, National Taiwan University Centers of Genomic and Precision Medicine | Shiou-Hwei Yeh, You-Yu Lin, Ya-Yun Lai, Chiao-Ling Li, Shan-Chwen Chang, Pei-Jer Chen, Sui-Yuan Chang |
| EPI_ISL_463010, EPI_ISL_463011, EPI_ISL_463012, EPI_ISL_463013, EPI_ISL_463014, EPI_ISL_463015, EPI_ISL_463016, EPI_ISL_463017, EPI_ISL_463018, EPI_ISL_463019, EPI_ISL_463020, EPI_ISL_463021, EPI_ISL_463022, EPI_ISL_463023, EPI_ISL_463024, EPI_ISL_463025, EPI_ISL_463026, EPI_ISL_463027, EPI_ISL_463028, EPI_ISL_463029, EPI_ISL_463030 | | | |
| see above | Institute of Life Sciences, Bhubaneswar | Immunogenomics lab, Institute of Life Sciences, Bhubaneswar | Sunil Raghav, Arup Ghosh, Atimukta Jha, Viplov K. Biswas, Swati Madhulika, Manasi Priyadarshini, Shuchi Smita, Kaushik Sen, Hiren G. Dodia, Deepak Singh, Jeky Chawla, Shamima Ansari, Rupesh Dash, Soma Chattopadhyay, Ghulam Hussain Syed, Shanti Senapati, Tushar K. Beuria, Rajeeb Swain, Punit Prasad, ILS COVID-19 TEAM, Orissa COVID-19 Study Group, DBT's PAN-INDIA 1000 SARS-CoV2 RNA genome sequencing consortium, Ajay Parida |
| EPI_ISL_463031, EPI_ISL_463032, EPI_ISL_463033, EPI_ISL_463034, EPI_ISL_463035, EPI_ISL_463036, EPI_ISL_463037, EPI_ISL_463038, EPI_ISL_463039, EPI_ISL_463040, EPI_ISL_463041, EPI_ISL_463042, EPI_ISL_463043, EPI_ISL_463044, EPI_ISL_463045, EPI_ISL_463046, EPI_ISL_463047, EPI_ISL_463048, EPI_ISL_463049, EPI_ISL_463050, EPI_ISL_463051 | | | |
| see above | Institute of Life Sciences, Bhubaneswar | Immunogenomics lab, Institute of Life Sciences, Bhubaneswar | Sunil Raghav, Arup Ghosh, Atimukta Jha, Viplov K. Biswas, Swati Madhulika, Manasi Priyadarshini, Shuchi Smita, O. P. Shriwas, Priyanka Mohapatra, Satya Ranjan Sahu, Aliva Minz, Debyashrita Barik, Rupesh Dash, Soma Chattopadhyay, Ghulam Hussain Syed, Shanti Senapati, Tushar K. Beuria, Rajeeb Swain, Punit Prasad, ILS COVID-19 TEAM, Orissa COVID-19 Study Group, DBT's PAN-INDIA 1000 SARS-CoV2 RNA genome sequencing consortium, Ajay Parida |
| EPI_ISL_463052, EPI_ISL_463053, EPI_ISL_463054, EPI_ISL_463055, EPI_ISL_463056, EPI_ISL_463057, EPI_ISL_463058, EPI_ISL_463059, EPI_ISL_463060, EPI_ISL_463061, EPI_ISL_463062, EPI_ISL_463063, EPI_ISL_463064, EPI_ISL_463065, EPI_ISL_463066, EPI_ISL_463067, EPI_ISL_463068, EPI_ISL_463069, EPI_ISL_463070, EPI_ISL_463071, EPI_ISL_463072 | | | |
| see above | Institute of Life Sciences, Bhubaneswar | Immunogenomics lab, Institute of Life Sciences, Bhubaneswar | Sunil Raghav, Arup Ghosh, Atimukta Jha, Viplov K. Biswas, Swati Madhulika, Manasi Priyadarshini, Shuchi Smita, Sifu Agarwal, Sanchari Chatterjee, Avula Kiran, Parej Nath, Supriya Suman, Rina Yadav, Rupesh Dash, Soma Chattopadhyay, Ghulam Hussain Syed, Shanti Senapati, Tushar K. Beuria, Rajeeb Swain, Punit Prasad, ILS COVID-19 TEAM, Orissa COVID-19 Study Group, DBT's PAN-INDIA 1000 SARS-CoV2 RNA genome sequencing consortium, Ajay Parida |
| EPI_ISL_463073, EPI_ISL_463074, EPI_ISL_463075, EPI_ISL_463076, EPI_ISL_463077, EPI_ISL_463078, EPI_ISL_463079, EPI_ISL_463080, EPI_ISL_463081, EPI_ISL_463082, EPI_ISL_463083, EPI_ISL_463084, EPI_ISL_463085, EPI_ISL_463086, EPI_ISL_463087, EPI_ISL_463088, EPI_ISL_463089, EPI_ISL_463090, EPI_ISL_463091, EPI_ISL_463092, EPI_ISL_463093 | | | |
| see above | Institute of Life Sciences, Bhubaneswar | Immunogenomics lab, Institute of Life Sciences, Bhubaneswar | Sunil Raghav, Arup Ghosh, Atimukta Jha, Viplov K. Biswas, Swati Madhulika, Manasi Priyadarshini, Shuchi Smita, Kautilya Kumar Jena, Sandhya Suranjika, Neha Singh, Eshna Laha, Saiket De, Rupesh Dash, Soma Chattopadhyay, Ghulam Hussain Syed, Shanti Senapati, Tushar K. Beuria, Rajeeb Swain, Punit Prasad, ILS COVID-19 TEAM, Orissa COVID-19 Study Group, DBT's PAN-INDIA 1000 SARS-CoV2 RNA genome sequencing consortium, Ajay Parida |
| EPI_ISL_463277, EPI_ISL_463278, EPI_ISL_463279, EPI_ISL_463280, EPI_ISL_463281, EPI_ISL_463282, EPI_ISL_463283, EPI_ISL_463284, EPI_ISL_463285, EPI_ISL_463286, EPI_ISL_463287, EPI_ISL_463288, EPI_ISL_463289, EPI_ISL_463290, EPI_ISL_463291, EPI_ISL_463292, EPI_ISL_463293, EPI_ISL_463294, EPI_ISL_463295, EPI_ISL_463296, EPI_ISL_463297, EPI_ISL_463298, EPI_ISL_463299, EPI_ISL_463300 | | | |
| see above | Ochsner Health | Bioinfoexperts, LLC | Susanna L. Lamers, David J. Nolan, Rebecca Rose, Sissy Cross, David Moraga Amador, Tong Yang, Luke Caruso, Wayra Navia, Lydia Von Borstel, Xiao Hui Zhou, Amy Feehan, Julia-Garcia-Diaz |
| EPI_ISL_464112, EPI_ISL_464113, EPI_ISL_464114, EPI_ISL_464115, EPI_ISL_464116, EPI_ISL_464117, EPI_ISL_464118, EPI_ISL_464119, EPI_ISL_464120, EPI_ISL_464121, EPI_ISL_464122, EPI_ISL_464123, EPI_ISL_464124, EPI_ISL_464125, EPI_ISL_464126, EPI_ISL_464127, EPI_ISL_464128, EPI_ISL_464129, EPI_ISL_464130, EPI_ISL_464131, EPI_ISL_464132, EPI_ISL_464133 | | | |
| see above | National Health Laboratory Service (NHLS), Tygerberg | Division of Medical Virology, Stellenbosch University and National Health Laboratory Service (NHLS) | Susan Engelbrecht, Kayla Delaney, Bronwyn Kleinhans, Houriiyah Tegally, Eduan Wilkindon, Gert van Zyl, Wolfgang Preiser, Tulio de Oliveira |
| EPI_ISL_464134 | National Health Laboratory Service (NHLS), Tygerberg | Stellenbosch University and NHLS | Susan Engelbrecht, Kayla Delaney, Bronwyn Kleinhans, Houriiyah Tegally, Eduan Wilkindon, Gert van Zyl, Wolfgang Preiser, Tulio de Oliveira |
| EPI_ISL_464135, EPI_ISL_464136, EPI_ISL_464137, EPI_ISL_464138, EPI_ISL_464139, EPI_ISL_464140, EPI_ISL_464141, EPI_ISL_464142, EPI_ISL_464143, EPI_ISL_464144, EPI_ISL_464145, EPI_ISL_464146, EPI_ISL_464147, EPI_ISL_464148, EPI_ISL_464149, EPI_ISL_464150, EPI_ISL_464151, EPI_ISL_464152, EPI_ISL_464153, EPI_ISL_464154, EPI_ISL_464155, EPI_ISL_464156, EPI_ISL_464157, EPI_ISL_464158 | | | |
| see above | National Health Laboratory Service (NHLS), Tygerberg | Division of Medical Virology, Stellenbosch University and National Health Laboratory Service (NHLS) | Susan Engelbrecht, Kayla Delaney, Bronwyn Kleinhans, Houriiyah Tegally, Eduan Wilkindon, Gert van Zyl, Wolfgang Preiser, Tulio de Oliveira |
| EPI_ISL_464163, EPI_ISL_464164 | National Institute of Laboratory Medicine and Referral Center | Genomic Research Lab, BCSIR | Tanjina Akhter Banu, Abu Sayeed Mohammad Mahmud, Mohammad Samir Uzzaman, Eshrar Osman, Md. Ahasan Habib, Shahina Akter, Md. Murshed Hasan Sarker, Barna Goswami, Iffat Jahan, Md. Saddam Hossain, Tasnim Nafisa, Md. Maruf Ahmed Molla, Mahmuda Yeasmin, Asish Kumar Ghosh, Arifa Akram, A. K. M. Shamsuzzaman, Sheikh Md. Selim Al Din, Utpal Chandra Ray, Salek Ahmed Sajib, Md. Salim Khan |
| EPI_ISL_466648 | Innovative Genomics Institute, UCB | Innovative Genomics Institute, UCB | Stacia Wyman, Haridha Shivram, Liana Lareau, Shana McDevitt, Justin Choi |
| EPI_ISL_467033 | GMERS Medical College and Hospital, Gandhinagar | Gujarat Biotechnology Research Centre | Tejas Shah, Ankit Hinsu, Pritesh Sabara, Apurvasinh Puvar, Janvi Raval, Zarna Patel, Monika Gandhi, Pinal Trivedi, Maharshi Pandya, Nidhi Patel, Nitin Savaliya, Raghawendra Kumar, Dinesh Kumar, Zuber Saiyed, Komal Patel, Labdhi Pandya, Snehal Bagatharia, Seema Bhatt, Gaurishankar Shrimali, Bhavesh Modi, Bharti Rajani, Priti Pandita, R D Dixit, A M Kadri, Harsh Bakshi, Chaitanya Joshi, Madhvi Joshi |
| EPI_ISL_467051 | B.J. Medical College and Civil hospital | Gujarat Biotechnology Research Centre | Snehal Bagatharia, Pranay Shah, Kamlesh J Upadhyay, Nirav Mungalpara, Tejas Shah, Ankit Hinsu, Pritesh Sabara, Apurvasinh Puvar, Janvi Raval, Zarna |

| | | | |
|---|---|---|---|
| | | | Patel, Monika Gandhi, Pinal Trivedi, Maharshi Pandya, Nidhi Patel, Nitin Savaliya, Raghawendra Kumar, Dinesh Kumar, Zuber Saiyed, Komal Patel, Labdhi Pandya, Bhavya Jindal, R D Dixit, A M Kadri, Harsh Bakshi, Chaitanya Joshi, Madhvi Joshi, |
| EPI_ISL_467092, EPI_ISL_467093, EPI_ISL_467094, EPI_ISL_467095, EPI_ISL_467096, EPI_ISL_467097, EPI_ISL_467098, EPI_ISL_467099, EPI_ISL_467100, EPI_ISL_467101, EPI_ISL_467102, EPI_ISL_467103, EPI_ISL_467104, EPI_ISL_467105, EPI_ISL_467106, EPI_ISL_467107, EPI_ISL_467108, EPI_ISL_467109, EPI_ISL_467110, EPI_ISL_467111, EPI_ISL_467112, EPI_ISL_467113, EPI_ISL_467114, EPI_ISL_467115, EPI_ISL_467116, EPI_ISL_467117, EPI_ISL_467118, EPI_ISL_467119, EPI_ISL_467120, EPI_ISL_467121, EPI_ISL_467122, EPI_ISL_467123, EPI_ISL_467124, EPI_ISL_467125, EPI_ISL_467126, EPI_ISL_467127, EPI_ISL_467128, EPI_ISL_467129, EPI_ISL_467130, EPI_ISL_467131, EPI_ISL_467132, EPI_ISL_467133, EPI_ISL_467134, EPI_ISL_467135, EPI_ISL_467136, EPI_ISL_467137, EPI_ISL_467138, EPI_ISL_467139, EPI_ISL_467140, EPI_ISL_467141, EPI_ISL_467142, EPI_ISL_467143, EPI_ISL_467144, EPI_ISL_467145, EPI_ISL_467146, EPI_ISL_467147, EPI_ISL_467148, EPI_ISL_467149, EPI_ISL_467150, EPI_ISL_467151, EPI_ISL_467152, EPI_ISL_467153, EPI_ISL_467154, EPI_ISL_467155, EPI_ISL_467156, EPI_ISL_467157, EPI_ISL_467158, EPI_ISL_467159, EPI_ISL_467160, EPI_ISL_467161, EPI_ISL_467162, EPI_ISL_467163, EPI_ISL_467164, EPI_ISL_467165, EPI_ISL_467166, EPI_ISL_467167, EPI_ISL_467168, EPI_ISL_467169, EPI_ISL_467170, EPI_ISL_467171, EPI_ISL_467172, EPI_ISL_467173, EPI_ISL_467174, EPI_ISL_467175, EPI_ISL_467176, EPI_ISL_467177, EPI_ISL_467178, EPI_ISL_467179, EPI_ISL_467180, EPI_ISL_467181, EPI_ISL_467182, EPI_ISL_467183 | | | |
| see above | Hospital Universitario Araba. Vitoria-Gasteiz | SeqCOVID-SPAIN consortium/IBV(CSIC) | Silvia Hernáez Crespo, Carmen Gómez González, Amaia Aguirre Quiñonero, Marina Fernández Torres, Mª Rosario Almela Ferrer, Mª Concepción Lecaroz Agara, Andrés Canut Blasco. and SeqCOVID-SPAIN consortium |
| EPI_ISL_467693, EPI_ISL_467694, EPI_ISL_467695, EPI_ISL_467696, EPI_ISL_467697, EPI_ISL_467698, EPI_ISL_467699, EPI_ISL_467700, EPI_ISL_467701, EPI_ISL_467702, EPI_ISL_467703, EPI_ISL_467704, EPI_ISL_467705, EPI_ISL_467706, EPI_ISL_467707, EPI_ISL_467708, EPI_ISL_467709, EPI_ISL_467710, EPI_ISL_467711, EPI_ISL_467712, EPI_ISL_467713, EPI_ISL_467714, EPI_ISL_467715, EPI_ISL_467716, EPI_ISL_467717, EPI_ISL_467718, EPI_ISL_467719 | | | |
| see above | PHE South West Regional Laboratory, National Infection Service | Wellcome Sanger Institute for the COVID-19 Genomics UK (COG-UK) consortium | Stephanie Hutchings, Hannah Pymont, Dr Peter Muir, Barry Vipond, Rich Hopes; and Alex Alderton, Roberto Amato, Sonia Goncalves, Ewan Harrison, David K. Jackson, Ian Johnston, Dominic Kwiatkowski, Cordelia Langford, John Sillitoe on behalf of the Wellcome Sanger Institute COVID-19 Surveillance Team (http://www.sanger.ac.uk/covid-team) |
| EPI_ISL_467774, EPI_ISL_467775 | Molecular diagnostic laboratory of Federal Budget Institution of Science "Central Research Institute of Epidemiology" of The Federal Service on Customers' Rights Protection and Human Well-being Surveillance | Group of Genomics and Postgenomic Technologies of Central Research Institute of Epidemiology | Speranskaya AS, Kaptelova VV, Samoilov AE, Korneenko EV, Sizova TV, Tivanova EV, Shipulina OY, Akimkin VG |
| EPI_ISL_467946, EPI_ISL_467947, EPI_ISL_467948, EPI_ISL_467949 | Innovative Genomics Institute, UC Berkeley | Innovative Genomics Institute, UC Berkeley | Stacia Wyman, Haridha Shivram, Liana Lareau, Shana McDevitt, Justin Choi |
| EPI_ISL_468718 | Environmental and Global Health | Environmental and Global Health | Stephenson,C.J., Subramaniam,K., Waltzek,T.B., Merck,L.H., Gibson,J.C., Morris,J.G. |
| EPI_ISL_468720 | University of Florida | University of Florida | Stephenson,C.J., Subramaniam,K., Waltzek,T.B., Lauzardo,M., Morris,J.G., Lednicky,J.A. |
| EPI_ISL_468721 | University of Florida | University of Florida | Stephenson,C.J., Subramaniam,K., Waltzek,T.B., Lauzardo,M., Gibson,J.C., Morris,J.G., Lednicky,J.A. |
| EPI_ISL_468723 | University of Florida | University of Florida | Stephenson,C.J., Subramaniam,K., Waltzek,T.B., Lauzardo,M., Morris,J.G., Lednicky,J.A. |
| EPI_ISL_469024 | B.J. Medical College and Civil hospital | Gujarat Biotechnology Research Centre | Tejas Shah, Ankit Hinsu, Pritesh Sabara, Apurvasinh Puvar, Janvi Raval, Zarna Patel, Monika Gandhi, Pinal Trivedi, Maharshi Pandya, Nidhi Patel, Nitin Savaliya, Raghawendra Kumar, Dinesh Kumar, Zuber Saiyed, Komal Patel, Labdhi Pandya, Snehal Bagatharia, Pranay Shah, Kamlesh J Upadhyay, Nirav Mungalpara, Priti Pandita, R D Dixit, A M Kadri, Harsh Bakshi, Chaitanya Joshi, Madhvi Joshi |
| EPI_ISL_469040 | GMERS Medical College & Hospital | Gujarat Biotechnology Research Centre | Snehal Bagatharia, Meenakshi Shah, Neena Doshi, Varsha Godbole, Tejas Shah, Ankit Hinsu, Pritesh Sabara, Apurvasinh Puvar, Janvi Raval, Zarna Patel, Monika Gandhi, Pinal Trivedi, Maharshi Pandya, Nidhi Patel, Nitin Savaliya, Raghawendra Kumar, Dinesh Kumar, Zuber Saiyed, Komal Patel, Labdhi Pandya, Fenil Patel, R D Dixit, A M Kadri, Harsh Bakshi, Chaitanya Joshi, Madhvi Joshi |
| EPI_ISL_469045 | Dr. N. D. Desai Medical College & Hospital | Gujarat Biotechnology Research Centre | Supreet Prabhu, Tejas Shah, Ankit Hinsu, Pritesh Sabara, Apurvasinh Puvar, Janvi Raval, Zarna Patel, Monika Gandhi, Pinal Trivedi, Maharshi Pandya, Nidhi Patel, Nitin Savaliya, Raghawendra Kumar, Dinesh Kumar, Zuber Saiyed, Komal Patel, Labdhi Pandya, Snehal Bagatharia, J G Buch, Jigar Gusani, Priyanka P Vatsa, R D Dixit, A M Kadri, Harsh Bakshi, Chaitanya Joshi, Madhvi Joshi |
| EPI_ISL_469046 | Dr. N. D. Desai Medical College & Hospital | Gujarat Biotechnology Research Centre | Tejas Shah, Ankit Hinsu, Pritesh Sabara, Apurvasinh Puvar, Janvi Raval, Zarna Patel, Monika Gandhi, Pinal Trivedi, Maharshi Pandya, Nidhi Patel, Nitin Savaliya, Raghawendra Kumar, Dinesh Kumar, Zuber Saiyed, Komal Patel, Labdhi Pandya, Snehal Bagatharia, J G Buch, Jigar Gusani, Supreet Prabhu, Pooja P Doshi, R D Dixit, A M Kadri, Harsh Bakshi, Chaitanya Joshi, Madhvi Joshi |
| EPI_ISL_469286 | National Institute of Laboratory Medicine and Referral Center | Genomic Research Lab, BCSIR | Tanjina Akhter Banu, Abu Sayeed Mohammad Mahmud, Mohammad Samir Uzzaman, Eshrar Osman, Md. Ahasan Habib, Shahina Akter, Md. Murshed Hasan Sarkar, Iffat Jahan, Barna Goswami, Md. Saddam Hossain, Tasnim Nafisa, Md. Maruf Ahmed Molla, Mahmuda Yeasmin, Asish Kumar Ghosh, Bayzid Bin Monir, A. K. M. Shamsuzzaman, Sheikh Md. Selim Al Din, Utpal Chandra Ray, Salek Ahmed Sajib, Md. Salim Khan |
| EPI_ISL_469345, EPI_ISL_469346, EPI_ISL_469347, EPI_ISL_469348, EPI_ISL_469349, EPI_ISL_469350, EPI_ISL_469351, EPI_ISL_469352, EPI_ISL_469353, EPI_ISL_469354, EPI_ISL_469355, EPI_ISL_469356, EPI_ISL_469357, EPI_ISL_469358, EPI_ISL_469359, EPI_ISL_469360, EPI_ISL_469361, EPI_ISL_469362, EPI_ISL_469363, EPI_ISL_469364, EPI_ISL_469365, EPI_ISL_469366, EPI_ISL_469367, EPI_ISL_469368, EPI_ISL_469369, EPI_ISL_469370, EPI_ISL_469371, EPI_ISL_469372, EPI_ISL_469373, EPI_ISL_469374, EPI_ISL_469375, EPI_ISL_469376, EPI_ISL_469377, EPI_ISL_469378, EPI_ISL_469379, EPI_ISL_469380, EPI_ISL_469381, EPI_ISL_469382, EPI_ISL_469383, EPI_ISL_469384, EPI_ISL_469385, EPI_ISL_469386, EPI_ISL_469387, EPI_ISL_469388, EPI_ISL_469389, EPI_ISL_469390, EPI_ISL_469391, EPI_ISL_469392, EPI_ISL_469393, EPI_ISL_469394, EPI_ISL_469395, EPI_ISL_469396, EPI_ISL_469397, EPI_ISL_469398, EPI_ISL_469399, EPI_ISL_469400, EPI_ISL_469401, EPI_ISL_469402, EPI_ISL_469403, EPI_ISL_469404, EPI_ISL_469405, EPI_ISL_469406, EPI_ISL_469407, EPI_ISL_469408, EPI_ISL_469409, EPI_ISL_469410, EPI_ISL_469411, EPI_ISL_469412, EPI_ISL_469413, EPI_ISL_469414, EPI_ISL_469415, EPI_ISL_469416, EPI_ISL_469417, EPI_ISL_469418, EPI_ISL_469419, EPI_ISL_469420, EPI_ISL_469421, EPI_ISL_469422, EPI_ISL_469423, EPI_ISL_469424, EPI_ISL_469425, EPI_ISL_469426, EPI_ISL_469427, EPI_ISL_469428, EPI_ISL_469429, EPI_ISL_469430, EPI_ISL_469431, EPI_ISL_469432, EPI_ISL_469433, EPI_ISL_469434, EPI_ISL_469435, EPI_ISL_469436, EPI_ISL_469437, EPI_ISL_469438, EPI_ISL_469439, EPI_ISL_469440, EPI_ISL_469441, EPI_ISL_469442, EPI_ISL_469443, EPI_ISL_469444, EPI_ISL_469445, EPI_ISL_469446, EPI_ISL_469447, EPI_ISL_469448, EPI_ISL_469449, EPI_ISL_469450, EPI_ISL_469451, EPI_ISL_469452, EPI_ISL_469453, EPI_ISL_469454, EPI_ISL_469455, EPI_ISL_469456, EPI_ISL_469457, EPI_ISL_469458, EPI_ISL_469459, EPI_ISL_469460, EPI_ISL_469461, EPI_ISL_469462, EPI_ISL_469463, EPI_ISL_469464, EPI_ISL_469465, EPI_ISL_469466, EPI_ISL_469467, EPI_ISL_469468, EPI_ISL_469469, EPI_ISL_469470, EPI_ISL_469471, EPI_ISL_469472, EPI_ISL_469473, EPI_ISL_469474, EPI_ISL_469475, EPI_ISL_469476, EPI_ISL_469477, EPI_ISL_469478, EPI_ISL_469479, EPI_ISL_469480, EPI_ISL_469481, EPI_ISL_469482, EPI_ISL_469483, EPI_ISL_469484, EPI_ISL_469485, EPI_ISL_469486, EPI_ISL_469487, EPI_ISL_469488, EPI_ISL_469489, EPI_ISL_469490, EPI_ISL_469491, EPI_ISL_469492, EPI_ISL_469493, EPI_ISL_469494, EPI_ISL_469495, EPI_ISL_469496, EPI_ISL_469497, EPI_ISL_469498, EPI_ISL_469499, EPI_ISL_469500, EPI_ISL_469501, EPI_ISL_469502, EPI_ISL_469503, EPI_ISL_469504, EPI_ISL_469505, EPI_ISL_469506, EPI_ISL_469507, EPI_ISL_469508, EPI_ISL_469509, EPI_ISL_469510, EPI_ISL_469511, EPI_ISL_469512, EPI_ISL_469513, EPI_ISL_469514, EPI_ISL_469515, EPI_ISL_469516, EPI_ISL_469517, EPI_ISL_469518, EPI_ISL_469519, EPI_ISL_469520, EPI_ISL_469521, EPI_ISL_469522, EPI_ISL_469523, EPI_ISL_469524, EPI_ISL_469525, EPI_ISL_469526, EPI_ISL_469527, EPI_ISL_469528, EPI_ISL_469529, EPI_ISL_469530, EPI_ISL_469531, EPI_ISL_469532, EPI_ISL_469533, EPI_ISL_469534, EPI_ISL_469535, EPI_ISL_469536, EPI_ISL_469537, EPI_ISL_469538, EPI_ISL_469539, EPI_ISL_469540, EPI_ISL_469541, EPI_ISL_469542, EPI_ISL_469543, EPI_ISL_469544, EPI_ISL_469545, EPI_ISL_469546, EPI_ISL_469547, EPI_ISL_469548, EPI_ISL_469549, EPI_ISL_469550, EPI_ISL_469551, EPI_ISL_469552, EPI_ISL_469553, EPI_ISL_469554, EPI_ISL_469555, EPI_ISL_469556, EPI_ISL_469557, EPI_ISL_469558, EPI_ISL_469559, EPI_ISL_469560, EPI_ISL_469561, EPI_ISL_469562, EPI_ISL_469563, EPI_ISL_469564, EPI_ISL_469565, EPI_ISL_469566, EPI_ISL_469567, EPI_ISL_469568, EPI_ISL_469569, EPI_ISL_469570, EPI_ISL_469571, EPI_ISL_469572, EPI_ISL_469573, EPI_ISL_469574, EPI_ISL_469575, EPI_ISL_469576, EPI_ISL_469577, EPI_ISL_469578, EPI_ISL_469579, EPI_ISL_469580, EPI_ISL_469581, EPI_ISL_469582, EPI_ISL_469583, EPI_ISL_469584, EPI_ISL_469585, EPI_ISL_469586, EPI_ISL_469587, EPI_ISL_469588, EPI_ISL_469589, EPI_ISL_469590, EPI_ISL_469591, EPI_ISL_469592, EPI_ISL_469593, EPI_ISL_469594, EPI_ISL_469595, EPI_ISL_469596, EPI_ISL_469597, EPI_ISL_469598, EPI_ISL_469599, EPI_ISL_469600, EPI_ISL_469601, EPI_ISL_469602, EPI_ISL_469603, EPI_ISL_469604, EPI_ISL_469605, EPI_ISL_469606, EPI_ISL_469607, EPI_ISL_469608, EPI_ISL_469609, EPI_ISL_469610, EPI_ISL_469611, EPI_ISL_469612, EPI_ISL_469613, EPI_ISL_469614, EPI_ISL_469615, EPI_ISL_469616, EPI_ISL_469617, EPI_ISL_469618, EPI_ISL_469619, EPI_ISL_469620, EPI_ISL_469621, EPI_ISL_469622, EPI_ISL_469623, EPI_ISL_469624, EPI_ISL_469625, EPI_ISL_469626, EPI_ISL_469627, EPI_ISL_469628, EPI_ISL_469629, EPI_ISL_469630, EPI_ISL_469631, EPI_ISL_469632, EPI_ISL_469633, EPI_ISL_469634, EPI_ISL_469635, EPI_ISL_469636, EPI_ISL_469637, EPI_ISL_469638, EPI_ISL_469639, EPI_ISL_469640, EPI_ISL_469641, EPI_ISL_469642, EPI_ISL_469643, EPI_ISL_469644, EPI_ISL_469645, EPI_ISL_469646, EPI_ISL_469647, EPI_ISL_469648, EPI_ISL_469649, EPI_ISL_469650, EPI_ISL_469651, EPI_ISL_469652, EPI_ISL_469653, EPI_ISL_469654, EPI_ISL_469655, EPI_ISL_469656, EPI_ISL_469657, EPI_ISL_469658, EPI_ISL_469659, EPI_ISL_469660, EPI_ISL_469661, EPI_ISL_469662, EPI_ISL_469663, EPI_ISL_469664, EPI_ISL_469665, EPI_ISL_469666, EPI_ISL_469667, EPI_ISL_469668, EPI_ISL_469669, EPI_ISL_469670, EPI_ISL_469671, EPI_ISL_469672, EPI_ISL_469673, EPI_ISL_469674, EPI_ISL_469675, EPI_ISL_469676, EPI_ISL_469677, EPI_ISL_469678, EPI_ISL_469679, EPI_ISL_469680, EPI_ISL_469681, EPI_ISL_469682, EPI_ISL_469683, EPI_ISL_469684, EPI_ISL_469685, EPI_ISL_469686, EPI_ISL_469687, EPI_ISL_469688, EPI_ISL_469689, EPI_ISL_469690, EPI_ISL_469691, EPI_ISL_469692, EPI_ISL_469693, EPI_ISL_469694, EPI_ISL_469695, EPI_ISL_469696, EPI_ISL_469697, EPI_ISL_469698, EPI_ISL_469699, EPI_ISL_469700, EPI_ISL_469701, EPI_ISL_469702, EPI_ISL_469703, EPI_ISL_469704, EPI_ISL_469705, EPI_ISL_469706, EPI_ISL_469707, EPI_ISL_469708, EPI_ISL_469709, EPI_ISL_469710, EPI_ISL_469711, EPI_ISL_469712, EPI_ISL_469713, EPI_ISL_469714, EPI_ISL_469715, EPI_ISL_469716, EPI_ISL_469717, EPI_ISL_469718, EPI_ISL_469719, EPI_ISL_469720, EPI_ISL_469721, EPI_ISL_469722, EPI_ISL_469723, EPI_ISL_469724, EPI_ISL_469725, EPI_ISL_469726, EPI_ISL_469727, EPI_ISL_469728, EPI_ISL_469729, EPI_ISL_469730, EPI_ISL_469731, EPI_ISL_469732, EPI_ISL_469733, EPI_ISL_469734, EPI_ISL_469735, EPI_ISL_469736, EPI_ISL_469737, EPI_ISL_469738, EPI_ISL_469739, EPI_ISL_469740, EPI_ISL_469741, EPI_ISL_469742, EPI_ISL_469743, EPI_ISL_469744, EPI_ISL_469745, EPI_ISL_469746, EPI_ISL_469747, EPI_ISL_469748, EPI_ISL_469749, EPI_ISL_469750, EPI_ISL_469751, EPI_ISL_469752, EPI_ISL_469753, EPI_ISL_469754, EPI_ISL_469755, EPI_ISL_469756, EPI_ISL_469757, EPI_ISL_469758, EPI_ISL_469759, EPI_ISL_469760, EPI_ISL_469761, EPI_ISL_469762, EPI_ISL_469763, EPI_ISL_469764, EPI_ISL_469765, EPI_ISL_469766, EPI_ISL_469767, EPI_ISL_469768, EPI_ISL_469769, EPI_ISL_469770, EPI_ISL_469771, EPI_ISL_469772, EPI_ISL_469773, EPI_ISL_469774, EPI_ISL_469775, EPI_ISL_469776, EPI_ISL_469777, EPI_ISL_469778, EPI_ISL_469779, EPI_ISL_469780, EPI_ISL_469781, EPI_ISL_469782, EPI_ISL_469783, EPI_ISL_469784, EPI_ISL_469785, EPI_ISL_469786, EPI_ISL_469787, EPI_ISL_469788, EPI_ISL_469789, EPI_ISL_469790, EPI_ISL_469791, EPI_ISL_469792, EPI_ISL_469793, EPI_ISL_469794, EPI_ISL_469795, EPI_ISL_469796, EPI_ISL_469797, EPI_ISL_469798, EPI_ISL_469799, EPI_ISL_469800, EPI_ISL_469801, EPI_ISL_469802, EPI_ISL_469803, EPI_ISL_469804, EPI_ISL_469806, EPI_ISL_469807, EPI_ISL_469808, EPI_ISL_469809, EPI_ISL_469810, EPI_ISL_469811, EPI_ISL_469812, EPI_ISL_469813, EPI_ISL_469814, EPI_ISL_469815, EPI_ISL_469816, EPI_ISL_469817, EPI_ISL_469818, EPI_ISL_469819, EPI_ISL_469820, EPI_ISL_469821, EPI_ISL_469822, EPI_ISL_469823, EPI_ISL_469824, EPI_ISL_469825, EPI_ISL_469826, EPI_ISL_469827, EPI_ISL_469828, EPI_ISL_469829, EPI_ISL_469830, EPI_ISL_469831, EPI_ISL_469832, EPI_ISL_469833, EPI_ISL_469834, EPI_ISL_469835, EPI_ISL_469836, EPI_ISL_469837, EPI_ISL_469838, EPI_ISL_469839, EPI_ISL_469840, EPI_ISL_469841, EPI_ISL_469842, EPI_ISL_469843, EPI_ISL_469914, EPI_ISL_469916, EPI_ISL_469917, EPI_ISL_470356, EPI_ISL_470357, EPI_ISL_470529, EPI_ISL_470530 | | | |
| see above | PHE South West Regional Laboratory, National Infection Service | Wellcome Sanger Institute for the COVID-19 Genomics UK (COG-UK) consortium | Stephanie Hutchings, Hannah Pymont, Dr Peter Muir, Barry Vipond, Rich Hopes; and Alex Alderton, Roberto Amato, Sonia Goncalves, Ewan Harrison, David K. Jackson, Ian Johnston, Dominic Kwiatkowski, Cordelia Langford, John Sillitoe on behalf of the Wellcome Sanger Institute COVID-19 Surveillance Team (http://www.sanger.ac.uk/covid-team) |
| EPI_ISL_470539 | Molecular diagnostic laboratory of Federal Budget Institution of Science "Central Research Institute of Epidemiology" of The Federal Service on Customers' Rights Protection and Human Well-being Surveillance | Group of Genomics and Postgenomic Technologies of Central Research Institute of Epidemiology | Speranskaya AS, Kaptelova VV, Samoilov AE, Korneenko EV, Sizova TV, Tivanova EV, Shipulina OY, Akimkin VG |
| EPI_ISL_471267 | Hospital IESS Babahoyo | Institute of Microbiology, Universidad San Francisco de Quito | Sully Márquez, Belén Prado-Vivar, Juan José Guadalupe, Bernardo Gutiérrez, Francisco Cordova, Ninfa Henriquez, Killen Briones-Zamora, Killen Briones-Claudette, Verónica Barragán, Patricio Rojas-Silva, Gabriel Trueba, Michelle Grunauer, Paúl Cárdenas |

| | | | |
|---|---|---|---|
| EPI_ISL_471269, EPI_ISL_471270, EPI_ISL_471271 | Hospital Oncológico Solca Núcleo de Quito | Institute of Microbiology, Universidad San Francisco de Quito | Sully Márquez, Belén Prado-Vivar, Juan José Guadalupe, Bernardo Gutiérrez, Marcos Di Stefano, Grace Salazar, Verónica Barragán, Patricio Rojas-Silva, Gabriel Trueba, Michelle Grunauer, Paúl Cárdenas |
| EPI_ISL_471607 | CSIR-Centre for Cellular and Molecular Biology | CSIR-Centre for Cellular and Molecular Biology | Sofia Banu, Payel Mukherjee, Priya Singh, Dhiviya Vedagiri, Divya Gupta, Vishal Sah, Santosh Kumar Kuncha, Krishnan Harinivas Harshan, Archana Bharadwaj Siva, Karthik Bharadwaj Tallapaka, Shagufta Khan, Lamuk Zaveri, Namami Gaur, Sakshi Shambhavi, Tulasi Nagabandi, Purushotham Vodnala, Deepak Kumar, Devi Prasad Vijayashankar, Disha Nanda, Divya Das, Jotin Gogoi, Manish Bhattacharjee, Rakesh K Mishra, Divya Tej Sowpati |
| EPI_ISL_471608 | CSIR-Centre for Cellular and Molecular Biology | CSIR-Centre for Cellular and Molecular Biology | Sofia Banu, Payel Mukherjee, Priya Singh, Dhiviya Vedagiri, Divya Gupta, Vishal Sah, Santosh Kumar Kuncha, Krishnan Harinivas Harshan, Archana Bharadwaj Siva, Karthik Bharadwaj Tallapaka, Shagufta Khan, Lamuk Zaveri, Namami Gaur, Sakshi Shambhavi, Tulasi Nagabandi, Purushotham Vodnala, Disha Nanda, Divya Das, Jotin Gogoi, Manish Bhattacharjee, Ravi Prasad Mukku, Rakesh K Mishra, Divya Tej Sowpati |
| EPI_ISL_471609 | CSIR-Centre for Cellular and Molecular Biology | CSIR-Centre for Cellular and Molecular Biology | Sofia Banu, Payel Mukherjee, Priya Singh, Dhiviya Vedagiri, Divya Gupta, Vishal Sah, Santosh Kumar Kuncha, Krishnan Harinivas Harshan, Archana Bharadwaj Siva, Karthik Bharadwaj Tallapaka, Shagufta Khan, Lamuk Zaveri, Namami Gaur, Sakshi Shambhavi, Tulasi Nagabandi, Purushotham Vodnala, Gokulan C G, Gunjan Purohit, Hanuman Tulashiram Kale, Pankaj Kumar, Prachand Issarapu, Rakesh K Mishra, Divya Tej Sowpati |
| EPI_ISL_471610 | CSIR-Centre for Cellular and Molecular Biology | CSIR-Centre for Cellular and Molecular Biology | Sofia Banu, Payel Mukherjee, Priya Singh, Dhiviya Vedagiri, Divya Gupta, Vishal Sah, Santosh Kumar Kuncha, Krishnan Harinivas Harshan, Archana Bharadwaj Siva, Karthik Bharadwaj Tallapaka, Shagufta Khan, Lamuk Zaveri, Namami Gaur, Sakshi Shambhavi, Tulasi Nagabandi, Purushotham Vodnala,Preethi Jampala, Sharada Ravi Iyer, Sulagana Mukherjee, Swetha Sundar, Peddapuvala Sai Uday Kiran, Rakesh K Mishra, Divya Tej Sowpati |
| EPI_ISL_471635 | CSIR-Centre for Cellular and Molecular Biology | CSIR-Centre for Cellular and Molecular Biology | Sofia Banu, Payel Mukherjee, Priya Singh, Dhiviya Vedagiri, Divya Gupta, Vishal Sah, Santosh Kumar Kuncha, Krishnan Harinivas Harshan, Archana Bharadwaj Siva, Karthik Bharadwaj Tallapaka, Shagufta Khan, Lamuk Zaveri, Namami Gaur, Sakshi Shambhavi, Tulasi Nagabandi, Purushotham Vodnala, Deepak Kumar, Devi Prasad Vijayashankar, Disha Nanda, Divya Das, Jotin Gogoi, Manish Bhattacharjee, Rakesh K Mishra, Divya Tej Sowpati |
| EPI_ISL_471636 | CSIR-Centre for Cellular and Molecular Biology | CSIR-Centre for Cellular and Molecular Biology | Sofia Banu, Payel Mukherjee, Priya Singh, Dhiviya Vedagiri, Divya Gupta, Vishal Sah, Santosh Kumar Kuncha, Krishnan Harinivas Harshan, Archana Bharadwaj Siva, Karthik Bharadwaj Tallapaka, Shagufta Khan, Lamuk Zaveri, Namami Gaur, Sakshi Shambhavi, Tulasi Nagabandi, Purushotham Vodnala, Disha Nanda, Divya Das, Jotin Gogoi, Manish Bhattacharjee, Ravi Prasad Mukku, Rakesh K Mishra, Divya Tej Sowpati |
| EPI_ISL_471637 | CSIR-Centre for Cellular and Molecular Biology | CSIR-Centre for Cellular and Molecular Biology | Sofia Banu, Payel Mukherjee, Priya Singh, Dhiviya Vedagiri, Divya Gupta, Vishal Sah, Santosh Kumar Kuncha, Krishnan Harinivas Harshan, Archana Bharadwaj Siva, Karthik Bharadwaj Tallapaka, Shagufta Khan, Lamuk Zaveri, Namami Gaur, Sakshi Shambhavi, Tulasi Nagabandi, Purushotham Vodnala, Gokulan C G, Gunjan Purohit, Hanuman Tulashiram Kale, Pankaj Kumar, Prachand Issarapu, Rakesh K Mishra, Divya Tej Sowpati |
| EPI_ISL_471638 | CSIR-Centre for Cellular and Molecular Biology | CSIR-Centre for Cellular and Molecular Biology | Sofia Banu, Payel Mukherjee, Priya Singh, Dhiviya Vedagiri, Divya Gupta, Vishal Sah, Santosh Kumar Kuncha, Krishnan Harinivas Harshan, Archana Bharadwaj Siva, Karthik Bharadwaj Tallapaka, Shagufta Khan, Lamuk Zaveri, Namami Gaur, Sakshi Shambhavi, Tulasi Nagabandi, Purushotham Vodnala,Preethi Jampala, Sharada Ravi Iyer, Sulagana Mukherjee, Swetha Sundar, Peddapuvala Sai Uday Kiran, Rakesh K Mishra, Divya Tej Sowpati |
| EPI_ISL_475026 | Banas Medical College and Research Institute | Gujarat Biotechnology Research Centre | Sunil R Joshi, Viren s Doshi, Pritesh Sabara, Apurvasinh Puvar, Janvi Raval, Zarna Patel, Monika Gandhi, Pinal Trivedi, Maharshi Pandya, Nidhi Patel, Nitin Savaliya, Raghawendra Kumar, Dinesh Kumar, Zuber Saiyed, Komal Patel, Labdhi Pandya, Snehal Bagatharia, Radhika Khara, Neha Rajpara, R D Dixit, A M Kadri, Harsh Bakshi, Chaitanya Joshi, Madhvi Joshi |
| EPI_ISL_475042 | Department of MicroBiology, Government Medical College, Surat | Gujarat Biotechnology Research Centre | Snehal Bagatharia, Naresh Chauhan, Summaiya Mullan, Amit gamit, Pritesh Sabara, Apurvasinh Puvar, Janvi Raval, Zarna Patel, Monika Gandhi, Pinal Trivedi, Maharshi Pandya, Nidhi Patel, Nitin Savaliya, Raghawendra Kumar, Dinesh Kumar, Zuber Saiyed, Komal Patel, Labdhi Pandya, Bhavya Jindal, R D Dixit, A M Kadri, Harsh Bakshi, Chaitanya Joshi, Madhvi Joshi |
| EPI_ISL_475044 | Department of MicroBiology, Government Medical College, Surat | Gujarat Biotechnology Research Centre | Summaiya Mullan, Amit gamit, Pritesh Sabara, Apurvasinh Puvar, Janvi Raval, Zarna Patel, Monika Gandhi, Pinal Trivedi, Maharshi Pandya, Nidhi Patel, Nitin Savaliya, Raghawendra Kumar, Dinesh Kumar, Zuber Saiyed, Komal Patel, Labdhi Pandya, Snehal Bagatharia, Naresh Chauhan, Pooja P Doshi, R D Dixit, A M Kadri, Harsh Bakshi, Chaitanya Joshi, Madhvi Joshi |
| EPI_ISL_475165 | National Institute of Laboratory Medicine and Referral Center | Genomic Research Lab, BCSIR | Shahina Akter, Abu Sayeed Mohammad Mahmud, Mohammad Samir Uzzaman, Eshrar Osman, Md. Ahasan Habib, Tanjina Akhter Banu, Md. Murshed Hasan Sarkar, Barna Goswami, Iffat Jahan, Md. Saddam Hossain, Tasnim Nafisa, Md. Maruf Ahmed Molla, Mahmuda Yeasmin, Asish Kumar Ghosh, Bayzid Bin Monir, A. K. M. Shamsuzzaman, Sheikh Md. Selim Al Din, Utpal Chandra Ray, Salek Ahmed Sajib, Md. Salim Khan |
| EPI_ISL_475166 | National Institute of Laboratory Medicine and Referral Center | Genomic Research Lab, BCSIR | Tanjina Akhter Banu, Abu Sayeed Mohammad Mahmud, Mohammad Samir Uzzaman, Eshrar Osman, Md. Ahasan Habib, Shahina Akter, Md. Murshed Hasan Sarkar, Barna Goswami, Iffat Jahan, Md. Saddam Hossain, Tasnim Nafisa, Md. Maruf Ahmed Molla, Mahmuda Yeasmin, Asish Kumar Ghosh, Bayzid Bin Monir, A. K. M. Shamsuzzaman, Sheikh Md. Selim Al Din, Utpal Chandra Ray, Salek Ahmed Sajib, Md. Salim Khan |
| EPI_ISL_475756 | National Institute of Laboratory Medicine and Referral Center | Genomic Research Lab, BCSIR | Tanjina Akhter Banu, Abu Sayeed Mohammad Mahmud, Mohammad Samir Uzzaman, Eshrar Osman, Md. Ahasan Habib, Shahina Akter, Md. Murshed Hasan Sarkar, Barna Goswami, Iffat Jahan, Md. Saddam Hossain, Tasnim Nafisa, Md. Maruf Ahmed Molla, Mahmuda Yeasmin, Asish Kumar Ghosh, Arifa Akram, A. K. M. Shamsuzzaman, Sheikh Md. Selim Al Din, Utpal Chandra Ray, Salek Ahmed Sajib, Md. Salim Khan |
| EPI_ISL_476022 | Defence Research & Development Establishment | Defence Research & Development Establishment | Shashi Sharma, Paban Kumar Dash, Jyoti S Kumar, Sushil Kumar Sharma, Ambuj Shrivastava |
| EPI_ISL_476023, EPI_ISL_476840, EPI_ISL_476842, EPI_ISL_476844, EPI_ISL_476846, EPI_ISL_476848, EPI_ISL_476849, EPI_ISL_476850, EPI_ISL_476852, EPI_ISL_476853, EPI_ISL_476854 | | | |
| see above | Defence Research & Development Establishment (DRDE) | Defence Research & Development Establishment (DRDE) | Shashi Sharma, Paban Kumar Dash, Sushil Kumar Sharma, Ambuj Shrivastava, Jyoti S. Kumar |
| EPI_ISL_476871 | Department of MicroBiology, Government Medical College, Surat | Gujarat Biotechnology Research Centre | Summaiya Mullan, Amit gamit, Apurvasinh Puvar, Janvi Raval, Zarna Patel, Monika Gandhi, Pinal Trivedi, Maharshi Pandya, Nidhi Patel, Nitin Savaliya, Raghawendra Kumar, Dinesh Kumar, Zuber Saiyed, Komal Patel, Labdhi Pandya, Afzal Ansari, Nikha Trivedi, Naresh Chauhan, R D Dixit, A M Kadri, Harsh Bakshi, Chaitanya Joshi, Madhvi Joshi |
| EPI_ISL_476883, EPI_ISL_476884, EPI_ISL_476885, EPI_ISL_476886, EPI_ISL_476887, EPI_ISL_476888, EPI_ISL_476889, EPI_ISL_476890, EPI_ISL_476891, EPI_ISL_476892, EPI_ISL_476893, EPI_ISL_476894, EPI_ISL_476895, EPI_ISL_476896 | | | |
| see above | Defence Research & Development Establishment (DRDE) | Defence Research & Development Establishment (DRDE) | Shashi Sharma, Paban Kumar Dash, Sushil Kumar Sharma, Ambuj Shrivastava, Jyoti S. Kumar |
| EPI_ISL_477015 | Institute of Microbiology, Universidad San Francisco de Quito | Institute of Microbiology, Universidad San Francisco de Quito | Sully Márquez, Belén Prado-Vivar, Juan José Guadalupe, Monica Becerra-Wong, Carla Torres, Bernardo Gutiérrez, Jorge Luis Velez, Verónica Barragán, Patricio Rojas-Silva, Gabriel Trueba, Michelle Grunauer, Paúl Cárdenas |
| EPI_ISL_477169 | Department for Virology, Molecular Biology and Genome Research, R. G. Lugar Center for Public Health Research, National Center for Disease Control and Public Health (NCDC) of Georgia. | Department for Virology, Molecular Biology and Genome Research, R. G. Lugar Center for Public Health Research, National Center for Disease Control and Public Health (NCDC) of Georgia. | Tata Imnadze, Giorgi Tomashvili, Meri Pantsulaia, Gvantsa Brachveli, Gvantsa Chanturia, Ann Machablishvili, Nato Kotaria, Marine Murtskhvaladze, Lela Sabadze, Mari Gavashelidze, Ana Papkiauri, Tamar Jashiashvili, Tea Tevdoradze, Ketevan Sidamonidze, Ekaterine Khmaladze, Ekaterine Zhghenti, Roena Sukhiashvili, Mariam Zakalashvili, Lela Urushadze, Magda Dgebuadze, Davit Tsaguria, Ekaterine Zangaladze, Nino Berishvili, Adam Kotorashvili, Maia Alkhazashvili, Irma Burjanadze, Anna Kasradze, Khatuna Zakhashvili, Paata Imnadze, Amiran Gamkrelidze. |
| EPI_ISL_478729, EPI_ISL_478731, EPI_ISL_478732, EPI_ISL_478734, EPI_ISL_478735, EPI_ISL_478736, EPI_ISL_478738, EPI_ISL_478739, EPI_ISL_478740, EPI_ISL_478741, EPI_ISL_478742, EPI_ISL_478743, EPI_ISL_478744, EPI_ISL_478746, EPI_ISL_478747, EPI_ISL_478748, EPI_ISL_478749, EPI_ISL_478751, EPI_ISL_478752, EPI_ISL_478753, EPI_ISL_478754, EPI_ISL_478756, EPI_ISL_478757, EPI_ISL_478758, EPI_ISL_478759, EPI_ISL_478761, EPI_ISL_478763, EPI_ISL_478764, EPI_ISL_478766, EPI_ISL_478767, EPI_ISL_478768, EPI_ISL_478769, EPI_ISL_478770, EPI_ISL_478771, EPI_ISL_478772, EPI_ISL_478773, EPI_ISL_478774, EPI_ISL_478775, EPI_ISL_478776, EPI_ISL_478777, EPI_ISL_478779, EPI_ISL_478780, EPI_ISL_478782, EPI_ISL_478783, EPI_ISL_478784, EPI_ISL_478785, EPI_ISL_478786, EPI_ISL_478787, EPI_ISL_478789, EPI_ISL_478790, EPI_ISL_478791, EPI_ISL_478792, EPI_ISL_478794, EPI_ISL_478795, EPI_ISL_478797, EPI_ISL_478798, EPI_ISL_478799, EPI_ISL_478801, EPI_ISL_478802, EPI_ISL_478803, EPI_ISL_478806, EPI_ISL_478807, EPI_ISL_478808, EPI_ISL_478809, EPI_ISL_478810, EPI_ISL_478811, EPI_ISL_478812, EPI_ISL_478813, EPI_ISL_478814, EPI_ISL_478815, EPI_ISL_478816, EPI_ISL_478819, EPI_ISL_478820, EPI_ISL_478822, EPI_ISL_478823, EPI_ISL_478824, EPI_ISL_478826, EPI_ISL_478827, EPI_ISL_478828, EPI_ISL_478829, EPI_ISL_478830, EPI_ISL_478832, EPI_ISL_478834, EPI_ISL_478835, EPI_ISL_478836, EPI_ISL_478837, EPI_ISL_478838, EPI_ISL_478839, EPI_ISL_478840, EPI_ISL_478841, EPI_ISL_478842, EPI_ISL_478843, EPI_ISL_478844, EPI_ISL_478845, EPI_ISL_478847, EPI_ISL_478848, EPI_ISL_478850, EPI_ISL_478851, EPI_ISL_478852, EPI_ISL_478854, EPI_ISL_478855, EPI_ISL_478856, EPI_ISL_478857, EPI_ISL_478858, EPI_ISL_478859, EPI_ISL_478860, EPI_ISL_478861, EPI_ISL_478863, EPI_ISL_478864, EPI_ISL_478865, EPI_ISL_478866, EPI_ISL_478867, EPI_ISL_478868, EPI_ISL_478869, EPI_ISL_478870, EPI_ISL_478871, EPI_ISL_478872, EPI_ISL_478873, EPI_ISL_478874, EPI_ISL_478875, EPI_ISL_478876, EPI_ISL_478877, EPI_ISL_478878, EPI_ISL_478879, EPI_ISL_478880, EPI_ISL_478881, EPI_ISL_478882, EPI_ISL_478884, EPI_ISL_478886, EPI_ISL_478887, EPI_ISL_478888, EPI_ISL_478889, EPI_ISL_478890, EPI_ISL_478891, EPI_ISL_478893, EPI_ISL_478894, EPI_ISL_478895, EPI_ISL_478897, EPI_ISL_478898, EPI_ISL_478900, EPI_ISL_478901, EPI_ISL_478903, EPI_ISL_478904, EPI_ISL_478905, EPI_ISL_478906, EPI_ISL_478907, EPI_ISL_478908, EPI_ISL_478909, EPI_ISL_478911, EPI_ISL_478912, EPI_ISL_478914, EPI_ISL_478915, EPI_ISL_478916, EPI_ISL_478917, EPI_ISL_478918, EPI_ISL_478919, EPI_ISL_478920, EPI_ISL_478922, EPI_ISL_478923, EPI_ISL_478924, EPI_ISL_478926, EPI_ISL_478927, EPI_ISL_478928, EPI_ISL_478929, EPI_ISL_478930, EPI_ISL_478931, EPI_ISL_478932, EPI_ISL_478933, EPI_ISL_478935, EPI_ISL_478936, EPI_ISL_478937, EPI_ISL_478938, EPI_ISL_478939, EPI_ISL_478940, EPI_ISL_478942, EPI_ISL_478943, EPI_ISL_478945, EPI_ISL_478946, EPI_ISL_478948, EPI_ISL_478949, EPI_ISL_478950, EPI_ISL_478951, EPI_ISL_478952, EPI_ISL_478953, EPI_ISL_478954, EPI_ISL_478956, EPI_ISL_478957, EPI_ISL_478959, EPI_ISL_478960, EPI_ISL_478962, EPI_ISL_478963, EPI_ISL_478964, EPI_ISL_478967, EPI_ISL_478968, EPI_ISL_478969, EPI_ISL_478971, EPI_ISL_478972, EPI_ISL_478974, EPI_ISL_478975, EPI_ISL_478976, EPI_ISL_478977, EPI_ISL_478979, EPI_ISL_478980, EPI_ISL_478982, EPI_ISL_478983, EPI_ISL_478984, EPI_ISL_478986, EPI_ISL_478987, EPI_ISL_478989, EPI_ISL_478991, EPI_ISL_478992, EPI_ISL_478994, EPI_ISL_478995, EPI_ISL_478997, EPI_ISL_478998, EPI_ISL_478999, EPI_ISL_479001, EPI_ISL_479003, EPI_ISL_479004, EPI_ISL_479006, EPI_ISL_479008, EPI_ISL_479010, EPI_ISL_479012, EPI_ISL_479014, EPI_ISL_479017, EPI_ISL_479020, EPI_ISL_479022, EPI_ISL_479026, EPI_ISL_479027, EPI_ISL_479028, EPI_ISL_479029, EPI_ISL_479030, EPI_ISL_479031, EPI_ISL_479033, EPI_ISL_479034, EPI_ISL_479035, EPI_ISL_479036, EPI_ISL_479037, EPI_ISL_479038, EPI_ISL_479041, EPI_ISL_479042, EPI_ISL_479043, EPI_ISL_479044, EPI_ISL_479045, EPI_ISL_479046, EPI_ISL_479047, EPI_ISL_479048, EPI_ISL_479049, EPI_ISL_479050, EPI_ISL_479051, EPI_ISL_479052, EPI_ISL_479053, EPI_ISL_479054, EPI_ISL_479056, EPI_ISL_479057, EPI_ISL_479058, EPI_ISL_479059, EPI_ISL_479060, EPI_ISL_479061, EPI_ISL_479062, EPI_ISL_479064, EPI_ISL_479066, EPI_ISL_479067, EPI_ISL_479068, EPI_ISL_479069, EPI_ISL_479071, EPI_ISL_479072, EPI_ISL_479073, EPI_ISL_479074, EPI_ISL_479075, EPI_ISL_479076, EPI_ISL_479078, EPI_ISL_479079, EPI_ISL_479080, EPI_ISL_479082, EPI_ISL_479083, EPI_ISL_479084, EPI_ISL_479085, EPI_ISL_479086, EPI_ISL_479087, EPI_ISL_479089, EPI_ISL_479090, EPI_ISL_479091, EPI_ISL_479093, EPI_ISL_479095, EPI_ISL_479096, EPI_ISL_479097, EPI_ISL_479099, EPI_ISL_479100, EPI_ISL_479101, EPI_ISL_479102, EPI_ISL_479103, EPI_ISL_479104, EPI_ISL_479108, EPI_ISL_479109, EPI_ISL_479110, EPI_ISL_479111, EPI_ISL_479112, EPI_ISL_479113, EPI_ISL_479114, EPI_ISL_479116, EPI_ISL_479117, EPI_ISL_479118, EPI_ISL_479119, EPI_ISL_479120, EPI_ISL_479121, EPI_ISL_479122, EPI_ISL_479123, EPI_ISL_479124, EPI_ISL_479125, EPI_ISL_479126, EPI_ISL_479127, EPI_ISL_479128, EPI_ISL_479129, EPI_ISL_479130, EPI_ISL_479131, EPI_ISL_479132, EPI_ISL_479133, EPI_ISL_479134, EPI_ISL_479135, EPI_ISL_479136, EPI_ISL_479137, EPI_ISL_479138, EPI_ISL_479139, | | | |

| | | | |
|---|---|---|---|
| EPI_ISL_479140, EPI_ISL_479141, EPI_ISL_479142, EPI_ISL_479143, EPI_ISL_479144, EPI_ISL_479145, EPI_ISL_479146, EPI_ISL_479147, EPI_ISL_479148, EPI_ISL_479149, EPI_ISL_479150, EPI_ISL_479151, EPI_ISL_479152, EPI_ISL_479153, EPI_ISL_479154, EPI_ISL_479156, EPI_ISL_479157, EPI_ISL_479158, EPI_ISL_479159, EPI_ISL_479160, EPI_ISL_479161, EPI_ISL_479162, EPI_ISL_479163, EPI_ISL_479164, EPI_ISL_479165, EPI_ISL_479166, EPI_ISL_479167, EPI_ISL_479169, EPI_ISL_479170, EPI_ISL_479171, EPI_ISL_479172, EPI_ISL_479173, EPI_ISL_479174 | | | |
| see above | Oxford Viromics, NDM, University of Oxford; Oxford University Hospitals; Basingstoke and North Hampshire Hospital | COVID-19 Genomics UK (COG-UK) Consortium | Tanya Golubchik, David Bonsall, George Macintyre, Amy Trebes, Mariateresa de Cesare, Catrin Moore, Alex Mobbs, Anita Justice, Robert Shaw, Monique Andersson, Timothy Peto, Emma Wise, Nathan Moore, Jessica Lynch, Nick Cortes, Matilde Mori, Stephen Kidd, David Buck, John Todd, Christophe Fraser |
| EPI_ISL_479620, EPI_ISL_479621, EPI_ISL_479622, EPI_ISL_479623, EPI_ISL_479624 | Molecular diagnostic laboratory of Federal Budget Institution of Science "Central Research Institute of Epidemiology" of The Federal Service on Customers' Rights Protection and Human Well-being Surveillance | Group of Genomics and Postgenomic Technologies of Central Research Institute of Epidemilology | Speranskaya AS, Kaptelova VV,Valdokhina AV, Bulanenko VP, Samoilov AE, Korneenko EV, Sizova TV, Tivanova EV, Shipulina OY, Akimkin VG |
| EPI_ISL_480426, EPI_ISL_480427 | National Institute of Laboratory Medicine and Referral Center | Bangladesh Council of Scientific and Industrial Research | Shahina Akter, Abu Sayeed Mohammad Mahmud, Mohammad Samir Uzzaman, Eshrar Osman, Md. Ahasan Habib, Tanjina Akhter Banu, Md. Murshed Hasan Sarkar, Barna Goswami, Iffat Jahan, Md. Saddam Hossain, Tasnim Nafisa, Md. Maruf Ahmed Molla, Mahmuda Yeasmin, Asish Kumar Ghosh, Shahjahan Siddike, A. K. M. Shamsuzzaman, Sheikh Md. Selim Al Din, Utpal Chandra Ray, Salek Ahmed Sajib, Md. Salim Khan |
| EPI_ISL_480439, EPI_ISL_480440 | National Institute of Laboratory Medicine and Referral Center | Bangladesh Council of Scientific and Industrial Research | Tanjina Akhter Banu, Abu Sayeed Mohammad Mahmud, Mohammad Samir Uzzaman, Eshrar Osman, Md. Ahasan Habib, Shahina Akter, Md. Murshed Hasan Sarkar, Barna Goswami, Iffat Jahan, Md. Saddam Hossain, Tasnim Nafisa, Md. Maruf Ahmed Molla, Mahmuda Yeasmin, Asish Kumar Ghosh, Shahjahan Siddike, A. K. M. Shamsuzzaman, Sheikh Md. Selim Al Din, Utpal Chandra Ray, Salek Ahmed Sajib, Md. Salim Khan |
| EPI_ISL_480601, EPI_ISL_480602 | National Health Laboratory, Timor-Leste | MDU-PHL | Soares da Silva, E., Dolores de Jesus da Costa, M., Salles de Sousa, A., Jayanti Pereira Tilman, A., Antonia da Costa, E., Barreto, I., Marr, I., Wapling, J., Francis, J., Ximenes, J., Canisia, D., Freeman, K., Dakh, F., Douglas, N., Baird, R., Caly, L., Seemann, T., Sait, M., Schultz, M., Sherry, N. |
| EPI_ISL_481110, EPI_ISL_481111, EPI_ISL_481112, EPI_ISL_481113, EPI_ISL_481114, EPI_ISL_481115, EPI_ISL_481116, EPI_ISL_481117, EPI_ISL_481118, EPI_ISL_481119, EPI_ISL_481120, EPI_ISL_481121, EPI_ISL_481122, EPI_ISL_481123, EPI_ISL_481124, EPI_ISL_481125, EPI_ISL_481126, EPI_ISL_481127, EPI_ISL_481128, EPI_ISL_481129, EPI_ISL_481130, EPI_ISL_481131, EPI_ISL_481132, EPI_ISL_481133 | | | |
| see above | Immunogenomics lab, Institute of Life Sciences, Bhubaneswar | Immunogenomics lab, Institute of Life Sciences, Bhubaneswar | Sunil Raghav, Arup Ghosh, Deepika Singh, Ankita Datey, P. Sushree Shyamli, Bharati Singh, Neha Singh, Atimukta Jha, Viplov K. Biswas, Swati Madhulika, Manasi Priyadarshini, Sneha Dutta, Auromira Khuntia, Rupesh Dash, Soma Chattopadhyay, Ghulam Hussain Syed, Shanti Senapati, Tushar K. Beuria, Rajeeb Swain, Punit Prasad, Orissa COVID-19 Study Group, DBT's PAN-INDIA 1000 SARS-CoV2 RNA genome sequencing consortium, Ajay Parida |
| EPI_ISL_481134, EPI_ISL_481135, EPI_ISL_481136, EPI_ISL_481137, EPI_ISL_481138, EPI_ISL_481139, EPI_ISL_481140, EPI_ISL_481141, EPI_ISL_481142, EPI_ISL_481143, EPI_ISL_481144, EPI_ISL_481145, EPI_ISL_481146, EPI_ISL_481147, EPI_ISL_481148, EPI_ISL_481149, EPI_ISL_481150, EPI_ISL_481151, EPI_ISL_481152, EPI_ISL_481153, EPI_ISL_481154, EPI_ISL_481155, EPI_ISL_481156, EPI_ISL_481157 | | | |
| see above | Immunogenomics lab, Institute of Life Sciences, Bhubaneswar | Immunogenomics lab, Institute of Life Sciences, Bhubaneswar | Sunil Raghav, Arup Ghosh, Ankita Datey, P. Sushree Shyamli, Bharati Singh, Neha Singh, Deepika Singh, Atimukta Jha, Viplov K. Biswas, Swati Madhulika, Manasi Priyadarshini, Aditi Chatterjee, Rahul Das, Soumyajit Ghosh, Rupesh Dash, Soma Chattopadhyay, Ghulam Hussain Syed, Shanti Senapati, Tushar K. Beuria, Rajeeb Swain, Punit Prasad, Amol Ratnakar Suryawanshi, Dileep Vasudeva, Orissa COVID-19 Study Group, DBT's PAN-INDIA 1000 SARS-CoV2 RNA genome sequencing consortium, Ajay Parida |
| EPI_ISL_481158, EPI_ISL_481159, EPI_ISL_481160, EPI_ISL_481161, EPI_ISL_481162, EPI_ISL_481163, EPI_ISL_481164, EPI_ISL_481165, EPI_ISL_481166, EPI_ISL_481167, EPI_ISL_481168, EPI_ISL_481169, EPI_ISL_481170, EPI_ISL_481171, EPI_ISL_481172, EPI_ISL_481173, EPI_ISL_481174, EPI_ISL_481175, EPI_ISL_481176, EPI_ISL_481177, EPI_ISL_481178, EPI_ISL_481179, EPI_ISL_481180, EPI_ISL_481181 | | | |
| see above | Immunogenomics lab, Institute of Life Sciences, Bhubaneswar | Immunogenomics lab, Institute of Life Sciences, Bhubaneswar | Sunil Raghav, Arup Ghosh, P. Sushree Shyamli, Bharati Singh, Neha Singh, Ankita Datey, Deepika Singh, Atimukta Jha, Viplov K. Biswas, Swati Madhulika, Manasi Priyadarshini, Tsheten Sheroa, Auromira Khuntia, Rupesh Dash, Soma Chattopadhyay, Ghulam Hussain Syed, Shanti Senapati, Tushar K. Beuria, Rajeeb Swain, Punit Prasad, Amol Ratnakar Suryawanshi, Dileep Vasudevan, Orissa COVID-19 Study Group, DBT's PAN-INDIA 1000 SARS-CoV2 RNA genome sequencing consortium, Ajay Parida |
| EPI_ISL_481182, EPI_ISL_481183, EPI_ISL_481184, EPI_ISL_481185, EPI_ISL_481186, EPI_ISL_481187, EPI_ISL_481188, EPI_ISL_481189, EPI_ISL_481190, EPI_ISL_481191, EPI_ISL_481192, EPI_ISL_481193, EPI_ISL_481194, EPI_ISL_481195, EPI_ISL_481196, EPI_ISL_481197, EPI_ISL_481198, EPI_ISL_481199, EPI_ISL_481200, EPI_ISL_481201, EPI_ISL_481202, EPI_ISL_481203, EPI_ISL_481204, EPI_ISL_481205 | | | |
| see above | Immunogenomics lab, Institute of Life Sciences, Bhubaneswar | Immunogenomics lab, Institute of Life Sciences, Bhubaneswar | Sunil Raghav, Arup Ghosh, Atimukta Jha, Viplov K. Biswas, Swati Madhulika, Manasi Priyadarshini, Ajit Singh, Sivaram Krishna, Naga Jogayya Kothakota, Rupesh Dash, Soma Chattopadhyay, Ghulam Hussain Syed, Shanti Senapati, Tushar K. Beuria, Rajeeb Swain, Punit Prasad, Amol Ratnakar Suryawanshi, Dileep Vasudevan, Orissa COVID-19 Study Group, DBT's PAN-INDIA 1000 SARS-CoV2 RNA genome sequencing consortium, Ajay Parida |
| EPI_ISL_481513, EPI_ISL_481514, EPI_ISL_481515, EPI_ISL_481516, EPI_ISL_481517, EPI_ISL_481518, EPI_ISL_481519, EPI_ISL_481520, EPI_ISL_481521, EPI_ISL_481522, EPI_ISL_481523, EPI_ISL_481524, EPI_ISL_481525, EPI_ISL_481526, EPI_ISL_481527, EPI_ISL_481528, EPI_ISL_481529, EPI_ISL_481530, EPI_ISL_481531, EPI_ISL_481532, EPI_ISL_481533, EPI_ISL_481534, EPI_ISL_481535, EPI_ISL_481536, EPI_ISL_481537, EPI_ISL_481538, EPI_ISL_481539, EPI_ISL_481540, EPI_ISL_481541, EPI_ISL_481542, EPI_ISL_481543, EPI_ISL_481544, EPI_ISL_481545, EPI_ISL_481546, EPI_ISL_481547, EPI_ISL_481548, EPI_ISL_481549, EPI_ISL_481550, EPI_ISL_481551, EPI_ISL_481552, EPI_ISL_481553, EPI_ISL_481554, EPI_ISL_481555, EPI_ISL_481556, EPI_ISL_481557, EPI_ISL_481558, EPI_ISL_481559, EPI_ISL_481560, EPI_ISL_481561, EPI_ISL_481562, EPI_ISL_481563, EPI_ISL_481564, EPI_ISL_481565, EPI_ISL_481566, EPI_ISL_481567, EPI_ISL_481568, EPI_ISL_481569, EPI_ISL_481570, EPI_ISL_481571, EPI_ISL_481572, EPI_ISL_481573, EPI_ISL_481574, EPI_ISL_481575, EPI_ISL_481576, EPI_ISL_481577, EPI_ISL_481578, EPI_ISL_481579, EPI_ISL_481580, EPI_ISL_481581, EPI_ISL_481582, EPI_ISL_481583, EPI_ISL_481584, EPI_ISL_481585, EPI_ISL_481586, EPI_ISL_481587, EPI_ISL_481588, EPI_ISL_481589, EPI_ISL_481590, EPI_ISL_481591, EPI_ISL_481592, EPI_ISL_481593, EPI_ISL_481594, EPI_ISL_481595, EPI_ISL_481596, EPI_ISL_481597, EPI_ISL_481598, EPI_ISL_481599, EPI_ISL_481600, EPI_ISL_481601, EPI_ISL_481602, EPI_ISL_481603, EPI_ISL_481604, EPI_ISL_481605, EPI_ISL_481606, EPI_ISL_481607, EPI_ISL_481608, EPI_ISL_481609, EPI_ISL_481610, EPI_ISL_481611, EPI_ISL_481612, EPI_ISL_481613, EPI_ISL_481614, EPI_ISL_481615, EPI_ISL_481616, EPI_ISL_481617, EPI_ISL_481618, EPI_ISL_481619, EPI_ISL_481620, EPI_ISL_481621, EPI_ISL_481622, EPI_ISL_481623, EPI_ISL_481624, EPI_ISL_481625, EPI_ISL_481626, EPI_ISL_481627, EPI_ISL_481628, EPI_ISL_481629, EPI_ISL_481630, EPI_ISL_481631, EPI_ISL_481632, EPI_ISL_481633, EPI_ISL_481634, EPI_ISL_481635, EPI_ISL_481636, EPI_ISL_481637, EPI_ISL_481638, EPI_ISL_481639, EPI_ISL_481640, EPI_ISL_481641, EPI_ISL_481642, EPI_ISL_481643, EPI_ISL_481644, EPI_ISL_481645, EPI_ISL_481646, EPI_ISL_481647, EPI_ISL_481648, EPI_ISL_481649, EPI_ISL_481650, EPI_ISL_481651, EPI_ISL_481652, EPI_ISL_481653, EPI_ISL_481654, EPI_ISL_481655, EPI_ISL_481656, EPI_ISL_481657, EPI_ISL_481658, EPI_ISL_481659, EPI_ISL_481660, EPI_ISL_481661, EPI_ISL_481662, EPI_ISL_481663, EPI_ISL_481664, EPI_ISL_481665, EPI_ISL_481666, EPI_ISL_481667, EPI_ISL_481668, EPI_ISL_481669, EPI_ISL_481670, EPI_ISL_481671, EPI_ISL_481672, EPI_ISL_481673, EPI_ISL_481674, EPI_ISL_481675, EPI_ISL_481676, EPI_ISL_481677, EPI_ISL_481678, EPI_ISL_481679, EPI_ISL_481680, EPI_ISL_481681, EPI_ISL_481682, EPI_ISL_481683, EPI_ISL_481684, EPI_ISL_481685, EPI_ISL_481686, EPI_ISL_481687, EPI_ISL_481688, EPI_ISL_481689, EPI_ISL_481690, EPI_ISL_481691, EPI_ISL_481692, EPI_ISL_481693, EPI_ISL_481694, EPI_ISL_481695, EPI_ISL_481696, EPI_ISL_481697, EPI_ISL_481698, EPI_ISL_481699, EPI_ISL_481700, EPI_ISL_481701, EPI_ISL_481702, EPI_ISL_481703, EPI_ISL_481704, EPI_ISL_481705, EPI_ISL_481706, EPI_ISL_481707, EPI_ISL_481708, EPI_ISL_481709, EPI_ISL_481710, EPI_ISL_481711, EPI_ISL_481712, EPI_ISL_481713, EPI_ISL_481714, EPI_ISL_481715, EPI_ISL_481717, EPI_ISL_481718, EPI_ISL_481719, EPI_ISL_481720, EPI_ISL_481721, EPI_ISL_481722, EPI_ISL_481723, EPI_ISL_481724, EPI_ISL_481725, EPI_ISL_481726, EPI_ISL_481727, EPI_ISL_481728, EPI_ISL_481729, EPI_ISL_481730, EPI_ISL_481731, EPI_ISL_481732, EPI_ISL_481733, EPI_ISL_481734, EPI_ISL_481735, EPI_ISL_481736, EPI_ISL_481737, EPI_ISL_481738, EPI_ISL_481739, EPI_ISL_481740 | | | |
| see above | Department of Virology and Immunology, University of Helsinki and Helsinki University Hospital, Huslab Finland | Department of Virology, Faculty of Medicine, University of Helsinki, Helsinki, Finland | Teemu Smura, Hannimari Kallio-Kokko, Jenni Virtanen, Maija Suvanto, Sari Hannula, Harri Kangas, Pekka Ellonen, Olli Vapalahti |
| EPI_ISL_481763, EPI_ISL_481764, EPI_ISL_481765, EPI_ISL_481766, EPI_ISL_481767, EPI_ISL_481768, EPI_ISL_481769, EPI_ISL_481770, EPI_ISL_481771, EPI_ISL_481772, EPI_ISL_481773, EPI_ISL_481774, EPI_ISL_481775, EPI_ISL_481776, EPI_ISL_481777, EPI_ISL_481778, EPI_ISL_481779, EPI_ISL_481780, EPI_ISL_481781, EPI_ISL_481782, EPI_ISL_481783, EPI_ISL_481784, EPI_ISL_481785, EPI_ISL_481786, EPI_ISL_481787, EPI_ISL_481788, EPI_ISL_481789, EPI_ISL_481790, EPI_ISL_481791, EPI_ISL_481792, EPI_ISL_481793, EPI_ISL_481794, EPI_ISL_481795, EPI_ISL_481796, EPI_ISL_481797, EPI_ISL_481798, EPI_ISL_481799, EPI_ISL_481800, EPI_ISL_481801, EPI_ISL_481802, EPI_ISL_481803, EPI_ISL_481804, EPI_ISL_481805, EPI_ISL_481806, EPI_ISL_481807, EPI_ISL_481808, EPI_ISL_481809, EPI_ISL_481810, EPI_ISL_481811, EPI_ISL_481812, EPI_ISL_481813, EPI_ISL_481814, EPI_ISL_481815, EPI_ISL_481816, EPI_ISL_481817, EPI_ISL_481818, EPI_ISL_481819, EPI_ISL_481820, EPI_ISL_481821, EPI_ISL_481822, EPI_ISL_481823, EPI_ISL_481824, EPI_ISL_481825, EPI_ISL_481826, EPI_ISL_481827, EPI_ISL_481828, EPI_ISL_481829, EPI_ISL_481830, EPI_ISL_481831, EPI_ISL_481832, EPI_ISL_481833, EPI_ISL_481834, EPI_ISL_481835, EPI_ISL_481836, EPI_ISL_481837, EPI_ISL_481838, EPI_ISL_481839, EPI_ISL_481840, EPI_ISL_481841, EPI_ISL_481842, EPI_ISL_481843, EPI_ISL_481844, EPI_ISL_481845, EPI_ISL_481846, EPI_ISL_481847, EPI_ISL_481848, EPI_ISL_481849, EPI_ISL_481850, EPI_ISL_481851, EPI_ISL_481852, EPI_ISL_481853, EPI_ISL_481854, EPI_ISL_481855, EPI_ISL_481856, EPI_ISL_481857, EPI_ISL_481858, EPI_ISL_481859, EPI_ISL_481860, EPI_ISL_481861, EPI_ISL_481862, EPI_ISL_481863, EPI_ISL_481864, EPI_ISL_481865, EPI_ISL_481866, EPI_ISL_481867, EPI_ISL_481868, EPI_ISL_481869, EPI_ISL_481870, EPI_ISL_481871, EPI_ISL_481872, EPI_ISL_481873, EPI_ISL_481874, EPI_ISL_481875, EPI_ISL_481876, EPI_ISL_481877, EPI_ISL_481878, EPI_ISL_481879, EPI_ISL_481880, EPI_ISL_481881, EPI_ISL_481882, EPI_ISL_481883, EPI_ISL_481884, EPI_ISL_481885, EPI_ISL_481886, EPI_ISL_481887, EPI_ISL_481888, EPI_ISL_481889, EPI_ISL_481890, EPI_ISL_481891, EPI_ISL_481892, EPI_ISL_481893, EPI_ISL_481894, EPI_ISL_481895, EPI_ISL_481896, EPI_ISL_481897, EPI_ISL_481898, EPI_ISL_481899, EPI_ISL_481900, EPI_ISL_481901, EPI_ISL_481902, EPI_ISL_481903, EPI_ISL_481904, EPI_ISL_481905, EPI_ISL_481906, EPI_ISL_481907, EPI_ISL_481908, EPI_ISL_481909, EPI_ISL_481910, EPI_ISL_481911, EPI_ISL_481912, EPI_ISL_481913, EPI_ISL_481914, EPI_ISL_481915, EPI_ISL_481916, EPI_ISL_481917, EPI_ISL_481918, EPI_ISL_481919, EPI_ISL_481920, EPI_ISL_481921, EPI_ISL_481922, EPI_ISL_481923, EPI_ISL_481924, EPI_ISL_481925, EPI_ISL_481926, EPI_ISL_481927, EPI_ISL_481928, EPI_ISL_481929, EPI_ISL_481930, EPI_ISL_481931, EPI_ISL_481932, EPI_ISL_481933, EPI_ISL_481934, EPI_ISL_481935, EPI_ISL_481936, EPI_ISL_481937, EPI_ISL_481938, EPI_ISL_481939, EPI_ISL_481940, EPI_ISL_481941, EPI_ISL_481942, EPI_ISL_481943, EPI_ISL_481944, EPI_ISL_481945, EPI_ISL_481946, EPI_ISL_481947, EPI_ISL_481948, EPI_ISL_481949, EPI_ISL_481950, EPI_ISL_481951, EPI_ISL_481952, EPI_ISL_481953, EPI_ISL_481954, EPI_ISL_481955, EPI_ISL_481956, EPI_ISL_481957, EPI_ISL_481958, EPI_ISL_481959, EPI_ISL_481960, EPI_ISL_481961, EPI_ISL_481962, EPI_ISL_481963, EPI_ISL_481964, EPI_ISL_481965, EPI_ISL_481966, EPI_ISL_481967, EPI_ISL_481968, EPI_ISL_481969, EPI_ISL_481970, EPI_ISL_481971, EPI_ISL_481972, EPI_ISL_481973, EPI_ISL_481974, EPI_ISL_481975, EPI_ISL_481976, EPI_ISL_481977, EPI_ISL_481978, EPI_ISL_481979, EPI_ISL_481980, EPI_ISL_481981, EPI_ISL_481982, EPI_ISL_481983, EPI_ISL_481984, EPI_ISL_481985, EPI_ISL_481986, EPI_ISL_481987, EPI_ISL_481988, EPI_ISL_481989, EPI_ISL_481990, EPI_ISL_481991, EPI_ISL_481992, EPI_ISL_481993, EPI_ISL_481994, EPI_ISL_481995, EPI_ISL_481996, EPI_ISL_481997, EPI_ISL_481998, EPI_ISL_481999, EPI_ISL_482000, EPI_ISL_482001, EPI_ISL_482002, EPI_ISL_482003, EPI_ISL_482004, EPI_ISL_482005, EPI_ISL_482006, EPI_ISL_482007, EPI_ISL_482008, EPI_ISL_482009, EPI_ISL_482010, EPI_ISL_482011, EPI_ISL_482012, EPI_ISL_482013, EPI_ISL_482014, EPI_ISL_482015, EPI_ISL_482016, EPI_ISL_482017, EPI_ISL_482018, EPI_ISL_482019, EPI_ISL_482020, EPI_ISL_482021, EPI_ISL_482022, EPI_ISL_482023, EPI_ISL_482024, EPI_ISL_482025, EPI_ISL_482026, EPI_ISL_482027, EPI_ISL_482029, EPI_ISL_482030, EPI_ISL_482031 | | | |
| see above | PHE South West Regional Laboratory, National Infection Service | Wellcome Sanger Institute for the COVID-19 Genomics UK (COG-UK) consortium | Stephanie Hutchings, Hannah Pymont, Dr Peter Muir, Barry Vipond, Rich Hopes; and Alex Alderton, Roberto Amato, Sonia Goncalves, Ewan Harrison, David K. Jackson, Ian Johnston, Dominic Kwiatkowski, Cordelia Langford, John Sillitoe on behalf of the Wellcome Sanger Institute COVID-19 Surveillance Team (http://www.sanger.ac.uk/covid-team) |
| EPI_ISL_482745 | Medical Microbiology, Leiden University Medical Center | Medical Microbiology, Leiden University Medical Center | Snijder,E.J., Ogando,N.S., Zevenhoven,J.C., Dalebout,T.J., de Vries,J.C. and Sidorov,I. |
| EPI_ISL_482746 | Medical Microbiology, Leiden University Medical Center | Medical Microbiology, Leiden University Medical Center | Snijder,E.J., Ogando,N.S., Zevenhoven,J.C., Dalebout,T.J., de Vries,J.J. and Sidorov,I. |
| EPI_ISL_483065 | Centro de Desenvolvimento Tecnologico em Saude, Fundacao Oswaldo Cruz | Centro de Desenvolvimento Tecnologico em Saude, Fundacao Oswaldo Cruz | Souza,T.M., Fintelman-Rodrigues,N., De Paula,A.D., Tschoeke,D., Barroso,S.P., Gregorio,M.L., Oliveira,J.S., Saraiva,F.B., Ferreira,M.A., Sacramento,C.Q. |
| EPI_ISL_483622, EPI_ISL_483623 | National Institute of Laboratory Medicine and Referral Center | Genomic Research Lab, BCSIR | Tasnim Nafisa, Abu Sayeed Mohammad Mahmud, Mohammad Samir Uzzaman, Eshrar Osman, Md. Ahasan Habib, Shahina Akter, Tanjina Akhter Banu, |

| | | | |
|---|---|---|---|
| | | | Md. Murshed Hasan Sarkar, Barna Goswami, Iffat Jahan, Md. Saddam Hossain, Md. Maruf Ahmed Molla, Mahmuda Yeasmin, Asish Kumar Ghosh, A. K. M. Shamsuzzaman, Sheikh Md. Selim Al Din, Utpal Chandra Ray, Salek Ahmed Sajib, Md. Salim Khan |
| EPI_ISL_483634 | National Institute of Laboratory Medicine and Referral Center | Genomic Research Lab, BCSIR | Shahina Akter, Abu Sayeed Mohammad Mahmud, Mohammad Samir Uzzaman, Eshrar Osman, Md. Ahasan Habib, Tanjina Akhter Banu, Md. Murshed Hasan Sarkar, Barna Goswami, Iffat Jahan, Md. Saddam Hossain, Tasnim Nafisa, Md. Maruf Ahmed Molla, Mahmuda Yeasmin, Asish Kumar Ghosh, A. K. M. Shamsuzzaman, Sheikh Md. Selim Al Din, Utpal Chandra Ray, Salek Ahmed Sajib, Md. Salim Khan |
| EPI_ISL_483635, EPI_ISL_483636 | National Institute of Laboratory Medicine and Referral Center | Genomic Research Lab, BCSIR | Tanjina Akhter Banu, Abu Sayeed Mohammad Mahmud, Mohammad Samir Uzzaman, Eshrar Osman, Md. Ahasan Habib, Shahina Akter, Md. Murshed Hasan Sarkar, Barna Goswami, Iffat Jahan, Md. Saddam Hossain, Tasnim Nafisa, Md. Maruf Ahmed Molla, Mahmuda Yeasmin, Asish Kumar Ghosh, A. K. M. Shamsuzzaman, Sheikh Md. Selim Al Din, Utpal Chandra Ray, Salek Ahmed Sajib, Md. Salim Khan |
| EPI_ISL_483859, EPI_ISL_483877 | Department of Microbiology, Government Medical College, Surat | Gujarat Biotechnology Research Centre | Summaiya Mullan, Amit gamit, Apurvasinh Puvar, Janvi Raval, Zarna Patel, Monika Gandhi, Pinal Trivedi, Maharshi Pandya, Nidhi Patel, Nitin Savaliya, Raghawendra Kumar, Dinesh Kumar, Zuber Saiyed, Komal Patel, Labdhi Pandya, Afzal Ansari, Nikha Trivedi, Naresh Chauhan, R D Dixit, A M Kadri, Harsh Bakshi, Chaitanya Joshi, Madhvi Joshi |
| EPI_ISL_485399 | Institute of Human Genetics, Polish Academy of Sciences | Institute of Human Genetics, Polish Academy of Sciences | Szymon Hryhorowicz, Adam Ustaszewski, Marta Kaczmarek-Ry, Emilia Lis, Ewa Zitkiewicz, Micha Witt, Andrzej Pawski |
| EPI_ISL_485400 | Institute of Human Genetics, Polish Academy of Sciences | Institute of Human Genetics, Polish Academy of Sciences, | Szymon Hryhorowicz, Adam Ustaszewski, Marta Kaczmarek-Ry, Emilia Lis, Ewa Zitkiewicz, Micha Witt, Andrzej Pawski |
| EPI_ISL_486815, EPI_ISL_486816, EPI_ISL_486817, EPI_ISL_486818, EPI_ISL_486819, EPI_ISL_486820, EPI_ISL_486821, EPI_ISL_486822, EPI_ISL_486823, EPI_ISL_486824, EPI_ISL_486825, EPI_ISL_486826, EPI_ISL_486827, EPI_ISL_486828, EPI_ISL_486829 | | | |
| see above | Molecular diagnostic laboratory of Federal Budget Institution of Science "Central Research Institute of Epidemiology" of The Federal Service on Customers' Rights Protection and Human Well-being Surveillance | Group of Genomics and Postgenomic Technologies of Central Research Institute of Epidemiology | Speranskaya AS, Kaptelova VV, Valdokhina AV, Bulanenko VP, Samoilov AE, Korneenko EV, Tivanova EV, Shipulina OY, Akimkin VG |
| EPI_ISL_487363, EPI_ISL_487364 | National Institute of Laboratory Medicine and Referral Center | Genomic Research Lab, BCSIR | Tanjina Akhter Banu, Abu Sayeed Mohammad Mahmud, Mohammad Samir Uzzaman, Eshrar Osman, Md. Ahasan Habib, Shahina Akter, Md. Murshed Hasan Sarkar, Barna Goswami, Iffat Jahan, Md. Saddam Hossain, Tasnim Nafisa, Md. Maruf Ahmed Molla, Mahmuda Yeasmin, Asish Kumar Ghosh, A. K. M. Shamsuzzaman, Sheikh Md. Selim Al Din, Utpal Chandra Ray, Salek Ahmed Sajib, Md. Salim Khan |
| EPI_ISL_487367 | National Institute of Laboratory Medicine and Referral Center | Genomic Research Lab, BCSIR | Shahina Akter, Abu Sayeed Mohammad Mahmud, Mohammad Samir Uzzaman, Eshrar Osman, Md. Ahasan Habib, Tanjina Akhter Banu, Md. Murshed Hasan Sarkar, Barna Goswami, Iffat Jahan, Md. Saddam Hossain, Tasnim Nafisa, Md. Maruf Ahmed Molla, Mahmuda Yeasmin, Asish Kumar Ghosh, A. K. M. Shamsuzzaman, Sheikh Md. Selim Al Din, Utpal Chandra Ray, Salek Ahmed Sajib, Md. Salim Khan |
| EPI_ISL_487524, EPI_ISL_487526, EPI_ISL_487527, EPI_ISL_487528, EPI_ISL_487530, EPI_ISL_487531, EPI_ISL_487532, EPI_ISL_487533, EPI_ISL_487535, EPI_ISL_487536, EPI_ISL_487538, EPI_ISL_487539, EPI_ISL_487541, EPI_ISL_487542, EPI_ISL_487543, EPI_ISL_487544, EPI_ISL_487545, EPI_ISL_487547, EPI_ISL_487548, EPI_ISL_487549, EPI_ISL_487551, EPI_ISL_487552, EPI_ISL_487553, EPI_ISL_487554, EPI_ISL_487555, EPI_ISL_487560, EPI_ISL_487561, EPI_ISL_487563, EPI_ISL_487565, EPI_ISL_487566, EPI_ISL_487567, EPI_ISL_487570, EPI_ISL_487571, EPI_ISL_487573, EPI_ISL_487574, EPI_ISL_487575, EPI_ISL_487576, EPI_ISL_487577, EPI_ISL_487579, EPI_ISL_487580, EPI_ISL_487583, EPI_ISL_487586, EPI_ISL_487587, EPI_ISL_487588, EPI_ISL_487590, EPI_ISL_487591, EPI_ISL_487592, EPI_ISL_487594, EPI_ISL_487596, EPI_ISL_487598, EPI_ISL_487599, EPI_ISL_487601, EPI_ISL_487602, EPI_ISL_487603, EPI_ISL_487604, EPI_ISL_487605, EPI_ISL_487608, EPI_ISL_487610, EPI_ISL_487611, EPI_ISL_487612, EPI_ISL_487613, EPI_ISL_487617, EPI_ISL_487618, EPI_ISL_487619, EPI_ISL_487620, EPI_ISL_487622, EPI_ISL_487623, EPI_ISL_487624, EPI_ISL_487629, EPI_ISL_487634, EPI_ISL_488187, EPI_ISL_488188, EPI_ISL_488189, EPI_ISL_488190, EPI_ISL_488191, EPI_ISL_488192, EPI_ISL_488193, EPI_ISL_488194, EPI_ISL_488195, EPI_ISL_488196, EPI_ISL_488197, EPI_ISL_488198, EPI_ISL_488199, EPI_ISL_488200, EPI_ISL_488201, EPI_ISL_488202, EPI_ISL_488203, EPI_ISL_488204, EPI_ISL_488205, EPI_ISL_488206, EPI_ISL_488207, EPI_ISL_488208, EPI_ISL_488209, EPI_ISL_488210, EPI_ISL_488211, EPI_ISL_488212, EPI_ISL_488213, EPI_ISL_488214, EPI_ISL_488215, EPI_ISL_488216, EPI_ISL_488217, EPI_ISL_488218, EPI_ISL_488219, EPI_ISL_488220, EPI_ISL_488221, EPI_ISL_488222, EPI_ISL_488223, EPI_ISL_488224, EPI_ISL_488225, EPI_ISL_488226, EPI_ISL_488227, EPI_ISL_488228, EPI_ISL_488229, EPI_ISL_488230, EPI_ISL_488231, EPI_ISL_488232, EPI_ISL_488233, EPI_ISL_488234, EPI_ISL_488235, EPI_ISL_488236, EPI_ISL_488237, EPI_ISL_488238, EPI_ISL_488239, EPI_ISL_488240, EPI_ISL_488241, EPI_ISL_488242, EPI_ISL_488243, EPI_ISL_488244, EPI_ISL_488245, EPI_ISL_488246, EPI_ISL_488247, EPI_ISL_488248, EPI_ISL_488249, EPI_ISL_488250, EPI_ISL_488251, EPI_ISL_488252, EPI_ISL_488253, EPI_ISL_488254, EPI_ISL_488255, EPI_ISL_488256, EPI_ISL_488257, EPI_ISL_488258, EPI_ISL_488259, EPI_ISL_488260, EPI_ISL_488261, EPI_ISL_488262, EPI_ISL_488263, EPI_ISL_488264, EPI_ISL_488265, EPI_ISL_488266, EPI_ISL_488267, EPI_ISL_488268, EPI_ISL_488269, EPI_ISL_488270, EPI_ISL_488271, EPI_ISL_488272, EPI_ISL_488273, EPI_ISL_488274, EPI_ISL_488275, EPI_ISL_488276, EPI_ISL_488277, EPI_ISL_488278, EPI_ISL_488279, EPI_ISL_488280, EPI_ISL_488281, EPI_ISL_488282, EPI_ISL_488283, EPI_ISL_488284, EPI_ISL_488285, EPI_ISL_488286, EPI_ISL_488287, EPI_ISL_488288, EPI_ISL_488289, EPI_ISL_488290, EPI_ISL_488291, EPI_ISL_488292, EPI_ISL_488293, EPI_ISL_488294, EPI_ISL_488295, EPI_ISL_488296, EPI_ISL_488297, EPI_ISL_488298, EPI_ISL_488299, EPI_ISL_488300, EPI_ISL_488301, EPI_ISL_488302, EPI_ISL_488303, EPI_ISL_488304, EPI_ISL_488305, EPI_ISL_488306, EPI_ISL_488307, EPI_ISL_488308, EPI_ISL_488309, EPI_ISL_488310, EPI_ISL_488311, EPI_ISL_488312, EPI_ISL_488313, EPI_ISL_488314, EPI_ISL_488315, EPI_ISL_488316, EPI_ISL_488317, EPI_ISL_488318, EPI_ISL_488319, EPI_ISL_488320, EPI_ISL_488321, EPI_ISL_488322, EPI_ISL_488323, EPI_ISL_488324, EPI_ISL_488325, EPI_ISL_488326, EPI_ISL_488327, EPI_ISL_488328, EPI_ISL_488329, EPI_ISL_488330, EPI_ISL_488331, EPI_ISL_488332, EPI_ISL_488333, EPI_ISL_488334, EPI_ISL_488335, EPI_ISL_488336, EPI_ISL_488337, EPI_ISL_488338, EPI_ISL_488339, EPI_ISL_488340, EPI_ISL_488341, EPI_ISL_488343, EPI_ISL_488344, EPI_ISL_488345, EPI_ISL_488346, EPI_ISL_488347, EPI_ISL_488348, EPI_ISL_488349, EPI_ISL_488350, EPI_ISL_488351, EPI_ISL_488352, EPI_ISL_488353, EPI_ISL_488354, EPI_ISL_488355, EPI_ISL_488356, EPI_ISL_488357, EPI_ISL_488358, EPI_ISL_488359, EPI_ISL_488360, EPI_ISL_488361, EPI_ISL_488362, EPI_ISL_488363, EPI_ISL_488364, EPI_ISL_488365, EPI_ISL_488366, EPI_ISL_488367, EPI_ISL_488368, EPI_ISL_488369, EPI_ISL_488370, EPI_ISL_488371, EPI_ISL_488372, EPI_ISL_488373, EPI_ISL_488374, EPI_ISL_488375, EPI_ISL_488376, EPI_ISL_488377, EPI_ISL_488378, EPI_ISL_488379, EPI_ISL_488380, EPI_ISL_488381, EPI_ISL_488382, EPI_ISL_488383, EPI_ISL_488384, EPI_ISL_488385, EPI_ISL_488386, EPI_ISL_488387, EPI_ISL_488388, EPI_ISL_488389, EPI_ISL_488390, EPI_ISL_488391, EPI_ISL_488392, EPI_ISL_488393, EPI_ISL_488394, EPI_ISL_488395, EPI_ISL_488396, EPI_ISL_488397, EPI_ISL_488398, EPI_ISL_488399, EPI_ISL_488400, EPI_ISL_488401, EPI_ISL_488402, EPI_ISL_488403, EPI_ISL_488404, EPI_ISL_488405, EPI_ISL_488406, EPI_ISL_488407, EPI_ISL_488408, EPI_ISL_488410, EPI_ISL_488412, EPI_ISL_488413, EPI_ISL_488414, EPI_ISL_488415, EPI_ISL_488416, EPI_ISL_488417, EPI_ISL_488418, EPI_ISL_488419, EPI_ISL_488420, EPI_ISL_488421, EPI_ISL_488422, EPI_ISL_488423, EPI_ISL_488424, EPI_ISL_488425, EPI_ISL_488426, EPI_ISL_488427, EPI_ISL_488428, EPI_ISL_488429, EPI_ISL_488430, EPI_ISL_488431, EPI_ISL_488432, EPI_ISL_488433, EPI_ISL_488434, EPI_ISL_488435, EPI_ISL_488436, EPI_ISL_488437, EPI_ISL_488438, EPI_ISL_488439, EPI_ISL_488440, EPI_ISL_488441, EPI_ISL_488442, EPI_ISL_488443, EPI_ISL_488444, EPI_ISL_488445, EPI_ISL_488446, EPI_ISL_488447, EPI_ISL_488448, EPI_ISL_488449, EPI_ISL_488450, EPI_ISL_488451, EPI_ISL_488452, EPI_ISL_488453, EPI_ISL_488454, EPI_ISL_488455, EPI_ISL_488877, EPI_ISL_488878, EPI_ISL_489120 | | | |
| see above | PHE South West Regional Laboratory, National Infection Service | Wellcome Sanger Institute for the COVID-19 Genomics UK (COG-UK) consortium | Stephanie Hutchings, Hannah Pymont, Dr Peter Muir, Barry Vipond, Rich Hopes; and Alex Alderton, Roberto Amato, Sonia Goncalves, Ewan Harrison, David K. Jackson, Ian Johnston, Dominic Kwiatkowski, Cordelia Langford, John Sillitoe on behalf of the Wellcome Sanger Institute COVID-19 Surveillance Team (http://www.sanger.ac.uk/covid-team) |
| EPI_ISL_489992, EPI_ISL_489993, EPI_ISL_489994, EPI_ISL_490014, EPI_ISL_490015, EPI_ISL_490016, EPI_ISL_490047, EPI_ISL_490048, EPI_ISL_490089, EPI_ISL_490090, EPI_ISL_490091, EPI_ISL_490092, EPI_ISL_490093, EPI_ISL_490094, EPI_ISL_490095, EPI_ISL_490096, EPI_ISL_490097, EPI_ISL_490098, EPI_ISL_490099, EPI_ISL_490100, EPI_ISL_490101, EPI_ISL_490102, EPI_ISL_490103 | | | |
| see above | Institute for Medical Research, Infectious Disease Research Centre, National Institutes of Health, Ministry of Health Malaysia | Institute for Medical Research, Infectious Disease Research Centre, National Institutes of Health, Ministry of Health Malaysia | Suppiah J, Mohd-Zawawi Z, Kamel K, Kalyanasundram J, Thayan R |
| EPI_ISL_490114 | National Institute of Laboratory Medicine and Referral Center | Genomic Research Lab, BCSIR | Tanjina Akhter Banu, Abu Sayeed Mohammad Mahmud, Mohammad Samir Uzzaman, Eshrar Osman, Md. Ahasan Habib, Shahina Akter, Md. Murshed Hasan Sarkar, Barna Goswami, Iffat Jahan, Md. Saddam Hossain, Tasnim Nafisa, Md. Maruf Ahmed Molla, Mahmuda Yeasmin, Asish Kumar Ghosh, A. K. M. Shamsuzzaman, Sheikh Md. Selim Al Din, Utpal Chandra Ray, Salek Ahmed Sajib, Md. Salim Khan |
| EPI_ISL_491265, EPI_ISL_491266, EPI_ISL_491267, EPI_ISL_491268, EPI_ISL_491269, EPI_ISL_491270, EPI_ISL_491271, EPI_ISL_491272, EPI_ISL_491273, EPI_ISL_491274, EPI_ISL_491275, EPI_ISL_491276, EPI_ISL_491277, EPI_ISL_491278, EPI_ISL_491279, EPI_ISL_491280, EPI_ISL_491281, EPI_ISL_491282, EPI_ISL_491283, EPI_ISL_491284, EPI_ISL_491285, EPI_ISL_491286, EPI_ISL_491287, EPI_ISL_491288, EPI_ISL_491289, EPI_ISL_491290, EPI_ISL_491291, EPI_ISL_491292, EPI_ISL_491293, EPI_ISL_491294, EPI_ISL_491295, EPI_ISL_491296, EPI_ISL_491297 | | | |
| see above | Instituto Gulbenkian de Ciência | Instituto Gulbenkian de Ciência | Susana Ladeiro, João Costa, Cathy Paulino, Joao Sobral, Ricardo Leite |
| EPI_ISL_492185, EPI_ISL_492186, EPI_ISL_492187, EPI_ISL_492188, EPI_ISL_492195, EPI_ISL_492197, EPI_ISL_492202, EPI_ISL_492203, EPI_ISL_492211, EPI_ISL_492212, EPI_ISL_492213, EPI_ISL_492214, EPI_ISL_492215, EPI_ISL_492216, EPI_ISL_492217, EPI_ISL_492218, EPI_ISL_492219, EPI_ISL_492220, EPI_ISL_492221, EPI_ISL_492222, EPI_ISL_492223, EPI_ISL_492224, EPI_ISL_492225, EPI_ISL_492226, EPI_ISL_492227, EPI_ISL_492228, EPI_ISL_492229, EPI_ISL_492230, EPI_ISL_492231, EPI_ISL_492232, EPI_ISL_492233, EPI_ISL_492234, EPI_ISL_492235, EPI_ISL_492236, EPI_ISL_492237, EPI_ISL_492238, EPI_ISL_492239, EPI_ISL_492240, EPI_ISL_492241, EPI_ISL_492242, EPI_ISL_492243, EPI_ISL_492244, EPI_ISL_492245, EPI_ISL_492246, EPI_ISL_492247, EPI_ISL_492248, EPI_ISL_492249, EPI_ISL_492250, EPI_ISL_492251, EPI_ISL_492252, EPI_ISL_492253, EPI_ISL_492254, EPI_ISL_492255, EPI_ISL_492256, EPI_ISL_492257, EPI_ISL_492258, EPI_ISL_492259, EPI_ISL_492260, EPI_ISL_492261, EPI_ISL_492262, EPI_ISL_492263, EPI_ISL_492264, EPI_ISL_492265, EPI_ISL_492266, EPI_ISL_492267, EPI_ISL_492268, EPI_ISL_492269, EPI_ISL_492270, EPI_ISL_492271, EPI_ISL_492272, EPI_ISL_492273, EPI_ISL_492274, EPI_ISL_492275, EPI_ISL_492276, EPI_ISL_492277, EPI_ISL_492278, EPI_ISL_492279, EPI_ISL_492280, EPI_ISL_492281, EPI_ISL_492282, EPI_ISL_492283, EPI_ISL_492284, EPI_ISL_492285, EPI_ISL_492286, EPI_ISL_492287, EPI_ISL_492288, EPI_ISL_492289, EPI_ISL_492290, EPI_ISL_492291, EPI_ISL_492292, EPI_ISL_492293, EPI_ISL_492294, EPI_ISL_492295, EPI_ISL_492296, EPI_ISL_492297, EPI_ISL_492298, EPI_ISL_492299, EPI_ISL_492300, EPI_ISL_492301, EPI_ISL_492302, EPI_ISL_492303, EPI_ISL_492304, EPI_ISL_492305, EPI_ISL_492306, EPI_ISL_492307, EPI_ISL_492308, EPI_ISL_492309, EPI_ISL_492310, EPI_ISL_492311, EPI_ISL_492312, EPI_ISL_492313, EPI_ISL_492314, EPI_ISL_492315, EPI_ISL_492316, EPI_ISL_492317, EPI_ISL_492318, EPI_ISL_492319, EPI_ISL_492320, EPI_ISL_492321, EPI_ISL_492322, EPI_ISL_492323, EPI_ISL_492324, EPI_ISL_492325, EPI_ISL_492326, EPI_ISL_492327, EPI_ISL_492328, EPI_ISL_492329, EPI_ISL_492330, EPI_ISL_492331, EPI_ISL_492332, EPI_ISL_492333, EPI_ISL_492334, EPI_ISL_492335, EPI_ISL_492336, EPI_ISL_492337, EPI_ISL_492338, EPI_ISL_492339, EPI_ISL_492340, EPI_ISL_492341, EPI_ISL_492342, EPI_ISL_492343, EPI_ISL_492344, EPI_ISL_492345, EPI_ISL_492346, EPI_ISL_492347, EPI_ISL_492348, EPI_ISL_492349, EPI_ISL_492350, EPI_ISL_492351, EPI_ISL_492352, EPI_ISL_492353, EPI_ISL_492354, EPI_ISL_492355, EPI_ISL_492356, EPI_ISL_492357, EPI_ISL_492358, EPI_ISL_492359, EPI_ISL_492360, EPI_ISL_492361, EPI_ISL_492362, EPI_ISL_492363, EPI_ISL_492364, EPI_ISL_492365, EPI_ISL_492366, EPI_ISL_492367, EPI_ISL_492368, EPI_ISL_492369, EPI_ISL_492370, EPI_ISL_492371, EPI_ISL_492372, EPI_ISL_492373, EPI_ISL_492374, EPI_ISL_492375, EPI_ISL_492376, EPI_ISL_492377, EPI_ISL_492378, EPI_ISL_492379, EPI_ISL_492380, EPI_ISL_492381, EPI_ISL_492382, EPI_ISL_492383, EPI_ISL_492384, EPI_ISL_492385, EPI_ISL_492386, EPI_ISL_492387, EPI_ISL_492388, EPI_ISL_492389, EPI_ISL_492390, EPI_ISL_492391, EPI_ISL_492392, EPI_ISL_492393, EPI_ISL_492394, EPI_ISL_492395, EPI_ISL_492396, EPI_ISL_492397, EPI_ISL_492398, EPI_ISL_492399, EPI_ISL_492400, EPI_ISL_492401, EPI_ISL_492402, EPI_ISL_492403, EPI_ISL_492404, EPI_ISL_492405, EPI_ISL_492406, EPI_ISL_492407, EPI_ISL_492408, EPI_ISL_492409, EPI_ISL_492410, EPI_ISL_492411, EPI_ISL_492412, EPI_ISL_492413, EPI_ISL_492414, EPI_ISL_492415, EPI_ISL_492416, EPI_ISL_492417, EPI_ISL_492418, EPI_ISL_492419, EPI_ISL_492420, EPI_ISL_492421, EPI_ISL_492422, EPI_ISL_492423, EPI_ISL_492424, EPI_ISL_492425, EPI_ISL_492426, EPI_ISL_492427, EPI_ISL_492428, EPI_ISL_492429, EPI_ISL_492430, EPI_ISL_492431, EPI_ISL_492432, EPI_ISL_492433, EPI_ISL_492434, EPI_ISL_492435, EPI_ISL_492436, EPI_ISL_492437, EPI_ISL_492438, EPI_ISL_492439, EPI_ISL_492440, EPI_ISL_492441, EPI_ISL_492442, EPI_ISL_492443, EPI_ISL_492444, EPI_ISL_492445, EPI_ISL_492446, EPI_ISL_492447, EPI_ISL_492448, EPI_ISL_492449, EPI_ISL_492505, EPI_ISL_492506, EPI_ISL_492507, EPI_ISL_492508, EPI_ISL_492509, EPI_ISL_492510, EPI_ISL_492511, EPI_ISL_492512, EPI_ISL_492513, EPI_ISL_492515, EPI_ISL_492516, EPI_ISL_492517, EPI_ISL_492518, EPI_ISL_492519, EPI_ISL_492520, EPI_ISL_492521, EPI_ISL_492522, EPI_ISL_492523, EPI_ISL_492524, EPI_ISL_492525, EPI_ISL_492526, EPI_ISL_492527, EPI_ISL_492528, EPI_ISL_492529, EPI_ISL_492530, EPI_ISL_492531, EPI_ISL_492532, EPI_ISL_492533, EPI_ISL_492534, EPI_ISL_492535, EPI_ISL_492536, EPI_ISL_492537, EPI_ISL_492538, EPI_ISL_492539, EPI_ISL_492540, EPI_ISL_492541, EPI_ISL_492542, EPI_ISL_492543, EPI_ISL_492544, EPI_ISL_492545, EPI_ISL_492546, EPI_ISL_492547, EPI_ISL_492548, EPI_ISL_492549, EPI_ISL_492550, EPI_ISL_492551, EPI_ISL_492552, EPI_ISL_492553, EPI_ISL_492554, EPI_ISL_492555, EPI_ISL_492556, EPI_ISL_492557, EPI_ISL_492558, EPI_ISL_492559, EPI_ISL_492560, EPI_ISL_492561, EPI_ISL_492562, EPI_ISL_492563, EPI_ISL_492564, EPI_ISL_492565, EPI_ISL_492566, EPI_ISL_492567, EPI_ISL_492568, EPI_ISL_492569, EPI_ISL_492570, EPI_ISL_492571, EPI_ISL_492572, EPI_ISL_492573, EPI_ISL_492574, EPI_ISL_492575, EPI_ISL_492576, EPI_ISL_492577, EPI_ISL_492578, EPI_ISL_492579, EPI_ISL_492580, EPI_ISL_492581, EPI_ISL_492582, EPI_ISL_492583, EPI_ISL_492584, EPI_ISL_492585, EPI_ISL_492586, EPI_ISL_492587, EPI_ISL_492588, EPI_ISL_492589, EPI_ISL_492590, EPI_ISL_492591, EPI_ISL_492592, EPI_ISL_492593, EPI_ISL_492595, EPI_ISL_492597, EPI_ISL_492598, EPI_ISL_492599, EPI_ISL_492600, EPI_ISL_492601, EPI_ISL_492602, EPI_ISL_492603, EPI_ISL_492604, EPI_ISL_492605, EPI_ISL_492606, EPI_ISL_492607, EPI_ISL_492608, EPI_ISL_492609, EPI_ISL_492610, EPI_ISL_492611, EPI_ISL_492612, EPI_ISL_492613, EPI_ISL_492614, EPI_ISL_492615, EPI_ISL_492616, EPI_ISL_492617, EPI_ISL_492618, EPI_ISL_492619, EPI_ISL_492620, EPI_ISL_492621, EPI_ISL_492622, EPI_ISL_492623, EPI_ISL_492624, EPI_ISL_492625, EPI_ISL_492626, EPI_ISL_492627, EPI_ISL_492628, EPI_ISL_492629, EPI_ISL_492630, EPI_ISL_492631, EPI_ISL_492632, EPI_ISL_492633, EPI_ISL_492634, EPI_ISL_492635, EPI_ISL_492636, EPI_ISL_492637, EPI_ISL_492638, EPI_ISL_492639, EPI_ISL_492640, EPI_ISL_492641, EPI_ISL_492642, EPI_ISL_492643, EPI_ISL_492644, EPI_ISL_492645, EPI_ISL_492646, EPI_ISL_492647, EPI_ISL_492648, EPI_ISL_492649, EPI_ISL_492650, EPI_ISL_492651, EPI_ISL_492652, EPI_ISL_492653, EPI_ISL_492654, EPI_ISL_492655, EPI_ISL_492656, | | | |

| | | | |
|---|---|---|---|
| EPI_ISL_492657, EPI_ISL_492658, EPI_ISL_492659, EPI_ISL_492660, EPI_ISL_492661, EPI_ISL_492662, EPI_ISL_492663, EPI_ISL_492664, EPI_ISL_492665, EPI_ISL_492666, EPI_ISL_492667, EPI_ISL_492668, EPI_ISL_492669, EPI_ISL_492670, EPI_ISL_492671, EPI_ISL_492672, EPI_ISL_492673, EPI_ISL_492674, EPI_ISL_492675, EPI_ISL_492676, EPI_ISL_492677, EPI_ISL_492678, EPI_ISL_492679, EPI_ISL_492680, EPI_ISL_492681, EPI_ISL_492682, EPI_ISL_492683, EPI_ISL_492684, EPI_ISL_492685, EPI_ISL_492686, EPI_ISL_492687, EPI_ISL_492688, EPI_ISL_492689, EPI_ISL_492690, EPI_ISL_492691, EPI_ISL_492692, EPI_ISL_492693, EPI_ISL_492694, EPI_ISL_492695, EPI_ISL_492696, EPI_ISL_492697, EPI_ISL_492698, EPI_ISL_492699, EPI_ISL_492700, EPI_ISL_492701, EPI_ISL_492702, EPI_ISL_492703, EPI_ISL_492704, EPI_ISL_492705, EPI_ISL_492706, EPI_ISL_492707, EPI_ISL_492708, EPI_ISL_492709, EPI_ISL_492710, EPI_ISL_492711, EPI_ISL_492712, EPI_ISL_492713, EPI_ISL_492714, EPI_ISL_492715, EPI_ISL_492716, EPI_ISL_492717, EPI_ISL_492718, EPI_ISL_492719, EPI_ISL_492720, EPI_ISL_492721, EPI_ISL_492722, EPI_ISL_492723, EPI_ISL_492724, EPI_ISL_492725, EPI_ISL_492726, EPI_ISL_492727, EPI_ISL_492728, EPI_ISL_492729, EPI_ISL_492730, EPI_ISL_492731, EPI_ISL_492732, EPI_ISL_492733, EPI_ISL_492734, EPI_ISL_492735, EPI_ISL_492736, EPI_ISL_492737, EPI_ISL_492738, EPI_ISL_492739, EPI_ISL_492740, EPI_ISL_492741, EPI_ISL_492742, EPI_ISL_492743, EPI_ISL_492744, EPI_ISL_492745, EPI_ISL_492764, EPI_ISL_492765, EPI_ISL_492766, EPI_ISL_492767, EPI_ISL_492768, EPI_ISL_492769, EPI_ISL_492770, EPI_ISL_492771, EPI_ISL_492772, EPI_ISL_492773, EPI_ISL_492774, EPI_ISL_492775, EPI_ISL_492776, EPI_ISL_492777, EPI_ISL_492778, EPI_ISL_492779, EPI_ISL_492780, EPI_ISL_492781, EPI_ISL_492782, EPI_ISL_492783, EPI_ISL_492784, EPI_ISL_492785, EPI_ISL_492786, EPI_ISL_492787, EPI_ISL_492788, EPI_ISL_492789, EPI_ISL_492790, EPI_ISL_492791, EPI_ISL_492792, EPI_ISL_492793, EPI_ISL_492794, EPI_ISL_492795, EPI_ISL_492796, EPI_ISL_492797, EPI_ISL_492798, EPI_ISL_492799, EPI_ISL_492800, EPI_ISL_492801, EPI_ISL_492802, EPI_ISL_492803, EPI_ISL_492804, EPI_ISL_492805, EPI_ISL_492806, EPI_ISL_492807, EPI_ISL_492808, EPI_ISL_492809, EPI_ISL_492810, EPI_ISL_492811, EPI_ISL_492812, EPI_ISL_492813, EPI_ISL_492814, EPI_ISL_492815, EPI_ISL_492816, EPI_ISL_492817, EPI_ISL_492818, EPI_ISL_492819, EPI_ISL_492820, EPI_ISL_492821, EPI_ISL_492822, EPI_ISL_492823, EPI_ISL_492824, EPI_ISL_492825, EPI_ISL_492826, EPI_ISL_492827, EPI_ISL_492828, EPI_ISL_492829, EPI_ISL_492830, EPI_ISL_492831, EPI_ISL_492832, EPI_ISL_492833, EPI_ISL_492834, EPI_ISL_492835 | | | |
| see above | PHE South West Regional Laboratory, National Infection Service | Wellcome Sanger Institute for the COVID-19 Genomics UK (COG-UK) consortium | Stephanie Hutchings, Hannah Pymont, Dr Peter Muir, Barry Vipond, Rich Hopes; and Alex Alderton, Roberto Amato, Sonia Goncalves, Ewan Harrison, David K. Jackson, Ian Johnston, Dominic Kwiatkowski, Cordelia Langford, John Sillitoe on behalf of the Wellcome Sanger Institute COVID-19 Surveillance Team (http://www.sanger.ac.uk/covid-team) |
| EPI_ISL_492837, EPI_ISL_492840, EPI_ISL_492843, EPI_ISL_492844, EPI_ISL_492845, EPI_ISL_492846, EPI_ISL_492847, EPI_ISL_492850, EPI_ISL_492851, EPI_ISL_492852, EPI_ISL_492856, EPI_ISL_492863, EPI_ISL_492864, EPI_ISL_492865, EPI_ISL_492868, EPI_ISL_492869, EPI_ISL_492872, EPI_ISL_492874, EPI_ISL_492876, EPI_ISL_492877, EPI_ISL_492879, EPI_ISL_492880, EPI_ISL_492882, EPI_ISL_492883, EPI_ISL_492885, EPI_ISL_492887, EPI_ISL_492888, EPI_ISL_492890, EPI_ISL_492894, EPI_ISL_492901, EPI_ISL_492902, EPI_ISL_492905, EPI_ISL_492906, EPI_ISL_492908, EPI_ISL_492911, EPI_ISL_492914 | | | |
| see above | Royal Free Hospital / Health Services Laboratories | Wellcome Sanger Institute for the COVID-19 Genomics UK (COG-UK) consortium | Tanzina Haque, Tabitha Mahungu, Dianne Irish, Cate Goodlad, Jenny Cross, Judith Heaney and Alex Alderton, Roberto Amato, Sonia Goncalves, Ewan Harrison, David K. Jackson, Ian Johnston, Dominic Kwiatkowski, Cordelia Langford, John Sillitoe on behalf of the Wellcome Sanger Institute COVID-19 Surveillance Team (http://www.sanger.ac.uk/covid-team) |
| EPI_ISL_495022 | Government Medical College, Bhavnagar | Gujarat Biotechnology Research Centre | Shirish Patel, Kairavi Desai, Saklin Malek, Dinesh Kumar, Zuber Saiyed, Komal Patel, Labdhi Pandya, Afzal Ansari, Nikha Trivedi, Apurvasinh Puvar, Janvi Raval, Zarna Patel, Monika Gandhi, Pinal Trivedi, Maharshi Pandya, Nidhi Patel, Nitin Savaliya, Raghawendra Kumar, R D Dixit, A M Kadri, Harsh Bakshi, Chaitanya Joshi, Madhvi Joshi |
| EPI_ISL_495069 | Dr. N. D. Desai Medical College & Hospital | Gujarat Biotechnology Research Centre | Supreet Prabhu, Apurvasinh Puvar, Janvi Raval, Zarna Patel, Monika Gandhi, Pinal Trivedi, Maharshi Pandya, Nidhi Patel, Nitin Savaliya, Raghawendra Kumar, Dinesh Kumar, Zuber Saiyed, Komal Patel, Labdhi Pandya, Afzal Ansari, Nikha Trivedi, J G Buch, Jigar Gusani, R D Dixit, A M Kadri, Harsh Bakshi, Chaitanya Joshi, Madhvi Joshi |
| EPI_ISL_495097, EPI_ISL_495098, EPI_ISL_495099, EPI_ISL_495100, EPI_ISL_495101, EPI_ISL_495102, EPI_ISL_495103, EPI_ISL_495104, EPI_ISL_495105, EPI_ISL_495106, EPI_ISL_495107, EPI_ISL_495108, EPI_ISL_495109, EPI_ISL_495110, EPI_ISL_495111, EPI_ISL_495112, EPI_ISL_495113, EPI_ISL_495114, EPI_ISL_495115, EPI_ISL_495116, EPI_ISL_495117, EPI_ISL_495118, EPI_ISL_495119, EPI_ISL_495120, EPI_ISL_495121 | | | |
| see above | PHE South West Regional Laboratory, National Infection Service | Wellcome Sanger Institute for the COVID-19 Genomics UK (COG-UK) consortium | Stephanie Hutchings, Hannah Pymont, Dr Peter Muir, Barry Vipond, Rich Hopes; and Alex Alderton, Roberto Amato, Sonia Goncalves, Ewan Harrison, David K. Jackson, Ian Johnston, Dominic Kwiatkowski, Cordelia Langford, John Sillitoe on behalf of the Wellcome Sanger Institute COVID-19 Surveillance Team (http://www.sanger.ac.uk/covid-team) |
| EPI_ISL_495122, EPI_ISL_495123, EPI_ISL_495124, EPI_ISL_495125, EPI_ISL_495126, EPI_ISL_495127, EPI_ISL_495128, EPI_ISL_495129, EPI_ISL_495130, EPI_ISL_495131, EPI_ISL_495132, EPI_ISL_495133, EPI_ISL_495134, EPI_ISL_495135, EPI_ISL_495136, EPI_ISL_495137, EPI_ISL_495138, EPI_ISL_495139, EPI_ISL_495140, EPI_ISL_495141, EPI_ISL_495142, EPI_ISL_495143, EPI_ISL_495144, EPI_ISL_495145, EPI_ISL_495146, EPI_ISL_495147, EPI_ISL_495148, EPI_ISL_495149, EPI_ISL_495150, EPI_ISL_495151, EPI_ISL_495152, EPI_ISL_495153, EPI_ISL_495154, EPI_ISL_495155, EPI_ISL_495156, EPI_ISL_495157, EPI_ISL_495158, EPI_ISL_495159, EPI_ISL_495160 | | | |
| see above | Innovative Genomics Institute, UC Berkeley | Innovative Genomics Institute, UC Berkeley | Stacia Wyman, Haridha Shivram, Liana Lareau, Shana McDevitt, Justin Choi |
| EPI_ISL_495168, EPI_ISL_495189 | CSIR-Centre for Cellular and Molecular Biology | CSIR-Centre for Cellular and Molecular Biology | Sofia Banu, Payel Mukherjee, Priya Singh,Onkar Kulkarni, Dhiviya Vedagiri, Divya Gupta, Vishal Sah, Santosh Kumar Kuncha, Krishnan Harinivas Harshan, Archana Bharadwaj Siva, Karthik Bharadwaj Tallapaka, Shagufta Khan, Lamuk Zaveri, Namami Gaur, Sakshi Shambhavi, Nikhil Hajirnis, M Soujanya Reddy, Pratheusa Maccha,Tulasi Nagabandi, Purushotham Vodnala, Deepak Kumar, Devi Prasad Vijayashankar, Disha Nanda, Divya Das, Jotin Gogoi, Manish Bhattacharjee, Rakesh K Mishra, Divya Tej Sowpati |
| EPI_ISL_495192 | CSIR-Centre for Cellular and Molecular Biology | CSIR-Centre for Cellular and Molecular Biology | Sofia Banu, Payel Mukherjee, Priya Singh,Onkar Kulkarni, Dhiviya Vedagiri, Divya Gupta, Vishal Sah, Santosh Kumar Kuncha, Krishnan Harinivas Harshan, Archana Bharadwaj Siva, Karthik Bharadwaj Tallapaka, Shagufta Khan, Lamuk Zaveri, Namami Gaur, Sakshi Shambhavi, Tulasi Nagabandi, Nikhil Hajirnis, M Soujanya Reddy, Pratheusa Maccha, Purushotham Vodnala, Gokulan C G, Gunjan Purohit, Hanuman Tulashiram Kale, Pankaj Kumar, Prachand Issarapu, Rakesh K Mishra, Divya Tej Sowpati |
| EPI_ISL_495194 | CSIR-Centre for Cellular and Molecular Biology | CSIR-Centre for Cellular and Molecular Biology | Sofia Banu, Payel Mukherjee, Priya Singh,Onkar Kulkarni, Dhiviya Vedagiri, Divya Gupta, Vishal Sah, Santosh Kumar Kuncha, Krishnan Harinivas Harshan, Archana Bharadwaj Siva, Karthik Bharadwaj Tallapaka, Shagufta Khan, Lamuk Zaveri, Namami Gaur, Sakshi Shambhavi, Nikhil Hajirnis, M Soujanya Reddy, Pratheusa Maccha,Tulasi Nagabandi, Purushotham Vodnala, Deepak Kumar, Devi Prasad Vijayashankar, Disha Nanda, Divya Das, Jotin Gogoi, Manish Bhattacharjee, Rakesh K Mishra, Divya Tej Sowpati |
| EPI_ISL_495197 | CSIR-Centre for Cellular and Molecular Biology | CSIR-Centre for Cellular and Molecular Biology | Sofia Banu, Payel Mukherjee, Priya Singh,Onkar Kulkarni, Dhiviya Vedagiri, Divya Gupta, Vishal Sah, Santosh Kumar Kuncha, Krishnan Harinivas Harshan, Archana Bharadwaj Siva, Karthik Bharadwaj Tallapaka, Shagufta Khan, Lamuk Zaveri, Namami Gaur, Sakshi Shambhavi, Nikhil Hajirnis, M Soujanya Reddy, Pratheusa Maccha, Tulasi Nagabandi, Purushotham Vodnala, Disha Nanda, Divya Das, Jotin Gogoi, Manish Bhattacharjee, Ravi Prasad Mukku, Rakesh K Mishra, Divya Tej Sowpati |
| EPI_ISL_495210 | CSIR-Centre for Cellular and Molecular Biology | CSIR-Centre for Cellular and Molecular Biology | Sofia Banu, Payel Mukherjee, Priya Singh,Onkar Kulkarni, Dhiviya Vedagiri, Divya Gupta, Vishal Sah, Santosh Kumar Kuncha, Krishnan Harinivas Harshan, Archana Bharadwaj Siva, Karthik Bharadwaj Tallapaka, Shagufta Khan, Lamuk Zaveri, Namami Gaur, Sakshi Shambhavi, Nikhil Hajirnis, M Soujanya Reddy, Pratheusa Maccha,Tulasi Nagabandi, Purushotham Vodnala, Deepak Kumar, Devi Prasad Vijayashankar, Disha Nanda, Divya Das, Jotin Gogoi, Manish Bhattacharjee, Rakesh K Mishra, Divya Tej Sowpati |
| EPI_ISL_495219, EPI_ISL_495221 | CSIR-Centre for Cellular and Molecular Biology | CSIR-Centre for Cellular and Molecular Biology | Sofia Banu, Payel Mukherjee, Priya Singh,Onkar Kulkarni, Dhiviya Vedagiri, Divya Gupta, Vishal Sah, Santosh Kumar Kuncha, Krishnan Harinivas Harshan, Archana Bharadwaj Siva, Karthik Bharadwaj Tallapaka, Shagufta Khan, Lamuk Zaveri, Namami Gaur, Sakshi Shambhavi, Nikhil Hajirnis, M Soujanya Reddy, Pratheusa Maccha, Tulasi Nagabandi, Purushotham Vodnala, Disha Nanda, Divya Das, Jotin Gogoi, Manish Bhattacharjee, Ravi Prasad Mukku, Rakesh K Mishra, Divya Tej Sowpati |
| EPI_ISL_495230 | CSIR-Centre for Cellular and Molecular Biology | CSIR-Centre for Cellular and Molecular Biology | Sofia Banu, Payel Mukherjee, Priya Singh,Onkar Kulkarni, Dhiviya Vedagiri, Divya Gupta, Vishal Sah, Santosh Kumar Kuncha, Krishnan Harinivas Harshan, Archana Bharadwaj Siva, Karthik Bharadwaj Tallapaka, Shagufta Khan, Lamuk Zaveri, Namami Gaur, Sakshi Shambhavi, Tulasi Nagabandi, Nikhil Hajirnis, M Soujanya Reddy, Pratheusa Maccha, Purushotham Vodnala, Gokulan C G, Gunjan Purohit, Hanuman Tulashiram Kale, Pankaj Kumar, Prachand Issarapu, Rakesh K Mishra, Divya Tej Sowpati |
| EPI_ISL_495232 | CSIR-Centre for Cellular and Molecular Biology | CSIR-Centre for Cellular and Molecular Biology | Sofia Banu, Payel Mukherjee, Priya Singh,Onkar Kulkarni, Dhiviya Vedagiri, Divya Gupta, Vishal Sah, Santosh Kumar Kuncha, Krishnan Harinivas Harshan, Archana Bharadwaj Siva, Karthik Bharadwaj Tallapaka, Shagufta Khan, Lamuk Zaveri, Namami Gaur, Sakshi Shambhavi, Nikhil Hajirnis, M Soujanya Reddy, Pratheusa Maccha, Tulasi Nagabandi, Purushotham Vodnala, Disha Nanda, Divya Das, Jotin Gogoi, Manish Bhattacharjee, Ravi Prasad Mukku, Rakesh K Mishra, Divya Tej Sowpati |
| EPI_ISL_495262 | CSIR-Centre for Cellular and Molecular Biology | CSIR-Centre for Cellular and Molecular Biology | Sofia Banu, Payel Mukherjee, Priya Singh,Onkar Kulkarni, Dhiviya Vedagiri, Divya Gupta, Vishal Sah, Santosh Kumar Kuncha, Krishnan Harinivas Harshan, Archana Bharadwaj Siva, Karthik Bharadwaj Tallapaka, Shagufta Khan, Lamuk Zaveri, Namami Gaur, Sakshi Shambhavi, Tulasi Nagabandi, Nikhil Hajirnis, M Soujanya Reddy, Pratheusa Maccha, Purushotham Vodnala, Gokulan C G, Gunjan Purohit, Hanuman Tulashiram Kale, Pankaj Kumar, Prachand Issarapu, Rakesh K Mishra, Divya Tej Sowpati |
| EPI_ISL_495274 | Osmania Medical College | CSIR-Centre for Cellular and Molecular Biology | Shashikala Reddy, Mahboob Khan, Lamuk Zaveri, Sofia Banu,Payel Mukherjee, Shagufta Khan,Priya Singh, Onkar Kulkarni, Dhiviya Vedagiri, Divya Gupta, Vishal Sah, Santosh Kumar Kuncha, Krishnan Harinivas Harshan, Archana Bharadwaj Siva, Karthik Bharadwaj Tallapaka,Nikhil Hajirnis, M Soujanya Reddy, Pratheusa Maccha, Namami Gaur, Sakshi Shambhavi, Tulasi Nagabandi, Purushotham Vodnala, Rakesh K Mishra, Divya Tej Sowpati |
| EPI_ISL_495275 | Osmania Medical College | CSIR-Centre for Cellular and Molecular Biology | Shashikala Reddy, Mahboob Khan, Onkar Kulkarni, Payel Mukherjee, Sofia Banu, Priya Singh, Dhiviya Vedagiri, Divya Gupta, Vishal Sah, Santosh Kumar Kuncha, Krishnan Harinivas Harshan, Archana Bharadwaj Siva, Karthik Bharadwaj Tallapaka, Shagufta Khan, Lamuk Zaveri, Nikhil Hajirnis, M Soujanya Reddy, Pratheusa Maccha,Namami Gaur, Sakshi Shambhavi, Tulasi Nagabandi, Purushotham Vodnala, Rakesh K Mishra, Divya Tej Sowpati |
| EPI_ISL_495276 | Osmania Medical College | CSIR-Centre for Cellular and Molecular Biology | Shashikala Reddy, Mahboob Khan,Payel Mukherjee, Sofia Banu, Priya Singh, Onkar Kulkarni, Dhiviya Vedagiri, Divya Gupta, Vishal Sah, Santosh Kumar Kuncha, Krishnan Harinivas Harshan, Archana Bharadwaj Siva, Karthik Bharadwaj Tallapaka, Shagufta Khan, Lamuk Zaveri, Namami Gaur, Sakshi Shambhavi, Nikhil Hajirnis, M Soujanya Reddy, Pratheusa Maccha,Tulasi Nagabandi, Purushotham Vodnala, Rakesh K Mishra, Divya Tej Sowpati |

| | | | |
|---|---|---|---|
| EPI_ISL_495277 | Osmania Medical College | CSIR-Centre for Cellular and Molecular Biology | Shashikala Reddy, Mahboob Khan, Namami Gaur, Lamuk Zaveri,Shagufta Khan, Nikhil Hajirnis, M Soujanya Reddy, Pratheusa Maccha, Sofia Banu, Onkar Kulkarni, Payel Mukherjee, Priya Singh, Dhiviya Vedagiri, Divya Gupta, Vishal Sah, Santosh Kumar Kuncha, Krishnan Harinivas Harshan, Archana Bharadwaj Siva, Karthik Bharadwaj Tallapaka,Sakshi Shambhavi, Tulasi Nagabandi, Purushotham Vodnala, Rakesh K Mishra, Divya Tej Sowpati |
| EPI_ISL_495278 | Osmania Medical College | CSIR-Centre for Cellular and Molecular Biology | Shashikala Reddy, Mahboob Khan,Tulasi Nagabandi, Namami Gaur, Lamuk Zaveri,Shagufta Khan,Nikhil Hajirnis, M Soujanya Reddy, Pratheusa Maccha, Sofia Banu,Payel Mukherjee, Priya Singh, Onkar Kulkarni, Dhiviya Vedagiri, Divya Gupta, Vishal Sah, Santosh Kumar Kuncha, Krishnan Harinivas Harshan, Archana Bharadwaj Siva, Karthik Bharadwaj Tallapaka,Sakshi Shambhavi, Purushotham Vodnala, Rakesh K Mishra, Divya Tej Sowpati |
| EPI_ISL_495279 | Osmania Medical College | CSIR-Centre for Cellular and Molecular Biology | Shashikala Reddy, Mahboob Khan,Sakshi Shambhavi,Tulasi Nagabandi, Namami Gaur, Lamuk Zaveri,Shagufta Khan, Nikhil Hajirnis, M Soujanya Reddy, Pratheusa Maccha,Sofia Banu,Payel Mukherjee, Priya Singh,Onkar Kulkarni, Dhiviya Vedagiri, Divya Gupta, Vishal Sah, Santosh Kumar Kuncha, Krishnan Harinivas Harshan, Archana Bharadwaj Siva, Karthik Bharadwaj Tallapaka,Sakshi Shambhavi, Purushotham Vodnala, Rakesh K Mishra, Divya Tej Sowpati |
| EPI_ISL_495280 | Osmania Medical College | CSIR-Centre for Cellular and Molecular Biology | Shashikala Reddy, Mahboob Khan,Payel Mukherjee, Sofia Banu, Priya Singh, Onkar Kulkarni, Dhiviya Vedagiri, Divya Gupta, Vishal Sah, Santosh Kumar Kuncha, Krishnan Harinivas Harshan, Archana Bharadwaj Siva, Karthik Bharadwaj Tallapaka, Shagufta Khan, Lamuk Zaveri, Nikhil Hajirnis, M Soujanya Reddy, Pratheusa Maccha,Namami Gaur, Sakshi Shambhavi, Tulasi Nagabandi, Purushotham Vodnala, Rakesh K Mishra, Divya Tej Sowpati |
| EPI_ISL_495281 | Osmania Medical College | CSIR-Centre for Cellular and Molecular Biology | Shashikala Reddy, Mahboob Khan, Sofia Banu,Payel Mukherjee, Priya Singh,Onkar Kulkarni, Dhiviya Vedagiri, Divya Gupta, Vishal Sah, Santosh Kumar Kuncha, Krishnan Harinivas Harshan, Archana Bharadwaj Siva, Karthik Bharadwaj Tallapaka, Shagufta Khan, Lamuk Zaveri, Namami Gaur, Nikhil Hajirnis, M Soujanya Reddy, Pratheusa Maccha,Sakshi Shambhavi, Tulasi Nagabandi, Purushotham Vodnala, Rakesh K Mishra, Divya Tej Sowpati |
| EPI_ISL_495282 | Osmania Medical College | CSIR-Centre for Cellular and Molecular Biology | Shashikala Reddy, Mahboob Khan,Payel Mukherjee, Sofia Banu, Priya Singh, Onkar Kulkarni, Dhiviya Vedagiri, Divya Gupta, Vishal Sah, Santosh Kumar Kuncha, Krishnan Harinivas Harshan, Archana Bharadwaj Siva, Karthik Bharadwaj Tallapaka, Shagufta Khan, Lamuk Zaveri, Nikhil Hajirnis, M Soujanya Reddy, Pratheusa Maccha,Namami Gaur, Sakshi Shambhavi, Tulasi Nagabandi, Purushotham Vodnala, Rakesh K Mishra, Divya Tej Sowpati |
| EPI_ISL_495283 | Osmania Medical College | CSIR-Centre for Cellular and Molecular Biology | Shashikala Reddy, Mahboob Khan, Namami Gaur, Lamuk Zaveri,Shagufta Khan, Sofia Banu, Onkar Kulkarni, Payel Mukherjee, Priya Singh, Dhiviya Vedagiri, Divya Gupta, Vishal Sah, Santosh Kumar Kuncha, Krishnan Harinivas Harshan, Archana Bharadwaj Siva, Karthik Bharadwaj Tallapaka,Sakshi Shambhavi, Nikhil Hajirnis, M Soujanya Reddy, Pratheusa Maccha, Tulasi Nagabandi, Purushotham Vodnala, Rakesh K Mishra, Divya Tej Sowpati |
| EPI_ISL_495284 | Osmania Medical College | CSIR-Centre for Cellular and Molecular Biology | Shashikala Reddy, Mahboob Khan,Tulasi Nagabandi, Namami Gaur, Lamuk Zaveri,Shagufta Khan, Sofia Banu,Payel Mukherjee, Priya Singh, Onkar Kulkarni, Dhiviya Vedagiri, Divya Gupta, Vishal Sah, Santosh Kumar Kuncha, Krishnan Harinivas Harshan, Archana Bharadwaj Siva, Karthik Bharadwaj Tallapaka,Sakshi Shambhavi,Nikhil Hajirnis, M Soujanya Reddy, Pratheusa Maccha, Purushotham Vodnala, Rakesh K Mishra, Divya Tej Sowpati |
| EPI_ISL_495285 | Osmania Medical College | CSIR-Centre for Cellular and Molecular Biology | Shashikala Reddy, Mahboob Khan,Shagufta Khan, Sofia Banu,Payel Mukherjee, Priya Singh,Onkar Kulkarni, Dhiviya Vedagiri, Divya Gupta, Vishal Sah, Santosh Kumar Kuncha, Krishnan Harinivas Harshan, Archana Bharadwaj Siva, Karthik Bharadwaj Tallapaka, Lamuk Zaveri, Nikhil Hajirnis, M Soujanya Reddy, Pratheusa Maccha, Namami Gaur, Sakshi Shambhavi, Tulasi Nagabandi, Purushotham Vodnala, Rakesh K Mishra, Divya Tej Sowpati |
| EPI_ISL_495286 | Osmania Medical College | CSIR-Centre for Cellular and Molecular Biology | Shashikala Reddy, Mahboob Khan, Sofia Banu,Payel Mukherjee, Priya Singh,Onkar Kulkarni, Dhiviya Vedagiri, Divya Gupta, Vishal Sah, Santosh Kumar Kuncha, Krishnan Harinivas Harshan, Archana Bharadwaj Siva, Karthik Bharadwaj Tallapaka, Shagufta Khan, Lamuk Zaveri, Namami Gaur, Sakshi Shambhavi, Nikhil Hajirnis, M Soujanya Reddy, Pratheusa Maccha,Tulasi Nagabandi, Purushotham Vodnala, Rakesh K Mishra, Divya Tej Sowpati |
| EPI_ISL_495287 | Osmania Medical College | CSIR-Centre for Cellular and Molecular Biology | Shashikala Reddy, Mahboob Khan,Onkar Kulkarni, Sofia Banu,Payel Mukherjee, Priya Singh, Dhiviya Vedagiri, Divya Gupta, Vishal Sah, Santosh Kumar Kuncha, Krishnan Harinivas Harshan, Archana Bharadwaj Siva, Karthik Bharadwaj Tallapaka, Shagufta Khan, Nikhil Hajirnis, M Soujanya Reddy, Pratheusa Maccha, Lamuk Zaveri, Namami Gaur, Sakshi Shambhavi, Tulasi Nagabandi, Purushotham Vodnala, Rakesh K Mishra, Divya Tej Sowpati |
| EPI_ISL_495288 | Osmania Medical College | CSIR-Centre for Cellular and Molecular Biology | Shashikala Reddy, Mahboob Khan, Lamuk Zaveri, Sofia Banu,Payel Mukherjee, Shagufta Khan,Priya Singh, Onkar Kulkarni, Dhiviya Vedagiri, Divya Gupta, Vishal Sah, Santosh Kumar Kuncha, Krishnan Harinivas Harshan, Archana Bharadwaj Siva, Karthik Bharadwaj Tallapaka, Namami Gaur, Nikhil Hajirnis, M Soujanya Reddy, Pratheusa Maccha,Sakshi Shambhavi, Tulasi Nagabandi, Purushotham Vodnala, Rakesh K Mishra, Divya Tej Sowpati |
| EPI_ISL_495289 | Osmania Medical College | CSIR-Centre for Cellular and Molecular Biology | Shashikala Reddy, Mahboob Khan, Namami Gaur, Lamuk Zaveri,Shagufta Khan, Sofia Banu, Onkar Kulkarni, Payel Mukherjee, Priya Singh, Dhiviya Vedagiri, Divya Gupta, Vishal Sah, Santosh Kumar Kuncha, Krishnan Harinivas Harshan, Archana Bharadwaj Siva, Karthik Bharadwaj Tallapaka,Sakshi Shambhavi, Nikhil Hajirnis, M Soujanya Reddy, Pratheusa Maccha, Tulasi Nagabandi, Purushotham Vodnala, Rakesh K Mishra, Divya Tej Sowpati |
| EPI_ISL_495290 | Osmania Medical College | CSIR-Centre for Cellular and Molecular Biology | Shashikala Reddy, Mahboob Khan,Tulasi Nagabandi, Namami Gaur, Lamuk Zaveri,Shagufta Khan, Sofia Banu,Payel Mukherjee, Priya Singh, Onkar Kulkarni, Dhiviya Vedagiri, Divya Gupta, Vishal Sah, Santosh Kumar Kuncha, Krishnan Harinivas Harshan, Archana Bharadwaj Siva, Karthik Bharadwaj Tallapaka,Sakshi Shambhavi, Nikhil Hajirnis, M Soujanya Reddy, Pratheusa Maccha, Purushotham Vodnala, Rakesh K Mishra, Divya Tej Sowpati |
| EPI_ISL_495291 | Osmania Medical College | CSIR-Centre for Cellular and Molecular Biology | Shashikala Reddy, Mahboob Khan,Onkar Kulkarni,Shagufta Khan, Sofia Banu,Payel Mukherjee, Priya Singh, Dhiviya Vedagiri, Divya Gupta, Vishal Sah, Santosh Kumar Kuncha, Krishnan Harinivas Harshan, Archana Bharadwaj Siva, Karthik Bharadwaj Tallapaka, Lamuk Zaveri, Nikhil Hajirnis, M Soujanya Reddy, Pratheusa Maccha, Namami Gaur, Sakshi Shambhavi, Tulasi Nagabandi, Purushotham Vodnala, Rakesh K Mishra, Divya Tej Sowpati |
| EPI_ISL_495292 | Osmania Medical College | CSIR-Centre for Cellular and Molecular Biology | Shashikala Reddy, Mahboob Khan,Shagufta Khan, Sofia Banu,Payel Mukherjee, Priya Singh,Onkar Kulkarni, Dhiviya Vedagiri, Divya Gupta, Vishal Sah, Santosh Kumar Kuncha, Krishnan Harinivas Harshan, Archana Bharadwaj Siva, Karthik Bharadwaj Tallapaka, Lamuk Zaveri, Namami Gaur, Sakshi Shambhavi,Nikhil Hajirnis, M Soujanya Reddy, Pratheusa Maccha, Tulasi Nagabandi, Purushotham Vodnala, Rakesh K Mishra, Divya Tej Sowpati |
| EPI_ISL_495293 | Osmania Medical College | CSIR-Centre for Cellular and Molecular Biology | Shashikala Reddy, Mahboob Khan, Lamuk Zaveri, Sofia Banu,Payel Mukherjee, Shagufta Khan,Priya Singh, Onkar Kulkarni, Dhiviya Vedagiri, Divya Gupta, Vishal Sah, Santosh Kumar Kuncha, Krishnan Harinivas Harshan, Archana Bharadwaj Siva, Karthik Bharadwaj Tallapaka, Namami Gaur, Sakshi Shambhavi, Nikhil Hajirnis, M Soujanya Reddy, Pratheusa Maccha, Tulasi Nagabandi, Purushotham Vodnala, Rakesh K Mishra, Divya Tej Sowpati |
| EPI_ISL_495294 | Osmania Medical College | CSIR-Centre for Cellular and Molecular Biology | Shashikala Reddy, Mahboob Khan,Onkar Kulkarni, Lamuk Zaveri, Sofia Banu,Payel Mukherjee, Shagufta Khan,Priya Singh, Dhiviya Vedagiri, Divya Gupta, Vishal Sah, Santosh Kumar Kuncha, Krishnan Harinivas Harshan, Archana Bharadwaj Siva, Karthik Bharadwaj Tallapaka, Namami Gaur, Sakshi Shambhavi, Nikhil Hajirnis, M Soujanya Reddy, Pratheusa Maccha, Tulasi Nagabandi, Purushotham Vodnala, Rakesh K Mishra, Divya Tej Sowpati |
| EPI_ISL_495295 | Osmania Medical College | CSIR-Centre for Cellular and Molecular Biology | Shashikala Reddy, Mahboob Khan, Sofia Banu,Payel Mukherjee, Priya Singh,Onkar Kulkarni, Dhiviya Vedagiri, Divya Gupta, Vishal Sah, Santosh Kumar Kuncha, Krishnan Harinivas Harshan, Archana Bharadwaj Siva, Karthik Bharadwaj Tallapaka, Shagufta Khan, Lamuk Zaveri,Nikhil Hajirnis, M Soujanya Reddy, Pratheusa Maccha, Namami Gaur, Sakshi Shambhavi, Tulasi Nagabandi, Purushotham Vodnala, Rakesh K Mishra, Divya Tej Sowpati |
| EPI_ISL_495296 | Osmania Medical College | CSIR-Centre for Cellular and Molecular Biology | Shashikala Reddy, Mahboob Khan,Shagufta Khan, Sofia Banu,Payel Mukherjee, Priya Singh,Onkar Kulkarni, Dhiviya Vedagiri, Divya Gupta, Vishal Sah, Santosh Kumar Kuncha, Krishnan Harinivas Harshan, Archana Bharadwaj Siva, Karthik Bharadwaj Tallapaka, Lamuk Zaveri, Nikhil Hajirnis, M Soujanya Reddy, Pratheusa Maccha, Namami Gaur, Sakshi Shambhavi, Tulasi Nagabandi, Purushotham Vodnala, Rakesh K Mishra, Divya Tej Sowpati |
| EPI_ISL_495297 | Osmania Medical College | CSIR-Centre for Cellular and Molecular Biology | Shashikala Reddy, Mahboob Khan,Sakshi Shambhavi,Tulasi Nagabandi, Namami Gaur, Lamuk Zaveri,Shagufta Khan, Sofia Banu,Payel Mukherjee, Priya Singh,Onkar Kulkarni, Dhiviya Vedagiri, Divya Gupta, Vishal Sah, Santosh Kumar Kuncha, Krishnan Harinivas Harshan, Archana Bharadwaj Siva, Karthik Bharadwaj Tallapaka,Sakshi Shambhavi, Nikhil Hajirnis, M Soujanya Reddy, Pratheusa Maccha,Purushotham Vodnala, Rakesh K Mishra, Divya Tej Sowpati |
| EPI_ISL_498228 | National Institute of Laboratory Medicine and Referral Center | Genomic Research Lab, BCSIR | Shahina Akter, Abu Sayeed Mohammad Mahmud, Mohammad Samir Uzzaman, Eshrar Osman, Md. Ahasan Habib, Tanjina Akhter Banu, Md. Murshed Hasan Sarkar, Barna Goswami, Iffat Jahan, Md. Saddam Hossain, Tasnim Nafisa, Md. Maruf Ahmed Molla, Mahmuda Yeasmin, Asish Kumar Ghosh, A. K. M. Shamsuzzaman, Sheikh Md. Selim Al Din, Utpal Chandra Ray, Salek Ahmed Sajib, Md. Salim Khan |
| EPI_ISL_498253, EPI_ISL_498254 | National Institute of Laboratory Medicine and Referral Center | Genomic Research Lab, BCSIR | Tanjina Akhter Banu, Abu Sayeed Mohammad Mahmud, Mohammad Samir Uzzaman, Eshrar Osman, Md. Ahasan Habib, Shahina Akter, Md. Murshed Hasan Sarkar, Barna Goswami, Iffat Jahan, Md. Saddam Hossain, Tasnim Nafisa, Md. Maruf Ahmed Molla, Mahmuda Yeasmin, Asish Kumar Ghosh, A. K. M. Shamsuzzaman, Sheikh Md. Selim Al Din, Utpal Chandra Ray, Salek Ahmed Sajib, Md. Salim Khan |
| EPI_ISL_498274, EPI_ISL_498417 | National Institute of Laboratory Medicine and Referral Center | Genomic Research Lab, BCSIR | Tasnim Nafisa, Abu Sayeed Mohammad Mahmud, Mohammad Samir Uzzaman, Eshrar Osman, Md. Ahasan Habib, Shahina Akter, Tanjina Akhter Banu, Md. Murshed Hasan Sarkar, Barna Goswami, Iffat Jahan, Md. Saddam Hossain, Md. Maruf Ahmed Molla, Mahmuda Yeasmin, Asish Kumar Ghosh, A. K. M. Shamsuzzaman, Sheikh Md. Selim Al Din, Utpal Chandra Ray, Salek Ahmed Sajib, Md. Salim Khan |

EPI_ISL_500228, EPI_ISL_500229, EPI_ISL_500230, EPI_ISL_500231, EPI_ISL_500232, EPI_ISL_500233, EPI_ISL_500234, EPI_ISL_500235, EPI_ISL_500236, EPI_ISL_500237, EPI_ISL_500238, EPI_ISL_500239, EPI_ISL_500240, EPI_ISL_500241, EPI_ISL_500242, EPI_ISL_500243, EPI_ISL_500244, EPI_ISL_500245, EPI_ISL_500246, EPI_ISL_500247, EPI_ISL_500248, EPI_ISL_500249, EPI_ISL_500250, EPI_ISL_500251, EPI_ISL_500252, EPI_ISL_500253, EPI_ISL_500254, EPI_ISL_500255, EPI_ISL_500256, EPI_ISL_500257, EPI_ISL_500258, EPI_ISL_500259, EPI_ISL_500260, EPI_ISL_500261, EPI_ISL_500262, EPI_ISL_500263,

| | | | |
|---|---|---|---|
| EPI_ISL_500264, EPI_ISL_500265, EPI_ISL_500266, EPI_ISL_500267, EPI_ISL_500268, EPI_ISL_500269, EPI_ISL_500270, EPI_ISL_500271, EPI_ISL_500272, EPI_ISL_500273, EPI_ISL_500274, EPI_ISL_500275, EPI_ISL_500276, EPI_ISL_500277, EPI_ISL_500278, EPI_ISL_500279, EPI_ISL_500280, EPI_ISL_500281, EPI_ISL_500282, EPI_ISL_500283, EPI_ISL_500284, EPI_ISL_500285, EPI_ISL_500286 | | | |
| see above | Hospital Universitario Araba. Vitoria-Gasteiz | SeqCOVID-SPAIN consortium/IBV(CSIC) | Silvia Hernáez Crespo, Carmen Gómez González, Amaia Aguirre Quiñonero, Marina Fernández Torres, Mª Rosario Almela Ferrer, Mª Concepción Lecaroz Agara, Andrés Canut Blasco and SeqCOVID-SPAIN consortium |
| EPI_ISL_501555, EPI_ISL_501556, EPI_ISL_501558, EPI_ISL_501559, EPI_ISL_501560, EPI_ISL_501561, EPI_ISL_501563, EPI_ISL_501564, EPI_ISL_501565, EPI_ISL_501566, EPI_ISL_501567, EPI_ISL_501568, EPI_ISL_501569, EPI_ISL_501570, EPI_ISL_501571, EPI_ISL_501572, EPI_ISL_501573, EPI_ISL_501574, EPI_ISL_501575, EPI_ISL_501576, EPI_ISL_501577, EPI_ISL_501578, EPI_ISL_501579, EPI_ISL_501580, EPI_ISL_501581, EPI_ISL_501582, EPI_ISL_501583, EPI_ISL_501584, EPI_ISL_501585, EPI_ISL_501586, EPI_ISL_501587, EPI_ISL_501588, EPI_ISL_501589, EPI_ISL_501590, EPI_ISL_501591, EPI_ISL_501592, EPI_ISL_501593, EPI_ISL_501594, EPI_ISL_501595, EPI_ISL_501596, EPI_ISL_501597, EPI_ISL_501598, EPI_ISL_501599, EPI_ISL_501601, EPI_ISL_501603, EPI_ISL_501604, EPI_ISL_501605, EPI_ISL_501606, EPI_ISL_501607, EPI_ISL_501608, EPI_ISL_501609, EPI_ISL_501610, EPI_ISL_501611, EPI_ISL_501612 | | | |
| see above | PHE South West Regional Laboratory, National Infection Service | Wellcome Sanger Institute for the COVID-19 Genomics UK (COG-UK) consortium | Stephanie Hutchings, Hannah Pymont, Dr Peter Muir, Barry Vipond, Rich Hopes; and Alex Alderton, Roberto Amato, Sonia Goncalves, Ewan Harrison, David K. Jackson, Ian Johnston, Dominic Kwiatkowski, Cordelia Langford, John Sillitoe on behalf of the Wellcome Sanger Institute COVID-19 Surveillance Team (http://www.sanger.ac.uk/covid-team) |
| EPI_ISL_507008 | National Veterinary Institute | National Veterinary Institute | Siamak Zohari |
| EPI_ISL_509430, EPI_ISL_509431, EPI_ISL_509432, EPI_ISL_509433, EPI_ISL_509434, EPI_ISL_509435 | Centro de Desenvolvimento Tecnologico em Saude, Fundacao Oswaldo Cruz | Centro de Desenvolvimento Tecnologico em Saude, Fundacao Oswaldo Cruz | Souza,T.M., Fintelman-Rodrigues,N., De Paula,A.D., Saraiva,F.B., Ferreira,M.A., Sacramento,C.Q., Medeiros,M.A. |
| EPI_ISL_509616, EPI_ISL_509617, EPI_ISL_509618 | Hospital Universitario Araba. Vitoria-Gasteiz | SeqCOVID-SPAIN consortium/IBV(CSIC) | Silvia Hernáez Crespo, Carmen Gómez González, Amaia Aguirre Quiñonero, Marina Fernández Torres, Mª Rosario Almela Ferrer, Mª Concepción Lecaroz Agara, Andrés Canut Blasco and SeqCOVID-SPAIN consortium |
| EPI_ISL_510529 | School of Veterinary Medicine, Disease Control | School of Veterinary Medicine, Disease Control | Simulundu,E., Kapata,N., Mupeta,F., Kapata,P.C., Saasa,N., Changula,K., Muleya,W., Chitanga,S., Chambaro,H., Mubemba,B., Masahiro,K., Chanda,D., Mulenga,L., Fwoloshi,S., Shibemba,A.L., Kapaya,F., Zulu,P., Musonda,K., Monze,M., Sinyange,N., Liwewe,M.M., Kapin'a,M., Chipimo,P.J., Ngosa,W., Morales,A.N., Kayeyi,N., Malama,K., Tembo,J., Bates,M., Sawa,H., Takada,A., Nalubamba,K.S., Mukonka,V., Chilufya,C. and Zumla,A. |
| EPI_ISL_510531 | Communicable Disease Laboratory, Public Health Directorate | Communicable Disease Laboratory, Public Health Directorate | Shehab,F., Altaif,Z., AlWasti,H., Zaed,A. |
| EPI_ISL_510535 | Molecular Virology, Instituto Carlos Chagas / Fiocruz Paraná | Universidade Federal do Parana (UFPR) | Suzukawa,A., Tscha,M., Zanluca,C., Raboni,S., Duarte dos Santos,C. |
| EPI_ISL_510536 | Centro de Desenvolvimento Tecnologico em Saude, Fundacao Oswaldo Cruz | Centro de Desenvolvimento Tecnologico em Saude, Fundacao Oswaldo Cruz | Souza,T.M., Fintelman-Rodrigues,N., De Paula,A.D., Saraiva,F.B., Ferreira,M.A., Sacramento,C.Q. and Medeiros,M.A. |
| EPI_ISL_510541 | Centro de Desenvolvimento Tecnologico em Saude, Fundacao Oswaldo Cruz | Centro de Desenvolvimento Tecnologico em Saude, Fundacao Oswaldo Cruz | Souza,T.M., Fintelman-Rodrigues,N., De Paula,A.D., Saraiva,F.B., Ferreira,M.A., Sacramento,C.Q. and Medeiros,M.A. |
| EPI_ISL_511805, EPI_ISL_511806, EPI_ISL_511807, EPI_ISL_511808, EPI_ISL_511809, EPI_ISL_511810, EPI_ISL_511811, EPI_ISL_511812, EPI_ISL_511813, EPI_ISL_511814, EPI_ISL_511815, EPI_ISL_511816, EPI_ISL_511817, EPI_ISL_511818, EPI_ISL_511819, EPI_ISL_511820, EPI_ISL_511821, EPI_ISL_511822, EPI_ISL_511823, EPI_ISL_511824, EPI_ISL_511825, EPI_ISL_511826, EPI_ISL_511827, EPI_ISL_511828, EPI_ISL_511829, EPI_ISL_511830, EPI_ISL_511831, EPI_ISL_511832, EPI_ISL_511833, EPI_ISL_511834, EPI_ISL_511835, EPI_ISL_511836, EPI_ISL_511837, EPI_ISL_511838, EPI_ISL_511839, EPI_ISL_511840, EPI_ISL_511841, EPI_ISL_511842, EPI_ISL_511843, EPI_ISL_511844, EPI_ISL_511845, EPI_ISL_511846, EPI_ISL_511847, EPI_ISL_511848, EPI_ISL_511849, EPI_ISL_511850, EPI_ISL_511851, EPI_ISL_511862 | | | |
| see above | Innovative Genomics Institute, UC Berkeley | Innovative Genomics Institute, UC Berkeley | Stacia Wyman, Haridha Shivram, Liana Lareau, Shana McDevitt, Justin Choi |
| EPI_ISL_511955, EPI_ISL_511956, EPI_ISL_511957, EPI_ISL_511958, EPI_ISL_511960, EPI_ISL_511961, EPI_ISL_511962, EPI_ISL_511963, EPI_ISL_511964, EPI_ISL_511965, EPI_ISL_511966, EPI_ISL_511967, EPI_ISL_511968, EPI_ISL_511969, EPI_ISL_511971, EPI_ISL_511972, EPI_ISL_511973, EPI_ISL_511974, EPI_ISL_511975, EPI_ISL_511976, EPI_ISL_511977, EPI_ISL_511978, EPI_ISL_511979, EPI_ISL_511980, EPI_ISL_511981 | | | |
| see above | PHE South West Regional Laboratory, National Infection Service | Wellcome Sanger Institute for the COVID-19 Genomics UK (COG-UK) consortium | Stephanie Hutchings, Hannah Pymont, Dr Peter Muir, Barry Vipond, Rich Hopes; and Alex Alderton, Roberto Amato, Sonia Goncalves, Ewan Harrison, David K. Jackson, Ian Johnston, Dominic Kwiatkowski, Cordelia Langford, John Sillitoe on behalf of the Wellcome Sanger Institute COVID-19 Surveillance Team (http://www.sanger.ac.uk/covid-team) |
| EPI_ISL_512071 | Department of MicroBiology, Government Medical College, Surat | Gujarat Biotechnology Research Centre | Summaiya Mullan, Amit gamit, Apurvasinh Puvar, Janvi Raval, Zarna Patel, Monika Gandhi, Pinal Trivedi, Maharshi Pandya, Nidhi Patel, Nitin Savaliya, Raghawendra Kumar, Dinesh Kumar, Zuber Saiyed, Komal Patel, Labdhi Pandya, Afzal Ansari, Nikha Trivedi, Naresh Chauhan, R D Dixit, A M Kadri, Harsh Bakshi, Chaitanya Joshi, Madhvi Joshi |
| EPI_ISL_512908, EPI_ISL_512909, EPI_ISL_512910, EPI_ISL_512911, EPI_ISL_512912, EPI_ISL_512913, EPI_ISL_512914, EPI_ISL_512915, EPI_ISL_512916, EPI_ISL_512917, EPI_ISL_513121, EPI_ISL_513122, EPI_ISL_513123, EPI_ISL_513124, EPI_ISL_513125, EPI_ISL_513126, EPI_ISL_513127, EPI_ISL_513128, EPI_ISL_513129, EPI_ISL_513130, EPI_ISL_513131, EPI_ISL_513132, EPI_ISL_513133, EPI_ISL_513134, EPI_ISL_513135, EPI_ISL_513136, EPI_ISL_513137, EPI_ISL_513138, EPI_ISL_513139, EPI_ISL_513140, EPI_ISL_513141, EPI_ISL_513142, EPI_ISL_513143, EPI_ISL_513144, EPI_ISL_513145, EPI_ISL_513146, EPI_ISL_513214, EPI_ISL_513215, EPI_ISL_513216, EPI_ISL_513217, EPI_ISL_513218, EPI_ISL_513219, EPI_ISL_513220, EPI_ISL_513221, EPI_ISL_513222, EPI_ISL_513223, EPI_ISL_513224, EPI_ISL_513225, EPI_ISL_513226 | | | |
| see above | Pathogen Genomics Lab King Abdullah University of Science and Technology(KAUST) | Pathogen Genomics Lab King Abdullah University of Science and Technology(KAUST) | Sharif Hala, Fadwa Alofi, Sara Mfarrej, Amit Kumar Subudhi, Rahul P Salunke, Fathia Ben Rached, Amanda Ooi, Luke Esau, Afrah Alsomali, Asim Khogeer, Jumana Taha, Abdulaziz Alahmadi, Kahled Algithami, Raeece Naeem, Anwar Hashem, Naif Almontashiri, Arnab Pain |
| EPI_ISL_513584 | PHE South West Regional Laboratory, National Infection Service | Wellcome Sanger Institute for the COVID-19 Genomics UK (COG-UK) consortium | Stephanie Hutchings, Hannah Pymont, Dr Peter Muir, Barry Vipond, Rich Hopes; and Alex Alderton, Roberto Amato, Sonia Goncalves, Ewan Harrison, David K. Jackson, Ian Johnston, Dominic Kwiatkowski, Cordelia Langford, John Sillitoe on behalf of the Wellcome Sanger Institute COVID-19 Surveillance Team (http://www.sanger.ac.uk/covid-team) |
| EPI_ISL_514231, EPI_ISL_514232 | National Institute of Laboratory Medicine and Referral Center | Genomic Research Lab, BCSIR | Shahina Akter, Abu Sayeed Mohammad Mahmud, Mohammad Samir Uzzaman, Eshrar Osman, Md. Ahashan Habib, Tanjina Akhter Banu, Md. Murshed Hasan Sarkar, Barna Goswami, Iffat Jahan, Md. Saddam Hossain, Tasnim Nafisa, Md. Maruf Ahmed Molla, Mahmuda Yeasmin, Asish Kumar Ghosh, A. K. M. Shamsuzzaman, Sheikh Md. Selim Al Din, Utpal Chandra Ray, Salek Ahmed Sajib, Md. Salim Khan |
| EPI_ISL_514242, EPI_ISL_514243, EPI_ISL_514244 | National Institute of Laboratory Medicine and Referral Center | Genomic Research Lab, BCSIR | Tanjina Akhter Banu, Abu Sayeed Mohammad Mahmud, Mohammad Samir Uzzaman, Eshrar Osman, Md. Ahashan Habib, Shahina Akter, Md. Murshed Hasan Sarkar, Barna Goswami, Iffat Jahan, Md. Saddam Hossain, Tasnim Nafisa, Md. Maruf Ahmed Molla, Mahmuda Yeasmin, Asish Kumar Ghosh, A. K. M. Shamsuzzaman, Sheikh Md. Selim Al Din, Utpal Chandra Ray, Salek Ahmed Sajib, Md. Salim Khan |
| EPI_ISL_524436 | Department of Immunology, The Scripps Research Institute | Department of Immunology, The Scripps Research Institute | Smither,A., Sabino-Santos,G., Snarski,P., Melnik,L., Bell,A., Genemaras,K., Drouin,A., Fusco,D., Garry,R., SEARCH Alliance,S.D. |
| EPI_ISL_524474 | Department of Infectious Diseases, Cantonal Hospital Baden | Institute of Medical Virology, University of Zurich | Stefan Schmutz, Maryam Zaheri, Verena Kufner, Gabriela Ziltener, Patrick Redli, Fiona Steiner, Jon Huder, Riccarda Capaul, Andrea Zbinden, Jürg Böni, Michael Huber, Alexandra Trkola |
| EPI_ISL_524475 | Division of Infectious Diseases, University Hospital Zürich | Institute of Medical Virology, University of Zurich | Stefan Schmutz, Maryam Zaheri, Verena Kufner, Gabriela Ziltener, Patrick Redli, Fiona Steiner, Jon Huder, Riccarda Capaul, Andrea Zbinden, Jürg Böni, Michael Huber, Alexandra Trkola |
| EPI_ISL_524476 | Bülach Hospital | Institute of Medical Virology, University of Zurich | Stefan Schmutz, Maryam Zaheri, Verena Kufner, Gabriela Ziltener, Patrick Redli, Fiona Steiner, Jon Huder, Riccarda Capaul, Andrea Zbinden, Jürg Böni, Michael Huber, Alexandra Trkola |
| EPI_ISL_524477 | Division of Infectious Diseases, University Hospital Zürich | Institute of Medical Virology, University of Zurich | Stefan Schmutz, Maryam Zaheri, Verena Kufner, Gabriela Ziltener, Patrick Redli, Fiona Steiner, Jon Huder, Riccarda Capaul, Andrea Zbinden, Jürg Böni, Michael Huber, Alexandra Trkola |
| EPI_ISL_524478 | Cantonal Hospital Winterthur | Institute of Medical Virology, University of Zurich | Stefan Schmutz, Maryam Zaheri, Verena Kufner, Gabriela Ziltener, Patrick Redli, Fiona Steiner, Jon Huder, Riccarda Capaul, Andrea Zbinden, Jürg Böni, Michael Huber, Alexandra Trkola |
| EPI_ISL_524479 | Klinik Hirslanden Zürich | Institute of Medical Virology, University of Zurich | Stefan Schmutz, Maryam Zaheri, Verena Kufner, Gabriela Ziltener, Patrick Redli, Fiona Steiner, Jon Huder, Riccarda Capaul, Andrea Zbinden, Jürg Böni, Michael Huber, Alexandra Trkola |
| EPI_ISL_524480, EPI_ISL_524481 | Department of Infectious Diseases, Cantonal Hospital Baden | Institute of Medical Virology, University of Zurich | Stefan Schmutz, Maryam Zaheri, Verena Kufner, Gabriela Ziltener, Patrick Redli, Fiona Steiner, Jon Huder, Riccarda Capaul, Andrea Zbinden, Jürg Böni, Michael Huber, Alexandra Trkola |
| EPI_ISL_524486, EPI_ISL_524487, EPI_ISL_524488, EPI_ISL_524489, EPI_ISL_524490, EPI_ISL_524491, EPI_ISL_524492 | PHE South West Regional Laboratory, National Infection Service | Wellcome Sanger Institute for the COVID-19 Genomics UK (COG-UK) consortium | Stephanie Hutchings, Hannah Pymont, Dr Peter Muir, Barry Vipond, Rich Hopes; and Alex Alderton, Roberto Amato, Sonia Goncalves, Ewan Harrison, David K. Jackson, Ian Johnston, Dominic Kwiatkowski, Cordelia Langford, John Sillitoe on behalf of the Wellcome Sanger Institute COVID-19 Surveillance Team (http://www.sanger.ac.uk/covid-team) |
| EPI_ISL_524782 | University Hospital Zürich | Institute of Medical Virology, University of Zurich | Stefan Schmutz, Maryam Zaheri, Verena Kufner, Gabriela Ziltener, Patrick Redli, Fiona Steiner, Jon Huder, Riccarda Capaul, Andrea Zbinden, Jürg Böni, Michael Huber, Alexandra Trkola |

| | | | |
|---|---|---|---|
| EPI_ISL_528828, EPI_ISL_528832, EPI_ISL_528848, EPI_ISL_528864 | CSIR-Centre for Cellular and Molecular Biology | CSIR-Centre for Cellular and Molecular Biology | Sofia Banu, Payel Mukherjee, Priya Singh,Onkar Kulkarni, Dhiviya Vedagiri, Divya Gupta, Vishal Sah, Santosh Kumar Kuncha, Krishnan Harinivas Harshan, Archana Bharadwaj Siva, Karthik Bharadwaj Tallapaka, Shagufta Khan, Lamuk Zaveri, Namami Gaur, Sakshi Shambhavi, Nikhil Hajirnis, M Soujanya Reddy, Pratheusa Maccha,Tulasi Nagabandi, Purushotham Vodnala, Deepak Kumar, Devi Prasad Vijayashankar, Disha Nanda, Divya Das, Jotin Gogoi, Manish Bhattacharjee, Rakesh K Mishra, Divya Tej Sowpati |
| EPI_ISL_529030 | Kingston Health Sciences Center | Queen's Genomics Lab at Ongwanada (Q-GLO) | Sjaarda CP, Rustom N, Huang D, Perez-Patrigeon S, Hudson ML, Wong H,Guan H, Ayub M, Soares CN, Colautti R, Evans GA, Sheth P |
| EPI_ISL_529139 | Centro de Desenvolvimento Tecnologico em Saude, Fundacao Oswaldo Cruz | Centro de Desenvolvimento Tecnologico em Saude, Fundacao Oswaldo Cruz | Souza,T.M., Fintelman-Rodrigues,N., De Paula,A.D., Saraiva,F.B., Ferreira,M.A., Sacramento,C.Q., Medeiros,M.A. |
| EPI_ISL_529140 | Centro de Desenvolvimento Tecnologico em Saude, Fundacao Oswaldo Cruz | Centro de Desenvolvimento Tecnologico em Saude, Fundacao Oswaldo Cruz | Souza,T.M., Fintelman-Rodrigues,N., De Paula,A.D., Saraiva,F.B., Ferreira,M.A., Sacramento,C.Q., Medeiros,M.A. |
| EPI_ISL_530225, EPI_ISL_530226, EPI_ISL_530227, EPI_ISL_530228, EPI_ISL_530229, EPI_ISL_530230, EPI_ISL_530231, EPI_ISL_530232, EPI_ISL_530233, EPI_ISL_530234, EPI_ISL_530235, EPI_ISL_530236, EPI_ISL_530237, EPI_ISL_530238, EPI_ISL_530239, EPI_ISL_530240, EPI_ISL_530241, EPI_ISL_530242, EPI_ISL_530243, EPI_ISL_530244, EPI_ISL_530245, EPI_ISL_530246, EPI_ISL_530247, EPI_ISL_530248, EPI_ISL_530249, EPI_ISL_530250, EPI_ISL_530251, EPI_ISL_530252, EPI_ISL_530253, EPI_ISL_530254, EPI_ISL_530255, EPI_ISL_530256, EPI_ISL_530257, EPI_ISL_530258, EPI_ISL_530259, EPI_ISL_530260, EPI_ISL_530261, EPI_ISL_530262, EPI_ISL_530263, EPI_ISL_530264, EPI_ISL_530265, EPI_ISL_530266, EPI_ISL_530267, EPI_ISL_530268, EPI_ISL_530269, EPI_ISL_530270, EPI_ISL_530271, EPI_ISL_530272, EPI_ISL_530273, EPI_ISL_530274, EPI_ISL_530275, EPI_ISL_530276, EPI_ISL_530277, EPI_ISL_530278, EPI_ISL_530279, EPI_ISL_530280, EPI_ISL_530281, EPI_ISL_530282, EPI_ISL_530283 | | | |
| see above | Queensland Health Forensic and Scientific Services, Public Health Virology | Public Health Virology Laboratory, Forensic and Scientific Services, Queensland Health | Son Nguyen et al |
| EPI_ISL_534336 | Department of Laboratory Medicine, National Taiwan University Hospital | Microbial Genomics Core Lab, National Taiwan University Centers of Genomic and Precision Medicine | Shiou-Hwei Yeh, You-Yu Lin, Ya-Yun Lai, Chiao-Ling Li, Shan-Chwen Chang, Pei-Jer Chen, Sui-Yuan Chang |
| EPI_ISL_534337, EPI_ISL_534338, EPI_ISL_534339, EPI_ISL_534340, EPI_ISL_534341, EPI_ISL_534342, EPI_ISL_534343, EPI_ISL_534344, EPI_ISL_534345 | Molecular diagnostic laboratory of Federal Budget Institution of Science "Central Research Institute of Epidemiology" of The Federal Service on Customers' Rights Protection and Human Well-being Surveillance | Group of Genomics and Postgenomic Technologies of Central Research Institute of Epidemilology | Speranskaya AS, Kaptelova VV, Samoilov AE, Valdokhina AV, Bulanenko VP, Bukharina A.Y., Tivanova EV, Shipulina OY, Akimkin VG |
| EPI_ISL_534346, EPI_ISL_534347, EPI_ISL_534348, EPI_ISL_534349 | Molecular diagnostic laboratory of Federal Budget Institution of Science "Central Research Institute of Epidemiology" of The Federal Service on Customers' Rights Protection and Human Well-being Surveillance | Group of Genomics and Postgenomic Technologies of Central Research Institute of Epidemilology | Speranskaya AS, Kaptelova VV, Valdokhina AV, Bulanenko VP, Samoilov AE, Korneenko EV, Tivanova EV, Shipulina OY, Akimkin VG |
| EPI_ISL_534763, EPI_ISL_534764, EPI_ISL_534765, EPI_ISL_534766, EPI_ISL_534767, EPI_ISL_534768, EPI_ISL_534769, EPI_ISL_534770, EPI_ISL_534771, EPI_ISL_534772, EPI_ISL_534773, EPI_ISL_534774, EPI_ISL_534775, EPI_ISL_534776, EPI_ISL_534777, EPI_ISL_534778, EPI_ISL_534779, EPI_ISL_534780, EPI_ISL_534781, EPI_ISL_534782, EPI_ISL_534783, EPI_ISL_534784, EPI_ISL_534785, EPI_ISL_534786, EPI_ISL_534787, EPI_ISL_534788, EPI_ISL_534789, EPI_ISL_534790, EPI_ISL_534791, EPI_ISL_534792, EPI_ISL_534793, EPI_ISL_534794, EPI_ISL_534795, EPI_ISL_534796, EPI_ISL_534797, EPI_ISL_534798, EPI_ISL_534799, EPI_ISL_534800, EPI_ISL_534801, EPI_ISL_534802, EPI_ISL_534803, EPI_ISL_534804, EPI_ISL_534805, EPI_ISL_534806, EPI_ISL_534807, EPI_ISL_534808, EPI_ISL_534809, EPI_ISL_534810, EPI_ISL_534811, EPI_ISL_534812, EPI_ISL_534813, EPI_ISL_534814, EPI_ISL_534815, EPI_ISL_534816, EPI_ISL_534817, EPI_ISL_534818, EPI_ISL_534819, EPI_ISL_534820, EPI_ISL_534821, EPI_ISL_534822, EPI_ISL_534823, EPI_ISL_534824, EPI_ISL_534825, EPI_ISL_534826, EPI_ISL_534827, EPI_ISL_534828, EPI_ISL_534829, EPI_ISL_534830, EPI_ISL_534831, EPI_ISL_534832, EPI_ISL_534833, EPI_ISL_534834, EPI_ISL_534835, EPI_ISL_534836, EPI_ISL_534837, EPI_ISL_534838, EPI_ISL_534839, EPI_ISL_534840, EPI_ISL_534841, EPI_ISL_534842, EPI_ISL_534843, EPI_ISL_534844, EPI_ISL_534845, EPI_ISL_534846, EPI_ISL_534847, EPI_ISL_534848, EPI_ISL_534849, EPI_ISL_534850, EPI_ISL_534851, EPI_ISL_534852, EPI_ISL_534853, EPI_ISL_534854, EPI_ISL_534855, EPI_ISL_534856, EPI_ISL_534857, EPI_ISL_534858, EPI_ISL_534859, EPI_ISL_534860, EPI_ISL_534861, EPI_ISL_534862, EPI_ISL_534863, EPI_ISL_534864, EPI_ISL_534865, EPI_ISL_534866, EPI_ISL_534867, EPI_ISL_534868, EPI_ISL_534869, EPI_ISL_534870, EPI_ISL_534871, EPI_ISL_534872, EPI_ISL_534873, EPI_ISL_534874, EPI_ISL_534875, EPI_ISL_534876, EPI_ISL_534877, EPI_ISL_534878, EPI_ISL_534879, EPI_ISL_534880, EPI_ISL_534881, EPI_ISL_534882, EPI_ISL_534883, EPI_ISL_534884, EPI_ISL_534885, EPI_ISL_534886, EPI_ISL_534887, EPI_ISL_534888, EPI_ISL_534889, EPI_ISL_534890, EPI_ISL_534891, EPI_ISL_534892, EPI_ISL_534893, EPI_ISL_534894, EPI_ISL_534895, EPI_ISL_534896, EPI_ISL_534897, EPI_ISL_534898, EPI_ISL_534899, EPI_ISL_534900, EPI_ISL_534901, EPI_ISL_534902, EPI_ISL_534903, EPI_ISL_534904, EPI_ISL_534905, EPI_ISL_534906, EPI_ISL_534907, EPI_ISL_534908, EPI_ISL_534909, EPI_ISL_534910, EPI_ISL_534911, EPI_ISL_534912, EPI_ISL_534913, EPI_ISL_534914, EPI_ISL_534915, EPI_ISL_534916, EPI_ISL_534917, EPI_ISL_534918, EPI_ISL_534919, EPI_ISL_534920, EPI_ISL_534921, EPI_ISL_534922, EPI_ISL_534923, EPI_ISL_534924, EPI_ISL_534925, EPI_ISL_534926, EPI_ISL_534927, EPI_ISL_534928, EPI_ISL_534929, EPI_ISL_534930, EPI_ISL_534931, EPI_ISL_534932, EPI_ISL_534933, EPI_ISL_534934, EPI_ISL_534935, EPI_ISL_534936, EPI_ISL_534937, EPI_ISL_534938, EPI_ISL_534939, EPI_ISL_534940, EPI_ISL_534941, EPI_ISL_534942, EPI_ISL_534943, EPI_ISL_534944, EPI_ISL_534945, EPI_ISL_534946, EPI_ISL_534947, EPI_ISL_534948, EPI_ISL_534949, EPI_ISL_534950, EPI_ISL_534951, EPI_ISL_534952, EPI_ISL_534953, EPI_ISL_534954, EPI_ISL_534955, EPI_ISL_534956, EPI_ISL_534957, EPI_ISL_534958, EPI_ISL_534959, EPI_ISL_534960, EPI_ISL_534961, EPI_ISL_534962, EPI_ISL_534963, EPI_ISL_534964, EPI_ISL_534965, EPI_ISL_534966, EPI_ISL_534967, EPI_ISL_534968, EPI_ISL_534969, EPI_ISL_534970, EPI_ISL_534971, EPI_ISL_534972, EPI_ISL_534973, EPI_ISL_534974, EPI_ISL_534975, EPI_ISL_534976, EPI_ISL_534977, EPI_ISL_534978, EPI_ISL_534979, EPI_ISL_534980, EPI_ISL_534981, EPI_ISL_534982, EPI_ISL_534983, EPI_ISL_534984, EPI_ISL_534985, EPI_ISL_534986, EPI_ISL_534987, EPI_ISL_534988, EPI_ISL_534989, EPI_ISL_534990, EPI_ISL_534991, EPI_ISL_534992, EPI_ISL_534993, EPI_ISL_534994, EPI_ISL_534995, EPI_ISL_534996, EPI_ISL_534997, EPI_ISL_534998, EPI_ISL_534999, EPI_ISL_535000, EPI_ISL_535001, EPI_ISL_535002, EPI_ISL_535003, EPI_ISL_535004, EPI_ISL_535005, EPI_ISL_535006, EPI_ISL_535007, EPI_ISL_535008, EPI_ISL_535009, EPI_ISL_535010, EPI_ISL_535011, EPI_ISL_535012, EPI_ISL_535013, EPI_ISL_535014, EPI_ISL_535015, EPI_ISL_535016, EPI_ISL_535017, EPI_ISL_535018, EPI_ISL_535019, EPI_ISL_535020, EPI_ISL_535021, EPI_ISL_535022, EPI_ISL_535023 | | | |
| see above | Oxford Viromics, NDM, University of Oxford; Oxford University Hospitals; Basingstoke and North Hampshire Hospital | COVID-19 Genomics UK (COG-UK) Consortium | Tanya Golubchik, David Bonsall, George Macintyre, Amy Trebes, Mariateresa de Cesare, Catrin Moore, Alex Mobbs, Anita Justice, Robert Shaw, Monique Andersson, Timothy Peto, Emma Wise, Nathan Moore, Jessica Lynch, Nick Cortes, Matilde Mori, Stephen Kidd, David Buck, John Todd, Christophe Fraser |
| EPI_ISL_537874, EPI_ISL_537875, EPI_ISL_537876, EPI_ISL_537877, EPI_ISL_537878, EPI_ISL_537879, EPI_ISL_537880, EPI_ISL_537881, EPI_ISL_537882, EPI_ISL_537883, EPI_ISL_537884, EPI_ISL_537885, EPI_ISL_537886, EPI_ISL_537887, EPI_ISL_537888, EPI_ISL_537889, EPI_ISL_537890, EPI_ISL_537891, EPI_ISL_537892, EPI_ISL_537893, EPI_ISL_537894, EPI_ISL_537895, EPI_ISL_537896, EPI_ISL_537897, EPI_ISL_537898, EPI_ISL_537899, EPI_ISL_537900, EPI_ISL_537901, EPI_ISL_537902, EPI_ISL_537903, EPI_ISL_537904, EPI_ISL_537905, EPI_ISL_537906, EPI_ISL_537907, EPI_ISL_537908, EPI_ISL_537909, EPI_ISL_537910, EPI_ISL_537911, EPI_ISL_537912, EPI_ISL_537913, EPI_ISL_537914, EPI_ISL_537915, EPI_ISL_537916, EPI_ISL_537917, EPI_ISL_537918, EPI_ISL_537919, EPI_ISL_537920, EPI_ISL_537921, EPI_ISL_537922, EPI_ISL_537923, EPI_ISL_537924, EPI_ISL_537925, EPI_ISL_537926, EPI_ISL_537927, EPI_ISL_537928, EPI_ISL_537929, EPI_ISL_537930, EPI_ISL_537931, EPI_ISL_537932, EPI_ISL_537933, EPI_ISL_537934, EPI_ISL_537935, EPI_ISL_537936, EPI_ISL_537937, EPI_ISL_537938, EPI_ISL_537939, EPI_ISL_537940, EPI_ISL_537941, EPI_ISL_537942, EPI_ISL_537943, EPI_ISL_537944, EPI_ISL_537945 | | | |
| see above | Hospital Universitario Araba. Vitoria-Gasteiz | SeqCOVID-SPAIN consortium/IBV(CSIC) | Silvia Hernáez Crespo, Carmen Gómez González, Amaia Aguirre Quiñonero, Marina Fernández Torres, Ma Rosario Almela Ferrer, Ma Concepción Lecaroz Agara, Andrés Canut Blasco and SeqCOVID-SPAIN consortium |
| EPI_ISL_538118, EPI_ISL_538231, EPI_ISL_538232 | Hospital Universitario Araba. Vitoria-Gasteiz | SeqCOVID-SPAIN consortium/IBV(CSIC) | Silvia Hernáez Crespo, Carmen Gómez González, Amaia Aguirre Quiñonero, Marina Fernández Torres, Mª Rosario Almela Ferrer, Mª Concepción Lecaroz Agara, Andrés Canut Blasco and SeqCOVID-SPAIN consortium |
| EPI_ISL_539299 | Molecular diagnostic laboratory of Federal Budget Institution of Science "Central Research Institute of Epidemiology" of The Federal Service on Customers' Rights Protection and Human Well-being Surveillance | Group of Genomics and Postgenomic Technologies of Central Research Institute of Epidemilology | Speranskaya AS, Kaptelova VV, Valdokhina AV, Bulanenko VP, Samoilov AE, Korneenko EV, Tivanova EV, Shipulina OY, Akimkin VG |
| EPI_ISL_539640 | CSIR-Centre for Cellular and Molecular Biology | CSIR-Centre for Cellular and Molecular Biology | Sofia Banu, Payel Mukherjee, Priya Singh,Onkar Kulkarni, Dhiviya Vedagiri, Divya Gupta, Vishal Sah, Santosh Kumar Kuncha, Krishnan Harinivas Harshan, Archana Bharadwaj Siva, Karthik Bharadwaj Tallapaka, Shagufta Khan, Lamuk Zaveri, Namami Gaur, Sakshi Shambhavi, Nikhil Hajirnis, M Soujanya Reddy, Pratheusa Maccha, Tulasi Nagabandi, Purushotham Vodnala, Disha Nanda, Divya Das, Jotin Gogoi, Manish Bhattacharjee, Ravi Prasad Mukku, Rakesh K Mishra, Divya Tej Sowpati |
| EPI_ISL_539641 | CSIR-Centre for Cellular and Molecular Biology | CSIR-Centre for Cellular and Molecular Biology | Sofia Banu, Payel Mukherjee, Priya Singh,Onkar Kulkarni, Dhiviya Vedagiri, Divya Gupta, Vishal Sah, Santosh Kumar Kuncha, Krishnan Harinivas Harshan, Archana Bharadwaj Siva, Karthik Bharadwaj Tallapaka, Shagufta Khan, Lamuk Zaveri, Namami Gaur, Sakshi Shambhavi, Nikhil Hajirnis, M Soujanya Reddy, Pratheusa Maccha,Tulasi Nagabandi, Purushotham Vodnala, Deepak Kumar, Devi Prasad Vijayashankar, Disha Nanda, Divya Das, Jotin Gogoi, Manish Bhattacharjee, Rakesh K Mishra, Divya Tej Sowpati |
| EPI_ISL_539642 | CSIR-Centre for Cellular and Molecular Biology | CSIR-Centre for Cellular and Molecular Biology | Sofia Banu, Payel Mukherjee, Priya Singh,Onkar Kulkarni, Dhiviya Vedagiri, Divya Gupta, Vishal Sah, Santosh Kumar Kuncha, Krishnan Harinivas Harshan, Archana Bharadwaj Siva, Karthik Bharadwaj Tallapaka, Shagufta Khan, Lamuk Zaveri, Namami Gaur, Sakshi Shambhavi, Tulasi Nagabandi, Nikhil Hajirnis, M Soujanya Reddy, Pratheusa Maccha, Purushotham Vodnala, Gokulan C G, Gunjan Purohit, Hanuman Tulashiram Kale, Pankaj Kumar, Prachand Issarapu, Rakesh K Mishra, Divya Tej Sowpati |
| EPI_ISL_539670 | CSIR-Centre for Cellular and Molecular Biology | CSIR-Centre for Cellular and Molecular Biology | Sofia Banu, Payel Mukherjee, Priya Singh,Onkar Kulkarni, Dhiviya Vedagiri, Divya Gupta, Vishal Sah, Santosh Kumar Kuncha, Krishnan Harinivas Harshan, Archana Bharadwaj Siva, Karthik Bharadwaj Tallapaka, Shagufta Khan, Lamuk Zaveri, Namami Gaur, Sakshi Shambhavi, Nikhil Hajirnis, M Soujanya Reddy, Pratheusa Maccha, Tulasi Nagabandi, Purushotham Vodnala, Disha Nanda, Divya Das, Jotin Gogoi, Manish Bhattacharjee, Ravi Prasad Mukku, Rakesh K Mishra, Divya Tej Sowpati |
| EPI_ISL_539671 | CSIR-Centre for Cellular and Molecular Biology | CSIR-Centre for Cellular and Molecular Biology | Sofia Banu, Payel Mukherjee, Priya Singh,Onkar Kulkarni, Dhiviya Vedagiri, Divya Gupta, Vishal Sah, Santosh Kumar Kuncha, Krishnan Harinivas Harshan, Archana Bharadwaj Siva, Karthik Bharadwaj Tallapaka, Shagufta Khan, Lamuk Zaveri, Namami Gaur, Sakshi Shambhavi, Nikhil Hajirnis, M Soujanya Reddy, Pratheusa Maccha,Tulasi Nagabandi, Purushotham Vodnala, Deepak Kumar, Devi Prasad Vijayashankar, Disha Nanda, Divya Das, Jotin Gogoi, Manish Bhattacharjee, Rakesh K Mishra, Divya Tej Sowpati |
| EPI_ISL_539672 | CSIR-Centre for Cellular and Molecular Biology | CSIR-Centre for Cellular and Molecular Biology | Sofia Banu, Payel Mukherjee, Priya Singh,Onkar Kulkarni, Dhiviya Vedagiri, Divya Gupta, Vishal Sah, Santosh Kumar Kuncha, Krishnan Harinivas Harshan, Archana Bharadwaj Siva, Karthik Bharadwaj Tallapaka, Shagufta Khan, Lamuk Zaveri, Namami Gaur, Sakshi Shambhavi, Tulasi Nagabandi, |

| | | | |
|---|---|---|---|
| | | | Nikhil Hajirnis, M Soujanya Reddy, Pratheusa Maccha, Purushotham Vodnala, Gokulan C G, Gunjan Purohit, Hanuman Tulashiram Kale, Pankaj Kumar, Prachand Issarapu, Rakesh K Mishra, Divya Tej Sowpati |
| EPI_ISL_539700 | CSIR-Centre for Cellular and Molecular Biology | CSIR-Centre for Cellular and Molecular Biology | Sofia Banu, Payel Mukherjee, Priya Singh,Onkar Kulkarni, Dhiviya Vedagiri, Divya Gupta, Vishal Sah, Santosh Kumar Kuncha, Krishnan Harinivas Harshan, Archana Bharadwaj Siva, Karthik Bharadwaj Tallapaka, Shagufta Khan, Lamuk Zaveri, Namami Gaur, Sakshi Shambhavi, Nikhil Hajirnis, M Soujanya Reddy, Pratheusa Maccha, Tulasi Nagabandi, Purushotham Vodnala, Disha Nanda, Divya Das, Jotin Gogoi, Manish Bhattacharjee, Ravi Prasad Mukku, Rakesh K Mishra, Divya Tej Sowpati |
| EPI_ISL_539701 | CSIR-Centre for Cellular and Molecular Biology | CSIR-Centre for Cellular and Molecular Biology | Sofia Banu, Payel Mukherjee, Priya Singh,Onkar Kulkarni, Dhiviya Vedagiri, Divya Gupta, Vishal Sah, Santosh Kumar Kuncha, Krishnan Harinivas Harshan, Archana Bharadwaj Siva, Karthik Bharadwaj Tallapaka, Shagufta Khan, Lamuk Zaveri, Namami Gaur, Sakshi Shambhavi, Nikhil Hajirnis, M Soujanya Reddy, Pratheusa Maccha,Tulasi Nagabandi, Purushotham Vodnala, Deepak Kumar, Devi Prasad Vijayashankar, Disha Nanda, Divya Das, Jotin Gogoi, Manish Bhattacharjee, Rakesh K Mishra, Divya Tej Sowpati |
| EPI_ISL_539702 | CSIR-Centre for Cellular and Molecular Biology | CSIR-Centre for Cellular and Molecular Biology | Sofia Banu, Payel Mukherjee, Priya Singh,Onkar Kulkarni, Dhiviya Vedagiri, Divya Gupta, Vishal Sah, Santosh Kumar Kuncha, Krishnan Harinivas Harshan, Archana Bharadwaj Siva, Karthik Bharadwaj Tallapaka, Shagufta Khan, Lamuk Zaveri, Namami Gaur, Sakshi Shambhavi, Tulasi Nagabandi, Nikhil Hajirnis, M Soujanya Reddy, Pratheusa Maccha, Purushotham Vodnala, Gokulan C G, Gunjan Purohit, Hanuman Tulashiram Kale, Pankaj Kumar, Prachand Issarapu, Rakesh K Mishra, Divya Tej Sowpati |
| EPI_ISL_539730 | CSIR-Centre for Cellular and Molecular Biology | CSIR-Centre for Cellular and Molecular Biology | Sofia Banu, Payel Mukherjee, Priya Singh,Onkar Kulkarni, Dhiviya Vedagiri, Divya Gupta, Vishal Sah, Santosh Kumar Kuncha, Krishnan Harinivas Harshan, Archana Bharadwaj Siva, Karthik Bharadwaj Tallapaka, Shagufta Khan, Lamuk Zaveri, Namami Gaur, Sakshi Shambhavi, Nikhil Hajirnis, M Soujanya Reddy, Pratheusa Maccha, Tulasi Nagabandi, Purushotham Vodnala, Disha Nanda, Divya Das, Jotin Gogoi, Manish Bhattacharjee, Ravi Prasad Mukku, Rakesh K Mishra, Divya Tej Sowpati |
| EPI_ISL_539731 | CSIR-Centre for Cellular and Molecular Biology | CSIR-Centre for Cellular and Molecular Biology | Sofia Banu, Payel Mukherjee, Priya Singh,Onkar Kulkarni, Dhiviya Vedagiri, Divya Gupta, Vishal Sah, Santosh Kumar Kuncha, Krishnan Harinivas Harshan, Archana Bharadwaj Siva, Karthik Bharadwaj Tallapaka, Shagufta Khan, Lamuk Zaveri, Namami Gaur, Sakshi Shambhavi, Nikhil Hajirnis, M Soujanya Reddy, Pratheusa Maccha,Tulasi Nagabandi, Purushotham Vodnala, Deepak Kumar, Devi Prasad Vijayashankar, Disha Nanda, Divya Das, Jotin Gogoi, Manish Bhattacharjee, Rakesh K Mishra, Divya Tej Sowpati |
| EPI_ISL_539732 | CSIR-Centre for Cellular and Molecular Biology | CSIR-Centre for Cellular and Molecular Biology | Sofia Banu, Payel Mukherjee, Priya Singh,Onkar Kulkarni, Dhiviya Vedagiri, Divya Gupta, Vishal Sah, Santosh Kumar Kuncha, Krishnan Harinivas Harshan, Archana Bharadwaj Siva, Karthik Bharadwaj Tallapaka, Shagufta Khan, Lamuk Zaveri, Namami Gaur, Sakshi Shambhavi, Tulasi Nagabandi, Nikhil Hajirnis, M Soujanya Reddy, Pratheusa Maccha, Purushotham Vodnala, Gokulan C G, Gunjan Purohit, Hanuman Tulashiram Kale, Pankaj Kumar, Prachand Issarapu, Rakesh K Mishra, Divya Tej Sowpati |
| EPI_ISL_539760 | CSIR-Centre for Cellular and Molecular Biology | CSIR-Centre for Cellular and Molecular Biology | Sofia Banu, Payel Mukherjee, Priya Singh,Onkar Kulkarni, Dhiviya Vedagiri, Divya Gupta, Vishal Sah, Santosh Kumar Kuncha, Krishnan Harinivas Harshan, Archana Bharadwaj Siva, Karthik Bharadwaj Tallapaka, Shagufta Khan, Lamuk Zaveri, Namami Gaur, Sakshi Shambhavi, Nikhil Hajirnis, M Soujanya Reddy, Pratheusa Maccha, Tulasi Nagabandi, Purushotham Vodnala, Disha Nanda, Divya Das, Jotin Gogoi, Manish Bhattacharjee, Ravi Prasad Mukku, Rakesh K Mishra, Divya Tej Sowpati |
| EPI_ISL_539761 | CSIR-Centre for Cellular and Molecular Biology | CSIR-Centre for Cellular and Molecular Biology | Sofia Banu, Payel Mukherjee, Priya Singh,Onkar Kulkarni, Dhiviya Vedagiri, Divya Gupta, Vishal Sah, Santosh Kumar Kuncha, Krishnan Harinivas Harshan, Archana Bharadwaj Siva, Karthik Bharadwaj Tallapaka, Shagufta Khan, Lamuk Zaveri, Namami Gaur, Sakshi Shambhavi, Nikhil Hajirnis, M Soujanya Reddy, Pratheusa Maccha,Tulasi Nagabandi, Purushotham Vodnala, Deepak Kumar, Devi Prasad Vijayashankar, Disha Nanda, Divya Das, Jotin Gogoi, Manish Bhattacharjee, Rakesh K Mishra, Divya Tej Sowpati |
| EPI_ISL_539762 | CSIR-Centre for Cellular and Molecular Biology | CSIR-Centre for Cellular and Molecular Biology | Sofia Banu, Payel Mukherjee, Priya Singh,Onkar Kulkarni, Dhiviya Vedagiri, Divya Gupta, Vishal Sah, Santosh Kumar Kuncha, Krishnan Harinivas Harshan, Archana Bharadwaj Siva, Karthik Bharadwaj Tallapaka, Shagufta Khan, Lamuk Zaveri, Namami Gaur, Sakshi Shambhavi, Tulasi Nagabandi, Nikhil Hajirnis, M Soujanya Reddy, Pratheusa Maccha, Purushotham Vodnala, Gokulan C G, Gunjan Purohit, Hanuman Tulashiram Kale, Pankaj Kumar, Prachand Issarapu, Rakesh K Mishra, Divya Tej Sowpati |
| EPI_ISL_539894, EPI_ISL_539896 | PHE South West Regional Laboratory, National Infection Service | Wellcome Sanger Institute for the COVID-19 Genomics UK (COG-UK) consortium | Stephanie Hutchings, Hannah Pymont, Dr Peter Muir, Barry Vipond, Rich Hopes; and Alex Alderton, Roberto Amato, Sonia Goncalves, Ewan Harrison, David K. Jackson, Ian Johnston, Dominic Kwiatkowski, Cordelia Langford, John Sillitoe on behalf of the Wellcome Sanger Institute COVID-19 Surveillance Team |
| EPI_ISL_541752, EPI_ISL_541754, EPI_ISL_541755, EPI_ISL_541757, EPI_ISL_541758, EPI_ISL_541759, EPI_ISL_541760, EPI_ISL_541761, EPI_ISL_541762, EPI_ISL_541763, EPI_ISL_541764, EPI_ISL_541765, EPI_ISL_541766, EPI_ISL_541768, EPI_ISL_541769, EPI_ISL_541770, EPI_ISL_541771, EPI_ISL_541772, EPI_ISL_541775, EPI_ISL_541776, EPI_ISL_541777, EPI_ISL_541778, EPI_ISL_541779, EPI_ISL_541780, EPI_ISL_541781, EPI_ISL_541782, EPI_ISL_541783 | | | |
| see above | Barts Health NHS Trust | Wellcome Sanger Institute for the COVID-19 Genomics UK (COG-UK) consortium | Teresa Cutino-Moguel, Mark Hopkins, Beatrix Kele, David Harrington and Alex Alderton, Roberto Amato, Sonia Goncalves, Ewan Harrison, David K. Jackson, Ian Johnston, Dominic Kwiatkowski, Cordelia Langford, John Sillitoe on behalf of the Wellcome Sanger Institute COVID-19 Surveillance Team |
| EPI_ISL_549329, EPI_ISL_549330, EPI_ISL_549331, EPI_ISL_549332, EPI_ISL_549333, EPI_ISL_549335, EPI_ISL_549337, EPI_ISL_549338, EPI_ISL_549339, EPI_ISL_549340, EPI_ISL_549342, EPI_ISL_549345, EPI_ISL_549346, EPI_ISL_549347, EPI_ISL_549348, EPI_ISL_549396, EPI_ISL_549397, EPI_ISL_549398, EPI_ISL_549399, EPI_ISL_549400, EPI_ISL_549401, EPI_ISL_549402, EPI_ISL_549403, EPI_ISL_549404, EPI_ISL_549405, EPI_ISL_549406, EPI_ISL_549407, EPI_ISL_549408, EPI_ISL_549409, EPI_ISL_549410, EPI_ISL_549411, EPI_ISL_549412, EPI_ISL_549413, EPI_ISL_549414, EPI_ISL_549415, EPI_ISL_549416, EPI_ISL_549417, EPI_ISL_549418, EPI_ISL_549419, EPI_ISL_549420, EPI_ISL_549421, EPI_ISL_549422, EPI_ISL_549423, EPI_ISL_549424, EPI_ISL_549425, EPI_ISL_549426, EPI_ISL_549427, EPI_ISL_549428, EPI_ISL_549429, EPI_ISL_549430, EPI_ISL_549431, EPI_ISL_549432, EPI_ISL_549433, EPI_ISL_549434, EPI_ISL_549435, EPI_ISL_549436, EPI_ISL_549437, EPI_ISL_549438, EPI_ISL_549439, EPI_ISL_549440, EPI_ISL_549441, EPI_ISL_549442, EPI_ISL_549443, EPI_ISL_549444, EPI_ISL_549445, EPI_ISL_549446, EPI_ISL_549447, EPI_ISL_549448, EPI_ISL_549449, EPI_ISL_549450, EPI_ISL_549451, EPI_ISL_549452, EPI_ISL_549453, EPI_ISL_549454, EPI_ISL_549455, EPI_ISL_549456, EPI_ISL_549457, EPI_ISL_549458, EPI_ISL_549459, EPI_ISL_549523, EPI_ISL_549524, EPI_ISL_549525, EPI_ISL_549526, EPI_ISL_549527, EPI_ISL_549528, EPI_ISL_549529, EPI_ISL_549530, EPI_ISL_549531, EPI_ISL_549532, EPI_ISL_549533, EPI_ISL_549534, EPI_ISL_549535, EPI_ISL_549536, EPI_ISL_549537, EPI_ISL_549538, EPI_ISL_549539, EPI_ISL_549540, EPI_ISL_549541, EPI_ISL_549542, EPI_ISL_549543, EPI_ISL_549544, EPI_ISL_549545, EPI_ISL_549546 | | | |
| see above | Oxford Viromics, NDM, University of Oxford; Oxford University Hospitals; Basingstoke and North Hampshire Hospital | COVID-19 Genomics UK (COG-UK) Consortium | Tanya Golubchik, David Bonsall, George Macintyre, Amy Trebes, Mariateresa de Cesare, Catrin Moore, Alex Mobbs, Anita Justice, Robert Shaw, Monique Andersson, Timothy Peto, Emma Wise, Nathan Moore, Jessica Lynch, Nick Cortes, Matilde Mori, Stephen Kidd, David Buck, John Todd, Christophe Fraser |
| EPI_ISL_551354, EPI_ISL_551355 | Lighthouse Lab in Alderley Park | Wellcome Sanger Institute for the COVID-19 Genomics UK (COG-UK) consortium | The Lighthouse Lab in Alderley Park and Alex Alderton, Roberto Amato, Sonia Goncalves, Ewan Harrison, David K. Jackson, Ian Johnston, Dominic Kwiatkowski, Cordelia Langford, John Sillitoe on behalf of the Wellcome Sanger Institute COVID-19 Surveillance Team |
| EPI_ISL_551356, EPI_ISL_551357, EPI_ISL_551359 | Lighthouse Lab in Alderley Park | Wellcome Sanger Institute for the COVID-19 Genomics UK (COG-UK) consortium | The Lighthouse Lab in Alderley Park and Alex Alderton, Roberto Amato, Sonia Goncalves, Ewan Harrison, David K. Jackson, Ian Johnston, Dominic Kwiatkowski, Cordelia Langford, John Sillitoe on behalf of the Wellcome Sanger Institute COVID-19 Surveillance Team (http://www.sanger.ac.uk/covid-team) |
| EPI_ISL_551360 | Lighthouse Lab in Alderley Park | Wellcome Sanger Institute for the COVID-19 Genomics UK (COG-UK) consortium | The Lighthouse Lab in Alderley Park and Alex Alderton, Roberto Amato, Sonia Goncalves, Ewan Harrison, David K. Jackson, Ian Johnston, Dominic Kwiatkowski, Cordelia Langford, John Sillitoe on behalf of the Wellcome Sanger Institute COVID-19 Surveillance Team |
| EPI_ISL_551361 | Lighthouse Lab in Alderley Park | Wellcome Sanger Institute for the COVID-19 Genomics UK (COG-UK) consortium | The Lighthouse Lab in Alderley Park and Alex Alderton, Roberto Amato, Sonia Goncalves, Ewan Harrison, David K. Jackson, Ian Johnston, Dominic Kwiatkowski, Cordelia Langford, John Sillitoe on behalf of the Wellcome Sanger Institute COVID-19 Surveillance Team (http://www.sanger.ac.uk/covid-team) |
| EPI_ISL_551362 | Lighthouse Lab in Alderley Park | Wellcome Sanger Institute for the COVID-19 Genomics UK (COG-UK) consortium | The Lighthouse Lab in Alderley Park and Alex Alderton, Roberto Amato, Sonia Goncalves, Ewan Harrison, David K. Jackson, Ian Johnston, Dominic Kwiatkowski, Cordelia Langford, John Sillitoe on behalf of the Wellcome Sanger Institute COVID-19 Surveillance Team |
| EPI_ISL_551363 | Lighthouse Lab in Alderley Park | Wellcome Sanger Institute for the COVID-19 Genomics UK (COG-UK) consortium | The Lighthouse Lab in Alderley Park and Alex Alderton, Roberto Amato, Sonia Goncalves, Ewan Harrison, David K. Jackson, Ian Johnston, Dominic Kwiatkowski, Cordelia Langford, John Sillitoe on behalf of the Wellcome Sanger Institute COVID-19 Surveillance Team (http://www.sanger.ac.uk/covid-team) |
| EPI_ISL_551364 | Lighthouse Lab in Alderley Park | Wellcome Sanger Institute for the COVID-19 Genomics UK (COG-UK) consortium | The Lighthouse Lab in Alderley Park and Alex Alderton, Roberto Amato, Sonia Goncalves, Ewan Harrison, David K. Jackson, Ian Johnston, Dominic Kwiatkowski, Cordelia Langford, John Sillitoe on behalf of the Wellcome Sanger Institute COVID-19 Surveillance Team |
| EPI_ISL_551365, EPI_ISL_551366, EPI_ISL_551370 | Lighthouse Lab in Alderley Park | Wellcome Sanger Institute for the COVID-19 Genomics UK (COG-UK) consortium | The Lighthouse Lab in Alderley Park and Alex Alderton, Roberto Amato, Sonia Goncalves, Ewan Harrison, David K. Jackson, Ian Johnston, Dominic Kwiatkowski, Cordelia Langford, John Sillitoe on behalf of the Wellcome Sanger Institute COVID-19 Surveillance Team (http://www.sanger.ac.uk/covid-team) |
| EPI_ISL_551371 | Lighthouse Lab in Alderley Park | Wellcome Sanger Institute for the COVID-19 Genomics UK (COG-UK) consortium | The Lighthouse Lab in Alderley Park and Alex Alderton, Roberto Amato, Sonia Goncalves, Ewan Harrison, David K. Jackson, Ian Johnston, Dominic Kwiatkowski, Cordelia Langford, John Sillitoe on behalf of the Wellcome Sanger Institute COVID-19 Surveillance Team |

| | | | |
|---|---|---|---|
| EPI_ISL_551372 | Lighthouse Lab in Alderley Park | Wellcome Sanger Institute for the COVID-19 Genomics UK (COG-UK) consortium | The Lighthouse Lab in Alderley Park and Alex Alderton, Roberto Amato, Sonia Goncalves, Ewan Harrison, David K. Jackson, Ian Johnston, Dominic Kwiatkowski, Cordelia Langford, John Sillitoe on behalf of the Wellcome Sanger Institute COVID-19 Surveillance Team (http://www.sanger.ac.uk/covid-team) |
| EPI_ISL_551373 | Lighthouse Lab in Alderley Park | Wellcome Sanger Institute for the COVID-19 Genomics UK (COG-UK) consortium | The Lighthouse Lab in Alderley Park and Alex Alderton, Roberto Amato, Sonia Goncalves, Ewan Harrison, David K. Jackson, Ian Johnston, Dominic Kwiatkowski, Cordelia Langford, John Sillitoe on behalf of the Wellcome Sanger Institute COVID-19 Surveillance Team |
| EPI_ISL_551376 | Lighthouse Lab in Alderley Park | Wellcome Sanger Institute for the COVID-19 Genomics UK (COG-UK) consortium | The Lighthouse Lab in Alderley Park and Alex Alderton, Roberto Amato, Sonia Goncalves, Ewan Harrison, David K. Jackson, Ian Johnston, Dominic Kwiatkowski, Cordelia Langford, John Sillitoe on behalf of the Wellcome Sanger Institute COVID-19 Surveillance Team (http://www.sanger.ac.uk/covid-team) |
| EPI_ISL_551377, EPI_ISL_551379 | Lighthouse Lab in Alderley Park | Wellcome Sanger Institute for the COVID-19 Genomics UK (COG-UK) consortium | The Lighthouse Lab in Alderley Park and Alex Alderton, Roberto Amato, Sonia Goncalves, Ewan Harrison, David K. Jackson, Ian Johnston, Dominic Kwiatkowski, Cordelia Langford, John Sillitoe on behalf of the Wellcome Sanger Institute COVID-19 Surveillance Team |
| EPI_ISL_551380, EPI_ISL_551381, EPI_ISL_551382, EPI_ISL_551383, EPI_ISL_551385 | Lighthouse Lab in Alderley Park | Wellcome Sanger Institute for the COVID-19 Genomics UK (COG-UK) consortium | The Lighthouse Lab in Alderley Park and Alex Alderton, Roberto Amato, Sonia Goncalves, Ewan Harrison, David K. Jackson, Ian Johnston, Dominic Kwiatkowski, Cordelia Langford, John Sillitoe on behalf of the Wellcome Sanger Institute COVID-19 Surveillance Team (http://www.sanger.ac.uk/covid-team) |
| EPI_ISL_551386, EPI_ISL_551387, EPI_ISL_551389, EPI_ISL_551390, EPI_ISL_551392 | Lighthouse Lab in Alderley Park | Wellcome Sanger Institute for the COVID-19 Genomics UK (COG-UK) consortium | The Lighthouse Lab in Alderley Park and Alex Alderton, Roberto Amato, Sonia Goncalves, Ewan Harrison, David K. Jackson, Ian Johnston, Dominic Kwiatkowski, Cordelia Langford, John Sillitoe on behalf of the Wellcome Sanger Institute COVID-19 Surveillance Team |
| EPI_ISL_551393, EPI_ISL_551394 | Lighthouse Lab in Alderley Park | Wellcome Sanger Institute for the COVID-19 Genomics UK (COG-UK) consortium | The Lighthouse Lab in Alderley Park and Alex Alderton, Roberto Amato, Sonia Goncalves, Ewan Harrison, David K. Jackson, Ian Johnston, Dominic Kwiatkowski, Cordelia Langford, John Sillitoe on behalf of the Wellcome Sanger Institute COVID-19 Surveillance Team (http://www.sanger.ac.uk/covid-team) |
| EPI_ISL_551395, EPI_ISL_551397 | Lighthouse Lab in Alderley Park | Wellcome Sanger Institute for the COVID-19 Genomics UK (COG-UK) consortium | The Lighthouse Lab in Alderley Park and Alex Alderton, Roberto Amato, Sonia Goncalves, Ewan Harrison, David K. Jackson, Ian Johnston, Dominic Kwiatkowski, Cordelia Langford, John Sillitoe on behalf of the Wellcome Sanger Institute COVID-19 Surveillance Team |
| EPI_ISL_551399 | Lighthouse Lab in Alderley Park | Wellcome Sanger Institute for the COVID-19 Genomics UK (COG-UK) consortium | The Lighthouse Lab in Alderley Park and Alex Alderton, Roberto Amato, Sonia Goncalves, Ewan Harrison, David K. Jackson, Ian Johnston, Dominic Kwiatkowski, Cordelia Langford, John Sillitoe on behalf of the Wellcome Sanger Institute COVID-19 Surveillance Team (http://www.sanger.ac.uk/covid-team) |
| EPI_ISL_551403, EPI_ISL_551408 | Lighthouse Lab in Alderley Park | Wellcome Sanger Institute for the COVID-19 Genomics UK (COG-UK) consortium | The Lighthouse Lab in Alderley Park and Alex Alderton, Roberto Amato, Sonia Goncalves, Ewan Harrison, David K. Jackson, Ian Johnston, Dominic Kwiatkowski, Cordelia Langford, John Sillitoe on behalf of the Wellcome Sanger Institute COVID-19 Surveillance Team |
| EPI_ISL_551409 | Lighthouse Lab in Alderley Park | Wellcome Sanger Institute for the COVID-19 Genomics UK (COG-UK) consortium | The Lighthouse Lab in Alderley Park and Alex Alderton, Roberto Amato, Sonia Goncalves, Ewan Harrison, David K. Jackson, Ian Johnston, Dominic Kwiatkowski, Cordelia Langford, John Sillitoe on behalf of the Wellcome Sanger Institute COVID-19 Surveillance Team (http://www.sanger.ac.uk/covid-team) |
| EPI_ISL_551411, EPI_ISL_551415, EPI_ISL_551416 | Lighthouse Lab in Alderley Park | Wellcome Sanger Institute for the COVID-19 Genomics UK (COG-UK) consortium | The Lighthouse Lab in Alderley Park and Alex Alderton, Roberto Amato, Sonia Goncalves, Ewan Harrison, David K. Jackson, Ian Johnston, Dominic Kwiatkowski, Cordelia Langford, John Sillitoe on behalf of the Wellcome Sanger Institute COVID-19 Surveillance Team |
| EPI_ISL_551417, EPI_ISL_551420 | Lighthouse Lab in Alderley Park | Wellcome Sanger Institute for the COVID-19 Genomics UK (COG-UK) consortium | The Lighthouse Lab in Alderley Park and Alex Alderton, Roberto Amato, Sonia Goncalves, Ewan Harrison, David K. Jackson, Ian Johnston, Dominic Kwiatkowski, Cordelia Langford, John Sillitoe on behalf of the Wellcome Sanger Institute COVID-19 Surveillance Team (http://www.sanger.ac.uk/covid-team) |
| EPI_ISL_551421 | Lighthouse Lab in Alderley Park | Wellcome Sanger Institute for the COVID-19 Genomics UK (COG-UK) consortium | The Lighthouse Lab in Alderley Park and Alex Alderton, Roberto Amato, Sonia Goncalves, Ewan Harrison, David K. Jackson, Ian Johnston, Dominic Kwiatkowski, Cordelia Langford, John Sillitoe on behalf of the Wellcome Sanger Institute COVID-19 Surveillance Team |
| EPI_ISL_551422 | Lighthouse Lab in Alderley Park | Wellcome Sanger Institute for the COVID-19 Genomics UK (COG-UK) consortium | The Lighthouse Lab in Alderley Park and Alex Alderton, Roberto Amato, Sonia Goncalves, Ewan Harrison, David K. Jackson, Ian Johnston, Dominic Kwiatkowski, Cordelia Langford, John Sillitoe on behalf of the Wellcome Sanger Institute COVID-19 Surveillance Team (http://www.sanger.ac.uk/covid-team) |
| EPI_ISL_551423, EPI_ISL_551424, EPI_ISL_551425 | Lighthouse Lab in Alderley Park | Wellcome Sanger Institute for the COVID-19 Genomics UK (COG-UK) consortium | The Lighthouse Lab in Alderley Park and Alex Alderton, Roberto Amato, Sonia Goncalves, Ewan Harrison, David K. Jackson, Ian Johnston, Dominic Kwiatkowski, Cordelia Langford, John Sillitoe on behalf of the Wellcome Sanger Institute COVID-19 Surveillance Team |
| EPI_ISL_551426 | Lighthouse Lab in Alderley Park | Wellcome Sanger Institute for the COVID-19 Genomics UK (COG-UK) consortium | The Lighthouse Lab in Alderley Park and Alex Alderton, Roberto Amato, Sonia Goncalves, Ewan Harrison, David K. Jackson, Ian Johnston, Dominic Kwiatkowski, Cordelia Langford, John Sillitoe on behalf of the Wellcome Sanger Institute COVID-19 Surveillance Team (http://www.sanger.ac.uk/covid-team) |
| EPI_ISL_551427 | Lighthouse Lab in Alderley Park | Wellcome Sanger Institute for the COVID-19 Genomics UK (COG-UK) consortium | The Lighthouse Lab in Alderley Park and Alex Alderton, Roberto Amato, Sonia Goncalves, Ewan Harrison, David K. Jackson, Ian Johnston, Dominic Kwiatkowski, Cordelia Langford, John Sillitoe on behalf of the Wellcome Sanger Institute COVID-19 Surveillance Team |
| EPI_ISL_551428 | Lighthouse Lab in Alderley Park | Wellcome Sanger Institute for the COVID-19 Genomics UK (COG-UK) consortium | The Lighthouse Lab in Alderley Park and Alex Alderton, Roberto Amato, Sonia Goncalves, Ewan Harrison, David K. Jackson, Ian Johnston, Dominic Kwiatkowski, Cordelia Langford, John Sillitoe on behalf of the Wellcome Sanger Institute COVID-19 Surveillance Team (http://www.sanger.ac.uk/covid-team) |
| EPI_ISL_551429 | Lighthouse Lab in Alderley Park | Wellcome Sanger Institute for the COVID-19 Genomics UK (COG-UK) consortium | The Lighthouse Lab in Alderley Park and Alex Alderton, Roberto Amato, Sonia Goncalves, Ewan Harrison, David K. Jackson, Ian Johnston, Dominic Kwiatkowski, Cordelia Langford, John Sillitoe on behalf of the Wellcome Sanger Institute COVID-19 Surveillance Team |
| EPI_ISL_551434, EPI_ISL_551435 | Lighthouse Lab in Alderley Park | Wellcome Sanger Institute for the COVID-19 Genomics UK (COG-UK) consortium | The Lighthouse Lab in Alderley Park and Alex Alderton, Roberto Amato, Sonia Goncalves, Ewan Harrison, David K. Jackson, Ian Johnston, Dominic Kwiatkowski, Cordelia Langford, John Sillitoe on behalf of the Wellcome Sanger Institute COVID-19 Surveillance Team (http://www.sanger.ac.uk/covid-team) |
| EPI_ISL_551436, EPI_ISL_551437, EPI_ISL_551438 | Lighthouse Lab in Alderley Park | Wellcome Sanger Institute for the COVID-19 Genomics UK (COG-UK) consortium | The Lighthouse Lab in Alderley Park and Alex Alderton, Roberto Amato, Sonia Goncalves, Ewan Harrison, David K. Jackson, Ian Johnston, Dominic Kwiatkowski, Cordelia Langford, John Sillitoe on behalf of the Wellcome Sanger Institute COVID-19 Surveillance Team |
| EPI_ISL_551439 | Lighthouse Lab in Alderley Park | Wellcome Sanger Institute for the COVID-19 Genomics UK (COG-UK) consortium | The Lighthouse Lab in Alderley Park and Alex Alderton, Roberto Amato, Sonia Goncalves, Ewan Harrison, David K. Jackson, Ian Johnston, Dominic Kwiatkowski, Cordelia Langford, John Sillitoe on behalf of the Wellcome Sanger Institute COVID-19 Surveillance Team (http://www.sanger.ac.uk/covid-team) |
| EPI_ISL_551440, EPI_ISL_551441, EPI_ISL_551443, EPI_ISL_551444 | Lighthouse Lab in Alderley Park | Wellcome Sanger Institute for the COVID-19 Genomics UK (COG-UK) consortium | The Lighthouse Lab in Alderley Park and Alex Alderton, Roberto Amato, Sonia Goncalves, Ewan Harrison, David K. Jackson, Ian Johnston, Dominic Kwiatkowski, Cordelia Langford, John Sillitoe on behalf of the Wellcome Sanger Institute COVID-19 Surveillance Team |
| EPI_ISL_551445, EPI_ISL_551446 | Lighthouse Lab in Alderley Park | Wellcome Sanger Institute for the COVID-19 Genomics UK (COG-UK) consortium | The Lighthouse Lab in Alderley Park and Alex Alderton, Roberto Amato, Sonia Goncalves, Ewan Harrison, David K. Jackson, Ian Johnston, Dominic Kwiatkowski, Cordelia Langford, John Sillitoe on behalf of the Wellcome Sanger Institute COVID-19 Surveillance Team (http://www.sanger.ac.uk/covid-team) |
| EPI_ISL_551447 | Lighthouse Lab in Alderley Park | Wellcome Sanger Institute for the COVID-19 Genomics UK (COG-UK) consortium | The Lighthouse Lab in Alderley Park and Alex Alderton, Roberto Amato, Sonia Goncalves, Ewan Harrison, David K. Jackson, Ian Johnston, Dominic Kwiatkowski, Cordelia Langford, John Sillitoe on behalf of the Wellcome Sanger Institute COVID-19 Surveillance Team |
| EPI_ISL_551448 | Lighthouse Lab in Alderley Park | Wellcome Sanger Institute for the COVID-19 Genomics UK (COG-UK) consortium | The Lighthouse Lab in Alderley Park and Alex Alderton, Roberto Amato, Sonia Goncalves, Ewan Harrison, David K. Jackson, Ian Johnston, Dominic Kwiatkowski, Cordelia Langford, John Sillitoe on behalf of the Wellcome Sanger Institute COVID-19 Surveillance Team (http://www.sanger.ac.uk/covid-team) |
| EPI_ISL_551449, EPI_ISL_551450, EPI_ISL_551451, EPI_ISL_551452, EPI_ISL_551454, EPI_ISL_551455, EPI_ISL_551456, EPI_ISL_551457 | Lighthouse Lab in Alderley Park | Wellcome Sanger Institute for the COVID-19 Genomics UK (COG-UK) consortium | The Lighthouse Lab in Alderley Park and Alex Alderton, Roberto Amato, Sonia Goncalves, Ewan Harrison, David K. Jackson, Ian Johnston, Dominic Kwiatkowski, Cordelia Langford, John Sillitoe on behalf of the Wellcome Sanger Institute COVID-19 Surveillance Team |

| | | | |
|---|---|---|---|
| EPI_ISL_551458, EPI_ISL_551459 | Lighthouse Lab in Alderley Park | Wellcome Sanger Institute for the COVID-19 Genomics UK (COG-UK) consortium | The Lighthouse Lab in Alderley Park and Alex Alderton, Roberto Amato, Sonia Goncalves, Ewan Harrison, David K. Jackson, Ian Johnston, Dominic Kwiatkowski, Cordelia Langford, John Sillitoe on behalf of the Wellcome Sanger Institute COVID-19 Surveillance Team (http://www.sanger.ac.uk/covid-team) |
| EPI_ISL_551460 | Lighthouse Lab in Alderley Park | Wellcome Sanger Institute for the COVID-19 Genomics UK (COG-UK) consortium | The Lighthouse Lab in Alderley Park and Alex Alderton, Roberto Amato, Sonia Goncalves, Ewan Harrison, David K. Jackson, Ian Johnston, Dominic Kwiatkowski, Cordelia Langford, John Sillitoe on behalf of the Wellcome Sanger Institute COVID-19 Surveillance Team |
| EPI_ISL_551461, EPI_ISL_551462, EPI_ISL_551463, EPI_ISL_551464, EPI_ISL_551465, EPI_ISL_551467, EPI_ISL_551468 | Lighthouse Lab in Alderley Park | Wellcome Sanger Institute for the COVID-19 Genomics UK (COG-UK) consortium | The Lighthouse Lab in Alderley Park and Alex Alderton, Roberto Amato, Sonia Goncalves, Ewan Harrison, David K. Jackson, Ian Johnston, Dominic Kwiatkowski, Cordelia Langford, John Sillitoe on behalf of the Wellcome Sanger Institute COVID-19 Surveillance Team (http://www.sanger.ac.uk/covid-team) |
| EPI_ISL_551470, EPI_ISL_551471 | Lighthouse Lab in Alderley Park | Wellcome Sanger Institute for the COVID-19 Genomics UK (COG-UK) consortium | The Lighthouse Lab in Alderley Park and Alex Alderton, Roberto Amato, Sonia Goncalves, Ewan Harrison, David K. Jackson, Ian Johnston, Dominic Kwiatkowski, Cordelia Langford, John Sillitoe on behalf of the Wellcome Sanger Institute COVID-19 Surveillance Team |
| EPI_ISL_551472, EPI_ISL_551473, EPI_ISL_551474, EPI_ISL_551475, EPI_ISL_551477 | Lighthouse Lab in Alderley Park | Wellcome Sanger Institute for the COVID-19 Genomics UK (COG-UK) consortium | The Lighthouse Lab in Alderley Park and Alex Alderton, Roberto Amato, Sonia Goncalves, Ewan Harrison, David K. Jackson, Ian Johnston, Dominic Kwiatkowski, Cordelia Langford, John Sillitoe on behalf of the Wellcome Sanger Institute COVID-19 Surveillance Team (http://www.sanger.ac.uk/covid-team) |
| EPI_ISL_551478 | Lighthouse Lab in Alderley Park | Wellcome Sanger Institute for the COVID-19 Genomics UK (COG-UK) consortium | The Lighthouse Lab in Alderley Park and Alex Alderton, Roberto Amato, Sonia Goncalves, Ewan Harrison, David K. Jackson, Ian Johnston, Dominic Kwiatkowski, Cordelia Langford, John Sillitoe on behalf of the Wellcome Sanger Institute COVID-19 Surveillance Team |
| EPI_ISL_551479 | Lighthouse Lab in Alderley Park | Wellcome Sanger Institute for the COVID-19 Genomics UK (COG-UK) consortium | The Lighthouse Lab in Alderley Park and Alex Alderton, Roberto Amato, Sonia Goncalves, Ewan Harrison, David K. Jackson, Ian Johnston, Dominic Kwiatkowski, Cordelia Langford, John Sillitoe on behalf of the Wellcome Sanger Institute COVID-19 Surveillance Team (http://www.sanger.ac.uk/covid-team) |
| EPI_ISL_551480, EPI_ISL_551481, EPI_ISL_551482 | Lighthouse Lab in Alderley Park | Wellcome Sanger Institute for the COVID-19 Genomics UK (COG-UK) consortium | The Lighthouse Lab in Alderley Park and Alex Alderton, Roberto Amato, Sonia Goncalves, Ewan Harrison, David K. Jackson, Ian Johnston, Dominic Kwiatkowski, Cordelia Langford, John Sillitoe on behalf of the Wellcome Sanger Institute COVID-19 Surveillance Team |
| EPI_ISL_551485 | Lighthouse Lab in Alderley Park | Wellcome Sanger Institute for the COVID-19 Genomics UK (COG-UK) consortium | The Lighthouse Lab in Alderley Park and Alex Alderton, Roberto Amato, Sonia Goncalves, Ewan Harrison, David K. Jackson, Ian Johnston, Dominic Kwiatkowski, Cordelia Langford, John Sillitoe on behalf of the Wellcome Sanger Institute COVID-19 Surveillance Team (http://www.sanger.ac.uk/covid-team) |
| EPI_ISL_551486, EPI_ISL_551487 | Lighthouse Lab in Alderley Park | Wellcome Sanger Institute for the COVID-19 Genomics UK (COG-UK) consortium | The Lighthouse Lab in Alderley Park and Alex Alderton, Roberto Amato, Sonia Goncalves, Ewan Harrison, David K. Jackson, Ian Johnston, Dominic Kwiatkowski, Cordelia Langford, John Sillitoe on behalf of the Wellcome Sanger Institute COVID-19 Surveillance Team |
| EPI_ISL_551488, EPI_ISL_551491, EPI_ISL_551492 | Lighthouse Lab in Alderley Park | Wellcome Sanger Institute for the COVID-19 Genomics UK (COG-UK) consortium | The Lighthouse Lab in Alderley Park and Alex Alderton, Roberto Amato, Sonia Goncalves, Ewan Harrison, David K. Jackson, Ian Johnston, Dominic Kwiatkowski, Cordelia Langford, John Sillitoe on behalf of the Wellcome Sanger Institute COVID-19 Surveillance Team (http://www.sanger.ac.uk/covid-team) |
| EPI_ISL_551494 | Lighthouse Lab in Alderley Park | Wellcome Sanger Institute for the COVID-19 Genomics UK (COG-UK) consortium | The Lighthouse Lab in Alderley Park and Alex Alderton, Roberto Amato, Sonia Goncalves, Ewan Harrison, David K. Jackson, Ian Johnston, Dominic Kwiatkowski, Cordelia Langford, John Sillitoe on behalf of the Wellcome Sanger Institute COVID-19 Surveillance Team |
| EPI_ISL_551497, EPI_ISL_551498 | Lighthouse Lab in Alderley Park | Wellcome Sanger Institute for the COVID-19 Genomics UK (COG-UK) consortium | The Lighthouse Lab in Alderley Park and Alex Alderton, Roberto Amato, Sonia Goncalves, Ewan Harrison, David K. Jackson, Ian Johnston, Dominic Kwiatkowski, Cordelia Langford, John Sillitoe on behalf of the Wellcome Sanger Institute COVID-19 Surveillance Team (http://www.sanger.ac.uk/covid-team) |
| EPI_ISL_551501, EPI_ISL_551503, EPI_ISL_551505 | Lighthouse Lab in Alderley Park | Wellcome Sanger Institute for the COVID-19 Genomics UK (COG-UK) consortium | The Lighthouse Lab in Alderley Park and Alex Alderton, Roberto Amato, Sonia Goncalves, Ewan Harrison, David K. Jackson, Ian Johnston, Dominic Kwiatkowski, Cordelia Langford, John Sillitoe on behalf of the Wellcome Sanger Institute COVID-19 Surveillance Team |
| EPI_ISL_551506 | Lighthouse Lab in Alderley Park | Wellcome Sanger Institute for the COVID-19 Genomics UK (COG-UK) consortium | The Lighthouse Lab in Alderley Park and Alex Alderton, Roberto Amato, Sonia Goncalves, Ewan Harrison, David K. Jackson, Ian Johnston, Dominic Kwiatkowski, Cordelia Langford, John Sillitoe on behalf of the Wellcome Sanger Institute COVID-19 Surveillance Team (http://www.sanger.ac.uk/covid-team) |
| EPI_ISL_551507, EPI_ISL_551508 | Lighthouse Lab in Alderley Park | Wellcome Sanger Institute for the COVID-19 Genomics UK (COG-UK) consortium | The Lighthouse Lab in Alderley Park and Alex Alderton, Roberto Amato, Sonia Goncalves, Ewan Harrison, David K. Jackson, Ian Johnston, Dominic Kwiatkowski, Cordelia Langford, John Sillitoe on behalf of the Wellcome Sanger Institute COVID-19 Surveillance Team |
| EPI_ISL_551509, EPI_ISL_551510, EPI_ISL_551512, EPI_ISL_551514, EPI_ISL_551516, EPI_ISL_551517 | Lighthouse Lab in Alderley Park | Wellcome Sanger Institute for the COVID-19 Genomics UK (COG-UK) consortium | The Lighthouse Lab in Alderley Park and Alex Alderton, Roberto Amato, Sonia Goncalves, Ewan Harrison, David K. Jackson, Ian Johnston, Dominic Kwiatkowski, Cordelia Langford, John Sillitoe on behalf of the Wellcome Sanger Institute COVID-19 Surveillance Team (http://www.sanger.ac.uk/covid-team) |
| EPI_ISL_551518, EPI_ISL_551520 | Lighthouse Lab in Alderley Park | Wellcome Sanger Institute for the COVID-19 Genomics UK (COG-UK) consortium | The Lighthouse Lab in Alderley Park and Alex Alderton, Roberto Amato, Sonia Goncalves, Ewan Harrison, David K. Jackson, Ian Johnston, Dominic Kwiatkowski, Cordelia Langford, John Sillitoe on behalf of the Wellcome Sanger Institute COVID-19 Surveillance Team |
| EPI_ISL_551521 | Lighthouse Lab in Alderley Park | Wellcome Sanger Institute for the COVID-19 Genomics UK (COG-UK) consortium | The Lighthouse Lab in Alderley Park and Alex Alderton, Roberto Amato, Sonia Goncalves, Ewan Harrison, David K. Jackson, Ian Johnston, Dominic Kwiatkowski, Cordelia Langford, John Sillitoe on behalf of the Wellcome Sanger Institute COVID-19 Surveillance Team (http://www.sanger.ac.uk/covid-team) |
| EPI_ISL_551522 | Lighthouse Lab in Alderley Park | Wellcome Sanger Institute for the COVID-19 Genomics UK (COG-UK) consortium | The Lighthouse Lab in Alderley Park and Alex Alderton, Roberto Amato, Sonia Goncalves, Ewan Harrison, David K. Jackson, Ian Johnston, Dominic Kwiatkowski, Cordelia Langford, John Sillitoe on behalf of the Wellcome Sanger Institute COVID-19 Surveillance Team |
| EPI_ISL_551524 | Lighthouse Lab in Alderley Park | Wellcome Sanger Institute for the COVID-19 Genomics UK (COG-UK) consortium | The Lighthouse Lab in Alderley Park and Alex Alderton, Roberto Amato, Sonia Goncalves, Ewan Harrison, David K. Jackson, Ian Johnston, Dominic Kwiatkowski, Cordelia Langford, John Sillitoe on behalf of the Wellcome Sanger Institute COVID-19 Surveillance Team (http://www.sanger.ac.uk/covid-team) |
| EPI_ISL_551525 | Lighthouse Lab in Alderley Park | Wellcome Sanger Institute for the COVID-19 Genomics UK (COG-UK) consortium | The Lighthouse Lab in Alderley Park and Alex Alderton, Roberto Amato, Sonia Goncalves, Ewan Harrison, David K. Jackson, Ian Johnston, Dominic Kwiatkowski, Cordelia Langford, John Sillitoe on behalf of the Wellcome Sanger Institute COVID-19 Surveillance Team |
| EPI_ISL_551529, EPI_ISL_551533 | Lighthouse Lab in Alderley Park | Wellcome Sanger Institute for the COVID-19 Genomics UK (COG-UK) consortium | The Lighthouse Lab in Alderley Park and Alex Alderton, Roberto Amato, Sonia Goncalves, Ewan Harrison, David K. Jackson, Ian Johnston, Dominic Kwiatkowski, Cordelia Langford, John Sillitoe on behalf of the Wellcome Sanger Institute COVID-19 Surveillance Team (http://www.sanger.ac.uk/covid-team) |
| EPI_ISL_551534 | Lighthouse Lab in Alderley Park | Wellcome Sanger Institute for the COVID-19 Genomics UK (COG-UK) consortium | The Lighthouse Lab in Alderley Park and Alex Alderton, Roberto Amato, Sonia Goncalves, Ewan Harrison, David K. Jackson, Ian Johnston, Dominic Kwiatkowski, Cordelia Langford, John Sillitoe on behalf of the Wellcome Sanger Institute COVID-19 Surveillance Team |
| EPI_ISL_551535 | Lighthouse Lab in Alderley Park | Wellcome Sanger Institute for the COVID-19 Genomics UK (COG-UK) consortium | The Lighthouse Lab in Alderley Park and Alex Alderton, Roberto Amato, Sonia Goncalves, Ewan Harrison, David K. Jackson, Ian Johnston, Dominic Kwiatkowski, Cordelia Langford, John Sillitoe on behalf of the Wellcome Sanger Institute COVID-19 Surveillance Team (http://www.sanger.ac.uk/covid-team) |
| EPI_ISL_551536 | Lighthouse Lab in Alderley Park | Wellcome Sanger Institute for the COVID-19 Genomics UK (COG-UK) consortium | The Lighthouse Lab in Alderley Park and Alex Alderton, Roberto Amato, Sonia Goncalves, Ewan Harrison, David K. Jackson, Ian Johnston, Dominic Kwiatkowski, Cordelia Langford, John Sillitoe on behalf of the Wellcome Sanger Institute COVID-19 Surveillance Team |
| EPI_ISL_551537 | Lighthouse Lab in Alderley Park | Wellcome Sanger Institute for the COVID-19 Genomics UK (COG-UK) consortium | The Lighthouse Lab in Alderley Park and Alex Alderton, Roberto Amato, Sonia Goncalves, Ewan Harrison, David K. Jackson, Ian Johnston, Dominic Kwiatkowski, Cordelia Langford, John Sillitoe on behalf of the Wellcome Sanger Institute COVID-19 Surveillance Team (http://www.sanger.ac.uk/covid-team) |
| EPI_ISL_551538 | Lighthouse Lab in Alderley Park | Wellcome Sanger Institute for the COVID-19 Genomics UK (COG-UK) consortium | The Lighthouse Lab in Alderley Park and Alex Alderton, Roberto Amato, Sonia Goncalves, Ewan Harrison, David K. Jackson, Ian Johnston, Dominic Kwiatkowski, Cordelia Langford, John Sillitoe on behalf of the Wellcome Sanger Institute COVID-19 Surveillance Team |
| EPI_ISL_551539 | Lighthouse Lab in Alderley Park | Wellcome Sanger Institute for the COVID-19 Genomics UK (COG-UK) consortium | The Lighthouse Lab in Alderley Park and Alex Alderton, Roberto Amato, Sonia Goncalves, Ewan Harrison, David K. Jackson, Ian Johnston, Dominic Kwiatkowski, Cordelia Langford, John Sillitoe on behalf of the Wellcome Sanger Institute COVID-19 Surveillance Team |

| | | | |
|---|---|---|---|
| | | | (http://www.sanger.ac.uk/covid-team) |
| EPI_ISL_551540 | Lighthouse Lab in Alderley Park | Wellcome Sanger Institute for the COVID-19 Genomics UK (COG-UK) consortium | The Lighthouse Lab in Alderley Park and Alex Alderton, Roberto Amato, Sonia Goncalves, Ewan Harrison, David K. Jackson, Ian Johnston, Dominic Kwiatkowski, Cordelia Langford, John Sillitoe on behalf of the Wellcome Sanger Institute COVID-19 Surveillance Team |
| EPI_ISL_551541 | Lighthouse Lab in Alderley Park | Wellcome Sanger Institute for the COVID-19 Genomics UK (COG-UK) consortium | The Lighthouse Lab in Alderley Park and Alex Alderton, Roberto Amato, Sonia Goncalves, Ewan Harrison, David K. Jackson, Ian Johnston, Dominic Kwiatkowski, Cordelia Langford, John Sillitoe on behalf of the Wellcome Sanger Institute COVID-19 Surveillance Team (http://www.sanger.ac.uk/covid-team) |
| EPI_ISL_551542 | Lighthouse Lab in Alderley Park | Wellcome Sanger Institute for the COVID-19 Genomics UK (COG-UK) consortium | The Lighthouse Lab in Alderley Park and Alex Alderton, Roberto Amato, Sonia Goncalves, Ewan Harrison, David K. Jackson, Ian Johnston, Dominic Kwiatkowski, Cordelia Langford, John Sillitoe on behalf of the Wellcome Sanger Institute COVID-19 Surveillance Team |
| EPI_ISL_551543 | Lighthouse Lab in Alderley Park | Wellcome Sanger Institute for the COVID-19 Genomics UK (COG-UK) consortium | The Lighthouse Lab in Alderley Park and Alex Alderton, Roberto Amato, Sonia Goncalves, Ewan Harrison, David K. Jackson, Ian Johnston, Dominic Kwiatkowski, Cordelia Langford, John Sillitoe on behalf of the Wellcome Sanger Institute COVID-19 Surveillance Team (http://www.sanger.ac.uk/covid-team) |
| EPI_ISL_551544, EPI_ISL_551545 | Lighthouse Lab in Alderley Park | Wellcome Sanger Institute for the COVID-19 Genomics UK (COG-UK) consortium | The Lighthouse Lab in Alderley Park and Alex Alderton, Roberto Amato, Sonia Goncalves, Ewan Harrison, David K. Jackson, Ian Johnston, Dominic Kwiatkowski, Cordelia Langford, John Sillitoe on behalf of the Wellcome Sanger Institute COVID-19 Surveillance Team |
| EPI_ISL_551546, EPI_ISL_551547 | Lighthouse Lab in Alderley Park | Wellcome Sanger Institute for the COVID-19 Genomics UK (COG-UK) consortium | The Lighthouse Lab in Alderley Park and Alex Alderton, Roberto Amato, Sonia Goncalves, Ewan Harrison, David K. Jackson, Ian Johnston, Dominic Kwiatkowski, Cordelia Langford, John Sillitoe on behalf of the Wellcome Sanger Institute COVID-19 Surveillance Team (http://www.sanger.ac.uk/covid-team) |
| EPI_ISL_551548 | Lighthouse Lab in Alderley Park | Wellcome Sanger Institute for the COVID-19 Genomics UK (COG-UK) consortium | The Lighthouse Lab in Alderley Park and Alex Alderton, Roberto Amato, Sonia Goncalves, Ewan Harrison, David K. Jackson, Ian Johnston, Dominic Kwiatkowski, Cordelia Langford, John Sillitoe on behalf of the Wellcome Sanger Institute COVID-19 Surveillance Team |
| EPI_ISL_551549, EPI_ISL_551551, EPI_ISL_551552 | Lighthouse Lab in Alderley Park | Wellcome Sanger Institute for the COVID-19 Genomics UK (COG-UK) consortium | The Lighthouse Lab in Alderley Park and Alex Alderton, Roberto Amato, Sonia Goncalves, Ewan Harrison, David K. Jackson, Ian Johnston, Dominic Kwiatkowski, Cordelia Langford, John Sillitoe on behalf of the Wellcome Sanger Institute COVID-19 Surveillance Team (http://www.sanger.ac.uk/covid-team) |
| EPI_ISL_551553 | Lighthouse Lab in Alderley Park | Wellcome Sanger Institute for the COVID-19 Genomics UK (COG-UK) consortium | The Lighthouse Lab in Alderley Park and Alex Alderton, Roberto Amato, Sonia Goncalves, Ewan Harrison, David K. Jackson, Ian Johnston, Dominic Kwiatkowski, Cordelia Langford, John Sillitoe on behalf of the Wellcome Sanger Institute COVID-19 Surveillance Team |
| EPI_ISL_551554, EPI_ISL_551556, EPI_ISL_551557, EPI_ISL_551558 | Lighthouse Lab in Alderley Park | Wellcome Sanger Institute for the COVID-19 Genomics UK (COG-UK) consortium | The Lighthouse Lab in Alderley Park and Alex Alderton, Roberto Amato, Sonia Goncalves, Ewan Harrison, David K. Jackson, Ian Johnston, Dominic Kwiatkowski, Cordelia Langford, John Sillitoe on behalf of the Wellcome Sanger Institute COVID-19 Surveillance Team (http://www.sanger.ac.uk/covid-team) |
| EPI_ISL_551559 | Lighthouse Lab in Alderley Park | Wellcome Sanger Institute for the COVID-19 Genomics UK (COG-UK) consortium | The Lighthouse Lab in Alderley Park and Alex Alderton, Roberto Amato, Sonia Goncalves, Ewan Harrison, David K. Jackson, Ian Johnston, Dominic Kwiatkowski, Cordelia Langford, John Sillitoe on behalf of the Wellcome Sanger Institute COVID-19 Surveillance Team |
| EPI_ISL_551560, EPI_ISL_551561, EPI_ISL_551563, EPI_ISL_551564 | Lighthouse Lab in Alderley Park | Wellcome Sanger Institute for the COVID-19 Genomics UK (COG-UK) consortium | The Lighthouse Lab in Alderley Park and Alex Alderton, Roberto Amato, Sonia Goncalves, Ewan Harrison, David K. Jackson, Ian Johnston, Dominic Kwiatkowski, Cordelia Langford, John Sillitoe on behalf of the Wellcome Sanger Institute COVID-19 Surveillance Team (http://www.sanger.ac.uk/covid-team) |
| EPI_ISL_551565, EPI_ISL_551566 | Lighthouse Lab in Alderley Park | Wellcome Sanger Institute for the COVID-19 Genomics UK (COG-UK) consortium | The Lighthouse Lab in Alderley Park and Alex Alderton, Roberto Amato, Sonia Goncalves, Ewan Harrison, David K. Jackson, Ian Johnston, Dominic Kwiatkowski, Cordelia Langford, John Sillitoe on behalf of the Wellcome Sanger Institute COVID-19 Surveillance Team |
| EPI_ISL_551569 | Lighthouse Lab in Alderley Park | Wellcome Sanger Institute for the COVID-19 Genomics UK (COG-UK) consortium | The Lighthouse Lab in Alderley Park and Alex Alderton, Roberto Amato, Sonia Goncalves, Ewan Harrison, David K. Jackson, Ian Johnston, Dominic Kwiatkowski, Cordelia Langford, John Sillitoe on behalf of the Wellcome Sanger Institute COVID-19 Surveillance Team (http://www.sanger.ac.uk/covid-team) |
| EPI_ISL_551570 | Lighthouse Lab in Alderley Park | Wellcome Sanger Institute for the COVID-19 Genomics UK (COG-UK) consortium | The Lighthouse Lab in Alderley Park and Alex Alderton, Roberto Amato, Sonia Goncalves, Ewan Harrison, David K. Jackson, Ian Johnston, Dominic Kwiatkowski, Cordelia Langford, John Sillitoe on behalf of the Wellcome Sanger Institute COVID-19 Surveillance Team |
| EPI_ISL_551572, EPI_ISL_551573, EPI_ISL_551574, EPI_ISL_551575, EPI_ISL_551576 | Lighthouse Lab in Alderley Park | Wellcome Sanger Institute for the COVID-19 Genomics UK (COG-UK) consortium | The Lighthouse Lab in Alderley Park and Alex Alderton, Roberto Amato, Sonia Goncalves, Ewan Harrison, David K. Jackson, Ian Johnston, Dominic Kwiatkowski, Cordelia Langford, John Sillitoe on behalf of the Wellcome Sanger Institute COVID-19 Surveillance Team (http://www.sanger.ac.uk/covid-team) |
| EPI_ISL_551735, EPI_ISL_551736 | Lighthouse Lab in Alderley Park | Wellcome Sanger Institute for the COVID-19 Genomics UK (COG-UK) consortium | The Lighthouse Lab in Alderley Park and Alex Alderton, Roberto Amato, Sonia Goncalves, Ewan Harrison, David K. Jackson, Ian Johnston, Dominic Kwiatkowski, Cordelia Langford, John Sillitoe on behalf of the Wellcome Sanger Institute COVID-19 Surveillance Team |
| EPI_ISL_551740, EPI_ISL_551742, EPI_ISL_551743, EPI_ISL_551749 | Lighthouse Lab in Alderley Park | Wellcome Sanger Institute for the COVID-19 Genomics UK (COG-UK) consortium | The Lighthouse Lab in Alderley Park and Alex Alderton, Roberto Amato, Sonia Goncalves, Ewan Harrison, David K. Jackson, Ian Johnston, Dominic Kwiatkowski, Cordelia Langford, John Sillitoe on behalf of the Wellcome Sanger Institute COVID-19 Surveillance Team (http://www.sanger.ac.uk/covid-team) |
| EPI_ISL_551751, EPI_ISL_551755, EPI_ISL_551756, EPI_ISL_551759, EPI_ISL_551761, EPI_ISL_551762 | Lighthouse Lab in Alderley Park | Wellcome Sanger Institute for the COVID-19 Genomics UK (COG-UK) consortium | The Lighthouse Lab in Alderley Park and Alex Alderton, Roberto Amato, Sonia Goncalves, Ewan Harrison, David K. Jackson, Ian Johnston, Dominic Kwiatkowski, Cordelia Langford, John Sillitoe on behalf of the Wellcome Sanger Institute COVID-19 Surveillance Team |
| EPI_ISL_551763 | Lighthouse Lab in Alderley Park | Wellcome Sanger Institute for the COVID-19 Genomics UK (COG-UK) consortium | The Lighthouse Lab in Alderley Park and Alex Alderton, Roberto Amato, Sonia Goncalves, Ewan Harrison, David K. Jackson, Ian Johnston, Dominic Kwiatkowski, Cordelia Langford, John Sillitoe on behalf of the Wellcome Sanger Institute COVID-19 Surveillance Team (http://www.sanger.ac.uk/covid-team) |
| EPI_ISL_551764, EPI_ISL_551769, EPI_ISL_551772, EPI_ISL_551776, EPI_ISL_551779, EPI_ISL_551780, EPI_ISL_551784, EPI_ISL_551785, EPI_ISL_551789, EPI_ISL_551795, EPI_ISL_551796, EPI_ISL_551798, EPI_ISL_551799 | | | |
| see above | Lighthouse Lab in Alderley Park | Wellcome Sanger Institute for the COVID-19 Genomics UK (COG-UK) consortium | The Lighthouse Lab in Alderley Park and Alex Alderton, Roberto Amato, Sonia Goncalves, Ewan Harrison, David K. Jackson, Ian Johnston, Dominic Kwiatkowski, Cordelia Langford, John Sillitoe on behalf of the Wellcome Sanger Institute COVID-19 Surveillance Team |
| EPI_ISL_551801 | Lighthouse Lab in Alderley Park | Wellcome Sanger Institute for the COVID-19 Genomics UK (COG-UK) consortium | The Lighthouse Lab in Alderley Park and Alex Alderton, Roberto Amato, Sonia Goncalves, Ewan Harrison, David K. Jackson, Ian Johnston, Dominic Kwiatkowski, Cordelia Langford, John Sillitoe on behalf of the Wellcome Sanger Institute COVID-19 Surveillance Team (http://www.sanger.ac.uk/covid-team) |
| EPI_ISL_551803, EPI_ISL_551804, EPI_ISL_551806, EPI_ISL_551807, EPI_ISL_551808, EPI_ISL_551810, EPI_ISL_551811, EPI_ISL_551812 | Lighthouse Lab in Alderley Park | Wellcome Sanger Institute for the COVID-19 Genomics UK (COG-UK) consortium | The Lighthouse Lab in Alderley Park and Alex Alderton, Roberto Amato, Sonia Goncalves, Ewan Harrison, David K. Jackson, Ian Johnston, Dominic Kwiatkowski, Cordelia Langford, John Sillitoe on behalf of the Wellcome Sanger Institute COVID-19 Surveillance Team |
| EPI_ISL_551815 | Lighthouse Lab in Alderley Park | Wellcome Sanger Institute for the COVID-19 Genomics UK (COG-UK) consortium | The Lighthouse Lab in Alderley Park and Alex Alderton, Roberto Amato, Sonia Goncalves, Ewan Harrison, David K. Jackson, Ian Johnston, Dominic Kwiatkowski, Cordelia Langford, John Sillitoe on behalf of the Wellcome Sanger Institute COVID-19 Surveillance Team (http://www.sanger.ac.uk/covid-team) |
| EPI_ISL_551818, EPI_ISL_551819, EPI_ISL_551823 | Lighthouse Lab in Alderley Park | Wellcome Sanger Institute for the COVID-19 Genomics UK (COG-UK) consortium | The Lighthouse Lab in Alderley Park and Alex Alderton, Roberto Amato, Sonia Goncalves, Ewan Harrison, David K. Jackson, Ian Johnston, Dominic Kwiatkowski, Cordelia Langford, John Sillitoe on behalf of the Wellcome Sanger Institute COVID-19 Surveillance Team |
| EPI_ISL_551825 | Lighthouse Lab in Alderley Park | Wellcome Sanger Institute for the COVID-19 Genomics UK (COG-UK) consortium | The Lighthouse Lab in Alderley Park and Alex Alderton, Roberto Amato, Sonia Goncalves, Ewan Harrison, David K. Jackson, Ian Johnston, Dominic Kwiatkowski, Cordelia Langford, John Sillitoe on behalf of the Wellcome Sanger Institute COVID-19 Surveillance Team (http://www.sanger.ac.uk/covid-team) |
| EPI_ISL_551826, EPI_ISL_551827 | Lighthouse Lab in Alderley Park | Wellcome Sanger Institute for the COVID-19 Genomics UK (COG-UK) consortium | The Lighthouse Lab in Alderley Park and Alex Alderton, Roberto Amato, Sonia Goncalves, Ewan Harrison, David K. Jackson, Ian Johnston, Dominic Kwiatkowski, Cordelia Langford, John Sillitoe on behalf of the Wellcome Sanger Institute COVID-19 Surveillance Team |
| EPI_ISL_551832 | Lighthouse Lab in Alderley Park | Wellcome Sanger Institute for the COVID-19 Genomics UK | The Lighthouse Lab in Alderley Park and Alex Alderton, Roberto Amato, Sonia Goncalves, Ewan Harrison, David K. Jackson, Ian Johnston, Dominic |

| | | | |
|---|---|---|---|
| | | (COG-UK) consortium | Kwiatkowski, Cordelia Langford, John Sillitoe on behalf of the Wellcome Sanger Institute COVID-19 Surveillance Team (http://www.sanger.ac.uk/covid-team) |
| EPI_ISL_551834 | Lighthouse Lab in Alderley Park | Wellcome Sanger Institute for the COVID-19 Genomics UK (COG-UK) consortium | The Lighthouse Lab in Alderley Park and Alex Alderton, Roberto Amato, Sonia Goncalves, Ewan Harrison, David K. Jackson, Ian Johnston, Dominic Kwiatkowski, Cordelia Langford, John Sillitoe on behalf of the Wellcome Sanger Institute COVID-19 Surveillance Team |
| EPI_ISL_552797, EPI_ISL_552798 | Lighthouse Lab in Alderley Park | Wellcome Sanger Institute for the COVID-19 Genomics UK (COG-UK) consortium | The Lighthouse Lab in Alderley Park and Alex Alderton, Roberto Amato, Sonia Goncalves, Ewan Harrison, David K. Jackson, Ian Johnston, Dominic Kwiatkowski, Cordelia Langford, John Sillitoe on behalf of the Wellcome Sanger Institute COVID-19 Surveillance Team (http://www.sanger.ac.uk/covid-team) |
| EPI_ISL_552800, EPI_ISL_552801, EPI_ISL_552803, EPI_ISL_552804, EPI_ISL_552805, EPI_ISL_552806, EPI_ISL_552807 | Lighthouse Lab in Alderley Park | Wellcome Sanger Institute for the COVID-19 Genomics UK (COG-UK) consortium | The Lighthouse Lab in Alderley Park and Alex Alderton, Roberto Amato, Sonia Goncalves, Ewan Harrison, David K. Jackson, Ian Johnston, Dominic Kwiatkowski, Cordelia Langford, John Sillitoe on behalf of the Wellcome Sanger Institute COVID-19 Surveillance Team |
| EPI_ISL_552808 | Lighthouse Lab in Alderley Park | Wellcome Sanger Institute for the COVID-19 Genomics UK (COG-UK) consortium | The Lighthouse Lab in Alderley Park and Alex Alderton, Roberto Amato, Sonia Goncalves, Ewan Harrison, David K. Jackson, Ian Johnston, Dominic Kwiatkowski, Cordelia Langford, John Sillitoe on behalf of the Wellcome Sanger Institute COVID-19 Surveillance Team (http://www.sanger.ac.uk/covid-team) |
| EPI_ISL_552812, EPI_ISL_552813, EPI_ISL_552815, EPI_ISL_552817, EPI_ISL_552819, EPI_ISL_552821, EPI_ISL_552822, EPI_ISL_552825, EPI_ISL_552827, EPI_ISL_552828, EPI_ISL_552829, EPI_ISL_552830, EPI_ISL_552831 | | | |
| see above | Lighthouse Lab in Alderley Park | Wellcome Sanger Institute for the COVID-19 Genomics UK (COG-UK) consortium | The Lighthouse Lab in Alderley Park and Alex Alderton, Roberto Amato, Sonia Goncalves, Ewan Harrison, David K. Jackson, Ian Johnston, Dominic Kwiatkowski, Cordelia Langford, John Sillitoe on behalf of the Wellcome Sanger Institute COVID-19 Surveillance Team |
| EPI_ISL_552832 | Lighthouse Lab in Alderley Park | Wellcome Sanger Institute for the COVID-19 Genomics UK (COG-UK) consortium | The Lighthouse Lab in Alderley Park and Alex Alderton, Roberto Amato, Sonia Goncalves, Ewan Harrison, David K. Jackson, Ian Johnston, Dominic Kwiatkowski, Cordelia Langford, John Sillitoe on behalf of the Wellcome Sanger Institute COVID-19 Surveillance Team (http://www.sanger.ac.uk/covid-team) |
| EPI_ISL_552833, EPI_ISL_552834, EPI_ISL_552835 | Lighthouse Lab in Alderley Park | Wellcome Sanger Institute for the COVID-19 Genomics UK (COG-UK) consortium | The Lighthouse Lab in Alderley Park and Alex Alderton, Roberto Amato, Sonia Goncalves, Ewan Harrison, David K. Jackson, Ian Johnston, Dominic Kwiatkowski, Cordelia Langford, John Sillitoe on behalf of the Wellcome Sanger Institute COVID-19 Surveillance Team |
| EPI_ISL_552836 | Lighthouse Lab in Alderley Park | Wellcome Sanger Institute for the COVID-19 Genomics UK (COG-UK) consortium | The Lighthouse Lab in Alderley Park and Alex Alderton, Roberto Amato, Sonia Goncalves, Ewan Harrison, David K. Jackson, Ian Johnston, Dominic Kwiatkowski, Cordelia Langford, John Sillitoe on behalf of the Wellcome Sanger Institute COVID-19 Surveillance Team (http://www.sanger.ac.uk/covid-team) |
| EPI_ISL_552837 | Lighthouse Lab in Alderley Park | Wellcome Sanger Institute for the COVID-19 Genomics UK (COG-UK) consortium | The Lighthouse Lab in Alderley Park and Alex Alderton, Roberto Amato, Sonia Goncalves, Ewan Harrison, David K. Jackson, Ian Johnston, Dominic Kwiatkowski, Cordelia Langford, John Sillitoe on behalf of the Wellcome Sanger Institute COVID-19 Surveillance Team |
| EPI_ISL_552839 | Lighthouse Lab in Alderley Park | Wellcome Sanger Institute for the COVID-19 Genomics UK (COG-UK) consortium | The Lighthouse Lab in Alderley Park and Alex Alderton, Roberto Amato, Sonia Goncalves, Ewan Harrison, David K. Jackson, Ian Johnston, Dominic Kwiatkowski, Cordelia Langford, John Sillitoe on behalf of the Wellcome Sanger Institute COVID-19 Surveillance Team (http://www.sanger.ac.uk/covid-team) |
| EPI_ISL_552840, EPI_ISL_552843, EPI_ISL_552844, EPI_ISL_552845, EPI_ISL_552846, EPI_ISL_552847, EPI_ISL_552848, EPI_ISL_552849 | Lighthouse Lab in Alderley Park | Wellcome Sanger Institute for the COVID-19 Genomics UK (COG-UK) consortium | The Lighthouse Lab in Alderley Park and Alex Alderton, Roberto Amato, Sonia Goncalves, Ewan Harrison, David K. Jackson, Ian Johnston, Dominic Kwiatkowski, Cordelia Langford, John Sillitoe on behalf of the Wellcome Sanger Institute COVID-19 Surveillance Team |
| EPI_ISL_552850 | Lighthouse Lab in Alderley Park | Wellcome Sanger Institute for the COVID-19 Genomics UK (COG-UK) consortium | The Lighthouse Lab in Alderley Park and Alex Alderton, Roberto Amato, Sonia Goncalves, Ewan Harrison, David K. Jackson, Ian Johnston, Dominic Kwiatkowski, Cordelia Langford, John Sillitoe on behalf of the Wellcome Sanger Institute COVID-19 Surveillance Team (http://www.sanger.ac.uk/covid-team) |
| EPI_ISL_552851, EPI_ISL_552853 | Lighthouse Lab in Alderley Park | Wellcome Sanger Institute for the COVID-19 Genomics UK (COG-UK) consortium | The Lighthouse Lab in Alderley Park and Alex Alderton, Roberto Amato, Sonia Goncalves, Ewan Harrison, David K. Jackson, Ian Johnston, Dominic Kwiatkowski, Cordelia Langford, John Sillitoe on behalf of the Wellcome Sanger Institute COVID-19 Surveillance Team |
| EPI_ISL_552855, EPI_ISL_552856 | Lighthouse Lab in Alderley Park | Wellcome Sanger Institute for the COVID-19 Genomics UK (COG-UK) consortium | The Lighthouse Lab in Alderley Park and Alex Alderton, Roberto Amato, Sonia Goncalves, Ewan Harrison, David K. Jackson, Ian Johnston, Dominic Kwiatkowski, Cordelia Langford, John Sillitoe on behalf of the Wellcome Sanger Institute COVID-19 Surveillance Team (http://www.sanger.ac.uk/covid-team) |
| EPI_ISL_552857 | Lighthouse Lab in Alderley Park | Wellcome Sanger Institute for the COVID-19 Genomics UK (COG-UK) consortium | The Lighthouse Lab in Alderley Park and Alex Alderton, Roberto Amato, Sonia Goncalves, Ewan Harrison, David K. Jackson, Ian Johnston, Dominic Kwiatkowski, Cordelia Langford, John Sillitoe on behalf of the Wellcome Sanger Institute COVID-19 Surveillance Team |
| EPI_ISL_552858 | Lighthouse Lab in Alderley Park | Wellcome Sanger Institute for the COVID-19 Genomics UK (COG-UK) consortium | The Lighthouse Lab in Alderley Park and Alex Alderton, Roberto Amato, Sonia Goncalves, Ewan Harrison, David K. Jackson, Ian Johnston, Dominic Kwiatkowski, Cordelia Langford, John Sillitoe on behalf of the Wellcome Sanger Institute COVID-19 Surveillance Team (http://www.sanger.ac.uk/covid-team) |
| EPI_ISL_552859, EPI_ISL_552860, EPI_ISL_552861, EPI_ISL_552862 | Lighthouse Lab in Alderley Park | Wellcome Sanger Institute for the COVID-19 Genomics UK (COG-UK) consortium | The Lighthouse Lab in Alderley Park and Alex Alderton, Roberto Amato, Sonia Goncalves, Ewan Harrison, David K. Jackson, Ian Johnston, Dominic Kwiatkowski, Cordelia Langford, John Sillitoe on behalf of the Wellcome Sanger Institute COVID-19 Surveillance Team |
| EPI_ISL_552863 | Lighthouse Lab in Alderley Park | Wellcome Sanger Institute for the COVID-19 Genomics UK (COG-UK) consortium | The Lighthouse Lab in Alderley Park and Alex Alderton, Roberto Amato, Sonia Goncalves, Ewan Harrison, David K. Jackson, Ian Johnston, Dominic Kwiatkowski, Cordelia Langford, John Sillitoe on behalf of the Wellcome Sanger Institute COVID-19 Surveillance Team (http://www.sanger.ac.uk/covid-team) |
| EPI_ISL_552864, EPI_ISL_552865, EPI_ISL_552866, EPI_ISL_552868 | Lighthouse Lab in Alderley Park | Wellcome Sanger Institute for the COVID-19 Genomics UK (COG-UK) consortium | The Lighthouse Lab in Alderley Park and Alex Alderton, Roberto Amato, Sonia Goncalves, Ewan Harrison, David K. Jackson, Ian Johnston, Dominic Kwiatkowski, Cordelia Langford, John Sillitoe on behalf of the Wellcome Sanger Institute COVID-19 Surveillance Team |
| EPI_ISL_552869, EPI_ISL_552870 | Lighthouse Lab in Alderley Park | Wellcome Sanger Institute for the COVID-19 Genomics UK (COG-UK) consortium | The Lighthouse Lab in Alderley Park and Alex Alderton, Roberto Amato, Sonia Goncalves, Ewan Harrison, David K. Jackson, Ian Johnston, Dominic Kwiatkowski, Cordelia Langford, John Sillitoe on behalf of the Wellcome Sanger Institute COVID-19 Surveillance Team (http://www.sanger.ac.uk/covid-team) |
| EPI_ISL_552871 | Lighthouse Lab in Alderley Park | Wellcome Sanger Institute for the COVID-19 Genomics UK (COG-UK) consortium | The Lighthouse Lab in Alderley Park and Alex Alderton, Roberto Amato, Sonia Goncalves, Ewan Harrison, David K. Jackson, Ian Johnston, Dominic Kwiatkowski, Cordelia Langford, John Sillitoe on behalf of the Wellcome Sanger Institute COVID-19 Surveillance Team |
| EPI_ISL_552872, EPI_ISL_552873, EPI_ISL_552876 | Lighthouse Lab in Alderley Park | Wellcome Sanger Institute for the COVID-19 Genomics UK (COG-UK) consortium | The Lighthouse Lab in Alderley Park and Alex Alderton, Roberto Amato, Sonia Goncalves, Ewan Harrison, David K. Jackson, Ian Johnston, Dominic Kwiatkowski, Cordelia Langford, John Sillitoe on behalf of the Wellcome Sanger Institute COVID-19 Surveillance Team (http://www.sanger.ac.uk/covid-team) |
| EPI_ISL_552878 | Lighthouse Lab in Alderley Park | Wellcome Sanger Institute for the COVID-19 Genomics UK (COG-UK) consortium | The Lighthouse Lab in Alderley Park and Alex Alderton, Roberto Amato, Sonia Goncalves, Ewan Harrison, David K. Jackson, Ian Johnston, Dominic Kwiatkowski, Cordelia Langford, John Sillitoe on behalf of the Wellcome Sanger Institute COVID-19 Surveillance Team |
| EPI_ISL_552881 | Lighthouse Lab in Alderley Park | Wellcome Sanger Institute for the COVID-19 Genomics UK (COG-UK) consortium | The Lighthouse Lab in Alderley Park and Alex Alderton, Roberto Amato, Sonia Goncalves, Ewan Harrison, David K. Jackson, Ian Johnston, Dominic Kwiatkowski, Cordelia Langford, John Sillitoe on behalf of the Wellcome Sanger Institute COVID-19 Surveillance Team (http://www.sanger.ac.uk/covid-team) |
| EPI_ISL_552882, EPI_ISL_552883 | Lighthouse Lab in Alderley Park | Wellcome Sanger Institute for the COVID-19 Genomics UK (COG-UK) consortium | The Lighthouse Lab in Alderley Park and Alex Alderton, Roberto Amato, Sonia Goncalves, Ewan Harrison, David K. Jackson, Ian Johnston, Dominic Kwiatkowski, Cordelia Langford, John Sillitoe on behalf of the Wellcome Sanger Institute COVID-19 Surveillance Team |
| EPI_ISL_552884 | Lighthouse Lab in Alderley Park | Wellcome Sanger Institute for the COVID-19 Genomics UK (COG-UK) consortium | The Lighthouse Lab in Alderley Park and Alex Alderton, Roberto Amato, Sonia Goncalves, Ewan Harrison, David K. Jackson, Ian Johnston, Dominic Kwiatkowski, Cordelia Langford, John Sillitoe on behalf of the Wellcome Sanger Institute COVID-19 Surveillance Team (http://www.sanger.ac.uk/covid-team) |
| EPI_ISL_552885 | Lighthouse Lab in Alderley Park | Wellcome Sanger Institute for the COVID-19 Genomics UK | The Lighthouse Lab in Alderley Park and Alex Alderton, Roberto Amato, Sonia Goncalves, Ewan Harrison, David K. Jackson, Ian Johnston, Dominic |

| | | | |
|---|---|---|---|
| | | (COG-UK) consortium | Kwiatkowski, Cordelia Langford, John Sillitoe on behalf of the Wellcome Sanger Institute COVID-19 Surveillance Team |
| EPI_ISL_552887 | Lighthouse Lab in Alderley Park | Wellcome Sanger Institute for the COVID-19 Genomics UK (COG-UK) consortium | The Lighthouse Lab in Alderley Park and Alex Alderton, Roberto Amato, Sonia Goncalves, Ewan Harrison, David K. Jackson, Ian Johnston, Dominic Kwiatkowski, Cordelia Langford, John Sillitoe on behalf of the Wellcome Sanger Institute COVID-19 Surveillance Team (http://www.sanger.ac.uk/covid-team) |
| EPI_ISL_552888 | Lighthouse Lab in Alderley Park | Wellcome Sanger Institute for the COVID-19 Genomics UK (COG-UK) consortium | The Lighthouse Lab in Alderley Park and Alex Alderton, Roberto Amato, Sonia Goncalves, Ewan Harrison, David K. Jackson, Ian Johnston, Dominic Kwiatkowski, Cordelia Langford, John Sillitoe on behalf of the Wellcome Sanger Institute COVID-19 Surveillance Team |
| EPI_ISL_552889 | Lighthouse Lab in Alderley Park | Wellcome Sanger Institute for the COVID-19 Genomics UK (COG-UK) consortium | The Lighthouse Lab in Alderley Park and Alex Alderton, Roberto Amato, Sonia Goncalves, Ewan Harrison, David K. Jackson, Ian Johnston, Dominic Kwiatkowski, Cordelia Langford, John Sillitoe on behalf of the Wellcome Sanger Institute COVID-19 Surveillance Team (http://www.sanger.ac.uk/covid-team) |
| EPI_ISL_552890, EPI_ISL_552891, EPI_ISL_552892 | Lighthouse Lab in Alderley Park | Wellcome Sanger Institute for the COVID-19 Genomics UK (COG-UK) consortium | The Lighthouse Lab in Alderley Park and Alex Alderton, Roberto Amato, Sonia Goncalves, Ewan Harrison, David K. Jackson, Ian Johnston, Dominic Kwiatkowski, Cordelia Langford, John Sillitoe on behalf of the Wellcome Sanger Institute COVID-19 Surveillance Team |
| EPI_ISL_552894, EPI_ISL_552895 | Lighthouse Lab in Alderley Park | Wellcome Sanger Institute for the COVID-19 Genomics UK (COG-UK) consortium | The Lighthouse Lab in Alderley Park and Alex Alderton, Roberto Amato, Sonia Goncalves, Ewan Harrison, David K. Jackson, Ian Johnston, Dominic Kwiatkowski, Cordelia Langford, John Sillitoe on behalf of the Wellcome Sanger Institute COVID-19 Surveillance Team (http://www.sanger.ac.uk/covid-team) |
| EPI_ISL_552896, EPI_ISL_552897, EPI_ISL_552899, EPI_ISL_552901, EPI_ISL_552902, EPI_ISL_552903, EPI_ISL_552904 | Lighthouse Lab in Alderley Park | Wellcome Sanger Institute for the COVID-19 Genomics UK (COG-UK) consortium | The Lighthouse Lab in Alderley Park and Alex Alderton, Roberto Amato, Sonia Goncalves, Ewan Harrison, David K. Jackson, Ian Johnston, Dominic Kwiatkowski, Cordelia Langford, John Sillitoe on behalf of the Wellcome Sanger Institute COVID-19 Surveillance Team |
| EPI_ISL_552906, EPI_ISL_552907, EPI_ISL_552908 | Lighthouse Lab in Alderley Park | Wellcome Sanger Institute for the COVID-19 Genomics UK (COG-UK) consortium | The Lighthouse Lab in Alderley Park and Alex Alderton, Roberto Amato, Sonia Goncalves, Ewan Harrison, David K. Jackson, Ian Johnston, Dominic Kwiatkowski, Cordelia Langford, John Sillitoe on behalf of the Wellcome Sanger Institute COVID-19 Surveillance Team (http://www.sanger.ac.uk/covid-team) |
| EPI_ISL_552909 | Lighthouse Lab in Alderley Park | Wellcome Sanger Institute for the COVID-19 Genomics UK (COG-UK) consortium | The Lighthouse Lab in Alderley Park and Alex Alderton, Roberto Amato, Sonia Goncalves, Ewan Harrison, David K. Jackson, Ian Johnston, Dominic Kwiatkowski, Cordelia Langford, John Sillitoe on behalf of the Wellcome Sanger Institute COVID-19 Surveillance Team |
| EPI_ISL_552911 | Lighthouse Lab in Alderley Park | Wellcome Sanger Institute for the COVID-19 Genomics UK (COG-UK) consortium | The Lighthouse Lab in Alderley Park and Alex Alderton, Roberto Amato, Sonia Goncalves, Ewan Harrison, David K. Jackson, Ian Johnston, Dominic Kwiatkowski, Cordelia Langford, John Sillitoe on behalf of the Wellcome Sanger Institute COVID-19 Surveillance Team (http://www.sanger.ac.uk/covid-team) |
| EPI_ISL_552912 | Lighthouse Lab in Alderley Park | Wellcome Sanger Institute for the COVID-19 Genomics UK (COG-UK) consortium | The Lighthouse Lab in Alderley Park and Alex Alderton, Roberto Amato, Sonia Goncalves, Ewan Harrison, David K. Jackson, Ian Johnston, Dominic Kwiatkowski, Cordelia Langford, John Sillitoe on behalf of the Wellcome Sanger Institute COVID-19 Surveillance Team |
| EPI_ISL_552913, EPI_ISL_552914 | Lighthouse Lab in Alderley Park | Wellcome Sanger Institute for the COVID-19 Genomics UK (COG-UK) consortium | The Lighthouse Lab in Alderley Park and Alex Alderton, Roberto Amato, Sonia Goncalves, Ewan Harrison, David K. Jackson, Ian Johnston, Dominic Kwiatkowski, Cordelia Langford, John Sillitoe on behalf of the Wellcome Sanger Institute COVID-19 Surveillance Team (http://www.sanger.ac.uk/covid-team) |
| EPI_ISL_552915, EPI_ISL_552917, EPI_ISL_552918, EPI_ISL_552919, EPI_ISL_552920, EPI_ISL_552921 | Lighthouse Lab in Alderley Park | Wellcome Sanger Institute for the COVID-19 Genomics UK (COG-UK) consortium | The Lighthouse Lab in Alderley Park and Alex Alderton, Roberto Amato, Sonia Goncalves, Ewan Harrison, David K. Jackson, Ian Johnston, Dominic Kwiatkowski, Cordelia Langford, John Sillitoe on behalf of the Wellcome Sanger Institute COVID-19 Surveillance Team |
| EPI_ISL_552922 | Lighthouse Lab in Alderley Park | Wellcome Sanger Institute for the COVID-19 Genomics UK (COG-UK) consortium | The Lighthouse Lab in Alderley Park and Alex Alderton, Roberto Amato, Sonia Goncalves, Ewan Harrison, David K. Jackson, Ian Johnston, Dominic Kwiatkowski, Cordelia Langford, John Sillitoe on behalf of the Wellcome Sanger Institute COVID-19 Surveillance Team (http://www.sanger.ac.uk/covid-team) |
| EPI_ISL_552923, EPI_ISL_552924, EPI_ISL_552925, EPI_ISL_552926, EPI_ISL_552927, EPI_ISL_552928, EPI_ISL_552929, EPI_ISL_552930, EPI_ISL_552931, EPI_ISL_552933, EPI_ISL_552934, EPI_ISL_552935, EPI_ISL_552937, EPI_ISL_552940, EPI_ISL_552941 | | | |
| see above | Lighthouse Lab in Alderley Park | Wellcome Sanger Institute for the COVID-19 Genomics UK (COG-UK) consortium | The Lighthouse Lab in Alderley Park and Alex Alderton, Roberto Amato, Sonia Goncalves, Ewan Harrison, David K. Jackson, Ian Johnston, Dominic Kwiatkowski, Cordelia Langford, John Sillitoe on behalf of the Wellcome Sanger Institute COVID-19 Surveillance Team |
| EPI_ISL_552942 | Lighthouse Lab in Alderley Park | Wellcome Sanger Institute for the COVID-19 Genomics UK (COG-UK) consortium | The Lighthouse Lab in Alderley Park and Alex Alderton, Roberto Amato, Sonia Goncalves, Ewan Harrison, David K. Jackson, Ian Johnston, Dominic Kwiatkowski, Cordelia Langford, John Sillitoe on behalf of the Wellcome Sanger Institute COVID-19 Surveillance Team (http://www.sanger.ac.uk/covid-team) |
| EPI_ISL_552944, EPI_ISL_552945, EPI_ISL_552946, EPI_ISL_552951, EPI_ISL_552954 | Lighthouse Lab in Alderley Park | Wellcome Sanger Institute for the COVID-19 Genomics UK (COG-UK) consortium | The Lighthouse Lab in Alderley Park and Alex Alderton, Roberto Amato, Sonia Goncalves, Ewan Harrison, David K. Jackson, Ian Johnston, Dominic Kwiatkowski, Cordelia Langford, John Sillitoe on behalf of the Wellcome Sanger Institute COVID-19 Surveillance Team |
| EPI_ISL_552955 | Lighthouse Lab in Alderley Park | Wellcome Sanger Institute for the COVID-19 Genomics UK (COG-UK) consortium | The Lighthouse Lab in Alderley Park and Alex Alderton, Roberto Amato, Sonia Goncalves, Ewan Harrison, David K. Jackson, Ian Johnston, Dominic Kwiatkowski, Cordelia Langford, John Sillitoe on behalf of the Wellcome Sanger Institute COVID-19 Surveillance Team (http://www.sanger.ac.uk/covid-team) |
| EPI_ISL_552956, EPI_ISL_552957, EPI_ISL_552958, EPI_ISL_552959, EPI_ISL_552962 | Lighthouse Lab in Alderley Park | Wellcome Sanger Institute for the COVID-19 Genomics UK (COG-UK) consortium | The Lighthouse Lab in Alderley Park and Alex Alderton, Roberto Amato, Sonia Goncalves, Ewan Harrison, David K. Jackson, Ian Johnston, Dominic Kwiatkowski, Cordelia Langford, John Sillitoe on behalf of the Wellcome Sanger Institute COVID-19 Surveillance Team |
| EPI_ISL_552964 | Lighthouse Lab in Alderley Park | Wellcome Sanger Institute for the COVID-19 Genomics UK (COG-UK) consortium | The Lighthouse Lab in Alderley Park and Alex Alderton, Roberto Amato, Sonia Goncalves, Ewan Harrison, David K. Jackson, Ian Johnston, Dominic Kwiatkowski, Cordelia Langford, John Sillitoe on behalf of the Wellcome Sanger Institute COVID-19 Surveillance Team (http://www.sanger.ac.uk/covid-team) |
| EPI_ISL_552965, EPI_ISL_552966, EPI_ISL_552968, EPI_ISL_552969, EPI_ISL_552971 | Lighthouse Lab in Alderley Park | Wellcome Sanger Institute for the COVID-19 Genomics UK (COG-UK) consortium | The Lighthouse Lab in Alderley Park and Alex Alderton, Roberto Amato, Sonia Goncalves, Ewan Harrison, David K. Jackson, Ian Johnston, Dominic Kwiatkowski, Cordelia Langford, John Sillitoe on behalf of the Wellcome Sanger Institute COVID-19 Surveillance Team |
| EPI_ISL_552976 | Lighthouse Lab in Alderley Park | Wellcome Sanger Institute for the COVID-19 Genomics UK (COG-UK) consortium | The Lighthouse Lab in Alderley Park and Alex Alderton, Roberto Amato, Sonia Goncalves, Ewan Harrison, David K. Jackson, Ian Johnston, Dominic Kwiatkowski, Cordelia Langford, John Sillitoe on behalf of the Wellcome Sanger Institute COVID-19 Surveillance Team (http://www.sanger.ac.uk/covid-team) |
| EPI_ISL_552977 | Lighthouse Lab in Alderley Park | Wellcome Sanger Institute for the COVID-19 Genomics UK (COG-UK) consortium | The Lighthouse Lab in Alderley Park and Alex Alderton, Roberto Amato, Sonia Goncalves, Ewan Harrison, David K. Jackson, Ian Johnston, Dominic Kwiatkowski, Cordelia Langford, John Sillitoe on behalf of the Wellcome Sanger Institute COVID-19 Surveillance Team |
| EPI_ISL_552978 | Lighthouse Lab in Alderley Park | Wellcome Sanger Institute for the COVID-19 Genomics UK (COG-UK) consortium | The Lighthouse Lab in Alderley Park and Alex Alderton, Roberto Amato, Sonia Goncalves, Ewan Harrison, David K. Jackson, Ian Johnston, Dominic Kwiatkowski, Cordelia Langford, John Sillitoe on behalf of the Wellcome Sanger Institute COVID-19 Surveillance Team (http://www.sanger.ac.uk/covid-team) |
| EPI_ISL_552980, EPI_ISL_552981 | Lighthouse Lab in Alderley Park | Wellcome Sanger Institute for the COVID-19 Genomics UK (COG-UK) consortium | The Lighthouse Lab in Alderley Park and Alex Alderton, Roberto Amato, Sonia Goncalves, Ewan Harrison, David K. Jackson, Ian Johnston, Dominic Kwiatkowski, Cordelia Langford, John Sillitoe on behalf of the Wellcome Sanger Institute COVID-19 Surveillance Team |
| EPI_ISL_552983 | Lighthouse Lab in Alderley Park | Wellcome Sanger Institute for the COVID-19 Genomics UK (COG-UK) consortium | The Lighthouse Lab in Alderley Park and Alex Alderton, Roberto Amato, Sonia Goncalves, Ewan Harrison, David K. Jackson, Ian Johnston, Dominic Kwiatkowski, Cordelia Langford, John Sillitoe on behalf of the Wellcome Sanger Institute COVID-19 Surveillance Team (http://www.sanger.ac.uk/covid-team) |
| EPI_ISL_552984, EPI_ISL_552985, EPI_ISL_552986, EPI_ISL_552988, EPI_ISL_552989, EPI_ISL_552991, | Lighthouse Lab in Alderley Park | Wellcome Sanger Institute for the COVID-19 Genomics UK (COG-UK) consortium | The Lighthouse Lab in Alderley Park and Alex Alderton, Roberto Amato, Sonia Goncalves, Ewan Harrison, David K. Jackson, Ian Johnston, Dominic Kwiatkowski, Cordelia Langford, John Sillitoe on behalf of the Wellcome Sanger Institute COVID-19 Surveillance Team |

| | | | |
|---|---|---|---|
| EPI_ISL_552992, EPI_ISL_552994 | | | |
| EPI_ISL_552996 | Lighthouse Lab in Alderley Park | Wellcome Sanger Institute for the COVID-19 Genomics UK (COG-UK) consortium | The Lighthouse Lab in Alderley Park and Alex Alderton, Roberto Amato, Sonia Goncalves, Ewan Harrison, David K. Jackson, Ian Johnston, Dominic Kwiatkowski, Cordelia Langford, John Sillitoe on behalf of the Wellcome Sanger Institute COVID-19 Surveillance Team (http://www.sanger.ac.uk/covid-team) |
| EPI_ISL_552998, EPI_ISL_552999, EPI_ISL_553000, EPI_ISL_553001, EPI_ISL_553002, EPI_ISL_553003, EPI_ISL_553004, EPI_ISL_553005 | Lighthouse Lab in Alderley Park | Wellcome Sanger Institute for the COVID-19 Genomics UK (COG-UK) consortium | The Lighthouse Lab in Alderley Park and Alex Alderton, Roberto Amato, Sonia Goncalves, Ewan Harrison, David K. Jackson, Ian Johnston, Dominic Kwiatkowski, Cordelia Langford, John Sillitoe on behalf of the Wellcome Sanger Institute COVID-19 Surveillance Team |
| EPI_ISL_553269, EPI_ISL_553271 | Lighthouse Lab in Alderley Park | Wellcome Sanger Institute for the COVID-19 Genomics UK (COG-UK) consortium | The Lighthouse Lab in Alderley Park and Alex Alderton, Roberto Amato, Sonia Goncalves, Ewan Harrison, David K. Jackson, Ian Johnston, Dominic Kwiatkowski, Cordelia Langford, John Sillitoe on behalf of the Wellcome Sanger Institute COVID-19 Surveillance Team (http://www.sanger.ac.uk/covid-team) |
| EPI_ISL_553273 | Lighthouse Lab in Alderley Park | Wellcome Sanger Institute for the COVID-19 Genomics UK (COG-UK) consortium | The Lighthouse Lab in Alderley Park and Alex Alderton, Roberto Amato, Sonia Goncalves, Ewan Harrison, David K. Jackson, Ian Johnston, Dominic Kwiatkowski, Cordelia Langford, John Sillitoe on behalf of the Wellcome Sanger Institute COVID-19 Surveillance Team |
| EPI_ISL_553274, EPI_ISL_553277 | Lighthouse Lab in Alderley Park | Wellcome Sanger Institute for the COVID-19 Genomics UK (COG-UK) consortium | The Lighthouse Lab in Alderley Park and Alex Alderton, Roberto Amato, Sonia Goncalves, Ewan Harrison, David K. Jackson, Ian Johnston, Dominic Kwiatkowski, Cordelia Langford, John Sillitoe on behalf of the Wellcome Sanger Institute COVID-19 Surveillance Team (http://www.sanger.ac.uk/covid-team) |
| EPI_ISL_553278 | Lighthouse Lab in Alderley Park | Wellcome Sanger Institute for the COVID-19 Genomics UK (COG-UK) consortium | The Lighthouse Lab in Alderley Park and Alex Alderton, Roberto Amato, Sonia Goncalves, Ewan Harrison, David K. Jackson, Ian Johnston, Dominic Kwiatkowski, Cordelia Langford, John Sillitoe on behalf of the Wellcome Sanger Institute COVID-19 Surveillance Team |
| EPI_ISL_553279, EPI_ISL_553280 | Lighthouse Lab in Alderley Park | Wellcome Sanger Institute for the COVID-19 Genomics UK (COG-UK) consortium | The Lighthouse Lab in Alderley Park and Alex Alderton, Roberto Amato, Sonia Goncalves, Ewan Harrison, David K. Jackson, Ian Johnston, Dominic Kwiatkowski, Cordelia Langford, John Sillitoe on behalf of the Wellcome Sanger Institute COVID-19 Surveillance Team (http://www.sanger.ac.uk/covid-team) |
| EPI_ISL_553281, EPI_ISL_553282, EPI_ISL_553283 | Lighthouse Lab in Alderley Park | Wellcome Sanger Institute for the COVID-19 Genomics UK (COG-UK) consortium | The Lighthouse Lab in Alderley Park and Alex Alderton, Roberto Amato, Sonia Goncalves, Ewan Harrison, David K. Jackson, Ian Johnston, Dominic Kwiatkowski, Cordelia Langford, John Sillitoe on behalf of the Wellcome Sanger Institute COVID-19 Surveillance Team |
| EPI_ISL_553284, EPI_ISL_553285 | Lighthouse Lab in Alderley Park | Wellcome Sanger Institute for the COVID-19 Genomics UK (COG-UK) consortium | The Lighthouse Lab in Alderley Park and Alex Alderton, Roberto Amato, Sonia Goncalves, Ewan Harrison, David K. Jackson, Ian Johnston, Dominic Kwiatkowski, Cordelia Langford, John Sillitoe on behalf of the Wellcome Sanger Institute COVID-19 Surveillance Team (http://www.sanger.ac.uk/covid-team) |
| EPI_ISL_553286 | Lighthouse Lab in Alderley Park | Wellcome Sanger Institute for the COVID-19 Genomics UK (COG-UK) consortium | The Lighthouse Lab in Alderley Park and Alex Alderton, Roberto Amato, Sonia Goncalves, Ewan Harrison, David K. Jackson, Ian Johnston, Dominic Kwiatkowski, Cordelia Langford, John Sillitoe on behalf of the Wellcome Sanger Institute COVID-19 Surveillance Team |
| EPI_ISL_553287 | Lighthouse Lab in Alderley Park | Wellcome Sanger Institute for the COVID-19 Genomics UK (COG-UK) consortium | The Lighthouse Lab in Alderley Park and Alex Alderton, Roberto Amato, Sonia Goncalves, Ewan Harrison, David K. Jackson, Ian Johnston, Dominic Kwiatkowski, Cordelia Langford, John Sillitoe on behalf of the Wellcome Sanger Institute COVID-19 Surveillance Team (http://www.sanger.ac.uk/covid-team) |
| EPI_ISL_553288 | Lighthouse Lab in Alderley Park | Wellcome Sanger Institute for the COVID-19 Genomics UK (COG-UK) consortium | The Lighthouse Lab in Alderley Park and Alex Alderton, Roberto Amato, Sonia Goncalves, Ewan Harrison, David K. Jackson, Ian Johnston, Dominic Kwiatkowski, Cordelia Langford, John Sillitoe on behalf of the Wellcome Sanger Institute COVID-19 Surveillance Team |
| EPI_ISL_553289, EPI_ISL_553290, EPI_ISL_553291, EPI_ISL_553292, EPI_ISL_553293, EPI_ISL_553294 | Lighthouse Lab in Alderley Park | Wellcome Sanger Institute for the COVID-19 Genomics UK (COG-UK) consortium | The Lighthouse Lab in Alderley Park and Alex Alderton, Roberto Amato, Sonia Goncalves, Ewan Harrison, David K. Jackson, Ian Johnston, Dominic Kwiatkowski, Cordelia Langford, John Sillitoe on behalf of the Wellcome Sanger Institute COVID-19 Surveillance Team (http://www.sanger.ac.uk/covid-team) |
| EPI_ISL_553296 | Lighthouse Lab in Alderley Park | Wellcome Sanger Institute for the COVID-19 Genomics UK (COG-UK) consortium | The Lighthouse Lab in Alderley Park and Alex Alderton, Roberto Amato, Sonia Goncalves, Ewan Harrison, David K. Jackson, Ian Johnston, Dominic Kwiatkowski, Cordelia Langford, John Sillitoe on behalf of the Wellcome Sanger Institute COVID-19 Surveillance Team |
| EPI_ISL_553297 | Lighthouse Lab in Alderley Park | Wellcome Sanger Institute for the COVID-19 Genomics UK (COG-UK) consortium | The Lighthouse Lab in Alderley Park and Alex Alderton, Roberto Amato, Sonia Goncalves, Ewan Harrison, David K. Jackson, Ian Johnston, Dominic Kwiatkowski, Cordelia Langford, John Sillitoe on behalf of the Wellcome Sanger Institute COVID-19 Surveillance Team (http://www.sanger.ac.uk/covid-team) |
| EPI_ISL_553298, EPI_ISL_553299 | Lighthouse Lab in Alderley Park | Wellcome Sanger Institute for the COVID-19 Genomics UK (COG-UK) consortium | The Lighthouse Lab in Alderley Park and Alex Alderton, Roberto Amato, Sonia Goncalves, Ewan Harrison, David K. Jackson, Ian Johnston, Dominic Kwiatkowski, Cordelia Langford, John Sillitoe on behalf of the Wellcome Sanger Institute COVID-19 Surveillance Team |
| EPI_ISL_553300, EPI_ISL_553301, EPI_ISL_553302, EPI_ISL_553303 | Lighthouse Lab in Alderley Park | Wellcome Sanger Institute for the COVID-19 Genomics UK (COG-UK) consortium | The Lighthouse Lab in Alderley Park and Alex Alderton, Roberto Amato, Sonia Goncalves, Ewan Harrison, David K. Jackson, Ian Johnston, Dominic Kwiatkowski, Cordelia Langford, John Sillitoe on behalf of the Wellcome Sanger Institute COVID-19 Surveillance Team (http://www.sanger.ac.uk/covid-team) |
| EPI_ISL_553304 | Lighthouse Lab in Alderley Park | Wellcome Sanger Institute for the COVID-19 Genomics UK (COG-UK) consortium | The Lighthouse Lab in Alderley Park and Alex Alderton, Roberto Amato, Sonia Goncalves, Ewan Harrison, David K. Jackson, Ian Johnston, Dominic Kwiatkowski, Cordelia Langford, John Sillitoe on behalf of the Wellcome Sanger Institute COVID-19 Surveillance Team |
| EPI_ISL_553305 | Lighthouse Lab in Alderley Park | Wellcome Sanger Institute for the COVID-19 Genomics UK (COG-UK) consortium | The Lighthouse Lab in Alderley Park and Alex Alderton, Roberto Amato, Sonia Goncalves, Ewan Harrison, David K. Jackson, Ian Johnston, Dominic Kwiatkowski, Cordelia Langford, John Sillitoe on behalf of the Wellcome Sanger Institute COVID-19 Surveillance Team (http://www.sanger.ac.uk/covid-team) |
| EPI_ISL_553306 | Lighthouse Lab in Alderley Park | Wellcome Sanger Institute for the COVID-19 Genomics UK (COG-UK) consortium | The Lighthouse Lab in Alderley Park and Alex Alderton, Roberto Amato, Sonia Goncalves, Ewan Harrison, David K. Jackson, Ian Johnston, Dominic Kwiatkowski, Cordelia Langford, John Sillitoe on behalf of the Wellcome Sanger Institute COVID-19 Surveillance Team |
| EPI_ISL_553307 | Lighthouse Lab in Alderley Park | Wellcome Sanger Institute for the COVID-19 Genomics UK (COG-UK) consortium | The Lighthouse Lab in Alderley Park and Alex Alderton, Roberto Amato, Sonia Goncalves, Ewan Harrison, David K. Jackson, Ian Johnston, Dominic Kwiatkowski, Cordelia Langford, John Sillitoe on behalf of the Wellcome Sanger Institute COVID-19 Surveillance Team (http://www.sanger.ac.uk/covid-team) |
| EPI_ISL_553308, EPI_ISL_553309 | Lighthouse Lab in Alderley Park | Wellcome Sanger Institute for the COVID-19 Genomics UK (COG-UK) consortium | The Lighthouse Lab in Alderley Park and Alex Alderton, Roberto Amato, Sonia Goncalves, Ewan Harrison, David K. Jackson, Ian Johnston, Dominic Kwiatkowski, Cordelia Langford, John Sillitoe on behalf of the Wellcome Sanger Institute COVID-19 Surveillance Team |
| EPI_ISL_553310, EPI_ISL_553311 | Lighthouse Lab in Alderley Park | Wellcome Sanger Institute for the COVID-19 Genomics UK (COG-UK) consortium | The Lighthouse Lab in Alderley Park and Alex Alderton, Roberto Amato, Sonia Goncalves, Ewan Harrison, David K. Jackson, Ian Johnston, Dominic Kwiatkowski, Cordelia Langford, John Sillitoe on behalf of the Wellcome Sanger Institute COVID-19 Surveillance Team (http://www.sanger.ac.uk/covid-team) |
| EPI_ISL_553312 | Lighthouse Lab in Alderley Park | Wellcome Sanger Institute for the COVID-19 Genomics UK (COG-UK) consortium | The Lighthouse Lab in Alderley Park and Alex Alderton, Roberto Amato, Sonia Goncalves, Ewan Harrison, David K. Jackson, Ian Johnston, Dominic Kwiatkowski, Cordelia Langford, John Sillitoe on behalf of the Wellcome Sanger Institute COVID-19 Surveillance Team |
| EPI_ISL_553313, EPI_ISL_553314 | Lighthouse Lab in Alderley Park | Wellcome Sanger Institute for the COVID-19 Genomics UK (COG-UK) consortium | The Lighthouse Lab in Alderley Park and Alex Alderton, Roberto Amato, Sonia Goncalves, Ewan Harrison, David K. Jackson, Ian Johnston, Dominic Kwiatkowski, Cordelia Langford, John Sillitoe on behalf of the Wellcome Sanger Institute COVID-19 Surveillance Team (http://www.sanger.ac.uk/covid-team) |
| EPI_ISL_553315 | Lighthouse Lab in Alderley Park | Wellcome Sanger Institute for the COVID-19 Genomics UK (COG-UK) consortium | The Lighthouse Lab in Alderley Park and Alex Alderton, Roberto Amato, Sonia Goncalves, Ewan Harrison, David K. Jackson, Ian Johnston, Dominic Kwiatkowski, Cordelia Langford, John Sillitoe on behalf of the Wellcome Sanger Institute COVID-19 Surveillance Team |
| EPI_ISL_553316, EPI_ISL_553317, EPI_ISL_553318 | Lighthouse Lab in Alderley Park | Wellcome Sanger Institute for the COVID-19 Genomics UK (COG-UK) consortium | The Lighthouse Lab in Alderley Park and Alex Alderton, Roberto Amato, Sonia Goncalves, Ewan Harrison, David K. Jackson, Ian Johnston, Dominic Kwiatkowski, Cordelia Langford, John Sillitoe on behalf of the Wellcome Sanger Institute COVID-19 Surveillance Team (http://www.sanger.ac.uk/covid-team) |
| EPI_ISL_553319, EPI_ISL_553320 | Lighthouse Lab in Alderley Park | Wellcome Sanger Institute for the COVID-19 Genomics UK (COG-UK) consortium | The Lighthouse Lab in Alderley Park and Alex Alderton, Roberto Amato, Sonia Goncalves, Ewan Harrison, David K. Jackson, Ian Johnston, Dominic Kwiatkowski, Cordelia Langford, John Sillitoe on behalf of the Wellcome Sanger Institute COVID-19 Surveillance Team |

| | | | |
|---|---|---|---|
| EPI_ISL_553321, EPI_ISL_553322, EPI_ISL_553324, EPI_ISL_553325, EPI_ISL_553326 | Lighthouse Lab in Alderley Park | Wellcome Sanger Institute for the COVID-19 Genomics UK (COG-UK) consortium | The Lighthouse Lab in Alderley Park and Alex Alderton, Roberto Amato, Sonia Goncalves, Ewan Harrison, David K. Jackson, Ian Johnston, Dominic Kwiatkowski, Cordelia Langford, John Sillitoe on behalf of the Wellcome Sanger Institute COVID-19 Surveillance Team (http://www.sanger.ac.uk/covid-team) |
| EPI_ISL_553328 | Lighthouse Lab in Alderley Park | Wellcome Sanger Institute for the COVID-19 Genomics UK (COG-UK) consortium | The Lighthouse Lab in Alderley Park and Alex Alderton, Roberto Amato, Sonia Goncalves, Ewan Harrison, David K. Jackson, Ian Johnston, Dominic Kwiatkowski, Cordelia Langford, John Sillitoe on behalf of the Wellcome Sanger Institute COVID-19 Surveillance Team |
| EPI_ISL_553329 | Lighthouse Lab in Alderley Park | Wellcome Sanger Institute for the COVID-19 Genomics UK (COG-UK) consortium | The Lighthouse Lab in Alderley Park and Alex Alderton, Roberto Amato, Sonia Goncalves, Ewan Harrison, David K. Jackson, Ian Johnston, Dominic Kwiatkowski, Cordelia Langford, John Sillitoe on behalf of the Wellcome Sanger Institute COVID-19 Surveillance Team (http://www.sanger.ac.uk/covid-team) |
| EPI_ISL_553331 | Lighthouse Lab in Alderley Park | Wellcome Sanger Institute for the COVID-19 Genomics UK (COG-UK) consortium | The Lighthouse Lab in Alderley Park and Alex Alderton, Roberto Amato, Sonia Goncalves, Ewan Harrison, David K. Jackson, Ian Johnston, Dominic Kwiatkowski, Cordelia Langford, John Sillitoe on behalf of the Wellcome Sanger Institute COVID-19 Surveillance Team |
| EPI_ISL_553332, EPI_ISL_553333 | Lighthouse Lab in Alderley Park | Wellcome Sanger Institute for the COVID-19 Genomics UK (COG-UK) consortium | The Lighthouse Lab in Alderley Park and Alex Alderton, Roberto Amato, Sonia Goncalves, Ewan Harrison, David K. Jackson, Ian Johnston, Dominic Kwiatkowski, Cordelia Langford, John Sillitoe on behalf of the Wellcome Sanger Institute COVID-19 Surveillance Team (http://www.sanger.ac.uk/covid-team) |
| EPI_ISL_553334 | Lighthouse Lab in Alderley Park | Wellcome Sanger Institute for the COVID-19 Genomics UK (COG-UK) consortium | The Lighthouse Lab in Alderley Park and Alex Alderton, Roberto Amato, Sonia Goncalves, Ewan Harrison, David K. Jackson, Ian Johnston, Dominic Kwiatkowski, Cordelia Langford, John Sillitoe on behalf of the Wellcome Sanger Institute COVID-19 Surveillance Team |
| EPI_ISL_553335, EPI_ISL_553336, EPI_ISL_553337 | Lighthouse Lab in Alderley Park | Wellcome Sanger Institute for the COVID-19 Genomics UK (COG-UK) consortium | The Lighthouse Lab in Alderley Park and Alex Alderton, Roberto Amato, Sonia Goncalves, Ewan Harrison, David K. Jackson, Ian Johnston, Dominic Kwiatkowski, Cordelia Langford, John Sillitoe on behalf of the Wellcome Sanger Institute COVID-19 Surveillance Team (http://www.sanger.ac.uk/covid-team) |
| EPI_ISL_553338, EPI_ISL_553339 | Lighthouse Lab in Alderley Park | Wellcome Sanger Institute for the COVID-19 Genomics UK (COG-UK) consortium | The Lighthouse Lab in Alderley Park and Alex Alderton, Roberto Amato, Sonia Goncalves, Ewan Harrison, David K. Jackson, Ian Johnston, Dominic Kwiatkowski, Cordelia Langford, John Sillitoe on behalf of the Wellcome Sanger Institute COVID-19 Surveillance Team |
| EPI_ISL_553340, EPI_ISL_553341, EPI_ISL_553342, EPI_ISL_553343 | Lighthouse Lab in Alderley Park | Wellcome Sanger Institute for the COVID-19 Genomics UK (COG-UK) consortium | The Lighthouse Lab in Alderley Park and Alex Alderton, Roberto Amato, Sonia Goncalves, Ewan Harrison, David K. Jackson, Ian Johnston, Dominic Kwiatkowski, Cordelia Langford, John Sillitoe on behalf of the Wellcome Sanger Institute COVID-19 Surveillance Team (http://www.sanger.ac.uk/covid-team) |
| EPI_ISL_553344 | Lighthouse Lab in Alderley Park | Wellcome Sanger Institute for the COVID-19 Genomics UK (COG-UK) consortium | The Lighthouse Lab in Alderley Park and Alex Alderton, Roberto Amato, Sonia Goncalves, Ewan Harrison, David K. Jackson, Ian Johnston, Dominic Kwiatkowski, Cordelia Langford, John Sillitoe on behalf of the Wellcome Sanger Institute COVID-19 Surveillance Team |
| EPI_ISL_553345 | Lighthouse Lab in Alderley Park | Wellcome Sanger Institute for the COVID-19 Genomics UK (COG-UK) consortium | The Lighthouse Lab in Alderley Park and Alex Alderton, Roberto Amato, Sonia Goncalves, Ewan Harrison, David K. Jackson, Ian Johnston, Dominic Kwiatkowski, Cordelia Langford, John Sillitoe on behalf of the Wellcome Sanger Institute COVID-19 Surveillance Team (http://www.sanger.ac.uk/covid-team) |
| EPI_ISL_553699, EPI_ISL_553700 | Lighthouse Lab in Alderley Park | Wellcome Sanger Institute for the COVID-19 Genomics UK (COG-UK) consortium | The Lighthouse Lab in Alderley Park and Alex Alderton, Roberto Amato, Sonia Goncalves, Ewan Harrison, David K. Jackson, Ian Johnston, Dominic Kwiatkowski, Cordelia Langford, John Sillitoe on behalf of the Wellcome Sanger Institute COVID-19 Surveillance Team |
| EPI_ISL_553701, EPI_ISL_553702, EPI_ISL_553703, EPI_ISL_553705 | Lighthouse Lab in Alderley Park | Wellcome Sanger Institute for the COVID-19 Genomics UK (COG-UK) consortium | The Lighthouse Lab in Alderley Park and Alex Alderton, Roberto Amato, Sonia Goncalves, Ewan Harrison, David K. Jackson, Ian Johnston, Dominic Kwiatkowski, Cordelia Langford, John Sillitoe on behalf of the Wellcome Sanger Institute COVID-19 Surveillance Team (http://www.sanger.ac.uk/covid-team) |
| EPI_ISL_553707 | Lighthouse Lab in Alderley Park | Wellcome Sanger Institute for the COVID-19 Genomics UK (COG-UK) consortium | The Lighthouse Lab in Alderley Park and Alex Alderton, Roberto Amato, Sonia Goncalves, Ewan Harrison, David K. Jackson, Ian Johnston, Dominic Kwiatkowski, Cordelia Langford, John Sillitoe on behalf of the Wellcome Sanger Institute COVID-19 Surveillance Team |
| EPI_ISL_553708 | Lighthouse Lab in Alderley Park | Wellcome Sanger Institute for the COVID-19 Genomics UK (COG-UK) consortium | The Lighthouse Lab in Alderley Park and Alex Alderton, Roberto Amato, Sonia Goncalves, Ewan Harrison, David K. Jackson, Ian Johnston, Dominic Kwiatkowski, Cordelia Langford, John Sillitoe on behalf of the Wellcome Sanger Institute COVID-19 Surveillance Team (http://www.sanger.ac.uk/covid-team) |
| EPI_ISL_553709 | Lighthouse Lab in Alderley Park | Wellcome Sanger Institute for the COVID-19 Genomics UK (COG-UK) consortium | The Lighthouse Lab in Alderley Park and Alex Alderton, Roberto Amato, Sonia Goncalves, Ewan Harrison, David K. Jackson, Ian Johnston, Dominic Kwiatkowski, Cordelia Langford, John Sillitoe on behalf of the Wellcome Sanger Institute COVID-19 Surveillance Team |
| EPI_ISL_553710 | Lighthouse Lab in Alderley Park | Wellcome Sanger Institute for the COVID-19 Genomics UK (COG-UK) consortium | The Lighthouse Lab in Alderley Park and Alex Alderton, Roberto Amato, Sonia Goncalves, Ewan Harrison, David K. Jackson, Ian Johnston, Dominic Kwiatkowski, Cordelia Langford, John Sillitoe on behalf of the Wellcome Sanger Institute COVID-19 Surveillance Team (http://www.sanger.ac.uk/covid-team) |
| EPI_ISL_553711 | Lighthouse Lab in Alderley Park | Wellcome Sanger Institute for the COVID-19 Genomics UK (COG-UK) consortium | The Lighthouse Lab in Alderley Park and Alex Alderton, Roberto Amato, Sonia Goncalves, Ewan Harrison, David K. Jackson, Ian Johnston, Dominic Kwiatkowski, Cordelia Langford, John Sillitoe on behalf of the Wellcome Sanger Institute COVID-19 Surveillance Team |
| EPI_ISL_553713 | Lighthouse Lab in Alderley Park | Wellcome Sanger Institute for the COVID-19 Genomics UK (COG-UK) consortium | The Lighthouse Lab in Alderley Park and Alex Alderton, Roberto Amato, Sonia Goncalves, Ewan Harrison, David K. Jackson, Ian Johnston, Dominic Kwiatkowski, Cordelia Langford, John Sillitoe on behalf of the Wellcome Sanger Institute COVID-19 Surveillance Team (http://www.sanger.ac.uk/covid-team) |
| EPI_ISL_553714, EPI_ISL_553715 | Lighthouse Lab in Alderley Park | Wellcome Sanger Institute for the COVID-19 Genomics UK (COG-UK) consortium | The Lighthouse Lab in Alderley Park and Alex Alderton, Roberto Amato, Sonia Goncalves, Ewan Harrison, David K. Jackson, Ian Johnston, Dominic Kwiatkowski, Cordelia Langford, John Sillitoe on behalf of the Wellcome Sanger Institute COVID-19 Surveillance Team |
| EPI_ISL_553716 | Lighthouse Lab in Alderley Park | Wellcome Sanger Institute for the COVID-19 Genomics UK (COG-UK) consortium | The Lighthouse Lab in Alderley Park and Alex Alderton, Roberto Amato, Sonia Goncalves, Ewan Harrison, David K. Jackson, Ian Johnston, Dominic Kwiatkowski, Cordelia Langford, John Sillitoe on behalf of the Wellcome Sanger Institute COVID-19 Surveillance Team (http://www.sanger.ac.uk/covid-team) |
| EPI_ISL_553717 | Lighthouse Lab in Alderley Park | Wellcome Sanger Institute for the COVID-19 Genomics UK (COG-UK) consortium | The Lighthouse Lab in Alderley Park and Alex Alderton, Roberto Amato, Sonia Goncalves, Ewan Harrison, David K. Jackson, Ian Johnston, Dominic Kwiatkowski, Cordelia Langford, John Sillitoe on behalf of the Wellcome Sanger Institute COVID-19 Surveillance Team |
| EPI_ISL_553718 | Lighthouse Lab in Alderley Park | Wellcome Sanger Institute for the COVID-19 Genomics UK (COG-UK) consortium | The Lighthouse Lab in Alderley Park and Alex Alderton, Roberto Amato, Sonia Goncalves, Ewan Harrison, David K. Jackson, Ian Johnston, Dominic Kwiatkowski, Cordelia Langford, John Sillitoe on behalf of the Wellcome Sanger Institute COVID-19 Surveillance Team (http://www.sanger.ac.uk/covid-team) |
| EPI_ISL_553719 | Lighthouse Lab in Alderley Park | Wellcome Sanger Institute for the COVID-19 Genomics UK (COG-UK) consortium | The Lighthouse Lab in Alderley Park and Alex Alderton, Roberto Amato, Sonia Goncalves, Ewan Harrison, David K. Jackson, Ian Johnston, Dominic Kwiatkowski, Cordelia Langford, John Sillitoe on behalf of the Wellcome Sanger Institute COVID-19 Surveillance Team |
| EPI_ISL_553720 | Lighthouse Lab in Alderley Park | Wellcome Sanger Institute for the COVID-19 Genomics UK (COG-UK) consortium | The Lighthouse Lab in Alderley Park and Alex Alderton, Roberto Amato, Sonia Goncalves, Ewan Harrison, David K. Jackson, Ian Johnston, Dominic Kwiatkowski, Cordelia Langford, John Sillitoe on behalf of the Wellcome Sanger Institute COVID-19 Surveillance Team (http://www.sanger.ac.uk/covid-team) |
| EPI_ISL_553721 | Lighthouse Lab in Alderley Park | Wellcome Sanger Institute for the COVID-19 Genomics UK (COG-UK) consortium | The Lighthouse Lab in Alderley Park and Alex Alderton, Roberto Amato, Sonia Goncalves, Ewan Harrison, David K. Jackson, Ian Johnston, Dominic Kwiatkowski, Cordelia Langford, John Sillitoe on behalf of the Wellcome Sanger Institute COVID-19 Surveillance Team |
| EPI_ISL_553722, EPI_ISL_553723 | Lighthouse Lab in Alderley Park | Wellcome Sanger Institute for the COVID-19 Genomics UK (COG-UK) consortium | The Lighthouse Lab in Alderley Park and Alex Alderton, Roberto Amato, Sonia Goncalves, Ewan Harrison, David K. Jackson, Ian Johnston, Dominic Kwiatkowski, Cordelia Langford, John Sillitoe on behalf of the Wellcome Sanger Institute COVID-19 Surveillance Team (http://www.sanger.ac.uk/covid-team) |
| EPI_ISL_553724 | Lighthouse Lab in Alderley Park | Wellcome Sanger Institute for the COVID-19 Genomics UK (COG-UK) consortium | The Lighthouse Lab in Alderley Park and Alex Alderton, Roberto Amato, Sonia Goncalves, Ewan Harrison, David K. Jackson, Ian Johnston, Dominic Kwiatkowski, Cordelia Langford, John Sillitoe on behalf of the Wellcome Sanger Institute COVID-19 Surveillance Team |
| EPI_ISL_553725, EPI_ISL_553726 | Lighthouse Lab in Alderley Park | Wellcome Sanger Institute for the COVID-19 Genomics UK (COG-UK) consortium | The Lighthouse Lab in Alderley Park and Alex Alderton, Roberto Amato, Sonia Goncalves, Ewan Harrison, David K. Jackson, Ian Johnston, Dominic Kwiatkowski, Cordelia Langford, John Sillitoe on behalf of the Wellcome Sanger Institute COVID-19 Surveillance Team (http://www.sanger.ac.uk/covid-team) |

| | | | |
|---|---|---|---|
| EPI_ISL_553728 | Lighthouse Lab in Alderley Park | Wellcome Sanger Institute for the COVID-19 Genomics UK (COG-UK) consortium | The Lighthouse Lab in Alderley Park and Alex Alderton, Roberto Amato, Sonia Goncalves, Ewan Harrison, David K. Jackson, Ian Johnston, Dominic Kwiatkowski, Cordelia Langford, John Sillitoe on behalf of the Wellcome Sanger Institute COVID-19 Surveillance Team |
| EPI_ISL_553729, EPI_ISL_553730 | Lighthouse Lab in Alderley Park | Wellcome Sanger Institute for the COVID-19 Genomics UK (COG-UK) consortium | The Lighthouse Lab in Alderley Park and Alex Alderton, Roberto Amato, Sonia Goncalves, Ewan Harrison, David K. Jackson, Ian Johnston, Dominic Kwiatkowski, Cordelia Langford, John Sillitoe on behalf of the Wellcome Sanger Institute COVID-19 Surveillance Team (http://www.sanger.ac.uk/covid-team) |
| EPI_ISL_553731, EPI_ISL_554616, EPI_ISL_554617, EPI_ISL_554618, EPI_ISL_554620, EPI_ISL_554624, EPI_ISL_554625, EPI_ISL_554626 | Lighthouse Lab in Alderley Park | Wellcome Sanger Institute for the COVID-19 Genomics UK (COG-UK) consortium | The Lighthouse Lab in Alderley Park and Alex Alderton, Roberto Amato, Sonia Goncalves, Ewan Harrison, David K. Jackson, Ian Johnston, Dominic Kwiatkowski, Cordelia Langford, John Sillitoe on behalf of the Wellcome Sanger Institute COVID-19 Surveillance Team |
| EPI_ISL_554630, EPI_ISL_554631 | Lighthouse Lab in Alderley Park | Wellcome Sanger Institute for the COVID-19 Genomics UK (COG-UK) consortium | The Lighthouse Lab in Alderley Park and Alex Alderton, Roberto Amato, Sonia Goncalves, Ewan Harrison, David K. Jackson, Ian Johnston, Dominic Kwiatkowski, Cordelia Langford, John Sillitoe on behalf of the Wellcome Sanger Institute COVID-19 Surveillance Team (http://www.sanger.ac.uk/covid-team) |
| EPI_ISL_554640, EPI_ISL_554651, EPI_ISL_554653, EPI_ISL_554654, EPI_ISL_554656, EPI_ISL_554657 | Lighthouse Lab in Alderley Park | Wellcome Sanger Institute for the COVID-19 Genomics UK (COG-UK) consortium | The Lighthouse Lab in Alderley Park and Alex Alderton, Roberto Amato, Sonia Goncalves, Ewan Harrison, David K. Jackson, Ian Johnston, Dominic Kwiatkowski, Cordelia Langford, John Sillitoe on behalf of the Wellcome Sanger Institute COVID-19 Surveillance Team |
| EPI_ISL_554663 | Lighthouse Lab in Alderley Park | Wellcome Sanger Institute for the COVID-19 Genomics UK (COG-UK) consortium | The Lighthouse Lab in Alderley Park and Alex Alderton, Roberto Amato, Sonia Goncalves, Ewan Harrison, David K. Jackson, Ian Johnston, Dominic Kwiatkowski, Cordelia Langford, John Sillitoe on behalf of the Wellcome Sanger Institute COVID-19 Surveillance Team (http://www.sanger.ac.uk/covid-team) |
| EPI_ISL_554664, EPI_ISL_554677, EPI_ISL_554680, EPI_ISL_554681, EPI_ISL_554686, EPI_ISL_554688 | Lighthouse Lab in Alderley Park | Wellcome Sanger Institute for the COVID-19 Genomics UK (COG-UK) consortium | The Lighthouse Lab in Alderley Park and Alex Alderton, Roberto Amato, Sonia Goncalves, Ewan Harrison, David K. Jackson, Ian Johnston, Dominic Kwiatkowski, Cordelia Langford, John Sillitoe on behalf of the Wellcome Sanger Institute COVID-19 Surveillance Team |
| EPI_ISL_554689 | Lighthouse Lab in Alderley Park | Wellcome Sanger Institute for the COVID-19 Genomics UK (COG-UK) consortium | The Lighthouse Lab in Alderley Park and Alex Alderton, Roberto Amato, Sonia Goncalves, Ewan Harrison, David K. Jackson, Ian Johnston, Dominic Kwiatkowski, Cordelia Langford, John Sillitoe on behalf of the Wellcome Sanger Institute COVID-19 Surveillance Team (http://www.sanger.ac.uk/covid-team) |
| EPI_ISL_554692 | Lighthouse Lab in Alderley Park | Wellcome Sanger Institute for the COVID-19 Genomics UK (COG-UK) consortium | The Lighthouse Lab in Alderley Park and Alex Alderton, Roberto Amato, Sonia Goncalves, Ewan Harrison, David K. Jackson, Ian Johnston, Dominic Kwiatkowski, Cordelia Langford, John Sillitoe on behalf of the Wellcome Sanger Institute COVID-19 Surveillance Team |
| EPI_ISL_554694 | Lighthouse Lab in Alderley Park | Wellcome Sanger Institute for the COVID-19 Genomics UK (COG-UK) consortium | The Lighthouse Lab in Alderley Park and Alex Alderton, Roberto Amato, Sonia Goncalves, Ewan Harrison, David K. Jackson, Ian Johnston, Dominic Kwiatkowski, Cordelia Langford, John Sillitoe on behalf of the Wellcome Sanger Institute COVID-19 Surveillance Team (http://www.sanger.ac.uk/covid-team) |
| EPI_ISL_554696, EPI_ISL_554697, EPI_ISL_554703, EPI_ISL_554708, EPI_ISL_554710, EPI_ISL_554716, EPI_ISL_554725 | Lighthouse Lab in Alderley Park | Wellcome Sanger Institute for the COVID-19 Genomics UK (COG-UK) consortium | The Lighthouse Lab in Alderley Park and Alex Alderton, Roberto Amato, Sonia Goncalves, Ewan Harrison, David K. Jackson, Ian Johnston, Dominic Kwiatkowski, Cordelia Langford, John Sillitoe on behalf of the Wellcome Sanger Institute COVID-19 Surveillance Team |
| EPI_ISL_554726 | Lighthouse Lab in Alderley Park | Wellcome Sanger Institute for the COVID-19 Genomics UK (COG-UK) consortium | The Lighthouse Lab in Alderley Park and Alex Alderton, Roberto Amato, Sonia Goncalves, Ewan Harrison, David K. Jackson, Ian Johnston, Dominic Kwiatkowski, Cordelia Langford, John Sillitoe on behalf of the Wellcome Sanger Institute COVID-19 Surveillance Team (http://www.sanger.ac.uk/covid-team) |
| EPI_ISL_554727, EPI_ISL_554730 | Lighthouse Lab in Alderley Park | Wellcome Sanger Institute for the COVID-19 Genomics UK (COG-UK) consortium | The Lighthouse Lab in Alderley Park and Alex Alderton, Roberto Amato, Sonia Goncalves, Ewan Harrison, David K. Jackson, Ian Johnston, Dominic Kwiatkowski, Cordelia Langford, John Sillitoe on behalf of the Wellcome Sanger Institute COVID-19 Surveillance Team |
| EPI_ISL_554735, EPI_ISL_554741, EPI_ISL_554743 | Lighthouse Lab in Alderley Park | Wellcome Sanger Institute for the COVID-19 Genomics UK (COG-UK) consortium | The Lighthouse Lab in Alderley Park and Alex Alderton, Roberto Amato, Sonia Goncalves, Ewan Harrison, David K. Jackson, Ian Johnston, Dominic Kwiatkowski, Cordelia Langford, John Sillitoe on behalf of the Wellcome Sanger Institute COVID-19 Surveillance Team (http://www.sanger.ac.uk/covid-team) |
| EPI_ISL_554754, EPI_ISL_554755, EPI_ISL_554756, EPI_ISL_554757, EPI_ISL_554759, EPI_ISL_554761 | Lighthouse Lab in Alderley Park | Wellcome Sanger Institute for the COVID-19 Genomics UK (COG-UK) consortium | The Lighthouse Lab in Alderley Park and Alex Alderton, Roberto Amato, Sonia Goncalves, Ewan Harrison, David K. Jackson, Ian Johnston, Dominic Kwiatkowski, Cordelia Langford, John Sillitoe on behalf of the Wellcome Sanger Institute COVID-19 Surveillance Team |
| EPI_ISL_554767 | Lighthouse Lab in Alderley Park | Wellcome Sanger Institute for the COVID-19 Genomics UK (COG-UK) consortium | The Lighthouse Lab in Alderley Park and Alex Alderton, Roberto Amato, Sonia Goncalves, Ewan Harrison, David K. Jackson, Ian Johnston, Dominic Kwiatkowski, Cordelia Langford, John Sillitoe on behalf of the Wellcome Sanger Institute COVID-19 Surveillance Team (http://www.sanger.ac.uk/covid-team) |
| EPI_ISL_554771, EPI_ISL_554772, EPI_ISL_554777, EPI_ISL_554778, EPI_ISL_554779 | Lighthouse Lab in Alderley Park | Wellcome Sanger Institute for the COVID-19 Genomics UK (COG-UK) consortium | The Lighthouse Lab in Alderley Park and Alex Alderton, Roberto Amato, Sonia Goncalves, Ewan Harrison, David K. Jackson, Ian Johnston, Dominic Kwiatkowski, Cordelia Langford, John Sillitoe on behalf of the Wellcome Sanger Institute COVID-19 Surveillance Team |
| EPI_ISL_554780 | Lighthouse Lab in Alderley Park | Wellcome Sanger Institute for the COVID-19 Genomics UK (COG-UK) consortium | The Lighthouse Lab in Alderley Park and Alex Alderton, Roberto Amato, Sonia Goncalves, Ewan Harrison, David K. Jackson, Ian Johnston, Dominic Kwiatkowski, Cordelia Langford, John Sillitoe on behalf of the Wellcome Sanger Institute COVID-19 Surveillance Team (http://www.sanger.ac.uk/covid-team) |
| EPI_ISL_554782, EPI_ISL_554792, EPI_ISL_554794, EPI_ISL_554795, EPI_ISL_554797, EPI_ISL_554799, EPI_ISL_554802, EPI_ISL_554804, EPI_ISL_554807, EPI_ISL_554808, EPI_ISL_554809, EPI_ISL_554810, EPI_ISL_554811, EPI_ISL_554812, EPI_ISL_554813, EPI_ISL_554815, EPI_ISL_554816, EPI_ISL_554820 | | | |
| see above | Lighthouse Lab in Alderley Park | Wellcome Sanger Institute for the COVID-19 Genomics UK (COG-UK) consortium | The Lighthouse Lab in Alderley Park and Alex Alderton, Roberto Amato, Sonia Goncalves, Ewan Harrison, David K. Jackson, Ian Johnston, Dominic Kwiatkowski, Cordelia Langford, John Sillitoe on behalf of the Wellcome Sanger Institute COVID-19 Surveillance Team |
| EPI_ISL_554827 | Lighthouse Lab in Alderley Park | Wellcome Sanger Institute for the COVID-19 Genomics UK (COG-UK) consortium | The Lighthouse Lab in Alderley Park and Alex Alderton, Roberto Amato, Sonia Goncalves, Ewan Harrison, David K. Jackson, Ian Johnston, Dominic Kwiatkowski, Cordelia Langford, John Sillitoe on behalf of the Wellcome Sanger Institute COVID-19 Surveillance Team (http://www.sanger.ac.uk/covid-team) |
| EPI_ISL_554830, EPI_ISL_554832, EPI_ISL_554833, EPI_ISL_554834, EPI_ISL_554836, EPI_ISL_554837, EPI_ISL_554841, EPI_ISL_554846, EPI_ISL_554847, EPI_ISL_554848, EPI_ISL_554850, EPI_ISL_554853, EPI_ISL_554856 | | | |
| see above | Lighthouse Lab in Alderley Park | Wellcome Sanger Institute for the COVID-19 Genomics UK (COG-UK) consortium | The Lighthouse Lab in Alderley Park and Alex Alderton, Roberto Amato, Sonia Goncalves, Ewan Harrison, David K. Jackson, Ian Johnston, Dominic Kwiatkowski, Cordelia Langford, John Sillitoe on behalf of the Wellcome Sanger Institute COVID-19 Surveillance Team |
| EPI_ISL_554899 | Lighthouse Lab in Alderley Park | Wellcome Sanger Institute for the COVID-19 Genomics UK (COG-UK) consortium | The Lighthouse Lab in Alderley Park and Alex Alderton, Roberto Amato, Sonia Goncalves, Ewan Harrison, David K. Jackson, Ian Johnston, Dominic Kwiatkowski, Cordelia Langford, John Sillitoe on behalf of the Wellcome Sanger Institute COVID-19 Surveillance Team (http://www.sanger.ac.uk/covid-team) |
| EPI_ISL_555507 | Lighthouse Lab in Alderley Park | Wellcome Sanger Institute for the COVID-19 Genomics UK (COG-UK) consortium | The Lighthouse Lab in Alderley Park and Alex Alderton, Roberto Amato, Sonia Goncalves, Ewan Harrison, David K. Jackson, Ian Johnston, Dominic Kwiatkowski, Cordelia Langford, John Sillitoe on behalf of the Wellcome Sanger Institute COVID-19 Surveillance Team |
| EPI_ISL_555509 | Lighthouse Lab in Alderley Park | Wellcome Sanger Institute for the COVID-19 Genomics UK (COG-UK) consortium | The Lighthouse Lab in Alderley Park and Alex Alderton, Roberto Amato, Sonia Goncalves, Ewan Harrison, David K. Jackson, Ian Johnston, Dominic Kwiatkowski, Cordelia Langford, John Sillitoe on behalf of the Wellcome Sanger Institute COVID-19 Surveillance Team (http://www.sanger.ac.uk/covid-team) |
| EPI_ISL_555512, EPI_ISL_555515, EPI_ISL_555521 | Lighthouse Lab in Alderley Park | Wellcome Sanger Institute for the COVID-19 Genomics UK (COG-UK) consortium | The Lighthouse Lab in Alderley Park and Alex Alderton, Roberto Amato, Sonia Goncalves, Ewan Harrison, David K. Jackson, Ian Johnston, Dominic Kwiatkowski, Cordelia Langford, John Sillitoe on behalf of the Wellcome Sanger Institute COVID-19 Surveillance Team |
| EPI_ISL_555522 | Lighthouse Lab in Alderley Park | Wellcome Sanger Institute for the COVID-19 Genomics UK (COG-UK) consortium | The Lighthouse Lab in Alderley Park and Alex Alderton, Roberto Amato, Sonia Goncalves, Ewan Harrison, David K. Jackson, Ian Johnston, Dominic Kwiatkowski, Cordelia Langford, John Sillitoe on behalf of the Wellcome Sanger Institute COVID-19 Surveillance Team |

| | | | |
|---|---|---|---|
| | | | (http://www.sanger.ac.uk/covid-team) |
| EPI_ISL_555525, EPI_ISL_555528, EPI_ISL_555530, EPI_ISL_555532 | Lighthouse Lab in Alderley Park | Wellcome Sanger Institute for the COVID-19 Genomics UK (COG-UK) consortium | The Lighthouse Lab in Alderley Park and Alex Alderton, Roberto Amato, Sonia Goncalves, Ewan Harrison, David K. Jackson, Ian Johnston, Dominic Kwiatkowski, Cordelia Langford, John Sillitoe on behalf of the Wellcome Sanger Institute COVID-19 Surveillance Team |
| EPI_ISL_555534, EPI_ISL_555536 | Lighthouse Lab in Alderley Park | Wellcome Sanger Institute for the COVID-19 Genomics UK (COG-UK) consortium | The Lighthouse Lab in Alderley Park and Alex Alderton, Roberto Amato, Sonia Goncalves, Ewan Harrison, David K. Jackson, Ian Johnston, Dominic Kwiatkowski, Cordelia Langford, John Sillitoe on behalf of the Wellcome Sanger Institute COVID-19 Surveillance Team (http://www.sanger.ac.uk/covid-team) |
| EPI_ISL_555538, EPI_ISL_555539, EPI_ISL_555545, EPI_ISL_555555, EPI_ISL_555556, EPI_ISL_555562, EPI_ISL_555569, EPI_ISL_555571 | Lighthouse Lab in Alderley Park | Wellcome Sanger Institute for the COVID-19 Genomics UK (COG-UK) consortium | The Lighthouse Lab in Alderley Park and Alex Alderton, Roberto Amato, Sonia Goncalves, Ewan Harrison, David K. Jackson, Ian Johnston, Dominic Kwiatkowski, Cordelia Langford, John Sillitoe on behalf of the Wellcome Sanger Institute COVID-19 Surveillance Team |
| EPI_ISL_555574 | Lighthouse Lab in Alderley Park | Wellcome Sanger Institute for the COVID-19 Genomics UK (COG-UK) consortium | The Lighthouse Lab in Alderley Park and Alex Alderton, Roberto Amato, Sonia Goncalves, Ewan Harrison, David K. Jackson, Ian Johnston, Dominic Kwiatkowski, Cordelia Langford, John Sillitoe on behalf of the Wellcome Sanger Institute COVID-19 Surveillance Team (http://www.sanger.ac.uk/covid-team) |
| EPI_ISL_555575, EPI_ISL_555577, EPI_ISL_555578, EPI_ISL_555586, EPI_ISL_555590, EPI_ISL_555591, EPI_ISL_555595, EPI_ISL_555599, EPI_ISL_555603, EPI_ISL_555605, EPI_ISL_555607, EPI_ISL_555609, EPI_ISL_555610, EPI_ISL_555611, EPI_ISL_555612, EPI_ISL_555618, EPI_ISL_555621, EPI_ISL_555624, EPI_ISL_555625 | | | |
| see above | Lighthouse Lab in Alderley Park | Wellcome Sanger Institute for the COVID-19 Genomics UK (COG-UK) consortium | The Lighthouse Lab in Alderley Park and Alex Alderton, Roberto Amato, Sonia Goncalves, Ewan Harrison, David K. Jackson, Ian Johnston, Dominic Kwiatkowski, Cordelia Langford, John Sillitoe on behalf of the Wellcome Sanger Institute COVID-19 Surveillance Team |
| EPI_ISL_555630, EPI_ISL_555634, EPI_ISL_555637, EPI_ISL_555638, EPI_ISL_555639 | Lighthouse Lab in Alderley Park | Wellcome Sanger Institute for the COVID-19 Genomics UK (COG-UK) consortium | The Lighthouse Lab in Alderley Park and Alex Alderton, Roberto Amato, Sonia Goncalves, Ewan Harrison, David K. Jackson, Ian Johnston, Dominic Kwiatkowski, Cordelia Langford, John Sillitoe on behalf of the Wellcome Sanger Institute COVID-19 Surveillance Team (http://www.sanger.ac.uk/covid-team) |
| EPI_ISL_555640 | Lighthouse Lab in Alderley Park | Wellcome Sanger Institute for the COVID-19 Genomics UK (COG-UK) consortium | The Lighthouse Lab in Alderley Park and Alex Alderton, Roberto Amato, Sonia Goncalves, Ewan Harrison, David K. Jackson, Ian Johnston, Dominic Kwiatkowski, Cordelia Langford, John Sillitoe on behalf of the Wellcome Sanger Institute COVID-19 Surveillance Team |
| EPI_ISL_555641 | Lighthouse Lab in Alderley Park | Wellcome Sanger Institute for the COVID-19 Genomics UK (COG-UK) consortium | The Lighthouse Lab in Alderley Park and Alex Alderton, Roberto Amato, Sonia Goncalves, Ewan Harrison, David K. Jackson, Ian Johnston, Dominic Kwiatkowski, Cordelia Langford, John Sillitoe on behalf of the Wellcome Sanger Institute COVID-19 Surveillance Team (http://www.sanger.ac.uk/covid-team) |
| EPI_ISL_555643, EPI_ISL_555644, EPI_ISL_555645, EPI_ISL_555646, EPI_ISL_555647, EPI_ISL_555648, EPI_ISL_555649 | Lighthouse Lab in Alderley Park | Wellcome Sanger Institute for the COVID-19 Genomics UK (COG-UK) consortium | The Lighthouse Lab in Alderley Park and Alex Alderton, Roberto Amato, Sonia Goncalves, Ewan Harrison, David K. Jackson, Ian Johnston, Dominic Kwiatkowski, Cordelia Langford, John Sillitoe on behalf of the Wellcome Sanger Institute COVID-19 Surveillance Team |
| EPI_ISL_555654 | Lighthouse Lab in Alderley Park | Wellcome Sanger Institute for the COVID-19 Genomics UK (COG-UK) consortium | The Lighthouse Lab in Alderley Park and Alex Alderton, Roberto Amato, Sonia Goncalves, Ewan Harrison, David K. Jackson, Ian Johnston, Dominic Kwiatkowski, Cordelia Langford, John Sillitoe on behalf of the Wellcome Sanger Institute COVID-19 Surveillance Team (http://www.sanger.ac.uk/covid-team) |
| EPI_ISL_555655, EPI_ISL_555661, EPI_ISL_555662, EPI_ISL_555665 | Lighthouse Lab in Alderley Park | Wellcome Sanger Institute for the COVID-19 Genomics UK (COG-UK) consortium | The Lighthouse Lab in Alderley Park and Alex Alderton, Roberto Amato, Sonia Goncalves, Ewan Harrison, David K. Jackson, Ian Johnston, Dominic Kwiatkowski, Cordelia Langford, John Sillitoe on behalf of the Wellcome Sanger Institute COVID-19 Surveillance Team |
| EPI_ISL_555674 | Lighthouse Lab in Alderley Park | Wellcome Sanger Institute for the COVID-19 Genomics UK (COG-UK) consortium | The Lighthouse Lab in Alderley Park and Alex Alderton, Roberto Amato, Sonia Goncalves, Ewan Harrison, David K. Jackson, Ian Johnston, Dominic Kwiatkowski, Cordelia Langford, John Sillitoe on behalf of the Wellcome Sanger Institute COVID-19 Surveillance Team (http://www.sanger.ac.uk/covid-team) |
| EPI_ISL_555675 | Lighthouse Lab in Alderley Park | Wellcome Sanger Institute for the COVID-19 Genomics UK (COG-UK) consortium | The Lighthouse Lab in Alderley Park and Alex Alderton, Roberto Amato, Sonia Goncalves, Ewan Harrison, David K. Jackson, Ian Johnston, Dominic Kwiatkowski, Cordelia Langford, John Sillitoe on behalf of the Wellcome Sanger Institute COVID-19 Surveillance Team |
| EPI_ISL_555676 | Lighthouse Lab in Alderley Park | Wellcome Sanger Institute for the COVID-19 Genomics UK (COG-UK) consortium | The Lighthouse Lab in Alderley Park and Alex Alderton, Roberto Amato, Sonia Goncalves, Ewan Harrison, David K. Jackson, Ian Johnston, Dominic Kwiatkowski, Cordelia Langford, John Sillitoe on behalf of the Wellcome Sanger Institute COVID-19 Surveillance Team (http://www.sanger.ac.uk/covid-team) |
| EPI_ISL_555684 | Lighthouse Lab in Alderley Park | Wellcome Sanger Institute for the COVID-19 Genomics UK (COG-UK) consortium | The Lighthouse Lab in Alderley Park and Alex Alderton, Roberto Amato, Sonia Goncalves, Ewan Harrison, David K. Jackson, Ian Johnston, Dominic Kwiatkowski, Cordelia Langford, John Sillitoe on behalf of the Wellcome Sanger Institute COVID-19 Surveillance Team |
| EPI_ISL_555686 | Lighthouse Lab in Alderley Park | Wellcome Sanger Institute for the COVID-19 Genomics UK (COG-UK) consortium | The Lighthouse Lab in Alderley Park and Alex Alderton, Roberto Amato, Sonia Goncalves, Ewan Harrison, David K. Jackson, Ian Johnston, Dominic Kwiatkowski, Cordelia Langford, John Sillitoe on behalf of the Wellcome Sanger Institute COVID-19 Surveillance Team (http://www.sanger.ac.uk/covid-team) |
| EPI_ISL_555687, EPI_ISL_555688, EPI_ISL_555690, EPI_ISL_555691, EPI_ISL_555692, EPI_ISL_555693, EPI_ISL_555697, EPI_ISL_555699, EPI_ISL_555701, EPI_ISL_555703, EPI_ISL_555704, EPI_ISL_555710, EPI_ISL_555720, EPI_ISL_555721, EPI_ISL_555722, EPI_ISL_555724, EPI_ISL_555730, EPI_ISL_555733, EPI_ISL_555734 | | | |
| see above | Lighthouse Lab in Alderley Park | Wellcome Sanger Institute for the COVID-19 Genomics UK (COG-UK) consortium | The Lighthouse Lab in Alderley Park and Alex Alderton, Roberto Amato, Sonia Goncalves, Ewan Harrison, David K. Jackson, Ian Johnston, Dominic Kwiatkowski, Cordelia Langford, John Sillitoe on behalf of the Wellcome Sanger Institute COVID-19 Surveillance Team |
| EPI_ISL_555736 | Lighthouse Lab in Alderley Park | Wellcome Sanger Institute for the COVID-19 Genomics UK (COG-UK) consortium | The Lighthouse Lab in Alderley Park and Alex Alderton, Roberto Amato, Sonia Goncalves, Ewan Harrison, David K. Jackson, Ian Johnston, Dominic Kwiatkowski, Cordelia Langford, John Sillitoe on behalf of the Wellcome Sanger Institute COVID-19 Surveillance Team (http://www.sanger.ac.uk/covid-team) |
| EPI_ISL_555737, EPI_ISL_555738 | Lighthouse Lab in Alderley Park | Wellcome Sanger Institute for the COVID-19 Genomics UK (COG-UK) consortium | The Lighthouse Lab in Alderley Park and Alex Alderton, Roberto Amato, Sonia Goncalves, Ewan Harrison, David K. Jackson, Ian Johnston, Dominic Kwiatkowski, Cordelia Langford, John Sillitoe on behalf of the Wellcome Sanger Institute COVID-19 Surveillance Team |
| EPI_ISL_555740, EPI_ISL_555743, EPI_ISL_555747 | Lighthouse Lab in Alderley Park | Wellcome Sanger Institute for the COVID-19 Genomics UK (COG-UK) consortium | The Lighthouse Lab in Alderley Park and Alex Alderton, Roberto Amato, Sonia Goncalves, Ewan Harrison, David K. Jackson, Ian Johnston, Dominic Kwiatkowski, Cordelia Langford, John Sillitoe on behalf of the Wellcome Sanger Institute COVID-19 Surveillance Team (http://www.sanger.ac.uk/covid-team) |
| EPI_ISL_555750 | Lighthouse Lab in Alderley Park | Wellcome Sanger Institute for the COVID-19 Genomics UK (COG-UK) consortium | The Lighthouse Lab in Alderley Park and Alex Alderton, Roberto Amato, Sonia Goncalves, Ewan Harrison, David K. Jackson, Ian Johnston, Dominic Kwiatkowski, Cordelia Langford, John Sillitoe on behalf of the Wellcome Sanger Institute COVID-19 Surveillance Team |
| EPI_ISL_555751 | Lighthouse Lab in Alderley Park | Wellcome Sanger Institute for the COVID-19 Genomics UK (COG-UK) consortium | The Lighthouse Lab in Alderley Park and Alex Alderton, Roberto Amato, Sonia Goncalves, Ewan Harrison, David K. Jackson, Ian Johnston, Dominic Kwiatkowski, Cordelia Langford, John Sillitoe on behalf of the Wellcome Sanger Institute COVID-19 Surveillance Team (http://www.sanger.ac.uk/covid-team) |
| EPI_ISL_555753, EPI_ISL_555755, EPI_ISL_555756, EPI_ISL_555757, EPI_ISL_555758 | Lighthouse Lab in Alderley Park | Wellcome Sanger Institute for the COVID-19 Genomics UK (COG-UK) consortium | The Lighthouse Lab in Alderley Park and Alex Alderton, Roberto Amato, Sonia Goncalves, Ewan Harrison, David K. Jackson, Ian Johnston, Dominic Kwiatkowski, Cordelia Langford, John Sillitoe on behalf of the Wellcome Sanger Institute COVID-19 Surveillance Team |
| EPI_ISL_556074 | Lighthouse Lab in Alderley Park | Wellcome Sanger Institute for the COVID-19 Genomics UK (COG-UK) consortium | The Lighthouse Lab in Alderley Park and Alex Alderton, Roberto Amato, Sonia Goncalves, Ewan Harrison, David K. Jackson, Ian Johnston, Dominic Kwiatkowski, Cordelia Langford, John Sillitoe on behalf of the Wellcome Sanger Institute COVID-19 Surveillance Team (http://www.sanger.ac.uk/covid-team) |
| EPI_ISL_556078 | Lighthouse Lab in Alderley Park | Wellcome Sanger Institute for the COVID-19 Genomics UK (COG-UK) consortium | The Lighthouse Lab in Alderley Park and Alex Alderton, Roberto Amato, Sonia Goncalves, Ewan Harrison, David K. Jackson, Ian Johnston, Dominic Kwiatkowski, Cordelia Langford, John Sillitoe on behalf of the Wellcome Sanger Institute COVID-19 Surveillance Team |
| EPI_ISL_556080, EPI_ISL_556081, | Lighthouse Lab in Alderley Park | Wellcome Sanger Institute for the COVID-19 Genomics UK | The Lighthouse Lab in Alderley Park and Alex Alderton, Roberto Amato, Sonia Goncalves, Ewan Harrison, David K. Jackson, Ian Johnston, Dominic |

| | | | |
|---|---|---|---|
| EPI_ISL_556082, EPI_ISL_556084 | | (COG-UK) consortium | Kwiatkowski, Cordelia Langford, John Sillitoe on behalf of the Wellcome Sanger Institute COVID-19 Surveillance Team (http://www.sanger.ac.uk/covid-team) |
| EPI_ISL_556086 | Lighthouse Lab in Alderley Park | Wellcome Sanger Institute for the COVID-19 Genomics UK (COG-UK) consortium | The Lighthouse Lab in Alderley Park and Alex Alderton, Roberto Amato, Sonia Goncalves, Ewan Harrison, David K. Jackson, Ian Johnston, Dominic Kwiatkowski, Cordelia Langford, John Sillitoe on behalf of the Wellcome Sanger Institute COVID-19 Surveillance Team |
| EPI_ISL_556089 | Lighthouse Lab in Alderley Park | Wellcome Sanger Institute for the COVID-19 Genomics UK (COG-UK) consortium | The Lighthouse Lab in Alderley Park and Alex Alderton, Roberto Amato, Sonia Goncalves, Ewan Harrison, David K. Jackson, Ian Johnston, Dominic Kwiatkowski, Cordelia Langford, John Sillitoe on behalf of the Wellcome Sanger Institute COVID-19 Surveillance Team (http://www.sanger.ac.uk/covid-team) |
| EPI_ISL_556096, EPI_ISL_556099, EPI_ISL_556100, EPI_ISL_556101, EPI_ISL_556103 | Lighthouse Lab in Alderley Park | Wellcome Sanger Institute for the COVID-19 Genomics UK (COG-UK) consortium | The Lighthouse Lab in Alderley Park and Alex Alderton, Roberto Amato, Sonia Goncalves, Ewan Harrison, David K. Jackson, Ian Johnston, Dominic Kwiatkowski, Cordelia Langford, John Sillitoe on behalf of the Wellcome Sanger Institute COVID-19 Surveillance Team |
| EPI_ISL_556105, EPI_ISL_556106, EPI_ISL_556107 | Lighthouse Lab in Alderley Park | Wellcome Sanger Institute for the COVID-19 Genomics UK (COG-UK) consortium | The Lighthouse Lab in Alderley Park and Alex Alderton, Roberto Amato, Sonia Goncalves, Ewan Harrison, David K. Jackson, Ian Johnston, Dominic Kwiatkowski, Cordelia Langford, John Sillitoe on behalf of the Wellcome Sanger Institute COVID-19 Surveillance Team (http://www.sanger.ac.uk/covid-team) |
| EPI_ISL_556108 | Lighthouse Lab in Alderley Park | Wellcome Sanger Institute for the COVID-19 Genomics UK (COG-UK) consortium | The Lighthouse Lab in Alderley Park and Alex Alderton, Roberto Amato, Sonia Goncalves, Ewan Harrison, David K. Jackson, Ian Johnston, Dominic Kwiatkowski, Cordelia Langford, John Sillitoe on behalf of the Wellcome Sanger Institute COVID-19 Surveillance Team |
| EPI_ISL_556113, EPI_ISL_556114, EPI_ISL_556117, EPI_ISL_556119, EPI_ISL_556120, EPI_ISL_556121, EPI_ISL_556122, EPI_ISL_556125, EPI_ISL_556127, EPI_ISL_556128, EPI_ISL_556129, EPI_ISL_556132, EPI_ISL_556135 | | | |
| see above | Lighthouse Lab in Alderley Park | Wellcome Sanger Institute for the COVID-19 Genomics UK (COG-UK) consortium | The Lighthouse Lab in Alderley Park and Alex Alderton, Roberto Amato, Sonia Goncalves, Ewan Harrison, David K. Jackson, Ian Johnston, Dominic Kwiatkowski, Cordelia Langford, John Sillitoe on behalf of the Wellcome Sanger Institute COVID-19 Surveillance Team (http://www.sanger.ac.uk/covid-team) |
| EPI_ISL_556142, EPI_ISL_556143, EPI_ISL_556145, EPI_ISL_556147 | Lighthouse Lab in Alderley Park | Wellcome Sanger Institute for the COVID-19 Genomics UK (COG-UK) consortium | The Lighthouse Lab in Alderley Park and Alex Alderton, Roberto Amato, Sonia Goncalves, Ewan Harrison, David K. Jackson, Ian Johnston, Dominic Kwiatkowski, Cordelia Langford, John Sillitoe on behalf of the Wellcome Sanger Institute COVID-19 Surveillance Team |
| EPI_ISL_556149 | Lighthouse Lab in Alderley Park | Wellcome Sanger Institute for the COVID-19 Genomics UK (COG-UK) consortium | The Lighthouse Lab in Alderley Park and Alex Alderton, Roberto Amato, Sonia Goncalves, Ewan Harrison, David K. Jackson, Ian Johnston, Dominic Kwiatkowski, Cordelia Langford, John Sillitoe on behalf of the Wellcome Sanger Institute COVID-19 Surveillance Team (http://www.sanger.ac.uk/covid-team) |
| EPI_ISL_556153 | Lighthouse Lab in Alderley Park | Wellcome Sanger Institute for the COVID-19 Genomics UK (COG-UK) consortium | The Lighthouse Lab in Alderley Park and Alex Alderton, Roberto Amato, Sonia Goncalves, Ewan Harrison, David K. Jackson, Ian Johnston, Dominic Kwiatkowski, Cordelia Langford, John Sillitoe on behalf of the Wellcome Sanger Institute COVID-19 Surveillance Team |
| EPI_ISL_556155, EPI_ISL_556156, EPI_ISL_556157 | Lighthouse Lab in Alderley Park | Wellcome Sanger Institute for the COVID-19 Genomics UK (COG-UK) consortium | The Lighthouse Lab in Alderley Park and Alex Alderton, Roberto Amato, Sonia Goncalves, Ewan Harrison, David K. Jackson, Ian Johnston, Dominic Kwiatkowski, Cordelia Langford, John Sillitoe on behalf of the Wellcome Sanger Institute COVID-19 Surveillance Team (http://www.sanger.ac.uk/covid-team) |
| EPI_ISL_556161, EPI_ISL_556162 | Lighthouse Lab in Alderley Park | Wellcome Sanger Institute for the COVID-19 Genomics UK (COG-UK) consortium | The Lighthouse Lab in Alderley Park and Alex Alderton, Roberto Amato, Sonia Goncalves, Ewan Harrison, David K. Jackson, Ian Johnston, Dominic Kwiatkowski, Cordelia Langford, John Sillitoe on behalf of the Wellcome Sanger Institute COVID-19 Surveillance Team |
| EPI_ISL_556167, EPI_ISL_556168 | Lighthouse Lab in Alderley Park | Wellcome Sanger Institute for the COVID-19 Genomics UK (COG-UK) consortium | The Lighthouse Lab in Alderley Park and Alex Alderton, Roberto Amato, Sonia Goncalves, Ewan Harrison, David K. Jackson, Ian Johnston, Dominic Kwiatkowski, Cordelia Langford, John Sillitoe on behalf of the Wellcome Sanger Institute COVID-19 Surveillance Team (http://www.sanger.ac.uk/covid-team) |
| EPI_ISL_556169 | Lighthouse Lab in Alderley Park | Wellcome Sanger Institute for the COVID-19 Genomics UK (COG-UK) consortium | The Lighthouse Lab in Alderley Park and Alex Alderton, Roberto Amato, Sonia Goncalves, Ewan Harrison, David K. Jackson, Ian Johnston, Dominic Kwiatkowski, Cordelia Langford, John Sillitoe on behalf of the Wellcome Sanger Institute COVID-19 Surveillance Team |
| EPI_ISL_556171, EPI_ISL_556174, EPI_ISL_556175, EPI_ISL_556179 | Lighthouse Lab in Alderley Park | Wellcome Sanger Institute for the COVID-19 Genomics UK (COG-UK) consortium | The Lighthouse Lab in Alderley Park and Alex Alderton, Roberto Amato, Sonia Goncalves, Ewan Harrison, David K. Jackson, Ian Johnston, Dominic Kwiatkowski, Cordelia Langford, John Sillitoe on behalf of the Wellcome Sanger Institute COVID-19 Surveillance Team (http://www.sanger.ac.uk/covid-team) |
| EPI_ISL_556181 | Lighthouse Lab in Alderley Park | Wellcome Sanger Institute for the COVID-19 Genomics UK (COG-UK) consortium | The Lighthouse Lab in Alderley Park and Alex Alderton, Roberto Amato, Sonia Goncalves, Ewan Harrison, David K. Jackson, Ian Johnston, Dominic Kwiatkowski, Cordelia Langford, John Sillitoe on behalf of the Wellcome Sanger Institute COVID-19 Surveillance Team |
| EPI_ISL_556185, EPI_ISL_556188 | Lighthouse Lab in Alderley Park | Wellcome Sanger Institute for the COVID-19 Genomics UK (COG-UK) consortium | The Lighthouse Lab in Alderley Park and Alex Alderton, Roberto Amato, Sonia Goncalves, Ewan Harrison, David K. Jackson, Ian Johnston, Dominic Kwiatkowski, Cordelia Langford, John Sillitoe on behalf of the Wellcome Sanger Institute COVID-19 Surveillance Team (http://www.sanger.ac.uk/covid-team) |
| EPI_ISL_556189 | Lighthouse Lab in Alderley Park | Wellcome Sanger Institute for the COVID-19 Genomics UK (COG-UK) consortium | The Lighthouse Lab in Alderley Park and Alex Alderton, Roberto Amato, Sonia Goncalves, Ewan Harrison, David K. Jackson, Ian Johnston, Dominic Kwiatkowski, Cordelia Langford, John Sillitoe on behalf of the Wellcome Sanger Institute COVID-19 Surveillance Team |
| EPI_ISL_556190 | Lighthouse Lab in Alderley Park | Wellcome Sanger Institute for the COVID-19 Genomics UK (COG-UK) consortium | The Lighthouse Lab in Alderley Park and Alex Alderton, Roberto Amato, Sonia Goncalves, Ewan Harrison, David K. Jackson, Ian Johnston, Dominic Kwiatkowski, Cordelia Langford, John Sillitoe on behalf of the Wellcome Sanger Institute COVID-19 Surveillance Team (http://www.sanger.ac.uk/covid-team) |
| EPI_ISL_556191 | Lighthouse Lab in Alderley Park | Wellcome Sanger Institute for the COVID-19 Genomics UK (COG-UK) consortium | The Lighthouse Lab in Alderley Park and Alex Alderton, Roberto Amato, Sonia Goncalves, Ewan Harrison, David K. Jackson, Ian Johnston, Dominic Kwiatkowski, Cordelia Langford, John Sillitoe on behalf of the Wellcome Sanger Institute COVID-19 Surveillance Team |
| EPI_ISL_556192, EPI_ISL_556193 | Lighthouse Lab in Alderley Park | Wellcome Sanger Institute for the COVID-19 Genomics UK (COG-UK) consortium | The Lighthouse Lab in Alderley Park and Alex Alderton, Roberto Amato, Sonia Goncalves, Ewan Harrison, David K. Jackson, Ian Johnston, Dominic Kwiatkowski, Cordelia Langford, John Sillitoe on behalf of the Wellcome Sanger Institute COVID-19 Surveillance Team (http://www.sanger.ac.uk/covid-team) |
| EPI_ISL_556194 | Lighthouse Lab in Alderley Park | Wellcome Sanger Institute for the COVID-19 Genomics UK (COG-UK) consortium | The Lighthouse Lab in Alderley Park and Alex Alderton, Roberto Amato, Sonia Goncalves, Ewan Harrison, David K. Jackson, Ian Johnston, Dominic Kwiatkowski, Cordelia Langford, John Sillitoe on behalf of the Wellcome Sanger Institute COVID-19 Surveillance Team |
| EPI_ISL_556196, EPI_ISL_556200, EPI_ISL_556202, EPI_ISL_556205, EPI_ISL_556206, EPI_ISL_556209, EPI_ISL_556210, EPI_ISL_556212, EPI_ISL_556215 | Lighthouse Lab in Alderley Park | Wellcome Sanger Institute for the COVID-19 Genomics UK (COG-UK) consortium | The Lighthouse Lab in Alderley Park and Alex Alderton, Roberto Amato, Sonia Goncalves, Ewan Harrison, David K. Jackson, Ian Johnston, Dominic Kwiatkowski, Cordelia Langford, John Sillitoe on behalf of the Wellcome Sanger Institute COVID-19 Surveillance Team (http://www.sanger.ac.uk/covid-team) |
| EPI_ISL_556216, EPI_ISL_556217, EPI_ISL_556219 | Lighthouse Lab in Alderley Park | Wellcome Sanger Institute for the COVID-19 Genomics UK (COG-UK) consortium | The Lighthouse Lab in Alderley Park and Alex Alderton, Roberto Amato, Sonia Goncalves, Ewan Harrison, David K. Jackson, Ian Johnston, Dominic Kwiatkowski, Cordelia Langford, John Sillitoe on behalf of the Wellcome Sanger Institute COVID-19 Surveillance Team |
| EPI_ISL_556220 | Lighthouse Lab in Alderley Park | Wellcome Sanger Institute for the COVID-19 Genomics UK (COG-UK) consortium | The Lighthouse Lab in Alderley Park and Alex Alderton, Roberto Amato, Sonia Goncalves, Ewan Harrison, David K. Jackson, Ian Johnston, Dominic Kwiatkowski, Cordelia Langford, John Sillitoe on behalf of the Wellcome Sanger Institute COVID-19 Surveillance Team (http://www.sanger.ac.uk/covid-team) |
| EPI_ISL_556222 | Lighthouse Lab in Alderley Park | Wellcome Sanger Institute for the COVID-19 Genomics UK (COG-UK) consortium | The Lighthouse Lab in Alderley Park and Alex Alderton, Roberto Amato, Sonia Goncalves, Ewan Harrison, David K. Jackson, Ian Johnston, Dominic Kwiatkowski, Cordelia Langford, John Sillitoe on behalf of the Wellcome Sanger Institute COVID-19 Surveillance Team |
| EPI_ISL_556224, EPI_ISL_556225 | Lighthouse Lab in Alderley Park | Wellcome Sanger Institute for the COVID-19 Genomics UK (COG-UK) consortium | The Lighthouse Lab in Alderley Park and Alex Alderton, Roberto Amato, Sonia Goncalves, Ewan Harrison, David K. Jackson, Ian Johnston, Dominic Kwiatkowski, Cordelia Langford, John Sillitoe on behalf of the Wellcome Sanger Institute COVID-19 Surveillance Team (http://www.sanger.ac.uk/covid-team) |
| EPI_ISL_556226, EPI_ISL_556228 | Lighthouse Lab in Alderley Park | Wellcome Sanger Institute for the COVID-19 Genomics UK (COG-UK) consortium | The Lighthouse Lab in Alderley Park and Alex Alderton, Roberto Amato, Sonia Goncalves, Ewan Harrison, David K. Jackson, Ian Johnston, Dominic Kwiatkowski, Cordelia Langford, John Sillitoe on behalf of the Wellcome Sanger Institute COVID-19 Surveillance Team |

| | | | |
|---|---|---|---|
| EPI_ISL_556229, EPI_ISL_556230 | Lighthouse Lab in Alderley Park | Wellcome Sanger Institute for the COVID-19 Genomics UK (COG-UK) consortium | The Lighthouse Lab in Alderley Park and Alex Alderton, Roberto Amato, Sonia Goncalves, Ewan Harrison, David K. Jackson, Ian Johnston, Dominic Kwiatkowski, Cordelia Langford, John Sillitoe on behalf of the Wellcome Sanger Institute COVID-19 Surveillance Team (http://www.sanger.ac.uk/covid-team) |
| EPI_ISL_556236 | Lighthouse Lab in Alderley Park | Wellcome Sanger Institute for the COVID-19 Genomics UK (COG-UK) consortium | The Lighthouse Lab in Alderley Park and Alex Alderton, Roberto Amato, Sonia Goncalves, Ewan Harrison, David K. Jackson, Ian Johnston, Dominic Kwiatkowski, Cordelia Langford, John Sillitoe on behalf of the Wellcome Sanger Institute COVID-19 Surveillance Team |
| EPI_ISL_556238, EPI_ISL_556239, EPI_ISL_556240 | Lighthouse Lab in Alderley Park | Wellcome Sanger Institute for the COVID-19 Genomics UK (COG-UK) consortium | The Lighthouse Lab in Alderley Park and Alex Alderton, Roberto Amato, Sonia Goncalves, Ewan Harrison, David K. Jackson, Ian Johnston, Dominic Kwiatkowski, Cordelia Langford, John Sillitoe on behalf of the Wellcome Sanger Institute COVID-19 Surveillance Team (http://www.sanger.ac.uk/covid-team) |
| EPI_ISL_556241 | Lighthouse Lab in Alderley Park | Wellcome Sanger Institute for the COVID-19 Genomics UK (COG-UK) consortium | The Lighthouse Lab in Alderley Park and Alex Alderton, Roberto Amato, Sonia Goncalves, Ewan Harrison, David K. Jackson, Ian Johnston, Dominic Kwiatkowski, Cordelia Langford, John Sillitoe on behalf of the Wellcome Sanger Institute COVID-19 Surveillance Team |
| EPI_ISL_556243, EPI_ISL_556245, EPI_ISL_556246 | Lighthouse Lab in Alderley Park | Wellcome Sanger Institute for the COVID-19 Genomics UK (COG-UK) consortium | The Lighthouse Lab in Alderley Park and Alex Alderton, Roberto Amato, Sonia Goncalves, Ewan Harrison, David K. Jackson, Ian Johnston, Dominic Kwiatkowski, Cordelia Langford, John Sillitoe on behalf of the Wellcome Sanger Institute COVID-19 Surveillance Team (http://www.sanger.ac.uk/covid-team) |
| EPI_ISL_556247, EPI_ISL_556249 | Lighthouse Lab in Alderley Park | Wellcome Sanger Institute for the COVID-19 Genomics UK (COG-UK) consortium | The Lighthouse Lab in Alderley Park and Alex Alderton, Roberto Amato, Sonia Goncalves, Ewan Harrison, David K. Jackson, Ian Johnston, Dominic Kwiatkowski, Cordelia Langford, John Sillitoe on behalf of the Wellcome Sanger Institute COVID-19 Surveillance Team |
| EPI_ISL_556250, EPI_ISL_556252, EPI_ISL_556254 | Lighthouse Lab in Alderley Park | Wellcome Sanger Institute for the COVID-19 Genomics UK (COG-UK) consortium | The Lighthouse Lab in Alderley Park and Alex Alderton, Roberto Amato, Sonia Goncalves, Ewan Harrison, David K. Jackson, Ian Johnston, Dominic Kwiatkowski, Cordelia Langford, John Sillitoe on behalf of the Wellcome Sanger Institute COVID-19 Surveillance Team (http://www.sanger.ac.uk/covid-team) |
| EPI_ISL_556255, EPI_ISL_556257 | Lighthouse Lab in Alderley Park | Wellcome Sanger Institute for the COVID-19 Genomics UK (COG-UK) consortium | The Lighthouse Lab in Alderley Park and Alex Alderton, Roberto Amato, Sonia Goncalves, Ewan Harrison, David K. Jackson, Ian Johnston, Dominic Kwiatkowski, Cordelia Langford, John Sillitoe on behalf of the Wellcome Sanger Institute COVID-19 Surveillance Team |
| EPI_ISL_556258, EPI_ISL_556260, EPI_ISL_556264, EPI_ISL_556265 | Lighthouse Lab in Alderley Park | Wellcome Sanger Institute for the COVID-19 Genomics UK (COG-UK) consortium | The Lighthouse Lab in Alderley Park and Alex Alderton, Roberto Amato, Sonia Goncalves, Ewan Harrison, David K. Jackson, Ian Johnston, Dominic Kwiatkowski, Cordelia Langford, John Sillitoe on behalf of the Wellcome Sanger Institute COVID-19 Surveillance Team (http://www.sanger.ac.uk/covid-team) |
| EPI_ISL_556268, EPI_ISL_556271 | Lighthouse Lab in Alderley Park | Wellcome Sanger Institute for the COVID-19 Genomics UK (COG-UK) consortium | The Lighthouse Lab in Alderley Park and Alex Alderton, Roberto Amato, Sonia Goncalves, Ewan Harrison, David K. Jackson, Ian Johnston, Dominic Kwiatkowski, Cordelia Langford, John Sillitoe on behalf of the Wellcome Sanger Institute COVID-19 Surveillance Team |
| EPI_ISL_556273, EPI_ISL_556274, EPI_ISL_556278, EPI_ISL_556279, EPI_ISL_556281, EPI_ISL_556284 | Lighthouse Lab in Alderley Park | Wellcome Sanger Institute for the COVID-19 Genomics UK (COG-UK) consortium | The Lighthouse Lab in Alderley Park and Alex Alderton, Roberto Amato, Sonia Goncalves, Ewan Harrison, David K. Jackson, Ian Johnston, Dominic Kwiatkowski, Cordelia Langford, John Sillitoe on behalf of the Wellcome Sanger Institute COVID-19 Surveillance Team (http://www.sanger.ac.uk/covid-team) |
| EPI_ISL_556286 | Lighthouse Lab in Alderley Park | Wellcome Sanger Institute for the COVID-19 Genomics UK (COG-UK) consortium | The Lighthouse Lab in Alderley Park and Alex Alderton, Roberto Amato, Sonia Goncalves, Ewan Harrison, David K. Jackson, Ian Johnston, Dominic Kwiatkowski, Cordelia Langford, John Sillitoe on behalf of the Wellcome Sanger Institute COVID-19 Surveillance Team |
| EPI_ISL_556288, EPI_ISL_556290, EPI_ISL_556292, EPI_ISL_556299, EPI_ISL_556301, EPI_ISL_556302, EPI_ISL_556303, EPI_ISL_556335, EPI_ISL_556339, EPI_ISL_556352, EPI_ISL_556394 | | | |
| see above | Lighthouse Lab in Alderley Park | Wellcome Sanger Institute for the COVID-19 Genomics UK (COG-UK) consortium | The Lighthouse Lab in Alderley Park and Alex Alderton, Roberto Amato, Sonia Goncalves, Ewan Harrison, David K. Jackson, Ian Johnston, Dominic Kwiatkowski, Cordelia Langford, John Sillitoe on behalf of the Wellcome Sanger Institute COVID-19 Surveillance Team (http://www.sanger.ac.uk/covid-team) |
| EPI_ISL_556414, EPI_ISL_556477, EPI_ISL_556487, EPI_ISL_556492, EPI_ISL_556503, EPI_ISL_556504 | Lighthouse Lab in Alderley Park | Wellcome Sanger Institute for the COVID-19 Genomics UK (COG-UK) consortium | The Lighthouse Lab in Alderley Park and Alex Alderton, Roberto Amato, Sonia Goncalves, Ewan Harrison, David K. Jackson, Ian Johnston, Dominic Kwiatkowski, Cordelia Langford, John Sillitoe on behalf of the Wellcome Sanger Institute COVID-19 Surveillance Team |
| EPI_ISL_556577, EPI_ISL_556578, EPI_ISL_556583, EPI_ISL_556585, EPI_ISL_556586, EPI_ISL_556587, EPI_ISL_556588, EPI_ISL_556589, EPI_ISL_556592, EPI_ISL_556596, EPI_ISL_556598, EPI_ISL_556601, EPI_ISL_556619, EPI_ISL_556620, EPI_ISL_556622, EPI_ISL_556623, EPI_ISL_556625, EPI_ISL_556627, EPI_ISL_556629, EPI_ISL_556631, EPI_ISL_556635, EPI_ISL_556985, EPI_ISL_556987, EPI_ISL_556989, EPI_ISL_556990, EPI_ISL_556991, EPI_ISL_556992, EPI_ISL_556994, EPI_ISL_556995, EPI_ISL_556996, EPI_ISL_556998, EPI_ISL_557000, EPI_ISL_557003, EPI_ISL_557004, EPI_ISL_557005, EPI_ISL_557006, EPI_ISL_557007, EPI_ISL_557008, EPI_ISL_557009, EPI_ISL_557010, EPI_ISL_557012, EPI_ISL_557013, EPI_ISL_557014, EPI_ISL_557015, EPI_ISL_557017, EPI_ISL_557018, EPI_ISL_557019, EPI_ISL_557020, EPI_ISL_557021, EPI_ISL_557022, EPI_ISL_557023, EPI_ISL_557024, EPI_ISL_557025, EPI_ISL_557026, EPI_ISL_557027, EPI_ISL_557029, EPI_ISL_557033, EPI_ISL_557034, EPI_ISL_557036, EPI_ISL_557037, EPI_ISL_557038, EPI_ISL_557040, EPI_ISL_557041, EPI_ISL_557042, EPI_ISL_557044, EPI_ISL_557045, EPI_ISL_557046, EPI_ISL_557047, EPI_ISL_557048, EPI_ISL_557050, EPI_ISL_557051, EPI_ISL_557052, EPI_ISL_557053, EPI_ISL_557054, EPI_ISL_557057, EPI_ISL_557058, EPI_ISL_557059, EPI_ISL_557060, EPI_ISL_557064, EPI_ISL_557066, EPI_ISL_557067, EPI_ISL_557068, EPI_ISL_557070, EPI_ISL_557072, EPI_ISL_557073, EPI_ISL_557074, EPI_ISL_557075, EPI_ISL_557076, EPI_ISL_557077, EPI_ISL_557078, EPI_ISL_557080, EPI_ISL_557084, EPI_ISL_557085, EPI_ISL_557088, EPI_ISL_557090, EPI_ISL_557091, EPI_ISL_557092, EPI_ISL_557095, EPI_ISL_557096, EPI_ISL_557097, EPI_ISL_557098, EPI_ISL_557099, EPI_ISL_557100, EPI_ISL_557101, EPI_ISL_557105, EPI_ISL_557106, EPI_ISL_557108, EPI_ISL_557111, EPI_ISL_557113, EPI_ISL_557114, EPI_ISL_557115, EPI_ISL_557119, EPI_ISL_557120, EPI_ISL_557121, EPI_ISL_557122, EPI_ISL_557124, EPI_ISL_557125, EPI_ISL_557127, EPI_ISL_557128, EPI_ISL_557132, EPI_ISL_557133, EPI_ISL_557134, EPI_ISL_557136, EPI_ISL_557137, EPI_ISL_557138, EPI_ISL_557139, EPI_ISL_557140, EPI_ISL_557143, EPI_ISL_557144, EPI_ISL_557145, EPI_ISL_557146, EPI_ISL_557147, EPI_ISL_557148, EPI_ISL_557150, EPI_ISL_557151, EPI_ISL_557152, EPI_ISL_557153, EPI_ISL_557155, EPI_ISL_557157, EPI_ISL_557158, EPI_ISL_557160, EPI_ISL_557162, EPI_ISL_557163, EPI_ISL_557165, EPI_ISL_557356, EPI_ISL_557359, EPI_ISL_557361, EPI_ISL_557363, EPI_ISL_557373, EPI_ISL_557374, EPI_ISL_557378, EPI_ISL_557380, EPI_ISL_557381, EPI_ISL_557390, EPI_ISL_557398, EPI_ISL_557399, EPI_ISL_557404, EPI_ISL_557408, EPI_ISL_557411, EPI_ISL_557415, EPI_ISL_557417, EPI_ISL_557419, EPI_ISL_557423, EPI_ISL_557428, EPI_ISL_557433, EPI_ISL_557434, EPI_ISL_557436, EPI_ISL_557439, EPI_ISL_557440, EPI_ISL_557446, EPI_ISL_557447, EPI_ISL_557449, EPI_ISL_557452, EPI_ISL_557453, EPI_ISL_557454, EPI_ISL_557468, EPI_ISL_557469, EPI_ISL_557471, EPI_ISL_557474, EPI_ISL_557476, EPI_ISL_557481, EPI_ISL_557482, EPI_ISL_557484, EPI_ISL_557485, EPI_ISL_557487, EPI_ISL_557488, EPI_ISL_557491, EPI_ISL_557493, EPI_ISL_557497, EPI_ISL_557500, EPI_ISL_557507, EPI_ISL_557511, EPI_ISL_557518, EPI_ISL_557523, EPI_ISL_557526, EPI_ISL_557527, EPI_ISL_557530, EPI_ISL_557531, EPI_ISL_557533, EPI_ISL_557537, EPI_ISL_557539, EPI_ISL_557541, EPI_ISL_557543, EPI_ISL_557544 | | | |
| see above | Lighthouse Lab in Alderley Park | Wellcome Sanger Institute for the COVID-19 Genomics UK (COG-UK) consortium | The Lighthouse Lab in Alderley Park and Alex Alderton, Roberto Amato, Sonia Goncalves, Ewan Harrison, David K. Jackson, Ian Johnston, Dominic Kwiatkowski, Cordelia Langford, John Sillitoe on behalf of the Wellcome Sanger Institute COVID-19 Surveillance Team (http://www.sanger.ac.uk/covid-team) |
| EPI_ISL_557551, EPI_ISL_557553, EPI_ISL_557554, EPI_ISL_557555, EPI_ISL_557556, EPI_ISL_557557, EPI_ISL_557558, EPI_ISL_557559, EPI_ISL_557560, EPI_ISL_557561, EPI_ISL_557562, EPI_ISL_557563, EPI_ISL_557564, EPI_ISL_557565, EPI_ISL_557566, EPI_ISL_557567, EPI_ISL_557568, EPI_ISL_557569 | | | |
| see above | Lighthouse Lab in Alderley Park | Wellcome Sanger Institute for the COVID-19 Genomics UK (COG-UK) consortium | The Lighthouse Lab in Alderley Park and Alex Alderton, Roberto Amato, Sonia Goncalves, Ewan Harrison, David K. Jackson, Ian Johnston, Dominic Kwiatkowski, Cordelia Langford, John Sillitoe on behalf of the Wellcome Sanger Institute COVID-19 Surveillance Team |
| EPI_ISL_557570 | Lighthouse Lab in Alderley Park | Wellcome Sanger Institute for the COVID-19 Genomics UK (COG-UK) consortium | The Lighthouse Lab in Alderley Park and Alex Alderton, Roberto Amato, Sonia Goncalves, Ewan Harrison, David K. Jackson, Ian Johnston, Dominic Kwiatkowski, Cordelia Langford, John Sillitoe on behalf of the Wellcome Sanger Institute COVID-19 Surveillance Team (http://www.sanger.ac.uk/covid-team) |
| EPI_ISL_557571, EPI_ISL_557572, EPI_ISL_557573, EPI_ISL_557574, EPI_ISL_557575, EPI_ISL_557576, EPI_ISL_557577, EPI_ISL_557578, EPI_ISL_557579, EPI_ISL_557580, EPI_ISL_557581, EPI_ISL_557582, EPI_ISL_557583, EPI_ISL_557584, EPI_ISL_557585, EPI_ISL_557586, EPI_ISL_557587, EPI_ISL_557588, EPI_ISL_557589, EPI_ISL_557591, EPI_ISL_557592, EPI_ISL_557593, EPI_ISL_557594, EPI_ISL_557595 | | | |
| see above | Lighthouse Lab in Alderley Park | Wellcome Sanger Institute for the COVID-19 Genomics UK (COG-UK) consortium | The Lighthouse Lab in Alderley Park and Alex Alderton, Roberto Amato, Sonia Goncalves, Ewan Harrison, David K. Jackson, Ian Johnston, Dominic Kwiatkowski, Cordelia Langford, John Sillitoe on behalf of the Wellcome Sanger Institute COVID-19 Surveillance Team |
| EPI_ISL_557596 | Lighthouse Lab in Alderley Park | Wellcome Sanger Institute for the COVID-19 Genomics UK (COG-UK) consortium | The Lighthouse Lab in Alderley Park and Alex Alderton, Roberto Amato, Sonia Goncalves, Ewan Harrison, David K. Jackson, Ian Johnston, Dominic Kwiatkowski, Cordelia Langford, John Sillitoe on behalf of the Wellcome Sanger Institute COVID-19 Surveillance Team (http://www.sanger.ac.uk/covid-team) |
| EPI_ISL_557597, EPI_ISL_557598, EPI_ISL_557599, EPI_ISL_557600, EPI_ISL_557601, EPI_ISL_557602, EPI_ISL_557603, EPI_ISL_557604, EPI_ISL_557605, EPI_ISL_557606, EPI_ISL_557608, EPI_ISL_557609, EPI_ISL_557610, EPI_ISL_557611 | | | |
| see above | Lighthouse Lab in Alderley Park | Wellcome Sanger Institute for the COVID-19 Genomics UK (COG-UK) consortium | The Lighthouse Lab in Alderley Park and Alex Alderton, Roberto Amato, Sonia Goncalves, Ewan Harrison, David K. Jackson, Ian Johnston, Dominic Kwiatkowski, Cordelia Langford, John Sillitoe on behalf of the Wellcome Sanger Institute COVID-19 Surveillance Team |
| EPI_ISL_557612 | Lighthouse Lab in Alderley Park | Wellcome Sanger Institute for the COVID-19 Genomics UK (COG-UK) consortium | The Lighthouse Lab in Alderley Park and Alex Alderton, Roberto Amato, Sonia Goncalves, Ewan Harrison, David K. Jackson, Ian Johnston, Dominic Kwiatkowski, Cordelia Langford, John Sillitoe on behalf of the Wellcome Sanger Institute COVID-19 Surveillance Team (http://www.sanger.ac.uk/covid-team) |
| EPI_ISL_557613, EPI_ISL_557614, EPI_ISL_557616, EPI_ISL_557617, EPI_ISL_557618, EPI_ISL_557619, EPI_ISL_557620, EPI_ISL_557621, EPI_ISL_557622, EPI_ISL_557623, EPI_ISL_557624, EPI_ISL_557626, EPI_ISL_557627, EPI_ISL_557628, EPI_ISL_557629, EPI_ISL_557631, EPI_ISL_557632, EPI_ISL_557633, EPI_ISL_557634, EPI_ISL_557635, EPI_ISL_557636, EPI_ISL_557637, EPI_ISL_557638, EPI_ISL_557639, EPI_ISL_557640, EPI_ISL_557641, EPI_ISL_557643, EPI_ISL_557644, EPI_ISL_557645, EPI_ISL_557646, EPI_ISL_557648, EPI_ISL_557649, EPI_ISL_557650, EPI_ISL_557651, EPI_ISL_557652, EPI_ISL_557653, EPI_ISL_557654, EPI_ISL_557656, EPI_ISL_557657 | | | |

| | | | |
|---|---|---|---|
| see above | Lighthouse Lab in Alderley Park | Wellcome Sanger Institute for the COVID-19 Genomics UK (COG-UK) consortium | The Lighthouse Lab in Alderley Park and Alex Alderton, Roberto Amato, Sonia Goncalves, Ewan Harrison, David K. Jackson, Ian Johnston, Dominic Kwiatkowski, Cordelia Langford, John Sillitoe on behalf of the Wellcome Sanger Institute COVID-19 Surveillance Team |
| EPI_ISL_557658 | Lighthouse Lab in Alderley Park | Wellcome Sanger Institute for the COVID-19 Genomics UK (COG-UK) consortium | The Lighthouse Lab in Alderley Park and Alex Alderton, Roberto Amato, Sonia Goncalves, Ewan Harrison, David K. Jackson, Ian Johnston, Dominic Kwiatkowski, Cordelia Langford, John Sillitoe on behalf of the Wellcome Sanger Institute COVID-19 Surveillance Team (http://www.sanger.ac.uk/covid-team) |
| EPI_ISL_557659, EPI_ISL_557660, EPI_ISL_557661, EPI_ISL_557662, EPI_ISL_557663, EPI_ISL_557664, EPI_ISL_557665, EPI_ISL_557666, EPI_ISL_557667, EPI_ISL_557669, EPI_ISL_557670, EPI_ISL_557671, EPI_ISL_557673, EPI_ISL_557674, EPI_ISL_557675, EPI_ISL_557677, EPI_ISL_557678, EPI_ISL_557679, EPI_ISL_557680, EPI_ISL_557682, EPI_ISL_557683, EPI_ISL_557684, EPI_ISL_557686, EPI_ISL_557687, EPI_ISL_557688, EPI_ISL_557689, EPI_ISL_557691, EPI_ISL_557692, EPI_ISL_557693, EPI_ISL_557694, EPI_ISL_557695, EPI_ISL_557696, EPI_ISL_557697, EPI_ISL_557698, EPI_ISL_557699 | | | |
| see above | Lighthouse Lab in Alderley Park | Wellcome Sanger Institute for the COVID-19 Genomics UK (COG-UK) consortium | The Lighthouse Lab in Alderley Park and Alex Alderton, Roberto Amato, Sonia Goncalves, Ewan Harrison, David K. Jackson, Ian Johnston, Dominic Kwiatkowski, Cordelia Langford, John Sillitoe on behalf of the Wellcome Sanger Institute COVID-19 Surveillance Team |
| EPI_ISL_557700 | Lighthouse Lab in Alderley Park | Wellcome Sanger Institute for the COVID-19 Genomics UK (COG-UK) consortium | The Lighthouse Lab in Alderley Park and Alex Alderton, Roberto Amato, Sonia Goncalves, Ewan Harrison, David K. Jackson, Ian Johnston, Dominic Kwiatkowski, Cordelia Langford, John Sillitoe on behalf of the Wellcome Sanger Institute COVID-19 Surveillance Team (http://www.sanger.ac.uk/covid-team) |
| EPI_ISL_557702, EPI_ISL_557703, EPI_ISL_557704, EPI_ISL_557705, EPI_ISL_557706, EPI_ISL_557707, EPI_ISL_557708, EPI_ISL_557710, EPI_ISL_557712, EPI_ISL_557714, EPI_ISL_557715, EPI_ISL_557716, EPI_ISL_557717, EPI_ISL_557718, EPI_ISL_557719, EPI_ISL_557720, EPI_ISL_557721, EPI_ISL_557722, EPI_ISL_557723, EPI_ISL_557724, EPI_ISL_557725, EPI_ISL_557726, EPI_ISL_557727, EPI_ISL_557729, EPI_ISL_557731, EPI_ISL_557732, EPI_ISL_557733, EPI_ISL_557734, EPI_ISL_557735, EPI_ISL_557736, EPI_ISL_557737, EPI_ISL_557738, EPI_ISL_557743, EPI_ISL_557744, EPI_ISL_557747, EPI_ISL_557749, EPI_ISL_557751, EPI_ISL_557754, EPI_ISL_557755, EPI_ISL_557756, EPI_ISL_557759, EPI_ISL_557761, EPI_ISL_557762, EPI_ISL_557763, EPI_ISL_557769, EPI_ISL_557770, EPI_ISL_557772, EPI_ISL_557773, EPI_ISL_557776, EPI_ISL_557779, EPI_ISL_557780, EPI_ISL_557785, EPI_ISL_557787, EPI_ISL_557790, EPI_ISL_557794, EPI_ISL_557797, EPI_ISL_557798, EPI_ISL_557799, EPI_ISL_557803, EPI_ISL_557804, EPI_ISL_557805, EPI_ISL_557806, EPI_ISL_557809, EPI_ISL_557810, EPI_ISL_557813, EPI_ISL_557814, EPI_ISL_557815, EPI_ISL_557818, EPI_ISL_557819, EPI_ISL_557820, EPI_ISL_557825, EPI_ISL_557826, EPI_ISL_557831, EPI_ISL_557833, EPI_ISL_557834, EPI_ISL_557835, EPI_ISL_557836, EPI_ISL_557837, EPI_ISL_557839, EPI_ISL_557841, EPI_ISL_557842, EPI_ISL_557844, EPI_ISL_557850, EPI_ISL_557851, EPI_ISL_557855, EPI_ISL_557856, EPI_ISL_557857, EPI_ISL_557860, EPI_ISL_557861, EPI_ISL_557862, EPI_ISL_557863, EPI_ISL_557872, EPI_ISL_557873, EPI_ISL_557874, EPI_ISL_557875, EPI_ISL_557883, EPI_ISL_557887, EPI_ISL_557888, EPI_ISL_557891, EPI_ISL_557892, EPI_ISL_557897, EPI_ISL_557901, EPI_ISL_557903, EPI_ISL_557904, EPI_ISL_557906, EPI_ISL_557912, EPI_ISL_557914, EPI_ISL_557916, EPI_ISL_557918, EPI_ISL_557920, EPI_ISL_557922, EPI_ISL_557923, EPI_ISL_557924, EPI_ISL_557928, EPI_ISL_557929, EPI_ISL_557930, EPI_ISL_557931, EPI_ISL_557932, EPI_ISL_557934, EPI_ISL_557935, EPI_ISL_557939, EPI_ISL_557943, EPI_ISL_557950, EPI_ISL_557951, EPI_ISL_557955, EPI_ISL_557960, EPI_ISL_557961 | | | |
| see above | Lighthouse Lab in Alderley Park | Wellcome Sanger Institute for the COVID-19 Genomics UK (COG-UK) consortium | The Lighthouse Lab in Alderley Park and Alex Alderton, Roberto Amato, Sonia Goncalves, Ewan Harrison, David K. Jackson, Ian Johnston, Dominic Kwiatkowski, Cordelia Langford, John Sillitoe on behalf of the Wellcome Sanger Institute COVID-19 Surveillance Team |
| EPI_ISL_557966 | Lighthouse Lab in Alderley Park | Wellcome Sanger Institute for the COVID-19 Genomics UK (COG-UK) consortium | The Lighthouse Lab in Alderley Park and Alex Alderton, Roberto Amato, Sonia Goncalves, Ewan Harrison, David K. Jackson, Ian Johnston, Dominic Kwiatkowski, Cordelia Langford, John Sillitoe on behalf of the Wellcome Sanger Institute COVID-19 Surveillance Team (http://www.sanger.ac.uk/covid-team) |
| EPI_ISL_557968, EPI_ISL_557971, EPI_ISL_557973, EPI_ISL_557975, EPI_ISL_557978, EPI_ISL_557980, EPI_ISL_557981, EPI_ISL_557982, EPI_ISL_557983, EPI_ISL_557985, EPI_ISL_557986, EPI_ISL_557987, EPI_ISL_557990, EPI_ISL_557991, EPI_ISL_557992, EPI_ISL_557993, EPI_ISL_557994, EPI_ISL_557995, EPI_ISL_557997, EPI_ISL_557998, EPI_ISL_558000, EPI_ISL_558003, EPI_ISL_558004, EPI_ISL_558005, EPI_ISL_558007, EPI_ISL_558008, EPI_ISL_558009, EPI_ISL_558010, EPI_ISL_558011, EPI_ISL_558012, EPI_ISL_558013, EPI_ISL_558014, EPI_ISL_558015, EPI_ISL_558018, EPI_ISL_558019, EPI_ISL_558022, EPI_ISL_558025, EPI_ISL_558027, EPI_ISL_558029, EPI_ISL_558032, EPI_ISL_558033, EPI_ISL_558034, EPI_ISL_558035, EPI_ISL_558036, EPI_ISL_558039, EPI_ISL_558040, EPI_ISL_558042, EPI_ISL_558043, EPI_ISL_558044, EPI_ISL_558045, EPI_ISL_558046, EPI_ISL_558047, EPI_ISL_558048, EPI_ISL_558049, EPI_ISL_558050, EPI_ISL_558051, EPI_ISL_558053, EPI_ISL_558054, EPI_ISL_558057, EPI_ISL_558059, EPI_ISL_558060, EPI_ISL_558061, EPI_ISL_558062, EPI_ISL_558063, EPI_ISL_558065, EPI_ISL_558066, EPI_ISL_558067, EPI_ISL_558069, EPI_ISL_558070, EPI_ISL_558072, EPI_ISL_558074, EPI_ISL_558075, EPI_ISL_558076, EPI_ISL_558077, EPI_ISL_558078, EPI_ISL_558080, EPI_ISL_558081, EPI_ISL_558082, EPI_ISL_558083, EPI_ISL_558084, EPI_ISL_558085, EPI_ISL_558086, EPI_ISL_558087, EPI_ISL_558088, EPI_ISL_558090, EPI_ISL_558091, EPI_ISL_558092, EPI_ISL_558093, EPI_ISL_558094, EPI_ISL_558095, EPI_ISL_558097, EPI_ISL_558098, EPI_ISL_558099, EPI_ISL_558100, EPI_ISL_558101, EPI_ISL_558102, EPI_ISL_558104, EPI_ISL_558105, EPI_ISL_558108, EPI_ISL_558110, EPI_ISL_558112, EPI_ISL_558114, EPI_ISL_558115, EPI_ISL_558116, EPI_ISL_558117, EPI_ISL_558118, EPI_ISL_558119, EPI_ISL_558120, EPI_ISL_558121, EPI_ISL_558122, EPI_ISL_558123, EPI_ISL_558124, EPI_ISL_558125, EPI_ISL_558128, EPI_ISL_558129, EPI_ISL_558130, EPI_ISL_558131, EPI_ISL_558132, EPI_ISL_558133, EPI_ISL_558134, EPI_ISL_558136, EPI_ISL_558137, EPI_ISL_558138, EPI_ISL_558139, EPI_ISL_558140 | | | |
| see above | Lighthouse Lab in Alderley Park | Wellcome Sanger Institute for the COVID-19 Genomics UK (COG-UK) consortium | The Lighthouse Lab in Alderley Park and Alex Alderton, Roberto Amato, Sonia Goncalves, Ewan Harrison, David K. Jackson, Ian Johnston, Dominic Kwiatkowski, Cordelia Langford, John Sillitoe on behalf of the Wellcome Sanger Institute COVID-19 Surveillance Team |
| EPI_ISL_558141 | Lighthouse Lab in Alderley Park | Wellcome Sanger Institute for the COVID-19 Genomics UK (COG-UK) consortium | The Lighthouse Lab in Alderley Park and Alex Alderton, Roberto Amato, Sonia Goncalves, Ewan Harrison, David K. Jackson, Ian Johnston, Dominic Kwiatkowski, Cordelia Langford, John Sillitoe on behalf of the Wellcome Sanger Institute COVID-19 Surveillance Team (http://www.sanger.ac.uk/covid-team) |
| EPI_ISL_558142, EPI_ISL_558144, EPI_ISL_558145, EPI_ISL_558146, EPI_ISL_558147, EPI_ISL_558148, EPI_ISL_558150, EPI_ISL_558151, EPI_ISL_558152, EPI_ISL_558153, EPI_ISL_558154, EPI_ISL_558156, EPI_ISL_558159, EPI_ISL_558160, EPI_ISL_558162, EPI_ISL_558164, EPI_ISL_558166, EPI_ISL_558167, EPI_ISL_558168, EPI_ISL_558169, EPI_ISL_558170, EPI_ISL_558171, EPI_ISL_558172, EPI_ISL_558173, EPI_ISL_558176 | | | |
| see above | Lighthouse Lab in Alderley Park | Wellcome Sanger Institute for the COVID-19 Genomics UK (COG-UK) consortium | The Lighthouse Lab in Alderley Park and Alex Alderton, Roberto Amato, Sonia Goncalves, Ewan Harrison, David K. Jackson, Ian Johnston, Dominic Kwiatkowski, Cordelia Langford, John Sillitoe on behalf of the Wellcome Sanger Institute COVID-19 Surveillance Team |
| EPI_ISL_558177, EPI_ISL_558178 | Lighthouse Lab in Alderley Park | Wellcome Sanger Institute for the COVID-19 Genomics UK (COG-UK) consortium | The Lighthouse Lab in Alderley Park and Alex Alderton, Roberto Amato, Sonia Goncalves, Ewan Harrison, David K. Jackson, Ian Johnston, Dominic Kwiatkowski, Cordelia Langford, John Sillitoe on behalf of the Wellcome Sanger Institute COVID-19 Surveillance Team (http://www.sanger.ac.uk/covid-team) |
| EPI_ISL_558180, EPI_ISL_558181, EPI_ISL_558182, EPI_ISL_558183, EPI_ISL_558184, EPI_ISL_558186, EPI_ISL_558187 | Lighthouse Lab in Alderley Park | Wellcome Sanger Institute for the COVID-19 Genomics UK (COG-UK) consortium | The Lighthouse Lab in Alderley Park and Alex Alderton, Roberto Amato, Sonia Goncalves, Ewan Harrison, David K. Jackson, Ian Johnston, Dominic Kwiatkowski, Cordelia Langford, John Sillitoe on behalf of the Wellcome Sanger Institute COVID-19 Surveillance Team |
| EPI_ISL_558188 | Lighthouse Lab in Alderley Park | Wellcome Sanger Institute for the COVID-19 Genomics UK (COG-UK) consortium | The Lighthouse Lab in Alderley Park and Alex Alderton, Roberto Amato, Sonia Goncalves, Ewan Harrison, David K. Jackson, Ian Johnston, Dominic Kwiatkowski, Cordelia Langford, John Sillitoe on behalf of the Wellcome Sanger Institute COVID-19 Surveillance Team (http://www.sanger.ac.uk/covid-team) |
| EPI_ISL_558189, EPI_ISL_558190, EPI_ISL_558191, EPI_ISL_558192, EPI_ISL_558193, EPI_ISL_558194, EPI_ISL_558196, EPI_ISL_558197, EPI_ISL_558198, EPI_ISL_558199, EPI_ISL_558200, EPI_ISL_558201, EPI_ISL_558202, EPI_ISL_558203, EPI_ISL_558204, EPI_ISL_558205, EPI_ISL_558206, EPI_ISL_558207, EPI_ISL_558208, EPI_ISL_558209, EPI_ISL_558210, EPI_ISL_558211, EPI_ISL_558212, EPI_ISL_558213 | | | |
| see above | Lighthouse Lab in Alderley Park | Wellcome Sanger Institute for the COVID-19 Genomics UK (COG-UK) consortium | The Lighthouse Lab in Alderley Park and Alex Alderton, Roberto Amato, Sonia Goncalves, Ewan Harrison, David K. Jackson, Ian Johnston, Dominic Kwiatkowski, Cordelia Langford, John Sillitoe on behalf of the Wellcome Sanger Institute COVID-19 Surveillance Team |
| EPI_ISL_558215 | Lighthouse Lab in Alderley Park | Wellcome Sanger Institute for the COVID-19 Genomics UK (COG-UK) consortium | The Lighthouse Lab in Alderley Park and Alex Alderton, Roberto Amato, Sonia Goncalves, Ewan Harrison, David K. Jackson, Ian Johnston, Dominic Kwiatkowski, Cordelia Langford, John Sillitoe on behalf of the Wellcome Sanger Institute COVID-19 Surveillance Team (http://www.sanger.ac.uk/covid-team) |
| EPI_ISL_558216, EPI_ISL_558217 | Lighthouse Lab in Alderley Park | Wellcome Sanger Institute for the COVID-19 Genomics UK (COG-UK) consortium | The Lighthouse Lab in Alderley Park and Alex Alderton, Roberto Amato, Sonia Goncalves, Ewan Harrison, David K. Jackson, Ian Johnston, Dominic Kwiatkowski, Cordelia Langford, John Sillitoe on behalf of the Wellcome Sanger Institute COVID-19 Surveillance Team |
| EPI_ISL_558218 | Lighthouse Lab in Alderley Park | Wellcome Sanger Institute for the COVID-19 Genomics UK (COG-UK) consortium | The Lighthouse Lab in Alderley Park and Alex Alderton, Roberto Amato, Sonia Goncalves, Ewan Harrison, David K. Jackson, Ian Johnston, Dominic Kwiatkowski, Cordelia Langford, John Sillitoe on behalf of the Wellcome Sanger Institute COVID-19 Surveillance Team (http://www.sanger.ac.uk/covid-team) |
| EPI_ISL_558219, EPI_ISL_558220, EPI_ISL_558221, EPI_ISL_558223, EPI_ISL_558225, EPI_ISL_558226, EPI_ISL_558227, EPI_ISL_558228, EPI_ISL_558229, EPI_ISL_558230, EPI_ISL_558231, EPI_ISL_558232, EPI_ISL_558233, EPI_ISL_558235, EPI_ISL_558236, EPI_ISL_558404, EPI_ISL_558405, EPI_ISL_558406, EPI_ISL_558407 | | | |
| see above | Lighthouse Lab in Alderley Park | Wellcome Sanger Institute for the COVID-19 Genomics UK (COG-UK) consortium | The Lighthouse Lab in Alderley Park and Alex Alderton, Roberto Amato, Sonia Goncalves, Ewan Harrison, David K. Jackson, Ian Johnston, Dominic Kwiatkowski, Cordelia Langford, John Sillitoe on behalf of the Wellcome Sanger Institute COVID-19 Surveillance Team |
| EPI_ISL_558408 | Lighthouse Lab in Alderley Park | Wellcome Sanger Institute for the COVID-19 Genomics UK (COG-UK) consortium | The Lighthouse Lab in Alderley Park and Alex Alderton, Roberto Amato, Sonia Goncalves, Ewan Harrison, David K. Jackson, Ian Johnston, Dominic Kwiatkowski, Cordelia Langford, John Sillitoe on behalf of the Wellcome Sanger Institute COVID-19 Surveillance Team (http://www.sanger.ac.uk/covid-team) |
| EPI_ISL_558409, EPI_ISL_558410, EPI_ISL_558411, EPI_ISL_558412, EPI_ISL_558413, EPI_ISL_558414, EPI_ISL_558416, EPI_ISL_558417, EPI_ISL_558418, EPI_ISL_558419, EPI_ISL_558420, EPI_ISL_558421, EPI_ISL_558423, EPI_ISL_558424, EPI_ISL_558425, EPI_ISL_558426, EPI_ISL_558427 | | | |
| see above | Lighthouse Lab in Alderley Park | Wellcome Sanger Institute for the COVID-19 Genomics UK (COG-UK) consortium | The Lighthouse Lab in Alderley Park and Alex Alderton, Roberto Amato, Sonia Goncalves, Ewan Harrison, David K. Jackson, Ian Johnston, Dominic Kwiatkowski, Cordelia Langford, John Sillitoe on behalf of the Wellcome Sanger Institute COVID-19 Surveillance Team |
| EPI_ISL_558428 | Lighthouse Lab in Alderley Park | Wellcome Sanger Institute for the COVID-19 Genomics UK | The Lighthouse Lab in Alderley Park and Alex Alderton, Roberto Amato, Sonia Goncalves, Ewan Harrison, David K. Jackson, Ian Johnston, Dominic |

| | | | |
|---|---|---|---|
| | | (COG-UK) consortium | Kwiatkowski, Cordelia Langford, John Sillitoe on behalf of the Wellcome Sanger Institute COVID-19 Surveillance Team (http://www.sanger.ac.uk/covid-team) |
| EPI_ISL_558429, EPI_ISL_558430, EPI_ISL_558431, EPI_ISL_558433, EPI_ISL_558434, EPI_ISL_558435, EPI_ISL_558436, EPI_ISL_558437, EPI_ISL_558438, EPI_ISL_558442, EPI_ISL_558443, EPI_ISL_558445, EPI_ISL_558447, EPI_ISL_558448, EPI_ISL_558449, EPI_ISL_558450, EPI_ISL_558451, EPI_ISL_558452, EPI_ISL_558453, EPI_ISL_558454, EPI_ISL_558455 | | | |
| see above | Lighthouse Lab in Alderley Park | Wellcome Sanger Institute for the COVID-19 Genomics UK (COG-UK) consortium | The Lighthouse Lab in Alderley Park and Alex Alderton, Roberto Amato, Sonia Goncalves, Ewan Harrison, David K. Jackson, Ian Johnston, Dominic Kwiatkowski, Cordelia Langford, John Sillitoe on behalf of the Wellcome Sanger Institute COVID-19 Surveillance Team |
| EPI_ISL_558456 | Lighthouse Lab in Alderley Park | Wellcome Sanger Institute for the COVID-19 Genomics UK (COG-UK) consortium | The Lighthouse Lab in Alderley Park and Alex Alderton, Roberto Amato, Sonia Goncalves, Ewan Harrison, David K. Jackson, Ian Johnston, Dominic Kwiatkowski, Cordelia Langford, John Sillitoe on behalf of the Wellcome Sanger Institute COVID-19 Surveillance Team (http://www.sanger.ac.uk/covid-team) |
| EPI_ISL_558457, EPI_ISL_558459, EPI_ISL_558460, EPI_ISL_558461, EPI_ISL_558462, EPI_ISL_558463, EPI_ISL_558464, EPI_ISL_558465 | Lighthouse Lab in Alderley Park | Wellcome Sanger Institute for the COVID-19 Genomics UK (COG-UK) consortium | The Lighthouse Lab in Alderley Park and Alex Alderton, Roberto Amato, Sonia Goncalves, Ewan Harrison, David K. Jackson, Ian Johnston, Dominic Kwiatkowski, Cordelia Langford, John Sillitoe on behalf of the Wellcome Sanger Institute COVID-19 Surveillance Team |
| EPI_ISL_558466 | Lighthouse Lab in Alderley Park | Wellcome Sanger Institute for the COVID-19 Genomics UK (COG-UK) consortium | The Lighthouse Lab in Alderley Park and Alex Alderton, Roberto Amato, Sonia Goncalves, Ewan Harrison, David K. Jackson, Ian Johnston, Dominic Kwiatkowski, Cordelia Langford, John Sillitoe on behalf of the Wellcome Sanger Institute COVID-19 Surveillance Team (http://www.sanger.ac.uk/covid-team) |
| EPI_ISL_558467, EPI_ISL_558468, EPI_ISL_558469, EPI_ISL_558470, EPI_ISL_558472, EPI_ISL_558473, EPI_ISL_558475, EPI_ISL_558476, EPI_ISL_558477, EPI_ISL_558479, EPI_ISL_558480, EPI_ISL_558481, EPI_ISL_558482, EPI_ISL_558483, EPI_ISL_558484, EPI_ISL_558486, EPI_ISL_558487, EPI_ISL_558488, EPI_ISL_558491, EPI_ISL_558492, EPI_ISL_558493, EPI_ISL_558495, EPI_ISL_558496, EPI_ISL_558497, EPI_ISL_558498, EPI_ISL_558499, EPI_ISL_558500, EPI_ISL_558501, EPI_ISL_558502, EPI_ISL_558503, EPI_ISL_558504, EPI_ISL_558505, EPI_ISL_558506, EPI_ISL_558507, EPI_ISL_558508, EPI_ISL_558510, EPI_ISL_558511, EPI_ISL_558513, EPI_ISL_558514, EPI_ISL_558515, EPI_ISL_558516, EPI_ISL_558518, EPI_ISL_558519, EPI_ISL_558520, EPI_ISL_558521, EPI_ISL_558522, EPI_ISL_558523, EPI_ISL_558524, EPI_ISL_558525, EPI_ISL_558526, EPI_ISL_558527, EPI_ISL_558528, EPI_ISL_558531, EPI_ISL_558532, EPI_ISL_558533, EPI_ISL_558534, EPI_ISL_558535, EPI_ISL_558536, EPI_ISL_558537, EPI_ISL_558538, EPI_ISL_558539, EPI_ISL_558540, EPI_ISL_558541, EPI_ISL_558542, EPI_ISL_558544, EPI_ISL_558546, EPI_ISL_558547, EPI_ISL_558548, EPI_ISL_558549, EPI_ISL_558550, EPI_ISL_558551, EPI_ISL_558552, EPI_ISL_558555, EPI_ISL_558556, EPI_ISL_558557, EPI_ISL_558558, EPI_ISL_558559, EPI_ISL_558560, EPI_ISL_558562, EPI_ISL_558563, EPI_ISL_558564, EPI_ISL_558565, EPI_ISL_558566, EPI_ISL_558568, EPI_ISL_558569, EPI_ISL_558570, EPI_ISL_558571, EPI_ISL_558572, EPI_ISL_558575, EPI_ISL_558576, EPI_ISL_558578, EPI_ISL_558580, EPI_ISL_558581, EPI_ISL_558582, EPI_ISL_558583, EPI_ISL_558584, EPI_ISL_558590, EPI_ISL_558591, EPI_ISL_558593, EPI_ISL_558595, EPI_ISL_558596, EPI_ISL_558597, EPI_ISL_558598, EPI_ISL_558599, EPI_ISL_558600, EPI_ISL_558602, EPI_ISL_558603, EPI_ISL_558604, EPI_ISL_558606, EPI_ISL_558607, EPI_ISL_558610, EPI_ISL_558611, EPI_ISL_558612, EPI_ISL_558613, EPI_ISL_558614, EPI_ISL_558615, EPI_ISL_558616 | | | |
| see above | Lighthouse Lab in Alderley Park | Wellcome Sanger Institute for the COVID-19 Genomics UK (COG-UK) consortium | The Lighthouse Lab in Alderley Park and Alex Alderton, Roberto Amato, Sonia Goncalves, Ewan Harrison, David K. Jackson, Ian Johnston, Dominic Kwiatkowski, Cordelia Langford, John Sillitoe on behalf of the Wellcome Sanger Institute COVID-19 Surveillance Team |
| EPI_ISL_559753, EPI_ISL_559754, EPI_ISL_559755, EPI_ISL_559756, EPI_ISL_559757, EPI_ISL_559758, EPI_ISL_559759, EPI_ISL_559760, EPI_ISL_559761, EPI_ISL_559762, EPI_ISL_559763, EPI_ISL_559764, EPI_ISL_559765, EPI_ISL_559766, EPI_ISL_559767, EPI_ISL_559768, EPI_ISL_559769, EPI_ISL_559770, EPI_ISL_559771, EPI_ISL_559772, EPI_ISL_559773, EPI_ISL_559774, EPI_ISL_559775, EPI_ISL_559776, EPI_ISL_559777, EPI_ISL_559778, EPI_ISL_559779, EPI_ISL_559780, EPI_ISL_559781, EPI_ISL_559782, EPI_ISL_559783, EPI_ISL_559784, EPI_ISL_559785, EPI_ISL_559786, EPI_ISL_559787, EPI_ISL_559788, EPI_ISL_559789, EPI_ISL_559790, EPI_ISL_559791, EPI_ISL_559792, EPI_ISL_559793, EPI_ISL_559794, EPI_ISL_559795, EPI_ISL_559796, EPI_ISL_559797, EPI_ISL_559798, EPI_ISL_559799, EPI_ISL_559800, EPI_ISL_559801, EPI_ISL_559802, EPI_ISL_559803, EPI_ISL_559804, EPI_ISL_559805, EPI_ISL_559806, EPI_ISL_559807, EPI_ISL_559808, EPI_ISL_559809, EPI_ISL_559810, EPI_ISL_559811, EPI_ISL_559812, EPI_ISL_559813, EPI_ISL_559814, EPI_ISL_559815, EPI_ISL_559816, EPI_ISL_559817, EPI_ISL_559818, EPI_ISL_559819, EPI_ISL_559820, EPI_ISL_559821, EPI_ISL_559822, EPI_ISL_559823, EPI_ISL_559824, EPI_ISL_559825, EPI_ISL_559826, EPI_ISL_559827, EPI_ISL_559828, EPI_ISL_559829, EPI_ISL_559830, EPI_ISL_559831, EPI_ISL_559832, EPI_ISL_559833, EPI_ISL_559834, EPI_ISL_559835, EPI_ISL_559836, EPI_ISL_559837, EPI_ISL_559838, EPI_ISL_559839, EPI_ISL_559840, EPI_ISL_559841, EPI_ISL_559842, EPI_ISL_559843, EPI_ISL_559844, EPI_ISL_559845, EPI_ISL_559846, EPI_ISL_559847, EPI_ISL_559848, EPI_ISL_559849, EPI_ISL_559850, EPI_ISL_559851, EPI_ISL_559852, EPI_ISL_559853, EPI_ISL_559854, EPI_ISL_559855, EPI_ISL_559856, EPI_ISL_559857, EPI_ISL_559858, EPI_ISL_559859, EPI_ISL_559860, EPI_ISL_559861, EPI_ISL_559862, EPI_ISL_559863, EPI_ISL_559864, EPI_ISL_559865, EPI_ISL_559866, EPI_ISL_559867, EPI_ISL_559868, EPI_ISL_559869, EPI_ISL_559870, EPI_ISL_559871, EPI_ISL_559872, EPI_ISL_559873, EPI_ISL_559874, EPI_ISL_559875, EPI_ISL_559876, EPI_ISL_559877, EPI_ISL_559878, EPI_ISL_559879, EPI_ISL_559880, EPI_ISL_559881, EPI_ISL_559882, EPI_ISL_559883, EPI_ISL_559884, EPI_ISL_559885, EPI_ISL_559886, EPI_ISL_559887, EPI_ISL_559888, EPI_ISL_559890, EPI_ISL_559891, EPI_ISL_559892, EPI_ISL_559893, EPI_ISL_559894, EPI_ISL_559895, EPI_ISL_559896, EPI_ISL_559897, EPI_ISL_559984, EPI_ISL_559985, EPI_ISL_559986, EPI_ISL_559987, EPI_ISL_559988, EPI_ISL_559989, EPI_ISL_559990, EPI_ISL_559991, EPI_ISL_559992, EPI_ISL_559993, EPI_ISL_559994, EPI_ISL_559995, EPI_ISL_559996, EPI_ISL_559997, EPI_ISL_559998, EPI_ISL_559999, EPI_ISL_560000, EPI_ISL_560001, EPI_ISL_560002, EPI_ISL_560003, EPI_ISL_560004, EPI_ISL_560005, EPI_ISL_560006, EPI_ISL_560007, EPI_ISL_560008, EPI_ISL_560009, EPI_ISL_560010, EPI_ISL_560011, EPI_ISL_560012, EPI_ISL_560013, EPI_ISL_560014, EPI_ISL_560015, EPI_ISL_560016, EPI_ISL_560017, EPI_ISL_560018, EPI_ISL_560019, EPI_ISL_560020, EPI_ISL_560021, EPI_ISL_560022, EPI_ISL_560023, EPI_ISL_560024, EPI_ISL_560025, EPI_ISL_560026, EPI_ISL_560027, EPI_ISL_560028, EPI_ISL_560029, EPI_ISL_560030, EPI_ISL_560031, EPI_ISL_560032, EPI_ISL_560033, EPI_ISL_560034, EPI_ISL_560035, EPI_ISL_560036, EPI_ISL_560037, EPI_ISL_560038, EPI_ISL_560039, EPI_ISL_560040, EPI_ISL_560041, EPI_ISL_560042, EPI_ISL_560043, EPI_ISL_560044, EPI_ISL_560045, EPI_ISL_560046, EPI_ISL_560047, EPI_ISL_560048, EPI_ISL_560049, EPI_ISL_560050, EPI_ISL_560051, EPI_ISL_560052, EPI_ISL_560053, EPI_ISL_560069, EPI_ISL_560070, EPI_ISL_560071, EPI_ISL_560072, EPI_ISL_560073, EPI_ISL_560074, EPI_ISL_560075, EPI_ISL_560076, EPI_ISL_560077, EPI_ISL_560078, EPI_ISL_560079, EPI_ISL_560080, EPI_ISL_560081, EPI_ISL_560082, EPI_ISL_560305, EPI_ISL_560306 | | | |
| see above | Oxford Viromics, NDM, University of Oxford; Oxford University Hospitals; Basingstoke and North Hampshire Hospital | COVID-19 Genomics UK (COG-UK) Consortium | Tanya Golubchik, David Bonsall, George Macintyre, Amy Trebes, Mariateresa de Cesare, Catrin Moore, Alex Mobbs, Anita Justice, Robert Shaw, Monique Andersson, Timothy Peto, Emma Wise, Nathan Moore, Jessica Lynch, Nick Cortes, Matilde Mori, Stephen Kidd, David Buck, John Todd, Christophe Fraser |
| EPI_ISL_560568, EPI_ISL_560569, EPI_ISL_560570, EPI_ISL_560571, EPI_ISL_560572, EPI_ISL_560573, EPI_ISL_560574, EPI_ISL_560575, EPI_ISL_560576, EPI_ISL_560577, EPI_ISL_560578, EPI_ISL_560579, EPI_ISL_560580 | | | |
| see above | hôpital | National Reference Center for Viruses of Respiratory Infections, Institut Pasteur, Paris | Sylvie Behillil, Fabiana Gambaro, Etienne Simon-Lorière, Vincent Enouf, Maud Vanpeene, Sylvie van der Werf |
| EPI_ISL_560581, EPI_ISL_560582, EPI_ISL_560583, EPI_ISL_560584, EPI_ISL_560585, EPI_ISL_560586, EPI_ISL_560587, EPI_ISL_560588, EPI_ISL_560589, EPI_ISL_560590 | Hopital | National Reference Center for Viruses of Respiratory Infections, Institut Pasteur, Paris | Sylvie Behillil, Fabiana Gambaro, Etienne Simon-Lorière, Vincent Enouf, Maud Vanpeene, Sylvie van der Werf |
| EPI_ISL_560591, EPI_ISL_560592, EPI_ISL_560593, EPI_ISL_560594, EPI_ISL_560595, EPI_ISL_560596, EPI_ISL_560597 | hopital | National Reference Center for Viruses of Respiratory Infections, Institut Pasteur, Paris | Sylvie Behillil, Fabiana Gambaro, Etienne Simon-Lorière, Vincent Enouf, Maud Vanpeene, Sylvie van der Werf |
| EPI_ISL_560598, EPI_ISL_560599, EPI_ISL_560600, EPI_ISL_560601, EPI_ISL_560602, EPI_ISL_560603, EPI_ISL_560604, EPI_ISL_560605, EPI_ISL_560606, EPI_ISL_560607, EPI_ISL_560608, EPI_ISL_560609, EPI_ISL_560610, EPI_ISL_560611, EPI_ISL_560612, EPI_ISL_560613, EPI_ISL_560614, EPI_ISL_560615, EPI_ISL_560616, EPI_ISL_560617, EPI_ISL_560618, EPI_ISL_560619, EPI_ISL_560620, EPI_ISL_560621, EPI_ISL_560622, EPI_ISL_560623, EPI_ISL_560624, EPI_ISL_560625, EPI_ISL_560626, EPI_ISL_560627, EPI_ISL_560628, EPI_ISL_560629, EPI_ISL_560630, EPI_ISL_560631, EPI_ISL_560632, EPI_ISL_560633, EPI_ISL_560634, EPI_ISL_560635, EPI_ISL_560636 | | | |
| see above | Hospital | National Reference Center for Viruses of Respiratory Infections, Institut Pasteur, Paris | Sylvie Behillil, Fabiana Gambaro, Etienne Simon-Lorière, Vincent Enouf, Maud Vanpeene, Sylvie van der Werf |
| EPI_ISL_560637, EPI_ISL_560638, EPI_ISL_560639, EPI_ISL_560640, EPI_ISL_560641, EPI_ISL_560642 | Labo Analyses Med | National Reference Center for Viruses of Respiratory Infections, Institut Pasteur, Paris | Sylvie Behillil, Fabiana Gambaro, Etienne Simon-Lorière, Vincent Enouf, Maud Vanpeene, Sylvie van der Werf |
| EPI_ISL_560643, EPI_ISL_560644, EPI_ISL_560645, EPI_ISL_560646 | Hospital | National Reference Center for Viruses of Respiratory Infections, Institut Pasteur, Paris | Sylvie Behillil, Fabiana Gambaro, Etienne Simon-Lorière, Vincent Enouf, Maud Vanpeene, Sylvie van der Werf |
| EPI_ISL_569664, EPI_ISL_569665, EPI_ISL_569666, EPI_ISL_569667, EPI_ISL_569668, EPI_ISL_569669, EPI_ISL_569670, EPI_ISL_569671, EPI_ISL_569672, EPI_ISL_569673, EPI_ISL_569674, EPI_ISL_569675, EPI_ISL_569676, EPI_ISL_569677, EPI_ISL_569678, EPI_ISL_569679, EPI_ISL_569680, EPI_ISL_569681, EPI_ISL_569682, EPI_ISL_569683, EPI_ISL_569684, EPI_ISL_569685, EPI_ISL_569686, EPI_ISL_569687, EPI_ISL_569688 | | | |
| see above | Lee Lab | Lee Lab | Sung Yong Park, Gina Faraci, Pamela M. Ward, Jane F. Emerson, and Ha Youn Lee |
| EPI_ISL_569859, EPI_ISL_569860, EPI_ISL_569861, EPI_ISL_569862, EPI_ISL_569863 | CSIR-Indian Institute of Chemical Biology, MEDICA Supercpecialty Hospital Kolkata | CSIR-Indian Institute of Chemical Biology, MEDICA Supercpecialty Hospital Kolkata | Sujay Krishna Maity, Priyanka Mallick, Debaleena Bhowmik, Abhishake Lahiri, Dr. Aviral Roy, Dr. Soumen Saha, Dr. Arpita Ghosh Mitra, Dr. Rajesh Pandey, Dr. Sandip Paul, Dr. Partha Chakrabarti, Dr. Saikat Chakrabarti |
| EPI_ISL_569887, EPI_ISL_569888, EPI_ISL_569889, EPI_ISL_569890, EPI_ISL_569891, EPI_ISL_569892, EPI_ISL_569893, EPI_ISL_569894, EPI_ISL_569895, EPI_ISL_569896, EPI_ISL_569897, EPI_ISL_569898, EPI_ISL_569899, EPI_ISL_569900, EPI_ISL_569901, EPI_ISL_569902, EPI_ISL_569903, EPI_ISL_569904, EPI_ISL_569905, EPI_ISL_569906, EPI_ISL_569907, EPI_ISL_569908, EPI_ISL_569909, EPI_ISL_569910, EPI_ISL_569911, EPI_ISL_569912, EPI_ISL_569913, EPI_ISL_569914, EPI_ISL_569915, EPI_ISL_569916, EPI_ISL_569917, EPI_ISL_569918, EPI_ISL_569919, EPI_ISL_569920, EPI_ISL_569921, EPI_ISL_569922, EPI_ISL_569923, EPI_ISL_569924, EPI_ISL_569925, EPI_ISL_569926, EPI_ISL_569927, EPI_ISL_569928, EPI_ISL_569929, EPI_ISL_569930, EPI_ISL_569931, EPI_ISL_569932, EPI_ISL_569933, EPI_ISL_569934, EPI_ISL_569935, EPI_ISL_569936, EPI_ISL_569937, EPI_ISL_569938, EPI_ISL_569939, EPI_ISL_569940, EPI_ISL_569941, EPI_ISL_569942, EPI_ISL_569943, EPI_ISL_569944, EPI_ISL_569945, EPI_ISL_569946, EPI_ISL_569947, EPI_ISL_569948 | | | |
| see above | Innovative Genomics Institute, UC Berkeley | Innovative Genomics Institute, UC Berkeley | Stacia Wyman, Haridha Shivram, Phil Frankino, Liana Lareau, Shana McDevitt, Justin Choi |
| EPI_ISL_572714, EPI_ISL_572723, EPI_ISL_572724, EPI_ISL_572726, EPI_ISL_572727, EPI_ISL_572728, EPI_ISL_572846, EPI_ISL_572849, EPI_ISL_572850, EPI_ISL_572857, EPI_ISL_572858, EPI_ISL_572859, EPI_ISL_572860, EPI_ISL_572861, EPI_ISL_572869, EPI_ISL_572870, EPI_ISL_572873, EPI_ISL_572891, EPI_ISL_572892, EPI_ISL_572893, EPI_ISL_572894, EPI_ISL_572895, EPI_ISL_572896, EPI_ISL_572897, EPI_ISL_572902, EPI_ISL_572903, EPI_ISL_572917, EPI_ISL_572918, EPI_ISL_572919, EPI_ISL_572920, EPI_ISL_572921, EPI_ISL_572922, EPI_ISL_572923, EPI_ISL_572924, EPI_ISL_572925, EPI_ISL_572926, EPI_ISL_572933, EPI_ISL_572934, EPI_ISL_572935, EPI_ISL_572936, EPI_ISL_572937, EPI_ISL_572947, EPI_ISL_572952, EPI_ISL_572953, EPI_ISL_572958, EPI_ISL_572960, EPI_ISL_572962, EPI_ISL_572963, EPI_ISL_572964, EPI_ISL_572965, EPI_ISL_572966, EPI_ISL_572967, EPI_ISL_572968, EPI_ISL_572969, | | | |

| | | | |
|---|---|---|---|
| EPI_ISL_572970, EPI_ISL_572971, EPI_ISL_572972, EPI_ISL_572973, EPI_ISL_572974, EPI_ISL_572975, EPI_ISL_572976, EPI_ISL_572977, EPI_ISL_572978, EPI_ISL_572979, EPI_ISL_572980, EPI_ISL_572981, EPI_ISL_572982, EPI_ISL_572983, EPI_ISL_572984, EPI_ISL_572985, EPI_ISL_572986, EPI_ISL_572987, EPI_ISL_572988, EPI_ISL_572989, EPI_ISL_572990, EPI_ISL_572991, EPI_ISL_572992, EPI_ISL_572993, EPI_ISL_572994, EPI_ISL_572995, EPI_ISL_572996, EPI_ISL_572997, EPI_ISL_572998, EPI_ISL_572999, EPI_ISL_573000, EPI_ISL_573001, EPI_ISL_573002, EPI_ISL_573003, EPI_ISL_573004, EPI_ISL_573005, EPI_ISL_573006, EPI_ISL_573007, EPI_ISL_573008, EPI_ISL_573009, EPI_ISL_573010, EPI_ISL_573011, EPI_ISL_573012, EPI_ISL_573013, EPI_ISL_573014, EPI_ISL_573015, EPI_ISL_573016, EPI_ISL_573017, EPI_ISL_573018, EPI_ISL_573019, EPI_ISL_573020, EPI_ISL_573021, EPI_ISL_573022, EPI_ISL_573023, EPI_ISL_573024, EPI_ISL_573025, EPI_ISL_573026, EPI_ISL_573027, EPI_ISL_573028, EPI_ISL_573029, EPI_ISL_573030, EPI_ISL_573031, EPI_ISL_573032, EPI_ISL_573033, EPI_ISL_573034, EPI_ISL_573035, EPI_ISL_573036, EPI_ISL_573037, EPI_ISL_573038, EPI_ISL_573039, EPI_ISL_573040, EPI_ISL_573041, EPI_ISL_573042, EPI_ISL_573043, EPI_ISL_573044, EPI_ISL_573045, EPI_ISL_573046, EPI_ISL_573047, EPI_ISL_573048, EPI_ISL_573049, EPI_ISL_573050, EPI_ISL_573051, EPI_ISL_573052, EPI_ISL_573053, EPI_ISL_573054, EPI_ISL_573055, EPI_ISL_573056, EPI_ISL_573057, EPI_ISL_573058, EPI_ISL_573059, EPI_ISL_573060, EPI_ISL_573061, EPI_ISL_573062, EPI_ISL_573063, EPI_ISL_573064, EPI_ISL_573065, EPI_ISL_573066, EPI_ISL_573067, EPI_ISL_573068, EPI_ISL_573069, EPI_ISL_573070, EPI_ISL_573071, EPI_ISL_573072, EPI_ISL_573073, EPI_ISL_573074, EPI_ISL_573075, EPI_ISL_573076, EPI_ISL_573077, EPI_ISL_573078, EPI_ISL_573079, EPI_ISL_573080, EPI_ISL_573081, EPI_ISL_573082, EPI_ISL_573083, EPI_ISL_573084, EPI_ISL_573154, EPI_ISL_573155, EPI_ISL_573156, EPI_ISL_573157, EPI_ISL_573158, EPI_ISL_573159, EPI_ISL_573160, EPI_ISL_573161, EPI_ISL_573162, EPI_ISL_573163, EPI_ISL_573164, EPI_ISL_573165, EPI_ISL_573166, EPI_ISL_573167, EPI_ISL_573168, EPI_ISL_573169, EPI_ISL_573170, EPI_ISL_573171, EPI_ISL_573172, EPI_ISL_573173, EPI_ISL_573174, EPI_ISL_573175, EPI_ISL_573176, EPI_ISL_573177, EPI_ISL_573178, EPI_ISL_573179, EPI_ISL_573180, EPI_ISL_573181, EPI_ISL_573182, EPI_ISL_573183, EPI_ISL_573184, EPI_ISL_573185, EPI_ISL_573186, EPI_ISL_573187, EPI_ISL_573188, EPI_ISL_573189, EPI_ISL_573190, EPI_ISL_573191, EPI_ISL_573192, EPI_ISL_573193, EPI_ISL_573194, EPI_ISL_573195, EPI_ISL_573196, EPI_ISL_573197, EPI_ISL_573198, EPI_ISL_573199, EPI_ISL_573200, EPI_ISL_573201, EPI_ISL_573202, EPI_ISL_573203, EPI_ISL_573204, EPI_ISL_573205, EPI_ISL_573206, EPI_ISL_573207, EPI_ISL_573208, EPI_ISL_573209, EPI_ISL_573210, EPI_ISL_573211, EPI_ISL_573212, EPI_ISL_573213, EPI_ISL_573214, EPI_ISL_573215, EPI_ISL_573216, EPI_ISL_573217, EPI_ISL_573218, EPI_ISL_573219, EPI_ISL_573220, EPI_ISL_573221, EPI_ISL_573222, EPI_ISL_573223, EPI_ISL_573224, EPI_ISL_573225, EPI_ISL_573226, EPI_ISL_573227, EPI_ISL_573228, EPI_ISL_573229, EPI_ISL_573230, EPI_ISL_573231, EPI_ISL_573232, EPI_ISL_573233, EPI_ISL_573234, EPI_ISL_573235, EPI_ISL_573566, EPI_ISL_573567, EPI_ISL_573568, EPI_ISL_573569, EPI_ISL_573570, EPI_ISL_573571, EPI_ISL_573572, EPI_ISL_573573, EPI_ISL_573574, EPI_ISL_573575, EPI_ISL_573576, EPI_ISL_573577, EPI_ISL_573578, EPI_ISL_573579, EPI_ISL_573580, EPI_ISL_573581, EPI_ISL_573582, EPI_ISL_573583, EPI_ISL_573584, EPI_ISL_573585, EPI_ISL_573586, EPI_ISL_573587, EPI_ISL_573588, EPI_ISL_573589, EPI_ISL_573590, EPI_ISL_573591, EPI_ISL_573592, EPI_ISL_573593, EPI_ISL_573594, EPI_ISL_573595, EPI_ISL_573596, EPI_ISL_573597, EPI_ISL_573598, EPI_ISL_573599, EPI_ISL_573600, EPI_ISL_573601, EPI_ISL_573602, EPI_ISL_573603, EPI_ISL_573604, EPI_ISL_573605, EPI_ISL_573606, EPI_ISL_573607, EPI_ISL_573608, EPI_ISL_573609, EPI_ISL_573610, EPI_ISL_573611, EPI_ISL_573612, EPI_ISL_573613, EPI_ISL_573614, EPI_ISL_573615, EPI_ISL_573616, EPI_ISL_573617, EPI_ISL_573618, EPI_ISL_573619, EPI_ISL_573620, EPI_ISL_573621, EPI_ISL_573622, EPI_ISL_573623, EPI_ISL_573624, EPI_ISL_573625, EPI_ISL_573626, EPI_ISL_573627, EPI_ISL_573628, EPI_ISL_573629, EPI_ISL_573630, EPI_ISL_573631, EPI_ISL_573632, EPI_ISL_573633, EPI_ISL_573634, EPI_ISL_573635, EPI_ISL_573636, EPI_ISL_573637, EPI_ISL_573638, EPI_ISL_573639, EPI_ISL_573640, EPI_ISL_573641, EPI_ISL_573642, EPI_ISL_573643, EPI_ISL_573644, EPI_ISL_573645, EPI_ISL_573646, EPI_ISL_573647, EPI_ISL_573648, EPI_ISL_573649, EPI_ISL_573650, EPI_ISL_573651, EPI_ISL_573759, EPI_ISL_573811, EPI_ISL_573812, EPI_ISL_573813, EPI_ISL_573814, EPI_ISL_573815, EPI_ISL_573816, EPI_ISL_573817, EPI_ISL_573818, EPI_ISL_573819, EPI_ISL_573820, EPI_ISL_573821, EPI_ISL_573822, EPI_ISL_573823, EPI_ISL_573824, EPI_ISL_573825, EPI_ISL_573826, EPI_ISL_573827, EPI_ISL_573828, EPI_ISL_573829, EPI_ISL_573830, EPI_ISL_573831, EPI_ISL_573832, EPI_ISL_573833, EPI_ISL_573834, EPI_ISL_573835, EPI_ISL_573836, EPI_ISL_573837, EPI_ISL_573838, EPI_ISL_573839, EPI_ISL_573840, EPI_ISL_573841, EPI_ISL_573842, EPI_ISL_573843, EPI_ISL_573844, EPI_ISL_573845, EPI_ISL_573846, EPI_ISL_573847, EPI_ISL_573848, EPI_ISL_573849, EPI_ISL_573850, EPI_ISL_573851, EPI_ISL_573852, EPI_ISL_573853, EPI_ISL_573854, EPI_ISL_573855, EPI_ISL_573866, EPI_ISL_573867, EPI_ISL_573868, EPI_ISL_573869, EPI_ISL_574257 | | | |
| see above | Oxford Viromics, NDM, University of Oxford; Oxford University Hospitals; Basingstoke and North Hampshire Hospital | COVID-19 Genomics UK (COG-UK) Consortium | Tanya Golubchik, David Bonsall, George Macintyre, Amy Trebes, Mariateresa de Cesare, Catrin Moore, Alex Mobbs, Anita Justice, Robert Shaw, Monique Andersson, Timothy Peto, Emma Wise, Nathan Moore, Jessica Lynch, Nick Cortes, Matilde Mori, Stephen Kidd, David Buck, John Todd, Christophe Fraser |
| EPI_ISL_576131, EPI_ISL_576132, EPI_ISL_576133, EPI_ISL_576134, EPI_ISL_576135, EPI_ISL_576136, EPI_ISL_576137, EPI_ISL_576138, EPI_ISL_576139, EPI_ISL_576140, EPI_ISL_576141, EPI_ISL_576142, EPI_ISL_576143, EPI_ISL_576144 | | | |
| see above | Public Health Virology Laboratory, Forensic and Scientific Services (PHV-FSS) | Public Health Virology Laboratory, Forensic and Scientific Services (PHV-FSS) | Son Nguyen et al |
| EPI_ISL_576518, EPI_ISL_576519, EPI_ISL_576520, EPI_ISL_576521, EPI_ISL_576522, EPI_ISL_576523, EPI_ISL_576524, EPI_ISL_576525, EPI_ISL_576526, EPI_ISL_576527, EPI_ISL_576528, EPI_ISL_576529, EPI_ISL_576530, EPI_ISL_576531, EPI_ISL_576532, EPI_ISL_576533, EPI_ISL_576534, EPI_ISL_576535, EPI_ISL_576536, EPI_ISL_576537, EPI_ISL_576538, EPI_ISL_576539, EPI_ISL_576540, EPI_ISL_576541, EPI_ISL_576542, EPI_ISL_576543, EPI_ISL_576544, EPI_ISL_576545, EPI_ISL_576546, EPI_ISL_576547, EPI_ISL_576548, EPI_ISL_576549, EPI_ISL_576550, EPI_ISL_576551, EPI_ISL_576552, EPI_ISL_576553, EPI_ISL_576554, EPI_ISL_576555, EPI_ISL_576556 | | | |
| see above | Innovative Genomics Institute, UC Berkeley | Innovative Genomics Institute, UC Berkeley | Stacia Wyman, Haridha Shivram, Phil Frankino, Liana Lareau, Shana McDevitt, Justin Choi |
| EPI_ISL_576569, EPI_ISL_576570, EPI_ISL_576571, EPI_ISL_576572, EPI_ISL_576573, EPI_ISL_576574, EPI_ISL_576575, EPI_ISL_576576, EPI_ISL_576577, EPI_ISL_576578, EPI_ISL_576579, EPI_ISL_576580, EPI_ISL_576581, EPI_ISL_576582, EPI_ISL_576583, EPI_ISL_576584, EPI_ISL_576585, EPI_ISL_576586, EPI_ISL_576587, EPI_ISL_576588, EPI_ISL_576589, EPI_ISL_576590, EPI_ISL_576591, EPI_ISL_576592, EPI_ISL_576593, EPI_ISL_576594, EPI_ISL_576595, EPI_ISL_576596, EPI_ISL_576597, EPI_ISL_576598, EPI_ISL_576599, EPI_ISL_576600, EPI_ISL_576601, EPI_ISL_576602, EPI_ISL_576603, EPI_ISL_576604, EPI_ISL_576605, EPI_ISL_576606, EPI_ISL_576607, EPI_ISL_576608, EPI_ISL_576609, EPI_ISL_576610, EPI_ISL_576611, EPI_ISL_576612, EPI_ISL_576613, EPI_ISL_576614, EPI_ISL_576615, EPI_ISL_576616, EPI_ISL_576617, EPI_ISL_576618, EPI_ISL_576619, EPI_ISL_576620, EPI_ISL_576621, EPI_ISL_576622, EPI_ISL_576623, EPI_ISL_576624, EPI_ISL_576625, EPI_ISL_576626, EPI_ISL_576627, EPI_ISL_576628, EPI_ISL_576629, EPI_ISL_576630, EPI_ISL_576631, EPI_ISL_576632, EPI_ISL_576633, EPI_ISL_576634, EPI_ISL_576635, EPI_ISL_576636, EPI_ISL_576637, EPI_ISL_576638, EPI_ISL_576639, EPI_ISL_576640, EPI_ISL_576641, EPI_ISL_576642, EPI_ISL_576643, EPI_ISL_576644, EPI_ISL_576645, EPI_ISL_576646, EPI_ISL_576647, EPI_ISL_576648, EPI_ISL_576649, EPI_ISL_576650, EPI_ISL_576651, EPI_ISL_576652, EPI_ISL_576653, EPI_ISL_576654, EPI_ISL_576655, EPI_ISL_576656, EPI_ISL_576657, EPI_ISL_576658, EPI_ISL_576659, EPI_ISL_576660, EPI_ISL_576661, EPI_ISL_576662, EPI_ISL_576663, EPI_ISL_576664, EPI_ISL_576665, EPI_ISL_576666, EPI_ISL_576667, EPI_ISL_576668, EPI_ISL_576669, EPI_ISL_576670, EPI_ISL_576671, EPI_ISL_576672, EPI_ISL_576673, EPI_ISL_576674, EPI_ISL_576675, EPI_ISL_576676, EPI_ISL_576677, EPI_ISL_576678, EPI_ISL_576679, EPI_ISL_576680, EPI_ISL_576681, EPI_ISL_576682, EPI_ISL_576683, EPI_ISL_576684, EPI_ISL_576685, EPI_ISL_576686, EPI_ISL_576687, EPI_ISL_576688, EPI_ISL_576689, EPI_ISL_576690, EPI_ISL_576691, EPI_ISL_576692, EPI_ISL_576693, EPI_ISL_576694, EPI_ISL_576695, EPI_ISL_576696, EPI_ISL_576697, EPI_ISL_576698, EPI_ISL_576699, EPI_ISL_576700, EPI_ISL_576701, EPI_ISL_576702, EPI_ISL_576703, EPI_ISL_576704, EPI_ISL_576705, EPI_ISL_576706, EPI_ISL_576707, EPI_ISL_576708, EPI_ISL_576709, EPI_ISL_576710, EPI_ISL_576711, EPI_ISL_576712, EPI_ISL_576713, EPI_ISL_576714, EPI_ISL_576715, EPI_ISL_576716, EPI_ISL_576717, EPI_ISL_576718, EPI_ISL_576719, EPI_ISL_576720, EPI_ISL_576721, EPI_ISL_576722, EPI_ISL_576723, EPI_ISL_576724, EPI_ISL_576725, EPI_ISL_576726, EPI_ISL_576727, EPI_ISL_576728, EPI_ISL_576729, EPI_ISL_576730, EPI_ISL_576731, EPI_ISL_576732, EPI_ISL_576733, EPI_ISL_576734, EPI_ISL_576735, EPI_ISL_576736, EPI_ISL_576737, EPI_ISL_576738, EPI_ISL_576739, EPI_ISL_576740, EPI_ISL_576741, EPI_ISL_576742, EPI_ISL_576743, EPI_ISL_576744, EPI_ISL_576745, EPI_ISL_576746, EPI_ISL_576747, EPI_ISL_576748, EPI_ISL_576749, EPI_ISL_576750, EPI_ISL_576751, EPI_ISL_576752, EPI_ISL_576753, EPI_ISL_576754, EPI_ISL_576755, EPI_ISL_576756, EPI_ISL_576757, EPI_ISL_576758, EPI_ISL_576759, EPI_ISL_576760, EPI_ISL_576761, EPI_ISL_576762, EPI_ISL_576763, EPI_ISL_576764, EPI_ISL_576765, EPI_ISL_576766, EPI_ISL_576767, EPI_ISL_576768, EPI_ISL_576769, EPI_ISL_576770, EPI_ISL_576771, EPI_ISL_576772, EPI_ISL_576773, EPI_ISL_576774, EPI_ISL_576775, EPI_ISL_576776, EPI_ISL_576777, EPI_ISL_576778, EPI_ISL_576779, EPI_ISL_576780, EPI_ISL_576781, EPI_ISL_576782, EPI_ISL_576783, EPI_ISL_576784, EPI_ISL_576785, EPI_ISL_576786, EPI_ISL_576787, EPI_ISL_576788, EPI_ISL_576789, EPI_ISL_576790, EPI_ISL_576791, EPI_ISL_576792, EPI_ISL_576793, EPI_ISL_576794, EPI_ISL_576795, EPI_ISL_576796, EPI_ISL_576797, EPI_ISL_576798, EPI_ISL_576799, EPI_ISL_576800, EPI_ISL_576801, EPI_ISL_576802, EPI_ISL_576803, EPI_ISL_576804, EPI_ISL_576805, EPI_ISL_576806, EPI_ISL_576807, EPI_ISL_576808, EPI_ISL_576809, EPI_ISL_576810, EPI_ISL_576811, EPI_ISL_576812, EPI_ISL_576813, EPI_ISL_576814, EPI_ISL_576815, EPI_ISL_576816, EPI_ISL_576817, EPI_ISL_576818, EPI_ISL_576819, EPI_ISL_576820, EPI_ISL_576821, EPI_ISL_576822, EPI_ISL_576823, EPI_ISL_576824, EPI_ISL_576825, EPI_ISL_576826, EPI_ISL_576827, EPI_ISL_576828, EPI_ISL_576829, EPI_ISL_576830, EPI_ISL_576831, EPI_ISL_576832, EPI_ISL_576833, EPI_ISL_576834, EPI_ISL_576835, EPI_ISL_576836, EPI_ISL_576837, EPI_ISL_576838, EPI_ISL_576839, EPI_ISL_576840, EPI_ISL_576841, EPI_ISL_576842, EPI_ISL_576843, EPI_ISL_576844, EPI_ISL_576845, EPI_ISL_576846, EPI_ISL_576847, EPI_ISL_576848, EPI_ISL_576849, EPI_ISL_576850, EPI_ISL_576851, EPI_ISL_576852, EPI_ISL_576853, EPI_ISL_576854, EPI_ISL_576855, EPI_ISL_576856, EPI_ISL_576857, EPI_ISL_576858, EPI_ISL_576859, EPI_ISL_576860, EPI_ISL_576861, EPI_ISL_576862, EPI_ISL_576863, EPI_ISL_576864, EPI_ISL_576865, EPI_ISL_576866, EPI_ISL_576867, EPI_ISL_576868, EPI_ISL_576869, EPI_ISL_576870, EPI_ISL_576871, EPI_ISL_576872, EPI_ISL_576873, EPI_ISL_576874, EPI_ISL_576875, EPI_ISL_576876, EPI_ISL_576920, EPI_ISL_576921, EPI_ISL_576922, EPI_ISL_576923, EPI_ISL_576924, EPI_ISL_576925, EPI_ISL_576926, EPI_ISL_576927, EPI_ISL_576928, EPI_ISL_576929, EPI_ISL_576930, EPI_ISL_576931, EPI_ISL_576932, EPI_ISL_576933, EPI_ISL_576934, EPI_ISL_576935, EPI_ISL_576936, EPI_ISL_576937, EPI_ISL_576938, EPI_ISL_576939, EPI_ISL_576940, EPI_ISL_576941, EPI_ISL_576942, EPI_ISL_576943, EPI_ISL_576944, EPI_ISL_576945, EPI_ISL_576946, EPI_ISL_576947, EPI_ISL_576948, EPI_ISL_576949, EPI_ISL_576950, EPI_ISL_576951, EPI_ISL_576952, EPI_ISL_576953, EPI_ISL_576954, EPI_ISL_576955, EPI_ISL_576956, EPI_ISL_576957, EPI_ISL_576958, EPI_ISL_576959, EPI_ISL_576960, EPI_ISL_577068, EPI_ISL_577069, EPI_ISL_577070, EPI_ISL_577071, EPI_ISL_577072, EPI_ISL_577073, EPI_ISL_577074, EPI_ISL_577075, EPI_ISL_577076, EPI_ISL_577077, EPI_ISL_577078, EPI_ISL_577079, EPI_ISL_577080, EPI_ISL_577081, EPI_ISL_577082, EPI_ISL_577083, EPI_ISL_577084, EPI_ISL_577085, EPI_ISL_577086, EPI_ISL_577087, EPI_ISL_577088, EPI_ISL_577089, EPI_ISL_577090, EPI_ISL_577091, EPI_ISL_577092, EPI_ISL_577093, EPI_ISL_577094, EPI_ISL_577095, EPI_ISL_577383, EPI_ISL_577384, EPI_ISL_577401, EPI_ISL_577402, EPI_ISL_577403, EPI_ISL_577404, EPI_ISL_577405, EPI_ISL_577406, EPI_ISL_577407, EPI_ISL_577408, EPI_ISL_577409, EPI_ISL_577410, EPI_ISL_577411, EPI_ISL_577412, EPI_ISL_577413, EPI_ISL_577414, EPI_ISL_577415, EPI_ISL_577416 | | | |
| see above | Oxford Viromics, NDM, University of Oxford; Oxford University Hospitals; Basingstoke and North Hampshire Hospital | COVID-19 Genomics UK (COG-UK) Consortium | Tanya Golubchik, David Bonsall, George Macintyre, Amy Trebes, Mariateresa de Cesare, Catrin Moore, Alex Mobbs, Anita Justice, Robert Shaw, Monique Andersson, Timothy Peto, Emma Wise, Nathan Moore, Jessica Lynch, Nick Cortes, Matilde Mori, Stephen Kidd, David Buck, John Todd, Christophe Fraser |
| EPI_ISL_578080 | CSIR-Indian Institute of Chemical Biology, MEDICA Supercpecialty Hospital Kolkata | CSIR-Indian Institute of Chemical Biology, MEDICA Supercpecialty Hospital Kolkata | Sujay Krishna Maity, Priyanka Mallick, Debaleena Bhowmik, Abhishake Lahiri, Dr. AviralRoy, Dr. Soumen Saha, Dr. Arpita Ghosh Mitra, Dr. Rajesh Pandey, Dr. Sandip Paul, Dr.Partha Chakrabarti, Dr. Saikat Chakrabarti |
| EPI_ISL_578081, EPI_ISL_578168, EPI_ISL_578175, EPI_ISL_578178, EPI_ISL_578179, EPI_ISL_578180, EPI_ISL_578181, EPI_ISL_578182, EPI_ISL_578183, EPI_ISL_578184, EPI_ISL_581449, EPI_ISL_581450, EPI_ISL_581451, EPI_ISL_581494, EPI_ISL_581495, EPI_ISL_581496, EPI_ISL_581497, EPI_ISL_581499, EPI_ISL_581500, EPI_ISL_581501, EPI_ISL_581502 | | | |
| see above | CSIR-Indian Institute of Chemical Biology, MEDICA Supercpecialty Hospital Kolkata | CSIR-Indian Institute of Chemical Biology, MEDICA Supercpecialty Hospital Kolkata | Sujay Krishna Maity, Priyanka Mallick, Debaleena Bhowmik, Abhishake Lahiri, Dr. Aviral Roy, Dr. Soumen Saha, Dr. Arpita Ghosh Mitra, Dr. Rajesh Pandey, Dr. Sandip Paul, Dr. Partha Chakrabarti, Dr. Saikat Chakrabarti |
| EPI_ISL_581503, EPI_ISL_581504 | CSIR-Indian Institute of Chemical Biology, MEDICA Supercpecialty Hospital Kolkata | CSIR-Indian Institute of Chemical Biology, MEDICA Supercpecialty Hospital Kolkata | Sujay Krishna Maity, Priyanka Mallick, Debaleena Bhowmik, Abhishake Lahiri, Dr. AviralRoy, Dr. Soumen Saha, Dr. Arpita Ghosh Mitra, Dr. Rajesh Pandey, Dr. Sandip Paul, Dr.Partha Chakrabarti, Dr. Saikat Chakrabarti |
| EPI_ISL_582028 | CSIR-Indian Institute of Chemical Biology, MEDICA Supercpecialty Hospital Kolkata | CSIR-Indian Institute of Chemical Biology, MEDICA Supercpecialty Hospital Kolkata | Sujay Krishna Maity, Priyanka Mallick, Debaleena Bhowmik, Abhishake Lahiri, Dr. Aviral Roy, Dr. Soumen Saha, Dr. Arpita Ghosh Mitra, Dr. Rajesh Pandey, Dr. Sandip Paul, Dr. Partha Chakrabarti, Dr. Saikat Chakrabarti |
| EPI_ISL_582031, EPI_ISL_582032 | Institute of Human Genetics, Polish Academy of Sciences | Institute of Human Genetics, Polish Academy of Sciences | Szymon Hryhorowicz, Adam Ustaszewski, Marta Kaczmarek-Ry, Emilia Lis, Ewa Zitkiewicz, Micha Witt, Andrzej Pawski |
| EPI_ISL_583426 | Garcia-Sastre Laboratory, Department of Microbiology, Icahn School of Medicine at Mount Sinai | van Bakel Laboratory, Genetics and Genomics Sciences, Icahn School of Medicine at Mount Sinai | Teresa Aydillo, Ana S. Gonzalez-Reiche, Sadaf Aslam, Adriana van de Guchte, Zenab Khan, Ajay Obla, Jayeeta Dutta, Harm van Bakel, Judith Aberg, Adolfo García-Sastre, Gunjan Shah, Tobias Hohl, Genovefa Papanicolaou, Miguel-Angel Perales, Kent Sepkowitz, Ngolela Esther Babady, and Mini Kamboj |
| EPI_ISL_583427, EPI_ISL_583428 | Memorial Sloan Kettering Cancer Center | van Bakel Laboratory, Genetics and Genomics Sciences, Icahn School of Medicine at Mount Sinai | Teresa Aydillo, Ana S. Gonzalez-Reiche, Sadaf Aslam, Adriana van de Guchte, Zenab Khan, Ajay Obla, Jayeeta Dutta, Harm van Bakel, Judith Aberg, Adolfo García-Sastre, Gunjan Shah, Tobias Hohl, Genovefa Papanicolaou, Miguel-Angel Perales, Kent Sepkowitz, Ngolela Esther Babady, and Mini Kamboj |
| EPI_ISL_583429 | Garcia-Sastre Laboratory, Department of Microbiology, Icahn School of Medicine at Mount Sinai | van Bakel Laboratory, Genetics and Genomics Sciences, Icahn School of Medicine at Mount Sinai | Teresa Aydillo, Ana S. Gonzalez-Reiche, Sadaf Aslam, Adriana van de Guchte, Zenab Khan, Ajay Obla, Jayeeta Dutta, Harm van Bakel, Judith Aberg, Adolfo García-Sastre, Gunjan Shah, Tobias Hohl, Genovefa Papanicolaou, Miguel-Angel Perales, Kent Sepkowitz, Ngolela Esther Babady, and Mini Kamboj |
| EPI_ISL_583430, EPI_ISL_583431, EPI_ISL_583432, EPI_ISL_583433 | Memorial Sloan Kettering Cancer Center | van Bakel Laboratory, Genetics and Genomics Sciences, Icahn School of Medicine at Mount Sinai | Teresa Aydillo, Ana S. Gonzalez-Reiche, Sadaf Aslam, Adriana van de Guchte, Zenab Khan, Ajay Obla, Jayeeta Dutta, Harm van Bakel, Judith Aberg, Adolfo García-Sastre, Gunjan Shah, Tobias Hohl, Genovefa Papanicolaou, Miguel-Angel Perales, Kent Sepkowitz, Ngolela Esther Babady, and Mini Kamboj |
| EPI_ISL_583434, EPI_ISL_583435, EPI_ISL_583436 | Garcia-Sastre Laboratory, Department of Microbiology, Icahn School of Medicine at Mount Sinai | van Bakel Laboratory, Genetics and Genomics Sciences, Icahn School of Medicine at Mount Sinai | Teresa Aydillo, Ana S. Gonzalez-Reiche, Sadaf Aslam, Adriana van de Guchte, Zenab Khan, Ajay Obla, Jayeeta Dutta, Harm van Bakel, Judith Aberg, Adolfo García-Sastre, Gunjan Shah, Tobias Hohl, Genovefa Papanicolaou, Miguel-Angel Perales, Kent Sepkowitz, Ngolela Esther Babady, and Mini Kamboj |

| | | | |
|---|---|---|---|
| EPI_ISL_583437, EPI_ISL_583438, EPI_ISL_583439, EPI_ISL_583440 | Memorial Sloan Kettering Cancer Center | van Bakel Laboratory, Genetics and Genomics Sciences, Icahn School of Medicine at Mount Sinai | Teresa Aydillo, Ana S. Gonzalez-Reiche, Sadaf Aslam, Adriana van de Guchte, Zenab Khan, Ajay Obla, Jayeeta Dutta, Harm van Bakel, Judith Aberg, Adolfo García-Sastre, Gunjan Shah, Tobias Hohl, Genovefa Papanicolaou, Miguel-Angel Perales, Kent Sepkowitz, Ngolela Esther Babady, and Mini Kamboj |
| EPI_ISL_583441, EPI_ISL_583442 | Garcia-Sastre Laboratory, Department of Microbiology, Icahn School of Medicine at Mount Sinai | van Bakel Laboratory, Genetics and Genomics Sciences, Icahn School of Medicine at Mount Sinai | Teresa Aydillo, Ana S. Gonzalez-Reiche, Sadaf Aslam, Adriana van de Guchte, Zenab Khan, Ajay Obla, Jayeeta Dutta, Harm van Bakel, Judith Aberg, Adolfo García-Sastre, Gunjan Shah, Tobias Hohl, Genovefa Papanicolaou, Miguel-Angel Perales, Kent Sepkowitz, Ngolela Esther Babady, and Mini Kamboj |
| EPI_ISL_583443, EPI_ISL_583444, EPI_ISL_583445, EPI_ISL_583446, EPI_ISL_583447, EPI_ISL_583448, EPI_ISL_583449, EPI_ISL_583450, EPI_ISL_583451 | Memorial Sloan Kettering Cancer Center | van Bakel Laboratory, Genetics and Genomics Sciences, Icahn School of Medicine at Mount Sinai | Teresa Aydillo, Ana S. Gonzalez-Reiche, Sadaf Aslam, Adriana van de Guchte, Zenab Khan, Ajay Obla, Jayeeta Dutta, Harm van Bakel, Judith Aberg, Adolfo García-Sastre, Gunjan Shah, Tobias Hohl, Genovefa Papanicolaou, Miguel-Angel Perales, Kent Sepkowitz, Ngolela Esther Babady, and Mini Kamboj |
| EPI_ISL_583452, EPI_ISL_583453, EPI_ISL_583454 | Garcia-Sastre Laboratory, Department of Microbiology, Icahn School of Medicine at Mount Sinai | van Bakel Laboratory, Genetics and Genomics Sciences, Icahn School of Medicine at Mount Sinai | Teresa Aydillo, Ana S. Gonzalez-Reiche, Sadaf Aslam, Adriana van de Guchte, Zenab Khan, Ajay Obla, Jayeeta Dutta, Harm van Bakel, Judith Aberg, Adolfo García-Sastre, Gunjan Shah, Tobias Hohl, Genovefa Papanicolaou, Miguel-Angel Perales, Kent Sepkowitz, Ngolela Esther Babady, and Mini Kamboj |
| EPI_ISL_583455, EPI_ISL_583456, EPI_ISL_583457 | Memorial Sloan Kettering Cancer Center | van Bakel Laboratory, Genetics and Genomics Sciences, Icahn School of Medicine at Mount Sinai | Teresa Aydillo, Ana S. Gonzalez-Reiche, Sadaf Aslam, Adriana van de Guchte, Zenab Khan, Ajay Obla, Jayeeta Dutta, Harm van Bakel, Judith Aberg, Adolfo García-Sastre, Gunjan Shah, Tobias Hohl, Genovefa Papanicolaou, Miguel-Angel Perales, Kent Sepkowitz, Ngolela Esther Babady, and Mini Kamboj |
| EPI_ISL_583458, EPI_ISL_583459 | Garcia-Sastre Laboratory, Department of Microbiology, Icahn School of Medicine at Mount Sinai | van Bakel Laboratory, Genetics and Genomics Sciences, Icahn School of Medicine at Mount Sinai | Teresa Aydillo, Ana S. Gonzalez-Reiche, Sadaf Aslam, Adriana van de Guchte, Zenab Khan, Ajay Obla, Jayeeta Dutta, Harm van Bakel, Judith Aberg, Adolfo García-Sastre, Gunjan Shah, Tobias Hohl, Genovefa Papanicolaou, Miguel-Angel Perales, Kent Sepkowitz, Ngolela Esther Babady, and Mini Kamboj |
| EPI_ISL_583460, EPI_ISL_583461, EPI_ISL_583462 | Memorial Sloan Kettering Cancer Center | van Bakel Laboratory, Genetics and Genomics Sciences, Icahn School of Medicine at Mount Sinai | Teresa Aydillo, Ana S. Gonzalez-Reiche, Sadaf Aslam, Adriana van de Guchte, Zenab Khan, Ajay Obla, Jayeeta Dutta, Harm van Bakel, Judith Aberg, Adolfo García-Sastre, Gunjan Shah, Tobias Hohl, Genovefa Papanicolaou, Miguel-Angel Perales, Kent Sepkowitz, Ngolela Esther Babady, and Mini Kamboj |
| EPI_ISL_583463, EPI_ISL_583464 | Garcia-Sastre Laboratory, Department of Microbiology, Icahn School of Medicine at Mount Sinai | van Bakel Laboratory, Genetics and Genomics Sciences, Icahn School of Medicine at Mount Sinai | Teresa Aydillo, Ana S. Gonzalez-Reiche, Sadaf Aslam, Adriana van de Guchte, Zenab Khan, Ajay Obla, Jayeeta Dutta, Harm van Bakel, Judith Aberg, Adolfo García-Sastre, Gunjan Shah, Tobias Hohl, Genovefa Papanicolaou, Miguel-Angel Perales, Kent Sepkowitz, Ngolela Esther Babady, and Mini Kamboj |
| EPI_ISL_583465, EPI_ISL_583466 | Memorial Sloan Kettering Cancer Center | van Bakel Laboratory, Genetics and Genomics Sciences, Icahn School of Medicine at Mount Sinai | Teresa Aydillo, Ana S. Gonzalez-Reiche, Sadaf Aslam, Adriana van de Guchte, Zenab Khan, Ajay Obla, Jayeeta Dutta, Harm van Bakel, Judith Aberg, Adolfo García-Sastre, Gunjan Shah, Tobias Hohl, Genovefa Papanicolaou, Miguel-Angel Perales, Kent Sepkowitz, Ngolela Esther Babady, and Mini Kamboj |
| EPI_ISL_583467 | Garcia-Sastre Laboratory, Department of Microbiology, Icahn School of Medicine at Mount Sinai | van Bakel Laboratory, Genetics and Genomics Sciences, Icahn School of Medicine at Mount Sinai | Teresa Aydillo, Ana S. Gonzalez-Reiche, Sadaf Aslam, Adriana van de Guchte, Zenab Khan, Ajay Obla, Jayeeta Dutta, Harm van Bakel, Judith Aberg, Adolfo García-Sastre, Gunjan Shah, Tobias Hohl, Genovefa Papanicolaou, Miguel-Angel Perales, Kent Sepkowitz, Ngolela Esther Babady, and Mini Kamboj |
| EPI_ISL_583468, EPI_ISL_583469, EPI_ISL_583470 | Memorial Sloan Kettering Cancer Center | van Bakel Laboratory, Genetics and Genomics Sciences, Icahn School of Medicine at Mount Sinai | Teresa Aydillo, Ana S. Gonzalez-Reiche, Sadaf Aslam, Adriana van de Guchte, Zenab Khan, Ajay Obla, Jayeeta Dutta, Harm van Bakel, Judith Aberg, Adolfo García-Sastre, Gunjan Shah, Tobias Hohl, Genovefa Papanicolaou, Miguel-Angel Perales, Kent Sepkowitz, Ngolela Esther Babady, and Mini Kamboj |
| EPI_ISL_583471 | Garcia-Sastre Laboratory, Department of Microbiology, Icahn School of Medicine at Mount Sinai | van Bakel Laboratory, Genetics and Genomics Sciences, Icahn School of Medicine at Mount Sinai | Teresa Aydillo, Ana S. Gonzalez-Reiche, Sadaf Aslam, Adriana van de Guchte, Zenab Khan, Ajay Obla, Jayeeta Dutta, Harm van Bakel, Judith Aberg, Adolfo García-Sastre, Gunjan Shah, Tobias Hohl, Genovefa Papanicolaou, Miguel-Angel Perales, Kent Sepkowitz, Ngolela Esther Babady, and Mini Kamboj |
| EPI_ISL_583472 | Memorial Sloan Kettering Cancer Center | van Bakel Laboratory, Genetics and Genomics Sciences, Icahn School of Medicine at Mount Sinai | Teresa Aydillo, Ana S. Gonzalez-Reiche, Sadaf Aslam, Adriana van de Guchte, Zenab Khan, Ajay Obla, Jayeeta Dutta, Harm van Bakel, Judith Aberg, Adolfo García-Sastre, Gunjan Shah, Tobias Hohl, Genovefa Papanicolaou, Miguel-Angel Perales, Kent Sepkowitz, Ngolela Esther Babady, and Mini Kamboj |
| EPI_ISL_583473 | Garcia-Sastre Laboratory, Department of Microbiology, Icahn School of Medicine at Mount Sinai | van Bakel Laboratory, Genetics and Genomics Sciences, Icahn School of Medicine at Mount Sinai | Teresa Aydillo, Ana S. Gonzalez-Reiche, Sadaf Aslam, Adriana van de Guchte, Zenab Khan, Ajay Obla, Jayeeta Dutta, Harm van Bakel, Judith Aberg, Adolfo García-Sastre, Gunjan Shah, Tobias Hohl, Genovefa Papanicolaou, Miguel-Angel Perales, Kent Sepkowitz, Ngolela Esther Babady, and Mini Kamboj |
| EPI_ISL_583474, EPI_ISL_583475, EPI_ISL_583476 | Memorial Sloan Kettering Cancer Center | van Bakel Laboratory, Genetics and Genomics Sciences, Icahn School of Medicine at Mount Sinai | Teresa Aydillo, Ana S. Gonzalez-Reiche, Sadaf Aslam, Adriana van de Guchte, Zenab Khan, Ajay Obla, Jayeeta Dutta, Harm van Bakel, Judith Aberg, Adolfo García-Sastre, Gunjan Shah, Tobias Hohl, Genovefa Papanicolaou, Miguel-Angel Perales, Kent Sepkowitz, Ngolela Esther Babady, and Mini Kamboj |
| EPI_ISL_583477 | Garcia-Sastre Laboratory, Department of Microbiology, Icahn School of Medicine at Mount Sinai | van Bakel Laboratory, Genetics and Genomics Sciences, Icahn School of Medicine at Mount Sinai | Teresa Aydillo, Ana S. Gonzalez-Reiche, Sadaf Aslam, Adriana van de Guchte, Zenab Khan, Ajay Obla, Jayeeta Dutta, Harm van Bakel, Judith Aberg, Adolfo García-Sastre, Gunjan Shah, Tobias Hohl, Genovefa Papanicolaou, Miguel-Angel Perales, Kent Sepkowitz, Ngolela Esther Babady, and Mini Kamboj |
| EPI_ISL_583478, EPI_ISL_583479, EPI_ISL_583480 | Memorial Sloan Kettering Cancer Center | van Bakel Laboratory, Genetics and Genomics Sciences, Icahn School of Medicine at Mount Sinai | Teresa Aydillo, Ana S. Gonzalez-Reiche, Sadaf Aslam, Adriana van de Guchte, Zenab Khan, Ajay Obla, Jayeeta Dutta, Harm van Bakel, Judith Aberg, Adolfo García-Sastre, Gunjan Shah, Tobias Hohl, Genovefa Papanicolaou, Miguel-Angel Perales, Kent Sepkowitz, Ngolela Esther Babady, and Mini Kamboj |
| EPI_ISL_584680 | Oxford Viromics, NDM, University of Oxford; Oxford University Hospitals; Basingstoke and North Hampshire Hospital | COVID-19 Genomics UK (COG-UK) Consortium | Tanya Golubchik, David Bonsall, George Macintyre, Amy Trebes, Mariateresa de Cesare, Catrin Moore, Alex Mobbs, Anita Justice, Robert Shaw, Monique Andersson, Timothy Peto, Emma Wise, Nathan Moore, Jessica Lynch, Nick Cortes, Matilde Mori, Stephen Kidd, David Buck, John Todd, Christophe Fraser |
| EPI_ISL_586569 | GCRI,Ahmedabad | Gujarat Biotechnology Research Centre | Shashank Pandya, Harsha Panchal, Apurvasinh Puvar, Janvi Raval, Zarna Patel, Monika Gandhi, Pinal Trivedi, Maharshi Pandya, Nidhi Patel, Nitin Savaliya, Raghawendra Kumar, Dinesh Kumar, Zuber Saiyed, Komal Patel, Labdhi Pandya, Afzal Ansari, Nikha Trivedi, R D Dixit, A M Kadri, Harsh Bakshi, Chaitanya Joshi, Madhvi Joshi |
| EPI_ISL_593597, EPI_ISL_593598, EPI_ISL_593599, EPI_ISL_593600, EPI_ISL_593601, EPI_ISL_593602, EPI_ISL_593603, EPI_ISL_593604, EPI_ISL_593605, EPI_ISL_593606, EPI_ISL_593607, EPI_ISL_593608, EPI_ISL_593609, EPI_ISL_593610, EPI_ISL_593611, EPI_ISL_593612, EPI_ISL_593613, EPI_ISL_593614, EPI_ISL_593615, EPI_ISL_593616, EPI_ISL_593617, EPI_ISL_593618, EPI_ISL_593619, EPI_ISL_593620, EPI_ISL_593621, EPI_ISL_593622, EPI_ISL_593623, EPI_ISL_593624, EPI_ISL_593625, EPI_ISL_593626, EPI_ISL_593627, EPI_ISL_593628, EPI_ISL_593629, EPI_ISL_593630, EPI_ISL_593631, EPI_ISL_593632, EPI_ISL_593633, EPI_ISL_593634, EPI_ISL_593635, EPI_ISL_593636, EPI_ISL_593637, EPI_ISL_593638, EPI_ISL_593639, EPI_ISL_593640, EPI_ISL_593641, EPI_ISL_593642, EPI_ISL_593643, EPI_ISL_593644, EPI_ISL_593645 | | | |
| see above | Public Health Virology Laboratory, Forensic and Scientific Services (PHV-FSS) | Public Health Virology Laboratory, Forensic and Scientific Services (PHV-FSS) | Son Nguyen et al |
| EPI_ISL_593902 | Sentinelles, Limay | National Reference Center for Viruses of Respiratory Infections, Institut Pasteur, Paris | Sylvie Behillil, Fabiana Gambaro, Etienne Simon-Lorière, Vincent Enouf, Maud Vanpeene, Sylvie van der Werf |
| EPI_ISL_593903 | Hospital, Argenteuil | National Reference Center for Viruses of Respiratory Infections, Institut Pasteur, Paris | Sylvie Behillil, Fabiana Gambaro, Etienne Simon-Lorière, Vincent Enouf, Maud Vanpeene, Sylvie van der Werf |
| EPI_ISL_593904, EPI_ISL_593905, EPI_ISL_593906, EPI_ISL_593907, EPI_ISL_593908, EPI_ISL_593909 | Labo Analyses Med, Sarcelles | National Reference Center for Viruses of Respiratory Infections, Institut Pasteur, Paris | Sylvie Behillil, Fabiana Gambaro, Etienne Simon-Lorière, Vincent Enouf, Maud Vanpeene, Sylvie van der Werf |
| EPI_ISL_593910 | Hospital, Boulogne-Billancourt | National Reference Center for Viruses of Respiratory Infections, Institut Pasteur, Paris | Sylvie Behillil, Fabiana Gambaro, Etienne Simon-Lorière, Vincent Enouf, Maud Vanpeene, Sylvie van der Werf |
| EPI_ISL_593911 | Hospital, Le Chesnay Cédex | National Reference Center for Viruses of Respiratory Infections, Institut Pasteur, Paris | Sylvie Behillil, Fabiana Gambaro, Etienne Simon-Lorière, Vincent Enouf, Maud Vanpeene, Sylvie van der Werf |

| | | | |
|---|---|---|---|
| EPI_ISL_593912 | Hospital, Marange Silvange | National Reference Center for Viruses of Respiratory Infections, Institut Pasteur, Paris | Sylvie Behillil, Fabiana Gambaro, Etienne Simon-Lorière, Vincent Enouf, Maud Vanpeene, Sylvie van der Werf |
| EPI_ISL_593913 | Hospital, Talange | National Reference Center for Viruses of Respiratory Infections, Institut Pasteur, Paris | Sylvie Behillil, Fabiana Gambaro, Etienne Simon-Lorière, Vincent Enouf, Maud Vanpeene, Sylvie van der Werf |
| EPI_ISL_593914 | Hospital, Metz | National Reference Center for Viruses of Respiratory Infections, Institut Pasteur, Paris | Sylvie Behillil, Fabiana Gambaro, Etienne Simon-Lorière, Vincent Enouf, Maud Vanpeene, Sylvie van der Werf |
| EPI_ISL_593915 | Hospital, Amneville | National Reference Center for Viruses of Respiratory Infections, Institut Pasteur, Paris | Sylvie Behillil, Fabiana Gambaro, Etienne Simon-Lorière, Vincent Enouf, Maud Vanpeene, Sylvie van der Werf |
| EPI_ISL_593916 | Hospital, Sarreguemines | National Reference Center for Viruses of Respiratory Infections, Institut Pasteur, Paris | Sylvie Behillil, Fabiana Gambaro, Etienne Simon-Lorière, Vincent Enouf, Maud Vanpeene, Sylvie van der Werf |
| EPI_ISL_593917 | Hospital, Montigny les Metz | National Reference Center for Viruses of Respiratory Infections, Institut Pasteur, Paris | Sylvie Behillil, Fabiana Gambaro, Etienne Simon-Lorière, Vincent Enouf, Maud Vanpeene, Sylvie van der Werf |
| EPI_ISL_593918 | Hospital, Joeuf | National Reference Center for Viruses of Respiratory Infections, Institut Pasteur, Paris | Sylvie Behillil, Fabiana Gambaro, Etienne Simon-Lorière, Vincent Enouf, Maud Vanpeene, Sylvie van der Werf |
| EPI_ISL_593919 | Hospital, Ottonville | National Reference Center for Viruses of Respiratory Infections, Institut Pasteur, Paris | Sylvie Behillil, Fabiana Gambaro, Etienne Simon-Lorière, Vincent Enouf, Maud Vanpeene, Sylvie van der Werf |
| EPI_ISL_593920 | Hospital, Metz | National Reference Center for Viruses of Respiratory Infections, Institut Pasteur, Paris | Sylvie Behillil, Fabiana Gambaro, Etienne Simon-Lorière, Vincent Enouf, Maud Vanpeene, Sylvie van der Werf |
| EPI_ISL_593921 | Hospital, Fameck | National Reference Center for Viruses of Respiratory Infections, Institut Pasteur, Paris | Sylvie Behillil, Fabiana Gambaro, Etienne Simon-Lorière, Vincent Enouf, Maud Vanpeene, Sylvie van der Werf |
| EPI_ISL_593922, EPI_ISL_593923 | Hospital, Maizières les Metz | National Reference Center for Viruses of Respiratory Infections, Institut Pasteur, Paris | Sylvie Behillil, Fabiana Gambaro, Etienne Simon-Lorière, Vincent Enouf, Maud Vanpeene, Sylvie van der Werf |
| EPI_ISL_593924, EPI_ISL_593925, EPI_ISL_593926, EPI_ISL_593927, EPI_ISL_593928, EPI_ISL_593929, EPI_ISL_593930, EPI_ISL_593931, EPI_ISL_593932, EPI_ISL_593933 | Labo Analyses Med, Puteaux | National Reference Center for Viruses of Respiratory Infections, Institut Pasteur, Paris | Sylvie Behillil, Fabiana Gambaro, Etienne Simon-Lorière, Vincent Enouf, Maud Vanpeene, Sylvie van der Werf |
| EPI_ISL_593934 | Sentinelles, Chanteloup-En-Brie | National Reference Center for Viruses of Respiratory Infections, Institut Pasteur, Paris | Sylvie Behillil, Fabiana Gambaro, Etienne Simon-Lorière, Vincent Enouf, Maud Vanpeene, Sylvie van der Werf |
| EPI_ISL_593935 | Sentinelles, Plessis-Trevise | National Reference Center for Viruses of Respiratory Infections, Institut Pasteur, Paris | Sylvie Behillil, Fabiana Gambaro, Etienne Simon-Lorière, Vincent Enouf, Maud Vanpeene, Sylvie van der Werf |
| EPI_ISL_593936 | Sentinelles, Fondettes | National Reference Center for Viruses of Respiratory Infections, Institut Pasteur, Paris | Sylvie Behillil, Fabiana Gambaro, Etienne Simon-Lorière, Vincent Enouf, Maud Vanpeene, Sylvie van der Werf |
| EPI_ISL_594163, EPI_ISL_594164, EPI_ISL_594165 | hopital | National Reference Center for Viruses of Respiratory Infections, Institut Pasteur, Paris | Sylvie Behillil, Fabiana Gambaro, Etienne Simon-Lorière, Vincent Enouf, Maud Vanpeene, Sylvie van der Werf |
| EPI_ISL_594467, EPI_ISL_594468, EPI_ISL_594469, EPI_ISL_594470, EPI_ISL_594471, EPI_ISL_594472, EPI_ISL_594473, EPI_ISL_594474, EPI_ISL_594475, EPI_ISL_594476, EPI_ISL_594477, EPI_ISL_594478, EPI_ISL_594479, EPI_ISL_594480, EPI_ISL_594481, EPI_ISL_594482, EPI_ISL_594483, EPI_ISL_594617, EPI_ISL_594618, EPI_ISL_594619, EPI_ISL_594620, EPI_ISL_594621, EPI_ISL_594622, EPI_ISL_594623, EPI_ISL_594624, EPI_ISL_594625, EPI_ISL_594626, EPI_ISL_594627, EPI_ISL_594628, EPI_ISL_594629, EPI_ISL_594630, EPI_ISL_594631, EPI_ISL_594632, EPI_ISL_594633, EPI_ISL_594634, EPI_ISL_594635, EPI_ISL_594636, EPI_ISL_594637, EPI_ISL_594638, EPI_ISL_594639, EPI_ISL_594640, EPI_ISL_594641, EPI_ISL_594642, EPI_ISL_594643, EPI_ISL_594644, EPI_ISL_594889, EPI_ISL_594890, EPI_ISL_594891, EPI_ISL_594892, EPI_ISL_594893, EPI_ISL_594894, EPI_ISL_594895, EPI_ISL_594896, EPI_ISL_594897, EPI_ISL_594898, EPI_ISL_594899, EPI_ISL_594900, EPI_ISL_594901, EPI_ISL_594902, EPI_ISL_594903, EPI_ISL_594904, EPI_ISL_594905, EPI_ISL_594906, EPI_ISL_594907, EPI_ISL_594908, EPI_ISL_594909, EPI_ISL_594910, EPI_ISL_594911, EPI_ISL_594912, EPI_ISL_594913, EPI_ISL_594914, EPI_ISL_594915, EPI_ISL_594916, EPI_ISL_594917, EPI_ISL_594918, EPI_ISL_594919, EPI_ISL_594920, EPI_ISL_594921, EPI_ISL_594922, EPI_ISL_595326, EPI_ISL_595327, EPI_ISL_595328, EPI_ISL_595617, EPI_ISL_595618, EPI_ISL_595619, EPI_ISL_595620, EPI_ISL_595621, EPI_ISL_595622, EPI_ISL_595623, EPI_ISL_595624, EPI_ISL_595625, EPI_ISL_595626, EPI_ISL_595627, EPI_ISL_595628, EPI_ISL_595629, EPI_ISL_595630, EPI_ISL_595631, EPI_ISL_595632, EPI_ISL_595633, EPI_ISL_595634, EPI_ISL_595635, EPI_ISL_595636, EPI_ISL_595637, EPI_ISL_595638, EPI_ISL_595639, EPI_ISL_595640, EPI_ISL_595641, EPI_ISL_595642, EPI_ISL_595643, EPI_ISL_595644, EPI_ISL_595645, EPI_ISL_595646, EPI_ISL_595647, EPI_ISL_595648, EPI_ISL_595649, EPI_ISL_595650, EPI_ISL_595651, EPI_ISL_595652, EPI_ISL_595653, EPI_ISL_595654, EPI_ISL_595655, EPI_ISL_595656, EPI_ISL_595657, EPI_ISL_595658, EPI_ISL_595659, EPI_ISL_595660, EPI_ISL_595661, EPI_ISL_595662, EPI_ISL_595663, EPI_ISL_595664, EPI_ISL_595665, EPI_ISL_595666, EPI_ISL_595667, EPI_ISL_595668, EPI_ISL_595669, EPI_ISL_595670, EPI_ISL_595671, EPI_ISL_595672, EPI_ISL_595673, EPI_ISL_595674, EPI_ISL_595675, EPI_ISL_595676, EPI_ISL_595677, EPI_ISL_595678, EPI_ISL_595679, EPI_ISL_595680, EPI_ISL_595681, EPI_ISL_595682, EPI_ISL_595683, EPI_ISL_595684, EPI_ISL_595685, EPI_ISL_595686, EPI_ISL_595687, EPI_ISL_595688, EPI_ISL_595689, EPI_ISL_595690, EPI_ISL_595691, EPI_ISL_595692, EPI_ISL_595693, EPI_ISL_595694, EPI_ISL_595695, EPI_ISL_595696, EPI_ISL_595697, EPI_ISL_595698, EPI_ISL_595699, EPI_ISL_595700, EPI_ISL_595701, EPI_ISL_595702, EPI_ISL_595703, EPI_ISL_595704, EPI_ISL_595705, EPI_ISL_595706, EPI_ISL_595707, EPI_ISL_595708, EPI_ISL_595709, EPI_ISL_595710, EPI_ISL_595711, EPI_ISL_595712, EPI_ISL_595713, EPI_ISL_595714, EPI_ISL_595715, EPI_ISL_595716, EPI_ISL_595717, EPI_ISL_595718, EPI_ISL_595719, EPI_ISL_595720, EPI_ISL_595721, EPI_ISL_595722, EPI_ISL_595723, EPI_ISL_595724, EPI_ISL_595725, EPI_ISL_595726, EPI_ISL_595727, EPI_ISL_595728, EPI_ISL_595729, EPI_ISL_595730, EPI_ISL_595731, EPI_ISL_595732, EPI_ISL_595733, EPI_ISL_595734, EPI_ISL_595735, EPI_ISL_595736, EPI_ISL_595737, EPI_ISL_595738, EPI_ISL_595739, EPI_ISL_595740, EPI_ISL_595741, EPI_ISL_595742, EPI_ISL_595743, EPI_ISL_595744, EPI_ISL_595745, EPI_ISL_595746, EPI_ISL_595747, EPI_ISL_595748, EPI_ISL_595749, EPI_ISL_595750, EPI_ISL_595751, EPI_ISL_595752, EPI_ISL_595753, EPI_ISL_595754, EPI_ISL_595755, EPI_ISL_595756, EPI_ISL_595757, EPI_ISL_595758, EPI_ISL_595759, EPI_ISL_595760, EPI_ISL_595761, EPI_ISL_595762, EPI_ISL_595763, EPI_ISL_595764, EPI_ISL_595765, EPI_ISL_595766, EPI_ISL_595767, EPI_ISL_595768, EPI_ISL_595769, EPI_ISL_595770, EPI_ISL_595771, EPI_ISL_595772, EPI_ISL_595773, EPI_ISL_595774, EPI_ISL_595775, EPI_ISL_595776, EPI_ISL_595777, EPI_ISL_595778, EPI_ISL_595779, EPI_ISL_595780, EPI_ISL_595781, EPI_ISL_595782, EPI_ISL_595783, EPI_ISL_595784, EPI_ISL_595785, EPI_ISL_595786, EPI_ISL_595787, EPI_ISL_595788, EPI_ISL_595789, EPI_ISL_595790, EPI_ISL_595791, EPI_ISL_595792, EPI_ISL_595793, EPI_ISL_595794, EPI_ISL_595795, EPI_ISL_595796, EPI_ISL_595797, EPI_ISL_595798 | | | |
| see above | Oxford Viromics, NDM, University of Oxford; Oxford University Hospitals; Basingstoke and North Hampshire Hospital | COVID-19 Genomics UK (COG-UK) Consortium | Tanya Golubchik, David Bonsall, George Macintyre, Amy Trebes, Mariateresa de Cesare, Catrin Moore, Alex Mobbs, Anita Justice, Robert Shaw, Monique Andersson, Timothy Peto, Emma Wise, Nathan Moore, Jessica Lynch, Nick Cortes, Matilde Mori, Stephen Kidd, David Buck, John Todd, Christophe Fraser |
| EPI_ISL_596449 | Institute for Medical Research, Infectious Disease Research Centre, National Institutes of Health, Ministry of Health Malaysia | Institute for Medical Research, Infectious Disease Research Centre, National Institutes of Health, Ministry of Health Malaysia | Suppiah J, Kamel K, Mohd-Zawawi Z, Thayan R |
| EPI_ISL_603240, EPI_ISL_603241 | National Institute of Laboratory Medicine and Referral Center | Genomic Research Lab, BCSIR | Tanjina Akhter Banu, Abu Sayeed Mohammad Mahmud, Mohammad Samir Uzzaman, Eshrar Osman, Md. Ahashan Habib, Shahina Akter, Md. Murshed Hasan Sarkar, Barna Goswami, Iffat Jahan, Md. Saddam Hossain, Tasnim Nafisa, Md. Maruf Ahmed Molla, Mahmuda Yeasmin, Asish Kumar Ghosh, A. K. M. Shamsuzzaman, Monira Parveen, Md. Masum Hossain Arif, Md. Salim Khan |
| EPI_ISL_610163, EPI_ISL_610164, EPI_ISL_610165, EPI_ISL_610166, EPI_ISL_610167, EPI_ISL_610168, EPI_ISL_610169, EPI_ISL_610170, EPI_ISL_610171, EPI_ISL_610172, EPI_ISL_610173, EPI_ISL_610174, EPI_ISL_610175, EPI_ISL_610176, EPI_ISL_610177, EPI_ISL_610178, EPI_ISL_610179, EPI_ISL_610180, EPI_ISL_610181, EPI_ISL_610182, EPI_ISL_610183, EPI_ISL_610184, EPI_ISL_610185, EPI_ISL_610186, EPI_ISL_610187, EPI_ISL_610188, EPI_ISL_610189, EPI_ISL_610190, EPI_ISL_610191, EPI_ISL_610192, EPI_ISL_610193, EPI_ISL_610194, EPI_ISL_610195, EPI_ISL_610196, EPI_ISL_610197, EPI_ISL_610198, EPI_ISL_610199, EPI_ISL_610200, EPI_ISL_610201, EPI_ISL_610202, EPI_ISL_610203, EPI_ISL_610204, EPI_ISL_610205, EPI_ISL_610206, EPI_ISL_610207, EPI_ISL_610208, EPI_ISL_610209, EPI_ISL_610210, EPI_ISL_610211, EPI_ISL_610212, EPI_ISL_610213, EPI_ISL_610214, EPI_ISL_610215, EPI_ISL_610216, EPI_ISL_610217, EPI_ISL_610218, EPI_ISL_610219, EPI_ISL_610220, EPI_ISL_610221, EPI_ISL_610222, EPI_ISL_610223, EPI_ISL_610224, EPI_ISL_610225, EPI_ISL_610226 | | | |
| see above | Department of Health Technology and Informatics, The Hong Kong Polytechnic University | Department of Health Technology and Informatics, The Hong Kong Polytechnic University | Siu,G.K.-H., Lee,L.-K., Leung,K.S.-S., Leung,J.S.-L., Ng,T.T.-L., Chan,C.T.-M., Tam,K.K.-G., Lao,H.-Y., Wu,A.K.-L., Yau,M.C.-Y., Lai,Y.W.-M., Fung,K.S.-C., Chau,S.K.-Y., Wong,B.K.-C., To,W.-K., Luk,K., Ho,A.Y.-M., Que,T.-L., Yip,K.-T., Yam,W.C., Shum,D.H.-K., Yip,S.P. |
| EPI_ISL_613563, EPI_ISL_613564, EPI_ISL_613707, EPI_ISL_613708 | Laboratory of Molecular Biology, Blood Center of Ribeirão Preto | Laboratory of Molecular Biology, Blood Center of Ribeirão Preto, Faculty of Medicine of Ribeirão Preto, University of São Paulo | Svetoslav N Slavov, Marta Giovanetti, Vagner Fonseca, Elaine V Santos, Evandra S Rodrigues, Talita Adelino, Joilson Xavier, Glauco de Carvalho Pereira, Aparecida Y Yamamoto, Diego Villa Clé, Rodrigo T Calado; Dimas T Covas, Luiz CJ Alcantara, Simone Kashima |
| EPI_ISL_613709, EPI_ISL_613711, EPI_ISL_613951, EPI_ISL_613954, EPI_ISL_613956, EPI_ISL_613961, EPI_ISL_613962, EPI_ISL_613964, EPI_ISL_613965, EPI_ISL_614011, EPI_ISL_614155, EPI_ISL_614156 | | | |
| see above | Laboratory of Molecular Biology, Blood Center of Ribeirão Preto, Faculty of Medicine of Ribeirão Preto, University of São Paulo | Laboratory of Molecular Biology, Blood Center of Ribeirão Preto, Faculty of Medicine of Ribeirão Preto, University of São Paulo | Svetoslav N Slavov, Marta Giovanetti, Vagner Fonseca, Elaine V Santos, Evandra S Rodrigues, Talita Adelino, Joilson Xavier, Glauco de Carvalho Pereira, Aparecida Y Yamamoto, Diego Villa Clé, Rodrigo T Calado; Dimas T Covas, Luiz CJ Alcantara, Simone Kashima |
| EPI_ISL_627079, EPI_ISL_627085, EPI_ISL_627250, EPI_ISL_627251, EPI_ISL_627255, EPI_ISL_627256, EPI_ISL_627257, EPI_ISL_627258, EPI_ISL_627259, EPI_ISL_627260, EPI_ISL_627261, EPI_ISL_627262, EPI_ISL_627263, EPI_ISL_627281, EPI_ISL_627282, EPI_ISL_627283, EPI_ISL_627284, EPI_ISL_627285, EPI_ISL_627286, EPI_ISL_627287, EPI_ISL_627288, EPI_ISL_627289, EPI_ISL_627290, EPI_ISL_627292, EPI_ISL_627294, EPI_ISL_627297, EPI_ISL_627299, EPI_ISL_627302, EPI_ISL_627303, EPI_ISL_627305, EPI_ISL_628334, EPI_ISL_628335, EPI_ISL_628336, EPI_ISL_628337, EPI_ISL_628338, EPI_ISL_628339, EPI_ISL_628340, EPI_ISL_628341, EPI_ISL_628342, EPI_ISL_628343, EPI_ISL_628344, EPI_ISL_628345, EPI_ISL_628346, EPI_ISL_628347, EPI_ISL_628348, EPI_ISL_628349, EPI_ISL_628350, EPI_ISL_628351, EPI_ISL_628352, EPI_ISL_628353, EPI_ISL_628354, EPI_ISL_628355, EPI_ISL_628356, EPI_ISL_628357, EPI_ISL_628358, EPI_ISL_628359, EPI_ISL_628360, EPI_ISL_628361, EPI_ISL_628362, EPI_ISL_628363, EPI_ISL_628364, EPI_ISL_628365, EPI_ISL_628366, EPI_ISL_628367, EPI_ISL_628368, EPI_ISL_628414, EPI_ISL_628415, EPI_ISL_628416, EPI_ISL_628417, EPI_ISL_628418, EPI_ISL_628419, EPI_ISL_628420, | | | |

EPI_ISL_628422, EPI_ISL_628423, EPI_ISL_628424, EPI_ISL_628425, EPI_ISL_628426, EPI_ISL_628427, EPI_ISL_628428, EPI_ISL_628429, EPI_ISL_628430, EPI_ISL_628431, EPI_ISL_628432, EPI_ISL_628433, EPI_ISL_628434, EPI_ISL_628435, EPI_ISL_628436, EPI_ISL_628437, EPI_ISL_628438, EPI_ISL_628439, EPI_ISL_628440, EPI_ISL_628441, EPI_ISL_628442, EPI_ISL_628443, EPI_ISL_628444, EPI_ISL_628445, EPI_ISL_628446, EPI_ISL_628447, EPI_ISL_628448, EPI_ISL_628449, EPI_ISL_628450, EPI_ISL_628451, EPI_ISL_628452, EPI_ISL_628453, EPI_ISL_628454, EPI_ISL_628455, EPI_ISL_628456, EPI_ISL_628457, EPI_ISL_628458, EPI_ISL_628459, EPI_ISL_628460, EPI_ISL_628461, EPI_ISL_628462, EPI_ISL_628463, EPI_ISL_628464, EPI_ISL_628465, EPI_ISL_628466, EPI_ISL_628467, EPI_ISL_628468, EPI_ISL_628469, EPI_ISL_628470, EPI_ISL_628471, EPI_ISL_628472, EPI_ISL_628473, EPI_ISL_628474, EPI_ISL_628475, EPI_ISL_628476, EPI_ISL_628477, EPI_ISL_628478, EPI_ISL_628479, EPI_ISL_628480, EPI_ISL_628481, EPI_ISL_628482, EPI_ISL_628483, EPI_ISL_628484, EPI_ISL_628485, EPI_ISL_628486, EPI_ISL_628487, EPI_ISL_628488, EPI_ISL_628489, EPI_ISL_628490, EPI_ISL_628491, EPI_ISL_628492, EPI_ISL_628493, EPI_ISL_628494, EPI_ISL_628495, EPI_ISL_628496, EPI_ISL_628497, EPI_ISL_628498, EPI_ISL_628499, EPI_ISL_628500, EPI_ISL_628501, EPI_ISL_628502, EPI_ISL_628503, EPI_ISL_628504, EPI_ISL_628505, EPI_ISL_628506, EPI_ISL_628507, EPI_ISL_628508, EPI_ISL_628509, EPI_ISL_628510, EPI_ISL_628511, EPI_ISL_628512, EPI_ISL_628513, EPI_ISL_628514, EPI_ISL_628515, EPI_ISL_628516, EPI_ISL_628517, EPI_ISL_628518, EPI_ISL_628519, EPI_ISL_628520, EPI_ISL_628521, EPI_ISL_628522, EPI_ISL_628523, EPI_ISL_628524, EPI_ISL_628525, EPI_ISL_628526, EPI_ISL_628527, EPI_ISL_628528, EPI_ISL_628529, EPI_ISL_628530, EPI_ISL_628531, EPI_ISL_628532, EPI_ISL_628533, EPI_ISL_628534, EPI_ISL_628535, EPI_ISL_628536, EPI_ISL_628537, EPI_ISL_628538, EPI_ISL_628539, EPI_ISL_628540, EPI_ISL_628541, EPI_ISL_628542, EPI_ISL_628543, EPI_ISL_628544, EPI_ISL_628545, EPI_ISL_628546, EPI_ISL_628547, EPI_ISL_628548, EPI_ISL_628549, EPI_ISL_628550, EPI_ISL_628551, EPI_ISL_628552, EPI_ISL_628553, EPI_ISL_628554, EPI_ISL_628555, EPI_ISL_628556, EPI_ISL_628557, EPI_ISL_628558, EPI_ISL_628559, EPI_ISL_628560, EPI_ISL_628561, EPI_ISL_628562, EPI_ISL_628563, EPI_ISL_628564, EPI_ISL_628565, EPI_ISL_628566, EPI_ISL_628567, EPI_ISL_628568, EPI_ISL_628569, EPI_ISL_628570, EPI_ISL_628571, EPI_ISL_628572, EPI_ISL_628573, EPI_ISL_628574, EPI_ISL_628575, EPI_ISL_628576, EPI_ISL_628577, EPI_ISL_628578, EPI_ISL_628579, EPI_ISL_628580, EPI_ISL_628581, EPI_ISL_628582, EPI_ISL_628583, EPI_ISL_628584, EPI_ISL_628585, EPI_ISL_628586, EPI_ISL_628587, EPI_ISL_628588, EPI_ISL_628589, EPI_ISL_628590, EPI_ISL_628591, EPI_ISL_628592, EPI_ISL_628593, EPI_ISL_628594, EPI_ISL_628595, EPI_ISL_628596, EPI_ISL_628597, EPI_ISL_628598, EPI_ISL_628599, EPI_ISL_628600, EPI_ISL_628601, EPI_ISL_628602, EPI_ISL_628603, EPI_ISL_628604, EPI_ISL_628605, EPI_ISL_628606, EPI_ISL_628607, EPI_ISL_628608, EPI_ISL_628609, EPI_ISL_628610, EPI_ISL_628611, EPI_ISL_628612, EPI_ISL_628613, EPI_ISL_628614, EPI_ISL_628615, EPI_ISL_628616, EPI_ISL_628617, EPI_ISL_628618, EPI_ISL_628619, EPI_ISL_628620, EPI_ISL_628621, EPI_ISL_628622, EPI_ISL_628623, EPI_ISL_628624, EPI_ISL_628625, EPI_ISL_628626, EPI_ISL_628627, EPI_ISL_628628, EPI_ISL_628629, EPI_ISL_628630, EPI_ISL_628631, EPI_ISL_628632, EPI_ISL_628633, EPI_ISL_628634, EPI_ISL_628635, EPI_ISL_628636, EPI_ISL_628637, EPI_ISL_628638, EPI_ISL_628639, EPI_ISL_628640, EPI_ISL_628641, EPI_ISL_628642, EPI_ISL_628643, EPI_ISL_628644, EPI_ISL_628645, EPI_ISL_628646, EPI_ISL_628647, EPI_ISL_628648, EPI_ISL_628649, EPI_ISL_628650, EPI_ISL_628651, EPI_ISL_628652, EPI_ISL_628653, EPI_ISL_628654, EPI_ISL_628655, EPI_ISL_628656, EPI_ISL_628657, EPI_ISL_628658, EPI_ISL_628659, EPI_ISL_628660, EPI_ISL_628661, EPI_ISL_628662, EPI_ISL_628663, EPI_ISL_628664, EPI_ISL_628665, EPI_ISL_628666, EPI_ISL_628667, EPI_ISL_628668, EPI_ISL_628669, EPI_ISL_628670, EPI_ISL_628671, EPI_ISL_628672, EPI_ISL_628673, EPI_ISL_628674, EPI_ISL_628675, EPI_ISL_628676, EPI_ISL_628677, EPI_ISL_628678, EPI_ISL_628679, EPI_ISL_628680, EPI_ISL_628681, EPI_ISL_628682, EPI_ISL_628683, EPI_ISL_628684, EPI_ISL_628685, EPI_ISL_628686, EPI_ISL_628687, EPI_ISL_628688, EPI_ISL_628689, EPI_ISL_628690, EPI_ISL_628691, EPI_ISL_628692, EPI_ISL_628693, EPI_ISL_628694, EPI_ISL_628695, EPI_ISL_628696, EPI_ISL_628697, EPI_ISL_628698, EPI_ISL_628699, EPI_ISL_628700

| | | | |
|---|---|---|---|
| see above | Oxford Viromics, NDM, University of Oxford; Oxford University Hospitals; Basingstoke and North Hampshire Hospital | COVID-19 Genomics UK (COG-UK) Consortium | Tanya Golubchik, David Bonsall, George Macintyre, Amy Trebes, Mariateresa de Cesare, Catrin Moore, Alex Mobbs, Anita Justice, Robert Shaw, Monique Andersson, Timothy Peto, Emma Wise, Nathan Moore, Jessica Lynch, Nick Cortes, Matilde Mori, Stephen Kidd, David Buck, John Todd, Christophe Fraser |

We gratefully acknowledge the following Authors from the Originating laboratories responsible for obtaining the specimens, as well as the Submitting laboratories where the genome data were generated and shared via GISAID, on which this research is based.

All Submitters of data may be contacted directly via www.gisaid.org

Authors are sorted alphabetically.

| Accession ID | Originating Laboratory | Submitting Laboratory | Authors |
|---|---|---|---|
| EPI_ISL_549808, EPI_ISL_549810, EPI_ISL_549811, EPI_ISL_549812, EPI_ISL_549814, EPI_ISL_549815, EPI_ISL_549816, EPI_ISL_549817, EPI_ISL_549818, EPI_ISL_549819, EPI_ISL_549820 | | | |
| see above | Lighthouse Lab in Milton Keynes | Wellcome Sanger Institute for the COVID-19 Genomics UK (COG-UK) consortium | The Lighthouse Lab in Milton Keynes and Alex Alderton, Roberto Amato, Sonia Goncalves, Ewan Harrison, David K. Jackson, Ian Johnston, Dominic Kwiatkowski, Cordelia Langford, John Sillitoe on behalf of the Wellcome Sanger Institute COVID-19 Surveillance Team (http://www.sanger.ac.uk/covid-team) |
| EPI_ISL_549821 | Lighthouse Lab in Milton Keynes | Wellcome Sanger Institute for the COVID-19 Genomics UK (COG-UK) consortium | The Lighthouse Lab in Alderley Park and Alex Alderton, Roberto Amato, Sonia Goncalves, Ewan Harrison, David K. Jackson, Ian Johnston, Dominic Kwiatkowski, Cordelia Langford, John Sillitoe on behalf of the Wellcome Sanger Institute COVID-19 Surveillance Team |
| EPI_ISL_549822, EPI_ISL_549823, EPI_ISL_549825, EPI_ISL_549826, EPI_ISL_549827, EPI_ISL_549828, EPI_ISL_549830, EPI_ISL_549831, EPI_ISL_549832, EPI_ISL_549833, EPI_ISL_549834, EPI_ISL_549835, EPI_ISL_549836, EPI_ISL_549837, EPI_ISL_549838, EPI_ISL_549839, EPI_ISL_549840, EPI_ISL_549841, EPI_ISL_549842, EPI_ISL_549843, EPI_ISL_549844, EPI_ISL_549845, EPI_ISL_549846, EPI_ISL_549847, EPI_ISL_549848, EPI_ISL_549849, EPI_ISL_549850, EPI_ISL_549852, EPI_ISL_549853, EPI_ISL_549854, EPI_ISL_549855, EPI_ISL_549856, EPI_ISL_549857, EPI_ISL_549858, EPI_ISL_549859, EPI_ISL_549860, EPI_ISL_549862 | | | |
| see above | Lighthouse Lab in Milton Keynes | Wellcome Sanger Institute for the COVID-19 Genomics UK (COG-UK) consortium | The Lighthouse Lab in Milton Keynes and Alex Alderton, Roberto Amato, Sonia Goncalves, Ewan Harrison, David K. Jackson, Ian Johnston, Dominic Kwiatkowski, Cordelia Langford, John Sillitoe on behalf of the Wellcome Sanger Institute COVID-19 Surveillance Team (http://www.sanger.ac.uk/covid-team) |
| EPI_ISL_549863 | Lighthouse Lab in Milton Keynes | Wellcome Sanger Institute for the COVID-19 Genomics UK (COG-UK) consortium | The Lighthouse Lab in Alderley Park and Alex Alderton, Roberto Amato, Sonia Goncalves, Ewan Harrison, David K. Jackson, Ian Johnston, Dominic Kwiatkowski, Cordelia Langford, John Sillitoe on behalf of the Wellcome Sanger Institute COVID-19 Surveillance Team |
| EPI_ISL_549864, EPI_ISL_549865, EPI_ISL_549866, EPI_ISL_549867, EPI_ISL_549868, EPI_ISL_549869, EPI_ISL_549870, EPI_ISL_549871, EPI_ISL_549872, EPI_ISL_549873, EPI_ISL_549875, EPI_ISL_549876, EPI_ISL_549877, EPI_ISL_549878, EPI_ISL_549879, EPI_ISL_549880, EPI_ISL_549881, EPI_ISL_549882, EPI_ISL_549883, EPI_ISL_549885, EPI_ISL_549886, EPI_ISL_549887, EPI_ISL_549888, EPI_ISL_549889, EPI_ISL_549890, EPI_ISL_549892, EPI_ISL_549893, EPI_ISL_549894, EPI_ISL_549895, EPI_ISL_549896, EPI_ISL_549897, EPI_ISL_549898, EPI_ISL_549899, EPI_ISL_549900, EPI_ISL_549901, EPI_ISL_549902, EPI_ISL_549903, EPI_ISL_549904, EPI_ISL_549905, EPI_ISL_549906, EPI_ISL_549908, EPI_ISL_549909, EPI_ISL_549910, EPI_ISL_549911, EPI_ISL_549912, EPI_ISL_549913, EPI_ISL_549914, EPI_ISL_549915, EPI_ISL_549916, EPI_ISL_549917, EPI_ISL_549918, EPI_ISL_549919, EPI_ISL_549920, EPI_ISL_549921, EPI_ISL_549922, EPI_ISL_549923, EPI_ISL_549924, EPI_ISL_549925, EPI_ISL_549926, EPI_ISL_549927, EPI_ISL_549928, EPI_ISL_549929, EPI_ISL_549930, EPI_ISL_549931, EPI_ISL_549932, EPI_ISL_549933, EPI_ISL_549934, EPI_ISL_549935, EPI_ISL_549936, EPI_ISL_549937, EPI_ISL_549939 | | | |
| see above | Lighthouse Lab in Milton Keynes | Wellcome Sanger Institute for the COVID-19 Genomics UK (COG-UK) consortium | The Lighthouse Lab in Milton Keynes and Alex Alderton, Roberto Amato, Sonia Goncalves, Ewan Harrison, David K. Jackson, Ian Johnston, Dominic Kwiatkowski, Cordelia Langford, John Sillitoe on behalf of the Wellcome Sanger Institute COVID-19 Surveillance Team (http://www.sanger.ac.uk/covid-team) |
| EPI_ISL_549940 | Lighthouse Lab in Milton Keynes | Wellcome Sanger Institute for the COVID-19 Genomics UK (COG-UK) consortium | The Lighthouse Lab in Alderley Park and Alex Alderton, Roberto Amato, Sonia Goncalves, Ewan Harrison, David K. Jackson, Ian Johnston, Dominic Kwiatkowski, Cordelia Langford, John Sillitoe on behalf of the Wellcome Sanger Institute COVID-19 Surveillance Team |
| EPI_ISL_549941, EPI_ISL_549942, EPI_ISL_549944, EPI_ISL_549945, EPI_ISL_549947, EPI_ISL_549948, EPI_ISL_549949, EPI_ISL_549950, EPI_ISL_549951, EPI_ISL_549952, EPI_ISL_549953, EPI_ISL_549954, EPI_ISL_549955 | | | |
| see above | Lighthouse Lab in Milton Keynes | Wellcome Sanger Institute for the COVID-19 Genomics UK (COG-UK) consortium | The Lighthouse Lab in Milton Keynes and Alex Alderton, Roberto Amato, Sonia Goncalves, Ewan Harrison, David K. Jackson, Ian Johnston, Dominic Kwiatkowski, Cordelia Langford, John Sillitoe on behalf of the Wellcome Sanger Institute COVID-19 Surveillance Team (http://www.sanger.ac.uk/covid-team) |
| EPI_ISL_549956 | Lighthouse Lab in Milton Keynes | Wellcome Sanger Institute for the COVID-19 Genomics UK (COG-UK) consortium | The Lighthouse Lab in Alderley Park and Alex Alderton, Roberto Amato, Sonia Goncalves, Ewan Harrison, David K. Jackson, Ian Johnston, Dominic Kwiatkowski, Cordelia Langford, John Sillitoe on behalf of the Wellcome Sanger Institute COVID-19 Surveillance Team |
| EPI_ISL_549957, EPI_ISL_549958, EPI_ISL_549959, EPI_ISL_549960, EPI_ISL_549961, EPI_ISL_549962, EPI_ISL_549963, EPI_ISL_549964, EPI_ISL_549965, EPI_ISL_549966, EPI_ISL_549967, EPI_ISL_549968, EPI_ISL_549969, EPI_ISL_549970, EPI_ISL_549971, EPI_ISL_549973, EPI_ISL_549974, EPI_ISL_549975, EPI_ISL_549976, EPI_ISL_549977, EPI_ISL_549978 | | | |
| see above | Lighthouse Lab in Milton Keynes | Wellcome Sanger Institute for the COVID-19 Genomics UK (COG-UK) consortium | The Lighthouse Lab in Milton Keynes and Alex Alderton, Roberto Amato, Sonia Goncalves, Ewan Harrison, David K. Jackson, Ian Johnston, Dominic Kwiatkowski, Cordelia Langford, John Sillitoe on behalf of the Wellcome Sanger Institute COVID-19 Surveillance Team (http://www.sanger.ac.uk/covid-team) |
| EPI_ISL_549979 | Lighthouse Lab in Milton Keynes | Wellcome Sanger Institute for the COVID-19 Genomics UK (COG-UK) consortium | The Lighthouse Lab in Alderley Park and Alex Alderton, Roberto Amato, Sonia Goncalves, Ewan Harrison, David K. Jackson, Ian Johnston, Dominic Kwiatkowski, Cordelia Langford, John Sillitoe on behalf of the Wellcome Sanger Institute COVID-19 Surveillance Team |
| EPI_ISL_549982, EPI_ISL_549983, EPI_ISL_549984, EPI_ISL_549985, EPI_ISL_549986, EPI_ISL_549987, EPI_ISL_549988, EPI_ISL_549989, EPI_ISL_549990, EPI_ISL_549991, EPI_ISL_549992, EPI_ISL_549993, EPI_ISL_549994, EPI_ISL_549995, EPI_ISL_549996, EPI_ISL_549997, EPI_ISL_549998, EPI_ISL_549999 | | | |
| see above | Lighthouse Lab in Milton Keynes | Wellcome Sanger Institute for the COVID-19 Genomics UK (COG-UK) consortium | The Lighthouse Lab in Milton Keynes and Alex Alderton, Roberto Amato, Sonia Goncalves, Ewan Harrison, David K. Jackson, Ian Johnston, Dominic Kwiatkowski, Cordelia Langford, John Sillitoe on behalf of the Wellcome Sanger Institute COVID-19 Surveillance Team (http://www.sanger.ac.uk/covid-team) |
| EPI_ISL_550000 | Lighthouse Lab in Milton Keynes | Wellcome Sanger Institute for the COVID-19 Genomics UK (COG-UK) consortium | The Lighthouse Lab in Alderley Park and Alex Alderton, Roberto Amato, Sonia Goncalves, Ewan Harrison, David K. Jackson, Ian Johnston, Dominic Kwiatkowski, Cordelia Langford, John Sillitoe on behalf of the Wellcome Sanger Institute COVID-19 Surveillance Team |
| EPI_ISL_550001, EPI_ISL_550002 | Lighthouse Lab in Milton Keynes | Wellcome Sanger Institute for the COVID-19 Genomics UK (COG-UK) consortium | The Lighthouse Lab in Milton Keynes and Alex Alderton, Roberto Amato, Sonia Goncalves, Ewan Harrison, David K. Jackson, Ian Johnston, Dominic Kwiatkowski, Cordelia Langford, John Sillitoe on behalf of the Wellcome Sanger Institute COVID-19 Surveillance Team (http://www.sanger.ac.uk/covid-team) |
| EPI_ISL_550003 | Lighthouse Lab in Milton Keynes | Wellcome Sanger Institute for the COVID-19 Genomics UK (COG-UK) consortium | The Lighthouse Lab in Alderley Park and Alex Alderton, Roberto Amato, Sonia Goncalves, Ewan Harrison, David K. Jackson, Ian Johnston, Dominic Kwiatkowski, Cordelia Langford, John Sillitoe on behalf of the Wellcome Sanger Institute COVID-19 Surveillance Team |
| EPI_ISL_550004, EPI_ISL_550005, EPI_ISL_550006, EPI_ISL_550007, EPI_ISL_550008, EPI_ISL_550009, EPI_ISL_550010, EPI_ISL_550011, EPI_ISL_550012, EPI_ISL_550013, EPI_ISL_550014, EPI_ISL_550015, EPI_ISL_550016 | | | |
| see above | Lighthouse Lab in Milton Keynes | Wellcome Sanger Institute for the COVID-19 Genomics UK (COG-UK) consortium | The Lighthouse Lab in Milton Keynes and Alex Alderton, Roberto Amato, Sonia Goncalves, Ewan Harrison, David K. Jackson, Ian Johnston, Dominic Kwiatkowski, Cordelia Langford, John Sillitoe on behalf of the Wellcome Sanger Institute COVID-19 Surveillance Team (http://www.sanger.ac.uk/covid-team) |
| EPI_ISL_550017 | Lighthouse Lab in Milton Keynes | Wellcome Sanger Institute for the COVID-19 Genomics UK (COG-UK) consortium | The Lighthouse Lab in Alderley Park and Alex Alderton, Roberto Amato, Sonia Goncalves, Ewan Harrison, David K. Jackson, Ian Johnston, Dominic Kwiatkowski, Cordelia Langford, John Sillitoe on behalf of the Wellcome Sanger Institute COVID-19 Surveillance Team |
| EPI_ISL_550018 | Lighthouse Lab in Milton Keynes | Wellcome Sanger Institute for the COVID-19 Genomics UK (COG-UK) consortium | The Lighthouse Lab in Milton Keynes and Alex Alderton, Roberto Amato, Sonia Goncalves, Ewan Harrison, David K. Jackson, Ian Johnston, Dominic Kwiatkowski, Cordelia Langford, John Sillitoe on behalf of the Wellcome Sanger Institute COVID-19 Surveillance Team (http://www.sanger.ac.uk/covid-team) |
| EPI_ISL_550019 | Lighthouse Lab in Milton Keynes | Wellcome Sanger Institute for the COVID-19 Genomics UK (COG-UK) consortium | The Lighthouse Lab in Alderley Park and Alex Alderton, Roberto Amato, Sonia Goncalves, Ewan Harrison, David K. Jackson, Ian Johnston, Dominic Kwiatkowski, Cordelia Langford, John Sillitoe on behalf of the Wellcome Sanger Institute COVID-19 Surveillance Team |
| EPI_ISL_550020, EPI_ISL_550022 | Lighthouse Lab in Milton Keynes | Wellcome Sanger Institute for the COVID-19 Genomics UK (COG-UK) consortium | The Lighthouse Lab in Milton Keynes and Alex Alderton, Roberto Amato, Sonia Goncalves, Ewan Harrison, David K. Jackson, Ian Johnston, Dominic Kwiatkowski, Cordelia Langford, John Sillitoe on behalf of the Wellcome Sanger Institute COVID-19 Surveillance Team (http://www.sanger.ac.uk/covid-team) |
| EPI_ISL_550023 | Lighthouse Lab in Milton Keynes | Wellcome Sanger Institute for the COVID-19 Genomics UK | The Lighthouse Lab in Alderley Park and Alex Alderton, Roberto Amato, Sonia Goncalves, Ewan Harrison, David K. Jackson, Ian Johnston, Dominic |

| | | | |
|---|---|---|---|
| | | (COG-UK) consortium | Kwiatkowski, Cordelia Langford, John Sillitoe on behalf of the Wellcome Sanger Institute COVID-19 Surveillance Team |
| EPI_ISL_550024, EPI_ISL_550025, EPI_ISL_550026, EPI_ISL_550027, EPI_ISL_550028, EPI_ISL_550029 | Lighthouse Lab in Milton Keynes | Wellcome Sanger Institute for the COVID-19 Genomics UK (COG-UK) consortium | The Lighthouse Lab in Milton Keynes and Alex Alderton, Roberto Amato, Sonia Goncalves, Ewan Harrison, David K. Jackson, Ian Johnston, Dominic Kwiatkowski, Cordelia Langford, John Sillitoe on behalf of the Wellcome Sanger Institute COVID-19 Surveillance Team (http://www.sanger.ac.uk/covid-team) |
| EPI_ISL_550030 | Lighthouse Lab in Milton Keynes | Wellcome Sanger Institute for the COVID-19 Genomics UK (COG-UK) consortium | The Lighthouse Lab in Alderley Park and Alex Alderton, Roberto Amato, Sonia Goncalves, Ewan Harrison, David K. Jackson, Ian Johnston, Dominic Kwiatkowski, Cordelia Langford, John Sillitoe on behalf of the Wellcome Sanger Institute COVID-19 Surveillance Team |
| EPI_ISL_550031, EPI_ISL_550032, EPI_ISL_550033, EPI_ISL_550034, EPI_ISL_550035 | Lighthouse Lab in Milton Keynes | Wellcome Sanger Institute for the COVID-19 Genomics UK (COG-UK) consortium | The Lighthouse Lab in Milton Keynes and Alex Alderton, Roberto Amato, Sonia Goncalves, Ewan Harrison, David K. Jackson, Ian Johnston, Dominic Kwiatkowski, Cordelia Langford, John Sillitoe on behalf of the Wellcome Sanger Institute COVID-19 Surveillance Team (http://www.sanger.ac.uk/covid-team) |
| EPI_ISL_550036, EPI_ISL_550037 | Lighthouse Lab in Milton Keynes | Wellcome Sanger Institute for the COVID-19 Genomics UK (COG-UK) consortium | The Lighthouse Lab in Alderley Park and Alex Alderton, Roberto Amato, Sonia Goncalves, Ewan Harrison, David K. Jackson, Ian Johnston, Dominic Kwiatkowski, Cordelia Langford, John Sillitoe on behalf of the Wellcome Sanger Institute COVID-19 Surveillance Team |
| EPI_ISL_550038, EPI_ISL_550040 | Lighthouse Lab in Milton Keynes | Wellcome Sanger Institute for the COVID-19 Genomics UK (COG-UK) consortium | The Lighthouse Lab in Milton Keynes and Alex Alderton, Roberto Amato, Sonia Goncalves, Ewan Harrison, David K. Jackson, Ian Johnston, Dominic Kwiatkowski, Cordelia Langford, John Sillitoe on behalf of the Wellcome Sanger Institute COVID-19 Surveillance Team (http://www.sanger.ac.uk/covid-team) |
| EPI_ISL_550041 | Lighthouse Lab in Milton Keynes | Wellcome Sanger Institute for the COVID-19 Genomics UK (COG-UK) consortium | The Lighthouse Lab in Alderley Park and Alex Alderton, Roberto Amato, Sonia Goncalves, Ewan Harrison, David K. Jackson, Ian Johnston, Dominic Kwiatkowski, Cordelia Langford, John Sillitoe on behalf of the Wellcome Sanger Institute COVID-19 Surveillance Team |
| EPI_ISL_550042, EPI_ISL_550043, EPI_ISL_550044, EPI_ISL_550045 | Lighthouse Lab in Milton Keynes | Wellcome Sanger Institute for the COVID-19 Genomics UK (COG-UK) consortium | The Lighthouse Lab in Milton Keynes and Alex Alderton, Roberto Amato, Sonia Goncalves, Ewan Harrison, David K. Jackson, Ian Johnston, Dominic Kwiatkowski, Cordelia Langford, John Sillitoe on behalf of the Wellcome Sanger Institute COVID-19 Surveillance Team (http://www.sanger.ac.uk/covid-team) |
| EPI_ISL_550046 | Lighthouse Lab in Milton Keynes | Wellcome Sanger Institute for the COVID-19 Genomics UK (COG-UK) consortium | The Lighthouse Lab in Alderley Park and Alex Alderton, Roberto Amato, Sonia Goncalves, Ewan Harrison, David K. Jackson, Ian Johnston, Dominic Kwiatkowski, Cordelia Langford, John Sillitoe on behalf of the Wellcome Sanger Institute COVID-19 Surveillance Team |
| EPI_ISL_550048, EPI_ISL_550049, EPI_ISL_550050, EPI_ISL_550052, EPI_ISL_550053, EPI_ISL_550054, EPI_ISL_550055, EPI_ISL_550056, EPI_ISL_550057, EPI_ISL_550058, EPI_ISL_550059, EPI_ISL_550060, EPI_ISL_550061, EPI_ISL_550062, EPI_ISL_550064, EPI_ISL_550065, EPI_ISL_550066, EPI_ISL_550067, EPI_ISL_550068, EPI_ISL_550070 | | | |
| see above | Lighthouse Lab in Milton Keynes | Wellcome Sanger Institute for the COVID-19 Genomics UK (COG-UK) consortium | The Lighthouse Lab in Milton Keynes and Alex Alderton, Roberto Amato, Sonia Goncalves, Ewan Harrison, David K. Jackson, Ian Johnston, Dominic Kwiatkowski, Cordelia Langford, John Sillitoe on behalf of the Wellcome Sanger Institute COVID-19 Surveillance Team (http://www.sanger.ac.uk/covid-team) |
| EPI_ISL_550071 | Lighthouse Lab in Milton Keynes | Wellcome Sanger Institute for the COVID-19 Genomics UK (COG-UK) consortium | The Lighthouse Lab in Alderley Park and Alex Alderton, Roberto Amato, Sonia Goncalves, Ewan Harrison, David K. Jackson, Ian Johnston, Dominic Kwiatkowski, Cordelia Langford, John Sillitoe on behalf of the Wellcome Sanger Institute COVID-19 Surveillance Team |
| EPI_ISL_550073, EPI_ISL_550074, EPI_ISL_550075, EPI_ISL_550077, EPI_ISL_550078, EPI_ISL_550079, EPI_ISL_550080, EPI_ISL_550081, EPI_ISL_550082, EPI_ISL_550083, EPI_ISL_550084, EPI_ISL_550085, EPI_ISL_550086, EPI_ISL_550087, EPI_ISL_550088, EPI_ISL_550089, EPI_ISL_550090, EPI_ISL_550091, EPI_ISL_550092, EPI_ISL_550093, EPI_ISL_550094, EPI_ISL_550095, EPI_ISL_550096, EPI_ISL_550097, EPI_ISL_550098, EPI_ISL_550100, EPI_ISL_550101, EPI_ISL_550102, EPI_ISL_550103, EPI_ISL_550104, EPI_ISL_550105, EPI_ISL_550106, EPI_ISL_550107, EPI_ISL_550108, EPI_ISL_550109, EPI_ISL_550110, EPI_ISL_550111, EPI_ISL_550112, EPI_ISL_550113, EPI_ISL_550114, EPI_ISL_550115, EPI_ISL_550117, EPI_ISL_550118, EPI_ISL_550119, EPI_ISL_550120, EPI_ISL_550121, EPI_ISL_550122, EPI_ISL_550123, EPI_ISL_550124, EPI_ISL_550127, EPI_ISL_550128, EPI_ISL_550129, EPI_ISL_550130, EPI_ISL_550131, EPI_ISL_550132, EPI_ISL_550133, EPI_ISL_550134, EPI_ISL_550135, EPI_ISL_550136, EPI_ISL_550137 | | | |
| see above | Lighthouse Lab in Milton Keynes | Wellcome Sanger Institute for the COVID-19 Genomics UK (COG-UK) consortium | The Lighthouse Lab in Milton Keynes and Alex Alderton, Roberto Amato, Sonia Goncalves, Ewan Harrison, David K. Jackson, Ian Johnston, Dominic Kwiatkowski, Cordelia Langford, John Sillitoe on behalf of the Wellcome Sanger Institute COVID-19 Surveillance Team (http://www.sanger.ac.uk/covid-team) |
| EPI_ISL_550138 | Lighthouse Lab in Milton Keynes | Wellcome Sanger Institute for the COVID-19 Genomics UK (COG-UK) consortium | The Lighthouse Lab in Alderley Park and Alex Alderton, Roberto Amato, Sonia Goncalves, Ewan Harrison, David K. Jackson, Ian Johnston, Dominic Kwiatkowski, Cordelia Langford, John Sillitoe on behalf of the Wellcome Sanger Institute COVID-19 Surveillance Team |
| EPI_ISL_550139, EPI_ISL_550140, EPI_ISL_550141, EPI_ISL_550142, EPI_ISL_550143 | Lighthouse Lab in Milton Keynes | Wellcome Sanger Institute for the COVID-19 Genomics UK (COG-UK) consortium | The Lighthouse Lab in Milton Keynes and Alex Alderton, Roberto Amato, Sonia Goncalves, Ewan Harrison, David K. Jackson, Ian Johnston, Dominic Kwiatkowski, Cordelia Langford, John Sillitoe on behalf of the Wellcome Sanger Institute COVID-19 Surveillance Team (http://www.sanger.ac.uk/covid-team) |
| EPI_ISL_550145, EPI_ISL_550146 | Lighthouse Lab in Milton Keynes | Wellcome Sanger Institute for the COVID-19 Genomics UK (COG-UK) consortium | The Lighthouse Lab in Alderley Park and Alex Alderton, Roberto Amato, Sonia Goncalves, Ewan Harrison, David K. Jackson, Ian Johnston, Dominic Kwiatkowski, Cordelia Langford, John Sillitoe on behalf of the Wellcome Sanger Institute COVID-19 Surveillance Team |
| EPI_ISL_550147, EPI_ISL_550148, EPI_ISL_550149, EPI_ISL_550150, EPI_ISL_550151, EPI_ISL_550152, EPI_ISL_550153, EPI_ISL_550155, EPI_ISL_550156, EPI_ISL_550157, EPI_ISL_550158 | | | |
| see above | Lighthouse Lab in Milton Keynes | Wellcome Sanger Institute for the COVID-19 Genomics UK (COG-UK) consortium | The Lighthouse Lab in Milton Keynes and Alex Alderton, Roberto Amato, Sonia Goncalves, Ewan Harrison, David K. Jackson, Ian Johnston, Dominic Kwiatkowski, Cordelia Langford, John Sillitoe on behalf of the Wellcome Sanger Institute COVID-19 Surveillance Team (http://www.sanger.ac.uk/covid-team) |
| EPI_ISL_550159 | Lighthouse Lab in Milton Keynes | Wellcome Sanger Institute for the COVID-19 Genomics UK (COG-UK) consortium | The Lighthouse Lab in Alderley Park and Alex Alderton, Roberto Amato, Sonia Goncalves, Ewan Harrison, David K. Jackson, Ian Johnston, Dominic Kwiatkowski, Cordelia Langford, John Sillitoe on behalf of the Wellcome Sanger Institute COVID-19 Surveillance Team |
| EPI_ISL_550160, EPI_ISL_550161 | Lighthouse Lab in Milton Keynes | Wellcome Sanger Institute for the COVID-19 Genomics UK (COG-UK) consortium | The Lighthouse Lab in Milton Keynes and Alex Alderton, Roberto Amato, Sonia Goncalves, Ewan Harrison, David K. Jackson, Ian Johnston, Dominic Kwiatkowski, Cordelia Langford, John Sillitoe on behalf of the Wellcome Sanger Institute COVID-19 Surveillance Team (http://www.sanger.ac.uk/covid-team) |
| EPI_ISL_550163 | Lighthouse Lab in Milton Keynes | Wellcome Sanger Institute for the COVID-19 Genomics UK (COG-UK) consortium | The Lighthouse Lab in Alderley Park and Alex Alderton, Roberto Amato, Sonia Goncalves, Ewan Harrison, David K. Jackson, Ian Johnston, Dominic Kwiatkowski, Cordelia Langford, John Sillitoe on behalf of the Wellcome Sanger Institute COVID-19 Surveillance Team |
| EPI_ISL_550164, EPI_ISL_550165, EPI_ISL_550166, EPI_ISL_550167, EPI_ISL_550169, EPI_ISL_550171, EPI_ISL_550172, EPI_ISL_550173, EPI_ISL_550174 | Lighthouse Lab in Milton Keynes | Wellcome Sanger Institute for the COVID-19 Genomics UK (COG-UK) consortium | The Lighthouse Lab in Milton Keynes and Alex Alderton, Roberto Amato, Sonia Goncalves, Ewan Harrison, David K. Jackson, Ian Johnston, Dominic Kwiatkowski, Cordelia Langford, John Sillitoe on behalf of the Wellcome Sanger Institute COVID-19 Surveillance Team (http://www.sanger.ac.uk/covid-team) |
| EPI_ISL_550175 | Lighthouse Lab in Milton Keynes | Wellcome Sanger Institute for the COVID-19 Genomics UK (COG-UK) consortium | The Lighthouse Lab in Alderley Park and Alex Alderton, Roberto Amato, Sonia Goncalves, Ewan Harrison, David K. Jackson, Ian Johnston, Dominic Kwiatkowski, Cordelia Langford, John Sillitoe on behalf of the Wellcome Sanger Institute COVID-19 Surveillance Team |
| EPI_ISL_550176, EPI_ISL_550177, EPI_ISL_550178, EPI_ISL_550180, EPI_ISL_550181, EPI_ISL_550182, EPI_ISL_550183, EPI_ISL_550184 | Lighthouse Lab in Milton Keynes | Wellcome Sanger Institute for the COVID-19 Genomics UK (COG-UK) consortium | The Lighthouse Lab in Milton Keynes and Alex Alderton, Roberto Amato, Sonia Goncalves, Ewan Harrison, David K. Jackson, Ian Johnston, Dominic Kwiatkowski, Cordelia Langford, John Sillitoe on behalf of the Wellcome Sanger Institute COVID-19 Surveillance Team (http://www.sanger.ac.uk/covid-team) |
| EPI_ISL_550185, EPI_ISL_550186, EPI_ISL_550187 | Lighthouse Lab in Milton Keynes | Wellcome Sanger Institute for the COVID-19 Genomics UK (COG-UK) consortium | The Lighthouse Lab in Alderley Park and Alex Alderton, Roberto Amato, Sonia Goncalves, Ewan Harrison, David K. Jackson, Ian Johnston, Dominic Kwiatkowski, Cordelia Langford, John Sillitoe on behalf of the Wellcome Sanger Institute COVID-19 Surveillance Team |
| EPI_ISL_550188, EPI_ISL_550189, EPI_ISL_550190, EPI_ISL_550192, EPI_ISL_550193, EPI_ISL_550194, EPI_ISL_550195, EPI_ISL_550196, EPI_ISL_550197, EPI_ISL_550198, EPI_ISL_550199, EPI_ISL_550200, EPI_ISL_550201, EPI_ISL_550202, EPI_ISL_550203, EPI_ISL_550204, EPI_ISL_550205, EPI_ISL_550206, EPI_ISL_550207, EPI_ISL_550208, EPI_ISL_550209, EPI_ISL_550210, EPI_ISL_550211, EPI_ISL_550212, EPI_ISL_550213, EPI_ISL_550214, EPI_ISL_550215, EPI_ISL_550216, EPI_ISL_550217, EPI_ISL_550218, EPI_ISL_550219, EPI_ISL_550220, EPI_ISL_550221, EPI_ISL_550222, EPI_ISL_550223, EPI_ISL_550224, EPI_ISL_550225, EPI_ISL_550226, EPI_ISL_550227, EPI_ISL_550228, EPI_ISL_550229, EPI_ISL_550230, EPI_ISL_550231, EPI_ISL_550232, EPI_ISL_550233, EPI_ISL_550234, EPI_ISL_550235, EPI_ISL_550236, EPI_ISL_550237, EPI_ISL_550238, EPI_ISL_550239, EPI_ISL_550240, EPI_ISL_550241, EPI_ISL_550242 | | | |
| see above | Lighthouse Lab in Milton Keynes | Wellcome Sanger Institute for the COVID-19 Genomics UK (COG-UK) consortium | The Lighthouse Lab in Milton Keynes and Alex Alderton, Roberto Amato, Sonia Goncalves, Ewan Harrison, David K. Jackson, Ian Johnston, Dominic Kwiatkowski, Cordelia Langford, John Sillitoe on behalf of the Wellcome Sanger Institute COVID-19 Surveillance Team (http://www.sanger.ac.uk/covid-team) |
| EPI_ISL_550244 | Lighthouse Lab in Milton Keynes | Wellcome Sanger Institute for the COVID-19 Genomics UK | The Lighthouse Lab in Alderley Park and Alex Alderton, Roberto Amato, Sonia Goncalves, Ewan Harrison, David K. Jackson, Ian Johnston, Dominic |

| | | | |
|---|---|---|---|
| | | (COG-UK) consortium | Kwiatkowski, Cordelia Langford, John Sillitoe on behalf of the Wellcome Sanger Institute COVID-19 Surveillance Team |
| EPI_ISL_550245, EPI_ISL_550246, EPI_ISL_550247, EPI_ISL_550248, EPI_ISL_550249, EPI_ISL_550250 | Lighthouse Lab in Milton Keynes | Wellcome Sanger Institute for the COVID-19 Genomics UK (COG-UK) consortium | The Lighthouse Lab in Milton Keynes and Alex Alderton, Roberto Amato, Sonia Goncalves, Ewan Harrison, David K. Jackson, Ian Johnston, Dominic Kwiatkowski, Cordelia Langford, John Sillitoe on behalf of the Wellcome Sanger Institute COVID-19 Surveillance Team (http://www.sanger.ac.uk/covid-team) |
| EPI_ISL_550251 | Lighthouse Lab in Milton Keynes | Wellcome Sanger Institute for the COVID-19 Genomics UK (COG-UK) consortium | The Lighthouse Lab in Alderley Park and Alex Alderton, Roberto Amato, Sonia Goncalves, Ewan Harrison, David K. Jackson, Ian Johnston, Dominic Kwiatkowski, Cordelia Langford, John Sillitoe on behalf of the Wellcome Sanger Institute COVID-19 Surveillance Team |
| EPI_ISL_550252, EPI_ISL_550253, EPI_ISL_550254, EPI_ISL_550255, EPI_ISL_550256 | Lighthouse Lab in Milton Keynes | Wellcome Sanger Institute for the COVID-19 Genomics UK (COG-UK) consortium | The Lighthouse Lab in Milton Keynes and Alex Alderton, Roberto Amato, Sonia Goncalves, Ewan Harrison, David K. Jackson, Ian Johnston, Dominic Kwiatkowski, Cordelia Langford, John Sillitoe on behalf of the Wellcome Sanger Institute COVID-19 Surveillance Team (http://www.sanger.ac.uk/covid-team) |
| EPI_ISL_550257 | Lighthouse Lab in Milton Keynes | Wellcome Sanger Institute for the COVID-19 Genomics UK (COG-UK) consortium | The Lighthouse Lab in Alderley Park and Alex Alderton, Roberto Amato, Sonia Goncalves, Ewan Harrison, David K. Jackson, Ian Johnston, Dominic Kwiatkowski, Cordelia Langford, John Sillitoe on behalf of the Wellcome Sanger Institute COVID-19 Surveillance Team |
| EPI_ISL_550258, EPI_ISL_550259, EPI_ISL_550260, EPI_ISL_550261, EPI_ISL_550262, EPI_ISL_550263, EPI_ISL_550264, EPI_ISL_550265, EPI_ISL_550266, EPI_ISL_550267, EPI_ISL_550268, EPI_ISL_550269, EPI_ISL_550270, EPI_ISL_550271, EPI_ISL_550272, EPI_ISL_550273, EPI_ISL_550274, EPI_ISL_550275, EPI_ISL_550276 | | | |
| see above | Lighthouse Lab in Milton Keynes | Wellcome Sanger Institute for the COVID-19 Genomics UK (COG-UK) consortium | The Lighthouse Lab in Milton Keynes and Alex Alderton, Roberto Amato, Sonia Goncalves, Ewan Harrison, David K. Jackson, Ian Johnston, Dominic Kwiatkowski, Cordelia Langford, John Sillitoe on behalf of the Wellcome Sanger Institute COVID-19 Surveillance Team (http://www.sanger.ac.uk/covid-team) |
| EPI_ISL_550277 | Lighthouse Lab in Milton Keynes | Wellcome Sanger Institute for the COVID-19 Genomics UK (COG-UK) consortium | The Lighthouse Lab in Alderley Park and Alex Alderton, Roberto Amato, Sonia Goncalves, Ewan Harrison, David K. Jackson, Ian Johnston, Dominic Kwiatkowski, Cordelia Langford, John Sillitoe on behalf of the Wellcome Sanger Institute COVID-19 Surveillance Team |
| EPI_ISL_550278, EPI_ISL_550279, EPI_ISL_550280, EPI_ISL_550281, EPI_ISL_550282, EPI_ISL_550283, EPI_ISL_550284, EPI_ISL_550285, EPI_ISL_550286, EPI_ISL_550287, EPI_ISL_550288, EPI_ISL_550289, EPI_ISL_550290, EPI_ISL_550291, EPI_ISL_550292, EPI_ISL_550293, EPI_ISL_550294, EPI_ISL_550295, EPI_ISL_550296, EPI_ISL_550297, EPI_ISL_550298, EPI_ISL_550301, EPI_ISL_550302, EPI_ISL_550303, EPI_ISL_550304, EPI_ISL_550305, EPI_ISL_550306, EPI_ISL_550307, EPI_ISL_550308, EPI_ISL_550309, EPI_ISL_550310, EPI_ISL_550311, EPI_ISL_550313, EPI_ISL_550314, EPI_ISL_550315, EPI_ISL_550316, EPI_ISL_550317, EPI_ISL_550318 | | | |
| see above | Lighthouse Lab in Milton Keynes | Wellcome Sanger Institute for the COVID-19 Genomics UK (COG-UK) consortium | The Lighthouse Lab in Milton Keynes and Alex Alderton, Roberto Amato, Sonia Goncalves, Ewan Harrison, David K. Jackson, Ian Johnston, Dominic Kwiatkowski, Cordelia Langford, John Sillitoe on behalf of the Wellcome Sanger Institute COVID-19 Surveillance Team (http://www.sanger.ac.uk/covid-team) |
| EPI_ISL_550319 | Lighthouse Lab in Milton Keynes | Wellcome Sanger Institute for the COVID-19 Genomics UK (COG-UK) consortium | The Lighthouse Lab in Alderley Park and Alex Alderton, Roberto Amato, Sonia Goncalves, Ewan Harrison, David K. Jackson, Ian Johnston, Dominic Kwiatkowski, Cordelia Langford, John Sillitoe on behalf of the Wellcome Sanger Institute COVID-19 Surveillance Team |
| EPI_ISL_550320, EPI_ISL_550321, EPI_ISL_550322, EPI_ISL_550323, EPI_ISL_550324, EPI_ISL_550326, EPI_ISL_550327 | Lighthouse Lab in Milton Keynes | Wellcome Sanger Institute for the COVID-19 Genomics UK (COG-UK) consortium | The Lighthouse Lab in Milton Keynes and Alex Alderton, Roberto Amato, Sonia Goncalves, Ewan Harrison, David K. Jackson, Ian Johnston, Dominic Kwiatkowski, Cordelia Langford, John Sillitoe on behalf of the Wellcome Sanger Institute COVID-19 Surveillance Team (http://www.sanger.ac.uk/covid-team) |
| EPI_ISL_550328 | Lighthouse Lab in Milton Keynes | Wellcome Sanger Institute for the COVID-19 Genomics UK (COG-UK) consortium | The Lighthouse Lab in Alderley Park and Alex Alderton, Roberto Amato, Sonia Goncalves, Ewan Harrison, David K. Jackson, Ian Johnston, Dominic Kwiatkowski, Cordelia Langford, John Sillitoe on behalf of the Wellcome Sanger Institute COVID-19 Surveillance Team |
| EPI_ISL_550329, EPI_ISL_550330, EPI_ISL_550331, EPI_ISL_550332, EPI_ISL_550333, EPI_ISL_550334, EPI_ISL_550335, EPI_ISL_550336, EPI_ISL_550337, EPI_ISL_550338, EPI_ISL_550339, EPI_ISL_550340, EPI_ISL_550342, EPI_ISL_550343, EPI_ISL_550344, EPI_ISL_550345, EPI_ISL_550347, EPI_ISL_550348, EPI_ISL_550349, EPI_ISL_550350, EPI_ISL_550351, EPI_ISL_550352, EPI_ISL_550354, EPI_ISL_550355, EPI_ISL_550356 | | | |
| see above | Lighthouse Lab in Milton Keynes | Wellcome Sanger Institute for the COVID-19 Genomics UK (COG-UK) consortium | The Lighthouse Lab in Milton Keynes and Alex Alderton, Roberto Amato, Sonia Goncalves, Ewan Harrison, David K. Jackson, Ian Johnston, Dominic Kwiatkowski, Cordelia Langford, John Sillitoe on behalf of the Wellcome Sanger Institute COVID-19 Surveillance Team (http://www.sanger.ac.uk/covid-team) |
| EPI_ISL_550357 | Lighthouse Lab in Milton Keynes | Wellcome Sanger Institute for the COVID-19 Genomics UK (COG-UK) consortium | The Lighthouse Lab in Alderley Park and Alex Alderton, Roberto Amato, Sonia Goncalves, Ewan Harrison, David K. Jackson, Ian Johnston, Dominic Kwiatkowski, Cordelia Langford, John Sillitoe on behalf of the Wellcome Sanger Institute COVID-19 Surveillance Team |
| EPI_ISL_550359, EPI_ISL_550360, EPI_ISL_550361, EPI_ISL_550362, EPI_ISL_550363 | Lighthouse Lab in Milton Keynes | Wellcome Sanger Institute for the COVID-19 Genomics UK (COG-UK) consortium | The Lighthouse Lab in Milton Keynes and Alex Alderton, Roberto Amato, Sonia Goncalves, Ewan Harrison, David K. Jackson, Ian Johnston, Dominic Kwiatkowski, Cordelia Langford, John Sillitoe on behalf of the Wellcome Sanger Institute COVID-19 Surveillance Team (http://www.sanger.ac.uk/covid-team) |
| EPI_ISL_550364 | Lighthouse Lab in Milton Keynes | Wellcome Sanger Institute for the COVID-19 Genomics UK (COG-UK) consortium | The Lighthouse Lab in Alderley Park and Alex Alderton, Roberto Amato, Sonia Goncalves, Ewan Harrison, David K. Jackson, Ian Johnston, Dominic Kwiatkowski, Cordelia Langford, John Sillitoe on behalf of the Wellcome Sanger Institute COVID-19 Surveillance Team |
| EPI_ISL_550365, EPI_ISL_550366, EPI_ISL_550367, EPI_ISL_550368, EPI_ISL_550369, EPI_ISL_550371, EPI_ISL_550372 | Lighthouse Lab in Milton Keynes | Wellcome Sanger Institute for the COVID-19 Genomics UK (COG-UK) consortium | The Lighthouse Lab in Milton Keynes and Alex Alderton, Roberto Amato, Sonia Goncalves, Ewan Harrison, David K. Jackson, Ian Johnston, Dominic Kwiatkowski, Cordelia Langford, John Sillitoe on behalf of the Wellcome Sanger Institute COVID-19 Surveillance Team (http://www.sanger.ac.uk/covid-team) |
| EPI_ISL_550373 | Lighthouse Lab in Milton Keynes | Wellcome Sanger Institute for the COVID-19 Genomics UK (COG-UK) consortium | The Lighthouse Lab in Alderley Park and Alex Alderton, Roberto Amato, Sonia Goncalves, Ewan Harrison, David K. Jackson, Ian Johnston, Dominic Kwiatkowski, Cordelia Langford, John Sillitoe on behalf of the Wellcome Sanger Institute COVID-19 Surveillance Team |
| EPI_ISL_550374, EPI_ISL_550375, EPI_ISL_550376, EPI_ISL_550377, EPI_ISL_550379, EPI_ISL_550380, EPI_ISL_550381, EPI_ISL_550382, EPI_ISL_550383, EPI_ISL_550384, EPI_ISL_550385, EPI_ISL_550386, EPI_ISL_550387, EPI_ISL_550388, EPI_ISL_550389, EPI_ISL_550390, EPI_ISL_550391, EPI_ISL_550392, EPI_ISL_550393, EPI_ISL_550394, EPI_ISL_550395 | | | |
| see above | Lighthouse Lab in Milton Keynes | Wellcome Sanger Institute for the COVID-19 Genomics UK (COG-UK) consortium | The Lighthouse Lab in Milton Keynes and Alex Alderton, Roberto Amato, Sonia Goncalves, Ewan Harrison, David K. Jackson, Ian Johnston, Dominic Kwiatkowski, Cordelia Langford, John Sillitoe on behalf of the Wellcome Sanger Institute COVID-19 Surveillance Team (http://www.sanger.ac.uk/covid-team) |
| EPI_ISL_550396, EPI_ISL_550397 | Lighthouse Lab in Milton Keynes | Wellcome Sanger Institute for the COVID-19 Genomics UK (COG-UK) consortium | The Lighthouse Lab in Alderley Park and Alex Alderton, Roberto Amato, Sonia Goncalves, Ewan Harrison, David K. Jackson, Ian Johnston, Dominic Kwiatkowski, Cordelia Langford, John Sillitoe on behalf of the Wellcome Sanger Institute COVID-19 Surveillance Team |
| EPI_ISL_550398, EPI_ISL_550399 | Lighthouse Lab in Milton Keynes | Wellcome Sanger Institute for the COVID-19 Genomics UK (COG-UK) consortium | The Lighthouse Lab in Milton Keynes and Alex Alderton, Roberto Amato, Sonia Goncalves, Ewan Harrison, David K. Jackson, Ian Johnston, Dominic Kwiatkowski, Cordelia Langford, John Sillitoe on behalf of the Wellcome Sanger Institute COVID-19 Surveillance Team (http://www.sanger.ac.uk/covid-team) |
| EPI_ISL_550400 | Lighthouse Lab in Milton Keynes | Wellcome Sanger Institute for the COVID-19 Genomics UK (COG-UK) consortium | The Lighthouse Lab in Alderley Park and Alex Alderton, Roberto Amato, Sonia Goncalves, Ewan Harrison, David K. Jackson, Ian Johnston, Dominic Kwiatkowski, Cordelia Langford, John Sillitoe on behalf of the Wellcome Sanger Institute COVID-19 Surveillance Team |
| EPI_ISL_550401, EPI_ISL_550402, EPI_ISL_550403, EPI_ISL_550404, EPI_ISL_550405, EPI_ISL_550406, EPI_ISL_550407, EPI_ISL_550408, EPI_ISL_550409, EPI_ISL_550410 | Lighthouse Lab in Milton Keynes | Wellcome Sanger Institute for the COVID-19 Genomics UK (COG-UK) consortium | The Lighthouse Lab in Milton Keynes and Alex Alderton, Roberto Amato, Sonia Goncalves, Ewan Harrison, David K. Jackson, Ian Johnston, Dominic Kwiatkowski, Cordelia Langford, John Sillitoe on behalf of the Wellcome Sanger Institute COVID-19 Surveillance Team (http://www.sanger.ac.uk/covid-team) |
| EPI_ISL_550411 | Lighthouse Lab in Milton Keynes | Wellcome Sanger Institute for the COVID-19 Genomics UK (COG-UK) consortium | The Lighthouse Lab in Alderley Park and Alex Alderton, Roberto Amato, Sonia Goncalves, Ewan Harrison, David K. Jackson, Ian Johnston, Dominic Kwiatkowski, Cordelia Langford, John Sillitoe on behalf of the Wellcome Sanger Institute COVID-19 Surveillance Team |
| EPI_ISL_550412, EPI_ISL_550413, EPI_ISL_550414, EPI_ISL_550415, EPI_ISL_550416, EPI_ISL_550417, EPI_ISL_550418, EPI_ISL_550419, EPI_ISL_550420 | Lighthouse Lab in Milton Keynes | Wellcome Sanger Institute for the COVID-19 Genomics UK (COG-UK) consortium | The Lighthouse Lab in Milton Keynes and Alex Alderton, Roberto Amato, Sonia Goncalves, Ewan Harrison, David K. Jackson, Ian Johnston, Dominic Kwiatkowski, Cordelia Langford, John Sillitoe on behalf of the Wellcome Sanger Institute COVID-19 Surveillance Team (http://www.sanger.ac.uk/covid-team) |

| | | | |
|---|---|---|---|
| EPI_ISL_550421 | Lighthouse Lab in Milton Keynes | Wellcome Sanger Institute for the COVID-19 Genomics UK (COG-UK) consortium | The Lighthouse Lab in Alderley Park and Alex Alderton, Roberto Amato, Sonia Goncalves, Ewan Harrison, David K. Jackson, Ian Johnston, Dominic Kwiatkowski, Cordelia Langford, John Sillitoe on behalf of the Wellcome Sanger Institute COVID-19 Surveillance Team |
| EPI_ISL_550422, EPI_ISL_550423, EPI_ISL_550424, EPI_ISL_550425, EPI_ISL_550426, EPI_ISL_550427, EPI_ISL_550428, EPI_ISL_550429, EPI_ISL_550430, EPI_ISL_550431, EPI_ISL_550432, EPI_ISL_550434, EPI_ISL_550435, EPI_ISL_550436, EPI_ISL_550437, EPI_ISL_550438, EPI_ISL_550439, EPI_ISL_550440, EPI_ISL_550441, EPI_ISL_550442, EPI_ISL_550443, EPI_ISL_550444, EPI_ISL_550445, EPI_ISL_550446, EPI_ISL_550447, EPI_ISL_550448, EPI_ISL_550449, EPI_ISL_550450, EPI_ISL_550451, EPI_ISL_550452, EPI_ISL_550453, EPI_ISL_550454 | | | |
| see above | Lighthouse Lab in Milton Keynes | Wellcome Sanger Institute for the COVID-19 Genomics UK (COG-UK) consortium | The Lighthouse Lab in Milton Keynes and Alex Alderton, Roberto Amato, Sonia Goncalves, Ewan Harrison, David K. Jackson, Ian Johnston, Dominic Kwiatkowski, Cordelia Langford, John Sillitoe on behalf of the Wellcome Sanger Institute COVID-19 Surveillance Team (http://www.sanger.ac.uk/covid-team) |
| EPI_ISL_550455 | Lighthouse Lab in Milton Keynes | Wellcome Sanger Institute for the COVID-19 Genomics UK (COG-UK) consortium | The Lighthouse Lab in Alderley Park and Alex Alderton, Roberto Amato, Sonia Goncalves, Ewan Harrison, David K. Jackson, Ian Johnston, Dominic Kwiatkowski, Cordelia Langford, John Sillitoe on behalf of the Wellcome Sanger Institute COVID-19 Surveillance Team |
| EPI_ISL_550456, EPI_ISL_550457 | Lighthouse Lab in Milton Keynes | Wellcome Sanger Institute for the COVID-19 Genomics UK (COG-UK) consortium | The Lighthouse Lab in Milton Keynes and Alex Alderton, Roberto Amato, Sonia Goncalves, Ewan Harrison, David K. Jackson, Ian Johnston, Dominic Kwiatkowski, Cordelia Langford, John Sillitoe on behalf of the Wellcome Sanger Institute COVID-19 Surveillance Team (http://www.sanger.ac.uk/covid-team) |
| EPI_ISL_550458 | Lighthouse Lab in Milton Keynes | Wellcome Sanger Institute for the COVID-19 Genomics UK (COG-UK) consortium | The Lighthouse Lab in Alderley Park and Alex Alderton, Roberto Amato, Sonia Goncalves, Ewan Harrison, David K. Jackson, Ian Johnston, Dominic Kwiatkowski, Cordelia Langford, John Sillitoe on behalf of the Wellcome Sanger Institute COVID-19 Surveillance Team |
| EPI_ISL_550459, EPI_ISL_550460, EPI_ISL_550461, EPI_ISL_550462, EPI_ISL_550463, EPI_ISL_550464, EPI_ISL_550465, EPI_ISL_550466, EPI_ISL_550467, EPI_ISL_550468, EPI_ISL_550469, EPI_ISL_550470, EPI_ISL_550471, EPI_ISL_550472, EPI_ISL_550473, EPI_ISL_550474, EPI_ISL_550475, EPI_ISL_550477, EPI_ISL_550478, EPI_ISL_550479, EPI_ISL_550481, EPI_ISL_550483, EPI_ISL_550484, EPI_ISL_550486, EPI_ISL_550487, EPI_ISL_550488, EPI_ISL_550489, EPI_ISL_550490, EPI_ISL_550491, EPI_ISL_550492, EPI_ISL_550494, EPI_ISL_550495, EPI_ISL_550496, EPI_ISL_550497, EPI_ISL_550498, EPI_ISL_550499, EPI_ISL_550500, EPI_ISL_550501, EPI_ISL_550502, EPI_ISL_550503, EPI_ISL_550504, EPI_ISL_550505, EPI_ISL_550506, EPI_ISL_550507, EPI_ISL_550508, EPI_ISL_550509, EPI_ISL_550510, EPI_ISL_550511, EPI_ISL_550513, EPI_ISL_550514, EPI_ISL_550515, EPI_ISL_550516, EPI_ISL_550517, EPI_ISL_550518, EPI_ISL_550519, EPI_ISL_550520, EPI_ISL_550521, EPI_ISL_550522, EPI_ISL_550523, EPI_ISL_550524, EPI_ISL_550525, EPI_ISL_550526, EPI_ISL_550527, EPI_ISL_550528, EPI_ISL_550529, EPI_ISL_550530, EPI_ISL_550531, EPI_ISL_550532, EPI_ISL_550533, EPI_ISL_550534 | | | |
| see above | Lighthouse Lab in Milton Keynes | Wellcome Sanger Institute for the COVID-19 Genomics UK (COG-UK) consortium | The Lighthouse Lab in Milton Keynes and Alex Alderton, Roberto Amato, Sonia Goncalves, Ewan Harrison, David K. Jackson, Ian Johnston, Dominic Kwiatkowski, Cordelia Langford, John Sillitoe on behalf of the Wellcome Sanger Institute COVID-19 Surveillance Team (http://www.sanger.ac.uk/covid-team) |
| EPI_ISL_550535, EPI_ISL_550536 | Lighthouse Lab in Milton Keynes | Wellcome Sanger Institute for the COVID-19 Genomics UK (COG-UK) consortium | The Lighthouse Lab in Alderley Park and Alex Alderton, Roberto Amato, Sonia Goncalves, Ewan Harrison, David K. Jackson, Ian Johnston, Dominic Kwiatkowski, Cordelia Langford, John Sillitoe on behalf of the Wellcome Sanger Institute COVID-19 Surveillance Team |
| EPI_ISL_550537, EPI_ISL_550539, EPI_ISL_550540, EPI_ISL_550541, EPI_ISL_550542, EPI_ISL_550543, EPI_ISL_550544, EPI_ISL_550545, EPI_ISL_550546, EPI_ISL_550547, EPI_ISL_550548, EPI_ISL_550549, EPI_ISL_550550 | | | |
| see above | Lighthouse Lab in Milton Keynes | Wellcome Sanger Institute for the COVID-19 Genomics UK (COG-UK) consortium | The Lighthouse Lab in Milton Keynes and Alex Alderton, Roberto Amato, Sonia Goncalves, Ewan Harrison, David K. Jackson, Ian Johnston, Dominic Kwiatkowski, Cordelia Langford, John Sillitoe on behalf of the Wellcome Sanger Institute COVID-19 Surveillance Team (http://www.sanger.ac.uk/covid-team) |
| EPI_ISL_550551, EPI_ISL_550552 | Lighthouse Lab in Milton Keynes | Wellcome Sanger Institute for the COVID-19 Genomics UK (COG-UK) consortium | The Lighthouse Lab in Alderley Park and Alex Alderton, Roberto Amato, Sonia Goncalves, Ewan Harrison, David K. Jackson, Ian Johnston, Dominic Kwiatkowski, Cordelia Langford, John Sillitoe on behalf of the Wellcome Sanger Institute COVID-19 Surveillance Team |
| EPI_ISL_550553, EPI_ISL_550554, EPI_ISL_550555, EPI_ISL_550556, EPI_ISL_550557, EPI_ISL_550558, EPI_ISL_550559, EPI_ISL_550560, EPI_ISL_550561, EPI_ISL_550563, EPI_ISL_550564, EPI_ISL_550565, EPI_ISL_550566, EPI_ISL_550567, EPI_ISL_550568, EPI_ISL_550569, EPI_ISL_550570, EPI_ISL_550571, EPI_ISL_550572, EPI_ISL_550573, EPI_ISL_550574 | | | |
| see above | Lighthouse Lab in Milton Keynes | Wellcome Sanger Institute for the COVID-19 Genomics UK (COG-UK) consortium | The Lighthouse Lab in Milton Keynes and Alex Alderton, Roberto Amato, Sonia Goncalves, Ewan Harrison, David K. Jackson, Ian Johnston, Dominic Kwiatkowski, Cordelia Langford, John Sillitoe on behalf of the Wellcome Sanger Institute COVID-19 Surveillance Team (http://www.sanger.ac.uk/covid-team) |
| EPI_ISL_550575 | Lighthouse Lab in Milton Keynes | Wellcome Sanger Institute for the COVID-19 Genomics UK (COG-UK) consortium | The Lighthouse Lab in Alderley Park and Alex Alderton, Roberto Amato, Sonia Goncalves, Ewan Harrison, David K. Jackson, Ian Johnston, Dominic Kwiatkowski, Cordelia Langford, John Sillitoe on behalf of the Wellcome Sanger Institute COVID-19 Surveillance Team |
| EPI_ISL_550576, EPI_ISL_550577, EPI_ISL_550578, EPI_ISL_550579, EPI_ISL_550580, EPI_ISL_550581, EPI_ISL_550582, EPI_ISL_550583, EPI_ISL_550584, EPI_ISL_550585, EPI_ISL_550586 | | | |
| see above | Lighthouse Lab in Milton Keynes | Wellcome Sanger Institute for the COVID-19 Genomics UK (COG-UK) consortium | The Lighthouse Lab in Milton Keynes and Alex Alderton, Roberto Amato, Sonia Goncalves, Ewan Harrison, David K. Jackson, Ian Johnston, Dominic Kwiatkowski, Cordelia Langford, John Sillitoe on behalf of the Wellcome Sanger Institute COVID-19 Surveillance Team (http://www.sanger.ac.uk/covid-team) |
| EPI_ISL_550587 | Lighthouse Lab in Milton Keynes | Wellcome Sanger Institute for the COVID-19 Genomics UK (COG-UK) consortium | The Lighthouse Lab in Alderley Park and Alex Alderton, Roberto Amato, Sonia Goncalves, Ewan Harrison, David K. Jackson, Ian Johnston, Dominic Kwiatkowski, Cordelia Langford, John Sillitoe on behalf of the Wellcome Sanger Institute COVID-19 Surveillance Team |
| EPI_ISL_550588, EPI_ISL_550589, EPI_ISL_550591, EPI_ISL_550592, EPI_ISL_550595, EPI_ISL_550596, EPI_ISL_550597, EPI_ISL_550598, EPI_ISL_550599, EPI_ISL_550600, EPI_ISL_550601, EPI_ISL_550602, EPI_ISL_550604, EPI_ISL_550605, EPI_ISL_550606, EPI_ISL_550607, EPI_ISL_550608, EPI_ISL_550609, EPI_ISL_550610, EPI_ISL_550611, EPI_ISL_550612, EPI_ISL_550613 | | | |
| see above | Lighthouse Lab in Milton Keynes | Wellcome Sanger Institute for the COVID-19 Genomics UK (COG-UK) consortium | The Lighthouse Lab in Milton Keynes and Alex Alderton, Roberto Amato, Sonia Goncalves, Ewan Harrison, David K. Jackson, Ian Johnston, Dominic Kwiatkowski, Cordelia Langford, John Sillitoe on behalf of the Wellcome Sanger Institute COVID-19 Surveillance Team (http://www.sanger.ac.uk/covid-team) |
| EPI_ISL_550614 | Lighthouse Lab in Milton Keynes | Wellcome Sanger Institute for the COVID-19 Genomics UK (COG-UK) consortium | The Lighthouse Lab in Alderley Park and Alex Alderton, Roberto Amato, Sonia Goncalves, Ewan Harrison, David K. Jackson, Ian Johnston, Dominic Kwiatkowski, Cordelia Langford, John Sillitoe on behalf of the Wellcome Sanger Institute COVID-19 Surveillance Team |
| EPI_ISL_550615, EPI_ISL_550616, EPI_ISL_550617, EPI_ISL_550618, EPI_ISL_550619, EPI_ISL_550620, EPI_ISL_550621, EPI_ISL_550622, EPI_ISL_550623, EPI_ISL_550624, EPI_ISL_550625, EPI_ISL_550626, EPI_ISL_550627, EPI_ISL_550628, EPI_ISL_550629, EPI_ISL_550630, EPI_ISL_550631, EPI_ISL_550632, EPI_ISL_550633, EPI_ISL_550634, EPI_ISL_550635, EPI_ISL_550636, EPI_ISL_550637 | | | |
| see above | Lighthouse Lab in Milton Keynes | Wellcome Sanger Institute for the COVID-19 Genomics UK (COG-UK) consortium | The Lighthouse Lab in Milton Keynes and Alex Alderton, Roberto Amato, Sonia Goncalves, Ewan Harrison, David K. Jackson, Ian Johnston, Dominic Kwiatkowski, Cordelia Langford, John Sillitoe on behalf of the Wellcome Sanger Institute COVID-19 Surveillance Team (http://www.sanger.ac.uk/covid-team) |
| EPI_ISL_550638 | Lighthouse Lab in Milton Keynes | Wellcome Sanger Institute for the COVID-19 Genomics UK (COG-UK) consortium | The Lighthouse Lab in Alderley Park and Alex Alderton, Roberto Amato, Sonia Goncalves, Ewan Harrison, David K. Jackson, Ian Johnston, Dominic Kwiatkowski, Cordelia Langford, John Sillitoe on behalf of the Wellcome Sanger Institute COVID-19 Surveillance Team |
| EPI_ISL_550639, EPI_ISL_550640, EPI_ISL_550642, EPI_ISL_550643, EPI_ISL_550645 | Lighthouse Lab in Milton Keynes | Wellcome Sanger Institute for the COVID-19 Genomics UK (COG-UK) consortium | The Lighthouse Lab in Milton Keynes and Alex Alderton, Roberto Amato, Sonia Goncalves, Ewan Harrison, David K. Jackson, Ian Johnston, Dominic Kwiatkowski, Cordelia Langford, John Sillitoe on behalf of the Wellcome Sanger Institute COVID-19 Surveillance Team (http://www.sanger.ac.uk/covid-team) |
| EPI_ISL_550646 | Lighthouse Lab in Milton Keynes | Wellcome Sanger Institute for the COVID-19 Genomics UK (COG-UK) consortium | The Lighthouse Lab in Alderley Park and Alex Alderton, Roberto Amato, Sonia Goncalves, Ewan Harrison, David K. Jackson, Ian Johnston, Dominic Kwiatkowski, Cordelia Langford, John Sillitoe on behalf of the Wellcome Sanger Institute COVID-19 Surveillance Team |
| EPI_ISL_550647, EPI_ISL_550648, EPI_ISL_550649, EPI_ISL_550650, EPI_ISL_550651, EPI_ISL_550652, EPI_ISL_550653, EPI_ISL_550654, EPI_ISL_550655, EPI_ISL_550656, EPI_ISL_550657, EPI_ISL_550658, EPI_ISL_550659, EPI_ISL_550661, EPI_ISL_550662, EPI_ISL_550663, EPI_ISL_550664 | | | |
| see above | Lighthouse Lab in Milton Keynes | Wellcome Sanger Institute for the COVID-19 Genomics UK (COG-UK) consortium | The Lighthouse Lab in Milton Keynes and Alex Alderton, Roberto Amato, Sonia Goncalves, Ewan Harrison, David K. Jackson, Ian Johnston, Dominic Kwiatkowski, Cordelia Langford, John Sillitoe on behalf of the Wellcome Sanger Institute COVID-19 Surveillance Team (http://www.sanger.ac.uk/covid-team) |
| EPI_ISL_550665 | Lighthouse Lab in Milton Keynes | Wellcome Sanger Institute for the COVID-19 Genomics UK (COG-UK) consortium | The Lighthouse Lab in Alderley Park and Alex Alderton, Roberto Amato, Sonia Goncalves, Ewan Harrison, David K. Jackson, Ian Johnston, Dominic Kwiatkowski, Cordelia Langford, John Sillitoe on behalf of the Wellcome Sanger Institute COVID-19 Surveillance Team |
| EPI_ISL_550666, EPI_ISL_550667, EPI_ISL_550668, EPI_ISL_550669, EPI_ISL_550670, EPI_ISL_550671, EPI_ISL_550672 | Lighthouse Lab in Milton Keynes | Wellcome Sanger Institute for the COVID-19 Genomics UK (COG-UK) consortium | The Lighthouse Lab in Milton Keynes and Alex Alderton, Roberto Amato, Sonia Goncalves, Ewan Harrison, David K. Jackson, Ian Johnston, Dominic Kwiatkowski, Cordelia Langford, John Sillitoe on behalf of the Wellcome Sanger Institute COVID-19 Surveillance Team (http://www.sanger.ac.uk/covid-team) |
| EPI_ISL_550673 | Lighthouse Lab in Milton Keynes | Wellcome Sanger Institute for the COVID-19 Genomics UK | The Lighthouse Lab in Alderley Park and Alex Alderton, Roberto Amato, Sonia Goncalves, Ewan Harrison, David K. Jackson, Ian Johnston, Dominic |

| | | | |
|---|---|---|---|
| | | (COG-UK) consortium | Kwiatkowski, Cordelia Langford, John Sillitoe on behalf of the Wellcome Sanger Institute COVID-19 Surveillance Team |
| EPI_ISL_550674 | Lighthouse Lab in Milton Keynes | Wellcome Sanger Institute for the COVID-19 Genomics UK (COG-UK) consortium | The Lighthouse Lab in Milton Keynes and Alex Alderton, Roberto Amato, Sonia Goncalves, Ewan Harrison, David K. Jackson, Ian Johnston, Dominic Kwiatkowski, Cordelia Langford, John Sillitoe on behalf of the Wellcome Sanger Institute COVID-19 Surveillance Team (http://www.sanger.ac.uk/covid-team) |
| EPI_ISL_550675 | Lighthouse Lab in Milton Keynes | Wellcome Sanger Institute for the COVID-19 Genomics UK (COG-UK) consortium | The Lighthouse Lab in Alderley Park and Alex Alderton, Roberto Amato, Sonia Goncalves, Ewan Harrison, David K. Jackson, Ian Johnston, Dominic Kwiatkowski, Cordelia Langford, John Sillitoe on behalf of the Wellcome Sanger Institute COVID-19 Surveillance Team |
| EPI_ISL_550676, EPI_ISL_550678, EPI_ISL_550679, EPI_ISL_550680, EPI_ISL_550681 | Lighthouse Lab in Milton Keynes | Wellcome Sanger Institute for the COVID-19 Genomics UK (COG-UK) consortium | The Lighthouse Lab in Milton Keynes and Alex Alderton, Roberto Amato, Sonia Goncalves, Ewan Harrison, David K. Jackson, Ian Johnston, Dominic Kwiatkowski, Cordelia Langford, John Sillitoe on behalf of the Wellcome Sanger Institute COVID-19 Surveillance Team (http://www.sanger.ac.uk/covid-team) |
| EPI_ISL_550682 | Lighthouse Lab in Milton Keynes | Wellcome Sanger Institute for the COVID-19 Genomics UK (COG-UK) consortium | The Lighthouse Lab in Alderley Park and Alex Alderton, Roberto Amato, Sonia Goncalves, Ewan Harrison, David K. Jackson, Ian Johnston, Dominic Kwiatkowski, Cordelia Langford, John Sillitoe on behalf of the Wellcome Sanger Institute COVID-19 Surveillance Team |
| EPI_ISL_550683, EPI_ISL_550684, EPI_ISL_550685, EPI_ISL_550686, EPI_ISL_550687, EPI_ISL_550688, EPI_ISL_550689, EPI_ISL_550690, EPI_ISL_550691, EPI_ISL_550692, EPI_ISL_550693, EPI_ISL_550694, EPI_ISL_550695, EPI_ISL_550696, EPI_ISL_550697, EPI_ISL_550698, EPI_ISL_550699, EPI_ISL_550700, EPI_ISL_550701, EPI_ISL_550702, EPI_ISL_550703, EPI_ISL_550704, EPI_ISL_550705, EPI_ISL_550706, EPI_ISL_550707, EPI_ISL_550708, EPI_ISL_550709, EPI_ISL_550710 | | | |
| see above | Lighthouse Lab in Milton Keynes | Wellcome Sanger Institute for the COVID-19 Genomics UK (COG-UK) consortium | The Lighthouse Lab in Milton Keynes and Alex Alderton, Roberto Amato, Sonia Goncalves, Ewan Harrison, David K. Jackson, Ian Johnston, Dominic Kwiatkowski, Cordelia Langford, John Sillitoe on behalf of the Wellcome Sanger Institute COVID-19 Surveillance Team (http://www.sanger.ac.uk/covid-team) |
| EPI_ISL_550711 | Lighthouse Lab in Milton Keynes | Wellcome Sanger Institute for the COVID-19 Genomics UK (COG-UK) consortium | The Lighthouse Lab in Alderley Park and Alex Alderton, Roberto Amato, Sonia Goncalves, Ewan Harrison, David K. Jackson, Ian Johnston, Dominic Kwiatkowski, Cordelia Langford, John Sillitoe on behalf of the Wellcome Sanger Institute COVID-19 Surveillance Team |
| EPI_ISL_550712, EPI_ISL_550713, EPI_ISL_550714, EPI_ISL_550715, EPI_ISL_550716, EPI_ISL_550717, EPI_ISL_550718, EPI_ISL_550719, EPI_ISL_550720, EPI_ISL_550721, EPI_ISL_550722, EPI_ISL_550723, EPI_ISL_550725 | | | |
| see above | Lighthouse Lab in Milton Keynes | Wellcome Sanger Institute for the COVID-19 Genomics UK (COG-UK) consortium | The Lighthouse Lab in Milton Keynes and Alex Alderton, Roberto Amato, Sonia Goncalves, Ewan Harrison, David K. Jackson, Ian Johnston, Dominic Kwiatkowski, Cordelia Langford, John Sillitoe on behalf of the Wellcome Sanger Institute COVID-19 Surveillance Team (http://www.sanger.ac.uk/covid-team) |
| EPI_ISL_550726 | Lighthouse Lab in Milton Keynes | Wellcome Sanger Institute for the COVID-19 Genomics UK (COG-UK) consortium | The Lighthouse Lab in Alderley Park and Alex Alderton, Roberto Amato, Sonia Goncalves, Ewan Harrison, David K. Jackson, Ian Johnston, Dominic Kwiatkowski, Cordelia Langford, John Sillitoe on behalf of the Wellcome Sanger Institute COVID-19 Surveillance Team |
| EPI_ISL_550727, EPI_ISL_550728 | Lighthouse Lab in Milton Keynes | Wellcome Sanger Institute for the COVID-19 Genomics UK (COG-UK) consortium | The Lighthouse Lab in Milton Keynes and Alex Alderton, Roberto Amato, Sonia Goncalves, Ewan Harrison, David K. Jackson, Ian Johnston, Dominic Kwiatkowski, Cordelia Langford, John Sillitoe on behalf of the Wellcome Sanger Institute COVID-19 Surveillance Team (http://www.sanger.ac.uk/covid-team) |
| EPI_ISL_550729 | Lighthouse Lab in Milton Keynes | Wellcome Sanger Institute for the COVID-19 Genomics UK (COG-UK) consortium | The Lighthouse Lab in Alderley Park and Alex Alderton, Roberto Amato, Sonia Goncalves, Ewan Harrison, David K. Jackson, Ian Johnston, Dominic Kwiatkowski, Cordelia Langford, John Sillitoe on behalf of the Wellcome Sanger Institute COVID-19 Surveillance Team |
| EPI_ISL_550730, EPI_ISL_550731, EPI_ISL_550732, EPI_ISL_550733, EPI_ISL_550734, EPI_ISL_550735, EPI_ISL_550736, EPI_ISL_550737, EPI_ISL_550738, EPI_ISL_550740, EPI_ISL_550741, EPI_ISL_550742, EPI_ISL_550743, EPI_ISL_550744, EPI_ISL_550747, EPI_ISL_550748, EPI_ISL_550749 | | | |
| see above | Lighthouse Lab in Milton Keynes | Wellcome Sanger Institute for the COVID-19 Genomics UK (COG-UK) consortium | The Lighthouse Lab in Milton Keynes and Alex Alderton, Roberto Amato, Sonia Goncalves, Ewan Harrison, David K. Jackson, Ian Johnston, Dominic Kwiatkowski, Cordelia Langford, John Sillitoe on behalf of the Wellcome Sanger Institute COVID-19 Surveillance Team (http://www.sanger.ac.uk/covid-team) |
| EPI_ISL_550750 | Lighthouse Lab in Milton Keynes | Wellcome Sanger Institute for the COVID-19 Genomics UK (COG-UK) consortium | The Lighthouse Lab in Alderley Park and Alex Alderton, Roberto Amato, Sonia Goncalves, Ewan Harrison, David K. Jackson, Ian Johnston, Dominic Kwiatkowski, Cordelia Langford, John Sillitoe on behalf of the Wellcome Sanger Institute COVID-19 Surveillance Team |
| EPI_ISL_550751, EPI_ISL_550752, EPI_ISL_550753, EPI_ISL_550754, EPI_ISL_550755, EPI_ISL_550756, EPI_ISL_550757, EPI_ISL_550758, EPI_ISL_550759, EPI_ISL_550761, EPI_ISL_550762, EPI_ISL_550763, EPI_ISL_550764, EPI_ISL_550765, EPI_ISL_550766 | | | |
| see above | Lighthouse Lab in Milton Keynes | Wellcome Sanger Institute for the COVID-19 Genomics UK (COG-UK) consortium | The Lighthouse Lab in Milton Keynes and Alex Alderton, Roberto Amato, Sonia Goncalves, Ewan Harrison, David K. Jackson, Ian Johnston, Dominic Kwiatkowski, Cordelia Langford, John Sillitoe on behalf of the Wellcome Sanger Institute COVID-19 Surveillance Team (http://www.sanger.ac.uk/covid-team) |
| EPI_ISL_550767 | Lighthouse Lab in Milton Keynes | Wellcome Sanger Institute for the COVID-19 Genomics UK (COG-UK) consortium | The Lighthouse Lab in Alderley Park and Alex Alderton, Roberto Amato, Sonia Goncalves, Ewan Harrison, David K. Jackson, Ian Johnston, Dominic Kwiatkowski, Cordelia Langford, John Sillitoe on behalf of the Wellcome Sanger Institute COVID-19 Surveillance Team |
| EPI_ISL_550768, EPI_ISL_550769 | Lighthouse Lab in Milton Keynes | Wellcome Sanger Institute for the COVID-19 Genomics UK (COG-UK) consortium | The Lighthouse Lab in Milton Keynes and Alex Alderton, Roberto Amato, Sonia Goncalves, Ewan Harrison, David K. Jackson, Ian Johnston, Dominic Kwiatkowski, Cordelia Langford, John Sillitoe on behalf of the Wellcome Sanger Institute COVID-19 Surveillance Team (http://www.sanger.ac.uk/covid-team) |
| EPI_ISL_550770, EPI_ISL_550771 | Lighthouse Lab in Milton Keynes | Wellcome Sanger Institute for the COVID-19 Genomics UK (COG-UK) consortium | The Lighthouse Lab in Alderley Park and Alex Alderton, Roberto Amato, Sonia Goncalves, Ewan Harrison, David K. Jackson, Ian Johnston, Dominic Kwiatkowski, Cordelia Langford, John Sillitoe on behalf of the Wellcome Sanger Institute COVID-19 Surveillance Team |
| EPI_ISL_550772, EPI_ISL_550773, EPI_ISL_550774, EPI_ISL_550775, EPI_ISL_550776, EPI_ISL_550777, EPI_ISL_550778, EPI_ISL_550779, EPI_ISL_550780, EPI_ISL_550781, EPI_ISL_550782, EPI_ISL_550783, EPI_ISL_550784, EPI_ISL_550785, EPI_ISL_550786, EPI_ISL_550787, EPI_ISL_550788, EPI_ISL_550789, EPI_ISL_550790, EPI_ISL_550791, EPI_ISL_550792, EPI_ISL_550793, EPI_ISL_550794, EPI_ISL_550795, EPI_ISL_550796, EPI_ISL_550797, EPI_ISL_550798, EPI_ISL_550799, EPI_ISL_550800, EPI_ISL_550801, EPI_ISL_550802, EPI_ISL_550803, EPI_ISL_550804, EPI_ISL_550805, EPI_ISL_550806, EPI_ISL_550807, EPI_ISL_550808, EPI_ISL_550809, EPI_ISL_550810 | | | |
| see above | Lighthouse Lab in Milton Keynes | Wellcome Sanger Institute for the COVID-19 Genomics UK (COG-UK) consortium | The Lighthouse Lab in Milton Keynes and Alex Alderton, Roberto Amato, Sonia Goncalves, Ewan Harrison, David K. Jackson, Ian Johnston, Dominic Kwiatkowski, Cordelia Langford, John Sillitoe on behalf of the Wellcome Sanger Institute COVID-19 Surveillance Team (http://www.sanger.ac.uk/covid-team) |
| EPI_ISL_550811 | Lighthouse Lab in Milton Keynes | Wellcome Sanger Institute for the COVID-19 Genomics UK (COG-UK) consortium | The Lighthouse Lab in Alderley Park and Alex Alderton, Roberto Amato, Sonia Goncalves, Ewan Harrison, David K. Jackson, Ian Johnston, Dominic Kwiatkowski, Cordelia Langford, John Sillitoe on behalf of the Wellcome Sanger Institute COVID-19 Surveillance Team |
| EPI_ISL_550812, EPI_ISL_550813, EPI_ISL_550814, EPI_ISL_550815, EPI_ISL_550816, EPI_ISL_550817, EPI_ISL_550818, EPI_ISL_550819, EPI_ISL_550820, EPI_ISL_550821, EPI_ISL_550822, EPI_ISL_550823, EPI_ISL_550824 | | | |
| see above | Lighthouse Lab in Milton Keynes | Wellcome Sanger Institute for the COVID-19 Genomics UK (COG-UK) consortium | The Lighthouse Lab in Milton Keynes and Alex Alderton, Roberto Amato, Sonia Goncalves, Ewan Harrison, David K. Jackson, Ian Johnston, Dominic Kwiatkowski, Cordelia Langford, John Sillitoe on behalf of the Wellcome Sanger Institute COVID-19 Surveillance Team (http://www.sanger.ac.uk/covid-team) |
| EPI_ISL_550825, EPI_ISL_550826 | Lighthouse Lab in Milton Keynes | Wellcome Sanger Institute for the COVID-19 Genomics UK (COG-UK) consortium | The Lighthouse Lab in Alderley Park and Alex Alderton, Roberto Amato, Sonia Goncalves, Ewan Harrison, David K. Jackson, Ian Johnston, Dominic Kwiatkowski, Cordelia Langford, John Sillitoe on behalf of the Wellcome Sanger Institute COVID-19 Surveillance Team |
| EPI_ISL_550827, EPI_ISL_550828, EPI_ISL_550829, EPI_ISL_550830, EPI_ISL_550831, EPI_ISL_550832, EPI_ISL_550833 | Lighthouse Lab in Milton Keynes | Wellcome Sanger Institute for the COVID-19 Genomics UK (COG-UK) consortium | The Lighthouse Lab in Milton Keynes and Alex Alderton, Roberto Amato, Sonia Goncalves, Ewan Harrison, David K. Jackson, Ian Johnston, Dominic Kwiatkowski, Cordelia Langford, John Sillitoe on behalf of the Wellcome Sanger Institute COVID-19 Surveillance Team (http://www.sanger.ac.uk/covid-team) |
| EPI_ISL_550834 | Lighthouse Lab in Milton Keynes | Wellcome Sanger Institute for the COVID-19 Genomics UK (COG-UK) consortium | The Lighthouse Lab in Alderley Park and Alex Alderton, Roberto Amato, Sonia Goncalves, Ewan Harrison, David K. Jackson, Ian Johnston, Dominic Kwiatkowski, Cordelia Langford, John Sillitoe on behalf of the Wellcome Sanger Institute COVID-19 Surveillance Team |
| EPI_ISL_550835, EPI_ISL_550836, EPI_ISL_550837, EPI_ISL_550839, EPI_ISL_550840, EPI_ISL_550842, EPI_ISL_550843, EPI_ISL_550844, EPI_ISL_550845 | Lighthouse Lab in Milton Keynes | Wellcome Sanger Institute for the COVID-19 Genomics UK (COG-UK) consortium | The Lighthouse Lab in Milton Keynes and Alex Alderton, Roberto Amato, Sonia Goncalves, Ewan Harrison, David K. Jackson, Ian Johnston, Dominic Kwiatkowski, Cordelia Langford, John Sillitoe on behalf of the Wellcome Sanger Institute COVID-19 Surveillance Team (http://www.sanger.ac.uk/covid-team) |
| EPI_ISL_550846 | Lighthouse Lab in Milton Keynes | Wellcome Sanger Institute for the COVID-19 Genomics UK (COG-UK) consortium | The Lighthouse Lab in Alderley Park and Alex Alderton, Roberto Amato, Sonia Goncalves, Ewan Harrison, David K. Jackson, Ian Johnston, Dominic Kwiatkowski, Cordelia Langford, John Sillitoe on behalf of the Wellcome Sanger Institute COVID-19 Surveillance Team |

| | | | |
|---|---|---|---|
| EPI_ISL_550847, EPI_ISL_550848 | Lighthouse Lab in Milton Keynes | Wellcome Sanger Institute for the COVID-19 Genomics UK (COG-UK) consortium | The Lighthouse Lab in Milton Keynes and Alex Alderton, Roberto Amato, Sonia Goncalves, Ewan Harrison, David K. Jackson, Ian Johnston, Dominic Kwiatkowski, Cordelia Langford, John Sillitoe on behalf of the Wellcome Sanger Institute COVID-19 Surveillance Team (http://www.sanger.ac.uk/covid-team) |
| EPI_ISL_550850 | Lighthouse Lab in Milton Keynes | Wellcome Sanger Institute for the COVID-19 Genomics UK (COG-UK) consortium | The Lighthouse Lab in Alderley Park and Alex Alderton, Roberto Amato, Sonia Goncalves, Ewan Harrison, David K. Jackson, Ian Johnston, Dominic Kwiatkowski, Cordelia Langford, John Sillitoe on behalf of the Wellcome Sanger Institute COVID-19 Surveillance Team |
| EPI_ISL_550851, EPI_ISL_550852, EPI_ISL_550853, EPI_ISL_550854, EPI_ISL_550855, EPI_ISL_550857, EPI_ISL_550858, EPI_ISL_550859, EPI_ISL_550860, EPI_ISL_550861, EPI_ISL_550862, EPI_ISL_550864, EPI_ISL_550865, EPI_ISL_550866, EPI_ISL_550867 | | | |
| see above | Lighthouse Lab in Milton Keynes | Wellcome Sanger Institute for the COVID-19 Genomics UK (COG-UK) consortium | The Lighthouse Lab in Milton Keynes and Alex Alderton, Roberto Amato, Sonia Goncalves, Ewan Harrison, David K. Jackson, Ian Johnston, Dominic Kwiatkowski, Cordelia Langford, John Sillitoe on behalf of the Wellcome Sanger Institute COVID-19 Surveillance Team (http://www.sanger.ac.uk/covid-team) |
| EPI_ISL_550870 | Lighthouse Lab in Milton Keynes | Wellcome Sanger Institute for the COVID-19 Genomics UK (COG-UK) consortium | The Lighthouse Lab in Alderley Park and Alex Alderton, Roberto Amato, Sonia Goncalves, Ewan Harrison, David K. Jackson, Ian Johnston, Dominic Kwiatkowski, Cordelia Langford, John Sillitoe on behalf of the Wellcome Sanger Institute COVID-19 Surveillance Team |
| EPI_ISL_550871, EPI_ISL_550872, EPI_ISL_550873, EPI_ISL_550874, EPI_ISL_550875, EPI_ISL_550876, EPI_ISL_550877, EPI_ISL_550878, EPI_ISL_550879, EPI_ISL_550880, EPI_ISL_550881, EPI_ISL_550882, EPI_ISL_550883, EPI_ISL_550884, EPI_ISL_550885, EPI_ISL_550886, EPI_ISL_550887, EPI_ISL_550888, EPI_ISL_550889, EPI_ISL_550890, EPI_ISL_550891, EPI_ISL_550892, EPI_ISL_550893, EPI_ISL_550894, EPI_ISL_550895, EPI_ISL_550896, EPI_ISL_550897, EPI_ISL_550898, EPI_ISL_550899, EPI_ISL_550900 | | | |
| see above | Lighthouse Lab in Milton Keynes | Wellcome Sanger Institute for the COVID-19 Genomics UK (COG-UK) consortium | The Lighthouse Lab in Milton Keynes and Alex Alderton, Roberto Amato, Sonia Goncalves, Ewan Harrison, David K. Jackson, Ian Johnston, Dominic Kwiatkowski, Cordelia Langford, John Sillitoe on behalf of the Wellcome Sanger Institute COVID-19 Surveillance Team (http://www.sanger.ac.uk/covid-team) |
| EPI_ISL_550901 | Lighthouse Lab in Milton Keynes | Wellcome Sanger Institute for the COVID-19 Genomics UK (COG-UK) consortium | The Lighthouse Lab in Alderley Park and Alex Alderton, Roberto Amato, Sonia Goncalves, Ewan Harrison, David K. Jackson, Ian Johnston, Dominic Kwiatkowski, Cordelia Langford, John Sillitoe on behalf of the Wellcome Sanger Institute COVID-19 Surveillance Team |
| EPI_ISL_550902, EPI_ISL_550903, EPI_ISL_550904, EPI_ISL_550905, EPI_ISL_550906, EPI_ISL_550907, EPI_ISL_550908, EPI_ISL_550909, EPI_ISL_550910, EPI_ISL_550911, EPI_ISL_550912, EPI_ISL_550913 | | | |
| see above | Lighthouse Lab in Milton Keynes | Wellcome Sanger Institute for the COVID-19 Genomics UK (COG-UK) consortium | The Lighthouse Lab in Milton Keynes and Alex Alderton, Roberto Amato, Sonia Goncalves, Ewan Harrison, David K. Jackson, Ian Johnston, Dominic Kwiatkowski, Cordelia Langford, John Sillitoe on behalf of the Wellcome Sanger Institute COVID-19 Surveillance Team (http://www.sanger.ac.uk/covid-team) |
| EPI_ISL_550914 | Lighthouse Lab in Milton Keynes | Wellcome Sanger Institute for the COVID-19 Genomics UK (COG-UK) consortium | The Lighthouse Lab in Alderley Park and Alex Alderton, Roberto Amato, Sonia Goncalves, Ewan Harrison, David K. Jackson, Ian Johnston, Dominic Kwiatkowski, Cordelia Langford, John Sillitoe on behalf of the Wellcome Sanger Institute COVID-19 Surveillance Team |
| EPI_ISL_550915, EPI_ISL_550916, EPI_ISL_550918, EPI_ISL_550920, EPI_ISL_550921, EPI_ISL_550922, EPI_ISL_550923, EPI_ISL_550924, EPI_ISL_550925, EPI_ISL_550926, EPI_ISL_550927, EPI_ISL_550928, EPI_ISL_550930, EPI_ISL_550931, EPI_ISL_550932, EPI_ISL_550933, EPI_ISL_550934, EPI_ISL_550935, EPI_ISL_550937, EPI_ISL_550938, EPI_ISL_550939, EPI_ISL_550940, EPI_ISL_550941, EPI_ISL_550942, EPI_ISL_550943, EPI_ISL_550944 | | | |
| see above | Lighthouse Lab in Milton Keynes | Wellcome Sanger Institute for the COVID-19 Genomics UK (COG-UK) consortium | The Lighthouse Lab in Milton Keynes and Alex Alderton, Roberto Amato, Sonia Goncalves, Ewan Harrison, David K. Jackson, Ian Johnston, Dominic Kwiatkowski, Cordelia Langford, John Sillitoe on behalf of the Wellcome Sanger Institute COVID-19 Surveillance Team (http://www.sanger.ac.uk/covid-team) |
| EPI_ISL_550945 | Lighthouse Lab in Milton Keynes | Wellcome Sanger Institute for the COVID-19 Genomics UK (COG-UK) consortium | The Lighthouse Lab in Alderley Park and Alex Alderton, Roberto Amato, Sonia Goncalves, Ewan Harrison, David K. Jackson, Ian Johnston, Dominic Kwiatkowski, Cordelia Langford, John Sillitoe on behalf of the Wellcome Sanger Institute COVID-19 Surveillance Team |
| EPI_ISL_550946, EPI_ISL_550947, EPI_ISL_550948, EPI_ISL_550949, EPI_ISL_550950, EPI_ISL_550951, EPI_ISL_550952, EPI_ISL_550953, EPI_ISL_550954, EPI_ISL_550955, EPI_ISL_550956, EPI_ISL_550957, EPI_ISL_550958, EPI_ISL_550959, EPI_ISL_550960, EPI_ISL_550961, EPI_ISL_550962, EPI_ISL_550963, EPI_ISL_550964, EPI_ISL_550965, EPI_ISL_550966, EPI_ISL_550967, EPI_ISL_550968, EPI_ISL_550969, EPI_ISL_550970, EPI_ISL_550971, EPI_ISL_550972, EPI_ISL_550973, EPI_ISL_550974, EPI_ISL_550975, EPI_ISL_550976, EPI_ISL_550977 | | | |
| see above | Lighthouse Lab in Milton Keynes | Wellcome Sanger Institute for the COVID-19 Genomics UK (COG-UK) consortium | The Lighthouse Lab in Milton Keynes and Alex Alderton, Roberto Amato, Sonia Goncalves, Ewan Harrison, David K. Jackson, Ian Johnston, Dominic Kwiatkowski, Cordelia Langford, John Sillitoe on behalf of the Wellcome Sanger Institute COVID-19 Surveillance Team (http://www.sanger.ac.uk/covid-team) |
| EPI_ISL_550978 | Lighthouse Lab in Milton Keynes | Wellcome Sanger Institute for the COVID-19 Genomics UK (COG-UK) consortium | The Lighthouse Lab in Alderley Park and Alex Alderton, Roberto Amato, Sonia Goncalves, Ewan Harrison, David K. Jackson, Ian Johnston, Dominic Kwiatkowski, Cordelia Langford, John Sillitoe on behalf of the Wellcome Sanger Institute COVID-19 Surveillance Team |
| EPI_ISL_550979, EPI_ISL_550980 | Lighthouse Lab in Milton Keynes | Wellcome Sanger Institute for the COVID-19 Genomics UK (COG-UK) consortium | The Lighthouse Lab in Milton Keynes and Alex Alderton, Roberto Amato, Sonia Goncalves, Ewan Harrison, David K. Jackson, Ian Johnston, Dominic Kwiatkowski, Cordelia Langford, John Sillitoe on behalf of the Wellcome Sanger Institute COVID-19 Surveillance Team (http://www.sanger.ac.uk/covid-team) |
| EPI_ISL_550981 | Lighthouse Lab in Milton Keynes | Wellcome Sanger Institute for the COVID-19 Genomics UK (COG-UK) consortium | The Lighthouse Lab in Alderley Park and Alex Alderton, Roberto Amato, Sonia Goncalves, Ewan Harrison, David K. Jackson, Ian Johnston, Dominic Kwiatkowski, Cordelia Langford, John Sillitoe on behalf of the Wellcome Sanger Institute COVID-19 Surveillance Team |
| EPI_ISL_550982, EPI_ISL_550984, EPI_ISL_550985, EPI_ISL_550986 | Lighthouse Lab in Milton Keynes | Wellcome Sanger Institute for the COVID-19 Genomics UK (COG-UK) consortium | The Lighthouse Lab in Milton Keynes and Alex Alderton, Roberto Amato, Sonia Goncalves, Ewan Harrison, David K. Jackson, Ian Johnston, Dominic Kwiatkowski, Cordelia Langford, John Sillitoe on behalf of the Wellcome Sanger Institute COVID-19 Surveillance Team (http://www.sanger.ac.uk/covid-team) |
| EPI_ISL_550987 | Lighthouse Lab in Milton Keynes | Wellcome Sanger Institute for the COVID-19 Genomics UK (COG-UK) consortium | The Lighthouse Lab in Alderley Park and Alex Alderton, Roberto Amato, Sonia Goncalves, Ewan Harrison, David K. Jackson, Ian Johnston, Dominic Kwiatkowski, Cordelia Langford, John Sillitoe on behalf of the Wellcome Sanger Institute COVID-19 Surveillance Team |
| EPI_ISL_550988, EPI_ISL_550989, EPI_ISL_550990, EPI_ISL_550991, EPI_ISL_550992, EPI_ISL_550994, EPI_ISL_550995, EPI_ISL_550996, EPI_ISL_550997, EPI_ISL_550998, EPI_ISL_550999, EPI_ISL_551000, EPI_ISL_551002, EPI_ISL_551003, EPI_ISL_551004 | | | |
| see above | Lighthouse Lab in Milton Keynes | Wellcome Sanger Institute for the COVID-19 Genomics UK (COG-UK) consortium | The Lighthouse Lab in Milton Keynes and Alex Alderton, Roberto Amato, Sonia Goncalves, Ewan Harrison, David K. Jackson, Ian Johnston, Dominic Kwiatkowski, Cordelia Langford, John Sillitoe on behalf of the Wellcome Sanger Institute COVID-19 Surveillance Team (http://www.sanger.ac.uk/covid-team) |
| EPI_ISL_551005 | Lighthouse Lab in Milton Keynes | Wellcome Sanger Institute for the COVID-19 Genomics UK (COG-UK) consortium | The Lighthouse Lab in Alderley Park and Alex Alderton, Roberto Amato, Sonia Goncalves, Ewan Harrison, David K. Jackson, Ian Johnston, Dominic Kwiatkowski, Cordelia Langford, John Sillitoe on behalf of the Wellcome Sanger Institute COVID-19 Surveillance Team |
| EPI_ISL_551006, EPI_ISL_551008, EPI_ISL_551009, EPI_ISL_551010, EPI_ISL_551011, EPI_ISL_551012, EPI_ISL_551014, EPI_ISL_551015, EPI_ISL_551016, EPI_ISL_551017, EPI_ISL_551018, EPI_ISL_551019, EPI_ISL_551020 | | | |
| see above | Lighthouse Lab in Milton Keynes | Wellcome Sanger Institute for the COVID-19 Genomics UK (COG-UK) consortium | The Lighthouse Lab in Milton Keynes and Alex Alderton, Roberto Amato, Sonia Goncalves, Ewan Harrison, David K. Jackson, Ian Johnston, Dominic Kwiatkowski, Cordelia Langford, John Sillitoe on behalf of the Wellcome Sanger Institute COVID-19 Surveillance Team (http://www.sanger.ac.uk/covid-team) |
| EPI_ISL_551021 | Lighthouse Lab in Milton Keynes | Wellcome Sanger Institute for the COVID-19 Genomics UK (COG-UK) consortium | The Lighthouse Lab in Alderley Park and Alex Alderton, Roberto Amato, Sonia Goncalves, Ewan Harrison, David K. Jackson, Ian Johnston, Dominic Kwiatkowski, Cordelia Langford, John Sillitoe on behalf of the Wellcome Sanger Institute COVID-19 Surveillance Team |
| EPI_ISL_551022, EPI_ISL_551023, EPI_ISL_551024, EPI_ISL_551025, EPI_ISL_551026, EPI_ISL_551027, EPI_ISL_551028, EPI_ISL_551029, EPI_ISL_551030, EPI_ISL_551031, EPI_ISL_551032, EPI_ISL_551033, EPI_ISL_551034, EPI_ISL_551035, EPI_ISL_551036, EPI_ISL_551037, EPI_ISL_551038, EPI_ISL_551039, EPI_ISL_551040, EPI_ISL_551041, EPI_ISL_551042, EPI_ISL_551043 | | | |
| see above | Lighthouse Lab in Milton Keynes | Wellcome Sanger Institute for the COVID-19 Genomics UK (COG-UK) consortium | The Lighthouse Lab in Milton Keynes and Alex Alderton, Roberto Amato, Sonia Goncalves, Ewan Harrison, David K. Jackson, Ian Johnston, Dominic Kwiatkowski, Cordelia Langford, John Sillitoe on behalf of the Wellcome Sanger Institute COVID-19 Surveillance Team (http://www.sanger.ac.uk/covid-team) |
| EPI_ISL_551044 | Lighthouse Lab in Milton Keynes | Wellcome Sanger Institute for the COVID-19 Genomics UK (COG-UK) consortium | The Lighthouse Lab in Alderley Park and Alex Alderton, Roberto Amato, Sonia Goncalves, Ewan Harrison, David K. Jackson, Ian Johnston, Dominic Kwiatkowski, Cordelia Langford, John Sillitoe on behalf of the Wellcome Sanger Institute COVID-19 Surveillance Team |
| EPI_ISL_551045, EPI_ISL_551046, EPI_ISL_551048, EPI_ISL_551049, EPI_ISL_551050, EPI_ISL_551052, EPI_ISL_551053, EPI_ISL_551054, EPI_ISL_551055, EPI_ISL_551056 | Lighthouse Lab in Milton Keynes | Wellcome Sanger Institute for the COVID-19 Genomics UK (COG-UK) consortium | The Lighthouse Lab in Milton Keynes and Alex Alderton, Roberto Amato, Sonia Goncalves, Ewan Harrison, David K. Jackson, Ian Johnston, Dominic Kwiatkowski, Cordelia Langford, John Sillitoe on behalf of the Wellcome Sanger Institute COVID-19 Surveillance Team (http://www.sanger.ac.uk/covid-team) |

| | | | |
|---|---|---|---|
| EPI_ISL_551057, EPI_ISL_551058 | Lighthouse Lab in Milton Keynes | Wellcome Sanger Institute for the COVID-19 Genomics UK (COG-UK) consortium | The Lighthouse Lab in Alderley Park and Alex Alderton, Roberto Amato, Sonia Goncalves, Ewan Harrison, David K. Jackson, Ian Johnston, Dominic Kwiatkowski, Cordelia Langford, John Sillitoe on behalf of the Wellcome Sanger Institute COVID-19 Surveillance Team |
| EPI_ISL_551059, EPI_ISL_551060, EPI_ISL_551061, EPI_ISL_551062, EPI_ISL_551063, EPI_ISL_551064, EPI_ISL_551065, EPI_ISL_551066, EPI_ISL_551067, EPI_ISL_551068, EPI_ISL_551069, EPI_ISL_551070, EPI_ISL_551071, EPI_ISL_551072, EPI_ISL_551073, EPI_ISL_551074, EPI_ISL_551075, EPI_ISL_551076, EPI_ISL_551077, EPI_ISL_551078, EPI_ISL_551079, EPI_ISL_551080, EPI_ISL_551081, EPI_ISL_551082, EPI_ISL_551083, EPI_ISL_551084, EPI_ISL_551085, EPI_ISL_551086, EPI_ISL_551087, EPI_ISL_551089, EPI_ISL_551091, EPI_ISL_551093, EPI_ISL_551094, EPI_ISL_551095, EPI_ISL_551096, EPI_ISL_551097, EPI_ISL_551098, EPI_ISL_551099, EPI_ISL_551100, EPI_ISL_551101, EPI_ISL_551102, EPI_ISL_551103, EPI_ISL_551104, EPI_ISL_551105, EPI_ISL_551106, EPI_ISL_551107, EPI_ISL_551109, EPI_ISL_551110, EPI_ISL_551111, EPI_ISL_551112, EPI_ISL_551113 | | | |
| see above | Lighthouse Lab in Milton Keynes | Wellcome Sanger Institute for the COVID-19 Genomics UK (COG-UK) consortium | The Lighthouse Lab in Milton Keynes and Alex Alderton, Roberto Amato, Sonia Goncalves, Ewan Harrison, David K. Jackson, Ian Johnston, Dominic Kwiatkowski, Cordelia Langford, John Sillitoe on behalf of the Wellcome Sanger Institute COVID-19 Surveillance Team (http://www.sanger.ac.uk/covid-team) |
| EPI_ISL_551114 | Lighthouse Lab in Milton Keynes | Wellcome Sanger Institute for the COVID-19 Genomics UK (COG-UK) consortium | The Lighthouse Lab in Alderley Park and Alex Alderton, Roberto Amato, Sonia Goncalves, Ewan Harrison, David K. Jackson, Ian Johnston, Dominic Kwiatkowski, Cordelia Langford, John Sillitoe on behalf of the Wellcome Sanger Institute COVID-19 Surveillance Team |
| EPI_ISL_551115, EPI_ISL_551116, EPI_ISL_551117, EPI_ISL_551118, EPI_ISL_551119, EPI_ISL_551120, EPI_ISL_551121, EPI_ISL_551122, EPI_ISL_551124, EPI_ISL_551126, EPI_ISL_551127, EPI_ISL_551128, EPI_ISL_551129, EPI_ISL_551130, EPI_ISL_551131, EPI_ISL_551132, EPI_ISL_551133, EPI_ISL_551134, EPI_ISL_551135, EPI_ISL_551136, EPI_ISL_551137, EPI_ISL_551138, EPI_ISL_551139, EPI_ISL_551140, EPI_ISL_551141, EPI_ISL_551142, EPI_ISL_551143, EPI_ISL_551144, EPI_ISL_551145, EPI_ISL_551146, EPI_ISL_551147, EPI_ISL_551148, EPI_ISL_551149, EPI_ISL_551150, EPI_ISL_551151, EPI_ISL_551152, EPI_ISL_551153, EPI_ISL_551154, EPI_ISL_551155, EPI_ISL_551156, EPI_ISL_551157, EPI_ISL_551158, EPI_ISL_551159, EPI_ISL_551160, EPI_ISL_551161, EPI_ISL_551162, EPI_ISL_551163, EPI_ISL_551164, EPI_ISL_551165 | | | |
| see above | Lighthouse Lab in Milton Keynes | Wellcome Sanger Institute for the COVID-19 Genomics UK (COG-UK) consortium | The Lighthouse Lab in Milton Keynes and Alex Alderton, Roberto Amato, Sonia Goncalves, Ewan Harrison, David K. Jackson, Ian Johnston, Dominic Kwiatkowski, Cordelia Langford, John Sillitoe on behalf of the Wellcome Sanger Institute COVID-19 Surveillance Team (http://www.sanger.ac.uk/covid-team) |
| EPI_ISL_551166 | Lighthouse Lab in Milton Keynes | Wellcome Sanger Institute for the COVID-19 Genomics UK (COG-UK) consortium | The Lighthouse Lab in Alderley Park and Alex Alderton, Roberto Amato, Sonia Goncalves, Ewan Harrison, David K. Jackson, Ian Johnston, Dominic Kwiatkowski, Cordelia Langford, John Sillitoe on behalf of the Wellcome Sanger Institute COVID-19 Surveillance Team |
| EPI_ISL_551167, EPI_ISL_551168, EPI_ISL_551169, EPI_ISL_551170, EPI_ISL_551171, EPI_ISL_551172, EPI_ISL_551173, EPI_ISL_551174, EPI_ISL_551175, EPI_ISL_551176, EPI_ISL_551178, EPI_ISL_551179, EPI_ISL_551180, EPI_ISL_551181, EPI_ISL_551182, EPI_ISL_551183, EPI_ISL_551184, EPI_ISL_551185, EPI_ISL_551186, EPI_ISL_551187, EPI_ISL_551188, EPI_ISL_551189, EPI_ISL_551190, EPI_ISL_551191, EPI_ISL_551192, EPI_ISL_551193, EPI_ISL_551194, EPI_ISL_551195, EPI_ISL_551196, EPI_ISL_551197, EPI_ISL_551198, EPI_ISL_551199, EPI_ISL_551201, EPI_ISL_551202, EPI_ISL_551203, EPI_ISL_551205, EPI_ISL_551206, EPI_ISL_551207, EPI_ISL_551208, EPI_ISL_551209, EPI_ISL_551210, EPI_ISL_551211, EPI_ISL_551212, EPI_ISL_551213, EPI_ISL_551214 | | | |
| see above | Lighthouse Lab in Milton Keynes | Wellcome Sanger Institute for the COVID-19 Genomics UK (COG-UK) consortium | The Lighthouse Lab in Milton Keynes and Alex Alderton, Roberto Amato, Sonia Goncalves, Ewan Harrison, David K. Jackson, Ian Johnston, Dominic Kwiatkowski, Cordelia Langford, John Sillitoe on behalf of the Wellcome Sanger Institute COVID-19 Surveillance Team (http://www.sanger.ac.uk/covid-team) |
| EPI_ISL_551215 | Lighthouse Lab in Milton Keynes | Wellcome Sanger Institute for the COVID-19 Genomics UK (COG-UK) consortium | The Lighthouse Lab in Alderley Park and Alex Alderton, Roberto Amato, Sonia Goncalves, Ewan Harrison, David K. Jackson, Ian Johnston, Dominic Kwiatkowski, Cordelia Langford, John Sillitoe on behalf of the Wellcome Sanger Institute COVID-19 Surveillance Team |
| EPI_ISL_551217, EPI_ISL_551219, EPI_ISL_551220, EPI_ISL_551221, EPI_ISL_551222, EPI_ISL_551223, EPI_ISL_551224, EPI_ISL_551225, EPI_ISL_551227, EPI_ISL_551228, EPI_ISL_551229, EPI_ISL_551230, EPI_ISL_551231, EPI_ISL_551232, EPI_ISL_551233, EPI_ISL_551234, EPI_ISL_551235, EPI_ISL_551236, EPI_ISL_551237, EPI_ISL_551238, EPI_ISL_551239, EPI_ISL_551240, EPI_ISL_551241, EPI_ISL_551242, EPI_ISL_551243 | | | |
| see above | Lighthouse Lab in Milton Keynes | Wellcome Sanger Institute for the COVID-19 Genomics UK (COG-UK) consortium | The Lighthouse Lab in Milton Keynes and Alex Alderton, Roberto Amato, Sonia Goncalves, Ewan Harrison, David K. Jackson, Ian Johnston, Dominic Kwiatkowski, Cordelia Langford, John Sillitoe on behalf of the Wellcome Sanger Institute COVID-19 Surveillance Team (http://www.sanger.ac.uk/covid-team) |
| EPI_ISL_551244 | Lighthouse Lab in Milton Keynes | Wellcome Sanger Institute for the COVID-19 Genomics UK (COG-UK) consortium | The Lighthouse Lab in Alderley Park and Alex Alderton, Roberto Amato, Sonia Goncalves, Ewan Harrison, David K. Jackson, Ian Johnston, Dominic Kwiatkowski, Cordelia Langford, John Sillitoe on behalf of the Wellcome Sanger Institute COVID-19 Surveillance Team |
| EPI_ISL_551245, EPI_ISL_551246, EPI_ISL_551247, EPI_ISL_551248, EPI_ISL_551249, EPI_ISL_551251, EPI_ISL_551252, EPI_ISL_551253, EPI_ISL_551255, EPI_ISL_551257, EPI_ISL_551258, EPI_ISL_551259, EPI_ISL_551260, EPI_ISL_551261, EPI_ISL_551262 | | | |
| see above | Lighthouse Lab in Milton Keynes | Wellcome Sanger Institute for the COVID-19 Genomics UK (COG-UK) consortium | The Lighthouse Lab in Milton Keynes and Alex Alderton, Roberto Amato, Sonia Goncalves, Ewan Harrison, David K. Jackson, Ian Johnston, Dominic Kwiatkowski, Cordelia Langford, John Sillitoe on behalf of the Wellcome Sanger Institute COVID-19 Surveillance Team (http://www.sanger.ac.uk/covid-team) |
| EPI_ISL_551263 | Lighthouse Lab in Milton Keynes | Wellcome Sanger Institute for the COVID-19 Genomics UK (COG-UK) consortium | The Lighthouse Lab in Alderley Park and Alex Alderton, Roberto Amato, Sonia Goncalves, Ewan Harrison, David K. Jackson, Ian Johnston, Dominic Kwiatkowski, Cordelia Langford, John Sillitoe on behalf of the Wellcome Sanger Institute COVID-19 Surveillance Team |
| EPI_ISL_551264, EPI_ISL_551265, EPI_ISL_551267, EPI_ISL_551268, EPI_ISL_551269, EPI_ISL_551270, EPI_ISL_551271, EPI_ISL_551273, EPI_ISL_551274, EPI_ISL_551275, EPI_ISL_551276, EPI_ISL_551277, EPI_ISL_551278, EPI_ISL_551279, EPI_ISL_551280, EPI_ISL_551281, EPI_ISL_551282, EPI_ISL_551284 | | | |
| see above | Lighthouse Lab in Milton Keynes | Wellcome Sanger Institute for the COVID-19 Genomics UK (COG-UK) consortium | The Lighthouse Lab in Milton Keynes and Alex Alderton, Roberto Amato, Sonia Goncalves, Ewan Harrison, David K. Jackson, Ian Johnston, Dominic Kwiatkowski, Cordelia Langford, John Sillitoe on behalf of the Wellcome Sanger Institute COVID-19 Surveillance Team (http://www.sanger.ac.uk/covid-team) |
| EPI_ISL_551285, EPI_ISL_551286, EPI_ISL_551287 | Lighthouse Lab in Alderley Park | Wellcome Sanger Institute for the COVID-19 Genomics UK (COG-UK) consortium | The Lighthouse Lab in Alderley Park and Alex Alderton, Roberto Amato, Sonia Goncalves, Ewan Harrison, David K. Jackson, Ian Johnston, Dominic Kwiatkowski, Cordelia Langford, John Sillitoe on behalf of the Wellcome Sanger Institute COVID-19 Surveillance Team |
| EPI_ISL_551288 | Lighthouse Lab in Alderley Park | Wellcome Sanger Institute for the COVID-19 Genomics UK (COG-UK) consortium | The Lighthouse Lab in Alderley Park and Alex Alderton, Roberto Amato, Sonia Goncalves, Ewan Harrison, David K. Jackson, Ian Johnston, Dominic Kwiatkowski, Cordelia Langford, John Sillitoe on behalf of the Wellcome Sanger Institute COVID-19 Surveillance Team (http://www.sanger.ac.uk/covid-team) |
| EPI_ISL_551289, EPI_ISL_551290, EPI_ISL_551291, EPI_ISL_551292, EPI_ISL_551293, EPI_ISL_551295, EPI_ISL_551296, EPI_ISL_551297, EPI_ISL_551298, EPI_ISL_551299, EPI_ISL_551300, EPI_ISL_551301, EPI_ISL_551302, EPI_ISL_551303, EPI_ISL_551304, EPI_ISL_551305, EPI_ISL_551306, EPI_ISL_551307, EPI_ISL_551308, EPI_ISL_551309, EPI_ISL_551310, EPI_ISL_551311, EPI_ISL_551312, EPI_ISL_551313 | | | |
| see above | Lighthouse Lab in Alderley Park | Wellcome Sanger Institute for the COVID-19 Genomics UK (COG-UK) consortium | The Lighthouse Lab in Alderley Park and Alex Alderton, Roberto Amato, Sonia Goncalves, Ewan Harrison, David K. Jackson, Ian Johnston, Dominic Kwiatkowski, Cordelia Langford, John Sillitoe on behalf of the Wellcome Sanger Institute COVID-19 Surveillance Team |
| EPI_ISL_551314 | Lighthouse Lab in Alderley Park | Wellcome Sanger Institute for the COVID-19 Genomics UK (COG-UK) consortium | The Lighthouse Lab in Alderley Park and Alex Alderton, Roberto Amato, Sonia Goncalves, Ewan Harrison, David K. Jackson, Ian Johnston, Dominic Kwiatkowski, Cordelia Langford, John Sillitoe on behalf of the Wellcome Sanger Institute COVID-19 Surveillance Team (http://www.sanger.ac.uk/covid-team) |
| EPI_ISL_551315, EPI_ISL_551316, EPI_ISL_551317, EPI_ISL_551318, EPI_ISL_551319, EPI_ISL_551320, EPI_ISL_551321, EPI_ISL_551322, EPI_ISL_551323, EPI_ISL_551324, EPI_ISL_551325, EPI_ISL_551326, EPI_ISL_551328, EPI_ISL_551329, EPI_ISL_551330, EPI_ISL_551331, EPI_ISL_551332, EPI_ISL_551333, EPI_ISL_551334, EPI_ISL_551335, EPI_ISL_551336, EPI_ISL_551337, EPI_ISL_551338, EPI_ISL_551339, EPI_ISL_551340, EPI_ISL_551341, EPI_ISL_551342, EPI_ISL_551343, EPI_ISL_551344, EPI_ISL_551345, EPI_ISL_551346, EPI_ISL_551347, EPI_ISL_551348 | | | |
| see above | Lighthouse Lab in Alderley Park | Wellcome Sanger Institute for the COVID-19 Genomics UK (COG-UK) consortium | The Lighthouse Lab in Alderley Park and Alex Alderton, Roberto Amato, Sonia Goncalves, Ewan Harrison, David K. Jackson, Ian Johnston, Dominic Kwiatkowski, Cordelia Langford, John Sillitoe on behalf of the Wellcome Sanger Institute COVID-19 Surveillance Team |
| EPI_ISL_551349 | Lighthouse Lab in Alderley Park | Wellcome Sanger Institute for the COVID-19 Genomics UK (COG-UK) consortium | The Lighthouse Lab in Alderley Park and Alex Alderton, Roberto Amato, Sonia Goncalves, Ewan Harrison, David K. Jackson, Ian Johnston, Dominic Kwiatkowski, Cordelia Langford, John Sillitoe on behalf of the Wellcome Sanger Institute COVID-19 Surveillance Team (http://www.sanger.ac.uk/covid-team) |
| EPI_ISL_551350, EPI_ISL_551351, EPI_ISL_551352, EPI_ISL_551353, EPI_ISL_551358 | Lighthouse Lab in Alderley Park | Wellcome Sanger Institute for the COVID-19 Genomics UK (COG-UK) consortium | The Lighthouse Lab in Alderley Park and Alex Alderton, Roberto Amato, Sonia Goncalves, Ewan Harrison, David K. Jackson, Ian Johnston, Dominic Kwiatkowski, Cordelia Langford, John Sillitoe on behalf of the Wellcome Sanger Institute COVID-19 Surveillance Team |
| EPI_ISL_551367, EPI_ISL_551368, EPI_ISL_551369, EPI_ISL_551374, EPI_ISL_551375, EPI_ISL_551378, EPI_ISL_551384, EPI_ISL_551388, EPI_ISL_551391, EPI_ISL_551396, EPI_ISL_551398, EPI_ISL_551400, EPI_ISL_551401, EPI_ISL_551402, EPI_ISL_551404, EPI_ISL_551405 | | | |
| see above | Lighthouse Lab in Alderley Park | Wellcome Sanger Institute for the COVID-19 Genomics UK (COG-UK) consortium | The Lighthouse Lab in Alderley Park and Alex Alderton, Roberto Amato, Sonia Goncalves, Ewan Harrison, David K. Jackson, Ian Johnston, Dominic Kwiatkowski, Cordelia Langford, John Sillitoe on behalf of the Wellcome Sanger Institute COVID-19 Surveillance Team (http://www.sanger.ac.uk/covid-team) |
| EPI_ISL_551406 | Lighthouse Lab in Alderley Park | Wellcome Sanger Institute for the COVID-19 Genomics UK (COG-UK) consortium | The Lighthouse Lab in Alderley Park and Alex Alderton, Roberto Amato, Sonia Goncalves, Ewan Harrison, David K. Jackson, Ian Johnston, Dominic Kwiatkowski, Cordelia Langford, John Sillitoe on behalf of the Wellcome Sanger Institute COVID-19 Surveillance Team |
| EPI_ISL_551407, EPI_ISL_551410, EPI_ISL_551413, EPI_ISL_551414, | Lighthouse Lab in Alderley Park | Wellcome Sanger Institute for the COVID-19 Genomics UK (COG-UK) consortium | The Lighthouse Lab in Alderley Park and Alex Alderton, Roberto Amato, Sonia Goncalves, Ewan Harrison, David K. Jackson, Ian Johnston, Dominic Kwiatkowski, Cordelia Langford, John Sillitoe on behalf of the Wellcome Sanger Institute COVID-19 Surveillance Team |

| | | | |
|---|---|---|---|
| EPI_ISL_551418, EPI_ISL_551419, EPI_ISL_551430, EPI_ISL_551431, EPI_ISL_551432, EPI_ISL_551433 | | | (http://www.sanger.ac.uk/covid-team) |
| EPI_ISL_551442 | Lighthouse Lab in Alderley Park | Wellcome Sanger Institute for the COVID-19 Genomics UK (COG-UK) consortium | The Lighthouse Lab in Alderley Park and Alex Alderton, Roberto Amato, Sonia Goncalves, Ewan Harrison, David K. Jackson, Ian Johnston, Dominic Kwiatkowski, Cordelia Langford, John Sillitoe on behalf of the Wellcome Sanger Institute COVID-19 Surveillance Team |
| EPI_ISL_551466, EPI_ISL_551469, EPI_ISL_551476, EPI_ISL_551483, EPI_ISL_551489, EPI_ISL_551490, EPI_ISL_551493, EPI_ISL_551495, EPI_ISL_551499, EPI_ISL_551500, EPI_ISL_551502, EPI_ISL_551504, EPI_ISL_551511 | | | |
| see above | Lighthouse Lab in Alderley Park | Wellcome Sanger Institute for the COVID-19 Genomics UK (COG-UK) consortium | The Lighthouse Lab in Alderley Park and Alex Alderton, Roberto Amato, Sonia Goncalves, Ewan Harrison, David K. Jackson, Ian Johnston, Dominic Kwiatkowski, Cordelia Langford, John Sillitoe on behalf of the Wellcome Sanger Institute COVID-19 Surveillance Team (http://www.sanger.ac.uk/covid-team) |
| EPI_ISL_551513 | Lighthouse Lab in Alderley Park | Wellcome Sanger Institute for the COVID-19 Genomics UK (COG-UK) consortium | The Lighthouse Lab in Alderley Park and Alex Alderton, Roberto Amato, Sonia Goncalves, Ewan Harrison, David K. Jackson, Ian Johnston, Dominic Kwiatkowski, Cordelia Langford, John Sillitoe on behalf of the Wellcome Sanger Institute COVID-19 Surveillance Team |
| EPI_ISL_551515 | Lighthouse Lab in Alderley Park | Wellcome Sanger Institute for the COVID-19 Genomics UK (COG-UK) consortium | The Lighthouse Lab in Alderley Park and Alex Alderton, Roberto Amato, Sonia Goncalves, Ewan Harrison, David K. Jackson, Ian Johnston, Dominic Kwiatkowski, Cordelia Langford, John Sillitoe on behalf of the Wellcome Sanger Institute COVID-19 Surveillance Team (http://www.sanger.ac.uk/covid-team) |
| EPI_ISL_551519 | Lighthouse Lab in Alderley Park | Wellcome Sanger Institute for the COVID-19 Genomics UK (COG-UK) consortium | The Lighthouse Lab in Alderley Park and Alex Alderton, Roberto Amato, Sonia Goncalves, Ewan Harrison, David K. Jackson, Ian Johnston, Dominic Kwiatkowski, Cordelia Langford, John Sillitoe on behalf of the Wellcome Sanger Institute COVID-19 Surveillance Team |
| EPI_ISL_551523, EPI_ISL_551526, EPI_ISL_551527, EPI_ISL_551528, EPI_ISL_551531 | Lighthouse Lab in Alderley Park | Wellcome Sanger Institute for the COVID-19 Genomics UK (COG-UK) consortium | The Lighthouse Lab in Alderley Park and Alex Alderton, Roberto Amato, Sonia Goncalves, Ewan Harrison, David K. Jackson, Ian Johnston, Dominic Kwiatkowski, Cordelia Langford, John Sillitoe on behalf of the Wellcome Sanger Institute COVID-19 Surveillance Team (http://www.sanger.ac.uk/covid-team) |
| EPI_ISL_551532 | Lighthouse Lab in Alderley Park | Wellcome Sanger Institute for the COVID-19 Genomics UK (COG-UK) consortium | The Lighthouse Lab in Alderley Park and Alex Alderton, Roberto Amato, Sonia Goncalves, Ewan Harrison, David K. Jackson, Ian Johnston, Dominic Kwiatkowski, Cordelia Langford, John Sillitoe on behalf of the Wellcome Sanger Institute COVID-19 Surveillance Team |
| EPI_ISL_551550, EPI_ISL_551555, EPI_ISL_551562, EPI_ISL_551567, EPI_ISL_551568, EPI_ISL_551571 | Lighthouse Lab in Alderley Park | Wellcome Sanger Institute for the COVID-19 Genomics UK (COG-UK) consortium | The Lighthouse Lab in Alderley Park and Alex Alderton, Roberto Amato, Sonia Goncalves, Ewan Harrison, David K. Jackson, Ian Johnston, Dominic Kwiatkowski, Cordelia Langford, John Sillitoe on behalf of the Wellcome Sanger Institute COVID-19 Surveillance Team (http://www.sanger.ac.uk/covid-team) |
| EPI_ISL_551577, EPI_ISL_551578, EPI_ISL_551579 | Lighthouse Lab in Milton Keynes | Wellcome Sanger Institute for the COVID-19 Genomics UK (COG-UK) consortium | The Lighthouse Lab in Milton Keynes and Alex Alderton, Roberto Amato, Sonia Goncalves, Ewan Harrison, David K. Jackson, Ian Johnston, Dominic Kwiatkowski, Cordelia Langford, John Sillitoe on behalf of the Wellcome Sanger Institute COVID-19 Surveillance Team (http://www.sanger.ac.uk/covid-team) |
| EPI_ISL_551580 | Lighthouse Lab in Milton Keynes | Wellcome Sanger Institute for the COVID-19 Genomics UK (COG-UK) consortium | The Lighthouse Lab in Alderley Park and Alex Alderton, Roberto Amato, Sonia Goncalves, Ewan Harrison, David K. Jackson, Ian Johnston, Dominic Kwiatkowski, Cordelia Langford, John Sillitoe on behalf of the Wellcome Sanger Institute COVID-19 Surveillance Team |
| EPI_ISL_551581, EPI_ISL_551582, EPI_ISL_551583 | Lighthouse Lab in Milton Keynes | Wellcome Sanger Institute for the COVID-19 Genomics UK (COG-UK) consortium | The Lighthouse Lab in Milton Keynes and Alex Alderton, Roberto Amato, Sonia Goncalves, Ewan Harrison, David K. Jackson, Ian Johnston, Dominic Kwiatkowski, Cordelia Langford, John Sillitoe on behalf of the Wellcome Sanger Institute COVID-19 Surveillance Team (http://www.sanger.ac.uk/covid-team) |
| EPI_ISL_551584, EPI_ISL_551585 | Lighthouse Lab in Milton Keynes | Wellcome Sanger Institute for the COVID-19 Genomics UK (COG-UK) consortium | The Lighthouse Lab in Alderley Park and Alex Alderton, Roberto Amato, Sonia Goncalves, Ewan Harrison, David K. Jackson, Ian Johnston, Dominic Kwiatkowski, Cordelia Langford, John Sillitoe on behalf of the Wellcome Sanger Institute COVID-19 Surveillance Team |
| EPI_ISL_551586, EPI_ISL_551587, EPI_ISL_551588, EPI_ISL_551589, EPI_ISL_551591, EPI_ISL_551592, EPI_ISL_551593, EPI_ISL_551594, EPI_ISL_551595, EPI_ISL_551596, EPI_ISL_551597, EPI_ISL_551598, EPI_ISL_551600, EPI_ISL_551601 | | | |
| see above | Lighthouse Lab in Milton Keynes | Wellcome Sanger Institute for the COVID-19 Genomics UK (COG-UK) consortium | The Lighthouse Lab in Milton Keynes and Alex Alderton, Roberto Amato, Sonia Goncalves, Ewan Harrison, David K. Jackson, Ian Johnston, Dominic Kwiatkowski, Cordelia Langford, John Sillitoe on behalf of the Wellcome Sanger Institute COVID-19 Surveillance Team (http://www.sanger.ac.uk/covid-team) |
| EPI_ISL_551603 | Lighthouse Lab in Milton Keynes | Wellcome Sanger Institute for the COVID-19 Genomics UK (COG-UK) consortium | The Lighthouse Lab in Alderley Park and Alex Alderton, Roberto Amato, Sonia Goncalves, Ewan Harrison, David K. Jackson, Ian Johnston, Dominic Kwiatkowski, Cordelia Langford, John Sillitoe on behalf of the Wellcome Sanger Institute COVID-19 Surveillance Team |
| EPI_ISL_551604, EPI_ISL_551605, EPI_ISL_551606, EPI_ISL_551607, EPI_ISL_551608, EPI_ISL_551609, EPI_ISL_551610, EPI_ISL_551611, EPI_ISL_551612, EPI_ISL_551613, EPI_ISL_551615, EPI_ISL_551616, EPI_ISL_551617, EPI_ISL_551618, EPI_ISL_551619, EPI_ISL_551620, EPI_ISL_551621, EPI_ISL_551622, EPI_ISL_551623 | | | |
| see above | Lighthouse Lab in Milton Keynes | Wellcome Sanger Institute for the COVID-19 Genomics UK (COG-UK) consortium | The Lighthouse Lab in Milton Keynes and Alex Alderton, Roberto Amato, Sonia Goncalves, Ewan Harrison, David K. Jackson, Ian Johnston, Dominic Kwiatkowski, Cordelia Langford, John Sillitoe on behalf of the Wellcome Sanger Institute COVID-19 Surveillance Team (http://www.sanger.ac.uk/covid-team) |
| EPI_ISL_551624 | Lighthouse Lab in Milton Keynes | Wellcome Sanger Institute for the COVID-19 Genomics UK (COG-UK) consortium | The Lighthouse Lab in Alderley Park and Alex Alderton, Roberto Amato, Sonia Goncalves, Ewan Harrison, David K. Jackson, Ian Johnston, Dominic Kwiatkowski, Cordelia Langford, John Sillitoe on behalf of the Wellcome Sanger Institute COVID-19 Surveillance Team |
| EPI_ISL_551625, EPI_ISL_551626 | Lighthouse Lab in Milton Keynes | Wellcome Sanger Institute for the COVID-19 Genomics UK (COG-UK) consortium | The Lighthouse Lab in Milton Keynes and Alex Alderton, Roberto Amato, Sonia Goncalves, Ewan Harrison, David K. Jackson, Ian Johnston, Dominic Kwiatkowski, Cordelia Langford, John Sillitoe on behalf of the Wellcome Sanger Institute COVID-19 Surveillance Team (http://www.sanger.ac.uk/covid-team) |
| EPI_ISL_551627 | Lighthouse Lab in Milton Keynes | Wellcome Sanger Institute for the COVID-19 Genomics UK (COG-UK) consortium | The Lighthouse Lab in Alderley Park and Alex Alderton, Roberto Amato, Sonia Goncalves, Ewan Harrison, David K. Jackson, Ian Johnston, Dominic Kwiatkowski, Cordelia Langford, John Sillitoe on behalf of the Wellcome Sanger Institute COVID-19 Surveillance Team |
| EPI_ISL_551628, EPI_ISL_551629, EPI_ISL_551630, EPI_ISL_551631, EPI_ISL_551632, EPI_ISL_551634, EPI_ISL_551636 | Lighthouse Lab in Milton Keynes | Wellcome Sanger Institute for the COVID-19 Genomics UK (COG-UK) consortium | The Lighthouse Lab in Milton Keynes and Alex Alderton, Roberto Amato, Sonia Goncalves, Ewan Harrison, David K. Jackson, Ian Johnston, Dominic Kwiatkowski, Cordelia Langford, John Sillitoe on behalf of the Wellcome Sanger Institute COVID-19 Surveillance Team (http://www.sanger.ac.uk/covid-team) |
| EPI_ISL_551637 | Lighthouse Lab in Milton Keynes | Wellcome Sanger Institute for the COVID-19 Genomics UK (COG-UK) consortium | The Lighthouse Lab in Alderley Park and Alex Alderton, Roberto Amato, Sonia Goncalves, Ewan Harrison, David K. Jackson, Ian Johnston, Dominic Kwiatkowski, Cordelia Langford, John Sillitoe on behalf of the Wellcome Sanger Institute COVID-19 Surveillance Team |
| EPI_ISL_551638, EPI_ISL_551639, EPI_ISL_551640, EPI_ISL_551641 | Lighthouse Lab in Milton Keynes | Wellcome Sanger Institute for the COVID-19 Genomics UK (COG-UK) consortium | The Lighthouse Lab in Milton Keynes and Alex Alderton, Roberto Amato, Sonia Goncalves, Ewan Harrison, David K. Jackson, Ian Johnston, Dominic Kwiatkowski, Cordelia Langford, John Sillitoe on behalf of the Wellcome Sanger Institute COVID-19 Surveillance Team (http://www.sanger.ac.uk/covid-team) |
| EPI_ISL_551642 | Lighthouse Lab in Milton Keynes | Wellcome Sanger Institute for the COVID-19 Genomics UK (COG-UK) consortium | The Lighthouse Lab in Alderley Park and Alex Alderton, Roberto Amato, Sonia Goncalves, Ewan Harrison, David K. Jackson, Ian Johnston, Dominic Kwiatkowski, Cordelia Langford, John Sillitoe on behalf of the Wellcome Sanger Institute COVID-19 Surveillance Team |
| EPI_ISL_551643, EPI_ISL_551644, EPI_ISL_551645, EPI_ISL_551646, EPI_ISL_551647, EPI_ISL_551648 | Lighthouse Lab in Milton Keynes | Wellcome Sanger Institute for the COVID-19 Genomics UK (COG-UK) consortium | The Lighthouse Lab in Milton Keynes and Alex Alderton, Roberto Amato, Sonia Goncalves, Ewan Harrison, David K. Jackson, Ian Johnston, Dominic Kwiatkowski, Cordelia Langford, John Sillitoe on behalf of the Wellcome Sanger Institute COVID-19 Surveillance Team (http://www.sanger.ac.uk/covid-team) |
| EPI_ISL_551649 | Lighthouse Lab in Milton Keynes | Wellcome Sanger Institute for the COVID-19 Genomics UK (COG-UK) consortium | The Lighthouse Lab in Alderley Park and Alex Alderton, Roberto Amato, Sonia Goncalves, Ewan Harrison, David K. Jackson, Ian Johnston, Dominic Kwiatkowski, Cordelia Langford, John Sillitoe on behalf of the Wellcome Sanger Institute COVID-19 Surveillance Team |
| EPI_ISL_551650, EPI_ISL_551651, EPI_ISL_551652, EPI_ISL_551653, EPI_ISL_551654, EPI_ISL_551655, EPI_ISL_551656, EPI_ISL_551658, EPI_ISL_551661, EPI_ISL_551662, EPI_ISL_551663, EPI_ISL_551665, EPI_ISL_551667, EPI_ISL_551669, EPI_ISL_551670, EPI_ISL_551671, EPI_ISL_551672, EPI_ISL_551673, EPI_ISL_551675, EPI_ISL_551676, EPI_ISL_551677, EPI_ISL_551678, EPI_ISL_551679, EPI_ISL_551680 | | | |
| see above | Lighthouse Lab in Milton Keynes | Wellcome Sanger Institute for the COVID-19 Genomics UK (COG-UK) consortium | The Lighthouse Lab in Milton Keynes and Alex Alderton, Roberto Amato, Sonia Goncalves, Ewan Harrison, David K. Jackson, Ian Johnston, Dominic Kwiatkowski, Cordelia Langford, John Sillitoe on behalf of the Wellcome Sanger Institute COVID-19 Surveillance Team (http://www.sanger.ac.uk/covid-team) |

| | | | |
|---|---|---|---|
| EPI_ISL_551681 | Lighthouse Lab in Milton Keynes | Wellcome Sanger Institute for the COVID-19 Genomics UK (COG-UK) consortium | The Lighthouse Lab in Alderley Park and Alex Alderton, Roberto Amato, Sonia Goncalves, Ewan Harrison, David K. Jackson, Ian Johnston, Dominic Kwiatkowski, Cordelia Langford, John Sillitoe on behalf of the Wellcome Sanger Institute COVID-19 Surveillance Team |
| EPI_ISL_551682, EPI_ISL_551683, EPI_ISL_551684, EPI_ISL_551685, EPI_ISL_551686, EPI_ISL_551688, EPI_ISL_551689, EPI_ISL_551690, EPI_ISL_551691 | Lighthouse Lab in Milton Keynes | Wellcome Sanger Institute for the COVID-19 Genomics UK (COG-UK) consortium | The Lighthouse Lab in Milton Keynes and Alex Alderton, Roberto Amato, Sonia Goncalves, Ewan Harrison, David K. Jackson, Ian Johnston, Dominic Kwiatkowski, Cordelia Langford, John Sillitoe on behalf of the Wellcome Sanger Institute COVID-19 Surveillance Team (http://www.sanger.ac.uk/covid-team) |
| EPI_ISL_551692 | Lighthouse Lab in Milton Keynes | Wellcome Sanger Institute for the COVID-19 Genomics UK (COG-UK) consortium | The Lighthouse Lab in Alderley Park and Alex Alderton, Roberto Amato, Sonia Goncalves, Ewan Harrison, David K. Jackson, Ian Johnston, Dominic Kwiatkowski, Cordelia Langford, John Sillitoe on behalf of the Wellcome Sanger Institute COVID-19 Surveillance Team |
| EPI_ISL_551693, EPI_ISL_551695 | Lighthouse Lab in Milton Keynes | Wellcome Sanger Institute for the COVID-19 Genomics UK (COG-UK) consortium | The Lighthouse Lab in Milton Keynes and Alex Alderton, Roberto Amato, Sonia Goncalves, Ewan Harrison, David K. Jackson, Ian Johnston, Dominic Kwiatkowski, Cordelia Langford, John Sillitoe on behalf of the Wellcome Sanger Institute COVID-19 Surveillance Team (http://www.sanger.ac.uk/covid-team) |
| EPI_ISL_551696 | Lighthouse Lab in Milton Keynes | Wellcome Sanger Institute for the COVID-19 Genomics UK (COG-UK) consortium | The Lighthouse Lab in Alderley Park and Alex Alderton, Roberto Amato, Sonia Goncalves, Ewan Harrison, David K. Jackson, Ian Johnston, Dominic Kwiatkowski, Cordelia Langford, John Sillitoe on behalf of the Wellcome Sanger Institute COVID-19 Surveillance Team |
| EPI_ISL_551697, EPI_ISL_551699, EPI_ISL_551700, EPI_ISL_551701, EPI_ISL_551702, EPI_ISL_551703, EPI_ISL_551704, EPI_ISL_551705, EPI_ISL_551706, EPI_ISL_551707, EPI_ISL_551708, EPI_ISL_551709, EPI_ISL_551710, EPI_ISL_551711, EPI_ISL_551712, EPI_ISL_551713, EPI_ISL_551714, EPI_ISL_551715 | | | |
| see above | Lighthouse Lab in Milton Keynes | Wellcome Sanger Institute for the COVID-19 Genomics UK (COG-UK) consortium | The Lighthouse Lab in Milton Keynes and Alex Alderton, Roberto Amato, Sonia Goncalves, Ewan Harrison, David K. Jackson, Ian Johnston, Dominic Kwiatkowski, Cordelia Langford, John Sillitoe on behalf of the Wellcome Sanger Institute COVID-19 Surveillance Team (http://www.sanger.ac.uk/covid-team) |
| EPI_ISL_551716 | Lighthouse Lab in Milton Keynes | Wellcome Sanger Institute for the COVID-19 Genomics UK (COG-UK) consortium | The Lighthouse Lab in Alderley Park and Alex Alderton, Roberto Amato, Sonia Goncalves, Ewan Harrison, David K. Jackson, Ian Johnston, Dominic Kwiatkowski, Cordelia Langford, John Sillitoe on behalf of the Wellcome Sanger Institute COVID-19 Surveillance Team |
| EPI_ISL_551717, EPI_ISL_551718, EPI_ISL_551719, EPI_ISL_551720, EPI_ISL_551721, EPI_ISL_551723, EPI_ISL_551724, EPI_ISL_551726, EPI_ISL_551727, EPI_ISL_551728, EPI_ISL_551729, EPI_ISL_551730, EPI_ISL_551731, EPI_ISL_551732, EPI_ISL_551733, EPI_ISL_551734, EPI_ISL_551737, EPI_ISL_551738, EPI_ISL_551739, EPI_ISL_551741, EPI_ISL_551744, EPI_ISL_551745, EPI_ISL_551746, EPI_ISL_551747, EPI_ISL_551748, EPI_ISL_551750, EPI_ISL_551752, EPI_ISL_551753, EPI_ISL_551754, EPI_ISL_551757, EPI_ISL_551758 | | | |
| see above | Lighthouse Lab in Milton Keynes | Wellcome Sanger Institute for the COVID-19 Genomics UK (COG-UK) consortium | The Lighthouse Lab in Milton Keynes and Alex Alderton, Roberto Amato, Sonia Goncalves, Ewan Harrison, David K. Jackson, Ian Johnston, Dominic Kwiatkowski, Cordelia Langford, John Sillitoe on behalf of the Wellcome Sanger Institute COVID-19 Surveillance Team (http://www.sanger.ac.uk/covid-team) |
| EPI_ISL_551760 | Lighthouse Lab in Alderley Park | Wellcome Sanger Institute for the COVID-19 Genomics UK (COG-UK) consortium | The Lighthouse Lab in Alderley Park and Alex Alderton, Roberto Amato, Sonia Goncalves, Ewan Harrison, David K. Jackson, Ian Johnston, Dominic Kwiatkowski, Cordelia Langford, John Sillitoe on behalf of the Wellcome Sanger Institute COVID-19 Surveillance Team |
| EPI_ISL_551765, EPI_ISL_551766, EPI_ISL_551767, EPI_ISL_551768, EPI_ISL_551773, EPI_ISL_551774, EPI_ISL_551775 | Lighthouse Lab in Milton Keynes | Wellcome Sanger Institute for the COVID-19 Genomics UK (COG-UK) consortium | The Lighthouse Lab in Milton Keynes and Alex Alderton, Roberto Amato, Sonia Goncalves, Ewan Harrison, David K. Jackson, Ian Johnston, Dominic Kwiatkowski, Cordelia Langford, John Sillitoe on behalf of the Wellcome Sanger Institute COVID-19 Surveillance Team (http://www.sanger.ac.uk/covid-team) |
| EPI_ISL_551777 | Lighthouse Lab in Milton Keynes | Wellcome Sanger Institute for the COVID-19 Genomics UK (COG-UK) consortium | The Lighthouse Lab in Alderley Park and Alex Alderton, Roberto Amato, Sonia Goncalves, Ewan Harrison, David K. Jackson, Ian Johnston, Dominic Kwiatkowski, Cordelia Langford, John Sillitoe on behalf of the Wellcome Sanger Institute COVID-19 Surveillance Team |
| EPI_ISL_551781, EPI_ISL_551782, EPI_ISL_551786, EPI_ISL_551787, EPI_ISL_551790, EPI_ISL_551791, EPI_ISL_551792, EPI_ISL_551793, EPI_ISL_551797, EPI_ISL_551800, EPI_ISL_551802, EPI_ISL_551809, EPI_ISL_551813, EPI_ISL_551814, EPI_ISL_551817, EPI_ISL_551820, EPI_ISL_551821, EPI_ISL_551822 | | | |
| see above | Lighthouse Lab in Milton Keynes | Wellcome Sanger Institute for the COVID-19 Genomics UK (COG-UK) consortium | The Lighthouse Lab in Milton Keynes and Alex Alderton, Roberto Amato, Sonia Goncalves, Ewan Harrison, David K. Jackson, Ian Johnston, Dominic Kwiatkowski, Cordelia Langford, John Sillitoe on behalf of the Wellcome Sanger Institute COVID-19 Surveillance Team (http://www.sanger.ac.uk/covid-team) |
| EPI_ISL_551824 | Lighthouse Lab in Alderley Park | Wellcome Sanger Institute for the COVID-19 Genomics UK (COG-UK) consortium | The Lighthouse Lab in Alderley Park and Alex Alderton, Roberto Amato, Sonia Goncalves, Ewan Harrison, David K. Jackson, Ian Johnston, Dominic Kwiatkowski, Cordelia Langford, John Sillitoe on behalf of the Wellcome Sanger Institute COVID-19 Surveillance Team |
| EPI_ISL_551828, EPI_ISL_551829, EPI_ISL_551830, EPI_ISL_551831, EPI_ISL_551833, EPI_ISL_551923, EPI_ISL_551924, EPI_ISL_551926 | Lighthouse Lab in Milton Keynes | Wellcome Sanger Institute for the COVID-19 Genomics UK (COG-UK) consortium | The Lighthouse Lab in Milton Keynes and Alex Alderton, Roberto Amato, Sonia Goncalves, Ewan Harrison, David K. Jackson, Ian Johnston, Dominic Kwiatkowski, Cordelia Langford, John Sillitoe on behalf of the Wellcome Sanger Institute COVID-19 Surveillance Team (http://www.sanger.ac.uk/covid-team) |
| EPI_ISL_551927 | Lighthouse Lab in Milton Keynes | Wellcome Sanger Institute for the COVID-19 Genomics UK (COG-UK) consortium | The Lighthouse Lab in Alderley Park and Alex Alderton, Roberto Amato, Sonia Goncalves, Ewan Harrison, David K. Jackson, Ian Johnston, Dominic Kwiatkowski, Cordelia Langford, John Sillitoe on behalf of the Wellcome Sanger Institute COVID-19 Surveillance Team |
| EPI_ISL_551928, EPI_ISL_551929, EPI_ISL_551930, EPI_ISL_551931, EPI_ISL_551933, EPI_ISL_551934, EPI_ISL_551935 | Lighthouse Lab in Milton Keynes | Wellcome Sanger Institute for the COVID-19 Genomics UK (COG-UK) consortium | The Lighthouse Lab in Milton Keynes and Alex Alderton, Roberto Amato, Sonia Goncalves, Ewan Harrison, David K. Jackson, Ian Johnston, Dominic Kwiatkowski, Cordelia Langford, John Sillitoe on behalf of the Wellcome Sanger Institute COVID-19 Surveillance Team (http://www.sanger.ac.uk/covid-team) |
| EPI_ISL_551938 | Lighthouse Lab in Milton Keynes | Wellcome Sanger Institute for the COVID-19 Genomics UK (COG-UK) consortium | The Lighthouse Lab in Alderley Park and Alex Alderton, Roberto Amato, Sonia Goncalves, Ewan Harrison, David K. Jackson, Ian Johnston, Dominic Kwiatkowski, Cordelia Langford, John Sillitoe on behalf of the Wellcome Sanger Institute COVID-19 Surveillance Team |
| EPI_ISL_551939, EPI_ISL_551940, EPI_ISL_551943, EPI_ISL_551944, EPI_ISL_551945, EPI_ISL_551946, EPI_ISL_551947, EPI_ISL_551948 | Lighthouse Lab in Milton Keynes | Wellcome Sanger Institute for the COVID-19 Genomics UK (COG-UK) consortium | The Lighthouse Lab in Milton Keynes and Alex Alderton, Roberto Amato, Sonia Goncalves, Ewan Harrison, David K. Jackson, Ian Johnston, Dominic Kwiatkowski, Cordelia Langford, John Sillitoe on behalf of the Wellcome Sanger Institute COVID-19 Surveillance Team (http://www.sanger.ac.uk/covid-team) |
| EPI_ISL_551949 | Lighthouse Lab in Milton Keynes | Wellcome Sanger Institute for the COVID-19 Genomics UK (COG-UK) consortium | The Lighthouse Lab in Alderley Park and Alex Alderton, Roberto Amato, Sonia Goncalves, Ewan Harrison, David K. Jackson, Ian Johnston, Dominic Kwiatkowski, Cordelia Langford, John Sillitoe on behalf of the Wellcome Sanger Institute COVID-19 Surveillance Team |
| EPI_ISL_551950, EPI_ISL_551952, EPI_ISL_551954, EPI_ISL_551955, EPI_ISL_551956, EPI_ISL_551958, EPI_ISL_551959, EPI_ISL_551960, EPI_ISL_551962, EPI_ISL_551963, EPI_ISL_551965, EPI_ISL_551966, EPI_ISL_551968, EPI_ISL_551969, EPI_ISL_551970, EPI_ISL_551971, EPI_ISL_551972, EPI_ISL_551973 | | | |
| see above | Lighthouse Lab in Milton Keynes | Wellcome Sanger Institute for the COVID-19 Genomics UK (COG-UK) consortium | The Lighthouse Lab in Milton Keynes and Alex Alderton, Roberto Amato, Sonia Goncalves, Ewan Harrison, David K. Jackson, Ian Johnston, Dominic Kwiatkowski, Cordelia Langford, John Sillitoe on behalf of the Wellcome Sanger Institute COVID-19 Surveillance Team (http://www.sanger.ac.uk/covid-team) |
| EPI_ISL_551974 | Lighthouse Lab in Milton Keynes | Wellcome Sanger Institute for the COVID-19 Genomics UK (COG-UK) consortium | The Lighthouse Lab in Alderley Park and Alex Alderton, Roberto Amato, Sonia Goncalves, Ewan Harrison, David K. Jackson, Ian Johnston, Dominic Kwiatkowski, Cordelia Langford, John Sillitoe on behalf of the Wellcome Sanger Institute COVID-19 Surveillance Team |
| EPI_ISL_551975, EPI_ISL_551976, EPI_ISL_551977, EPI_ISL_551978, EPI_ISL_551979, EPI_ISL_551981, EPI_ISL_551982, EPI_ISL_551984, EPI_ISL_551985, EPI_ISL_551986, EPI_ISL_551987, EPI_ISL_551988, EPI_ISL_551989, EPI_ISL_551992, EPI_ISL_551995, EPI_ISL_551996, EPI_ISL_551998, EPI_ISL_551999, EPI_ISL_552000 | | | |
| see above | Lighthouse Lab in Milton Keynes | Wellcome Sanger Institute for the COVID-19 Genomics UK (COG-UK) consortium | The Lighthouse Lab in Milton Keynes and Alex Alderton, Roberto Amato, Sonia Goncalves, Ewan Harrison, David K. Jackson, Ian Johnston, Dominic Kwiatkowski, Cordelia Langford, John Sillitoe on behalf of the Wellcome Sanger Institute COVID-19 Surveillance Team (http://www.sanger.ac.uk/covid-team) |
| EPI_ISL_552002, EPI_ISL_552003 | Lighthouse Lab in Milton Keynes | Wellcome Sanger Institute for the COVID-19 Genomics UK (COG-UK) consortium | The Lighthouse Lab in Alderley Park and Alex Alderton, Roberto Amato, Sonia Goncalves, Ewan Harrison, David K. Jackson, Ian Johnston, Dominic Kwiatkowski, Cordelia Langford, John Sillitoe on behalf of the Wellcome Sanger Institute COVID-19 Surveillance Team |
| EPI_ISL_552004, EPI_ISL_552007, EPI_ISL_552011, EPI_ISL_552012 | Lighthouse Lab in Milton Keynes | Wellcome Sanger Institute for the COVID-19 Genomics UK (COG-UK) consortium | The Lighthouse Lab in Milton Keynes and Alex Alderton, Roberto Amato, Sonia Goncalves, Ewan Harrison, David K. Jackson, Ian Johnston, Dominic Kwiatkowski, Cordelia Langford, John Sillitoe on behalf of the Wellcome Sanger Institute COVID-19 Surveillance Team (http://www.sanger.ac.uk/covid-team) |

| | | | |
|---|---|---|---|
| EPI_ISL_552013 | Lighthouse Lab in Milton Keynes | Wellcome Sanger Institute for the COVID-19 Genomics UK (COG-UK) consortium | The Lighthouse Lab in Alderley Park and Alex Alderton, Roberto Amato, Sonia Goncalves, Ewan Harrison, David K. Jackson, Ian Johnston, Dominic Kwiatkowski, Cordelia Langford, John Sillitoe on behalf of the Wellcome Sanger Institute COVID-19 Surveillance Team |
| EPI_ISL_552016, EPI_ISL_552018, EPI_ISL_552019, EPI_ISL_552022, EPI_ISL_552023, EPI_ISL_552024, EPI_ISL_552025, EPI_ISL_552026, EPI_ISL_552027, EPI_ISL_552028, EPI_ISL_552029, EPI_ISL_552031, EPI_ISL_552032, EPI_ISL_552033, EPI_ISL_552034, EPI_ISL_552035, EPI_ISL_552036, EPI_ISL_552038, EPI_ISL_552039 | | | |
| see above | Lighthouse Lab in Milton Keynes | Wellcome Sanger Institute for the COVID-19 Genomics UK (COG-UK) consortium | The Lighthouse Lab in Milton Keynes and Alex Alderton, Roberto Amato, Sonia Goncalves, Ewan Harrison, David K. Jackson, Ian Johnston, Dominic Kwiatkowski, Cordelia Langford, John Sillitoe on behalf of the Wellcome Sanger Institute COVID-19 Surveillance Team (http://www.sanger.ac.uk/covid-team) |
| EPI_ISL_552040 | Lighthouse Lab in Milton Keynes | Wellcome Sanger Institute for the COVID-19 Genomics UK (COG-UK) consortium | The Lighthouse Lab in Alderley Park and Alex Alderton, Roberto Amato, Sonia Goncalves, Ewan Harrison, David K. Jackson, Ian Johnston, Dominic Kwiatkowski, Cordelia Langford, John Sillitoe on behalf of the Wellcome Sanger Institute COVID-19 Surveillance Team |
| EPI_ISL_552041, EPI_ISL_552042, EPI_ISL_552043 | Lighthouse Lab in Milton Keynes | Wellcome Sanger Institute for the COVID-19 Genomics UK (COG-UK) consortium | The Lighthouse Lab in Milton Keynes and Alex Alderton, Roberto Amato, Sonia Goncalves, Ewan Harrison, David K. Jackson, Ian Johnston, Dominic Kwiatkowski, Cordelia Langford, John Sillitoe on behalf of the Wellcome Sanger Institute COVID-19 Surveillance Team (http://www.sanger.ac.uk/covid-team) |
| EPI_ISL_552045 | Lighthouse Lab in Milton Keynes | Wellcome Sanger Institute for the COVID-19 Genomics UK (COG-UK) consortium | The Lighthouse Lab in Alderley Park and Alex Alderton, Roberto Amato, Sonia Goncalves, Ewan Harrison, David K. Jackson, Ian Johnston, Dominic Kwiatkowski, Cordelia Langford, John Sillitoe on behalf of the Wellcome Sanger Institute COVID-19 Surveillance Team |
| EPI_ISL_552047, EPI_ISL_552048, EPI_ISL_552049, EPI_ISL_552050, EPI_ISL_552052 | Lighthouse Lab in Milton Keynes | Wellcome Sanger Institute for the COVID-19 Genomics UK (COG-UK) consortium | The Lighthouse Lab in Milton Keynes and Alex Alderton, Roberto Amato, Sonia Goncalves, Ewan Harrison, David K. Jackson, Ian Johnston, Dominic Kwiatkowski, Cordelia Langford, John Sillitoe on behalf of the Wellcome Sanger Institute COVID-19 Surveillance Team (http://www.sanger.ac.uk/covid-team) |
| EPI_ISL_552053 | Lighthouse Lab in Milton Keynes | Wellcome Sanger Institute for the COVID-19 Genomics UK (COG-UK) consortium | The Lighthouse Lab in Alderley Park and Alex Alderton, Roberto Amato, Sonia Goncalves, Ewan Harrison, David K. Jackson, Ian Johnston, Dominic Kwiatkowski, Cordelia Langford, John Sillitoe on behalf of the Wellcome Sanger Institute COVID-19 Surveillance Team |
| EPI_ISL_552054, EPI_ISL_552055 | Lighthouse Lab in Milton Keynes | Wellcome Sanger Institute for the COVID-19 Genomics UK (COG-UK) consortium | The Lighthouse Lab in Milton Keynes and Alex Alderton, Roberto Amato, Sonia Goncalves, Ewan Harrison, David K. Jackson, Ian Johnston, Dominic Kwiatkowski, Cordelia Langford, John Sillitoe on behalf of the Wellcome Sanger Institute COVID-19 Surveillance Team (http://www.sanger.ac.uk/covid-team) |
| EPI_ISL_552056 | Lighthouse Lab in Milton Keynes | Wellcome Sanger Institute for the COVID-19 Genomics UK (COG-UK) consortium | The Lighthouse Lab in Alderley Park and Alex Alderton, Roberto Amato, Sonia Goncalves, Ewan Harrison, David K. Jackson, Ian Johnston, Dominic Kwiatkowski, Cordelia Langford, John Sillitoe on behalf of the Wellcome Sanger Institute COVID-19 Surveillance Team |
| EPI_ISL_552057 | Lighthouse Lab in Milton Keynes | Wellcome Sanger Institute for the COVID-19 Genomics UK (COG-UK) consortium | The Lighthouse Lab in Milton Keynes and Alex Alderton, Roberto Amato, Sonia Goncalves, Ewan Harrison, David K. Jackson, Ian Johnston, Dominic Kwiatkowski, Cordelia Langford, John Sillitoe on behalf of the Wellcome Sanger Institute COVID-19 Surveillance Team (http://www.sanger.ac.uk/covid-team) |
| EPI_ISL_552058 | Lighthouse Lab in Milton Keynes | Wellcome Sanger Institute for the COVID-19 Genomics UK (COG-UK) consortium | The Lighthouse Lab in Alderley Park and Alex Alderton, Roberto Amato, Sonia Goncalves, Ewan Harrison, David K. Jackson, Ian Johnston, Dominic Kwiatkowski, Cordelia Langford, John Sillitoe on behalf of the Wellcome Sanger Institute COVID-19 Surveillance Team |
| EPI_ISL_552059, EPI_ISL_552060 | Lighthouse Lab in Milton Keynes | Wellcome Sanger Institute for the COVID-19 Genomics UK (COG-UK) consortium | The Lighthouse Lab in Milton Keynes and Alex Alderton, Roberto Amato, Sonia Goncalves, Ewan Harrison, David K. Jackson, Ian Johnston, Dominic Kwiatkowski, Cordelia Langford, John Sillitoe on behalf of the Wellcome Sanger Institute COVID-19 Surveillance Team (http://www.sanger.ac.uk/covid-team) |
| EPI_ISL_552062 | Lighthouse Lab in Milton Keynes | Wellcome Sanger Institute for the COVID-19 Genomics UK (COG-UK) consortium | The Lighthouse Lab in Alderley Park and Alex Alderton, Roberto Amato, Sonia Goncalves, Ewan Harrison, David K. Jackson, Ian Johnston, Dominic Kwiatkowski, Cordelia Langford, John Sillitoe on behalf of the Wellcome Sanger Institute COVID-19 Surveillance Team |
| EPI_ISL_552063, EPI_ISL_552064 | Lighthouse Lab in Milton Keynes | Wellcome Sanger Institute for the COVID-19 Genomics UK (COG-UK) consortium | The Lighthouse Lab in Milton Keynes and Alex Alderton, Roberto Amato, Sonia Goncalves, Ewan Harrison, David K. Jackson, Ian Johnston, Dominic Kwiatkowski, Cordelia Langford, John Sillitoe on behalf of the Wellcome Sanger Institute COVID-19 Surveillance Team (http://www.sanger.ac.uk/covid-team) |
| EPI_ISL_552066 | Lighthouse Lab in Milton Keynes | Wellcome Sanger Institute for the COVID-19 Genomics UK (COG-UK) consortium | The Lighthouse Lab in Alderley Park and Alex Alderton, Roberto Amato, Sonia Goncalves, Ewan Harrison, David K. Jackson, Ian Johnston, Dominic Kwiatkowski, Cordelia Langford, John Sillitoe on behalf of the Wellcome Sanger Institute COVID-19 Surveillance Team |
| EPI_ISL_552067, EPI_ISL_552068, EPI_ISL_552069, EPI_ISL_552070, EPI_ISL_552071, EPI_ISL_552072, EPI_ISL_552074, EPI_ISL_552075, EPI_ISL_552076, EPI_ISL_552077, EPI_ISL_552078, EPI_ISL_552079, EPI_ISL_552080 | | | |
| see above | Lighthouse Lab in Milton Keynes | Wellcome Sanger Institute for the COVID-19 Genomics UK (COG-UK) consortium | The Lighthouse Lab in Milton Keynes and Alex Alderton, Roberto Amato, Sonia Goncalves, Ewan Harrison, David K. Jackson, Ian Johnston, Dominic Kwiatkowski, Cordelia Langford, John Sillitoe on behalf of the Wellcome Sanger Institute COVID-19 Surveillance Team (http://www.sanger.ac.uk/covid-team) |
| EPI_ISL_552081 | Lighthouse Lab in Milton Keynes | Wellcome Sanger Institute for the COVID-19 Genomics UK (COG-UK) consortium | The Lighthouse Lab in Alderley Park and Alex Alderton, Roberto Amato, Sonia Goncalves, Ewan Harrison, David K. Jackson, Ian Johnston, Dominic Kwiatkowski, Cordelia Langford, John Sillitoe on behalf of the Wellcome Sanger Institute COVID-19 Surveillance Team |
| EPI_ISL_552084, EPI_ISL_552085, EPI_ISL_552086, EPI_ISL_552087, EPI_ISL_552088, EPI_ISL_552089, EPI_ISL_552090, EPI_ISL_552092, EPI_ISL_552093, EPI_ISL_552096, EPI_ISL_552097, EPI_ISL_552099, EPI_ISL_552100, EPI_ISL_552101, EPI_ISL_552102, EPI_ISL_552103, EPI_ISL_552104, EPI_ISL_552107, EPI_ISL_552108, EPI_ISL_552109, EPI_ISL_552110, EPI_ISL_552112, EPI_ISL_552113, EPI_ISL_552114, EPI_ISL_552115, EPI_ISL_552116 | | | |
| see above | Lighthouse Lab in Milton Keynes | Wellcome Sanger Institute for the COVID-19 Genomics UK (COG-UK) consortium | The Lighthouse Lab in Milton Keynes and Alex Alderton, Roberto Amato, Sonia Goncalves, Ewan Harrison, David K. Jackson, Ian Johnston, Dominic Kwiatkowski, Cordelia Langford, John Sillitoe on behalf of the Wellcome Sanger Institute COVID-19 Surveillance Team (http://www.sanger.ac.uk/covid-team) |
| EPI_ISL_552117 | Lighthouse Lab in Milton Keynes | Wellcome Sanger Institute for the COVID-19 Genomics UK (COG-UK) consortium | The Lighthouse Lab in Alderley Park and Alex Alderton, Roberto Amato, Sonia Goncalves, Ewan Harrison, David K. Jackson, Ian Johnston, Dominic Kwiatkowski, Cordelia Langford, John Sillitoe on behalf of the Wellcome Sanger Institute COVID-19 Surveillance Team |
| EPI_ISL_552118, EPI_ISL_552119, EPI_ISL_552120, EPI_ISL_552121, EPI_ISL_552122, EPI_ISL_552123, EPI_ISL_552125, EPI_ISL_552127, EPI_ISL_552128, EPI_ISL_552129, EPI_ISL_552130, EPI_ISL_552131 | | | |
| see above | Lighthouse Lab in Milton Keynes | Wellcome Sanger Institute for the COVID-19 Genomics UK (COG-UK) consortium | The Lighthouse Lab in Milton Keynes and Alex Alderton, Roberto Amato, Sonia Goncalves, Ewan Harrison, David K. Jackson, Ian Johnston, Dominic Kwiatkowski, Cordelia Langford, John Sillitoe on behalf of the Wellcome Sanger Institute COVID-19 Surveillance Team (http://www.sanger.ac.uk/covid-team) |
| EPI_ISL_552132 | Lighthouse Lab in Milton Keynes | Wellcome Sanger Institute for the COVID-19 Genomics UK (COG-UK) consortium | The Lighthouse Lab in Alderley Park and Alex Alderton, Roberto Amato, Sonia Goncalves, Ewan Harrison, David K. Jackson, Ian Johnston, Dominic Kwiatkowski, Cordelia Langford, John Sillitoe on behalf of the Wellcome Sanger Institute COVID-19 Surveillance Team |
| EPI_ISL_552133, EPI_ISL_552134, EPI_ISL_552135, EPI_ISL_552136, EPI_ISL_552137, EPI_ISL_552139, EPI_ISL_552140, EPI_ISL_552141, EPI_ISL_552142, EPI_ISL_552143 | Lighthouse Lab in Milton Keynes | Wellcome Sanger Institute for the COVID-19 Genomics UK (COG-UK) consortium | The Lighthouse Lab in Milton Keynes and Alex Alderton, Roberto Amato, Sonia Goncalves, Ewan Harrison, David K. Jackson, Ian Johnston, Dominic Kwiatkowski, Cordelia Langford, John Sillitoe on behalf of the Wellcome Sanger Institute COVID-19 Surveillance Team (http://www.sanger.ac.uk/covid-team) |
| EPI_ISL_552144, EPI_ISL_552145 | Lighthouse Lab in Milton Keynes | Wellcome Sanger Institute for the COVID-19 Genomics UK (COG-UK) consortium | The Lighthouse Lab in Alderley Park and Alex Alderton, Roberto Amato, Sonia Goncalves, Ewan Harrison, David K. Jackson, Ian Johnston, Dominic Kwiatkowski, Cordelia Langford, John Sillitoe on behalf of the Wellcome Sanger Institute COVID-19 Surveillance Team |
| EPI_ISL_552146, EPI_ISL_552147, EPI_ISL_552149, EPI_ISL_552150, EPI_ISL_552152, EPI_ISL_552153, EPI_ISL_552154, EPI_ISL_552155, EPI_ISL_552156, EPI_ISL_552158, EPI_ISL_552159 | | | |
| see above | Lighthouse Lab in Milton Keynes | Wellcome Sanger Institute for the COVID-19 Genomics UK (COG-UK) consortium | The Lighthouse Lab in Milton Keynes and Alex Alderton, Roberto Amato, Sonia Goncalves, Ewan Harrison, David K. Jackson, Ian Johnston, Dominic Kwiatkowski, Cordelia Langford, John Sillitoe on behalf of the Wellcome Sanger Institute COVID-19 Surveillance Team (http://www.sanger.ac.uk/covid-team) |
| EPI_ISL_552162 | Lighthouse Lab in Milton Keynes | Wellcome Sanger Institute for the COVID-19 Genomics UK (COG-UK) consortium | The Lighthouse Lab in Alderley Park and Alex Alderton, Roberto Amato, Sonia Goncalves, Ewan Harrison, David K. Jackson, Ian Johnston, Dominic Kwiatkowski, Cordelia Langford, John Sillitoe on behalf of the Wellcome Sanger Institute COVID-19 Surveillance Team |
| EPI_ISL_552163 | Lighthouse Lab in Milton Keynes | Wellcome Sanger Institute for the COVID-19 Genomics UK (COG-UK) consortium | The Lighthouse Lab in Milton Keynes and Alex Alderton, Roberto Amato, Sonia Goncalves, Ewan Harrison, David K. Jackson, Ian Johnston, Dominic Kwiatkowski, Cordelia Langford, John Sillitoe on behalf of the Wellcome Sanger Institute COVID-19 Surveillance Team |

| | | | |
|---|---|---|---|
| | | | (http://www.sanger.ac.uk/covid-team) |
| EPI_ISL_552164 | Lighthouse Lab in Milton Keynes | Wellcome Sanger Institute for the COVID-19 Genomics UK (COG-UK) consortium | The Lighthouse Lab in Alderley Park and Alex Alderton, Roberto Amato, Sonia Goncalves, Ewan Harrison, David K. Jackson, Ian Johnston, Dominic Kwiatkowski, Cordelia Langford, John Sillitoe on behalf of the Wellcome Sanger Institute COVID-19 Surveillance Team |
| EPI_ISL_552165, EPI_ISL_552167, EPI_ISL_552168, EPI_ISL_552169, EPI_ISL_552171, EPI_ISL_552172, EPI_ISL_552174, EPI_ISL_552175, EPI_ISL_552176, EPI_ISL_552177, EPI_ISL_552178, EPI_ISL_552179, EPI_ISL_552181 | | | |
| see above | Lighthouse Lab in Milton Keynes | Wellcome Sanger Institute for the COVID-19 Genomics UK (COG-UK) consortium | The Lighthouse Lab in Milton Keynes and Alex Alderton, Roberto Amato, Sonia Goncalves, Ewan Harrison, David K. Jackson, Ian Johnston, Dominic Kwiatkowski, Cordelia Langford, John Sillitoe on behalf of the Wellcome Sanger Institute COVID-19 Surveillance Team (http://www.sanger.ac.uk/covid-team) |
| EPI_ISL_552182 | Lighthouse Lab in Milton Keynes | Wellcome Sanger Institute for the COVID-19 Genomics UK (COG-UK) consortium | The Lighthouse Lab in Alderley Park and Alex Alderton, Roberto Amato, Sonia Goncalves, Ewan Harrison, David K. Jackson, Ian Johnston, Dominic Kwiatkowski, Cordelia Langford, John Sillitoe on behalf of the Wellcome Sanger Institute COVID-19 Surveillance Team |
| EPI_ISL_552183, EPI_ISL_552184, EPI_ISL_552186, EPI_ISL_552187, EPI_ISL_552188, EPI_ISL_552189, EPI_ISL_552190 | Lighthouse Lab in Milton Keynes | Wellcome Sanger Institute for the COVID-19 Genomics UK (COG-UK) consortium | The Lighthouse Lab in Milton Keynes and Alex Alderton, Roberto Amato, Sonia Goncalves, Ewan Harrison, David K. Jackson, Ian Johnston, Dominic Kwiatkowski, Cordelia Langford, John Sillitoe on behalf of the Wellcome Sanger Institute COVID-19 Surveillance Team (http://www.sanger.ac.uk/covid-team) |
| EPI_ISL_552191 | Lighthouse Lab in Milton Keynes | Wellcome Sanger Institute for the COVID-19 Genomics UK (COG-UK) consortium | The Lighthouse Lab in Alderley Park and Alex Alderton, Roberto Amato, Sonia Goncalves, Ewan Harrison, David K. Jackson, Ian Johnston, Dominic Kwiatkowski, Cordelia Langford, John Sillitoe on behalf of the Wellcome Sanger Institute COVID-19 Surveillance Team |
| EPI_ISL_552192, EPI_ISL_552193, EPI_ISL_552194, EPI_ISL_552195, EPI_ISL_552196, EPI_ISL_552197, EPI_ISL_552198, EPI_ISL_552199, EPI_ISL_552200, EPI_ISL_552201, EPI_ISL_552203 | | | |
| see above | Lighthouse Lab in Milton Keynes | Wellcome Sanger Institute for the COVID-19 Genomics UK (COG-UK) consortium | The Lighthouse Lab in Milton Keynes and Alex Alderton, Roberto Amato, Sonia Goncalves, Ewan Harrison, David K. Jackson, Ian Johnston, Dominic Kwiatkowski, Cordelia Langford, John Sillitoe on behalf of the Wellcome Sanger Institute COVID-19 Surveillance Team (http://www.sanger.ac.uk/covid-team) |
| EPI_ISL_552205 | Lighthouse Lab in Milton Keynes | Wellcome Sanger Institute for the COVID-19 Genomics UK (COG-UK) consortium | The Lighthouse Lab in Alderley Park and Alex Alderton, Roberto Amato, Sonia Goncalves, Ewan Harrison, David K. Jackson, Ian Johnston, Dominic Kwiatkowski, Cordelia Langford, John Sillitoe on behalf of the Wellcome Sanger Institute COVID-19 Surveillance Team |
| EPI_ISL_552206, EPI_ISL_552208, EPI_ISL_552211 | Lighthouse Lab in Milton Keynes | Wellcome Sanger Institute for the COVID-19 Genomics UK (COG-UK) consortium | The Lighthouse Lab in Milton Keynes and Alex Alderton, Roberto Amato, Sonia Goncalves, Ewan Harrison, David K. Jackson, Ian Johnston, Dominic Kwiatkowski, Cordelia Langford, John Sillitoe on behalf of the Wellcome Sanger Institute COVID-19 Surveillance Team (http://www.sanger.ac.uk/covid-team) |
| EPI_ISL_552212 | Lighthouse Lab in Milton Keynes | Wellcome Sanger Institute for the COVID-19 Genomics UK (COG-UK) consortium | The Lighthouse Lab in Alderley Park and Alex Alderton, Roberto Amato, Sonia Goncalves, Ewan Harrison, David K. Jackson, Ian Johnston, Dominic Kwiatkowski, Cordelia Langford, John Sillitoe on behalf of the Wellcome Sanger Institute COVID-19 Surveillance Team |
| EPI_ISL_552213, EPI_ISL_552214, EPI_ISL_552215, EPI_ISL_552216, EPI_ISL_552217, EPI_ISL_552218, EPI_ISL_552219, EPI_ISL_552220, EPI_ISL_552221, EPI_ISL_552222, EPI_ISL_552223, EPI_ISL_552224, EPI_ISL_552225, EPI_ISL_552226, EPI_ISL_552227, EPI_ISL_552228, EPI_ISL_552229, EPI_ISL_552230, EPI_ISL_552231, EPI_ISL_552232, EPI_ISL_552234, EPI_ISL_552235, EPI_ISL_552236, EPI_ISL_552237, EPI_ISL_552238, EPI_ISL_552241, EPI_ISL_552242, EPI_ISL_552244, EPI_ISL_552245, EPI_ISL_552247, EPI_ISL_552248, EPI_ISL_552249, EPI_ISL_552250, EPI_ISL_552252, EPI_ISL_552253, EPI_ISL_552254, EPI_ISL_552255, EPI_ISL_552256, EPI_ISL_552257, EPI_ISL_552258, EPI_ISL_552260, EPI_ISL_552261, EPI_ISL_552262, EPI_ISL_552263, EPI_ISL_552264, EPI_ISL_552265, EPI_ISL_552266, EPI_ISL_552267, EPI_ISL_552268, EPI_ISL_552269, EPI_ISL_552270, EPI_ISL_552271, EPI_ISL_552272, EPI_ISL_552273, EPI_ISL_552275, EPI_ISL_552276, EPI_ISL_552277 | | | |
| see above | Lighthouse Lab in Milton Keynes | Wellcome Sanger Institute for the COVID-19 Genomics UK (COG-UK) consortium | The Lighthouse Lab in Milton Keynes and Alex Alderton, Roberto Amato, Sonia Goncalves, Ewan Harrison, David K. Jackson, Ian Johnston, Dominic Kwiatkowski, Cordelia Langford, John Sillitoe on behalf of the Wellcome Sanger Institute COVID-19 Surveillance Team (http://www.sanger.ac.uk/covid-team) |
| EPI_ISL_552278 | Lighthouse Lab in Milton Keynes | Wellcome Sanger Institute for the COVID-19 Genomics UK (COG-UK) consortium | The Lighthouse Lab in Alderley Park and Alex Alderton, Roberto Amato, Sonia Goncalves, Ewan Harrison, David K. Jackson, Ian Johnston, Dominic Kwiatkowski, Cordelia Langford, John Sillitoe on behalf of the Wellcome Sanger Institute COVID-19 Surveillance Team |
| EPI_ISL_552279, EPI_ISL_552280, EPI_ISL_552281, EPI_ISL_552282, EPI_ISL_552283, EPI_ISL_552284, EPI_ISL_552285, EPI_ISL_552286, EPI_ISL_552287, EPI_ISL_552291, EPI_ISL_552293, EPI_ISL_552295, EPI_ISL_552296, EPI_ISL_552297, EPI_ISL_552298, EPI_ISL_552299, EPI_ISL_552300, EPI_ISL_552301, EPI_ISL_552302, EPI_ISL_552303, EPI_ISL_552305, EPI_ISL_552308, EPI_ISL_552309, EPI_ISL_552310, EPI_ISL_552311, EPI_ISL_552312, EPI_ISL_552313, EPI_ISL_552315, EPI_ISL_552316, EPI_ISL_552317, EPI_ISL_552318, EPI_ISL_552320, EPI_ISL_552321, EPI_ISL_552322, EPI_ISL_552323, EPI_ISL_552324, EPI_ISL_552325, EPI_ISL_552326, EPI_ISL_552327, EPI_ISL_552329, EPI_ISL_552330, EPI_ISL_552332 | | | |
| see above | Lighthouse Lab in Milton Keynes | Wellcome Sanger Institute for the COVID-19 Genomics UK (COG-UK) consortium | The Lighthouse Lab in Milton Keynes and Alex Alderton, Roberto Amato, Sonia Goncalves, Ewan Harrison, David K. Jackson, Ian Johnston, Dominic Kwiatkowski, Cordelia Langford, John Sillitoe on behalf of the Wellcome Sanger Institute COVID-19 Surveillance Team (http://www.sanger.ac.uk/covid-team) |
| EPI_ISL_552334 | Lighthouse Lab in Milton Keynes | Wellcome Sanger Institute for the COVID-19 Genomics UK (COG-UK) consortium | The Lighthouse Lab in Alderley Park and Alex Alderton, Roberto Amato, Sonia Goncalves, Ewan Harrison, David K. Jackson, Ian Johnston, Dominic Kwiatkowski, Cordelia Langford, John Sillitoe on behalf of the Wellcome Sanger Institute COVID-19 Surveillance Team |
| EPI_ISL_552335, EPI_ISL_552336, EPI_ISL_552337, EPI_ISL_552338, EPI_ISL_552339, EPI_ISL_552340, EPI_ISL_552341, EPI_ISL_552343, EPI_ISL_552344, EPI_ISL_552345, EPI_ISL_552347, EPI_ISL_552349, EPI_ISL_552350, EPI_ISL_552351, EPI_ISL_552352, EPI_ISL_552353, EPI_ISL_552355, EPI_ISL_552356, EPI_ISL_552357 | | | |
| see above | Lighthouse Lab in Milton Keynes | Wellcome Sanger Institute for the COVID-19 Genomics UK (COG-UK) consortium | The Lighthouse Lab in Milton Keynes and Alex Alderton, Roberto Amato, Sonia Goncalves, Ewan Harrison, David K. Jackson, Ian Johnston, Dominic Kwiatkowski, Cordelia Langford, John Sillitoe on behalf of the Wellcome Sanger Institute COVID-19 Surveillance Team (http://www.sanger.ac.uk/covid-team) |
| EPI_ISL_552358, EPI_ISL_552359 | Lighthouse Lab in Milton Keynes | Wellcome Sanger Institute for the COVID-19 Genomics UK (COG-UK) consortium | The Lighthouse Lab in Alderley Park and Alex Alderton, Roberto Amato, Sonia Goncalves, Ewan Harrison, David K. Jackson, Ian Johnston, Dominic Kwiatkowski, Cordelia Langford, John Sillitoe on behalf of the Wellcome Sanger Institute COVID-19 Surveillance Team |
| EPI_ISL_552360, EPI_ISL_552361, EPI_ISL_552362, EPI_ISL_552363, EPI_ISL_552364, EPI_ISL_552365, EPI_ISL_552367, EPI_ISL_552368, EPI_ISL_552369, EPI_ISL_552370, EPI_ISL_552371, EPI_ISL_552372, EPI_ISL_552373, EPI_ISL_552375, EPI_ISL_552376, EPI_ISL_552377, EPI_ISL_552378, EPI_ISL_552380, EPI_ISL_552381, EPI_ISL_552382, EPI_ISL_552383, EPI_ISL_552384 | | | |
| see above | Lighthouse Lab in Milton Keynes | Wellcome Sanger Institute for the COVID-19 Genomics UK (COG-UK) consortium | The Lighthouse Lab in Milton Keynes and Alex Alderton, Roberto Amato, Sonia Goncalves, Ewan Harrison, David K. Jackson, Ian Johnston, Dominic Kwiatkowski, Cordelia Langford, John Sillitoe on behalf of the Wellcome Sanger Institute COVID-19 Surveillance Team (http://www.sanger.ac.uk/covid-team) |
| EPI_ISL_552385 | Lighthouse Lab in Milton Keynes | Wellcome Sanger Institute for the COVID-19 Genomics UK (COG-UK) consortium | The Lighthouse Lab in Alderley Park and Alex Alderton, Roberto Amato, Sonia Goncalves, Ewan Harrison, David K. Jackson, Ian Johnston, Dominic Kwiatkowski, Cordelia Langford, John Sillitoe on behalf of the Wellcome Sanger Institute COVID-19 Surveillance Team |
| EPI_ISL_552386, EPI_ISL_552387, EPI_ISL_552389, EPI_ISL_552391, EPI_ISL_552393, EPI_ISL_552394, EPI_ISL_552395, EPI_ISL_552396, EPI_ISL_552397, EPI_ISL_552398 | Lighthouse Lab in Milton Keynes | Wellcome Sanger Institute for the COVID-19 Genomics UK (COG-UK) consortium | The Lighthouse Lab in Milton Keynes and Alex Alderton, Roberto Amato, Sonia Goncalves, Ewan Harrison, David K. Jackson, Ian Johnston, Dominic Kwiatkowski, Cordelia Langford, John Sillitoe on behalf of the Wellcome Sanger Institute COVID-19 Surveillance Team (http://www.sanger.ac.uk/covid-team) |
| EPI_ISL_552399 | Lighthouse Lab in Milton Keynes | Wellcome Sanger Institute for the COVID-19 Genomics UK (COG-UK) consortium | The Lighthouse Lab in Alderley Park and Alex Alderton, Roberto Amato, Sonia Goncalves, Ewan Harrison, David K. Jackson, Ian Johnston, Dominic Kwiatkowski, Cordelia Langford, John Sillitoe on behalf of the Wellcome Sanger Institute COVID-19 Surveillance Team |
| EPI_ISL_552400, EPI_ISL_552401, EPI_ISL_552402, EPI_ISL_552403, EPI_ISL_552405, EPI_ISL_552406, EPI_ISL_552407 | Lighthouse Lab in Milton Keynes | Wellcome Sanger Institute for the COVID-19 Genomics UK (COG-UK) consortium | The Lighthouse Lab in Milton Keynes and Alex Alderton, Roberto Amato, Sonia Goncalves, Ewan Harrison, David K. Jackson, Ian Johnston, Dominic Kwiatkowski, Cordelia Langford, John Sillitoe on behalf of the Wellcome Sanger Institute COVID-19 Surveillance Team (http://www.sanger.ac.uk/covid-team) |
| EPI_ISL_552408 | Lighthouse Lab in Milton Keynes | Wellcome Sanger Institute for the COVID-19 Genomics UK (COG-UK) consortium | The Lighthouse Lab in Alderley Park and Alex Alderton, Roberto Amato, Sonia Goncalves, Ewan Harrison, David K. Jackson, Ian Johnston, Dominic Kwiatkowski, Cordelia Langford, John Sillitoe on behalf of the Wellcome Sanger Institute COVID-19 Surveillance Team |
| EPI_ISL_552409, EPI_ISL_552410, EPI_ISL_552411, EPI_ISL_552412 | Lighthouse Lab in Milton Keynes | Wellcome Sanger Institute for the COVID-19 Genomics UK (COG-UK) consortium | The Lighthouse Lab in Milton Keynes and Alex Alderton, Roberto Amato, Sonia Goncalves, Ewan Harrison, David K. Jackson, Ian Johnston, Dominic Kwiatkowski, Cordelia Langford, John Sillitoe on behalf of the Wellcome Sanger Institute COVID-19 Surveillance Team (http://www.sanger.ac.uk/covid-team) |

| | | | |
|---|---|---|---|
| EPI_ISL_552414 | Lighthouse Lab in Milton Keynes | Wellcome Sanger Institute for the COVID-19 Genomics UK (COG-UK) consortium | The Lighthouse Lab in Alderley Park and Alex Alderton, Roberto Amato, Sonia Goncalves, Ewan Harrison, David K. Jackson, Ian Johnston, Dominic Kwiatkowski, Cordelia Langford, John Sillitoe on behalf of the Wellcome Sanger Institute COVID-19 Surveillance Team |
| EPI_ISL_552415 | Lighthouse Lab in Milton Keynes | Wellcome Sanger Institute for the COVID-19 Genomics UK (COG-UK) consortium | The Lighthouse Lab in Milton Keynes and Alex Alderton, Roberto Amato, Sonia Goncalves, Ewan Harrison, David K. Jackson, Ian Johnston, Dominic Kwiatkowski, Cordelia Langford, John Sillitoe on behalf of the Wellcome Sanger Institute COVID-19 Surveillance Team (http://www.sanger.ac.uk/covid-team) |
| EPI_ISL_552417 | Lighthouse Lab in Milton Keynes | Wellcome Sanger Institute for the COVID-19 Genomics UK (COG-UK) consortium | The Lighthouse Lab in Alderley Park and Alex Alderton, Roberto Amato, Sonia Goncalves, Ewan Harrison, David K. Jackson, Ian Johnston, Dominic Kwiatkowski, Cordelia Langford, John Sillitoe on behalf of the Wellcome Sanger Institute COVID-19 Surveillance Team |
| EPI_ISL_552418, EPI_ISL_552419, EPI_ISL_552420, EPI_ISL_552421, EPI_ISL_552422, EPI_ISL_552423, EPI_ISL_552426, EPI_ISL_552427, EPI_ISL_552428, EPI_ISL_552429, EPI_ISL_552430, EPI_ISL_552431, EPI_ISL_552432, EPI_ISL_552433, EPI_ISL_552435, EPI_ISL_552436, EPI_ISL_552438, EPI_ISL_552439, EPI_ISL_552440 | | | |
| see above | Lighthouse Lab in Milton Keynes | Wellcome Sanger Institute for the COVID-19 Genomics UK (COG-UK) consortium | The Lighthouse Lab in Milton Keynes and Alex Alderton, Roberto Amato, Sonia Goncalves, Ewan Harrison, David K. Jackson, Ian Johnston, Dominic Kwiatkowski, Cordelia Langford, John Sillitoe on behalf of the Wellcome Sanger Institute COVID-19 Surveillance Team (http://www.sanger.ac.uk/covid-team) |
| EPI_ISL_552441 | Lighthouse Lab in Milton Keynes | Wellcome Sanger Institute for the COVID-19 Genomics UK (COG-UK) consortium | The Lighthouse Lab in Alderley Park and Alex Alderton, Roberto Amato, Sonia Goncalves, Ewan Harrison, David K. Jackson, Ian Johnston, Dominic Kwiatkowski, Cordelia Langford, John Sillitoe on behalf of the Wellcome Sanger Institute COVID-19 Surveillance Team |
| EPI_ISL_552442, EPI_ISL_552443, EPI_ISL_552444, EPI_ISL_552445 | Lighthouse Lab in Milton Keynes | Wellcome Sanger Institute for the COVID-19 Genomics UK (COG-UK) consortium | The Lighthouse Lab in Milton Keynes and Alex Alderton, Roberto Amato, Sonia Goncalves, Ewan Harrison, David K. Jackson, Ian Johnston, Dominic Kwiatkowski, Cordelia Langford, John Sillitoe on behalf of the Wellcome Sanger Institute COVID-19 Surveillance Team (http://www.sanger.ac.uk/covid-team) |
| EPI_ISL_552446 | Lighthouse Lab in Alderley Park | Wellcome Sanger Institute for the COVID-19 Genomics UK (COG-UK) consortium | The Lighthouse Lab in Alderley Park and Alex Alderton, Roberto Amato, Sonia Goncalves, Ewan Harrison, David K. Jackson, Ian Johnston, Dominic Kwiatkowski, Cordelia Langford, John Sillitoe on behalf of the Wellcome Sanger Institute COVID-19 Surveillance Team |
| EPI_ISL_552447, EPI_ISL_552448 | Lighthouse Lab in Milton Keynes | Wellcome Sanger Institute for the COVID-19 Genomics UK (COG-UK) consortium | The Lighthouse Lab in Milton Keynes and Alex Alderton, Roberto Amato, Sonia Goncalves, Ewan Harrison, David K. Jackson, Ian Johnston, Dominic Kwiatkowski, Cordelia Langford, John Sillitoe on behalf of the Wellcome Sanger Institute COVID-19 Surveillance Team (http://www.sanger.ac.uk/covid-team) |
| EPI_ISL_552449 | Lighthouse Lab in Alderley Park | Wellcome Sanger Institute for the COVID-19 Genomics UK (COG-UK) consortium | The Lighthouse Lab in Alderley Park and Alex Alderton, Roberto Amato, Sonia Goncalves, Ewan Harrison, David K. Jackson, Ian Johnston, Dominic Kwiatkowski, Cordelia Langford, John Sillitoe on behalf of the Wellcome Sanger Institute COVID-19 Surveillance Team |
| EPI_ISL_552450, EPI_ISL_552451, EPI_ISL_552452, EPI_ISL_552453, EPI_ISL_552454, EPI_ISL_552455, EPI_ISL_552456, EPI_ISL_552459, EPI_ISL_552460, EPI_ISL_552461, EPI_ISL_552462 | | | |
| see above | Lighthouse Lab in Milton Keynes | Wellcome Sanger Institute for the COVID-19 Genomics UK (COG-UK) consortium | The Lighthouse Lab in Milton Keynes and Alex Alderton, Roberto Amato, Sonia Goncalves, Ewan Harrison, David K. Jackson, Ian Johnston, Dominic Kwiatkowski, Cordelia Langford, John Sillitoe on behalf of the Wellcome Sanger Institute COVID-19 Surveillance Team (http://www.sanger.ac.uk/covid-team) |
| EPI_ISL_552463 | Lighthouse Lab in Alderley Park | Wellcome Sanger Institute for the COVID-19 Genomics UK (COG-UK) consortium | The Lighthouse Lab in Alderley Park and Alex Alderton, Roberto Amato, Sonia Goncalves, Ewan Harrison, David K. Jackson, Ian Johnston, Dominic Kwiatkowski, Cordelia Langford, John Sillitoe on behalf of the Wellcome Sanger Institute COVID-19 Surveillance Team (http://www.sanger.ac.uk/covid-team) |
| EPI_ISL_552464 | Lighthouse Lab in Alderley Park | Wellcome Sanger Institute for the COVID-19 Genomics UK (COG-UK) consortium | The Lighthouse Lab in Alderley Park and Alex Alderton, Roberto Amato, Sonia Goncalves, Ewan Harrison, David K. Jackson, Ian Johnston, Dominic Kwiatkowski, Cordelia Langford, John Sillitoe on behalf of the Wellcome Sanger Institute COVID-19 Surveillance Team |
| EPI_ISL_552465 | Lighthouse Lab in Milton Keynes | Wellcome Sanger Institute for the COVID-19 Genomics UK (COG-UK) consortium | The Lighthouse Lab in Milton Keynes and Alex Alderton, Roberto Amato, Sonia Goncalves, Ewan Harrison, David K. Jackson, Ian Johnston, Dominic Kwiatkowski, Cordelia Langford, John Sillitoe on behalf of the Wellcome Sanger Institute COVID-19 Surveillance Team (http://www.sanger.ac.uk/covid-team) |
| EPI_ISL_552466 | Lighthouse Lab in Alderley Park | Wellcome Sanger Institute for the COVID-19 Genomics UK (COG-UK) consortium | The Lighthouse Lab in Alderley Park and Alex Alderton, Roberto Amato, Sonia Goncalves, Ewan Harrison, David K. Jackson, Ian Johnston, Dominic Kwiatkowski, Cordelia Langford, John Sillitoe on behalf of the Wellcome Sanger Institute COVID-19 Surveillance Team |
| EPI_ISL_552467 | Lighthouse Lab in Milton Keynes | Wellcome Sanger Institute for the COVID-19 Genomics UK (COG-UK) consortium | The Lighthouse Lab in Milton Keynes and Alex Alderton, Roberto Amato, Sonia Goncalves, Ewan Harrison, David K. Jackson, Ian Johnston, Dominic Kwiatkowski, Cordelia Langford, John Sillitoe on behalf of the Wellcome Sanger Institute COVID-19 Surveillance Team (http://www.sanger.ac.uk/covid-team) |
| EPI_ISL_552468 | Lighthouse Lab in Alderley Park | Wellcome Sanger Institute for the COVID-19 Genomics UK (COG-UK) consortium | The Lighthouse Lab in Alderley Park and Alex Alderton, Roberto Amato, Sonia Goncalves, Ewan Harrison, David K. Jackson, Ian Johnston, Dominic Kwiatkowski, Cordelia Langford, John Sillitoe on behalf of the Wellcome Sanger Institute COVID-19 Surveillance Team |
| EPI_ISL_552469, EPI_ISL_552470, EPI_ISL_552471, EPI_ISL_552472 | Lighthouse Lab in Milton Keynes | Wellcome Sanger Institute for the COVID-19 Genomics UK (COG-UK) consortium | The Lighthouse Lab in Milton Keynes and Alex Alderton, Roberto Amato, Sonia Goncalves, Ewan Harrison, David K. Jackson, Ian Johnston, Dominic Kwiatkowski, Cordelia Langford, John Sillitoe on behalf of the Wellcome Sanger Institute COVID-19 Surveillance Team (http://www.sanger.ac.uk/covid-team) |
| EPI_ISL_552473 | Lighthouse Lab in Alderley Park | Wellcome Sanger Institute for the COVID-19 Genomics UK (COG-UK) consortium | The Lighthouse Lab in Alderley Park and Alex Alderton, Roberto Amato, Sonia Goncalves, Ewan Harrison, David K. Jackson, Ian Johnston, Dominic Kwiatkowski, Cordelia Langford, John Sillitoe on behalf of the Wellcome Sanger Institute COVID-19 Surveillance Team |
| EPI_ISL_552474, EPI_ISL_552475 | Lighthouse Lab in Milton Keynes | Wellcome Sanger Institute for the COVID-19 Genomics UK (COG-UK) consortium | The Lighthouse Lab in Milton Keynes and Alex Alderton, Roberto Amato, Sonia Goncalves, Ewan Harrison, David K. Jackson, Ian Johnston, Dominic Kwiatkowski, Cordelia Langford, John Sillitoe on behalf of the Wellcome Sanger Institute COVID-19 Surveillance Team (http://www.sanger.ac.uk/covid-team) |
| EPI_ISL_552476 | Lighthouse Lab in Alderley Park | Wellcome Sanger Institute for the COVID-19 Genomics UK (COG-UK) consortium | The Lighthouse Lab in Alderley Park and Alex Alderton, Roberto Amato, Sonia Goncalves, Ewan Harrison, David K. Jackson, Ian Johnston, Dominic Kwiatkowski, Cordelia Langford, John Sillitoe on behalf of the Wellcome Sanger Institute COVID-19 Surveillance Team |
| EPI_ISL_552477, EPI_ISL_552478 | Lighthouse Lab in Milton Keynes | Wellcome Sanger Institute for the COVID-19 Genomics UK (COG-UK) consortium | The Lighthouse Lab in Milton Keynes and Alex Alderton, Roberto Amato, Sonia Goncalves, Ewan Harrison, David K. Jackson, Ian Johnston, Dominic Kwiatkowski, Cordelia Langford, John Sillitoe on behalf of the Wellcome Sanger Institute COVID-19 Surveillance Team (http://www.sanger.ac.uk/covid-team) |
| EPI_ISL_552479 | Lighthouse Lab in Alderley Park | Wellcome Sanger Institute for the COVID-19 Genomics UK (COG-UK) consortium | The Lighthouse Lab in Alderley Park and Alex Alderton, Roberto Amato, Sonia Goncalves, Ewan Harrison, David K. Jackson, Ian Johnston, Dominic Kwiatkowski, Cordelia Langford, John Sillitoe on behalf of the Wellcome Sanger Institute COVID-19 Surveillance Team |
| EPI_ISL_552480 | Lighthouse Lab in Milton Keynes | Wellcome Sanger Institute for the COVID-19 Genomics UK (COG-UK) consortium | The Lighthouse Lab in Milton Keynes and Alex Alderton, Roberto Amato, Sonia Goncalves, Ewan Harrison, David K. Jackson, Ian Johnston, Dominic Kwiatkowski, Cordelia Langford, John Sillitoe on behalf of the Wellcome Sanger Institute COVID-19 Surveillance Team (http://www.sanger.ac.uk/covid-team) |
| EPI_ISL_552481 | Lighthouse Lab in Alderley Park | Wellcome Sanger Institute for the COVID-19 Genomics UK (COG-UK) consortium | The Lighthouse Lab in Alderley Park and Alex Alderton, Roberto Amato, Sonia Goncalves, Ewan Harrison, David K. Jackson, Ian Johnston, Dominic Kwiatkowski, Cordelia Langford, John Sillitoe on behalf of the Wellcome Sanger Institute COVID-19 Surveillance Team |
| EPI_ISL_552482, EPI_ISL_552484 | Lighthouse Lab in Milton Keynes | Wellcome Sanger Institute for the COVID-19 Genomics UK (COG-UK) consortium | The Lighthouse Lab in Milton Keynes and Alex Alderton, Roberto Amato, Sonia Goncalves, Ewan Harrison, David K. Jackson, Ian Johnston, Dominic Kwiatkowski, Cordelia Langford, John Sillitoe on behalf of the Wellcome Sanger Institute COVID-19 Surveillance Team (http://www.sanger.ac.uk/covid-team) |
| EPI_ISL_552486 | Lighthouse Lab in Alderley Park | Wellcome Sanger Institute for the COVID-19 Genomics UK (COG-UK) consortium | The Lighthouse Lab in Alderley Park and Alex Alderton, Roberto Amato, Sonia Goncalves, Ewan Harrison, David K. Jackson, Ian Johnston, Dominic Kwiatkowski, Cordelia Langford, John Sillitoe on behalf of the Wellcome Sanger Institute COVID-19 Surveillance Team |
| EPI_ISL_552487 | Lighthouse Lab in Milton Keynes | Wellcome Sanger Institute for the COVID-19 Genomics UK (COG-UK) consortium | The Lighthouse Lab in Milton Keynes and Alex Alderton, Roberto Amato, Sonia Goncalves, Ewan Harrison, David K. Jackson, Ian Johnston, Dominic Kwiatkowski, Cordelia Langford, John Sillitoe on behalf of the Wellcome Sanger Institute COVID-19 Surveillance Team (http://www.sanger.ac.uk/covid-team) |
| EPI_ISL_552488 | Lighthouse Lab in Alderley Park | Wellcome Sanger Institute for the COVID-19 Genomics UK (COG-UK) consortium | The Lighthouse Lab in Alderley Park and Alex Alderton, Roberto Amato, Sonia Goncalves, Ewan Harrison, David K. Jackson, Ian Johnston, Dominic Kwiatkowski, Cordelia Langford, John Sillitoe on behalf of the Wellcome Sanger Institute COVID-19 Surveillance Team |

| | | | |
|---|---|---|---|
| | | | (http://www.sanger.ac.uk/covid-team) |
| EPI_ISL_552489, EPI_ISL_552490, EPI_ISL_552491, EPI_ISL_552494, EPI_ISL_552495 | Lighthouse Lab in Milton Keynes | Wellcome Sanger Institute for the COVID-19 Genomics UK (COG-UK) consortium | The Lighthouse Lab in Milton Keynes and Alex Alderton, Roberto Amato, Sonia Goncalves, Ewan Harrison, David K. Jackson, Ian Johnston, Dominic Kwiatkowski, Cordelia Langford, John Sillitoe on behalf of the Wellcome Sanger Institute COVID-19 Surveillance Team (http://www.sanger.ac.uk/covid-team) |
| EPI_ISL_552496 | Lighthouse Lab in Milton Keynes | Wellcome Sanger Institute for the COVID-19 Genomics UK (COG-UK) consortium | The Lighthouse Lab in Alderley Park and Alex Alderton, Roberto Amato, Sonia Goncalves, Ewan Harrison, David K. Jackson, Ian Johnston, Dominic Kwiatkowski, Cordelia Langford, John Sillitoe on behalf of the Wellcome Sanger Institute COVID-19 Surveillance Team |
| EPI_ISL_552497, EPI_ISL_552498, EPI_ISL_552499, EPI_ISL_552500, EPI_ISL_552501, EPI_ISL_552502, EPI_ISL_552503, EPI_ISL_552504, EPI_ISL_552505 | Lighthouse Lab in Milton Keynes | Wellcome Sanger Institute for the COVID-19 Genomics UK (COG-UK) consortium | The Lighthouse Lab in Milton Keynes and Alex Alderton, Roberto Amato, Sonia Goncalves, Ewan Harrison, David K. Jackson, Ian Johnston, Dominic Kwiatkowski, Cordelia Langford, John Sillitoe on behalf of the Wellcome Sanger Institute COVID-19 Surveillance Team (http://www.sanger.ac.uk/covid-team) |
| EPI_ISL_552506, EPI_ISL_552507, EPI_ISL_552508 | Lighthouse Lab in Alderley Park | Wellcome Sanger Institute for the COVID-19 Genomics UK (COG-UK) consortium | The Lighthouse Lab in Alderley Park and Alex Alderton, Roberto Amato, Sonia Goncalves, Ewan Harrison, David K. Jackson, Ian Johnston, Dominic Kwiatkowski, Cordelia Langford, John Sillitoe on behalf of the Wellcome Sanger Institute COVID-19 Surveillance Team |
| EPI_ISL_552509, EPI_ISL_552510 | Lighthouse Lab in Milton Keynes | Wellcome Sanger Institute for the COVID-19 Genomics UK (COG-UK) consortium | The Lighthouse Lab in Milton Keynes and Alex Alderton, Roberto Amato, Sonia Goncalves, Ewan Harrison, David K. Jackson, Ian Johnston, Dominic Kwiatkowski, Cordelia Langford, John Sillitoe on behalf of the Wellcome Sanger Institute COVID-19 Surveillance Team (http://www.sanger.ac.uk/covid-team) |
| EPI_ISL_552512 | Lighthouse Lab in Alderley Park | Wellcome Sanger Institute for the COVID-19 Genomics UK (COG-UK) consortium | The Lighthouse Lab in Alderley Park and Alex Alderton, Roberto Amato, Sonia Goncalves, Ewan Harrison, David K. Jackson, Ian Johnston, Dominic Kwiatkowski, Cordelia Langford, John Sillitoe on behalf of the Wellcome Sanger Institute COVID-19 Surveillance Team |
| EPI_ISL_552513, EPI_ISL_552514 | Lighthouse Lab in Milton Keynes | Wellcome Sanger Institute for the COVID-19 Genomics UK (COG-UK) consortium | The Lighthouse Lab in Milton Keynes and Alex Alderton, Roberto Amato, Sonia Goncalves, Ewan Harrison, David K. Jackson, Ian Johnston, Dominic Kwiatkowski, Cordelia Langford, John Sillitoe on behalf of the Wellcome Sanger Institute COVID-19 Surveillance Team (http://www.sanger.ac.uk/covid-team) |
| EPI_ISL_552515, EPI_ISL_552516 | Lighthouse Lab in Alderley Park | Wellcome Sanger Institute for the COVID-19 Genomics UK (COG-UK) consortium | The Lighthouse Lab in Alderley Park and Alex Alderton, Roberto Amato, Sonia Goncalves, Ewan Harrison, David K. Jackson, Ian Johnston, Dominic Kwiatkowski, Cordelia Langford, John Sillitoe on behalf of the Wellcome Sanger Institute COVID-19 Surveillance Team |
| EPI_ISL_552517, EPI_ISL_552518, EPI_ISL_552520, EPI_ISL_552521, EPI_ISL_552522 | Lighthouse Lab in Milton Keynes | Wellcome Sanger Institute for the COVID-19 Genomics UK (COG-UK) consortium | The Lighthouse Lab in Milton Keynes and Alex Alderton, Roberto Amato, Sonia Goncalves, Ewan Harrison, David K. Jackson, Ian Johnston, Dominic Kwiatkowski, Cordelia Langford, John Sillitoe on behalf of the Wellcome Sanger Institute COVID-19 Surveillance Team (http://www.sanger.ac.uk/covid-team) |
| EPI_ISL_552523 | Lighthouse Lab in Alderley Park | Wellcome Sanger Institute for the COVID-19 Genomics UK (COG-UK) consortium | The Lighthouse Lab in Alderley Park and Alex Alderton, Roberto Amato, Sonia Goncalves, Ewan Harrison, David K. Jackson, Ian Johnston, Dominic Kwiatkowski, Cordelia Langford, John Sillitoe on behalf of the Wellcome Sanger Institute COVID-19 Surveillance Team |
| EPI_ISL_552525 | Lighthouse Lab in Milton Keynes | Wellcome Sanger Institute for the COVID-19 Genomics UK (COG-UK) consortium | The Lighthouse Lab in Milton Keynes and Alex Alderton, Roberto Amato, Sonia Goncalves, Ewan Harrison, David K. Jackson, Ian Johnston, Dominic Kwiatkowski, Cordelia Langford, John Sillitoe on behalf of the Wellcome Sanger Institute COVID-19 Surveillance Team (http://www.sanger.ac.uk/covid-team) |
| EPI_ISL_552526, EPI_ISL_552527 | Lighthouse Lab in Alderley Park | Wellcome Sanger Institute for the COVID-19 Genomics UK (COG-UK) consortium | The Lighthouse Lab in Alderley Park and Alex Alderton, Roberto Amato, Sonia Goncalves, Ewan Harrison, David K. Jackson, Ian Johnston, Dominic Kwiatkowski, Cordelia Langford, John Sillitoe on behalf of the Wellcome Sanger Institute COVID-19 Surveillance Team |
| EPI_ISL_552528, EPI_ISL_552529 | Lighthouse Lab in Milton Keynes | Wellcome Sanger Institute for the COVID-19 Genomics UK (COG-UK) consortium | The Lighthouse Lab in Milton Keynes and Alex Alderton, Roberto Amato, Sonia Goncalves, Ewan Harrison, David K. Jackson, Ian Johnston, Dominic Kwiatkowski, Cordelia Langford, John Sillitoe on behalf of the Wellcome Sanger Institute COVID-19 Surveillance Team (http://www.sanger.ac.uk/covid-team) |
| EPI_ISL_552530 | Lighthouse Lab in Alderley Park | Wellcome Sanger Institute for the COVID-19 Genomics UK (COG-UK) consortium | The Lighthouse Lab in Alderley Park and Alex Alderton, Roberto Amato, Sonia Goncalves, Ewan Harrison, David K. Jackson, Ian Johnston, Dominic Kwiatkowski, Cordelia Langford, John Sillitoe on behalf of the Wellcome Sanger Institute COVID-19 Surveillance Team |
| EPI_ISL_552531 | Lighthouse Lab in Milton Keynes | Wellcome Sanger Institute for the COVID-19 Genomics UK (COG-UK) consortium | The Lighthouse Lab in Milton Keynes and Alex Alderton, Roberto Amato, Sonia Goncalves, Ewan Harrison, David K. Jackson, Ian Johnston, Dominic Kwiatkowski, Cordelia Langford, John Sillitoe on behalf of the Wellcome Sanger Institute COVID-19 Surveillance Team (http://www.sanger.ac.uk/covid-team) |
| EPI_ISL_552533 | Lighthouse Lab in Alderley Park | Wellcome Sanger Institute for the COVID-19 Genomics UK (COG-UK) consortium | The Lighthouse Lab in Alderley Park and Alex Alderton, Roberto Amato, Sonia Goncalves, Ewan Harrison, David K. Jackson, Ian Johnston, Dominic Kwiatkowski, Cordelia Langford, John Sillitoe on behalf of the Wellcome Sanger Institute COVID-19 Surveillance Team |
| EPI_ISL_552534 | Lighthouse Lab in Alderley Park | Wellcome Sanger Institute for the COVID-19 Genomics UK (COG-UK) consortium | The Lighthouse Lab in Alderley Park and Alex Alderton, Roberto Amato, Sonia Goncalves, Ewan Harrison, David K. Jackson, Ian Johnston, Dominic Kwiatkowski, Cordelia Langford, John Sillitoe on behalf of the Wellcome Sanger Institute COVID-19 Surveillance Team (http://www.sanger.ac.uk/covid-team) |
| EPI_ISL_552535, EPI_ISL_552536, EPI_ISL_552538 | Lighthouse Lab in Milton Keynes | Wellcome Sanger Institute for the COVID-19 Genomics UK (COG-UK) consortium | The Lighthouse Lab in Milton Keynes and Alex Alderton, Roberto Amato, Sonia Goncalves, Ewan Harrison, David K. Jackson, Ian Johnston, Dominic Kwiatkowski, Cordelia Langford, John Sillitoe on behalf of the Wellcome Sanger Institute COVID-19 Surveillance Team (http://www.sanger.ac.uk/covid-team) |
| EPI_ISL_552539, EPI_ISL_552540 | Lighthouse Lab in Alderley Park | Wellcome Sanger Institute for the COVID-19 Genomics UK (COG-UK) consortium | The Lighthouse Lab in Alderley Park and Alex Alderton, Roberto Amato, Sonia Goncalves, Ewan Harrison, David K. Jackson, Ian Johnston, Dominic Kwiatkowski, Cordelia Langford, John Sillitoe on behalf of the Wellcome Sanger Institute COVID-19 Surveillance Team |
| EPI_ISL_552541, EPI_ISL_552542, EPI_ISL_552544, EPI_ISL_552545 | Lighthouse Lab in Milton Keynes | Wellcome Sanger Institute for the COVID-19 Genomics UK (COG-UK) consortium | The Lighthouse Lab in Milton Keynes and Alex Alderton, Roberto Amato, Sonia Goncalves, Ewan Harrison, David K. Jackson, Ian Johnston, Dominic Kwiatkowski, Cordelia Langford, John Sillitoe on behalf of the Wellcome Sanger Institute COVID-19 Surveillance Team (http://www.sanger.ac.uk/covid-team) |
| EPI_ISL_552547 | Lighthouse Lab in Alderley Park | Wellcome Sanger Institute for the COVID-19 Genomics UK (COG-UK) consortium | The Lighthouse Lab in Alderley Park and Alex Alderton, Roberto Amato, Sonia Goncalves, Ewan Harrison, David K. Jackson, Ian Johnston, Dominic Kwiatkowski, Cordelia Langford, John Sillitoe on behalf of the Wellcome Sanger Institute COVID-19 Surveillance Team |
| EPI_ISL_552548, EPI_ISL_552549, EPI_ISL_552550, EPI_ISL_552551, EPI_ISL_552553, EPI_ISL_552555 | Lighthouse Lab in Milton Keynes | Wellcome Sanger Institute for the COVID-19 Genomics UK (COG-UK) consortium | The Lighthouse Lab in Milton Keynes and Alex Alderton, Roberto Amato, Sonia Goncalves, Ewan Harrison, David K. Jackson, Ian Johnston, Dominic Kwiatkowski, Cordelia Langford, John Sillitoe on behalf of the Wellcome Sanger Institute COVID-19 Surveillance Team (http://www.sanger.ac.uk/covid-team) |
| EPI_ISL_552556 | Lighthouse Lab in Alderley Park | Wellcome Sanger Institute for the COVID-19 Genomics UK (COG-UK) consortium | The Lighthouse Lab in Alderley Park and Alex Alderton, Roberto Amato, Sonia Goncalves, Ewan Harrison, David K. Jackson, Ian Johnston, Dominic Kwiatkowski, Cordelia Langford, John Sillitoe on behalf of the Wellcome Sanger Institute COVID-19 Surveillance Team |
| EPI_ISL_552557, EPI_ISL_552558, EPI_ISL_552559, EPI_ISL_552561 | Lighthouse Lab in Milton Keynes | Wellcome Sanger Institute for the COVID-19 Genomics UK (COG-UK) consortium | The Lighthouse Lab in Milton Keynes and Alex Alderton, Roberto Amato, Sonia Goncalves, Ewan Harrison, David K. Jackson, Ian Johnston, Dominic Kwiatkowski, Cordelia Langford, John Sillitoe on behalf of the Wellcome Sanger Institute COVID-19 Surveillance Team (http://www.sanger.ac.uk/covid-team) |
| EPI_ISL_552562 | Lighthouse Lab in Alderley Park | Wellcome Sanger Institute for the COVID-19 Genomics UK (COG-UK) consortium | The Lighthouse Lab in Alderley Park and Alex Alderton, Roberto Amato, Sonia Goncalves, Ewan Harrison, David K. Jackson, Ian Johnston, Dominic Kwiatkowski, Cordelia Langford, John Sillitoe on behalf of the Wellcome Sanger Institute COVID-19 Surveillance Team |
| EPI_ISL_552563, EPI_ISL_552564 | Lighthouse Lab in Milton Keynes | Wellcome Sanger Institute for the COVID-19 Genomics UK (COG-UK) consortium | The Lighthouse Lab in Milton Keynes and Alex Alderton, Roberto Amato, Sonia Goncalves, Ewan Harrison, David K. Jackson, Ian Johnston, Dominic Kwiatkowski, Cordelia Langford, John Sillitoe on behalf of the Wellcome Sanger Institute COVID-19 Surveillance Team (http://www.sanger.ac.uk/covid-team) |
| EPI_ISL_552565, EPI_ISL_552566 | Lighthouse Lab in Alderley Park | Wellcome Sanger Institute for the COVID-19 Genomics UK (COG-UK) consortium | The Lighthouse Lab in Alderley Park and Alex Alderton, Roberto Amato, Sonia Goncalves, Ewan Harrison, David K. Jackson, Ian Johnston, Dominic Kwiatkowski, Cordelia Langford, John Sillitoe on behalf of the Wellcome Sanger Institute COVID-19 Surveillance Team |
| EPI_ISL_552567, EPI_ISL_552568, EPI_ISL_552569, EPI_ISL_552571 | Lighthouse Lab in Milton Keynes | Wellcome Sanger Institute for the COVID-19 Genomics UK (COG-UK) consortium | The Lighthouse Lab in Milton Keynes and Alex Alderton, Roberto Amato, Sonia Goncalves, Ewan Harrison, David K. Jackson, Ian Johnston, Dominic Kwiatkowski, Cordelia Langford, John Sillitoe on behalf of the Wellcome Sanger Institute COVID-19 Surveillance Team |

| | | | |
|---|---|---|---|
| | | | (http://www.sanger.ac.uk/covid-team) |
| EPI_ISL_552572 | Lighthouse Lab in Alderley Park | Wellcome Sanger Institute for the COVID-19 Genomics UK (COG-UK) consortium | The Lighthouse Lab in Alderley Park and Alex Alderton, Roberto Amato, Sonia Goncalves, Ewan Harrison, David K. Jackson, Ian Johnston, Dominic Kwiatkowski, Cordelia Langford, John Sillitoe on behalf of the Wellcome Sanger Institute COVID-19 Surveillance Team |
| EPI_ISL_552574 | Lighthouse Lab in Alderley Park | Wellcome Sanger Institute for the COVID-19 Genomics UK (COG-UK) consortium | The Lighthouse Lab in Alderley Park and Alex Alderton, Roberto Amato, Sonia Goncalves, Ewan Harrison, David K. Jackson, Ian Johnston, Dominic Kwiatkowski, Cordelia Langford, John Sillitoe on behalf of the Wellcome Sanger Institute COVID-19 Surveillance Team (http://www.sanger.ac.uk/covid-team) |
| EPI_ISL_552576, EPI_ISL_552577 | Lighthouse Lab in Alderley Park | Wellcome Sanger Institute for the COVID-19 Genomics UK (COG-UK) consortium | The Lighthouse Lab in Alderley Park and Alex Alderton, Roberto Amato, Sonia Goncalves, Ewan Harrison, David K. Jackson, Ian Johnston, Dominic Kwiatkowski, Cordelia Langford, John Sillitoe on behalf of the Wellcome Sanger Institute COVID-19 Surveillance Team |
| EPI_ISL_552578 | Lighthouse Lab in Milton Keynes | Wellcome Sanger Institute for the COVID-19 Genomics UK (COG-UK) consortium | The Lighthouse Lab in Milton Keynes and Alex Alderton, Roberto Amato, Sonia Goncalves, Ewan Harrison, David K. Jackson, Ian Johnston, Dominic Kwiatkowski, Cordelia Langford, John Sillitoe on behalf of the Wellcome Sanger Institute COVID-19 Surveillance Team (http://www.sanger.ac.uk/covid-team) |
| EPI_ISL_552579, EPI_ISL_552580 | Lighthouse Lab in Alderley Park | Wellcome Sanger Institute for the COVID-19 Genomics UK (COG-UK) consortium | The Lighthouse Lab in Alderley Park and Alex Alderton, Roberto Amato, Sonia Goncalves, Ewan Harrison, David K. Jackson, Ian Johnston, Dominic Kwiatkowski, Cordelia Langford, John Sillitoe on behalf of the Wellcome Sanger Institute COVID-19 Surveillance Team |
| EPI_ISL_552581, EPI_ISL_552582 | Lighthouse Lab in Milton Keynes | Wellcome Sanger Institute for the COVID-19 Genomics UK (COG-UK) consortium | The Lighthouse Lab in Milton Keynes and Alex Alderton, Roberto Amato, Sonia Goncalves, Ewan Harrison, David K. Jackson, Ian Johnston, Dominic Kwiatkowski, Cordelia Langford, John Sillitoe on behalf of the Wellcome Sanger Institute COVID-19 Surveillance Team (http://www.sanger.ac.uk/covid-team) |
| EPI_ISL_552583 | Lighthouse Lab in Alderley Park | Wellcome Sanger Institute for the COVID-19 Genomics UK (COG-UK) consortium | The Lighthouse Lab in Alderley Park and Alex Alderton, Roberto Amato, Sonia Goncalves, Ewan Harrison, David K. Jackson, Ian Johnston, Dominic Kwiatkowski, Cordelia Langford, John Sillitoe on behalf of the Wellcome Sanger Institute COVID-19 Surveillance Team |
| EPI_ISL_552584, EPI_ISL_552585, EPI_ISL_552586 | Lighthouse Lab in Milton Keynes | Wellcome Sanger Institute for the COVID-19 Genomics UK (COG-UK) consortium | The Lighthouse Lab in Milton Keynes and Alex Alderton, Roberto Amato, Sonia Goncalves, Ewan Harrison, David K. Jackson, Ian Johnston, Dominic Kwiatkowski, Cordelia Langford, John Sillitoe on behalf of the Wellcome Sanger Institute COVID-19 Surveillance Team (http://www.sanger.ac.uk/covid-team) |
| EPI_ISL_552587 | Lighthouse Lab in Alderley Park | Wellcome Sanger Institute for the COVID-19 Genomics UK (COG-UK) consortium | The Lighthouse Lab in Alderley Park and Alex Alderton, Roberto Amato, Sonia Goncalves, Ewan Harrison, David K. Jackson, Ian Johnston, Dominic Kwiatkowski, Cordelia Langford, John Sillitoe on behalf of the Wellcome Sanger Institute COVID-19 Surveillance Team |
| EPI_ISL_552588, EPI_ISL_552589 | Lighthouse Lab in Milton Keynes | Wellcome Sanger Institute for the COVID-19 Genomics UK (COG-UK) consortium | The Lighthouse Lab in Milton Keynes and Alex Alderton, Roberto Amato, Sonia Goncalves, Ewan Harrison, David K. Jackson, Ian Johnston, Dominic Kwiatkowski, Cordelia Langford, John Sillitoe on behalf of the Wellcome Sanger Institute COVID-19 Surveillance Team (http://www.sanger.ac.uk/covid-team) |
| EPI_ISL_552590, EPI_ISL_552591 | Lighthouse Lab in Alderley Park | Wellcome Sanger Institute for the COVID-19 Genomics UK (COG-UK) consortium | The Lighthouse Lab in Alderley Park and Alex Alderton, Roberto Amato, Sonia Goncalves, Ewan Harrison, David K. Jackson, Ian Johnston, Dominic Kwiatkowski, Cordelia Langford, John Sillitoe on behalf of the Wellcome Sanger Institute COVID-19 Surveillance Team |
| EPI_ISL_552594 | Lighthouse Lab in Milton Keynes | Wellcome Sanger Institute for the COVID-19 Genomics UK (COG-UK) consortium | The Lighthouse Lab in Milton Keynes and Alex Alderton, Roberto Amato, Sonia Goncalves, Ewan Harrison, David K. Jackson, Ian Johnston, Dominic Kwiatkowski, Cordelia Langford, John Sillitoe on behalf of the Wellcome Sanger Institute COVID-19 Surveillance Team (http://www.sanger.ac.uk/covid-team) |
| EPI_ISL_552595, EPI_ISL_552596 | Lighthouse Lab in Alderley Park | Wellcome Sanger Institute for the COVID-19 Genomics UK (COG-UK) consortium | The Lighthouse Lab in Alderley Park and Alex Alderton, Roberto Amato, Sonia Goncalves, Ewan Harrison, David K. Jackson, Ian Johnston, Dominic Kwiatkowski, Cordelia Langford, John Sillitoe on behalf of the Wellcome Sanger Institute COVID-19 Surveillance Team |
| EPI_ISL_552597 | Lighthouse Lab in Milton Keynes | Wellcome Sanger Institute for the COVID-19 Genomics UK (COG-UK) consortium | The Lighthouse Lab in Milton Keynes and Alex Alderton, Roberto Amato, Sonia Goncalves, Ewan Harrison, David K. Jackson, Ian Johnston, Dominic Kwiatkowski, Cordelia Langford, John Sillitoe on behalf of the Wellcome Sanger Institute COVID-19 Surveillance Team (http://www.sanger.ac.uk/covid-team) |
| EPI_ISL_552598 | Lighthouse Lab in Alderley Park | Wellcome Sanger Institute for the COVID-19 Genomics UK (COG-UK) consortium | The Lighthouse Lab in Alderley Park and Alex Alderton, Roberto Amato, Sonia Goncalves, Ewan Harrison, David K. Jackson, Ian Johnston, Dominic Kwiatkowski, Cordelia Langford, John Sillitoe on behalf of the Wellcome Sanger Institute COVID-19 Surveillance Team |
| EPI_ISL_552599, EPI_ISL_552600, EPI_ISL_552601, EPI_ISL_552602, EPI_ISL_552603, EPI_ISL_552604, EPI_ISL_552606 | Lighthouse Lab in Milton Keynes | Wellcome Sanger Institute for the COVID-19 Genomics UK (COG-UK) consortium | The Lighthouse Lab in Milton Keynes and Alex Alderton, Roberto Amato, Sonia Goncalves, Ewan Harrison, David K. Jackson, Ian Johnston, Dominic Kwiatkowski, Cordelia Langford, John Sillitoe on behalf of the Wellcome Sanger Institute COVID-19 Surveillance Team (http://www.sanger.ac.uk/covid-team) |
| EPI_ISL_552607, EPI_ISL_552608 | Lighthouse Lab in Alderley Park | Wellcome Sanger Institute for the COVID-19 Genomics UK (COG-UK) consortium | The Lighthouse Lab in Alderley Park and Alex Alderton, Roberto Amato, Sonia Goncalves, Ewan Harrison, David K. Jackson, Ian Johnston, Dominic Kwiatkowski, Cordelia Langford, John Sillitoe on behalf of the Wellcome Sanger Institute COVID-19 Surveillance Team |
| EPI_ISL_552609, EPI_ISL_552610, EPI_ISL_552611, EPI_ISL_552612, EPI_ISL_552613 | Lighthouse Lab in Milton Keynes | Wellcome Sanger Institute for the COVID-19 Genomics UK (COG-UK) consortium | The Lighthouse Lab in Milton Keynes and Alex Alderton, Roberto Amato, Sonia Goncalves, Ewan Harrison, David K. Jackson, Ian Johnston, Dominic Kwiatkowski, Cordelia Langford, John Sillitoe on behalf of the Wellcome Sanger Institute COVID-19 Surveillance Team (http://www.sanger.ac.uk/covid-team) |
| EPI_ISL_552614 | Lighthouse Lab in Alderley Park | Wellcome Sanger Institute for the COVID-19 Genomics UK (COG-UK) consortium | The Lighthouse Lab in Alderley Park and Alex Alderton, Roberto Amato, Sonia Goncalves, Ewan Harrison, David K. Jackson, Ian Johnston, Dominic Kwiatkowski, Cordelia Langford, John Sillitoe on behalf of the Wellcome Sanger Institute COVID-19 Surveillance Team |
| EPI_ISL_552616 | Lighthouse Lab in Milton Keynes | Wellcome Sanger Institute for the COVID-19 Genomics UK (COG-UK) consortium | The Lighthouse Lab in Milton Keynes and Alex Alderton, Roberto Amato, Sonia Goncalves, Ewan Harrison, David K. Jackson, Ian Johnston, Dominic Kwiatkowski, Cordelia Langford, John Sillitoe on behalf of the Wellcome Sanger Institute COVID-19 Surveillance Team (http://www.sanger.ac.uk/covid-team) |
| EPI_ISL_552617, EPI_ISL_552618, EPI_ISL_552620 | Lighthouse Lab in Alderley Park | Wellcome Sanger Institute for the COVID-19 Genomics UK (COG-UK) consortium | The Lighthouse Lab in Alderley Park and Alex Alderton, Roberto Amato, Sonia Goncalves, Ewan Harrison, David K. Jackson, Ian Johnston, Dominic Kwiatkowski, Cordelia Langford, John Sillitoe on behalf of the Wellcome Sanger Institute COVID-19 Surveillance Team |
| EPI_ISL_552621 | Lighthouse Lab in Milton Keynes | Wellcome Sanger Institute for the COVID-19 Genomics UK (COG-UK) consortium | The Lighthouse Lab in Milton Keynes and Alex Alderton, Roberto Amato, Sonia Goncalves, Ewan Harrison, David K. Jackson, Ian Johnston, Dominic Kwiatkowski, Cordelia Langford, John Sillitoe on behalf of the Wellcome Sanger Institute COVID-19 Surveillance Team (http://www.sanger.ac.uk/covid-team) |
| EPI_ISL_552622 | Lighthouse Lab in Alderley Park | Wellcome Sanger Institute for the COVID-19 Genomics UK (COG-UK) consortium | The Lighthouse Lab in Alderley Park and Alex Alderton, Roberto Amato, Sonia Goncalves, Ewan Harrison, David K. Jackson, Ian Johnston, Dominic Kwiatkowski, Cordelia Langford, John Sillitoe on behalf of the Wellcome Sanger Institute COVID-19 Surveillance Team |
| EPI_ISL_552623, EPI_ISL_552624, EPI_ISL_552625, EPI_ISL_552626, EPI_ISL_552627, EPI_ISL_552628 | Lighthouse Lab in Milton Keynes | Wellcome Sanger Institute for the COVID-19 Genomics UK (COG-UK) consortium | The Lighthouse Lab in Milton Keynes and Alex Alderton, Roberto Amato, Sonia Goncalves, Ewan Harrison, David K. Jackson, Ian Johnston, Dominic Kwiatkowski, Cordelia Langford, John Sillitoe on behalf of the Wellcome Sanger Institute COVID-19 Surveillance Team (http://www.sanger.ac.uk/covid-team) |
| EPI_ISL_552629 | Lighthouse Lab in Alderley Park | Wellcome Sanger Institute for the COVID-19 Genomics UK (COG-UK) consortium | The Lighthouse Lab in Alderley Park and Alex Alderton, Roberto Amato, Sonia Goncalves, Ewan Harrison, David K. Jackson, Ian Johnston, Dominic Kwiatkowski, Cordelia Langford, John Sillitoe on behalf of the Wellcome Sanger Institute COVID-19 Surveillance Team |
| EPI_ISL_552630 | Lighthouse Lab in Milton Keynes | Wellcome Sanger Institute for the COVID-19 Genomics UK (COG-UK) consortium | The Lighthouse Lab in Alderley Park and Alex Alderton, Roberto Amato, Sonia Goncalves, Ewan Harrison, David K. Jackson, Ian Johnston, Dominic Kwiatkowski, Cordelia Langford, John Sillitoe on behalf of the Wellcome Sanger Institute COVID-19 Surveillance Team |
| EPI_ISL_552631, EPI_ISL_552632, EPI_ISL_552633, EPI_ISL_552634 | Lighthouse Lab in Milton Keynes | Wellcome Sanger Institute for the COVID-19 Genomics UK (COG-UK) consortium | The Lighthouse Lab in Milton Keynes and Alex Alderton, Roberto Amato, Sonia Goncalves, Ewan Harrison, David K. Jackson, Ian Johnston, Dominic Kwiatkowski, Cordelia Langford, John Sillitoe on behalf of the Wellcome Sanger Institute COVID-19 Surveillance Team (http://www.sanger.ac.uk/covid-team) |
| EPI_ISL_552635 | Lighthouse Lab in Milton Keynes | Wellcome Sanger Institute for the COVID-19 Genomics UK (COG-UK) consortium | The Lighthouse Lab in Alderley Park and Alex Alderton, Roberto Amato, Sonia Goncalves, Ewan Harrison, David K. Jackson, Ian Johnston, Dominic Kwiatkowski, Cordelia Langford, John Sillitoe on behalf of the Wellcome Sanger Institute COVID-19 Surveillance Team |
| EPI_ISL_552636, EPI_ISL_552637, EPI_ISL_552638 | Lighthouse Lab in Milton Keynes | Wellcome Sanger Institute for the COVID-19 Genomics UK (COG-UK) consortium | The Lighthouse Lab in Milton Keynes and Alex Alderton, Roberto Amato, Sonia Goncalves, Ewan Harrison, David K. Jackson, Ian Johnston, Dominic Kwiatkowski, Cordelia Langford, John Sillitoe on behalf of the Wellcome Sanger Institute COVID-19 Surveillance Team |

| | | | |
|---|---|---|---|
| | | | (http://www.sanger.ac.uk/covid-team) |
| EPI_ISL_552639 | Lighthouse Lab in Alderley Park | Wellcome Sanger Institute for the COVID-19 Genomics UK (COG-UK) consortium | The Lighthouse Lab in Alderley Park and Alex Alderton, Roberto Amato, Sonia Goncalves, Ewan Harrison, David K. Jackson, Ian Johnston, Dominic Kwiatkowski, Cordelia Langford, John Sillitoe on behalf of the Wellcome Sanger Institute COVID-19 Surveillance Team (http://www.sanger.ac.uk/covid-team) |
| EPI_ISL_552640, EPI_ISL_552641 | Lighthouse Lab in Milton Keynes | Wellcome Sanger Institute for the COVID-19 Genomics UK (COG-UK) consortium | The Lighthouse Lab in Milton Keynes and Alex Alderton, Roberto Amato, Sonia Goncalves, Ewan Harrison, David K. Jackson, Ian Johnston, Dominic Kwiatkowski, Cordelia Langford, John Sillitoe on behalf of the Wellcome Sanger Institute COVID-19 Surveillance Team (http://www.sanger.ac.uk/covid-team) |
| EPI_ISL_552642 | Lighthouse Lab in Alderley Park | Wellcome Sanger Institute for the COVID-19 Genomics UK (COG-UK) consortium | The Lighthouse Lab in Alderley Park and Alex Alderton, Roberto Amato, Sonia Goncalves, Ewan Harrison, David K. Jackson, Ian Johnston, Dominic Kwiatkowski, Cordelia Langford, John Sillitoe on behalf of the Wellcome Sanger Institute COVID-19 Surveillance Team |
| EPI_ISL_552643 | Lighthouse Lab in Milton Keynes | Wellcome Sanger Institute for the COVID-19 Genomics UK (COG-UK) consortium | The Lighthouse Lab in Milton Keynes and Alex Alderton, Roberto Amato, Sonia Goncalves, Ewan Harrison, David K. Jackson, Ian Johnston, Dominic Kwiatkowski, Cordelia Langford, John Sillitoe on behalf of the Wellcome Sanger Institute COVID-19 Surveillance Team (http://www.sanger.ac.uk/covid-team) |
| EPI_ISL_552644 | Lighthouse Lab in Alderley Park | Wellcome Sanger Institute for the COVID-19 Genomics UK (COG-UK) consortium | The Lighthouse Lab in Alderley Park and Alex Alderton, Roberto Amato, Sonia Goncalves, Ewan Harrison, David K. Jackson, Ian Johnston, Dominic Kwiatkowski, Cordelia Langford, John Sillitoe on behalf of the Wellcome Sanger Institute COVID-19 Surveillance Team |
| EPI_ISL_552645 | Lighthouse Lab in Milton Keynes | Wellcome Sanger Institute for the COVID-19 Genomics UK (COG-UK) consortium | The Lighthouse Lab in Milton Keynes and Alex Alderton, Roberto Amato, Sonia Goncalves, Ewan Harrison, David K. Jackson, Ian Johnston, Dominic Kwiatkowski, Cordelia Langford, John Sillitoe on behalf of the Wellcome Sanger Institute COVID-19 Surveillance Team (http://www.sanger.ac.uk/covid-team) |
| EPI_ISL_552646 | Lighthouse Lab in Alderley Park | Wellcome Sanger Institute for the COVID-19 Genomics UK (COG-UK) consortium | The Lighthouse Lab in Alderley Park and Alex Alderton, Roberto Amato, Sonia Goncalves, Ewan Harrison, David K. Jackson, Ian Johnston, Dominic Kwiatkowski, Cordelia Langford, John Sillitoe on behalf of the Wellcome Sanger Institute COVID-19 Surveillance Team |
| EPI_ISL_552647, EPI_ISL_552648, EPI_ISL_552650, EPI_ISL_552651, EPI_ISL_552652, EPI_ISL_552653, EPI_ISL_552654 | Lighthouse Lab in Milton Keynes | Wellcome Sanger Institute for the COVID-19 Genomics UK (COG-UK) consortium | The Lighthouse Lab in Milton Keynes and Alex Alderton, Roberto Amato, Sonia Goncalves, Ewan Harrison, David K. Jackson, Ian Johnston, Dominic Kwiatkowski, Cordelia Langford, John Sillitoe on behalf of the Wellcome Sanger Institute COVID-19 Surveillance Team (http://www.sanger.ac.uk/covid-team) |
| EPI_ISL_552655 | Lighthouse Lab in Alderley Park | Wellcome Sanger Institute for the COVID-19 Genomics UK (COG-UK) consortium | The Lighthouse Lab in Alderley Park and Alex Alderton, Roberto Amato, Sonia Goncalves, Ewan Harrison, David K. Jackson, Ian Johnston, Dominic Kwiatkowski, Cordelia Langford, John Sillitoe on behalf of the Wellcome Sanger Institute COVID-19 Surveillance Team |
| EPI_ISL_552656, EPI_ISL_552657, EPI_ISL_552658 | Lighthouse Lab in Milton Keynes | Wellcome Sanger Institute for the COVID-19 Genomics UK (COG-UK) consortium | The Lighthouse Lab in Milton Keynes and Alex Alderton, Roberto Amato, Sonia Goncalves, Ewan Harrison, David K. Jackson, Ian Johnston, Dominic Kwiatkowski, Cordelia Langford, John Sillitoe on behalf of the Wellcome Sanger Institute COVID-19 Surveillance Team (http://www.sanger.ac.uk/covid-team) |
| EPI_ISL_552659 | Lighthouse Lab in Alderley Park | Wellcome Sanger Institute for the COVID-19 Genomics UK (COG-UK) consortium | The Lighthouse Lab in Alderley Park and Alex Alderton, Roberto Amato, Sonia Goncalves, Ewan Harrison, David K. Jackson, Ian Johnston, Dominic Kwiatkowski, Cordelia Langford, John Sillitoe on behalf of the Wellcome Sanger Institute COVID-19 Surveillance Team |
| EPI_ISL_552660, EPI_ISL_552661, EPI_ISL_552662 | Lighthouse Lab in Milton Keynes | Wellcome Sanger Institute for the COVID-19 Genomics UK (COG-UK) consortium | The Lighthouse Lab in Milton Keynes and Alex Alderton, Roberto Amato, Sonia Goncalves, Ewan Harrison, David K. Jackson, Ian Johnston, Dominic Kwiatkowski, Cordelia Langford, John Sillitoe on behalf of the Wellcome Sanger Institute COVID-19 Surveillance Team (http://www.sanger.ac.uk/covid-team) |
| EPI_ISL_552663 | Lighthouse Lab in Milton Keynes | Wellcome Sanger Institute for the COVID-19 Genomics UK (COG-UK) consortium | The Lighthouse Lab in Alderley Park and Alex Alderton, Roberto Amato, Sonia Goncalves, Ewan Harrison, David K. Jackson, Ian Johnston, Dominic Kwiatkowski, Cordelia Langford, John Sillitoe on behalf of the Wellcome Sanger Institute COVID-19 Surveillance Team |
| EPI_ISL_552664 | Lighthouse Lab in Milton Keynes | Wellcome Sanger Institute for the COVID-19 Genomics UK (COG-UK) consortium | The Lighthouse Lab in Milton Keynes and Alex Alderton, Roberto Amato, Sonia Goncalves, Ewan Harrison, David K. Jackson, Ian Johnston, Dominic Kwiatkowski, Cordelia Langford, John Sillitoe on behalf of the Wellcome Sanger Institute COVID-19 Surveillance Team (http://www.sanger.ac.uk/covid-team) |
| EPI_ISL_552665, EPI_ISL_552666 | Lighthouse Lab in Alderley Park | Wellcome Sanger Institute for the COVID-19 Genomics UK (COG-UK) consortium | The Lighthouse Lab in Alderley Park and Alex Alderton, Roberto Amato, Sonia Goncalves, Ewan Harrison, David K. Jackson, Ian Johnston, Dominic Kwiatkowski, Cordelia Langford, John Sillitoe on behalf of the Wellcome Sanger Institute COVID-19 Surveillance Team |
| EPI_ISL_552667 | Lighthouse Lab in Milton Keynes | Wellcome Sanger Institute for the COVID-19 Genomics UK (COG-UK) consortium | The Lighthouse Lab in Milton Keynes and Alex Alderton, Roberto Amato, Sonia Goncalves, Ewan Harrison, David K. Jackson, Ian Johnston, Dominic Kwiatkowski, Cordelia Langford, John Sillitoe on behalf of the Wellcome Sanger Institute COVID-19 Surveillance Team (http://www.sanger.ac.uk/covid-team) |
| EPI_ISL_552668 | Lighthouse Lab in Milton Keynes | Wellcome Sanger Institute for the COVID-19 Genomics UK (COG-UK) consortium | The Lighthouse Lab in Alderley Park and Alex Alderton, Roberto Amato, Sonia Goncalves, Ewan Harrison, David K. Jackson, Ian Johnston, Dominic Kwiatkowski, Cordelia Langford, John Sillitoe on behalf of the Wellcome Sanger Institute COVID-19 Surveillance Team |
| EPI_ISL_552669 | Lighthouse Lab in Alderley Park | Wellcome Sanger Institute for the COVID-19 Genomics UK (COG-UK) consortium | The Lighthouse Lab in Alderley Park and Alex Alderton, Roberto Amato, Sonia Goncalves, Ewan Harrison, David K. Jackson, Ian Johnston, Dominic Kwiatkowski, Cordelia Langford, John Sillitoe on behalf of the Wellcome Sanger Institute COVID-19 Surveillance Team |
| EPI_ISL_552670 | Lighthouse Lab in Milton Keynes | Wellcome Sanger Institute for the COVID-19 Genomics UK (COG-UK) consortium | The Lighthouse Lab in Milton Keynes and Alex Alderton, Roberto Amato, Sonia Goncalves, Ewan Harrison, David K. Jackson, Ian Johnston, Dominic Kwiatkowski, Cordelia Langford, John Sillitoe on behalf of the Wellcome Sanger Institute COVID-19 Surveillance Team (http://www.sanger.ac.uk/covid-team) |
| EPI_ISL_552671 | Lighthouse Lab in Alderley Park | Wellcome Sanger Institute for the COVID-19 Genomics UK (COG-UK) consortium | The Lighthouse Lab in Alderley Park and Alex Alderton, Roberto Amato, Sonia Goncalves, Ewan Harrison, David K. Jackson, Ian Johnston, Dominic Kwiatkowski, Cordelia Langford, John Sillitoe on behalf of the Wellcome Sanger Institute COVID-19 Surveillance Team |
| EPI_ISL_552672, EPI_ISL_552673, EPI_ISL_552674, EPI_ISL_552675, EPI_ISL_552676, EPI_ISL_552677, EPI_ISL_552678 | Lighthouse Lab in Milton Keynes | Wellcome Sanger Institute for the COVID-19 Genomics UK (COG-UK) consortium | The Lighthouse Lab in Milton Keynes and Alex Alderton, Roberto Amato, Sonia Goncalves, Ewan Harrison, David K. Jackson, Ian Johnston, Dominic Kwiatkowski, Cordelia Langford, John Sillitoe on behalf of the Wellcome Sanger Institute COVID-19 Surveillance Team (http://www.sanger.ac.uk/covid-team) |
| EPI_ISL_552679 | Lighthouse Lab in Alderley Park | Wellcome Sanger Institute for the COVID-19 Genomics UK (COG-UK) consortium | The Lighthouse Lab in Alderley Park and Alex Alderton, Roberto Amato, Sonia Goncalves, Ewan Harrison, David K. Jackson, Ian Johnston, Dominic Kwiatkowski, Cordelia Langford, John Sillitoe on behalf of the Wellcome Sanger Institute COVID-19 Surveillance Team |
| EPI_ISL_552680 | Lighthouse Lab in Milton Keynes | Wellcome Sanger Institute for the COVID-19 Genomics UK (COG-UK) consortium | The Lighthouse Lab in Milton Keynes and Alex Alderton, Roberto Amato, Sonia Goncalves, Ewan Harrison, David K. Jackson, Ian Johnston, Dominic Kwiatkowski, Cordelia Langford, John Sillitoe on behalf of the Wellcome Sanger Institute COVID-19 Surveillance Team (http://www.sanger.ac.uk/covid-team) |
| EPI_ISL_552681 | Lighthouse Lab in Alderley Park | Wellcome Sanger Institute for the COVID-19 Genomics UK (COG-UK) consortium | The Lighthouse Lab in Alderley Park and Alex Alderton, Roberto Amato, Sonia Goncalves, Ewan Harrison, David K. Jackson, Ian Johnston, Dominic Kwiatkowski, Cordelia Langford, John Sillitoe on behalf of the Wellcome Sanger Institute COVID-19 Surveillance Team |
| EPI_ISL_552682, EPI_ISL_552683, EPI_ISL_552684, EPI_ISL_552685 | Lighthouse Lab in Milton Keynes | Wellcome Sanger Institute for the COVID-19 Genomics UK (COG-UK) consortium | The Lighthouse Lab in Milton Keynes and Alex Alderton, Roberto Amato, Sonia Goncalves, Ewan Harrison, David K. Jackson, Ian Johnston, Dominic Kwiatkowski, Cordelia Langford, John Sillitoe on behalf of the Wellcome Sanger Institute COVID-19 Surveillance Team (http://www.sanger.ac.uk/covid-team) |
| EPI_ISL_552686 | Lighthouse Lab in Alderley Park | Wellcome Sanger Institute for the COVID-19 Genomics UK (COG-UK) consortium | The Lighthouse Lab in Alderley Park and Alex Alderton, Roberto Amato, Sonia Goncalves, Ewan Harrison, David K. Jackson, Ian Johnston, Dominic Kwiatkowski, Cordelia Langford, John Sillitoe on behalf of the Wellcome Sanger Institute COVID-19 Surveillance Team |
| EPI_ISL_552688 | Lighthouse Lab in Milton Keynes | Wellcome Sanger Institute for the COVID-19 Genomics UK (COG-UK) consortium | The Lighthouse Lab in Milton Keynes and Alex Alderton, Roberto Amato, Sonia Goncalves, Ewan Harrison, David K. Jackson, Ian Johnston, Dominic Kwiatkowski, Cordelia Langford, John Sillitoe on behalf of the Wellcome Sanger Institute COVID-19 Surveillance Team (http://www.sanger.ac.uk/covid-team) |
| EPI_ISL_552689 | Lighthouse Lab in Alderley Park | Wellcome Sanger Institute for the COVID-19 Genomics UK (COG-UK) consortium | The Lighthouse Lab in Alderley Park and Alex Alderton, Roberto Amato, Sonia Goncalves, Ewan Harrison, David K. Jackson, Ian Johnston, Dominic Kwiatkowski, Cordelia Langford, John Sillitoe on behalf of the Wellcome Sanger Institute COVID-19 Surveillance Team |

| | | | |
|---|---|---|---|
| | | | (http://www.sanger.ac.uk/covid-team) |
| EPI_ISL_552690, EPI_ISL_552691, EPI_ISL_552692, EPI_ISL_552693, EPI_ISL_552694, EPI_ISL_552695, EPI_ISL_552696, EPI_ISL_552697, EPI_ISL_552698 | Lighthouse Lab in Milton Keynes | Wellcome Sanger Institute for the COVID-19 Genomics UK (COG-UK) consortium | The Lighthouse Lab in Milton Keynes and Alex Alderton, Roberto Amato, Sonia Goncalves, Ewan Harrison, David K. Jackson, Ian Johnston, Dominic Kwiatkowski, Cordelia Langford, John Sillitoe on behalf of the Wellcome Sanger Institute COVID-19 Surveillance Team (http://www.sanger.ac.uk/covid-team) |
| EPI_ISL_552699 | Lighthouse Lab in Alderley Park | Wellcome Sanger Institute for the COVID-19 Genomics UK (COG-UK) consortium | The Lighthouse Lab in Alderley Park and Alex Alderton, Roberto Amato, Sonia Goncalves, Ewan Harrison, David K. Jackson, Ian Johnston, Dominic Kwiatkowski, Cordelia Langford, John Sillitoe on behalf of the Wellcome Sanger Institute COVID-19 Surveillance Team |
| EPI_ISL_552700, EPI_ISL_552701, EPI_ISL_552702 | Lighthouse Lab in Milton Keynes | Wellcome Sanger Institute for the COVID-19 Genomics UK (COG-UK) consortium | The Lighthouse Lab in Milton Keynes and Alex Alderton, Roberto Amato, Sonia Goncalves, Ewan Harrison, David K. Jackson, Ian Johnston, Dominic Kwiatkowski, Cordelia Langford, John Sillitoe on behalf of the Wellcome Sanger Institute COVID-19 Surveillance Team (http://www.sanger.ac.uk/covid-team) |
| EPI_ISL_552703 | Lighthouse Lab in Alderley Park | Wellcome Sanger Institute for the COVID-19 Genomics UK (COG-UK) consortium | The Lighthouse Lab in Alderley Park and Alex Alderton, Roberto Amato, Sonia Goncalves, Ewan Harrison, David K. Jackson, Ian Johnston, Dominic Kwiatkowski, Cordelia Langford, John Sillitoe on behalf of the Wellcome Sanger Institute COVID-19 Surveillance Team |
| EPI_ISL_552704, EPI_ISL_552705 | Lighthouse Lab in Milton Keynes | Wellcome Sanger Institute for the COVID-19 Genomics UK (COG-UK) consortium | The Lighthouse Lab in Milton Keynes and Alex Alderton, Roberto Amato, Sonia Goncalves, Ewan Harrison, David K. Jackson, Ian Johnston, Dominic Kwiatkowski, Cordelia Langford, John Sillitoe on behalf of the Wellcome Sanger Institute COVID-19 Surveillance Team (http://www.sanger.ac.uk/covid-team) |
| EPI_ISL_552706, EPI_ISL_552707 | Lighthouse Lab in Alderley Park | Wellcome Sanger Institute for the COVID-19 Genomics UK (COG-UK) consortium | The Lighthouse Lab in Alderley Park and Alex Alderton, Roberto Amato, Sonia Goncalves, Ewan Harrison, David K. Jackson, Ian Johnston, Dominic Kwiatkowski, Cordelia Langford, John Sillitoe on behalf of the Wellcome Sanger Institute COVID-19 Surveillance Team |
| EPI_ISL_552708, EPI_ISL_552709, EPI_ISL_552710, EPI_ISL_552711 | Lighthouse Lab in Milton Keynes | Wellcome Sanger Institute for the COVID-19 Genomics UK (COG-UK) consortium | The Lighthouse Lab in Milton Keynes and Alex Alderton, Roberto Amato, Sonia Goncalves, Ewan Harrison, David K. Jackson, Ian Johnston, Dominic Kwiatkowski, Cordelia Langford, John Sillitoe on behalf of the Wellcome Sanger Institute COVID-19 Surveillance Team (http://www.sanger.ac.uk/covid-team) |
| EPI_ISL_552712, EPI_ISL_552713 | Lighthouse Lab in Alderley Park | Wellcome Sanger Institute for the COVID-19 Genomics UK (COG-UK) consortium | The Lighthouse Lab in Alderley Park and Alex Alderton, Roberto Amato, Sonia Goncalves, Ewan Harrison, David K. Jackson, Ian Johnston, Dominic Kwiatkowski, Cordelia Langford, John Sillitoe on behalf of the Wellcome Sanger Institute COVID-19 Surveillance Team |
| EPI_ISL_552715, EPI_ISL_552716 | Lighthouse Lab in Milton Keynes | Wellcome Sanger Institute for the COVID-19 Genomics UK (COG-UK) consortium | The Lighthouse Lab in Milton Keynes and Alex Alderton, Roberto Amato, Sonia Goncalves, Ewan Harrison, David K. Jackson, Ian Johnston, Dominic Kwiatkowski, Cordelia Langford, John Sillitoe on behalf of the Wellcome Sanger Institute COVID-19 Surveillance Team (http://www.sanger.ac.uk/covid-team) |
| EPI_ISL_552717 | Lighthouse Lab in Alderley Park | Wellcome Sanger Institute for the COVID-19 Genomics UK (COG-UK) consortium | The Lighthouse Lab in Alderley Park and Alex Alderton, Roberto Amato, Sonia Goncalves, Ewan Harrison, David K. Jackson, Ian Johnston, Dominic Kwiatkowski, Cordelia Langford, John Sillitoe on behalf of the Wellcome Sanger Institute COVID-19 Surveillance Team |
| EPI_ISL_552718 | Lighthouse Lab in Milton Keynes | Wellcome Sanger Institute for the COVID-19 Genomics UK (COG-UK) consortium | The Lighthouse Lab in Milton Keynes and Alex Alderton, Roberto Amato, Sonia Goncalves, Ewan Harrison, David K. Jackson, Ian Johnston, Dominic Kwiatkowski, Cordelia Langford, John Sillitoe on behalf of the Wellcome Sanger Institute COVID-19 Surveillance Team (http://www.sanger.ac.uk/covid-team) |
| EPI_ISL_552719, EPI_ISL_552720, EPI_ISL_552721, EPI_ISL_552722, EPI_ISL_552723, EPI_ISL_552724 | Lighthouse Lab in Alderley Park | Wellcome Sanger Institute for the COVID-19 Genomics UK (COG-UK) consortium | The Lighthouse Lab in Alderley Park and Alex Alderton, Roberto Amato, Sonia Goncalves, Ewan Harrison, David K. Jackson, Ian Johnston, Dominic Kwiatkowski, Cordelia Langford, John Sillitoe on behalf of the Wellcome Sanger Institute COVID-19 Surveillance Team |
| EPI_ISL_552725 | Lighthouse Lab in Milton Keynes | Wellcome Sanger Institute for the COVID-19 Genomics UK (COG-UK) consortium | The Lighthouse Lab in Alderley Park and Alex Alderton, Roberto Amato, Sonia Goncalves, Ewan Harrison, David K. Jackson, Ian Johnston, Dominic Kwiatkowski, Cordelia Langford, John Sillitoe on behalf of the Wellcome Sanger Institute COVID-19 Surveillance Team |
| EPI_ISL_552726 | Lighthouse Lab in Alderley Park | Wellcome Sanger Institute for the COVID-19 Genomics UK (COG-UK) consortium | The Lighthouse Lab in Alderley Park and Alex Alderton, Roberto Amato, Sonia Goncalves, Ewan Harrison, David K. Jackson, Ian Johnston, Dominic Kwiatkowski, Cordelia Langford, John Sillitoe on behalf of the Wellcome Sanger Institute COVID-19 Surveillance Team |
| EPI_ISL_552727 | Lighthouse Lab in Milton Keynes | Wellcome Sanger Institute for the COVID-19 Genomics UK (COG-UK) consortium | The Lighthouse Lab in Milton Keynes and Alex Alderton, Roberto Amato, Sonia Goncalves, Ewan Harrison, David K. Jackson, Ian Johnston, Dominic Kwiatkowski, Cordelia Langford, John Sillitoe on behalf of the Wellcome Sanger Institute COVID-19 Surveillance Team (http://www.sanger.ac.uk/covid-team) |
| EPI_ISL_552728, EPI_ISL_552729 | Lighthouse Lab in Alderley Park | Wellcome Sanger Institute for the COVID-19 Genomics UK (COG-UK) consortium | The Lighthouse Lab in Alderley Park and Alex Alderton, Roberto Amato, Sonia Goncalves, Ewan Harrison, David K. Jackson, Ian Johnston, Dominic Kwiatkowski, Cordelia Langford, John Sillitoe on behalf of the Wellcome Sanger Institute COVID-19 Surveillance Team |
| EPI_ISL_552730 | Lighthouse Lab in Alderley Park | Wellcome Sanger Institute for the COVID-19 Genomics UK (COG-UK) consortium | The Lighthouse Lab in Alderley Park and Alex Alderton, Roberto Amato, Sonia Goncalves, Ewan Harrison, David K. Jackson, Ian Johnston, Dominic Kwiatkowski, Cordelia Langford, John Sillitoe on behalf of the Wellcome Sanger Institute COVID-19 Surveillance Team (http://www.sanger.ac.uk/covid-team) |
| EPI_ISL_552731, EPI_ISL_552732 | Lighthouse Lab in Alderley Park | Wellcome Sanger Institute for the COVID-19 Genomics UK (COG-UK) consortium | The Lighthouse Lab in Alderley Park and Alex Alderton, Roberto Amato, Sonia Goncalves, Ewan Harrison, David K. Jackson, Ian Johnston, Dominic Kwiatkowski, Cordelia Langford, John Sillitoe on behalf of the Wellcome Sanger Institute COVID-19 Surveillance Team |
| EPI_ISL_552733, EPI_ISL_552734 | Lighthouse Lab in Milton Keynes | Wellcome Sanger Institute for the COVID-19 Genomics UK (COG-UK) consortium | The Lighthouse Lab in Milton Keynes and Alex Alderton, Roberto Amato, Sonia Goncalves, Ewan Harrison, David K. Jackson, Ian Johnston, Dominic Kwiatkowski, Cordelia Langford, John Sillitoe on behalf of the Wellcome Sanger Institute COVID-19 Surveillance Team (http://www.sanger.ac.uk/covid-team) |
| EPI_ISL_552735 | Lighthouse Lab in Alderley Park | Wellcome Sanger Institute for the COVID-19 Genomics UK (COG-UK) consortium | The Lighthouse Lab in Alderley Park and Alex Alderton, Roberto Amato, Sonia Goncalves, Ewan Harrison, David K. Jackson, Ian Johnston, Dominic Kwiatkowski, Cordelia Langford, John Sillitoe on behalf of the Wellcome Sanger Institute COVID-19 Surveillance Team |
| EPI_ISL_552736 | Lighthouse Lab in Milton Keynes | Wellcome Sanger Institute for the COVID-19 Genomics UK (COG-UK) consortium | The Lighthouse Lab in Milton Keynes and Alex Alderton, Roberto Amato, Sonia Goncalves, Ewan Harrison, David K. Jackson, Ian Johnston, Dominic Kwiatkowski, Cordelia Langford, John Sillitoe on behalf of the Wellcome Sanger Institute COVID-19 Surveillance Team (http://www.sanger.ac.uk/covid-team) |
| EPI_ISL_552737 | Lighthouse Lab in Milton Keynes | Wellcome Sanger Institute for the COVID-19 Genomics UK (COG-UK) consortium | The Lighthouse Lab in Alderley Park and Alex Alderton, Roberto Amato, Sonia Goncalves, Ewan Harrison, David K. Jackson, Ian Johnston, Dominic Kwiatkowski, Cordelia Langford, John Sillitoe on behalf of the Wellcome Sanger Institute COVID-19 Surveillance Team |
| EPI_ISL_552738, EPI_ISL_552739 | Lighthouse Lab in Milton Keynes | Wellcome Sanger Institute for the COVID-19 Genomics UK (COG-UK) consortium | The Lighthouse Lab in Milton Keynes and Alex Alderton, Roberto Amato, Sonia Goncalves, Ewan Harrison, David K. Jackson, Ian Johnston, Dominic Kwiatkowski, Cordelia Langford, John Sillitoe on behalf of the Wellcome Sanger Institute COVID-19 Surveillance Team (http://www.sanger.ac.uk/covid-team) |
| EPI_ISL_552740 | Lighthouse Lab in Milton Keynes | Wellcome Sanger Institute for the COVID-19 Genomics UK (COG-UK) consortium | The Lighthouse Lab in Alderley Park and Alex Alderton, Roberto Amato, Sonia Goncalves, Ewan Harrison, David K. Jackson, Ian Johnston, Dominic Kwiatkowski, Cordelia Langford, John Sillitoe on behalf of the Wellcome Sanger Institute COVID-19 Surveillance Team |
| EPI_ISL_552741, EPI_ISL_552742 | Lighthouse Lab in Alderley Park | Wellcome Sanger Institute for the COVID-19 Genomics UK (COG-UK) consortium | The Lighthouse Lab in Alderley Park and Alex Alderton, Roberto Amato, Sonia Goncalves, Ewan Harrison, David K. Jackson, Ian Johnston, Dominic Kwiatkowski, Cordelia Langford, John Sillitoe on behalf of the Wellcome Sanger Institute COVID-19 Surveillance Team (http://www.sanger.ac.uk/covid-team) |
| EPI_ISL_552743, EPI_ISL_552744 | Lighthouse Lab in Milton Keynes | Wellcome Sanger Institute for the COVID-19 Genomics UK (COG-UK) consortium | The Lighthouse Lab in Milton Keynes and Alex Alderton, Roberto Amato, Sonia Goncalves, Ewan Harrison, David K. Jackson, Ian Johnston, Dominic Kwiatkowski, Cordelia Langford, John Sillitoe on behalf of the Wellcome Sanger Institute COVID-19 Surveillance Team (http://www.sanger.ac.uk/covid-team) |
| EPI_ISL_552745, EPI_ISL_552746 | Lighthouse Lab in Alderley Park | Wellcome Sanger Institute for the COVID-19 Genomics UK (COG-UK) consortium | The Lighthouse Lab in Alderley Park and Alex Alderton, Roberto Amato, Sonia Goncalves, Ewan Harrison, David K. Jackson, Ian Johnston, Dominic Kwiatkowski, Cordelia Langford, John Sillitoe on behalf of the Wellcome Sanger Institute COVID-19 Surveillance Team (http://www.sanger.ac.uk/covid-team) |
| EPI_ISL_552747 | Lighthouse Lab in Milton Keynes | Wellcome Sanger Institute for the COVID-19 Genomics UK | The Lighthouse Lab in Milton Keynes and Alex Alderton, Roberto Amato, Sonia Goncalves, Ewan Harrison, David K. Jackson, Ian Johnston, Dominic |

| | | | |
|---|---|---|---|
| | | (COG-UK) consortium | Kwiatkowski, Cordelia Langford, John Sillitoe on behalf of the Wellcome Sanger Institute COVID-19 Surveillance Team (http://www.sanger.ac.uk/covid-team) |
| EPI_ISL_552748, EPI_ISL_552749 | Lighthouse Lab in Alderley Park | Wellcome Sanger Institute for the COVID-19 Genomics UK (COG-UK) consortium | The Lighthouse Lab in Alderley Park and Alex Alderton, Roberto Amato, Sonia Goncalves, Ewan Harrison, David K. Jackson, Ian Johnston, Dominic Kwiatkowski, Cordelia Langford, John Sillitoe on behalf of the Wellcome Sanger Institute COVID-19 Surveillance Team (http://www.sanger.ac.uk/covid-team) |
| EPI_ISL_552750 | Lighthouse Lab in Alderley Park | Wellcome Sanger Institute for the COVID-19 Genomics UK (COG-UK) consortium | The Lighthouse Lab in Alderley Park and Alex Alderton, Roberto Amato, Sonia Goncalves, Ewan Harrison, David K. Jackson, Ian Johnston, Dominic Kwiatkowski, Cordelia Langford, John Sillitoe on behalf of the Wellcome Sanger Institute COVID-19 Surveillance Team |
| EPI_ISL_552751 | Lighthouse Lab in Milton Keynes | Wellcome Sanger Institute for the COVID-19 Genomics UK (COG-UK) consortium | The Lighthouse Lab in Milton Keynes and Alex Alderton, Roberto Amato, Sonia Goncalves, Ewan Harrison, David K. Jackson, Ian Johnston, Dominic Kwiatkowski, Cordelia Langford, John Sillitoe on behalf of the Wellcome Sanger Institute COVID-19 Surveillance Team (http://www.sanger.ac.uk/covid-team) |
| EPI_ISL_552752 | Lighthouse Lab in Milton Keynes | Wellcome Sanger Institute for the COVID-19 Genomics UK (COG-UK) consortium | The Lighthouse Lab in Alderley Park and Alex Alderton, Roberto Amato, Sonia Goncalves, Ewan Harrison, David K. Jackson, Ian Johnston, Dominic Kwiatkowski, Cordelia Langford, John Sillitoe on behalf of the Wellcome Sanger Institute COVID-19 Surveillance Team |
| EPI_ISL_552753, EPI_ISL_552755, EPI_ISL_552756 | Lighthouse Lab in Milton Keynes | Wellcome Sanger Institute for the COVID-19 Genomics UK (COG-UK) consortium | The Lighthouse Lab in Milton Keynes and Alex Alderton, Roberto Amato, Sonia Goncalves, Ewan Harrison, David K. Jackson, Ian Johnston, Dominic Kwiatkowski, Cordelia Langford, John Sillitoe on behalf of the Wellcome Sanger Institute COVID-19 Surveillance Team (http://www.sanger.ac.uk/covid-team) |
| EPI_ISL_552759 | Lighthouse Lab in Milton Keynes | Wellcome Sanger Institute for the COVID-19 Genomics UK (COG-UK) consortium | The Lighthouse Lab in Alderley Park and Alex Alderton, Roberto Amato, Sonia Goncalves, Ewan Harrison, David K. Jackson, Ian Johnston, Dominic Kwiatkowski, Cordelia Langford, John Sillitoe on behalf of the Wellcome Sanger Institute COVID-19 Surveillance Team |
| EPI_ISL_552760, EPI_ISL_552761 | Lighthouse Lab in Alderley Park | Wellcome Sanger Institute for the COVID-19 Genomics UK (COG-UK) consortium | The Lighthouse Lab in Alderley Park and Alex Alderton, Roberto Amato, Sonia Goncalves, Ewan Harrison, David K. Jackson, Ian Johnston, Dominic Kwiatkowski, Cordelia Langford, John Sillitoe on behalf of the Wellcome Sanger Institute COVID-19 Surveillance Team |
| EPI_ISL_552762, EPI_ISL_552763, EPI_ISL_552764 | Lighthouse Lab in Milton Keynes | Wellcome Sanger Institute for the COVID-19 Genomics UK (COG-UK) consortium | The Lighthouse Lab in Milton Keynes and Alex Alderton, Roberto Amato, Sonia Goncalves, Ewan Harrison, David K. Jackson, Ian Johnston, Dominic Kwiatkowski, Cordelia Langford, John Sillitoe on behalf of the Wellcome Sanger Institute COVID-19 Surveillance Team (http://www.sanger.ac.uk/covid-team) |
| EPI_ISL_552765, EPI_ISL_552766 | Lighthouse Lab in Milton Keynes | Wellcome Sanger Institute for the COVID-19 Genomics UK (COG-UK) consortium | The Lighthouse Lab in Alderley Park and Alex Alderton, Roberto Amato, Sonia Goncalves, Ewan Harrison, David K. Jackson, Ian Johnston, Dominic Kwiatkowski, Cordelia Langford, John Sillitoe on behalf of the Wellcome Sanger Institute COVID-19 Surveillance Team |
| EPI_ISL_552767 | Lighthouse Lab in Milton Keynes | Wellcome Sanger Institute for the COVID-19 Genomics UK (COG-UK) consortium | The Lighthouse Lab in Milton Keynes and Alex Alderton, Roberto Amato, Sonia Goncalves, Ewan Harrison, David K. Jackson, Ian Johnston, Dominic Kwiatkowski, Cordelia Langford, John Sillitoe on behalf of the Wellcome Sanger Institute COVID-19 Surveillance Team (http://www.sanger.ac.uk/covid-team) |
| EPI_ISL_552768 | Lighthouse Lab in Alderley Park | Wellcome Sanger Institute for the COVID-19 Genomics UK (COG-UK) consortium | The Lighthouse Lab in Alderley Park and Alex Alderton, Roberto Amato, Sonia Goncalves, Ewan Harrison, David K. Jackson, Ian Johnston, Dominic Kwiatkowski, Cordelia Langford, John Sillitoe on behalf of the Wellcome Sanger Institute COVID-19 Surveillance Team |
| EPI_ISL_552769 | Lighthouse Lab in Milton Keynes | Wellcome Sanger Institute for the COVID-19 Genomics UK (COG-UK) consortium | The Lighthouse Lab in Alderley Park and Alex Alderton, Roberto Amato, Sonia Goncalves, Ewan Harrison, David K. Jackson, Ian Johnston, Dominic Kwiatkowski, Cordelia Langford, John Sillitoe on behalf of the Wellcome Sanger Institute COVID-19 Surveillance Team |
| EPI_ISL_552770, EPI_ISL_552771 | Lighthouse Lab in Alderley Park | Wellcome Sanger Institute for the COVID-19 Genomics UK (COG-UK) consortium | The Lighthouse Lab in Alderley Park and Alex Alderton, Roberto Amato, Sonia Goncalves, Ewan Harrison, David K. Jackson, Ian Johnston, Dominic Kwiatkowski, Cordelia Langford, John Sillitoe on behalf of the Wellcome Sanger Institute COVID-19 Surveillance Team |
| EPI_ISL_552772 | Lighthouse Lab in Milton Keynes | Wellcome Sanger Institute for the COVID-19 Genomics UK (COG-UK) consortium | The Lighthouse Lab in Milton Keynes and Alex Alderton, Roberto Amato, Sonia Goncalves, Ewan Harrison, David K. Jackson, Ian Johnston, Dominic Kwiatkowski, Cordelia Langford, John Sillitoe on behalf of the Wellcome Sanger Institute COVID-19 Surveillance Team (http://www.sanger.ac.uk/covid-team) |
| EPI_ISL_552773 | Lighthouse Lab in Alderley Park | Wellcome Sanger Institute for the COVID-19 Genomics UK (COG-UK) consortium | The Lighthouse Lab in Alderley Park and Alex Alderton, Roberto Amato, Sonia Goncalves, Ewan Harrison, David K. Jackson, Ian Johnston, Dominic Kwiatkowski, Cordelia Langford, John Sillitoe on behalf of the Wellcome Sanger Institute COVID-19 Surveillance Team |
| EPI_ISL_552774 | Lighthouse Lab in Alderley Park | Wellcome Sanger Institute for the COVID-19 Genomics UK (COG-UK) consortium | The Lighthouse Lab in Alderley Park and Alex Alderton, Roberto Amato, Sonia Goncalves, Ewan Harrison, David K. Jackson, Ian Johnston, Dominic Kwiatkowski, Cordelia Langford, John Sillitoe on behalf of the Wellcome Sanger Institute COVID-19 Surveillance Team (http://www.sanger.ac.uk/covid-team) |
| EPI_ISL_552775, EPI_ISL_552776, EPI_ISL_552777, EPI_ISL_552779, EPI_ISL_552780 | Lighthouse Lab in Alderley Park | Wellcome Sanger Institute for the COVID-19 Genomics UK (COG-UK) consortium | The Lighthouse Lab in Alderley Park and Alex Alderton, Roberto Amato, Sonia Goncalves, Ewan Harrison, David K. Jackson, Ian Johnston, Dominic Kwiatkowski, Cordelia Langford, John Sillitoe on behalf of the Wellcome Sanger Institute COVID-19 Surveillance Team |
| EPI_ISL_552781 | Lighthouse Lab in Milton Keynes | Wellcome Sanger Institute for the COVID-19 Genomics UK (COG-UK) consortium | The Lighthouse Lab in Milton Keynes and Alex Alderton, Roberto Amato, Sonia Goncalves, Ewan Harrison, David K. Jackson, Ian Johnston, Dominic Kwiatkowski, Cordelia Langford, John Sillitoe on behalf of the Wellcome Sanger Institute COVID-19 Surveillance Team (http://www.sanger.ac.uk/covid-team) |
| EPI_ISL_552782, EPI_ISL_552783 | Lighthouse Lab in Alderley Park | Wellcome Sanger Institute for the COVID-19 Genomics UK (COG-UK) consortium | The Lighthouse Lab in Alderley Park and Alex Alderton, Roberto Amato, Sonia Goncalves, Ewan Harrison, David K. Jackson, Ian Johnston, Dominic Kwiatkowski, Cordelia Langford, John Sillitoe on behalf of the Wellcome Sanger Institute COVID-19 Surveillance Team |
| EPI_ISL_552784 | Lighthouse Lab in Milton Keynes | Wellcome Sanger Institute for the COVID-19 Genomics UK (COG-UK) consortium | The Lighthouse Lab in Milton Keynes and Alex Alderton, Roberto Amato, Sonia Goncalves, Ewan Harrison, David K. Jackson, Ian Johnston, Dominic Kwiatkowski, Cordelia Langford, John Sillitoe on behalf of the Wellcome Sanger Institute COVID-19 Surveillance Team (http://www.sanger.ac.uk/covid-team) |
| EPI_ISL_552785 | Lighthouse Lab in Alderley Park | Wellcome Sanger Institute for the COVID-19 Genomics UK (COG-UK) consortium | The Lighthouse Lab in Alderley Park and Alex Alderton, Roberto Amato, Sonia Goncalves, Ewan Harrison, David K. Jackson, Ian Johnston, Dominic Kwiatkowski, Cordelia Langford, John Sillitoe on behalf of the Wellcome Sanger Institute COVID-19 Surveillance Team |
| EPI_ISL_552787, EPI_ISL_552788, EPI_ISL_552789, EPI_ISL_552790, EPI_ISL_552791 | Lighthouse Lab in Milton Keynes | Wellcome Sanger Institute for the COVID-19 Genomics UK (COG-UK) consortium | The Lighthouse Lab in Milton Keynes and Alex Alderton, Roberto Amato, Sonia Goncalves, Ewan Harrison, David K. Jackson, Ian Johnston, Dominic Kwiatkowski, Cordelia Langford, John Sillitoe on behalf of the Wellcome Sanger Institute COVID-19 Surveillance Team (http://www.sanger.ac.uk/covid-team) |
| EPI_ISL_552792, EPI_ISL_552793, EPI_ISL_552794 | Lighthouse Lab in Alderley Park | Wellcome Sanger Institute for the COVID-19 Genomics UK (COG-UK) consortium | The Lighthouse Lab in Alderley Park and Alex Alderton, Roberto Amato, Sonia Goncalves, Ewan Harrison, David K. Jackson, Ian Johnston, Dominic Kwiatkowski, Cordelia Langford, John Sillitoe on behalf of the Wellcome Sanger Institute COVID-19 Surveillance Team |
| EPI_ISL_552795 | Lighthouse Lab in Milton Keynes | Wellcome Sanger Institute for the COVID-19 Genomics UK (COG-UK) consortium | The Lighthouse Lab in Milton Keynes and Alex Alderton, Roberto Amato, Sonia Goncalves, Ewan Harrison, David K. Jackson, Ian Johnston, Dominic Kwiatkowski, Cordelia Langford, John Sillitoe on behalf of the Wellcome Sanger Institute COVID-19 Surveillance Team (http://www.sanger.ac.uk/covid-team) |
| EPI_ISL_552796 | Lighthouse Lab in Alderley Park | Wellcome Sanger Institute for the COVID-19 Genomics UK (COG-UK) consortium | The Lighthouse Lab in Alderley Park and Alex Alderton, Roberto Amato, Sonia Goncalves, Ewan Harrison, David K. Jackson, Ian Johnston, Dominic Kwiatkowski, Cordelia Langford, John Sillitoe on behalf of the Wellcome Sanger Institute COVID-19 Surveillance Team |
| EPI_ISL_552799, EPI_ISL_552802, EPI_ISL_552809, EPI_ISL_552810, EPI_ISL_552811, EPI_ISL_552814 | Lighthouse Lab in Milton Keynes | Wellcome Sanger Institute for the COVID-19 Genomics UK (COG-UK) consortium | The Lighthouse Lab in Milton Keynes and Alex Alderton, Roberto Amato, Sonia Goncalves, Ewan Harrison, David K. Jackson, Ian Johnston, Dominic Kwiatkowski, Cordelia Langford, John Sillitoe on behalf of the Wellcome Sanger Institute COVID-19 Surveillance Team (http://www.sanger.ac.uk/covid-team) |
| EPI_ISL_552816 | Lighthouse Lab in Alderley Park | Wellcome Sanger Institute for the COVID-19 Genomics UK (COG-UK) consortium | The Lighthouse Lab in Alderley Park and Alex Alderton, Roberto Amato, Sonia Goncalves, Ewan Harrison, David K. Jackson, Ian Johnston, Dominic Kwiatkowski, Cordelia Langford, John Sillitoe on behalf of the Wellcome Sanger Institute COVID-19 Surveillance Team |
| EPI_ISL_552820, EPI_ISL_552823 | Lighthouse Lab in Milton Keynes | Wellcome Sanger Institute for the COVID-19 Genomics UK (COG-UK) consortium | The Lighthouse Lab in Milton Keynes and Alex Alderton, Roberto Amato, Sonia Goncalves, Ewan Harrison, David K. Jackson, Ian Johnston, Dominic Kwiatkowski, Cordelia Langford, John Sillitoe on behalf of the Wellcome Sanger Institute COVID-19 Surveillance Team (http://www.sanger.ac.uk/covid-team) |
| EPI_ISL_552824, EPI_ISL_552826, | Lighthouse Lab in Alderley Park | Wellcome Sanger Institute for the COVID-19 Genomics UK | The Lighthouse Lab in Alderley Park and Alex Alderton, Roberto Amato, Sonia Goncalves, Ewan Harrison, David K. Jackson, Ian Johnston, Dominic |

| | | | |
|---|---|---|---|
| EPI_ISL_552838 | | (COG-UK) consortium | Kwiatkowski, Cordelia Langford, John Sillitoe on behalf of the Wellcome Sanger Institute COVID-19 Surveillance Team |
| EPI_ISL_552842, EPI_ISL_552852 | Lighthouse Lab in Milton Keynes | Wellcome Sanger Institute for the COVID-19 Genomics UK (COG-UK) consortium | The Lighthouse Lab in Milton Keynes and Alex Alderton, Roberto Amato, Sonia Goncalves, Ewan Harrison, David K. Jackson, Ian Johnston, Dominic Kwiatkowski, Cordelia Langford, John Sillitoe on behalf of the Wellcome Sanger Institute COVID-19 Surveillance Team (http://www.sanger.ac.uk/covid-team) |
| EPI_ISL_552854 | Lighthouse Lab in Alderley Park | Wellcome Sanger Institute for the COVID-19 Genomics UK (COG-UK) consortium | The Lighthouse Lab in Alderley Park and Alex Alderton, Roberto Amato, Sonia Goncalves, Ewan Harrison, David K. Jackson, Ian Johnston, Dominic Kwiatkowski, Cordelia Langford, John Sillitoe on behalf of the Wellcome Sanger Institute COVID-19 Surveillance Team (http://www.sanger.ac.uk/covid-team) |
| EPI_ISL_552867, EPI_ISL_552874, EPI_ISL_552875 | Lighthouse Lab in Milton Keynes | Wellcome Sanger Institute for the COVID-19 Genomics UK (COG-UK) consortium | The Lighthouse Lab in Milton Keynes and Alex Alderton, Roberto Amato, Sonia Goncalves, Ewan Harrison, David K. Jackson, Ian Johnston, Dominic Kwiatkowski, Cordelia Langford, John Sillitoe on behalf of the Wellcome Sanger Institute COVID-19 Surveillance Team (http://www.sanger.ac.uk/covid-team) |
| EPI_ISL_552877 | Lighthouse Lab in Alderley Park | Wellcome Sanger Institute for the COVID-19 Genomics UK (COG-UK) consortium | The Lighthouse Lab in Alderley Park and Alex Alderton, Roberto Amato, Sonia Goncalves, Ewan Harrison, David K. Jackson, Ian Johnston, Dominic Kwiatkowski, Cordelia Langford, John Sillitoe on behalf of the Wellcome Sanger Institute COVID-19 Surveillance Team |
| EPI_ISL_552879 | Lighthouse Lab in Milton Keynes | Wellcome Sanger Institute for the COVID-19 Genomics UK (COG-UK) consortium | The Lighthouse Lab in Milton Keynes and Alex Alderton, Roberto Amato, Sonia Goncalves, Ewan Harrison, David K. Jackson, Ian Johnston, Dominic Kwiatkowski, Cordelia Langford, John Sillitoe on behalf of the Wellcome Sanger Institute COVID-19 Surveillance Team (http://www.sanger.ac.uk/covid-team) |
| EPI_ISL_552886, EPI_ISL_552893, EPI_ISL_552898 | Lighthouse Lab in Alderley Park | Wellcome Sanger Institute for the COVID-19 Genomics UK (COG-UK) consortium | The Lighthouse Lab in Alderley Park and Alex Alderton, Roberto Amato, Sonia Goncalves, Ewan Harrison, David K. Jackson, Ian Johnston, Dominic Kwiatkowski, Cordelia Langford, John Sillitoe on behalf of the Wellcome Sanger Institute COVID-19 Surveillance Team |
| EPI_ISL_552900 | Lighthouse Lab in Alderley Park | Wellcome Sanger Institute for the COVID-19 Genomics UK (COG-UK) consortium | The Lighthouse Lab in Alderley Park and Alex Alderton, Roberto Amato, Sonia Goncalves, Ewan Harrison, David K. Jackson, Ian Johnston, Dominic Kwiatkowski, Cordelia Langford, John Sillitoe on behalf of the Wellcome Sanger Institute COVID-19 Surveillance Team (http://www.sanger.ac.uk/covid-team) |
| EPI_ISL_552905, EPI_ISL_552910, EPI_ISL_552916 | Lighthouse Lab in Milton Keynes | Wellcome Sanger Institute for the COVID-19 Genomics UK (COG-UK) consortium | The Lighthouse Lab in Milton Keynes and Alex Alderton, Roberto Amato, Sonia Goncalves, Ewan Harrison, David K. Jackson, Ian Johnston, Dominic Kwiatkowski, Cordelia Langford, John Sillitoe on behalf of the Wellcome Sanger Institute COVID-19 Surveillance Team (http://www.sanger.ac.uk/covid-team) |
| EPI_ISL_552932 | Lighthouse Lab in Alderley Park | Wellcome Sanger Institute for the COVID-19 Genomics UK (COG-UK) consortium | The Lighthouse Lab in Alderley Park and Alex Alderton, Roberto Amato, Sonia Goncalves, Ewan Harrison, David K. Jackson, Ian Johnston, Dominic Kwiatkowski, Cordelia Langford, John Sillitoe on behalf of the Wellcome Sanger Institute COVID-19 Surveillance Team |
| EPI_ISL_552936, EPI_ISL_552938, EPI_ISL_552939 | Lighthouse Lab in Milton Keynes | Wellcome Sanger Institute for the COVID-19 Genomics UK (COG-UK) consortium | The Lighthouse Lab in Milton Keynes and Alex Alderton, Roberto Amato, Sonia Goncalves, Ewan Harrison, David K. Jackson, Ian Johnston, Dominic Kwiatkowski, Cordelia Langford, John Sillitoe on behalf of the Wellcome Sanger Institute COVID-19 Surveillance Team (http://www.sanger.ac.uk/covid-team) |
| EPI_ISL_552943 | Lighthouse Lab in Alderley Park | Wellcome Sanger Institute for the COVID-19 Genomics UK (COG-UK) consortium | The Lighthouse Lab in Alderley Park and Alex Alderton, Roberto Amato, Sonia Goncalves, Ewan Harrison, David K. Jackson, Ian Johnston, Dominic Kwiatkowski, Cordelia Langford, John Sillitoe on behalf of the Wellcome Sanger Institute COVID-19 Surveillance Team |
| EPI_ISL_552947, EPI_ISL_552948, EPI_ISL_552949, EPI_ISL_552950 | Lighthouse Lab in Milton Keynes | Wellcome Sanger Institute for the COVID-19 Genomics UK (COG-UK) consortium | The Lighthouse Lab in Milton Keynes and Alex Alderton, Roberto Amato, Sonia Goncalves, Ewan Harrison, David K. Jackson, Ian Johnston, Dominic Kwiatkowski, Cordelia Langford, John Sillitoe on behalf of the Wellcome Sanger Institute COVID-19 Surveillance Team (http://www.sanger.ac.uk/covid-team) |
| EPI_ISL_552952 | Lighthouse Lab in Alderley Park | Wellcome Sanger Institute for the COVID-19 Genomics UK (COG-UK) consortium | The Lighthouse Lab in Alderley Park and Alex Alderton, Roberto Amato, Sonia Goncalves, Ewan Harrison, David K. Jackson, Ian Johnston, Dominic Kwiatkowski, Cordelia Langford, John Sillitoe on behalf of the Wellcome Sanger Institute COVID-19 Surveillance Team |
| EPI_ISL_552953 | Lighthouse Lab in Milton Keynes | Wellcome Sanger Institute for the COVID-19 Genomics UK (COG-UK) consortium | The Lighthouse Lab in Alderley Park and Alex Alderton, Roberto Amato, Sonia Goncalves, Ewan Harrison, David K. Jackson, Ian Johnston, Dominic Kwiatkowski, Cordelia Langford, John Sillitoe on behalf of the Wellcome Sanger Institute COVID-19 Surveillance Team |
| EPI_ISL_552960, EPI_ISL_552970 | Lighthouse Lab in Alderley Park | Wellcome Sanger Institute for the COVID-19 Genomics UK (COG-UK) consortium | The Lighthouse Lab in Alderley Park and Alex Alderton, Roberto Amato, Sonia Goncalves, Ewan Harrison, David K. Jackson, Ian Johnston, Dominic Kwiatkowski, Cordelia Langford, John Sillitoe on behalf of the Wellcome Sanger Institute COVID-19 Surveillance Team |
| EPI_ISL_552974 | Lighthouse Lab in Alderley Park | Wellcome Sanger Institute for the COVID-19 Genomics UK (COG-UK) consortium | The Lighthouse Lab in Alderley Park and Alex Alderton, Roberto Amato, Sonia Goncalves, Ewan Harrison, David K. Jackson, Ian Johnston, Dominic Kwiatkowski, Cordelia Langford, John Sillitoe on behalf of the Wellcome Sanger Institute COVID-19 Surveillance Team (http://www.sanger.ac.uk/covid-team) |
| EPI_ISL_552979 | Lighthouse Lab in Milton Keynes | Wellcome Sanger Institute for the COVID-19 Genomics UK (COG-UK) consortium | The Lighthouse Lab in Milton Keynes and Alex Alderton, Roberto Amato, Sonia Goncalves, Ewan Harrison, David K. Jackson, Ian Johnston, Dominic Kwiatkowski, Cordelia Langford, John Sillitoe on behalf of the Wellcome Sanger Institute COVID-19 Surveillance Team (http://www.sanger.ac.uk/covid-team) |
| EPI_ISL_552982 | Lighthouse Lab in Alderley Park | Wellcome Sanger Institute for the COVID-19 Genomics UK (COG-UK) consortium | The Lighthouse Lab in Alderley Park and Alex Alderton, Roberto Amato, Sonia Goncalves, Ewan Harrison, David K. Jackson, Ian Johnston, Dominic Kwiatkowski, Cordelia Langford, John Sillitoe on behalf of the Wellcome Sanger Institute COVID-19 Surveillance Team (http://www.sanger.ac.uk/covid-team) |
| EPI_ISL_552987, EPI_ISL_552990 | Lighthouse Lab in Milton Keynes | Wellcome Sanger Institute for the COVID-19 Genomics UK (COG-UK) consortium | The Lighthouse Lab in Milton Keynes and Alex Alderton, Roberto Amato, Sonia Goncalves, Ewan Harrison, David K. Jackson, Ian Johnston, Dominic Kwiatkowski, Cordelia Langford, John Sillitoe on behalf of the Wellcome Sanger Institute COVID-19 Surveillance Team (http://www.sanger.ac.uk/covid-team) |
| EPI_ISL_552993 | Lighthouse Lab in Alderley Park | Wellcome Sanger Institute for the COVID-19 Genomics UK (COG-UK) consortium | The Lighthouse Lab in Alderley Park and Alex Alderton, Roberto Amato, Sonia Goncalves, Ewan Harrison, David K. Jackson, Ian Johnston, Dominic Kwiatkowski, Cordelia Langford, John Sillitoe on behalf of the Wellcome Sanger Institute COVID-19 Surveillance Team |
| EPI_ISL_552995, EPI_ISL_552997 | Lighthouse Lab in Milton Keynes | Wellcome Sanger Institute for the COVID-19 Genomics UK (COG-UK) consortium | The Lighthouse Lab in Milton Keynes and Alex Alderton, Roberto Amato, Sonia Goncalves, Ewan Harrison, David K. Jackson, Ian Johnston, Dominic Kwiatkowski, Cordelia Langford, John Sillitoe on behalf of the Wellcome Sanger Institute COVID-19 Surveillance Team (http://www.sanger.ac.uk/covid-team) |
| EPI_ISL_553006 | Lighthouse Lab in Alderley Park | Wellcome Sanger Institute for the COVID-19 Genomics UK (COG-UK) consortium | The Lighthouse Lab in Alderley Park and Alex Alderton, Roberto Amato, Sonia Goncalves, Ewan Harrison, David K. Jackson, Ian Johnston, Dominic Kwiatkowski, Cordelia Langford, John Sillitoe on behalf of the Wellcome Sanger Institute COVID-19 Surveillance Team (http://www.sanger.ac.uk/covid-team) |
| EPI_ISL_553008 | Lighthouse Lab in Milton Keynes | Wellcome Sanger Institute for the COVID-19 Genomics UK (COG-UK) consortium | The Lighthouse Lab in Alderley Park and Alex Alderton, Roberto Amato, Sonia Goncalves, Ewan Harrison, David K. Jackson, Ian Johnston, Dominic Kwiatkowski, Cordelia Langford, John Sillitoe on behalf of the Wellcome Sanger Institute COVID-19 Surveillance Team |
| EPI_ISL_553009, EPI_ISL_553010, EPI_ISL_553011, EPI_ISL_553012, EPI_ISL_553013, EPI_ISL_553014 | Lighthouse Lab in Milton Keynes | Wellcome Sanger Institute for the COVID-19 Genomics UK (COG-UK) consortium | The Lighthouse Lab in Milton Keynes and Alex Alderton, Roberto Amato, Sonia Goncalves, Ewan Harrison, David K. Jackson, Ian Johnston, Dominic Kwiatkowski, Cordelia Langford, John Sillitoe on behalf of the Wellcome Sanger Institute COVID-19 Surveillance Team (http://www.sanger.ac.uk/covid-team) |
| EPI_ISL_553015, EPI_ISL_553016 | Lighthouse Lab in Milton Keynes | Wellcome Sanger Institute for the COVID-19 Genomics UK (COG-UK) consortium | The Lighthouse Lab in Alderley Park and Alex Alderton, Roberto Amato, Sonia Goncalves, Ewan Harrison, David K. Jackson, Ian Johnston, Dominic Kwiatkowski, Cordelia Langford, John Sillitoe on behalf of the Wellcome Sanger Institute COVID-19 Surveillance Team |
| EPI_ISL_553017, EPI_ISL_553018, EPI_ISL_553019, EPI_ISL_553020, EPI_ISL_553021 | Lighthouse Lab in Milton Keynes | Wellcome Sanger Institute for the COVID-19 Genomics UK (COG-UK) consortium | The Lighthouse Lab in Milton Keynes and Alex Alderton, Roberto Amato, Sonia Goncalves, Ewan Harrison, David K. Jackson, Ian Johnston, Dominic Kwiatkowski, Cordelia Langford, John Sillitoe on behalf of the Wellcome Sanger Institute COVID-19 Surveillance Team (http://www.sanger.ac.uk/covid-team) |
| EPI_ISL_553023 | Lighthouse Lab in Milton Keynes | Wellcome Sanger Institute for the COVID-19 Genomics UK (COG-UK) consortium | The Lighthouse Lab in Alderley Park and Alex Alderton, Roberto Amato, Sonia Goncalves, Ewan Harrison, David K. Jackson, Ian Johnston, Dominic Kwiatkowski, Cordelia Langford, John Sillitoe on behalf of the Wellcome Sanger Institute COVID-19 Surveillance Team |
| EPI_ISL_553025, EPI_ISL_553026, EPI_ISL_553027, EPI_ISL_553028 | Lighthouse Lab in Milton Keynes | Wellcome Sanger Institute for the COVID-19 Genomics UK (COG-UK) consortium | The Lighthouse Lab in Milton Keynes and Alex Alderton, Roberto Amato, Sonia Goncalves, Ewan Harrison, David K. Jackson, Ian Johnston, Dominic Kwiatkowski, Cordelia Langford, John Sillitoe on behalf of the Wellcome Sanger Institute COVID-19 Surveillance Team |

| | | | |
|---|---|---|---|
| | | | (http://www.sanger.ac.uk/covid-team) |
| EPI_ISL_553029 | Lighthouse Lab in Milton Keynes | Wellcome Sanger Institute for the COVID-19 Genomics UK (COG-UK) consortium | The Lighthouse Lab in Alderley Park and Alex Alderton, Roberto Amato, Sonia Goncalves, Ewan Harrison, David K. Jackson, Ian Johnston, Dominic Kwiatkowski, Cordelia Langford, John Sillitoe on behalf of the Wellcome Sanger Institute COVID-19 Surveillance Team |
| EPI_ISL_553031, EPI_ISL_553032, EPI_ISL_553033, EPI_ISL_553034, EPI_ISL_553035, EPI_ISL_553036, EPI_ISL_553040, EPI_ISL_553041, EPI_ISL_553043, EPI_ISL_553045, EPI_ISL_553046, EPI_ISL_553047, EPI_ISL_553048, EPI_ISL_553049, EPI_ISL_553051, EPI_ISL_553053, EPI_ISL_553056, EPI_ISL_553057, EPI_ISL_553059, EPI_ISL_553061, EPI_ISL_553062, EPI_ISL_553063, EPI_ISL_553064, EPI_ISL_553065, EPI_ISL_553066, EPI_ISL_553067, EPI_ISL_553069, EPI_ISL_553070, EPI_ISL_553072, EPI_ISL_553073, EPI_ISL_553074, EPI_ISL_553075, EPI_ISL_553076, EPI_ISL_553077, EPI_ISL_553078, EPI_ISL_553080, EPI_ISL_553081, EPI_ISL_553082, EPI_ISL_553083, EPI_ISL_553086 | | | |
| see above | Lighthouse Lab in Milton Keynes | Wellcome Sanger Institute for the COVID-19 Genomics UK (COG-UK) consortium | The Lighthouse Lab in Milton Keynes and Alex Alderton, Roberto Amato, Sonia Goncalves, Ewan Harrison, David K. Jackson, Ian Johnston, Dominic Kwiatkowski, Cordelia Langford, John Sillitoe on behalf of the Wellcome Sanger Institute COVID-19 Surveillance Team (http://www.sanger.ac.uk/covid-team) |
| EPI_ISL_553087 | Lighthouse Lab in Milton Keynes | Wellcome Sanger Institute for the COVID-19 Genomics UK (COG-UK) consortium | The Lighthouse Lab in Alderley Park and Alex Alderton, Roberto Amato, Sonia Goncalves, Ewan Harrison, David K. Jackson, Ian Johnston, Dominic Kwiatkowski, Cordelia Langford, John Sillitoe on behalf of the Wellcome Sanger Institute COVID-19 Surveillance Team |
| EPI_ISL_553088, EPI_ISL_553089, EPI_ISL_553091, EPI_ISL_553092, EPI_ISL_553094, EPI_ISL_553095, EPI_ISL_553096, EPI_ISL_553097, EPI_ISL_553098, EPI_ISL_553099, EPI_ISL_553101, EPI_ISL_553102 | | | |
| see above | Lighthouse Lab in Milton Keynes | Wellcome Sanger Institute for the COVID-19 Genomics UK (COG-UK) consortium | The Lighthouse Lab in Milton Keynes and Alex Alderton, Roberto Amato, Sonia Goncalves, Ewan Harrison, David K. Jackson, Ian Johnston, Dominic Kwiatkowski, Cordelia Langford, John Sillitoe on behalf of the Wellcome Sanger Institute COVID-19 Surveillance Team (http://www.sanger.ac.uk/covid-team) |
| EPI_ISL_553103 | Lighthouse Lab in Milton Keynes | Wellcome Sanger Institute for the COVID-19 Genomics UK (COG-UK) consortium | The Lighthouse Lab in Alderley Park and Alex Alderton, Roberto Amato, Sonia Goncalves, Ewan Harrison, David K. Jackson, Ian Johnston, Dominic Kwiatkowski, Cordelia Langford, John Sillitoe on behalf of the Wellcome Sanger Institute COVID-19 Surveillance Team |
| EPI_ISL_553104 | Lighthouse Lab in Milton Keynes | Wellcome Sanger Institute for the COVID-19 Genomics UK (COG-UK) consortium | The Lighthouse Lab in Milton Keynes and Alex Alderton, Roberto Amato, Sonia Goncalves, Ewan Harrison, David K. Jackson, Ian Johnston, Dominic Kwiatkowski, Cordelia Langford, John Sillitoe on behalf of the Wellcome Sanger Institute COVID-19 Surveillance Team (http://www.sanger.ac.uk/covid-team) |
| EPI_ISL_553105 | Lighthouse Lab in Milton Keynes | Wellcome Sanger Institute for the COVID-19 Genomics UK (COG-UK) consortium | The Lighthouse Lab in Alderley Park and Alex Alderton, Roberto Amato, Sonia Goncalves, Ewan Harrison, David K. Jackson, Ian Johnston, Dominic Kwiatkowski, Cordelia Langford, John Sillitoe on behalf of the Wellcome Sanger Institute COVID-19 Surveillance Team |
| EPI_ISL_553106, EPI_ISL_553107, EPI_ISL_553111, EPI_ISL_553112 | Lighthouse Lab in Milton Keynes | Wellcome Sanger Institute for the COVID-19 Genomics UK (COG-UK) consortium | The Lighthouse Lab in Milton Keynes and Alex Alderton, Roberto Amato, Sonia Goncalves, Ewan Harrison, David K. Jackson, Ian Johnston, Dominic Kwiatkowski, Cordelia Langford, John Sillitoe on behalf of the Wellcome Sanger Institute COVID-19 Surveillance Team (http://www.sanger.ac.uk/covid-team) |
| EPI_ISL_553113 | Lighthouse Lab in Milton Keynes | Wellcome Sanger Institute for the COVID-19 Genomics UK (COG-UK) consortium | The Lighthouse Lab in Alderley Park and Alex Alderton, Roberto Amato, Sonia Goncalves, Ewan Harrison, David K. Jackson, Ian Johnston, Dominic Kwiatkowski, Cordelia Langford, John Sillitoe on behalf of the Wellcome Sanger Institute COVID-19 Surveillance Team |
| EPI_ISL_553114, EPI_ISL_553115, EPI_ISL_553116, EPI_ISL_553117, EPI_ISL_553118, EPI_ISL_553119, EPI_ISL_553120, EPI_ISL_553121, EPI_ISL_553122, EPI_ISL_553123, EPI_ISL_553124 | | | |
| see above | Lighthouse Lab in Milton Keynes | Wellcome Sanger Institute for the COVID-19 Genomics UK (COG-UK) consortium | The Lighthouse Lab in Milton Keynes and Alex Alderton, Roberto Amato, Sonia Goncalves, Ewan Harrison, David K. Jackson, Ian Johnston, Dominic Kwiatkowski, Cordelia Langford, John Sillitoe on behalf of the Wellcome Sanger Institute COVID-19 Surveillance Team (http://www.sanger.ac.uk/covid-team) |
| EPI_ISL_553125 | Lighthouse Lab in Milton Keynes | Wellcome Sanger Institute for the COVID-19 Genomics UK (COG-UK) consortium | The Lighthouse Lab in Alderley Park and Alex Alderton, Roberto Amato, Sonia Goncalves, Ewan Harrison, David K. Jackson, Ian Johnston, Dominic Kwiatkowski, Cordelia Langford, John Sillitoe on behalf of the Wellcome Sanger Institute COVID-19 Surveillance Team |
| EPI_ISL_553126 | Lighthouse Lab in Milton Keynes | Wellcome Sanger Institute for the COVID-19 Genomics UK (COG-UK) consortium | The Lighthouse Lab in Milton Keynes and Alex Alderton, Roberto Amato, Sonia Goncalves, Ewan Harrison, David K. Jackson, Ian Johnston, Dominic Kwiatkowski, Cordelia Langford, John Sillitoe on behalf of the Wellcome Sanger Institute COVID-19 Surveillance Team (http://www.sanger.ac.uk/covid-team) |
| EPI_ISL_553127 | Lighthouse Lab in Milton Keynes | Wellcome Sanger Institute for the COVID-19 Genomics UK (COG-UK) consortium | The Lighthouse Lab in Alderley Park and Alex Alderton, Roberto Amato, Sonia Goncalves, Ewan Harrison, David K. Jackson, Ian Johnston, Dominic Kwiatkowski, Cordelia Langford, John Sillitoe on behalf of the Wellcome Sanger Institute COVID-19 Surveillance Team |
| EPI_ISL_553128, EPI_ISL_553130 | Lighthouse Lab in Milton Keynes | Wellcome Sanger Institute for the COVID-19 Genomics UK (COG-UK) consortium | The Lighthouse Lab in Milton Keynes and Alex Alderton, Roberto Amato, Sonia Goncalves, Ewan Harrison, David K. Jackson, Ian Johnston, Dominic Kwiatkowski, Cordelia Langford, John Sillitoe on behalf of the Wellcome Sanger Institute COVID-19 Surveillance Team (http://www.sanger.ac.uk/covid-team) |
| EPI_ISL_553131 | Lighthouse Lab in Milton Keynes | Wellcome Sanger Institute for the COVID-19 Genomics UK (COG-UK) consortium | The Lighthouse Lab in Alderley Park and Alex Alderton, Roberto Amato, Sonia Goncalves, Ewan Harrison, David K. Jackson, Ian Johnston, Dominic Kwiatkowski, Cordelia Langford, John Sillitoe on behalf of the Wellcome Sanger Institute COVID-19 Surveillance Team |
| EPI_ISL_553132, EPI_ISL_553133, EPI_ISL_553134, EPI_ISL_553135, EPI_ISL_553137, EPI_ISL_553138, EPI_ISL_553140 | Lighthouse Lab in Milton Keynes | Wellcome Sanger Institute for the COVID-19 Genomics UK (COG-UK) consortium | The Lighthouse Lab in Milton Keynes and Alex Alderton, Roberto Amato, Sonia Goncalves, Ewan Harrison, David K. Jackson, Ian Johnston, Dominic Kwiatkowski, Cordelia Langford, John Sillitoe on behalf of the Wellcome Sanger Institute COVID-19 Surveillance Team (http://www.sanger.ac.uk/covid-team) |
| EPI_ISL_553141 | Lighthouse Lab in Milton Keynes | Wellcome Sanger Institute for the COVID-19 Genomics UK (COG-UK) consortium | The Lighthouse Lab in Alderley Park and Alex Alderton, Roberto Amato, Sonia Goncalves, Ewan Harrison, David K. Jackson, Ian Johnston, Dominic Kwiatkowski, Cordelia Langford, John Sillitoe on behalf of the Wellcome Sanger Institute COVID-19 Surveillance Team |
| EPI_ISL_553142, EPI_ISL_553143, EPI_ISL_553144, EPI_ISL_553145, EPI_ISL_553147, EPI_ISL_553150, EPI_ISL_553151, EPI_ISL_553152, EPI_ISL_553153, EPI_ISL_553154, EPI_ISL_553155, EPI_ISL_553156, EPI_ISL_553157, EPI_ISL_553158, EPI_ISL_553159, EPI_ISL_553160, EPI_ISL_553162, EPI_ISL_553163, EPI_ISL_553164, EPI_ISL_553165 | | | |
| see above | Lighthouse Lab in Milton Keynes | Wellcome Sanger Institute for the COVID-19 Genomics UK (COG-UK) consortium | The Lighthouse Lab in Milton Keynes and Alex Alderton, Roberto Amato, Sonia Goncalves, Ewan Harrison, David K. Jackson, Ian Johnston, Dominic Kwiatkowski, Cordelia Langford, John Sillitoe on behalf of the Wellcome Sanger Institute COVID-19 Surveillance Team (http://www.sanger.ac.uk/covid-team) |
| EPI_ISL_553166 | Lighthouse Lab in Milton Keynes | Wellcome Sanger Institute for the COVID-19 Genomics UK (COG-UK) consortium | The Lighthouse Lab in Alderley Park and Alex Alderton, Roberto Amato, Sonia Goncalves, Ewan Harrison, David K. Jackson, Ian Johnston, Dominic Kwiatkowski, Cordelia Langford, John Sillitoe on behalf of the Wellcome Sanger Institute COVID-19 Surveillance Team |
| EPI_ISL_553167, EPI_ISL_553168, EPI_ISL_553169, EPI_ISL_553170, EPI_ISL_553171, EPI_ISL_553172, EPI_ISL_553174, EPI_ISL_553175, EPI_ISL_553176, EPI_ISL_553177, EPI_ISL_553178, EPI_ISL_553179, EPI_ISL_553180, EPI_ISL_553181, EPI_ISL_553183, EPI_ISL_553184, EPI_ISL_553185, EPI_ISL_553188, EPI_ISL_553189, EPI_ISL_553190, EPI_ISL_553191, EPI_ISL_553192, EPI_ISL_553194 | | | |
| see above | Lighthouse Lab in Milton Keynes | Wellcome Sanger Institute for the COVID-19 Genomics UK (COG-UK) consortium | The Lighthouse Lab in Milton Keynes and Alex Alderton, Roberto Amato, Sonia Goncalves, Ewan Harrison, David K. Jackson, Ian Johnston, Dominic Kwiatkowski, Cordelia Langford, John Sillitoe on behalf of the Wellcome Sanger Institute COVID-19 Surveillance Team (http://www.sanger.ac.uk/covid-team) |
| EPI_ISL_553195 | Lighthouse Lab in Milton Keynes | Wellcome Sanger Institute for the COVID-19 Genomics UK (COG-UK) consortium | The Lighthouse Lab in Alderley Park and Alex Alderton, Roberto Amato, Sonia Goncalves, Ewan Harrison, David K. Jackson, Ian Johnston, Dominic Kwiatkowski, Cordelia Langford, John Sillitoe on behalf of the Wellcome Sanger Institute COVID-19 Surveillance Team |
| EPI_ISL_553196, EPI_ISL_553197, EPI_ISL_553198, EPI_ISL_553199, EPI_ISL_553200, EPI_ISL_553201, EPI_ISL_553203, EPI_ISL_553204, EPI_ISL_553205, EPI_ISL_553206, EPI_ISL_553207, EPI_ISL_553208, EPI_ISL_553209, EPI_ISL_553210, EPI_ISL_553211 | | | |
| see above | Lighthouse Lab in Milton Keynes | Wellcome Sanger Institute for the COVID-19 Genomics UK (COG-UK) consortium | The Lighthouse Lab in Milton Keynes and Alex Alderton, Roberto Amato, Sonia Goncalves, Ewan Harrison, David K. Jackson, Ian Johnston, Dominic Kwiatkowski, Cordelia Langford, John Sillitoe on behalf of the Wellcome Sanger Institute COVID-19 Surveillance Team (http://www.sanger.ac.uk/covid-team) |
| EPI_ISL_553212 | Lighthouse Lab in Milton Keynes | Wellcome Sanger Institute for the COVID-19 Genomics UK (COG-UK) consortium | The Lighthouse Lab in Alderley Park and Alex Alderton, Roberto Amato, Sonia Goncalves, Ewan Harrison, David K. Jackson, Ian Johnston, Dominic Kwiatkowski, Cordelia Langford, John Sillitoe on behalf of the Wellcome Sanger Institute COVID-19 Surveillance Team |
| EPI_ISL_553213, EPI_ISL_553214, EPI_ISL_553215, EPI_ISL_553216, EPI_ISL_553217, EPI_ISL_553218 | Lighthouse Lab in Milton Keynes | Wellcome Sanger Institute for the COVID-19 Genomics UK (COG-UK) consortium | The Lighthouse Lab in Milton Keynes and Alex Alderton, Roberto Amato, Sonia Goncalves, Ewan Harrison, David K. Jackson, Ian Johnston, Dominic Kwiatkowski, Cordelia Langford, John Sillitoe on behalf of the Wellcome Sanger Institute COVID-19 Surveillance Team (http://www.sanger.ac.uk/covid-team) |

| | | | |
|---|---|---|---|
| EPI_ISL_553219 | Lighthouse Lab in Milton Keynes | Wellcome Sanger Institute for the COVID-19 Genomics UK (COG-UK) consortium | The Lighthouse Lab in Alderley Park and Alex Alderton, Roberto Amato, Sonia Goncalves, Ewan Harrison, David K. Jackson, Ian Johnston, Dominic Kwiatkowski, Cordelia Langford, John Sillitoe on behalf of the Wellcome Sanger Institute COVID-19 Surveillance Team |
| EPI_ISL_553220 | Lighthouse Lab in Milton Keynes | Wellcome Sanger Institute for the COVID-19 Genomics UK (COG-UK) consortium | The Lighthouse Lab in Milton Keynes and Alex Alderton, Roberto Amato, Sonia Goncalves, Ewan Harrison, David K. Jackson, Ian Johnston, Dominic Kwiatkowski, Cordelia Langford, John Sillitoe on behalf of the Wellcome Sanger Institute COVID-19 Surveillance Team (http://www.sanger.ac.uk/covid-team) |
| EPI_ISL_553221 | Lighthouse Lab in Milton Keynes | Wellcome Sanger Institute for the COVID-19 Genomics UK (COG-UK) consortium | The Lighthouse Lab in Alderley Park and Alex Alderton, Roberto Amato, Sonia Goncalves, Ewan Harrison, David K. Jackson, Ian Johnston, Dominic Kwiatkowski, Cordelia Langford, John Sillitoe on behalf of the Wellcome Sanger Institute COVID-19 Surveillance Team |
| EPI_ISL_553222 | Lighthouse Lab in Milton Keynes | Wellcome Sanger Institute for the COVID-19 Genomics UK (COG-UK) consortium | The Lighthouse Lab in Milton Keynes and Alex Alderton, Roberto Amato, Sonia Goncalves, Ewan Harrison, David K. Jackson, Ian Johnston, Dominic Kwiatkowski, Cordelia Langford, John Sillitoe on behalf of the Wellcome Sanger Institute COVID-19 Surveillance Team (http://www.sanger.ac.uk/covid-team) |
| EPI_ISL_553223 | Lighthouse Lab in Milton Keynes | Wellcome Sanger Institute for the COVID-19 Genomics UK (COG-UK) consortium | The Lighthouse Lab in Alderley Park and Alex Alderton, Roberto Amato, Sonia Goncalves, Ewan Harrison, David K. Jackson, Ian Johnston, Dominic Kwiatkowski, Cordelia Langford, John Sillitoe on behalf of the Wellcome Sanger Institute COVID-19 Surveillance Team |
| EPI_ISL_553224, EPI_ISL_553225, EPI_ISL_553226, EPI_ISL_553227, EPI_ISL_553228, EPI_ISL_553229, EPI_ISL_553230, EPI_ISL_553232, EPI_ISL_553233 | Lighthouse Lab in Milton Keynes | Wellcome Sanger Institute for the COVID-19 Genomics UK (COG-UK) consortium | The Lighthouse Lab in Milton Keynes and Alex Alderton, Roberto Amato, Sonia Goncalves, Ewan Harrison, David K. Jackson, Ian Johnston, Dominic Kwiatkowski, Cordelia Langford, John Sillitoe on behalf of the Wellcome Sanger Institute COVID-19 Surveillance Team (http://www.sanger.ac.uk/covid-team) |
| EPI_ISL_553234 | Lighthouse Lab in Milton Keynes | Wellcome Sanger Institute for the COVID-19 Genomics UK (COG-UK) consortium | The Lighthouse Lab in Alderley Park and Alex Alderton, Roberto Amato, Sonia Goncalves, Ewan Harrison, David K. Jackson, Ian Johnston, Dominic Kwiatkowski, Cordelia Langford, John Sillitoe on behalf of the Wellcome Sanger Institute COVID-19 Surveillance Team |
| EPI_ISL_553236, EPI_ISL_553237, EPI_ISL_553238 | Lighthouse Lab in Milton Keynes | Wellcome Sanger Institute for the COVID-19 Genomics UK (COG-UK) consortium | The Lighthouse Lab in Milton Keynes and Alex Alderton, Roberto Amato, Sonia Goncalves, Ewan Harrison, David K. Jackson, Ian Johnston, Dominic Kwiatkowski, Cordelia Langford, John Sillitoe on behalf of the Wellcome Sanger Institute COVID-19 Surveillance Team (http://www.sanger.ac.uk/covid-team) |
| EPI_ISL_553239 | Lighthouse Lab in Milton Keynes | Wellcome Sanger Institute for the COVID-19 Genomics UK (COG-UK) consortium | The Lighthouse Lab in Milton Keynes and Alex Alderton, Roberto Amato, Sonia Goncalves, Ewan Harrison, David K. Jackson, Ian Johnston, Dominic Kwiatkowski, Cordelia Langford, John Sillitoe on behalf of the Wellcome Sanger Institute COVID-19 Surveillance Team |
| EPI_ISL_553243, EPI_ISL_553244, EPI_ISL_553245, EPI_ISL_553247, EPI_ISL_553249, EPI_ISL_553250, EPI_ISL_553253, EPI_ISL_553254, EPI_ISL_553255, EPI_ISL_553256, EPI_ISL_553257, EPI_ISL_553258, EPI_ISL_553259 | | | |
| see above | Lighthouse Lab in Milton Keynes | Wellcome Sanger Institute for the COVID-19 Genomics UK (COG-UK) consortium | The Lighthouse Lab in Milton Keynes and Alex Alderton, Roberto Amato, Sonia Goncalves, Ewan Harrison, David K. Jackson, Ian Johnston, Dominic Kwiatkowski, Cordelia Langford, John Sillitoe on behalf of the Wellcome Sanger Institute COVID-19 Surveillance Team (http://www.sanger.ac.uk/covid-team) |
| EPI_ISL_553260 | Lighthouse Lab in Milton Keynes | Wellcome Sanger Institute for the COVID-19 Genomics UK (COG-UK) consortium | The Lighthouse Lab in Alderley Park and Alex Alderton, Roberto Amato, Sonia Goncalves, Ewan Harrison, David K. Jackson, Ian Johnston, Dominic Kwiatkowski, Cordelia Langford, John Sillitoe on behalf of the Wellcome Sanger Institute COVID-19 Surveillance Team |
| EPI_ISL_553261, EPI_ISL_553262, EPI_ISL_553263, EPI_ISL_553264, EPI_ISL_553265, EPI_ISL_553266, EPI_ISL_553267, EPI_ISL_553268, EPI_ISL_553270 | Lighthouse Lab in Milton Keynes | Wellcome Sanger Institute for the COVID-19 Genomics UK (COG-UK) consortium | The Lighthouse Lab in Milton Keynes and Alex Alderton, Roberto Amato, Sonia Goncalves, Ewan Harrison, David K. Jackson, Ian Johnston, Dominic Kwiatkowski, Cordelia Langford, John Sillitoe on behalf of the Wellcome Sanger Institute COVID-19 Surveillance Team (http://www.sanger.ac.uk/covid-team) |
| EPI_ISL_553272 | Lighthouse Lab in Alderley Park | Wellcome Sanger Institute for the COVID-19 Genomics UK (COG-UK) consortium | The Lighthouse Lab in Alderley Park and Alex Alderton, Roberto Amato, Sonia Goncalves, Ewan Harrison, David K. Jackson, Ian Johnston, Dominic Kwiatkowski, Cordelia Langford, John Sillitoe on behalf of the Wellcome Sanger Institute COVID-19 Surveillance Team (http://www.sanger.ac.uk/covid-team) |
| EPI_ISL_553275, EPI_ISL_553276, EPI_ISL_553295, EPI_ISL_553323 | Lighthouse Lab in Milton Keynes | Wellcome Sanger Institute for the COVID-19 Genomics UK (COG-UK) consortium | The Lighthouse Lab in Milton Keynes and Alex Alderton, Roberto Amato, Sonia Goncalves, Ewan Harrison, David K. Jackson, Ian Johnston, Dominic Kwiatkowski, Cordelia Langford, John Sillitoe on behalf of the Wellcome Sanger Institute COVID-19 Surveillance Team (http://www.sanger.ac.uk/covid-team) |
| EPI_ISL_553327, EPI_ISL_553330, EPI_ISL_553346, EPI_ISL_553347 | Lighthouse Lab in Alderley Park | Wellcome Sanger Institute for the COVID-19 Genomics UK (COG-UK) consortium | The Lighthouse Lab in Alderley Park and Alex Alderton, Roberto Amato, Sonia Goncalves, Ewan Harrison, David K. Jackson, Ian Johnston, Dominic Kwiatkowski, Cordelia Langford, John Sillitoe on behalf of the Wellcome Sanger Institute COVID-19 Surveillance Team (http://www.sanger.ac.uk/covid-team) |
| EPI_ISL_553348, EPI_ISL_553349, EPI_ISL_553350 | Lighthouse Lab in Alderley Park | Wellcome Sanger Institute for the COVID-19 Genomics UK (COG-UK) consortium | The Lighthouse Lab in Alderley Park and Alex Alderton, Roberto Amato, Sonia Goncalves, Ewan Harrison, David K. Jackson, Ian Johnston, Dominic Kwiatkowski, Cordelia Langford, John Sillitoe on behalf of the Wellcome Sanger Institute COVID-19 Surveillance Team |
| EPI_ISL_553351 | Lighthouse Lab in Alderley Park | Wellcome Sanger Institute for the COVID-19 Genomics UK (COG-UK) consortium | The Lighthouse Lab in Alderley Park and Alex Alderton, Roberto Amato, Sonia Goncalves, Ewan Harrison, David K. Jackson, Ian Johnston, Dominic Kwiatkowski, Cordelia Langford, John Sillitoe on behalf of the Wellcome Sanger Institute COVID-19 Surveillance Team (http://www.sanger.ac.uk/covid-team) |
| EPI_ISL_553352, EPI_ISL_553353 | Lighthouse Lab in Alderley Park | Wellcome Sanger Institute for the COVID-19 Genomics UK (COG-UK) consortium | The Lighthouse Lab in Alderley Park and Alex Alderton, Roberto Amato, Sonia Goncalves, Ewan Harrison, David K. Jackson, Ian Johnston, Dominic Kwiatkowski, Cordelia Langford, John Sillitoe on behalf of the Wellcome Sanger Institute COVID-19 Surveillance Team |
| EPI_ISL_553354, EPI_ISL_553355, EPI_ISL_553356 | Lighthouse Lab in Alderley Park | Wellcome Sanger Institute for the COVID-19 Genomics UK (COG-UK) consortium | The Lighthouse Lab in Alderley Park and Alex Alderton, Roberto Amato, Sonia Goncalves, Ewan Harrison, David K. Jackson, Ian Johnston, Dominic Kwiatkowski, Cordelia Langford, John Sillitoe on behalf of the Wellcome Sanger Institute COVID-19 Surveillance Team (http://www.sanger.ac.uk/covid-team) |
| EPI_ISL_553357 | Lighthouse Lab in Alderley Park | Wellcome Sanger Institute for the COVID-19 Genomics UK (COG-UK) consortium | The Lighthouse Lab in Alderley Park and Alex Alderton, Roberto Amato, Sonia Goncalves, Ewan Harrison, David K. Jackson, Ian Johnston, Dominic Kwiatkowski, Cordelia Langford, John Sillitoe on behalf of the Wellcome Sanger Institute COVID-19 Surveillance Team |
| EPI_ISL_553358 | Lighthouse Lab in Alderley Park | Wellcome Sanger Institute for the COVID-19 Genomics UK (COG-UK) consortium | The Lighthouse Lab in Alderley Park and Alex Alderton, Roberto Amato, Sonia Goncalves, Ewan Harrison, David K. Jackson, Ian Johnston, Dominic Kwiatkowski, Cordelia Langford, John Sillitoe on behalf of the Wellcome Sanger Institute COVID-19 Surveillance Team (http://www.sanger.ac.uk/covid-team) |
| EPI_ISL_553359, EPI_ISL_553360, EPI_ISL_553361, EPI_ISL_553362 | Lighthouse Lab in Alderley Park | Wellcome Sanger Institute for the COVID-19 Genomics UK (COG-UK) consortium | The Lighthouse Lab in Alderley Park and Alex Alderton, Roberto Amato, Sonia Goncalves, Ewan Harrison, David K. Jackson, Ian Johnston, Dominic Kwiatkowski, Cordelia Langford, John Sillitoe on behalf of the Wellcome Sanger Institute COVID-19 Surveillance Team |
| EPI_ISL_553363 | Lighthouse Lab in Alderley Park | Wellcome Sanger Institute for the COVID-19 Genomics UK (COG-UK) consortium | The Lighthouse Lab in Alderley Park and Alex Alderton, Roberto Amato, Sonia Goncalves, Ewan Harrison, David K. Jackson, Ian Johnston, Dominic Kwiatkowski, Cordelia Langford, John Sillitoe on behalf of the Wellcome Sanger Institute COVID-19 Surveillance Team (http://www.sanger.ac.uk/covid-team) |
| EPI_ISL_553364, EPI_ISL_553365 | Lighthouse Lab in Alderley Park | Wellcome Sanger Institute for the COVID-19 Genomics UK (COG-UK) consortium | The Lighthouse Lab in Alderley Park and Alex Alderton, Roberto Amato, Sonia Goncalves, Ewan Harrison, David K. Jackson, Ian Johnston, Dominic Kwiatkowski, Cordelia Langford, John Sillitoe on behalf of the Wellcome Sanger Institute COVID-19 Surveillance Team |
| EPI_ISL_553366 | Lighthouse Lab in Alderley Park | Wellcome Sanger Institute for the COVID-19 Genomics UK (COG-UK) consortium | The Lighthouse Lab in Alderley Park and Alex Alderton, Roberto Amato, Sonia Goncalves, Ewan Harrison, David K. Jackson, Ian Johnston, Dominic Kwiatkowski, Cordelia Langford, John Sillitoe on behalf of the Wellcome Sanger Institute COVID-19 Surveillance Team (http://www.sanger.ac.uk/covid-team) |
| EPI_ISL_553367 | Lighthouse Lab in Alderley Park | Wellcome Sanger Institute for the COVID-19 Genomics UK (COG-UK) consortium | The Lighthouse Lab in Alderley Park and Alex Alderton, Roberto Amato, Sonia Goncalves, Ewan Harrison, David K. Jackson, Ian Johnston, Dominic Kwiatkowski, Cordelia Langford, John Sillitoe on behalf of the Wellcome Sanger Institute COVID-19 Surveillance Team |
| EPI_ISL_553368 | Lighthouse Lab in Alderley Park | Wellcome Sanger Institute for the COVID-19 Genomics UK (COG-UK) consortium | The Lighthouse Lab in Alderley Park and Alex Alderton, Roberto Amato, Sonia Goncalves, Ewan Harrison, David K. Jackson, Ian Johnston, Dominic Kwiatkowski, Cordelia Langford, John Sillitoe on behalf of the Wellcome Sanger Institute COVID-19 Surveillance Team (http://www.sanger.ac.uk/covid-team) |

| | | | |
|---|---|---|---|
| EPI_ISL_553369, EPI_ISL_553370, EPI_ISL_553371 | Lighthouse Lab in Alderley Park | Wellcome Sanger Institute for the COVID-19 Genomics UK (COG-UK) consortium | The Lighthouse Lab in Alderley Park and Alex Alderton, Roberto Amato, Sonia Goncalves, Ewan Harrison, David K. Jackson, Ian Johnston, Dominic Kwiatkowski, Cordelia Langford, John Sillitoe on behalf of the Wellcome Sanger Institute COVID-19 Surveillance Team |
| EPI_ISL_553372 | Lighthouse Lab in Alderley Park | Wellcome Sanger Institute for the COVID-19 Genomics UK (COG-UK) consortium | The Lighthouse Lab in Alderley Park and Alex Alderton, Roberto Amato, Sonia Goncalves, Ewan Harrison, David K. Jackson, Ian Johnston, Dominic Kwiatkowski, Cordelia Langford, John Sillitoe on behalf of the Wellcome Sanger Institute COVID-19 Surveillance Team (http://www.sanger.ac.uk/covid-team) |
| EPI_ISL_553373 | Lighthouse Lab in Alderley Park | Wellcome Sanger Institute for the COVID-19 Genomics UK (COG-UK) consortium | The Lighthouse Lab in Alderley Park and Alex Alderton, Roberto Amato, Sonia Goncalves, Ewan Harrison, David K. Jackson, Ian Johnston, Dominic Kwiatkowski, Cordelia Langford, John Sillitoe on behalf of the Wellcome Sanger Institute COVID-19 Surveillance Team |
| EPI_ISL_553374, EPI_ISL_553375 | Lighthouse Lab in Alderley Park | Wellcome Sanger Institute for the COVID-19 Genomics UK (COG-UK) consortium | The Lighthouse Lab in Alderley Park and Alex Alderton, Roberto Amato, Sonia Goncalves, Ewan Harrison, David K. Jackson, Ian Johnston, Dominic Kwiatkowski, Cordelia Langford, John Sillitoe on behalf of the Wellcome Sanger Institute COVID-19 Surveillance Team (http://www.sanger.ac.uk/covid-team) |
| EPI_ISL_553376, EPI_ISL_553377, EPI_ISL_553378 | Lighthouse Lab in Alderley Park | Wellcome Sanger Institute for the COVID-19 Genomics UK (COG-UK) consortium | The Lighthouse Lab in Alderley Park and Alex Alderton, Roberto Amato, Sonia Goncalves, Ewan Harrison, David K. Jackson, Ian Johnston, Dominic Kwiatkowski, Cordelia Langford, John Sillitoe on behalf of the Wellcome Sanger Institute COVID-19 Surveillance Team |
| EPI_ISL_553379, EPI_ISL_553380 | Lighthouse Lab in Alderley Park | Wellcome Sanger Institute for the COVID-19 Genomics UK (COG-UK) consortium | The Lighthouse Lab in Alderley Park and Alex Alderton, Roberto Amato, Sonia Goncalves, Ewan Harrison, David K. Jackson, Ian Johnston, Dominic Kwiatkowski, Cordelia Langford, John Sillitoe on behalf of the Wellcome Sanger Institute COVID-19 Surveillance Team (http://www.sanger.ac.uk/covid-team) |
| EPI_ISL_553381, EPI_ISL_553382, EPI_ISL_553383 | Lighthouse Lab in Alderley Park | Wellcome Sanger Institute for the COVID-19 Genomics UK (COG-UK) consortium | The Lighthouse Lab in Alderley Park and Alex Alderton, Roberto Amato, Sonia Goncalves, Ewan Harrison, David K. Jackson, Ian Johnston, Dominic Kwiatkowski, Cordelia Langford, John Sillitoe on behalf of the Wellcome Sanger Institute COVID-19 Surveillance Team |
| EPI_ISL_553384, EPI_ISL_553385 | Lighthouse Lab in Alderley Park | Wellcome Sanger Institute for the COVID-19 Genomics UK (COG-UK) consortium | The Lighthouse Lab in Alderley Park and Alex Alderton, Roberto Amato, Sonia Goncalves, Ewan Harrison, David K. Jackson, Ian Johnston, Dominic Kwiatkowski, Cordelia Langford, John Sillitoe on behalf of the Wellcome Sanger Institute COVID-19 Surveillance Team (http://www.sanger.ac.uk/covid-team) |
| EPI_ISL_553386, EPI_ISL_553387, EPI_ISL_553388 | Lighthouse Lab in Alderley Park | Wellcome Sanger Institute for the COVID-19 Genomics UK (COG-UK) consortium | The Lighthouse Lab in Alderley Park and Alex Alderton, Roberto Amato, Sonia Goncalves, Ewan Harrison, David K. Jackson, Ian Johnston, Dominic Kwiatkowski, Cordelia Langford, John Sillitoe on behalf of the Wellcome Sanger Institute COVID-19 Surveillance Team |
| EPI_ISL_553389, EPI_ISL_553390 | Lighthouse Lab in Alderley Park | Wellcome Sanger Institute for the COVID-19 Genomics UK (COG-UK) consortium | The Lighthouse Lab in Alderley Park and Alex Alderton, Roberto Amato, Sonia Goncalves, Ewan Harrison, David K. Jackson, Ian Johnston, Dominic Kwiatkowski, Cordelia Langford, John Sillitoe on behalf of the Wellcome Sanger Institute COVID-19 Surveillance Team (http://www.sanger.ac.uk/covid-team) |
| EPI_ISL_553391 | Lighthouse Lab in Alderley Park | Wellcome Sanger Institute for the COVID-19 Genomics UK (COG-UK) consortium | The Lighthouse Lab in Alderley Park and Alex Alderton, Roberto Amato, Sonia Goncalves, Ewan Harrison, David K. Jackson, Ian Johnston, Dominic Kwiatkowski, Cordelia Langford, John Sillitoe on behalf of the Wellcome Sanger Institute COVID-19 Surveillance Team |
| EPI_ISL_553392, EPI_ISL_553393, EPI_ISL_553394 | Lighthouse Lab in Alderley Park | Wellcome Sanger Institute for the COVID-19 Genomics UK (COG-UK) consortium | The Lighthouse Lab in Alderley Park and Alex Alderton, Roberto Amato, Sonia Goncalves, Ewan Harrison, David K. Jackson, Ian Johnston, Dominic Kwiatkowski, Cordelia Langford, John Sillitoe on behalf of the Wellcome Sanger Institute COVID-19 Surveillance Team (http://www.sanger.ac.uk/covid-team) |
| EPI_ISL_553395 | Lighthouse Lab in Alderley Park | Wellcome Sanger Institute for the COVID-19 Genomics UK (COG-UK) consortium | The Lighthouse Lab in Alderley Park and Alex Alderton, Roberto Amato, Sonia Goncalves, Ewan Harrison, David K. Jackson, Ian Johnston, Dominic Kwiatkowski, Cordelia Langford, John Sillitoe on behalf of the Wellcome Sanger Institute COVID-19 Surveillance Team |
| EPI_ISL_553396 | Lighthouse Lab in Alderley Park | Wellcome Sanger Institute for the COVID-19 Genomics UK (COG-UK) consortium | The Lighthouse Lab in Alderley Park and Alex Alderton, Roberto Amato, Sonia Goncalves, Ewan Harrison, David K. Jackson, Ian Johnston, Dominic Kwiatkowski, Cordelia Langford, John Sillitoe on behalf of the Wellcome Sanger Institute COVID-19 Surveillance Team (http://www.sanger.ac.uk/covid-team) |
| EPI_ISL_553397, EPI_ISL_553398 | Lighthouse Lab in Alderley Park | Wellcome Sanger Institute for the COVID-19 Genomics UK (COG-UK) consortium | The Lighthouse Lab in Alderley Park and Alex Alderton, Roberto Amato, Sonia Goncalves, Ewan Harrison, David K. Jackson, Ian Johnston, Dominic Kwiatkowski, Cordelia Langford, John Sillitoe on behalf of the Wellcome Sanger Institute COVID-19 Surveillance Team |
| EPI_ISL_553399 | Lighthouse Lab in Alderley Park | Wellcome Sanger Institute for the COVID-19 Genomics UK (COG-UK) consortium | The Lighthouse Lab in Alderley Park and Alex Alderton, Roberto Amato, Sonia Goncalves, Ewan Harrison, David K. Jackson, Ian Johnston, Dominic Kwiatkowski, Cordelia Langford, John Sillitoe on behalf of the Wellcome Sanger Institute COVID-19 Surveillance Team (http://www.sanger.ac.uk/covid-team) |
| EPI_ISL_553400, EPI_ISL_553401, EPI_ISL_553402 | Lighthouse Lab in Alderley Park | Wellcome Sanger Institute for the COVID-19 Genomics UK (COG-UK) consortium | The Lighthouse Lab in Alderley Park and Alex Alderton, Roberto Amato, Sonia Goncalves, Ewan Harrison, David K. Jackson, Ian Johnston, Dominic Kwiatkowski, Cordelia Langford, John Sillitoe on behalf of the Wellcome Sanger Institute COVID-19 Surveillance Team |
| EPI_ISL_553403 | Lighthouse Lab in Alderley Park | Wellcome Sanger Institute for the COVID-19 Genomics UK (COG-UK) consortium | The Lighthouse Lab in Alderley Park and Alex Alderton, Roberto Amato, Sonia Goncalves, Ewan Harrison, David K. Jackson, Ian Johnston, Dominic Kwiatkowski, Cordelia Langford, John Sillitoe on behalf of the Wellcome Sanger Institute COVID-19 Surveillance Team (http://www.sanger.ac.uk/covid-team) |
| EPI_ISL_553404, EPI_ISL_553405, EPI_ISL_553406, EPI_ISL_553407, EPI_ISL_553408, EPI_ISL_553409 | Lighthouse Lab in Alderley Park | Wellcome Sanger Institute for the COVID-19 Genomics UK (COG-UK) consortium | The Lighthouse Lab in Alderley Park and Alex Alderton, Roberto Amato, Sonia Goncalves, Ewan Harrison, David K. Jackson, Ian Johnston, Dominic Kwiatkowski, Cordelia Langford, John Sillitoe on behalf of the Wellcome Sanger Institute COVID-19 Surveillance Team |
| EPI_ISL_553410 | Lighthouse Lab in Alderley Park | Wellcome Sanger Institute for the COVID-19 Genomics UK (COG-UK) consortium | The Lighthouse Lab in Alderley Park and Alex Alderton, Roberto Amato, Sonia Goncalves, Ewan Harrison, David K. Jackson, Ian Johnston, Dominic Kwiatkowski, Cordelia Langford, John Sillitoe on behalf of the Wellcome Sanger Institute COVID-19 Surveillance Team (http://www.sanger.ac.uk/covid-team) |
| EPI_ISL_553411 | Lighthouse Lab in Alderley Park | Wellcome Sanger Institute for the COVID-19 Genomics UK (COG-UK) consortium | The Lighthouse Lab in Alderley Park and Alex Alderton, Roberto Amato, Sonia Goncalves, Ewan Harrison, David K. Jackson, Ian Johnston, Dominic Kwiatkowski, Cordelia Langford, John Sillitoe on behalf of the Wellcome Sanger Institute COVID-19 Surveillance Team |
| EPI_ISL_553412 | Lighthouse Lab in Alderley Park | Wellcome Sanger Institute for the COVID-19 Genomics UK (COG-UK) consortium | The Lighthouse Lab in Alderley Park and Alex Alderton, Roberto Amato, Sonia Goncalves, Ewan Harrison, David K. Jackson, Ian Johnston, Dominic Kwiatkowski, Cordelia Langford, John Sillitoe on behalf of the Wellcome Sanger Institute COVID-19 Surveillance Team (http://www.sanger.ac.uk/covid-team) |
| EPI_ISL_553413 | Lighthouse Lab in Alderley Park | Wellcome Sanger Institute for the COVID-19 Genomics UK (COG-UK) consortium | The Lighthouse Lab in Alderley Park and Alex Alderton, Roberto Amato, Sonia Goncalves, Ewan Harrison, David K. Jackson, Ian Johnston, Dominic Kwiatkowski, Cordelia Langford, John Sillitoe on behalf of the Wellcome Sanger Institute COVID-19 Surveillance Team |
| EPI_ISL_553414, EPI_ISL_553415 | Lighthouse Lab in Alderley Park | Wellcome Sanger Institute for the COVID-19 Genomics UK (COG-UK) consortium | The Lighthouse Lab in Alderley Park and Alex Alderton, Roberto Amato, Sonia Goncalves, Ewan Harrison, David K. Jackson, Ian Johnston, Dominic Kwiatkowski, Cordelia Langford, John Sillitoe on behalf of the Wellcome Sanger Institute COVID-19 Surveillance Team (http://www.sanger.ac.uk/covid-team) |
| EPI_ISL_553416 | Lighthouse Lab in Alderley Park | Wellcome Sanger Institute for the COVID-19 Genomics UK (COG-UK) consortium | The Lighthouse Lab in Alderley Park and Alex Alderton, Roberto Amato, Sonia Goncalves, Ewan Harrison, David K. Jackson, Ian Johnston, Dominic Kwiatkowski, Cordelia Langford, John Sillitoe on behalf of the Wellcome Sanger Institute COVID-19 Surveillance Team |
| EPI_ISL_553417 | Lighthouse Lab in Alderley Park | Wellcome Sanger Institute for the COVID-19 Genomics UK (COG-UK) consortium | The Lighthouse Lab in Alderley Park and Alex Alderton, Roberto Amato, Sonia Goncalves, Ewan Harrison, David K. Jackson, Ian Johnston, Dominic Kwiatkowski, Cordelia Langford, John Sillitoe on behalf of the Wellcome Sanger Institute COVID-19 Surveillance Team (http://www.sanger.ac.uk/covid-team) |
| EPI_ISL_553418 | Lighthouse Lab in Alderley Park | Wellcome Sanger Institute for the COVID-19 Genomics UK (COG-UK) consortium | The Lighthouse Lab in Alderley Park and Alex Alderton, Roberto Amato, Sonia Goncalves, Ewan Harrison, David K. Jackson, Ian Johnston, Dominic Kwiatkowski, Cordelia Langford, John Sillitoe on behalf of the Wellcome Sanger Institute COVID-19 Surveillance Team |
| EPI_ISL_553419, EPI_ISL_553420 | Lighthouse Lab in Alderley Park | Wellcome Sanger Institute for the COVID-19 Genomics UK (COG-UK) consortium | The Lighthouse Lab in Alderley Park and Alex Alderton, Roberto Amato, Sonia Goncalves, Ewan Harrison, David K. Jackson, Ian Johnston, Dominic Kwiatkowski, Cordelia Langford, John Sillitoe on behalf of the Wellcome Sanger Institute COVID-19 Surveillance Team (http://www.sanger.ac.uk/covid-team) |
| EPI_ISL_553422, EPI_ISL_553423 | Lighthouse Lab in Alderley Park | Wellcome Sanger Institute for the COVID-19 Genomics UK (COG-UK) consortium | The Lighthouse Lab in Alderley Park and Alex Alderton, Roberto Amato, Sonia Goncalves, Ewan Harrison, David K. Jackson, Ian Johnston, Dominic Kwiatkowski, Cordelia Langford, John Sillitoe on behalf of the Wellcome Sanger Institute COVID-19 Surveillance Team |

| | | | |
|---|---|---|---|
| EPI_ISL_553424 | Lighthouse Lab in Alderley Park | Wellcome Sanger Institute for the COVID-19 Genomics UK (COG-UK) consortium | The Lighthouse Lab in Alderley Park and Alex Alderton, Roberto Amato, Sonia Goncalves, Ewan Harrison, David K. Jackson, Ian Johnston, Dominic Kwiatkowski, Cordelia Langford, John Sillitoe on behalf of the Wellcome Sanger Institute COVID-19 Surveillance Team (http://www.sanger.ac.uk/covid-team) |
| EPI_ISL_553425 | Lighthouse Lab in Alderley Park | Wellcome Sanger Institute for the COVID-19 Genomics UK (COG-UK) consortium | The Lighthouse Lab in Alderley Park and Alex Alderton, Roberto Amato, Sonia Goncalves, Ewan Harrison, David K. Jackson, Ian Johnston, Dominic Kwiatkowski, Cordelia Langford, John Sillitoe on behalf of the Wellcome Sanger Institute COVID-19 Surveillance Team |
| EPI_ISL_553426 | Lighthouse Lab in Alderley Park | Wellcome Sanger Institute for the COVID-19 Genomics UK (COG-UK) consortium | The Lighthouse Lab in Alderley Park and Alex Alderton, Roberto Amato, Sonia Goncalves, Ewan Harrison, David K. Jackson, Ian Johnston, Dominic Kwiatkowski, Cordelia Langford, John Sillitoe on behalf of the Wellcome Sanger Institute COVID-19 Surveillance Team (http://www.sanger.ac.uk/covid-team) |
| EPI_ISL_553427, EPI_ISL_553428 | Lighthouse Lab in Alderley Park | Wellcome Sanger Institute for the COVID-19 Genomics UK (COG-UK) consortium | The Lighthouse Lab in Alderley Park and Alex Alderton, Roberto Amato, Sonia Goncalves, Ewan Harrison, David K. Jackson, Ian Johnston, Dominic Kwiatkowski, Cordelia Langford, John Sillitoe on behalf of the Wellcome Sanger Institute COVID-19 Surveillance Team |
| EPI_ISL_553429 | Lighthouse Lab in Alderley Park | Wellcome Sanger Institute for the COVID-19 Genomics UK (COG-UK) consortium | The Lighthouse Lab in Alderley Park and Alex Alderton, Roberto Amato, Sonia Goncalves, Ewan Harrison, David K. Jackson, Ian Johnston, Dominic Kwiatkowski, Cordelia Langford, John Sillitoe on behalf of the Wellcome Sanger Institute COVID-19 Surveillance Team (http://www.sanger.ac.uk/covid-team) |
| EPI_ISL_553430, EPI_ISL_553431, EPI_ISL_553432 | Lighthouse Lab in Alderley Park | Wellcome Sanger Institute for the COVID-19 Genomics UK (COG-UK) consortium | The Lighthouse Lab in Alderley Park and Alex Alderton, Roberto Amato, Sonia Goncalves, Ewan Harrison, David K. Jackson, Ian Johnston, Dominic Kwiatkowski, Cordelia Langford, John Sillitoe on behalf of the Wellcome Sanger Institute COVID-19 Surveillance Team |
| EPI_ISL_553433 | Lighthouse Lab in Alderley Park | Wellcome Sanger Institute for the COVID-19 Genomics UK (COG-UK) consortium | The Lighthouse Lab in Alderley Park and Alex Alderton, Roberto Amato, Sonia Goncalves, Ewan Harrison, David K. Jackson, Ian Johnston, Dominic Kwiatkowski, Cordelia Langford, John Sillitoe on behalf of the Wellcome Sanger Institute COVID-19 Surveillance Team (http://www.sanger.ac.uk/covid-team) |
| EPI_ISL_553434 | Lighthouse Lab in Alderley Park | Wellcome Sanger Institute for the COVID-19 Genomics UK (COG-UK) consortium | The Lighthouse Lab in Alderley Park and Alex Alderton, Roberto Amato, Sonia Goncalves, Ewan Harrison, David K. Jackson, Ian Johnston, Dominic Kwiatkowski, Cordelia Langford, John Sillitoe on behalf of the Wellcome Sanger Institute COVID-19 Surveillance Team |
| EPI_ISL_553435 | Lighthouse Lab in Alderley Park | Wellcome Sanger Institute for the COVID-19 Genomics UK (COG-UK) consortium | The Lighthouse Lab in Alderley Park and Alex Alderton, Roberto Amato, Sonia Goncalves, Ewan Harrison, David K. Jackson, Ian Johnston, Dominic Kwiatkowski, Cordelia Langford, John Sillitoe on behalf of the Wellcome Sanger Institute COVID-19 Surveillance Team (http://www.sanger.ac.uk/covid-team) |
| EPI_ISL_553436 | Lighthouse Lab in Alderley Park | Wellcome Sanger Institute for the COVID-19 Genomics UK (COG-UK) consortium | The Lighthouse Lab in Alderley Park and Alex Alderton, Roberto Amato, Sonia Goncalves, Ewan Harrison, David K. Jackson, Ian Johnston, Dominic Kwiatkowski, Cordelia Langford, John Sillitoe on behalf of the Wellcome Sanger Institute COVID-19 Surveillance Team |
| EPI_ISL_553437, EPI_ISL_553438, EPI_ISL_553439, EPI_ISL_553440, EPI_ISL_553441 | Lighthouse Lab in Alderley Park | Wellcome Sanger Institute for the COVID-19 Genomics UK (COG-UK) consortium | The Lighthouse Lab in Alderley Park and Alex Alderton, Roberto Amato, Sonia Goncalves, Ewan Harrison, David K. Jackson, Ian Johnston, Dominic Kwiatkowski, Cordelia Langford, John Sillitoe on behalf of the Wellcome Sanger Institute COVID-19 Surveillance Team (http://www.sanger.ac.uk/covid-team) |
| EPI_ISL_553442 | Lighthouse Lab in Alderley Park | Wellcome Sanger Institute for the COVID-19 Genomics UK (COG-UK) consortium | The Lighthouse Lab in Alderley Park and Alex Alderton, Roberto Amato, Sonia Goncalves, Ewan Harrison, David K. Jackson, Ian Johnston, Dominic Kwiatkowski, Cordelia Langford, John Sillitoe on behalf of the Wellcome Sanger Institute COVID-19 Surveillance Team |
| EPI_ISL_553443, EPI_ISL_553444 | Lighthouse Lab in Alderley Park | Wellcome Sanger Institute for the COVID-19 Genomics UK (COG-UK) consortium | The Lighthouse Lab in Alderley Park and Alex Alderton, Roberto Amato, Sonia Goncalves, Ewan Harrison, David K. Jackson, Ian Johnston, Dominic Kwiatkowski, Cordelia Langford, John Sillitoe on behalf of the Wellcome Sanger Institute COVID-19 Surveillance Team (http://www.sanger.ac.uk/covid-team) |
| EPI_ISL_553445, EPI_ISL_553446 | Lighthouse Lab in Alderley Park | Wellcome Sanger Institute for the COVID-19 Genomics UK (COG-UK) consortium | The Lighthouse Lab in Alderley Park and Alex Alderton, Roberto Amato, Sonia Goncalves, Ewan Harrison, David K. Jackson, Ian Johnston, Dominic Kwiatkowski, Cordelia Langford, John Sillitoe on behalf of the Wellcome Sanger Institute COVID-19 Surveillance Team |
| EPI_ISL_553447 | Lighthouse Lab in Alderley Park | Wellcome Sanger Institute for the COVID-19 Genomics UK (COG-UK) consortium | The Lighthouse Lab in Alderley Park and Alex Alderton, Roberto Amato, Sonia Goncalves, Ewan Harrison, David K. Jackson, Ian Johnston, Dominic Kwiatkowski, Cordelia Langford, John Sillitoe on behalf of the Wellcome Sanger Institute COVID-19 Surveillance Team (http://www.sanger.ac.uk/covid-team) |
| EPI_ISL_553448, EPI_ISL_553449 | Lighthouse Lab in Alderley Park | Wellcome Sanger Institute for the COVID-19 Genomics UK (COG-UK) consortium | The Lighthouse Lab in Alderley Park and Alex Alderton, Roberto Amato, Sonia Goncalves, Ewan Harrison, David K. Jackson, Ian Johnston, Dominic Kwiatkowski, Cordelia Langford, John Sillitoe on behalf of the Wellcome Sanger Institute COVID-19 Surveillance Team |
| EPI_ISL_553450 | Lighthouse Lab in Alderley Park | Wellcome Sanger Institute for the COVID-19 Genomics UK (COG-UK) consortium | The Lighthouse Lab in Alderley Park and Alex Alderton, Roberto Amato, Sonia Goncalves, Ewan Harrison, David K. Jackson, Ian Johnston, Dominic Kwiatkowski, Cordelia Langford, John Sillitoe on behalf of the Wellcome Sanger Institute COVID-19 Surveillance Team (http://www.sanger.ac.uk/covid-team) |
| EPI_ISL_553451, EPI_ISL_553452, EPI_ISL_553453, EPI_ISL_553454, EPI_ISL_553455, EPI_ISL_553456, EPI_ISL_553457, EPI_ISL_553458, EPI_ISL_553459, EPI_ISL_553460, EPI_ISL_553461, EPI_ISL_553462 | | | |
| see above | Lighthouse Lab in Alderley Park | Wellcome Sanger Institute for the COVID-19 Genomics UK (COG-UK) consortium | The Lighthouse Lab in Alderley Park and Alex Alderton, Roberto Amato, Sonia Goncalves, Ewan Harrison, David K. Jackson, Ian Johnston, Dominic Kwiatkowski, Cordelia Langford, John Sillitoe on behalf of the Wellcome Sanger Institute COVID-19 Surveillance Team |
| EPI_ISL_553463 | Lighthouse Lab in Alderley Park | Wellcome Sanger Institute for the COVID-19 Genomics UK (COG-UK) consortium | The Lighthouse Lab in Alderley Park and Alex Alderton, Roberto Amato, Sonia Goncalves, Ewan Harrison, David K. Jackson, Ian Johnston, Dominic Kwiatkowski, Cordelia Langford, John Sillitoe on behalf of the Wellcome Sanger Institute COVID-19 Surveillance Team (http://www.sanger.ac.uk/covid-team) |
| EPI_ISL_553464, EPI_ISL_553465 | Lighthouse Lab in Alderley Park | Wellcome Sanger Institute for the COVID-19 Genomics UK (COG-UK) consortium | The Lighthouse Lab in Alderley Park and Alex Alderton, Roberto Amato, Sonia Goncalves, Ewan Harrison, David K. Jackson, Ian Johnston, Dominic Kwiatkowski, Cordelia Langford, John Sillitoe on behalf of the Wellcome Sanger Institute COVID-19 Surveillance Team |
| EPI_ISL_553466 | Lighthouse Lab in Alderley Park | Wellcome Sanger Institute for the COVID-19 Genomics UK (COG-UK) consortium | The Lighthouse Lab in Alderley Park and Alex Alderton, Roberto Amato, Sonia Goncalves, Ewan Harrison, David K. Jackson, Ian Johnston, Dominic Kwiatkowski, Cordelia Langford, John Sillitoe on behalf of the Wellcome Sanger Institute COVID-19 Surveillance Team (http://www.sanger.ac.uk/covid-team) |
| EPI_ISL_553467 | Lighthouse Lab in Alderley Park | Wellcome Sanger Institute for the COVID-19 Genomics UK (COG-UK) consortium | The Lighthouse Lab in Alderley Park and Alex Alderton, Roberto Amato, Sonia Goncalves, Ewan Harrison, David K. Jackson, Ian Johnston, Dominic Kwiatkowski, Cordelia Langford, John Sillitoe on behalf of the Wellcome Sanger Institute COVID-19 Surveillance Team |
| EPI_ISL_553468 | Lighthouse Lab in Alderley Park | Wellcome Sanger Institute for the COVID-19 Genomics UK (COG-UK) consortium | The Lighthouse Lab in Alderley Park and Alex Alderton, Roberto Amato, Sonia Goncalves, Ewan Harrison, David K. Jackson, Ian Johnston, Dominic Kwiatkowski, Cordelia Langford, John Sillitoe on behalf of the Wellcome Sanger Institute COVID-19 Surveillance Team (http://www.sanger.ac.uk/covid-team) |
| EPI_ISL_553469, EPI_ISL_553470 | Lighthouse Lab in Alderley Park | Wellcome Sanger Institute for the COVID-19 Genomics UK (COG-UK) consortium | The Lighthouse Lab in Alderley Park and Alex Alderton, Roberto Amato, Sonia Goncalves, Ewan Harrison, David K. Jackson, Ian Johnston, Dominic Kwiatkowski, Cordelia Langford, John Sillitoe on behalf of the Wellcome Sanger Institute COVID-19 Surveillance Team |
| EPI_ISL_553471 | Lighthouse Lab in Alderley Park | Wellcome Sanger Institute for the COVID-19 Genomics UK (COG-UK) consortium | The Lighthouse Lab in Alderley Park and Alex Alderton, Roberto Amato, Sonia Goncalves, Ewan Harrison, David K. Jackson, Ian Johnston, Dominic Kwiatkowski, Cordelia Langford, John Sillitoe on behalf of the Wellcome Sanger Institute COVID-19 Surveillance Team (http://www.sanger.ac.uk/covid-team) |
| EPI_ISL_553472, EPI_ISL_553473 | Lighthouse Lab in Alderley Park | Wellcome Sanger Institute for the COVID-19 Genomics UK (COG-UK) consortium | The Lighthouse Lab in Alderley Park and Alex Alderton, Roberto Amato, Sonia Goncalves, Ewan Harrison, David K. Jackson, Ian Johnston, Dominic Kwiatkowski, Cordelia Langford, John Sillitoe on behalf of the Wellcome Sanger Institute COVID-19 Surveillance Team |
| EPI_ISL_553474, EPI_ISL_553475, EPI_ISL_553476 | Lighthouse Lab in Alderley Park | Wellcome Sanger Institute for the COVID-19 Genomics UK (COG-UK) consortium | The Lighthouse Lab in Alderley Park and Alex Alderton, Roberto Amato, Sonia Goncalves, Ewan Harrison, David K. Jackson, Ian Johnston, Dominic Kwiatkowski, Cordelia Langford, John Sillitoe on behalf of the Wellcome Sanger Institute COVID-19 Surveillance Team (http://www.sanger.ac.uk/covid-team) |
| EPI_ISL_553477 | Lighthouse Lab in Alderley Park | Wellcome Sanger Institute for the COVID-19 Genomics UK (COG-UK) consortium | The Lighthouse Lab in Alderley Park and Alex Alderton, Roberto Amato, Sonia Goncalves, Ewan Harrison, David K. Jackson, Ian Johnston, Dominic Kwiatkowski, Cordelia Langford, John Sillitoe on behalf of the Wellcome Sanger Institute COVID-19 Surveillance Team |
| EPI_ISL_553478 | Lighthouse Lab in Alderley Park | Wellcome Sanger Institute for the COVID-19 Genomics UK (COG-UK) consortium | The Lighthouse Lab in Alderley Park and Alex Alderton, Roberto Amato, Sonia Goncalves, Ewan Harrison, David K. Jackson, Ian Johnston, Dominic Kwiatkowski, Cordelia Langford, John Sillitoe on behalf of the Wellcome Sanger Institute COVID-19 Surveillance Team |

| | | | |
|---|---|---|---|
| | | | (http://www.sanger.ac.uk/covid-team) |
| EPI_ISL_553479, EPI_ISL_553480, EPI_ISL_553481 | Lighthouse Lab in Alderley Park | Wellcome Sanger Institute for the COVID-19 Genomics UK (COG-UK) consortium | The Lighthouse Lab in Alderley Park and Alex Alderton, Roberto Amato, Sonia Goncalves, Ewan Harrison, David K. Jackson, Ian Johnston, Dominic Kwiatkowski, Cordelia Langford, John Sillitoe on behalf of the Wellcome Sanger Institute COVID-19 Surveillance Team |
| EPI_ISL_553482 | Lighthouse Lab in Alderley Park | Wellcome Sanger Institute for the COVID-19 Genomics UK (COG-UK) consortium | The Lighthouse Lab in Alderley Park and Alex Alderton, Roberto Amato, Sonia Goncalves, Ewan Harrison, David K. Jackson, Ian Johnston, Dominic Kwiatkowski, Cordelia Langford, John Sillitoe on behalf of the Wellcome Sanger Institute COVID-19 Surveillance Team (http://www.sanger.ac.uk/covid-team) |
| EPI_ISL_553483, EPI_ISL_553484, EPI_ISL_553485 | Lighthouse Lab in Alderley Park | Wellcome Sanger Institute for the COVID-19 Genomics UK (COG-UK) consortium | The Lighthouse Lab in Alderley Park and Alex Alderton, Roberto Amato, Sonia Goncalves, Ewan Harrison, David K. Jackson, Ian Johnston, Dominic Kwiatkowski, Cordelia Langford, John Sillitoe on behalf of the Wellcome Sanger Institute COVID-19 Surveillance Team |
| EPI_ISL_553486 | Lighthouse Lab in Alderley Park | Wellcome Sanger Institute for the COVID-19 Genomics UK (COG-UK) consortium | The Lighthouse Lab in Alderley Park and Alex Alderton, Roberto Amato, Sonia Goncalves, Ewan Harrison, David K. Jackson, Ian Johnston, Dominic Kwiatkowski, Cordelia Langford, John Sillitoe on behalf of the Wellcome Sanger Institute COVID-19 Surveillance Team (http://www.sanger.ac.uk/covid-team) |
| EPI_ISL_553487, EPI_ISL_553488, EPI_ISL_553489 | Lighthouse Lab in Alderley Park | Wellcome Sanger Institute for the COVID-19 Genomics UK (COG-UK) consortium | The Lighthouse Lab in Alderley Park and Alex Alderton, Roberto Amato, Sonia Goncalves, Ewan Harrison, David K. Jackson, Ian Johnston, Dominic Kwiatkowski, Cordelia Langford, John Sillitoe on behalf of the Wellcome Sanger Institute COVID-19 Surveillance Team |
| EPI_ISL_553490, EPI_ISL_553491 | Lighthouse Lab in Alderley Park | Wellcome Sanger Institute for the COVID-19 Genomics UK (COG-UK) consortium | The Lighthouse Lab in Alderley Park and Alex Alderton, Roberto Amato, Sonia Goncalves, Ewan Harrison, David K. Jackson, Ian Johnston, Dominic Kwiatkowski, Cordelia Langford, John Sillitoe on behalf of the Wellcome Sanger Institute COVID-19 Surveillance Team (http://www.sanger.ac.uk/covid-team) |
| EPI_ISL_553492, EPI_ISL_553493 | Lighthouse Lab in Alderley Park | Wellcome Sanger Institute for the COVID-19 Genomics UK (COG-UK) consortium | The Lighthouse Lab in Alderley Park and Alex Alderton, Roberto Amato, Sonia Goncalves, Ewan Harrison, David K. Jackson, Ian Johnston, Dominic Kwiatkowski, Cordelia Langford, John Sillitoe on behalf of the Wellcome Sanger Institute COVID-19 Surveillance Team |
| EPI_ISL_553494 | Lighthouse Lab in Alderley Park | Wellcome Sanger Institute for the COVID-19 Genomics UK (COG-UK) consortium | The Lighthouse Lab in Alderley Park and Alex Alderton, Roberto Amato, Sonia Goncalves, Ewan Harrison, David K. Jackson, Ian Johnston, Dominic Kwiatkowski, Cordelia Langford, John Sillitoe on behalf of the Wellcome Sanger Institute COVID-19 Surveillance Team (http://www.sanger.ac.uk/covid-team) |
| EPI_ISL_553495 | Lighthouse Lab in Alderley Park | Wellcome Sanger Institute for the COVID-19 Genomics UK (COG-UK) consortium | The Lighthouse Lab in Alderley Park and Alex Alderton, Roberto Amato, Sonia Goncalves, Ewan Harrison, David K. Jackson, Ian Johnston, Dominic Kwiatkowski, Cordelia Langford, John Sillitoe on behalf of the Wellcome Sanger Institute COVID-19 Surveillance Team |
| EPI_ISL_553496 | Lighthouse Lab in Alderley Park | Wellcome Sanger Institute for the COVID-19 Genomics UK (COG-UK) consortium | The Lighthouse Lab in Alderley Park and Alex Alderton, Roberto Amato, Sonia Goncalves, Ewan Harrison, David K. Jackson, Ian Johnston, Dominic Kwiatkowski, Cordelia Langford, John Sillitoe on behalf of the Wellcome Sanger Institute COVID-19 Surveillance Team (http://www.sanger.ac.uk/covid-team) |
| EPI_ISL_553497, EPI_ISL_553498 | Lighthouse Lab in Alderley Park | Wellcome Sanger Institute for the COVID-19 Genomics UK (COG-UK) consortium | The Lighthouse Lab in Alderley Park and Alex Alderton, Roberto Amato, Sonia Goncalves, Ewan Harrison, David K. Jackson, Ian Johnston, Dominic Kwiatkowski, Cordelia Langford, John Sillitoe on behalf of the Wellcome Sanger Institute COVID-19 Surveillance Team |
| EPI_ISL_553499 | Lighthouse Lab in Alderley Park | Wellcome Sanger Institute for the COVID-19 Genomics UK (COG-UK) consortium | The Lighthouse Lab in Alderley Park and Alex Alderton, Roberto Amato, Sonia Goncalves, Ewan Harrison, David K. Jackson, Ian Johnston, Dominic Kwiatkowski, Cordelia Langford, John Sillitoe on behalf of the Wellcome Sanger Institute COVID-19 Surveillance Team (http://www.sanger.ac.uk/covid-team) |
| EPI_ISL_553500, EPI_ISL_553501 | Lighthouse Lab in Alderley Park | Wellcome Sanger Institute for the COVID-19 Genomics UK (COG-UK) consortium | The Lighthouse Lab in Alderley Park and Alex Alderton, Roberto Amato, Sonia Goncalves, Ewan Harrison, David K. Jackson, Ian Johnston, Dominic Kwiatkowski, Cordelia Langford, John Sillitoe on behalf of the Wellcome Sanger Institute COVID-19 Surveillance Team |
| EPI_ISL_553502 | Lighthouse Lab in Alderley Park | Wellcome Sanger Institute for the COVID-19 Genomics UK (COG-UK) consortium | The Lighthouse Lab in Alderley Park and Alex Alderton, Roberto Amato, Sonia Goncalves, Ewan Harrison, David K. Jackson, Ian Johnston, Dominic Kwiatkowski, Cordelia Langford, John Sillitoe on behalf of the Wellcome Sanger Institute COVID-19 Surveillance Team (http://www.sanger.ac.uk/covid-team) |
| EPI_ISL_553503 | Lighthouse Lab in Alderley Park | Wellcome Sanger Institute for the COVID-19 Genomics UK (COG-UK) consortium | The Lighthouse Lab in Alderley Park and Alex Alderton, Roberto Amato, Sonia Goncalves, Ewan Harrison, David K. Jackson, Ian Johnston, Dominic Kwiatkowski, Cordelia Langford, John Sillitoe on behalf of the Wellcome Sanger Institute COVID-19 Surveillance Team |
| EPI_ISL_553504, EPI_ISL_553505, EPI_ISL_553506 | Lighthouse Lab in Alderley Park | Wellcome Sanger Institute for the COVID-19 Genomics UK (COG-UK) consortium | The Lighthouse Lab in Alderley Park and Alex Alderton, Roberto Amato, Sonia Goncalves, Ewan Harrison, David K. Jackson, Ian Johnston, Dominic Kwiatkowski, Cordelia Langford, John Sillitoe on behalf of the Wellcome Sanger Institute COVID-19 Surveillance Team (http://www.sanger.ac.uk/covid-team) |
| EPI_ISL_553507 | Lighthouse Lab in Alderley Park | Wellcome Sanger Institute for the COVID-19 Genomics UK (COG-UK) consortium | The Lighthouse Lab in Alderley Park and Alex Alderton, Roberto Amato, Sonia Goncalves, Ewan Harrison, David K. Jackson, Ian Johnston, Dominic Kwiatkowski, Cordelia Langford, John Sillitoe on behalf of the Wellcome Sanger Institute COVID-19 Surveillance Team |
| EPI_ISL_553508, EPI_ISL_553509 | Lighthouse Lab in Alderley Park | Wellcome Sanger Institute for the COVID-19 Genomics UK (COG-UK) consortium | The Lighthouse Lab in Alderley Park and Alex Alderton, Roberto Amato, Sonia Goncalves, Ewan Harrison, David K. Jackson, Ian Johnston, Dominic Kwiatkowski, Cordelia Langford, John Sillitoe on behalf of the Wellcome Sanger Institute COVID-19 Surveillance Team (http://www.sanger.ac.uk/covid-team) |
| EPI_ISL_553510 | Lighthouse Lab in Alderley Park | Wellcome Sanger Institute for the COVID-19 Genomics UK (COG-UK) consortium | The Lighthouse Lab in Alderley Park and Alex Alderton, Roberto Amato, Sonia Goncalves, Ewan Harrison, David K. Jackson, Ian Johnston, Dominic Kwiatkowski, Cordelia Langford, John Sillitoe on behalf of the Wellcome Sanger Institute COVID-19 Surveillance Team |
| EPI_ISL_553511, EPI_ISL_553512 | Lighthouse Lab in Alderley Park | Wellcome Sanger Institute for the COVID-19 Genomics UK (COG-UK) consortium | The Lighthouse Lab in Alderley Park and Alex Alderton, Roberto Amato, Sonia Goncalves, Ewan Harrison, David K. Jackson, Ian Johnston, Dominic Kwiatkowski, Cordelia Langford, John Sillitoe on behalf of the Wellcome Sanger Institute COVID-19 Surveillance Team (http://www.sanger.ac.uk/covid-team) |
| EPI_ISL_553513, EPI_ISL_553514 | Lighthouse Lab in Alderley Park | Wellcome Sanger Institute for the COVID-19 Genomics UK (COG-UK) consortium | The Lighthouse Lab in Alderley Park and Alex Alderton, Roberto Amato, Sonia Goncalves, Ewan Harrison, David K. Jackson, Ian Johnston, Dominic Kwiatkowski, Cordelia Langford, John Sillitoe on behalf of the Wellcome Sanger Institute COVID-19 Surveillance Team |
| EPI_ISL_553515, EPI_ISL_553516 | Lighthouse Lab in Alderley Park | Wellcome Sanger Institute for the COVID-19 Genomics UK (COG-UK) consortium | The Lighthouse Lab in Alderley Park and Alex Alderton, Roberto Amato, Sonia Goncalves, Ewan Harrison, David K. Jackson, Ian Johnston, Dominic Kwiatkowski, Cordelia Langford, John Sillitoe on behalf of the Wellcome Sanger Institute COVID-19 Surveillance Team (http://www.sanger.ac.uk/covid-team) |
| EPI_ISL_553517 | Lighthouse Lab in Alderley Park | Wellcome Sanger Institute for the COVID-19 Genomics UK (COG-UK) consortium | The Lighthouse Lab in Alderley Park and Alex Alderton, Roberto Amato, Sonia Goncalves, Ewan Harrison, David K. Jackson, Ian Johnston, Dominic Kwiatkowski, Cordelia Langford, John Sillitoe on behalf of the Wellcome Sanger Institute COVID-19 Surveillance Team |
| EPI_ISL_553518 | Lighthouse Lab in Alderley Park | Wellcome Sanger Institute for the COVID-19 Genomics UK (COG-UK) consortium | The Lighthouse Lab in Alderley Park and Alex Alderton, Roberto Amato, Sonia Goncalves, Ewan Harrison, David K. Jackson, Ian Johnston, Dominic Kwiatkowski, Cordelia Langford, John Sillitoe on behalf of the Wellcome Sanger Institute COVID-19 Surveillance Team (http://www.sanger.ac.uk/covid-team) |
| EPI_ISL_553519, EPI_ISL_553520, EPI_ISL_553521 | Lighthouse Lab in Alderley Park | Wellcome Sanger Institute for the COVID-19 Genomics UK (COG-UK) consortium | The Lighthouse Lab in Alderley Park and Alex Alderton, Roberto Amato, Sonia Goncalves, Ewan Harrison, David K. Jackson, Ian Johnston, Dominic Kwiatkowski, Cordelia Langford, John Sillitoe on behalf of the Wellcome Sanger Institute COVID-19 Surveillance Team |
| EPI_ISL_553522, EPI_ISL_553523, EPI_ISL_553524 | Lighthouse Lab in Alderley Park | Wellcome Sanger Institute for the COVID-19 Genomics UK (COG-UK) consortium | The Lighthouse Lab in Alderley Park and Alex Alderton, Roberto Amato, Sonia Goncalves, Ewan Harrison, David K. Jackson, Ian Johnston, Dominic Kwiatkowski, Cordelia Langford, John Sillitoe on behalf of the Wellcome Sanger Institute COVID-19 Surveillance Team (http://www.sanger.ac.uk/covid-team) |
| EPI_ISL_553525, EPI_ISL_553526, EPI_ISL_553527, EPI_ISL_553528 | Lighthouse Lab in Alderley Park | Wellcome Sanger Institute for the COVID-19 Genomics UK (COG-UK) consortium | The Lighthouse Lab in Alderley Park and Alex Alderton, Roberto Amato, Sonia Goncalves, Ewan Harrison, David K. Jackson, Ian Johnston, Dominic Kwiatkowski, Cordelia Langford, John Sillitoe on behalf of the Wellcome Sanger Institute COVID-19 Surveillance Team |
| EPI_ISL_553529 | Lighthouse Lab in Alderley Park | Wellcome Sanger Institute for the COVID-19 Genomics UK (COG-UK) consortium | The Lighthouse Lab in Alderley Park and Alex Alderton, Roberto Amato, Sonia Goncalves, Ewan Harrison, David K. Jackson, Ian Johnston, Dominic Kwiatkowski, Cordelia Langford, John Sillitoe on behalf of the Wellcome Sanger Institute COVID-19 Surveillance Team (http://www.sanger.ac.uk/covid-team) |
| EPI_ISL_553530, EPI_ISL_553531, EPI_ISL_553532, EPI_ISL_553533, | Lighthouse Lab in Alderley Park | Wellcome Sanger Institute for the COVID-19 Genomics UK (COG-UK) consortium | The Lighthouse Lab in Alderley Park and Alex Alderton, Roberto Amato, Sonia Goncalves, Ewan Harrison, David K. Jackson, Ian Johnston, Dominic Kwiatkowski, Cordelia Langford, John Sillitoe on behalf of the Wellcome Sanger Institute COVID-19 Surveillance Team |

| | | | |
|---|---|---|---|
| EPI_ISL_553534 | | | |
| EPI_ISL_553535, EPI_ISL_553536 | Lighthouse Lab in Alderley Park | Wellcome Sanger Institute for the COVID-19 Genomics UK (COG-UK) consortium | The Lighthouse Lab in Alderley Park and Alex Alderton, Roberto Amato, Sonia Goncalves, Ewan Harrison, David K. Jackson, Ian Johnston, Dominic Kwiatkowski, Cordelia Langford, John Sillitoe on behalf of the Wellcome Sanger Institute COVID-19 Surveillance Team (http://www.sanger.ac.uk/covid-team) |
| EPI_ISL_553537, EPI_ISL_553538, EPI_ISL_553539 | Lighthouse Lab in Alderley Park | Wellcome Sanger Institute for the COVID-19 Genomics UK (COG-UK) consortium | The Lighthouse Lab in Alderley Park and Alex Alderton, Roberto Amato, Sonia Goncalves, Ewan Harrison, David K. Jackson, Ian Johnston, Dominic Kwiatkowski, Cordelia Langford, John Sillitoe on behalf of the Wellcome Sanger Institute COVID-19 Surveillance Team |
| EPI_ISL_553540 | Lighthouse Lab in Alderley Park | Wellcome Sanger Institute for the COVID-19 Genomics UK (COG-UK) consortium | The Lighthouse Lab in Alderley Park and Alex Alderton, Roberto Amato, Sonia Goncalves, Ewan Harrison, David K. Jackson, Ian Johnston, Dominic Kwiatkowski, Cordelia Langford, John Sillitoe on behalf of the Wellcome Sanger Institute COVID-19 Surveillance Team (http://www.sanger.ac.uk/covid-team) |
| EPI_ISL_553541, EPI_ISL_553542, EPI_ISL_553543, EPI_ISL_553544, EPI_ISL_553545, EPI_ISL_553546, EPI_ISL_553547 | Lighthouse Lab in Alderley Park | Wellcome Sanger Institute for the COVID-19 Genomics UK (COG-UK) consortium | The Lighthouse Lab in Alderley Park and Alex Alderton, Roberto Amato, Sonia Goncalves, Ewan Harrison, David K. Jackson, Ian Johnston, Dominic Kwiatkowski, Cordelia Langford, John Sillitoe on behalf of the Wellcome Sanger Institute COVID-19 Surveillance Team |
| EPI_ISL_553548, EPI_ISL_553549 | Lighthouse Lab in Alderley Park | Wellcome Sanger Institute for the COVID-19 Genomics UK (COG-UK) consortium | The Lighthouse Lab in Alderley Park and Alex Alderton, Roberto Amato, Sonia Goncalves, Ewan Harrison, David K. Jackson, Ian Johnston, Dominic Kwiatkowski, Cordelia Langford, John Sillitoe on behalf of the Wellcome Sanger Institute COVID-19 Surveillance Team (http://www.sanger.ac.uk/covid-team) |
| EPI_ISL_553550 | Lighthouse Lab in Alderley Park | Wellcome Sanger Institute for the COVID-19 Genomics UK (COG-UK) consortium | The Lighthouse Lab in Alderley Park and Alex Alderton, Roberto Amato, Sonia Goncalves, Ewan Harrison, David K. Jackson, Ian Johnston, Dominic Kwiatkowski, Cordelia Langford, John Sillitoe on behalf of the Wellcome Sanger Institute COVID-19 Surveillance Team |
| EPI_ISL_553551, EPI_ISL_553552 | Lighthouse Lab in Alderley Park | Wellcome Sanger Institute for the COVID-19 Genomics UK (COG-UK) consortium | The Lighthouse Lab in Alderley Park and Alex Alderton, Roberto Amato, Sonia Goncalves, Ewan Harrison, David K. Jackson, Ian Johnston, Dominic Kwiatkowski, Cordelia Langford, John Sillitoe on behalf of the Wellcome Sanger Institute COVID-19 Surveillance Team (http://www.sanger.ac.uk/covid-team) |
| EPI_ISL_553553 | Lighthouse Lab in Alderley Park | Wellcome Sanger Institute for the COVID-19 Genomics UK (COG-UK) consortium | The Lighthouse Lab in Alderley Park and Alex Alderton, Roberto Amato, Sonia Goncalves, Ewan Harrison, David K. Jackson, Ian Johnston, Dominic Kwiatkowski, Cordelia Langford, John Sillitoe on behalf of the Wellcome Sanger Institute COVID-19 Surveillance Team |
| EPI_ISL_553554 | Lighthouse Lab in Alderley Park | Wellcome Sanger Institute for the COVID-19 Genomics UK (COG-UK) consortium | The Lighthouse Lab in Alderley Park and Alex Alderton, Roberto Amato, Sonia Goncalves, Ewan Harrison, David K. Jackson, Ian Johnston, Dominic Kwiatkowski, Cordelia Langford, John Sillitoe on behalf of the Wellcome Sanger Institute COVID-19 Surveillance Team (http://www.sanger.ac.uk/covid-team) |
| EPI_ISL_553555, EPI_ISL_553556, EPI_ISL_553557, EPI_ISL_553558 | Lighthouse Lab in Alderley Park | Wellcome Sanger Institute for the COVID-19 Genomics UK (COG-UK) consortium | The Lighthouse Lab in Alderley Park and Alex Alderton, Roberto Amato, Sonia Goncalves, Ewan Harrison, David K. Jackson, Ian Johnston, Dominic Kwiatkowski, Cordelia Langford, John Sillitoe on behalf of the Wellcome Sanger Institute COVID-19 Surveillance Team |
| EPI_ISL_553559, EPI_ISL_553560, EPI_ISL_553561 | Lighthouse Lab in Alderley Park | Wellcome Sanger Institute for the COVID-19 Genomics UK (COG-UK) consortium | The Lighthouse Lab in Alderley Park and Alex Alderton, Roberto Amato, Sonia Goncalves, Ewan Harrison, David K. Jackson, Ian Johnston, Dominic Kwiatkowski, Cordelia Langford, John Sillitoe on behalf of the Wellcome Sanger Institute COVID-19 Surveillance Team (http://www.sanger.ac.uk/covid-team) |
| EPI_ISL_553562 | Lighthouse Lab in Alderley Park | Wellcome Sanger Institute for the COVID-19 Genomics UK (COG-UK) consortium | The Lighthouse Lab in Alderley Park and Alex Alderton, Roberto Amato, Sonia Goncalves, Ewan Harrison, David K. Jackson, Ian Johnston, Dominic Kwiatkowski, Cordelia Langford, John Sillitoe on behalf of the Wellcome Sanger Institute COVID-19 Surveillance Team |
| EPI_ISL_553563, EPI_ISL_553564 | Lighthouse Lab in Alderley Park | Wellcome Sanger Institute for the COVID-19 Genomics UK (COG-UK) consortium | The Lighthouse Lab in Alderley Park and Alex Alderton, Roberto Amato, Sonia Goncalves, Ewan Harrison, David K. Jackson, Ian Johnston, Dominic Kwiatkowski, Cordelia Langford, John Sillitoe on behalf of the Wellcome Sanger Institute COVID-19 Surveillance Team (http://www.sanger.ac.uk/covid-team) |
| EPI_ISL_553565, EPI_ISL_553566 | Lighthouse Lab in Alderley Park | Wellcome Sanger Institute for the COVID-19 Genomics UK (COG-UK) consortium | The Lighthouse Lab in Alderley Park and Alex Alderton, Roberto Amato, Sonia Goncalves, Ewan Harrison, David K. Jackson, Ian Johnston, Dominic Kwiatkowski, Cordelia Langford, John Sillitoe on behalf of the Wellcome Sanger Institute COVID-19 Surveillance Team |
| EPI_ISL_553567 | Lighthouse Lab in Alderley Park | Wellcome Sanger Institute for the COVID-19 Genomics UK (COG-UK) consortium | The Lighthouse Lab in Alderley Park and Alex Alderton, Roberto Amato, Sonia Goncalves, Ewan Harrison, David K. Jackson, Ian Johnston, Dominic Kwiatkowski, Cordelia Langford, John Sillitoe on behalf of the Wellcome Sanger Institute COVID-19 Surveillance Team (http://www.sanger.ac.uk/covid-team) |
| EPI_ISL_553569 | Lighthouse Lab in Alderley Park | Wellcome Sanger Institute for the COVID-19 Genomics UK (COG-UK) consortium | The Lighthouse Lab in Alderley Park and Alex Alderton, Roberto Amato, Sonia Goncalves, Ewan Harrison, David K. Jackson, Ian Johnston, Dominic Kwiatkowski, Cordelia Langford, John Sillitoe on behalf of the Wellcome Sanger Institute COVID-19 Surveillance Team |
| EPI_ISL_553570 | Lighthouse Lab in Alderley Park | Wellcome Sanger Institute for the COVID-19 Genomics UK (COG-UK) consortium | The Lighthouse Lab in Alderley Park and Alex Alderton, Roberto Amato, Sonia Goncalves, Ewan Harrison, David K. Jackson, Ian Johnston, Dominic Kwiatkowski, Cordelia Langford, John Sillitoe on behalf of the Wellcome Sanger Institute COVID-19 Surveillance Team (http://www.sanger.ac.uk/covid-team) |
| EPI_ISL_553571 | Lighthouse Lab in Alderley Park | Wellcome Sanger Institute for the COVID-19 Genomics UK (COG-UK) consortium | The Lighthouse Lab in Alderley Park and Alex Alderton, Roberto Amato, Sonia Goncalves, Ewan Harrison, David K. Jackson, Ian Johnston, Dominic Kwiatkowski, Cordelia Langford, John Sillitoe on behalf of the Wellcome Sanger Institute COVID-19 Surveillance Team |
| EPI_ISL_553572, EPI_ISL_553573, EPI_ISL_553574, EPI_ISL_553575, EPI_ISL_553576 | Lighthouse Lab in Alderley Park | Wellcome Sanger Institute for the COVID-19 Genomics UK (COG-UK) consortium | The Lighthouse Lab in Alderley Park and Alex Alderton, Roberto Amato, Sonia Goncalves, Ewan Harrison, David K. Jackson, Ian Johnston, Dominic Kwiatkowski, Cordelia Langford, John Sillitoe on behalf of the Wellcome Sanger Institute COVID-19 Surveillance Team (http://www.sanger.ac.uk/covid-team) |
| EPI_ISL_553577, EPI_ISL_553578 | Lighthouse Lab in Alderley Park | Wellcome Sanger Institute for the COVID-19 Genomics UK (COG-UK) consortium | The Lighthouse Lab in Alderley Park and Alex Alderton, Roberto Amato, Sonia Goncalves, Ewan Harrison, David K. Jackson, Ian Johnston, Dominic Kwiatkowski, Cordelia Langford, John Sillitoe on behalf of the Wellcome Sanger Institute COVID-19 Surveillance Team |
| EPI_ISL_553579, EPI_ISL_553580 | Lighthouse Lab in Alderley Park | Wellcome Sanger Institute for the COVID-19 Genomics UK (COG-UK) consortium | The Lighthouse Lab in Alderley Park and Alex Alderton, Roberto Amato, Sonia Goncalves, Ewan Harrison, David K. Jackson, Ian Johnston, Dominic Kwiatkowski, Cordelia Langford, John Sillitoe on behalf of the Wellcome Sanger Institute COVID-19 Surveillance Team (http://www.sanger.ac.uk/covid-team) |
| EPI_ISL_553581 | Lighthouse Lab in Alderley Park | Wellcome Sanger Institute for the COVID-19 Genomics UK (COG-UK) consortium | The Lighthouse Lab in Alderley Park and Alex Alderton, Roberto Amato, Sonia Goncalves, Ewan Harrison, David K. Jackson, Ian Johnston, Dominic Kwiatkowski, Cordelia Langford, John Sillitoe on behalf of the Wellcome Sanger Institute COVID-19 Surveillance Team |
| EPI_ISL_553582 | Lighthouse Lab in Alderley Park | Wellcome Sanger Institute for the COVID-19 Genomics UK (COG-UK) consortium | The Lighthouse Lab in Alderley Park and Alex Alderton, Roberto Amato, Sonia Goncalves, Ewan Harrison, David K. Jackson, Ian Johnston, Dominic Kwiatkowski, Cordelia Langford, John Sillitoe on behalf of the Wellcome Sanger Institute COVID-19 Surveillance Team (http://www.sanger.ac.uk/covid-team) |
| EPI_ISL_553583 | Lighthouse Lab in Alderley Park | Wellcome Sanger Institute for the COVID-19 Genomics UK (COG-UK) consortium | The Lighthouse Lab in Alderley Park and Alex Alderton, Roberto Amato, Sonia Goncalves, Ewan Harrison, David K. Jackson, Ian Johnston, Dominic Kwiatkowski, Cordelia Langford, John Sillitoe on behalf of the Wellcome Sanger Institute COVID-19 Surveillance Team |
| EPI_ISL_553584 | Lighthouse Lab in Alderley Park | Wellcome Sanger Institute for the COVID-19 Genomics UK (COG-UK) consortium | The Lighthouse Lab in Alderley Park and Alex Alderton, Roberto Amato, Sonia Goncalves, Ewan Harrison, David K. Jackson, Ian Johnston, Dominic Kwiatkowski, Cordelia Langford, John Sillitoe on behalf of the Wellcome Sanger Institute COVID-19 Surveillance Team (http://www.sanger.ac.uk/covid-team) |
| EPI_ISL_553585, EPI_ISL_553586 | Lighthouse Lab in Alderley Park | Wellcome Sanger Institute for the COVID-19 Genomics UK (COG-UK) consortium | The Lighthouse Lab in Alderley Park and Alex Alderton, Roberto Amato, Sonia Goncalves, Ewan Harrison, David K. Jackson, Ian Johnston, Dominic Kwiatkowski, Cordelia Langford, John Sillitoe on behalf of the Wellcome Sanger Institute COVID-19 Surveillance Team |
| EPI_ISL_553587 | Lighthouse Lab in Alderley Park | Wellcome Sanger Institute for the COVID-19 Genomics UK (COG-UK) consortium | The Lighthouse Lab in Alderley Park and Alex Alderton, Roberto Amato, Sonia Goncalves, Ewan Harrison, David K. Jackson, Ian Johnston, Dominic Kwiatkowski, Cordelia Langford, John Sillitoe on behalf of the Wellcome Sanger Institute COVID-19 Surveillance Team (http://www.sanger.ac.uk/covid-team) |
| EPI_ISL_553588 | Lighthouse Lab in Alderley Park | Wellcome Sanger Institute for the COVID-19 Genomics UK (COG-UK) consortium | The Lighthouse Lab in Alderley Park and Alex Alderton, Roberto Amato, Sonia Goncalves, Ewan Harrison, David K. Jackson, Ian Johnston, Dominic Kwiatkowski, Cordelia Langford, John Sillitoe on behalf of the Wellcome Sanger Institute COVID-19 Surveillance Team |

| | | | |
|---|---|---|---|
| EPI_ISL_553589 | Lighthouse Lab in Alderley Park | Wellcome Sanger Institute for the COVID-19 Genomics UK (COG-UK) consortium | The Lighthouse Lab in Alderley Park and Alex Alderton, Roberto Amato, Sonia Goncalves, Ewan Harrison, David K. Jackson, Ian Johnston, Dominic Kwiatkowski, Cordelia Langford, John Sillitoe on behalf of the Wellcome Sanger Institute COVID-19 Surveillance Team (http://www.sanger.ac.uk/covid-team) |
| EPI_ISL_553590, EPI_ISL_553591 | Lighthouse Lab in Alderley Park | Wellcome Sanger Institute for the COVID-19 Genomics UK (COG-UK) consortium | The Lighthouse Lab in Alderley Park and Alex Alderton, Roberto Amato, Sonia Goncalves, Ewan Harrison, David K. Jackson, Ian Johnston, Dominic Kwiatkowski, Cordelia Langford, John Sillitoe on behalf of the Wellcome Sanger Institute COVID-19 Surveillance Team |
| EPI_ISL_553592 | Lighthouse Lab in Alderley Park | Wellcome Sanger Institute for the COVID-19 Genomics UK (COG-UK) consortium | The Lighthouse Lab in Alderley Park and Alex Alderton, Roberto Amato, Sonia Goncalves, Ewan Harrison, David K. Jackson, Ian Johnston, Dominic Kwiatkowski, Cordelia Langford, John Sillitoe on behalf of the Wellcome Sanger Institute COVID-19 Surveillance Team (http://www.sanger.ac.uk/covid-team) |
| EPI_ISL_553593, EPI_ISL_553594, EPI_ISL_553595, EPI_ISL_553596, EPI_ISL_553597, EPI_ISL_553598 | Lighthouse Lab in Alderley Park | Wellcome Sanger Institute for the COVID-19 Genomics UK (COG-UK) consortium | The Lighthouse Lab in Alderley Park and Alex Alderton, Roberto Amato, Sonia Goncalves, Ewan Harrison, David K. Jackson, Ian Johnston, Dominic Kwiatkowski, Cordelia Langford, John Sillitoe on behalf of the Wellcome Sanger Institute COVID-19 Surveillance Team |
| EPI_ISL_553599 | Lighthouse Lab in Alderley Park | Wellcome Sanger Institute for the COVID-19 Genomics UK (COG-UK) consortium | The Lighthouse Lab in Alderley Park and Alex Alderton, Roberto Amato, Sonia Goncalves, Ewan Harrison, David K. Jackson, Ian Johnston, Dominic Kwiatkowski, Cordelia Langford, John Sillitoe on behalf of the Wellcome Sanger Institute COVID-19 Surveillance Team (http://www.sanger.ac.uk/covid-team) |
| EPI_ISL_553600, EPI_ISL_553601, EPI_ISL_553602, EPI_ISL_553603 | Lighthouse Lab in Alderley Park | Wellcome Sanger Institute for the COVID-19 Genomics UK (COG-UK) consortium | The Lighthouse Lab in Alderley Park and Alex Alderton, Roberto Amato, Sonia Goncalves, Ewan Harrison, David K. Jackson, Ian Johnston, Dominic Kwiatkowski, Cordelia Langford, John Sillitoe on behalf of the Wellcome Sanger Institute COVID-19 Surveillance Team |
| EPI_ISL_553604 | Lighthouse Lab in Alderley Park | Wellcome Sanger Institute for the COVID-19 Genomics UK (COG-UK) consortium | The Lighthouse Lab in Alderley Park and Alex Alderton, Roberto Amato, Sonia Goncalves, Ewan Harrison, David K. Jackson, Ian Johnston, Dominic Kwiatkowski, Cordelia Langford, John Sillitoe on behalf of the Wellcome Sanger Institute COVID-19 Surveillance Team (http://www.sanger.ac.uk/covid-team) |
| EPI_ISL_553605 | Lighthouse Lab in Alderley Park | Wellcome Sanger Institute for the COVID-19 Genomics UK (COG-UK) consortium | The Lighthouse Lab in Alderley Park and Alex Alderton, Roberto Amato, Sonia Goncalves, Ewan Harrison, David K. Jackson, Ian Johnston, Dominic Kwiatkowski, Cordelia Langford, John Sillitoe on behalf of the Wellcome Sanger Institute COVID-19 Surveillance Team |
| EPI_ISL_553606 | Lighthouse Lab in Alderley Park | Wellcome Sanger Institute for the COVID-19 Genomics UK (COG-UK) consortium | The Lighthouse Lab in Alderley Park and Alex Alderton, Roberto Amato, Sonia Goncalves, Ewan Harrison, David K. Jackson, Ian Johnston, Dominic Kwiatkowski, Cordelia Langford, John Sillitoe on behalf of the Wellcome Sanger Institute COVID-19 Surveillance Team (http://www.sanger.ac.uk/covid-team) |
| EPI_ISL_553607, EPI_ISL_553608, EPI_ISL_553609 | Lighthouse Lab in Alderley Park | Wellcome Sanger Institute for the COVID-19 Genomics UK (COG-UK) consortium | The Lighthouse Lab in Alderley Park and Alex Alderton, Roberto Amato, Sonia Goncalves, Ewan Harrison, David K. Jackson, Ian Johnston, Dominic Kwiatkowski, Cordelia Langford, John Sillitoe on behalf of the Wellcome Sanger Institute COVID-19 Surveillance Team |
| EPI_ISL_553610, EPI_ISL_553611, EPI_ISL_553612, EPI_ISL_553613 | Lighthouse Lab in Alderley Park | Wellcome Sanger Institute for the COVID-19 Genomics UK (COG-UK) consortium | The Lighthouse Lab in Alderley Park and Alex Alderton, Roberto Amato, Sonia Goncalves, Ewan Harrison, David K. Jackson, Ian Johnston, Dominic Kwiatkowski, Cordelia Langford, John Sillitoe on behalf of the Wellcome Sanger Institute COVID-19 Surveillance Team (http://www.sanger.ac.uk/covid-team) |
| EPI_ISL_553614, EPI_ISL_553615 | Lighthouse Lab in Alderley Park | Wellcome Sanger Institute for the COVID-19 Genomics UK (COG-UK) consortium | The Lighthouse Lab in Alderley Park and Alex Alderton, Roberto Amato, Sonia Goncalves, Ewan Harrison, David K. Jackson, Ian Johnston, Dominic Kwiatkowski, Cordelia Langford, John Sillitoe on behalf of the Wellcome Sanger Institute COVID-19 Surveillance Team |
| EPI_ISL_553616, EPI_ISL_553617 | Lighthouse Lab in Alderley Park | Wellcome Sanger Institute for the COVID-19 Genomics UK (COG-UK) consortium | The Lighthouse Lab in Alderley Park and Alex Alderton, Roberto Amato, Sonia Goncalves, Ewan Harrison, David K. Jackson, Ian Johnston, Dominic Kwiatkowski, Cordelia Langford, John Sillitoe on behalf of the Wellcome Sanger Institute COVID-19 Surveillance Team (http://www.sanger.ac.uk/covid-team) |
| EPI_ISL_553618, EPI_ISL_553619, EPI_ISL_553620, EPI_ISL_553621, EPI_ISL_553622, EPI_ISL_553623, EPI_ISL_553624 | Lighthouse Lab in Alderley Park | Wellcome Sanger Institute for the COVID-19 Genomics UK (COG-UK) consortium | The Lighthouse Lab in Alderley Park and Alex Alderton, Roberto Amato, Sonia Goncalves, Ewan Harrison, David K. Jackson, Ian Johnston, Dominic Kwiatkowski, Cordelia Langford, John Sillitoe on behalf of the Wellcome Sanger Institute COVID-19 Surveillance Team |
| EPI_ISL_553625 | Lighthouse Lab in Alderley Park | Wellcome Sanger Institute for the COVID-19 Genomics UK (COG-UK) consortium | The Lighthouse Lab in Alderley Park and Alex Alderton, Roberto Amato, Sonia Goncalves, Ewan Harrison, David K. Jackson, Ian Johnston, Dominic Kwiatkowski, Cordelia Langford, John Sillitoe on behalf of the Wellcome Sanger Institute COVID-19 Surveillance Team (http://www.sanger.ac.uk/covid-team) |
| EPI_ISL_553626 | Lighthouse Lab in Alderley Park | Wellcome Sanger Institute for the COVID-19 Genomics UK (COG-UK) consortium | The Lighthouse Lab in Alderley Park and Alex Alderton, Roberto Amato, Sonia Goncalves, Ewan Harrison, David K. Jackson, Ian Johnston, Dominic Kwiatkowski, Cordelia Langford, John Sillitoe on behalf of the Wellcome Sanger Institute COVID-19 Surveillance Team |
| EPI_ISL_553627 | Lighthouse Lab in Alderley Park | Wellcome Sanger Institute for the COVID-19 Genomics UK (COG-UK) consortium | The Lighthouse Lab in Alderley Park and Alex Alderton, Roberto Amato, Sonia Goncalves, Ewan Harrison, David K. Jackson, Ian Johnston, Dominic Kwiatkowski, Cordelia Langford, John Sillitoe on behalf of the Wellcome Sanger Institute COVID-19 Surveillance Team (http://www.sanger.ac.uk/covid-team) |
| EPI_ISL_553628, EPI_ISL_553629 | Lighthouse Lab in Alderley Park | Wellcome Sanger Institute for the COVID-19 Genomics UK (COG-UK) consortium | The Lighthouse Lab in Alderley Park and Alex Alderton, Roberto Amato, Sonia Goncalves, Ewan Harrison, David K. Jackson, Ian Johnston, Dominic Kwiatkowski, Cordelia Langford, John Sillitoe on behalf of the Wellcome Sanger Institute COVID-19 Surveillance Team |
| EPI_ISL_553630, EPI_ISL_553631, EPI_ISL_553632, EPI_ISL_553633, EPI_ISL_553634, EPI_ISL_553635, EPI_ISL_553636, EPI_ISL_553637, EPI_ISL_553638, EPI_ISL_553639, EPI_ISL_553640 | | | |
| see above | Lighthouse Lab in Alderley Park | Wellcome Sanger Institute for the COVID-19 Genomics UK (COG-UK) consortium | The Lighthouse Lab in Alderley Park and Alex Alderton, Roberto Amato, Sonia Goncalves, Ewan Harrison, David K. Jackson, Ian Johnston, Dominic Kwiatkowski, Cordelia Langford, John Sillitoe on behalf of the Wellcome Sanger Institute COVID-19 Surveillance Team (http://www.sanger.ac.uk/covid-team) |
| EPI_ISL_553641, EPI_ISL_553642, EPI_ISL_553643, EPI_ISL_553644, EPI_ISL_553645, EPI_ISL_553646, EPI_ISL_553647, EPI_ISL_553648, EPI_ISL_553649, EPI_ISL_553650, EPI_ISL_553651, EPI_ISL_553652, EPI_ISL_553653, EPI_ISL_553654, EPI_ISL_553655, EPI_ISL_553656, EPI_ISL_553657, EPI_ISL_553658, EPI_ISL_553659, EPI_ISL_553660, EPI_ISL_553661, EPI_ISL_553662 | | | |
| see above | Lighthouse Lab in Alderley Park | Wellcome Sanger Institute for the COVID-19 Genomics UK (COG-UK) consortium | The Lighthouse Lab in Alderley Park and Alex Alderton, Roberto Amato, Sonia Goncalves, Ewan Harrison, David K. Jackson, Ian Johnston, Dominic Kwiatkowski, Cordelia Langford, John Sillitoe on behalf of the Wellcome Sanger Institute COVID-19 Surveillance Team |
| EPI_ISL_553663 | Lighthouse Lab in Alderley Park | Wellcome Sanger Institute for the COVID-19 Genomics UK (COG-UK) consortium | The Lighthouse Lab in Alderley Park and Alex Alderton, Roberto Amato, Sonia Goncalves, Ewan Harrison, David K. Jackson, Ian Johnston, Dominic Kwiatkowski, Cordelia Langford, John Sillitoe on behalf of the Wellcome Sanger Institute COVID-19 Surveillance Team (http://www.sanger.ac.uk/covid-team) |
| EPI_ISL_553664, EPI_ISL_553665, EPI_ISL_553666, EPI_ISL_553667, EPI_ISL_553668, EPI_ISL_553669, EPI_ISL_553670, EPI_ISL_553671, EPI_ISL_553672 | Lighthouse Lab in Alderley Park | Wellcome Sanger Institute for the COVID-19 Genomics UK (COG-UK) consortium | The Lighthouse Lab in Alderley Park and Alex Alderton, Roberto Amato, Sonia Goncalves, Ewan Harrison, David K. Jackson, Ian Johnston, Dominic Kwiatkowski, Cordelia Langford, John Sillitoe on behalf of the Wellcome Sanger Institute COVID-19 Surveillance Team |
| EPI_ISL_553673, EPI_ISL_553674 | Lighthouse Lab in Alderley Park | Wellcome Sanger Institute for the COVID-19 Genomics UK (COG-UK) consortium | The Lighthouse Lab in Alderley Park and Alex Alderton, Roberto Amato, Sonia Goncalves, Ewan Harrison, David K. Jackson, Ian Johnston, Dominic Kwiatkowski, Cordelia Langford, John Sillitoe on behalf of the Wellcome Sanger Institute COVID-19 Surveillance Team (http://www.sanger.ac.uk/covid-team) |
| EPI_ISL_553675, EPI_ISL_553676, EPI_ISL_553677, EPI_ISL_553678 | Lighthouse Lab in Alderley Park | Wellcome Sanger Institute for the COVID-19 Genomics UK (COG-UK) consortium | The Lighthouse Lab in Alderley Park and Alex Alderton, Roberto Amato, Sonia Goncalves, Ewan Harrison, David K. Jackson, Ian Johnston, Dominic Kwiatkowski, Cordelia Langford, John Sillitoe on behalf of the Wellcome Sanger Institute COVID-19 Surveillance Team |
| EPI_ISL_553679 | Lighthouse Lab in Alderley Park | Wellcome Sanger Institute for the COVID-19 Genomics UK (COG-UK) consortium | The Lighthouse Lab in Alderley Park and Alex Alderton, Roberto Amato, Sonia Goncalves, Ewan Harrison, David K. Jackson, Ian Johnston, Dominic Kwiatkowski, Cordelia Langford, John Sillitoe on behalf of the Wellcome Sanger Institute COVID-19 Surveillance Team (http://www.sanger.ac.uk/covid-team) |
| EPI_ISL_553680, EPI_ISL_553681, EPI_ISL_553682, EPI_ISL_553683, EPI_ISL_553684, EPI_ISL_553685, EPI_ISL_553686, EPI_ISL_553687, EPI_ISL_553688, EPI_ISL_553689, EPI_ISL_553690, EPI_ISL_553691, EPI_ISL_553692, EPI_ISL_553693, EPI_ISL_553694, EPI_ISL_553695 | | | |

| | | | |
|---|---|---|---|
| see above | Lighthouse Lab in Alderley Park | Wellcome Sanger Institute for the COVID-19 Genomics UK (COG-UK) consortium | The Lighthouse Lab in Alderley Park and Alex Alderton, Roberto Amato, Sonia Goncalves, Ewan Harrison, David K. Jackson, Ian Johnston, Dominic Kwiatkowski, Cordelia Langford, John Sillitoe on behalf of the Wellcome Sanger Institute COVID-19 Surveillance Team |
| EPI_ISL_553696 | Lighthouse Lab in Alderley Park | Wellcome Sanger Institute for the COVID-19 Genomics UK (COG-UK) consortium | The Lighthouse Lab in Alderley Park and Alex Alderton, Roberto Amato, Sonia Goncalves, Ewan Harrison, David K. Jackson, Ian Johnston, Dominic Kwiatkowski, Cordelia Langford, John Sillitoe on behalf of the Wellcome Sanger Institute COVID-19 Surveillance Team (http://www.sanger.ac.uk/covid-team) |
| EPI_ISL_553697, EPI_ISL_553698, EPI_ISL_553704 | Lighthouse Lab in Alderley Park | Wellcome Sanger Institute for the COVID-19 Genomics UK (COG-UK) consortium | The Lighthouse Lab in Alderley Park and Alex Alderton, Roberto Amato, Sonia Goncalves, Ewan Harrison, David K. Jackson, Ian Johnston, Dominic Kwiatkowski, Cordelia Langford, John Sillitoe on behalf of the Wellcome Sanger Institute COVID-19 Surveillance Team |
| EPI_ISL_553706, EPI_ISL_553712, EPI_ISL_553727 | Lighthouse Lab in Alderley Park | Wellcome Sanger Institute for the COVID-19 Genomics UK (COG-UK) consortium | The Lighthouse Lab in Alderley Park and Alex Alderton, Roberto Amato, Sonia Goncalves, Ewan Harrison, David K. Jackson, Ian Johnston, Dominic Kwiatkowski, Cordelia Langford, John Sillitoe on behalf of the Wellcome Sanger Institute COVID-19 Surveillance Team (http://www.sanger.ac.uk/covid-team) |
| EPI_ISL_553733, EPI_ISL_553734, EPI_ISL_553735, EPI_ISL_553736, EPI_ISL_553737, EPI_ISL_553738, EPI_ISL_553739 | Lighthouse Lab in Alderley Park | Wellcome Sanger Institute for the COVID-19 Genomics UK (COG-UK) consortium | The Lighthouse Lab in Alderley Park and Alex Alderton, Roberto Amato, Sonia Goncalves, Ewan Harrison, David K. Jackson, Ian Johnston, Dominic Kwiatkowski, Cordelia Langford, John Sillitoe on behalf of the Wellcome Sanger Institute COVID-19 Surveillance Team |
| EPI_ISL_553740, EPI_ISL_553741, EPI_ISL_553742, EPI_ISL_553743, EPI_ISL_553744, EPI_ISL_553745, EPI_ISL_553746, EPI_ISL_553747, EPI_ISL_553748, EPI_ISL_553749 | Lighthouse Lab in Alderley Park | Wellcome Sanger Institute for the COVID-19 Genomics UK (COG-UK) consortium | The Lighthouse Lab in Alderley Park and Alex Alderton, Roberto Amato, Sonia Goncalves, Ewan Harrison, David K. Jackson, Ian Johnston, Dominic Kwiatkowski, Cordelia Langford, John Sillitoe on behalf of the Wellcome Sanger Institute COVID-19 Surveillance Team (http://www.sanger.ac.uk/covid-team) |
| EPI_ISL_553750, EPI_ISL_553751, EPI_ISL_553752, EPI_ISL_553753, EPI_ISL_553754, EPI_ISL_553755, EPI_ISL_553756, EPI_ISL_553757, EPI_ISL_553758, EPI_ISL_553759, EPI_ISL_553760, EPI_ISL_553761, EPI_ISL_553762 | | | |
| see above | Lighthouse Lab in Alderley Park | Wellcome Sanger Institute for the COVID-19 Genomics UK (COG-UK) consortium | The Lighthouse Lab in Alderley Park and Alex Alderton, Roberto Amato, Sonia Goncalves, Ewan Harrison, David K. Jackson, Ian Johnston, Dominic Kwiatkowski, Cordelia Langford, John Sillitoe on behalf of the Wellcome Sanger Institute COVID-19 Surveillance Team |
| EPI_ISL_553763 | Lighthouse Lab in Alderley Park | Wellcome Sanger Institute for the COVID-19 Genomics UK (COG-UK) consortium | The Lighthouse Lab in Alderley Park and Alex Alderton, Roberto Amato, Sonia Goncalves, Ewan Harrison, David K. Jackson, Ian Johnston, Dominic Kwiatkowski, Cordelia Langford, John Sillitoe on behalf of the Wellcome Sanger Institute COVID-19 Surveillance Team (http://www.sanger.ac.uk/covid-team) |
| EPI_ISL_553764, EPI_ISL_553765, EPI_ISL_553766, EPI_ISL_553767, EPI_ISL_553768, EPI_ISL_553769, EPI_ISL_553770, EPI_ISL_553771, EPI_ISL_553772, EPI_ISL_553773 | Lighthouse Lab in Alderley Park | Wellcome Sanger Institute for the COVID-19 Genomics UK (COG-UK) consortium | The Lighthouse Lab in Alderley Park and Alex Alderton, Roberto Amato, Sonia Goncalves, Ewan Harrison, David K. Jackson, Ian Johnston, Dominic Kwiatkowski, Cordelia Langford, John Sillitoe on behalf of the Wellcome Sanger Institute COVID-19 Surveillance Team |
| EPI_ISL_553774 | Lighthouse Lab in Alderley Park | Wellcome Sanger Institute for the COVID-19 Genomics UK (COG-UK) consortium | The Lighthouse Lab in Alderley Park and Alex Alderton, Roberto Amato, Sonia Goncalves, Ewan Harrison, David K. Jackson, Ian Johnston, Dominic Kwiatkowski, Cordelia Langford, John Sillitoe on behalf of the Wellcome Sanger Institute COVID-19 Surveillance Team (http://www.sanger.ac.uk/covid-team) |
| EPI_ISL_553775, EPI_ISL_553776, EPI_ISL_553777, EPI_ISL_553779, EPI_ISL_553780, EPI_ISL_553781, EPI_ISL_553782, EPI_ISL_553783, EPI_ISL_553785, EPI_ISL_553786, EPI_ISL_553787, EPI_ISL_553788, EPI_ISL_553789, EPI_ISL_553790, EPI_ISL_553791, EPI_ISL_553792, EPI_ISL_553793, EPI_ISL_553794 | | | |
| see above | Lighthouse Lab in Alderley Park | Wellcome Sanger Institute for the COVID-19 Genomics UK (COG-UK) consortium | The Lighthouse Lab in Alderley Park and Alex Alderton, Roberto Amato, Sonia Goncalves, Ewan Harrison, David K. Jackson, Ian Johnston, Dominic Kwiatkowski, Cordelia Langford, John Sillitoe on behalf of the Wellcome Sanger Institute COVID-19 Surveillance Team |
| EPI_ISL_553795 | Lighthouse Lab in Alderley Park | Wellcome Sanger Institute for the COVID-19 Genomics UK (COG-UK) consortium | The Lighthouse Lab in Alderley Park and Alex Alderton, Roberto Amato, Sonia Goncalves, Ewan Harrison, David K. Jackson, Ian Johnston, Dominic Kwiatkowski, Cordelia Langford, John Sillitoe on behalf of the Wellcome Sanger Institute COVID-19 Surveillance Team (http://www.sanger.ac.uk/covid-team) |
| EPI_ISL_553796, EPI_ISL_553797, EPI_ISL_553798, EPI_ISL_553799, EPI_ISL_553800, EPI_ISL_553801, EPI_ISL_553802, EPI_ISL_553803, EPI_ISL_553804, EPI_ISL_553805, EPI_ISL_553806, EPI_ISL_553807, EPI_ISL_553808, EPI_ISL_553809, EPI_ISL_553810, EPI_ISL_553811, EPI_ISL_553813, EPI_ISL_553814, EPI_ISL_553815, EPI_ISL_553816, EPI_ISL_553817, EPI_ISL_553818, EPI_ISL_553819, EPI_ISL_553820, EPI_ISL_553821, EPI_ISL_553822, EPI_ISL_553823, EPI_ISL_553824, EPI_ISL_553825, EPI_ISL_553826, EPI_ISL_553827, EPI_ISL_553828, EPI_ISL_553829, EPI_ISL_553830, EPI_ISL_553831, EPI_ISL_553832, EPI_ISL_553833, EPI_ISL_553834, EPI_ISL_553835, EPI_ISL_553836, EPI_ISL_553837, EPI_ISL_553838, EPI_ISL_553839, EPI_ISL_553840, EPI_ISL_553841, EPI_ISL_553842, EPI_ISL_553843, EPI_ISL_553844, EPI_ISL_553845, EPI_ISL_553847, EPI_ISL_553848, EPI_ISL_553849 | | | |
| see above | Lighthouse Lab in Alderley Park | Wellcome Sanger Institute for the COVID-19 Genomics UK (COG-UK) consortium | The Lighthouse Lab in Alderley Park and Alex Alderton, Roberto Amato, Sonia Goncalves, Ewan Harrison, David K. Jackson, Ian Johnston, Dominic Kwiatkowski, Cordelia Langford, John Sillitoe on behalf of the Wellcome Sanger Institute COVID-19 Surveillance Team |
| EPI_ISL_553850 | Lighthouse Lab in Alderley Park | Wellcome Sanger Institute for the COVID-19 Genomics UK (COG-UK) consortium | The Lighthouse Lab in Alderley Park and Alex Alderton, Roberto Amato, Sonia Goncalves, Ewan Harrison, David K. Jackson, Ian Johnston, Dominic Kwiatkowski, Cordelia Langford, John Sillitoe on behalf of the Wellcome Sanger Institute COVID-19 Surveillance Team (http://www.sanger.ac.uk/covid-team) |
| EPI_ISL_553851, EPI_ISL_553852, EPI_ISL_553853, EPI_ISL_553854, EPI_ISL_553855 | Lighthouse Lab in Alderley Park | Wellcome Sanger Institute for the COVID-19 Genomics UK (COG-UK) consortium | The Lighthouse Lab in Alderley Park and Alex Alderton, Roberto Amato, Sonia Goncalves, Ewan Harrison, David K. Jackson, Ian Johnston, Dominic Kwiatkowski, Cordelia Langford, John Sillitoe on behalf of the Wellcome Sanger Institute COVID-19 Surveillance Team |
| EPI_ISL_553856 | Lighthouse Lab in Alderley Park | Wellcome Sanger Institute for the COVID-19 Genomics UK (COG-UK) consortium | The Lighthouse Lab in Alderley Park and Alex Alderton, Roberto Amato, Sonia Goncalves, Ewan Harrison, David K. Jackson, Ian Johnston, Dominic Kwiatkowski, Cordelia Langford, John Sillitoe on behalf of the Wellcome Sanger Institute COVID-19 Surveillance Team (http://www.sanger.ac.uk/covid-team) |
| EPI_ISL_553857, EPI_ISL_553858, EPI_ISL_553859, EPI_ISL_553860, EPI_ISL_553861, EPI_ISL_553862 | Lighthouse Lab in Alderley Park | Wellcome Sanger Institute for the COVID-19 Genomics UK (COG-UK) consortium | The Lighthouse Lab in Alderley Park and Alex Alderton, Roberto Amato, Sonia Goncalves, Ewan Harrison, David K. Jackson, Ian Johnston, Dominic Kwiatkowski, Cordelia Langford, John Sillitoe on behalf of the Wellcome Sanger Institute COVID-19 Surveillance Team |
| EPI_ISL_553863 | Lighthouse Lab in Alderley Park | Wellcome Sanger Institute for the COVID-19 Genomics UK (COG-UK) consortium | The Lighthouse Lab in Alderley Park and Alex Alderton, Roberto Amato, Sonia Goncalves, Ewan Harrison, David K. Jackson, Ian Johnston, Dominic Kwiatkowski, Cordelia Langford, John Sillitoe on behalf of the Wellcome Sanger Institute COVID-19 Surveillance Team (http://www.sanger.ac.uk/covid-team) |
| EPI_ISL_553864, EPI_ISL_553865, EPI_ISL_553867, EPI_ISL_553868, EPI_ISL_553869 | Lighthouse Lab in Alderley Park | Wellcome Sanger Institute for the COVID-19 Genomics UK (COG-UK) consortium | The Lighthouse Lab in Alderley Park and Alex Alderton, Roberto Amato, Sonia Goncalves, Ewan Harrison, David K. Jackson, Ian Johnston, Dominic Kwiatkowski, Cordelia Langford, John Sillitoe on behalf of the Wellcome Sanger Institute COVID-19 Surveillance Team |
| EPI_ISL_553870 | Lighthouse Lab in Alderley Park | Wellcome Sanger Institute for the COVID-19 Genomics UK (COG-UK) consortium | The Lighthouse Lab in Alderley Park and Alex Alderton, Roberto Amato, Sonia Goncalves, Ewan Harrison, David K. Jackson, Ian Johnston, Dominic Kwiatkowski, Cordelia Langford, John Sillitoe on behalf of the Wellcome Sanger Institute COVID-19 Surveillance Team (http://www.sanger.ac.uk/covid-team) |
| EPI_ISL_553872, EPI_ISL_553873 | Lighthouse Lab in Alderley Park | Wellcome Sanger Institute for the COVID-19 Genomics UK (COG-UK) consortium | The Lighthouse Lab in Alderley Park and Alex Alderton, Roberto Amato, Sonia Goncalves, Ewan Harrison, David K. Jackson, Ian Johnston, Dominic Kwiatkowski, Cordelia Langford, John Sillitoe on behalf of the Wellcome Sanger Institute COVID-19 Surveillance Team |
| EPI_ISL_553874, EPI_ISL_553875 | Lighthouse Lab in Alderley Park | Wellcome Sanger Institute for the COVID-19 Genomics UK (COG-UK) consortium | The Lighthouse Lab in Alderley Park and Alex Alderton, Roberto Amato, Sonia Goncalves, Ewan Harrison, David K. Jackson, Ian Johnston, Dominic Kwiatkowski, Cordelia Langford, John Sillitoe on behalf of the Wellcome Sanger Institute COVID-19 Surveillance Team (http://www.sanger.ac.uk/covid-team) |
| EPI_ISL_553876, EPI_ISL_553877, EPI_ISL_553878 | Lighthouse Lab in Alderley Park | Wellcome Sanger Institute for the COVID-19 Genomics UK (COG-UK) consortium | The Lighthouse Lab in Alderley Park and Alex Alderton, Roberto Amato, Sonia Goncalves, Ewan Harrison, David K. Jackson, Ian Johnston, Dominic Kwiatkowski, Cordelia Langford, John Sillitoe on behalf of the Wellcome Sanger Institute COVID-19 Surveillance Team |
| EPI_ISL_553879, EPI_ISL_553880 | Lighthouse Lab in Milton Keynes | Wellcome Sanger Institute for the COVID-19 Genomics UK (COG-UK) consortium | The Lighthouse Lab in Milton Keynes and Alex Alderton, Roberto Amato, Sonia Goncalves, Ewan Harrison, David K. Jackson, Ian Johnston, Dominic Kwiatkowski, Cordelia Langford, John Sillitoe on behalf of the Wellcome Sanger Institute COVID-19 Surveillance Team |

| | | | |
|---|---|---|---|
| | | | (http://www.sanger.ac.uk/covid-team) |
| EPI_ISL_553881, EPI_ISL_553882, EPI_ISL_553883, EPI_ISL_553884, EPI_ISL_553886 | Lighthouse Lab in Alderley Park | Wellcome Sanger Institute for the COVID-19 Genomics UK (COG-UK) consortium | The Lighthouse Lab in Alderley Park and Alex Alderton, Roberto Amato, Sonia Goncalves, Ewan Harrison, David K. Jackson, Ian Johnston, Dominic Kwiatkowski, Cordelia Langford, John Sillitoe on behalf of the Wellcome Sanger Institute COVID-19 Surveillance Team |
| EPI_ISL_553887 | Lighthouse Lab in Alderley Park | Wellcome Sanger Institute for the COVID-19 Genomics UK (COG-UK) consortium | The Lighthouse Lab in Alderley Park and Alex Alderton, Roberto Amato, Sonia Goncalves, Ewan Harrison, David K. Jackson, Ian Johnston, Dominic Kwiatkowski, Cordelia Langford, John Sillitoe on behalf of the Wellcome Sanger Institute COVID-19 Surveillance Team (http://www.sanger.ac.uk/covid-team) |
| EPI_ISL_553888 | Lighthouse Lab in Alderley Park | Wellcome Sanger Institute for the COVID-19 Genomics UK (COG-UK) consortium | The Lighthouse Lab in Alderley Park and Alex Alderton, Roberto Amato, Sonia Goncalves, Ewan Harrison, David K. Jackson, Ian Johnston, Dominic Kwiatkowski, Cordelia Langford, John Sillitoe on behalf of the Wellcome Sanger Institute COVID-19 Surveillance Team |
| EPI_ISL_553889 | Lighthouse Lab in Milton Keynes | Wellcome Sanger Institute for the COVID-19 Genomics UK (COG-UK) consortium | The Lighthouse Lab in Milton Keynes and Alex Alderton, Roberto Amato, Sonia Goncalves, Ewan Harrison, David K. Jackson, Ian Johnston, Dominic Kwiatkowski, Cordelia Langford, John Sillitoe on behalf of the Wellcome Sanger Institute COVID-19 Surveillance Team (http://www.sanger.ac.uk/covid-team) |
| EPI_ISL_553890 | Lighthouse Lab in Alderley Park | Wellcome Sanger Institute for the COVID-19 Genomics UK (COG-UK) consortium | The Lighthouse Lab in Alderley Park and Alex Alderton, Roberto Amato, Sonia Goncalves, Ewan Harrison, David K. Jackson, Ian Johnston, Dominic Kwiatkowski, Cordelia Langford, John Sillitoe on behalf of the Wellcome Sanger Institute COVID-19 Surveillance Team (http://www.sanger.ac.uk/covid-team) |
| EPI_ISL_553891, EPI_ISL_553892 | Lighthouse Lab in Alderley Park | Wellcome Sanger Institute for the COVID-19 Genomics UK (COG-UK) consortium | The Lighthouse Lab in Alderley Park and Alex Alderton, Roberto Amato, Sonia Goncalves, Ewan Harrison, David K. Jackson, Ian Johnston, Dominic Kwiatkowski, Cordelia Langford, John Sillitoe on behalf of the Wellcome Sanger Institute COVID-19 Surveillance Team |
| EPI_ISL_553893, EPI_ISL_553894 | Lighthouse Lab in Milton Keynes | Wellcome Sanger Institute for the COVID-19 Genomics UK (COG-UK) consortium | The Lighthouse Lab in Milton Keynes and Alex Alderton, Roberto Amato, Sonia Goncalves, Ewan Harrison, David K. Jackson, Ian Johnston, Dominic Kwiatkowski, Cordelia Langford, John Sillitoe on behalf of the Wellcome Sanger Institute COVID-19 Surveillance Team (http://www.sanger.ac.uk/covid-team) |
| EPI_ISL_553895 | Lighthouse Lab in Alderley Park | Wellcome Sanger Institute for the COVID-19 Genomics UK (COG-UK) consortium | The Lighthouse Lab in Alderley Park and Alex Alderton, Roberto Amato, Sonia Goncalves, Ewan Harrison, David K. Jackson, Ian Johnston, Dominic Kwiatkowski, Cordelia Langford, John Sillitoe on behalf of the Wellcome Sanger Institute COVID-19 Surveillance Team |
| EPI_ISL_553896, EPI_ISL_553897, EPI_ISL_553898 | Lighthouse Lab in Milton Keynes | Wellcome Sanger Institute for the COVID-19 Genomics UK (COG-UK) consortium | The Lighthouse Lab in Milton Keynes and Alex Alderton, Roberto Amato, Sonia Goncalves, Ewan Harrison, David K. Jackson, Ian Johnston, Dominic Kwiatkowski, Cordelia Langford, John Sillitoe on behalf of the Wellcome Sanger Institute COVID-19 Surveillance Team (http://www.sanger.ac.uk/covid-team) |
| EPI_ISL_553899, EPI_ISL_553900, EPI_ISL_553901 | Lighthouse Lab in Alderley Park | Wellcome Sanger Institute for the COVID-19 Genomics UK (COG-UK) consortium | The Lighthouse Lab in Alderley Park and Alex Alderton, Roberto Amato, Sonia Goncalves, Ewan Harrison, David K. Jackson, Ian Johnston, Dominic Kwiatkowski, Cordelia Langford, John Sillitoe on behalf of the Wellcome Sanger Institute COVID-19 Surveillance Team |
| EPI_ISL_553902 | Lighthouse Lab in Alderley Park | Wellcome Sanger Institute for the COVID-19 Genomics UK (COG-UK) consortium | The Lighthouse Lab in Alderley Park and Alex Alderton, Roberto Amato, Sonia Goncalves, Ewan Harrison, David K. Jackson, Ian Johnston, Dominic Kwiatkowski, Cordelia Langford, John Sillitoe on behalf of the Wellcome Sanger Institute COVID-19 Surveillance Team (http://www.sanger.ac.uk/covid-team) |
| EPI_ISL_553903, EPI_ISL_553904 | Lighthouse Lab in Alderley Park | Wellcome Sanger Institute for the COVID-19 Genomics UK (COG-UK) consortium | The Lighthouse Lab in Alderley Park and Alex Alderton, Roberto Amato, Sonia Goncalves, Ewan Harrison, David K. Jackson, Ian Johnston, Dominic Kwiatkowski, Cordelia Langford, John Sillitoe on behalf of the Wellcome Sanger Institute COVID-19 Surveillance Team |
| EPI_ISL_553905 | Lighthouse Lab in Milton Keynes | Wellcome Sanger Institute for the COVID-19 Genomics UK (COG-UK) consortium | The Lighthouse Lab in Milton Keynes and Alex Alderton, Roberto Amato, Sonia Goncalves, Ewan Harrison, David K. Jackson, Ian Johnston, Dominic Kwiatkowski, Cordelia Langford, John Sillitoe on behalf of the Wellcome Sanger Institute COVID-19 Surveillance Team (http://www.sanger.ac.uk/covid-team) |
| EPI_ISL_553906, EPI_ISL_553908, EPI_ISL_553909, EPI_ISL_553910, EPI_ISL_553911, EPI_ISL_553912, EPI_ISL_553913, EPI_ISL_553914, EPI_ISL_553915, EPI_ISL_553916 | Lighthouse Lab in Alderley Park | Wellcome Sanger Institute for the COVID-19 Genomics UK (COG-UK) consortium | The Lighthouse Lab in Alderley Park and Alex Alderton, Roberto Amato, Sonia Goncalves, Ewan Harrison, David K. Jackson, Ian Johnston, Dominic Kwiatkowski, Cordelia Langford, John Sillitoe on behalf of the Wellcome Sanger Institute COVID-19 Surveillance Team |
| EPI_ISL_553917 | Lighthouse Lab in Alderley Park | Wellcome Sanger Institute for the COVID-19 Genomics UK (COG-UK) consortium | The Lighthouse Lab in Alderley Park and Alex Alderton, Roberto Amato, Sonia Goncalves, Ewan Harrison, David K. Jackson, Ian Johnston, Dominic Kwiatkowski, Cordelia Langford, John Sillitoe on behalf of the Wellcome Sanger Institute COVID-19 Surveillance Team (http://www.sanger.ac.uk/covid-team) |
| EPI_ISL_553918, EPI_ISL_553919, EPI_ISL_553920, EPI_ISL_553921 | Lighthouse Lab in Alderley Park | Wellcome Sanger Institute for the COVID-19 Genomics UK (COG-UK) consortium | The Lighthouse Lab in Alderley Park and Alex Alderton, Roberto Amato, Sonia Goncalves, Ewan Harrison, David K. Jackson, Ian Johnston, Dominic Kwiatkowski, Cordelia Langford, John Sillitoe on behalf of the Wellcome Sanger Institute COVID-19 Surveillance Team |
| EPI_ISL_553922 | Lighthouse Lab in Alderley Park | Wellcome Sanger Institute for the COVID-19 Genomics UK (COG-UK) consortium | The Lighthouse Lab in Alderley Park and Alex Alderton, Roberto Amato, Sonia Goncalves, Ewan Harrison, David K. Jackson, Ian Johnston, Dominic Kwiatkowski, Cordelia Langford, John Sillitoe on behalf of the Wellcome Sanger Institute COVID-19 Surveillance Team (http://www.sanger.ac.uk/covid-team) |
| EPI_ISL_553923, EPI_ISL_553924 | Lighthouse Lab in Alderley Park | Wellcome Sanger Institute for the COVID-19 Genomics UK (COG-UK) consortium | The Lighthouse Lab in Alderley Park and Alex Alderton, Roberto Amato, Sonia Goncalves, Ewan Harrison, David K. Jackson, Ian Johnston, Dominic Kwiatkowski, Cordelia Langford, John Sillitoe on behalf of the Wellcome Sanger Institute COVID-19 Surveillance Team |
| EPI_ISL_553925 | Lighthouse Lab in Alderley Park | Wellcome Sanger Institute for the COVID-19 Genomics UK (COG-UK) consortium | The Lighthouse Lab in Alderley Park and Alex Alderton, Roberto Amato, Sonia Goncalves, Ewan Harrison, David K. Jackson, Ian Johnston, Dominic Kwiatkowski, Cordelia Langford, John Sillitoe on behalf of the Wellcome Sanger Institute COVID-19 Surveillance Team (http://www.sanger.ac.uk/covid-team) |
| EPI_ISL_553926, EPI_ISL_553927 | Lighthouse Lab in Alderley Park | Wellcome Sanger Institute for the COVID-19 Genomics UK (COG-UK) consortium | The Lighthouse Lab in Alderley Park and Alex Alderton, Roberto Amato, Sonia Goncalves, Ewan Harrison, David K. Jackson, Ian Johnston, Dominic Kwiatkowski, Cordelia Langford, John Sillitoe on behalf of the Wellcome Sanger Institute COVID-19 Surveillance Team |
| EPI_ISL_553928 | Lighthouse Lab in Alderley Park | Wellcome Sanger Institute for the COVID-19 Genomics UK (COG-UK) consortium | The Lighthouse Lab in Alderley Park and Alex Alderton, Roberto Amato, Sonia Goncalves, Ewan Harrison, David K. Jackson, Ian Johnston, Dominic Kwiatkowski, Cordelia Langford, John Sillitoe on behalf of the Wellcome Sanger Institute COVID-19 Surveillance Team (http://www.sanger.ac.uk/covid-team) |
| EPI_ISL_553929 | Lighthouse Lab in Milton Keynes | Wellcome Sanger Institute for the COVID-19 Genomics UK (COG-UK) consortium | The Lighthouse Lab in Alderley Park and Alex Alderton, Roberto Amato, Sonia Goncalves, Ewan Harrison, David K. Jackson, Ian Johnston, Dominic Kwiatkowski, Cordelia Langford, John Sillitoe on behalf of the Wellcome Sanger Institute COVID-19 Surveillance Team |
| EPI_ISL_553930, EPI_ISL_553931 | Lighthouse Lab in Alderley Park | Wellcome Sanger Institute for the COVID-19 Genomics UK (COG-UK) consortium | The Lighthouse Lab in Alderley Park and Alex Alderton, Roberto Amato, Sonia Goncalves, Ewan Harrison, David K. Jackson, Ian Johnston, Dominic Kwiatkowski, Cordelia Langford, John Sillitoe on behalf of the Wellcome Sanger Institute COVID-19 Surveillance Team |
| EPI_ISL_553932 | Lighthouse Lab in Alderley Park | Wellcome Sanger Institute for the COVID-19 Genomics UK (COG-UK) consortium | The Lighthouse Lab in Alderley Park and Alex Alderton, Roberto Amato, Sonia Goncalves, Ewan Harrison, David K. Jackson, Ian Johnston, Dominic Kwiatkowski, Cordelia Langford, John Sillitoe on behalf of the Wellcome Sanger Institute COVID-19 Surveillance Team (http://www.sanger.ac.uk/covid-team) |
| EPI_ISL_553933 | Lighthouse Lab in Alderley Park | Wellcome Sanger Institute for the COVID-19 Genomics UK (COG-UK) consortium | The Lighthouse Lab in Alderley Park and Alex Alderton, Roberto Amato, Sonia Goncalves, Ewan Harrison, David K. Jackson, Ian Johnston, Dominic Kwiatkowski, Cordelia Langford, John Sillitoe on behalf of the Wellcome Sanger Institute COVID-19 Surveillance Team |
| EPI_ISL_553934 | Lighthouse Lab in Milton Keynes | Wellcome Sanger Institute for the COVID-19 Genomics UK (COG-UK) consortium | The Lighthouse Lab in Milton Keynes and Alex Alderton, Roberto Amato, Sonia Goncalves, Ewan Harrison, David K. Jackson, Ian Johnston, Dominic Kwiatkowski, Cordelia Langford, John Sillitoe on behalf of the Wellcome Sanger Institute COVID-19 Surveillance Team (http://www.sanger.ac.uk/covid-team) |
| EPI_ISL_553935 | Lighthouse Lab in Alderley Park | Wellcome Sanger Institute for the COVID-19 Genomics UK (COG-UK) consortium | The Lighthouse Lab in Alderley Park and Alex Alderton, Roberto Amato, Sonia Goncalves, Ewan Harrison, David K. Jackson, Ian Johnston, Dominic Kwiatkowski, Cordelia Langford, John Sillitoe on behalf of the Wellcome Sanger Institute COVID-19 Surveillance Team |
| EPI_ISL_553936, EPI_ISL_553937 | Lighthouse Lab in Alderley Park | Wellcome Sanger Institute for the COVID-19 Genomics UK | The Lighthouse Lab in Alderley Park and Alex Alderton, Roberto Amato, Sonia Goncalves, Ewan Harrison, David K. Jackson, Ian Johnston, Dominic |

| | | | |
|---|---|---|---|
| | | (COG-UK) consortium | Kwiatkowski, Cordelia Langford, John Sillitoe on behalf of the Wellcome Sanger Institute COVID-19 Surveillance Team (http://www.sanger.ac.uk/covid-team) |
| EPI_ISL_553938 | Lighthouse Lab in Alderley Park | Wellcome Sanger Institute for the COVID-19 Genomics UK (COG-UK) consortium | The Lighthouse Lab in Alderley Park and Alex Alderton, Roberto Amato, Sonia Goncalves, Ewan Harrison, David K. Jackson, Ian Johnston, Dominic Kwiatkowski, Cordelia Langford, John Sillitoe on behalf of the Wellcome Sanger Institute COVID-19 Surveillance Team |
| EPI_ISL_553939 | Lighthouse Lab in Milton Keynes | Wellcome Sanger Institute for the COVID-19 Genomics UK (COG-UK) consortium | The Lighthouse Lab in Milton Keynes and Alex Alderton, Roberto Amato, Sonia Goncalves, Ewan Harrison, David K. Jackson, Ian Johnston, Dominic Kwiatkowski, Cordelia Langford, John Sillitoe on behalf of the Wellcome Sanger Institute COVID-19 Surveillance Team (http://www.sanger.ac.uk/covid-team) |
| EPI_ISL_553940, EPI_ISL_553941, EPI_ISL_553942 | Lighthouse Lab in Alderley Park | Wellcome Sanger Institute for the COVID-19 Genomics UK (COG-UK) consortium | The Lighthouse Lab in Alderley Park and Alex Alderton, Roberto Amato, Sonia Goncalves, Ewan Harrison, David K. Jackson, Ian Johnston, Dominic Kwiatkowski, Cordelia Langford, John Sillitoe on behalf of the Wellcome Sanger Institute COVID-19 Surveillance Team |
| EPI_ISL_553943 | Lighthouse Lab in Alderley Park | Wellcome Sanger Institute for the COVID-19 Genomics UK (COG-UK) consortium | The Lighthouse Lab in Alderley Park and Alex Alderton, Roberto Amato, Sonia Goncalves, Ewan Harrison, David K. Jackson, Ian Johnston, Dominic Kwiatkowski, Cordelia Langford, John Sillitoe on behalf of the Wellcome Sanger Institute COVID-19 Surveillance Team (http://www.sanger.ac.uk/covid-team) |
| EPI_ISL_553944, EPI_ISL_553945, EPI_ISL_553947, EPI_ISL_553948, EPI_ISL_553949, EPI_ISL_553950, EPI_ISL_553951 | Lighthouse Lab in Alderley Park | Wellcome Sanger Institute for the COVID-19 Genomics UK (COG-UK) consortium | The Lighthouse Lab in Alderley Park and Alex Alderton, Roberto Amato, Sonia Goncalves, Ewan Harrison, David K. Jackson, Ian Johnston, Dominic Kwiatkowski, Cordelia Langford, John Sillitoe on behalf of the Wellcome Sanger Institute COVID-19 Surveillance Team |
| EPI_ISL_553952 | Lighthouse Lab in Milton Keynes | Wellcome Sanger Institute for the COVID-19 Genomics UK (COG-UK) consortium | The Lighthouse Lab in Milton Keynes and Alex Alderton, Roberto Amato, Sonia Goncalves, Ewan Harrison, David K. Jackson, Ian Johnston, Dominic Kwiatkowski, Cordelia Langford, John Sillitoe on behalf of the Wellcome Sanger Institute COVID-19 Surveillance Team (http://www.sanger.ac.uk/covid-team) |
| EPI_ISL_553953 | Lighthouse Lab in Alderley Park | Wellcome Sanger Institute for the COVID-19 Genomics UK (COG-UK) consortium | The Lighthouse Lab in Alderley Park and Alex Alderton, Roberto Amato, Sonia Goncalves, Ewan Harrison, David K. Jackson, Ian Johnston, Dominic Kwiatkowski, Cordelia Langford, John Sillitoe on behalf of the Wellcome Sanger Institute COVID-19 Surveillance Team |
| EPI_ISL_553954, EPI_ISL_553955, EPI_ISL_553957 | Lighthouse Lab in Milton Keynes | Wellcome Sanger Institute for the COVID-19 Genomics UK (COG-UK) consortium | The Lighthouse Lab in Milton Keynes and Alex Alderton, Roberto Amato, Sonia Goncalves, Ewan Harrison, David K. Jackson, Ian Johnston, Dominic Kwiatkowski, Cordelia Langford, John Sillitoe on behalf of the Wellcome Sanger Institute COVID-19 Surveillance Team (http://www.sanger.ac.uk/covid-team) |
| EPI_ISL_553958, EPI_ISL_553959, EPI_ISL_553960, EPI_ISL_553961, EPI_ISL_553962 | Lighthouse Lab in Alderley Park | Wellcome Sanger Institute for the COVID-19 Genomics UK (COG-UK) consortium | The Lighthouse Lab in Alderley Park and Alex Alderton, Roberto Amato, Sonia Goncalves, Ewan Harrison, David K. Jackson, Ian Johnston, Dominic Kwiatkowski, Cordelia Langford, John Sillitoe on behalf of the Wellcome Sanger Institute COVID-19 Surveillance Team |
| EPI_ISL_553963 | Lighthouse Lab in Alderley Park | Wellcome Sanger Institute for the COVID-19 Genomics UK (COG-UK) consortium | The Lighthouse Lab in Alderley Park and Alex Alderton, Roberto Amato, Sonia Goncalves, Ewan Harrison, David K. Jackson, Ian Johnston, Dominic Kwiatkowski, Cordelia Langford, John Sillitoe on behalf of the Wellcome Sanger Institute COVID-19 Surveillance Team (http://www.sanger.ac.uk/covid-team) |
| EPI_ISL_553964, EPI_ISL_553966, EPI_ISL_553967, EPI_ISL_553968, EPI_ISL_553969, EPI_ISL_553970, EPI_ISL_553971 | Lighthouse Lab in Alderley Park | Wellcome Sanger Institute for the COVID-19 Genomics UK (COG-UK) consortium | The Lighthouse Lab in Alderley Park and Alex Alderton, Roberto Amato, Sonia Goncalves, Ewan Harrison, David K. Jackson, Ian Johnston, Dominic Kwiatkowski, Cordelia Langford, John Sillitoe on behalf of the Wellcome Sanger Institute COVID-19 Surveillance Team |
| EPI_ISL_553972 | Lighthouse Lab in Milton Keynes | Wellcome Sanger Institute for the COVID-19 Genomics UK (COG-UK) consortium | The Lighthouse Lab in Milton Keynes and Alex Alderton, Roberto Amato, Sonia Goncalves, Ewan Harrison, David K. Jackson, Ian Johnston, Dominic Kwiatkowski, Cordelia Langford, John Sillitoe on behalf of the Wellcome Sanger Institute COVID-19 Surveillance Team (http://www.sanger.ac.uk/covid-team) |
| EPI_ISL_553973 | Lighthouse Lab in Alderley Park | Wellcome Sanger Institute for the COVID-19 Genomics UK (COG-UK) consortium | The Lighthouse Lab in Alderley Park and Alex Alderton, Roberto Amato, Sonia Goncalves, Ewan Harrison, David K. Jackson, Ian Johnston, Dominic Kwiatkowski, Cordelia Langford, John Sillitoe on behalf of the Wellcome Sanger Institute COVID-19 Surveillance Team |
| EPI_ISL_553974 | Lighthouse Lab in Alderley Park | Wellcome Sanger Institute for the COVID-19 Genomics UK (COG-UK) consortium | The Lighthouse Lab in Alderley Park and Alex Alderton, Roberto Amato, Sonia Goncalves, Ewan Harrison, David K. Jackson, Ian Johnston, Dominic Kwiatkowski, Cordelia Langford, John Sillitoe on behalf of the Wellcome Sanger Institute COVID-19 Surveillance Team (http://www.sanger.ac.uk/covid-team) |
| EPI_ISL_553975, EPI_ISL_553976, EPI_ISL_553977, EPI_ISL_553978, EPI_ISL_553979, EPI_ISL_553980, EPI_ISL_553981, EPI_ISL_553982 | Lighthouse Lab in Alderley Park | Wellcome Sanger Institute for the COVID-19 Genomics UK (COG-UK) consortium | The Lighthouse Lab in Alderley Park and Alex Alderton, Roberto Amato, Sonia Goncalves, Ewan Harrison, David K. Jackson, Ian Johnston, Dominic Kwiatkowski, Cordelia Langford, John Sillitoe on behalf of the Wellcome Sanger Institute COVID-19 Surveillance Team |
| EPI_ISL_553983 | Lighthouse Lab in Milton Keynes | Wellcome Sanger Institute for the COVID-19 Genomics UK (COG-UK) consortium | The Lighthouse Lab in Milton Keynes and Alex Alderton, Roberto Amato, Sonia Goncalves, Ewan Harrison, David K. Jackson, Ian Johnston, Dominic Kwiatkowski, Cordelia Langford, John Sillitoe on behalf of the Wellcome Sanger Institute COVID-19 Surveillance Team (http://www.sanger.ac.uk/covid-team) |
| EPI_ISL_553984, EPI_ISL_553986 | Lighthouse Lab in Alderley Park | Wellcome Sanger Institute for the COVID-19 Genomics UK (COG-UK) consortium | The Lighthouse Lab in Alderley Park and Alex Alderton, Roberto Amato, Sonia Goncalves, Ewan Harrison, David K. Jackson, Ian Johnston, Dominic Kwiatkowski, Cordelia Langford, John Sillitoe on behalf of the Wellcome Sanger Institute COVID-19 Surveillance Team |
| EPI_ISL_553987 | Lighthouse Lab in Milton Keynes | Wellcome Sanger Institute for the COVID-19 Genomics UK (COG-UK) consortium | The Lighthouse Lab in Milton Keynes and Alex Alderton, Roberto Amato, Sonia Goncalves, Ewan Harrison, David K. Jackson, Ian Johnston, Dominic Kwiatkowski, Cordelia Langford, John Sillitoe on behalf of the Wellcome Sanger Institute COVID-19 Surveillance Team (http://www.sanger.ac.uk/covid-team) |
| EPI_ISL_553988 | Lighthouse Lab in Alderley Park | Wellcome Sanger Institute for the COVID-19 Genomics UK (COG-UK) consortium | The Lighthouse Lab in Alderley Park and Alex Alderton, Roberto Amato, Sonia Goncalves, Ewan Harrison, David K. Jackson, Ian Johnston, Dominic Kwiatkowski, Cordelia Langford, John Sillitoe on behalf of the Wellcome Sanger Institute COVID-19 Surveillance Team |
| EPI_ISL_553989 | Lighthouse Lab in Milton Keynes | Wellcome Sanger Institute for the COVID-19 Genomics UK (COG-UK) consortium | The Lighthouse Lab in Milton Keynes and Alex Alderton, Roberto Amato, Sonia Goncalves, Ewan Harrison, David K. Jackson, Ian Johnston, Dominic Kwiatkowski, Cordelia Langford, John Sillitoe on behalf of the Wellcome Sanger Institute COVID-19 Surveillance Team (http://www.sanger.ac.uk/covid-team) |
| EPI_ISL_553990, EPI_ISL_553991, EPI_ISL_553992 | Lighthouse Lab in Alderley Park | Wellcome Sanger Institute for the COVID-19 Genomics UK (COG-UK) consortium | The Lighthouse Lab in Alderley Park and Alex Alderton, Roberto Amato, Sonia Goncalves, Ewan Harrison, David K. Jackson, Ian Johnston, Dominic Kwiatkowski, Cordelia Langford, John Sillitoe on behalf of the Wellcome Sanger Institute COVID-19 Surveillance Team |
| EPI_ISL_553993, EPI_ISL_553994 | Lighthouse Lab in Milton Keynes | Wellcome Sanger Institute for the COVID-19 Genomics UK (COG-UK) consortium | The Lighthouse Lab in Milton Keynes and Alex Alderton, Roberto Amato, Sonia Goncalves, Ewan Harrison, David K. Jackson, Ian Johnston, Dominic Kwiatkowski, Cordelia Langford, John Sillitoe on behalf of the Wellcome Sanger Institute COVID-19 Surveillance Team (http://www.sanger.ac.uk/covid-team) |
| EPI_ISL_553995, EPI_ISL_553996, EPI_ISL_553997, EPI_ISL_553998, EPI_ISL_553999, EPI_ISL_554000, EPI_ISL_554001 | Lighthouse Lab in Alderley Park | Wellcome Sanger Institute for the COVID-19 Genomics UK (COG-UK) consortium | The Lighthouse Lab in Alderley Park and Alex Alderton, Roberto Amato, Sonia Goncalves, Ewan Harrison, David K. Jackson, Ian Johnston, Dominic Kwiatkowski, Cordelia Langford, John Sillitoe on behalf of the Wellcome Sanger Institute COVID-19 Surveillance Team |
| EPI_ISL_554002 | Lighthouse Lab in Alderley Park | Wellcome Sanger Institute for the COVID-19 Genomics UK (COG-UK) consortium | The Lighthouse Lab in Alderley Park and Alex Alderton, Roberto Amato, Sonia Goncalves, Ewan Harrison, David K. Jackson, Ian Johnston, Dominic Kwiatkowski, Cordelia Langford, John Sillitoe on behalf of the Wellcome Sanger Institute COVID-19 Surveillance Team (http://www.sanger.ac.uk/covid-team) |
| EPI_ISL_554004, EPI_ISL_554006 | Lighthouse Lab in Alderley Park | Wellcome Sanger Institute for the COVID-19 Genomics UK (COG-UK) consortium | The Lighthouse Lab in Alderley Park and Alex Alderton, Roberto Amato, Sonia Goncalves, Ewan Harrison, David K. Jackson, Ian Johnston, Dominic Kwiatkowski, Cordelia Langford, John Sillitoe on behalf of the Wellcome Sanger Institute COVID-19 Surveillance Team |
| EPI_ISL_554007 | Lighthouse Lab in Alderley Park | Wellcome Sanger Institute for the COVID-19 Genomics UK | The Lighthouse Lab in Alderley Park and Alex Alderton, Roberto Amato, Sonia Goncalves, Ewan Harrison, David K. Jackson, Ian Johnston, Dominic |

| | | | |
|---|---|---|---|
| | | (COG-UK) consortium | Kwiatkowski, Cordelia Langford, John Sillitoe on behalf of the Wellcome Sanger Institute COVID-19 Surveillance Team (http://www.sanger.ac.uk/covid-team) |
| EPI_ISL_554008 | Lighthouse Lab in Alderley Park | Wellcome Sanger Institute for the COVID-19 Genomics UK (COG-UK) consortium | The Lighthouse Lab in Alderley Park and Alex Alderton, Roberto Amato, Sonia Goncalves, Ewan Harrison, David K. Jackson, Ian Johnston, Dominic Kwiatkowski, Cordelia Langford, John Sillitoe on behalf of the Wellcome Sanger Institute COVID-19 Surveillance Team |
| EPI_ISL_554009 | Lighthouse Lab in Milton Keynes | Wellcome Sanger Institute for the COVID-19 Genomics UK (COG-UK) consortium | The Lighthouse Lab in Milton Keynes and Alex Alderton, Roberto Amato, Sonia Goncalves, Ewan Harrison, David K. Jackson, Ian Johnston, Dominic Kwiatkowski, Cordelia Langford, John Sillitoe on behalf of the Wellcome Sanger Institute COVID-19 Surveillance Team (http://www.sanger.ac.uk/covid-team) |
| EPI_ISL_554010, EPI_ISL_554011, EPI_ISL_554012, EPI_ISL_554013 | Lighthouse Lab in Alderley Park | Wellcome Sanger Institute for the COVID-19 Genomics UK (COG-UK) consortium | The Lighthouse Lab in Alderley Park and Alex Alderton, Roberto Amato, Sonia Goncalves, Ewan Harrison, David K. Jackson, Ian Johnston, Dominic Kwiatkowski, Cordelia Langford, John Sillitoe on behalf of the Wellcome Sanger Institute COVID-19 Surveillance Team |
| EPI_ISL_554014 | Lighthouse Lab in Alderley Park | Wellcome Sanger Institute for the COVID-19 Genomics UK (COG-UK) consortium | The Lighthouse Lab in Alderley Park and Alex Alderton, Roberto Amato, Sonia Goncalves, Ewan Harrison, David K. Jackson, Ian Johnston, Dominic Kwiatkowski, Cordelia Langford, John Sillitoe on behalf of the Wellcome Sanger Institute COVID-19 Surveillance Team (http://www.sanger.ac.uk/covid-team) |
| EPI_ISL_554015 | Lighthouse Lab in Milton Keynes | Wellcome Sanger Institute for the COVID-19 Genomics UK (COG-UK) consortium | The Lighthouse Lab in Milton Keynes and Alex Alderton, Roberto Amato, Sonia Goncalves, Ewan Harrison, David K. Jackson, Ian Johnston, Dominic Kwiatkowski, Cordelia Langford, John Sillitoe on behalf of the Wellcome Sanger Institute COVID-19 Surveillance Team (http://www.sanger.ac.uk/covid-team) |
| EPI_ISL_554016, EPI_ISL_554017, EPI_ISL_554018, EPI_ISL_554019, EPI_ISL_554020 | Lighthouse Lab in Alderley Park | Wellcome Sanger Institute for the COVID-19 Genomics UK (COG-UK) consortium | The Lighthouse Lab in Alderley Park and Alex Alderton, Roberto Amato, Sonia Goncalves, Ewan Harrison, David K. Jackson, Ian Johnston, Dominic Kwiatkowski, Cordelia Langford, John Sillitoe on behalf of the Wellcome Sanger Institute COVID-19 Surveillance Team |
| EPI_ISL_554021 | Lighthouse Lab in Alderley Park | Wellcome Sanger Institute for the COVID-19 Genomics UK (COG-UK) consortium | The Lighthouse Lab in Alderley Park and Alex Alderton, Roberto Amato, Sonia Goncalves, Ewan Harrison, David K. Jackson, Ian Johnston, Dominic Kwiatkowski, Cordelia Langford, John Sillitoe on behalf of the Wellcome Sanger Institute COVID-19 Surveillance Team (http://www.sanger.ac.uk/covid-team) |
| EPI_ISL_554022, EPI_ISL_554023 | Lighthouse Lab in Alderley Park | Wellcome Sanger Institute for the COVID-19 Genomics UK (COG-UK) consortium | The Lighthouse Lab in Alderley Park and Alex Alderton, Roberto Amato, Sonia Goncalves, Ewan Harrison, David K. Jackson, Ian Johnston, Dominic Kwiatkowski, Cordelia Langford, John Sillitoe on behalf of the Wellcome Sanger Institute COVID-19 Surveillance Team |
| EPI_ISL_554024, EPI_ISL_554025, EPI_ISL_554026 | Lighthouse Lab in Milton Keynes | Wellcome Sanger Institute for the COVID-19 Genomics UK (COG-UK) consortium | The Lighthouse Lab in Milton Keynes and Alex Alderton, Roberto Amato, Sonia Goncalves, Ewan Harrison, David K. Jackson, Ian Johnston, Dominic Kwiatkowski, Cordelia Langford, John Sillitoe on behalf of the Wellcome Sanger Institute COVID-19 Surveillance Team (http://www.sanger.ac.uk/covid-team) |
| EPI_ISL_554027 | Lighthouse Lab in Alderley Park | Wellcome Sanger Institute for the COVID-19 Genomics UK (COG-UK) consortium | The Lighthouse Lab in Alderley Park and Alex Alderton, Roberto Amato, Sonia Goncalves, Ewan Harrison, David K. Jackson, Ian Johnston, Dominic Kwiatkowski, Cordelia Langford, John Sillitoe on behalf of the Wellcome Sanger Institute COVID-19 Surveillance Team |
| EPI_ISL_554028 | Lighthouse Lab in Milton Keynes | Wellcome Sanger Institute for the COVID-19 Genomics UK (COG-UK) consortium | The Lighthouse Lab in Milton Keynes and Alex Alderton, Roberto Amato, Sonia Goncalves, Ewan Harrison, David K. Jackson, Ian Johnston, Dominic Kwiatkowski, Cordelia Langford, John Sillitoe on behalf of the Wellcome Sanger Institute COVID-19 Surveillance Team (http://www.sanger.ac.uk/covid-team) |
| EPI_ISL_554029, EPI_ISL_554030, EPI_ISL_554031, EPI_ISL_554032, EPI_ISL_554033, EPI_ISL_554034, EPI_ISL_554035, EPI_ISL_554036, EPI_ISL_554037, EPI_ISL_554038, EPI_ISL_554039, EPI_ISL_554040, EPI_ISL_554041 | | | |
| see above | Lighthouse Lab in Alderley Park | Wellcome Sanger Institute for the COVID-19 Genomics UK (COG-UK) consortium | The Lighthouse Lab in Alderley Park and Alex Alderton, Roberto Amato, Sonia Goncalves, Ewan Harrison, David K. Jackson, Ian Johnston, Dominic Kwiatkowski, Cordelia Langford, John Sillitoe on behalf of the Wellcome Sanger Institute COVID-19 Surveillance Team |
| EPI_ISL_554042 | Lighthouse Lab in Milton Keynes | Wellcome Sanger Institute for the COVID-19 Genomics UK (COG-UK) consortium | The Lighthouse Lab in Alderley Park and Alex Alderton, Roberto Amato, Sonia Goncalves, Ewan Harrison, David K. Jackson, Ian Johnston, Dominic Kwiatkowski, Cordelia Langford, John Sillitoe on behalf of the Wellcome Sanger Institute COVID-19 Surveillance Team |
| EPI_ISL_554043, EPI_ISL_554044, EPI_ISL_554045, EPI_ISL_554046, EPI_ISL_554047 | Lighthouse Lab in Alderley Park | Wellcome Sanger Institute for the COVID-19 Genomics UK (COG-UK) consortium | The Lighthouse Lab in Alderley Park and Alex Alderton, Roberto Amato, Sonia Goncalves, Ewan Harrison, David K. Jackson, Ian Johnston, Dominic Kwiatkowski, Cordelia Langford, John Sillitoe on behalf of the Wellcome Sanger Institute COVID-19 Surveillance Team |
| EPI_ISL_554048 | Lighthouse Lab in Milton Keynes | Wellcome Sanger Institute for the COVID-19 Genomics UK (COG-UK) consortium | The Lighthouse Lab in Milton Keynes and Alex Alderton, Roberto Amato, Sonia Goncalves, Ewan Harrison, David K. Jackson, Ian Johnston, Dominic Kwiatkowski, Cordelia Langford, John Sillitoe on behalf of the Wellcome Sanger Institute COVID-19 Surveillance Team (http://www.sanger.ac.uk/covid-team) |
| EPI_ISL_554049, EPI_ISL_554050, EPI_ISL_554051, EPI_ISL_554052, EPI_ISL_554053 | Lighthouse Lab in Alderley Park | Wellcome Sanger Institute for the COVID-19 Genomics UK (COG-UK) consortium | The Lighthouse Lab in Alderley Park and Alex Alderton, Roberto Amato, Sonia Goncalves, Ewan Harrison, David K. Jackson, Ian Johnston, Dominic Kwiatkowski, Cordelia Langford, John Sillitoe on behalf of the Wellcome Sanger Institute COVID-19 Surveillance Team |
| EPI_ISL_554054 | Lighthouse Lab in Milton Keynes | Wellcome Sanger Institute for the COVID-19 Genomics UK (COG-UK) consortium | The Lighthouse Lab in Milton Keynes and Alex Alderton, Roberto Amato, Sonia Goncalves, Ewan Harrison, David K. Jackson, Ian Johnston, Dominic Kwiatkowski, Cordelia Langford, John Sillitoe on behalf of the Wellcome Sanger Institute COVID-19 Surveillance Team (http://www.sanger.ac.uk/covid-team) |
| EPI_ISL_554055, EPI_ISL_554056 | Lighthouse Lab in Alderley Park | Wellcome Sanger Institute for the COVID-19 Genomics UK (COG-UK) consortium | The Lighthouse Lab in Alderley Park and Alex Alderton, Roberto Amato, Sonia Goncalves, Ewan Harrison, David K. Jackson, Ian Johnston, Dominic Kwiatkowski, Cordelia Langford, John Sillitoe on behalf of the Wellcome Sanger Institute COVID-19 Surveillance Team (http://www.sanger.ac.uk/covid-team) |
| EPI_ISL_554057, EPI_ISL_554058 | Lighthouse Lab in Milton Keynes | Wellcome Sanger Institute for the COVID-19 Genomics UK (COG-UK) consortium | The Lighthouse Lab in Milton Keynes and Alex Alderton, Roberto Amato, Sonia Goncalves, Ewan Harrison, David K. Jackson, Ian Johnston, Dominic Kwiatkowski, Cordelia Langford, John Sillitoe on behalf of the Wellcome Sanger Institute COVID-19 Surveillance Team (http://www.sanger.ac.uk/covid-team) |
| EPI_ISL_554059, EPI_ISL_554060, EPI_ISL_554061, EPI_ISL_554062 | Lighthouse Lab in Alderley Park | Wellcome Sanger Institute for the COVID-19 Genomics UK (COG-UK) consortium | The Lighthouse Lab in Alderley Park and Alex Alderton, Roberto Amato, Sonia Goncalves, Ewan Harrison, David K. Jackson, Ian Johnston, Dominic Kwiatkowski, Cordelia Langford, John Sillitoe on behalf of the Wellcome Sanger Institute COVID-19 Surveillance Team |
| EPI_ISL_554063 | Lighthouse Lab in Alderley Park | Wellcome Sanger Institute for the COVID-19 Genomics UK (COG-UK) consortium | The Lighthouse Lab in Alderley Park and Alex Alderton, Roberto Amato, Sonia Goncalves, Ewan Harrison, David K. Jackson, Ian Johnston, Dominic Kwiatkowski, Cordelia Langford, John Sillitoe on behalf of the Wellcome Sanger Institute COVID-19 Surveillance Team (http://www.sanger.ac.uk/covid-team) |
| EPI_ISL_554064 | Lighthouse Lab in Alderley Park | Wellcome Sanger Institute for the COVID-19 Genomics UK (COG-UK) consortium | The Lighthouse Lab in Alderley Park and Alex Alderton, Roberto Amato, Sonia Goncalves, Ewan Harrison, David K. Jackson, Ian Johnston, Dominic Kwiatkowski, Cordelia Langford, John Sillitoe on behalf of the Wellcome Sanger Institute COVID-19 Surveillance Team |
| EPI_ISL_554065 | Lighthouse Lab in Milton Keynes | Wellcome Sanger Institute for the COVID-19 Genomics UK (COG-UK) consortium | The Lighthouse Lab in Milton Keynes and Alex Alderton, Roberto Amato, Sonia Goncalves, Ewan Harrison, David K. Jackson, Ian Johnston, Dominic Kwiatkowski, Cordelia Langford, John Sillitoe on behalf of the Wellcome Sanger Institute COVID-19 Surveillance Team (http://www.sanger.ac.uk/covid-team) |
| EPI_ISL_554066, EPI_ISL_554067, EPI_ISL_554068 | Lighthouse Lab in Alderley Park | Wellcome Sanger Institute for the COVID-19 Genomics UK (COG-UK) consortium | The Lighthouse Lab in Alderley Park and Alex Alderton, Roberto Amato, Sonia Goncalves, Ewan Harrison, David K. Jackson, Ian Johnston, Dominic Kwiatkowski, Cordelia Langford, John Sillitoe on behalf of the Wellcome Sanger Institute COVID-19 Surveillance Team |
| EPI_ISL_554069 | Lighthouse Lab in Alderley Park | Wellcome Sanger Institute for the COVID-19 Genomics UK (COG-UK) consortium | The Lighthouse Lab in Alderley Park and Alex Alderton, Roberto Amato, Sonia Goncalves, Ewan Harrison, David K. Jackson, Ian Johnston, Dominic Kwiatkowski, Cordelia Langford, John Sillitoe on behalf of the Wellcome Sanger Institute COVID-19 Surveillance Team (http://www.sanger.ac.uk/covid-team) |
| EPI_ISL_554070, EPI_ISL_554071, EPI_ISL_554072, EPI_ISL_554073 | Lighthouse Lab in Alderley Park | Wellcome Sanger Institute for the COVID-19 Genomics UK (COG-UK) consortium | The Lighthouse Lab in Alderley Park and Alex Alderton, Roberto Amato, Sonia Goncalves, Ewan Harrison, David K. Jackson, Ian Johnston, Dominic Kwiatkowski, Cordelia Langford, John Sillitoe on behalf of the Wellcome Sanger Institute COVID-19 Surveillance Team |
| EPI_ISL_554074 | Lighthouse Lab in Alderley Park | Wellcome Sanger Institute for the COVID-19 Genomics UK (COG-UK) consortium | The Lighthouse Lab in Alderley Park and Alex Alderton, Roberto Amato, Sonia Goncalves, Ewan Harrison, David K. Jackson, Ian Johnston, Dominic Kwiatkowski, Cordelia Langford, John Sillitoe on behalf of the Wellcome Sanger Institute COVID-19 Surveillance Team |

| | | | |
|---|---|---|---|
| | | | (http://www.sanger.ac.uk/covid-team) |
| EPI_ISL_554075, EPI_ISL_554076, EPI_ISL_554077, EPI_ISL_554078, EPI_ISL_554079 | Lighthouse Lab in Alderley Park | Wellcome Sanger Institute for the COVID-19 Genomics UK (COG-UK) consortium | The Lighthouse Lab in Alderley Park and Alex Alderton, Roberto Amato, Sonia Goncalves, Ewan Harrison, David K. Jackson, Ian Johnston, Dominic Kwiatkowski, Cordelia Langford, John Sillitoe on behalf of the Wellcome Sanger Institute COVID-19 Surveillance Team |
| EPI_ISL_554080 | Lighthouse Lab in Milton Keynes | Wellcome Sanger Institute for the COVID-19 Genomics UK (COG-UK) consortium | The Lighthouse Lab in Milton Keynes and Alex Alderton, Roberto Amato, Sonia Goncalves, Ewan Harrison, David K. Jackson, Ian Johnston, Dominic Kwiatkowski, Cordelia Langford, John Sillitoe on behalf of the Wellcome Sanger Institute COVID-19 Surveillance Team (http://www.sanger.ac.uk/covid-team) |
| EPI_ISL_554081, EPI_ISL_554082, EPI_ISL_554083, EPI_ISL_554084, EPI_ISL_554085, EPI_ISL_554086, EPI_ISL_554087, EPI_ISL_554088, EPI_ISL_554089, EPI_ISL_554090, EPI_ISL_554091, EPI_ISL_554092, EPI_ISL_554093, EPI_ISL_554094, EPI_ISL_554095, EPI_ISL_554096 | | | |
| see above | Lighthouse Lab in Alderley Park | Wellcome Sanger Institute for the COVID-19 Genomics UK (COG-UK) consortium | The Lighthouse Lab in Alderley Park and Alex Alderton, Roberto Amato, Sonia Goncalves, Ewan Harrison, David K. Jackson, Ian Johnston, Dominic Kwiatkowski, Cordelia Langford, John Sillitoe on behalf of the Wellcome Sanger Institute COVID-19 Surveillance Team |
| EPI_ISL_554097 | Lighthouse Lab in Milton Keynes | Wellcome Sanger Institute for the COVID-19 Genomics UK (COG-UK) consortium | The Lighthouse Lab in Milton Keynes and Alex Alderton, Roberto Amato, Sonia Goncalves, Ewan Harrison, David K. Jackson, Ian Johnston, Dominic Kwiatkowski, Cordelia Langford, John Sillitoe on behalf of the Wellcome Sanger Institute COVID-19 Surveillance Team (http://www.sanger.ac.uk/covid-team) |
| EPI_ISL_554098 | Lighthouse Lab in Alderley Park | Wellcome Sanger Institute for the COVID-19 Genomics UK (COG-UK) consortium | The Lighthouse Lab in Alderley Park and Alex Alderton, Roberto Amato, Sonia Goncalves, Ewan Harrison, David K. Jackson, Ian Johnston, Dominic Kwiatkowski, Cordelia Langford, John Sillitoe on behalf of the Wellcome Sanger Institute COVID-19 Surveillance Team |
| EPI_ISL_554099, EPI_ISL_554100, EPI_ISL_554101 | Lighthouse Lab in Milton Keynes | Wellcome Sanger Institute for the COVID-19 Genomics UK (COG-UK) consortium | The Lighthouse Lab in Milton Keynes and Alex Alderton, Roberto Amato, Sonia Goncalves, Ewan Harrison, David K. Jackson, Ian Johnston, Dominic Kwiatkowski, Cordelia Langford, John Sillitoe on behalf of the Wellcome Sanger Institute COVID-19 Surveillance Team (http://www.sanger.ac.uk/covid-team) |
| EPI_ISL_554102, EPI_ISL_554103 | Lighthouse Lab in Alderley Park | Wellcome Sanger Institute for the COVID-19 Genomics UK (COG-UK) consortium | The Lighthouse Lab in Alderley Park and Alex Alderton, Roberto Amato, Sonia Goncalves, Ewan Harrison, David K. Jackson, Ian Johnston, Dominic Kwiatkowski, Cordelia Langford, John Sillitoe on behalf of the Wellcome Sanger Institute COVID-19 Surveillance Team |
| EPI_ISL_554104, EPI_ISL_554105 | Lighthouse Lab in Alderley Park | Wellcome Sanger Institute for the COVID-19 Genomics UK (COG-UK) consortium | The Lighthouse Lab in Alderley Park and Alex Alderton, Roberto Amato, Sonia Goncalves, Ewan Harrison, David K. Jackson, Ian Johnston, Dominic Kwiatkowski, Cordelia Langford, John Sillitoe on behalf of the Wellcome Sanger Institute COVID-19 Surveillance Team (http://www.sanger.ac.uk/covid-team) |
| EPI_ISL_554106, EPI_ISL_554107, EPI_ISL_554108, EPI_ISL_554109, EPI_ISL_554110, EPI_ISL_554111 | Lighthouse Lab in Alderley Park | Wellcome Sanger Institute for the COVID-19 Genomics UK (COG-UK) consortium | The Lighthouse Lab in Alderley Park and Alex Alderton, Roberto Amato, Sonia Goncalves, Ewan Harrison, David K. Jackson, Ian Johnston, Dominic Kwiatkowski, Cordelia Langford, John Sillitoe on behalf of the Wellcome Sanger Institute COVID-19 Surveillance Team |
| EPI_ISL_554112 | Lighthouse Lab in Alderley Park | Wellcome Sanger Institute for the COVID-19 Genomics UK (COG-UK) consortium | The Lighthouse Lab in Alderley Park and Alex Alderton, Roberto Amato, Sonia Goncalves, Ewan Harrison, David K. Jackson, Ian Johnston, Dominic Kwiatkowski, Cordelia Langford, John Sillitoe on behalf of the Wellcome Sanger Institute COVID-19 Surveillance Team (http://www.sanger.ac.uk/covid-team) |
| EPI_ISL_554113, EPI_ISL_554114, EPI_ISL_554115, EPI_ISL_554116, EPI_ISL_554117 | Lighthouse Lab in Alderley Park | Wellcome Sanger Institute for the COVID-19 Genomics UK (COG-UK) consortium | The Lighthouse Lab in Alderley Park and Alex Alderton, Roberto Amato, Sonia Goncalves, Ewan Harrison, David K. Jackson, Ian Johnston, Dominic Kwiatkowski, Cordelia Langford, John Sillitoe on behalf of the Wellcome Sanger Institute COVID-19 Surveillance Team |
| EPI_ISL_554118 | Lighthouse Lab in Milton Keynes | Wellcome Sanger Institute for the COVID-19 Genomics UK (COG-UK) consortium | The Lighthouse Lab in Milton Keynes and Alex Alderton, Roberto Amato, Sonia Goncalves, Ewan Harrison, David K. Jackson, Ian Johnston, Dominic Kwiatkowski, Cordelia Langford, John Sillitoe on behalf of the Wellcome Sanger Institute COVID-19 Surveillance Team (http://www.sanger.ac.uk/covid-team) |
| EPI_ISL_554120 | Lighthouse Lab in Alderley Park | Wellcome Sanger Institute for the COVID-19 Genomics UK (COG-UK) consortium | The Lighthouse Lab in Alderley Park and Alex Alderton, Roberto Amato, Sonia Goncalves, Ewan Harrison, David K. Jackson, Ian Johnston, Dominic Kwiatkowski, Cordelia Langford, John Sillitoe on behalf of the Wellcome Sanger Institute COVID-19 Surveillance Team |
| EPI_ISL_554121 | Lighthouse Lab in Milton Keynes | Wellcome Sanger Institute for the COVID-19 Genomics UK (COG-UK) consortium | The Lighthouse Lab in Milton Keynes and Alex Alderton, Roberto Amato, Sonia Goncalves, Ewan Harrison, David K. Jackson, Ian Johnston, Dominic Kwiatkowski, Cordelia Langford, John Sillitoe on behalf of the Wellcome Sanger Institute COVID-19 Surveillance Team (http://www.sanger.ac.uk/covid-team) |
| EPI_ISL_554122 | Lighthouse Lab in Alderley Park | Wellcome Sanger Institute for the COVID-19 Genomics UK (COG-UK) consortium | The Lighthouse Lab in Alderley Park and Alex Alderton, Roberto Amato, Sonia Goncalves, Ewan Harrison, David K. Jackson, Ian Johnston, Dominic Kwiatkowski, Cordelia Langford, John Sillitoe on behalf of the Wellcome Sanger Institute COVID-19 Surveillance Team |
| EPI_ISL_554123, EPI_ISL_554124 | Lighthouse Lab in Milton Keynes | Wellcome Sanger Institute for the COVID-19 Genomics UK (COG-UK) consortium | The Lighthouse Lab in Milton Keynes and Alex Alderton, Roberto Amato, Sonia Goncalves, Ewan Harrison, David K. Jackson, Ian Johnston, Dominic Kwiatkowski, Cordelia Langford, John Sillitoe on behalf of the Wellcome Sanger Institute COVID-19 Surveillance Team (http://www.sanger.ac.uk/covid-team) |
| EPI_ISL_554126 | Lighthouse Lab in Alderley Park | Wellcome Sanger Institute for the COVID-19 Genomics UK (COG-UK) consortium | The Lighthouse Lab in Alderley Park and Alex Alderton, Roberto Amato, Sonia Goncalves, Ewan Harrison, David K. Jackson, Ian Johnston, Dominic Kwiatkowski, Cordelia Langford, John Sillitoe on behalf of the Wellcome Sanger Institute COVID-19 Surveillance Team |
| EPI_ISL_554127 | Lighthouse Lab in Milton Keynes | Wellcome Sanger Institute for the COVID-19 Genomics UK (COG-UK) consortium | The Lighthouse Lab in Milton Keynes and Alex Alderton, Roberto Amato, Sonia Goncalves, Ewan Harrison, David K. Jackson, Ian Johnston, Dominic Kwiatkowski, Cordelia Langford, John Sillitoe on behalf of the Wellcome Sanger Institute COVID-19 Surveillance Team (http://www.sanger.ac.uk/covid-team) |
| EPI_ISL_554129, EPI_ISL_554130, EPI_ISL_554131 | Lighthouse Lab in Alderley Park | Wellcome Sanger Institute for the COVID-19 Genomics UK (COG-UK) consortium | The Lighthouse Lab in Alderley Park and Alex Alderton, Roberto Amato, Sonia Goncalves, Ewan Harrison, David K. Jackson, Ian Johnston, Dominic Kwiatkowski, Cordelia Langford, John Sillitoe on behalf of the Wellcome Sanger Institute COVID-19 Surveillance Team |
| EPI_ISL_554132, EPI_ISL_554133, EPI_ISL_554134 | Lighthouse Lab in Milton Keynes | Wellcome Sanger Institute for the COVID-19 Genomics UK (COG-UK) consortium | The Lighthouse Lab in Milton Keynes and Alex Alderton, Roberto Amato, Sonia Goncalves, Ewan Harrison, David K. Jackson, Ian Johnston, Dominic Kwiatkowski, Cordelia Langford, John Sillitoe on behalf of the Wellcome Sanger Institute COVID-19 Surveillance Team (http://www.sanger.ac.uk/covid-team) |
| EPI_ISL_554136 | Lighthouse Lab in Alderley Park | Wellcome Sanger Institute for the COVID-19 Genomics UK (COG-UK) consortium | The Lighthouse Lab in Alderley Park and Alex Alderton, Roberto Amato, Sonia Goncalves, Ewan Harrison, David K. Jackson, Ian Johnston, Dominic Kwiatkowski, Cordelia Langford, John Sillitoe on behalf of the Wellcome Sanger Institute COVID-19 Surveillance Team |
| EPI_ISL_554138, EPI_ISL_554139 | Lighthouse Lab in Milton Keynes | Wellcome Sanger Institute for the COVID-19 Genomics UK (COG-UK) consortium | The Lighthouse Lab in Milton Keynes and Alex Alderton, Roberto Amato, Sonia Goncalves, Ewan Harrison, David K. Jackson, Ian Johnston, Dominic Kwiatkowski, Cordelia Langford, John Sillitoe on behalf of the Wellcome Sanger Institute COVID-19 Surveillance Team (http://www.sanger.ac.uk/covid-team) |
| EPI_ISL_554140 | Lighthouse Lab in Alderley Park | Wellcome Sanger Institute for the COVID-19 Genomics UK (COG-UK) consortium | The Lighthouse Lab in Alderley Park and Alex Alderton, Roberto Amato, Sonia Goncalves, Ewan Harrison, David K. Jackson, Ian Johnston, Dominic Kwiatkowski, Cordelia Langford, John Sillitoe on behalf of the Wellcome Sanger Institute COVID-19 Surveillance Team |
| EPI_ISL_554142 | Lighthouse Lab in Milton Keynes | Wellcome Sanger Institute for the COVID-19 Genomics UK (COG-UK) consortium | The Lighthouse Lab in Alderley Park and Alex Alderton, Roberto Amato, Sonia Goncalves, Ewan Harrison, David K. Jackson, Ian Johnston, Dominic Kwiatkowski, Cordelia Langford, John Sillitoe on behalf of the Wellcome Sanger Institute COVID-19 Surveillance Team |
| EPI_ISL_554143, EPI_ISL_554144, EPI_ISL_554146, EPI_ISL_554147, EPI_ISL_554148 | Lighthouse Lab in Milton Keynes | Wellcome Sanger Institute for the COVID-19 Genomics UK (COG-UK) consortium | The Lighthouse Lab in Milton Keynes and Alex Alderton, Roberto Amato, Sonia Goncalves, Ewan Harrison, David K. Jackson, Ian Johnston, Dominic Kwiatkowski, Cordelia Langford, John Sillitoe on behalf of the Wellcome Sanger Institute COVID-19 Surveillance Team (http://www.sanger.ac.uk/covid-team) |
| EPI_ISL_554149 | Lighthouse Lab in Alderley Park | Wellcome Sanger Institute for the COVID-19 Genomics UK (COG-UK) consortium | The Lighthouse Lab in Alderley Park and Alex Alderton, Roberto Amato, Sonia Goncalves, Ewan Harrison, David K. Jackson, Ian Johnston, Dominic Kwiatkowski, Cordelia Langford, John Sillitoe on behalf of the Wellcome Sanger Institute COVID-19 Surveillance Team |
| EPI_ISL_554150 | Lighthouse Lab in Milton Keynes | Wellcome Sanger Institute for the COVID-19 Genomics UK (COG-UK) consortium | The Lighthouse Lab in Milton Keynes and Alex Alderton, Roberto Amato, Sonia Goncalves, Ewan Harrison, David K. Jackson, Ian Johnston, Dominic Kwiatkowski, Cordelia Langford, John Sillitoe on behalf of the Wellcome Sanger Institute COVID-19 Surveillance Team (http://www.sanger.ac.uk/covid-team) |
| EPI_ISL_554151 | Lighthouse Lab in Alderley Park | Wellcome Sanger Institute for the COVID-19 Genomics UK | The Lighthouse Lab in Alderley Park and Alex Alderton, Roberto Amato, Sonia Goncalves, Ewan Harrison, David K. Jackson, Ian Johnston, Dominic |

| | | | |
|---|---|---|---|
| | | (COG-UK) consortium | Kwiatkowski, Cordelia Langford, John Sillitoe on behalf of the Wellcome Sanger Institute COVID-19 Surveillance Team |
| EPI_ISL_554152, EPI_ISL_554153, EPI_ISL_554154 | Lighthouse Lab in Milton Keynes | Wellcome Sanger Institute for the COVID-19 Genomics UK (COG-UK) consortium | The Lighthouse Lab in Milton Keynes and Alex Alderton, Roberto Amato, Sonia Goncalves, Ewan Harrison, David K. Jackson, Ian Johnston, Dominic Kwiatkowski, Cordelia Langford, John Sillitoe on behalf of the Wellcome Sanger Institute COVID-19 Surveillance Team (http://www.sanger.ac.uk/covid-team) |
| EPI_ISL_554155, EPI_ISL_554156, EPI_ISL_554157, EPI_ISL_554158 | Lighthouse Lab in Alderley Park | Wellcome Sanger Institute for the COVID-19 Genomics UK (COG-UK) consortium | The Lighthouse Lab in Alderley Park and Alex Alderton, Roberto Amato, Sonia Goncalves, Ewan Harrison, David K. Jackson, Ian Johnston, Dominic Kwiatkowski, Cordelia Langford, John Sillitoe on behalf of the Wellcome Sanger Institute COVID-19 Surveillance Team |
| EPI_ISL_554159, EPI_ISL_554160, EPI_ISL_554163 | Lighthouse Lab in Milton Keynes | Wellcome Sanger Institute for the COVID-19 Genomics UK (COG-UK) consortium | The Lighthouse Lab in Milton Keynes and Alex Alderton, Roberto Amato, Sonia Goncalves, Ewan Harrison, David K. Jackson, Ian Johnston, Dominic Kwiatkowski, Cordelia Langford, John Sillitoe on behalf of the Wellcome Sanger Institute COVID-19 Surveillance Team (http://www.sanger.ac.uk/covid-team) |
| EPI_ISL_554164 | Lighthouse Lab in Alderley Park | Wellcome Sanger Institute for the COVID-19 Genomics UK (COG-UK) consortium | The Lighthouse Lab in Alderley Park and Alex Alderton, Roberto Amato, Sonia Goncalves, Ewan Harrison, David K. Jackson, Ian Johnston, Dominic Kwiatkowski, Cordelia Langford, John Sillitoe on behalf of the Wellcome Sanger Institute COVID-19 Surveillance Team (http://www.sanger.ac.uk/covid-team) |
| EPI_ISL_554165 | Lighthouse Lab in Alderley Park | Wellcome Sanger Institute for the COVID-19 Genomics UK (COG-UK) consortium | The Lighthouse Lab in Alderley Park and Alex Alderton, Roberto Amato, Sonia Goncalves, Ewan Harrison, David K. Jackson, Ian Johnston, Dominic Kwiatkowski, Cordelia Langford, John Sillitoe on behalf of the Wellcome Sanger Institute COVID-19 Surveillance Team |
| EPI_ISL_554166, EPI_ISL_554167, EPI_ISL_554168 | Lighthouse Lab in Milton Keynes | Wellcome Sanger Institute for the COVID-19 Genomics UK (COG-UK) consortium | The Lighthouse Lab in Milton Keynes and Alex Alderton, Roberto Amato, Sonia Goncalves, Ewan Harrison, David K. Jackson, Ian Johnston, Dominic Kwiatkowski, Cordelia Langford, John Sillitoe on behalf of the Wellcome Sanger Institute COVID-19 Surveillance Team (http://www.sanger.ac.uk/covid-team) |
| EPI_ISL_554170, EPI_ISL_554171, EPI_ISL_554172 | Lighthouse Lab in Alderley Park | Wellcome Sanger Institute for the COVID-19 Genomics UK (COG-UK) consortium | The Lighthouse Lab in Alderley Park and Alex Alderton, Roberto Amato, Sonia Goncalves, Ewan Harrison, David K. Jackson, Ian Johnston, Dominic Kwiatkowski, Cordelia Langford, John Sillitoe on behalf of the Wellcome Sanger Institute COVID-19 Surveillance Team |
| EPI_ISL_554173 | Lighthouse Lab in Milton Keynes | Wellcome Sanger Institute for the COVID-19 Genomics UK (COG-UK) consortium | The Lighthouse Lab in Milton Keynes and Alex Alderton, Roberto Amato, Sonia Goncalves, Ewan Harrison, David K. Jackson, Ian Johnston, Dominic Kwiatkowski, Cordelia Langford, John Sillitoe on behalf of the Wellcome Sanger Institute COVID-19 Surveillance Team (http://www.sanger.ac.uk/covid-team) |
| EPI_ISL_554174 | Lighthouse Lab in Alderley Park | Wellcome Sanger Institute for the COVID-19 Genomics UK (COG-UK) consortium | The Lighthouse Lab in Alderley Park and Alex Alderton, Roberto Amato, Sonia Goncalves, Ewan Harrison, David K. Jackson, Ian Johnston, Dominic Kwiatkowski, Cordelia Langford, John Sillitoe on behalf of the Wellcome Sanger Institute COVID-19 Surveillance Team (http://www.sanger.ac.uk/covid-team) |
| EPI_ISL_554175, EPI_ISL_554176 | Lighthouse Lab in Alderley Park | Wellcome Sanger Institute for the COVID-19 Genomics UK (COG-UK) consortium | The Lighthouse Lab in Alderley Park and Alex Alderton, Roberto Amato, Sonia Goncalves, Ewan Harrison, David K. Jackson, Ian Johnston, Dominic Kwiatkowski, Cordelia Langford, John Sillitoe on behalf of the Wellcome Sanger Institute COVID-19 Surveillance Team |
| EPI_ISL_554177, EPI_ISL_554178, EPI_ISL_554179, EPI_ISL_554180, EPI_ISL_554181, EPI_ISL_554182 | Lighthouse Lab in Milton Keynes | Wellcome Sanger Institute for the COVID-19 Genomics UK (COG-UK) consortium | The Lighthouse Lab in Milton Keynes and Alex Alderton, Roberto Amato, Sonia Goncalves, Ewan Harrison, David K. Jackson, Ian Johnston, Dominic Kwiatkowski, Cordelia Langford, John Sillitoe on behalf of the Wellcome Sanger Institute COVID-19 Surveillance Team (http://www.sanger.ac.uk/covid-team) |
| EPI_ISL_554183 | Lighthouse Lab in Alderley Park | Wellcome Sanger Institute for the COVID-19 Genomics UK (COG-UK) consortium | The Lighthouse Lab in Alderley Park and Alex Alderton, Roberto Amato, Sonia Goncalves, Ewan Harrison, David K. Jackson, Ian Johnston, Dominic Kwiatkowski, Cordelia Langford, John Sillitoe on behalf of the Wellcome Sanger Institute COVID-19 Surveillance Team |
| EPI_ISL_554184, EPI_ISL_554185 | Lighthouse Lab in Milton Keynes | Wellcome Sanger Institute for the COVID-19 Genomics UK (COG-UK) consortium | The Lighthouse Lab in Milton Keynes and Alex Alderton, Roberto Amato, Sonia Goncalves, Ewan Harrison, David K. Jackson, Ian Johnston, Dominic Kwiatkowski, Cordelia Langford, John Sillitoe on behalf of the Wellcome Sanger Institute COVID-19 Surveillance Team (http://www.sanger.ac.uk/covid-team) |
| EPI_ISL_554186, EPI_ISL_554187, EPI_ISL_554188, EPI_ISL_554189, EPI_ISL_554190 | Lighthouse Lab in Alderley Park | Wellcome Sanger Institute for the COVID-19 Genomics UK (COG-UK) consortium | The Lighthouse Lab in Alderley Park and Alex Alderton, Roberto Amato, Sonia Goncalves, Ewan Harrison, David K. Jackson, Ian Johnston, Dominic Kwiatkowski, Cordelia Langford, John Sillitoe on behalf of the Wellcome Sanger Institute COVID-19 Surveillance Team |
| EPI_ISL_554191 | Lighthouse Lab in Milton Keynes | Wellcome Sanger Institute for the COVID-19 Genomics UK (COG-UK) consortium | The Lighthouse Lab in Milton Keynes and Alex Alderton, Roberto Amato, Sonia Goncalves, Ewan Harrison, David K. Jackson, Ian Johnston, Dominic Kwiatkowski, Cordelia Langford, John Sillitoe on behalf of the Wellcome Sanger Institute COVID-19 Surveillance Team (http://www.sanger.ac.uk/covid-team) |
| EPI_ISL_554192, EPI_ISL_554193 | Lighthouse Lab in Alderley Park | Wellcome Sanger Institute for the COVID-19 Genomics UK (COG-UK) consortium | The Lighthouse Lab in Alderley Park and Alex Alderton, Roberto Amato, Sonia Goncalves, Ewan Harrison, David K. Jackson, Ian Johnston, Dominic Kwiatkowski, Cordelia Langford, John Sillitoe on behalf of the Wellcome Sanger Institute COVID-19 Surveillance Team (http://www.sanger.ac.uk/covid-team) |
| EPI_ISL_554194, EPI_ISL_554196, EPI_ISL_554197, EPI_ISL_554198, EPI_ISL_554199 | Lighthouse Lab in Alderley Park | Wellcome Sanger Institute for the COVID-19 Genomics UK (COG-UK) consortium | The Lighthouse Lab in Alderley Park and Alex Alderton, Roberto Amato, Sonia Goncalves, Ewan Harrison, David K. Jackson, Ian Johnston, Dominic Kwiatkowski, Cordelia Langford, John Sillitoe on behalf of the Wellcome Sanger Institute COVID-19 Surveillance Team |
| EPI_ISL_554200, EPI_ISL_554201 | Lighthouse Lab in Milton Keynes | Wellcome Sanger Institute for the COVID-19 Genomics UK (COG-UK) consortium | The Lighthouse Lab in Milton Keynes and Alex Alderton, Roberto Amato, Sonia Goncalves, Ewan Harrison, David K. Jackson, Ian Johnston, Dominic Kwiatkowski, Cordelia Langford, John Sillitoe on behalf of the Wellcome Sanger Institute COVID-19 Surveillance Team (http://www.sanger.ac.uk/covid-team) |
| EPI_ISL_554202, EPI_ISL_554203 | Lighthouse Lab in Alderley Park | Wellcome Sanger Institute for the COVID-19 Genomics UK (COG-UK) consortium | The Lighthouse Lab in Alderley Park and Alex Alderton, Roberto Amato, Sonia Goncalves, Ewan Harrison, David K. Jackson, Ian Johnston, Dominic Kwiatkowski, Cordelia Langford, John Sillitoe on behalf of the Wellcome Sanger Institute COVID-19 Surveillance Team |
| EPI_ISL_554204, EPI_ISL_554205 | Lighthouse Lab in Milton Keynes | Wellcome Sanger Institute for the COVID-19 Genomics UK (COG-UK) consortium | The Lighthouse Lab in Milton Keynes and Alex Alderton, Roberto Amato, Sonia Goncalves, Ewan Harrison, David K. Jackson, Ian Johnston, Dominic Kwiatkowski, Cordelia Langford, John Sillitoe on behalf of the Wellcome Sanger Institute COVID-19 Surveillance Team (http://www.sanger.ac.uk/covid-team) |
| EPI_ISL_554206, EPI_ISL_554207 | Lighthouse Lab in Alderley Park | Wellcome Sanger Institute for the COVID-19 Genomics UK (COG-UK) consortium | The Lighthouse Lab in Alderley Park and Alex Alderton, Roberto Amato, Sonia Goncalves, Ewan Harrison, David K. Jackson, Ian Johnston, Dominic Kwiatkowski, Cordelia Langford, John Sillitoe on behalf of the Wellcome Sanger Institute COVID-19 Surveillance Team |
| EPI_ISL_554208 | Lighthouse Lab in Milton Keynes | Wellcome Sanger Institute for the COVID-19 Genomics UK (COG-UK) consortium | The Lighthouse Lab in Milton Keynes and Alex Alderton, Roberto Amato, Sonia Goncalves, Ewan Harrison, David K. Jackson, Ian Johnston, Dominic Kwiatkowski, Cordelia Langford, John Sillitoe on behalf of the Wellcome Sanger Institute COVID-19 Surveillance Team (http://www.sanger.ac.uk/covid-team) |
| EPI_ISL_554209 | Lighthouse Lab in Alderley Park | Wellcome Sanger Institute for the COVID-19 Genomics UK (COG-UK) consortium | The Lighthouse Lab in Alderley Park and Alex Alderton, Roberto Amato, Sonia Goncalves, Ewan Harrison, David K. Jackson, Ian Johnston, Dominic Kwiatkowski, Cordelia Langford, John Sillitoe on behalf of the Wellcome Sanger Institute COVID-19 Surveillance Team |
| EPI_ISL_554211 | Lighthouse Lab in Milton Keynes | Wellcome Sanger Institute for the COVID-19 Genomics UK (COG-UK) consortium | The Lighthouse Lab in Milton Keynes and Alex Alderton, Roberto Amato, Sonia Goncalves, Ewan Harrison, David K. Jackson, Ian Johnston, Dominic Kwiatkowski, Cordelia Langford, John Sillitoe on behalf of the Wellcome Sanger Institute COVID-19 Surveillance Team (http://www.sanger.ac.uk/covid-team) |
| EPI_ISL_554212 | Lighthouse Lab in Alderley Park | Wellcome Sanger Institute for the COVID-19 Genomics UK (COG-UK) consortium | The Lighthouse Lab in Alderley Park and Alex Alderton, Roberto Amato, Sonia Goncalves, Ewan Harrison, David K. Jackson, Ian Johnston, Dominic Kwiatkowski, Cordelia Langford, John Sillitoe on behalf of the Wellcome Sanger Institute COVID-19 Surveillance Team |
| EPI_ISL_554214 | Lighthouse Lab in Milton Keynes | Wellcome Sanger Institute for the COVID-19 Genomics UK (COG-UK) consortium | The Lighthouse Lab in Milton Keynes and Alex Alderton, Roberto Amato, Sonia Goncalves, Ewan Harrison, David K. Jackson, Ian Johnston, Dominic Kwiatkowski, Cordelia Langford, John Sillitoe on behalf of the Wellcome Sanger Institute COVID-19 Surveillance Team (http://www.sanger.ac.uk/covid-team) |
| EPI_ISL_554215 | Lighthouse Lab in Alderley Park | Wellcome Sanger Institute for the COVID-19 Genomics UK (COG-UK) consortium | The Lighthouse Lab in Alderley Park and Alex Alderton, Roberto Amato, Sonia Goncalves, Ewan Harrison, David K. Jackson, Ian Johnston, Dominic Kwiatkowski, Cordelia Langford, John Sillitoe on behalf of the Wellcome Sanger Institute COVID-19 Surveillance Team |
| EPI_ISL_554216, EPI_ISL_554217, | Lighthouse Lab in Milton Keynes | Wellcome Sanger Institute for the COVID-19 Genomics UK | The Lighthouse Lab in Milton Keynes and Alex Alderton, Roberto Amato, Sonia Goncalves, Ewan Harrison, David K. Jackson, Ian Johnston, Dominic |

| | | | |
|---|---|---|---|
| EPI_ISL_554218 | | (COG-UK) consortium | Kwiatkowski, Cordelia Langford, John Sillitoe on behalf of the Wellcome Sanger Institute COVID-19 Surveillance Team (http://www.sanger.ac.uk/covid-team) |
| EPI_ISL_554219 | Lighthouse Lab in Alderley Park | Wellcome Sanger Institute for the COVID-19 Genomics UK (COG-UK) consortium | The Lighthouse Lab in Alderley Park and Alex Alderton, Roberto Amato, Sonia Goncalves, Ewan Harrison, David K. Jackson, Ian Johnston, Dominic Kwiatkowski, Cordelia Langford, John Sillitoe on behalf of the Wellcome Sanger Institute COVID-19 Surveillance Team |
| EPI_ISL_554220, EPI_ISL_554221 | Lighthouse Lab in Milton Keynes | Wellcome Sanger Institute for the COVID-19 Genomics UK (COG-UK) consortium | The Lighthouse Lab in Milton Keynes and Alex Alderton, Roberto Amato, Sonia Goncalves, Ewan Harrison, David K. Jackson, Ian Johnston, Dominic Kwiatkowski, Cordelia Langford, John Sillitoe on behalf of the Wellcome Sanger Institute COVID-19 Surveillance Team (http://www.sanger.ac.uk/covid-team) |
| EPI_ISL_554222 | Lighthouse Lab in Alderley Park | Wellcome Sanger Institute for the COVID-19 Genomics UK (COG-UK) consortium | The Lighthouse Lab in Alderley Park and Alex Alderton, Roberto Amato, Sonia Goncalves, Ewan Harrison, David K. Jackson, Ian Johnston, Dominic Kwiatkowski, Cordelia Langford, John Sillitoe on behalf of the Wellcome Sanger Institute COVID-19 Surveillance Team (http://www.sanger.ac.uk/covid-team) |
| EPI_ISL_554223, EPI_ISL_554224 | Lighthouse Lab in Milton Keynes | Wellcome Sanger Institute for the COVID-19 Genomics UK (COG-UK) consortium | The Lighthouse Lab in Milton Keynes and Alex Alderton, Roberto Amato, Sonia Goncalves, Ewan Harrison, David K. Jackson, Ian Johnston, Dominic Kwiatkowski, Cordelia Langford, John Sillitoe on behalf of the Wellcome Sanger Institute COVID-19 Surveillance Team (http://www.sanger.ac.uk/covid-team) |
| EPI_ISL_554225 | Lighthouse Lab in Alderley Park | Wellcome Sanger Institute for the COVID-19 Genomics UK (COG-UK) consortium | The Lighthouse Lab in Alderley Park and Alex Alderton, Roberto Amato, Sonia Goncalves, Ewan Harrison, David K. Jackson, Ian Johnston, Dominic Kwiatkowski, Cordelia Langford, John Sillitoe on behalf of the Wellcome Sanger Institute COVID-19 Surveillance Team (http://www.sanger.ac.uk/covid-team) |
| EPI_ISL_554226 | Lighthouse Lab in Milton Keynes | Wellcome Sanger Institute for the COVID-19 Genomics UK (COG-UK) consortium | The Lighthouse Lab in Milton Keynes and Alex Alderton, Roberto Amato, Sonia Goncalves, Ewan Harrison, David K. Jackson, Ian Johnston, Dominic Kwiatkowski, Cordelia Langford, John Sillitoe on behalf of the Wellcome Sanger Institute COVID-19 Surveillance Team (http://www.sanger.ac.uk/covid-team) |
| EPI_ISL_554227 | Lighthouse Lab in Alderley Park | Wellcome Sanger Institute for the COVID-19 Genomics UK (COG-UK) consortium | The Lighthouse Lab in Alderley Park and Alex Alderton, Roberto Amato, Sonia Goncalves, Ewan Harrison, David K. Jackson, Ian Johnston, Dominic Kwiatkowski, Cordelia Langford, John Sillitoe on behalf of the Wellcome Sanger Institute COVID-19 Surveillance Team |
| EPI_ISL_554228 | Lighthouse Lab in Milton Keynes | Wellcome Sanger Institute for the COVID-19 Genomics UK (COG-UK) consortium | The Lighthouse Lab in Milton Keynes and Alex Alderton, Roberto Amato, Sonia Goncalves, Ewan Harrison, David K. Jackson, Ian Johnston, Dominic Kwiatkowski, Cordelia Langford, John Sillitoe on behalf of the Wellcome Sanger Institute COVID-19 Surveillance Team (http://www.sanger.ac.uk/covid-team) |
| EPI_ISL_554229, EPI_ISL_554230 | Lighthouse Lab in Alderley Park | Wellcome Sanger Institute for the COVID-19 Genomics UK (COG-UK) consortium | The Lighthouse Lab in Alderley Park and Alex Alderton, Roberto Amato, Sonia Goncalves, Ewan Harrison, David K. Jackson, Ian Johnston, Dominic Kwiatkowski, Cordelia Langford, John Sillitoe on behalf of the Wellcome Sanger Institute COVID-19 Surveillance Team |
| EPI_ISL_554231, EPI_ISL_554232 | Lighthouse Lab in Milton Keynes | Wellcome Sanger Institute for the COVID-19 Genomics UK (COG-UK) consortium | The Lighthouse Lab in Milton Keynes and Alex Alderton, Roberto Amato, Sonia Goncalves, Ewan Harrison, David K. Jackson, Ian Johnston, Dominic Kwiatkowski, Cordelia Langford, John Sillitoe on behalf of the Wellcome Sanger Institute COVID-19 Surveillance Team (http://www.sanger.ac.uk/covid-team) |
| EPI_ISL_554233, EPI_ISL_554235 | Lighthouse Lab in Alderley Park | Wellcome Sanger Institute for the COVID-19 Genomics UK (COG-UK) consortium | The Lighthouse Lab in Alderley Park and Alex Alderton, Roberto Amato, Sonia Goncalves, Ewan Harrison, David K. Jackson, Ian Johnston, Dominic Kwiatkowski, Cordelia Langford, John Sillitoe on behalf of the Wellcome Sanger Institute COVID-19 Surveillance Team |
| EPI_ISL_554237 | Lighthouse Lab in Milton Keynes | Wellcome Sanger Institute for the COVID-19 Genomics UK (COG-UK) consortium | The Lighthouse Lab in Milton Keynes and Alex Alderton, Roberto Amato, Sonia Goncalves, Ewan Harrison, David K. Jackson, Ian Johnston, Dominic Kwiatkowski, Cordelia Langford, John Sillitoe on behalf of the Wellcome Sanger Institute COVID-19 Surveillance Team (http://www.sanger.ac.uk/covid-team) |
| EPI_ISL_554238 | Lighthouse Lab in Alderley Park | Wellcome Sanger Institute for the COVID-19 Genomics UK (COG-UK) consortium | The Lighthouse Lab in Alderley Park and Alex Alderton, Roberto Amato, Sonia Goncalves, Ewan Harrison, David K. Jackson, Ian Johnston, Dominic Kwiatkowski, Cordelia Langford, John Sillitoe on behalf of the Wellcome Sanger Institute COVID-19 Surveillance Team (http://www.sanger.ac.uk/covid-team) |
| EPI_ISL_554239, EPI_ISL_554240 | Lighthouse Lab in Milton Keynes | Wellcome Sanger Institute for the COVID-19 Genomics UK (COG-UK) consortium | The Lighthouse Lab in Milton Keynes and Alex Alderton, Roberto Amato, Sonia Goncalves, Ewan Harrison, David K. Jackson, Ian Johnston, Dominic Kwiatkowski, Cordelia Langford, John Sillitoe on behalf of the Wellcome Sanger Institute COVID-19 Surveillance Team (http://www.sanger.ac.uk/covid-team) |
| EPI_ISL_554241 | Lighthouse Lab in Alderley Park | Wellcome Sanger Institute for the COVID-19 Genomics UK (COG-UK) consortium | The Lighthouse Lab in Alderley Park and Alex Alderton, Roberto Amato, Sonia Goncalves, Ewan Harrison, David K. Jackson, Ian Johnston, Dominic Kwiatkowski, Cordelia Langford, John Sillitoe on behalf of the Wellcome Sanger Institute COVID-19 Surveillance Team |
| EPI_ISL_554242, EPI_ISL_554243, EPI_ISL_554244, EPI_ISL_554245, EPI_ISL_554246, EPI_ISL_554247, EPI_ISL_554248, EPI_ISL_554249 | Lighthouse Lab in Milton Keynes | Wellcome Sanger Institute for the COVID-19 Genomics UK (COG-UK) consortium | The Lighthouse Lab in Milton Keynes and Alex Alderton, Roberto Amato, Sonia Goncalves, Ewan Harrison, David K. Jackson, Ian Johnston, Dominic Kwiatkowski, Cordelia Langford, John Sillitoe on behalf of the Wellcome Sanger Institute COVID-19 Surveillance Team (http://www.sanger.ac.uk/covid-team) |
| EPI_ISL_554250 | Lighthouse Lab in Alderley Park | Wellcome Sanger Institute for the COVID-19 Genomics UK (COG-UK) consortium | The Lighthouse Lab in Alderley Park and Alex Alderton, Roberto Amato, Sonia Goncalves, Ewan Harrison, David K. Jackson, Ian Johnston, Dominic Kwiatkowski, Cordelia Langford, John Sillitoe on behalf of the Wellcome Sanger Institute COVID-19 Surveillance Team |
| EPI_ISL_554251 | Lighthouse Lab in Milton Keynes | Wellcome Sanger Institute for the COVID-19 Genomics UK (COG-UK) consortium | The Lighthouse Lab in Milton Keynes and Alex Alderton, Roberto Amato, Sonia Goncalves, Ewan Harrison, David K. Jackson, Ian Johnston, Dominic Kwiatkowski, Cordelia Langford, John Sillitoe on behalf of the Wellcome Sanger Institute COVID-19 Surveillance Team (http://www.sanger.ac.uk/covid-team) |
| EPI_ISL_554252 | Lighthouse Lab in Alderley Park | Wellcome Sanger Institute for the COVID-19 Genomics UK (COG-UK) consortium | The Lighthouse Lab in Alderley Park and Alex Alderton, Roberto Amato, Sonia Goncalves, Ewan Harrison, David K. Jackson, Ian Johnston, Dominic Kwiatkowski, Cordelia Langford, John Sillitoe on behalf of the Wellcome Sanger Institute COVID-19 Surveillance Team (http://www.sanger.ac.uk/covid-team) |
| EPI_ISL_554253, EPI_ISL_554254, EPI_ISL_554255, EPI_ISL_554256 | Lighthouse Lab in Milton Keynes | Wellcome Sanger Institute for the COVID-19 Genomics UK (COG-UK) consortium | The Lighthouse Lab in Milton Keynes and Alex Alderton, Roberto Amato, Sonia Goncalves, Ewan Harrison, David K. Jackson, Ian Johnston, Dominic Kwiatkowski, Cordelia Langford, John Sillitoe on behalf of the Wellcome Sanger Institute COVID-19 Surveillance Team (http://www.sanger.ac.uk/covid-team) |
| EPI_ISL_554257 | Lighthouse Lab in Alderley Park | Wellcome Sanger Institute for the COVID-19 Genomics UK (COG-UK) consortium | The Lighthouse Lab in Alderley Park and Alex Alderton, Roberto Amato, Sonia Goncalves, Ewan Harrison, David K. Jackson, Ian Johnston, Dominic Kwiatkowski, Cordelia Langford, John Sillitoe on behalf of the Wellcome Sanger Institute COVID-19 Surveillance Team |
| EPI_ISL_554258 | Lighthouse Lab in Milton Keynes | Wellcome Sanger Institute for the COVID-19 Genomics UK (COG-UK) consortium | The Lighthouse Lab in Milton Keynes and Alex Alderton, Roberto Amato, Sonia Goncalves, Ewan Harrison, David K. Jackson, Ian Johnston, Dominic Kwiatkowski, Cordelia Langford, John Sillitoe on behalf of the Wellcome Sanger Institute COVID-19 Surveillance Team (http://www.sanger.ac.uk/covid-team) |
| EPI_ISL_554260 | Lighthouse Lab in Alderley Park | Wellcome Sanger Institute for the COVID-19 Genomics UK (COG-UK) consortium | The Lighthouse Lab in Alderley Park and Alex Alderton, Roberto Amato, Sonia Goncalves, Ewan Harrison, David K. Jackson, Ian Johnston, Dominic Kwiatkowski, Cordelia Langford, John Sillitoe on behalf of the Wellcome Sanger Institute COVID-19 Surveillance Team (http://www.sanger.ac.uk/covid-team) |
| EPI_ISL_554261, EPI_ISL_554262 | Lighthouse Lab in Milton Keynes | Wellcome Sanger Institute for the COVID-19 Genomics UK (COG-UK) consortium | The Lighthouse Lab in Milton Keynes and Alex Alderton, Roberto Amato, Sonia Goncalves, Ewan Harrison, David K. Jackson, Ian Johnston, Dominic Kwiatkowski, Cordelia Langford, John Sillitoe on behalf of the Wellcome Sanger Institute COVID-19 Surveillance Team (http://www.sanger.ac.uk/covid-team) |
| EPI_ISL_554263 | Lighthouse Lab in Alderley Park | Wellcome Sanger Institute for the COVID-19 Genomics UK (COG-UK) consortium | The Lighthouse Lab in Alderley Park and Alex Alderton, Roberto Amato, Sonia Goncalves, Ewan Harrison, David K. Jackson, Ian Johnston, Dominic Kwiatkowski, Cordelia Langford, John Sillitoe on behalf of the Wellcome Sanger Institute COVID-19 Surveillance Team (http://www.sanger.ac.uk/covid-team) |
| EPI_ISL_554264 | Lighthouse Lab in Milton Keynes | Wellcome Sanger Institute for the COVID-19 Genomics UK (COG-UK) consortium | The Lighthouse Lab in Milton Keynes and Alex Alderton, Roberto Amato, Sonia Goncalves, Ewan Harrison, David K. Jackson, Ian Johnston, Dominic Kwiatkowski, Cordelia Langford, John Sillitoe on behalf of the Wellcome Sanger Institute COVID-19 Surveillance Team (http://www.sanger.ac.uk/covid-team) |

| | | | |
|---|---|---|---|
| EPI_ISL_554265, EPI_ISL_554266 | Lighthouse Lab in Alderley Park | Wellcome Sanger Institute for the COVID-19 Genomics UK (COG-UK) consortium | The Lighthouse Lab in Alderley Park and Alex Alderton, Roberto Amato, Sonia Goncalves, Ewan Harrison, David K. Jackson, Ian Johnston, Dominic Kwiatkowski, Cordelia Langford, John Sillitoe on behalf of the Wellcome Sanger Institute COVID-19 Surveillance Team |
| EPI_ISL_554267 | Lighthouse Lab in Milton Keynes | Wellcome Sanger Institute for the COVID-19 Genomics UK (COG-UK) consortium | The Lighthouse Lab in Milton Keynes and Alex Alderton, Roberto Amato, Sonia Goncalves, Ewan Harrison, David K. Jackson, Ian Johnston, Dominic Kwiatkowski, Cordelia Langford, John Sillitoe on behalf of the Wellcome Sanger Institute COVID-19 Surveillance Team (http://www.sanger.ac.uk/covid-team) |
| EPI_ISL_554268 | Lighthouse Lab in Alderley Park | Wellcome Sanger Institute for the COVID-19 Genomics UK (COG-UK) consortium | The Lighthouse Lab in Alderley Park and Alex Alderton, Roberto Amato, Sonia Goncalves, Ewan Harrison, David K. Jackson, Ian Johnston, Dominic Kwiatkowski, Cordelia Langford, John Sillitoe on behalf of the Wellcome Sanger Institute COVID-19 Surveillance Team |
| EPI_ISL_554269 | Lighthouse Lab in Milton Keynes | Wellcome Sanger Institute for the COVID-19 Genomics UK (COG-UK) consortium | The Lighthouse Lab in Milton Keynes and Alex Alderton, Roberto Amato, Sonia Goncalves, Ewan Harrison, David K. Jackson, Ian Johnston, Dominic Kwiatkowski, Cordelia Langford, John Sillitoe on behalf of the Wellcome Sanger Institute COVID-19 Surveillance Team (http://www.sanger.ac.uk/covid-team) |
| EPI_ISL_554270 | Lighthouse Lab in Alderley Park | Wellcome Sanger Institute for the COVID-19 Genomics UK (COG-UK) consortium | The Lighthouse Lab in Alderley Park and Alex Alderton, Roberto Amato, Sonia Goncalves, Ewan Harrison, David K. Jackson, Ian Johnston, Dominic Kwiatkowski, Cordelia Langford, John Sillitoe on behalf of the Wellcome Sanger Institute COVID-19 Surveillance Team |
| EPI_ISL_554271, EPI_ISL_554272, EPI_ISL_554274 | Lighthouse Lab in Milton Keynes | Wellcome Sanger Institute for the COVID-19 Genomics UK (COG-UK) consortium | The Lighthouse Lab in Milton Keynes and Alex Alderton, Roberto Amato, Sonia Goncalves, Ewan Harrison, David K. Jackson, Ian Johnston, Dominic Kwiatkowski, Cordelia Langford, John Sillitoe on behalf of the Wellcome Sanger Institute COVID-19 Surveillance Team (http://www.sanger.ac.uk/covid-team) |
| EPI_ISL_554275 | Lighthouse Lab in Alderley Park | Wellcome Sanger Institute for the COVID-19 Genomics UK (COG-UK) consortium | The Lighthouse Lab in Alderley Park and Alex Alderton, Roberto Amato, Sonia Goncalves, Ewan Harrison, David K. Jackson, Ian Johnston, Dominic Kwiatkowski, Cordelia Langford, John Sillitoe on behalf of the Wellcome Sanger Institute COVID-19 Surveillance Team |
| EPI_ISL_554276, EPI_ISL_554277 | Lighthouse Lab in Milton Keynes | Wellcome Sanger Institute for the COVID-19 Genomics UK (COG-UK) consortium | The Lighthouse Lab in Milton Keynes and Alex Alderton, Roberto Amato, Sonia Goncalves, Ewan Harrison, David K. Jackson, Ian Johnston, Dominic Kwiatkowski, Cordelia Langford, John Sillitoe on behalf of the Wellcome Sanger Institute COVID-19 Surveillance Team (http://www.sanger.ac.uk/covid-team) |
| EPI_ISL_554278, EPI_ISL_554279, EPI_ISL_554280, EPI_ISL_554282, EPI_ISL_554283 | Lighthouse Lab in Alderley Park | Wellcome Sanger Institute for the COVID-19 Genomics UK (COG-UK) consortium | The Lighthouse Lab in Alderley Park and Alex Alderton, Roberto Amato, Sonia Goncalves, Ewan Harrison, David K. Jackson, Ian Johnston, Dominic Kwiatkowski, Cordelia Langford, John Sillitoe on behalf of the Wellcome Sanger Institute COVID-19 Surveillance Team |
| EPI_ISL_554284, EPI_ISL_554285, EPI_ISL_554286, EPI_ISL_554287 | Lighthouse Lab in Milton Keynes | Wellcome Sanger Institute for the COVID-19 Genomics UK (COG-UK) consortium | The Lighthouse Lab in Milton Keynes and Alex Alderton, Roberto Amato, Sonia Goncalves, Ewan Harrison, David K. Jackson, Ian Johnston, Dominic Kwiatkowski, Cordelia Langford, John Sillitoe on behalf of the Wellcome Sanger Institute COVID-19 Surveillance Team (http://www.sanger.ac.uk/covid-team) |
| EPI_ISL_554288 | Lighthouse Lab in Alderley Park | Wellcome Sanger Institute for the COVID-19 Genomics UK (COG-UK) consortium | The Lighthouse Lab in Alderley Park and Alex Alderton, Roberto Amato, Sonia Goncalves, Ewan Harrison, David K. Jackson, Ian Johnston, Dominic Kwiatkowski, Cordelia Langford, John Sillitoe on behalf of the Wellcome Sanger Institute COVID-19 Surveillance Team (http://www.sanger.ac.uk/covid-team) |
| EPI_ISL_554289, EPI_ISL_554290, EPI_ISL_554291 | Lighthouse Lab in Milton Keynes | Wellcome Sanger Institute for the COVID-19 Genomics UK (COG-UK) consortium | The Lighthouse Lab in Milton Keynes and Alex Alderton, Roberto Amato, Sonia Goncalves, Ewan Harrison, David K. Jackson, Ian Johnston, Dominic Kwiatkowski, Cordelia Langford, John Sillitoe on behalf of the Wellcome Sanger Institute COVID-19 Surveillance Team (http://www.sanger.ac.uk/covid-team) |
| EPI_ISL_554292 | Lighthouse Lab in Alderley Park | Wellcome Sanger Institute for the COVID-19 Genomics UK (COG-UK) consortium | The Lighthouse Lab in Alderley Park and Alex Alderton, Roberto Amato, Sonia Goncalves, Ewan Harrison, David K. Jackson, Ian Johnston, Dominic Kwiatkowski, Cordelia Langford, John Sillitoe on behalf of the Wellcome Sanger Institute COVID-19 Surveillance Team |
| EPI_ISL_554293, EPI_ISL_554294, EPI_ISL_554295 | Lighthouse Lab in Milton Keynes | Wellcome Sanger Institute for the COVID-19 Genomics UK (COG-UK) consortium | The Lighthouse Lab in Milton Keynes and Alex Alderton, Roberto Amato, Sonia Goncalves, Ewan Harrison, David K. Jackson, Ian Johnston, Dominic Kwiatkowski, Cordelia Langford, John Sillitoe on behalf of the Wellcome Sanger Institute COVID-19 Surveillance Team (http://www.sanger.ac.uk/covid-team) |
| EPI_ISL_554296 | Lighthouse Lab in Alderley Park | Wellcome Sanger Institute for the COVID-19 Genomics UK (COG-UK) consortium | The Lighthouse Lab in Alderley Park and Alex Alderton, Roberto Amato, Sonia Goncalves, Ewan Harrison, David K. Jackson, Ian Johnston, Dominic Kwiatkowski, Cordelia Langford, John Sillitoe on behalf of the Wellcome Sanger Institute COVID-19 Surveillance Team |
| EPI_ISL_554297 | Lighthouse Lab in Milton Keynes | Wellcome Sanger Institute for the COVID-19 Genomics UK (COG-UK) consortium | The Lighthouse Lab in Milton Keynes and Alex Alderton, Roberto Amato, Sonia Goncalves, Ewan Harrison, David K. Jackson, Ian Johnston, Dominic Kwiatkowski, Cordelia Langford, John Sillitoe on behalf of the Wellcome Sanger Institute COVID-19 Surveillance Team (http://www.sanger.ac.uk/covid-team) |
| EPI_ISL_554298, EPI_ISL_554299 | Lighthouse Lab in Alderley Park | Wellcome Sanger Institute for the COVID-19 Genomics UK (COG-UK) consortium | The Lighthouse Lab in Alderley Park and Alex Alderton, Roberto Amato, Sonia Goncalves, Ewan Harrison, David K. Jackson, Ian Johnston, Dominic Kwiatkowski, Cordelia Langford, John Sillitoe on behalf of the Wellcome Sanger Institute COVID-19 Surveillance Team |
| EPI_ISL_554300 | Lighthouse Lab in Milton Keynes | Wellcome Sanger Institute for the COVID-19 Genomics UK (COG-UK) consortium | The Lighthouse Lab in Milton Keynes and Alex Alderton, Roberto Amato, Sonia Goncalves, Ewan Harrison, David K. Jackson, Ian Johnston, Dominic Kwiatkowski, Cordelia Langford, John Sillitoe on behalf of the Wellcome Sanger Institute COVID-19 Surveillance Team (http://www.sanger.ac.uk/covid-team) |
| EPI_ISL_554301 | Lighthouse Lab in Alderley Park | Wellcome Sanger Institute for the COVID-19 Genomics UK (COG-UK) consortium | The Lighthouse Lab in Alderley Park and Alex Alderton, Roberto Amato, Sonia Goncalves, Ewan Harrison, David K. Jackson, Ian Johnston, Dominic Kwiatkowski, Cordelia Langford, John Sillitoe on behalf of the Wellcome Sanger Institute COVID-19 Surveillance Team |
| EPI_ISL_554302, EPI_ISL_554304 | Lighthouse Lab in Milton Keynes | Wellcome Sanger Institute for the COVID-19 Genomics UK (COG-UK) consortium | The Lighthouse Lab in Milton Keynes and Alex Alderton, Roberto Amato, Sonia Goncalves, Ewan Harrison, David K. Jackson, Ian Johnston, Dominic Kwiatkowski, Cordelia Langford, John Sillitoe on behalf of the Wellcome Sanger Institute COVID-19 Surveillance Team (http://www.sanger.ac.uk/covid-team) |
| EPI_ISL_554305 | Lighthouse Lab in Alderley Park | Wellcome Sanger Institute for the COVID-19 Genomics UK (COG-UK) consortium | The Lighthouse Lab in Alderley Park and Alex Alderton, Roberto Amato, Sonia Goncalves, Ewan Harrison, David K. Jackson, Ian Johnston, Dominic Kwiatkowski, Cordelia Langford, John Sillitoe on behalf of the Wellcome Sanger Institute COVID-19 Surveillance Team |
| EPI_ISL_554306, EPI_ISL_554308 | Lighthouse Lab in Milton Keynes | Wellcome Sanger Institute for the COVID-19 Genomics UK (COG-UK) consortium | The Lighthouse Lab in Milton Keynes and Alex Alderton, Roberto Amato, Sonia Goncalves, Ewan Harrison, David K. Jackson, Ian Johnston, Dominic Kwiatkowski, Cordelia Langford, John Sillitoe on behalf of the Wellcome Sanger Institute COVID-19 Surveillance Team (http://www.sanger.ac.uk/covid-team) |
| EPI_ISL_554309 | Lighthouse Lab in Alderley Park | Wellcome Sanger Institute for the COVID-19 Genomics UK (COG-UK) consortium | The Lighthouse Lab in Alderley Park and Alex Alderton, Roberto Amato, Sonia Goncalves, Ewan Harrison, David K. Jackson, Ian Johnston, Dominic Kwiatkowski, Cordelia Langford, John Sillitoe on behalf of the Wellcome Sanger Institute COVID-19 Surveillance Team |
| EPI_ISL_554310, EPI_ISL_554312 | Lighthouse Lab in Milton Keynes | Wellcome Sanger Institute for the COVID-19 Genomics UK (COG-UK) consortium | The Lighthouse Lab in Milton Keynes and Alex Alderton, Roberto Amato, Sonia Goncalves, Ewan Harrison, David K. Jackson, Ian Johnston, Dominic Kwiatkowski, Cordelia Langford, John Sillitoe on behalf of the Wellcome Sanger Institute COVID-19 Surveillance Team (http://www.sanger.ac.uk/covid-team) |
| EPI_ISL_554313 | Lighthouse Lab in Milton Keynes | Wellcome Sanger Institute for the COVID-19 Genomics UK (COG-UK) consortium | The Lighthouse Lab in Alderley Park and Alex Alderton, Roberto Amato, Sonia Goncalves, Ewan Harrison, David K. Jackson, Ian Johnston, Dominic Kwiatkowski, Cordelia Langford, John Sillitoe on behalf of the Wellcome Sanger Institute COVID-19 Surveillance Team |
| EPI_ISL_554314, EPI_ISL_554315 | Lighthouse Lab in Milton Keynes | Wellcome Sanger Institute for the COVID-19 Genomics UK (COG-UK) consortium | The Lighthouse Lab in Milton Keynes and Alex Alderton, Roberto Amato, Sonia Goncalves, Ewan Harrison, David K. Jackson, Ian Johnston, Dominic Kwiatkowski, Cordelia Langford, John Sillitoe on behalf of the Wellcome Sanger Institute COVID-19 Surveillance Team (http://www.sanger.ac.uk/covid-team) |
| EPI_ISL_554318 | Lighthouse Lab in Alderley Park | Wellcome Sanger Institute for the COVID-19 Genomics UK (COG-UK) consortium | The Lighthouse Lab in Alderley Park and Alex Alderton, Roberto Amato, Sonia Goncalves, Ewan Harrison, David K. Jackson, Ian Johnston, Dominic Kwiatkowski, Cordelia Langford, John Sillitoe on behalf of the Wellcome Sanger Institute COVID-19 Surveillance Team |
| EPI_ISL_554319 | Lighthouse Lab in Milton Keynes | Wellcome Sanger Institute for the COVID-19 Genomics UK (COG-UK) consortium | The Lighthouse Lab in Milton Keynes and Alex Alderton, Roberto Amato, Sonia Goncalves, Ewan Harrison, David K. Jackson, Ian Johnston, Dominic Kwiatkowski, Cordelia Langford, John Sillitoe on behalf of the Wellcome Sanger Institute COVID-19 Surveillance Team (http://www.sanger.ac.uk/covid-team) |
| EPI_ISL_554320, EPI_ISL_554321, | Lighthouse Lab in Alderley Park | Wellcome Sanger Institute for the COVID-19 Genomics UK | The Lighthouse Lab in Alderley Park and Alex Alderton, Roberto Amato, Sonia Goncalves, Ewan Harrison, David K. Jackson, Ian Johnston, Dominic |

| | | | |
|---|---|---|---|
| EPI_ISL_554322, EPI_ISL_554323, EPI_ISL_554324, EPI_ISL_554325, EPI_ISL_554326, EPI_ISL_554327 | | (COG-UK) consortium | Kwiatkowski, Cordelia Langford, John Sillitoe on behalf of the Wellcome Sanger Institute COVID-19 Surveillance Team |
| EPI_ISL_554328, EPI_ISL_554329 | Lighthouse Lab in Alderley Park | Wellcome Sanger Institute for the COVID-19 Genomics UK (COG-UK) consortium | The Lighthouse Lab in Alderley Park and Alex Alderton, Roberto Amato, Sonia Goncalves, Ewan Harrison, David K. Jackson, Ian Johnston, Dominic Kwiatkowski, Cordelia Langford, John Sillitoe on behalf of the Wellcome Sanger Institute COVID-19 Surveillance Team (http://www.sanger.ac.uk/covid-team) |
| EPI_ISL_554330, EPI_ISL_554331, EPI_ISL_554332, EPI_ISL_554333, EPI_ISL_554334, EPI_ISL_554335, EPI_ISL_554336, EPI_ISL_554337, EPI_ISL_554338, EPI_ISL_554339, EPI_ISL_554340, EPI_ISL_554341, EPI_ISL_554342, EPI_ISL_554343, EPI_ISL_554344, EPI_ISL_554345, EPI_ISL_554346, EPI_ISL_554347, EPI_ISL_554348, EPI_ISL_554349, EPI_ISL_554350, EPI_ISL_554351, EPI_ISL_554352, EPI_ISL_554353, EPI_ISL_554354, EPI_ISL_554355, EPI_ISL_554356, EPI_ISL_554357, EPI_ISL_554358, EPI_ISL_554359, EPI_ISL_554360, EPI_ISL_554361, EPI_ISL_554362 | | | |
| see above | Lighthouse Lab in Alderley Park | Wellcome Sanger Institute for the COVID-19 Genomics UK (COG-UK) consortium | The Lighthouse Lab in Alderley Park and Alex Alderton, Roberto Amato, Sonia Goncalves, Ewan Harrison, David K. Jackson, Ian Johnston, Dominic Kwiatkowski, Cordelia Langford, John Sillitoe on behalf of the Wellcome Sanger Institute COVID-19 Surveillance Team |
| EPI_ISL_554363, EPI_ISL_554364 | Lighthouse Lab in Alderley Park | Wellcome Sanger Institute for the COVID-19 Genomics UK (COG-UK) consortium | The Lighthouse Lab in Alderley Park and Alex Alderton, Roberto Amato, Sonia Goncalves, Ewan Harrison, David K. Jackson, Ian Johnston, Dominic Kwiatkowski, Cordelia Langford, John Sillitoe on behalf of the Wellcome Sanger Institute COVID-19 Surveillance Team (http://www.sanger.ac.uk/covid-team) |
| EPI_ISL_554365, EPI_ISL_554366, EPI_ISL_554367 | Lighthouse Lab in Alderley Park | Wellcome Sanger Institute for the COVID-19 Genomics UK (COG-UK) consortium | The Lighthouse Lab in Alderley Park and Alex Alderton, Roberto Amato, Sonia Goncalves, Ewan Harrison, David K. Jackson, Ian Johnston, Dominic Kwiatkowski, Cordelia Langford, John Sillitoe on behalf of the Wellcome Sanger Institute COVID-19 Surveillance Team |
| EPI_ISL_554368 | Lighthouse Lab in Alderley Park | Wellcome Sanger Institute for the COVID-19 Genomics UK (COG-UK) consortium | The Lighthouse Lab in Alderley Park and Alex Alderton, Roberto Amato, Sonia Goncalves, Ewan Harrison, David K. Jackson, Ian Johnston, Dominic Kwiatkowski, Cordelia Langford, John Sillitoe on behalf of the Wellcome Sanger Institute COVID-19 Surveillance Team (http://www.sanger.ac.uk/covid-team) |
| EPI_ISL_554369, EPI_ISL_554370, EPI_ISL_554371, EPI_ISL_554372, EPI_ISL_554373 | Lighthouse Lab in Alderley Park | Wellcome Sanger Institute for the COVID-19 Genomics UK (COG-UK) consortium | The Lighthouse Lab in Alderley Park and Alex Alderton, Roberto Amato, Sonia Goncalves, Ewan Harrison, David K. Jackson, Ian Johnston, Dominic Kwiatkowski, Cordelia Langford, John Sillitoe on behalf of the Wellcome Sanger Institute COVID-19 Surveillance Team |
| EPI_ISL_554374 | Lighthouse Lab in Alderley Park | Wellcome Sanger Institute for the COVID-19 Genomics UK (COG-UK) consortium | The Lighthouse Lab in Alderley Park and Alex Alderton, Roberto Amato, Sonia Goncalves, Ewan Harrison, David K. Jackson, Ian Johnston, Dominic Kwiatkowski, Cordelia Langford, John Sillitoe on behalf of the Wellcome Sanger Institute COVID-19 Surveillance Team (http://www.sanger.ac.uk/covid-team) |
| EPI_ISL_554375, EPI_ISL_554376, EPI_ISL_554377, EPI_ISL_554378, EPI_ISL_554379, EPI_ISL_554380, EPI_ISL_554381, EPI_ISL_554382, EPI_ISL_554383 | Lighthouse Lab in Alderley Park | Wellcome Sanger Institute for the COVID-19 Genomics UK (COG-UK) consortium | The Lighthouse Lab in Alderley Park and Alex Alderton, Roberto Amato, Sonia Goncalves, Ewan Harrison, David K. Jackson, Ian Johnston, Dominic Kwiatkowski, Cordelia Langford, John Sillitoe on behalf of the Wellcome Sanger Institute COVID-19 Surveillance Team |
| EPI_ISL_554384 | Lighthouse Lab in Alderley Park | Wellcome Sanger Institute for the COVID-19 Genomics UK (COG-UK) consortium | The Lighthouse Lab in Alderley Park and Alex Alderton, Roberto Amato, Sonia Goncalves, Ewan Harrison, David K. Jackson, Ian Johnston, Dominic Kwiatkowski, Cordelia Langford, John Sillitoe on behalf of the Wellcome Sanger Institute COVID-19 Surveillance Team (http://www.sanger.ac.uk/covid-team) |
| EPI_ISL_554385, EPI_ISL_554386, EPI_ISL_554387, EPI_ISL_554388, EPI_ISL_554389, EPI_ISL_554390, EPI_ISL_554391, EPI_ISL_554392, EPI_ISL_554393, EPI_ISL_554394, EPI_ISL_554395, EPI_ISL_554396, EPI_ISL_554397, EPI_ISL_554398, EPI_ISL_554399, EPI_ISL_554400, EPI_ISL_554401, EPI_ISL_554402 | | | |
| see above | Lighthouse Lab in Alderley Park | Wellcome Sanger Institute for the COVID-19 Genomics UK (COG-UK) consortium | The Lighthouse Lab in Alderley Park and Alex Alderton, Roberto Amato, Sonia Goncalves, Ewan Harrison, David K. Jackson, Ian Johnston, Dominic Kwiatkowski, Cordelia Langford, John Sillitoe on behalf of the Wellcome Sanger Institute COVID-19 Surveillance Team |
| EPI_ISL_554403, EPI_ISL_554404 | Lighthouse Lab in Alderley Park | Wellcome Sanger Institute for the COVID-19 Genomics UK (COG-UK) consortium | The Lighthouse Lab in Alderley Park and Alex Alderton, Roberto Amato, Sonia Goncalves, Ewan Harrison, David K. Jackson, Ian Johnston, Dominic Kwiatkowski, Cordelia Langford, John Sillitoe on behalf of the Wellcome Sanger Institute COVID-19 Surveillance Team (http://www.sanger.ac.uk/covid-team) |
| EPI_ISL_554405, EPI_ISL_554406, EPI_ISL_554407, EPI_ISL_554408, EPI_ISL_554409, EPI_ISL_554410, EPI_ISL_554411, EPI_ISL_554412 | Lighthouse Lab in Alderley Park | Wellcome Sanger Institute for the COVID-19 Genomics UK (COG-UK) consortium | The Lighthouse Lab in Alderley Park and Alex Alderton, Roberto Amato, Sonia Goncalves, Ewan Harrison, David K. Jackson, Ian Johnston, Dominic Kwiatkowski, Cordelia Langford, John Sillitoe on behalf of the Wellcome Sanger Institute COVID-19 Surveillance Team |
| EPI_ISL_554413 | Lighthouse Lab in Alderley Park | Wellcome Sanger Institute for the COVID-19 Genomics UK (COG-UK) consortium | The Lighthouse Lab in Alderley Park and Alex Alderton, Roberto Amato, Sonia Goncalves, Ewan Harrison, David K. Jackson, Ian Johnston, Dominic Kwiatkowski, Cordelia Langford, John Sillitoe on behalf of the Wellcome Sanger Institute COVID-19 Surveillance Team (http://www.sanger.ac.uk/covid-team) |
| EPI_ISL_554414, EPI_ISL_554415, EPI_ISL_554416, EPI_ISL_554417, EPI_ISL_554418, EPI_ISL_554419, EPI_ISL_554420, EPI_ISL_554421, EPI_ISL_554422, EPI_ISL_554423, EPI_ISL_554424, EPI_ISL_554425 | | | |
| see above | Lighthouse Lab in Alderley Park | Wellcome Sanger Institute for the COVID-19 Genomics UK (COG-UK) consortium | The Lighthouse Lab in Alderley Park and Alex Alderton, Roberto Amato, Sonia Goncalves, Ewan Harrison, David K. Jackson, Ian Johnston, Dominic Kwiatkowski, Cordelia Langford, John Sillitoe on behalf of the Wellcome Sanger Institute COVID-19 Surveillance Team |
| EPI_ISL_554426 | Lighthouse Lab in Alderley Park | Wellcome Sanger Institute for the COVID-19 Genomics UK (COG-UK) consortium | The Lighthouse Lab in Alderley Park and Alex Alderton, Roberto Amato, Sonia Goncalves, Ewan Harrison, David K. Jackson, Ian Johnston, Dominic Kwiatkowski, Cordelia Langford, John Sillitoe on behalf of the Wellcome Sanger Institute COVID-19 Surveillance Team (http://www.sanger.ac.uk/covid-team) |
| EPI_ISL_554427, EPI_ISL_554428, EPI_ISL_554429, EPI_ISL_554430, EPI_ISL_554431, EPI_ISL_554432, EPI_ISL_554433, EPI_ISL_554434, EPI_ISL_554435, EPI_ISL_554436, EPI_ISL_554437, EPI_ISL_554438, EPI_ISL_554439 | | | |
| see above | Lighthouse Lab in Alderley Park | Wellcome Sanger Institute for the COVID-19 Genomics UK (COG-UK) consortium | The Lighthouse Lab in Alderley Park and Alex Alderton, Roberto Amato, Sonia Goncalves, Ewan Harrison, David K. Jackson, Ian Johnston, Dominic Kwiatkowski, Cordelia Langford, John Sillitoe on behalf of the Wellcome Sanger Institute COVID-19 Surveillance Team |
| EPI_ISL_554440 | Lighthouse Lab in Alderley Park | Wellcome Sanger Institute for the COVID-19 Genomics UK (COG-UK) consortium | The Lighthouse Lab in Alderley Park and Alex Alderton, Roberto Amato, Sonia Goncalves, Ewan Harrison, David K. Jackson, Ian Johnston, Dominic Kwiatkowski, Cordelia Langford, John Sillitoe on behalf of the Wellcome Sanger Institute COVID-19 Surveillance Team (http://www.sanger.ac.uk/covid-team) |
| EPI_ISL_554441, EPI_ISL_554442 | Lighthouse Lab in Alderley Park | Wellcome Sanger Institute for the COVID-19 Genomics UK (COG-UK) consortium | The Lighthouse Lab in Alderley Park and Alex Alderton, Roberto Amato, Sonia Goncalves, Ewan Harrison, David K. Jackson, Ian Johnston, Dominic Kwiatkowski, Cordelia Langford, John Sillitoe on behalf of the Wellcome Sanger Institute COVID-19 Surveillance Team |
| EPI_ISL_554443 | Lighthouse Lab in Alderley Park | Wellcome Sanger Institute for the COVID-19 Genomics UK (COG-UK) consortium | The Lighthouse Lab in Alderley Park and Alex Alderton, Roberto Amato, Sonia Goncalves, Ewan Harrison, David K. Jackson, Ian Johnston, Dominic Kwiatkowski, Cordelia Langford, John Sillitoe on behalf of the Wellcome Sanger Institute COVID-19 Surveillance Team (http://www.sanger.ac.uk/covid-team) |
| EPI_ISL_554444, EPI_ISL_554445, EPI_ISL_554446, EPI_ISL_554447, EPI_ISL_554448 | Lighthouse Lab in Alderley Park | Wellcome Sanger Institute for the COVID-19 Genomics UK (COG-UK) consortium | The Lighthouse Lab in Alderley Park and Alex Alderton, Roberto Amato, Sonia Goncalves, Ewan Harrison, David K. Jackson, Ian Johnston, Dominic Kwiatkowski, Cordelia Langford, John Sillitoe on behalf of the Wellcome Sanger Institute COVID-19 Surveillance Team |
| EPI_ISL_554449 | Lighthouse Lab in Alderley Park | Wellcome Sanger Institute for the COVID-19 Genomics UK (COG-UK) consortium | The Lighthouse Lab in Alderley Park and Alex Alderton, Roberto Amato, Sonia Goncalves, Ewan Harrison, David K. Jackson, Ian Johnston, Dominic Kwiatkowski, Cordelia Langford, John Sillitoe on behalf of the Wellcome Sanger Institute COVID-19 Surveillance Team (http://www.sanger.ac.uk/covid-team) |
| EPI_ISL_554450, EPI_ISL_554451 | Lighthouse Lab in Alderley Park | Wellcome Sanger Institute for the COVID-19 Genomics UK (COG-UK) consortium | The Lighthouse Lab in Alderley Park and Alex Alderton, Roberto Amato, Sonia Goncalves, Ewan Harrison, David K. Jackson, Ian Johnston, Dominic Kwiatkowski, Cordelia Langford, John Sillitoe on behalf of the Wellcome Sanger Institute COVID-19 Surveillance Team |
| EPI_ISL_554452 | Lighthouse Lab in Alderley Park | Wellcome Sanger Institute for the COVID-19 Genomics UK (COG-UK) consortium | The Lighthouse Lab in Alderley Park and Alex Alderton, Roberto Amato, Sonia Goncalves, Ewan Harrison, David K. Jackson, Ian Johnston, Dominic Kwiatkowski, Cordelia Langford, John Sillitoe on behalf of the Wellcome Sanger Institute COVID-19 Surveillance Team (http://www.sanger.ac.uk/covid-team) |

| | | | |
|---|---|---|---|
| EPI_ISL_554453, EPI_ISL_554454, EPI_ISL_554455, EPI_ISL_554456, EPI_ISL_554457, EPI_ISL_554458, EPI_ISL_554459, EPI_ISL_554460, EPI_ISL_554461 | Lighthouse Lab in Alderley Park | Wellcome Sanger Institute for the COVID-19 Genomics UK (COG-UK) consortium | The Lighthouse Lab in Alderley Park and Alex Alderton, Roberto Amato, Sonia Goncalves, Ewan Harrison, David K. Jackson, Ian Johnston, Dominic Kwiatkowski, Cordelia Langford, John Sillitoe on behalf of the Wellcome Sanger Institute COVID-19 Surveillance Team |
| EPI_ISL_554462, EPI_ISL_554463 | Lighthouse Lab in Alderley Park | Wellcome Sanger Institute for the COVID-19 Genomics UK (COG-UK) consortium | The Lighthouse Lab in Alderley Park and Alex Alderton, Roberto Amato, Sonia Goncalves, Ewan Harrison, David K. Jackson, Ian Johnston, Dominic Kwiatkowski, Cordelia Langford, John Sillitoe on behalf of the Wellcome Sanger Institute COVID-19 Surveillance Team (http://www.sanger.ac.uk/covid-team) |
| EPI_ISL_554464, EPI_ISL_554465, EPI_ISL_554466, EPI_ISL_554467, EPI_ISL_554468, EPI_ISL_554469, EPI_ISL_554470, EPI_ISL_554471, EPI_ISL_554472, EPI_ISL_554473 | Lighthouse Lab in Alderley Park | Wellcome Sanger Institute for the COVID-19 Genomics UK (COG-UK) consortium | The Lighthouse Lab in Alderley Park and Alex Alderton, Roberto Amato, Sonia Goncalves, Ewan Harrison, David K. Jackson, Ian Johnston, Dominic Kwiatkowski, Cordelia Langford, John Sillitoe on behalf of the Wellcome Sanger Institute COVID-19 Surveillance Team |
| EPI_ISL_554474 | Lighthouse Lab in Alderley Park | Wellcome Sanger Institute for the COVID-19 Genomics UK (COG-UK) consortium | The Lighthouse Lab in Alderley Park and Alex Alderton, Roberto Amato, Sonia Goncalves, Ewan Harrison, David K. Jackson, Ian Johnston, Dominic Kwiatkowski, Cordelia Langford, John Sillitoe on behalf of the Wellcome Sanger Institute COVID-19 Surveillance Team (http://www.sanger.ac.uk/covid-team) |
| EPI_ISL_554475, EPI_ISL_554476, EPI_ISL_554477, EPI_ISL_554479, EPI_ISL_554480, EPI_ISL_554481, EPI_ISL_554482, EPI_ISL_554483, EPI_ISL_554485, EPI_ISL_554486, EPI_ISL_554487, EPI_ISL_554488, EPI_ISL_554489 | | | |
| see above | Lighthouse Lab in Alderley Park | Wellcome Sanger Institute for the COVID-19 Genomics UK (COG-UK) consortium | The Lighthouse Lab in Alderley Park and Alex Alderton, Roberto Amato, Sonia Goncalves, Ewan Harrison, David K. Jackson, Ian Johnston, Dominic Kwiatkowski, Cordelia Langford, John Sillitoe on behalf of the Wellcome Sanger Institute COVID-19 Surveillance Team |
| EPI_ISL_554490, EPI_ISL_554491 | Lighthouse Lab in Alderley Park | Wellcome Sanger Institute for the COVID-19 Genomics UK (COG-UK) consortium | The Lighthouse Lab in Alderley Park and Alex Alderton, Roberto Amato, Sonia Goncalves, Ewan Harrison, David K. Jackson, Ian Johnston, Dominic Kwiatkowski, Cordelia Langford, John Sillitoe on behalf of the Wellcome Sanger Institute COVID-19 Surveillance Team (http://www.sanger.ac.uk/covid-team) |
| EPI_ISL_554492, EPI_ISL_554493, EPI_ISL_554494, EPI_ISL_554495 | Lighthouse Lab in Alderley Park | Wellcome Sanger Institute for the COVID-19 Genomics UK (COG-UK) consortium | The Lighthouse Lab in Alderley Park and Alex Alderton, Roberto Amato, Sonia Goncalves, Ewan Harrison, David K. Jackson, Ian Johnston, Dominic Kwiatkowski, Cordelia Langford, John Sillitoe on behalf of the Wellcome Sanger Institute COVID-19 Surveillance Team |
| EPI_ISL_554496 | Lighthouse Lab in Alderley Park | Wellcome Sanger Institute for the COVID-19 Genomics UK (COG-UK) consortium | The Lighthouse Lab in Alderley Park and Alex Alderton, Roberto Amato, Sonia Goncalves, Ewan Harrison, David K. Jackson, Ian Johnston, Dominic Kwiatkowski, Cordelia Langford, John Sillitoe on behalf of the Wellcome Sanger Institute COVID-19 Surveillance Team (http://www.sanger.ac.uk/covid-team) |
| EPI_ISL_554498, EPI_ISL_554499, EPI_ISL_554500, EPI_ISL_554501, EPI_ISL_554502, EPI_ISL_554503 | Lighthouse Lab in Alderley Park | Wellcome Sanger Institute for the COVID-19 Genomics UK (COG-UK) consortium | The Lighthouse Lab in Alderley Park and Alex Alderton, Roberto Amato, Sonia Goncalves, Ewan Harrison, David K. Jackson, Ian Johnston, Dominic Kwiatkowski, Cordelia Langford, John Sillitoe on behalf of the Wellcome Sanger Institute COVID-19 Surveillance Team |
| EPI_ISL_554504 | Lighthouse Lab in Alderley Park | Wellcome Sanger Institute for the COVID-19 Genomics UK (COG-UK) consortium | The Lighthouse Lab in Alderley Park and Alex Alderton, Roberto Amato, Sonia Goncalves, Ewan Harrison, David K. Jackson, Ian Johnston, Dominic Kwiatkowski, Cordelia Langford, John Sillitoe on behalf of the Wellcome Sanger Institute COVID-19 Surveillance Team (http://www.sanger.ac.uk/covid-team) |
| EPI_ISL_554505, EPI_ISL_554506, EPI_ISL_554507, EPI_ISL_554508, EPI_ISL_554509, EPI_ISL_554510, EPI_ISL_554511, EPI_ISL_554512 | Lighthouse Lab in Alderley Park | Wellcome Sanger Institute for the COVID-19 Genomics UK (COG-UK) consortium | The Lighthouse Lab in Alderley Park and Alex Alderton, Roberto Amato, Sonia Goncalves, Ewan Harrison, David K. Jackson, Ian Johnston, Dominic Kwiatkowski, Cordelia Langford, John Sillitoe on behalf of the Wellcome Sanger Institute COVID-19 Surveillance Team |
| EPI_ISL_554513 | Lighthouse Lab in Alderley Park | Wellcome Sanger Institute for the COVID-19 Genomics UK (COG-UK) consortium | The Lighthouse Lab in Alderley Park and Alex Alderton, Roberto Amato, Sonia Goncalves, Ewan Harrison, David K. Jackson, Ian Johnston, Dominic Kwiatkowski, Cordelia Langford, John Sillitoe on behalf of the Wellcome Sanger Institute COVID-19 Surveillance Team (http://www.sanger.ac.uk/covid-team) |
| EPI_ISL_554514, EPI_ISL_554515, EPI_ISL_554516, EPI_ISL_554517 | Lighthouse Lab in Alderley Park | Wellcome Sanger Institute for the COVID-19 Genomics UK (COG-UK) consortium | The Lighthouse Lab in Alderley Park and Alex Alderton, Roberto Amato, Sonia Goncalves, Ewan Harrison, David K. Jackson, Ian Johnston, Dominic Kwiatkowski, Cordelia Langford, John Sillitoe on behalf of the Wellcome Sanger Institute COVID-19 Surveillance Team |
| EPI_ISL_554518 | Lighthouse Lab in Alderley Park | Wellcome Sanger Institute for the COVID-19 Genomics UK (COG-UK) consortium | The Lighthouse Lab in Alderley Park and Alex Alderton, Roberto Amato, Sonia Goncalves, Ewan Harrison, David K. Jackson, Ian Johnston, Dominic Kwiatkowski, Cordelia Langford, John Sillitoe on behalf of the Wellcome Sanger Institute COVID-19 Surveillance Team (http://www.sanger.ac.uk/covid-team) |
| EPI_ISL_554519, EPI_ISL_554520, EPI_ISL_554521, EPI_ISL_554522, EPI_ISL_554523, EPI_ISL_554524, EPI_ISL_554525, EPI_ISL_554526 | Lighthouse Lab in Alderley Park | Wellcome Sanger Institute for the COVID-19 Genomics UK (COG-UK) consortium | The Lighthouse Lab in Alderley Park and Alex Alderton, Roberto Amato, Sonia Goncalves, Ewan Harrison, David K. Jackson, Ian Johnston, Dominic Kwiatkowski, Cordelia Langford, John Sillitoe on behalf of the Wellcome Sanger Institute COVID-19 Surveillance Team |
| EPI_ISL_554527, EPI_ISL_554528 | Lighthouse Lab in Alderley Park | Wellcome Sanger Institute for the COVID-19 Genomics UK (COG-UK) consortium | The Lighthouse Lab in Alderley Park and Alex Alderton, Roberto Amato, Sonia Goncalves, Ewan Harrison, David K. Jackson, Ian Johnston, Dominic Kwiatkowski, Cordelia Langford, John Sillitoe on behalf of the Wellcome Sanger Institute COVID-19 Surveillance Team (http://www.sanger.ac.uk/covid-team) |
| EPI_ISL_554529, EPI_ISL_554530, EPI_ISL_554531 | Lighthouse Lab in Alderley Park | Wellcome Sanger Institute for the COVID-19 Genomics UK (COG-UK) consortium | The Lighthouse Lab in Alderley Park and Alex Alderton, Roberto Amato, Sonia Goncalves, Ewan Harrison, David K. Jackson, Ian Johnston, Dominic Kwiatkowski, Cordelia Langford, John Sillitoe on behalf of the Wellcome Sanger Institute COVID-19 Surveillance Team |
| EPI_ISL_554532 | Lighthouse Lab in Alderley Park | Wellcome Sanger Institute for the COVID-19 Genomics UK (COG-UK) consortium | The Lighthouse Lab in Alderley Park and Alex Alderton, Roberto Amato, Sonia Goncalves, Ewan Harrison, David K. Jackson, Ian Johnston, Dominic Kwiatkowski, Cordelia Langford, John Sillitoe on behalf of the Wellcome Sanger Institute COVID-19 Surveillance Team (http://www.sanger.ac.uk/covid-team) |
| EPI_ISL_554533, EPI_ISL_554534, EPI_ISL_554535, EPI_ISL_554536, EPI_ISL_554538, EPI_ISL_554539 | Lighthouse Lab in Alderley Park | Wellcome Sanger Institute for the COVID-19 Genomics UK (COG-UK) consortium | The Lighthouse Lab in Alderley Park and Alex Alderton, Roberto Amato, Sonia Goncalves, Ewan Harrison, David K. Jackson, Ian Johnston, Dominic Kwiatkowski, Cordelia Langford, John Sillitoe on behalf of the Wellcome Sanger Institute COVID-19 Surveillance Team |
| EPI_ISL_554540 | Lighthouse Lab in Alderley Park | Wellcome Sanger Institute for the COVID-19 Genomics UK (COG-UK) consortium | The Lighthouse Lab in Alderley Park and Alex Alderton, Roberto Amato, Sonia Goncalves, Ewan Harrison, David K. Jackson, Ian Johnston, Dominic Kwiatkowski, Cordelia Langford, John Sillitoe on behalf of the Wellcome Sanger Institute COVID-19 Surveillance Team (http://www.sanger.ac.uk/covid-team) |
| EPI_ISL_554541, EPI_ISL_554542, EPI_ISL_554543, EPI_ISL_554544, EPI_ISL_554545, EPI_ISL_554546, EPI_ISL_554547, EPI_ISL_554548, EPI_ISL_554549, EPI_ISL_554550, EPI_ISL_554551, EPI_ISL_554552, EPI_ISL_554553, EPI_ISL_554554, EPI_ISL_554555, EPI_ISL_554556, EPI_ISL_554557, EPI_ISL_554558 | | | |
| see above | Lighthouse Lab in Alderley Park | Wellcome Sanger Institute for the COVID-19 Genomics UK (COG-UK) consortium | The Lighthouse Lab in Alderley Park and Alex Alderton, Roberto Amato, Sonia Goncalves, Ewan Harrison, David K. Jackson, Ian Johnston, Dominic Kwiatkowski, Cordelia Langford, John Sillitoe on behalf of the Wellcome Sanger Institute COVID-19 Surveillance Team |
| EPI_ISL_554559 | Lighthouse Lab in Alderley Park | Wellcome Sanger Institute for the COVID-19 Genomics UK (COG-UK) consortium | The Lighthouse Lab in Alderley Park and Alex Alderton, Roberto Amato, Sonia Goncalves, Ewan Harrison, David K. Jackson, Ian Johnston, Dominic Kwiatkowski, Cordelia Langford, John Sillitoe on behalf of the Wellcome Sanger Institute COVID-19 Surveillance Team (http://www.sanger.ac.uk/covid-team) |
| EPI_ISL_554560, EPI_ISL_554561, EPI_ISL_554562 | Lighthouse Lab in Alderley Park | Wellcome Sanger Institute for the COVID-19 Genomics UK (COG-UK) consortium | The Lighthouse Lab in Alderley Park and Alex Alderton, Roberto Amato, Sonia Goncalves, Ewan Harrison, David K. Jackson, Ian Johnston, Dominic Kwiatkowski, Cordelia Langford, John Sillitoe on behalf of the Wellcome Sanger Institute COVID-19 Surveillance Team |
| EPI_ISL_554563 | Lighthouse Lab in Alderley Park | Wellcome Sanger Institute for the COVID-19 Genomics UK (COG-UK) consortium | The Lighthouse Lab in Alderley Park and Alex Alderton, Roberto Amato, Sonia Goncalves, Ewan Harrison, David K. Jackson, Ian Johnston, Dominic Kwiatkowski, Cordelia Langford, John Sillitoe on behalf of the Wellcome Sanger Institute COVID-19 Surveillance Team (http://www.sanger.ac.uk/covid-team) |

| | | | |
|---|---|---|---|
| EPI_ISL_554564, EPI_ISL_554565, EPI_ISL_554566, EPI_ISL_554567, EPI_ISL_554568, EPI_ISL_554569, EPI_ISL_554570, EPI_ISL_554571, EPI_ISL_554572, EPI_ISL_554573 | Lighthouse Lab in Alderley Park | Wellcome Sanger Institute for the COVID-19 Genomics UK (COG-UK) consortium | The Lighthouse Lab in Alderley Park and Alex Alderton, Roberto Amato, Sonia Goncalves, Ewan Harrison, David K. Jackson, Ian Johnston, Dominic Kwiatkowski, Cordelia Langford, John Sillitoe on behalf of the Wellcome Sanger Institute COVID-19 Surveillance Team |
| EPI_ISL_554574 | Lighthouse Lab in Alderley Park | Wellcome Sanger Institute for the COVID-19 Genomics UK (COG-UK) consortium | The Lighthouse Lab in Alderley Park and Alex Alderton, Roberto Amato, Sonia Goncalves, Ewan Harrison, David K. Jackson, Ian Johnston, Dominic Kwiatkowski, Cordelia Langford, John Sillitoe on behalf of the Wellcome Sanger Institute COVID-19 Surveillance Team (http://www.sanger.ac.uk/covid-team) |
| EPI_ISL_554575, EPI_ISL_554576, EPI_ISL_554577, EPI_ISL_554578, EPI_ISL_554579, EPI_ISL_554580, EPI_ISL_554581, EPI_ISL_554582, EPI_ISL_554583, EPI_ISL_554584, EPI_ISL_554585, EPI_ISL_554586, EPI_ISL_554587, EPI_ISL_554588, EPI_ISL_554589, EPI_ISL_554590, EPI_ISL_554591, EPI_ISL_554592, EPI_ISL_554593, EPI_ISL_554594, EPI_ISL_554595, EPI_ISL_554596, EPI_ISL_554597, EPI_ISL_554598, EPI_ISL_554599, EPI_ISL_554600, EPI_ISL_554602, EPI_ISL_554603, EPI_ISL_554604, EPI_ISL_554605, EPI_ISL_554606, EPI_ISL_554607, EPI_ISL_554608, EPI_ISL_554609, EPI_ISL_554610, EPI_ISL_554611 | | | |
| see above | Lighthouse Lab in Alderley Park | Wellcome Sanger Institute for the COVID-19 Genomics UK (COG-UK) consortium | The Lighthouse Lab in Alderley Park and Alex Alderton, Roberto Amato, Sonia Goncalves, Ewan Harrison, David K. Jackson, Ian Johnston, Dominic Kwiatkowski, Cordelia Langford, John Sillitoe on behalf of the Wellcome Sanger Institute COVID-19 Surveillance Team |
| EPI_ISL_554612 | Lighthouse Lab in Alderley Park | Wellcome Sanger Institute for the COVID-19 Genomics UK (COG-UK) consortium | The Lighthouse Lab in Alderley Park and Alex Alderton, Roberto Amato, Sonia Goncalves, Ewan Harrison, David K. Jackson, Ian Johnston, Dominic Kwiatkowski, Cordelia Langford, John Sillitoe on behalf of the Wellcome Sanger Institute COVID-19 Surveillance Team (http://www.sanger.ac.uk/covid-team) |
| EPI_ISL_554613, EPI_ISL_554614, EPI_ISL_554615 | Lighthouse Lab in Alderley Park | Wellcome Sanger Institute for the COVID-19 Genomics UK (COG-UK) consortium | The Lighthouse Lab in Alderley Park and Alex Alderton, Roberto Amato, Sonia Goncalves, Ewan Harrison, David K. Jackson, Ian Johnston, Dominic Kwiatkowski, Cordelia Langford, John Sillitoe on behalf of the Wellcome Sanger Institute COVID-19 Surveillance Team |
| EPI_ISL_554621 | Lighthouse Lab in Milton Keynes | Wellcome Sanger Institute for the COVID-19 Genomics UK (COG-UK) consortium | The Lighthouse Lab in Alderley Park and Alex Alderton, Roberto Amato, Sonia Goncalves, Ewan Harrison, David K. Jackson, Ian Johnston, Dominic Kwiatkowski, Cordelia Langford, John Sillitoe on behalf of the Wellcome Sanger Institute COVID-19 Surveillance Team |
| EPI_ISL_554622 | Lighthouse Lab in Milton Keynes | Wellcome Sanger Institute for the COVID-19 Genomics UK (COG-UK) consortium | The Lighthouse Lab in Milton Keynes and Alex Alderton, Roberto Amato, Sonia Goncalves, Ewan Harrison, David K. Jackson, Ian Johnston, Dominic Kwiatkowski, Cordelia Langford, John Sillitoe on behalf of the Wellcome Sanger Institute COVID-19 Surveillance Team (http://www.sanger.ac.uk/covid-team) |
| EPI_ISL_554623 | Lighthouse Lab in Alderley Park | Wellcome Sanger Institute for the COVID-19 Genomics UK (COG-UK) consortium | The Lighthouse Lab in Alderley Park and Alex Alderton, Roberto Amato, Sonia Goncalves, Ewan Harrison, David K. Jackson, Ian Johnston, Dominic Kwiatkowski, Cordelia Langford, John Sillitoe on behalf of the Wellcome Sanger Institute COVID-19 Surveillance Team |
| EPI_ISL_554628, EPI_ISL_554629 | Lighthouse Lab in Milton Keynes | Wellcome Sanger Institute for the COVID-19 Genomics UK (COG-UK) consortium | The Lighthouse Lab in Milton Keynes and Alex Alderton, Roberto Amato, Sonia Goncalves, Ewan Harrison, David K. Jackson, Ian Johnston, Dominic Kwiatkowski, Cordelia Langford, John Sillitoe on behalf of the Wellcome Sanger Institute COVID-19 Surveillance Team (http://www.sanger.ac.uk/covid-team) |
| EPI_ISL_554632 | Lighthouse Lab in Alderley Park | Wellcome Sanger Institute for the COVID-19 Genomics UK (COG-UK) consortium | The Lighthouse Lab in Alderley Park and Alex Alderton, Roberto Amato, Sonia Goncalves, Ewan Harrison, David K. Jackson, Ian Johnston, Dominic Kwiatkowski, Cordelia Langford, John Sillitoe on behalf of the Wellcome Sanger Institute COVID-19 Surveillance Team (http://www.sanger.ac.uk/covid-team) |
| EPI_ISL_554633 | Lighthouse Lab in Milton Keynes | Wellcome Sanger Institute for the COVID-19 Genomics UK (COG-UK) consortium | The Lighthouse Lab in Alderley Park and Alex Alderton, Roberto Amato, Sonia Goncalves, Ewan Harrison, David K. Jackson, Ian Johnston, Dominic Kwiatkowski, Cordelia Langford, John Sillitoe on behalf of the Wellcome Sanger Institute COVID-19 Surveillance Team |
| EPI_ISL_554634, EPI_ISL_554635, EPI_ISL_554636, EPI_ISL_554637, EPI_ISL_554639, EPI_ISL_554641 | Lighthouse Lab in Milton Keynes | Wellcome Sanger Institute for the COVID-19 Genomics UK (COG-UK) consortium | The Lighthouse Lab in Milton Keynes and Alex Alderton, Roberto Amato, Sonia Goncalves, Ewan Harrison, David K. Jackson, Ian Johnston, Dominic Kwiatkowski, Cordelia Langford, John Sillitoe on behalf of the Wellcome Sanger Institute COVID-19 Surveillance Team (http://www.sanger.ac.uk/covid-team) |
| EPI_ISL_554642 | Lighthouse Lab in Alderley Park | Wellcome Sanger Institute for the COVID-19 Genomics UK (COG-UK) consortium | The Lighthouse Lab in Alderley Park and Alex Alderton, Roberto Amato, Sonia Goncalves, Ewan Harrison, David K. Jackson, Ian Johnston, Dominic Kwiatkowski, Cordelia Langford, John Sillitoe on behalf of the Wellcome Sanger Institute COVID-19 Surveillance Team |
| EPI_ISL_554643, EPI_ISL_554644, EPI_ISL_554645 | Lighthouse Lab in Milton Keynes | Wellcome Sanger Institute for the COVID-19 Genomics UK (COG-UK) consortium | The Lighthouse Lab in Milton Keynes and Alex Alderton, Roberto Amato, Sonia Goncalves, Ewan Harrison, David K. Jackson, Ian Johnston, Dominic Kwiatkowski, Cordelia Langford, John Sillitoe on behalf of the Wellcome Sanger Institute COVID-19 Surveillance Team (http://www.sanger.ac.uk/covid-team) |
| EPI_ISL_554646 | Lighthouse Lab in Alderley Park | Wellcome Sanger Institute for the COVID-19 Genomics UK (COG-UK) consortium | The Lighthouse Lab in Alderley Park and Alex Alderton, Roberto Amato, Sonia Goncalves, Ewan Harrison, David K. Jackson, Ian Johnston, Dominic Kwiatkowski, Cordelia Langford, John Sillitoe on behalf of the Wellcome Sanger Institute COVID-19 Surveillance Team |
| EPI_ISL_554647, EPI_ISL_554648, EPI_ISL_554649, EPI_ISL_554652, EPI_ISL_554658, EPI_ISL_554659, EPI_ISL_554660 | Lighthouse Lab in Milton Keynes | Wellcome Sanger Institute for the COVID-19 Genomics UK (COG-UK) consortium | The Lighthouse Lab in Milton Keynes and Alex Alderton, Roberto Amato, Sonia Goncalves, Ewan Harrison, David K. Jackson, Ian Johnston, Dominic Kwiatkowski, Cordelia Langford, John Sillitoe on behalf of the Wellcome Sanger Institute COVID-19 Surveillance Team (http://www.sanger.ac.uk/covid-team) |
| EPI_ISL_554661, EPI_ISL_554666 | Lighthouse Lab in Alderley Park | Wellcome Sanger Institute for the COVID-19 Genomics UK (COG-UK) consortium | The Lighthouse Lab in Alderley Park and Alex Alderton, Roberto Amato, Sonia Goncalves, Ewan Harrison, David K. Jackson, Ian Johnston, Dominic Kwiatkowski, Cordelia Langford, John Sillitoe on behalf of the Wellcome Sanger Institute COVID-19 Surveillance Team |
| EPI_ISL_554668, EPI_ISL_554670, EPI_ISL_554671, EPI_ISL_554672, EPI_ISL_554673, EPI_ISL_554674, EPI_ISL_554675, EPI_ISL_554676, EPI_ISL_554678, EPI_ISL_554679, EPI_ISL_554682, EPI_ISL_554683, EPI_ISL_554684, EPI_ISL_554685, EPI_ISL_554687, EPI_ISL_554690, EPI_ISL_554691, EPI_ISL_554695, EPI_ISL_554698, EPI_ISL_554699, EPI_ISL_554700, EPI_ISL_554701, EPI_ISL_554702, EPI_ISL_554704, EPI_ISL_554705, EPI_ISL_554706, EPI_ISL_554707, EPI_ISL_554711, EPI_ISL_554712, EPI_ISL_554713, EPI_ISL_554714, EPI_ISL_554715, EPI_ISL_554717, EPI_ISL_554718, EPI_ISL_554719, EPI_ISL_554720, EPI_ISL_554721, EPI_ISL_554722, EPI_ISL_554723, EPI_ISL_554724, EPI_ISL_554729, EPI_ISL_554731 | | | |
| see above | Lighthouse Lab in Milton Keynes | Wellcome Sanger Institute for the COVID-19 Genomics UK (COG-UK) consortium | The Lighthouse Lab in Milton Keynes and Alex Alderton, Roberto Amato, Sonia Goncalves, Ewan Harrison, David K. Jackson, Ian Johnston, Dominic Kwiatkowski, Cordelia Langford, John Sillitoe on behalf of the Wellcome Sanger Institute COVID-19 Surveillance Team (http://www.sanger.ac.uk/covid-team) |
| EPI_ISL_554732 | Lighthouse Lab in Alderley Park | Wellcome Sanger Institute for the COVID-19 Genomics UK (COG-UK) consortium | The Lighthouse Lab in Alderley Park and Alex Alderton, Roberto Amato, Sonia Goncalves, Ewan Harrison, David K. Jackson, Ian Johnston, Dominic Kwiatkowski, Cordelia Langford, John Sillitoe on behalf of the Wellcome Sanger Institute COVID-19 Surveillance Team |
| EPI_ISL_554733 | Lighthouse Lab in Milton Keynes | Wellcome Sanger Institute for the COVID-19 Genomics UK (COG-UK) consortium | The Lighthouse Lab in Milton Keynes and Alex Alderton, Roberto Amato, Sonia Goncalves, Ewan Harrison, David K. Jackson, Ian Johnston, Dominic Kwiatkowski, Cordelia Langford, John Sillitoe on behalf of the Wellcome Sanger Institute COVID-19 Surveillance Team (http://www.sanger.ac.uk/covid-team) |
| EPI_ISL_554734 | Lighthouse Lab in Milton Keynes | Wellcome Sanger Institute for the COVID-19 Genomics UK (COG-UK) consortium | The Lighthouse Lab in Alderley Park and Alex Alderton, Roberto Amato, Sonia Goncalves, Ewan Harrison, David K. Jackson, Ian Johnston, Dominic Kwiatkowski, Cordelia Langford, John Sillitoe on behalf of the Wellcome Sanger Institute COVID-19 Surveillance Team |
| EPI_ISL_554737, EPI_ISL_554738, EPI_ISL_554739, EPI_ISL_554740, EPI_ISL_554745, EPI_ISL_554746 | Lighthouse Lab in Milton Keynes | Wellcome Sanger Institute for the COVID-19 Genomics UK (COG-UK) consortium | The Lighthouse Lab in Milton Keynes and Alex Alderton, Roberto Amato, Sonia Goncalves, Ewan Harrison, David K. Jackson, Ian Johnston, Dominic Kwiatkowski, Cordelia Langford, John Sillitoe on behalf of the Wellcome Sanger Institute COVID-19 Surveillance Team (http://www.sanger.ac.uk/covid-team) |
| EPI_ISL_554747 | Lighthouse Lab in Milton Keynes | Wellcome Sanger Institute for the COVID-19 Genomics UK (COG-UK) consortium | The Lighthouse Lab in Alderley Park and Alex Alderton, Roberto Amato, Sonia Goncalves, Ewan Harrison, David K. Jackson, Ian Johnston, Dominic Kwiatkowski, Cordelia Langford, John Sillitoe on behalf of the Wellcome Sanger Institute COVID-19 Surveillance Team |
| EPI_ISL_554749, EPI_ISL_554751, EPI_ISL_554752, EPI_ISL_554753, EPI_ISL_554758, EPI_ISL_554760, EPI_ISL_554762, EPI_ISL_554765, EPI_ISL_554766, EPI_ISL_554768, EPI_ISL_554769, EPI_ISL_554770, EPI_ISL_554773 | | | |
| see above | Lighthouse Lab in Milton Keynes | Wellcome Sanger Institute for the COVID-19 Genomics UK (COG-UK) consortium | The Lighthouse Lab in Milton Keynes and Alex Alderton, Roberto Amato, Sonia Goncalves, Ewan Harrison, David K. Jackson, Ian Johnston, Dominic Kwiatkowski, Cordelia Langford, John Sillitoe on behalf of the Wellcome Sanger Institute COVID-19 Surveillance Team (http://www.sanger.ac.uk/covid-team) |
| EPI_ISL_554774 | Lighthouse Lab in Milton Keynes | Wellcome Sanger Institute for the COVID-19 Genomics UK (COG-UK) consortium | The Lighthouse Lab in Alderley Park and Alex Alderton, Roberto Amato, Sonia Goncalves, Ewan Harrison, David K. Jackson, Ian Johnston, Dominic Kwiatkowski, Cordelia Langford, John Sillitoe on behalf of the Wellcome Sanger Institute COVID-19 Surveillance Team |
| EPI_ISL_554775 | Lighthouse Lab in Milton Keynes | Wellcome Sanger Institute for the COVID-19 Genomics UK | The Lighthouse Lab in Milton Keynes and Alex Alderton, Roberto Amato, Sonia Goncalves, Ewan Harrison, David K. Jackson, Ian Johnston, Dominic |

| | | | |
|---|---|---|---|
| | | (COG-UK) consortium | Kwiatkowski, Cordelia Langford, John Sillitoe on behalf of the Wellcome Sanger Institute COVID-19 Surveillance Team (http://www.sanger.ac.uk/covid-team) |
| EPI_ISL_554776 | Lighthouse Lab in Alderley Park | Wellcome Sanger Institute for the COVID-19 Genomics UK (COG-UK) consortium | The Lighthouse Lab in Alderley Park and Alex Alderton, Roberto Amato, Sonia Goncalves, Ewan Harrison, David K. Jackson, Ian Johnston, Dominic Kwiatkowski, Cordelia Langford, John Sillitoe on behalf of the Wellcome Sanger Institute COVID-19 Surveillance Team |
| EPI_ISL_554783 | Lighthouse Lab in Milton Keynes | Wellcome Sanger Institute for the COVID-19 Genomics UK (COG-UK) consortium | The Lighthouse Lab in Alderley Park and Alex Alderton, Roberto Amato, Sonia Goncalves, Ewan Harrison, David K. Jackson, Ian Johnston, Dominic Kwiatkowski, Cordelia Langford, John Sillitoe on behalf of the Wellcome Sanger Institute COVID-19 Surveillance Team |
| EPI_ISL_554784, EPI_ISL_554785, EPI_ISL_554786, EPI_ISL_554787, EPI_ISL_554788, EPI_ISL_554789, EPI_ISL_554790, EPI_ISL_554791, EPI_ISL_554793, EPI_ISL_554796, EPI_ISL_554801, EPI_ISL_554805, EPI_ISL_554806, EPI_ISL_554814, EPI_ISL_554817, EPI_ISL_554818, EPI_ISL_554819, EPI_ISL_554821, EPI_ISL_554822, EPI_ISL_554823, EPI_ISL_554824, EPI_ISL_554825, EPI_ISL_554826, EPI_ISL_554828, EPI_ISL_554829, EPI_ISL_554831, EPI_ISL_554835, EPI_ISL_554838, EPI_ISL_554839 | | | |
| see above | Lighthouse Lab in Milton Keynes | Wellcome Sanger Institute for the COVID-19 Genomics UK (COG-UK) consortium | The Lighthouse Lab in Milton Keynes and Alex Alderton, Roberto Amato, Sonia Goncalves, Ewan Harrison, David K. Jackson, Ian Johnston, Dominic Kwiatkowski, Cordelia Langford, John Sillitoe on behalf of the Wellcome Sanger Institute COVID-19 Surveillance Team (http://www.sanger.ac.uk/covid-team) |
| EPI_ISL_554840 | Lighthouse Lab in Milton Keynes | Wellcome Sanger Institute for the COVID-19 Genomics UK (COG-UK) consortium | The Lighthouse Lab in Alderley Park and Alex Alderton, Roberto Amato, Sonia Goncalves, Ewan Harrison, David K. Jackson, Ian Johnston, Dominic Kwiatkowski, Cordelia Langford, John Sillitoe on behalf of the Wellcome Sanger Institute COVID-19 Surveillance Team |
| EPI_ISL_554842, EPI_ISL_554843, EPI_ISL_554844 | Lighthouse Lab in Milton Keynes | Wellcome Sanger Institute for the COVID-19 Genomics UK (COG-UK) consortium | The Lighthouse Lab in Milton Keynes and Alex Alderton, Roberto Amato, Sonia Goncalves, Ewan Harrison, David K. Jackson, Ian Johnston, Dominic Kwiatkowski, Cordelia Langford, John Sillitoe on behalf of the Wellcome Sanger Institute COVID-19 Surveillance Team (http://www.sanger.ac.uk/covid-team) |
| EPI_ISL_554849 | Lighthouse Lab in Milton Keynes | Wellcome Sanger Institute for the COVID-19 Genomics UK (COG-UK) consortium | The Lighthouse Lab in Alderley Park and Alex Alderton, Roberto Amato, Sonia Goncalves, Ewan Harrison, David K. Jackson, Ian Johnston, Dominic Kwiatkowski, Cordelia Langford, John Sillitoe on behalf of the Wellcome Sanger Institute COVID-19 Surveillance Team |
| EPI_ISL_554851, EPI_ISL_554852, EPI_ISL_554854, EPI_ISL_554855, EPI_ISL_554857, EPI_ISL_554860 | Lighthouse Lab in Milton Keynes | Wellcome Sanger Institute for the COVID-19 Genomics UK (COG-UK) consortium | The Lighthouse Lab in Milton Keynes and Alex Alderton, Roberto Amato, Sonia Goncalves, Ewan Harrison, David K. Jackson, Ian Johnston, Dominic Kwiatkowski, Cordelia Langford, John Sillitoe on behalf of the Wellcome Sanger Institute COVID-19 Surveillance Team (http://www.sanger.ac.uk/covid-team) |
| EPI_ISL_554861 | Lighthouse Lab in Alderley Park | Wellcome Sanger Institute for the COVID-19 Genomics UK (COG-UK) consortium | The Lighthouse Lab in Alderley Park and Alex Alderton, Roberto Amato, Sonia Goncalves, Ewan Harrison, David K. Jackson, Ian Johnston, Dominic Kwiatkowski, Cordelia Langford, John Sillitoe on behalf of the Wellcome Sanger Institute COVID-19 Surveillance Team |
| EPI_ISL_554862, EPI_ISL_554863 | Lighthouse Lab in Alderley Park | Wellcome Sanger Institute for the COVID-19 Genomics UK (COG-UK) consortium | The Lighthouse Lab in Alderley Park and Alex Alderton, Roberto Amato, Sonia Goncalves, Ewan Harrison, David K. Jackson, Ian Johnston, Dominic Kwiatkowski, Cordelia Langford, John Sillitoe on behalf of the Wellcome Sanger Institute COVID-19 Surveillance Team (http://www.sanger.ac.uk/covid-team) |
| EPI_ISL_554864, EPI_ISL_554865 | Lighthouse Lab in Alderley Park | Wellcome Sanger Institute for the COVID-19 Genomics UK (COG-UK) consortium | The Lighthouse Lab in Alderley Park and Alex Alderton, Roberto Amato, Sonia Goncalves, Ewan Harrison, David K. Jackson, Ian Johnston, Dominic Kwiatkowski, Cordelia Langford, John Sillitoe on behalf of the Wellcome Sanger Institute COVID-19 Surveillance Team |
| EPI_ISL_554866, EPI_ISL_554867, EPI_ISL_554869, EPI_ISL_554870, EPI_ISL_554871, EPI_ISL_554872, EPI_ISL_554873, EPI_ISL_554874, EPI_ISL_554875, EPI_ISL_554876, EPI_ISL_554877, EPI_ISL_554878, EPI_ISL_554879, EPI_ISL_554880, EPI_ISL_554881, EPI_ISL_554882, EPI_ISL_554883, EPI_ISL_554884, EPI_ISL_554885, EPI_ISL_554886, EPI_ISL_554887, EPI_ISL_554888, EPI_ISL_554889, EPI_ISL_554890, EPI_ISL_554891, EPI_ISL_554892, EPI_ISL_554893, EPI_ISL_554894, EPI_ISL_554895, EPI_ISL_554896, EPI_ISL_554897, EPI_ISL_554898, EPI_ISL_554900, EPI_ISL_554901 | | | |
| see above | Lighthouse Lab in Alderley Park | Wellcome Sanger Institute for the COVID-19 Genomics UK (COG-UK) consortium | The Lighthouse Lab in Alderley Park and Alex Alderton, Roberto Amato, Sonia Goncalves, Ewan Harrison, David K. Jackson, Ian Johnston, Dominic Kwiatkowski, Cordelia Langford, John Sillitoe on behalf of the Wellcome Sanger Institute COVID-19 Surveillance Team (http://www.sanger.ac.uk/covid-team) |
| EPI_ISL_554902 | Lighthouse Lab in Alderley Park | Wellcome Sanger Institute for the COVID-19 Genomics UK (COG-UK) consortium | The Lighthouse Lab in Alderley Park and Alex Alderton, Roberto Amato, Sonia Goncalves, Ewan Harrison, David K. Jackson, Ian Johnston, Dominic Kwiatkowski, Cordelia Langford, John Sillitoe on behalf of the Wellcome Sanger Institute COVID-19 Surveillance Team |
| EPI_ISL_554903, EPI_ISL_554904, EPI_ISL_554905, EPI_ISL_554906, EPI_ISL_554907, EPI_ISL_554908, EPI_ISL_554909, EPI_ISL_554910, EPI_ISL_554911, EPI_ISL_554912, EPI_ISL_554913, EPI_ISL_554914, EPI_ISL_554915, EPI_ISL_554916, EPI_ISL_554917, EPI_ISL_554918, EPI_ISL_554919, EPI_ISL_554920, EPI_ISL_554921, EPI_ISL_554922, EPI_ISL_554923 | | | |
| see above | Lighthouse Lab in Alderley Park | Wellcome Sanger Institute for the COVID-19 Genomics UK (COG-UK) consortium | The Lighthouse Lab in Alderley Park and Alex Alderton, Roberto Amato, Sonia Goncalves, Ewan Harrison, David K. Jackson, Ian Johnston, Dominic Kwiatkowski, Cordelia Langford, John Sillitoe on behalf of the Wellcome Sanger Institute COVID-19 Surveillance Team (http://www.sanger.ac.uk/covid-team) |
| EPI_ISL_554924 | Lighthouse Lab in Alderley Park | Wellcome Sanger Institute for the COVID-19 Genomics UK (COG-UK) consortium | The Lighthouse Lab in Alderley Park and Alex Alderton, Roberto Amato, Sonia Goncalves, Ewan Harrison, David K. Jackson, Ian Johnston, Dominic Kwiatkowski, Cordelia Langford, John Sillitoe on behalf of the Wellcome Sanger Institute COVID-19 Surveillance Team |
| EPI_ISL_554925, EPI_ISL_554926, EPI_ISL_554927 | Lighthouse Lab in Alderley Park | Wellcome Sanger Institute for the COVID-19 Genomics UK (COG-UK) consortium | The Lighthouse Lab in Alderley Park and Alex Alderton, Roberto Amato, Sonia Goncalves, Ewan Harrison, David K. Jackson, Ian Johnston, Dominic Kwiatkowski, Cordelia Langford, John Sillitoe on behalf of the Wellcome Sanger Institute COVID-19 Surveillance Team (http://www.sanger.ac.uk/covid-team) |
| EPI_ISL_554928 | Lighthouse Lab in Alderley Park | Wellcome Sanger Institute for the COVID-19 Genomics UK (COG-UK) consortium | The Lighthouse Lab in Alderley Park and Alex Alderton, Roberto Amato, Sonia Goncalves, Ewan Harrison, David K. Jackson, Ian Johnston, Dominic Kwiatkowski, Cordelia Langford, John Sillitoe on behalf of the Wellcome Sanger Institute COVID-19 Surveillance Team |
| EPI_ISL_554930, EPI_ISL_554931, EPI_ISL_554932, EPI_ISL_554933, EPI_ISL_554934 | Lighthouse Lab in Alderley Park | Wellcome Sanger Institute for the COVID-19 Genomics UK (COG-UK) consortium | The Lighthouse Lab in Alderley Park and Alex Alderton, Roberto Amato, Sonia Goncalves, Ewan Harrison, David K. Jackson, Ian Johnston, Dominic Kwiatkowski, Cordelia Langford, John Sillitoe on behalf of the Wellcome Sanger Institute COVID-19 Surveillance Team (http://www.sanger.ac.uk/covid-team) |
| EPI_ISL_554935, EPI_ISL_554936, EPI_ISL_554937 | Lighthouse Lab in Alderley Park | Wellcome Sanger Institute for the COVID-19 Genomics UK (COG-UK) consortium | The Lighthouse Lab in Alderley Park and Alex Alderton, Roberto Amato, Sonia Goncalves, Ewan Harrison, David K. Jackson, Ian Johnston, Dominic Kwiatkowski, Cordelia Langford, John Sillitoe on behalf of the Wellcome Sanger Institute COVID-19 Surveillance Team |
| EPI_ISL_554938, EPI_ISL_554939, EPI_ISL_554940, EPI_ISL_554941, EPI_ISL_554942, EPI_ISL_554943 | Lighthouse Lab in Alderley Park | Wellcome Sanger Institute for the COVID-19 Genomics UK (COG-UK) consortium | The Lighthouse Lab in Alderley Park and Alex Alderton, Roberto Amato, Sonia Goncalves, Ewan Harrison, David K. Jackson, Ian Johnston, Dominic Kwiatkowski, Cordelia Langford, John Sillitoe on behalf of the Wellcome Sanger Institute COVID-19 Surveillance Team (http://www.sanger.ac.uk/covid-team) |
| EPI_ISL_554944 | Lighthouse Lab in Alderley Park | Wellcome Sanger Institute for the COVID-19 Genomics UK (COG-UK) consortium | The Lighthouse Lab in Alderley Park and Alex Alderton, Roberto Amato, Sonia Goncalves, Ewan Harrison, David K. Jackson, Ian Johnston, Dominic Kwiatkowski, Cordelia Langford, John Sillitoe on behalf of the Wellcome Sanger Institute COVID-19 Surveillance Team |
| EPI_ISL_554945, EPI_ISL_554946, EPI_ISL_554947, EPI_ISL_554948, EPI_ISL_554949, EPI_ISL_554950, EPI_ISL_554951, EPI_ISL_554952, EPI_ISL_554953, EPI_ISL_554954, EPI_ISL_554955, EPI_ISL_554956, EPI_ISL_554957, EPI_ISL_554958, EPI_ISL_554959, EPI_ISL_554960 | | | |
| see above | Lighthouse Lab in Alderley Park | Wellcome Sanger Institute for the COVID-19 Genomics UK (COG-UK) consortium | The Lighthouse Lab in Alderley Park and Alex Alderton, Roberto Amato, Sonia Goncalves, Ewan Harrison, David K. Jackson, Ian Johnston, Dominic Kwiatkowski, Cordelia Langford, John Sillitoe on behalf of the Wellcome Sanger Institute COVID-19 Surveillance Team (http://www.sanger.ac.uk/covid-team) |
| EPI_ISL_554961 | Lighthouse Lab in Alderley Park | Wellcome Sanger Institute for the COVID-19 Genomics UK (COG-UK) consortium | The Lighthouse Lab in Alderley Park and Alex Alderton, Roberto Amato, Sonia Goncalves, Ewan Harrison, David K. Jackson, Ian Johnston, Dominic Kwiatkowski, Cordelia Langford, John Sillitoe on behalf of the Wellcome Sanger Institute COVID-19 Surveillance Team |
| EPI_ISL_554962, EPI_ISL_554963, EPI_ISL_554964, EPI_ISL_554965, EPI_ISL_554966, EPI_ISL_554967 | Lighthouse Lab in Alderley Park | Wellcome Sanger Institute for the COVID-19 Genomics UK (COG-UK) consortium | The Lighthouse Lab in Alderley Park and Alex Alderton, Roberto Amato, Sonia Goncalves, Ewan Harrison, David K. Jackson, Ian Johnston, Dominic Kwiatkowski, Cordelia Langford, John Sillitoe on behalf of the Wellcome Sanger Institute COVID-19 Surveillance Team (http://www.sanger.ac.uk/covid-team) |
| EPI_ISL_554968, EPI_ISL_554969 | Lighthouse Lab in Alderley Park | Wellcome Sanger Institute for the COVID-19 Genomics UK (COG-UK) consortium | The Lighthouse Lab in Alderley Park and Alex Alderton, Roberto Amato, Sonia Goncalves, Ewan Harrison, David K. Jackson, Ian Johnston, Dominic Kwiatkowski, Cordelia Langford, John Sillitoe on behalf of the Wellcome Sanger Institute COVID-19 Surveillance Team |
| EPI_ISL_554970, EPI_ISL_554971, EPI_ISL_554972, EPI_ISL_554973, EPI_ISL_554974, EPI_ISL_554975, EPI_ISL_554976, EPI_ISL_554977 | Lighthouse Lab in Alderley Park | Wellcome Sanger Institute for the COVID-19 Genomics UK (COG-UK) consortium | The Lighthouse Lab in Alderley Park and Alex Alderton, Roberto Amato, Sonia Goncalves, Ewan Harrison, David K. Jackson, Ian Johnston, Dominic Kwiatkowski, Cordelia Langford, John Sillitoe on behalf of the Wellcome Sanger Institute COVID-19 Surveillance Team (http://www.sanger.ac.uk/covid-team) |
| EPI_ISL_554978 | Lighthouse Lab in Alderley Park | Wellcome Sanger Institute for the COVID-19 Genomics UK | The Lighthouse Lab in Alderley Park and Alex Alderton, Roberto Amato, Sonia Goncalves, Ewan Harrison, David K. Jackson, Ian Johnston, Dominic |

| | | | |
|---|---|---|---|
| | | (COG-UK) consortium | Kwiatkowski, Cordelia Langford, John Sillitoe on behalf of the Wellcome Sanger Institute COVID-19 Surveillance Team |
| EPI_ISL_554979, EPI_ISL_554980, EPI_ISL_554981, EPI_ISL_554982, EPI_ISL_554983, EPI_ISL_554984, EPI_ISL_554985, EPI_ISL_554986, EPI_ISL_554987, EPI_ISL_554988 | Lighthouse Lab in Alderley Park | Wellcome Sanger Institute for the COVID-19 Genomics UK (COG-UK) consortium | The Lighthouse Lab in Alderley Park and Alex Alderton, Roberto Amato, Sonia Goncalves, Ewan Harrison, David K. Jackson, Ian Johnston, Dominic Kwiatkowski, Cordelia Langford, John Sillitoe on behalf of the Wellcome Sanger Institute COVID-19 Surveillance Team (http://www.sanger.ac.uk/covid-team) |
| EPI_ISL_554989 | Lighthouse Lab in Alderley Park | Wellcome Sanger Institute for the COVID-19 Genomics UK (COG-UK) consortium | The Lighthouse Lab in Alderley Park and Alex Alderton, Roberto Amato, Sonia Goncalves, Ewan Harrison, David K. Jackson, Ian Johnston, Dominic Kwiatkowski, Cordelia Langford, John Sillitoe on behalf of the Wellcome Sanger Institute COVID-19 Surveillance Team |
| EPI_ISL_554990, EPI_ISL_554991, EPI_ISL_554992, EPI_ISL_554993, EPI_ISL_554994, EPI_ISL_554995 | Lighthouse Lab in Alderley Park | Wellcome Sanger Institute for the COVID-19 Genomics UK (COG-UK) consortium | The Lighthouse Lab in Alderley Park and Alex Alderton, Roberto Amato, Sonia Goncalves, Ewan Harrison, David K. Jackson, Ian Johnston, Dominic Kwiatkowski, Cordelia Langford, John Sillitoe on behalf of the Wellcome Sanger Institute COVID-19 Surveillance Team (http://www.sanger.ac.uk/covid-team) |
| EPI_ISL_554996 | Lighthouse Lab in Alderley Park | Wellcome Sanger Institute for the COVID-19 Genomics UK (COG-UK) consortium | The Lighthouse Lab in Alderley Park and Alex Alderton, Roberto Amato, Sonia Goncalves, Ewan Harrison, David K. Jackson, Ian Johnston, Dominic Kwiatkowski, Cordelia Langford, John Sillitoe on behalf of the Wellcome Sanger Institute COVID-19 Surveillance Team |
| EPI_ISL_554997, EPI_ISL_554998, EPI_ISL_554999 | Lighthouse Lab in Alderley Park | Wellcome Sanger Institute for the COVID-19 Genomics UK (COG-UK) consortium | The Lighthouse Lab in Alderley Park and Alex Alderton, Roberto Amato, Sonia Goncalves, Ewan Harrison, David K. Jackson, Ian Johnston, Dominic Kwiatkowski, Cordelia Langford, John Sillitoe on behalf of the Wellcome Sanger Institute COVID-19 Surveillance Team (http://www.sanger.ac.uk/covid-team) |
| EPI_ISL_555000 | Lighthouse Lab in Alderley Park | Wellcome Sanger Institute for the COVID-19 Genomics UK (COG-UK) consortium | The Lighthouse Lab in Alderley Park and Alex Alderton, Roberto Amato, Sonia Goncalves, Ewan Harrison, David K. Jackson, Ian Johnston, Dominic Kwiatkowski, Cordelia Langford, John Sillitoe on behalf of the Wellcome Sanger Institute COVID-19 Surveillance Team |
| EPI_ISL_555001, EPI_ISL_555002 | Lighthouse Lab in Alderley Park | Wellcome Sanger Institute for the COVID-19 Genomics UK (COG-UK) consortium | The Lighthouse Lab in Alderley Park and Alex Alderton, Roberto Amato, Sonia Goncalves, Ewan Harrison, David K. Jackson, Ian Johnston, Dominic Kwiatkowski, Cordelia Langford, John Sillitoe on behalf of the Wellcome Sanger Institute COVID-19 Surveillance Team (http://www.sanger.ac.uk/covid-team) |
| EPI_ISL_555003 | Lighthouse Lab in Alderley Park | Wellcome Sanger Institute for the COVID-19 Genomics UK (COG-UK) consortium | The Lighthouse Lab in Alderley Park and Alex Alderton, Roberto Amato, Sonia Goncalves, Ewan Harrison, David K. Jackson, Ian Johnston, Dominic Kwiatkowski, Cordelia Langford, John Sillitoe on behalf of the Wellcome Sanger Institute COVID-19 Surveillance Team |
| EPI_ISL_555004, EPI_ISL_555005, EPI_ISL_555006, EPI_ISL_555007, EPI_ISL_555008, EPI_ISL_555009, EPI_ISL_555010, EPI_ISL_555011 | Lighthouse Lab in Alderley Park | Wellcome Sanger Institute for the COVID-19 Genomics UK (COG-UK) consortium | The Lighthouse Lab in Alderley Park and Alex Alderton, Roberto Amato, Sonia Goncalves, Ewan Harrison, David K. Jackson, Ian Johnston, Dominic Kwiatkowski, Cordelia Langford, John Sillitoe on behalf of the Wellcome Sanger Institute COVID-19 Surveillance Team (http://www.sanger.ac.uk/covid-team) |
| EPI_ISL_555012 | Lighthouse Lab in Alderley Park | Wellcome Sanger Institute for the COVID-19 Genomics UK (COG-UK) consortium | The Lighthouse Lab in Alderley Park and Alex Alderton, Roberto Amato, Sonia Goncalves, Ewan Harrison, David K. Jackson, Ian Johnston, Dominic Kwiatkowski, Cordelia Langford, John Sillitoe on behalf of the Wellcome Sanger Institute COVID-19 Surveillance Team |
| EPI_ISL_555013, EPI_ISL_555014, EPI_ISL_555015, EPI_ISL_555016, EPI_ISL_555017 | Lighthouse Lab in Alderley Park | Wellcome Sanger Institute for the COVID-19 Genomics UK (COG-UK) consortium | The Lighthouse Lab in Alderley Park and Alex Alderton, Roberto Amato, Sonia Goncalves, Ewan Harrison, David K. Jackson, Ian Johnston, Dominic Kwiatkowski, Cordelia Langford, John Sillitoe on behalf of the Wellcome Sanger Institute COVID-19 Surveillance Team (http://www.sanger.ac.uk/covid-team) |
| EPI_ISL_555018 | Lighthouse Lab in Alderley Park | Wellcome Sanger Institute for the COVID-19 Genomics UK (COG-UK) consortium | The Lighthouse Lab in Alderley Park and Alex Alderton, Roberto Amato, Sonia Goncalves, Ewan Harrison, David K. Jackson, Ian Johnston, Dominic Kwiatkowski, Cordelia Langford, John Sillitoe on behalf of the Wellcome Sanger Institute COVID-19 Surveillance Team |
| EPI_ISL_555020, EPI_ISL_555021, EPI_ISL_555022 | Lighthouse Lab in Alderley Park | Wellcome Sanger Institute for the COVID-19 Genomics UK (COG-UK) consortium | The Lighthouse Lab in Alderley Park and Alex Alderton, Roberto Amato, Sonia Goncalves, Ewan Harrison, David K. Jackson, Ian Johnston, Dominic Kwiatkowski, Cordelia Langford, John Sillitoe on behalf of the Wellcome Sanger Institute COVID-19 Surveillance Team (http://www.sanger.ac.uk/covid-team) |
| EPI_ISL_555023, EPI_ISL_555024 | Lighthouse Lab in Alderley Park | Wellcome Sanger Institute for the COVID-19 Genomics UK (COG-UK) consortium | The Lighthouse Lab in Alderley Park and Alex Alderton, Roberto Amato, Sonia Goncalves, Ewan Harrison, David K. Jackson, Ian Johnston, Dominic Kwiatkowski, Cordelia Langford, John Sillitoe on behalf of the Wellcome Sanger Institute COVID-19 Surveillance Team |
| EPI_ISL_555025, EPI_ISL_555026, EPI_ISL_555027, EPI_ISL_555028 | Lighthouse Lab in Alderley Park | Wellcome Sanger Institute for the COVID-19 Genomics UK (COG-UK) consortium | The Lighthouse Lab in Alderley Park and Alex Alderton, Roberto Amato, Sonia Goncalves, Ewan Harrison, David K. Jackson, Ian Johnston, Dominic Kwiatkowski, Cordelia Langford, John Sillitoe on behalf of the Wellcome Sanger Institute COVID-19 Surveillance Team (http://www.sanger.ac.uk/covid-team) |
| EPI_ISL_555029 | Lighthouse Lab in Alderley Park | Wellcome Sanger Institute for the COVID-19 Genomics UK (COG-UK) consortium | The Lighthouse Lab in Alderley Park and Alex Alderton, Roberto Amato, Sonia Goncalves, Ewan Harrison, David K. Jackson, Ian Johnston, Dominic Kwiatkowski, Cordelia Langford, John Sillitoe on behalf of the Wellcome Sanger Institute COVID-19 Surveillance Team |
| EPI_ISL_555030, EPI_ISL_555031, EPI_ISL_555032, EPI_ISL_555033, EPI_ISL_555034, EPI_ISL_555035, EPI_ISL_555036, EPI_ISL_555037, EPI_ISL_555038 | Lighthouse Lab in Alderley Park | Wellcome Sanger Institute for the COVID-19 Genomics UK (COG-UK) consortium | The Lighthouse Lab in Alderley Park and Alex Alderton, Roberto Amato, Sonia Goncalves, Ewan Harrison, David K. Jackson, Ian Johnston, Dominic Kwiatkowski, Cordelia Langford, John Sillitoe on behalf of the Wellcome Sanger Institute COVID-19 Surveillance Team (http://www.sanger.ac.uk/covid-team) |
| EPI_ISL_555039 | Lighthouse Lab in Alderley Park | Wellcome Sanger Institute for the COVID-19 Genomics UK (COG-UK) consortium | The Lighthouse Lab in Alderley Park and Alex Alderton, Roberto Amato, Sonia Goncalves, Ewan Harrison, David K. Jackson, Ian Johnston, Dominic Kwiatkowski, Cordelia Langford, John Sillitoe on behalf of the Wellcome Sanger Institute COVID-19 Surveillance Team |
| EPI_ISL_555040, EPI_ISL_555042 | Lighthouse Lab in Alderley Park | Wellcome Sanger Institute for the COVID-19 Genomics UK (COG-UK) consortium | The Lighthouse Lab in Alderley Park and Alex Alderton, Roberto Amato, Sonia Goncalves, Ewan Harrison, David K. Jackson, Ian Johnston, Dominic Kwiatkowski, Cordelia Langford, John Sillitoe on behalf of the Wellcome Sanger Institute COVID-19 Surveillance Team (http://www.sanger.ac.uk/covid-team) |
| EPI_ISL_555043 | Lighthouse Lab in Alderley Park | Wellcome Sanger Institute for the COVID-19 Genomics UK (COG-UK) consortium | The Lighthouse Lab in Alderley Park and Alex Alderton, Roberto Amato, Sonia Goncalves, Ewan Harrison, David K. Jackson, Ian Johnston, Dominic Kwiatkowski, Cordelia Langford, John Sillitoe on behalf of the Wellcome Sanger Institute COVID-19 Surveillance Team |
| EPI_ISL_555044, EPI_ISL_555045, EPI_ISL_555046, EPI_ISL_555047 | Lighthouse Lab in Alderley Park | Wellcome Sanger Institute for the COVID-19 Genomics UK (COG-UK) consortium | The Lighthouse Lab in Alderley Park and Alex Alderton, Roberto Amato, Sonia Goncalves, Ewan Harrison, David K. Jackson, Ian Johnston, Dominic Kwiatkowski, Cordelia Langford, John Sillitoe on behalf of the Wellcome Sanger Institute COVID-19 Surveillance Team (http://www.sanger.ac.uk/covid-team) |
| EPI_ISL_555048, EPI_ISL_555049 | Lighthouse Lab in Alderley Park | Wellcome Sanger Institute for the COVID-19 Genomics UK (COG-UK) consortium | The Lighthouse Lab in Alderley Park and Alex Alderton, Roberto Amato, Sonia Goncalves, Ewan Harrison, David K. Jackson, Ian Johnston, Dominic Kwiatkowski, Cordelia Langford, John Sillitoe on behalf of the Wellcome Sanger Institute COVID-19 Surveillance Team |
| EPI_ISL_555050, EPI_ISL_555051, EPI_ISL_555052 | Lighthouse Lab in Alderley Park | Wellcome Sanger Institute for the COVID-19 Genomics UK (COG-UK) consortium | The Lighthouse Lab in Alderley Park and Alex Alderton, Roberto Amato, Sonia Goncalves, Ewan Harrison, David K. Jackson, Ian Johnston, Dominic Kwiatkowski, Cordelia Langford, John Sillitoe on behalf of the Wellcome Sanger Institute COVID-19 Surveillance Team (http://www.sanger.ac.uk/covid-team) |
| EPI_ISL_555053 | Lighthouse Lab in Alderley Park | Wellcome Sanger Institute for the COVID-19 Genomics UK (COG-UK) consortium | The Lighthouse Lab in Alderley Park and Alex Alderton, Roberto Amato, Sonia Goncalves, Ewan Harrison, David K. Jackson, Ian Johnston, Dominic Kwiatkowski, Cordelia Langford, John Sillitoe on behalf of the Wellcome Sanger Institute COVID-19 Surveillance Team |
| EPI_ISL_555054, EPI_ISL_555055, EPI_ISL_555056, EPI_ISL_555057, EPI_ISL_555058, EPI_ISL_555059 | Lighthouse Lab in Alderley Park | Wellcome Sanger Institute for the COVID-19 Genomics UK (COG-UK) consortium | The Lighthouse Lab in Alderley Park and Alex Alderton, Roberto Amato, Sonia Goncalves, Ewan Harrison, David K. Jackson, Ian Johnston, Dominic Kwiatkowski, Cordelia Langford, John Sillitoe on behalf of the Wellcome Sanger Institute COVID-19 Surveillance Team (http://www.sanger.ac.uk/covid-team) |
| EPI_ISL_555060 | Lighthouse Lab in Alderley Park | Wellcome Sanger Institute for the COVID-19 Genomics UK (COG-UK) consortium | The Lighthouse Lab in Alderley Park and Alex Alderton, Roberto Amato, Sonia Goncalves, Ewan Harrison, David K. Jackson, Ian Johnston, Dominic Kwiatkowski, Cordelia Langford, John Sillitoe on behalf of the Wellcome Sanger Institute COVID-19 Surveillance Team |
| EPI_ISL_555061 | Lighthouse Lab in Alderley Park | Wellcome Sanger Institute for the COVID-19 Genomics UK (COG-UK) consortium | The Lighthouse Lab in Alderley Park and Alex Alderton, Roberto Amato, Sonia Goncalves, Ewan Harrison, David K. Jackson, Ian Johnston, Dominic Kwiatkowski, Cordelia Langford, John Sillitoe on behalf of the Wellcome Sanger Institute COVID-19 Surveillance Team (http://www.sanger.ac.uk/covid-team) |

| | | | |
|---|---|---|---|
| EPI_ISL_555062 | Lighthouse Lab in Alderley Park | Wellcome Sanger Institute for the COVID-19 Genomics UK (COG-UK) consortium | The Lighthouse Lab in Alderley Park and Alex Alderton, Roberto Amato, Sonia Goncalves, Ewan Harrison, David K. Jackson, Ian Johnston, Dominic Kwiatkowski, Cordelia Langford, John Sillitoe on behalf of the Wellcome Sanger Institute COVID-19 Surveillance Team |
| EPI_ISL_555063, EPI_ISL_555064, EPI_ISL_555065 | Lighthouse Lab in Alderley Park | Wellcome Sanger Institute for the COVID-19 Genomics UK (COG-UK) consortium | The Lighthouse Lab in Alderley Park and Alex Alderton, Roberto Amato, Sonia Goncalves, Ewan Harrison, David K. Jackson, Ian Johnston, Dominic Kwiatkowski, Cordelia Langford, John Sillitoe on behalf of the Wellcome Sanger Institute COVID-19 Surveillance Team (http://www.sanger.ac.uk/covid-team) |
| EPI_ISL_555066 | Lighthouse Lab in Alderley Park | Wellcome Sanger Institute for the COVID-19 Genomics UK (COG-UK) consortium | The Lighthouse Lab in Alderley Park and Alex Alderton, Roberto Amato, Sonia Goncalves, Ewan Harrison, David K. Jackson, Ian Johnston, Dominic Kwiatkowski, Cordelia Langford, John Sillitoe on behalf of the Wellcome Sanger Institute COVID-19 Surveillance Team |
| EPI_ISL_555067, EPI_ISL_555068 | Lighthouse Lab in Alderley Park | Wellcome Sanger Institute for the COVID-19 Genomics UK (COG-UK) consortium | The Lighthouse Lab in Alderley Park and Alex Alderton, Roberto Amato, Sonia Goncalves, Ewan Harrison, David K. Jackson, Ian Johnston, Dominic Kwiatkowski, Cordelia Langford, John Sillitoe on behalf of the Wellcome Sanger Institute COVID-19 Surveillance Team (http://www.sanger.ac.uk/covid-team) |
| EPI_ISL_555069, EPI_ISL_555070 | Lighthouse Lab in Alderley Park | Wellcome Sanger Institute for the COVID-19 Genomics UK (COG-UK) consortium | The Lighthouse Lab in Alderley Park and Alex Alderton, Roberto Amato, Sonia Goncalves, Ewan Harrison, David K. Jackson, Ian Johnston, Dominic Kwiatkowski, Cordelia Langford, John Sillitoe on behalf of the Wellcome Sanger Institute COVID-19 Surveillance Team |
| EPI_ISL_555071, EPI_ISL_555072, EPI_ISL_555073, EPI_ISL_555074, EPI_ISL_555075, EPI_ISL_555076, EPI_ISL_555077, EPI_ISL_555078, EPI_ISL_555079, EPI_ISL_555080, EPI_ISL_555081, EPI_ISL_555082, EPI_ISL_555083, EPI_ISL_555084 | | | |
| see above | Lighthouse Lab in Alderley Park | Wellcome Sanger Institute for the COVID-19 Genomics UK (COG-UK) consortium | The Lighthouse Lab in Alderley Park and Alex Alderton, Roberto Amato, Sonia Goncalves, Ewan Harrison, David K. Jackson, Ian Johnston, Dominic Kwiatkowski, Cordelia Langford, John Sillitoe on behalf of the Wellcome Sanger Institute COVID-19 Surveillance Team (http://www.sanger.ac.uk/covid-team) |
| EPI_ISL_555085 | Lighthouse Lab in Alderley Park | Wellcome Sanger Institute for the COVID-19 Genomics UK (COG-UK) consortium | The Lighthouse Lab in Alderley Park and Alex Alderton, Roberto Amato, Sonia Goncalves, Ewan Harrison, David K. Jackson, Ian Johnston, Dominic Kwiatkowski, Cordelia Langford, John Sillitoe on behalf of the Wellcome Sanger Institute COVID-19 Surveillance Team |
| EPI_ISL_555086, EPI_ISL_555087, EPI_ISL_555088 | Lighthouse Lab in Alderley Park | Wellcome Sanger Institute for the COVID-19 Genomics UK (COG-UK) consortium | The Lighthouse Lab in Alderley Park and Alex Alderton, Roberto Amato, Sonia Goncalves, Ewan Harrison, David K. Jackson, Ian Johnston, Dominic Kwiatkowski, Cordelia Langford, John Sillitoe on behalf of the Wellcome Sanger Institute COVID-19 Surveillance Team (http://www.sanger.ac.uk/covid-team) |
| EPI_ISL_555089 | Lighthouse Lab in Alderley Park | Wellcome Sanger Institute for the COVID-19 Genomics UK (COG-UK) consortium | The Lighthouse Lab in Alderley Park and Alex Alderton, Roberto Amato, Sonia Goncalves, Ewan Harrison, David K. Jackson, Ian Johnston, Dominic Kwiatkowski, Cordelia Langford, John Sillitoe on behalf of the Wellcome Sanger Institute COVID-19 Surveillance Team |
| EPI_ISL_555090 | Lighthouse Lab in Alderley Park | Wellcome Sanger Institute for the COVID-19 Genomics UK (COG-UK) consortium | The Lighthouse Lab in Alderley Park and Alex Alderton, Roberto Amato, Sonia Goncalves, Ewan Harrison, David K. Jackson, Ian Johnston, Dominic Kwiatkowski, Cordelia Langford, John Sillitoe on behalf of the Wellcome Sanger Institute COVID-19 Surveillance Team (http://www.sanger.ac.uk/covid-team) |
| EPI_ISL_555091 | Lighthouse Lab in Alderley Park | Wellcome Sanger Institute for the COVID-19 Genomics UK (COG-UK) consortium | The Lighthouse Lab in Alderley Park and Alex Alderton, Roberto Amato, Sonia Goncalves, Ewan Harrison, David K. Jackson, Ian Johnston, Dominic Kwiatkowski, Cordelia Langford, John Sillitoe on behalf of the Wellcome Sanger Institute COVID-19 Surveillance Team |
| EPI_ISL_555092, EPI_ISL_555093, EPI_ISL_555094, EPI_ISL_555095, EPI_ISL_555096, EPI_ISL_555097, EPI_ISL_555098, EPI_ISL_555099, EPI_ISL_555100, EPI_ISL_555101, EPI_ISL_555102, EPI_ISL_555103 | | | |
| see above | Lighthouse Lab in Alderley Park | Wellcome Sanger Institute for the COVID-19 Genomics UK (COG-UK) consortium | The Lighthouse Lab in Alderley Park and Alex Alderton, Roberto Amato, Sonia Goncalves, Ewan Harrison, David K. Jackson, Ian Johnston, Dominic Kwiatkowski, Cordelia Langford, John Sillitoe on behalf of the Wellcome Sanger Institute COVID-19 Surveillance Team (http://www.sanger.ac.uk/covid-team) |
| EPI_ISL_555104, EPI_ISL_555105 | Lighthouse Lab in Alderley Park | Wellcome Sanger Institute for the COVID-19 Genomics UK (COG-UK) consortium | The Lighthouse Lab in Alderley Park and Alex Alderton, Roberto Amato, Sonia Goncalves, Ewan Harrison, David K. Jackson, Ian Johnston, Dominic Kwiatkowski, Cordelia Langford, John Sillitoe on behalf of the Wellcome Sanger Institute COVID-19 Surveillance Team |
| EPI_ISL_555106, EPI_ISL_555107, EPI_ISL_555108, EPI_ISL_555109, EPI_ISL_555110 | Lighthouse Lab in Alderley Park | Wellcome Sanger Institute for the COVID-19 Genomics UK (COG-UK) consortium | The Lighthouse Lab in Alderley Park and Alex Alderton, Roberto Amato, Sonia Goncalves, Ewan Harrison, David K. Jackson, Ian Johnston, Dominic Kwiatkowski, Cordelia Langford, John Sillitoe on behalf of the Wellcome Sanger Institute COVID-19 Surveillance Team (http://www.sanger.ac.uk/covid-team) |
| EPI_ISL_555111 | Lighthouse Lab in Alderley Park | Wellcome Sanger Institute for the COVID-19 Genomics UK (COG-UK) consortium | The Lighthouse Lab in Alderley Park and Alex Alderton, Roberto Amato, Sonia Goncalves, Ewan Harrison, David K. Jackson, Ian Johnston, Dominic Kwiatkowski, Cordelia Langford, John Sillitoe on behalf of the Wellcome Sanger Institute COVID-19 Surveillance Team |
| EPI_ISL_555112, EPI_ISL_555113, EPI_ISL_555114, EPI_ISL_555115, EPI_ISL_555116 | Lighthouse Lab in Alderley Park | Wellcome Sanger Institute for the COVID-19 Genomics UK (COG-UK) consortium | The Lighthouse Lab in Alderley Park and Alex Alderton, Roberto Amato, Sonia Goncalves, Ewan Harrison, David K. Jackson, Ian Johnston, Dominic Kwiatkowski, Cordelia Langford, John Sillitoe on behalf of the Wellcome Sanger Institute COVID-19 Surveillance Team (http://www.sanger.ac.uk/covid-team) |
| EPI_ISL_555117 | Lighthouse Lab in Alderley Park | Wellcome Sanger Institute for the COVID-19 Genomics UK (COG-UK) consortium | The Lighthouse Lab in Alderley Park and Alex Alderton, Roberto Amato, Sonia Goncalves, Ewan Harrison, David K. Jackson, Ian Johnston, Dominic Kwiatkowski, Cordelia Langford, John Sillitoe on behalf of the Wellcome Sanger Institute COVID-19 Surveillance Team |
| EPI_ISL_555118, EPI_ISL_555119, EPI_ISL_555120, EPI_ISL_555121 | Lighthouse Lab in Alderley Park | Wellcome Sanger Institute for the COVID-19 Genomics UK (COG-UK) consortium | The Lighthouse Lab in Alderley Park and Alex Alderton, Roberto Amato, Sonia Goncalves, Ewan Harrison, David K. Jackson, Ian Johnston, Dominic Kwiatkowski, Cordelia Langford, John Sillitoe on behalf of the Wellcome Sanger Institute COVID-19 Surveillance Team (http://www.sanger.ac.uk/covid-team) |
| EPI_ISL_555122 | Lighthouse Lab in Alderley Park | Wellcome Sanger Institute for the COVID-19 Genomics UK (COG-UK) consortium | The Lighthouse Lab in Alderley Park and Alex Alderton, Roberto Amato, Sonia Goncalves, Ewan Harrison, David K. Jackson, Ian Johnston, Dominic Kwiatkowski, Cordelia Langford, John Sillitoe on behalf of the Wellcome Sanger Institute COVID-19 Surveillance Team |
| EPI_ISL_555123, EPI_ISL_555124 | Lighthouse Lab in Alderley Park | Wellcome Sanger Institute for the COVID-19 Genomics UK (COG-UK) consortium | The Lighthouse Lab in Alderley Park and Alex Alderton, Roberto Amato, Sonia Goncalves, Ewan Harrison, David K. Jackson, Ian Johnston, Dominic Kwiatkowski, Cordelia Langford, John Sillitoe on behalf of the Wellcome Sanger Institute COVID-19 Surveillance Team (http://www.sanger.ac.uk/covid-team) |
| EPI_ISL_555125 | Lighthouse Lab in Alderley Park | Wellcome Sanger Institute for the COVID-19 Genomics UK (COG-UK) consortium | The Lighthouse Lab in Alderley Park and Alex Alderton, Roberto Amato, Sonia Goncalves, Ewan Harrison, David K. Jackson, Ian Johnston, Dominic Kwiatkowski, Cordelia Langford, John Sillitoe on behalf of the Wellcome Sanger Institute COVID-19 Surveillance Team |
| EPI_ISL_555126, EPI_ISL_555128 | Lighthouse Lab in Alderley Park | Wellcome Sanger Institute for the COVID-19 Genomics UK (COG-UK) consortium | The Lighthouse Lab in Alderley Park and Alex Alderton, Roberto Amato, Sonia Goncalves, Ewan Harrison, David K. Jackson, Ian Johnston, Dominic Kwiatkowski, Cordelia Langford, John Sillitoe on behalf of the Wellcome Sanger Institute COVID-19 Surveillance Team (http://www.sanger.ac.uk/covid-team) |
| EPI_ISL_555130, EPI_ISL_555131, EPI_ISL_555132, EPI_ISL_555133, EPI_ISL_555134, EPI_ISL_555135, EPI_ISL_555136, EPI_ISL_555137, EPI_ISL_555138, EPI_ISL_555139 | Lighthouse Lab in Alderley Park | Wellcome Sanger Institute for the COVID-19 Genomics UK (COG-UK) consortium | The Lighthouse Lab in Alderley Park and Alex Alderton, Roberto Amato, Sonia Goncalves, Ewan Harrison, David K. Jackson, Ian Johnston, Dominic Kwiatkowski, Cordelia Langford, John Sillitoe on behalf of the Wellcome Sanger Institute COVID-19 Surveillance Team |
| EPI_ISL_555140 | Lighthouse Lab in Alderley Park | Wellcome Sanger Institute for the COVID-19 Genomics UK (COG-UK) consortium | The Lighthouse Lab in Alderley Park and Alex Alderton, Roberto Amato, Sonia Goncalves, Ewan Harrison, David K. Jackson, Ian Johnston, Dominic Kwiatkowski, Cordelia Langford, John Sillitoe on behalf of the Wellcome Sanger Institute COVID-19 Surveillance Team (http://www.sanger.ac.uk/covid-team) |
| EPI_ISL_555141 | Lighthouse Lab in Alderley Park | Wellcome Sanger Institute for the COVID-19 Genomics UK (COG-UK) consortium | The Lighthouse Lab in Alderley Park and Alex Alderton, Roberto Amato, Sonia Goncalves, Ewan Harrison, David K. Jackson, Ian Johnston, Dominic Kwiatkowski, Cordelia Langford, John Sillitoe on behalf of the Wellcome Sanger Institute COVID-19 Surveillance Team |
| EPI_ISL_555142 | Lighthouse Lab in Alderley Park | Wellcome Sanger Institute for the COVID-19 Genomics UK (COG-UK) consortium | The Lighthouse Lab in Alderley Park and Alex Alderton, Roberto Amato, Sonia Goncalves, Ewan Harrison, David K. Jackson, Ian Johnston, Dominic Kwiatkowski, Cordelia Langford, John Sillitoe on behalf of the Wellcome Sanger Institute COVID-19 Surveillance Team (http://www.sanger.ac.uk/covid-team) |
| EPI_ISL_555143 | Lighthouse Lab in Alderley Park | Wellcome Sanger Institute for the COVID-19 Genomics UK (COG-UK) consortium | The Lighthouse Lab in Alderley Park and Alex Alderton, Roberto Amato, Sonia Goncalves, Ewan Harrison, David K. Jackson, Ian Johnston, Dominic Kwiatkowski, Cordelia Langford, John Sillitoe on behalf of the Wellcome Sanger Institute COVID-19 Surveillance Team |
| EPI_ISL_555144, EPI_ISL_555145 | Lighthouse Lab in Alderley Park | Wellcome Sanger Institute for the COVID-19 Genomics UK | The Lighthouse Lab in Alderley Park and Alex Alderton, Roberto Amato, Sonia Goncalves, Ewan Harrison, David K. Jackson, Ian Johnston, Dominic |

| | | | |
|---|---|---|---|
| | | (COG-UK) consortium | Kwiatkowski, Cordelia Langford, John Sillitoe on behalf of the Wellcome Sanger Institute COVID-19 Surveillance Team (http://www.sanger.ac.uk/covid-team) |
| EPI_ISL_555146 | Lighthouse Lab in Alderley Park | Wellcome Sanger Institute for the COVID-19 Genomics UK (COG-UK) consortium | The Lighthouse Lab in Alderley Park and Alex Alderton, Roberto Amato, Sonia Goncalves, Ewan Harrison, David K. Jackson, Ian Johnston, Dominic Kwiatkowski, Cordelia Langford, John Sillitoe on behalf of the Wellcome Sanger Institute COVID-19 Surveillance Team |
| EPI_ISL_555147 | Lighthouse Lab in Alderley Park | Wellcome Sanger Institute for the COVID-19 Genomics UK (COG-UK) consortium | The Lighthouse Lab in Alderley Park and Alex Alderton, Roberto Amato, Sonia Goncalves, Ewan Harrison, David K. Jackson, Ian Johnston, Dominic Kwiatkowski, Cordelia Langford, John Sillitoe on behalf of the Wellcome Sanger Institute COVID-19 Surveillance Team (http://www.sanger.ac.uk/covid-team) |
| EPI_ISL_555149, EPI_ISL_555150, EPI_ISL_555151, EPI_ISL_555152, EPI_ISL_555153, EPI_ISL_555154 | Lighthouse Lab in Alderley Park | Wellcome Sanger Institute for the COVID-19 Genomics UK (COG-UK) consortium | The Lighthouse Lab in Alderley Park and Alex Alderton, Roberto Amato, Sonia Goncalves, Ewan Harrison, David K. Jackson, Ian Johnston, Dominic Kwiatkowski, Cordelia Langford, John Sillitoe on behalf of the Wellcome Sanger Institute COVID-19 Surveillance Team |
| EPI_ISL_555155 | Lighthouse Lab in Alderley Park | Wellcome Sanger Institute for the COVID-19 Genomics UK (COG-UK) consortium | The Lighthouse Lab in Alderley Park and Alex Alderton, Roberto Amato, Sonia Goncalves, Ewan Harrison, David K. Jackson, Ian Johnston, Dominic Kwiatkowski, Cordelia Langford, John Sillitoe on behalf of the Wellcome Sanger Institute COVID-19 Surveillance Team (http://www.sanger.ac.uk/covid-team) |
| EPI_ISL_555156, EPI_ISL_555157, EPI_ISL_555158 | Lighthouse Lab in Alderley Park | Wellcome Sanger Institute for the COVID-19 Genomics UK (COG-UK) consortium | The Lighthouse Lab in Alderley Park and Alex Alderton, Roberto Amato, Sonia Goncalves, Ewan Harrison, David K. Jackson, Ian Johnston, Dominic Kwiatkowski, Cordelia Langford, John Sillitoe on behalf of the Wellcome Sanger Institute COVID-19 Surveillance Team |
| EPI_ISL_555159 | Lighthouse Lab in Alderley Park | Wellcome Sanger Institute for the COVID-19 Genomics UK (COG-UK) consortium | The Lighthouse Lab in Alderley Park and Alex Alderton, Roberto Amato, Sonia Goncalves, Ewan Harrison, David K. Jackson, Ian Johnston, Dominic Kwiatkowski, Cordelia Langford, John Sillitoe on behalf of the Wellcome Sanger Institute COVID-19 Surveillance Team (http://www.sanger.ac.uk/covid-team) |
| EPI_ISL_555160, EPI_ISL_555161, EPI_ISL_555162 | Lighthouse Lab in Alderley Park | Wellcome Sanger Institute for the COVID-19 Genomics UK (COG-UK) consortium | The Lighthouse Lab in Alderley Park and Alex Alderton, Roberto Amato, Sonia Goncalves, Ewan Harrison, David K. Jackson, Ian Johnston, Dominic Kwiatkowski, Cordelia Langford, John Sillitoe on behalf of the Wellcome Sanger Institute COVID-19 Surveillance Team |
| EPI_ISL_555163 | Lighthouse Lab in Alderley Park | Wellcome Sanger Institute for the COVID-19 Genomics UK (COG-UK) consortium | The Lighthouse Lab in Alderley Park and Alex Alderton, Roberto Amato, Sonia Goncalves, Ewan Harrison, David K. Jackson, Ian Johnston, Dominic Kwiatkowski, Cordelia Langford, John Sillitoe on behalf of the Wellcome Sanger Institute COVID-19 Surveillance Team (http://www.sanger.ac.uk/covid-team) |
| EPI_ISL_555164 | Lighthouse Lab in Alderley Park | Wellcome Sanger Institute for the COVID-19 Genomics UK (COG-UK) consortium | The Lighthouse Lab in Alderley Park and Alex Alderton, Roberto Amato, Sonia Goncalves, Ewan Harrison, David K. Jackson, Ian Johnston, Dominic Kwiatkowski, Cordelia Langford, John Sillitoe on behalf of the Wellcome Sanger Institute COVID-19 Surveillance Team |
| EPI_ISL_555165 | Lighthouse Lab in Alderley Park | Wellcome Sanger Institute for the COVID-19 Genomics UK (COG-UK) consortium | The Lighthouse Lab in Alderley Park and Alex Alderton, Roberto Amato, Sonia Goncalves, Ewan Harrison, David K. Jackson, Ian Johnston, Dominic Kwiatkowski, Cordelia Langford, John Sillitoe on behalf of the Wellcome Sanger Institute COVID-19 Surveillance Team (http://www.sanger.ac.uk/covid-team) |
| EPI_ISL_555166, EPI_ISL_555167 | Lighthouse Lab in Alderley Park | Wellcome Sanger Institute for the COVID-19 Genomics UK (COG-UK) consortium | The Lighthouse Lab in Alderley Park and Alex Alderton, Roberto Amato, Sonia Goncalves, Ewan Harrison, David K. Jackson, Ian Johnston, Dominic Kwiatkowski, Cordelia Langford, John Sillitoe on behalf of the Wellcome Sanger Institute COVID-19 Surveillance Team |
| EPI_ISL_555168 | Lighthouse Lab in Alderley Park | Wellcome Sanger Institute for the COVID-19 Genomics UK (COG-UK) consortium | The Lighthouse Lab in Alderley Park and Alex Alderton, Roberto Amato, Sonia Goncalves, Ewan Harrison, David K. Jackson, Ian Johnston, Dominic Kwiatkowski, Cordelia Langford, John Sillitoe on behalf of the Wellcome Sanger Institute COVID-19 Surveillance Team (http://www.sanger.ac.uk/covid-team) |
| EPI_ISL_555169, EPI_ISL_555170, EPI_ISL_555171, EPI_ISL_555172, EPI_ISL_555173 | Lighthouse Lab in Alderley Park | Wellcome Sanger Institute for the COVID-19 Genomics UK (COG-UK) consortium | The Lighthouse Lab in Alderley Park and Alex Alderton, Roberto Amato, Sonia Goncalves, Ewan Harrison, David K. Jackson, Ian Johnston, Dominic Kwiatkowski, Cordelia Langford, John Sillitoe on behalf of the Wellcome Sanger Institute COVID-19 Surveillance Team |
| EPI_ISL_555174 | Lighthouse Lab in Alderley Park | Wellcome Sanger Institute for the COVID-19 Genomics UK (COG-UK) consortium | The Lighthouse Lab in Alderley Park and Alex Alderton, Roberto Amato, Sonia Goncalves, Ewan Harrison, David K. Jackson, Ian Johnston, Dominic Kwiatkowski, Cordelia Langford, John Sillitoe on behalf of the Wellcome Sanger Institute COVID-19 Surveillance Team (http://www.sanger.ac.uk/covid-team) |
| EPI_ISL_555175, EPI_ISL_555176, EPI_ISL_555177 | Lighthouse Lab in Alderley Park | Wellcome Sanger Institute for the COVID-19 Genomics UK (COG-UK) consortium | The Lighthouse Lab in Alderley Park and Alex Alderton, Roberto Amato, Sonia Goncalves, Ewan Harrison, David K. Jackson, Ian Johnston, Dominic Kwiatkowski, Cordelia Langford, John Sillitoe on behalf of the Wellcome Sanger Institute COVID-19 Surveillance Team |
| EPI_ISL_555178, EPI_ISL_555179 | Lighthouse Lab in Alderley Park | Wellcome Sanger Institute for the COVID-19 Genomics UK (COG-UK) consortium | The Lighthouse Lab in Alderley Park and Alex Alderton, Roberto Amato, Sonia Goncalves, Ewan Harrison, David K. Jackson, Ian Johnston, Dominic Kwiatkowski, Cordelia Langford, John Sillitoe on behalf of the Wellcome Sanger Institute COVID-19 Surveillance Team (http://www.sanger.ac.uk/covid-team) |
| EPI_ISL_555180, EPI_ISL_555181, EPI_ISL_555182, EPI_ISL_555183, EPI_ISL_555184, EPI_ISL_555185 | Lighthouse Lab in Alderley Park | Wellcome Sanger Institute for the COVID-19 Genomics UK (COG-UK) consortium | The Lighthouse Lab in Alderley Park and Alex Alderton, Roberto Amato, Sonia Goncalves, Ewan Harrison, David K. Jackson, Ian Johnston, Dominic Kwiatkowski, Cordelia Langford, John Sillitoe on behalf of the Wellcome Sanger Institute COVID-19 Surveillance Team |
| EPI_ISL_555186, EPI_ISL_555187 | Lighthouse Lab in Alderley Park | Wellcome Sanger Institute for the COVID-19 Genomics UK (COG-UK) consortium | The Lighthouse Lab in Alderley Park and Alex Alderton, Roberto Amato, Sonia Goncalves, Ewan Harrison, David K. Jackson, Ian Johnston, Dominic Kwiatkowski, Cordelia Langford, John Sillitoe on behalf of the Wellcome Sanger Institute COVID-19 Surveillance Team (http://www.sanger.ac.uk/covid-team) |
| EPI_ISL_555188, EPI_ISL_555189, EPI_ISL_555190, EPI_ISL_555191 | Lighthouse Lab in Alderley Park | Wellcome Sanger Institute for the COVID-19 Genomics UK (COG-UK) consortium | The Lighthouse Lab in Alderley Park and Alex Alderton, Roberto Amato, Sonia Goncalves, Ewan Harrison, David K. Jackson, Ian Johnston, Dominic Kwiatkowski, Cordelia Langford, John Sillitoe on behalf of the Wellcome Sanger Institute COVID-19 Surveillance Team |
| EPI_ISL_555192 | Lighthouse Lab in Alderley Park | Wellcome Sanger Institute for the COVID-19 Genomics UK (COG-UK) consortium | The Lighthouse Lab in Alderley Park and Alex Alderton, Roberto Amato, Sonia Goncalves, Ewan Harrison, David K. Jackson, Ian Johnston, Dominic Kwiatkowski, Cordelia Langford, John Sillitoe on behalf of the Wellcome Sanger Institute COVID-19 Surveillance Team (http://www.sanger.ac.uk/covid-team) |
| EPI_ISL_555193, EPI_ISL_555194, EPI_ISL_555195 | Lighthouse Lab in Alderley Park | Wellcome Sanger Institute for the COVID-19 Genomics UK (COG-UK) consortium | The Lighthouse Lab in Alderley Park and Alex Alderton, Roberto Amato, Sonia Goncalves, Ewan Harrison, David K. Jackson, Ian Johnston, Dominic Kwiatkowski, Cordelia Langford, John Sillitoe on behalf of the Wellcome Sanger Institute COVID-19 Surveillance Team |
| EPI_ISL_555196 | Lighthouse Lab in Alderley Park | Wellcome Sanger Institute for the COVID-19 Genomics UK (COG-UK) consortium | The Lighthouse Lab in Alderley Park and Alex Alderton, Roberto Amato, Sonia Goncalves, Ewan Harrison, David K. Jackson, Ian Johnston, Dominic Kwiatkowski, Cordelia Langford, John Sillitoe on behalf of the Wellcome Sanger Institute COVID-19 Surveillance Team (http://www.sanger.ac.uk/covid-team) |
| EPI_ISL_555197, EPI_ISL_555198 | Lighthouse Lab in Alderley Park | Wellcome Sanger Institute for the COVID-19 Genomics UK (COG-UK) consortium | The Lighthouse Lab in Alderley Park and Alex Alderton, Roberto Amato, Sonia Goncalves, Ewan Harrison, David K. Jackson, Ian Johnston, Dominic Kwiatkowski, Cordelia Langford, John Sillitoe on behalf of the Wellcome Sanger Institute COVID-19 Surveillance Team |
| EPI_ISL_555199, EPI_ISL_555200 | Lighthouse Lab in Alderley Park | Wellcome Sanger Institute for the COVID-19 Genomics UK (COG-UK) consortium | The Lighthouse Lab in Alderley Park and Alex Alderton, Roberto Amato, Sonia Goncalves, Ewan Harrison, David K. Jackson, Ian Johnston, Dominic Kwiatkowski, Cordelia Langford, John Sillitoe on behalf of the Wellcome Sanger Institute COVID-19 Surveillance Team (http://www.sanger.ac.uk/covid-team) |
| EPI_ISL_555201, EPI_ISL_555202, EPI_ISL_555203 | Lighthouse Lab in Alderley Park | Wellcome Sanger Institute for the COVID-19 Genomics UK (COG-UK) consortium | The Lighthouse Lab in Alderley Park and Alex Alderton, Roberto Amato, Sonia Goncalves, Ewan Harrison, David K. Jackson, Ian Johnston, Dominic Kwiatkowski, Cordelia Langford, John Sillitoe on behalf of the Wellcome Sanger Institute COVID-19 Surveillance Team |
| EPI_ISL_555204 | Lighthouse Lab in Alderley Park | Wellcome Sanger Institute for the COVID-19 Genomics UK (COG-UK) consortium | The Lighthouse Lab in Alderley Park and Alex Alderton, Roberto Amato, Sonia Goncalves, Ewan Harrison, David K. Jackson, Ian Johnston, Dominic Kwiatkowski, Cordelia Langford, John Sillitoe on behalf of the Wellcome Sanger Institute COVID-19 Surveillance Team (http://www.sanger.ac.uk/covid-team) |
| EPI_ISL_555205, EPI_ISL_555206, EPI_ISL_555207 | Lighthouse Lab in Alderley Park | Wellcome Sanger Institute for the COVID-19 Genomics UK (COG-UK) consortium | The Lighthouse Lab in Alderley Park and Alex Alderton, Roberto Amato, Sonia Goncalves, Ewan Harrison, David K. Jackson, Ian Johnston, Dominic Kwiatkowski, Cordelia Langford, John Sillitoe on behalf of the Wellcome Sanger Institute COVID-19 Surveillance Team |
| EPI_ISL_555208 | Lighthouse Lab in Alderley Park | Wellcome Sanger Institute for the COVID-19 Genomics UK | The Lighthouse Lab in Alderley Park and Alex Alderton, Roberto Amato, Sonia Goncalves, Ewan Harrison, David K. Jackson, Ian Johnston, Dominic |

| | | | |
|---|---|---|---|
| | | (COG-UK) consortium | Kwiatkowski, Cordelia Langford, John Sillitoe on behalf of the Wellcome Sanger Institute COVID-19 Surveillance Team (http://www.sanger.ac.uk/covid-team) |
| EPI_ISL_555209 | Lighthouse Lab in Alderley Park | Wellcome Sanger Institute for the COVID-19 Genomics UK (COG-UK) consortium | The Lighthouse Lab in Alderley Park and Alex Alderton, Roberto Amato, Sonia Goncalves, Ewan Harrison, David K. Jackson, Ian Johnston, Dominic Kwiatkowski, Cordelia Langford, John Sillitoe on behalf of the Wellcome Sanger Institute COVID-19 Surveillance Team |
| EPI_ISL_555210 | Lighthouse Lab in Alderley Park | Wellcome Sanger Institute for the COVID-19 Genomics UK (COG-UK) consortium | The Lighthouse Lab in Alderley Park and Alex Alderton, Roberto Amato, Sonia Goncalves, Ewan Harrison, David K. Jackson, Ian Johnston, Dominic Kwiatkowski, Cordelia Langford, John Sillitoe on behalf of the Wellcome Sanger Institute COVID-19 Surveillance Team (http://www.sanger.ac.uk/covid-team) |
| EPI_ISL_555211, EPI_ISL_555212 | Lighthouse Lab in Alderley Park | Wellcome Sanger Institute for the COVID-19 Genomics UK (COG-UK) consortium | The Lighthouse Lab in Alderley Park and Alex Alderton, Roberto Amato, Sonia Goncalves, Ewan Harrison, David K. Jackson, Ian Johnston, Dominic Kwiatkowski, Cordelia Langford, John Sillitoe on behalf of the Wellcome Sanger Institute COVID-19 Surveillance Team |
| EPI_ISL_555213 | Lighthouse Lab in Alderley Park | Wellcome Sanger Institute for the COVID-19 Genomics UK (COG-UK) consortium | The Lighthouse Lab in Alderley Park and Alex Alderton, Roberto Amato, Sonia Goncalves, Ewan Harrison, David K. Jackson, Ian Johnston, Dominic Kwiatkowski, Cordelia Langford, John Sillitoe on behalf of the Wellcome Sanger Institute COVID-19 Surveillance Team (http://www.sanger.ac.uk/covid-team) |
| EPI_ISL_555214, EPI_ISL_555215 | Lighthouse Lab in Alderley Park | Wellcome Sanger Institute for the COVID-19 Genomics UK (COG-UK) consortium | The Lighthouse Lab in Alderley Park and Alex Alderton, Roberto Amato, Sonia Goncalves, Ewan Harrison, David K. Jackson, Ian Johnston, Dominic Kwiatkowski, Cordelia Langford, John Sillitoe on behalf of the Wellcome Sanger Institute COVID-19 Surveillance Team |
| EPI_ISL_555216 | Lighthouse Lab in Alderley Park | Wellcome Sanger Institute for the COVID-19 Genomics UK (COG-UK) consortium | The Lighthouse Lab in Alderley Park and Alex Alderton, Roberto Amato, Sonia Goncalves, Ewan Harrison, David K. Jackson, Ian Johnston, Dominic Kwiatkowski, Cordelia Langford, John Sillitoe on behalf of the Wellcome Sanger Institute COVID-19 Surveillance Team (http://www.sanger.ac.uk/covid-team) |
| EPI_ISL_555217, EPI_ISL_555218, EPI_ISL_555219 | Lighthouse Lab in Alderley Park | Wellcome Sanger Institute for the COVID-19 Genomics UK (COG-UK) consortium | The Lighthouse Lab in Alderley Park and Alex Alderton, Roberto Amato, Sonia Goncalves, Ewan Harrison, David K. Jackson, Ian Johnston, Dominic Kwiatkowski, Cordelia Langford, John Sillitoe on behalf of the Wellcome Sanger Institute COVID-19 Surveillance Team |
| EPI_ISL_555220, EPI_ISL_555221 | Lighthouse Lab in Alderley Park | Wellcome Sanger Institute for the COVID-19 Genomics UK (COG-UK) consortium | The Lighthouse Lab in Alderley Park and Alex Alderton, Roberto Amato, Sonia Goncalves, Ewan Harrison, David K. Jackson, Ian Johnston, Dominic Kwiatkowski, Cordelia Langford, John Sillitoe on behalf of the Wellcome Sanger Institute COVID-19 Surveillance Team (http://www.sanger.ac.uk/covid-team) |
| EPI_ISL_555222 | Lighthouse Lab in Alderley Park | Wellcome Sanger Institute for the COVID-19 Genomics UK (COG-UK) consortium | The Lighthouse Lab in Alderley Park and Alex Alderton, Roberto Amato, Sonia Goncalves, Ewan Harrison, David K. Jackson, Ian Johnston, Dominic Kwiatkowski, Cordelia Langford, John Sillitoe on behalf of the Wellcome Sanger Institute COVID-19 Surveillance Team |
| EPI_ISL_555223 | Lighthouse Lab in Alderley Park | Wellcome Sanger Institute for the COVID-19 Genomics UK (COG-UK) consortium | The Lighthouse Lab in Alderley Park and Alex Alderton, Roberto Amato, Sonia Goncalves, Ewan Harrison, David K. Jackson, Ian Johnston, Dominic Kwiatkowski, Cordelia Langford, John Sillitoe on behalf of the Wellcome Sanger Institute COVID-19 Surveillance Team (http://www.sanger.ac.uk/covid-team) |
| EPI_ISL_555224, EPI_ISL_555225, EPI_ISL_555226, EPI_ISL_555227 | Lighthouse Lab in Alderley Park | Wellcome Sanger Institute for the COVID-19 Genomics UK (COG-UK) consortium | The Lighthouse Lab in Alderley Park and Alex Alderton, Roberto Amato, Sonia Goncalves, Ewan Harrison, David K. Jackson, Ian Johnston, Dominic Kwiatkowski, Cordelia Langford, John Sillitoe on behalf of the Wellcome Sanger Institute COVID-19 Surveillance Team |
| EPI_ISL_555228 | Lighthouse Lab in Alderley Park | Wellcome Sanger Institute for the COVID-19 Genomics UK (COG-UK) consortium | The Lighthouse Lab in Alderley Park and Alex Alderton, Roberto Amato, Sonia Goncalves, Ewan Harrison, David K. Jackson, Ian Johnston, Dominic Kwiatkowski, Cordelia Langford, John Sillitoe on behalf of the Wellcome Sanger Institute COVID-19 Surveillance Team (http://www.sanger.ac.uk/covid-team) |
| EPI_ISL_555229 | Lighthouse Lab in Alderley Park | Wellcome Sanger Institute for the COVID-19 Genomics UK (COG-UK) consortium | The Lighthouse Lab in Alderley Park and Alex Alderton, Roberto Amato, Sonia Goncalves, Ewan Harrison, David K. Jackson, Ian Johnston, Dominic Kwiatkowski, Cordelia Langford, John Sillitoe on behalf of the Wellcome Sanger Institute COVID-19 Surveillance Team |
| EPI_ISL_555230, EPI_ISL_555231, EPI_ISL_555232 | Lighthouse Lab in Alderley Park | Wellcome Sanger Institute for the COVID-19 Genomics UK (COG-UK) consortium | The Lighthouse Lab in Alderley Park and Alex Alderton, Roberto Amato, Sonia Goncalves, Ewan Harrison, David K. Jackson, Ian Johnston, Dominic Kwiatkowski, Cordelia Langford, John Sillitoe on behalf of the Wellcome Sanger Institute COVID-19 Surveillance Team (http://www.sanger.ac.uk/covid-team) |
| EPI_ISL_555233 | Lighthouse Lab in Alderley Park | Wellcome Sanger Institute for the COVID-19 Genomics UK (COG-UK) consortium | The Lighthouse Lab in Alderley Park and Alex Alderton, Roberto Amato, Sonia Goncalves, Ewan Harrison, David K. Jackson, Ian Johnston, Dominic Kwiatkowski, Cordelia Langford, John Sillitoe on behalf of the Wellcome Sanger Institute COVID-19 Surveillance Team |
| EPI_ISL_555234, EPI_ISL_555235 | Lighthouse Lab in Alderley Park | Wellcome Sanger Institute for the COVID-19 Genomics UK (COG-UK) consortium | The Lighthouse Lab in Alderley Park and Alex Alderton, Roberto Amato, Sonia Goncalves, Ewan Harrison, David K. Jackson, Ian Johnston, Dominic Kwiatkowski, Cordelia Langford, John Sillitoe on behalf of the Wellcome Sanger Institute COVID-19 Surveillance Team (http://www.sanger.ac.uk/covid-team) |
| EPI_ISL_555236, EPI_ISL_555237, EPI_ISL_555238, EPI_ISL_555239, EPI_ISL_555240, EPI_ISL_555241, EPI_ISL_555242, EPI_ISL_555243, EPI_ISL_555244, EPI_ISL_555245, EPI_ISL_555246 | | | |
| see above | Lighthouse Lab in Alderley Park | Wellcome Sanger Institute for the COVID-19 Genomics UK (COG-UK) consortium | The Lighthouse Lab in Alderley Park and Alex Alderton, Roberto Amato, Sonia Goncalves, Ewan Harrison, David K. Jackson, Ian Johnston, Dominic Kwiatkowski, Cordelia Langford, John Sillitoe on behalf of the Wellcome Sanger Institute COVID-19 Surveillance Team |
| EPI_ISL_555247 | Lighthouse Lab in Alderley Park | Wellcome Sanger Institute for the COVID-19 Genomics UK (COG-UK) consortium | The Lighthouse Lab in Alderley Park and Alex Alderton, Roberto Amato, Sonia Goncalves, Ewan Harrison, David K. Jackson, Ian Johnston, Dominic Kwiatkowski, Cordelia Langford, John Sillitoe on behalf of the Wellcome Sanger Institute COVID-19 Surveillance Team (http://www.sanger.ac.uk/covid-team) |
| EPI_ISL_555248, EPI_ISL_555249 | Lighthouse Lab in Alderley Park | Wellcome Sanger Institute for the COVID-19 Genomics UK (COG-UK) consortium | The Lighthouse Lab in Alderley Park and Alex Alderton, Roberto Amato, Sonia Goncalves, Ewan Harrison, David K. Jackson, Ian Johnston, Dominic Kwiatkowski, Cordelia Langford, John Sillitoe on behalf of the Wellcome Sanger Institute COVID-19 Surveillance Team |
| EPI_ISL_555250, EPI_ISL_555251 | Lighthouse Lab in Alderley Park | Wellcome Sanger Institute for the COVID-19 Genomics UK (COG-UK) consortium | The Lighthouse Lab in Alderley Park and Alex Alderton, Roberto Amato, Sonia Goncalves, Ewan Harrison, David K. Jackson, Ian Johnston, Dominic Kwiatkowski, Cordelia Langford, John Sillitoe on behalf of the Wellcome Sanger Institute COVID-19 Surveillance Team (http://www.sanger.ac.uk/covid-team) |
| EPI_ISL_555252 | Lighthouse Lab in Alderley Park | Wellcome Sanger Institute for the COVID-19 Genomics UK (COG-UK) consortium | The Lighthouse Lab in Alderley Park and Alex Alderton, Roberto Amato, Sonia Goncalves, Ewan Harrison, David K. Jackson, Ian Johnston, Dominic Kwiatkowski, Cordelia Langford, John Sillitoe on behalf of the Wellcome Sanger Institute COVID-19 Surveillance Team |
| EPI_ISL_555253 | Lighthouse Lab in Alderley Park | Wellcome Sanger Institute for the COVID-19 Genomics UK (COG-UK) consortium | The Lighthouse Lab in Alderley Park and Alex Alderton, Roberto Amato, Sonia Goncalves, Ewan Harrison, David K. Jackson, Ian Johnston, Dominic Kwiatkowski, Cordelia Langford, John Sillitoe on behalf of the Wellcome Sanger Institute COVID-19 Surveillance Team (http://www.sanger.ac.uk/covid-team) |
| EPI_ISL_555254 | Lighthouse Lab in Alderley Park | Wellcome Sanger Institute for the COVID-19 Genomics UK (COG-UK) consortium | The Lighthouse Lab in Alderley Park and Alex Alderton, Roberto Amato, Sonia Goncalves, Ewan Harrison, David K. Jackson, Ian Johnston, Dominic Kwiatkowski, Cordelia Langford, John Sillitoe on behalf of the Wellcome Sanger Institute COVID-19 Surveillance Team |
| EPI_ISL_555255 | Lighthouse Lab in Alderley Park | Wellcome Sanger Institute for the COVID-19 Genomics UK (COG-UK) consortium | The Lighthouse Lab in Alderley Park and Alex Alderton, Roberto Amato, Sonia Goncalves, Ewan Harrison, David K. Jackson, Ian Johnston, Dominic Kwiatkowski, Cordelia Langford, John Sillitoe on behalf of the Wellcome Sanger Institute COVID-19 Surveillance Team (http://www.sanger.ac.uk/covid-team) |
| EPI_ISL_555256, EPI_ISL_555257 | Lighthouse Lab in Alderley Park | Wellcome Sanger Institute for the COVID-19 Genomics UK (COG-UK) consortium | The Lighthouse Lab in Alderley Park and Alex Alderton, Roberto Amato, Sonia Goncalves, Ewan Harrison, David K. Jackson, Ian Johnston, Dominic Kwiatkowski, Cordelia Langford, John Sillitoe on behalf of the Wellcome Sanger Institute COVID-19 Surveillance Team |
| EPI_ISL_555258 | Lighthouse Lab in Alderley Park | Wellcome Sanger Institute for the COVID-19 Genomics UK (COG-UK) consortium | The Lighthouse Lab in Alderley Park and Alex Alderton, Roberto Amato, Sonia Goncalves, Ewan Harrison, David K. Jackson, Ian Johnston, Dominic Kwiatkowski, Cordelia Langford, John Sillitoe on behalf of the Wellcome Sanger Institute COVID-19 Surveillance Team (http://www.sanger.ac.uk/covid-team) |
| EPI_ISL_555260, EPI_ISL_555261, EPI_ISL_555262 | Lighthouse Lab in Alderley Park | Wellcome Sanger Institute for the COVID-19 Genomics UK (COG-UK) consortium | The Lighthouse Lab in Alderley Park and Alex Alderton, Roberto Amato, Sonia Goncalves, Ewan Harrison, David K. Jackson, Ian Johnston, Dominic Kwiatkowski, Cordelia Langford, John Sillitoe on behalf of the Wellcome Sanger Institute COVID-19 Surveillance Team |
| EPI_ISL_555263, EPI_ISL_555264 | Lighthouse Lab in Alderley Park | Wellcome Sanger Institute for the COVID-19 Genomics UK (COG-UK) consortium | The Lighthouse Lab in Alderley Park and Alex Alderton, Roberto Amato, Sonia Goncalves, Ewan Harrison, David K. Jackson, Ian Johnston, Dominic Kwiatkowski, Cordelia Langford, John Sillitoe on behalf of the Wellcome Sanger Institute COVID-19 Surveillance Team (http://www.sanger.ac.uk/covid-team) |

| | | | |
|---|---|---|---|
| EPI_ISL_555265, EPI_ISL_555266, EPI_ISL_555267, EPI_ISL_555268, EPI_ISL_555269, EPI_ISL_555270, EPI_ISL_555271, EPI_ISL_555272, EPI_ISL_555273 | Lighthouse Lab in Alderley Park | Wellcome Sanger Institute for the COVID-19 Genomics UK (COG-UK) consortium | The Lighthouse Lab in Alderley Park and Alex Alderton, Roberto Amato, Sonia Goncalves, Ewan Harrison, David K. Jackson, Ian Johnston, Dominic Kwiatkowski, Cordelia Langford, John Sillitoe on behalf of the Wellcome Sanger Institute COVID-19 Surveillance Team |
| EPI_ISL_555274 | Lighthouse Lab in Alderley Park | Wellcome Sanger Institute for the COVID-19 Genomics UK (COG-UK) consortium | The Lighthouse Lab in Alderley Park and Alex Alderton, Roberto Amato, Sonia Goncalves, Ewan Harrison, David K. Jackson, Ian Johnston, Dominic Kwiatkowski, Cordelia Langford, John Sillitoe on behalf of the Wellcome Sanger Institute COVID-19 Surveillance Team (http://www.sanger.ac.uk/covid-team) |
| EPI_ISL_555275, EPI_ISL_555276 | Lighthouse Lab in Alderley Park | Wellcome Sanger Institute for the COVID-19 Genomics UK (COG-UK) consortium | The Lighthouse Lab in Alderley Park and Alex Alderton, Roberto Amato, Sonia Goncalves, Ewan Harrison, David K. Jackson, Ian Johnston, Dominic Kwiatkowski, Cordelia Langford, John Sillitoe on behalf of the Wellcome Sanger Institute COVID-19 Surveillance Team |
| EPI_ISL_555277, EPI_ISL_555278 | Lighthouse Lab in Alderley Park | Wellcome Sanger Institute for the COVID-19 Genomics UK (COG-UK) consortium | The Lighthouse Lab in Alderley Park and Alex Alderton, Roberto Amato, Sonia Goncalves, Ewan Harrison, David K. Jackson, Ian Johnston, Dominic Kwiatkowski, Cordelia Langford, John Sillitoe on behalf of the Wellcome Sanger Institute COVID-19 Surveillance Team (http://www.sanger.ac.uk/covid-team) |
| EPI_ISL_555279, EPI_ISL_555280 | Lighthouse Lab in Alderley Park | Wellcome Sanger Institute for the COVID-19 Genomics UK (COG-UK) consortium | The Lighthouse Lab in Alderley Park and Alex Alderton, Roberto Amato, Sonia Goncalves, Ewan Harrison, David K. Jackson, Ian Johnston, Dominic Kwiatkowski, Cordelia Langford, John Sillitoe on behalf of the Wellcome Sanger Institute COVID-19 Surveillance Team |
| EPI_ISL_555281 | Lighthouse Lab in Alderley Park | Wellcome Sanger Institute for the COVID-19 Genomics UK (COG-UK) consortium | The Lighthouse Lab in Alderley Park and Alex Alderton, Roberto Amato, Sonia Goncalves, Ewan Harrison, David K. Jackson, Ian Johnston, Dominic Kwiatkowski, Cordelia Langford, John Sillitoe on behalf of the Wellcome Sanger Institute COVID-19 Surveillance Team (http://www.sanger.ac.uk/covid-team) |
| EPI_ISL_555282, EPI_ISL_555283 | Lighthouse Lab in Alderley Park | Wellcome Sanger Institute for the COVID-19 Genomics UK (COG-UK) consortium | The Lighthouse Lab in Alderley Park and Alex Alderton, Roberto Amato, Sonia Goncalves, Ewan Harrison, David K. Jackson, Ian Johnston, Dominic Kwiatkowski, Cordelia Langford, John Sillitoe on behalf of the Wellcome Sanger Institute COVID-19 Surveillance Team |
| EPI_ISL_555284 | Lighthouse Lab in Alderley Park | Wellcome Sanger Institute for the COVID-19 Genomics UK (COG-UK) consortium | The Lighthouse Lab in Alderley Park and Alex Alderton, Roberto Amato, Sonia Goncalves, Ewan Harrison, David K. Jackson, Ian Johnston, Dominic Kwiatkowski, Cordelia Langford, John Sillitoe on behalf of the Wellcome Sanger Institute COVID-19 Surveillance Team (http://www.sanger.ac.uk/covid-team) |
| EPI_ISL_555285, EPI_ISL_555286, EPI_ISL_555287, EPI_ISL_555288, EPI_ISL_555289, EPI_ISL_555290, EPI_ISL_555291, EPI_ISL_555292, EPI_ISL_555293, EPI_ISL_555294, EPI_ISL_555295 | | | |
| see above | Lighthouse Lab in Alderley Park | Wellcome Sanger Institute for the COVID-19 Genomics UK (COG-UK) consortium | The Lighthouse Lab in Alderley Park and Alex Alderton, Roberto Amato, Sonia Goncalves, Ewan Harrison, David K. Jackson, Ian Johnston, Dominic Kwiatkowski, Cordelia Langford, John Sillitoe on behalf of the Wellcome Sanger Institute COVID-19 Surveillance Team |
| EPI_ISL_555296 | Lighthouse Lab in Alderley Park | Wellcome Sanger Institute for the COVID-19 Genomics UK (COG-UK) consortium | The Lighthouse Lab in Alderley Park and Alex Alderton, Roberto Amato, Sonia Goncalves, Ewan Harrison, David K. Jackson, Ian Johnston, Dominic Kwiatkowski, Cordelia Langford, John Sillitoe on behalf of the Wellcome Sanger Institute COVID-19 Surveillance Team (http://www.sanger.ac.uk/covid-team) |
| EPI_ISL_555297, EPI_ISL_555298, EPI_ISL_555299, EPI_ISL_555300, EPI_ISL_555301, EPI_ISL_555302 | Lighthouse Lab in Alderley Park | Wellcome Sanger Institute for the COVID-19 Genomics UK (COG-UK) consortium | The Lighthouse Lab in Alderley Park and Alex Alderton, Roberto Amato, Sonia Goncalves, Ewan Harrison, David K. Jackson, Ian Johnston, Dominic Kwiatkowski, Cordelia Langford, John Sillitoe on behalf of the Wellcome Sanger Institute COVID-19 Surveillance Team |
| EPI_ISL_555305 | Lighthouse Lab in Alderley Park | Wellcome Sanger Institute for the COVID-19 Genomics UK (COG-UK) consortium | The Lighthouse Lab in Alderley Park and Alex Alderton, Roberto Amato, Sonia Goncalves, Ewan Harrison, David K. Jackson, Ian Johnston, Dominic Kwiatkowski, Cordelia Langford, John Sillitoe on behalf of the Wellcome Sanger Institute COVID-19 Surveillance Team (http://www.sanger.ac.uk/covid-team) |
| EPI_ISL_555306, EPI_ISL_555307, EPI_ISL_555308 | Lighthouse Lab in Alderley Park | Wellcome Sanger Institute for the COVID-19 Genomics UK (COG-UK) consortium | The Lighthouse Lab in Alderley Park and Alex Alderton, Roberto Amato, Sonia Goncalves, Ewan Harrison, David K. Jackson, Ian Johnston, Dominic Kwiatkowski, Cordelia Langford, John Sillitoe on behalf of the Wellcome Sanger Institute COVID-19 Surveillance Team |
| EPI_ISL_555309, EPI_ISL_555310 | Lighthouse Lab in Alderley Park | Wellcome Sanger Institute for the COVID-19 Genomics UK (COG-UK) consortium | The Lighthouse Lab in Alderley Park and Alex Alderton, Roberto Amato, Sonia Goncalves, Ewan Harrison, David K. Jackson, Ian Johnston, Dominic Kwiatkowski, Cordelia Langford, John Sillitoe on behalf of the Wellcome Sanger Institute COVID-19 Surveillance Team (http://www.sanger.ac.uk/covid-team) |
| EPI_ISL_555311 | Lighthouse Lab in Alderley Park | Wellcome Sanger Institute for the COVID-19 Genomics UK (COG-UK) consortium | The Lighthouse Lab in Alderley Park and Alex Alderton, Roberto Amato, Sonia Goncalves, Ewan Harrison, David K. Jackson, Ian Johnston, Dominic Kwiatkowski, Cordelia Langford, John Sillitoe on behalf of the Wellcome Sanger Institute COVID-19 Surveillance Team |
| EPI_ISL_555312 | Lighthouse Lab in Alderley Park | Wellcome Sanger Institute for the COVID-19 Genomics UK (COG-UK) consortium | The Lighthouse Lab in Alderley Park and Alex Alderton, Roberto Amato, Sonia Goncalves, Ewan Harrison, David K. Jackson, Ian Johnston, Dominic Kwiatkowski, Cordelia Langford, John Sillitoe on behalf of the Wellcome Sanger Institute COVID-19 Surveillance Team (http://www.sanger.ac.uk/covid-team) |
| EPI_ISL_555313, EPI_ISL_555314, EPI_ISL_555315, EPI_ISL_555316, EPI_ISL_555317, EPI_ISL_555318, EPI_ISL_555319, EPI_ISL_555320, EPI_ISL_555321, EPI_ISL_555322, EPI_ISL_555323, EPI_ISL_555324, EPI_ISL_555325, EPI_ISL_555326, EPI_ISL_555327, EPI_ISL_555328, EPI_ISL_555329, EPI_ISL_555330, EPI_ISL_555331, EPI_ISL_555332, EPI_ISL_555333, EPI_ISL_555334, EPI_ISL_555335, EPI_ISL_555336 | | | |
| see above | Lighthouse Lab in Alderley Park | Wellcome Sanger Institute for the COVID-19 Genomics UK (COG-UK) consortium | The Lighthouse Lab in Alderley Park and Alex Alderton, Roberto Amato, Sonia Goncalves, Ewan Harrison, David K. Jackson, Ian Johnston, Dominic Kwiatkowski, Cordelia Langford, John Sillitoe on behalf of the Wellcome Sanger Institute COVID-19 Surveillance Team |
| EPI_ISL_555337 | Lighthouse Lab in Alderley Park | Wellcome Sanger Institute for the COVID-19 Genomics UK (COG-UK) consortium | The Lighthouse Lab in Alderley Park and Alex Alderton, Roberto Amato, Sonia Goncalves, Ewan Harrison, David K. Jackson, Ian Johnston, Dominic Kwiatkowski, Cordelia Langford, John Sillitoe on behalf of the Wellcome Sanger Institute COVID-19 Surveillance Team (http://www.sanger.ac.uk/covid-team) |
| EPI_ISL_555338, EPI_ISL_555339, EPI_ISL_555340, EPI_ISL_555341, EPI_ISL_555342, EPI_ISL_555343, EPI_ISL_555344, EPI_ISL_555345, EPI_ISL_555346, EPI_ISL_555347 | Lighthouse Lab in Alderley Park | Wellcome Sanger Institute for the COVID-19 Genomics UK (COG-UK) consortium | The Lighthouse Lab in Alderley Park and Alex Alderton, Roberto Amato, Sonia Goncalves, Ewan Harrison, David K. Jackson, Ian Johnston, Dominic Kwiatkowski, Cordelia Langford, John Sillitoe on behalf of the Wellcome Sanger Institute COVID-19 Surveillance Team |
| EPI_ISL_555348 | Lighthouse Lab in Alderley Park | Wellcome Sanger Institute for the COVID-19 Genomics UK (COG-UK) consortium | The Lighthouse Lab in Alderley Park and Alex Alderton, Roberto Amato, Sonia Goncalves, Ewan Harrison, David K. Jackson, Ian Johnston, Dominic Kwiatkowski, Cordelia Langford, John Sillitoe on behalf of the Wellcome Sanger Institute COVID-19 Surveillance Team (http://www.sanger.ac.uk/covid-team) |
| EPI_ISL_555349, EPI_ISL_555350, EPI_ISL_555351, EPI_ISL_555352, EPI_ISL_555353, EPI_ISL_555354 | Lighthouse Lab in Alderley Park | Wellcome Sanger Institute for the COVID-19 Genomics UK (COG-UK) consortium | The Lighthouse Lab in Alderley Park and Alex Alderton, Roberto Amato, Sonia Goncalves, Ewan Harrison, David K. Jackson, Ian Johnston, Dominic Kwiatkowski, Cordelia Langford, John Sillitoe on behalf of the Wellcome Sanger Institute COVID-19 Surveillance Team |
| EPI_ISL_555355, EPI_ISL_555356 | Lighthouse Lab in Alderley Park | Wellcome Sanger Institute for the COVID-19 Genomics UK (COG-UK) consortium | The Lighthouse Lab in Alderley Park and Alex Alderton, Roberto Amato, Sonia Goncalves, Ewan Harrison, David K. Jackson, Ian Johnston, Dominic Kwiatkowski, Cordelia Langford, John Sillitoe on behalf of the Wellcome Sanger Institute COVID-19 Surveillance Team (http://www.sanger.ac.uk/covid-team) |
| EPI_ISL_555357, EPI_ISL_555358, EPI_ISL_555359 | Lighthouse Lab in Alderley Park | Wellcome Sanger Institute for the COVID-19 Genomics UK (COG-UK) consortium | The Lighthouse Lab in Alderley Park and Alex Alderton, Roberto Amato, Sonia Goncalves, Ewan Harrison, David K. Jackson, Ian Johnston, Dominic Kwiatkowski, Cordelia Langford, John Sillitoe on behalf of the Wellcome Sanger Institute COVID-19 Surveillance Team |
| EPI_ISL_555360 | Lighthouse Lab in Alderley Park | Wellcome Sanger Institute for the COVID-19 Genomics UK (COG-UK) consortium | The Lighthouse Lab in Alderley Park and Alex Alderton, Roberto Amato, Sonia Goncalves, Ewan Harrison, David K. Jackson, Ian Johnston, Dominic Kwiatkowski, Cordelia Langford, John Sillitoe on behalf of the Wellcome Sanger Institute COVID-19 Surveillance Team (http://www.sanger.ac.uk/covid-team) |
| EPI_ISL_555361 | Lighthouse Lab in Alderley Park | Wellcome Sanger Institute for the COVID-19 Genomics UK (COG-UK) consortium | The Lighthouse Lab in Alderley Park and Alex Alderton, Roberto Amato, Sonia Goncalves, Ewan Harrison, David K. Jackson, Ian Johnston, Dominic Kwiatkowski, Cordelia Langford, John Sillitoe on behalf of the Wellcome Sanger Institute COVID-19 Surveillance Team |
| EPI_ISL_555362, EPI_ISL_555363 | Lighthouse Lab in Alderley Park | Wellcome Sanger Institute for the COVID-19 Genomics UK | The Lighthouse Lab in Alderley Park and Alex Alderton, Roberto Amato, Sonia Goncalves, Ewan Harrison, David K. Jackson, Ian Johnston, Dominic |

| | | | |
|---|---|---|---|
| | | (COG-UK) consortium | Kwiatkowski, Cordelia Langford, John Sillitoe on behalf of the Wellcome Sanger Institute COVID-19 Surveillance Team (http://www.sanger.ac.uk/covid-team) |
| EPI_ISL_555364, EPI_ISL_555365, EPI_ISL_555366, EPI_ISL_555367, EPI_ISL_555368, EPI_ISL_555369, EPI_ISL_555370, EPI_ISL_555371 | Lighthouse Lab in Alderley Park | Wellcome Sanger Institute for the COVID-19 Genomics UK (COG-UK) consortium | The Lighthouse Lab in Alderley Park and Alex Alderton, Roberto Amato, Sonia Goncalves, Ewan Harrison, David K. Jackson, Ian Johnston, Dominic Kwiatkowski, Cordelia Langford, John Sillitoe on behalf of the Wellcome Sanger Institute COVID-19 Surveillance Team |
| EPI_ISL_555372, EPI_ISL_555373, EPI_ISL_555374 | Lighthouse Lab in Alderley Park | Wellcome Sanger Institute for the COVID-19 Genomics UK (COG-UK) consortium | The Lighthouse Lab in Alderley Park and Alex Alderton, Roberto Amato, Sonia Goncalves, Ewan Harrison, David K. Jackson, Ian Johnston, Dominic Kwiatkowski, Cordelia Langford, John Sillitoe on behalf of the Wellcome Sanger Institute COVID-19 Surveillance Team (http://www.sanger.ac.uk/covid-team) |
| EPI_ISL_555375, EPI_ISL_555376, EPI_ISL_555377, EPI_ISL_555378, EPI_ISL_555379, EPI_ISL_555380, EPI_ISL_555381, EPI_ISL_555382 | Lighthouse Lab in Alderley Park | Wellcome Sanger Institute for the COVID-19 Genomics UK (COG-UK) consortium | The Lighthouse Lab in Alderley Park and Alex Alderton, Roberto Amato, Sonia Goncalves, Ewan Harrison, David K. Jackson, Ian Johnston, Dominic Kwiatkowski, Cordelia Langford, John Sillitoe on behalf of the Wellcome Sanger Institute COVID-19 Surveillance Team |
| EPI_ISL_555383 | Lighthouse Lab in Alderley Park | Wellcome Sanger Institute for the COVID-19 Genomics UK (COG-UK) consortium | The Lighthouse Lab in Alderley Park and Alex Alderton, Roberto Amato, Sonia Goncalves, Ewan Harrison, David K. Jackson, Ian Johnston, Dominic Kwiatkowski, Cordelia Langford, John Sillitoe on behalf of the Wellcome Sanger Institute COVID-19 Surveillance Team (http://www.sanger.ac.uk/covid-team) |
| EPI_ISL_555384, EPI_ISL_555385, EPI_ISL_555386, EPI_ISL_555387, EPI_ISL_555388, EPI_ISL_555389, EPI_ISL_555390, EPI_ISL_555392 | Lighthouse Lab in Alderley Park | Wellcome Sanger Institute for the COVID-19 Genomics UK (COG-UK) consortium | The Lighthouse Lab in Alderley Park and Alex Alderton, Roberto Amato, Sonia Goncalves, Ewan Harrison, David K. Jackson, Ian Johnston, Dominic Kwiatkowski, Cordelia Langford, John Sillitoe on behalf of the Wellcome Sanger Institute COVID-19 Surveillance Team |
| EPI_ISL_555393 | Lighthouse Lab in Alderley Park | Wellcome Sanger Institute for the COVID-19 Genomics UK (COG-UK) consortium | The Lighthouse Lab in Alderley Park and Alex Alderton, Roberto Amato, Sonia Goncalves, Ewan Harrison, David K. Jackson, Ian Johnston, Dominic Kwiatkowski, Cordelia Langford, John Sillitoe on behalf of the Wellcome Sanger Institute COVID-19 Surveillance Team (http://www.sanger.ac.uk/covid-team) |
| EPI_ISL_555394, EPI_ISL_555395 | Lighthouse Lab in Alderley Park | Wellcome Sanger Institute for the COVID-19 Genomics UK (COG-UK) consortium | The Lighthouse Lab in Alderley Park and Alex Alderton, Roberto Amato, Sonia Goncalves, Ewan Harrison, David K. Jackson, Ian Johnston, Dominic Kwiatkowski, Cordelia Langford, John Sillitoe on behalf of the Wellcome Sanger Institute COVID-19 Surveillance Team |
| EPI_ISL_555396 | Lighthouse Lab in Alderley Park | Wellcome Sanger Institute for the COVID-19 Genomics UK (COG-UK) consortium | The Lighthouse Lab in Alderley Park and Alex Alderton, Roberto Amato, Sonia Goncalves, Ewan Harrison, David K. Jackson, Ian Johnston, Dominic Kwiatkowski, Cordelia Langford, John Sillitoe on behalf of the Wellcome Sanger Institute COVID-19 Surveillance Team (http://www.sanger.ac.uk/covid-team) |
| EPI_ISL_555397, EPI_ISL_555399, EPI_ISL_555400, EPI_ISL_555401, EPI_ISL_555402, EPI_ISL_555403 | Lighthouse Lab in Alderley Park | Wellcome Sanger Institute for the COVID-19 Genomics UK (COG-UK) consortium | The Lighthouse Lab in Alderley Park and Alex Alderton, Roberto Amato, Sonia Goncalves, Ewan Harrison, David K. Jackson, Ian Johnston, Dominic Kwiatkowski, Cordelia Langford, John Sillitoe on behalf of the Wellcome Sanger Institute COVID-19 Surveillance Team |
| EPI_ISL_555404 | Lighthouse Lab in Alderley Park | Wellcome Sanger Institute for the COVID-19 Genomics UK (COG-UK) consortium | The Lighthouse Lab in Alderley Park and Alex Alderton, Roberto Amato, Sonia Goncalves, Ewan Harrison, David K. Jackson, Ian Johnston, Dominic Kwiatkowski, Cordelia Langford, John Sillitoe on behalf of the Wellcome Sanger Institute COVID-19 Surveillance Team (http://www.sanger.ac.uk/covid-team) |
| EPI_ISL_555405, EPI_ISL_555406, EPI_ISL_555407, EPI_ISL_555408, EPI_ISL_555409, EPI_ISL_555410, EPI_ISL_555411, EPI_ISL_555412, EPI_ISL_555413, EPI_ISL_555414, EPI_ISL_555415, EPI_ISL_555416, EPI_ISL_555417, EPI_ISL_555418, EPI_ISL_555419, EPI_ISL_555420, EPI_ISL_555421 | | | |
| see above | Lighthouse Lab in Alderley Park | Wellcome Sanger Institute for the COVID-19 Genomics UK (COG-UK) consortium | The Lighthouse Lab in Alderley Park and Alex Alderton, Roberto Amato, Sonia Goncalves, Ewan Harrison, David K. Jackson, Ian Johnston, Dominic Kwiatkowski, Cordelia Langford, John Sillitoe on behalf of the Wellcome Sanger Institute COVID-19 Surveillance Team |
| EPI_ISL_555422 | Lighthouse Lab in Alderley Park | Wellcome Sanger Institute for the COVID-19 Genomics UK (COG-UK) consortium | The Lighthouse Lab in Alderley Park and Alex Alderton, Roberto Amato, Sonia Goncalves, Ewan Harrison, David K. Jackson, Ian Johnston, Dominic Kwiatkowski, Cordelia Langford, John Sillitoe on behalf of the Wellcome Sanger Institute COVID-19 Surveillance Team (http://www.sanger.ac.uk/covid-team) |
| EPI_ISL_555423, EPI_ISL_555424, EPI_ISL_555425, EPI_ISL_555426, EPI_ISL_555427, EPI_ISL_555428, EPI_ISL_555429, EPI_ISL_555430, EPI_ISL_555431 | Lighthouse Lab in Alderley Park | Wellcome Sanger Institute for the COVID-19 Genomics UK (COG-UK) consortium | The Lighthouse Lab in Alderley Park and Alex Alderton, Roberto Amato, Sonia Goncalves, Ewan Harrison, David K. Jackson, Ian Johnston, Dominic Kwiatkowski, Cordelia Langford, John Sillitoe on behalf of the Wellcome Sanger Institute COVID-19 Surveillance Team |
| EPI_ISL_555434 | Lighthouse Lab in Alderley Park | Wellcome Sanger Institute for the COVID-19 Genomics UK (COG-UK) consortium | The Lighthouse Lab in Alderley Park and Alex Alderton, Roberto Amato, Sonia Goncalves, Ewan Harrison, David K. Jackson, Ian Johnston, Dominic Kwiatkowski, Cordelia Langford, John Sillitoe on behalf of the Wellcome Sanger Institute COVID-19 Surveillance Team (http://www.sanger.ac.uk/covid-team) |
| EPI_ISL_555435, EPI_ISL_555436, EPI_ISL_555437, EPI_ISL_555438, EPI_ISL_555439, EPI_ISL_555440 | Lighthouse Lab in Alderley Park | Wellcome Sanger Institute for the COVID-19 Genomics UK (COG-UK) consortium | The Lighthouse Lab in Alderley Park and Alex Alderton, Roberto Amato, Sonia Goncalves, Ewan Harrison, David K. Jackson, Ian Johnston, Dominic Kwiatkowski, Cordelia Langford, John Sillitoe on behalf of the Wellcome Sanger Institute COVID-19 Surveillance Team |
| EPI_ISL_555441 | Lighthouse Lab in Alderley Park | Wellcome Sanger Institute for the COVID-19 Genomics UK (COG-UK) consortium | The Lighthouse Lab in Alderley Park and Alex Alderton, Roberto Amato, Sonia Goncalves, Ewan Harrison, David K. Jackson, Ian Johnston, Dominic Kwiatkowski, Cordelia Langford, John Sillitoe on behalf of the Wellcome Sanger Institute COVID-19 Surveillance Team (http://www.sanger.ac.uk/covid-team) |
| EPI_ISL_555442, EPI_ISL_555443, EPI_ISL_555444, EPI_ISL_555445, EPI_ISL_555446, EPI_ISL_555447, EPI_ISL_555448, EPI_ISL_555449, EPI_ISL_555450, EPI_ISL_555451, EPI_ISL_555452, EPI_ISL_555453, EPI_ISL_555454, EPI_ISL_555455, EPI_ISL_555456, EPI_ISL_555457, EPI_ISL_555458, EPI_ISL_555459, EPI_ISL_555460, EPI_ISL_555461 | | | |
| see above | Lighthouse Lab in Alderley Park | Wellcome Sanger Institute for the COVID-19 Genomics UK (COG-UK) consortium | The Lighthouse Lab in Alderley Park and Alex Alderton, Roberto Amato, Sonia Goncalves, Ewan Harrison, David K. Jackson, Ian Johnston, Dominic Kwiatkowski, Cordelia Langford, John Sillitoe on behalf of the Wellcome Sanger Institute COVID-19 Surveillance Team |
| EPI_ISL_555462, EPI_ISL_555463 | Lighthouse Lab in Alderley Park | Wellcome Sanger Institute for the COVID-19 Genomics UK (COG-UK) consortium | The Lighthouse Lab in Alderley Park and Alex Alderton, Roberto Amato, Sonia Goncalves, Ewan Harrison, David K. Jackson, Ian Johnston, Dominic Kwiatkowski, Cordelia Langford, John Sillitoe on behalf of the Wellcome Sanger Institute COVID-19 Surveillance Team (http://www.sanger.ac.uk/covid-team) |
| EPI_ISL_555464, EPI_ISL_555465, EPI_ISL_555466, EPI_ISL_555467, EPI_ISL_555468, EPI_ISL_555469, EPI_ISL_555470, EPI_ISL_555471 | Lighthouse Lab in Alderley Park | Wellcome Sanger Institute for the COVID-19 Genomics UK (COG-UK) consortium | The Lighthouse Lab in Alderley Park and Alex Alderton, Roberto Amato, Sonia Goncalves, Ewan Harrison, David K. Jackson, Ian Johnston, Dominic Kwiatkowski, Cordelia Langford, John Sillitoe on behalf of the Wellcome Sanger Institute COVID-19 Surveillance Team |
| EPI_ISL_555472 | Lighthouse Lab in Alderley Park | Wellcome Sanger Institute for the COVID-19 Genomics UK (COG-UK) consortium | The Lighthouse Lab in Alderley Park and Alex Alderton, Roberto Amato, Sonia Goncalves, Ewan Harrison, David K. Jackson, Ian Johnston, Dominic Kwiatkowski, Cordelia Langford, John Sillitoe on behalf of the Wellcome Sanger Institute COVID-19 Surveillance Team (http://www.sanger.ac.uk/covid-team) |
| EPI_ISL_555473 | Lighthouse Lab in Alderley Park | Wellcome Sanger Institute for the COVID-19 Genomics UK (COG-UK) consortium | The Lighthouse Lab in Alderley Park and Alex Alderton, Roberto Amato, Sonia Goncalves, Ewan Harrison, David K. Jackson, Ian Johnston, Dominic Kwiatkowski, Cordelia Langford, John Sillitoe on behalf of the Wellcome Sanger Institute COVID-19 Surveillance Team |
| EPI_ISL_555474 | Lighthouse Lab in Alderley Park | Wellcome Sanger Institute for the COVID-19 Genomics UK (COG-UK) consortium | The Lighthouse Lab in Alderley Park and Alex Alderton, Roberto Amato, Sonia Goncalves, Ewan Harrison, David K. Jackson, Ian Johnston, Dominic Kwiatkowski, Cordelia Langford, John Sillitoe on behalf of the Wellcome Sanger Institute COVID-19 Surveillance Team (http://www.sanger.ac.uk/covid-team) |
| EPI_ISL_555475, EPI_ISL_555476, EPI_ISL_555477, EPI_ISL_555478 | Lighthouse Lab in Alderley Park | Wellcome Sanger Institute for the COVID-19 Genomics UK (COG-UK) consortium | The Lighthouse Lab in Alderley Park and Alex Alderton, Roberto Amato, Sonia Goncalves, Ewan Harrison, David K. Jackson, Ian Johnston, Dominic Kwiatkowski, Cordelia Langford, John Sillitoe on behalf of the Wellcome Sanger Institute COVID-19 Surveillance Team |

| | | | |
|---|---|---|---|
| EPI_ISL_555479 | Lighthouse Lab in Alderley Park | Wellcome Sanger Institute for the COVID-19 Genomics UK (COG-UK) consortium | The Lighthouse Lab in Alderley Park and Alex Alderton, Roberto Amato, Sonia Goncalves, Ewan Harrison, David K. Jackson, Ian Johnston, Dominic Kwiatkowski, Cordelia Langford, John Sillitoe on behalf of the Wellcome Sanger Institute COVID-19 Surveillance Team (http://www.sanger.ac.uk/covid-team) |
| EPI_ISL_555480 | Lighthouse Lab in Alderley Park | Wellcome Sanger Institute for the COVID-19 Genomics UK (COG-UK) consortium | The Lighthouse Lab in Alderley Park and Alex Alderton, Roberto Amato, Sonia Goncalves, Ewan Harrison, David K. Jackson, Ian Johnston, Dominic Kwiatkowski, Cordelia Langford, John Sillitoe on behalf of the Wellcome Sanger Institute COVID-19 Surveillance Team |
| EPI_ISL_555481 | Lighthouse Lab in Alderley Park | Wellcome Sanger Institute for the COVID-19 Genomics UK (COG-UK) consortium | The Lighthouse Lab in Alderley Park and Alex Alderton, Roberto Amato, Sonia Goncalves, Ewan Harrison, David K. Jackson, Ian Johnston, Dominic Kwiatkowski, Cordelia Langford, John Sillitoe on behalf of the Wellcome Sanger Institute COVID-19 Surveillance Team (http://www.sanger.ac.uk/covid-team) |
| EPI_ISL_555482, EPI_ISL_555483, EPI_ISL_555484, EPI_ISL_555485, EPI_ISL_555486, EPI_ISL_555487, EPI_ISL_555488, EPI_ISL_555489, EPI_ISL_555490, EPI_ISL_555491, EPI_ISL_555492, EPI_ISL_555493, EPI_ISL_555494 | | | |
| see above | Lighthouse Lab in Alderley Park | Wellcome Sanger Institute for the COVID-19 Genomics UK (COG-UK) consortium | The Lighthouse Lab in Alderley Park and Alex Alderton, Roberto Amato, Sonia Goncalves, Ewan Harrison, David K. Jackson, Ian Johnston, Dominic Kwiatkowski, Cordelia Langford, John Sillitoe on behalf of the Wellcome Sanger Institute COVID-19 Surveillance Team |
| EPI_ISL_555495 | Lighthouse Lab in Alderley Park | Wellcome Sanger Institute for the COVID-19 Genomics UK (COG-UK) consortium | The Lighthouse Lab in Alderley Park and Alex Alderton, Roberto Amato, Sonia Goncalves, Ewan Harrison, David K. Jackson, Ian Johnston, Dominic Kwiatkowski, Cordelia Langford, John Sillitoe on behalf of the Wellcome Sanger Institute COVID-19 Surveillance Team (http://www.sanger.ac.uk/covid-team) |
| EPI_ISL_555496, EPI_ISL_555497, EPI_ISL_555498, EPI_ISL_555499 | Lighthouse Lab in Alderley Park | Wellcome Sanger Institute for the COVID-19 Genomics UK (COG-UK) consortium | The Lighthouse Lab in Alderley Park and Alex Alderton, Roberto Amato, Sonia Goncalves, Ewan Harrison, David K. Jackson, Ian Johnston, Dominic Kwiatkowski, Cordelia Langford, John Sillitoe on behalf of the Wellcome Sanger Institute COVID-19 Surveillance Team |
| EPI_ISL_555500 | Lighthouse Lab in Alderley Park | Wellcome Sanger Institute for the COVID-19 Genomics UK (COG-UK) consortium | The Lighthouse Lab in Alderley Park and Alex Alderton, Roberto Amato, Sonia Goncalves, Ewan Harrison, David K. Jackson, Ian Johnston, Dominic Kwiatkowski, Cordelia Langford, John Sillitoe on behalf of the Wellcome Sanger Institute COVID-19 Surveillance Team (http://www.sanger.ac.uk/covid-team) |
| EPI_ISL_555501 | Lighthouse Lab in Alderley Park | Wellcome Sanger Institute for the COVID-19 Genomics UK (COG-UK) consortium | The Lighthouse Lab in Alderley Park and Alex Alderton, Roberto Amato, Sonia Goncalves, Ewan Harrison, David K. Jackson, Ian Johnston, Dominic Kwiatkowski, Cordelia Langford, John Sillitoe on behalf of the Wellcome Sanger Institute COVID-19 Surveillance Team |
| EPI_ISL_555503 | Lighthouse Lab in Alderley Park | Wellcome Sanger Institute for the COVID-19 Genomics UK (COG-UK) consortium | The Lighthouse Lab in Alderley Park and Alex Alderton, Roberto Amato, Sonia Goncalves, Ewan Harrison, David K. Jackson, Ian Johnston, Dominic Kwiatkowski, Cordelia Langford, John Sillitoe on behalf of the Wellcome Sanger Institute COVID-19 Surveillance Team (http://www.sanger.ac.uk/covid-team) |
| EPI_ISL_555504, EPI_ISL_555505 | Lighthouse Lab in Alderley Park | Wellcome Sanger Institute for the COVID-19 Genomics UK (COG-UK) consortium | The Lighthouse Lab in Alderley Park and Alex Alderton, Roberto Amato, Sonia Goncalves, Ewan Harrison, David K. Jackson, Ian Johnston, Dominic Kwiatkowski, Cordelia Langford, John Sillitoe on behalf of the Wellcome Sanger Institute COVID-19 Surveillance Team |
| EPI_ISL_555506, EPI_ISL_555508, EPI_ISL_555513 | Lighthouse Lab in Alderley Park | Wellcome Sanger Institute for the COVID-19 Genomics UK (COG-UK) consortium | The Lighthouse Lab in Alderley Park and Alex Alderton, Roberto Amato, Sonia Goncalves, Ewan Harrison, David K. Jackson, Ian Johnston, Dominic Kwiatkowski, Cordelia Langford, John Sillitoe on behalf of the Wellcome Sanger Institute COVID-19 Surveillance Team (http://www.sanger.ac.uk/covid-team) |
| EPI_ISL_555514 | Lighthouse Lab in Alderley Park | Wellcome Sanger Institute for the COVID-19 Genomics UK (COG-UK) consortium | The Lighthouse Lab in Alderley Park and Alex Alderton, Roberto Amato, Sonia Goncalves, Ewan Harrison, David K. Jackson, Ian Johnston, Dominic Kwiatkowski, Cordelia Langford, John Sillitoe on behalf of the Wellcome Sanger Institute COVID-19 Surveillance Team |
| EPI_ISL_555516, EPI_ISL_555517 | Lighthouse Lab in Alderley Park | Wellcome Sanger Institute for the COVID-19 Genomics UK (COG-UK) consortium | The Lighthouse Lab in Alderley Park and Alex Alderton, Roberto Amato, Sonia Goncalves, Ewan Harrison, David K. Jackson, Ian Johnston, Dominic Kwiatkowski, Cordelia Langford, John Sillitoe on behalf of the Wellcome Sanger Institute COVID-19 Surveillance Team (http://www.sanger.ac.uk/covid-team) |
| EPI_ISL_555518, EPI_ISL_555519 | Lighthouse Lab in Alderley Park | Wellcome Sanger Institute for the COVID-19 Genomics UK (COG-UK) consortium | The Lighthouse Lab in Alderley Park and Alex Alderton, Roberto Amato, Sonia Goncalves, Ewan Harrison, David K. Jackson, Ian Johnston, Dominic Kwiatkowski, Cordelia Langford, John Sillitoe on behalf of the Wellcome Sanger Institute COVID-19 Surveillance Team |
| EPI_ISL_555520, EPI_ISL_555523, EPI_ISL_555524 | Lighthouse Lab in Alderley Park | Wellcome Sanger Institute for the COVID-19 Genomics UK (COG-UK) consortium | The Lighthouse Lab in Alderley Park and Alex Alderton, Roberto Amato, Sonia Goncalves, Ewan Harrison, David K. Jackson, Ian Johnston, Dominic Kwiatkowski, Cordelia Langford, John Sillitoe on behalf of the Wellcome Sanger Institute COVID-19 Surveillance Team (http://www.sanger.ac.uk/covid-team) |
| EPI_ISL_555526, EPI_ISL_555527 | Lighthouse Lab in Alderley Park | Wellcome Sanger Institute for the COVID-19 Genomics UK (COG-UK) consortium | The Lighthouse Lab in Alderley Park and Alex Alderton, Roberto Amato, Sonia Goncalves, Ewan Harrison, David K. Jackson, Ian Johnston, Dominic Kwiatkowski, Cordelia Langford, John Sillitoe on behalf of the Wellcome Sanger Institute COVID-19 Surveillance Team |
| EPI_ISL_555529 | Lighthouse Lab in Alderley Park | Wellcome Sanger Institute for the COVID-19 Genomics UK (COG-UK) consortium | The Lighthouse Lab in Alderley Park and Alex Alderton, Roberto Amato, Sonia Goncalves, Ewan Harrison, David K. Jackson, Ian Johnston, Dominic Kwiatkowski, Cordelia Langford, John Sillitoe on behalf of the Wellcome Sanger Institute COVID-19 Surveillance Team (http://www.sanger.ac.uk/covid-team) |
| EPI_ISL_555531, EPI_ISL_555533, EPI_ISL_555535 | Lighthouse Lab in Alderley Park | Wellcome Sanger Institute for the COVID-19 Genomics UK (COG-UK) consortium | The Lighthouse Lab in Alderley Park and Alex Alderton, Roberto Amato, Sonia Goncalves, Ewan Harrison, David K. Jackson, Ian Johnston, Dominic Kwiatkowski, Cordelia Langford, John Sillitoe on behalf of the Wellcome Sanger Institute COVID-19 Surveillance Team |
| EPI_ISL_555537 | Lighthouse Lab in Alderley Park | Wellcome Sanger Institute for the COVID-19 Genomics UK (COG-UK) consortium | The Lighthouse Lab in Alderley Park and Alex Alderton, Roberto Amato, Sonia Goncalves, Ewan Harrison, David K. Jackson, Ian Johnston, Dominic Kwiatkowski, Cordelia Langford, John Sillitoe on behalf of the Wellcome Sanger Institute COVID-19 Surveillance Team (http://www.sanger.ac.uk/covid-team) |
| EPI_ISL_555540 | Lighthouse Lab in Alderley Park | Wellcome Sanger Institute for the COVID-19 Genomics UK (COG-UK) consortium | The Lighthouse Lab in Alderley Park and Alex Alderton, Roberto Amato, Sonia Goncalves, Ewan Harrison, David K. Jackson, Ian Johnston, Dominic Kwiatkowski, Cordelia Langford, John Sillitoe on behalf of the Wellcome Sanger Institute COVID-19 Surveillance Team |
| EPI_ISL_555541, EPI_ISL_555542, EPI_ISL_555543, EPI_ISL_555544, EPI_ISL_555546 | Lighthouse Lab in Alderley Park | Wellcome Sanger Institute for the COVID-19 Genomics UK (COG-UK) consortium | The Lighthouse Lab in Alderley Park and Alex Alderton, Roberto Amato, Sonia Goncalves, Ewan Harrison, David K. Jackson, Ian Johnston, Dominic Kwiatkowski, Cordelia Langford, John Sillitoe on behalf of the Wellcome Sanger Institute COVID-19 Surveillance Team (http://www.sanger.ac.uk/covid-team) |
| EPI_ISL_555547 | Lighthouse Lab in Alderley Park | Wellcome Sanger Institute for the COVID-19 Genomics UK (COG-UK) consortium | The Lighthouse Lab in Alderley Park and Alex Alderton, Roberto Amato, Sonia Goncalves, Ewan Harrison, David K. Jackson, Ian Johnston, Dominic Kwiatkowski, Cordelia Langford, John Sillitoe on behalf of the Wellcome Sanger Institute COVID-19 Surveillance Team |
| EPI_ISL_555548 | Lighthouse Lab in Alderley Park | Wellcome Sanger Institute for the COVID-19 Genomics UK (COG-UK) consortium | The Lighthouse Lab in Alderley Park and Alex Alderton, Roberto Amato, Sonia Goncalves, Ewan Harrison, David K. Jackson, Ian Johnston, Dominic Kwiatkowski, Cordelia Langford, John Sillitoe on behalf of the Wellcome Sanger Institute COVID-19 Surveillance Team (http://www.sanger.ac.uk/covid-team) |
| EPI_ISL_555549, EPI_ISL_555550 | Lighthouse Lab in Alderley Park | Wellcome Sanger Institute for the COVID-19 Genomics UK (COG-UK) consortium | The Lighthouse Lab in Alderley Park and Alex Alderton, Roberto Amato, Sonia Goncalves, Ewan Harrison, David K. Jackson, Ian Johnston, Dominic Kwiatkowski, Cordelia Langford, John Sillitoe on behalf of the Wellcome Sanger Institute COVID-19 Surveillance Team |
| EPI_ISL_555551, EPI_ISL_555552, EPI_ISL_555553 | Lighthouse Lab in Alderley Park | Wellcome Sanger Institute for the COVID-19 Genomics UK (COG-UK) consortium | The Lighthouse Lab in Alderley Park and Alex Alderton, Roberto Amato, Sonia Goncalves, Ewan Harrison, David K. Jackson, Ian Johnston, Dominic Kwiatkowski, Cordelia Langford, John Sillitoe on behalf of the Wellcome Sanger Institute COVID-19 Surveillance Team (http://www.sanger.ac.uk/covid-team) |
| EPI_ISL_555554 | Lighthouse Lab in Alderley Park | Wellcome Sanger Institute for the COVID-19 Genomics UK (COG-UK) consortium | The Lighthouse Lab in Alderley Park and Alex Alderton, Roberto Amato, Sonia Goncalves, Ewan Harrison, David K. Jackson, Ian Johnston, Dominic Kwiatkowski, Cordelia Langford, John Sillitoe on behalf of the Wellcome Sanger Institute COVID-19 Surveillance Team |
| EPI_ISL_555557 | Lighthouse Lab in Alderley Park | Wellcome Sanger Institute for the COVID-19 Genomics UK (COG-UK) consortium | The Lighthouse Lab in Alderley Park and Alex Alderton, Roberto Amato, Sonia Goncalves, Ewan Harrison, David K. Jackson, Ian Johnston, Dominic Kwiatkowski, Cordelia Langford, John Sillitoe on behalf of the Wellcome Sanger Institute COVID-19 Surveillance Team (http://www.sanger.ac.uk/covid-team) |
| EPI_ISL_555558 | Lighthouse Lab in Milton Keynes | Wellcome Sanger Institute for the COVID-19 Genomics UK (COG-UK) consortium | The Lighthouse Lab in Milton Keynes and Alex Alderton, Roberto Amato, Sonia Goncalves, Ewan Harrison, David K. Jackson, Ian Johnston, Dominic Kwiatkowski, Cordelia Langford, John Sillitoe on behalf of the Wellcome Sanger Institute COVID-19 Surveillance Team (http://www.sanger.ac.uk/covid-team) |
| EPI_ISL_555559 | Lighthouse Lab in Alderley Park | Wellcome Sanger Institute for the COVID-19 Genomics UK | The Lighthouse Lab in Alderley Park and Alex Alderton, Roberto Amato, Sonia Goncalves, Ewan Harrison, David K. Jackson, Ian Johnston, Dominic |

| | | | |
|---|---|---|---|
| | | (COG-UK) consortium | Kwiatkowski, Cordelia Langford, John Sillitoe on behalf of the Wellcome Sanger Institute COVID-19 Surveillance Team |
| EPI_ISL_555560, EPI_ISL_555561, EPI_ISL_555563, EPI_ISL_555564, EPI_ISL_555565, EPI_ISL_555566 | Lighthouse Lab in Alderley Park | Wellcome Sanger Institute for the COVID-19 Genomics UK (COG-UK) consortium | The Lighthouse Lab in Alderley Park and Alex Alderton, Roberto Amato, Sonia Goncalves, Ewan Harrison, David K. Jackson, Ian Johnston, Dominic Kwiatkowski, Cordelia Langford, John Sillitoe on behalf of the Wellcome Sanger Institute COVID-19 Surveillance Team (http://www.sanger.ac.uk/covid-team) |
| EPI_ISL_555567 | Lighthouse Lab in Alderley Park | Wellcome Sanger Institute for the COVID-19 Genomics UK (COG-UK) consortium | The Lighthouse Lab in Alderley Park and Alex Alderton, Roberto Amato, Sonia Goncalves, Ewan Harrison, David K. Jackson, Ian Johnston, Dominic Kwiatkowski, Cordelia Langford, John Sillitoe on behalf of the Wellcome Sanger Institute COVID-19 Surveillance Team |
| EPI_ISL_555568, EPI_ISL_555570, EPI_ISL_555572, EPI_ISL_555573, EPI_ISL_555576, EPI_ISL_555579 | Lighthouse Lab in Alderley Park | Wellcome Sanger Institute for the COVID-19 Genomics UK (COG-UK) consortium | The Lighthouse Lab in Alderley Park and Alex Alderton, Roberto Amato, Sonia Goncalves, Ewan Harrison, David K. Jackson, Ian Johnston, Dominic Kwiatkowski, Cordelia Langford, John Sillitoe on behalf of the Wellcome Sanger Institute COVID-19 Surveillance Team (http://www.sanger.ac.uk/covid-team) |
| EPI_ISL_555580, EPI_ISL_555581 | Lighthouse Lab in Alderley Park | Wellcome Sanger Institute for the COVID-19 Genomics UK (COG-UK) consortium | The Lighthouse Lab in Alderley Park and Alex Alderton, Roberto Amato, Sonia Goncalves, Ewan Harrison, David K. Jackson, Ian Johnston, Dominic Kwiatkowski, Cordelia Langford, John Sillitoe on behalf of the Wellcome Sanger Institute COVID-19 Surveillance Team |
| EPI_ISL_555582 | Lighthouse Lab in Alderley Park | Wellcome Sanger Institute for the COVID-19 Genomics UK (COG-UK) consortium | The Lighthouse Lab in Alderley Park and Alex Alderton, Roberto Amato, Sonia Goncalves, Ewan Harrison, David K. Jackson, Ian Johnston, Dominic Kwiatkowski, Cordelia Langford, John Sillitoe on behalf of the Wellcome Sanger Institute COVID-19 Surveillance Team (http://www.sanger.ac.uk/covid-team) |
| EPI_ISL_555583 | Lighthouse Lab in Alderley Park | Wellcome Sanger Institute for the COVID-19 Genomics UK (COG-UK) consortium | The Lighthouse Lab in Alderley Park and Alex Alderton, Roberto Amato, Sonia Goncalves, Ewan Harrison, David K. Jackson, Ian Johnston, Dominic Kwiatkowski, Cordelia Langford, John Sillitoe on behalf of the Wellcome Sanger Institute COVID-19 Surveillance Team |
| EPI_ISL_555584, EPI_ISL_555585, EPI_ISL_555587 | Lighthouse Lab in Alderley Park | Wellcome Sanger Institute for the COVID-19 Genomics UK (COG-UK) consortium | The Lighthouse Lab in Alderley Park and Alex Alderton, Roberto Amato, Sonia Goncalves, Ewan Harrison, David K. Jackson, Ian Johnston, Dominic Kwiatkowski, Cordelia Langford, John Sillitoe on behalf of the Wellcome Sanger Institute COVID-19 Surveillance Team (http://www.sanger.ac.uk/covid-team) |
| EPI_ISL_555588, EPI_ISL_555589 | Lighthouse Lab in Alderley Park | Wellcome Sanger Institute for the COVID-19 Genomics UK (COG-UK) consortium | The Lighthouse Lab in Alderley Park and Alex Alderton, Roberto Amato, Sonia Goncalves, Ewan Harrison, David K. Jackson, Ian Johnston, Dominic Kwiatkowski, Cordelia Langford, John Sillitoe on behalf of the Wellcome Sanger Institute COVID-19 Surveillance Team |
| EPI_ISL_555592 | Lighthouse Lab in Alderley Park | Wellcome Sanger Institute for the COVID-19 Genomics UK (COG-UK) consortium | The Lighthouse Lab in Alderley Park and Alex Alderton, Roberto Amato, Sonia Goncalves, Ewan Harrison, David K. Jackson, Ian Johnston, Dominic Kwiatkowski, Cordelia Langford, John Sillitoe on behalf of the Wellcome Sanger Institute COVID-19 Surveillance Team (http://www.sanger.ac.uk/covid-team) |
| EPI_ISL_555593, EPI_ISL_555594, EPI_ISL_555596, EPI_ISL_555597 | Lighthouse Lab in Alderley Park | Wellcome Sanger Institute for the COVID-19 Genomics UK (COG-UK) consortium | The Lighthouse Lab in Alderley Park and Alex Alderton, Roberto Amato, Sonia Goncalves, Ewan Harrison, David K. Jackson, Ian Johnston, Dominic Kwiatkowski, Cordelia Langford, John Sillitoe on behalf of the Wellcome Sanger Institute COVID-19 Surveillance Team |
| EPI_ISL_555598 | Lighthouse Lab in Alderley Park | Wellcome Sanger Institute for the COVID-19 Genomics UK (COG-UK) consortium | The Lighthouse Lab in Alderley Park and Alex Alderton, Roberto Amato, Sonia Goncalves, Ewan Harrison, David K. Jackson, Ian Johnston, Dominic Kwiatkowski, Cordelia Langford, John Sillitoe on behalf of the Wellcome Sanger Institute COVID-19 Surveillance Team (http://www.sanger.ac.uk/covid-team) |
| EPI_ISL_555600 | Lighthouse Lab in Alderley Park | Wellcome Sanger Institute for the COVID-19 Genomics UK (COG-UK) consortium | The Lighthouse Lab in Alderley Park and Alex Alderton, Roberto Amato, Sonia Goncalves, Ewan Harrison, David K. Jackson, Ian Johnston, Dominic Kwiatkowski, Cordelia Langford, John Sillitoe on behalf of the Wellcome Sanger Institute COVID-19 Surveillance Team |
| EPI_ISL_555601 | Lighthouse Lab in Alderley Park | Wellcome Sanger Institute for the COVID-19 Genomics UK (COG-UK) consortium | The Lighthouse Lab in Alderley Park and Alex Alderton, Roberto Amato, Sonia Goncalves, Ewan Harrison, David K. Jackson, Ian Johnston, Dominic Kwiatkowski, Cordelia Langford, John Sillitoe on behalf of the Wellcome Sanger Institute COVID-19 Surveillance Team (http://www.sanger.ac.uk/covid-team) |
| EPI_ISL_555602 | Lighthouse Lab in Alderley Park | Wellcome Sanger Institute for the COVID-19 Genomics UK (COG-UK) consortium | The Lighthouse Lab in Alderley Park and Alex Alderton, Roberto Amato, Sonia Goncalves, Ewan Harrison, David K. Jackson, Ian Johnston, Dominic Kwiatkowski, Cordelia Langford, John Sillitoe on behalf of the Wellcome Sanger Institute COVID-19 Surveillance Team |
| EPI_ISL_555604, EPI_ISL_555606 | Lighthouse Lab in Alderley Park | Wellcome Sanger Institute for the COVID-19 Genomics UK (COG-UK) consortium | The Lighthouse Lab in Alderley Park and Alex Alderton, Roberto Amato, Sonia Goncalves, Ewan Harrison, David K. Jackson, Ian Johnston, Dominic Kwiatkowski, Cordelia Langford, John Sillitoe on behalf of the Wellcome Sanger Institute COVID-19 Surveillance Team (http://www.sanger.ac.uk/covid-team) |
| EPI_ISL_555608 | Lighthouse Lab in Alderley Park | Wellcome Sanger Institute for the COVID-19 Genomics UK (COG-UK) consortium | The Lighthouse Lab in Alderley Park and Alex Alderton, Roberto Amato, Sonia Goncalves, Ewan Harrison, David K. Jackson, Ian Johnston, Dominic Kwiatkowski, Cordelia Langford, John Sillitoe on behalf of the Wellcome Sanger Institute COVID-19 Surveillance Team |
| EPI_ISL_555613 | Lighthouse Lab in Alderley Park | Wellcome Sanger Institute for the COVID-19 Genomics UK (COG-UK) consortium | The Lighthouse Lab in Alderley Park and Alex Alderton, Roberto Amato, Sonia Goncalves, Ewan Harrison, David K. Jackson, Ian Johnston, Dominic Kwiatkowski, Cordelia Langford, John Sillitoe on behalf of the Wellcome Sanger Institute COVID-19 Surveillance Team (http://www.sanger.ac.uk/covid-team) |
| EPI_ISL_555614 | Lighthouse Lab in Alderley Park | Wellcome Sanger Institute for the COVID-19 Genomics UK (COG-UK) consortium | The Lighthouse Lab in Alderley Park and Alex Alderton, Roberto Amato, Sonia Goncalves, Ewan Harrison, David K. Jackson, Ian Johnston, Dominic Kwiatkowski, Cordelia Langford, John Sillitoe on behalf of the Wellcome Sanger Institute COVID-19 Surveillance Team |
| EPI_ISL_555615 | Lighthouse Lab in Alderley Park | Wellcome Sanger Institute for the COVID-19 Genomics UK (COG-UK) consortium | The Lighthouse Lab in Alderley Park and Alex Alderton, Roberto Amato, Sonia Goncalves, Ewan Harrison, David K. Jackson, Ian Johnston, Dominic Kwiatkowski, Cordelia Langford, John Sillitoe on behalf of the Wellcome Sanger Institute COVID-19 Surveillance Team (http://www.sanger.ac.uk/covid-team) |
| EPI_ISL_555616 | Lighthouse Lab in Milton Keynes | Wellcome Sanger Institute for the COVID-19 Genomics UK (COG-UK) consortium | The Lighthouse Lab in Milton Keynes and Alex Alderton, Roberto Amato, Sonia Goncalves, Ewan Harrison, David K. Jackson, Ian Johnston, Dominic Kwiatkowski, Cordelia Langford, John Sillitoe on behalf of the Wellcome Sanger Institute COVID-19 Surveillance Team (http://www.sanger.ac.uk/covid-team) |
| EPI_ISL_555617 | Lighthouse Lab in Alderley Park | Wellcome Sanger Institute for the COVID-19 Genomics UK (COG-UK) consortium | The Lighthouse Lab in Alderley Park and Alex Alderton, Roberto Amato, Sonia Goncalves, Ewan Harrison, David K. Jackson, Ian Johnston, Dominic Kwiatkowski, Cordelia Langford, John Sillitoe on behalf of the Wellcome Sanger Institute COVID-19 Surveillance Team (http://www.sanger.ac.uk/covid-team) |
| EPI_ISL_555619, EPI_ISL_555620, EPI_ISL_555622 | Lighthouse Lab in Alderley Park | Wellcome Sanger Institute for the COVID-19 Genomics UK (COG-UK) consortium | The Lighthouse Lab in Alderley Park and Alex Alderton, Roberto Amato, Sonia Goncalves, Ewan Harrison, David K. Jackson, Ian Johnston, Dominic Kwiatkowski, Cordelia Langford, John Sillitoe on behalf of the Wellcome Sanger Institute COVID-19 Surveillance Team |
| EPI_ISL_555623, EPI_ISL_555626, EPI_ISL_555627, EPI_ISL_555628, EPI_ISL_555629, EPI_ISL_555631, EPI_ISL_555632, EPI_ISL_555633, EPI_ISL_555635, EPI_ISL_555636 | Lighthouse Lab in Alderley Park | Wellcome Sanger Institute for the COVID-19 Genomics UK (COG-UK) consortium | The Lighthouse Lab in Alderley Park and Alex Alderton, Roberto Amato, Sonia Goncalves, Ewan Harrison, David K. Jackson, Ian Johnston, Dominic Kwiatkowski, Cordelia Langford, John Sillitoe on behalf of the Wellcome Sanger Institute COVID-19 Surveillance Team (http://www.sanger.ac.uk/covid-team) |
| EPI_ISL_555642, EPI_ISL_555652 | Lighthouse Lab in Alderley Park | Wellcome Sanger Institute for the COVID-19 Genomics UK (COG-UK) consortium | The Lighthouse Lab in Alderley Park and Alex Alderton, Roberto Amato, Sonia Goncalves, Ewan Harrison, David K. Jackson, Ian Johnston, Dominic Kwiatkowski, Cordelia Langford, John Sillitoe on behalf of the Wellcome Sanger Institute COVID-19 Surveillance Team |
| EPI_ISL_555653, EPI_ISL_555656, EPI_ISL_555657 | Lighthouse Lab in Alderley Park | Wellcome Sanger Institute for the COVID-19 Genomics UK (COG-UK) consortium | The Lighthouse Lab in Alderley Park and Alex Alderton, Roberto Amato, Sonia Goncalves, Ewan Harrison, David K. Jackson, Ian Johnston, Dominic Kwiatkowski, Cordelia Langford, John Sillitoe on behalf of the Wellcome Sanger Institute COVID-19 Surveillance Team (http://www.sanger.ac.uk/covid-team) |
| EPI_ISL_555658 | Lighthouse Lab in Alderley Park | Wellcome Sanger Institute for the COVID-19 Genomics UK (COG-UK) consortium | The Lighthouse Lab in Alderley Park and Alex Alderton, Roberto Amato, Sonia Goncalves, Ewan Harrison, David K. Jackson, Ian Johnston, Dominic Kwiatkowski, Cordelia Langford, John Sillitoe on behalf of the Wellcome Sanger Institute COVID-19 Surveillance Team |
| EPI_ISL_555659, EPI_ISL_555663 | Lighthouse Lab in Alderley Park | Wellcome Sanger Institute for the COVID-19 Genomics UK (COG-UK) consortium | The Lighthouse Lab in Alderley Park and Alex Alderton, Roberto Amato, Sonia Goncalves, Ewan Harrison, David K. Jackson, Ian Johnston, Dominic Kwiatkowski, Cordelia Langford, John Sillitoe on behalf of the Wellcome Sanger Institute COVID-19 Surveillance Team (http://www.sanger.ac.uk/covid-team) |
| EPI_ISL_555664 | Lighthouse Lab in Alderley Park | Wellcome Sanger Institute for the COVID-19 Genomics UK | The Lighthouse Lab in Alderley Park and Alex Alderton, Roberto Amato, Sonia Goncalves, Ewan Harrison, David K. Jackson, Ian Johnston, Dominic |

| | | | |
|---|---|---|---|
| | | (COG-UK) consortium | Kwiatkowski, Cordelia Langford, John Sillitoe on behalf of the Wellcome Sanger Institute COVID-19 Surveillance Team |
| EPI_ISL_555666, EPI_ISL_555667, EPI_ISL_555668 | Lighthouse Lab in Alderley Park | Wellcome Sanger Institute for the COVID-19 Genomics UK (COG-UK) consortium | The Lighthouse Lab in Alderley Park and Alex Alderton, Roberto Amato, Sonia Goncalves, Ewan Harrison, David K. Jackson, Ian Johnston, Dominic Kwiatkowski, Cordelia Langford, John Sillitoe on behalf of the Wellcome Sanger Institute COVID-19 Surveillance Team (http://www.sanger.ac.uk/covid-team) |
| EPI_ISL_555669 | Lighthouse Lab in Alderley Park | Wellcome Sanger Institute for the COVID-19 Genomics UK (COG-UK) consortium | The Lighthouse Lab in Alderley Park and Alex Alderton, Roberto Amato, Sonia Goncalves, Ewan Harrison, David K. Jackson, Ian Johnston, Dominic Kwiatkowski, Cordelia Langford, John Sillitoe on behalf of the Wellcome Sanger Institute COVID-19 Surveillance Team |
| EPI_ISL_555670 | Lighthouse Lab in Alderley Park | Wellcome Sanger Institute for the COVID-19 Genomics UK (COG-UK) consortium | The Lighthouse Lab in Alderley Park and Alex Alderton, Roberto Amato, Sonia Goncalves, Ewan Harrison, David K. Jackson, Ian Johnston, Dominic Kwiatkowski, Cordelia Langford, John Sillitoe on behalf of the Wellcome Sanger Institute COVID-19 Surveillance Team (http://www.sanger.ac.uk/covid-team) |
| EPI_ISL_555671 | Lighthouse Lab in Alderley Park | Wellcome Sanger Institute for the COVID-19 Genomics UK (COG-UK) consortium | The Lighthouse Lab in Alderley Park and Alex Alderton, Roberto Amato, Sonia Goncalves, Ewan Harrison, David K. Jackson, Ian Johnston, Dominic Kwiatkowski, Cordelia Langford, John Sillitoe on behalf of the Wellcome Sanger Institute COVID-19 Surveillance Team |
| EPI_ISL_555672, EPI_ISL_555673, EPI_ISL_555677, EPI_ISL_555678 | Lighthouse Lab in Alderley Park | Wellcome Sanger Institute for the COVID-19 Genomics UK (COG-UK) consortium | The Lighthouse Lab in Alderley Park and Alex Alderton, Roberto Amato, Sonia Goncalves, Ewan Harrison, David K. Jackson, Ian Johnston, Dominic Kwiatkowski, Cordelia Langford, John Sillitoe on behalf of the Wellcome Sanger Institute COVID-19 Surveillance Team (http://www.sanger.ac.uk/covid-team) |
| EPI_ISL_555679 | Lighthouse Lab in Alderley Park | Wellcome Sanger Institute for the COVID-19 Genomics UK (COG-UK) consortium | The Lighthouse Lab in Alderley Park and Alex Alderton, Roberto Amato, Sonia Goncalves, Ewan Harrison, David K. Jackson, Ian Johnston, Dominic Kwiatkowski, Cordelia Langford, John Sillitoe on behalf of the Wellcome Sanger Institute COVID-19 Surveillance Team |
| EPI_ISL_555680 | Lighthouse Lab in Alderley Park | Wellcome Sanger Institute for the COVID-19 Genomics UK (COG-UK) consortium | The Lighthouse Lab in Alderley Park and Alex Alderton, Roberto Amato, Sonia Goncalves, Ewan Harrison, David K. Jackson, Ian Johnston, Dominic Kwiatkowski, Cordelia Langford, John Sillitoe on behalf of the Wellcome Sanger Institute COVID-19 Surveillance Team (http://www.sanger.ac.uk/covid-team) |
| EPI_ISL_555681 | Lighthouse Lab in Alderley Park | Wellcome Sanger Institute for the COVID-19 Genomics UK (COG-UK) consortium | The Lighthouse Lab in Alderley Park and Alex Alderton, Roberto Amato, Sonia Goncalves, Ewan Harrison, David K. Jackson, Ian Johnston, Dominic Kwiatkowski, Cordelia Langford, John Sillitoe on behalf of the Wellcome Sanger Institute COVID-19 Surveillance Team |
| EPI_ISL_555682 | Lighthouse Lab in Alderley Park | Wellcome Sanger Institute for the COVID-19 Genomics UK (COG-UK) consortium | The Lighthouse Lab in Alderley Park and Alex Alderton, Roberto Amato, Sonia Goncalves, Ewan Harrison, David K. Jackson, Ian Johnston, Dominic Kwiatkowski, Cordelia Langford, John Sillitoe on behalf of the Wellcome Sanger Institute COVID-19 Surveillance Team (http://www.sanger.ac.uk/covid-team) |
| EPI_ISL_555683 | Lighthouse Lab in Alderley Park | Wellcome Sanger Institute for the COVID-19 Genomics UK (COG-UK) consortium | The Lighthouse Lab in Alderley Park and Alex Alderton, Roberto Amato, Sonia Goncalves, Ewan Harrison, David K. Jackson, Ian Johnston, Dominic Kwiatkowski, Cordelia Langford, John Sillitoe on behalf of the Wellcome Sanger Institute COVID-19 Surveillance Team |
| EPI_ISL_555685 | Lighthouse Lab in Alderley Park | Wellcome Sanger Institute for the COVID-19 Genomics UK (COG-UK) consortium | The Lighthouse Lab in Alderley Park and Alex Alderton, Roberto Amato, Sonia Goncalves, Ewan Harrison, David K. Jackson, Ian Johnston, Dominic Kwiatkowski, Cordelia Langford, John Sillitoe on behalf of the Wellcome Sanger Institute COVID-19 Surveillance Team (http://www.sanger.ac.uk/covid-team) |
| EPI_ISL_555689 | Lighthouse Lab in Alderley Park | Wellcome Sanger Institute for the COVID-19 Genomics UK (COG-UK) consortium | The Lighthouse Lab in Alderley Park and Alex Alderton, Roberto Amato, Sonia Goncalves, Ewan Harrison, David K. Jackson, Ian Johnston, Dominic Kwiatkowski, Cordelia Langford, John Sillitoe on behalf of the Wellcome Sanger Institute COVID-19 Surveillance Team |
| EPI_ISL_555694 | Lighthouse Lab in Alderley Park | Wellcome Sanger Institute for the COVID-19 Genomics UK (COG-UK) consortium | The Lighthouse Lab in Alderley Park and Alex Alderton, Roberto Amato, Sonia Goncalves, Ewan Harrison, David K. Jackson, Ian Johnston, Dominic Kwiatkowski, Cordelia Langford, John Sillitoe on behalf of the Wellcome Sanger Institute COVID-19 Surveillance Team (http://www.sanger.ac.uk/covid-team) |
| EPI_ISL_555695 | Lighthouse Lab in Alderley Park | Wellcome Sanger Institute for the COVID-19 Genomics UK (COG-UK) consortium | The Lighthouse Lab in Alderley Park and Alex Alderton, Roberto Amato, Sonia Goncalves, Ewan Harrison, David K. Jackson, Ian Johnston, Dominic Kwiatkowski, Cordelia Langford, John Sillitoe on behalf of the Wellcome Sanger Institute COVID-19 Surveillance Team |
| EPI_ISL_555696 | Lighthouse Lab in Alderley Park | Wellcome Sanger Institute for the COVID-19 Genomics UK (COG-UK) consortium | The Lighthouse Lab in Alderley Park and Alex Alderton, Roberto Amato, Sonia Goncalves, Ewan Harrison, David K. Jackson, Ian Johnston, Dominic Kwiatkowski, Cordelia Langford, John Sillitoe on behalf of the Wellcome Sanger Institute COVID-19 Surveillance Team (http://www.sanger.ac.uk/covid-team) |
| EPI_ISL_555698, EPI_ISL_555700 | Lighthouse Lab in Alderley Park | Wellcome Sanger Institute for the COVID-19 Genomics UK (COG-UK) consortium | The Lighthouse Lab in Alderley Park and Alex Alderton, Roberto Amato, Sonia Goncalves, Ewan Harrison, David K. Jackson, Ian Johnston, Dominic Kwiatkowski, Cordelia Langford, John Sillitoe on behalf of the Wellcome Sanger Institute COVID-19 Surveillance Team |
| EPI_ISL_555702 | Lighthouse Lab in Alderley Park | Wellcome Sanger Institute for the COVID-19 Genomics UK (COG-UK) consortium | The Lighthouse Lab in Alderley Park and Alex Alderton, Roberto Amato, Sonia Goncalves, Ewan Harrison, David K. Jackson, Ian Johnston, Dominic Kwiatkowski, Cordelia Langford, John Sillitoe on behalf of the Wellcome Sanger Institute COVID-19 Surveillance Team (http://www.sanger.ac.uk/covid-team) |
| EPI_ISL_555705 | Lighthouse Lab in Alderley Park | Wellcome Sanger Institute for the COVID-19 Genomics UK (COG-UK) consortium | The Lighthouse Lab in Alderley Park and Alex Alderton, Roberto Amato, Sonia Goncalves, Ewan Harrison, David K. Jackson, Ian Johnston, Dominic Kwiatkowski, Cordelia Langford, John Sillitoe on behalf of the Wellcome Sanger Institute COVID-19 Surveillance Team |
| EPI_ISL_555706 | Lighthouse Lab in Alderley Park | Wellcome Sanger Institute for the COVID-19 Genomics UK (COG-UK) consortium | The Lighthouse Lab in Alderley Park and Alex Alderton, Roberto Amato, Sonia Goncalves, Ewan Harrison, David K. Jackson, Ian Johnston, Dominic Kwiatkowski, Cordelia Langford, John Sillitoe on behalf of the Wellcome Sanger Institute COVID-19 Surveillance Team (http://www.sanger.ac.uk/covid-team) |
| EPI_ISL_555707 | Lighthouse Lab in Alderley Park | Wellcome Sanger Institute for the COVID-19 Genomics UK (COG-UK) consortium | The Lighthouse Lab in Alderley Park and Alex Alderton, Roberto Amato, Sonia Goncalves, Ewan Harrison, David K. Jackson, Ian Johnston, Dominic Kwiatkowski, Cordelia Langford, John Sillitoe on behalf of the Wellcome Sanger Institute COVID-19 Surveillance Team |
| EPI_ISL_555708 | Lighthouse Lab in Alderley Park | Wellcome Sanger Institute for the COVID-19 Genomics UK (COG-UK) consortium | The Lighthouse Lab in Alderley Park and Alex Alderton, Roberto Amato, Sonia Goncalves, Ewan Harrison, David K. Jackson, Ian Johnston, Dominic Kwiatkowski, Cordelia Langford, John Sillitoe on behalf of the Wellcome Sanger Institute COVID-19 Surveillance Team (http://www.sanger.ac.uk/covid-team) |
| EPI_ISL_555709, EPI_ISL_555711 | Lighthouse Lab in Alderley Park | Wellcome Sanger Institute for the COVID-19 Genomics UK (COG-UK) consortium | The Lighthouse Lab in Alderley Park and Alex Alderton, Roberto Amato, Sonia Goncalves, Ewan Harrison, David K. Jackson, Ian Johnston, Dominic Kwiatkowski, Cordelia Langford, John Sillitoe on behalf of the Wellcome Sanger Institute COVID-19 Surveillance Team |
| EPI_ISL_555712, EPI_ISL_555713 | Lighthouse Lab in Alderley Park | Wellcome Sanger Institute for the COVID-19 Genomics UK (COG-UK) consortium | The Lighthouse Lab in Alderley Park and Alex Alderton, Roberto Amato, Sonia Goncalves, Ewan Harrison, David K. Jackson, Ian Johnston, Dominic Kwiatkowski, Cordelia Langford, John Sillitoe on behalf of the Wellcome Sanger Institute COVID-19 Surveillance Team (http://www.sanger.ac.uk/covid-team) |
| EPI_ISL_555714 | Lighthouse Lab in Alderley Park | Wellcome Sanger Institute for the COVID-19 Genomics UK (COG-UK) consortium | The Lighthouse Lab in Alderley Park and Alex Alderton, Roberto Amato, Sonia Goncalves, Ewan Harrison, David K. Jackson, Ian Johnston, Dominic Kwiatkowski, Cordelia Langford, John Sillitoe on behalf of the Wellcome Sanger Institute COVID-19 Surveillance Team |
| EPI_ISL_555715 | Lighthouse Lab in Alderley Park | Wellcome Sanger Institute for the COVID-19 Genomics UK (COG-UK) consortium | The Lighthouse Lab in Alderley Park and Alex Alderton, Roberto Amato, Sonia Goncalves, Ewan Harrison, David K. Jackson, Ian Johnston, Dominic Kwiatkowski, Cordelia Langford, John Sillitoe on behalf of the Wellcome Sanger Institute COVID-19 Surveillance Team (http://www.sanger.ac.uk/covid-team) |
| EPI_ISL_555716, EPI_ISL_555717 | Lighthouse Lab in Alderley Park | Wellcome Sanger Institute for the COVID-19 Genomics UK (COG-UK) consortium | The Lighthouse Lab in Alderley Park and Alex Alderton, Roberto Amato, Sonia Goncalves, Ewan Harrison, David K. Jackson, Ian Johnston, Dominic Kwiatkowski, Cordelia Langford, John Sillitoe on behalf of the Wellcome Sanger Institute COVID-19 Surveillance Team |
| EPI_ISL_555718, EPI_ISL_555719 | Lighthouse Lab in Alderley Park | Wellcome Sanger Institute for the COVID-19 Genomics UK (COG-UK) consortium | The Lighthouse Lab in Alderley Park and Alex Alderton, Roberto Amato, Sonia Goncalves, Ewan Harrison, David K. Jackson, Ian Johnston, Dominic Kwiatkowski, Cordelia Langford, John Sillitoe on behalf of the Wellcome Sanger Institute COVID-19 Surveillance Team (http://www.sanger.ac.uk/covid-team) |
| EPI_ISL_555723 | Lighthouse Lab in Alderley Park | Wellcome Sanger Institute for the COVID-19 Genomics UK (COG-UK) consortium | The Lighthouse Lab in Alderley Park and Alex Alderton, Roberto Amato, Sonia Goncalves, Ewan Harrison, David K. Jackson, Ian Johnston, Dominic Kwiatkowski, Cordelia Langford, John Sillitoe on behalf of the Wellcome Sanger Institute COVID-19 Surveillance Team |
| EPI_ISL_555725, EPI_ISL_555726, EPI_ISL_555727, EPI_ISL_555728 | Lighthouse Lab in Alderley Park | Wellcome Sanger Institute for the COVID-19 Genomics UK (COG-UK) consortium | The Lighthouse Lab in Alderley Park and Alex Alderton, Roberto Amato, Sonia Goncalves, Ewan Harrison, David K. Jackson, Ian Johnston, Dominic Kwiatkowski, Cordelia Langford, John Sillitoe on behalf of the Wellcome Sanger Institute COVID-19 Surveillance Team |

| | | | |
|---|---|---|---|
| | | | (http://www.sanger.ac.uk/covid-team) |
| EPI_ISL_555729 | Lighthouse Lab in Alderley Park | Wellcome Sanger Institute for the COVID-19 Genomics UK (COG-UK) consortium | The Lighthouse Lab in Alderley Park and Alex Alderton, Roberto Amato, Sonia Goncalves, Ewan Harrison, David K. Jackson, Ian Johnston, Dominic Kwiatkowski, Cordelia Langford, John Sillitoe on behalf of the Wellcome Sanger Institute COVID-19 Surveillance Team |
| EPI_ISL_555731, EPI_ISL_555732 | Lighthouse Lab in Alderley Park | Wellcome Sanger Institute for the COVID-19 Genomics UK (COG-UK) consortium | The Lighthouse Lab in Alderley Park and Alex Alderton, Roberto Amato, Sonia Goncalves, Ewan Harrison, David K. Jackson, Ian Johnston, Dominic Kwiatkowski, Cordelia Langford, John Sillitoe on behalf of the Wellcome Sanger Institute COVID-19 Surveillance Team (http://www.sanger.ac.uk/covid-team) |
| EPI_ISL_555735 | Lighthouse Lab in Alderley Park | Wellcome Sanger Institute for the COVID-19 Genomics UK (COG-UK) consortium | The Lighthouse Lab in Alderley Park and Alex Alderton, Roberto Amato, Sonia Goncalves, Ewan Harrison, David K. Jackson, Ian Johnston, Dominic Kwiatkowski, Cordelia Langford, John Sillitoe on behalf of the Wellcome Sanger Institute COVID-19 Surveillance Team |
| EPI_ISL_555739, EPI_ISL_555741, EPI_ISL_555742, EPI_ISL_555744, EPI_ISL_555745, EPI_ISL_555746, EPI_ISL_555748, EPI_ISL_555749, EPI_ISL_555752, EPI_ISL_555754, EPI_ISL_555759 | | | |
| see above | Lighthouse Lab in Alderley Park | Wellcome Sanger Institute for the COVID-19 Genomics UK (COG-UK) consortium | The Lighthouse Lab in Alderley Park and Alex Alderton, Roberto Amato, Sonia Goncalves, Ewan Harrison, David K. Jackson, Ian Johnston, Dominic Kwiatkowski, Cordelia Langford, John Sillitoe on behalf of the Wellcome Sanger Institute COVID-19 Surveillance Team (http://www.sanger.ac.uk/covid-team) |
| EPI_ISL_555760 | Lighthouse Lab in Alderley Park | Wellcome Sanger Institute for the COVID-19 Genomics UK (COG-UK) consortium | The Lighthouse Lab in Alderley Park and Alex Alderton, Roberto Amato, Sonia Goncalves, Ewan Harrison, David K. Jackson, Ian Johnston, Dominic Kwiatkowski, Cordelia Langford, John Sillitoe on behalf of the Wellcome Sanger Institute COVID-19 Surveillance Team |
| EPI_ISL_555761, EPI_ISL_555762, EPI_ISL_555763 | Lighthouse Lab in Alderley Park | Wellcome Sanger Institute for the COVID-19 Genomics UK (COG-UK) consortium | The Lighthouse Lab in Alderley Park and Alex Alderton, Roberto Amato, Sonia Goncalves, Ewan Harrison, David K. Jackson, Ian Johnston, Dominic Kwiatkowski, Cordelia Langford, John Sillitoe on behalf of the Wellcome Sanger Institute COVID-19 Surveillance Team (http://www.sanger.ac.uk/covid-team) |
| EPI_ISL_555764, EPI_ISL_555765, EPI_ISL_555766, EPI_ISL_555767, EPI_ISL_555768, EPI_ISL_555769, EPI_ISL_555770, EPI_ISL_555771, EPI_ISL_555772, EPI_ISL_555773 | Lighthouse Lab in Alderley Park | Wellcome Sanger Institute for the COVID-19 Genomics UK (COG-UK) consortium | The Lighthouse Lab in Alderley Park and Alex Alderton, Roberto Amato, Sonia Goncalves, Ewan Harrison, David K. Jackson, Ian Johnston, Dominic Kwiatkowski, Cordelia Langford, John Sillitoe on behalf of the Wellcome Sanger Institute COVID-19 Surveillance Team |
| EPI_ISL_555774 | Lighthouse Lab in Alderley Park | Wellcome Sanger Institute for the COVID-19 Genomics UK (COG-UK) consortium | The Lighthouse Lab in Alderley Park and Alex Alderton, Roberto Amato, Sonia Goncalves, Ewan Harrison, David K. Jackson, Ian Johnston, Dominic Kwiatkowski, Cordelia Langford, John Sillitoe on behalf of the Wellcome Sanger Institute COVID-19 Surveillance Team (http://www.sanger.ac.uk/covid-team) |
| EPI_ISL_555775, EPI_ISL_555776, EPI_ISL_555777, EPI_ISL_555778, EPI_ISL_555779, EPI_ISL_555780, EPI_ISL_555781, EPI_ISL_555782, EPI_ISL_555783, EPI_ISL_555784 | Lighthouse Lab in Alderley Park | Wellcome Sanger Institute for the COVID-19 Genomics UK (COG-UK) consortium | The Lighthouse Lab in Alderley Park and Alex Alderton, Roberto Amato, Sonia Goncalves, Ewan Harrison, David K. Jackson, Ian Johnston, Dominic Kwiatkowski, Cordelia Langford, John Sillitoe on behalf of the Wellcome Sanger Institute COVID-19 Surveillance Team |
| EPI_ISL_555785 | Lighthouse Lab in Alderley Park | Wellcome Sanger Institute for the COVID-19 Genomics UK (COG-UK) consortium | The Lighthouse Lab in Alderley Park and Alex Alderton, Roberto Amato, Sonia Goncalves, Ewan Harrison, David K. Jackson, Ian Johnston, Dominic Kwiatkowski, Cordelia Langford, John Sillitoe on behalf of the Wellcome Sanger Institute COVID-19 Surveillance Team (http://www.sanger.ac.uk/covid-team) |
| EPI_ISL_555786, EPI_ISL_555787 | Lighthouse Lab in Alderley Park | Wellcome Sanger Institute for the COVID-19 Genomics UK (COG-UK) consortium | The Lighthouse Lab in Alderley Park and Alex Alderton, Roberto Amato, Sonia Goncalves, Ewan Harrison, David K. Jackson, Ian Johnston, Dominic Kwiatkowski, Cordelia Langford, John Sillitoe on behalf of the Wellcome Sanger Institute COVID-19 Surveillance Team |
| EPI_ISL_555788 | Lighthouse Lab in Alderley Park | Wellcome Sanger Institute for the COVID-19 Genomics UK (COG-UK) consortium | The Lighthouse Lab in Alderley Park and Alex Alderton, Roberto Amato, Sonia Goncalves, Ewan Harrison, David K. Jackson, Ian Johnston, Dominic Kwiatkowski, Cordelia Langford, John Sillitoe on behalf of the Wellcome Sanger Institute COVID-19 Surveillance Team (http://www.sanger.ac.uk/covid-team) |
| EPI_ISL_555789, EPI_ISL_555790, EPI_ISL_555791, EPI_ISL_555792, EPI_ISL_555793 | Lighthouse Lab in Alderley Park | Wellcome Sanger Institute for the COVID-19 Genomics UK (COG-UK) consortium | The Lighthouse Lab in Alderley Park and Alex Alderton, Roberto Amato, Sonia Goncalves, Ewan Harrison, David K. Jackson, Ian Johnston, Dominic Kwiatkowski, Cordelia Langford, John Sillitoe on behalf of the Wellcome Sanger Institute COVID-19 Surveillance Team |
| EPI_ISL_555795 | Lighthouse Lab in Alderley Park | Wellcome Sanger Institute for the COVID-19 Genomics UK (COG-UK) consortium | The Lighthouse Lab in Alderley Park and Alex Alderton, Roberto Amato, Sonia Goncalves, Ewan Harrison, David K. Jackson, Ian Johnston, Dominic Kwiatkowski, Cordelia Langford, John Sillitoe on behalf of the Wellcome Sanger Institute COVID-19 Surveillance Team (http://www.sanger.ac.uk/covid-team) |
| EPI_ISL_555796, EPI_ISL_555797, EPI_ISL_555798, EPI_ISL_555799, EPI_ISL_555800, EPI_ISL_555801, EPI_ISL_555802, EPI_ISL_555803 | Lighthouse Lab in Alderley Park | Wellcome Sanger Institute for the COVID-19 Genomics UK (COG-UK) consortium | The Lighthouse Lab in Alderley Park and Alex Alderton, Roberto Amato, Sonia Goncalves, Ewan Harrison, David K. Jackson, Ian Johnston, Dominic Kwiatkowski, Cordelia Langford, John Sillitoe on behalf of the Wellcome Sanger Institute COVID-19 Surveillance Team |
| EPI_ISL_555804 | Lighthouse Lab in Alderley Park | Wellcome Sanger Institute for the COVID-19 Genomics UK (COG-UK) consortium | The Lighthouse Lab in Alderley Park and Alex Alderton, Roberto Amato, Sonia Goncalves, Ewan Harrison, David K. Jackson, Ian Johnston, Dominic Kwiatkowski, Cordelia Langford, John Sillitoe on behalf of the Wellcome Sanger Institute COVID-19 Surveillance Team (http://www.sanger.ac.uk/covid-team) |
| EPI_ISL_555805, EPI_ISL_555806, EPI_ISL_555808, EPI_ISL_555809 | Lighthouse Lab in Alderley Park | Wellcome Sanger Institute for the COVID-19 Genomics UK (COG-UK) consortium | The Lighthouse Lab in Alderley Park and Alex Alderton, Roberto Amato, Sonia Goncalves, Ewan Harrison, David K. Jackson, Ian Johnston, Dominic Kwiatkowski, Cordelia Langford, John Sillitoe on behalf of the Wellcome Sanger Institute COVID-19 Surveillance Team |
| EPI_ISL_555810 | Lighthouse Lab in Alderley Park | Wellcome Sanger Institute for the COVID-19 Genomics UK (COG-UK) consortium | The Lighthouse Lab in Alderley Park and Alex Alderton, Roberto Amato, Sonia Goncalves, Ewan Harrison, David K. Jackson, Ian Johnston, Dominic Kwiatkowski, Cordelia Langford, John Sillitoe on behalf of the Wellcome Sanger Institute COVID-19 Surveillance Team (http://www.sanger.ac.uk/covid-team) |
| EPI_ISL_555811, EPI_ISL_555812, EPI_ISL_555813, EPI_ISL_555814, EPI_ISL_555816, EPI_ISL_555817, EPI_ISL_555818, EPI_ISL_555819, EPI_ISL_555820, EPI_ISL_555821, EPI_ISL_555822, EPI_ISL_555823, EPI_ISL_555824, EPI_ISL_555825, EPI_ISL_555826, EPI_ISL_555827, EPI_ISL_555828 | | | |
| see above | Lighthouse Lab in Alderley Park | Wellcome Sanger Institute for the COVID-19 Genomics UK (COG-UK) consortium | The Lighthouse Lab in Alderley Park and Alex Alderton, Roberto Amato, Sonia Goncalves, Ewan Harrison, David K. Jackson, Ian Johnston, Dominic Kwiatkowski, Cordelia Langford, John Sillitoe on behalf of the Wellcome Sanger Institute COVID-19 Surveillance Team |
| EPI_ISL_555829 | Lighthouse Lab in Alderley Park | Wellcome Sanger Institute for the COVID-19 Genomics UK (COG-UK) consortium | The Lighthouse Lab in Alderley Park and Alex Alderton, Roberto Amato, Sonia Goncalves, Ewan Harrison, David K. Jackson, Ian Johnston, Dominic Kwiatkowski, Cordelia Langford, John Sillitoe on behalf of the Wellcome Sanger Institute COVID-19 Surveillance Team (http://www.sanger.ac.uk/covid-team) |
| EPI_ISL_555830, EPI_ISL_555831 | Lighthouse Lab in Alderley Park | Wellcome Sanger Institute for the COVID-19 Genomics UK (COG-UK) consortium | The Lighthouse Lab in Alderley Park and Alex Alderton, Roberto Amato, Sonia Goncalves, Ewan Harrison, David K. Jackson, Ian Johnston, Dominic Kwiatkowski, Cordelia Langford, John Sillitoe on behalf of the Wellcome Sanger Institute COVID-19 Surveillance Team |
| EPI_ISL_555832 | Lighthouse Lab in Alderley Park | Wellcome Sanger Institute for the COVID-19 Genomics UK (COG-UK) consortium | The Lighthouse Lab in Alderley Park and Alex Alderton, Roberto Amato, Sonia Goncalves, Ewan Harrison, David K. Jackson, Ian Johnston, Dominic Kwiatkowski, Cordelia Langford, John Sillitoe on behalf of the Wellcome Sanger Institute COVID-19 Surveillance Team (http://www.sanger.ac.uk/covid-team) |
| EPI_ISL_555833 | Lighthouse Lab in Alderley Park | Wellcome Sanger Institute for the COVID-19 Genomics UK (COG-UK) consortium | The Lighthouse Lab in Alderley Park and Alex Alderton, Roberto Amato, Sonia Goncalves, Ewan Harrison, David K. Jackson, Ian Johnston, Dominic Kwiatkowski, Cordelia Langford, John Sillitoe on behalf of the Wellcome Sanger Institute COVID-19 Surveillance Team |
| EPI_ISL_555834, EPI_ISL_555835, EPI_ISL_555836, EPI_ISL_555837 | Lighthouse Lab in Alderley Park | Wellcome Sanger Institute for the COVID-19 Genomics UK (COG-UK) consortium | The Lighthouse Lab in Alderley Park and Alex Alderton, Roberto Amato, Sonia Goncalves, Ewan Harrison, David K. Jackson, Ian Johnston, Dominic Kwiatkowski, Cordelia Langford, John Sillitoe on behalf of the Wellcome Sanger Institute COVID-19 Surveillance Team (http://www.sanger.ac.uk/covid-team) |
| EPI_ISL_555838 | Lighthouse Lab in Alderley Park | Wellcome Sanger Institute for the COVID-19 Genomics UK (COG-UK) consortium | The Lighthouse Lab in Alderley Park and Alex Alderton, Roberto Amato, Sonia Goncalves, Ewan Harrison, David K. Jackson, Ian Johnston, Dominic Kwiatkowski, Cordelia Langford, John Sillitoe on behalf of the Wellcome Sanger Institute COVID-19 Surveillance Team |

| | | | |
|---|---|---|---|
| EPI_ISL_555839, EPI_ISL_555840, EPI_ISL_555841 | Lighthouse Lab in Alderley Park | Wellcome Sanger Institute for the COVID-19 Genomics UK (COG-UK) consortium | The Lighthouse Lab in Alderley Park and Alex Alderton, Roberto Amato, Sonia Goncalves, Ewan Harrison, David K. Jackson, Ian Johnston, Dominic Kwiatkowski, Cordelia Langford, John Sillitoe on behalf of the Wellcome Sanger Institute COVID-19 Surveillance Team (http://www.sanger.ac.uk/covid-team) |
| EPI_ISL_555842 | Lighthouse Lab in Alderley Park | Wellcome Sanger Institute for the COVID-19 Genomics UK (COG-UK) consortium | The Lighthouse Lab in Alderley Park and Alex Alderton, Roberto Amato, Sonia Goncalves, Ewan Harrison, David K. Jackson, Ian Johnston, Dominic Kwiatkowski, Cordelia Langford, John Sillitoe on behalf of the Wellcome Sanger Institute COVID-19 Surveillance Team |
| EPI_ISL_555845 | Lighthouse Lab in Alderley Park | Wellcome Sanger Institute for the COVID-19 Genomics UK (COG-UK) consortium | The Lighthouse Lab in Alderley Park and Alex Alderton, Roberto Amato, Sonia Goncalves, Ewan Harrison, David K. Jackson, Ian Johnston, Dominic Kwiatkowski, Cordelia Langford, John Sillitoe on behalf of the Wellcome Sanger Institute COVID-19 Surveillance Team (http://www.sanger.ac.uk/covid-team) |
| EPI_ISL_555846 | Lighthouse Lab in Alderley Park | Wellcome Sanger Institute for the COVID-19 Genomics UK (COG-UK) consortium | The Lighthouse Lab in Alderley Park and Alex Alderton, Roberto Amato, Sonia Goncalves, Ewan Harrison, David K. Jackson, Ian Johnston, Dominic Kwiatkowski, Cordelia Langford, John Sillitoe on behalf of the Wellcome Sanger Institute COVID-19 Surveillance Team |
| EPI_ISL_555847, EPI_ISL_555848, EPI_ISL_555849, EPI_ISL_555850, EPI_ISL_555851, EPI_ISL_555852, EPI_ISL_555853, EPI_ISL_555854 | Lighthouse Lab in Alderley Park | Wellcome Sanger Institute for the COVID-19 Genomics UK (COG-UK) consortium | The Lighthouse Lab in Alderley Park and Alex Alderton, Roberto Amato, Sonia Goncalves, Ewan Harrison, David K. Jackson, Ian Johnston, Dominic Kwiatkowski, Cordelia Langford, John Sillitoe on behalf of the Wellcome Sanger Institute COVID-19 Surveillance Team (http://www.sanger.ac.uk/covid-team) |
| EPI_ISL_555855 | Lighthouse Lab in Alderley Park | Wellcome Sanger Institute for the COVID-19 Genomics UK (COG-UK) consortium | The Lighthouse Lab in Alderley Park and Alex Alderton, Roberto Amato, Sonia Goncalves, Ewan Harrison, David K. Jackson, Ian Johnston, Dominic Kwiatkowski, Cordelia Langford, John Sillitoe on behalf of the Wellcome Sanger Institute COVID-19 Surveillance Team |
| EPI_ISL_555856, EPI_ISL_555857, EPI_ISL_555858 | Lighthouse Lab in Alderley Park | Wellcome Sanger Institute for the COVID-19 Genomics UK (COG-UK) consortium | The Lighthouse Lab in Alderley Park and Alex Alderton, Roberto Amato, Sonia Goncalves, Ewan Harrison, David K. Jackson, Ian Johnston, Dominic Kwiatkowski, Cordelia Langford, John Sillitoe on behalf of the Wellcome Sanger Institute COVID-19 Surveillance Team (http://www.sanger.ac.uk/covid-team) |
| EPI_ISL_555859 | Lighthouse Lab in Alderley Park | Wellcome Sanger Institute for the COVID-19 Genomics UK (COG-UK) consortium | The Lighthouse Lab in Alderley Park and Alex Alderton, Roberto Amato, Sonia Goncalves, Ewan Harrison, David K. Jackson, Ian Johnston, Dominic Kwiatkowski, Cordelia Langford, John Sillitoe on behalf of the Wellcome Sanger Institute COVID-19 Surveillance Team |
| EPI_ISL_555860, EPI_ISL_555861, EPI_ISL_555862, EPI_ISL_555863, EPI_ISL_555865, EPI_ISL_555866, EPI_ISL_555867 | Lighthouse Lab in Alderley Park | Wellcome Sanger Institute for the COVID-19 Genomics UK (COG-UK) consortium | The Lighthouse Lab in Alderley Park and Alex Alderton, Roberto Amato, Sonia Goncalves, Ewan Harrison, David K. Jackson, Ian Johnston, Dominic Kwiatkowski, Cordelia Langford, John Sillitoe on behalf of the Wellcome Sanger Institute COVID-19 Surveillance Team (http://www.sanger.ac.uk/covid-team) |
| EPI_ISL_555869, EPI_ISL_555870 | Lighthouse Lab in Alderley Park | Wellcome Sanger Institute for the COVID-19 Genomics UK (COG-UK) consortium | The Lighthouse Lab in Alderley Park and Alex Alderton, Roberto Amato, Sonia Goncalves, Ewan Harrison, David K. Jackson, Ian Johnston, Dominic Kwiatkowski, Cordelia Langford, John Sillitoe on behalf of the Wellcome Sanger Institute COVID-19 Surveillance Team |
| EPI_ISL_555871, EPI_ISL_555872 | Lighthouse Lab in Alderley Park | Wellcome Sanger Institute for the COVID-19 Genomics UK (COG-UK) consortium | The Lighthouse Lab in Alderley Park and Alex Alderton, Roberto Amato, Sonia Goncalves, Ewan Harrison, David K. Jackson, Ian Johnston, Dominic Kwiatkowski, Cordelia Langford, John Sillitoe on behalf of the Wellcome Sanger Institute COVID-19 Surveillance Team (http://www.sanger.ac.uk/covid-team) |
| EPI_ISL_555873 | Lighthouse Lab in Alderley Park | Wellcome Sanger Institute for the COVID-19 Genomics UK (COG-UK) consortium | The Lighthouse Lab in Alderley Park and Alex Alderton, Roberto Amato, Sonia Goncalves, Ewan Harrison, David K. Jackson, Ian Johnston, Dominic Kwiatkowski, Cordelia Langford, John Sillitoe on behalf of the Wellcome Sanger Institute COVID-19 Surveillance Team |
| EPI_ISL_555874, EPI_ISL_555876, EPI_ISL_555877, EPI_ISL_555878, EPI_ISL_555879, EPI_ISL_555880 | Lighthouse Lab in Alderley Park | Wellcome Sanger Institute for the COVID-19 Genomics UK (COG-UK) consortium | The Lighthouse Lab in Alderley Park and Alex Alderton, Roberto Amato, Sonia Goncalves, Ewan Harrison, David K. Jackson, Ian Johnston, Dominic Kwiatkowski, Cordelia Langford, John Sillitoe on behalf of the Wellcome Sanger Institute COVID-19 Surveillance Team (http://www.sanger.ac.uk/covid-team) |
| EPI_ISL_555881 | Lighthouse Lab in Alderley Park | Wellcome Sanger Institute for the COVID-19 Genomics UK (COG-UK) consortium | The Lighthouse Lab in Alderley Park and Alex Alderton, Roberto Amato, Sonia Goncalves, Ewan Harrison, David K. Jackson, Ian Johnston, Dominic Kwiatkowski, Cordelia Langford, John Sillitoe on behalf of the Wellcome Sanger Institute COVID-19 Surveillance Team |
| EPI_ISL_555882, EPI_ISL_555883 | Lighthouse Lab in Alderley Park | Wellcome Sanger Institute for the COVID-19 Genomics UK (COG-UK) consortium | The Lighthouse Lab in Alderley Park and Alex Alderton, Roberto Amato, Sonia Goncalves, Ewan Harrison, David K. Jackson, Ian Johnston, Dominic Kwiatkowski, Cordelia Langford, John Sillitoe on behalf of the Wellcome Sanger Institute COVID-19 Surveillance Team (http://www.sanger.ac.uk/covid-team) |
| EPI_ISL_555884 | Lighthouse Lab in Alderley Park | Wellcome Sanger Institute for the COVID-19 Genomics UK (COG-UK) consortium | The Lighthouse Lab in Alderley Park and Alex Alderton, Roberto Amato, Sonia Goncalves, Ewan Harrison, David K. Jackson, Ian Johnston, Dominic Kwiatkowski, Cordelia Langford, John Sillitoe on behalf of the Wellcome Sanger Institute COVID-19 Surveillance Team |
| EPI_ISL_555885, EPI_ISL_555886, EPI_ISL_555888, EPI_ISL_555889, EPI_ISL_555890, EPI_ISL_555891 | Lighthouse Lab in Alderley Park | Wellcome Sanger Institute for the COVID-19 Genomics UK (COG-UK) consortium | The Lighthouse Lab in Alderley Park and Alex Alderton, Roberto Amato, Sonia Goncalves, Ewan Harrison, David K. Jackson, Ian Johnston, Dominic Kwiatkowski, Cordelia Langford, John Sillitoe on behalf of the Wellcome Sanger Institute COVID-19 Surveillance Team (http://www.sanger.ac.uk/covid-team) |
| EPI_ISL_555892 | Lighthouse Lab in Alderley Park | Wellcome Sanger Institute for the COVID-19 Genomics UK (COG-UK) consortium | The Lighthouse Lab in Alderley Park and Alex Alderton, Roberto Amato, Sonia Goncalves, Ewan Harrison, David K. Jackson, Ian Johnston, Dominic Kwiatkowski, Cordelia Langford, John Sillitoe on behalf of the Wellcome Sanger Institute COVID-19 Surveillance Team |
| EPI_ISL_555893 | Lighthouse Lab in Alderley Park | Wellcome Sanger Institute for the COVID-19 Genomics UK (COG-UK) consortium | The Lighthouse Lab in Alderley Park and Alex Alderton, Roberto Amato, Sonia Goncalves, Ewan Harrison, David K. Jackson, Ian Johnston, Dominic Kwiatkowski, Cordelia Langford, John Sillitoe on behalf of the Wellcome Sanger Institute COVID-19 Surveillance Team (http://www.sanger.ac.uk/covid-team) |
| EPI_ISL_555894, EPI_ISL_555895 | Lighthouse Lab in Alderley Park | Wellcome Sanger Institute for the COVID-19 Genomics UK (COG-UK) consortium | The Lighthouse Lab in Alderley Park and Alex Alderton, Roberto Amato, Sonia Goncalves, Ewan Harrison, David K. Jackson, Ian Johnston, Dominic Kwiatkowski, Cordelia Langford, John Sillitoe on behalf of the Wellcome Sanger Institute COVID-19 Surveillance Team |
| EPI_ISL_555896, EPI_ISL_555897, EPI_ISL_555898, EPI_ISL_555899, EPI_ISL_555900, EPI_ISL_555901, EPI_ISL_555902, EPI_ISL_555903, EPI_ISL_555904 | Lighthouse Lab in Alderley Park | Wellcome Sanger Institute for the COVID-19 Genomics UK (COG-UK) consortium | The Lighthouse Lab in Alderley Park and Alex Alderton, Roberto Amato, Sonia Goncalves, Ewan Harrison, David K. Jackson, Ian Johnston, Dominic Kwiatkowski, Cordelia Langford, John Sillitoe on behalf of the Wellcome Sanger Institute COVID-19 Surveillance Team (http://www.sanger.ac.uk/covid-team) |
| EPI_ISL_555905 | Lighthouse Lab in Alderley Park | Wellcome Sanger Institute for the COVID-19 Genomics UK (COG-UK) consortium | The Lighthouse Lab in Alderley Park and Alex Alderton, Roberto Amato, Sonia Goncalves, Ewan Harrison, David K. Jackson, Ian Johnston, Dominic Kwiatkowski, Cordelia Langford, John Sillitoe on behalf of the Wellcome Sanger Institute COVID-19 Surveillance Team |
| EPI_ISL_555906, EPI_ISL_555907, EPI_ISL_555908, EPI_ISL_555909, EPI_ISL_555911, EPI_ISL_555912, EPI_ISL_555913, EPI_ISL_555914, EPI_ISL_555915 | Lighthouse Lab in Alderley Park | Wellcome Sanger Institute for the COVID-19 Genomics UK (COG-UK) consortium | The Lighthouse Lab in Alderley Park and Alex Alderton, Roberto Amato, Sonia Goncalves, Ewan Harrison, David K. Jackson, Ian Johnston, Dominic Kwiatkowski, Cordelia Langford, John Sillitoe on behalf of the Wellcome Sanger Institute COVID-19 Surveillance Team (http://www.sanger.ac.uk/covid-team) |
| EPI_ISL_555916 | Lighthouse Lab in Alderley Park | Wellcome Sanger Institute for the COVID-19 Genomics UK (COG-UK) consortium | The Lighthouse Lab in Alderley Park and Alex Alderton, Roberto Amato, Sonia Goncalves, Ewan Harrison, David K. Jackson, Ian Johnston, Dominic Kwiatkowski, Cordelia Langford, John Sillitoe on behalf of the Wellcome Sanger Institute COVID-19 Surveillance Team |
| EPI_ISL_555917, EPI_ISL_555918, EPI_ISL_555919, EPI_ISL_555920, EPI_ISL_555921, EPI_ISL_555922, EPI_ISL_555923, EPI_ISL_555924, EPI_ISL_555925, EPI_ISL_555926, EPI_ISL_555927, EPI_ISL_555928 | | | |
| see above | Lighthouse Lab in Alderley Park | Wellcome Sanger Institute for the COVID-19 Genomics UK (COG-UK) consortium | The Lighthouse Lab in Alderley Park and Alex Alderton, Roberto Amato, Sonia Goncalves, Ewan Harrison, David K. Jackson, Ian Johnston, Dominic Kwiatkowski, Cordelia Langford, John Sillitoe on behalf of the Wellcome Sanger Institute COVID-19 Surveillance Team (http://www.sanger.ac.uk/covid-team) |
| EPI_ISL_555929 | Lighthouse Lab in Alderley Park | Wellcome Sanger Institute for the COVID-19 Genomics UK (COG-UK) consortium | The Lighthouse Lab in Alderley Park and Alex Alderton, Roberto Amato, Sonia Goncalves, Ewan Harrison, David K. Jackson, Ian Johnston, Dominic Kwiatkowski, Cordelia Langford, John Sillitoe on behalf of the Wellcome Sanger Institute COVID-19 Surveillance Team |
| EPI_ISL_555930, EPI_ISL_555931, EPI_ISL_555932, EPI_ISL_555933 | Lighthouse Lab in Alderley Park | Wellcome Sanger Institute for the COVID-19 Genomics UK (COG-UK) consortium | The Lighthouse Lab in Alderley Park and Alex Alderton, Roberto Amato, Sonia Goncalves, Ewan Harrison, David K. Jackson, Ian Johnston, Dominic Kwiatkowski, Cordelia Langford, John Sillitoe on behalf of the Wellcome Sanger Institute COVID-19 Surveillance Team |

| | | | |
|---|---|---|---|
| | | | (http://www.sanger.ac.uk/covid-team) |
| EPI_ISL_555934 | Lighthouse Lab in Alderley Park | Wellcome Sanger Institute for the COVID-19 Genomics UK (COG-UK) consortium | The Lighthouse Lab in Alderley Park and Alex Alderton, Roberto Amato, Sonia Goncalves, Ewan Harrison, David K. Jackson, Ian Johnston, Dominic Kwiatkowski, Cordelia Langford, John Sillitoe on behalf of the Wellcome Sanger Institute COVID-19 Surveillance Team |
| EPI_ISL_555935, EPI_ISL_555936, EPI_ISL_555937, EPI_ISL_555938, EPI_ISL_555940, EPI_ISL_555941, EPI_ISL_555942, EPI_ISL_555943, EPI_ISL_555944, EPI_ISL_555946, EPI_ISL_555949, EPI_ISL_555950, EPI_ISL_555951, EPI_ISL_555952 | | | |
| see above | Lighthouse Lab in Alderley Park | Wellcome Sanger Institute for the COVID-19 Genomics UK (COG-UK) consortium | The Lighthouse Lab in Alderley Park and Alex Alderton, Roberto Amato, Sonia Goncalves, Ewan Harrison, David K. Jackson, Ian Johnston, Dominic Kwiatkowski, Cordelia Langford, John Sillitoe on behalf of the Wellcome Sanger Institute COVID-19 Surveillance Team (http://www.sanger.ac.uk/covid-team) |
| EPI_ISL_555953 | Lighthouse Lab in Alderley Park | Wellcome Sanger Institute for the COVID-19 Genomics UK (COG-UK) consortium | The Lighthouse Lab in Alderley Park and Alex Alderton, Roberto Amato, Sonia Goncalves, Ewan Harrison, David K. Jackson, Ian Johnston, Dominic Kwiatkowski, Cordelia Langford, John Sillitoe on behalf of the Wellcome Sanger Institute COVID-19 Surveillance Team |
| EPI_ISL_555954, EPI_ISL_555956, EPI_ISL_555957, EPI_ISL_555958, EPI_ISL_555959 | Lighthouse Lab in Alderley Park | Wellcome Sanger Institute for the COVID-19 Genomics UK (COG-UK) consortium | The Lighthouse Lab in Alderley Park and Alex Alderton, Roberto Amato, Sonia Goncalves, Ewan Harrison, David K. Jackson, Ian Johnston, Dominic Kwiatkowski, Cordelia Langford, John Sillitoe on behalf of the Wellcome Sanger Institute COVID-19 Surveillance Team (http://www.sanger.ac.uk/covid-team) |
| EPI_ISL_555960, EPI_ISL_555961 | Lighthouse Lab in Alderley Park | Wellcome Sanger Institute for the COVID-19 Genomics UK (COG-UK) consortium | The Lighthouse Lab in Alderley Park and Alex Alderton, Roberto Amato, Sonia Goncalves, Ewan Harrison, David K. Jackson, Ian Johnston, Dominic Kwiatkowski, Cordelia Langford, John Sillitoe on behalf of the Wellcome Sanger Institute COVID-19 Surveillance Team |
| EPI_ISL_555962, EPI_ISL_555963, EPI_ISL_555964, EPI_ISL_555965, EPI_ISL_555966, EPI_ISL_555967, EPI_ISL_555968 | Lighthouse Lab in Alderley Park | Wellcome Sanger Institute for the COVID-19 Genomics UK (COG-UK) consortium | The Lighthouse Lab in Alderley Park and Alex Alderton, Roberto Amato, Sonia Goncalves, Ewan Harrison, David K. Jackson, Ian Johnston, Dominic Kwiatkowski, Cordelia Langford, John Sillitoe on behalf of the Wellcome Sanger Institute COVID-19 Surveillance Team (http://www.sanger.ac.uk/covid-team) |
| EPI_ISL_555969 | Lighthouse Lab in Alderley Park | Wellcome Sanger Institute for the COVID-19 Genomics UK (COG-UK) consortium | The Lighthouse Lab in Alderley Park and Alex Alderton, Roberto Amato, Sonia Goncalves, Ewan Harrison, David K. Jackson, Ian Johnston, Dominic Kwiatkowski, Cordelia Langford, John Sillitoe on behalf of the Wellcome Sanger Institute COVID-19 Surveillance Team |
| EPI_ISL_555970 | Lighthouse Lab in Alderley Park | Wellcome Sanger Institute for the COVID-19 Genomics UK (COG-UK) consortium | The Lighthouse Lab in Alderley Park and Alex Alderton, Roberto Amato, Sonia Goncalves, Ewan Harrison, David K. Jackson, Ian Johnston, Dominic Kwiatkowski, Cordelia Langford, John Sillitoe on behalf of the Wellcome Sanger Institute COVID-19 Surveillance Team (http://www.sanger.ac.uk/covid-team) |
| EPI_ISL_555971, EPI_ISL_555972 | Lighthouse Lab in Alderley Park | Wellcome Sanger Institute for the COVID-19 Genomics UK (COG-UK) consortium | The Lighthouse Lab in Alderley Park and Alex Alderton, Roberto Amato, Sonia Goncalves, Ewan Harrison, David K. Jackson, Ian Johnston, Dominic Kwiatkowski, Cordelia Langford, John Sillitoe on behalf of the Wellcome Sanger Institute COVID-19 Surveillance Team |
| EPI_ISL_555974, EPI_ISL_555975 | Lighthouse Lab in Alderley Park | Wellcome Sanger Institute for the COVID-19 Genomics UK (COG-UK) consortium | The Lighthouse Lab in Alderley Park and Alex Alderton, Roberto Amato, Sonia Goncalves, Ewan Harrison, David K. Jackson, Ian Johnston, Dominic Kwiatkowski, Cordelia Langford, John Sillitoe on behalf of the Wellcome Sanger Institute COVID-19 Surveillance Team (http://www.sanger.ac.uk/covid-team) |
| EPI_ISL_555977 | Lighthouse Lab in Alderley Park | Wellcome Sanger Institute for the COVID-19 Genomics UK (COG-UK) consortium | The Lighthouse Lab in Alderley Park and Alex Alderton, Roberto Amato, Sonia Goncalves, Ewan Harrison, David K. Jackson, Ian Johnston, Dominic Kwiatkowski, Cordelia Langford, John Sillitoe on behalf of the Wellcome Sanger Institute COVID-19 Surveillance Team |
| EPI_ISL_555978 | Lighthouse Lab in Alderley Park | Wellcome Sanger Institute for the COVID-19 Genomics UK (COG-UK) consortium | The Lighthouse Lab in Alderley Park and Alex Alderton, Roberto Amato, Sonia Goncalves, Ewan Harrison, David K. Jackson, Ian Johnston, Dominic Kwiatkowski, Cordelia Langford, John Sillitoe on behalf of the Wellcome Sanger Institute COVID-19 Surveillance Team (http://www.sanger.ac.uk/covid-team) |
| EPI_ISL_555979 | Lighthouse Lab in Alderley Park | Wellcome Sanger Institute for the COVID-19 Genomics UK (COG-UK) consortium | The Lighthouse Lab in Alderley Park and Alex Alderton, Roberto Amato, Sonia Goncalves, Ewan Harrison, David K. Jackson, Ian Johnston, Dominic Kwiatkowski, Cordelia Langford, John Sillitoe on behalf of the Wellcome Sanger Institute COVID-19 Surveillance Team |
| EPI_ISL_555980, EPI_ISL_555981, EPI_ISL_555982, EPI_ISL_555983 | Lighthouse Lab in Alderley Park | Wellcome Sanger Institute for the COVID-19 Genomics UK (COG-UK) consortium | The Lighthouse Lab in Alderley Park and Alex Alderton, Roberto Amato, Sonia Goncalves, Ewan Harrison, David K. Jackson, Ian Johnston, Dominic Kwiatkowski, Cordelia Langford, John Sillitoe on behalf of the Wellcome Sanger Institute COVID-19 Surveillance Team (http://www.sanger.ac.uk/covid-team) |
| EPI_ISL_555984 | Lighthouse Lab in Alderley Park | Wellcome Sanger Institute for the COVID-19 Genomics UK (COG-UK) consortium | The Lighthouse Lab in Alderley Park and Alex Alderton, Roberto Amato, Sonia Goncalves, Ewan Harrison, David K. Jackson, Ian Johnston, Dominic Kwiatkowski, Cordelia Langford, John Sillitoe on behalf of the Wellcome Sanger Institute COVID-19 Surveillance Team |
| EPI_ISL_555985, EPI_ISL_555987, EPI_ISL_555988 | Lighthouse Lab in Alderley Park | Wellcome Sanger Institute for the COVID-19 Genomics UK (COG-UK) consortium | The Lighthouse Lab in Alderley Park and Alex Alderton, Roberto Amato, Sonia Goncalves, Ewan Harrison, David K. Jackson, Ian Johnston, Dominic Kwiatkowski, Cordelia Langford, John Sillitoe on behalf of the Wellcome Sanger Institute COVID-19 Surveillance Team (http://www.sanger.ac.uk/covid-team) |
| EPI_ISL_555989 | Lighthouse Lab in Alderley Park | Wellcome Sanger Institute for the COVID-19 Genomics UK (COG-UK) consortium | The Lighthouse Lab in Alderley Park and Alex Alderton, Roberto Amato, Sonia Goncalves, Ewan Harrison, David K. Jackson, Ian Johnston, Dominic Kwiatkowski, Cordelia Langford, John Sillitoe on behalf of the Wellcome Sanger Institute COVID-19 Surveillance Team |
| EPI_ISL_555991 | Lighthouse Lab in Alderley Park | Wellcome Sanger Institute for the COVID-19 Genomics UK (COG-UK) consortium | The Lighthouse Lab in Alderley Park and Alex Alderton, Roberto Amato, Sonia Goncalves, Ewan Harrison, David K. Jackson, Ian Johnston, Dominic Kwiatkowski, Cordelia Langford, John Sillitoe on behalf of the Wellcome Sanger Institute COVID-19 Surveillance Team (http://www.sanger.ac.uk/covid-team) |
| EPI_ISL_555992, EPI_ISL_555994, EPI_ISL_555995 | Lighthouse Lab in Alderley Park | Wellcome Sanger Institute for the COVID-19 Genomics UK (COG-UK) consortium | The Lighthouse Lab in Alderley Park and Alex Alderton, Roberto Amato, Sonia Goncalves, Ewan Harrison, David K. Jackson, Ian Johnston, Dominic Kwiatkowski, Cordelia Langford, John Sillitoe on behalf of the Wellcome Sanger Institute COVID-19 Surveillance Team |
| EPI_ISL_555996 | Lighthouse Lab in Alderley Park | Wellcome Sanger Institute for the COVID-19 Genomics UK (COG-UK) consortium | The Lighthouse Lab in Alderley Park and Alex Alderton, Roberto Amato, Sonia Goncalves, Ewan Harrison, David K. Jackson, Ian Johnston, Dominic Kwiatkowski, Cordelia Langford, John Sillitoe on behalf of the Wellcome Sanger Institute COVID-19 Surveillance Team (http://www.sanger.ac.uk/covid-team) |
| EPI_ISL_555997, EPI_ISL_555998 | Lighthouse Lab in Alderley Park | Wellcome Sanger Institute for the COVID-19 Genomics UK (COG-UK) consortium | The Lighthouse Lab in Alderley Park and Alex Alderton, Roberto Amato, Sonia Goncalves, Ewan Harrison, David K. Jackson, Ian Johnston, Dominic Kwiatkowski, Cordelia Langford, John Sillitoe on behalf of the Wellcome Sanger Institute COVID-19 Surveillance Team |
| EPI_ISL_555999, EPI_ISL_556001, EPI_ISL_556002 | Lighthouse Lab in Alderley Park | Wellcome Sanger Institute for the COVID-19 Genomics UK (COG-UK) consortium | The Lighthouse Lab in Alderley Park and Alex Alderton, Roberto Amato, Sonia Goncalves, Ewan Harrison, David K. Jackson, Ian Johnston, Dominic Kwiatkowski, Cordelia Langford, John Sillitoe on behalf of the Wellcome Sanger Institute COVID-19 Surveillance Team (http://www.sanger.ac.uk/covid-team) |
| EPI_ISL_556003 | Lighthouse Lab in Alderley Park | Wellcome Sanger Institute for the COVID-19 Genomics UK (COG-UK) consortium | The Lighthouse Lab in Alderley Park and Alex Alderton, Roberto Amato, Sonia Goncalves, Ewan Harrison, David K. Jackson, Ian Johnston, Dominic Kwiatkowski, Cordelia Langford, John Sillitoe on behalf of the Wellcome Sanger Institute COVID-19 Surveillance Team |
| EPI_ISL_556004 | Lighthouse Lab in Alderley Park | Wellcome Sanger Institute for the COVID-19 Genomics UK (COG-UK) consortium | The Lighthouse Lab in Alderley Park and Alex Alderton, Roberto Amato, Sonia Goncalves, Ewan Harrison, David K. Jackson, Ian Johnston, Dominic Kwiatkowski, Cordelia Langford, John Sillitoe on behalf of the Wellcome Sanger Institute COVID-19 Surveillance Team (http://www.sanger.ac.uk/covid-team) |
| EPI_ISL_556005, EPI_ISL_556006 | Lighthouse Lab in Alderley Park | Wellcome Sanger Institute for the COVID-19 Genomics UK (COG-UK) consortium | The Lighthouse Lab in Alderley Park and Alex Alderton, Roberto Amato, Sonia Goncalves, Ewan Harrison, David K. Jackson, Ian Johnston, Dominic Kwiatkowski, Cordelia Langford, John Sillitoe on behalf of the Wellcome Sanger Institute COVID-19 Surveillance Team |
| EPI_ISL_556007, EPI_ISL_556008, EPI_ISL_556009 | Lighthouse Lab in Alderley Park | Wellcome Sanger Institute for the COVID-19 Genomics UK (COG-UK) consortium | The Lighthouse Lab in Alderley Park and Alex Alderton, Roberto Amato, Sonia Goncalves, Ewan Harrison, David K. Jackson, Ian Johnston, Dominic Kwiatkowski, Cordelia Langford, John Sillitoe on behalf of the Wellcome Sanger Institute COVID-19 Surveillance Team (http://www.sanger.ac.uk/covid-team) |
| EPI_ISL_556010 | Lighthouse Lab in Alderley Park | Wellcome Sanger Institute for the COVID-19 Genomics UK (COG-UK) consortium | The Lighthouse Lab in Alderley Park and Alex Alderton, Roberto Amato, Sonia Goncalves, Ewan Harrison, David K. Jackson, Ian Johnston, Dominic Kwiatkowski, Cordelia Langford, John Sillitoe on behalf of the Wellcome Sanger Institute COVID-19 Surveillance Team |
| EPI_ISL_556011, EPI_ISL_556013, EPI_ISL_556014 | Lighthouse Lab in Alderley Park | Wellcome Sanger Institute for the COVID-19 Genomics UK (COG-UK) consortium | The Lighthouse Lab in Alderley Park and Alex Alderton, Roberto Amato, Sonia Goncalves, Ewan Harrison, David K. Jackson, Ian Johnston, Dominic Kwiatkowski, Cordelia Langford, John Sillitoe on behalf of the Wellcome Sanger Institute COVID-19 Surveillance Team (http://www.sanger.ac.uk/covid-team) |

| | | | |
|---|---|---|---|
| EPI_ISL_556015 | Lighthouse Lab in Alderley Park | Wellcome Sanger Institute for the COVID-19 Genomics UK (COG-UK) consortium | The Lighthouse Lab in Alderley Park and Alex Alderton, Roberto Amato, Sonia Goncalves, Ewan Harrison, David K. Jackson, Ian Johnston, Dominic Kwiatkowski, Cordelia Langford, John Sillitoe on behalf of the Wellcome Sanger Institute COVID-19 Surveillance Team |
| EPI_ISL_556016, EPI_ISL_556017 | Lighthouse Lab in Alderley Park | Wellcome Sanger Institute for the COVID-19 Genomics UK (COG-UK) consortium | The Lighthouse Lab in Alderley Park and Alex Alderton, Roberto Amato, Sonia Goncalves, Ewan Harrison, David K. Jackson, Ian Johnston, Dominic Kwiatkowski, Cordelia Langford, John Sillitoe on behalf of the Wellcome Sanger Institute COVID-19 Surveillance Team (http://www.sanger.ac.uk/covid-team) |
| EPI_ISL_556018, EPI_ISL_556019, EPI_ISL_556020 | Lighthouse Lab in Alderley Park | Wellcome Sanger Institute for the COVID-19 Genomics UK (COG-UK) consortium | The Lighthouse Lab in Alderley Park and Alex Alderton, Roberto Amato, Sonia Goncalves, Ewan Harrison, David K. Jackson, Ian Johnston, Dominic Kwiatkowski, Cordelia Langford, John Sillitoe on behalf of the Wellcome Sanger Institute COVID-19 Surveillance Team |
| EPI_ISL_556023, EPI_ISL_556024 | Lighthouse Lab in Alderley Park | Wellcome Sanger Institute for the COVID-19 Genomics UK (COG-UK) consortium | The Lighthouse Lab in Alderley Park and Alex Alderton, Roberto Amato, Sonia Goncalves, Ewan Harrison, David K. Jackson, Ian Johnston, Dominic Kwiatkowski, Cordelia Langford, John Sillitoe on behalf of the Wellcome Sanger Institute COVID-19 Surveillance Team (http://www.sanger.ac.uk/covid-team) |
| EPI_ISL_556025 | Lighthouse Lab in Alderley Park | Wellcome Sanger Institute for the COVID-19 Genomics UK (COG-UK) consortium | The Lighthouse Lab in Alderley Park and Alex Alderton, Roberto Amato, Sonia Goncalves, Ewan Harrison, David K. Jackson, Ian Johnston, Dominic Kwiatkowski, Cordelia Langford, John Sillitoe on behalf of the Wellcome Sanger Institute COVID-19 Surveillance Team |
| EPI_ISL_556027, EPI_ISL_556028, EPI_ISL_556029, EPI_ISL_556030 | Lighthouse Lab in Alderley Park | Wellcome Sanger Institute for the COVID-19 Genomics UK (COG-UK) consortium | The Lighthouse Lab in Alderley Park and Alex Alderton, Roberto Amato, Sonia Goncalves, Ewan Harrison, David K. Jackson, Ian Johnston, Dominic Kwiatkowski, Cordelia Langford, John Sillitoe on behalf of the Wellcome Sanger Institute COVID-19 Surveillance Team (http://www.sanger.ac.uk/covid-team) |
| EPI_ISL_556031 | Lighthouse Lab in Alderley Park | Wellcome Sanger Institute for the COVID-19 Genomics UK (COG-UK) consortium | The Lighthouse Lab in Alderley Park and Alex Alderton, Roberto Amato, Sonia Goncalves, Ewan Harrison, David K. Jackson, Ian Johnston, Dominic Kwiatkowski, Cordelia Langford, John Sillitoe on behalf of the Wellcome Sanger Institute COVID-19 Surveillance Team |
| EPI_ISL_556032, EPI_ISL_556033, EPI_ISL_556034, EPI_ISL_556035, EPI_ISL_556036, EPI_ISL_556037, EPI_ISL_556038, EPI_ISL_556040, EPI_ISL_556042, EPI_ISL_556043, EPI_ISL_556044 | | | |
| see above | Lighthouse Lab in Alderley Park | Wellcome Sanger Institute for the COVID-19 Genomics UK (COG-UK) consortium | The Lighthouse Lab in Alderley Park and Alex Alderton, Roberto Amato, Sonia Goncalves, Ewan Harrison, David K. Jackson, Ian Johnston, Dominic Kwiatkowski, Cordelia Langford, John Sillitoe on behalf of the Wellcome Sanger Institute COVID-19 Surveillance Team (http://www.sanger.ac.uk/covid-team) |
| EPI_ISL_556045 | Lighthouse Lab in Alderley Park | Wellcome Sanger Institute for the COVID-19 Genomics UK (COG-UK) consortium | The Lighthouse Lab in Alderley Park and Alex Alderton, Roberto Amato, Sonia Goncalves, Ewan Harrison, David K. Jackson, Ian Johnston, Dominic Kwiatkowski, Cordelia Langford, John Sillitoe on behalf of the Wellcome Sanger Institute COVID-19 Surveillance Team |
| EPI_ISL_556046, EPI_ISL_556048, EPI_ISL_556049, EPI_ISL_556050, EPI_ISL_556051, EPI_ISL_556052, EPI_ISL_556053, EPI_ISL_556054, EPI_ISL_556055, EPI_ISL_556056, EPI_ISL_556057, EPI_ISL_556059 | | | |
| see above | Lighthouse Lab in Alderley Park | Wellcome Sanger Institute for the COVID-19 Genomics UK (COG-UK) consortium | The Lighthouse Lab in Alderley Park and Alex Alderton, Roberto Amato, Sonia Goncalves, Ewan Harrison, David K. Jackson, Ian Johnston, Dominic Kwiatkowski, Cordelia Langford, John Sillitoe on behalf of the Wellcome Sanger Institute COVID-19 Surveillance Team (http://www.sanger.ac.uk/covid-team) |
| EPI_ISL_556060 | Lighthouse Lab in Alderley Park | Wellcome Sanger Institute for the COVID-19 Genomics UK (COG-UK) consortium | The Lighthouse Lab in Alderley Park and Alex Alderton, Roberto Amato, Sonia Goncalves, Ewan Harrison, David K. Jackson, Ian Johnston, Dominic Kwiatkowski, Cordelia Langford, John Sillitoe on behalf of the Wellcome Sanger Institute COVID-19 Surveillance Team |
| EPI_ISL_556061, EPI_ISL_556062 | Lighthouse Lab in Alderley Park | Wellcome Sanger Institute for the COVID-19 Genomics UK (COG-UK) consortium | The Lighthouse Lab in Alderley Park and Alex Alderton, Roberto Amato, Sonia Goncalves, Ewan Harrison, David K. Jackson, Ian Johnston, Dominic Kwiatkowski, Cordelia Langford, John Sillitoe on behalf of the Wellcome Sanger Institute COVID-19 Surveillance Team (http://www.sanger.ac.uk/covid-team) |
| EPI_ISL_556064 | Lighthouse Lab in Alderley Park | Wellcome Sanger Institute for the COVID-19 Genomics UK (COG-UK) consortium | The Lighthouse Lab in Alderley Park and Alex Alderton, Roberto Amato, Sonia Goncalves, Ewan Harrison, David K. Jackson, Ian Johnston, Dominic Kwiatkowski, Cordelia Langford, John Sillitoe on behalf of the Wellcome Sanger Institute COVID-19 Surveillance Team |
| EPI_ISL_556065, EPI_ISL_556067, EPI_ISL_556068, EPI_ISL_556070, EPI_ISL_556071, EPI_ISL_556072, EPI_ISL_556073 | Lighthouse Lab in Alderley Park | Wellcome Sanger Institute for the COVID-19 Genomics UK (COG-UK) consortium | The Lighthouse Lab in Alderley Park and Alex Alderton, Roberto Amato, Sonia Goncalves, Ewan Harrison, David K. Jackson, Ian Johnston, Dominic Kwiatkowski, Cordelia Langford, John Sillitoe on behalf of the Wellcome Sanger Institute COVID-19 Surveillance Team (http://www.sanger.ac.uk/covid-team) |
| EPI_ISL_556075, EPI_ISL_556076, EPI_ISL_556077, EPI_ISL_556079 | Lighthouse Lab in Milton Keynes | Wellcome Sanger Institute for the COVID-19 Genomics UK (COG-UK) consortium | The Lighthouse Lab in Milton Keynes and Alex Alderton, Roberto Amato, Sonia Goncalves, Ewan Harrison, David K. Jackson, Ian Johnston, Dominic Kwiatkowski, Cordelia Langford, John Sillitoe on behalf of the Wellcome Sanger Institute COVID-19 Surveillance Team (http://www.sanger.ac.uk/covid-team) |
| EPI_ISL_556083 | Lighthouse Lab in Milton Keynes | Wellcome Sanger Institute for the COVID-19 Genomics UK (COG-UK) consortium | The Lighthouse Lab in Alderley Park and Alex Alderton, Roberto Amato, Sonia Goncalves, Ewan Harrison, David K. Jackson, Ian Johnston, Dominic Kwiatkowski, Cordelia Langford, John Sillitoe on behalf of the Wellcome Sanger Institute COVID-19 Surveillance Team |
| EPI_ISL_556085 | Lighthouse Lab in Alderley Park | Wellcome Sanger Institute for the COVID-19 Genomics UK (COG-UK) consortium | The Lighthouse Lab in Alderley Park and Alex Alderton, Roberto Amato, Sonia Goncalves, Ewan Harrison, David K. Jackson, Ian Johnston, Dominic Kwiatkowski, Cordelia Langford, John Sillitoe on behalf of the Wellcome Sanger Institute COVID-19 Surveillance Team |
| EPI_ISL_556087 | Lighthouse Lab in Milton Keynes | Wellcome Sanger Institute for the COVID-19 Genomics UK (COG-UK) consortium | The Lighthouse Lab in Milton Keynes and Alex Alderton, Roberto Amato, Sonia Goncalves, Ewan Harrison, David K. Jackson, Ian Johnston, Dominic Kwiatkowski, Cordelia Langford, John Sillitoe on behalf of the Wellcome Sanger Institute COVID-19 Surveillance Team (http://www.sanger.ac.uk/covid-team) |
| EPI_ISL_556088 | Lighthouse Lab in Alderley Park | Wellcome Sanger Institute for the COVID-19 Genomics UK (COG-UK) consortium | The Lighthouse Lab in Alderley Park and Alex Alderton, Roberto Amato, Sonia Goncalves, Ewan Harrison, David K. Jackson, Ian Johnston, Dominic Kwiatkowski, Cordelia Langford, John Sillitoe on behalf of the Wellcome Sanger Institute COVID-19 Surveillance Team (http://www.sanger.ac.uk/covid-team) |
| EPI_ISL_556090, EPI_ISL_556092, EPI_ISL_556093, EPI_ISL_556095, EPI_ISL_556097, EPI_ISL_556098, EPI_ISL_556102, EPI_ISL_556104, EPI_ISL_556109 | Lighthouse Lab in Milton Keynes | Wellcome Sanger Institute for the COVID-19 Genomics UK (COG-UK) consortium | The Lighthouse Lab in Milton Keynes and Alex Alderton, Roberto Amato, Sonia Goncalves, Ewan Harrison, David K. Jackson, Ian Johnston, Dominic Kwiatkowski, Cordelia Langford, John Sillitoe on behalf of the Wellcome Sanger Institute COVID-19 Surveillance Team (http://www.sanger.ac.uk/covid-team) |
| EPI_ISL_556110 | Lighthouse Lab in Alderley Park | Wellcome Sanger Institute for the COVID-19 Genomics UK (COG-UK) consortium | The Lighthouse Lab in Alderley Park and Alex Alderton, Roberto Amato, Sonia Goncalves, Ewan Harrison, David K. Jackson, Ian Johnston, Dominic Kwiatkowski, Cordelia Langford, John Sillitoe on behalf of the Wellcome Sanger Institute COVID-19 Surveillance Team (http://www.sanger.ac.uk/covid-team) |
| EPI_ISL_556111 | Lighthouse Lab in Alderley Park | Wellcome Sanger Institute for the COVID-19 Genomics UK (COG-UK) consortium | The Lighthouse Lab in Alderley Park and Alex Alderton, Roberto Amato, Sonia Goncalves, Ewan Harrison, David K. Jackson, Ian Johnston, Dominic Kwiatkowski, Cordelia Langford, John Sillitoe on behalf of the Wellcome Sanger Institute COVID-19 Surveillance Team |
| EPI_ISL_556112, EPI_ISL_556115, EPI_ISL_556116 | Lighthouse Lab in Milton Keynes | Wellcome Sanger Institute for the COVID-19 Genomics UK (COG-UK) consortium | The Lighthouse Lab in Milton Keynes and Alex Alderton, Roberto Amato, Sonia Goncalves, Ewan Harrison, David K. Jackson, Ian Johnston, Dominic Kwiatkowski, Cordelia Langford, John Sillitoe on behalf of the Wellcome Sanger Institute COVID-19 Surveillance Team (http://www.sanger.ac.uk/covid-team) |
| EPI_ISL_556118 | Lighthouse Lab in Alderley Park | Wellcome Sanger Institute for the COVID-19 Genomics UK (COG-UK) consortium | The Lighthouse Lab in Alderley Park and Alex Alderton, Roberto Amato, Sonia Goncalves, Ewan Harrison, David K. Jackson, Ian Johnston, Dominic Kwiatkowski, Cordelia Langford, John Sillitoe on behalf of the Wellcome Sanger Institute COVID-19 Surveillance Team |
| EPI_ISL_556123 | Lighthouse Lab in Milton Keynes | Wellcome Sanger Institute for the COVID-19 Genomics UK (COG-UK) consortium | The Lighthouse Lab in Milton Keynes and Alex Alderton, Roberto Amato, Sonia Goncalves, Ewan Harrison, David K. Jackson, Ian Johnston, Dominic Kwiatkowski, Cordelia Langford, John Sillitoe on behalf of the Wellcome Sanger Institute COVID-19 Surveillance Team (http://www.sanger.ac.uk/covid-team) |
| EPI_ISL_556124 | Lighthouse Lab in Alderley Park | Wellcome Sanger Institute for the COVID-19 Genomics UK (COG-UK) consortium | The Lighthouse Lab in Alderley Park and Alex Alderton, Roberto Amato, Sonia Goncalves, Ewan Harrison, David K. Jackson, Ian Johnston, Dominic Kwiatkowski, Cordelia Langford, John Sillitoe on behalf of the Wellcome Sanger Institute COVID-19 Surveillance Team (http://www.sanger.ac.uk/covid-team) |
| EPI_ISL_556126 | Lighthouse Lab in Milton Keynes | Wellcome Sanger Institute for the COVID-19 Genomics UK | The Lighthouse Lab in Milton Keynes and Alex Alderton, Roberto Amato, Sonia Goncalves, Ewan Harrison, David K. Jackson, Ian Johnston, Dominic |

| | | | |
|---|---|---|---|
| | | (COG-UK) consortium | Kwiatkowski, Cordelia Langford, John Sillitoe on behalf of the Wellcome Sanger Institute COVID-19 Surveillance Team (http://www.sanger.ac.uk/covid-team) |
| EPI_ISL_556130, EPI_ISL_556131, EPI_ISL_556136 | Lighthouse Lab in Alderley Park | Wellcome Sanger Institute for the COVID-19 Genomics UK (COG-UK) consortium | The Lighthouse Lab in Alderley Park and Alex Alderton, Roberto Amato, Sonia Goncalves, Ewan Harrison, David K. Jackson, Ian Johnston, Dominic Kwiatkowski, Cordelia Langford, John Sillitoe on behalf of the Wellcome Sanger Institute COVID-19 Surveillance Team |
| EPI_ISL_556137 | Lighthouse Lab in Alderley Park | Wellcome Sanger Institute for the COVID-19 Genomics UK (COG-UK) consortium | The Lighthouse Lab in Alderley Park and Alex Alderton, Roberto Amato, Sonia Goncalves, Ewan Harrison, David K. Jackson, Ian Johnston, Dominic Kwiatkowski, Cordelia Langford, John Sillitoe on behalf of the Wellcome Sanger Institute COVID-19 Surveillance Team (http://www.sanger.ac.uk/covid-team) |
| EPI_ISL_556138, EPI_ISL_556139 | Lighthouse Lab in Milton Keynes | Wellcome Sanger Institute for the COVID-19 Genomics UK (COG-UK) consortium | The Lighthouse Lab in Milton Keynes and Alex Alderton, Roberto Amato, Sonia Goncalves, Ewan Harrison, David K. Jackson, Ian Johnston, Dominic Kwiatkowski, Cordelia Langford, John Sillitoe on behalf of the Wellcome Sanger Institute COVID-19 Surveillance Team (http://www.sanger.ac.uk/covid-team) |
| EPI_ISL_556140, EPI_ISL_556144 | Lighthouse Lab in Alderley Park | Wellcome Sanger Institute for the COVID-19 Genomics UK (COG-UK) consortium | The Lighthouse Lab in Alderley Park and Alex Alderton, Roberto Amato, Sonia Goncalves, Ewan Harrison, David K. Jackson, Ian Johnston, Dominic Kwiatkowski, Cordelia Langford, John Sillitoe on behalf of the Wellcome Sanger Institute COVID-19 Surveillance Team (http://www.sanger.ac.uk/covid-team) |
| EPI_ISL_556146, EPI_ISL_556148, EPI_ISL_556150 | Lighthouse Lab in Milton Keynes | Wellcome Sanger Institute for the COVID-19 Genomics UK (COG-UK) consortium | The Lighthouse Lab in Milton Keynes and Alex Alderton, Roberto Amato, Sonia Goncalves, Ewan Harrison, David K. Jackson, Ian Johnston, Dominic Kwiatkowski, Cordelia Langford, John Sillitoe on behalf of the Wellcome Sanger Institute COVID-19 Surveillance Team (http://www.sanger.ac.uk/covid-team) |
| EPI_ISL_556151 | Lighthouse Lab in Alderley Park | Wellcome Sanger Institute for the COVID-19 Genomics UK (COG-UK) consortium | The Lighthouse Lab in Alderley Park and Alex Alderton, Roberto Amato, Sonia Goncalves, Ewan Harrison, David K. Jackson, Ian Johnston, Dominic Kwiatkowski, Cordelia Langford, John Sillitoe on behalf of the Wellcome Sanger Institute COVID-19 Surveillance Team |
| EPI_ISL_556152 | Lighthouse Lab in Alderley Park | Wellcome Sanger Institute for the COVID-19 Genomics UK (COG-UK) consortium | The Lighthouse Lab in Alderley Park and Alex Alderton, Roberto Amato, Sonia Goncalves, Ewan Harrison, David K. Jackson, Ian Johnston, Dominic Kwiatkowski, Cordelia Langford, John Sillitoe on behalf of the Wellcome Sanger Institute COVID-19 Surveillance Team (http://www.sanger.ac.uk/covid-team) |
| EPI_ISL_556154 | Lighthouse Lab in Alderley Park | Wellcome Sanger Institute for the COVID-19 Genomics UK (COG-UK) consortium | The Lighthouse Lab in Alderley Park and Alex Alderton, Roberto Amato, Sonia Goncalves, Ewan Harrison, David K. Jackson, Ian Johnston, Dominic Kwiatkowski, Cordelia Langford, John Sillitoe on behalf of the Wellcome Sanger Institute COVID-19 Surveillance Team |
| EPI_ISL_556158 | Lighthouse Lab in Milton Keynes | Wellcome Sanger Institute for the COVID-19 Genomics UK (COG-UK) consortium | The Lighthouse Lab in Milton Keynes and Alex Alderton, Roberto Amato, Sonia Goncalves, Ewan Harrison, David K. Jackson, Ian Johnston, Dominic Kwiatkowski, Cordelia Langford, John Sillitoe on behalf of the Wellcome Sanger Institute COVID-19 Surveillance Team (http://www.sanger.ac.uk/covid-team) |
| EPI_ISL_556159 | Lighthouse Lab in Alderley Park | Wellcome Sanger Institute for the COVID-19 Genomics UK (COG-UK) consortium | The Lighthouse Lab in Alderley Park and Alex Alderton, Roberto Amato, Sonia Goncalves, Ewan Harrison, David K. Jackson, Ian Johnston, Dominic Kwiatkowski, Cordelia Langford, John Sillitoe on behalf of the Wellcome Sanger Institute COVID-19 Surveillance Team (http://www.sanger.ac.uk/covid-team) |
| EPI_ISL_556160 | Lighthouse Lab in Alderley Park | Wellcome Sanger Institute for the COVID-19 Genomics UK (COG-UK) consortium | The Lighthouse Lab in Alderley Park and Alex Alderton, Roberto Amato, Sonia Goncalves, Ewan Harrison, David K. Jackson, Ian Johnston, Dominic Kwiatkowski, Cordelia Langford, John Sillitoe on behalf of the Wellcome Sanger Institute COVID-19 Surveillance Team |
| EPI_ISL_556163, EPI_ISL_556166, EPI_ISL_556170, EPI_ISL_556172 | Lighthouse Lab in Milton Keynes | Wellcome Sanger Institute for the COVID-19 Genomics UK (COG-UK) consortium | The Lighthouse Lab in Milton Keynes and Alex Alderton, Roberto Amato, Sonia Goncalves, Ewan Harrison, David K. Jackson, Ian Johnston, Dominic Kwiatkowski, Cordelia Langford, John Sillitoe on behalf of the Wellcome Sanger Institute COVID-19 Surveillance Team (http://www.sanger.ac.uk/covid-team) |
| EPI_ISL_556173 | Lighthouse Lab in Alderley Park | Wellcome Sanger Institute for the COVID-19 Genomics UK (COG-UK) consortium | The Lighthouse Lab in Alderley Park and Alex Alderton, Roberto Amato, Sonia Goncalves, Ewan Harrison, David K. Jackson, Ian Johnston, Dominic Kwiatkowski, Cordelia Langford, John Sillitoe on behalf of the Wellcome Sanger Institute COVID-19 Surveillance Team (http://www.sanger.ac.uk/covid-team) |
| EPI_ISL_556176 | Lighthouse Lab in Milton Keynes | Wellcome Sanger Institute for the COVID-19 Genomics UK (COG-UK) consortium | The Lighthouse Lab in Milton Keynes and Alex Alderton, Roberto Amato, Sonia Goncalves, Ewan Harrison, David K. Jackson, Ian Johnston, Dominic Kwiatkowski, Cordelia Langford, John Sillitoe on behalf of the Wellcome Sanger Institute COVID-19 Surveillance Team (http://www.sanger.ac.uk/covid-team) |
| EPI_ISL_556177 | Lighthouse Lab in Alderley Park | Wellcome Sanger Institute for the COVID-19 Genomics UK (COG-UK) consortium | The Lighthouse Lab in Alderley Park and Alex Alderton, Roberto Amato, Sonia Goncalves, Ewan Harrison, David K. Jackson, Ian Johnston, Dominic Kwiatkowski, Cordelia Langford, John Sillitoe on behalf of the Wellcome Sanger Institute COVID-19 Surveillance Team (http://www.sanger.ac.uk/covid-team) |
| EPI_ISL_556178 | Lighthouse Lab in Alderley Park | Wellcome Sanger Institute for the COVID-19 Genomics UK (COG-UK) consortium | The Lighthouse Lab in Alderley Park and Alex Alderton, Roberto Amato, Sonia Goncalves, Ewan Harrison, David K. Jackson, Ian Johnston, Dominic Kwiatkowski, Cordelia Langford, John Sillitoe on behalf of the Wellcome Sanger Institute COVID-19 Surveillance Team |
| EPI_ISL_556180, EPI_ISL_556182 | Lighthouse Lab in Alderley Park | Wellcome Sanger Institute for the COVID-19 Genomics UK (COG-UK) consortium | The Lighthouse Lab in Alderley Park and Alex Alderton, Roberto Amato, Sonia Goncalves, Ewan Harrison, David K. Jackson, Ian Johnston, Dominic Kwiatkowski, Cordelia Langford, John Sillitoe on behalf of the Wellcome Sanger Institute COVID-19 Surveillance Team (http://www.sanger.ac.uk/covid-team) |
| EPI_ISL_556183 | Lighthouse Lab in Milton Keynes | Wellcome Sanger Institute for the COVID-19 Genomics UK (COG-UK) consortium | The Lighthouse Lab in Milton Keynes and Alex Alderton, Roberto Amato, Sonia Goncalves, Ewan Harrison, David K. Jackson, Ian Johnston, Dominic Kwiatkowski, Cordelia Langford, John Sillitoe on behalf of the Wellcome Sanger Institute COVID-19 Surveillance Team (http://www.sanger.ac.uk/covid-team) |
| EPI_ISL_556184 | Lighthouse Lab in Alderley Park | Wellcome Sanger Institute for the COVID-19 Genomics UK (COG-UK) consortium | The Lighthouse Lab in Alderley Park and Alex Alderton, Roberto Amato, Sonia Goncalves, Ewan Harrison, David K. Jackson, Ian Johnston, Dominic Kwiatkowski, Cordelia Langford, John Sillitoe on behalf of the Wellcome Sanger Institute COVID-19 Surveillance Team |
| EPI_ISL_556187, EPI_ISL_556195 | Lighthouse Lab in Milton Keynes | Wellcome Sanger Institute for the COVID-19 Genomics UK (COG-UK) consortium | The Lighthouse Lab in Milton Keynes and Alex Alderton, Roberto Amato, Sonia Goncalves, Ewan Harrison, David K. Jackson, Ian Johnston, Dominic Kwiatkowski, Cordelia Langford, John Sillitoe on behalf of the Wellcome Sanger Institute COVID-19 Surveillance Team (http://www.sanger.ac.uk/covid-team) |
| EPI_ISL_556197, EPI_ISL_556198 | Lighthouse Lab in Milton Keynes | Wellcome Sanger Institute for the COVID-19 Genomics UK (COG-UK) consortium | The Lighthouse Lab in Alderley Park and Alex Alderton, Roberto Amato, Sonia Goncalves, Ewan Harrison, David K. Jackson, Ian Johnston, Dominic Kwiatkowski, Cordelia Langford, John Sillitoe on behalf of the Wellcome Sanger Institute COVID-19 Surveillance Team |
| EPI_ISL_556199, EPI_ISL_556201 | Lighthouse Lab in Milton Keynes | Wellcome Sanger Institute for the COVID-19 Genomics UK (COG-UK) consortium | The Lighthouse Lab in Milton Keynes and Alex Alderton, Roberto Amato, Sonia Goncalves, Ewan Harrison, David K. Jackson, Ian Johnston, Dominic Kwiatkowski, Cordelia Langford, John Sillitoe on behalf of the Wellcome Sanger Institute COVID-19 Surveillance Team (http://www.sanger.ac.uk/covid-team) |
| EPI_ISL_556203 | Lighthouse Lab in Alderley Park | Wellcome Sanger Institute for the COVID-19 Genomics UK (COG-UK) consortium | The Lighthouse Lab in Alderley Park and Alex Alderton, Roberto Amato, Sonia Goncalves, Ewan Harrison, David K. Jackson, Ian Johnston, Dominic Kwiatkowski, Cordelia Langford, John Sillitoe on behalf of the Wellcome Sanger Institute COVID-19 Surveillance Team |
| EPI_ISL_556204, EPI_ISL_556208, EPI_ISL_556211, EPI_ISL_556213, EPI_ISL_556214 | Lighthouse Lab in Milton Keynes | Wellcome Sanger Institute for the COVID-19 Genomics UK (COG-UK) consortium | The Lighthouse Lab in Milton Keynes and Alex Alderton, Roberto Amato, Sonia Goncalves, Ewan Harrison, David K. Jackson, Ian Johnston, Dominic Kwiatkowski, Cordelia Langford, John Sillitoe on behalf of the Wellcome Sanger Institute COVID-19 Surveillance Team (http://www.sanger.ac.uk/covid-team) |
| EPI_ISL_556221 | Lighthouse Lab in Alderley Park | Wellcome Sanger Institute for the COVID-19 Genomics UK (COG-UK) consortium | The Lighthouse Lab in Alderley Park and Alex Alderton, Roberto Amato, Sonia Goncalves, Ewan Harrison, David K. Jackson, Ian Johnston, Dominic Kwiatkowski, Cordelia Langford, John Sillitoe on behalf of the Wellcome Sanger Institute COVID-19 Surveillance Team (http://www.sanger.ac.uk/covid-team) |
| EPI_ISL_556223 | Lighthouse Lab in Milton Keynes | Wellcome Sanger Institute for the COVID-19 Genomics UK (COG-UK) consortium | The Lighthouse Lab in Milton Keynes and Alex Alderton, Roberto Amato, Sonia Goncalves, Ewan Harrison, David K. Jackson, Ian Johnston, Dominic Kwiatkowski, Cordelia Langford, John Sillitoe on behalf of the Wellcome Sanger Institute COVID-19 Surveillance Team (http://www.sanger.ac.uk/covid-team) |
| EPI_ISL_556227 | Lighthouse Lab in Alderley Park | Wellcome Sanger Institute for the COVID-19 Genomics UK (COG-UK) consortium | The Lighthouse Lab in Alderley Park and Alex Alderton, Roberto Amato, Sonia Goncalves, Ewan Harrison, David K. Jackson, Ian Johnston, Dominic Kwiatkowski, Cordelia Langford, John Sillitoe on behalf of the Wellcome Sanger Institute COVID-19 Surveillance Team |

| | | | |
|---|---|---|---|
| | | | (http://www.sanger.ac.uk/covid-team) |
| EPI_ISL_556231 | Lighthouse Lab in Alderley Park | Wellcome Sanger Institute for the COVID-19 Genomics UK (COG-UK) consortium | The Lighthouse Lab in Alderley Park and Alex Alderton, Roberto Amato, Sonia Goncalves, Ewan Harrison, David K. Jackson, Ian Johnston, Dominic Kwiatkowski, Cordelia Langford, John Sillitoe on behalf of the Wellcome Sanger Institute COVID-19 Surveillance Team |
| EPI_ISL_556232 | Lighthouse Lab in Milton Keynes | Wellcome Sanger Institute for the COVID-19 Genomics UK (COG-UK) consortium | The Lighthouse Lab in Milton Keynes and Alex Alderton, Roberto Amato, Sonia Goncalves, Ewan Harrison, David K. Jackson, Ian Johnston, Dominic Kwiatkowski, Cordelia Langford, John Sillitoe on behalf of the Wellcome Sanger Institute COVID-19 Surveillance Team (http://www.sanger.ac.uk/covid-team) |
| EPI_ISL_556233 | Lighthouse Lab in Milton Keynes | Wellcome Sanger Institute for the COVID-19 Genomics UK (COG-UK) consortium | The Lighthouse Lab in Milton Keynes and Alex Alderton, Roberto Amato, Sonia Goncalves, Ewan Harrison, David K. Jackson, Ian Johnston, Dominic Kwiatkowski, Cordelia Langford, John Sillitoe on behalf of the Wellcome Sanger Institute COVID-19 Surveillance Team |
| EPI_ISL_556234 | Lighthouse Lab in Alderley Park | Wellcome Sanger Institute for the COVID-19 Genomics UK (COG-UK) consortium | The Lighthouse Lab in Alderley Park and Alex Alderton, Roberto Amato, Sonia Goncalves, Ewan Harrison, David K. Jackson, Ian Johnston, Dominic Kwiatkowski, Cordelia Langford, John Sillitoe on behalf of the Wellcome Sanger Institute COVID-19 Surveillance Team |
| EPI_ISL_556235 | Lighthouse Lab in Milton Keynes | Wellcome Sanger Institute for the COVID-19 Genomics UK (COG-UK) consortium | The Lighthouse Lab in Alderley Park and Alex Alderton, Roberto Amato, Sonia Goncalves, Ewan Harrison, David K. Jackson, Ian Johnston, Dominic Kwiatkowski, Cordelia Langford, John Sillitoe on behalf of the Wellcome Sanger Institute COVID-19 Surveillance Team |
| EPI_ISL_556237, EPI_ISL_556244, EPI_ISL_556248, EPI_ISL_556251, EPI_ISL_556253 | Lighthouse Lab in Milton Keynes | Wellcome Sanger Institute for the COVID-19 Genomics UK (COG-UK) consortium | The Lighthouse Lab in Milton Keynes and Alex Alderton, Roberto Amato, Sonia Goncalves, Ewan Harrison, David K. Jackson, Ian Johnston, Dominic Kwiatkowski, Cordelia Langford, John Sillitoe on behalf of the Wellcome Sanger Institute COVID-19 Surveillance Team (http://www.sanger.ac.uk/covid-team) |
| EPI_ISL_556256 | Lighthouse Lab in Alderley Park | Wellcome Sanger Institute for the COVID-19 Genomics UK (COG-UK) consortium | The Lighthouse Lab in Alderley Park and Alex Alderton, Roberto Amato, Sonia Goncalves, Ewan Harrison, David K. Jackson, Ian Johnston, Dominic Kwiatkowski, Cordelia Langford, John Sillitoe on behalf of the Wellcome Sanger Institute COVID-19 Surveillance Team |
| EPI_ISL_556259, EPI_ISL_556261 | Lighthouse Lab in Milton Keynes | Wellcome Sanger Institute for the COVID-19 Genomics UK (COG-UK) consortium | The Lighthouse Lab in Milton Keynes and Alex Alderton, Roberto Amato, Sonia Goncalves, Ewan Harrison, David K. Jackson, Ian Johnston, Dominic Kwiatkowski, Cordelia Langford, John Sillitoe on behalf of the Wellcome Sanger Institute COVID-19 Surveillance Team (http://www.sanger.ac.uk/covid-team) |
| EPI_ISL_556262 | Lighthouse Lab in Alderley Park | Wellcome Sanger Institute for the COVID-19 Genomics UK (COG-UK) consortium | The Lighthouse Lab in Alderley Park and Alex Alderton, Roberto Amato, Sonia Goncalves, Ewan Harrison, David K. Jackson, Ian Johnston, Dominic Kwiatkowski, Cordelia Langford, John Sillitoe on behalf of the Wellcome Sanger Institute COVID-19 Surveillance Team |
| EPI_ISL_556263, EPI_ISL_556267, EPI_ISL_556269 | Lighthouse Lab in Milton Keynes | Wellcome Sanger Institute for the COVID-19 Genomics UK (COG-UK) consortium | The Lighthouse Lab in Milton Keynes and Alex Alderton, Roberto Amato, Sonia Goncalves, Ewan Harrison, David K. Jackson, Ian Johnston, Dominic Kwiatkowski, Cordelia Langford, John Sillitoe on behalf of the Wellcome Sanger Institute COVID-19 Surveillance Team (http://www.sanger.ac.uk/covid-team) |
| EPI_ISL_556270, EPI_ISL_556272 | Lighthouse Lab in Alderley Park | Wellcome Sanger Institute for the COVID-19 Genomics UK (COG-UK) consortium | The Lighthouse Lab in Alderley Park and Alex Alderton, Roberto Amato, Sonia Goncalves, Ewan Harrison, David K. Jackson, Ian Johnston, Dominic Kwiatkowski, Cordelia Langford, John Sillitoe on behalf of the Wellcome Sanger Institute COVID-19 Surveillance Team (http://www.sanger.ac.uk/covid-team) |
| EPI_ISL_556275, EPI_ISL_556276 | Lighthouse Lab in Milton Keynes | Wellcome Sanger Institute for the COVID-19 Genomics UK (COG-UK) consortium | The Lighthouse Lab in Milton Keynes and Alex Alderton, Roberto Amato, Sonia Goncalves, Ewan Harrison, David K. Jackson, Ian Johnston, Dominic Kwiatkowski, Cordelia Langford, John Sillitoe on behalf of the Wellcome Sanger Institute COVID-19 Surveillance Team (http://www.sanger.ac.uk/covid-team) |
| EPI_ISL_556277 | Lighthouse Lab in Alderley Park | Wellcome Sanger Institute for the COVID-19 Genomics UK (COG-UK) consortium | The Lighthouse Lab in Alderley Park and Alex Alderton, Roberto Amato, Sonia Goncalves, Ewan Harrison, David K. Jackson, Ian Johnston, Dominic Kwiatkowski, Cordelia Langford, John Sillitoe on behalf of the Wellcome Sanger Institute COVID-19 Surveillance Team (http://www.sanger.ac.uk/covid-team) |
| EPI_ISL_556280, EPI_ISL_556283 | Lighthouse Lab in Alderley Park | Wellcome Sanger Institute for the COVID-19 Genomics UK (COG-UK) consortium | The Lighthouse Lab in Alderley Park and Alex Alderton, Roberto Amato, Sonia Goncalves, Ewan Harrison, David K. Jackson, Ian Johnston, Dominic Kwiatkowski, Cordelia Langford, John Sillitoe on behalf of the Wellcome Sanger Institute COVID-19 Surveillance Team |
| EPI_ISL_556285, EPI_ISL_556289, EPI_ISL_556293, EPI_ISL_556294 | Lighthouse Lab in Milton Keynes | Wellcome Sanger Institute for the COVID-19 Genomics UK (COG-UK) consortium | The Lighthouse Lab in Milton Keynes and Alex Alderton, Roberto Amato, Sonia Goncalves, Ewan Harrison, David K. Jackson, Ian Johnston, Dominic Kwiatkowski, Cordelia Langford, John Sillitoe on behalf of the Wellcome Sanger Institute COVID-19 Surveillance Team (http://www.sanger.ac.uk/covid-team) |
| EPI_ISL_556295, EPI_ISL_556296 | Lighthouse Lab in Alderley Park | Wellcome Sanger Institute for the COVID-19 Genomics UK (COG-UK) consortium | The Lighthouse Lab in Alderley Park and Alex Alderton, Roberto Amato, Sonia Goncalves, Ewan Harrison, David K. Jackson, Ian Johnston, Dominic Kwiatkowski, Cordelia Langford, John Sillitoe on behalf of the Wellcome Sanger Institute COVID-19 Surveillance Team (http://www.sanger.ac.uk/covid-team) |
| EPI_ISL_556297 | Lighthouse Lab in Alderley Park | Wellcome Sanger Institute for the COVID-19 Genomics UK (COG-UK) consortium | The Lighthouse Lab in Alderley Park and Alex Alderton, Roberto Amato, Sonia Goncalves, Ewan Harrison, David K. Jackson, Ian Johnston, Dominic Kwiatkowski, Cordelia Langford, John Sillitoe on behalf of the Wellcome Sanger Institute COVID-19 Surveillance Team |
| EPI_ISL_556298, EPI_ISL_556300 | Lighthouse Lab in Milton Keynes | Wellcome Sanger Institute for the COVID-19 Genomics UK (COG-UK) consortium | The Lighthouse Lab in Milton Keynes and Alex Alderton, Roberto Amato, Sonia Goncalves, Ewan Harrison, David K. Jackson, Ian Johnston, Dominic Kwiatkowski, Cordelia Langford, John Sillitoe on behalf of the Wellcome Sanger Institute COVID-19 Surveillance Team (http://www.sanger.ac.uk/covid-team) |
| EPI_ISL_556304 | Lighthouse Lab in Milton Keynes | Wellcome Sanger Institute for the COVID-19 Genomics UK (COG-UK) consortium | The Lighthouse Lab in Alderley Park and Alex Alderton, Roberto Amato, Sonia Goncalves, Ewan Harrison, David K. Jackson, Ian Johnston, Dominic Kwiatkowski, Cordelia Langford, John Sillitoe on behalf of the Wellcome Sanger Institute COVID-19 Surveillance Team |
| EPI_ISL_556306, EPI_ISL_556307, EPI_ISL_556308, EPI_ISL_556310, EPI_ISL_556311, EPI_ISL_556312, EPI_ISL_556313, EPI_ISL_556314, EPI_ISL_556315, EPI_ISL_556316, EPI_ISL_556319, EPI_ISL_556320, EPI_ISL_556322, EPI_ISL_556323, EPI_ISL_556325, EPI_ISL_556326, EPI_ISL_556327, EPI_ISL_556328, EPI_ISL_556329, EPI_ISL_556330, EPI_ISL_556331, EPI_ISL_556332, EPI_ISL_556333, EPI_ISL_556334, EPI_ISL_556336, EPI_ISL_556337, EPI_ISL_556340, EPI_ISL_556341, EPI_ISL_556342, EPI_ISL_556343, EPI_ISL_556345, EPI_ISL_556346, EPI_ISL_556347, EPI_ISL_556353, EPI_ISL_556354, EPI_ISL_556355, EPI_ISL_556356, EPI_ISL_556358, EPI_ISL_556359, EPI_ISL_556360, EPI_ISL_556361, EPI_ISL_556362, EPI_ISL_556363, EPI_ISL_556364, EPI_ISL_556365, EPI_ISL_556366, EPI_ISL_556367, EPI_ISL_556368, EPI_ISL_556369, EPI_ISL_556370, EPI_ISL_556371, EPI_ISL_556372, EPI_ISL_556373, EPI_ISL_556374, EPI_ISL_556375, EPI_ISL_556376, EPI_ISL_556377, EPI_ISL_556378, EPI_ISL_556379, EPI_ISL_556380, EPI_ISL_556381, EPI_ISL_556382, EPI_ISL_556383, EPI_ISL_556384, EPI_ISL_556385, EPI_ISL_556386, EPI_ISL_556387, EPI_ISL_556388, EPI_ISL_556389, EPI_ISL_556390, EPI_ISL_556391, EPI_ISL_556392, EPI_ISL_556393, EPI_ISL_556395, EPI_ISL_556396, EPI_ISL_556397, EPI_ISL_556398, EPI_ISL_556399, EPI_ISL_556400 | | | |
| see above | Lighthouse Lab in Alderley Park | Wellcome Sanger Institute for the COVID-19 Genomics UK (COG-UK) consortium | The Lighthouse Lab in Alderley Park and Alex Alderton, Roberto Amato, Sonia Goncalves, Ewan Harrison, David K. Jackson, Ian Johnston, Dominic Kwiatkowski, Cordelia Langford, John Sillitoe on behalf of the Wellcome Sanger Institute COVID-19 Surveillance Team (http://www.sanger.ac.uk/covid-team) |
| EPI_ISL_556401 | Lighthouse Lab in Alderley Park | Wellcome Sanger Institute for the COVID-19 Genomics UK (COG-UK) consortium | The Lighthouse Lab in Alderley Park and Alex Alderton, Roberto Amato, Sonia Goncalves, Ewan Harrison, David K. Jackson, Ian Johnston, Dominic Kwiatkowski, Cordelia Langford, John Sillitoe on behalf of the Wellcome Sanger Institute COVID-19 Surveillance Team |
| EPI_ISL_556402 | Lighthouse Lab in Alderley Park | Wellcome Sanger Institute for the COVID-19 Genomics UK (COG-UK) consortium | The Lighthouse Lab in Alderley Park and Alex Alderton, Roberto Amato, Sonia Goncalves, Ewan Harrison, David K. Jackson, Ian Johnston, Dominic Kwiatkowski, Cordelia Langford, John Sillitoe on behalf of the Wellcome Sanger Institute COVID-19 Surveillance Team (http://www.sanger.ac.uk/covid-team) |
| EPI_ISL_556403, EPI_ISL_556404, EPI_ISL_556405, EPI_ISL_556406, EPI_ISL_556407, EPI_ISL_556408, EPI_ISL_556409, EPI_ISL_556410, EPI_ISL_556412, EPI_ISL_556413, EPI_ISL_556415 | | | |
| see above | Lighthouse Lab in Alderley Park | Wellcome Sanger Institute for the COVID-19 Genomics UK (COG-UK) consortium | The Lighthouse Lab in Alderley Park and Alex Alderton, Roberto Amato, Sonia Goncalves, Ewan Harrison, David K. Jackson, Ian Johnston, Dominic Kwiatkowski, Cordelia Langford, John Sillitoe on behalf of the Wellcome Sanger Institute COVID-19 Surveillance Team |
| EPI_ISL_556416 | Lighthouse Lab in Alderley Park | Wellcome Sanger Institute for the COVID-19 Genomics UK (COG-UK) consortium | The Lighthouse Lab in Alderley Park and Alex Alderton, Roberto Amato, Sonia Goncalves, Ewan Harrison, David K. Jackson, Ian Johnston, Dominic Kwiatkowski, Cordelia Langford, John Sillitoe on behalf of the Wellcome Sanger Institute COVID-19 Surveillance Team (http://www.sanger.ac.uk/covid-team) |
| EPI_ISL_556417, EPI_ISL_556418, EPI_ISL_556419 | Lighthouse Lab in Alderley Park | Wellcome Sanger Institute for the COVID-19 Genomics UK (COG-UK) consortium | The Lighthouse Lab in Alderley Park and Alex Alderton, Roberto Amato, Sonia Goncalves, Ewan Harrison, David K. Jackson, Ian Johnston, Dominic Kwiatkowski, Cordelia Langford, John Sillitoe on behalf of the Wellcome Sanger Institute COVID-19 Surveillance Team |
| EPI_ISL_556420 | Lighthouse Lab in Alderley Park | Wellcome Sanger Institute for the COVID-19 Genomics UK (COG-UK) consortium | The Lighthouse Lab in Alderley Park and Alex Alderton, Roberto Amato, Sonia Goncalves, Ewan Harrison, David K. Jackson, Ian Johnston, Dominic Kwiatkowski, Cordelia Langford, John Sillitoe on behalf of the Wellcome Sanger Institute COVID-19 Surveillance Team (http://www.sanger.ac.uk/covid-team) |

| | | | |
|---|---|---|---|
| EPI_ISL_556421 | Lighthouse Lab in Alderley Park | Wellcome Sanger Institute for the COVID-19 Genomics UK (COG-UK) consortium | The Lighthouse Lab in Alderley Park and Alex Alderton, Roberto Amato, Sonia Goncalves, Ewan Harrison, David K. Jackson, Ian Johnston, Dominic Kwiatkowski, Cordelia Langford, John Sillitoe on behalf of the Wellcome Sanger Institute COVID-19 Surveillance Team |
| EPI_ISL_556422 | Lighthouse Lab in Alderley Park | Wellcome Sanger Institute for the COVID-19 Genomics UK (COG-UK) consortium | The Lighthouse Lab in Alderley Park and Alex Alderton, Roberto Amato, Sonia Goncalves, Ewan Harrison, David K. Jackson, Ian Johnston, Dominic Kwiatkowski, Cordelia Langford, John Sillitoe on behalf of the Wellcome Sanger Institute COVID-19 Surveillance Team (http://www.sanger.ac.uk/covid-team) |
| EPI_ISL_556423, EPI_ISL_556424, EPI_ISL_556426, EPI_ISL_556427, EPI_ISL_556429, EPI_ISL_556430, EPI_ISL_556431, EPI_ISL_556432, EPI_ISL_556436, EPI_ISL_556437, EPI_ISL_556438, EPI_ISL_556439, EPI_ISL_556440 | | | |
| see above | Lighthouse Lab in Alderley Park | Wellcome Sanger Institute for the COVID-19 Genomics UK (COG-UK) consortium | The Lighthouse Lab in Alderley Park and Alex Alderton, Roberto Amato, Sonia Goncalves, Ewan Harrison, David K. Jackson, Ian Johnston, Dominic Kwiatkowski, Cordelia Langford, John Sillitoe on behalf of the Wellcome Sanger Institute COVID-19 Surveillance Team |
| EPI_ISL_556441, EPI_ISL_556442, EPI_ISL_556443 | Lighthouse Lab in Alderley Park | Wellcome Sanger Institute for the COVID-19 Genomics UK (COG-UK) consortium | The Lighthouse Lab in Alderley Park and Alex Alderton, Roberto Amato, Sonia Goncalves, Ewan Harrison, David K. Jackson, Ian Johnston, Dominic Kwiatkowski, Cordelia Langford, John Sillitoe on behalf of the Wellcome Sanger Institute COVID-19 Surveillance Team (http://www.sanger.ac.uk/covid-team) |
| EPI_ISL_556444, EPI_ISL_556445, EPI_ISL_556447, EPI_ISL_556448, EPI_ISL_556449, EPI_ISL_556450 | Lighthouse Lab in Alderley Park | Wellcome Sanger Institute for the COVID-19 Genomics UK (COG-UK) consortium | The Lighthouse Lab in Alderley Park and Alex Alderton, Roberto Amato, Sonia Goncalves, Ewan Harrison, David K. Jackson, Ian Johnston, Dominic Kwiatkowski, Cordelia Langford, John Sillitoe on behalf of the Wellcome Sanger Institute COVID-19 Surveillance Team |
| EPI_ISL_556451 | Lighthouse Lab in Alderley Park | Wellcome Sanger Institute for the COVID-19 Genomics UK (COG-UK) consortium | The Lighthouse Lab in Alderley Park and Alex Alderton, Roberto Amato, Sonia Goncalves, Ewan Harrison, David K. Jackson, Ian Johnston, Dominic Kwiatkowski, Cordelia Langford, John Sillitoe on behalf of the Wellcome Sanger Institute COVID-19 Surveillance Team (http://www.sanger.ac.uk/covid-team) |
| EPI_ISL_556452, EPI_ISL_556453, EPI_ISL_556454 | Lighthouse Lab in Alderley Park | Wellcome Sanger Institute for the COVID-19 Genomics UK (COG-UK) consortium | The Lighthouse Lab in Alderley Park and Alex Alderton, Roberto Amato, Sonia Goncalves, Ewan Harrison, David K. Jackson, Ian Johnston, Dominic Kwiatkowski, Cordelia Langford, John Sillitoe on behalf of the Wellcome Sanger Institute COVID-19 Surveillance Team |
| EPI_ISL_556455 | Lighthouse Lab in Alderley Park | Wellcome Sanger Institute for the COVID-19 Genomics UK (COG-UK) consortium | The Lighthouse Lab in Alderley Park and Alex Alderton, Roberto Amato, Sonia Goncalves, Ewan Harrison, David K. Jackson, Ian Johnston, Dominic Kwiatkowski, Cordelia Langford, John Sillitoe on behalf of the Wellcome Sanger Institute COVID-19 Surveillance Team (http://www.sanger.ac.uk/covid-team) |
| EPI_ISL_556456, EPI_ISL_556457, EPI_ISL_556458, EPI_ISL_556460, EPI_ISL_556461, EPI_ISL_556462 | Lighthouse Lab in Alderley Park | Wellcome Sanger Institute for the COVID-19 Genomics UK (COG-UK) consortium | The Lighthouse Lab in Alderley Park and Alex Alderton, Roberto Amato, Sonia Goncalves, Ewan Harrison, David K. Jackson, Ian Johnston, Dominic Kwiatkowski, Cordelia Langford, John Sillitoe on behalf of the Wellcome Sanger Institute COVID-19 Surveillance Team |
| EPI_ISL_556463, EPI_ISL_556464 | Lighthouse Lab in Alderley Park | Wellcome Sanger Institute for the COVID-19 Genomics UK (COG-UK) consortium | The Lighthouse Lab in Alderley Park and Alex Alderton, Roberto Amato, Sonia Goncalves, Ewan Harrison, David K. Jackson, Ian Johnston, Dominic Kwiatkowski, Cordelia Langford, John Sillitoe on behalf of the Wellcome Sanger Institute COVID-19 Surveillance Team (http://www.sanger.ac.uk/covid-team) |
| EPI_ISL_556465, EPI_ISL_556466 | Lighthouse Lab in Alderley Park | Wellcome Sanger Institute for the COVID-19 Genomics UK (COG-UK) consortium | The Lighthouse Lab in Alderley Park and Alex Alderton, Roberto Amato, Sonia Goncalves, Ewan Harrison, David K. Jackson, Ian Johnston, Dominic Kwiatkowski, Cordelia Langford, John Sillitoe on behalf of the Wellcome Sanger Institute COVID-19 Surveillance Team |
| EPI_ISL_556467 | Lighthouse Lab in Alderley Park | Wellcome Sanger Institute for the COVID-19 Genomics UK (COG-UK) consortium | The Lighthouse Lab in Alderley Park and Alex Alderton, Roberto Amato, Sonia Goncalves, Ewan Harrison, David K. Jackson, Ian Johnston, Dominic Kwiatkowski, Cordelia Langford, John Sillitoe on behalf of the Wellcome Sanger Institute COVID-19 Surveillance Team (http://www.sanger.ac.uk/covid-team) |
| EPI_ISL_556468, EPI_ISL_556469, EPI_ISL_556470, EPI_ISL_556472 | Lighthouse Lab in Alderley Park | Wellcome Sanger Institute for the COVID-19 Genomics UK (COG-UK) consortium | The Lighthouse Lab in Alderley Park and Alex Alderton, Roberto Amato, Sonia Goncalves, Ewan Harrison, David K. Jackson, Ian Johnston, Dominic Kwiatkowski, Cordelia Langford, John Sillitoe on behalf of the Wellcome Sanger Institute COVID-19 Surveillance Team |
| EPI_ISL_556473, EPI_ISL_556474 | Lighthouse Lab in Alderley Park | Wellcome Sanger Institute for the COVID-19 Genomics UK (COG-UK) consortium | The Lighthouse Lab in Alderley Park and Alex Alderton, Roberto Amato, Sonia Goncalves, Ewan Harrison, David K. Jackson, Ian Johnston, Dominic Kwiatkowski, Cordelia Langford, John Sillitoe on behalf of the Wellcome Sanger Institute COVID-19 Surveillance Team (http://www.sanger.ac.uk/covid-team) |
| EPI_ISL_556475 | Lighthouse Lab in Alderley Park | Wellcome Sanger Institute for the COVID-19 Genomics UK (COG-UK) consortium | The Lighthouse Lab in Alderley Park and Alex Alderton, Roberto Amato, Sonia Goncalves, Ewan Harrison, David K. Jackson, Ian Johnston, Dominic Kwiatkowski, Cordelia Langford, John Sillitoe on behalf of the Wellcome Sanger Institute COVID-19 Surveillance Team |
| EPI_ISL_556476 | Lighthouse Lab in Alderley Park | Wellcome Sanger Institute for the COVID-19 Genomics UK (COG-UK) consortium | The Lighthouse Lab in Alderley Park and Alex Alderton, Roberto Amato, Sonia Goncalves, Ewan Harrison, David K. Jackson, Ian Johnston, Dominic Kwiatkowski, Cordelia Langford, John Sillitoe on behalf of the Wellcome Sanger Institute COVID-19 Surveillance Team (http://www.sanger.ac.uk/covid-team) |
| EPI_ISL_556478 | Lighthouse Lab in Alderley Park | Wellcome Sanger Institute for the COVID-19 Genomics UK (COG-UK) consortium | The Lighthouse Lab in Alderley Park and Alex Alderton, Roberto Amato, Sonia Goncalves, Ewan Harrison, David K. Jackson, Ian Johnston, Dominic Kwiatkowski, Cordelia Langford, John Sillitoe on behalf of the Wellcome Sanger Institute COVID-19 Surveillance Team |
| EPI_ISL_556479 | Lighthouse Lab in Alderley Park | Wellcome Sanger Institute for the COVID-19 Genomics UK (COG-UK) consortium | The Lighthouse Lab in Alderley Park and Alex Alderton, Roberto Amato, Sonia Goncalves, Ewan Harrison, David K. Jackson, Ian Johnston, Dominic Kwiatkowski, Cordelia Langford, John Sillitoe on behalf of the Wellcome Sanger Institute COVID-19 Surveillance Team (http://www.sanger.ac.uk/covid-team) |
| EPI_ISL_556480, EPI_ISL_556481, EPI_ISL_556483, EPI_ISL_556484 | Lighthouse Lab in Alderley Park | Wellcome Sanger Institute for the COVID-19 Genomics UK (COG-UK) consortium | The Lighthouse Lab in Alderley Park and Alex Alderton, Roberto Amato, Sonia Goncalves, Ewan Harrison, David K. Jackson, Ian Johnston, Dominic Kwiatkowski, Cordelia Langford, John Sillitoe on behalf of the Wellcome Sanger Institute COVID-19 Surveillance Team |
| EPI_ISL_556485 | Lighthouse Lab in Alderley Park | Wellcome Sanger Institute for the COVID-19 Genomics UK (COG-UK) consortium | The Lighthouse Lab in Alderley Park and Alex Alderton, Roberto Amato, Sonia Goncalves, Ewan Harrison, David K. Jackson, Ian Johnston, Dominic Kwiatkowski, Cordelia Langford, John Sillitoe on behalf of the Wellcome Sanger Institute COVID-19 Surveillance Team (http://www.sanger.ac.uk/covid-team) |
| EPI_ISL_556486, EPI_ISL_556489, EPI_ISL_556490, EPI_ISL_556491 | Lighthouse Lab in Alderley Park | Wellcome Sanger Institute for the COVID-19 Genomics UK (COG-UK) consortium | The Lighthouse Lab in Alderley Park and Alex Alderton, Roberto Amato, Sonia Goncalves, Ewan Harrison, David K. Jackson, Ian Johnston, Dominic Kwiatkowski, Cordelia Langford, John Sillitoe on behalf of the Wellcome Sanger Institute COVID-19 Surveillance Team |
| EPI_ISL_556493, EPI_ISL_556495 | Lighthouse Lab in Alderley Park | Wellcome Sanger Institute for the COVID-19 Genomics UK (COG-UK) consortium | The Lighthouse Lab in Alderley Park and Alex Alderton, Roberto Amato, Sonia Goncalves, Ewan Harrison, David K. Jackson, Ian Johnston, Dominic Kwiatkowski, Cordelia Langford, John Sillitoe on behalf of the Wellcome Sanger Institute COVID-19 Surveillance Team (http://www.sanger.ac.uk/covid-team) |
| EPI_ISL_556496, EPI_ISL_556497 | Lighthouse Lab in Alderley Park | Wellcome Sanger Institute for the COVID-19 Genomics UK (COG-UK) consortium | The Lighthouse Lab in Alderley Park and Alex Alderton, Roberto Amato, Sonia Goncalves, Ewan Harrison, David K. Jackson, Ian Johnston, Dominic Kwiatkowski, Cordelia Langford, John Sillitoe on behalf of the Wellcome Sanger Institute COVID-19 Surveillance Team |
| EPI_ISL_556498, EPI_ISL_556499 | Lighthouse Lab in Alderley Park | Wellcome Sanger Institute for the COVID-19 Genomics UK (COG-UK) consortium | The Lighthouse Lab in Alderley Park and Alex Alderton, Roberto Amato, Sonia Goncalves, Ewan Harrison, David K. Jackson, Ian Johnston, Dominic Kwiatkowski, Cordelia Langford, John Sillitoe on behalf of the Wellcome Sanger Institute COVID-19 Surveillance Team (http://www.sanger.ac.uk/covid-team) |
| EPI_ISL_556501, EPI_ISL_556502, EPI_ISL_556506, EPI_ISL_556507, EPI_ISL_556511, EPI_ISL_556512, EPI_ISL_556513 | Lighthouse Lab in Alderley Park | Wellcome Sanger Institute for the COVID-19 Genomics UK (COG-UK) consortium | The Lighthouse Lab in Alderley Park and Alex Alderton, Roberto Amato, Sonia Goncalves, Ewan Harrison, David K. Jackson, Ian Johnston, Dominic Kwiatkowski, Cordelia Langford, John Sillitoe on behalf of the Wellcome Sanger Institute COVID-19 Surveillance Team |
| EPI_ISL_556514 | Lighthouse Lab in Alderley Park | Wellcome Sanger Institute for the COVID-19 Genomics UK (COG-UK) consortium | The Lighthouse Lab in Alderley Park and Alex Alderton, Roberto Amato, Sonia Goncalves, Ewan Harrison, David K. Jackson, Ian Johnston, Dominic Kwiatkowski, Cordelia Langford, John Sillitoe on behalf of the Wellcome Sanger Institute COVID-19 Surveillance Team (http://www.sanger.ac.uk/covid-team) |
| EPI_ISL_556515 | Lighthouse Lab in Alderley Park | Wellcome Sanger Institute for the COVID-19 Genomics UK (COG-UK) consortium | The Lighthouse Lab in Alderley Park and Alex Alderton, Roberto Amato, Sonia Goncalves, Ewan Harrison, David K. Jackson, Ian Johnston, Dominic Kwiatkowski, Cordelia Langford, John Sillitoe on behalf of the Wellcome Sanger Institute COVID-19 Surveillance Team |
| EPI_ISL_556576, EPI_ISL_556579, EPI_ISL_556580, EPI_ISL_556581, EPI_ISL_556582, EPI_ISL_556584, EPI_ISL_556590, EPI_ISL_556591, EPI_ISL_556593, EPI_ISL_556594, EPI_ISL_556595, EPI_ISL_556597, EPI_ISL_556599, EPI_ISL_556600, EPI_ISL_556602, EPI_ISL_556603, EPI_ISL_556604, EPI_ISL_556605, | | | |

| | | | |
|---|---|---|---|
| EPI_ISL_556606, EPI_ISL_556607, EPI_ISL_556608, EPI_ISL_556609, EPI_ISL_556610, EPI_ISL_556611, EPI_ISL_556612, EPI_ISL_556613, EPI_ISL_556614, EPI_ISL_556615, EPI_ISL_556616, EPI_ISL_556617, EPI_ISL_556618, EPI_ISL_556621, EPI_ISL_556624, EPI_ISL_556626, EPI_ISL_556628, EPI_ISL_556630, EPI_ISL_556632, EPI_ISL_556633, EPI_ISL_556634, EPI_ISL_556636, EPI_ISL_556637, EPI_ISL_556638, EPI_ISL_556986, EPI_ISL_556988, EPI_ISL_556993, EPI_ISL_556997, EPI_ISL_556999, EPI_ISL_557001, EPI_ISL_557011, EPI_ISL_557016, EPI_ISL_557028, EPI_ISL_557030, EPI_ISL_557031, EPI_ISL_557032, EPI_ISL_557035, EPI_ISL_557039, EPI_ISL_557049, EPI_ISL_557055, EPI_ISL_557056, EPI_ISL_557061, EPI_ISL_557062, EPI_ISL_557063, EPI_ISL_557065, EPI_ISL_557069, EPI_ISL_557071, EPI_ISL_557079, EPI_ISL_557081, EPI_ISL_557082, EPI_ISL_557083, EPI_ISL_557086, EPI_ISL_557087, EPI_ISL_557089, EPI_ISL_557093, EPI_ISL_557094, EPI_ISL_557102, EPI_ISL_557103, EPI_ISL_557104, EPI_ISL_557107, EPI_ISL_557109, EPI_ISL_557112, EPI_ISL_557116, EPI_ISL_557117, EPI_ISL_557118, EPI_ISL_557123, EPI_ISL_557126, EPI_ISL_557129, EPI_ISL_557130, EPI_ISL_557131, EPI_ISL_557135, EPI_ISL_557141, EPI_ISL_557149, EPI_ISL_557154, EPI_ISL_557156, EPI_ISL_557159, EPI_ISL_557161, EPI_ISL_557164 | | | |
| see above | Lighthouse Lab in Alderley Park | Wellcome Sanger Institute for the COVID-19 Genomics UK (COG-UK) consortium | The Lighthouse Lab in Alderley Park and Alex Alderton, Roberto Amato, Sonia Goncalves, Ewan Harrison, David K. Jackson, Ian Johnston, Dominic Kwiatkowski, Cordelia Langford, John Sillitoe on behalf of the Wellcome Sanger Institute COVID-19 Surveillance Team (http://www.sanger.ac.uk/covid-team) |
| EPI_ISL_557167, EPI_ISL_557168, EPI_ISL_557170, EPI_ISL_557171, EPI_ISL_557172, EPI_ISL_557173, EPI_ISL_557174, EPI_ISL_557175, EPI_ISL_557177, EPI_ISL_557178, EPI_ISL_557179, EPI_ISL_557181, EPI_ISL_557182, EPI_ISL_557183, EPI_ISL_557184, EPI_ISL_557185, EPI_ISL_557188, EPI_ISL_557190, EPI_ISL_557191, EPI_ISL_557194, EPI_ISL_557195, EPI_ISL_557196, EPI_ISL_557197, EPI_ISL_557198, EPI_ISL_557202, EPI_ISL_557203, EPI_ISL_557204, EPI_ISL_557205, EPI_ISL_557206, EPI_ISL_557208, EPI_ISL_557209, EPI_ISL_557210, EPI_ISL_557211, EPI_ISL_557212, EPI_ISL_557214, EPI_ISL_557215, EPI_ISL_557216, EPI_ISL_557217, EPI_ISL_557218, EPI_ISL_557219, EPI_ISL_557220, EPI_ISL_557221, EPI_ISL_557222, EPI_ISL_557223, EPI_ISL_557224, EPI_ISL_557226, EPI_ISL_557228, EPI_ISL_557230, EPI_ISL_557231, EPI_ISL_557233, EPI_ISL_557234, EPI_ISL_557235, EPI_ISL_557236, EPI_ISL_557239, EPI_ISL_557240, EPI_ISL_557241, EPI_ISL_557242, EPI_ISL_557243, EPI_ISL_557244, EPI_ISL_557246, EPI_ISL_557247, EPI_ISL_557248, EPI_ISL_557249, EPI_ISL_557251, EPI_ISL_557252, EPI_ISL_557253, EPI_ISL_557256, EPI_ISL_557257, EPI_ISL_557258, EPI_ISL_557259, EPI_ISL_557260, EPI_ISL_557261, EPI_ISL_557262, EPI_ISL_557263, EPI_ISL_557264, EPI_ISL_557266, EPI_ISL_557267, EPI_ISL_557268, EPI_ISL_557269, EPI_ISL_557270, EPI_ISL_557273, EPI_ISL_557275, EPI_ISL_557276, EPI_ISL_557277, EPI_ISL_557278, EPI_ISL_557279, EPI_ISL_557280, EPI_ISL_557282, EPI_ISL_557285, EPI_ISL_557286, EPI_ISL_557287, EPI_ISL_557288, EPI_ISL_557289, EPI_ISL_557290, EPI_ISL_557291, EPI_ISL_557292, EPI_ISL_557296, EPI_ISL_557297, EPI_ISL_557298, EPI_ISL_557300, EPI_ISL_557302, EPI_ISL_557303, EPI_ISL_557305, EPI_ISL_557309, EPI_ISL_557310, EPI_ISL_557311, EPI_ISL_557312, EPI_ISL_557314, EPI_ISL_557315, EPI_ISL_557316, EPI_ISL_557317, EPI_ISL_557318, EPI_ISL_557319, EPI_ISL_557321, EPI_ISL_557322, EPI_ISL_557323, EPI_ISL_557326, EPI_ISL_557327, EPI_ISL_557328, EPI_ISL_557329, EPI_ISL_557331, EPI_ISL_557332, EPI_ISL_557333, EPI_ISL_557334, EPI_ISL_557336 | | | |
| see above | Lighthouse Lab in Milton Keynes | Wellcome Sanger Institute for the COVID-19 Genomics UK (COG-UK) consortium | The Lighthouse Lab in Milton Keynes and Alex Alderton, Roberto Amato, Sonia Goncalves, Ewan Harrison, David K. Jackson, Ian Johnston, Dominic Kwiatkowski, Cordelia Langford, John Sillitoe on behalf of the Wellcome Sanger Institute COVID-19 Surveillance Team (http://www.sanger.ac.uk/covid-team) |
| EPI_ISL_557337 | Lighthouse Lab in Milton Keynes | Wellcome Sanger Institute for the COVID-19 Genomics UK (COG-UK) consortium | The Lighthouse Lab in Alderley Park and Alex Alderton, Roberto Amato, Sonia Goncalves, Ewan Harrison, David K. Jackson, Ian Johnston, Dominic Kwiatkowski, Cordelia Langford, John Sillitoe on behalf of the Wellcome Sanger Institute COVID-19 Surveillance Team |
| EPI_ISL_557339, EPI_ISL_557341, EPI_ISL_557342, EPI_ISL_557343, EPI_ISL_557344, EPI_ISL_557345, EPI_ISL_557346, EPI_ISL_557347, EPI_ISL_557348, EPI_ISL_557349, EPI_ISL_557350 | | | |
| see above | Lighthouse Lab in Milton Keynes | Wellcome Sanger Institute for the COVID-19 Genomics UK (COG-UK) consortium | The Lighthouse Lab in Milton Keynes and Alex Alderton, Roberto Amato, Sonia Goncalves, Ewan Harrison, David K. Jackson, Ian Johnston, Dominic Kwiatkowski, Cordelia Langford, John Sillitoe on behalf of the Wellcome Sanger Institute COVID-19 Surveillance Team (http://www.sanger.ac.uk/covid-team) |
| EPI_ISL_557351 | Lighthouse Lab in Alderley Park | Wellcome Sanger Institute for the COVID-19 Genomics UK (COG-UK) consortium | The Lighthouse Lab in Alderley Park and Alex Alderton, Roberto Amato, Sonia Goncalves, Ewan Harrison, David K. Jackson, Ian Johnston, Dominic Kwiatkowski, Cordelia Langford, John Sillitoe on behalf of the Wellcome Sanger Institute COVID-19 Surveillance Team (http://www.sanger.ac.uk/covid-team) |
| EPI_ISL_557352 | Lighthouse Lab in Milton Keynes | Wellcome Sanger Institute for the COVID-19 Genomics UK (COG-UK) consortium | The Lighthouse Lab in Milton Keynes and Alex Alderton, Roberto Amato, Sonia Goncalves, Ewan Harrison, David K. Jackson, Ian Johnston, Dominic Kwiatkowski, Cordelia Langford, John Sillitoe on behalf of the Wellcome Sanger Institute COVID-19 Surveillance Team (http://www.sanger.ac.uk/covid-team) |
| EPI_ISL_557354 | Lighthouse Lab in Alderley Park | Wellcome Sanger Institute for the COVID-19 Genomics UK (COG-UK) consortium | The Lighthouse Lab in Alderley Park and Alex Alderton, Roberto Amato, Sonia Goncalves, Ewan Harrison, David K. Jackson, Ian Johnston, Dominic Kwiatkowski, Cordelia Langford, John Sillitoe on behalf of the Wellcome Sanger Institute COVID-19 Surveillance Team (http://www.sanger.ac.uk/covid-team) |
| EPI_ISL_557357, EPI_ISL_557360, EPI_ISL_557362, EPI_ISL_557364, EPI_ISL_557365 | Lighthouse Lab in Milton Keynes | Wellcome Sanger Institute for the COVID-19 Genomics UK (COG-UK) consortium | The Lighthouse Lab in Milton Keynes and Alex Alderton, Roberto Amato, Sonia Goncalves, Ewan Harrison, David K. Jackson, Ian Johnston, Dominic Kwiatkowski, Cordelia Langford, John Sillitoe on behalf of the Wellcome Sanger Institute COVID-19 Surveillance Team (http://www.sanger.ac.uk/covid-team) |
| EPI_ISL_557366 | Lighthouse Lab in Alderley Park | Wellcome Sanger Institute for the COVID-19 Genomics UK (COG-UK) consortium | The Lighthouse Lab in Alderley Park and Alex Alderton, Roberto Amato, Sonia Goncalves, Ewan Harrison, David K. Jackson, Ian Johnston, Dominic Kwiatkowski, Cordelia Langford, John Sillitoe on behalf of the Wellcome Sanger Institute COVID-19 Surveillance Team (http://www.sanger.ac.uk/covid-team) |
| EPI_ISL_557367, EPI_ISL_557369 | Lighthouse Lab in Milton Keynes | Wellcome Sanger Institute for the COVID-19 Genomics UK (COG-UK) consortium | The Lighthouse Lab in Milton Keynes and Alex Alderton, Roberto Amato, Sonia Goncalves, Ewan Harrison, David K. Jackson, Ian Johnston, Dominic Kwiatkowski, Cordelia Langford, John Sillitoe on behalf of the Wellcome Sanger Institute COVID-19 Surveillance Team (http://www.sanger.ac.uk/covid-team) |
| EPI_ISL_557371 | Lighthouse Lab in Alderley Park | Wellcome Sanger Institute for the COVID-19 Genomics UK (COG-UK) consortium | The Lighthouse Lab in Alderley Park and Alex Alderton, Roberto Amato, Sonia Goncalves, Ewan Harrison, David K. Jackson, Ian Johnston, Dominic Kwiatkowski, Cordelia Langford, John Sillitoe on behalf of the Wellcome Sanger Institute COVID-19 Surveillance Team (http://www.sanger.ac.uk/covid-team) |
| EPI_ISL_557372, EPI_ISL_557375, EPI_ISL_557376, EPI_ISL_557377, EPI_ISL_557382 | Lighthouse Lab in Milton Keynes | Wellcome Sanger Institute for the COVID-19 Genomics UK (COG-UK) consortium | The Lighthouse Lab in Milton Keynes and Alex Alderton, Roberto Amato, Sonia Goncalves, Ewan Harrison, David K. Jackson, Ian Johnston, Dominic Kwiatkowski, Cordelia Langford, John Sillitoe on behalf of the Wellcome Sanger Institute COVID-19 Surveillance Team (http://www.sanger.ac.uk/covid-team) |
| EPI_ISL_557383 | Lighthouse Lab in Alderley Park | Wellcome Sanger Institute for the COVID-19 Genomics UK (COG-UK) consortium | The Lighthouse Lab in Alderley Park and Alex Alderton, Roberto Amato, Sonia Goncalves, Ewan Harrison, David K. Jackson, Ian Johnston, Dominic Kwiatkowski, Cordelia Langford, John Sillitoe on behalf of the Wellcome Sanger Institute COVID-19 Surveillance Team (http://www.sanger.ac.uk/covid-team) |
| EPI_ISL_557384, EPI_ISL_557386, EPI_ISL_557387 | Lighthouse Lab in Milton Keynes | Wellcome Sanger Institute for the COVID-19 Genomics UK (COG-UK) consortium | The Lighthouse Lab in Milton Keynes and Alex Alderton, Roberto Amato, Sonia Goncalves, Ewan Harrison, David K. Jackson, Ian Johnston, Dominic Kwiatkowski, Cordelia Langford, John Sillitoe on behalf of the Wellcome Sanger Institute COVID-19 Surveillance Team (http://www.sanger.ac.uk/covid-team) |
| EPI_ISL_557388 | Lighthouse Lab in Alderley Park | Wellcome Sanger Institute for the COVID-19 Genomics UK (COG-UK) consortium | The Lighthouse Lab in Alderley Park and Alex Alderton, Roberto Amato, Sonia Goncalves, Ewan Harrison, David K. Jackson, Ian Johnston, Dominic Kwiatkowski, Cordelia Langford, John Sillitoe on behalf of the Wellcome Sanger Institute COVID-19 Surveillance Team (http://www.sanger.ac.uk/covid-team) |
| EPI_ISL_557391, EPI_ISL_557393, EPI_ISL_557394, EPI_ISL_557395, EPI_ISL_557396, EPI_ISL_557397, EPI_ISL_557400, EPI_ISL_557401 | Lighthouse Lab in Milton Keynes | Wellcome Sanger Institute for the COVID-19 Genomics UK (COG-UK) consortium | The Lighthouse Lab in Milton Keynes and Alex Alderton, Roberto Amato, Sonia Goncalves, Ewan Harrison, David K. Jackson, Ian Johnston, Dominic Kwiatkowski, Cordelia Langford, John Sillitoe on behalf of the Wellcome Sanger Institute COVID-19 Surveillance Team (http://www.sanger.ac.uk/covid-team) |
| EPI_ISL_557405, EPI_ISL_557406 | Lighthouse Lab in Alderley Park | Wellcome Sanger Institute for the COVID-19 Genomics UK (COG-UK) consortium | The Lighthouse Lab in Alderley Park and Alex Alderton, Roberto Amato, Sonia Goncalves, Ewan Harrison, David K. Jackson, Ian Johnston, Dominic Kwiatkowski, Cordelia Langford, John Sillitoe on behalf of the Wellcome Sanger Institute COVID-19 Surveillance Team (http://www.sanger.ac.uk/covid-team) |
| EPI_ISL_557407, EPI_ISL_557410 | Lighthouse Lab in Milton Keynes | Wellcome Sanger Institute for the COVID-19 Genomics UK (COG-UK) consortium | The Lighthouse Lab in Milton Keynes and Alex Alderton, Roberto Amato, Sonia Goncalves, Ewan Harrison, David K. Jackson, Ian Johnston, Dominic Kwiatkowski, Cordelia Langford, John Sillitoe on behalf of the Wellcome Sanger Institute COVID-19 Surveillance Team (http://www.sanger.ac.uk/covid-team) |
| EPI_ISL_557412 | Lighthouse Lab in Alderley Park | Wellcome Sanger Institute for the COVID-19 Genomics UK (COG-UK) consortium | The Lighthouse Lab in Alderley Park and Alex Alderton, Roberto Amato, Sonia Goncalves, Ewan Harrison, David K. Jackson, Ian Johnston, Dominic Kwiatkowski, Cordelia Langford, John Sillitoe on behalf of the Wellcome Sanger Institute COVID-19 Surveillance Team (http://www.sanger.ac.uk/covid-team) |
| EPI_ISL_557413 | Lighthouse Lab in Milton Keynes | Wellcome Sanger Institute for the COVID-19 Genomics UK (COG-UK) consortium | The Lighthouse Lab in Milton Keynes and Alex Alderton, Roberto Amato, Sonia Goncalves, Ewan Harrison, David K. Jackson, Ian Johnston, Dominic Kwiatkowski, Cordelia Langford, John Sillitoe on behalf of the Wellcome Sanger Institute COVID-19 Surveillance Team (http://www.sanger.ac.uk/covid-team) |
| EPI_ISL_557414 | Lighthouse Lab in Alderley Park | Wellcome Sanger Institute for the COVID-19 Genomics UK (COG-UK) consortium | The Lighthouse Lab in Alderley Park and Alex Alderton, Roberto Amato, Sonia Goncalves, Ewan Harrison, David K. Jackson, Ian Johnston, Dominic Kwiatkowski, Cordelia Langford, John Sillitoe on behalf of the Wellcome Sanger Institute COVID-19 Surveillance Team (http://www.sanger.ac.uk/covid-team) |

| | | | |
|---|---|---|---|
| EPI_ISL_557416, EPI_ISL_557418, EPI_ISL_557421 | Lighthouse Lab in Milton Keynes | Wellcome Sanger Institute for the COVID-19 Genomics UK (COG-UK) consortium | The Lighthouse Lab in Milton Keynes and Alex Alderton, Roberto Amato, Sonia Goncalves, Ewan Harrison, David K. Jackson, Ian Johnston, Dominic Kwiatkowski, Cordelia Langford, John Sillitoe on behalf of the Wellcome Sanger Institute COVID-19 Surveillance Team (http://www.sanger.ac.uk/covid-team) |
| EPI_ISL_557425 | Lighthouse Lab in Alderley Park | Wellcome Sanger Institute for the COVID-19 Genomics UK (COG-UK) consortium | The Lighthouse Lab in Alderley Park and Alex Alderton, Roberto Amato, Sonia Goncalves, Ewan Harrison, David K. Jackson, Ian Johnston, Dominic Kwiatkowski, Cordelia Langford, John Sillitoe on behalf of the Wellcome Sanger Institute COVID-19 Surveillance Team (http://www.sanger.ac.uk/covid-team) |
| EPI_ISL_557426, EPI_ISL_557427, EPI_ISL_557429 | Lighthouse Lab in Milton Keynes | Wellcome Sanger Institute for the COVID-19 Genomics UK (COG-UK) consortium | The Lighthouse Lab in Milton Keynes and Alex Alderton, Roberto Amato, Sonia Goncalves, Ewan Harrison, David K. Jackson, Ian Johnston, Dominic Kwiatkowski, Cordelia Langford, John Sillitoe on behalf of the Wellcome Sanger Institute COVID-19 Surveillance Team (http://www.sanger.ac.uk/covid-team) |
| EPI_ISL_557430 | Lighthouse Lab in Alderley Park | Wellcome Sanger Institute for the COVID-19 Genomics UK (COG-UK) consortium | The Lighthouse Lab in Alderley Park and Alex Alderton, Roberto Amato, Sonia Goncalves, Ewan Harrison, David K. Jackson, Ian Johnston, Dominic Kwiatkowski, Cordelia Langford, John Sillitoe on behalf of the Wellcome Sanger Institute COVID-19 Surveillance Team (http://www.sanger.ac.uk/covid-team) |
| EPI_ISL_557431, EPI_ISL_557432, EPI_ISL_557435, EPI_ISL_557437, EPI_ISL_557441, EPI_ISL_557442, EPI_ISL_557443, EPI_ISL_557444, EPI_ISL_557445, EPI_ISL_557450, EPI_ISL_557451, EPI_ISL_557455, EPI_ISL_557456, EPI_ISL_557457, EPI_ISL_557458 | | | |
| see above | Lighthouse Lab in Milton Keynes | Wellcome Sanger Institute for the COVID-19 Genomics UK (COG-UK) consortium | The Lighthouse Lab in Milton Keynes and Alex Alderton, Roberto Amato, Sonia Goncalves, Ewan Harrison, David K. Jackson, Ian Johnston, Dominic Kwiatkowski, Cordelia Langford, John Sillitoe on behalf of the Wellcome Sanger Institute COVID-19 Surveillance Team (http://www.sanger.ac.uk/covid-team) |
| EPI_ISL_557459 | Lighthouse Lab in Milton Keynes | Wellcome Sanger Institute for the COVID-19 Genomics UK (COG-UK) consortium | The Lighthouse Lab in Milton Keynes and Alex Alderton, Roberto Amato, Sonia Goncalves, Ewan Harrison, David K. Jackson, Ian Johnston, Dominic Kwiatkowski, Cordelia Langford, John Sillitoe on behalf of the Wellcome Sanger Institute COVID-19 Surveillance Team |
| EPI_ISL_557460, EPI_ISL_557461, EPI_ISL_557463, EPI_ISL_557464, EPI_ISL_557465 | Lighthouse Lab in Milton Keynes | Wellcome Sanger Institute for the COVID-19 Genomics UK (COG-UK) consortium | The Lighthouse Lab in Milton Keynes and Alex Alderton, Roberto Amato, Sonia Goncalves, Ewan Harrison, David K. Jackson, Ian Johnston, Dominic Kwiatkowski, Cordelia Langford, John Sillitoe on behalf of the Wellcome Sanger Institute COVID-19 Surveillance Team (http://www.sanger.ac.uk/covid-team) |
| EPI_ISL_557466 | Lighthouse Lab in Alderley Park | Wellcome Sanger Institute for the COVID-19 Genomics UK (COG-UK) consortium | The Lighthouse Lab in Alderley Park and Alex Alderton, Roberto Amato, Sonia Goncalves, Ewan Harrison, David K. Jackson, Ian Johnston, Dominic Kwiatkowski, Cordelia Langford, John Sillitoe on behalf of the Wellcome Sanger Institute COVID-19 Surveillance Team (http://www.sanger.ac.uk/covid-team) |
| EPI_ISL_557467, EPI_ISL_557470, EPI_ISL_557473, EPI_ISL_557475, EPI_ISL_557479, EPI_ISL_557483, EPI_ISL_557486, EPI_ISL_557489, EPI_ISL_557490, EPI_ISL_557492, EPI_ISL_557495, EPI_ISL_557496, EPI_ISL_557498 | | | |
| see above | Lighthouse Lab in Milton Keynes | Wellcome Sanger Institute for the COVID-19 Genomics UK (COG-UK) consortium | The Lighthouse Lab in Milton Keynes and Alex Alderton, Roberto Amato, Sonia Goncalves, Ewan Harrison, David K. Jackson, Ian Johnston, Dominic Kwiatkowski, Cordelia Langford, John Sillitoe on behalf of the Wellcome Sanger Institute COVID-19 Surveillance Team (http://www.sanger.ac.uk/covid-team) |
| EPI_ISL_557499 | Lighthouse Lab in Alderley Park | Wellcome Sanger Institute for the COVID-19 Genomics UK (COG-UK) consortium | The Lighthouse Lab in Alderley Park and Alex Alderton, Roberto Amato, Sonia Goncalves, Ewan Harrison, David K. Jackson, Ian Johnston, Dominic Kwiatkowski, Cordelia Langford, John Sillitoe on behalf of the Wellcome Sanger Institute COVID-19 Surveillance Team (http://www.sanger.ac.uk/covid-team) |
| EPI_ISL_557505, EPI_ISL_557506, EPI_ISL_557508, EPI_ISL_557510, EPI_ISL_557512, EPI_ISL_557513, EPI_ISL_557514, EPI_ISL_557515, EPI_ISL_557517, EPI_ISL_557519, EPI_ISL_557520, EPI_ISL_557521 | | | |
| see above | Lighthouse Lab in Milton Keynes | Wellcome Sanger Institute for the COVID-19 Genomics UK (COG-UK) consortium | The Lighthouse Lab in Milton Keynes and Alex Alderton, Roberto Amato, Sonia Goncalves, Ewan Harrison, David K. Jackson, Ian Johnston, Dominic Kwiatkowski, Cordelia Langford, John Sillitoe on behalf of the Wellcome Sanger Institute COVID-19 Surveillance Team (http://www.sanger.ac.uk/covid-team) |
| EPI_ISL_557522 | Lighthouse Lab in Alderley Park | Wellcome Sanger Institute for the COVID-19 Genomics UK (COG-UK) consortium | The Lighthouse Lab in Alderley Park and Alex Alderton, Roberto Amato, Sonia Goncalves, Ewan Harrison, David K. Jackson, Ian Johnston, Dominic Kwiatkowski, Cordelia Langford, John Sillitoe on behalf of the Wellcome Sanger Institute COVID-19 Surveillance Team (http://www.sanger.ac.uk/covid-team) |
| EPI_ISL_557525 | Lighthouse Lab in Milton Keynes | Wellcome Sanger Institute for the COVID-19 Genomics UK (COG-UK) consortium | The Lighthouse Lab in Milton Keynes and Alex Alderton, Roberto Amato, Sonia Goncalves, Ewan Harrison, David K. Jackson, Ian Johnston, Dominic Kwiatkowski, Cordelia Langford, John Sillitoe on behalf of the Wellcome Sanger Institute COVID-19 Surveillance Team (http://www.sanger.ac.uk/covid-team) |
| EPI_ISL_557528 | Lighthouse Lab in Alderley Park | Wellcome Sanger Institute for the COVID-19 Genomics UK (COG-UK) consortium | The Lighthouse Lab in Alderley Park and Alex Alderton, Roberto Amato, Sonia Goncalves, Ewan Harrison, David K. Jackson, Ian Johnston, Dominic Kwiatkowski, Cordelia Langford, John Sillitoe on behalf of the Wellcome Sanger Institute COVID-19 Surveillance Team (http://www.sanger.ac.uk/covid-team) |
| EPI_ISL_557529, EPI_ISL_557532, EPI_ISL_557534, EPI_ISL_557536, EPI_ISL_557538, EPI_ISL_557540, EPI_ISL_557542, EPI_ISL_557545, EPI_ISL_557546, EPI_ISL_557547, EPI_ISL_557548, EPI_ISL_557550, EPI_ISL_557552 | | | |
| see above | Lighthouse Lab in Milton Keynes | Wellcome Sanger Institute for the COVID-19 Genomics UK (COG-UK) consortium | The Lighthouse Lab in Milton Keynes and Alex Alderton, Roberto Amato, Sonia Goncalves, Ewan Harrison, David K. Jackson, Ian Johnston, Dominic Kwiatkowski, Cordelia Langford, John Sillitoe on behalf of the Wellcome Sanger Institute COVID-19 Surveillance Team (http://www.sanger.ac.uk/covid-team) |
| EPI_ISL_557590 | Lighthouse Lab in Alderley Park | Wellcome Sanger Institute for the COVID-19 Genomics UK (COG-UK) consortium | The Lighthouse Lab in Alderley Park and Alex Alderton, Roberto Amato, Sonia Goncalves, Ewan Harrison, David K. Jackson, Ian Johnston, Dominic Kwiatkowski, Cordelia Langford, John Sillitoe on behalf of the Wellcome Sanger Institute COVID-19 Surveillance Team |
| EPI_ISL_557607 | Lighthouse Lab in Alderley Park | Wellcome Sanger Institute for the COVID-19 Genomics UK (COG-UK) consortium | The Lighthouse Lab in Alderley Park and Alex Alderton, Roberto Amato, Sonia Goncalves, Ewan Harrison, David K. Jackson, Ian Johnston, Dominic Kwiatkowski, Cordelia Langford, John Sillitoe on behalf of the Wellcome Sanger Institute COVID-19 Surveillance Team (http://www.sanger.ac.uk/covid-team) |
| EPI_ISL_557615, EPI_ISL_557630, EPI_ISL_557642, EPI_ISL_557647, EPI_ISL_557655, EPI_ISL_557668 | Lighthouse Lab in Alderley Park | Wellcome Sanger Institute for the COVID-19 Genomics UK (COG-UK) consortium | The Lighthouse Lab in Alderley Park and Alex Alderton, Roberto Amato, Sonia Goncalves, Ewan Harrison, David K. Jackson, Ian Johnston, Dominic Kwiatkowski, Cordelia Langford, John Sillitoe on behalf of the Wellcome Sanger Institute COVID-19 Surveillance Team |
| EPI_ISL_557672 | Lighthouse Lab in Alderley Park | Wellcome Sanger Institute for the COVID-19 Genomics UK (COG-UK) consortium | The Lighthouse Lab in Alderley Park and Alex Alderton, Roberto Amato, Sonia Goncalves, Ewan Harrison, David K. Jackson, Ian Johnston, Dominic Kwiatkowski, Cordelia Langford, John Sillitoe on behalf of the Wellcome Sanger Institute COVID-19 Surveillance Team (http://www.sanger.ac.uk/covid-team) |
| EPI_ISL_557676, EPI_ISL_557681 | Lighthouse Lab in Alderley Park | Wellcome Sanger Institute for the COVID-19 Genomics UK (COG-UK) consortium | The Lighthouse Lab in Alderley Park and Alex Alderton, Roberto Amato, Sonia Goncalves, Ewan Harrison, David K. Jackson, Ian Johnston, Dominic Kwiatkowski, Cordelia Langford, John Sillitoe on behalf of the Wellcome Sanger Institute COVID-19 Surveillance Team |
| EPI_ISL_557685 | Lighthouse Lab in Alderley Park | Wellcome Sanger Institute for the COVID-19 Genomics UK (COG-UK) consortium | The Lighthouse Lab in Alderley Park and Alex Alderton, Roberto Amato, Sonia Goncalves, Ewan Harrison, David K. Jackson, Ian Johnston, Dominic Kwiatkowski, Cordelia Langford, John Sillitoe on behalf of the Wellcome Sanger Institute COVID-19 Surveillance Team (http://www.sanger.ac.uk/covid-team) |
| EPI_ISL_557690 | Lighthouse Lab in Alderley Park | Wellcome Sanger Institute for the COVID-19 Genomics UK (COG-UK) consortium | The Lighthouse Lab in Alderley Park and Alex Alderton, Roberto Amato, Sonia Goncalves, Ewan Harrison, David K. Jackson, Ian Johnston, Dominic Kwiatkowski, Cordelia Langford, John Sillitoe on behalf of the Wellcome Sanger Institute COVID-19 Surveillance Team |
| EPI_ISL_557709, EPI_ISL_557713 | Lighthouse Lab in Alderley Park | Wellcome Sanger Institute for the COVID-19 Genomics UK (COG-UK) consortium | The Lighthouse Lab in Alderley Park and Alex Alderton, Roberto Amato, Sonia Goncalves, Ewan Harrison, David K. Jackson, Ian Johnston, Dominic Kwiatkowski, Cordelia Langford, John Sillitoe on behalf of the Wellcome Sanger Institute COVID-19 Surveillance Team (http://www.sanger.ac.uk/covid-team) |
| EPI_ISL_557740, EPI_ISL_557742 | Lighthouse Lab in Milton Keynes | Wellcome Sanger Institute for the COVID-19 Genomics UK (COG-UK) consortium | The Lighthouse Lab in Milton Keynes and Alex Alderton, Roberto Amato, Sonia Goncalves, Ewan Harrison, David K. Jackson, Ian Johnston, Dominic Kwiatkowski, Cordelia Langford, John Sillitoe on behalf of the Wellcome Sanger Institute COVID-19 Surveillance Team (http://www.sanger.ac.uk/covid-team) |
| EPI_ISL_557746 | Lighthouse Lab in Alderley Park | Wellcome Sanger Institute for the COVID-19 Genomics UK | The Lighthouse Lab in Alderley Park and Alex Alderton, Roberto Amato, Sonia Goncalves, Ewan Harrison, David K. Jackson, Ian Johnston, Dominic |

| | | | |
|---|---|---|---|
| | | (COG-UK) consortium | Kwiatkowski, Cordelia Langford, John Sillitoe on behalf of the Wellcome Sanger Institute COVID-19 Surveillance Team |
| EPI_ISL_557748 | Lighthouse Lab in Milton Keynes | Wellcome Sanger Institute for the COVID-19 Genomics UK (COG-UK) consortium | The Lighthouse Lab in Milton Keynes and Alex Alderton, Roberto Amato, Sonia Goncalves, Ewan Harrison, David K. Jackson, Ian Johnston, Dominic Kwiatkowski, Cordelia Langford, John Sillitoe on behalf of the Wellcome Sanger Institute COVID-19 Surveillance Team (http://www.sanger.ac.uk/covid-team) |
| EPI_ISL_557752 | Lighthouse Lab in Alderley Park | Wellcome Sanger Institute for the COVID-19 Genomics UK (COG-UK) consortium | The Lighthouse Lab in Alderley Park and Alex Alderton, Roberto Amato, Sonia Goncalves, Ewan Harrison, David K. Jackson, Ian Johnston, Dominic Kwiatkowski, Cordelia Langford, John Sillitoe on behalf of the Wellcome Sanger Institute COVID-19 Surveillance Team |
| EPI_ISL_557753, EPI_ISL_557757, EPI_ISL_557765, EPI_ISL_557766 | Lighthouse Lab in Milton Keynes | Wellcome Sanger Institute for the COVID-19 Genomics UK (COG-UK) consortium | The Lighthouse Lab in Milton Keynes and Alex Alderton, Roberto Amato, Sonia Goncalves, Ewan Harrison, David K. Jackson, Ian Johnston, Dominic Kwiatkowski, Cordelia Langford, John Sillitoe on behalf of the Wellcome Sanger Institute COVID-19 Surveillance Team (http://www.sanger.ac.uk/covid-team) |
| EPI_ISL_557767 | Lighthouse Lab in Alderley Park | Wellcome Sanger Institute for the COVID-19 Genomics UK (COG-UK) consortium | The Lighthouse Lab in Alderley Park and Alex Alderton, Roberto Amato, Sonia Goncalves, Ewan Harrison, David K. Jackson, Ian Johnston, Dominic Kwiatkowski, Cordelia Langford, John Sillitoe on behalf of the Wellcome Sanger Institute COVID-19 Surveillance Team |
| EPI_ISL_557774 | Lighthouse Lab in Alderley Park | Wellcome Sanger Institute for the COVID-19 Genomics UK (COG-UK) consortium | The Lighthouse Lab in Alderley Park and Alex Alderton, Roberto Amato, Sonia Goncalves, Ewan Harrison, David K. Jackson, Ian Johnston, Dominic Kwiatkowski, Cordelia Langford, John Sillitoe on behalf of the Wellcome Sanger Institute COVID-19 Surveillance Team (http://www.sanger.ac.uk/covid-team) |
| EPI_ISL_557777 | Lighthouse Lab in Alderley Park | Wellcome Sanger Institute for the COVID-19 Genomics UK (COG-UK) consortium | The Lighthouse Lab in Alderley Park and Alex Alderton, Roberto Amato, Sonia Goncalves, Ewan Harrison, David K. Jackson, Ian Johnston, Dominic Kwiatkowski, Cordelia Langford, John Sillitoe on behalf of the Wellcome Sanger Institute COVID-19 Surveillance Team |
| EPI_ISL_557778, EPI_ISL_557781 | Lighthouse Lab in Milton Keynes | Wellcome Sanger Institute for the COVID-19 Genomics UK (COG-UK) consortium | The Lighthouse Lab in Milton Keynes and Alex Alderton, Roberto Amato, Sonia Goncalves, Ewan Harrison, David K. Jackson, Ian Johnston, Dominic Kwiatkowski, Cordelia Langford, John Sillitoe on behalf of the Wellcome Sanger Institute COVID-19 Surveillance Team (http://www.sanger.ac.uk/covid-team) |
| EPI_ISL_557783 | Lighthouse Lab in Alderley Park | Wellcome Sanger Institute for the COVID-19 Genomics UK (COG-UK) consortium | The Lighthouse Lab in Alderley Park and Alex Alderton, Roberto Amato, Sonia Goncalves, Ewan Harrison, David K. Jackson, Ian Johnston, Dominic Kwiatkowski, Cordelia Langford, John Sillitoe on behalf of the Wellcome Sanger Institute COVID-19 Surveillance Team |
| EPI_ISL_557784 | Lighthouse Lab in Milton Keynes | Wellcome Sanger Institute for the COVID-19 Genomics UK (COG-UK) consortium | The Lighthouse Lab in Milton Keynes and Alex Alderton, Roberto Amato, Sonia Goncalves, Ewan Harrison, David K. Jackson, Ian Johnston, Dominic Kwiatkowski, Cordelia Langford, John Sillitoe on behalf of the Wellcome Sanger Institute COVID-19 Surveillance Team (http://www.sanger.ac.uk/covid-team) |
| EPI_ISL_557788 | Lighthouse Lab in Alderley Park | Wellcome Sanger Institute for the COVID-19 Genomics UK (COG-UK) consortium | The Lighthouse Lab in Alderley Park and Alex Alderton, Roberto Amato, Sonia Goncalves, Ewan Harrison, David K. Jackson, Ian Johnston, Dominic Kwiatkowski, Cordelia Langford, John Sillitoe on behalf of the Wellcome Sanger Institute COVID-19 Surveillance Team |
| EPI_ISL_557789 | Lighthouse Lab in Milton Keynes | Wellcome Sanger Institute for the COVID-19 Genomics UK (COG-UK) consortium | The Lighthouse Lab in Milton Keynes and Alex Alderton, Roberto Amato, Sonia Goncalves, Ewan Harrison, David K. Jackson, Ian Johnston, Dominic Kwiatkowski, Cordelia Langford, John Sillitoe on behalf of the Wellcome Sanger Institute COVID-19 Surveillance Team (http://www.sanger.ac.uk/covid-team) |
| EPI_ISL_557792 | Lighthouse Lab in Alderley Park | Wellcome Sanger Institute for the COVID-19 Genomics UK (COG-UK) consortium | The Lighthouse Lab in Alderley Park and Alex Alderton, Roberto Amato, Sonia Goncalves, Ewan Harrison, David K. Jackson, Ian Johnston, Dominic Kwiatkowski, Cordelia Langford, John Sillitoe on behalf of the Wellcome Sanger Institute COVID-19 Surveillance Team |
| EPI_ISL_557795 | Lighthouse Lab in Milton Keynes | Wellcome Sanger Institute for the COVID-19 Genomics UK (COG-UK) consortium | The Lighthouse Lab in Milton Keynes and Alex Alderton, Roberto Amato, Sonia Goncalves, Ewan Harrison, David K. Jackson, Ian Johnston, Dominic Kwiatkowski, Cordelia Langford, John Sillitoe on behalf of the Wellcome Sanger Institute COVID-19 Surveillance Team (http://www.sanger.ac.uk/covid-team) |
| EPI_ISL_557796 | Lighthouse Lab in Milton Keynes | Wellcome Sanger Institute for the COVID-19 Genomics UK (COG-UK) consortium | The Lighthouse Lab in Alderley Park and Alex Alderton, Roberto Amato, Sonia Goncalves, Ewan Harrison, David K. Jackson, Ian Johnston, Dominic Kwiatkowski, Cordelia Langford, John Sillitoe on behalf of the Wellcome Sanger Institute COVID-19 Surveillance Team |
| EPI_ISL_557800 | Lighthouse Lab in Milton Keynes | Wellcome Sanger Institute for the COVID-19 Genomics UK (COG-UK) consortium | The Lighthouse Lab in Milton Keynes and Alex Alderton, Roberto Amato, Sonia Goncalves, Ewan Harrison, David K. Jackson, Ian Johnston, Dominic Kwiatkowski, Cordelia Langford, John Sillitoe on behalf of the Wellcome Sanger Institute COVID-19 Surveillance Team (http://www.sanger.ac.uk/covid-team) |
| EPI_ISL_557801 | Lighthouse Lab in Alderley Park | Wellcome Sanger Institute for the COVID-19 Genomics UK (COG-UK) consortium | The Lighthouse Lab in Alderley Park and Alex Alderton, Roberto Amato, Sonia Goncalves, Ewan Harrison, David K. Jackson, Ian Johnston, Dominic Kwiatkowski, Cordelia Langford, John Sillitoe on behalf of the Wellcome Sanger Institute COVID-19 Surveillance Team (http://www.sanger.ac.uk/covid-team) |
| EPI_ISL_557802, EPI_ISL_557812, EPI_ISL_557821, EPI_ISL_557822, EPI_ISL_557823, EPI_ISL_557824, EPI_ISL_557827, EPI_ISL_557829, EPI_ISL_557830, EPI_ISL_557845 | Lighthouse Lab in Milton Keynes | Wellcome Sanger Institute for the COVID-19 Genomics UK (COG-UK) consortium | The Lighthouse Lab in Milton Keynes and Alex Alderton, Roberto Amato, Sonia Goncalves, Ewan Harrison, David K. Jackson, Ian Johnston, Dominic Kwiatkowski, Cordelia Langford, John Sillitoe on behalf of the Wellcome Sanger Institute COVID-19 Surveillance Team (http://www.sanger.ac.uk/covid-team) |
| EPI_ISL_557846 | Lighthouse Lab in Alderley Park | Wellcome Sanger Institute for the COVID-19 Genomics UK (COG-UK) consortium | The Lighthouse Lab in Alderley Park and Alex Alderton, Roberto Amato, Sonia Goncalves, Ewan Harrison, David K. Jackson, Ian Johnston, Dominic Kwiatkowski, Cordelia Langford, John Sillitoe on behalf of the Wellcome Sanger Institute COVID-19 Surveillance Team |
| EPI_ISL_557847, EPI_ISL_557848, EPI_ISL_557849, EPI_ISL_557853, EPI_ISL_557854, EPI_ISL_557858 | Lighthouse Lab in Milton Keynes | Wellcome Sanger Institute for the COVID-19 Genomics UK (COG-UK) consortium | The Lighthouse Lab in Milton Keynes and Alex Alderton, Roberto Amato, Sonia Goncalves, Ewan Harrison, David K. Jackson, Ian Johnston, Dominic Kwiatkowski, Cordelia Langford, John Sillitoe on behalf of the Wellcome Sanger Institute COVID-19 Surveillance Team (http://www.sanger.ac.uk/covid-team) |
| EPI_ISL_557859 | Lighthouse Lab in Alderley Park | Wellcome Sanger Institute for the COVID-19 Genomics UK (COG-UK) consortium | The Lighthouse Lab in Alderley Park and Alex Alderton, Roberto Amato, Sonia Goncalves, Ewan Harrison, David K. Jackson, Ian Johnston, Dominic Kwiatkowski, Cordelia Langford, John Sillitoe on behalf of the Wellcome Sanger Institute COVID-19 Surveillance Team |
| EPI_ISL_557864 | Lighthouse Lab in Alderley Park | Wellcome Sanger Institute for the COVID-19 Genomics UK (COG-UK) consortium | The Lighthouse Lab in Alderley Park and Alex Alderton, Roberto Amato, Sonia Goncalves, Ewan Harrison, David K. Jackson, Ian Johnston, Dominic Kwiatkowski, Cordelia Langford, John Sillitoe on behalf of the Wellcome Sanger Institute COVID-19 Surveillance Team (http://www.sanger.ac.uk/covid-team) |
| EPI_ISL_557866, EPI_ISL_557867, EPI_ISL_557869, EPI_ISL_557870 | Lighthouse Lab in Milton Keynes | Wellcome Sanger Institute for the COVID-19 Genomics UK (COG-UK) consortium | The Lighthouse Lab in Milton Keynes and Alex Alderton, Roberto Amato, Sonia Goncalves, Ewan Harrison, David K. Jackson, Ian Johnston, Dominic Kwiatkowski, Cordelia Langford, John Sillitoe on behalf of the Wellcome Sanger Institute COVID-19 Surveillance Team (http://www.sanger.ac.uk/covid-team) |
| EPI_ISL_557871 | Lighthouse Lab in Alderley Park | Wellcome Sanger Institute for the COVID-19 Genomics UK (COG-UK) consortium | The Lighthouse Lab in Alderley Park and Alex Alderton, Roberto Amato, Sonia Goncalves, Ewan Harrison, David K. Jackson, Ian Johnston, Dominic Kwiatkowski, Cordelia Langford, John Sillitoe on behalf of the Wellcome Sanger Institute COVID-19 Surveillance Team |
| EPI_ISL_557876, EPI_ISL_557877, EPI_ISL_557878, EPI_ISL_557879 | Lighthouse Lab in Milton Keynes | Wellcome Sanger Institute for the COVID-19 Genomics UK (COG-UK) consortium | The Lighthouse Lab in Milton Keynes and Alex Alderton, Roberto Amato, Sonia Goncalves, Ewan Harrison, David K. Jackson, Ian Johnston, Dominic Kwiatkowski, Cordelia Langford, John Sillitoe on behalf of the Wellcome Sanger Institute COVID-19 Surveillance Team (http://www.sanger.ac.uk/covid-team) |
| EPI_ISL_557881, EPI_ISL_557882, EPI_ISL_557886 | Lighthouse Lab in Alderley Park | Wellcome Sanger Institute for the COVID-19 Genomics UK (COG-UK) consortium | The Lighthouse Lab in Alderley Park and Alex Alderton, Roberto Amato, Sonia Goncalves, Ewan Harrison, David K. Jackson, Ian Johnston, Dominic Kwiatkowski, Cordelia Langford, John Sillitoe on behalf of the Wellcome Sanger Institute COVID-19 Surveillance Team |
| EPI_ISL_557890, EPI_ISL_557893, EPI_ISL_557894, EPI_ISL_557895 | Lighthouse Lab in Milton Keynes | Wellcome Sanger Institute for the COVID-19 Genomics UK (COG-UK) consortium | The Lighthouse Lab in Milton Keynes and Alex Alderton, Roberto Amato, Sonia Goncalves, Ewan Harrison, David K. Jackson, Ian Johnston, Dominic Kwiatkowski, Cordelia Langford, John Sillitoe on behalf of the Wellcome Sanger Institute COVID-19 Surveillance Team (http://www.sanger.ac.uk/covid-team) |
| EPI_ISL_557899 | Lighthouse Lab in Alderley Park | Wellcome Sanger Institute for the COVID-19 Genomics UK (COG-UK) consortium | The Lighthouse Lab in Alderley Park and Alex Alderton, Roberto Amato, Sonia Goncalves, Ewan Harrison, David K. Jackson, Ian Johnston, Dominic Kwiatkowski, Cordelia Langford, John Sillitoe on behalf of the Wellcome Sanger Institute COVID-19 Surveillance Team |
| EPI_ISL_557900 | Lighthouse Lab in Alderley Park | Wellcome Sanger Institute for the COVID-19 Genomics UK | The Lighthouse Lab in Alderley Park and Alex Alderton, Roberto Amato, Sonia Goncalves, Ewan Harrison, David K. Jackson, Ian Johnston, Dominic |

| | | | |
|---|---|---|---|
| | | (COG-UK) consortium | Kwiatkowski, Cordelia Langford, John Sillitoe on behalf of the Wellcome Sanger Institute COVID-19 Surveillance Team (http://www.sanger.ac.uk/covid-team) |
| EPI_ISL_557902 | Lighthouse Lab in Alderley Park | Wellcome Sanger Institute for the COVID-19 Genomics UK (COG-UK) consortium | The Lighthouse Lab in Alderley Park and Alex Alderton, Roberto Amato, Sonia Goncalves, Ewan Harrison, David K. Jackson, Ian Johnston, Dominic Kwiatkowski, Cordelia Langford, John Sillitoe on behalf of the Wellcome Sanger Institute COVID-19 Surveillance Team |
| EPI_ISL_557905, EPI_ISL_557907 | Lighthouse Lab in Milton Keynes | Wellcome Sanger Institute for the COVID-19 Genomics UK (COG-UK) consortium | The Lighthouse Lab in Milton Keynes and Alex Alderton, Roberto Amato, Sonia Goncalves, Ewan Harrison, David K. Jackson, Ian Johnston, Dominic Kwiatkowski, Cordelia Langford, John Sillitoe on behalf of the Wellcome Sanger Institute COVID-19 Surveillance Team (http://www.sanger.ac.uk/covid-team) |
| EPI_ISL_557910 | Lighthouse Lab in Alderley Park | Wellcome Sanger Institute for the COVID-19 Genomics UK (COG-UK) consortium | The Lighthouse Lab in Alderley Park and Alex Alderton, Roberto Amato, Sonia Goncalves, Ewan Harrison, David K. Jackson, Ian Johnston, Dominic Kwiatkowski, Cordelia Langford, John Sillitoe on behalf of the Wellcome Sanger Institute COVID-19 Surveillance Team (http://www.sanger.ac.uk/covid-team) |
| EPI_ISL_557911 | Lighthouse Lab in Milton Keynes | Wellcome Sanger Institute for the COVID-19 Genomics UK (COG-UK) consortium | The Lighthouse Lab in Milton Keynes and Alex Alderton, Roberto Amato, Sonia Goncalves, Ewan Harrison, David K. Jackson, Ian Johnston, Dominic Kwiatkowski, Cordelia Langford, John Sillitoe on behalf of the Wellcome Sanger Institute COVID-19 Surveillance Team (http://www.sanger.ac.uk/covid-team) |
| EPI_ISL_557913 | Lighthouse Lab in Alderley Park | Wellcome Sanger Institute for the COVID-19 Genomics UK (COG-UK) consortium | The Lighthouse Lab in Alderley Park and Alex Alderton, Roberto Amato, Sonia Goncalves, Ewan Harrison, David K. Jackson, Ian Johnston, Dominic Kwiatkowski, Cordelia Langford, John Sillitoe on behalf of the Wellcome Sanger Institute COVID-19 Surveillance Team |
| EPI_ISL_557919, EPI_ISL_557925, EPI_ISL_557926, EPI_ISL_557927 | Lighthouse Lab in Milton Keynes | Wellcome Sanger Institute for the COVID-19 Genomics UK (COG-UK) consortium | The Lighthouse Lab in Milton Keynes and Alex Alderton, Roberto Amato, Sonia Goncalves, Ewan Harrison, David K. Jackson, Ian Johnston, Dominic Kwiatkowski, Cordelia Langford, John Sillitoe on behalf of the Wellcome Sanger Institute COVID-19 Surveillance Team (http://www.sanger.ac.uk/covid-team) |
| EPI_ISL_557936 | Lighthouse Lab in Alderley Park | Wellcome Sanger Institute for the COVID-19 Genomics UK (COG-UK) consortium | The Lighthouse Lab in Alderley Park and Alex Alderton, Roberto Amato, Sonia Goncalves, Ewan Harrison, David K. Jackson, Ian Johnston, Dominic Kwiatkowski, Cordelia Langford, John Sillitoe on behalf of the Wellcome Sanger Institute COVID-19 Surveillance Team (http://www.sanger.ac.uk/covid-team) |
| EPI_ISL_557938, EPI_ISL_557940, EPI_ISL_557941, EPI_ISL_557942, EPI_ISL_557944 | Lighthouse Lab in Milton Keynes | Wellcome Sanger Institute for the COVID-19 Genomics UK (COG-UK) consortium | The Lighthouse Lab in Milton Keynes and Alex Alderton, Roberto Amato, Sonia Goncalves, Ewan Harrison, David K. Jackson, Ian Johnston, Dominic Kwiatkowski, Cordelia Langford, John Sillitoe on behalf of the Wellcome Sanger Institute COVID-19 Surveillance Team (http://www.sanger.ac.uk/covid-team) |
| EPI_ISL_557945, EPI_ISL_557946 | Lighthouse Lab in Alderley Park | Wellcome Sanger Institute for the COVID-19 Genomics UK (COG-UK) consortium | The Lighthouse Lab in Alderley Park and Alex Alderton, Roberto Amato, Sonia Goncalves, Ewan Harrison, David K. Jackson, Ian Johnston, Dominic Kwiatkowski, Cordelia Langford, John Sillitoe on behalf of the Wellcome Sanger Institute COVID-19 Surveillance Team |
| EPI_ISL_557947, EPI_ISL_557949, EPI_ISL_557953, EPI_ISL_557954, EPI_ISL_557956 | Lighthouse Lab in Milton Keynes | Wellcome Sanger Institute for the COVID-19 Genomics UK (COG-UK) consortium | The Lighthouse Lab in Milton Keynes and Alex Alderton, Roberto Amato, Sonia Goncalves, Ewan Harrison, David K. Jackson, Ian Johnston, Dominic Kwiatkowski, Cordelia Langford, John Sillitoe on behalf of the Wellcome Sanger Institute COVID-19 Surveillance Team (http://www.sanger.ac.uk/covid-team) |
| EPI_ISL_557957 | Lighthouse Lab in Alderley Park | Wellcome Sanger Institute for the COVID-19 Genomics UK (COG-UK) consortium | The Lighthouse Lab in Alderley Park and Alex Alderton, Roberto Amato, Sonia Goncalves, Ewan Harrison, David K. Jackson, Ian Johnston, Dominic Kwiatkowski, Cordelia Langford, John Sillitoe on behalf of the Wellcome Sanger Institute COVID-19 Surveillance Team |
| EPI_ISL_557959 | Lighthouse Lab in Milton Keynes | Wellcome Sanger Institute for the COVID-19 Genomics UK (COG-UK) consortium | The Lighthouse Lab in Milton Keynes and Alex Alderton, Roberto Amato, Sonia Goncalves, Ewan Harrison, David K. Jackson, Ian Johnston, Dominic Kwiatkowski, Cordelia Langford, John Sillitoe on behalf of the Wellcome Sanger Institute COVID-19 Surveillance Team (http://www.sanger.ac.uk/covid-team) |
| EPI_ISL_557962 | Lighthouse Lab in Alderley Park | Wellcome Sanger Institute for the COVID-19 Genomics UK (COG-UK) consortium | The Lighthouse Lab in Alderley Park and Alex Alderton, Roberto Amato, Sonia Goncalves, Ewan Harrison, David K. Jackson, Ian Johnston, Dominic Kwiatkowski, Cordelia Langford, John Sillitoe on behalf of the Wellcome Sanger Institute COVID-19 Surveillance Team |
| EPI_ISL_557964, EPI_ISL_557965 | Lighthouse Lab in Milton Keynes | Wellcome Sanger Institute for the COVID-19 Genomics UK (COG-UK) consortium | The Lighthouse Lab in Milton Keynes and Alex Alderton, Roberto Amato, Sonia Goncalves, Ewan Harrison, David K. Jackson, Ian Johnston, Dominic Kwiatkowski, Cordelia Langford, John Sillitoe on behalf of the Wellcome Sanger Institute COVID-19 Surveillance Team (http://www.sanger.ac.uk/covid-team) |
| EPI_ISL_557969 | Lighthouse Lab in Alderley Park | Wellcome Sanger Institute for the COVID-19 Genomics UK (COG-UK) consortium | The Lighthouse Lab in Alderley Park and Alex Alderton, Roberto Amato, Sonia Goncalves, Ewan Harrison, David K. Jackson, Ian Johnston, Dominic Kwiatkowski, Cordelia Langford, John Sillitoe on behalf of the Wellcome Sanger Institute COVID-19 Surveillance Team |
| EPI_ISL_557972, EPI_ISL_557974 | Lighthouse Lab in Milton Keynes | Wellcome Sanger Institute for the COVID-19 Genomics UK (COG-UK) consortium | The Lighthouse Lab in Milton Keynes and Alex Alderton, Roberto Amato, Sonia Goncalves, Ewan Harrison, David K. Jackson, Ian Johnston, Dominic Kwiatkowski, Cordelia Langford, John Sillitoe on behalf of the Wellcome Sanger Institute COVID-19 Surveillance Team (http://www.sanger.ac.uk/covid-team) |
| EPI_ISL_557977 | Lighthouse Lab in Milton Keynes | Wellcome Sanger Institute for the COVID-19 Genomics UK (COG-UK) consortium | The Lighthouse Lab in Alderley Park and Alex Alderton, Roberto Amato, Sonia Goncalves, Ewan Harrison, David K. Jackson, Ian Johnston, Dominic Kwiatkowski, Cordelia Langford, John Sillitoe on behalf of the Wellcome Sanger Institute COVID-19 Surveillance Team |
| EPI_ISL_557979, EPI_ISL_557984 | Lighthouse Lab in Milton Keynes | Wellcome Sanger Institute for the COVID-19 Genomics UK (COG-UK) consortium | The Lighthouse Lab in Milton Keynes and Alex Alderton, Roberto Amato, Sonia Goncalves, Ewan Harrison, David K. Jackson, Ian Johnston, Dominic Kwiatkowski, Cordelia Langford, John Sillitoe on behalf of the Wellcome Sanger Institute COVID-19 Surveillance Team (http://www.sanger.ac.uk/covid-team) |
| EPI_ISL_557988 | Lighthouse Lab in Alderley Park | Wellcome Sanger Institute for the COVID-19 Genomics UK (COG-UK) consortium | The Lighthouse Lab in Alderley Park and Alex Alderton, Roberto Amato, Sonia Goncalves, Ewan Harrison, David K. Jackson, Ian Johnston, Dominic Kwiatkowski, Cordelia Langford, John Sillitoe on behalf of the Wellcome Sanger Institute COVID-19 Surveillance Team (http://www.sanger.ac.uk/covid-team) |
| EPI_ISL_557989 | Lighthouse Lab in Milton Keynes | Wellcome Sanger Institute for the COVID-19 Genomics UK (COG-UK) consortium | The Lighthouse Lab in Milton Keynes and Alex Alderton, Roberto Amato, Sonia Goncalves, Ewan Harrison, David K. Jackson, Ian Johnston, Dominic Kwiatkowski, Cordelia Langford, John Sillitoe on behalf of the Wellcome Sanger Institute COVID-19 Surveillance Team (http://www.sanger.ac.uk/covid-team) |
| EPI_ISL_557996, EPI_ISL_558002, EPI_ISL_558006 | Lighthouse Lab in Alderley Park | Wellcome Sanger Institute for the COVID-19 Genomics UK (COG-UK) consortium | The Lighthouse Lab in Alderley Park and Alex Alderton, Roberto Amato, Sonia Goncalves, Ewan Harrison, David K. Jackson, Ian Johnston, Dominic Kwiatkowski, Cordelia Langford, John Sillitoe on behalf of the Wellcome Sanger Institute COVID-19 Surveillance Team |
| EPI_ISL_558016, EPI_ISL_558017 | Lighthouse Lab in Alderley Park | Wellcome Sanger Institute for the COVID-19 Genomics UK (COG-UK) consortium | The Lighthouse Lab in Alderley Park and Alex Alderton, Roberto Amato, Sonia Goncalves, Ewan Harrison, David K. Jackson, Ian Johnston, Dominic Kwiatkowski, Cordelia Langford, John Sillitoe on behalf of the Wellcome Sanger Institute COVID-19 Surveillance Team (http://www.sanger.ac.uk/covid-team) |
| EPI_ISL_558020 | Lighthouse Lab in Alderley Park | Wellcome Sanger Institute for the COVID-19 Genomics UK (COG-UK) consortium | The Lighthouse Lab in Alderley Park and Alex Alderton, Roberto Amato, Sonia Goncalves, Ewan Harrison, David K. Jackson, Ian Johnston, Dominic Kwiatkowski, Cordelia Langford, John Sillitoe on behalf of the Wellcome Sanger Institute COVID-19 Surveillance Team |
| EPI_ISL_558021 | Lighthouse Lab in Milton Keynes | Wellcome Sanger Institute for the COVID-19 Genomics UK (COG-UK) consortium | The Lighthouse Lab in Milton Keynes and Alex Alderton, Roberto Amato, Sonia Goncalves, Ewan Harrison, David K. Jackson, Ian Johnston, Dominic Kwiatkowski, Cordelia Langford, John Sillitoe on behalf of the Wellcome Sanger Institute COVID-19 Surveillance Team (http://www.sanger.ac.uk/covid-team) |
| EPI_ISL_558023 | Lighthouse Lab in Alderley Park | Wellcome Sanger Institute for the COVID-19 Genomics UK (COG-UK) consortium | The Lighthouse Lab in Alderley Park and Alex Alderton, Roberto Amato, Sonia Goncalves, Ewan Harrison, David K. Jackson, Ian Johnston, Dominic Kwiatkowski, Cordelia Langford, John Sillitoe on behalf of the Wellcome Sanger Institute COVID-19 Surveillance Team (http://www.sanger.ac.uk/covid-team) |
| EPI_ISL_558026, EPI_ISL_558028, EPI_ISL_558030 | Lighthouse Lab in Milton Keynes | Wellcome Sanger Institute for the COVID-19 Genomics UK (COG-UK) consortium | The Lighthouse Lab in Milton Keynes and Alex Alderton, Roberto Amato, Sonia Goncalves, Ewan Harrison, David K. Jackson, Ian Johnston, Dominic Kwiatkowski, Cordelia Langford, John Sillitoe on behalf of the Wellcome Sanger Institute COVID-19 Surveillance Team (http://www.sanger.ac.uk/covid-team) |
| EPI_ISL_558031 | Lighthouse Lab in Alderley Park | Wellcome Sanger Institute for the COVID-19 Genomics UK (COG-UK) consortium | The Lighthouse Lab in Alderley Park and Alex Alderton, Roberto Amato, Sonia Goncalves, Ewan Harrison, David K. Jackson, Ian Johnston, Dominic Kwiatkowski, Cordelia Langford, John Sillitoe on behalf of the Wellcome Sanger Institute COVID-19 Surveillance Team |

| | | | |
|---|---|---|---|
| EPI_ISL_558037 | Lighthouse Lab in Milton Keynes | Wellcome Sanger Institute for the COVID-19 Genomics UK (COG-UK) consortium | The Lighthouse Lab in Milton Keynes and Alex Alderton, Roberto Amato, Sonia Goncalves, Ewan Harrison, David K. Jackson, Ian Johnston, Dominic Kwiatkowski, Cordelia Langford, John Sillitoe on behalf of the Wellcome Sanger Institute COVID-19 Surveillance Team (http://www.sanger.ac.uk/covid-team) |
| EPI_ISL_558038 | Lighthouse Lab in Alderley Park | Wellcome Sanger Institute for the COVID-19 Genomics UK (COG-UK) consortium | The Lighthouse Lab in Alderley Park and Alex Alderton, Roberto Amato, Sonia Goncalves, Ewan Harrison, David K. Jackson, Ian Johnston, Dominic Kwiatkowski, Cordelia Langford, John Sillitoe on behalf of the Wellcome Sanger Institute COVID-19 Surveillance Team |
| EPI_ISL_558041 | Lighthouse Lab in Milton Keynes | Wellcome Sanger Institute for the COVID-19 Genomics UK (COG-UK) consortium | The Lighthouse Lab in Milton Keynes and Alex Alderton, Roberto Amato, Sonia Goncalves, Ewan Harrison, David K. Jackson, Ian Johnston, Dominic Kwiatkowski, Cordelia Langford, John Sillitoe on behalf of the Wellcome Sanger Institute COVID-19 Surveillance Team (http://www.sanger.ac.uk/covid-team) |
| EPI_ISL_558052 | Lighthouse Lab in Alderley Park | Wellcome Sanger Institute for the COVID-19 Genomics UK (COG-UK) consortium | The Lighthouse Lab in Alderley Park and Alex Alderton, Roberto Amato, Sonia Goncalves, Ewan Harrison, David K. Jackson, Ian Johnston, Dominic Kwiatkowski, Cordelia Langford, John Sillitoe on behalf of the Wellcome Sanger Institute COVID-19 Surveillance Team (http://www.sanger.ac.uk/covid-team) |
| EPI_ISL_558055 | Lighthouse Lab in Alderley Park | Wellcome Sanger Institute for the COVID-19 Genomics UK (COG-UK) consortium | The Lighthouse Lab in Alderley Park and Alex Alderton, Roberto Amato, Sonia Goncalves, Ewan Harrison, David K. Jackson, Ian Johnston, Dominic Kwiatkowski, Cordelia Langford, John Sillitoe on behalf of the Wellcome Sanger Institute COVID-19 Surveillance Team |
| EPI_ISL_558056 | Lighthouse Lab in Alderley Park | Wellcome Sanger Institute for the COVID-19 Genomics UK (COG-UK) consortium | The Lighthouse Lab in Alderley Park and Alex Alderton, Roberto Amato, Sonia Goncalves, Ewan Harrison, David K. Jackson, Ian Johnston, Dominic Kwiatkowski, Cordelia Langford, John Sillitoe on behalf of the Wellcome Sanger Institute COVID-19 Surveillance Team (http://www.sanger.ac.uk/covid-team) |
| EPI_ISL_558058 | Lighthouse Lab in Milton Keynes | Wellcome Sanger Institute for the COVID-19 Genomics UK (COG-UK) consortium | The Lighthouse Lab in Milton Keynes and Alex Alderton, Roberto Amato, Sonia Goncalves, Ewan Harrison, David K. Jackson, Ian Johnston, Dominic Kwiatkowski, Cordelia Langford, John Sillitoe on behalf of the Wellcome Sanger Institute COVID-19 Surveillance Team (http://www.sanger.ac.uk/covid-team) |
| EPI_ISL_558064 | Lighthouse Lab in Alderley Park | Wellcome Sanger Institute for the COVID-19 Genomics UK (COG-UK) consortium | The Lighthouse Lab in Alderley Park and Alex Alderton, Roberto Amato, Sonia Goncalves, Ewan Harrison, David K. Jackson, Ian Johnston, Dominic Kwiatkowski, Cordelia Langford, John Sillitoe on behalf of the Wellcome Sanger Institute COVID-19 Surveillance Team (http://www.sanger.ac.uk/covid-team) |
| EPI_ISL_558068 | Lighthouse Lab in Milton Keynes | Wellcome Sanger Institute for the COVID-19 Genomics UK (COG-UK) consortium | The Lighthouse Lab in Milton Keynes and Alex Alderton, Roberto Amato, Sonia Goncalves, Ewan Harrison, David K. Jackson, Ian Johnston, Dominic Kwiatkowski, Cordelia Langford, John Sillitoe on behalf of the Wellcome Sanger Institute COVID-19 Surveillance Team (http://www.sanger.ac.uk/covid-team) |
| EPI_ISL_558071 | Lighthouse Lab in Alderley Park | Wellcome Sanger Institute for the COVID-19 Genomics UK (COG-UK) consortium | The Lighthouse Lab in Alderley Park and Alex Alderton, Roberto Amato, Sonia Goncalves, Ewan Harrison, David K. Jackson, Ian Johnston, Dominic Kwiatkowski, Cordelia Langford, John Sillitoe on behalf of the Wellcome Sanger Institute COVID-19 Surveillance Team |
| EPI_ISL_558073 | Lighthouse Lab in Alderley Park | Wellcome Sanger Institute for the COVID-19 Genomics UK (COG-UK) consortium | The Lighthouse Lab in Alderley Park and Alex Alderton, Roberto Amato, Sonia Goncalves, Ewan Harrison, David K. Jackson, Ian Johnston, Dominic Kwiatkowski, Cordelia Langford, John Sillitoe on behalf of the Wellcome Sanger Institute COVID-19 Surveillance Team (http://www.sanger.ac.uk/covid-team) |
| EPI_ISL_558079 | Lighthouse Lab in Alderley Park | Wellcome Sanger Institute for the COVID-19 Genomics UK (COG-UK) consortium | The Lighthouse Lab in Alderley Park and Alex Alderton, Roberto Amato, Sonia Goncalves, Ewan Harrison, David K. Jackson, Ian Johnston, Dominic Kwiatkowski, Cordelia Langford, John Sillitoe on behalf of the Wellcome Sanger Institute COVID-19 Surveillance Team |
| EPI_ISL_558089, EPI_ISL_558096, EPI_ISL_558103 | Lighthouse Lab in Milton Keynes | Wellcome Sanger Institute for the COVID-19 Genomics UK (COG-UK) consortium | The Lighthouse Lab in Milton Keynes and Alex Alderton, Roberto Amato, Sonia Goncalves, Ewan Harrison, David K. Jackson, Ian Johnston, Dominic Kwiatkowski, Cordelia Langford, John Sillitoe on behalf of the Wellcome Sanger Institute COVID-19 Surveillance Team (http://www.sanger.ac.uk/covid-team) |
| EPI_ISL_558107 | Lighthouse Lab in Alderley Park | Wellcome Sanger Institute for the COVID-19 Genomics UK (COG-UK) consortium | The Lighthouse Lab in Alderley Park and Alex Alderton, Roberto Amato, Sonia Goncalves, Ewan Harrison, David K. Jackson, Ian Johnston, Dominic Kwiatkowski, Cordelia Langford, John Sillitoe on behalf of the Wellcome Sanger Institute COVID-19 Surveillance Team (http://www.sanger.ac.uk/covid-team) |
| EPI_ISL_558109, EPI_ISL_558111, EPI_ISL_558113, EPI_ISL_558135, EPI_ISL_558143, EPI_ISL_558149 | Lighthouse Lab in Alderley Park | Wellcome Sanger Institute for the COVID-19 Genomics UK (COG-UK) consortium | The Lighthouse Lab in Alderley Park and Alex Alderton, Roberto Amato, Sonia Goncalves, Ewan Harrison, David K. Jackson, Ian Johnston, Dominic Kwiatkowski, Cordelia Langford, John Sillitoe on behalf of the Wellcome Sanger Institute COVID-19 Surveillance Team |
| EPI_ISL_558155 | Lighthouse Lab in Milton Keynes | Wellcome Sanger Institute for the COVID-19 Genomics UK (COG-UK) consortium | The Lighthouse Lab in Milton Keynes and Alex Alderton, Roberto Amato, Sonia Goncalves, Ewan Harrison, David K. Jackson, Ian Johnston, Dominic Kwiatkowski, Cordelia Langford, John Sillitoe on behalf of the Wellcome Sanger Institute COVID-19 Surveillance Team (http://www.sanger.ac.uk/covid-team) |
| EPI_ISL_558157, EPI_ISL_558158, EPI_ISL_558161, EPI_ISL_558163 | Lighthouse Lab in Alderley Park | Wellcome Sanger Institute for the COVID-19 Genomics UK (COG-UK) consortium | The Lighthouse Lab in Alderley Park and Alex Alderton, Roberto Amato, Sonia Goncalves, Ewan Harrison, David K. Jackson, Ian Johnston, Dominic Kwiatkowski, Cordelia Langford, John Sillitoe on behalf of the Wellcome Sanger Institute COVID-19 Surveillance Team |
| EPI_ISL_558165 | Lighthouse Lab in Alderley Park | Wellcome Sanger Institute for the COVID-19 Genomics UK (COG-UK) consortium | The Lighthouse Lab in Alderley Park and Alex Alderton, Roberto Amato, Sonia Goncalves, Ewan Harrison, David K. Jackson, Ian Johnston, Dominic Kwiatkowski, Cordelia Langford, John Sillitoe on behalf of the Wellcome Sanger Institute COVID-19 Surveillance Team (http://www.sanger.ac.uk/covid-team) |
| EPI_ISL_558174, EPI_ISL_558175, EPI_ISL_558179 | Lighthouse Lab in Alderley Park | Wellcome Sanger Institute for the COVID-19 Genomics UK (COG-UK) consortium | The Lighthouse Lab in Alderley Park and Alex Alderton, Roberto Amato, Sonia Goncalves, Ewan Harrison, David K. Jackson, Ian Johnston, Dominic Kwiatkowski, Cordelia Langford, John Sillitoe on behalf of the Wellcome Sanger Institute COVID-19 Surveillance Team |
| EPI_ISL_558195 | Lighthouse Lab in Milton Keynes | Wellcome Sanger Institute for the COVID-19 Genomics UK (COG-UK) consortium | The Lighthouse Lab in Milton Keynes and Alex Alderton, Roberto Amato, Sonia Goncalves, Ewan Harrison, David K. Jackson, Ian Johnston, Dominic Kwiatkowski, Cordelia Langford, John Sillitoe on behalf of the Wellcome Sanger Institute COVID-19 Surveillance Team (http://www.sanger.ac.uk/covid-team) |
| EPI_ISL_558214, EPI_ISL_558224, EPI_ISL_558238, EPI_ISL_558240 | Lighthouse Lab in Alderley Park | Wellcome Sanger Institute for the COVID-19 Genomics UK (COG-UK) consortium | The Lighthouse Lab in Alderley Park and Alex Alderton, Roberto Amato, Sonia Goncalves, Ewan Harrison, David K. Jackson, Ian Johnston, Dominic Kwiatkowski, Cordelia Langford, John Sillitoe on behalf of the Wellcome Sanger Institute COVID-19 Surveillance Team |
| EPI_ISL_558241 | Lighthouse Lab in Milton Keynes | Wellcome Sanger Institute for the COVID-19 Genomics UK (COG-UK) consortium | The Lighthouse Lab in Milton Keynes and Alex Alderton, Roberto Amato, Sonia Goncalves, Ewan Harrison, David K. Jackson, Ian Johnston, Dominic Kwiatkowski, Cordelia Langford, John Sillitoe on behalf of the Wellcome Sanger Institute COVID-19 Surveillance Team (http://www.sanger.ac.uk/covid-team) |
| EPI_ISL_558242 | Lighthouse Lab in Milton Keynes | Wellcome Sanger Institute for the COVID-19 Genomics UK (COG-UK) consortium | The Lighthouse Lab in Alderley Park and Alex Alderton, Roberto Amato, Sonia Goncalves, Ewan Harrison, David K. Jackson, Ian Johnston, Dominic Kwiatkowski, Cordelia Langford, John Sillitoe on behalf of the Wellcome Sanger Institute COVID-19 Surveillance Team |
| EPI_ISL_558243 | Lighthouse Lab in Milton Keynes | Wellcome Sanger Institute for the COVID-19 Genomics UK (COG-UK) consortium | The Lighthouse Lab in Milton Keynes and Alex Alderton, Roberto Amato, Sonia Goncalves, Ewan Harrison, David K. Jackson, Ian Johnston, Dominic Kwiatkowski, Cordelia Langford, John Sillitoe on behalf of the Wellcome Sanger Institute COVID-19 Surveillance Team (http://www.sanger.ac.uk/covid-team) |
| EPI_ISL_558244 | Lighthouse Lab in Alderley Park | Wellcome Sanger Institute for the COVID-19 Genomics UK (COG-UK) consortium | The Lighthouse Lab in Alderley Park and Alex Alderton, Roberto Amato, Sonia Goncalves, Ewan Harrison, David K. Jackson, Ian Johnston, Dominic Kwiatkowski, Cordelia Langford, John Sillitoe on behalf of the Wellcome Sanger Institute COVID-19 Surveillance Team |
| EPI_ISL_558245, EPI_ISL_558246, EPI_ISL_558247, EPI_ISL_558248, EPI_ISL_558249, EPI_ISL_558251, EPI_ISL_558253, EPI_ISL_558254, EPI_ISL_558255 | Lighthouse Lab in Milton Keynes | Wellcome Sanger Institute for the COVID-19 Genomics UK (COG-UK) consortium | The Lighthouse Lab in Milton Keynes and Alex Alderton, Roberto Amato, Sonia Goncalves, Ewan Harrison, David K. Jackson, Ian Johnston, Dominic Kwiatkowski, Cordelia Langford, John Sillitoe on behalf of the Wellcome Sanger Institute COVID-19 Surveillance Team (http://www.sanger.ac.uk/covid-team) |
| EPI_ISL_558256 | Lighthouse Lab in Alderley Park | Wellcome Sanger Institute for the COVID-19 Genomics UK (COG-UK) consortium | The Lighthouse Lab in Alderley Park and Alex Alderton, Roberto Amato, Sonia Goncalves, Ewan Harrison, David K. Jackson, Ian Johnston, Dominic Kwiatkowski, Cordelia Langford, John Sillitoe on behalf of the Wellcome Sanger Institute COVID-19 Surveillance Team |
| EPI_ISL_558257, EPI_ISL_558258 | Lighthouse Lab in Milton Keynes | Wellcome Sanger Institute for the COVID-19 Genomics UK | The Lighthouse Lab in Milton Keynes and Alex Alderton, Roberto Amato, Sonia Goncalves, Ewan Harrison, David K. Jackson, Ian Johnston, Dominic |

| | | | |
|---|---|---|---|
| | | (COG-UK) consortium | Kwiatkowski, Cordelia Langford, John Sillitoe on behalf of the Wellcome Sanger Institute COVID-19 Surveillance Team (http://www.sanger.ac.uk/covid-team) |
| EPI_ISL_558259, EPI_ISL_558260 | Lighthouse Lab in Alderley Park | Wellcome Sanger Institute for the COVID-19 Genomics UK (COG-UK) consortium | The Lighthouse Lab in Alderley Park and Alex Alderton, Roberto Amato, Sonia Goncalves, Ewan Harrison, David K. Jackson, Ian Johnston, Dominic Kwiatkowski, Cordelia Langford, John Sillitoe on behalf of the Wellcome Sanger Institute COVID-19 Surveillance Team |
| EPI_ISL_558262 | Lighthouse Lab in Milton Keynes | Wellcome Sanger Institute for the COVID-19 Genomics UK (COG-UK) consortium | The Lighthouse Lab in Milton Keynes and Alex Alderton, Roberto Amato, Sonia Goncalves, Ewan Harrison, David K. Jackson, Ian Johnston, Dominic Kwiatkowski, Cordelia Langford, John Sillitoe on behalf of the Wellcome Sanger Institute COVID-19 Surveillance Team (http://www.sanger.ac.uk/covid-team) |
| EPI_ISL_558263 | Lighthouse Lab in Alderley Park | Wellcome Sanger Institute for the COVID-19 Genomics UK (COG-UK) consortium | The Lighthouse Lab in Alderley Park and Alex Alderton, Roberto Amato, Sonia Goncalves, Ewan Harrison, David K. Jackson, Ian Johnston, Dominic Kwiatkowski, Cordelia Langford, John Sillitoe on behalf of the Wellcome Sanger Institute COVID-19 Surveillance Team |
| EPI_ISL_558264, EPI_ISL_558265, EPI_ISL_558266, EPI_ISL_558267 | Lighthouse Lab in Milton Keynes | Wellcome Sanger Institute for the COVID-19 Genomics UK (COG-UK) consortium | The Lighthouse Lab in Milton Keynes and Alex Alderton, Roberto Amato, Sonia Goncalves, Ewan Harrison, David K. Jackson, Ian Johnston, Dominic Kwiatkowski, Cordelia Langford, John Sillitoe on behalf of the Wellcome Sanger Institute COVID-19 Surveillance Team (http://www.sanger.ac.uk/covid-team) |
| EPI_ISL_558268, EPI_ISL_558269, EPI_ISL_558270 | Lighthouse Lab in Alderley Park | Wellcome Sanger Institute for the COVID-19 Genomics UK (COG-UK) consortium | The Lighthouse Lab in Alderley Park and Alex Alderton, Roberto Amato, Sonia Goncalves, Ewan Harrison, David K. Jackson, Ian Johnston, Dominic Kwiatkowski, Cordelia Langford, John Sillitoe on behalf of the Wellcome Sanger Institute COVID-19 Surveillance Team |
| EPI_ISL_558272, EPI_ISL_558273 | Lighthouse Lab in Milton Keynes | Wellcome Sanger Institute for the COVID-19 Genomics UK (COG-UK) consortium | The Lighthouse Lab in Milton Keynes and Alex Alderton, Roberto Amato, Sonia Goncalves, Ewan Harrison, David K. Jackson, Ian Johnston, Dominic Kwiatkowski, Cordelia Langford, John Sillitoe on behalf of the Wellcome Sanger Institute COVID-19 Surveillance Team (http://www.sanger.ac.uk/covid-team) |
| EPI_ISL_558274 | Lighthouse Lab in Alderley Park | Wellcome Sanger Institute for the COVID-19 Genomics UK (COG-UK) consortium | The Lighthouse Lab in Alderley Park and Alex Alderton, Roberto Amato, Sonia Goncalves, Ewan Harrison, David K. Jackson, Ian Johnston, Dominic Kwiatkowski, Cordelia Langford, John Sillitoe on behalf of the Wellcome Sanger Institute COVID-19 Surveillance Team |
| EPI_ISL_558275 | Lighthouse Lab in Milton Keynes | Wellcome Sanger Institute for the COVID-19 Genomics UK (COG-UK) consortium | The Lighthouse Lab in Milton Keynes and Alex Alderton, Roberto Amato, Sonia Goncalves, Ewan Harrison, David K. Jackson, Ian Johnston, Dominic Kwiatkowski, Cordelia Langford, John Sillitoe on behalf of the Wellcome Sanger Institute COVID-19 Surveillance Team (http://www.sanger.ac.uk/covid-team) |
| EPI_ISL_558276, EPI_ISL_558277 | Lighthouse Lab in Alderley Park | Wellcome Sanger Institute for the COVID-19 Genomics UK (COG-UK) consortium | The Lighthouse Lab in Alderley Park and Alex Alderton, Roberto Amato, Sonia Goncalves, Ewan Harrison, David K. Jackson, Ian Johnston, Dominic Kwiatkowski, Cordelia Langford, John Sillitoe on behalf of the Wellcome Sanger Institute COVID-19 Surveillance Team |
| EPI_ISL_558278, EPI_ISL_558279, EPI_ISL_558280, EPI_ISL_558281, EPI_ISL_558282, EPI_ISL_558284, EPI_ISL_558286, EPI_ISL_558287 | Lighthouse Lab in Milton Keynes | Wellcome Sanger Institute for the COVID-19 Genomics UK (COG-UK) consortium | The Lighthouse Lab in Milton Keynes and Alex Alderton, Roberto Amato, Sonia Goncalves, Ewan Harrison, David K. Jackson, Ian Johnston, Dominic Kwiatkowski, Cordelia Langford, John Sillitoe on behalf of the Wellcome Sanger Institute COVID-19 Surveillance Team (http://www.sanger.ac.uk/covid-team) |
| EPI_ISL_558288, EPI_ISL_558289 | Lighthouse Lab in Alderley Park | Wellcome Sanger Institute for the COVID-19 Genomics UK (COG-UK) consortium | The Lighthouse Lab in Alderley Park and Alex Alderton, Roberto Amato, Sonia Goncalves, Ewan Harrison, David K. Jackson, Ian Johnston, Dominic Kwiatkowski, Cordelia Langford, John Sillitoe on behalf of the Wellcome Sanger Institute COVID-19 Surveillance Team |
| EPI_ISL_558290 | Lighthouse Lab in Milton Keynes | Wellcome Sanger Institute for the COVID-19 Genomics UK (COG-UK) consortium | The Lighthouse Lab in Milton Keynes and Alex Alderton, Roberto Amato, Sonia Goncalves, Ewan Harrison, David K. Jackson, Ian Johnston, Dominic Kwiatkowski, Cordelia Langford, John Sillitoe on behalf of the Wellcome Sanger Institute COVID-19 Surveillance Team (http://www.sanger.ac.uk/covid-team) |
| EPI_ISL_558291 | Lighthouse Lab in Alderley Park | Wellcome Sanger Institute for the COVID-19 Genomics UK (COG-UK) consortium | The Lighthouse Lab in Alderley Park and Alex Alderton, Roberto Amato, Sonia Goncalves, Ewan Harrison, David K. Jackson, Ian Johnston, Dominic Kwiatkowski, Cordelia Langford, John Sillitoe on behalf of the Wellcome Sanger Institute COVID-19 Surveillance Team |
| EPI_ISL_558292, EPI_ISL_558293, EPI_ISL_558294, EPI_ISL_558295 | Lighthouse Lab in Milton Keynes | Wellcome Sanger Institute for the COVID-19 Genomics UK (COG-UK) consortium | The Lighthouse Lab in Milton Keynes and Alex Alderton, Roberto Amato, Sonia Goncalves, Ewan Harrison, David K. Jackson, Ian Johnston, Dominic Kwiatkowski, Cordelia Langford, John Sillitoe on behalf of the Wellcome Sanger Institute COVID-19 Surveillance Team (http://www.sanger.ac.uk/covid-team) |
| EPI_ISL_558296 | Lighthouse Lab in Milton Keynes | Wellcome Sanger Institute for the COVID-19 Genomics UK (COG-UK) consortium | The Lighthouse Lab in Alderley Park and Alex Alderton, Roberto Amato, Sonia Goncalves, Ewan Harrison, David K. Jackson, Ian Johnston, Dominic Kwiatkowski, Cordelia Langford, John Sillitoe on behalf of the Wellcome Sanger Institute COVID-19 Surveillance Team |
| EPI_ISL_558297, EPI_ISL_558298, EPI_ISL_558299, EPI_ISL_558300, EPI_ISL_558302, EPI_ISL_558303, EPI_ISL_558304 | Lighthouse Lab in Milton Keynes | Wellcome Sanger Institute for the COVID-19 Genomics UK (COG-UK) consortium | The Lighthouse Lab in Milton Keynes and Alex Alderton, Roberto Amato, Sonia Goncalves, Ewan Harrison, David K. Jackson, Ian Johnston, Dominic Kwiatkowski, Cordelia Langford, John Sillitoe on behalf of the Wellcome Sanger Institute COVID-19 Surveillance Team (http://www.sanger.ac.uk/covid-team) |
| EPI_ISL_558307 | Lighthouse Lab in Alderley Park | Wellcome Sanger Institute for the COVID-19 Genomics UK (COG-UK) consortium | The Lighthouse Lab in Alderley Park and Alex Alderton, Roberto Amato, Sonia Goncalves, Ewan Harrison, David K. Jackson, Ian Johnston, Dominic Kwiatkowski, Cordelia Langford, John Sillitoe on behalf of the Wellcome Sanger Institute COVID-19 Surveillance Team |
| EPI_ISL_558309, EPI_ISL_558310 | Lighthouse Lab in Milton Keynes | Wellcome Sanger Institute for the COVID-19 Genomics UK (COG-UK) consortium | The Lighthouse Lab in Milton Keynes and Alex Alderton, Roberto Amato, Sonia Goncalves, Ewan Harrison, David K. Jackson, Ian Johnston, Dominic Kwiatkowski, Cordelia Langford, John Sillitoe on behalf of the Wellcome Sanger Institute COVID-19 Surveillance Team (http://www.sanger.ac.uk/covid-team) |
| EPI_ISL_558312 | Lighthouse Lab in Alderley Park | Wellcome Sanger Institute for the COVID-19 Genomics UK (COG-UK) consortium | The Lighthouse Lab in Alderley Park and Alex Alderton, Roberto Amato, Sonia Goncalves, Ewan Harrison, David K. Jackson, Ian Johnston, Dominic Kwiatkowski, Cordelia Langford, John Sillitoe on behalf of the Wellcome Sanger Institute COVID-19 Surveillance Team |
| EPI_ISL_558313, EPI_ISL_558314, EPI_ISL_558315 | Lighthouse Lab in Milton Keynes | Wellcome Sanger Institute for the COVID-19 Genomics UK (COG-UK) consortium | The Lighthouse Lab in Milton Keynes and Alex Alderton, Roberto Amato, Sonia Goncalves, Ewan Harrison, David K. Jackson, Ian Johnston, Dominic Kwiatkowski, Cordelia Langford, John Sillitoe on behalf of the Wellcome Sanger Institute COVID-19 Surveillance Team (http://www.sanger.ac.uk/covid-team) |
| EPI_ISL_558316 | Lighthouse Lab in Alderley Park | Wellcome Sanger Institute for the COVID-19 Genomics UK (COG-UK) consortium | The Lighthouse Lab in Alderley Park and Alex Alderton, Roberto Amato, Sonia Goncalves, Ewan Harrison, David K. Jackson, Ian Johnston, Dominic Kwiatkowski, Cordelia Langford, John Sillitoe on behalf of the Wellcome Sanger Institute COVID-19 Surveillance Team |
| EPI_ISL_558317, EPI_ISL_558318, EPI_ISL_558319, EPI_ISL_558320 | Lighthouse Lab in Milton Keynes | Wellcome Sanger Institute for the COVID-19 Genomics UK (COG-UK) consortium | The Lighthouse Lab in Milton Keynes and Alex Alderton, Roberto Amato, Sonia Goncalves, Ewan Harrison, David K. Jackson, Ian Johnston, Dominic Kwiatkowski, Cordelia Langford, John Sillitoe on behalf of the Wellcome Sanger Institute COVID-19 Surveillance Team (http://www.sanger.ac.uk/covid-team) |
| EPI_ISL_558321 | Lighthouse Lab in Alderley Park | Wellcome Sanger Institute for the COVID-19 Genomics UK (COG-UK) consortium | The Lighthouse Lab in Alderley Park and Alex Alderton, Roberto Amato, Sonia Goncalves, Ewan Harrison, David K. Jackson, Ian Johnston, Dominic Kwiatkowski, Cordelia Langford, John Sillitoe on behalf of the Wellcome Sanger Institute COVID-19 Surveillance Team |
| EPI_ISL_558322, EPI_ISL_558323 | Lighthouse Lab in Milton Keynes | Wellcome Sanger Institute for the COVID-19 Genomics UK (COG-UK) consortium | The Lighthouse Lab in Milton Keynes and Alex Alderton, Roberto Amato, Sonia Goncalves, Ewan Harrison, David K. Jackson, Ian Johnston, Dominic Kwiatkowski, Cordelia Langford, John Sillitoe on behalf of the Wellcome Sanger Institute COVID-19 Surveillance Team (http://www.sanger.ac.uk/covid-team) |
| EPI_ISL_558324 | Lighthouse Lab in Alderley Park | Wellcome Sanger Institute for the COVID-19 Genomics UK (COG-UK) consortium | The Lighthouse Lab in Alderley Park and Alex Alderton, Roberto Amato, Sonia Goncalves, Ewan Harrison, David K. Jackson, Ian Johnston, Dominic Kwiatkowski, Cordelia Langford, John Sillitoe on behalf of the Wellcome Sanger Institute COVID-19 Surveillance Team |
| EPI_ISL_558325, EPI_ISL_558326, EPI_ISL_558327 | Lighthouse Lab in Milton Keynes | Wellcome Sanger Institute for the COVID-19 Genomics UK (COG-UK) consortium | The Lighthouse Lab in Milton Keynes and Alex Alderton, Roberto Amato, Sonia Goncalves, Ewan Harrison, David K. Jackson, Ian Johnston, Dominic Kwiatkowski, Cordelia Langford, John Sillitoe on behalf of the Wellcome Sanger Institute COVID-19 Surveillance Team (http://www.sanger.ac.uk/covid-team) |
| EPI_ISL_558329 | Lighthouse Lab in Alderley Park | Wellcome Sanger Institute for the COVID-19 Genomics UK (COG-UK) consortium | The Lighthouse Lab in Alderley Park and Alex Alderton, Roberto Amato, Sonia Goncalves, Ewan Harrison, David K. Jackson, Ian Johnston, Dominic Kwiatkowski, Cordelia Langford, John Sillitoe on behalf of the Wellcome Sanger Institute COVID-19 Surveillance Team |
| EPI_ISL_558331 | Lighthouse Lab in Milton Keynes | Wellcome Sanger Institute for the COVID-19 Genomics UK (COG-UK) consortium | The Lighthouse Lab in Milton Keynes and Alex Alderton, Roberto Amato, Sonia Goncalves, Ewan Harrison, David K. Jackson, Ian Johnston, Dominic Kwiatkowski, Cordelia Langford, John Sillitoe on behalf of the Wellcome Sanger Institute COVID-19 Surveillance Team |

| | | | |
|---|---|---|---|
| | | | (http://www.sanger.ac.uk/covid-team) |
| EPI_ISL_558332 | Lighthouse Lab in Alderley Park | Wellcome Sanger Institute for the COVID-19 Genomics UK (COG-UK) consortium | The Lighthouse Lab in Alderley Park and Alex Alderton, Roberto Amato, Sonia Goncalves, Ewan Harrison, David K. Jackson, Ian Johnston, Dominic Kwiatkowski, Cordelia Langford, John Sillitoe on behalf of the Wellcome Sanger Institute COVID-19 Surveillance Team |
| EPI_ISL_558333 | Lighthouse Lab in Milton Keynes | Wellcome Sanger Institute for the COVID-19 Genomics UK (COG-UK) consortium | The Lighthouse Lab in Alderley Park and Alex Alderton, Roberto Amato, Sonia Goncalves, Ewan Harrison, David K. Jackson, Ian Johnston, Dominic Kwiatkowski, Cordelia Langford, John Sillitoe on behalf of the Wellcome Sanger Institute COVID-19 Surveillance Team |
| EPI_ISL_558334 | Lighthouse Lab in Milton Keynes | Wellcome Sanger Institute for the COVID-19 Genomics UK (COG-UK) consortium | The Lighthouse Lab in Milton Keynes and Alex Alderton, Roberto Amato, Sonia Goncalves, Ewan Harrison, David K. Jackson, Ian Johnston, Dominic Kwiatkowski, Cordelia Langford, John Sillitoe on behalf of the Wellcome Sanger Institute COVID-19 Surveillance Team (http://www.sanger.ac.uk/covid-team) |
| EPI_ISL_558335 | Lighthouse Lab in Alderley Park | Wellcome Sanger Institute for the COVID-19 Genomics UK (COG-UK) consortium | The Lighthouse Lab in Alderley Park and Alex Alderton, Roberto Amato, Sonia Goncalves, Ewan Harrison, David K. Jackson, Ian Johnston, Dominic Kwiatkowski, Cordelia Langford, John Sillitoe on behalf of the Wellcome Sanger Institute COVID-19 Surveillance Team |
| EPI_ISL_558336, EPI_ISL_558337 | Lighthouse Lab in Milton Keynes | Wellcome Sanger Institute for the COVID-19 Genomics UK (COG-UK) consortium | The Lighthouse Lab in Milton Keynes and Alex Alderton, Roberto Amato, Sonia Goncalves, Ewan Harrison, David K. Jackson, Ian Johnston, Dominic Kwiatkowski, Cordelia Langford, John Sillitoe on behalf of the Wellcome Sanger Institute COVID-19 Surveillance Team (http://www.sanger.ac.uk/covid-team) |
| EPI_ISL_558338 | Lighthouse Lab in Alderley Park | Wellcome Sanger Institute for the COVID-19 Genomics UK (COG-UK) consortium | The Lighthouse Lab in Alderley Park and Alex Alderton, Roberto Amato, Sonia Goncalves, Ewan Harrison, David K. Jackson, Ian Johnston, Dominic Kwiatkowski, Cordelia Langford, John Sillitoe on behalf of the Wellcome Sanger Institute COVID-19 Surveillance Team |
| EPI_ISL_558340, EPI_ISL_558341, EPI_ISL_558342 | Lighthouse Lab in Milton Keynes | Wellcome Sanger Institute for the COVID-19 Genomics UK (COG-UK) consortium | The Lighthouse Lab in Milton Keynes and Alex Alderton, Roberto Amato, Sonia Goncalves, Ewan Harrison, David K. Jackson, Ian Johnston, Dominic Kwiatkowski, Cordelia Langford, John Sillitoe on behalf of the Wellcome Sanger Institute COVID-19 Surveillance Team (http://www.sanger.ac.uk/covid-team) |
| EPI_ISL_558343 | Lighthouse Lab in Alderley Park | Wellcome Sanger Institute for the COVID-19 Genomics UK (COG-UK) consortium | The Lighthouse Lab in Alderley Park and Alex Alderton, Roberto Amato, Sonia Goncalves, Ewan Harrison, David K. Jackson, Ian Johnston, Dominic Kwiatkowski, Cordelia Langford, John Sillitoe on behalf of the Wellcome Sanger Institute COVID-19 Surveillance Team |
| EPI_ISL_558344 | Lighthouse Lab in Milton Keynes | Wellcome Sanger Institute for the COVID-19 Genomics UK (COG-UK) consortium | The Lighthouse Lab in Milton Keynes and Alex Alderton, Roberto Amato, Sonia Goncalves, Ewan Harrison, David K. Jackson, Ian Johnston, Dominic Kwiatkowski, Cordelia Langford, John Sillitoe on behalf of the Wellcome Sanger Institute COVID-19 Surveillance Team (http://www.sanger.ac.uk/covid-team) |
| EPI_ISL_558345 | Lighthouse Lab in Alderley Park | Wellcome Sanger Institute for the COVID-19 Genomics UK (COG-UK) consortium | The Lighthouse Lab in Alderley Park and Alex Alderton, Roberto Amato, Sonia Goncalves, Ewan Harrison, David K. Jackson, Ian Johnston, Dominic Kwiatkowski, Cordelia Langford, John Sillitoe on behalf of the Wellcome Sanger Institute COVID-19 Surveillance Team |
| EPI_ISL_558346, EPI_ISL_558347 | Lighthouse Lab in Milton Keynes | Wellcome Sanger Institute for the COVID-19 Genomics UK (COG-UK) consortium | The Lighthouse Lab in Milton Keynes and Alex Alderton, Roberto Amato, Sonia Goncalves, Ewan Harrison, David K. Jackson, Ian Johnston, Dominic Kwiatkowski, Cordelia Langford, John Sillitoe on behalf of the Wellcome Sanger Institute COVID-19 Surveillance Team (http://www.sanger.ac.uk/covid-team) |
| EPI_ISL_558348 | Lighthouse Lab in Alderley Park | Wellcome Sanger Institute for the COVID-19 Genomics UK (COG-UK) consortium | The Lighthouse Lab in Alderley Park and Alex Alderton, Roberto Amato, Sonia Goncalves, Ewan Harrison, David K. Jackson, Ian Johnston, Dominic Kwiatkowski, Cordelia Langford, John Sillitoe on behalf of the Wellcome Sanger Institute COVID-19 Surveillance Team |
| EPI_ISL_558349 | Lighthouse Lab in Milton Keynes | Wellcome Sanger Institute for the COVID-19 Genomics UK (COG-UK) consortium | The Lighthouse Lab in Milton Keynes and Alex Alderton, Roberto Amato, Sonia Goncalves, Ewan Harrison, David K. Jackson, Ian Johnston, Dominic Kwiatkowski, Cordelia Langford, John Sillitoe on behalf of the Wellcome Sanger Institute COVID-19 Surveillance Team (http://www.sanger.ac.uk/covid-team) |
| EPI_ISL_558350 | Lighthouse Lab in Alderley Park | Wellcome Sanger Institute for the COVID-19 Genomics UK (COG-UK) consortium | The Lighthouse Lab in Alderley Park and Alex Alderton, Roberto Amato, Sonia Goncalves, Ewan Harrison, David K. Jackson, Ian Johnston, Dominic Kwiatkowski, Cordelia Langford, John Sillitoe on behalf of the Wellcome Sanger Institute COVID-19 Surveillance Team |
| EPI_ISL_558351, EPI_ISL_558352, EPI_ISL_558353 | Lighthouse Lab in Milton Keynes | Wellcome Sanger Institute for the COVID-19 Genomics UK (COG-UK) consortium | The Lighthouse Lab in Milton Keynes and Alex Alderton, Roberto Amato, Sonia Goncalves, Ewan Harrison, David K. Jackson, Ian Johnston, Dominic Kwiatkowski, Cordelia Langford, John Sillitoe on behalf of the Wellcome Sanger Institute COVID-19 Surveillance Team (http://www.sanger.ac.uk/covid-team) |
| EPI_ISL_558354 | Lighthouse Lab in Alderley Park | Wellcome Sanger Institute for the COVID-19 Genomics UK (COG-UK) consortium | The Lighthouse Lab in Alderley Park and Alex Alderton, Roberto Amato, Sonia Goncalves, Ewan Harrison, David K. Jackson, Ian Johnston, Dominic Kwiatkowski, Cordelia Langford, John Sillitoe on behalf of the Wellcome Sanger Institute COVID-19 Surveillance Team (http://www.sanger.ac.uk/covid-team) |
| EPI_ISL_558355 | Lighthouse Lab in Alderley Park | Wellcome Sanger Institute for the COVID-19 Genomics UK (COG-UK) consortium | The Lighthouse Lab in Alderley Park and Alex Alderton, Roberto Amato, Sonia Goncalves, Ewan Harrison, David K. Jackson, Ian Johnston, Dominic Kwiatkowski, Cordelia Langford, John Sillitoe on behalf of the Wellcome Sanger Institute COVID-19 Surveillance Team |
| EPI_ISL_558357, EPI_ISL_558358 | Lighthouse Lab in Milton Keynes | Wellcome Sanger Institute for the COVID-19 Genomics UK (COG-UK) consortium | The Lighthouse Lab in Milton Keynes and Alex Alderton, Roberto Amato, Sonia Goncalves, Ewan Harrison, David K. Jackson, Ian Johnston, Dominic Kwiatkowski, Cordelia Langford, John Sillitoe on behalf of the Wellcome Sanger Institute COVID-19 Surveillance Team (http://www.sanger.ac.uk/covid-team) |
| EPI_ISL_558359 | Lighthouse Lab in Milton Keynes | Wellcome Sanger Institute for the COVID-19 Genomics UK (COG-UK) consortium | The Lighthouse Lab in Alderley Park and Alex Alderton, Roberto Amato, Sonia Goncalves, Ewan Harrison, David K. Jackson, Ian Johnston, Dominic Kwiatkowski, Cordelia Langford, John Sillitoe on behalf of the Wellcome Sanger Institute COVID-19 Surveillance Team |
| EPI_ISL_558360, EPI_ISL_558361, EPI_ISL_558363 | Lighthouse Lab in Milton Keynes | Wellcome Sanger Institute for the COVID-19 Genomics UK (COG-UK) consortium | The Lighthouse Lab in Milton Keynes and Alex Alderton, Roberto Amato, Sonia Goncalves, Ewan Harrison, David K. Jackson, Ian Johnston, Dominic Kwiatkowski, Cordelia Langford, John Sillitoe on behalf of the Wellcome Sanger Institute COVID-19 Surveillance Team (http://www.sanger.ac.uk/covid-team) |
| EPI_ISL_558364 | Lighthouse Lab in Alderley Park | Wellcome Sanger Institute for the COVID-19 Genomics UK (COG-UK) consortium | The Lighthouse Lab in Alderley Park and Alex Alderton, Roberto Amato, Sonia Goncalves, Ewan Harrison, David K. Jackson, Ian Johnston, Dominic Kwiatkowski, Cordelia Langford, John Sillitoe on behalf of the Wellcome Sanger Institute COVID-19 Surveillance Team |
| EPI_ISL_558365, EPI_ISL_558366, EPI_ISL_558367 | Lighthouse Lab in Milton Keynes | Wellcome Sanger Institute for the COVID-19 Genomics UK (COG-UK) consortium | The Lighthouse Lab in Milton Keynes and Alex Alderton, Roberto Amato, Sonia Goncalves, Ewan Harrison, David K. Jackson, Ian Johnston, Dominic Kwiatkowski, Cordelia Langford, John Sillitoe on behalf of the Wellcome Sanger Institute COVID-19 Surveillance Team (http://www.sanger.ac.uk/covid-team) |
| EPI_ISL_558368 | Lighthouse Lab in Alderley Park | Wellcome Sanger Institute for the COVID-19 Genomics UK (COG-UK) consortium | The Lighthouse Lab in Alderley Park and Alex Alderton, Roberto Amato, Sonia Goncalves, Ewan Harrison, David K. Jackson, Ian Johnston, Dominic Kwiatkowski, Cordelia Langford, John Sillitoe on behalf of the Wellcome Sanger Institute COVID-19 Surveillance Team (http://www.sanger.ac.uk/covid-team) |
| EPI_ISL_558369, EPI_ISL_558371, EPI_ISL_558372, EPI_ISL_558373, EPI_ISL_558374, EPI_ISL_558375, EPI_ISL_558376, EPI_ISL_558377, EPI_ISL_558378 | Lighthouse Lab in Milton Keynes | Wellcome Sanger Institute for the COVID-19 Genomics UK (COG-UK) consortium | The Lighthouse Lab in Milton Keynes and Alex Alderton, Roberto Amato, Sonia Goncalves, Ewan Harrison, David K. Jackson, Ian Johnston, Dominic Kwiatkowski, Cordelia Langford, John Sillitoe on behalf of the Wellcome Sanger Institute COVID-19 Surveillance Team (http://www.sanger.ac.uk/covid-team) |
| EPI_ISL_558379 | Lighthouse Lab in Alderley Park | Wellcome Sanger Institute for the COVID-19 Genomics UK (COG-UK) consortium | The Lighthouse Lab in Alderley Park and Alex Alderton, Roberto Amato, Sonia Goncalves, Ewan Harrison, David K. Jackson, Ian Johnston, Dominic Kwiatkowski, Cordelia Langford, John Sillitoe on behalf of the Wellcome Sanger Institute COVID-19 Surveillance Team |
| EPI_ISL_558380, EPI_ISL_558381, EPI_ISL_558382, EPI_ISL_558383, EPI_ISL_558384, EPI_ISL_558385, EPI_ISL_558386, EPI_ISL_558388 | Lighthouse Lab in Milton Keynes | Wellcome Sanger Institute for the COVID-19 Genomics UK (COG-UK) consortium | The Lighthouse Lab in Milton Keynes and Alex Alderton, Roberto Amato, Sonia Goncalves, Ewan Harrison, David K. Jackson, Ian Johnston, Dominic Kwiatkowski, Cordelia Langford, John Sillitoe on behalf of the Wellcome Sanger Institute COVID-19 Surveillance Team (http://www.sanger.ac.uk/covid-team) |
| EPI_ISL_558389 | Lighthouse Lab in Alderley Park | Wellcome Sanger Institute for the COVID-19 Genomics UK (COG-UK) consortium | The Lighthouse Lab in Alderley Park and Alex Alderton, Roberto Amato, Sonia Goncalves, Ewan Harrison, David K. Jackson, Ian Johnston, Dominic Kwiatkowski, Cordelia Langford, John Sillitoe on behalf of the Wellcome Sanger Institute COVID-19 Surveillance Team |
| EPI_ISL_558390, EPI_ISL_558391, | Lighthouse Lab in Milton Keynes | Wellcome Sanger Institute for the COVID-19 Genomics UK | The Lighthouse Lab in Milton Keynes and Alex Alderton, Roberto Amato, Sonia Goncalves, Ewan Harrison, David K. Jackson, Ian Johnston, Dominic |

| | | | |
|---|---|---|---|
| EPI_ISL_558393 | | (COG-UK) consortium | Kwiatkowski, Cordelia Langford, John Sillitoe on behalf of the Wellcome Sanger Institute COVID-19 Surveillance Team (http://www.sanger.ac.uk/covid-team) |
| EPI_ISL_558394, EPI_ISL_558395, EPI_ISL_558396 | Lighthouse Lab in Alderley Park | Wellcome Sanger Institute for the COVID-19 Genomics UK (COG-UK) consortium | The Lighthouse Lab in Alderley Park and Alex Alderton, Roberto Amato, Sonia Goncalves, Ewan Harrison, David K. Jackson, Ian Johnston, Dominic Kwiatkowski, Cordelia Langford, John Sillitoe on behalf of the Wellcome Sanger Institute COVID-19 Surveillance Team |
| EPI_ISL_558397, EPI_ISL_558399, EPI_ISL_558400, EPI_ISL_558401, EPI_ISL_558403 | Lighthouse Lab in Milton Keynes | Wellcome Sanger Institute for the COVID-19 Genomics UK (COG-UK) consortium | The Lighthouse Lab in Milton Keynes and Alex Alderton, Roberto Amato, Sonia Goncalves, Ewan Harrison, David K. Jackson, Ian Johnston, Dominic Kwiatkowski, Cordelia Langford, John Sillitoe on behalf of the Wellcome Sanger Institute COVID-19 Surveillance Team (http://www.sanger.ac.uk/covid-team) |
| EPI_ISL_558415, EPI_ISL_558439, EPI_ISL_558440, EPI_ISL_558441, EPI_ISL_558444, EPI_ISL_558446, EPI_ISL_558458, EPI_ISL_558471, EPI_ISL_558474, EPI_ISL_558478, EPI_ISL_558485, EPI_ISL_558489, EPI_ISL_558490, EPI_ISL_558494, EPI_ISL_558509, EPI_ISL_558512, EPI_ISL_558517, EPI_ISL_558529, EPI_ISL_558530 | | | |
| see above | Lighthouse Lab in Alderley Park | Wellcome Sanger Institute for the COVID-19 Genomics UK (COG-UK) consortium | The Lighthouse Lab in Alderley Park and Alex Alderton, Roberto Amato, Sonia Goncalves, Ewan Harrison, David K. Jackson, Ian Johnston, Dominic Kwiatkowski, Cordelia Langford, John Sillitoe on behalf of the Wellcome Sanger Institute COVID-19 Surveillance Team |
| EPI_ISL_558545 | Lighthouse Lab in Alderley Park | Wellcome Sanger Institute for the COVID-19 Genomics UK (COG-UK) consortium | The Lighthouse Lab in Alderley Park and Alex Alderton, Roberto Amato, Sonia Goncalves, Ewan Harrison, David K. Jackson, Ian Johnston, Dominic Kwiatkowski, Cordelia Langford, John Sillitoe on behalf of the Wellcome Sanger Institute COVID-19 Surveillance Team (http://www.sanger.ac.uk/covid-team) |
| EPI_ISL_558553, EPI_ISL_558554, EPI_ISL_558561, EPI_ISL_558567, EPI_ISL_558573, EPI_ISL_558574 | Lighthouse Lab in Alderley Park | Wellcome Sanger Institute for the COVID-19 Genomics UK (COG-UK) consortium | The Lighthouse Lab in Alderley Park and Alex Alderton, Roberto Amato, Sonia Goncalves, Ewan Harrison, David K. Jackson, Ian Johnston, Dominic Kwiatkowski, Cordelia Langford, John Sillitoe on behalf of the Wellcome Sanger Institute COVID-19 Surveillance Team |
| EPI_ISL_558577 | Lighthouse Lab in Alderley Park | Wellcome Sanger Institute for the COVID-19 Genomics UK (COG-UK) consortium | The Lighthouse Lab in Alderley Park and Alex Alderton, Roberto Amato, Sonia Goncalves, Ewan Harrison, David K. Jackson, Ian Johnston, Dominic Kwiatkowski, Cordelia Langford, John Sillitoe on behalf of the Wellcome Sanger Institute COVID-19 Surveillance Team (http://www.sanger.ac.uk/covid-team) |
| EPI_ISL_558579, EPI_ISL_558585, EPI_ISL_558586, EPI_ISL_558587 | Lighthouse Lab in Alderley Park | Wellcome Sanger Institute for the COVID-19 Genomics UK (COG-UK) consortium | The Lighthouse Lab in Alderley Park and Alex Alderton, Roberto Amato, Sonia Goncalves, Ewan Harrison, David K. Jackson, Ian Johnston, Dominic Kwiatkowski, Cordelia Langford, John Sillitoe on behalf of the Wellcome Sanger Institute COVID-19 Surveillance Team |
| EPI_ISL_558588 | Lighthouse Lab in Alderley Park | Wellcome Sanger Institute for the COVID-19 Genomics UK (COG-UK) consortium | The Lighthouse Lab in Alderley Park and Alex Alderton, Roberto Amato, Sonia Goncalves, Ewan Harrison, David K. Jackson, Ian Johnston, Dominic Kwiatkowski, Cordelia Langford, John Sillitoe on behalf of the Wellcome Sanger Institute COVID-19 Surveillance Team (http://www.sanger.ac.uk/covid-team) |
| EPI_ISL_558589 | Lighthouse Lab in Alderley Park | Wellcome Sanger Institute for the COVID-19 Genomics UK (COG-UK) consortium | The Lighthouse Lab in Alderley Park and Alex Alderton, Roberto Amato, Sonia Goncalves, Ewan Harrison, David K. Jackson, Ian Johnston, Dominic Kwiatkowski, Cordelia Langford, John Sillitoe on behalf of the Wellcome Sanger Institute COVID-19 Surveillance Team |
| EPI_ISL_558592 | Lighthouse Lab in Alderley Park | Wellcome Sanger Institute for the COVID-19 Genomics UK (COG-UK) consortium | The Lighthouse Lab in Alderley Park and Alex Alderton, Roberto Amato, Sonia Goncalves, Ewan Harrison, David K. Jackson, Ian Johnston, Dominic Kwiatkowski, Cordelia Langford, John Sillitoe on behalf of the Wellcome Sanger Institute COVID-19 Surveillance Team (http://www.sanger.ac.uk/covid-team) |
| EPI_ISL_558594 | Lighthouse Lab in Alderley Park | Wellcome Sanger Institute for the COVID-19 Genomics UK (COG-UK) consortium | The Lighthouse Lab in Alderley Park and Alex Alderton, Roberto Amato, Sonia Goncalves, Ewan Harrison, David K. Jackson, Ian Johnston, Dominic Kwiatkowski, Cordelia Langford, John Sillitoe on behalf of the Wellcome Sanger Institute COVID-19 Surveillance Team |
| EPI_ISL_558601 | Lighthouse Lab in Alderley Park | Wellcome Sanger Institute for the COVID-19 Genomics UK (COG-UK) consortium | The Lighthouse Lab in Alderley Park and Alex Alderton, Roberto Amato, Sonia Goncalves, Ewan Harrison, David K. Jackson, Ian Johnston, Dominic Kwiatkowski, Cordelia Langford, John Sillitoe on behalf of the Wellcome Sanger Institute COVID-19 Surveillance Team (http://www.sanger.ac.uk/covid-team) |
| EPI_ISL_558608 | Lighthouse Lab in Alderley Park | Wellcome Sanger Institute for the COVID-19 Genomics UK (COG-UK) consortium | The Lighthouse Lab in Alderley Park and Alex Alderton, Roberto Amato, Sonia Goncalves, Ewan Harrison, David K. Jackson, Ian Johnston, Dominic Kwiatkowski, Cordelia Langford, John Sillitoe on behalf of the Wellcome Sanger Institute COVID-19 Surveillance Team |
| EPI_ISL_558609 | Lighthouse Lab in Alderley Park | Wellcome Sanger Institute for the COVID-19 Genomics UK (COG-UK) consortium | The Lighthouse Lab in Alderley Park and Alex Alderton, Roberto Amato, Sonia Goncalves, Ewan Harrison, David K. Jackson, Ian Johnston, Dominic Kwiatkowski, Cordelia Langford, John Sillitoe on behalf of the Wellcome Sanger Institute COVID-19 Surveillance Team (http://www.sanger.ac.uk/covid-team) |
| EPI_ISL_558618, EPI_ISL_558619, EPI_ISL_558620, EPI_ISL_558621, EPI_ISL_558622, EPI_ISL_558623, EPI_ISL_558624, EPI_ISL_558625, EPI_ISL_558626, EPI_ISL_558627, EPI_ISL_558628, EPI_ISL_558629, EPI_ISL_558630, EPI_ISL_558631, EPI_ISL_558632, EPI_ISL_558633, EPI_ISL_558634, EPI_ISL_558635, EPI_ISL_558636, EPI_ISL_558637, EPI_ISL_558639, EPI_ISL_558640, EPI_ISL_558641 | | | |
| see above | Lighthouse Lab in Alderley Park | Wellcome Sanger Institute for the COVID-19 Genomics UK (COG-UK) consortium | The Lighthouse Lab in Alderley Park and Alex Alderton, Roberto Amato, Sonia Goncalves, Ewan Harrison, David K. Jackson, Ian Johnston, Dominic Kwiatkowski, Cordelia Langford, John Sillitoe on behalf of the Wellcome Sanger Institute COVID-19 Surveillance Team |
| EPI_ISL_558642 | Lighthouse Lab in Alderley Park | Wellcome Sanger Institute for the COVID-19 Genomics UK (COG-UK) consortium | The Lighthouse Lab in Alderley Park and Alex Alderton, Roberto Amato, Sonia Goncalves, Ewan Harrison, David K. Jackson, Ian Johnston, Dominic Kwiatkowski, Cordelia Langford, John Sillitoe on behalf of the Wellcome Sanger Institute COVID-19 Surveillance Team (http://www.sanger.ac.uk/covid-team) |
| EPI_ISL_558643, EPI_ISL_558644, EPI_ISL_558645, EPI_ISL_558646, EPI_ISL_558647, EPI_ISL_558648, EPI_ISL_558649, EPI_ISL_558650, EPI_ISL_558651, EPI_ISL_558652, EPI_ISL_558653, EPI_ISL_558654, EPI_ISL_558655, EPI_ISL_558656, EPI_ISL_558657, EPI_ISL_558658, EPI_ISL_558659, EPI_ISL_558660, EPI_ISL_558661, EPI_ISL_558662, EPI_ISL_558663, EPI_ISL_558664, EPI_ISL_558665, EPI_ISL_558666, EPI_ISL_558667, EPI_ISL_558668, EPI_ISL_558669, EPI_ISL_558670, EPI_ISL_558671, EPI_ISL_558672, EPI_ISL_558673, EPI_ISL_558674, EPI_ISL_558675, EPI_ISL_558676, EPI_ISL_558677, EPI_ISL_558678, EPI_ISL_558679, EPI_ISL_558680, EPI_ISL_558681, EPI_ISL_558682, EPI_ISL_558683, EPI_ISL_558684, EPI_ISL_558685, EPI_ISL_558686, EPI_ISL_558687, EPI_ISL_558688, EPI_ISL_558689, EPI_ISL_558690, EPI_ISL_558691, EPI_ISL_558692, EPI_ISL_558693, EPI_ISL_558694, EPI_ISL_558695, EPI_ISL_558696, EPI_ISL_558697, EPI_ISL_558698, EPI_ISL_558699, EPI_ISL_558700 | | | |
| see above | Lighthouse Lab in Alderley Park | Wellcome Sanger Institute for the COVID-19 Genomics UK (COG-UK) consortium | The Lighthouse Lab in Alderley Park and Alex Alderton, Roberto Amato, Sonia Goncalves, Ewan Harrison, David K. Jackson, Ian Johnston, Dominic Kwiatkowski, Cordelia Langford, John Sillitoe on behalf of the Wellcome Sanger Institute COVID-19 Surveillance Team |
| EPI_ISL_558701 | Lighthouse Lab in Alderley Park | Wellcome Sanger Institute for the COVID-19 Genomics UK (COG-UK) consortium | The Lighthouse Lab in Alderley Park and Alex Alderton, Roberto Amato, Sonia Goncalves, Ewan Harrison, David K. Jackson, Ian Johnston, Dominic Kwiatkowski, Cordelia Langford, John Sillitoe on behalf of the Wellcome Sanger Institute COVID-19 Surveillance Team (http://www.sanger.ac.uk/covid-team) |
| EPI_ISL_558702, EPI_ISL_558703, EPI_ISL_558704, EPI_ISL_558705, EPI_ISL_558706, EPI_ISL_558707, EPI_ISL_558708, EPI_ISL_558709, EPI_ISL_558710, EPI_ISL_558711, EPI_ISL_558712, EPI_ISL_558713, EPI_ISL_558714, EPI_ISL_558715, EPI_ISL_558716, EPI_ISL_558717, EPI_ISL_558718, EPI_ISL_558719, EPI_ISL_558720, EPI_ISL_558721, EPI_ISL_558722, EPI_ISL_558723, EPI_ISL_558724, EPI_ISL_558725, EPI_ISL_558726, EPI_ISL_558727, EPI_ISL_558728, EPI_ISL_558729, EPI_ISL_558730 | | | |
| see above | Lighthouse Lab in Alderley Park | Wellcome Sanger Institute for the COVID-19 Genomics UK (COG-UK) consortium | The Lighthouse Lab in Alderley Park and Alex Alderton, Roberto Amato, Sonia Goncalves, Ewan Harrison, David K. Jackson, Ian Johnston, Dominic Kwiatkowski, Cordelia Langford, John Sillitoe on behalf of the Wellcome Sanger Institute COVID-19 Surveillance Team |
| EPI_ISL_558731 | Lighthouse Lab in Milton Keynes | Wellcome Sanger Institute for the COVID-19 Genomics UK (COG-UK) consortium | The Lighthouse Lab in Milton Keynes and Alex Alderton, Roberto Amato, Sonia Goncalves, Ewan Harrison, David K. Jackson, Ian Johnston, Dominic Kwiatkowski, Cordelia Langford, John Sillitoe on behalf of the Wellcome Sanger Institute COVID-19 Surveillance Team (http://www.sanger.ac.uk/covid-team) |
| EPI_ISL_558732, EPI_ISL_558733, EPI_ISL_558734, EPI_ISL_558735, EPI_ISL_558736, EPI_ISL_558737, EPI_ISL_558738, EPI_ISL_558739, EPI_ISL_558740, EPI_ISL_558741, EPI_ISL_558742, EPI_ISL_558743, EPI_ISL_558744, EPI_ISL_558745, EPI_ISL_558746, EPI_ISL_558747, EPI_ISL_558748, EPI_ISL_558749, EPI_ISL_558750, EPI_ISL_558751, EPI_ISL_558752, EPI_ISL_558753, EPI_ISL_558754, EPI_ISL_558755 | | | |
| see above | Lighthouse Lab in Alderley Park | Wellcome Sanger Institute for the COVID-19 Genomics UK (COG-UK) consortium | The Lighthouse Lab in Alderley Park and Alex Alderton, Roberto Amato, Sonia Goncalves, Ewan Harrison, David K. Jackson, Ian Johnston, Dominic Kwiatkowski, Cordelia Langford, John Sillitoe on behalf of the Wellcome Sanger Institute COVID-19 Surveillance Team |
| EPI_ISL_558756 | Lighthouse Lab in Alderley Park | Wellcome Sanger Institute for the COVID-19 Genomics UK (COG-UK) consortium | The Lighthouse Lab in Alderley Park and Alex Alderton, Roberto Amato, Sonia Goncalves, Ewan Harrison, David K. Jackson, Ian Johnston, Dominic Kwiatkowski, Cordelia Langford, John Sillitoe on behalf of the Wellcome Sanger Institute COVID-19 Surveillance Team (http://www.sanger.ac.uk/covid-team) |
| EPI_ISL_558757, EPI_ISL_558758, EPI_ISL_558759, EPI_ISL_558760, EPI_ISL_558761, EPI_ISL_558762, EPI_ISL_558763, EPI_ISL_558764, EPI_ISL_558765, EPI_ISL_558766, EPI_ISL_558767, EPI_ISL_558768, EPI_ISL_558769, EPI_ISL_558770, EPI_ISL_558771, EPI_ISL_558772, EPI_ISL_558773, EPI_ISL_558774, EPI_ISL_558775, EPI_ISL_558776, EPI_ISL_558777, EPI_ISL_558778, EPI_ISL_558779, EPI_ISL_558780, EPI_ISL_558781, EPI_ISL_558782, EPI_ISL_558783, EPI_ISL_558784, EPI_ISL_558785, EPI_ISL_558786, EPI_ISL_558787, EPI_ISL_558788, EPI_ISL_558789 | | | |
| see above | Lighthouse Lab in Alderley Park | Wellcome Sanger Institute for the COVID-19 Genomics UK | The Lighthouse Lab in Alderley Park and Alex Alderton, Roberto Amato, Sonia Goncalves, Ewan Harrison, David K. Jackson, Ian Johnston, Dominic |

| | | | |
|---|---|---|---|
| | | (COG-UK) consortium | Kwiatkowski, Cordelia Langford, John Sillitoe on behalf of the Wellcome Sanger Institute COVID-19 Surveillance Team |
| EPI_ISL_558790 | Lighthouse Lab in Alderley Park | Wellcome Sanger Institute for the COVID-19 Genomics UK (COG-UK) consortium | The Lighthouse Lab in Alderley Park and Alex Alderton, Roberto Amato, Sonia Goncalves, Ewan Harrison, David K. Jackson, Ian Johnston, Dominic Kwiatkowski, Cordelia Langford, John Sillitoe on behalf of the Wellcome Sanger Institute COVID-19 Surveillance Team (http://www.sanger.ac.uk/covid-team) |
| EPI_ISL_558791, EPI_ISL_558792, EPI_ISL_558793 | Lighthouse Lab in Alderley Park | Wellcome Sanger Institute for the COVID-19 Genomics UK (COG-UK) consortium | The Lighthouse Lab in Alderley Park and Alex Alderton, Roberto Amato, Sonia Goncalves, Ewan Harrison, David K. Jackson, Ian Johnston, Dominic Kwiatkowski, Cordelia Langford, John Sillitoe on behalf of the Wellcome Sanger Institute COVID-19 Surveillance Team |
| EPI_ISL_558794 | Lighthouse Lab in Alderley Park | Wellcome Sanger Institute for the COVID-19 Genomics UK (COG-UK) consortium | The Lighthouse Lab in Alderley Park and Alex Alderton, Roberto Amato, Sonia Goncalves, Ewan Harrison, David K. Jackson, Ian Johnston, Dominic Kwiatkowski, Cordelia Langford, John Sillitoe on behalf of the Wellcome Sanger Institute COVID-19 Surveillance Team (http://www.sanger.ac.uk/covid-team) |
| EPI_ISL_558795, EPI_ISL_558797, EPI_ISL_558798, EPI_ISL_558799, EPI_ISL_558800, EPI_ISL_558801, EPI_ISL_558802, EPI_ISL_558803, EPI_ISL_558804, EPI_ISL_558805, EPI_ISL_558806, EPI_ISL_558807, EPI_ISL_558808, EPI_ISL_558809, EPI_ISL_558810, EPI_ISL_558811, EPI_ISL_558812, EPI_ISL_558813, EPI_ISL_558814, EPI_ISL_558815, EPI_ISL_558816, EPI_ISL_558817, EPI_ISL_558818, EPI_ISL_558819, EPI_ISL_558820, EPI_ISL_558821, EPI_ISL_558822, EPI_ISL_558823, EPI_ISL_558824, EPI_ISL_558825, EPI_ISL_558826, EPI_ISL_558827, EPI_ISL_558828, EPI_ISL_558829, EPI_ISL_558830, EPI_ISL_558831, EPI_ISL_558832, EPI_ISL_558833, EPI_ISL_558834, EPI_ISL_558835, EPI_ISL_558836, EPI_ISL_558837, EPI_ISL_558838, EPI_ISL_558839, EPI_ISL_558840, EPI_ISL_558841, EPI_ISL_558842, EPI_ISL_558843, EPI_ISL_558844, EPI_ISL_558845, EPI_ISL_558847, EPI_ISL_558848, EPI_ISL_558849, EPI_ISL_558850, EPI_ISL_558851, EPI_ISL_558852, EPI_ISL_558853, EPI_ISL_558854, EPI_ISL_558855, EPI_ISL_558856, EPI_ISL_558857, EPI_ISL_558858, EPI_ISL_558859, EPI_ISL_558860 | | | |
| see above | Lighthouse Lab in Alderley Park | Wellcome Sanger Institute for the COVID-19 Genomics UK (COG-UK) consortium | The Lighthouse Lab in Alderley Park and Alex Alderton, Roberto Amato, Sonia Goncalves, Ewan Harrison, David K. Jackson, Ian Johnston, Dominic Kwiatkowski, Cordelia Langford, John Sillitoe on behalf of the Wellcome Sanger Institute COVID-19 Surveillance Team |
| EPI_ISL_558861 | Lighthouse Lab in Alderley Park | Wellcome Sanger Institute for the COVID-19 Genomics UK (COG-UK) consortium | The Lighthouse Lab in Alderley Park and Alex Alderton, Roberto Amato, Sonia Goncalves, Ewan Harrison, David K. Jackson, Ian Johnston, Dominic Kwiatkowski, Cordelia Langford, John Sillitoe on behalf of the Wellcome Sanger Institute COVID-19 Surveillance Team (http://www.sanger.ac.uk/covid-team) |
| EPI_ISL_558862, EPI_ISL_558863, EPI_ISL_558864, EPI_ISL_558865, EPI_ISL_558866, EPI_ISL_558867, EPI_ISL_558868, EPI_ISL_558869, EPI_ISL_558870, EPI_ISL_558871, EPI_ISL_558873, EPI_ISL_558874, EPI_ISL_558875, EPI_ISL_558876, EPI_ISL_558878, EPI_ISL_558879, EPI_ISL_558880, EPI_ISL_558881, EPI_ISL_558882, EPI_ISL_558883, EPI_ISL_558884, EPI_ISL_558885, EPI_ISL_558886, EPI_ISL_558887, EPI_ISL_558889, EPI_ISL_558890, EPI_ISL_558891, EPI_ISL_558892, EPI_ISL_558893 | | | |
| see above | Lighthouse Lab in Alderley Park | Wellcome Sanger Institute for the COVID-19 Genomics UK (COG-UK) consortium | The Lighthouse Lab in Alderley Park and Alex Alderton, Roberto Amato, Sonia Goncalves, Ewan Harrison, David K. Jackson, Ian Johnston, Dominic Kwiatkowski, Cordelia Langford, John Sillitoe on behalf of the Wellcome Sanger Institute COVID-19 Surveillance Team |
| EPI_ISL_558894 | Lighthouse Lab in Alderley Park | Wellcome Sanger Institute for the COVID-19 Genomics UK (COG-UK) consortium | The Lighthouse Lab in Alderley Park and Alex Alderton, Roberto Amato, Sonia Goncalves, Ewan Harrison, David K. Jackson, Ian Johnston, Dominic Kwiatkowski, Cordelia Langford, John Sillitoe on behalf of the Wellcome Sanger Institute COVID-19 Surveillance Team (http://www.sanger.ac.uk/covid-team) |
| EPI_ISL_558895, EPI_ISL_558896, EPI_ISL_558897, EPI_ISL_558898, EPI_ISL_558899, EPI_ISL_558900, EPI_ISL_558901, EPI_ISL_558902 | Lighthouse Lab in Alderley Park | Wellcome Sanger Institute for the COVID-19 Genomics UK (COG-UK) consortium | The Lighthouse Lab in Alderley Park and Alex Alderton, Roberto Amato, Sonia Goncalves, Ewan Harrison, David K. Jackson, Ian Johnston, Dominic Kwiatkowski, Cordelia Langford, John Sillitoe on behalf of the Wellcome Sanger Institute COVID-19 Surveillance Team |
| EPI_ISL_558903 | Lighthouse Lab in Milton Keynes | Wellcome Sanger Institute for the COVID-19 Genomics UK (COG-UK) consortium | The Lighthouse Lab in Milton Keynes and Alex Alderton, Roberto Amato, Sonia Goncalves, Ewan Harrison, David K. Jackson, Ian Johnston, Dominic Kwiatkowski, Cordelia Langford, John Sillitoe on behalf of the Wellcome Sanger Institute COVID-19 Surveillance Team (http://www.sanger.ac.uk/covid-team) |
| EPI_ISL_558904, EPI_ISL_558905, EPI_ISL_558906, EPI_ISL_558907, EPI_ISL_558908, EPI_ISL_558909, EPI_ISL_558910, EPI_ISL_558911, EPI_ISL_558912 | Lighthouse Lab in Alderley Park | Wellcome Sanger Institute for the COVID-19 Genomics UK (COG-UK) consortium | The Lighthouse Lab in Alderley Park and Alex Alderton, Roberto Amato, Sonia Goncalves, Ewan Harrison, David K. Jackson, Ian Johnston, Dominic Kwiatkowski, Cordelia Langford, John Sillitoe on behalf of the Wellcome Sanger Institute COVID-19 Surveillance Team |
| EPI_ISL_558913 | Lighthouse Lab in Alderley Park | Wellcome Sanger Institute for the COVID-19 Genomics UK (COG-UK) consortium | The Lighthouse Lab in Alderley Park and Alex Alderton, Roberto Amato, Sonia Goncalves, Ewan Harrison, David K. Jackson, Ian Johnston, Dominic Kwiatkowski, Cordelia Langford, John Sillitoe on behalf of the Wellcome Sanger Institute COVID-19 Surveillance Team (http://www.sanger.ac.uk/covid-team) |
| EPI_ISL_558914 | Lighthouse Lab in Milton Keynes | Wellcome Sanger Institute for the COVID-19 Genomics UK (COG-UK) consortium | The Lighthouse Lab in Milton Keynes and Alex Alderton, Roberto Amato, Sonia Goncalves, Ewan Harrison, David K. Jackson, Ian Johnston, Dominic Kwiatkowski, Cordelia Langford, John Sillitoe on behalf of the Wellcome Sanger Institute COVID-19 Surveillance Team (http://www.sanger.ac.uk/covid-team) |
| EPI_ISL_558915, EPI_ISL_558916, EPI_ISL_558917, EPI_ISL_558918, EPI_ISL_558919 | Lighthouse Lab in Alderley Park | Wellcome Sanger Institute for the COVID-19 Genomics UK (COG-UK) consortium | The Lighthouse Lab in Alderley Park and Alex Alderton, Roberto Amato, Sonia Goncalves, Ewan Harrison, David K. Jackson, Ian Johnston, Dominic Kwiatkowski, Cordelia Langford, John Sillitoe on behalf of the Wellcome Sanger Institute COVID-19 Surveillance Team |
| EPI_ISL_558920 | Lighthouse Lab in Milton Keynes | Wellcome Sanger Institute for the COVID-19 Genomics UK (COG-UK) consortium | The Lighthouse Lab in Milton Keynes and Alex Alderton, Roberto Amato, Sonia Goncalves, Ewan Harrison, David K. Jackson, Ian Johnston, Dominic Kwiatkowski, Cordelia Langford, John Sillitoe on behalf of the Wellcome Sanger Institute COVID-19 Surveillance Team (http://www.sanger.ac.uk/covid-team) |
| EPI_ISL_558922 | Lighthouse Lab in Alderley Park | Wellcome Sanger Institute for the COVID-19 Genomics UK (COG-UK) consortium | The Lighthouse Lab in Alderley Park and Alex Alderton, Roberto Amato, Sonia Goncalves, Ewan Harrison, David K. Jackson, Ian Johnston, Dominic Kwiatkowski, Cordelia Langford, John Sillitoe on behalf of the Wellcome Sanger Institute COVID-19 Surveillance Team |
| EPI_ISL_558923 | Lighthouse Lab in Milton Keynes | Wellcome Sanger Institute for the COVID-19 Genomics UK (COG-UK) consortium | The Lighthouse Lab in Milton Keynes and Alex Alderton, Roberto Amato, Sonia Goncalves, Ewan Harrison, David K. Jackson, Ian Johnston, Dominic Kwiatkowski, Cordelia Langford, John Sillitoe on behalf of the Wellcome Sanger Institute COVID-19 Surveillance Team (http://www.sanger.ac.uk/covid-team) |
| EPI_ISL_558924 | Lighthouse Lab in Alderley Park | Wellcome Sanger Institute for the COVID-19 Genomics UK (COG-UK) consortium | The Lighthouse Lab in Alderley Park and Alex Alderton, Roberto Amato, Sonia Goncalves, Ewan Harrison, David K. Jackson, Ian Johnston, Dominic Kwiatkowski, Cordelia Langford, John Sillitoe on behalf of the Wellcome Sanger Institute COVID-19 Surveillance Team |
| EPI_ISL_558925 | Lighthouse Lab in Milton Keynes | Wellcome Sanger Institute for the COVID-19 Genomics UK (COG-UK) consortium | The Lighthouse Lab in Milton Keynes and Alex Alderton, Roberto Amato, Sonia Goncalves, Ewan Harrison, David K. Jackson, Ian Johnston, Dominic Kwiatkowski, Cordelia Langford, John Sillitoe on behalf of the Wellcome Sanger Institute COVID-19 Surveillance Team (http://www.sanger.ac.uk/covid-team) |
| EPI_ISL_558927 | Lighthouse Lab in Alderley Park | Wellcome Sanger Institute for the COVID-19 Genomics UK (COG-UK) consortium | The Lighthouse Lab in Alderley Park and Alex Alderton, Roberto Amato, Sonia Goncalves, Ewan Harrison, David K. Jackson, Ian Johnston, Dominic Kwiatkowski, Cordelia Langford, John Sillitoe on behalf of the Wellcome Sanger Institute COVID-19 Surveillance Team |
| EPI_ISL_558928 | Lighthouse Lab in Milton Keynes | Wellcome Sanger Institute for the COVID-19 Genomics UK (COG-UK) consortium | The Lighthouse Lab in Milton Keynes and Alex Alderton, Roberto Amato, Sonia Goncalves, Ewan Harrison, David K. Jackson, Ian Johnston, Dominic Kwiatkowski, Cordelia Langford, John Sillitoe on behalf of the Wellcome Sanger Institute COVID-19 Surveillance Team (http://www.sanger.ac.uk/covid-team) |
| EPI_ISL_558929, EPI_ISL_558930, EPI_ISL_558931, EPI_ISL_558933 | Lighthouse Lab in Alderley Park | Wellcome Sanger Institute for the COVID-19 Genomics UK (COG-UK) consortium | The Lighthouse Lab in Alderley Park and Alex Alderton, Roberto Amato, Sonia Goncalves, Ewan Harrison, David K. Jackson, Ian Johnston, Dominic Kwiatkowski, Cordelia Langford, John Sillitoe on behalf of the Wellcome Sanger Institute COVID-19 Surveillance Team |
| EPI_ISL_558934 | Lighthouse Lab in Milton Keynes | Wellcome Sanger Institute for the COVID-19 Genomics UK (COG-UK) consortium | The Lighthouse Lab in Milton Keynes and Alex Alderton, Roberto Amato, Sonia Goncalves, Ewan Harrison, David K. Jackson, Ian Johnston, Dominic Kwiatkowski, Cordelia Langford, John Sillitoe on behalf of the Wellcome Sanger Institute COVID-19 Surveillance Team (http://www.sanger.ac.uk/covid-team) |
| EPI_ISL_558935 | Lighthouse Lab in Alderley Park | Wellcome Sanger Institute for the COVID-19 Genomics UK (COG-UK) consortium | The Lighthouse Lab in Alderley Park and Alex Alderton, Roberto Amato, Sonia Goncalves, Ewan Harrison, David K. Jackson, Ian Johnston, Dominic Kwiatkowski, Cordelia Langford, John Sillitoe on behalf of the Wellcome Sanger Institute COVID-19 Surveillance Team (http://www.sanger.ac.uk/covid-team) |
| EPI_ISL_558936 | Lighthouse Lab in Alderley Park | Wellcome Sanger Institute for the COVID-19 Genomics UK (COG-UK) consortium | The Lighthouse Lab in Alderley Park and Alex Alderton, Roberto Amato, Sonia Goncalves, Ewan Harrison, David K. Jackson, Ian Johnston, Dominic Kwiatkowski, Cordelia Langford, John Sillitoe on behalf of the Wellcome Sanger Institute COVID-19 Surveillance Team |

| | | | |
|---|---|---|---|
| EPI_ISL_558937 | Lighthouse Lab in Milton Keynes | Wellcome Sanger Institute for the COVID-19 Genomics UK (COG-UK) consortium | The Lighthouse Lab in Milton Keynes and Alex Alderton, Roberto Amato, Sonia Goncalves, Ewan Harrison, David K. Jackson, Ian Johnston, Dominic Kwiatkowski, Cordelia Langford, John Sillitoe on behalf of the Wellcome Sanger Institute COVID-19 Surveillance Team (http://www.sanger.ac.uk/covid-team) |
| EPI_ISL_558938, EPI_ISL_558939, EPI_ISL_558940, EPI_ISL_558941, EPI_ISL_558942, EPI_ISL_558943, EPI_ISL_558944, EPI_ISL_558945, EPI_ISL_558946, EPI_ISL_558947, EPI_ISL_558948 | | | |
| see above | Lighthouse Lab in Alderley Park | Wellcome Sanger Institute for the COVID-19 Genomics UK (COG-UK) consortium | The Lighthouse Lab in Alderley Park and Alex Alderton, Roberto Amato, Sonia Goncalves, Ewan Harrison, David K. Jackson, Ian Johnston, Dominic Kwiatkowski, Cordelia Langford, John Sillitoe on behalf of the Wellcome Sanger Institute COVID-19 Surveillance Team |
| EPI_ISL_558949, EPI_ISL_558951 | Lighthouse Lab in Milton Keynes | Wellcome Sanger Institute for the COVID-19 Genomics UK (COG-UK) consortium | The Lighthouse Lab in Milton Keynes and Alex Alderton, Roberto Amato, Sonia Goncalves, Ewan Harrison, David K. Jackson, Ian Johnston, Dominic Kwiatkowski, Cordelia Langford, John Sillitoe on behalf of the Wellcome Sanger Institute COVID-19 Surveillance Team (http://www.sanger.ac.uk/covid-team) |
| EPI_ISL_558952, EPI_ISL_558953, EPI_ISL_558954 | Lighthouse Lab in Alderley Park | Wellcome Sanger Institute for the COVID-19 Genomics UK (COG-UK) consortium | The Lighthouse Lab in Alderley Park and Alex Alderton, Roberto Amato, Sonia Goncalves, Ewan Harrison, David K. Jackson, Ian Johnston, Dominic Kwiatkowski, Cordelia Langford, John Sillitoe on behalf of the Wellcome Sanger Institute COVID-19 Surveillance Team |
| EPI_ISL_558955 | Lighthouse Lab in Alderley Park | Wellcome Sanger Institute for the COVID-19 Genomics UK (COG-UK) consortium | The Lighthouse Lab in Alderley Park and Alex Alderton, Roberto Amato, Sonia Goncalves, Ewan Harrison, David K. Jackson, Ian Johnston, Dominic Kwiatkowski, Cordelia Langford, John Sillitoe on behalf of the Wellcome Sanger Institute COVID-19 Surveillance Team (http://www.sanger.ac.uk/covid-team) |
| EPI_ISL_558956, EPI_ISL_558957, EPI_ISL_558958 | Lighthouse Lab in Alderley Park | Wellcome Sanger Institute for the COVID-19 Genomics UK (COG-UK) consortium | The Lighthouse Lab in Alderley Park and Alex Alderton, Roberto Amato, Sonia Goncalves, Ewan Harrison, David K. Jackson, Ian Johnston, Dominic Kwiatkowski, Cordelia Langford, John Sillitoe on behalf of the Wellcome Sanger Institute COVID-19 Surveillance Team |
| EPI_ISL_558959 | Lighthouse Lab in Milton Keynes | Wellcome Sanger Institute for the COVID-19 Genomics UK (COG-UK) consortium | The Lighthouse Lab in Milton Keynes and Alex Alderton, Roberto Amato, Sonia Goncalves, Ewan Harrison, David K. Jackson, Ian Johnston, Dominic Kwiatkowski, Cordelia Langford, John Sillitoe on behalf of the Wellcome Sanger Institute COVID-19 Surveillance Team (http://www.sanger.ac.uk/covid-team) |
| EPI_ISL_558960, EPI_ISL_558961, EPI_ISL_558962, EPI_ISL_558963, EPI_ISL_558964 | Lighthouse Lab in Alderley Park | Wellcome Sanger Institute for the COVID-19 Genomics UK (COG-UK) consortium | The Lighthouse Lab in Alderley Park and Alex Alderton, Roberto Amato, Sonia Goncalves, Ewan Harrison, David K. Jackson, Ian Johnston, Dominic Kwiatkowski, Cordelia Langford, John Sillitoe on behalf of the Wellcome Sanger Institute COVID-19 Surveillance Team |
| EPI_ISL_558965 | Lighthouse Lab in Milton Keynes | Wellcome Sanger Institute for the COVID-19 Genomics UK (COG-UK) consortium | The Lighthouse Lab in Milton Keynes and Alex Alderton, Roberto Amato, Sonia Goncalves, Ewan Harrison, David K. Jackson, Ian Johnston, Dominic Kwiatkowski, Cordelia Langford, John Sillitoe on behalf of the Wellcome Sanger Institute COVID-19 Surveillance Team (http://www.sanger.ac.uk/covid-team) |
| EPI_ISL_558966, EPI_ISL_558967, EPI_ISL_558968, EPI_ISL_558969 | Lighthouse Lab in Alderley Park | Wellcome Sanger Institute for the COVID-19 Genomics UK (COG-UK) consortium | The Lighthouse Lab in Alderley Park and Alex Alderton, Roberto Amato, Sonia Goncalves, Ewan Harrison, David K. Jackson, Ian Johnston, Dominic Kwiatkowski, Cordelia Langford, John Sillitoe on behalf of the Wellcome Sanger Institute COVID-19 Surveillance Team |
| EPI_ISL_558970 | Lighthouse Lab in Milton Keynes | Wellcome Sanger Institute for the COVID-19 Genomics UK (COG-UK) consortium | The Lighthouse Lab in Milton Keynes and Alex Alderton, Roberto Amato, Sonia Goncalves, Ewan Harrison, David K. Jackson, Ian Johnston, Dominic Kwiatkowski, Cordelia Langford, John Sillitoe on behalf of the Wellcome Sanger Institute COVID-19 Surveillance Team (http://www.sanger.ac.uk/covid-team) |
| EPI_ISL_558971, EPI_ISL_558972 | Lighthouse Lab in Alderley Park | Wellcome Sanger Institute for the COVID-19 Genomics UK (COG-UK) consortium | The Lighthouse Lab in Alderley Park and Alex Alderton, Roberto Amato, Sonia Goncalves, Ewan Harrison, David K. Jackson, Ian Johnston, Dominic Kwiatkowski, Cordelia Langford, John Sillitoe on behalf of the Wellcome Sanger Institute COVID-19 Surveillance Team |
| EPI_ISL_558973 | Lighthouse Lab in Milton Keynes | Wellcome Sanger Institute for the COVID-19 Genomics UK (COG-UK) consortium | The Lighthouse Lab in Milton Keynes and Alex Alderton, Roberto Amato, Sonia Goncalves, Ewan Harrison, David K. Jackson, Ian Johnston, Dominic Kwiatkowski, Cordelia Langford, John Sillitoe on behalf of the Wellcome Sanger Institute COVID-19 Surveillance Team (http://www.sanger.ac.uk/covid-team) |
| EPI_ISL_558974, EPI_ISL_558976, EPI_ISL_558977, EPI_ISL_558978, EPI_ISL_558979 | Lighthouse Lab in Alderley Park | Wellcome Sanger Institute for the COVID-19 Genomics UK (COG-UK) consortium | The Lighthouse Lab in Alderley Park and Alex Alderton, Roberto Amato, Sonia Goncalves, Ewan Harrison, David K. Jackson, Ian Johnston, Dominic Kwiatkowski, Cordelia Langford, John Sillitoe on behalf of the Wellcome Sanger Institute COVID-19 Surveillance Team |
| EPI_ISL_558980, EPI_ISL_558981 | Lighthouse Lab in Milton Keynes | Wellcome Sanger Institute for the COVID-19 Genomics UK (COG-UK) consortium | The Lighthouse Lab in Milton Keynes and Alex Alderton, Roberto Amato, Sonia Goncalves, Ewan Harrison, David K. Jackson, Ian Johnston, Dominic Kwiatkowski, Cordelia Langford, John Sillitoe on behalf of the Wellcome Sanger Institute COVID-19 Surveillance Team (http://www.sanger.ac.uk/covid-team) |
| EPI_ISL_558982, EPI_ISL_558983, EPI_ISL_558984, EPI_ISL_558985, EPI_ISL_558986 | Lighthouse Lab in Alderley Park | Wellcome Sanger Institute for the COVID-19 Genomics UK (COG-UK) consortium | The Lighthouse Lab in Alderley Park and Alex Alderton, Roberto Amato, Sonia Goncalves, Ewan Harrison, David K. Jackson, Ian Johnston, Dominic Kwiatkowski, Cordelia Langford, John Sillitoe on behalf of the Wellcome Sanger Institute COVID-19 Surveillance Team |
| EPI_ISL_558987 | Lighthouse Lab in Milton Keynes | Wellcome Sanger Institute for the COVID-19 Genomics UK (COG-UK) consortium | The Lighthouse Lab in Milton Keynes and Alex Alderton, Roberto Amato, Sonia Goncalves, Ewan Harrison, David K. Jackson, Ian Johnston, Dominic Kwiatkowski, Cordelia Langford, John Sillitoe on behalf of the Wellcome Sanger Institute COVID-19 Surveillance Team (http://www.sanger.ac.uk/covid-team) |
| EPI_ISL_558988, EPI_ISL_558989, EPI_ISL_558990, EPI_ISL_558991, EPI_ISL_558992 | Lighthouse Lab in Alderley Park | Wellcome Sanger Institute for the COVID-19 Genomics UK (COG-UK) consortium | The Lighthouse Lab in Alderley Park and Alex Alderton, Roberto Amato, Sonia Goncalves, Ewan Harrison, David K. Jackson, Ian Johnston, Dominic Kwiatkowski, Cordelia Langford, John Sillitoe on behalf of the Wellcome Sanger Institute COVID-19 Surveillance Team |
| EPI_ISL_558995 | Lighthouse Lab in Alderley Park | Wellcome Sanger Institute for the COVID-19 Genomics UK (COG-UK) consortium | The Lighthouse Lab in Alderley Park and Alex Alderton, Roberto Amato, Sonia Goncalves, Ewan Harrison, David K. Jackson, Ian Johnston, Dominic Kwiatkowski, Cordelia Langford, John Sillitoe on behalf of the Wellcome Sanger Institute COVID-19 Surveillance Team (http://www.sanger.ac.uk/covid-team) |
| EPI_ISL_558996, EPI_ISL_558997, EPI_ISL_558998, EPI_ISL_558999 | Lighthouse Lab in Alderley Park | Wellcome Sanger Institute for the COVID-19 Genomics UK (COG-UK) consortium | The Lighthouse Lab in Alderley Park and Alex Alderton, Roberto Amato, Sonia Goncalves, Ewan Harrison, David K. Jackson, Ian Johnston, Dominic Kwiatkowski, Cordelia Langford, John Sillitoe on behalf of the Wellcome Sanger Institute COVID-19 Surveillance Team |
| EPI_ISL_559000 | Lighthouse Lab in Milton Keynes | Wellcome Sanger Institute for the COVID-19 Genomics UK (COG-UK) consortium | The Lighthouse Lab in Milton Keynes and Alex Alderton, Roberto Amato, Sonia Goncalves, Ewan Harrison, David K. Jackson, Ian Johnston, Dominic Kwiatkowski, Cordelia Langford, John Sillitoe on behalf of the Wellcome Sanger Institute COVID-19 Surveillance Team (http://www.sanger.ac.uk/covid-team) |
| EPI_ISL_559001, EPI_ISL_559002, EPI_ISL_559003, EPI_ISL_559004 | Lighthouse Lab in Alderley Park | Wellcome Sanger Institute for the COVID-19 Genomics UK (COG-UK) consortium | The Lighthouse Lab in Alderley Park and Alex Alderton, Roberto Amato, Sonia Goncalves, Ewan Harrison, David K. Jackson, Ian Johnston, Dominic Kwiatkowski, Cordelia Langford, John Sillitoe on behalf of the Wellcome Sanger Institute COVID-19 Surveillance Team |
| EPI_ISL_559005 | Lighthouse Lab in Milton Keynes | Wellcome Sanger Institute for the COVID-19 Genomics UK (COG-UK) consortium | The Lighthouse Lab in Alderley Park and Alex Alderton, Roberto Amato, Sonia Goncalves, Ewan Harrison, David K. Jackson, Ian Johnston, Dominic Kwiatkowski, Cordelia Langford, John Sillitoe on behalf of the Wellcome Sanger Institute COVID-19 Surveillance Team |
| EPI_ISL_559006, EPI_ISL_559007, EPI_ISL_559008 | Lighthouse Lab in Alderley Park | Wellcome Sanger Institute for the COVID-19 Genomics UK (COG-UK) consortium | The Lighthouse Lab in Alderley Park and Alex Alderton, Roberto Amato, Sonia Goncalves, Ewan Harrison, David K. Jackson, Ian Johnston, Dominic Kwiatkowski, Cordelia Langford, John Sillitoe on behalf of the Wellcome Sanger Institute COVID-19 Surveillance Team |
| EPI_ISL_559009 | Lighthouse Lab in Milton Keynes | Wellcome Sanger Institute for the COVID-19 Genomics UK (COG-UK) consortium | The Lighthouse Lab in Alderley Park and Alex Alderton, Roberto Amato, Sonia Goncalves, Ewan Harrison, David K. Jackson, Ian Johnston, Dominic Kwiatkowski, Cordelia Langford, John Sillitoe on behalf of the Wellcome Sanger Institute COVID-19 Surveillance Team |
| EPI_ISL_559010 | Lighthouse Lab in Alderley Park | Wellcome Sanger Institute for the COVID-19 Genomics UK (COG-UK) consortium | The Lighthouse Lab in Alderley Park and Alex Alderton, Roberto Amato, Sonia Goncalves, Ewan Harrison, David K. Jackson, Ian Johnston, Dominic Kwiatkowski, Cordelia Langford, John Sillitoe on behalf of the Wellcome Sanger Institute COVID-19 Surveillance Team |
| EPI_ISL_559011 | Lighthouse Lab in Milton Keynes | Wellcome Sanger Institute for the COVID-19 Genomics UK (COG-UK) consortium | The Lighthouse Lab in Milton Keynes and Alex Alderton, Roberto Amato, Sonia Goncalves, Ewan Harrison, David K. Jackson, Ian Johnston, Dominic Kwiatkowski, Cordelia Langford, John Sillitoe on behalf of the Wellcome Sanger Institute COVID-19 Surveillance Team (http://www.sanger.ac.uk/covid-team) |
| EPI_ISL_559012, EPI_ISL_559013, EPI_ISL_559014, EPI_ISL_559015, | Lighthouse Lab in Alderley Park | Wellcome Sanger Institute for the COVID-19 Genomics UK (COG-UK) consortium | The Lighthouse Lab in Alderley Park and Alex Alderton, Roberto Amato, Sonia Goncalves, Ewan Harrison, David K. Jackson, Ian Johnston, Dominic Kwiatkowski, Cordelia Langford, John Sillitoe on behalf of the Wellcome Sanger Institute COVID-19 Surveillance Team |

| | | | |
|---|---|---|---|
| EPI_ISL_559016, EPI_ISL_559018, EPI_ISL_559020, EPI_ISL_559021 | | | |
| EPI_ISL_559022 | Lighthouse Lab in Milton Keynes | Wellcome Sanger Institute for the COVID-19 Genomics UK (COG-UK) consortium | The Lighthouse Lab in Milton Keynes and Alex Alderton, Roberto Amato, Sonia Goncalves, Ewan Harrison, David K. Jackson, Ian Johnston, Dominic Kwiatkowski, Cordelia Langford, John Sillitoe on behalf of the Wellcome Sanger Institute COVID-19 Surveillance Team (http://www.sanger.ac.uk/covid-team) |
| EPI_ISL_559023, EPI_ISL_559024, EPI_ISL_559025, EPI_ISL_559027 | Lighthouse Lab in Alderley Park | Wellcome Sanger Institute for the COVID-19 Genomics UK (COG-UK) consortium | The Lighthouse Lab in Alderley Park and Alex Alderton, Roberto Amato, Sonia Goncalves, Ewan Harrison, David K. Jackson, Ian Johnston, Dominic Kwiatkowski, Cordelia Langford, John Sillitoe on behalf of the Wellcome Sanger Institute COVID-19 Surveillance Team |
| EPI_ISL_559028 | Lighthouse Lab in Milton Keynes | Wellcome Sanger Institute for the COVID-19 Genomics UK (COG-UK) consortium | The Lighthouse Lab in Milton Keynes and Alex Alderton, Roberto Amato, Sonia Goncalves, Ewan Harrison, David K. Jackson, Ian Johnston, Dominic Kwiatkowski, Cordelia Langford, John Sillitoe on behalf of the Wellcome Sanger Institute COVID-19 Surveillance Team (http://www.sanger.ac.uk/covid-team) |
| EPI_ISL_559029, EPI_ISL_559030, EPI_ISL_559031, EPI_ISL_559032, EPI_ISL_559033, EPI_ISL_559034, EPI_ISL_559035, EPI_ISL_559036, EPI_ISL_559037, EPI_ISL_559038, EPI_ISL_559039, EPI_ISL_559040, EPI_ISL_559041 | | | |
| see above | Lighthouse Lab in Alderley Park | Wellcome Sanger Institute for the COVID-19 Genomics UK (COG-UK) consortium | The Lighthouse Lab in Alderley Park and Alex Alderton, Roberto Amato, Sonia Goncalves, Ewan Harrison, David K. Jackson, Ian Johnston, Dominic Kwiatkowski, Cordelia Langford, John Sillitoe on behalf of the Wellcome Sanger Institute COVID-19 Surveillance Team |
| EPI_ISL_559043 | Lighthouse Lab in Milton Keynes | Wellcome Sanger Institute for the COVID-19 Genomics UK (COG-UK) consortium | The Lighthouse Lab in Milton Keynes and Alex Alderton, Roberto Amato, Sonia Goncalves, Ewan Harrison, David K. Jackson, Ian Johnston, Dominic Kwiatkowski, Cordelia Langford, John Sillitoe on behalf of the Wellcome Sanger Institute COVID-19 Surveillance Team (http://www.sanger.ac.uk/covid-team) |
| EPI_ISL_559044, EPI_ISL_559045 | Lighthouse Lab in Alderley Park | Wellcome Sanger Institute for the COVID-19 Genomics UK (COG-UK) consortium | The Lighthouse Lab in Alderley Park and Alex Alderton, Roberto Amato, Sonia Goncalves, Ewan Harrison, David K. Jackson, Ian Johnston, Dominic Kwiatkowski, Cordelia Langford, John Sillitoe on behalf of the Wellcome Sanger Institute COVID-19 Surveillance Team |
| EPI_ISL_559047 | Lighthouse Lab in Milton Keynes | Wellcome Sanger Institute for the COVID-19 Genomics UK (COG-UK) consortium | The Lighthouse Lab in Milton Keynes and Alex Alderton, Roberto Amato, Sonia Goncalves, Ewan Harrison, David K. Jackson, Ian Johnston, Dominic Kwiatkowski, Cordelia Langford, John Sillitoe on behalf of the Wellcome Sanger Institute COVID-19 Surveillance Team (http://www.sanger.ac.uk/covid-team) |
| EPI_ISL_559048 | Lighthouse Lab in Alderley Park | Wellcome Sanger Institute for the COVID-19 Genomics UK (COG-UK) consortium | The Lighthouse Lab in Alderley Park and Alex Alderton, Roberto Amato, Sonia Goncalves, Ewan Harrison, David K. Jackson, Ian Johnston, Dominic Kwiatkowski, Cordelia Langford, John Sillitoe on behalf of the Wellcome Sanger Institute COVID-19 Surveillance Team |
| EPI_ISL_559049 | Lighthouse Lab in Milton Keynes | Wellcome Sanger Institute for the COVID-19 Genomics UK (COG-UK) consortium | The Lighthouse Lab in Alderley Park and Alex Alderton, Roberto Amato, Sonia Goncalves, Ewan Harrison, David K. Jackson, Ian Johnston, Dominic Kwiatkowski, Cordelia Langford, John Sillitoe on behalf of the Wellcome Sanger Institute COVID-19 Surveillance Team |
| EPI_ISL_559050, EPI_ISL_559051, EPI_ISL_559052, EPI_ISL_559053, EPI_ISL_559054, EPI_ISL_559055, EPI_ISL_559056 | Lighthouse Lab in Alderley Park | Wellcome Sanger Institute for the COVID-19 Genomics UK (COG-UK) consortium | The Lighthouse Lab in Alderley Park and Alex Alderton, Roberto Amato, Sonia Goncalves, Ewan Harrison, David K. Jackson, Ian Johnston, Dominic Kwiatkowski, Cordelia Langford, John Sillitoe on behalf of the Wellcome Sanger Institute COVID-19 Surveillance Team |
| EPI_ISL_559057 | Lighthouse Lab in Milton Keynes | Wellcome Sanger Institute for the COVID-19 Genomics UK (COG-UK) consortium | The Lighthouse Lab in Milton Keynes and Alex Alderton, Roberto Amato, Sonia Goncalves, Ewan Harrison, David K. Jackson, Ian Johnston, Dominic Kwiatkowski, Cordelia Langford, John Sillitoe on behalf of the Wellcome Sanger Institute COVID-19 Surveillance Team (http://www.sanger.ac.uk/covid-team) |
| EPI_ISL_559058, EPI_ISL_559059, EPI_ISL_559060 | Lighthouse Lab in Alderley Park | Wellcome Sanger Institute for the COVID-19 Genomics UK (COG-UK) consortium | The Lighthouse Lab in Alderley Park and Alex Alderton, Roberto Amato, Sonia Goncalves, Ewan Harrison, David K. Jackson, Ian Johnston, Dominic Kwiatkowski, Cordelia Langford, John Sillitoe on behalf of the Wellcome Sanger Institute COVID-19 Surveillance Team |
| EPI_ISL_559061 | Lighthouse Lab in Milton Keynes | Wellcome Sanger Institute for the COVID-19 Genomics UK (COG-UK) consortium | The Lighthouse Lab in Milton Keynes and Alex Alderton, Roberto Amato, Sonia Goncalves, Ewan Harrison, David K. Jackson, Ian Johnston, Dominic Kwiatkowski, Cordelia Langford, John Sillitoe on behalf of the Wellcome Sanger Institute COVID-19 Surveillance Team (http://www.sanger.ac.uk/covid-team) |
| EPI_ISL_559062 | Lighthouse Lab in Alderley Park | Wellcome Sanger Institute for the COVID-19 Genomics UK (COG-UK) consortium | The Lighthouse Lab in Alderley Park and Alex Alderton, Roberto Amato, Sonia Goncalves, Ewan Harrison, David K. Jackson, Ian Johnston, Dominic Kwiatkowski, Cordelia Langford, John Sillitoe on behalf of the Wellcome Sanger Institute COVID-19 Surveillance Team |
| EPI_ISL_559063 | Lighthouse Lab in Milton Keynes | Wellcome Sanger Institute for the COVID-19 Genomics UK (COG-UK) consortium | The Lighthouse Lab in Milton Keynes and Alex Alderton, Roberto Amato, Sonia Goncalves, Ewan Harrison, David K. Jackson, Ian Johnston, Dominic Kwiatkowski, Cordelia Langford, John Sillitoe on behalf of the Wellcome Sanger Institute COVID-19 Surveillance Team (http://www.sanger.ac.uk/covid-team) |
| EPI_ISL_559064, EPI_ISL_559065, EPI_ISL_559066, EPI_ISL_559067, EPI_ISL_559068, EPI_ISL_559069, EPI_ISL_559070, EPI_ISL_559072, EPI_ISL_559073, EPI_ISL_559074, EPI_ISL_559075, EPI_ISL_559076, EPI_ISL_559077, EPI_ISL_559078, EPI_ISL_559079 | | | |
| see above | Lighthouse Lab in Alderley Park | Wellcome Sanger Institute for the COVID-19 Genomics UK (COG-UK) consortium | The Lighthouse Lab in Alderley Park and Alex Alderton, Roberto Amato, Sonia Goncalves, Ewan Harrison, David K. Jackson, Ian Johnston, Dominic Kwiatkowski, Cordelia Langford, John Sillitoe on behalf of the Wellcome Sanger Institute COVID-19 Surveillance Team |
| EPI_ISL_559080, EPI_ISL_559081 | Lighthouse Lab in Milton Keynes | Wellcome Sanger Institute for the COVID-19 Genomics UK (COG-UK) consortium | The Lighthouse Lab in Milton Keynes and Alex Alderton, Roberto Amato, Sonia Goncalves, Ewan Harrison, David K. Jackson, Ian Johnston, Dominic Kwiatkowski, Cordelia Langford, John Sillitoe on behalf of the Wellcome Sanger Institute COVID-19 Surveillance Team (http://www.sanger.ac.uk/covid-team) |
| EPI_ISL_559082 | Lighthouse Lab in Alderley Park | Wellcome Sanger Institute for the COVID-19 Genomics UK (COG-UK) consortium | The Lighthouse Lab in Alderley Park and Alex Alderton, Roberto Amato, Sonia Goncalves, Ewan Harrison, David K. Jackson, Ian Johnston, Dominic Kwiatkowski, Cordelia Langford, John Sillitoe on behalf of the Wellcome Sanger Institute COVID-19 Surveillance Team |
| EPI_ISL_559084 | Lighthouse Lab in Milton Keynes | Wellcome Sanger Institute for the COVID-19 Genomics UK (COG-UK) consortium | The Lighthouse Lab in Milton Keynes and Alex Alderton, Roberto Amato, Sonia Goncalves, Ewan Harrison, David K. Jackson, Ian Johnston, Dominic Kwiatkowski, Cordelia Langford, John Sillitoe on behalf of the Wellcome Sanger Institute COVID-19 Surveillance Team (http://www.sanger.ac.uk/covid-team) |
| EPI_ISL_559085, EPI_ISL_559086, EPI_ISL_559087 | Lighthouse Lab in Alderley Park | Wellcome Sanger Institute for the COVID-19 Genomics UK (COG-UK) consortium | The Lighthouse Lab in Alderley Park and Alex Alderton, Roberto Amato, Sonia Goncalves, Ewan Harrison, David K. Jackson, Ian Johnston, Dominic Kwiatkowski, Cordelia Langford, John Sillitoe on behalf of the Wellcome Sanger Institute COVID-19 Surveillance Team |
| EPI_ISL_559088 | Lighthouse Lab in Milton Keynes | Wellcome Sanger Institute for the COVID-19 Genomics UK (COG-UK) consortium | The Lighthouse Lab in Milton Keynes and Alex Alderton, Roberto Amato, Sonia Goncalves, Ewan Harrison, David K. Jackson, Ian Johnston, Dominic Kwiatkowski, Cordelia Langford, John Sillitoe on behalf of the Wellcome Sanger Institute COVID-19 Surveillance Team (http://www.sanger.ac.uk/covid-team) |
| EPI_ISL_559089 | Lighthouse Lab in Alderley Park | Wellcome Sanger Institute for the COVID-19 Genomics UK (COG-UK) consortium | The Lighthouse Lab in Alderley Park and Alex Alderton, Roberto Amato, Sonia Goncalves, Ewan Harrison, David K. Jackson, Ian Johnston, Dominic Kwiatkowski, Cordelia Langford, John Sillitoe on behalf of the Wellcome Sanger Institute COVID-19 Surveillance Team (http://www.sanger.ac.uk/covid-team) |
| EPI_ISL_559090, EPI_ISL_559091, EPI_ISL_559092, EPI_ISL_559093, EPI_ISL_559094, EPI_ISL_559095, EPI_ISL_559096, EPI_ISL_559097, EPI_ISL_559098, EPI_ISL_559099 | Lighthouse Lab in Alderley Park | Wellcome Sanger Institute for the COVID-19 Genomics UK (COG-UK) consortium | The Lighthouse Lab in Alderley Park and Alex Alderton, Roberto Amato, Sonia Goncalves, Ewan Harrison, David K. Jackson, Ian Johnston, Dominic Kwiatkowski, Cordelia Langford, John Sillitoe on behalf of the Wellcome Sanger Institute COVID-19 Surveillance Team |
| EPI_ISL_559100 | Lighthouse Lab in Milton Keynes | Wellcome Sanger Institute for the COVID-19 Genomics UK (COG-UK) consortium | The Lighthouse Lab in Milton Keynes and Alex Alderton, Roberto Amato, Sonia Goncalves, Ewan Harrison, David K. Jackson, Ian Johnston, Dominic Kwiatkowski, Cordelia Langford, John Sillitoe on behalf of the Wellcome Sanger Institute COVID-19 Surveillance Team (http://www.sanger.ac.uk/covid-team) |
| EPI_ISL_559101, EPI_ISL_559102 | Lighthouse Lab in Alderley Park | Wellcome Sanger Institute for the COVID-19 Genomics UK (COG-UK) consortium | The Lighthouse Lab in Alderley Park and Alex Alderton, Roberto Amato, Sonia Goncalves, Ewan Harrison, David K. Jackson, Ian Johnston, Dominic Kwiatkowski, Cordelia Langford, John Sillitoe on behalf of the Wellcome Sanger Institute COVID-19 Surveillance Team |
| EPI_ISL_559103 | Lighthouse Lab in Alderley Park | Wellcome Sanger Institute for the COVID-19 Genomics UK (COG-UK) consortium | The Lighthouse Lab in Alderley Park and Alex Alderton, Roberto Amato, Sonia Goncalves, Ewan Harrison, David K. Jackson, Ian Johnston, Dominic Kwiatkowski, Cordelia Langford, John Sillitoe on behalf of the Wellcome Sanger Institute COVID-19 Surveillance Team |

| | | | |
|---|---|---|---|
| | | | (http://www.sanger.ac.uk/covid-team) |
| EPI_ISL_559104 | Lighthouse Lab in Milton Keynes | Wellcome Sanger Institute for the COVID-19 Genomics UK (COG-UK) consortium | The Lighthouse Lab in Milton Keynes and Alex Alderton, Roberto Amato, Sonia Goncalves, Ewan Harrison, David K. Jackson, Ian Johnston, Dominic Kwiatkowski, Cordelia Langford, John Sillitoe on behalf of the Wellcome Sanger Institute COVID-19 Surveillance Team (http://www.sanger.ac.uk/covid-team) |
| EPI_ISL_559105 | Lighthouse Lab in Alderley Park | Wellcome Sanger Institute for the COVID-19 Genomics UK (COG-UK) consortium | The Lighthouse Lab in Alderley Park and Alex Alderton, Roberto Amato, Sonia Goncalves, Ewan Harrison, David K. Jackson, Ian Johnston, Dominic Kwiatkowski, Cordelia Langford, John Sillitoe on behalf of the Wellcome Sanger Institute COVID-19 Surveillance Team |
| EPI_ISL_559106 | Lighthouse Lab in Milton Keynes | Wellcome Sanger Institute for the COVID-19 Genomics UK (COG-UK) consortium | The Lighthouse Lab in Milton Keynes and Alex Alderton, Roberto Amato, Sonia Goncalves, Ewan Harrison, David K. Jackson, Ian Johnston, Dominic Kwiatkowski, Cordelia Langford, John Sillitoe on behalf of the Wellcome Sanger Institute COVID-19 Surveillance Team (http://www.sanger.ac.uk/covid-team) |
| EPI_ISL_559107, EPI_ISL_559108, EPI_ISL_559109, EPI_ISL_559110, EPI_ISL_559111, EPI_ISL_559112, EPI_ISL_559113, EPI_ISL_559114, EPI_ISL_559115, EPI_ISL_559116 | Lighthouse Lab in Alderley Park | Wellcome Sanger Institute for the COVID-19 Genomics UK (COG-UK) consortium | The Lighthouse Lab in Alderley Park and Alex Alderton, Roberto Amato, Sonia Goncalves, Ewan Harrison, David K. Jackson, Ian Johnston, Dominic Kwiatkowski, Cordelia Langford, John Sillitoe on behalf of the Wellcome Sanger Institute COVID-19 Surveillance Team |
| EPI_ISL_559117 | Lighthouse Lab in Milton Keynes | Wellcome Sanger Institute for the COVID-19 Genomics UK (COG-UK) consortium | The Lighthouse Lab in Milton Keynes and Alex Alderton, Roberto Amato, Sonia Goncalves, Ewan Harrison, David K. Jackson, Ian Johnston, Dominic Kwiatkowski, Cordelia Langford, John Sillitoe on behalf of the Wellcome Sanger Institute COVID-19 Surveillance Team (http://www.sanger.ac.uk/covid-team) |
| EPI_ISL_559118 | Lighthouse Lab in Alderley Park | Wellcome Sanger Institute for the COVID-19 Genomics UK (COG-UK) consortium | The Lighthouse Lab in Alderley Park and Alex Alderton, Roberto Amato, Sonia Goncalves, Ewan Harrison, David K. Jackson, Ian Johnston, Dominic Kwiatkowski, Cordelia Langford, John Sillitoe on behalf of the Wellcome Sanger Institute COVID-19 Surveillance Team |
| EPI_ISL_559119 | Lighthouse Lab in Milton Keynes | Wellcome Sanger Institute for the COVID-19 Genomics UK (COG-UK) consortium | The Lighthouse Lab in Milton Keynes and Alex Alderton, Roberto Amato, Sonia Goncalves, Ewan Harrison, David K. Jackson, Ian Johnston, Dominic Kwiatkowski, Cordelia Langford, John Sillitoe on behalf of the Wellcome Sanger Institute COVID-19 Surveillance Team (http://www.sanger.ac.uk/covid-team) |
| EPI_ISL_559121, EPI_ISL_559122, EPI_ISL_559123, EPI_ISL_559124 | Lighthouse Lab in Alderley Park | Wellcome Sanger Institute for the COVID-19 Genomics UK (COG-UK) consortium | The Lighthouse Lab in Alderley Park and Alex Alderton, Roberto Amato, Sonia Goncalves, Ewan Harrison, David K. Jackson, Ian Johnston, Dominic Kwiatkowski, Cordelia Langford, John Sillitoe on behalf of the Wellcome Sanger Institute COVID-19 Surveillance Team |
| EPI_ISL_559126 | Lighthouse Lab in Milton Keynes | Wellcome Sanger Institute for the COVID-19 Genomics UK (COG-UK) consortium | The Lighthouse Lab in Alderley Park and Alex Alderton, Roberto Amato, Sonia Goncalves, Ewan Harrison, David K. Jackson, Ian Johnston, Dominic Kwiatkowski, Cordelia Langford, John Sillitoe on behalf of the Wellcome Sanger Institute COVID-19 Surveillance Team |
| EPI_ISL_559127, EPI_ISL_559128, EPI_ISL_559130 | Lighthouse Lab in Alderley Park | Wellcome Sanger Institute for the COVID-19 Genomics UK (COG-UK) consortium | The Lighthouse Lab in Alderley Park and Alex Alderton, Roberto Amato, Sonia Goncalves, Ewan Harrison, David K. Jackson, Ian Johnston, Dominic Kwiatkowski, Cordelia Langford, John Sillitoe on behalf of the Wellcome Sanger Institute COVID-19 Surveillance Team |
| EPI_ISL_559131, EPI_ISL_559132 | Lighthouse Lab in Milton Keynes | Wellcome Sanger Institute for the COVID-19 Genomics UK (COG-UK) consortium | The Lighthouse Lab in Milton Keynes and Alex Alderton, Roberto Amato, Sonia Goncalves, Ewan Harrison, David K. Jackson, Ian Johnston, Dominic Kwiatkowski, Cordelia Langford, John Sillitoe on behalf of the Wellcome Sanger Institute COVID-19 Surveillance Team (http://www.sanger.ac.uk/covid-team) |
| EPI_ISL_559134, EPI_ISL_559135 | Lighthouse Lab in Alderley Park | Wellcome Sanger Institute for the COVID-19 Genomics UK (COG-UK) consortium | The Lighthouse Lab in Alderley Park and Alex Alderton, Roberto Amato, Sonia Goncalves, Ewan Harrison, David K. Jackson, Ian Johnston, Dominic Kwiatkowski, Cordelia Langford, John Sillitoe on behalf of the Wellcome Sanger Institute COVID-19 Surveillance Team |
| EPI_ISL_559136 | Lighthouse Lab in Milton Keynes | Wellcome Sanger Institute for the COVID-19 Genomics UK (COG-UK) consortium | The Lighthouse Lab in Milton Keynes and Alex Alderton, Roberto Amato, Sonia Goncalves, Ewan Harrison, David K. Jackson, Ian Johnston, Dominic Kwiatkowski, Cordelia Langford, John Sillitoe on behalf of the Wellcome Sanger Institute COVID-19 Surveillance Team (http://www.sanger.ac.uk/covid-team) |
| EPI_ISL_559137, EPI_ISL_559138, EPI_ISL_559139, EPI_ISL_559140, EPI_ISL_559141, EPI_ISL_559142, EPI_ISL_559143, EPI_ISL_559144, EPI_ISL_559145, EPI_ISL_559146 | Lighthouse Lab in Alderley Park | Wellcome Sanger Institute for the COVID-19 Genomics UK (COG-UK) consortium | The Lighthouse Lab in Alderley Park and Alex Alderton, Roberto Amato, Sonia Goncalves, Ewan Harrison, David K. Jackson, Ian Johnston, Dominic Kwiatkowski, Cordelia Langford, John Sillitoe on behalf of the Wellcome Sanger Institute COVID-19 Surveillance Team |
| EPI_ISL_559147 | Lighthouse Lab in Milton Keynes | Wellcome Sanger Institute for the COVID-19 Genomics UK (COG-UK) consortium | The Lighthouse Lab in Milton Keynes and Alex Alderton, Roberto Amato, Sonia Goncalves, Ewan Harrison, David K. Jackson, Ian Johnston, Dominic Kwiatkowski, Cordelia Langford, John Sillitoe on behalf of the Wellcome Sanger Institute COVID-19 Surveillance Team (http://www.sanger.ac.uk/covid-team) |
| EPI_ISL_559148 | Lighthouse Lab in Alderley Park | Wellcome Sanger Institute for the COVID-19 Genomics UK (COG-UK) consortium | The Lighthouse Lab in Alderley Park and Alex Alderton, Roberto Amato, Sonia Goncalves, Ewan Harrison, David K. Jackson, Ian Johnston, Dominic Kwiatkowski, Cordelia Langford, John Sillitoe on behalf of the Wellcome Sanger Institute COVID-19 Surveillance Team |
| EPI_ISL_559149 | Lighthouse Lab in Milton Keynes | Wellcome Sanger Institute for the COVID-19 Genomics UK (COG-UK) consortium | The Lighthouse Lab in Milton Keynes and Alex Alderton, Roberto Amato, Sonia Goncalves, Ewan Harrison, David K. Jackson, Ian Johnston, Dominic Kwiatkowski, Cordelia Langford, John Sillitoe on behalf of the Wellcome Sanger Institute COVID-19 Surveillance Team (http://www.sanger.ac.uk/covid-team) |
| EPI_ISL_559150, EPI_ISL_559151, EPI_ISL_559152, EPI_ISL_559153 | Lighthouse Lab in Alderley Park | Wellcome Sanger Institute for the COVID-19 Genomics UK (COG-UK) consortium | The Lighthouse Lab in Alderley Park and Alex Alderton, Roberto Amato, Sonia Goncalves, Ewan Harrison, David K. Jackson, Ian Johnston, Dominic Kwiatkowski, Cordelia Langford, John Sillitoe on behalf of the Wellcome Sanger Institute COVID-19 Surveillance Team |
| EPI_ISL_559154 | Lighthouse Lab in Milton Keynes | Wellcome Sanger Institute for the COVID-19 Genomics UK (COG-UK) consortium | The Lighthouse Lab in Milton Keynes and Alex Alderton, Roberto Amato, Sonia Goncalves, Ewan Harrison, David K. Jackson, Ian Johnston, Dominic Kwiatkowski, Cordelia Langford, John Sillitoe on behalf of the Wellcome Sanger Institute COVID-19 Surveillance Team (http://www.sanger.ac.uk/covid-team) |
| EPI_ISL_559155, EPI_ISL_559156 | Lighthouse Lab in Alderley Park | Wellcome Sanger Institute for the COVID-19 Genomics UK (COG-UK) consortium | The Lighthouse Lab in Alderley Park and Alex Alderton, Roberto Amato, Sonia Goncalves, Ewan Harrison, David K. Jackson, Ian Johnston, Dominic Kwiatkowski, Cordelia Langford, John Sillitoe on behalf of the Wellcome Sanger Institute COVID-19 Surveillance Team |
| EPI_ISL_559157, EPI_ISL_559158 | Lighthouse Lab in Milton Keynes | Wellcome Sanger Institute for the COVID-19 Genomics UK (COG-UK) consortium | The Lighthouse Lab in Milton Keynes and Alex Alderton, Roberto Amato, Sonia Goncalves, Ewan Harrison, David K. Jackson, Ian Johnston, Dominic Kwiatkowski, Cordelia Langford, John Sillitoe on behalf of the Wellcome Sanger Institute COVID-19 Surveillance Team (http://www.sanger.ac.uk/covid-team) |
| EPI_ISL_559159, EPI_ISL_559160, EPI_ISL_559161, EPI_ISL_559162 | Lighthouse Lab in Alderley Park | Wellcome Sanger Institute for the COVID-19 Genomics UK (COG-UK) consortium | The Lighthouse Lab in Alderley Park and Alex Alderton, Roberto Amato, Sonia Goncalves, Ewan Harrison, David K. Jackson, Ian Johnston, Dominic Kwiatkowski, Cordelia Langford, John Sillitoe on behalf of the Wellcome Sanger Institute COVID-19 Surveillance Team |
| EPI_ISL_559163, EPI_ISL_559164 | Lighthouse Lab in Milton Keynes | Wellcome Sanger Institute for the COVID-19 Genomics UK (COG-UK) consortium | The Lighthouse Lab in Milton Keynes and Alex Alderton, Roberto Amato, Sonia Goncalves, Ewan Harrison, David K. Jackson, Ian Johnston, Dominic Kwiatkowski, Cordelia Langford, John Sillitoe on behalf of the Wellcome Sanger Institute COVID-19 Surveillance Team (http://www.sanger.ac.uk/covid-team) |
| EPI_ISL_559165 | Lighthouse Lab in Alderley Park | Wellcome Sanger Institute for the COVID-19 Genomics UK (COG-UK) consortium | The Lighthouse Lab in Alderley Park and Alex Alderton, Roberto Amato, Sonia Goncalves, Ewan Harrison, David K. Jackson, Ian Johnston, Dominic Kwiatkowski, Cordelia Langford, John Sillitoe on behalf of the Wellcome Sanger Institute COVID-19 Surveillance Team |
| EPI_ISL_559167 | Lighthouse Lab in Milton Keynes | Wellcome Sanger Institute for the COVID-19 Genomics UK (COG-UK) consortium | The Lighthouse Lab in Milton Keynes and Alex Alderton, Roberto Amato, Sonia Goncalves, Ewan Harrison, David K. Jackson, Ian Johnston, Dominic Kwiatkowski, Cordelia Langford, John Sillitoe on behalf of the Wellcome Sanger Institute COVID-19 Surveillance Team (http://www.sanger.ac.uk/covid-team) |
| EPI_ISL_559168, EPI_ISL_559169, EPI_ISL_559170, EPI_ISL_559172, EPI_ISL_559173 | Lighthouse Lab in Alderley Park | Wellcome Sanger Institute for the COVID-19 Genomics UK (COG-UK) consortium | The Lighthouse Lab in Alderley Park and Alex Alderton, Roberto Amato, Sonia Goncalves, Ewan Harrison, David K. Jackson, Ian Johnston, Dominic Kwiatkowski, Cordelia Langford, John Sillitoe on behalf of the Wellcome Sanger Institute COVID-19 Surveillance Team |
| EPI_ISL_559174 | Lighthouse Lab in Milton Keynes | Wellcome Sanger Institute for the COVID-19 Genomics UK (COG-UK) consortium | The Lighthouse Lab in Milton Keynes and Alex Alderton, Roberto Amato, Sonia Goncalves, Ewan Harrison, David K. Jackson, Ian Johnston, Dominic Kwiatkowski, Cordelia Langford, John Sillitoe on behalf of the Wellcome Sanger Institute COVID-19 Surveillance Team |

| | | | |
|---|---|---|---|
| | | | (http://www.sanger.ac.uk/covid-team) |
| EPI_ISL_559175 | Lighthouse Lab in Alderley Park | Wellcome Sanger Institute for the COVID-19 Genomics UK (COG-UK) consortium | The Lighthouse Lab in Alderley Park and Alex Alderton, Roberto Amato, Sonia Goncalves, Ewan Harrison, David K. Jackson, Ian Johnston, Dominic Kwiatkowski, Cordelia Langford, John Sillitoe on behalf of the Wellcome Sanger Institute COVID-19 Surveillance Team |
| EPI_ISL_559176 | Lighthouse Lab in Alderley Park | Wellcome Sanger Institute for the COVID-19 Genomics UK (COG-UK) consortium | The Lighthouse Lab in Alderley Park and Alex Alderton, Roberto Amato, Sonia Goncalves, Ewan Harrison, David K. Jackson, Ian Johnston, Dominic Kwiatkowski, Cordelia Langford, John Sillitoe on behalf of the Wellcome Sanger Institute COVID-19 Surveillance Team (http://www.sanger.ac.uk/covid-team) |
| EPI_ISL_559177, EPI_ISL_559178, EPI_ISL_559179, EPI_ISL_559180 | Lighthouse Lab in Alderley Park | Wellcome Sanger Institute for the COVID-19 Genomics UK (COG-UK) consortium | The Lighthouse Lab in Alderley Park and Alex Alderton, Roberto Amato, Sonia Goncalves, Ewan Harrison, David K. Jackson, Ian Johnston, Dominic Kwiatkowski, Cordelia Langford, John Sillitoe on behalf of the Wellcome Sanger Institute COVID-19 Surveillance Team |
| EPI_ISL_559181 | Lighthouse Lab in Milton Keynes | Wellcome Sanger Institute for the COVID-19 Genomics UK (COG-UK) consortium | The Lighthouse Lab in Milton Keynes and Alex Alderton, Roberto Amato, Sonia Goncalves, Ewan Harrison, David K. Jackson, Ian Johnston, Dominic Kwiatkowski, Cordelia Langford, John Sillitoe on behalf of the Wellcome Sanger Institute COVID-19 Surveillance Team (http://www.sanger.ac.uk/covid-team) |
| EPI_ISL_559182, EPI_ISL_559183, EPI_ISL_559184, EPI_ISL_559185, EPI_ISL_559186 | Lighthouse Lab in Alderley Park | Wellcome Sanger Institute for the COVID-19 Genomics UK (COG-UK) consortium | The Lighthouse Lab in Alderley Park and Alex Alderton, Roberto Amato, Sonia Goncalves, Ewan Harrison, David K. Jackson, Ian Johnston, Dominic Kwiatkowski, Cordelia Langford, John Sillitoe on behalf of the Wellcome Sanger Institute COVID-19 Surveillance Team |
| EPI_ISL_559187 | Lighthouse Lab in Milton Keynes | Wellcome Sanger Institute for the COVID-19 Genomics UK (COG-UK) consortium | The Lighthouse Lab in Milton Keynes and Alex Alderton, Roberto Amato, Sonia Goncalves, Ewan Harrison, David K. Jackson, Ian Johnston, Dominic Kwiatkowski, Cordelia Langford, John Sillitoe on behalf of the Wellcome Sanger Institute COVID-19 Surveillance Team (http://www.sanger.ac.uk/covid-team) |
| EPI_ISL_559188, EPI_ISL_559189 | Lighthouse Lab in Alderley Park | Wellcome Sanger Institute for the COVID-19 Genomics UK (COG-UK) consortium | The Lighthouse Lab in Alderley Park and Alex Alderton, Roberto Amato, Sonia Goncalves, Ewan Harrison, David K. Jackson, Ian Johnston, Dominic Kwiatkowski, Cordelia Langford, John Sillitoe on behalf of the Wellcome Sanger Institute COVID-19 Surveillance Team |
| EPI_ISL_559190 | Lighthouse Lab in Alderley Park | Wellcome Sanger Institute for the COVID-19 Genomics UK (COG-UK) consortium | The Lighthouse Lab in Alderley Park and Alex Alderton, Roberto Amato, Sonia Goncalves, Ewan Harrison, David K. Jackson, Ian Johnston, Dominic Kwiatkowski, Cordelia Langford, John Sillitoe on behalf of the Wellcome Sanger Institute COVID-19 Surveillance Team (http://www.sanger.ac.uk/covid-team) |
| EPI_ISL_559191 | Lighthouse Lab in Alderley Park | Wellcome Sanger Institute for the COVID-19 Genomics UK (COG-UK) consortium | The Lighthouse Lab in Alderley Park and Alex Alderton, Roberto Amato, Sonia Goncalves, Ewan Harrison, David K. Jackson, Ian Johnston, Dominic Kwiatkowski, Cordelia Langford, John Sillitoe on behalf of the Wellcome Sanger Institute COVID-19 Surveillance Team |
| EPI_ISL_559192 | Lighthouse Lab in Alderley Park | Wellcome Sanger Institute for the COVID-19 Genomics UK (COG-UK) consortium | The Lighthouse Lab in Alderley Park and Alex Alderton, Roberto Amato, Sonia Goncalves, Ewan Harrison, David K. Jackson, Ian Johnston, Dominic Kwiatkowski, Cordelia Langford, John Sillitoe on behalf of the Wellcome Sanger Institute COVID-19 Surveillance Team (http://www.sanger.ac.uk/covid-team) |
| EPI_ISL_559193 | Lighthouse Lab in Alderley Park | Wellcome Sanger Institute for the COVID-19 Genomics UK (COG-UK) consortium | The Lighthouse Lab in Alderley Park and Alex Alderton, Roberto Amato, Sonia Goncalves, Ewan Harrison, David K. Jackson, Ian Johnston, Dominic Kwiatkowski, Cordelia Langford, John Sillitoe on behalf of the Wellcome Sanger Institute COVID-19 Surveillance Team |
| EPI_ISL_559194, EPI_ISL_559195 | Lighthouse Lab in Milton Keynes | Wellcome Sanger Institute for the COVID-19 Genomics UK (COG-UK) consortium | The Lighthouse Lab in Milton Keynes and Alex Alderton, Roberto Amato, Sonia Goncalves, Ewan Harrison, David K. Jackson, Ian Johnston, Dominic Kwiatkowski, Cordelia Langford, John Sillitoe on behalf of the Wellcome Sanger Institute COVID-19 Surveillance Team (http://www.sanger.ac.uk/covid-team) |
| EPI_ISL_559196, EPI_ISL_559197, EPI_ISL_559198, EPI_ISL_559199, EPI_ISL_559200 | Lighthouse Lab in Alderley Park | Wellcome Sanger Institute for the COVID-19 Genomics UK (COG-UK) consortium | The Lighthouse Lab in Alderley Park and Alex Alderton, Roberto Amato, Sonia Goncalves, Ewan Harrison, David K. Jackson, Ian Johnston, Dominic Kwiatkowski, Cordelia Langford, John Sillitoe on behalf of the Wellcome Sanger Institute COVID-19 Surveillance Team |
| EPI_ISL_559201 | Lighthouse Lab in Milton Keynes | Wellcome Sanger Institute for the COVID-19 Genomics UK (COG-UK) consortium | The Lighthouse Lab in Milton Keynes and Alex Alderton, Roberto Amato, Sonia Goncalves, Ewan Harrison, David K. Jackson, Ian Johnston, Dominic Kwiatkowski, Cordelia Langford, John Sillitoe on behalf of the Wellcome Sanger Institute COVID-19 Surveillance Team (http://www.sanger.ac.uk/covid-team) |
| EPI_ISL_559202, EPI_ISL_559203, EPI_ISL_559205 | Lighthouse Lab in Alderley Park | Wellcome Sanger Institute for the COVID-19 Genomics UK (COG-UK) consortium | The Lighthouse Lab in Alderley Park and Alex Alderton, Roberto Amato, Sonia Goncalves, Ewan Harrison, David K. Jackson, Ian Johnston, Dominic Kwiatkowski, Cordelia Langford, John Sillitoe on behalf of the Wellcome Sanger Institute COVID-19 Surveillance Team |
| EPI_ISL_559206 | Lighthouse Lab in Alderley Park | Wellcome Sanger Institute for the COVID-19 Genomics UK (COG-UK) consortium | The Lighthouse Lab in Alderley Park and Alex Alderton, Roberto Amato, Sonia Goncalves, Ewan Harrison, David K. Jackson, Ian Johnston, Dominic Kwiatkowski, Cordelia Langford, John Sillitoe on behalf of the Wellcome Sanger Institute COVID-19 Surveillance Team (http://www.sanger.ac.uk/covid-team) |
| EPI_ISL_559207 | Lighthouse Lab in Milton Keynes | Wellcome Sanger Institute for the COVID-19 Genomics UK (COG-UK) consortium | The Lighthouse Lab in Milton Keynes and Alex Alderton, Roberto Amato, Sonia Goncalves, Ewan Harrison, David K. Jackson, Ian Johnston, Dominic Kwiatkowski, Cordelia Langford, John Sillitoe on behalf of the Wellcome Sanger Institute COVID-19 Surveillance Team (http://www.sanger.ac.uk/covid-team) |
| EPI_ISL_559208, EPI_ISL_559209, EPI_ISL_559210, EPI_ISL_559211, EPI_ISL_559212, EPI_ISL_559214, EPI_ISL_559216, EPI_ISL_559217 | Lighthouse Lab in Alderley Park | Wellcome Sanger Institute for the COVID-19 Genomics UK (COG-UK) consortium | The Lighthouse Lab in Alderley Park and Alex Alderton, Roberto Amato, Sonia Goncalves, Ewan Harrison, David K. Jackson, Ian Johnston, Dominic Kwiatkowski, Cordelia Langford, John Sillitoe on behalf of the Wellcome Sanger Institute COVID-19 Surveillance Team |
| EPI_ISL_559218 | Lighthouse Lab in Milton Keynes | Wellcome Sanger Institute for the COVID-19 Genomics UK (COG-UK) consortium | The Lighthouse Lab in Milton Keynes and Alex Alderton, Roberto Amato, Sonia Goncalves, Ewan Harrison, David K. Jackson, Ian Johnston, Dominic Kwiatkowski, Cordelia Langford, John Sillitoe on behalf of the Wellcome Sanger Institute COVID-19 Surveillance Team (http://www.sanger.ac.uk/covid-team) |
| EPI_ISL_559219, EPI_ISL_559220 | Lighthouse Lab in Alderley Park | Wellcome Sanger Institute for the COVID-19 Genomics UK (COG-UK) consortium | The Lighthouse Lab in Alderley Park and Alex Alderton, Roberto Amato, Sonia Goncalves, Ewan Harrison, David K. Jackson, Ian Johnston, Dominic Kwiatkowski, Cordelia Langford, John Sillitoe on behalf of the Wellcome Sanger Institute COVID-19 Surveillance Team |
| EPI_ISL_559221 | Lighthouse Lab in Alderley Park | Wellcome Sanger Institute for the COVID-19 Genomics UK (COG-UK) consortium | The Lighthouse Lab in Alderley Park and Alex Alderton, Roberto Amato, Sonia Goncalves, Ewan Harrison, David K. Jackson, Ian Johnston, Dominic Kwiatkowski, Cordelia Langford, John Sillitoe on behalf of the Wellcome Sanger Institute COVID-19 Surveillance Team (http://www.sanger.ac.uk/covid-team) |
| EPI_ISL_559222 | Lighthouse Lab in Alderley Park | Wellcome Sanger Institute for the COVID-19 Genomics UK (COG-UK) consortium | The Lighthouse Lab in Alderley Park and Alex Alderton, Roberto Amato, Sonia Goncalves, Ewan Harrison, David K. Jackson, Ian Johnston, Dominic Kwiatkowski, Cordelia Langford, John Sillitoe on behalf of the Wellcome Sanger Institute COVID-19 Surveillance Team |
| EPI_ISL_559223 | Lighthouse Lab in Milton Keynes | Wellcome Sanger Institute for the COVID-19 Genomics UK (COG-UK) consortium | The Lighthouse Lab in Milton Keynes and Alex Alderton, Roberto Amato, Sonia Goncalves, Ewan Harrison, David K. Jackson, Ian Johnston, Dominic Kwiatkowski, Cordelia Langford, John Sillitoe on behalf of the Wellcome Sanger Institute COVID-19 Surveillance Team (http://www.sanger.ac.uk/covid-team) |
| EPI_ISL_559224, EPI_ISL_559225, EPI_ISL_559226, EPI_ISL_559227, EPI_ISL_559229, EPI_ISL_559230, EPI_ISL_559231, EPI_ISL_559233, EPI_ISL_559234 | Lighthouse Lab in Alderley Park | Wellcome Sanger Institute for the COVID-19 Genomics UK (COG-UK) consortium | The Lighthouse Lab in Alderley Park and Alex Alderton, Roberto Amato, Sonia Goncalves, Ewan Harrison, David K. Jackson, Ian Johnston, Dominic Kwiatkowski, Cordelia Langford, John Sillitoe on behalf of the Wellcome Sanger Institute COVID-19 Surveillance Team |
| EPI_ISL_559235, EPI_ISL_559236 | Lighthouse Lab in Milton Keynes | Wellcome Sanger Institute for the COVID-19 Genomics UK (COG-UK) consortium | The Lighthouse Lab in Milton Keynes and Alex Alderton, Roberto Amato, Sonia Goncalves, Ewan Harrison, David K. Jackson, Ian Johnston, Dominic Kwiatkowski, Cordelia Langford, John Sillitoe on behalf of the Wellcome Sanger Institute COVID-19 Surveillance Team (http://www.sanger.ac.uk/covid-team) |
| EPI_ISL_559237, EPI_ISL_559238, EPI_ISL_559239 | Lighthouse Lab in Alderley Park | Wellcome Sanger Institute for the COVID-19 Genomics UK (COG-UK) consortium | The Lighthouse Lab in Alderley Park and Alex Alderton, Roberto Amato, Sonia Goncalves, Ewan Harrison, David K. Jackson, Ian Johnston, Dominic Kwiatkowski, Cordelia Langford, John Sillitoe on behalf of the Wellcome Sanger Institute COVID-19 Surveillance Team |
| EPI_ISL_559240 | Lighthouse Lab in Alderley Park | Wellcome Sanger Institute for the COVID-19 Genomics UK | The Lighthouse Lab in Alderley Park and Alex Alderton, Roberto Amato, Sonia Goncalves, Ewan Harrison, David K. Jackson, Ian Johnston, Dominic |

| | | | |
|---|---|---|---|
| | | (COG-UK) consortium | Kwiatkowski, Cordelia Langford, John Sillitoe on behalf of the Wellcome Sanger Institute COVID-19 Surveillance Team (http://www.sanger.ac.uk/covid-team) |
| EPI_ISL_559241 | Lighthouse Lab in Milton Keynes | Wellcome Sanger Institute for the COVID-19 Genomics UK (COG-UK) consortium | The Lighthouse Lab in Milton Keynes and Alex Alderton, Roberto Amato, Sonia Goncalves, Ewan Harrison, David K. Jackson, Ian Johnston, Dominic Kwiatkowski, Cordelia Langford, John Sillitoe on behalf of the Wellcome Sanger Institute COVID-19 Surveillance Team (http://www.sanger.ac.uk/covid-team) |
| EPI_ISL_559242, EPI_ISL_559243 | Lighthouse Lab in Alderley Park | Wellcome Sanger Institute for the COVID-19 Genomics UK (COG-UK) consortium | The Lighthouse Lab in Alderley Park and Alex Alderton, Roberto Amato, Sonia Goncalves, Ewan Harrison, David K. Jackson, Ian Johnston, Dominic Kwiatkowski, Cordelia Langford, John Sillitoe on behalf of the Wellcome Sanger Institute COVID-19 Surveillance Team |
| EPI_ISL_559244 | Lighthouse Lab in Milton Keynes | Wellcome Sanger Institute for the COVID-19 Genomics UK (COG-UK) consortium | The Lighthouse Lab in Milton Keynes and Alex Alderton, Roberto Amato, Sonia Goncalves, Ewan Harrison, David K. Jackson, Ian Johnston, Dominic Kwiatkowski, Cordelia Langford, John Sillitoe on behalf of the Wellcome Sanger Institute COVID-19 Surveillance Team (http://www.sanger.ac.uk/covid-team) |
| EPI_ISL_559245 | Lighthouse Lab in Alderley Park | Wellcome Sanger Institute for the COVID-19 Genomics UK (COG-UK) consortium | The Lighthouse Lab in Alderley Park and Alex Alderton, Roberto Amato, Sonia Goncalves, Ewan Harrison, David K. Jackson, Ian Johnston, Dominic Kwiatkowski, Cordelia Langford, John Sillitoe on behalf of the Wellcome Sanger Institute COVID-19 Surveillance Team |
| EPI_ISL_559246, EPI_ISL_559247 | Lighthouse Lab in Milton Keynes | Wellcome Sanger Institute for the COVID-19 Genomics UK (COG-UK) consortium | The Lighthouse Lab in Milton Keynes and Alex Alderton, Roberto Amato, Sonia Goncalves, Ewan Harrison, David K. Jackson, Ian Johnston, Dominic Kwiatkowski, Cordelia Langford, John Sillitoe on behalf of the Wellcome Sanger Institute COVID-19 Surveillance Team (http://www.sanger.ac.uk/covid-team) |
| EPI_ISL_559248, EPI_ISL_559249, EPI_ISL_559250, EPI_ISL_559251, EPI_ISL_559252 | Lighthouse Lab in Alderley Park | Wellcome Sanger Institute for the COVID-19 Genomics UK (COG-UK) consortium | The Lighthouse Lab in Alderley Park and Alex Alderton, Roberto Amato, Sonia Goncalves, Ewan Harrison, David K. Jackson, Ian Johnston, Dominic Kwiatkowski, Cordelia Langford, John Sillitoe on behalf of the Wellcome Sanger Institute COVID-19 Surveillance Team |
| EPI_ISL_559253, EPI_ISL_559254 | Lighthouse Lab in Milton Keynes | Wellcome Sanger Institute for the COVID-19 Genomics UK (COG-UK) consortium | The Lighthouse Lab in Milton Keynes and Alex Alderton, Roberto Amato, Sonia Goncalves, Ewan Harrison, David K. Jackson, Ian Johnston, Dominic Kwiatkowski, Cordelia Langford, John Sillitoe on behalf of the Wellcome Sanger Institute COVID-19 Surveillance Team (http://www.sanger.ac.uk/covid-team) |
| EPI_ISL_559255 | Lighthouse Lab in Alderley Park | Wellcome Sanger Institute for the COVID-19 Genomics UK (COG-UK) consortium | The Lighthouse Lab in Alderley Park and Alex Alderton, Roberto Amato, Sonia Goncalves, Ewan Harrison, David K. Jackson, Ian Johnston, Dominic Kwiatkowski, Cordelia Langford, John Sillitoe on behalf of the Wellcome Sanger Institute COVID-19 Surveillance Team |
| EPI_ISL_559256 | Lighthouse Lab in Milton Keynes | Wellcome Sanger Institute for the COVID-19 Genomics UK (COG-UK) consortium | The Lighthouse Lab in Milton Keynes and Alex Alderton, Roberto Amato, Sonia Goncalves, Ewan Harrison, David K. Jackson, Ian Johnston, Dominic Kwiatkowski, Cordelia Langford, John Sillitoe on behalf of the Wellcome Sanger Institute COVID-19 Surveillance Team (http://www.sanger.ac.uk/covid-team) |
| EPI_ISL_559257, EPI_ISL_559258 | Lighthouse Lab in Alderley Park | Wellcome Sanger Institute for the COVID-19 Genomics UK (COG-UK) consortium | The Lighthouse Lab in Alderley Park and Alex Alderton, Roberto Amato, Sonia Goncalves, Ewan Harrison, David K. Jackson, Ian Johnston, Dominic Kwiatkowski, Cordelia Langford, John Sillitoe on behalf of the Wellcome Sanger Institute COVID-19 Surveillance Team |
| EPI_ISL_559259 | Lighthouse Lab in Milton Keynes | Wellcome Sanger Institute for the COVID-19 Genomics UK (COG-UK) consortium | The Lighthouse Lab in Milton Keynes and Alex Alderton, Roberto Amato, Sonia Goncalves, Ewan Harrison, David K. Jackson, Ian Johnston, Dominic Kwiatkowski, Cordelia Langford, John Sillitoe on behalf of the Wellcome Sanger Institute COVID-19 Surveillance Team (http://www.sanger.ac.uk/covid-team) |
| EPI_ISL_559260, EPI_ISL_559262 | Lighthouse Lab in Alderley Park | Wellcome Sanger Institute for the COVID-19 Genomics UK (COG-UK) consortium | The Lighthouse Lab in Alderley Park and Alex Alderton, Roberto Amato, Sonia Goncalves, Ewan Harrison, David K. Jackson, Ian Johnston, Dominic Kwiatkowski, Cordelia Langford, John Sillitoe on behalf of the Wellcome Sanger Institute COVID-19 Surveillance Team |
| EPI_ISL_559263 | Lighthouse Lab in Milton Keynes | Wellcome Sanger Institute for the COVID-19 Genomics UK (COG-UK) consortium | The Lighthouse Lab in Milton Keynes and Alex Alderton, Roberto Amato, Sonia Goncalves, Ewan Harrison, David K. Jackson, Ian Johnston, Dominic Kwiatkowski, Cordelia Langford, John Sillitoe on behalf of the Wellcome Sanger Institute COVID-19 Surveillance Team (http://www.sanger.ac.uk/covid-team) |
| EPI_ISL_559265 | Lighthouse Lab in Alderley Park | Wellcome Sanger Institute for the COVID-19 Genomics UK (COG-UK) consortium | The Lighthouse Lab in Alderley Park and Alex Alderton, Roberto Amato, Sonia Goncalves, Ewan Harrison, David K. Jackson, Ian Johnston, Dominic Kwiatkowski, Cordelia Langford, John Sillitoe on behalf of the Wellcome Sanger Institute COVID-19 Surveillance Team (http://www.sanger.ac.uk/covid-team) |
| EPI_ISL_559266, EPI_ISL_559267, EPI_ISL_559268, EPI_ISL_559269, EPI_ISL_559270, EPI_ISL_559271, EPI_ISL_559272, EPI_ISL_559273, EPI_ISL_559274, EPI_ISL_559276 | Lighthouse Lab in Alderley Park | Wellcome Sanger Institute for the COVID-19 Genomics UK (COG-UK) consortium | The Lighthouse Lab in Alderley Park and Alex Alderton, Roberto Amato, Sonia Goncalves, Ewan Harrison, David K. Jackson, Ian Johnston, Dominic Kwiatkowski, Cordelia Langford, John Sillitoe on behalf of the Wellcome Sanger Institute COVID-19 Surveillance Team |
| EPI_ISL_559277, EPI_ISL_559278 | Lighthouse Lab in Milton Keynes | Wellcome Sanger Institute for the COVID-19 Genomics UK (COG-UK) consortium | The Lighthouse Lab in Milton Keynes and Alex Alderton, Roberto Amato, Sonia Goncalves, Ewan Harrison, David K. Jackson, Ian Johnston, Dominic Kwiatkowski, Cordelia Langford, John Sillitoe on behalf of the Wellcome Sanger Institute COVID-19 Surveillance Team (http://www.sanger.ac.uk/covid-team) |
| EPI_ISL_559279, EPI_ISL_559280, EPI_ISL_559281, EPI_ISL_559282 | Lighthouse Lab in Alderley Park | Wellcome Sanger Institute for the COVID-19 Genomics UK (COG-UK) consortium | The Lighthouse Lab in Alderley Park and Alex Alderton, Roberto Amato, Sonia Goncalves, Ewan Harrison, David K. Jackson, Ian Johnston, Dominic Kwiatkowski, Cordelia Langford, John Sillitoe on behalf of the Wellcome Sanger Institute COVID-19 Surveillance Team |
| EPI_ISL_559283, EPI_ISL_559284 | Lighthouse Lab in Milton Keynes | Wellcome Sanger Institute for the COVID-19 Genomics UK (COG-UK) consortium | The Lighthouse Lab in Milton Keynes and Alex Alderton, Roberto Amato, Sonia Goncalves, Ewan Harrison, David K. Jackson, Ian Johnston, Dominic Kwiatkowski, Cordelia Langford, John Sillitoe on behalf of the Wellcome Sanger Institute COVID-19 Surveillance Team (http://www.sanger.ac.uk/covid-team) |
| EPI_ISL_559285, EPI_ISL_559286, EPI_ISL_559288, EPI_ISL_559289 | Lighthouse Lab in Alderley Park | Wellcome Sanger Institute for the COVID-19 Genomics UK (COG-UK) consortium | The Lighthouse Lab in Alderley Park and Alex Alderton, Roberto Amato, Sonia Goncalves, Ewan Harrison, David K. Jackson, Ian Johnston, Dominic Kwiatkowski, Cordelia Langford, John Sillitoe on behalf of the Wellcome Sanger Institute COVID-19 Surveillance Team |
| EPI_ISL_559290 | Lighthouse Lab in Milton Keynes | Wellcome Sanger Institute for the COVID-19 Genomics UK (COG-UK) consortium | The Lighthouse Lab in Milton Keynes and Alex Alderton, Roberto Amato, Sonia Goncalves, Ewan Harrison, David K. Jackson, Ian Johnston, Dominic Kwiatkowski, Cordelia Langford, John Sillitoe on behalf of the Wellcome Sanger Institute COVID-19 Surveillance Team (http://www.sanger.ac.uk/covid-team) |
| EPI_ISL_559291, EPI_ISL_559292, EPI_ISL_559293, EPI_ISL_559294, EPI_ISL_559295 | Lighthouse Lab in Alderley Park | Wellcome Sanger Institute for the COVID-19 Genomics UK (COG-UK) consortium | The Lighthouse Lab in Alderley Park and Alex Alderton, Roberto Amato, Sonia Goncalves, Ewan Harrison, David K. Jackson, Ian Johnston, Dominic Kwiatkowski, Cordelia Langford, John Sillitoe on behalf of the Wellcome Sanger Institute COVID-19 Surveillance Team |
| EPI_ISL_559296 | Lighthouse Lab in Milton Keynes | Wellcome Sanger Institute for the COVID-19 Genomics UK (COG-UK) consortium | The Lighthouse Lab in Milton Keynes and Alex Alderton, Roberto Amato, Sonia Goncalves, Ewan Harrison, David K. Jackson, Ian Johnston, Dominic Kwiatkowski, Cordelia Langford, John Sillitoe on behalf of the Wellcome Sanger Institute COVID-19 Surveillance Team (http://www.sanger.ac.uk/covid-team) |
| EPI_ISL_559297, EPI_ISL_559298, EPI_ISL_559299 | Lighthouse Lab in Alderley Park | Wellcome Sanger Institute for the COVID-19 Genomics UK (COG-UK) consortium | The Lighthouse Lab in Alderley Park and Alex Alderton, Roberto Amato, Sonia Goncalves, Ewan Harrison, David K. Jackson, Ian Johnston, Dominic Kwiatkowski, Cordelia Langford, John Sillitoe on behalf of the Wellcome Sanger Institute COVID-19 Surveillance Team |
| EPI_ISL_559300 | Lighthouse Lab in Milton Keynes | Wellcome Sanger Institute for the COVID-19 Genomics UK (COG-UK) consortium | The Lighthouse Lab in Milton Keynes and Alex Alderton, Roberto Amato, Sonia Goncalves, Ewan Harrison, David K. Jackson, Ian Johnston, Dominic Kwiatkowski, Cordelia Langford, John Sillitoe on behalf of the Wellcome Sanger Institute COVID-19 Surveillance Team (http://www.sanger.ac.uk/covid-team) |
| EPI_ISL_559301, EPI_ISL_559302, EPI_ISL_559303, EPI_ISL_559304, EPI_ISL_559305, EPI_ISL_559306, EPI_ISL_559307, EPI_ISL_559308, EPI_ISL_559309, EPI_ISL_559310, EPI_ISL_559311 | | | |
| see above | Lighthouse Lab in Alderley Park | Wellcome Sanger Institute for the COVID-19 Genomics UK (COG-UK) consortium | The Lighthouse Lab in Alderley Park and Alex Alderton, Roberto Amato, Sonia Goncalves, Ewan Harrison, David K. Jackson, Ian Johnston, Dominic Kwiatkowski, Cordelia Langford, John Sillitoe on behalf of the Wellcome Sanger Institute COVID-19 Surveillance Team |
| EPI_ISL_559312 | Lighthouse Lab in Milton Keynes | Wellcome Sanger Institute for the COVID-19 Genomics UK (COG-UK) consortium | The Lighthouse Lab in Milton Keynes and Alex Alderton, Roberto Amato, Sonia Goncalves, Ewan Harrison, David K. Jackson, Ian Johnston, Dominic Kwiatkowski, Cordelia Langford, John Sillitoe on behalf of the Wellcome Sanger Institute COVID-19 Surveillance Team (http://www.sanger.ac.uk/covid-team) |

| | | | |
|---|---|---|---|
| EPI_ISL_559313 | Lighthouse Lab in Alderley Park | Wellcome Sanger Institute for the COVID-19 Genomics UK (COG-UK) consortium | The Lighthouse Lab in Alderley Park and Alex Alderton, Roberto Amato, Sonia Goncalves, Ewan Harrison, David K. Jackson, Ian Johnston, Dominic Kwiatkowski, Cordelia Langford, John Sillitoe on behalf of the Wellcome Sanger Institute COVID-19 Surveillance Team |
| EPI_ISL_559314 | Lighthouse Lab in Milton Keynes | Wellcome Sanger Institute for the COVID-19 Genomics UK (COG-UK) consortium | The Lighthouse Lab in Milton Keynes and Alex Alderton, Roberto Amato, Sonia Goncalves, Ewan Harrison, David K. Jackson, Ian Johnston, Dominic Kwiatkowski, Cordelia Langford, John Sillitoe on behalf of the Wellcome Sanger Institute COVID-19 Surveillance Team (http://www.sanger.ac.uk/covid-team) |
| EPI_ISL_559315, EPI_ISL_559316, EPI_ISL_559317, EPI_ISL_559318 | Lighthouse Lab in Alderley Park | Wellcome Sanger Institute for the COVID-19 Genomics UK (COG-UK) consortium | The Lighthouse Lab in Alderley Park and Alex Alderton, Roberto Amato, Sonia Goncalves, Ewan Harrison, David K. Jackson, Ian Johnston, Dominic Kwiatkowski, Cordelia Langford, John Sillitoe on behalf of the Wellcome Sanger Institute COVID-19 Surveillance Team |
| EPI_ISL_559319, EPI_ISL_559320 | Lighthouse Lab in Milton Keynes | Wellcome Sanger Institute for the COVID-19 Genomics UK (COG-UK) consortium | The Lighthouse Lab in Milton Keynes and Alex Alderton, Roberto Amato, Sonia Goncalves, Ewan Harrison, David K. Jackson, Ian Johnston, Dominic Kwiatkowski, Cordelia Langford, John Sillitoe on behalf of the Wellcome Sanger Institute COVID-19 Surveillance Team (http://www.sanger.ac.uk/covid-team) |
| EPI_ISL_559321, EPI_ISL_559322, EPI_ISL_559323, EPI_ISL_559324, EPI_ISL_559325 | Lighthouse Lab in Alderley Park | Wellcome Sanger Institute for the COVID-19 Genomics UK (COG-UK) consortium | The Lighthouse Lab in Alderley Park and Alex Alderton, Roberto Amato, Sonia Goncalves, Ewan Harrison, David K. Jackson, Ian Johnston, Dominic Kwiatkowski, Cordelia Langford, John Sillitoe on behalf of the Wellcome Sanger Institute COVID-19 Surveillance Team |
| EPI_ISL_559326 | Lighthouse Lab in Milton Keynes | Wellcome Sanger Institute for the COVID-19 Genomics UK (COG-UK) consortium | The Lighthouse Lab in Milton Keynes and Alex Alderton, Roberto Amato, Sonia Goncalves, Ewan Harrison, David K. Jackson, Ian Johnston, Dominic Kwiatkowski, Cordelia Langford, John Sillitoe on behalf of the Wellcome Sanger Institute COVID-19 Surveillance Team (http://www.sanger.ac.uk/covid-team) |
| EPI_ISL_559327, EPI_ISL_559328, EPI_ISL_559329, EPI_ISL_559330 | Lighthouse Lab in Alderley Park | Wellcome Sanger Institute for the COVID-19 Genomics UK (COG-UK) consortium | The Lighthouse Lab in Alderley Park and Alex Alderton, Roberto Amato, Sonia Goncalves, Ewan Harrison, David K. Jackson, Ian Johnston, Dominic Kwiatkowski, Cordelia Langford, John Sillitoe on behalf of the Wellcome Sanger Institute COVID-19 Surveillance Team |
| EPI_ISL_559331 | Lighthouse Lab in Milton Keynes | Wellcome Sanger Institute for the COVID-19 Genomics UK (COG-UK) consortium | The Lighthouse Lab in Milton Keynes and Alex Alderton, Roberto Amato, Sonia Goncalves, Ewan Harrison, David K. Jackson, Ian Johnston, Dominic Kwiatkowski, Cordelia Langford, John Sillitoe on behalf of the Wellcome Sanger Institute COVID-19 Surveillance Team (http://www.sanger.ac.uk/covid-team) |
| EPI_ISL_559332, EPI_ISL_559333, EPI_ISL_559334 | Lighthouse Lab in Alderley Park | Wellcome Sanger Institute for the COVID-19 Genomics UK (COG-UK) consortium | The Lighthouse Lab in Alderley Park and Alex Alderton, Roberto Amato, Sonia Goncalves, Ewan Harrison, David K. Jackson, Ian Johnston, Dominic Kwiatkowski, Cordelia Langford, John Sillitoe on behalf of the Wellcome Sanger Institute COVID-19 Surveillance Team |
| EPI_ISL_559335 | Lighthouse Lab in Milton Keynes | Wellcome Sanger Institute for the COVID-19 Genomics UK (COG-UK) consortium | The Lighthouse Lab in Milton Keynes and Alex Alderton, Roberto Amato, Sonia Goncalves, Ewan Harrison, David K. Jackson, Ian Johnston, Dominic Kwiatkowski, Cordelia Langford, John Sillitoe on behalf of the Wellcome Sanger Institute COVID-19 Surveillance Team (http://www.sanger.ac.uk/covid-team) |
| EPI_ISL_559336, EPI_ISL_559338, EPI_ISL_559339, EPI_ISL_559340, EPI_ISL_559341, EPI_ISL_559342, EPI_ISL_559343, EPI_ISL_559344, EPI_ISL_559345, EPI_ISL_559346, EPI_ISL_559347, EPI_ISL_559348, EPI_ISL_559349, EPI_ISL_559350, EPI_ISL_559351 | | | |
| see above | Lighthouse Lab in Alderley Park | Wellcome Sanger Institute for the COVID-19 Genomics UK (COG-UK) consortium | The Lighthouse Lab in Alderley Park and Alex Alderton, Roberto Amato, Sonia Goncalves, Ewan Harrison, David K. Jackson, Ian Johnston, Dominic Kwiatkowski, Cordelia Langford, John Sillitoe on behalf of the Wellcome Sanger Institute COVID-19 Surveillance Team |
| EPI_ISL_559352, EPI_ISL_559354, EPI_ISL_559355 | Lighthouse Lab in Milton Keynes | Wellcome Sanger Institute for the COVID-19 Genomics UK (COG-UK) consortium | The Lighthouse Lab in Milton Keynes and Alex Alderton, Roberto Amato, Sonia Goncalves, Ewan Harrison, David K. Jackson, Ian Johnston, Dominic Kwiatkowski, Cordelia Langford, John Sillitoe on behalf of the Wellcome Sanger Institute COVID-19 Surveillance Team (http://www.sanger.ac.uk/covid-team) |
| EPI_ISL_559356, EPI_ISL_559357, EPI_ISL_559358, EPI_ISL_559359, EPI_ISL_559360, EPI_ISL_559361, EPI_ISL_559362, EPI_ISL_559363, EPI_ISL_559364, EPI_ISL_559365, EPI_ISL_559366, EPI_ISL_559367 | | | |
| see above | Lighthouse Lab in Alderley Park | Wellcome Sanger Institute for the COVID-19 Genomics UK (COG-UK) consortium | The Lighthouse Lab in Alderley Park and Alex Alderton, Roberto Amato, Sonia Goncalves, Ewan Harrison, David K. Jackson, Ian Johnston, Dominic Kwiatkowski, Cordelia Langford, John Sillitoe on behalf of the Wellcome Sanger Institute COVID-19 Surveillance Team |
| EPI_ISL_559368 | Lighthouse Lab in Milton Keynes | Wellcome Sanger Institute for the COVID-19 Genomics UK (COG-UK) consortium | The Lighthouse Lab in Milton Keynes and Alex Alderton, Roberto Amato, Sonia Goncalves, Ewan Harrison, David K. Jackson, Ian Johnston, Dominic Kwiatkowski, Cordelia Langford, John Sillitoe on behalf of the Wellcome Sanger Institute COVID-19 Surveillance Team (http://www.sanger.ac.uk/covid-team) |
| EPI_ISL_559369, EPI_ISL_559370, EPI_ISL_559371, EPI_ISL_559372, EPI_ISL_559373, EPI_ISL_559374, EPI_ISL_559375, EPI_ISL_559376, EPI_ISL_559377, EPI_ISL_559378, EPI_ISL_559379, EPI_ISL_559380, EPI_ISL_559381, EPI_ISL_559382, EPI_ISL_559383, EPI_ISL_559384, EPI_ISL_559385, EPI_ISL_559386, EPI_ISL_559387, EPI_ISL_559388, EPI_ISL_559389, EPI_ISL_559390, EPI_ISL_559391, EPI_ISL_559392, EPI_ISL_559393, EPI_ISL_559394, EPI_ISL_559395, EPI_ISL_559396, EPI_ISL_559397, EPI_ISL_559398, EPI_ISL_559399, EPI_ISL_559400, EPI_ISL_559401, EPI_ISL_559402, EPI_ISL_559403, EPI_ISL_559404, EPI_ISL_559405, EPI_ISL_559406, EPI_ISL_559407, EPI_ISL_559408, EPI_ISL_559409, EPI_ISL_559410, EPI_ISL_559411, EPI_ISL_559412, EPI_ISL_559413, EPI_ISL_559414, EPI_ISL_559415, EPI_ISL_559416, EPI_ISL_559417 | | | |
| see above | Lighthouse Lab in Alderley Park | Wellcome Sanger Institute for the COVID-19 Genomics UK (COG-UK) consortium | The Lighthouse Lab in Alderley Park and Alex Alderton, Roberto Amato, Sonia Goncalves, Ewan Harrison, David K. Jackson, Ian Johnston, Dominic Kwiatkowski, Cordelia Langford, John Sillitoe on behalf of the Wellcome Sanger Institute COVID-19 Surveillance Team |
| EPI_ISL_559418 | Lighthouse Lab in Alderley Park | Wellcome Sanger Institute for the COVID-19 Genomics UK (COG-UK) consortium | The Lighthouse Lab in Alderley Park and Alex Alderton, Roberto Amato, Sonia Goncalves, Ewan Harrison, David K. Jackson, Ian Johnston, Dominic Kwiatkowski, Cordelia Langford, John Sillitoe on behalf of the Wellcome Sanger Institute COVID-19 Surveillance Team (http://www.sanger.ac.uk/covid-team) |
| EPI_ISL_559419, EPI_ISL_559420, EPI_ISL_559421, EPI_ISL_559422, EPI_ISL_559423, EPI_ISL_559424, EPI_ISL_559425, EPI_ISL_559426, EPI_ISL_559427, EPI_ISL_559428, EPI_ISL_559429, EPI_ISL_559430, EPI_ISL_559431, EPI_ISL_559432, EPI_ISL_559433, EPI_ISL_559434, EPI_ISL_559435, EPI_ISL_559436, EPI_ISL_559437, EPI_ISL_559438, EPI_ISL_559439, EPI_ISL_559440, EPI_ISL_559441, EPI_ISL_559442, EPI_ISL_559443, EPI_ISL_559444, EPI_ISL_559445, EPI_ISL_559446, EPI_ISL_559448, EPI_ISL_559449, EPI_ISL_559450, EPI_ISL_559451, EPI_ISL_559452, EPI_ISL_559453, EPI_ISL_559454, EPI_ISL_559455, EPI_ISL_559456, EPI_ISL_559457, EPI_ISL_559458, EPI_ISL_559459, EPI_ISL_559460, EPI_ISL_559461, EPI_ISL_559462, EPI_ISL_559463, EPI_ISL_559464 | | | |
| see above | Lighthouse Lab in Alderley Park | Wellcome Sanger Institute for the COVID-19 Genomics UK (COG-UK) consortium | The Lighthouse Lab in Alderley Park and Alex Alderton, Roberto Amato, Sonia Goncalves, Ewan Harrison, David K. Jackson, Ian Johnston, Dominic Kwiatkowski, Cordelia Langford, John Sillitoe on behalf of the Wellcome Sanger Institute COVID-19 Surveillance Team |
| EPI_ISL_559465 | Lighthouse Lab in Alderley Park | Wellcome Sanger Institute for the COVID-19 Genomics UK (COG-UK) consortium | The Lighthouse Lab in Alderley Park and Alex Alderton, Roberto Amato, Sonia Goncalves, Ewan Harrison, David K. Jackson, Ian Johnston, Dominic Kwiatkowski, Cordelia Langford, John Sillitoe on behalf of the Wellcome Sanger Institute COVID-19 Surveillance Team (http://www.sanger.ac.uk/covid-team) |
| EPI_ISL_559466, EPI_ISL_559467, EPI_ISL_559468, EPI_ISL_559469, EPI_ISL_559470, EPI_ISL_559471, EPI_ISL_559472, EPI_ISL_559473, EPI_ISL_559474, EPI_ISL_559475, EPI_ISL_559476, EPI_ISL_559477, EPI_ISL_559478, EPI_ISL_559479, EPI_ISL_559480, EPI_ISL_559481, EPI_ISL_559482, EPI_ISL_559483, EPI_ISL_559484, EPI_ISL_559485, EPI_ISL_559486, EPI_ISL_559487, EPI_ISL_559488, EPI_ISL_559489, EPI_ISL_559490, EPI_ISL_559491, EPI_ISL_559492, EPI_ISL_559493 | | | |
| see above | Lighthouse Lab in Alderley Park | Wellcome Sanger Institute for the COVID-19 Genomics UK (COG-UK) consortium | The Lighthouse Lab in Alderley Park and Alex Alderton, Roberto Amato, Sonia Goncalves, Ewan Harrison, David K. Jackson, Ian Johnston, Dominic Kwiatkowski, Cordelia Langford, John Sillitoe on behalf of the Wellcome Sanger Institute COVID-19 Surveillance Team |
| EPI_ISL_559494 | Lighthouse Lab in Alderley Park | Wellcome Sanger Institute for the COVID-19 Genomics UK (COG-UK) consortium | The Lighthouse Lab in Alderley Park and Alex Alderton, Roberto Amato, Sonia Goncalves, Ewan Harrison, David K. Jackson, Ian Johnston, Dominic Kwiatkowski, Cordelia Langford, John Sillitoe on behalf of the Wellcome Sanger Institute COVID-19 Surveillance Team (http://www.sanger.ac.uk/covid-team) |
| EPI_ISL_559495, EPI_ISL_559496, EPI_ISL_559497, EPI_ISL_559498, EPI_ISL_559499, EPI_ISL_559500, EPI_ISL_559501, EPI_ISL_559502, EPI_ISL_559503, EPI_ISL_559504, EPI_ISL_559505, EPI_ISL_559506, EPI_ISL_559507, EPI_ISL_559508, EPI_ISL_559509, EPI_ISL_559510, EPI_ISL_559511, EPI_ISL_559512, EPI_ISL_559513, EPI_ISL_559514, EPI_ISL_559515, EPI_ISL_559516, EPI_ISL_559517, EPI_ISL_559518, EPI_ISL_559519, EPI_ISL_559520, EPI_ISL_559521, EPI_ISL_559522, EPI_ISL_559523 | | | |
| see above | Lighthouse Lab in Alderley Park | Wellcome Sanger Institute for the COVID-19 Genomics UK (COG-UK) consortium | The Lighthouse Lab in Alderley Park and Alex Alderton, Roberto Amato, Sonia Goncalves, Ewan Harrison, David K. Jackson, Ian Johnston, Dominic Kwiatkowski, Cordelia Langford, John Sillitoe on behalf of the Wellcome Sanger Institute COVID-19 Surveillance Team |
| EPI_ISL_559524 | Lighthouse Lab in Alderley Park | Wellcome Sanger Institute for the COVID-19 Genomics UK (COG-UK) consortium | The Lighthouse Lab in Alderley Park and Alex Alderton, Roberto Amato, Sonia Goncalves, Ewan Harrison, David K. Jackson, Ian Johnston, Dominic Kwiatkowski, Cordelia Langford, John Sillitoe on behalf of the Wellcome Sanger Institute COVID-19 Surveillance Team (http://www.sanger.ac.uk/covid-team) |
| EPI_ISL_559525, EPI_ISL_559526, EPI_ISL_559527, EPI_ISL_559528, EPI_ISL_559529, EPI_ISL_559530, EPI_ISL_559531, EPI_ISL_559532, EPI_ISL_559533, EPI_ISL_559534, EPI_ISL_559535, EPI_ISL_559536, EPI_ISL_559537, EPI_ISL_559538, EPI_ISL_559539, EPI_ISL_559540, EPI_ISL_559541, EPI_ISL_559542, EPI_ISL_559543, EPI_ISL_559544 | | | |
| see above | Lighthouse Lab in Alderley Park | Wellcome Sanger Institute for the COVID-19 Genomics UK (COG-UK) consortium | The Lighthouse Lab in Alderley Park and Alex Alderton, Roberto Amato, Sonia Goncalves, Ewan Harrison, David K. Jackson, Ian Johnston, Dominic Kwiatkowski, Cordelia Langford, John Sillitoe on behalf of the Wellcome Sanger Institute COVID-19 Surveillance Team |

| | | | |
|---|---|---|---|
| EPI_ISL_559545 | Lighthouse Lab in Alderley Park | Wellcome Sanger Institute for the COVID-19 Genomics UK (COG-UK) consortium | The Lighthouse Lab in Alderley Park and Alex Alderton, Roberto Amato, Sonia Goncalves, Ewan Harrison, David K. Jackson, Ian Johnston, Dominic Kwiatkowski, Cordelia Langford, John Sillitoe on behalf of the Wellcome Sanger Institute COVID-19 Surveillance Team (http://www.sanger.ac.uk/covid-team) |
| EPI_ISL_559546, EPI_ISL_559547 | Lighthouse Lab in Alderley Park | Wellcome Sanger Institute for the COVID-19 Genomics UK (COG-UK) consortium | The Lighthouse Lab in Alderley Park and Alex Alderton, Roberto Amato, Sonia Goncalves, Ewan Harrison, David K. Jackson, Ian Johnston, Dominic Kwiatkowski, Cordelia Langford, John Sillitoe on behalf of the Wellcome Sanger Institute COVID-19 Surveillance Team |
| EPI_ISL_559548 | Lighthouse Lab in Alderley Park | Wellcome Sanger Institute for the COVID-19 Genomics UK (COG-UK) consortium | The Lighthouse Lab in Alderley Park and Alex Alderton, Roberto Amato, Sonia Goncalves, Ewan Harrison, David K. Jackson, Ian Johnston, Dominic Kwiatkowski, Cordelia Langford, John Sillitoe on behalf of the Wellcome Sanger Institute COVID-19 Surveillance Team (http://www.sanger.ac.uk/covid-team) |
| EPI_ISL_559549 | Lighthouse Lab in Alderley Park | Wellcome Sanger Institute for the COVID-19 Genomics UK (COG-UK) consortium | The Lighthouse Lab in Alderley Park and Alex Alderton, Roberto Amato, Sonia Goncalves, Ewan Harrison, David K. Jackson, Ian Johnston, Dominic Kwiatkowski, Cordelia Langford, John Sillitoe on behalf of the Wellcome Sanger Institute COVID-19 Surveillance Team |
| EPI_ISL_559550, EPI_ISL_559551, EPI_ISL_559552 | Lighthouse Lab in Alderley Park | Wellcome Sanger Institute for the COVID-19 Genomics UK (COG-UK) consortium | The Lighthouse Lab in Alderley Park and Alex Alderton, Roberto Amato, Sonia Goncalves, Ewan Harrison, David K. Jackson, Ian Johnston, Dominic Kwiatkowski, Cordelia Langford, John Sillitoe on behalf of the Wellcome Sanger Institute COVID-19 Surveillance Team (http://www.sanger.ac.uk/covid-team) |
| EPI_ISL_559553 | Lighthouse Lab in Alderley Park | Wellcome Sanger Institute for the COVID-19 Genomics UK (COG-UK) consortium | The Lighthouse Lab in Alderley Park and Alex Alderton, Roberto Amato, Sonia Goncalves, Ewan Harrison, David K. Jackson, Ian Johnston, Dominic Kwiatkowski, Cordelia Langford, John Sillitoe on behalf of the Wellcome Sanger Institute COVID-19 Surveillance Team |
| EPI_ISL_559554, EPI_ISL_559555, EPI_ISL_559556, EPI_ISL_559557, EPI_ISL_559558 | Lighthouse Lab in Alderley Park | Wellcome Sanger Institute for the COVID-19 Genomics UK (COG-UK) consortium | The Lighthouse Lab in Alderley Park and Alex Alderton, Roberto Amato, Sonia Goncalves, Ewan Harrison, David K. Jackson, Ian Johnston, Dominic Kwiatkowski, Cordelia Langford, John Sillitoe on behalf of the Wellcome Sanger Institute COVID-19 Surveillance Team (http://www.sanger.ac.uk/covid-team) |
| EPI_ISL_559559 | Lighthouse Lab in Alderley Park | Wellcome Sanger Institute for the COVID-19 Genomics UK (COG-UK) consortium | The Lighthouse Lab in Alderley Park and Alex Alderton, Roberto Amato, Sonia Goncalves, Ewan Harrison, David K. Jackson, Ian Johnston, Dominic Kwiatkowski, Cordelia Langford, John Sillitoe on behalf of the Wellcome Sanger Institute COVID-19 Surveillance Team |
| EPI_ISL_559560, EPI_ISL_559561, EPI_ISL_559562, EPI_ISL_559563, EPI_ISL_559564, EPI_ISL_559565, EPI_ISL_559566, EPI_ISL_559567, EPI_ISL_559568, EPI_ISL_559569 | Lighthouse Lab in Alderley Park | Wellcome Sanger Institute for the COVID-19 Genomics UK (COG-UK) consortium | The Lighthouse Lab in Alderley Park and Alex Alderton, Roberto Amato, Sonia Goncalves, Ewan Harrison, David K. Jackson, Ian Johnston, Dominic Kwiatkowski, Cordelia Langford, John Sillitoe on behalf of the Wellcome Sanger Institute COVID-19 Surveillance Team (http://www.sanger.ac.uk/covid-team) |
| EPI_ISL_559570, EPI_ISL_559571 | Lighthouse Lab in Alderley Park | Wellcome Sanger Institute for the COVID-19 Genomics UK (COG-UK) consortium | The Lighthouse Lab in Alderley Park and Alex Alderton, Roberto Amato, Sonia Goncalves, Ewan Harrison, David K. Jackson, Ian Johnston, Dominic Kwiatkowski, Cordelia Langford, John Sillitoe on behalf of the Wellcome Sanger Institute COVID-19 Surveillance Team |
| EPI_ISL_559572, EPI_ISL_559573, EPI_ISL_559574 | Lighthouse Lab in Alderley Park | Wellcome Sanger Institute for the COVID-19 Genomics UK (COG-UK) consortium | The Lighthouse Lab in Alderley Park and Alex Alderton, Roberto Amato, Sonia Goncalves, Ewan Harrison, David K. Jackson, Ian Johnston, Dominic Kwiatkowski, Cordelia Langford, John Sillitoe on behalf of the Wellcome Sanger Institute COVID-19 Surveillance Team (http://www.sanger.ac.uk/covid-team) |
| EPI_ISL_559575 | Lighthouse Lab in Alderley Park | Wellcome Sanger Institute for the COVID-19 Genomics UK (COG-UK) consortium | The Lighthouse Lab in Alderley Park and Alex Alderton, Roberto Amato, Sonia Goncalves, Ewan Harrison, David K. Jackson, Ian Johnston, Dominic Kwiatkowski, Cordelia Langford, John Sillitoe on behalf of the Wellcome Sanger Institute COVID-19 Surveillance Team |
| EPI_ISL_559577 | Lighthouse Lab in Alderley Park | Wellcome Sanger Institute for the COVID-19 Genomics UK (COG-UK) consortium | The Lighthouse Lab in Alderley Park and Alex Alderton, Roberto Amato, Sonia Goncalves, Ewan Harrison, David K. Jackson, Ian Johnston, Dominic Kwiatkowski, Cordelia Langford, John Sillitoe on behalf of the Wellcome Sanger Institute COVID-19 Surveillance Team (http://www.sanger.ac.uk/covid-team) |
| EPI_ISL_559578 | Lighthouse Lab in Alderley Park | Wellcome Sanger Institute for the COVID-19 Genomics UK (COG-UK) consortium | The Lighthouse Lab in Alderley Park and Alex Alderton, Roberto Amato, Sonia Goncalves, Ewan Harrison, David K. Jackson, Ian Johnston, Dominic Kwiatkowski, Cordelia Langford, John Sillitoe on behalf of the Wellcome Sanger Institute COVID-19 Surveillance Team |
| EPI_ISL_559579, EPI_ISL_559580, EPI_ISL_559581, EPI_ISL_559582, EPI_ISL_559583, EPI_ISL_559584, EPI_ISL_559585, EPI_ISL_559586, EPI_ISL_559587, EPI_ISL_559588, EPI_ISL_559589, EPI_ISL_559590, EPI_ISL_559592 | | | |
| see above | Lighthouse Lab in Alderley Park | Wellcome Sanger Institute for the COVID-19 Genomics UK (COG-UK) consortium | The Lighthouse Lab in Alderley Park and Alex Alderton, Roberto Amato, Sonia Goncalves, Ewan Harrison, David K. Jackson, Ian Johnston, Dominic Kwiatkowski, Cordelia Langford, John Sillitoe on behalf of the Wellcome Sanger Institute COVID-19 Surveillance Team (http://www.sanger.ac.uk/covid-team) |
| EPI_ISL_559593 | Lighthouse Lab in Alderley Park | Wellcome Sanger Institute for the COVID-19 Genomics UK (COG-UK) consortium | The Lighthouse Lab in Alderley Park and Alex Alderton, Roberto Amato, Sonia Goncalves, Ewan Harrison, David K. Jackson, Ian Johnston, Dominic Kwiatkowski, Cordelia Langford, John Sillitoe on behalf of the Wellcome Sanger Institute COVID-19 Surveillance Team |
| EPI_ISL_559594, EPI_ISL_559595, EPI_ISL_559596, EPI_ISL_559597, EPI_ISL_559598, EPI_ISL_559599 | Lighthouse Lab in Alderley Park | Wellcome Sanger Institute for the COVID-19 Genomics UK (COG-UK) consortium | The Lighthouse Lab in Alderley Park and Alex Alderton, Roberto Amato, Sonia Goncalves, Ewan Harrison, David K. Jackson, Ian Johnston, Dominic Kwiatkowski, Cordelia Langford, John Sillitoe on behalf of the Wellcome Sanger Institute COVID-19 Surveillance Team (http://www.sanger.ac.uk/covid-team) |
| EPI_ISL_559600 | Lighthouse Lab in Alderley Park | Wellcome Sanger Institute for the COVID-19 Genomics UK (COG-UK) consortium | The Lighthouse Lab in Alderley Park and Alex Alderton, Roberto Amato, Sonia Goncalves, Ewan Harrison, David K. Jackson, Ian Johnston, Dominic Kwiatkowski, Cordelia Langford, John Sillitoe on behalf of the Wellcome Sanger Institute COVID-19 Surveillance Team |
| EPI_ISL_559601, EPI_ISL_559602, EPI_ISL_559603 | Lighthouse Lab in Alderley Park | Wellcome Sanger Institute for the COVID-19 Genomics UK (COG-UK) consortium | The Lighthouse Lab in Alderley Park and Alex Alderton, Roberto Amato, Sonia Goncalves, Ewan Harrison, David K. Jackson, Ian Johnston, Dominic Kwiatkowski, Cordelia Langford, John Sillitoe on behalf of the Wellcome Sanger Institute COVID-19 Surveillance Team (http://www.sanger.ac.uk/covid-team) |
| EPI_ISL_559604 | Lighthouse Lab in Alderley Park | Wellcome Sanger Institute for the COVID-19 Genomics UK (COG-UK) consortium | The Lighthouse Lab in Alderley Park and Alex Alderton, Roberto Amato, Sonia Goncalves, Ewan Harrison, David K. Jackson, Ian Johnston, Dominic Kwiatkowski, Cordelia Langford, John Sillitoe on behalf of the Wellcome Sanger Institute COVID-19 Surveillance Team |
| EPI_ISL_559605, EPI_ISL_559606, EPI_ISL_559607, EPI_ISL_559608 | Lighthouse Lab in Alderley Park | Wellcome Sanger Institute for the COVID-19 Genomics UK (COG-UK) consortium | The Lighthouse Lab in Alderley Park and Alex Alderton, Roberto Amato, Sonia Goncalves, Ewan Harrison, David K. Jackson, Ian Johnston, Dominic Kwiatkowski, Cordelia Langford, John Sillitoe on behalf of the Wellcome Sanger Institute COVID-19 Surveillance Team (http://www.sanger.ac.uk/covid-team) |
| EPI_ISL_559609 | Lighthouse Lab in Alderley Park | Wellcome Sanger Institute for the COVID-19 Genomics UK (COG-UK) consortium | The Lighthouse Lab in Alderley Park and Alex Alderton, Roberto Amato, Sonia Goncalves, Ewan Harrison, David K. Jackson, Ian Johnston, Dominic Kwiatkowski, Cordelia Langford, John Sillitoe on behalf of the Wellcome Sanger Institute COVID-19 Surveillance Team |
| EPI_ISL_559610, EPI_ISL_559611, EPI_ISL_559612, EPI_ISL_559613, EPI_ISL_559614, EPI_ISL_559615, EPI_ISL_559617, EPI_ISL_559618 | Lighthouse Lab in Alderley Park | Wellcome Sanger Institute for the COVID-19 Genomics UK (COG-UK) consortium | The Lighthouse Lab in Alderley Park and Alex Alderton, Roberto Amato, Sonia Goncalves, Ewan Harrison, David K. Jackson, Ian Johnston, Dominic Kwiatkowski, Cordelia Langford, John Sillitoe on behalf of the Wellcome Sanger Institute COVID-19 Surveillance Team (http://www.sanger.ac.uk/covid-team) |
| EPI_ISL_559619 | Lighthouse Lab in Alderley Park | Wellcome Sanger Institute for the COVID-19 Genomics UK (COG-UK) consortium | The Lighthouse Lab in Alderley Park and Alex Alderton, Roberto Amato, Sonia Goncalves, Ewan Harrison, David K. Jackson, Ian Johnston, Dominic Kwiatkowski, Cordelia Langford, John Sillitoe on behalf of the Wellcome Sanger Institute COVID-19 Surveillance Team |
| EPI_ISL_559620, EPI_ISL_559621, EPI_ISL_559622 | Lighthouse Lab in Alderley Park | Wellcome Sanger Institute for the COVID-19 Genomics UK (COG-UK) consortium | The Lighthouse Lab in Alderley Park and Alex Alderton, Roberto Amato, Sonia Goncalves, Ewan Harrison, David K. Jackson, Ian Johnston, Dominic Kwiatkowski, Cordelia Langford, John Sillitoe on behalf of the Wellcome Sanger Institute COVID-19 Surveillance Team (http://www.sanger.ac.uk/covid-team) |
| EPI_ISL_559623 | Lighthouse Lab in Alderley Park | Wellcome Sanger Institute for the COVID-19 Genomics UK (COG-UK) consortium | The Lighthouse Lab in Alderley Park and Alex Alderton, Roberto Amato, Sonia Goncalves, Ewan Harrison, David K. Jackson, Ian Johnston, Dominic Kwiatkowski, Cordelia Langford, John Sillitoe on behalf of the Wellcome Sanger Institute COVID-19 Surveillance Team |
| EPI_ISL_559624, EPI_ISL_559625, EPI_ISL_559626, EPI_ISL_559627 | Lighthouse Lab in Alderley Park | Wellcome Sanger Institute for the COVID-19 Genomics UK (COG-UK) consortium | The Lighthouse Lab in Alderley Park and Alex Alderton, Roberto Amato, Sonia Goncalves, Ewan Harrison, David K. Jackson, Ian Johnston, Dominic Kwiatkowski, Cordelia Langford, John Sillitoe on behalf of the Wellcome Sanger Institute COVID-19 Surveillance Team (http://www.sanger.ac.uk/covid-team) |
| EPI_ISL_559628 | Lighthouse Lab in Alderley Park | Wellcome Sanger Institute for the COVID-19 Genomics UK | The Lighthouse Lab in Alderley Park and Alex Alderton, Roberto Amato, Sonia Goncalves, Ewan Harrison, David K. Jackson, Ian Johnston, Dominic |

| | | | |
|---|---|---|---|
| | | (COG-UK) consortium | Kwiatkowski, Cordelia Langford, John Sillitoe on behalf of the Wellcome Sanger Institute COVID-19 Surveillance Team |
| EPI_ISL_559629, EPI_ISL_559630, EPI_ISL_559631, EPI_ISL_559632, EPI_ISL_559633, EPI_ISL_559634, EPI_ISL_559635, EPI_ISL_559636, EPI_ISL_559637, EPI_ISL_559638, EPI_ISL_559639, EPI_ISL_559640, EPI_ISL_559641, EPI_ISL_559642, EPI_ISL_559643, EPI_ISL_559645, EPI_ISL_559646, EPI_ISL_559647, EPI_ISL_559648 | | | |
| see above | Lighthouse Lab in Alderley Park | Wellcome Sanger Institute for the COVID-19 Genomics UK (COG-UK) consortium | The Lighthouse Lab in Alderley Park and Alex Alderton, Roberto Amato, Sonia Goncalves, Ewan Harrison, David K. Jackson, Ian Johnston, Dominic Kwiatkowski, Cordelia Langford, John Sillitoe on behalf of the Wellcome Sanger Institute COVID-19 Surveillance Team (http://www.sanger.ac.uk/covid-team) |
| EPI_ISL_559649 | Lighthouse Lab in Alderley Park | Wellcome Sanger Institute for the COVID-19 Genomics UK (COG-UK) consortium | The Lighthouse Lab in Alderley Park and Alex Alderton, Roberto Amato, Sonia Goncalves, Ewan Harrison, David K. Jackson, Ian Johnston, Dominic Kwiatkowski, Cordelia Langford, John Sillitoe on behalf of the Wellcome Sanger Institute COVID-19 Surveillance Team |
| EPI_ISL_559650, EPI_ISL_559651, EPI_ISL_559652, EPI_ISL_559653, EPI_ISL_559654, EPI_ISL_559655, EPI_ISL_559656, EPI_ISL_559657 | Lighthouse Lab in Alderley Park | Wellcome Sanger Institute for the COVID-19 Genomics UK (COG-UK) consortium | The Lighthouse Lab in Alderley Park and Alex Alderton, Roberto Amato, Sonia Goncalves, Ewan Harrison, David K. Jackson, Ian Johnston, Dominic Kwiatkowski, Cordelia Langford, John Sillitoe on behalf of the Wellcome Sanger Institute COVID-19 Surveillance Team (http://www.sanger.ac.uk/covid-team) |
| EPI_ISL_559658 | Lighthouse Lab in Alderley Park | Wellcome Sanger Institute for the COVID-19 Genomics UK (COG-UK) consortium | The Lighthouse Lab in Alderley Park and Alex Alderton, Roberto Amato, Sonia Goncalves, Ewan Harrison, David K. Jackson, Ian Johnston, Dominic Kwiatkowski, Cordelia Langford, John Sillitoe on behalf of the Wellcome Sanger Institute COVID-19 Surveillance Team |
| EPI_ISL_559659, EPI_ISL_559660 | Lighthouse Lab in Alderley Park | Wellcome Sanger Institute for the COVID-19 Genomics UK (COG-UK) consortium | The Lighthouse Lab in Alderley Park and Alex Alderton, Roberto Amato, Sonia Goncalves, Ewan Harrison, David K. Jackson, Ian Johnston, Dominic Kwiatkowski, Cordelia Langford, John Sillitoe on behalf of the Wellcome Sanger Institute COVID-19 Surveillance Team (http://www.sanger.ac.uk/covid-team) |
| EPI_ISL_559661 | Lighthouse Lab in Alderley Park | Wellcome Sanger Institute for the COVID-19 Genomics UK (COG-UK) consortium | The Lighthouse Lab in Alderley Park and Alex Alderton, Roberto Amato, Sonia Goncalves, Ewan Harrison, David K. Jackson, Ian Johnston, Dominic Kwiatkowski, Cordelia Langford, John Sillitoe on behalf of the Wellcome Sanger Institute COVID-19 Surveillance Team |
| EPI_ISL_559662, EPI_ISL_559663, EPI_ISL_559664, EPI_ISL_559665, EPI_ISL_559666, EPI_ISL_559667, EPI_ISL_559668, EPI_ISL_559669, EPI_ISL_559671, EPI_ISL_559672, EPI_ISL_559675, EPI_ISL_559676, EPI_ISL_559677, EPI_ISL_559678, EPI_ISL_559679, EPI_ISL_559680, EPI_ISL_559681, EPI_ISL_559682, EPI_ISL_559683, EPI_ISL_559684, EPI_ISL_559685, EPI_ISL_559686, EPI_ISL_559689, EPI_ISL_559690, EPI_ISL_559691, EPI_ISL_559692, EPI_ISL_559693, EPI_ISL_559694, EPI_ISL_559695, EPI_ISL_559696, EPI_ISL_559697, EPI_ISL_559698, EPI_ISL_559701, EPI_ISL_559702 | | | |
| see above | Lighthouse Lab in Milton Keynes | Wellcome Sanger Institute for the COVID-19 Genomics UK (COG-UK) consortium | The Lighthouse Lab in Milton Keynes and Alex Alderton, Roberto Amato, Sonia Goncalves, Ewan Harrison, David K. Jackson, Ian Johnston, Dominic Kwiatkowski, Cordelia Langford, John Sillitoe on behalf of the Wellcome Sanger Institute COVID-19 Surveillance Team (http://www.sanger.ac.uk/covid-team) |
| EPI_ISL_559703 | Lighthouse Lab in Milton Keynes | Wellcome Sanger Institute for the COVID-19 Genomics UK (COG-UK) consortium | The Lighthouse Lab in Milton Keynes and Alex Alderton, Roberto Amato, Sonia Goncalves, Ewan Harrison, David K. Jackson, Ian Johnston, Dominic Kwiatkowski, Cordelia Langford, John Sillitoe on behalf of the Wellcome Sanger Institute COVID-19 Surveillance Team |
| EPI_ISL_559704 | Lighthouse Lab in Milton Keynes | Wellcome Sanger Institute for the COVID-19 Genomics UK (COG-UK) consortium | The Lighthouse Lab in Milton Keynes and Alex Alderton, Roberto Amato, Sonia Goncalves, Ewan Harrison, David K. Jackson, Ian Johnston, Dominic Kwiatkowski, Cordelia Langford, John Sillitoe on behalf of the Wellcome Sanger Institute COVID-19 Surveillance Team (http://www.sanger.ac.uk/covid-team) |
| EPI_ISL_559705, EPI_ISL_559707 | Lighthouse Lab in Milton Keynes | Wellcome Sanger Institute for the COVID-19 Genomics UK (COG-UK) consortium | The Lighthouse Lab in Milton Keynes and Alex Alderton, Roberto Amato, Sonia Goncalves, Ewan Harrison, David K. Jackson, Ian Johnston, Dominic Kwiatkowski, Cordelia Langford, John Sillitoe on behalf of the Wellcome Sanger Institute COVID-19 Surveillance Team |
| EPI_ISL_559708 | Lighthouse Lab in Milton Keynes | Wellcome Sanger Institute for the COVID-19 Genomics UK (COG-UK) consortium | The Lighthouse Lab in Milton Keynes and Alex Alderton, Roberto Amato, Sonia Goncalves, Ewan Harrison, David K. Jackson, Ian Johnston, Dominic Kwiatkowski, Cordelia Langford, John Sillitoe on behalf of the Wellcome Sanger Institute COVID-19 Surveillance Team (http://www.sanger.ac.uk/covid-team) |
| EPI_ISL_559710, EPI_ISL_559711, EPI_ISL_559712 | Lighthouse Lab in Milton Keynes | Wellcome Sanger Institute for the COVID-19 Genomics UK (COG-UK) consortium | The Lighthouse Lab in Milton Keynes and Alex Alderton, Roberto Amato, Sonia Goncalves, Ewan Harrison, David K. Jackson, Ian Johnston, Dominic Kwiatkowski, Cordelia Langford, John Sillitoe on behalf of the Wellcome Sanger Institute COVID-19 Surveillance Team |
| EPI_ISL_559713 | Lighthouse Lab in Milton Keynes | Wellcome Sanger Institute for the COVID-19 Genomics UK (COG-UK) consortium | The Lighthouse Lab in Milton Keynes and Alex Alderton, Roberto Amato, Sonia Goncalves, Ewan Harrison, David K. Jackson, Ian Johnston, Dominic Kwiatkowski, Cordelia Langford, John Sillitoe on behalf of the Wellcome Sanger Institute COVID-19 Surveillance Team (http://www.sanger.ac.uk/covid-team) |
| EPI_ISL_559714 | Lighthouse Lab in Milton Keynes | Wellcome Sanger Institute for the COVID-19 Genomics UK (COG-UK) consortium | The Lighthouse Lab in Milton Keynes and Alex Alderton, Roberto Amato, Sonia Goncalves, Ewan Harrison, David K. Jackson, Ian Johnston, Dominic Kwiatkowski, Cordelia Langford, John Sillitoe on behalf of the Wellcome Sanger Institute COVID-19 Surveillance Team |
| EPI_ISL_559716, EPI_ISL_559717 | Lighthouse Lab in Milton Keynes | Wellcome Sanger Institute for the COVID-19 Genomics UK (COG-UK) consortium | The Lighthouse Lab in Milton Keynes and Alex Alderton, Roberto Amato, Sonia Goncalves, Ewan Harrison, David K. Jackson, Ian Johnston, Dominic Kwiatkowski, Cordelia Langford, John Sillitoe on behalf of the Wellcome Sanger Institute COVID-19 Surveillance Team (http://www.sanger.ac.uk/covid-team) |
| EPI_ISL_559718, EPI_ISL_559719, EPI_ISL_559720, EPI_ISL_559721, EPI_ISL_559722, EPI_ISL_559723, EPI_ISL_559725, EPI_ISL_559726, EPI_ISL_559727, EPI_ISL_559728, EPI_ISL_559729 | | | |
| see above | Lighthouse Lab in Milton Keynes | Wellcome Sanger Institute for the COVID-19 Genomics UK (COG-UK) consortium | The Lighthouse Lab in Milton Keynes and Alex Alderton, Roberto Amato, Sonia Goncalves, Ewan Harrison, David K. Jackson, Ian Johnston, Dominic Kwiatkowski, Cordelia Langford, John Sillitoe on behalf of the Wellcome Sanger Institute COVID-19 Surveillance Team |
| EPI_ISL_559730 | Lighthouse Lab in Milton Keynes | Wellcome Sanger Institute for the COVID-19 Genomics UK (COG-UK) consortium | The Lighthouse Lab in Milton Keynes and Alex Alderton, Roberto Amato, Sonia Goncalves, Ewan Harrison, David K. Jackson, Ian Johnston, Dominic Kwiatkowski, Cordelia Langford, John Sillitoe on behalf of the Wellcome Sanger Institute COVID-19 Surveillance Team (http://www.sanger.ac.uk/covid-team) |
| EPI_ISL_559731 | Lighthouse Lab in Milton Keynes | Wellcome Sanger Institute for the COVID-19 Genomics UK (COG-UK) consortium | The Lighthouse Lab in Milton Keynes and Alex Alderton, Roberto Amato, Sonia Goncalves, Ewan Harrison, David K. Jackson, Ian Johnston, Dominic Kwiatkowski, Cordelia Langford, John Sillitoe on behalf of the Wellcome Sanger Institute COVID-19 Surveillance Team |
| EPI_ISL_559732 | Lighthouse Lab in Milton Keynes | Wellcome Sanger Institute for the COVID-19 Genomics UK (COG-UK) consortium | The Lighthouse Lab in Milton Keynes and Alex Alderton, Roberto Amato, Sonia Goncalves, Ewan Harrison, David K. Jackson, Ian Johnston, Dominic Kwiatkowski, Cordelia Langford, John Sillitoe on behalf of the Wellcome Sanger Institute COVID-19 Surveillance Team (http://www.sanger.ac.uk/covid-team) |
| EPI_ISL_559735 | Lighthouse Lab in Milton Keynes | Wellcome Sanger Institute for the COVID-19 Genomics UK (COG-UK) consortium | The Lighthouse Lab in Milton Keynes and Alex Alderton, Roberto Amato, Sonia Goncalves, Ewan Harrison, David K. Jackson, Ian Johnston, Dominic Kwiatkowski, Cordelia Langford, John Sillitoe on behalf of the Wellcome Sanger Institute COVID-19 Surveillance Team |
| EPI_ISL_559736 | Lighthouse Lab in Milton Keynes | Wellcome Sanger Institute for the COVID-19 Genomics UK (COG-UK) consortium | The Lighthouse Lab in Milton Keynes and Alex Alderton, Roberto Amato, Sonia Goncalves, Ewan Harrison, David K. Jackson, Ian Johnston, Dominic Kwiatkowski, Cordelia Langford, John Sillitoe on behalf of the Wellcome Sanger Institute COVID-19 Surveillance Team (http://www.sanger.ac.uk/covid-team) |
| EPI_ISL_559737, EPI_ISL_559738, EPI_ISL_559739, EPI_ISL_559741, EPI_ISL_559742 | Lighthouse Lab in Milton Keynes | Wellcome Sanger Institute for the COVID-19 Genomics UK (COG-UK) consortium | The Lighthouse Lab in Milton Keynes and Alex Alderton, Roberto Amato, Sonia Goncalves, Ewan Harrison, David K. Jackson, Ian Johnston, Dominic Kwiatkowski, Cordelia Langford, John Sillitoe on behalf of the Wellcome Sanger Institute COVID-19 Surveillance Team |
| EPI_ISL_559743 | Lighthouse Lab in Milton Keynes | Wellcome Sanger Institute for the COVID-19 Genomics UK (COG-UK) consortium | The Lighthouse Lab in Alderley Park and Alex Alderton, Roberto Amato, Sonia Goncalves, Ewan Harrison, David K. Jackson, Ian Johnston, Dominic Kwiatkowski, Cordelia Langford, John Sillitoe on behalf of the Wellcome Sanger Institute COVID-19 Surveillance Team |
| EPI_ISL_559744, EPI_ISL_559745 | Lighthouse Lab in Milton Keynes | Wellcome Sanger Institute for the COVID-19 Genomics UK (COG-UK) consortium | The Lighthouse Lab in Milton Keynes and Alex Alderton, Roberto Amato, Sonia Goncalves, Ewan Harrison, David K. Jackson, Ian Johnston, Dominic Kwiatkowski, Cordelia Langford, John Sillitoe on behalf of the Wellcome Sanger Institute COVID-19 Surveillance Team (http://www.sanger.ac.uk/covid-team) |
| EPI_ISL_559746, EPI_ISL_559747, EPI_ISL_559748, EPI_ISL_559749, EPI_ISL_559750, EPI_ISL_559751, EPI_ISL_566421, EPI_ISL_566422, EPI_ISL_566424, EPI_ISL_566425, EPI_ISL_566426, EPI_ISL_566427, EPI_ISL_566430, EPI_ISL_566431, EPI_ISL_566432, EPI_ISL_566433, EPI_ISL_566434, EPI_ISL_566436, EPI_ISL_566437, EPI_ISL_566438, EPI_ISL_566439, EPI_ISL_566440, EPI_ISL_566441, EPI_ISL_566442, EPI_ISL_566443, EPI_ISL_566447, EPI_ISL_566448, EPI_ISL_566449, EPI_ISL_566450, EPI_ISL_566451, EPI_ISL_566452, EPI_ISL_566454, EPI_ISL_566458, EPI_ISL_566460, EPI_ISL_566461, EPI_ISL_566462, EPI_ISL_566463, EPI_ISL_566464, EPI_ISL_566465, EPI_ISL_566466, EPI_ISL_566469, EPI_ISL_566470, EPI_ISL_566471, EPI_ISL_566475, EPI_ISL_566476, EPI_ISL_566477, EPI_ISL_566478, EPI_ISL_566479, EPI_ISL_566480, EPI_ISL_566481, EPI_ISL_566482, EPI_ISL_566483, EPI_ISL_566484, EPI_ISL_566485, EPI_ISL_566486, EPI_ISL_566488, EPI_ISL_566489, EPI_ISL_566490, EPI_ISL_566491, EPI_ISL_566494, EPI_ISL_566496, EPI_ISL_566497, EPI_ISL_566498, EPI_ISL_566499, EPI_ISL_566501, EPI_ISL_566502, EPI_ISL_566503, EPI_ISL_566504, EPI_ISL_566505, EPI_ISL_566506, EPI_ISL_566507, EPI_ISL_566509, EPI_ISL_566510, EPI_ISL_566511, EPI_ISL_566512, EPI_ISL_566514, EPI_ISL_566515, EPI_ISL_566516, EPI_ISL_566517, EPI_ISL_566520, EPI_ISL_566521, EPI_ISL_566522, EPI_ISL_566523, EPI_ISL_566525, EPI_ISL_566526, EPI_ISL_566527, EPI_ISL_566529, EPI_ISL_566530, EPI_ISL_566531, EPI_ISL_566532, | | | |

EPI_ISL_566533, EPI_ISL_566535, EPI_ISL_566536, EPI_ISL_566538, EPI_ISL_566539, EPI_ISL_566540, EPI_ISL_566541, EPI_ISL_566545, EPI_ISL_566546, EPI_ISL_566547, EPI_ISL_566548, EPI_ISL_566549, EPI_ISL_566550, EPI_ISL_566551, EPI_ISL_566552, EPI_ISL_566553, EPI_ISL_566554, EPI_ISL_566555, EPI_ISL_566556, EPI_ISL_566557, EPI_ISL_566558, EPI_ISL_566559, EPI_ISL_566560, EPI_ISL_566561, EPI_ISL_566562, EPI_ISL_566563, EPI_ISL_566564, EPI_ISL_566565, EPI_ISL_566566, EPI_ISL_566568, EPI_ISL_566569, EPI_ISL_566570, EPI_ISL_566571, EPI_ISL_566572, EPI_ISL_566573, EPI_ISL_566574, EPI_ISL_566575, EPI_ISL_566577, EPI_ISL_566578, EPI_ISL_566579, EPI_ISL_566580, EPI_ISL_566581, EPI_ISL_566582, EPI_ISL_566583, EPI_ISL_566584, EPI_ISL_566585, EPI_ISL_566587, EPI_ISL_566588, EPI_ISL_566590, EPI_ISL_566591, EPI_ISL_566592, EPI_ISL_566594, EPI_ISL_566595, EPI_ISL_566596, EPI_ISL_566597, EPI_ISL_566598, EPI_ISL_566599, EPI_ISL_566601, EPI_ISL_566602, EPI_ISL_566604, EPI_ISL_566605, EPI_ISL_566606, EPI_ISL_566607, EPI_ISL_566608, EPI_ISL_566609, EPI_ISL_566612, EPI_ISL_566613, EPI_ISL_566614, EPI_ISL_566615, EPI_ISL_566616, EPI_ISL_566617, EPI_ISL_566618, EPI_ISL_566619, EPI_ISL_566620, EPI_ISL_566621, EPI_ISL_566622, EPI_ISL_566623, EPI_ISL_566624, EPI_ISL_566626, EPI_ISL_566627, EPI_ISL_566628, EPI_ISL_566629, EPI_ISL_566630, EPI_ISL_566632, EPI_ISL_566633, EPI_ISL_566634, EPI_ISL_566635, EPI_ISL_566636, EPI_ISL_566640, EPI_ISL_566641, EPI_ISL_566642, EPI_ISL_566643, EPI_ISL_566644, EPI_ISL_566645, EPI_ISL_566646, EPI_ISL_566647, EPI_ISL_566648, EPI_ISL_566649, EPI_ISL_566650, EPI_ISL_566651, EPI_ISL_566652, EPI_ISL_566653, EPI_ISL_566654, EPI_ISL_566655, EPI_ISL_566656, EPI_ISL_566657, EPI_ISL_566658, EPI_ISL_566660, EPI_ISL_566663, EPI_ISL_566664, EPI_ISL_566665, EPI_ISL_566667, EPI_ISL_566668, EPI_ISL_566669, EPI_ISL_566670, EPI_ISL_566671, EPI_ISL_566673, EPI_ISL_566675, EPI_ISL_566676, EPI_ISL_566677, EPI_ISL_566678, EPI_ISL_566679, EPI_ISL_566680, EPI_ISL_566681, EPI_ISL_566682, EPI_ISL_566683, EPI_ISL_566684, EPI_ISL_566685, EPI_ISL_566686, EPI_ISL_566687, EPI_ISL_566688, EPI_ISL_566689, EPI_ISL_566690, EPI_ISL_566691, EPI_ISL_566692, EPI_ISL_566693, EPI_ISL_566694, EPI_ISL_566695, EPI_ISL_566696, EPI_ISL_566697, EPI_ISL_566698, EPI_ISL_566699, EPI_ISL_566700, EPI_ISL_566701, EPI_ISL_566702, EPI_ISL_566704, EPI_ISL_566705, EPI_ISL_566706, EPI_ISL_566708, EPI_ISL_566709, EPI_ISL_566710, EPI_ISL_566711, EPI_ISL_566712, EPI_ISL_566713, EPI_ISL_566714, EPI_ISL_566715, EPI_ISL_566716, EPI_ISL_566717, EPI_ISL_566718, EPI_ISL_566719, EPI_ISL_566720, EPI_ISL_566721, EPI_ISL_566722, EPI_ISL_566723, EPI_ISL_566725, EPI_ISL_566726, EPI_ISL_566727, EPI_ISL_566728, EPI_ISL_566729, EPI_ISL_566730, EPI_ISL_566731, EPI_ISL_566732, EPI_ISL_566733, EPI_ISL_566734, EPI_ISL_566735, EPI_ISL_566736, EPI_ISL_566737, EPI_ISL_566739, EPI_ISL_566740, EPI_ISL_566741, EPI_ISL_566742, EPI_ISL_566743, EPI_ISL_566744, EPI_ISL_566745, EPI_ISL_566746, EPI_ISL_566747, EPI_ISL_566748, EPI_ISL_566749, EPI_ISL_566750, EPI_ISL_566751, EPI_ISL_566752, EPI_ISL_566753, EPI_ISL_566754, EPI_ISL_566755, EPI_ISL_566756, EPI_ISL_566757, EPI_ISL_566758, EPI_ISL_566759, EPI_ISL_566760, EPI_ISL_566761, EPI_ISL_566762, EPI_ISL_566763, EPI_ISL_566764, EPI_ISL_566765, EPI_ISL_566766, EPI_ISL_566767, EPI_ISL_566768, EPI_ISL_566769, EPI_ISL_566770, EPI_ISL_566771, EPI_ISL_566772, EPI_ISL_566773, EPI_ISL_566774, EPI_ISL_566775, EPI_ISL_566776, EPI_ISL_566777, EPI_ISL_566778, EPI_ISL_566779, EPI_ISL_566780, EPI_ISL_566781, EPI_ISL_566782, EPI_ISL_566783, EPI_ISL_566784, EPI_ISL_566785, EPI_ISL_566786, EPI_ISL_566787, EPI_ISL_566788, EPI_ISL_566789, EPI_ISL_566790, EPI_ISL_566791, EPI_ISL_566792, EPI_ISL_566794, EPI_ISL_566795, EPI_ISL_566796, EPI_ISL_566797, EPI_ISL_566798, EPI_ISL_566799, EPI_ISL_566800, EPI_ISL_566801, EPI_ISL_566803, EPI_ISL_566804, EPI_ISL_566805, EPI_ISL_566806, EPI_ISL_566807, EPI_ISL_566808, EPI_ISL_566809, EPI_ISL_566810, EPI_ISL_566811, EPI_ISL_566812, EPI_ISL_566813, EPI_ISL_566814, EPI_ISL_566815, EPI_ISL_566816, EPI_ISL_566817, EPI_ISL_566818, EPI_ISL_566819, EPI_ISL_566820, EPI_ISL_566821, EPI_ISL_566822, EPI_ISL_566823, EPI_ISL_566824, EPI_ISL_566825, EPI_ISL_566826, EPI_ISL_566827, EPI_ISL_566828, EPI_ISL_566829, EPI_ISL_566830, EPI_ISL_566831, EPI_ISL_566832, EPI_ISL_566833, EPI_ISL_566834, EPI_ISL_566835, EPI_ISL_566836, EPI_ISL_566837, EPI_ISL_566838, EPI_ISL_566839, EPI_ISL_566840, EPI_ISL_566841, EPI_ISL_566842, EPI_ISL_566843, EPI_ISL_566844, EPI_ISL_566845, EPI_ISL_566846, EPI_ISL_566847, EPI_ISL_566848, EPI_ISL_566849, EPI_ISL_566850, EPI_ISL_566851, EPI_ISL_566852, EPI_ISL_566853, EPI_ISL_566854, EPI_ISL_566855, EPI_ISL_566856, EPI_ISL_566857, EPI_ISL_566858, EPI_ISL_566859, EPI_ISL_566860, EPI_ISL_566861, EPI_ISL_566862, EPI_ISL_566863, EPI_ISL_566864, EPI_ISL_566865, EPI_ISL_566866, EPI_ISL_566867, EPI_ISL_566868, EPI_ISL_566869, EPI_ISL_566870, EPI_ISL_566871, EPI_ISL_566872, EPI_ISL_566873, EPI_ISL_566874, EPI_ISL_566875, EPI_ISL_566876, EPI_ISL_566877, EPI_ISL_566878, EPI_ISL_566879, EPI_ISL_566880, EPI_ISL_566881, EPI_ISL_566882, EPI_ISL_566883, EPI_ISL_566884, EPI_ISL_566885, EPI_ISL_566886, EPI_ISL_566887, EPI_ISL_566888, EPI_ISL_566889, EPI_ISL_566890, EPI_ISL_566891, EPI_ISL_566892, EPI_ISL_566893, EPI_ISL_566894, EPI_ISL_566895, EPI_ISL_566896, EPI_ISL_566897, EPI_ISL_566898, EPI_ISL_566899, EPI_ISL_566900, EPI_ISL_566901, EPI_ISL_566902, EPI_ISL_566903, EPI_ISL_566904, EPI_ISL_566905, EPI_ISL_566906, EPI_ISL_566907, EPI_ISL_566909, EPI_ISL_566910, EPI_ISL_566911, EPI_ISL_566912, EPI_ISL_566913, EPI_ISL_566914, EPI_ISL_566915, EPI_ISL_566916, EPI_ISL_566917, EPI_ISL_566918, EPI_ISL_566919, EPI_ISL_566920, EPI_ISL_566921, EPI_ISL_566923, EPI_ISL_566924, EPI_ISL_566925, EPI_ISL_566926, EPI_ISL_566927, EPI_ISL_566928, EPI_ISL_566929, EPI_ISL_566930, EPI_ISL_566931, EPI_ISL_566933, EPI_ISL_566934, EPI_ISL_566935, EPI_ISL_566936, EPI_ISL_566937, EPI_ISL_566938, EPI_ISL_566939, EPI_ISL_566940, EPI_ISL_566941, EPI_ISL_566942, EPI_ISL_566943, EPI_ISL_566944, EPI_ISL_566945, EPI_ISL_566946, EPI_ISL_566947, EPI_ISL_566948, EPI_ISL_566949, EPI_ISL_566950, EPI_ISL_566952, EPI_ISL_566953, EPI_ISL_566954, EPI_ISL_566955, EPI_ISL_566957, EPI_ISL_566958, EPI_ISL_566959, EPI_ISL_566960, EPI_ISL_566961, EPI_ISL_566962, EPI_ISL_566963, EPI_ISL_566964, EPI_ISL_566965, EPI_ISL_566966, EPI_ISL_566967, EPI_ISL_566968, EPI_ISL_566969, EPI_ISL_566970, EPI_ISL_566971, EPI_ISL_566972, EPI_ISL_566974, EPI_ISL_566975, EPI_ISL_566976, EPI_ISL_566977, EPI_ISL_566978, EPI_ISL_566979, EPI_ISL_566980, EPI_ISL_566981, EPI_ISL_566982, EPI_ISL_566983, EPI_ISL_566984, EPI_ISL_566985, EPI_ISL_566987, EPI_ISL_566988, EPI_ISL_566989, EPI_ISL_566990, EPI_ISL_566991, EPI_ISL_566992, EPI_ISL_566993, EPI_ISL_566995, EPI_ISL_566996, EPI_ISL_566997, EPI_ISL_568289, EPI_ISL_568290, EPI_ISL_568364, EPI_ISL_568469, EPI_ISL_575761, EPI_ISL_575762, EPI_ISL_575763, EPI_ISL_575765, EPI_ISL_575766, EPI_ISL_575767, EPI_ISL_575768, EPI_ISL_575769, EPI_ISL_575770, EPI_ISL_575771, EPI_ISL_575772, EPI_ISL_575773, EPI_ISL_575774, EPI_ISL_575775, EPI_ISL_575776, EPI_ISL_575777, EPI_ISL_575778, EPI_ISL_575779, EPI_ISL_575780, EPI_ISL_575781, EPI_ISL_575782, EPI_ISL_575783, EPI_ISL_575784, EPI_ISL_575785, EPI_ISL_575786, EPI_ISL_575787, EPI_ISL_575789, EPI_ISL_575790, EPI_ISL_575791, EPI_ISL_575792, EPI_ISL_575793, EPI_ISL_575794, EPI_ISL_575796, EPI_ISL_575797, EPI_ISL_575798, EPI_ISL_575800, EPI_ISL_575801, EPI_ISL_575803, EPI_ISL_575806, EPI_ISL_575807, EPI_ISL_575808, EPI_ISL_575809, EPI_ISL_575810, EPI_ISL_575811, EPI_ISL_575812, EPI_ISL_575813, EPI_ISL_575814, EPI_ISL_575815, EPI_ISL_575816, EPI_ISL_575817, EPI_ISL_575818, EPI_ISL_575819, EPI_ISL_575820, EPI_ISL_575821, EPI_ISL_575822, EPI_ISL_575823, EPI_ISL_575824, EPI_ISL_575825, EPI_ISL_575826, EPI_ISL_575827, EPI_ISL_575828, EPI_ISL_575829, EPI_ISL_575830, EPI_ISL_575831, EPI_ISL_575832, EPI_ISL_575833, EPI_ISL_575834, EPI_ISL_575835, EPI_ISL_575836, EPI_ISL_575837, EPI_ISL_575838, EPI_ISL_575839, EPI_ISL_575840, EPI_ISL_575842, EPI_ISL_575843, EPI_ISL_575844, EPI_ISL_575845, EPI_ISL_575847, EPI_ISL_575848, EPI_ISL_575849, EPI_ISL_575850, EPI_ISL_575851, EPI_ISL_575852, EPI_ISL_575853, EPI_ISL_575854, EPI_ISL_575855, EPI_ISL_575856, EPI_ISL_575857, EPI_ISL_575858, EPI_ISL_575859, EPI_ISL_575860, EPI_ISL_575861, EPI_ISL_575862, EPI_ISL_575863, EPI_ISL_575864, EPI_ISL_575865, EPI_ISL_575866, EPI_ISL_575867, EPI_ISL_575868, EPI_ISL_575869, EPI_ISL_575870, EPI_ISL_575872, EPI_ISL_575873, EPI_ISL_575874, EPI_ISL_575875, EPI_ISL_575876, EPI_ISL_575877, EPI_ISL_575878, EPI_ISL_575879, EPI_ISL_575880, EPI_ISL_575881, EPI_ISL_575882, EPI_ISL_575883, EPI_ISL_575884, EPI_ISL_575885, EPI_ISL_575886, EPI_ISL_575887, EPI_ISL_575888, EPI_ISL_575889, EPI_ISL_575890, EPI_ISL_575891, EPI_ISL_575892, EPI_ISL_575893, EPI_ISL_575894, EPI_ISL_575896, EPI_ISL_575897, EPI_ISL_575898, EPI_ISL_575899, EPI_ISL_575900, EPI_ISL_575901, EPI_ISL_575902, EPI_ISL_575903, EPI_ISL_575904, EPI_ISL_575905, EPI_ISL_575906, EPI_ISL_575907, EPI_ISL_575908, EPI_ISL_575909, EPI_ISL_575910, EPI_ISL_575911, EPI_ISL_575913, EPI_ISL_575914, EPI_ISL_575915, EPI_ISL_575916, EPI_ISL_575917, EPI_ISL_575918, EPI_ISL_575919, EPI_ISL_575920, EPI_ISL_575921, EPI_ISL_575922, EPI_ISL_575923, EPI_ISL_575924, EPI_ISL_575925, EPI_ISL_575926, EPI_ISL_575927, EPI_ISL_575928, EPI_ISL_575930, EPI_ISL_575931, EPI_ISL_575932, EPI_ISL_575933, EPI_ISL_575934, EPI_ISL_575935, EPI_ISL_575936, EPI_ISL_575937, EPI_ISL_575938, EPI_ISL_575939, EPI_ISL_575940, EPI_ISL_575941, EPI_ISL_575942, EPI_ISL_575944, EPI_ISL_575945, EPI_ISL_575946, EPI_ISL_575947, EPI_ISL_575948, EPI_ISL_575949, EPI_ISL_575950, EPI_ISL_575951, EPI_ISL_575952, EPI_ISL_575953, EPI_ISL_575955, EPI_ISL_575956, EPI_ISL_575958, EPI_ISL_575959, EPI_ISL_575961, EPI_ISL_575962, EPI_ISL_575963, EPI_ISL_575964, EPI_ISL_575965, EPI_ISL_575966, EPI_ISL_575967, EPI_ISL_575968, EPI_ISL_575969, EPI_ISL_575970, EPI_ISL_575971, EPI_ISL_575972, EPI_ISL_575973, EPI_ISL_575974, EPI_ISL_575975, EPI_ISL_575977

| | | | |
|---|---|---|---|
| see above | Lighthouse Lab in Milton Keynes | Wellcome Sanger Institute for the COVID-19 Genomics UK (COG-UK) consortium | The Lighthouse Lab in Milton Keynes and Alex Alderton, Roberto Amato, Sonia Goncalves, Ewan Harrison, David K. Jackson, Ian Johnston, Dominic Kwiatkowski, Cordelia Langford, John Sillitoe on behalf of the Wellcome Sanger Institute COVID-19 Surveillance Team |
| EPI_ISL_575978, EPI_ISL_575980, EPI_ISL_575981, EPI_ISL_576094 | Lighthouse Lab in Milton Keynes | Wellcome Sanger Institute for the COVID-19 Genomics UK (COG-UK) consortium | The Lighthouse Lab in Milton Keynes and Alex Alderton, Roberto Amato, Sonia Goncalves, Ewan Harrison, David K. Jackson, Ian Johnston, Dominic Kwiatkowski, Cordelia Langford, John Sillitoe on behalf of the Wellcome Sanger Institute COVID-19 Surveillance Team (http://www.sanger.ac.uk/covid-team) |

EPI_ISL_580637, EPI_ISL_580646, EPI_ISL_580647, EPI_ISL_580656, EPI_ISL_580666, EPI_ISL_580671, EPI_ISL_580698, EPI_ISL_580700, EPI_ISL_580711, EPI_ISL_580712, EPI_ISL_580718, EPI_ISL_580719, EPI_ISL_580726, EPI_ISL_580728, EPI_ISL_580732, EPI_ISL_580733, EPI_ISL_580736, EPI_ISL_580743, EPI_ISL_580745, EPI_ISL_580746, EPI_ISL_580750, EPI_ISL_580752, EPI_ISL_580753, EPI_ISL_580766, EPI_ISL_580768, EPI_ISL_580772, EPI_ISL_580773, EPI_ISL_580776, EPI_ISL_580797, EPI_ISL_580805, EPI_ISL_580812, EPI_ISL_580813, EPI_ISL_580819, EPI_ISL_580830, EPI_ISL_580841, EPI_ISL_580847, EPI_ISL_580848, EPI_ISL_580850, EPI_ISL_580855, EPI_ISL_580856, EPI_ISL_580861, EPI_ISL_580864, EPI_ISL_580873, EPI_ISL_580875, EPI_ISL_580877, EPI_ISL_580882, EPI_ISL_580896, EPI_ISL_580898, EPI_ISL_580900, EPI_ISL_580902, EPI_ISL_580904, EPI_ISL_580921, EPI_ISL_580929, EPI_ISL_580935, EPI_ISL_580948, EPI_ISL_580961, EPI_ISL_580979, EPI_ISL_580980, EPI_ISL_580990, EPI_ISL_580993, EPI_ISL_581000, EPI_ISL_581029, EPI_ISL_581038, EPI_ISL_581046, EPI_ISL_581054, EPI_ISL_581058, EPI_ISL_581064, EPI_ISL_581067, EPI_ISL_581074, EPI_ISL_581076, EPI_ISL_581078, EPI_ISL_581084, EPI_ISL_581086, EPI_ISL_581093, EPI_ISL_581106, EPI_ISL_581120, EPI_ISL_581122, EPI_ISL_581126, EPI_ISL_581128, EPI_ISL_581134, EPI_ISL_581142, EPI_ISL_581143, EPI_ISL_581152, EPI_ISL_581153, EPI_ISL_581159, EPI_ISL_581161, EPI_ISL_581168, EPI_ISL_581173, EPI_ISL_581178, EPI_ISL_581188, EPI_ISL_581198, EPI_ISL_581201, EPI_ISL_581202, EPI_ISL_581209, EPI_ISL_581212, EPI_ISL_581222, EPI_ISL_581225, EPI_ISL_581232, EPI_ISL_581234, EPI_ISL_581243, EPI_ISL_581250, EPI_ISL_581254, EPI_ISL_581259, EPI_ISL_581263, EPI_ISL_581272, EPI_ISL_581275, EPI_ISL_581278, EPI_ISL_581282, EPI_ISL_581283, EPI_ISL_581284, EPI_ISL_581287, EPI_ISL_581306, EPI_ISL_581311, EPI_ISL_581314, EPI_ISL_581315, EPI_ISL_581325, EPI_ISL_581326, EPI_ISL_581328, EPI_ISL_581335, EPI_ISL_581339, EPI_ISL_581343, EPI_ISL_581361, EPI_ISL_581368, EPI_ISL_581369, EPI_ISL_581370, EPI_ISL_581371, EPI_ISL_581375, EPI_ISL_581376, EPI_ISL_581382, EPI_ISL_581383, EPI_ISL_581385, EPI_ISL_581386, EPI_ISL_581389, EPI_ISL_581390, EPI_ISL_581397, EPI_ISL_581398, EPI_ISL_581399, EPI_ISL_581400, EPI_ISL_581401, EPI_ISL_581415, EPI_ISL_581421, EPI_ISL_581428, EPI_ISL_581434, EPI_ISL_581438, EPI_ISL_581447, EPI_ISL_589067, EPI_ISL_589070, EPI_ISL_589086, EPI_ISL_589161, EPI_ISL_589174, EPI_ISL_589179, EPI_ISL_589196, EPI_ISL_589200, EPI_ISL_589253, EPI_ISL_589259

| | | | |
|---|---|---|---|
| see above | Lighthouse Lab in Milton Keynes | Wellcome Sanger Institute for the COVID-19 Genomics UK (COG-UK) consortium | The Lighthouse Lab in Milton Keynes and Alex Alderton, Roberto Amato, Sonia Goncalves, Ewan Harrison, David K. Jackson, Ian Johnston, Dominic Kwiatkowski, Cordelia Langford, John Sillitoe on behalf of the Wellcome Sanger Institute COVID-19 Surveillance Team |

EPI_ISL_589577, EPI_ISL_589580, EPI_ISL_589584, EPI_ISL_589607, EPI_ISL_589644, EPI_ISL_589646, EPI_ISL_589647, EPI_ISL_589668, EPI_ISL_589676, EPI_ISL_589681, EPI_ISL_589699, EPI_ISL_589715, EPI_ISL_589724, EPI_ISL_589758, EPI_ISL_589763, EPI_ISL_589764, EPI_ISL_589766, EPI_ISL_589767, EPI_ISL_589768, EPI_ISL_589769, EPI_ISL_589770, EPI_ISL_589771, EPI_ISL_589772, EPI_ISL_589773, EPI_ISL_589787, EPI_ISL_589789, EPI_ISL_589791, EPI_ISL_589792, EPI_ISL_589794, EPI_ISL_589795, EPI_ISL_589797, EPI_ISL_589798, EPI_ISL_589799, EPI_ISL_589800, EPI_ISL_589801, EPI_ISL_590172, EPI_ISL_590174, EPI_ISL_590175, EPI_ISL_598256, EPI_ISL_598266, EPI_ISL_598271, EPI_ISL_598278, EPI_ISL_598279, EPI_ISL_598281, EPI_ISL_598282, EPI_ISL_598297, EPI_ISL_598310, EPI_ISL_598317, EPI_ISL_598321, EPI_ISL_598322, EPI_ISL_598325, EPI_ISL_598327, EPI_ISL_598328, EPI_ISL_598329, EPI_ISL_598352, EPI_ISL_598355, EPI_ISL_598370, EPI_ISL_598376, EPI_ISL_598384, EPI_ISL_598387, EPI_ISL_598390, EPI_ISL_598391, EPI_ISL_598394, EPI_ISL_598395, EPI_ISL_598400, EPI_ISL_598406, EPI_ISL_598411, EPI_ISL_598416, EPI_ISL_598417, EPI_ISL_598422, EPI_ISL_598423, EPI_ISL_598440, EPI_ISL_598446, EPI_ISL_598454, EPI_ISL_598456, EPI_ISL_598457, EPI_ISL_598460, EPI_ISL_598464, EPI_ISL_598473, EPI_ISL_598477, EPI_ISL_598480, EPI_ISL_598490, EPI_ISL_598492, EPI_ISL_598495, EPI_ISL_598499, EPI_ISL_598508, EPI_ISL_598515, EPI_ISL_598525, EPI_ISL_598526, EPI_ISL_598527, EPI_ISL_598528, EPI_ISL_598529, EPI_ISL_598530, EPI_ISL_598533, EPI_ISL_598534, EPI_ISL_598535, EPI_ISL_598536, EPI_ISL_598537, EPI_ISL_598538, EPI_ISL_598539, EPI_ISL_598541, EPI_ISL_598542, EPI_ISL_598543, EPI_ISL_598544, EPI_ISL_598545, EPI_ISL_598546, EPI_ISL_598549, EPI_ISL_598550, EPI_ISL_598553, EPI_ISL_598554, EPI_ISL_598555, EPI_ISL_598556, EPI_ISL_598558, EPI_ISL_598560, EPI_ISL_598561, EPI_ISL_598562, EPI_ISL_598563, EPI_ISL_598565, EPI_ISL_598568, EPI_ISL_598569, EPI_ISL_598572, EPI_ISL_598573, EPI_ISL_598574, EPI_ISL_598576, EPI_ISL_598577, EPI_ISL_598579, EPI_ISL_598580, EPI_ISL_598581, EPI_ISL_598582, EPI_ISL_598583, EPI_ISL_598584, EPI_ISL_598585, EPI_ISL_598586, EPI_ISL_598587, EPI_ISL_598588, EPI_ISL_598589, EPI_ISL_598591, EPI_ISL_598593, EPI_ISL_598595, EPI_ISL_598597, EPI_ISL_598598, EPI_ISL_598599, EPI_ISL_598601, EPI_ISL_598602, EPI_ISL_598604, EPI_ISL_598605, EPI_ISL_598607, EPI_ISL_598608, EPI_ISL_598610, EPI_ISL_598611, EPI_ISL_598614, EPI_ISL_598615, EPI_ISL_598616, EPI_ISL_598617, EPI_ISL_598618, EPI_ISL_598619, EPI_ISL_598620, EPI_ISL_598621, EPI_ISL_598624, EPI_ISL_598625, EPI_ISL_598626, EPI_ISL_598627, EPI_ISL_598628, EPI_ISL_598629, EPI_ISL_598630, EPI_ISL_598632, EPI_ISL_598633, EPI_ISL_598635, EPI_ISL_598639, EPI_ISL_598641, EPI_ISL_598642, EPI_ISL_598643, EPI_ISL_598644, EPI_ISL_598645, EPI_ISL_598646, EPI_ISL_598648, EPI_ISL_598649, EPI_ISL_598651, EPI_ISL_598652, EPI_ISL_598653, EPI_ISL_598654, EPI_ISL_598657, EPI_ISL_598661, EPI_ISL_598662, EPI_ISL_598664, EPI_ISL_598665, EPI_ISL_598666, EPI_ISL_598668, EPI_ISL_598669, EPI_ISL_598670, EPI_ISL_598671, EPI_ISL_598672, EPI_ISL_598673, EPI_ISL_598674, EPI_ISL_598675, EPI_ISL_598676, EPI_ISL_598677, EPI_ISL_598679, EPI_ISL_598681, EPI_ISL_598682, EPI_ISL_598683, EPI_ISL_598684, EPI_ISL_598687, EPI_ISL_598688, EPI_ISL_598689, EPI_ISL_598690, EPI_ISL_598691, EPI_ISL_598692, EPI_ISL_598694, EPI_ISL_598696, EPI_ISL_598697, EPI_ISL_598698, EPI_ISL_598699, EPI_ISL_598700, EPI_ISL_598701, EPI_ISL_598702, EPI_ISL_598703, EPI_ISL_598704, EPI_ISL_598705, EPI_ISL_598707, EPI_ISL_598708, EPI_ISL_598709, EPI_ISL_598710, EPI_ISL_598712, EPI_ISL_598713, EPI_ISL_598714, EPI_ISL_598715, EPI_ISL_598716, EPI_ISL_598717, EPI_ISL_598718, EPI_ISL_598719, EPI_ISL_598720, EPI_ISL_598723, EPI_ISL_598724, EPI_ISL_598727, EPI_ISL_598729, EPI_ISL_598730, EPI_ISL_598731, EPI_ISL_598732, EPI_ISL_598736, EPI_ISL_598738, EPI_ISL_598740, EPI_ISL_598742, EPI_ISL_598743, EPI_ISL_598744, EPI_ISL_598745, EPI_ISL_598746, EPI_ISL_598751, EPI_ISL_598752, EPI_ISL_598753, EPI_ISL_598754, EPI_ISL_598756, EPI_ISL_598757, EPI_ISL_598759, EPI_ISL_598760, EPI_ISL_598761, EPI_ISL_598762, EPI_ISL_598763, EPI_ISL_598764, EPI_ISL_598766, EPI_ISL_598767, EPI_ISL_598768, EPI_ISL_598769, EPI_ISL_598770, EPI_ISL_598771, EPI_ISL_598772, EPI_ISL_598773, EPI_ISL_598774, EPI_ISL_598775, EPI_ISL_598776, EPI_ISL_598777, EPI_ISL_598778, EPI_ISL_598779, EPI_ISL_598780, EPI_ISL_598783, EPI_ISL_598784, EPI_ISL_598785, EPI_ISL_598788, EPI_ISL_598790, EPI_ISL_598791, EPI_ISL_598792, EPI_ISL_598793, EPI_ISL_598794, EPI_ISL_598795, EPI_ISL_598796, EPI_ISL_598797, EPI_ISL_598798, EPI_ISL_598802, EPI_ISL_598803, EPI_ISL_598804, EPI_ISL_598805, EPI_ISL_598806, EPI_ISL_598807, EPI_ISL_598808, EPI_ISL_598809, EPI_ISL_598811, EPI_ISL_598812, EPI_ISL_598813, EPI_ISL_598814, EPI_ISL_598816, EPI_ISL_598822, EPI_ISL_599830, EPI_ISL_599831, EPI_ISL_599835, EPI_ISL_599839, EPI_ISL_599843, EPI_ISL_599847, EPI_ISL_599850, EPI_ISL_599851, EPI_ISL_599878, EPI_ISL_599882, EPI_ISL_599884, EPI_ISL_599886, EPI_ISL_599895, EPI_ISL_599897, EPI_ISL_599907, EPI_ISL_599910, EPI_ISL_599946, EPI_ISL_599947, EPI_ISL_599950, EPI_ISL_599952, EPI_ISL_599953, EPI_ISL_599955, EPI_ISL_599966, EPI_ISL_599971, EPI_ISL_599974, EPI_ISL_599982, EPI_ISL_599985, EPI_ISL_599988, EPI_ISL_599991, EPI_ISL_599994, EPI_ISL_599996, EPI_ISL_599997, EPI_ISL_600006, EPI_ISL_600019, EPI_ISL_600027, EPI_ISL_600031, EPI_ISL_600043, EPI_ISL_600044, EPI_ISL_600050, EPI_ISL_600064, EPI_ISL_600065, EPI_ISL_600067, EPI_ISL_600071, EPI_ISL_600077, EPI_ISL_600079, EPI_ISL_600081, EPI_ISL_600088, EPI_ISL_600108, EPI_ISL_600123, EPI_ISL_600124, EPI_ISL_600126, EPI_ISL_600892, EPI_ISL_600901, EPI_ISL_600912, EPI_ISL_600913, EPI_ISL_600915, EPI_ISL_600923, EPI_ISL_600925, EPI_ISL_600926, EPI_ISL_600927, EPI_ISL_600928, EPI_ISL_600929, EPI_ISL_600930, EPI_ISL_601243, EPI_ISL_601263, EPI_ISL_601264, EPI_ISL_601273, EPI_ISL_601277, EPI_ISL_601283, EPI_ISL_601289, EPI_ISL_601292, EPI_ISL_601293, EPI_ISL_601297, EPI_ISL_601306, EPI_ISL_601310, EPI_ISL_601311, EPI_ISL_601312, EPI_ISL_601316, EPI_ISL_601321, EPI_ISL_601322, EPI_ISL_601337, EPI_ISL_601346, EPI_ISL_601350, EPI_ISL_601367, EPI_ISL_601378, EPI_ISL_601381, EPI_ISL_601382, EPI_ISL_601385, EPI_ISL_601396, EPI_ISL_601397, EPI_ISL_601398, EPI_ISL_601415, EPI_ISL_601419, EPI_ISL_601422, EPI_ISL_601428, EPI_ISL_601440, EPI_ISL_601441, EPI_ISL_601457, EPI_ISL_601458, EPI_ISL_601472, EPI_ISL_601475, EPI_ISL_601485, EPI_ISL_601487, EPI_ISL_601490, EPI_ISL_601494, EPI_ISL_601496, EPI_ISL_601497, EPI_ISL_601498, EPI_ISL_601500, EPI_ISL_601503, EPI_ISL_601504, EPI_ISL_601508, EPI_ISL_601509, EPI_ISL_601510, EPI_ISL_601512, EPI_ISL_601516, EPI_ISL_601517, EPI_ISL_601519, EPI_ISL_601527, EPI_ISL_601529, EPI_ISL_601534, EPI_ISL_601537, EPI_ISL_601538, EPI_ISL_601542, EPI_ISL_601544, EPI_ISL_601551, EPI_ISL_601553, EPI_ISL_601559, EPI_ISL_601560, EPI_ISL_601564, EPI_ISL_601565, EPI_ISL_601566, EPI_ISL_601568, EPI_ISL_601572, EPI_ISL_601576, EPI_ISL_601592, EPI_ISL_601593, EPI_ISL_601597, EPI_ISL_601601, EPI_ISL_601602, EPI_ISL_601605, EPI_ISL_601607, EPI_ISL_601608, EPI_ISL_601609, EPI_ISL_601612, EPI_ISL_601613, EPI_ISL_601620, EPI_ISL_601621, EPI_ISL_601623, EPI_ISL_601625, EPI_ISL_601626, EPI_ISL_601629, EPI_ISL_601630, EPI_ISL_601635, EPI_ISL_601636, EPI_ISL_601637, EPI_ISL_601638, EPI_ISL_601640, EPI_ISL_601643, EPI_ISL_601645, EPI_ISL_601646, EPI_ISL_601654, EPI_ISL_601655, EPI_ISL_601661, EPI_ISL_601662, EPI_ISL_601665, EPI_ISL_601669, EPI_ISL_601672, EPI_ISL_601674, EPI_ISL_601675, EPI_ISL_601678, EPI_ISL_601679, EPI_ISL_601681, EPI_ISL_601683, EPI_ISL_601684, EPI_ISL_601686, EPI_ISL_601687, EPI_ISL_601689, EPI_ISL_601693, EPI_ISL_601694, EPI_ISL_601695, EPI_ISL_601696, EPI_ISL_601697, EPI_ISL_601703, EPI_ISL_601705, EPI_ISL_601707, EPI_ISL_601712, EPI_ISL_601713, EPI_ISL_601714, EPI_ISL_601716, EPI_ISL_601718, EPI_ISL_601719, EPI_ISL_601720, EPI_ISL_601722, EPI_ISL_601724, EPI_ISL_601726, EPI_ISL_601727, EPI_ISL_601729, EPI_ISL_601735, EPI_ISL_601736, EPI_ISL_601737, EPI_ISL_601741, EPI_ISL_601742, EPI_ISL_601745, EPI_ISL_601746, EPI_ISL_601750, EPI_ISL_601751, EPI_ISL_601752, EPI_ISL_601754, EPI_ISL_601758, EPI_ISL_601759, EPI_ISL_601760, EPI_ISL_601762, EPI_ISL_601767, EPI_ISL_601769, EPI_ISL_601770, EPI_ISL_601772, EPI_ISL_601775, EPI_ISL_601781, EPI_ISL_601782, EPI_ISL_601784, EPI_ISL_601786, EPI_ISL_602027, EPI_ISL_602028, EPI_ISL_602029, EPI_ISL_602145, EPI_ISL_602146, EPI_ISL_602147, EPI_ISL_602148, EPI_ISL_602149, EPI_ISL_602150, EPI_ISL_602151, EPI_ISL_602152, EPI_ISL_602153

| | | | |
|---|---|---|---|
| see above | Lighthouse Lab in Milton Keynes | Wellcome Sanger Institute for the COVID-19 Genomics UK (COG-UK) consortium | The Lighthouse Lab in Milton Keynes and Alex Alderton, Roberto Amato, Sonia Goncalves, Ewan Harrison, David K. Jackson, Ian Johnston, Dominic Kwiatkowski, Cordelia Langford, John Sillitoe on behalf of the Wellcome Sanger Institute COVID-19 Surveillance Team (http://www.sanger.ac.uk/covid-team) |

EPI_ISL_606296, EPI_ISL_606297, EPI_ISL_606298, EPI_ISL_606299, EPI_ISL_606300, EPI_ISL_606301, EPI_ISL_606302, EPI_ISL_606303, EPI_ISL_606304, EPI_ISL_606305, EPI_ISL_606306, EPI_ISL_606307, EPI_ISL_606308, EPI_ISL_606309, EPI_ISL_606310, EPI_ISL_606312, EPI_ISL_606313, EPI_ISL_606315, EPI_ISL_606316, EPI_ISL_606317, EPI_ISL_606318, EPI_ISL_606319, EPI_ISL_606320, EPI_ISL_606322, EPI_ISL_606323, EPI_ISL_606324, EPI_ISL_606325, EPI_ISL_606326, EPI_ISL_606327, EPI_ISL_606329, EPI_ISL_606330, EPI_ISL_606331, EPI_ISL_606332, EPI_ISL_606333, EPI_ISL_606334, EPI_ISL_606335, EPI_ISL_606336, EPI_ISL_606337, EPI_ISL_606338, EPI_ISL_606339, EPI_ISL_606340, EPI_ISL_606341, EPI_ISL_606343, EPI_ISL_606344, EPI_ISL_606345, EPI_ISL_606346, EPI_ISL_606347, EPI_ISL_606348, EPI_ISL_606349, EPI_ISL_606350, EPI_ISL_606351, EPI_ISL_606352, EPI_ISL_606354, EPI_ISL_606355, EPI_ISL_606356, EPI_ISL_606357, EPI_ISL_606358, EPI_ISL_606359, EPI_ISL_606360, EPI_ISL_606361, EPI_ISL_606363, EPI_ISL_606364, EPI_ISL_606365, EPI_ISL_606366, EPI_ISL_606367, EPI_ISL_606368, EPI_ISL_606369, EPI_ISL_606370, EPI_ISL_606371, EPI_ISL_606372, EPI_ISL_606373, EPI_ISL_606374, EPI_ISL_606375, EPI_ISL_606376, EPI_ISL_606377, EPI_ISL_606378, EPI_ISL_606380, EPI_ISL_606382, EPI_ISL_606383, EPI_ISL_606384, EPI_ISL_606385, EPI_ISL_606386, EPI_ISL_606387, EPI_ISL_606388, EPI_ISL_606389, EPI_ISL_606390, EPI_ISL_606391, EPI_ISL_606392, EPI_ISL_606393, EPI_ISL_606394, EPI_ISL_606395, EPI_ISL_606396, EPI_ISL_606397, EPI_ISL_606398, EPI_ISL_606399, EPI_ISL_606400, EPI_ISL_606403, EPI_ISL_606404, EPI_ISL_606405, EPI_ISL_606406, EPI_ISL_606407, EPI_ISL_606408, EPI_ISL_606409, EPI_ISL_606410, EPI_ISL_606411, EPI_ISL_606412, EPI_ISL_606413, EPI_ISL_606414, EPI_ISL_606416, EPI_ISL_606417, EPI_ISL_606418, EPI_ISL_606419, EPI_ISL_606420, EPI_ISL_606421, EPI_ISL_606422, EPI_ISL_606424, EPI_ISL_606425, EPI_ISL_606426, EPI_ISL_606427, EPI_ISL_606428, EPI_ISL_606429, EPI_ISL_606430, EPI_ISL_606431, EPI_ISL_606432, EPI_ISL_606433, EPI_ISL_606434, EPI_ISL_606435, EPI_ISL_606436, EPI_ISL_606437, EPI_ISL_606438, EPI_ISL_606439, EPI_ISL_606440, EPI_ISL_606441, EPI_ISL_606442, EPI_ISL_606443, EPI_ISL_606444, EPI_ISL_606445, EPI_ISL_606446, EPI_ISL_606447, EPI_ISL_606448, EPI_ISL_606449, EPI_ISL_606450, EPI_ISL_606451, EPI_ISL_606452, EPI_ISL_606453, EPI_ISL_606454, EPI_ISL_606455, EPI_ISL_606456, EPI_ISL_606457, EPI_ISL_606458, EPI_ISL_606459, EPI_ISL_606460, EPI_ISL_606461, EPI_ISL_606463, EPI_ISL_606464, EPI_ISL_606465, EPI_ISL_606466, EPI_ISL_606467, EPI_ISL_606468, EPI_ISL_606469, EPI_ISL_606470, EPI_ISL_606471, EPI_ISL_606472, EPI_ISL_606473, EPI_ISL_606474, EPI_ISL_606475, EPI_ISL_606476, EPI_ISL_606477, EPI_ISL_606478, EPI_ISL_606479, EPI_ISL_606480, EPI_ISL_606482, EPI_ISL_606484, EPI_ISL_606485, EPI_ISL_606486, EPI_ISL_606487, EPI_ISL_606489, EPI_ISL_606490, EPI_ISL_606491, EPI_ISL_606492, EPI_ISL_606493, EPI_ISL_606494, EPI_ISL_606495, EPI_ISL_606496, EPI_ISL_606497, EPI_ISL_606498, EPI_ISL_606499, EPI_ISL_606500, EPI_ISL_606501, EPI_ISL_606502, EPI_ISL_606504, EPI_ISL_606505, EPI_ISL_606506, EPI_ISL_606507, EPI_ISL_606508, EPI_ISL_606509, EPI_ISL_606510, EPI_ISL_606511, EPI_ISL_606514, EPI_ISL_606515, EPI_ISL_606517, EPI_ISL_606518, EPI_ISL_606519, EPI_ISL_606520, EPI_ISL_606521, EPI_ISL_606522, EPI_ISL_606525, EPI_ISL_606526, EPI_ISL_606527, EPI_ISL_606528, EPI_ISL_606529, EPI_ISL_606530, EPI_ISL_606531, EPI_ISL_606532, EPI_ISL_606533, EPI_ISL_606534, EPI_ISL_606535, EPI_ISL_606536, EPI_ISL_606537, EPI_ISL_606538, EPI_ISL_606539, EPI_ISL_606540, EPI_ISL_606541, EPI_ISL_606542, EPI_ISL_606543, EPI_ISL_606544, EPI_ISL_606545, EPI_ISL_606546, EPI_ISL_606548, EPI_ISL_606549, EPI_ISL_606550, EPI_ISL_606551, EPI_ISL_606552, EPI_ISL_606553, EPI_ISL_606555, EPI_ISL_606556, EPI_ISL_606557, EPI_ISL_606558, EPI_ISL_606559, EPI_ISL_606560, EPI_ISL_606561, EPI_ISL_606562, EPI_ISL_606563, EPI_ISL_606564, EPI_ISL_606565, EPI_ISL_606566, EPI_ISL_606567, EPI_ISL_606568, EPI_ISL_606569, EPI_ISL_606570, EPI_ISL_606571, EPI_ISL_606572, EPI_ISL_606573, EPI_ISL_606574, EPI_ISL_606575, EPI_ISL_606576, EPI_ISL_606577, EPI_ISL_606578, EPI_ISL_606579, EPI_ISL_606580, EPI_ISL_606581, EPI_ISL_606582, EPI_ISL_606583, EPI_ISL_606584, EPI_ISL_606586, EPI_ISL_606587, EPI_ISL_606589, EPI_ISL_606590, EPI_ISL_606591, EPI_ISL_606592, EPI_ISL_606593, EPI_ISL_606594, EPI_ISL_606595, EPI_ISL_606596, EPI_ISL_606597, EPI_ISL_606598, EPI_ISL_606599, EPI_ISL_606600, EPI_ISL_606601, EPI_ISL_606602, EPI_ISL_606603, EPI_ISL_606604, EPI_ISL_606607, EPI_ISL_606609, EPI_ISL_606610, EPI_ISL_606611, EPI_ISL_606612, EPI_ISL_606613, EPI_ISL_606616, EPI_ISL_606617, EPI_ISL_606618, EPI_ISL_606621, EPI_ISL_606623, EPI_ISL_606626, EPI_ISL_606627, EPI_ISL_606628, EPI_ISL_606629, EPI_ISL_606630, EPI_ISL_606633, EPI_ISL_606636, EPI_ISL_606637, EPI_ISL_606638, EPI_ISL_606639, EPI_ISL_606640, EPI_ISL_606641, EPI_ISL_606642, EPI_ISL_606643, EPI_ISL_606644, EPI_ISL_606645, EPI_ISL_606646, EPI_ISL_606647, EPI_ISL_606648, EPI_ISL_606649, EPI_ISL_606650, EPI_ISL_606651, EPI_ISL_606653, EPI_ISL_606655, EPI_ISL_606656, EPI_ISL_606657, EPI_ISL_606658, EPI_ISL_606659, EPI_ISL_606660, EPI_ISL_606664, EPI_ISL_606665, EPI_ISL_606666, EPI_ISL_606667, EPI_ISL_606668, EPI_ISL_606669, EPI_ISL_606670, EPI_ISL_606671, EPI_ISL_606673, EPI_ISL_606674, EPI_ISL_606675, EPI_ISL_606678, EPI_ISL_606679, EPI_ISL_606681, EPI_ISL_606682, EPI_ISL_606687, EPI_ISL_606688, EPI_ISL_606689, EPI_ISL_606690, EPI_ISL_606691, EPI_ISL_606693, EPI_ISL_606694, EPI_ISL_606695, EPI_ISL_606696, EPI_ISL_606697, EPI_ISL_606699, EPI_ISL_606700, EPI_ISL_606702, EPI_ISL_606705, EPI_ISL_606706, EPI_ISL_606707, EPI_ISL_606708, EPI_ISL_606709, EPI_ISL_606711, EPI_ISL_606712, EPI_ISL_606713, EPI_ISL_606714, EPI_ISL_606716, EPI_ISL_606718, EPI_ISL_606719, EPI_ISL_606720, EPI_ISL_606722, EPI_ISL_606723, EPI_ISL_606724, EPI_ISL_606725, EPI_ISL_606726, EPI_ISL_606727, EPI_ISL_606728, EPI_ISL_606729, EPI_ISL_606730, EPI_ISL_606731, EPI_ISL_606734, EPI_ISL_606735, EPI_ISL_606736, EPI_ISL_606741, EPI_ISL_606743, EPI_ISL_606744, EPI_ISL_606746, EPI_ISL_606748, EPI_ISL_606749, EPI_ISL_606750, EPI_ISL_606751, EPI_ISL_606752, EPI_ISL_606753, EPI_ISL_606754, EPI_ISL_606755, EPI_ISL_606756, EPI_ISL_606758, EPI_ISL_606761, EPI_ISL_606764, EPI_ISL_606765, EPI_ISL_606767, EPI_ISL_606768, EPI_ISL_606769, EPI_ISL_606770, EPI_ISL_606771, EPI_ISL_606774, EPI_ISL_606775, EPI_ISL_606777, EPI_ISL_606778, EPI_ISL_606782, EPI_ISL_606784, EPI_ISL_606786, EPI_ISL_606787, EPI_ISL_606789, EPI_ISL_606790, EPI_ISL_606792, EPI_ISL_606793, EPI_ISL_606794, EPI_ISL_606795, EPI_ISL_606796, EPI_ISL_606798, EPI_ISL_606800, EPI_ISL_606801, EPI_ISL_606802, EPI_ISL_606803, EPI_ISL_606804, EPI_ISL_606805, EPI_ISL_606806, EPI_ISL_606807, EPI_ISL_606808, EPI_ISL_606809, EPI_ISL_606812, EPI_ISL_606813, EPI_ISL_606814, EPI_ISL_606815, EPI_ISL_606819, EPI_ISL_606820, EPI_ISL_606822, EPI_ISL_606823, EPI_ISL_606826, EPI_ISL_606828, EPI_ISL_606829, EPI_ISL_606830, EPI_ISL_606831, EPI_ISL_606832, EPI_ISL_606833, EPI_ISL_606834, EPI_ISL_606835, EPI_ISL_606836, EPI_ISL_606837, EPI_ISL_606838, EPI_ISL_606843, EPI_ISL_606844, EPI_ISL_606845, EPI_ISL_606848, EPI_ISL_606849, EPI_ISL_606850, EPI_ISL_606851, EPI_ISL_606853, EPI_ISL_606854, EPI_ISL_606855, EPI_ISL_606856, EPI_ISL_606857, EPI_ISL_606858, EPI_ISL_606859, EPI_ISL_606861, EPI_ISL_606862, EPI_ISL_606863, EPI_ISL_606866, EPI_ISL_606868, EPI_ISL_606869, EPI_ISL_606870, EPI_ISL_606872, EPI_ISL_606873, EPI_ISL_606874, EPI_ISL_606876, EPI_ISL_606877, EPI_ISL_606878, EPI_ISL_606880, EPI_ISL_606881, EPI_ISL_606882, EPI_ISL_606883, EPI_ISL_606884, EPI_ISL_606885, EPI_ISL_606886, EPI_ISL_606887, EPI_ISL_606888, EPI_ISL_606890, EPI_ISL_606894, EPI_ISL_606895, EPI_ISL_606896, EPI_ISL_606897, EPI_ISL_606899, EPI_ISL_606900, EPI_ISL_606902, EPI_ISL_606904, EPI_ISL_606905, EPI_ISL_606906, EPI_ISL_606907, EPI_ISL_606908, EPI_ISL_606912, EPI_ISL_606913, EPI_ISL_606914, EPI_ISL_606915, EPI_ISL_606917, EPI_ISL_606918, EPI_ISL_606919, EPI_ISL_606920, EPI_ISL_606921, EPI_ISL_606922, EPI_ISL_606923, EPI_ISL_607291, EPI_ISL_607292, EPI_ISL_607293, EPI_ISL_607294, EPI_ISL_607295, EPI_ISL_607296, EPI_ISL_607297, EPI_ISL_607299, EPI_ISL_607300, EPI_ISL_607301, EPI_ISL_607303, EPI_ISL_607306, EPI_ISL_607308, EPI_ISL_607309, EPI_ISL_607310, EPI_ISL_607311, EPI_ISL_607312, EPI_ISL_607314, EPI_ISL_607315, EPI_ISL_607316, EPI_ISL_607317, EPI_ISL_607318, EPI_ISL_607320, EPI_ISL_607321, EPI_ISL_607324, EPI_ISL_607325, EPI_ISL_607328, EPI_ISL_607330, EPI_ISL_607331, EPI_ISL_607332, EPI_ISL_607333, EPI_ISL_607335, EPI_ISL_607336, EPI_ISL_607337, EPI_ISL_607339, EPI_ISL_607342, EPI_ISL_607343, EPI_ISL_607344, EPI_ISL_607345, EPI_ISL_607346, EPI_ISL_607348, EPI_ISL_607350, EPI_ISL_607351, EPI_ISL_607352, EPI_ISL_607353, EPI_ISL_607354, EPI_ISL_607355, EPI_ISL_607356, EPI_ISL_607357, EPI_ISL_607358, EPI_ISL_607359, EPI_ISL_607360, EPI_ISL_607361, EPI_ISL_607362, EPI_ISL_607363, EPI_ISL_607364, EPI_ISL_607365, EPI_ISL_607366, EPI_ISL_607367, EPI_ISL_607369, EPI_ISL_607370, EPI_ISL_607371, EPI_ISL_607372, EPI_ISL_607373, EPI_ISL_607374, EPI_ISL_607375, EPI_ISL_607376, EPI_ISL_607378, EPI_ISL_607379, EPI_ISL_607380, EPI_ISL_607381, EPI_ISL_607382, EPI_ISL_607383, EPI_ISL_607384, EPI_ISL_607385, EPI_ISL_607386, EPI_ISL_607388, EPI_ISL_607389, EPI_ISL_607390, EPI_ISL_607393, EPI_ISL_607394, EPI_ISL_607395, EPI_ISL_607396, EPI_ISL_607397, EPI_ISL_607398, EPI_ISL_607399, EPI_ISL_607400, EPI_ISL_607401, EPI_ISL_607402, EPI_ISL_607403, EPI_ISL_607404, EPI_ISL_607405, EPI_ISL_607406, EPI_ISL_607407, EPI_ISL_607409, EPI_ISL_607411, EPI_ISL_607412, EPI_ISL_607413, EPI_ISL_607415, EPI_ISL_607417, EPI_ISL_607419, EPI_ISL_607421, EPI_ISL_607422, EPI_ISL_607423, EPI_ISL_607424, EPI_ISL_607425, EPI_ISL_607426, EPI_ISL_607427, EPI_ISL_607428, EPI_ISL_607429, EPI_ISL_607430, EPI_ISL_607431, EPI_ISL_607432, EPI_ISL_607433, EPI_ISL_607434, EPI_ISL_607435, EPI_ISL_607436, EPI_ISL_607437, EPI_ISL_607438, EPI_ISL_607439, EPI_ISL_607441, EPI_ISL_607442, EPI_ISL_607443, EPI_ISL_607444, EPI_ISL_607445, EPI_ISL_607446, EPI_ISL_607447, EPI_ISL_607449, EPI_ISL_607450, EPI_ISL_607451, EPI_ISL_607452, EPI_ISL_607453, EPI_ISL_607454, EPI_ISL_607456, EPI_ISL_607457, EPI_ISL_607458, EPI_ISL_607459, EPI_ISL_607460, EPI_ISL_607462, EPI_ISL_607463, EPI_ISL_607464, EPI_ISL_607466, EPI_ISL_607467, EPI_ISL_607468, EPI_ISL_607469, EPI_ISL_607470, EPI_ISL_607471, EPI_ISL_607472, EPI_ISL_607473, EPI_ISL_607474, EPI_ISL_607476, EPI_ISL_607477, EPI_ISL_607478, EPI_ISL_607479, EPI_ISL_607480, EPI_ISL_607481, EPI_ISL_607482, EPI_ISL_607483, EPI_ISL_607484, EPI_ISL_607485, EPI_ISL_607486, EPI_ISL_607487, EPI_ISL_607488, EPI_ISL_607489, EPI_ISL_607490, EPI_ISL_607491, EPI_ISL_607492, EPI_ISL_607493, EPI_ISL_607494, EPI_ISL_607495, EPI_ISL_607496, EPI_ISL_607497, EPI_ISL_607498, EPI_ISL_607499, EPI_ISL_607500, EPI_ISL_607501, EPI_ISL_607502, EPI_ISL_607503, EPI_ISL_607504, EPI_ISL_607506, EPI_ISL_607507, EPI_ISL_607508, EPI_ISL_607510, EPI_ISL_607511, EPI_ISL_607512, EPI_ISL_607513, EPI_ISL_607514, EPI_ISL_607515, EPI_ISL_607517, EPI_ISL_607518, EPI_ISL_607519, EPI_ISL_607520, EPI_ISL_607521, EPI_ISL_607522, EPI_ISL_607524, EPI_ISL_607525, EPI_ISL_607526, EPI_ISL_607527, EPI_ISL_607528, EPI_ISL_607529, EPI_ISL_607530, EPI_ISL_607531, EPI_ISL_607532, EPI_ISL_607533, EPI_ISL_607534, EPI_ISL_607535, EPI_ISL_607536, EPI_ISL_607537, EPI_ISL_607538, EPI_ISL_607539, EPI_ISL_607540, EPI_ISL_607541, EPI_ISL_607542, EPI_ISL_607543, EPI_ISL_607546, EPI_ISL_607547, EPI_ISL_607548, EPI_ISL_607549, EPI_ISL_607551, EPI_ISL_607552, EPI_ISL_607553, EPI_ISL_607554, EPI_ISL_607555, EPI_ISL_607556, EPI_ISL_607557, EPI_ISL_607558, EPI_ISL_607559, EPI_ISL_607560, EPI_ISL_607561, EPI_ISL_607562, EPI_ISL_607563, EPI_ISL_607564, EPI_ISL_607565, EPI_ISL_607567, EPI_ISL_607568, EPI_ISL_607569, EPI_ISL_607570, EPI_ISL_607571, EPI_ISL_607574, EPI_ISL_607576, EPI_ISL_607577, EPI_ISL_607578, EPI_ISL_607580, EPI_ISL_607582, EPI_ISL_607584, EPI_ISL_607588, EPI_ISL_607589, EPI_ISL_607590, EPI_ISL_607591, EPI_ISL_607592, EPI_ISL_607593, EPI_ISL_607594, EPI_ISL_607595, EPI_ISL_607596, EPI_ISL_607597, EPI_ISL_607598, EPI_ISL_607599, EPI_ISL_607600, EPI_ISL_607602, EPI_ISL_607603, EPI_ISL_607604, EPI_ISL_607605, EPI_ISL_607606, EPI_ISL_607607, EPI_ISL_607609, EPI_ISL_607610, EPI_ISL_607611, EPI_ISL_607612, EPI_ISL_607613, EPI_ISL_607615, EPI_ISL_607616, EPI_ISL_607617, EPI_ISL_607618, EPI_ISL_607619, EPI_ISL_607620, EPI_ISL_607621, EPI_ISL_607622, EPI_ISL_608698, EPI_ISL_608704, EPI_ISL_608722, EPI_ISL_608745, EPI_ISL_608752, EPI_ISL_608766, EPI_ISL_608799, EPI_ISL_608829, EPI_ISL_608870, EPI_ISL_608901, EPI_ISL_608951, EPI_ISL_609026, EPI_ISL_609044, EPI_ISL_609139, EPI_ISL_609219, EPI_ISL_609220, EPI_ISL_609222, EPI_ISL_609225, EPI_ISL_609228, EPI_ISL_609229, EPI_ISL_609232, EPI_ISL_609233, EPI_ISL_609234, EPI_ISL_609235, EPI_ISL_609237, EPI_ISL_609238, EPI_ISL_609239, EPI_ISL_609241, EPI_ISL_609246, EPI_ISL_609247, EPI_ISL_609248, EPI_ISL_609249, EPI_ISL_609250, EPI_ISL_609251, EPI_ISL_609252, EPI_ISL_609254, EPI_ISL_609256, EPI_ISL_609257, EPI_ISL_609258, EPI_ISL_609260, EPI_ISL_609261, EPI_ISL_609262, EPI_ISL_609263, EPI_ISL_609264, EPI_ISL_609268, EPI_ISL_609269, EPI_ISL_609270, EPI_ISL_609271, EPI_ISL_609276, EPI_ISL_609277, EPI_ISL_609280, EPI_ISL_609281, EPI_ISL_609282, EPI_ISL_609283, EPI_ISL_609284, EPI_ISL_609285, EPI_ISL_609286, EPI_ISL_609287, EPI_ISL_609288, EPI_ISL_609289, EPI_ISL_609290, EPI_ISL_609291, EPI_ISL_609292, EPI_ISL_609293, EPI_ISL_609294, EPI_ISL_609295, EPI_ISL_609296, EPI_ISL_609297, EPI_ISL_609298, EPI_ISL_609299, EPI_ISL_609300, EPI_ISL_609301, EPI_ISL_609303, EPI_ISL_609304, EPI_ISL_609305, EPI_ISL_609306, EPI_ISL_609307

| | | | |
|---|---|---|---|
| see above | Lighthouse Lab in Milton Keynes | Wellcome Sanger Institute for the COVID-19 Genomics UK (COG-UK) consortium | The Lighthouse Lab in Milton Keynes and Alex Alderton, Roberto Amato, Sonia Goncalves, Ewan Harrison, David K. Jackson, Ian Johnston, Dominic Kwiatkowski, Cordelia Langford, John Sillitoe on behalf of the Wellcome Sanger Institute COVID-19 Surveillance Team |
| EPI_ISL_623218, EPI_ISL_623262, EPI_ISL_623486 | Lighthouse Lab in Milton Keynes | Wellcome Sanger Institute for the COVID-19 Genomics UK (COG-UK) consortium | The Lighthouse Lab in Milton Keynes and Alex Alderton, Roberto Amato, Sonia Goncalves, Ewan Harrison, David K. Jackson, Ian Johnston, Dominic Kwiatkowski, Cordelia Langford, John Sillitoe on behalf of the Wellcome Sanger Institute COVID-19 Surveillance Team (http://www.sanger.ac.uk/covid-team) |

EPI_ISL_629109, EPI_ISL_629110, EPI_ISL_629112, EPI_ISL_629114, EPI_ISL_629115, EPI_ISL_629116, EPI_ISL_629117, EPI_ISL_629118, EPI_ISL_629119, EPI_ISL_629120, EPI_ISL_629121, EPI_ISL_629124, EPI_ISL_629126, EPI_ISL_629127, EPI_ISL_629130, EPI_ISL_629131, EPI_ISL_629132, EPI_ISL_629133, EPI_ISL_629134, EPI_ISL_629137, EPI_ISL_629139, EPI_ISL_629141, EPI_ISL_629142, EPI_ISL_629146, EPI_ISL_629148, EPI_ISL_629149, EPI_ISL_629150, EPI_ISL_629152, EPI_ISL_629154, EPI_ISL_629155, EPI_ISL_629156, EPI_ISL_629157, EPI_ISL_629158, EPI_ISL_629159, EPI_ISL_629161, EPI_ISL_629163, EPI_ISL_629164, EPI_ISL_629167, EPI_ISL_629169, EPI_ISL_629171, EPI_ISL_629172, EPI_ISL_629174, EPI_ISL_629175, EPI_ISL_629176, EPI_ISL_629181, EPI_ISL_629183, EPI_ISL_629185, EPI_ISL_629186, EPI_ISL_629187, EPI_ISL_629188, EPI_ISL_629190, EPI_ISL_629191, EPI_ISL_629192, EPI_ISL_629193, EPI_ISL_629195, EPI_ISL_629196, EPI_ISL_629200, EPI_ISL_629201, EPI_ISL_629203, EPI_ISL_629204, EPI_ISL_629205, EPI_ISL_629207, EPI_ISL_629208, EPI_ISL_629210, EPI_ISL_629211, EPI_ISL_629212, EPI_ISL_629214, EPI_ISL_629215, EPI_ISL_629216, EPI_ISL_629218, EPI_ISL_629220, EPI_ISL_629221, EPI_ISL_629222, EPI_ISL_629223, EPI_ISL_629224, EPI_ISL_629225, EPI_ISL_629226, EPI_ISL_629227, EPI_ISL_629228, EPI_ISL_629229, EPI_ISL_629230, EPI_ISL_629231, EPI_ISL_629234, EPI_ISL_629235, EPI_ISL_629236, EPI_ISL_629237, EPI_ISL_629238, EPI_ISL_629239, EPI_ISL_629240, EPI_ISL_629241, EPI_ISL_629242, EPI_ISL_629244, EPI_ISL_629248, EPI_ISL_629250, EPI_ISL_629252, EPI_ISL_629253, EPI_ISL_629255, EPI_ISL_629256, EPI_ISL_629257, EPI_ISL_629258, EPI_ISL_629259, EPI_ISL_629260, EPI_ISL_629262, EPI_ISL_629263, EPI_ISL_629264, EPI_ISL_629267, EPI_ISL_629268, EPI_ISL_629269, EPI_ISL_629272, EPI_ISL_629273, EPI_ISL_629275, EPI_ISL_629280, EPI_ISL_629283, EPI_ISL_629284, EPI_ISL_629285, EPI_ISL_629290, EPI_ISL_629294, EPI_ISL_629296, EPI_ISL_629297, EPI_ISL_629298, EPI_ISL_629299, EPI_ISL_629300, EPI_ISL_629301, EPI_ISL_629302, EPI_ISL_629303, EPI_ISL_629306, EPI_ISL_629307, EPI_ISL_629310, EPI_ISL_629312, EPI_ISL_629313, EPI_ISL_629314, EPI_ISL_629315, EPI_ISL_629316, EPI_ISL_629317, EPI_ISL_629318, EPI_ISL_629319, EPI_ISL_629320, EPI_ISL_629321, EPI_ISL_629322, EPI_ISL_629323, EPI_ISL_629324, EPI_ISL_629326, EPI_ISL_629327, EPI_ISL_629329, EPI_ISL_629331, EPI_ISL_629332, EPI_ISL_629333, EPI_ISL_629334, EPI_ISL_629336, EPI_ISL_629337, EPI_ISL_629338, EPI_ISL_629339, EPI_ISL_629341, EPI_ISL_629344, EPI_ISL_629346, EPI_ISL_629347, EPI_ISL_629349, EPI_ISL_629350, EPI_ISL_629351, EPI_ISL_629352, EPI_ISL_629353, EPI_ISL_629355, EPI_ISL_629356, EPI_ISL_629358, EPI_ISL_629362, EPI_ISL_629363, EPI_ISL_629364, EPI_ISL_629365, EPI_ISL_629367, EPI_ISL_629368, EPI_ISL_629371, EPI_ISL_629372, EPI_ISL_629373, EPI_ISL_629374, EPI_ISL_629375, EPI_ISL_629376, EPI_ISL_629377, EPI_ISL_629378, EPI_ISL_629381, EPI_ISL_629382, EPI_ISL_629384, EPI_ISL_629385, EPI_ISL_629386, EPI_ISL_629387, EPI_ISL_629388, EPI_ISL_629393, EPI_ISL_629394, EPI_ISL_629395, EPI_ISL_629396, EPI_ISL_629397, EPI_ISL_629399, EPI_ISL_629400, EPI_ISL_629401, EPI_ISL_629402, EPI_ISL_629403, EPI_ISL_629404, EPI_ISL_629405, EPI_ISL_629406, EPI_ISL_629407, EPI_ISL_629408, EPI_ISL_629410, EPI_ISL_629412, EPI_ISL_629415, EPI_ISL_629416, EPI_ISL_629417, EPI_ISL_629418, EPI_ISL_629419, EPI_ISL_629420, EPI_ISL_629421, EPI_ISL_629422, EPI_ISL_629423, EPI_ISL_629424, EPI_ISL_629425, EPI_ISL_629426, EPI_ISL_629427, EPI_ISL_629428, EPI_ISL_629429, EPI_ISL_629430, EPI_ISL_629434, EPI_ISL_629435, EPI_ISL_629437, EPI_ISL_629438, EPI_ISL_629440, EPI_ISL_629444, EPI_ISL_629447, EPI_ISL_629450, EPI_ISL_629451, EPI_ISL_629457, EPI_ISL_629458, EPI_ISL_629462, EPI_ISL_629469, EPI_ISL_629472, EPI_ISL_629473, EPI_ISL_629475, EPI_ISL_629476, EPI_ISL_629477, EPI_ISL_629481, EPI_ISL_629483, EPI_ISL_629486, EPI_ISL_629488, EPI_ISL_629489, EPI_ISL_629493, EPI_ISL_629494, EPI_ISL_629496, EPI_ISL_629497, EPI_ISL_629498, EPI_ISL_629500, EPI_ISL_629502, EPI_ISL_629503, EPI_ISL_629508, EPI_ISL_629509, EPI_ISL_629511, EPI_ISL_629513, EPI_ISL_629515, EPI_ISL_629518, EPI_ISL_629520, EPI_ISL_629521, EPI_ISL_629525, EPI_ISL_629529, EPI_ISL_629530, EPI_ISL_629532, EPI_ISL_629533, EPI_ISL_629538, EPI_ISL_629542, EPI_ISL_629543, EPI_ISL_629546, EPI_ISL_629550, EPI_ISL_629551, EPI_ISL_629552, EPI_ISL_629553, EPI_ISL_629556, EPI_ISL_629557, EPI_ISL_629561, EPI_ISL_629563, EPI_ISL_629570, EPI_ISL_629572, EPI_ISL_629574, EPI_ISL_629577, EPI_ISL_629578, EPI_ISL_629579, EPI_ISL_629582, EPI_ISL_629586, EPI_ISL_629588, EPI_ISL_629589, EPI_ISL_629594, EPI_ISL_629596, EPI_ISL_629598, EPI_ISL_629600, EPI_ISL_629602, EPI_ISL_629606, EPI_ISL_629609, EPI_ISL_629610, EPI_ISL_629613, EPI_ISL_629616, EPI_ISL_629618, EPI_ISL_629619, EPI_ISL_629621, EPI_ISL_629622, EPI_ISL_629623, EPI_ISL_629624, EPI_ISL_629626, EPI_ISL_629627, EPI_ISL_629633, EPI_ISL_629636, EPI_ISL_629637, EPI_ISL_629638, EPI_ISL_629639, EPI_ISL_629643, EPI_ISL_629646, EPI_ISL_629649, EPI_ISL_629652, EPI_ISL_629654, EPI_ISL_629655, EPI_ISL_629656, EPI_ISL_629657, EPI_ISL_629658, EPI_ISL_629662, EPI_ISL_629664, EPI_ISL_629666, EPI_ISL_629667, EPI_ISL_629670, EPI_ISL_629677, EPI_ISL_629678, EPI_ISL_629680, EPI_ISL_629681, EPI_ISL_629682, EPI_ISL_629684, EPI_ISL_629686, EPI_ISL_629687, EPI_ISL_629691, EPI_ISL_629693, EPI_ISL_629694, EPI_ISL_629695, EPI_ISL_629697, EPI_ISL_629698, EPI_ISL_629699, EPI_ISL_629700, EPI_ISL_629707, EPI_ISL_629711, EPI_ISL_629712, EPI_ISL_629714, EPI_ISL_629715, EPI_ISL_629718, EPI_ISL_629719, EPI_ISL_629720, EPI_ISL_629726, EPI_ISL_629732, EPI_ISL_629734, EPI_ISL_629736, EPI_ISL_629741, EPI_ISL_629742, EPI_ISL_629744, EPI_ISL_629748, EPI_ISL_629751, EPI_ISL_629753, EPI_ISL_630007, EPI_ISL_630008, EPI_ISL_630009, EPI_ISL_630011, EPI_ISL_630013, EPI_ISL_630016, EPI_ISL_630017, EPI_ISL_630018, EPI_ISL_630019, EPI_ISL_630020, EPI_ISL_630021, EPI_ISL_630022, EPI_ISL_630024, EPI_ISL_630025, EPI_ISL_630026, EPI_ISL_630027, EPI_ISL_630028, EPI_ISL_630029, EPI_ISL_630030, EPI_ISL_630031, EPI_ISL_630033, EPI_ISL_630036, EPI_ISL_630037, EPI_ISL_630038, EPI_ISL_630039, EPI_ISL_630040, EPI_ISL_630041, EPI_ISL_630042, EPI_ISL_630043, EPI_ISL_630044, EPI_ISL_630045, EPI_ISL_630046, EPI_ISL_630047, EPI_ISL_630048, EPI_ISL_630049, EPI_ISL_630050, EPI_ISL_630052, EPI_ISL_630054, EPI_ISL_630056, EPI_ISL_630057, EPI_ISL_630059, EPI_ISL_630060, EPI_ISL_630061, EPI_ISL_630062, EPI_ISL_630064, EPI_ISL_630069, EPI_ISL_630071, EPI_ISL_630072, EPI_ISL_630073, EPI_ISL_630074, EPI_ISL_630075, EPI_ISL_630077, EPI_ISL_630079, EPI_ISL_630081, EPI_ISL_630083, EPI_ISL_630084, EPI_ISL_630085, EPI_ISL_630086, EPI_ISL_630088, EPI_ISL_630090, EPI_ISL_630091, EPI_ISL_630093, EPI_ISL_630096, EPI_ISL_630097, EPI_ISL_630100, EPI_ISL_630101, EPI_ISL_630102, EPI_ISL_630104, EPI_ISL_630106, EPI_ISL_630107, EPI_ISL_630108, EPI_ISL_630109, EPI_ISL_630110, EPI_ISL_630112, EPI_ISL_630113, EPI_ISL_630116, EPI_ISL_630117, EPI_ISL_630118, EPI_ISL_630119, EPI_ISL_630120, EPI_ISL_630121, EPI_ISL_630122, EPI_ISL_630123, EPI_ISL_630124, EPI_ISL_630125, EPI_ISL_630126, EPI_ISL_630129, EPI_ISL_630130, EPI_ISL_630131, EPI_ISL_630133, EPI_ISL_630134, EPI_ISL_630136, EPI_ISL_630138, EPI_ISL_630139, EPI_ISL_630141, EPI_ISL_630144, EPI_ISL_630146, EPI_ISL_630148, EPI_ISL_630149, EPI_ISL_630150, EPI_ISL_630152, EPI_ISL_630153, EPI_ISL_630154, EPI_ISL_630156, EPI_ISL_630158, EPI_ISL_630159, EPI_ISL_630160, EPI_ISL_630161, EPI_ISL_630163, EPI_ISL_630165, EPI_ISL_630166, EPI_ISL_630167, EPI_ISL_630168, EPI_ISL_630169, EPI_ISL_630172, EPI_ISL_630173, EPI_ISL_630174, EPI_ISL_630175, EPI_ISL_630176, EPI_ISL_630178, EPI_ISL_630180, EPI_ISL_630181, EPI_ISL_630182, EPI_ISL_630183, EPI_ISL_630185, EPI_ISL_630186, EPI_ISL_630187, EPI_ISL_630188, EPI_ISL_630189, EPI_ISL_630192, EPI_ISL_630193, EPI_ISL_630194, EPI_ISL_630196, EPI_ISL_630197, EPI_ISL_630198, EPI_ISL_630199, EPI_ISL_630202, EPI_ISL_630204, EPI_ISL_630205, EPI_ISL_630206, EPI_ISL_630207, EPI_ISL_630208, EPI_ISL_630209, EPI_ISL_630210, EPI_ISL_630212, EPI_ISL_630215, EPI_ISL_630216, EPI_ISL_630217, EPI_ISL_630220, EPI_ISL_630221, EPI_ISL_630222, EPI_ISL_630223, EPI_ISL_630224, EPI_ISL_630225, EPI_ISL_630226, EPI_ISL_630229, EPI_ISL_630230, EPI_ISL_630231, EPI_ISL_630232, EPI_ISL_630234, EPI_ISL_630235, EPI_ISL_630236, EPI_ISL_630237, EPI_ISL_630238, EPI_ISL_630241, EPI_ISL_630242, EPI_ISL_630243, EPI_ISL_630245, EPI_ISL_630246, EPI_ISL_630247, EPI_ISL_630248, EPI_ISL_630250, EPI_ISL_630253, EPI_ISL_630254, EPI_ISL_630256, EPI_ISL_630257, EPI_ISL_630258, EPI_ISL_630260, EPI_ISL_630261, EPI_ISL_630262, EPI_ISL_630264, EPI_ISL_630265, EPI_ISL_630267, EPI_ISL_630270, EPI_ISL_630271, EPI_ISL_630272, EPI_ISL_630273, EPI_ISL_630275, EPI_ISL_630277, EPI_ISL_630278, EPI_ISL_630279, EPI_ISL_630281, EPI_ISL_630282, EPI_ISL_630283, EPI_ISL_630285, EPI_ISL_630288, EPI_ISL_630289, EPI_ISL_630290, EPI_ISL_630291, EPI_ISL_630292, EPI_ISL_630293, EPI_ISL_630295, EPI_ISL_630296,

EPI_ISL_630297, EPI_ISL_630298, EPI_ISL_630300, EPI_ISL_630301, EPI_ISL_630302, EPI_ISL_630303, EPI_ISL_630304, EPI_ISL_630307, EPI_ISL_630308, EPI_ISL_630309, EPI_ISL_630311, EPI_ISL_630312, EPI_ISL_630314, EPI_ISL_630316, EPI_ISL_630318, EPI_ISL_630319, EPI_ISL_630321, EPI_ISL_630322, EPI_ISL_630323, EPI_ISL_630324, EPI_ISL_630325, EPI_ISL_630327, EPI_ISL_630328, EPI_ISL_630329, EPI_ISL_630330, EPI_ISL_630332, EPI_ISL_630333, EPI_ISL_630334, EPI_ISL_630335, EPI_ISL_630336, EPI_ISL_630337, EPI_ISL_630338, EPI_ISL_630339, EPI_ISL_630340, EPI_ISL_630341, EPI_ISL_630342, EPI_ISL_630343, EPI_ISL_630345, EPI_ISL_630346, EPI_ISL_630347, EPI_ISL_630974, EPI_ISL_631043, EPI_ISL_631054, EPI_ISL_631072, EPI_ISL_631103, EPI_ISL_631121, EPI_ISL_631167, EPI_ISL_631207

| | | | |
|---|---|---|---|
| see above | Lighthouse Lab in Milton Keynes | Wellcome Sanger Institute for the COVID-19 Genomics UK (COG-UK) consortium | The Lighthouse Lab in Milton Keynes and Alex Alderton, Roberto Amato, Sonia Goncalves, Ewan Harrison, David K. Jackson, Ian Johnston, Dominic Kwiatkowski, Cordelia Langford, John Sillitoe on behalf of the Wellcome Sanger Institute COVID-19 Surveillance Team |

We gratefully acknowledge the following Authors from the Originating laboratories responsible for obtaining the specimens, as well as the Submitting laboratories where the genome data were generated and shared via GISAID, on which this research is based.

All Submitters of data may be contacted directly via www.gisaid.org

Authors are sorted alphabetically.

| Accession ID | Originating Laboratory | Submitting Laboratory | Authors |
|---|---|---|---|
| EPI_ISL_402119 | National Institute for Viral Disease Control and Prevention, China CDC | National Institute for Viral Disease Control and Prevention, China CDC | Wenjie TanXiang ZhaoWenling WangXuejun MaYongzhong JiangRoujian Lu, Ji Wang, Weimin ZhouPeihua NiuPeipei LiuFaxian ZhanWeifeng ShiBaoying HuangJun LiuLi ZhaoYao MengXiaozhou HeFei YeNa ZhuYang LiJing ChenWenbo XuGeorge F. GaoGuizhen Wu |
| EPI_ISL_402120 | National Institute for Viral Disease Control and Prevention, China CDC | National Institute for Viral Disease Control and Prevention, China CDC | Wenjie TanXiang ZhaoWenling WangXuejun MaYongzhong JiangRoujian LuJi WangWeimin ZhouPeihua NiuPeipei LiuFaxian ZhanWeifeng ShiBaoying HuangJun LiuLi ZhaoYao MengXiaozhou HeFei YeNa ZhuYang LiJing ChenWenbo XuGeorge F. GaoGuizhen Wu |
| EPI_ISL_402121 | National Institute for Viral Disease Control and Prevention, China CDC | National Institute for Viral Disease Control and Prevention, China CDC | Wenjie TanXuejun MaXiang ZhaoWenling WangYongzhong JiangRoujian LuJi WangPeihua Niu, Weimin Zhou, Faxian ZhanWeifeng ShiBaoying HuangJun LiuLi ZhaoYao MengFei YeNa Zhu, Xiaozhou HePeipei Liu, Yang LiJing ChenWenbo XuGeorge F. GaoGuizhen Wu |
| EPI_ISL_406798, EPI_ISL_406800, EPI_ISL_406801 | General Hospital of Central Theater Command of People's Liberation Army of China | BGI & Institute of Microbiology, Chinese Academy of Sciences & Shandong First Medical University & Shandong Academy of Medical Sciences & General Hospital of Central Theater Command of People's Liberation Army of China | Weijun Chen, Yuhai Bi, Weifeng Shi and Zhenhong Hu |
| EPI_ISL_406862 | Charité Universitätsmedizin Berlin, Institute of Virology; Institut für Mikrobiologie der Bundeswehr, Munich | Charité Universitätsmedizin Berlin, Institute of Virology | Victor M Corman, Julia Schneider, Talitha Veith, Barbara Mühlemann, Markus Antwerpen, Christian Drosten, Roman Wölfel |
| EPI_ISL_407084 | Department of Virology III, National Institute of Infectious Diseases | Pathogen Genomics Center, National Institute of Infectious Diseases | Tsuyoshi Sekizuka, Shutoku Matsuyama, Naganori Nao, Kazuya Shirato, Shinji Watanabe, Makoto Takeda, Makoto Kuroda |
| EPI_ISL_408480 | National Institute for Viral Disease Control and Prevention, China CDC | National Institute for Viral Disease Control & Prevention, CCDC | Wenjie TanXiaoqing FuXiang ZhaoWenling Wang Peihua NiuRoujian Lu,Yanhong SunBaoying HuangLi ZhaoFei YeWenbo XuGeorge F. GaoGuizhen Wu |
| EPI_ISL_408481 | National Institute for Viral Disease Control and Prevention, China CDC | National Institute for Viral Disease Control & Prevention, CCDC | Wenjie Tan, Hengqin Wang, Xiang Zhao, Wenling Wang, Peihua Niu, Roujian Lu, Sheng Ye, Baoying Huang, Li Zhao, Fei Ye, Wenbo Xu, George F. Gao, Guizhen Wu |
| EPI_ISL_408482 | National Institute for Viral Disease Control and Prevention, China CDC | National Institute for Viral Disease Control & Prevention, CCDC | Wenjie Tan, Zhaoguo Wang, Xiang Zhao, Wenling Wang, Peihua Niu, Roujian Lu, Ti Liu, Baoying Huang, Li Zhao, Fei Ye, Wenbo Xu, George F. Gao, Guizhen Wu |
| EPI_ISL_408483 | National Institute for Viral Disease Control and Prevention, China CDC | National Institute for Viral Disease Control & Prevention, CCDC | Wenjie TanZhen Teng,Xiang ZhaoWenling Wang Peihua NiuRoujian Lu,Chongshan Li,Baoying HuangLi ZhaoFei YeWenbo XuGeorge F. GaoGuizhen Wu |
| EPI_ISL_408484 | National Institute for Viral Disease Control and Prevention, China CDC | National Institute for Viral Disease Control & Prevention, CCDC | Wenjie Tan, Jianan Xu, Wenling Wang, Peihua Niu, Roujian Lu, Huiping Yang, Xiang Zhao, Baoying Huang, Li Zhao, Fei Ye, Wenbo Xu, George F. Gao, Guizhen Wu |
| EPI_ISL_408485 | National Institute for Viral Disease Control and Prevention, China CDC | National Institute for Viral Disease Control & Prevention, CCDC | Wenjie Tan,Quanyi Wang,Wenling Wang, Peihua Niu,Roujian Lu,Yang Pan,Xiang Zhao,Baoying Huang,Li Zhao,Fei Ye,Wenbo Xu,George F. Gao,Guizhen Wu |
| EPI_ISL_408486 | National Institute for Viral Disease Control and Prevention, China CDC | National Institute for Viral Disease Control & Prevention, CCDC | Wenjie Tan, Yong Shi, Wenling Wang, Peihua Niu, Roujian Lu, Jianxiong Li, Xiang Zhao, Baoying Huang, Li Zhao, Fei Ye, Wenbo Xu, George F. Gao, Guizhen Wu |
| EPI_ISL_408487 | National Institute for Viral Disease Control and Prevention, China CDC | National Institute for Viral Disease Control & Prevention, China CDC | Wenjie Tan, Jin Xu, Wenling Wang, Peihua Niu, Roujian Lu, Xueyong Huang, Xiang Zhao, Baoying Huang, Li Zhao, Fei Ye, Wenbo Xu, George F. Gao, Guizhen Wu |
| EPI_ISL_408488 | National Institute for Viral Disease Control and Prevention, China CDC | National Institute for Viral Disease Control & Prevention, CCDC | Wenjie Tan, Shenjiao Wang, Wenling Wang, Peihua Niu, Roujian Lu, Kangchen Zhao, Xiang Zhao, Baoying Huang, Li Zhao, Fei Ye, Wenbo Xu, George F. Gao, Guizhen Wu |
| EPI_ISL_408665, EPI_ISL_408666, EPI_ISL_408667 | Dept. of Virology III, National Institute of Infectious Diseases | Pathogen Genomics Center, National Institute of Infectious Diseases | Tsuyoshi Sekizuka, Shutoku Matsuyama, Naganori Nao, Kazuya Shirato, Makoto Takeda, Makoto Kuroda |
| EPI_ISL_408668 | National Influenza Center - National Institute of Hygiene and Epidemiology (NIHE) | National Influenza Center - National Institute of Hygiene and Epidemiology (NIHE) | Ung Thi Hong Trang, Hoang Vu Mai Phuong, Nguyen Le Khanh Hang, Nguyen Vu Son, Le Thi Thanh, Vuong Duc Cuong, Nguyen Phuong Anh, Pham Thi Hien, Tran Thu Huong, Le Thi Quynh Mai, |
| EPI_ISL_408669 | Dept. of Virology III, National Institute of Infectious Diseases | Pathogen Genomics Center, National Institute of Infectious Diseases | Tsuyoshi Sekizuka, Shutoku Matsuyama, Naganori Nao, Kazuya Shirato, Makoto Takeda, Makoto Kuroda |
| EPI_ISL_408978 | Wuhan Fourth Hospital | Beijing Genomics Institute (BGI) | Weijun Chen |
| EPI_ISL_410531, EPI_ISL_410532 | Dept. of Pathology, National Institute of Infectious Diseases | Pathogen Genomics Center, National Institute of Infectious Diseases | Tsuyoshi Sekizuka, Harutaka Katano, Shutoku Matsuyama, Naganori Nao, Kazuya Shirato, Motoi Suzuki, Hideki Hasegawa, Takaji Wakita, Makoto Takeda, Tadaki Suzuki, Makoto Kuroda |
| EPI_ISL_412912 | State Health Office Baden-Württemberg | Charite Universitatsmedizin Berlin, Institute of Virology | Victor M Corman, Julia Schneider, Barbara Muhlemann, Talitha Veith, Jörn Beheim-Schwarzbach, Terry Jones, Rainer Oehme, Silke Fischer, Christian Drosten |
| EPI_ISL_413459 | Department of Pathology, Toshima Hospital | Pathogen Genomics Center, National Institute of Infectious Diseases | Tsuyoshi Sekizuka, Kentaro Itokawa, Takuya Adachi, Masahiro Sano, Jun Yamazaki, Ippei Miyamoto, Haruka Nishioka, Ja-Mun Chong, Noriko Nakajima, Yuko Sato, Minoru Tobiume, Harutaka Katano, Tadaki Suzuki, Makoto Kuroda |
| EPI_ISL_413485 | Department of microbiology laboratory,Anhui Provincial Center for Disease Control and Prevention | Department of microbiology laboratory,Anhui Provincial Center for Disease Control and Prevention | Weiwei Li,Jun He,Yong Sun,Junling Yu,Qingqing Chen,Yuan Yuan,Yonglin Shi,Zhuhui Zhang,Yinglu Ge,Weidong Li,Bin Su,Zhirong Liu |
| EPI_ISL_413557, EPI_ISL_413558, EPI_ISL_413559, EPI_ISL_413561 | California Department of Public Health | Chiu Laboratory, University of California, San Francisco | Xianding Deng, Scot Federman, Chao-Yang Pan, Hugo Guevara,Wei Gu, Debra A. Wadford, and Charles Y. Chiu |
| EPI_ISL_413599 | Centre for Infectious Diseases and Microbiology - Public Health | NSW Health Pathology - Institute of Clinical Pathology and Medical Research; Westmead Hospital; University of Sydney | Timms, V, Eden J-S, Lam C, Gray K, Rockett R, Gall, M, Carter I, Rahman H, Holmes EC, O'Sullivan MV, Sintchenko V, Chen SC, Maddocks S, Kok J and Dwyer DE for the 2019-nCoV Study Group* |
| EPI_ISL_413922, EPI_ISL_413924, EPI_ISL_413925, EPI_ISL_413926, EPI_ISL_413928, EPI_ISL_413931 | California Department of Public Health | Chiu Laboratory, University of California, San Francisco | Xianding Deng, Scot Federman, Chao-Yang Pan, Hugo Guevara,Wei Gu, Debra A. Wadford, and Charles Y. Chiu |
| EPI_ISL_414500, EPI_ISL_414501 | Virology Department, Sheffield Teaching Hospitals NHS Foundation Trust | Department of Infection, Immunity and Cardiovascular Disease, The Florey Institute, The Medical School, University of Sheffield | Thushan de Silva, Matthew Parker, Matthew Wyles, Mehmet Yavuz, Mohammad Raza, Cariad Evans |
| EPI_ISL_414934, EPI_ISL_414935, EPI_ISL_414936, EPI_ISL_414937, EPI_ISL_414938, EPI_ISL_414939, EPI_ISL_414940, EPI_ISL_414941 | Shandong Provincial Center for Disease Control and Prevention | Beijing Institute of Microbiology and Epidemiology | Xiao-Lin Jiang, Xiao-Li Zhang, Xiang-Na Zhao, Cun-Bao Li, Jie Lei, Zeng-Qiang Kou, Wen-Kui Sun, Yang Hang, Feng Gao, Sheng-Xiang Ji, Can-Fang Lin, Bo Pang, Ming-Xiao Yao, Guo-Lin Wang, Lin Yao, Li-Jun Duan, Xiao Wei, Dian-Ming Kang, Mai-Juan Ma |

| | | | |
|---|---|---|---|
| EPI_ISL_416538 | Wellington Hospital | Institute of Environmental Science and Research (ESR) | Wellington SCL, Wellington Hospital, Riddiford Street, Newtown, Wellington 6021, New Zealand |
| EPI_ISL_416565, EPI_ISL_416566, EPI_ISL_416567, EPI_ISL_416568, EPI_ISL_416569, EPI_ISL_416570, EPI_ISL_416571, EPI_ISL_416572, EPI_ISL_416573, EPI_ISL_416574, EPI_ISL_416575, EPI_ISL_416576, EPI_ISL_416577, EPI_ISL_416578, EPI_ISL_416579, EPI_ISL_416580, EPI_ISL_416581, EPI_ISL_416582, EPI_ISL_416583, EPI_ISL_416584, EPI_ISL_416585, EPI_ISL_416586, EPI_ISL_416587, EPI_ISL_416588, EPI_ISL_416589, EPI_ISL_416590, EPI_ISL_416591, EPI_ISL_416592, EPI_ISL_416593, EPI_ISL_416594, EPI_ISL_416595, EPI_ISL_416596, EPI_ISL_416597, EPI_ISL_416598, EPI_ISL_416599, EPI_ISL_416600, EPI_ISL_416601, EPI_ISL_416602, EPI_ISL_416603, EPI_ISL_416604, EPI_ISL_416605, EPI_ISL_416606, EPI_ISL_416607, EPI_ISL_416608, EPI_ISL_416609, EPI_ISL_416610, EPI_ISL_416611, EPI_ISL_416612, EPI_ISL_416613, EPI_ISL_416614, EPI_ISL_416615, EPI_ISL_416616, EPI_ISL_416617, EPI_ISL_416618, EPI_ISL_416619, EPI_ISL_416620, EPI_ISL_416621, EPI_ISL_416622, EPI_ISL_416623, EPI_ISL_416624, EPI_ISL_416625, EPI_ISL_416626, EPI_ISL_416627, EPI_ISL_416628, EPI_ISL_416629, EPI_ISL_416630, EPI_ISL_416631, EPI_ISL_416632, EPI_ISL_416633, EPI_ISL_416634 | | | |
| see above | Japanese Quarantine Stations | Pathogen Genomics Center, National Institute of Infectious Diseases | Tsuyoshi Sekizuka, Kentaro Itokawa, Rina Tanaka, Masanori Hashino, Tsutomu Kageyama, Shinji Saito, Ikuyo Takayama, Hideki Hasegawa, Takuri Takahashi, Hajime Kamiya, Takuya Yamagishi, Motoi Suzuki, Takaji Wakita, Makoto Kuroda |
| EPI_ISL_416730, EPI_ISL_416731, EPI_ISL_416732, EPI_ISL_416733, EPI_ISL_416734, EPI_ISL_416735, EPI_ISL_416736, EPI_ISL_416737, EPI_ISL_416738, EPI_ISL_416739, EPI_ISL_416740 | | | |
| see above | Virology Department, Sheffield Teaching Hospitals NHS Foundation Trust | Department of Infection, Immunity and Cardiovascular Disease, The Florey Institute, The Medical School, University of Sheffield | Thushan de Silva, Matthew Parker, Adri Angyal, Rebecca Brown, Matthew Wyles, Mehmet Yavuz, Mohammad Raza, Cariad Evans |
| EPI_ISL_416742, EPI_ISL_416743 | NRL for Influenza, Centrum Epidemiology and Microbiology of National Institute of Public Health, Czech Republic | Charite Universitaetsmedizin Berlin, Institute of Virology | Victor M Corman, Julia Schneider, Jorn Beheim-Schwarzbach, Talitha Veith, Barbara Muehlemann, Terry Jones, Akexander Nagy, Jaromira Vecerova, Dusan Trnka, Ludmila Novakova, Helena Jirincova, Christian Drosten |
| EPI_ISL_417034 | Laboratorio de Ecologia de Doencas Transmissiveis na Amazonia, Instituto Leonidas e Maria Deane - Fiocruz Amazonia | Laboratorio de Ecologia de Doencas Transmissiveis na Amazonia, Instituto Leonidas e Maria Deane - Fiocruz Amazonia | Valdinete Nascimento, André Corado, Fernanda Nascimento, Ágatha Costa, Debora Duarte, Luciana Gonçalves, Michele Jesus, Sérgio Luz, Felipe Naveca |
| EPI_ISL_417316 | Chiu Laboratory, University of California, San Francisco | Chiu Laboratory, University of California, San Francisco | Xianding Deng, Scot Federman, Wei Gu, and Charles Y. Chiu |
| EPI_ISL_417317, EPI_ISL_417318, EPI_ISL_417319, EPI_ISL_417320 | Santa Clara County Public Health Department | Chiu Laboratory, University of California, San Francisco | Xianding Deng, Scot Federman, Wei Gu, Elsa Villarino, Brandon Bonin, Debra A. Wadford, and Charles Y. Chiu |
| EPI_ISL_417323, EPI_ISL_417324, EPI_ISL_417325, EPI_ISL_417327 | California Department of Public Health | Chiu Laboratory, University of California, San Francisco | Xianding Deng, Scot Federman, Chao-Yang Pan, Hugo Guevara,Wei Gu, Debra A. Wadford, and Charles Y. Chiu |
| EPI_ISL_417330, EPI_ISL_417332 | Chiu Laboratory, University of California, San Francisco | Chiu Laboratory, University of California, San Francisco | Xianding Deng, Scot Federman, Wei Gu, and Charles Y. Chiu |
| EPI_ISL_417387, EPI_ISL_417404 | Centre for Infectious Diseases and Microbiology Public Health | NSW Health Pathology - Institute of Clinical Pathology and Medical Research; Westmead Hospital; University of Sydney | Timms V, Gall M, Arnott A, Sadsad R, Carter I, Rahman H, Holmes EC, O'Sullivan MV, Sintchenko V, Chen SC, Maddocks S, Kok J, Dwyer DE, Rockett R, Eden J-S, Lam C and Gray K for the 2019-nCoV Study Group |
| EPI_ISL_418270 | KU Leuven, Clinical and Epidemiological Virology | KU Leuven, Clinical and Epidemiological Virology | Tony Wawina, Joan Marti-Carreras, Bert Vanmechelen, Piet Maes |
| EPI_ISL_418286, EPI_ISL_418287, EPI_ISL_418288, EPI_ISL_418289, EPI_ISL_418290, EPI_ISL_418291, EPI_ISL_418292, EPI_ISL_418293, EPI_ISL_418294, EPI_ISL_418295, EPI_ISL_418296, EPI_ISL_418297, EPI_ISL_418298, EPI_ISL_418299, EPI_ISL_418300, EPI_ISL_418301, EPI_ISL_418302, EPI_ISL_418303, EPI_ISL_418304, EPI_ISL_418305, EPI_ISL_418306, EPI_ISL_418307, EPI_ISL_418308, EPI_ISL_418309, EPI_ISL_418310, EPI_ISL_418311, EPI_ISL_418312, EPI_ISL_418313, EPI_ISL_418314, EPI_ISL_418315, EPI_ISL_418316, EPI_ISL_418317, EPI_ISL_418318, EPI_ISL_418319, EPI_ISL_418320, EPI_ISL_418321 | | | |
| see above | Virology Department, Sheffield Teaching Hospitals NHS Foundation Trust | Department of Infection, Immunity and Cardiovascular Disease, The Florey Institute, The Medical School, University of Sheffield | Thushan de Silva, Matthew Parker, Adri Angyal, Rebecca Brown, Rachel Tucker, Paul Parsons, Danielle Groves, Alex Keeley, Dave Partridge, Matthew Wyles, Benjamin Lindsey, Mehmet Yavuz, Mohammad Raza, Cariad Evans |
| EPI_ISL_418864, EPI_ISL_418956, EPI_ISL_418957, EPI_ISL_418958, EPI_ISL_419256, EPI_ISL_419257, EPI_ISL_419258, EPI_ISL_419260, EPI_ISL_419261, EPI_ISL_419262, EPI_ISL_419263 | | | |
| see above | Virginia DCLS | Virginia DCLS | Virginia DCLS |
| EPI_ISL_419296 | Kochi Prefectural Institute of Public Health | Pathogen Genomics Center, National Institute of Infectious Diseases | Tsuyoshi Sekizuka, Akihiko Tokaji, Kentaro Itokawa, Rina Tanaka, Masanori Hashino, Hajime Kamiya, Motoi Suzuki, Makoto Kuroda |
| EPI_ISL_419297, EPI_ISL_419298 | Chiba Prefectural Institute of Public Health | Pathogen Genomics Center, National Institute of Infectious Diseases | Tsuyoshi Sekizuka, Masakatsu Taira, Yushi Hachisu, Kentaro Itokawa, Rina Tanaka, Masanori Hashino, Hajime Kamiya, Motoi Suzuki, Makoto Kuroda |
| EPI_ISL_419299, EPI_ISL_419300 | Ishikawa Prefectural Institute of Public Health and Environmental Science | Pathogen Genomics Center, National Institute of Infectious Diseases | Tsuyoshi Sekizuka, Sanae Kuramoto, Eri Nariai, Kentaro Itokawa, Rina Tanaka, Masanori Hashino, Hajime Kamiya, Motoi Suzuki, Makoto Kuroda |
| EPI_ISL_419301, EPI_ISL_419302, EPI_ISL_419303, EPI_ISL_419304, EPI_ISL_419305, EPI_ISL_419306, EPI_ISL_419307, EPI_ISL_419308 | Saitama Prefectural Institute of Public Health | Pathogen Genomics Center, National Institute of Infectious Diseases | Tsuyoshi Sekizuka, Michiyo Shinohara, Tsuyoshi Kishimoto, Kentaro Itokawa, Rina Tanaka, Masanori Hashino, Hajime Kamiya, Motoi Suzuki, Makoto Kuroda |
| EPI_ISL_419309, EPI_ISL_419310, EPI_ISL_419311 | Chiba Prefectural Institute of Public Health | Pathogen Genomics Center, National Institute of Infectious Diseases | Tsuyoshi Sekizuka, Masakatsu Taira, Yushi Hachisu, Kentaro Itokawa, Rina Tanaka, Masanori Hashino, Hajime Kamiya, Motoi Suzuki, Makoto Kuroda |
| EPI_ISL_419706, EPI_ISL_419708, EPI_ISL_419710, EPI_ISL_419711, EPI_ISL_419712, EPI_ISL_419713, EPI_ISL_420018, EPI_ISL_420019, EPI_ISL_420020, EPI_ISL_420021, EPI_ISL_420022, EPI_ISL_420023, EPI_ISL_420024, EPI_ISL_420025, EPI_ISL_420026, EPI_ISL_420027, EPI_ISL_420028, EPI_ISL_420029 | | | |
| see above | Virginia DCLS | Virginia DCLS | Virginia DCLS |
| EPI_ISL_420121 | Servicio de Microbiología. Consorcio Hospital General Universitario de Valencia | Sequencing and Bioinformatics Service and Molecular Epidemiology Research Group. FISABIO-Public Health | Vicente Soriano Chirona, Maria Alma Bracho, Griselda De Marco, Beatriz Beamud, Lidia Ruiz Roldan, Marta Pla Diaz,Neris Garcia-Gonzalez, Loreto Ferrús Abad, Inma Galán Vendrell, Paula Ruiz-Hueso, Mariana Reyes-Prieto, Maria Dolores Ocete, Llúcia Martínez-Priego, Concepcion Gimeno, Giuseppe D'Auria, Fernando Gonzalez-Candelas |
| EPI_ISL_420154, EPI_ISL_420155, EPI_ISL_420156, EPI_ISL_420157, EPI_ISL_420158, EPI_ISL_420159, EPI_ISL_420160, EPI_ISL_420161, EPI_ISL_420162, EPI_ISL_420163, EPI_ISL_420164, EPI_ISL_420165, EPI_ISL_420166, EPI_ISL_420167, EPI_ISL_420168, EPI_ISL_420169, EPI_ISL_420170, EPI_ISL_420171, EPI_ISL_420172, EPI_ISL_420173, EPI_ISL_420174, EPI_ISL_420175, EPI_ISL_420176, EPI_ISL_420177, EPI_ISL_420178, EPI_ISL_420179, EPI_ISL_420180, EPI_ISL_420181, EPI_ISL_420182, EPI_ISL_420183, EPI_ISL_420184, EPI_ISL_420185, EPI_ISL_420186, EPI_ISL_420187, EPI_ISL_420188, EPI_ISL_420189, EPI_ISL_420190, EPI_ISL_420191, EPI_ISL_420192, EPI_ISL_420193, EPI_ISL_420194, EPI_ISL_420195, EPI_ISL_420196, EPI_ISL_420197, EPI_ISL_420198, EPI_ISL_420199, EPI_ISL_420200, EPI_ISL_420201, EPI_ISL_420202, EPI_ISL_420203, EPI_ISL_420204, EPI_ISL_420205, EPI_ISL_420206, EPI_ISL_420207, EPI_ISL_420208, EPI_ISL_420209, EPI_ISL_420210, EPI_ISL_420211, EPI_ISL_420212, EPI_ISL_420213, EPI_ISL_420214, EPI_ISL_420215, EPI_ISL_420216, EPI_ISL_420217, EPI_ISL_420218, EPI_ISL_420219, EPI_ISL_420220, EPI_ISL_420221, EPI_ISL_420222, EPI_ISL_420223, EPI_ISL_420224, EPI_ISL_420225, EPI_ISL_420226, EPI_ISL_420227, EPI_ISL_420228, EPI_ISL_420229, EPI_ISL_420230, EPI_ISL_420231, EPI_ISL_420232, EPI_ISL_420233, EPI_ISL_420234, EPI_ISL_420235, EPI_ISL_420236, EPI_ISL_420237, EPI_ISL_420238, EPI_ISL_420239, EPI_ISL_420240, EPI_ISL_420241, EPI_ISL_420242, EPI_ISL_420243, EPI_ISL_420244, EPI_ISL_420245, EPI_ISL_420246, EPI_ISL_420247, EPI_ISL_420248, EPI_ISL_420249, EPI_ISL_420250, EPI_ISL_420251, EPI_ISL_420252, EPI_ISL_420253, EPI_ISL_420254, EPI_ISL_420255, EPI_ISL_420256, EPI_ISL_420257, EPI_ISL_420258, EPI_ISL_420259, EPI_ISL_420260, EPI_ISL_420261, EPI_ISL_420262, EPI_ISL_420263, EPI_ISL_420264, EPI_ISL_420265, EPI_ISL_420266, EPI_ISL_420267, EPI_ISL_420268, EPI_ISL_420269, EPI_ISL_420270, EPI_ISL_420271, EPI_ISL_420272, EPI_ISL_420273, EPI_ISL_420274, EPI_ISL_420275, EPI_ISL_420276, EPI_ISL_420277, EPI_ISL_420278, EPI_ISL_420279, EPI_ISL_420280, EPI_ISL_420281, EPI_ISL_420282, EPI_ISL_420283, EPI_ISL_420284, EPI_ISL_420285, EPI_ISL_420286, EPI_ISL_420287, EPI_ISL_420288, EPI_ISL_420289, EPI_ISL_420290, EPI_ISL_420291, EPI_ISL_420292 | | | |
| see above | Virology Department, Sheffield Teaching Hospitals NHS Foundation Trust | Department of Infection, Immunity and Cardiovascular Disease, The Florey Institute, The Medical School, University of Sheffield | Thushan de Silva, Matthew Parker, Adri Angyal, Rebecca Brown, Rachel Tucker, Paul Parsons, Luke Green, Danielle Groves, Alex Keeley, Dave Partridge, Matthew Wyles, Benjamin Lindsey, Mehmet Yavuz, Mohammad Raza, Cariad Evans |
| EPI_ISL_420541 | Institute of Microbiology and Immunology, Faculty of Medicine, University of Ljubljana | Institute of Microbiology and Immunology, Faculty of Medicine, University of Ljubljana | Tomaž Mark Zorec, Samo Zakotnik, Lucijan Skubic, Miša Korva, Tatjana Avši - Županc, Mario Poljak |
| EPI_ISL_420628, EPI_ISL_420629, EPI_ISL_420630 | Virginia DCLS | Virginia DCLS | Virginia DCLS |
| EPI_ISL_421279, EPI_ISL_421281 | Clinical Diagnostics Laboratory, Diagnostic & Experimental Pathology, Lilly Research Laboratories | Clinical Diagnostics Laboratory, Diagnostic & Experimental Pathology, Lilly Research Laboratories | Tim Holzer, Mayuri Vaidya, Angie Fulford, Sam McNeely, Rachael Redmond, Phil Ebert, John Calley, Leslie O'Neill Reising, Pat Finnegan, Erin Wray, John McElwee, Jeff Fill, Joe Oakley, Andrew Schade |
| EPI_ISL_425190 | Servicio de Microbiología. Consorcio Hospital General Universitario de Valencia | Sequencing and Bioinformatics Service and Molecular Epidemiology Research Group. FISABIO-Public Health | Vicente Soriano Chirona, Ivan Ansari, David Navarro, Maria Alma Bracho, Griselda De Marco, Beatriz Beamud, Lidia Ruiz Roldan, Marta Pla Diaz, Neris Garcia-Gonzalez, Inma Galán Vendrell, Sandra Carbo, Loreto Ferrús Abad, Paula Ruiz-Hueso, Mariana Reyes-Prieto, Llúcia Martínez-Priego, Giuseppe D'Auria, Fernando Gonzalez-Candelas |
| EPI_ISL_425202 | Servicio de Microbiologia. Hospital Clinico Universitario de Valencia | Sequencing and Bioinformatics Service and Molecular Epidemiology Research Group. FISABIO-Public Health | Vicente Soriano Chirona, Maria Alma Bracho, Griselda De Marco, Beatriz Beamud, Lidia Ruiz Roldan, Marta Pla Diaz,Neris Garcia-Gonzalez, Loreto Ferrús Abad, Maria Dolores Ocete, Inma Galán Vendrell, Paula Ruiz-Hueso, Mariana Reyes-Prieto, Llúcia Martínez-Priego, Concepcion Gimeno, Giuseppe D'Auria, Fernando Gonzalez-Candelas |

| | | | |
|---|---|---|---|
| EPI_ISL_426454, EPI_ISL_426455, EPI_ISL_426456, EPI_ISL_426457, EPI_ISL_426458, EPI_ISL_426459, EPI_ISL_426460, EPI_ISL_426461, EPI_ISL_426462, EPI_ISL_426463, EPI_ISL_426464, EPI_ISL_426465, EPI_ISL_426466, EPI_ISL_426467, EPI_ISL_426468, EPI_ISL_426469, EPI_ISL_426470, EPI_ISL_426471, EPI_ISL_426472, EPI_ISL_426473, EPI_ISL_426474, EPI_ISL_426475 | | | |
| see above | Virginia DCLS | Virginia DCLS | Virginia DCLS |
| EPI_ISL_427643, EPI_ISL_427646, EPI_ISL_427649, EPI_ISL_427663, EPI_ISL_427664, EPI_ISL_427665, EPI_ISL_427678, EPI_ISL_427682, EPI_ISL_427692, EPI_ISL_427695, EPI_ISL_427698, EPI_ISL_427704 | | | |
| see above | Centre for Infectious Diseases and Microbiology Public Health | NSW Health Pathology - Institute of Clinical Pathology and Medical Research; Westmead Hospital; University of Sydney | Timms V, Gall M, Arnott A, Sadsad R, Draper J, Sim E, Bachmann N, Rockett R, Lam C, Gray K, Carter I, Holmes EC, O'Sullivan MV, Byun R, Sintchenko V, Chen SC, Eden JS, Maddocks S, Kok J, Propenko M, Sorrell T, Chang S, Basile K, Dwyer DE for the 2019-nCoV Study Group |
| EPI_ISL_427719 | ACT Pathology, The Canberra Hospital | NSW Health Pathology - Institute of Clinical Pathology and Medical Research; Westmead Hospital; University of Sydney | Timms V, Gall M, Arnott A, Sadsad R, Draper J, Sim E, Bachmann N, Rockett R, Lam C, Gray K, Carter I, Holmes EC, O'Sullivan MV, Byun R, Sintchenko V, Chen SC, Eden JS, Maddocks S, Kok J, Propenko M, Sorrell T, Chang S, Basile K, Dwyer DE for the 2019-nCoV Study Group |
| EPI_ISL_427725, EPI_ISL_427734, EPI_ISL_427748, EPI_ISL_427771, EPI_ISL_427794, EPI_ISL_427803, EPI_ISL_427806 | Centre for Infectious Diseases and Microbiology Public Health | NSW Health Pathology - Institute of Clinical Pathology and Medical Research; Westmead Hospital; University of Sydney | Timms V, Gall M, Arnott A, Sadsad R, Draper J, Sim E, Bachmann N, Rockett R, Lam C, Gray K, Carter I, Holmes EC, O'Sullivan MV, Byun R, Sintchenko V, Chen SC, Eden JS, Maddocks S, Kok J, Propenko M, Sorrell T, Chang S, Basile K, Dwyer DE for the 2019-nCoV Study Group |
| EPI_ISL_428202, EPI_ISL_428203, EPI_ISL_428204, EPI_ISL_428205, EPI_ISL_428206 | Nebraska Public Health Laboratory | UNMC COVID-19 Response Team | UNMC COVID-19 Response Team |
| EPI_ISL_428925, EPI_ISL_428926, EPI_ISL_428927, EPI_ISL_428928, EPI_ISL_428929, EPI_ISL_428930, EPI_ISL_428931, EPI_ISL_428932 | ViroGenetics - BSL3 Laboratory of Virology; Human Genome Variation Research Group & Genomics Centre MCB; Bioinformatics Research Group; Wojewódzka Stacja Sanitarno-Epidemiologiczna w Krakowie | ViroGenetics - BSL3 Laboratory of Virology; Human Genome Variation Research Group & Genomics Centre MCB; Bioinformatics Research Group | Wojciech Branicki, Ewelina Popiech, Micha Kowalski, Agnieszka Starowicz, Adrianna Klajmon, Aleksandra Pisarek, Danuta Piniewska-Róg, Kamila Marszaek, Tomasz Gromowski, Katarzyna Kopera, Katarzyna Dudek, Inga Drebot, Katarzyna Gua, Magda Pachota, Aleksandra Synowiec, Marek Sanak, Jarosaw Foremny, Pawe P abaj, Krzysztof Pyr |
| EPI_ISL_429126, EPI_ISL_429127 | Unilabs Skovde | The Public Health Agency of Sweden | Tobias Kollberg, Helena Enroth, Olov Svartstrom, Maria Lind Karlberg, Anna-Malin Linde, Oskar Karlsson Lindsjo, Anna Risberg, Shaman Muradrasoli, Karin Tegmark-Wisell |
| EPI_ISL_429852, EPI_ISL_429853, EPI_ISL_429854, EPI_ISL_429855 | Centers for Disease Control and Prevention of Lishui | Department of InspectionCenters for Disease Control and Prevention of Lishui | Wang Xiaoguang,Ji Qiaoying,Ji Jiansong,Ye Bifeng,Ye Ling |
| EPI_ISL_429876 | California Department of Public Health | Chiu Laboratory, University of California, San Francisco | Xianding Deng, Scot Federman, Chao-Yang Pan, Hugo Guevara,Wei Gu, Debra A. Wadford, and Charles Y. Chiu |
| EPI_ISL_429877, EPI_ISL_429878 | Chiu Laboratory, University of California, San Francisco | Chiu Laboratory, University of California, San Francisco | Xianding Deng, Scot Federman, Wei Gu, and Charles Y. Chiu |
| EPI_ISL_429879, EPI_ISL_429880 | Santa Clara County Public Health Department | Chiu Laboratory, University of California, San Francisco | Xianding Deng, Scot Federman, Wei Gu, Elsa Villarino, Brandon Bonin, Debra A. Wadford, and Charles Y. Chiu |
| EPI_ISL_429881 | California Department of Public Health | Chiu Laboratory, University of California, San Francisco | Xianding Deng, Scot Federman, Wei Gu, and Charles Y. Chiu |
| EPI_ISL_429969, EPI_ISL_429970, EPI_ISL_429971, EPI_ISL_429972, EPI_ISL_429973, EPI_ISL_429974, EPI_ISL_429975, EPI_ISL_429976, EPI_ISL_429977, EPI_ISL_429978, EPI_ISL_429979, EPI_ISL_429980, EPI_ISL_429981, EPI_ISL_429982, EPI_ISL_429983, EPI_ISL_429984, EPI_ISL_429985, EPI_ISL_429986, EPI_ISL_429987, EPI_ISL_429988, EPI_ISL_429989 | | | |
| see above | Virginia DCLS | Virginia DCLS | Virginia DCLS |
| EPI_ISL_430469 | Hellenic Pasteur Institute, Public Health Laboratories | Hellenic Pasteur Institute, Public Health Laboratories, Unit of Bioinformatics and Applied Genomics | Vasiliki Pogka, Timokratis Karamitros, Athanasios Kossyvakis, Antonios Kalliaropoulos, Horefti Elina, Evangelidou Maria, Androniki Voulgari-Kokota, Aspasia Kontou, Andreas Mentis |
| EPI_ISL_431117 | Department of Microbiology, Gandhi Medical College and Hospital, Secendrabad, Hyderabad, India | Department of Microbiology, Gandhi Medical College and Hospital, Secendrabad, Hyderabad, India | Thrilok Chander B, Muttineni Radhakrishna, Nagamani K, Raja Rao M, Kalyani Putty, Ravikumar P, Sunitha P, Pankaj Singh D, Anand Kumar K, Amit A. Upadhyay, Steven E.Bosinger, Rama Amara |
| EPI_ISL_432451, EPI_ISL_432452, EPI_ISL_432453, EPI_ISL_432454, EPI_ISL_432455, EPI_ISL_432456, EPI_ISL_432457, EPI_ISL_432458, EPI_ISL_432459, EPI_ISL_432460, EPI_ISL_432461, EPI_ISL_432462, EPI_ISL_432463, EPI_ISL_432464, EPI_ISL_432465, EPI_ISL_432466, EPI_ISL_432467, EPI_ISL_432468, EPI_ISL_432469, EPI_ISL_432470, EPI_ISL_432471, EPI_ISL_432472, EPI_ISL_432473, EPI_ISL_432474, EPI_ISL_432475, EPI_ISL_432476, EPI_ISL_432477, EPI_ISL_432478, EPI_ISL_432479, EPI_ISL_432480, EPI_ISL_432481, EPI_ISL_432482, EPI_ISL_432483, EPI_ISL_432484, EPI_ISL_432485, EPI_ISL_432486, EPI_ISL_432487, EPI_ISL_432488, EPI_ISL_432489, EPI_ISL_432490, EPI_ISL_432491, EPI_ISL_432492, EPI_ISL_432493, EPI_ISL_432494, EPI_ISL_432495, EPI_ISL_432496, EPI_ISL_432497, EPI_ISL_432498, EPI_ISL_432499, EPI_ISL_432500, EPI_ISL_432501, EPI_ISL_432502, EPI_ISL_432503, EPI_ISL_432504, EPI_ISL_432505, EPI_ISL_432506, EPI_ISL_432507, EPI_ISL_432508, EPI_ISL_432509, EPI_ISL_432510, EPI_ISL_432511, EPI_ISL_432512, EPI_ISL_432513, EPI_ISL_432514, EPI_ISL_432515, EPI_ISL_432516, EPI_ISL_432517, EPI_ISL_432518, EPI_ISL_432519, EPI_ISL_432520, EPI_ISL_432521, EPI_ISL_432522, EPI_ISL_432523, EPI_ISL_432524, EPI_ISL_432525, EPI_ISL_432526, EPI_ISL_432527, EPI_ISL_432528, EPI_ISL_432529, EPI_ISL_432530, EPI_ISL_432531, EPI_ISL_432532, EPI_ISL_432533, EPI_ISL_432534, EPI_ISL_432535, EPI_ISL_432536, EPI_ISL_432537, EPI_ISL_432538, EPI_ISL_432539, EPI_ISL_432540, EPI_ISL_432541, EPI_ISL_432542, EPI_ISL_432543, EPI_ISL_432544, EPI_ISL_432545, EPI_ISL_432546, EPI_ISL_432547, EPI_ISL_432548, EPI_ISL_432549, EPI_ISL_432550, EPI_ISL_432551, EPI_ISL_432552, EPI_ISL_432553, EPI_ISL_432554, EPI_ISL_432555, EPI_ISL_432556, EPI_ISL_432557, EPI_ISL_432558, EPI_ISL_432559, EPI_ISL_432560, EPI_ISL_432561, EPI_ISL_432562, EPI_ISL_432563, EPI_ISL_432564, EPI_ISL_432565, EPI_ISL_432566, EPI_ISL_432567, EPI_ISL_432568, EPI_ISL_432569, EPI_ISL_432570, EPI_ISL_432571, EPI_ISL_432572, EPI_ISL_432573, EPI_ISL_432574, EPI_ISL_432575, EPI_ISL_432576, EPI_ISL_432577, EPI_ISL_432578, EPI_ISL_432579, EPI_ISL_432580, EPI_ISL_432581, EPI_ISL_432582, EPI_ISL_432583, EPI_ISL_432584, EPI_ISL_432585, EPI_ISL_432586, EPI_ISL_432587, EPI_ISL_432588, EPI_ISL_432589, EPI_ISL_432590, EPI_ISL_432591, EPI_ISL_432592, EPI_ISL_432593, EPI_ISL_432594, EPI_ISL_432595, EPI_ISL_432596, EPI_ISL_432597, EPI_ISL_432598, EPI_ISL_432599, EPI_ISL_432600, EPI_ISL_432601, EPI_ISL_432602, EPI_ISL_432603, EPI_ISL_432604, EPI_ISL_432605, EPI_ISL_432606, EPI_ISL_432607, EPI_ISL_432608, EPI_ISL_432609, EPI_ISL_432610, EPI_ISL_432611, EPI_ISL_432612, EPI_ISL_432613, EPI_ISL_432614, EPI_ISL_432615, EPI_ISL_432616, EPI_ISL_432617, EPI_ISL_432618, EPI_ISL_432619, EPI_ISL_432620, EPI_ISL_432621, EPI_ISL_432622, EPI_ISL_432623, EPI_ISL_432624, EPI_ISL_432625, EPI_ISL_432626, EPI_ISL_432627, EPI_ISL_432628, EPI_ISL_432629, EPI_ISL_432630, EPI_ISL_432631, EPI_ISL_432632, EPI_ISL_432633, EPI_ISL_432634, EPI_ISL_432635, EPI_ISL_432636, EPI_ISL_432637, EPI_ISL_432638, EPI_ISL_432639, EPI_ISL_432640, EPI_ISL_432641, EPI_ISL_432642, EPI_ISL_432643, EPI_ISL_432644, EPI_ISL_432645, EPI_ISL_432646, EPI_ISL_432647, EPI_ISL_432648, EPI_ISL_432649, EPI_ISL_432650, EPI_ISL_432651, EPI_ISL_432652, EPI_ISL_432653, EPI_ISL_432654, EPI_ISL_432655, EPI_ISL_432656, EPI_ISL_432657, EPI_ISL_432658, EPI_ISL_432659, EPI_ISL_432660, EPI_ISL_432661, EPI_ISL_432662, EPI_ISL_432663, EPI_ISL_432664, EPI_ISL_432665, EPI_ISL_432666, EPI_ISL_432667, EPI_ISL_432710, EPI_ISL_432711, EPI_ISL_432712, EPI_ISL_432713, EPI_ISL_432714, EPI_ISL_432715, EPI_ISL_432716, EPI_ISL_432717, EPI_ISL_432718, EPI_ISL_432719, EPI_ISL_432720, EPI_ISL_432721, EPI_ISL_432722, EPI_ISL_432723, EPI_ISL_432724, EPI_ISL_432725, EPI_ISL_432726, EPI_ISL_432727, EPI_ISL_432728, EPI_ISL_432729, EPI_ISL_432730, EPI_ISL_432731, EPI_ISL_432732, EPI_ISL_432733, EPI_ISL_432734, EPI_ISL_432735, EPI_ISL_432736, EPI_ISL_432737, EPI_ISL_432738, EPI_ISL_432739, EPI_ISL_432740, EPI_ISL_432741, EPI_ISL_432742, EPI_ISL_432743, EPI_ISL_432744, EPI_ISL_432745, EPI_ISL_432746, EPI_ISL_432747, EPI_ISL_432748, EPI_ISL_432749, EPI_ISL_432750, EPI_ISL_432751, EPI_ISL_432752, EPI_ISL_432753, EPI_ISL_432754, EPI_ISL_432755, EPI_ISL_432756, EPI_ISL_432757, EPI_ISL_432758, EPI_ISL_432759, EPI_ISL_432760, EPI_ISL_432761, EPI_ISL_432762, EPI_ISL_432763, EPI_ISL_432764, EPI_ISL_432765, EPI_ISL_432766, EPI_ISL_432767, EPI_ISL_432768, EPI_ISL_432769, EPI_ISL_432770, EPI_ISL_432771, EPI_ISL_432772, EPI_ISL_432773, EPI_ISL_432774, EPI_ISL_432775, EPI_ISL_432776, EPI_ISL_432777, EPI_ISL_432778, EPI_ISL_432779, EPI_ISL_432780, EPI_ISL_432781, EPI_ISL_432782, EPI_ISL_432783, EPI_ISL_432784, EPI_ISL_432785, EPI_ISL_432786, EPI_ISL_432787, EPI_ISL_432788, EPI_ISL_432789, EPI_ISL_432790, EPI_ISL_432791, EPI_ISL_432792, EPI_ISL_432793, EPI_ISL_432794, EPI_ISL_432795, EPI_ISL_432796, EPI_ISL_432797, EPI_ISL_432798, EPI_ISL_432799, EPI_ISL_432800, EPI_ISL_432801, EPI_ISL_432802, EPI_ISL_432803, EPI_ISL_432804, EPI_ISL_432805, EPI_ISL_432806, EPI_ISL_432807, EPI_ISL_432808, EPI_ISL_432809, EPI_ISL_432810, EPI_ISL_432811, EPI_ISL_432812, EPI_ISL_432813, EPI_ISL_432814, EPI_ISL_432815, EPI_ISL_432816, EPI_ISL_432817, EPI_ISL_432818, EPI_ISL_432819, EPI_ISL_432820, EPI_ISL_432821, EPI_ISL_432822, EPI_ISL_432823, EPI_ISL_432824, EPI_ISL_432825, EPI_ISL_432826, EPI_ISL_432827, EPI_ISL_432828, EPI_ISL_432829, EPI_ISL_432830, EPI_ISL_432831, EPI_ISL_432832, EPI_ISL_432833, EPI_ISL_432834, EPI_ISL_432835, EPI_ISL_432836, EPI_ISL_432837, EPI_ISL_432838, EPI_ISL_432839, EPI_ISL_432840, EPI_ISL_432841, EPI_ISL_432842, EPI_ISL_432843, EPI_ISL_432844, EPI_ISL_432845, EPI_ISL_432846, EPI_ISL_432847, EPI_ISL_432848, EPI_ISL_432849, EPI_ISL_432850, EPI_ISL_432851, EPI_ISL_432852, EPI_ISL_432853, EPI_ISL_432854, EPI_ISL_432855, EPI_ISL_432856, EPI_ISL_432857, EPI_ISL_432858, EPI_ISL_432859, EPI_ISL_432860, EPI_ISL_432861, EPI_ISL_432862, EPI_ISL_432863, EPI_ISL_432864, EPI_ISL_432865, EPI_ISL_432866, EPI_ISL_432867 | | | |
| see above | Virology Department, Sheffield Teaching Hospitals NHS Foundation Trust / Virology Department, Sheffield Teaching Hospitals NHS Foundation Trust | COVID-19 Genomics UK (COG-UK) Consortium | Thushan de Silva, Matthew Parker,Adri Angyal, Rebecca Brown, Luke Green, Rachel Tucker, Paul Parsons, Danielle Groves, Alex Keeley, Dave Partridge, Matthew Wyles, Benjamin Lindsey, Mehmet Yavuz, Mohammad Raza, Cariad Evans |
| EPI_ISL_434534 | National Institute for Viral Disease Control and Prevention, China CDC | National Institute for Viral Disease Control and Prevention, China CDC, Yunnan Provincial CDC | Wenjie Tan, Roujian Lu, Wenling Wang, Peihua Niu, Huijuan Wang, Baoying Huang, Li Zhao, Fei Ye, Guizhen Wu |
| EPI_ISL_434563, EPI_ISL_434564, EPI_ISL_434565, EPI_ISL_434566, EPI_ISL_434567, EPI_ISL_434568, EPI_ISL_434569 | unknown | Microbiology, The University of Hong Kong | To,K.K.W. and Yuen,K.-Y. |
| EPI_ISL_434570 | unknown | Microbiology | To,K.K.W. and Yuen,K.-Y. |
| EPI_ISL_434590, EPI_ISL_434591, EPI_ISL_434592, EPI_ISL_434593, EPI_ISL_434594, EPI_ISL_434595, EPI_ISL_434596, EPI_ISL_434597, EPI_ISL_434598, EPI_ISL_434599, EPI_ISL_434600, EPI_ISL_434601, EPI_ISL_434602, EPI_ISL_434603, EPI_ISL_434604, EPI_ISL_434605, EPI_ISL_434606 | | | |
| see above | Virginia DCLS | Virginia DCLS | Virginia DCLS |
| EPI_ISL_434645, EPI_ISL_434649 | Svardsjo VC | The Public Health Agency of Sweden | Tommy Janers, Oskar Karlsson Lindsjo, Maria Lind Karlberg, Anna-Malin Linde, Olov Svartstrom, Anna Risberg, Theresa Enkirch, Mia Brytting, Karin Tegmark-Wisell |
| EPI_ISL_435580, EPI_ISL_435581, EPI_ISL_435582, EPI_ISL_435583, EPI_ISL_435584, EPI_ISL_435585, EPI_ISL_435586, EPI_ISL_435587, EPI_ISL_435588, EPI_ISL_435589, EPI_ISL_435590, EPI_ISL_435591, EPI_ISL_435592, EPI_ISL_435594, EPI_ISL_435595, EPI_ISL_435596, EPI_ISL_435597, EPI_ISL_435598, EPI_ISL_435599, EPI_ISL_435600, EPI_ISL_435601, EPI_ISL_435602, EPI_ISL_435603, EPI_ISL_435604, EPI_ISL_435605, EPI_ISL_435606, EPI_ISL_435607, EPI_ISL_435608, EPI_ISL_435609, EPI_ISL_435610, EPI_ISL_435611, EPI_ISL_435612, EPI_ISL_435613, EPI_ISL_435615, EPI_ISL_435616, EPI_ISL_435617, EPI_ISL_435618, EPI_ISL_435619, EPI_ISL_435620, EPI_ISL_435621, EPI_ISL_435622, EPI_ISL_435624, EPI_ISL_435625, EPI_ISL_435626, EPI_ISL_435627, EPI_ISL_435628, EPI_ISL_435629, EPI_ISL_435630, EPI_ISL_435631, EPI_ISL_435632, EPI_ISL_435633, EPI_ISL_435634, EPI_ISL_435635, EPI_ISL_435636, EPI_ISL_435637, EPI_ISL_435638, EPI_ISL_435639, EPI_ISL_435640, EPI_ISL_435641, EPI_ISL_435643, EPI_ISL_435644, EPI_ISL_435645, EPI_ISL_435646, EPI_ISL_435647, EPI_ISL_435649, EPI_ISL_435650, EPI_ISL_435651, EPI_ISL_435652, EPI_ISL_435653, EPI_ISL_435654, EPI_ISL_435655, EPI_ISL_435656, EPI_ISL_435657, EPI_ISL_435658, EPI_ISL_435659, EPI_ISL_435660, EPI_ISL_435661, EPI_ISL_435662, EPI_ISL_435663, EPI_ISL_435664, EPI_ISL_435665, EPI_ISL_435666, EPI_ISL_435667, EPI_ISL_435668, EPI_ISL_435669, EPI_ISL_435670, EPI_ISL_435671, EPI_ISL_435673 | | | |

| | | | |
|---|---|---|---|
| see above | Santa Clara County Public Health Department | Chiu Laboratory, University of California, San Francisco | Xianding Deng, Scot Federman, Wei Gu, Elsa Villarino, Brandon Bonin, Debra A. Wadford, and Charles Y. Chiu |
| EPI_ISL_436289, EPI_ISL_436309, EPI_ISL_436329, EPI_ISL_436349, EPI_ISL_436400 | Servicio de Microbiologia. Hospital Clinico Universitario de Valencia | Sequencing and Bioinformatics Service and Molecular Epidemiology Research Group. FISABIO-Public Health | Vicente Soriano Chirona, Ivan Ansari, David Navarro, Maria Alma Bracho, Griselda De Marco, Beatriz Beamud, Lidia Ruiz Roldan, Marta Pla Diaz, Neris Garcia-Gonzalez, Inma Galán Vendrell, Sandra Carbo, Loreto Ferrús Abad, Paula Ruiz-Hueso, Mariana Reyes-Prieto, Llúcia Martínez-Priego, Giuseppe D'Auria, Fernando Gonzalez-Candelas |
| EPI_ISL_437387, EPI_ISL_437388, EPI_ISL_437389, EPI_ISL_437390, EPI_ISL_437391, EPI_ISL_437392, EPI_ISL_437393, EPI_ISL_437394, EPI_ISL_437395, EPI_ISL_437396, EPI_ISL_437397, EPI_ISL_437398, EPI_ISL_437399, EPI_ISL_437400, EPI_ISL_437401, EPI_ISL_437402, EPI_ISL_437403, EPI_ISL_437404, EPI_ISL_437405, EPI_ISL_437406, EPI_ISL_437407, EPI_ISL_437408, EPI_ISL_437409, EPI_ISL_437410, EPI_ISL_437411, EPI_ISL_437412, EPI_ISL_437413, EPI_ISL_437414, EPI_ISL_437415, EPI_ISL_437416, EPI_ISL_437417, EPI_ISL_437418, EPI_ISL_437419, EPI_ISL_437420, EPI_ISL_437421, EPI_ISL_437422, EPI_ISL_437423, EPI_ISL_437424, EPI_ISL_437425, EPI_ISL_437426, EPI_ISL_437427, EPI_ISL_437428, EPI_ISL_437429, EPI_ISL_437430, EPI_ISL_437431, EPI_ISL_437432 | | | |
| see above | Virginia DCLS | Virginia DCLS | Virginia DCLS |
| EPI_ISL_437455, EPI_ISL_437456, EPI_ISL_437457, EPI_ISL_437458 | Clinical Diagnostics Laboratory, Diagnostic & Experimental Pathology, Lilly Research Laboratories | Clinical Diagnostics Laboratory, Diagnostic & Experimental Pathology, Lilly Research Laboratories | Tim Holzer, Mayuri Vaidya, Angie Fulford, Sam McNeely, Rachael Redmond, Phil Ebert, John Calley, Leslie ONeill Reising, Pat Finnegan, Erin Wray, John McElwee, Jeff Fill, Joe Oakley, Andrew Schade |
| EPI_ISL_437689 | Laboratory for Urgent Response to Biological Threats | Institut Pasteur CIBU / ERI | V. Caro, A. Kwasiborski, V. Hourdel, C. Balière, J. Vanhomwegen, C. Batéjat, JC. Manuguerra |
| EPI_ISL_437690 | Laboratory for Urgent Response to Biological Threats | Institut Pasteur CIBU /ERI | V. Caro, A. Kwasiborski, H. Hourdel, C. Balière, J. Vanhomwegen, C. Batéjat, JC. Manuguerra |
| EPI_ISL_437763, EPI_ISL_437764, EPI_ISL_437765, EPI_ISL_437766, EPI_ISL_437767, EPI_ISL_437768, EPI_ISL_437769, EPI_ISL_437770, EPI_ISL_437771, EPI_ISL_437772, EPI_ISL_437773, EPI_ISL_437774, EPI_ISL_437775, EPI_ISL_437776, EPI_ISL_437777, EPI_ISL_437778, EPI_ISL_437779, EPI_ISL_437780, EPI_ISL_437781, EPI_ISL_437782, EPI_ISL_437783, EPI_ISL_437784, EPI_ISL_437785, EPI_ISL_437786, EPI_ISL_437787, EPI_ISL_437788, EPI_ISL_437789, EPI_ISL_437790, EPI_ISL_437791, EPI_ISL_437792, EPI_ISL_437793, EPI_ISL_437794, EPI_ISL_437795, EPI_ISL_437796, EPI_ISL_437797, EPI_ISL_437798, EPI_ISL_437799, EPI_ISL_437800, EPI_ISL_437801, EPI_ISL_437802 | | | |
| see above | Virginia DCLS | Virginia DCLS | Virginia DCLS |
| EPI_ISL_441898, EPI_ISL_441899, EPI_ISL_441900, EPI_ISL_441901, EPI_ISL_441902, EPI_ISL_441903, EPI_ISL_441904, EPI_ISL_441905, EPI_ISL_441906, EPI_ISL_441907, EPI_ISL_441908, EPI_ISL_441909, EPI_ISL_441910, EPI_ISL_441911, EPI_ISL_441912, EPI_ISL_441913, EPI_ISL_441914, EPI_ISL_441915, EPI_ISL_441916, EPI_ISL_441917, EPI_ISL_441918, EPI_ISL_441919, EPI_ISL_441920, EPI_ISL_441921, EPI_ISL_441922, EPI_ISL_441923, EPI_ISL_441924, EPI_ISL_441925, EPI_ISL_441926, EPI_ISL_441927, EPI_ISL_441928, EPI_ISL_441929, EPI_ISL_441930, EPI_ISL_441931, EPI_ISL_441932, EPI_ISL_441933, EPI_ISL_441934, EPI_ISL_441935, EPI_ISL_441936, EPI_ISL_441937, EPI_ISL_441938, EPI_ISL_441939, EPI_ISL_441940, EPI_ISL_441941, EPI_ISL_441942, EPI_ISL_441943, EPI_ISL_441944, EPI_ISL_441945, EPI_ISL_441946, EPI_ISL_441947, EPI_ISL_441948, EPI_ISL_441949, EPI_ISL_441950, EPI_ISL_441951, EPI_ISL_441952, EPI_ISL_441953, EPI_ISL_441954, EPI_ISL_441955, EPI_ISL_441956, EPI_ISL_441957, EPI_ISL_441958, EPI_ISL_441959, EPI_ISL_441960, EPI_ISL_441961, EPI_ISL_441962, EPI_ISL_441963, EPI_ISL_441964, EPI_ISL_441965, EPI_ISL_441966, EPI_ISL_441967, EPI_ISL_441968, EPI_ISL_441969, EPI_ISL_441970, EPI_ISL_441971, EPI_ISL_441972, EPI_ISL_441973, EPI_ISL_441974, EPI_ISL_441975, EPI_ISL_441976, EPI_ISL_441977, EPI_ISL_441978, EPI_ISL_441979, EPI_ISL_441980, EPI_ISL_441981, EPI_ISL_441982, EPI_ISL_441983, EPI_ISL_441984, EPI_ISL_441985, EPI_ISL_441986, EPI_ISL_441987, EPI_ISL_441988, EPI_ISL_441989, EPI_ISL_441990, EPI_ISL_441991, EPI_ISL_441992, EPI_ISL_441993, EPI_ISL_441994, EPI_ISL_441995, EPI_ISL_441996, EPI_ISL_441997, EPI_ISL_441998, EPI_ISL_441999, EPI_ISL_442000, EPI_ISL_442001, EPI_ISL_442002, EPI_ISL_442003, EPI_ISL_442004, EPI_ISL_442005, EPI_ISL_442006, EPI_ISL_442007, EPI_ISL_442008, EPI_ISL_442009, EPI_ISL_442010, EPI_ISL_442011, EPI_ISL_442012, EPI_ISL_442013, EPI_ISL_442014, EPI_ISL_442015, EPI_ISL_442016, EPI_ISL_442017, EPI_ISL_442018, EPI_ISL_442019, EPI_ISL_442020, EPI_ISL_442021, EPI_ISL_442022, EPI_ISL_442023, EPI_ISL_442024, EPI_ISL_442025, EPI_ISL_442026, EPI_ISL_442027, EPI_ISL_442028, EPI_ISL_442029, EPI_ISL_442030, EPI_ISL_442031, EPI_ISL_442032, EPI_ISL_442033, EPI_ISL_442034, EPI_ISL_442035, EPI_ISL_442036, EPI_ISL_442037, EPI_ISL_442038, EPI_ISL_442039, EPI_ISL_442040, EPI_ISL_442041, EPI_ISL_442042, EPI_ISL_442043, EPI_ISL_442344, EPI_ISL_442345, EPI_ISL_442346, EPI_ISL_442347, EPI_ISL_442348, EPI_ISL_442349, EPI_ISL_442350, EPI_ISL_442351, EPI_ISL_442352, EPI_ISL_442353, EPI_ISL_442354, EPI_ISL_442355, EPI_ISL_442356, EPI_ISL_442357, EPI_ISL_442358, EPI_ISL_442359, EPI_ISL_442360, EPI_ISL_442361, EPI_ISL_442362, EPI_ISL_442363, EPI_ISL_442364, EPI_ISL_442365, EPI_ISL_442366, EPI_ISL_442367, EPI_ISL_442368, EPI_ISL_442369, EPI_ISL_442370, EPI_ISL_442371, EPI_ISL_442372, EPI_ISL_442373, EPI_ISL_442374, EPI_ISL_442375, EPI_ISL_442376, EPI_ISL_442377, EPI_ISL_442378, EPI_ISL_442379, EPI_ISL_442380, EPI_ISL_442381, EPI_ISL_442382, EPI_ISL_442383, EPI_ISL_442384, EPI_ISL_442385, EPI_ISL_442386, EPI_ISL_442387, EPI_ISL_442388, EPI_ISL_442389, EPI_ISL_442390, EPI_ISL_442391, EPI_ISL_442392, EPI_ISL_442393, EPI_ISL_442394, EPI_ISL_442395, EPI_ISL_442396, EPI_ISL_442397, EPI_ISL_442398, EPI_ISL_442399, EPI_ISL_442400, EPI_ISL_442401, EPI_ISL_442402, EPI_ISL_442403, EPI_ISL_442404, EPI_ISL_442405, EPI_ISL_442406, EPI_ISL_442407, EPI_ISL_442408, EPI_ISL_442409, EPI_ISL_442410, EPI_ISL_442411, EPI_ISL_442412, EPI_ISL_442413, EPI_ISL_442414, EPI_ISL_442415, EPI_ISL_442416, EPI_ISL_442417, EPI_ISL_442418, EPI_ISL_442419, EPI_ISL_442420, EPI_ISL_442421, EPI_ISL_442422, EPI_ISL_442423, EPI_ISL_442424, EPI_ISL_442425, EPI_ISL_442426, EPI_ISL_442427, EPI_ISL_442428, EPI_ISL_442429, EPI_ISL_442430, EPI_ISL_442431, EPI_ISL_442432, EPI_ISL_442433, EPI_ISL_442434, EPI_ISL_442435, EPI_ISL_442436, EPI_ISL_442437, EPI_ISL_442438, EPI_ISL_442439, EPI_ISL_442440, EPI_ISL_442441, EPI_ISL_442442, EPI_ISL_442443, EPI_ISL_442444, EPI_ISL_442445, EPI_ISL_442446, EPI_ISL_442447, EPI_ISL_442448, EPI_ISL_442449, EPI_ISL_442450, EPI_ISL_442451, EPI_ISL_442452, EPI_ISL_442453, EPI_ISL_442454, EPI_ISL_442455, EPI_ISL_442456, EPI_ISL_442457, EPI_ISL_442458, EPI_ISL_442459, EPI_ISL_442460, EPI_ISL_442461, EPI_ISL_442462, EPI_ISL_442463, EPI_ISL_442464, EPI_ISL_442465, EPI_ISL_442466, EPI_ISL_442467, EPI_ISL_442468, EPI_ISL_442469, EPI_ISL_442470, EPI_ISL_442471, EPI_ISL_442472, EPI_ISL_442473, EPI_ISL_442474, EPI_ISL_442475, EPI_ISL_442476, EPI_ISL_442477, EPI_ISL_442478, EPI_ISL_442479, EPI_ISL_442480, EPI_ISL_442481, EPI_ISL_442482, EPI_ISL_442483, EPI_ISL_442484, EPI_ISL_442485, EPI_ISL_442486, EPI_ISL_442487, EPI_ISL_442488, EPI_ISL_442489, EPI_ISL_442490, EPI_ISL_442491, EPI_ISL_442492, EPI_ISL_442493, EPI_ISL_442494, EPI_ISL_442495, EPI_ISL_442496, EPI_ISL_442497, EPI_ISL_442498, EPI_ISL_442499, EPI_ISL_442500, EPI_ISL_442501, EPI_ISL_442502, EPI_ISL_442503, EPI_ISL_442504, EPI_ISL_442505, EPI_ISL_442506, EPI_ISL_442507, EPI_ISL_442508, EPI_ISL_442509, EPI_ISL_442510, EPI_ISL_442511, EPI_ISL_442512, EPI_ISL_442513, EPI_ISL_442514, EPI_ISL_442515, EPI_ISL_442516, EPI_ISL_442517, EPI_ISL_442518, EPI_ISL_442519, EPI_ISL_442520, EPI_ISL_442521, EPI_ISL_442522 | | | |
| see above | Virology Department, Sheffield Teaching Hospitals NHS Foundation Trust/Department of Infection, Immunity and Cardiovascular Disease, The Medical School, University of Sheffield | COVID-19 Genomics UK (COG-UK) Consortium | Thushan de Silva, Matthew Parker, Nikki Smith, Adri Angyal, Rebecca Brown, Luke Green, Rachel Tucker, Paul Parsons, Danielle Groves, Katie Johnson, Laura Carrilero, Alex Keeley, Dave Partridge, Matthew Wyles, Benjamin Lindsey, Mehmet Yavuz, Mohammad Raza, Cariad Evans |
| EPI_ISL_444022, EPI_ISL_445078, EPI_ISL_445079, EPI_ISL_445080, EPI_ISL_445081, EPI_ISL_445082, EPI_ISL_445083, EPI_ISL_445084 | Baylor College of Medicine | Baylor College of Medicine: HGSC | Vasanthi Avadhanula, Erin Nicholson, David Henke, Pedro Piedra, Harsha Doddapaneni, Donna Muzny, Qingchang Meng, Hsu Chao, Zeineen Momin, Hua Shen, George Weissenberger, Kavya Kottapalli, Yimiti Meiheerguli, Sejal Salvi, Ginger Metcalf, Vipin Menon, Sara J.J. Cregeen, Matthew C. Ross, Tulin Ayvaz, Richard Sucgang, Kristi L. Hoffman, Matthew Wong, Joseph F. Petrosino |
| EPI_ISL_445086 | Laboratory Diagnostic, Veterinary Specialized Institute Kraljevo | Laboratory Diagnostic, Veterinary Specialized Institute Kraljevo | Vidanovic,D., Tesovic,B., Sekler,M., Dmitric,M., Debeljak,Z., Matovic,K., Vaskovic,N., Petrovic,T., Volkening,J. and Alfonso,C.L. |
| EPI_ISL_445087 | Laboratory Diagnostic, Veterinary Specialized Institute Kraljevo | Laboratory Diagnostic, Veterinary Specialized Institute Kraljevo | Vidanovic,D., Tesovic,B., Sekler,M., Dmitric,M., Debeljak,Z., Matovic,K., Vaskovic,N., Petrovic,T., Volkening,J. and Alfonso,C. |
| EPI_ISL_447481, EPI_ISL_447495, EPI_ISL_447509, EPI_ISL_447522 | Servicio de Microbiologia. Hospital Clinico Universitario de Valencia | Sequencing and Bioinformatics Service and Molecular Epidemiology Research Group. FISABIO-Public Health | Vicente Soriano Chirona, Ivan Ansari, Llúcia Martínez-Priego, Giuseppe 'Auria, David Navarro, Eliseo Albert, Maria Alma Bracho, Lidia Ruiz Roldan, Neris Garcia-Gonzalez, Inma Galán Vendrell, Sandra Carbo, Loreto Ferrús Abad, Paula Ruiz-Hueso, Mariana Reyes-Prieto, Fernando Gonzalez-Candelas |
| EPI_ISL_447579, EPI_ISL_447580, EPI_ISL_447582, EPI_ISL_447583, EPI_ISL_447860, EPI_ISL_447861 | CSIR-Centre for Cellular and Molecular Biology | CSIR-Centre for Cellular and Molecular Biology | Tulasi Nagabandi, Namami Gaur, Sakshi Shambhavi, Lamuk Zaveri, Shagufta Khan, Purushotham Vodnala, Payel Mukherjee, Sofia Banu, Priya Singh, Dhiviya Vedagiri, Divya Gupta, Vishal Sah, Santosh Kumar Kuncha, Krishnan Harinivas Harshan, Archana Bharadwaj Siva, Karthik Bharadwaj Tallapaka, Rakesh K Mishra, Divya Tej Sowpati |
| EPI_ISL_448826, EPI_ISL_448827, EPI_ISL_448828, EPI_ISL_448829, EPI_ISL_448830, EPI_ISL_448831, EPI_ISL_448832, EPI_ISL_448833, EPI_ISL_448834, EPI_ISL_448835, EPI_ISL_448836, EPI_ISL_448837, EPI_ISL_448838, EPI_ISL_448839, EPI_ISL_448840, EPI_ISL_448841, EPI_ISL_448842, EPI_ISL_448843 | | | |
| see above | Virology Department, Sheffield Teaching Hospitals NHS Foundation Trust/Department of Infection, Immunity and Cardiovascular Disease, The Medical School, University of Sheffield | COVID-19 Genomics UK (COG-UK) Consortium | Thushan de Silva, Matthew Parker, Nikki Smith, Adri Angyal, Rebecca Brown, Luke Green, Rachel Tucker, Paul Parsons, Danielle Groves, Katie Johnson, Laura Carrilero, Alex Keeley, Dave Partridge, Matthew Wyles, Benjamin Lindsey, Mehmet Yavuz, Mohammad Raza, Cariad Evans |
| EPI_ISL_448844 | Virology Laboratory, Castle Hill Hospital, Hull University Teaching Hospitals NHS Trust/Department of Infection, Immunity and Cardiovascular Disease, The Medical School, University of Sheffield | COVID-19 Genomics UK (COG-UK) Consortium | Thushan de Silva, Matthew Parker, Nikki Smith, Adri Angyal, Rebecca Brown, Luke Green, Rachel Tucker, Paul Parsons, Danielle Groves, Katie Johnson, Laura Carrilero, Alex Keeley, Dave Partridge, Matthew Wyles, Benjamin Lindsey, Mehmet Yavuz, Mohammad Raza, Cariad Evans |
| EPI_ISL_448845, EPI_ISL_448846, EPI_ISL_448847, EPI_ISL_448848, EPI_ISL_448849 | Virology Department, Sheffield Teaching Hospitals NHS Foundation Trust/Department of Infection, Immunity and Cardiovascular Disease, The Medical School, University of Sheffield | COVID-19 Genomics UK (COG-UK) Consortium | Thushan de Silva, Matthew Parker, Nikki Smith, Adri Angyal, Rebecca Brown, Luke Green, Rachel Tucker, Paul Parsons, Danielle Groves, Katie Johnson, Laura Carrilero, Alex Keeley, Dave Partridge, Matthew Wyles, Benjamin Lindsey, Mehmet Yavuz, Mohammad Raza, Cariad Evans |
| EPI_ISL_448850 | Virology Laboratory, Castle Hill Hospital, Hull University Teaching Hospitals NHS Trust/Department of Infection, Immunity and Cardiovascular Disease, The Medical School, University of Sheffield | COVID-19 Genomics UK (COG-UK) Consortium | Thushan de Silva, Matthew Parker, Nikki Smith, Adri Angyal, Rebecca Brown, Luke Green, Rachel Tucker, Paul Parsons, Danielle Groves, Katie Johnson, Laura Carrilero, Alex Keeley, Dave Partridge, Matthew Wyles, Benjamin Lindsey, Mehmet Yavuz, Mohammad Raza, Cariad Evans |
| EPI_ISL_448851, EPI_ISL_448852, EPI_ISL_448853, EPI_ISL_448854, EPI_ISL_448855, EPI_ISL_448856, EPI_ISL_448857, EPI_ISL_448858, EPI_ISL_448859, EPI_ISL_448860, EPI_ISL_448861, EPI_ISL_448862, EPI_ISL_448863 | | | |
| see above | Virology Department, Sheffield Teaching Hospitals NHS Foundation Trust/Department of Infection, Immunity and Cardiovascular Disease, The Medical School, University of Sheffield | COVID-19 Genomics UK (COG-UK) Consortium | Thushan de Silva, Matthew Parker, Nikki Smith, Adri Angyal, Rebecca Brown, Luke Green, Rachel Tucker, Paul Parsons, Danielle Groves, Katie Johnson, Laura Carrilero, Alex Keeley, Dave Partridge, Matthew Wyles, Benjamin Lindsey, Mehmet Yavuz, Mohammad Raza, Cariad Evans |
| EPI_ISL_448864 | Virology Laboratory, Castle Hill Hospital, Hull University Teaching Hospitals NHS Trust/Department of Infection, | COVID-19 Genomics UK (COG-UK) Consortium | Thushan de Silva, Matthew Parker, Nikki Smith, Adri Angyal, Rebecca Brown, Luke Green, Rachel Tucker, Paul Parsons, Danielle Groves, Katie Johnson, Laura Carrilero, Alex Keeley, Dave Partridge, Matthew Wyles, Benjamin Lindsey, Mehmet Yavuz, Mohammad Raza, Cariad Evans |

| | | | |
|---|---|---|---|
| | Immunity and Cardiovascular Disease, The Medical School, University of Sheffield | | |
| EPI_ISL_448865 | Virology Department, Sheffield Teaching Hospitals NHS Foundation Trust/Department of Infection, Immunity and Cardiovascular Disease, The Medical School, University of Sheffield | COVID-19 Genomics UK (COG-UK) Consortium | Thushan de Silva, Matthew Parker, Nikki Smith, Adri Angyal, Rebecca Brown, Luke Green, Rachel Tucker, Paul Parsons, Danielle Groves, Katie Johnson, Laura Carrilero, Alex Keeley, Dave Partridge, Matthew Wyles, Benjamin Lindsey, Mehmet Yavuz, Mohammad Raza, Cariad Evans |
| EPI_ISL_448866 | Virology Laboratory, Castle Hill Hospital, Hull University Teaching Hospitals NHS Trust/Department of Infection, Immunity and Cardiovascular Disease, The Medical School, University of Sheffield | COVID-19 Genomics UK (COG-UK) Consortium | Thushan de Silva, Matthew Parker, Nikki Smith, Adri Angyal, Rebecca Brown, Luke Green, Rachel Tucker, Paul Parsons, Danielle Groves, Katie Johnson, Laura Carrilero, Alex Keeley, Dave Partridge, Matthew Wyles, Benjamin Lindsey, Mehmet Yavuz, Mohammad Raza, Cariad Evans |
| EPI_ISL_448867, EPI_ISL_448868, EPI_ISL_448869, EPI_ISL_448870, EPI_ISL_448871, EPI_ISL_448872, EPI_ISL_448873, EPI_ISL_448874, EPI_ISL_448875, EPI_ISL_448876, EPI_ISL_448877, EPI_ISL_448878, EPI_ISL_448879, EPI_ISL_448880, EPI_ISL_448881, EPI_ISL_448882, EPI_ISL_448883, EPI_ISL_448884, EPI_ISL_448885, EPI_ISL_448886, EPI_ISL_448887, EPI_ISL_448888, EPI_ISL_448889, EPI_ISL_448890, EPI_ISL_448891, EPI_ISL_448892, EPI_ISL_448893 | | | |
| see above | Virology Department, Sheffield Teaching Hospitals NHS Foundation Trust/Department of Infection, Immunity and Cardiovascular Disease, The Medical School, University of Sheffield | COVID-19 Genomics UK (COG-UK) Consortium | Thushan de Silva, Matthew Parker, Nikki Smith, Adri Angyal, Rebecca Brown, Luke Green, Rachel Tucker, Paul Parsons, Danielle Groves, Katie Johnson, Laura Carrilero, Alex Keeley, Dave Partridge, Matthew Wyles, Benjamin Lindsey, Mehmet Yavuz, Mohammad Raza, Cariad Evans |
| EPI_ISL_448894 | Virology Laboratory, Castle Hill Hospital, Hull University Teaching Hospitals NHS Trust/Department of Infection, Immunity and Cardiovascular Disease, The Medical School, University of Sheffield | COVID-19 Genomics UK (COG-UK) Consortium | Thushan de Silva, Matthew Parker, Nikki Smith, Adri Angyal, Rebecca Brown, Luke Green, Rachel Tucker, Paul Parsons, Danielle Groves, Katie Johnson, Laura Carrilero, Alex Keeley, Dave Partridge, Matthew Wyles, Benjamin Lindsey, Mehmet Yavuz, Mohammad Raza, Cariad Evans |
| EPI_ISL_448895 | Virology Department, Sheffield Teaching Hospitals NHS Foundation Trust/Department of Infection, Immunity and Cardiovascular Disease, The Medical School, University of Sheffield | COVID-19 Genomics UK (COG-UK) Consortium | Thushan de Silva, Matthew Parker, Nikki Smith, Adri Angyal, Rebecca Brown, Luke Green, Rachel Tucker, Paul Parsons, Danielle Groves, Katie Johnson, Laura Carrilero, Alex Keeley, Dave Partridge, Matthew Wyles, Benjamin Lindsey, Mehmet Yavuz, Mohammad Raza, Cariad Evans |
| EPI_ISL_448896 | Virology Laboratory, Castle Hill Hospital, Hull University Teaching Hospitals NHS Trust/Department of Infection, Immunity and Cardiovascular Disease, The Medical School, University of Sheffield | COVID-19 Genomics UK (COG-UK) Consortium | Thushan de Silva, Matthew Parker, Nikki Smith, Adri Angyal, Rebecca Brown, Luke Green, Rachel Tucker, Paul Parsons, Danielle Groves, Katie Johnson, Laura Carrilero, Alex Keeley, Dave Partridge, Matthew Wyles, Benjamin Lindsey, Mehmet Yavuz, Mohammad Raza, Cariad Evans |
| EPI_ISL_448897, EPI_ISL_448898, EPI_ISL_448899, EPI_ISL_448900, EPI_ISL_448901 | Virology Department, Sheffield Teaching Hospitals NHS Foundation Trust/Department of Infection, Immunity and Cardiovascular Disease, The Medical School, University of Sheffield | COVID-19 Genomics UK (COG-UK) Consortium | Thushan de Silva, Matthew Parker, Nikki Smith, Adri Angyal, Rebecca Brown, Luke Green, Rachel Tucker, Paul Parsons, Danielle Groves, Katie Johnson, Laura Carrilero, Alex Keeley, Dave Partridge, Matthew Wyles, Benjamin Lindsey, Mehmet Yavuz, Mohammad Raza, Cariad Evans |
| EPI_ISL_448902 | Virology Laboratory, Castle Hill Hospital, Hull University Teaching Hospitals NHS Trust/Department of Infection, Immunity and Cardiovascular Disease, The Medical School, University of Sheffield | COVID-19 Genomics UK (COG-UK) Consortium | Thushan de Silva, Matthew Parker, Nikki Smith, Adri Angyal, Rebecca Brown, Luke Green, Rachel Tucker, Paul Parsons, Danielle Groves, Katie Johnson, Laura Carrilero, Alex Keeley, Dave Partridge, Matthew Wyles, Benjamin Lindsey, Mehmet Yavuz, Mohammad Raza, Cariad Evans |
| EPI_ISL_448903, EPI_ISL_448904, EPI_ISL_448905 | Virology Department, Sheffield Teaching Hospitals NHS Foundation Trust/Department of Infection, Immunity and Cardiovascular Disease, The Medical School, University of Sheffield | COVID-19 Genomics UK (COG-UK) Consortium | Thushan de Silva, Matthew Parker, Nikki Smith, Adri Angyal, Rebecca Brown, Luke Green, Rachel Tucker, Paul Parsons, Danielle Groves, Katie Johnson, Laura Carrilero, Alex Keeley, Dave Partridge, Matthew Wyles, Benjamin Lindsey, Mehmet Yavuz, Mohammad Raza, Cariad Evans |
| EPI_ISL_448906 | Virology Laboratory, Castle Hill Hospital, Hull University Teaching Hospitals NHS Trust/Department of Infection, Immunity and Cardiovascular Disease, The Medical School, University of Sheffield | COVID-19 Genomics UK (COG-UK) Consortium | Thushan de Silva, Matthew Parker, Nikki Smith, Adri Angyal, Rebecca Brown, Luke Green, Rachel Tucker, Paul Parsons, Danielle Groves, Katie Johnson, Laura Carrilero, Alex Keeley, Dave Partridge, Matthew Wyles, Benjamin Lindsey, Mehmet Yavuz, Mohammad Raza, Cariad Evans |
| EPI_ISL_448907, EPI_ISL_448908, EPI_ISL_448909, EPI_ISL_448910, EPI_ISL_448911, EPI_ISL_448912, EPI_ISL_448913, EPI_ISL_448914, EPI_ISL_448915, EPI_ISL_448916, EPI_ISL_448917 | | | |
| see above | Virology Department, Sheffield Teaching Hospitals NHS Foundation Trust/Department of Infection, Immunity and Cardiovascular Disease, The Medical School, University of Sheffield | COVID-19 Genomics UK (COG-UK) Consortium | Thushan de Silva, Matthew Parker, Nikki Smith, Adri Angyal, Rebecca Brown, Luke Green, Rachel Tucker, Paul Parsons, Danielle Groves, Katie Johnson, Laura Carrilero, Alex Keeley, Dave Partridge, Matthew Wyles, Benjamin Lindsey, Mehmet Yavuz, Mohammad Raza, Cariad Evans |
| EPI_ISL_449476, EPI_ISL_449477, EPI_ISL_449478, EPI_ISL_449479, EPI_ISL_449480, EPI_ISL_449481, EPI_ISL_449482, EPI_ISL_449483, EPI_ISL_449484, EPI_ISL_449485, EPI_ISL_449487 | | | |
| see above | unknown | Department of Respiratory and Critical Care | Wang,X., Zhou,Q., He,Y., Liu,L., Ma,X., Wei,X., Jiang,N., Liang,L., Zheng,Y., Ma,L., Xu,Y., Yang,D., Zhang,J., Yang,B., Jiang,N., Zheng,Y., Ma,L., Xu,Y., Yang,D., Zhang,J., Yang,B., Jiang,N., Deng,T., Zhai,B., Gao,Y., Liu,W., Bai,X., Pan,T., Wang,G., Chang,Y., Zhang,Z., Shi,H., Ma,W.L. and Gao,Z. |
| EPI_ISL_450232, EPI_ISL_450233, EPI_ISL_450234, EPI_ISL_450235, EPI_ISL_450236, EPI_ISL_450237, EPI_ISL_450238, EPI_ISL_450239, EPI_ISL_450240 | UCSF Clinical Microbiology Laboratory | Chiu Laboratory, University of California, San Francisco | Xianding Deng, Scot Federman, Wei Gu, and Charles Y. Chiu |
| EPI_ISL_450331 | CSIR-Centre for Cellular and Molecular Biology | CSIR-Centre for Cellular and Molecular Biology | Tulasi Nagabandi, Namami Gaur, Sakshi Shambhavi, Lamuk Zaveri, Shagufta Khan, Purushotham Vodnala, Payel Mukherjee, Sofia Banu, Priya Singh, Dhiviya Vedagiri, Divya Gupta, Vishal Sah, Santosh Kumar Kuncha, Krishnan Harinivas Harshan, Archana Bharadwaj Siva, Karthik Bharadwaj Tallapaka,G. Aditya Kumar, Koushick Sivakumar, Pooja Ramesh Gupta, Rajan Kumar Jha, Shraddha Vijay Lahoti, Deepak Kumar, Devi Prasad Vijayashankara, Disha Nanda, Divya Das, Jotin Gogoi, Manish Bhattacharjee, Rakesh K Mishra, Divya Tej Sowpati |
| EPI_ISL_450408, EPI_ISL_450409, EPI_ISL_450410, EPI_ISL_450411, EPI_ISL_450412 | unknown | Microbiology | To,K.K.W., Yuen,K.-Y. |
| EPI_ISL_450835 | Unilabs Skovde | The Public Health Agency of Sweden | Tobias Kollberg, Helena Enroth, Anna-Malin Linde, Maria Lind Karlberg, Oskar Karlsson Lindsjo, Olov Svartstrom, Anna Risberg, Theresa Enkirch, Mia Brytting, Karin Tegmark-Wisell |
| EPI_ISL_451310, EPI_ISL_451311 | Hellenic Pasteur Institute, National Influenza Reference laboratory of Southern Greece & Unit of Bioinformatics and Applied Genomics | Hellenic Pasteur Institute, National Influenza Reference laboratory of Southern Greece & Unit of Bioinformatics and Applied Genomics | Vasiliki Pogka, Timokratis Karamitros, Athanasios Kossyvakis, Antonios Kalliaropoulos, Horefti Elina, Evangelidou Maria, Androniki Voulgari-Kokota, Aspasia Kontou, Andreas Mentis |
| EPI_ISL_452218, EPI_ISL_452219, EPI_ISL_452220, EPI_ISL_452221, EPI_ISL_452222, EPI_ISL_452223 | Goethe University Hospital Frankfurt | Institute for Medical Virology, Goethe University Hospital Frankfurt | Tuna Toptan, Sebastian Hoehl, Sandra Westhaus, Denisa Bojkova, Annemarie Berger, Björn Rotter, Klaus Hoffmeier, Jindrich Cinatl, Sandra Ciesek, and Marek Widera |
| EPI_ISL_452796, EPI_ISL_452797, EPI_ISL_452798, EPI_ISL_452799, EPI_ISL_452800, EPI_ISL_452801, EPI_ISL_452802, EPI_ISL_452803, EPI_ISL_452804, EPI_ISL_452805, EPI_ISL_452806, EPI_ISL_452807, EPI_ISL_452808, EPI_ISL_452809, EPI_ISL_452810, EPI_ISL_452811, EPI_ISL_452812, EPI_ISL_452813, EPI_ISL_452814, EPI_ISL_452815, EPI_ISL_452816, EPI_ISL_452817, EPI_ISL_452818, EPI_ISL_452819, EPI_ISL_452820, EPI_ISL_452821, EPI_ISL_452822, EPI_ISL_452823, EPI_ISL_452824, EPI_ISL_452825, EPI_ISL_452826, EPI_ISL_452827, EPI_ISL_452828, EPI_ISL_452829, EPI_ISL_452830, EPI_ISL_452831, EPI_ISL_452832, EPI_ISL_452833, EPI_ISL_452834, EPI_ISL_452835, EPI_ISL_452836, EPI_ISL_452837, EPI_ISL_452838, EPI_ISL_452839, EPI_ISL_452840, EPI_ISL_452841, EPI_ISL_452842, EPI_ISL_452843, EPI_ISL_452844, EPI_ISL_452845, EPI_ISL_452846, EPI_ISL_452847, EPI_ISL_452848, EPI_ISL_452849, EPI_ISL_452850, EPI_ISL_452851, EPI_ISL_452852, EPI_ISL_452853, EPI_ISL_452854, EPI_ISL_452855, EPI_ISL_452856 | | | |

| | | | |
|---|---|---|---|
| see above | Virginia DCLS | Virginia DCLS | Virginia DCLS |
| EPI_ISL_453696, EPI_ISL_453697, EPI_ISL_453698, EPI_ISL_453699, EPI_ISL_453700, EPI_ISL_453701, EPI_ISL_453702, EPI_ISL_453703, EPI_ISL_453704, EPI_ISL_453705, EPI_ISL_453706, EPI_ISL_453707, EPI_ISL_453708, EPI_ISL_453709, EPI_ISL_453710, EPI_ISL_453711, EPI_ISL_453712, EPI_ISL_453713, EPI_ISL_453714, EPI_ISL_453715, EPI_ISL_453716, EPI_ISL_453717, EPI_ISL_453718, EPI_ISL_453719, EPI_ISL_453720, EPI_ISL_453721, EPI_ISL_453722, EPI_ISL_453723, EPI_ISL_453724, EPI_ISL_453725, EPI_ISL_453726, EPI_ISL_453727, EPI_ISL_453728, EPI_ISL_453729, EPI_ISL_453730, EPI_ISL_453731, EPI_ISL_453732, EPI_ISL_453733, EPI_ISL_453734, EPI_ISL_453735, EPI_ISL_453736, EPI_ISL_453737, EPI_ISL_453738, EPI_ISL_453739, EPI_ISL_453740, EPI_ISL_453741, EPI_ISL_453742, EPI_ISL_453743, EPI_ISL_453744, EPI_ISL_453745, EPI_ISL_453746, EPI_ISL_453747, EPI_ISL_453748, EPI_ISL_453749, EPI_ISL_453750, EPI_ISL_453751, EPI_ISL_453752, EPI_ISL_453753, EPI_ISL_453754, EPI_ISL_453755, EPI_ISL_453756, EPI_ISL_453757, EPI_ISL_453758, EPI_ISL_453759, EPI_ISL_453760, EPI_ISL_453761, EPI_ISL_453762, EPI_ISL_453763, EPI_ISL_453764, EPI_ISL_453765, EPI_ISL_453766, EPI_ISL_453767, EPI_ISL_453768, EPI_ISL_453769, EPI_ISL_453770, EPI_ISL_453771, EPI_ISL_453772, EPI_ISL_453773, EPI_ISL_453774, EPI_ISL_453775, EPI_ISL_453776, EPI_ISL_453777, EPI_ISL_453778 | | | |
| see above | Virology Department, Sheffield Teaching Hospitals NHS Foundation Trust/Department of Infection, Immunity and Cardiovascular Disease, The Medical School, University of Sheffield | COVID-19 Genomics UK (COG-UK) Consortium | Thushan de Silva, Matthew Parker, Nikki Smith, Adri Angyal, Rebecca Brown, Luke Green, Rachel Tucker, Paul Parsons, Danielle Groves, Katie Johnson, Laura Carrilero, Alex Keeley, Dave Partridge, Matthew Wyles, Benjamin Lindsey, Mehmet Yavuz, Mohammad Raza, Cariad Evans |
| EPI_ISL_454749 | Japanese Quarantine Stations | Pathogen Genomics Center, National Institute of Infectious Diseases | Tsuyoshi Sekizuka, Kentaro Itokawa, Rina Tanaka, Masanori Hashino, Tsutomu Kageyama, Shinji Saito, Ikuyo Takayama, Hideki Hasegawa, Takuri Takahashi, Hajime Kamiya, Takuya Yamagishi, Motoi Suzuki, Takaji Wakita, Makoto Kuroda |
| EPI_ISL_454795 | Veterinary Specialized Institute Kraljevo | Veterinary Specialized Institute Kraljevo | Vidanovic,D., Tesovic,B., Sekler,M., Dmitric,M., Debeljak,Z., Matovic,K., Vaskovic,N., Petrovic,T., Volkening,J. and Alfonso,C.L. |
| EPI_ISL_455480 | Veterinary Specialized Institute Kraljevo | Veterinary Specialized Institute Kraljevo | Vidanovic,D., Tesovic,B., Banovic Djeri,B., Sekler,M., Dmitric,M., Debeljak,Z., Matovic,K., Vaskovic,N., Petrovic,T., Volkening,J. and Alfonso,C.L. |
| EPI_ISL_455723 | Servicio de Microbiologia. Hospital Clinico Universitario de Valencia | Sequencing and Bioinformatics Service and Molecular Epidemiology Research Group. FISABIO-Public Health, and SeqCOVID-Spain Consortium | Vicente Soriano Chirona, Ivan Ansari, Llúcia Martínez-Priego, Giuseppe 'Auria, David Navarro, Eliseo Albert, Maria Alma Bracho, Lidia Ruiz Roldan, Neris Garcia-Gonzalez, Inma Galán Vendrell, Sandra Carbo, Loreto Ferrús Abad, Paula Ruiz-Hueso, Mariana Reyes-Prieto, Fernando Gonzalez-Candelas |
| EPI_ISL_455735, EPI_ISL_455736 | Servicio de Microbiologia. Hospital_Arnau_de_Vilanova | Sequencing and Bioinformatics Service and Molecular Epidemiology Research Group. FISABIO-Public Health, and SeqCOVID-Spain Consortium | Victoria Dominguez, Maria Alma Bracho, Griselda De Marco, Lidia Ruiz Roldan, Neris Garcia-Gonzalez, Inma Galán Vendrell, Sandra Carbo, Loreto Ferrús Abad, Paula Ruiz-Hueso, Mariana Reyes-Prieto, Vicente Soriano Chirona, Ivan Ansari, Llúcia Martínez-Priego, Giuseppe 'Auria, Fernando Gonzalez-Candelas |
| EPI_ISL_457581, EPI_ISL_457582, EPI_ISL_457583, EPI_ISL_457584, EPI_ISL_457585, EPI_ISL_457586, EPI_ISL_457587, EPI_ISL_457588, EPI_ISL_457589, EPI_ISL_457590, EPI_ISL_457591, EPI_ISL_457592, EPI_ISL_457593, EPI_ISL_457594, EPI_ISL_457595, EPI_ISL_457596, EPI_ISL_457597, EPI_ISL_457598, EPI_ISL_457599, EPI_ISL_457600, EPI_ISL_457601, EPI_ISL_457602, EPI_ISL_457603, EPI_ISL_457604, EPI_ISL_457605, EPI_ISL_457606, EPI_ISL_457607, EPI_ISL_457608, EPI_ISL_457609, EPI_ISL_457610, EPI_ISL_457611, EPI_ISL_457612, EPI_ISL_457613, EPI_ISL_457614, EPI_ISL_457615, EPI_ISL_457616, EPI_ISL_457617, EPI_ISL_457618, EPI_ISL_457619, EPI_ISL_457620, EPI_ISL_457621, EPI_ISL_457622, EPI_ISL_457623, EPI_ISL_457624, EPI_ISL_457625, EPI_ISL_457626, EPI_ISL_457627, EPI_ISL_457628, EPI_ISL_457629, EPI_ISL_457630, EPI_ISL_457631, EPI_ISL_457632, EPI_ISL_457633, EPI_ISL_457634, EPI_ISL_457635, EPI_ISL_457636, EPI_ISL_457637, EPI_ISL_457638, EPI_ISL_457639, EPI_ISL_457640, EPI_ISL_457641, EPI_ISL_457642, EPI_ISL_457643, EPI_ISL_457644, EPI_ISL_457645, EPI_ISL_457646, EPI_ISL_457647, EPI_ISL_457648, EPI_ISL_457649, EPI_ISL_457650, EPI_ISL_457651, EPI_ISL_457652, EPI_ISL_457653, EPI_ISL_457654, EPI_ISL_457655, EPI_ISL_457656, EPI_ISL_457657, EPI_ISL_457658, EPI_ISL_457659, EPI_ISL_457660, EPI_ISL_457661, EPI_ISL_457662, EPI_ISL_457663, EPI_ISL_457664, EPI_ISL_457665, EPI_ISL_457666, EPI_ISL_457667, EPI_ISL_457668, EPI_ISL_457669, EPI_ISL_457670, EPI_ISL_457671, EPI_ISL_457672, EPI_ISL_457673, EPI_ISL_457674, EPI_ISL_457675, EPI_ISL_457676, EPI_ISL_457677, EPI_ISL_457678, EPI_ISL_457679, EPI_ISL_457680, EPI_ISL_457681, EPI_ISL_457682, EPI_ISL_457683, EPI_ISL_457684, EPI_ISL_457685, EPI_ISL_457686 | | | |
| see above | Virology Department, Sheffield Teaching Hospitals NHS Foundation Trust/Department of Infection, Immunity and Cardiovascular Disease, The Medical School, University of Sheffield | COVID-19 Genomics UK (COG-UK) Consortium | Thushan de Silva, Matthew Parker, Nikki Smith, Adri Angyal, Rebecca Brown, Luke Green, Rachel Tucker, Paul Parsons, Danielle Groves, Katie Johnson, Laura Carrilero, Alex Keeley, Dave Partridge, Matthew Wyles, Benjamin Lindsey, Mehmet Yavuz, Mohammad Raza, Cariad Evans |
| EPI_ISL_458050 | CSIR-Centre for Cellular and Molecular Biology | CSIR-Centre for Cellular and Molecular Biology | Tulasi Nagabandi, Namami Gaur, Sakshi Shambhavi, Lamuk Zaveri, Shagufta Khan, Purushotham Vodnala, Payel Mukherjee, Sofia Banu, Priya Singh, Dhiviya Vedagiri, Divya Gupta, Vishal Sah, Santosh Kumar Kuncha, Krishnan Harinivas Harshan, Archana Bharadwaj Siva, Karthik Bharadwaj Tallapaka,Kezia J Ann, Radhika Khandelwal, Roshan Maku Venkata, Shemin Mansuri, Sonu Uday, Rakesh K Mishra, Divya Tej Sowpati |
| EPI_ISL_458056 | CSIR-Centre for Cellular and Molecular Biology | CSIR-Centre for Cellular and Molecular Biology | Tulasi Nagabandi, Namami Gaur, Sakshi Shambhavi, Lamuk Zaveri, Shagufta Khan, Purushotham Vodnala, Payel Mukherjee, Sofia Banu, Priya Singh, Dhiviya Vedagiri, Divya Gupta, Vishal Sah, Santosh Kumar Kuncha, Krishnan Harinivas Harshan, Archana Bharadwaj Siva, Karthik Bharadwaj Tallapaka,G. Aditya Kumar, Koushick Sivakumar, Pooja Ramesh Gupta, Rajan Kumar Jha, Shraddha Vijay Lahoti, Rakesh K Mishra, Divya Tej Sowpati |
| EPI_ISL_458156, EPI_ISL_458157, EPI_ISL_458158, EPI_ISL_458159, EPI_ISL_458160, EPI_ISL_458161, EPI_ISL_458162, EPI_ISL_458163, EPI_ISL_458164, EPI_ISL_458165, EPI_ISL_458166, EPI_ISL_458167, EPI_ISL_458168, EPI_ISL_458169, EPI_ISL_458170, EPI_ISL_458171, EPI_ISL_458172, EPI_ISL_458173, EPI_ISL_458174, EPI_ISL_458175, EPI_ISL_458176, EPI_ISL_458177, EPI_ISL_458178, EPI_ISL_458179, EPI_ISL_458180, EPI_ISL_458181, EPI_ISL_458182, EPI_ISL_458183, EPI_ISL_458184, EPI_ISL_458185, EPI_ISL_458186, EPI_ISL_458187, EPI_ISL_458188, EPI_ISL_458189, EPI_ISL_458190, EPI_ISL_458191, EPI_ISL_458192, EPI_ISL_458193, EPI_ISL_458194, EPI_ISL_458195, EPI_ISL_458196, EPI_ISL_458197, EPI_ISL_458198, EPI_ISL_458199, EPI_ISL_458200, EPI_ISL_458201, EPI_ISL_458202, EPI_ISL_458203, EPI_ISL_458204, EPI_ISL_458205, EPI_ISL_458206, EPI_ISL_458207, EPI_ISL_458208, EPI_ISL_458209, EPI_ISL_458210, EPI_ISL_458211, EPI_ISL_458212, EPI_ISL_458213, EPI_ISL_458214, EPI_ISL_458215, EPI_ISL_458216, EPI_ISL_458217, EPI_ISL_458218, EPI_ISL_458219, EPI_ISL_458220, EPI_ISL_458221, EPI_ISL_458222, EPI_ISL_458223, EPI_ISL_458224, EPI_ISL_458225, EPI_ISL_458226, EPI_ISL_458227, EPI_ISL_458228, EPI_ISL_458229, EPI_ISL_458230, EPI_ISL_458231, EPI_ISL_458232, EPI_ISL_458233, EPI_ISL_458234, EPI_ISL_458235 | | | |
| see above | KU Leuven, Rega Institute, Clinical and Epidemiological Virology | KU Leuven, Rega Institute, Clinical and Epidemiological Virology | Tony Wawina-Bokalanga, Bert Vanmechelen, Joan Marti-Carerras, Piet Maes |
| EPI_ISL_461999, EPI_ISL_462000, EPI_ISL_462001, EPI_ISL_462002, EPI_ISL_462003, EPI_ISL_462004, EPI_ISL_462005, EPI_ISL_462006, EPI_ISL_462007, EPI_ISL_462008, EPI_ISL_462009, EPI_ISL_462010, EPI_ISL_462011, EPI_ISL_462012, EPI_ISL_462013, EPI_ISL_462014, EPI_ISL_462015, EPI_ISL_462016, EPI_ISL_462017, EPI_ISL_462018, EPI_ISL_462019, EPI_ISL_462020, EPI_ISL_462021, EPI_ISL_462022, EPI_ISL_462023, EPI_ISL_462024, EPI_ISL_462025, EPI_ISL_462026, EPI_ISL_462027, EPI_ISL_462028, EPI_ISL_462029, EPI_ISL_462030, EPI_ISL_462031, EPI_ISL_462032, EPI_ISL_462033, EPI_ISL_462034, EPI_ISL_462035, EPI_ISL_462036, EPI_ISL_462037, EPI_ISL_462038, EPI_ISL_462039, EPI_ISL_462040, EPI_ISL_462041, EPI_ISL_462042, EPI_ISL_462043, EPI_ISL_462044, EPI_ISL_462045, EPI_ISL_462046, EPI_ISL_462047, EPI_ISL_462048, EPI_ISL_462049, EPI_ISL_462050, EPI_ISL_462051, EPI_ISL_462052, EPI_ISL_462053, EPI_ISL_462054, EPI_ISL_462055, EPI_ISL_462056, EPI_ISL_462057, EPI_ISL_462058, EPI_ISL_462059, EPI_ISL_462060, EPI_ISL_462061, EPI_ISL_462062, EPI_ISL_462063, EPI_ISL_462064, EPI_ISL_462065, EPI_ISL_462066, EPI_ISL_462067, EPI_ISL_462068, EPI_ISL_462069, EPI_ISL_462070, EPI_ISL_462071, EPI_ISL_462072, EPI_ISL_462073, EPI_ISL_462074, EPI_ISL_462075, EPI_ISL_462076, EPI_ISL_462077, EPI_ISL_462078, EPI_ISL_462079, EPI_ISL_462080, EPI_ISL_462081, EPI_ISL_462082, EPI_ISL_462083, EPI_ISL_462084 | | | |
| see above | Virology Department, Sheffield Teaching Hospitals NHS Foundation Trust/Department of Infection, Immunity and Cardiovascular Disease, The Medical School, University of Sheffield | COVID-19 Genomics UK (COG-UK) Consortium | Thushan de Silva, Matthew Parker, Nikki Smith, Adri Angyal, Rebecca Brown, Luke Green, Rachel Tucker, Paul Parsons, Danielle Groves, Katie Johnson, Laura Carrilero, Alex Keeley, Dave Partridge, Matthew Wyles, Benjamin Lindsey, Mehmet Yavuz, Mohammad Raza, Cariad Evans |
| EPI_ISL_462151, EPI_ISL_462152, EPI_ISL_462153, EPI_ISL_462154, EPI_ISL_462155, EPI_ISL_462156, EPI_ISL_462157, EPI_ISL_462158, EPI_ISL_462159, EPI_ISL_462160, EPI_ISL_462161, EPI_ISL_462162, EPI_ISL_462163, EPI_ISL_462164, EPI_ISL_462165, EPI_ISL_462166, EPI_ISL_462167, EPI_ISL_462168, EPI_ISL_462169, EPI_ISL_462170, EPI_ISL_462171, EPI_ISL_462172, EPI_ISL_462173, EPI_ISL_462174, EPI_ISL_462175, EPI_ISL_462176, EPI_ISL_462177, EPI_ISL_462178, EPI_ISL_462179, EPI_ISL_462180, EPI_ISL_462181, EPI_ISL_462182, EPI_ISL_462183, EPI_ISL_462184, EPI_ISL_462185, EPI_ISL_462186, EPI_ISL_462187, EPI_ISL_462188, EPI_ISL_462189, EPI_ISL_462190, EPI_ISL_462191, EPI_ISL_462192, EPI_ISL_462193, EPI_ISL_462194, EPI_ISL_462195, EPI_ISL_462196, EPI_ISL_462197, EPI_ISL_462198, EPI_ISL_462199, EPI_ISL_462200, EPI_ISL_462201, EPI_ISL_462202, EPI_ISL_462203, EPI_ISL_462204, EPI_ISL_462205, EPI_ISL_462206, EPI_ISL_462207, EPI_ISL_462208, EPI_ISL_462209, EPI_ISL_462210, EPI_ISL_462211, EPI_ISL_462212, EPI_ISL_462213, EPI_ISL_462214, EPI_ISL_462215, EPI_ISL_462216, EPI_ISL_462217, EPI_ISL_462218, EPI_ISL_462219, EPI_ISL_462220, EPI_ISL_462221, EPI_ISL_462222, EPI_ISL_462223, EPI_ISL_462224, EPI_ISL_462225, EPI_ISL_462226, EPI_ISL_462227, EPI_ISL_462228, EPI_ISL_462229, EPI_ISL_462230, EPI_ISL_462231, EPI_ISL_462232, EPI_ISL_462233, EPI_ISL_462234, EPI_ISL_462235, EPI_ISL_462236, EPI_ISL_462237, EPI_ISL_462238, EPI_ISL_462239, EPI_ISL_462240, EPI_ISL_462241, EPI_ISL_462242, EPI_ISL_462243, EPI_ISL_462244, EPI_ISL_462245, EPI_ISL_462246, EPI_ISL_462247, EPI_ISL_462248, EPI_ISL_462249, EPI_ISL_462250, EPI_ISL_462251, EPI_ISL_462252, EPI_ISL_462253, EPI_ISL_462254, EPI_ISL_462255, EPI_ISL_462256, EPI_ISL_462257, EPI_ISL_462258, EPI_ISL_462259, EPI_ISL_462260, EPI_ISL_462261, EPI_ISL_462262, EPI_ISL_462263, EPI_ISL_462264, EPI_ISL_462265, EPI_ISL_462266, EPI_ISL_462267, EPI_ISL_462268, EPI_ISL_462269, EPI_ISL_462270, EPI_ISL_462271, EPI_ISL_462272, EPI_ISL_462273, EPI_ISL_462274, EPI_ISL_462275 | | | |
| see above | KU Leuven, Rega Institute, Clinical and Epidemiological Virology | KU Leuven, Rega Institute, Clinical and Epidemiological Virology | Tony Wawina-Bokalanga, Bert Vanmechelen, Joan Marti-Carerras, Piet Maes |
| EPI_ISL_462434, EPI_ISL_462435, EPI_ISL_462436, EPI_ISL_462437, EPI_ISL_462438 | unknown | Laboratory Diagnostic | Vidanovic,D., Tesovic,B., Banovic Djeri,B., Knezevic,A., Jankovic,M., Sekler,M., Dmitric,M., Petrovic,T., Volkening,J., Afonso,C.L. |
| EPI_ISL_462450, EPI_ISL_462451, EPI_ISL_462452, EPI_ISL_462453, EPI_ISL_462454, EPI_ISL_462455, EPI_ISL_462456, EPI_ISL_462457, EPI_ISL_462458, EPI_ISL_462459, EPI_ISL_462460, EPI_ISL_462461, EPI_ISL_462462, EPI_ISL_462463, EPI_ISL_462464, EPI_ISL_462465, EPI_ISL_462466, EPI_ISL_462467, EPI_ISL_462468, EPI_ISL_462469, EPI_ISL_462470, EPI_ISL_462471, EPI_ISL_462472, EPI_ISL_462473, EPI_ISL_462474, EPI_ISL_462475, EPI_ISL_462476 | | | |
| see above | Clinical Center, University of Sarajevo | Charite Universitatsmedizin Berlin, Institute of Virology | Victor M Corman, Jorn Beheim-Schwarzbach, Barbara Muehlemann, Talitha Veith, Julia Schneider, Terry Jones, Amela Dedeic-Ljubovic, Irma Salimovic-Besic, Suzana Arapcic, Almedina Hadzihasanovic-Moro, Selma Mutevelic, Christian Drosten |
| EPI_ISL_463094, EPI_ISL_463095, EPI_ISL_463096, EPI_ISL_463097, EPI_ISL_463098, EPI_ISL_463099, EPI_ISL_463100, EPI_ISL_463101, EPI_ISL_463102, EPI_ISL_463103, EPI_ISL_463104, EPI_ISL_463105, EPI_ISL_463106, EPI_ISL_463107, EPI_ISL_463108, EPI_ISL_463109, EPI_ISL_463110, EPI_ISL_463111, EPI_ISL_463112, EPI_ISL_463113, EPI_ISL_463114, EPI_ISL_463115, EPI_ISL_463116, EPI_ISL_463117, EPI_ISL_463118, EPI_ISL_463119, EPI_ISL_463120, EPI_ISL_463121, EPI_ISL_463122, EPI_ISL_463123, EPI_ISL_463124, EPI_ISL_463125, EPI_ISL_463126, EPI_ISL_463127, EPI_ISL_463128, EPI_ISL_463129, EPI_ISL_463130, EPI_ISL_463131, EPI_ISL_463132, EPI_ISL_463133, EPI_ISL_463134, EPI_ISL_463135, EPI_ISL_463136, EPI_ISL_463137 | | | |
| see above | Virginia DCLS | Virginia DCLS | Virginia DCLS |
| EPI_ISL_464065, EPI_ISL_464066, EPI_ISL_464067, EPI_ISL_464068, EPI_ISL_464069, EPI_ISL_464070, EPI_ISL_464071, EPI_ISL_464072, EPI_ISL_464073, EPI_ISL_464074, EPI_ISL_464075, EPI_ISL_464076, EPI_ISL_464077, EPI_ISL_464078, EPI_ISL_464079, EPI_ISL_464080, EPI_ISL_464081, EPI_ISL_464082, EPI_ISL_464083, EPI_ISL_464084, EPI_ISL_464085, EPI_ISL_464086, EPI_ISL_464087, EPI_ISL_464088, EPI_ISL_464089, EPI_ISL_464090 | | | |
| see above | KU Leuven, Rega Institute, Clinical and Epidemiological | KU Leuven, Rega Institute, Clinical and Epidemiological | Tony Wawina-Bokalanga, Bert Vanmechelen, Joan Marti-Carerras, Piet Maes |

| | | | |
|---|---|---|---|
| | Virology | Virology | |
| EPI_ISL_466652, EPI_ISL_466653, EPI_ISL_466654, EPI_ISL_466655, EPI_ISL_466656, EPI_ISL_466657, EPI_ISL_466658, EPI_ISL_466659, EPI_ISL_466660, EPI_ISL_466661, EPI_ISL_466662, EPI_ISL_466663, EPI_ISL_466664, EPI_ISL_466665, EPI_ISL_466666, EPI_ISL_466667, EPI_ISL_466668, EPI_ISL_466669, EPI_ISL_466670, EPI_ISL_466671, EPI_ISL_466672, EPI_ISL_466673, EPI_ISL_466674, EPI_ISL_466675, EPI_ISL_466676, EPI_ISL_466677, EPI_ISL_466678, EPI_ISL_466679, EPI_ISL_466680, EPI_ISL_466681, EPI_ISL_466682, EPI_ISL_466683, EPI_ISL_467298 | | | |
| see above | Nebraska Public Health Laboratory | UNMC COVID-19 Response Team | UNMC COVID-19 Response Team |
| EPI_ISL_467778, EPI_ISL_467779, EPI_ISL_467780, EPI_ISL_467781 | National Influenza Centre Romania | Charite Universitatsmedizin Berlin, Institute of Virology | Victor M Corman, Jorn Beheim-Schwarzbach, Barbara Muehlemann, Talitha Veith, Julia Schneider, Terry Jones, L. Ustea, N. Paraschiv, M. Lazar, Christian Drosten |
| EPI_ISL_467782, EPI_ISL_467783, EPI_ISL_467784, EPI_ISL_467785, EPI_ISL_467786, EPI_ISL_467787, EPI_ISL_467788, EPI_ISL_467789, EPI_ISL_467790, EPI_ISL_467791, EPI_ISL_467792, EPI_ISL_467793, EPI_ISL_467794, EPI_ISL_467795, EPI_ISL_467796, EPI_ISL_467797, EPI_ISL_467798, EPI_ISL_467799, EPI_ISL_467800, EPI_ISL_467801, EPI_ISL_467802, EPI_ISL_467803, EPI_ISL_467804, EPI_ISL_467805, EPI_ISL_467806, EPI_ISL_467807, EPI_ISL_467808 | | | |
| see above | Virginia DCLS | Virginia DCLS | Virginia DCLS |
| EPI_ISL_467809 | Cedars-Sinai Medical Center, Department of Pathology & Laboratory Medicine, Molecular Pathology Laboratory | Cedars-Sinai Medical Center, Molecular Pathology Laboratory of Department of Pathology & Laboratory Medicine and Genomic Core | Wenjuan Zhang, John Paul Govindavari, Brian Davis, Stephanie Chen, Jong Taek Kim, Jianbo Song, Jean Lopategui, Jasmine T Plummer, Eric Vail |
| EPI_ISL_467928, EPI_ISL_467929, EPI_ISL_467930, EPI_ISL_467931, EPI_ISL_467932, EPI_ISL_467933, EPI_ISL_467934, EPI_ISL_467935, EPI_ISL_467936, EPI_ISL_467937, EPI_ISL_467938, EPI_ISL_467939, EPI_ISL_467940, EPI_ISL_467941, EPI_ISL_467942, EPI_ISL_467943, EPI_ISL_467944 | | | |
| see above | Virginia DCLS | Virginia DCLS | Virginia DCLS |
| EPI_ISL_469254 | National Institute for Viral Disease Control and Prevention, China CDC | Institute of Viral Disease Control and Prevention, China CDC | Wenjie Tan, Lijuan Chen, Peihua NiuBaoying Huang, Li Zhao, Yubai Bi, Wenling Wang, Roujian Lu, Dayan Wang, Wenbo Xu, George Fu Gao, Chun Huang, Guizhen Wu |
| EPI_ISL_469255 | National Institute for Viral Disease Control and Prevention, China CDC | Institute of Viral Disease Control and Prevention, China CDC | Xiang ZhaoLijuan Chen, Dayan Wang, Yong Zhang, Yao MengZhixiao ChenYuchao Wu, Jun Han, Weifeng Shi, Yanhai Wang, William J. Liu, Shiwen Wang, George F. Gao, Wenbo Xu, Chun Huang, Guizhen Wu |
| EPI_ISL_469918, EPI_ISL_469919, EPI_ISL_469920, EPI_ISL_469921, EPI_ISL_469922, EPI_ISL_469923, EPI_ISL_469924, EPI_ISL_469925, EPI_ISL_469926, EPI_ISL_469927, EPI_ISL_469928, EPI_ISL_469929 | | | |
| see above | Virology Department, Sheffield Teaching Hospitals NHS Foundation Trust / Department of Infection, Immunity and Cardiovascular Disease, The Medical School, University of Sheffield | Wellcome Sanger Institute for the COVID-19 Genomics UK (COG-UK) consortium | Thushan de Silva, Matthew Parker,Adri Angyal, Rebecca Brown, Luke Green, Rachel Tucker, Paul Parsons, Danielle Groves, Alex Keeley, Dave Partridge, Matthew Wyles, Benjamin Lindsey, Mehmet Yavuz, Mohammad Raza, Cariad Evans and Alex Alderton, Roberto Amato, Sonia Goncalves, Ewan Harrison, David K. Jackson, Ian Johnston, Dominic Kwiatkowski, Cordelia Langford, John Sillitoe on behalf of the Wellcome Sanger Institute COVID-19 Surveillance Team (http://www.sanger.ac.uk/covid-team) |
| EPI_ISL_471611 | CSIR-Centre for Cellular and Molecular Biology | CSIR-Centre for Cellular and Molecular Biology | Tulasi Nagabandi, Namami Gaur, Sakshi Shambhavi, Lamuk Zaveri, Shagufta Khan, Purushotham Vodnala, Payel Mukherjee, Sofia Banu, Priya Singh, Dhiviya Vedagiri, Divya Gupta, Vishal Sah, Santosh Kumar Kuncha, Krishnan Harinivas Harshan, Archana Bharadwaj Siva, Karthik Bharadwaj Tallapaka,G. Aditya Kumar, Koushick Sivakumar, Pooja Ramesh Gupta, Rajan Kumar Jha, Shraddha Vijay Lahoti, Rakesh K Mishra, Divya Tej Sowpati |
| EPI_ISL_471612 | CSIR-Centre for Cellular and Molecular Biology | CSIR-Centre for Cellular and Molecular Biology | Tulasi Nagabandi, Namami Gaur, Sakshi Shambhavi, Lamuk Zaveri, Shagufta Khan, Purushotham Vodnala, Payel Mukherjee, Sofia Banu, Priya Singh, Dhiviya Vedagiri, Divya Gupta, Vishal Sah, Santosh Kumar Kuncha, Krishnan Harinivas Harshan, Archana Bharadwaj Siva, Karthik Bharadwaj Tallapaka,Kezia J Ann, Radhika Khandelwal, Roshan Maku Venkata, Shemin Mansuri, Sonu Uday, Rakesh K Mishra, Divya Tej Sowpati |
| EPI_ISL_471613 | CSIR-Centre for Cellular and Molecular Biology | CSIR-Centre for Cellular and Molecular Biology | Tulasi Nagabandi, Namami Gaur, Sakshi Shambhavi, Lamuk Zaveri, Shagufta Khan, Purushotham Vodnala, Payel Mukherjee, Sofia Banu, Priya Singh, Dhiviya Vedagiri, Divya Gupta, Vishal Sah, Santosh Kumar Kuncha, Krishnan Harinivas Harshan, Archana Bharadwaj Siva, Karthik Bharadwaj Tallapaka,G. Aditya Kumar, Koushick Sivakumar, Pooja Ramesh Gupta, Rajan Kumar Jha, Shraddha Vijay Lahoti, Rakesh K Mishra, Divya Tej Sowpati |
| EPI_ISL_471614 | CSIR-Centre for Cellular and Molecular Biology | CSIR-Centre for Cellular and Molecular Biology | Tulasi Nagabandi, Namami Gaur, Sakshi Shambhavi, Lamuk Zaveri, Shagufta Khan, Purushotham Vodnala, Payel Mukherjee, Sofia Banu, Priya Singh, Dhiviya Vedagiri, Divya Gupta, Vishal Sah, Santosh Kumar Kuncha, Krishnan Harinivas Harshan, Archana Bharadwaj Siva, Karthik Bharadwaj Tallapaka,Kezia J Ann, Radhika Khandelwal, Roshan Maku Venkata, Shemin Mansuri, Sonu Uday, Rakesh K Mishra, Divya Tej Sowpati |
| EPI_ISL_471639 | CSIR-Centre for Cellular and Molecular Biology | CSIR-Centre for Cellular and Molecular Biology | Tulasi Nagabandi, Namami Gaur, Sakshi Shambhavi, Lamuk Zaveri, Shagufta Khan, Purushotham Vodnala, Payel Mukherjee, Sofia Banu, Priya Singh, Dhiviya Vedagiri, Divya Gupta, Vishal Sah, Santosh Kumar Kuncha, Krishnan Harinivas Harshan, Archana Bharadwaj Siva, Karthik Bharadwaj Tallapaka,G. Aditya Kumar, Koushick Sivakumar, Pooja Ramesh Gupta, Rajan Kumar Jha, Shraddha Vijay Lahoti, Rakesh K Mishra, Divya Tej Sowpati |
| EPI_ISL_471640 | CSIR-Centre for Cellular and Molecular Biology | CSIR-Centre for Cellular and Molecular Biology | Tulasi Nagabandi, Namami Gaur, Sakshi Shambhavi, Lamuk Zaveri, Shagufta Khan, Purushotham Vodnala, Payel Mukherjee, Sofia Banu, Priya Singh, Dhiviya Vedagiri, Divya Gupta, Vishal Sah, Santosh Kumar Kuncha, Krishnan Harinivas Harshan, Archana Bharadwaj Siva, Karthik Bharadwaj Tallapaka,Kezia J Ann, Radhika Khandelwal, Roshan Maku Venkata, Shemin Mansuri, Sonu Uday, Rakesh K Mishra, Divya Tej Sowpati |
| EPI_ISL_471643 | CSIR-Centre for Cellular and Molecular Biology | CSIR-Centre for Cellular and Molecular Biology | Tulasi Nagabandi, Namami Gaur, Sakshi Shambhavi, Lamuk Zaveri, Shagufta Khan, Purushotham Vodnala, Payel Mukherjee, Sofia Banu, Priya Singh, Dhiviya Vedagiri, Divya Gupta, Vishal Sah, Santosh Kumar Kuncha, Krishnan Harinivas Harshan, Archana Bharadwaj Siva, Karthik Bharadwaj Tallapaka,G. Aditya Kumar, Koushick Sivakumar, Pooja Ramesh Gupta, Rajan Kumar Jha, Shraddha Vijay Lahoti, Rakesh K Mishra, Divya Tej Sowpati |
| EPI_ISL_471644 | CSIR-Centre for Cellular and Molecular Biology | CSIR-Centre for Cellular and Molecular Biology | Tulasi Nagabandi, Namami Gaur, Sakshi Shambhavi, Lamuk Zaveri, Shagufta Khan, Purushotham Vodnala, Payel Mukherjee, Sofia Banu, Priya Singh, Dhiviya Vedagiri, Divya Gupta, Vishal Sah, Santosh Kumar Kuncha, Krishnan Harinivas Harshan, Archana Bharadwaj Siva, Karthik Bharadwaj Tallapaka,Kezia J Ann, Radhika Khandelwal, Roshan Maku Venkata, Shemin Mansuri, Sonu Uday, Rakesh K Mishra, Divya Tej Sowpati |
| EPI_ISL_475027 | Banas Medical College and Research Institute | Gujarat Biotechnology Research Centre | Viren s Doshi, Pritesh Sabara, Apurvasinh Puvar, Janvi Raval, Zarna Patel, Monika Gandhi, Pinal Trivedi, Maharshi Pandya, Nidhi Patel, Nitin Savaliya, Raghawendra Kumar, Dinesh Kumar, Zuber Saiyed, Komal Patel, Labdhi Pandya, Snehal Bagatharia, Radhika Khara, Sunil R Joshi, Afzal Ansari, R D Dixit, A M Kadri, Harsh Bakshi, Chaitanya Joshi, Madhvi Joshi |
| EPI_ISL_475174, EPI_ISL_475175, EPI_ISL_475176, EPI_ISL_475177, EPI_ISL_475178, EPI_ISL_475179, EPI_ISL_475180, EPI_ISL_475181, EPI_ISL_475182, EPI_ISL_475183, EPI_ISL_475184, EPI_ISL_475185, EPI_ISL_475186, EPI_ISL_475187, EPI_ISL_475188, EPI_ISL_475189, EPI_ISL_475190, EPI_ISL_475191, EPI_ISL_475192, EPI_ISL_475193, EPI_ISL_475194, EPI_ISL_475195, EPI_ISL_475196, EPI_ISL_475197, EPI_ISL_475198, EPI_ISL_475199, EPI_ISL_475200, EPI_ISL_475201, EPI_ISL_475202, EPI_ISL_475203, EPI_ISL_475204, EPI_ISL_475205, EPI_ISL_475206, EPI_ISL_475207, EPI_ISL_475208, EPI_ISL_475209, EPI_ISL_475210, EPI_ISL_475211, EPI_ISL_475212, EPI_ISL_475213, EPI_ISL_475214, EPI_ISL_475215, EPI_ISL_475216, EPI_ISL_475217, EPI_ISL_475218, EPI_ISL_475219, EPI_ISL_475220, EPI_ISL_475221, EPI_ISL_475222, EPI_ISL_475223, EPI_ISL_475224, EPI_ISL_475225, EPI_ISL_475226, EPI_ISL_475227, EPI_ISL_475228, EPI_ISL_475229, EPI_ISL_475230, EPI_ISL_475231, EPI_ISL_475232, EPI_ISL_475233, EPI_ISL_475234, EPI_ISL_475235, EPI_ISL_475236, EPI_ISL_475237 | | | |
| see above | Nebraska Public Health Laboratory | UNMC COVID-19 Response Team | UNMC COVID-19 Response Team |
| EPI_ISL_475342, EPI_ISL_475343, EPI_ISL_475344, EPI_ISL_475345, EPI_ISL_475346, EPI_ISL_475347, EPI_ISL_475348, EPI_ISL_475349, EPI_ISL_475350, EPI_ISL_475351, EPI_ISL_475352, EPI_ISL_475353, EPI_ISL_475354, EPI_ISL_475355, EPI_ISL_475356, EPI_ISL_475357, EPI_ISL_475358, EPI_ISL_475359, EPI_ISL_475360, EPI_ISL_475361, EPI_ISL_475362, EPI_ISL_475363, EPI_ISL_475364, EPI_ISL_475365, EPI_ISL_475366, EPI_ISL_475367, EPI_ISL_475368, EPI_ISL_475369, EPI_ISL_475370, EPI_ISL_475371, EPI_ISL_475372, EPI_ISL_475373, EPI_ISL_475374, EPI_ISL_475375, EPI_ISL_475376, EPI_ISL_475377, EPI_ISL_475378, EPI_ISL_475379, EPI_ISL_475380, EPI_ISL_475381, EPI_ISL_475382, EPI_ISL_475383, EPI_ISL_475384, EPI_ISL_475385, EPI_ISL_475386, EPI_ISL_475387, EPI_ISL_475388, EPI_ISL_475389, EPI_ISL_475390, EPI_ISL_475391, EPI_ISL_475392, EPI_ISL_475393, EPI_ISL_475394, EPI_ISL_475395, EPI_ISL_475396, EPI_ISL_475397, EPI_ISL_475398, EPI_ISL_475399, EPI_ISL_475400, EPI_ISL_475401, EPI_ISL_475402, EPI_ISL_475403, EPI_ISL_475404, EPI_ISL_475405, EPI_ISL_475406, EPI_ISL_475407, EPI_ISL_475408, EPI_ISL_475409, EPI_ISL_475410, EPI_ISL_475411, EPI_ISL_475412, EPI_ISL_475413, EPI_ISL_475414, EPI_ISL_475415, EPI_ISL_475416, EPI_ISL_475417, EPI_ISL_475418, EPI_ISL_475419, EPI_ISL_475420, EPI_ISL_475421, EPI_ISL_475422, EPI_ISL_475423, EPI_ISL_475424, EPI_ISL_475425, EPI_ISL_475426, EPI_ISL_475427, EPI_ISL_475428, EPI_ISL_475429, EPI_ISL_475430, EPI_ISL_475431, EPI_ISL_475432, EPI_ISL_475433, EPI_ISL_475434, EPI_ISL_475435, EPI_ISL_475436, EPI_ISL_475437, EPI_ISL_475438, EPI_ISL_475439, EPI_ISL_475440, EPI_ISL_475441, EPI_ISL_475442, EPI_ISL_475443, EPI_ISL_475444, EPI_ISL_475445, EPI_ISL_475446, EPI_ISL_475447, EPI_ISL_475448, EPI_ISL_475449, EPI_ISL_475450, EPI_ISL_475451, EPI_ISL_475452, EPI_ISL_475453, EPI_ISL_475454, EPI_ISL_475455, EPI_ISL_475456, EPI_ISL_475457, EPI_ISL_475458, EPI_ISL_475459, EPI_ISL_475460, EPI_ISL_475461, EPI_ISL_475462, EPI_ISL_475463, EPI_ISL_475464, EPI_ISL_475465, EPI_ISL_475466, EPI_ISL_475467, EPI_ISL_475468, EPI_ISL_475469, EPI_ISL_475470, EPI_ISL_475471, EPI_ISL_475472, EPI_ISL_475473, EPI_ISL_475474, EPI_ISL_475475, EPI_ISL_475476, EPI_ISL_475477, EPI_ISL_475478, EPI_ISL_475479, EPI_ISL_475480, EPI_ISL_475481, EPI_ISL_475482, EPI_ISL_475483, EPI_ISL_475484, EPI_ISL_475485, EPI_ISL_475486, EPI_ISL_475487, EPI_ISL_475488, EPI_ISL_475489, EPI_ISL_475490, EPI_ISL_475491, EPI_ISL_475492, EPI_ISL_475493, EPI_ISL_475494, EPI_ISL_475495, EPI_ISL_475496, EPI_ISL_475497, EPI_ISL_475498, EPI_ISL_475499, EPI_ISL_475500, EPI_ISL_475501, EPI_ISL_475502, EPI_ISL_475503, EPI_ISL_475504, EPI_ISL_475505, EPI_ISL_475506, EPI_ISL_475507, EPI_ISL_475508, EPI_ISL_475509, EPI_ISL_475510 | | | |
| see above | Virology Department, Sheffield Teaching Hospitals NHS Foundation Trust/Department of Infection, Immunity and Cardiovascular Disease, The Medical School, University of Sheffield | COVID-19 Genomics UK (COG-UK) Consortium | Thushan de Silva, Matthew Parker, Nikki Smith, Adri Angyal, Rebecca Brown, Luke Green, Rachel Tucker, Paul Parsons, Danielle Groves, Katie Johnson, Laura Carrilero, Alex Keeley, Dave Partridge, Matthew Wyles, Benjamin Lindsey, Mehmet Yavuz, Mohammad Raza, Cariad Evans |
| EPI_ISL_475574, EPI_ISL_475575, EPI_ISL_475576, EPI_ISL_475577, EPI_ISL_475578, EPI_ISL_475579, EPI_ISL_475580, EPI_ISL_475581, EPI_ISL_475582, EPI_ISL_475583, EPI_ISL_475584, EPI_ISL_475585, EPI_ISL_475586, EPI_ISL_475587, EPI_ISL_475588, EPI_ISL_475589, EPI_ISL_475590, EPI_ISL_475591, EPI_ISL_475592, EPI_ISL_475593, EPI_ISL_475594, EPI_ISL_475595, EPI_ISL_475596, EPI_ISL_475597, EPI_ISL_475598, EPI_ISL_475599, EPI_ISL_475600, EPI_ISL_475601, EPI_ISL_475602, EPI_ISL_475603, EPI_ISL_475604, EPI_ISL_475605, EPI_ISL_475606, EPI_ISL_475607, EPI_ISL_475608, EPI_ISL_475609, EPI_ISL_475610, EPI_ISL_475611, EPI_ISL_475612, EPI_ISL_475613, EPI_ISL_475614, EPI_ISL_475615, EPI_ISL_475616, EPI_ISL_475617, EPI_ISL_475618, EPI_ISL_475619, EPI_ISL_475620, EPI_ISL_475621, EPI_ISL_475622, EPI_ISL_475623, EPI_ISL_475624, EPI_ISL_475625, EPI_ISL_475626, EPI_ISL_475627, EPI_ISL_475628, EPI_ISL_475629, EPI_ISL_475630, EPI_ISL_475631, EPI_ISL_475632, EPI_ISL_475633, EPI_ISL_475634, EPI_ISL_475635, EPI_ISL_475636, EPI_ISL_475637, EPI_ISL_475638, EPI_ISL_475639, EPI_ISL_475640, EPI_ISL_475641, EPI_ISL_475642, EPI_ISL_475643, EPI_ISL_475644, EPI_ISL_475645, EPI_ISL_475646, EPI_ISL_475647, EPI_ISL_475648, EPI_ISL_475649, EPI_ISL_475650, EPI_ISL_475651, EPI_ISL_475652, EPI_ISL_475653, EPI_ISL_475654, EPI_ISL_475655, EPI_ISL_475656, EPI_ISL_475657, EPI_ISL_475658, EPI_ISL_475659, EPI_ISL_475660, EPI_ISL_475661, EPI_ISL_475662, EPI_ISL_475663, | | | |

| | | | |
|---|---|---|---|
| EPI_ISL_475664, EPI_ISL_475665, EPI_ISL_475666, EPI_ISL_475667, EPI_ISL_475668, EPI_ISL_475669, EPI_ISL_475670, EPI_ISL_475671, EPI_ISL_475672, EPI_ISL_475673, EPI_ISL_475674, EPI_ISL_475675, EPI_ISL_475676, EPI_ISL_475677, EPI_ISL_475678, EPI_ISL_475679, EPI_ISL_475680, EPI_ISL_475681, EPI_ISL_475682, EPI_ISL_475683, EPI_ISL_475684, EPI_ISL_475685, EPI_ISL_475686, EPI_ISL_475687, EPI_ISL_475688, EPI_ISL_475689, EPI_ISL_475690, EPI_ISL_475691, EPI_ISL_475692, EPI_ISL_475693, EPI_ISL_475694, EPI_ISL_475695, EPI_ISL_475696, EPI_ISL_475697, EPI_ISL_475698, EPI_ISL_475699, EPI_ISL_475700, EPI_ISL_475701, EPI_ISL_475702, EPI_ISL_475703, EPI_ISL_475704, EPI_ISL_475705, EPI_ISL_475706, EPI_ISL_475707, EPI_ISL_475708, EPI_ISL_475709, EPI_ISL_475710, EPI_ISL_475711, EPI_ISL_475712, EPI_ISL_475713, EPI_ISL_475714, EPI_ISL_475715, EPI_ISL_475716 | | | |
| see above | Cedars-Sinai Medical Center, Department of Pathology & Laboratory Medicine, Molecular Pathology Laboratory | Cedars-Sinai Medical Center, Molecular Pathology Laboratory of Department of Pathology & Laboratory Medicine and Genomic Core | Wenjuan Zhang, John Paul Govindavari, Brian Davis, Stephanie Chen, Jong Taek Kim, Jianbo Song, Jean Lopategui, Jasmine T Plummer, Eric Vail |
| EPI_ISL_476705 | Labor Kneißler GmbH & Co. KG | Heinrich Pette Institute, Leibniz Institute for Experimental Virology | Thomas Günther, Adam Grundhoff, Manja Czech-Sioli, Nicole Fischer, Matthias Ottinger, Melanie M. Brinkmann |
| EPI_ISL_476941, EPI_ISL_476942, EPI_ISL_476943, EPI_ISL_476944, EPI_ISL_476945, EPI_ISL_476946, EPI_ISL_476947, EPI_ISL_476948, EPI_ISL_476949, EPI_ISL_476950, EPI_ISL_476951, EPI_ISL_476952, EPI_ISL_476953, EPI_ISL_476954, EPI_ISL_476955, EPI_ISL_476956, EPI_ISL_476957, EPI_ISL_476958, EPI_ISL_476959, EPI_ISL_476960, EPI_ISL_476961, EPI_ISL_476962, EPI_ISL_476963, EPI_ISL_476964, EPI_ISL_476965, EPI_ISL_476966, EPI_ISL_476967, EPI_ISL_476968, EPI_ISL_476969, EPI_ISL_476970, EPI_ISL_476971, EPI_ISL_476972, EPI_ISL_476973, EPI_ISL_476974, EPI_ISL_476975, EPI_ISL_476976, EPI_ISL_476977, EPI_ISL_476978, EPI_ISL_476979, EPI_ISL_476980, EPI_ISL_476981, EPI_ISL_476982, EPI_ISL_476983, EPI_ISL_476984, EPI_ISL_476985, EPI_ISL_476986, EPI_ISL_476987, EPI_ISL_476988, EPI_ISL_476989, EPI_ISL_476990, EPI_ISL_476991, EPI_ISL_476992, EPI_ISL_476993, EPI_ISL_476994, EPI_ISL_476995, EPI_ISL_476996, EPI_ISL_476997, EPI_ISL_476998, EPI_ISL_476999, EPI_ISL_477000, EPI_ISL_477001, EPI_ISL_477002, EPI_ISL_477003, EPI_ISL_477004, EPI_ISL_477005, EPI_ISL_477006, EPI_ISL_477007 | | | |
| see above | KU Leuven, Rega Institute, Clinical and Epidemiological Virology | KU Leuven, Rega Institute, Clinical and Epidemiological Virology | Tony Wawina-Bokalanga, Joan Marti-Carerras, Bert Vanmechelen, Piet Maes |
| EPI_ISL_477204 | Prof. Massimo Zollo CEINGE TASK-FORCE COVID19 - Regione Campania | Prof. Massimo Zollo CEINGE TASK-FORCE COVID19 - Regione Campania | Veronica Ferrucci1,2, Dae young Kong8, Fatemeh asadzadeh1,2, Laura Marrone1,2, Roberto Siciliano1,2, Rino Cerino3, Giovanna Fusco3, Marika Comegna1,2, Angelo Boccia2, Maurizio Viscardi3, Giorgia Borriello3, Sergio Brandi3, Claudia Tiberio4, Luigi Atripaldi4, Giovanni Paolella1,2, Giuseppe Castaldo1,2, Stefano Pascarella4, Martina Bianchi4, Lorenzo Chiariotti1,2, Jae Myun Lee5, Jae Ho Jung6, Kyong Seop Yun7, Hong Yeoul Kim 7,8* and Massimo Zollo1,2* 1 CEINGE Biotecnologie Avanzate, Naples, Italia 2 Dipartimento di Medicina Molecolare e Biotecnologie Mediche DMMBM University of Naples Federico II, Italia 3 Istituto Zooprofilattico Sperimentale del Mezzogiorno, Naples, Italia 4 -U.O.C. di Patologia Clinica Ospedale D. Cotugno, Azienda Sanitaria Ospedali dei Colli, Naples, Italy. 5 Università La Sapienza di Roma, Italia 6 Department of Microbiology, Yonsei University College of Medicine, Seoul, Korea 7 Department of Surgery, Yonsei University College of Medicine, Seoul, Korea 8 Haim bio co., Ltd, , Indust |
| EPI_ISL_477631, EPI_ISL_477632, EPI_ISL_477633, EPI_ISL_477634, EPI_ISL_477635, EPI_ISL_477636, EPI_ISL_477637, EPI_ISL_477638, EPI_ISL_477639, EPI_ISL_477640, EPI_ISL_477641, EPI_ISL_477642, EPI_ISL_477643, EPI_ISL_477644, EPI_ISL_477645, EPI_ISL_477646, EPI_ISL_477647, EPI_ISL_477648, EPI_ISL_477649, EPI_ISL_477650, EPI_ISL_477651, EPI_ISL_477652, EPI_ISL_477653, EPI_ISL_477654, EPI_ISL_477655, EPI_ISL_477656, EPI_ISL_477657, EPI_ISL_477658, EPI_ISL_477659, EPI_ISL_477660, EPI_ISL_477661, EPI_ISL_477662, EPI_ISL_477663, EPI_ISL_477664, EPI_ISL_477665, EPI_ISL_477666, EPI_ISL_477667, EPI_ISL_477668, EPI_ISL_477669, EPI_ISL_477670, EPI_ISL_477671, EPI_ISL_477672 | | | |
| see above | Virginia DCLS | Virginia DCLS | Virginia DCLS |
| EPI_ISL_479195, EPI_ISL_479196, EPI_ISL_479197, EPI_ISL_479198, EPI_ISL_479199, EPI_ISL_479200, EPI_ISL_479201, EPI_ISL_479202, EPI_ISL_479203, EPI_ISL_479204, EPI_ISL_479205, EPI_ISL_479206, EPI_ISL_479207, EPI_ISL_479208, EPI_ISL_479209, EPI_ISL_479210, EPI_ISL_479211, EPI_ISL_479212, EPI_ISL_479213, EPI_ISL_479214, EPI_ISL_479215, EPI_ISL_479216, EPI_ISL_479217, EPI_ISL_479218, EPI_ISL_479219, EPI_ISL_479220, EPI_ISL_479221, EPI_ISL_479222, EPI_ISL_479223, EPI_ISL_479224, EPI_ISL_479225, EPI_ISL_479226, EPI_ISL_479227, EPI_ISL_479228, EPI_ISL_479229, EPI_ISL_479230, EPI_ISL_479231, EPI_ISL_479232, EPI_ISL_479233, EPI_ISL_479234, EPI_ISL_479235, EPI_ISL_479236, EPI_ISL_479237, EPI_ISL_479238, EPI_ISL_479239, EPI_ISL_479240, EPI_ISL_479241, EPI_ISL_479242, EPI_ISL_479243, EPI_ISL_479244, EPI_ISL_479245, EPI_ISL_479246, EPI_ISL_479247, EPI_ISL_479248, EPI_ISL_479249, EPI_ISL_479250, EPI_ISL_479251, EPI_ISL_479252, EPI_ISL_479253, EPI_ISL_479254, EPI_ISL_479255, EPI_ISL_479256, EPI_ISL_479257, EPI_ISL_479258, EPI_ISL_479259, EPI_ISL_479260, EPI_ISL_479261, EPI_ISL_479262, EPI_ISL_479263, EPI_ISL_479264, EPI_ISL_479265, EPI_ISL_479266, EPI_ISL_479267, EPI_ISL_479268, EPI_ISL_479269, EPI_ISL_479270, EPI_ISL_479271, EPI_ISL_479272, EPI_ISL_479273, EPI_ISL_479274, EPI_ISL_479275, EPI_ISL_479276, EPI_ISL_479277, EPI_ISL_479278, EPI_ISL_479279, EPI_ISL_479280, EPI_ISL_479281, EPI_ISL_479282, EPI_ISL_479283 | | | |
| see above | Virology Department, Sheffield Teaching Hospitals NHS Foundation Trust/Department of Infection, Immunity and Cardiovascular Disease, The Medical School, University of Sheffield | COVID-19 Genomics UK (COG-UK) Consortium | Thushan de Silva, Matthew Parker, Nikki Smith, Adri Angyal, Rebecca Brown, Luke Green, Rachel Tucker, Paul Parsons, Danielle Groves, Katie Johnson, Laura Carrilero, Alex Keeley, Dave Partridge, Matthew Wyles, Benjamin Lindsey, Mehmet Yavuz, Mohammad Raza, Cariad Evans |
| EPI_ISL_479797, EPI_ISL_479798 | Sagamihara City Public Health Research Institute | Pathogen Genomics Center, National Institute of Infectious Diseases | Tsuyoshi Sekizuka, Hiroshi Nakamura, Kentaro Itokawa, Rina Tanaka, Masanori Hashino, Hajime Kamiya, Motoi Suzuki, Makoto Kuroda |
| EPI_ISL_479799, EPI_ISL_479800 | Sapporo City Institute of Public Health | Pathogen Genomics Center, National Institute of Infectious Diseases | Tsuyoshi Sekizuka, Asami Ohnishi, Kentaro Itokawa, Rina Tanaka, Masanori Hashino, Hajime Kamiya, Motoi Suzuki, Makoto Kuroda |
| EPI_ISL_479802, EPI_ISL_479803, EPI_ISL_479804 | Sagamihara City Public Health Research Institute | Pathogen Genomics Center, National Institute of Infectious Diseases | Tsuyoshi Sekizuka, Hiroshi Nakamura, Kentaro Itokawa, Rina Tanaka, Masanori Hashino, Hajime Kamiya, Motoi Suzuki, Makoto Kuroda |
| EPI_ISL_479805, EPI_ISL_479806, EPI_ISL_479807, EPI_ISL_479808 | Saitama Prefectural Institute of Public Health | Pathogen Genomics Center, National Institute of Infectious Diseases | Tsuyoshi Sekizuka, Hayato Ehara, Kentaro Itokawa, Rina Tanaka, Masanori Hashino, Hajime Kamiya, Motoi Suzuki, Makoto Kuroda |
| EPI_ISL_479812, EPI_ISL_479813, EPI_ISL_479814, EPI_ISL_479815, EPI_ISL_479816, EPI_ISL_479817, EPI_ISL_479818, EPI_ISL_479819, EPI_ISL_479820 | Hokkaido Institute of Public Health | Pathogen Genomics Center, National Institute of Infectious Diseases | Tsuyoshi Sekizuka, Rika Komagome, Kentaro Itokawa, Rina Tanaka, Masanori Hashino, Hajime Kamiya, Motoi Suzuki, Makoto Kuroda |
| EPI_ISL_479821, EPI_ISL_479822 | Department of Infectious Diseases, Kobe Institute of Health | Pathogen Genomics Center, National Institute of Infectious Diseases | Tsuyoshi Sekizuka, Ryohei Nomoto, Kentaro Itokawa, Rina Tanaka, Masanori Hashino, Hajime Kamiya, Motoi Suzuki, Makoto Kuroda |
| EPI_ISL_479823 | Kochi Prefectural Institute of Public Health | Pathogen Genomics Center, National Institute of Infectious Diseases | Tsuyoshi Sekizuka, Akihiko Tokaji, Kentaro Itokawa, Rina Tanaka, Masanori Hashino, Hajime Kamiya, Motoi Suzuki, Makoto Kuroda |
| EPI_ISL_479824 | Kumamoto PrefecturalInstituteof Public-Health and Environmental Science | Pathogen Genomics Center, National Institute of Infectious Diseases | Tsuyoshi Sekizuka, Shunsuke Yahiro, Kentaro Itokawa, Rina Tanaka, Masanori Hashino, Hajime Kamiya, Motoi Suzuki, Makoto Kuroda |
| EPI_ISL_479826, EPI_ISL_479827, EPI_ISL_479828, EPI_ISL_479829, EPI_ISL_479830, EPI_ISL_479831, EPI_ISL_479832, EPI_ISL_479833, EPI_ISL_479834, EPI_ISL_479835, EPI_ISL_479836, EPI_ISL_479837, EPI_ISL_479838, EPI_ISL_479839, EPI_ISL_479840, EPI_ISL_479841, EPI_ISL_479842, EPI_ISL_479843, EPI_ISL_479844, EPI_ISL_479845, EPI_ISL_479846, EPI_ISL_479847, EPI_ISL_479848, EPI_ISL_479849 | | | |
| see above | Sapporo City Institute of Public Health | Pathogen Genomics Center, National Institute of Infectious Diseases | Tsuyoshi Sekizuka, Asami Ohnishi, Kentaro Itokawa, Rina Tanaka, Masanori Hashino, Hajime Kamiya, Motoi Suzuki, Makoto Kuroda |
| EPI_ISL_479850, EPI_ISL_479851, EPI_ISL_479852, EPI_ISL_479853, EPI_ISL_479854 | Gunma Prefectural Institute of Public Health and Environmental Sciences | Pathogen Genomics Center, National Institute of Infectious Diseases | Tsuyoshi Sekizuka, Hiroyuki Tsukagoshi, Kentaro Itokawa, Rina Tanaka, Masanori Hashino, Hajime Kamiya, Motoi Suzuki, Makoto Kuroda |
| EPI_ISL_479855, EPI_ISL_479856, EPI_ISL_479857, EPI_ISL_479858, EPI_ISL_479859, EPI_ISL_479860, EPI_ISL_479861 | Department of Infectious Diseases, Kobe Institute of Health | Pathogen Genomics Center, National Institute of Infectious Diseases | Tsuyoshi Sekizuka, Ryohei Nomoto, Kentaro Itokawa, Rina Tanaka, Masanori Hashino, Hajime Kamiya, Motoi Suzuki, Makoto Kuroda |
| EPI_ISL_479862, EPI_ISL_479863, EPI_ISL_479864 | Wakayama Prefectural Research Center of Environment and Public Health | Pathogen Genomics Center, National Institute of Infectious Diseases | Tsuyoshi Sekizuka, Fumio Terasoma, Yosuke Hamajima, Kentaro Itokawa, Rina Tanaka, Masanori Hashino, Hajime Kamiya, Motoi Suzuki, Makoto Kuroda |
| EPI_ISL_479868 | Department of Infectious Diseases, Kobe Institute of Health | Pathogen Genomics Center, National Institute of Infectious Diseases | Tsuyoshi Sekizuka, Ryohei Nomoto, Kentaro Itokawa, Rina Tanaka, Masanori Hashino, Hajime Kamiya, Motoi Suzuki, Makoto Kuroda |
| EPI_ISL_479869 | Niigata Prefectural Institute of Public Health and Environmental Sciences | Pathogen Genomics Center, National Institute of Infectious Diseases | Tsuyoshi Sekizuka, Reiko Arai, Kentaro Itokawa, Rina Tanaka, Masanori Hashino, Hajime Kamiya, Motoi Suzuki, Makoto Kuroda |
| EPI_ISL_479870, EPI_ISL_479871 | Sagamihara City Public Health Research Institute | Pathogen Genomics Center, National Institute of Infectious Diseases | Tsuyoshi Sekizuka, Hiroshi Nakamura, Kentaro Itokawa, Rina Tanaka, Masanori Hashino, Hajime Kamiya, Motoi Suzuki, Makoto Kuroda |
| EPI_ISL_479872, EPI_ISL_479873, EPI_ISL_479874, EPI_ISL_479875, EPI_ISL_479876, EPI_ISL_479877, EPI_ISL_479878, EPI_ISL_479879, EPI_ISL_479880, EPI_ISL_479881, EPI_ISL_479882, EPI_ISL_479883, EPI_ISL_479884, EPI_ISL_479885 | | | |
| see above | Sapporo City Institute of Public Health | Pathogen Genomics Center, National Institute of Infectious | Tsuyoshi Sekizuka, Asami Ohnishi, Kentaro Itokawa, Rina Tanaka, Masanori Hashino, Hajime Kamiya, Motoi Suzuki, Makoto Kuroda |

| | | | |
|---|---|---|---|
| | | Diseases | |
| EPI_ISL_479886, EPI_ISL_479887, EPI_ISL_479888, EPI_ISL_479889, EPI_ISL_479890, EPI_ISL_479891, EPI_ISL_479892, EPI_ISL_479893, EPI_ISL_479894, EPI_ISL_479895 | Tokyo Metropolitan Institute of Public Health | Pathogen Genomics Center, National Institute of Infectious Diseases | Tsuyoshi Sekizuka, Kenji Sadamasu, Takashi Chiba, Mami Nagashima, Kentaro Itokawa, Rina Tanaka, Masanori Hashino, Hajime Kamiya, Motoi Suzuki, Makoto Kuroda |
| EPI_ISL_479896, EPI_ISL_479897, EPI_ISL_479898, EPI_ISL_479899, EPI_ISL_479900, EPI_ISL_479901 | Gunma Prefectural Institute of Public Health and Environmental Sciences | Pathogen Genomics Center, National Institute of Infectious Diseases | Tsuyoshi Sekizuka, Hiroyuki Tsukagoshi, Kentaro Itokawa, Rina Tanaka, Masanori Hashino, Hajime Kamiya, Motoi Suzuki, Makoto Kuroda |
| EPI_ISL_479902 | Niigata Prefectural Institute of Public Health and Environmental Sciences | Pathogen Genomics Center, National Institute of Infectious Diseases | Tsuyoshi Sekizuka, Reiko Arai, Kentaro Itokawa, Rina Tanaka, Masanori Hashino, Hajime Kamiya, Motoi Suzuki, Makoto Kuroda |
| EPI_ISL_479903, EPI_ISL_479904, EPI_ISL_479905, EPI_ISL_479906, EPI_ISL_479907, EPI_ISL_479908, EPI_ISL_479909, EPI_ISL_479910, EPI_ISL_479911, EPI_ISL_479912 | Himeji City Institute of Environment and Health | Pathogen Genomics Center, National Institute of Infectious Diseases | Tsuyoshi Sekizuka, Kentaro Itokawa, Rina Tanaka, Masanori Hashino, Hajime Kamiya, Motoi Suzuki, Makoto Kuroda |
| EPI_ISL_479913, EPI_ISL_479914, EPI_ISL_479915, EPI_ISL_479916, EPI_ISL_479917, EPI_ISL_479918, EPI_ISL_479919, EPI_ISL_479920, EPI_ISL_479921, EPI_ISL_479922, EPI_ISL_479923, EPI_ISL_479924 | | | |
| see above | Niigata City Public Health Research Institute | Pathogen Genomics Center, National Institute of Infectious Diseases | Tsuyoshi Sekizuka, Yurie Takahashi, Kentaro Itokawa, Rina Tanaka, Masanori Hashino, Hajime Kamiya, Motoi Suzuki, Makoto Kuroda |
| EPI_ISL_479925, EPI_ISL_479926, EPI_ISL_479927 | Sakai City Institute of Public Health | Pathogen Genomics Center, National Institute of Infectious Diseases | Tsuyoshi Sekizuka, Tatsuya Miyoshi, Kentaro Itokawa, Rina Tanaka, Masanori Hashino, Hajime Kamiya, Motoi Suzuki, Makoto Kuroda |
| EPI_ISL_479928, EPI_ISL_479929, EPI_ISL_479930, EPI_ISL_479931, EPI_ISL_479932, EPI_ISL_479933, EPI_ISL_479934, EPI_ISL_479935 | Saitama Prefectural Institute of Public Health | Pathogen Genomics Center, National Institute of Infectious Diseases | Tsuyoshi Sekizuka, Hayato Ehara, Kentaro Itokawa, Rina Tanaka, Masanori Hashino, Hajime Kamiya, Motoi Suzuki, Makoto Kuroda |
| EPI_ISL_479936, EPI_ISL_479937, EPI_ISL_479938, EPI_ISL_479939, EPI_ISL_479940, EPI_ISL_479941, EPI_ISL_479942, EPI_ISL_479943 | Ibaraki Prefectural Institute of Public Health | Pathogen Genomics Center, National Institute of Infectious Diseases | Tsuyoshi Sekizuka, Keiko Goto, Kentaro Itokawa, Rina Tanaka, Masanori Hashino, Hajime Kamiya, Motoi Suzuki, Makoto Kuroda |
| EPI_ISL_479959, EPI_ISL_479960, EPI_ISL_479961, EPI_ISL_479962, EPI_ISL_479963, EPI_ISL_479964, EPI_ISL_479965 | Tokyo Metropolitan Institute of Public Health | Pathogen Genomics Center, National Institute of Infectious Diseases | Tsuyoshi Sekizuka, Kenji Sadamasu, Takashi Chiba, Mami Nagashima, Kentaro Itokawa, Rina Tanaka, Masanori Hashino, Hajime Kamiya, Motoi Suzuki, Makoto Kuroda |
| EPI_ISL_479967, EPI_ISL_479968, EPI_ISL_479969, EPI_ISL_479970, EPI_ISL_479971, EPI_ISL_479972, EPI_ISL_479973, EPI_ISL_479974, EPI_ISL_479975, EPI_ISL_479976, EPI_ISL_479977, EPI_ISL_479978 | | | |
| see above | Fukui Prefectural Institute of Public Health and Environmental Science | Pathogen Genomics Center, National Institute of Infectious Diseases | Tsuyoshi Sekizuka, Miho Toho, Kentaro Itokawa, Rina Tanaka, Masanori Hashino, Hajime Kamiya, Motoi Suzuki, Makoto Kuroda |
| EPI_ISL_479979, EPI_ISL_479980, EPI_ISL_479981, EPI_ISL_479982, EPI_ISL_479983, EPI_ISL_479984 | Oita Prefectural Institute of Public Health and Environmental Science | Pathogen Genomics Center, National Institute of Infectious Diseases | Tsuyoshi Sekizuka, Mari Sasaki, Kentaro Itokawa, Rina Tanaka, Masanori Hashino, Hajime Kamiya, Motoi Suzuki, Makoto Kuroda |
| EPI_ISL_479985 | Ibaraki Prefectural Institute of Public Health | Pathogen Genomics Center, National Institute of Infectious Diseases | Tsuyoshi Sekizuka, Keiko Goto, Kentaro Itokawa, Rina Tanaka, Masanori Hashino, Hajime Kamiya, Motoi Suzuki, Makoto Kuroda |
| EPI_ISL_479986, EPI_ISL_479987, EPI_ISL_479988, EPI_ISL_479989 | Department of Infectious Diseases, Kobe Institute of Health | Pathogen Genomics Center, National Institute of Infectious Diseases | Tsuyoshi Sekizuka, Ryohei Nomoto, Kentaro Itokawa, Rina Tanaka, Masanori Hashino, Hajime Kamiya, Motoi Suzuki, Makoto Kuroda |
| EPI_ISL_479990 | Kitakyushu City Institute of Health and Environmental Sciences | Pathogen Genomics Center, National Institute of Infectious Diseases | Tsuyoshi Sekizuka, Katsuya Obata, Asuka Kikuchi Kentaro Itokawa, Rina Tanaka, Masanori Hashino, Hajime Kamiya, Motoi Suzuki, Makoto Kuroda |
| EPI_ISL_479991, EPI_ISL_479992, EPI_ISL_479993, EPI_ISL_479994, EPI_ISL_479995, EPI_ISL_479996 | Kumamoto City Public Health Research Institute | Pathogen Genomics Center, National Institute of Infectious Diseases | Tsuyoshi Sekizuka, Kaori Tashiro, Kentaro Itokawa, Rina Tanaka, Masanori Hashino, Hajime Kamiya, Motoi Suzuki, Makoto Kuroda |
| EPI_ISL_479997, EPI_ISL_479998, EPI_ISL_479999, EPI_ISL_480000, EPI_ISL_480001 | Nagano Environmental Conservation Research Institute | Pathogen Genomics Center, National Institute of Infectious Diseases | Tsuyoshi Sekizuka, Naoko Shimodaira, Kentaro Itokawa, Rina Tanaka, Masanori Hashino, Hajime Kamiya, Motoi Suzuki, Makoto Kuroda |
| EPI_ISL_480002, EPI_ISL_480003 | Nagasaki Prefectural Institute for Environmental Research and Public Health | Pathogen Genomics Center, National Institute of Infectious Diseases | Tsuyoshi Sekizuka, Fumiaki Matsumoto, Kentaro Itokawa, Rina Tanaka, Masanori Hashino, Hajime Kamiya, Motoi Suzuki, Makoto Kuroda |
| EPI_ISL_480004, EPI_ISL_480005, EPI_ISL_480006, EPI_ISL_480007, EPI_ISL_480008, EPI_ISL_480009, EPI_ISL_480010, EPI_ISL_480011, EPI_ISL_480012, EPI_ISL_480013, EPI_ISL_480014 | | | |
| see above | Chiba Prefectural Institute of Public Health | Pathogen Genomics Center, National Institute of Infectious Diseases | Tsuyoshi Sekizuka, Masakatsu Taira, Kentaro Itokawa, Rina Tanaka, Masanori Hashino, Hajime Kamiya, Motoi Suzuki, Makoto Kuroda |
| EPI_ISL_480015, EPI_ISL_480016, EPI_ISL_480017, EPI_ISL_480018, EPI_ISL_480019, EPI_ISL_480020 | Gunma Prefectural Institute of Public Health and Environmental Sciences | Pathogen Genomics Center, National Institute of Infectious Diseases | Tsuyoshi Sekizuka, Hiroyuki Tsukagoshi, Kentaro Itokawa, Rina Tanaka, Masanori Hashino, Hajime Kamiya, Motoi Suzuki, Makoto Kuroda |
| EPI_ISL_480021, EPI_ISL_480022, EPI_ISL_480023, EPI_ISL_480024, EPI_ISL_480025, EPI_ISL_480026, EPI_ISL_480027, EPI_ISL_480028, EPI_ISL_480029 | Ibaraki Prefectural Institute of Public Health | Pathogen Genomics Center, National Institute of Infectious Diseases | Tsuyoshi Sekizuka, Keiko Goto, Kentaro Itokawa, Rina Tanaka, Masanori Hashino, Hajime Kamiya, Motoi Suzuki, Makoto Kuroda |
| EPI_ISL_480030, EPI_ISL_480031, EPI_ISL_480032, EPI_ISL_480033, EPI_ISL_480034, EPI_ISL_480035, EPI_ISL_480036, EPI_ISL_480037, EPI_ISL_480038, EPI_ISL_480039, EPI_ISL_480040, EPI_ISL_480041 | | | |
| see above | Tochigi Prefectural Institute of Public Health and Environmental Science | Pathogen Genomics Center, National Institute of Infectious Diseases | Tsuyoshi Sekizuka, Ako Nakajima, Kentaro Itokawa, Rina Tanaka, Masanori Hashino, Hajime Kamiya, Motoi Suzuki, Makoto Kuroda |
| EPI_ISL_480042, EPI_ISL_480043, EPI_ISL_480044, EPI_ISL_480045, EPI_ISL_480046, EPI_ISL_480047, EPI_ISL_480048, EPI_ISL_480049, EPI_ISL_480050, EPI_ISL_480051, EPI_ISL_480052, EPI_ISL_480053, EPI_ISL_480054, EPI_ISL_480055, EPI_ISL_480056, EPI_ISL_480057, EPI_ISL_480058, EPI_ISL_480059, EPI_ISL_480060, EPI_ISL_480061, EPI_ISL_480062, EPI_ISL_480063, EPI_ISL_480064 | | | |
| see above | Nagoya City Public Health Research Institute | Pathogen Genomics Center, National Institute of Infectious Diseases | Tsuyoshi Sekizuka, Takuya Miki, Shinichiro Shibata, Kentaro Itokawa, Rina Tanaka, Masanori Hashino, Hajime Kamiya, Motoi Suzuki, Makoto Kuroda |
| EPI_ISL_480065, EPI_ISL_480066, EPI_ISL_480067, EPI_ISL_480068, | Sakai City Institute of Public Health | Pathogen Genomics Center, National Institute of Infectious Diseases | Tsuyoshi Sekizuka, Tatsuya Miyoshi, Kentaro Itokawa, Rina Tanaka, Masanori Hashino, Hajime Kamiya, Motoi Suzuki, Makoto Kuroda |

| | | | |
|---|---|---|---|
| EPI_ISL_480069, EPI_ISL_480070, EPI_ISL_480071, EPI_ISL_480072 | | | |
| EPI_ISL_480073 | Tochigi Prefectural Institute of Public Health and Environmental Science | Pathogen Genomics Center, National Institute of Infectious Diseases | Tsuyoshi Sekizuka, Ako Nakajima, Kentaro Itokawa, Rina Tanaka, Masanori Hashino, Hajime Kamiya, Motoi Suzuki, Makoto Kuroda |
| EPI_ISL_480074, EPI_ISL_480075, EPI_ISL_480076, EPI_ISL_480077, EPI_ISL_480078, EPI_ISL_480079, EPI_ISL_480080, EPI_ISL_480081, EPI_ISL_480082 | Shizuoka City Institute of Environmental Sciences and Public Health | Pathogen Genomics Center, National Institute of Infectious Diseases | Tsuyoshi Sekizuka, Takaharu Maehata,Sou Okamura,Yuji Kanazawa,Kenji Yagi, Kentaro Itokawa, Rina Tanaka, Masanori Hashino, Hajime Kamiya, Motoi Suzuki, Makoto Kuroda |
| EPI_ISL_480083, EPI_ISL_480084, EPI_ISL_480085, EPI_ISL_480086, EPI_ISL_480087, EPI_ISL_480088, EPI_ISL_480089 | Gifu Prefectural Institute of Public Health and Environmental Sciences | Pathogen Genomics Center, National Institute of Infectious Diseases | Tsuyoshi Sekizuka, Yoshihiko Kameyama, Kentaro Itokawa, Rina Tanaka, Masanori Hashino, Hajime Kamiya, Motoi Suzuki, Makoto Kuroda |
| EPI_ISL_480090, EPI_ISL_480091, EPI_ISL_480092, EPI_ISL_480093, EPI_ISL_480094, EPI_ISL_480095, EPI_ISL_480096, EPI_ISL_480097, EPI_ISL_480098, EPI_ISL_480099, EPI_ISL_480100, EPI_ISL_480101, EPI_ISL_480102 | | | |
| see above | Department of Infectious Diseases, Kobe Institute of Health | Pathogen Genomics Center, National Institute of Infectious Diseases | Tsuyoshi Sekizuka, Ryohei Nomoto, Kentaro Itokawa, Rina Tanaka, Masanori Hashino, Hajime Kamiya, Motoi Suzuki, Makoto Kuroda |
| EPI_ISL_480103, EPI_ISL_480104, EPI_ISL_480105, EPI_ISL_480106, EPI_ISL_480107, EPI_ISL_480108 | Koshigaya City Public Health Center | Pathogen Genomics Center, National Institute of Infectious Diseases | Tsuyoshi Sekizuka, Yuka Furui, Aya Tamura, Kyohei Sakata, Takumi Daimon, Yoko Togawa, Yoshiko Hamada, Kentaro Itokawa, Rina Tanaka, Masanori Hashino, Hajime Kamiya, Motoi Suzuki, Makoto Kuroda |
| EPI_ISL_480109, EPI_ISL_480110, EPI_ISL_480111, EPI_ISL_480112, EPI_ISL_480113, EPI_ISL_480114, EPI_ISL_480115, EPI_ISL_480116, EPI_ISL_480117, EPI_ISL_480118, EPI_ISL_480119 | | | |
| see above | Oita Prefectural Institute of Public Health and Environmental Science | Pathogen Genomics Center, National Institute of Infectious Diseases | Tsuyoshi Sekizuka, Mari Sasaki, Kentaro Itokawa, Rina Tanaka, Masanori Hashino, Hajime Kamiya, Motoi Suzuki, Makoto Kuroda |
| EPI_ISL_480120, EPI_ISL_480121, EPI_ISL_480122, EPI_ISL_480123, EPI_ISL_480124, EPI_ISL_480125, EPI_ISL_480126, EPI_ISL_480127, EPI_ISL_480128, EPI_ISL_480129, EPI_ISL_480130, EPI_ISL_480131, EPI_ISL_480132, EPI_ISL_480133, EPI_ISL_480134, EPI_ISL_480135, EPI_ISL_480136, EPI_ISL_480137, EPI_ISL_480138, EPI_ISL_480139, EPI_ISL_480140, EPI_ISL_480141, EPI_ISL_480142, EPI_ISL_480143, EPI_ISL_480144, EPI_ISL_480145, EPI_ISL_480146, EPI_ISL_480147, EPI_ISL_480148, EPI_ISL_480149, EPI_ISL_480150, EPI_ISL_480151, EPI_ISL_480152, EPI_ISL_480153, EPI_ISL_480154, EPI_ISL_480155, EPI_ISL_480156, EPI_ISL_480157, EPI_ISL_480158, EPI_ISL_480159, EPI_ISL_480160, EPI_ISL_480161, EPI_ISL_480162, EPI_ISL_480163, EPI_ISL_480164, EPI_ISL_480165, EPI_ISL_480166, EPI_ISL_480167, EPI_ISL_480168 | | | |
| see above | Fukui Prefectural Institute of Public Health and Environmental Science | Pathogen Genomics Center, National Institute of Infectious Diseases | Tsuyoshi Sekizuka, Miho Toho, Kentaro Itokawa, Rina Tanaka, Masanori Hashino, Hajime Kamiya, Motoi Suzuki, Makoto Kuroda |
| EPI_ISL_480169, EPI_ISL_480170, EPI_ISL_480171, EPI_ISL_480172, EPI_ISL_480173, EPI_ISL_480174, EPI_ISL_480175, EPI_ISL_480176, EPI_ISL_480177 | Gunma Prefectural Institute of Public Health and Environmental Sciences | Pathogen Genomics Center, National Institute of Infectious Diseases | Tsuyoshi Sekizuka, Hiroyuki Tsukagoshi, Kentaro Itokawa, Rina Tanaka, Masanori Hashino, Hajime Kamiya, Motoi Suzuki, Makoto Kuroda |
| EPI_ISL_480178, EPI_ISL_480179 | Hiroshima City Institute of Public Health | Pathogen Genomics Center, National Institute of Infectious Diseases | Tsuyoshi Sekizuka, Kota Noritsune, Kentaro Itokawa, Rina Tanaka, Masanori Hashino, Hajime Kamiya, Motoi Suzuki, Makoto Kuroda |
| EPI_ISL_480180, EPI_ISL_480181, EPI_ISL_480182, EPI_ISL_480183, EPI_ISL_480184, EPI_ISL_480185, EPI_ISL_480186, EPI_ISL_480187, EPI_ISL_480188, EPI_ISL_480189 | Ibaraki Prefectural Institute of Public Health | Pathogen Genomics Center, National Institute of Infectious Diseases | Tsuyoshi Sekizuka, Keiko Goto, Kentaro Itokawa, Rina Tanaka, Masanori Hashino, Hajime Kamiya, Motoi Suzuki, Makoto Kuroda |
| EPI_ISL_480196, EPI_ISL_480197, EPI_ISL_480198, EPI_ISL_480199, EPI_ISL_480200, EPI_ISL_480201, EPI_ISL_480202, EPI_ISL_480203 | Toyama Institute of Health | Pathogen Genomics Center, National Institute of Infectious Diseases | Tsuyoshi Sekizuka, Masae Itamochi, Kazunori Oishi, Kentaro Itokawa, Rina Tanaka, Masanori Hashino, Hajime Kamiya, Motoi Suzuki, Makoto Kuroda |
| EPI_ISL_480204 | Akita City Public Health Center | Pathogen Genomics Center, National Institute of Infectious Diseases | Tsuyoshi Sekizuka, Koichi Ito, Kentaro Itokawa, Rina Tanaka, Masanori Hashino, Hajime Kamiya, Motoi Suzuki, Makoto Kuroda |
| EPI_ISL_480205, EPI_ISL_480206, EPI_ISL_480207 | Department of Infectious Diseases, Kobe Institute of Health | Pathogen Genomics Center, National Institute of Infectious Diseases | Tsuyoshi Sekizuka, Ryohei Nomoto, Kentaro Itokawa, Rina Tanaka, Masanori Hashino, Hajime Kamiya, Motoi Suzuki, Makoto Kuroda |
| EPI_ISL_480208 | Pathogen Genomics Center, National Institute of Infectious Diseases | Pathogen Genomics Center, National Institute of Infectious Diseases | Tsuyoshi Sekizuka, Ryohei Nomoto, Kentaro Itokawa, Rina Tanaka, Masanori Hashino, Hajime Kamiya, Motoi Suzuki, Makoto Kuroda |
| EPI_ISL_480209, EPI_ISL_480210, EPI_ISL_480211, EPI_ISL_480212, EPI_ISL_480213, EPI_ISL_480214, EPI_ISL_480215, EPI_ISL_480216, EPI_ISL_480217, EPI_ISL_480218, EPI_ISL_480219, EPI_ISL_480220 | | | |
| see above | Department of Infectious Diseases, Kobe Institute of Health | Pathogen Genomics Center, National Institute of Infectious Diseases | Tsuyoshi Sekizuka, Ryohei Nomoto, Kentaro Itokawa, Rina Tanaka, Masanori Hashino, Hajime Kamiya, Motoi Suzuki, Makoto Kuroda |
| EPI_ISL_480221, EPI_ISL_480222, EPI_ISL_480223 | Koshigaya City Public Health Center | Pathogen Genomics Center, National Institute of Infectious Diseases | Tsuyoshi Sekizuka, Yuka Furui, Aya Tamura, Kyohei Sakata, Takumi Daimon, Yoko Togawa, Yoshiko Hamada, Kentaro Itokawa, Rina Tanaka, Masanori Hashino, Hajime Kamiya, Motoi Suzuki, Makoto Kuroda |
| EPI_ISL_480225 | Fukui Prefectural Institute of Public Health and Environmental Science | Pathogen Genomics Center, National Institute of Infectious Diseases | Tsuyoshi Sekizuka, Miho Toho, Kentaro Itokawa, Rina Tanaka, Masanori Hashino, Hajime Kamiya, Motoi Suzuki, Makoto Kuroda |
| EPI_ISL_480226 | Niigata Prefectural Institute of Public Health and Environmental Sciences | Pathogen Genomics Center, National Institute of Infectious Diseases | Tsuyoshi Sekizuka, Reiko Arai, Kentaro Itokawa, Rina Tanaka, Masanori Hashino, Hajime Kamiya, Motoi Suzuki, Makoto Kuroda |
| EPI_ISL_480227 | Tokyo Metropolitan Institute of Public Health | Pathogen Genomics Center, National Institute of Infectious Diseases | Tsuyoshi Sekizuka, Kenji Sadamasu, Takashi Chiba, Mami Nagashima, Kentaro Itokawa, Rina Tanaka, Masanori Hashino, Hajime Kamiya, Motoi Suzuki, Makoto Kuroda |
| EPI_ISL_482744 | Laboratory Diagnostic, Veterinary Specialized Institute Kraljevo | Laboratory Diagnostic, Veterinary Specialized Institute Kraljevo | Vidanovic,D., Tesovic,B., Banovic Djeri,B., Knezevic,A., Vidanovic,D., Tesovic,B., Banovic Djeri,B., Knezevic,A., Afonso,C. |
| EPI_ISL_484433, EPI_ISL_484434, EPI_ISL_484435, EPI_ISL_484436, EPI_ISL_484437, EPI_ISL_484438, EPI_ISL_484439, EPI_ISL_484440, EPI_ISL_484441, EPI_ISL_484442, EPI_ISL_484443, EPI_ISL_484444, EPI_ISL_484445, EPI_ISL_484446, EPI_ISL_484447, EPI_ISL_484448, EPI_ISL_484449, EPI_ISL_484450, EPI_ISL_484451, EPI_ISL_484452, EPI_ISL_484453, EPI_ISL_484454, EPI_ISL_484455, EPI_ISL_484456, EPI_ISL_484457, EPI_ISL_484458, EPI_ISL_484459, EPI_ISL_484460, EPI_ISL_484461, EPI_ISL_484462, EPI_ISL_484463, EPI_ISL_484464, EPI_ISL_484465, EPI_ISL_484466, EPI_ISL_484467, EPI_ISL_484468, EPI_ISL_484469, EPI_ISL_484470, EPI_ISL_484471, EPI_ISL_484472, EPI_ISL_484473, EPI_ISL_484474, EPI_ISL_484475, EPI_ISL_484476, EPI_ISL_484477, EPI_ISL_484478, EPI_ISL_484479, EPI_ISL_484480, EPI_ISL_484481, EPI_ISL_484482, EPI_ISL_484483, EPI_ISL_484484, EPI_ISL_484485, EPI_ISL_484486, EPI_ISL_484487, EPI_ISL_484488, EPI_ISL_484489, EPI_ISL_484490, EPI_ISL_484491, EPI_ISL_484492, EPI_ISL_484493, EPI_ISL_484494, EPI_ISL_484495, EPI_ISL_484496, EPI_ISL_484497, EPI_ISL_484498, EPI_ISL_484499, EPI_ISL_484500, EPI_ISL_484501, EPI_ISL_484502, EPI_ISL_484503, EPI_ISL_484504, EPI_ISL_484505, EPI_ISL_484506, EPI_ISL_484507, EPI_ISL_484508, EPI_ISL_484509, EPI_ISL_484510, EPI_ISL_484511, EPI_ISL_484512, EPI_ISL_484513, EPI_ISL_484514, EPI_ISL_484515, EPI_ISL_484516, EPI_ISL_484517, EPI_ISL_484518 | | | |
| see above | Virology Department, Sheffield Teaching Hospitals NHS Foundation Trust/Department of Infection, Immunity and Cardiovascular Disease, The Medical School, University of Sheffield | COVID-19 Genomics UK (COG-UK) Consortium | Thushan de Silva, Matthew Parker, Nikki Smith, Adri Angyal, Rebecca Brown, Luke Green, Rachel Tucker, Paul Parsons, Danielle Groves, Katie Johnson, Laura Carrilero, Alex Keeley, Dave Partridge, Matthew Wyles, Benjamin Lindsey, Mehmet Yavuz, Mohammad Raza, Cariad Evans |
| EPI_ISL_484709, EPI_ISL_484710, EPI_ISL_484711, EPI_ISL_484712, EPI_ISL_484713, EPI_ISL_484714, EPI_ISL_484715, EPI_ISL_484716, EPI_ISL_484717, EPI_ISL_484718, EPI_ISL_484719, EPI_ISL_484720, EPI_ISL_484721, EPI_ISL_484722, EPI_ISL_484723, EPI_ISL_484724, EPI_ISL_484725, EPI_ISL_484726, EPI_ISL_484727, EPI_ISL_484728, EPI_ISL_484729, EPI_ISL_484730, EPI_ISL_484731, EPI_ISL_484732, EPI_ISL_484733, EPI_ISL_484734, EPI_ISL_484735, EPI_ISL_484736, EPI_ISL_484737, EPI_ISL_484738, EPI_ISL_484739, EPI_ISL_484740, EPI_ISL_484741, EPI_ISL_484742, EPI_ISL_484743, EPI_ISL_484744, EPI_ISL_484745, EPI_ISL_484746, EPI_ISL_484747, EPI_ISL_484748, EPI_ISL_484749, EPI_ISL_484750, EPI_ISL_484751, EPI_ISL_484752, EPI_ISL_484753, EPI_ISL_484754, EPI_ISL_484755, EPI_ISL_484756, EPI_ISL_484757, EPI_ISL_484758, EPI_ISL_484759, EPI_ISL_484760, EPI_ISL_484761, EPI_ISL_484762, | | | |

| | | | |
|---|---|---|---|
| EPI_ISL_484763, EPI_ISL_484764, EPI_ISL_484765, EPI_ISL_484766, EPI_ISL_484767, EPI_ISL_484768, EPI_ISL_484769, EPI_ISL_484770, EPI_ISL_484771, EPI_ISL_484772, EPI_ISL_484773, EPI_ISL_484774, EPI_ISL_484775, EPI_ISL_484776, EPI_ISL_484777, EPI_ISL_484778, EPI_ISL_484779, EPI_ISL_484780, EPI_ISL_484781, EPI_ISL_484782, EPI_ISL_484783, EPI_ISL_484784, EPI_ISL_484785, EPI_ISL_484786, EPI_ISL_484787, EPI_ISL_484788, EPI_ISL_484789, EPI_ISL_484790, EPI_ISL_484791, EPI_ISL_484792, EPI_ISL_484793, EPI_ISL_484794, EPI_ISL_484795, EPI_ISL_484796, EPI_ISL_484797, EPI_ISL_484798, EPI_ISL_484799, EPI_ISL_484800, EPI_ISL_484801, EPI_ISL_484802, EPI_ISL_484803, EPI_ISL_484804, EPI_ISL_484805, EPI_ISL_484806 | | | |
| see above | University of Michigan Clinical Microbiology Laboratory | Lauring Lab, University of Michigan, Department of Microbiology and Immunology | Valesano et al. |
| EPI_ISL_485814, EPI_ISL_485815, EPI_ISL_485816, EPI_ISL_485817, EPI_ISL_485818, EPI_ISL_485819, EPI_ISL_485820, EPI_ISL_485821, EPI_ISL_485822, EPI_ISL_485823, EPI_ISL_485824, EPI_ISL_485825, EPI_ISL_485826, EPI_ISL_485827, EPI_ISL_485828, EPI_ISL_485829, EPI_ISL_485830, EPI_ISL_485831, EPI_ISL_485832, EPI_ISL_485833, EPI_ISL_485834, EPI_ISL_485835, EPI_ISL_485836, EPI_ISL_485837, EPI_ISL_485838, EPI_ISL_485839, EPI_ISL_485840, EPI_ISL_485841, EPI_ISL_485842, EPI_ISL_485843, EPI_ISL_485844, EPI_ISL_485845, EPI_ISL_485846, EPI_ISL_485847, EPI_ISL_485848, EPI_ISL_485849, EPI_ISL_485850, EPI_ISL_485851, EPI_ISL_485852, EPI_ISL_485853, EPI_ISL_485854, EPI_ISL_485855, EPI_ISL_485856, EPI_ISL_485857, EPI_ISL_485858, EPI_ISL_485859, EPI_ISL_485860, EPI_ISL_485861, EPI_ISL_485862, EPI_ISL_485863, EPI_ISL_485864, EPI_ISL_485865, EPI_ISL_485866, EPI_ISL_485867, EPI_ISL_485868, EPI_ISL_485869, EPI_ISL_485870, EPI_ISL_485871 | | | |
| see above | Virginia DCLS | Virginia DCLS | Virginia DCLS |
| EPI_ISL_487229, EPI_ISL_487230 | University of Michigan Clinical Microbiology Laboratory | Lauring Lab, University of Michigan, Department of Microbiology and Immunology | Valesano et al. |
| EPI_ISL_487370, EPI_ISL_487377, EPI_ISL_487379, EPI_ISL_487381 | Hellenic Pasteur Institute, National Influenza Reference laboratory of Southern Greece & Unit of Bioinformatics and Applied Genomics | Hellenic Pasteur Institute, National Influenza Reference laboratory of Southern Greece & Unit of Bioinformatics and Applied Genomics | Vasiliki Pogka, Timokratis Karamitros, Athanasios Kossyvakis, Antonios Kalliaropoulos, Horefti Elina, Evangelidou Maria, Androniki Voulgari-Kokota, Aspasia Kontou, Andreas Mentis |
| EPI_ISL_487398, EPI_ISL_487399, EPI_ISL_487400, EPI_ISL_487401, EPI_ISL_487402, EPI_ISL_487403, EPI_ISL_487404, EPI_ISL_487405, EPI_ISL_487406, EPI_ISL_487407, EPI_ISL_487408, EPI_ISL_487409, EPI_ISL_487410, EPI_ISL_487411, EPI_ISL_487412, EPI_ISL_487413, EPI_ISL_487414, EPI_ISL_487415, EPI_ISL_487416, EPI_ISL_487417, EPI_ISL_487418, EPI_ISL_487419, EPI_ISL_487420, EPI_ISL_487421, EPI_ISL_487422, EPI_ISL_487423, EPI_ISL_487424, EPI_ISL_487425, EPI_ISL_487426, EPI_ISL_487427, EPI_ISL_487428, EPI_ISL_487429, EPI_ISL_487430, EPI_ISL_487431 | | | |
| see above | Labor Kneißler GmbH & Co. KG | Heinrich Pette Institute, Leibniz Institute for Experimental Virology | Thomas Günther, Adam Grundhoff, Manja Czech-Sioli, Nicole Fischer, Matthias Ottinger, Melanie M. Brinkmann |
| EPI_ISL_490210, EPI_ISL_490211, EPI_ISL_490212, EPI_ISL_490213, EPI_ISL_490214, EPI_ISL_490215, EPI_ISL_490216, EPI_ISL_490217, EPI_ISL_490218, EPI_ISL_490219, EPI_ISL_490220, EPI_ISL_490221, EPI_ISL_490222, EPI_ISL_490223 | | | |
| see above | Quest Diagnostics | Q Squared Solutions - QRTP facility | Victor J Weigman |
| EPI_ISL_490581, EPI_ISL_490582, EPI_ISL_490583, EPI_ISL_490584, EPI_ISL_490585, EPI_ISL_490586, EPI_ISL_490587, EPI_ISL_490588, EPI_ISL_490589, EPI_ISL_490590, EPI_ISL_490591, EPI_ISL_490592, EPI_ISL_490593, EPI_ISL_490594, EPI_ISL_490595, EPI_ISL_490596, EPI_ISL_490597, EPI_ISL_490598, EPI_ISL_490599, EPI_ISL_490600, EPI_ISL_490601, EPI_ISL_490602, EPI_ISL_490603, EPI_ISL_490604, EPI_ISL_490605, EPI_ISL_490606, EPI_ISL_490607, EPI_ISL_490608, EPI_ISL_490609, EPI_ISL_490610, EPI_ISL_490611, EPI_ISL_490612, EPI_ISL_490613, EPI_ISL_490614, EPI_ISL_490615, EPI_ISL_490616, EPI_ISL_490617, EPI_ISL_490618, EPI_ISL_490619, EPI_ISL_490620, EPI_ISL_490621, EPI_ISL_490622, EPI_ISL_490623, EPI_ISL_490624, EPI_ISL_490625, EPI_ISL_490626, EPI_ISL_490627, EPI_ISL_490628, EPI_ISL_490629, EPI_ISL_490630, EPI_ISL_490631, EPI_ISL_490632, EPI_ISL_490633, EPI_ISL_490634, EPI_ISL_490635, EPI_ISL_490636, EPI_ISL_490637, EPI_ISL_490638, EPI_ISL_490639, EPI_ISL_490640, EPI_ISL_490641, EPI_ISL_490642, EPI_ISL_490643, EPI_ISL_490644, EPI_ISL_490645 | | | |
| see above | Virology Department, Sheffield Teaching Hospitals NHS Foundation Trust/Department of Infection, Immunity and Cardiovascular Disease, The Medical School, University of Sheffield | COVID-19 Genomics UK (COG-UK) Consortium | Thushan de Silva, Matthew Parker, Nikki Smith, Adri Angyal, Rebecca Brown, Luke Green, Rachel Tucker, Paul Parsons, Danielle Groves, Katie Johnson, Laura Carrilero, Alex Keeley, Dave Partridge, Matthew Wyles, Benjamin Lindsey, Mehmet Yavuz, Mohammad Raza, Cariad Evans |
| EPI_ISL_491298 | Rural Health Unit - Calauan, Laguna | Research Institute for Tropical Medicine | Tujan, M.A.A., Onza,O.J.T., Polotan,F.G.M., Medado,I.A.P., Bautista,C.T., Brunker, K., Mercado,E.S., Manalo, D.L., Demetria, C.S. |
| EPI_ISL_492075, EPI_ISL_492076, EPI_ISL_492077, EPI_ISL_492078, EPI_ISL_492079, EPI_ISL_492080, EPI_ISL_492081, EPI_ISL_492082, EPI_ISL_492083, EPI_ISL_492084, EPI_ISL_492085, EPI_ISL_492086 | | | |
| see above | Institute for Public Health of the Republic of North Macedonia | Charite Universitatsmedizin Berlin, Institute of Virology | Victor M Corman, Joern Beheim-Schwarzbach, Barbara Muhlemann, Talitha Veith, Julia Schneider, Elizabeta Jancheska, Maja Kuzmanovska, Golubinka Bosevska, Terry Jones, Christian Drosten |
| EPI_ISL_493666, EPI_ISL_493667, EPI_ISL_493668, EPI_ISL_493669, EPI_ISL_493670, EPI_ISL_493671, EPI_ISL_493672, EPI_ISL_493673, EPI_ISL_493674, EPI_ISL_493675, EPI_ISL_493676, EPI_ISL_493677, EPI_ISL_493678, EPI_ISL_493679, EPI_ISL_493680, EPI_ISL_493681, EPI_ISL_493682, EPI_ISL_493683, EPI_ISL_493684, EPI_ISL_493685, EPI_ISL_493686, EPI_ISL_493687, EPI_ISL_493688, EPI_ISL_493689, EPI_ISL_493690, EPI_ISL_493691, EPI_ISL_493692, EPI_ISL_493693, EPI_ISL_493694, EPI_ISL_493695, EPI_ISL_493696, EPI_ISL_493697, EPI_ISL_493698, EPI_ISL_493699, EPI_ISL_493700, EPI_ISL_493701, EPI_ISL_493702, EPI_ISL_493703, EPI_ISL_493704, EPI_ISL_493705, EPI_ISL_493706, EPI_ISL_493707, EPI_ISL_493708, EPI_ISL_493709, EPI_ISL_493710, EPI_ISL_493711, EPI_ISL_493712, EPI_ISL_493713, EPI_ISL_493714, EPI_ISL_493715, EPI_ISL_493716, EPI_ISL_493717, EPI_ISL_493718, EPI_ISL_493719, EPI_ISL_493720, EPI_ISL_493721, EPI_ISL_493722, EPI_ISL_493723, EPI_ISL_493724, EPI_ISL_493725, EPI_ISL_493726, EPI_ISL_493727, EPI_ISL_493728, EPI_ISL_493729, EPI_ISL_493730, EPI_ISL_493731, EPI_ISL_493732, EPI_ISL_493733, EPI_ISL_493734, EPI_ISL_493735, EPI_ISL_493736, EPI_ISL_493737, EPI_ISL_493738, EPI_ISL_493739, EPI_ISL_493740 | | | |
| see above | Virology Department, Sheffield Teaching Hospitals NHS Foundation Trust/Department of Infection, Immunity and Cardiovascular Disease, The Medical School, University of Sheffield | COVID-19 Genomics UK (COG-UK) Consortium | Thushan de Silva, Matthew Parker, Nikki Smith, Adri Angyal, Rebecca Brown, Luke Green, Rachel Tucker, Paul Parsons, Danielle Groves, Katie Johnson, Laura Carrilero, Alex Keeley, Dave Partridge, Matthew Wyles, Benjamin Lindsey, Mehmet Yavuz, Mohammad Raza, Cariad Evans |
| EPI_ISL_495032, EPI_ISL_495050 | GMERS Medical College & Hospital, Gotri, Vadodara | Gujarat Biotechnology Research Centre | Varsha Godbole, Apurvasinh Puvar, Janvi Raval, Zarna Patel, Monika Gandhi, Pinal Trivedi, Maharshi Pandya, Nidhi Patel, Nitin Savaliya, Raghawendra Kumar, Dinesh Kumar, Zuber Saiyed, Komal Patel, Labdhi Pandya, Afzal Ansari, Nikha Trivedi, Meenakshi Shah, Neena Doshi, R D Dixit, A M Kadri, Harsh Bakshi, Chaitanya Joshi, Madhvi Joshi |
| EPI_ISL_495176 | CSIR-Centre for Cellular and Molecular Biology | CSIR-Centre for Cellular and Molecular Biology | Tulasi Nagabandi, Namami Gaur, Sakshi Shambhavi, Lamuk Zaveri, Shagufta Khan, Nikhil Hajirnis, M Soujanya Reddy, Pratheusa Maccha, Purushotham Vodnala, Payel Mukherjee, Sofia Banu, Priya Singh, Onkar Kulkarni, Dhiviya Vedagiri, Divya Gupta, Vishal Sah, Santosh Kumar Kuncha, Krishnan Harinivas Harshan, Archana Bharadwaj Siva, Karthik Bharadwaj Tallapaka,G. Aditya Kumar, Koushick Sivakumar, Pooja Ramesh Gupta, Rajan Kumar Jha, Shraddha Vijay Lahoti, Rakesh K Mishra, Divya Tej Sowpati |
| EPI_ISL_495198 | CSIR-Centre for Cellular and Molecular Biology | CSIR-Centre for Cellular and Molecular Biology | Tulasi Nagabandi, Namami Gaur, Sakshi Shambhavi, Lamuk Zaveri, Shagufta Khan, Nikhil Hajirnis, M Soujanya Reddy, Pratheusa Maccha, Purushotham Vodnala, Payel Mukherjee, Sofia Banu, Priya Singh,Onkar Kulkarni, Dhiviya Vedagiri, Divya Gupta, Vishal Sah, Santosh Kumar Kuncha, Krishnan Harinivas Harshan, Archana Bharadwaj Siva, Karthik Bharadwaj Tallapaka,Kezia J Ann, Radhika Khandelwal, Roshan Maku Venkata, Shemin Mansuri, Sonu Uday, Rakesh K Mishra, Divya Tej Sowpati |
| EPI_ISL_495203, EPI_ISL_495214 | CSIR-Centre for Cellular and Molecular Biology | CSIR-Centre for Cellular and Molecular Biology | Tulasi Nagabandi, Namami Gaur, Sakshi Shambhavi, Lamuk Zaveri, Shagufta Khan, Nikhil Hajirnis, M Soujanya Reddy, Pratheusa Maccha, Purushotham Vodnala, Payel Mukherjee, Sofia Banu, Priya Singh,Onkar Kulkarni , Dhiviya Vedagiri, Divya Gupta, Vishal Sah, Santosh Kumar Kuncha, Krishnan Harinivas Harshan, Archana Bharadwaj Siva, Karthik Bharadwaj Tallapaka,G. Aditya Kumar, Koushick Sivakumar, Pooja Ramesh Gupta, Rajan Kumar Jha, Shraddha Vijay Lahoti, Rakesh K Mishra, Divya Tej Sowpati |
| EPI_ISL_495235 | CSIR-Centre for Cellular and Molecular Biology | CSIR-Centre for Cellular and Molecular Biology | Tulasi Nagabandi, Namami Gaur, Sakshi Shambhavi, Lamuk Zaveri, Shagufta Khan, Nikhil Hajirnis, M Soujanya Reddy, Pratheusa Maccha, Purushotham Vodnala, Payel Mukherjee, Sofia Banu, Priya Singh, Onkar Kulkarni, Dhiviya Vedagiri, Divya Gupta, Vishal Sah, Santosh Kumar Kuncha, Krishnan Harinivas Harshan, Archana Bharadwaj Siva, Karthik Bharadwaj Tallapaka,G. Aditya Kumar, Koushick Sivakumar, Pooja Ramesh Gupta, Rajan Kumar Jha, Shraddha Vijay Lahoti, Rakesh K Mishra, Divya Tej Sowpati |
| EPI_ISL_495242 | CSIR-Centre for Cellular and Molecular Biology | CSIR-Centre for Cellular and Molecular Biology | Tulasi Nagabandi, Namami Gaur, Sakshi Shambhavi, Lamuk Zaveri, Shagufta Khan, Nikhil Hajirnis, M Soujanya Reddy, Pratheusa Maccha, Purushotham Vodnala, Payel Mukherjee, Sofia Banu, Priya Singh,Onkar Kulkarni, Dhiviya Vedagiri, Divya Gupta, Vishal Sah, Santosh Kumar Kuncha, Krishnan Harinivas Harshan, Archana Bharadwaj Siva, Karthik Bharadwaj Tallapaka,Kezia J Ann, Radhika Khandelwal, Roshan Maku Venkata, Shemin Mansuri, Sonu Uday, Rakesh K Mishra, Divya Tej Sowpati |
| EPI_ISL_495255, EPI_ISL_495263 | CSIR-Centre for Cellular and Molecular Biology | CSIR-Centre for Cellular and Molecular Biology | Tulasi Nagabandi, Namami Gaur, Sakshi Shambhavi, Lamuk Zaveri, Shagufta Khan, Nikhil Hajirnis, M Soujanya Reddy, Pratheusa Maccha, Purushotham Vodnala, Payel Mukherjee, Sofia Banu, Priya Singh, Onkar Kulkarni, Dhiviya Vedagiri, Divya Gupta, Vishal Sah, Santosh Kumar Kuncha, Krishnan Harinivas Harshan, Archana Bharadwaj Siva, Karthik Bharadwaj Tallapaka,G. Aditya Kumar, Koushick Sivakumar, Pooja Ramesh Gupta, Rajan Kumar Jha, Shraddha Vijay Lahoti, Rakesh K Mishra, Divya Tej Sowpati |
| EPI_ISL_495267 | CSIR-Centre for Cellular and Molecular Biology | CSIR-Centre for Cellular and Molecular Biology | Tulasi Nagabandi, Namami Gaur, Sakshi Shambhavi, Lamuk Zaveri, Shagufta Khan, Nikhil Hajirnis, M Soujanya Reddy, Pratheusa Maccha, Purushotham Vodnala, Payel Mukherjee, Sofia Banu, Priya Singh,Onkar Kulkarni, Dhiviya Vedagiri, Divya Gupta, Vishal Sah, Santosh Kumar Kuncha, Krishnan Harinivas Harshan, Archana Bharadwaj Siva, Karthik Bharadwaj Tallapaka,Kezia J Ann, Radhika Khandelwal, Roshan Maku Venkata, Shemin Mansuri, Sonu Uday, Rakesh K Mishra, Divya Tej Sowpati |
| EPI_ISL_495268 | CSIR-Centre for Cellular and Molecular Biology | CSIR-Centre for Cellular and Molecular Biology | Tulasi Nagabandi, Namami Gaur, Sakshi Shambhavi, Lamuk Zaveri, Shagufta Khan, Nikhil Hajirnis, M Soujanya Reddy, Pratheusa Maccha, Purushotham Vodnala, Payel Mukherjee, Sofia Banu, Priya Singh,Onkar Kulkarni , Dhiviya Vedagiri, Divya Gupta, Vishal Sah, Santosh Kumar Kuncha, Krishnan |

| | | | |
|---|---|---|---|
| | | | Harinivas Harshan, Archana Bharadwaj Siva, Karthik Bharadwaj Tallapaka,G. Aditya Kumar, Koushick Sivakumar, Pooja Ramesh Gupta, Rajan Kumar Jha, Shraddha Vijay Lahoti, Rakesh K Mishra, Divya Tej Sowpati |
| EPI_ISL_495298, EPI_ISL_495299, EPI_ISL_495300, EPI_ISL_495301, EPI_ISL_495302, EPI_ISL_495303, EPI_ISL_495304, EPI_ISL_495305, EPI_ISL_495306, EPI_ISL_495307, EPI_ISL_495308, EPI_ISL_495309, EPI_ISL_495310, EPI_ISL_495311, EPI_ISL_495312, EPI_ISL_495313, EPI_ISL_495314, EPI_ISL_495315, EPI_ISL_495316, EPI_ISL_495317, EPI_ISL_495318, EPI_ISL_495319, EPI_ISL_495320, EPI_ISL_495321, EPI_ISL_495322, EPI_ISL_495323, EPI_ISL_495324, EPI_ISL_495325, EPI_ISL_495326, EPI_ISL_495327, EPI_ISL_495328, EPI_ISL_495329 | | | |
| see above | Virginia DCLS | Virginia DCLS | Virginia DCLS |
| EPI_ISL_495459 | Centers for Disease Control and Prevention of Lishui | Department of InspectionCenters for Disease Control and Prevention of Lishui | Wang Xiaoguang,Ji Qiaoying,Ji Jiansong,Ye Bifeng,Ye Ling |
| EPI_ISL_495563, EPI_ISL_495564, EPI_ISL_495565, EPI_ISL_495566, EPI_ISL_495567, EPI_ISL_495568, EPI_ISL_495569, EPI_ISL_495570, EPI_ISL_495571, EPI_ISL_495572, EPI_ISL_495573, EPI_ISL_495574, EPI_ISL_495575, EPI_ISL_495576, EPI_ISL_495577, EPI_ISL_495578, EPI_ISL_495579, EPI_ISL_495580, EPI_ISL_495581, EPI_ISL_495582, EPI_ISL_495583, EPI_ISL_495584, EPI_ISL_495585, EPI_ISL_495586, EPI_ISL_495587, EPI_ISL_495588, EPI_ISL_495589, EPI_ISL_495590, EPI_ISL_495591, EPI_ISL_495592, EPI_ISL_495593, EPI_ISL_495594, EPI_ISL_495595, EPI_ISL_495596 | | | |
| see above | University of Michigan Clinical Microbiology Laboratory | Lauring Lab, University of Michigan, Department of Microbiology and Immunology | Valesano et al. |
| EPI_ISL_498691, EPI_ISL_498692, EPI_ISL_498693, EPI_ISL_498694 | National Institute for Viral Disease Control and Prevention, China CDC | National Institute for Viral Disease Control and Prevention, China CDC | Xiang Zhao,LingLing Mao,Yao Meng,Zhixiao Chen,Yuchao Wu,Yong ZhangBo ZhijianJianqun Zhang,Yang Song,Dayan Wang,WenQing YaoWenbo Xu |
| EPI_ISL_499330, EPI_ISL_499331, EPI_ISL_499332, EPI_ISL_499333, EPI_ISL_499334, EPI_ISL_499335, EPI_ISL_499336, EPI_ISL_499337, EPI_ISL_499338, EPI_ISL_499339, EPI_ISL_499340, EPI_ISL_499341, EPI_ISL_499342, EPI_ISL_499343, EPI_ISL_499344, EPI_ISL_499345, EPI_ISL_499346, EPI_ISL_499347, EPI_ISL_499348, EPI_ISL_499349, EPI_ISL_499350, EPI_ISL_499351 | | | |
| see above | Virology Department, Sheffield Teaching Hospitals NHS Foundation Trust/Department of Infection, Immunity and Cardiovascular Disease, The Medical School, University of Sheffield | COVID-19 Genomics UK (COG-UK) Consortium | Thushan de Silva, Matthew Parker, Nikki Smith, Adri Angyal, Rebecca Brown, Luke Green, Rachel Tucker, Paul Parsons, Danielle Groves, Katie Johnson, Laura Carrilero, Alex Keeley, Dave Partridge, Matthew Wyles, Benjamin Lindsey, Mehmet Yavuz, Mohammad Raza, Cariad Evans |
| EPI_ISL_500831, EPI_ISL_500832, EPI_ISL_500833, EPI_ISL_500834, EPI_ISL_500835, EPI_ISL_500836, EPI_ISL_500837, EPI_ISL_500838, EPI_ISL_500839, EPI_ISL_500840, EPI_ISL_500841, EPI_ISL_500842, EPI_ISL_500843, EPI_ISL_500844, EPI_ISL_500845, EPI_ISL_500846, EPI_ISL_500847, EPI_ISL_500848, EPI_ISL_500849, EPI_ISL_500850, EPI_ISL_500851, EPI_ISL_500852, EPI_ISL_500853, EPI_ISL_500854, EPI_ISL_500855, EPI_ISL_500856, EPI_ISL_500857, EPI_ISL_500858, EPI_ISL_500859, EPI_ISL_500860, EPI_ISL_500861, EPI_ISL_500862, EPI_ISL_500863 | | | |
| see above | Virginia DCLS | Virginia DCLS | Virginia DCLS |
| EPI_ISL_501167, EPI_ISL_501168, EPI_ISL_501169, EPI_ISL_501171, EPI_ISL_501172, EPI_ISL_501173, EPI_ISL_501174 | Baylor College of Medicine | Baylor College of Medicine: HGSC | Vasanthi Avadhanula, Erin Nicholson, David Henke, Pedro Piedra, Harsha Doddapaneni, Donna Muzny, Qingchang Meng, Hsu Chao, Zeineen Momin, Hua Shen, George Weissenberger, Kavya Kottapalli, Yimiti Meiheerguli, Sejal Salvi, Ginger Metcalf, Vipin Menon, Sara J.J. Cregeen, Matthew C. Ross, Tulin Ayvaz, Richard Sucgang, Kristi L. Hoffman, Matthew Wong, Joseph F. Petrosino |
| EPI_ISL_501230 | Hellenic Pasteur Institute, Public Health Laboratories | Hellenic Pasteur Institute, National Influenza Reference laboratory of Southern Greece & Unit of Bioinformatics and Applied Genomics | Vasiliki Pogka, Timokratis Karamitros, Athanasios Kossyvakis, Antonios Kalliaropoulos, Horefti Elina, Evangelidou Maria, Androniki Voulgari-Kokota, Aspasia Kontou, Andreas Mentis |
| EPI_ISL_501231, EPI_ISL_501233, EPI_ISL_501234, EPI_ISL_501235, EPI_ISL_501236, EPI_ISL_501237, EPI_ISL_501238, EPI_ISL_501241, EPI_ISL_501242, EPI_ISL_501248, EPI_ISL_501249, EPI_ISL_501250, EPI_ISL_501251 | | | |
| see above | Hellenic Pasteur Institute, National Influenza Reference laboratory of Southern Greece & Unit of Bioinformatics and Applied Genomics | Hellenic Pasteur Institute, National Influenza Reference laboratory of Southern Greece & Unit of Bioinformatics and Applied Genomics | Vasiliki Pogka, Timokratis Karamitros, Athanasios Kossyvakis, Antonios Kalliaropoulos, Horefti Elina, Evangelidou Maria, Androniki Voulgari-Kokota, Aspasia Kontou, Andreas Mentis |
| EPI_ISL_502779 | LACEN/PE | LABBE, Federal University of Pernambuco | WILSON JOSE DA SILVA JUNIOR, HEIDI LACERDA ALVES DA CRUZ, MARCOS DA SILVEIRA REGUEIRA NETO, BRUNO SAMPAIO, SERGIO DE SA LEITAO PAIVA JUNIOR, ZILDENE DE SOUSA SILVEIRA, MAIRA GALDINO DA ROCHA PITTA, MICHELLY CRISTINY PEREIRA, MARCOS ANTONIO DE MORAIS JUNIOR, ANTONIO CARLOS DE FREITAS, VALDIR DE QUEIROZ BALBINO. |
| EPI_ISL_502875 | LACEN/PE | LABBE, Federal University of Pernambuco | WILSON JOSE DA SILVA JUNIOR, HEIDI LACERDA ALVES DA CRUZ, MARCOS DA SILVEIRA REGUEIRA NETO, BRUNO SAMPAIO, SERGIO DE SA LEITAO PAIVA JUNIOR, ZILDENE DE SOUSA SILVEIRA, MAIRA GALDINO DA ROCHA PITTA, MICHELLY CRISTINY PEREIRA, REGINALDO GONCALVES DE LIMA NETO, MARCOS ANTONIO DE MORAIS JUNIOR, ANTONIO CARLOS DE FREITAS, VALDIR DE QUEIROZ BALBINO. |
| EPI_ISL_507160, EPI_ISL_507161, EPI_ISL_507162, EPI_ISL_507163, EPI_ISL_507164, EPI_ISL_507165, EPI_ISL_507166, EPI_ISL_507167, EPI_ISL_507168, EPI_ISL_507169, EPI_ISL_507170, EPI_ISL_507171, EPI_ISL_507172, EPI_ISL_507173, EPI_ISL_507174, EPI_ISL_507175, EPI_ISL_507176, EPI_ISL_507177, EPI_ISL_507178, EPI_ISL_507179 | | | |
| see above | Virology Department, Sheffield Teaching Hospitals NHS Foundation Trust/Department of Infection, Immunity and Cardiovascular Disease, The Medical School, University of Sheffield | COVID-19 Genomics UK (COG-UK) Consortium | Thushan de Silva, Matthew Parker, Nikki Smith, Adri Angyal, Rebecca Brown, Luke Green, Rachel Tucker, Paul Parsons, Danielle Groves, Katie Johnson, Laura Carrilero, Alex Keeley, Dave Partridge, Matthew Wyles, Benjamin Lindsey, Mehmet Yavuz, Mohammad Raza, Cariad Evans |
| EPI_ISL_508864, EPI_ISL_508866, EPI_ISL_508868, EPI_ISL_508870 | Division of Infectious Diseases and Hospital Epidemiology, University Hospital Zürich | Institute of Medical Virology, University of Zurich | Verena Kufner, Maryam Zaheri, Dana Weissberg, Jürg Böni, Silvana K. Rampini, Peter W. Schreiber, Irene A. Abela, Hugo Sax, Aline Wolfensberger, Michael Huber |
| EPI_ISL_509424, EPI_ISL_509425, EPI_ISL_509426, EPI_ISL_509427, EPI_ISL_509428, EPI_ISL_509429 | Microbiology and Immunology, University of South Alabama | Microbiology and Immunology, University of South Alabama | Wood,R.R., Roberts,R.A., Houserova,D., Borchert,G.M., Fouty,B., Rayner,J.O. |
| EPI_ISL_513294 | Baylor College of Medicine | Baylor College of Medicine: HGSC | Vasanthi Avadhanula, Erin Nicholson, David Henke, Pedro Piedra, Harsha Doddapaneni, Donna Muzny, Qingchang Meng, Hsu Chao, Zeineen Momin, Hua Shen, George Weissenberger, Kavya Kottapalli, Yimiti Meiheerguli, Sejal Salvi, Ginger Metcalf, Vipin Menon, Sara J.J. Cregeen, Matthew C. Ross, Tulin Ayvaz, Richard Sucgang, Kristi L. Hoffman, Matthew Wong, Joseph F. Petrosino |
| EPI_ISL_514425, EPI_ISL_514426, EPI_ISL_514427, EPI_ISL_514428, EPI_ISL_514429, EPI_ISL_514430, EPI_ISL_514431 | Laboratory Diagnostic, Veterinary Specialized Institute Kraljevo | Laboratory Diagnostic, Veterinary Specialized Institute Kraljevo | Vidanovic,D., Tesovic,B., Knezevic,A., Jankovic,M., Sekler,M., Banovic Djeri,B., Volkening,J., Afonso,C., Petrovic,T. |
| EPI_ISL_514432 | Prof. Massimo Zollo CEINGE TASK-FORCE COVID19 - Regione Campania | Prof. Massimo Zollo CEINGE TASK-FORCE COVID19 - Regione Campania | Veronica Ferrucci, Dae young Kong, Fatemeh asadzadeh, Laura Marrone, Roberto Siciliano, Rino Cerino, Giovanna Fusco, Marika Comegna, Angelo Boccia, Maurizio Viscardi, Giorgia Borriello, Sergio Brandi, Claudia Tiberio, Luigi Atripaldi, Giovanni Paolella, Giuseppe Castaldo, Stefano Pascarella, Martina Bianchi, Lorenzo Chiariotti, Jae Myun Lee, Jae Ho Jung, Kyong Seop Yun, Hong Yeoul Kim and Massimo Zollo |
| EPI_ISL_516629, EPI_ISL_516630, EPI_ISL_516631, EPI_ISL_516632, EPI_ISL_516633, EPI_ISL_516634, EPI_ISL_516635, EPI_ISL_516636, EPI_ISL_516637, EPI_ISL_516638, EPI_ISL_516639, EPI_ISL_516640, EPI_ISL_516641, EPI_ISL_516642, EPI_ISL_516643, EPI_ISL_516644, EPI_ISL_516645 | | | |
| see above | Charite Universitatsmedizin Berlin, Institut fur Virologie/Labor Berlin | Charite Universitatsmedizin Berlin, Institut fur Virologie/Labor Berlin | Victor M Corman, Barbara Muhlemann, Jörn Beheim-Schwarzbach, Julia Schneider, Talitha Veith, Terry Jones, Christian Drosten |
| EPI_ISL_516655, EPI_ISL_516656, EPI_ISL_516657, EPI_ISL_516658, EPI_ISL_516659, EPI_ISL_516660, EPI_ISL_516661, EPI_ISL_516662, EPI_ISL_516663, EPI_ISL_516664, EPI_ISL_516665, EPI_ISL_516666, EPI_ISL_516667, EPI_ISL_516668, EPI_ISL_516669, EPI_ISL_516670, EPI_ISL_516671, EPI_ISL_516672, EPI_ISL_516673, EPI_ISL_516674, EPI_ISL_516675, EPI_ISL_516676, EPI_ISL_516677, EPI_ISL_516678, EPI_ISL_516679, EPI_ISL_516680, EPI_ISL_516681, EPI_ISL_516682, EPI_ISL_516683, EPI_ISL_516684, EPI_ISL_516687, EPI_ISL_516688, EPI_ISL_516689, EPI_ISL_516690, EPI_ISL_516691, EPI_ISL_516692, EPI_ISL_516693, EPI_ISL_516694, EPI_ISL_516695, EPI_ISL_516696, EPI_ISL_516697, EPI_ISL_516698, EPI_ISL_516699, EPI_ISL_516700, EPI_ISL_516701, EPI_ISL_516702, EPI_ISL_516703, EPI_ISL_516704, EPI_ISL_516705, EPI_ISL_516706, EPI_ISL_516707, EPI_ISL_516708, EPI_ISL_516709, EPI_ISL_516710, EPI_ISL_516711, EPI_ISL_516712, EPI_ISL_516713, EPI_ISL_516714, EPI_ISL_516715, EPI_ISL_516716, EPI_ISL_516717, EPI_ISL_516718, EPI_ISL_516719, EPI_ISL_516720 | | | |
| see above | Virginia DCLS | Virginia DCLS | Virginia DCLS |
| EPI_ISL_516922, EPI_ISL_516923, EPI_ISL_516924, EPI_ISL_516925, EPI_ISL_516926, EPI_ISL_516927, EPI_ISL_516928, EPI_ISL_516929, EPI_ISL_516930, EPI_ISL_516931, EPI_ISL_516932, EPI_ISL_516933 | | | |
| see above | Department for Molecular Diagnostics, Centre for Medical Microbiology, Institute of Public Health of Montenegro | Charite Universitatsmedizin Berlin, Institut fur Virologie | Victor M Corman, Terry Jones, Jörn Beheim-Schwarzbach, Barbara Muehlemann, Talitha Veith, Julia Schneider, Marija Govedarica and Danijela Vujoševi, Christian Drosten |
| EPI_ISL_516934, EPI_ISL_516935, EPI_ISL_516936, EPI_ISL_516938 | Nicolae Testemitanu State University of Medicine and Pharmacy | International Centre for Genetic Engineering and Biotechnology (ICGEB) and ARGO Open Lab Platform for | Ulinici M, Licastro D, Dal Monego S, Rajasekharan S, Marcello A |

| | | | |
|---|---|---|---|
| EPI_ISL_517543, EPI_ISL_517544, EPI_ISL_517545, EPI_ISL_517546, EPI_ISL_517547, EPI_ISL_517548, EPI_ISL_517549, EPI_ISL_517550, EPI_ISL_517551, EPI_ISL_517552, EPI_ISL_517553, EPI_ISL_517554, EPI_ISL_517555, EPI_ISL_517556, EPI_ISL_517557, EPI_ISL_517558, EPI_ISL_517559, EPI_ISL_517560, EPI_ISL_517561, EPI_ISL_517562, EPI_ISL_517563, EPI_ISL_517564, EPI_ISL_517565, EPI_ISL_517566, EPI_ISL_517567, EPI_ISL_517568, EPI_ISL_517569, EPI_ISL_517570, EPI_ISL_517571, EPI_ISL_517572, EPI_ISL_517573, EPI_ISL_517574, EPI_ISL_517575, EPI_ISL_517576, EPI_ISL_517577, EPI_ISL_517578, EPI_ISL_517579 | | | |
| see above | Virology Department, Sheffield Teaching Hospitals NHS Foundation Trust/Department of Infection, Immunity and Cardiovascular Disease, The Medical School, University of Sheffield | COVID-19 Genomics UK (COG-UK) Consortium | Thushan de Silva, Matthew Parker, Nikki Smith, Adri Angyal, Rebecca Brown, Luke Green, Rachel Tucker, Paul Parsons, Danielle Groves, Katie Johnson, Laura Carrilero, Alex Keeley, Dave Partridge, Matthew Wyles, Benjamin Lindsey, Mehmet Yavuz, Mohammad Raza, Cariad Evans |
| EPI_ISL_522349, EPI_ISL_522350 | KU Leuven, Rega Institute, Clinical and Epidemiological Virology | KU Leuven, Rega Institute, Clinical and Epidemiological Virology | Tony Wawina-Bokalanga, Joan Marti-Carerras, Bert Vanmechelen, Piet Maes |
| EPI_ISL_522777, EPI_ISL_522778, EPI_ISL_522779, EPI_ISL_522780, EPI_ISL_522781, EPI_ISL_522782, EPI_ISL_522783, EPI_ISL_522784, EPI_ISL_522785, EPI_ISL_522786, EPI_ISL_522787, EPI_ISL_522788, EPI_ISL_522789, EPI_ISL_522790, EPI_ISL_522791, EPI_ISL_522792, EPI_ISL_522793, EPI_ISL_522794, EPI_ISL_522795, EPI_ISL_522796, EPI_ISL_522797, EPI_ISL_522798, EPI_ISL_522799, EPI_ISL_522800, EPI_ISL_522801, EPI_ISL_522802, EPI_ISL_522803, EPI_ISL_522804, EPI_ISL_522805, EPI_ISL_522806, EPI_ISL_522807, EPI_ISL_522808, EPI_ISL_522809, EPI_ISL_522810, EPI_ISL_522811, EPI_ISL_522812, EPI_ISL_522813, EPI_ISL_522814, EPI_ISL_522815, EPI_ISL_522816, EPI_ISL_522817, EPI_ISL_522818, EPI_ISL_522819, EPI_ISL_522820, EPI_ISL_522821, EPI_ISL_522822, EPI_ISL_522823 | | | |
| see above | Virginia DCLS | Virginia DCLS | Virginia DCLS |
| EPI_ISL_523960 | National Agency for Public Health, Republic of Moldova | Charite Universitatsmedizin Berlin, Institute of Virology | Victor M Corman, Jorn Beheim-Schwarzbach, Barbara Mühlemann, Talitha Veith, Julia Schneider, Ala Halacu, Mariana Apostol, Terry Jones, Christian Drosten |
| EPI_ISL_523962 | National Agency for Public Health, Republic of Moldova | Charite Universitatsmedizin Berlin, Institute of Virology | Victor M Corman, Joern Beheim-Schwarzbach, Barbara Muhlemann, Talitha Veith, Julia Schneider, Elizabeta Jancheska, Maja Kuzmanovska, Golubinka Bosevska, Terry Jones, Christian Drosten |
| EPI_ISL_523964, EPI_ISL_523966, EPI_ISL_523968 | National Agency for Public Health, Republic of Moldova | Charite Universitatsmedizin Berlin, Institute of Virology | Victor M Corman, Jorn Beheim-Schwarzbach, Barbara Mühlemann, Talitha Veith, Julia Schneider, Ala Halacu, Mariana Apostol, Terry Jones, Christian Drosten |
| EPI_ISL_524482 | University Hospital Zürich | Institute of Medical Virology, University of Zurich | Verena Kufner, Maryam Zaheri, Gabriela Ziltener, Stefan Schmutz, Patrick Redli, Fiona Steiner, Jon Huder, Riccarda Capaul, Andrea Zbinden, Jürg Böni, Michael Huber, Alexandra Trkola |
| EPI_ISL_524483 | Division of Infectious Diseases, University Hospital Zürich | Institute of Medical Virology, University of Zurich | Verena Kufner, Maryam Zaheri, Gabriela Ziltener, Stefan Schmutz, Patrick Redli, Fiona Steiner, Jon Huder, Riccarda Capaul, Andrea Zbinden, Jürg Böni, Michael Huber, Alexandra Trkola |
| EPI_ISL_524484 | Hospital Schwyz | Institute of Medical Virology, University of Zurich | Verena Kufner, Maryam Zaheri, Gabriela Ziltener, Stefan Schmutz, Patrick Redli, Fiona Steiner, Jon Huder, Riccarda Capaul, Andrea Zbinden, Jürg Böni, Michael Huber, Alexandra Trkola |
| EPI_ISL_524485 | Division of Infectious Diseases, University Hospital Zürich | Institute of Medical Virology, University of Zurich | Verena Kufner, Maryam Zaheri, Gabriela Ziltener, Stefan Schmutz, Patrick Redli, Fiona Steiner, Jon Huder, Riccarda Capaul, Andrea Zbinden, Jürg Böni, Michael Huber, Alexandra Trkola |
| EPI_ISL_525189, EPI_ISL_525190, EPI_ISL_525191, EPI_ISL_525192, EPI_ISL_525193, EPI_ISL_525194, EPI_ISL_525195, EPI_ISL_525196, EPI_ISL_525197, EPI_ISL_525198, EPI_ISL_525199, EPI_ISL_525200, EPI_ISL_525201, EPI_ISL_525202, EPI_ISL_525203, EPI_ISL_525204, EPI_ISL_525205 | | | |
| see above | Virginia DCLS | Virginia DCLS | Virginia DCLS |
| EPI_ISL_526437, EPI_ISL_526438, EPI_ISL_526439, EPI_ISL_526440, EPI_ISL_526441, EPI_ISL_526442, EPI_ISL_526443, EPI_ISL_526444, EPI_ISL_526445, EPI_ISL_526446, EPI_ISL_526447, EPI_ISL_526448, EPI_ISL_526449, EPI_ISL_526450 | | | |
| see above | Virology Department, Sheffield Teaching Hospitals NHS Foundation Trust/Department of Infection, Immunity and Cardiovascular Disease, The Medical School, University of Sheffield | COVID-19 Genomics UK (COG-UK) Consortium | Thushan de Silva, Matthew Parker, Nikki Smith, Adri Angyal, Rebecca Brown, Luke Green, Rachel Tucker, Paul Parsons, Danielle Groves, Katie Johnson, Laura Carrilero, Alex Keeley, Dave Partridge, Matthew Wyles, Benjamin Lindsey, Mehmet Yavuz, Mohammad Raza, Cariad Evans |
| EPI_ISL_526775, EPI_ISL_526776, EPI_ISL_526777, EPI_ISL_526778, EPI_ISL_526779, EPI_ISL_526780, EPI_ISL_526781, EPI_ISL_526782, EPI_ISL_526783, EPI_ISL_526784, EPI_ISL_526785, EPI_ISL_526786, EPI_ISL_526787, EPI_ISL_526788, EPI_ISL_526789, EPI_ISL_526790, EPI_ISL_526791, EPI_ISL_526792, EPI_ISL_526793, EPI_ISL_526794, EPI_ISL_526795, EPI_ISL_526796, EPI_ISL_526797, EPI_ISL_526798, EPI_ISL_526799, EPI_ISL_526800, EPI_ISL_526801, EPI_ISL_526802, EPI_ISL_526803, EPI_ISL_526804, EPI_ISL_526805, EPI_ISL_526806, EPI_ISL_526807, EPI_ISL_526808, EPI_ISL_526809, EPI_ISL_526810, EPI_ISL_526811, EPI_ISL_526812, EPI_ISL_526813, EPI_ISL_526814, EPI_ISL_526815, EPI_ISL_526816, EPI_ISL_526817, EPI_ISL_526818, EPI_ISL_526819, EPI_ISL_526820, EPI_ISL_526821, EPI_ISL_526822, EPI_ISL_526823, EPI_ISL_526824, EPI_ISL_526825, EPI_ISL_526826, EPI_ISL_526827, EPI_ISL_526828, EPI_ISL_526829, EPI_ISL_526830, EPI_ISL_526831, EPI_ISL_526832, EPI_ISL_526833, EPI_ISL_526834, EPI_ISL_526835, EPI_ISL_526836, EPI_ISL_526837, EPI_ISL_526838, EPI_ISL_526839, EPI_ISL_526840, EPI_ISL_526841, EPI_ISL_526842, EPI_ISL_526843, EPI_ISL_526844, EPI_ISL_526845, EPI_ISL_526846, EPI_ISL_526847, EPI_ISL_526848, EPI_ISL_526849, EPI_ISL_526850, EPI_ISL_526851, EPI_ISL_526852, EPI_ISL_526853, EPI_ISL_526854, EPI_ISL_526855, EPI_ISL_526856, EPI_ISL_526857, EPI_ISL_526858, EPI_ISL_526859, EPI_ISL_526860, EPI_ISL_526861, EPI_ISL_526862, EPI_ISL_526863, EPI_ISL_526864, EPI_ISL_526865, EPI_ISL_526866, EPI_ISL_526867, EPI_ISL_526868, EPI_ISL_526869, EPI_ISL_526870, EPI_ISL_526871, EPI_ISL_526872, EPI_ISL_526873, EPI_ISL_526874, EPI_ISL_526875, EPI_ISL_526876, EPI_ISL_526877, EPI_ISL_526878, EPI_ISL_526879, EPI_ISL_526880, EPI_ISL_526881, EPI_ISL_526882, EPI_ISL_526883, EPI_ISL_526884, EPI_ISL_526885, EPI_ISL_526886, EPI_ISL_526887, EPI_ISL_526888, EPI_ISL_526889, EPI_ISL_526890, EPI_ISL_526891, EPI_ISL_526892, EPI_ISL_526893, EPI_ISL_526894, EPI_ISL_526895, EPI_ISL_526896, EPI_ISL_526897, EPI_ISL_526898, EPI_ISL_526899, EPI_ISL_526900, EPI_ISL_526901, EPI_ISL_526902, EPI_ISL_526903, EPI_ISL_526904, EPI_ISL_526905, EPI_ISL_526906, EPI_ISL_526907, EPI_ISL_526908, EPI_ISL_526909, EPI_ISL_526910, EPI_ISL_526911, EPI_ISL_526912, EPI_ISL_526913, EPI_ISL_526914, EPI_ISL_526915, EPI_ISL_526916, EPI_ISL_526917, EPI_ISL_526918, EPI_ISL_526919, EPI_ISL_526920, EPI_ISL_526921, EPI_ISL_526922, EPI_ISL_526923, EPI_ISL_526924, EPI_ISL_526925, EPI_ISL_526926, EPI_ISL_526927, EPI_ISL_526928, EPI_ISL_526929, EPI_ISL_526930, EPI_ISL_526931 | | | |
| see above | Virginia DCLS | Virginia DCLS | Virginia DCLS |
| EPI_ISL_528573, EPI_ISL_528574, EPI_ISL_528575, EPI_ISL_528576, EPI_ISL_528577, EPI_ISL_528578, EPI_ISL_528579, EPI_ISL_528580 | National Genomics Core-Center for DNA Fingerprinting and Diagnostics | National Genomics Core- Center for DNA Fingerprinting and Diagnostics (NGC-CDFD)- DBT's PAN-INDIA-1000 Genome consortium | Vinay Donipadi, G Shashikanth, Heena Shah, Bala Pratyusha, Parveen Kumar, Sandip Patra, Mugdha Singh, Reelina Basu, Dhanraj Adey, Bharath Kumar, Bhavani Sontam, Shaik Nasar Vali, R Harinarayanan, Rashna Bhandari, Murali Dharan Bashyam, Debashish Mitra, Divya Vashisht, Ashwin Dalal |
| EPI_ISL_528639, EPI_ISL_528640, EPI_ISL_528641, EPI_ISL_528642, EPI_ISL_528643, EPI_ISL_528644, EPI_ISL_528645, EPI_ISL_528646, EPI_ISL_528647, EPI_ISL_528648, EPI_ISL_528649, EPI_ISL_528650, EPI_ISL_528651, EPI_ISL_528652, EPI_ISL_528653, EPI_ISL_528654, EPI_ISL_528655, EPI_ISL_528656, EPI_ISL_528657, EPI_ISL_528658, EPI_ISL_528659, EPI_ISL_528660, EPI_ISL_528661, EPI_ISL_528662 | | | |
| see above | Virginia DCLS | Virginia DCLS | Virginia DCLS |
| EPI_ISL_528663, EPI_ISL_528664, EPI_ISL_528665, EPI_ISL_528666, EPI_ISL_528667, EPI_ISL_528668, EPI_ISL_528669, EPI_ISL_528670, EPI_ISL_528671, EPI_ISL_528672, EPI_ISL_528673, EPI_ISL_528674, EPI_ISL_528675, EPI_ISL_528676, EPI_ISL_528677, EPI_ISL_528678, EPI_ISL_528679, EPI_ISL_528680, EPI_ISL_528681, EPI_ISL_528682 | | | |
| see above | Virginia Division of Consolidated Laboratory Services (DCLS) | Virginia Division of Consolidated Laboratory Services (DCLS) | Virginia DCLS |
| EPI_ISL_528683, EPI_ISL_528684 | NCDC Institute of Genomics and Integrative Biology | NCDC Institute of Genomics and Integrative Biology | Vivekanand A, Mahesh S. Dhar, Bharathram Uppili, Akshay Kanakan, Simmi Tiwari, RadhaKrishnan VS, Robin Marwal, Azka Khan, Ajit Shewale, Pooja Sharma, Tushar Nale, Rajesh Pandey, Sandhya Kabra, Mohammed Faruq, Sujeet Singh, Anurag Agrawal, Partha Rakshit |
| EPI_ISL_528685 | IIP Institute of Genomics and Integrative Biology | IIP Institute of Genomics and Integrative Biology | Vivekanand A, Mahesh S. Dhar, Bharathram Uppili, Akshay Kanakan, Simmi Tiwari, RadhaKrishnan VS, Robin Marwal, Azka Khan, Ajit Shewale, Pooja Sharma, Tushar Nale, Rajesh Pandey, Sandhya Kabra, Mohammed Faruq, Sujeet Singh, Anurag Agrawal, Partha Rakshit |
| EPI_ISL_528810, EPI_ISL_528811, EPI_ISL_528812 | Department of Medicine, Gandhi hospital, Hyderabad | CSIR-Centre for Cellular and Molecular Biology | Thrilok Chander Bingi,Rajarao Mesipogu ,Vinayasekhar Aedula,Lamuk Zaveri, Shagufta Khan, Namami Gaur, Sakshi Shambhavi, Nikhil Hajirnis, M Soujanya Reddy, Pratheusa Maccha, Tulasi Nagabandi, Purushotham Vodnala, Payel Mukherjee, Sofia Banu, Priya Singh, Onkar Kulkarni, Dhiviya Vedagiri, Divya Gupta, Vishal Sah, Santosh Kumar Kuncha, Krishnan Harinivas Harshan, Archana Bharadwaj Siva, Karthik Bharadwaj Tallapaka, Renu Sudhakar, Somesh Gorde, Gangumala Srinivas Reddy, Sujoy Deb, Swati Bayyana, Rakesh K Mishra, Divya Tej Sowpati |
| EPI_ISL_528813, EPI_ISL_528815 | Department of Medicine, Gandhi hospital, Hyderabad | CSIR-Centre for Cellular and Molecular Biology | Vinayasekhar Aedula,Thrilok Chander Bingi, Rajarao Mesipogu, Shagufta Khan, Lamuk Zaveri, Namami Gaur, Sakshi Shambhavi, Nikhil Hajirnis, M Soujanya Reddy, Pratheusa Maccha,Tulasi Nagabandi, Purushotham Vodnala, Payel Mukherjee, Sofia Banu, Priya Singh, Onkar Kulkarni, Dhiviya Vedagiri, Divya Gupta, Vishal Sah, Santosh Kumar Kuncha, Krishnan Harinivas Harshan, Archana Bharadwaj Siva, Karthik Bharadwaj Tallapaka,Umesh Kumar, Unis Ahmad Bhat, Ajay Sarawagi, Priyanka Pant, Rajkanwar Nathawat, Rakesh K Mishra, Divya Tej Sowpati |
| EPI_ISL_528817 | Department of Medicine, Gandhi hospital, Hyderabad | CSIR-Centre for Cellular and Molecular Biology | Thrilok Chander Bingi,Rajarao Mesipogu ,Vinayasekhar Aedula,Lamuk Zaveri, Shagufta Khan, Namami Gaur, Sakshi Shambhavi, Nikhil Hajirnis, M Soujanya Reddy, Pratheusa Maccha, Tulasi Nagabandi, Purushotham Vodnala, Payel Mukherjee, Sofia Banu, Priya Singh, Onkar Kulkarni, Dhiviya Vedagiri, Divya Gupta, Vishal Sah, Santosh Kumar Kuncha, Krishnan Harinivas Harshan, Archana Bharadwaj Siva, Karthik Bharadwaj Tallapaka, Renu Sudhakar, Somesh Gorde, Gangumala Srinivas Reddy, Sujoy Deb, Swati Bayyana, Rakesh K Mishra, Divya Tej Sowpati |
| EPI_ISL_528818 | Department of Medicine, Gandhi hospital, Hyderabad | CSIR-Centre for Cellular and Molecular Biology | Vinayasekhar Aedula,Thrilok Chander Bingi, Rajarao Mesipogu, Shagufta Khan, Lamuk Zaveri, Namami Gaur, Sakshi Shambhavi, Nikhil Hajirnis, M Soujanya Reddy, Pratheusa Maccha,Tulasi Nagabandi, Purushotham Vodnala, Payel Mukherjee, Sofia Banu, Priya Singh, Onkar Kulkarni, Dhiviya |

| | | | |
|---|---|---|---|
| | | | Vedagiri, Divya Gupta, Vishal Sah, Santosh Kumar Kuncha, Krishnan Harinivas Harshan, Archana Bharadwaj Siva, Karthik Bharadwaj Tallapaka,Umesh Kumar, Unis Ahmad Bhat, Ajay Sarawagi, Priyanka Pant, Rajkanwar Nathawat, Rakesh K Mishra, Divya Tej Sowpati |
| EPI_ISL_528821 | Department of Medicine, Gandhi hospital, Hyderabad | CSIR-Centre for Cellular and Molecular Biology | Thrilok Chander Bingi,Rajarao Mesipogu ,Vinayasekhar Aedula,Lamuk Zaveri, Shagufta Khan, Namami Gaur, Sakshi Shambhavi, Nikhil Hajirnis, M Soujanya Reddy, Pratheusa Maccha, Tulasi Nagabandi, Purushotham Vodnala, Payel Mukherjee, Sofia Banu, Priya Singh, Onkar Kulkarni, Dhiviya Vedagiri, Divya Gupta, Vishal Sah, Santosh Kumar Kuncha, Krishnan Harinivas Harshan, Archana Bharadwaj Siva, Karthik Bharadwaj Tallapaka, Renu Sudhakar, Somesh Gorde, Gangumala Srinivas Reddy, Sujoy Deb, Swati Bayyana, Rakesh K Mishra, Divya Tej Sowpati |
| EPI_ISL_528822 | Department of Medicine, Gandhi hospital, Hyderabad | CSIR-Centre for Cellular and Molecular Biology | Vinayasekhar Aedula,Thrilok Chander Bingi, Rajarao Mesipogu, Shagufta Khan, Lamuk Zaveri, Namami Gaur, Sakshi Shambhavi, Nikhil Hajirnis, M Soujanya Reddy, Pratheusa Maccha,Tulasi Nagabandi, Purushotham Vodnala, Payel Mukherjee, Sofia Banu, Priya Singh, Onkar Kulkarni, Dhiviya Vedagiri, Divya Gupta, Vishal Sah, Santosh Kumar Kuncha, Krishnan Harinivas Harshan, Archana Bharadwaj Siva, Karthik Bharadwaj Tallapaka,Umesh Kumar, Unis Ahmad Bhat, Ajay Sarawagi, Priyanka Pant, Rajkanwar Nathawat, Rakesh K Mishra, Divya Tej Sowpati |
| EPI_ISL_528824, EPI_ISL_528830, EPI_ISL_528836, EPI_ISL_528840, EPI_ISL_528853, EPI_ISL_528860 | CSIR-Centre for Cellular and Molecular Biology | CSIR-Centre for Cellular and Molecular Biology | Tulasi Nagabandi, Namami Gaur, Sakshi Shambhavi, Lamuk Zaveri, Shagufta Khan, Nikhil Hajirnis, M Soujanya Reddy, Pratheusa Maccha, Purushotham Vodnala, Payel Mukherjee, Sofia Banu, Priya Singh, Onkar Kulkarni, Dhiviya Vedagiri, Divya Gupta, Vishal Sah, Santosh Kumar Kuncha, Krishnan Harinivas Harshan, Archana Bharadwaj Siva, Karthik Bharadwaj Tallapaka,G. Aditya Kumar, Koushick Sivakumar, Pooja Ramesh Gupta, Rajan Kumar Jha, Shraddha Vijay Lahoti, Rakesh K Mishra, Divya Tej Sowpati |
| EPI_ISL_529135 | Department of Virology III, National Institute of Infectious Diseases | Pathogen Genomics Center, National Institute of Infectious Diseases | Tsuyoshi Sekizuka, Shutoku Matsuyama, Kentaro Itokawa, Rina Tanaka, Masanori Hashino, Makoto Takeda, Takaji Wakita, Makoto Kuroda |
| EPI_ISL_529301, EPI_ISL_529353, EPI_ISL_529354, EPI_ISL_529355, EPI_ISL_529356, EPI_ISL_529382, EPI_ISL_529407, EPI_ISL_529419, EPI_ISL_529433, EPI_ISL_529446, EPI_ISL_529449, EPI_ISL_529500, EPI_ISL_529501, EPI_ISL_529502, EPI_ISL_529503, EPI_ISL_529521, EPI_ISL_529675, EPI_ISL_529676, EPI_ISL_529677, EPI_ISL_529678, EPI_ISL_529679, EPI_ISL_529680, EPI_ISL_529681, EPI_ISL_529682, EPI_ISL_529683 | | | |
| see above | Virology Department, Sheffield Teaching Hospitals NHS Foundation Trust/Department of Infection, Immunity and Cardiovascular Disease, The Medical School, University of Sheffield | COVID-19 Genomics UK (COG-UK) Consortium | Thushan de Silva, Matthew Parker, Nikki Smith, Adri Angyal, Rebecca Brown, Luke Green, Rachel Tucker, Paul Parsons, Danielle Groves, Katie Johnson, Laura Carrilero, Alex Keeley, Dave Partridge, Matthew Wyles, Benjamin Lindsey, Mehmet Yavuz, Mohammad Raza, Cariad Evans |
| EPI_ISL_529908, EPI_ISL_529909, EPI_ISL_529910, EPI_ISL_529911, EPI_ISL_529912, EPI_ISL_529913, EPI_ISL_529914, EPI_ISL_529915, EPI_ISL_529916, EPI_ISL_529917, EPI_ISL_529918, EPI_ISL_529919, EPI_ISL_529920, EPI_ISL_529921, EPI_ISL_529922, EPI_ISL_529923, EPI_ISL_529924, EPI_ISL_529925, EPI_ISL_529926, EPI_ISL_529927, EPI_ISL_529928, EPI_ISL_529929, EPI_ISL_529930, EPI_ISL_529931, EPI_ISL_529932, EPI_ISL_529933, EPI_ISL_529934, EPI_ISL_529935, EPI_ISL_529936 | | | |
| see above | Virginia Division of Consolidated Laboratory Services | Virginia Division of Consolidated Laboratory Services | Virginia DCLS |
| EPI_ISL_529937, EPI_ISL_529938, EPI_ISL_529939, EPI_ISL_529940, EPI_ISL_529941, EPI_ISL_529942, EPI_ISL_529943, EPI_ISL_529944, EPI_ISL_529945, EPI_ISL_529946, EPI_ISL_529947, EPI_ISL_529948, EPI_ISL_529949, EPI_ISL_529950, EPI_ISL_529951, EPI_ISL_529952, EPI_ISL_529953, EPI_ISL_529954, EPI_ISL_529955, EPI_ISL_529956, EPI_ISL_529957, EPI_ISL_529958, EPI_ISL_529959, EPI_ISL_529960 | | | |
| see above | Virginia DCLS | Virginia DCLS | Virginia DCLS |
| EPI_ISL_535043, EPI_ISL_535044, EPI_ISL_535045, EPI_ISL_535046, EPI_ISL_535047, EPI_ISL_535048, EPI_ISL_535049, EPI_ISL_535050, EPI_ISL_535051, EPI_ISL_535052, EPI_ISL_535053, EPI_ISL_535054, EPI_ISL_535055, EPI_ISL_535056, EPI_ISL_535057, EPI_ISL_535058, EPI_ISL_535059, EPI_ISL_535060, EPI_ISL_535061, EPI_ISL_535062, EPI_ISL_535063, EPI_ISL_535064, EPI_ISL_535065, EPI_ISL_535066, EPI_ISL_535067, EPI_ISL_535068, EPI_ISL_535069, EPI_ISL_535070, EPI_ISL_535071, EPI_ISL_535072, EPI_ISL_535073, EPI_ISL_535074, EPI_ISL_535075, EPI_ISL_535076, EPI_ISL_535077, EPI_ISL_535078, EPI_ISL_535079, EPI_ISL_535080, EPI_ISL_535081, EPI_ISL_535082, EPI_ISL_535083, EPI_ISL_535084, EPI_ISL_535085, EPI_ISL_535086, EPI_ISL_535087, EPI_ISL_535088, EPI_ISL_535089, EPI_ISL_535090, EPI_ISL_535091, EPI_ISL_535092, EPI_ISL_535093, EPI_ISL_535094, EPI_ISL_535095, EPI_ISL_535096, EPI_ISL_535097, EPI_ISL_535098, EPI_ISL_535099, EPI_ISL_535100, EPI_ISL_535101 | | | |
| see above | Virology Department, Sheffield Teaching Hospitals NHS Foundation Trust/Department of Infection, Immunity and Cardiovascular Disease, The Medical School, University of Sheffield | COVID-19 Genomics UK (COG-UK) Consortium | Thushan de Silva, Matthew Parker, Nikki Smith, Adri Angyal, Rebecca Brown, Luke Green, Rachel Tucker, Paul Parsons, Danielle Groves, Katie Johnson, Laura Carrilero, Alex Keeley, Dave Partridge, Matthew Wyles, Benjamin Lindsey, Mehmet Yavuz, Mohammad Raza, Cariad Evans |
| EPI_ISL_536399 | Laboratory of Immunovirology. Universidad de Antioquia | Instituto Nacional de Salud - Unidad de Secuenciación y Genómica | Wbeimar Aguilar-Jimenez, Lizdany Flórez, Francisco J. Díaz, Katherine Laiton-Donato, Carlos Franco-Muñoz, Diego Álvarez-Diaz and Marcela Mercado-Reyes |
| EPI_ISL_536788 | Southern Community Labs Dunedin | Institute of Environmental Science and Research (ESR) | Xiaoyun Ren, Matt Storey, Nikki Freed, Muhammad Faisal, Jing Wang, Hermes Perez, Anja Werno, Antje van der Linden, Arlo Upton, Chris Mansell, David Hammer, Dragana Drinkovic, Gary McAuliffe, Hana Sofia Andersson, James Ussher, Jill Sherwood, Josh Freeman, Julia Howard, Juliet Elvy, Mary DeAlmeida, Matt Blakiston, Matthew Rogers, Max Bloomfield, Michael Addidle, Michelle Balm, Sally Roberts, Sarah Jefferies, Sharmini Muttaiyah, Susan Morpeth, Susan Taylor, Timothy Blackmore, Vani Sathyendran, Veronica Playle, Virginia Hope, Erasmus Smit, Lauren Jelly, Olin Silander, Joep de Ligt |
| EPI_ISL_536790, EPI_ISL_536791 | Waikato Hospital | Institute of Environmental Science and Research (ESR) | Xiaoyun Ren, Matt Storey, Nikki Freed, Muhammad Faisal, Jing Wang, Hermes Perez, Anja Werno, Antje van der Linden, Arlo Upton, Chris Mansell, David Hammer, Dragana Drinkovic, Gary McAuliffe, Hana Sofia Andersson, James Ussher, Jill Sherwood, Josh Freeman, Julia Howard, Juliet Elvy, Mary DeAlmeida, Matt Blakiston, Matthew Rogers, Max Bloomfield, Michael Addidle, Michelle Balm, Sally Roberts, Sarah Jefferies, Sharmini Muttaiyah, Susan Morpeth, Susan Taylor, Timothy Blackmore, Vani Sathyendran, Veronica Playle, Virginia Hope, Erasmus Smit, Lauren Jelly, Olin Silander, Joep de Ligt |
| EPI_ISL_536792 | Canterbury Health Laboratories | Institute of Environmental Science and Research (ESR) | Xiaoyun Ren, Matt Storey, Nikki Freed, Muhammad Faisal, Jing Wang, Hermes Perez, Anja Werno, Antje van der Linden, Arlo Upton, Chris Mansell, David Hammer, Dragana Drinkovic, Gary McAuliffe, Hana Sofia Andersson, James Ussher, Jill Sherwood, Josh Freeman, Julia Howard, Juliet Elvy, Mary DeAlmeida, Matt Blakiston, Matthew Rogers, Max Bloomfield, Michael Addidle, Michelle Balm, Sally Roberts, Sarah Jefferies, Sharmini Muttaiyah, Susan Morpeth, Susan Taylor, Timothy Blackmore, Vani Sathyendran, Veronica Playle, Virginia Hope, Erasmus Smit, Lauren Jelly, Olin Silander, Joep de Ligt |
| EPI_ISL_537206, EPI_ISL_537207, EPI_ISL_537208, EPI_ISL_537209, EPI_ISL_537210, EPI_ISL_537211, EPI_ISL_537212, EPI_ISL_537214, EPI_ISL_537215, EPI_ISL_537216, EPI_ISL_537217, EPI_ISL_537218, EPI_ISL_537219, EPI_ISL_537220, EPI_ISL_537221, EPI_ISL_537222, EPI_ISL_537223, EPI_ISL_537224, EPI_ISL_537225, EPI_ISL_537226, EPI_ISL_537227, EPI_ISL_537228, EPI_ISL_537229, EPI_ISL_537230, EPI_ISL_537231, EPI_ISL_537232, EPI_ISL_537233, EPI_ISL_537234, EPI_ISL_537235, EPI_ISL_537236, EPI_ISL_537237, EPI_ISL_537238, EPI_ISL_537239, EPI_ISL_537240, EPI_ISL_537241, EPI_ISL_537242, EPI_ISL_537243, EPI_ISL_537244, EPI_ISL_537245, EPI_ISL_537246, EPI_ISL_537247, EPI_ISL_537248, EPI_ISL_537249, EPI_ISL_537250, EPI_ISL_537251, EPI_ISL_537252, EPI_ISL_537253, EPI_ISL_537255, EPI_ISL_537256, EPI_ISL_537257, EPI_ISL_537258, EPI_ISL_537259, EPI_ISL_537260, EPI_ISL_537261, EPI_ISL_537262, EPI_ISL_537263, EPI_ISL_537264, EPI_ISL_537265, EPI_ISL_537266, EPI_ISL_537267, EPI_ISL_537268, EPI_ISL_537269, EPI_ISL_537270, EPI_ISL_537271, EPI_ISL_537272, EPI_ISL_537273, EPI_ISL_537274, EPI_ISL_537275, EPI_ISL_537276, EPI_ISL_537277, EPI_ISL_537278, EPI_ISL_537280, EPI_ISL_537281, EPI_ISL_537282, EPI_ISL_537283, EPI_ISL_537284 | | | |
| see above | Virology Department, Sheffield Teaching Hospitals NHS Foundation Trust / Department of Infection, Immunity and Cardiovascular Disease, The Medical School, University of Sheffield | Wellcome Sanger Institute for the COVID-19 Genomics UK (COG-UK) consortium | Thushan de Silva, Matthew Parker,Adri Angyal, Rebecca Brown, Luke Green, Rachel Tucker, Paul Parsons, Danielle Groves, Alex Keeley, Dave Partridge, Matthew Wyles, Benjamin Lindsey, Mehmet Yavuz, Mohammad Raza, Cariad Evans and Alex Alderton, Roberto Amato, Sonia Goncalves, Ewan Harrison, David K. Jackson, Ian Johnston, Dominic Kwiatkowski, Cordelia Langford, John Sillitoe on behalf of the Wellcome Sanger Institute COVID-19 Surveillance Team |
| EPI_ISL_537996, EPI_ISL_537997, EPI_ISL_537998, EPI_ISL_537999, EPI_ISL_538000, EPI_ISL_538001, EPI_ISL_538002 | Servicio de Microbiologia. Hospital Arnau de Vilanova | SeqCOVID-SPAIN consortium/IBV(CSIC) | Victoria Dominguez, Maria Alma Bracho, Griselda De Marco, Lidia Ruiz Roldan, Neris Garcia-Gonzalez, Inma Galán Vendrell, Sandra Carbo, Loreto Ferrús Abad, Paula Ruiz-Hueso, Mariana Reyes-Prieto, Vicente Soriano Chirona, Ivan Ansari, Llúcia Martínez-Priego, Giuseppe D'Auria, Fernando Gonzalez-Candelas and SeqCOVID-SPAIN consortium |
| EPI_ISL_539643 | CSIR-Centre for Cellular and Molecular Biology | CSIR-Centre for Cellular and Molecular Biology | Tulasi Nagabandi, Namami Gaur, Sakshi Shambhavi, Lamuk Zaveri, Shagufta Khan, Nikhil Hajirnis, M Soujanya Reddy, Pratheusa Maccha, Purushotham Vodnala, Payel Mukherjee, Sofia Banu, Priya Singh, Onkar Kulkarni, Dhiviya Vedagiri, Divya Gupta, Vishal Sah, Santosh Kumar Kuncha, Krishnan Harinivas Harshan, Archana Bharadwaj Siva, Karthik Bharadwaj Tallapaka,G. Aditya Kumar, Koushick Sivakumar, Pooja Ramesh Gupta, Rajan Kumar Jha, Shraddha Vijay Lahoti, Rakesh K Mishra, Divya Tej Sowpati |
| EPI_ISL_539644 | CSIR-Centre for Cellular and Molecular Biology | CSIR-Centre for Cellular and Molecular Biology | Tulasi Nagabandi, Namami Gaur, Sakshi Shambhavi, Lamuk Zaveri, Shagufta Khan, Nikhil Hajirnis, M Soujanya Reddy, Pratheusa Maccha, Purushotham Vodnala, Payel Mukherjee, Sofia Banu, Priya Singh,Onkar Kulkarni , Dhiviya Vedagiri, Divya Gupta, Vishal Sah, Santosh Kumar Kuncha, Krishnan Harinivas Harshan, Archana Bharadwaj Siva, Karthik Bharadwaj Tallapaka,G. Aditya Kumar, Koushick Sivakumar, Pooja Ramesh Gupta, Rajan Kumar Jha, Shraddha Vijay Lahoti, Rakesh K Mishra, Divya Tej Sowpati |
| EPI_ISL_539645 | CSIR-Centre for Cellular and Molecular Biology | CSIR-Centre for Cellular and Molecular Biology | Tulasi Nagabandi, Namami Gaur, Sakshi Shambhavi, Lamuk Zaveri, Shagufta Khan, Nikhil Hajirnis, M Soujanya Reddy, Pratheusa Maccha, Purushotham Vodnala, Payel Mukherjee, Sofia Banu, Priya Singh,Onkar Kulkarni, Dhiviya Vedagiri, Divya Gupta, Vishal Sah, Santosh Kumar Kuncha, Krishnan Harinivas Harshan, Archana Bharadwaj Siva, Karthik Bharadwaj Tallapaka,Kezia J Ann, Radhika Khandelwal, Roshan Maku Venkata, Shemin Mansuri, Sonu Uday, Rakesh K Mishra, Divya Tej Sowpati |
| EPI_ISL_539673 | CSIR-Centre for Cellular and Molecular Biology | CSIR-Centre for Cellular and Molecular Biology | Tulasi Nagabandi, Namami Gaur, Sakshi Shambhavi, Lamuk Zaveri, Shagufta Khan, Nikhil Hajirnis, M Soujanya Reddy, Pratheusa Maccha, Purushotham |

| | | | |
|---|---|---|---|
| | | | Vodnala, Payel Mukherjee, Sofia Banu, Priya Singh, Onkar Kulkarni, Dhiviya Vedagiri, Divya Gupta, Vishal Sah, Santosh Kumar Kuncha, Krishnan Harinivas Harshan, Archana Bharadwaj Siva, Karthik Bharadwaj Tallapaka,G. Aditya Kumar, Koushick Sivakumar, Pooja Ramesh Gupta, Rajan Kumar Jha, Shraddha Vijay Lahoti, Rakesh K Mishra, Divya Tej Sowpati |
| EPI_ISL_539674 | CSIR-Centre for Cellular and Molecular Biology | CSIR-Centre for Cellular and Molecular Biology | Tulasi Nagabandi, Namami Gaur, Sakshi Shambhavi, Lamuk Zaveri, Shagufta Khan, Nikhil Hajirnis, M Soujanya Reddy, Pratheusa Maccha, Purushotham Vodnala, Payel Mukherjee, Sofia Banu, Priya Singh,Onkar Kulkarni , Dhiviya Vedagiri, Divya Gupta, Vishal Sah, Santosh Kumar Kuncha, Krishnan Harinivas Harshan, Archana Bharadwaj Siva, Karthik Bharadwaj Tallapaka,G. Aditya Kumar, Koushick Sivakumar, Pooja Ramesh Gupta, Rajan Kumar Jha, Shraddha Vijay Lahoti, Rakesh K Mishra, Divya Tej Sowpati |
| EPI_ISL_539675 | CSIR-Centre for Cellular and Molecular Biology | CSIR-Centre for Cellular and Molecular Biology | Tulasi Nagabandi, Namami Gaur, Sakshi Shambhavi, Lamuk Zaveri, Shagufta Khan, Nikhil Hajirnis, M Soujanya Reddy, Pratheusa Maccha, Purushotham Vodnala, Payel Mukherjee, Sofia Banu, Priya Singh, Onkar Kulkarni, Dhiviya Vedagiri, Divya Gupta, Vishal Sah, Santosh Kumar Kuncha, Krishnan Harinivas Harshan, Archana Bharadwaj Siva, Karthik Bharadwaj Tallapaka,Kezia J Ann, Radhika Khandelwal, Roshan Maku Venkata, Shemin Mansuri, Sonu Uday, Rakesh K Mishra, Divya Tej Sowpati |
| EPI_ISL_539703 | CSIR-Centre for Cellular and Molecular Biology | CSIR-Centre for Cellular and Molecular Biology | Tulasi Nagabandi, Namami Gaur, Sakshi Shambhavi, Lamuk Zaveri, Shagufta Khan, Nikhil Hajirnis, M Soujanya Reddy, Pratheusa Maccha, Purushotham Vodnala, Payel Mukherjee, Sofia Banu, Priya Singh, Onkar Kulkarni, Dhiviya Vedagiri, Divya Gupta, Vishal Sah, Santosh Kumar Kuncha, Krishnan Harinivas Harshan, Archana Bharadwaj Siva, Karthik Bharadwaj Tallapaka,G. Aditya Kumar, Koushick Sivakumar, Pooja Ramesh Gupta, Rajan Kumar Jha, Shraddha Vijay Lahoti, Rakesh K Mishra, Divya Tej Sowpati |
| EPI_ISL_539704 | CSIR-Centre for Cellular and Molecular Biology | CSIR-Centre for Cellular and Molecular Biology | Tulasi Nagabandi, Namami Gaur, Sakshi Shambhavi, Lamuk Zaveri, Shagufta Khan, Nikhil Hajirnis, M Soujanya Reddy, Pratheusa Maccha, Purushotham Vodnala, Payel Mukherjee, Sofia Banu, Priya Singh,Onkar Kulkarni , Dhiviya Vedagiri, Divya Gupta, Vishal Sah, Santosh Kumar Kuncha, Krishnan Harinivas Harshan, Archana Bharadwaj Siva, Karthik Bharadwaj Tallapaka,G. Aditya Kumar, Koushick Sivakumar, Pooja Ramesh Gupta, Rajan Kumar Jha, Shraddha Vijay Lahoti, Rakesh K Mishra, Divya Tej Sowpati |
| EPI_ISL_539705 | CSIR-Centre for Cellular and Molecular Biology | CSIR-Centre for Cellular and Molecular Biology | Tulasi Nagabandi, Namami Gaur, Sakshi Shambhavi, Lamuk Zaveri, Shagufta Khan, Nikhil Hajirnis, M Soujanya Reddy, Pratheusa Maccha, Purushotham Vodnala, Payel Mukherjee, Sofia Banu, Priya Singh, Onkar Kulkarni, Dhiviya Vedagiri, Divya Gupta, Vishal Sah, Santosh Kumar Kuncha, Krishnan Harinivas Harshan, Archana Bharadwaj Siva, Karthik Bharadwaj Tallapaka,Kezia J Ann, Radhika Khandelwal, Roshan Maku Venkata, Shemin Mansuri, Sonu Uday, Rakesh K Mishra, Divya Tej Sowpati |
| EPI_ISL_539733 | CSIR-Centre for Cellular and Molecular Biology | CSIR-Centre for Cellular and Molecular Biology | Tulasi Nagabandi, Namami Gaur, Sakshi Shambhavi, Lamuk Zaveri, Shagufta Khan, Nikhil Hajirnis, M Soujanya Reddy, Pratheusa Maccha, Purushotham Vodnala, Payel Mukherjee, Sofia Banu, Priya Singh, Onkar Kulkarni, Dhiviya Vedagiri, Divya Gupta, Vishal Sah, Santosh Kumar Kuncha, Krishnan Harinivas Harshan, Archana Bharadwaj Siva, Karthik Bharadwaj Tallapaka,G. Aditya Kumar, Koushick Sivakumar, Pooja Ramesh Gupta, Rajan Kumar Jha, Shraddha Vijay Lahoti, Rakesh K Mishra, Divya Tej Sowpati |
| EPI_ISL_539734 | CSIR-Centre for Cellular and Molecular Biology | CSIR-Centre for Cellular and Molecular Biology | Tulasi Nagabandi, Namami Gaur, Sakshi Shambhavi, Lamuk Zaveri, Shagufta Khan, Nikhil Hajirnis, M Soujanya Reddy, Pratheusa Maccha, Purushotham Vodnala, Payel Mukherjee, Sofia Banu, Priya Singh,Onkar Kulkarni , Dhiviya Vedagiri, Divya Gupta, Vishal Sah, Santosh Kumar Kuncha, Krishnan Harinivas Harshan, Archana Bharadwaj Siva, Karthik Bharadwaj Tallapaka,G. Aditya Kumar, Koushick Sivakumar, Pooja Ramesh Gupta, Rajan Kumar Jha, Shraddha Vijay Lahoti, Rakesh K Mishra, Divya Tej Sowpati |
| EPI_ISL_539735 | CSIR-Centre for Cellular and Molecular Biology | CSIR-Centre for Cellular and Molecular Biology | Tulasi Nagabandi, Namami Gaur, Sakshi Shambhavi, Lamuk Zaveri, Shagufta Khan, Nikhil Hajirnis, M Soujanya Reddy, Pratheusa Maccha, Purushotham Vodnala, Payel Mukherjee, Sofia Banu, Priya Singh,Onkar Kulkarni, Dhiviya Vedagiri, Divya Gupta, Vishal Sah, Santosh Kumar Kuncha, Krishnan Harinivas Harshan, Archana Bharadwaj Siva, Karthik Bharadwaj Tallapaka,Kezia J Ann, Radhika Khandelwal, Roshan Maku Venkata, Shemin Mansuri, Sonu Uday, Rakesh K Mishra, Divya Tej Sowpati |
| EPI_ISL_539763 | CSIR-Centre for Cellular and Molecular Biology | CSIR-Centre for Cellular and Molecular Biology | Tulasi Nagabandi, Namami Gaur, Sakshi Shambhavi, Lamuk Zaveri, Shagufta Khan, Nikhil Hajirnis, M Soujanya Reddy, Pratheusa Maccha, Purushotham Vodnala, Payel Mukherjee, Sofia Banu, Priya Singh, Onkar Kulkarni, Dhiviya Vedagiri, Divya Gupta, Vishal Sah, Santosh Kumar Kuncha, Krishnan Harinivas Harshan, Archana Bharadwaj Siva, Karthik Bharadwaj Tallapaka,G. Aditya Kumar, Koushick Sivakumar, Pooja Ramesh Gupta, Rajan Kumar Jha, Shraddha Vijay Lahoti, Rakesh K Mishra, Divya Tej Sowpati |
| EPI_ISL_539764 | CSIR-Centre for Cellular and Molecular Biology | CSIR-Centre for Cellular and Molecular Biology | Tulasi Nagabandi, Namami Gaur, Sakshi Shambhavi, Lamuk Zaveri, Shagufta Khan, Nikhil Hajirnis, M Soujanya Reddy, Pratheusa Maccha, Purushotham Vodnala, Payel Mukherjee, Sofia Banu, Priya Singh,Onkar Kulkarni , Dhiviya Vedagiri, Divya Gupta, Vishal Sah, Santosh Kumar Kuncha, Krishnan Harinivas Harshan, Archana Bharadwaj Siva, Karthik Bharadwaj Tallapaka,G. Aditya Kumar, Koushick Sivakumar, Pooja Ramesh Gupta, Rajan Kumar Jha, Shraddha Vijay Lahoti, Rakesh K Mishra, Divya Tej Sowpati |
| EPI_ISL_539765 | CSIR-Centre for Cellular and Molecular Biology | CSIR-Centre for Cellular and Molecular Biology | Tulasi Nagabandi, Namami Gaur, Sakshi Shambhavi, Lamuk Zaveri, Shagufta Khan, Nikhil Hajirnis, M Soujanya Reddy, Pratheusa Maccha, Purushotham Vodnala, Payel Mukherjee, Sofia Banu, Priya Singh, Onkar Kulkarni, Dhiviya Vedagiri, Divya Gupta, Vishal Sah, Santosh Kumar Kuncha, Krishnan Harinivas Harshan, Archana Bharadwaj Siva, Karthik Bharadwaj Tallapaka,Kezia J Ann, Radhika Khandelwal, Roshan Maku Venkata, Shemin Mansuri, Sonu Uday, Rakesh K Mishra, Divya Tej Sowpati |
| EPI_ISL_539766 | CSIR-Centre for Cellular and Molecular Biology | CSIR-Centre for Cellular and Molecular Biology | Tulasi Nagabandi, Namami Gaur, Sakshi Shambhavi, Lamuk Zaveri, Shagufta Khan, Nikhil Hajirnis, M Soujanya Reddy, Pratheusa Maccha, Purushotham Vodnala, Payel Mukherjee, Sofia Banu, Priya Singh, Onkar Kulkarni, Dhiviya Vedagiri, Divya Gupta, Vishal Sah, Santosh Kumar Kuncha, Krishnan Harinivas Harshan, Archana Bharadwaj Siva, Karthik Bharadwaj Tallapaka,G. Aditya Kumar, Koushick Sivakumar, Pooja Ramesh Gupta, Rajan Kumar Jha, Shraddha Vijay Lahoti, Rakesh K Mishra, Divya Tej Sowpati |
| EPI_ISL_539767 | CSIR-Centre for Cellular and Molecular Biology | CSIR-Centre for Cellular and Molecular Biology | Tulasi Nagabandi, Namami Gaur, Sakshi Shambhavi, Lamuk Zaveri, Shagufta Khan, Nikhil Hajirnis, M Soujanya Reddy, Pratheusa Maccha, Purushotham Vodnala, Payel Mukherjee, Sofia Banu, Priya Singh,Onkar Kulkarni , Dhiviya Vedagiri, Divya Gupta, Vishal Sah, Santosh Kumar Kuncha, Krishnan Harinivas Harshan, Archana Bharadwaj Siva, Karthik Bharadwaj Tallapaka,G. Aditya Kumar, Koushick Sivakumar, Pooja Ramesh Gupta, Rajan Kumar Jha, Shraddha Vijay Lahoti, Rakesh K Mishra, Divya Tej Sowpati |
| EPI_ISL_539768 | CSIR-Centre for Cellular and Molecular Biology | CSIR-Centre for Cellular and Molecular Biology | Tulasi Nagabandi, Namami Gaur, Sakshi Shambhavi, Lamuk Zaveri, Shagufta Khan, Nikhil Hajirnis, M Soujanya Reddy, Pratheusa Maccha, Purushotham Vodnala, Payel Mukherjee, Sofia Banu, Priya Singh, Onkar Kulkarni, Dhiviya Vedagiri, Divya Gupta, Vishal Sah, Santosh Kumar Kuncha, Krishnan Harinivas Harshan, Archana Bharadwaj Siva, Karthik Bharadwaj Tallapaka,Kezia J Ann, Radhika Khandelwal, Roshan Maku Venkata, Shemin Mansuri, Sonu Uday, Rakesh K Mishra, Divya Tej Sowpati |
| EPI_ISL_540719, EPI_ISL_540720, EPI_ISL_540721, EPI_ISL_540722, EPI_ISL_540723, EPI_ISL_540724, EPI_ISL_540725, EPI_ISL_540726, EPI_ISL_540727, EPI_ISL_540728, EPI_ISL_540729, EPI_ISL_540730, EPI_ISL_540731, EPI_ISL_540732, EPI_ISL_540733, EPI_ISL_540734, EPI_ISL_540735, EPI_ISL_540736, EPI_ISL_540737, EPI_ISL_540738, EPI_ISL_540739, EPI_ISL_540740, EPI_ISL_540741, EPI_ISL_540742, EPI_ISL_540743, EPI_ISL_540744, EPI_ISL_540745, EPI_ISL_540746, EPI_ISL_540747, EPI_ISL_540748, EPI_ISL_540749, EPI_ISL_540750, EPI_ISL_540751, EPI_ISL_540752, EPI_ISL_540753, EPI_ISL_540754, EPI_ISL_540755, EPI_ISL_540756, EPI_ISL_540757, EPI_ISL_540758, EPI_ISL_540759, EPI_ISL_540760, EPI_ISL_540761, EPI_ISL_540762, EPI_ISL_540763, EPI_ISL_540764, EPI_ISL_540765, EPI_ISL_540766, EPI_ISL_540767, EPI_ISL_540768, EPI_ISL_540769, EPI_ISL_540770, EPI_ISL_540771, EPI_ISL_540772, EPI_ISL_540773, EPI_ISL_540774 | | | |
| see above | Virology Department, Sheffield Teaching Hospitals NHS Foundation Trust/Department of Infection, Immunity and Cardiovascular Disease, The Medical School, University of Sheffield | COVID-19 Genomics UK (COG-UK) Consortium | Thushan de Silva, Matthew Parker, Nikki Smith, Adri Angyal, Rebecca Brown, Luke Green, Rachel Tucker, Paul Parsons, Danielle Groves, Katie Johnson, Laura Carrilero, Alex Keeley, Dave Partridge, Matthew Wyles, Benjamin Lindsey, Mehmet Yavuz, Mohammad Raza, Cariad Evans |
| EPI_ISL_541649, EPI_ISL_541650, EPI_ISL_541651, EPI_ISL_541652, EPI_ISL_541653, EPI_ISL_541654, EPI_ISL_541655 | Laboratory Diagnostic, Veterinary Specialized Institute Kraljevo | Laboratory Diagnostic, Veterinary Specialized Institute Kraljevo | Vidanovic,D., Tesovic,B., Knezevic,A., Jovanovic,T., Jankovic,M., Sekler,M., Banovic Djeri,B., Volkening,J., Afonso,C., Petrovic,T. |
| EPI_ISL_541656 | Laboratory Diagnostic, Veterinary Specialized Institute Kraljevo | Laboratory Diagnostic, Veterinary Specialized Institute Kraljevo | Vidanovic,D., Tesovic,B., Knezevic,A., Jovanovic,T., Jankovic,M., Sekler,M., Banovic Djeri,B., Volkening,J., Afonso,C., Petrovic,T. |
| EPI_ISL_541657, EPI_ISL_541658, EPI_ISL_541659, EPI_ISL_541660, EPI_ISL_541661, EPI_ISL_541662 | Laboratory Diagnostic, Veterinary Specialized Institute Kraljevo | Laboratory Diagnostic, Veterinary Specialized Institute Kraljevo | Vidanovic,D., Tesovic,B., Knezevic,A., Jovanovic,T., Jankovic,M., Sekler,M., Banovic Djeri,B., Volkening,J., Afonso,C., Petrovic,T. |
| EPI_ISL_548038, EPI_ISL_548039 | Middlemore Hospital | Institute of Environmental Science and Research (ESR) | Xiaoyun Ren, Matt Storey, Nikki Freed, Muhammad Faisal, Jing Wang, Hermes Perez, Anja Werno, Antje van der Linden, Arlo Upton, Chris Mansell, David |

| | | | |
|---|---|---|---|
| | | | Hammer, Dragana Drinkovic, Gary McAuliffe, Hana Sofia Andersson, James Ussher, Jill Sherwood, Josh Freeman, Julia Howard, Juliet Elvy, Mary DeAlmeida, Matt Blakiston, Matthew Rogers, Max Bloomfield, Michael Addidle, Michelle Balm, Sally Roberts, Sarah Jefferies, Sharmini Muttaiyah, Susan Morpeth, Susan Taylor, Timothy Blackmore, Vani Sathyendran, Veronica Playle, Virginia Hope, Erasmus Smit, Lauren Jelly, Olin Silander, Joep de Ligt |
| EPI_ISL_548051 | LabPLUS | Institute of Environmental Science and Research (ESR) | Xiaoyun Ren, Matt Storey, Nikki Freed, Muhammad Faisal, Jing Wang, Hermes Perez, Anja Werno, Antje van der Linden, Arlo Upton, Chris Mansell, David Hammer, Dragana Drinkovic, Gary McAuliffe, Hana Sofia Andersson, James Ussher, Jill Sherwood, Josh Freeman, Julia Howard, Juliet Elvy, Mary DeAlmeida, Matt Blakiston, Matthew Rogers, Max Bloomfield, Michael Addidle, Michelle Balm, Sally Roberts, Sarah Jefferies, Sharmini Muttaiyah, Susan Morpeth, Susan Taylor, Timothy Blackmore, Vani Sathyendran, Veronica Playle, Virginia Hope, Erasmus Smit, Lauren Jelly, Olin Silander, Joep de Ligt |
| EPI_ISL_548052, EPI_ISL_548053, EPI_ISL_548054, EPI_ISL_548055, EPI_ISL_548056 | Middlemore Hospital | Institute of Environmental Science and Research (ESR) | Xiaoyun Ren, Matt Storey, Nikki Freed, Muhammad Faisal, Jing Wang, Hermes Perez, Anja Werno, Antje van der Linden, Arlo Upton, Chris Mansell, David Hammer, Dragana Drinkovic, Gary McAuliffe, Hana Sofia Andersson, James Ussher, Jill Sherwood, Josh Freeman, Julia Howard, Juliet Elvy, Mary DeAlmeida, Matt Blakiston, Matthew Rogers, Max Bloomfield, Michael Addidle, Michelle Balm, Sally Roberts, Sarah Jefferies, Sharmini Muttaiyah, Susan Morpeth, Susan Taylor, Timothy Blackmore, Vani Sathyendran, Veronica Playle, Virginia Hope, Erasmus Smit, Lauren Jelly, Olin Silander, Joep de Ligt |
| EPI_ISL_548058, EPI_ISL_548059, EPI_ISL_548060, EPI_ISL_548062, EPI_ISL_548063, EPI_ISL_548065, EPI_ISL_548066 | North Shore Hospital | Institute of Environmental Science and Research (ESR) | Xiaoyun Ren, Matt Storey, Nikki Freed, Muhammad Faisal, Jing Wang, Hermes Perez, Anja Werno, Antje van der Linden, Arlo Upton, Chris Mansell, David Hammer, Dragana Drinkovic, Gary McAuliffe, Hana Sofia Andersson, James Ussher, Jill Sherwood, Josh Freeman, Julia Howard, Juliet Elvy, Mary DeAlmeida, Matt Blakiston, Matthew Rogers, Max Bloomfield, Michael Addidle, Michelle Balm, Sally Roberts, Sarah Jefferies, Sharmini Muttaiyah, Susan Morpeth, Susan Taylor, Timothy Blackmore, Vani Sathyendran, Veronica Playle, Virginia Hope, Erasmus Smit, Lauren Jelly, Olin Silander, Joep de Ligt |
| EPI_ISL_548068 | LabPLUS | Institute of Environmental Science and Research (ESR) | Xiaoyun Ren, Matt Storey, Nikki Freed, Muhammad Faisal, Jing Wang, Hermes Perez, Anja Werno, Antje van der Linden, Arlo Upton, Chris Mansell, David Hammer, Dragana Drinkovic, Gary McAuliffe, Hana Sofia Andersson, James Ussher, Jill Sherwood, Josh Freeman, Julia Howard, Juliet Elvy, Mary DeAlmeida, Matt Blakiston, Matthew Rogers, Max Bloomfield, Michael Addidle, Michelle Balm, Sally Roberts, Sarah Jefferies, Sharmini Muttaiyah, Susan Morpeth, Susan Taylor, Timothy Blackmore, Vani Sathyendran, Veronica Playle, Virginia Hope, Erasmus Smit, Lauren Jelly, Olin Silander, Joep de Ligt |
| EPI_ISL_548074 | Middlemore Hospital | Institute of Environmental Science and Research (ESR) | Xiaoyun Ren, Matt Storey, Nikki Freed, Muhammad Faisal, Jing Wang, Hermes Perez, Anja Werno, Antje van der Linden, Arlo Upton, Chris Mansell, David Hammer, Dragana Drinkovic, Gary McAuliffe, Hana Sofia Andersson, James Ussher, Jill Sherwood, Josh Freeman, Julia Howard, Juliet Elvy, Mary DeAlmeida, Matt Blakiston, Matthew Rogers, Max Bloomfield, Michael Addidle, Michelle Balm, Sally Roberts, Sarah Jefferies, Sharmini Muttaiyah, Susan Morpeth, Susan Taylor, Timothy Blackmore, Vani Sathyendran, Veronica Playle, Virginia Hope, Erasmus Smit, Lauren Jelly, Olin Silander, Joep de Ligt |
| EPI_ISL_548080, EPI_ISL_548083 | LabPLUS | Institute of Environmental Science and Research (ESR) | Xiaoyun Ren, Matt Storey, Nikki Freed, Muhammad Faisal, Jing Wang, Hermes Perez, Anja Werno, Antje van der Linden, Arlo Upton, Chris Mansell, David Hammer, Dragana Drinkovic, Gary McAuliffe, Hana Sofia Andersson, James Ussher, Jill Sherwood, Josh Freeman, Julia Howard, Juliet Elvy, Mary DeAlmeida, Matt Blakiston, Matthew Rogers, Max Bloomfield, Michael Addidle, Michelle Balm, Sally Roberts, Sarah Jefferies, Sharmini Muttaiyah, Susan Morpeth, Susan Taylor, Timothy Blackmore, Vani Sathyendran, Veronica Playle, Virginia Hope, Erasmus Smit, Lauren Jelly, Olin Silander, Joep de Ligt |
| EPI_ISL_548094, EPI_ISL_548095, EPI_ISL_548096, EPI_ISL_548097 | Middlemore Hospital | Institute of Environmental Science and Research (ESR) | Xiaoyun Ren, Matt Storey, Nikki Freed, Muhammad Faisal, Jing Wang, Hermes Perez, Anja Werno, Antje van der Linden, Arlo Upton, Chris Mansell, David Hammer, Dragana Drinkovic, Gary McAuliffe, Hana Sofia Andersson, James Ussher, Jill Sherwood, Josh Freeman, Julia Howard, Juliet Elvy, Mary DeAlmeida, Matt Blakiston, Matthew Rogers, Max Bloomfield, Michael Addidle, Michelle Balm, Sally Roberts, Sarah Jefferies, Sharmini Muttaiyah, Susan Morpeth, Susan Taylor, Timothy Blackmore, Vani Sathyendran, Veronica Playle, Virginia Hope, Erasmus Smit, Lauren Jelly, Olin Silander, Joep de Ligt |
| EPI_ISL_548102 | Waikato Hospital | Institute of Environmental Science and Research (ESR) | Xiaoyun Ren, Matt Storey, Nikki Freed, Muhammad Faisal, Jing Wang, Hermes Perez, Anja Werno, Antje van der Linden, Arlo Upton, Chris Mansell, David Hammer, Dragana Drinkovic, Gary McAuliffe, Hana Sofia Andersson, James Ussher, Jill Sherwood, Josh Freeman, Julia Howard, Juliet Elvy, Mary DeAlmeida, Matt Blakiston, Matthew Rogers, Max Bloomfield, Michael Addidle, Michelle Balm, Sally Roberts, Sarah Jefferies, Sharmini Muttaiyah, Susan Morpeth, Susan Taylor, Timothy Blackmore, Vani Sathyendran, Veronica Playle, Virginia Hope, Erasmus Smit, Lauren Jelly, Olin Silander, Joep de Ligt |
| EPI_ISL_548103 | Middlemore Hospital | Institute of Environmental Science and Research (ESR) | Xiaoyun Ren, Matt Storey, Nikki Freed, Muhammad Faisal, Jing Wang, Hermes Perez, Anja Werno, Antje van der Linden, Arlo Upton, Chris Mansell, David Hammer, Dragana Drinkovic, Gary McAuliffe, Hana Sofia Andersson, James Ussher, Jill Sherwood, Josh Freeman, Julia Howard, Juliet Elvy, Mary DeAlmeida, Matt Blakiston, Matthew Rogers, Max Bloomfield, Michael Addidle, Michelle Balm, Sally Roberts, Sarah Jefferies, Sharmini Muttaiyah, Susan Morpeth, Susan Taylor, Timothy Blackmore, Vani Sathyendran, Veronica Playle, Virginia Hope, Erasmus Smit, Lauren Jelly, Olin Silander, Joep de Ligt |
| EPI_ISL_548109 | LabPLUS | Institute of Environmental Science and Research (ESR) | Xiaoyun Ren, Matt Storey, Nikki Freed, Muhammad Faisal, Jing Wang, Hermes Perez, Anja Werno, Antje van der Linden, Arlo Upton, Chris Mansell, David Hammer, Dragana Drinkovic, Gary McAuliffe, Hana Sofia Andersson, James Ussher, Jill Sherwood, Josh Freeman, Julia Howard, Juliet Elvy, Mary DeAlmeida, Matt Blakiston, Matthew Rogers, Max Bloomfield, Michael Addidle, Michelle Balm, Sally Roberts, Sarah Jefferies, Sharmini Muttaiyah, Susan Morpeth, Susan Taylor, Timothy Blackmore, Vani Sathyendran, Veronica Playle, Virginia Hope, Erasmus Smit, Lauren Jelly, Olin Silander, Joep de Ligt |
| EPI_ISL_548112, EPI_ISL_548113, EPI_ISL_548114, EPI_ISL_548115 | Middlemore Hospital | Institute of Environmental Science and Research (ESR) | Xiaoyun Ren, Matt Storey, Nikki Freed, Muhammad Faisal, Jing Wang, Hermes Perez, Anja Werno, Antje van der Linden, Arlo Upton, Chris Mansell, David Hammer, Dragana Drinkovic, Gary McAuliffe, Hana Sofia Andersson, James Ussher, Jill Sherwood, Josh Freeman, Julia Howard, Juliet Elvy, Mary DeAlmeida, Matt Blakiston, Matthew Rogers, Max Bloomfield, Michael Addidle, Michelle Balm, Sally Roberts, Sarah Jefferies, Sharmini Muttaiyah, Susan Morpeth, Susan Taylor, Timothy Blackmore, Vani Sathyendran, Veronica Playle, Virginia Hope, Erasmus Smit, Lauren Jelly, Olin Silander, Joep de Ligt |
| EPI_ISL_548116, EPI_ISL_548117, EPI_ISL_548118 | Canterbury Health Laboratories | Institute of Environmental Science and Research (ESR) | Xiaoyun Ren, Matt Storey, Nikki Freed, Muhammad Faisal, Jing Wang, Hermes Perez, Anja Werno, Antje van der Linden, Arlo Upton, Chris Mansell, David Hammer, Dragana Drinkovic, Gary McAuliffe, Hana Sofia Andersson, James Ussher, Jill Sherwood, Josh Freeman, Julia Howard, Juliet Elvy, Mary DeAlmeida, Matt Blakiston, Matthew Rogers, Max Bloomfield, Michael Addidle, Michelle Balm, Sally Roberts, Sarah Jefferies, Sharmini Muttaiyah, Susan Morpeth, Susan Taylor, Timothy Blackmore, Vani Sathyendran, Veronica Playle, Virginia Hope, Erasmus Smit, Lauren Jelly, Olin Silander, Joep de Ligt |
| EPI_ISL_548119, EPI_ISL_548122 | LabPLUS | Institute of Environmental Science and Research (ESR) | Xiaoyun Ren, Matt Storey, Nikki Freed, Muhammad Faisal, Jing Wang, Hermes Perez, Anja Werno, Antje van der Linden, Arlo Upton, Chris Mansell, David Hammer, Dragana Drinkovic, Gary McAuliffe, Hana Sofia Andersson, James Ussher, Jill Sherwood, Josh Freeman, Julia Howard, Juliet Elvy, Mary DeAlmeida, Matt Blakiston, Matthew Rogers, Max Bloomfield, Michael Addidle, Michelle Balm, Sally Roberts, Sarah Jefferies, Sharmini Muttaiyah, Susan Morpeth, Susan Taylor, Timothy Blackmore, Vani Sathyendran, Veronica Playle, Virginia Hope, Erasmus Smit, Lauren Jelly, Olin Silander, Joep de Ligt |
| EPI_ISL_548125, EPI_ISL_548127, EPI_ISL_548130, EPI_ISL_548133 | Middlemore Hospital | Institute of Environmental Science and Research (ESR) | Xiaoyun Ren, Matt Storey, Nikki Freed, Muhammad Faisal, Jing Wang, Hermes Perez, Anja Werno, Antje van der Linden, Arlo Upton, Chris Mansell, David Hammer, Dragana Drinkovic, Gary McAuliffe, Hana Sofia Andersson, James Ussher, Jill Sherwood, Josh Freeman, Julia Howard, Juliet Elvy, Mary DeAlmeida, Matt Blakiston, Matthew Rogers, Max Bloomfield, Michael Addidle, Michelle Balm, Sally Roberts, Sarah Jefferies, Sharmini Muttaiyah, Susan Morpeth, Susan Taylor, Timothy Blackmore, Vani Sathyendran, Veronica Playle, Virginia Hope, Erasmus Smit, Lauren Jelly, Olin Silander, Joep de Ligt |
| EPI_ISL_548134, EPI_ISL_548138 | Canterbury Health Laboratories | Institute of Environmental Science and Research (ESR) | Xiaoyun Ren, Matt Storey, Nikki Freed, Muhammad Faisal, Jing Wang, Hermes Perez, Anja Werno, Antje van der Linden, Arlo Upton, Chris Mansell, David Hammer, Dragana Drinkovic, Gary McAuliffe, Hana Sofia Andersson, James Ussher, Jill Sherwood, Josh Freeman, Julia Howard, Juliet Elvy, Mary DeAlmeida, Matt Blakiston, Matthew Rogers, Max Bloomfield, Michael Addidle, Michelle Balm, Sally Roberts, Sarah Jefferies, Sharmini Muttaiyah, Susan Morpeth, Susan Taylor, Timothy Blackmore, Vani Sathyendran, Veronica Playle, Virginia Hope, Erasmus Smit, Lauren Jelly, Olin Silander, Joep de Ligt |
| EPI_ISL_548144, EPI_ISL_548145, EPI_ISL_548146 | Middlemore Hospital | Institute of Environmental Science and Research (ESR) | Xiaoyun Ren, Matt Storey, Nikki Freed, Muhammad Faisal, Jing Wang, Hermes Perez, Anja Werno, Antje van der Linden, Arlo Upton, Chris Mansell, David Hammer, Dragana Drinkovic, Gary McAuliffe, Hana Sofia Andersson, James Ussher, Jill Sherwood, Josh Freeman, Julia Howard, Juliet Elvy, Mary DeAlmeida, Matt Blakiston, Matthew Rogers, Max Bloomfield, Michael Addidle, Michelle Balm, Sally Roberts, Sarah Jefferies, Sharmini Muttaiyah, Susan Morpeth, Susan Taylor, Timothy Blackmore, Vani Sathyendran, Veronica Playle, Virginia Hope, Erasmus Smit, Lauren Jelly, Olin Silander, Joep de Ligt |
| EPI_ISL_548966, EPI_ISL_548967, EPI_ISL_548968, EPI_ISL_548969, EPI_ISL_548970, EPI_ISL_548971 | Expo2020 Emergency Center | Agiomix | Walaa Allam, Cherif Ben Hamada, Cengiz Yakicier, Walid Dridi, Rashid Mohammed, Tamer Degheidy |
| EPI_ISL_549809, EPI_ISL_549813, EPI_ISL_549824, EPI_ISL_549851, EPI_ISL_549874, EPI_ISL_549884, EPI_ISL_549891, EPI_ISL_549907, EPI_ISL_549938, EPI_ISL_549943, EPI_ISL_549946, EPI_ISL_549972, EPI_ISL_549980, EPI_ISL_549981, EPI_ISL_550021, EPI_ISL_550039, EPI_ISL_550047, EPI_ISL_550051, EPI_ISL_550063, EPI_ISL_550072, EPI_ISL_550076, EPI_ISL_550099, EPI_ISL_550116, EPI_ISL_550125, EPI_ISL_550126, EPI_ISL_550144, EPI_ISL_550154, EPI_ISL_550162, EPI_ISL_550168, EPI_ISL_550170, EPI_ISL_550179, EPI_ISL_550191, EPI_ISL_550243 | | | |
| see above | Lighthouse Lab in Milton Keynes | Wellcome Sanger Institute for the COVID-19 Genomics UK (COG-UK) consortium | The Lighthouse Lab in Milton Keynes and Alex Alderton, Roberto Amato, Sonia Goncalves, Ewan Harrison, David K. Jackson, Ian Johnston, Dominic Kwiatkowski, Cordelia Langford, John Sillitoe on behalf of the Wellcome Sanger Institute COVID-19 Surveillance Team (http://www.sanger.ac.uk/covid-team) |
| EPI_ISL_550299 | Lighthouse Lab in Milton Keynes | Wellcome Sanger Institute for the COVID-19 Genomics UK (COG-UK) consortium | The Lighthouse Lab in Milton Keynes and Alex Alderton, Roberto Amato, Sonia Goncalves, Ewan Harrison, David K. Jackson, Ian Johnston, Dominic Kwiatkowski, Cordelia Langford, John Sillitoe on behalf of the Wellcome Sanger Institute COVID-19 Surveillance Team |

| | | | |
|---|---|---|---|
| EPI_ISL_550300, EPI_ISL_550312, EPI_ISL_550325, EPI_ISL_550341, EPI_ISL_550346, EPI_ISL_550353, EPI_ISL_550358, EPI_ISL_550370, EPI_ISL_550378, EPI_ISL_550433, EPI_ISL_550476, EPI_ISL_550480, EPI_ISL_550482, EPI_ISL_550493, EPI_ISL_550512, EPI_ISL_550538, EPI_ISL_550562, EPI_ISL_550590, EPI_ISL_550593, EPI_ISL_550594, EPI_ISL_550603, EPI_ISL_550641, EPI_ISL_550644, EPI_ISL_550660, EPI_ISL_550677, EPI_ISL_550724, EPI_ISL_550739, EPI_ISL_550745, EPI_ISL_550746, EPI_ISL_550760, EPI_ISL_550838, EPI_ISL_550841, EPI_ISL_550856, EPI_ISL_550863, EPI_ISL_550868, EPI_ISL_550869, EPI_ISL_550917, EPI_ISL_550919, EPI_ISL_550929, EPI_ISL_550936, EPI_ISL_550993, EPI_ISL_551047, EPI_ISL_551051, EPI_ISL_551088, EPI_ISL_551090, EPI_ISL_551108, EPI_ISL_551123, EPI_ISL_551125, EPI_ISL_551177, EPI_ISL_551200, EPI_ISL_551204, EPI_ISL_551216, EPI_ISL_551226, EPI_ISL_551250, EPI_ISL_551254, EPI_ISL_551256, EPI_ISL_551266, EPI_ISL_551272, EPI_ISL_551283, EPI_ISL_551590, EPI_ISL_551599, EPI_ISL_551602, EPI_ISL_551614, EPI_ISL_551633, EPI_ISL_551635, EPI_ISL_551657, EPI_ISL_551659, EPI_ISL_551660, EPI_ISL_551664, EPI_ISL_551666, EPI_ISL_551668, EPI_ISL_551674, EPI_ISL_551687, EPI_ISL_551694, EPI_ISL_551698, EPI_ISL_551722, EPI_ISL_551725, EPI_ISL_551770, EPI_ISL_551771, EPI_ISL_551778, EPI_ISL_551783, EPI_ISL_551788, EPI_ISL_551794, EPI_ISL_551805, EPI_ISL_551816, EPI_ISL_551922, EPI_ISL_551936, EPI_ISL_551937, EPI_ISL_551941, EPI_ISL_551942, EPI_ISL_551957, EPI_ISL_551961, EPI_ISL_551964, EPI_ISL_551991, EPI_ISL_551993, EPI_ISL_551997, EPI_ISL_552006, EPI_ISL_552014, EPI_ISL_552015, EPI_ISL_552020, EPI_ISL_552021, EPI_ISL_552030, EPI_ISL_552037, EPI_ISL_552044, EPI_ISL_552046, EPI_ISL_552061, EPI_ISL_552082, EPI_ISL_552083, EPI_ISL_552095, EPI_ISL_552111, EPI_ISL_552126, EPI_ISL_552138, EPI_ISL_552148, EPI_ISL_552160, EPI_ISL_552161, EPI_ISL_552202, EPI_ISL_552204, EPI_ISL_552207, EPI_ISL_552209, EPI_ISL_552210, EPI_ISL_552233, EPI_ISL_552239, EPI_ISL_552243, EPI_ISL_552246, EPI_ISL_552251, EPI_ISL_552259, EPI_ISL_552274, EPI_ISL_552288, EPI_ISL_552289, EPI_ISL_552290, EPI_ISL_552292, EPI_ISL_552294, EPI_ISL_552304, EPI_ISL_552307, EPI_ISL_552314, EPI_ISL_552319, EPI_ISL_552328, EPI_ISL_552331, EPI_ISL_552333, EPI_ISL_552342, EPI_ISL_552346, EPI_ISL_552348, EPI_ISL_552354, EPI_ISL_552366, EPI_ISL_552374, EPI_ISL_552379, EPI_ISL_552388, EPI_ISL_552390, EPI_ISL_552392, EPI_ISL_552404, EPI_ISL_552413, EPI_ISL_552416, EPI_ISL_552424, EPI_ISL_552425, EPI_ISL_552434, EPI_ISL_552437, EPI_ISL_552457, EPI_ISL_552483, EPI_ISL_552485, EPI_ISL_552493, EPI_ISL_552511, EPI_ISL_552519, EPI_ISL_552524, EPI_ISL_552532, EPI_ISL_552537, EPI_ISL_552543, EPI_ISL_552546, EPI_ISL_552552, EPI_ISL_552554, EPI_ISL_552560, EPI_ISL_552570, EPI_ISL_552573, EPI_ISL_552575, EPI_ISL_552592, EPI_ISL_552593, EPI_ISL_552605, EPI_ISL_552615, EPI_ISL_552619, EPI_ISL_552649, EPI_ISL_552687, EPI_ISL_552714, EPI_ISL_552754, EPI_ISL_552757, EPI_ISL_552758, EPI_ISL_552778, EPI_ISL_552786, EPI_ISL_552961, EPI_ISL_552963, EPI_ISL_552967, EPI_ISL_552975, EPI_ISL_553007, EPI_ISL_553022, EPI_ISL_553024, EPI_ISL_553030, EPI_ISL_553037, EPI_ISL_553038, EPI_ISL_553039, EPI_ISL_553042, EPI_ISL_553044, EPI_ISL_553050, EPI_ISL_553052, EPI_ISL_553054, EPI_ISL_553055, EPI_ISL_553058, EPI_ISL_553060, EPI_ISL_553068, EPI_ISL_553071, EPI_ISL_553079, EPI_ISL_553085, EPI_ISL_553090, EPI_ISL_553093, EPI_ISL_553100, EPI_ISL_553108, EPI_ISL_553109, EPI_ISL_553110, EPI_ISL_553129, EPI_ISL_553136, EPI_ISL_553139, EPI_ISL_553146, EPI_ISL_553148, EPI_ISL_553149, EPI_ISL_553161, EPI_ISL_553173, EPI_ISL_553182, EPI_ISL_553186, EPI_ISL_553187, EPI_ISL_553193, EPI_ISL_553202, EPI_ISL_553231, EPI_ISL_553235, EPI_ISL_553240, EPI_ISL_553241, EPI_ISL_553242, EPI_ISL_553246, EPI_ISL_553248, EPI_ISL_553251, EPI_ISL_553252, EPI_ISL_553871, EPI_ISL_553885, EPI_ISL_553907, EPI_ISL_553956, EPI_ISL_553965, EPI_ISL_553985, EPI_ISL_554003, EPI_ISL_554005, EPI_ISL_554119, EPI_ISL_554125, EPI_ISL_554128, EPI_ISL_554135, EPI_ISL_554137, EPI_ISL_554141, EPI_ISL_554145, EPI_ISL_554161, EPI_ISL_554162, EPI_ISL_554169, EPI_ISL_554210, EPI_ISL_554234, EPI_ISL_554236, EPI_ISL_554259, EPI_ISL_554303, EPI_ISL_554307, EPI_ISL_554311, EPI_ISL_554316, EPI_ISL_554317, EPI_ISL_554619, EPI_ISL_554627, EPI_ISL_554638, EPI_ISL_554650, EPI_ISL_554655, EPI_ISL_554665, EPI_ISL_554669, EPI_ISL_554693, EPI_ISL_554728, EPI_ISL_554736, EPI_ISL_554742, EPI_ISL_554744, EPI_ISL_554748, EPI_ISL_554750, EPI_ISL_554764, EPI_ISL_554781, EPI_ISL_554800, EPI_ISL_554803, EPI_ISL_554858, EPI_ISL_554859, EPI_ISL_556091, EPI_ISL_556094, EPI_ISL_556133, EPI_ISL_556134, EPI_ISL_556164, EPI_ISL_556165, EPI_ISL_556186, EPI_ISL_556218, EPI_ISL_556242, EPI_ISL_556266, EPI_ISL_556282, EPI_ISL_556287, EPI_ISL_556291, EPI_ISL_557166, EPI_ISL_557169, EPI_ISL_557176, EPI_ISL_557180, EPI_ISL_557187, EPI_ISL_557192, EPI_ISL_557193, EPI_ISL_557199, EPI_ISL_557200, EPI_ISL_557201, EPI_ISL_557207, EPI_ISL_557225, EPI_ISL_557227, EPI_ISL_557229, EPI_ISL_557232, EPI_ISL_557237, EPI_ISL_557245, EPI_ISL_557250, EPI_ISL_557254, EPI_ISL_557255, EPI_ISL_557265, EPI_ISL_557271, EPI_ISL_557272, EPI_ISL_557274, EPI_ISL_557281, EPI_ISL_557283, EPI_ISL_557284, EPI_ISL_557293, EPI_ISL_557294, EPI_ISL_557295, EPI_ISL_557299, EPI_ISL_557301, EPI_ISL_557304, EPI_ISL_557306, EPI_ISL_557307, EPI_ISL_557308, EPI_ISL_557313, EPI_ISL_557320, EPI_ISL_557324, EPI_ISL_557325, EPI_ISL_557330, EPI_ISL_557335, EPI_ISL_557338, EPI_ISL_557340, EPI_ISL_557353, EPI_ISL_557358, EPI_ISL_557368, EPI_ISL_557370, EPI_ISL_557385, EPI_ISL_557389, EPI_ISL_557392, EPI_ISL_557402, EPI_ISL_557403, EPI_ISL_557409, EPI_ISL_557420, EPI_ISL_557422, EPI_ISL_557424, EPI_ISL_557438, EPI_ISL_557448, EPI_ISL_557462, EPI_ISL_557472, EPI_ISL_557477, EPI_ISL_557478, EPI_ISL_557480, EPI_ISL_557494, EPI_ISL_557501, EPI_ISL_557502, EPI_ISL_557503, EPI_ISL_557509, EPI_ISL_557516, EPI_ISL_557524, EPI_ISL_557535, EPI_ISL_557549, EPI_ISL_557701, EPI_ISL_557741, EPI_ISL_557745, EPI_ISL_557750, EPI_ISL_557758, EPI_ISL_557764, EPI_ISL_557768, EPI_ISL_557771, EPI_ISL_557775, EPI_ISL_557782, EPI_ISL_557786, EPI_ISL_557791, EPI_ISL_557793, EPI_ISL_557807, EPI_ISL_557808, EPI_ISL_557811, EPI_ISL_557816, EPI_ISL_557817, EPI_ISL_557828, EPI_ISL_557832, EPI_ISL_557838, EPI_ISL_557865, EPI_ISL_557868, EPI_ISL_557880, EPI_ISL_557884, EPI_ISL_557889, EPI_ISL_557898, EPI_ISL_557908, EPI_ISL_557909, EPI_ISL_557915, EPI_ISL_557917, EPI_ISL_557921, EPI_ISL_557933, EPI_ISL_557937, EPI_ISL_557948, EPI_ISL_557958, EPI_ISL_557963, EPI_ISL_557967, EPI_ISL_557970, EPI_ISL_557976, EPI_ISL_558001, EPI_ISL_558106, EPI_ISL_558127, EPI_ISL_558234, EPI_ISL_558239, EPI_ISL_558250, EPI_ISL_558252, EPI_ISL_558271, EPI_ISL_558285, EPI_ISL_558301, EPI_ISL_558305, EPI_ISL_558306, EPI_ISL_558308, EPI_ISL_558330, EPI_ISL_558339, EPI_ISL_558356, EPI_ISL_558370, EPI_ISL_558392, EPI_ISL_558398, EPI_ISL_558402, EPI_ISL_558638, EPI_ISL_558926, EPI_ISL_558932, EPI_ISL_558975, EPI_ISL_558993, EPI_ISL_559026, EPI_ISL_559071, EPI_ISL_559083, EPI_ISL_559125, EPI_ISL_559133, EPI_ISL_559166, EPI_ISL_559171, EPI_ISL_559204, EPI_ISL_559215, EPI_ISL_559232, EPI_ISL_559673, EPI_ISL_559674, EPI_ISL_559687, EPI_ISL_559688, EPI_ISL_559699, EPI_ISL_559700 | | | |
| see above | Lighthouse Lab in Milton Keynes | Wellcome Sanger Institute for the COVID-19 Genomics UK (COG-UK) consortium | The Lighthouse Lab in Milton Keynes and Alex Alderton, Roberto Amato, Sonia Goncalves, Ewan Harrison, David K. Jackson, Ian Johnston, Dominic Kwiatkowski, Cordelia Langford, John Sillitoe on behalf of the Wellcome Sanger Institute COVID-19 Surveillance Team (http://www.sanger.ac.uk/covid-team) |
| EPI_ISL_559709, EPI_ISL_559715, EPI_ISL_559752 | Lighthouse Lab in Milton Keynes | Wellcome Sanger Institute for the COVID-19 Genomics UK (COG-UK) consortium | The Lighthouse Lab in Milton Keynes and Alex Alderton, Roberto Amato, Sonia Goncalves, Ewan Harrison, David K. Jackson, Ian Johnston, Dominic Kwiatkowski, Cordelia Langford, John Sillitoe on behalf of the Wellcome Sanger Institute COVID-19 Surveillance Team |
| EPI_ISL_559898, EPI_ISL_559899, EPI_ISL_559900, EPI_ISL_559901, EPI_ISL_559902, EPI_ISL_559903, EPI_ISL_559904, EPI_ISL_559905, EPI_ISL_559906, EPI_ISL_559907, EPI_ISL_559908, EPI_ISL_559909, EPI_ISL_559910, EPI_ISL_559911, EPI_ISL_559912, EPI_ISL_559913, EPI_ISL_559914, EPI_ISL_559915, EPI_ISL_559916, EPI_ISL_559917, EPI_ISL_559918, EPI_ISL_559919, EPI_ISL_559920, EPI_ISL_559921, EPI_ISL_559922, EPI_ISL_559923, EPI_ISL_559924, EPI_ISL_559925, EPI_ISL_559926, EPI_ISL_559927, EPI_ISL_559928, EPI_ISL_559929, EPI_ISL_559930, EPI_ISL_559931, EPI_ISL_559932, EPI_ISL_559933, EPI_ISL_559934, EPI_ISL_559935, EPI_ISL_559936, EPI_ISL_559937, EPI_ISL_559938, EPI_ISL_559939, EPI_ISL_559940, EPI_ISL_559941, EPI_ISL_559942, EPI_ISL_559943, EPI_ISL_559944, EPI_ISL_559945, EPI_ISL_559946, EPI_ISL_560054, EPI_ISL_560055, EPI_ISL_560056, EPI_ISL_560057, EPI_ISL_560058, EPI_ISL_560059, EPI_ISL_560060, EPI_ISL_560061, EPI_ISL_560062, EPI_ISL_560063, EPI_ISL_560064, EPI_ISL_560065, EPI_ISL_560066, EPI_ISL_560067, EPI_ISL_560068 | | | |
| see above | Virology Department, Sheffield Teaching Hospitals NHS Foundation Trust/Department of Infection, Immunity and Cardiovascular Disease, The Medical School, University of Sheffield | COVID-19 Genomics UK (COG-UK) Consortium | Thushan de Silva, Matthew Parker, Nikki Smith, Adri Angyal, Rebecca Brown, Luke Green, Rachel Tucker, Paul Parsons, Danielle Groves, Katie Johnson, Laura Carrilero, Alex Keeley, Dave Partridge, Matthew Wyles, Benjamin Lindsey, Mehmet Yavuz, Mohammad Raza, Cariad Evans |
| EPI_ISL_566435, EPI_ISL_566446, EPI_ISL_566459, EPI_ISL_566468, EPI_ISL_566472, EPI_ISL_566487, EPI_ISL_566513, EPI_ISL_566542, EPI_ISL_566593, EPI_ISL_566600, EPI_ISL_566603, EPI_ISL_566637, EPI_ISL_566639, EPI_ISL_566703, EPI_ISL_566707, EPI_ISL_566724, EPI_ISL_566738, EPI_ISL_566793, EPI_ISL_566802, EPI_ISL_566908, EPI_ISL_566922, EPI_ISL_566932, EPI_ISL_566956, EPI_ISL_566973, EPI_ISL_566994, EPI_ISL_568365, EPI_ISL_568386, EPI_ISL_568387, EPI_ISL_568395, EPI_ISL_568466, EPI_ISL_568470, EPI_ISL_568471, EPI_ISL_568476 | | | |
| see above | Lighthouse Lab in Milton Keynes | Wellcome Sanger Institute for the COVID-19 Genomics UK (COG-UK) consortium | The Lighthouse Lab in Milton Keynes and Alex Alderton, Roberto Amato, Sonia Goncalves, Ewan Harrison, David K. Jackson, Ian Johnston, Dominic Kwiatkowski, Cordelia Langford, John Sillitoe on behalf of the Wellcome Sanger Institute COVID-19 Surveillance Team |
| EPI_ISL_572195, EPI_ISL_572196, EPI_ISL_572197, EPI_ISL_572198, EPI_ISL_572199, EPI_ISL_572200, EPI_ISL_572201, EPI_ISL_572202, EPI_ISL_572203, EPI_ISL_572204, EPI_ISL_572205, EPI_ISL_572206, EPI_ISL_572207, EPI_ISL_572208, EPI_ISL_572209, EPI_ISL_572210, EPI_ISL_572211, EPI_ISL_572212, EPI_ISL_572213, EPI_ISL_572214, EPI_ISL_572215, EPI_ISL_572216, EPI_ISL_572217, EPI_ISL_572218, EPI_ISL_572219, EPI_ISL_572220, EPI_ISL_572221, EPI_ISL_572222, EPI_ISL_572223, EPI_ISL_572224, EPI_ISL_572225, EPI_ISL_572226, EPI_ISL_572227, EPI_ISL_572228, EPI_ISL_572229, EPI_ISL_572230, EPI_ISL_572231, EPI_ISL_572232, EPI_ISL_572233, EPI_ISL_572234, EPI_ISL_572235, EPI_ISL_572236, EPI_ISL_572237, EPI_ISL_572238, EPI_ISL_572239, EPI_ISL_572240, EPI_ISL_572241, EPI_ISL_572242, EPI_ISL_572243, EPI_ISL_572244, EPI_ISL_572245, EPI_ISL_572246, EPI_ISL_572247, EPI_ISL_572248, EPI_ISL_572249, EPI_ISL_572250, EPI_ISL_572251, EPI_ISL_572252, EPI_ISL_572253, EPI_ISL_572254, EPI_ISL_572255, EPI_ISL_572256, EPI_ISL_572257, EPI_ISL_572258, EPI_ISL_572259, EPI_ISL_572260, EPI_ISL_572261, EPI_ISL_572262, EPI_ISL_572263, EPI_ISL_572264, EPI_ISL_572265, EPI_ISL_572266, EPI_ISL_572267, EPI_ISL_572268, EPI_ISL_572269, EPI_ISL_572270, EPI_ISL_572271, EPI_ISL_572272, EPI_ISL_572273, EPI_ISL_572274, EPI_ISL_572275, EPI_ISL_572276, EPI_ISL_572277, EPI_ISL_572278, EPI_ISL_572279, EPI_ISL_572280, EPI_ISL_572281, EPI_ISL_572282, EPI_ISL_572283, EPI_ISL_572284, EPI_ISL_572285, EPI_ISL_572286, EPI_ISL_572287, EPI_ISL_572288, EPI_ISL_572289, EPI_ISL_572290, EPI_ISL_572291, EPI_ISL_572292, EPI_ISL_572293, EPI_ISL_572294, EPI_ISL_572295, EPI_ISL_572296, EPI_ISL_572297, EPI_ISL_572298, EPI_ISL_572299, EPI_ISL_572300, EPI_ISL_572301, EPI_ISL_572302, EPI_ISL_572303, EPI_ISL_572304, EPI_ISL_572305, EPI_ISL_572306, EPI_ISL_572307, EPI_ISL_572308, EPI_ISL_572309, EPI_ISL_572310, EPI_ISL_572311, EPI_ISL_572312, EPI_ISL_572313, EPI_ISL_572314, EPI_ISL_572315, EPI_ISL_572316, EPI_ISL_572317, EPI_ISL_572318 | | | |
| see above | Virginia DCLS | Virginia DCLS | Virginia DCLS |
| EPI_ISL_572417, EPI_ISL_572418, EPI_ISL_572420, EPI_ISL_572422, EPI_ISL_572428, EPI_ISL_572432, EPI_ISL_572433, EPI_ISL_572457, EPI_ISL_572458, EPI_ISL_572467, EPI_ISL_572481, EPI_ISL_572482, EPI_ISL_572516, EPI_ISL_572522, EPI_ISL_572529, EPI_ISL_572567, EPI_ISL_572587, EPI_ISL_572589, EPI_ISL_572609, EPI_ISL_572613, EPI_ISL_572618, EPI_ISL_572619, EPI_ISL_572678, EPI_ISL_572722, EPI_ISL_572743, EPI_ISL_572752, EPI_ISL_572757, EPI_ISL_572767, EPI_ISL_572796, EPI_ISL_572820, EPI_ISL_572841, EPI_ISL_572848, EPI_ISL_572912, EPI_ISL_572913, EPI_ISL_572914, EPI_ISL_572915, EPI_ISL_572916, EPI_ISL_572927, EPI_ISL_572928, EPI_ISL_572929, EPI_ISL_572930, EPI_ISL_572945, EPI_ISL_572946, EPI_ISL_572948, EPI_ISL_572949, EPI_ISL_572956, EPI_ISL_572959, EPI_ISL_572961, EPI_ISL_573526, EPI_ISL_573527, EPI_ISL_573528, EPI_ISL_573529, EPI_ISL_573530, EPI_ISL_573531, EPI_ISL_573532, EPI_ISL_573533, EPI_ISL_573534, EPI_ISL_573535, EPI_ISL_573536, EPI_ISL_573537, EPI_ISL_573538, EPI_ISL_573539, EPI_ISL_573540, EPI_ISL_573541, EPI_ISL_573542, EPI_ISL_573543, EPI_ISL_573544, EPI_ISL_573545, EPI_ISL_573546, EPI_ISL_573547, EPI_ISL_573548, EPI_ISL_573549, EPI_ISL_573550, EPI_ISL_573551, EPI_ISL_573552, EPI_ISL_573553, EPI_ISL_573554, EPI_ISL_573555, EPI_ISL_573556, EPI_ISL_573557, EPI_ISL_573558, EPI_ISL_573559, EPI_ISL_573560, EPI_ISL_573561, EPI_ISL_573562, EPI_ISL_573563, EPI_ISL_573564, EPI_ISL_573565, EPI_ISL_573686, EPI_ISL_573687, EPI_ISL_573688, EPI_ISL_573689, EPI_ISL_573690, EPI_ISL_573691, EPI_ISL_573692, EPI_ISL_573693, EPI_ISL_573694, EPI_ISL_573695, EPI_ISL_573696, EPI_ISL_573697, EPI_ISL_573698, EPI_ISL_573699, EPI_ISL_573700, EPI_ISL_573701, EPI_ISL_573702, EPI_ISL_573703, EPI_ISL_573704, EPI_ISL_573705, EPI_ISL_573706, EPI_ISL_573707, EPI_ISL_573708, EPI_ISL_573709, EPI_ISL_573710, EPI_ISL_573711, EPI_ISL_573712, EPI_ISL_573713, EPI_ISL_573714, EPI_ISL_573715, EPI_ISL_573716, EPI_ISL_573717, EPI_ISL_573718, EPI_ISL_573719, EPI_ISL_573720, EPI_ISL_573721, EPI_ISL_573722, EPI_ISL_573723, EPI_ISL_573724, EPI_ISL_573725, EPI_ISL_573726, EPI_ISL_573727, EPI_ISL_573728, EPI_ISL_573729, EPI_ISL_573730, EPI_ISL_573731, EPI_ISL_573732, EPI_ISL_573733, EPI_ISL_573734, EPI_ISL_573735, EPI_ISL_573736, EPI_ISL_573737, EPI_ISL_573738, EPI_ISL_573739, EPI_ISL_573740, EPI_ISL_573741, EPI_ISL_573742, EPI_ISL_573743, EPI_ISL_573744, EPI_ISL_573745, EPI_ISL_573746, EPI_ISL_573747, EPI_ISL_573748, EPI_ISL_573749, EPI_ISL_573750, EPI_ISL_573751, EPI_ISL_573752, EPI_ISL_573753, EPI_ISL_573754, EPI_ISL_573755, EPI_ISL_573756, EPI_ISL_573757, EPI_ISL_573758, EPI_ISL_573762, EPI_ISL_573763, EPI_ISL_573806, EPI_ISL_573807, EPI_ISL_573808, EPI_ISL_573809, EPI_ISL_573810 | | | |
| see above | Virology Department, Sheffield Teaching Hospitals NHS Foundation Trust/Department of Infection, Immunity and Cardiovascular Disease, The Medical School, University of Sheffield | COVID-19 Genomics UK (COG-UK) Consortium | Thushan de Silva, Matthew Parker, Nikki Smith, Adri Angyal, Rebecca Brown, Luke Green, Rachel Tucker, Paul Parsons, Danielle Groves, Katie Johnson, Laura Carrilero, Alex Keeley, Dave Partridge, Matthew Wyles, Benjamin Lindsey, Mehmet Yavuz, Mohammad Raza, Cariad Evans |
| EPI_ISL_574479, EPI_ISL_574480, EPI_ISL_574481, EPI_ISL_574482, EPI_ISL_574483, EPI_ISL_574484, EPI_ISL_574485, EPI_ISL_574486, EPI_ISL_574487, EPI_ISL_574488, EPI_ISL_574489, EPI_ISL_574490, EPI_ISL_574491, EPI_ISL_574492 | | | |
| see above | Programme in Emerging Infectious Diseases, Duke-NUS Medical School | National Public Health Laboratory, National Centre for Infectious Diseases | Tze Minn Mak, Sophie Octavia, Zhenyang Zhou, Danielle E Anderson, Adrian Eng Zheng Kang, Lin Cui, Raymond Tzer Pin Lin |
| EPI_ISL_574493, EPI_ISL_574494, EPI_ISL_574495, EPI_ISL_574496, EPI_ISL_574497, EPI_ISL_574498, EPI_ISL_574499, EPI_ISL_574500, EPI_ISL_574501, EPI_ISL_574502, EPI_ISL_574503, EPI_ISL_574504, EPI_ISL_574505, EPI_ISL_574506, EPI_ISL_574507, EPI_ISL_574508, EPI_ISL_574509, EPI_ISL_574510, EPI_ISL_574511, EPI_ISL_574512, EPI_ISL_574513, EPI_ISL_574514, EPI_ISL_574515, EPI_ISL_574516, EPI_ISL_574517, EPI_ISL_574518, EPI_ISL_574519, EPI_ISL_574520, EPI_ISL_574521, EPI_ISL_574522, EPI_ISL_574523, EPI_ISL_574524, EPI_ISL_574525, EPI_ISL_574526, EPI_ISL_574527, EPI_ISL_574528, EPI_ISL_574529, EPI_ISL_574530, EPI_ISL_574531, EPI_ISL_574532, EPI_ISL_574533, EPI_ISL_574534, EPI_ISL_574535, EPI_ISL_574536, EPI_ISL_574537, EPI_ISL_574538, EPI_ISL_574539, EPI_ISL_574540, EPI_ISL_574541, EPI_ISL_574542 | | | |
| see above | National Public Health Laboratory, National Centre for Infectious Diseases | National Public Health Laboratory, National Centre for Infectious Diseases | Tze Minn Mak, Sophie Octavia, Zhenyang Zhou, Lin Cui, Raymond Tzer Pin Lin |
| EPI_ISL_575367, EPI_ISL_575764, EPI_ISL_575788, EPI_ISL_575795, EPI_ISL_575799, EPI_ISL_575802, EPI_ISL_575804, EPI_ISL_575805, EPI_ISL_575841, EPI_ISL_575846, EPI_ISL_575871, EPI_ISL_575895, EPI_ISL_575912, EPI_ISL_575929, EPI_ISL_575943, EPI_ISL_575954, EPI_ISL_575957, EPI_ISL_575960 | | | |
| see above | Lighthouse Lab in Milton Keynes | Wellcome Sanger Institute for the COVID-19 Genomics UK (COG-UK) consortium | The Lighthouse Lab in Milton Keynes and Alex Alderton, Roberto Amato, Sonia Goncalves, Ewan Harrison, David K. Jackson, Ian Johnston, Dominic Kwiatkowski, Cordelia Langford, John Sillitoe on behalf of the Wellcome Sanger Institute COVID-19 Surveillance Team |
| EPI_ISL_575979, EPI_ISL_575984, EPI_ISL_575985, EPI_ISL_575986, EPI_ISL_575987, EPI_ISL_575988, EPI_ISL_575989, EPI_ISL_575990, EPI_ISL_575991, EPI_ISL_575992, EPI_ISL_575993, EPI_ISL_575994 | | | |
| see above | Lighthouse Lab in Milton Keynes | Wellcome Sanger Institute for the COVID-19 Genomics UK | The Lighthouse Lab in Milton Keynes and Alex Alderton, Roberto Amato, Sonia Goncalves, Ewan Harrison, David K. Jackson, Ian Johnston, Dominic |

| | | | |
|---|---|---|---|
| | | (COG-UK) consortium | Kwiatkowski, Cordelia Langford, John Sillitoe on behalf of the Wellcome Sanger Institute COVID-19 Surveillance Team (http://www.sanger.ac.uk/covid-team) |
| EPI_ISL_577734, EPI_ISL_577736, EPI_ISL_577738 | Institute of Virology, Biomedical Research Center of the Slovak Academy of Sciences, Bratislava | Faculty of Natural Sciences, Comenius University, Bratislava | Viktória Hodorová, Kristína Boršová, Broa Brejová, Viktória abanová, Dominika Friová, Sabina Fumaová Havlíková, Juraj Kopáek, Martina Liková, ubomíra Lukáiková, Martina Neboháová, Monika Sláviková, Edita Staroová, Elena Tichá, Tomáš Vina, Jozef Nosek, Boris Klempa |
| EPI_ISL_578082, EPI_ISL_578083, EPI_ISL_578084, EPI_ISL_578085, EPI_ISL_578086, EPI_ISL_578087, EPI_ISL_578088, EPI_ISL_578089, EPI_ISL_578090, EPI_ISL_578091, EPI_ISL_578092, EPI_ISL_578093, EPI_ISL_578094, EPI_ISL_578095, EPI_ISL_578096, EPI_ISL_578097, EPI_ISL_578098, EPI_ISL_578099, EPI_ISL_578100, EPI_ISL_578101, EPI_ISL_578102, EPI_ISL_578103, EPI_ISL_578104, EPI_ISL_578105, EPI_ISL_578106, EPI_ISL_578107, EPI_ISL_578108, EPI_ISL_578109, EPI_ISL_578110, EPI_ISL_578111, EPI_ISL_578112, EPI_ISL_578113, EPI_ISL_578114, EPI_ISL_578115, EPI_ISL_578116, EPI_ISL_578117, EPI_ISL_578118, EPI_ISL_578119, EPI_ISL_578120, EPI_ISL_578121, EPI_ISL_578122, EPI_ISL_578123, EPI_ISL_578124, EPI_ISL_578125, EPI_ISL_578126, EPI_ISL_578127, EPI_ISL_578128, EPI_ISL_578129, EPI_ISL_578130, EPI_ISL_578131, EPI_ISL_578132, EPI_ISL_578133, EPI_ISL_578134, EPI_ISL_578135, EPI_ISL_578136, EPI_ISL_578137, EPI_ISL_578138, EPI_ISL_578139, EPI_ISL_578140, EPI_ISL_578141, EPI_ISL_578142, EPI_ISL_578143, EPI_ISL_578144, EPI_ISL_578145, EPI_ISL_578146, EPI_ISL_578147, EPI_ISL_578148, EPI_ISL_578149, EPI_ISL_578150, EPI_ISL_578151, EPI_ISL_578152, EPI_ISL_578153, EPI_ISL_578154, EPI_ISL_578155, EPI_ISL_578156, EPI_ISL_578157, EPI_ISL_578158, EPI_ISL_578159, EPI_ISL_578160, EPI_ISL_578161, EPI_ISL_578162, EPI_ISL_578163, EPI_ISL_578164, EPI_ISL_578165, EPI_ISL_578166, EPI_ISL_578167, EPI_ISL_578169, EPI_ISL_578170, EPI_ISL_578171, EPI_ISL_578172, EPI_ISL_578173, EPI_ISL_578174, EPI_ISL_578185 | | | |
| see above | University of Michigan Clinical Microbiology Laboratory | Lauring Lab, University of Michigan, Department of Microbiology and Immunology | Valesano |
| EPI_ISL_579076, EPI_ISL_579087, EPI_ISL_579088, EPI_ISL_579089, EPI_ISL_579090, EPI_ISL_579091, EPI_ISL_579092 | Canterbury Health Laboratories | Institute of Environmental Science and Research (ESR) | Xiaoyun Ren, Matt Storey, Nikki Freed, Muhammad Faisal, Jing Wang, Hermes Perez, Anja Werno, Antje van der Linden, Arlo Upton, Chris Mansell, David Hammer, Dragana Drinkovic, Gary McAuliffe, Hana Sofia Andersson, James Ussher, Jill Sherwood, Josh Freeman, Julia Howard, Juliet Elvy, Mary DeAlmeida, Matt Blakiston, Matthew Rogers, Max Bloomfield, Michael Addidle, Michelle Balm, Sally Roberts, Sarah Jefferies, Sharmini Muttaiyah, Susan Morpeth, Susan Taylor, Timothy Blackmore, Vani Sathyendran, Veronica Playle, Virginia Hope, Erasmus Smit, Lauren Jelly, Olin Silander, Joep de Ligt |
| EPI_ISL_579093, EPI_ISL_579094, EPI_ISL_579095, EPI_ISL_579096 | North Shore Hospital | Institute of Environmental Science and Research (ESR) | Xiaoyun Ren, Matt Storey, Nikki Freed, Muhammad Faisal, Jing Wang, Hermes Perez, Anja Werno, Antje van der Linden, Arlo Upton, Chris Mansell, David Hammer, Dragana Drinkovic, Gary McAuliffe, Hana Sofia Andersson, James Ussher, Jill Sherwood, Josh Freeman, Julia Howard, Juliet Elvy, Mary DeAlmeida, Matt Blakiston, Matthew Rogers, Max Bloomfield, Michael Addidle, Michelle Balm, Sally Roberts, Sarah Jefferies, Sharmini Muttaiyah, Susan Morpeth, Susan Taylor, Timothy Blackmore, Vani Sathyendran, Veronica Playle, Virginia Hope, Erasmus Smit, Lauren Jelly, Olin Silander, Joep de Ligt |
| EPI_ISL_579097, EPI_ISL_579098 | LabPLUS | Institute of Environmental Science and Research (ESR) | Xiaoyun Ren, Matt Storey, Nikki Freed, Muhammad Faisal, Jing Wang, Hermes Perez, Anja Werno, Antje van der Linden, Arlo Upton, Chris Mansell, David Hammer, Dragana Drinkovic, Gary McAuliffe, Hana Sofia Andersson, James Ussher, Jill Sherwood, Josh Freeman, Julia Howard, Juliet Elvy, Mary DeAlmeida, Matt Blakiston, Matthew Rogers, Max Bloomfield, Michael Addidle, Michelle Balm, Sally Roberts, Sarah Jefferies, Sharmini Muttaiyah, Susan Morpeth, Susan Taylor, Timothy Blackmore, Vani Sathyendran, Veronica Playle, Virginia Hope, Erasmus Smit, Lauren Jelly, Olin Silander, Joep de Ligt |
| EPI_ISL_579100 | Middlemore Hospital | Institute of Environmental Science and Research (ESR) | Xiaoyun Ren, Matt Storey, Nikki Freed, Muhammad Faisal, Jing Wang, Hermes Perez, Anja Werno, Antje van der Linden, Arlo Upton, Chris Mansell, David Hammer, Dragana Drinkovic, Gary McAuliffe, Hana Sofia Andersson, James Ussher, Jill Sherwood, Josh Freeman, Julia Howard, Juliet Elvy, Mary DeAlmeida, Matt Blakiston, Matthew Rogers, Max Bloomfield, Michael Addidle, Michelle Balm, Sally Roberts, Sarah Jefferies, Sharmini Muttaiyah, Susan Morpeth, Susan Taylor, Timothy Blackmore, Vani Sathyendran, Veronica Playle, Virginia Hope, Erasmus Smit, Lauren Jelly, Olin Silander, Joep de Ligt |
| EPI_ISL_579108, EPI_ISL_579109, EPI_ISL_579110, EPI_ISL_579112, EPI_ISL_579120, EPI_ISL_579123 | LabPLUS | Institute of Environmental Science and Research (ESR) | Xiaoyun Ren, Matt Storey, Nikki Freed, Muhammad Faisal, Jing Wang, Hermes Perez, Anja Werno, Antje van der Linden, Arlo Upton, Chris Mansell, David Hammer, Dragana Drinkovic, Gary McAuliffe, Hana Sofia Andersson, James Ussher, Jill Sherwood, Josh Freeman, Julia Howard, Juliet Elvy, Mary DeAlmeida, Matt Blakiston, Matthew Rogers, Max Bloomfield, Michael Addidle, Michelle Balm, Sally Roberts, Sarah Jefferies, Sharmini Muttaiyah, Susan Morpeth, Susan Taylor, Timothy Blackmore, Vani Sathyendran, Veronica Playle, Virginia Hope, Erasmus Smit, Lauren Jelly, Olin Silander, Joep de Ligt |
| EPI_ISL_579124, EPI_ISL_579125, EPI_ISL_579127 | Southern Community Labs Dunedin | Institute of Environmental Science and Research (ESR) | Xiaoyun Ren, Matt Storey, Nikki Freed, Muhammad Faisal, Jing Wang, Hermes Perez, Anja Werno, Antje van der Linden, Arlo Upton, Chris Mansell, David Hammer, Dragana Drinkovic, Gary McAuliffe, Hana Sofia Andersson, James Ussher, Jill Sherwood, Josh Freeman, Julia Howard, Juliet Elvy, Mary DeAlmeida, Matt Blakiston, Matthew Rogers, Max Bloomfield, Michael Addidle, Michelle Balm, Sally Roberts, Sarah Jefferies, Sharmini Muttaiyah, Susan Morpeth, Susan Taylor, Timothy Blackmore, Vani Sathyendran, Veronica Playle, Virginia Hope, Erasmus Smit, Lauren Jelly, Olin Silander, Joep de Ligt |
| EPI_ISL_579132 | Middlemore Hospital | Institute of Environmental Science and Research (ESR) | Xiaoyun Ren, Matt Storey, Nikki Freed, Muhammad Faisal, Jing Wang, Hermes Perez, Anja Werno, Antje van der Linden, Arlo Upton, Chris Mansell, David Hammer, Dragana Drinkovic, Gary McAuliffe, Hana Sofia Andersson, James Ussher, Jill Sherwood, Josh Freeman, Julia Howard, Juliet Elvy, Mary DeAlmeida, Matt Blakiston, Matthew Rogers, Max Bloomfield, Michael Addidle, Michelle Balm, Sally Roberts, Sarah Jefferies, Sharmini Muttaiyah, Susan Morpeth, Susan Taylor, Timothy Blackmore, Vani Sathyendran, Veronica Playle, Virginia Hope, Erasmus Smit, Lauren Jelly, Olin Silander, Joep de Ligt |
| EPI_ISL_579133, EPI_ISL_579149, EPI_ISL_579162, EPI_ISL_579166, EPI_ISL_579168, EPI_ISL_579172, EPI_ISL_579175, EPI_ISL_579182, EPI_ISL_579185, EPI_ISL_579219 | Southern Community Labs Dunedin | Institute of Environmental Science and Research (ESR) | Xiaoyun Ren, Matt Storey, Nikki Freed, Muhammad Faisal, Jing Wang, Hermes Perez, Anja Werno, Antje van der Linden, Arlo Upton, Chris Mansell, David Hammer, Dragana Drinkovic, Gary McAuliffe, Hana Sofia Andersson, James Ussher, Jill Sherwood, Josh Freeman, Julia Howard, Juliet Elvy, Mary DeAlmeida, Matt Blakiston, Matthew Rogers, Max Bloomfield, Michael Addidle, Michelle Balm, Sally Roberts, Sarah Jefferies, Sharmini Muttaiyah, Susan Morpeth, Susan Taylor, Timothy Blackmore, Vani Sathyendran, Veronica Playle, Virginia Hope, Erasmus Smit, Lauren Jelly, Olin Silander, Joep de Ligt |
| EPI_ISL_579220, EPI_ISL_579223, EPI_ISL_579224, EPI_ISL_579225, EPI_ISL_579226 | Canterbury Health Laboratories | Institute of Environmental Science and Research (ESR) | Xiaoyun Ren, Matt Storey, Nikki Freed, Muhammad Faisal, Jing Wang, Hermes Perez, Anja Werno, Antje van der Linden, Arlo Upton, Chris Mansell, David Hammer, Dragana Drinkovic, Gary McAuliffe, Hana Sofia Andersson, James Ussher, Jill Sherwood, Josh Freeman, Julia Howard, Juliet Elvy, Mary DeAlmeida, Matt Blakiston, Matthew Rogers, Max Bloomfield, Michael Addidle, Michelle Balm, Sally Roberts, Sarah Jefferies, Sharmini Muttaiyah, Susan Morpeth, Susan Taylor, Timothy Blackmore, Vani Sathyendran, Veronica Playle, Virginia Hope, Erasmus Smit, Lauren Jelly, Olin Silander, Joep de Ligt |
| EPI_ISL_579228, EPI_ISL_579232, EPI_ISL_579303, EPI_ISL_579306, EPI_ISL_579308, EPI_ISL_579309, EPI_ISL_579311, EPI_ISL_579312, EPI_ISL_579313, EPI_ISL_579314 | Middlemore Hospital | Institute of Environmental Science and Research (ESR) | Xiaoyun Ren, Matt Storey, Nikki Freed, Muhammad Faisal, Jing Wang, Hermes Perez, Anja Werno, Antje van der Linden, Arlo Upton, Chris Mansell, David Hammer, Dragana Drinkovic, Gary McAuliffe, Hana Sofia Andersson, James Ussher, Jill Sherwood, Josh Freeman, Julia Howard, Juliet Elvy, Mary DeAlmeida, Matt Blakiston, Matthew Rogers, Max Bloomfield, Michael Addidle, Michelle Balm, Sally Roberts, Sarah Jefferies, Sharmini Muttaiyah, Susan Morpeth, Susan Taylor, Timothy Blackmore, Vani Sathyendran, Veronica Playle, Virginia Hope, Erasmus Smit, Lauren Jelly, Olin Silander, Joep de Ligt |
| EPI_ISL_579403 | LabPLUS | Institute of Environmental Science and Research (ESR) | Xiaoyun Ren, Matt Storey, Nikki Freed, Muhammad Faisal, Jing Wang, Hermes Perez, Anja Werno, Antje van der Linden, Arlo Upton, Chris Mansell, David Hammer, Dragana Drinkovic, Gary McAuliffe, Hana Sofia Andersson, James Ussher, Jill Sherwood, Josh Freeman, Julia Howard, Juliet Elvy, Mary DeAlmeida, Matt Blakiston, Matthew Rogers, Max Bloomfield, Michael Addidle, Michelle Balm, Sally Roberts, Sarah Jefferies, Sharmini Muttaiyah, Susan Morpeth, Susan Taylor, Timothy Blackmore, Vani Sathyendran, Veronica Playle, Virginia Hope, Erasmus Smit, Lauren Jelly, Olin Silander, Joep de Ligt |
| EPI_ISL_579404 | Middlemore Hospital | Institute of Environmental Science and Research (ESR) | Xiaoyun Ren, Matt Storey, Nikki Freed, Muhammad Faisal, Jing Wang, Hermes Perez, Anja Werno, Antje van der Linden, Arlo Upton, Chris Mansell, David Hammer, Dragana Drinkovic, Gary McAuliffe, Hana Sofia Andersson, James Ussher, Jill Sherwood, Josh Freeman, Julia Howard, Juliet Elvy, Mary DeAlmeida, Matt Blakiston, Matthew Rogers, Max Bloomfield, Michael Addidle, Michelle Balm, Sally Roberts, Sarah Jefferies, Sharmini Muttaiyah, Susan Morpeth, Susan Taylor, Timothy Blackmore, Vani Sathyendran, Veronica Playle, Virginia Hope, Erasmus Smit, Lauren Jelly, Olin Silander, Joep de Ligt |
| EPI_ISL_579405 | LabPLUS | Institute of Environmental Science and Research (ESR) | Xiaoyun Ren, Matt Storey, Nikki Freed, Muhammad Faisal, Jing Wang, Hermes Perez, Anja Werno, Antje van der Linden, Arlo Upton, Chris Mansell, David Hammer, Dragana Drinkovic, Gary McAuliffe, Hana Sofia Andersson, James Ussher, Jill Sherwood, Josh Freeman, Julia Howard, Juliet Elvy, Mary DeAlmeida, Matt Blakiston, Matthew Rogers, Max Bloomfield, Michael Addidle, Michelle Balm, Sally Roberts, Sarah Jefferies, Sharmini Muttaiyah, Susan Morpeth, Susan Taylor, Timothy Blackmore, Vani Sathyendran, Veronica Playle, Virginia Hope, Erasmus Smit, Lauren Jelly, Olin Silander, Joep de Ligt |
| EPI_ISL_579406, EPI_ISL_579407 | North Shore Hospital | Institute of Environmental Science and Research (ESR) | Xiaoyun Ren, Matt Storey, Nikki Freed, Muhammad Faisal, Jing Wang, Hermes Perez, Anja Werno, Antje van der Linden, Arlo Upton, Chris Mansell, David Hammer, Dragana Drinkovic, Gary McAuliffe, Hana Sofia Andersson, James Ussher, Jill Sherwood, Josh Freeman, Julia Howard, Juliet Elvy, Mary DeAlmeida, Matt Blakiston, Matthew Rogers, Max Bloomfield, Michael Addidle, Michelle Balm, Sally Roberts, Sarah Jefferies, Sharmini Muttaiyah, Susan Morpeth, Susan Taylor, Timothy Blackmore, Vani Sathyendran, Veronica Playle, Virginia Hope, Erasmus Smit, Lauren Jelly, Olin Silander, Joep de Ligt |
| EPI_ISL_579418, EPI_ISL_579420 | Wellington SCL (WN) | Institute of Environmental Science and Research (ESR) | Xiaoyun Ren, Matt Storey, Nikki Freed, Muhammad Faisal, Jing Wang, Hermes Perez, Anja Werno, Antje van der Linden, Arlo Upton, Chris Mansell, David Hammer, Dragana Drinkovic, Gary McAuliffe, Hana Sofia Andersson, James Ussher, Jill Sherwood, Josh Freeman, Julia Howard, Juliet Elvy, Mary DeAlmeida, Matt Blakiston, Matthew Rogers, Max Bloomfield, Michael Addidle, Michelle Balm, Sally Roberts, Sarah Jefferies, Sharmini Muttaiyah, Susan Morpeth, Susan Taylor, Timothy Blackmore, Vani Sathyendran, Veronica Playle, Virginia Hope, Erasmus Smit, Lauren Jelly, Olin Silander, Joep de Ligt |
| EPI_ISL_579421, EPI_ISL_579422, EPI_ISL_579423, EPI_ISL_579424 | LabTests | Institute of Environmental Science and Research (ESR) | Xiaoyun Ren, Matt Storey, Nikki Freed, Muhammad Faisal, Jing Wang, Hermes Perez, Anja Werno, Antje van der Linden, Arlo Upton, Chris Mansell, David Hammer, Dragana Drinkovic, Gary McAuliffe, Hana Sofia Andersson, James Ussher, Jill Sherwood, Josh Freeman, Julia Howard, Juliet Elvy, Mary DeAlmeida, Matt Blakiston, Matthew Rogers, Max Bloomfield, Michael Addidle, Michelle Balm, Sally Roberts, Sarah Jefferies, Sharmini Muttaiyah, Susan Morpeth, Susan Taylor, Timothy Blackmore, Vani Sathyendran, Veronica Playle, Virginia Hope, Erasmus Smit, Lauren Jelly, Olin Silander, Joep de Ligt |

| | | | |
|---|---|---|---|
| EPI_ISL_579425 | LabPLUS | Institute of Environmental Science and Research (ESR) | Xiaoyun Ren, Matt Storey, Nikki Freed, Muhammad Faisal, Jing Wang, Hermes Perez, Anja Werno, Antje van der Linden, Arlo Upton, Chris Mansell, David Hammer, Dragana Drinkovic, Gary McAuliffe, Hana Sofia Andersson, James Ussher, Jill Sherwood, Josh Freeman, Julia Howard, Juliet Elvy, Mary DeAlmeida, Matt Blakiston, Matthew Rogers, Max Bloomfield, Michael Addidle, Michelle Balm, Sally Roberts, Sarah Jefferies, Sharmini Muttaiyah, Susan Morpeth, Susan Taylor, Timothy Blackmore, Vani Sathyendran, Veronica Playle, Virginia Hope, Erasmus Smit, Lauren Jelly, Olin Silander, Joep de Ligt |
| EPI_ISL_579426, EPI_ISL_579427, EPI_ISL_579428, EPI_ISL_579429, EPI_ISL_579430, EPI_ISL_579431, EPI_ISL_579432, EPI_ISL_579433, EPI_ISL_579434, EPI_ISL_579435, EPI_ISL_579436, EPI_ISL_579437, EPI_ISL_579438, EPI_ISL_579439, EPI_ISL_579440, EPI_ISL_579442, EPI_ISL_579443, EPI_ISL_579446, EPI_ISL_579447, EPI_ISL_579449, EPI_ISL_579450, EPI_ISL_579451, EPI_ISL_579452, EPI_ISL_579455, EPI_ISL_579457, EPI_ISL_579460, EPI_ISL_579467, EPI_ISL_579476, EPI_ISL_579477, EPI_ISL_579479, EPI_ISL_579483, EPI_ISL_579484, EPI_ISL_579486, EPI_ISL_579489, EPI_ISL_579490, EPI_ISL_579491, EPI_ISL_579492, EPI_ISL_579502 | | | |
| see above | Canterbury Health Laboratories | Institute of Environmental Science and Research (ESR) | Xiaoyun Ren, Matt Storey, Nikki Freed, Muhammad Faisal, Jing Wang, Hermes Perez, Anja Werno, Antje van der Linden, Arlo Upton, Chris Mansell, David Hammer, Dragana Drinkovic, Gary McAuliffe, Hana Sofia Andersson, James Ussher, Jill Sherwood, Josh Freeman, Julia Howard, Juliet Elvy, Mary DeAlmeida, Matt Blakiston, Matthew Rogers, Max Bloomfield, Michael Addidle, Michelle Balm, Sally Roberts, Sarah Jefferies, Sharmini Muttaiyah, Susan Morpeth, Susan Taylor, Timothy Blackmore, Vani Sathyendran, Veronica Playle, Virginia Hope, Erasmus Smit, Lauren Jelly, Olin Silander, Joep de Ligt |
| EPI_ISL_580631, EPI_ISL_580632, EPI_ISL_580634, EPI_ISL_580638, EPI_ISL_580639, EPI_ISL_580641, EPI_ISL_580642, EPI_ISL_580643, EPI_ISL_580644, EPI_ISL_580649, EPI_ISL_580652, EPI_ISL_580653, EPI_ISL_580654, EPI_ISL_580657, EPI_ISL_580658, EPI_ISL_580659, EPI_ISL_580661, EPI_ISL_580662, EPI_ISL_580664, EPI_ISL_580665, EPI_ISL_580667, EPI_ISL_580668, EPI_ISL_580669, EPI_ISL_580670, EPI_ISL_580672, EPI_ISL_580673, EPI_ISL_580676, EPI_ISL_580683, EPI_ISL_580684, EPI_ISL_580685, EPI_ISL_580687, EPI_ISL_580688, EPI_ISL_580693, EPI_ISL_580694, EPI_ISL_580701, EPI_ISL_580702, EPI_ISL_580703, EPI_ISL_580704, EPI_ISL_580707, EPI_ISL_580709, EPI_ISL_580713, EPI_ISL_580714, EPI_ISL_580715, EPI_ISL_580716, EPI_ISL_580717, EPI_ISL_580721, EPI_ISL_580724, EPI_ISL_580725, EPI_ISL_580727, EPI_ISL_580729, EPI_ISL_580731, EPI_ISL_580735, EPI_ISL_580739, EPI_ISL_580741, EPI_ISL_580742, EPI_ISL_580744, EPI_ISL_580747, EPI_ISL_580748, EPI_ISL_580749, EPI_ISL_580751, EPI_ISL_580755, EPI_ISL_580756, EPI_ISL_580757, EPI_ISL_580758, EPI_ISL_580759, EPI_ISL_580760, EPI_ISL_580761, EPI_ISL_580762, EPI_ISL_580765, EPI_ISL_580767, EPI_ISL_580770, EPI_ISL_580774, EPI_ISL_580775, EPI_ISL_580777, EPI_ISL_580778, EPI_ISL_580779, EPI_ISL_580781, EPI_ISL_580782, EPI_ISL_580783, EPI_ISL_580785, EPI_ISL_580786, EPI_ISL_580790, EPI_ISL_580792, EPI_ISL_580793, EPI_ISL_580794, EPI_ISL_580795, EPI_ISL_580796, EPI_ISL_580798, EPI_ISL_580799, EPI_ISL_580800, EPI_ISL_580801, EPI_ISL_580802, EPI_ISL_580803, EPI_ISL_580804, EPI_ISL_580806, EPI_ISL_580807, EPI_ISL_580808, EPI_ISL_580809, EPI_ISL_580810, EPI_ISL_580811, EPI_ISL_580814, EPI_ISL_580815, EPI_ISL_580816, EPI_ISL_580817, EPI_ISL_580818, EPI_ISL_580820, EPI_ISL_580826, EPI_ISL_580827, EPI_ISL_580829, EPI_ISL_580831, EPI_ISL_580832, EPI_ISL_580834, EPI_ISL_580836, EPI_ISL_580837, EPI_ISL_580838, EPI_ISL_580839, EPI_ISL_580842, EPI_ISL_580843, EPI_ISL_580844, EPI_ISL_580845, EPI_ISL_580846, EPI_ISL_580852, EPI_ISL_580854, EPI_ISL_580857, EPI_ISL_580885, EPI_ISL_580892, EPI_ISL_580897, EPI_ISL_580903, EPI_ISL_580910, EPI_ISL_580913, EPI_ISL_580915, EPI_ISL_580917, EPI_ISL_580918, EPI_ISL_580940, EPI_ISL_580950, EPI_ISL_580951, EPI_ISL_580959, EPI_ISL_580976, EPI_ISL_580991, EPI_ISL_580995, EPI_ISL_580996, EPI_ISL_580998, EPI_ISL_581004, EPI_ISL_581008, EPI_ISL_581009, EPI_ISL_581018, EPI_ISL_581023, EPI_ISL_581025, EPI_ISL_581026, EPI_ISL_581028, EPI_ISL_581032, EPI_ISL_581035, EPI_ISL_581040, EPI_ISL_581041, EPI_ISL_581056, EPI_ISL_581065, EPI_ISL_581069, EPI_ISL_581097, EPI_ISL_581099, EPI_ISL_581103, EPI_ISL_581107, EPI_ISL_581111, EPI_ISL_581113, EPI_ISL_581114, EPI_ISL_581115, EPI_ISL_581116, EPI_ISL_581117, EPI_ISL_581118, EPI_ISL_581119, EPI_ISL_581121, EPI_ISL_581123, EPI_ISL_581124, EPI_ISL_581125, EPI_ISL_581129, EPI_ISL_581130, EPI_ISL_581131, EPI_ISL_581135, EPI_ISL_581136, EPI_ISL_581137, EPI_ISL_581138, EPI_ISL_581139, EPI_ISL_581140, EPI_ISL_581144, EPI_ISL_581145, EPI_ISL_581147, EPI_ISL_581148, EPI_ISL_581150, EPI_ISL_581151, EPI_ISL_581155, EPI_ISL_581156, EPI_ISL_581157, EPI_ISL_581158, EPI_ISL_581160, EPI_ISL_581162, EPI_ISL_581163, EPI_ISL_581164, EPI_ISL_581165, EPI_ISL_581167, EPI_ISL_581169, EPI_ISL_581170, EPI_ISL_581171, EPI_ISL_581172, EPI_ISL_581175, EPI_ISL_581176, EPI_ISL_581177, EPI_ISL_581179, EPI_ISL_581180, EPI_ISL_581181, EPI_ISL_581184, EPI_ISL_581185, EPI_ISL_581186, EPI_ISL_581187, EPI_ISL_581190, EPI_ISL_581191, EPI_ISL_581192, EPI_ISL_581195, EPI_ISL_581197, EPI_ISL_581199, EPI_ISL_581200, EPI_ISL_581203, EPI_ISL_581204, EPI_ISL_581205, EPI_ISL_581206, EPI_ISL_581207, EPI_ISL_581208, EPI_ISL_581210, EPI_ISL_581211, EPI_ISL_581213, EPI_ISL_581215, EPI_ISL_581216, EPI_ISL_581218, EPI_ISL_581219, EPI_ISL_581220, EPI_ISL_581221, EPI_ISL_581224, EPI_ISL_581227, EPI_ISL_581228, EPI_ISL_581231, EPI_ISL_581233, EPI_ISL_581235, EPI_ISL_581236, EPI_ISL_581237, EPI_ISL_581238, EPI_ISL_581239, EPI_ISL_581240, EPI_ISL_581241, EPI_ISL_581242, EPI_ISL_581244, EPI_ISL_581246, EPI_ISL_581248, EPI_ISL_581249, EPI_ISL_581251, EPI_ISL_581252, EPI_ISL_581253, EPI_ISL_581255, EPI_ISL_581256, EPI_ISL_581257, EPI_ISL_581258, EPI_ISL_581260, EPI_ISL_581264, EPI_ISL_581265, EPI_ISL_581266, EPI_ISL_581268, EPI_ISL_581269, EPI_ISL_581270, EPI_ISL_581271, EPI_ISL_581273, EPI_ISL_581274, EPI_ISL_581276, EPI_ISL_581277, EPI_ISL_581279, EPI_ISL_581280, EPI_ISL_581281, EPI_ISL_581286, EPI_ISL_581288, EPI_ISL_581289, EPI_ISL_581290, EPI_ISL_581292, EPI_ISL_581293, EPI_ISL_581294, EPI_ISL_581295, EPI_ISL_581296, EPI_ISL_581297, EPI_ISL_581298, EPI_ISL_581299, EPI_ISL_581300, EPI_ISL_581301, EPI_ISL_581302, EPI_ISL_581303, EPI_ISL_581304, EPI_ISL_581305, EPI_ISL_581307, EPI_ISL_581308, EPI_ISL_581309, EPI_ISL_581312, EPI_ISL_581313, EPI_ISL_581316, EPI_ISL_581317, EPI_ISL_581318, EPI_ISL_581319, EPI_ISL_581320, EPI_ISL_581321, EPI_ISL_581322, EPI_ISL_581327, EPI_ISL_581330, EPI_ISL_581331, EPI_ISL_581332, EPI_ISL_581333, EPI_ISL_581336, EPI_ISL_581338, EPI_ISL_581340, EPI_ISL_581342, EPI_ISL_581345, EPI_ISL_581347, EPI_ISL_581348, EPI_ISL_581349, EPI_ISL_581351, EPI_ISL_581352, EPI_ISL_581353, EPI_ISL_581355, EPI_ISL_581356, EPI_ISL_581357, EPI_ISL_581360, EPI_ISL_581362, EPI_ISL_581363, EPI_ISL_581364, EPI_ISL_581365, EPI_ISL_581372, EPI_ISL_581373, EPI_ISL_581374, EPI_ISL_581388, EPI_ISL_581393, EPI_ISL_581422, EPI_ISL_581440 | | | |
| see above | Lighthouse Lab in Milton Keynes | Wellcome Sanger Institute for the COVID-19 Genomics UK (COG-UK) consortium | The Lighthouse Lab in Milton Keynes and Alex Alderton, Roberto Amato, Sonia Goncalves, Ewan Harrison, David K. Jackson, Ian Johnston, Dominic Kwiatkowski, Cordelia Langford, John Sillitoe on behalf of the Wellcome Sanger Institute COVID-19 Surveillance Team |
| EPI_ISL_581505, EPI_ISL_581506, EPI_ISL_581507 | NCDC/IGIB | NCDC/IGIB | Vivekanand A, Mahesh S. Dhar, Bharathram Uppili, Nishu Tyagi, Pooja Sharma, Akshay Kanakan, Simmi Tiwari, RadhaKrishnan VS, Robin Marwal, Azka Khan, Ajit Shewale, Tushar Nale, Rajesh Pandey, Sandhya Kabra, Mohammed Faruq, Sujeet Singh, Anurag Agrawal, Partha Rakshit |
| EPI_ISL_581508, EPI_ISL_581509, EPI_ISL_581510, EPI_ISL_581511, EPI_ISL_581512, EPI_ISL_581513, EPI_ISL_581514, EPI_ISL_581515, EPI_ISL_581516, EPI_ISL_581517, EPI_ISL_581518, EPI_ISL_581520, EPI_ISL_581522, EPI_ISL_581523, EPI_ISL_581524, EPI_ISL_581525, EPI_ISL_581526, EPI_ISL_581527, EPI_ISL_581528, EPI_ISL_581529, EPI_ISL_581530, EPI_ISL_581531, EPI_ISL_581532, EPI_ISL_581533, EPI_ISL_581534, EPI_ISL_581535, EPI_ISL_581536, EPI_ISL_581537, EPI_ISL_581538, EPI_ISL_581539, EPI_ISL_581540, EPI_ISL_581541, EPI_ISL_581542, EPI_ISL_581543, EPI_ISL_581544, EPI_ISL_581545, EPI_ISL_581546, EPI_ISL_581547, EPI_ISL_581548, EPI_ISL_581549, EPI_ISL_581550, EPI_ISL_581551, EPI_ISL_581552, EPI_ISL_581553, EPI_ISL_581554, EPI_ISL_581555, EPI_ISL_581556, EPI_ISL_581557, EPI_ISL_581558, EPI_ISL_581559, EPI_ISL_581560, EPI_ISL_581561, EPI_ISL_581562, EPI_ISL_581563, EPI_ISL_581564, EPI_ISL_581565, EPI_ISL_581566, EPI_ISL_581567, EPI_ISL_581568, EPI_ISL_581569, EPI_ISL_581570, EPI_ISL_581571, EPI_ISL_581572, EPI_ISL_581573, EPI_ISL_581574 | | | |
| see above | Virginia DCLS | Virginia DCLS | Virginia DCLS |
| EPI_ISL_582134 | Protzer Lab | Protzer Lab | Ulrike Protzer, Dieter Hoffmann, Eva Schulte, Andrea Theumer, Oliver Drechsel, Max von Kleist, Alexander Karollus, Julien Gagneur |
| EPI_ISL_582524, EPI_ISL_582525, EPI_ISL_582526, EPI_ISL_582527, EPI_ISL_582528, EPI_ISL_582529 | Veterinary Specialized Institute "Kraljevo", Serbia | Veterinary Specialized Institute "Kraljevo", Serbia | Vidanovic,D., Tesovic,B., Knezevic,A., Jovanovic,T., Jankovic,M., Sekler,M., Banovic Djeri,B., Petrovic,T., Volkening,J., Afonso,C. |
| EPI_ISL_582530 | Laboratory Diagnostic, Veterinary Specialized Institute Kraljevo | Laboratory Diagnostic, Veterinary Specialized Institute Kraljevo | Vidanovic,D., Tesovic,B., Knezevic,A., Jovanovic,T., Jankovic,M., Sekler,M., Banovic Djeri,B., Petrovic,T., Volkening,J., Afonso,C. |
| EPI_ISL_582531, EPI_ISL_582532, EPI_ISL_582533, EPI_ISL_582534 | Veterinary Specialized Institute "Kraljevo", Serbia | Veterinary Specialized Institute "Kraljevo", Serbia | Vidanovic,D., Tesovic,B., Knezevic,A., Jovanovic,T., Jankovic,M., Sekler,M., Banovic Djeri,B., Petrovic,T., Volkening,J., Afonso,C. |
| EPI_ISL_583249, EPI_ISL_583250, EPI_ISL_583251, EPI_ISL_583252, EPI_ISL_583253, EPI_ISL_583254, EPI_ISL_583255, EPI_ISL_583256, EPI_ISL_583257, EPI_ISL_583258, EPI_ISL_583259, EPI_ISL_583260, EPI_ISL_583261, EPI_ISL_583262, EPI_ISL_583263, EPI_ISL_583264, EPI_ISL_583265, EPI_ISL_583266, EPI_ISL_583267, EPI_ISL_583268, EPI_ISL_583269, EPI_ISL_583270, EPI_ISL_583271, EPI_ISL_583272, EPI_ISL_583273, EPI_ISL_583274, EPI_ISL_583275, EPI_ISL_583276, EPI_ISL_583277, EPI_ISL_583278, EPI_ISL_583279, EPI_ISL_583280, EPI_ISL_583281, EPI_ISL_583282, EPI_ISL_583283, EPI_ISL_583284, EPI_ISL_583285, EPI_ISL_583286, EPI_ISL_583287, EPI_ISL_583288, EPI_ISL_583289, EPI_ISL_583290, EPI_ISL_583291, EPI_ISL_583292, EPI_ISL_583293, EPI_ISL_583294, EPI_ISL_583295, EPI_ISL_583296, EPI_ISL_583297, EPI_ISL_583298, EPI_ISL_583299, EPI_ISL_583300, EPI_ISL_583301, EPI_ISL_583302, EPI_ISL_583303, EPI_ISL_583304, EPI_ISL_583305, EPI_ISL_583306, EPI_ISL_583307, EPI_ISL_583308, EPI_ISL_583309, EPI_ISL_583310, EPI_ISL_583311, EPI_ISL_583312, EPI_ISL_583313, EPI_ISL_583314, EPI_ISL_583315, EPI_ISL_583316, EPI_ISL_583317, EPI_ISL_583318, EPI_ISL_583319, EPI_ISL_583320, EPI_ISL_583321, EPI_ISL_583322, EPI_ISL_583323, EPI_ISL_583324, EPI_ISL_583325, EPI_ISL_583326, EPI_ISL_583327, EPI_ISL_583328, EPI_ISL_583329, EPI_ISL_583330, EPI_ISL_583331, EPI_ISL_583332, EPI_ISL_583333, EPI_ISL_583334, EPI_ISL_583335, EPI_ISL_583336, EPI_ISL_583337, EPI_ISL_583338, EPI_ISL_583339, EPI_ISL_583340, EPI_ISL_583341, EPI_ISL_583342, EPI_ISL_583343, EPI_ISL_583344, EPI_ISL_583345, EPI_ISL_583346, EPI_ISL_583347, EPI_ISL_583348, EPI_ISL_583349, EPI_ISL_583350, EPI_ISL_583351, EPI_ISL_583352, EPI_ISL_583353, EPI_ISL_583354, EPI_ISL_583355, EPI_ISL_583356, EPI_ISL_583357, EPI_ISL_583358, EPI_ISL_583359, EPI_ISL_583360, EPI_ISL_583361, EPI_ISL_583362, EPI_ISL_583363, EPI_ISL_583364, EPI_ISL_583365, EPI_ISL_583366, EPI_ISL_583367, EPI_ISL_583368, EPI_ISL_583369, EPI_ISL_583370, EPI_ISL_583371, EPI_ISL_583372, EPI_ISL_583373, EPI_ISL_583374, EPI_ISL_583375, EPI_ISL_583376, EPI_ISL_583377, EPI_ISL_583378, EPI_ISL_583379, EPI_ISL_583380, EPI_ISL_583381, EPI_ISL_583382, EPI_ISL_583383, EPI_ISL_583384, EPI_ISL_583385, EPI_ISL_583386, EPI_ISL_583387, EPI_ISL_583388, EPI_ISL_583389, EPI_ISL_583390, EPI_ISL_583391, EPI_ISL_583392, EPI_ISL_583393, EPI_ISL_583394, EPI_ISL_583395, EPI_ISL_583396, EPI_ISL_583397, EPI_ISL_583398, EPI_ISL_583399, EPI_ISL_583400, EPI_ISL_583401, EPI_ISL_583402, EPI_ISL_583403, EPI_ISL_583404, EPI_ISL_583405, EPI_ISL_583406, EPI_ISL_583407, EPI_ISL_583408, EPI_ISL_583409, EPI_ISL_583410, EPI_ISL_583411, EPI_ISL_583412, EPI_ISL_583413, EPI_ISL_583414, EPI_ISL_583415, EPI_ISL_583416, EPI_ISL_583417, EPI_ISL_583418, EPI_ISL_583419, EPI_ISL_583420, EPI_ISL_583421, EPI_ISL_583422, EPI_ISL_583423, EPI_ISL_583424, EPI_ISL_583425 | | | |
| see above | University of Michigan Clinical Microbiology Laboratory | Lauring Lab, University of Michigan, Department of Microbiology and Immunology | Valesano |
| EPI_ISL_583481, EPI_ISL_583483, EPI_ISL_583485, EPI_ISL_583489 | Institute of Virology, Biomedical Research Center of the Slovak Academy of Sciences, Bratislava | Faculty of Natural Sciences, Comenius University, Bratislava | Viktória Hodorová, Kristína Boršová, Broa Brejová, Viktória abanová, Dominika Friová, Sabina Fumaová Havlíková, Juraj Kopáek, Martina Liková, ubomíra Lukáiková, Martina Neboháová, Monika Sláviková, Edita Staroová, Elena Tichá, Tomáš Vina, Jozef Nosek, Boris Klempa |
| EPI_ISL_584095, EPI_ISL_584096, EPI_ISL_584097 | University of Michigan Clinical Microbiology Laboratory | Lauring Lab, University of Michigan, Department of Microbiology and Immunology | Valesano |
| EPI_ISL_584167, EPI_ISL_584188, EPI_ISL_584189, EPI_ISL_584190, EPI_ISL_584191, EPI_ISL_584192, EPI_ISL_584196, EPI_ISL_584197, EPI_ISL_584198, EPI_ISL_584199, EPI_ISL_584207, EPI_ISL_584208, EPI_ISL_584209, EPI_ISL_584210, EPI_ISL_584211, EPI_ISL_584217, EPI_ISL_584218, EPI_ISL_584219, EPI_ISL_584220, EPI_ISL_584221, EPI_ISL_584222, EPI_ISL_584223, EPI_ISL_584224, EPI_ISL_584225, EPI_ISL_584226, EPI_ISL_584227, EPI_ISL_584228, EPI_ISL_584229, EPI_ISL_584230, EPI_ISL_584231, EPI_ISL_584232, EPI_ISL_584233, EPI_ISL_584234, EPI_ISL_584235, EPI_ISL_584236, EPI_ISL_584237, EPI_ISL_584238, EPI_ISL_584239, EPI_ISL_584240, EPI_ISL_584241, EPI_ISL_584242, EPI_ISL_584243, EPI_ISL_584244, EPI_ISL_584245, EPI_ISL_584246, EPI_ISL_584247, EPI_ISL_584248, EPI_ISL_584249, EPI_ISL_584250, EPI_ISL_584251, EPI_ISL_584252, EPI_ISL_584253, EPI_ISL_584254, EPI_ISL_584255, EPI_ISL_584256, EPI_ISL_584257, EPI_ISL_584258, EPI_ISL_584259, EPI_ISL_584260, EPI_ISL_584261, EPI_ISL_584262, EPI_ISL_584263, EPI_ISL_584264, EPI_ISL_584265, EPI_ISL_584266, EPI_ISL_584267, EPI_ISL_584268, EPI_ISL_584269, EPI_ISL_584270, EPI_ISL_584271, EPI_ISL_584272, EPI_ISL_584273, EPI_ISL_584274, EPI_ISL_584275, EPI_ISL_584276, EPI_ISL_584277, EPI_ISL_584278, EPI_ISL_584279, EPI_ISL_584280, EPI_ISL_584281, EPI_ISL_584282, EPI_ISL_584324, EPI_ISL_584325, EPI_ISL_584326, EPI_ISL_584327, EPI_ISL_584328, EPI_ISL_584329, EPI_ISL_584330, EPI_ISL_584331, EPI_ISL_584332, EPI_ISL_584333, EPI_ISL_584334, EPI_ISL_584335, EPI_ISL_584336, EPI_ISL_584337, EPI_ISL_584338, EPI_ISL_584339, EPI_ISL_584340, EPI_ISL_584341, EPI_ISL_584342, EPI_ISL_584343, EPI_ISL_584344, EPI_ISL_584345, EPI_ISL_584346, EPI_ISL_584347, EPI_ISL_584348, EPI_ISL_584349, EPI_ISL_584350, EPI_ISL_584351, EPI_ISL_584352, EPI_ISL_584353, EPI_ISL_584354, EPI_ISL_584355, EPI_ISL_584356, EPI_ISL_584357, EPI_ISL_584358, EPI_ISL_584359, EPI_ISL_584360, EPI_ISL_584361, EPI_ISL_584362, EPI_ISL_584363, EPI_ISL_584364, EPI_ISL_584365, EPI_ISL_584366, EPI_ISL_584367, EPI_ISL_584368, EPI_ISL_584369, EPI_ISL_584370, EPI_ISL_584371, EPI_ISL_584372, EPI_ISL_584373, EPI_ISL_584374, EPI_ISL_584375, EPI_ISL_584376, EPI_ISL_584377, EPI_ISL_584378, EPI_ISL_584379, EPI_ISL_584380, EPI_ISL_585009, EPI_ISL_585010, EPI_ISL_585011, EPI_ISL_585012, EPI_ISL_585013, EPI_ISL_585014, EPI_ISL_585015, EPI_ISL_585016, EPI_ISL_585017, EPI_ISL_585018, EPI_ISL_585019, EPI_ISL_585020, EPI_ISL_585021, EPI_ISL_585022, EPI_ISL_585023, EPI_ISL_585024, EPI_ISL_585025, EPI_ISL_585026, EPI_ISL_585027, EPI_ISL_585028, EPI_ISL_585029, EPI_ISL_585030, EPI_ISL_585031, EPI_ISL_585032, EPI_ISL_585033, EPI_ISL_585034, EPI_ISL_585035, EPI_ISL_585036, EPI_ISL_585037, EPI_ISL_585038, EPI_ISL_585039, EPI_ISL_585040, EPI_ISL_585041, EPI_ISL_585042, EPI_ISL_585043, EPI_ISL_585044, EPI_ISL_585045, EPI_ISL_585046, EPI_ISL_585047, EPI_ISL_585048, EPI_ISL_585049, EPI_ISL_585050, EPI_ISL_585051, EPI_ISL_585052, EPI_ISL_585053, EPI_ISL_585054, EPI_ISL_585055, EPI_ISL_585056, EPI_ISL_585057, EPI_ISL_585058, EPI_ISL_585059, EPI_ISL_585060, EPI_ISL_585061, EPI_ISL_585062, EPI_ISL_585063, EPI_ISL_585064, EPI_ISL_585065, EPI_ISL_585066, EPI_ISL_585067, EPI_ISL_585068, EPI_ISL_585069, EPI_ISL_585070, EPI_ISL_585071, EPI_ISL_585072, EPI_ISL_585073, EPI_ISL_585074, EPI_ISL_585075, EPI_ISL_585076, EPI_ISL_585077, EPI_ISL_585078, EPI_ISL_585079, EPI_ISL_585080, EPI_ISL_585081, EPI_ISL_585082, EPI_ISL_585083, EPI_ISL_585084, EPI_ISL_585085, EPI_ISL_585086, EPI_ISL_585087, EPI_ISL_585088, EPI_ISL_585089, EPI_ISL_585090, EPI_ISL_585091, EPI_ISL_585620, EPI_ISL_585621, EPI_ISL_585622, EPI_ISL_585623, EPI_ISL_585624, EPI_ISL_585625, EPI_ISL_585626 | | | |
| see above | Virology Department, Sheffield Teaching Hospitals NHS Foundation Trust/Department of Infection, Immunity and Cardiovascular Disease, The Medical School, University of | COVID-19 Genomics UK (COG-UK) Consortium | Thushan de Silva, Matthew Parker, Nikki Smith, Adri Angyal, Rebecca Brown, Luke Green, Rachel Tucker, Paul Parsons, Danielle Groves, Katie Johnson, Laura Carrilero, Alex Keeley, Dave Partridge, Matthew Wyles, Benjamin Lindsey, Mehmet Yavuz, Mohammad Raza, Cariad Evans |

| | | | |
|---|---|---|---|
| | Sheffield | | |
| EPI_ISL_589017, EPI_ISL_589018, EPI_ISL_589019, EPI_ISL_589021, EPI_ISL_589024, EPI_ISL_589026, EPI_ISL_589028, EPI_ISL_589029, EPI_ISL_589030, EPI_ISL_589034, EPI_ISL_589039, EPI_ISL_589040, EPI_ISL_589041, EPI_ISL_589043, EPI_ISL_589051, EPI_ISL_589057, EPI_ISL_589063, EPI_ISL_589074, EPI_ISL_589075, EPI_ISL_589077, EPI_ISL_589079, EPI_ISL_589080, EPI_ISL_589081, EPI_ISL_589083, EPI_ISL_589090, EPI_ISL_589091, EPI_ISL_589095, EPI_ISL_589097, EPI_ISL_589098, EPI_ISL_589099, EPI_ISL_589101, EPI_ISL_589103, EPI_ISL_589104, EPI_ISL_589108, EPI_ISL_589109, EPI_ISL_589110, EPI_ISL_589111, EPI_ISL_589114, EPI_ISL_589115, EPI_ISL_589117, EPI_ISL_589118, EPI_ISL_589121, EPI_ISL_589122, EPI_ISL_589123, EPI_ISL_589127, EPI_ISL_589133, EPI_ISL_589135, EPI_ISL_589137, EPI_ISL_589147, EPI_ISL_589153, EPI_ISL_589157, EPI_ISL_589160, EPI_ISL_589166, EPI_ISL_589167, EPI_ISL_589168, EPI_ISL_589169, EPI_ISL_589170, EPI_ISL_589171, EPI_ISL_589172, EPI_ISL_589173, EPI_ISL_589176, EPI_ISL_589182, EPI_ISL_589183, EPI_ISL_589185, EPI_ISL_589187, EPI_ISL_589195, EPI_ISL_589198, EPI_ISL_589199, EPI_ISL_589205, EPI_ISL_589211, EPI_ISL_589212, EPI_ISL_589221, EPI_ISL_589224, EPI_ISL_589225, EPI_ISL_589226, EPI_ISL_589227, EPI_ISL_589229, EPI_ISL_589231, EPI_ISL_589233, EPI_ISL_589235, EPI_ISL_589237, EPI_ISL_589242, EPI_ISL_589243, EPI_ISL_589247, EPI_ISL_589251, EPI_ISL_589256, EPI_ISL_589260, EPI_ISL_589261, EPI_ISL_589264, EPI_ISL_589265, EPI_ISL_589266, EPI_ISL_589297, EPI_ISL_589304, EPI_ISL_589335, EPI_ISL_589355, EPI_ISL_589377, EPI_ISL_589407, EPI_ISL_589417, EPI_ISL_589423, EPI_ISL_589430, EPI_ISL_589440, EPI_ISL_589493, EPI_ISL_589519, EPI_ISL_589520, EPI_ISL_589521, EPI_ISL_589526 | | | |
| see above | Lighthouse Lab in Milton Keynes | Wellcome Sanger Institute for the COVID-19 Genomics UK (COG-UK) consortium | The Lighthouse Lab in Milton Keynes and Alex Alderton, Roberto Amato, Sonia Goncalves, Ewan Harrison, David K. Jackson, Ian Johnston, Dominic Kwiatkowski, Cordelia Langford, John Sillitoe on behalf of the Wellcome Sanger Institute COVID-19 Surveillance Team |
| EPI_ISL_589572, EPI_ISL_589573, EPI_ISL_589574, EPI_ISL_589575, EPI_ISL_589576, EPI_ISL_589578, EPI_ISL_589579, EPI_ISL_589581, EPI_ISL_589582, EPI_ISL_589583, EPI_ISL_589585, EPI_ISL_589586, EPI_ISL_589587, EPI_ISL_589588, EPI_ISL_589589, EPI_ISL_589590, EPI_ISL_589591, EPI_ISL_589592, EPI_ISL_589593, EPI_ISL_589594, EPI_ISL_589595, EPI_ISL_589596, EPI_ISL_589597, EPI_ISL_589598, EPI_ISL_589599, EPI_ISL_589600, EPI_ISL_589601, EPI_ISL_589602, EPI_ISL_589603, EPI_ISL_589604, EPI_ISL_589605, EPI_ISL_589606, EPI_ISL_589608, EPI_ISL_589609, EPI_ISL_589610, EPI_ISL_589611, EPI_ISL_589612, EPI_ISL_589613, EPI_ISL_589614, EPI_ISL_589615, EPI_ISL_589616, EPI_ISL_589617, EPI_ISL_589618, EPI_ISL_589619, EPI_ISL_589620, EPI_ISL_589621, EPI_ISL_589622, EPI_ISL_589623, EPI_ISL_589624, EPI_ISL_589625, EPI_ISL_589626, EPI_ISL_589627, EPI_ISL_589628, EPI_ISL_589629, EPI_ISL_589630, EPI_ISL_589631, EPI_ISL_589632, EPI_ISL_589633, EPI_ISL_589634, EPI_ISL_589635, EPI_ISL_589636, EPI_ISL_589637, EPI_ISL_589638, EPI_ISL_589639, EPI_ISL_589640, EPI_ISL_589641, EPI_ISL_589642, EPI_ISL_589643, EPI_ISL_589645, EPI_ISL_589648, EPI_ISL_589649, EPI_ISL_589650, EPI_ISL_589651, EPI_ISL_589652, EPI_ISL_589653, EPI_ISL_589654, EPI_ISL_589655, EPI_ISL_589656, EPI_ISL_589657, EPI_ISL_589658, EPI_ISL_589659, EPI_ISL_589660, EPI_ISL_589661, EPI_ISL_589662, EPI_ISL_589663, EPI_ISL_589664, EPI_ISL_589665, EPI_ISL_589666, EPI_ISL_589667, EPI_ISL_589669, EPI_ISL_589670, EPI_ISL_589671, EPI_ISL_589672, EPI_ISL_589673, EPI_ISL_589674, EPI_ISL_589675, EPI_ISL_589677, EPI_ISL_589678, EPI_ISL_589679, EPI_ISL_589680, EPI_ISL_589682, EPI_ISL_589683, EPI_ISL_589684, EPI_ISL_589685, EPI_ISL_589686, EPI_ISL_589687, EPI_ISL_589688, EPI_ISL_589689, EPI_ISL_589690, EPI_ISL_589691, EPI_ISL_589692, EPI_ISL_589693, EPI_ISL_589694, EPI_ISL_589695, EPI_ISL_589696, EPI_ISL_589697, EPI_ISL_589698, EPI_ISL_589700, EPI_ISL_589701, EPI_ISL_589702, EPI_ISL_589703, EPI_ISL_589704, EPI_ISL_589705, EPI_ISL_589706, EPI_ISL_589707, EPI_ISL_589708, EPI_ISL_589709, EPI_ISL_589710, EPI_ISL_589711, EPI_ISL_589712, EPI_ISL_589713, EPI_ISL_589714, EPI_ISL_589716, EPI_ISL_589717, EPI_ISL_589718, EPI_ISL_589719, EPI_ISL_589720, EPI_ISL_589721, EPI_ISL_589722, EPI_ISL_589723, EPI_ISL_589725, EPI_ISL_589726, EPI_ISL_589727, EPI_ISL_589728, EPI_ISL_589729, EPI_ISL_589730, EPI_ISL_589731, EPI_ISL_589732, EPI_ISL_589733, EPI_ISL_589734, EPI_ISL_589735, EPI_ISL_589736, EPI_ISL_589737, EPI_ISL_589738, EPI_ISL_589739, EPI_ISL_589740, EPI_ISL_589741, EPI_ISL_589742, EPI_ISL_589743, EPI_ISL_589744, EPI_ISL_589745, EPI_ISL_589746, EPI_ISL_589747, EPI_ISL_589748, EPI_ISL_589749, EPI_ISL_589750, EPI_ISL_589751, EPI_ISL_589752, EPI_ISL_589753, EPI_ISL_589754, EPI_ISL_589755, EPI_ISL_589756, EPI_ISL_589757, EPI_ISL_589759, EPI_ISL_589760, EPI_ISL_589761, EPI_ISL_589796 | | | |
| see above | Lighthouse Lab in Milton Keynes | Wellcome Sanger Institute for the COVID-19 Genomics UK (COG-UK) consortium | The Lighthouse Lab in Milton Keynes and Alex Alderton, Roberto Amato, Sonia Goncalves, Ewan Harrison, David K. Jackson, Ian Johnston, Dominic Kwiatkowski, Cordelia Langford, John Sillitoe on behalf of the Wellcome Sanger Institute COVID-19 Surveillance Team (http://www.sanger.ac.uk/covid-team) |
| EPI_ISL_590699, EPI_ISL_590700, EPI_ISL_590701, EPI_ISL_590702, EPI_ISL_590703, EPI_ISL_590704, EPI_ISL_590705, EPI_ISL_590706, EPI_ISL_590707, EPI_ISL_590708, EPI_ISL_590709, EPI_ISL_590710, EPI_ISL_590711, EPI_ISL_590712, EPI_ISL_590713, EPI_ISL_590714, EPI_ISL_590715, EPI_ISL_590716, EPI_ISL_590717, EPI_ISL_590718, EPI_ISL_590719, EPI_ISL_590720, EPI_ISL_590721, EPI_ISL_590722, EPI_ISL_590723, EPI_ISL_590724, EPI_ISL_590725, EPI_ISL_590726, EPI_ISL_590727, EPI_ISL_590728, EPI_ISL_590729, EPI_ISL_590730, EPI_ISL_590731, EPI_ISL_590732, EPI_ISL_590733, EPI_ISL_590734, EPI_ISL_590735, EPI_ISL_590736, EPI_ISL_590737, EPI_ISL_590738, EPI_ISL_590739, EPI_ISL_590740, EPI_ISL_590741, EPI_ISL_590742, EPI_ISL_590743, EPI_ISL_590744, EPI_ISL_590745, EPI_ISL_590746, EPI_ISL_590747, EPI_ISL_590748, EPI_ISL_590749, EPI_ISL_590750, EPI_ISL_590751, EPI_ISL_590752 | | | |
| see above | University of Michigan Clinical Microbiology Laboratory | Lauring Lab, University of Michigan, Department of Microbiology and Immunology | Valesano |
| EPI_ISL_591054, EPI_ISL_591055, EPI_ISL_591056 | Virus Ecology, Rocky Mountain Laboratories, National Institutes of Health | Virus Ecology, Rocky Mountain Laboratories, National Institutes of Health | Van Doremalen,N., Holbrook,M.G., Barbian,K.D., Bushmaker,C., Bushmaker,T., Martens,C.A., Munster,V.J. |
| EPI_ISL_591531, EPI_ISL_591532, EPI_ISL_591533, EPI_ISL_591534 | Laboratorio de Infectologia y virologia molecular | Center for Mathematical Modeling and Center for Genome Regulation. Santiago, Chile | Valiente F, Gaete A, Travisany D, Palma R, Urra C, Varas M, Allende ML, Maass A, González M, Ferres M. |
| EPI_ISL_595802, EPI_ISL_595803, EPI_ISL_595804, EPI_ISL_595805, EPI_ISL_595806, EPI_ISL_595807, EPI_ISL_595808, EPI_ISL_595809, EPI_ISL_595810, EPI_ISL_595811, EPI_ISL_595812, EPI_ISL_595813, EPI_ISL_595814, EPI_ISL_595815, EPI_ISL_595816, EPI_ISL_595817, EPI_ISL_595818, EPI_ISL_595819, EPI_ISL_595820, EPI_ISL_595821, EPI_ISL_595823, EPI_ISL_595824, EPI_ISL_595825, EPI_ISL_595826, EPI_ISL_595827, EPI_ISL_595828, EPI_ISL_595829, EPI_ISL_595830, EPI_ISL_595831, EPI_ISL_595832, EPI_ISL_595833, EPI_ISL_595834, EPI_ISL_595835, EPI_ISL_595836, EPI_ISL_596191, EPI_ISL_596192, EPI_ISL_596193, EPI_ISL_596194, EPI_ISL_596195, EPI_ISL_596196, EPI_ISL_596197, EPI_ISL_596198, EPI_ISL_596199, EPI_ISL_596200, EPI_ISL_596201, EPI_ISL_596202, EPI_ISL_596203, EPI_ISL_596204, EPI_ISL_596205, EPI_ISL_596206, EPI_ISL_596207, EPI_ISL_596208, EPI_ISL_596209, EPI_ISL_596210, EPI_ISL_596211, EPI_ISL_596212, EPI_ISL_596213, EPI_ISL_596214, EPI_ISL_596215, EPI_ISL_596216, EPI_ISL_596217, EPI_ISL_596218, EPI_ISL_596219, EPI_ISL_596220, EPI_ISL_596221, EPI_ISL_596222, EPI_ISL_596223, EPI_ISL_596224, EPI_ISL_596225, EPI_ISL_596226, EPI_ISL_596227 | | | |
| see above | Virology Department, Sheffield Teaching Hospitals NHS Foundation Trust/Department of Infection, Immunity and Cardiovascular Disease, The Medical School, University of Sheffield | COVID-19 Genomics UK (COG-UK) Consortium | Thushan de Silva, Matthew Parker, Nikki Smith, Adri Angyal, Rebecca Brown, Luke Green, Rachel Tucker, Paul Parsons, Danielle Groves, Katie Johnson, Laura Carrilero, Alex Keeley, Dave Partridge, Matthew Wyles, Benjamin Lindsey, Mehmet Yavuz, Mohammad Raza, Cariad Evans |
| EPI_ISL_596456, EPI_ISL_596457, EPI_ISL_596458, EPI_ISL_596459, EPI_ISL_596460, EPI_ISL_596461, EPI_ISL_596462, EPI_ISL_596463, EPI_ISL_596464, EPI_ISL_596465, EPI_ISL_596466, EPI_ISL_596467, EPI_ISL_596468, EPI_ISL_596469, EPI_ISL_596470, EPI_ISL_596471, EPI_ISL_596472, EPI_ISL_596473, EPI_ISL_596474, EPI_ISL_596475, EPI_ISL_596476, EPI_ISL_596477, EPI_ISL_596478, EPI_ISL_596479, EPI_ISL_596480, EPI_ISL_596481, EPI_ISL_596482, EPI_ISL_596483, EPI_ISL_596484, EPI_ISL_596485, EPI_ISL_596486, EPI_ISL_596487, EPI_ISL_596488, EPI_ISL_596489, EPI_ISL_596490, EPI_ISL_596491, EPI_ISL_596492, EPI_ISL_596493, EPI_ISL_596494, EPI_ISL_596495, EPI_ISL_596496, EPI_ISL_596497 | | | |
| see above | National Public Health Laboratory, National Centre for Infectious Diseases | National Public Health Laboratory, National Centre for Infectious Diseases | Tze Minn Mak, Sophie Octavia, Zhenyang Zhou, Lin Cui, Raymond Tzer Pin Lin |
| EPI_ISL_596569, EPI_ISL_596570, EPI_ISL_596571, EPI_ISL_596572, EPI_ISL_596573, EPI_ISL_596574, EPI_ISL_596575, EPI_ISL_596576, EPI_ISL_596577, EPI_ISL_596578, EPI_ISL_596579, EPI_ISL_596580, EPI_ISL_596581, EPI_ISL_596582, EPI_ISL_596583, EPI_ISL_596584, EPI_ISL_596585, EPI_ISL_596586, EPI_ISL_596587, EPI_ISL_596588, EPI_ISL_596589, EPI_ISL_596590, EPI_ISL_596591, EPI_ISL_596592, EPI_ISL_596593, EPI_ISL_596594, EPI_ISL_596595, EPI_ISL_596596, EPI_ISL_596597, EPI_ISL_596598, EPI_ISL_596599, EPI_ISL_596600, EPI_ISL_596601, EPI_ISL_596602, EPI_ISL_596603, EPI_ISL_596604, EPI_ISL_596605, EPI_ISL_596606, EPI_ISL_596607, EPI_ISL_596608, EPI_ISL_596609, EPI_ISL_596610, EPI_ISL_596611, EPI_ISL_596612, EPI_ISL_596613, EPI_ISL_596614, EPI_ISL_596615, EPI_ISL_596616, EPI_ISL_596617, EPI_ISL_596618, EPI_ISL_596619, EPI_ISL_596620, EPI_ISL_596621, EPI_ISL_596622, EPI_ISL_596623, EPI_ISL_596624, EPI_ISL_596625, EPI_ISL_596626 | | | |
| see above | University of Michigan Clinical Microbiology Laboratory | Lauring Lab, University of Michigan, Department of Microbiology and Immunology | Valesano |
| EPI_ISL_598250, EPI_ISL_598251, EPI_ISL_598252, EPI_ISL_598253, EPI_ISL_598254, EPI_ISL_598255, EPI_ISL_598257, EPI_ISL_598258, EPI_ISL_598259, EPI_ISL_598260, EPI_ISL_598261, EPI_ISL_598262, EPI_ISL_598263, EPI_ISL_598264, EPI_ISL_598265, EPI_ISL_598267, EPI_ISL_598268, EPI_ISL_598269, EPI_ISL_598270, EPI_ISL_598272, EPI_ISL_598273, EPI_ISL_598274, EPI_ISL_598275, EPI_ISL_598276, EPI_ISL_598277, EPI_ISL_598280, EPI_ISL_598283, EPI_ISL_598284, EPI_ISL_598285, EPI_ISL_598286, EPI_ISL_598287, EPI_ISL_598288, EPI_ISL_598289, EPI_ISL_598290, EPI_ISL_598291, EPI_ISL_598292, EPI_ISL_598293, EPI_ISL_598294, EPI_ISL_598295, EPI_ISL_598296, EPI_ISL_598298, EPI_ISL_598299, EPI_ISL_598300, EPI_ISL_598301, EPI_ISL_598302, EPI_ISL_598303, EPI_ISL_598304, EPI_ISL_598306, EPI_ISL_598307, EPI_ISL_598308, EPI_ISL_598309, EPI_ISL_598311, EPI_ISL_598312, EPI_ISL_598313, EPI_ISL_598314, EPI_ISL_598315, EPI_ISL_598316, EPI_ISL_598318, EPI_ISL_598319, EPI_ISL_598320, EPI_ISL_598323, EPI_ISL_598324, EPI_ISL_598326, EPI_ISL_598332, EPI_ISL_598333, EPI_ISL_598334, EPI_ISL_598335, EPI_ISL_598336, EPI_ISL_598337, EPI_ISL_598338, EPI_ISL_598339, EPI_ISL_598340, EPI_ISL_598341, EPI_ISL_598342, EPI_ISL_598343, EPI_ISL_598344, EPI_ISL_598345, EPI_ISL_598346, EPI_ISL_598347, EPI_ISL_598348, EPI_ISL_598349, EPI_ISL_598350, EPI_ISL_598351, EPI_ISL_598353, EPI_ISL_598354, EPI_ISL_598356, EPI_ISL_598357, EPI_ISL_598358, EPI_ISL_598359, EPI_ISL_598360, EPI_ISL_598362, EPI_ISL_598363, EPI_ISL_598364, EPI_ISL_598365, EPI_ISL_598366, EPI_ISL_598367, EPI_ISL_598368, EPI_ISL_598369, EPI_ISL_598371, EPI_ISL_598372, EPI_ISL_598373, EPI_ISL_598374, EPI_ISL_598375, EPI_ISL_598377, EPI_ISL_598378, EPI_ISL_598379, EPI_ISL_598380, EPI_ISL_598381, EPI_ISL_598382, EPI_ISL_598383, EPI_ISL_598385, EPI_ISL_598386, EPI_ISL_598388, EPI_ISL_598389, EPI_ISL_598392, EPI_ISL_598393, EPI_ISL_598396, EPI_ISL_598397, EPI_ISL_598398, EPI_ISL_598399, EPI_ISL_598401, EPI_ISL_598402, EPI_ISL_598403, EPI_ISL_598404, EPI_ISL_598405, EPI_ISL_598407, EPI_ISL_598409, EPI_ISL_598410, EPI_ISL_598412, EPI_ISL_598413, EPI_ISL_598414, EPI_ISL_598415, EPI_ISL_598418, EPI_ISL_598419, EPI_ISL_598420, EPI_ISL_598421, EPI_ISL_598424, EPI_ISL_598425, EPI_ISL_598426, EPI_ISL_598427, EPI_ISL_598428, EPI_ISL_598429, EPI_ISL_598430, EPI_ISL_598431, EPI_ISL_598432, EPI_ISL_598434, EPI_ISL_598435, EPI_ISL_598437, EPI_ISL_598438, EPI_ISL_598439, EPI_ISL_598441, EPI_ISL_598442, EPI_ISL_598443, EPI_ISL_598444, EPI_ISL_598445, EPI_ISL_598447, EPI_ISL_598448, EPI_ISL_598449, EPI_ISL_598450, EPI_ISL_598451, EPI_ISL_598452, EPI_ISL_598453, EPI_ISL_598455, EPI_ISL_598458, EPI_ISL_598459, EPI_ISL_598461, EPI_ISL_598462, EPI_ISL_598463, EPI_ISL_598465, EPI_ISL_598466, EPI_ISL_598467, EPI_ISL_598468, EPI_ISL_598469, EPI_ISL_598470, EPI_ISL_598471, EPI_ISL_598472, EPI_ISL_598474, EPI_ISL_598475, EPI_ISL_598476, EPI_ISL_598478, EPI_ISL_598479, EPI_ISL_598481, EPI_ISL_598482, EPI_ISL_598483, EPI_ISL_598484, EPI_ISL_598485, EPI_ISL_598486, EPI_ISL_598487, EPI_ISL_598488, EPI_ISL_598489, EPI_ISL_598491, EPI_ISL_598493, EPI_ISL_598494, EPI_ISL_598496, EPI_ISL_598497, EPI_ISL_598498, EPI_ISL_598500, EPI_ISL_598501, EPI_ISL_598502, EPI_ISL_598503, EPI_ISL_598504, EPI_ISL_598505, EPI_ISL_598506, EPI_ISL_598507, EPI_ISL_598509, EPI_ISL_598510, EPI_ISL_598511, EPI_ISL_598512, EPI_ISL_598513, EPI_ISL_598514, EPI_ISL_598516, EPI_ISL_598517, EPI_ISL_598518, EPI_ISL_598519, EPI_ISL_598520, EPI_ISL_598521, EPI_ISL_598522, EPI_ISL_598523, EPI_ISL_598524, EPI_ISL_598531, EPI_ISL_598532, EPI_ISL_598540, EPI_ISL_598547, EPI_ISL_598548, EPI_ISL_598551, EPI_ISL_598552, EPI_ISL_598557, EPI_ISL_598559, EPI_ISL_598564, EPI_ISL_598566, EPI_ISL_598567, EPI_ISL_598570, EPI_ISL_598571, EPI_ISL_598575, EPI_ISL_598578, EPI_ISL_598590, EPI_ISL_598592, EPI_ISL_598594, EPI_ISL_598596, EPI_ISL_598600, EPI_ISL_598603, EPI_ISL_598609, EPI_ISL_598612, EPI_ISL_598613, EPI_ISL_598622, EPI_ISL_598623, EPI_ISL_598631, EPI_ISL_598634, EPI_ISL_598636, EPI_ISL_598637, EPI_ISL_598640, EPI_ISL_598647, EPI_ISL_598650, EPI_ISL_598655, EPI_ISL_598656, EPI_ISL_598658, EPI_ISL_598659, EPI_ISL_598660, EPI_ISL_598663, EPI_ISL_598678, EPI_ISL_598680, EPI_ISL_598685, EPI_ISL_598686, EPI_ISL_598693, EPI_ISL_598695, EPI_ISL_598706, EPI_ISL_598711, EPI_ISL_598721, EPI_ISL_598722, EPI_ISL_598725, EPI_ISL_598726, EPI_ISL_598728, EPI_ISL_598733, EPI_ISL_598734, EPI_ISL_598735, EPI_ISL_598737, EPI_ISL_598739, EPI_ISL_598741, EPI_ISL_598747, EPI_ISL_598748, EPI_ISL_598749, EPI_ISL_598750, EPI_ISL_598755, EPI_ISL_598758, EPI_ISL_598765, EPI_ISL_598781, EPI_ISL_598782, EPI_ISL_598786, EPI_ISL_598787, EPI_ISL_598789, EPI_ISL_598799, EPI_ISL_598800, EPI_ISL_598801, EPI_ISL_598810, EPI_ISL_598815, EPI_ISL_598817, EPI_ISL_598818, EPI_ISL_598819, EPI_ISL_598820, EPI_ISL_598823, EPI_ISL_599818, EPI_ISL_599819, EPI_ISL_599820, EPI_ISL_599821, EPI_ISL_599822, EPI_ISL_599823, EPI_ISL_599824, EPI_ISL_599825, EPI_ISL_599826, EPI_ISL_599827, EPI_ISL_599828, EPI_ISL_599829, EPI_ISL_599832, EPI_ISL_599833, EPI_ISL_599834, EPI_ISL_599836, EPI_ISL_599837, EPI_ISL_599838, EPI_ISL_599840, EPI_ISL_599841, EPI_ISL_599842, EPI_ISL_599844, EPI_ISL_599846, EPI_ISL_599848, EPI_ISL_599849, EPI_ISL_599852, EPI_ISL_599853, EPI_ISL_599854, EPI_ISL_599855, EPI_ISL_599856, EPI_ISL_599857, EPI_ISL_599858, EPI_ISL_599859, EPI_ISL_599860, EPI_ISL_599861, EPI_ISL_599862, EPI_ISL_599863, EPI_ISL_599864, EPI_ISL_599865, EPI_ISL_599866, EPI_ISL_599867, EPI_ISL_599868, EPI_ISL_599869, EPI_ISL_599870, EPI_ISL_599871, EPI_ISL_599872, EPI_ISL_599873, EPI_ISL_599874, EPI_ISL_599875, EPI_ISL_599876, EPI_ISL_599877, EPI_ISL_599879, EPI_ISL_599880, EPI_ISL_599881, EPI_ISL_599883, EPI_ISL_599885, EPI_ISL_599887, EPI_ISL_599889, EPI_ISL_599890, EPI_ISL_599891, EPI_ISL_599892, EPI_ISL_599893, EPI_ISL_599894, EPI_ISL_599896, EPI_ISL_599898, EPI_ISL_599899, EPI_ISL_599900, EPI_ISL_599901, EPI_ISL_599902, EPI_ISL_599903, EPI_ISL_599904, EPI_ISL_599905, EPI_ISL_599906, EPI_ISL_599908, EPI_ISL_599909, EPI_ISL_599911, EPI_ISL_599912, EPI_ISL_599913, EPI_ISL_599914, EPI_ISL_599915, EPI_ISL_599916, EPI_ISL_599917, EPI_ISL_599918, EPI_ISL_599919, EPI_ISL_599920, EPI_ISL_599921, EPI_ISL_599922, EPI_ISL_599923, EPI_ISL_599924, EPI_ISL_599925, EPI_ISL_599926, EPI_ISL_599927, EPI_ISL_599928, EPI_ISL_599929, EPI_ISL_599930, EPI_ISL_599931, EPI_ISL_599932, EPI_ISL_599933, EPI_ISL_599934, EPI_ISL_599935, EPI_ISL_599936, EPI_ISL_599937, EPI_ISL_599938, EPI_ISL_599939, EPI_ISL_599940, EPI_ISL_599941, EPI_ISL_599942, EPI_ISL_599943, EPI_ISL_599944, EPI_ISL_599945, EPI_ISL_599949, EPI_ISL_599951, EPI_ISL_599954, EPI_ISL_599956, EPI_ISL_599957, EPI_ISL_599958, EPI_ISL_599959, EPI_ISL_599961, EPI_ISL_599962, EPI_ISL_599963, EPI_ISL_599964, EPI_ISL_599965, EPI_ISL_599967, EPI_ISL_599968, EPI_ISL_599969, EPI_ISL_599970, EPI_ISL_599972, EPI_ISL_599973, EPI_ISL_599975, EPI_ISL_599976, EPI_ISL_599977, EPI_ISL_599978, EPI_ISL_599979, EPI_ISL_599980, EPI_ISL_599981, EPI_ISL_599983, EPI_ISL_599984, EPI_ISL_599986, EPI_ISL_599987, EPI_ISL_599989, EPI_ISL_599990, EPI_ISL_599992, EPI_ISL_599993, EPI_ISL_599995, EPI_ISL_599998, EPI_ISL_599999, EPI_ISL_600000, EPI_ISL_600001, EPI_ISL_600002, EPI_ISL_600003, EPI_ISL_600004, EPI_ISL_600005, EPI_ISL_600007, EPI_ISL_600008, EPI_ISL_600009, EPI_ISL_600010, EPI_ISL_600011, EPI_ISL_600012, EPI_ISL_600013, EPI_ISL_600014, EPI_ISL_600015, EPI_ISL_600016, EPI_ISL_600017, EPI_ISL_600018, EPI_ISL_600020, EPI_ISL_600021, EPI_ISL_600022, EPI_ISL_600023, EPI_ISL_600024, EPI_ISL_600025, EPI_ISL_600026, EPI_ISL_600028, EPI_ISL_600029, EPI_ISL_600030, EPI_ISL_600032, EPI_ISL_600033, EPI_ISL_600034, EPI_ISL_600035, EPI_ISL_600036, EPI_ISL_600037, EPI_ISL_600038, EPI_ISL_600039, EPI_ISL_600040, EPI_ISL_600041, EPI_ISL_600042, EPI_ISL_600045, EPI_ISL_600046, EPI_ISL_600047, EPI_ISL_600048, EPI_ISL_600049, EPI_ISL_600051, EPI_ISL_600052, EPI_ISL_600053, EPI_ISL_600054, EPI_ISL_600055, EPI_ISL_600057, EPI_ISL_600058, EPI_ISL_600059, EPI_ISL_600060, EPI_ISL_600061, EPI_ISL_600062, EPI_ISL_600063, EPI_ISL_600066, EPI_ISL_600068, EPI_ISL_600069, EPI_ISL_600070, EPI_ISL_600072, EPI_ISL_600073, EPI_ISL_600074, EPI_ISL_600075, EPI_ISL_600076, EPI_ISL_600078, EPI_ISL_600080, EPI_ISL_600082, EPI_ISL_600083, EPI_ISL_600084, EPI_ISL_600085, EPI_ISL_600086, EPI_ISL_600087, EPI_ISL_600089, EPI_ISL_600090, EPI_ISL_600091, EPI_ISL_600092, EPI_ISL_600093, EPI_ISL_600094, EPI_ISL_600095, EPI_ISL_600096, EPI_ISL_600097, EPI_ISL_600098, EPI_ISL_600099, EPI_ISL_600100, EPI_ISL_600101, EPI_ISL_600102, EPI_ISL_600103, EPI_ISL_600104, EPI_ISL_600105, EPI_ISL_600106, EPI_ISL_600107, EPI_ISL_600109, EPI_ISL_600110, | | | |

| | | | |
|---|---|---|---|
| EPI_ISL_600111, EPI_ISL_600112, EPI_ISL_600113, EPI_ISL_600114, EPI_ISL_600115, EPI_ISL_600116, EPI_ISL_600117, EPI_ISL_600118, EPI_ISL_600119, EPI_ISL_600120, EPI_ISL_600121, EPI_ISL_600122, EPI_ISL_600125, EPI_ISL_600127, EPI_ISL_600931, EPI_ISL_601224, EPI_ISL_601234, EPI_ISL_601235, EPI_ISL_601239, EPI_ISL_601240, EPI_ISL_601242, EPI_ISL_601247, EPI_ISL_601248, EPI_ISL_601249, EPI_ISL_601256, EPI_ISL_601258, EPI_ISL_601267, EPI_ISL_601268, EPI_ISL_601270, EPI_ISL_601272, EPI_ISL_601275, EPI_ISL_601278, EPI_ISL_601280, EPI_ISL_601281, EPI_ISL_601282, EPI_ISL_601287, EPI_ISL_601291, EPI_ISL_601294, EPI_ISL_601298, EPI_ISL_601299, EPI_ISL_601301, EPI_ISL_601302, EPI_ISL_601303, EPI_ISL_601304, EPI_ISL_601305, EPI_ISL_601307, EPI_ISL_601313, EPI_ISL_601314, EPI_ISL_601317, EPI_ISL_601320, EPI_ISL_601324, EPI_ISL_601325, EPI_ISL_601326, EPI_ISL_601327, EPI_ISL_601329, EPI_ISL_601330, EPI_ISL_601338, EPI_ISL_601340, EPI_ISL_601343, EPI_ISL_601345, EPI_ISL_601349, EPI_ISL_601354, EPI_ISL_601359, EPI_ISL_601360, EPI_ISL_601363, EPI_ISL_601365, EPI_ISL_601368, EPI_ISL_601369, EPI_ISL_601370, EPI_ISL_601371, EPI_ISL_601373, EPI_ISL_601375, EPI_ISL_601376, EPI_ISL_601379, EPI_ISL_601383, EPI_ISL_601388, EPI_ISL_601390, EPI_ISL_601392, EPI_ISL_601394, EPI_ISL_601395, EPI_ISL_601404, EPI_ISL_601406, EPI_ISL_601409, EPI_ISL_601410, EPI_ISL_601412, EPI_ISL_601417, EPI_ISL_601418, EPI_ISL_601420, EPI_ISL_601427, EPI_ISL_601431, EPI_ISL_601434, EPI_ISL_601437, EPI_ISL_601438, EPI_ISL_601439, EPI_ISL_601442, EPI_ISL_601443, EPI_ISL_601444, EPI_ISL_601447, EPI_ISL_601448, EPI_ISL_601449, EPI_ISL_601450, EPI_ISL_601451, EPI_ISL_601452, EPI_ISL_601455, EPI_ISL_601456, EPI_ISL_601463, EPI_ISL_601464, EPI_ISL_601465, EPI_ISL_601468, EPI_ISL_601470, EPI_ISL_601473, EPI_ISL_601474, EPI_ISL_601476, EPI_ISL_601478, EPI_ISL_601480, EPI_ISL_601484, EPI_ISL_601491, EPI_ISL_601492, EPI_ISL_601502, EPI_ISL_601505, EPI_ISL_601507, EPI_ISL_601511, EPI_ISL_601513, EPI_ISL_601514, EPI_ISL_601518, EPI_ISL_601520, EPI_ISL_601521, EPI_ISL_601523, EPI_ISL_601524, EPI_ISL_601525, EPI_ISL_601526, EPI_ISL_601528, EPI_ISL_601531, EPI_ISL_601532, EPI_ISL_601533, EPI_ISL_601535, EPI_ISL_601536, EPI_ISL_601539, EPI_ISL_601540, EPI_ISL_601543, EPI_ISL_601545, EPI_ISL_601546, EPI_ISL_601548, EPI_ISL_601552, EPI_ISL_601554, EPI_ISL_601556, EPI_ISL_601557, EPI_ISL_601558, EPI_ISL_601561, EPI_ISL_601562, EPI_ISL_601563, EPI_ISL_601570, EPI_ISL_601573, EPI_ISL_601574, EPI_ISL_601575, EPI_ISL_601577, EPI_ISL_601581, EPI_ISL_601582, EPI_ISL_601583, EPI_ISL_601584, EPI_ISL_601586, EPI_ISL_601587, EPI_ISL_601595, EPI_ISL_601596, EPI_ISL_601598, EPI_ISL_601599, EPI_ISL_601600, EPI_ISL_601603, EPI_ISL_601610, EPI_ISL_601614, EPI_ISL_601616, EPI_ISL_601631, EPI_ISL_601632, EPI_ISL_601633, EPI_ISL_601642, EPI_ISL_601644, EPI_ISL_601647, EPI_ISL_601649, EPI_ISL_601651, EPI_ISL_601653, EPI_ISL_601656, EPI_ISL_601658, EPI_ISL_601659, EPI_ISL_601660, EPI_ISL_601670, EPI_ISL_601671, EPI_ISL_601676, EPI_ISL_601677, EPI_ISL_601680, EPI_ISL_601688, EPI_ISL_601692, EPI_ISL_601698, EPI_ISL_601699, EPI_ISL_601701, EPI_ISL_601702, EPI_ISL_601706, EPI_ISL_601708, EPI_ISL_601710, EPI_ISL_601711, EPI_ISL_601717, EPI_ISL_601721, EPI_ISL_601723, EPI_ISL_601725, EPI_ISL_601731, EPI_ISL_601732, EPI_ISL_601734, EPI_ISL_601738, EPI_ISL_601739, EPI_ISL_601743, EPI_ISL_601744, EPI_ISL_601749, EPI_ISL_601753, EPI_ISL_601755, EPI_ISL_601756, EPI_ISL_601757, EPI_ISL_601761, EPI_ISL_601763, EPI_ISL_601773, EPI_ISL_601774, EPI_ISL_601776, EPI_ISL_601777, EPI_ISL_601779, EPI_ISL_601780 | | | |
| see above | Lighthouse Lab in Milton Keynes | Wellcome Sanger Institute for the COVID-19 Genomics UK (COG-UK) consortium | The Lighthouse Lab in Milton Keynes and Alex Alderton, Roberto Amato, Sonia Goncalves, Ewan Harrison, David K. Jackson, Ian Johnston, Dominic Kwiatkowski, Cordelia Langford, John Sillitoe on behalf of the Wellcome Sanger Institute COVID-19 Surveillance Team (http://www.sanger.ac.uk/covid-team) |
| EPI_ISL_605817, EPI_ISL_605818, EPI_ISL_605819, EPI_ISL_605820, EPI_ISL_605821, EPI_ISL_605822, EPI_ISL_605823, EPI_ISL_605824 | National Public Health Laboratory, National Centre for Infectious Diseases | National Public Health Laboratory, National Centre for Infectious Diseases | Tze Minn Mak, Sophie Octavia, Zhenyang Zhou, Lin Cui, Raymond Tzer Pin Lin |
| EPI_ISL_606321, EPI_ISL_606328, EPI_ISL_606342, EPI_ISL_606362, EPI_ISL_606379, EPI_ISL_606381, EPI_ISL_606401, EPI_ISL_606423, EPI_ISL_606481, EPI_ISL_606483, EPI_ISL_606488, EPI_ISL_606503, EPI_ISL_606512, EPI_ISL_606513, EPI_ISL_606523, EPI_ISL_606547, EPI_ISL_606554, EPI_ISL_606585, EPI_ISL_606605, EPI_ISL_606606, EPI_ISL_606614, EPI_ISL_606615, EPI_ISL_606619, EPI_ISL_606620, EPI_ISL_606622, EPI_ISL_606624, EPI_ISL_606625, EPI_ISL_606631, EPI_ISL_606632, EPI_ISL_606634, EPI_ISL_606635, EPI_ISL_606652, EPI_ISL_606661, EPI_ISL_606662, EPI_ISL_606663, EPI_ISL_606672, EPI_ISL_606676, EPI_ISL_606677, EPI_ISL_606680, EPI_ISL_606683, EPI_ISL_606684, EPI_ISL_606685, EPI_ISL_606686, EPI_ISL_606692, EPI_ISL_606698, EPI_ISL_606701, EPI_ISL_606703, EPI_ISL_606704, EPI_ISL_606710, EPI_ISL_606715, EPI_ISL_606717, EPI_ISL_606721, EPI_ISL_606732, EPI_ISL_606733, EPI_ISL_606737, EPI_ISL_606738, EPI_ISL_606739, EPI_ISL_606740, EPI_ISL_606742, EPI_ISL_606745, EPI_ISL_606747, EPI_ISL_606757, EPI_ISL_606759, EPI_ISL_606760, EPI_ISL_606762, EPI_ISL_606763, EPI_ISL_606766, EPI_ISL_606772, EPI_ISL_606773, EPI_ISL_606776, EPI_ISL_606779, EPI_ISL_606780, EPI_ISL_606781, EPI_ISL_606783, EPI_ISL_606785, EPI_ISL_606788, EPI_ISL_606791, EPI_ISL_606799, EPI_ISL_606810, EPI_ISL_606811, EPI_ISL_606816, EPI_ISL_606817, EPI_ISL_606818, EPI_ISL_606821, EPI_ISL_606824, EPI_ISL_606825, EPI_ISL_606827, EPI_ISL_606839, EPI_ISL_606840, EPI_ISL_606841, EPI_ISL_606842, EPI_ISL_606846, EPI_ISL_606847, EPI_ISL_606852, EPI_ISL_606860, EPI_ISL_606864, EPI_ISL_606865, EPI_ISL_606867, EPI_ISL_606871, EPI_ISL_606875, EPI_ISL_606879, EPI_ISL_606889, EPI_ISL_606891, EPI_ISL_606892, EPI_ISL_606893, EPI_ISL_606898, EPI_ISL_606901, EPI_ISL_606903, EPI_ISL_606909, EPI_ISL_606910, EPI_ISL_606911, EPI_ISL_606916, EPI_ISL_607298, EPI_ISL_607302, EPI_ISL_607304, EPI_ISL_607305, EPI_ISL_607307, EPI_ISL_607313, EPI_ISL_607319, EPI_ISL_607322, EPI_ISL_607323, EPI_ISL_607326, EPI_ISL_607327, EPI_ISL_607329, EPI_ISL_607334, EPI_ISL_607338, EPI_ISL_607340, EPI_ISL_607341, EPI_ISL_607347, EPI_ISL_607349, EPI_ISL_607368, EPI_ISL_607377, EPI_ISL_607387, EPI_ISL_607391, EPI_ISL_607392, EPI_ISL_607408, EPI_ISL_607410, EPI_ISL_607414, EPI_ISL_607416, EPI_ISL_607418, EPI_ISL_607420, EPI_ISL_607440, EPI_ISL_607455, EPI_ISL_607465, EPI_ISL_607475, EPI_ISL_607505, EPI_ISL_607509, EPI_ISL_607523, EPI_ISL_607544, EPI_ISL_607545, EPI_ISL_607566, EPI_ISL_607573, EPI_ISL_607575, EPI_ISL_607581, EPI_ISL_607585, EPI_ISL_607586, EPI_ISL_607601, EPI_ISL_607608, EPI_ISL_607614, EPI_ISL_608697, EPI_ISL_608699, EPI_ISL_608700, EPI_ISL_608701, EPI_ISL_608702, EPI_ISL_608703, EPI_ISL_608705, EPI_ISL_608706, EPI_ISL_608707, EPI_ISL_608708, EPI_ISL_608709, EPI_ISL_608710, EPI_ISL_608711, EPI_ISL_608712, EPI_ISL_608713, EPI_ISL_608714, EPI_ISL_608715, EPI_ISL_608716, EPI_ISL_608717, EPI_ISL_608718, EPI_ISL_608719, EPI_ISL_608720, EPI_ISL_608721, EPI_ISL_608723, EPI_ISL_608724, EPI_ISL_608725, EPI_ISL_608726, EPI_ISL_608727, EPI_ISL_608728, EPI_ISL_608729, EPI_ISL_608730, EPI_ISL_608731, EPI_ISL_608732, EPI_ISL_608733, EPI_ISL_608734, EPI_ISL_608735, EPI_ISL_608736, EPI_ISL_608737, EPI_ISL_608738, EPI_ISL_608739, EPI_ISL_608740, EPI_ISL_608741, EPI_ISL_608742, EPI_ISL_608744, EPI_ISL_608746, EPI_ISL_608747, EPI_ISL_608749, EPI_ISL_608750, EPI_ISL_608751, EPI_ISL_608753, EPI_ISL_608754, EPI_ISL_608755, EPI_ISL_608756, EPI_ISL_608757, EPI_ISL_608758, EPI_ISL_608759, EPI_ISL_608760, EPI_ISL_608761, EPI_ISL_608762, EPI_ISL_608763, EPI_ISL_608764, EPI_ISL_608765, EPI_ISL_608767, EPI_ISL_608768, EPI_ISL_608769, EPI_ISL_608770, EPI_ISL_608771, EPI_ISL_608772, EPI_ISL_608773, EPI_ISL_608774, EPI_ISL_608775, EPI_ISL_608776, EPI_ISL_608777, EPI_ISL_608778, EPI_ISL_608779, EPI_ISL_608780, EPI_ISL_608781, EPI_ISL_608782, EPI_ISL_608783, EPI_ISL_608784, EPI_ISL_608785, EPI_ISL_608786, EPI_ISL_608787, EPI_ISL_608788, EPI_ISL_608789, EPI_ISL_608790, EPI_ISL_608791, EPI_ISL_608792, EPI_ISL_608793, EPI_ISL_608794, EPI_ISL_608795, EPI_ISL_608796, EPI_ISL_608797, EPI_ISL_608798, EPI_ISL_608800, EPI_ISL_608801, EPI_ISL_608802, EPI_ISL_608803, EPI_ISL_608804, EPI_ISL_608805, EPI_ISL_608806, EPI_ISL_608807, EPI_ISL_608808, EPI_ISL_608809, EPI_ISL_608810, EPI_ISL_608811, EPI_ISL_608812, EPI_ISL_608813, EPI_ISL_608814, EPI_ISL_608815, EPI_ISL_608816, EPI_ISL_608817, EPI_ISL_608818, EPI_ISL_608819, EPI_ISL_608820, EPI_ISL_608821, EPI_ISL_608822, EPI_ISL_608823, EPI_ISL_608824, EPI_ISL_608825, EPI_ISL_608826, EPI_ISL_608827, EPI_ISL_608828, EPI_ISL_608830, EPI_ISL_608831, EPI_ISL_608832, EPI_ISL_608833, EPI_ISL_608835, EPI_ISL_608836, EPI_ISL_608837, EPI_ISL_608838, EPI_ISL_608839, EPI_ISL_608840, EPI_ISL_608841, EPI_ISL_608842, EPI_ISL_608843, EPI_ISL_608844, EPI_ISL_608846, EPI_ISL_608848, EPI_ISL_608850, EPI_ISL_608854, EPI_ISL_608857, EPI_ISL_608859, EPI_ISL_608861, EPI_ISL_608871, EPI_ISL_608882, EPI_ISL_608893, EPI_ISL_608894, EPI_ISL_608905, EPI_ISL_608936, EPI_ISL_608940, EPI_ISL_608954, EPI_ISL_608975, EPI_ISL_608977, EPI_ISL_608978, EPI_ISL_608986, EPI_ISL_609001, EPI_ISL_609006, EPI_ISL_609008, EPI_ISL_609015, EPI_ISL_609017, EPI_ISL_609021, EPI_ISL_609024, EPI_ISL_609027, EPI_ISL_609029, EPI_ISL_609030, EPI_ISL_609036, EPI_ISL_609038, EPI_ISL_609042, EPI_ISL_609046, EPI_ISL_609049, EPI_ISL_609058, EPI_ISL_609059, EPI_ISL_609062, EPI_ISL_609063, EPI_ISL_609066, EPI_ISL_609071, EPI_ISL_609076, EPI_ISL_609084, EPI_ISL_609108, EPI_ISL_609110, EPI_ISL_609115, EPI_ISL_609116, EPI_ISL_609117, EPI_ISL_609126, EPI_ISL_609128, EPI_ISL_609136, EPI_ISL_609137, EPI_ISL_609138, EPI_ISL_609140, EPI_ISL_609149, EPI_ISL_609151, EPI_ISL_609155, EPI_ISL_609221, EPI_ISL_609223, EPI_ISL_609224, EPI_ISL_609226, EPI_ISL_609227, EPI_ISL_609231, EPI_ISL_609236, EPI_ISL_609240, EPI_ISL_609242, EPI_ISL_609243, EPI_ISL_609244, EPI_ISL_609245, EPI_ISL_609253, EPI_ISL_609255, EPI_ISL_609259, EPI_ISL_609265, EPI_ISL_609266, EPI_ISL_609272, EPI_ISL_609273, EPI_ISL_609274, EPI_ISL_609275, EPI_ISL_609278, EPI_ISL_609279 | | | |
| see above | Lighthouse Lab in Milton Keynes | Wellcome Sanger Institute for the COVID-19 Genomics UK (COG-UK) consortium | The Lighthouse Lab in Milton Keynes and Alex Alderton, Roberto Amato, Sonia Goncalves, Ewan Harrison, David K. Jackson, Ian Johnston, Dominic Kwiatkowski, Cordelia Langford, John Sillitoe on behalf of the Wellcome Sanger Institute COVID-19 Surveillance Team |
| EPI_ISL_609971, EPI_ISL_609972, EPI_ISL_609973, EPI_ISL_609974, EPI_ISL_609975, EPI_ISL_609976, EPI_ISL_609977, EPI_ISL_609978, EPI_ISL_609979, EPI_ISL_609980, EPI_ISL_609981, EPI_ISL_609982, EPI_ISL_609983, EPI_ISL_609984, EPI_ISL_609985, EPI_ISL_609986, EPI_ISL_609987, EPI_ISL_609988 | | | |
| see above | Virginia DCLS | Virginia DCLS | Virginia DCLS |
| EPI_ISL_610048, EPI_ISL_610049, EPI_ISL_610050, EPI_ISL_610051, EPI_ISL_610052, EPI_ISL_610053, EPI_ISL_610054, EPI_ISL_610055, EPI_ISL_610056, EPI_ISL_610057, EPI_ISL_610058, EPI_ISL_610059, EPI_ISL_610060, EPI_ISL_610061, EPI_ISL_610062, EPI_ISL_610063, EPI_ISL_610064, EPI_ISL_610065, EPI_ISL_610066, EPI_ISL_610067, EPI_ISL_610068, EPI_ISL_610069, EPI_ISL_610070, EPI_ISL_610071, EPI_ISL_610072, EPI_ISL_610073, EPI_ISL_610074, EPI_ISL_610075, EPI_ISL_610076, EPI_ISL_610077, EPI_ISL_610078, EPI_ISL_610079, EPI_ISL_610080, EPI_ISL_610081, EPI_ISL_610082, EPI_ISL_610083, EPI_ISL_610084, EPI_ISL_610085, EPI_ISL_610086, EPI_ISL_610087, EPI_ISL_610088, EPI_ISL_610089, EPI_ISL_610090, EPI_ISL_610091, EPI_ISL_610092, EPI_ISL_610093, EPI_ISL_610094, EPI_ISL_610095, EPI_ISL_610096, EPI_ISL_610097, EPI_ISL_610098, EPI_ISL_610099, EPI_ISL_610100, EPI_ISL_610101, EPI_ISL_610102, EPI_ISL_610103, EPI_ISL_610104, EPI_ISL_610105, EPI_ISL_610106, EPI_ISL_610107, EPI_ISL_610108, EPI_ISL_610109, EPI_ISL_610110, EPI_ISL_610111, EPI_ISL_610112, EPI_ISL_610113, EPI_ISL_610114, EPI_ISL_610115, EPI_ISL_610116, EPI_ISL_610117 | | | |
| see above | University of Michigan Clinical Microbiology Laboratory | Lauring Lab, University of Michigan, Department of Microbiology and Immunology | Valesano |
| EPI_ISL_610118, EPI_ISL_610119, EPI_ISL_610120, EPI_ISL_610121, EPI_ISL_610122, EPI_ISL_610123, EPI_ISL_610124, EPI_ISL_610125, EPI_ISL_610126, EPI_ISL_610127, EPI_ISL_610128, EPI_ISL_610129, EPI_ISL_610130, EPI_ISL_610131, EPI_ISL_610132, EPI_ISL_610133, EPI_ISL_610134, EPI_ISL_610135, EPI_ISL_610136, EPI_ISL_610137, EPI_ISL_610138, EPI_ISL_610139, EPI_ISL_610140, EPI_ISL_610141, EPI_ISL_610142, EPI_ISL_610143, EPI_ISL_610144, EPI_ISL_610145, EPI_ISL_610146, EPI_ISL_610147, EPI_ISL_610148, EPI_ISL_610149, EPI_ISL_610150, EPI_ISL_610151 | | | |
| see above | Virginia DCLS | Virginia DCLS | Virginia DCLS |
| EPI_ISL_610246, EPI_ISL_610247, EPI_ISL_610248 | University of Michigan Clinical Microbiology Laboratory | Lauring Lab, University of Michigan, Department of Microbiology and Immunology | Valesano |
| EPI_ISL_611520, EPI_ISL_611525, EPI_ISL_611528, EPI_ISL_611536, EPI_ISL_611541, EPI_ISL_611545, EPI_ISL_611553, EPI_ISL_611583, EPI_ISL_611584, EPI_ISL_611587, EPI_ISL_611594, EPI_ISL_611598, EPI_ISL_611602, EPI_ISL_611606, EPI_ISL_611650, EPI_ISL_611651, EPI_ISL_611655, EPI_ISL_611656, EPI_ISL_611661, EPI_ISL_611669, EPI_ISL_611678, EPI_ISL_611679, EPI_ISL_611681, EPI_ISL_611682, EPI_ISL_611690, EPI_ISL_611698, EPI_ISL_611699, EPI_ISL_611733, EPI_ISL_611744, EPI_ISL_611745, EPI_ISL_611774, EPI_ISL_611775, EPI_ISL_611777, EPI_ISL_611778, EPI_ISL_611779, EPI_ISL_611780, EPI_ISL_611781, EPI_ISL_611782, EPI_ISL_611783, EPI_ISL_611787, EPI_ISL_611788, EPI_ISL_611789, EPI_ISL_611793, EPI_ISL_611794, EPI_ISL_611795, EPI_ISL_611797, EPI_ISL_611798, EPI_ISL_611799, EPI_ISL_611800, EPI_ISL_611801, EPI_ISL_611802, EPI_ISL_611803, EPI_ISL_611804, EPI_ISL_611805, EPI_ISL_611806, EPI_ISL_611816, EPI_ISL_611817, EPI_ISL_611829, EPI_ISL_611830, EPI_ISL_611861, EPI_ISL_611862, EPI_ISL_611869, EPI_ISL_611920, EPI_ISL_611936, EPI_ISL_611979, EPI_ISL_611988, EPI_ISL_611997, EPI_ISL_612005, EPI_ISL_612020, EPI_ISL_612022, EPI_ISL_612023, EPI_ISL_612033, EPI_ISL_612050, EPI_ISL_612055, EPI_ISL_612058, EPI_ISL_612062, EPI_ISL_612067, EPI_ISL_612084, EPI_ISL_612102, EPI_ISL_612196, EPI_ISL_613356, EPI_ISL_613357, EPI_ISL_613358, EPI_ISL_613359, EPI_ISL_613360, EPI_ISL_613361, EPI_ISL_613362, EPI_ISL_613363, EPI_ISL_613364, EPI_ISL_613365, EPI_ISL_613366, EPI_ISL_613367, EPI_ISL_613368, EPI_ISL_613369, EPI_ISL_613370, EPI_ISL_613371, EPI_ISL_613372, EPI_ISL_613373, EPI_ISL_613374, EPI_ISL_613375, EPI_ISL_613376, EPI_ISL_613377, EPI_ISL_613378, EPI_ISL_613379, EPI_ISL_613380, EPI_ISL_613381, EPI_ISL_613382, EPI_ISL_613383, EPI_ISL_613384, EPI_ISL_613385, EPI_ISL_613386, EPI_ISL_613387, EPI_ISL_613388, EPI_ISL_613389, EPI_ISL_613390, EPI_ISL_613391, EPI_ISL_613392, EPI_ISL_613393, EPI_ISL_613394, EPI_ISL_613395, EPI_ISL_613396, EPI_ISL_613397, EPI_ISL_613398, EPI_ISL_613399, EPI_ISL_613400, EPI_ISL_613401, EPI_ISL_613402, EPI_ISL_613403, EPI_ISL_613404, EPI_ISL_613405, EPI_ISL_613406, EPI_ISL_613407, EPI_ISL_613408, EPI_ISL_613409 | | | |
| see above | Virology Department, Sheffield Teaching Hospitals NHS Foundation Trust/Department of Infection, Immunity and Cardiovascular Disease, The Medical School, University of Sheffield | COVID-19 Genomics UK (COG-UK) Consortium | Thushan de Silva, Matthew Parker, Nikki Smith, Adri Angyal, Rebecca Brown, Luke Green, Rachel Tucker, Paul Parsons, Danielle Groves, Katie Johnson, Laura Carrilero, Alex Keeley, Dave Partridge, Matthew Wyles, Benjamin Lindsey, Mehmet Yavuz, Mohammad Raza, Cariad Evans |
| EPI_ISL_614012, EPI_ISL_614013, EPI_ISL_614014, EPI_ISL_614015, EPI_ISL_614016, EPI_ISL_614017, EPI_ISL_614018, EPI_ISL_614019, EPI_ISL_614020, EPI_ISL_614021, EPI_ISL_614022, EPI_ISL_614023, EPI_ISL_614024, EPI_ISL_614025, EPI_ISL_614026, EPI_ISL_614027, EPI_ISL_614028, EPI_ISL_614029, EPI_ISL_614030, EPI_ISL_614031, EPI_ISL_614032, EPI_ISL_614033, EPI_ISL_614034, EPI_ISL_614035, EPI_ISL_614036, EPI_ISL_614037, EPI_ISL_614038, EPI_ISL_614039, EPI_ISL_614040, EPI_ISL_614041, EPI_ISL_614042, EPI_ISL_614043, EPI_ISL_614044, EPI_ISL_614045, EPI_ISL_614046, EPI_ISL_614047, EPI_ISL_614048, EPI_ISL_614049, EPI_ISL_614050, EPI_ISL_614051, EPI_ISL_614052, EPI_ISL_614053, EPI_ISL_614054, EPI_ISL_614055, EPI_ISL_614056, EPI_ISL_614057, EPI_ISL_614058, EPI_ISL_614059, EPI_ISL_614060, EPI_ISL_614061, EPI_ISL_614062, EPI_ISL_614063, EPI_ISL_614064, EPI_ISL_614065, EPI_ISL_614066, EPI_ISL_614067, EPI_ISL_614068, EPI_ISL_614069, EPI_ISL_614070, EPI_ISL_614071, EPI_ISL_614072, EPI_ISL_614073, EPI_ISL_614074, EPI_ISL_614075, EPI_ISL_614076, EPI_ISL_614077, EPI_ISL_614078, EPI_ISL_614079, EPI_ISL_614080, EPI_ISL_614081, EPI_ISL_614082, EPI_ISL_614083, EPI_ISL_614084, EPI_ISL_614085, EPI_ISL_614086, EPI_ISL_614087, EPI_ISL_614088, EPI_ISL_614089, EPI_ISL_614090, EPI_ISL_614091, EPI_ISL_614092, EPI_ISL_614093, EPI_ISL_614094, EPI_ISL_614096, EPI_ISL_614097, EPI_ISL_614098, EPI_ISL_614099, EPI_ISL_614100, EPI_ISL_614101, EPI_ISL_614102, EPI_ISL_614103, EPI_ISL_614104, EPI_ISL_614105, EPI_ISL_614106, EPI_ISL_614107, EPI_ISL_614108, EPI_ISL_614109, EPI_ISL_614110, EPI_ISL_614111, EPI_ISL_614112, EPI_ISL_614113, EPI_ISL_614114, EPI_ISL_614115, EPI_ISL_614116, EPI_ISL_614117, EPI_ISL_614118, EPI_ISL_614119, EPI_ISL_614120, EPI_ISL_614121, EPI_ISL_614122, EPI_ISL_614123, EPI_ISL_614124, EPI_ISL_614125, EPI_ISL_614126, EPI_ISL_614127, EPI_ISL_614128, EPI_ISL_614129, EPI_ISL_614130, EPI_ISL_614131, EPI_ISL_614132, EPI_ISL_614133, EPI_ISL_614134, EPI_ISL_614135, EPI_ISL_614136, EPI_ISL_614137, EPI_ISL_614138, EPI_ISL_614139, EPI_ISL_614140, EPI_ISL_614141, EPI_ISL_614142, EPI_ISL_614143, EPI_ISL_614144, EPI_ISL_614145, EPI_ISL_614146, EPI_ISL_614147, EPI_ISL_614148, EPI_ISL_614149, EPI_ISL_614150, EPI_ISL_614151, EPI_ISL_614152, EPI_ISL_614153, EPI_ISL_614154 | | | |
| see above | Virginia DCLS | Virginia DCLS | Virginia DCLS |
| EPI_ISL_614157 | NCDC/IGIB | NCDC/IGIB | Vivekanand A, Mahesh S. Dhar, Bharathram Uppili, Akshay Kanakan, Simmi Tiwari, RadhaKrishnan VS, Robin Marwal, Azka Khan, Ajit Shewale, Pooja Sharma, Tushar Nale, Rajesh Pandey, Sandhya Kabra, Mohammed Faruq, Sujeet Singh, Anurag Agrawal, Partha Rakshit |

| | | | |
|---|---|---|---|
| EPI_ISL_622767 | Laboratory Services Section, Texas Department of State Health Services-SARS-CoV-2 submission group | Laboratory Services Section, Texas Department of State Health Services-SARS-CoV-2 submission group | Tuladhar,R., Oh,B., Zhang,J., Rahman,M., Pokharel,A., Koag,M.,Wang,C., Lee,R., Kubin,G. and Pedrueza,M. |
| EPI_ISL_622768 | Canterbury Health Laboratories | Institute of Environmental Science and Research (ESR) | Xiaoyun Ren, Matt Storey, Nikki Freed, Muhammad Faisal, Jing Wang, Hermes Perez, Anja Werno, Antje van der Linden, Arlo Upton, Chris Mansell, David Hammer, Dragana Drinkovic, Gary McAuliffe, Hana Sofia Andersson, James Ussher, Jill Sherwood, Josh Freeman, Julia Howard, Juliet Elvy, Mary DeAlmeida, Matt Blakiston, Matthew Rogers, Max Bloomfield, Michael Addidle, Michelle Balm, Sally Roberts, Sarah Jefferies, Sharmini Muttaiyah, Susan Morpeth, Susan Taylor, Timothy Blackmore, Vani Sathyendran, Veronica Playle, Virginia Hope, Erasmus Smit, Lauren Jelly, Olin Silander, Joep de Ligt |
| EPI_ISL_622773 | LabPLUS | Institute of Environmental Science and Research (ESR) | Xiaoyun Ren, Matt Storey, Nikki Freed, Muhammad Faisal, Jing Wang, Hermes Perez, Anja Werno, Antje van der Linden, Arlo Upton, Chris Mansell, David Hammer, Dragana Drinkovic, Gary McAuliffe, Hana Sofia Andersson, James Ussher, Jill Sherwood, Josh Freeman, Julia Howard, Juliet Elvy, Mary DeAlmeida, Matt Blakiston, Matthew Rogers, Max Bloomfield, Michael Addidle, Michelle Balm, Sally Roberts, Sarah Jefferies, Sharmini Muttaiyah, Susan Morpeth, Susan Taylor, Timothy Blackmore, Vani Sathyendran, Veronica Playle, Virginia Hope, Erasmus Smit, Lauren Jelly, Olin Silander, Joep de Ligt |
| EPI_ISL_622776 | Middlemore Hospital | Institute of Environmental Science and Research (ESR) | Xiaoyun Ren, Matt Storey, Nikki Freed, Muhammad Faisal, Jing Wang, Hermes Perez, Anja Werno, Antje van der Linden, Arlo Upton, Chris Mansell, David Hammer, Dragana Drinkovic, Gary McAuliffe, Hana Sofia Andersson, James Ussher, Jill Sherwood, Josh Freeman, Julia Howard, Juliet Elvy, Mary DeAlmeida, Matt Blakiston, Matthew Rogers, Max Bloomfield, Michael Addidle, Michelle Balm, Sally Roberts, Sarah Jefferies, Sharmini Muttaiyah, Susan Morpeth, Susan Taylor, Timothy Blackmore, Vani Sathyendran, Veronica Playle, Virginia Hope, Erasmus Smit, Lauren Jelly, Olin Silander, Joep de Ligt |
| EPI_ISL_622777 | LabPLUS | Institute of Environmental Science and Research (ESR) | Xiaoyun Ren, Matt Storey, Nikki Freed, Muhammad Faisal, Jing Wang, Hermes Perez, Anja Werno, Antje van der Linden, Arlo Upton, Chris Mansell, David Hammer, Dragana Drinkovic, Gary McAuliffe, Hana Sofia Andersson, James Ussher, Jill Sherwood, Josh Freeman, Julia Howard, Juliet Elvy, Mary DeAlmeida, Matt Blakiston, Matthew Rogers, Max Bloomfield, Michael Addidle, Michelle Balm, Sally Roberts, Sarah Jefferies, Sharmini Muttaiyah, Susan Morpeth, Susan Taylor, Timothy Blackmore, Vani Sathyendran, Veronica Playle, Virginia Hope, Erasmus Smit, Lauren Jelly, Olin Silander, Joep de Ligt |
| EPI_ISL_622780 | Canterbury Health Laboratories | Institute of Environmental Science and Research (ESR) | Xiaoyun Ren, Matt Storey, Nikki Freed, Muhammad Faisal, Jing Wang, Hermes Perez, Anja Werno, Antje van der Linden, Arlo Upton, Chris Mansell, David Hammer, Dragana Drinkovic, Gary McAuliffe, Hana Sofia Andersson, James Ussher, Jill Sherwood, Josh Freeman, Julia Howard, Juliet Elvy, Mary DeAlmeida, Matt Blakiston, Matthew Rogers, Max Bloomfield, Michael Addidle, Michelle Balm, Sally Roberts, Sarah Jefferies, Sharmini Muttaiyah, Susan Morpeth, Susan Taylor, Timothy Blackmore, Vani Sathyendran, Veronica Playle, Virginia Hope, Erasmus Smit, Lauren Jelly, Olin Silander, Joep de Ligt |
| EPI_ISL_622783 | Middlemore Hospital | Institute of Environmental Science and Research (ESR) | Xiaoyun Ren, Matt Storey, Nikki Freed, Muhammad Faisal, Jing Wang, Hermes Perez, Anja Werno, Antje van der Linden, Arlo Upton, Chris Mansell, David Hammer, Dragana Drinkovic, Gary McAuliffe, Hana Sofia Andersson, James Ussher, Jill Sherwood, Josh Freeman, Julia Howard, Juliet Elvy, Mary DeAlmeida, Matt Blakiston, Matthew Rogers, Max Bloomfield, Michael Addidle, Michelle Balm, Sally Roberts, Sarah Jefferies, Sharmini Muttaiyah, Susan Morpeth, Susan Taylor, Timothy Blackmore, Vani Sathyendran, Veronica Playle, Virginia Hope, Erasmus Smit, Lauren Jelly, Olin Silander, Joep de Ligt |
| EPI_ISL_622788, EPI_ISL_622790, EPI_ISL_622800 | Canterbury Health Laboratories | Institute of Environmental Science and Research (ESR) | Xiaoyun Ren, Matt Storey, Nikki Freed, Muhammad Faisal, Jing Wang, Hermes Perez, Anja Werno, Antje van der Linden, Arlo Upton, Chris Mansell, David Hammer, Dragana Drinkovic, Gary McAuliffe, Hana Sofia Andersson, James Ussher, Jill Sherwood, Josh Freeman, Julia Howard, Juliet Elvy, Mary DeAlmeida, Matt Blakiston, Matthew Rogers, Max Bloomfield, Michael Addidle, Michelle Balm, Sally Roberts, Sarah Jefferies, Sharmini Muttaiyah, Susan Morpeth, Susan Taylor, Timothy Blackmore, Vani Sathyendran, Veronica Playle, Virginia Hope, Erasmus Smit, Lauren Jelly, Olin Silander, Joep de Ligt |
| EPI_ISL_622819 | Middlemore Hospital | Institute of Environmental Science and Research (ESR) | Xiaoyun Ren, Matt Storey, Nikki Freed, Muhammad Faisal, Jing Wang, Hermes Perez, Anja Werno, Antje van der Linden, Arlo Upton, Chris Mansell, David Hammer, Dragana Drinkovic, Gary McAuliffe, Hana Sofia Andersson, James Ussher, Jill Sherwood, Josh Freeman, Julia Howard, Juliet Elvy, Mary DeAlmeida, Matt Blakiston, Matthew Rogers, Max Bloomfield, Michael Addidle, Michelle Balm, Sally Roberts, Sarah Jefferies, Sharmini Muttaiyah, Susan Morpeth, Susan Taylor, Timothy Blackmore, Vani Sathyendran, Veronica Playle, Virginia Hope, Erasmus Smit, Lauren Jelly, Olin Silander, Joep de Ligt |
| EPI_ISL_623211, EPI_ISL_623212, EPI_ISL_623213, EPI_ISL_623215, EPI_ISL_623216, EPI_ISL_623217, EPI_ISL_623219, EPI_ISL_623221, EPI_ISL_623223, EPI_ISL_623224, EPI_ISL_623225, EPI_ISL_623226, EPI_ISL_623227, EPI_ISL_623228, EPI_ISL_623229, EPI_ISL_623230, EPI_ISL_623231, EPI_ISL_623232, EPI_ISL_623233, EPI_ISL_623235, EPI_ISL_623236, EPI_ISL_623237, EPI_ISL_623238, EPI_ISL_623239, EPI_ISL_623240, EPI_ISL_623241, EPI_ISL_623242, EPI_ISL_623243, EPI_ISL_623244, EPI_ISL_623246, EPI_ISL_623249, EPI_ISL_623250, EPI_ISL_623251, EPI_ISL_623252, EPI_ISL_623253, EPI_ISL_623254, EPI_ISL_623255, EPI_ISL_623257, EPI_ISL_623258, EPI_ISL_623260, EPI_ISL_623261, EPI_ISL_623263, EPI_ISL_623264, EPI_ISL_623265, EPI_ISL_623266, EPI_ISL_623267, EPI_ISL_623269, EPI_ISL_623270, EPI_ISL_623271, EPI_ISL_623272, EPI_ISL_623274, EPI_ISL_623275, EPI_ISL_623277, EPI_ISL_623278, EPI_ISL_623279, EPI_ISL_623280, EPI_ISL_623282, EPI_ISL_623283, EPI_ISL_623284, EPI_ISL_623285, EPI_ISL_623287, EPI_ISL_623289, EPI_ISL_623290, EPI_ISL_623291, EPI_ISL_623292, EPI_ISL_623293, EPI_ISL_623294, EPI_ISL_623295, EPI_ISL_623296, EPI_ISL_623297, EPI_ISL_623299, EPI_ISL_623300, EPI_ISL_623302, EPI_ISL_623303, EPI_ISL_623304, EPI_ISL_623305, EPI_ISL_623306, EPI_ISL_623307, EPI_ISL_623309, EPI_ISL_623310, EPI_ISL_623314, EPI_ISL_623315, EPI_ISL_623316, EPI_ISL_623317, EPI_ISL_623318, EPI_ISL_623319, EPI_ISL_623320, EPI_ISL_623322, EPI_ISL_623324, EPI_ISL_623325, EPI_ISL_623326, EPI_ISL_623327, EPI_ISL_623328, EPI_ISL_623329, EPI_ISL_623331, EPI_ISL_623332, EPI_ISL_623335, EPI_ISL_623336, EPI_ISL_623338, EPI_ISL_623339, EPI_ISL_623340, EPI_ISL_623341, EPI_ISL_623342, EPI_ISL_623348, EPI_ISL_623349, EPI_ISL_623350, EPI_ISL_623351, EPI_ISL_623352, EPI_ISL_623353, EPI_ISL_623354, EPI_ISL_623355, EPI_ISL_623356, EPI_ISL_623357, EPI_ISL_623358, EPI_ISL_623361, EPI_ISL_623362, EPI_ISL_623365, EPI_ISL_623367, EPI_ISL_623368, EPI_ISL_623369, EPI_ISL_623371, EPI_ISL_623372, EPI_ISL_623373, EPI_ISL_623374, EPI_ISL_623375, EPI_ISL_623376, EPI_ISL_623377, EPI_ISL_623378, EPI_ISL_623379, EPI_ISL_623380, EPI_ISL_623381, EPI_ISL_623382, EPI_ISL_623383, EPI_ISL_623384, EPI_ISL_623387, EPI_ISL_623388, EPI_ISL_623389, EPI_ISL_623391, EPI_ISL_623394, EPI_ISL_623395, EPI_ISL_623396, EPI_ISL_623399, EPI_ISL_623400, EPI_ISL_623401, EPI_ISL_623402, EPI_ISL_623403, EPI_ISL_623405, EPI_ISL_623406, EPI_ISL_623407, EPI_ISL_623408, EPI_ISL_623410, EPI_ISL_623412, EPI_ISL_623413, EPI_ISL_623414, EPI_ISL_623415, EPI_ISL_623416, EPI_ISL_623417, EPI_ISL_623418, EPI_ISL_623419, EPI_ISL_623420, EPI_ISL_623421, EPI_ISL_623422, EPI_ISL_623423, EPI_ISL_623429, EPI_ISL_623430, EPI_ISL_623431, EPI_ISL_623432, EPI_ISL_623434, EPI_ISL_623435, EPI_ISL_623436, EPI_ISL_623438, EPI_ISL_623439, EPI_ISL_623441, EPI_ISL_623442, EPI_ISL_623443, EPI_ISL_623446, EPI_ISL_623447, EPI_ISL_623450, EPI_ISL_623452, EPI_ISL_623453, EPI_ISL_623454, EPI_ISL_623455, EPI_ISL_623457, EPI_ISL_623458, EPI_ISL_623459, EPI_ISL_623460, EPI_ISL_623461, EPI_ISL_623462, EPI_ISL_623463, EPI_ISL_623464, EPI_ISL_623465, EPI_ISL_623466, EPI_ISL_623467, EPI_ISL_623468, EPI_ISL_623469, EPI_ISL_623470, EPI_ISL_623471, EPI_ISL_623472, EPI_ISL_623473, EPI_ISL_623476, EPI_ISL_623478, EPI_ISL_623479, EPI_ISL_623480, EPI_ISL_623481, EPI_ISL_623483, EPI_ISL_623487, EPI_ISL_623491, EPI_ISL_623492, EPI_ISL_623493, EPI_ISL_623496, EPI_ISL_623497, EPI_ISL_623498, EPI_ISL_623499, EPI_ISL_623500, EPI_ISL_623501, EPI_ISL_623502, EPI_ISL_623503, EPI_ISL_623504, EPI_ISL_623505, EPI_ISL_623506, EPI_ISL_623507, EPI_ISL_623508, EPI_ISL_623509, EPI_ISL_623510, EPI_ISL_623511, EPI_ISL_623512, EPI_ISL_623513, EPI_ISL_623514, EPI_ISL_623516, EPI_ISL_623517, EPI_ISL_623518, EPI_ISL_623519, EPI_ISL_623521, EPI_ISL_623523, EPI_ISL_623524, EPI_ISL_623525, EPI_ISL_623526, EPI_ISL_623529, EPI_ISL_623530, EPI_ISL_623533, EPI_ISL_623534, EPI_ISL_623535, EPI_ISL_623536, EPI_ISL_623538, EPI_ISL_623540, EPI_ISL_623541, EPI_ISL_623542, EPI_ISL_623543, EPI_ISL_623546, EPI_ISL_623549, EPI_ISL_623552, EPI_ISL_623553, EPI_ISL_623554, EPI_ISL_623561, EPI_ISL_623563, EPI_ISL_623564, EPI_ISL_623565, EPI_ISL_623566, EPI_ISL_623571, EPI_ISL_623573, EPI_ISL_623577, EPI_ISL_623579, EPI_ISL_623580, EPI_ISL_623582, EPI_ISL_623583, EPI_ISL_623586, EPI_ISL_623588, EPI_ISL_623589, EPI_ISL_623590, EPI_ISL_623591, EPI_ISL_623594, EPI_ISL_623596, EPI_ISL_623597, EPI_ISL_623598, EPI_ISL_623599, EPI_ISL_623601, EPI_ISL_623603, EPI_ISL_623604, EPI_ISL_623606, EPI_ISL_623607, EPI_ISL_623610, EPI_ISL_623613, EPI_ISL_623614, EPI_ISL_623615, EPI_ISL_623618, EPI_ISL_623620, EPI_ISL_623621, EPI_ISL_623622, EPI_ISL_623623, EPI_ISL_623624, EPI_ISL_623625, EPI_ISL_623627, EPI_ISL_623628, EPI_ISL_623629, EPI_ISL_623631, EPI_ISL_623632, EPI_ISL_623634, EPI_ISL_623635, EPI_ISL_623636, EPI_ISL_623637, EPI_ISL_623638, EPI_ISL_623642, EPI_ISL_623646, EPI_ISL_623648, EPI_ISL_623649, EPI_ISL_623656, EPI_ISL_623659, EPI_ISL_623660, EPI_ISL_623668, EPI_ISL_623669, EPI_ISL_623670, EPI_ISL_623671, EPI_ISL_623672, EPI_ISL_623673, EPI_ISL_623676, EPI_ISL_623677, EPI_ISL_623678, EPI_ISL_623681, EPI_ISL_623682, EPI_ISL_623684, EPI_ISL_623685, EPI_ISL_623686, EPI_ISL_623687, EPI_ISL_623689, EPI_ISL_623691, EPI_ISL_623692, EPI_ISL_623693, EPI_ISL_623695, EPI_ISL_623696, EPI_ISL_623697, EPI_ISL_623698, EPI_ISL_623699, EPI_ISL_623701, EPI_ISL_623704, EPI_ISL_623705, EPI_ISL_623706, EPI_ISL_623707, EPI_ISL_623708, EPI_ISL_623709, EPI_ISL_623711, EPI_ISL_623713, EPI_ISL_623714, EPI_ISL_623715, EPI_ISL_623718, EPI_ISL_623719, EPI_ISL_623720, EPI_ISL_623722, EPI_ISL_623723, EPI_ISL_623727, EPI_ISL_623728, EPI_ISL_623729, EPI_ISL_623730, EPI_ISL_623732, EPI_ISL_623734, EPI_ISL_623737, EPI_ISL_623738, EPI_ISL_623739, EPI_ISL_623743, EPI_ISL_623744, EPI_ISL_623745, EPI_ISL_623747, EPI_ISL_623748, EPI_ISL_623752, EPI_ISL_623753, EPI_ISL_623755, EPI_ISL_623756, EPI_ISL_623757, EPI_ISL_623760, EPI_ISL_623761, EPI_ISL_623765, EPI_ISL_623766, EPI_ISL_623767, EPI_ISL_623768, EPI_ISL_623769, EPI_ISL_623770, EPI_ISL_623771, EPI_ISL_623773, EPI_ISL_623777, EPI_ISL_623778, EPI_ISL_623779, EPI_ISL_623780, EPI_ISL_623782, EPI_ISL_623783, EPI_ISL_623784, EPI_ISL_623786, EPI_ISL_623787, EPI_ISL_623789, EPI_ISL_623792, EPI_ISL_623795, EPI_ISL_623801, EPI_ISL_623803, EPI_ISL_623804, EPI_ISL_623805, EPI_ISL_623807, EPI_ISL_623813, EPI_ISL_623814, EPI_ISL_623817, EPI_ISL_623818, EPI_ISL_623819, EPI_ISL_623820, EPI_ISL_623821, EPI_ISL_623824, EPI_ISL_623825, EPI_ISL_623826, EPI_ISL_623827, EPI_ISL_623830, EPI_ISL_623835, EPI_ISL_623836, EPI_ISL_623837, EPI_ISL_623839, EPI_ISL_623842, EPI_ISL_623845, EPI_ISL_623850, EPI_ISL_623851, EPI_ISL_623852, EPI_ISL_623853, EPI_ISL_625405, EPI_ISL_625406, EPI_ISL_625407, EPI_ISL_625408, EPI_ISL_625409, EPI_ISL_625410, EPI_ISL_625411, EPI_ISL_625412, EPI_ISL_625413, EPI_ISL_625414, EPI_ISL_625415, EPI_ISL_625416, EPI_ISL_625417, EPI_ISL_625418, EPI_ISL_625419, EPI_ISL_625420, EPI_ISL_625421, EPI_ISL_625424, EPI_ISL_625426, EPI_ISL_625428, EPI_ISL_625430 | | | |
| see above | Lighthouse Lab in Milton Keynes | Wellcome Sanger Institute for the COVID-19 Genomics UK (COG-UK) consortium | The Lighthouse Lab in Milton Keynes and Alex Alderton, Roberto Amato, Sonia Goncalves, Ewan Harrison, David K. Jackson, Ian Johnston, Dominic Kwiatkowski, Cordelia Langford, John Sillitoe on behalf of the Wellcome Sanger Institute COVID-19 Surveillance Team (http://www.sanger.ac.uk/covid-team) |
| EPI_ISL_626628, EPI_ISL_626629, EPI_ISL_626630, EPI_ISL_626631, EPI_ISL_626632, EPI_ISL_626633, EPI_ISL_626634, EPI_ISL_626635, EPI_ISL_626636, EPI_ISL_626637, EPI_ISL_626638, EPI_ISL_626639, EPI_ISL_626640, EPI_ISL_626641, EPI_ISL_626642, EPI_ISL_626643, EPI_ISL_626644, EPI_ISL_626645, EPI_ISL_626646 | | | |
| see above | National Public Health Laboratory, National Centre for Infectious Diseases | National Public Health Laboratory, National Centre for Infectious Diseases | Tze Minn Mak, Sophie Octavia, Zhenyang Zhou, Lin Cui, Raymond Tzer Pin Lin |
| EPI_ISL_626664, EPI_ISL_626679, EPI_ISL_626680, EPI_ISL_626710, EPI_ISL_626734, EPI_ISL_626735, EPI_ISL_626739, EPI_ISL_626743, EPI_ISL_626747, EPI_ISL_626752, EPI_ISL_626754, EPI_ISL_626757, EPI_ISL_626771, EPI_ISL_626781, EPI_ISL_626786, EPI_ISL_626791, EPI_ISL_626809, EPI_ISL_626817, EPI_ISL_626821, EPI_ISL_626823, EPI_ISL_626824, EPI_ISL_626827, EPI_ISL_626830, EPI_ISL_626852, EPI_ISL_626896, EPI_ISL_626898, EPI_ISL_626899, EPI_ISL_626900, EPI_ISL_626901, EPI_ISL_626902, EPI_ISL_626903, EPI_ISL_626906, EPI_ISL_626907, EPI_ISL_626908, EPI_ISL_626909, EPI_ISL_626911, EPI_ISL_626912, EPI_ISL_626913, EPI_ISL_626915, EPI_ISL_626916, EPI_ISL_626918, EPI_ISL_626922, EPI_ISL_626923, EPI_ISL_626924, EPI_ISL_626929, EPI_ISL_626932, EPI_ISL_626938, EPI_ISL_626947, EPI_ISL_626948, EPI_ISL_626949, EPI_ISL_626950, EPI_ISL_626951, EPI_ISL_627019, EPI_ISL_627047, EPI_ISL_627103, EPI_ISL_627130, EPI_ISL_627164, EPI_ISL_627184, EPI_ISL_627191, EPI_ISL_627252, EPI_ISL_627253, EPI_ISL_627254, EPI_ISL_627264, EPI_ISL_627280, EPI_ISL_627298, EPI_ISL_627313, EPI_ISL_627492, EPI_ISL_627493, EPI_ISL_627494, EPI_ISL_627495, EPI_ISL_627496, EPI_ISL_627497, EPI_ISL_627498, EPI_ISL_627499, EPI_ISL_627500, EPI_ISL_627501, EPI_ISL_627502, EPI_ISL_627503, EPI_ISL_627504, EPI_ISL_627505, EPI_ISL_627506, EPI_ISL_627507, EPI_ISL_627508, EPI_ISL_627509, EPI_ISL_627510, EPI_ISL_627511, EPI_ISL_627512, EPI_ISL_627513, EPI_ISL_627514, EPI_ISL_627515, EPI_ISL_627516, EPI_ISL_627517, EPI_ISL_628369, EPI_ISL_628370, EPI_ISL_628371, EPI_ISL_628372, EPI_ISL_628373, EPI_ISL_628374, EPI_ISL_628375, EPI_ISL_628376, EPI_ISL_628377, EPI_ISL_628378, EPI_ISL_628379, EPI_ISL_628380, EPI_ISL_628381, EPI_ISL_628382, EPI_ISL_628383, EPI_ISL_628384, EPI_ISL_628385, EPI_ISL_628386, EPI_ISL_628387, EPI_ISL_628388, EPI_ISL_628389, EPI_ISL_628390, EPI_ISL_628391, EPI_ISL_628392, EPI_ISL_628393, EPI_ISL_628394, EPI_ISL_628395, EPI_ISL_628396, EPI_ISL_628397, EPI_ISL_628398, EPI_ISL_628399, EPI_ISL_628400, EPI_ISL_628401, EPI_ISL_628402, EPI_ISL_628403, EPI_ISL_628404, EPI_ISL_628405, EPI_ISL_628406, EPI_ISL_628407, EPI_ISL_628408, EPI_ISL_628409, EPI_ISL_628410, EPI_ISL_628411, EPI_ISL_628412, EPI_ISL_628413, EPI_ISL_628421 | | | |
| see above | Virology Department, Sheffield Teaching Hospitals NHS Foundation Trust/Department of Infection, Immunity and Cardiovascular Disease, The Medical School, University of Sheffield | COVID-19 Genomics UK (COG-UK) Consortium | Thushan de Silva, Matthew Parker, Nikki Smith, Adri Angyal, Rebecca Brown, Luke Green, Rachel Tucker, Paul Parsons, Danielle Groves, Katie Johnson, Laura Carrilero, Alex Keeley, Dave Partridge, Matthew Wyles, Benjamin Lindsey, Mehmet Yavuz, Mohammad Raza, Cariad Evans |
| EPI_ISL_628804, EPI_ISL_628805, EPI_ISL_628806, EPI_ISL_628807, EPI_ISL_628808, EPI_ISL_628809, EPI_ISL_628810, EPI_ISL_628811, EPI_ISL_628812, EPI_ISL_628813, EPI_ISL_628814, EPI_ISL_628815, EPI_ISL_628816, EPI_ISL_628817, EPI_ISL_628818, EPI_ISL_628819, EPI_ISL_628820, EPI_ISL_628821, EPI_ISL_628822, EPI_ISL_628823, EPI_ISL_628824, EPI_ISL_628825, EPI_ISL_628826, EPI_ISL_628827, EPI_ISL_628828, EPI_ISL_628829, EPI_ISL_628830, EPI_ISL_628831, EPI_ISL_628832, EPI_ISL_628833, EPI_ISL_628834, EPI_ISL_628835, EPI_ISL_628836, EPI_ISL_628837, EPI_ISL_628838, EPI_ISL_628839, | | | |

| | | | |
|---|---|---|---|
| EPI_ISL_628840, EPI_ISL_628841, EPI_ISL_628842, EPI_ISL_628843, EPI_ISL_628844, EPI_ISL_628845, EPI_ISL_628846, EPI_ISL_628847, EPI_ISL_628848, EPI_ISL_628849, EPI_ISL_628850, EPI_ISL_628851, EPI_ISL_628852, EPI_ISL_628853, EPI_ISL_628854, EPI_ISL_628855, EPI_ISL_628856, EPI_ISL_628857, EPI_ISL_628858, EPI_ISL_628859, EPI_ISL_628860, EPI_ISL_628861, EPI_ISL_628862, EPI_ISL_628863, EPI_ISL_628864, EPI_ISL_628865, EPI_ISL_628866, EPI_ISL_628867, EPI_ISL_628868, EPI_ISL_628869, EPI_ISL_628870, EPI_ISL_628871, EPI_ISL_628872, EPI_ISL_628873, EPI_ISL_628874, EPI_ISL_628875, EPI_ISL_628876, EPI_ISL_628877, EPI_ISL_628878, EPI_ISL_628879, EPI_ISL_628880, EPI_ISL_628881, EPI_ISL_628882, EPI_ISL_628883, EPI_ISL_628884, EPI_ISL_628885, EPI_ISL_628886 | | | |
| see above | University of Michigan Clinical Microbiology Laboratory | Lauring Lab, University of Michigan, Department of Microbiology and Immunology | Valesano |
| EPI_ISL_629111, EPI_ISL_629113, EPI_ISL_629122, EPI_ISL_629123, EPI_ISL_629125, EPI_ISL_629128, EPI_ISL_629129, EPI_ISL_629135, EPI_ISL_629136, EPI_ISL_629140, EPI_ISL_629143, EPI_ISL_629144, EPI_ISL_629145, EPI_ISL_629147, EPI_ISL_629151, EPI_ISL_629153, EPI_ISL_629160, EPI_ISL_629162, EPI_ISL_629165, EPI_ISL_629166, EPI_ISL_629168, EPI_ISL_629170, EPI_ISL_629173, EPI_ISL_629177, EPI_ISL_629178, EPI_ISL_629179, EPI_ISL_629180, EPI_ISL_629182, EPI_ISL_629184, EPI_ISL_629189, EPI_ISL_629194, EPI_ISL_629197, EPI_ISL_629198, EPI_ISL_629202, EPI_ISL_629206, EPI_ISL_629209, EPI_ISL_629213, EPI_ISL_629217, EPI_ISL_629232, EPI_ISL_629233, EPI_ISL_629243, EPI_ISL_629246, EPI_ISL_629247, EPI_ISL_629249, EPI_ISL_629251, EPI_ISL_629254, EPI_ISL_629261, EPI_ISL_629265, EPI_ISL_629266, EPI_ISL_629270, EPI_ISL_629271, EPI_ISL_629274, EPI_ISL_629276, EPI_ISL_629277, EPI_ISL_629278, EPI_ISL_629279, EPI_ISL_629281, EPI_ISL_629282, EPI_ISL_629286, EPI_ISL_629287, EPI_ISL_629288, EPI_ISL_629289, EPI_ISL_629291, EPI_ISL_629292, EPI_ISL_629293, EPI_ISL_629295, EPI_ISL_629304, EPI_ISL_629305, EPI_ISL_629308, EPI_ISL_629309, EPI_ISL_629311, EPI_ISL_629325, EPI_ISL_629328, EPI_ISL_629330, EPI_ISL_629335, EPI_ISL_629340, EPI_ISL_629342, EPI_ISL_629343, EPI_ISL_629345, EPI_ISL_629348, EPI_ISL_629354, EPI_ISL_629357, EPI_ISL_629359, EPI_ISL_629360, EPI_ISL_629361, EPI_ISL_629366, EPI_ISL_629369, EPI_ISL_629370, EPI_ISL_629379, EPI_ISL_629380, EPI_ISL_629383, EPI_ISL_629389, EPI_ISL_629390, EPI_ISL_629391, EPI_ISL_629392, EPI_ISL_629398, EPI_ISL_629411, EPI_ISL_629413, EPI_ISL_629414, EPI_ISL_629431, EPI_ISL_629432, EPI_ISL_629433, EPI_ISL_629436, EPI_ISL_629439, EPI_ISL_629441, EPI_ISL_629442, EPI_ISL_629443, EPI_ISL_629445, EPI_ISL_629446, EPI_ISL_629448, EPI_ISL_629449, EPI_ISL_629452, EPI_ISL_629453, EPI_ISL_629454, EPI_ISL_629455, EPI_ISL_629456, EPI_ISL_629459, EPI_ISL_629460, EPI_ISL_629461, EPI_ISL_629463, EPI_ISL_629464, EPI_ISL_629465, EPI_ISL_629466, EPI_ISL_629467, EPI_ISL_629468, EPI_ISL_629470, EPI_ISL_629471, EPI_ISL_629474, EPI_ISL_629478, EPI_ISL_629479, EPI_ISL_629480, EPI_ISL_629482, EPI_ISL_629484, EPI_ISL_629485, EPI_ISL_629487, EPI_ISL_629491, EPI_ISL_629492, EPI_ISL_629495, EPI_ISL_629499, EPI_ISL_629501, EPI_ISL_629504, EPI_ISL_629505, EPI_ISL_629506, EPI_ISL_629507, EPI_ISL_629510, EPI_ISL_629514, EPI_ISL_629516, EPI_ISL_629517, EPI_ISL_629519, EPI_ISL_629522, EPI_ISL_629523, EPI_ISL_629524, EPI_ISL_629527, EPI_ISL_629528, EPI_ISL_629531, EPI_ISL_629534, EPI_ISL_629535, EPI_ISL_629536, EPI_ISL_629537, EPI_ISL_629539, EPI_ISL_629540, EPI_ISL_629541, EPI_ISL_629544, EPI_ISL_629545, EPI_ISL_629547, EPI_ISL_629548, EPI_ISL_629549, EPI_ISL_629554, EPI_ISL_629555, EPI_ISL_629558, EPI_ISL_629559, EPI_ISL_629560, EPI_ISL_629562, EPI_ISL_629564, EPI_ISL_629565, EPI_ISL_629566, EPI_ISL_629567, EPI_ISL_629568, EPI_ISL_629569, EPI_ISL_629571, EPI_ISL_629573, EPI_ISL_629575, EPI_ISL_629576, EPI_ISL_629581, EPI_ISL_629583, EPI_ISL_629584, EPI_ISL_629585, EPI_ISL_629587, EPI_ISL_629591, EPI_ISL_629593, EPI_ISL_629595, EPI_ISL_629597, EPI_ISL_629599, EPI_ISL_629601, EPI_ISL_629603, EPI_ISL_629604, EPI_ISL_629605, EPI_ISL_629607, EPI_ISL_629608, EPI_ISL_629611, EPI_ISL_629612, EPI_ISL_629614, EPI_ISL_629615, EPI_ISL_629617, EPI_ISL_629620, EPI_ISL_629625, EPI_ISL_629628, EPI_ISL_629629, EPI_ISL_629630, EPI_ISL_629631, EPI_ISL_629632, EPI_ISL_629634, EPI_ISL_629635, EPI_ISL_629640, EPI_ISL_629641, EPI_ISL_629642, EPI_ISL_629644, EPI_ISL_629645, EPI_ISL_629647, EPI_ISL_629648, EPI_ISL_629650, EPI_ISL_629651, EPI_ISL_629653, EPI_ISL_629659, EPI_ISL_629660, EPI_ISL_629661, EPI_ISL_629663, EPI_ISL_629665, EPI_ISL_629668, EPI_ISL_629669, EPI_ISL_629671, EPI_ISL_629672, EPI_ISL_629673, EPI_ISL_629674, EPI_ISL_629675, EPI_ISL_629676, EPI_ISL_629679, EPI_ISL_629683, EPI_ISL_629685, EPI_ISL_629688, EPI_ISL_629689, EPI_ISL_629690, EPI_ISL_629692, EPI_ISL_629696, EPI_ISL_629701, EPI_ISL_629702, EPI_ISL_629703, EPI_ISL_629704, EPI_ISL_629705, EPI_ISL_629706, EPI_ISL_629708, EPI_ISL_629709, EPI_ISL_629710, EPI_ISL_629713, EPI_ISL_629716, EPI_ISL_629717, EPI_ISL_629721, EPI_ISL_629722, EPI_ISL_629723, EPI_ISL_629724, EPI_ISL_629725, EPI_ISL_629728, EPI_ISL_629729, EPI_ISL_629730, EPI_ISL_629731, EPI_ISL_629733, EPI_ISL_629735, EPI_ISL_629737, EPI_ISL_629738, EPI_ISL_629739, EPI_ISL_629740, EPI_ISL_629743, EPI_ISL_629745, EPI_ISL_629746, EPI_ISL_629747, EPI_ISL_629749, EPI_ISL_629750, EPI_ISL_629752, EPI_ISL_630010, EPI_ISL_630012, EPI_ISL_630014, EPI_ISL_630015, EPI_ISL_630023, EPI_ISL_630032, EPI_ISL_630034, EPI_ISL_630035, EPI_ISL_630051, EPI_ISL_630053, EPI_ISL_630055, EPI_ISL_630058, EPI_ISL_630063, EPI_ISL_630065, EPI_ISL_630066, EPI_ISL_630067, EPI_ISL_630068, EPI_ISL_630070, EPI_ISL_630076, EPI_ISL_630078, EPI_ISL_630080, EPI_ISL_630082, EPI_ISL_630087, EPI_ISL_630089, EPI_ISL_630092, EPI_ISL_630094, EPI_ISL_630095, EPI_ISL_630098, EPI_ISL_630099, EPI_ISL_630103, EPI_ISL_630105, EPI_ISL_630111, EPI_ISL_630114, EPI_ISL_630115, EPI_ISL_630127, EPI_ISL_630128, EPI_ISL_630132, EPI_ISL_630135, EPI_ISL_630137, EPI_ISL_630140, EPI_ISL_630142, EPI_ISL_630143, EPI_ISL_630145, EPI_ISL_630147, EPI_ISL_630151, EPI_ISL_630155, EPI_ISL_630157, EPI_ISL_630162, EPI_ISL_630164, EPI_ISL_630170, EPI_ISL_630171, EPI_ISL_630177, EPI_ISL_630179, EPI_ISL_630184, EPI_ISL_630190, EPI_ISL_630195, EPI_ISL_630200, EPI_ISL_630201, EPI_ISL_630203, EPI_ISL_630211, EPI_ISL_630213, EPI_ISL_630214, EPI_ISL_630218, EPI_ISL_630227, EPI_ISL_630228, EPI_ISL_630233, EPI_ISL_630239, EPI_ISL_630240, EPI_ISL_630244, EPI_ISL_630251, EPI_ISL_630252, EPI_ISL_630255, EPI_ISL_630259, EPI_ISL_630263, EPI_ISL_630266, EPI_ISL_630268, EPI_ISL_630269, EPI_ISL_630274, EPI_ISL_630276, EPI_ISL_630280, EPI_ISL_630284, EPI_ISL_630286, EPI_ISL_630287, EPI_ISL_630294, EPI_ISL_630299, EPI_ISL_630305, EPI_ISL_630306, EPI_ISL_630310, EPI_ISL_630313, EPI_ISL_630315, EPI_ISL_630317, EPI_ISL_630320, EPI_ISL_630326, EPI_ISL_630331, EPI_ISL_630344, EPI_ISL_630973, EPI_ISL_630978, EPI_ISL_630980, EPI_ISL_630981, EPI_ISL_630986, EPI_ISL_630987, EPI_ISL_630990, EPI_ISL_630991, EPI_ISL_630993, EPI_ISL_630995, EPI_ISL_630998, EPI_ISL_631000, EPI_ISL_631002, EPI_ISL_631004, EPI_ISL_631006, EPI_ISL_631007, EPI_ISL_631009, EPI_ISL_631010, EPI_ISL_631013, EPI_ISL_631015, EPI_ISL_631019, EPI_ISL_631020, EPI_ISL_631021, EPI_ISL_631022, EPI_ISL_631024, EPI_ISL_631026, EPI_ISL_631029, EPI_ISL_631030, EPI_ISL_631032, EPI_ISL_631034, EPI_ISL_631036, EPI_ISL_631037, EPI_ISL_631040, EPI_ISL_631042, EPI_ISL_631047, EPI_ISL_631048, EPI_ISL_631049, EPI_ISL_631051, EPI_ISL_631052, EPI_ISL_631058, EPI_ISL_631061, EPI_ISL_631064, EPI_ISL_631065, EPI_ISL_631066, EPI_ISL_631067, EPI_ISL_631068, EPI_ISL_631070, EPI_ISL_631071, EPI_ISL_631073, EPI_ISL_631075, EPI_ISL_631076, EPI_ISL_631077, EPI_ISL_631079, EPI_ISL_631082, EPI_ISL_631083, EPI_ISL_631085, EPI_ISL_631086, EPI_ISL_631087, EPI_ISL_631088, EPI_ISL_631091, EPI_ISL_631092, EPI_ISL_631095, EPI_ISL_631097, EPI_ISL_631098, EPI_ISL_631100, EPI_ISL_631104, EPI_ISL_631106, EPI_ISL_631108, EPI_ISL_631110, EPI_ISL_631112, EPI_ISL_631114, EPI_ISL_631115, EPI_ISL_631119, EPI_ISL_631120, EPI_ISL_631122, EPI_ISL_631124, EPI_ISL_631125, EPI_ISL_631126, EPI_ISL_631130, EPI_ISL_631134, EPI_ISL_631136, EPI_ISL_631138, EPI_ISL_631139, EPI_ISL_631140, EPI_ISL_631141, EPI_ISL_631143, EPI_ISL_631145, EPI_ISL_631146, EPI_ISL_631147, EPI_ISL_631151, EPI_ISL_631152, EPI_ISL_631153, EPI_ISL_631156, EPI_ISL_631157, EPI_ISL_631158, EPI_ISL_631161, EPI_ISL_631162, EPI_ISL_631166, EPI_ISL_631168, EPI_ISL_631169, EPI_ISL_631170, EPI_ISL_631172, EPI_ISL_631176, EPI_ISL_631178, EPI_ISL_631179, EPI_ISL_631182, EPI_ISL_631183, EPI_ISL_631186, EPI_ISL_631187, EPI_ISL_631188, EPI_ISL_631193, EPI_ISL_631195, EPI_ISL_631196, EPI_ISL_631197, EPI_ISL_631198, EPI_ISL_631200, EPI_ISL_631201, EPI_ISL_631203, EPI_ISL_631205, EPI_ISL_631206, EPI_ISL_631209, EPI_ISL_631210, EPI_ISL_631211, EPI_ISL_631213, EPI_ISL_631214, EPI_ISL_631216, EPI_ISL_631218, EPI_ISL_631223, EPI_ISL_631226, EPI_ISL_631227, EPI_ISL_631229, EPI_ISL_631230, EPI_ISL_631232, EPI_ISL_631233, EPI_ISL_631234, EPI_ISL_631236, EPI_ISL_631238, EPI_ISL_631239, EPI_ISL_631240, EPI_ISL_631242, EPI_ISL_631243, EPI_ISL_631244, EPI_ISL_631246, EPI_ISL_631247, EPI_ISL_631248, EPI_ISL_631249, EPI_ISL_631251, EPI_ISL_631252, EPI_ISL_631253, EPI_ISL_631255, EPI_ISL_631256, EPI_ISL_631258, EPI_ISL_631259, EPI_ISL_631260, EPI_ISL_631261, EPI_ISL_631263, EPI_ISL_631264, EPI_ISL_631267, EPI_ISL_631275 | | | |
| see above | Lighthouse Lab in Milton Keynes | Wellcome Sanger Institute for the COVID-19 Genomics UK (COG-UK) consortium | The Lighthouse Lab in Milton Keynes and Alex Alderton, Roberto Amato, Sonia Goncalves, Ewan Harrison, David K. Jackson, Ian Johnston, Dominic Kwiatkowski, Cordelia Langford, John Sillitoe on behalf of the Wellcome Sanger Institute COVID-19 Surveillance Team |
| EPI_ISL_632286 | Laboratory Services Section, Texas Department of State Health Services | Laboratory Services Section, Texas Department of State Health Services | Tuladhar,R., Oh,B., Zhang,J., Rahman,M., Pokharel,A., Pedrueza,M.,Bonser,J., Koag,M., Wang,C., Lee,R. and Kubin,G. |

We gratefully acknowledge the following Authors from the Originating laboratories responsible for obtaining the specimens, as well as the Submitting laboratories where the genome data were generated and shared via GISAID, on which this research is based.

All Submitters of data may be contacted directly via www.gisaid.org

Authors are sorted alphabetically.

| Accession ID | Originating Laboratory | Submitting Laboratory | Authors |
|---|---|---|---|
| EPI_ISL_402125 | National Institute for Communicable Disease Control and Prevention (ICDC) Chinese Center for Disease Control and Prevention (China CDC) | National Institute for Communicable Disease Control and Prevention (ICDC) Chinese Center for Disease Control and Prevention (China CDC) | Zhang,Y.-Z., Wu,F., Chen,Y.-M., Pei,Y.-Y., Xu,L., Wang,W., Zhao,S., Yu,B., Hu,Y., Tao,Z.-W., Song,Z.-G., Tian,J.-H., Zhang,Y.-L., Liu,Y., Zheng,J.-J., Dai,F.-H., Wang,Q.-M., She,J.-L. and Zhu,T.-Y. |
| EPI_ISL_404227 | Zhejiang Provincial Center for Disease Control and Prevention | Department of Microbiology, Zhejiang Provincial Center for Disease Control and Prevention | Yin Chen, Yanjun Zhang, Haiyan Mao, Junhang Pan, Xiuyu Lou, Yiyu Lu, Juying Yan, Hanping Zhu, Jian Gao, Yan Feng, Yi Sun, Hao Yan, Zhen Li, Yisheng Sun, Liming Gong, Qiong Ge, Wen Shi, Xinying Wang, Wenwu Yao, Zhangnv Yang, Fang Xu, Chen Chen, Enfu Chen, Zhen Wang, Zhiping Chen, Jianmin Jiang, Chonggao Hu |
| EPI_ISL_404228 | Zhejiang Provincial Center for Disease Control and Prevention | Department of Microbiology, Zhejiang Provincial Center for Disease Control and Prevention | Yanjun Zhang, Yin Chen, Haiyan Mao, Junhang Pan, Xiuyu Lou, Yiyu Lu, Juying Yan, Hanping Zhu, Jian Gao, Yan Feng, Yi Sun, Hao Yan, Zhen Li, Yisheng Sun, Liming Gong, Qiong Ge, Wen Shi, Xinying Wang, Wenwu Yao, Zhangnv Yang, Fang Xu, Chen Chen, Enfu Chen, Zhen Wang, Zhiping Chen, Jianmin Jiang, Chonggao Hu |
| EPI_ISL_404253 | IL Department of Public Health Chicago Laboratory | Pathogen Discovery, Respiratory Viruses Branch, Division of Viral Diseases, Centers for Dieases Control and Prevention | Ying Tao, Krista Queen, Clinton R. Paden, Jing Zhang, Yan Li, Anna Uehara, Xiaoyan Lu, Brian Lynch, Senthil Kumar K. Sakthivel, Brett L. Whitaker, Shifaq Kamili, Lijuan Wang, Janna' R. Murray, Susan I. Gerber, Stephen Lindstrom, Suxiang Tong |
| EPI_ISL_406223 | Arizona Department of Health Services | Pathogen Discovery, Respiratory Viruses Branch, Division of Viral Diseases, Centers for Disease Control and Prevention | Ying Tao, Clinton R. Paden, Krista Queen, Anna Uehara, Yan Li, Jing Zhang, Xiaoyan Lu, Brian Lynch, Senthil Kumar K. Sakthivel, Brett L. Whitaker, Shifaq Kamili, Lijuan Wang, Janna' R. Murray, Susan I. Gerber, Stephen Lindstrom, Suxiang Tong |
| EPI_ISL_406592 | Shenzhen Third People's Hospital | Shenzhen Key Laboratory of Pathogen and Immunity, National Clinical Research Center for Infectious Disease,Shenzhen Third People's Hospital | Yang Yang, Chenguang Shen, Li Xing, Zhixiang Xu, Haixia Zheng, Yingxia Liu |
| EPI_ISL_406593, EPI_ISL_406594, EPI_ISL_406595 | Shenzhen Key Laboratory of Pathogen and Immunity, National Clinical Research Center for Infectious Disease, Shenzhen Third People's Hospital | Shenzhen Key Laboratory of Pathogen and Immunity, National Clinical Research Center for Infectious Disease, Shenzhen Third People's Hospital | Yang Yang, Chenguang Shen, Li Xing, Zhixiang Xu, Haixia Zheng, Yingxia Liu |
| EPI_ISL_406970 | Hangzhou Center for Disease and Control Microbiology Lab | Hangzhou Center for Disease and Control Microbiology Lab | Yu Hua, Wang Haoqiu, Li Jun, Yu Xinfeng |
| EPI_ISL_408010 | California Department of Health | Pathogen Discovery, Respiratory Viruses Branch, Division of Viral Diseases, Centers for Dieases Control and Prevention | Ying Tao, Krista Queen, Jing Zhang, Yan Li, Anna Uehara, Clinton Paden, Xiaoyan Lu, Brian Lynch, Senthil Kumar K. Sakthivel, Brett L. Whitaker, Shifaq Kamili, Lijuan Wang, Janna' R. Murray, Susan I. Gerber, Stephen Lindstrom, Suxiang Tong |
| EPI_ISL_408478 | Yongchuan District Center for Disease Control and Prevention | Chongqing Municipal Center for Disease Control and Prevention | Ye Sheng, Tang Yun, Ling Hua,Yu zhen,Chen Shuang,Tan ZhangPing, Su Kun, Li Qing, Tang Wenge, Rong Rong |
| EPI_ISL_408479 | Zhongxian Center for Disease Control and Prevention | Chongqing Municipal Center for Disease Control and Prevention | Ye Sheng, Tang Yun, Ling Hua, Zhang Hong, Yu zhen,Chen Shuang,Tan ZhangPing, Su Kun, Li Qin, Tang Wenge, Rong Rong |
| EPI_ISL_410045 | IL Department of Public Health Chicago Laboratory | Pathogen Discovery, Respiratory Viruses Branch, Division of Viral Diseases, Centers for Dieases Control and Prevention | Yan Li, Jing Zhang, Krista Queen, Ying Tao, Anna Uehara, Clinton R. Paden, Xiaoyan Lu, Brian Lynch, Senthil Kumar K. Sakthivel, Brett L. Whitaker, Shifaq Kamili, Lijuan Wang, Janna' R. Murray, Susan I. Gerber, Stephen Lindstrom, Suxiang Tong |
| EPI_ISL_412386 | Beijing Ditan Hospital, Capital Medical University | National Institute for Communicable Disease Control and Prevention, Chinese Center for Disease Control and Prevention | Xinmin Xu, Xin Lu, Pan Xiang, Haijian Zhou, Biao Kan, Yajie Wang, Jingyuan Liu, Yanwen Xiong, Huizhu Wang, Ruihong Li, Fangfang Jin, Jie Gong, Xiaoping Chen, Lili Gao, Haofeng Xiong, Lin Pu, Chuansheng Li, Ming Zhang, Jianbo Tan, Yao Sun, Yufeng Liu, Hebing Guo, Jingjing Hao |
| EPI_ISL_413612, EPI_ISL_413613, EPI_ISL_413614, EPI_ISL_413615, EPI_ISL_413616, EPI_ISL_413617 | unknown | Pathogen Discovery, Respiratory Viruses Branch, Division of Viral Diseases, Centers for Dieases Control and Prevention | Ying Tao, Clinton R. Paden, Krista Queen, Anna Uehara, Jing Zhang, Yan Li, Haibin Wang, Shifaq Kamili, Xiaoyan Lu, Brian Lynch, Senthil Kumar K. Sakthivel, Brett L. Whitaker, Lijuan Wang, Janna' R. Murray, Jasmine Padilla, Justin Lee, Susan I. Gerber, Stephen Lindstrom, Suxiang Tong |
| EPI_ISL_413869 | Guangdong Provincial Institution of Public Health, Guangdong Provinical Center for Disease Control and Prevention | Guangdong Provincial Institution of Public Health | Jing Lu, Louis du Plessis, Liu Zhe, Jiufeng Sun, Sarah François, Huifang Lin, Moritz Kraemer, Jingju Peng, Qianlin Xiong, Runyu Yuan, Lilian Zeng, Pingping Zhou, Chuming Liang, Tao Liu, Wei Li, Juan Su, Huanying Zheng, Kang Min, Song Tie, Bo Peng, Shisong Fang, Wenzhe Su, Kuibiao Li, Ruilin Sun, Ru bai, Xi Tang, Minfeng Liang, Nuno Faria, Josh Quick, Andrew Rambaut, Verity Hill, Wenjun Ma, Nick Loman, Oliver Pybus, Changwen Ke |
| EPI_ISL_414479, EPI_ISL_414480, EPI_ISL_414481 | unknown | Pathogen Discovery, Respiratory Viruses Branch, Division of Viral Diseases, Centers for Dieases Control and Prevention | Ying Tao, Krista Queen, Clinton R. Paden, Anna Uehara, Jing Zhang, Yan Li, Mary S. Keckler, Alison S. Laufer Halpin, Haibin Wang, Jasmine Padilla, Justin Lee, Christopher A. Elkins, Susan I. Gerber, Suxiang Tong |
| EPI_ISL_414510 | Key Laboratory of Medical Molecular Virology (MOE/NHC/CAMS), School of Basic Medicine, Shanghai Medical College, Fudan University | Key Laboratory of Medical Molecular Virology (MOE/NHC/CAMS), School of Basic Medicine, Shanghai Medical College, Fudan University | Zhang,R., Yi,Z., Wang,Y., Teng,Z., Xu,W., Song,W., Cai,X., Sun,Z., Gu,C., Zhou,Y., Chen,H., Ye,R., Han,W., Zhu,Y., Feng,F., Fang,F., Li,C., Zhang,X., Qu,D., Fu,C., Xie,Y. and Yuan,Z. |
| EPI_ISL_414663, EPI_ISL_414686 | State Key Laboratory of Respiratory Disease, National Clinical Research Center for Respiratory Disease, Guangzhou Institute of Respiratory Health, the First Affiliated Hospital of Guangzhou Medical University | The First Affiliated Hospital of Guangzhou Medical University & BGI-Shenzhen | Zhao et al |
| EPI_ISL_414687 | State Key Laboratory of Respiratory Disease, National Clinical Research Center for Respiratory Disease, Guangzhou Institute of Respiratory Health, the First Affiliated Hospital of Guangzhou Medical University | the First Affiliated Hospital of Guangzhou Medical University & BGI-Shenzhen | Zhao et al |
| EPI_ISL_414688, EPI_ISL_414689, EPI_ISL_414690, EPI_ISL_414691, EPI_ISL_414692 | State Key Laboratory of Respiratory Disease, National Clinical Research Center for Respiratory Disease, Guangzhou Institute of Respiratory Health, the First Affiliated Hospital of Guangzhou Medical University | The First Affiliated Hospital of Guangzhou Medical University & BGI-Shenzhen | Zhao et al |
| EPI_ISL_416314, EPI_ISL_416315 | Department of Microbiology, Faculty of Medicine, The Chinese University of Hong Kong, Hong Kong SAR, China | Department of Microbiology, Faculty of Medicine, Chinese University of Hong Kong, Hong Kong SAR, China | Zigui Chen, Paul KS Chan |
| EPI_ISL_417917 | Department of Medical Microbiology, University Malaya Medical Centre | Department of Medical Microbiology | Yoong Min CHONG, Sasheela PONNAMPALAVANAR, Sharifah Faridah SYED OMAR, Adeeba KAMARULZAMAN,Vijayan MUNUSAMY, Chee Kuan WONG, Cindy Shuan Ju TEH, I-Ching SAM, Yoke Fun Chan, University Malaya Medical Centre COVID Team |
| EPI_ISL_417918 | Department of Medical Microbiology, University Malaya Medical Centre | Department of Medical Microbiology, Faculty of Medicine, University of Malaya | Yoong Min CHONG, Sasheela PONNAMPALAVANAR, Sharifah Faridah SYED OMAR, Adeeba KAMARULZAMAN,Vijayan MUNUSAMY, Chee Kuan WONG, Cindy Shuan Ju TEH, I-Ching SAM, Yoke Fun Chan, University Malaya Medical Centre COVID Team |
| EPI_ISL_417919 | Department of Medical Microbiology, University Malaya Medical Centre | Department of Medical Microbiology, Faculty of Medicine, University of Malaya | Yoong Min CHONG, Sasheela PONNAMPALAVANAR, Sharifah Faridah SYED OMAR, Adeeba KAMARULZAMAN,Vijayan MUNUSAMY, Chee Kuan WONG, Fadhil Hadi JAMALUDDIN, Han Ming GAN, Cindy Shuan Ju TEH, I-Ching SAM, Yoke Fun CHAN, University Malaya Medical Centre COVID Team |
| EPI_ISL_417920 | Department of Medical Microbiology, University Malaya | Department of Medical Microbiology, Faculty of Medicine, | Yoong Min CHONG, Sasheela PONNAMPALAVANAR, Sharifah Faridah SYED OMAR, Adeeba KAMARULZAMAN,Vijayan MUNUSAMY, Chee Kuan |

| | | | |
|---|---|---|---|
| | Medical Centre | University of Malaya | WONG, Fadhil Hadi JAMALUDDIN, Cindy Shuan Ju TEH, I-Ching SAM, Yoke Fun Chan, University Malaya Medical Centre COVID Team |
| EPI_ISL_418441, EPI_ISL_418442, EPI_ISL_418502, EPI_ISL_418503, EPI_ISL_418504 | Hangzhou Center for Disease Control and Prevention | Inspection Center of Hangzhou Center for Disease Control and Prevention | Yu hua, Wang haoqiu, Li jun, Yu xinfeng, Pan jingcao |
| EPI_ISL_418506 | Hangzhou Center for Disease Control and Prevention | Inspection Center of Hanghzou Center for Disease Control and Prevention | Yu hua, Wang haoqiu, Li jun, Yu xinfeng, Pan jingcao |
| EPI_ISL_418507, EPI_ISL_418508, EPI_ISL_418509 | Hangzhou Center for Disease Control and Prevention | Inspection Center of Hangzhou Center for Disease Control and Prevention | Yu hua, Wang haoqiu, Li jun, Yu xinfeng, Pan jingcao |
| EPI_ISL_418510 | Hangzhou Center for Disease Control and Prevention | Insepction Center of Hangzhou Center for Disease Control and Prevention | Yu hua, Wang haoqiu, Li jun, Yu xinfeng, Pan jingcao |
| EPI_ISL_418511, EPI_ISL_418512, EPI_ISL_418513, EPI_ISL_418514, EPI_ISL_418515 | Hangzhou Center for Disease Control and Prevention | Inspection Center of Hangzhou Center for Disease Control and Prevention | Yu hua, Wang haoqiu, Li jun, Yu xinfeng, Pan jingcao |
| EPI_ISL_418778, EPI_ISL_418779, EPI_ISL_418780, EPI_ISL_418781, EPI_ISL_418782, EPI_ISL_418783, EPI_ISL_418784, EPI_ISL_418785, EPI_ISL_418786, EPI_ISL_418787, EPI_ISL_418788, EPI_ISL_418789, EPI_ISL_418790, EPI_ISL_418791 | | | |
| see above | WA State Department of Health | Pathogen Discovery, Respiratory Viruses Branch, Division of Viral Diseases, Centers for Disease Control and Prevention | Ying Tao, Jing Zhang, Clinton R. Paden, Krista Queen, Anna Uehara, Yan Li, Haibin Wang, Jesica Jacobs, Denny Russell, Brian Hiatt, Jessica Gant, Suxiang Tong |
| EPI_ISL_419553 | RI State Health Laboratories | Pathogen Discovery, Respiratory Viruses Branch, Division of Viral Diseases, Centers for Disease Control and Prevention | Ying Tao, Jing Zhang, Krista Queen, Anna Uehara, Clinton R. Paden, Yan Li, Haibin Wang, Jasmine Padilla, Justin Lee, Suxiang Tong |
| EPI_ISL_419554 | California Department of Public Health | Pathogen Discovery, Respiratory Viruses Branch, Division of Viral Diseases, Centers for Disease Control and Prevention | Ying Tao, Jing Zhang, Krista Queen, Anna Uehara, Clinton R. Paden, Yan Li, Haibin Wang, Jasmine Padilla, Justin Lee, Suxiang Tong |
| EPI_ISL_419555 | WA State Department of Health | Pathogen Discovery, Respiratory Viruses Branch, Division of Viral Diseases, Centers for Disease Control and Prevention | Ying Tao, Jing Zhang, Krista Queen, Anna Uehara, Clinton R. Paden, Yan Li, Haibin Wang, Jasmine Padilla, Justin Lee, Suxiang Tong |
| EPI_ISL_419556, EPI_ISL_419557 | GA Department of Public Health Laboratory | Pathogen Discovery, Respiratory Viruses Branch, Division of Viral Diseases, Centers for Disease Control and Prevention | Ying Tao, Jing Zhang, Krista Queen, Anna Uehara, Clinton R. Paden, Yan Li, Haibin Wang, Jasmine Padilla, Justin Lee, Suxiang Tong |
| EPI_ISL_419558 | OR State PHL-Virology/Immunology Section | Pathogen Discovery, Respiratory Viruses Branch, Division of Viral Diseases, Centers for Disease Control and Prevention | Ying Tao, Jing Zhang, Krista Queen, Anna Uehara, Clinton R. Paden, Yan Li, Haibin Wang, Jasmine Padilla, Justin Lee, Suxiang Tong |
| EPI_ISL_422425 | Zhejiang Provincial Center for Disease Control and Prevention | Zhejiang Provincial Center for Disease Control and Prevention | Yanjun Zhang, Yi Sun |
| EPI_ISL_424349 | Human Genetic Research Center, Kawsar Biotech Company | Human Genetic Research Center, Kawsar Biotech Company | Zeinali,S., Khosravi,M.A., Abbasalipour Bashash,M., Mostafavi Jabbari,S., Firoozi,M., Pourtavakoli,S., Khateri,E., Zeinali,R. and Hoseini,F. |
| EPI_ISL_424841, EPI_ISL_424842 | SC Dept of Health and Env. Control-Bureau of Laboratories | Pathogen Discovery, Respiratory Viruses Branch, Division of Viral Diseases, Centers for Disease Control and Prevention | Yan Li, Krista Queen, Clinton R. Paden, Rachel Marine, Anna Uehara, Ying Tao, Jing Zhang, Haibin Wang, Mary S. Keckler, Alison S. Laufer Halpin, Christopher A. Elkins, Suxiang Tong |
| EPI_ISL_424843, EPI_ISL_424844, EPI_ISL_424845, EPI_ISL_424846, EPI_ISL_424847 | MA State Public Health Laboratory | Pathogen Discovery, Respiratory Viruses Branch, Division of Viral Diseases, Centers for Disease Control and Prevention | Yan Li, Krista Queen, Clinton R. Paden, Rachel Marine, Anna Uehara, Ying Tao, Jing Zhang, Haibin Wang, Mary S. Keckler, Alison S. Laufer Halpin, Christopher A. Elkins, Suxiang Tong |
| EPI_ISL_424848, EPI_ISL_424849 | AZ SPHL, Arizona Department of Health Services | Pathogen Discovery, Respiratory Viruses Branch, Division of Viral Diseases, Centers for Disease Control and Prevention | Yan Li, Krista Queen, Clinton R. Paden, Rachel Marine, Anna Uehara, Ying Tao, Jing Zhang, Haibin Wang, Mary S. Keckler, Alison S. Laufer Halpin, Christopher A. Elkins, Suxiang Tong |
| EPI_ISL_424850, EPI_ISL_424851 | IL Department of Public Health Chicago Laboratory | Pathogen Discovery, Respiratory Viruses Branch, Division of Viral Diseases, Centers for Disease Control and Prevention | Yan Li, Krista Queen, Clinton R. Paden, Rachel Marine, Anna Uehara, Ying Tao, Jing Zhang, Haibin Wang, Mary S. Keckler, Alison S. Laufer Halpin, Christopher A. Elkins, Suxiang Tong |
| EPI_ISL_424852 | DC Public Health Lab/ Dept. of Forensic Sciences | Pathogen Discovery, Respiratory Viruses Branch, Division of Viral Diseases, Centers for Disease Control and Prevention | Yan Li, Krista Queen, Clinton R. Paden, Rachel Marine, Anna Uehara, Ying Tao, Jing Zhang, Haibin Wang, Mary S. Keckler, Alison S. Laufer Halpin, Christopher A. Elkins, Suxiang Tong |
| EPI_ISL_424853, EPI_ISL_424854 | FL Bureau of Public Health Laboratories-Miami | Pathogen Discovery, Respiratory Viruses Branch, Division of Viral Diseases, Centers for Disease Control and Prevention | Yan Li, Krista Queen, Clinton R. Paden, Rachel Marine, Anna Uehara, Ying Tao, Jing Zhang, Haibin Wang, Mary S. Keckler, Alison S. Laufer Halpin, Christopher A. Elkins, Suxiang Tong |
| EPI_ISL_424855 | FL Bureau of Public Health Laboratories-Tampa | Pathogen Discovery, Respiratory Viruses Branch, Division of Viral Diseases, Centers for Disease Control and Prevention | Yan Li, Krista Queen, Clinton R. Paden, Rachel Marine, Anna Uehara, Ying Tao, Jing Zhang, Haibin Wang, Mary S. Keckler, Alison S. Laufer Halpin, Christopher A. Elkins, Suxiang Tong |
| EPI_ISL_424856, EPI_ISL_424857 | FL Bur. of Public Health Laboratories-Jacksonville | Pathogen Discovery, Respiratory Viruses Branch, Division of Viral Diseases, Centers for Disease Control and Prevention | Yan Li, Krista Queen, Clinton R. Paden, Rachel Marine, Anna Uehara, Ying Tao, Jing Zhang, Haibin Wang, Mary S. Keckler, Alison S. Laufer Halpin, Christopher A. Elkins, Suxiang Tong |
| EPI_ISL_424858, EPI_ISL_424859, EPI_ISL_424861, EPI_ISL_424864 | GA Department of Public Health Laboratory | Pathogen Discovery, Respiratory Viruses Branch, Division of Viral Diseases, Centers for Disease Control and Prevention | Yan Li, Krista Queen, Clinton R. Paden, Rachel Marine, Anna Uehara, Ying Tao, Jing Zhang, Haibin Wang, Mary S. Keckler, Alison S. Laufer Halpin, Christopher A. Elkins, Suxiang Tong |
| EPI_ISL_424865 | IA State Hygienic Laboratory | Pathogen Discovery, Respiratory Viruses Branch, Division of Viral Diseases, Centers for Disease Control and Prevention | Yan Li, Krista Queen, Clinton R. Paden, Rachel Marine, Anna Uehara, Ying Tao, Jing Zhang, Haibin Wang, Mary S. Keckler, Alison S. Laufer Halpin, Christopher A. Elkins, Suxiang Tong |
| EPI_ISL_424866 | IN State Department of Health Laboratory Services | Pathogen Discovery, Respiratory Viruses Branch, Division of Viral Diseases, Centers for Disease Control and Prevention | Yan Li, Krista Queen, Clinton R. Paden, Rachel Marine, Anna Uehara, Ying Tao, Jing Zhang, Haibin Wang, Mary S. Keckler, Alison S. Laufer Halpin, Christopher A. Elkins, Suxiang Tong |
| EPI_ISL_424867 | KS Health and Environmental Laboratories | Pathogen Discovery, Respiratory Viruses Branch, Division of Viral Diseases, Centers for Disease Control and Prevention | Yan Li, Krista Queen, Clinton R. Paden, Rachel Marine, Anna Uehara, Ying Tao, Jing Zhang, Haibin Wang, Mary S. Keckler, Alison S. Laufer Halpin, Christopher A. Elkins, Suxiang Tong |
| EPI_ISL_424868 | LA Office of Public Health Laboratories | Pathogen Discovery, Respiratory Viruses Branch, Division of Viral Diseases, Centers for Disease Control and Prevention | Yan Li, Krista Queen, Clinton R. Paden, Rachel Marine, Anna Uehara, Ying Tao, Jing Zhang, Haibin Wang, Mary S. Keckler, Alison S. Laufer Halpin, Christopher A. Elkins, Suxiang Tong |
| EPI_ISL_424869 | MD DOH Laboratories Administration | Pathogen Discovery, Respiratory Viruses Branch, Division of Viral Diseases, Centers for Disease Control and Prevention | Yan Li, Krista Queen, Clinton R. Paden, Rachel Marine, Anna Uehara, Ying Tao, Jing Zhang, Haibin Wang, Mary S. Keckler, Alison S. Laufer Halpin, Christopher A. Elkins, Suxiang Tong |
| EPI_ISL_424870 | MO State Public Health Laboratory | Pathogen Discovery, Respiratory Viruses Branch, Division of Viral Diseases, Centers for Disease Control and Prevention | Yan Li, Krista Queen, Clinton R. Paden, Rachel Marine, Anna Uehara, Ying Tao, Jing Zhang, Haibin Wang, Mary S. Keckler, Alison S. Laufer Halpin, Christopher A. Elkins, Suxiang Tong |
| EPI_ISL_424871, EPI_ISL_424872, EPI_ISL_424873 | NC State Laboratory of Public Health | Pathogen Discovery, Respiratory Viruses Branch, Division of Viral Diseases, Centers for Disease Control and Prevention | Yan Li, Krista Queen, Clinton R. Paden, Rachel Marine, Anna Uehara, Ying Tao, Jing Zhang, Haibin Wang, Mary S. Keckler, Alison S. Laufer Halpin, Christopher A. Elkins, Suxiang Tong |
| EPI_ISL_424874, EPI_ISL_424875 | NE Public Health Laboratory | Pathogen Discovery, Respiratory Viruses Branch, Division of Viral Diseases, Centers for Disease Control and Prevention | Yan Li, Krista Queen, Clinton R. Paden, Rachel Marine, Anna Uehara, Ying Tao, Jing Zhang, Haibin Wang, Mary S. Keckler, Alison S. Laufer Halpin, Christopher A. Elkins, Suxiang Tong |
| EPI_ISL_424876, EPI_ISL_424877 | NH Dept. of Health and Human Services Public Health Labs | Pathogen Discovery, Respiratory Viruses Branch, Division of Viral Diseases, Centers for Disease Control and Prevention | Yan Li, Krista Queen, Clinton R. Paden, Rachel Marine, Anna Uehara, Ying Tao, Jing Zhang, Haibin Wang, Mary S. Keckler, Alison S. Laufer Halpin, Christopher A. Elkins, Suxiang Tong |
| EPI_ISL_424878 | NJ Public Health and Environmental Laboratories | Pathogen Discovery, Respiratory Viruses Branch, Division of Viral Diseases, Centers for Disease Control and Prevention | Yan Li, Krista Queen, Clinton R. Paden, Rachel Marine, Anna Uehara, Ying Tao, Jing Zhang, Haibin Wang, Mary S. Keckler, Alison S. Laufer Halpin, Christopher A. Elkins, Suxiang Tong |
| EPI_ISL_424879 | NV-Southern Nevada Public Health Laboratory | Pathogen Discovery, Respiratory Viruses Branch, Division of Viral Diseases, Centers for Disease Control and Prevention | Yan Li, Krista Queen, Clinton R. Paden, Rachel Marine, Anna Uehara, Ying Tao, Jing Zhang, Haibin Wang, Mary S. Keckler, Alison S. Laufer Halpin, Christopher A. Elkins, Suxiang Tong |
| EPI_ISL_424880 | OH Department of Health Laboratory | Pathogen Discovery, Respiratory Viruses Branch, Division of | Yan Li, Krista Queen, Clinton R. Paden, Rachel Marine, Anna Uehara, Ying Tao, Jing Zhang, Haibin Wang, Mary S. Keckler, Alison S. Laufer Halpin, |

| | | | |
|---|---|---|---|
| | | Viral Diseases, Centers for Disease Control and Prevention | Christopher A. Elkins, Suxiang Tong |
| EPI_ISL_424881, EPI_ISL_424882, EPI_ISL_424883, EPI_ISL_424884, EPI_ISL_424885, EPI_ISL_424886 | PA Department of Health, Bureau of Laboratories | Pathogen Discovery, Respiratory Viruses Branch, Division of Viral Diseases, Centers for Disease Control and Prevention | Yan Li, Krista Queen, Clinton R. Paden, Rachel Marine, Anna Uehara, Ying Tao, Jing Zhang, Haibin Wang, Mary S. Keckler, Alison S. Laufer Halpin, Christopher A. Elkins, Suxiang Tong |
| EPI_ISL_424887 | RI State Health Laboratories | Pathogen Discovery, Respiratory Viruses Branch, Division of Viral Diseases, Centers for Disease Control and Prevention | Yan Li, Krista Queen, Clinton R. Paden, Rachel Marine, Anna Uehara, Ying Tao, Jing Zhang, Haibin Wang, Mary S. Keckler, Alison S. Laufer Halpin, Christopher A. Elkins, Suxiang Tong |
| EPI_ISL_424888 | SC Dept of Health and Env. Control-Bureau of Laboratories | Pathogen Discovery, Respiratory Viruses Branch, Division of Viral Diseases, Centers for Disease Control and Prevention | Yan Li, Krista Queen, Clinton R. Paden, Rachel Marine, Anna Uehara, Ying Tao, Jing Zhang, Haibin Wang, Mary S. Keckler, Alison S. Laufer Halpin, Christopher A. Elkins, Suxiang Tong |
| EPI_ISL_424889, EPI_ISL_424890 | UT-Unified State Labs: Public Health Utah DOH | Pathogen Discovery, Respiratory Viruses Branch, Division of Viral Diseases, Centers for Disease Control and Prevention | Yan Li, Krista Queen, Clinton R. Paden, Rachel Marine, Anna Uehara, Ying Tao, Jing Zhang, Haibin Wang, Mary S. Keckler, Alison S. Laufer Halpin, Christopher A. Elkins, Suxiang Tong |
| EPI_ISL_424891, EPI_ISL_424892, EPI_ISL_424893, EPI_ISL_424894 | VA-Division of Consolidated Laboratory Services | Pathogen Discovery, Respiratory Viruses Branch, Division of Viral Diseases, Centers for Disease Control and Prevention | Yan Li, Krista Queen, Clinton R. Paden, Rachel Marine, Anna Uehara, Ying Tao, Jing Zhang, Haibin Wang, Mary S. Keckler, Alison S. Laufer Halpin, Christopher A. Elkins, Suxiang Tong |
| EPI_ISL_424895, EPI_ISL_424896, EPI_ISL_424897, EPI_ISL_424898, EPI_ISL_424899, EPI_ISL_424900 | IA State Hygienic Laboratory | Pathogen Discovery, Respiratory Viruses Branch, Division of Viral Diseases, Centers for Disease Control and Prevention | Ying Tao, Clinton R. Paden, Jing Zhang, Krista Queen, Anna Uehara, Yan Li, Haibin Wang, Mary S. Keckler, Alison S. Laufer Halpin, Christopher A. Elkins, Suxiang Tong |
| EPI_ISL_424901 | NJ Public Health and Environmental Laboratories | Pathogen Discovery, Respiratory Viruses Branch, Division of Viral Diseases, Centers for Disease Control and Prevention | Ying Tao, Clinton R. Paden, Jing Zhang, Krista Queen, Anna Uehara, Yan Li, Haibin Wang, Mary S. Keckler, Alison S. Laufer Halpin, Christopher A. Elkins, Suxiang Tong |
| EPI_ISL_424902, EPI_ISL_424903, EPI_ISL_424904, EPI_ISL_424905 | SC Dept of Health and Env. Control-Bureau of Laboratories | Pathogen Discovery, Respiratory Viruses Branch, Division of Viral Diseases, Centers for Disease Control and Prevention | Ying Tao, Clinton R. Paden, Jing Zhang, Krista Queen, Anna Uehara, Yan Li, Haibin Wang, Mary S. Keckler, Alison S. Laufer Halpin, Christopher A. Elkins, Suxiang Tong |
| EPI_ISL_424906 | NYC Department of Health and Mental Hygiene | Pathogen Discovery, Respiratory Viruses Branch, Division of Viral Diseases, Centers for Disease Control and Prevention | Ying Tao, Clinton R. Paden, Jing Zhang, Krista Queen, Anna Uehara, Yan Li, Haibin Wang, Mary S. Keckler, Alison S. Laufer Halpin, Christopher A. Elkins, Suxiang Tong |
| EPI_ISL_424907 | VA-Division of Consolidated Laboratory Services | Pathogen Discovery, Respiratory Viruses Branch, Division of Viral Diseases, Centers for Disease Control and Prevention | Ying Tao, Clinton R. Paden, Jing Zhang, Krista Queen, Anna Uehara, Yan Li, Haibin Wang, Mary S. Keckler, Alison S. Laufer Halpin, Christopher A. Elkins, Suxiang Tong |
| EPI_ISL_424908, EPI_ISL_424909, EPI_ISL_424910, EPI_ISL_424911, EPI_ISL_424912, EPI_ISL_424913, EPI_ISL_424914, EPI_ISL_424915, EPI_ISL_424916, EPI_ISL_424917, EPI_ISL_424918, EPI_ISL_424919, EPI_ISL_424920 | | | |
| see above | MA State Public Health Laboratory | Pathogen Discovery, Respiratory Viruses Branch, Division of Viral Diseases, Centers for Disease Control and Prevention | Ying Tao, Clinton R. Paden, Jing Zhang, Krista Queen, Anna Uehara, Yan Li, Haibin Wang, Mary S. Keckler, Alison S. Laufer Halpin, Christopher A. Elkins, Suxiang Tong |
| EPI_ISL_426437, EPI_ISL_426440, EPI_ISL_426441, EPI_ISL_426443, EPI_ISL_426444, EPI_ISL_426446, EPI_ISL_426447, EPI_ISL_426448, EPI_ISL_426449, EPI_ISL_426451, EPI_ISL_426452, EPI_ISL_426453 | | | |
| see above | WA State Department of Health | Pathogen Discovery, Respiratory Viruses Branch, Division of Viral Diseases, Centers for Disease Control and Prevention | Ying Tao, Jing Zhang, Clinton R. Paden, Krista Queen, Anna Uehara, Yan Li, Haibin Wang, Jesica Jacobs, Denny Russell, Brian Hiatt, Jessica Gant, Suxiang Tong |
| EPI_ISL_429074, EPI_ISL_429075, EPI_ISL_429077, EPI_ISL_429082, EPI_ISL_429083, EPI_ISL_429085, EPI_ISL_429088, EPI_ISL_429089, EPI_ISL_429090, EPI_ISL_429091, EPI_ISL_429092, EPI_ISL_429093, EPI_ISL_429096, EPI_ISL_429097, EPI_ISL_429098 | | | |
| see above | The First Affiliated Hospital of Guangzhou Medical University | BGI-shenzhen & The First Affiliated Hospital of Guangzhou Medical University | Yanqun Wang, Daxi Wang, Lu Zhang, Wanying Sun, Zhaoyong Zhang et al. |
| EPI_ISL_429239 | Department of Clinical Laboratory, the First People's Hospital of Yunnan Province | Department of Clinical Laboratory, the First People's Hospital of Yunnan Province | Yi Sun,Ziqin Dian,Ya Xu,Guiqian Zhang,Xin Fan,Yu Zhang |
| EPI_ISL_430847 | HS mikrobiologi virus | The Public Health Agency of Sweden | Zhibing Yun, Oskar Karlsson Lindsjo, Maria Lind Karlberg, Anna-Malin Linde, Olov Svartstrom, Anna Risberg, Shaman Muradrasoli, Karin Tegmark-Wisell |
| EPI_ISL_436040, EPI_ISL_436041, EPI_ISL_436042, EPI_ISL_436043 | DC Public Health Lab Dept of Forensic Science | Pathogen Discovery, Respiratory Viruses Branch, Division of Viral Diseases, Centers for Disease Control and Prevention | Ying Tao, Jing Zhang, Krista Queen, Yan Li, Anna Uehara, Clinton R. Paden, Haibin Wang, Zachary Weiner, Bettina Bankamp, Suxiang Tong |
| EPI_ISL_436044 | Louisiana Office of Public Health Laboratories | Pathogen Discovery, Respiratory Viruses Branch, Division of Viral Diseases, Centers for Disease Control and Prevention | Ying Tao, Jing Zhang, Krista Queen, Yan Li, Anna Uehara, Clinton R. Paden, Haibin Wang, Zachary Weiner, Bettina Bankamp, Suxiang Tong |
| EPI_ISL_436045, EPI_ISL_436046 | US VI Department of Health | Pathogen Discovery, Respiratory Viruses Branch, Division of Viral Diseases, Centers for Disease Control and Prevention | Ying Tao, Jing Zhang, Krista Queen, Yan Li, Anna Uehara, Clinton R. Paden, Haibin Wang, Zachary Weiner, Bettina Bankamp, Suxiang Tong |
| EPI_ISL_436047, EPI_ISL_436048, EPI_ISL_436049, EPI_ISL_436050, EPI_ISL_436051, EPI_ISL_436052, EPI_ISL_436053, EPI_ISL_436054, EPI_ISL_436055, EPI_ISL_436056, EPI_ISL_436057, EPI_ISL_436058, EPI_ISL_436059, EPI_ISL_436060, EPI_ISL_436061, EPI_ISL_436062, EPI_ISL_436063, EPI_ISL_436064, EPI_ISL_436065, EPI_ISL_436066, EPI_ISL_436067, EPI_ISL_436068, EPI_ISL_436069, EPI_ISL_436070, EPI_ISL_436071, EPI_ISL_436072, EPI_ISL_436073, EPI_ISL_436074, EPI_ISL_436075, EPI_ISL_436076, EPI_ISL_436077, EPI_ISL_436078, EPI_ISL_436079, EPI_ISL_436080, EPI_ISL_436081, EPI_ISL_436082 | | | |
| see above | NYC Department of Health and Mental Hygiene | Pathogen Discovery, Respiratory Viruses Branch, Division of Viral Diseases, Centers for Disease Control and Prevention | Ying Tao, Krista Queen, Christy Harrison, Jennifer Rakeman, Clinton R. Paden, Jing Zhang, Anna Uehara, Yan Li, Haibin Wang, Jasmine Padilla, Justin Lee, Bettina Bankamp, Zachary Weiner, Suxiang Tong |
| EPI_ISL_437443 | Department of MicroBiology, Government Medical College, Surat | Gujarat Biotechnology Research Centre | Zuber Saiyed, Dipa Kinariwala, Disha Patel, Binita Aring, Neeta Khandelwal, Geeta Vaghela, Sonia Barve, Bhavesh Modi, Kairavi Joshi, Gaurishankar Shrimali, Nidhi Sood, Pranay Shah, R D Dixit, Snehal Bagatharia, Kamlesh J Upadhyay, Ramesh Pandit, Tejas Shah, Ankit Hinsu, Pritesh Sabara, Apurvasinh Puvar, Janvi Raval, Monika Gandhi, Pinal Trivedi, Maharshi Pandya, Amit Kanani, Akanksha Verma, Nitin Savaliya, Raghawendra Kumar, Dinesh Kumar, Sharmistha Majumdar, Chaitanya Joshi, Madhvi Joshi |
| EPI_ISL_444473 | B.J. Medical College and Civil hospital | Gujarat Biotechnology Research Centre | Zuber Saiyed, Dipa Kinariwala, Disha Patel, Binita Aring, Neeta Khandelwal, Geeta Vaghela, Sonia Barve, Bhavesh Modi, Kairavi Joshi, Gaurishankar Shrimali, Nidhi Sood, Pranay Shah, R D Dixit, Snehal Bagatharia, Kamlesh J Upadhyay, Ramesh Pandit, Tejas Shah, Ankit Hinsu, Pritesh Sabara, Apurvasinh Puvar, Janvi Raval, Monika Gandhi, Pinal Trivedi, Maharshi Pandya, Amit Kanani, Akanksha Verma, Nitin Savaliya, Raghawendra Kumar, Dinesh Kumar, Armi Chaudhari, Chaitanya Joshi, Madhvi Joshi |
| EPI_ISL_447045 | B.J. Medical College and Civil hospital | Gujarat Biotechnology Research Centre | Zuber Saiyed, Dipa Kinariwala, Disha Patel, Binita Aring, Neeta Khandelwal, Geeta Vaghela, Sonia Barve, Bhavesh Modi, Kairavi Joshi, Gaurishankar Shrimali, Nidhi Sood, Pranay Shah, R D Dixit, Snehal Bagatharia, Kamlesh J Upadhyay, Ramesh Pandit, Tejas Shah, Ankit Hinsu, Pritesh Sabara, Apurvasinh Puvar, Janvi Raval, Monika Gandhi, Pinal Trivedi, Maharshi Pandya, Amit Kanani, Akanksha Verma, Nitin Savaliya, Raghawendra Kumar, Dinesh Kumar, Neha Rajpara, Chaitanya Joshi, Madhvi Joshi |
| EPI_ISL_447554 | GMERS Medical College and Hospital, Gandhinagar | Gujarat Biotechnology Research Centre | Zuber Saiyed, Dipa Kinariwala, Disha Patel, Binita Aring, Neeta Khandelwal, Geeta Vaghela, Sonia Barve, Bhavesh Modi, Kairavi Joshi, Gaurishankar Shrimali, Nidhi Sood, Pranay Shah, R D Dixit, Snehal Bagatharia, Kamlesh J Upadhyay, Ramesh Pandit, Tejas Shah, Ankit Hinsu, Pritesh Sabara, Apurvasinh Puvar, Janvi Raval, Monika Gandhi, Pinal Trivedi, Maharshi Pandya, Amit Kanani, Akanksha Verma, Nitin Savaliya, Raghawendra Kumar, Dinesh Kumar, Sharmistha Majumdar, Chaitanya Joshi, Madhvi Joshi |
| EPI_ISL_447886 | unknown | Pathogen Discovery | Ying Tao, Yan Li, Jing Zhang, Clinton R. Paden, Krista Queen, Anna Uehara, Haibin Wang, Julu Bhatnagar, Suxiang Tong |
| EPI_ISL_450444 | 20 Dongda Street, Fengtai District, Beijing, Beijing 100071, China | Dept. OPA, Beijing Institute of Microbiology and Epidemiology | Zhang,X.A., Fan,H., Qi,R.Z., Zheng,W., Zheng,K., Gong,J.H., Fang,L.Q. and Liu,W. |
| EPI_ISL_450784 | Government Medical College-Bhavnagar | Gujarat Biotechnology Research Centre | Zarna Patel, Ramesh Pandit, Tejas Shah, Ankit Hinsu, Pritesh Sabara, Apurvasinh Puvar, Janvi Raval, Monika Gandhi, Pinal Trivedi, Maharshi Pandya, Amit Kanani, Nidhi Patel, Nitin Savaliya, Raghawendra Kumar, Dinesh Kumar, Zuber Saiyed, Komal Patel, Labdhi Pandya, Snehal Bagatharia, Kairavi Desai, Saklain Malek, Shirish Patel, Bhavesh Modi, Gaurishankar Shrimali, R D Dixit, A M Kadri, Afzal Ansari, Chaitanya Joshi, Madhvi Joshi |
| EPI_ISL_450788 | Pandit Deendayal Upadhyay Government Medical College, Rajkot | Gujarat Biotechnology Research Centre | Zarna Patel, Tejas Shah, Ankit Hinsu, Pritesh Sabara, Apurvasinh Puvar, Janvi Raval, Monika Gandhi, Pinal Trivedi, Maharshi Pandya, Amit Kanani, Nidhi Patel, Nitin Savaliya, Raghawendra Kumar, Dinesh Kumar, Zuber Saiyed, Komal Patel, Labdhi Pandya, Snehal Bagatharia, Prakash Modi, Sejul Antala, Manish Pattani, Ramesh Pandit, Bhavesh Modi, Gaurishankar Shrimali, R D Dixit, A M Kadri, Camellia Chakraborty, Chaitanya Joshi, Madhvi Joshi |
| EPI_ISL_450790 | Pandit Deendayal Upadhyay Government Medical College, Rajkot | Gujarat Biotechnology Research Centre | Zarna Patel, Pritesh Sabara, Apurvasinh Puvar, Janvi Raval, Monika Gandhi, Pinal Trivedi, Maharshi Pandya, Amit Kanani, Nidhi Patel, Nitin Savaliya, Raghawendra Kumar, Dinesh Kumar, Zuber Saiyed, Komal Patel, Labdhi Pandya, Snehal Bagatharia, Prakash Modi, Sejul Antala, Manish Pattani, |

| | | | |
|---|---|---|---|
| | | | Ramesh Pandit, Tejas Shah, Ankit Hinsu, Bhavesh Modi, Gaurishankar Shrimali, R D Dixit, A M Kadri, Sharmistha Majumdar, Chaitanya Joshi, Madhvi Joshi |
| EPI_ISL_450791 | Pandit Deendayal Upadhyay Government Medical College, Rajkot | Gujarat Biotechnology Research Centre | Zarna Patel, Apurvasinh Puvar, Janvi Raval, Monika Gandhi, Pinal Trivedi, Maharshi Pandya, Amit Kanani, Nidhi Patel, Nitin Savaliya, Raghawendra Kumar, Dinesh Kumar, Zuber Saiyed, Komal Patel, Labdhi Pandya, Snehal Bagatharia, Prakash Modi, Sejul Antala, Manish Pattani, Ramesh Pandit, Tejas Shah, Ankit Hinsu, Pritesh Sabara, Bhavesh Modi, Gaurishankar Shrimali, R D Dixit, A M Kadri, Pooja P Doshi, Chaitanya Joshi, Madhvi Joshi |
| EPI_ISL_450793, EPI_ISL_450794, EPI_ISL_450795, EPI_ISL_450796, EPI_ISL_450797, EPI_ISL_450799 | Jamaica Ministry of Health and Wellness | Pathogen Discovery, Respiratory Viruses Branch, Division of Viral Diseases, Centers for Disease Control and Prevention | Yan Li, Anna Montmayeur, Ying Tao, Krista Queen, Jing Zhang, Anna Uehara, Clinton R. Paden, Rachel Marine, Haibin Wang, Zachary Weiner, Bettina Bankamp, Suxiang Tong |
| EPI_ISL_450800 | AR Dept. of Health-Public Health Lab | Pathogen Discovery, Respiratory Viruses Branch, Division of Viral Diseases, Centers for Disease Control and Prevention | Yan Li, Anna Montmayeur, Ying Tao, Krista Queen, Jing Zhang, Anna Uehara, Clinton R. Paden, Rachel Marine, Haibin Wang, Zachary Weiner, Bettina Bankamp, Suxiang Tong |
| EPI_ISL_450801 | Georgia Department of Health | Pathogen Discovery, Respiratory Viruses Branch, Division of Viral Diseases, Centers for Disease Control and Prevention | Yan Li, Anna Montmayeur, Ying Tao, Krista Queen, Jing Zhang, Anna Uehara, Clinton R. Paden, Rachel Marine, Haibin Wang, Zachary Weiner, Bettina Bankamp, Suxiang Tong |
| EPI_ISL_450802, EPI_ISL_450803 | PA Department of Health, Bureau of Laboratories | Pathogen Discovery, Respiratory Viruses Branch, Division of Viral Diseases, Centers for Disease Control and Prevention | Yan Li, Anna Montmayeur, Ying Tao, Krista Queen, Jing Zhang, Anna Uehara, Clinton R. Paden, Rachel Marine, Haibin Wang, Zachary Weiner, Bettina Bankamp, Suxiang Tong |
| EPI_ISL_450804, EPI_ISL_450806 | VI-US Virgin Islands Department of Health | Pathogen Discovery, Respiratory Viruses Branch, Division of Viral Diseases, Centers for Disease Control and Prevention | Yan Li, Anna Montmayeur, Ying Tao, Krista Queen, Jing Zhang, Anna Uehara, Clinton R. Paden, Rachel Marine, Haibin Wang, Zachary Weiner, Bettina Bankamp, Suxiang Tong |
| EPI_ISL_451150 | M.P Shah Government Medocal college Jamnagar | Gujarat Biotechnology Research Centre | Zarna Patel, Monika Gandhi, Pinal Trivedi, Maharshi Pandya, Amit Kanani, Nidhi Patel, Nitin Savaliya, Raghawendra Kumar, Dinesh Kumar, Zuber Saiyed, Komal Patel, Labdhi Pandya, Snehal Bagatharia, Ramesh Pandit, Tejas Shah, Ankit Hinsu, Pritesh Sabara, Apurvasinh Puvar, Binita Aring, Janvi Raval, Bhavesh Modi, Gaurishankar Shrimali, R D Dixit, A M Kadri, Pragya Sharma, Chaitanya Joshi, Madhvi Joshi, |
| EPI_ISL_451161 | Government Medical College, Vadodara | Gujarat Biotechnology Research Centre | Zuber Saiyed, Komal Patel, Labdhi Pandya, Snehal Bagatharia, Ramesh Pandit, Tejas Shah, Ankit Hinsu, Pritesh Sabara, Apurvasinh Puvar, Janvi Raval, Zarna Patel, Monika Gandhi, Pinal Trivedi, Maharshi Pandya, Manish Pattani, Tanuja Javadekar , Amit Kanani, Nidhi Patel, Nitin Savaliya, Raghawendra Kumar, Dinesh Kumar, Bhavesh Modi, Gaurishankar Shrimali, R D Dixit, A M Kadri, Akanksha Verma, Chaitanya Joshi, Madhvi Joshi |
| EPI_ISL_451934 | Research Unit, University Hospital for Infectious Diseases "Dr. Fran Mihaljevi" | Cicin Sain lab, Helmholtz Centre for Infection Research | Zeeshan Chaudhry, Kathrin Eschke, Željka Maak Šafranko, Ivan-Christian Kurolt |
| EPI_ISL_452105 | Texas DSHS Lab Services | Pathogen Discovery, Respiratory Viruses Branch, Division of Viral Diseases, Centers for Disease Control and Prevention | Yan Li, Anna Montmayeur, Ying Tao, Krista Queen, Jing Zhang, Anna Uehara, Clinton R. Paden, Rachel Marine, Mary S. Keckler, Alison S. Laufer Halpin, Haibin Wang, Christopher A. Elkins, Zachary Weiner, Suxiang Tong |
| EPI_ISL_452106, EPI_ISL_452107 | LA Office of Public Health Laboratories | Pathogen Discovery, Respiratory Viruses Branch, Division of Viral Diseases, Centers for Disease Control and Prevention | Yan Li, Anna Montmayeur, Ying Tao, Krista Queen, Jing Zhang, Anna Uehara, Clinton R. Paden, Rachel Marine, Mary S. Keckler, Alison S. Laufer Halpin, Haibin Wang, Christopher A. Elkins, Zachary Weiner, Suxiang Tong |
| EPI_ISL_452108 | MN Department of Health | Pathogen Discovery, Respiratory Viruses Branch, Division of Viral Diseases, Centers for Disease Control and Prevention | Yan Li, Anna Montmayeur, Ying Tao, Krista Queen, Jing Zhang, Anna Uehara, Clinton R. Paden, Rachel Marine, Mary S. Keckler, Alison S. Laufer Halpin, Haibin Wang, Christopher A. Elkins, Zachary Weiner, Suxiang Tong |
| EPI_ISL_452109 | NM Department of Health | Pathogen Discovery, Respiratory Viruses Branch, Division of Viral Diseases, Centers for Disease Control and Prevention | Yan Li, Anna Montmayeur, Ying Tao, Krista Queen, Jing Zhang, Anna Uehara, Clinton R. Paden, Rachel Marine, Mary S. Keckler, Alison S. Laufer Halpin, Haibin Wang, Christopher A. Elkins, Zachary Weiner, Suxiang Tong |
| EPI_ISL_452110, EPI_ISL_452111 | FL Bureau of Public Health Laboratories | Pathogen Discovery, Respiratory Viruses Branch, Division of Viral Diseases, Centers for Disease Control and Prevention | Yan Li, Anna Montmayeur, Ying Tao, Krista Queen, Jing Zhang, Anna Uehara, Clinton R. Paden, Rachel Marine, Mary S. Keckler, Alison S. Laufer Halpin, Haibin Wang, Christopher A. Elkins, Zachary Weiner, Suxiang Tong |
| EPI_ISL_452112 | Texas DSHS Lab Services | Pathogen Discovery, Respiratory Viruses Branch, Division of Viral Diseases, Centers for Disease Control and Prevention | Yan Li, Anna Montmayeur, Ying Tao, Krista Queen, Jing Zhang, Anna Uehara, Clinton R. Paden, Rachel Marine, Mary S. Keckler, Alison S. Laufer Halpin, Haibin Wang, Christopher A. Elkins, Zachary Weiner, Suxiang Tong |
| EPI_ISL_452113 | MN Department of Health | Pathogen Discovery, Respiratory Viruses Branch, Division of Viral Diseases, Centers for Disease Control and Prevention | Yan Li, Anna Montmayeur, Ying Tao, Krista Queen, Jing Zhang, Anna Uehara, Clinton R. Paden, Rachel Marine, Mary S. Keckler, Alison S. Laufer Halpin, Haibin Wang, Christopher A. Elkins, Zachary Weiner, Suxiang Tong |
| EPI_ISL_452114 | KS Health and Environmental Laboratories | Pathogen Discovery, Respiratory Viruses Branch, Division of Viral Diseases, Centers for Disease Control and Prevention | Yan Li, Anna Montmayeur, Ying Tao, Krista Queen, Jing Zhang, Anna Uehara, Clinton R. Paden, Rachel Marine, Mary S. Keckler, Alison S. Laufer Halpin, Haibin Wang, Christopher A. Elkins, Zachary Weiner, Suxiang Tong |
| EPI_ISL_452115 | Texas DSHS Lab Services | Pathogen Discovery, Respiratory Viruses Branch, Division of Viral Diseases, Centers for Disease Control and Prevention | Yan Li, Anna Montmayeur, Ying Tao, Krista Queen, Jing Zhang, Anna Uehara, Clinton R. Paden, Rachel Marine, Mary S. Keckler, Alison S. Laufer Halpin, Haibin Wang, Christopher A. Elkins, Zachary Weiner, Suxiang Tong |
| EPI_ISL_452116, EPI_ISL_452117 | NJ Public Health and Environmental Laboratories | Pathogen Discovery, Respiratory Viruses Branch, Division of Viral Diseases, Centers for Disease Control and Prevention | Yan Li, Anna Montmayeur, Ying Tao, Krista Queen, Jing Zhang, Anna Uehara, Clinton R. Paden, Rachel Marine, Mary S. Keckler, Alison S. Laufer Halpin, Haibin Wang, Christopher A. Elkins, Zachary Weiner, Suxiang Tong |
| EPI_ISL_452118 | AZ SPHL, Arizona Department of Health Services | Pathogen Discovery, Respiratory Viruses Branch, Division of Viral Diseases, Centers for Disease Control and Prevention | Yan Li, Anna Montmayeur, Ying Tao, Krista Queen, Jing Zhang, Anna Uehara, Clinton R. Paden, Rachel Marine, Mary S. Keckler, Alison S. Laufer Halpin, Haibin Wang, Christopher A. Elkins, Zachary Weiner, Suxiang Tong |
| EPI_ISL_452119 | VI-US Virgin Islands Department of Health | Pathogen Discovery, Respiratory Viruses Branch, Division of Viral Diseases, Centers for Disease Control and Prevention | Yan Li, Anna Montmayeur, Ying Tao, Krista Queen, Jing Zhang, Anna Uehara, Clinton R. Paden, Rachel Marine, Mary S. Keckler, Alison S. Laufer Halpin, Haibin Wang, Christopher A. Elkins, Zachary Weiner, Suxiang Tong |
| EPI_ISL_454646 | City of El Paso Department of Public Health Laboratory | Pathogen Discovery, Respiratory Viruses Branch, Division of Viral Diseases, Centers for Disease Control and Prevention | Ying Tao, Clinton R. Paden, Jing Zhang, Anna Uehara, Krista Queen, Yan Li, Haibin Wang, Zachary Weiner, Michael Bowen, Suxiang Tong |
| EPI_ISL_454647, EPI_ISL_454648, EPI_ISL_454649, EPI_ISL_454650, EPI_ISL_454651, EPI_ISL_454652 | VI-US Virgin Islands Department of Health | Pathogen Discovery, Respiratory Viruses Branch, Division of Viral Diseases, Centers for Disease Control and Prevention | Ying Tao, Clinton R. Paden, Jing Zhang, Anna Uehara, Krista Queen, Yan Li, Haibin Wang, Zachary Weiner, Bettina Bankamp, Suxiang Tong |
| EPI_ISL_455025 | Government Medical College, Vadodara | Gujarat Biotechnology Research Centre | Zarna Patel, Monika Gandhi, Pinal Trivedi, Maharshi Pandya, Amit Kanani, Nidhi Patel, Nitin Savaliya, Raghawendra Kumar, Dinesh Kumar, Zuber Saiyed, Komal Patel, Labdhi Pandya, Snehal Bagatharia, Tanuja Javadekar , R N Daveshwar, Ramesh Pandit, Tejas Shah, Ankit Hinsu, Pritesh Sabara, Apurvasinh Puvar, Janvi Raval, Akanksha Verma, Bhavesh Modi, Gaurishankar Shrimali, R D Dixit, A M Kadri, Umang Mishra, Chaitanya Joshi, Madhvi Joshi, |
| EPI_ISL_457687, EPI_ISL_457688, EPI_ISL_457689, EPI_ISL_457690, EPI_ISL_457691, EPI_ISL_457692, EPI_ISL_457693, EPI_ISL_457694, EPI_ISL_457695, EPI_ISL_457696, EPI_ISL_457697, EPI_ISL_457698 | | | |
| see above | The First Affiliated Hospital of Guangzhou Medical University, Guangzhou, China | BGI-shenzhen & The First Affiliated Hospital of Guangzhou Medical University | Yanqun Wang, Daxi Wang, Lu Zhang, Wanying Sun, Zhaoyong Zhang et al. |
| EPI_ISL_458097 | B.J. Medical College and Civil hospital | Gujarat Biotechnology Research Centre | Zuber Saiyed, Komal Patel, Labdhi Pandya, Snehal Bagatharia, Dhaval Vaghela, Ramesh Patel, Pranay Shah, Kamlesh J Upadhyay, Ramesh Pandit, Tejas Shah, Ankit Hinsu, Pritesh Sabara, Apurvasinh Puvar, Janvi Raval, Zarna Patel, Monika Gandhi, Pinal Trivedi, Maharshi Pandya, Amit Kanani, Nidhi Patel, Nitin Savaliya, Raghawendra Kumar, Dinesh Kumar, Priti Pandita, Bhavesh Modi, Gaurishankar Shrimali, R D Dixit, A M Kadri, Umang Mishra, Chaitanya Joshi, Madhvi Joshi |
| EPI_ISL_458109 | Gujarat Biotechnology Research Centre | Gujarat Biotechnology Research Centre | Zarna Patel, Monika Gandhi, Pinal Trivedi, Maharshi Pandya, Amit Kanani, Nidhi Patel, Nitin Savaliya, Raghawendra Kumar, Dinesh Kumar, Zuber Saiyed, Komal Patel, Labdhi Pandya, Snehal Bagatharia, Ramesh Pandit, Tejas Shah, Ankit Hinsu, Pritesh Sabara, Apurvasinh Puvar, Janvi Raval, Akanksha Verma, Bhavesh Modi, Gaurishankar Shrimali, R D Dixit, A M Kadri, Umang Mishra, Chaitanya Joshi, Madhvi Joshi, , , , , |
| EPI_ISL_459909 | Zoonotic and Exotic infection Diseases Division, Harbin Veterinary Research Institute, CAAS | Zoonotic and Exotic infection Diseases Division, Harbin Veterinary Research Institute, CAAS | Zhigao Bu, Jinliang Wang |
| EPI_ISL_467038 | GMERS Medical College and Hospital, Gandhinagar | Gujarat Biotechnology Research Centre | Zarna Patel, Monika Gandhi, Pinal Trivedi, Maharshi Pandya, Nidhi Patel, Nitin Savaliya, Raghawendra Kumar, Dinesh Kumar, Zuber Saiyed, Komal Patel, Labdhi Pandya, Snehal Bagatharia, Seema Bhatt, Gaurishankar Shrimali, Bhavesh Modi, Bharti Rajani, Tejas Shah, Ankit Hinsu, Pritesh Sabara, Apurvasinh Puvar, Janvi Raval, Neelam Nathani, R D Dixit, A M Kadri, Harsh Bakshi, Chaitanya Joshi, Madhvi Joshi |

| | | | |
|---|---|---|---|
| EPI_ISL_467048 | B.J. Medical College and Civil hospital | Gujarat Biotechnology Research Centre | Zuber Saiyed, Komal Patel, Labdhi Pandya, Snehal Bagatharia, Pranay Shah, Kamlesh J Upadhyay, Nirav Mungalpara, Tejas Shah, Ankit Hinsu, Pritesh Sabara, Apurvasinh Puvar, Janvi Raval, Zarna Patel, Monika Gandhi, Pinal Trivedi, Maharshi Pandya, Nidhi Patel, Nitin Savaliya, Raghawendra Kumar, Dinesh Kumar, Fenil Patel, R D Dixit, A M Kadri, Harsh Bakshi, Chaitanya Joshi, Madhvi Joshi, |
| EPI_ISL_468044, EPI_ISL_468045, EPI_ISL_468046, EPI_ISL_468047, EPI_ISL_468048, EPI_ISL_468049, EPI_ISL_468050, EPI_ISL_468051, EPI_ISL_468052, EPI_ISL_468053, EPI_ISL_468054, EPI_ISL_468055 | | | |
| see above | Egyptian National Cancer Institute (ENCI) | Egyptian National Cancer Institute (ENCI) | Zekri, Abdel Rahman N, Amer,K.E., Ahmed,O.S., Soliman,H.K., Hafez,M.M., Bahnassy,A.A., Abdelhamid,W., Gad,A., Ali,M., Hassan,W., Samir,M., Raouf,A., Hamdy,M.S., Soliman,M.S., Elsissy,M.H., Elkhateeb,S.M., Ezzelarab,M.H., Abouelhoda, Mohamed |
| EPI_ISL_468056 | Egyptian National Cancer Institute (ENCI) | Egyptian National Cancer Institute (ENCI) | Zekri, Abdel Rahman N., Amer,K.E., Ahmed,O.S., Soliman,H.K., Ali,M.A., Hassan,W.A., Mahmoud,A.A., Khattab,A.A., Hafez,M.M., Abouelhoda, Mohamed |
| EPI_ISL_468057, EPI_ISL_468058, EPI_ISL_468059 | Egyptian National Cancer Institute (ENCI) | Egyptian National Cancer Institute (ENCI) | Zekri, Abdel Rahman N, Amer,K.E., Ahmed,O.S., Soliman,H.K., Hafez,M.M., Bahnassy,A.A., Abdelhamid,W., Gad,A., Ali,M., Hassan,W., Samir,M., Raouf,A., Hamdy,M.S., Soliman,M.S., Elsissy,M.H., Elkhateeb,S.M., Ezzelarab,M.H., Abouelhoda, Mohamed |
| EPI_ISL_468060, EPI_ISL_468062 | Egyptian National Cancer Institute (ENCI) | Egyptian National Cancer Institute (ENCI) | Zekri, Abdel Rahman N., Amer,K.E., Ahmed,O.S., Soliman,H.K., Ali,M.A., Hassan,W.A., Mahmoud,A.A., Khattab,A.A., Hafez,M.M., Abouelhoda, Mohamed |
| EPI_ISL_469029 | Government Medical College, Vadodara | Gujarat Biotechnology Research Centre | Zarna Patel, Monika Gandhi, Pinal Trivedi, Maharshi Pandya, Nidhi Patel, Nitin Savaliya, Raghawendra Kumar, Dinesh Kumar, Zuber Saiyed, Komal Patel, Labdhi Pandya, Snehal Bagatharia, Meenakshi Shah, Neena Doshi, Varsha Godbole, Tejas Shah, Ankit Hinsu, Pritesh Sabara, Apurvasinh Puvar, Janvi Raval, Neelam Nathani, R D Dixit, A M Kadri, Harsh Bakshi, Chaitanya Joshi, Madhvi Joshi |
| EPI_ISL_469037 | GMERS Medical College & Hospital | Gujarat Biotechnology Research Centre | Zuber Saiyed, Komal Patel, Labdhi Pandya, Snehal Bagatharia, Meenakshi Shah, Neena Doshi, Varsha Godbole, Tejas Shah, Ankit Hinsu, Pritesh Sabara, Apurvasinh Puvar, Janvi Raval, Zarna Patel, Monika Gandhi, Pinal Trivedi, Maharshi Pandya, Nidhi Patel, Nitin Savaliya, Raghawendra Kumar, Dinesh Kumar, Pragya Sharma, R D Dixit, A M Kadri, Harsh Bakshi, Chaitanya Joshi, Madhvi Joshi |
| EPI_ISL_469275 | Egyptian National Cancer Institute (ENCI) | Human Genome Center | Zekri, Abdel Rahman N, Amer,K.E., Ahmed,O.S., Soliman,H.K., Hafez,M.M., Bahnassy,A.A., Abdelhamid,W., Gad,A., Ali,M., Hassan,W., Samir,M., Raouf,A., Hamdy,M.S., Soliman,M.S., Elsissy,M.H., Elkhateeb,S.M., Ezzelarab,M.H., Abouelhoda, Mohamed |
| EPI_ISL_475031 | Department of MicroBiology, Government Medical College, Surat | Gujarat Biotechnology Research Centre | Zarna Patel, Monika Gandhi, Pinal Trivedi, Maharshi Pandya, Nidhi Patel, Nitin Savaliya, Raghawendra Kumar, Dinesh Kumar, Zuber Saiyed, Komal Patel, Labdhi Pandya, Snehal Bagatharia, Naresh Chauhan, Summaiya Mullan, Amit gamit, Pritesh Sabara, Apurvasinh Puvar, Janvi Raval, Bhavya Jindal, R D Dixit, A M Kadri, Harsh Bakshi, Chaitanya Joshi, Madhvi Joshi |
| EPI_ISL_475039 | Department of MicroBiology, Government Medical College, Surat | Gujarat Biotechnology Research Centre | Zuber Saiyed, Komal Patel, Labdhi Pandya, Snehal Bagatharia, Naresh Chauhan, Summaiya Mullan, Amit gamit, Pritesh Sabara, Apurvasinh Puvar, Janvi Raval, Zarna Patel, Monika Gandhi, Pinal Trivedi, Maharshi Pandya, Nidhi Patel, Nitin Savaliya, Raghawendra Kumar, Dinesh Kumar, Fenil Patel, R D Dixit, A M Kadri, Harsh Bakshi, Chaitanya Joshi, Madhvi Joshi |
| EPI_ISL_475049 | GMERS Medical College & Hospital | Gujarat Biotechnology Research Centre | Zarna Patel, Monika Gandhi, Pinal Trivedi, Maharshi Pandya, Nidhi Patel, Nitin Savaliya, Raghawendra Kumar, Dinesh Kumar, Zuber Saiyed, Komal Patel, Labdhi Pandya, Snehal Bagatharia, Meenakshi Shah, Neena Doshi, Varsha Godbole, Pritesh Sabara, Apurvasinh Puvar, Janvi Raval, Afzal Ansari, R D Dixit, A M Kadri, Harsh Bakshi, Chaitanya Joshi, Madhvi Joshi |
| EPI_ISL_475057 | Dr. N. D. Desai Medical College & Hospital | Gujarat Biotechnology Research Centre | Zuber Saiyed, Komal Patel, Labdhi Pandya, Supreet Prabhu, Snehal Bagatharia, Jigar Gusani, J G Buch, Pritesh Sabara, Apurvasinh Puvar, Janvi Raval, Zarna Patel, Monika Gandhi, Pinal Trivedi, Maharshi Pandya, Nidhi Patel, Nitin Savaliya, Raghawendra Kumar, Dinesh Kumar, Priti Pandita, R D Dixit, A M Kadri, Harsh Bakshi, Chaitanya Joshi, Madhvi Joshi |
| EPI_ISL_475722, EPI_ISL_475723, EPI_ISL_475724 | Egyptian National Cancer Institute (ENCI) | Egyptian National Cancer Institute (ENCI) | Zekri, Abdel Rahman N, Amer,K.E., Ahmed,O.S., Soliman,H.K., Hafez,M.M., Bahnassy,A.A., Abdelhamid,W., Gad,A., Ali,M., Hassan,W., Samir,M., Raouf,A., Hamdy,M.S., Soliman,M.S., Elsissy,M.H., Elkhateeb,S.M., Ezzelarab,M.H., Abouelhoda, Mohamed |
| EPI_ISL_476822, EPI_ISL_476823, EPI_ISL_476824, EPI_ISL_476825, EPI_ISL_476826, EPI_ISL_476827, EPI_ISL_476828, EPI_ISL_476829, EPI_ISL_476830, EPI_ISL_476831, EPI_ISL_476833, EPI_ISL_476834 | | | |
| see above | Laboratoire des Fièvres Hémorragiques Virales du Benin | Charité-Universitätsmedizin Berlin | Yadouleton, Anges; Sander Anna-Lena; Moreira-Soto Andres; Drexler, Jan Felix |
| EPI_ISL_476857 | GMERS Medical College & Hospital, Gotri, Vadodara | Gujarat Biotechnology Research Centre | Zarna Patel, Monika Gandhi, Pinal Trivedi, Maharshi Pandya, Nidhi Patel, Nitin Savaliya, Raghawendra Kumar, Dinesh Kumar, Zuber Saiyed, Komal Patel, Labdhi Pandya, Afzal Ansari, Nikha Trivedi, Meenakshi Shah, Neena Doshi, Varsha Godbole, Apurvasinh Puvar, Janvi Raval, R D Dixit, A M Kadri, Harsh Bakshi, Chaitanya Joshi, Madhvi Joshi |
| EPI_ISL_476865 | GMERS Medical College and Hospital, Gandhinagar | Gujarat Biotechnology Research Centre | Zuber Saiyed, Komal Patel, Labdhi Pandya, Afzal Ansari, Nikha Trivedi, Seema Bhatt, Gaurishankar Shrimali, Bhavesh Modi, Bharti Rajani, Apurvasinh Puvar, Janvi Raval, Zarna Patel, Monika Gandhi, Pinal Trivedi, Maharshi Pandya, Nidhi Patel, Nitin Savaliya, Raghawendra Kumar, Dinesh Kumar, R D Dixit, A M Kadri, Harsh Bakshi, Chaitanya Joshi, Madhvi Joshi |
| EPI_ISL_476875 | Department of MicroBiology, Government Medical College, Surat | Gujarat Biotechnology Research Centre | Zarna Patel, Monika Gandhi, Pinal Trivedi, Maharshi Pandya, Nidhi Patel, Nitin Savaliya, Raghawendra Kumar, Dinesh Kumar, Zuber Saiyed, Komal Patel, Labdhi Pandya, Afzal Ansari, Nikha Trivedi, Naresh Chauhan, Summaiya Mullan, Amit gamit, Apurvasinh Puvar, Janvi Raval, R D Dixit, A M Kadri, Harsh Bakshi, Chaitanya Joshi, Madhvi Joshi |
| EPI_ISL_476882 | Department of MicroBiology, Government Medical College, Surat | Gujarat Biotechnology Research Centre | Zuber Saiyed, Komal Patel, Labdhi Pandya, Afzal Ansari, Nikha Trivedi, Naresh Chauhan, Summaiya Mullan, Amit gamit, Apurvasinh Puvar, Janvi Raval, Zarna Patel, Monika Gandhi, Pinal Trivedi, Maharshi Pandya, Nidhi Patel, Nitin Savaliya, Raghawendra Kumar, Dinesh Kumar, R D Dixit, A M Kadri, Harsh Bakshi, Chaitanya Joshi, Madhvi Joshi |
| EPI_ISL_477161, EPI_ISL_478672, EPI_ISL_479686, EPI_ISL_479687, EPI_ISL_479688, EPI_ISL_479689, EPI_ISL_479690, EPI_ISL_479691, EPI_ISL_479692, EPI_ISL_479693, EPI_ISL_479694, EPI_ISL_479695, EPI_ISL_479696, EPI_ISL_479697, EPI_ISL_479698, EPI_ISL_479699, EPI_ISL_479700, EPI_ISL_479701, EPI_ISL_479702, EPI_ISL_479703, EPI_ISL_479704, EPI_ISL_479705, EPI_ISL_479706, EPI_ISL_479707, EPI_ISL_479708, EPI_ISL_479709, EPI_ISL_479710, EPI_ISL_479711, EPI_ISL_479712, EPI_ISL_479713, EPI_ISL_479714, EPI_ISL_479715, EPI_ISL_479716, EPI_ISL_479717, EPI_ISL_479718, EPI_ISL_479719, EPI_ISL_479720, EPI_ISL_479721, EPI_ISL_479722, EPI_ISL_479723, EPI_ISL_479724, EPI_ISL_479725, EPI_ISL_479726, EPI_ISL_479727 | | | |
| see above | Egyptian National Cancer Institute (ENCI) | Egyptian National Cancer Institute (ENCI) | Zekri, Abdel Rahman N, Amer,K.E., Ahmed,O.S., Soliman,H.K., Hafez,M.M., Bahnassy,A.A., Abdelhamid,W., Gad,A., Ali,M., Hassan,W., Samir,M., Raouf,A., Hamdy,M.S., Soliman,M.S., Elsissy,M.H., Elkhateeb,S.M., Ezzelarab,M.H., Abouelhoda, Mohamed |
| EPI_ISL_479728 | Egyptian National Cancer Institute (ENCI) | Egyptian National Cancer Institute (ENCI) | Zekri,A.N., Amer,K.E., Ahmed,O.S., Soliman,H.K., Bahnassy,A.A., Ali,M., Abdelhamid,W., Gad,A., Hassan,W., Samir,M., Raouf,A., Hamdy,M.S., Soliman,M.S., Elsissy,M.H., Elkhateeb,S.M., Ezzelarab,M.H., Abouelhoda,M. |
| EPI_ISL_479729, EPI_ISL_479730, EPI_ISL_479731, EPI_ISL_479732, EPI_ISL_479733, EPI_ISL_479734, EPI_ISL_479735 | Egyptian National Cancer Institute (ENCI) | Egyptian National Cancer Institute (ENCI) | Zekri, Abdel Rahman N, Amer,K.E., Ahmed,O.S., Soliman,H.K., Hafez,M.M., Bahnassy,A.A., Abdelhamid,W., Gad,A., Ali,M., Hassan,W., Samir,M., Raouf,A., Hamdy,M.S., Soliman,M.S., Elsissy,M.H., Elkhateeb,S.M., Ezzelarab,M.H., Abouelhoda, Mohamed |
| EPI_ISL_483828 | GMERS Medical College Himmatnagar | Gujarat Biotechnology Research Centre | Zarna Patel, Monika Gandhi, Pinal Trivedi, Maharshi Pandya, Nidhi Patel, Nitin Savaliya, Raghawendra Kumar, Dinesh Kumar, Zuber Saiyed, Komal Patel, Labdhi Pandya, Afzal Ansari, Nikha Trivedi, Himanshu Khatri, Mayur Gandhi, Apurvasinh Puvar, Janvi Raval, R D Dixit, A M Kadri, Harsh Bakshi, Chaitanya Joshi, Madhvi Joshi |
| EPI_ISL_483836 | Pandit Deendayal Upadhyay Government Medical College, Rajkot | Gujarat Biotechnology Research Centre | Zuber Saiyed, Komal Patel, Labdhi Pandya, Afzal Ansari, Nikha Trivedi, Gauravi Dhruv, Arti Trivedi, Apurvasinh Puvar, Janvi Raval, Zarna Patel, Monika Gandhi, Pinal Trivedi, Maharshi Pandya, Nidhi Patel, Nitin Savaliya, Raghawendra Kumar, Dinesh Kumar, R D Dixit, A M Kadri, Harsh Bakshi, Chaitanya Joshi, Madhvi Joshi |
| EPI_ISL_483845 | Pandit Deendayal Upadhyay Government Medical College, Rajkot | Gujarat Biotechnology Research Centre | Zarna Patel, Monika Gandhi, Pinal Trivedi, Maharshi Pandya, Nidhi Patel, Nitin Savaliya, Raghawendra Kumar, Dinesh Kumar, Zuber Saiyed, Komal Patel, Labdhi Pandya, Afzal Ansari, Nikha Trivedi, Gauravi Dhruv, Arti Trivedi, Apurvasinh Puvar, Janvi Raval, R D Dixit, A M Kadri, Harsh Bakshi, Chaitanya Joshi, Madhvi Joshi |
| EPI_ISL_483853 | Department of Microbiology, Government Medical College, Surat | Gujarat Biotechnology Research Centre | Zuber Saiyed, Komal Patel, Labdhi Pandya, Afzal Ansari, Nikha Trivedi, Naresh Chauhan, Summaiya Mullan, Amit gamit, Apurvasinh Puvar, Janvi Raval, Zarna Patel, Monika Gandhi, Pinal Trivedi, Maharshi Pandya, Nidhi Patel, Nitin Savaliya, Raghawendra Kumar, Dinesh Kumar, R D Dixit, A M Kadri, Harsh Bakshi, Chaitanya Joshi, Madhvi Joshi |
| EPI_ISL_483863 | Department of Microbiology, Government Medical College, Surat | Gujarat Biotechnology Research Centre | Zarna Patel, Monika Gandhi, Pinal Trivedi, Maharshi Pandya, Nidhi Patel, Nitin Savaliya, Raghawendra Kumar, Dinesh Kumar, Zuber Saiyed, Komal Patel, Labdhi Pandya, Afzal Ansari, Nikha Trivedi, Naresh Chauhan, Summaiya Mullan, Amit gamit, Apurvasinh Puvar, Janvi Raval, R D Dixit, A M Kadri, Harsh Bakshi, Chaitanya Joshi, Madhvi Joshi |
| EPI_ISL_483871 | Department of Microbiology, Government Medical College, | Gujarat Biotechnology Research Centre | Zuber Saiyed, Komal Patel, Labdhi Pandya, Afzal Ansari, Nikha Trivedi, Naresh Chauhan, Summaiya Mullan, Amit gamit, Apurvasinh Puvar, Janvi Raval, |

| | | | |
|---|---|---|---|
| | Surat | | Zarna Patel, Monika Gandhi, Pinal Trivedi, Maharshi Pandya, Nidhi Patel, Nitin Savaliya, Raghawendra Kumar, Dinesh Kumar, R D Dixit, A M Kadri, Harsh Bakshi, Chaitanya Joshi, Madhvi Joshi |
| EPI_ISL_485401 | Communicable Disease Laboratory, Public Health Directorate | Communicable Disease Laboratory, Public Health Directorate | Zaed,A., Al-Wasti,H., Al-Taif,Z. and Shehab,F. |
| EPI_ISL_486887 | National Influenza Center, Bahrain | National Influenza Center, Bahrain | Zaed,A., Altaif,Z., Shehab,F., AlWasti,H. |
| EPI_ISL_487273 | unknown | Communicable Disease Laboratory, Public Health Directorate | Zaed,A., Shehab,F., AlWasti,H., Altaif,Z. |
| EPI_ISL_495015 | B.J. Medical College and Civil hospital | Gujarat Biotechnology Research Centre | Zarna Patel, Monika Gandhi, Pinal Trivedi, Maharshi Pandya, Nidhi Patel, Nitin Savaliya, Raghawendra Kumar, Dinesh Kumar, Zuber Saiyed, Komal Patel, Labdhi Pandya, Afzal Ansari, Nikha Trivedi, Pranay Shah, Kamlesh J Upadhyay, Sanjay Kapadia, Apurvasinh Puvar, Janvi Raval, R D Dixit, A M Kadri, Harsh Bakshi, Chaitanya Joshi, Madhvi Joshi |
| EPI_ISL_495035 | GMERS Medical College & Hospital, Gotri, Vadodara | Gujarat Biotechnology Research Centre | Zarna Patel, Monika Gandhi, Pinal Trivedi, Maharshi Pandya, Nidhi Patel, Nitin Savaliya, Raghawendra Kumar, Dinesh Kumar, Zuber Saiyed, Komal Patel, Labdhi Pandya, Afzal Ansari, Nikha Trivedi, Meenakshi Shah, Neena Doshi, Varsha Godbole, Apurvasinh Puvar, Janvi Raval, R D Dixit, A M Kadri, Harsh Bakshi, Chaitanya Joshi, Madhvi Joshi |
| EPI_ISL_495043 | GMERS Medical College & Hospital, Gotri, Vadodara | Gujarat Biotechnology Research Centre | Zuber Saiyed, Komal Patel, Labdhi Pandya, Afzal Ansari, Nikha Trivedi, Meenakshi Shah, Neena Doshi, Varsha Godbole, Apurvasinh Puvar, Janvi Raval, Zarna Patel, Monika Gandhi, Pinal Trivedi, Maharshi Pandya, Nidhi Patel, Nitin Savaliya, Raghawendra Kumar, Dinesh Kumar, R D Dixit, A M Kadri, Harsh Bakshi, Chaitanya Joshi, Madhvi Joshi |
| EPI_ISL_495053 | GMERS Medical College & Hospital, Gotri, Vadodara | Gujarat Biotechnology Research Centre | Zarna Patel, Monika Gandhi, Pinal Trivedi, Maharshi Pandya, Nidhi Patel, Nitin Savaliya, Raghawendra Kumar, Dinesh Kumar, Zuber Saiyed, Komal Patel, Labdhi Pandya, Afzal Ansari, Nikha Trivedi, Meenakshi Shah, Neena Doshi, Varsha Godbole, Apurvasinh Puvar, Janvi Raval, R D Dixit, A M Kadri, Harsh Bakshi, Chaitanya Joshi, Madhvi Joshi |
| EPI_ISL_495061 | GMERS Medical College & Hospital, Gotri, Vadodara | Gujarat Biotechnology Research Centre | Zuber Saiyed, Komal Patel, Labdhi Pandya, Afzal Ansari, Nikha Trivedi, Meenakshi Shah, Neena Doshi, Varsha Godbole, Apurvasinh Puvar, Janvi Raval, Zarna Patel, Monika Gandhi, Pinal Trivedi, Maharshi Pandya, Nidhi Patel, Nitin Savaliya, Raghawendra Kumar, Dinesh Kumar, R D Dixit, A M Kadri, Harsh Bakshi, Chaitanya Joshi, Madhvi Joshi |
| EPI_ISL_495072 | Department of MicroBiology, Government Medical College, Surat | Gujarat Biotechnology Research Centre | Zarna Patel, Monika Gandhi, Pinal Trivedi, Maharshi Pandya, Nidhi Patel, Nitin Savaliya, Raghawendra Kumar, Dinesh Kumar, Zuber Saiyed, Komal Patel, Labdhi Pandya, Afzal Ansari, Nikha Trivedi, Naresh Chauhan, Summaiya Mullan, Amit gamit, Apurvasinh Puvar, Janvi Raval, R D Dixit, A M Kadri, Harsh Bakshi, Chaitanya Joshi, Madhvi Joshi |
| EPI_ISL_497950 | Shaoxing CDC | Zhejiang Provincial Center for Disease Control and Prevention | Yin Chen, Yanjun Zhang, Haiyan Mao, Junhang Pan, Xiuyu Lou, Yi Sun, Hao Yan, Zhen Li, Wen Shi |
| EPI_ISL_500946 | GMERS Medical College & Hospital, Gotri, Vadodara | Gujarat Biotechnology Research Centre | Zuber Saiyed, Komal Patel, Labdhi Pandya, Afzal Ansari, Nikha Trivedi, Meenakshi Shah, Neena Doshi, Varsha Godbole, Apurvasinh Puvar, Janvi Raval, Zarna Patel, Monika Gandhi, Pinal Trivedi, Maharshi Pandya, Nidhi Patel, Nitin Savaliya, Raghawendra Kumar, Dinesh Kumar, R D Dixit, A M Kadri, Harsh Bakshi, Chaitanya Joshi, Madhvi Joshi |
| EPI_ISL_500948 | Department of MicroBiology, Government Medical College, Surat | Gujarat Biotechnology Research Centre | Zuber Saiyed, Komal Patel, Labdhi Pandya, Afzal Ansari, Nikha Trivedi, Naresh Chauhan, Summaiya Mullan, Amit gamit, Apurvasinh Puvar, Janvi Raval, Zarna Patel, Monika Gandhi, Pinal Trivedi, Maharshi Pandya, Nidhi Patel, Nitin Savaliya, Raghawendra Kumar, Dinesh Kumar, R D Dixit, A M Kadri, Harsh Bakshi, Chaitanya Joshi, Madhvi Joshi |
| EPI_ISL_501176, EPI_ISL_501177, EPI_ISL_501178, EPI_ISL_501179, EPI_ISL_501180, EPI_ISL_501181, EPI_ISL_501182, EPI_ISL_501183, EPI_ISL_501184, EPI_ISL_501185, EPI_ISL_501186, EPI_ISL_501187, EPI_ISL_501188, EPI_ISL_501189, EPI_ISL_501190, EPI_ISL_501191, EPI_ISL_501192, EPI_ISL_501193, EPI_ISL_501194, EPI_ISL_501195, EPI_ISL_501196, EPI_ISL_501197, EPI_ISL_501198, EPI_ISL_501199, EPI_ISL_501200, EPI_ISL_501201, EPI_ISL_501202, EPI_ISL_501203, EPI_ISL_501204, EPI_ISL_501205, EPI_ISL_501206, EPI_ISL_501207, EPI_ISL_501208, EPI_ISL_501209, EPI_ISL_501210, EPI_ISL_501211, EPI_ISL_501212, EPI_ISL_501213, EPI_ISL_501214, EPI_ISL_501215, EPI_ISL_501216, EPI_ISL_501217, EPI_ISL_501218, EPI_ISL_501219, EPI_ISL_501220, EPI_ISL_501221, EPI_ISL_501222, EPI_ISL_501223, EPI_ISL_501224, EPI_ISL_501225, EPI_ISL_501226, EPI_ISL_501227, EPI_ISL_501228, EPI_ISL_506996, EPI_ISL_506997, EPI_ISL_506998, EPI_ISL_506999, EPI_ISL_507000 | | | |
| see above | Department of Medical Microbiology, University Malaya Medical Centre | Department of Medical Microbiology, Faculty of Medicine, University of Malaya | Yoong Min CHONG, Jennifer Chong, I-Ching SAM, Yoke Fun CHAN, University Malaya Medical Centre COVID Team |
| EPI_ISL_509662 | Georgia Department of Health | Pathogen Discovery, Respiratory Viruses Branch, Division of Viral Diseases, Centers for Disease Control and Prevention | Yan Li, Anna Montmayeur, Jing Zhang, Krista Queen, Anna Uehara, Ying Tao, Rachel Marine, Clinton R. Paden, Haibin Wang, Suxiang Tong |
| EPI_ISL_509663, EPI_ISL_509664, EPI_ISL_509665, EPI_ISL_509666, EPI_ISL_509667, EPI_ISL_509668, EPI_ISL_509670, EPI_ISL_509671, EPI_ISL_509672, EPI_ISL_509673, EPI_ISL_509674, EPI_ISL_509675, EPI_ISL_509676, EPI_ISL_509677, EPI_ISL_509678, EPI_ISL_509679, EPI_ISL_509680, EPI_ISL_509681, EPI_ISL_509682 | | | |
| see above | AR Dept. of Health-Public Health Lab | Pathogen Discovery, Respiratory Viruses Branch, Division of Viral Diseases, Centers for Disease Control and Prevention | Yan Li, Anna Montmayeur, Jing Zhang, Krista Queen, Anna Uehara, Ying Tao, Rachel Marine, Clinton R. Paden, Haibin Wang, Suxiang Tong |
| EPI_ISL_509688 | Alabama Department of Public Health Bureau of Clinical Laboratories | Pathogen Discovery, Respiratory Viruses Branch, Division of Viral Diseases, Centers for Disease Control and Prevention | Ying Tao, Jing Zhang, Krista Queen, Anna Uehara, Yan Li, Clinton Paden, Haibin Wang, Suxiang Tong |
| EPI_ISL_509689, EPI_ISL_509690, EPI_ISL_509691, EPI_ISL_509692, EPI_ISL_509693, EPI_ISL_509694 | Utah Public Health Laboratory | Pathogen Discovery, Respiratory Viruses Branch, Division of Viral Diseases, Centers for Disease Control and Prevention | Ying Tao, Jing Zhang, Krista Queen, Anna Uehara, Yan Li, Clinton Paden, Haibin Wang, Suxiang Tong |
| EPI_ISL_509695, EPI_ISL_509696, EPI_ISL_509697, EPI_ISL_509698, EPI_ISL_509699, EPI_ISL_509700, EPI_ISL_509701, EPI_ISL_509702, EPI_ISL_509703 | Guatemala Ministry of Public Health | Pathogen Discovery, Respiratory Viruses Branch, Division of Viral Diseases, Centers for Disease Control and Prevention | Ying Tao, Jing Zhang, Krista Queen, Anna Uehara, Yan Li, Clinton Paden, Haibin Wang, Suxiang Tong |
| EPI_ISL_509704, EPI_ISL_509705, EPI_ISL_509706 | Wisconsin Department of Health Services | Pathogen Discovery, Respiratory Viruses Branch, Division of Viral Diseases, Centers for Disease Control and Prevention | Ying Tao, Jing Zhang, Krista Queen, Anna Uehara, Yan Li, Clinton Paden, Haibin Wang, Suxiang Tong |
| EPI_ISL_512065 | B.J. Medical College and Civil hospital, Ahmedabad | Gujarat Biotechnology Research Centre | Zuber Saiyed, Komal Patel, Labdhi Pandya, Afzal Ansari, Nikha Trivedi, Pranay Shah, Kamlesh J Upadhyay, Sanjay Kapadia, Apurvasinh Puvar, Janvi Raval, Zarna Patel, Monika Gandhi, Pinal Trivedi, Maharshi Pandya, Nidhi Patel, Nitin Savaliya, Raghawendra Kumar, Dinesh Kumar, R D Dixit, A M Kadri, Harsh Bakshi, Chaitanya Joshi, Madhvi Joshi |
| EPI_ISL_514062, EPI_ISL_514063, EPI_ISL_514064, EPI_ISL_514066, EPI_ISL_514069, EPI_ISL_514070, EPI_ISL_514073, EPI_ISL_514074, EPI_ISL_514077, EPI_ISL_514078, EPI_ISL_514079, EPI_ISL_514084, EPI_ISL_514088, EPI_ISL_514090, EPI_ISL_514091, EPI_ISL_514092 | | | |
| see above | Navy and Marine Corps Public Health Center | Pathogen Discovery, Respiratory Viruses Branch, Division of Viral Diseases, Centers for Disease Control and Prevention | Yan Li, Anna Montmayeur, Krista Queen, Jing Zhang, Ying Tao, Anna Uehara, Rachel Marine, Clinton R. Paden, Haibin Wang, Suxiang Tong |
| EPI_ISL_514436 | Dr. RSS Hospital, Modasa | Gujarat Biotechnology Research Centre | Zarna Patel, Monika Gandhi, Pinal Trivedi, Maharshi Pandya, Nidhi Patel, Nitin Savaliya, Raghawendra Kumar, Dinesh Kumar, Zuber Saiyed, Komal Patel, Labdhi Pandya, Afzal Ansari, Nikha Trivedi, Harsh Chaudhari, Apurvasinh Puvar, Janvi Raval, R D Dixit, A M Kadri, Harsh Bakshi, Chaitanya Joshi, Madhvi Joshi |
| EPI_ISL_514584 | B.J. Medical College and Civil hospital, Ahmedabad | Gujarat Biotechnology Research Centre | Zuber Saiyed, Komal Patel, Labdhi Pandya, Afzal Ansari, Nikha Trivedi, Pranay Shah, Kamlesh J Upadhyay, Sanjay Kapadia, Apurvasinh Puvar, Janvi Raval, Zarna Patel, Monika Gandhi, Pinal Trivedi, Maharshi Pandya, Nidhi Patel, Nitin Savaliya, Raghawendra Kumar, Dinesh Kumar, R D Dixit, A M Kadri, Harsh Bakshi, Chaitanya Joshi, Madhvi Joshi |
| EPI_ISL_514594 | B.J. Medical College and Civil hospital, Ahmedabad | Gujarat Biotechnology Research Centre | Zarna Patel, Monika Gandhi, Pinal Trivedi, Maharshi Pandya, Nidhi Patel, Nitin Savaliya, Raghawendra Kumar, Dinesh Kumar, Zuber Saiyed, Komal Patel, Labdhi Pandya, Afzal Ansari, Nikha Trivedi, Pranay Shah, Kamlesh J Upadhyay, Sanjay Kapadia, Apurvasinh Puvar, Janvi Raval, R D Dixit, A M Kadri, Harsh Bakshi, Chaitanya Joshi, Madhvi Joshi |
| EPI_ISL_514602 | B.J. Medical College and Civil hospital, Ahmedabad | Gujarat Biotechnology Research Centre | Zuber Saiyed, Komal Patel, Labdhi Pandya, Afzal Ansari, Nikha Trivedi, Pranay Shah, Kamlesh J Upadhyay, Sanjay Kapadia, Apurvasinh Puvar, Janvi Raval, Zarna Patel, Monika Gandhi, Pinal Trivedi, Maharshi Pandya, Nidhi Patel, Nitin Savaliya, Raghawendra Kumar, Dinesh Kumar, R D Dixit, A M Kadri, Harsh Bakshi, Chaitanya Joshi, Madhvi Joshi |
| EPI_ISL_514751 | CoronaNet Lab- TaskForce Regione Campania, CEINGE | CoronaNet Lab- TaskForce Regione Campania, CEINGE | Zollo,M., Ferrucci,V., Kong,Dy., Asadzadeh,F., Marrone,L.,Siciliano,R., Cerino,R., Fusco,G., Comegna,M., Boccia,A.,Viscardi,M., Borriello,G., Brandi,S., |

| | | | |
|---|---|---|---|
| | Biotecnologie Avanzate, Via G. Salvatore | Biotecnologie Avanzate, Via G. Salvatore | Tiberio,C., Atripaldi,L.,Paolella,G., Castaldo,G., Pascarella,S., Bianchi,M., Chiariotti,L.,Lee,J.M., Jung,J.H., Yun,K.S. and Kim,H.Y. |
| EPI_ISL_524426 | Egyptian National Cancer Institute (ENCI) | Egyptian National Cancer Institute (ENCI) | Zekri, Abdel Rahman N., Amer,K.E., Ahmed,O.S., Soliman,H.K., Ali,M.A., Hassan,W.A., Mahmoud,A.A., Khattab,A.A., Hafez,M.M., Abouelhoda, Mohamed |
| EPI_ISL_524427 | Egyptian National Cancer Institute (ENCI) | Egyptian National Cancer Institute (ENCI) | Zekri, Abdel Rahman N, Amer,K.E., Ahmed,O.S., Soliman,H.K., Hafez,M.M., Bahnassy,A.A., Abdelhamid,W., Gad,A., Ali,M., Hassan,W., Samir,M., Raouf,A., Hamdy,M.S., Soliman,M.S., Elsissy,M.H., Elkhateeb,S.M., Ezzelarab,M.H., Abouelhoda, Mohamed |
| EPI_ISL_524715 | B.J. Medical College and Civil hospital, Ahmedabad | Gujarat Biotechnology Research Centre | Zarna Patel, Monika Gandhi, Pinal Trivedi, Maharshi Pandya, Nidhi Patel, Nitin Savaliya, Raghawendra Kumar, Dinesh Kumar, Zuber Saiyed, Komal Patel, Labdhi Pandya, Afzal Ansari, Nikha Trivedi, Pranay Shah, Kamlesh J Upadhyay, Sanjay Kapadia, Apurvasinh Puvar, Janvi Raval, R D Dixit, A M Kadri, Harsh Bakshi, Chaitanya Joshi, Madhvi Joshi |
| EPI_ISL_524723 | B.J. Medical College and Civil hospital, Ahmedabad | Gujarat Biotechnology Research Centre | Zuber Saiyed, Komal Patel, Labdhi Pandya, Afzal Ansari, Nikha Trivedi, Pranay Shah, Kamlesh J Upadhyay, Sanjay Kapadia, Apurvasinh Puvar, Janvi Raval, Zarna Patel, Monika Gandhi, Pinal Trivedi, Maharshi Pandya, Nidhi Patel, Nitin Savaliya, Raghawendra Kumar, Dinesh Kumar, R D Dixit, A M Kadri, Harsh Bakshi, Chaitanya Joshi, Madhvi Joshi |
| EPI_ISL_524737 | GMERS Medical College and Hospital, Gandhinagar | Gujarat Biotechnology Research Centre | Zarna Patel, Monika Gandhi, Pinal Trivedi, Maharshi Pandya, Nidhi Patel, Nitin Savaliya, Raghawendra Kumar, Dinesh Kumar, Zuber Saiyed, Komal Patel, Labdhi Pandya, Afzal Ansari, Nikha Trivedi, Seema Bhatt, Gaurishankar Shrimali, Bhavesh Modi, Bharti Rajani, Apurvasinh Puvar, Janvi Raval, R D Dixit, A M Kadri, Harsh Bakshi, Chaitanya Joshi, Madhvi Joshi |
| EPI_ISL_524743 | GMERS Medical College and Hospital, Gandhinagar | Gujarat Biotechnology Research Centre | Zuber Saiyed, Komal Patel, Labdhi Pandya, Afzal Ansari, Nikha Trivedi, Seema Bhatt, Gaurishankar Shrimali, Bhavesh Modi, Bharti Rajani, Apurvasinh Puvar, Janvi Raval, Zarna Patel, Monika Gandhi, Pinal Trivedi, Maharshi Pandya, Nidhi Patel, Nitin Savaliya, Raghawendra Kumar, Dinesh Kumar, R D Dixit, A M Kadri, Harsh Bakshi, Chaitanya Joshi, Madhvi Joshi |
| EPI_ISL_524751 | Sir M P Shah Government Medical College, Jamnagar | Gujarat Biotechnology Research Centre | Zarna Patel, Monika Gandhi, Pinal Trivedi, Maharshi Pandya, Nidhi Patel, Nitin Savaliya, Raghawendra Kumar, Dinesh Kumar, Zuber Saiyed, Komal Patel, Labdhi Pandya, Afzal Ansari, Nikha Trivedi, Binita Aring, Apurvasinh Puvar, Janvi Raval, R D Dixit, A M Kadri, Harsh Bakshi, Chaitanya Joshi, Madhvi Joshi |
| EPI_ISL_524759 | GMERS Medical College and Hospital, Dharpur, Patan | Gujarat Biotechnology Research Centre | Zuber Saiyed, Komal Patel, Labdhi Pandya, Afzal Ansari, Nikha Trivedi, A N Parmar, Apurvasinh Puvar, Janvi Raval, Zarna Patel, Monika Gandhi, Pinal Trivedi, Maharshi Pandya, Nidhi Patel, Nitin Savaliya, Raghawendra Kumar, Dinesh Kumar, R D Dixit, A M Kadri, Harsh Bakshi, Chaitanya Joshi, Madhvi Joshi |
| EPI_ISL_525421 | B.J. Medical College and Civil hospital, Ahmedabad | Gujarat Biotechnology Research Centre | Zarna Patel, Monika Gandhi, Pinal Trivedi, Maharshi Pandya, Nidhi Patel, Nitin Savaliya, Raghawendra Kumar, Dinesh Kumar, Zuber Saiyed, Komal Patel, Labdhi Pandya, Afzal Ansari, Nikha Trivedi, Pranay Shah, Kamlesh J Upadhyay, Sanjay Kapadia, Apurvasinh Puvar, Janvi Raval, R D Dixit, A M Kadri, Harsh Bakshi, Chaitanya Joshi, Madhvi Joshi |
| EPI_ISL_527632, EPI_ISL_527633, EPI_ISL_527634, EPI_ISL_527635, EPI_ISL_527636, EPI_ISL_527637, EPI_ISL_527638, EPI_ISL_527639, EPI_ISL_527640, EPI_ISL_527641, EPI_ISL_527642, EPI_ISL_527643, EPI_ISL_527644, EPI_ISL_527645, EPI_ISL_527646, EPI_ISL_527647, EPI_ISL_527648, EPI_ISL_527649, EPI_ISL_527650, EPI_ISL_527651, EPI_ISL_527652, EPI_ISL_527653, EPI_ISL_527654, EPI_ISL_527655, EPI_ISL_527656, EPI_ISL_527657 | | | |
| see above | AR Dept. of Health-Public Health Lab | Pathogen Discovery, Respiratory Viruses Branch, Division of Viral Diseases, Centers for Disease Control and Prevention | Ying Tao, Jing Zhang, Yan Li, Krista Queen, Anna Uehara, Clinton Paden, Haibin Wang, Suxiang Tong |
| EPI_ISL_527658, EPI_ISL_527659, EPI_ISL_527660, EPI_ISL_527661, EPI_ISL_527662, EPI_ISL_527663, EPI_ISL_527664, EPI_ISL_527665 | GA Department of Public Health Laboratory | Pathogen Discovery, Respiratory Viruses Branch, Division of Viral Diseases, Centers for Disease Control and Prevention | Yan Li, Anna Montmayeur, Jing Zhang, Krista Queen, Ying Tao, Anna Uehara, Rachel Marine, Clinton R. Paden, Haibin Wang, Suxiang Tong |
| EPI_ISL_527667, EPI_ISL_527668, EPI_ISL_527669, EPI_ISL_527670, EPI_ISL_527671, EPI_ISL_527672, EPI_ISL_527673, EPI_ISL_527674, EPI_ISL_527675, EPI_ISL_527676, EPI_ISL_527677, EPI_ISL_527678, EPI_ISL_527679, EPI_ISL_527680, EPI_ISL_527681, EPI_ISL_527682, EPI_ISL_527683, EPI_ISL_527684, EPI_ISL_527685, EPI_ISL_527686, EPI_ISL_527687, EPI_ISL_527688, EPI_ISL_527689, EPI_ISL_527690, EPI_ISL_527691, EPI_ISL_527693, EPI_ISL_527694, EPI_ISL_527695, EPI_ISL_527696, EPI_ISL_527697, EPI_ISL_527698, EPI_ISL_527700, EPI_ISL_527702, EPI_ISL_527703, EPI_ISL_527704, EPI_ISL_527705, EPI_ISL_527706, EPI_ISL_527707, EPI_ISL_527708, EPI_ISL_527709, EPI_ISL_527710, EPI_ISL_527711, EPI_ISL_527712, EPI_ISL_527713, EPI_ISL_527715, EPI_ISL_527718, EPI_ISL_527719, EPI_ISL_527721, EPI_ISL_527722, EPI_ISL_527723, EPI_ISL_527725, EPI_ISL_527726, EPI_ISL_527727, EPI_ISL_527728, EPI_ISL_527729, EPI_ISL_527730, EPI_ISL_527732, EPI_ISL_527733, EPI_ISL_527736 | | | |
| see above | MN PHL Division, Minnesota Department of Health | Pathogen Discovery, Respiratory Viruses Branch, Division of Viral Diseases, Centers for Disease Control and Prevention | Yan Li, Anna Montmayeur, Jing Zhang, Krista Queen, Ying Tao, Anna Uehara, Rachel Marine, Clinton R. Paden, Haibin Wang, Suxiang Tong |
| EPI_ISL_527737 | WI State Laboratory of Hygiene | Pathogen Discovery, Respiratory Viruses Branch, Division of Viral Diseases, Centers for Disease Control and Prevention | Ying Tao, Jing Zhang, Yan Li, Krista Queen, Anna Uehara, Clinton Paden, Haibin Wang, Suxiang Tong |
| EPI_ISL_527871 | Institute of Medical Biology, Chinese Academy of Medical Sciences and Peking Union Medical College | Institute of Medical Biology, Chinese Academy of Medical Sciences and Peking Union Medical College | Xu,X., Liao,Y., Wang,L., Zhou,X., Xie,Z., Chen,H., Fan,S., Liu,L.,Zheng,H., Jiang,G. and Li,Q. |
| EPI_ISL_528746 | Immanuel Hospital | Molecular Genetics Laboratory-Faculty of Medicine-Universitas Padjadjaran; School of Life Sciences and Technology & School of Pharmacy-Institut Teknologi Bandung; Laboratorium Kesehatan Provinsi Jawa Barat | Yunia Sribudiani,Tri Hanggono Achmad, Mas Rizky A.A. Syamsunarno, Fensi Amalina, Catur Riani, Azzania Fibriani, Husna Nugrahapraja, Marselina Irasonia Tan, Tarwadi, Ema Rahmawati, Savira Ekawardhani, Hesti Lina Wiraswati, Ryan Bayusantika Ristandi, Rifky Waluyajati Rachman, Cut Nur Cinthia Alamanda, Lia Faridah, Hammam Riza, Sony Solistia Wirawan, Agung Eru Wibowo, Irvan Faizal |
| EPI_ISL_528751 | Santo Borromeus Hospital | Molecular Genetics Laboratory-Faculty of Medicine-Universitas Padjadjaran; School of Life Sciences and Technology & School of Pharmacy-Institut Teknologi Bandung; Laboratorium Kesehatan Provinsi Jawa Barat | Yunia Sribudiani, Tri Hanggono Achmad, Mas Rizky A.A. Syamsunarno, Fensi Amalina, Catur Riani, Azzania Fibriani, Husna Nugrahapraja, Marselina Irasonia Tan, Tarwadi, Ema Rahmawati, Savira Ekawardhani, Hesti Lina Wiraswati, Ryan Bayusantika Ristandi, Rifky Waluyajati Rachman, Cut Nur Cinthia Alamanda, Lia Faridah, Hammam Riza, Sony Solistia Wirawan, Agung Eru Wibowo, Irvan Faizal |
| EPI_ISL_529141, EPI_ISL_529142, EPI_ISL_529143, EPI_ISL_529144, EPI_ISL_529145 | Egyptian National Cancer Institute (ENCI) | Egyptian National Cancer Institute (ENCI) | Zekri, Abdel Rahman N., Amer,K.E., Ahmed,O.S., Soliman,H.K., Ali,M.A., Hassan,W.A., Mahmoud,A.A., Khattab,A.A., Hafez,M.M., Abouelhoda, Mohamed |
| EPI_ISL_535305, EPI_ISL_535306, EPI_ISL_535307, EPI_ISL_535308, EPI_ISL_535309, EPI_ISL_535310, EPI_ISL_535311, EPI_ISL_535312, EPI_ISL_535313, EPI_ISL_535314, EPI_ISL_535315, EPI_ISL_535316, EPI_ISL_535317, EPI_ISL_535318, EPI_ISL_535319, EPI_ISL_535320, EPI_ISL_535321, EPI_ISL_535322, EPI_ISL_535323, EPI_ISL_535324, EPI_ISL_535325, EPI_ISL_535326, EPI_ISL_535327, EPI_ISL_535328, EPI_ISL_535329, EPI_ISL_535330, EPI_ISL_535331, EPI_ISL_535332, EPI_ISL_535333, EPI_ISL_535334, EPI_ISL_535335, EPI_ISL_535336, EPI_ISL_535337, EPI_ISL_535338, EPI_ISL_535339, EPI_ISL_535340, EPI_ISL_535343, EPI_ISL_535347, EPI_ISL_535349, EPI_ISL_535350, EPI_ISL_535352, EPI_ISL_535353, EPI_ISL_535354, EPI_ISL_535355, EPI_ISL_535356, EPI_ISL_535358 | | | |
| see above | LA Office of Public Health Laboratories | Pathogen Discovery, Respiratory Viruses Branch, Division of Viral Diseases, Centers for Disease Control and Prevention | Ying Tao, Jing Zhang, Yan Li, Krista Queen, Anna Uehara, Clinton Paden, Haibin Wang, Suxiang Tong |
| EPI_ISL_535651, EPI_ISL_535653, EPI_ISL_535655, EPI_ISL_535656, EPI_ISL_535657, EPI_ISL_535660 | AR Dept. of Health-Public Health Lab | Pathogen Discovery, Respiratory Viruses Branch, Division of Viral Diseases, Centers for Disease Control and Prevention | Yan Li, Jing Zhang, Ying Tao, Krista Queen, Brian Lynch, Anna Uehara, Clinton R. Paden, Rachel Marine, Haibin Wang, Suxiang Tong |
| EPI_ISL_535662, EPI_ISL_535663, EPI_ISL_535664, EPI_ISL_535665, EPI_ISL_535666, EPI_ISL_535667, EPI_ISL_535668, EPI_ISL_535669, EPI_ISL_535670, EPI_ISL_535671, EPI_ISL_535672, EPI_ISL_535673, EPI_ISL_535674, EPI_ISL_535676, EPI_ISL_535677, EPI_ISL_535678, EPI_ISL_535679, EPI_ISL_535680, EPI_ISL_535681, EPI_ISL_535682, EPI_ISL_535683, EPI_ISL_535684, EPI_ISL_535685, EPI_ISL_535686, EPI_ISL_535687, EPI_ISL_535688, EPI_ISL_535689, EPI_ISL_535690, EPI_ISL_535691, EPI_ISL_535692, EPI_ISL_535693, EPI_ISL_535694, EPI_ISL_535695, EPI_ISL_535696, EPI_ISL_535697, EPI_ISL_535698, EPI_ISL_535699, EPI_ISL_535700, EPI_ISL_535702 | | | |
| see above | CDPH, Microbial Diseases Laboratory | Pathogen Discovery, Respiratory Viruses Branch, Division of Viral Diseases, Centers for Disease Control and Prevention | Yan Li, Jing Zhang, Ying Tao, Krista Queen, Brian Lynch, Anna Uehara, Clinton R. Paden, Rachel Marine, Haibin Wang, Suxiang Tong |
| EPI_ISL_535703 | CDPH, Microbial Diseases Laboratory | Pathogen Discovery, Respiratory Viruses Branch, Division of Viral Diseases, Centers for Disease Control and Prevention | Ying Tao, Jing Zhang, Yan Li, Krista Queen, Anna Uehara, Clinton R. Paden, Haibin Wang, Suxiang Tong |
| EPI_ISL_535704, EPI_ISL_535705 | CDPH, Microbial Diseases Laboratory | Pathogen Discovery, Respiratory Viruses Branch, Division of Viral Diseases, Centers for Disease Control and Prevention | Yan Li, Jing Zhang, Ying Tao, Krista Queen, Brian Lynch, Anna Uehara, Clinton R. Paden, Rachel Marine, Haibin Wang, Suxiang Tong |
| EPI_ISL_535706 | CDPH, Microbial Diseases Laboratory | Pathogen Discovery, Respiratory Viruses Branch, Division of Viral Diseases, Centers for Disease Control and Prevention | Ying Tao, Jing Zhang, Yan Li, Krista Queen, Anna Uehara, Clinton R. Paden, Haibin Wang, Suxiang Tong |

| | | | |
|---|---|---|---|
| EPI_ISL_535707, EPI_ISL_535709, EPI_ISL_535710, EPI_ISL_535711, EPI_ISL_535712, EPI_ISL_535713, EPI_ISL_535714, EPI_ISL_535715 | CDPH, Microbial Diseases Laboratory | Pathogen Discovery, Respiratory Viruses Branch, Division of Viral Diseases, Centers for Disease Control and Prevention | Yan Li, Jing Zhang, Ying Tao, Krista Queen, Brian Lynch, Anna Uehara, Clinton R. Paden, Rachel Marine, Haibin Wang, Suxiang Tong |
| EPI_ISL_536789 | Southern Community Labs Dunedin | Institute of Environmental Science and Research (ESR) | Xiaoyun Ren, Matt Storey, Nikki Freed, Muhammad Faisal, Jing Wang, Hermes Perez, Anja Werno, Antje van der Linden, Arlo Upton, Chris Mansell, David Hammer, Dragana Drinkovic, Gary McAuliffe, Hana Sofia Andersson, Jill Sherwood, Josh Freeman, Julia Howard, Mary DeAlmeida, Matt Blakiston, Matthew Rogers, Max Bloomfield, Michael Addidle, Michelle Balm, Sally Roberts, Sarah Jefferies, Sharmini Muttaiyah, Susan Morpeth, Susan Taylor, Timothy Blackmore, Vani Sathyendran, Veronica Playle, Virginia Hope, Erasmus Smit, Lauren Jelly, Olin Silander, Joep de Ligt |
| EPI_ISL_540431 | IN State Department of Health Laboratory Services | Pathogen Discovery, Respiratory Viruses Branch, Division of Viral Diseases, Centers for Disease Control and Prevention | Yan Li, Jing Zhang, Anna Montmayeur, Krista Queen,Ying Tao, Anna Uehara, Clinton R. Paden, Rachel Marine, Haibin Wang, Suxiang Tong |
| EPI_ISL_540432, EPI_ISL_540433 | TN Division of Laboratory Services | Pathogen Discovery, Respiratory Viruses Branch, Division of Viral Diseases, Centers for Disease Control and Prevention | Yan Li, Jing Zhang, Anna Montmayeur, Krista Queen,Ying Tao, Anna Uehara, Clinton R. Paden, Rachel Marine, Haibin Wang, Suxiang Tong |
| EPI_ISL_540434 | TN Division of Laboratory Services | Pathogen Discovery, Respiratory Viruses Branch, Division of Viral Diseases, Centers for Disease Control and Prevention | Ying Tao, Yan Li, Jing Zhang, Krista Queen, Anna Uehara, Clinton R. Paden, Haibin Wang, Suxiang Tong |
| EPI_ISL_540436, EPI_ISL_540437, EPI_ISL_540438 | WI State Laboratory of Hygiene | Pathogen Discovery, Respiratory Viruses Branch, Division of Viral Diseases, Centers for Disease Control and Prevention | Yan Li, Jing Zhang, Anna Montmayeur, Krista Queen,Ying Tao, Anna Uehara, Clinton R. Paden, Rachel Marine, Haibin Wang, Suxiang Tong |
| EPI_ISL_540439 | WVDHHR - Office of Laboratory Services | Pathogen Discovery, Respiratory Viruses Branch, Division of Viral Diseases, Centers for Disease Control and Prevention | Ying Tao, Yan Li, Jing Zhang, Krista Queen, Anna Uehara, Clinton R. Paden, Haibin Wang, Suxiang Tong |
| EPI_ISL_540441 | NC State Laboratory of Public Health | Pathogen Discovery, Respiratory Viruses Branch, Division of Viral Diseases, Centers for Disease Control and Prevention | Yan Li, Jing Zhang, Anna Montmayeur, Krista Queen,Ying Tao, Anna Uehara, Clinton R. Paden, Rachel Marine, Haibin Wang, Suxiang Tong |
| EPI_ISL_547969 | LabPLUS | Institute of Environmental Science and Research (ESR) | Xiaoyun Ren, Matt Storey, Nikki Freed, Muhammad Faisal, Jing Wang, Hermes Perez, Anja Werno, Antje van der Linden, Arlo Upton, Chris Mansell, David Hammer, Dragana Drinkovic, Gary McAuliffe, Hana Sofia Andersson, James Ussher, Jill Sherwood, Josh Freeman, Julia Howard, Juliet Elvy, Mary DeAlmeida, Matt Blakiston, Matthew Rogers, Max Bloomfield, Michael Addidle, Michelle Balm, Sally Roberts, Sarah Jefferies, Sharmini Muttaiyah, Susan Morpeth, Susan Taylor, Timothy Blackmore, Vani Sathyendran, Veronica Playle, Virginia Hope, Erasmus Smit, Lauren Jelly, Olin Silander, Joep de Ligt |
| EPI_ISL_547970, EPI_ISL_547971, EPI_ISL_547972, EPI_ISL_547973, EPI_ISL_547974, EPI_ISL_547975 | LabTests | Institute of Environmental Science and Research (ESR) | Xiaoyun Ren, Matt Storey, Nikki Freed, Muhammad Faisal, Jing Wang, Hermes Perez, Anja Werno, Antje van der Linden, Arlo Upton, Chris Mansell, David Hammer, Dragana Drinkovic, Gary McAuliffe, Hana Sofia Andersson, James Ussher, Jill Sherwood, Josh Freeman, Julia Howard, Juliet Elvy, Mary DeAlmeida, Matt Blakiston, Matthew Rogers, Max Bloomfield, Michael Addidle, Michelle Balm, Sally Roberts, Sarah Jefferies, Sharmini Muttaiyah, Susan Morpeth, Susan Taylor, Timothy Blackmore, Vani Sathyendran, Veronica Playle, Virginia Hope, Erasmus Smit, Lauren Jelly, Olin Silander, Joep de Ligt |
| EPI_ISL_547976 | LabPLUS | Institute of Environmental Science and Research (ESR) | Xiaoyun Ren, Matt Storey, Nikki Freed, Muhammad Faisal, Jing Wang, Hermes Perez, Anja Werno, Antje van der Linden, Arlo Upton, Chris Mansell, David Hammer, Dragana Drinkovic, Gary McAuliffe, Hana Sofia Andersson, James Ussher, Jill Sherwood, Josh Freeman, Julia Howard, Juliet Elvy, Mary DeAlmeida, Matt Blakiston, Matthew Rogers, Max Bloomfield, Michael Addidle, Michelle Balm, Sally Roberts, Sarah Jefferies, Sharmini Muttaiyah, Susan Morpeth, Susan Taylor, Timothy Blackmore, Vani Sathyendran, Veronica Playle, Virginia Hope, Erasmus Smit, Lauren Jelly, Olin Silander, Joep de Ligt |
| EPI_ISL_547977 | LabTests | Institute of Environmental Science and Research (ESR) | Xiaoyun Ren, Matt Storey, Nikki Freed, Muhammad Faisal, Jing Wang, Hermes Perez, Anja Werno, Antje van der Linden, Arlo Upton, Chris Mansell, David Hammer, Dragana Drinkovic, Gary McAuliffe, Hana Sofia Andersson, James Ussher, Jill Sherwood, Josh Freeman, Julia Howard, Juliet Elvy, Mary DeAlmeida, Matt Blakiston, Matthew Rogers, Max Bloomfield, Michael Addidle, Michelle Balm, Sally Roberts, Sarah Jefferies, Sharmini Muttaiyah, Susan Morpeth, Susan Taylor, Timothy Blackmore, Vani Sathyendran, Veronica Playle, Virginia Hope, Erasmus Smit, Lauren Jelly, Olin Silander, Joep de Ligt |
| EPI_ISL_547978, EPI_ISL_547979 | LabPLUS | Institute of Environmental Science and Research (ESR) | Xiaoyun Ren, Matt Storey, Nikki Freed, Muhammad Faisal, Jing Wang, Hermes Perez, Anja Werno, Antje van der Linden, Arlo Upton, Chris Mansell, David Hammer, Dragana Drinkovic, Gary McAuliffe, Hana Sofia Andersson, James Ussher, Jill Sherwood, Josh Freeman, Julia Howard, Juliet Elvy, Mary DeAlmeida, Matt Blakiston, Matthew Rogers, Max Bloomfield, Michael Addidle, Michelle Balm, Sally Roberts, Sarah Jefferies, Sharmini Muttaiyah, Susan Morpeth, Susan Taylor, Timothy Blackmore, Vani Sathyendran, Veronica Playle, Virginia Hope, Erasmus Smit, Lauren Jelly, Olin Silander, Joep de Ligt |
| EPI_ISL_547980, EPI_ISL_547981, EPI_ISL_547982, EPI_ISL_547983, EPI_ISL_547984, EPI_ISL_547985 | Canterbury Health Laboratories | Institute of Environmental Science and Research (ESR) | Xiaoyun Ren, Matt Storey, Nikki Freed, Muhammad Faisal, Jing Wang, Hermes Perez, Anja Werno, Antje van der Linden, Arlo Upton, Chris Mansell, David Hammer, Dragana Drinkovic, Gary McAuliffe, Hana Sofia Andersson, James Ussher, Jill Sherwood, Josh Freeman, Julia Howard, Juliet Elvy, Mary DeAlmeida, Matt Blakiston, Matthew Rogers, Max Bloomfield, Michael Addidle, Michelle Balm, Sally Roberts, Sarah Jefferies, Sharmini Muttaiyah, Susan Morpeth, Susan Taylor, Timothy Blackmore, Vani Sathyendran, Veronica Playle, Virginia Hope, Erasmus Smit, Lauren Jelly, Olin Silander, Joep de Ligt |
| EPI_ISL_547986, EPI_ISL_547987, EPI_ISL_547988, EPI_ISL_547989 | LabTests | Institute of Environmental Science and Research (ESR) | Xiaoyun Ren, Matt Storey, Nikki Freed, Muhammad Faisal, Jing Wang, Hermes Perez, Anja Werno, Antje van der Linden, Arlo Upton, Chris Mansell, David Hammer, Dragana Drinkovic, Gary McAuliffe, Hana Sofia Andersson, James Ussher, Jill Sherwood, Josh Freeman, Julia Howard, Juliet Elvy, Mary DeAlmeida, Matt Blakiston, Matthew Rogers, Max Bloomfield, Michael Addidle, Michelle Balm, Sally Roberts, Sarah Jefferies, Sharmini Muttaiyah, Susan Morpeth, Susan Taylor, Timothy Blackmore, Vani Sathyendran, Veronica Playle, Virginia Hope, Erasmus Smit, Lauren Jelly, Olin Silander, Joep de Ligt |
| EPI_ISL_547990, EPI_ISL_547991, EPI_ISL_547992 | LabPLUS | Institute of Environmental Science and Research (ESR) | Xiaoyun Ren, Matt Storey, Nikki Freed, Muhammad Faisal, Jing Wang, Hermes Perez, Anja Werno, Antje van der Linden, Arlo Upton, Chris Mansell, David Hammer, Dragana Drinkovic, Gary McAuliffe, Hana Sofia Andersson, James Ussher, Jill Sherwood, Josh Freeman, Julia Howard, Juliet Elvy, Mary DeAlmeida, Matt Blakiston, Matthew Rogers, Max Bloomfield, Michael Addidle, Michelle Balm, Sally Roberts, Sarah Jefferies, Sharmini Muttaiyah, Susan Morpeth, Susan Taylor, Timothy Blackmore, Vani Sathyendran, Veronica Playle, Virginia Hope, Erasmus Smit, Lauren Jelly, Olin Silander, Joep de Ligt |
| EPI_ISL_547993, EPI_ISL_547994, EPI_ISL_547995 | LabTests | Institute of Environmental Science and Research (ESR) | Xiaoyun Ren, Matt Storey, Nikki Freed, Muhammad Faisal, Jing Wang, Hermes Perez, Anja Werno, Antje van der Linden, Arlo Upton, Chris Mansell, David Hammer, Dragana Drinkovic, Gary McAuliffe, Hana Sofia Andersson, James Ussher, Jill Sherwood, Josh Freeman, Julia Howard, Juliet Elvy, Mary DeAlmeida, Matt Blakiston, Matthew Rogers, Max Bloomfield, Michael Addidle, Michelle Balm, Sally Roberts, Sarah Jefferies, Sharmini Muttaiyah, Susan Morpeth, Susan Taylor, Timothy Blackmore, Vani Sathyendran, Veronica Playle, Virginia Hope, Erasmus Smit, Lauren Jelly, Olin Silander, Joep de Ligt |
| EPI_ISL_548001, EPI_ISL_548002, EPI_ISL_548003, EPI_ISL_548004, EPI_ISL_548005, EPI_ISL_548006, EPI_ISL_548007, EPI_ISL_548008, EPI_ISL_548009, EPI_ISL_548010, EPI_ISL_548011, EPI_ISL_548012 | | | |
| see above | Middlemore Hospital | Institute of Environmental Science and Research (ESR) | Xiaoyun Ren, Matt Storey, Nikki Freed, Muhammad Faisal, Jing Wang, Hermes Perez, Anja Werno, Antje van der Linden, Arlo Upton, Chris Mansell, David Hammer, Dragana Drinkovic, Gary McAuliffe, Hana Sofia Andersson, James Ussher, Jill Sherwood, Josh Freeman, Julia Howard, Juliet Elvy, Mary DeAlmeida, Matt Blakiston, Matthew Rogers, Max Bloomfield, Michael Addidle, Michelle Balm, Sally Roberts, Sarah Jefferies, Sharmini Muttaiyah, Susan Morpeth, Susan Taylor, Timothy Blackmore, Vani Sathyendran, Veronica Playle, Virginia Hope, Erasmus Smit, Lauren Jelly, Olin Silander, Joep de Ligt |
| EPI_ISL_548013, EPI_ISL_548014, EPI_ISL_548015, EPI_ISL_548016 | LabTests | Institute of Environmental Science and Research (ESR) | Xiaoyun Ren, Matt Storey, Nikki Freed, Muhammad Faisal, Jing Wang, Hermes Perez, Anja Werno, Antje van der Linden, Arlo Upton, Chris Mansell, David Hammer, Dragana Drinkovic, Gary McAuliffe, Hana Sofia Andersson, James Ussher, Jill Sherwood, Josh Freeman, Julia Howard, Juliet Elvy, Mary DeAlmeida, Matt Blakiston, Matthew Rogers, Max Bloomfield, Michael Addidle, Michelle Balm, Sally Roberts, Sarah Jefferies, Sharmini Muttaiyah, Susan Morpeth, Susan Taylor, Timothy Blackmore, Vani Sathyendran, Veronica Playle, Virginia Hope, Erasmus Smit, Lauren Jelly, Olin Silander, Joep de Ligt |
| EPI_ISL_548017 | LabPLUS | Institute of Environmental Science and Research (ESR) | Xiaoyun Ren, Matt Storey, Nikki Freed, Muhammad Faisal, Jing Wang, Hermes Perez, Anja Werno, Antje van der Linden, Arlo Upton, Chris Mansell, David Hammer, Dragana Drinkovic, Gary McAuliffe, Hana Sofia Andersson, James Ussher, Jill Sherwood, Josh Freeman, Julia Howard, Juliet Elvy, Mary DeAlmeida, Matt Blakiston, Matthew Rogers, Max Bloomfield, Michael Addidle, Michelle Balm, Sally Roberts, Sarah Jefferies, Sharmini Muttaiyah, Susan Morpeth, Susan Taylor, Timothy Blackmore, Vani Sathyendran, Veronica Playle, Virginia Hope, Erasmus Smit, Lauren Jelly, Olin Silander, Joep de Ligt |
| EPI_ISL_548018, EPI_ISL_548019, EPI_ISL_548020 | North Shore Hospital | Institute of Environmental Science and Research (ESR) | Xiaoyun Ren, Matt Storey, Nikki Freed, Muhammad Faisal, Jing Wang, Hermes Perez, Anja Werno, Antje van der Linden, Arlo Upton, Chris Mansell, David Hammer, Dragana Drinkovic, Gary McAuliffe, Hana Sofia Andersson, James Ussher, Jill Sherwood, Josh Freeman, Julia Howard, Juliet Elvy, Mary DeAlmeida, Matt Blakiston, Matthew Rogers, Max Bloomfield, Michael Addidle, Michelle Balm, Sally Roberts, Sarah Jefferies, Sharmini Muttaiyah, Susan Morpeth, Susan Taylor, Timothy Blackmore, Vani Sathyendran, Veronica Playle, Virginia Hope, Erasmus Smit, Lauren Jelly, Olin Silander, Joep de Ligt |
| EPI_ISL_548021, EPI_ISL_548022, EPI_ISL_548023, EPI_ISL_548024, EPI_ISL_548025, EPI_ISL_548026, | LabPLUS | Institute of Environmental Science and Research (ESR) | Xiaoyun Ren, Matt Storey, Nikki Freed, Muhammad Faisal, Jing Wang, Hermes Perez, Anja Werno, Antje van der Linden, Arlo Upton, Chris Mansell, David Hammer, Dragana Drinkovic, Gary McAuliffe, Hana Sofia Andersson, James Ussher, Jill Sherwood, Josh Freeman, Julia Howard, Juliet Elvy, Mary DeAlmeida, Matt Blakiston, Matthew Rogers, Max Bloomfield, Michael Addidle, Michelle Balm, Sally Roberts, Sarah Jefferies, Sharmini Muttaiyah, Susan |

| | | | |
|---|---|---|---|
| EPI_ISL_548027 | | | Morpeth, Susan Taylor, Timothy Blackmore, Vani Sathyendran, Veronica Playle, Virginia Hope, Erasmus Smit, Lauren Jelly, Olin Silander, Joep de Ligt |
| EPI_ISL_548028, EPI_ISL_548029, EPI_ISL_548030 | North Shore Hospital | Institute of Environmental Science and Research (ESR) | Xiaoyun Ren, Matt Storey, Nikki Freed, Muhammad Faisal, Jing Wang, Hermes Perez, Anja Werno, Antje van der Linden, Arlo Upton, Chris Mansell, David Hammer, Dragana Drinkovic, Gary McAuliffe, Hana Sofia Andersson, James Ussher, Jill Sherwood, Josh Freeman, Julia Howard, Juliet Elvy, Mary DeAlmeida, Matt Blakiston, Matthew Rogers, Max Bloomfield, Michael Addidle, Michelle Balm, Sally Roberts, Sarah Jefferies, Sharmini Muttaiyah, Susan Morpeth, Susan Taylor, Timothy Blackmore, Vani Sathyendran, Veronica Playle, Virginia Hope, Erasmus Smit, Lauren Jelly, Olin Silander, Joep de Ligt |
| EPI_ISL_548031, EPI_ISL_548032, EPI_ISL_548033, EPI_ISL_548034, EPI_ISL_548035, EPI_ISL_548036, EPI_ISL_548037 | Middlemore Hospital | Institute of Environmental Science and Research (ESR) | Xiaoyun Ren, Matt Storey, Nikki Freed, Muhammad Faisal, Jing Wang, Hermes Perez, Anja Werno, Antje van der Linden, Arlo Upton, Chris Mansell, David Hammer, Dragana Drinkovic, Gary McAuliffe, Hana Sofia Andersson, James Ussher, Jill Sherwood, Josh Freeman, Julia Howard, Juliet Elvy, Mary DeAlmeida, Matt Blakiston, Matthew Rogers, Max Bloomfield, Michael Addidle, Michelle Balm, Sally Roberts, Sarah Jefferies, Sharmini Muttaiyah, Susan Morpeth, Susan Taylor, Timothy Blackmore, Vani Sathyendran, Veronica Playle, Virginia Hope, Erasmus Smit, Lauren Jelly, Olin Silander, Joep de Ligt |
| EPI_ISL_548040, EPI_ISL_548041 | LabPLUS | Institute of Environmental Science and Research (ESR) | Xiaoyun Ren, Matt Storey, Nikki Freed, Muhammad Faisal, Jing Wang, Hermes Perez, Anja Werno, Antje van der Linden, Arlo Upton, Chris Mansell, David Hammer, Dragana Drinkovic, Gary McAuliffe, Hana Sofia Andersson, James Ussher, Jill Sherwood, Josh Freeman, Julia Howard, Juliet Elvy, Mary DeAlmeida, Matt Blakiston, Matthew Rogers, Max Bloomfield, Michael Addidle, Michelle Balm, Sally Roberts, Sarah Jefferies, Sharmini Muttaiyah, Susan Morpeth, Susan Taylor, Timothy Blackmore, Vani Sathyendran, Veronica Playle, Virginia Hope, Erasmus Smit, Lauren Jelly, Olin Silander, Joep de Ligt |
| EPI_ISL_548042, EPI_ISL_548043, EPI_ISL_548044 | North Shore Hospital | Institute of Environmental Science and Research (ESR) | Xiaoyun Ren, Matt Storey, Nikki Freed, Muhammad Faisal, Jing Wang, Hermes Perez, Anja Werno, Antje van der Linden, Arlo Upton, Chris Mansell, David Hammer, Dragana Drinkovic, Gary McAuliffe, Hana Sofia Andersson, James Ussher, Jill Sherwood, Josh Freeman, Julia Howard, Juliet Elvy, Mary DeAlmeida, Matt Blakiston, Matthew Rogers, Max Bloomfield, Michael Addidle, Michelle Balm, Sally Roberts, Sarah Jefferies, Sharmini Muttaiyah, Susan Morpeth, Susan Taylor, Timothy Blackmore, Vani Sathyendran, Veronica Playle, Virginia Hope, Erasmus Smit, Lauren Jelly, Olin Silander, Joep de Ligt |
| EPI_ISL_548046 | LabTests | Institute of Environmental Science and Research (ESR) | Xiaoyun Ren, Matt Storey, Nikki Freed, Muhammad Faisal, Jing Wang, Hermes Perez, Anja Werno, Antje van der Linden, Arlo Upton, Chris Mansell, David Hammer, Dragana Drinkovic, Gary McAuliffe, Hana Sofia Andersson, James Ussher, Jill Sherwood, Josh Freeman, Julia Howard, Juliet Elvy, Mary DeAlmeida, Matt Blakiston, Matthew Rogers, Max Bloomfield, Michael Addidle, Michelle Balm, Sally Roberts, Sarah Jefferies, Sharmini Muttaiyah, Susan Morpeth, Susan Taylor, Timothy Blackmore, Vani Sathyendran, Veronica Playle, Virginia Hope, Erasmus Smit, Lauren Jelly, Olin Silander, Joep de Ligt |
| EPI_ISL_548047, EPI_ISL_548048 | North Shore Hospital | Institute of Environmental Science and Research (ESR) | Xiaoyun Ren, Matt Storey, Nikki Freed, Muhammad Faisal, Jing Wang, Hermes Perez, Anja Werno, Antje van der Linden, Arlo Upton, Chris Mansell, David Hammer, Dragana Drinkovic, Gary McAuliffe, Hana Sofia Andersson, James Ussher, Jill Sherwood, Josh Freeman, Julia Howard, Juliet Elvy, Mary DeAlmeida, Matt Blakiston, Matthew Rogers, Max Bloomfield, Michael Addidle, Michelle Balm, Sally Roberts, Sarah Jefferies, Sharmini Muttaiyah, Susan Morpeth, Susan Taylor, Timothy Blackmore, Vani Sathyendran, Veronica Playle, Virginia Hope, Erasmus Smit, Lauren Jelly, Olin Silander, Joep de Ligt |
| EPI_ISL_548049, EPI_ISL_548050 | LabPLUS | Institute of Environmental Science and Research (ESR) | Xiaoyun Ren, Matt Storey, Nikki Freed, Muhammad Faisal, Jing Wang, Hermes Perez, Anja Werno, Antje van der Linden, Arlo Upton, Chris Mansell, David Hammer, Dragana Drinkovic, Gary McAuliffe, Hana Sofia Andersson, James Ussher, Jill Sherwood, Josh Freeman, Julia Howard, Juliet Elvy, Mary DeAlmeida, Matt Blakiston, Matthew Rogers, Max Bloomfield, Michael Addidle, Michelle Balm, Sally Roberts, Sarah Jefferies, Sharmini Muttaiyah, Susan Morpeth, Susan Taylor, Timothy Blackmore, Vani Sathyendran, Veronica Playle, Virginia Hope, Erasmus Smit, Lauren Jelly, Olin Silander, Joep de Ligt |
| EPI_ISL_548057, EPI_ISL_548061 | North Shore Hospital | Institute of Environmental Science and Research (ESR) | Xiaoyun Ren, Matt Storey, Nikki Freed, Muhammad Faisal, Jing Wang, Hermes Perez, Anja Werno, Antje van der Linden, Arlo Upton, Chris Mansell, David Hammer, Dragana Drinkovic, Gary McAuliffe, Hana Sofia Andersson, James Ussher, Jill Sherwood, Josh Freeman, Julia Howard, Juliet Elvy, Mary DeAlmeida, Matt Blakiston, Matthew Rogers, Max Bloomfield, Michael Addidle, Michelle Balm, Sally Roberts, Sarah Jefferies, Sharmini Muttaiyah, Susan Morpeth, Susan Taylor, Timothy Blackmore, Vani Sathyendran, Veronica Playle, Virginia Hope, Erasmus Smit, Lauren Jelly, Olin Silander, Joep de Ligt |
| EPI_ISL_548064 | Waikato Hospital | Institute of Environmental Science and Research (ESR) | Xiaoyun Ren, Matt Storey, Nikki Freed, Muhammad Faisal, Jing Wang, Hermes Perez, Anja Werno, Antje van der Linden, Arlo Upton, Chris Mansell, David Hammer, Dragana Drinkovic, Gary McAuliffe, Hana Sofia Andersson, James Ussher, Jill Sherwood, Josh Freeman, Julia Howard, Juliet Elvy, Mary DeAlmeida, Matt Blakiston, Matthew Rogers, Max Bloomfield, Michael Addidle, Michelle Balm, Sally Roberts, Sarah Jefferies, Sharmini Muttaiyah, Susan Morpeth, Susan Taylor, Timothy Blackmore, Vani Sathyendran, Veronica Playle, Virginia Hope, Erasmus Smit, Lauren Jelly, Olin Silander, Joep de Ligt |
| EPI_ISL_548067 | North Shore Hospital | Institute of Environmental Science and Research (ESR) | Xiaoyun Ren, Matt Storey, Nikki Freed, Muhammad Faisal, Jing Wang, Hermes Perez, Anja Werno, Antje van der Linden, Arlo Upton, Chris Mansell, David Hammer, Dragana Drinkovic, Gary McAuliffe, Hana Sofia Andersson, James Ussher, Jill Sherwood, Josh Freeman, Julia Howard, Juliet Elvy, Mary DeAlmeida, Matt Blakiston, Matthew Rogers, Max Bloomfield, Michael Addidle, Michelle Balm, Sally Roberts, Sarah Jefferies, Sharmini Muttaiyah, Susan Morpeth, Susan Taylor, Timothy Blackmore, Vani Sathyendran, Veronica Playle, Virginia Hope, Erasmus Smit, Lauren Jelly, Olin Silander, Joep de Ligt |
| EPI_ISL_548069, EPI_ISL_548070, EPI_ISL_548071, EPI_ISL_548072, EPI_ISL_548073 | LabPLUS | Institute of Environmental Science and Research (ESR) | Xiaoyun Ren, Matt Storey, Nikki Freed, Muhammad Faisal, Jing Wang, Hermes Perez, Anja Werno, Antje van der Linden, Arlo Upton, Chris Mansell, David Hammer, Dragana Drinkovic, Gary McAuliffe, Hana Sofia Andersson, James Ussher, Jill Sherwood, Josh Freeman, Julia Howard, Juliet Elvy, Mary DeAlmeida, Matt Blakiston, Matthew Rogers, Max Bloomfield, Michael Addidle, Michelle Balm, Sally Roberts, Sarah Jefferies, Sharmini Muttaiyah, Susan Morpeth, Susan Taylor, Timothy Blackmore, Vani Sathyendran, Veronica Playle, Virginia Hope, Erasmus Smit, Lauren Jelly, Olin Silander, Joep de Ligt |
| EPI_ISL_548075, EPI_ISL_548076 | North Shore Hospital | Institute of Environmental Science and Research (ESR) | Xiaoyun Ren, Matt Storey, Nikki Freed, Muhammad Faisal, Jing Wang, Hermes Perez, Anja Werno, Antje van der Linden, Arlo Upton, Chris Mansell, David Hammer, Dragana Drinkovic, Gary McAuliffe, Hana Sofia Andersson, James Ussher, Jill Sherwood, Josh Freeman, Julia Howard, Juliet Elvy, Mary DeAlmeida, Matt Blakiston, Matthew Rogers, Max Bloomfield, Michael Addidle, Michelle Balm, Sally Roberts, Sarah Jefferies, Sharmini Muttaiyah, Susan Morpeth, Susan Taylor, Timothy Blackmore, Vani Sathyendran, Veronica Playle, Virginia Hope, Erasmus Smit, Lauren Jelly, Olin Silander, Joep de Ligt |
| EPI_ISL_548077, EPI_ISL_548078, EPI_ISL_548079 | LabPLUS | Institute of Environmental Science and Research (ESR) | Xiaoyun Ren, Matt Storey, Nikki Freed, Muhammad Faisal, Jing Wang, Hermes Perez, Anja Werno, Antje van der Linden, Arlo Upton, Chris Mansell, David Hammer, Dragana Drinkovic, Gary McAuliffe, Hana Sofia Andersson, James Ussher, Jill Sherwood, Josh Freeman, Julia Howard, Juliet Elvy, Mary DeAlmeida, Matt Blakiston, Matthew Rogers, Max Bloomfield, Michael Addidle, Michelle Balm, Sally Roberts, Sarah Jefferies, Sharmini Muttaiyah, Susan Morpeth, Susan Taylor, Timothy Blackmore, Vani Sathyendran, Veronica Playle, Virginia Hope, Erasmus Smit, Lauren Jelly, Olin Silander, Joep de Ligt |
| EPI_ISL_548081, EPI_ISL_548082 | Middlemore Hospital | Institute of Environmental Science and Research (ESR) | Xiaoyun Ren, Matt Storey, Nikki Freed, Muhammad Faisal, Jing Wang, Hermes Perez, Anja Werno, Antje van der Linden, Arlo Upton, Chris Mansell, David Hammer, Dragana Drinkovic, Gary McAuliffe, Hana Sofia Andersson, James Ussher, Jill Sherwood, Josh Freeman, Julia Howard, Juliet Elvy, Mary DeAlmeida, Matt Blakiston, Matthew Rogers, Max Bloomfield, Michael Addidle, Michelle Balm, Sally Roberts, Sarah Jefferies, Sharmini Muttaiyah, Susan Morpeth, Susan Taylor, Timothy Blackmore, Vani Sathyendran, Veronica Playle, Virginia Hope, Erasmus Smit, Lauren Jelly, Olin Silander, Joep de Ligt |
| EPI_ISL_548084 | LabPLUS | Institute of Environmental Science and Research (ESR) | Xiaoyun Ren, Matt Storey, Nikki Freed, Muhammad Faisal, Jing Wang, Hermes Perez, Anja Werno, Antje van der Linden, Arlo Upton, Chris Mansell, David Hammer, Dragana Drinkovic, Gary McAuliffe, Hana Sofia Andersson, James Ussher, Jill Sherwood, Josh Freeman, Julia Howard, Juliet Elvy, Mary DeAlmeida, Matt Blakiston, Matthew Rogers, Max Bloomfield, Michael Addidle, Michelle Balm, Sally Roberts, Sarah Jefferies, Sharmini Muttaiyah, Susan Morpeth, Susan Taylor, Timothy Blackmore, Vani Sathyendran, Veronica Playle, Virginia Hope, Erasmus Smit, Lauren Jelly, Olin Silander, Joep de Ligt |
| EPI_ISL_548101 | Waikato Hospital | Institute of Environmental Science and Research (ESR) | Xiaoyun Ren, Matt Storey, Nikki Freed, Muhammad Faisal, Jing Wang, Hermes Perez, Anja Werno, Antje van der Linden, Arlo Upton, Chris Mansell, David Hammer, Dragana Drinkovic, Gary McAuliffe, Hana Sofia Andersson, James Ussher, Jill Sherwood, Josh Freeman, Julia Howard, Juliet Elvy, Mary DeAlmeida, Matt Blakiston, Matthew Rogers, Max Bloomfield, Michael Addidle, Michelle Balm, Sally Roberts, Sarah Jefferies, Sharmini Muttaiyah, Susan Morpeth, Susan Taylor, Timothy Blackmore, Vani Sathyendran, Veronica Playle, Virginia Hope, Erasmus Smit, Lauren Jelly, Olin Silander, Joep de Ligt |
| EPI_ISL_548104, EPI_ISL_548105 | LabTests | Institute of Environmental Science and Research (ESR) | Xiaoyun Ren, Matt Storey, Nikki Freed, Muhammad Faisal, Jing Wang, Hermes Perez, Anja Werno, Antje van der Linden, Arlo Upton, Chris Mansell, David Hammer, Dragana Drinkovic, Gary McAuliffe, Hana Sofia Andersson, James Ussher, Jill Sherwood, Josh Freeman, Julia Howard, Juliet Elvy, Mary DeAlmeida, Matt Blakiston, Matthew Rogers, Max Bloomfield, Michael Addidle, Michelle Balm, Sally Roberts, Sarah Jefferies, Sharmini Muttaiyah, Susan Morpeth, Susan Taylor, Timothy Blackmore, Vani Sathyendran, Veronica Playle, Virginia Hope, Erasmus Smit, Lauren Jelly, Olin Silander, Joep de Ligt |
| EPI_ISL_548106, EPI_ISL_548108, EPI_ISL_548110, EPI_ISL_548111, EPI_ISL_548120 | LabPLUS | Institute of Environmental Science and Research (ESR) | Xiaoyun Ren, Matt Storey, Nikki Freed, Muhammad Faisal, Jing Wang, Hermes Perez, Anja Werno, Antje van der Linden, Arlo Upton, Chris Mansell, David Hammer, Dragana Drinkovic, Gary McAuliffe, Hana Sofia Andersson, James Ussher, Jill Sherwood, Josh Freeman, Julia Howard, Juliet Elvy, Mary DeAlmeida, Matt Blakiston, Matthew Rogers, Max Bloomfield, Michael Addidle, Michelle Balm, Sally Roberts, Sarah Jefferies, Sharmini Muttaiyah, Susan Morpeth, Susan Taylor, Timothy Blackmore, Vani Sathyendran, Veronica Playle, Virginia Hope, Erasmus Smit, Lauren Jelly, Olin Silander, Joep de Ligt |
| EPI_ISL_548121, EPI_ISL_548123, EPI_ISL_548124 | Middlemore Hospital | Institute of Environmental Science and Research (ESR) | Xiaoyun Ren, Matt Storey, Nikki Freed, Muhammad Faisal, Jing Wang, Hermes Perez, Anja Werno, Antje van der Linden, Arlo Upton, Chris Mansell, David Hammer, Dragana Drinkovic, Gary McAuliffe, Hana Sofia Andersson, James Ussher, Jill Sherwood, Josh Freeman, Julia Howard, Juliet Elvy, Mary DeAlmeida, Matt Blakiston, Matthew Rogers, Max Bloomfield, Michael Addidle, Michelle Balm, Sally Roberts, Sarah Jefferies, Sharmini Muttaiyah, Susan Morpeth, Susan Taylor, Timothy Blackmore, Vani Sathyendran, Veronica Playle, Virginia Hope, Erasmus Smit, Lauren Jelly, Olin Silander, Joep de Ligt |

| | | | |
|---|---|---|---|
| EPI_ISL_548129 | Canterbury Health Laboratories | Institute of Environmental Science and Research (ESR) | Xiaoyun Ren, Matt Storey, Nikki Freed, Muhammad Faisal, Jing Wang, Hermes Perez, Anja Werno, Antje van der Linden, Arlo Upton, Chris Mansell, David Hammer, Dragana Drinkovic, Gary McAuliffe, Hana Sofia Andersson, James Ussher, Jill Sherwood, Josh Freeman, Julia Howard, Juliet Elvy, Mary DeAlmeida, Matt Blakiston, Matthew Rogers, Max Bloomfield, Michael Addidle, Michelle Balm, Sally Roberts, Sarah Jefferies, Sharmini Muttaiyah, Susan Morpeth, Susan Taylor, Timothy Blackmore, Vani Sathyendran, Veronica Playle, Virginia Hope, Erasmus Smit, Lauren Jelly, Olin Silander, Joep de Ligt |
| EPI_ISL_548132 | Middlemore Hospital | Institute of Environmental Science and Research (ESR) | Xiaoyun Ren, Matt Storey, Nikki Freed, Muhammad Faisal, Jing Wang, Hermes Perez, Anja Werno, Antje van der Linden, Arlo Upton, Chris Mansell, David Hammer, Dragana Drinkovic, Gary McAuliffe, Hana Sofia Andersson, James Ussher, Jill Sherwood, Josh Freeman, Julia Howard, Juliet Elvy, Mary DeAlmeida, Matt Blakiston, Matthew Rogers, Max Bloomfield, Michael Addidle, Michelle Balm, Sally Roberts, Sarah Jefferies, Sharmini Muttaiyah, Susan Morpeth, Susan Taylor, Timothy Blackmore, Vani Sathyendran, Veronica Playle, Virginia Hope, Erasmus Smit, Lauren Jelly, Olin Silander, Joep de Ligt |
| EPI_ISL_548139, EPI_ISL_548140 | Canterbury Health Laboratories | Institute of Environmental Science and Research (ESR) | Xiaoyun Ren, Matt Storey, Nikki Freed, Muhammad Faisal, Jing Wang, Hermes Perez, Anja Werno, Antje van der Linden, Arlo Upton, Chris Mansell, David Hammer, Dragana Drinkovic, Gary McAuliffe, Hana Sofia Andersson, James Ussher, Jill Sherwood, Josh Freeman, Julia Howard, Juliet Elvy, Mary DeAlmeida, Matt Blakiston, Matthew Rogers, Max Bloomfield, Michael Addidle, Michelle Balm, Sally Roberts, Sarah Jefferies, Sharmini Muttaiyah, Susan Morpeth, Susan Taylor, Timothy Blackmore, Vani Sathyendran, Veronica Playle, Virginia Hope, Erasmus Smit, Lauren Jelly, Olin Silander, Joep de Ligt |
| EPI_ISL_548942, EPI_ISL_548943, EPI_ISL_548944, EPI_ISL_548945, EPI_ISL_548946 | Institute of Microbiology, University of Veterinary and Animal sciences | Institute of Microbiology, University of Veterinary and Animal sciences | Yaqub,T., Nawaz,M., Ali,M.A., Altaf,I., Raza,S., Shabbir,M.A., Ashraf,M.A., Aziz,S.Z., Cheema,S.Q., Shah,M.B., Hassan,S., Rafique,S., Sardar,N., Mehmood,A., Aziz,M.W., Fazal,S., Khan,N., Khan,M.T., Attique,M.M., Asif,A., Anwar,M., Awan,N.A., Younis,M.U., Bhatti,M.A., Tahir,Z., Mukhtar,N., Sarwar,H., Rana,M.S., Shabbir,M.Z. |
| EPI_ISL_568556, EPI_ISL_568557, EPI_ISL_568558, EPI_ISL_568559, EPI_ISL_568560, EPI_ISL_568561, EPI_ISL_568562, EPI_ISL_568563, EPI_ISL_568564, EPI_ISL_568565, EPI_ISL_568566, EPI_ISL_568567, EPI_ISL_568568, EPI_ISL_568569, EPI_ISL_568570, EPI_ISL_568571, EPI_ISL_568572, EPI_ISL_568573, EPI_ISL_568574, EPI_ISL_568575, EPI_ISL_568576 | | | |
| see above | Department of Infectious Diseases and Immunology, National Hospital Organization Nagoya Medical Center | Clinical Research Center, National Hospital Organization Nagoya Medical Center | Yoshihiro Nakata, Hirotaka Ode, Mai Kubota, Masakazu Matsuda, Kazuhiro Matsuoka, Nakasuji Miho, Mikiko Mori, Mayumi Imahashi, Yoshiyuki Yokomaku, Yasumasa Iwatani |
| EPI_ISL_576170, EPI_ISL_576171 | PA Department of Health, Bureau of Laboratories | Pathogen Discovery, Respiratory Viruses Branch, Division of Viral Diseases, Centers for Disease Control and Prevention | Yan Li, Anna Montmayeur, Brian Lynch, Jing Zhang, Krista Queen, Ying Tao, Anna Uehara, Rachel Marine, Clinton R. Paden, Peter Cook, Haibin Wang, Suxiang Tong |
| EPI_ISL_576172 | TN Division of Laboratory Services | Pathogen Discovery, Respiratory Viruses Branch, Division of Viral Diseases, Centers for Disease Control and Prevention | Yan Li, Anna Montmayeur, Brian Lynch, Jing Zhang, Krista Queen, Ying Tao, Anna Uehara, Rachel Marine, Clinton R. Paden, Peter Cook, Haibin Wang, Suxiang Tong |
| EPI_ISL_576173 | FL Department of Health-Bureau of Epidemiology | Pathogen Discovery, Respiratory Viruses Branch, Division of Viral Diseases, Centers for Disease Control and Prevention | Ying Tao, Jing Zhang, Brian Lynch, Yan Li, Krista Queen, Anna Uehara, Clinton R. Paden, Peter Cook, Haibin Wang, Suxiang Tong |
| EPI_ISL_576174 | VA-Division of Consolidated Laboratory Services | Pathogen Discovery, Respiratory Viruses Branch, Division of Viral Diseases, Centers for Disease Control and Prevention | Ying Tao, Jing Zhang, Brian Lynch, Yan Li, Krista Queen, Anna Uehara, Clinton R. Paden, Peter Cook, Haibin Wang, Suxiang Tong |
| EPI_ISL_576175, EPI_ISL_576176 | TN Division of Laboratory Services | Pathogen Discovery, Respiratory Viruses Branch, Division of Viral Diseases, Centers for Disease Control and Prevention | Yan Li, Anna Montmayeur, Brian Lynch, Jing Zhang, Krista Queen, Ying Tao, Anna Uehara, Rachel Marine, Clinton R. Paden, Peter Cook, Haibin Wang, Suxiang Tong |
| EPI_ISL_576177 | CA, CDPH, Viral and Rickettsial Disease Laboratory | Pathogen Discovery, Respiratory Viruses Branch, Division of Viral Diseases, Centers for Disease Control and Prevention | Ying Tao, Jing Zhang, Brian Lynch, Yan Li, Krista Queen, Anna Uehara, Clinton R. Paden, Peter Cook, Haibin Wang, Suxiang Tong |
| EPI_ISL_576178 | DC Public Health Lab/ Dept. of Forensic Sciences | Pathogen Discovery, Respiratory Viruses Branch, Division of Viral Diseases, Centers for Disease Control and Prevention | Ying Tao, Jing Zhang, Brian Lynch, Yan Li, Krista Queen, Anna Uehara, Clinton R. Paden, Peter Cook, Haibin Wang, Suxiang Tong |
| EPI_ISL_576179, EPI_ISL_576180, EPI_ISL_576181, EPI_ISL_576182, EPI_ISL_576183, EPI_ISL_576184, EPI_ISL_576185, EPI_ISL_576186, EPI_ISL_576187, EPI_ISL_576188 | WI State Laboratory of Hygiene | Pathogen Discovery, Respiratory Viruses Branch, Division of Viral Diseases, Centers for Disease Control and Prevention | Ying Tao, Jing Zhang, Brian Lynch, Yan Li, Krista Queen, Anna Uehara, Clinton R. Paden, Peter Cook, Haibin Wang, Suxiang Tong |
| EPI_ISL_576189, EPI_ISL_576190, EPI_ISL_576191, EPI_ISL_576192, EPI_ISL_576193, EPI_ISL_576194, EPI_ISL_576195 | GA Department of Public Health Laboratory | Pathogen Discovery, Respiratory Viruses Branch, Division of Viral Diseases, Centers for Disease Control and Prevention | Ying Tao, Jing Zhang, Brian Lynch, Yan Li, Krista Queen, Anna Uehara, Clinton R. Paden, Peter Cook, Haibin Wang, Suxiang Tong |
| EPI_ISL_576196, EPI_ISL_576197 | AR Dept. of Health-Public Health Lab | Pathogen Discovery, Respiratory Viruses Branch, Division of Viral Diseases, Centers for Disease Control and Prevention | Ying Tao, Jing Zhang, Brian Lynch, Yan Li, Krista Queen, Anna Uehara, Clinton R. Paden, Peter Cook, Haibin Wang, Suxiang Tong |
| EPI_ISL_576363, EPI_ISL_576364, EPI_ISL_576365, EPI_ISL_576366, EPI_ISL_576367, EPI_ISL_576368, EPI_ISL_576369, EPI_ISL_576370 | Utah Public Health Laboratory, Utah Public Health Laboratory Infectious Disease submission group | Utah Public Health Laboratory, Utah Public Health Laboratory Infectious Disease submission group | Young,E.L., Oakeson,K., Sangster,A., Hirschi,B., Butz,H. |
| EPI_ISL_576371, EPI_ISL_576372, EPI_ISL_576373 | Cancer Biology Department, National Cancer Institute | Cancer Biology Department, National Cancer Institute | Zekri,A.N., Soliman,H.K., Ahmed,O.S., Hafez,M.M., Hamdy,M.S., Abouelhoda,M. |
| EPI_ISL_577598 | Utah Public Health Laboratory, Utah Public Health Laboratory Infectious Disease submission group | Utah Public Health Laboratory | Young,E.L., Oakeson,K., Sangster,A., Hirschi,B., Butz,H. |
| EPI_ISL_577599 | Utah Public Health Laboratory, Utah Public Health Laboratory Infectious Disease submission group | Utah Public Health Laboratory, Utah Public Health Laboratory Infectious Disease submission group | Young,E.L., Oakeson,K., Sangster,A., Hirschi,B., Butz,H. |
| EPI_ISL_577600 | Utah Public Health Laboratory, Utah Public Health Laboratory Infectious Disease submission group | Utah Public Health Laboratory | Young,E.L., Oakeson,K., Sangster,A., Hirschi,B., Butz,H. |
| EPI_ISL_577601 | Utah Public Health Laboratory, Utah Public Health Laboratory Infectious Disease submission group | Utah Public Health Laboratory, Utah Public Health Laboratory Infectious Disease submission group | Young,E.L., Oakeson,K., Sangster,A., Hirschi,B., Butz,H. |
| EPI_ISL_579058 | LabTests | Institute of Environmental Science and Research (ESR) | Xiaoyun Ren, Matt Storey, Nikki Freed, Muhammad Faisal, Jing Wang, Hermes Perez, Anja Werno, Antje van der Linden, Arlo Upton, Chris Mansell, David Hammer, Dragana Drinkovic, Gary McAuliffe, Hana Sofia Andersson, James Ussher, Jill Sherwood, Josh Freeman, Julia Howard, Juliet Elvy, Mary DeAlmeida, Matt Blakiston, Matthew Rogers, Max Bloomfield, Michael Addidle, Michelle Balm, Sally Roberts, Sarah Jefferies, Sharmini Muttaiyah, Susan Morpeth, Susan Taylor, Timothy Blackmore, Vani Sathyendran, Veronica Playle, Virginia Hope, Erasmus Smit, Lauren Jelly, Olin Silander, Joep de Ligt |
| EPI_ISL_579059 | LabPLUS | Institute of Environmental Science and Research (ESR) | Xiaoyun Ren, Matt Storey, Nikki Freed, Muhammad Faisal, Jing Wang, Hermes Perez, Anja Werno, Antje van der Linden, Arlo Upton, Chris Mansell, David Hammer, Dragana Drinkovic, Gary McAuliffe, Hana Sofia Andersson, James Ussher, Jill Sherwood, Josh Freeman, Julia Howard, Juliet Elvy, Mary DeAlmeida, Matt Blakiston, Matthew Rogers, Max Bloomfield, Michael Addidle, Michelle Balm, Sally Roberts, Sarah Jefferies, Sharmini Muttaiyah, Susan Morpeth, Susan Taylor, Timothy Blackmore, Vani Sathyendran, Veronica Playle, Virginia Hope, Erasmus Smit, Lauren Jelly, Olin Silander, Joep de Ligt |
| EPI_ISL_579060, EPI_ISL_579061, EPI_ISL_579062, EPI_ISL_579063, EPI_ISL_579064, EPI_ISL_579065, EPI_ISL_579066, EPI_ISL_579067, EPI_ISL_579068, EPI_ISL_579069, EPI_ISL_579070, EPI_ISL_579071, EPI_ISL_579072, EPI_ISL_579073, EPI_ISL_579074, EPI_ISL_579075, EPI_ISL_579077, EPI_ISL_579078, EPI_ISL_579079, EPI_ISL_579080, EPI_ISL_579081, EPI_ISL_579082, EPI_ISL_579083, EPI_ISL_579084, EPI_ISL_579085, EPI_ISL_579086 | | | |
| see above | Canterbury Health Laboratories | Institute of Environmental Science and Research (ESR) | Xiaoyun Ren, Matt Storey, Nikki Freed, Muhammad Faisal, Jing Wang, Hermes Perez, Anja Werno, Antje van der Linden, Arlo Upton, Chris Mansell, David Hammer, Dragana Drinkovic, Gary McAuliffe, Hana Sofia Andersson, James Ussher, Jill Sherwood, Josh Freeman, Julia Howard, Juliet Elvy, Mary DeAlmeida, Matt Blakiston, Matthew Rogers, Max Bloomfield, Michael Addidle, Michelle Balm, Sally Roberts, Sarah Jefferies, Sharmini Muttaiyah, Susan Morpeth, Susan Taylor, Timothy Blackmore, Vani Sathyendran, Veronica Playle, Virginia Hope, Erasmus Smit, Lauren Jelly, Olin Silander, Joep de Ligt |
| EPI_ISL_579099 | LabTests | Institute of Environmental Science and Research (ESR) | Xiaoyun Ren, Matt Storey, Nikki Freed, Muhammad Faisal, Jing Wang, Hermes Perez, Anja Werno, Antje van der Linden, Arlo Upton, Chris Mansell, David Hammer, Dragana Drinkovic, Gary McAuliffe, Hana Sofia Andersson, James Ussher, Jill Sherwood, Josh Freeman, Julia Howard, Juliet Elvy, Mary |

| | | | |
|---|---|---|---|
| | | | DeAlmeida, Matt Blakiston, Matthew Rogers, Max Bloomfield, Michael Addidle, Michelle Balm, Sally Roberts, Sarah Jefferies, Sharmini Muttaiyah, Susan Morpeth, Susan Taylor, Timothy Blackmore, Vani Sathyendran, Veronica Playle, Virginia Hope, Erasmus Smit, Lauren Jelly, Olin Silander, Joep de Ligt |
| EPI_ISL_579101, EPI_ISL_579102, EPI_ISL_579103, EPI_ISL_579104 | Middlemore Hospital | Institute of Environmental Science and Research (ESR) | Xiaoyun Ren, Matt Storey, Nikki Freed, Muhammad Faisal, Jing Wang, Hermes Perez, Anja Werno, Antje van der Linden, Arlo Upton, Chris Mansell, David Hammer, Dragana Drinkovic, Gary McAuliffe, Hana Sofia Andersson, James Ussher, Jill Sherwood, Josh Freeman, Julia Howard, Juliet Elvy, Mary DeAlmeida, Matt Blakiston, Matthew Rogers, Max Bloomfield, Michael Addidle, Michelle Balm, Sally Roberts, Sarah Jefferies, Sharmini Muttaiyah, Susan Morpeth, Susan Taylor, Timothy Blackmore, Vani Sathyendran, Veronica Playle, Virginia Hope, Erasmus Smit, Lauren Jelly, Olin Silander, Joep de Ligt |
| EPI_ISL_579105, EPI_ISL_579106, EPI_ISL_579107, EPI_ISL_579111, EPI_ISL_579113, EPI_ISL_579114, EPI_ISL_579115, EPI_ISL_579116, EPI_ISL_579117 | LabPLUS | Institute of Environmental Science and Research (ESR) | Xiaoyun Ren, Matt Storey, Nikki Freed, Muhammad Faisal, Jing Wang, Hermes Perez, Anja Werno, Antje van der Linden, Arlo Upton, Chris Mansell, David Hammer, Dragana Drinkovic, Gary McAuliffe, Hana Sofia Andersson, James Ussher, Jill Sherwood, Josh Freeman, Julia Howard, Juliet Elvy, Mary DeAlmeida, Matt Blakiston, Matthew Rogers, Max Bloomfield, Michael Addidle, Michelle Balm, Sally Roberts, Sarah Jefferies, Sharmini Muttaiyah, Susan Morpeth, Susan Taylor, Timothy Blackmore, Vani Sathyendran, Veronica Playle, Virginia Hope, Erasmus Smit, Lauren Jelly, Olin Silander, Joep de Ligt |
| EPI_ISL_579118 | Waikato Hospital | Institute of Environmental Science and Research (ESR) | Xiaoyun Ren, Matt Storey, Nikki Freed, Muhammad Faisal, Jing Wang, Hermes Perez, Anja Werno, Antje van der Linden, Arlo Upton, Chris Mansell, David Hammer, Dragana Drinkovic, Gary McAuliffe, Hana Sofia Andersson, James Ussher, Jill Sherwood, Josh Freeman, Julia Howard, Juliet Elvy, Mary DeAlmeida, Matt Blakiston, Matthew Rogers, Max Bloomfield, Michael Addidle, Michelle Balm, Sally Roberts, Sarah Jefferies, Sharmini Muttaiyah, Susan Morpeth, Susan Taylor, Timothy Blackmore, Vani Sathyendran, Veronica Playle, Virginia Hope, Erasmus Smit, Lauren Jelly, Olin Silander, Joep de Ligt |
| EPI_ISL_579119 | LabPLUS | Institute of Environmental Science and Research (ESR) | Xiaoyun Ren, Matt Storey, Nikki Freed, Muhammad Faisal, Jing Wang, Hermes Perez, Anja Werno, Antje van der Linden, Arlo Upton, Chris Mansell, David Hammer, Dragana Drinkovic, Gary McAuliffe, Hana Sofia Andersson, James Ussher, Jill Sherwood, Josh Freeman, Julia Howard, Juliet Elvy, Mary DeAlmeida, Matt Blakiston, Matthew Rogers, Max Bloomfield, Michael Addidle, Michelle Balm, Sally Roberts, Sarah Jefferies, Sharmini Muttaiyah, Susan Morpeth, Susan Taylor, Timothy Blackmore, Vani Sathyendran, Veronica Playle, Virginia Hope, Erasmus Smit, Lauren Jelly, Olin Silander, Joep de Ligt |
| EPI_ISL_579121 | Waikato Hospital | Institute of Environmental Science and Research (ESR) | Xiaoyun Ren, Matt Storey, Nikki Freed, Muhammad Faisal, Jing Wang, Hermes Perez, Anja Werno, Antje van der Linden, Arlo Upton, Chris Mansell, David Hammer, Dragana Drinkovic, Gary McAuliffe, Hana Sofia Andersson, James Ussher, Jill Sherwood, Josh Freeman, Julia Howard, Juliet Elvy, Mary DeAlmeida, Matt Blakiston, Matthew Rogers, Max Bloomfield, Michael Addidle, Michelle Balm, Sally Roberts, Sarah Jefferies, Sharmini Muttaiyah, Susan Morpeth, Susan Taylor, Timothy Blackmore, Vani Sathyendran, Veronica Playle, Virginia Hope, Erasmus Smit, Lauren Jelly, Olin Silander, Joep de Ligt |
| EPI_ISL_579122 | LabPLUS | Institute of Environmental Science and Research (ESR) | Xiaoyun Ren, Matt Storey, Nikki Freed, Muhammad Faisal, Jing Wang, Hermes Perez, Anja Werno, Antje van der Linden, Arlo Upton, Chris Mansell, David Hammer, Dragana Drinkovic, Gary McAuliffe, Hana Sofia Andersson, James Ussher, Jill Sherwood, Josh Freeman, Julia Howard, Juliet Elvy, Mary DeAlmeida, Matt Blakiston, Matthew Rogers, Max Bloomfield, Michael Addidle, Michelle Balm, Sally Roberts, Sarah Jefferies, Sharmini Muttaiyah, Susan Morpeth, Susan Taylor, Timothy Blackmore, Vani Sathyendran, Veronica Playle, Virginia Hope, Erasmus Smit, Lauren Jelly, Olin Silander, Joep de Ligt |
| EPI_ISL_579126, EPI_ISL_579128 | Southern Community Labs Dunedin | Institute of Environmental Science and Research (ESR) | Xiaoyun Ren, Matt Storey, Nikki Freed, Muhammad Faisal, Jing Wang, Hermes Perez, Anja Werno, Antje van der Linden, Arlo Upton, Chris Mansell, David Hammer, Dragana Drinkovic, Gary McAuliffe, Hana Sofia Andersson, James Ussher, Jill Sherwood, Josh Freeman, Julia Howard, Juliet Elvy, Mary DeAlmeida, Matt Blakiston, Matthew Rogers, Max Bloomfield, Michael Addidle, Michelle Balm, Sally Roberts, Sarah Jefferies, Sharmini Muttaiyah, Susan Morpeth, Susan Taylor, Timothy Blackmore, Vani Sathyendran, Veronica Playle, Virginia Hope, Erasmus Smit, Lauren Jelly, Olin Silander, Joep de Ligt |
| EPI_ISL_579129, EPI_ISL_579130, EPI_ISL_579131 | Middlemore Hospital | Institute of Environmental Science and Research (ESR) | Xiaoyun Ren, Matt Storey, Nikki Freed, Muhammad Faisal, Jing Wang, Hermes Perez, Anja Werno, Antje van der Linden, Arlo Upton, Chris Mansell, David Hammer, Dragana Drinkovic, Gary McAuliffe, Hana Sofia Andersson, James Ussher, Jill Sherwood, Josh Freeman, Julia Howard, Juliet Elvy, Mary DeAlmeida, Matt Blakiston, Matthew Rogers, Max Bloomfield, Michael Addidle, Michelle Balm, Sally Roberts, Sarah Jefferies, Sharmini Muttaiyah, Susan Morpeth, Susan Taylor, Timothy Blackmore, Vani Sathyendran, Veronica Playle, Virginia Hope, Erasmus Smit, Lauren Jelly, Olin Silander, Joep de Ligt |
| EPI_ISL_579134, EPI_ISL_579135, EPI_ISL_579136, EPI_ISL_579137, EPI_ISL_579138, EPI_ISL_579139, EPI_ISL_579140, EPI_ISL_579141, EPI_ISL_579142, EPI_ISL_579143, EPI_ISL_579144, EPI_ISL_579145, EPI_ISL_579146, EPI_ISL_579147, EPI_ISL_579148, EPI_ISL_579150, EPI_ISL_579151, EPI_ISL_579152, EPI_ISL_579153, EPI_ISL_579154, EPI_ISL_579155 | | | |
| see above | Southern Community Labs Dunedin | Institute of Environmental Science and Research (ESR) | Xiaoyun Ren, Matt Storey, Nikki Freed, Muhammad Faisal, Jing Wang, Hermes Perez, Anja Werno, Antje van der Linden, Arlo Upton, Chris Mansell, David Hammer, Dragana Drinkovic, Gary McAuliffe, Hana Sofia Andersson, James Ussher, Jill Sherwood, Josh Freeman, Julia Howard, Juliet Elvy, Mary DeAlmeida, Matt Blakiston, Matthew Rogers, Max Bloomfield, Michael Addidle, Michelle Balm, Sally Roberts, Sarah Jefferies, Sharmini Muttaiyah, Susan Morpeth, Susan Taylor, Timothy Blackmore, Vani Sathyendran, Veronica Playle, Virginia Hope, Erasmus Smit, Lauren Jelly, Olin Silander, Joep de Ligt |
| EPI_ISL_579156 | LabPLUS | Institute of Environmental Science and Research (ESR) | Xiaoyun Ren, Matt Storey, Nikki Freed, Muhammad Faisal, Jing Wang, Hermes Perez, Anja Werno, Antje van der Linden, Arlo Upton, Chris Mansell, David Hammer, Dragana Drinkovic, Gary McAuliffe, Hana Sofia Andersson, James Ussher, Jill Sherwood, Josh Freeman, Julia Howard, Juliet Elvy, Mary DeAlmeida, Matt Blakiston, Matthew Rogers, Max Bloomfield, Michael Addidle, Michelle Balm, Sally Roberts, Sarah Jefferies, Sharmini Muttaiyah, Susan Morpeth, Susan Taylor, Timothy Blackmore, Vani Sathyendran, Veronica Playle, Virginia Hope, Erasmus Smit, Lauren Jelly, Olin Silander, Joep de Ligt |
| EPI_ISL_579157, EPI_ISL_579158, EPI_ISL_579159, EPI_ISL_579160, EPI_ISL_579161, EPI_ISL_579163, EPI_ISL_579164, EPI_ISL_579165, EPI_ISL_579167, EPI_ISL_579169, EPI_ISL_579170, EPI_ISL_579171, EPI_ISL_579173, EPI_ISL_579174, EPI_ISL_579176, EPI_ISL_579177, EPI_ISL_579178, EPI_ISL_579179, EPI_ISL_579180, EPI_ISL_579181, EPI_ISL_579183, EPI_ISL_579184, EPI_ISL_579186, EPI_ISL_579187, EPI_ISL_579188, EPI_ISL_579189 | | | |
| see above | Southern Community Labs Dunedin | Institute of Environmental Science and Research (ESR) | Xiaoyun Ren, Matt Storey, Nikki Freed, Muhammad Faisal, Jing Wang, Hermes Perez, Anja Werno, Antje van der Linden, Arlo Upton, Chris Mansell, David Hammer, Dragana Drinkovic, Gary McAuliffe, Hana Sofia Andersson, James Ussher, Jill Sherwood, Josh Freeman, Julia Howard, Juliet Elvy, Mary DeAlmeida, Matt Blakiston, Matthew Rogers, Max Bloomfield, Michael Addidle, Michelle Balm, Sally Roberts, Sarah Jefferies, Sharmini Muttaiyah, Susan Morpeth, Susan Taylor, Timothy Blackmore, Vani Sathyendran, Veronica Playle, Virginia Hope, Erasmus Smit, Lauren Jelly, Olin Silander, Joep de Ligt |
| EPI_ISL_579221, EPI_ISL_579222, EPI_ISL_579227 | Canterbury Health Laboratories | Institute of Environmental Science and Research (ESR) | Xiaoyun Ren, Matt Storey, Nikki Freed, Muhammad Faisal, Jing Wang, Hermes Perez, Anja Werno, Antje van der Linden, Arlo Upton, Chris Mansell, David Hammer, Dragana Drinkovic, Gary McAuliffe, Hana Sofia Andersson, James Ussher, Jill Sherwood, Josh Freeman, Julia Howard, Juliet Elvy, Mary DeAlmeida, Matt Blakiston, Matthew Rogers, Max Bloomfield, Michael Addidle, Michelle Balm, Sally Roberts, Sarah Jefferies, Sharmini Muttaiyah, Susan Morpeth, Susan Taylor, Timothy Blackmore, Vani Sathyendran, Veronica Playle, Virginia Hope, Erasmus Smit, Lauren Jelly, Olin Silander, Joep de Ligt |
| EPI_ISL_579229, EPI_ISL_579230, EPI_ISL_579231, EPI_ISL_579301, EPI_ISL_579302, EPI_ISL_579304, EPI_ISL_579305, EPI_ISL_579307, EPI_ISL_579310 | Middlemore Hospital | Institute of Environmental Science and Research (ESR) | Xiaoyun Ren, Matt Storey, Nikki Freed, Muhammad Faisal, Jing Wang, Hermes Perez, Anja Werno, Antje van der Linden, Arlo Upton, Chris Mansell, David Hammer, Dragana Drinkovic, Gary McAuliffe, Hana Sofia Andersson, James Ussher, Jill Sherwood, Josh Freeman, Julia Howard, Juliet Elvy, Mary DeAlmeida, Matt Blakiston, Matthew Rogers, Max Bloomfield, Michael Addidle, Michelle Balm, Sally Roberts, Sarah Jefferies, Sharmini Muttaiyah, Susan Morpeth, Susan Taylor, Timothy Blackmore, Vani Sathyendran, Veronica Playle, Virginia Hope, Erasmus Smit, Lauren Jelly, Olin Silander, Joep de Ligt |
| EPI_ISL_579441, EPI_ISL_579444, EPI_ISL_579445, EPI_ISL_579448, EPI_ISL_579454, EPI_ISL_579456, EPI_ISL_579458, EPI_ISL_579459, EPI_ISL_579461, EPI_ISL_579462, EPI_ISL_579463, EPI_ISL_579464, EPI_ISL_579465, EPI_ISL_579466, EPI_ISL_579468, EPI_ISL_579469, EPI_ISL_579470, EPI_ISL_579471, EPI_ISL_579473, EPI_ISL_579474, EPI_ISL_579475, EPI_ISL_579478, EPI_ISL_579480, EPI_ISL_579481, EPI_ISL_579482, EPI_ISL_579485, EPI_ISL_579487, EPI_ISL_579488, EPI_ISL_579493, EPI_ISL_579494, EPI_ISL_579495, EPI_ISL_579496, EPI_ISL_579497, EPI_ISL_579498, EPI_ISL_579499, EPI_ISL_579500, EPI_ISL_579501, EPI_ISL_579503, EPI_ISL_579504, EPI_ISL_579505, EPI_ISL_579506 | | | |
| see above | Canterbury Health Laboratories | Institute of Environmental Science and Research (ESR) | Xiaoyun Ren, Matt Storey, Nikki Freed, Muhammad Faisal, Jing Wang, Hermes Perez, Anja Werno, Antje van der Linden, Arlo Upton, Chris Mansell, David Hammer, Dragana Drinkovic, Gary McAuliffe, Hana Sofia Andersson, James Ussher, Jill Sherwood, Josh Freeman, Julia Howard, Juliet Elvy, Mary DeAlmeida, Matt Blakiston, Matthew Rogers, Max Bloomfield, Michael Addidle, Michelle Balm, Sally Roberts, Sarah Jefferies, Sharmini Muttaiyah, Susan Morpeth, Susan Taylor, Timothy Blackmore, Vani Sathyendran, Veronica Playle, Virginia Hope, Erasmus Smit, Lauren Jelly, Olin Silander, Joep de Ligt |
| EPI_ISL_582017 | LabPLUS | Institute of Environmental Science and Research (ESR) | Xiaoyun Ren, Matt Storey, Nikki Freed, Muhammad Faisal, Jing Wang, Hermes Perez, Anja Werno, Antje van der Linden, Arlo Upton, Chris Mansell, David Hammer, Dragana Drinkovic, Gary McAuliffe, Hana Sofia Andersson, James Ussher, Jill Sherwood, Josh Freeman, Julia Howard, Juliet Elvy, Mary DeAlmeida, Matt Blakiston, Matthew Rogers, Max Bloomfield, Michael Addidle, Michelle Balm, Sally Roberts, Sarah Jefferies, Sharmini Muttaiyah, Susan Morpeth, Susan Taylor, Timothy Blackmore, Vani Sathyendran, Veronica Playle, Virginia Hope, Erasmus Smit, Lauren Jelly, Olin Silander, Joep de Ligt |
| EPI_ISL_582018, EPI_ISL_582019, EPI_ISL_582020 | PathLab Bay of Plenty | Institute of Environmental Science and Research (ESR) | Xiaoyun Ren, Matt Storey, Nikki Freed, Muhammad Faisal, Jing Wang, Hermes Perez, Anja Werno, Antje van der Linden, Arlo Upton, Chris Mansell, David Hammer, Dragana Drinkovic, Gary McAuliffe, Hana Sofia Andersson, James Ussher, Jill Sherwood, Josh Freeman, Julia Howard, Juliet Elvy, Mary DeAlmeida, Matt Blakiston, Matthew Rogers, Max Bloomfield, Michael Addidle, Michelle Balm, Sally Roberts, Sarah Jefferies, Sharmini Muttaiyah, Susan Morpeth, Susan Taylor, Timothy Blackmore, Vani Sathyendran, Veronica Playle, Virginia Hope, Erasmus Smit, Lauren Jelly, Olin Silander, Joep de Ligt |
| EPI_ISL_582022, EPI_ISL_582023 | Pathlab Lakes | Institute of Environmental Science and Research (ESR) | Xiaoyun Ren, Matt Storey, Nikki Freed, Muhammad Faisal, Jing Wang, Hermes Perez, Anja Werno, Antje van der Linden, Arlo Upton, Chris Mansell, David Hammer, Dragana Drinkovic, Gary McAuliffe, Hana Sofia Andersson, James Ussher, Jill Sherwood, Josh Freeman, Julia Howard, Juliet Elvy, Mary |

| | | | |
|---|---|---|---|
| | | | DeAlmeida, Matt Blakiston, Matthew Rogers, Max Bloomfield, Michael Addidle, Michelle Balm, Sally Roberts, Sarah Jefferies, Sharmini Muttaiyah, Susan Morpeth, Susan Taylor, Timothy Blackmore, Vani Sathyendran, Veronica Playle, Virginia Hope, Erasmus Smit, Lauren Jelly, Olin Silander, Joep de Ligt |
| EPI_ISL_582033 | Department of Pathology, School of Medicine, Imam Khomeini Hospital, Tehran University of Medical Sciences | Genetics Research Center. University Of Social Welfare And Rehabilitation Sciences | Zohreh Fattahi, Marzieh Mohseni, Khadijeh Jalalvand, Azam Ghaziasadi, Seyedeh elham Mortazavi, Ali Jafarpour, Azar Hadadi, Alireza Abdollahi, Ali Jafarpour, Azam Ghaziasad, Seyedeh elham Mortazavi, Saber Soltani, Reza Najafipour , Kimia Kahrizi, Seyed Mohammad Jazayeri, Hossein Najmabadi |
| EPI_ISL_586519 | B.J. Medical College and Civil hospital, Ahmedabad | Gujarat Biotechnology Research Centre | Zuber Saiyed, Komal Patel, Labdhi Pandya, Afzal Ansari, Nikha Trivedi, Pranay Shah, Kamlesh J Upadhyay, Sanjay Kapadia, Apurvasinh Puvar, Janvi Raval, Zarna Patel, Monika Gandhi, Pinal Trivedi, Maharshi Pandya, Nidhi Patel, Nitin Savaliya, Raghawendra Kumar, Dinesh Kumar, R D Dixit, A M Kadri, Harsh Bakshi, Chaitanya Joshi, Madhvi Joshi |
| EPI_ISL_586528 | B.J. Medical College and Civil hospital, Ahmedabad | Gujarat Biotechnology Research Centre | Zarna Patel, Monika Gandhi, Pinal Trivedi, Maharshi Pandya, Nidhi Patel, Nitin Savaliya, Raghawendra Kumar, Dinesh Kumar, Zuber Saiyed, Komal Patel, Labdhi Pandya, Afzal Ansari, Nikha Trivedi, Pranay Shah, Kamlesh J Upadhyay, Sanjay Kapadia, Apurvasinh Puvar, Janvi Raval, R D Dixit, A M Kadri, Harsh Bakshi, Chaitanya Joshi, Madhvi Joshi |
| EPI_ISL_586531 | General Hospital,Chhotaudepur | Gujarat Biotechnology Research Centre | Yogesh Parmar, Arti Thakur, Dharak Pandya, Maharshi Pandya, Nidhi Patel, Nitin Savaliya, Raghawendra Kumar, Dinesh Kumar, Zuber Saiyed, Komal Patel, Labdhi Pandya, Afzal Ansari, Nikha Trivedi, Apurvasinh Puvar, Janvi Raval, Zarna Patel, Monika Gandhi, Pinal Trivedi, R D Dixit, A M Kadri, Harsh Bakshi, Chaitanya Joshi, Madhvi Joshi |
| EPI_ISL_586541 | Zydus Medical Hospital,Dahod | Gujarat Biotechnology Research Centre | Zuber Saiyed, Komal Patel, Labdhi Pandya, Afzal Ansari, Nikha Trivedi, Apurvasinh Puvar, Janvi Raval, Zarna Patel, Monika Gandhi, Pinal Trivedi, Maharshi Pandya, Nidhi Patel, Kalpesh Labana, Mohit Desai, Nitin Savaliya, Raghawendra Kumar, Dinesh Kumar, R D Dixit, A M Kadri, Harsh Bakshi, Chaitanya Joshi, Madhvi Joshi |
| EPI_ISL_586558 | CCC,Veraval | Gujarat Biotechnology Research Centre | Zuber Saiyed, Komal Patel, Labdhi Pandya, Afzal Ansari, Nikha Trivedi, Jignesh Parmar, Jitendra Bamrotia, Apurvasinh Puvar, Janvi Raval, Zarna Patel, Monika Gandhi, Pinal Trivedi, Maharshi Pandya, Nidhi Patel, Nitin Savaliya, Raghawendra Kumar, Dinesh Kumar, R D Dixit, A M Kadri, Harsh Bakshi, Chaitanya Joshi, Madhvi Joshi |
| EPI_ISL_586567, EPI_ISL_586568 | Saikrishna Hospital,Mehsana | Gujarat Biotechnology Research Centre | Zarna Patel, Monika Gandhi, Pinal Trivedi, Maharshi Pandya, Nidhi Patel, Nitin Savaliya, Raghawendra Kumar, Dinesh Kumar, Zuber Saiyed, Komal Patel, Labdhi Pandya, Afzal Ansari, Nikha Trivedi, Harshadbhai Parmar, Apurvasinh Puvar, Janvi Raval, R D Dixit, A M Kadri, Harsh Bakshi, Chaitanya Joshi, Madhvi Joshi |
| EPI_ISL_590692 | Government Hospital,Veraval | Gujarat Biotechnology Research Centre | Zarna Patel, Monika Gandhi, Pinal Trivedi, Maharshi Pandya, Nidhi Patel, Nitin Savaliya, Raghawendra Kumar, Dinesh Kumar, Zuber Saiyed, Komal Patel, Labdhi Pandya, Afzal Ansari, Nikha Trivedi, Jignesh Parmar, Jitendra Bamrotia, Apurvasinh Puvar, Janvi Raval, R D Dixit, A M Kadri, Harsh Bakshi, Chaitanya Joshi, Madhvi Joshi |
| EPI_ISL_594185, EPI_ISL_594186, EPI_ISL_594187, EPI_ISL_594188 | Department of Pathology, School of Medicine, Imam Khomeini Hospital, Tehran University of Medical Sciences | Genetics Research Center, University of Social Welfare and Rehabilitation Sciences | Zohreh Fattahi, Marzieh Mohseni, Khadijeh Jalalvand, Azam Ghaziasadi, Seyedeh elham Mortazavi, Ali Jafarpour, Azar Hadadi, Alireza Abdollahi, Ali Jafarpour, Azam Ghaziasad, Seyedeh elham Mortazavi, Saber Soltani, Reza Najafipour, Kimia Kahrizi, Seyed Mohammad Jazayeri, Hossein Najmabadi |
| EPI_ISL_594447, EPI_ISL_594448, EPI_ISL_594449, EPI_ISL_594450, EPI_ISL_594451, EPI_ISL_594452, EPI_ISL_594453, EPI_ISL_594454 | Washington State Public Health Laboratories | Pathogen Discovery, Respiratory Viruses Branch, Division of Viral Diseases, Centers for Disease Control and Prevention | Ying Tao, Yan Li, Clinton Paden, Jing Zhang, Krista Queen, Anna Uehara, Haibin Wang, Julu Bhatnagar, Suxiang Tong |
| EPI_ISL_594455 | Texas Department of State Health Services Public Health Laboratory | Pathogen Discovery, Respiratory Viruses Branch, Division of Viral Diseases, Centers for Disease Control and Prevention | Ying Tao, Yan Li, Clinton Paden, Jing Zhang, Krista Queen, Anna Uehara, Haibin Wang, Julu Bhatnagar, Suxiang Tong |
| EPI_ISL_594456 | Georgia Public Health Laboratory | Pathogen Discovery, Respiratory Viruses Branch, Division of Viral Diseases, Centers for Disease Control and Prevention | Ying Tao, Yan Li, Clinton Paden, Jing Zhang, Krista Queen, Anna Uehara, Haibin Wang, Julu Bhatnagar, Suxiang Tong |
| EPI_ISL_594457 | NYS Department of Health | Pathogen Discovery, Respiratory Viruses Branch, Division of Viral Diseases, Centers for Disease Control and Prevention | Ying Tao, Yan Li, Clinton Paden, Jing Zhang, Krista Queen, Anna Uehara, Haibin Wang, Julu Bhatnagar, Suxiang Tong |
| EPI_ISL_594458 | California Department of Public Health | Pathogen Discovery, Respiratory Viruses Branch, Division of Viral Diseases, Centers for Disease Control and Prevention | Ying Tao, Yan Li, Clinton Paden, Jing Zhang, Krista Queen, Anna Uehara, Haibin Wang, Julu Bhatnagar, Suxiang Tong |
| EPI_ISL_594459 | Washington State Public Health Laboratories | Pathogen Discovery, Respiratory Viruses Branch, Division of Viral Diseases, Centers for Disease Control and Prevention | Ying Tao, Yan Li, Clinton Paden, Jing Zhang, Krista Queen, Anna Uehara, Haibin Wang, Julu Bhatnagar, Suxiang Tong |
| EPI_ISL_594460 | DC Department of Forensic Sciences | Pathogen Discovery, Respiratory Viruses Branch, Division of Viral Diseases, Centers for Disease Control and Prevention | Ying Tao, Yan Li, Clinton Paden, Jing Zhang, Krista Queen, Anna Uehara, Haibin Wang, Julu Bhatnagar, Suxiang Tong |
| EPI_ISL_594461 | FL Bureau of Public Health Laboratories | Pathogen Discovery, Respiratory Viruses Branch, Division of Viral Diseases, Centers for Disease Control and Prevention | Ying Tao, Yan Li, Clinton Paden, Jing Zhang, Krista Queen, Anna Uehara, Haibin Wang, Julu Bhatnagar, Suxiang Tong |
| EPI_ISL_594462 | NM Department of Health | Pathogen Discovery, Respiratory Viruses Branch, Division of Viral Diseases, Centers for Disease Control and Prevention | Ying Tao, Yan Li, Clinton Paden, Jing Zhang, Krista Queen, Anna Uehara, Haibin Wang, Julu Bhatnagar, Suxiang Tong |
| EPI_ISL_594463 | MN Department of Health | Pathogen Discovery, Respiratory Viruses Branch, Division of Viral Diseases, Centers for Disease Control and Prevention | Ying Tao, Yan Li, Clinton Paden, Jing Zhang, Krista Queen, Anna Uehara, Haibin Wang, Julu Bhatnagar, Suxiang Tong |
| EPI_ISL_594464 | Texas Department of State Health Services Public Health Laboratory | Pathogen Discovery, Respiratory Viruses Branch, Division of Viral Diseases, Centers for Disease Control and Prevention | Ying Tao, Yan Li, Clinton Paden, Jing Zhang, Krista Queen, Anna Uehara, Haibin Wang, Julu Bhatnagar, Suxiang Tong |
| EPI_ISL_594465 | FL Bureau of Public Health Laboratories | Pathogen Discovery, Respiratory Viruses Branch, Division of Viral Diseases, Centers for Disease Control and Prevention | Ying Tao, Yan Li, Clinton Paden, Jing Zhang, Krista Queen, Anna Uehara, Haibin Wang, Julu Bhatnagar, Suxiang Tong |
| EPI_ISL_596451, EPI_ISL_596452 | Department of Pathology, School of Medicine, Imam Khomeini Hospital, Tehran University of Medical Sciences | Genetics Research Center, University of Social Welfare and Rehabilitation Sciences | Zohreh Fattahi, Marzieh Mohseni, Khadijeh Jalalvand, Azam Ghaziasadi, Seyedeh elham Mortazavi, Ali Jafarpour, Azar Hadadi, Alireza Abdollahi, Ali Jafarpour, Azam Ghaziasad, Seyedeh elham Mortazavi, Saber Soltani, Reza Najafipour, Kimia Kahrizi, Seyed Mohammad Jazayeri, Hossein Najmabadi |
| EPI_ISL_596453 | Booali laboratory, Qom, Iran. Department of Virology, School of Public Health, Tehran University of Medical Sciences, Tehran, Iran. | Genetics Research Center, University of Social Welfare and Rehabilitation Sciences | Zohreh Fattahi, Marzieh Mohseni, Khadijeh Jalalvand, Azam Ghaziasadi, Seyedeh elham Mortazavi, Ali Jafarpour, Mohammad Khazeni, Seyed Amir Momeni, Kimia Kahrizi, Seyed Mohammad Jazayeri, Hossein Najmbadi |
| EPI_ISL_596454 | Infectious Disease and Tropical Medicine Research Center, Resistant Tuberculosis Institute, Zahedan University of Medical Sciences, Zahedan, Iran. | Genetics Research Center, University of Social Welfare and Rehabilitation Sciences | Zohreh Fattahi, Marzieh Mohseni, Khadijeh Jalalvand, Azam Ghaziasadi, Seyedeh elham Mortazavi, Ali Jafarpour, Ebrahim Kord, Seyed Mohammad Hashemi-Shahri, Kimia Kahrizi, Seyed Mohammad Jazayeri, Hossein Najmabadi |
| EPI_ISL_596455 | Department of Pathology, School of Medicine, Imam Khomeini Hospital, Tehran University of Medical Sciences | Genetics Research Center, University of Social Welfare and Rehabilitation Sciences | Zohreh Fattahi, Marzieh Mohseni, Khadijeh Jalalvand, Azam Ghaziasadi, Seyedeh elham Mortazavi, Ali Jafarpour, Azar Hadadi, Alireza Abdollahi, Ali Jafarpour, Azam Ghaziasad, Seyedeh elham Mortazavi, Saber Soltani, Reza Najafipour, Kimia Kahrizi, Seyed Mohammad Jazayeri, Hossein Najmabadi |
| EPI_ISL_602556, EPI_ISL_602558, EPI_ISL_603049, EPI_ISL_603050, EPI_ISL_603051, EPI_ISL_603052, EPI_ISL_603053, EPI_ISL_603054, EPI_ISL_603055, EPI_ISL_603058, EPI_ISL_603059, EPI_ISL_603061, EPI_ISL_603064, EPI_ISL_603065, EPI_ISL_603066, EPI_ISL_603067, EPI_ISL_603068, EPI_ISL_603069, EPI_ISL_603070, EPI_ISL_603072, EPI_ISL_603073, EPI_ISL_603074, EPI_ISL_603075, EPI_ISL_603078, EPI_ISL_603079 | | | |
| see above | Utah Public Health Laboratory, Utah Public Health Laboratory Infectious Disease submission group | Utah Public Health Laboratory, Utah Public Health Laboratory Infectious Disease submission group | Young,E.L., Oakeson,K. |
| EPI_ISL_605931 | Utah Public Health Laboratory, Utah Public Health Laboratory Infectious Disease submission group | Utah Public Health Laboratory, Utah Public Health Laboratory Infectious Disease submission group | Young,E.L. and Oakeson,K. |
| EPI_ISL_620407, EPI_ISL_620408, EPI_ISL_620410 | Utah Public Health Laboratory, Utah Public Health Laboratory Infectious Disease submission group | Utah Public Health Laboratory, Utah Public Health Laboratory Infectious Disease submission group | Young,E.L., Oakeson,K.F. |
| EPI_ISL_622769, EPI_ISL_622770, EPI_ISL_622771 | Canterbury Health Laboratories | Institute of Environmental Science and Research (ESR) | Xiaoyun Ren, Matt Storey, Nikki Freed, Muhammad Faisal, Jing Wang, Hermes Perez, Anja Werno, Antje van der Linden, Arlo Upton, Chris Mansell, David Hammer, Dragana Drinkovic, Gary McAuliffe, Hana Sofia Andersson, James Ussher, Jill Sherwood, Josh Freeman, Julia Howard, Juliet Elvy, Mary |

| | | | |
|---|---|---|---|
| | | | DeAlmeida, Matt Blakiston, Matthew Rogers, Max Bloomfield, Michael Addidle, Michelle Balm, Sally Roberts, Sarah Jefferies, Sharmini Muttaiyah, Susan Morpeth, Susan Taylor, Timothy Blackmore, Vani Sathyendran, Veronica Playle, Virginia Hope, Erasmus Smit, Lauren Jelly, Olin Silander, Joep de Ligt |
| EPI_ISL_622772 | Middlemore Hospital | Institute of Environmental Science and Research (ESR) | Xiaoyun Ren, Matt Storey, Nikki Freed, Muhammad Faisal, Jing Wang, Hermes Perez, Anja Werno, Antje van der Linden, Arlo Upton, Chris Mansell, David Hammer, Dragana Drinkovic, Gary McAuliffe, Hana Sofia Andersson, James Ussher, Jill Sherwood, Josh Freeman, Julia Howard, Juliet Elvy, Mary DeAlmeida, Matt Blakiston, Matthew Rogers, Max Bloomfield, Michael Addidle, Michelle Balm, Sally Roberts, Sarah Jefferies, Sharmini Muttaiyah, Susan Morpeth, Susan Taylor, Timothy Blackmore, Vani Sathyendran, Veronica Playle, Virginia Hope, Erasmus Smit, Lauren Jelly, Olin Silander, Joep de Ligt |
| EPI_ISL_622774 | LabPLUS | Institute of Environmental Science and Research (ESR) | Xiaoyun Ren, Matt Storey, Nikki Freed, Muhammad Faisal, Jing Wang, Hermes Perez, Anja Werno, Antje van der Linden, Arlo Upton, Chris Mansell, David Hammer, Dragana Drinkovic, Gary McAuliffe, Hana Sofia Andersson, James Ussher, Jill Sherwood, Josh Freeman, Julia Howard, Juliet Elvy, Mary DeAlmeida, Matt Blakiston, Matthew Rogers, Max Bloomfield, Michael Addidle, Michelle Balm, Sally Roberts, Sarah Jefferies, Sharmini Muttaiyah, Susan Morpeth, Susan Taylor, Timothy Blackmore, Vani Sathyendran, Veronica Playle, Virginia Hope, Erasmus Smit, Lauren Jelly, Olin Silander, Joep de Ligt |
| EPI_ISL_622775 | Canterbury Health Laboratories | Institute of Environmental Science and Research (ESR) | Xiaoyun Ren, Matt Storey, Nikki Freed, Muhammad Faisal, Jing Wang, Hermes Perez, Anja Werno, Antje van der Linden, Arlo Upton, Chris Mansell, David Hammer, Dragana Drinkovic, Gary McAuliffe, Hana Sofia Andersson, James Ussher, Jill Sherwood, Josh Freeman, Julia Howard, Juliet Elvy, Mary DeAlmeida, Matt Blakiston, Matthew Rogers, Max Bloomfield, Michael Addidle, Michelle Balm, Sally Roberts, Sarah Jefferies, Sharmini Muttaiyah, Susan Morpeth, Susan Taylor, Timothy Blackmore, Vani Sathyendran, Veronica Playle, Virginia Hope, Erasmus Smit, Lauren Jelly, Olin Silander, Joep de Ligt |
| EPI_ISL_622778 | LabTests | Institute of Environmental Science and Research (ESR) | Xiaoyun Ren, Matt Storey, Nikki Freed, Muhammad Faisal, Jing Wang, Hermes Perez, Anja Werno, Antje van der Linden, Arlo Upton, Chris Mansell, David Hammer, Dragana Drinkovic, Gary McAuliffe, Hana Sofia Andersson, James Ussher, Jill Sherwood, Josh Freeman, Julia Howard, Juliet Elvy, Mary DeAlmeida, Matt Blakiston, Matthew Rogers, Max Bloomfield, Michael Addidle, Michelle Balm, Sally Roberts, Sarah Jefferies, Sharmini Muttaiyah, Susan Morpeth, Susan Taylor, Timothy Blackmore, Vani Sathyendran, Veronica Playle, Virginia Hope, Erasmus Smit, Lauren Jelly, Olin Silander, Joep de Ligt |
| EPI_ISL_622779 | LabPLUS | Institute of Environmental Science and Research (ESR) | Xiaoyun Ren, Matt Storey, Nikki Freed, Muhammad Faisal, Jing Wang, Hermes Perez, Anja Werno, Antje van der Linden, Arlo Upton, Chris Mansell, David Hammer, Dragana Drinkovic, Gary McAuliffe, Hana Sofia Andersson, James Ussher, Jill Sherwood, Josh Freeman, Julia Howard, Juliet Elvy, Mary DeAlmeida, Matt Blakiston, Matthew Rogers, Max Bloomfield, Michael Addidle, Michelle Balm, Sally Roberts, Sarah Jefferies, Sharmini Muttaiyah, Susan Morpeth, Susan Taylor, Timothy Blackmore, Vani Sathyendran, Veronica Playle, Virginia Hope, Erasmus Smit, Lauren Jelly, Olin Silander, Joep de Ligt |
| EPI_ISL_622781, EPI_ISL_622782, EPI_ISL_622784 | Canterbury Health Laboratories | Institute of Environmental Science and Research (ESR) | Xiaoyun Ren, Matt Storey, Nikki Freed, Muhammad Faisal, Jing Wang, Hermes Perez, Anja Werno, Antje van der Linden, Arlo Upton, Chris Mansell, David Hammer, Dragana Drinkovic, Gary McAuliffe, Hana Sofia Andersson, James Ussher, Jill Sherwood, Josh Freeman, Julia Howard, Juliet Elvy, Mary DeAlmeida, Matt Blakiston, Matthew Rogers, Max Bloomfield, Michael Addidle, Michelle Balm, Sally Roberts, Sarah Jefferies, Sharmini Muttaiyah, Susan Morpeth, Susan Taylor, Timothy Blackmore, Vani Sathyendran, Veronica Playle, Virginia Hope, Erasmus Smit, Lauren Jelly, Olin Silander, Joep de Ligt |
| EPI_ISL_622785 | Waikato Hospital | Institute of Environmental Science and Research (ESR) | Xiaoyun Ren, Matt Storey, Nikki Freed, Muhammad Faisal, Jing Wang, Hermes Perez, Anja Werno, Antje van der Linden, Arlo Upton, Chris Mansell, David Hammer, Dragana Drinkovic, Gary McAuliffe, Hana Sofia Andersson, James Ussher, Jill Sherwood, Josh Freeman, Julia Howard, Juliet Elvy, Mary DeAlmeida, Matt Blakiston, Matthew Rogers, Max Bloomfield, Michael Addidle, Michelle Balm, Sally Roberts, Sarah Jefferies, Sharmini Muttaiyah, Susan Morpeth, Susan Taylor, Timothy Blackmore, Vani Sathyendran, Veronica Playle, Virginia Hope, Erasmus Smit, Lauren Jelly, Olin Silander, Joep de Ligt |
| EPI_ISL_622786 | Canterbury Health Laboratories | Institute of Environmental Science and Research (ESR) | Xiaoyun Ren, Matt Storey, Nikki Freed, Muhammad Faisal, Jing Wang, Hermes Perez, Anja Werno, Antje van der Linden, Arlo Upton, Chris Mansell, David Hammer, Dragana Drinkovic, Gary McAuliffe, Hana Sofia Andersson, James Ussher, Jill Sherwood, Josh Freeman, Julia Howard, Juliet Elvy, Mary DeAlmeida, Matt Blakiston, Matthew Rogers, Max Bloomfield, Michael Addidle, Michelle Balm, Sally Roberts, Sarah Jefferies, Sharmini Muttaiyah, Susan Morpeth, Susan Taylor, Timothy Blackmore, Vani Sathyendran, Veronica Playle, Virginia Hope, Erasmus Smit, Lauren Jelly, Olin Silander, Joep de Ligt |
| EPI_ISL_622787 | Middlemore Hospital | Institute of Environmental Science and Research (ESR) | Xiaoyun Ren, Matt Storey, Nikki Freed, Muhammad Faisal, Jing Wang, Hermes Perez, Anja Werno, Antje van der Linden, Arlo Upton, Chris Mansell, David Hammer, Dragana Drinkovic, Gary McAuliffe, Hana Sofia Andersson, James Ussher, Jill Sherwood, Josh Freeman, Julia Howard, Juliet Elvy, Mary DeAlmeida, Matt Blakiston, Matthew Rogers, Max Bloomfield, Michael Addidle, Michelle Balm, Sally Roberts, Sarah Jefferies, Sharmini Muttaiyah, Susan Morpeth, Susan Taylor, Timothy Blackmore, Vani Sathyendran, Veronica Playle, Virginia Hope, Erasmus Smit, Lauren Jelly, Olin Silander, Joep de Ligt |
| EPI_ISL_622789, EPI_ISL_622791 | Canterbury Health Laboratories | Institute of Environmental Science and Research (ESR) | Xiaoyun Ren, Matt Storey, Nikki Freed, Muhammad Faisal, Jing Wang, Hermes Perez, Anja Werno, Antje van der Linden, Arlo Upton, Chris Mansell, David Hammer, Dragana Drinkovic, Gary McAuliffe, Hana Sofia Andersson, James Ussher, Jill Sherwood, Josh Freeman, Julia Howard, Juliet Elvy, Mary DeAlmeida, Matt Blakiston, Matthew Rogers, Max Bloomfield, Michael Addidle, Michelle Balm, Sally Roberts, Sarah Jefferies, Sharmini Muttaiyah, Susan Morpeth, Susan Taylor, Timothy Blackmore, Vani Sathyendran, Veronica Playle, Virginia Hope, Erasmus Smit, Lauren Jelly, Olin Silander, Joep de Ligt |
| EPI_ISL_622792 | Middlemore Hospital | Institute of Environmental Science and Research (ESR) | Xiaoyun Ren, Matt Storey, Nikki Freed, Muhammad Faisal, Jing Wang, Hermes Perez, Anja Werno, Antje van der Linden, Arlo Upton, Chris Mansell, David Hammer, Dragana Drinkovic, Gary McAuliffe, Hana Sofia Andersson, James Ussher, Jill Sherwood, Josh Freeman, Julia Howard, Juliet Elvy, Mary DeAlmeida, Matt Blakiston, Matthew Rogers, Max Bloomfield, Michael Addidle, Michelle Balm, Sally Roberts, Sarah Jefferies, Sharmini Muttaiyah, Susan Morpeth, Susan Taylor, Timothy Blackmore, Vani Sathyendran, Veronica Playle, Virginia Hope, Erasmus Smit, Lauren Jelly, Olin Silander, Joep de Ligt |
| EPI_ISL_622793 | LabTests | Institute of Environmental Science and Research (ESR) | Xiaoyun Ren, Matt Storey, Nikki Freed, Muhammad Faisal, Jing Wang, Hermes Perez, Anja Werno, Antje van der Linden, Arlo Upton, Chris Mansell, David Hammer, Dragana Drinkovic, Gary McAuliffe, Hana Sofia Andersson, James Ussher, Jill Sherwood, Josh Freeman, Julia Howard, Juliet Elvy, Mary DeAlmeida, Matt Blakiston, Matthew Rogers, Max Bloomfield, Michael Addidle, Michelle Balm, Sally Roberts, Sarah Jefferies, Sharmini Muttaiyah, Susan Morpeth, Susan Taylor, Timothy Blackmore, Vani Sathyendran, Veronica Playle, Virginia Hope, Erasmus Smit, Lauren Jelly, Olin Silander, Joep de Ligt |
| EPI_ISL_622794, EPI_ISL_622795, EPI_ISL_622796, EPI_ISL_622797, EPI_ISL_622798, EPI_ISL_622799, EPI_ISL_622801 | Canterbury Health Laboratories | Institute of Environmental Science and Research (ESR) | Xiaoyun Ren, Matt Storey, Nikki Freed, Muhammad Faisal, Jing Wang, Hermes Perez, Anja Werno, Antje van der Linden, Arlo Upton, Chris Mansell, David Hammer, Dragana Drinkovic, Gary McAuliffe, Hana Sofia Andersson, James Ussher, Jill Sherwood, Josh Freeman, Julia Howard, Juliet Elvy, Mary DeAlmeida, Matt Blakiston, Matthew Rogers, Max Bloomfield, Michael Addidle, Michelle Balm, Sally Roberts, Sarah Jefferies, Sharmini Muttaiyah, Susan Morpeth, Susan Taylor, Timothy Blackmore, Vani Sathyendran, Veronica Playle, Virginia Hope, Erasmus Smit, Lauren Jelly, Olin Silander, Joep de Ligt |
| EPI_ISL_622802 | Middlemore Hospital | Institute of Environmental Science and Research (ESR) | Xiaoyun Ren, Matt Storey, Nikki Freed, Muhammad Faisal, Jing Wang, Hermes Perez, Anja Werno, Antje van der Linden, Arlo Upton, Chris Mansell, David Hammer, Dragana Drinkovic, Gary McAuliffe, Hana Sofia Andersson, James Ussher, Jill Sherwood, Josh Freeman, Julia Howard, Juliet Elvy, Mary DeAlmeida, Matt Blakiston, Matthew Rogers, Max Bloomfield, Michael Addidle, Michelle Balm, Sally Roberts, Sarah Jefferies, Sharmini Muttaiyah, Susan Morpeth, Susan Taylor, Timothy Blackmore, Vani Sathyendran, Veronica Playle, Virginia Hope, Erasmus Smit, Lauren Jelly, Olin Silander, Joep de Ligt |
| EPI_ISL_622803 | LabPLUS | Institute of Environmental Science and Research (ESR) | Xiaoyun Ren, Matt Storey, Nikki Freed, Muhammad Faisal, Jing Wang, Hermes Perez, Anja Werno, Antje van der Linden, Arlo Upton, Chris Mansell, David Hammer, Dragana Drinkovic, Gary McAuliffe, Hana Sofia Andersson, James Ussher, Jill Sherwood, Josh Freeman, Julia Howard, Juliet Elvy, Mary DeAlmeida, Matt Blakiston, Matthew Rogers, Max Bloomfield, Michael Addidle, Michelle Balm, Sally Roberts, Sarah Jefferies, Sharmini Muttaiyah, Susan Morpeth, Susan Taylor, Timothy Blackmore, Vani Sathyendran, Veronica Playle, Virginia Hope, Erasmus Smit, Lauren Jelly, Olin Silander, Joep de Ligt |
| EPI_ISL_622805, EPI_ISL_622806 | PathLab Bay of Plenty | Institute of Environmental Science and Research (ESR) | Xiaoyun Ren, Matt Storey, Nikki Freed, Muhammad Faisal, Jing Wang, Hermes Perez, Anja Werno, Antje van der Linden, Arlo Upton, Chris Mansell, David Hammer, Dragana Drinkovic, Gary McAuliffe, Hana Sofia Andersson, James Ussher, Jill Sherwood, Josh Freeman, Julia Howard, Juliet Elvy, Mary DeAlmeida, Matt Blakiston, Matthew Rogers, Max Bloomfield, Michael Addidle, Michelle Balm, Sally Roberts, Sarah Jefferies, Sharmini Muttaiyah, Susan Morpeth, Susan Taylor, Timothy Blackmore, Vani Sathyendran, Veronica Playle, Virginia Hope, Erasmus Smit, Lauren Jelly, Olin Silander, Joep de Ligt |
| EPI_ISL_622807 | LabPLUS | Institute of Environmental Science and Research (ESR) | Xiaoyun Ren, Matt Storey, Nikki Freed, Muhammad Faisal, Jing Wang, Hermes Perez, Anja Werno, Antje van der Linden, Arlo Upton, Chris Mansell, David Hammer, Dragana Drinkovic, Gary McAuliffe, Hana Sofia Andersson, James Ussher, Jill Sherwood, Josh Freeman, Julia Howard, Juliet Elvy, Mary DeAlmeida, Matt Blakiston, Matthew Rogers, Max Bloomfield, Michael Addidle, Michelle Balm, Sally Roberts, Sarah Jefferies, Sharmini Muttaiyah, Susan Morpeth, Susan Taylor, Timothy Blackmore, Vani Sathyendran, Veronica Playle, Virginia Hope, Erasmus Smit, Lauren Jelly, Olin Silander, Joep de Ligt |
| EPI_ISL_622808, EPI_ISL_622809, EPI_ISL_622810, EPI_ISL_622811, EPI_ISL_622812, EPI_ISL_622813 | Canterbury Health Laboratories | Institute of Environmental Science and Research (ESR) | Xiaoyun Ren, Matt Storey, Nikki Freed, Muhammad Faisal, Jing Wang, Hermes Perez, Anja Werno, Antje van der Linden, Arlo Upton, Chris Mansell, David Hammer, Dragana Drinkovic, Gary McAuliffe, Hana Sofia Andersson, James Ussher, Jill Sherwood, Josh Freeman, Julia Howard, Juliet Elvy, Mary DeAlmeida, Matt Blakiston, Matthew Rogers, Max Bloomfield, Michael Addidle, Michelle Balm, Sally Roberts, Sarah Jefferies, Sharmini Muttaiyah, Susan Morpeth, Susan Taylor, Timothy Blackmore, Vani Sathyendran, Veronica Playle, Virginia Hope, Erasmus Smit, Lauren Jelly, Olin Silander, Joep de Ligt |
| EPI_ISL_622814, EPI_ISL_622815, EPI_ISL_622816 | LabPLUS | Institute of Environmental Science and Research (ESR) | Xiaoyun Ren, Matt Storey, Nikki Freed, Muhammad Faisal, Jing Wang, Hermes Perez, Anja Werno, Antje van der Linden, Arlo Upton, Chris Mansell, David Hammer, Dragana Drinkovic, Gary McAuliffe, Hana Sofia Andersson, James Ussher, Jill Sherwood, Josh Freeman, Julia Howard, Juliet Elvy, Mary DeAlmeida, Matt Blakiston, Matthew Rogers, Max Bloomfield, Michael Addidle, Michelle Balm, Sally Roberts, Sarah Jefferies, Sharmini Muttaiyah, Susan Morpeth, Susan Taylor, Timothy Blackmore, Vani Sathyendran, Veronica Playle, Virginia Hope, Erasmus Smit, Lauren Jelly, Olin Silander, Joep de Ligt |

| | | | |
|---|---|---|---|
| EPI_ISL_622817, EPI_ISL_622818, EPI_ISL_622820 | Middlemore Hospital | Institute of Environmental Science and Research (ESR) | Xiaoyun Ren, Matt Storey, Nikki Freed, Muhammad Faisal, Jing Wang, Hermes Perez, Anja Werno, Antje van der Linden, Arlo Upton, Chris Mansell, David Hammer, Dragana Drinkovic, Gary McAuliffe, Hana Sofia Andersson, James Ussher, Jill Sherwood, Josh Freeman, Julia Howard, Juliet Elvy, Mary DeAlmeida, Matt Blakiston, Matthew Rogers, Max Bloomfield, Michael Addidle, Michelle Balm, Sally Roberts, Sarah Jefferies, Sharmini Muttaiyah, Susan Morpeth, Susan Taylor, Timothy Blackmore, Vani Sathyendran, Veronica Playle, Virginia Hope, Erasmus Smit, Lauren Jelly, Olin Silander, Joep de Ligt |
| EPI_ISL_622821, EPI_ISL_622822, EPI_ISL_622823 | LabPLUS | Institute of Environmental Science and Research (ESR) | Xiaoyun Ren, Matt Storey, Nikki Freed, Muhammad Faisal, Jing Wang, Hermes Perez, Anja Werno, Antje van der Linden, Arlo Upton, Chris Mansell, David Hammer, Dragana Drinkovic, Gary McAuliffe, Hana Sofia Andersson, James Ussher, Jill Sherwood, Josh Freeman, Julia Howard, Juliet Elvy, Mary DeAlmeida, Matt Blakiston, Matthew Rogers, Max Bloomfield, Michael Addidle, Michelle Balm, Sally Roberts, Sarah Jefferies, Sharmini Muttaiyah, Susan Morpeth, Susan Taylor, Timothy Blackmore, Vani Sathyendran, Veronica Playle, Virginia Hope, Erasmus Smit, Lauren Jelly, Olin Silander, Joep de Ligt |
| EPI_ISL_622824, EPI_ISL_622825, EPI_ISL_622826, EPI_ISL_622827, EPI_ISL_622828, EPI_ISL_622829, EPI_ISL_622830 | Canterbury Health Laboratories | Institute of Environmental Science and Research (ESR) | Xiaoyun Ren, Matt Storey, Nikki Freed, Muhammad Faisal, Jing Wang, Hermes Perez, Anja Werno, Antje van der Linden, Arlo Upton, Chris Mansell, David Hammer, Dragana Drinkovic, Gary McAuliffe, Hana Sofia Andersson, James Ussher, Jill Sherwood, Josh Freeman, Julia Howard, Juliet Elvy, Mary DeAlmeida, Matt Blakiston, Matthew Rogers, Max Bloomfield, Michael Addidle, Michelle Balm, Sally Roberts, Sarah Jefferies, Sharmini Muttaiyah, Susan Morpeth, Susan Taylor, Timothy Blackmore, Vani Sathyendran, Veronica Playle, Virginia Hope, Erasmus Smit, Lauren Jelly, Olin Silander, Joep de Ligt |
| EPI_ISL_622831, EPI_ISL_622832 | Middlemore Hospital | Institute of Environmental Science and Research (ESR) | Xiaoyun Ren, Matt Storey, Nikki Freed, Muhammad Faisal, Jing Wang, Hermes Perez, Anja Werno, Antje van der Linden, Arlo Upton, Chris Mansell, David Hammer, Dragana Drinkovic, Gary McAuliffe, Hana Sofia Andersson, James Ussher, Jill Sherwood, Josh Freeman, Julia Howard, Juliet Elvy, Mary DeAlmeida, Matt Blakiston, Matthew Rogers, Max Bloomfield, Michael Addidle, Michelle Balm, Sally Roberts, Sarah Jefferies, Sharmini Muttaiyah, Susan Morpeth, Susan Taylor, Timothy Blackmore, Vani Sathyendran, Veronica Playle, Virginia Hope, Erasmus Smit, Lauren Jelly, Olin Silander, Joep de Ligt |
| EPI_ISL_625683 | National Reference Laboratory for COVID-19, Pasteur Institute of Iran | National Reference Laboratory for COVID-19, Pasteur Institute of Iran | Zahra Ahmadi, Zahra Fereydouni, Tahmineh Jalali, Mohammad Hassan Pouriayevali, Mahsa Tavakkoli, Marzieh Sajjadi, Setareh Kashanian, Sanam Azad-Manjiri, Heasam Nemati, Tahereh Mohammadi, Kayhan Azadmanesh, Zabihollah Shoja, Sana Eybpoosh, Ahmad Ghasemi, Parastoo Yekta, Sepideh Gerdooei, Farideh Niknam, Mostafa Salehi-Vaziri |
| EPI_ISL_626352, EPI_ISL_626353, EPI_ISL_626354, EPI_ISL_626355, EPI_ISL_626356 | LabPLUS | Institute of Environmental Science and Research (ESR) | Xiaoyun Ren, Matt Storey, Nikki Freed, Muhammad Faisal, Jing Wang, Hermes Perez, Anja Werno, Antje van der Linden, Arlo Upton, Chris Mansell, David Hammer, Dragana Drinkovic, Gary McAuliffe, Hana Sofia Andersson, James Ussher, Jill Sherwood, Josh Freeman, Julia Howard, Juliet Elvy, Mary DeAlmeida, Matt Blakiston, Matthew Rogers, Max Bloomfield, Michael Addidle, Michelle Balm, Sally Roberts, Sarah Jefferies, Sharmini Muttaiyah, Susan Morpeth, Susan Taylor, Timothy Blackmore, Vani Sathyendran, Veronica Playle, Virginia Hope, Erasmus Smit, Lauren Jelly, Olin Silander, Joep de Ligt |
